# A Bayesian neural network for toxicity prediction

**DOI:** 10.1101/2020.04.28.065532

**Authors:** Elizaveta Semenova, Dominic P. Williams, Avid M. Afzal, Stanley E. Lazic

## Abstract

Predicting the toxicity of a compound preclinically enables better decision making, thereby reducing development costs and increasing patient safety. It is a complex issue, but *in vitro* assays and physico-chemical properties of compounds can be used to predict clinical toxicity. Neural networks (NNs) are a popular predictive tool due to their flexibility and ability to model non-linearities, but they are prone to overfitting and therefore are not recommended for small data sets. Furthermore, they don’t quantify uncertainty in the predictions. Bayesian neural networks (BNNs) are able to avoid these pitfalls by using prior distributions on the parameters of a NN model and representing uncertainty about the predictions in the form of a distribution. We model the severity of drug-induced liver injury (DILI) to provide an example of a BNN performing better than a traditional but less flexible proportional odds logistic regression (POLR) model. We use appropriate metrics to evaluate predictions of the ordinal data type. To demonstrate the effect of a hierarchical prior for BNNs as an alternative to hyperparameter optimisation for NNs, we compare the performance of a BNN against NNs with dropout or penalty regularisation. We reduce the task to multiclass classification in order to be able to perform this comparison. A BNN trained for the multiclass classification produces poorer results than a BNN that captures the order. The current work lays a foundation for more complex models built on larger datasets, but can already be adopted by safety pharmacologists for risk quantification.

## Introduction

Drug-induced liver injury (DILI) is the most frequent cause of acute liver failure in most Western countries [1] and may require discontinuation of treatment or hospitalisation. As a consequence, early drug development focuses on characterising the safety profiles of compounds to avoid adverse effects in humans and costly drug withdrawal at the post-marketing stage.

Predicting clinical hepatotoxicity is difficult due to its multi-mechanistic nature: drugs are taken up from circulation into the liver, where they can be metabolised by cytochrome P450-mediated enzymatic reactions. These reactions can produce reactive metabolites that can bind to and alter the function of proteins, or form haptens, which can initiate an immune response. Both of these processes can contribute to liver injury. Additionally, some drugs or their metabolites can block the export of bile from hepatocytes, leading to intracellular damage from bile. Finally, drug-induced cellular changes can lead to hepatocyte death or promote an immune response, leading to liver injury. Chemical properties of the drug and unknown contributors within the biology of patients also play a role.

A range of *in vivo* [2] and *in silico* [3, 4] approaches have been developed to estimate human DILI risk. However, retrospective analyses have shown that preclinical animal studies fail to make correct predictions in about 45% of clinical trials [5, 6]. In turn, classical statistical and machine learning models require a sufficient amount of data for adequate use, and collecting information about toxicological properties of compounds is resource consuming, e.g. some adverse effects become known only after a long period of a drug being on the market. As a result, the datasets available for analysis contain few observations. Traditional models, which have been applied for DILI prediction, such as linear discriminant analysis, artificial neural networks [7], and support vector machine [8], may struggle to analyse small amounts of data reliably, and they provide no information about the uncertainty of the predictions. A Bayesian approach to model formulation and analysis offers a toolset to resolve these shortcomings and provides considerable flexibility: a broad range of models can be formalised with the help of probabilistic programming languages, prior knowledge can be taken into account, and multiple sources of uncertainty can be incorporated and propagated into the estimates and predictions. Bayesian models have been used to predict the presence or absence of DILI, but few studies distinguish between different levels of severity [9, 10]. In our previous work [10] we used a Bayesian proportional odds logistic regression (POLR) model that included pair-wise interactions of predictors to allow for more complex relationships. It would be possible to add higher order interactions but the number of parameters grows quickly and overfitting becomes more of a problem.

Deep neural networks (DNNs) have recently been achieving state of the art results in many fields and have been used to predict toxicity for both a single target and multiple targets (multitask learning) [7, 11, 12]. NNs are popular due to their flexibility, but are prone to overfitting [13] and do not capture uncertainty. This might lead to overconfident predictions even when they are erroneous. Bayesian neural networks (BNNs) use priors to avoid overfitting and provide uncertainty in the predictions [14, 15]. They represent each estimated parameter as a distribution, rather than as a single point. Unlike some other Bayesian models where prior information about individual parameters can be used explicitly, the role of priors for BNNs is in regularisation. Bayesian Deep Learning – a field following the Bayes approach for neural network models – is rapidly developing [14, 16, 17, 18]. Inference methodology for BNNs is being actively researched, but computational limitations remain a bottleneck, and real-life applications are rare. Due to the computational difficulties, deep BNNs are often inferred using variational inference – an approximate method that makes strong assumptions about the posterior (that it is a multivariate Gaussian). Our model uses Markov Chain Monte Carlo sampling as we have a small dataset and a small network (it is not deep and has only one hidden layer). By separating model and inference from each other, modern probabilistic programming languages (PPLs) make inference for Bayesian models straightforward [19]. They provide an intuitive syntax to define a model, and a set of sampling algorithms to run the inference. Hence, only a generative model needs to be defined by a user to estimate parameters and make predictions. In the context of probabilistic modelling, the Julia language [20] provides a fertile ground: several PPLs have been recently developed on its basis - Turing, Gen, Stheno, SOSS, Omega. The current model was implemented in Turing – a PPL embedded in Julia. The advantage of Julia and Turing combination is that packages in Julia can be seamlessly combined and any generative model, formulated using Julia tools, can be plugged into a PPL for inference without modifications. For instance, a neural network architecture can be defined using Flux.jl and used both for classical and Bayesian inference, when combined with a probabilistic modelling language, such as Turing; probability distributions implemented in the Distributions.jl package can be used to formulate the model. For comparison, Python-based PPLs, such as PyMC3 [21], Edward2 [22] or TensorFlow Probability [23], would require re-writing generative models using Theano or TensorFlow tensors and distributions implemented directly in the corresponding PPLs. Same applies to Stan [24], which represents a stand-alone PPL with multiple interfaces.

In the current paper we propose a Bayesian neural network to predict clinical severity of liver injury from preclinical *in vitro* assay results and physicochemical properties of compounds. Based only on these data, along with the clinical dose, the model enables drug safety scientists to better predict liver toxicity. In this way, we extend the recently proposed POLR model [10] and model different levels of severity by accounting for potential non-linear relationships. We use performance measures appropriate for imbalanced and ordinal data to evaluate the models and employ bootstrapping to asses variation in the metrics.

In addition, we compare the performance of traditional and Bayesian NNs on the given dataset for the multiclass classification task. Our results suggest that hierarchical priors provide a natural tool to work with hyper-parameters: for standard NNs hyper-parameter tuning is required, but BNNs learn hyper-parameters as part of the model fitting.

## Materials and methods

### Data

Data used for the analysis are described in [9], where information on the compounds is provided, including labels, assay results and physicochemical properties as detailed below. The dataset contains 237 labelled compounds and the DILI severity labels are: “no-DILI”, “less-DILI”, “most-DILI”, “ambiguous” and “clinical development”. Records belonging to the first three categories (overall 184 compounds) were used to train and evaluate the model; records with labels “ambiguous”and “clinical development” were excluded from the analysis. The DILI severity labelling was created by encoding the verbally expressed categories into numerical ordinal data: compounds of the most DILI concern were attributed to category 3, compounds of moderate concern were attributed to category 2, and safe compounds were attributed to category 1. The split into training and test sets was created to match the proportions of each of the three categories to avoid imbalances. The training and test sets contained 147 and 37 compounds, respectively. The data, classified as ambiguous, was not used. Assay data included IC50 values for bile salt export pump (BSEP) inhibition, cytotoxicity (THP1, THLE), HepG2 Glucose cytotoxicity, HepG2 Glucose-Galactose cytotoxicity ratio (mitochondrial toxicity) and carbon bond saturation (Fsp^3^), i.e. the number of sp^3^ hybridized carbons divided by the total carbon count. The assay readouts were available as continuous censored values. Lipophilicity (cLogP) as well as the maximum plasma concentration in humans (total *C*_max_) were available as non-censored continuous variables. A graphical description of the predictors and severity of liver injury is shown in Figures 1 and 2.

**Figure 1:**
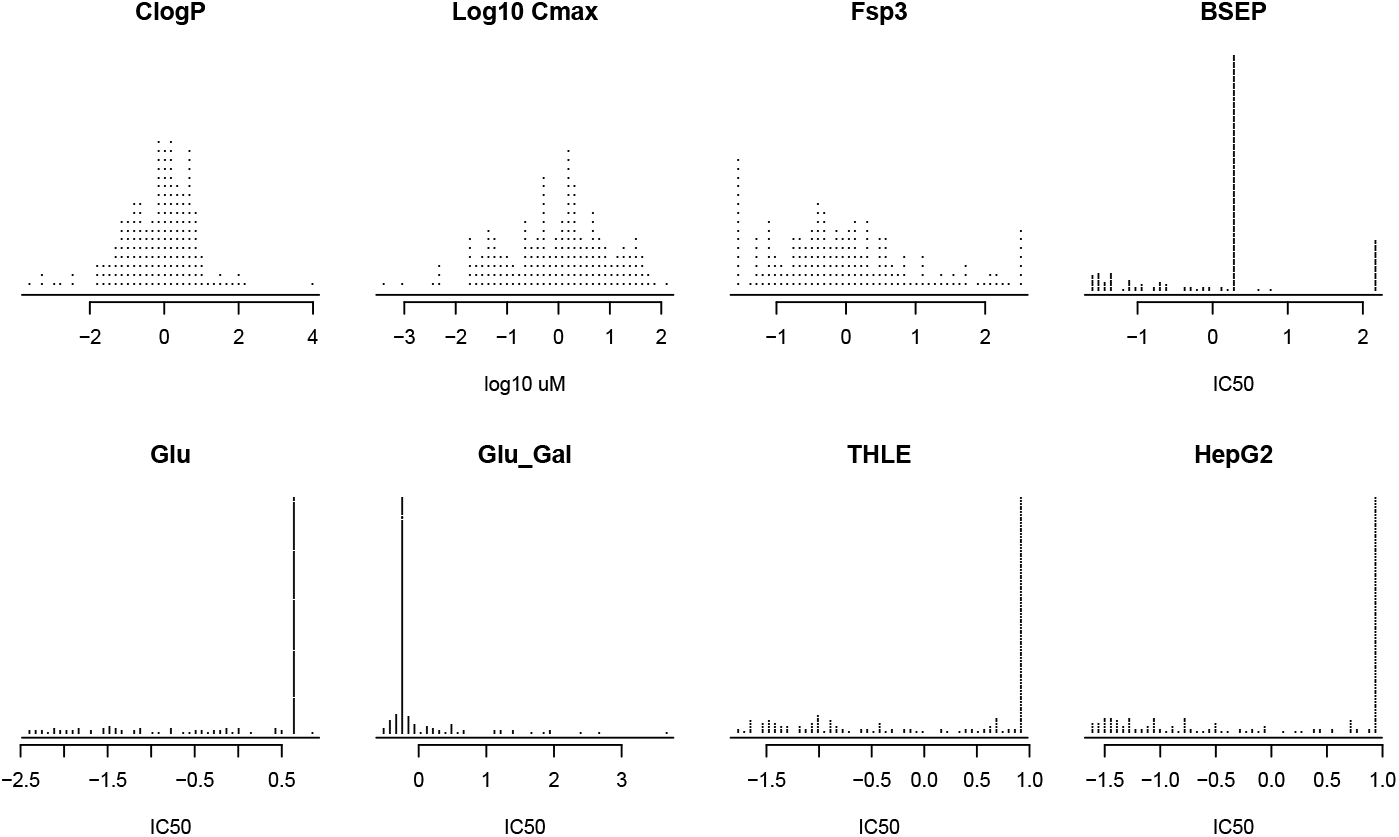
Predictors: assay data, physicochemical properties, and *C*_max_.

**Figure 2:**
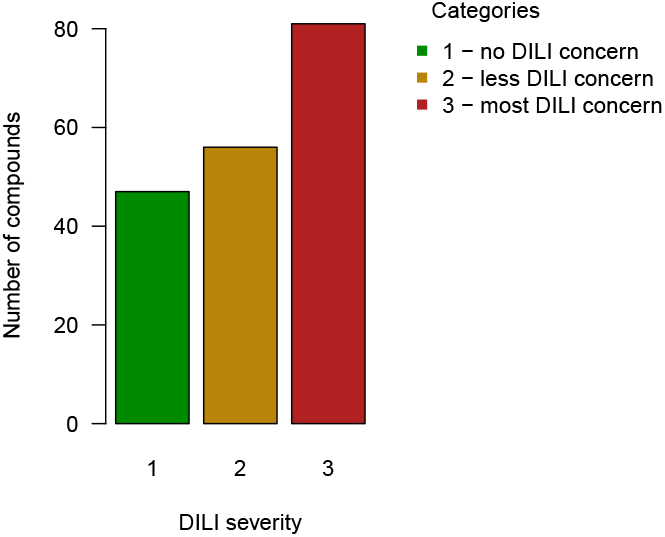
Severity score distribution of the data.

### Models

We propose a BNN to predict toxicity and compare it to our previous POLR model, which is used as a baseline. The input of both models is the assay readouts, physicochemical properties, and *C*_max_. These covariates (and their interactions in case of the POLR model) comprise a design matrix, i.e. the list of predictors. Both models can be viewed as networks: POLR has no hidden layer, only an input and output layer (Figure 3), while the BNN has an additional intermediate hidden layer (Figure 14). Each node of the hidden layer is connected with all of the input covariates and with the output node. Due to the Bayesian formulation of the model, any uncertainty present in the predictions of the latent layer is propagated into the uncertainty of the output.

**Figure 3:**
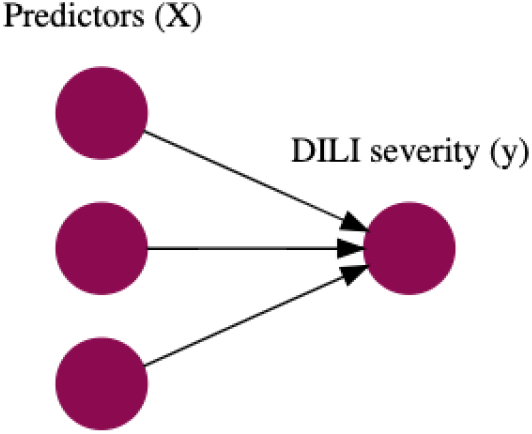
POLR model structure.

The ordering of DILI categories induces a specific structure; neighbouring categories need to be more highly correlated than distant categories. This structure is achieved on the level of the latent continuous variable *η*: it is computed by the models from the design matrix and is then thresholded to yield the ordinal classes. The severity of liver injury *y* is governed by such an unobserved continuous variable called the linear predictor. Three severity classes are defined using two thresholds *c*_1_,*c*_2_, which are estimated from the data. Consequently, the outcome *y* is modelled via a proportional odds logistic regression, also known as a cumulative logit or ordered logit model: the predicted value equals 1 if the unbounded latent variable is smaller than *c*_1_, 3 if the latent variable is larger than *c*_2_, and 2 otherwise. If needed, both the linear predictor and the thresholds can be mapped to the probability scale by the inverse logit transformation.

The Bayesian model formulation consists of two parts: a likelihood and preliminary information, expressed as prior distributions of parameters. The likelihood reflects the assumptions about the data generating process and allows for evaluation of the model against observed data. The inference is made by updating the distributions of the parameters according to how well the model and the data match. We have used zero-mean Gaussian priors for the weights which is a common approach [25, 26, 27, 28]. The POLR and BNN models are described in more detail below.

### Proportional odds logistic regression (POLR)

Under the POLR, model associations between the predictors and the outcome are estimated directly (Fig. 3). All 8 main effects and 21 pair-wise interactions (excluding interactions with *C*_max_) were used as predictors. Interactions with *C*_max_ were excluded following [10]: since *C*_max_ is not experimentally measured when making predictions, it is estimated from a pharmacokinetic model. There is a lot of uncertainty in *C*_max_ values and we want to minimise its influence in the model.

The hierarchical Bayesian POLR model is therefore formulated as

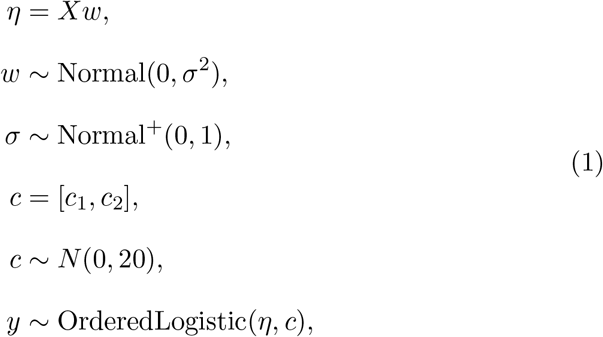

where *X* denotes the design matrix (147 × 29 for the training and 37 × 29 for the test sets), *η* denotes the linear predictor, *w* denotes the set of coefficients with shared variance *σ*^2^, which is a hyper-parameter of the model. All of the parameters - *w, σ* and *c* - are inferred from data.

### Bayesian Neural Network (BNN)

Neural networks (NNs) are built by including hidden layers between input and output layers. Each hidden layer consists of latent nodes applying a predefined computation on the input value to pass the result forward to the next layers. If there is more than one hidden layer in the network, it is considered to be deep. The number of nodes and activation functions need to be defined to complete the NN specification. A NN is governed by a set of parameters: weights and biases. In traditional NNs, all parameters are defined as single numbers which need to be estimated. Training is done via optimisation, i.e. a learning algorithm aims to find a single optimal set of parameters. In the Bayesian setting each parameter is described by a distribution and the task of Bayesian inference is to learn that distribution from data given some initial assumption about the distribution. The parameter space is explored by a learning algorithm and results are obtained via marginalization. The hierarchical BNN model is formulated as presented in Figure 14 and described by formula (2). The thresholds separating the classes *c*, the variance hyper-parameter *σ*^2^, and the weights *w*, are estimated from data.

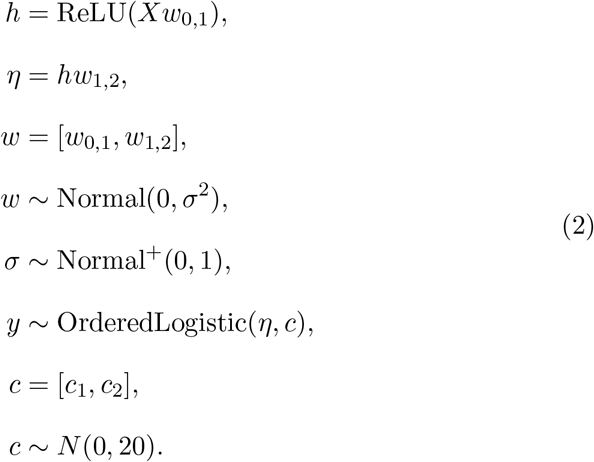

Here, ReLU is the rectified linear unit function, *w*_0,1_ is a set of parameters connecting the input and the hidden layer, and *w*_1,2_ is a set of parameters connecting the hidden and the output layer. The design matrix *X* has dimensions 147 × 8 for the training and 37 × 8 for the test sets.

### Evaluation measures

To evaluate the models, several of its properties need to be taken into account: Bayesian model formulation, ordered outcome, class imbalances, and the continuous nature of the latent prediction. A set of metrics that address these properties were used to compare the baseline and the proposed models: Watanabe-Akaike Information criterion, balanced accuracy, ordered Brier Score, and Brier Skill Score.

*Watanabe-Akaike Information criterion* (WAIC) [29] serves as a tool for Bayesian model selection. It is applicable to models with non-normal posteriors and models with smaller WAIC are preferred.

#### Accuracy

Calculated as the ratio of the number of correct predictions to the number of all examples, is frequently used to evaluate performance of predictive models: if *TP*, *TN*, *N*, *P* are the numbers of true positives, true negatives, all negatives and all positives, correspondingly, the expression for accuracy is (*TP* + *TN*)/(*P* + *N*). Applications of this measure are limited when the observed data is imbalanced and/or ordered. For imbalanced data high accuracy can be easily achieved when the model, potentially erroneously, frequently predicts the dominant class. Sensitivity and specificity suffer from similar issues. A better solution, accounting for imbalances, is given by the *balanced accuracy* (BA) [30]: in case of three classes, given the confusion matrix in Table 1, it is calculated as shown in Formula (3).

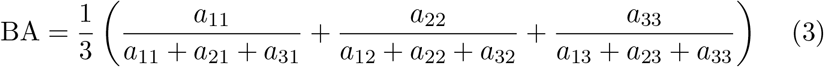

**Table 1:** Confusion matrix for an ordinal variable with three classes.

|  |  | observed |  |  |
| --- | --- | --- | --- | --- |
|  |  | 1 | 2 | 3 |
| predicted | 1 | $a_{11}$ | $a_{12}$ | $a_{13}$ |
| | 2 | $a_{21}$ | $a_{22}$ | $a_{23}$ |
| | 3 | $a_{31}$ | $a_{32}$ | $a_{33}$ |

Balanced accuracy addresses imbalances in data but is not able to capture the ordered nature of the data: predicting the safety class as 1 or 2 should be penalised differently, if the observed class is 3. To take the ordering into account we apply the *ordered Brier Score* (OBS) [31], which measures the distance from the predicted probability to the true class, accounting for the ordered nature of the data; this measure is more suitable than balanced accuracy for ordered outcomes. Models with smaller OBS are preferred. OBS is calculated as

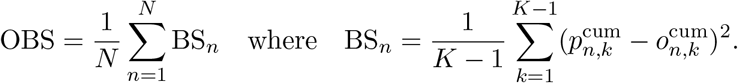

Here 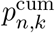 is the *cumulative* predicted probability for the *n*-th observation to be in the *k*-th class (i.e. it is the probability to lie within the first *k* classes), 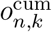 is the *cumulative* true outcome (0 or 1), i.e. it is the outcome of the event “to lie within the first *k* classes”. Since each of the summands lies between 0 and 1, so will the OBS. An example of the OBS calculation is given in the Supplement. OBS is a multiclass generalisation of the Brier score [32] for binary outcomes 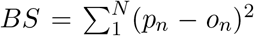, where *o_n_* is the 0 or 1 outcome and *p_n_* is the predicted probability of *o_n_*. The smaller the value of the BS or its cumulative version, the closer is the model’s prediction to the observed data. In the Bayesian setting the posterior distribution of the probabilities 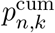 can be used to derive the distribution of the BS.

*Brier Skill Score* (BSS) [32] is an intuitive metric to compare a model in question to a baseline model and models with larger BSS are preferred. The baseline model can be, for instance, set to predict the observed frequencies of classes and yield the Brier score BS^b^. If BS^m^ is the score given by the proposed model, then the BSS can be computed as 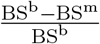. Closeness of the BSS to 0 means that the proposed model is not much better than the baseline, and closeness to 1 means that predictions made by the model are good.

Such measures as accuracy and BA are overly sensitive on small datasets and we advocate for the continuous measures, such as OBS and BSS, to be used as primary evaluation metrics, whenever appropriate.

To assess the variation in metrics, we generated 20 bootstrap datasets from the training data. We used bootstrapping instead of cross-validation because the training set was small. Since bootstrapping samples with replacement, the bootstrapped datasets are the same size as the original, making better use of the limited data [33]. For each bootstrapped data set, we sampled the 147 training compounds with replacement and trained the models on these “in-sample” compounds. Compounds not selected formed the “out-of-sample” data, and typically 2/3 of the compounds are in the insample set and 1/3 in the out-of-sample set. Evaluation metrics were then computed on the out-of-sample and test data (see Appendix E). We expect the performance on the test data to deteriorate when training on the insample data compared with the full training data, but such results are still useful since they can show variation and trends in metrics for the POLR and BNN models. We used the bootstrap experiments to assess how the performance metrics change as distance to the training data increases. For each bootstrap dataset we find the centroid of in-sample data, and then calculate the mean Euclidean distance of the test data to the centroid.

We have performed several additional tests to study the POLR and BNN models: one more way to demonstrate that a model is not overfitting is to apply a random permutation to the labels and fit the model on this new data with broken correlations between inputs and outputs. The model should not be able to predict better than chance.

### Comparing Bayesian and non-Bayesian neural networks

Inference for NNs with ordered outcomes is not a standard task and does not have out-of-the-box tools, and so we reduced the task to multiclass classification, i.e. we ignore the order in the outcome variable. We compare two models with the same architecture, i.e. a multilayered perceptron with the classification task. The Bayesian Neural Network uses a hierarchical prior to prevent overfitting. The NN is initially trained without regularisation, and then we explore two regularisation techniques: dropout [34] and quadratic penalty. The two problems are formulated as follows:

NN, inference via optimisation :

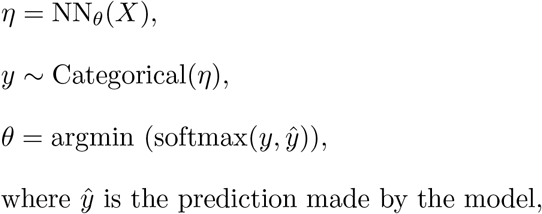
BNN, inference via marginalization :

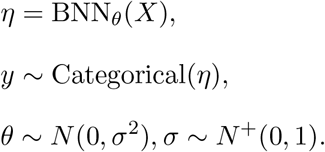
The BNN was tested both under common priors for all weights and biases in the model, i.e.

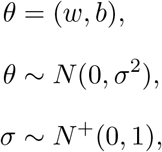

as well as with separate priors for weights and biases, i.e.

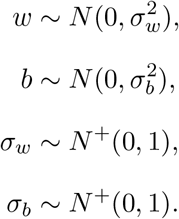
We have also tried separating weights of different layers, i.e.

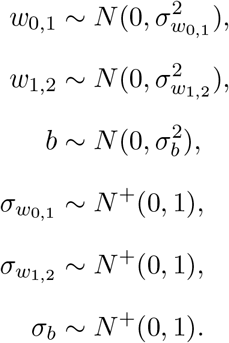

The neural network was trained using the ADAM optimizer. We started with a NN without any regularisation. Consequently, we have added a Dropout layer and tested a grid of dropout probabilities from 0.1 to 0.9 with step 0.1. Finally, we have explored the effect of the *L*_2_ penalty for a set of values of the regularising parameter. For the ADAM algorithm, we have explored several learning rates (*α* =0.0001, 0.001, 0.01 and 0.1). The presented results are for *α* = 0.001 since it has shown the best evaluation measures. The number of epochs was chosen as when results have converged to stable estimates. To account for random effects in the process of NN optimisation (such as the Glorot, a.k.a. Xavier, initialisation of the starting values), we have repeated each of the NN experiments 100 times and report average metrics.

The computational workflow was implemented in Julia [20] and we used the No-U-Turn sampler in Turing [19] for Bayesian inference. The Flux.jl package was used to specify the architecture of neural networks. Trace plots and R-hat statistics were used to evaluate model convergence.

## Results

Table 2 presents the metrics for the BNN and POLR models: WAIC is smaller for BNN, which indicates that it should generalise better to unseen examples. Mean and median test and train OBS are smaller for BNN showing that the error made on the continuous scale is smaller for BNN. Moreover, mean and median train and test BSS are larger for BNN showing that BNN is performing better than POLR when both models are compared to the model predicting class frequencies. BA is higher for BNN for train and test sets, reflecting the quality of class-specific predictions.

**Table 2:** Model comparison according to performance measures

| Model | WAIC | Mean OBS<br>Train / Test | Median OBS<br>Train / Test | Mean BSS<br>Train / Test | Median BSS<br>Train / Test | BA<br>Train / Test |
| --- | --- | --- | --- | --- | --- | --- |
| POLR | 267.3 | 0.14 / 0.16 | 0.11 / 0.12 | 0.24 / 0.20 | 0.36 / 0.36 | 0.61 / 0.61 |
| BNN | 252.8 | 0.12 / 0.14 | 0.08 / 0.10 | 0.37 / 0.31 | 0.46 / 0.39 | 0.70 / 0.67 |

Predictions, together with the uncertainty profiles, were computed for each compound. For example, Figure 5 displays results for a safe compound (Folic acid) by the two models: each of the panels – (a) and (b) – show the continuous predictor on the left and the posterior predictive distribution on the right. Ideally, we would like the most probable class to coincide with the true value, since this would mean that our model has made a correct prediction. More uncertainty in the continuous predictor translates into higher uncertainty of the predicted class. Posterior predictive distributions visualise the uncertainty of the predictions. For this example, we can see that the POLR prediction is less certain than the BNN even though they make the same prediction. The ordered Brier score is sensitive to these differences. Figure 6 compares the profiles of three compounds that are structurally similar but have different DILI severities: Rosiglitazone and Pioglitazone are class 2 and Troglitazone is class 3. Figure 7 shows the posterior predictive distributions for the same three compounds. The BNN is able to better separate these structurally similar compounds: the continuous predictors of the three compounds obtained by POLR are closer to each other than those obtained by BNN (Fig. 6). POLR misclassifies Rosiglitazone and Pioglitazone, while BNN misclassifies only the latter. Misclassification would be reflected in the balanced accuracy, and the confidence of this misclassification will be reflected in the OBS. The overview of all predictions made by the two models is graphically presented in Figure 8. Predicted severity on the continuous probability scale (the point estimate is obtained as the median of the posterior distribution) on the y-axis is plotted against the true class on the x-axis. The horizontal dashed lines display the estimated thresholds. The BNN displays sharper separation between categories. Figure 9 presents the posterior distribution of the ordered Brier score for training and test sets. All distributions are skewed towards zero, which indicates the model’s good predictive ability. Despite the similarity of the OBS plots for POLR and BNN on the training data, the OBS on test data has overall lower values for BNN. Similarly, the BSS is shifted towards 1 for the BNN (Figure 10).

**Figure 4:**
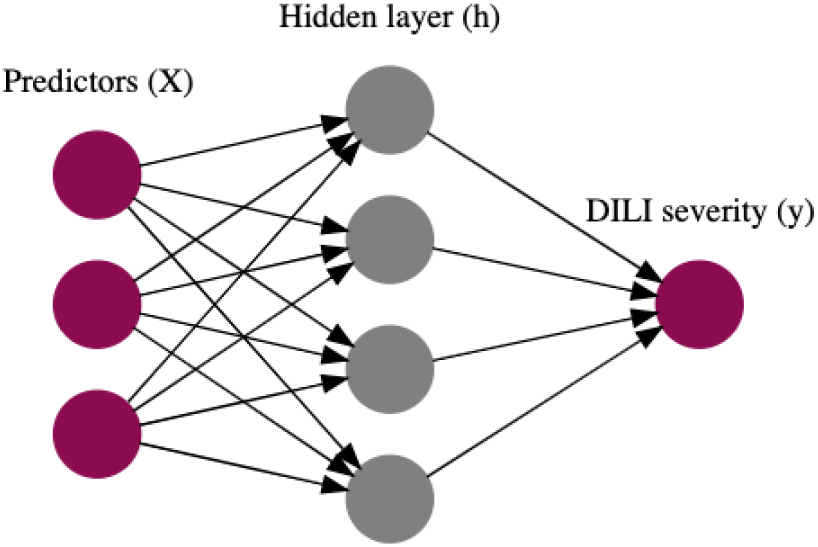
BNN model structure: purple nodes are observed and grey nodes are hidden.

**Figure 5:**
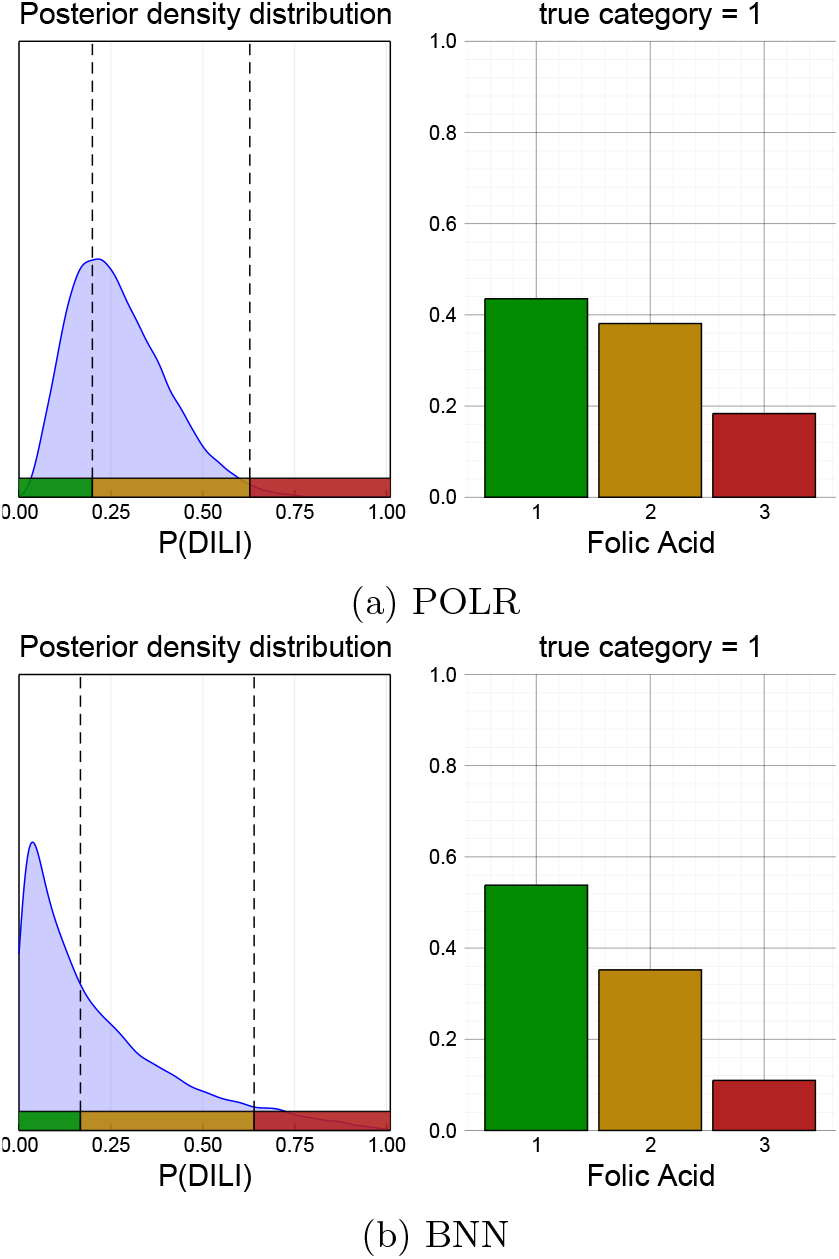
Comparison of predictions by POLR (a) and BNN (b) models for one compound: continuous latent prediction on the left and posterior predictive distribution on the right

**Figure 6:**
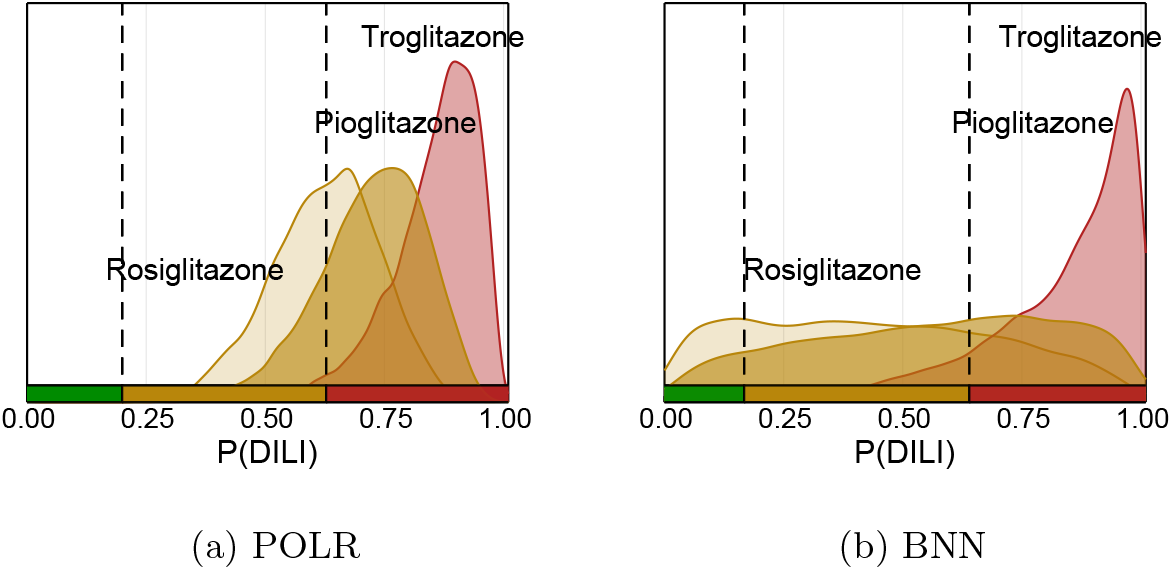
Profiles for three compounds with similar chemical structure but different toxicity classes.

**Figure 7:**
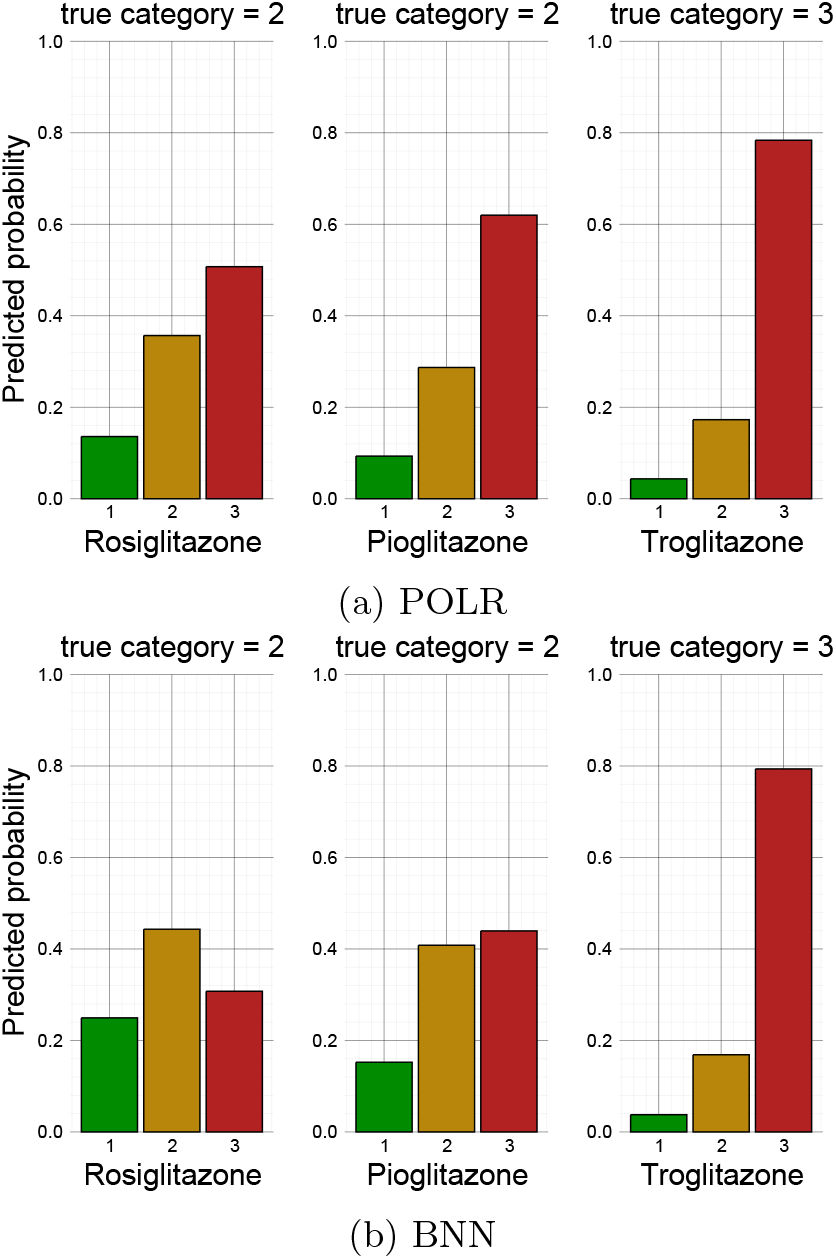
Posterior predictive distributions for three compounds with similar chemical structure but different toxicity classes.

**Figure 8:**
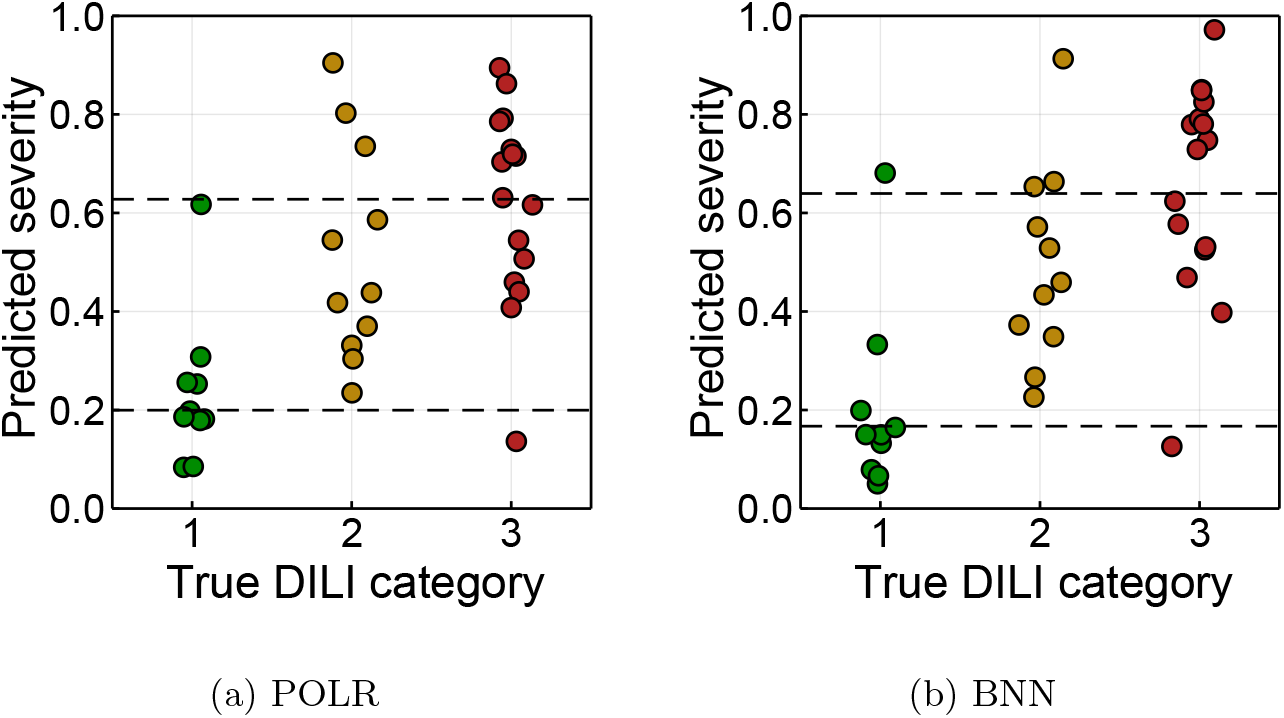
Overview of predictions made by both models on test data. Values are posterior means.

**Figure 9:**
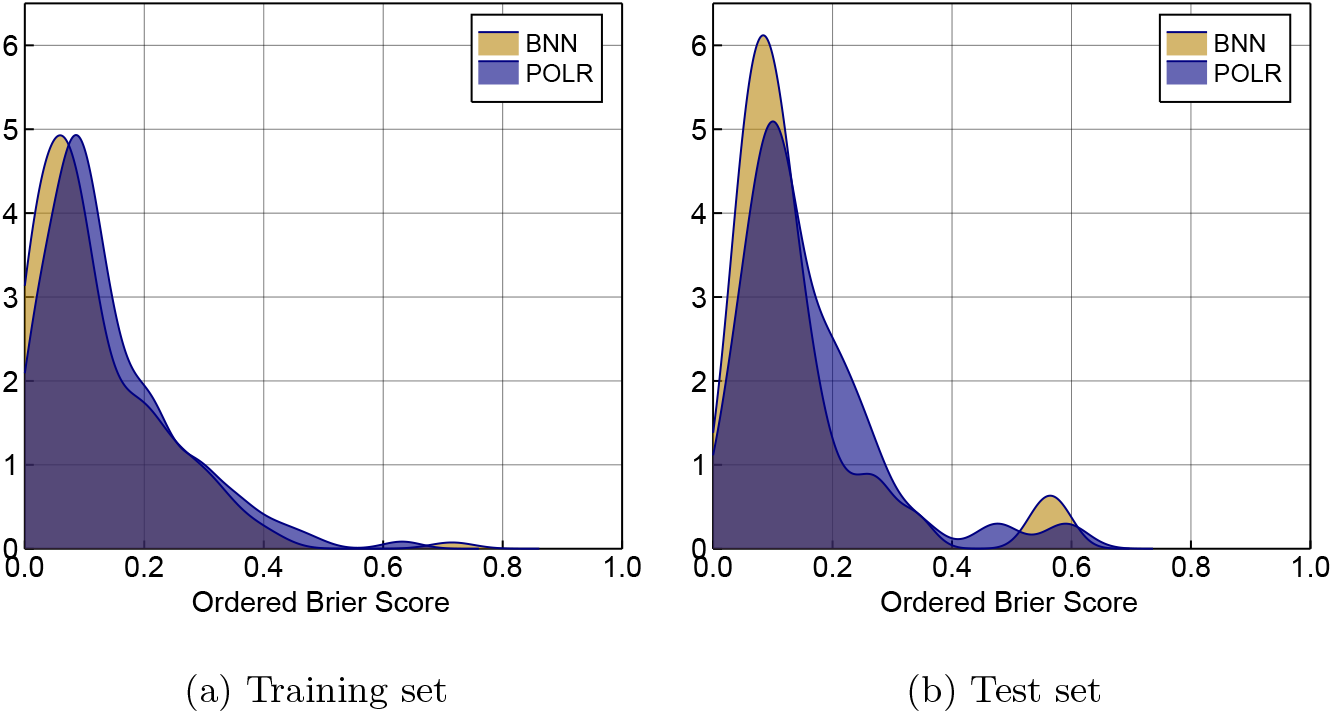
Ordered Brier score for training and test sets for both models.

**Figure 10:**
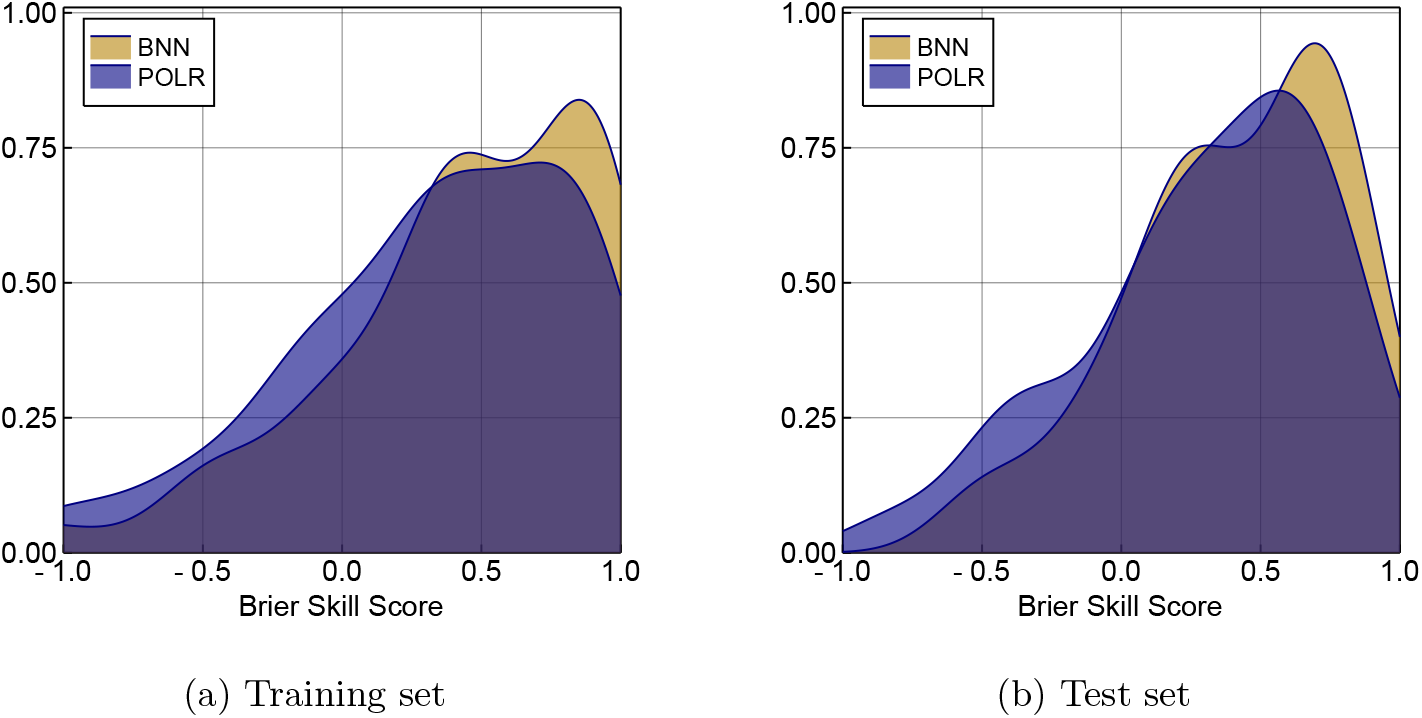
Brier Skill score for training and test sets for both models.

Table 3 shows the medians and standard deviations of the metrics based on the bootstrapping experiments. Trends in OBS, BSS and BA measures as dependence on the distance from the centroid of the corresponding in-sample data set are shown on Figure 11: scatter plots display individual bootstrap iterations and straight lines show trends, estimated by linear models.

**Table 3:** Results of bootstrapping experiments

| Model | Median (sd) OBS | Median (sd) BSS | BA (sd) |
| --- | --- | --- | --- |
|  | Out-of-sample / Test | Out-of-sample / Test | Out-of-sample / Test |
| POLR | 0.12 (0.02) / 0.12 (0.02) | 0.34 (0.11) / 0.35 (0.08) | 0.53 (0.05) / 0.56 (0.04) |
| BNN | 0.08 (0.02) / 0.08 (0.02) | 0.57 (0.13) / 0.59 (0.13) | 0.61 (0.05) / 0.60 (0.04) |

**Figure 11:**
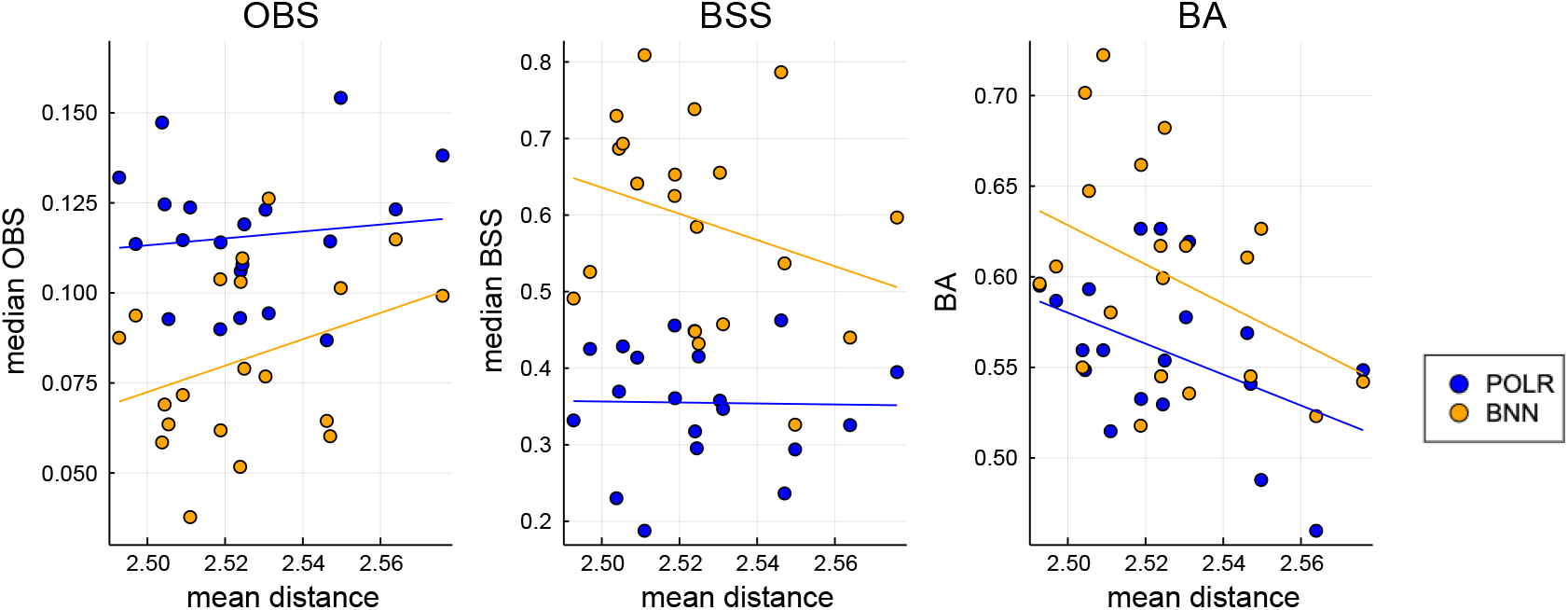
Trends in OBS, BSS and BA measures as dependence on the distance from the centroid of the corresponding in-sample data set.

Table 4 compares the BNN and NN models for the multiclass classification task. Separating the priors for the weights and biases for the BNN has not led to improved results compared with common priors. BNN trained for multiclass classification task produces poorer results than the BNN model capturing the order. Test balanced accuracy is the highest for BNN with common priors for weights and biases. Among the NN models the best test balanced accuracy was achieved by the NN via penalty regularisation. However, it does not exceed the BNN result and it required a grid search for the hyper-parameter value, which might have been not an optimal strategy. BNNs take longer to train, but hyper-parameters are inferred as a part of model fitting.

**Table 4:** Accuracy of the models with an unordered multiclass outcome with 15 hidden nodes.

| Model | Train accuracy | Test accuracy | Train BA | Test BA |
| --- | --- | --- | --- | --- |
| BNN | 0.69 | **0.63** | 0.68 | **0.62** |
| BNN, separate priors $w, b$ | 0.70 | **0.62** | 0.69 | **0.62** |
| NN without regularisation | 0.99 | 0.51 | 0.99 | 0.53 |
| NN Dropout p=0.1 | 0.79 | 0.55 | 0.79 | 0.54 |
| NN Dropout p=0.2 | 0.76 | 0.58 | 0.76 | 0.57 |
| NN Dropout p=0.3 | 0.74 | 0.58 | 0.73 | 0.58 |
| NN Dropout p=0.4 | 0.70 | 0.58 | 0.70 | 0.56 |
| NN Dropout p=0.5 | 0.68 | 0.59 | 0.66 | 0.56 |
| NN Dropout p=0.6 | 0.65 | 0.58 | 0.63 | 0.56 |
| NN Dropout p=0.7 | 0.61 | 0.57 | 0.58 | 0.54 |
| NN Penalty l=0.1 | 0.54 | 0.53 | 0.47 | 0.45 |
| NN Penalty l=0.05 | 0.62 | 0.57 | 0.59 | 0.57 |
| NN Penalty l=0.01 | 0.76 | 0.61 | 0.76 | 0.60 |
| NN Penalty l=0.007 | 0.80 | 0.60 | 0.80 | 0.60 |
| NN Penalty l=0.001 | 0.89 | 0.55 | 0.90 | 0.56 |

Under permutation of labels, both POLR and BNN models do not show signs of overfitting (Appendix C). Calibration is a useful way to evaluate models [35], and we have performed the assessment for POLR and BNN (Appendix D).

Due to the higher number of parameters, BNN requires longer run-times to converge: for POLR model 2000 iterations were enough to achieve convergence and took 4 minutes, while for BNN we sampled for 7000 iterations, which took 35 minutes of elapsed CPU time as measured with the help of the CPUTime.jl package on a laptop with 2.3 GHz Intel Core i5 and 8 GB Memory.

## Discussion and future work

Pharmaceutical companies need to prevent attrition due to adverse drug effects at the earliest possible stage of drug development. We have proposed a Bayesian neural network model to predict toxicity from assay data and physicochemical properties of compounds. The BNN with ordered outcome was able to make more accurate predictions on the test set as compared to a traditional but less flexible POLR model, and multiclass BNN showed better performabnce than non-Bayesian multiclass NNs with the same architecture. Futhermore, the BNN does not overfit on a relatively small number of compounds (147).

When compared to a non-bayesian NN, the BNN showed more flexibility by being able to model ordered outcomes with the off-the-shelf modelling tools. Approaches for fitting ordered outcomes do exist in the NN literature but are not implemented in common deep learning software packages [36]. In the multi-class classification task, the target vector is set via the one-hot encoding to *t* = (0, ‥, 0, 1, 0, 0, …, 0) and the goal is to obtain a probability distribution vector (*o*_1_, *o*_2_, …, *o_k_*, …, *o_K_*) with *o_k_* being closer to 1 and other elements being closer to zero. There is a constraint 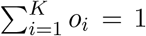 and the soft-max function 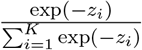 is used to compute the probabilities. For ordered outcomes, the target is being re-coded as *t* = (1, 1, ‥, 1, 1, 0, 0, 0, 0) where the components *t_i_*(1 ≤ *i* ≤ *k*) are set to one and other elements to zero. The goal is to compute the (non-normalised) cumulative probability distribution (*o*_1_, *o*_2_, …, *o_k_*, …, *o_K_*) where *o_i_*(*i ≤ k*) is close to 1 and *o_i_*(*i ≥ k*) is close to 0. Each probability *o_i_* is now being predicted by a sigmoid function 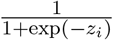. The drawback of using unordered categories is that it does not guarantee a monotonic relation *o*_1_ ≤ *o*_2_ ≤ … ≤ *o_K_*, which is desirable for making predictions. Hyper-parameter search needs to be performed for NNs with cross-validation, while in the Bayesian framework it is done as a part of model fitting. Inference is being performed via marginalisation, which can be thought of as an ensemble of models.

Compounds that are chemically similar might have different toxicity properties [37]. An example is the group of Rosiglitazone, Pioglitazone and Troglitazone: the first two compounds are moderately toxic, while the toxicity of the last one is more severe. BNN model was better able to distingush the profiles than the POLR model. Better understanding of the issue could be gained by including chemical representation of molecules into the model.

Well established performance metrics such as accuracy, sensitivity, and specificity, are limited for uncertainty quantification. Accuracy is ignorant of the amount of error made in case of an incorrect prediction and discards the level of certainty for correct predictions, which is particularly dangerous for small data sets. Brier scores address the issue well since they use continuous predictions. The version for binary outcomes has been used, for cardiotoxicity [38], and we have used the ordered version to handle the ordered data and predictions. Qualitatively, balanced accuracy and Brier score agree well in terms of selecting the best model.

Some continuous variables were censored, but we did not perform modelling of the censored data. Bayesian formulation could include such step as a part of the model, while in the classical context one would need to separate the procedure into two steps: first, the censored data needs to be imputed without regard of the whole model, and then the NN model can be applied.

The number of nodes in the hidden layer was chosen empirically, i.e. we have explored both the compression (e.g. 5 nodes) and expansion (e.g. 25 nodes) of data and the best results were given by the hidden layer with 15 nodes. Results for the multiclass classification task with a small number of nodes (5) can be found in the Supplement (Table 5). The combination of empirically chosen number of nodes and a shrinkage prior (hierarchical normal) have produced good results. Stronger shrinkage priors, such as horseshoe or regularised horseshoe, are known to be able to handle even highly over-estimated number of nodes well [25].

We acknowledge that non-linearities can also be modelled in alternative ways. The options include Gaussian Processes (GPs) and splines, which are both more complex than the proposed model. The connection between GPs and infinitely wide NNs has long been known: a single-layer fully-connected neural network with independent identically distributed priors over its parameters is equivalent to a GP [28]. The result has also been extended to more than one-layer networks [39] and further researched [40, 41]. But since in the presented model, we do not scale the priors by the number of nodes, our model is not a subclass of those representable by GPs. For the decision between a BNN and a GP we used the following heuristic: as long as the largest weight matrix has both the number of rows and columns smaller than the number of the observations in the dataset, it is reasonable to chose a BNN: if *n* is the number of observations, complexity of a GP is *O*(*n*^3^); complexity of a BNN is *O*(*nn*_1_(*n*_0_ + *n*_2_)), where *n*_0_ is the number of input features, *n*_1_ is the number of nodes in the hidden layer and *n*_2_ is the number of output nodes. Consequently, as long as *O*(*nn*_1_(*n*_0_ + *n*_2_)) < *O*(*n*^3^), BNN is computationally more efficient. In our case *n* = 147, *n*_0_ = 8, *n*_1_ = 15, *n*_2_ = 1.

There is a lot of discussion around Bayesian neural networks with the main conclusion being that due to bad scalability there have been “no publicized deployments of Bayesian neural networks in industrial practice” [42] by early 2020. Our work demonstrates the usefulness of BNNs for applied questions, such as toxicology. Scalability for BNNs is an issue, but research is being done to overcome it [14, 16, 17, 18, 43]. To our knowledge, the current work provides the first application of a Bayesian neural network with an ordinal outcome – a flexible predictive model able to capture non-linearities and which generalises well to a new data and yields information about the degree of uncertainty – to DILI prediction from *in vitro* data.

## Supplementary

Supplementary File BNN_Julia.zip contains data and code in Julia and Turing.

## Acknowledgements

We are thankful to Martin Trapp, Mohamed Tarek and Tor Fjelde for their help in understanding Turing.

## Author Contributions

Elizaveta Semenova: Conceptualization, Formal analysis, Methodology, Software, Original draft, Review & editing; Dominic P. Williams: Funding acquisition, Supervision, Review & editing; Avid M. Afzal: Supervision, Review & editing; Stanley E. Lazic: Conceptualization, Funding acquisition, Supervision, Review & editing.

## Supplementary material

### A. Ordered Brier score - an example

**Model 1.**

| Class | 1 | 2 | 3 |
| --- | --- | --- | --- |
| predicted probability | 0 | 2/3 | 1/3 |
| observed outcome | 0 | 1 | 0 |
| cumulative predicted probability | 0 | 2/3 | 3/3 |
| cumulative outcome | 0 | 1 | 1 |

**Model 2.**

Model 1
| Class | 1 | 2 | 3 |
| --- | --- | --- | --- |
| predicted probability | 0 | 0 | 3/3 |
| observed outcome | 1 | 0 | 0 |
| cumulative predicted probability | 0 | 0 | 3/3 |
| cumulative outcome | 1 | 1 | 1 |

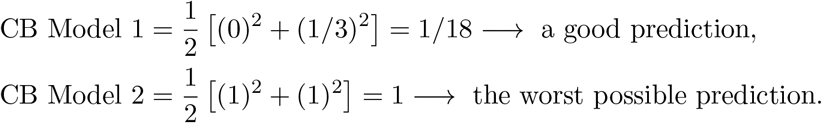

We compared hierarchical and non-hierarchical BNNs with an ordinal outcome. The hierarchical model shows better results than the non-hierarchical model on the training data, but comparable results on the test data. We conclude that the non-hierarchical model is overfitting.

### B. Comparison of frequentist optimisation and Bayesian inference for NNs with multinomial outcome and 5 hidden nodes in the hidden layer

We have tried different number of nodes in the hidden layer: numbers larger than the input dimension (8) to expand the information from the inputs in the hidden layer, and numbers smaller than the input dimension to condense the information. Here we present results of a model with 5 hidden nodes.

**Table 5:** Accuracy of the models with multiclass outcome, 5 hidden nodes.

| model | train accuracy | test accuracy | train BA | test BA |
| --- | --- | --- | --- | --- |
| BNN | 0.70 | **0.59** | 0.70 | **0.59** |
| BNN, separate priors $w, b$ | 0.72 | **0.59** | 0.71 | **0.59** |
| NN without regularization | 0.73 | 0.57 | 0.72 | 0.57 |
| NN Dropout p=0.1 | 0.68 | 0.58 | 0.68 | 0.56 |
| NN Dropout p=0.2 | 0.65 | 0.58 | 0.64 | 0.57 |
| NN Dropout p=0.3 | 0.62 | 0.56 | 0.60 | 0.54 |
| NN Dropout p=0.4 | 0.60 | 0.56 | 0.57 | 0.53 |
| NN Dropout p=0.5 | 0.57 | 0.53 | 0.53 | 0.51 |
| NN Dropout p=0.6 | 0.55 | 0.54 | 0.50 | 0.48 |
| NN Dropout p=0.7 | 0.52 | 0.50 | 0.45 | 0.45 |
| NN Penalty | 0.44 | 0.42 | 0.33 | 0.33 |

### C. Results on outcomes with permutation

**Figure 12:**
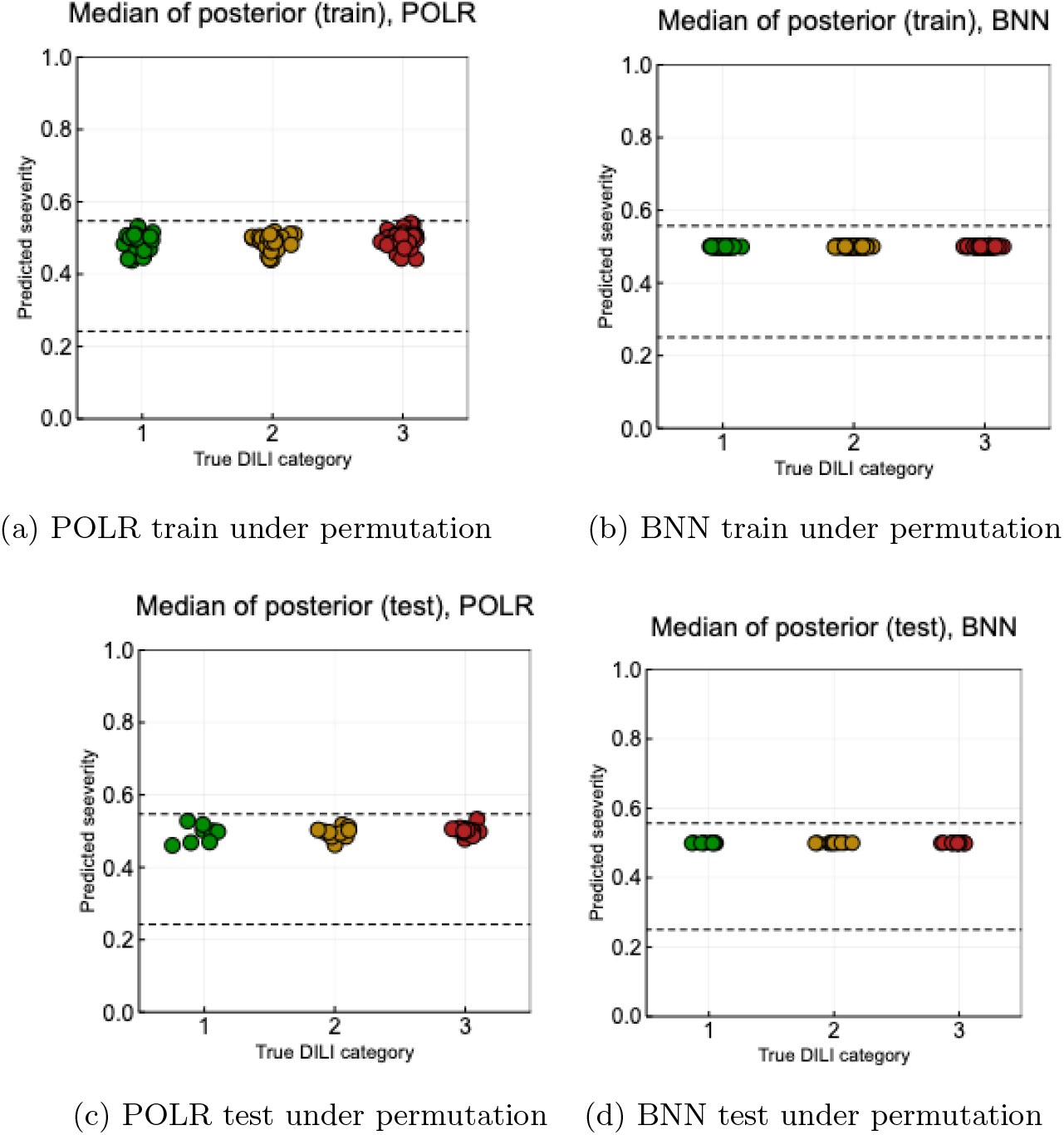
Results on the outcomes with permutation

An additional way to demonstrate that a model isn’t overfitting is to apply a random permutation to the labels and fit the model on this new data with broken correlations between inputs and outputs. The model shouldn’t be able to predict better than chance. Both models do not show signs of overfitting.

### D. Calibration

Calibration performance procedures are well established for binary outcomes, but this is not the case for ordered outcomes. That is why we compared category 1 vs 2 and 3, and categories 1 and 2 vs category 3. The calibration curve for both models deviated from the diagonal line for the 1 vs 2+3 comparison, but this was due to a few compounds that were difficult to predict and the low number of compounds in category 1. We do not see a difference in results for POLR and BNN.

**Figure 13:**
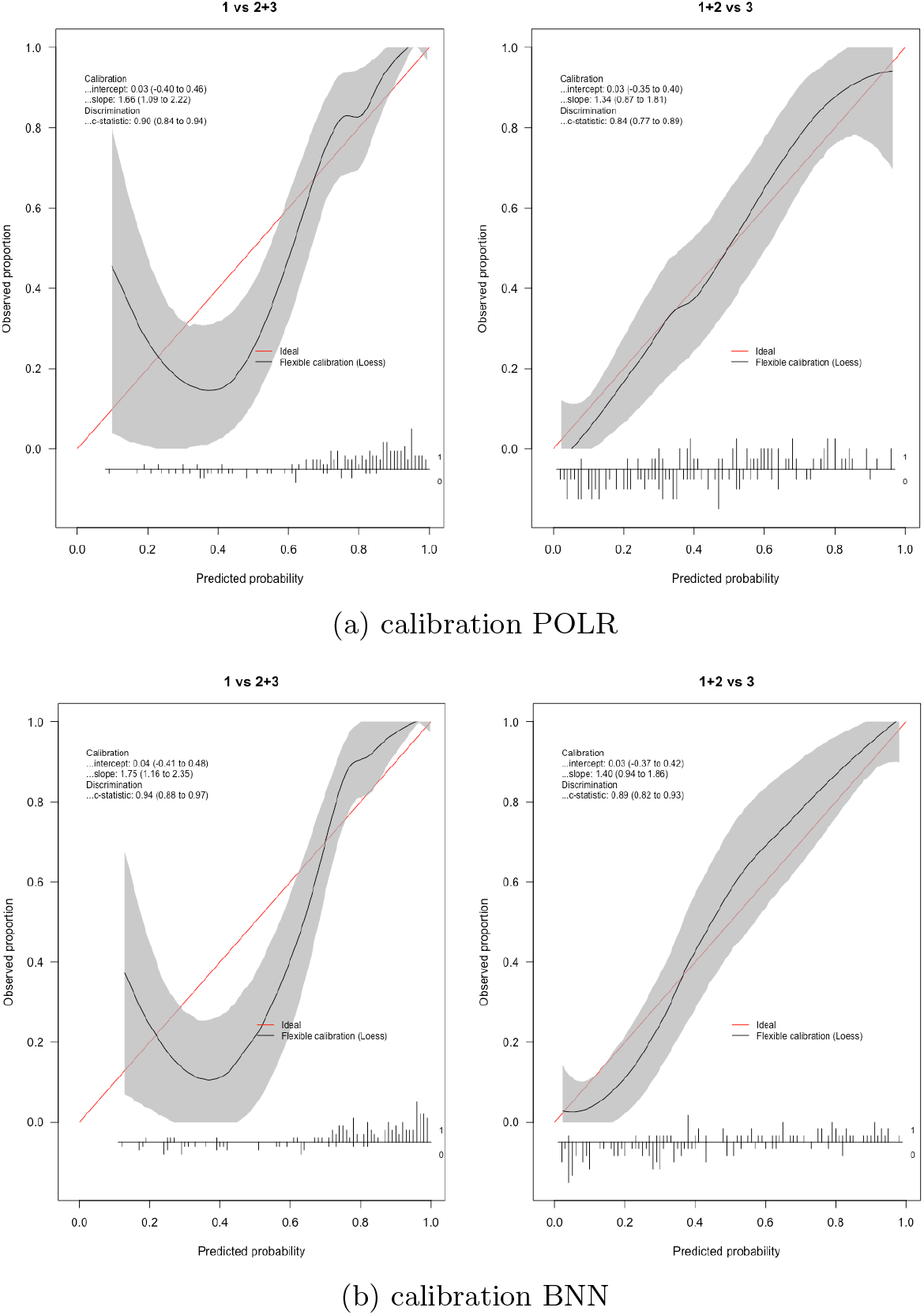
Calibration results

### E. Bootstrapping procedure

**Figure 14:**
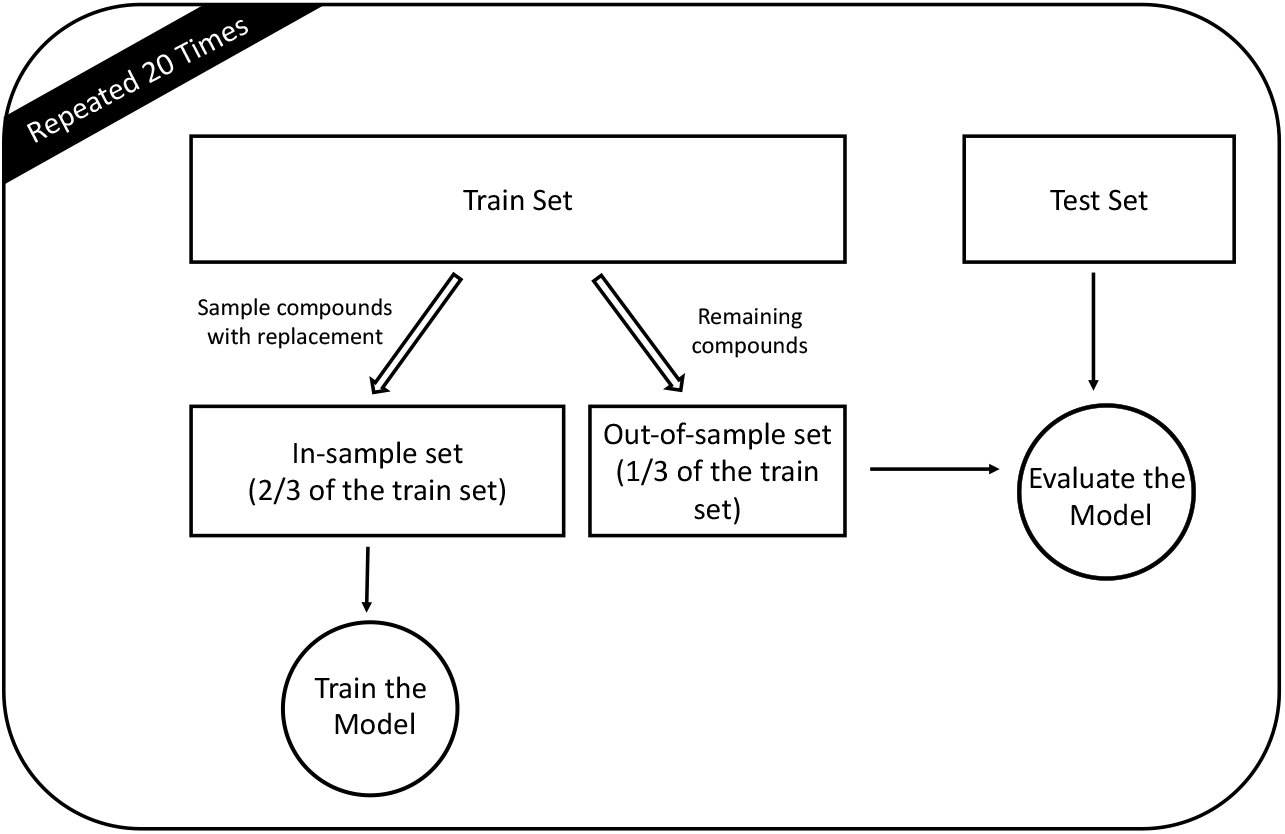
Flow of data withing the bootstrapping procedure.

## References

[1] J. H. Hoofnagle and E. S. Björnsson, “Drug-induced liver injury—types and phenotypes,” New England Journal of Medicine, vol. 381, no. 3, pp. 264–273, 2019.

[2] L. A. Vernetti, A. Vogt, A. Gough, and D. L. Taylor, “Evolution of experimental models of the liver to predict human drug hepatotoxicity and efficacy,” Clinics in liver disease, vol. 21, no. 1, pp. 197–214, 2017.

[3] H. Zhang, L. Ding, Y. Zou, S.-Q. Hu, H.-G. Huang, W.-B. Kong, and J. Zhang, “Predicting drug-induced liver injury in human with naïve bayes classifier approach,” Journal of computer-aided molecular design, vol. 30, no. 10, pp. 889–898, 2016.

[4] M. Cruz-Monteagudo, M. N. D. Cordeiro, and F. Borges, “Computational chemistry approach for the early detection of drug-induced idiosyncratic liver toxicity,” Journal of computational chemistry, vol. 29, no. 4, pp. 533–549, 2008.

[5] H. Olson, G. Betton, D. Robinson, K. Thomas, A. Monro, G. Kolaja, P. Lilly, J. Sanders, G. Sipes, W. Bracken, et al., “Concordance of the toxicity of pharmaceuticals in humans and in animals,” Regulatory Toxicology and Pharmacology, vol. 32, no. 1, pp. 56–67, 2000.

[6] C. Alden, A. Lynn, A. Bourdeau, D. Morton, F. Sistare, V. Kadambi, and L. Silverman, “A critical review of the effectiveness of rodent pharmaceutical carcinogenesis testing in predicting for human risk,” Veterinary pathology, vol. 48, no. 3, pp. 772–784, 2011.

[7] Y. Xu, Z. Dai, F. Chen, S. Gao, J. Pei, and L. Lai, “Deep learning for drug-induced liver injury,” Journal of chemical information and modeling, vol. 55, no. 10, pp. 2085–2093, 2015.

[8] D. Fourches, J. C. Barnes, N. C. Day, P. Bradley, J. Z. Reed, and A. Tropsha, “Cheminformatics analysis of assertions mined from literature that describe drug-induced liver injury in different species,” Chemical research in toxicology, vol. 23, no. 1, pp. 171–183, 2009.

[9] M. D. Aleo, F. Shah, S. Allen, H. A. Barton, C. Costales, S. Lazzaro, L. Leung, A. Nilson, R. S. Obach, A. D. Rodrigues, et al., “Moving beyond binary predictions of human drug-induced liver injury (dili) towards contrasting relative risk potential,” Chemical research in toxicology, 2019.

[10] D. P. Williams, S. Lazic, A. J. Foster, E. Semenova, and P. Morgan, “Predicting drug-induced liver injury with bayesian machine learning,” Chemical research in toxicology, vol. 33, no. 1, pp. 239–248, 2019.

[11] S. Sosnin, D. Karlov, I. V. Tetko, and M. V. Fedorov, “Comparative study of multitask toxicity modeling on a broad chemical space,” Journal of chemical information and modeling, vol. 59, no. 3, pp. 1062–1072, 2018.

[12] M. Kawaguchi, T. Nukaga, S. Sekine, A. Takemura, T. Susukida, S. Oeda, A. Kodama, M. Hirota, H. Kouzuki, and K. Ito, “Mechanism-based integrated assay systems for the prediction of drug-induced liver injury,” Toxicology and Applied Pharmacology, p. 114958, 2020.

[13] S. Salman and X. Liu, “Overfitting mechanism and avoidance in deep neural networks,” arXiv preprint arXiv:1901.06566, 2019.

[14] C. Blundell, J. Cornebise, K. Kavukcuoglu, and D. Wierstra, “Weight uncertainty in neural networks,” arXiv preprint arXiv:1505.05424, 2015.

[15] A.-L. Popkes, H. Overweg, A. Ercole, Y. Li, J. M. Hernández-Lobato, Y. Zaykov, and C. Zhang, “Interpretable outcome prediction with sparse bayesian neural networks in intensive care,” arXiv preprint arXiv:1905.02599, 2019.

[16] Y. Gal and Z. Ghahramani, “Dropout as a bayesian approximation: Representing model uncertainty in deep learning,” in international conference on machine learning, pp. 1050–1059, 2016.

[17] A. Kristiadi, M. Hein, and P. Hennig, “Being bayesian, even just a bit, fixes overconfidence in relu networks,” arXiv preprint arXiv:2002.10118, 2020.

[18] M. Welling and Y. W. Teh, “Bayesian learning via stochastic gradient langevin dynamics,” in Proceedings of the 28th international conference on machine learning (ICML-11), pp. 681–688, 2011.

[19] H. Ge, K. Xu, and Z. Ghahramani, “Turing: A language for flexible probabilistic inference,” in International Conference on Artificial Intelligence and Statistics, pp. 1682–1690, 2018.

[20] J. Bezanson, A. Edelman, S. Karpinski, and V. B. Shah, “Julia: A fresh approach to numerical computing,” SIAM review, vol. 59, no. 1, pp. 65–98, 2017.

[21] J. Salvatier, T. V. Wiecki, and C. Fonnesbeck, “Probabilistic programming in python using PyMC3,” PeerJ Computer Science, vol. 2, p. e55, apr 2016.

[22] D. Tran, M. D. Hoffman, D. Moore, C. Suter, S. Vasudevan, A. Radul, M. Johnson, and R. A. Saurous, “Simple, distributed, and accelerated probabilistic programming,” in Neural Information Processing Systems, 2018.

[23] J. V. Dillon, I. Langmore, D. Tran, E. Brevdo, S. Vasudevan, D. Moore, B. Patton, A. Alemi, M. Hoffman, and R. A. Saurous, “Tensorflow distributions,” arXiv preprint arXiv:1711.10604, 2017.

[24] B. Carpenter, A. Gelman, M. D. Hoffman, D. Lee, B. Goodrich, M. Betancourt, M. Brubaker, J. Guo, P. Li, and A. Riddell, “Stan: A probabilistic programming language,” Journal of statistical software, vol. 76, no. 1, 2017.

[25] S. Ghosh, J. Yao, and F. Doshi-Velez, “Model selection in bayesian neural networks via horseshoe priors,” Journal of Machine Learning Research, vol. 20, no. 182, pp. 1–46, 2019.

[26] D. J. MacKay, “A practical bayesian framework for backpropagation networks,” Neural computation, vol. 4, no. 3, pp. 448–472, 1992.

[27] R. M. Neal, “Bayesian learning via stochastic dynamics,” in Advances in neural information processing systems, pp. 475–482, 1993.

[28] R. M. Neal, Bayesian learning for neural networks, vol. 118. Springer Science & Business Media, 2012.

[29] S. Watanabe, “A widely applicable bayesian information criterion,” Journal of Machine Learning Research, vol. 14, no. Mar, pp. 867–897, 2013.

[30] K. H. Brodersen, C. S. Ong, K. E. Stephan, and J. M. Buhmann, “The balanced accuracy and its posterior distribution,” in 2010-20th International Conference on Pattern Recognition, pp. 3121–3124, IEEE, 2010.

[31] E. C. Merkle, “Weighted brier score decompositions for topically heterogenous forecasting tournaments,” 2018.

[32] G. W. Brier, “Verification of forecasts expressed in terms of probability,” Monthly weather review, vol. 78, no. 1, pp. 1–3, 1950.

[33] E. W. Steyerberg et al., Clinical prediction models. Springer, 2019.

[34] N. Srivastava, G. Hinton, A. Krizhevsky, I. Sutskever, and R. Salakhutdinov, “Dropout: a simple way to prevent neural networks from overfitting,” The journal of machine learning research, vol. 15, no. 1, pp. 1929–1958, 2014.

[35] B. Van Calster, D. J. McLernon, M. van Smeden, L. Wynants, and E. W. Steyerberg, “Calibration: the achilles heel of predictive analytics,” BMC medicine, vol. 17, no. 1, pp. 1–7, 2019.

[36] J. Cheng, Z. Wang, and G. Pollastri, “A neural network approach to ordinal regression,” in 2008 IEEE International Joint Conference on Neural Networks (IEEE World Congress on Computational Intelligence), pp. 1279–1284, IEEE, 2008.

[37] M. Chen, J. Borlak, and W. Tong, “A model to predict severity of drug-induced liver injury in humans,” Hepatology, vol. 64, no. 3, pp. 931–940, 2016.

[38] S. E. Lazic, N. Edmunds, and C. E. Pollard, “Predicting drug safety and communicating risk: benefits of a bayesian approach,” Toxicological Sciences, vol. 162, no. 1, pp. 89–98, 2017.

[39] A. G. d. G. Matthews, M. Rowland, J. Hron, R. E. Turner, and Z. Ghahramani, “Gaussian process behaviour in wide deep neural networks,” arXiv preprint arXiv:1804.11271, 2018.

[40] J. Lee, Y. Bahri, R. Novak, S. S. Schoenholz, J. Pennington, and J. Sohl-Dickstein, “Deep neural networks as gaussian processes,” arXiv preprint arXiv:1711.00165, 2017.

[41] A. Jacot, F. Gabriel, and C. Hongler, “Neural tangent kernel: Convergence and generalization in neural networks,” in Advances in neural information processing systems, pp. 8571–8580, 2018.

[42] F. Wenzel, K. Roth, B. S. Veeling, J. Światkowski, L. Tran, S. Mandt, J. Snoek, T. Salimans, R. Jenatton, and S. Nowozin, “How good is the bayes posterior in deep neural networks really?,” arXiv preprint arXiv:2002.02405, 2020.

[43] J. M. Hernández-Lobato and R. Adams, “Probabilistic backpropagation for scalable learning of bayesian neural networks,” in International Conference on Machine Learning, pp. 1861–1869, 2015.

