## Supplementary material for "A Bayesian neural network for toxicity prediction": 1_POLR.html

1\_POLR\_submit


In [108]:

```
## ---------------------------------------------------------
## Import libraries and functions
## ---------------------------------------------------------

using CSV
using DataFrames
using Turing, Flux, Plots, Random
using StatsBase
using StatsFuns: logistic
using MLBase
using StatsModels
using FreqTables
using KernelDensity
using JLD
using CPUTime
using MLPreprocessing

Turing.turnprogress(true);
Turing.setadbackend(:reverse_diff)

include("functions.jl")
```

```
┌ Info: [Turing]: global PROGRESS is set as true
└ @ Turing /Users/kcft114/.julia/packages/Turing/m05p3/src/Turing.jl:24
```

Out[108]:

```
euc_dist (generic function with 1 method)
```

In [109]:

```
## ---------------------------------------------------------
## Read and prepare data
## ---------------------------------------------------------

# read in data
df_train = CSV.read("data/Aleo_train_match.csv")
df_test = CSV.read("data/Aleo_test_match.csv")
df_ambig = CSV.read("data/Aleo_ambig.csv");
```

In [110]:

```
println(names(df_test))
println(size(df_train))
println(size(df_test))
println(size(df_ambig))

# select predictor variables
df_Xtrain = df_train[!, [:ClogP, :BSEP, :Glu, :Glu_Gal, :THLE, :HepG2, :Fsp3, :log10cmax]]
df_Xtest = df_test[!, [:ClogP, :BSEP, :Glu, :Glu_Gal, :THLE, :HepG2, :Fsp3, :log10cmax]]
df_Xambig = df_ambig[!, [:ClogP, :BSEP, :Glu, :Glu_Gal, :THLE, :HepG2, :Fsp3, :log10cmax]]

# select outcome variable
df_y_train = df_train[!, :dili_sev]
df_y_test = df_test[!, :dili_sev];
df_y_ambig = df_ambig[!, :dili_sev];

# combine X and y in one column
df_yXtrain = hcat(df_y_train, df_Xtrain)
df_yXtest = hcat(df_y_test, df_Xtest)
df_yXambig = hcat(df_y_ambig, df_Xambig)

# formula for design matrix
f = @formula(x1 ~  0 + ClogP + BSEP + Glu + Glu_Gal + THLE + HepG2+ Fsp3  +  log10cmax +
                      ClogP * (BSEP + Glu + Glu_Gal + THLE + HepG2+ Fsp3) +
                              BSEP * (Glu + Glu_Gal + THLE + HepG2+ Fsp3) +
                                      Glu * (Glu_Gal + THLE + HepG2+ Fsp3) +
                                             Glu_Gal * (THLE + HepG2+ Fsp3)+
                                                        THLE * (HepG2+ Fsp3)+
                                                                HepG2 * Fsp3);

# model frame
mf_train = ModelFrame(f, df_yXtrain)
mf_test = ModelFrame(f, df_yXtest)
mf_ambig = ModelFrame(f, df_yXambig)
coefnames(mf_test)

# model matrix
mm_ambig = modelmatrix(mf_ambig)
mm_test = modelmatrix(mf_test)
mm_train = modelmatrix(mf_train)

# model matrix as DataFrame (for scaling)
mm_train_df = convert(DataFrame, mm_train)
mm_test_df = convert(DataFrame, mm_test)
mm_ambig_df = convert(DataFrame, mm_ambig)

# standardize train, test and ambig data
scaler = fit(StandardScaler, mm_train_df)
transform!(mm_train_df, scaler)
transform!(mm_test_df, scaler)
transform!(mm_ambig_df, scaler)

# outcome variable as Int
y_train = Int.(convert(Array, df_y_train))
y_test = Int.(convert(Array, df_y_test))

println(sort(countmap(y_train)))
println(sort(countmap(y_test)))

# convert to matrix
X_train = convert(Matrix, mm_train_df)
X_test = convert(Matrix, mm_test_df);
X_ambig = convert(Matrix, mm_ambig_df);

X_train = slicematrix(X_train)
X_test = slicematrix(X_test)
X_ambig = slicematrix(X_ambig)

drug_names_train = df_train[:,:Drug];
drug_names_test = df_test[:,:Drug];
drug_names_ambig = df_ambig[:,:Drug];

#number of features
size(X_train[1], 1)
```

```
Symbol[:Column1, :Drug, :ClogP, :BSEP, :Glu, :Glu_Gal, :THLE, :HepG2, :vDILIConcern, :Fsp3, :log10cmax, :dili_sev]
(147, 12)
(37, 12)
(53, 12)
OrderedCollections.OrderedDict(1=>37,2=>45,3=>65)
OrderedCollections.OrderedDict(1=>10,2=>11,3=>16)
```

Out[110]:

```
29
```

In [111]:

```
## ---------------------------------------------------------
## Network architechture: nodes_out, nodes_in, activation_function, include_bias_or_not
## ---------------------------------------------------------

network_shape = [(1, size(X_train[1], 1), :identity , 0)]

num_params = sum([i * o + i * b for (o, i, _, b) in network_shape])
```

Out[111]:

```
29
```

In [112]:

```
## ---------------------------------------------------------
## Inference
## ---------------------------------------------------------

# Uncomment this part to perform inference

# one chain:
#num_samples = 2_000
#@time @CPUtime ch2 = sample(bayes_nn(hcat(X_train...), y_train, network_shape, num_params), NUTS(num_samples, 0.65));

# for multiple chains run the following:
#num_chains = 4
#num_samples = 20_000
#chains = mapreduce(c -> sample(bayes_nn(hcat(X_train...), y_train, network_shape, num_params), NUTS(num_samples, 0.65)), chainscat, 1:num_chains)
```

In [113]:

```
## ---------------------------------------------------------
## Save/read results
## ---------------------------------------------------------

#write("chains/POLR.jls", ch2)

ch2 = read("chains/POLR.jls", Chains);
```

In [114]:

```
## ---------------------------------------------------------
## Check convergence
## ---------------------------------------------------------
```

In [115]:

```
# should be true
flag = !(sum(isnan.(summarystats(ch2)[:, :r_hat]))>0) & !(sum(abs.(summarystats(ch2)[:, :r_hat] .- 1) .> 0.1) > 0) & !(sum(summarystats(ch2)[:, :std] .< 1e-13) > 0)
```

Out[115]:

```
true
```

In [116]:

```
show(ch2)
```

```
Object of type Chains, with data of type 1000×42×1 Array{Union{Missing, Float64},3}

Log evidence      = 0.0
Iterations        = 1:1000
Thinning interval = 1
Chains            = 1
Samples per chain = 1000
internals         = eval_num, lp, acceptance_rate, hamiltonian_energy, is_accept, log_density, n_steps, numerical_error, step_size, tree_depth
parameters        = sig, c1, θ[22], θ[21], log_diff_c, θ[7], θ[10], θ[4], θ[15], θ[16], θ[11], θ[20], θ[19], θ[25], θ[8], θ[18], θ[13], θ[9], θ[1], θ[3], θ[5], θ[29], θ[24], θ[23], θ[12], θ[26], θ[14], θ[28], θ[17], θ[2], θ[6], θ[27]

2-element Array{ChainDataFrame,1}

Summary Statistics

│ Row │ parameters │ mean       │ std      │ naive_se   │ mcse       │ ess     │ r_hat    │
│     │ Symbol     │ Float64    │ Float64  │ Float64    │ Float64    │ Any     │ Any      │
├─────┼────────────┼────────────┼──────────┼────────────┼────────────┼─────────┼──────────┤
│ 1   │ c1         │ -1.51631   │ 0.250206 │ 0.00791222 │ 0.0110676  │ 457.908 │ 1.00883  │
│ 2   │ log_diff_c │ 0.622026   │ 0.141897 │ 0.00448718 │ 0.00637713 │ 293.977 │ 1.00925  │
│ 3   │ sig        │ 0.398623   │ 0.128601 │ 0.00406673 │ 0.00943822 │ 106.109 │ 1.02368  │
│ 4   │ θ[1]       │ 0.174228   │ 0.314214 │ 0.00993632 │ 0.00884999 │ 1000.0  │ 1.00298  │
│ 5   │ θ[2]       │ -0.333348  │ 0.340799 │ 0.010777   │ 0.0106398  │ 1000.0  │ 1.00349  │
│ 6   │ θ[3]       │ -0.100295  │ 0.334754 │ 0.0105859  │ 0.00955085 │ 1000.0  │ 1.0011   │
│ 7   │ θ[4]       │ 0.311787   │ 0.321626 │ 0.0101707  │ 0.0129817  │ 1000.0  │ 1.00141  │
│ 8   │ θ[5]       │ -0.208475  │ 0.359275 │ 0.0113613  │ 0.0103426  │ 1000.0  │ 0.999    │
│ 9   │ θ[6]       │ 0.0035606  │ 0.349758 │ 0.0110603  │ 0.0117165  │ 1000.0  │ 0.999735 │
│ 10  │ θ[7]       │ -0.0645731 │ 0.316255 │ 0.0100009  │ 0.0133307  │ 1000.0  │ 1.00048  │
│ 11  │ θ[8]       │ 0.781618   │ 0.229677 │ 0.00726303 │ 0.0143487  │ 191.953 │ 1.01679  │
│ 12  │ θ[9]       │ -0.327009  │ 0.326212 │ 0.0103157  │ 0.0098815  │ 486.091 │ 1.00059  │
│ 13  │ θ[10]      │ -0.0922668 │ 0.353909 │ 0.0111916  │ 0.0115269  │ 976.049 │ 1.00107  │
│ 14  │ θ[11]      │ 0.221084   │ 0.322151 │ 0.0101873  │ 0.00834804 │ 1000.0  │ 1.0005   │
│ 15  │ θ[12]      │ 0.0610853  │ 0.323421 │ 0.0102275  │ 0.00767863 │ 1000.0  │ 1.00039  │
│ 16  │ θ[13]      │ 0.0631842  │ 0.326433 │ 0.0103227  │ 0.00900093 │ 1000.0  │ 0.999116 │
│ 17  │ θ[14]      │ 0.231444   │ 0.275897 │ 0.00872462 │ 0.00866775 │ 587.267 │ 1.0014   │
│ 18  │ θ[15]      │ 0.0917991  │ 0.345014 │ 0.0109103  │ 0.0127968  │ 642.093 │ 0.999766 │
│ 19  │ θ[16]      │ 0.0664886  │ 0.30729  │ 0.00971737 │ 0.00959291 │ 1000.0  │ 1.00194  │
│ 20  │ θ[17]      │ -0.116357  │ 0.337102 │ 0.0106601  │ 0.00573573 │ 1000.0  │ 1.00043  │
│ 21  │ θ[18]      │ -0.0207566 │ 0.313759 │ 0.00992192 │ 0.0091549  │ 1000.0  │ 0.999048 │
│ 22  │ θ[19]      │ -0.201793  │ 0.32504  │ 0.0102787  │ 0.00916672 │ 1000.0  │ 0.999019 │
│ 23  │ θ[20]      │ 0.166389   │ 0.309653 │ 0.00979208 │ 0.00954789 │ 1000.0  │ 1.0037   │
│ 24  │ θ[21]      │ -0.15539   │ 0.361001 │ 0.0114158  │ 0.00970179 │ 1000.0  │ 1.0001   │
│ 25  │ θ[22]      │ 0.150197   │ 0.370881 │ 0.0117283  │ 0.0135806  │ 1000.0  │ 1.00497  │
│ 26  │ θ[23]      │ -0.0284688 │ 0.349177 │ 0.0110419  │ 0.00947799 │ 1000.0  │ 1.00058  │
│ 27  │ θ[24]      │ 0.0122302  │ 0.313799 │ 0.0099232  │ 0.00902763 │ 1000.0  │ 1.00007  │
│ 28  │ θ[25]      │ 0.084383   │ 0.314911 │ 0.00995835 │ 0.0070303  │ 1000.0  │ 0.9995   │
│ 29  │ θ[26]      │ -0.310066  │ 0.34076  │ 0.0107758  │ 0.0169711  │ 409.236 │ 1.01417  │
│ 30  │ θ[27]      │ -0.0206747 │ 0.351772 │ 0.011124   │ 0.00684168 │ 1000.0  │ 0.999019 │
│ 31  │ θ[28]      │ -0.154879  │ 0.371753 │ 0.0117559  │ 0.00636816 │ 1000.0  │ 0.999011 │
│ 32  │ θ[29]      │ 0.195169   │ 0.360692 │ 0.0114061  │ 0.0135983  │ 799.407 │ 1.00586  │

Quantiles

│ Row │ parameters │ 2.5%      │ 25.0%      │ 50.0%      │ 75.0%       │ 97.5%    │
│     │ Symbol     │ Float64   │ Float64    │ Float64    │ Float64     │ Float64  │
├─────┼────────────┼───────────┼────────────┼────────────┼─────────────┼──────────┤
│ 1   │ c1         │ -2.05821  │ -1.6776    │ -1.50431   │ -1.3386     │ -1.07011 │
│ 2   │ log_diff_c │ 0.321232  │ 0.536029   │ 0.624959   │ 0.720521    │ 0.868845 │
│ 3   │ sig        │ 0.204174  │ 0.304126   │ 0.381058   │ 0.479435    │ 0.71096  │
│ 4   │ θ[1]       │ -0.486344 │ -0.0158455 │ 0.162369   │ 0.366411    │ 0.813453 │
│ 5   │ θ[2]       │ -1.04927  │ -0.538486  │ -0.318841  │ -0.112634   │ 0.299022 │
│ 6   │ θ[3]       │ -0.799946 │ -0.305725  │ -0.0671733 │ 0.129255    │ 0.512122 │
│ 7   │ θ[4]       │ -0.297435 │ 0.117651   │ 0.294879   │ 0.489299    │ 1.00611  │
│ 8   │ θ[5]       │ -0.953189 │ -0.409154  │ -0.19576   │ 0.00673349  │ 0.483649 │
│ 9   │ θ[6]       │ -0.750124 │ -0.201413  │ 0.00144777 │ 0.22118     │ 0.698231 │
│ 10  │ θ[7]       │ -0.685393 │ -0.258925  │ -0.0727976 │ 0.120087    │ 0.611735 │
│ 11  │ θ[8]       │ 0.335953  │ 0.62249    │ 0.773373   │ 0.933537    │ 1.25224  │
│ 12  │ θ[9]       │ -1.03773  │ -0.527351  │ -0.297035  │ -0.12064    │ 0.247477 │
│ 13  │ θ[10]      │ -0.867953 │ -0.299063  │ -0.0777428 │ 0.13335     │ 0.580586 │
│ 14  │ θ[11]      │ -0.362481 │ 0.00688649 │ 0.208324   │ 0.417516    │ 0.925899 │
│ 15  │ θ[12]      │ -0.58545  │ -0.142399  │ 0.0551449  │ 0.273453    │ 0.715024 │
│ 16  │ θ[13]      │ -0.549687 │ -0.140006  │ 0.0606497  │ 0.251841    │ 0.711364 │
│ 17  │ θ[14]      │ -0.261324 │ 0.0397951  │ 0.217476   │ 0.404085    │ 0.820533 │
│ 18  │ θ[15]      │ -0.49241  │ -0.134857  │ 0.0637515  │ 0.2969      │ 0.839173 │
│ 19  │ θ[16]      │ -0.498147 │ -0.141649  │ 0.0548788  │ 0.251395    │ 0.720584 │
│ 20  │ θ[17]      │ -0.804604 │ -0.317677  │ -0.11482   │ 0.10128     │ 0.599577 │
│ 21  │ θ[18]      │ -0.675511 │ -0.207608  │ -0.0127926 │ 0.179702    │ 0.571838 │
│ 22  │ θ[19]      │ -0.875647 │ -0.395834  │ -0.203133  │ -0.00713561 │ 0.439627 │
│ 23  │ θ[20]      │ -0.421626 │ -0.0355168 │ 0.171454   │ 0.365174    │ 0.80005  │
│ 24  │ θ[21]      │ -0.920409 │ -0.378871  │ -0.137186  │ 0.0766129   │ 0.523043 │
│ 25  │ θ[22]      │ -0.567285 │ -0.0842463 │ 0.127285   │ 0.380206    │ 0.934306 │
│ 26  │ θ[23]      │ -0.72865  │ -0.246843  │ -0.0203735 │ 0.184442    │ 0.656964 │
│ 27  │ θ[24]      │ -0.600971 │ -0.165864  │ 0.0172164  │ 0.18588     │ 0.728538 │
│ 28  │ θ[25]      │ -0.546788 │ -0.106798  │ 0.0906885  │ 0.278761    │ 0.720571 │
│ 29  │ θ[26]      │ -1.07987  │ -0.517123  │ -0.280811  │ -0.0770069  │ 0.2892   │
│ 30  │ θ[27]      │ -0.694816 │ -0.242077  │ -0.0105075 │ 0.201806    │ 0.678199 │
│ 31  │ θ[28]      │ -0.93882  │ -0.378858  │ -0.154766  │ 0.0739203   │ 0.603205 │
│ 32  │ θ[29]      │ -0.441183 │ -0.0416351 │ 0.162595   │ 0.415851    │ 0.976743 │
```

In [117]:

```
# Extract the θ parameters from the sampled chain.
params2 = ch2[:θ].value.data

# cutpoints
e_log_diff_c = exp.(ch2[:log_diff_c].value.data)[:,1,1]
c1_est = ch2[:c1].value.data[:,1,1]
c2_est = c1_est + e_log_diff_c;
```

In [97]:

```
n_samps = size(ch2, 1)
```

Out[97]:

```
1000
```

In [98]:

```
## ---------------------------------------------------------
## predict for training data
## ---------------------------------------------------------

probs_mat_train, y_pred_samps_train, logpdf_mat_train, y_pred_train, eta_post_train = predict(X_train, y_train, n_samps);

## ---------------------------------------------------------
## metrics for training data
## ---------------------------------------------------------

waic = WAIC_logpfd(logpdf_mat_train)
tbl = freqtable(y_train)
ptbl = prop(tbl)
ptbl = convert(Array, ptbl)
v = vcat(fill(ptbl, length(y_train)))
probs_mat_freq = convert(Array{Float64,2}, hcat(v...)')

# model-based Brier score
cBrier_train = cumBrier(probs_mat_train, y_train)

# baseline Brier score
cBrier_BB_train = cumBrier(probs_mat_freq, y_train)

# Brier Skill score
BSS_train = (cBrier_BB_train .- cBrier_train) ./ cBrier_BB_train

y_pred = convert(Array{Int64,1}, y_pred_train);
y_train = convert(Array{Int64,1}, y_train);
C = confusmat(3, y_train, y_pred)
acc = (C[1,1] + C[2,2] + C[3,3]) / sum(C)
bacc = 1/3 *(C[1,1] / sum(C[1, :]) + C[2,2] / sum(C[2, :]) + C[3,3] / sum(C[3, :]))

println("WAIC = ", round(waic, digits=1))
println("mean cumBrier = ", round(mean(cBrier_train), digits=2))
println("median cumBrier = ", round(median(cBrier_train), digits=2))
println("mean BSS = ", round(mean(BSS_train), digits=2))
println("median BSS = ", round(median(BSS_train), digits=2))
println("Balanced accuracy = ", round(bacc, digits=2))
println("====== Confision Matrix:")
C
```

```
WAIC = 267.3
mean cumBrier = 0.14
median cumBrier = 0.11
mean BSS = 0.24
median BSS = 0.36
Balanced accuracy = 0.61
====== Confision Matrix:
```

Out[98]:

```
3×3 Array{Int64,2}:
 25   7   5
  4  17  24
  3  12  50
```

In [99]:

```
# Rosiglitazone
probsplot(drug_names_train, y_pred_samps_train, 38, y_train)
```

xml version="1.0" encoding="utf-8"?


1

2

3

0.0

0.2

0.4

0.6

0.8

1.0


Rosiglitazone, true category = 2

DILI class

Predicted probability

In [100]:

```
## extract predicted values and convert to 0-1 scale
post = logistic.(eta_post_train)
post_train = post

# cutpoints
c1 = logistic(mean(c1_est))
c2 = logistic(mean(c2_est))
```

Out[100]:

```
0.590224293131391
```

In [101]:

```
postplot(drug_names_train, post_train, 38, y_train)
```

Out[101]:

xml version="1.0" encoding="utf-8"?


0.00

0.25

0.50

0.75

1.00


Rosiglitazone

P(DILI)

In [102]:

```
probsplot(drug_names_train, y_pred_samps_train, 107, y_train)
```

xml version="1.0" encoding="utf-8"?


1

2

3

0.0

0.2

0.4

0.6

0.8

1.0


Troglitazone, true category = 3

DILI class

Predicted probability

In [103]:

```
postplot(drug_names_train, post_train, 107, y_train)
```

Out[103]:

xml version="1.0" encoding="utf-8"?


0.00

0.25

0.50

0.75

1.00


Troglitazone

P(DILI)

In [118]:

```
## ---------------------------------------------------------
## Predict for test data
## ---------------------------------------------------------

probs_mat_test, y_pred_samps_test, logpdf_mat_test, y_pred_test, eta_post_test = predict(X_test,y_test, n_samps);

## ---------------------------------------------------------
## metrics for test data
## ---------------------------------------------------------

#dic = DIC_logpfd(logpdf_mat)
waic = WAIC_logpfd(logpdf_mat_test)

tbl = freqtable(y_test)
ptbl = prop(tbl)
ptbl = convert(Array, ptbl)
v = vcat(fill(ptbl, length(y_test)))
probs_mat_freq = convert(Array{Float64,2}, hcat(v...)')

# model-based Brier score
cBrier_test = cumBrier(probs_mat_test, y_test)

# baseline Brier score
cBrier_BB_test = cumBrier(probs_mat_freq, y_test)

# Brier Skill score
BSS_test = (cBrier_BB_test .- cBrier_test) ./ cBrier_BB_test


y_pred = convert(Array{Int64,1}, y_pred_test);
y_test = convert(Array{Int64,1}, y_test);
C = confusmat(3, y_test, y_pred)
acc = (C[1,1] + C[2,2] + C[3,3]) / sum(C)
bacc = 1/3 *(C[1,1] / sum(C[1, :]) + C[2,2] / sum(C[2, :]) + C[3,3] / sum(C[3, :]))

#println("DIC = ", round(dic, digits=1))
#println("WAIC = ", round(waic, digits=1))
println("mean cumBrier = ", round(mean(cBrier_test), digits=2))
println("median cumBrier = ", round(median(cBrier_test), digits=2))
println("mean BSS = ", round(mean(BSS_test), digits=2))
println("median BSS = ", round(median(BSS_test), digits=2))
#println("Accuracy = ", round(acc, digits=2))
println("Balanced accuracy = ", round(bacc, digits=2))
println("====== Confision Matrix:")
C
```

```
mean cumBrier = 0.16
median cumBrier = 0.12
mean BSS = 0.2
median BSS = 0.36
Balanced accuracy = 0.61
====== Confision Matrix:
```

Out[118]:

```
3×3 Array{Int64,2}:
 8  1   1
 2  3   6
 1  3  12
```

In [147]:

```
sum(C)
```

Out[147]:

```
37
```

In [148]:

```
save("chains/polr_cBrier.jld", "polr_cBrier_train", cBrier_train, "polr_cBrier_test", cBrier_test, 
                               "polr_BSS_train", BSS_train, "polr_BSS_test", BSS_test )
```

In [149]:

```
## ---------------------------------------------------------
## Plots
## ---------------------------------------------------------

k1 = kde(cBrier_train)
k2 = kde(cBrier_test)
p1 = plot(k2.x, k2.density,  
          color = :navy,
          fill = (0, 0.6, :navy), 
          label="test data", 
          yticks = true, 
          legend = :topright,
          #legend = false,
          #ylim=(0, 1.05 * maximum(vcat(k1.density, k2.density))),
          framestyle = :box,
          #title="Ordered Brier Score.",
          #titlefont = font(10, "Calibri"),
          size=[350,350],
          xlabel="Ordered Brier Score",
          xguidefontsize=font(7, "Calibri"),
          xlim = (-0.3, 1.2), 
          ylim = (0, 6.5),
          grid = :xy,
          gridopacity = 0.5)
plot!(k1.x, k1.density, color = :purple, fill = (0, 0.6, :purple), label="training data")
```

Out[149]:

xml version="1.0" encoding="utf-8"?


-0.25

0.00

0.25

0.50

0.75

1.00

0

1

2

3

4

5

6


Ordered Brier Score


test data


training data

In [56]:

```
#savefig("figs/OBS_OL.pdf")
```

In [150]:

```
# Brier Skill score

k1 = kde(BSS_train)
k2 = kde(BSS_test)
p1 = plot(k2.x, k2.density,  
          color = :navy,
          fill = (0, 0.6, :navy), 
          label="test data", 
          yticks = true, 
          legend = :topleft,
          #legend = false,
          #ylim=(0, 1.05 * maximum(vcat(k1.density, k2.density))),
          framestyle = :box,
          #title="Ordered Brier Score.",
          #titlefont = font(10, "Calibri"),
          size=[350,350],
          xlabel="Brier Skill Score",
          xguidefontsize=font(7, "Calibri"),
          xlim = (-2, 2),
          ylim = (0, 1.01),
          grid = :xy,
          gridopacity = 0.5)
plot!(k1.x, k1.density, color = :purple, fill = (0, 0.6, :purple), label="training data")
```

Out[150]:

xml version="1.0" encoding="utf-8"?


-2

-1

0

1

2

0.00

0.25

0.50

0.75

1.00


Brier Skill Score


test data


training data

In [58]:

```
#savefig("figs/BSS_OL.pdf")
```

In [151]:

```
probsplot(drug_names_test, y_pred_samps_test, 6, y_test)
```

xml version="1.0" encoding="utf-8"?


1

2

3

0.0

0.2

0.4

0.6

0.8

1.0


Pioglitazone, true category = 2

DILI class

Predicted probability

In [152]:

```
## extract predicted values and convert to 0-1 scale
post = logistic.(eta_post_test)
post_test = post

# cutpoints
c1 = logistic(mean(c1_est))
c2 = logistic(mean(c2_est))
```

Out[152]:

```
0.5913546502643376
```

In [153]:

```
postplot(drug_names_test, post, 6, y_test)
```

Out[153]:

xml version="1.0" encoding="utf-8"?


0.00

0.25

0.50

0.75

1.00


Pioglitazone

P(DILI)

In [154]:

```
## ---------------------------------------------------------
## dot plot
## ---------------------------------------------------------

post_med = median(post, dims=1)';

col = []
for i in 1:length(y_test)
    if y_test[i] == 1
        col = vcat(col, :green4)
    elseif y_test[i] == 2
        col = vcat(col, :darkgoldenrod)
    else
        col = vcat(col, :firebrick)
    end
end 

scatter(y_test + randn(length(y_test))/15, post_med, 
        alpha=1.0, 
        xlim=(0.5,3.5), 
        ylim=(0,1), 
        size=(250,250),
        legend = false,
        xlabel="True DILI category" ,
        xguidefontsize=font(11, "Calibri"), 
        xticks = 1:1:3,
        ylabel="Predicted severity", 
        yguidefontsize=font(11, "Calibri"),
        #fillcolor = [:navy, :purple, :magenta],
        color = col,
        framestyle = :box,
        tickfont=font(10))
hline!([c1, c2], color = :black, linestyle = :dash)
```

Out[154]:

xml version="1.0" encoding="utf-8"?


1

2

3

0.0

0.2

0.4

0.6

0.8

1.0


True DILI category

Predicted severity

In [34]:

```
#savefig("figs/preds_dot_OL.pdf")
```

In [155]:

```
## ---------------------------------------------------------
## increasing human DILI - posteriors
## ---------------------------------------------------------
   
    # cutpoints
    c1 = logistic(mean(c1_est))
    c2 = logistic(mean(c2_est))

#---- Rosiglitazone ---------------------
    post = post_train

    ind = 38

    post_ind = post[:,ind]
    
    post_ind = vcat(0, post_ind, 1)
    
    kde_npoints = 2048
    #dens = kde(post_ind, npoints=kde_npoints, bandwidth=0.1)
    dens = kde(post_ind, npoints=kde_npoints)

    p1 = plot(dens.x, dens.density,  
              fill = (0, 0.2, :darkgoldenrod), 
              color = :darkgoldenrod, 
              #title = drug_names_train[ind], 
              xlabel="P(DILI)", 
              xlims = (0,1.01),
              ylims = (0,6),
              legend=false,
              yticks = false,
              framestyle = :box,
              size=[250,250],
              titlefont = font(14, "Calibri"),
              xguidefontsize=font(14, "Calibri"))
    vline!([c1, c2], color = :black, linestyle = :dash)

    # define a function that returns a Plots.Shape
    rectangle(w, h, x, y) = Shape(x .+ [0,w,w,0], y .+ [0,0,h,h])

    plot!(rectangle(c1+0.05, 0.25, -0.05,0), color = :green4, alpha = 0.9)
    plot!(rectangle(c2-c1,0.25,c1,0), color = :darkgoldenrod, alpha = 0.9)
    plot!(rectangle(1.05-c2,0.25,c2,0), color = :firebrick, alpha = 0.9)

#---- Troglitazone ---------------------
    post = post_train

    ind = 107

    post_ind = post[:,ind]
    
    post_ind = vcat(0, post_ind, 1)
    
    kde_npoints = 2048
    #dens = kde(post_ind, npoints=kde_npoints, bandwidth=0.1)
    dens = kde(post_ind, npoints=kde_npoints)

    plot!(dens.x, dens.density,  
              fill = (0, 0.4, :firebrick), 
              color = :firebrick, 
              #title = drug_names_train[ind], 
              xlabel="P(DILI)", 
              xlims = (0,1.01),
              ylims = (0,6),
              legend=false,
              yticks = false,
              framestyle = :box,
              size=[250,250],
              titlefont = font(14, "Calibri"),
              xguidefontsize=font(14, "Calibri"))
    vline!([c1, c2], color = :black, linestyle = :dash)

    # define a function that returns a Plots.Shape
    rectangle(w, h, x, y) = Shape(x .+ [0,w,w,0], y .+ [0,0,h,h])

    plot!(rectangle(c1+0.05, 0.25, -0.05,0), color = :green4, alpha = 0.9)
    plot!(rectangle(c2-c1,0.25,c1,0), color = :darkgoldenrod, alpha = 0.9)
    plot!(rectangle(1.05-c2,0.25,c2,0), color = :firebrick, alpha = 0.9)

#---- Pioglitazone ---------------------
    post = post_test

    ind = 6

    post_ind = post[:,ind]
    
    post_ind = vcat(0, post_ind, 1)
    
    kde_npoints = 2048
    #dens = kde(post_ind, npoints=kde_npoints, bandwidth=0.1)
    dens = kde(post_ind, npoints=kde_npoints)

    plot!(dens.x, dens.density,  
              fill = (0, 0.6, :darkgoldenrod), 
              color = :darkgoldenrod, 
             #title = drug_names_test[ind], 
              xlabel="P(DILI)", 
              xlims = (0,1.01),
              ylims = (0,6),
              legend=false,
              yticks = false,
              framestyle = :box,
              size=[250,250],
              titlefont = font(14, "Calibri"),
              xguidefontsize=font(10, "Calibri"))
    vline!([c1, c2], color = :black, linestyle = :dash)

    # define a function that returns a Plots.Shape
    rectangle(w, h, x, y) = Shape(x .+ [0,w,w,0], y .+ [0,0,h,h])

    plot!(rectangle(c1+0.05, 0.25, -0.05,0), color = :green4, alpha = 0.9)
    plot!(rectangle(c2-c1,0.25,c1,0), color = :darkgoldenrod, alpha = 0.9)
    plot!(rectangle(1.05-c2,0.25,c2,0), color = :firebrick, alpha = 0.9)

#------- annotations ------------
    annotate!(0.87, 5.5, text("Troglitazone", :balck, 9))
    annotate!(0.77, 4.2, text("Pioglitazone", :balck, 9))
    annotate!(0.35, 2, text("Rosiglitazone", :balck, 9))
```

Out[155]:

xml version="1.0" encoding="utf-8"?


0.00

0.25

0.50

0.75

1.00


P(DILI)


Troglitazone

Pioglitazone

Rosiglitazone

In [44]:

```
#savefig("figs/POLR_rosi_pio.pdf")
```

In [156]:

```
## ---------------------------------------------------------
## increasing human DILI - posterior predictive distributions
## ---------------------------------------------------------

#---- Rosiglitazone ---------------------
    y_pred_samps = y_pred_samps_train
    y = y_train
    drug_names = drug_names_train

    j = 38

    tbl = freqtable(y_pred_samps[:,j])
    ptbl = prop(tbl)
    b1 = bar( ptbl, 
             labels="10 sin values", 
             #size=[150,250],
              xlabel=drug_names[j], 
              ylabel="Predicted probability", 
              xticks = 1:1:3,
              legend=false,
              yticks = true,
              framestyle = :grid,
              ylimit = (0,1),
              title = "true category = " * string(y[j]) , 
              titlefont = font(14, "Calibri"),
              xguidefontsize=font(14, "Calibri"),
              yguidefontsize=font(14, "Calibri"),
              #fillcolor = [:navy, :purple, :magenta],
              fillcolor = [:green4, :darkgoldenrod, :firebrick],
              #alpha= [0.9, 0.9,0.6],  
              minorgrid = true,
              #gridcolor = :black,
              gridopacity = 0.5,
              grid = :xy,
              bar_width = 0.9);

#---- Troglitazone ---------------------
    y_pred_samps = y_pred_samps_train
    y = y_train   
    drug_names = drug_names_train

    j = 107

    tbl = freqtable(y_pred_samps[:,j])
    ptbl = prop(tbl)
    b2 = bar( ptbl, 
             labels="10 sin values", 
             #size=[150,250],
              xlabel=drug_names[j], 
              #ylabel="Predicted probability", 
              xticks = 1:1:3,
              legend=false,
              yticks = true,
              framestyle = :grid,
              ylimit = (0,1),
              title = "true category = " * string(y[j]) , 
              titlefont = font(14, "Calibri"),
              xguidefontsize=font(14, "Calibri"),
              yguidefontsize=font(14, "Calibri"),
              #fillcolor = [:navy, :purple, :magenta],
              fillcolor = [:green4, :darkgoldenrod, :firebrick],
              #alpha= [0.9, 0.9,0.6],  
              minorgrid = true,
              #gridcolor = :black,
              gridopacity = 0.5,
              grid = :xy,
              bar_width = 0.9);
#---- Pioglitazone ---------------------
    y_pred_samps = y_pred_samps_test
    y = y_test
    drug_names = drug_names_test

    j = 6

    tbl = freqtable(y_pred_samps[:,j])
    ptbl = prop(tbl)
    b3 = bar( ptbl, 
             labels="10 sin values", 
             #size=[150,250],
              xlabel=drug_names[j], 
              #ylabel="Predicted probability", 
              xticks = 1:1:3,
              legend=false,
              #yticks = true,
              #yticks = false,
              framestyle = :grid,
              ylimit = (0,1),
              title = "true category = " * string(y[j]) , 
              titlefont = font(14, "Calibri"),
              xguidefontsize=font(14, "Calibri"),
              #yguidefontsize=font(14, "Calibri"),
              #fillcolor = [:navy, :purple, :magenta],
              fillcolor = [:green4, :darkgoldenrod, :firebrick],
              #alpha= [0.9, 0.9,0.6],  
              minorgrid = true,
              #gridcolor = :black,
              gridopacity = 0.5,
              grid = :xy,
              bar_width = 0.9);

plot(b1, b3, b2, layout=(1, 3), label="")
```

Out[156]:

xml version="1.0" encoding="utf-8"?


1

2

3

0.0

0.2

0.4

0.6

0.8

1.0

true category = 2

Rosiglitazone

Predicted probability


1

2

3

0.0

0.2

0.4

0.6

0.8

1.0

true category = 2

Pioglitazone


1

2

3

0.0

0.2

0.4

0.6

0.8

1.0

true category = 3

Troglitazone

In [46]:

```
#savefig("figs/POLR_rosi_pio_pred.pdf")
```

In [157]:

```
## ---------------------------------------------------------
## Predict for ambiguous-category data
## ---------------------------------------------------------

probs_mat_ambig, y_pred_samps_ambig, logpdf_mat_ambig, y_pred_ambig, eta_post_ambig = predict(X_ambig, false, n_samps);
```

In [158]:

```
## extract predicted values and convert to 0-1 scale
post = logistic.(eta_post_ambig)
post_ambig = post

# cutpoints
c1 = logistic(mean(c1_est))
c2 = logistic(mean(c2_est))

postplot(drug_names_ambig, post_ambig, 2)
```

Out[158]:

xml version="1.0" encoding="utf-8"?


0.00

0.25

0.50

0.75

1.00


Ambrisentan

P(DILI)

In [159]:

```
postplot(drug_names_ambig, post, 3)
```

Out[159]:

xml version="1.0" encoding="utf-8"?


0.00

0.25

0.50

0.75

1.00


Buspirone

P(DILI)

In [160]:

```
probsplot(drug_names_ambig, y_pred_samps_ambig, 2, false)
```

xml version="1.0" encoding="utf-8"?


1

2

3

0.0

0.2

0.4

0.6

0.8

1.0


DILI class

Predicted probability

In [161]:

```
probsplot(drug_names_ambig, y_pred_samps_ambig, 3, false)
```

xml version="1.0" encoding="utf-8"?


1

2

3

0.0

0.2

0.4

0.6

0.8

1.0


DILI class

Predicted probability

In [162]:

```
#println(drug_names_train)
println(drug_names_train[85])
println(drug_names_ambig[3])
```

```
Nefazodone
Buspirone
```

In [163]:

```
## ---------------------------------------------------------
## increasing human DILI - posteriors
## ---------------------------------------------------------
   
    # cutpoints
    c1 = logistic(mean(c1_est))
    c2 = logistic(mean(c2_est))

#---- Nefazodone ---------------------
    # ---------inputs----------
    post = post_train
    drug_names = drug_names_train
    #ind = 112
    ind = 85
    
    # ---------function---------
    post_ind = post[:,ind]
    
    post_ind = vcat(0, post_ind, 1)
    
    kde_npoints = 2048
    #dens = kde(post_ind, npoints=kde_npoints, bandwidth=0.1)
    dens = kde(post_ind, npoints=kde_npoints)

    p1 = plot(dens.x, dens.density,  
              fill = (0, 0.2, :firebrick), 
              color = :firebrick, 
              title = drug_names[ind], 
              xlabel="P(DILI)", 
              xlims = (0,1.01),
              ylims = (0,6),
              legend=false,
              yticks = false,
              framestyle = :box,
              size=[250,250],
              titlefont = font(7, "Calibri"),
              xguidefontsize=font(7, "Calibri"))
    vline!([c1, c2], color = :black, linestyle = :dash)

    # define a function that returns a Plots.Shape
    rectangle(w, h, x, y) = Shape(x .+ [0,w,w,0], y .+ [0,0,h,h])

    plot!(rectangle(c1+0.05, 0.25, -0.05,0), color = :green4, alpha = 0.9)
    plot!(rectangle(c2-c1,0.25,c1,0), color = :darkgoldenrod, alpha = 0.9)
    plot!(rectangle(1.05-c2,0.25,c2,0), color = :firebrick, alpha = 0.9)

#---- Buspirone ---------------------
    # ---------inputs----------
    post = post_ambig
    drug_names = drug_names_ambig
    ind = 3
    
    # ---------function---------
    post_ind = post[:,ind]
    
    post_ind = vcat(0, post_ind, 1)
    
    kde_npoints = 2048
    #dens = kde(post_ind, npoints=kde_npoints, bandwidth=0.1)
    dens = kde(post_ind, npoints=kde_npoints)

    plot!(dens.x, dens.density,  
              fill = (0, 0.2, :blue), 
              color = :blue, 
              title = drug_names[ind], 
              xlabel="P(DILI)", 
              xlims = (0,1.01),
              ylims = (0,6),
              legend=false,
              yticks = false,
              framestyle = :box,
              size=[250,250],
              titlefont = font(7, "Calibri"),
              xguidefontsize=font(7, "Calibri"))
```

Out[163]:

xml version="1.0" encoding="utf-8"?


0.00

0.25

0.50

0.75

1.00


Buspirone

P(DILI)

In [164]:

```
println(drug_names_train[58])
println(drug_names_ambig[2])
```

```
Bosentan
Ambrisentan
```

In [165]:

```
#---- Bosentan ---------------------
    # ---------inputs----------
    post = post_train
    drug_names = drug_names_train
    #ind = 123
    ind = 58
    
    # ---------function---------
    post_ind = post[:,ind]
    
    post_ind = vcat(0, post_ind, 1)
    
    kde_npoints = 2048
    #dens = kde(post_ind, npoints=kde_npoints, bandwidth=0.1)
    dens = kde(post_ind, npoints=kde_npoints)

    p2 = plot(dens.x, dens.density,  
              fill = (0, 0.2, :firebrick), 
              color = :firebrick, 
              title = drug_names[ind], 
              xlabel="P(DILI)", 
              xlims = (0,1.01),
              ylims = (0,6),
              legend=false,
              yticks = false,
              framestyle = :box,
              #size=[250,250],
              titlefont = font(7, "Calibri"),
              xguidefontsize=font(7, "Calibri"))
    vline!([c1, c2], color = :black, linestyle = :dash)

    # define a function that returns a Plots.Shape
    rectangle(w, h, x, y) = Shape(x .+ [0,w,w,0], y .+ [0,0,h,h])

    plot!(rectangle(c1+0.05, 0.25, -0.05,0), color = :green4, alpha = 0.9)
    plot!(rectangle(c2-c1,0.25,c1,0), color = :darkgoldenrod, alpha = 0.9)
    plot!(rectangle(1.05-c2,0.25,c2,0), color = :firebrick, alpha = 0.9)

#---- Ambrisentan ---------------------
    # ---------inputs----------
    post = post_ambig
    drug_names = drug_names_ambig
    ind = 2
    
    # ---------function---------
    post_ind = post[:,ind]
    
    post_ind = vcat(0, post_ind, 1)
    
    kde_npoints = 2048
    #dens = kde(post_ind, npoints=kde_npoints, bandwidth=0.1)
    dens = kde(post_ind, npoints=kde_npoints)

    plot!(dens.x, dens.density,  
              fill = (0, 0.2, :blue), 
              color = :blue, 
              title = drug_names[ind], 
              xlabel="P(DILI)", 
              xlims = (0,1.01),
              ylims = (0,6),
              legend=false,
              yticks = false,
              framestyle = :box,
              size=[250,250],
              titlefont = font(7, "Calibri"),
              xguidefontsize=font(7, "Calibri"))
```

Out[165]:

xml version="1.0" encoding="utf-8"?


0.00

0.25

0.50

0.75

1.00


Ambrisentan

P(DILI)

In [166]:

```
println(drug_names_train[58])
println(drug_names_ambig[2])
```

```
Bosentan
Ambrisentan
```

In [167]:

```
#---- Bosentan ---------------------
    # ---------inputs----------
    post = post_train
    drug_names = drug_names_train
    ind = 58
    
    # ---------function---------
    post_ind = post[:,ind]
    
    post_ind = vcat(0, post_ind, 1)
    
    kde_npoints = 2048
    #dens = kde(post_ind, npoints=kde_npoints, bandwidth=0.1)
    dens = kde(post_ind, npoints=kde_npoints)

    p2 = plot(dens.x, dens.density,  
              fill = (0, 0.2, :firebrick), 
              color = :firebrick, 
              title = drug_names[ind], 
              xlabel="P(DILI)", 
              xlims = (0,1.01),
              ylims = (0,6),
              legend=false,
              yticks = false,
              framestyle = :box,
              #size=[250,250],
              titlefont = font(7, "Calibri"),
              xguidefontsize=font(7, "Calibri"))
    vline!([c1, c2], color = :black, linestyle = :dash)

    # define a function that returns a Plots.Shape
    rectangle(w, h, x, y) = Shape(x .+ [0,w,w,0], y .+ [0,0,h,h])

    plot!(rectangle(c1+0.05, 0.25, -0.05,0), color = :green4, alpha = 0.9)
    plot!(rectangle(c2-c1,0.25,c1,0), color = :darkgoldenrod, alpha = 0.9)
    plot!(rectangle(1.05-c2,0.25,c2,0), color = :firebrick, alpha = 0.9)

#---- Ambrisentan ---------------------
    # ---------inputs----------
    post = post_ambig
    drug_names = drug_names_ambig
    ind = 2
    
    # ---------function---------
    post_ind = post[:,ind]
    
    post_ind = vcat(0, post_ind, 1)
    
    kde_npoints = 2048
    #dens = kde(post_ind, npoints=kde_npoints, bandwidth=0.1)
    dens = kde(post_ind, npoints=kde_npoints)

    plot!(dens.x, dens.density,  
              fill = (0, 0.2, :blue), 
              color = :blue, 
              title = drug_names[ind], 
              xlabel="P(DILI)", 
              xlims = (0,1.01),
              ylims = (0,6),
              legend=false,
              yticks = false,
              framestyle = :box,
              size=[250,250],
              titlefont = font(7, "Calibri"),
              xguidefontsize=font(7, "Calibri"))
```

Out[167]:

xml version="1.0" encoding="utf-8"?


0.00

0.25

0.50

0.75

1.00


Ambrisentan

P(DILI)

In [168]:

```
println(drug_names_train[85])
println(drug_names_ambig[3])
```

```
Nefazodone
Buspirone
```

In [169]:

```
#---- Nefazodone ---------------------
    y_pred_samps = y_pred_samps_train
    y = y_train
    drug_names = drug_names_train

    #j = 112
    j = 85

    tbl = freqtable(y_pred_samps[:,j])
    ptbl = prop(tbl)
    b1 = bar( ptbl, 
             labels="10 sin values", 
             #size=[150,250],
              xlabel=drug_names[j], 
              ylabel="Predicted probability", 
              xticks = 1:1:3,
              legend=false,
              yticks = true,
              framestyle = :grid,
              ylimit = (0,1),
              title = "true category = " * string(y[j]) , 
              titlefont = font(7, "Calibri"),
              xguidefontsize=font(7, "Calibri"),
              yguidefontsize=font(7, "Calibri"),
              #fillcolor = [:navy, :purple, :magenta],
              fillcolor = [:green4, :darkgoldenrod, :firebrick],
              #alpha= [0.9, 0.9,0.6],  
              minorgrid = true,
              #gridcolor = :black,
              gridopacity = 0.5,
              grid = :xy,
              bar_width = 0.9);

#---- Buspirone ---------------------
    y_pred_samps = y_pred_samps_ambig
#    y = y_train   
    drug_names = drug_names_ambig

    j = 3

    tbl = freqtable(y_pred_samps[:,j])
    ptbl = prop(tbl)
    b2 = bar( ptbl, 
             labels="10 sin values", 
             #size=[150,250],
              xlabel=drug_names[j], 
              #ylabel="Predicted probability", 
              xticks = 1:1:3,
              legend=false,
              yticks = true,
              framestyle = :grid,
              ylimit = (0,1),
              #title = "true category = " * string(y[j]) , 
              titlefont = font(7, "Calibri"),
              xguidefontsize=font(7, "Calibri"),
              yguidefontsize=font(7, "Calibri"),
              #fillcolor = [:navy, :purple, :magenta],
              fillcolor = [:green4, :darkgoldenrod, :firebrick],
              #alpha= [0.9, 0.9,0.6],  
              minorgrid = true,
              #gridcolor = :black,
              gridopacity = 0.5,
              grid = :xy,
              bar_width = 0.9);

plot(b2, b1, layout=(1, 2), label="")
```

Out[169]:

xml version="1.0" encoding="utf-8"?


1

2

3

0.0

0.2

0.4

0.6

0.8

1.0

Buspirone


1

2

3

0.0

0.2

0.4

0.6

0.8

1.0

true category = 3

Nefazodone

Predicted probability

In [77]:

```
#---- Nefazodone ---------------------
    y_pred_samps = y_pred_samps_train
    y = y_train
    drug_names = drug_names_train

    #j = 112
    j = 85

    tbl = freqtable(y_pred_samps[:,j])
    ptbl = prop(tbl)
    b1 = bar( ptbl, 
             labels="10 sin values", 
             #size=[150,250],
              xlabel=drug_names[j], 
              ylabel="Predicted probability", 
              xticks = 1:1:3,
              legend=false,
              yticks = true,
              framestyle = :grid,
              ylimit = (0,1),
              title = "true category = " * string(y[j]) , 
              titlefont = font(7, "Calibri"),
              xguidefontsize=font(7, "Calibri"),
              yguidefontsize=font(7, "Calibri"),
              #fillcolor = [:navy, :purple, :magenta],
              fillcolor = [:green4, :darkgoldenrod, :firebrick],
              #alpha= [0.9, 0.9,0.6],  
              minorgrid = true,
              #gridcolor = :black,
              gridopacity = 0.5,
              grid = :xy,
              bar_width = 0.9);

#---- Buspirone ---------------------
    y_pred_samps = y_pred_samps_ambig
#    y = y_train   
    drug_names = drug_names_ambig

    j = 3

    tbl = freqtable(y_pred_samps[:,j])
    ptbl = prop(tbl)
    b2 = bar( ptbl, 
             labels="10 sin values", 
             #size=[150,250],
              xlabel=drug_names[j], 
              #ylabel="Predicted probability", 
              xticks = 1:1:3,
              legend=false,
              yticks = true,
              framestyle = :grid,
              ylimit = (0,1),
              #title = "true category = " * string(y[j]) , 
              titlefont = font(7, "Calibri"),
              xguidefontsize=font(7, "Calibri"),
              yguidefontsize=font(7, "Calibri"),
              #fillcolor = [:navy, :purple, :magenta],
              fillcolor = [:green4, :darkgoldenrod, :firebrick],
              #alpha= [0.9, 0.9,0.6],  
              minorgrid = true,
              #gridcolor = :black,
              gridopacity = 0.5,
              grid = :xy,
              bar_width = 0.9);

plot(b2, b1, layout=(1, 2), label="")
```

Out[77]:

xml version="1.0" encoding="utf-8"?


1

2

3

0.0

0.2

0.4

0.6

0.8

1.0

Buspirone


1

2

3

0.0

0.2

0.4

0.6

0.8

1.0

true category = 3

Nefazodone

Predicted probability

In [170]:

```
#---- Bozentan ---------------------
    y_pred_samps = y_pred_samps_train
    y = y_train
    drug_names = drug_names_train

    j = 58

    tbl = freqtable(y_pred_samps[:,j])
    ptbl = prop(tbl)
    b1 = bar( ptbl, 
             labels="10 sin values", 
             #size=[150,250],
              xlabel=drug_names[j], 
              ylabel="Predicted probability", 
              xticks = 1:1:3,
              legend=false,
              yticks = true,
              framestyle = :grid,
              ylimit = (0,1),
              title = "true category = " * string(y[j]) , 
              titlefont = font(14, "Calibri"),
              xguidefontsize=font(14, "Calibri"),
              yguidefontsize=font(14, "Calibri"),
              #fillcolor = [:navy, :purple, :magenta],
              fillcolor = [:green4, :darkgoldenrod, :firebrick],
              #alpha= [0.9, 0.9,0.6],  
              minorgrid = true,
              #gridcolor = :black,
              gridopacity = 0.5,
              grid = :xy,
              bar_width = 0.9);

#---- Buspirone ---------------------
    y_pred_samps = y_pred_samps_ambig
#    y = y_train   
    drug_names = drug_names_ambig

    j = 3

    tbl = freqtable(y_pred_samps[:,j])
    ptbl = prop(tbl)
    b2 = bar( ptbl, 
             labels="10 sin values", 
             #size=[150,250],
              xlabel=drug_names[j], 
              #ylabel="Predicted probability", 
              xticks = 1:1:3,
              legend=false,
              yticks = true,
              framestyle = :grid,
              ylimit = (0,1),
              #title = "true category = " * string(y[j]) , 
              titlefont = font(14, "Calibri"),
              xguidefontsize=font(14, "Calibri"),
              yguidefontsize=font(14, "Calibri"),
              #fillcolor = [:navy, :purple, :magenta],
              fillcolor = [:green4, :darkgoldenrod, :firebrick],
              #alpha= [0.9, 0.9,0.6],  
              minorgrid = true,
              #gridcolor = :black,
              gridopacity = 0.5,
              grid = :xy,
              bar_width = 0.9);

plot(b2, b1, layout=(1, 2), label="")
```

Out[170]:

xml version="1.0" encoding="utf-8"?


1

2

3

0.0

0.2

0.4

0.6

0.8

1.0

Buspirone


1

2

3

0.0

0.2

0.4

0.6

0.8

1.0

true category = 3

Bosentan

Predicted probability

In [171]:

```
#---- Folic acid ---------------------
   # cutpoints
    c1 = logistic(mean(c1_est))
    c2 = logistic(mean(c2_est))

    post = post_test
    drug_names = drug_names_test

    ind = 30

    post_ind = post[:,ind]
  
    post_ind = vcat(0, post_ind, 1)

    kde_npoints = 2048
    dens = kde(post_ind, npoints=kde_npoints)

    p1 = plot(dens.x, dens.density,  
              fill = (0, 0.2, :blue), 
              color = :blue, 
              #title = drug_names[ind], 
              title = "Posterior density distribution", 
              xlabel="P(DILI)", 
              xlims = (0,1.01),
              ylims = (0,6),
              legend=false,
              yticks = false,
              framestyle = :box,
              size=[250,250],
              titlefont = font(14, "Calibri"),
              xguidefontsize=font(14, "Calibri"),
              tickfont=font(9))
   vline!([c1, c2], color = :black, linestyle = :dash)

    # define a function that returns a Plots.Shape
    rectangle(w, h, x, y) = Shape(x .+ [0,w,w,0], y .+ [0,0,h,h])

    plot!(rectangle(c1+0.05, 0.25, -0.05,0), color = :green4, alpha = 0.9)
    plot!(rectangle(c2-c1,0.25,c1,0), color = :darkgoldenrod, alpha = 0.9)
    plot!(rectangle(1.05-c2,0.25,c2,0), color = :firebrick, alpha = 0.9)

#---- Folic acid ---------------------
    y_pred_samps = y_pred_samps_test
    y = y_test

    j = 30

    tbl = freqtable(y_pred_samps[:,j])
    ptbl = prop(tbl)
    p2 = bar( ptbl, 
             labels="10 sin values", 
             #size=[150,250],
              xlabel=drug_names[j], 
              #ylabel="Predicted probability", 
              xticks = 1:1:3,
              legend=false,
              #yticks = true,
            #yticks = false,
              framestyle = :grid,
              ylimit = (0,1),
              title = "true category = " * string(y[j]) , 
              titlefont = font(14, "Calibri"),
              xguidefontsize=font(14, "Calibri"),
              #yguidefontsize=font(7, "Calibri"),
              #fillcolor = [:navy, :purple, :magenta],
              fillcolor = [:green4, :darkgoldenrod, :firebrick],
              #alpha= [0.9, 0.9,0.6],  
              minorgrid = true,
              #gridcolor = :black,
              gridopacity = 0.5,
              grid = :xy,
              bar_width = 0.9,
              tickfont=font(9));

plot(p1, p2, layout=(1, 2), label="")
```

Out[171]:

xml version="1.0" encoding="utf-8"?


0.00

0.25

0.50

0.75

1.00


Posterior density distribution

P(DILI)


1

2

3

0.0

0.2

0.4

0.6

0.8

1.0

true category = 1

Folic Acid

In [85]:

```
#savefig("figs/POLR_FolicAcid.pdf")
```

In [142]:

```
using Pkg
Pkg.status()
```

```
    Status `~/Box Sync/27_BNN_DILI/revision_correct_submit/Project.toml`
  [336ed68f] CSV v0.5.9
  [a93c6f00] DataFrames v0.19.1
  [587475ba] Flux v0.8.3
  [da1fdf0e] FreqTables v0.3.1
  [4138dd39] JLD v0.9.2
  [5ab0869b] KernelDensity v0.5.1
  [f0e99cf1] MLBase v0.8.0
  [91a5bcdd] Plots v0.25.3
  [2913bbd2] StatsBase v0.32.0
  [4c63d2b9] StatsFuns v0.8.0
  [3eaba693] StatsModels v0.6.2
  [fce5fe82] Turing v0.6.23 #master (https://github.com/TuringLang/Turing.jl.git)
  [9a3f8284] Random
```

In [44]:

```
# Pkg.add(Pkg.PackageSpec(;name="Turing", version="0.6.23"))
```
