## Supplementary material for "A Bayesian neural network for toxicity prediction": 2_BNN.html

2\_BNN\_submit


In [153]:

```
## ---------------------------------------------------------
## Import libraries and functions
## ---------------------------------------------------------

using CSV
using DataFrames
using Turing, Flux, Plots, Random
using StatsBase
using StatsFuns: logistic
using MLBase
using StatsModels
using FreqTables
using KernelDensity
using JLD
using CPUTime
using MLPreprocessing

Turing.turnprogress(true);
Turing.setadbackend(:reverse_diff)

include("functions.jl")
```

```
┌ Info: [Turing]: global PROGRESS is set as true
└ @ Turing /Users/kcft114/.julia/packages/Turing/m05p3/src/Turing.jl:24
```

Out[153]:

```
euc_dist (generic function with 1 method)
```

In [154]:

```
## ---------------------------------------------------------
## Read and prepare data
## ---------------------------------------------------------

# read in data
df_ambig = CSV.read("data/Aleo_ambig.csv");
df_train = CSV.read("data/Aleo_train_match.csv")
df_test = CSV.read("data/Aleo_test_match.csv")

println(size(df_train))
println(size(df_test))
println(size(df_ambig))

# select predictor variables
df_Xtrain = df_train[!, [:ClogP, :BSEP, :Glu, :Glu_Gal, :THLE, :HepG2, :Fsp3, :log10cmax]]
df_Xtest = df_test[!, [:ClogP, :BSEP, :Glu, :Glu_Gal, :THLE, :HepG2, :Fsp3, :log10cmax]]
df_Xambig = df_ambig[!, [:ClogP, :BSEP, :Glu, :Glu_Gal, :THLE, :HepG2, :Fsp3, :log10cmax]]

# standardize train, test and ambig data
scaler = fit(StandardScaler, df_Xtrain)
transform!(df_Xtrain, scaler)
transform!(df_Xambig, scaler)
transform!(df_Xtest, scaler);

# select outcome variable
df_y_train = df_train[!, :dili_sev]
df_y_test = df_test[!, :dili_sev];
df_y_ambig = df_ambig[!, :dili_sev];

# combine X and y in one column
df_yXtrain = hcat(df_y_train, df_Xtrain)
df_yXtest = hcat(df_y_test, df_Xtest)
df_yXambig = hcat(df_y_ambig, df_Xambig)

# formula for design matrix
f = @formula(x1 ~  0 + ClogP + BSEP + Glu + Glu_Gal + THLE + HepG2+ Fsp3  +  log10cmax);

# model frame
mf_train = ModelFrame(f, df_yXtrain)
mf_test = ModelFrame(f, df_yXtest)
mf_ambig = ModelFrame(f, df_yXambig)
coefnames(mf_test)

# model matrix
X_train = modelmatrix(mf_train)
X_test = modelmatrix(mf_test)
X_ambig = modelmatrix(mf_ambig);

# outcome variable as Int
y_train = Int.(convert(Array, df_y_train))
y_test = Int.(convert(Array, df_y_test))

println(sort(countmap(y_train)))
println(sort(countmap(y_test)))

X_train = slicematrix(X_train)
X_test = slicematrix(X_test)
X_ambig = slicematrix(X_ambig)

drug_names_train = df_train[:,:Drug];
drug_names_test = df_test[:,:Drug];
drug_names_ambig = df_ambig[:,:Drug];

#number of features
size(X_train[1], 1)
```

```
(147, 12)
(37, 12)
(53, 12)
OrderedCollections.OrderedDict(1=>37,2=>45,3=>65)
OrderedCollections.OrderedDict(1=>10,2=>11,3=>16)
```

Out[154]:

```
8
```

In [155]:

```
## ---------------------------------------------------------
## Network architechture: nodes_out, nodes_in, activation_function, include_bias_or_not
## ---------------------------------------------------------

network_shape = [
    (15, size(X_train[1], 1), :relu, 1),
    (1,15, :identity, 0)]

num_params = sum([i * o + b * o for (o, i, _, b) in network_shape])
```

Out[155]:

```
150
```

In [167]:

```
## ---------------------------------------------------------
## Inference
## ---------------------------------------------------------

# Uncomment this part to perform inference

# one chain:
#num_samples = 7_000

#@time @CPUtime ch2 = sample(bayes_nn(hcat(X_train...), y_train, network_shape, num_params), NUTS(num_samples, 0.65));

# for multiple chains run the following:
#num_chains = 4
#num_samples = 7_000
#@time @CPUtime chains = mapreduce(c -> sample(bayes_nn(hcat(X...), y, network_shape, num_params), NUTS(num_samples, 0.65)), chainscat, 1:num_chains)
```

```
┌ Warning: The current proposal will be rejected due to numerical error(s).
│   isfiniteθ = true
│   isfiniter = true
│   isfiniteℓπ = false
│   isfiniteℓκ = true
└ @ AdvancedHMC /Users/kcft114/.julia/packages/AdvancedHMC/YWXfk/src/hamiltonian.jl:36
┌ Warning: The current proposal will be rejected due to numerical error(s).
│   isfiniteθ = false
│   isfiniter = false
│   isfiniteℓπ = false
│   isfiniteℓκ = false
└ @ AdvancedHMC /Users/kcft114/.julia/packages/AdvancedHMC/YWXfk/src/hamiltonian.jl:36
┌ Warning: The current proposal will be rejected due to numerical error(s).
│   isfiniteθ = false
│   isfiniter = false
│   isfiniteℓπ = false
│   isfiniteℓκ = false
└ @ AdvancedHMC /Users/kcft114/.julia/packages/AdvancedHMC/YWXfk/src/hamiltonian.jl:36
┌ Warning: The current proposal will be rejected due to numerical error(s).
│   isfiniteθ = false
│   isfiniter = false
│   isfiniteℓπ = false
│   isfiniteℓκ = false
└ @ AdvancedHMC /Users/kcft114/.julia/packages/AdvancedHMC/YWXfk/src/hamiltonian.jl:36
┌ Info: Found initial step size
│   init_ϵ = 0.07500000000000001
└ @ Turing.Inference /Users/kcft114/.julia/packages/Turing/m05p3/src/inference/hmc.jl:365
┌ Warning: The current proposal will be rejected due to numerical error(s).
│   isfiniteθ = true
│   isfiniter = true
│   isfiniteℓπ = false
│   isfiniteℓκ = true
└ @ AdvancedHMC /Users/kcft114/.julia/packages/AdvancedHMC/YWXfk/src/hamiltonian.jl:36
┌ Warning: The current proposal will be rejected due to numerical error(s).
│   isfiniteθ = false
│   isfiniter = false
│   isfiniteℓπ = false
│   isfiniteℓκ = false
└ @ AdvancedHMC /Users/kcft114/.julia/packages/AdvancedHMC/YWXfk/src/hamiltonian.jl:36
┌ Warning: Incorrect ϵ = NaN; ϵ_previous = 0.07500000000000001 is used instead.
└ @ AdvancedHMC.Adaptation /Users/kcft114/.julia/packages/AdvancedHMC/YWXfk/src/adaptation/stepsize.jl:103
┌ Warning: The current proposal will be rejected due to numerical error(s).
│   isfiniteθ = true
│   isfiniter = true
│   isfiniteℓπ = false
│   isfiniteℓκ = true
└ @ AdvancedHMC /Users/kcft114/.julia/packages/AdvancedHMC/YWXfk/src/hamiltonian.jl:36
┌ Warning: The current proposal will be rejected due to numerical error(s).
│   isfiniteθ = false
│   isfiniter = false
│   isfiniteℓπ = false
│   isfiniteℓκ = false
└ @ AdvancedHMC /Users/kcft114/.julia/packages/AdvancedHMC/YWXfk/src/hamiltonian.jl:36
┌ Warning: Incorrect ϵ = NaN; ϵ_previous = 0.07500000000000001 is used instead.
└ @ AdvancedHMC.Adaptation /Users/kcft114/.julia/packages/AdvancedHMC/YWXfk/src/adaptation/stepsize.jl:103
┌ Warning: The current proposal will be rejected due to numerical error(s).
│   isfiniteθ = true
│   isfiniter = true
│   isfiniteℓπ = false
│   isfiniteℓκ = true
└ @ AdvancedHMC /Users/kcft114/.julia/packages/AdvancedHMC/YWXfk/src/hamiltonian.jl:36
┌ Warning: The current proposal will be rejected due to numerical error(s).
│   isfiniteθ = false
│   isfiniter = false
│   isfiniteℓπ = false
│   isfiniteℓκ = false
└ @ AdvancedHMC /Users/kcft114/.julia/packages/AdvancedHMC/YWXfk/src/hamiltonian.jl:36
┌ Warning: Incorrect ϵ = NaN; ϵ_previous = 0.07500000000000001 is used instead.
└ @ AdvancedHMC.Adaptation /Users/kcft114/.julia/packages/AdvancedHMC/YWXfk/src/adaptation/stepsize.jl:103
┌ Warning: The current proposal will be rejected due to numerical error(s).
│   isfiniteθ = true
│   isfiniter = true
│   isfiniteℓπ = false
│   isfiniteℓκ = true
└ @ AdvancedHMC /Users/kcft114/.julia/packages/AdvancedHMC/YWXfk/src/hamiltonian.jl:36
┌ Warning: The current proposal will be rejected due to numerical error(s).
│   isfiniteθ = false
│   isfiniter = false
│   isfiniteℓπ = false
│   isfiniteℓκ = false
└ @ AdvancedHMC /Users/kcft114/.julia/packages/AdvancedHMC/YWXfk/src/hamiltonian.jl:36
┌ Warning: Incorrect ϵ = NaN; ϵ_previous = 0.07500000000000001 is used instead.
└ @ AdvancedHMC.Adaptation /Users/kcft114/.julia/packages/AdvancedHMC/YWXfk/src/adaptation/stepsize.jl:103
┌ Warning: The current proposal will be rejected due to numerical error(s).
│   isfiniteθ = true
│   isfiniter = true
│   isfiniteℓπ = false
│   isfiniteℓκ = true
└ @ AdvancedHMC /Users/kcft114/.julia/packages/AdvancedHMC/YWXfk/src/hamiltonian.jl:36
┌ Warning: The current proposal will be rejected due to numerical error(s).
│   isfiniteθ = false
│   isfiniter = false
│   isfiniteℓπ = false
│   isfiniteℓκ = false
└ @ AdvancedHMC /Users/kcft114/.julia/packages/AdvancedHMC/YWXfk/src/hamiltonian.jl:36
┌ Warning: Incorrect ϵ = NaN; ϵ_previous = 0.07500000000000001 is used instead.
└ @ AdvancedHMC.Adaptation /Users/kcft114/.julia/packages/AdvancedHMC/YWXfk/src/adaptation/stepsize.jl:103
┌ Warning: The current proposal will be rejected due to numerical error(s).
│   isfiniteθ = true
│   isfiniter = true
│   isfiniteℓπ = false
│   isfiniteℓκ = true
└ @ AdvancedHMC /Users/kcft114/.julia/packages/AdvancedHMC/YWXfk/src/hamiltonian.jl:36
┌ Warning: The current proposal will be rejected due to numerical error(s).
│   isfiniteθ = false
│   isfiniter = false
│   isfiniteℓπ = false
│   isfiniteℓκ = false
└ @ AdvancedHMC /Users/kcft114/.julia/packages/AdvancedHMC/YWXfk/src/hamiltonian.jl:36
┌ Warning: Incorrect ϵ = NaN; ϵ_previous = 0.07500000000000001 is used instead.
└ @ AdvancedHMC.Adaptation /Users/kcft114/.julia/packages/AdvancedHMC/YWXfk/src/adaptation/stepsize.jl:103
┌ Warning: The current proposal will be rejected due to numerical error(s).
│   isfiniteθ = true
│   isfiniter = true
│   isfiniteℓπ = false
│   isfiniteℓκ = true
└ @ AdvancedHMC /Users/kcft114/.julia/packages/AdvancedHMC/YWXfk/src/hamiltonian.jl:36
┌ Warning: The current proposal will be rejected due to numerical error(s).
│   isfiniteθ = false
│   isfiniter = false
│   isfiniteℓπ = false
│   isfiniteℓκ = false
└ @ AdvancedHMC /Users/kcft114/.julia/packages/AdvancedHMC/YWXfk/src/hamiltonian.jl:36
┌ Warning: Incorrect ϵ = NaN; ϵ_previous = 0.07500000000000001 is used instead.
└ @ AdvancedHMC.Adaptation /Users/kcft114/.julia/packages/AdvancedHMC/YWXfk/src/adaptation/stepsize.jl:103
┌ Warning: The current proposal will be rejected due to numerical error(s).
│   isfiniteθ = true
│   isfiniter = true
│   isfiniteℓπ = false
│   isfiniteℓκ = true
└ @ AdvancedHMC /Users/kcft114/.julia/packages/AdvancedHMC/YWXfk/src/hamiltonian.jl:36
┌ Warning: The current proposal will be rejected due to numerical error(s).
│   isfiniteθ = false
│   isfiniter = false
│   isfiniteℓπ = false
│   isfiniteℓκ = false
└ @ AdvancedHMC /Users/kcft114/.julia/packages/AdvancedHMC/YWXfk/src/hamiltonian.jl:36
┌ Warning: Incorrect ϵ = NaN; ϵ_previous = 0.07500000000000001 is used instead.
└ @ AdvancedHMC.Adaptation /Users/kcft114/.julia/packages/AdvancedHMC/YWXfk/src/adaptation/stepsize.jl:103
┌ Warning: The current proposal will be rejected due to numerical error(s).
│   isfiniteθ = true
│   isfiniter = true
│   isfiniteℓπ = false
│   isfiniteℓκ = true
└ @ AdvancedHMC /Users/kcft114/.julia/packages/AdvancedHMC/YWXfk/src/hamiltonian.jl:36
┌ Warning: The current proposal will be rejected due to numerical error(s).
│   isfiniteθ = false
│   isfiniter = false
│   isfiniteℓπ = false
│   isfiniteℓκ = false
└ @ AdvancedHMC /Users/kcft114/.julia/packages/AdvancedHMC/YWXfk/src/hamiltonian.jl:36
┌ Warning: Incorrect ϵ = NaN; ϵ_previous = 0.07500000000000001 is used instead.
└ @ AdvancedHMC.Adaptation /Users/kcft114/.julia/packages/AdvancedHMC/YWXfk/src/adaptation/stepsize.jl:103
┌ Warning: The current proposal will be rejected due to numerical error(s).
│   isfiniteθ = true
│   isfiniter = true
│   isfiniteℓπ = false
│   isfiniteℓκ = true
└ @ AdvancedHMC /Users/kcft114/.julia/packages/AdvancedHMC/YWXfk/src/hamiltonian.jl:36
┌ Warning: The current proposal will be rejected due to numerical error(s).
│   isfiniteθ = false
│   isfiniter = false
│   isfiniteℓπ = false
│   isfiniteℓκ = false
└ @ AdvancedHMC /Users/kcft114/.julia/packages/AdvancedHMC/YWXfk/src/hamiltonian.jl:36
┌ Warning: Incorrect ϵ = NaN; ϵ_previous = 0.07500000000000001 is used instead.
└ @ AdvancedHMC.Adaptation /Users/kcft114/.julia/packages/AdvancedHMC/YWXfk/src/adaptation/stepsize.jl:103
┌ Warning: The current proposal will be rejected due to numerical error(s).
│   isfiniteθ = true
│   isfiniter = true
│   isfiniteℓπ = false
│   isfiniteℓκ = true
└ @ AdvancedHMC /Users/kcft114/.julia/packages/AdvancedHMC/YWXfk/src/hamiltonian.jl:36
┌ Warning: The current proposal will be rejected due to numerical error(s).
│   isfiniteθ = false
│   isfiniter = false
│   isfiniteℓπ = false
│   isfiniteℓκ = false
└ @ AdvancedHMC /Users/kcft114/.julia/packages/AdvancedHMC/YWXfk/src/hamiltonian.jl:36
┌ Warning: Incorrect ϵ = NaN; ϵ_previous = 0.07500000000000001 is used instead.
└ @ AdvancedHMC.Adaptation /Users/kcft114/.julia/packages/AdvancedHMC/YWXfk/src/adaptation/stepsize.jl:103
┌ Warning: The current proposal will be rejected due to numerical error(s).
│   isfiniteθ = true
│   isfiniter = true
│   isfiniteℓπ = false
│   isfiniteℓκ = true
└ @ AdvancedHMC /Users/kcft114/.julia/packages/AdvancedHMC/YWXfk/src/hamiltonian.jl:36
┌ Warning: The current proposal will be rejected due to numerical error(s).
│   isfiniteθ = false
│   isfiniter = false
│   isfiniteℓπ = false
│   isfiniteℓκ = false
└ @ AdvancedHMC /Users/kcft114/.julia/packages/AdvancedHMC/YWXfk/src/hamiltonian.jl:36
┌ Warning: Incorrect ϵ = NaN; ϵ_previous = 0.07500000000000001 is used instead.
└ @ AdvancedHMC.Adaptation /Users/kcft114/.julia/packages/AdvancedHMC/YWXfk/src/adaptation/stepsize.jl:103
┌ Warning: The current proposal will be rejected due to numerical error(s).
│   isfiniteθ = true
│   isfiniter = true
│   isfiniteℓπ = false
│   isfiniteℓκ = true
└ @ AdvancedHMC /Users/kcft114/.julia/packages/AdvancedHMC/YWXfk/src/hamiltonian.jl:36
┌ Warning: The current proposal will be rejected due to numerical error(s).
│   isfiniteθ = false
│   isfiniter = false
│   isfiniteℓπ = false
│   isfiniteℓκ = false
└ @ AdvancedHMC /Users/kcft114/.julia/packages/AdvancedHMC/YWXfk/src/hamiltonian.jl:36
┌ Warning: Incorrect ϵ = NaN; ϵ_previous = 0.07500000000000001 is used instead.
└ @ AdvancedHMC.Adaptation /Users/kcft114/.julia/packages/AdvancedHMC/YWXfk/src/adaptation/stepsize.jl:103
┌ Warning: The current proposal will be rejected due to numerical error(s).
│   isfiniteθ = true
│   isfiniter = true
│   isfiniteℓπ = false
│   isfiniteℓκ = true
└ @ AdvancedHMC /Users/kcft114/.julia/packages/AdvancedHMC/YWXfk/src/hamiltonian.jl:36
┌ Warning: The current proposal will be rejected due to numerical error(s).
│   isfiniteθ = false
│   isfiniter = false
│   isfiniteℓπ = false
│   isfiniteℓκ = false
└ @ AdvancedHMC /Users/kcft114/.julia/packages/AdvancedHMC/YWXfk/src/hamiltonian.jl:36
┌ Warning: Incorrect ϵ = NaN; ϵ_previous = 0.07500000000000001 is used instead.
└ @ AdvancedHMC.Adaptation /Users/kcft114/.julia/packages/AdvancedHMC/YWXfk/src/adaptation/stepsize.jl:103
┌ Warning: The current proposal will be rejected due to numerical error(s).
│   isfiniteθ = true
│   isfiniter = true
│   isfiniteℓπ = false
│   isfiniteℓκ = true
└ @ AdvancedHMC /Users/kcft114/.julia/packages/AdvancedHMC/YWXfk/src/hamiltonian.jl:36
┌ Warning: The current proposal will be rejected due to numerical error(s).
│   isfiniteθ = false
│   isfiniter = false
│   isfiniteℓπ = false
│   isfiniteℓκ = false
└ @ AdvancedHMC /Users/kcft114/.julia/packages/AdvancedHMC/YWXfk/src/hamiltonian.jl:36
┌ Warning: Incorrect ϵ = NaN; ϵ_previous = 0.07500000000000001 is used instead.
└ @ AdvancedHMC.Adaptation /Users/kcft114/.julia/packages/AdvancedHMC/YWXfk/src/adaptation/stepsize.jl:103
┌ Warning: The current proposal will be rejected due to numerical error(s).
│   isfiniteθ = true
│   isfiniter = true
│   isfiniteℓπ = false
│   isfiniteℓκ = true
└ @ AdvancedHMC /Users/kcft114/.julia/packages/AdvancedHMC/YWXfk/src/hamiltonian.jl:36
┌ Warning: The current proposal will be rejected due to numerical error(s).
│   isfiniteθ = false
│   isfiniter = false
│   isfiniteℓπ = false
│   isfiniteℓκ = false
└ @ AdvancedHMC /Users/kcft114/.julia/packages/AdvancedHMC/YWXfk/src/hamiltonian.jl:36
┌ Warning: Incorrect ϵ = NaN; ϵ_previous = 0.07500000000000001 is used instead.
└ @ AdvancedHMC.Adaptation /Users/kcft114/.julia/packages/AdvancedHMC/YWXfk/src/adaptation/stepsize.jl:103
┌ Warning: The current proposal will be rejected due to numerical error(s).
│   isfiniteθ = true
│   isfiniter = true
│   isfiniteℓπ = false
│   isfiniteℓκ = true
└ @ AdvancedHMC /Users/kcft114/.julia/packages/AdvancedHMC/YWXfk/src/hamiltonian.jl:36
┌ Warning: The current proposal will be rejected due to numerical error(s).
│   isfiniteθ = false
│   isfiniter = false
│   isfiniteℓπ = false
│   isfiniteℓκ = false
└ @ AdvancedHMC /Users/kcft114/.julia/packages/AdvancedHMC/YWXfk/src/hamiltonian.jl:36
┌ Warning: Incorrect ϵ = NaN; ϵ_previous = 0.07500000000000001 is used instead.
└ @ AdvancedHMC.Adaptation /Users/kcft114/.julia/packages/AdvancedHMC/YWXfk/src/adaptation/stepsize.jl:103
┌ Warning: The current proposal will be rejected due to numerical error(s).
│   isfiniteθ = true
│   isfiniter = true
│   isfiniteℓπ = false
│   isfiniteℓκ = true
└ @ AdvancedHMC /Users/kcft114/.julia/packages/AdvancedHMC/YWXfk/src/hamiltonian.jl:36
┌ Warning: The current proposal will be rejected due to numerical error(s).
│   isfiniteθ = false
│   isfiniter = false
│   isfiniteℓπ = false
│   isfiniteℓκ = false
└ @ AdvancedHMC /Users/kcft114/.julia/packages/AdvancedHMC/YWXfk/src/hamiltonian.jl:36
┌ Warning: Incorrect ϵ = NaN; ϵ_previous = 0.07500000000000001 is used instead.
└ @ AdvancedHMC.Adaptation /Users/kcft114/.julia/packages/AdvancedHMC/YWXfk/src/adaptation/stepsize.jl:103
┌ Warning: The current proposal will be rejected due to numerical error(s).
│   isfiniteθ = true
│   isfiniter = true
│   isfiniteℓπ = false
│   isfiniteℓκ = true
└ @ AdvancedHMC /Users/kcft114/.julia/packages/AdvancedHMC/YWXfk/src/hamiltonian.jl:36
┌ Warning: The current proposal will be rejected due to numerical error(s).
│   isfiniteθ = false
│   isfiniter = false
│   isfiniteℓπ = false
│   isfiniteℓκ = false
└ @ AdvancedHMC /Users/kcft114/.julia/packages/AdvancedHMC/YWXfk/src/hamiltonian.jl:36
┌ Warning: Incorrect ϵ = NaN; ϵ_previous = 0.07500000000000001 is used instead.
└ @ AdvancedHMC.Adaptation /Users/kcft114/.julia/packages/AdvancedHMC/YWXfk/src/adaptation/stepsize.jl:103
┌ Warning: The current proposal will be rejected due to numerical error(s).
│   isfiniteθ = true
│   isfiniter = true
│   isfiniteℓπ = false
│   isfiniteℓκ = true
└ @ AdvancedHMC /Users/kcft114/.julia/packages/AdvancedHMC/YWXfk/src/hamiltonian.jl:36
┌ Warning: The current proposal will be rejected due to numerical error(s).
│   isfiniteθ = false
│   isfiniter = false
│   isfiniteℓπ = false
│   isfiniteℓκ = false
└ @ AdvancedHMC /Users/kcft114/.julia/packages/AdvancedHMC/YWXfk/src/hamiltonian.jl:36
┌ Warning: Incorrect ϵ = NaN; ϵ_previous = 0.07500000000000001 is used instead.
└ @ AdvancedHMC.Adaptation /Users/kcft114/.julia/packages/AdvancedHMC/YWXfk/src/adaptation/stepsize.jl:103
┌ Warning: The current proposal will be rejected due to numerical error(s).
│   isfiniteθ = true
│   isfiniter = true
│   isfiniteℓπ = false
│   isfiniteℓκ = true
└ @ AdvancedHMC /Users/kcft114/.julia/packages/AdvancedHMC/YWXfk/src/hamiltonian.jl:36
┌ Warning: The current proposal will be rejected due to numerical error(s).
│   isfiniteθ = false
│   isfiniter = false
│   isfiniteℓπ = false
│   isfiniteℓκ = false
└ @ AdvancedHMC /Users/kcft114/.julia/packages/AdvancedHMC/YWXfk/src/hamiltonian.jl:36
┌ Warning: Incorrect ϵ = NaN; ϵ_previous = 0.07500000000000001 is used instead.
└ @ AdvancedHMC.Adaptation /Users/kcft114/.julia/packages/AdvancedHMC/YWXfk/src/adaptation/stepsize.jl:103
┌ Warning: The current proposal will be rejected due to numerical error(s).
│   isfiniteθ = true
│   isfiniter = true
│   isfiniteℓπ = false
│   isfiniteℓκ = true
└ @ AdvancedHMC /Users/kcft114/.julia/packages/AdvancedHMC/YWXfk/src/hamiltonian.jl:36
┌ Warning: The current proposal will be rejected due to numerical error(s).
│   isfiniteθ = false
│   isfiniter = false
│   isfiniteℓπ = false
│   isfiniteℓκ = false
└ @ AdvancedHMC /Users/kcft114/.julia/packages/AdvancedHMC/YWXfk/src/hamiltonian.jl:36
┌ Warning: Incorrect ϵ = NaN; ϵ_previous = 0.07500000000000001 is used instead.
└ @ AdvancedHMC.Adaptation /Users/kcft114/.julia/packages/AdvancedHMC/YWXfk/src/adaptation/stepsize.jl:103
┌ Warning: The current proposal will be rejected due to numerical error(s).
│   isfiniteθ = true
│   isfiniter = true
│   isfiniteℓπ = false
│   isfiniteℓκ = true
└ @ AdvancedHMC /Users/kcft114/.julia/packages/AdvancedHMC/YWXfk/src/hamiltonian.jl:36
┌ Warning: The current proposal will be rejected due to numerical error(s).
│   isfiniteθ = false
│   isfiniter = false
│   isfiniteℓπ = false
│   isfiniteℓκ = false
└ @ AdvancedHMC /Users/kcft114/.julia/packages/AdvancedHMC/YWXfk/src/hamiltonian.jl:36
┌ Warning: Incorrect ϵ = NaN; ϵ_previous = 0.07500000000000001 is used instead.
└ @ AdvancedHMC.Adaptation /Users/kcft114/.julia/packages/AdvancedHMC/YWXfk/src/adaptation/stepsize.jl:103
┌ Warning: The current proposal will be rejected due to numerical error(s).
│   isfiniteθ = true
│   isfiniter = true
│   isfiniteℓπ = false
│   isfiniteℓκ = true
└ @ AdvancedHMC /Users/kcft114/.julia/packages/AdvancedHMC/YWXfk/src/hamiltonian.jl:36
┌ Warning: The current proposal will be rejected due to numerical error(s).
│   isfiniteθ = false
│   isfiniter = false
│   isfiniteℓπ = false
│   isfiniteℓκ = false
└ @ AdvancedHMC /Users/kcft114/.julia/packages/AdvancedHMC/YWXfk/src/hamiltonian.jl:36
┌ Warning: Incorrect ϵ = NaN; ϵ_previous = 0.07500000000000001 is used instead.
└ @ AdvancedHMC.Adaptation /Users/kcft114/.julia/packages/AdvancedHMC/YWXfk/src/adaptation/stepsize.jl:103
┌ Warning: The current proposal will be rejected due to numerical error(s).
│   isfiniteθ = true
│   isfiniter = true
│   isfiniteℓπ = false
│   isfiniteℓκ = true
└ @ AdvancedHMC /Users/kcft114/.julia/packages/AdvancedHMC/YWXfk/src/hamiltonian.jl:36
┌ Warning: The current proposal will be rejected due to numerical error(s).
│   isfiniteθ = false
│   isfiniter = false
│   isfiniteℓπ = false
│   isfiniteℓκ = false
└ @ AdvancedHMC /Users/kcft114/.julia/packages/AdvancedHMC/YWXfk/src/hamiltonian.jl:36
┌ Warning: Incorrect ϵ = NaN; ϵ_previous = 0.07500000000000001 is used instead.
└ @ AdvancedHMC.Adaptation /Users/kcft114/.julia/packages/AdvancedHMC/YWXfk/src/adaptation/stepsize.jl:103
┌ Warning: The current proposal will be rejected due to numerical error(s).
│   isfiniteθ = true
│   isfiniter = true
│   isfiniteℓπ = false
│   isfiniteℓκ = true
└ @ AdvancedHMC /Users/kcft114/.julia/packages/AdvancedHMC/YWXfk/src/hamiltonian.jl:36
┌ Warning: The current proposal will be rejected due to numerical error(s).
│   isfiniteθ = false
│   isfiniter = false
│   isfiniteℓπ = false
│   isfiniteℓκ = false
└ @ AdvancedHMC /Users/kcft114/.julia/packages/AdvancedHMC/YWXfk/src/hamiltonian.jl:36
┌ Warning: Incorrect ϵ = NaN; ϵ_previous = 0.07500000000000001 is used instead.
└ @ AdvancedHMC.Adaptation /Users/kcft114/.julia/packages/AdvancedHMC/YWXfk/src/adaptation/stepsize.jl:103
┌ Warning: The current proposal will be rejected due to numerical error(s).
│   isfiniteθ = true
│   isfiniter = true
│   isfiniteℓπ = false
│   isfiniteℓκ = true
└ @ AdvancedHMC /Users/kcft114/.julia/packages/AdvancedHMC/YWXfk/src/hamiltonian.jl:36
┌ Warning: The current proposal will be rejected due to numerical error(s).
│   isfiniteθ = false
│   isfiniter = false
│   isfiniteℓπ = false
│   isfiniteℓκ = false
└ @ AdvancedHMC /Users/kcft114/.julia/packages/AdvancedHMC/YWXfk/src/hamiltonian.jl:36
┌ Warning: Incorrect ϵ = NaN; ϵ_previous = 0.07500000000000001 is used instead.
└ @ AdvancedHMC.Adaptation /Users/kcft114/.julia/packages/AdvancedHMC/YWXfk/src/adaptation/stepsize.jl:103
┌ Warning: The current proposal will be rejected due to numerical error(s).
│   isfiniteθ = true
│   isfiniter = true
│   isfiniteℓπ = false
│   isfiniteℓκ = true
└ @ AdvancedHMC /Users/kcft114/.julia/packages/AdvancedHMC/YWXfk/src/hamiltonian.jl:36
┌ Warning: The current proposal will be rejected due to numerical error(s).
│   isfiniteθ = false
│   isfiniter = false
│   isfiniteℓπ = false
│   isfiniteℓκ = false
└ @ AdvancedHMC /Users/kcft114/.julia/packages/AdvancedHMC/YWXfk/src/hamiltonian.jl:36
┌ Warning: Incorrect ϵ = NaN; ϵ_previous = 0.07500000000000001 is used instead.
└ @ AdvancedHMC.Adaptation /Users/kcft114/.julia/packages/AdvancedHMC/YWXfk/src/adaptation/stepsize.jl:103
┌ Warning: The current proposal will be rejected due to numerical error(s).
│   isfiniteθ = true
│   isfiniter = true
│   isfiniteℓπ = false
│   isfiniteℓκ = true
└ @ AdvancedHMC /Users/kcft114/.julia/packages/AdvancedHMC/YWXfk/src/hamiltonian.jl:36
┌ Warning: The current proposal will be rejected due to numerical error(s).
│   isfiniteθ = false
│   isfiniter = false
│   isfiniteℓπ = false
│   isfiniteℓκ = false
└ @ AdvancedHMC /Users/kcft114/.julia/packages/AdvancedHMC/YWXfk/src/hamiltonian.jl:36
┌ Warning: Incorrect ϵ = NaN; ϵ_previous = 0.07500000000000001 is used instead.
└ @ AdvancedHMC.Adaptation /Users/kcft114/.julia/packages/AdvancedHMC/YWXfk/src/adaptation/stepsize.jl:103
┌ Warning: The current proposal will be rejected due to numerical error(s).
│   isfiniteθ = true
│   isfiniter = true
│   isfiniteℓπ = false
│   isfiniteℓκ = true
└ @ AdvancedHMC /Users/kcft114/.julia/packages/AdvancedHMC/YWXfk/src/hamiltonian.jl:36
┌ Warning: The current proposal will be rejected due to numerical error(s).
│   isfiniteθ = false
│   isfiniter = false
│   isfiniteℓπ = false
│   isfiniteℓκ = false
└ @ AdvancedHMC /Users/kcft114/.julia/packages/AdvancedHMC/YWXfk/src/hamiltonian.jl:36
┌ Warning: Incorrect ϵ = NaN; ϵ_previous = 0.07500000000000001 is used instead.
└ @ AdvancedHMC.Adaptation /Users/kcft114/.julia/packages/AdvancedHMC/YWXfk/src/adaptation/stepsize.jl:103
┌ Warning: The current proposal will be rejected due to numerical error(s).
│   isfiniteθ = true
│   isfiniter = true
│   isfiniteℓπ = false
│   isfiniteℓκ = true
└ @ AdvancedHMC /Users/kcft114/.julia/packages/AdvancedHMC/YWXfk/src/hamiltonian.jl:36
┌ Warning: The current proposal will be rejected due to numerical error(s).
│   isfiniteθ = false
│   isfiniter = false
│   isfiniteℓπ = false
│   isfiniteℓκ = false
└ @ AdvancedHMC /Users/kcft114/.julia/packages/AdvancedHMC/YWXfk/src/hamiltonian.jl:36
┌ Warning: Incorrect ϵ = NaN; ϵ_previous = 0.07500000000000001 is used instead.
└ @ AdvancedHMC.Adaptation /Users/kcft114/.julia/packages/AdvancedHMC/YWXfk/src/adaptation/stepsize.jl:103
┌ Warning: The current proposal will be rejected due to numerical error(s).
│   isfiniteθ = true
│   isfiniter = true
│   isfiniteℓπ = false
│   isfiniteℓκ = true
└ @ AdvancedHMC /Users/kcft114/.julia/packages/AdvancedHMC/YWXfk/src/hamiltonian.jl:36
┌ Warning: The current proposal will be rejected due to numerical error(s).
│   isfiniteθ = false
│   isfiniter = false
│   isfiniteℓπ = false
│   isfiniteℓκ = false
└ @ AdvancedHMC /Users/kcft114/.julia/packages/AdvancedHMC/YWXfk/src/hamiltonian.jl:36
┌ Warning: Incorrect ϵ = NaN; ϵ_previous = 0.07500000000000001 is used instead.
└ @ AdvancedHMC.Adaptation /Users/kcft114/.julia/packages/AdvancedHMC/YWXfk/src/adaptation/stepsize.jl:103
┌ Warning: The current proposal will be rejected due to numerical error(s).
│   isfiniteθ = true
│   isfiniter = true
│   isfiniteℓπ = false
│   isfiniteℓκ = true
└ @ AdvancedHMC /Users/kcft114/.julia/packages/AdvancedHMC/YWXfk/src/hamiltonian.jl:36
┌ Warning: The current proposal will be rejected due to numerical error(s).
│   isfiniteθ = false
│   isfiniter = false
│   isfiniteℓπ = false
│   isfiniteℓκ = false
└ @ AdvancedHMC /Users/kcft114/.julia/packages/AdvancedHMC/YWXfk/src/hamiltonian.jl:36
┌ Warning: Incorrect ϵ = NaN; ϵ_previous = 0.07500000000000001 is used instead.
└ @ AdvancedHMC.Adaptation /Users/kcft114/.julia/packages/AdvancedHMC/YWXfk/src/adaptation/stepsize.jl:103
┌ Warning: The current proposal will be rejected due to numerical error(s).
│   isfiniteθ = true
│   isfiniter = true
│   isfiniteℓπ = false
│   isfiniteℓκ = true
└ @ AdvancedHMC /Users/kcft114/.julia/packages/AdvancedHMC/YWXfk/src/hamiltonian.jl:36
┌ Warning: The current proposal will be rejected due to numerical error(s).
│   isfiniteθ = false
│   isfiniter = false
│   isfiniteℓπ = false
│   isfiniteℓκ = false
└ @ AdvancedHMC /Users/kcft114/.julia/packages/AdvancedHMC/YWXfk/src/hamiltonian.jl:36
┌ Warning: Incorrect ϵ = NaN; ϵ_previous = 0.07500000000000001 is used instead.
└ @ AdvancedHMC.Adaptation /Users/kcft114/.julia/packages/AdvancedHMC/YWXfk/src/adaptation/stepsize.jl:103
┌ Warning: The current proposal will be rejected due to numerical error(s).
│   isfiniteθ = true
│   isfiniter = true
│   isfiniteℓπ = false
│   isfiniteℓκ = true
└ @ AdvancedHMC /Users/kcft114/.julia/packages/AdvancedHMC/YWXfk/src/hamiltonian.jl:36
┌ Warning: The current proposal will be rejected due to numerical error(s).
│   isfiniteθ = false
│   isfiniter = false
│   isfiniteℓπ = false
│   isfiniteℓκ = false
└ @ AdvancedHMC /Users/kcft114/.julia/packages/AdvancedHMC/YWXfk/src/hamiltonian.jl:36
┌ Warning: Incorrect ϵ = NaN; ϵ_previous = 0.07500000000000001 is used instead.
└ @ AdvancedHMC.Adaptation /Users/kcft114/.julia/packages/AdvancedHMC/YWXfk/src/adaptation/stepsize.jl:103
┌ Warning: The current proposal will be rejected due to numerical error(s).
│   isfiniteθ = true
│   isfiniter = true
│   isfiniteℓπ = false
│   isfiniteℓκ = true
└ @ AdvancedHMC /Users/kcft114/.julia/packages/AdvancedHMC/YWXfk/src/hamiltonian.jl:36
┌ Warning: The current proposal will be rejected due to numerical error(s).
│   isfiniteθ = false
│   isfiniter = false
│   isfiniteℓπ = false
│   isfiniteℓκ = false
└ @ AdvancedHMC /Users/kcft114/.julia/packages/AdvancedHMC/YWXfk/src/hamiltonian.jl:36
┌ Warning: Incorrect ϵ = NaN; ϵ_previous = 0.07500000000000001 is used instead.
└ @ AdvancedHMC.Adaptation /Users/kcft114/.julia/packages/AdvancedHMC/YWXfk/src/adaptation/stepsize.jl:103
┌ Warning: The current proposal will be rejected due to numerical error(s).
│   isfiniteθ = true
│   isfiniter = true
│   isfiniteℓπ = false
│   isfiniteℓκ = true
└ @ AdvancedHMC /Users/kcft114/.julia/packages/AdvancedHMC/YWXfk/src/hamiltonian.jl:36
┌ Warning: The current proposal will be rejected due to numerical error(s).
│   isfiniteθ = false
│   isfiniter = false
│   isfiniteℓπ = false
│   isfiniteℓκ = false
└ @ AdvancedHMC /Users/kcft114/.julia/packages/AdvancedHMC/YWXfk/src/hamiltonian.jl:36
┌ Warning: Incorrect ϵ = NaN; ϵ_previous = 0.07500000000000001 is used instead.
└ @ AdvancedHMC.Adaptation /Users/kcft114/.julia/packages/AdvancedHMC/YWXfk/src/adaptation/stepsize.jl:103
┌ Warning: The current proposal will be rejected due to numerical error(s).
│   isfiniteθ = true
│   isfiniter = true
│   isfiniteℓπ = false
│   isfiniteℓκ = true
└ @ AdvancedHMC /Users/kcft114/.julia/packages/AdvancedHMC/YWXfk/src/hamiltonian.jl:36
┌ Warning: The current proposal will be rejected due to numerical error(s).
│   isfiniteθ = false
│   isfiniter = false
│   isfiniteℓπ = false
│   isfiniteℓκ = false
└ @ AdvancedHMC /Users/kcft114/.julia/packages/AdvancedHMC/YWXfk/src/hamiltonian.jl:36
┌ Warning: Incorrect ϵ = NaN; ϵ_previous = 0.07500000000000001 is used instead.
└ @ AdvancedHMC.Adaptation /Users/kcft114/.julia/packages/AdvancedHMC/YWXfk/src/adaptation/stepsize.jl:103
┌ Warning: The current proposal will be rejected due to numerical error(s).
│   isfiniteθ = true
│   isfiniter = true
│   isfiniteℓπ = false
│   isfiniteℓκ = true
└ @ AdvancedHMC /Users/kcft114/.julia/packages/AdvancedHMC/YWXfk/src/hamiltonian.jl:36
┌ Warning: The current proposal will be rejected due to numerical error(s).
│   isfiniteθ = false
│   isfiniter = false
│   isfiniteℓπ = false
│   isfiniteℓκ = false
└ @ AdvancedHMC /Users/kcft114/.julia/packages/AdvancedHMC/YWXfk/src/hamiltonian.jl:36
┌ Warning: Incorrect ϵ = NaN; ϵ_previous = 0.07500000000000001 is used instead.
└ @ AdvancedHMC.Adaptation /Users/kcft114/.julia/packages/AdvancedHMC/YWXfk/src/adaptation/stepsize.jl:103
┌ Warning: The current proposal will be rejected due to numerical error(s).
│   isfiniteθ = true
│   isfiniter = true
│   isfiniteℓπ = false
│   isfiniteℓκ = true
└ @ AdvancedHMC /Users/kcft114/.julia/packages/AdvancedHMC/YWXfk/src/hamiltonian.jl:36
┌ Warning: The current proposal will be rejected due to numerical error(s).
│   isfiniteθ = false
│   isfiniter = false
│   isfiniteℓπ = false
│   isfiniteℓκ = false
└ @ AdvancedHMC /Users/kcft114/.julia/packages/AdvancedHMC/YWXfk/src/hamiltonian.jl:36
┌ Warning: Incorrect ϵ = NaN; ϵ_previous = 0.07500000000000001 is used instead.
└ @ AdvancedHMC.Adaptation /Users/kcft114/.julia/packages/AdvancedHMC/YWXfk/src/adaptation/stepsize.jl:103
┌ Warning: The current proposal will be rejected due to numerical error(s).
│   isfiniteθ = true
│   isfiniter = true
│   isfiniteℓπ = false
│   isfiniteℓκ = true
└ @ AdvancedHMC /Users/kcft114/.julia/packages/AdvancedHMC/YWXfk/src/hamiltonian.jl:36
┌ Warning: The current proposal will be rejected due to numerical error(s).
│   isfiniteθ = false
│   isfiniter = false
│   isfiniteℓπ = false
│   isfiniteℓκ = false
└ @ AdvancedHMC /Users/kcft114/.julia/packages/AdvancedHMC/YWXfk/src/hamiltonian.jl:36
┌ Warning: Incorrect ϵ = NaN; ϵ_previous = 0.07500000000000001 is used instead.
└ @ AdvancedHMC.Adaptation /Users/kcft114/.julia/packages/AdvancedHMC/YWXfk/src/adaptation/stepsize.jl:103
┌ Warning: The current proposal will be rejected due to numerical error(s).
│   isfiniteθ = true
│   isfiniter = true
│   isfiniteℓπ = false
│   isfiniteℓκ = true
└ @ AdvancedHMC /Users/kcft114/.julia/packages/AdvancedHMC/YWXfk/src/hamiltonian.jl:36
┌ Warning: The current proposal will be rejected due to numerical error(s).
│   isfiniteθ = false
│   isfiniter = false
│   isfiniteℓπ = false
│   isfiniteℓκ = false
└ @ AdvancedHMC /Users/kcft114/.julia/packages/AdvancedHMC/YWXfk/src/hamiltonian.jl:36
┌ Warning: Incorrect ϵ = NaN; ϵ_previous = 0.07500000000000001 is used instead.
└ @ AdvancedHMC.Adaptation /Users/kcft114/.julia/packages/AdvancedHMC/YWXfk/src/adaptation/stepsize.jl:103
┌ Warning: The current proposal will be rejected due to numerical error(s).
│   isfiniteθ = true
│   isfiniter = true
│   isfiniteℓπ = false
│   isfiniteℓκ = true
└ @ AdvancedHMC /Users/kcft114/.julia/packages/AdvancedHMC/YWXfk/src/hamiltonian.jl:36
┌ Warning: The current proposal will be rejected due to numerical error(s).
│   isfiniteθ = false
│   isfiniter = false
│   isfiniteℓπ = false
│   isfiniteℓκ = false
└ @ AdvancedHMC /Users/kcft114/.julia/packages/AdvancedHMC/YWXfk/src/hamiltonian.jl:36
┌ Warning: Incorrect ϵ = NaN; ϵ_previous = 0.07500000000000001 is used instead.
└ @ AdvancedHMC.Adaptation /Users/kcft114/.julia/packages/AdvancedHMC/YWXfk/src/adaptation/stepsize.jl:103
┌ Warning: The current proposal will be rejected due to numerical error(s).
│   isfiniteθ = true
│   isfiniter = true
│   isfiniteℓπ = false
│   isfiniteℓκ = true
└ @ AdvancedHMC /Users/kcft114/.julia/packages/AdvancedHMC/YWXfk/src/hamiltonian.jl:36
┌ Warning: The current proposal will be rejected due to numerical error(s).
│   isfiniteθ = false
│   isfiniter = false
│   isfiniteℓπ = false
│   isfiniteℓκ = false
└ @ AdvancedHMC /Users/kcft114/.julia/packages/AdvancedHMC/YWXfk/src/hamiltonian.jl:36
┌ Warning: Incorrect ϵ = NaN; ϵ_previous = 0.07500000000000001 is used instead.
└ @ AdvancedHMC.Adaptation /Users/kcft114/.julia/packages/AdvancedHMC/YWXfk/src/adaptation/stepsize.jl:103
┌ Warning: The current proposal will be rejected due to numerical error(s).
│   isfiniteθ = true
│   isfiniter = true
│   isfiniteℓπ = false
│   isfiniteℓκ = true
└ @ AdvancedHMC /Users/kcft114/.julia/packages/AdvancedHMC/YWXfk/src/hamiltonian.jl:36
┌ Warning: The current proposal will be rejected due to numerical error(s).
│   isfiniteθ = false
│   isfiniter = false
│   isfiniteℓπ = false
│   isfiniteℓκ = false
└ @ AdvancedHMC /Users/kcft114/.julia/packages/AdvancedHMC/YWXfk/src/hamiltonian.jl:36
┌ Warning: Incorrect ϵ = NaN; ϵ_previous = 0.07500000000000001 is used instead.
└ @ AdvancedHMC.Adaptation /Users/kcft114/.julia/packages/AdvancedHMC/YWXfk/src/adaptation/stepsize.jl:103
┌ Warning: The current proposal will be rejected due to numerical error(s).
│   isfiniteθ = true
│   isfiniter = true
│   isfiniteℓπ = false
│   isfiniteℓκ = true
└ @ AdvancedHMC /Users/kcft114/.julia/packages/AdvancedHMC/YWXfk/src/hamiltonian.jl:36
┌ Warning: The current proposal will be rejected due to numerical error(s).
│   isfiniteθ = false
│   isfiniter = false
│   isfiniteℓπ = false
│   isfiniteℓκ = false
└ @ AdvancedHMC /Users/kcft114/.julia/packages/AdvancedHMC/YWXfk/src/hamiltonian.jl:36
┌ Warning: Incorrect ϵ = NaN; ϵ_previous = 0.07500000000000001 is used instead.
└ @ AdvancedHMC.Adaptation /Users/kcft114/.julia/packages/AdvancedHMC/YWXfk/src/adaptation/stepsize.jl:103
┌ Warning: The current proposal will be rejected due to numerical error(s).
│   isfiniteθ = true
│   isfiniter = true
│   isfiniteℓπ = false
│   isfiniteℓκ = true
└ @ AdvancedHMC /Users/kcft114/.julia/packages/AdvancedHMC/YWXfk/src/hamiltonian.jl:36
┌ Warning: The current proposal will be rejected due to numerical error(s).
│   isfiniteθ = false
│   isfiniter = false
│   isfiniteℓπ = false
│   isfiniteℓκ = false
└ @ AdvancedHMC /Users/kcft114/.julia/packages/AdvancedHMC/YWXfk/src/hamiltonian.jl:36
┌ Warning: Incorrect ϵ = NaN; ϵ_previous = 0.07500000000000001 is used instead.
└ @ AdvancedHMC.Adaptation /Users/kcft114/.julia/packages/AdvancedHMC/YWXfk/src/adaptation/stepsize.jl:103
┌ Warning: The current proposal will be rejected due to numerical error(s).
│   isfiniteθ = true
│   isfiniter = true
│   isfiniteℓπ = false
│   isfiniteℓκ = true
└ @ AdvancedHMC /Users/kcft114/.julia/packages/AdvancedHMC/YWXfk/src/hamiltonian.jl:36
┌ Warning: The current proposal will be rejected due to numerical error(s).
│   isfiniteθ = false
│   isfiniter = false
│   isfiniteℓπ = false
│   isfiniteℓκ = false
└ @ AdvancedHMC /Users/kcft114/.julia/packages/AdvancedHMC/YWXfk/src/hamiltonian.jl:36
┌ Warning: Incorrect ϵ = NaN; ϵ_previous = 0.07500000000000001 is used instead.
└ @ AdvancedHMC.Adaptation /Users/kcft114/.julia/packages/AdvancedHMC/YWXfk/src/adaptation/stepsize.jl:103
┌ Warning: The current proposal will be rejected due to numerical error(s).
│   isfiniteθ = true
│   isfiniter = true
│   isfiniteℓπ = false
│   isfiniteℓκ = true
└ @ AdvancedHMC /Users/kcft114/.julia/packages/AdvancedHMC/YWXfk/src/hamiltonian.jl:36
┌ Warning: The current proposal will be rejected due to numerical error(s).
│   isfiniteθ = false
│   isfiniter = false
│   isfiniteℓπ = false
│   isfiniteℓκ = false
└ @ AdvancedHMC /Users/kcft114/.julia/packages/AdvancedHMC/YWXfk/src/hamiltonian.jl:36
┌ Warning: Incorrect ϵ = NaN; ϵ_previous = 0.07500000000000001 is used instead.
└ @ AdvancedHMC.Adaptation /Users/kcft114/.julia/packages/AdvancedHMC/YWXfk/src/adaptation/stepsize.jl:103
┌ Warning: The current proposal will be rejected due to numerical error(s).
│   isfiniteθ = true
│   isfiniter = true
│   isfiniteℓπ = false
│   isfiniteℓκ = true
└ @ AdvancedHMC /Users/kcft114/.julia/packages/AdvancedHMC/YWXfk/src/hamiltonian.jl:36
┌ Warning: The current proposal will be rejected due to numerical error(s).
│   isfiniteθ = false
│   isfiniter = false
│   isfiniteℓπ = false
│   isfiniteℓκ = false
└ @ AdvancedHMC /Users/kcft114/.julia/packages/AdvancedHMC/YWXfk/src/hamiltonian.jl:36
┌ Warning: Incorrect ϵ = NaN; ϵ_previous = 0.07500000000000001 is used instead.
└ @ AdvancedHMC.Adaptation /Users/kcft114/.julia/packages/AdvancedHMC/YWXfk/src/adaptation/stepsize.jl:103
┌ Warning: The current proposal will be rejected due to numerical error(s).
│   isfiniteθ = true
│   isfiniter = true
│   isfiniteℓπ = false
│   isfiniteℓκ = true
└ @ AdvancedHMC /Users/kcft114/.julia/packages/AdvancedHMC/YWXfk/src/hamiltonian.jl:36
┌ Warning: The current proposal will be rejected due to numerical error(s).
│   isfiniteθ = false
│   isfiniter = false
│   isfiniteℓπ = false
│   isfiniteℓκ = false
└ @ AdvancedHMC /Users/kcft114/.julia/packages/AdvancedHMC/YWXfk/src/hamiltonian.jl:36
┌ Warning: Incorrect ϵ = NaN; ϵ_previous = 0.07500000000000001 is used instead.
└ @ AdvancedHMC.Adaptation /Users/kcft114/.julia/packages/AdvancedHMC/YWXfk/src/adaptation/stepsize.jl:103
┌ Warning: The current proposal will be rejected due to numerical error(s).
│   isfiniteθ = true
│   isfiniter = true
│   isfiniteℓπ = false
│   isfiniteℓκ = true
└ @ AdvancedHMC /Users/kcft114/.julia/packages/AdvancedHMC/YWXfk/src/hamiltonian.jl:36
┌ Warning: The current proposal will be rejected due to numerical error(s).
│   isfiniteθ = false
│   isfiniter = false
│   isfiniteℓπ = false
│   isfiniteℓκ = false
└ @ AdvancedHMC /Users/kcft114/.julia/packages/AdvancedHMC/YWXfk/src/hamiltonian.jl:36
┌ Warning: Incorrect ϵ = NaN; ϵ_previous = 0.07500000000000001 is used instead.
└ @ AdvancedHMC.Adaptation /Users/kcft114/.julia/packages/AdvancedHMC/YWXfk/src/adaptation/stepsize.jl:103
┌ Warning: The current proposal will be rejected due to numerical error(s).
│   isfiniteθ = true
│   isfiniter = true
│   isfiniteℓπ = false
│   isfiniteℓκ = true
└ @ AdvancedHMC /Users/kcft114/.julia/packages/AdvancedHMC/YWXfk/src/hamiltonian.jl:36
┌ Warning: The current proposal will be rejected due to numerical error(s).
│   isfiniteθ = false
│   isfiniter = false
│   isfiniteℓπ = false
│   isfiniteℓκ = false
└ @ AdvancedHMC /Users/kcft114/.julia/packages/AdvancedHMC/YWXfk/src/hamiltonian.jl:36
┌ Warning: Incorrect ϵ = NaN; ϵ_previous = 0.07500000000000001 is used instead.
└ @ AdvancedHMC.Adaptation /Users/kcft114/.julia/packages/AdvancedHMC/YWXfk/src/adaptation/stepsize.jl:103
┌ Warning: The current proposal will be rejected due to numerical error(s).
│   isfiniteθ = true
│   isfiniter = true
│   isfiniteℓπ = false
│   isfiniteℓκ = true
└ @ AdvancedHMC /Users/kcft114/.julia/packages/AdvancedHMC/YWXfk/src/hamiltonian.jl:36
┌ Warning: The current proposal will be rejected due to numerical error(s).
│   isfiniteθ = false
│   isfiniter = false
│   isfiniteℓπ = false
│   isfiniteℓκ = false
└ @ AdvancedHMC /Users/kcft114/.julia/packages/AdvancedHMC/YWXfk/src/hamiltonian.jl:36
┌ Warning: Incorrect ϵ = NaN; ϵ_previous = 0.07500000000000001 is used instead.
└ @ AdvancedHMC.Adaptation /Users/kcft114/.julia/packages/AdvancedHMC/YWXfk/src/adaptation/stepsize.jl:103
┌ Warning: The current proposal will be rejected due to numerical error(s).
│   isfiniteθ = true
│   isfiniter = true
│   isfiniteℓπ = false
│   isfiniteℓκ = true
└ @ AdvancedHMC /Users/kcft114/.julia/packages/AdvancedHMC/YWXfk/src/hamiltonian.jl:36
┌ Warning: The current proposal will be rejected due to numerical error(s).
│   isfiniteθ = false
│   isfiniter = false
│   isfiniteℓπ = false
│   isfiniteℓκ = false
└ @ AdvancedHMC /Users/kcft114/.julia/packages/AdvancedHMC/YWXfk/src/hamiltonian.jl:36
┌ Warning: Incorrect ϵ = NaN; ϵ_previous = 0.07500000000000001 is used instead.
└ @ AdvancedHMC.Adaptation /Users/kcft114/.julia/packages/AdvancedHMC/YWXfk/src/adaptation/stepsize.jl:103
┌ Warning: The current proposal will be rejected due to numerical error(s).
│   isfiniteθ = true
│   isfiniter = true
│   isfiniteℓπ = false
│   isfiniteℓκ = true
└ @ AdvancedHMC /Users/kcft114/.julia/packages/AdvancedHMC/YWXfk/src/hamiltonian.jl:36
┌ Warning: The current proposal will be rejected due to numerical error(s).
│   isfiniteθ = false
│   isfiniter = false
│   isfiniteℓπ = false
│   isfiniteℓκ = false
└ @ AdvancedHMC /Users/kcft114/.julia/packages/AdvancedHMC/YWXfk/src/hamiltonian.jl:36
┌ Warning: Incorrect ϵ = NaN; ϵ_previous = 0.07500000000000001 is used instead.
└ @ AdvancedHMC.Adaptation /Users/kcft114/.julia/packages/AdvancedHMC/YWXfk/src/adaptation/stepsize.jl:103
┌ Warning: The current proposal will be rejected due to numerical error(s).
│   isfiniteθ = true
│   isfiniter = true
│   isfiniteℓπ = false
│   isfiniteℓκ = true
└ @ AdvancedHMC /Users/kcft114/.julia/packages/AdvancedHMC/YWXfk/src/hamiltonian.jl:36
┌ Warning: The current proposal will be rejected due to numerical error(s).
│   isfiniteθ = false
│   isfiniter = false
│   isfiniteℓπ = false
│   isfiniteℓκ = false
└ @ AdvancedHMC /Users/kcft114/.julia/packages/AdvancedHMC/YWXfk/src/hamiltonian.jl:36
┌ Warning: Incorrect ϵ = NaN; ϵ_previous = 0.07500000000000001 is used instead.
└ @ AdvancedHMC.Adaptation /Users/kcft114/.julia/packages/AdvancedHMC/YWXfk/src/adaptation/stepsize.jl:103
┌ Warning: The current proposal will be rejected due to numerical error(s).
│   isfiniteθ = true
│   isfiniter = true
│   isfiniteℓπ = false
│   isfiniteℓκ = true
└ @ AdvancedHMC /Users/kcft114/.julia/packages/AdvancedHMC/YWXfk/src/hamiltonian.jl:36
┌ Warning: The current proposal will be rejected due to numerical error(s).
│   isfiniteθ = false
│   isfiniter = false
│   isfiniteℓπ = false
│   isfiniteℓκ = false
└ @ AdvancedHMC /Users/kcft114/.julia/packages/AdvancedHMC/YWXfk/src/hamiltonian.jl:36
┌ Warning: Incorrect ϵ = NaN; ϵ_previous = 0.07500000000000001 is used instead.
└ @ AdvancedHMC.Adaptation /Users/kcft114/.julia/packages/AdvancedHMC/YWXfk/src/adaptation/stepsize.jl:103
┌ Warning: The current proposal will be rejected due to numerical error(s).
│   isfiniteθ = true
│   isfiniter = true
│   isfiniteℓπ = false
│   isfiniteℓκ = true
└ @ AdvancedHMC /Users/kcft114/.julia/packages/AdvancedHMC/YWXfk/src/hamiltonian.jl:36
┌ Warning: The current proposal will be rejected due to numerical error(s).
│   isfiniteθ = false
│   isfiniter = false
│   isfiniteℓπ = false
│   isfiniteℓκ = false
└ @ AdvancedHMC /Users/kcft114/.julia/packages/AdvancedHMC/YWXfk/src/hamiltonian.jl:36
┌ Warning: Incorrect ϵ = NaN; ϵ_previous = 0.07500000000000001 is used instead.
└ @ AdvancedHMC.Adaptation /Users/kcft114/.julia/packages/AdvancedHMC/YWXfk/src/adaptation/stepsize.jl:103
┌ Warning: The current proposal will be rejected due to numerical error(s).
│   isfiniteθ = true
│   isfiniter = true
│   isfiniteℓπ = false
│   isfiniteℓκ = true
└ @ AdvancedHMC /Users/kcft114/.julia/packages/AdvancedHMC/YWXfk/src/hamiltonian.jl:36
┌ Warning: The current proposal will be rejected due to numerical error(s).
│   isfiniteθ = false
│   isfiniter = false
│   isfiniteℓπ = false
│   isfiniteℓκ = false
└ @ AdvancedHMC /Users/kcft114/.julia/packages/AdvancedHMC/YWXfk/src/hamiltonian.jl:36
┌ Warning: Incorrect ϵ = NaN; ϵ_previous = 0.07500000000000001 is used instead.
└ @ AdvancedHMC.Adaptation /Users/kcft114/.julia/packages/AdvancedHMC/YWXfk/src/adaptation/stepsize.jl:103
┌ Warning: The current proposal will be rejected due to numerical error(s).
│   isfiniteθ = true
│   isfiniter = true
│   isfiniteℓπ = false
│   isfiniteℓκ = true
└ @ AdvancedHMC /Users/kcft114/.julia/packages/AdvancedHMC/YWXfk/src/hamiltonian.jl:36
┌ Warning: The current proposal will be rejected due to numerical error(s).
│   isfiniteθ = false
│   isfiniter = false
│   isfiniteℓπ = false
│   isfiniteℓκ = false
└ @ AdvancedHMC /Users/kcft114/.julia/packages/AdvancedHMC/YWXfk/src/hamiltonian.jl:36
┌ Warning: Incorrect ϵ = NaN; ϵ_previous = 0.07500000000000001 is used instead.
└ @ AdvancedHMC.Adaptation /Users/kcft114/.julia/packages/AdvancedHMC/YWXfk/src/adaptation/stepsize.jl:103
┌ Warning: The current proposal will be rejected due to numerical error(s).
│   isfiniteθ = true
│   isfiniter = true
│   isfiniteℓπ = false
│   isfiniteℓκ = true
└ @ AdvancedHMC /Users/kcft114/.julia/packages/AdvancedHMC/YWXfk/src/hamiltonian.jl:36
┌ Warning: The current proposal will be rejected due to numerical error(s).
│   isfiniteθ = false
│   isfiniter = false
│   isfiniteℓπ = false
│   isfiniteℓκ = false
└ @ AdvancedHMC /Users/kcft114/.julia/packages/AdvancedHMC/YWXfk/src/hamiltonian.jl:36
┌ Warning: Incorrect ϵ = NaN; ϵ_previous = 0.07500000000000001 is used instead.
└ @ AdvancedHMC.Adaptation /Users/kcft114/.julia/packages/AdvancedHMC/YWXfk/src/adaptation/stepsize.jl:103
┌ Warning: The current proposal will be rejected due to numerical error(s).
│   isfiniteθ = true
│   isfiniter = true
│   isfiniteℓπ = false
│   isfiniteℓκ = true
└ @ AdvancedHMC /Users/kcft114/.julia/packages/AdvancedHMC/YWXfk/src/hamiltonian.jl:36
┌ Warning: The current proposal will be rejected due to numerical error(s).
│   isfiniteθ = false
│   isfiniter = false
│   isfiniteℓπ = false
│   isfiniteℓκ = false
└ @ AdvancedHMC /Users/kcft114/.julia/packages/AdvancedHMC/YWXfk/src/hamiltonian.jl:36
┌ Warning: Incorrect ϵ = NaN; ϵ_previous = 0.07500000000000001 is used instead.
└ @ AdvancedHMC.Adaptation /Users/kcft114/.julia/packages/AdvancedHMC/YWXfk/src/adaptation/stepsize.jl:103
┌ Warning: The current proposal will be rejected due to numerical error(s).
│   isfiniteθ = true
│   isfiniter = true
│   isfiniteℓπ = false
│   isfiniteℓκ = true
└ @ AdvancedHMC /Users/kcft114/.julia/packages/AdvancedHMC/YWXfk/src/hamiltonian.jl:36
┌ Warning: The current proposal will be rejected due to numerical error(s).
│   isfiniteθ = false
│   isfiniter = false
│   isfiniteℓπ = false
│   isfiniteℓκ = false
└ @ AdvancedHMC /Users/kcft114/.julia/packages/AdvancedHMC/YWXfk/src/hamiltonian.jl:36
┌ Warning: Incorrect ϵ = NaN; ϵ_previous = 0.07500000000000001 is used instead.
└ @ AdvancedHMC.Adaptation /Users/kcft114/.julia/packages/AdvancedHMC/YWXfk/src/adaptation/stepsize.jl:103
┌ Warning: The current proposal will be rejected due to numerical error(s).
│   isfiniteθ = true
│   isfiniter = true
│   isfiniteℓπ = false
│   isfiniteℓκ = true
└ @ AdvancedHMC /Users/kcft114/.julia/packages/AdvancedHMC/YWXfk/src/hamiltonian.jl:36
┌ Warning: The current proposal will be rejected due to numerical error(s).
│   isfiniteθ = false
│   isfiniter = false
│   isfiniteℓπ = false
│   isfiniteℓκ = false
└ @ AdvancedHMC /Users/kcft114/.julia/packages/AdvancedHMC/YWXfk/src/hamiltonian.jl:36
┌ Warning: Incorrect ϵ = NaN; ϵ_previous = 0.07500000000000001 is used instead.
└ @ AdvancedHMC.Adaptation /Users/kcft114/.julia/packages/AdvancedHMC/YWXfk/src/adaptation/stepsize.jl:103
┌ Warning: The current proposal will be rejected due to numerical error(s).
│   isfiniteθ = true
│   isfiniter = true
│   isfiniteℓπ = false
│   isfiniteℓκ = true
└ @ AdvancedHMC /Users/kcft114/.julia/packages/AdvancedHMC/YWXfk/src/hamiltonian.jl:36
┌ Warning: The current proposal will be rejected due to numerical error(s).
│   isfiniteθ = false
│   isfiniter = false
│   isfiniteℓπ = false
│   isfiniteℓκ = false
└ @ AdvancedHMC /Users/kcft114/.julia/packages/AdvancedHMC/YWXfk/src/hamiltonian.jl:36
┌ Warning: Incorrect ϵ = NaN; ϵ_previous = 0.07500000000000001 is used instead.
└ @ AdvancedHMC.Adaptation /Users/kcft114/.julia/packages/AdvancedHMC/YWXfk/src/adaptation/stepsize.jl:103
┌ Warning: The current proposal will be rejected due to numerical error(s).
│   isfiniteθ = true
│   isfiniter = true
│   isfiniteℓπ = false
│   isfiniteℓκ = true
└ @ AdvancedHMC /Users/kcft114/.julia/packages/AdvancedHMC/YWXfk/src/hamiltonian.jl:36
┌ Warning: The current proposal will be rejected due to numerical error(s).
│   isfiniteθ = false
│   isfiniter = false
│   isfiniteℓπ = false
│   isfiniteℓκ = false
└ @ AdvancedHMC /Users/kcft114/.julia/packages/AdvancedHMC/YWXfk/src/hamiltonian.jl:36
┌ Warning: Incorrect ϵ = NaN; ϵ_previous = 0.07500000000000001 is used instead.
└ @ AdvancedHMC.Adaptation /Users/kcft114/.julia/packages/AdvancedHMC/YWXfk/src/adaptation/stepsize.jl:103
┌ Warning: The current proposal will be rejected due to numerical error(s).
│   isfiniteθ = true
│   isfiniter = true
│   isfiniteℓπ = false
│   isfiniteℓκ = true
└ @ AdvancedHMC /Users/kcft114/.julia/packages/AdvancedHMC/YWXfk/src/hamiltonian.jl:36
┌ Warning: The current proposal will be rejected due to numerical error(s).
│   isfiniteθ = false
│   isfiniter = false
│   isfiniteℓπ = false
│   isfiniteℓκ = false
└ @ AdvancedHMC /Users/kcft114/.julia/packages/AdvancedHMC/YWXfk/src/hamiltonian.jl:36
┌ Warning: Incorrect ϵ = NaN; ϵ_previous = 0.07500000000000001 is used instead.
└ @ AdvancedHMC.Adaptation /Users/kcft114/.julia/packages/AdvancedHMC/YWXfk/src/adaptation/stepsize.jl:103
┌ Warning: The current proposal will be rejected due to numerical error(s).
│   isfiniteθ = true
│   isfiniter = true
│   isfiniteℓπ = false
│   isfiniteℓκ = true
└ @ AdvancedHMC /Users/kcft114/.julia/packages/AdvancedHMC/YWXfk/src/hamiltonian.jl:36
┌ Warning: The current proposal will be rejected due to numerical error(s).
│   isfiniteθ = false
│   isfiniter = false
│   isfiniteℓπ = false
│   isfiniteℓκ = false
└ @ AdvancedHMC /Users/kcft114/.julia/packages/AdvancedHMC/YWXfk/src/hamiltonian.jl:36
┌ Warning: Incorrect ϵ = NaN; ϵ_previous = 0.07500000000000001 is used instead.
└ @ AdvancedHMC.Adaptation /Users/kcft114/.julia/packages/AdvancedHMC/YWXfk/src/adaptation/stepsize.jl:103
┌ Warning: The current proposal will be rejected due to numerical error(s).
│   isfiniteθ = true
│   isfiniter = true
│   isfiniteℓπ = false
│   isfiniteℓκ = true
└ @ AdvancedHMC /Users/kcft114/.julia/packages/AdvancedHMC/YWXfk/src/hamiltonian.jl:36
┌ Warning: The current proposal will be rejected due to numerical error(s).
│   isfiniteθ = false
│   isfiniter = false
│   isfiniteℓπ = false
│   isfiniteℓκ = false
└ @ AdvancedHMC /Users/kcft114/.julia/packages/AdvancedHMC/YWXfk/src/hamiltonian.jl:36
┌ Warning: Incorrect ϵ = NaN; ϵ_previous = 0.07500000000000001 is used instead.
└ @ AdvancedHMC.Adaptation /Users/kcft114/.julia/packages/AdvancedHMC/YWXfk/src/adaptation/stepsize.jl:103
┌ Warning: The current proposal will be rejected due to numerical error(s).
│   isfiniteθ = true
│   isfiniter = true
│   isfiniteℓπ = false
│   isfiniteℓκ = true
└ @ AdvancedHMC /Users/kcft114/.julia/packages/AdvancedHMC/YWXfk/src/hamiltonian.jl:36
┌ Warning: The current proposal will be rejected due to numerical error(s).
│   isfiniteθ = false
│   isfiniter = false
│   isfiniteℓπ = false
│   isfiniteℓκ = false
└ @ AdvancedHMC /Users/kcft114/.julia/packages/AdvancedHMC/YWXfk/src/hamiltonian.jl:36
┌ Warning: Incorrect ϵ = NaN; ϵ_previous = 0.07500000000000001 is used instead.
└ @ AdvancedHMC.Adaptation /Users/kcft114/.julia/packages/AdvancedHMC/YWXfk/src/adaptation/stepsize.jl:103
┌ Warning: The current proposal will be rejected due to numerical error(s).
│   isfiniteθ = true
│   isfiniter = true
│   isfiniteℓπ = false
│   isfiniteℓκ = true
└ @ AdvancedHMC /Users/kcft114/.julia/packages/AdvancedHMC/YWXfk/src/hamiltonian.jl:36
┌ Warning: The current proposal will be rejected due to numerical error(s).
│   isfiniteθ = false
│   isfiniter = false
│   isfiniteℓπ = false
│   isfiniteℓκ = false
└ @ AdvancedHMC /Users/kcft114/.julia/packages/AdvancedHMC/YWXfk/src/hamiltonian.jl:36
┌ Warning: Incorrect ϵ = NaN; ϵ_previous = 0.07500000000000001 is used instead.
└ @ AdvancedHMC.Adaptation /Users/kcft114/.julia/packages/AdvancedHMC/YWXfk/src/adaptation/stepsize.jl:103
┌ Warning: The current proposal will be rejected due to numerical error(s).
│   isfiniteθ = true
│   isfiniter = true
│   isfiniteℓπ = false
│   isfiniteℓκ = true
└ @ AdvancedHMC /Users/kcft114/.julia/packages/AdvancedHMC/YWXfk/src/hamiltonian.jl:36
┌ Warning: The current proposal will be rejected due to numerical error(s).
│   isfiniteθ = false
│   isfiniter = false
│   isfiniteℓπ = false
│   isfiniteℓκ = false
└ @ AdvancedHMC /Users/kcft114/.julia/packages/AdvancedHMC/YWXfk/src/hamiltonian.jl:36
┌ Warning: Incorrect ϵ = NaN; ϵ_previous = 0.07500000000000001 is used instead.
└ @ AdvancedHMC.Adaptation /Users/kcft114/.julia/packages/AdvancedHMC/YWXfk/src/adaptation/stepsize.jl:103
┌ Warning: The current proposal will be rejected due to numerical error(s).
│   isfiniteθ = true
│   isfiniter = true
│   isfiniteℓπ = false
│   isfiniteℓκ = true
└ @ AdvancedHMC /Users/kcft114/.julia/packages/AdvancedHMC/YWXfk/src/hamiltonian.jl:36
┌ Warning: The current proposal will be rejected due to numerical error(s).
│   isfiniteθ = false
│   isfiniter = false
│   isfiniteℓπ = false
│   isfiniteℓκ = false
└ @ AdvancedHMC /Users/kcft114/.julia/packages/AdvancedHMC/YWXfk/src/hamiltonian.jl:36
┌ Warning: Incorrect ϵ = NaN; ϵ_previous = 0.07500000000000001 is used instead.
└ @ AdvancedHMC.Adaptation /Users/kcft114/.julia/packages/AdvancedHMC/YWXfk/src/adaptation/stepsize.jl:103
┌ Warning: The current proposal will be rejected due to numerical error(s).
│   isfiniteθ = true
│   isfiniter = true
│   isfiniteℓπ = false
│   isfiniteℓκ = true
└ @ AdvancedHMC /Users/kcft114/.julia/packages/AdvancedHMC/YWXfk/src/hamiltonian.jl:36
┌ Warning: The current proposal will be rejected due to numerical error(s).
│   isfiniteθ = false
│   isfiniter = false
│   isfiniteℓπ = false
│   isfiniteℓκ = false
└ @ AdvancedHMC /Users/kcft114/.julia/packages/AdvancedHMC/YWXfk/src/hamiltonian.jl:36
┌ Warning: Incorrect ϵ = NaN; ϵ_previous = 0.07500000000000001 is used instead.
└ @ AdvancedHMC.Adaptation /Users/kcft114/.julia/packages/AdvancedHMC/YWXfk/src/adaptation/stepsize.jl:103
┌ Warning: The current proposal will be rejected due to numerical error(s).
│   isfiniteθ = true
│   isfiniter = true
│   isfiniteℓπ = false
│   isfiniteℓκ = true
└ @ AdvancedHMC /Users/kcft114/.julia/packages/AdvancedHMC/YWXfk/src/hamiltonian.jl:36
┌ Warning: The current proposal will be rejected due to numerical error(s).
│   isfiniteθ = false
│   isfiniter = false
│   isfiniteℓπ = false
│   isfiniteℓκ = false
└ @ AdvancedHMC /Users/kcft114/.julia/packages/AdvancedHMC/YWXfk/src/hamiltonian.jl:36
┌ Warning: Incorrect ϵ = NaN; ϵ_previous = 0.07500000000000001 is used instead.
└ @ AdvancedHMC.Adaptation /Users/kcft114/.julia/packages/AdvancedHMC/YWXfk/src/adaptation/stepsize.jl:103
┌ Warning: The current proposal will be rejected due to numerical error(s).
│   isfiniteθ = true
│   isfiniter = true
│   isfiniteℓπ = false
│   isfiniteℓκ = true
└ @ AdvancedHMC /Users/kcft114/.julia/packages/AdvancedHMC/YWXfk/src/hamiltonian.jl:36
┌ Warning: The current proposal will be rejected due to numerical error(s).
│   isfiniteθ = false
│   isfiniter = false
│   isfiniteℓπ = false
│   isfiniteℓκ = false
└ @ AdvancedHMC /Users/kcft114/.julia/packages/AdvancedHMC/YWXfk/src/hamiltonian.jl:36
┌ Warning: Incorrect ϵ = NaN; ϵ_previous = 0.07500000000000001 is used instead.
└ @ AdvancedHMC.Adaptation /Users/kcft114/.julia/packages/AdvancedHMC/YWXfk/src/adaptation/stepsize.jl:103
┌ Warning: The current proposal will be rejected due to numerical error(s).
│   isfiniteθ = true
│   isfiniter = true
│   isfiniteℓπ = false
│   isfiniteℓκ = true
└ @ AdvancedHMC /Users/kcft114/.julia/packages/AdvancedHMC/YWXfk/src/hamiltonian.jl:36
┌ Warning: The current proposal will be rejected due to numerical error(s).
│   isfiniteθ = false
│   isfiniter = false
│   isfiniteℓπ = false
│   isfiniteℓκ = false
└ @ AdvancedHMC /Users/kcft114/.julia/packages/AdvancedHMC/YWXfk/src/hamiltonian.jl:36
┌ Warning: Incorrect ϵ = NaN; ϵ_previous = 0.07500000000000001 is used instead.
└ @ AdvancedHMC.Adaptation /Users/kcft114/.julia/packages/AdvancedHMC/YWXfk/src/adaptation/stepsize.jl:103
┌ Warning: The current proposal will be rejected due to numerical error(s).
│   isfiniteθ = true
│   isfiniter = true
│   isfiniteℓπ = false
│   isfiniteℓκ = true
└ @ AdvancedHMC /Users/kcft114/.julia/packages/AdvancedHMC/YWXfk/src/hamiltonian.jl:36
┌ Warning: The current proposal will be rejected due to numerical error(s).
│   isfiniteθ = false
│   isfiniter = false
│   isfiniteℓπ = false
│   isfiniteℓκ = false
└ @ AdvancedHMC /Users/kcft114/.julia/packages/AdvancedHMC/YWXfk/src/hamiltonian.jl:36
┌ Warning: Incorrect ϵ = NaN; ϵ_previous = 0.07500000000000001 is used instead.
└ @ AdvancedHMC.Adaptation /Users/kcft114/.julia/packages/AdvancedHMC/YWXfk/src/adaptation/stepsize.jl:103
┌ Warning: The current proposal will be rejected due to numerical error(s).
│   isfiniteθ = true
│   isfiniter = true
│   isfiniteℓπ = false
│   isfiniteℓκ = true
└ @ AdvancedHMC /Users/kcft114/.julia/packages/AdvancedHMC/YWXfk/src/hamiltonian.jl:36
┌ Warning: The current proposal will be rejected due to numerical error(s).
│   isfiniteθ = false
│   isfiniter = false
│   isfiniteℓπ = false
│   isfiniteℓκ = false
└ @ AdvancedHMC /Users/kcft114/.julia/packages/AdvancedHMC/YWXfk/src/hamiltonian.jl:36
┌ Warning: Incorrect ϵ = NaN; ϵ_previous = 0.07500000000000001 is used instead.
└ @ AdvancedHMC.Adaptation /Users/kcft114/.julia/packages/AdvancedHMC/YWXfk/src/adaptation/stepsize.jl:103
┌ Warning: The current proposal will be rejected due to numerical error(s).
│   isfiniteθ = true
│   isfiniter = true
│   isfiniteℓπ = false
│   isfiniteℓκ = true
└ @ AdvancedHMC /Users/kcft114/.julia/packages/AdvancedHMC/YWXfk/src/hamiltonian.jl:36
┌ Warning: The current proposal will be rejected due to numerical error(s).
│   isfiniteθ = false
│   isfiniter = false
│   isfiniteℓπ = false
│   isfiniteℓκ = false
└ @ AdvancedHMC /Users/kcft114/.julia/packages/AdvancedHMC/YWXfk/src/hamiltonian.jl:36
┌ Warning: Incorrect ϵ = NaN; ϵ_previous = 0.07500000000000001 is used instead.
└ @ AdvancedHMC.Adaptation /Users/kcft114/.julia/packages/AdvancedHMC/YWXfk/src/adaptation/stepsize.jl:103
┌ Warning: The current proposal will be rejected due to numerical error(s).
│   isfiniteθ = true
│   isfiniter = true
│   isfiniteℓπ = false
│   isfiniteℓκ = true
└ @ AdvancedHMC /Users/kcft114/.julia/packages/AdvancedHMC/YWXfk/src/hamiltonian.jl:36
┌ Warning: The current proposal will be rejected due to numerical error(s).
│   isfiniteθ = false
│   isfiniter = false
│   isfiniteℓπ = false
│   isfiniteℓκ = false
└ @ AdvancedHMC /Users/kcft114/.julia/packages/AdvancedHMC/YWXfk/src/hamiltonian.jl:36
┌ Warning: Incorrect ϵ = NaN; ϵ_previous = 0.07500000000000001 is used instead.
└ @ AdvancedHMC.Adaptation /Users/kcft114/.julia/packages/AdvancedHMC/YWXfk/src/adaptation/stepsize.jl:103
┌ Warning: The current proposal will be rejected due to numerical error(s).
│   isfiniteθ = true
│   isfiniter = true
│   isfiniteℓπ = false
│   isfiniteℓκ = true
└ @ AdvancedHMC /Users/kcft114/.julia/packages/AdvancedHMC/YWXfk/src/hamiltonian.jl:36
┌ Warning: The current proposal will be rejected due to numerical error(s).
│   isfiniteθ = false
│   isfiniter = false
│   isfiniteℓπ = false
│   isfiniteℓκ = false
└ @ AdvancedHMC /Users/kcft114/.julia/packages/AdvancedHMC/YWXfk/src/hamiltonian.jl:36
┌ Warning: Incorrect ϵ = NaN; ϵ_previous = 0.07500000000000001 is used instead.
└ @ AdvancedHMC.Adaptation /Users/kcft114/.julia/packages/AdvancedHMC/YWXfk/src/adaptation/stepsize.jl:103
┌ Warning: The current proposal will be rejected due to numerical error(s).
│   isfiniteθ = true
│   isfiniter = true
│   isfiniteℓπ = false
│   isfiniteℓκ = true
└ @ AdvancedHMC /Users/kcft114/.julia/packages/AdvancedHMC/YWXfk/src/hamiltonian.jl:36
┌ Warning: The current proposal will be rejected due to numerical error(s).
│   isfiniteθ = false
│   isfiniter = false
│   isfiniteℓπ = false
│   isfiniteℓκ = false
└ @ AdvancedHMC /Users/kcft114/.julia/packages/AdvancedHMC/YWXfk/src/hamiltonian.jl:36
┌ Warning: Incorrect ϵ = NaN; ϵ_previous = 0.07500000000000001 is used instead.
└ @ AdvancedHMC.Adaptation /Users/kcft114/.julia/packages/AdvancedHMC/YWXfk/src/adaptation/stepsize.jl:103
┌ Warning: The current proposal will be rejected due to numerical error(s).
│   isfiniteθ = true
│   isfiniter = true
│   isfiniteℓπ = false
│   isfiniteℓκ = true
└ @ AdvancedHMC /Users/kcft114/.julia/packages/AdvancedHMC/YWXfk/src/hamiltonian.jl:36
┌ Warning: The current proposal will be rejected due to numerical error(s).
│   isfiniteθ = false
│   isfiniter = false
│   isfiniteℓπ = false
│   isfiniteℓκ = false
└ @ AdvancedHMC /Users/kcft114/.julia/packages/AdvancedHMC/YWXfk/src/hamiltonian.jl:36
┌ Warning: Incorrect ϵ = NaN; ϵ_previous = 0.07500000000000001 is used instead.
└ @ AdvancedHMC.Adaptation /Users/kcft114/.julia/packages/AdvancedHMC/YWXfk/src/adaptation/stepsize.jl:103
┌ Warning: The current proposal will be rejected due to numerical error(s).
│   isfiniteθ = true
│   isfiniter = true
│   isfiniteℓπ = false
│   isfiniteℓκ = true
└ @ AdvancedHMC /Users/kcft114/.julia/packages/AdvancedHMC/YWXfk/src/hamiltonian.jl:36
┌ Warning: The current proposal will be rejected due to numerical error(s).
│   isfiniteθ = false
│   isfiniter = false
│   isfiniteℓπ = false
│   isfiniteℓκ = false
└ @ AdvancedHMC /Users/kcft114/.julia/packages/AdvancedHMC/YWXfk/src/hamiltonian.jl:36
┌ Warning: Incorrect ϵ = NaN; ϵ_previous = 0.07500000000000001 is used instead.
└ @ AdvancedHMC.Adaptation /Users/kcft114/.julia/packages/AdvancedHMC/YWXfk/src/adaptation/stepsize.jl:103
┌ Warning: The current proposal will be rejected due to numerical error(s).
│   isfiniteθ = true
│   isfiniter = true
│   isfiniteℓπ = false
│   isfiniteℓκ = true
└ @ AdvancedHMC /Users/kcft114/.julia/packages/AdvancedHMC/YWXfk/src/hamiltonian.jl:36
┌ Warning: The current proposal will be rejected due to numerical error(s).
│   isfiniteθ = false
│   isfiniter = false
│   isfiniteℓπ = false
│   isfiniteℓκ = false
└ @ AdvancedHMC /Users/kcft114/.julia/packages/AdvancedHMC/YWXfk/src/hamiltonian.jl:36
┌ Warning: Incorrect ϵ = NaN; ϵ_previous = 0.07500000000000001 is used instead.
└ @ AdvancedHMC.Adaptation /Users/kcft114/.julia/packages/AdvancedHMC/YWXfk/src/adaptation/stepsize.jl:103
┌ Warning: The current proposal will be rejected due to numerical error(s).
│   isfiniteθ = true
│   isfiniter = true
│   isfiniteℓπ = false
│   isfiniteℓκ = true
└ @ AdvancedHMC /Users/kcft114/.julia/packages/AdvancedHMC/YWXfk/src/hamiltonian.jl:36
┌ Warning: The current proposal will be rejected due to numerical error(s).
│   isfiniteθ = false
│   isfiniter = false
│   isfiniteℓπ = false
│   isfiniteℓκ = false
└ @ AdvancedHMC /Users/kcft114/.julia/packages/AdvancedHMC/YWXfk/src/hamiltonian.jl:36
┌ Warning: Incorrect ϵ = NaN; ϵ_previous = 0.07500000000000001 is used instead.
└ @ AdvancedHMC.Adaptation /Users/kcft114/.julia/packages/AdvancedHMC/YWXfk/src/adaptation/stepsize.jl:103
┌ Warning: The current proposal will be rejected due to numerical error(s).
│   isfiniteθ = true
│   isfiniter = true
│   isfiniteℓπ = false
│   isfiniteℓκ = true
└ @ AdvancedHMC /Users/kcft114/.julia/packages/AdvancedHMC/YWXfk/src/hamiltonian.jl:36
┌ Warning: The current proposal will be rejected due to numerical error(s).
│   isfiniteθ = false
│   isfiniter = false
│   isfiniteℓπ = false
│   isfiniteℓκ = false
└ @ AdvancedHMC /Users/kcft114/.julia/packages/AdvancedHMC/YWXfk/src/hamiltonian.jl:36
┌ Warning: Incorrect ϵ = NaN; ϵ_previous = 0.07500000000000001 is used instead.
└ @ AdvancedHMC.Adaptation /Users/kcft114/.julia/packages/AdvancedHMC/YWXfk/src/adaptation/stepsize.jl:103
┌ Warning: The current proposal will be rejected due to numerical error(s).
│   isfiniteθ = true
│   isfiniter = true
│   isfiniteℓπ = false
│   isfiniteℓκ = true
└ @ AdvancedHMC /Users/kcft114/.julia/packages/AdvancedHMC/YWXfk/src/hamiltonian.jl:36
┌ Warning: The current proposal will be rejected due to numerical error(s).
│   isfiniteθ = false
│   isfiniter = false
│   isfiniteℓπ = false
│   isfiniteℓκ = false
└ @ AdvancedHMC /Users/kcft114/.julia/packages/AdvancedHMC/YWXfk/src/hamiltonian.jl:36
┌ Warning: Incorrect ϵ = NaN; ϵ_previous = 0.07500000000000001 is used instead.
└ @ AdvancedHMC.Adaptation /Users/kcft114/.julia/packages/AdvancedHMC/YWXfk/src/adaptation/stepsize.jl:103
┌ Warning: The current proposal will be rejected due to numerical error(s).
│   isfiniteθ = true
│   isfiniter = true
│   isfiniteℓπ = false
│   isfiniteℓκ = true
└ @ AdvancedHMC /Users/kcft114/.julia/packages/AdvancedHMC/YWXfk/src/hamiltonian.jl:36
┌ Warning: The current proposal will be rejected due to numerical error(s).
│   isfiniteθ = false
│   isfiniter = false
│   isfiniteℓπ = false
│   isfiniteℓκ = false
└ @ AdvancedHMC /Users/kcft114/.julia/packages/AdvancedHMC/YWXfk/src/hamiltonian.jl:36
┌ Warning: Incorrect ϵ = NaN; ϵ_previous = 0.07500000000000001 is used instead.
└ @ AdvancedHMC.Adaptation /Users/kcft114/.julia/packages/AdvancedHMC/YWXfk/src/adaptation/stepsize.jl:103
┌ Warning: The current proposal will be rejected due to numerical error(s).
│   isfiniteθ = true
│   isfiniter = true
│   isfiniteℓπ = false
│   isfiniteℓκ = true
└ @ AdvancedHMC /Users/kcft114/.julia/packages/AdvancedHMC/YWXfk/src/hamiltonian.jl:36
┌ Warning: The current proposal will be rejected due to numerical error(s).
│   isfiniteθ = false
│   isfiniter = false
│   isfiniteℓπ = false
│   isfiniteℓκ = false
└ @ AdvancedHMC /Users/kcft114/.julia/packages/AdvancedHMC/YWXfk/src/hamiltonian.jl:36
┌ Warning: Incorrect ϵ = NaN; ϵ_previous = 0.07500000000000001 is used instead.
└ @ AdvancedHMC.Adaptation /Users/kcft114/.julia/packages/AdvancedHMC/YWXfk/src/adaptation/stepsize.jl:103
┌ Warning: The current proposal will be rejected due to numerical error(s).
│   isfiniteθ = true
│   isfiniter = true
│   isfiniteℓπ = false
│   isfiniteℓκ = true
└ @ AdvancedHMC /Users/kcft114/.julia/packages/AdvancedHMC/YWXfk/src/hamiltonian.jl:36
┌ Warning: The current proposal will be rejected due to numerical error(s).
│   isfiniteθ = false
│   isfiniter = false
│   isfiniteℓπ = false
│   isfiniteℓκ = false
└ @ AdvancedHMC /Users/kcft114/.julia/packages/AdvancedHMC/YWXfk/src/hamiltonian.jl:36
┌ Warning: Incorrect ϵ = NaN; ϵ_previous = 0.07500000000000001 is used instead.
└ @ AdvancedHMC.Adaptation /Users/kcft114/.julia/packages/AdvancedHMC/YWXfk/src/adaptation/stepsize.jl:103
┌ Warning: The current proposal will be rejected due to numerical error(s).
│   isfiniteθ = true
│   isfiniter = true
│   isfiniteℓπ = false
│   isfiniteℓκ = true
└ @ AdvancedHMC /Users/kcft114/.julia/packages/AdvancedHMC/YWXfk/src/hamiltonian.jl:36
┌ Warning: The current proposal will be rejected due to numerical error(s).
│   isfiniteθ = false
│   isfiniter = false
│   isfiniteℓπ = false
│   isfiniteℓκ = false
└ @ AdvancedHMC /Users/kcft114/.julia/packages/AdvancedHMC/YWXfk/src/hamiltonian.jl:36
┌ Warning: Incorrect ϵ = NaN; ϵ_previous = 0.07500000000000001 is used instead.
└ @ AdvancedHMC.Adaptation /Users/kcft114/.julia/packages/AdvancedHMC/YWXfk/src/adaptation/stepsize.jl:103
┌ Warning: The current proposal will be rejected due to numerical error(s).
│   isfiniteθ = true
│   isfiniter = true
│   isfiniteℓπ = false
│   isfiniteℓκ = true
└ @ AdvancedHMC /Users/kcft114/.julia/packages/AdvancedHMC/YWXfk/src/hamiltonian.jl:36
┌ Warning: The current proposal will be rejected due to numerical error(s).
│   isfiniteθ = false
│   isfiniter = false
│   isfiniteℓπ = false
│   isfiniteℓκ = false
└ @ AdvancedHMC /Users/kcft114/.julia/packages/AdvancedHMC/YWXfk/src/hamiltonian.jl:36
┌ Warning: Incorrect ϵ = NaN; ϵ_previous = 0.07500000000000001 is used instead.
└ @ AdvancedHMC.Adaptation /Users/kcft114/.julia/packages/AdvancedHMC/YWXfk/src/adaptation/stepsize.jl:103
┌ Warning: The current proposal will be rejected due to numerical error(s).
│   isfiniteθ = true
│   isfiniter = true
│   isfiniteℓπ = false
│   isfiniteℓκ = true
└ @ AdvancedHMC /Users/kcft114/.julia/packages/AdvancedHMC/YWXfk/src/hamiltonian.jl:36
┌ Warning: The current proposal will be rejected due to numerical error(s).
│   isfiniteθ = false
│   isfiniter = false
│   isfiniteℓπ = false
│   isfiniteℓκ = false
└ @ AdvancedHMC /Users/kcft114/.julia/packages/AdvancedHMC/YWXfk/src/hamiltonian.jl:36
┌ Warning: Incorrect ϵ = NaN; ϵ_previous = 0.07500000000000001 is used instead.
└ @ AdvancedHMC.Adaptation /Users/kcft114/.julia/packages/AdvancedHMC/YWXfk/src/adaptation/stepsize.jl:103
┌ Warning: The current proposal will be rejected due to numerical error(s).
│   isfiniteθ = true
│   isfiniter = true
│   isfiniteℓπ = false
│   isfiniteℓκ = true
└ @ AdvancedHMC /Users/kcft114/.julia/packages/AdvancedHMC/YWXfk/src/hamiltonian.jl:36
┌ Warning: The current proposal will be rejected due to numerical error(s).
│   isfiniteθ = false
│   isfiniter = false
│   isfiniteℓπ = false
│   isfiniteℓκ = false
└ @ AdvancedHMC /Users/kcft114/.julia/packages/AdvancedHMC/YWXfk/src/hamiltonian.jl:36
┌ Warning: Incorrect ϵ = NaN; ϵ_previous = 0.07500000000000001 is used instead.
└ @ AdvancedHMC.Adaptation /Users/kcft114/.julia/packages/AdvancedHMC/YWXfk/src/adaptation/stepsize.jl:103
┌ Warning: The current proposal will be rejected due to numerical error(s).
│   isfiniteθ = true
│   isfiniter = true
│   isfiniteℓπ = false
│   isfiniteℓκ = true
└ @ AdvancedHMC /Users/kcft114/.julia/packages/AdvancedHMC/YWXfk/src/hamiltonian.jl:36
┌ Warning: The current proposal will be rejected due to numerical error(s).
│   isfiniteθ = false
│   isfiniter = false
│   isfiniteℓπ = false
│   isfiniteℓκ = false
└ @ AdvancedHMC /Users/kcft114/.julia/packages/AdvancedHMC/YWXfk/src/hamiltonian.jl:36
┌ Warning: Incorrect ϵ = NaN; ϵ_previous = 0.07500000000000001 is used instead.
└ @ AdvancedHMC.Adaptation /Users/kcft114/.julia/packages/AdvancedHMC/YWXfk/src/adaptation/stepsize.jl:103
┌ Warning: The current proposal will be rejected due to numerical error(s).
│   isfiniteθ = true
│   isfiniter = true
│   isfiniteℓπ = false
│   isfiniteℓκ = true
└ @ AdvancedHMC /Users/kcft114/.julia/packages/AdvancedHMC/YWXfk/src/hamiltonian.jl:36
┌ Warning: The current proposal will be rejected due to numerical error(s).
│   isfiniteθ = false
│   isfiniter = false
│   isfiniteℓπ = false
│   isfiniteℓκ = false
└ @ AdvancedHMC /Users/kcft114/.julia/packages/AdvancedHMC/YWXfk/src/hamiltonian.jl:36
┌ Warning: Incorrect ϵ = NaN; ϵ_previous = 0.07500000000000001 is used instead.
└ @ AdvancedHMC.Adaptation /Users/kcft114/.julia/packages/AdvancedHMC/YWXfk/src/adaptation/stepsize.jl:103
┌ Warning: The current proposal will be rejected due to numerical error(s).
│   isfiniteθ = true
│   isfiniter = true
│   isfiniteℓπ = false
│   isfiniteℓκ = true
└ @ AdvancedHMC /Users/kcft114/.julia/packages/AdvancedHMC/YWXfk/src/hamiltonian.jl:36
┌ Warning: The current proposal will be rejected due to numerical error(s).
│   isfiniteθ = false
│   isfiniter = false
│   isfiniteℓπ = false
│   isfiniteℓκ = false
└ @ AdvancedHMC /Users/kcft114/.julia/packages/AdvancedHMC/YWXfk/src/hamiltonian.jl:36
┌ Warning: Incorrect ϵ = NaN; ϵ_previous = 0.07500000000000001 is used instead.
└ @ AdvancedHMC.Adaptation /Users/kcft114/.julia/packages/AdvancedHMC/YWXfk/src/adaptation/stepsize.jl:103
┌ Warning: The current proposal will be rejected due to numerical error(s).
│   isfiniteθ = true
│   isfiniter = true
│   isfiniteℓπ = false
│   isfiniteℓκ = true
└ @ AdvancedHMC /Users/kcft114/.julia/packages/AdvancedHMC/YWXfk/src/hamiltonian.jl:36
┌ Warning: The current proposal will be rejected due to numerical error(s).
│   isfiniteθ = false
│   isfiniter = false
│   isfiniteℓπ = false
│   isfiniteℓκ = false
└ @ AdvancedHMC /Users/kcft114/.julia/packages/AdvancedHMC/YWXfk/src/hamiltonian.jl:36
┌ Warning: Incorrect ϵ = NaN; ϵ_previous = 0.07500000000000001 is used instead.
└ @ AdvancedHMC.Adaptation /Users/kcft114/.julia/packages/AdvancedHMC/YWXfk/src/adaptation/stepsize.jl:103
┌ Warning: The current proposal will be rejected due to numerical error(s).
│   isfiniteθ = true
│   isfiniter = true
│   isfiniteℓπ = false
│   isfiniteℓκ = true
└ @ AdvancedHMC /Users/kcft114/.julia/packages/AdvancedHMC/YWXfk/src/hamiltonian.jl:36
┌ Warning: The current proposal will be rejected due to numerical error(s).
│   isfiniteθ = false
│   isfiniter = false
│   isfiniteℓπ = false
│   isfiniteℓκ = false
└ @ AdvancedHMC /Users/kcft114/.julia/packages/AdvancedHMC/YWXfk/src/hamiltonian.jl:36
┌ Warning: Incorrect ϵ = NaN; ϵ_previous = 0.07500000000000001 is used instead.
└ @ AdvancedHMC.Adaptation /Users/kcft114/.julia/packages/AdvancedHMC/YWXfk/src/adaptation/stepsize.jl:103
┌ Warning: The current proposal will be rejected due to numerical error(s).
│   isfiniteθ = true
│   isfiniter = true
│   isfiniteℓπ = false
│   isfiniteℓκ = true
└ @ AdvancedHMC /Users/kcft114/.julia/packages/AdvancedHMC/YWXfk/src/hamiltonian.jl:36
┌ Warning: The current proposal will be rejected due to numerical error(s).
│   isfiniteθ = false
│   isfiniter = false
│   isfiniteℓπ = false
│   isfiniteℓκ = false
└ @ AdvancedHMC /Users/kcft114/.julia/packages/AdvancedHMC/YWXfk/src/hamiltonian.jl:36
┌ Warning: Incorrect ϵ = NaN; ϵ_previous = 0.07500000000000001 is used instead.
└ @ AdvancedHMC.Adaptation /Users/kcft114/.julia/packages/AdvancedHMC/YWXfk/src/adaptation/stepsize.jl:103
┌ Warning: The current proposal will be rejected due to numerical error(s).
│   isfiniteθ = true
│   isfiniter = true
│   isfiniteℓπ = false
│   isfiniteℓκ = true
└ @ AdvancedHMC /Users/kcft114/.julia/packages/AdvancedHMC/YWXfk/src/hamiltonian.jl:36
┌ Warning: The current proposal will be rejected due to numerical error(s).
│   isfiniteθ = false
│   isfiniter = false
│   isfiniteℓπ = false
│   isfiniteℓκ = false
└ @ AdvancedHMC /Users/kcft114/.julia/packages/AdvancedHMC/YWXfk/src/hamiltonian.jl:36
┌ Warning: Incorrect ϵ = NaN; ϵ_previous = 0.07500000000000001 is used instead.
└ @ AdvancedHMC.Adaptation /Users/kcft114/.julia/packages/AdvancedHMC/YWXfk/src/adaptation/stepsize.jl:103
┌ Warning: The current proposal will be rejected due to numerical error(s).
│   isfiniteθ = true
│   isfiniter = true
│   isfiniteℓπ = false
│   isfiniteℓκ = true
└ @ AdvancedHMC /Users/kcft114/.julia/packages/AdvancedHMC/YWXfk/src/hamiltonian.jl:36
┌ Warning: The current proposal will be rejected due to numerical error(s).
│   isfiniteθ = false
│   isfiniter = false
│   isfiniteℓπ = false
│   isfiniteℓκ = false
└ @ AdvancedHMC /Users/kcft114/.julia/packages/AdvancedHMC/YWXfk/src/hamiltonian.jl:36
┌ Warning: Incorrect ϵ = NaN; ϵ_previous = 0.07500000000000001 is used instead.
└ @ AdvancedHMC.Adaptation /Users/kcft114/.julia/packages/AdvancedHMC/YWXfk/src/adaptation/stepsize.jl:103
┌ Warning: The current proposal will be rejected due to numerical error(s).
│   isfiniteθ = true
│   isfiniter = true
│   isfiniteℓπ = false
│   isfiniteℓκ = true
└ @ AdvancedHMC /Users/kcft114/.julia/packages/AdvancedHMC/YWXfk/src/hamiltonian.jl:36
┌ Warning: The current proposal will be rejected due to numerical error(s).
│   isfiniteθ = false
│   isfiniter = false
│   isfiniteℓπ = false
│   isfiniteℓκ = false
└ @ AdvancedHMC /Users/kcft114/.julia/packages/AdvancedHMC/YWXfk/src/hamiltonian.jl:36
┌ Warning: Incorrect ϵ = NaN; ϵ_previous = 0.07500000000000001 is used instead.
└ @ AdvancedHMC.Adaptation /Users/kcft114/.julia/packages/AdvancedHMC/YWXfk/src/adaptation/stepsize.jl:103
┌ Warning: The current proposal will be rejected due to numerical error(s).
│   isfiniteθ = true
│   isfiniter = true
│   isfiniteℓπ = false
│   isfiniteℓκ = true
└ @ AdvancedHMC /Users/kcft114/.julia/packages/AdvancedHMC/YWXfk/src/hamiltonian.jl:36
┌ Warning: The current proposal will be rejected due to numerical error(s).
│   isfiniteθ = false
│   isfiniter = false
│   isfiniteℓπ = false
│   isfiniteℓκ = false
└ @ AdvancedHMC /Users/kcft114/.julia/packages/AdvancedHMC/YWXfk/src/hamiltonian.jl:36
┌ Warning: Incorrect ϵ = NaN; ϵ_previous = 0.07500000000000001 is used instead.
└ @ AdvancedHMC.Adaptation /Users/kcft114/.julia/packages/AdvancedHMC/YWXfk/src/adaptation/stepsize.jl:103
┌ Warning: The current proposal will be rejected due to numerical error(s).
│   isfiniteθ = true
│   isfiniter = true
│   isfiniteℓπ = false
│   isfiniteℓκ = true
└ @ AdvancedHMC /Users/kcft114/.julia/packages/AdvancedHMC/YWXfk/src/hamiltonian.jl:36
┌ Warning: The current proposal will be rejected due to numerical error(s).
│   isfiniteθ = false
│   isfiniter = false
│   isfiniteℓπ = false
│   isfiniteℓκ = false
└ @ AdvancedHMC /Users/kcft114/.julia/packages/AdvancedHMC/YWXfk/src/hamiltonian.jl:36
┌ Warning: Incorrect ϵ = NaN; ϵ_previous = 0.07500000000000001 is used instead.
└ @ AdvancedHMC.Adaptation /Users/kcft114/.julia/packages/AdvancedHMC/YWXfk/src/adaptation/stepsize.jl:103
┌ Warning: The current proposal will be rejected due to numerical error(s).
│   isfiniteθ = true
│   isfiniter = true
│   isfiniteℓπ = false
│   isfiniteℓκ = true
└ @ AdvancedHMC /Users/kcft114/.julia/packages/AdvancedHMC/YWXfk/src/hamiltonian.jl:36
┌ Warning: The current proposal will be rejected due to numerical error(s).
│   isfiniteθ = false
│   isfiniter = false
│   isfiniteℓπ = false
│   isfiniteℓκ = false
└ @ AdvancedHMC /Users/kcft114/.julia/packages/AdvancedHMC/YWXfk/src/hamiltonian.jl:36
┌ Warning: Incorrect ϵ = NaN; ϵ_previous = 0.07500000000000001 is used instead.
└ @ AdvancedHMC.Adaptation /Users/kcft114/.julia/packages/AdvancedHMC/YWXfk/src/adaptation/stepsize.jl:103
┌ Warning: The current proposal will be rejected due to numerical error(s).
│   isfiniteθ = true
│   isfiniter = true
│   isfiniteℓπ = false
│   isfiniteℓκ = true
└ @ AdvancedHMC /Users/kcft114/.julia/packages/AdvancedHMC/YWXfk/src/hamiltonian.jl:36
┌ Warning: The current proposal will be rejected due to numerical error(s).
│   isfiniteθ = false
│   isfiniter = false
│   isfiniteℓπ = false
│   isfiniteℓκ = false
└ @ AdvancedHMC /Users/kcft114/.julia/packages/AdvancedHMC/YWXfk/src/hamiltonian.jl:36
┌ Warning: Incorrect ϵ = NaN; ϵ_previous = 0.07500000000000001 is used instead.
└ @ AdvancedHMC.Adaptation /Users/kcft114/.julia/packages/AdvancedHMC/YWXfk/src/adaptation/stepsize.jl:103
┌ Warning: The current proposal will be rejected due to numerical error(s).
│   isfiniteθ = true
│   isfiniter = true
│   isfiniteℓπ = false
│   isfiniteℓκ = true
└ @ AdvancedHMC /Users/kcft114/.julia/packages/AdvancedHMC/YWXfk/src/hamiltonian.jl:36
┌ Warning: The current proposal will be rejected due to numerical error(s).
│   isfiniteθ = false
│   isfiniter = false
│   isfiniteℓπ = false
│   isfiniteℓκ = false
└ @ AdvancedHMC /Users/kcft114/.julia/packages/AdvancedHMC/YWXfk/src/hamiltonian.jl:36
┌ Warning: Incorrect ϵ = NaN; ϵ_previous = 0.07500000000000001 is used instead.
└ @ AdvancedHMC.Adaptation /Users/kcft114/.julia/packages/AdvancedHMC/YWXfk/src/adaptation/stepsize.jl:103
┌ Warning: The current proposal will be rejected due to numerical error(s).
│   isfiniteθ = true
│   isfiniter = true
│   isfiniteℓπ = false
│   isfiniteℓκ = true
└ @ AdvancedHMC /Users/kcft114/.julia/packages/AdvancedHMC/YWXfk/src/hamiltonian.jl:36
┌ Warning: The current proposal will be rejected due to numerical error(s).
│   isfiniteθ = false
│   isfiniter = false
│   isfiniteℓπ = false
│   isfiniteℓκ = false
└ @ AdvancedHMC /Users/kcft114/.julia/packages/AdvancedHMC/YWXfk/src/hamiltonian.jl:36
┌ Warning: Incorrect ϵ = NaN; ϵ_previous = 0.07500000000000001 is used instead.
└ @ AdvancedHMC.Adaptation /Users/kcft114/.julia/packages/AdvancedHMC/YWXfk/src/adaptation/stepsize.jl:103
┌ Warning: The current proposal will be rejected due to numerical error(s).
│   isfiniteθ = true
│   isfiniter = true
│   isfiniteℓπ = false
│   isfiniteℓκ = true
└ @ AdvancedHMC /Users/kcft114/.julia/packages/AdvancedHMC/YWXfk/src/hamiltonian.jl:36
┌ Warning: The current proposal will be rejected due to numerical error(s).
│   isfiniteθ = false
│   isfiniter = false
│   isfiniteℓπ = false
│   isfiniteℓκ = false
└ @ AdvancedHMC /Users/kcft114/.julia/packages/AdvancedHMC/YWXfk/src/hamiltonian.jl:36
┌ Warning: Incorrect ϵ = NaN; ϵ_previous = 0.07500000000000001 is used instead.
└ @ AdvancedHMC.Adaptation /Users/kcft114/.julia/packages/AdvancedHMC/YWXfk/src/adaptation/stepsize.jl:103
┌ Warning: The current proposal will be rejected due to numerical error(s).
│   isfiniteθ = true
│   isfiniter = true
│   isfiniteℓπ = false
│   isfiniteℓκ = true
└ @ AdvancedHMC /Users/kcft114/.julia/packages/AdvancedHMC/YWXfk/src/hamiltonian.jl:36
┌ Warning: The current proposal will be rejected due to numerical error(s).
│   isfiniteθ = false
│   isfiniter = false
│   isfiniteℓπ = false
│   isfiniteℓκ = false
└ @ AdvancedHMC /Users/kcft114/.julia/packages/AdvancedHMC/YWXfk/src/hamiltonian.jl:36
┌ Warning: Incorrect ϵ = NaN; ϵ_previous = 0.07500000000000001 is used instead.
└ @ AdvancedHMC.Adaptation /Users/kcft114/.julia/packages/AdvancedHMC/YWXfk/src/adaptation/stepsize.jl:103
┌ Warning: The current proposal will be rejected due to numerical error(s).
│   isfiniteθ = true
│   isfiniter = true
│   isfiniteℓπ = false
│   isfiniteℓκ = true
└ @ AdvancedHMC /Users/kcft114/.julia/packages/AdvancedHMC/YWXfk/src/hamiltonian.jl:36
┌ Warning: The current proposal will be rejected due to numerical error(s).
│   isfiniteθ = false
│   isfiniter = false
│   isfiniteℓπ = false
│   isfiniteℓκ = false
└ @ AdvancedHMC /Users/kcft114/.julia/packages/AdvancedHMC/YWXfk/src/hamiltonian.jl:36
┌ Warning: Incorrect ϵ = NaN; ϵ_previous = 0.07500000000000001 is used instead.
└ @ AdvancedHMC.Adaptation /Users/kcft114/.julia/packages/AdvancedHMC/YWXfk/src/adaptation/stepsize.jl:103
┌ Warning: The current proposal will be rejected due to numerical error(s).
│   isfiniteθ = true
│   isfiniter = true
│   isfiniteℓπ = false
│   isfiniteℓκ = true
└ @ AdvancedHMC /Users/kcft114/.julia/packages/AdvancedHMC/YWXfk/src/hamiltonian.jl:36
┌ Warning: The current proposal will be rejected due to numerical error(s).
│   isfiniteθ = false
│   isfiniter = false
│   isfiniteℓπ = false
│   isfiniteℓκ = false
└ @ AdvancedHMC /Users/kcft114/.julia/packages/AdvancedHMC/YWXfk/src/hamiltonian.jl:36
┌ Warning: Incorrect ϵ = NaN; ϵ_previous = 0.07500000000000001 is used instead.
└ @ AdvancedHMC.Adaptation /Users/kcft114/.julia/packages/AdvancedHMC/YWXfk/src/adaptation/stepsize.jl:103
┌ Warning: The current proposal will be rejected due to numerical error(s).
│   isfiniteθ = true
│   isfiniter = true
│   isfiniteℓπ = false
│   isfiniteℓκ = true
└ @ AdvancedHMC /Users/kcft114/.julia/packages/AdvancedHMC/YWXfk/src/hamiltonian.jl:36
┌ Warning: The current proposal will be rejected due to numerical error(s).
│   isfiniteθ = false
│   isfiniter = false
│   isfiniteℓπ = false
│   isfiniteℓκ = false
└ @ AdvancedHMC /Users/kcft114/.julia/packages/AdvancedHMC/YWXfk/src/hamiltonian.jl:36
┌ Warning: Incorrect ϵ = NaN; ϵ_previous = 0.07500000000000001 is used instead.
└ @ AdvancedHMC.Adaptation /Users/kcft114/.julia/packages/AdvancedHMC/YWXfk/src/adaptation/stepsize.jl:103
┌ Warning: The current proposal will be rejected due to numerical error(s).
│   isfiniteθ = true
│   isfiniter = true
│   isfiniteℓπ = false
│   isfiniteℓκ = true
└ @ AdvancedHMC /Users/kcft114/.julia/packages/AdvancedHMC/YWXfk/src/hamiltonian.jl:36
┌ Warning: The current proposal will be rejected due to numerical error(s).
│   isfiniteθ = false
│   isfiniter = false
│   isfiniteℓπ = false
│   isfiniteℓκ = false
└ @ AdvancedHMC /Users/kcft114/.julia/packages/AdvancedHMC/YWXfk/src/hamiltonian.jl:36
┌ Warning: Incorrect ϵ = NaN; ϵ_previous = 0.07500000000000001 is used instead.
└ @ AdvancedHMC.Adaptation /Users/kcft114/.julia/packages/AdvancedHMC/YWXfk/src/adaptation/stepsize.jl:103
┌ Warning: The current proposal will be rejected due to numerical error(s).
│   isfiniteθ = true
│   isfiniter = true
│   isfiniteℓπ = false
│   isfiniteℓκ = true
└ @ AdvancedHMC /Users/kcft114/.julia/packages/AdvancedHMC/YWXfk/src/hamiltonian.jl:36
┌ Warning: The current proposal will be rejected due to numerical error(s).
│   isfiniteθ = false
│   isfiniter = false
│   isfiniteℓπ = false
│   isfiniteℓκ = false
└ @ AdvancedHMC /Users/kcft114/.julia/packages/AdvancedHMC/YWXfk/src/hamiltonian.jl:36
┌ Warning: Incorrect ϵ = NaN; ϵ_previous = 0.07500000000000001 is used instead.
└ @ AdvancedHMC.Adaptation /Users/kcft114/.julia/packages/AdvancedHMC/YWXfk/src/adaptation/stepsize.jl:103
┌ Warning: The current proposal will be rejected due to numerical error(s).
│   isfiniteθ = true
│   isfiniter = true
│   isfiniteℓπ = false
│   isfiniteℓκ = true
└ @ AdvancedHMC /Users/kcft114/.julia/packages/AdvancedHMC/YWXfk/src/hamiltonian.jl:36
┌ Warning: The current proposal will be rejected due to numerical error(s).
│   isfiniteθ = false
│   isfiniter = false
│   isfiniteℓπ = false
│   isfiniteℓκ = false
└ @ AdvancedHMC /Users/kcft114/.julia/packages/AdvancedHMC/YWXfk/src/hamiltonian.jl:36
┌ Warning: Incorrect ϵ = NaN; ϵ_previous = 0.07500000000000001 is used instead.
└ @ AdvancedHMC.Adaptation /Users/kcft114/.julia/packages/AdvancedHMC/YWXfk/src/adaptation/stepsize.jl:103
┌ Warning: The current proposal will be rejected due to numerical error(s).
│   isfiniteθ = true
│   isfiniter = true
│   isfiniteℓπ = false
│   isfiniteℓκ = true
└ @ AdvancedHMC /Users/kcft114/.julia/packages/AdvancedHMC/YWXfk/src/hamiltonian.jl:36
┌ Warning: The current proposal will be rejected due to numerical error(s).
│   isfiniteθ = false
│   isfiniter = false
│   isfiniteℓπ = false
│   isfiniteℓκ = false
└ @ AdvancedHMC /Users/kcft114/.julia/packages/AdvancedHMC/YWXfk/src/hamiltonian.jl:36
┌ Warning: Incorrect ϵ = NaN; ϵ_previous = 0.07500000000000001 is used instead.
└ @ AdvancedHMC.Adaptation /Users/kcft114/.julia/packages/AdvancedHMC/YWXfk/src/adaptation/stepsize.jl:103
┌ Warning: The current proposal will be rejected due to numerical error(s).
│   isfiniteθ = true
│   isfiniter = true
│   isfiniteℓπ = false
│   isfiniteℓκ = true
└ @ AdvancedHMC /Users/kcft114/.julia/packages/AdvancedHMC/YWXfk/src/hamiltonian.jl:36
┌ Warning: The current proposal will be rejected due to numerical error(s).
│   isfiniteθ = false
│   isfiniter = false
│   isfiniteℓπ = false
│   isfiniteℓκ = false
└ @ AdvancedHMC /Users/kcft114/.julia/packages/AdvancedHMC/YWXfk/src/hamiltonian.jl:36
┌ Warning: Incorrect ϵ = NaN; ϵ_previous = 0.07500000000000001 is used instead.
└ @ AdvancedHMC.Adaptation /Users/kcft114/.julia/packages/AdvancedHMC/YWXfk/src/adaptation/stepsize.jl:103
┌ Warning: The current proposal will be rejected due to numerical error(s).
│   isfiniteθ = true
│   isfiniter = true
│   isfiniteℓπ = false
│   isfiniteℓκ = true
└ @ AdvancedHMC /Users/kcft114/.julia/packages/AdvancedHMC/YWXfk/src/hamiltonian.jl:36
┌ Warning: The current proposal will be rejected due to numerical error(s).
│   isfiniteθ = false
│   isfiniter = false
│   isfiniteℓπ = false
│   isfiniteℓκ = false
└ @ AdvancedHMC /Users/kcft114/.julia/packages/AdvancedHMC/YWXfk/src/hamiltonian.jl:36
┌ Warning: Incorrect ϵ = NaN; ϵ_previous = 0.07500000000000001 is used instead.
└ @ AdvancedHMC.Adaptation /Users/kcft114/.julia/packages/AdvancedHMC/YWXfk/src/adaptation/stepsize.jl:103
┌ Warning: The current proposal will be rejected due to numerical error(s).
│   isfiniteθ = true
│   isfiniter = true
│   isfiniteℓπ = false
│   isfiniteℓκ = true
└ @ AdvancedHMC /Users/kcft114/.julia/packages/AdvancedHMC/YWXfk/src/hamiltonian.jl:36
┌ Warning: The current proposal will be rejected due to numerical error(s).
│   isfiniteθ = false
│   isfiniter = false
│   isfiniteℓπ = false
│   isfiniteℓκ = false
└ @ AdvancedHMC /Users/kcft114/.julia/packages/AdvancedHMC/YWXfk/src/hamiltonian.jl:36
┌ Warning: Incorrect ϵ = NaN; ϵ_previous = 0.07500000000000001 is used instead.
└ @ AdvancedHMC.Adaptation /Users/kcft114/.julia/packages/AdvancedHMC/YWXfk/src/adaptation/stepsize.jl:103
┌ Warning: The current proposal will be rejected due to numerical error(s).
│   isfiniteθ = true
│   isfiniter = true
│   isfiniteℓπ = false
│   isfiniteℓκ = true
└ @ AdvancedHMC /Users/kcft114/.julia/packages/AdvancedHMC/YWXfk/src/hamiltonian.jl:36
┌ Warning: The current proposal will be rejected due to numerical error(s).
│   isfiniteθ = false
│   isfiniter = false
│   isfiniteℓπ = false
│   isfiniteℓκ = false
└ @ AdvancedHMC /Users/kcft114/.julia/packages/AdvancedHMC/YWXfk/src/hamiltonian.jl:36
┌ Warning: Incorrect ϵ = NaN; ϵ_previous = 0.07500000000000001 is used instead.
└ @ AdvancedHMC.Adaptation /Users/kcft114/.julia/packages/AdvancedHMC/YWXfk/src/adaptation/stepsize.jl:103
┌ Warning: The current proposal will be rejected due to numerical error(s).
│   isfiniteθ = true
│   isfiniter = true
│   isfiniteℓπ = false
│   isfiniteℓκ = true
└ @ AdvancedHMC /Users/kcft114/.julia/packages/AdvancedHMC/YWXfk/src/hamiltonian.jl:36
┌ Warning: The current proposal will be rejected due to numerical error(s).
│   isfiniteθ = false
│   isfiniter = false
│   isfiniteℓπ = false
│   isfiniteℓκ = false
└ @ AdvancedHMC /Users/kcft114/.julia/packages/AdvancedHMC/YWXfk/src/hamiltonian.jl:36
┌ Warning: Incorrect ϵ = NaN; ϵ_previous = 0.07500000000000001 is used instead.
└ @ AdvancedHMC.Adaptation /Users/kcft114/.julia/packages/AdvancedHMC/YWXfk/src/adaptation/stepsize.jl:103
┌ Warning: The current proposal will be rejected due to numerical error(s).
│   isfiniteθ = true
│   isfiniter = true
│   isfiniteℓπ = false
│   isfiniteℓκ = true
└ @ AdvancedHMC /Users/kcft114/.julia/packages/AdvancedHMC/YWXfk/src/hamiltonian.jl:36
┌ Warning: The current proposal will be rejected due to numerical error(s).
│   isfiniteθ = false
│   isfiniter = false
│   isfiniteℓπ = false
│   isfiniteℓκ = false
└ @ AdvancedHMC /Users/kcft114/.julia/packages/AdvancedHMC/YWXfk/src/hamiltonian.jl:36
┌ Warning: Incorrect ϵ = NaN; ϵ_previous = 0.07500000000000001 is used instead.
└ @ AdvancedHMC.Adaptation /Users/kcft114/.julia/packages/AdvancedHMC/YWXfk/src/adaptation/stepsize.jl:103
┌ Warning: The current proposal will be rejected due to numerical error(s).
│   isfiniteθ = true
│   isfiniter = true
│   isfiniteℓπ = false
│   isfiniteℓκ = true
└ @ AdvancedHMC /Users/kcft114/.julia/packages/AdvancedHMC/YWXfk/src/hamiltonian.jl:36
┌ Warning: The current proposal will be rejected due to numerical error(s).
│   isfiniteθ = false
│   isfiniter = false
│   isfiniteℓπ = false
│   isfiniteℓκ = false
└ @ AdvancedHMC /Users/kcft114/.julia/packages/AdvancedHMC/YWXfk/src/hamiltonian.jl:36
┌ Warning: Incorrect ϵ = NaN; ϵ_previous = 0.07500000000000001 is used instead.
└ @ AdvancedHMC.Adaptation /Users/kcft114/.julia/packages/AdvancedHMC/YWXfk/src/adaptation/stepsize.jl:103
┌ Warning: The current proposal will be rejected due to numerical error(s).
│   isfiniteθ = true
│   isfiniter = true
│   isfiniteℓπ = false
│   isfiniteℓκ = true
└ @ AdvancedHMC /Users/kcft114/.julia/packages/AdvancedHMC/YWXfk/src/hamiltonian.jl:36
┌ Warning: The current proposal will be rejected due to numerical error(s).
│   isfiniteθ = false
│   isfiniter = false
│   isfiniteℓπ = false
│   isfiniteℓκ = false
└ @ AdvancedHMC /Users/kcft114/.julia/packages/AdvancedHMC/YWXfk/src/hamiltonian.jl:36
┌ Warning: Incorrect ϵ = NaN; ϵ_previous = 0.07500000000000001 is used instead.
└ @ AdvancedHMC.Adaptation /Users/kcft114/.julia/packages/AdvancedHMC/YWXfk/src/adaptation/stepsize.jl:103
┌ Warning: The current proposal will be rejected due to numerical error(s).
│   isfiniteθ = true
│   isfiniter = true
│   isfiniteℓπ = false
│   isfiniteℓκ = true
└ @ AdvancedHMC /Users/kcft114/.julia/packages/AdvancedHMC/YWXfk/src/hamiltonian.jl:36
┌ Warning: The current proposal will be rejected due to numerical error(s).
│   isfiniteθ = false
│   isfiniter = false
│   isfiniteℓπ = false
│   isfiniteℓκ = false
└ @ AdvancedHMC /Users/kcft114/.julia/packages/AdvancedHMC/YWXfk/src/hamiltonian.jl:36
┌ Warning: Incorrect ϵ = NaN; ϵ_previous = 0.07500000000000001 is used instead.
└ @ AdvancedHMC.Adaptation /Users/kcft114/.julia/packages/AdvancedHMC/YWXfk/src/adaptation/stepsize.jl:103
┌ Warning: The current proposal will be rejected due to numerical error(s).
│   isfiniteθ = true
│   isfiniter = true
│   isfiniteℓπ = false
│   isfiniteℓκ = true
└ @ AdvancedHMC /Users/kcft114/.julia/packages/AdvancedHMC/YWXfk/src/hamiltonian.jl:36
┌ Warning: The current proposal will be rejected due to numerical error(s).
│   isfiniteθ = false
│   isfiniter = false
│   isfiniteℓπ = false
│   isfiniteℓκ = false
└ @ AdvancedHMC /Users/kcft114/.julia/packages/AdvancedHMC/YWXfk/src/hamiltonian.jl:36
┌ Warning: Incorrect ϵ = NaN; ϵ_previous = 0.07500000000000001 is used instead.
└ @ AdvancedHMC.Adaptation /Users/kcft114/.julia/packages/AdvancedHMC/YWXfk/src/adaptation/stepsize.jl:103
┌ Warning: The current proposal will be rejected due to numerical error(s).
│   isfiniteθ = true
│   isfiniter = true
│   isfiniteℓπ = false
│   isfiniteℓκ = true
└ @ AdvancedHMC /Users/kcft114/.julia/packages/AdvancedHMC/YWXfk/src/hamiltonian.jl:36
┌ Warning: The current proposal will be rejected due to numerical error(s).
│   isfiniteθ = false
│   isfiniter = false
│   isfiniteℓπ = false
│   isfiniteℓκ = false
└ @ AdvancedHMC /Users/kcft114/.julia/packages/AdvancedHMC/YWXfk/src/hamiltonian.jl:36
┌ Warning: Incorrect ϵ = NaN; ϵ_previous = 0.07500000000000001 is used instead.
└ @ AdvancedHMC.Adaptation /Users/kcft114/.julia/packages/AdvancedHMC/YWXfk/src/adaptation/stepsize.jl:103
┌ Warning: The current proposal will be rejected due to numerical error(s).
│   isfiniteθ = true
│   isfiniter = true
│   isfiniteℓπ = false
│   isfiniteℓκ = true
└ @ AdvancedHMC /Users/kcft114/.julia/packages/AdvancedHMC/YWXfk/src/hamiltonian.jl:36
┌ Warning: The current proposal will be rejected due to numerical error(s).
│   isfiniteθ = false
│   isfiniter = false
│   isfiniteℓπ = false
│   isfiniteℓκ = false
└ @ AdvancedHMC /Users/kcft114/.julia/packages/AdvancedHMC/YWXfk/src/hamiltonian.jl:36
┌ Warning: Incorrect ϵ = NaN; ϵ_previous = 0.07500000000000001 is used instead.
└ @ AdvancedHMC.Adaptation /Users/kcft114/.julia/packages/AdvancedHMC/YWXfk/src/adaptation/stepsize.jl:103
┌ Warning: The current proposal will be rejected due to numerical error(s).
│   isfiniteθ = true
│   isfiniter = true
│   isfiniteℓπ = false
│   isfiniteℓκ = true
└ @ AdvancedHMC /Users/kcft114/.julia/packages/AdvancedHMC/YWXfk/src/hamiltonian.jl:36
┌ Warning: The current proposal will be rejected due to numerical error(s).
│   isfiniteθ = false
│   isfiniter = false
│   isfiniteℓπ = false
│   isfiniteℓκ = false
└ @ AdvancedHMC /Users/kcft114/.julia/packages/AdvancedHMC/YWXfk/src/hamiltonian.jl:36
┌ Warning: Incorrect ϵ = NaN; ϵ_previous = 0.07500000000000001 is used instead.
└ @ AdvancedHMC.Adaptation /Users/kcft114/.julia/packages/AdvancedHMC/YWXfk/src/adaptation/stepsize.jl:103
┌ Warning: The current proposal will be rejected due to numerical error(s).
│   isfiniteθ = true
│   isfiniter = true
│   isfiniteℓπ = false
│   isfiniteℓκ = true
└ @ AdvancedHMC /Users/kcft114/.julia/packages/AdvancedHMC/YWXfk/src/hamiltonian.jl:36
┌ Warning: The current proposal will be rejected due to numerical error(s).
│   isfiniteθ = false
│   isfiniter = false
│   isfiniteℓπ = false
│   isfiniteℓκ = false
└ @ AdvancedHMC /Users/kcft114/.julia/packages/AdvancedHMC/YWXfk/src/hamiltonian.jl:36
┌ Warning: Incorrect ϵ = NaN; ϵ_previous = 0.07500000000000001 is used instead.
└ @ AdvancedHMC.Adaptation /Users/kcft114/.julia/packages/AdvancedHMC/YWXfk/src/adaptation/stepsize.jl:103
┌ Warning: The current proposal will be rejected due to numerical error(s).
│   isfiniteθ = true
│   isfiniter = true
│   isfiniteℓπ = false
│   isfiniteℓκ = true
└ @ AdvancedHMC /Users/kcft114/.julia/packages/AdvancedHMC/YWXfk/src/hamiltonian.jl:36
┌ Warning: The current proposal will be rejected due to numerical error(s).
│   isfiniteθ = false
│   isfiniter = false
│   isfiniteℓπ = false
│   isfiniteℓκ = false
└ @ AdvancedHMC /Users/kcft114/.julia/packages/AdvancedHMC/YWXfk/src/hamiltonian.jl:36
┌ Warning: Incorrect ϵ = NaN; ϵ_previous = 0.07500000000000001 is used instead.
└ @ AdvancedHMC.Adaptation /Users/kcft114/.julia/packages/AdvancedHMC/YWXfk/src/adaptation/stepsize.jl:103
┌ Warning: The current proposal will be rejected due to numerical error(s).
│   isfiniteθ = true
│   isfiniter = true
│   isfiniteℓπ = false
│   isfiniteℓκ = true
└ @ AdvancedHMC /Users/kcft114/.julia/packages/AdvancedHMC/YWXfk/src/hamiltonian.jl:36
┌ Warning: The current proposal will be rejected due to numerical error(s).
│   isfiniteθ = false
│   isfiniter = false
│   isfiniteℓπ = false
│   isfiniteℓκ = false
└ @ AdvancedHMC /Users/kcft114/.julia/packages/AdvancedHMC/YWXfk/src/hamiltonian.jl:36
┌ Warning: Incorrect ϵ = NaN; ϵ_previous = 0.07500000000000001 is used instead.
└ @ AdvancedHMC.Adaptation /Users/kcft114/.julia/packages/AdvancedHMC/YWXfk/src/adaptation/stepsize.jl:103
┌ Warning: The current proposal will be rejected due to numerical error(s).
│   isfiniteθ = true
│   isfiniter = true
│   isfiniteℓπ = false
│   isfiniteℓκ = true
└ @ AdvancedHMC /Users/kcft114/.julia/packages/AdvancedHMC/YWXfk/src/hamiltonian.jl:36
┌ Warning: The current proposal will be rejected due to numerical error(s).
│   isfiniteθ = false
│   isfiniter = false
│   isfiniteℓπ = false
│   isfiniteℓκ = false
└ @ AdvancedHMC /Users/kcft114/.julia/packages/AdvancedHMC/YWXfk/src/hamiltonian.jl:36
┌ Warning: Incorrect ϵ = NaN; ϵ_previous = 0.07500000000000001 is used instead.
└ @ AdvancedHMC.Adaptation /Users/kcft114/.julia/packages/AdvancedHMC/YWXfk/src/adaptation/stepsize.jl:103
┌ Warning: The current proposal will be rejected due to numerical error(s).
│   isfiniteθ = true
│   isfiniter = true
│   isfiniteℓπ = false
│   isfiniteℓκ = true
└ @ AdvancedHMC /Users/kcft114/.julia/packages/AdvancedHMC/YWXfk/src/hamiltonian.jl:36
┌ Warning: The current proposal will be rejected due to numerical error(s).
│   isfiniteθ = false
│   isfiniter = false
│   isfiniteℓπ = false
│   isfiniteℓκ = false
└ @ AdvancedHMC /Users/kcft114/.julia/packages/AdvancedHMC/YWXfk/src/hamiltonian.jl:36
┌ Warning: Incorrect ϵ = NaN; ϵ_previous = 0.07500000000000001 is used instead.
└ @ AdvancedHMC.Adaptation /Users/kcft114/.julia/packages/AdvancedHMC/YWXfk/src/adaptation/stepsize.jl:103
┌ Warning: The current proposal will be rejected due to numerical error(s).
│   isfiniteθ = true
│   isfiniter = true
│   isfiniteℓπ = false
│   isfiniteℓκ = true
└ @ AdvancedHMC /Users/kcft114/.julia/packages/AdvancedHMC/YWXfk/src/hamiltonian.jl:36
┌ Warning: The current proposal will be rejected due to numerical error(s).
│   isfiniteθ = false
│   isfiniter = false
│   isfiniteℓπ = false
│   isfiniteℓκ = false
└ @ AdvancedHMC /Users/kcft114/.julia/packages/AdvancedHMC/YWXfk/src/hamiltonian.jl:36
┌ Warning: Incorrect ϵ = NaN; ϵ_previous = 0.07500000000000001 is used instead.
└ @ AdvancedHMC.Adaptation /Users/kcft114/.julia/packages/AdvancedHMC/YWXfk/src/adaptation/stepsize.jl:103
┌ Warning: The current proposal will be rejected due to numerical error(s).
│   isfiniteθ = true
│   isfiniter = true
│   isfiniteℓπ = false
│   isfiniteℓκ = true
└ @ AdvancedHMC /Users/kcft114/.julia/packages/AdvancedHMC/YWXfk/src/hamiltonian.jl:36
┌ Warning: The current proposal will be rejected due to numerical error(s).
│   isfiniteθ = false
│   isfiniter = false
│   isfiniteℓπ = false
│   isfiniteℓκ = false
└ @ AdvancedHMC /Users/kcft114/.julia/packages/AdvancedHMC/YWXfk/src/hamiltonian.jl:36
┌ Warning: Incorrect ϵ = NaN; ϵ_previous = 0.07500000000000001 is used instead.
└ @ AdvancedHMC.Adaptation /Users/kcft114/.julia/packages/AdvancedHMC/YWXfk/src/adaptation/stepsize.jl:103
┌ Warning: The current proposal will be rejected due to numerical error(s).
│   isfiniteθ = true
│   isfiniter = true
│   isfiniteℓπ = false
│   isfiniteℓκ = true
└ @ AdvancedHMC /Users/kcft114/.julia/packages/AdvancedHMC/YWXfk/src/hamiltonian.jl:36
┌ Warning: The current proposal will be rejected due to numerical error(s).
│   isfiniteθ = false
│   isfiniter = false
│   isfiniteℓπ = false
│   isfiniteℓκ = false
└ @ AdvancedHMC /Users/kcft114/.julia/packages/AdvancedHMC/YWXfk/src/hamiltonian.jl:36
┌ Warning: Incorrect ϵ = NaN; ϵ_previous = 0.07500000000000001 is used instead.
└ @ AdvancedHMC.Adaptation /Users/kcft114/.julia/packages/AdvancedHMC/YWXfk/src/adaptation/stepsize.jl:103
┌ Warning: The current proposal will be rejected due to numerical error(s).
│   isfiniteθ = true
│   isfiniter = true
│   isfiniteℓπ = false
│   isfiniteℓκ = true
└ @ AdvancedHMC /Users/kcft114/.julia/packages/AdvancedHMC/YWXfk/src/hamiltonian.jl:36
┌ Warning: The current proposal will be rejected due to numerical error(s).
│   isfiniteθ = false
│   isfiniter = false
│   isfiniteℓπ = false
│   isfiniteℓκ = false
└ @ AdvancedHMC /Users/kcft114/.julia/packages/AdvancedHMC/YWXfk/src/hamiltonian.jl:36
┌ Warning: Incorrect ϵ = NaN; ϵ_previous = 0.07500000000000001 is used instead.
└ @ AdvancedHMC.Adaptation /Users/kcft114/.julia/packages/AdvancedHMC/YWXfk/src/adaptation/stepsize.jl:103
┌ Warning: The current proposal will be rejected due to numerical error(s).
│   isfiniteθ = true
│   isfiniter = true
│   isfiniteℓπ = false
│   isfiniteℓκ = true
└ @ AdvancedHMC /Users/kcft114/.julia/packages/AdvancedHMC/YWXfk/src/hamiltonian.jl:36
┌ Warning: The current proposal will be rejected due to numerical error(s).
│   isfiniteθ = false
│   isfiniter = false
│   isfiniteℓπ = false
│   isfiniteℓκ = false
└ @ AdvancedHMC /Users/kcft114/.julia/packages/AdvancedHMC/YWXfk/src/hamiltonian.jl:36
┌ Warning: Incorrect ϵ = NaN; ϵ_previous = 0.07500000000000001 is used instead.
└ @ AdvancedHMC.Adaptation /Users/kcft114/.julia/packages/AdvancedHMC/YWXfk/src/adaptation/stepsize.jl:103
┌ Warning: The current proposal will be rejected due to numerical error(s).
│   isfiniteθ = true
│   isfiniter = true
│   isfiniteℓπ = false
│   isfiniteℓκ = true
└ @ AdvancedHMC /Users/kcft114/.julia/packages/AdvancedHMC/YWXfk/src/hamiltonian.jl:36
┌ Warning: The current proposal will be rejected due to numerical error(s).
│   isfiniteθ = false
│   isfiniter = false
│   isfiniteℓπ = false
│   isfiniteℓκ = false
└ @ AdvancedHMC /Users/kcft114/.julia/packages/AdvancedHMC/YWXfk/src/hamiltonian.jl:36
┌ Warning: Incorrect ϵ = NaN; ϵ_previous = 0.07500000000000001 is used instead.
└ @ AdvancedHMC.Adaptation /Users/kcft114/.julia/packages/AdvancedHMC/YWXfk/src/adaptation/stepsize.jl:103
┌ Warning: The current proposal will be rejected due to numerical error(s).
│   isfiniteθ = true
│   isfiniter = true
│   isfiniteℓπ = false
│   isfiniteℓκ = true
└ @ AdvancedHMC /Users/kcft114/.julia/packages/AdvancedHMC/YWXfk/src/hamiltonian.jl:36
┌ Warning: The current proposal will be rejected due to numerical error(s).
│   isfiniteθ = false
│   isfiniter = false
│   isfiniteℓπ = false
│   isfiniteℓκ = false
└ @ AdvancedHMC /Users/kcft114/.julia/packages/AdvancedHMC/YWXfk/src/hamiltonian.jl:36
┌ Warning: Incorrect ϵ = NaN; ϵ_previous = 0.07500000000000001 is used instead.
└ @ AdvancedHMC.Adaptation /Users/kcft114/.julia/packages/AdvancedHMC/YWXfk/src/adaptation/stepsize.jl:103
┌ Warning: The current proposal will be rejected due to numerical error(s).
│   isfiniteθ = true
│   isfiniter = true
│   isfiniteℓπ = false
│   isfiniteℓκ = true
└ @ AdvancedHMC /Users/kcft114/.julia/packages/AdvancedHMC/YWXfk/src/hamiltonian.jl:36
┌ Warning: The current proposal will be rejected due to numerical error(s).
│   isfiniteθ = false
│   isfiniter = false
│   isfiniteℓπ = false
│   isfiniteℓκ = false
└ @ AdvancedHMC /Users/kcft114/.julia/packages/AdvancedHMC/YWXfk/src/hamiltonian.jl:36
┌ Warning: Incorrect ϵ = NaN; ϵ_previous = 0.07500000000000001 is used instead.
└ @ AdvancedHMC.Adaptation /Users/kcft114/.julia/packages/AdvancedHMC/YWXfk/src/adaptation/stepsize.jl:103
┌ Warning: The current proposal will be rejected due to numerical error(s).
│   isfiniteθ = true
│   isfiniter = true
│   isfiniteℓπ = false
│   isfiniteℓκ = true
└ @ AdvancedHMC /Users/kcft114/.julia/packages/AdvancedHMC/YWXfk/src/hamiltonian.jl:36
┌ Warning: The current proposal will be rejected due to numerical error(s).
│   isfiniteθ = false
│   isfiniter = false
│   isfiniteℓπ = false
│   isfiniteℓκ = false
└ @ AdvancedHMC /Users/kcft114/.julia/packages/AdvancedHMC/YWXfk/src/hamiltonian.jl:36
┌ Warning: Incorrect ϵ = NaN; ϵ_previous = 0.07500000000000001 is used instead.
└ @ AdvancedHMC.Adaptation /Users/kcft114/.julia/packages/AdvancedHMC/YWXfk/src/adaptation/stepsize.jl:103
┌ Warning: The current proposal will be rejected due to numerical error(s).
│   isfiniteθ = true
│   isfiniter = true
│   isfiniteℓπ = false
│   isfiniteℓκ = true
└ @ AdvancedHMC /Users/kcft114/.julia/packages/AdvancedHMC/YWXfk/src/hamiltonian.jl:36
┌ Warning: The current proposal will be rejected due to numerical error(s).
│   isfiniteθ = false
│   isfiniter = false
│   isfiniteℓπ = false
│   isfiniteℓκ = false
└ @ AdvancedHMC /Users/kcft114/.julia/packages/AdvancedHMC/YWXfk/src/hamiltonian.jl:36
┌ Warning: Incorrect ϵ = NaN; ϵ_previous = 0.07500000000000001 is used instead.
└ @ AdvancedHMC.Adaptation /Users/kcft114/.julia/packages/AdvancedHMC/YWXfk/src/adaptation/stepsize.jl:103
┌ Warning: The current proposal will be rejected due to numerical error(s).
│   isfiniteθ = true
│   isfiniter = true
│   isfiniteℓπ = false
│   isfiniteℓκ = true
└ @ AdvancedHMC /Users/kcft114/.julia/packages/AdvancedHMC/YWXfk/src/hamiltonian.jl:36
┌ Warning: The current proposal will be rejected due to numerical error(s).
│   isfiniteθ = false
│   isfiniter = false
│   isfiniteℓπ = false
│   isfiniteℓκ = false
└ @ AdvancedHMC /Users/kcft114/.julia/packages/AdvancedHMC/YWXfk/src/hamiltonian.jl:36
┌ Warning: Incorrect ϵ = NaN; ϵ_previous = 0.07500000000000001 is used instead.
└ @ AdvancedHMC.Adaptation /Users/kcft114/.julia/packages/AdvancedHMC/YWXfk/src/adaptation/stepsize.jl:103
┌ Warning: The current proposal will be rejected due to numerical error(s).
│   isfiniteθ = true
│   isfiniter = true
│   isfiniteℓπ = false
│   isfiniteℓκ = true
└ @ AdvancedHMC /Users/kcft114/.julia/packages/AdvancedHMC/YWXfk/src/hamiltonian.jl:36
┌ Warning: The current proposal will be rejected due to numerical error(s).
│   isfiniteθ = false
│   isfiniter = false
│   isfiniteℓπ = false
│   isfiniteℓκ = false
└ @ AdvancedHMC /Users/kcft114/.julia/packages/AdvancedHMC/YWXfk/src/hamiltonian.jl:36
┌ Warning: Incorrect ϵ = NaN; ϵ_previous = 0.07500000000000001 is used instead.
└ @ AdvancedHMC.Adaptation /Users/kcft114/.julia/packages/AdvancedHMC/YWXfk/src/adaptation/stepsize.jl:103
┌ Warning: The current proposal will be rejected due to numerical error(s).
│   isfiniteθ = true
│   isfiniter = true
│   isfiniteℓπ = false
│   isfiniteℓκ = true
└ @ AdvancedHMC /Users/kcft114/.julia/packages/AdvancedHMC/YWXfk/src/hamiltonian.jl:36
┌ Warning: The current proposal will be rejected due to numerical error(s).
│   isfiniteθ = false
│   isfiniter = false
│   isfiniteℓπ = false
│   isfiniteℓκ = false
└ @ AdvancedHMC /Users/kcft114/.julia/packages/AdvancedHMC/YWXfk/src/hamiltonian.jl:36
┌ Warning: Incorrect ϵ = NaN; ϵ_previous = 0.07500000000000001 is used instead.
└ @ AdvancedHMC.Adaptation /Users/kcft114/.julia/packages/AdvancedHMC/YWXfk/src/adaptation/stepsize.jl:103
┌ Warning: The current proposal will be rejected due to numerical error(s).
│   isfiniteθ = true
│   isfiniter = true
│   isfiniteℓπ = false
│   isfiniteℓκ = true
└ @ AdvancedHMC /Users/kcft114/.julia/packages/AdvancedHMC/YWXfk/src/hamiltonian.jl:36
┌ Warning: The current proposal will be rejected due to numerical error(s).
│   isfiniteθ = false
│   isfiniter = false
│   isfiniteℓπ = false
│   isfiniteℓκ = false
└ @ AdvancedHMC /Users/kcft114/.julia/packages/AdvancedHMC/YWXfk/src/hamiltonian.jl:36
┌ Warning: Incorrect ϵ = NaN; ϵ_previous = 0.07500000000000001 is used instead.
└ @ AdvancedHMC.Adaptation /Users/kcft114/.julia/packages/AdvancedHMC/YWXfk/src/adaptation/stepsize.jl:103
┌ Warning: The current proposal will be rejected due to numerical error(s).
│   isfiniteθ = true
│   isfiniter = true
│   isfiniteℓπ = false
│   isfiniteℓκ = true
└ @ AdvancedHMC /Users/kcft114/.julia/packages/AdvancedHMC/YWXfk/src/hamiltonian.jl:36
┌ Warning: The current proposal will be rejected due to numerical error(s).
│   isfiniteθ = false
│   isfiniter = false
│   isfiniteℓπ = false
│   isfiniteℓκ = false
└ @ AdvancedHMC /Users/kcft114/.julia/packages/AdvancedHMC/YWXfk/src/hamiltonian.jl:36
┌ Warning: Incorrect ϵ = NaN; ϵ_previous = 0.07500000000000001 is used instead.
└ @ AdvancedHMC.Adaptation /Users/kcft114/.julia/packages/AdvancedHMC/YWXfk/src/adaptation/stepsize.jl:103
┌ Warning: The current proposal will be rejected due to numerical error(s).
│   isfiniteθ = true
│   isfiniter = true
│   isfiniteℓπ = false
│   isfiniteℓκ = true
└ @ AdvancedHMC /Users/kcft114/.julia/packages/AdvancedHMC/YWXfk/src/hamiltonian.jl:36
┌ Warning: The current proposal will be rejected due to numerical error(s).
│   isfiniteθ = false
│   isfiniter = false
│   isfiniteℓπ = false
│   isfiniteℓκ = false
└ @ AdvancedHMC /Users/kcft114/.julia/packages/AdvancedHMC/YWXfk/src/hamiltonian.jl:36
┌ Warning: Incorrect ϵ = NaN; ϵ_previous = 0.07500000000000001 is used instead.
└ @ AdvancedHMC.Adaptation /Users/kcft114/.julia/packages/AdvancedHMC/YWXfk/src/adaptation/stepsize.jl:103
┌ Warning: The current proposal will be rejected due to numerical error(s).
│   isfiniteθ = true
│   isfiniter = true
│   isfiniteℓπ = false
│   isfiniteℓκ = true
└ @ AdvancedHMC /Users/kcft114/.julia/packages/AdvancedHMC/YWXfk/src/hamiltonian.jl:36
┌ Warning: The current proposal will be rejected due to numerical error(s).
│   isfiniteθ = false
│   isfiniter = false
│   isfiniteℓπ = false
│   isfiniteℓκ = false
└ @ AdvancedHMC /Users/kcft114/.julia/packages/AdvancedHMC/YWXfk/src/hamiltonian.jl:36
┌ Warning: Incorrect ϵ = NaN; ϵ_previous = 0.07500000000000001 is used instead.
└ @ AdvancedHMC.Adaptation /Users/kcft114/.julia/packages/AdvancedHMC/YWXfk/src/adaptation/stepsize.jl:103
┌ Warning: The current proposal will be rejected due to numerical error(s).
│   isfiniteθ = true
│   isfiniter = true
│   isfiniteℓπ = false
│   isfiniteℓκ = true
└ @ AdvancedHMC /Users/kcft114/.julia/packages/AdvancedHMC/YWXfk/src/hamiltonian.jl:36
┌ Warning: The current proposal will be rejected due to numerical error(s).
│   isfiniteθ = false
│   isfiniter = false
│   isfiniteℓπ = false
│   isfiniteℓκ = false
└ @ AdvancedHMC /Users/kcft114/.julia/packages/AdvancedHMC/YWXfk/src/hamiltonian.jl:36
┌ Warning: Incorrect ϵ = NaN; ϵ_previous = 0.07500000000000001 is used instead.
└ @ AdvancedHMC.Adaptation /Users/kcft114/.julia/packages/AdvancedHMC/YWXfk/src/adaptation/stepsize.jl:103
┌ Warning: The current proposal will be rejected due to numerical error(s).
│   isfiniteθ = true
│   isfiniter = true
│   isfiniteℓπ = false
│   isfiniteℓκ = true
└ @ AdvancedHMC /Users/kcft114/.julia/packages/AdvancedHMC/YWXfk/src/hamiltonian.jl:36
┌ Warning: The current proposal will be rejected due to numerical error(s).
│   isfiniteθ = false
│   isfiniter = false
│   isfiniteℓπ = false
│   isfiniteℓκ = false
└ @ AdvancedHMC /Users/kcft114/.julia/packages/AdvancedHMC/YWXfk/src/hamiltonian.jl:36
┌ Warning: Incorrect ϵ = NaN; ϵ_previous = 0.07500000000000001 is used instead.
└ @ AdvancedHMC.Adaptation /Users/kcft114/.julia/packages/AdvancedHMC/YWXfk/src/adaptation/stepsize.jl:103
┌ Warning: The current proposal will be rejected due to numerical error(s).
│   isfiniteθ = true
│   isfiniter = true
│   isfiniteℓπ = false
│   isfiniteℓκ = true
└ @ AdvancedHMC /Users/kcft114/.julia/packages/AdvancedHMC/YWXfk/src/hamiltonian.jl:36
┌ Warning: The current proposal will be rejected due to numerical error(s).
│   isfiniteθ = false
│   isfiniter = false
│   isfiniteℓπ = false
│   isfiniteℓκ = false
└ @ AdvancedHMC /Users/kcft114/.julia/packages/AdvancedHMC/YWXfk/src/hamiltonian.jl:36
┌ Warning: Incorrect ϵ = NaN; ϵ_previous = 0.07500000000000001 is used instead.
└ @ AdvancedHMC.Adaptation /Users/kcft114/.julia/packages/AdvancedHMC/YWXfk/src/adaptation/stepsize.jl:103
┌ Warning: The current proposal will be rejected due to numerical error(s).
│   isfiniteθ = true
│   isfiniter = true
│   isfiniteℓπ = false
│   isfiniteℓκ = true
└ @ AdvancedHMC /Users/kcft114/.julia/packages/AdvancedHMC/YWXfk/src/hamiltonian.jl:36
┌ Warning: The current proposal will be rejected due to numerical error(s).
│   isfiniteθ = false
│   isfiniter = false
│   isfiniteℓπ = false
│   isfiniteℓκ = false
└ @ AdvancedHMC /Users/kcft114/.julia/packages/AdvancedHMC/YWXfk/src/hamiltonian.jl:36
┌ Warning: Incorrect ϵ = NaN; ϵ_previous = 0.07500000000000001 is used instead.
└ @ AdvancedHMC.Adaptation /Users/kcft114/.julia/packages/AdvancedHMC/YWXfk/src/adaptation/stepsize.jl:103
┌ Warning: The current proposal will be rejected due to numerical error(s).
│   isfiniteθ = true
│   isfiniter = true
│   isfiniteℓπ = false
│   isfiniteℓκ = true
└ @ AdvancedHMC /Users/kcft114/.julia/packages/AdvancedHMC/YWXfk/src/hamiltonian.jl:36
┌ Warning: The current proposal will be rejected due to numerical error(s).
│   isfiniteθ = false
│   isfiniter = false
│   isfiniteℓπ = false
│   isfiniteℓκ = false
└ @ AdvancedHMC /Users/kcft114/.julia/packages/AdvancedHMC/YWXfk/src/hamiltonian.jl:36
┌ Warning: Incorrect ϵ = NaN; ϵ_previous = 0.07500000000000001 is used instead.
└ @ AdvancedHMC.Adaptation /Users/kcft114/.julia/packages/AdvancedHMC/YWXfk/src/adaptation/stepsize.jl:103
┌ Warning: The current proposal will be rejected due to numerical error(s).
│   isfiniteθ = true
│   isfiniter = true
│   isfiniteℓπ = false
│   isfiniteℓκ = true
└ @ AdvancedHMC /Users/kcft114/.julia/packages/AdvancedHMC/YWXfk/src/hamiltonian.jl:36
┌ Warning: The current proposal will be rejected due to numerical error(s).
│   isfiniteθ = false
│   isfiniter = false
│   isfiniteℓπ = false
│   isfiniteℓκ = false
└ @ AdvancedHMC /Users/kcft114/.julia/packages/AdvancedHMC/YWXfk/src/hamiltonian.jl:36
┌ Warning: Incorrect ϵ = NaN; ϵ_previous = 0.07500000000000001 is used instead.
└ @ AdvancedHMC.Adaptation /Users/kcft114/.julia/packages/AdvancedHMC/YWXfk/src/adaptation/stepsize.jl:103
┌ Warning: The current proposal will be rejected due to numerical error(s).
│   isfiniteθ = true
│   isfiniter = true
│   isfiniteℓπ = false
│   isfiniteℓκ = true
└ @ AdvancedHMC /Users/kcft114/.julia/packages/AdvancedHMC/YWXfk/src/hamiltonian.jl:36
┌ Warning: The current proposal will be rejected due to numerical error(s).
│   isfiniteθ = false
│   isfiniter = false
│   isfiniteℓπ = false
│   isfiniteℓκ = false
└ @ AdvancedHMC /Users/kcft114/.julia/packages/AdvancedHMC/YWXfk/src/hamiltonian.jl:36
┌ Warning: Incorrect ϵ = NaN; ϵ_previous = 0.07500000000000001 is used instead.
└ @ AdvancedHMC.Adaptation /Users/kcft114/.julia/packages/AdvancedHMC/YWXfk/src/adaptation/stepsize.jl:103
┌ Warning: The current proposal will be rejected due to numerical error(s).
│   isfiniteθ = true
│   isfiniter = true
│   isfiniteℓπ = false
│   isfiniteℓκ = true
└ @ AdvancedHMC /Users/kcft114/.julia/packages/AdvancedHMC/YWXfk/src/hamiltonian.jl:36
┌ Warning: The current proposal will be rejected due to numerical error(s).
│   isfiniteθ = false
│   isfiniter = false
│   isfiniteℓπ = false
│   isfiniteℓκ = false
└ @ AdvancedHMC /Users/kcft114/.julia/packages/AdvancedHMC/YWXfk/src/hamiltonian.jl:36
┌ Warning: Incorrect ϵ = NaN; ϵ_previous = 0.07500000000000001 is used instead.
└ @ AdvancedHMC.Adaptation /Users/kcft114/.julia/packages/AdvancedHMC/YWXfk/src/adaptation/stepsize.jl:103
┌ Warning: The current proposal will be rejected due to numerical error(s).
│   isfiniteθ = true
│   isfiniter = true
│   isfiniteℓπ = false
│   isfiniteℓκ = true
└ @ AdvancedHMC /Users/kcft114/.julia/packages/AdvancedHMC/YWXfk/src/hamiltonian.jl:36
┌ Warning: The current proposal will be rejected due to numerical error(s).
│   isfiniteθ = false
│   isfiniter = false
│   isfiniteℓπ = false
│   isfiniteℓκ = false
└ @ AdvancedHMC /Users/kcft114/.julia/packages/AdvancedHMC/YWXfk/src/hamiltonian.jl:36
┌ Warning: Incorrect ϵ = NaN; ϵ_previous = 0.07500000000000001 is used instead.
└ @ AdvancedHMC.Adaptation /Users/kcft114/.julia/packages/AdvancedHMC/YWXfk/src/adaptation/stepsize.jl:103
┌ Warning: The current proposal will be rejected due to numerical error(s).
│   isfiniteθ = true
│   isfiniter = true
│   isfiniteℓπ = false
│   isfiniteℓκ = true
└ @ AdvancedHMC /Users/kcft114/.julia/packages/AdvancedHMC/YWXfk/src/hamiltonian.jl:36
┌ Warning: The current proposal will be rejected due to numerical error(s).
│   isfiniteθ = false
│   isfiniter = false
│   isfiniteℓπ = false
│   isfiniteℓκ = false
└ @ AdvancedHMC /Users/kcft114/.julia/packages/AdvancedHMC/YWXfk/src/hamiltonian.jl:36
┌ Warning: Incorrect ϵ = NaN; ϵ_previous = 0.07500000000000001 is used instead.
└ @ AdvancedHMC.Adaptation /Users/kcft114/.julia/packages/AdvancedHMC/YWXfk/src/adaptation/stepsize.jl:103
┌ Warning: The current proposal will be rejected due to numerical error(s).
│   isfiniteθ = true
│   isfiniter = true
│   isfiniteℓπ = false
│   isfiniteℓκ = true
└ @ AdvancedHMC /Users/kcft114/.julia/packages/AdvancedHMC/YWXfk/src/hamiltonian.jl:36
┌ Warning: The current proposal will be rejected due to numerical error(s).
│   isfiniteθ = false
│   isfiniter = false
│   isfiniteℓπ = false
│   isfiniteℓκ = false
└ @ AdvancedHMC /Users/kcft114/.julia/packages/AdvancedHMC/YWXfk/src/hamiltonian.jl:36
┌ Warning: Incorrect ϵ = NaN; ϵ_previous = 0.07500000000000001 is used instead.
└ @ AdvancedHMC.Adaptation /Users/kcft114/.julia/packages/AdvancedHMC/YWXfk/src/adaptation/stepsize.jl:103
┌ Warning: The current proposal will be rejected due to numerical error(s).
│   isfiniteθ = true
│   isfiniter = true
│   isfiniteℓπ = false
│   isfiniteℓκ = true
└ @ AdvancedHMC /Users/kcft114/.julia/packages/AdvancedHMC/YWXfk/src/hamiltonian.jl:36
┌ Warning: The current proposal will be rejected due to numerical error(s).
│   isfiniteθ = false
│   isfiniter = false
│   isfiniteℓπ = false
│   isfiniteℓκ = false
└ @ AdvancedHMC /Users/kcft114/.julia/packages/AdvancedHMC/YWXfk/src/hamiltonian.jl:36
┌ Warning: Incorrect ϵ = NaN; ϵ_previous = 0.07500000000000001 is used instead.
└ @ AdvancedHMC.Adaptation /Users/kcft114/.julia/packages/AdvancedHMC/YWXfk/src/adaptation/stepsize.jl:103
┌ Warning: The current proposal will be rejected due to numerical error(s).
│   isfiniteθ = true
│   isfiniter = true
│   isfiniteℓπ = false
│   isfiniteℓκ = true
└ @ AdvancedHMC /Users/kcft114/.julia/packages/AdvancedHMC/YWXfk/src/hamiltonian.jl:36
┌ Warning: The current proposal will be rejected due to numerical error(s).
│   isfiniteθ = false
│   isfiniter = false
│   isfiniteℓπ = false
│   isfiniteℓκ = false
└ @ AdvancedHMC /Users/kcft114/.julia/packages/AdvancedHMC/YWXfk/src/hamiltonian.jl:36
┌ Warning: Incorrect ϵ = NaN; ϵ_previous = 0.07500000000000001 is used instead.
└ @ AdvancedHMC.Adaptation /Users/kcft114/.julia/packages/AdvancedHMC/YWXfk/src/adaptation/stepsize.jl:103
┌ Warning: The current proposal will be rejected due to numerical error(s).
│   isfiniteθ = true
│   isfiniter = true
│   isfiniteℓπ = false
│   isfiniteℓκ = true
└ @ AdvancedHMC /Users/kcft114/.julia/packages/AdvancedHMC/YWXfk/src/hamiltonian.jl:36
┌ Warning: The current proposal will be rejected due to numerical error(s).
│   isfiniteθ = false
│   isfiniter = false
│   isfiniteℓπ = false
│   isfiniteℓκ = false
└ @ AdvancedHMC /Users/kcft114/.julia/packages/AdvancedHMC/YWXfk/src/hamiltonian.jl:36
┌ Warning: Incorrect ϵ = NaN; ϵ_previous = 0.07500000000000001 is used instead.
└ @ AdvancedHMC.Adaptation /Users/kcft114/.julia/packages/AdvancedHMC/YWXfk/src/adaptation/stepsize.jl:103
┌ Warning: The current proposal will be rejected due to numerical error(s).
│   isfiniteθ = true
│   isfiniter = true
│   isfiniteℓπ = false
│   isfiniteℓκ = true
└ @ AdvancedHMC /Users/kcft114/.julia/packages/AdvancedHMC/YWXfk/src/hamiltonian.jl:36
┌ Warning: The current proposal will be rejected due to numerical error(s).
│   isfiniteθ = false
│   isfiniter = false
│   isfiniteℓπ = false
│   isfiniteℓκ = false
└ @ AdvancedHMC /Users/kcft114/.julia/packages/AdvancedHMC/YWXfk/src/hamiltonian.jl:36
┌ Warning: Incorrect ϵ = NaN; ϵ_previous = 0.07500000000000001 is used instead.
└ @ AdvancedHMC.Adaptation /Users/kcft114/.julia/packages/AdvancedHMC/YWXfk/src/adaptation/stepsize.jl:103
┌ Warning: The current proposal will be rejected due to numerical error(s).
│   isfiniteθ = true
│   isfiniter = true
│   isfiniteℓπ = false
│   isfiniteℓκ = true
└ @ AdvancedHMC /Users/kcft114/.julia/packages/AdvancedHMC/YWXfk/src/hamiltonian.jl:36
┌ Warning: The current proposal will be rejected due to numerical error(s).
│   isfiniteθ = false
│   isfiniter = false
│   isfiniteℓπ = false
│   isfiniteℓκ = false
└ @ AdvancedHMC /Users/kcft114/.julia/packages/AdvancedHMC/YWXfk/src/hamiltonian.jl:36
┌ Warning: Incorrect ϵ = NaN; ϵ_previous = 0.07500000000000001 is used instead.
└ @ AdvancedHMC.Adaptation /Users/kcft114/.julia/packages/AdvancedHMC/YWXfk/src/adaptation/stepsize.jl:103
┌ Warning: The current proposal will be rejected due to numerical error(s).
│   isfiniteθ = true
│   isfiniter = true
│   isfiniteℓπ = false
│   isfiniteℓκ = true
└ @ AdvancedHMC /Users/kcft114/.julia/packages/AdvancedHMC/YWXfk/src/hamiltonian.jl:36
┌ Warning: The current proposal will be rejected due to numerical error(s).
│   isfiniteθ = false
│   isfiniter = false
│   isfiniteℓπ = false
│   isfiniteℓκ = false
└ @ AdvancedHMC /Users/kcft114/.julia/packages/AdvancedHMC/YWXfk/src/hamiltonian.jl:36
┌ Warning: Incorrect ϵ = NaN; ϵ_previous = 0.07500000000000001 is used instead.
└ @ AdvancedHMC.Adaptation /Users/kcft114/.julia/packages/AdvancedHMC/YWXfk/src/adaptation/stepsize.jl:103
┌ Warning: The current proposal will be rejected due to numerical error(s).
│   isfiniteθ = true
│   isfiniter = true
│   isfiniteℓπ = false
│   isfiniteℓκ = true
└ @ AdvancedHMC /Users/kcft114/.julia/packages/AdvancedHMC/YWXfk/src/hamiltonian.jl:36
┌ Warning: The current proposal will be rejected due to numerical error(s).
│   isfiniteθ = false
│   isfiniter = false
│   isfiniteℓπ = false
│   isfiniteℓκ = false
└ @ AdvancedHMC /Users/kcft114/.julia/packages/AdvancedHMC/YWXfk/src/hamiltonian.jl:36
┌ Warning: Incorrect ϵ = NaN; ϵ_previous = 0.07500000000000001 is used instead.
└ @ AdvancedHMC.Adaptation /Users/kcft114/.julia/packages/AdvancedHMC/YWXfk/src/adaptation/stepsize.jl:103
┌ Warning: The current proposal will be rejected due to numerical error(s).
│   isfiniteθ = true
│   isfiniter = true
│   isfiniteℓπ = false
│   isfiniteℓκ = true
└ @ AdvancedHMC /Users/kcft114/.julia/packages/AdvancedHMC/YWXfk/src/hamiltonian.jl:36
┌ Warning: The current proposal will be rejected due to numerical error(s).
│   isfiniteθ = false
│   isfiniter = false
│   isfiniteℓπ = false
│   isfiniteℓκ = false
└ @ AdvancedHMC /Users/kcft114/.julia/packages/AdvancedHMC/YWXfk/src/hamiltonian.jl:36
┌ Warning: Incorrect ϵ = NaN; ϵ_previous = 0.07500000000000001 is used instead.
└ @ AdvancedHMC.Adaptation /Users/kcft114/.julia/packages/AdvancedHMC/YWXfk/src/adaptation/stepsize.jl:103
┌ Warning: The current proposal will be rejected due to numerical error(s).
│   isfiniteθ = true
│   isfiniter = true
│   isfiniteℓπ = false
│   isfiniteℓκ = true
└ @ AdvancedHMC /Users/kcft114/.julia/packages/AdvancedHMC/YWXfk/src/hamiltonian.jl:36
┌ Warning: The current proposal will be rejected due to numerical error(s).
│   isfiniteθ = false
│   isfiniter = false
│   isfiniteℓπ = false
│   isfiniteℓκ = false
└ @ AdvancedHMC /Users/kcft114/.julia/packages/AdvancedHMC/YWXfk/src/hamiltonian.jl:36
┌ Warning: Incorrect ϵ = NaN; ϵ_previous = 0.07500000000000001 is used instead.
└ @ AdvancedHMC.Adaptation /Users/kcft114/.julia/packages/AdvancedHMC/YWXfk/src/adaptation/stepsize.jl:103
┌ Warning: The current proposal will be rejected due to numerical error(s).
│   isfiniteθ = true
│   isfiniter = true
│   isfiniteℓπ = false
│   isfiniteℓκ = true
└ @ AdvancedHMC /Users/kcft114/.julia/packages/AdvancedHMC/YWXfk/src/hamiltonian.jl:36
┌ Warning: The current proposal will be rejected due to numerical error(s).
│   isfiniteθ = false
│   isfiniter = false
│   isfiniteℓπ = false
│   isfiniteℓκ = false
└ @ AdvancedHMC /Users/kcft114/.julia/packages/AdvancedHMC/YWXfk/src/hamiltonian.jl:36
┌ Warning: Incorrect ϵ = NaN; ϵ_previous = 0.07500000000000001 is used instead.
└ @ AdvancedHMC.Adaptation /Users/kcft114/.julia/packages/AdvancedHMC/YWXfk/src/adaptation/stepsize.jl:103
┌ Warning: The current proposal will be rejected due to numerical error(s).
│   isfiniteθ = true
│   isfiniter = true
│   isfiniteℓπ = false
│   isfiniteℓκ = true
└ @ AdvancedHMC /Users/kcft114/.julia/packages/AdvancedHMC/YWXfk/src/hamiltonian.jl:36
┌ Warning: The current proposal will be rejected due to numerical error(s).
│   isfiniteθ = false
│   isfiniter = false
│   isfiniteℓπ = false
│   isfiniteℓκ = false
└ @ AdvancedHMC /Users/kcft114/.julia/packages/AdvancedHMC/YWXfk/src/hamiltonian.jl:36
┌ Warning: Incorrect ϵ = NaN; ϵ_previous = 0.07500000000000001 is used instead.
└ @ AdvancedHMC.Adaptation /Users/kcft114/.julia/packages/AdvancedHMC/YWXfk/src/adaptation/stepsize.jl:103
┌ Warning: The current proposal will be rejected due to numerical error(s).
│   isfiniteθ = true
│   isfiniter = true
│   isfiniteℓπ = false
│   isfiniteℓκ = true
└ @ AdvancedHMC /Users/kcft114/.julia/packages/AdvancedHMC/YWXfk/src/hamiltonian.jl:36
┌ Warning: The current proposal will be rejected due to numerical error(s).
│   isfiniteθ = false
│   isfiniter = false
│   isfiniteℓπ = false
│   isfiniteℓκ = false
└ @ AdvancedHMC /Users/kcft114/.julia/packages/AdvancedHMC/YWXfk/src/hamiltonian.jl:36
┌ Warning: Incorrect ϵ = NaN; ϵ_previous = 0.07500000000000001 is used instead.
└ @ AdvancedHMC.Adaptation /Users/kcft114/.julia/packages/AdvancedHMC/YWXfk/src/adaptation/stepsize.jl:103
┌ Warning: The current proposal will be rejected due to numerical error(s).
│   isfiniteθ = true
│   isfiniter = true
│   isfiniteℓπ = false
│   isfiniteℓκ = true
└ @ AdvancedHMC /Users/kcft114/.julia/packages/AdvancedHMC/YWXfk/src/hamiltonian.jl:36
┌ Warning: The current proposal will be rejected due to numerical error(s).
│   isfiniteθ = false
│   isfiniter = false
│   isfiniteℓπ = false
│   isfiniteℓκ = false
└ @ AdvancedHMC /Users/kcft114/.julia/packages/AdvancedHMC/YWXfk/src/hamiltonian.jl:36
┌ Warning: Incorrect ϵ = NaN; ϵ_previous = 0.07500000000000001 is used instead.
└ @ AdvancedHMC.Adaptation /Users/kcft114/.julia/packages/AdvancedHMC/YWXfk/src/adaptation/stepsize.jl:103
┌ Warning: The current proposal will be rejected due to numerical error(s).
│   isfiniteθ = true
│   isfiniter = true
│   isfiniteℓπ = false
│   isfiniteℓκ = true
└ @ AdvancedHMC /Users/kcft114/.julia/packages/AdvancedHMC/YWXfk/src/hamiltonian.jl:36
┌ Warning: The current proposal will be rejected due to numerical error(s).
│   isfiniteθ = false
│   isfiniter = false
│   isfiniteℓπ = false
│   isfiniteℓκ = false
└ @ AdvancedHMC /Users/kcft114/.julia/packages/AdvancedHMC/YWXfk/src/hamiltonian.jl:36
┌ Warning: Incorrect ϵ = NaN; ϵ_previous = 0.07500000000000001 is used instead.
└ @ AdvancedHMC.Adaptation /Users/kcft114/.julia/packages/AdvancedHMC/YWXfk/src/adaptation/stepsize.jl:103
┌ Warning: The current proposal will be rejected due to numerical error(s).
│   isfiniteθ = true
│   isfiniter = true
│   isfiniteℓπ = false
│   isfiniteℓκ = true
└ @ AdvancedHMC /Users/kcft114/.julia/packages/AdvancedHMC/YWXfk/src/hamiltonian.jl:36
┌ Warning: The current proposal will be rejected due to numerical error(s).
│   isfiniteθ = false
│   isfiniter = false
│   isfiniteℓπ = false
│   isfiniteℓκ = false
└ @ AdvancedHMC /Users/kcft114/.julia/packages/AdvancedHMC/YWXfk/src/hamiltonian.jl:36
┌ Warning: Incorrect ϵ = NaN; ϵ_previous = 0.07500000000000001 is used instead.
└ @ AdvancedHMC.Adaptation /Users/kcft114/.julia/packages/AdvancedHMC/YWXfk/src/adaptation/stepsize.jl:103
┌ Warning: The current proposal will be rejected due to numerical error(s).
│   isfiniteθ = true
│   isfiniter = true
│   isfiniteℓπ = false
│   isfiniteℓκ = true
└ @ AdvancedHMC /Users/kcft114/.julia/packages/AdvancedHMC/YWXfk/src/hamiltonian.jl:36
┌ Warning: The current proposal will be rejected due to numerical error(s).
│   isfiniteθ = false
│   isfiniter = false
│   isfiniteℓπ = false
│   isfiniteℓκ = false
└ @ AdvancedHMC /Users/kcft114/.julia/packages/AdvancedHMC/YWXfk/src/hamiltonian.jl:36
┌ Warning: Incorrect ϵ = NaN; ϵ_previous = 0.07500000000000001 is used instead.
└ @ AdvancedHMC.Adaptation /Users/kcft114/.julia/packages/AdvancedHMC/YWXfk/src/adaptation/stepsize.jl:103
┌ Warning: The current proposal will be rejected due to numerical error(s).
│   isfiniteθ = true
│   isfiniter = true
│   isfiniteℓπ = false
│   isfiniteℓκ = true
└ @ AdvancedHMC /Users/kcft114/.julia/packages/AdvancedHMC/YWXfk/src/hamiltonian.jl:36
┌ Warning: The current proposal will be rejected due to numerical error(s).
│   isfiniteθ = false
│   isfiniter = false
│   isfiniteℓπ = false
│   isfiniteℓκ = false
└ @ AdvancedHMC /Users/kcft114/.julia/packages/AdvancedHMC/YWXfk/src/hamiltonian.jl:36
┌ Warning: Incorrect ϵ = NaN; ϵ_previous = 0.07500000000000001 is used instead.
└ @ AdvancedHMC.Adaptation /Users/kcft114/.julia/packages/AdvancedHMC/YWXfk/src/adaptation/stepsize.jl:103
┌ Warning: The current proposal will be rejected due to numerical error(s).
│   isfiniteθ = true
│   isfiniter = true
│   isfiniteℓπ = false
│   isfiniteℓκ = true
└ @ AdvancedHMC /Users/kcft114/.julia/packages/AdvancedHMC/YWXfk/src/hamiltonian.jl:36
┌ Warning: The current proposal will be rejected due to numerical error(s).
│   isfiniteθ = false
│   isfiniter = false
│   isfiniteℓπ = false
│   isfiniteℓκ = false
└ @ AdvancedHMC /Users/kcft114/.julia/packages/AdvancedHMC/YWXfk/src/hamiltonian.jl:36
┌ Warning: Incorrect ϵ = NaN; ϵ_previous = 0.07500000000000001 is used instead.
└ @ AdvancedHMC.Adaptation /Users/kcft114/.julia/packages/AdvancedHMC/YWXfk/src/adaptation/stepsize.jl:103
┌ Warning: The current proposal will be rejected due to numerical error(s).
│   isfiniteθ = true
│   isfiniter = true
│   isfiniteℓπ = false
│   isfiniteℓκ = true
└ @ AdvancedHMC /Users/kcft114/.julia/packages/AdvancedHMC/YWXfk/src/hamiltonian.jl:36
┌ Warning: The current proposal will be rejected due to numerical error(s).
│   isfiniteθ = false
│   isfiniter = false
│   isfiniteℓπ = false
│   isfiniteℓκ = false
└ @ AdvancedHMC /Users/kcft114/.julia/packages/AdvancedHMC/YWXfk/src/hamiltonian.jl:36
┌ Warning: Incorrect ϵ = NaN; ϵ_previous = 0.07500000000000001 is used instead.
└ @ AdvancedHMC.Adaptation /Users/kcft114/.julia/packages/AdvancedHMC/YWXfk/src/adaptation/stepsize.jl:103
┌ Warning: The current proposal will be rejected due to numerical error(s).
│   isfiniteθ = true
│   isfiniter = true
│   isfiniteℓπ = false
│   isfiniteℓκ = true
└ @ AdvancedHMC /Users/kcft114/.julia/packages/AdvancedHMC/YWXfk/src/hamiltonian.jl:36
┌ Warning: The current proposal will be rejected due to numerical error(s).
│   isfiniteθ = false
│   isfiniter = false
│   isfiniteℓπ = false
│   isfiniteℓκ = false
└ @ AdvancedHMC /Users/kcft114/.julia/packages/AdvancedHMC/YWXfk/src/hamiltonian.jl:36
┌ Warning: Incorrect ϵ = NaN; ϵ_previous = 0.07500000000000001 is used instead.
└ @ AdvancedHMC.Adaptation /Users/kcft114/.julia/packages/AdvancedHMC/YWXfk/src/adaptation/stepsize.jl:103
┌ Warning: The current proposal will be rejected due to numerical error(s).
│   isfiniteθ = true
│   isfiniter = true
│   isfiniteℓπ = false
│   isfiniteℓκ = true
└ @ AdvancedHMC /Users/kcft114/.julia/packages/AdvancedHMC/YWXfk/src/hamiltonian.jl:36
┌ Warning: The current proposal will be rejected due to numerical error(s).
│   isfiniteθ = false
│   isfiniter = false
│   isfiniteℓπ = false
│   isfiniteℓκ = false
└ @ AdvancedHMC /Users/kcft114/.julia/packages/AdvancedHMC/YWXfk/src/hamiltonian.jl:36
┌ Warning: Incorrect ϵ = NaN; ϵ_previous = 0.07500000000000001 is used instead.
└ @ AdvancedHMC.Adaptation /Users/kcft114/.julia/packages/AdvancedHMC/YWXfk/src/adaptation/stepsize.jl:103
┌ Warning: The current proposal will be rejected due to numerical error(s).
│   isfiniteθ = true
│   isfiniter = true
│   isfiniteℓπ = false
│   isfiniteℓκ = true
└ @ AdvancedHMC /Users/kcft114/.julia/packages/AdvancedHMC/YWXfk/src/hamiltonian.jl:36
┌ Warning: The current proposal will be rejected due to numerical error(s).
│   isfiniteθ = false
│   isfiniter = false
│   isfiniteℓπ = false
│   isfiniteℓκ = false
└ @ AdvancedHMC /Users/kcft114/.julia/packages/AdvancedHMC/YWXfk/src/hamiltonian.jl:36
┌ Warning: Incorrect ϵ = NaN; ϵ_previous = 0.07500000000000001 is used instead.
└ @ AdvancedHMC.Adaptation /Users/kcft114/.julia/packages/AdvancedHMC/YWXfk/src/adaptation/stepsize.jl:103
┌ Warning: The current proposal will be rejected due to numerical error(s).
│   isfiniteθ = true
│   isfiniter = true
│   isfiniteℓπ = false
│   isfiniteℓκ = true
└ @ AdvancedHMC /Users/kcft114/.julia/packages/AdvancedHMC/YWXfk/src/hamiltonian.jl:36
┌ Warning: The current proposal will be rejected due to numerical error(s).
│   isfiniteθ = false
│   isfiniter = false
│   isfiniteℓπ = false
│   isfiniteℓκ = false
└ @ AdvancedHMC /Users/kcft114/.julia/packages/AdvancedHMC/YWXfk/src/hamiltonian.jl:36
┌ Warning: Incorrect ϵ = NaN; ϵ_previous = 0.07500000000000001 is used instead.
└ @ AdvancedHMC.Adaptation /Users/kcft114/.julia/packages/AdvancedHMC/YWXfk/src/adaptation/stepsize.jl:103
┌ Warning: The current proposal will be rejected due to numerical error(s).
│   isfiniteθ = true
│   isfiniter = true
│   isfiniteℓπ = false
│   isfiniteℓκ = true
└ @ AdvancedHMC /Users/kcft114/.julia/packages/AdvancedHMC/YWXfk/src/hamiltonian.jl:36
┌ Warning: The current proposal will be rejected due to numerical error(s).
│   isfiniteθ = false
│   isfiniter = false
│   isfiniteℓπ = false
│   isfiniteℓκ = false
└ @ AdvancedHMC /Users/kcft114/.julia/packages/AdvancedHMC/YWXfk/src/hamiltonian.jl:36
┌ Warning: Incorrect ϵ = NaN; ϵ_previous = 0.07500000000000001 is used instead.
└ @ AdvancedHMC.Adaptation /Users/kcft114/.julia/packages/AdvancedHMC/YWXfk/src/adaptation/stepsize.jl:103
┌ Warning: The current proposal will be rejected due to numerical error(s).
│   isfiniteθ = true
│   isfiniter = true
│   isfiniteℓπ = false
│   isfiniteℓκ = true
└ @ AdvancedHMC /Users/kcft114/.julia/packages/AdvancedHMC/YWXfk/src/hamiltonian.jl:36
┌ Warning: The current proposal will be rejected due to numerical error(s).
│   isfiniteθ = false
│   isfiniter = false
│   isfiniteℓπ = false
│   isfiniteℓκ = false
└ @ AdvancedHMC /Users/kcft114/.julia/packages/AdvancedHMC/YWXfk/src/hamiltonian.jl:36
┌ Warning: Incorrect ϵ = NaN; ϵ_previous = 0.07500000000000001 is used instead.
└ @ AdvancedHMC.Adaptation /Users/kcft114/.julia/packages/AdvancedHMC/YWXfk/src/adaptation/stepsize.jl:103
┌ Warning: The current proposal will be rejected due to numerical error(s).
│   isfiniteθ = true
│   isfiniter = true
│   isfiniteℓπ = false
│   isfiniteℓκ = true
└ @ AdvancedHMC /Users/kcft114/.julia/packages/AdvancedHMC/YWXfk/src/hamiltonian.jl:36
┌ Warning: The current proposal will be rejected due to numerical error(s).
│   isfiniteθ = false
│   isfiniter = false
│   isfiniteℓπ = false
│   isfiniteℓκ = false
└ @ AdvancedHMC /Users/kcft114/.julia/packages/AdvancedHMC/YWXfk/src/hamiltonian.jl:36
┌ Warning: Incorrect ϵ = NaN; ϵ_previous = 0.07500000000000001 is used instead.
└ @ AdvancedHMC.Adaptation /Users/kcft114/.julia/packages/AdvancedHMC/YWXfk/src/adaptation/stepsize.jl:103
┌ Warning: The current proposal will be rejected due to numerical error(s).
│   isfiniteθ = true
│   isfiniter = true
│   isfiniteℓπ = false
│   isfiniteℓκ = true
└ @ AdvancedHMC /Users/kcft114/.julia/packages/AdvancedHMC/YWXfk/src/hamiltonian.jl:36
┌ Warning: The current proposal will be rejected due to numerical error(s).
│   isfiniteθ = false
│   isfiniter = false
│   isfiniteℓπ = false
│   isfiniteℓκ = false
└ @ AdvancedHMC /Users/kcft114/.julia/packages/AdvancedHMC/YWXfk/src/hamiltonian.jl:36
┌ Warning: Incorrect ϵ = NaN; ϵ_previous = 0.07500000000000001 is used instead.
└ @ AdvancedHMC.Adaptation /Users/kcft114/.julia/packages/AdvancedHMC/YWXfk/src/adaptation/stepsize.jl:103
┌ Warning: The current proposal will be rejected due to numerical error(s).
│   isfiniteθ = true
│   isfiniter = true
│   isfiniteℓπ = false
│   isfiniteℓκ = true
└ @ AdvancedHMC /Users/kcft114/.julia/packages/AdvancedHMC/YWXfk/src/hamiltonian.jl:36
┌ Warning: The current proposal will be rejected due to numerical error(s).
│   isfiniteθ = false
│   isfiniter = false
│   isfiniteℓπ = false
│   isfiniteℓκ = false
└ @ AdvancedHMC /Users/kcft114/.julia/packages/AdvancedHMC/YWXfk/src/hamiltonian.jl:36
┌ Warning: Incorrect ϵ = NaN; ϵ_previous = 0.07500000000000001 is used instead.
└ @ AdvancedHMC.Adaptation /Users/kcft114/.julia/packages/AdvancedHMC/YWXfk/src/adaptation/stepsize.jl:103
┌ Warning: The current proposal will be rejected due to numerical error(s).
│   isfiniteθ = true
│   isfiniter = true
│   isfiniteℓπ = false
│   isfiniteℓκ = true
└ @ AdvancedHMC /Users/kcft114/.julia/packages/AdvancedHMC/YWXfk/src/hamiltonian.jl:36
┌ Warning: The current proposal will be rejected due to numerical error(s).
│   isfiniteθ = false
│   isfiniter = false
│   isfiniteℓπ = false
│   isfiniteℓκ = false
└ @ AdvancedHMC /Users/kcft114/.julia/packages/AdvancedHMC/YWXfk/src/hamiltonian.jl:36
┌ Warning: Incorrect ϵ = NaN; ϵ_previous = 0.07500000000000001 is used instead.
└ @ AdvancedHMC.Adaptation /Users/kcft114/.julia/packages/AdvancedHMC/YWXfk/src/adaptation/stepsize.jl:103
┌ Warning: The current proposal will be rejected due to numerical error(s).
│   isfiniteθ = true
│   isfiniter = true
│   isfiniteℓπ = false
│   isfiniteℓκ = true
└ @ AdvancedHMC /Users/kcft114/.julia/packages/AdvancedHMC/YWXfk/src/hamiltonian.jl:36
┌ Warning: The current proposal will be rejected due to numerical error(s).
│   isfiniteθ = false
│   isfiniter = false
│   isfiniteℓπ = false
│   isfiniteℓκ = false
└ @ AdvancedHMC /Users/kcft114/.julia/packages/AdvancedHMC/YWXfk/src/hamiltonian.jl:36
┌ Warning: Incorrect ϵ = NaN; ϵ_previous = 0.07500000000000001 is used instead.
└ @ AdvancedHMC.Adaptation /Users/kcft114/.julia/packages/AdvancedHMC/YWXfk/src/adaptation/stepsize.jl:103
┌ Warning: The current proposal will be rejected due to numerical error(s).
│   isfiniteθ = true
│   isfiniter = true
│   isfiniteℓπ = false
│   isfiniteℓκ = true
└ @ AdvancedHMC /Users/kcft114/.julia/packages/AdvancedHMC/YWXfk/src/hamiltonian.jl:36
┌ Warning: The current proposal will be rejected due to numerical error(s).
│   isfiniteθ = false
│   isfiniter = false
│   isfiniteℓπ = false
│   isfiniteℓκ = false
└ @ AdvancedHMC /Users/kcft114/.julia/packages/AdvancedHMC/YWXfk/src/hamiltonian.jl:36
┌ Warning: Incorrect ϵ = NaN; ϵ_previous = 0.07500000000000001 is used instead.
└ @ AdvancedHMC.Adaptation /Users/kcft114/.julia/packages/AdvancedHMC/YWXfk/src/adaptation/stepsize.jl:103
┌ Warning: The current proposal will be rejected due to numerical error(s).
│   isfiniteθ = true
│   isfiniter = true
│   isfiniteℓπ = false
│   isfiniteℓκ = true
└ @ AdvancedHMC /Users/kcft114/.julia/packages/AdvancedHMC/YWXfk/src/hamiltonian.jl:36
┌ Warning: The current proposal will be rejected due to numerical error(s).
│   isfiniteθ = false
│   isfiniter = false
│   isfiniteℓπ = false
│   isfiniteℓκ = false
└ @ AdvancedHMC /Users/kcft114/.julia/packages/AdvancedHMC/YWXfk/src/hamiltonian.jl:36
┌ Warning: Incorrect ϵ = NaN; ϵ_previous = 0.07500000000000001 is used instead.
└ @ AdvancedHMC.Adaptation /Users/kcft114/.julia/packages/AdvancedHMC/YWXfk/src/adaptation/stepsize.jl:103
┌ Warning: The current proposal will be rejected due to numerical error(s).
│   isfiniteθ = true
│   isfiniter = true
│   isfiniteℓπ = false
│   isfiniteℓκ = true
└ @ AdvancedHMC /Users/kcft114/.julia/packages/AdvancedHMC/YWXfk/src/hamiltonian.jl:36
┌ Warning: The current proposal will be rejected due to numerical error(s).
│   isfiniteθ = false
│   isfiniter = false
│   isfiniteℓπ = false
│   isfiniteℓκ = false
└ @ AdvancedHMC /Users/kcft114/.julia/packages/AdvancedHMC/YWXfk/src/hamiltonian.jl:36
┌ Warning: Incorrect ϵ = NaN; ϵ_previous = 0.07500000000000001 is used instead.
└ @ AdvancedHMC.Adaptation /Users/kcft114/.julia/packages/AdvancedHMC/YWXfk/src/adaptation/stepsize.jl:103
┌ Warning: The current proposal will be rejected due to numerical error(s).
│   isfiniteθ = true
│   isfiniter = true
│   isfiniteℓπ = false
│   isfiniteℓκ = true
└ @ AdvancedHMC /Users/kcft114/.julia/packages/AdvancedHMC/YWXfk/src/hamiltonian.jl:36
┌ Warning: The current proposal will be rejected due to numerical error(s).
│   isfiniteθ = false
│   isfiniter = false
│   isfiniteℓπ = false
│   isfiniteℓκ = false
└ @ AdvancedHMC /Users/kcft114/.julia/packages/AdvancedHMC/YWXfk/src/hamiltonian.jl:36
┌ Warning: Incorrect ϵ = NaN; ϵ_previous = 0.07500000000000001 is used instead.
└ @ AdvancedHMC.Adaptation /Users/kcft114/.julia/packages/AdvancedHMC/YWXfk/src/adaptation/stepsize.jl:103
┌ Warning: The current proposal will be rejected due to numerical error(s).
│   isfiniteθ = true
│   isfiniter = true
│   isfiniteℓπ = false
│   isfiniteℓκ = true
└ @ AdvancedHMC /Users/kcft114/.julia/packages/AdvancedHMC/YWXfk/src/hamiltonian.jl:36
┌ Warning: The current proposal will be rejected due to numerical error(s).
│   isfiniteθ = false
│   isfiniter = false
│   isfiniteℓπ = false
│   isfiniteℓκ = false
└ @ AdvancedHMC /Users/kcft114/.julia/packages/AdvancedHMC/YWXfk/src/hamiltonian.jl:36
┌ Warning: Incorrect ϵ = NaN; ϵ_previous = 0.07500000000000001 is used instead.
└ @ AdvancedHMC.Adaptation /Users/kcft114/.julia/packages/AdvancedHMC/YWXfk/src/adaptation/stepsize.jl:103
┌ Warning: The current proposal will be rejected due to numerical error(s).
│   isfiniteθ = true
│   isfiniter = true
│   isfiniteℓπ = false
│   isfiniteℓκ = true
└ @ AdvancedHMC /Users/kcft114/.julia/packages/AdvancedHMC/YWXfk/src/hamiltonian.jl:36
┌ Warning: The current proposal will be rejected due to numerical error(s).
│   isfiniteθ = false
│   isfiniter = false
│   isfiniteℓπ = false
│   isfiniteℓκ = false
└ @ AdvancedHMC /Users/kcft114/.julia/packages/AdvancedHMC/YWXfk/src/hamiltonian.jl:36
┌ Warning: Incorrect ϵ = NaN; ϵ_previous = 0.07500000000000001 is used instead.
└ @ AdvancedHMC.Adaptation /Users/kcft114/.julia/packages/AdvancedHMC/YWXfk/src/adaptation/stepsize.jl:103
┌ Warning: The current proposal will be rejected due to numerical error(s).
│   isfiniteθ = true
│   isfiniter = true
│   isfiniteℓπ = false
│   isfiniteℓκ = true
└ @ AdvancedHMC /Users/kcft114/.julia/packages/AdvancedHMC/YWXfk/src/hamiltonian.jl:36
┌ Warning: The current proposal will be rejected due to numerical error(s).
│   isfiniteθ = false
│   isfiniter = false
│   isfiniteℓπ = false
│   isfiniteℓκ = false
└ @ AdvancedHMC /Users/kcft114/.julia/packages/AdvancedHMC/YWXfk/src/hamiltonian.jl:36
┌ Warning: Incorrect ϵ = NaN; ϵ_previous = 0.07500000000000001 is used instead.
└ @ AdvancedHMC.Adaptation /Users/kcft114/.julia/packages/AdvancedHMC/YWXfk/src/adaptation/stepsize.jl:103
┌ Warning: The current proposal will be rejected due to numerical error(s).
│   isfiniteθ = true
│   isfiniter = true
│   isfiniteℓπ = false
│   isfiniteℓκ = true
└ @ AdvancedHMC /Users/kcft114/.julia/packages/AdvancedHMC/YWXfk/src/hamiltonian.jl:36
┌ Warning: The current proposal will be rejected due to numerical error(s).
│   isfiniteθ = false
│   isfiniter = false
│   isfiniteℓπ = false
│   isfiniteℓκ = false
└ @ AdvancedHMC /Users/kcft114/.julia/packages/AdvancedHMC/YWXfk/src/hamiltonian.jl:36
┌ Warning: Incorrect ϵ = NaN; ϵ_previous = 0.07500000000000001 is used instead.
└ @ AdvancedHMC.Adaptation /Users/kcft114/.julia/packages/AdvancedHMC/YWXfk/src/adaptation/stepsize.jl:103
┌ Warning: The current proposal will be rejected due to numerical error(s).
│   isfiniteθ = true
│   isfiniter = true
│   isfiniteℓπ = false
│   isfiniteℓκ = true
└ @ AdvancedHMC /Users/kcft114/.julia/packages/AdvancedHMC/YWXfk/src/hamiltonian.jl:36
┌ Warning: The current proposal will be rejected due to numerical error(s).
│   isfiniteθ = false
│   isfiniter = false
│   isfiniteℓπ = false
│   isfiniteℓκ = false
└ @ AdvancedHMC /Users/kcft114/.julia/packages/AdvancedHMC/YWXfk/src/hamiltonian.jl:36
┌ Warning: Incorrect ϵ = NaN; ϵ_previous = 0.07500000000000001 is used instead.
└ @ AdvancedHMC.Adaptation /Users/kcft114/.julia/packages/AdvancedHMC/YWXfk/src/adaptation/stepsize.jl:103
┌ Warning: The current proposal will be rejected due to numerical error(s).
│   isfiniteθ = true
│   isfiniter = true
│   isfiniteℓπ = false
│   isfiniteℓκ = true
└ @ AdvancedHMC /Users/kcft114/.julia/packages/AdvancedHMC/YWXfk/src/hamiltonian.jl:36
┌ Warning: The current proposal will be rejected due to numerical error(s).
│   isfiniteθ = false
│   isfiniter = false
│   isfiniteℓπ = false
│   isfiniteℓκ = false
└ @ AdvancedHMC /Users/kcft114/.julia/packages/AdvancedHMC/YWXfk/src/hamiltonian.jl:36
┌ Warning: Incorrect ϵ = NaN; ϵ_previous = 0.07500000000000001 is used instead.
└ @ AdvancedHMC.Adaptation /Users/kcft114/.julia/packages/AdvancedHMC/YWXfk/src/adaptation/stepsize.jl:103
┌ Warning: The current proposal will be rejected due to numerical error(s).
│   isfiniteθ = true
│   isfiniter = true
│   isfiniteℓπ = false
│   isfiniteℓκ = true
└ @ AdvancedHMC /Users/kcft114/.julia/packages/AdvancedHMC/YWXfk/src/hamiltonian.jl:36
┌ Warning: The current proposal will be rejected due to numerical error(s).
│   isfiniteθ = false
│   isfiniter = false
│   isfiniteℓπ = false
│   isfiniteℓκ = false
└ @ AdvancedHMC /Users/kcft114/.julia/packages/AdvancedHMC/YWXfk/src/hamiltonian.jl:36
┌ Warning: Incorrect ϵ = NaN; ϵ_previous = 0.07500000000000001 is used instead.
└ @ AdvancedHMC.Adaptation /Users/kcft114/.julia/packages/AdvancedHMC/YWXfk/src/adaptation/stepsize.jl:103
┌ Warning: The current proposal will be rejected due to numerical error(s).
│   isfiniteθ = true
│   isfiniter = true
│   isfiniteℓπ = false
│   isfiniteℓκ = true
└ @ AdvancedHMC /Users/kcft114/.julia/packages/AdvancedHMC/YWXfk/src/hamiltonian.jl:36
┌ Warning: The current proposal will be rejected due to numerical error(s).
│   isfiniteθ = false
│   isfiniter = false
│   isfiniteℓπ = false
│   isfiniteℓκ = false
└ @ AdvancedHMC /Users/kcft114/.julia/packages/AdvancedHMC/YWXfk/src/hamiltonian.jl:36
┌ Warning: Incorrect ϵ = NaN; ϵ_previous = 0.07500000000000001 is used instead.
└ @ AdvancedHMC.Adaptation /Users/kcft114/.julia/packages/AdvancedHMC/YWXfk/src/adaptation/stepsize.jl:103
┌ Warning: The current proposal will be rejected due to numerical error(s).
│   isfiniteθ = true
│   isfiniter = true
│   isfiniteℓπ = false
│   isfiniteℓκ = true
└ @ AdvancedHMC /Users/kcft114/.julia/packages/AdvancedHMC/YWXfk/src/hamiltonian.jl:36
┌ Warning: The current proposal will be rejected due to numerical error(s).
│   isfiniteθ = false
│   isfiniter = false
│   isfiniteℓπ = false
│   isfiniteℓκ = false
└ @ AdvancedHMC /Users/kcft114/.julia/packages/AdvancedHMC/YWXfk/src/hamiltonian.jl:36
┌ Warning: Incorrect ϵ = NaN; ϵ_previous = 0.07500000000000001 is used instead.
└ @ AdvancedHMC.Adaptation /Users/kcft114/.julia/packages/AdvancedHMC/YWXfk/src/adaptation/stepsize.jl:103
┌ Warning: The current proposal will be rejected due to numerical error(s).
│   isfiniteθ = true
│   isfiniter = true
│   isfiniteℓπ = false
│   isfiniteℓκ = true
└ @ AdvancedHMC /Users/kcft114/.julia/packages/AdvancedHMC/YWXfk/src/hamiltonian.jl:36
┌ Warning: The current proposal will be rejected due to numerical error(s).
│   isfiniteθ = false
│   isfiniter = false
│   isfiniteℓπ = false
│   isfiniteℓκ = false
└ @ AdvancedHMC /Users/kcft114/.julia/packages/AdvancedHMC/YWXfk/src/hamiltonian.jl:36
┌ Warning: Incorrect ϵ = NaN; ϵ_previous = 0.07500000000000001 is used instead.
└ @ AdvancedHMC.Adaptation /Users/kcft114/.julia/packages/AdvancedHMC/YWXfk/src/adaptation/stepsize.jl:103
┌ Warning: The current proposal will be rejected due to numerical error(s).
│   isfiniteθ = true
│   isfiniter = true
│   isfiniteℓπ = false
│   isfiniteℓκ = true
└ @ AdvancedHMC /Users/kcft114/.julia/packages/AdvancedHMC/YWXfk/src/hamiltonian.jl:36
┌ Warning: The current proposal will be rejected due to numerical error(s).
│   isfiniteθ = false
│   isfiniter = false
│   isfiniteℓπ = false
│   isfiniteℓκ = false
└ @ AdvancedHMC /Users/kcft114/.julia/packages/AdvancedHMC/YWXfk/src/hamiltonian.jl:36
┌ Warning: Incorrect ϵ = NaN; ϵ_previous = 0.07500000000000001 is used instead.
└ @ AdvancedHMC.Adaptation /Users/kcft114/.julia/packages/AdvancedHMC/YWXfk/src/adaptation/stepsize.jl:103
┌ Warning: The current proposal will be rejected due to numerical error(s).
│   isfiniteθ = true
│   isfiniter = true
│   isfiniteℓπ = false
│   isfiniteℓκ = true
└ @ AdvancedHMC /Users/kcft114/.julia/packages/AdvancedHMC/YWXfk/src/hamiltonian.jl:36
┌ Warning: The current proposal will be rejected due to numerical error(s).
│   isfiniteθ = false
│   isfiniter = false
│   isfiniteℓπ = false
│   isfiniteℓκ = false
└ @ AdvancedHMC /Users/kcft114/.julia/packages/AdvancedHMC/YWXfk/src/hamiltonian.jl:36
┌ Warning: Incorrect ϵ = NaN; ϵ_previous = 0.07500000000000001 is used instead.
└ @ AdvancedHMC.Adaptation /Users/kcft114/.julia/packages/AdvancedHMC/YWXfk/src/adaptation/stepsize.jl:103
┌ Warning: The current proposal will be rejected due to numerical error(s).
│   isfiniteθ = true
│   isfiniter = true
│   isfiniteℓπ = false
│   isfiniteℓκ = true
└ @ AdvancedHMC /Users/kcft114/.julia/packages/AdvancedHMC/YWXfk/src/hamiltonian.jl:36
┌ Warning: The current proposal will be rejected due to numerical error(s).
│   isfiniteθ = false
│   isfiniter = false
│   isfiniteℓπ = false
│   isfiniteℓκ = false
└ @ AdvancedHMC /Users/kcft114/.julia/packages/AdvancedHMC/YWXfk/src/hamiltonian.jl:36
┌ Warning: Incorrect ϵ = NaN; ϵ_previous = 0.07500000000000001 is used instead.
└ @ AdvancedHMC.Adaptation /Users/kcft114/.julia/packages/AdvancedHMC/YWXfk/src/adaptation/stepsize.jl:103
┌ Warning: The current proposal will be rejected due to numerical error(s).
│   isfiniteθ = true
│   isfiniter = true
│   isfiniteℓπ = false
│   isfiniteℓκ = true
└ @ AdvancedHMC /Users/kcft114/.julia/packages/AdvancedHMC/YWXfk/src/hamiltonian.jl:36
┌ Warning: The current proposal will be rejected due to numerical error(s).
│   isfiniteθ = false
│   isfiniter = false
│   isfiniteℓπ = false
│   isfiniteℓκ = false
└ @ AdvancedHMC /Users/kcft114/.julia/packages/AdvancedHMC/YWXfk/src/hamiltonian.jl:36
┌ Warning: Incorrect ϵ = NaN; ϵ_previous = 0.07500000000000001 is used instead.
└ @ AdvancedHMC.Adaptation /Users/kcft114/.julia/packages/AdvancedHMC/YWXfk/src/adaptation/stepsize.jl:103
┌ Warning: The current proposal will be rejected due to numerical error(s).
│   isfiniteθ = true
│   isfiniter = true
│   isfiniteℓπ = false
│   isfiniteℓκ = true
└ @ AdvancedHMC /Users/kcft114/.julia/packages/AdvancedHMC/YWXfk/src/hamiltonian.jl:36
┌ Warning: The current proposal will be rejected due to numerical error(s).
│   isfiniteθ = false
│   isfiniter = false
│   isfiniteℓπ = false
│   isfiniteℓκ = false
└ @ AdvancedHMC /Users/kcft114/.julia/packages/AdvancedHMC/YWXfk/src/hamiltonian.jl:36
┌ Warning: Incorrect ϵ = NaN; ϵ_previous = 0.07500000000000001 is used instead.
└ @ AdvancedHMC.Adaptation /Users/kcft114/.julia/packages/AdvancedHMC/YWXfk/src/adaptation/stepsize.jl:103
┌ Warning: The current proposal will be rejected due to numerical error(s).
│   isfiniteθ = true
│   isfiniter = true
│   isfiniteℓπ = false
│   isfiniteℓκ = true
└ @ AdvancedHMC /Users/kcft114/.julia/packages/AdvancedHMC/YWXfk/src/hamiltonian.jl:36
┌ Warning: The current proposal will be rejected due to numerical error(s).
│   isfiniteθ = false
│   isfiniter = false
│   isfiniteℓπ = false
│   isfiniteℓκ = false
└ @ AdvancedHMC /Users/kcft114/.julia/packages/AdvancedHMC/YWXfk/src/hamiltonian.jl:36
┌ Warning: Incorrect ϵ = NaN; ϵ_previous = 0.07500000000000001 is used instead.
└ @ AdvancedHMC.Adaptation /Users/kcft114/.julia/packages/AdvancedHMC/YWXfk/src/adaptation/stepsize.jl:103
┌ Warning: The current proposal will be rejected due to numerical error(s).
│   isfiniteθ = true
│   isfiniter = true
│   isfiniteℓπ = false
│   isfiniteℓκ = true
└ @ AdvancedHMC /Users/kcft114/.julia/packages/AdvancedHMC/YWXfk/src/hamiltonian.jl:36
┌ Warning: The current proposal will be rejected due to numerical error(s).
│   isfiniteθ = false
│   isfiniter = false
│   isfiniteℓπ = false
│   isfiniteℓκ = false
└ @ AdvancedHMC /Users/kcft114/.julia/packages/AdvancedHMC/YWXfk/src/hamiltonian.jl:36
┌ Warning: Incorrect ϵ = NaN; ϵ_previous = 0.07500000000000001 is used instead.
└ @ AdvancedHMC.Adaptation /Users/kcft114/.julia/packages/AdvancedHMC/YWXfk/src/adaptation/stepsize.jl:103
┌ Warning: The current proposal will be rejected due to numerical error(s).
│   isfiniteθ = true
│   isfiniter = true
│   isfiniteℓπ = false
│   isfiniteℓκ = true
└ @ AdvancedHMC /Users/kcft114/.julia/packages/AdvancedHMC/YWXfk/src/hamiltonian.jl:36
┌ Warning: The current proposal will be rejected due to numerical error(s).
│   isfiniteθ = false
│   isfiniter = false
│   isfiniteℓπ = false
│   isfiniteℓκ = false
└ @ AdvancedHMC /Users/kcft114/.julia/packages/AdvancedHMC/YWXfk/src/hamiltonian.jl:36
┌ Warning: Incorrect ϵ = NaN; ϵ_previous = 0.07500000000000001 is used instead.
└ @ AdvancedHMC.Adaptation /Users/kcft114/.julia/packages/AdvancedHMC/YWXfk/src/adaptation/stepsize.jl:103
┌ Warning: The current proposal will be rejected due to numerical error(s).
│   isfiniteθ = true
│   isfiniter = true
│   isfiniteℓπ = false
│   isfiniteℓκ = true
└ @ AdvancedHMC /Users/kcft114/.julia/packages/AdvancedHMC/YWXfk/src/hamiltonian.jl:36
┌ Warning: The current proposal will be rejected due to numerical error(s).
│   isfiniteθ = false
│   isfiniter = false
│   isfiniteℓπ = false
│   isfiniteℓκ = false
└ @ AdvancedHMC /Users/kcft114/.julia/packages/AdvancedHMC/YWXfk/src/hamiltonian.jl:36
┌ Warning: Incorrect ϵ = NaN; ϵ_previous = 0.07500000000000001 is used instead.
└ @ AdvancedHMC.Adaptation /Users/kcft114/.julia/packages/AdvancedHMC/YWXfk/src/adaptation/stepsize.jl:103
┌ Warning: The current proposal will be rejected due to numerical error(s).
│   isfiniteθ = true
│   isfiniter = true
│   isfiniteℓπ = false
│   isfiniteℓκ = true
└ @ AdvancedHMC /Users/kcft114/.julia/packages/AdvancedHMC/YWXfk/src/hamiltonian.jl:36
┌ Warning: The current proposal will be rejected due to numerical error(s).
│   isfiniteθ = false
│   isfiniter = false
│   isfiniteℓπ = false
│   isfiniteℓκ = false
└ @ AdvancedHMC /Users/kcft114/.julia/packages/AdvancedHMC/YWXfk/src/hamiltonian.jl:36
┌ Warning: Incorrect ϵ = NaN; ϵ_previous = 0.07500000000000001 is used instead.
└ @ AdvancedHMC.Adaptation /Users/kcft114/.julia/packages/AdvancedHMC/YWXfk/src/adaptation/stepsize.jl:103
┌ Warning: The current proposal will be rejected due to numerical error(s).
│   isfiniteθ = true
│   isfiniter = true
│   isfiniteℓπ = false
│   isfiniteℓκ = true
└ @ AdvancedHMC /Users/kcft114/.julia/packages/AdvancedHMC/YWXfk/src/hamiltonian.jl:36
┌ Warning: The current proposal will be rejected due to numerical error(s).
│   isfiniteθ = false
│   isfiniter = false
│   isfiniteℓπ = false
│   isfiniteℓκ = false
└ @ AdvancedHMC /Users/kcft114/.julia/packages/AdvancedHMC/YWXfk/src/hamiltonian.jl:36
┌ Warning: Incorrect ϵ = NaN; ϵ_previous = 0.07500000000000001 is used instead.
└ @ AdvancedHMC.Adaptation /Users/kcft114/.julia/packages/AdvancedHMC/YWXfk/src/adaptation/stepsize.jl:103
┌ Warning: The current proposal will be rejected due to numerical error(s).
│   isfiniteθ = true
│   isfiniter = true
│   isfiniteℓπ = false
│   isfiniteℓκ = true
└ @ AdvancedHMC /Users/kcft114/.julia/packages/AdvancedHMC/YWXfk/src/hamiltonian.jl:36
┌ Warning: The current proposal will be rejected due to numerical error(s).
│   isfiniteθ = false
│   isfiniter = false
│   isfiniteℓπ = false
│   isfiniteℓκ = false
└ @ AdvancedHMC /Users/kcft114/.julia/packages/AdvancedHMC/YWXfk/src/hamiltonian.jl:36
┌ Warning: Incorrect ϵ = NaN; ϵ_previous = 0.07500000000000001 is used instead.
└ @ AdvancedHMC.Adaptation /Users/kcft114/.julia/packages/AdvancedHMC/YWXfk/src/adaptation/stepsize.jl:103
┌ Warning: The current proposal will be rejected due to numerical error(s).
│   isfiniteθ = true
│   isfiniter = true
│   isfiniteℓπ = false
│   isfiniteℓκ = true
└ @ AdvancedHMC /Users/kcft114/.julia/packages/AdvancedHMC/YWXfk/src/hamiltonian.jl:36
┌ Warning: The current proposal will be rejected due to numerical error(s).
│   isfiniteθ = false
│   isfiniter = false
│   isfiniteℓπ = false
│   isfiniteℓκ = false
└ @ AdvancedHMC /Users/kcft114/.julia/packages/AdvancedHMC/YWXfk/src/hamiltonian.jl:36
┌ Warning: Incorrect ϵ = NaN; ϵ_previous = 0.07500000000000001 is used instead.
└ @ AdvancedHMC.Adaptation /Users/kcft114/.julia/packages/AdvancedHMC/YWXfk/src/adaptation/stepsize.jl:103
┌ Warning: The current proposal will be rejected due to numerical error(s).
│   isfiniteθ = true
│   isfiniter = true
│   isfiniteℓπ = false
│   isfiniteℓκ = true
└ @ AdvancedHMC /Users/kcft114/.julia/packages/AdvancedHMC/YWXfk/src/hamiltonian.jl:36
┌ Warning: The current proposal will be rejected due to numerical error(s).
│   isfiniteθ = false
│   isfiniter = false
│   isfiniteℓπ = false
│   isfiniteℓκ = false
└ @ AdvancedHMC /Users/kcft114/.julia/packages/AdvancedHMC/YWXfk/src/hamiltonian.jl:36
┌ Warning: Incorrect ϵ = NaN; ϵ_previous = 0.07500000000000001 is used instead.
└ @ AdvancedHMC.Adaptation /Users/kcft114/.julia/packages/AdvancedHMC/YWXfk/src/adaptation/stepsize.jl:103
┌ Warning: The current proposal will be rejected due to numerical error(s).
│   isfiniteθ = true
│   isfiniter = true
│   isfiniteℓπ = false
│   isfiniteℓκ = true
└ @ AdvancedHMC /Users/kcft114/.julia/packages/AdvancedHMC/YWXfk/src/hamiltonian.jl:36
┌ Warning: The current proposal will be rejected due to numerical error(s).
│   isfiniteθ = false
│   isfiniter = false
│   isfiniteℓπ = false
│   isfiniteℓκ = false
└ @ AdvancedHMC /Users/kcft114/.julia/packages/AdvancedHMC/YWXfk/src/hamiltonian.jl:36
┌ Warning: Incorrect ϵ = NaN; ϵ_previous = 0.07500000000000001 is used instead.
└ @ AdvancedHMC.Adaptation /Users/kcft114/.julia/packages/AdvancedHMC/YWXfk/src/adaptation/stepsize.jl:103
┌ Warning: The current proposal will be rejected due to numerical error(s).
│   isfiniteθ = true
│   isfiniter = true
│   isfiniteℓπ = false
│   isfiniteℓκ = true
└ @ AdvancedHMC /Users/kcft114/.julia/packages/AdvancedHMC/YWXfk/src/hamiltonian.jl:36
┌ Warning: The current proposal will be rejected due to numerical error(s).
│   isfiniteθ = false
│   isfiniter = false
│   isfiniteℓπ = false
│   isfiniteℓκ = false
└ @ AdvancedHMC /Users/kcft114/.julia/packages/AdvancedHMC/YWXfk/src/hamiltonian.jl:36
┌ Warning: Incorrect ϵ = NaN; ϵ_previous = 0.07500000000000001 is used instead.
└ @ AdvancedHMC.Adaptation /Users/kcft114/.julia/packages/AdvancedHMC/YWXfk/src/adaptation/stepsize.jl:103
┌ Warning: The current proposal will be rejected due to numerical error(s).
│   isfiniteθ = true
│   isfiniter = true
│   isfiniteℓπ = false
│   isfiniteℓκ = true
└ @ AdvancedHMC /Users/kcft114/.julia/packages/AdvancedHMC/YWXfk/src/hamiltonian.jl:36
┌ Warning: The current proposal will be rejected due to numerical error(s).
│   isfiniteθ = false
│   isfiniter = false
│   isfiniteℓπ = false
│   isfiniteℓκ = false
└ @ AdvancedHMC /Users/kcft114/.julia/packages/AdvancedHMC/YWXfk/src/hamiltonian.jl:36
┌ Warning: Incorrect ϵ = NaN; ϵ_previous = 0.07500000000000001 is used instead.
└ @ AdvancedHMC.Adaptation /Users/kcft114/.julia/packages/AdvancedHMC/YWXfk/src/adaptation/stepsize.jl:103
┌ Warning: The current proposal will be rejected due to numerical error(s).
│   isfiniteθ = true
│   isfiniter = true
│   isfiniteℓπ = false
│   isfiniteℓκ = true
└ @ AdvancedHMC /Users/kcft114/.julia/packages/AdvancedHMC/YWXfk/src/hamiltonian.jl:36
┌ Warning: The current proposal will be rejected due to numerical error(s).
│   isfiniteθ = false
│   isfiniter = false
│   isfiniteℓπ = false
│   isfiniteℓκ = false
└ @ AdvancedHMC /Users/kcft114/.julia/packages/AdvancedHMC/YWXfk/src/hamiltonian.jl:36
┌ Warning: Incorrect ϵ = NaN; ϵ_previous = 0.07500000000000001 is used instead.
└ @ AdvancedHMC.Adaptation /Users/kcft114/.julia/packages/AdvancedHMC/YWXfk/src/adaptation/stepsize.jl:103
┌ Warning: The current proposal will be rejected due to numerical error(s).
│   isfiniteθ = true
│   isfiniter = true
│   isfiniteℓπ = false
│   isfiniteℓκ = true
└ @ AdvancedHMC /Users/kcft114/.julia/packages/AdvancedHMC/YWXfk/src/hamiltonian.jl:36
┌ Warning: The current proposal will be rejected due to numerical error(s).
│   isfiniteθ = false
│   isfiniter = false
│   isfiniteℓπ = false
│   isfiniteℓκ = false
└ @ AdvancedHMC /Users/kcft114/.julia/packages/AdvancedHMC/YWXfk/src/hamiltonian.jl:36
┌ Warning: Incorrect ϵ = NaN; ϵ_previous = 0.07500000000000001 is used instead.
└ @ AdvancedHMC.Adaptation /Users/kcft114/.julia/packages/AdvancedHMC/YWXfk/src/adaptation/stepsize.jl:103
┌ Warning: The current proposal will be rejected due to numerical error(s).
│   isfiniteθ = true
│   isfiniter = true
│   isfiniteℓπ = false
│   isfiniteℓκ = true
└ @ AdvancedHMC /Users/kcft114/.julia/packages/AdvancedHMC/YWXfk/src/hamiltonian.jl:36
┌ Warning: The current proposal will be rejected due to numerical error(s).
│   isfiniteθ = false
│   isfiniter = false
│   isfiniteℓπ = false
│   isfiniteℓκ = false
└ @ AdvancedHMC /Users/kcft114/.julia/packages/AdvancedHMC/YWXfk/src/hamiltonian.jl:36
┌ Warning: Incorrect ϵ = NaN; ϵ_previous = 0.07500000000000001 is used instead.
└ @ AdvancedHMC.Adaptation /Users/kcft114/.julia/packages/AdvancedHMC/YWXfk/src/adaptation/stepsize.jl:103
┌ Warning: The current proposal will be rejected due to numerical error(s).
│   isfiniteθ = true
│   isfiniter = true
│   isfiniteℓπ = false
│   isfiniteℓκ = true
└ @ AdvancedHMC /Users/kcft114/.julia/packages/AdvancedHMC/YWXfk/src/hamiltonian.jl:36
┌ Warning: The current proposal will be rejected due to numerical error(s).
│   isfiniteθ = false
│   isfiniter = false
│   isfiniteℓπ = false
│   isfiniteℓκ = false
└ @ AdvancedHMC /Users/kcft114/.julia/packages/AdvancedHMC/YWXfk/src/hamiltonian.jl:36
┌ Warning: Incorrect ϵ = NaN; ϵ_previous = 0.07500000000000001 is used instead.
└ @ AdvancedHMC.Adaptation /Users/kcft114/.julia/packages/AdvancedHMC/YWXfk/src/adaptation/stepsize.jl:103
┌ Warning: The current proposal will be rejected due to numerical error(s).
│   isfiniteθ = true
│   isfiniter = true
│   isfiniteℓπ = false
│   isfiniteℓκ = true
└ @ AdvancedHMC /Users/kcft114/.julia/packages/AdvancedHMC/YWXfk/src/hamiltonian.jl:36
┌ Warning: The current proposal will be rejected due to numerical error(s).
│   isfiniteθ = false
│   isfiniter = false
│   isfiniteℓπ = false
│   isfiniteℓκ = false
└ @ AdvancedHMC /Users/kcft114/.julia/packages/AdvancedHMC/YWXfk/src/hamiltonian.jl:36
┌ Warning: Incorrect ϵ = NaN; ϵ_previous = 0.07500000000000001 is used instead.
└ @ AdvancedHMC.Adaptation /Users/kcft114/.julia/packages/AdvancedHMC/YWXfk/src/adaptation/stepsize.jl:103
┌ Warning: The current proposal will be rejected due to numerical error(s).
│   isfiniteθ = true
│   isfiniter = true
│   isfiniteℓπ = false
│   isfiniteℓκ = true
└ @ AdvancedHMC /Users/kcft114/.julia/packages/AdvancedHMC/YWXfk/src/hamiltonian.jl:36
┌ Warning: The current proposal will be rejected due to numerical error(s).
│   isfiniteθ = false
│   isfiniter = false
│   isfiniteℓπ = false
│   isfiniteℓκ = false
└ @ AdvancedHMC /Users/kcft114/.julia/packages/AdvancedHMC/YWXfk/src/hamiltonian.jl:36
┌ Warning: Incorrect ϵ = NaN; ϵ_previous = 0.07500000000000001 is used instead.
└ @ AdvancedHMC.Adaptation /Users/kcft114/.julia/packages/AdvancedHMC/YWXfk/src/adaptation/stepsize.jl:103
┌ Warning: The current proposal will be rejected due to numerical error(s).
│   isfiniteθ = true
│   isfiniter = true
│   isfiniteℓπ = false
│   isfiniteℓκ = true
└ @ AdvancedHMC /Users/kcft114/.julia/packages/AdvancedHMC/YWXfk/src/hamiltonian.jl:36
┌ Warning: The current proposal will be rejected due to numerical error(s).
│   isfiniteθ = false
│   isfiniter = false
│   isfiniteℓπ = false
│   isfiniteℓκ = false
└ @ AdvancedHMC /Users/kcft114/.julia/packages/AdvancedHMC/YWXfk/src/hamiltonian.jl:36
┌ Warning: Incorrect ϵ = NaN; ϵ_previous = 0.07500000000000001 is used instead.
└ @ AdvancedHMC.Adaptation /Users/kcft114/.julia/packages/AdvancedHMC/YWXfk/src/adaptation/stepsize.jl:103
┌ Warning: The current proposal will be rejected due to numerical error(s).
│   isfiniteθ = true
│   isfiniter = true
│   isfiniteℓπ = false
│   isfiniteℓκ = true
└ @ AdvancedHMC /Users/kcft114/.julia/packages/AdvancedHMC/YWXfk/src/hamiltonian.jl:36
┌ Warning: The current proposal will be rejected due to numerical error(s).
│   isfiniteθ = false
│   isfiniter = false
│   isfiniteℓπ = false
│   isfiniteℓκ = false
└ @ AdvancedHMC /Users/kcft114/.julia/packages/AdvancedHMC/YWXfk/src/hamiltonian.jl:36
┌ Warning: Incorrect ϵ = NaN; ϵ_previous = 0.07500000000000001 is used instead.
└ @ AdvancedHMC.Adaptation /Users/kcft114/.julia/packages/AdvancedHMC/YWXfk/src/adaptation/stepsize.jl:103
┌ Warning: The current proposal will be rejected due to numerical error(s).
│   isfiniteθ = true
│   isfiniter = true
│   isfiniteℓπ = false
│   isfiniteℓκ = true
└ @ AdvancedHMC /Users/kcft114/.julia/packages/AdvancedHMC/YWXfk/src/hamiltonian.jl:36
┌ Warning: The current proposal will be rejected due to numerical error(s).
│   isfiniteθ = false
│   isfiniter = false
│   isfiniteℓπ = false
│   isfiniteℓκ = false
└ @ AdvancedHMC /Users/kcft114/.julia/packages/AdvancedHMC/YWXfk/src/hamiltonian.jl:36
┌ Warning: Incorrect ϵ = NaN; ϵ_previous = 0.07500000000000001 is used instead.
└ @ AdvancedHMC.Adaptation /Users/kcft114/.julia/packages/AdvancedHMC/YWXfk/src/adaptation/stepsize.jl:103
┌ Warning: The current proposal will be rejected due to numerical error(s).
│   isfiniteθ = true
│   isfiniter = true
│   isfiniteℓπ = false
│   isfiniteℓκ = true
└ @ AdvancedHMC /Users/kcft114/.julia/packages/AdvancedHMC/YWXfk/src/hamiltonian.jl:36
┌ Warning: The current proposal will be rejected due to numerical error(s).
│   isfiniteθ = false
│   isfiniter = false
│   isfiniteℓπ = false
│   isfiniteℓκ = false
└ @ AdvancedHMC /Users/kcft114/.julia/packages/AdvancedHMC/YWXfk/src/hamiltonian.jl:36
┌ Warning: Incorrect ϵ = NaN; ϵ_previous = 0.07500000000000001 is used instead.
└ @ AdvancedHMC.Adaptation /Users/kcft114/.julia/packages/AdvancedHMC/YWXfk/src/adaptation/stepsize.jl:103
┌ Warning: The current proposal will be rejected due to numerical error(s).
│   isfiniteθ = true
│   isfiniter = true
│   isfiniteℓπ = false
│   isfiniteℓκ = true
└ @ AdvancedHMC /Users/kcft114/.julia/packages/AdvancedHMC/YWXfk/src/hamiltonian.jl:36
┌ Warning: The current proposal will be rejected due to numerical error(s).
│   isfiniteθ = false
│   isfiniter = false
│   isfiniteℓπ = false
│   isfiniteℓκ = false
└ @ AdvancedHMC /Users/kcft114/.julia/packages/AdvancedHMC/YWXfk/src/hamiltonian.jl:36
┌ Warning: Incorrect ϵ = NaN; ϵ_previous = 0.07500000000000001 is used instead.
└ @ AdvancedHMC.Adaptation /Users/kcft114/.julia/packages/AdvancedHMC/YWXfk/src/adaptation/stepsize.jl:103
┌ Warning: The current proposal will be rejected due to numerical error(s).
│   isfiniteθ = true
│   isfiniter = true
│   isfiniteℓπ = false
│   isfiniteℓκ = true
└ @ AdvancedHMC /Users/kcft114/.julia/packages/AdvancedHMC/YWXfk/src/hamiltonian.jl:36
┌ Warning: The current proposal will be rejected due to numerical error(s).
│   isfiniteθ = false
│   isfiniter = false
│   isfiniteℓπ = false
│   isfiniteℓκ = false
└ @ AdvancedHMC /Users/kcft114/.julia/packages/AdvancedHMC/YWXfk/src/hamiltonian.jl:36
┌ Warning: Incorrect ϵ = NaN; ϵ_previous = 0.07500000000000001 is used instead.
└ @ AdvancedHMC.Adaptation /Users/kcft114/.julia/packages/AdvancedHMC/YWXfk/src/adaptation/stepsize.jl:103
┌ Warning: The current proposal will be rejected due to numerical error(s).
│   isfiniteθ = true
│   isfiniter = true
│   isfiniteℓπ = false
│   isfiniteℓκ = true
└ @ AdvancedHMC /Users/kcft114/.julia/packages/AdvancedHMC/YWXfk/src/hamiltonian.jl:36
┌ Warning: The current proposal will be rejected due to numerical error(s).
│   isfiniteθ = false
│   isfiniter = false
│   isfiniteℓπ = false
│   isfiniteℓκ = false
└ @ AdvancedHMC /Users/kcft114/.julia/packages/AdvancedHMC/YWXfk/src/hamiltonian.jl:36
┌ Warning: Incorrect ϵ = NaN; ϵ_previous = 0.07500000000000001 is used instead.
└ @ AdvancedHMC.Adaptation /Users/kcft114/.julia/packages/AdvancedHMC/YWXfk/src/adaptation/stepsize.jl:103
┌ Warning: The current proposal will be rejected due to numerical error(s).
│   isfiniteθ = true
│   isfiniter = true
│   isfiniteℓπ = false
│   isfiniteℓκ = true
└ @ AdvancedHMC /Users/kcft114/.julia/packages/AdvancedHMC/YWXfk/src/hamiltonian.jl:36
┌ Warning: The current proposal will be rejected due to numerical error(s).
│   isfiniteθ = false
│   isfiniter = false
│   isfiniteℓπ = false
│   isfiniteℓκ = false
└ @ AdvancedHMC /Users/kcft114/.julia/packages/AdvancedHMC/YWXfk/src/hamiltonian.jl:36
┌ Warning: Incorrect ϵ = NaN; ϵ_previous = 0.07500000000000001 is used instead.
└ @ AdvancedHMC.Adaptation /Users/kcft114/.julia/packages/AdvancedHMC/YWXfk/src/adaptation/stepsize.jl:103
┌ Warning: The current proposal will be rejected due to numerical error(s).
│   isfiniteθ = true
│   isfiniter = true
│   isfiniteℓπ = false
│   isfiniteℓκ = true
└ @ AdvancedHMC /Users/kcft114/.julia/packages/AdvancedHMC/YWXfk/src/hamiltonian.jl:36
┌ Warning: The current proposal will be rejected due to numerical error(s).
│   isfiniteθ = false
│   isfiniter = false
│   isfiniteℓπ = false
│   isfiniteℓκ = false
└ @ AdvancedHMC /Users/kcft114/.julia/packages/AdvancedHMC/YWXfk/src/hamiltonian.jl:36
┌ Warning: Incorrect ϵ = NaN; ϵ_previous = 0.07500000000000001 is used instead.
└ @ AdvancedHMC.Adaptation /Users/kcft114/.julia/packages/AdvancedHMC/YWXfk/src/adaptation/stepsize.jl:103
┌ Warning: The current proposal will be rejected due to numerical error(s).
│   isfiniteθ = true
│   isfiniter = true
│   isfiniteℓπ = false
│   isfiniteℓκ = true
└ @ AdvancedHMC /Users/kcft114/.julia/packages/AdvancedHMC/YWXfk/src/hamiltonian.jl:36
┌ Warning: The current proposal will be rejected due to numerical error(s).
│   isfiniteθ = false
│   isfiniter = false
│   isfiniteℓπ = false
│   isfiniteℓκ = false
└ @ AdvancedHMC /Users/kcft114/.julia/packages/AdvancedHMC/YWXfk/src/hamiltonian.jl:36
┌ Warning: Incorrect ϵ = NaN; ϵ_previous = 0.07500000000000001 is used instead.
└ @ AdvancedHMC.Adaptation /Users/kcft114/.julia/packages/AdvancedHMC/YWXfk/src/adaptation/stepsize.jl:103
┌ Warning: The current proposal will be rejected due to numerical error(s).
│   isfiniteθ = true
│   isfiniter = true
│   isfiniteℓπ = false
│   isfiniteℓκ = true
└ @ AdvancedHMC /Users/kcft114/.julia/packages/AdvancedHMC/YWXfk/src/hamiltonian.jl:36
┌ Warning: The current proposal will be rejected due to numerical error(s).
│   isfiniteθ = false
│   isfiniter = false
│   isfiniteℓπ = false
│   isfiniteℓκ = false
└ @ AdvancedHMC /Users/kcft114/.julia/packages/AdvancedHMC/YWXfk/src/hamiltonian.jl:36
┌ Warning: Incorrect ϵ = NaN; ϵ_previous = 0.07500000000000001 is used instead.
└ @ AdvancedHMC.Adaptation /Users/kcft114/.julia/packages/AdvancedHMC/YWXfk/src/adaptation/stepsize.jl:103
┌ Warning: The current proposal will be rejected due to numerical error(s).
│   isfiniteθ = true
│   isfiniter = true
│   isfiniteℓπ = false
│   isfiniteℓκ = true
└ @ AdvancedHMC /Users/kcft114/.julia/packages/AdvancedHMC/YWXfk/src/hamiltonian.jl:36
┌ Warning: The current proposal will be rejected due to numerical error(s).
│   isfiniteθ = false
│   isfiniter = false
│   isfiniteℓπ = false
│   isfiniteℓκ = false
└ @ AdvancedHMC /Users/kcft114/.julia/packages/AdvancedHMC/YWXfk/src/hamiltonian.jl:36
┌ Warning: Incorrect ϵ = NaN; ϵ_previous = 0.07500000000000001 is used instead.
└ @ AdvancedHMC.Adaptation /Users/kcft114/.julia/packages/AdvancedHMC/YWXfk/src/adaptation/stepsize.jl:103
┌ Warning: The current proposal will be rejected due to numerical error(s).
│   isfiniteθ = true
│   isfiniter = true
│   isfiniteℓπ = false
│   isfiniteℓκ = true
└ @ AdvancedHMC /Users/kcft114/.julia/packages/AdvancedHMC/YWXfk/src/hamiltonian.jl:36
┌ Warning: The current proposal will be rejected due to numerical error(s).
│   isfiniteθ = false
│   isfiniter = false
│   isfiniteℓπ = false
│   isfiniteℓκ = false
└ @ AdvancedHMC /Users/kcft114/.julia/packages/AdvancedHMC/YWXfk/src/hamiltonian.jl:36
┌ Warning: Incorrect ϵ = NaN; ϵ_previous = 0.07500000000000001 is used instead.
└ @ AdvancedHMC.Adaptation /Users/kcft114/.julia/packages/AdvancedHMC/YWXfk/src/adaptation/stepsize.jl:103
┌ Warning: The current proposal will be rejected due to numerical error(s).
│   isfiniteθ = true
│   isfiniter = true
│   isfiniteℓπ = false
│   isfiniteℓκ = true
└ @ AdvancedHMC /Users/kcft114/.julia/packages/AdvancedHMC/YWXfk/src/hamiltonian.jl:36
┌ Warning: The current proposal will be rejected due to numerical error(s).
│   isfiniteθ = false
│   isfiniter = false
│   isfiniteℓπ = false
│   isfiniteℓκ = false
└ @ AdvancedHMC /Users/kcft114/.julia/packages/AdvancedHMC/YWXfk/src/hamiltonian.jl:36
┌ Warning: Incorrect ϵ = NaN; ϵ_previous = 0.07500000000000001 is used instead.
└ @ AdvancedHMC.Adaptation /Users/kcft114/.julia/packages/AdvancedHMC/YWXfk/src/adaptation/stepsize.jl:103
┌ Warning: The current proposal will be rejected due to numerical error(s).
│   isfiniteθ = true
│   isfiniter = true
│   isfiniteℓπ = false
│   isfiniteℓκ = true
└ @ AdvancedHMC /Users/kcft114/.julia/packages/AdvancedHMC/YWXfk/src/hamiltonian.jl:36
┌ Warning: The current proposal will be rejected due to numerical error(s).
│   isfiniteθ = false
│   isfiniter = false
│   isfiniteℓπ = false
│   isfiniteℓκ = false
└ @ AdvancedHMC /Users/kcft114/.julia/packages/AdvancedHMC/YWXfk/src/hamiltonian.jl:36
┌ Warning: Incorrect ϵ = NaN; ϵ_previous = 0.07500000000000001 is used instead.
└ @ AdvancedHMC.Adaptation /Users/kcft114/.julia/packages/AdvancedHMC/YWXfk/src/adaptation/stepsize.jl:103
┌ Warning: The current proposal will be rejected due to numerical error(s).
│   isfiniteθ = true
│   isfiniter = true
│   isfiniteℓπ = false
│   isfiniteℓκ = true
└ @ AdvancedHMC /Users/kcft114/.julia/packages/AdvancedHMC/YWXfk/src/hamiltonian.jl:36
┌ Warning: The current proposal will be rejected due to numerical error(s).
│   isfiniteθ = false
│   isfiniter = false
│   isfiniteℓπ = false
│   isfiniteℓκ = false
└ @ AdvancedHMC /Users/kcft114/.julia/packages/AdvancedHMC/YWXfk/src/hamiltonian.jl:36
┌ Warning: Incorrect ϵ = NaN; ϵ_previous = 0.07500000000000001 is used instead.
└ @ AdvancedHMC.Adaptation /Users/kcft114/.julia/packages/AdvancedHMC/YWXfk/src/adaptation/stepsize.jl:103
┌ Warning: The current proposal will be rejected due to numerical error(s).
│   isfiniteθ = true
│   isfiniter = true
│   isfiniteℓπ = false
│   isfiniteℓκ = true
└ @ AdvancedHMC /Users/kcft114/.julia/packages/AdvancedHMC/YWXfk/src/hamiltonian.jl:36
┌ Warning: The current proposal will be rejected due to numerical error(s).
│   isfiniteθ = false
│   isfiniter = false
│   isfiniteℓπ = false
│   isfiniteℓκ = false
└ @ AdvancedHMC /Users/kcft114/.julia/packages/AdvancedHMC/YWXfk/src/hamiltonian.jl:36
┌ Warning: Incorrect ϵ = NaN; ϵ_previous = 0.07500000000000001 is used instead.
└ @ AdvancedHMC.Adaptation /Users/kcft114/.julia/packages/AdvancedHMC/YWXfk/src/adaptation/stepsize.jl:103
┌ Warning: The current proposal will be rejected due to numerical error(s).
│   isfiniteθ = true
│   isfiniter = true
│   isfiniteℓπ = false
│   isfiniteℓκ = true
└ @ AdvancedHMC /Users/kcft114/.julia/packages/AdvancedHMC/YWXfk/src/hamiltonian.jl:36
┌ Warning: The current proposal will be rejected due to numerical error(s).
│   isfiniteθ = false
│   isfiniter = false
│   isfiniteℓπ = false
│   isfiniteℓκ = false
└ @ AdvancedHMC /Users/kcft114/.julia/packages/AdvancedHMC/YWXfk/src/hamiltonian.jl:36
┌ Warning: Incorrect ϵ = NaN; ϵ_previous = 0.07500000000000001 is used instead.
└ @ AdvancedHMC.Adaptation /Users/kcft114/.julia/packages/AdvancedHMC/YWXfk/src/adaptation/stepsize.jl:103
┌ Warning: The current proposal will be rejected due to numerical error(s).
│   isfiniteθ = true
│   isfiniter = true
│   isfiniteℓπ = false
│   isfiniteℓκ = true
└ @ AdvancedHMC /Users/kcft114/.julia/packages/AdvancedHMC/YWXfk/src/hamiltonian.jl:36
┌ Warning: The current proposal will be rejected due to numerical error(s).
│   isfiniteθ = false
│   isfiniter = false
│   isfiniteℓπ = false
│   isfiniteℓκ = false
└ @ AdvancedHMC /Users/kcft114/.julia/packages/AdvancedHMC/YWXfk/src/hamiltonian.jl:36
┌ Warning: Incorrect ϵ = NaN; ϵ_previous = 0.07500000000000001 is used instead.
└ @ AdvancedHMC.Adaptation /Users/kcft114/.julia/packages/AdvancedHMC/YWXfk/src/adaptation/stepsize.jl:103
┌ Warning: The current proposal will be rejected due to numerical error(s).
│   isfiniteθ = true
│   isfiniter = true
│   isfiniteℓπ = false
│   isfiniteℓκ = true
└ @ AdvancedHMC /Users/kcft114/.julia/packages/AdvancedHMC/YWXfk/src/hamiltonian.jl:36
┌ Warning: The current proposal will be rejected due to numerical error(s).
│   isfiniteθ = false
│   isfiniter = false
│   isfiniteℓπ = false
│   isfiniteℓκ = false
└ @ AdvancedHMC /Users/kcft114/.julia/packages/AdvancedHMC/YWXfk/src/hamiltonian.jl:36
┌ Warning: Incorrect ϵ = NaN; ϵ_previous = 0.07500000000000001 is used instead.
└ @ AdvancedHMC.Adaptation /Users/kcft114/.julia/packages/AdvancedHMC/YWXfk/src/adaptation/stepsize.jl:103
┌ Warning: The current proposal will be rejected due to numerical error(s).
│   isfiniteθ = true
│   isfiniter = true
│   isfiniteℓπ = false
│   isfiniteℓκ = true
└ @ AdvancedHMC /Users/kcft114/.julia/packages/AdvancedHMC/YWXfk/src/hamiltonian.jl:36
┌ Warning: The current proposal will be rejected due to numerical error(s).
│   isfiniteθ = false
│   isfiniter = false
│   isfiniteℓπ = false
│   isfiniteℓκ = false
└ @ AdvancedHMC /Users/kcft114/.julia/packages/AdvancedHMC/YWXfk/src/hamiltonian.jl:36
┌ Warning: Incorrect ϵ = NaN; ϵ_previous = 0.07500000000000001 is used instead.
└ @ AdvancedHMC.Adaptation /Users/kcft114/.julia/packages/AdvancedHMC/YWXfk/src/adaptation/stepsize.jl:103
┌ Warning: The current proposal will be rejected due to numerical error(s).
│   isfiniteθ = true
│   isfiniter = true
│   isfiniteℓπ = false
│   isfiniteℓκ = true
└ @ AdvancedHMC /Users/kcft114/.julia/packages/AdvancedHMC/YWXfk/src/hamiltonian.jl:36
┌ Warning: The current proposal will be rejected due to numerical error(s).
│   isfiniteθ = false
│   isfiniter = false
│   isfiniteℓπ = false
│   isfiniteℓκ = false
└ @ AdvancedHMC /Users/kcft114/.julia/packages/AdvancedHMC/YWXfk/src/hamiltonian.jl:36
┌ Warning: Incorrect ϵ = NaN; ϵ_previous = 0.07500000000000001 is used instead.
└ @ AdvancedHMC.Adaptation /Users/kcft114/.julia/packages/AdvancedHMC/YWXfk/src/adaptation/stepsize.jl:103
┌ Warning: The current proposal will be rejected due to numerical error(s).
│   isfiniteθ = true
│   isfiniter = true
│   isfiniteℓπ = false
│   isfiniteℓκ = true
└ @ AdvancedHMC /Users/kcft114/.julia/packages/AdvancedHMC/YWXfk/src/hamiltonian.jl:36
┌ Warning: The current proposal will be rejected due to numerical error(s).
│   isfiniteθ = false
│   isfiniter = false
│   isfiniteℓπ = false
│   isfiniteℓκ = false
└ @ AdvancedHMC /Users/kcft114/.julia/packages/AdvancedHMC/YWXfk/src/hamiltonian.jl:36
┌ Warning: Incorrect ϵ = NaN; ϵ_previous = 0.07500000000000001 is used instead.
└ @ AdvancedHMC.Adaptation /Users/kcft114/.julia/packages/AdvancedHMC/YWXfk/src/adaptation/stepsize.jl:103
┌ Warning: The current proposal will be rejected due to numerical error(s).
│   isfiniteθ = true
│   isfiniter = true
│   isfiniteℓπ = false
│   isfiniteℓκ = true
└ @ AdvancedHMC /Users/kcft114/.julia/packages/AdvancedHMC/YWXfk/src/hamiltonian.jl:36
┌ Warning: The current proposal will be rejected due to numerical error(s).
│   isfiniteθ = false
│   isfiniter = false
│   isfiniteℓπ = false
│   isfiniteℓκ = false
└ @ AdvancedHMC /Users/kcft114/.julia/packages/AdvancedHMC/YWXfk/src/hamiltonian.jl:36
┌ Warning: Incorrect ϵ = NaN; ϵ_previous = 0.07500000000000001 is used instead.
└ @ AdvancedHMC.Adaptation /Users/kcft114/.julia/packages/AdvancedHMC/YWXfk/src/adaptation/stepsize.jl:103
┌ Warning: The current proposal will be rejected due to numerical error(s).
│   isfiniteθ = true
│   isfiniter = true
│   isfiniteℓπ = false
│   isfiniteℓκ = true
└ @ AdvancedHMC /Users/kcft114/.julia/packages/AdvancedHMC/YWXfk/src/hamiltonian.jl:36
┌ Warning: The current proposal will be rejected due to numerical error(s).
│   isfiniteθ = false
│   isfiniter = false
│   isfiniteℓπ = false
│   isfiniteℓκ = false
└ @ AdvancedHMC /Users/kcft114/.julia/packages/AdvancedHMC/YWXfk/src/hamiltonian.jl:36
┌ Warning: Incorrect ϵ = NaN; ϵ_previous = 0.07500000000000001 is used instead.
└ @ AdvancedHMC.Adaptation /Users/kcft114/.julia/packages/AdvancedHMC/YWXfk/src/adaptation/stepsize.jl:103
┌ Warning: The current proposal will be rejected due to numerical error(s).
│   isfiniteθ = true
│   isfiniter = true
│   isfiniteℓπ = false
│   isfiniteℓκ = true
└ @ AdvancedHMC /Users/kcft114/.julia/packages/AdvancedHMC/YWXfk/src/hamiltonian.jl:36
┌ Warning: The current proposal will be rejected due to numerical error(s).
│   isfiniteθ = false
│   isfiniter = false
│   isfiniteℓπ = false
│   isfiniteℓκ = false
└ @ AdvancedHMC /Users/kcft114/.julia/packages/AdvancedHMC/YWXfk/src/hamiltonian.jl:36
┌ Warning: Incorrect ϵ = NaN; ϵ_previous = 0.07500000000000001 is used instead.
└ @ AdvancedHMC.Adaptation /Users/kcft114/.julia/packages/AdvancedHMC/YWXfk/src/adaptation/stepsize.jl:103
┌ Warning: The current proposal will be rejected due to numerical error(s).
│   isfiniteθ = true
│   isfiniter = true
│   isfiniteℓπ = false
│   isfiniteℓκ = true
└ @ AdvancedHMC /Users/kcft114/.julia/packages/AdvancedHMC/YWXfk/src/hamiltonian.jl:36
┌ Warning: The current proposal will be rejected due to numerical error(s).
│   isfiniteθ = false
│   isfiniter = false
│   isfiniteℓπ = false
│   isfiniteℓκ = false
└ @ AdvancedHMC /Users/kcft114/.julia/packages/AdvancedHMC/YWXfk/src/hamiltonian.jl:36
┌ Warning: Incorrect ϵ = NaN; ϵ_previous = 0.07500000000000001 is used instead.
└ @ AdvancedHMC.Adaptation /Users/kcft114/.julia/packages/AdvancedHMC/YWXfk/src/adaptation/stepsize.jl:103
┌ Warning: The current proposal will be rejected due to numerical error(s).
│   isfiniteθ = true
│   isfiniter = true
│   isfiniteℓπ = false
│   isfiniteℓκ = true
└ @ AdvancedHMC /Users/kcft114/.julia/packages/AdvancedHMC/YWXfk/src/hamiltonian.jl:36
┌ Warning: The current proposal will be rejected due to numerical error(s).
│   isfiniteθ = false
│   isfiniter = false
│   isfiniteℓπ = false
│   isfiniteℓκ = false
└ @ AdvancedHMC /Users/kcft114/.julia/packages/AdvancedHMC/YWXfk/src/hamiltonian.jl:36
┌ Warning: Incorrect ϵ = NaN; ϵ_previous = 0.07500000000000001 is used instead.
└ @ AdvancedHMC.Adaptation /Users/kcft114/.julia/packages/AdvancedHMC/YWXfk/src/adaptation/stepsize.jl:103
┌ Warning: The current proposal will be rejected due to numerical error(s).
│   isfiniteθ = true
│   isfiniter = true
│   isfiniteℓπ = false
│   isfiniteℓκ = true
└ @ AdvancedHMC /Users/kcft114/.julia/packages/AdvancedHMC/YWXfk/src/hamiltonian.jl:36
┌ Warning: The current proposal will be rejected due to numerical error(s).
│   isfiniteθ = false
│   isfiniter = false
│   isfiniteℓπ = false
│   isfiniteℓκ = false
└ @ AdvancedHMC /Users/kcft114/.julia/packages/AdvancedHMC/YWXfk/src/hamiltonian.jl:36
┌ Warning: Incorrect ϵ = NaN; ϵ_previous = 0.07500000000000001 is used instead.
└ @ AdvancedHMC.Adaptation /Users/kcft114/.julia/packages/AdvancedHMC/YWXfk/src/adaptation/stepsize.jl:103
┌ Warning: The current proposal will be rejected due to numerical error(s).
│   isfiniteθ = true
│   isfiniter = true
│   isfiniteℓπ = false
│   isfiniteℓκ = true
└ @ AdvancedHMC /Users/kcft114/.julia/packages/AdvancedHMC/YWXfk/src/hamiltonian.jl:36
┌ Warning: The current proposal will be rejected due to numerical error(s).
│   isfiniteθ = false
│   isfiniter = false
│   isfiniteℓπ = false
│   isfiniteℓκ = false
└ @ AdvancedHMC /Users/kcft114/.julia/packages/AdvancedHMC/YWXfk/src/hamiltonian.jl:36
┌ Warning: Incorrect ϵ = NaN; ϵ_previous = 0.07500000000000001 is used instead.
└ @ AdvancedHMC.Adaptation /Users/kcft114/.julia/packages/AdvancedHMC/YWXfk/src/adaptation/stepsize.jl:103
┌ Warning: The current proposal will be rejected due to numerical error(s).
│   isfiniteθ = true
│   isfiniter = true
│   isfiniteℓπ = false
│   isfiniteℓκ = true
└ @ AdvancedHMC /Users/kcft114/.julia/packages/AdvancedHMC/YWXfk/src/hamiltonian.jl:36
┌ Warning: The current proposal will be rejected due to numerical error(s).
│   isfiniteθ = false
│   isfiniter = false
│   isfiniteℓπ = false
│   isfiniteℓκ = false
└ @ AdvancedHMC /Users/kcft114/.julia/packages/AdvancedHMC/YWXfk/src/hamiltonian.jl:36
┌ Warning: Incorrect ϵ = NaN; ϵ_previous = 0.07500000000000001 is used instead.
└ @ AdvancedHMC.Adaptation /Users/kcft114/.julia/packages/AdvancedHMC/YWXfk/src/adaptation/stepsize.jl:103
┌ Warning: The current proposal will be rejected due to numerical error(s).
│   isfiniteθ = true
│   isfiniter = true
│   isfiniteℓπ = false
│   isfiniteℓκ = true
└ @ AdvancedHMC /Users/kcft114/.julia/packages/AdvancedHMC/YWXfk/src/hamiltonian.jl:36
┌ Warning: The current proposal will be rejected due to numerical error(s).
│   isfiniteθ = false
│   isfiniter = false
│   isfiniteℓπ = false
│   isfiniteℓκ = false
└ @ AdvancedHMC /Users/kcft114/.julia/packages/AdvancedHMC/YWXfk/src/hamiltonian.jl:36
┌ Warning: Incorrect ϵ = NaN; ϵ_previous = 0.07500000000000001 is used instead.
└ @ AdvancedHMC.Adaptation /Users/kcft114/.julia/packages/AdvancedHMC/YWXfk/src/adaptation/stepsize.jl:103
┌ Warning: The current proposal will be rejected due to numerical error(s).
│   isfiniteθ = true
│   isfiniter = true
│   isfiniteℓπ = false
│   isfiniteℓκ = true
└ @ AdvancedHMC /Users/kcft114/.julia/packages/AdvancedHMC/YWXfk/src/hamiltonian.jl:36
┌ Warning: The current proposal will be rejected due to numerical error(s).
│   isfiniteθ = false
│   isfiniter = false
│   isfiniteℓπ = false
│   isfiniteℓκ = false
└ @ AdvancedHMC /Users/kcft114/.julia/packages/AdvancedHMC/YWXfk/src/hamiltonian.jl:36
┌ Warning: Incorrect ϵ = NaN; ϵ_previous = 0.07500000000000001 is used instead.
└ @ AdvancedHMC.Adaptation /Users/kcft114/.julia/packages/AdvancedHMC/YWXfk/src/adaptation/stepsize.jl:103
┌ Warning: The current proposal will be rejected due to numerical error(s).
│   isfiniteθ = true
│   isfiniter = true
│   isfiniteℓπ = false
│   isfiniteℓκ = true
└ @ AdvancedHMC /Users/kcft114/.julia/packages/AdvancedHMC/YWXfk/src/hamiltonian.jl:36
┌ Warning: The current proposal will be rejected due to numerical error(s).
│   isfiniteθ = false
│   isfiniter = false
│   isfiniteℓπ = false
│   isfiniteℓκ = false
└ @ AdvancedHMC /Users/kcft114/.julia/packages/AdvancedHMC/YWXfk/src/hamiltonian.jl:36
┌ Warning: Incorrect ϵ = NaN; ϵ_previous = 0.07500000000000001 is used instead.
└ @ AdvancedHMC.Adaptation /Users/kcft114/.julia/packages/AdvancedHMC/YWXfk/src/adaptation/stepsize.jl:103
┌ Warning: The current proposal will be rejected due to numerical error(s).
│   isfiniteθ = true
│   isfiniter = true
│   isfiniteℓπ = false
│   isfiniteℓκ = true
└ @ AdvancedHMC /Users/kcft114/.julia/packages/AdvancedHMC/YWXfk/src/hamiltonian.jl:36
┌ Warning: The current proposal will be rejected due to numerical error(s).
│   isfiniteθ = false
│   isfiniter = false
│   isfiniteℓπ = false
│   isfiniteℓκ = false
└ @ AdvancedHMC /Users/kcft114/.julia/packages/AdvancedHMC/YWXfk/src/hamiltonian.jl:36
┌ Warning: Incorrect ϵ = NaN; ϵ_previous = 0.07500000000000001 is used instead.
└ @ AdvancedHMC.Adaptation /Users/kcft114/.julia/packages/AdvancedHMC/YWXfk/src/adaptation/stepsize.jl:103
┌ Warning: The current proposal will be rejected due to numerical error(s).
│   isfiniteθ = true
│   isfiniter = true
│   isfiniteℓπ = false
│   isfiniteℓκ = true
└ @ AdvancedHMC /Users/kcft114/.julia/packages/AdvancedHMC/YWXfk/src/hamiltonian.jl:36
┌ Warning: The current proposal will be rejected due to numerical error(s).
│   isfiniteθ = false
│   isfiniter = false
│   isfiniteℓπ = false
│   isfiniteℓκ = false
└ @ AdvancedHMC /Users/kcft114/.julia/packages/AdvancedHMC/YWXfk/src/hamiltonian.jl:36
┌ Warning: Incorrect ϵ = NaN; ϵ_previous = 0.07500000000000001 is used instead.
└ @ AdvancedHMC.Adaptation /Users/kcft114/.julia/packages/AdvancedHMC/YWXfk/src/adaptation/stepsize.jl:103
┌ Warning: The current proposal will be rejected due to numerical error(s).
│   isfiniteθ = true
│   isfiniter = true
│   isfiniteℓπ = false
│   isfiniteℓκ = true
└ @ AdvancedHMC /Users/kcft114/.julia/packages/AdvancedHMC/YWXfk/src/hamiltonian.jl:36
┌ Warning: The current proposal will be rejected due to numerical error(s).
│   isfiniteθ = false
│   isfiniter = false
│   isfiniteℓπ = false
│   isfiniteℓκ = false
└ @ AdvancedHMC /Users/kcft114/.julia/packages/AdvancedHMC/YWXfk/src/hamiltonian.jl:36
┌ Warning: Incorrect ϵ = NaN; ϵ_previous = 0.07500000000000001 is used instead.
└ @ AdvancedHMC.Adaptation /Users/kcft114/.julia/packages/AdvancedHMC/YWXfk/src/adaptation/stepsize.jl:103
┌ Warning: The current proposal will be rejected due to numerical error(s).
│   isfiniteθ = true
│   isfiniter = true
│   isfiniteℓπ = false
│   isfiniteℓκ = true
└ @ AdvancedHMC /Users/kcft114/.julia/packages/AdvancedHMC/YWXfk/src/hamiltonian.jl:36
┌ Warning: The current proposal will be rejected due to numerical error(s).
│   isfiniteθ = false
│   isfiniter = false
│   isfiniteℓπ = false
│   isfiniteℓκ = false
└ @ AdvancedHMC /Users/kcft114/.julia/packages/AdvancedHMC/YWXfk/src/hamiltonian.jl:36
┌ Warning: Incorrect ϵ = NaN; ϵ_previous = 0.07500000000000001 is used instead.
└ @ AdvancedHMC.Adaptation /Users/kcft114/.julia/packages/AdvancedHMC/YWXfk/src/adaptation/stepsize.jl:103
┌ Warning: The current proposal will be rejected due to numerical error(s).
│   isfiniteθ = true
│   isfiniter = true
│   isfiniteℓπ = false
│   isfiniteℓκ = true
└ @ AdvancedHMC /Users/kcft114/.julia/packages/AdvancedHMC/YWXfk/src/hamiltonian.jl:36
┌ Warning: The current proposal will be rejected due to numerical error(s).
│   isfiniteθ = false
│   isfiniter = false
│   isfiniteℓπ = false
│   isfiniteℓκ = false
└ @ AdvancedHMC /Users/kcft114/.julia/packages/AdvancedHMC/YWXfk/src/hamiltonian.jl:36
┌ Warning: Incorrect ϵ = NaN; ϵ_previous = 0.07500000000000001 is used instead.
└ @ AdvancedHMC.Adaptation /Users/kcft114/.julia/packages/AdvancedHMC/YWXfk/src/adaptation/stepsize.jl:103
┌ Warning: The current proposal will be rejected due to numerical error(s).
│   isfiniteθ = true
│   isfiniter = true
│   isfiniteℓπ = false
│   isfiniteℓκ = true
└ @ AdvancedHMC /Users/kcft114/.julia/packages/AdvancedHMC/YWXfk/src/hamiltonian.jl:36
┌ Warning: The current proposal will be rejected due to numerical error(s).
│   isfiniteθ = false
│   isfiniter = false
│   isfiniteℓπ = false
│   isfiniteℓκ = false
└ @ AdvancedHMC /Users/kcft114/.julia/packages/AdvancedHMC/YWXfk/src/hamiltonian.jl:36
┌ Warning: Incorrect ϵ = NaN; ϵ_previous = 0.07500000000000001 is used instead.
└ @ AdvancedHMC.Adaptation /Users/kcft114/.julia/packages/AdvancedHMC/YWXfk/src/adaptation/stepsize.jl:103
┌ Warning: The current proposal will be rejected due to numerical error(s).
│   isfiniteθ = true
│   isfiniter = true
│   isfiniteℓπ = false
│   isfiniteℓκ = true
└ @ AdvancedHMC /Users/kcft114/.julia/packages/AdvancedHMC/YWXfk/src/hamiltonian.jl:36
┌ Warning: The current proposal will be rejected due to numerical error(s).
│   isfiniteθ = false
│   isfiniter = false
│   isfiniteℓπ = false
│   isfiniteℓκ = false
└ @ AdvancedHMC /Users/kcft114/.julia/packages/AdvancedHMC/YWXfk/src/hamiltonian.jl:36
┌ Warning: Incorrect ϵ = NaN; ϵ_previous = 0.07500000000000001 is used instead.
└ @ AdvancedHMC.Adaptation /Users/kcft114/.julia/packages/AdvancedHMC/YWXfk/src/adaptation/stepsize.jl:103
┌ Warning: The current proposal will be rejected due to numerical error(s).
│   isfiniteθ = true
│   isfiniter = true
│   isfiniteℓπ = false
│   isfiniteℓκ = true
└ @ AdvancedHMC /Users/kcft114/.julia/packages/AdvancedHMC/YWXfk/src/hamiltonian.jl:36
┌ Warning: The current proposal will be rejected due to numerical error(s).
│   isfiniteθ = false
│   isfiniter = false
│   isfiniteℓπ = false
│   isfiniteℓκ = false
└ @ AdvancedHMC /Users/kcft114/.julia/packages/AdvancedHMC/YWXfk/src/hamiltonian.jl:36
┌ Warning: Incorrect ϵ = NaN; ϵ_previous = 0.07500000000000001 is used instead.
└ @ AdvancedHMC.Adaptation /Users/kcft114/.julia/packages/AdvancedHMC/YWXfk/src/adaptation/stepsize.jl:103
┌ Warning: The current proposal will be rejected due to numerical error(s).
│   isfiniteθ = true
│   isfiniter = true
│   isfiniteℓπ = false
│   isfiniteℓκ = true
└ @ AdvancedHMC /Users/kcft114/.julia/packages/AdvancedHMC/YWXfk/src/hamiltonian.jl:36
┌ Warning: The current proposal will be rejected due to numerical error(s).
│   isfiniteθ = false
│   isfiniter = false
│   isfiniteℓπ = false
│   isfiniteℓκ = false
└ @ AdvancedHMC /Users/kcft114/.julia/packages/AdvancedHMC/YWXfk/src/hamiltonian.jl:36
┌ Warning: Incorrect ϵ = NaN; ϵ_previous = 0.07500000000000001 is used instead.
└ @ AdvancedHMC.Adaptation /Users/kcft114/.julia/packages/AdvancedHMC/YWXfk/src/adaptation/stepsize.jl:103
┌ Warning: The current proposal will be rejected due to numerical error(s).
│   isfiniteθ = true
│   isfiniter = true
│   isfiniteℓπ = false
│   isfiniteℓκ = true
└ @ AdvancedHMC /Users/kcft114/.julia/packages/AdvancedHMC/YWXfk/src/hamiltonian.jl:36
┌ Warning: The current proposal will be rejected due to numerical error(s).
│   isfiniteθ = false
│   isfiniter = false
│   isfiniteℓπ = false
│   isfiniteℓκ = false
└ @ AdvancedHMC /Users/kcft114/.julia/packages/AdvancedHMC/YWXfk/src/hamiltonian.jl:36
┌ Warning: Incorrect ϵ = NaN; ϵ_previous = 0.07500000000000001 is used instead.
└ @ AdvancedHMC.Adaptation /Users/kcft114/.julia/packages/AdvancedHMC/YWXfk/src/adaptation/stepsize.jl:103
┌ Warning: The current proposal will be rejected due to numerical error(s).
│   isfiniteθ = true
│   isfiniter = true
│   isfiniteℓπ = false
│   isfiniteℓκ = true
└ @ AdvancedHMC /Users/kcft114/.julia/packages/AdvancedHMC/YWXfk/src/hamiltonian.jl:36
┌ Warning: The current proposal will be rejected due to numerical error(s).
│   isfiniteθ = false
│   isfiniter = false
│   isfiniteℓπ = false
│   isfiniteℓκ = false
└ @ AdvancedHMC /Users/kcft114/.julia/packages/AdvancedHMC/YWXfk/src/hamiltonian.jl:36
┌ Warning: Incorrect ϵ = NaN; ϵ_previous = 0.07500000000000001 is used instead.
└ @ AdvancedHMC.Adaptation /Users/kcft114/.julia/packages/AdvancedHMC/YWXfk/src/adaptation/stepsize.jl:103
┌ Warning: The current proposal will be rejected due to numerical error(s).
│   isfiniteθ = true
│   isfiniter = true
│   isfiniteℓπ = false
│   isfiniteℓκ = true
└ @ AdvancedHMC /Users/kcft114/.julia/packages/AdvancedHMC/YWXfk/src/hamiltonian.jl:36
┌ Warning: The current proposal will be rejected due to numerical error(s).
│   isfiniteθ = false
│   isfiniter = false
│   isfiniteℓπ = false
│   isfiniteℓκ = false
└ @ AdvancedHMC /Users/kcft114/.julia/packages/AdvancedHMC/YWXfk/src/hamiltonian.jl:36
┌ Warning: Incorrect ϵ = NaN; ϵ_previous = 0.07500000000000001 is used instead.
└ @ AdvancedHMC.Adaptation /Users/kcft114/.julia/packages/AdvancedHMC/YWXfk/src/adaptation/stepsize.jl:103
┌ Warning: The current proposal will be rejected due to numerical error(s).
│   isfiniteθ = true
│   isfiniter = true
│   isfiniteℓπ = false
│   isfiniteℓκ = true
└ @ AdvancedHMC /Users/kcft114/.julia/packages/AdvancedHMC/YWXfk/src/hamiltonian.jl:36
┌ Warning: The current proposal will be rejected due to numerical error(s).
│   isfiniteθ = false
│   isfiniter = false
│   isfiniteℓπ = false
│   isfiniteℓκ = false
└ @ AdvancedHMC /Users/kcft114/.julia/packages/AdvancedHMC/YWXfk/src/hamiltonian.jl:36
┌ Warning: Incorrect ϵ = NaN; ϵ_previous = 0.07500000000000001 is used instead.
└ @ AdvancedHMC.Adaptation /Users/kcft114/.julia/packages/AdvancedHMC/YWXfk/src/adaptation/stepsize.jl:103
┌ Warning: The current proposal will be rejected due to numerical error(s).
│   isfiniteθ = true
│   isfiniter = true
│   isfiniteℓπ = false
│   isfiniteℓκ = true
└ @ AdvancedHMC /Users/kcft114/.julia/packages/AdvancedHMC/YWXfk/src/hamiltonian.jl:36
┌ Warning: The current proposal will be rejected due to numerical error(s).
│   isfiniteθ = false
│   isfiniter = false
│   isfiniteℓπ = false
│   isfiniteℓκ = false
└ @ AdvancedHMC /Users/kcft114/.julia/packages/AdvancedHMC/YWXfk/src/hamiltonian.jl:36
┌ Warning: Incorrect ϵ = NaN; ϵ_previous = 0.07500000000000001 is used instead.
└ @ AdvancedHMC.Adaptation /Users/kcft114/.julia/packages/AdvancedHMC/YWXfk/src/adaptation/stepsize.jl:103
┌ Warning: The current proposal will be rejected due to numerical error(s).
│   isfiniteθ = true
│   isfiniter = true
│   isfiniteℓπ = false
│   isfiniteℓκ = true
└ @ AdvancedHMC /Users/kcft114/.julia/packages/AdvancedHMC/YWXfk/src/hamiltonian.jl:36
┌ Warning: The current proposal will be rejected due to numerical error(s).
│   isfiniteθ = false
│   isfiniter = false
│   isfiniteℓπ = false
│   isfiniteℓκ = false
└ @ AdvancedHMC /Users/kcft114/.julia/packages/AdvancedHMC/YWXfk/src/hamiltonian.jl:36
┌ Warning: Incorrect ϵ = NaN; ϵ_previous = 0.07500000000000001 is used instead.
└ @ AdvancedHMC.Adaptation /Users/kcft114/.julia/packages/AdvancedHMC/YWXfk/src/adaptation/stepsize.jl:103
┌ Warning: The current proposal will be rejected due to numerical error(s).
│   isfiniteθ = true
│   isfiniter = true
│   isfiniteℓπ = false
│   isfiniteℓκ = true
└ @ AdvancedHMC /Users/kcft114/.julia/packages/AdvancedHMC/YWXfk/src/hamiltonian.jl:36
┌ Warning: The current proposal will be rejected due to numerical error(s).
│   isfiniteθ = false
│   isfiniter = false
│   isfiniteℓπ = false
│   isfiniteℓκ = false
└ @ AdvancedHMC /Users/kcft114/.julia/packages/AdvancedHMC/YWXfk/src/hamiltonian.jl:36
┌ Warning: Incorrect ϵ = NaN; ϵ_previous = 0.07500000000000001 is used instead.
└ @ AdvancedHMC.Adaptation /Users/kcft114/.julia/packages/AdvancedHMC/YWXfk/src/adaptation/stepsize.jl:103
┌ Warning: The current proposal will be rejected due to numerical error(s).
│   isfiniteθ = true
│   isfiniter = true
│   isfiniteℓπ = false
│   isfiniteℓκ = true
└ @ AdvancedHMC /Users/kcft114/.julia/packages/AdvancedHMC/YWXfk/src/hamiltonian.jl:36
┌ Warning: The current proposal will be rejected due to numerical error(s).
│   isfiniteθ = false
│   isfiniter = false
│   isfiniteℓπ = false
│   isfiniteℓκ = false
└ @ AdvancedHMC /Users/kcft114/.julia/packages/AdvancedHMC/YWXfk/src/hamiltonian.jl:36
┌ Warning: Incorrect ϵ = NaN; ϵ_previous = 0.07500000000000001 is used instead.
└ @ AdvancedHMC.Adaptation /Users/kcft114/.julia/packages/AdvancedHMC/YWXfk/src/adaptation/stepsize.jl:103
┌ Warning: The current proposal will be rejected due to numerical error(s).
│   isfiniteθ = true
│   isfiniter = true
│   isfiniteℓπ = false
│   isfiniteℓκ = true
└ @ AdvancedHMC /Users/kcft114/.julia/packages/AdvancedHMC/YWXfk/src/hamiltonian.jl:36
┌ Warning: The current proposal will be rejected due to numerical error(s).
│   isfiniteθ = false
│   isfiniter = false
│   isfiniteℓπ = false
│   isfiniteℓκ = false
└ @ AdvancedHMC /Users/kcft114/.julia/packages/AdvancedHMC/YWXfk/src/hamiltonian.jl:36
┌ Warning: Incorrect ϵ = NaN; ϵ_previous = 0.07500000000000001 is used instead.
└ @ AdvancedHMC.Adaptation /Users/kcft114/.julia/packages/AdvancedHMC/YWXfk/src/adaptation/stepsize.jl:103
┌ Warning: The current proposal will be rejected due to numerical error(s).
│   isfiniteθ = true
│   isfiniter = true
│   isfiniteℓπ = false
│   isfiniteℓκ = true
└ @ AdvancedHMC /Users/kcft114/.julia/packages/AdvancedHMC/YWXfk/src/hamiltonian.jl:36
┌ Warning: The current proposal will be rejected due to numerical error(s).
│   isfiniteθ = false
│   isfiniter = false
│   isfiniteℓπ = false
│   isfiniteℓκ = false
└ @ AdvancedHMC /Users/kcft114/.julia/packages/AdvancedHMC/YWXfk/src/hamiltonian.jl:36
┌ Warning: Incorrect ϵ = NaN; ϵ_previous = 0.07500000000000001 is used instead.
└ @ AdvancedHMC.Adaptation /Users/kcft114/.julia/packages/AdvancedHMC/YWXfk/src/adaptation/stepsize.jl:103
┌ Warning: The current proposal will be rejected due to numerical error(s).
│   isfiniteθ = true
│   isfiniter = true
│   isfiniteℓπ = false
│   isfiniteℓκ = true
└ @ AdvancedHMC /Users/kcft114/.julia/packages/AdvancedHMC/YWXfk/src/hamiltonian.jl:36
┌ Warning: The current proposal will be rejected due to numerical error(s).
│   isfiniteθ = false
│   isfiniter = false
│   isfiniteℓπ = false
│   isfiniteℓκ = false
└ @ AdvancedHMC /Users/kcft114/.julia/packages/AdvancedHMC/YWXfk/src/hamiltonian.jl:36
┌ Warning: Incorrect ϵ = NaN; ϵ_previous = 0.07500000000000001 is used instead.
└ @ AdvancedHMC.Adaptation /Users/kcft114/.julia/packages/AdvancedHMC/YWXfk/src/adaptation/stepsize.jl:103
┌ Warning: The current proposal will be rejected due to numerical error(s).
│   isfiniteθ = true
│   isfiniter = true
│   isfiniteℓπ = false
│   isfiniteℓκ = true
└ @ AdvancedHMC /Users/kcft114/.julia/packages/AdvancedHMC/YWXfk/src/hamiltonian.jl:36
┌ Warning: The current proposal will be rejected due to numerical error(s).
│   isfiniteθ = false
│   isfiniter = false
│   isfiniteℓπ = false
│   isfiniteℓκ = false
└ @ AdvancedHMC /Users/kcft114/.julia/packages/AdvancedHMC/YWXfk/src/hamiltonian.jl:36
┌ Warning: Incorrect ϵ = NaN; ϵ_previous = 0.07500000000000001 is used instead.
└ @ AdvancedHMC.Adaptation /Users/kcft114/.julia/packages/AdvancedHMC/YWXfk/src/adaptation/stepsize.jl:103
┌ Warning: The current proposal will be rejected due to numerical error(s).
│   isfiniteθ = true
│   isfiniter = true
│   isfiniteℓπ = false
│   isfiniteℓκ = true
└ @ AdvancedHMC /Users/kcft114/.julia/packages/AdvancedHMC/YWXfk/src/hamiltonian.jl:36
┌ Warning: The current proposal will be rejected due to numerical error(s).
│   isfiniteθ = false
│   isfiniter = false
│   isfiniteℓπ = false
│   isfiniteℓκ = false
└ @ AdvancedHMC /Users/kcft114/.julia/packages/AdvancedHMC/YWXfk/src/hamiltonian.jl:36
┌ Warning: Incorrect ϵ = NaN; ϵ_previous = 0.07500000000000001 is used instead.
└ @ AdvancedHMC.Adaptation /Users/kcft114/.julia/packages/AdvancedHMC/YWXfk/src/adaptation/stepsize.jl:103
┌ Warning: The current proposal will be rejected due to numerical error(s).
│   isfiniteθ = true
│   isfiniter = true
│   isfiniteℓπ = false
│   isfiniteℓκ = true
└ @ AdvancedHMC /Users/kcft114/.julia/packages/AdvancedHMC/YWXfk/src/hamiltonian.jl:36
┌ Warning: The current proposal will be rejected due to numerical error(s).
│   isfiniteθ = false
│   isfiniter = false
│   isfiniteℓπ = false
│   isfiniteℓκ = false
└ @ AdvancedHMC /Users/kcft114/.julia/packages/AdvancedHMC/YWXfk/src/hamiltonian.jl:36
┌ Warning: Incorrect ϵ = NaN; ϵ_previous = 0.07500000000000001 is used instead.
└ @ AdvancedHMC.Adaptation /Users/kcft114/.julia/packages/AdvancedHMC/YWXfk/src/adaptation/stepsize.jl:103
┌ Warning: The current proposal will be rejected due to numerical error(s).
│   isfiniteθ = true
│   isfiniter = true
│   isfiniteℓπ = false
│   isfiniteℓκ = true
└ @ AdvancedHMC /Users/kcft114/.julia/packages/AdvancedHMC/YWXfk/src/hamiltonian.jl:36
┌ Warning: The current proposal will be rejected due to numerical error(s).
│   isfiniteθ = false
│   isfiniter = false
│   isfiniteℓπ = false
│   isfiniteℓκ = false
└ @ AdvancedHMC /Users/kcft114/.julia/packages/AdvancedHMC/YWXfk/src/hamiltonian.jl:36
┌ Warning: Incorrect ϵ = NaN; ϵ_previous = 0.07500000000000001 is used instead.
└ @ AdvancedHMC.Adaptation /Users/kcft114/.julia/packages/AdvancedHMC/YWXfk/src/adaptation/stepsize.jl:103
┌ Warning: The current proposal will be rejected due to numerical error(s).
│   isfiniteθ = true
│   isfiniter = true
│   isfiniteℓπ = false
│   isfiniteℓκ = true
└ @ AdvancedHMC /Users/kcft114/.julia/packages/AdvancedHMC/YWXfk/src/hamiltonian.jl:36
┌ Warning: The current proposal will be rejected due to numerical error(s).
│   isfiniteθ = false
│   isfiniter = false
│   isfiniteℓπ = false
│   isfiniteℓκ = false
└ @ AdvancedHMC /Users/kcft114/.julia/packages/AdvancedHMC/YWXfk/src/hamiltonian.jl:36
┌ Warning: Incorrect ϵ = NaN; ϵ_previous = 0.07500000000000001 is used instead.
└ @ AdvancedHMC.Adaptation /Users/kcft114/.julia/packages/AdvancedHMC/YWXfk/src/adaptation/stepsize.jl:103
┌ Warning: The current proposal will be rejected due to numerical error(s).
│   isfiniteθ = true
│   isfiniter = true
│   isfiniteℓπ = false
│   isfiniteℓκ = true
└ @ AdvancedHMC /Users/kcft114/.julia/packages/AdvancedHMC/YWXfk/src/hamiltonian.jl:36
┌ Warning: The current proposal will be rejected due to numerical error(s).
│   isfiniteθ = false
│   isfiniter = false
│   isfiniteℓπ = false
│   isfiniteℓκ = false
└ @ AdvancedHMC /Users/kcft114/.julia/packages/AdvancedHMC/YWXfk/src/hamiltonian.jl:36
┌ Warning: Incorrect ϵ = NaN; ϵ_previous = 0.07500000000000001 is used instead.
└ @ AdvancedHMC.Adaptation /Users/kcft114/.julia/packages/AdvancedHMC/YWXfk/src/adaptation/stepsize.jl:103
┌ Warning: The current proposal will be rejected due to numerical error(s).
│   isfiniteθ = true
│   isfiniter = true
│   isfiniteℓπ = false
│   isfiniteℓκ = true
└ @ AdvancedHMC /Users/kcft114/.julia/packages/AdvancedHMC/YWXfk/src/hamiltonian.jl:36
┌ Warning: The current proposal will be rejected due to numerical error(s).
│   isfiniteθ = false
│   isfiniter = false
│   isfiniteℓπ = false
│   isfiniteℓκ = false
└ @ AdvancedHMC /Users/kcft114/.julia/packages/AdvancedHMC/YWXfk/src/hamiltonian.jl:36
┌ Warning: Incorrect ϵ = NaN; ϵ_previous = 0.07500000000000001 is used instead.
└ @ AdvancedHMC.Adaptation /Users/kcft114/.julia/packages/AdvancedHMC/YWXfk/src/adaptation/stepsize.jl:103
┌ Warning: The current proposal will be rejected due to numerical error(s).
│   isfiniteθ = true
│   isfiniter = true
│   isfiniteℓπ = false
│   isfiniteℓκ = true
└ @ AdvancedHMC /Users/kcft114/.julia/packages/AdvancedHMC/YWXfk/src/hamiltonian.jl:36
┌ Warning: The current proposal will be rejected due to numerical error(s).
│   isfiniteθ = false
│   isfiniter = false
│   isfiniteℓπ = false
│   isfiniteℓκ = false
└ @ AdvancedHMC /Users/kcft114/.julia/packages/AdvancedHMC/YWXfk/src/hamiltonian.jl:36
┌ Warning: Incorrect ϵ = NaN; ϵ_previous = 0.07500000000000001 is used instead.
└ @ AdvancedHMC.Adaptation /Users/kcft114/.julia/packages/AdvancedHMC/YWXfk/src/adaptation/stepsize.jl:103
┌ Warning: The current proposal will be rejected due to numerical error(s).
│   isfiniteθ = true
│   isfiniter = true
│   isfiniteℓπ = false
│   isfiniteℓκ = true
└ @ AdvancedHMC /Users/kcft114/.julia/packages/AdvancedHMC/YWXfk/src/hamiltonian.jl:36
┌ Warning: The current proposal will be rejected due to numerical error(s).
│   isfiniteθ = false
│   isfiniter = false
│   isfiniteℓπ = false
│   isfiniteℓκ = false
└ @ AdvancedHMC /Users/kcft114/.julia/packages/AdvancedHMC/YWXfk/src/hamiltonian.jl:36
┌ Warning: Incorrect ϵ = NaN; ϵ_previous = 0.07500000000000001 is used instead.
└ @ AdvancedHMC.Adaptation /Users/kcft114/.julia/packages/AdvancedHMC/YWXfk/src/adaptation/stepsize.jl:103
┌ Warning: The current proposal will be rejected due to numerical error(s).
│   isfiniteθ = true
│   isfiniter = true
│   isfiniteℓπ = false
│   isfiniteℓκ = true
└ @ AdvancedHMC /Users/kcft114/.julia/packages/AdvancedHMC/YWXfk/src/hamiltonian.jl:36
┌ Warning: The current proposal will be rejected due to numerical error(s).
│   isfiniteθ = false
│   isfiniter = false
│   isfiniteℓπ = false
│   isfiniteℓκ = false
└ @ AdvancedHMC /Users/kcft114/.julia/packages/AdvancedHMC/YWXfk/src/hamiltonian.jl:36
┌ Warning: Incorrect ϵ = NaN; ϵ_previous = 0.07500000000000001 is used instead.
└ @ AdvancedHMC.Adaptation /Users/kcft114/.julia/packages/AdvancedHMC/YWXfk/src/adaptation/stepsize.jl:103
┌ Warning: The current proposal will be rejected due to numerical error(s).
│   isfiniteθ = true
│   isfiniter = true
│   isfiniteℓπ = false
│   isfiniteℓκ = true
└ @ AdvancedHMC /Users/kcft114/.julia/packages/AdvancedHMC/YWXfk/src/hamiltonian.jl:36
┌ Warning: The current proposal will be rejected due to numerical error(s).
│   isfiniteθ = false
│   isfiniter = false
│   isfiniteℓπ = false
│   isfiniteℓκ = false
└ @ AdvancedHMC /Users/kcft114/.julia/packages/AdvancedHMC/YWXfk/src/hamiltonian.jl:36
┌ Warning: Incorrect ϵ = NaN; ϵ_previous = 0.07500000000000001 is used instead.
└ @ AdvancedHMC.Adaptation /Users/kcft114/.julia/packages/AdvancedHMC/YWXfk/src/adaptation/stepsize.jl:103
┌ Warning: The current proposal will be rejected due to numerical error(s).
│   isfiniteθ = true
│   isfiniter = true
│   isfiniteℓπ = false
│   isfiniteℓκ = true
└ @ AdvancedHMC /Users/kcft114/.julia/packages/AdvancedHMC/YWXfk/src/hamiltonian.jl:36
┌ Warning: The current proposal will be rejected due to numerical error(s).
│   isfiniteθ = false
│   isfiniter = false
│   isfiniteℓπ = false
│   isfiniteℓκ = false
└ @ AdvancedHMC /Users/kcft114/.julia/packages/AdvancedHMC/YWXfk/src/hamiltonian.jl:36
┌ Warning: Incorrect ϵ = NaN; ϵ_previous = 0.07500000000000001 is used instead.
└ @ AdvancedHMC.Adaptation /Users/kcft114/.julia/packages/AdvancedHMC/YWXfk/src/adaptation/stepsize.jl:103
┌ Warning: The current proposal will be rejected due to numerical error(s).
│   isfiniteθ = true
│   isfiniter = true
│   isfiniteℓπ = false
│   isfiniteℓκ = true
└ @ AdvancedHMC /Users/kcft114/.julia/packages/AdvancedHMC/YWXfk/src/hamiltonian.jl:36
┌ Warning: The current proposal will be rejected due to numerical error(s).
│   isfiniteθ = false
│   isfiniter = false
│   isfiniteℓπ = false
│   isfiniteℓκ = false
└ @ AdvancedHMC /Users/kcft114/.julia/packages/AdvancedHMC/YWXfk/src/hamiltonian.jl:36
┌ Warning: Incorrect ϵ = NaN; ϵ_previous = 0.07500000000000001 is used instead.
└ @ AdvancedHMC.Adaptation /Users/kcft114/.julia/packages/AdvancedHMC/YWXfk/src/adaptation/stepsize.jl:103
┌ Warning: The current proposal will be rejected due to numerical error(s).
│   isfiniteθ = true
│   isfiniter = true
│   isfiniteℓπ = false
│   isfiniteℓκ = true
└ @ AdvancedHMC /Users/kcft114/.julia/packages/AdvancedHMC/YWXfk/src/hamiltonian.jl:36
┌ Warning: The current proposal will be rejected due to numerical error(s).
│   isfiniteθ = false
│   isfiniter = false
│   isfiniteℓπ = false
│   isfiniteℓκ = false
└ @ AdvancedHMC /Users/kcft114/.julia/packages/AdvancedHMC/YWXfk/src/hamiltonian.jl:36
┌ Warning: Incorrect ϵ = NaN; ϵ_previous = 0.07500000000000001 is used instead.
└ @ AdvancedHMC.Adaptation /Users/kcft114/.julia/packages/AdvancedHMC/YWXfk/src/adaptation/stepsize.jl:103
┌ Warning: The current proposal will be rejected due to numerical error(s).
│   isfiniteθ = true
│   isfiniter = true
│   isfiniteℓπ = false
│   isfiniteℓκ = true
└ @ AdvancedHMC /Users/kcft114/.julia/packages/AdvancedHMC/YWXfk/src/hamiltonian.jl:36
┌ Warning: The current proposal will be rejected due to numerical error(s).
│   isfiniteθ = false
│   isfiniter = false
│   isfiniteℓπ = false
│   isfiniteℓκ = false
└ @ AdvancedHMC /Users/kcft114/.julia/packages/AdvancedHMC/YWXfk/src/hamiltonian.jl:36
┌ Warning: Incorrect ϵ = NaN; ϵ_previous = 0.07500000000000001 is used instead.
└ @ AdvancedHMC.Adaptation /Users/kcft114/.julia/packages/AdvancedHMC/YWXfk/src/adaptation/stepsize.jl:103
┌ Warning: The current proposal will be rejected due to numerical error(s).
│   isfiniteθ = true
│   isfiniter = true
│   isfiniteℓπ = false
│   isfiniteℓκ = true
└ @ AdvancedHMC /Users/kcft114/.julia/packages/AdvancedHMC/YWXfk/src/hamiltonian.jl:36
┌ Warning: The current proposal will be rejected due to numerical error(s).
│   isfiniteθ = false
│   isfiniter = false
│   isfiniteℓπ = false
│   isfiniteℓκ = false
└ @ AdvancedHMC /Users/kcft114/.julia/packages/AdvancedHMC/YWXfk/src/hamiltonian.jl:36
┌ Warning: Incorrect ϵ = NaN; ϵ_previous = 0.07500000000000001 is used instead.
└ @ AdvancedHMC.Adaptation /Users/kcft114/.julia/packages/AdvancedHMC/YWXfk/src/adaptation/stepsize.jl:103
┌ Warning: The current proposal will be rejected due to numerical error(s).
│   isfiniteθ = true
│   isfiniter = true
│   isfiniteℓπ = false
│   isfiniteℓκ = true
└ @ AdvancedHMC /Users/kcft114/.julia/packages/AdvancedHMC/YWXfk/src/hamiltonian.jl:36
┌ Warning: The current proposal will be rejected due to numerical error(s).
│   isfiniteθ = false
│   isfiniter = false
│   isfiniteℓπ = false
│   isfiniteℓκ = false
└ @ AdvancedHMC /Users/kcft114/.julia/packages/AdvancedHMC/YWXfk/src/hamiltonian.jl:36
┌ Warning: Incorrect ϵ = NaN; ϵ_previous = 0.07500000000000001 is used instead.
└ @ AdvancedHMC.Adaptation /Users/kcft114/.julia/packages/AdvancedHMC/YWXfk/src/adaptation/stepsize.jl:103
┌ Warning: The current proposal will be rejected due to numerical error(s).
│   isfiniteθ = true
│   isfiniter = true
│   isfiniteℓπ = false
│   isfiniteℓκ = true
└ @ AdvancedHMC /Users/kcft114/.julia/packages/AdvancedHMC/YWXfk/src/hamiltonian.jl:36
┌ Warning: The current proposal will be rejected due to numerical error(s).
│   isfiniteθ = false
│   isfiniter = false
│   isfiniteℓπ = false
│   isfiniteℓκ = false
└ @ AdvancedHMC /Users/kcft114/.julia/packages/AdvancedHMC/YWXfk/src/hamiltonian.jl:36
┌ Warning: Incorrect ϵ = NaN; ϵ_previous = 0.07500000000000001 is used instead.
└ @ AdvancedHMC.Adaptation /Users/kcft114/.julia/packages/AdvancedHMC/YWXfk/src/adaptation/stepsize.jl:103
┌ Warning: The current proposal will be rejected due to numerical error(s).
│   isfiniteθ = true
│   isfiniter = true
│   isfiniteℓπ = false
│   isfiniteℓκ = true
└ @ AdvancedHMC /Users/kcft114/.julia/packages/AdvancedHMC/YWXfk/src/hamiltonian.jl:36
┌ Warning: The current proposal will be rejected due to numerical error(s).
│   isfiniteθ = false
│   isfiniter = false
│   isfiniteℓπ = false
│   isfiniteℓκ = false
└ @ AdvancedHMC /Users/kcft114/.julia/packages/AdvancedHMC/YWXfk/src/hamiltonian.jl:36
┌ Warning: Incorrect ϵ = NaN; ϵ_previous = 0.07500000000000001 is used instead.
└ @ AdvancedHMC.Adaptation /Users/kcft114/.julia/packages/AdvancedHMC/YWXfk/src/adaptation/stepsize.jl:103
┌ Warning: The current proposal will be rejected due to numerical error(s).
│   isfiniteθ = true
│   isfiniter = true
│   isfiniteℓπ = false
│   isfiniteℓκ = true
└ @ AdvancedHMC /Users/kcft114/.julia/packages/AdvancedHMC/YWXfk/src/hamiltonian.jl:36
┌ Warning: The current proposal will be rejected due to numerical error(s).
│   isfiniteθ = false
│   isfiniter = false
│   isfiniteℓπ = false
│   isfiniteℓκ = false
└ @ AdvancedHMC /Users/kcft114/.julia/packages/AdvancedHMC/YWXfk/src/hamiltonian.jl:36
┌ Warning: Incorrect ϵ = NaN; ϵ_previous = 0.07500000000000001 is used instead.
└ @ AdvancedHMC.Adaptation /Users/kcft114/.julia/packages/AdvancedHMC/YWXfk/src/adaptation/stepsize.jl:103
┌ Warning: The current proposal will be rejected due to numerical error(s).
│   isfiniteθ = true
│   isfiniter = true
│   isfiniteℓπ = false
│   isfiniteℓκ = true
└ @ AdvancedHMC /Users/kcft114/.julia/packages/AdvancedHMC/YWXfk/src/hamiltonian.jl:36
┌ Warning: The current proposal will be rejected due to numerical error(s).
│   isfiniteθ = false
│   isfiniter = false
│   isfiniteℓπ = false
│   isfiniteℓκ = false
└ @ AdvancedHMC /Users/kcft114/.julia/packages/AdvancedHMC/YWXfk/src/hamiltonian.jl:36
┌ Warning: Incorrect ϵ = NaN; ϵ_previous = 0.07500000000000001 is used instead.
└ @ AdvancedHMC.Adaptation /Users/kcft114/.julia/packages/AdvancedHMC/YWXfk/src/adaptation/stepsize.jl:103
┌ Warning: The current proposal will be rejected due to numerical error(s).
│   isfiniteθ = true
│   isfiniter = true
│   isfiniteℓπ = false
│   isfiniteℓκ = true
└ @ AdvancedHMC /Users/kcft114/.julia/packages/AdvancedHMC/YWXfk/src/hamiltonian.jl:36
┌ Warning: The current proposal will be rejected due to numerical error(s).
│   isfiniteθ = false
│   isfiniter = false
│   isfiniteℓπ = false
│   isfiniteℓκ = false
└ @ AdvancedHMC /Users/kcft114/.julia/packages/AdvancedHMC/YWXfk/src/hamiltonian.jl:36
┌ Warning: Incorrect ϵ = NaN; ϵ_previous = 0.07500000000000001 is used instead.
└ @ AdvancedHMC.Adaptation /Users/kcft114/.julia/packages/AdvancedHMC/YWXfk/src/adaptation/stepsize.jl:103
┌ Warning: The current proposal will be rejected due to numerical error(s).
│   isfiniteθ = true
│   isfiniter = true
│   isfiniteℓπ = false
│   isfiniteℓκ = true
└ @ AdvancedHMC /Users/kcft114/.julia/packages/AdvancedHMC/YWXfk/src/hamiltonian.jl:36
┌ Warning: The current proposal will be rejected due to numerical error(s).
│   isfiniteθ = false
│   isfiniter = false
│   isfiniteℓπ = false
│   isfiniteℓκ = false
└ @ AdvancedHMC /Users/kcft114/.julia/packages/AdvancedHMC/YWXfk/src/hamiltonian.jl:36
┌ Warning: Incorrect ϵ = NaN; ϵ_previous = 0.07500000000000001 is used instead.
└ @ AdvancedHMC.Adaptation /Users/kcft114/.julia/packages/AdvancedHMC/YWXfk/src/adaptation/stepsize.jl:103
┌ Warning: The current proposal will be rejected due to numerical error(s).
│   isfiniteθ = true
│   isfiniter = true
│   isfiniteℓπ = false
│   isfiniteℓκ = true
└ @ AdvancedHMC /Users/kcft114/.julia/packages/AdvancedHMC/YWXfk/src/hamiltonian.jl:36
┌ Warning: The current proposal will be rejected due to numerical error(s).
│   isfiniteθ = false
│   isfiniter = false
│   isfiniteℓπ = false
│   isfiniteℓκ = false
└ @ AdvancedHMC /Users/kcft114/.julia/packages/AdvancedHMC/YWXfk/src/hamiltonian.jl:36
┌ Warning: Incorrect ϵ = NaN; ϵ_previous = 0.07500000000000001 is used instead.
└ @ AdvancedHMC.Adaptation /Users/kcft114/.julia/packages/AdvancedHMC/YWXfk/src/adaptation/stepsize.jl:103
┌ Warning: The current proposal will be rejected due to numerical error(s).
│   isfiniteθ = true
│   isfiniter = true
│   isfiniteℓπ = false
│   isfiniteℓκ = true
└ @ AdvancedHMC /Users/kcft114/.julia/packages/AdvancedHMC/YWXfk/src/hamiltonian.jl:36
┌ Warning: The current proposal will be rejected due to numerical error(s).
│   isfiniteθ = false
│   isfiniter = false
│   isfiniteℓπ = false
│   isfiniteℓκ = false
└ @ AdvancedHMC /Users/kcft114/.julia/packages/AdvancedHMC/YWXfk/src/hamiltonian.jl:36
┌ Warning: Incorrect ϵ = NaN; ϵ_previous = 0.07500000000000001 is used instead.
└ @ AdvancedHMC.Adaptation /Users/kcft114/.julia/packages/AdvancedHMC/YWXfk/src/adaptation/stepsize.jl:103
┌ Warning: The current proposal will be rejected due to numerical error(s).
│   isfiniteθ = true
│   isfiniter = true
│   isfiniteℓπ = false
│   isfiniteℓκ = true
└ @ AdvancedHMC /Users/kcft114/.julia/packages/AdvancedHMC/YWXfk/src/hamiltonian.jl:36
┌ Warning: The current proposal will be rejected due to numerical error(s).
│   isfiniteθ = false
│   isfiniter = false
│   isfiniteℓπ = false
│   isfiniteℓκ = false
└ @ AdvancedHMC /Users/kcft114/.julia/packages/AdvancedHMC/YWXfk/src/hamiltonian.jl:36
┌ Warning: Incorrect ϵ = NaN; ϵ_previous = 0.07500000000000001 is used instead.
└ @ AdvancedHMC.Adaptation /Users/kcft114/.julia/packages/AdvancedHMC/YWXfk/src/adaptation/stepsize.jl:103
┌ Warning: The current proposal will be rejected due to numerical error(s).
│   isfiniteθ = true
│   isfiniter = true
│   isfiniteℓπ = false
│   isfiniteℓκ = true
└ @ AdvancedHMC /Users/kcft114/.julia/packages/AdvancedHMC/YWXfk/src/hamiltonian.jl:36
┌ Warning: The current proposal will be rejected due to numerical error(s).
│   isfiniteθ = false
│   isfiniter = false
│   isfiniteℓπ = false
│   isfiniteℓκ = false
└ @ AdvancedHMC /Users/kcft114/.julia/packages/AdvancedHMC/YWXfk/src/hamiltonian.jl:36
┌ Warning: Incorrect ϵ = NaN; ϵ_previous = 0.07500000000000001 is used instead.
└ @ AdvancedHMC.Adaptation /Users/kcft114/.julia/packages/AdvancedHMC/YWXfk/src/adaptation/stepsize.jl:103
┌ Warning: The current proposal will be rejected due to numerical error(s).
│   isfiniteθ = true
│   isfiniter = true
│   isfiniteℓπ = false
│   isfiniteℓκ = true
└ @ AdvancedHMC /Users/kcft114/.julia/packages/AdvancedHMC/YWXfk/src/hamiltonian.jl:36
┌ Warning: The current proposal will be rejected due to numerical error(s).
│   isfiniteθ = false
│   isfiniter = false
│   isfiniteℓπ = false
│   isfiniteℓκ = false
└ @ AdvancedHMC /Users/kcft114/.julia/packages/AdvancedHMC/YWXfk/src/hamiltonian.jl:36
┌ Warning: Incorrect ϵ = NaN; ϵ_previous = 0.07500000000000001 is used instead.
└ @ AdvancedHMC.Adaptation /Users/kcft114/.julia/packages/AdvancedHMC/YWXfk/src/adaptation/stepsize.jl:103
┌ Warning: The current proposal will be rejected due to numerical error(s).
│   isfiniteθ = true
│   isfiniter = true
│   isfiniteℓπ = false
│   isfiniteℓκ = true
└ @ AdvancedHMC /Users/kcft114/.julia/packages/AdvancedHMC/YWXfk/src/hamiltonian.jl:36
┌ Warning: The current proposal will be rejected due to numerical error(s).
│   isfiniteθ = false
│   isfiniter = false
│   isfiniteℓπ = false
│   isfiniteℓκ = false
└ @ AdvancedHMC /Users/kcft114/.julia/packages/AdvancedHMC/YWXfk/src/hamiltonian.jl:36
┌ Warning: Incorrect ϵ = NaN; ϵ_previous = 0.07500000000000001 is used instead.
└ @ AdvancedHMC.Adaptation /Users/kcft114/.julia/packages/AdvancedHMC/YWXfk/src/adaptation/stepsize.jl:103
┌ Warning: The current proposal will be rejected due to numerical error(s).
│   isfiniteθ = true
│   isfiniter = true
│   isfiniteℓπ = false
│   isfiniteℓκ = true
└ @ AdvancedHMC /Users/kcft114/.julia/packages/AdvancedHMC/YWXfk/src/hamiltonian.jl:36
┌ Warning: The current proposal will be rejected due to numerical error(s).
│   isfiniteθ = false
│   isfiniter = false
│   isfiniteℓπ = false
│   isfiniteℓκ = false
└ @ AdvancedHMC /Users/kcft114/.julia/packages/AdvancedHMC/YWXfk/src/hamiltonian.jl:36
┌ Warning: Incorrect ϵ = NaN; ϵ_previous = 0.07500000000000001 is used instead.
└ @ AdvancedHMC.Adaptation /Users/kcft114/.julia/packages/AdvancedHMC/YWXfk/src/adaptation/stepsize.jl:103
┌ Warning: The current proposal will be rejected due to numerical error(s).
│   isfiniteθ = true
│   isfiniter = true
│   isfiniteℓπ = false
│   isfiniteℓκ = true
└ @ AdvancedHMC /Users/kcft114/.julia/packages/AdvancedHMC/YWXfk/src/hamiltonian.jl:36
┌ Warning: The current proposal will be rejected due to numerical error(s).
│   isfiniteθ = false
│   isfiniter = false
│   isfiniteℓπ = false
│   isfiniteℓκ = false
└ @ AdvancedHMC /Users/kcft114/.julia/packages/AdvancedHMC/YWXfk/src/hamiltonian.jl:36
┌ Warning: Incorrect ϵ = NaN; ϵ_previous = 0.07500000000000001 is used instead.
└ @ AdvancedHMC.Adaptation /Users/kcft114/.julia/packages/AdvancedHMC/YWXfk/src/adaptation/stepsize.jl:103
┌ Warning: The current proposal will be rejected due to numerical error(s).
│   isfiniteθ = true
│   isfiniter = true
│   isfiniteℓπ = false
│   isfiniteℓκ = true
└ @ AdvancedHMC /Users/kcft114/.julia/packages/AdvancedHMC/YWXfk/src/hamiltonian.jl:36
┌ Warning: The current proposal will be rejected due to numerical error(s).
│   isfiniteθ = false
│   isfiniter = false
│   isfiniteℓπ = false
│   isfiniteℓκ = false
└ @ AdvancedHMC /Users/kcft114/.julia/packages/AdvancedHMC/YWXfk/src/hamiltonian.jl:36
┌ Warning: Incorrect ϵ = NaN; ϵ_previous = 0.07500000000000001 is used instead.
└ @ AdvancedHMC.Adaptation /Users/kcft114/.julia/packages/AdvancedHMC/YWXfk/src/adaptation/stepsize.jl:103
┌ Warning: The current proposal will be rejected due to numerical error(s).
│   isfiniteθ = true
│   isfiniter = true
│   isfiniteℓπ = false
│   isfiniteℓκ = true
└ @ AdvancedHMC /Users/kcft114/.julia/packages/AdvancedHMC/YWXfk/src/hamiltonian.jl:36
┌ Warning: The current proposal will be rejected due to numerical error(s).
│   isfiniteθ = false
│   isfiniter = false
│   isfiniteℓπ = false
│   isfiniteℓκ = false
└ @ AdvancedHMC /Users/kcft114/.julia/packages/AdvancedHMC/YWXfk/src/hamiltonian.jl:36
┌ Warning: Incorrect ϵ = NaN; ϵ_previous = 0.07500000000000001 is used instead.
└ @ AdvancedHMC.Adaptation /Users/kcft114/.julia/packages/AdvancedHMC/YWXfk/src/adaptation/stepsize.jl:103
┌ Warning: The current proposal will be rejected due to numerical error(s).
│   isfiniteθ = true
│   isfiniter = true
│   isfiniteℓπ = false
│   isfiniteℓκ = true
└ @ AdvancedHMC /Users/kcft114/.julia/packages/AdvancedHMC/YWXfk/src/hamiltonian.jl:36
┌ Warning: The current proposal will be rejected due to numerical error(s).
│   isfiniteθ = false
│   isfiniter = false
│   isfiniteℓπ = false
│   isfiniteℓκ = false
└ @ AdvancedHMC /Users/kcft114/.julia/packages/AdvancedHMC/YWXfk/src/hamiltonian.jl:36
┌ Warning: Incorrect ϵ = NaN; ϵ_previous = 0.07500000000000001 is used instead.
└ @ AdvancedHMC.Adaptation /Users/kcft114/.julia/packages/AdvancedHMC/YWXfk/src/adaptation/stepsize.jl:103
┌ Warning: The current proposal will be rejected due to numerical error(s).
│   isfiniteθ = true
│   isfiniter = true
│   isfiniteℓπ = false
│   isfiniteℓκ = true
└ @ AdvancedHMC /Users/kcft114/.julia/packages/AdvancedHMC/YWXfk/src/hamiltonian.jl:36
┌ Warning: The current proposal will be rejected due to numerical error(s).
│   isfiniteθ = false
│   isfiniter = false
│   isfiniteℓπ = false
│   isfiniteℓκ = false
└ @ AdvancedHMC /Users/kcft114/.julia/packages/AdvancedHMC/YWXfk/src/hamiltonian.jl:36
┌ Warning: Incorrect ϵ = NaN; ϵ_previous = 0.07500000000000001 is used instead.
└ @ AdvancedHMC.Adaptation /Users/kcft114/.julia/packages/AdvancedHMC/YWXfk/src/adaptation/stepsize.jl:103
┌ Warning: The current proposal will be rejected due to numerical error(s).
│   isfiniteθ = true
│   isfiniter = true
│   isfiniteℓπ = false
│   isfiniteℓκ = true
└ @ AdvancedHMC /Users/kcft114/.julia/packages/AdvancedHMC/YWXfk/src/hamiltonian.jl:36
┌ Warning: The current proposal will be rejected due to numerical error(s).
│   isfiniteθ = false
│   isfiniter = false
│   isfiniteℓπ = false
│   isfiniteℓκ = false
└ @ AdvancedHMC /Users/kcft114/.julia/packages/AdvancedHMC/YWXfk/src/hamiltonian.jl:36
┌ Warning: Incorrect ϵ = NaN; ϵ_previous = 0.07500000000000001 is used instead.
└ @ AdvancedHMC.Adaptation /Users/kcft114/.julia/packages/AdvancedHMC/YWXfk/src/adaptation/stepsize.jl:103
┌ Warning: The current proposal will be rejected due to numerical error(s).
│   isfiniteθ = true
│   isfiniter = true
│   isfiniteℓπ = false
│   isfiniteℓκ = true
└ @ AdvancedHMC /Users/kcft114/.julia/packages/AdvancedHMC/YWXfk/src/hamiltonian.jl:36
┌ Warning: The current proposal will be rejected due to numerical error(s).
│   isfiniteθ = false
│   isfiniter = false
│   isfiniteℓπ = false
│   isfiniteℓκ = false
└ @ AdvancedHMC /Users/kcft114/.julia/packages/AdvancedHMC/YWXfk/src/hamiltonian.jl:36
┌ Warning: Incorrect ϵ = NaN; ϵ_previous = 0.07500000000000001 is used instead.
└ @ AdvancedHMC.Adaptation /Users/kcft114/.julia/packages/AdvancedHMC/YWXfk/src/adaptation/stepsize.jl:103
┌ Warning: The current proposal will be rejected due to numerical error(s).
│   isfiniteθ = true
│   isfiniter = true
│   isfiniteℓπ = false
│   isfiniteℓκ = true
└ @ AdvancedHMC /Users/kcft114/.julia/packages/AdvancedHMC/YWXfk/src/hamiltonian.jl:36
┌ Warning: The current proposal will be rejected due to numerical error(s).
│   isfiniteθ = false
│   isfiniter = false
│   isfiniteℓπ = false
│   isfiniteℓκ = false
└ @ AdvancedHMC /Users/kcft114/.julia/packages/AdvancedHMC/YWXfk/src/hamiltonian.jl:36
┌ Warning: Incorrect ϵ = NaN; ϵ_previous = 0.07500000000000001 is used instead.
└ @ AdvancedHMC.Adaptation /Users/kcft114/.julia/packages/AdvancedHMC/YWXfk/src/adaptation/stepsize.jl:103
┌ Warning: The current proposal will be rejected due to numerical error(s).
│   isfiniteθ = true
│   isfiniter = true
│   isfiniteℓπ = false
│   isfiniteℓκ = true
└ @ AdvancedHMC /Users/kcft114/.julia/packages/AdvancedHMC/YWXfk/src/hamiltonian.jl:36
┌ Warning: The current proposal will be rejected due to numerical error(s).
│   isfiniteθ = false
│   isfiniter = false
│   isfiniteℓπ = false
│   isfiniteℓκ = false
└ @ AdvancedHMC /Users/kcft114/.julia/packages/AdvancedHMC/YWXfk/src/hamiltonian.jl:36
┌ Warning: Incorrect ϵ = NaN; ϵ_previous = 0.07500000000000001 is used instead.
└ @ AdvancedHMC.Adaptation /Users/kcft114/.julia/packages/AdvancedHMC/YWXfk/src/adaptation/stepsize.jl:103
┌ Warning: The current proposal will be rejected due to numerical error(s).
│   isfiniteθ = true
│   isfiniter = true
│   isfiniteℓπ = false
│   isfiniteℓκ = true
└ @ AdvancedHMC /Users/kcft114/.julia/packages/AdvancedHMC/YWXfk/src/hamiltonian.jl:36
┌ Warning: The current proposal will be rejected due to numerical error(s).
│   isfiniteθ = false
│   isfiniter = false
│   isfiniteℓπ = false
│   isfiniteℓκ = false
└ @ AdvancedHMC /Users/kcft114/.julia/packages/AdvancedHMC/YWXfk/src/hamiltonian.jl:36
┌ Warning: Incorrect ϵ = NaN; ϵ_previous = 0.07500000000000001 is used instead.
└ @ AdvancedHMC.Adaptation /Users/kcft114/.julia/packages/AdvancedHMC/YWXfk/src/adaptation/stepsize.jl:103
┌ Warning: The current proposal will be rejected due to numerical error(s).
│   isfiniteθ = true
│   isfiniter = true
│   isfiniteℓπ = false
│   isfiniteℓκ = true
└ @ AdvancedHMC /Users/kcft114/.julia/packages/AdvancedHMC/YWXfk/src/hamiltonian.jl:36
┌ Warning: The current proposal will be rejected due to numerical error(s).
│   isfiniteθ = false
│   isfiniter = false
│   isfiniteℓπ = false
│   isfiniteℓκ = false
└ @ AdvancedHMC /Users/kcft114/.julia/packages/AdvancedHMC/YWXfk/src/hamiltonian.jl:36
┌ Warning: Incorrect ϵ = NaN; ϵ_previous = 0.07500000000000001 is used instead.
└ @ AdvancedHMC.Adaptation /Users/kcft114/.julia/packages/AdvancedHMC/YWXfk/src/adaptation/stepsize.jl:103
┌ Warning: The current proposal will be rejected due to numerical error(s).
│   isfiniteθ = true
│   isfiniter = true
│   isfiniteℓπ = false
│   isfiniteℓκ = true
└ @ AdvancedHMC /Users/kcft114/.julia/packages/AdvancedHMC/YWXfk/src/hamiltonian.jl:36
┌ Warning: The current proposal will be rejected due to numerical error(s).
│   isfiniteθ = false
│   isfiniter = false
│   isfiniteℓπ = false
│   isfiniteℓκ = false
└ @ AdvancedHMC /Users/kcft114/.julia/packages/AdvancedHMC/YWXfk/src/hamiltonian.jl:36
┌ Warning: Incorrect ϵ = NaN; ϵ_previous = 0.07500000000000001 is used instead.
└ @ AdvancedHMC.Adaptation /Users/kcft114/.julia/packages/AdvancedHMC/YWXfk/src/adaptation/stepsize.jl:103
┌ Warning: The current proposal will be rejected due to numerical error(s).
│   isfiniteθ = true
│   isfiniter = true
│   isfiniteℓπ = false
│   isfiniteℓκ = true
└ @ AdvancedHMC /Users/kcft114/.julia/packages/AdvancedHMC/YWXfk/src/hamiltonian.jl:36
┌ Warning: The current proposal will be rejected due to numerical error(s).
│   isfiniteθ = false
│   isfiniter = false
│   isfiniteℓπ = false
│   isfiniteℓκ = false
└ @ AdvancedHMC /Users/kcft114/.julia/packages/AdvancedHMC/YWXfk/src/hamiltonian.jl:36
┌ Warning: Incorrect ϵ = NaN; ϵ_previous = 0.07500000000000001 is used instead.
└ @ AdvancedHMC.Adaptation /Users/kcft114/.julia/packages/AdvancedHMC/YWXfk/src/adaptation/stepsize.jl:103
┌ Warning: The current proposal will be rejected due to numerical error(s).
│   isfiniteθ = true
│   isfiniter = true
│   isfiniteℓπ = false
│   isfiniteℓκ = true
└ @ AdvancedHMC /Users/kcft114/.julia/packages/AdvancedHMC/YWXfk/src/hamiltonian.jl:36
┌ Warning: The current proposal will be rejected due to numerical error(s).
│   isfiniteθ = false
│   isfiniter = false
│   isfiniteℓπ = false
│   isfiniteℓκ = false
└ @ AdvancedHMC /Users/kcft114/.julia/packages/AdvancedHMC/YWXfk/src/hamiltonian.jl:36
┌ Warning: Incorrect ϵ = NaN; ϵ_previous = 0.07500000000000001 is used instead.
└ @ AdvancedHMC.Adaptation /Users/kcft114/.julia/packages/AdvancedHMC/YWXfk/src/adaptation/stepsize.jl:103
┌ Warning: The current proposal will be rejected due to numerical error(s).
│   isfiniteθ = true
│   isfiniter = true
│   isfiniteℓπ = false
│   isfiniteℓκ = true
└ @ AdvancedHMC /Users/kcft114/.julia/packages/AdvancedHMC/YWXfk/src/hamiltonian.jl:36
┌ Warning: The current proposal will be rejected due to numerical error(s).
│   isfiniteθ = false
│   isfiniter = false
│   isfiniteℓπ = false
│   isfiniteℓκ = false
└ @ AdvancedHMC /Users/kcft114/.julia/packages/AdvancedHMC/YWXfk/src/hamiltonian.jl:36
┌ Warning: Incorrect ϵ = NaN; ϵ_previous = 0.07500000000000001 is used instead.
└ @ AdvancedHMC.Adaptation /Users/kcft114/.julia/packages/AdvancedHMC/YWXfk/src/adaptation/stepsize.jl:103
┌ Warning: The current proposal will be rejected due to numerical error(s).
│   isfiniteθ = true
│   isfiniter = true
│   isfiniteℓπ = false
│   isfiniteℓκ = true
└ @ AdvancedHMC /Users/kcft114/.julia/packages/AdvancedHMC/YWXfk/src/hamiltonian.jl:36
┌ Warning: The current proposal will be rejected due to numerical error(s).
│   isfiniteθ = false
│   isfiniter = false
│   isfiniteℓπ = false
│   isfiniteℓκ = false
└ @ AdvancedHMC /Users/kcft114/.julia/packages/AdvancedHMC/YWXfk/src/hamiltonian.jl:36
┌ Warning: Incorrect ϵ = NaN; ϵ_previous = 0.07500000000000001 is used instead.
└ @ AdvancedHMC.Adaptation /Users/kcft114/.julia/packages/AdvancedHMC/YWXfk/src/adaptation/stepsize.jl:103
┌ Warning: The current proposal will be rejected due to numerical error(s).
│   isfiniteθ = true
│   isfiniter = true
│   isfiniteℓπ = false
│   isfiniteℓκ = true
└ @ AdvancedHMC /Users/kcft114/.julia/packages/AdvancedHMC/YWXfk/src/hamiltonian.jl:36
┌ Warning: The current proposal will be rejected due to numerical error(s).
│   isfiniteθ = false
│   isfiniter = false
│   isfiniteℓπ = false
│   isfiniteℓκ = false
└ @ AdvancedHMC /Users/kcft114/.julia/packages/AdvancedHMC/YWXfk/src/hamiltonian.jl:36
┌ Warning: Incorrect ϵ = NaN; ϵ_previous = 0.07500000000000001 is used instead.
└ @ AdvancedHMC.Adaptation /Users/kcft114/.julia/packages/AdvancedHMC/YWXfk/src/adaptation/stepsize.jl:103
┌ Warning: The current proposal will be rejected due to numerical error(s).
│   isfiniteθ = true
│   isfiniter = true
│   isfiniteℓπ = false
│   isfiniteℓκ = true
└ @ AdvancedHMC /Users/kcft114/.julia/packages/AdvancedHMC/YWXfk/src/hamiltonian.jl:36
┌ Warning: The current proposal will be rejected due to numerical error(s).
│   isfiniteθ = false
│   isfiniter = false
│   isfiniteℓπ = false
│   isfiniteℓκ = false
└ @ AdvancedHMC /Users/kcft114/.julia/packages/AdvancedHMC/YWXfk/src/hamiltonian.jl:36
┌ Warning: Incorrect ϵ = NaN; ϵ_previous = 0.07500000000000001 is used instead.
└ @ AdvancedHMC.Adaptation /Users/kcft114/.julia/packages/AdvancedHMC/YWXfk/src/adaptation/stepsize.jl:103
┌ Warning: The current proposal will be rejected due to numerical error(s).
│   isfiniteθ = true
│   isfiniter = true
│   isfiniteℓπ = false
│   isfiniteℓκ = true
└ @ AdvancedHMC /Users/kcft114/.julia/packages/AdvancedHMC/YWXfk/src/hamiltonian.jl:36
┌ Warning: The current proposal will be rejected due to numerical error(s).
│   isfiniteθ = false
│   isfiniter = false
│   isfiniteℓπ = false
│   isfiniteℓκ = false
└ @ AdvancedHMC /Users/kcft114/.julia/packages/AdvancedHMC/YWXfk/src/hamiltonian.jl:36
┌ Warning: Incorrect ϵ = NaN; ϵ_previous = 0.07500000000000001 is used instead.
└ @ AdvancedHMC.Adaptation /Users/kcft114/.julia/packages/AdvancedHMC/YWXfk/src/adaptation/stepsize.jl:103
┌ Warning: The current proposal will be rejected due to numerical error(s).
│   isfiniteθ = true
│   isfiniter = true
│   isfiniteℓπ = false
│   isfiniteℓκ = true
└ @ AdvancedHMC /Users/kcft114/.julia/packages/AdvancedHMC/YWXfk/src/hamiltonian.jl:36
┌ Warning: The current proposal will be rejected due to numerical error(s).
│   isfiniteθ = false
│   isfiniter = false
│   isfiniteℓπ = false
│   isfiniteℓκ = false
└ @ AdvancedHMC /Users/kcft114/.julia/packages/AdvancedHMC/YWXfk/src/hamiltonian.jl:36
┌ Warning: Incorrect ϵ = NaN; ϵ_previous = 0.07500000000000001 is used instead.
└ @ AdvancedHMC.Adaptation /Users/kcft114/.julia/packages/AdvancedHMC/YWXfk/src/adaptation/stepsize.jl:103
┌ Warning: The current proposal will be rejected due to numerical error(s).
│   isfiniteθ = true
│   isfiniter = true
│   isfiniteℓπ = false
│   isfiniteℓκ = true
└ @ AdvancedHMC /Users/kcft114/.julia/packages/AdvancedHMC/YWXfk/src/hamiltonian.jl:36
┌ Warning: The current proposal will be rejected due to numerical error(s).
│   isfiniteθ = false
│   isfiniter = false
│   isfiniteℓπ = false
│   isfiniteℓκ = false
└ @ AdvancedHMC /Users/kcft114/.julia/packages/AdvancedHMC/YWXfk/src/hamiltonian.jl:36
┌ Warning: Incorrect ϵ = NaN; ϵ_previous = 0.07500000000000001 is used instead.
└ @ AdvancedHMC.Adaptation /Users/kcft114/.julia/packages/AdvancedHMC/YWXfk/src/adaptation/stepsize.jl:103
┌ Warning: The current proposal will be rejected due to numerical error(s).
│   isfiniteθ = true
│   isfiniter = true
│   isfiniteℓπ = false
│   isfiniteℓκ = true
└ @ AdvancedHMC /Users/kcft114/.julia/packages/AdvancedHMC/YWXfk/src/hamiltonian.jl:36
┌ Warning: The current proposal will be rejected due to numerical error(s).
│   isfiniteθ = false
│   isfiniter = false
│   isfiniteℓπ = false
│   isfiniteℓκ = false
└ @ AdvancedHMC /Users/kcft114/.julia/packages/AdvancedHMC/YWXfk/src/hamiltonian.jl:36
┌ Warning: Incorrect ϵ = NaN; ϵ_previous = 0.07500000000000001 is used instead.
└ @ AdvancedHMC.Adaptation /Users/kcft114/.julia/packages/AdvancedHMC/YWXfk/src/adaptation/stepsize.jl:103
┌ Warning: The current proposal will be rejected due to numerical error(s).
│   isfiniteθ = true
│   isfiniter = true
│   isfiniteℓπ = false
│   isfiniteℓκ = true
└ @ AdvancedHMC /Users/kcft114/.julia/packages/AdvancedHMC/YWXfk/src/hamiltonian.jl:36
┌ Warning: The current proposal will be rejected due to numerical error(s).
│   isfiniteθ = false
│   isfiniter = false
│   isfiniteℓπ = false
│   isfiniteℓκ = false
└ @ AdvancedHMC /Users/kcft114/.julia/packages/AdvancedHMC/YWXfk/src/hamiltonian.jl:36
┌ Warning: Incorrect ϵ = NaN; ϵ_previous = 0.07500000000000001 is used instead.
└ @ AdvancedHMC.Adaptation /Users/kcft114/.julia/packages/AdvancedHMC/YWXfk/src/adaptation/stepsize.jl:103
┌ Warning: The current proposal will be rejected due to numerical error(s).
│   isfiniteθ = true
│   isfiniter = true
│   isfiniteℓπ = false
│   isfiniteℓκ = true
└ @ AdvancedHMC /Users/kcft114/.julia/packages/AdvancedHMC/YWXfk/src/hamiltonian.jl:36
┌ Warning: The current proposal will be rejected due to numerical error(s).
│   isfiniteθ = false
│   isfiniter = false
│   isfiniteℓπ = false
│   isfiniteℓκ = false
└ @ AdvancedHMC /Users/kcft114/.julia/packages/AdvancedHMC/YWXfk/src/hamiltonian.jl:36
┌ Warning: Incorrect ϵ = NaN; ϵ_previous = 0.07500000000000001 is used instead.
└ @ AdvancedHMC.Adaptation /Users/kcft114/.julia/packages/AdvancedHMC/YWXfk/src/adaptation/stepsize.jl:103
┌ Warning: The current proposal will be rejected due to numerical error(s).
│   isfiniteθ = true
│   isfiniter = true
│   isfiniteℓπ = false
│   isfiniteℓκ = true
└ @ AdvancedHMC /Users/kcft114/.julia/packages/AdvancedHMC/YWXfk/src/hamiltonian.jl:36
┌ Warning: The current proposal will be rejected due to numerical error(s).
│   isfiniteθ = false
│   isfiniter = false
│   isfiniteℓπ = false
│   isfiniteℓκ = false
└ @ AdvancedHMC /Users/kcft114/.julia/packages/AdvancedHMC/YWXfk/src/hamiltonian.jl:36
┌ Warning: Incorrect ϵ = NaN; ϵ_previous = 0.07500000000000001 is used instead.
└ @ AdvancedHMC.Adaptation /Users/kcft114/.julia/packages/AdvancedHMC/YWXfk/src/adaptation/stepsize.jl:103
┌ Warning: The current proposal will be rejected due to numerical error(s).
│   isfiniteθ = true
│   isfiniter = true
│   isfiniteℓπ = false
│   isfiniteℓκ = true
└ @ AdvancedHMC /Users/kcft114/.julia/packages/AdvancedHMC/YWXfk/src/hamiltonian.jl:36
┌ Warning: The current proposal will be rejected due to numerical error(s).
│   isfiniteθ = false
│   isfiniter = false
│   isfiniteℓπ = false
│   isfiniteℓκ = false
└ @ AdvancedHMC /Users/kcft114/.julia/packages/AdvancedHMC/YWXfk/src/hamiltonian.jl:36
┌ Warning: Incorrect ϵ = NaN; ϵ_previous = 0.07500000000000001 is used instead.
└ @ AdvancedHMC.Adaptation /Users/kcft114/.julia/packages/AdvancedHMC/YWXfk/src/adaptation/stepsize.jl:103
┌ Warning: The current proposal will be rejected due to numerical error(s).
│   isfiniteθ = true
│   isfiniter = true
│   isfiniteℓπ = false
│   isfiniteℓκ = true
└ @ AdvancedHMC /Users/kcft114/.julia/packages/AdvancedHMC/YWXfk/src/hamiltonian.jl:36
┌ Warning: The current proposal will be rejected due to numerical error(s).
│   isfiniteθ = false
│   isfiniter = false
│   isfiniteℓπ = false
│   isfiniteℓκ = false
└ @ AdvancedHMC /Users/kcft114/.julia/packages/AdvancedHMC/YWXfk/src/hamiltonian.jl:36
┌ Warning: Incorrect ϵ = NaN; ϵ_previous = 0.07500000000000001 is used instead.
└ @ AdvancedHMC.Adaptation /Users/kcft114/.julia/packages/AdvancedHMC/YWXfk/src/adaptation/stepsize.jl:103
┌ Warning: The current proposal will be rejected due to numerical error(s).
│   isfiniteθ = true
│   isfiniter = true
│   isfiniteℓπ = false
│   isfiniteℓκ = true
└ @ AdvancedHMC /Users/kcft114/.julia/packages/AdvancedHMC/YWXfk/src/hamiltonian.jl:36
┌ Warning: The current proposal will be rejected due to numerical error(s).
│   isfiniteθ = false
│   isfiniter = false
│   isfiniteℓπ = false
│   isfiniteℓκ = false
└ @ AdvancedHMC /Users/kcft114/.julia/packages/AdvancedHMC/YWXfk/src/hamiltonian.jl:36
┌ Warning: Incorrect ϵ = NaN; ϵ_previous = 0.07500000000000001 is used instead.
└ @ AdvancedHMC.Adaptation /Users/kcft114/.julia/packages/AdvancedHMC/YWXfk/src/adaptation/stepsize.jl:103
┌ Warning: The current proposal will be rejected due to numerical error(s).
│   isfiniteθ = true
│   isfiniter = true
│   isfiniteℓπ = false
│   isfiniteℓκ = true
└ @ AdvancedHMC /Users/kcft114/.julia/packages/AdvancedHMC/YWXfk/src/hamiltonian.jl:36
┌ Warning: The current proposal will be rejected due to numerical error(s).
│   isfiniteθ = false
│   isfiniter = false
│   isfiniteℓπ = false
│   isfiniteℓκ = false
└ @ AdvancedHMC /Users/kcft114/.julia/packages/AdvancedHMC/YWXfk/src/hamiltonian.jl:36
┌ Warning: Incorrect ϵ = NaN; ϵ_previous = 0.07500000000000001 is used instead.
└ @ AdvancedHMC.Adaptation /Users/kcft114/.julia/packages/AdvancedHMC/YWXfk/src/adaptation/stepsize.jl:103
┌ Warning: The current proposal will be rejected due to numerical error(s).
│   isfiniteθ = true
│   isfiniter = true
│   isfiniteℓπ = false
│   isfiniteℓκ = true
└ @ AdvancedHMC /Users/kcft114/.julia/packages/AdvancedHMC/YWXfk/src/hamiltonian.jl:36
┌ Warning: The current proposal will be rejected due to numerical error(s).
│   isfiniteθ = false
│   isfiniter = false
│   isfiniteℓπ = false
│   isfiniteℓκ = false
└ @ AdvancedHMC /Users/kcft114/.julia/packages/AdvancedHMC/YWXfk/src/hamiltonian.jl:36
┌ Warning: Incorrect ϵ = NaN; ϵ_previous = 0.07500000000000001 is used instead.
└ @ AdvancedHMC.Adaptation /Users/kcft114/.julia/packages/AdvancedHMC/YWXfk/src/adaptation/stepsize.jl:103
┌ Warning: The current proposal will be rejected due to numerical error(s).
│   isfiniteθ = true
│   isfiniter = true
│   isfiniteℓπ = false
│   isfiniteℓκ = true
└ @ AdvancedHMC /Users/kcft114/.julia/packages/AdvancedHMC/YWXfk/src/hamiltonian.jl:36
┌ Warning: The current proposal will be rejected due to numerical error(s).
│   isfiniteθ = false
│   isfiniter = false
│   isfiniteℓπ = false
│   isfiniteℓκ = false
└ @ AdvancedHMC /Users/kcft114/.julia/packages/AdvancedHMC/YWXfk/src/hamiltonian.jl:36
┌ Warning: Incorrect ϵ = NaN; ϵ_previous = 0.07500000000000001 is used instead.
└ @ AdvancedHMC.Adaptation /Users/kcft114/.julia/packages/AdvancedHMC/YWXfk/src/adaptation/stepsize.jl:103
┌ Warning: The current proposal will be rejected due to numerical error(s).
│   isfiniteθ = true
│   isfiniter = true
│   isfiniteℓπ = false
│   isfiniteℓκ = true
└ @ AdvancedHMC /Users/kcft114/.julia/packages/AdvancedHMC/YWXfk/src/hamiltonian.jl:36
┌ Warning: The current proposal will be rejected due to numerical error(s).
│   isfiniteθ = false
│   isfiniter = false
│   isfiniteℓπ = false
│   isfiniteℓκ = false
└ @ AdvancedHMC /Users/kcft114/.julia/packages/AdvancedHMC/YWXfk/src/hamiltonian.jl:36
┌ Warning: Incorrect ϵ = NaN; ϵ_previous = 0.07500000000000001 is used instead.
└ @ AdvancedHMC.Adaptation /Users/kcft114/.julia/packages/AdvancedHMC/YWXfk/src/adaptation/stepsize.jl:103
┌ Warning: The current proposal will be rejected due to numerical error(s).
│   isfiniteθ = true
│   isfiniter = true
│   isfiniteℓπ = false
│   isfiniteℓκ = true
└ @ AdvancedHMC /Users/kcft114/.julia/packages/AdvancedHMC/YWXfk/src/hamiltonian.jl:36
┌ Warning: The current proposal will be rejected due to numerical error(s).
│   isfiniteθ = false
│   isfiniter = false
│   isfiniteℓπ = false
│   isfiniteℓκ = false
└ @ AdvancedHMC /Users/kcft114/.julia/packages/AdvancedHMC/YWXfk/src/hamiltonian.jl:36
┌ Warning: Incorrect ϵ = NaN; ϵ_previous = 0.07500000000000001 is used instead.
└ @ AdvancedHMC.Adaptation /Users/kcft114/.julia/packages/AdvancedHMC/YWXfk/src/adaptation/stepsize.jl:103
┌ Warning: The current proposal will be rejected due to numerical error(s).
│   isfiniteθ = true
│   isfiniter = true
│   isfiniteℓπ = false
│   isfiniteℓκ = true
└ @ AdvancedHMC /Users/kcft114/.julia/packages/AdvancedHMC/YWXfk/src/hamiltonian.jl:36
┌ Warning: The current proposal will be rejected due to numerical error(s).
│   isfiniteθ = false
│   isfiniter = false
│   isfiniteℓπ = false
│   isfiniteℓκ = false
└ @ AdvancedHMC /Users/kcft114/.julia/packages/AdvancedHMC/YWXfk/src/hamiltonian.jl:36
┌ Warning: Incorrect ϵ = NaN; ϵ_previous = 0.07500000000000001 is used instead.
└ @ AdvancedHMC.Adaptation /Users/kcft114/.julia/packages/AdvancedHMC/YWXfk/src/adaptation/stepsize.jl:103
┌ Warning: The current proposal will be rejected due to numerical error(s).
│   isfiniteθ = true
│   isfiniter = true
│   isfiniteℓπ = false
│   isfiniteℓκ = true
└ @ AdvancedHMC /Users/kcft114/.julia/packages/AdvancedHMC/YWXfk/src/hamiltonian.jl:36
┌ Warning: The current proposal will be rejected due to numerical error(s).
│   isfiniteθ = false
│   isfiniter = false
│   isfiniteℓπ = false
│   isfiniteℓκ = false
└ @ AdvancedHMC /Users/kcft114/.julia/packages/AdvancedHMC/YWXfk/src/hamiltonian.jl:36
┌ Warning: Incorrect ϵ = NaN; ϵ_previous = 0.07500000000000001 is used instead.
└ @ AdvancedHMC.Adaptation /Users/kcft114/.julia/packages/AdvancedHMC/YWXfk/src/adaptation/stepsize.jl:103
┌ Warning: The current proposal will be rejected due to numerical error(s).
│   isfiniteθ = true
│   isfiniter = true
│   isfiniteℓπ = false
│   isfiniteℓκ = true
└ @ AdvancedHMC /Users/kcft114/.julia/packages/AdvancedHMC/YWXfk/src/hamiltonian.jl:36
┌ Warning: The current proposal will be rejected due to numerical error(s).
│   isfiniteθ = false
│   isfiniter = false
│   isfiniteℓπ = false
│   isfiniteℓκ = false
└ @ AdvancedHMC /Users/kcft114/.julia/packages/AdvancedHMC/YWXfk/src/hamiltonian.jl:36
┌ Warning: Incorrect ϵ = NaN; ϵ_previous = 0.07500000000000001 is used instead.
└ @ AdvancedHMC.Adaptation /Users/kcft114/.julia/packages/AdvancedHMC/YWXfk/src/adaptation/stepsize.jl:103
┌ Warning: The current proposal will be rejected due to numerical error(s).
│   isfiniteθ = true
│   isfiniter = true
│   isfiniteℓπ = false
│   isfiniteℓκ = true
└ @ AdvancedHMC /Users/kcft114/.julia/packages/AdvancedHMC/YWXfk/src/hamiltonian.jl:36
┌ Warning: The current proposal will be rejected due to numerical error(s).
│   isfiniteθ = false
│   isfiniter = false
│   isfiniteℓπ = false
│   isfiniteℓκ = false
└ @ AdvancedHMC /Users/kcft114/.julia/packages/AdvancedHMC/YWXfk/src/hamiltonian.jl:36
┌ Warning: Incorrect ϵ = NaN; ϵ_previous = 0.07500000000000001 is used instead.
└ @ AdvancedHMC.Adaptation /Users/kcft114/.julia/packages/AdvancedHMC/YWXfk/src/adaptation/stepsize.jl:103
┌ Warning: The current proposal will be rejected due to numerical error(s).
│   isfiniteθ = true
│   isfiniter = true
│   isfiniteℓπ = false
│   isfiniteℓκ = true
└ @ AdvancedHMC /Users/kcft114/.julia/packages/AdvancedHMC/YWXfk/src/hamiltonian.jl:36
┌ Warning: The current proposal will be rejected due to numerical error(s).
│   isfiniteθ = false
│   isfiniter = false
│   isfiniteℓπ = false
│   isfiniteℓκ = false
└ @ AdvancedHMC /Users/kcft114/.julia/packages/AdvancedHMC/YWXfk/src/hamiltonian.jl:36
┌ Warning: Incorrect ϵ = NaN; ϵ_previous = 0.07500000000000001 is used instead.
└ @ AdvancedHMC.Adaptation /Users/kcft114/.julia/packages/AdvancedHMC/YWXfk/src/adaptation/stepsize.jl:103
┌ Warning: The current proposal will be rejected due to numerical error(s).
│   isfiniteθ = true
│   isfiniter = true
│   isfiniteℓπ = false
│   isfiniteℓκ = true
└ @ AdvancedHMC /Users/kcft114/.julia/packages/AdvancedHMC/YWXfk/src/hamiltonian.jl:36
┌ Warning: The current proposal will be rejected due to numerical error(s).
│   isfiniteθ = false
│   isfiniter = false
│   isfiniteℓπ = false
│   isfiniteℓκ = false
└ @ AdvancedHMC /Users/kcft114/.julia/packages/AdvancedHMC/YWXfk/src/hamiltonian.jl:36
┌ Warning: Incorrect ϵ = NaN; ϵ_previous = 0.07500000000000001 is used instead.
└ @ AdvancedHMC.Adaptation /Users/kcft114/.julia/packages/AdvancedHMC/YWXfk/src/adaptation/stepsize.jl:103
┌ Warning: The current proposal will be rejected due to numerical error(s).
│   isfiniteθ = true
│   isfiniter = true
│   isfiniteℓπ = false
│   isfiniteℓκ = true
└ @ AdvancedHMC /Users/kcft114/.julia/packages/AdvancedHMC/YWXfk/src/hamiltonian.jl:36
┌ Warning: The current proposal will be rejected due to numerical error(s).
│   isfiniteθ = false
│   isfiniter = false
│   isfiniteℓπ = false
│   isfiniteℓκ = false
└ @ AdvancedHMC /Users/kcft114/.julia/packages/AdvancedHMC/YWXfk/src/hamiltonian.jl:36
┌ Warning: Incorrect ϵ = NaN; ϵ_previous = 0.07500000000000001 is used instead.
└ @ AdvancedHMC.Adaptation /Users/kcft114/.julia/packages/AdvancedHMC/YWXfk/src/adaptation/stepsize.jl:103
┌ Warning: The current proposal will be rejected due to numerical error(s).
│   isfiniteθ = true
│   isfiniter = true
│   isfiniteℓπ = false
│   isfiniteℓκ = true
└ @ AdvancedHMC /Users/kcft114/.julia/packages/AdvancedHMC/YWXfk/src/hamiltonian.jl:36
┌ Warning: The current proposal will be rejected due to numerical error(s).
│   isfiniteθ = false
│   isfiniter = false
│   isfiniteℓπ = false
│   isfiniteℓκ = false
└ @ AdvancedHMC /Users/kcft114/.julia/packages/AdvancedHMC/YWXfk/src/hamiltonian.jl:36
┌ Warning: Incorrect ϵ = NaN; ϵ_previous = 0.07500000000000001 is used instead.
└ @ AdvancedHMC.Adaptation /Users/kcft114/.julia/packages/AdvancedHMC/YWXfk/src/adaptation/stepsize.jl:103
┌ Warning: The current proposal will be rejected due to numerical error(s).
│   isfiniteθ = true
│   isfiniter = true
│   isfiniteℓπ = false
│   isfiniteℓκ = true
└ @ AdvancedHMC /Users/kcft114/.julia/packages/AdvancedHMC/YWXfk/src/hamiltonian.jl:36
┌ Warning: The current proposal will be rejected due to numerical error(s).
│   isfiniteθ = false
│   isfiniter = false
│   isfiniteℓπ = false
│   isfiniteℓκ = false
└ @ AdvancedHMC /Users/kcft114/.julia/packages/AdvancedHMC/YWXfk/src/hamiltonian.jl:36
┌ Warning: Incorrect ϵ = NaN; ϵ_previous = 0.07500000000000001 is used instead.
└ @ AdvancedHMC.Adaptation /Users/kcft114/.julia/packages/AdvancedHMC/YWXfk/src/adaptation/stepsize.jl:103
┌ Warning: The current proposal will be rejected due to numerical error(s).
│   isfiniteθ = true
│   isfiniter = true
│   isfiniteℓπ = false
│   isfiniteℓκ = true
└ @ AdvancedHMC /Users/kcft114/.julia/packages/AdvancedHMC/YWXfk/src/hamiltonian.jl:36
┌ Warning: The current proposal will be rejected due to numerical error(s).
│   isfiniteθ = false
│   isfiniter = false
│   isfiniteℓπ = false
│   isfiniteℓκ = false
└ @ AdvancedHMC /Users/kcft114/.julia/packages/AdvancedHMC/YWXfk/src/hamiltonian.jl:36
┌ Warning: Incorrect ϵ = NaN; ϵ_previous = 0.07500000000000001 is used instead.
└ @ AdvancedHMC.Adaptation /Users/kcft114/.julia/packages/AdvancedHMC/YWXfk/src/adaptation/stepsize.jl:103
┌ Warning: The current proposal will be rejected due to numerical error(s).
│   isfiniteθ = true
│   isfiniter = true
│   isfiniteℓπ = false
│   isfiniteℓκ = true
└ @ AdvancedHMC /Users/kcft114/.julia/packages/AdvancedHMC/YWXfk/src/hamiltonian.jl:36
┌ Warning: The current proposal will be rejected due to numerical error(s).
│   isfiniteθ = false
│   isfiniter = false
│   isfiniteℓπ = false
│   isfiniteℓκ = false
└ @ AdvancedHMC /Users/kcft114/.julia/packages/AdvancedHMC/YWXfk/src/hamiltonian.jl:36
┌ Warning: Incorrect ϵ = NaN; ϵ_previous = 0.07500000000000001 is used instead.
└ @ AdvancedHMC.Adaptation /Users/kcft114/.julia/packages/AdvancedHMC/YWXfk/src/adaptation/stepsize.jl:103
┌ Warning: The current proposal will be rejected due to numerical error(s).
│   isfiniteθ = true
│   isfiniter = true
│   isfiniteℓπ = false
│   isfiniteℓκ = true
└ @ AdvancedHMC /Users/kcft114/.julia/packages/AdvancedHMC/YWXfk/src/hamiltonian.jl:36
┌ Warning: The current proposal will be rejected due to numerical error(s).
│   isfiniteθ = false
│   isfiniter = false
│   isfiniteℓπ = false
│   isfiniteℓκ = false
└ @ AdvancedHMC /Users/kcft114/.julia/packages/AdvancedHMC/YWXfk/src/hamiltonian.jl:36
┌ Warning: Incorrect ϵ = NaN; ϵ_previous = 0.07500000000000001 is used instead.
└ @ AdvancedHMC.Adaptation /Users/kcft114/.julia/packages/AdvancedHMC/YWXfk/src/adaptation/stepsize.jl:103
┌ Warning: The current proposal will be rejected due to numerical error(s).
│   isfiniteθ = true
│   isfiniter = true
│   isfiniteℓπ = false
│   isfiniteℓκ = true
└ @ AdvancedHMC /Users/kcft114/.julia/packages/AdvancedHMC/YWXfk/src/hamiltonian.jl:36
┌ Warning: The current proposal will be rejected due to numerical error(s).
│   isfiniteθ = false
│   isfiniter = false
│   isfiniteℓπ = false
│   isfiniteℓκ = false
└ @ AdvancedHMC /Users/kcft114/.julia/packages/AdvancedHMC/YWXfk/src/hamiltonian.jl:36
┌ Warning: Incorrect ϵ = NaN; ϵ_previous = 0.07500000000000001 is used instead.
└ @ AdvancedHMC.Adaptation /Users/kcft114/.julia/packages/AdvancedHMC/YWXfk/src/adaptation/stepsize.jl:103
┌ Warning: The current proposal will be rejected due to numerical error(s).
│   isfiniteθ = true
│   isfiniter = true
│   isfiniteℓπ = false
│   isfiniteℓκ = true
└ @ AdvancedHMC /Users/kcft114/.julia/packages/AdvancedHMC/YWXfk/src/hamiltonian.jl:36
┌ Warning: The current proposal will be rejected due to numerical error(s).
│   isfiniteθ = false
│   isfiniter = false
│   isfiniteℓπ = false
│   isfiniteℓκ = false
└ @ AdvancedHMC /Users/kcft114/.julia/packages/AdvancedHMC/YWXfk/src/hamiltonian.jl:36
┌ Warning: Incorrect ϵ = NaN; ϵ_previous = 0.07500000000000001 is used instead.
└ @ AdvancedHMC.Adaptation /Users/kcft114/.julia/packages/AdvancedHMC/YWXfk/src/adaptation/stepsize.jl:103
┌ Warning: The current proposal will be rejected due to numerical error(s).
│   isfiniteθ = true
│   isfiniter = true
│   isfiniteℓπ = false
│   isfiniteℓκ = true
└ @ AdvancedHMC /Users/kcft114/.julia/packages/AdvancedHMC/YWXfk/src/hamiltonian.jl:36
┌ Warning: The current proposal will be rejected due to numerical error(s).
│   isfiniteθ = false
│   isfiniter = false
│   isfiniteℓπ = false
│   isfiniteℓκ = false
└ @ AdvancedHMC /Users/kcft114/.julia/packages/AdvancedHMC/YWXfk/src/hamiltonian.jl:36
┌ Warning: Incorrect ϵ = NaN; ϵ_previous = 0.07500000000000001 is used instead.
└ @ AdvancedHMC.Adaptation /Users/kcft114/.julia/packages/AdvancedHMC/YWXfk/src/adaptation/stepsize.jl:103
┌ Warning: The current proposal will be rejected due to numerical error(s).
│   isfiniteθ = true
│   isfiniter = true
│   isfiniteℓπ = false
│   isfiniteℓκ = true
└ @ AdvancedHMC /Users/kcft114/.julia/packages/AdvancedHMC/YWXfk/src/hamiltonian.jl:36
┌ Warning: The current proposal will be rejected due to numerical error(s).
│   isfiniteθ = false
│   isfiniter = false
│   isfiniteℓπ = false
│   isfiniteℓκ = false
└ @ AdvancedHMC /Users/kcft114/.julia/packages/AdvancedHMC/YWXfk/src/hamiltonian.jl:36
┌ Warning: Incorrect ϵ = NaN; ϵ_previous = 0.07500000000000001 is used instead.
└ @ AdvancedHMC.Adaptation /Users/kcft114/.julia/packages/AdvancedHMC/YWXfk/src/adaptation/stepsize.jl:103
┌ Warning: The current proposal will be rejected due to numerical error(s).
│   isfiniteθ = true
│   isfiniter = true
│   isfiniteℓπ = false
│   isfiniteℓκ = true
└ @ AdvancedHMC /Users/kcft114/.julia/packages/AdvancedHMC/YWXfk/src/hamiltonian.jl:36
┌ Warning: The current proposal will be rejected due to numerical error(s).
│   isfiniteθ = false
│   isfiniter = false
│   isfiniteℓπ = false
│   isfiniteℓκ = false
└ @ AdvancedHMC /Users/kcft114/.julia/packages/AdvancedHMC/YWXfk/src/hamiltonian.jl:36
┌ Warning: Incorrect ϵ = NaN; ϵ_previous = 0.07500000000000001 is used instead.
└ @ AdvancedHMC.Adaptation /Users/kcft114/.julia/packages/AdvancedHMC/YWXfk/src/adaptation/stepsize.jl:103
┌ Warning: The current proposal will be rejected due to numerical error(s).
│   isfiniteθ = true
│   isfiniter = true
│   isfiniteℓπ = false
│   isfiniteℓκ = true
└ @ AdvancedHMC /Users/kcft114/.julia/packages/AdvancedHMC/YWXfk/src/hamiltonian.jl:36
┌ Warning: The current proposal will be rejected due to numerical error(s).
│   isfiniteθ = false
│   isfiniter = false
│   isfiniteℓπ = false
│   isfiniteℓκ = false
└ @ AdvancedHMC /Users/kcft114/.julia/packages/AdvancedHMC/YWXfk/src/hamiltonian.jl:36
┌ Warning: Incorrect ϵ = NaN; ϵ_previous = 0.07500000000000001 is used instead.
└ @ AdvancedHMC.Adaptation /Users/kcft114/.julia/packages/AdvancedHMC/YWXfk/src/adaptation/stepsize.jl:103
┌ Warning: The current proposal will be rejected due to numerical error(s).
│   isfiniteθ = true
│   isfiniter = true
│   isfiniteℓπ = false
│   isfiniteℓκ = true
└ @ AdvancedHMC /Users/kcft114/.julia/packages/AdvancedHMC/YWXfk/src/hamiltonian.jl:36
┌ Warning: The current proposal will be rejected due to numerical error(s).
│   isfiniteθ = false
│   isfiniter = false
│   isfiniteℓπ = false
│   isfiniteℓκ = false
└ @ AdvancedHMC /Users/kcft114/.julia/packages/AdvancedHMC/YWXfk/src/hamiltonian.jl:36
┌ Warning: Incorrect ϵ = NaN; ϵ_previous = 0.07500000000000001 is used instead.
└ @ AdvancedHMC.Adaptation /Users/kcft114/.julia/packages/AdvancedHMC/YWXfk/src/adaptation/stepsize.jl:103
┌ Warning: The current proposal will be rejected due to numerical error(s).
│   isfiniteθ = true
│   isfiniter = true
│   isfiniteℓπ = false
│   isfiniteℓκ = true
└ @ AdvancedHMC /Users/kcft114/.julia/packages/AdvancedHMC/YWXfk/src/hamiltonian.jl:36
┌ Warning: The current proposal will be rejected due to numerical error(s).
│   isfiniteθ = false
│   isfiniter = false
│   isfiniteℓπ = false
│   isfiniteℓκ = false
└ @ AdvancedHMC /Users/kcft114/.julia/packages/AdvancedHMC/YWXfk/src/hamiltonian.jl:36
┌ Warning: Incorrect ϵ = NaN; ϵ_previous = 0.07500000000000001 is used instead.
└ @ AdvancedHMC.Adaptation /Users/kcft114/.julia/packages/AdvancedHMC/YWXfk/src/adaptation/stepsize.jl:103
┌ Warning: The current proposal will be rejected due to numerical error(s).
│   isfiniteθ = true
│   isfiniter = true
│   isfiniteℓπ = false
│   isfiniteℓκ = true
└ @ AdvancedHMC /Users/kcft114/.julia/packages/AdvancedHMC/YWXfk/src/hamiltonian.jl:36
┌ Warning: The current proposal will be rejected due to numerical error(s).
│   isfiniteθ = false
│   isfiniter = false
│   isfiniteℓπ = false
│   isfiniteℓκ = false
└ @ AdvancedHMC /Users/kcft114/.julia/packages/AdvancedHMC/YWXfk/src/hamiltonian.jl:36
┌ Warning: Incorrect ϵ = NaN; ϵ_previous = 0.07500000000000001 is used instead.
└ @ AdvancedHMC.Adaptation /Users/kcft114/.julia/packages/AdvancedHMC/YWXfk/src/adaptation/stepsize.jl:103
┌ Warning: The current proposal will be rejected due to numerical error(s).
│   isfiniteθ = true
│   isfiniter = true
│   isfiniteℓπ = false
│   isfiniteℓκ = true
└ @ AdvancedHMC /Users/kcft114/.julia/packages/AdvancedHMC/YWXfk/src/hamiltonian.jl:36
┌ Warning: The current proposal will be rejected due to numerical error(s).
│   isfiniteθ = false
│   isfiniter = false
│   isfiniteℓπ = false
│   isfiniteℓκ = false
└ @ AdvancedHMC /Users/kcft114/.julia/packages/AdvancedHMC/YWXfk/src/hamiltonian.jl:36
┌ Warning: Incorrect ϵ = NaN; ϵ_previous = 0.07500000000000001 is used instead.
└ @ AdvancedHMC.Adaptation /Users/kcft114/.julia/packages/AdvancedHMC/YWXfk/src/adaptation/stepsize.jl:103
┌ Warning: The current proposal will be rejected due to numerical error(s).
│   isfiniteθ = true
│   isfiniter = true
│   isfiniteℓπ = false
│   isfiniteℓκ = true
└ @ AdvancedHMC /Users/kcft114/.julia/packages/AdvancedHMC/YWXfk/src/hamiltonian.jl:36
┌ Warning: The current proposal will be rejected due to numerical error(s).
│   isfiniteθ = false
│   isfiniter = false
│   isfiniteℓπ = false
│   isfiniteℓκ = false
└ @ AdvancedHMC /Users/kcft114/.julia/packages/AdvancedHMC/YWXfk/src/hamiltonian.jl:36
┌ Warning: Incorrect ϵ = NaN; ϵ_previous = 0.07500000000000001 is used instead.
└ @ AdvancedHMC.Adaptation /Users/kcft114/.julia/packages/AdvancedHMC/YWXfk/src/adaptation/stepsize.jl:103
┌ Warning: The current proposal will be rejected due to numerical error(s).
│   isfiniteθ = true
│   isfiniter = true
│   isfiniteℓπ = false
│   isfiniteℓκ = true
└ @ AdvancedHMC /Users/kcft114/.julia/packages/AdvancedHMC/YWXfk/src/hamiltonian.jl:36
┌ Warning: The current proposal will be rejected due to numerical error(s).
│   isfiniteθ = false
│   isfiniter = false
│   isfiniteℓπ = false
│   isfiniteℓκ = false
└ @ AdvancedHMC /Users/kcft114/.julia/packages/AdvancedHMC/YWXfk/src/hamiltonian.jl:36
┌ Warning: Incorrect ϵ = NaN; ϵ_previous = 0.07500000000000001 is used instead.
└ @ AdvancedHMC.Adaptation /Users/kcft114/.julia/packages/AdvancedHMC/YWXfk/src/adaptation/stepsize.jl:103
┌ Warning: The current proposal will be rejected due to numerical error(s).
│   isfiniteθ = true
│   isfiniter = true
│   isfiniteℓπ = false
│   isfiniteℓκ = true
└ @ AdvancedHMC /Users/kcft114/.julia/packages/AdvancedHMC/YWXfk/src/hamiltonian.jl:36
┌ Warning: The current proposal will be rejected due to numerical error(s).
│   isfiniteθ = false
│   isfiniter = false
│   isfiniteℓπ = false
│   isfiniteℓκ = false
└ @ AdvancedHMC /Users/kcft114/.julia/packages/AdvancedHMC/YWXfk/src/hamiltonian.jl:36
┌ Warning: Incorrect ϵ = NaN; ϵ_previous = 0.07500000000000001 is used instead.
└ @ AdvancedHMC.Adaptation /Users/kcft114/.julia/packages/AdvancedHMC/YWXfk/src/adaptation/stepsize.jl:103
┌ Warning: The current proposal will be rejected due to numerical error(s).
│   isfiniteθ = true
│   isfiniter = true
│   isfiniteℓπ = false
│   isfiniteℓκ = true
└ @ AdvancedHMC /Users/kcft114/.julia/packages/AdvancedHMC/YWXfk/src/hamiltonian.jl:36
┌ Warning: The current proposal will be rejected due to numerical error(s).
│   isfiniteθ = false
│   isfiniter = false
│   isfiniteℓπ = false
│   isfiniteℓκ = false
└ @ AdvancedHMC /Users/kcft114/.julia/packages/AdvancedHMC/YWXfk/src/hamiltonian.jl:36
┌ Warning: Incorrect ϵ = NaN; ϵ_previous = 0.07500000000000001 is used instead.
└ @ AdvancedHMC.Adaptation /Users/kcft114/.julia/packages/AdvancedHMC/YWXfk/src/adaptation/stepsize.jl:103
┌ Warning: The current proposal will be rejected due to numerical error(s).
│   isfiniteθ = true
│   isfiniter = true
│   isfiniteℓπ = false
│   isfiniteℓκ = true
└ @ AdvancedHMC /Users/kcft114/.julia/packages/AdvancedHMC/YWXfk/src/hamiltonian.jl:36
┌ Warning: The current proposal will be rejected due to numerical error(s).
│   isfiniteθ = false
│   isfiniter = false
│   isfiniteℓπ = false
│   isfiniteℓκ = false
└ @ AdvancedHMC /Users/kcft114/.julia/packages/AdvancedHMC/YWXfk/src/hamiltonian.jl:36
┌ Warning: Incorrect ϵ = NaN; ϵ_previous = 0.07500000000000001 is used instead.
└ @ AdvancedHMC.Adaptation /Users/kcft114/.julia/packages/AdvancedHMC/YWXfk/src/adaptation/stepsize.jl:103
┌ Warning: The current proposal will be rejected due to numerical error(s).
│   isfiniteθ = true
│   isfiniter = true
│   isfiniteℓπ = false
│   isfiniteℓκ = true
└ @ AdvancedHMC /Users/kcft114/.julia/packages/AdvancedHMC/YWXfk/src/hamiltonian.jl:36
┌ Warning: The current proposal will be rejected due to numerical error(s).
│   isfiniteθ = false
│   isfiniter = false
│   isfiniteℓπ = false
│   isfiniteℓκ = false
└ @ AdvancedHMC /Users/kcft114/.julia/packages/AdvancedHMC/YWXfk/src/hamiltonian.jl:36
┌ Warning: Incorrect ϵ = NaN; ϵ_previous = 0.07500000000000001 is used instead.
└ @ AdvancedHMC.Adaptation /Users/kcft114/.julia/packages/AdvancedHMC/YWXfk/src/adaptation/stepsize.jl:103
┌ Warning: The current proposal will be rejected due to numerical error(s).
│   isfiniteθ = true
│   isfiniter = true
│   isfiniteℓπ = false
│   isfiniteℓκ = true
└ @ AdvancedHMC /Users/kcft114/.julia/packages/AdvancedHMC/YWXfk/src/hamiltonian.jl:36
┌ Warning: The current proposal will be rejected due to numerical error(s).
│   isfiniteθ = false
│   isfiniter = false
│   isfiniteℓπ = false
│   isfiniteℓκ = false
└ @ AdvancedHMC /Users/kcft114/.julia/packages/AdvancedHMC/YWXfk/src/hamiltonian.jl:36
┌ Warning: Incorrect ϵ = NaN; ϵ_previous = 0.07500000000000001 is used instead.
└ @ AdvancedHMC.Adaptation /Users/kcft114/.julia/packages/AdvancedHMC/YWXfk/src/adaptation/stepsize.jl:103
┌ Warning: The current proposal will be rejected due to numerical error(s).
│   isfiniteθ = true
│   isfiniter = true
│   isfiniteℓπ = false
│   isfiniteℓκ = true
└ @ AdvancedHMC /Users/kcft114/.julia/packages/AdvancedHMC/YWXfk/src/hamiltonian.jl:36
┌ Warning: The current proposal will be rejected due to numerical error(s).
│   isfiniteθ = false
│   isfiniter = false
│   isfiniteℓπ = false
│   isfiniteℓκ = false
└ @ AdvancedHMC /Users/kcft114/.julia/packages/AdvancedHMC/YWXfk/src/hamiltonian.jl:36
┌ Warning: Incorrect ϵ = NaN; ϵ_previous = 0.07500000000000001 is used instead.
└ @ AdvancedHMC.Adaptation /Users/kcft114/.julia/packages/AdvancedHMC/YWXfk/src/adaptation/stepsize.jl:103
┌ Warning: The current proposal will be rejected due to numerical error(s).
│   isfiniteθ = true
│   isfiniter = true
│   isfiniteℓπ = false
│   isfiniteℓκ = true
└ @ AdvancedHMC /Users/kcft114/.julia/packages/AdvancedHMC/YWXfk/src/hamiltonian.jl:36
┌ Warning: The current proposal will be rejected due to numerical error(s).
│   isfiniteθ = false
│   isfiniter = false
│   isfiniteℓπ = false
│   isfiniteℓκ = false
└ @ AdvancedHMC /Users/kcft114/.julia/packages/AdvancedHMC/YWXfk/src/hamiltonian.jl:36
┌ Warning: Incorrect ϵ = NaN; ϵ_previous = 0.07500000000000001 is used instead.
└ @ AdvancedHMC.Adaptation /Users/kcft114/.julia/packages/AdvancedHMC/YWXfk/src/adaptation/stepsize.jl:103
┌ Warning: The current proposal will be rejected due to numerical error(s).
│   isfiniteθ = true
│   isfiniter = true
│   isfiniteℓπ = false
│   isfiniteℓκ = true
└ @ AdvancedHMC /Users/kcft114/.julia/packages/AdvancedHMC/YWXfk/src/hamiltonian.jl:36
┌ Warning: The current proposal will be rejected due to numerical error(s).
│   isfiniteθ = false
│   isfiniter = false
│   isfiniteℓπ = false
│   isfiniteℓκ = false
└ @ AdvancedHMC /Users/kcft114/.julia/packages/AdvancedHMC/YWXfk/src/hamiltonian.jl:36
┌ Warning: Incorrect ϵ = NaN; ϵ_previous = 0.07500000000000001 is used instead.
└ @ AdvancedHMC.Adaptation /Users/kcft114/.julia/packages/AdvancedHMC/YWXfk/src/adaptation/stepsize.jl:103
┌ Warning: The current proposal will be rejected due to numerical error(s).
│   isfiniteθ = true
│   isfiniter = true
│   isfiniteℓπ = false
│   isfiniteℓκ = true
└ @ AdvancedHMC /Users/kcft114/.julia/packages/AdvancedHMC/YWXfk/src/hamiltonian.jl:36
┌ Warning: The current proposal will be rejected due to numerical error(s).
│   isfiniteθ = false
│   isfiniter = false
│   isfiniteℓπ = false
│   isfiniteℓκ = false
└ @ AdvancedHMC /Users/kcft114/.julia/packages/AdvancedHMC/YWXfk/src/hamiltonian.jl:36
┌ Warning: Incorrect ϵ = NaN; ϵ_previous = 0.07500000000000001 is used instead.
└ @ AdvancedHMC.Adaptation /Users/kcft114/.julia/packages/AdvancedHMC/YWXfk/src/adaptation/stepsize.jl:103
┌ Warning: The current proposal will be rejected due to numerical error(s).
│   isfiniteθ = true
│   isfiniter = true
│   isfiniteℓπ = false
│   isfiniteℓκ = true
└ @ AdvancedHMC /Users/kcft114/.julia/packages/AdvancedHMC/YWXfk/src/hamiltonian.jl:36
┌ Warning: The current proposal will be rejected due to numerical error(s).
│   isfiniteθ = false
│   isfiniter = false
│   isfiniteℓπ = false
│   isfiniteℓκ = false
└ @ AdvancedHMC /Users/kcft114/.julia/packages/AdvancedHMC/YWXfk/src/hamiltonian.jl:36
┌ Warning: Incorrect ϵ = NaN; ϵ_previous = 0.07500000000000001 is used instead.
└ @ AdvancedHMC.Adaptation /Users/kcft114/.julia/packages/AdvancedHMC/YWXfk/src/adaptation/stepsize.jl:103
┌ Warning: The current proposal will be rejected due to numerical error(s).
│   isfiniteθ = true
│   isfiniter = true
│   isfiniteℓπ = false
│   isfiniteℓκ = true
└ @ AdvancedHMC /Users/kcft114/.julia/packages/AdvancedHMC/YWXfk/src/hamiltonian.jl:36
┌ Warning: The current proposal will be rejected due to numerical error(s).
│   isfiniteθ = false
│   isfiniter = false
│   isfiniteℓπ = false
│   isfiniteℓκ = false
└ @ AdvancedHMC /Users/kcft114/.julia/packages/AdvancedHMC/YWXfk/src/hamiltonian.jl:36
┌ Warning: Incorrect ϵ = NaN; ϵ_previous = 0.07500000000000001 is used instead.
└ @ AdvancedHMC.Adaptation /Users/kcft114/.julia/packages/AdvancedHMC/YWXfk/src/adaptation/stepsize.jl:103
┌ Warning: The current proposal will be rejected due to numerical error(s).
│   isfiniteθ = true
│   isfiniter = true
│   isfiniteℓπ = false
│   isfiniteℓκ = true
└ @ AdvancedHMC /Users/kcft114/.julia/packages/AdvancedHMC/YWXfk/src/hamiltonian.jl:36
┌ Warning: The current proposal will be rejected due to numerical error(s).
│   isfiniteθ = false
│   isfiniter = false
│   isfiniteℓπ = false
│   isfiniteℓκ = false
└ @ AdvancedHMC /Users/kcft114/.julia/packages/AdvancedHMC/YWXfk/src/hamiltonian.jl:36
┌ Warning: Incorrect ϵ = NaN; ϵ_previous = 0.07500000000000001 is used instead.
└ @ AdvancedHMC.Adaptation /Users/kcft114/.julia/packages/AdvancedHMC/YWXfk/src/adaptation/stepsize.jl:103
┌ Warning: The current proposal will be rejected due to numerical error(s).
│   isfiniteθ = true
│   isfiniter = true
│   isfiniteℓπ = false
│   isfiniteℓκ = true
└ @ AdvancedHMC /Users/kcft114/.julia/packages/AdvancedHMC/YWXfk/src/hamiltonian.jl:36
┌ Warning: The current proposal will be rejected due to numerical error(s).
│   isfiniteθ = false
│   isfiniter = false
│   isfiniteℓπ = false
│   isfiniteℓκ = false
└ @ AdvancedHMC /Users/kcft114/.julia/packages/AdvancedHMC/YWXfk/src/hamiltonian.jl:36
┌ Warning: Incorrect ϵ = NaN; ϵ_previous = 0.07500000000000001 is used instead.
└ @ AdvancedHMC.Adaptation /Users/kcft114/.julia/packages/AdvancedHMC/YWXfk/src/adaptation/stepsize.jl:103
┌ Warning: The current proposal will be rejected due to numerical error(s).
│   isfiniteθ = true
│   isfiniter = true
│   isfiniteℓπ = false
│   isfiniteℓκ = true
└ @ AdvancedHMC /Users/kcft114/.julia/packages/AdvancedHMC/YWXfk/src/hamiltonian.jl:36
┌ Warning: The current proposal will be rejected due to numerical error(s).
│   isfiniteθ = false
│   isfiniter = false
│   isfiniteℓπ = false
│   isfiniteℓκ = false
└ @ AdvancedHMC /Users/kcft114/.julia/packages/AdvancedHMC/YWXfk/src/hamiltonian.jl:36
┌ Warning: Incorrect ϵ = NaN; ϵ_previous = 0.07500000000000001 is used instead.
└ @ AdvancedHMC.Adaptation /Users/kcft114/.julia/packages/AdvancedHMC/YWXfk/src/adaptation/stepsize.jl:103
┌ Warning: The current proposal will be rejected due to numerical error(s).
│   isfiniteθ = true
│   isfiniter = true
│   isfiniteℓπ = false
│   isfiniteℓκ = true
└ @ AdvancedHMC /Users/kcft114/.julia/packages/AdvancedHMC/YWXfk/src/hamiltonian.jl:36
┌ Warning: The current proposal will be rejected due to numerical error(s).
│   isfiniteθ = false
│   isfiniter = false
│   isfiniteℓπ = false
│   isfiniteℓκ = false
└ @ AdvancedHMC /Users/kcft114/.julia/packages/AdvancedHMC/YWXfk/src/hamiltonian.jl:36
┌ Warning: Incorrect ϵ = NaN; ϵ_previous = 0.07500000000000001 is used instead.
└ @ AdvancedHMC.Adaptation /Users/kcft114/.julia/packages/AdvancedHMC/YWXfk/src/adaptation/stepsize.jl:103
┌ Warning: The current proposal will be rejected due to numerical error(s).
│   isfiniteθ = true
│   isfiniter = true
│   isfiniteℓπ = false
│   isfiniteℓκ = true
└ @ AdvancedHMC /Users/kcft114/.julia/packages/AdvancedHMC/YWXfk/src/hamiltonian.jl:36
┌ Warning: The current proposal will be rejected due to numerical error(s).
│   isfiniteθ = false
│   isfiniter = false
│   isfiniteℓπ = false
│   isfiniteℓκ = false
└ @ AdvancedHMC /Users/kcft114/.julia/packages/AdvancedHMC/YWXfk/src/hamiltonian.jl:36
┌ Warning: Incorrect ϵ = NaN; ϵ_previous = 0.07500000000000001 is used instead.
└ @ AdvancedHMC.Adaptation /Users/kcft114/.julia/packages/AdvancedHMC/YWXfk/src/adaptation/stepsize.jl:103
┌ Warning: The current proposal will be rejected due to numerical error(s).
│   isfiniteθ = true
│   isfiniter = true
│   isfiniteℓπ = false
│   isfiniteℓκ = true
└ @ AdvancedHMC /Users/kcft114/.julia/packages/AdvancedHMC/YWXfk/src/hamiltonian.jl:36
┌ Warning: The current proposal will be rejected due to numerical error(s).
│   isfiniteθ = false
│   isfiniter = false
│   isfiniteℓπ = false
│   isfiniteℓκ = false
└ @ AdvancedHMC /Users/kcft114/.julia/packages/AdvancedHMC/YWXfk/src/hamiltonian.jl:36
┌ Warning: Incorrect ϵ = NaN; ϵ_previous = 0.07500000000000001 is used instead.
└ @ AdvancedHMC.Adaptation /Users/kcft114/.julia/packages/AdvancedHMC/YWXfk/src/adaptation/stepsize.jl:103
┌ Warning: The current proposal will be rejected due to numerical error(s).
│   isfiniteθ = true
│   isfiniter = true
│   isfiniteℓπ = false
│   isfiniteℓκ = true
└ @ AdvancedHMC /Users/kcft114/.julia/packages/AdvancedHMC/YWXfk/src/hamiltonian.jl:36
┌ Warning: The current proposal will be rejected due to numerical error(s).
│   isfiniteθ = false
│   isfiniter = false
│   isfiniteℓπ = false
│   isfiniteℓκ = false
└ @ AdvancedHMC /Users/kcft114/.julia/packages/AdvancedHMC/YWXfk/src/hamiltonian.jl:36
┌ Warning: Incorrect ϵ = NaN; ϵ_previous = 0.07500000000000001 is used instead.
└ @ AdvancedHMC.Adaptation /Users/kcft114/.julia/packages/AdvancedHMC/YWXfk/src/adaptation/stepsize.jl:103
┌ Warning: The current proposal will be rejected due to numerical error(s).
│   isfiniteθ = true
│   isfiniter = true
│   isfiniteℓπ = false
│   isfiniteℓκ = true
└ @ AdvancedHMC /Users/kcft114/.julia/packages/AdvancedHMC/YWXfk/src/hamiltonian.jl:36
┌ Warning: The current proposal will be rejected due to numerical error(s).
│   isfiniteθ = false
│   isfiniter = false
│   isfiniteℓπ = false
│   isfiniteℓκ = false
└ @ AdvancedHMC /Users/kcft114/.julia/packages/AdvancedHMC/YWXfk/src/hamiltonian.jl:36
┌ Warning: Incorrect ϵ = NaN; ϵ_previous = 0.07500000000000001 is used instead.
└ @ AdvancedHMC.Adaptation /Users/kcft114/.julia/packages/AdvancedHMC/YWXfk/src/adaptation/stepsize.jl:103
┌ Warning: The current proposal will be rejected due to numerical error(s).
│   isfiniteθ = true
│   isfiniter = true
│   isfiniteℓπ = false
│   isfiniteℓκ = true
└ @ AdvancedHMC /Users/kcft114/.julia/packages/AdvancedHMC/YWXfk/src/hamiltonian.jl:36
┌ Warning: The current proposal will be rejected due to numerical error(s).
│   isfiniteθ = false
│   isfiniter = false
│   isfiniteℓπ = false
│   isfiniteℓκ = false
└ @ AdvancedHMC /Users/kcft114/.julia/packages/AdvancedHMC/YWXfk/src/hamiltonian.jl:36
┌ Warning: Incorrect ϵ = NaN; ϵ_previous = 0.07500000000000001 is used instead.
└ @ AdvancedHMC.Adaptation /Users/kcft114/.julia/packages/AdvancedHMC/YWXfk/src/adaptation/stepsize.jl:103
┌ Warning: The current proposal will be rejected due to numerical error(s).
│   isfiniteθ = true
│   isfiniter = true
│   isfiniteℓπ = false
│   isfiniteℓκ = true
└ @ AdvancedHMC /Users/kcft114/.julia/packages/AdvancedHMC/YWXfk/src/hamiltonian.jl:36
┌ Warning: The current proposal will be rejected due to numerical error(s).
│   isfiniteθ = false
│   isfiniter = false
│   isfiniteℓπ = false
│   isfiniteℓκ = false
└ @ AdvancedHMC /Users/kcft114/.julia/packages/AdvancedHMC/YWXfk/src/hamiltonian.jl:36
┌ Warning: Incorrect ϵ = NaN; ϵ_previous = 0.07500000000000001 is used instead.
└ @ AdvancedHMC.Adaptation /Users/kcft114/.julia/packages/AdvancedHMC/YWXfk/src/adaptation/stepsize.jl:103
┌ Warning: The current proposal will be rejected due to numerical error(s).
│   isfiniteθ = true
│   isfiniter = true
│   isfiniteℓπ = false
│   isfiniteℓκ = true
└ @ AdvancedHMC /Users/kcft114/.julia/packages/AdvancedHMC/YWXfk/src/hamiltonian.jl:36
┌ Warning: The current proposal will be rejected due to numerical error(s).
│   isfiniteθ = false
│   isfiniter = false
│   isfiniteℓπ = false
│   isfiniteℓκ = false
└ @ AdvancedHMC /Users/kcft114/.julia/packages/AdvancedHMC/YWXfk/src/hamiltonian.jl:36
┌ Warning: Incorrect ϵ = NaN; ϵ_previous = 0.07500000000000001 is used instead.
└ @ AdvancedHMC.Adaptation /Users/kcft114/.julia/packages/AdvancedHMC/YWXfk/src/adaptation/stepsize.jl:103
┌ Warning: The current proposal will be rejected due to numerical error(s).
│   isfiniteθ = true
│   isfiniter = true
│   isfiniteℓπ = false
│   isfiniteℓκ = true
└ @ AdvancedHMC /Users/kcft114/.julia/packages/AdvancedHMC/YWXfk/src/hamiltonian.jl:36
┌ Warning: The current proposal will be rejected due to numerical error(s).
│   isfiniteθ = false
│   isfiniter = false
│   isfiniteℓπ = false
│   isfiniteℓκ = false
└ @ AdvancedHMC /Users/kcft114/.julia/packages/AdvancedHMC/YWXfk/src/hamiltonian.jl:36
┌ Warning: Incorrect ϵ = NaN; ϵ_previous = 0.07500000000000001 is used instead.
└ @ AdvancedHMC.Adaptation /Users/kcft114/.julia/packages/AdvancedHMC/YWXfk/src/adaptation/stepsize.jl:103
┌ Warning: The current proposal will be rejected due to numerical error(s).
│   isfiniteθ = true
│   isfiniter = true
│   isfiniteℓπ = false
│   isfiniteℓκ = true
└ @ AdvancedHMC /Users/kcft114/.julia/packages/AdvancedHMC/YWXfk/src/hamiltonian.jl:36
┌ Warning: The current proposal will be rejected due to numerical error(s).
│   isfiniteθ = false
│   isfiniter = false
│   isfiniteℓπ = false
│   isfiniteℓκ = false
└ @ AdvancedHMC /Users/kcft114/.julia/packages/AdvancedHMC/YWXfk/src/hamiltonian.jl:36
┌ Warning: Incorrect ϵ = NaN; ϵ_previous = 0.07500000000000001 is used instead.
└ @ AdvancedHMC.Adaptation /Users/kcft114/.julia/packages/AdvancedHMC/YWXfk/src/adaptation/stepsize.jl:103
┌ Warning: The current proposal will be rejected due to numerical error(s).
│   isfiniteθ = true
│   isfiniter = true
│   isfiniteℓπ = false
│   isfiniteℓκ = true
└ @ AdvancedHMC /Users/kcft114/.julia/packages/AdvancedHMC/YWXfk/src/hamiltonian.jl:36
┌ Warning: The current proposal will be rejected due to numerical error(s).
│   isfiniteθ = false
│   isfiniter = false
│   isfiniteℓπ = false
│   isfiniteℓκ = false
└ @ AdvancedHMC /Users/kcft114/.julia/packages/AdvancedHMC/YWXfk/src/hamiltonian.jl:36
┌ Warning: Incorrect ϵ = NaN; ϵ_previous = 0.07500000000000001 is used instead.
└ @ AdvancedHMC.Adaptation /Users/kcft114/.julia/packages/AdvancedHMC/YWXfk/src/adaptation/stepsize.jl:103
┌ Warning: The current proposal will be rejected due to numerical error(s).
│   isfiniteθ = true
│   isfiniter = true
│   isfiniteℓπ = false
│   isfiniteℓκ = true
└ @ AdvancedHMC /Users/kcft114/.julia/packages/AdvancedHMC/YWXfk/src/hamiltonian.jl:36
┌ Warning: The current proposal will be rejected due to numerical error(s).
│   isfiniteθ = false
│   isfiniter = false
│   isfiniteℓπ = false
│   isfiniteℓκ = false
└ @ AdvancedHMC /Users/kcft114/.julia/packages/AdvancedHMC/YWXfk/src/hamiltonian.jl:36
┌ Warning: Incorrect ϵ = NaN; ϵ_previous = 0.07500000000000001 is used instead.
└ @ AdvancedHMC.Adaptation /Users/kcft114/.julia/packages/AdvancedHMC/YWXfk/src/adaptation/stepsize.jl:103
┌ Warning: The current proposal will be rejected due to numerical error(s).
│   isfiniteθ = true
│   isfiniter = true
│   isfiniteℓπ = false
│   isfiniteℓκ = true
└ @ AdvancedHMC /Users/kcft114/.julia/packages/AdvancedHMC/YWXfk/src/hamiltonian.jl:36
┌ Warning: The current proposal will be rejected due to numerical error(s).
│   isfiniteθ = false
│   isfiniter = false
│   isfiniteℓπ = false
│   isfiniteℓκ = false
└ @ AdvancedHMC /Users/kcft114/.julia/packages/AdvancedHMC/YWXfk/src/hamiltonian.jl:36
┌ Warning: Incorrect ϵ = NaN; ϵ_previous = 0.07500000000000001 is used instead.
└ @ AdvancedHMC.Adaptation /Users/kcft114/.julia/packages/AdvancedHMC/YWXfk/src/adaptation/stepsize.jl:103
┌ Warning: The current proposal will be rejected due to numerical error(s).
│   isfiniteθ = true
│   isfiniter = true
│   isfiniteℓπ = false
│   isfiniteℓκ = true
└ @ AdvancedHMC /Users/kcft114/.julia/packages/AdvancedHMC/YWXfk/src/hamiltonian.jl:36
┌ Warning: The current proposal will be rejected due to numerical error(s).
│   isfiniteθ = false
│   isfiniter = false
│   isfiniteℓπ = false
│   isfiniteℓκ = false
└ @ AdvancedHMC /Users/kcft114/.julia/packages/AdvancedHMC/YWXfk/src/hamiltonian.jl:36
┌ Warning: Incorrect ϵ = NaN; ϵ_previous = 0.07500000000000001 is used instead.
└ @ AdvancedHMC.Adaptation /Users/kcft114/.julia/packages/AdvancedHMC/YWXfk/src/adaptation/stepsize.jl:103
┌ Warning: The current proposal will be rejected due to numerical error(s).
│   isfiniteθ = true
│   isfiniter = true
│   isfiniteℓπ = false
│   isfiniteℓκ = true
└ @ AdvancedHMC /Users/kcft114/.julia/packages/AdvancedHMC/YWXfk/src/hamiltonian.jl:36
┌ Warning: The current proposal will be rejected due to numerical error(s).
│   isfiniteθ = false
│   isfiniter = false
│   isfiniteℓπ = false
│   isfiniteℓκ = false
└ @ AdvancedHMC /Users/kcft114/.julia/packages/AdvancedHMC/YWXfk/src/hamiltonian.jl:36
┌ Warning: Incorrect ϵ = NaN; ϵ_previous = 0.07500000000000001 is used instead.
└ @ AdvancedHMC.Adaptation /Users/kcft114/.julia/packages/AdvancedHMC/YWXfk/src/adaptation/stepsize.jl:103
┌ Warning: The current proposal will be rejected due to numerical error(s).
│   isfiniteθ = true
│   isfiniter = true
│   isfiniteℓπ = false
│   isfiniteℓκ = true
└ @ AdvancedHMC /Users/kcft114/.julia/packages/AdvancedHMC/YWXfk/src/hamiltonian.jl:36
┌ Warning: The current proposal will be rejected due to numerical error(s).
│   isfiniteθ = false
│   isfiniter = false
│   isfiniteℓπ = false
│   isfiniteℓκ = false
└ @ AdvancedHMC /Users/kcft114/.julia/packages/AdvancedHMC/YWXfk/src/hamiltonian.jl:36
┌ Warning: Incorrect ϵ = NaN; ϵ_previous = 0.07500000000000001 is used instead.
└ @ AdvancedHMC.Adaptation /Users/kcft114/.julia/packages/AdvancedHMC/YWXfk/src/adaptation/stepsize.jl:103
┌ Warning: The current proposal will be rejected due to numerical error(s).
│   isfiniteθ = true
│   isfiniter = true
│   isfiniteℓπ = false
│   isfiniteℓκ = true
└ @ AdvancedHMC /Users/kcft114/.julia/packages/AdvancedHMC/YWXfk/src/hamiltonian.jl:36
┌ Warning: The current proposal will be rejected due to numerical error(s).
│   isfiniteθ = false
│   isfiniter = false
│   isfiniteℓπ = false
│   isfiniteℓκ = false
└ @ AdvancedHMC /Users/kcft114/.julia/packages/AdvancedHMC/YWXfk/src/hamiltonian.jl:36
┌ Warning: Incorrect ϵ = NaN; ϵ_previous = 0.07500000000000001 is used instead.
└ @ AdvancedHMC.Adaptation /Users/kcft114/.julia/packages/AdvancedHMC/YWXfk/src/adaptation/stepsize.jl:103
┌ Warning: The current proposal will be rejected due to numerical error(s).
│   isfiniteθ = true
│   isfiniter = true
│   isfiniteℓπ = false
│   isfiniteℓκ = true
└ @ AdvancedHMC /Users/kcft114/.julia/packages/AdvancedHMC/YWXfk/src/hamiltonian.jl:36
┌ Warning: The current proposal will be rejected due to numerical error(s).
│   isfiniteθ = false
│   isfiniter = false
│   isfiniteℓπ = false
│   isfiniteℓκ = false
└ @ AdvancedHMC /Users/kcft114/.julia/packages/AdvancedHMC/YWXfk/src/hamiltonian.jl:36
┌ Warning: Incorrect ϵ = NaN; ϵ_previous = 0.07500000000000001 is used instead.
└ @ AdvancedHMC.Adaptation /Users/kcft114/.julia/packages/AdvancedHMC/YWXfk/src/adaptation/stepsize.jl:103
┌ Warning: The current proposal will be rejected due to numerical error(s).
│   isfiniteθ = true
│   isfiniter = true
│   isfiniteℓπ = false
│   isfiniteℓκ = true
└ @ AdvancedHMC /Users/kcft114/.julia/packages/AdvancedHMC/YWXfk/src/hamiltonian.jl:36
┌ Warning: The current proposal will be rejected due to numerical error(s).
│   isfiniteθ = false
│   isfiniter = false
│   isfiniteℓπ = false
│   isfiniteℓκ = false
└ @ AdvancedHMC /Users/kcft114/.julia/packages/AdvancedHMC/YWXfk/src/hamiltonian.jl:36
┌ Warning: Incorrect ϵ = NaN; ϵ_previous = 0.07500000000000001 is used instead.
└ @ AdvancedHMC.Adaptation /Users/kcft114/.julia/packages/AdvancedHMC/YWXfk/src/adaptation/stepsize.jl:103
┌ Warning: The current proposal will be rejected due to numerical error(s).
│   isfiniteθ = true
│   isfiniter = true
│   isfiniteℓπ = false
│   isfiniteℓκ = true
└ @ AdvancedHMC /Users/kcft114/.julia/packages/AdvancedHMC/YWXfk/src/hamiltonian.jl:36
┌ Warning: The current proposal will be rejected due to numerical error(s).
│   isfiniteθ = false
│   isfiniter = false
│   isfiniteℓπ = false
│   isfiniteℓκ = false
└ @ AdvancedHMC /Users/kcft114/.julia/packages/AdvancedHMC/YWXfk/src/hamiltonian.jl:36
┌ Warning: Incorrect ϵ = NaN; ϵ_previous = 0.07500000000000001 is used instead.
└ @ AdvancedHMC.Adaptation /Users/kcft114/.julia/packages/AdvancedHMC/YWXfk/src/adaptation/stepsize.jl:103
┌ Warning: The current proposal will be rejected due to numerical error(s).
│   isfiniteθ = true
│   isfiniter = true
│   isfiniteℓπ = false
│   isfiniteℓκ = true
└ @ AdvancedHMC /Users/kcft114/.julia/packages/AdvancedHMC/YWXfk/src/hamiltonian.jl:36
┌ Warning: The current proposal will be rejected due to numerical error(s).
│   isfiniteθ = false
│   isfiniter = false
│   isfiniteℓπ = false
│   isfiniteℓκ = false
└ @ AdvancedHMC /Users/kcft114/.julia/packages/AdvancedHMC/YWXfk/src/hamiltonian.jl:36
┌ Warning: Incorrect ϵ = NaN; ϵ_previous = 0.07500000000000001 is used instead.
└ @ AdvancedHMC.Adaptation /Users/kcft114/.julia/packages/AdvancedHMC/YWXfk/src/adaptation/stepsize.jl:103
┌ Warning: The current proposal will be rejected due to numerical error(s).
│   isfiniteθ = true
│   isfiniter = true
│   isfiniteℓπ = false
│   isfiniteℓκ = true
└ @ AdvancedHMC /Users/kcft114/.julia/packages/AdvancedHMC/YWXfk/src/hamiltonian.jl:36
┌ Warning: The current proposal will be rejected due to numerical error(s).
│   isfiniteθ = false
│   isfiniter = false
│   isfiniteℓπ = false
│   isfiniteℓκ = false
└ @ AdvancedHMC /Users/kcft114/.julia/packages/AdvancedHMC/YWXfk/src/hamiltonian.jl:36
┌ Warning: Incorrect ϵ = NaN; ϵ_previous = 0.07500000000000001 is used instead.
└ @ AdvancedHMC.Adaptation /Users/kcft114/.julia/packages/AdvancedHMC/YWXfk/src/adaptation/stepsize.jl:103
┌ Warning: The current proposal will be rejected due to numerical error(s).
│   isfiniteθ = true
│   isfiniter = true
│   isfiniteℓπ = false
│   isfiniteℓκ = true
└ @ AdvancedHMC /Users/kcft114/.julia/packages/AdvancedHMC/YWXfk/src/hamiltonian.jl:36
┌ Warning: The current proposal will be rejected due to numerical error(s).
│   isfiniteθ = false
│   isfiniter = false
│   isfiniteℓπ = false
│   isfiniteℓκ = false
└ @ AdvancedHMC /Users/kcft114/.julia/packages/AdvancedHMC/YWXfk/src/hamiltonian.jl:36
┌ Warning: Incorrect ϵ = NaN; ϵ_previous = 0.07500000000000001 is used instead.
└ @ AdvancedHMC.Adaptation /Users/kcft114/.julia/packages/AdvancedHMC/YWXfk/src/adaptation/stepsize.jl:103
┌ Warning: The current proposal will be rejected due to numerical error(s).
│   isfiniteθ = true
│   isfiniter = true
│   isfiniteℓπ = false
│   isfiniteℓκ = true
└ @ AdvancedHMC /Users/kcft114/.julia/packages/AdvancedHMC/YWXfk/src/hamiltonian.jl:36
┌ Warning: The current proposal will be rejected due to numerical error(s).
│   isfiniteθ = false
│   isfiniter = false
│   isfiniteℓπ = false
│   isfiniteℓκ = false
└ @ AdvancedHMC /Users/kcft114/.julia/packages/AdvancedHMC/YWXfk/src/hamiltonian.jl:36
┌ Warning: Incorrect ϵ = NaN; ϵ_previous = 0.07500000000000001 is used instead.
└ @ AdvancedHMC.Adaptation /Users/kcft114/.julia/packages/AdvancedHMC/YWXfk/src/adaptation/stepsize.jl:103
┌ Warning: The current proposal will be rejected due to numerical error(s).
│   isfiniteθ = true
│   isfiniter = true
│   isfiniteℓπ = false
│   isfiniteℓκ = true
└ @ AdvancedHMC /Users/kcft114/.julia/packages/AdvancedHMC/YWXfk/src/hamiltonian.jl:36
┌ Warning: The current proposal will be rejected due to numerical error(s).
│   isfiniteθ = false
│   isfiniter = false
│   isfiniteℓπ = false
│   isfiniteℓκ = false
└ @ AdvancedHMC /Users/kcft114/.julia/packages/AdvancedHMC/YWXfk/src/hamiltonian.jl:36
┌ Warning: Incorrect ϵ = NaN; ϵ_previous = 0.07500000000000001 is used instead.
└ @ AdvancedHMC.Adaptation /Users/kcft114/.julia/packages/AdvancedHMC/YWXfk/src/adaptation/stepsize.jl:103
┌ Warning: The current proposal will be rejected due to numerical error(s).
│   isfiniteθ = true
│   isfiniter = true
│   isfiniteℓπ = false
│   isfiniteℓκ = true
└ @ AdvancedHMC /Users/kcft114/.julia/packages/AdvancedHMC/YWXfk/src/hamiltonian.jl:36
┌ Warning: The current proposal will be rejected due to numerical error(s).
│   isfiniteθ = false
│   isfiniter = false
│   isfiniteℓπ = false
│   isfiniteℓκ = false
└ @ AdvancedHMC /Users/kcft114/.julia/packages/AdvancedHMC/YWXfk/src/hamiltonian.jl:36
┌ Warning: Incorrect ϵ = NaN; ϵ_previous = 0.07500000000000001 is used instead.
└ @ AdvancedHMC.Adaptation /Users/kcft114/.julia/packages/AdvancedHMC/YWXfk/src/adaptation/stepsize.jl:103
┌ Warning: The current proposal will be rejected due to numerical error(s).
│   isfiniteθ = true
│   isfiniter = true
│   isfiniteℓπ = false
│   isfiniteℓκ = true
└ @ AdvancedHMC /Users/kcft114/.julia/packages/AdvancedHMC/YWXfk/src/hamiltonian.jl:36
┌ Warning: The current proposal will be rejected due to numerical error(s).
│   isfiniteθ = false
│   isfiniter = false
│   isfiniteℓπ = false
│   isfiniteℓκ = false
└ @ AdvancedHMC /Users/kcft114/.julia/packages/AdvancedHMC/YWXfk/src/hamiltonian.jl:36
┌ Warning: Incorrect ϵ = NaN; ϵ_previous = 0.07500000000000001 is used instead.
└ @ AdvancedHMC.Adaptation /Users/kcft114/.julia/packages/AdvancedHMC/YWXfk/src/adaptation/stepsize.jl:103
┌ Warning: The current proposal will be rejected due to numerical error(s).
│   isfiniteθ = true
│   isfiniter = true
│   isfiniteℓπ = false
│   isfiniteℓκ = true
└ @ AdvancedHMC /Users/kcft114/.julia/packages/AdvancedHMC/YWXfk/src/hamiltonian.jl:36
┌ Warning: The current proposal will be rejected due to numerical error(s).
│   isfiniteθ = false
│   isfiniter = false
│   isfiniteℓπ = false
│   isfiniteℓκ = false
└ @ AdvancedHMC /Users/kcft114/.julia/packages/AdvancedHMC/YWXfk/src/hamiltonian.jl:36
┌ Warning: Incorrect ϵ = NaN; ϵ_previous = 0.07500000000000001 is used instead.
└ @ AdvancedHMC.Adaptation /Users/kcft114/.julia/packages/AdvancedHMC/YWXfk/src/adaptation/stepsize.jl:103
┌ Warning: The current proposal will be rejected due to numerical error(s).
│   isfiniteθ = true
│   isfiniter = true
│   isfiniteℓπ = false
│   isfiniteℓκ = true
└ @ AdvancedHMC /Users/kcft114/.julia/packages/AdvancedHMC/YWXfk/src/hamiltonian.jl:36
┌ Warning: The current proposal will be rejected due to numerical error(s).
│   isfiniteθ = false
│   isfiniter = false
│   isfiniteℓπ = false
│   isfiniteℓκ = false
└ @ AdvancedHMC /Users/kcft114/.julia/packages/AdvancedHMC/YWXfk/src/hamiltonian.jl:36
┌ Warning: Incorrect ϵ = NaN; ϵ_previous = 0.07500000000000001 is used instead.
└ @ AdvancedHMC.Adaptation /Users/kcft114/.julia/packages/AdvancedHMC/YWXfk/src/adaptation/stepsize.jl:103
┌ Warning: The current proposal will be rejected due to numerical error(s).
│   isfiniteθ = true
│   isfiniter = true
│   isfiniteℓπ = false
│   isfiniteℓκ = true
└ @ AdvancedHMC /Users/kcft114/.julia/packages/AdvancedHMC/YWXfk/src/hamiltonian.jl:36
┌ Warning: The current proposal will be rejected due to numerical error(s).
│   isfiniteθ = false
│   isfiniter = false
│   isfiniteℓπ = false
│   isfiniteℓκ = false
└ @ AdvancedHMC /Users/kcft114/.julia/packages/AdvancedHMC/YWXfk/src/hamiltonian.jl:36
┌ Warning: Incorrect ϵ = NaN; ϵ_previous = 0.07500000000000001 is used instead.
└ @ AdvancedHMC.Adaptation /Users/kcft114/.julia/packages/AdvancedHMC/YWXfk/src/adaptation/stepsize.jl:103
┌ Warning: The current proposal will be rejected due to numerical error(s).
│   isfiniteθ = true
│   isfiniter = true
│   isfiniteℓπ = false
│   isfiniteℓκ = true
└ @ AdvancedHMC /Users/kcft114/.julia/packages/AdvancedHMC/YWXfk/src/hamiltonian.jl:36
┌ Warning: The current proposal will be rejected due to numerical error(s).
│   isfiniteθ = false
│   isfiniter = false
│   isfiniteℓπ = false
│   isfiniteℓκ = false
└ @ AdvancedHMC /Users/kcft114/.julia/packages/AdvancedHMC/YWXfk/src/hamiltonian.jl:36
┌ Warning: Incorrect ϵ = NaN; ϵ_previous = 0.07500000000000001 is used instead.
└ @ AdvancedHMC.Adaptation /Users/kcft114/.julia/packages/AdvancedHMC/YWXfk/src/adaptation/stepsize.jl:103
┌ Warning: The current proposal will be rejected due to numerical error(s).
│   isfiniteθ = true
│   isfiniter = true
│   isfiniteℓπ = false
│   isfiniteℓκ = true
└ @ AdvancedHMC /Users/kcft114/.julia/packages/AdvancedHMC/YWXfk/src/hamiltonian.jl:36
┌ Warning: The current proposal will be rejected due to numerical error(s).
│   isfiniteθ = false
│   isfiniter = false
│   isfiniteℓπ = false
│   isfiniteℓκ = false
└ @ AdvancedHMC /Users/kcft114/.julia/packages/AdvancedHMC/YWXfk/src/hamiltonian.jl:36
┌ Warning: Incorrect ϵ = NaN; ϵ_previous = 0.07500000000000001 is used instead.
└ @ AdvancedHMC.Adaptation /Users/kcft114/.julia/packages/AdvancedHMC/YWXfk/src/adaptation/stepsize.jl:103
┌ Warning: The current proposal will be rejected due to numerical error(s).
│   isfiniteθ = true
│   isfiniter = true
│   isfiniteℓπ = false
│   isfiniteℓκ = true
└ @ AdvancedHMC /Users/kcft114/.julia/packages/AdvancedHMC/YWXfk/src/hamiltonian.jl:36
┌ Warning: The current proposal will be rejected due to numerical error(s).
│   isfiniteθ = false
│   isfiniter = false
│   isfiniteℓπ = false
│   isfiniteℓκ = false
└ @ AdvancedHMC /Users/kcft114/.julia/packages/AdvancedHMC/YWXfk/src/hamiltonian.jl:36
┌ Warning: Incorrect ϵ = NaN; ϵ_previous = 0.07500000000000001 is used instead.
└ @ AdvancedHMC.Adaptation /Users/kcft114/.julia/packages/AdvancedHMC/YWXfk/src/adaptation/stepsize.jl:103
┌ Warning: The current proposal will be rejected due to numerical error(s).
│   isfiniteθ = true
│   isfiniter = true
│   isfiniteℓπ = false
│   isfiniteℓκ = true
└ @ AdvancedHMC /Users/kcft114/.julia/packages/AdvancedHMC/YWXfk/src/hamiltonian.jl:36
┌ Warning: The current proposal will be rejected due to numerical error(s).
│   isfiniteθ = false
│   isfiniter = false
│   isfiniteℓπ = false
│   isfiniteℓκ = false
└ @ AdvancedHMC /Users/kcft114/.julia/packages/AdvancedHMC/YWXfk/src/hamiltonian.jl:36
┌ Warning: Incorrect ϵ = NaN; ϵ_previous = 0.07500000000000001 is used instead.
└ @ AdvancedHMC.Adaptation /Users/kcft114/.julia/packages/AdvancedHMC/YWXfk/src/adaptation/stepsize.jl:103
┌ Warning: The current proposal will be rejected due to numerical error(s).
│   isfiniteθ = true
│   isfiniter = true
│   isfiniteℓπ = false
│   isfiniteℓκ = true
└ @ AdvancedHMC /Users/kcft114/.julia/packages/AdvancedHMC/YWXfk/src/hamiltonian.jl:36
┌ Warning: The current proposal will be rejected due to numerical error(s).
│   isfiniteθ = false
│   isfiniter = false
│   isfiniteℓπ = false
│   isfiniteℓκ = false
└ @ AdvancedHMC /Users/kcft114/.julia/packages/AdvancedHMC/YWXfk/src/hamiltonian.jl:36
┌ Warning: Incorrect ϵ = NaN; ϵ_previous = 0.07500000000000001 is used instead.
└ @ AdvancedHMC.Adaptation /Users/kcft114/.julia/packages/AdvancedHMC/YWXfk/src/adaptation/stepsize.jl:103
┌ Warning: The current proposal will be rejected due to numerical error(s).
│   isfiniteθ = true
│   isfiniter = true
│   isfiniteℓπ = false
│   isfiniteℓκ = true
└ @ AdvancedHMC /Users/kcft114/.julia/packages/AdvancedHMC/YWXfk/src/hamiltonian.jl:36
┌ Warning: The current proposal will be rejected due to numerical error(s).
│   isfiniteθ = false
│   isfiniter = false
│   isfiniteℓπ = false
│   isfiniteℓκ = false
└ @ AdvancedHMC /Users/kcft114/.julia/packages/AdvancedHMC/YWXfk/src/hamiltonian.jl:36
┌ Warning: Incorrect ϵ = NaN; ϵ_previous = 0.07500000000000001 is used instead.
└ @ AdvancedHMC.Adaptation /Users/kcft114/.julia/packages/AdvancedHMC/YWXfk/src/adaptation/stepsize.jl:103
┌ Warning: The current proposal will be rejected due to numerical error(s).
│   isfiniteθ = true
│   isfiniter = true
│   isfiniteℓπ = false
│   isfiniteℓκ = true
└ @ AdvancedHMC /Users/kcft114/.julia/packages/AdvancedHMC/YWXfk/src/hamiltonian.jl:36
┌ Warning: The current proposal will be rejected due to numerical error(s).
│   isfiniteθ = false
│   isfiniter = false
│   isfiniteℓπ = false
│   isfiniteℓκ = false
└ @ AdvancedHMC /Users/kcft114/.julia/packages/AdvancedHMC/YWXfk/src/hamiltonian.jl:36
┌ Warning: Incorrect ϵ = NaN; ϵ_previous = 0.07500000000000001 is used instead.
└ @ AdvancedHMC.Adaptation /Users/kcft114/.julia/packages/AdvancedHMC/YWXfk/src/adaptation/stepsize.jl:103
┌ Warning: The current proposal will be rejected due to numerical error(s).
│   isfiniteθ = true
│   isfiniter = true
│   isfiniteℓπ = false
│   isfiniteℓκ = true
└ @ AdvancedHMC /Users/kcft114/.julia/packages/AdvancedHMC/YWXfk/src/hamiltonian.jl:36
┌ Warning: The current proposal will be rejected due to numerical error(s).
│   isfiniteθ = false
│   isfiniter = false
│   isfiniteℓπ = false
│   isfiniteℓκ = false
└ @ AdvancedHMC /Users/kcft114/.julia/packages/AdvancedHMC/YWXfk/src/hamiltonian.jl:36
┌ Warning: Incorrect ϵ = NaN; ϵ_previous = 0.07500000000000001 is used instead.
└ @ AdvancedHMC.Adaptation /Users/kcft114/.julia/packages/AdvancedHMC/YWXfk/src/adaptation/stepsize.jl:103
┌ Warning: The current proposal will be rejected due to numerical error(s).
│   isfiniteθ = true
│   isfiniter = true
│   isfiniteℓπ = false
│   isfiniteℓκ = true
└ @ AdvancedHMC /Users/kcft114/.julia/packages/AdvancedHMC/YWXfk/src/hamiltonian.jl:36
┌ Warning: The current proposal will be rejected due to numerical error(s).
│   isfiniteθ = false
│   isfiniter = false
│   isfiniteℓπ = false
│   isfiniteℓκ = false
└ @ AdvancedHMC /Users/kcft114/.julia/packages/AdvancedHMC/YWXfk/src/hamiltonian.jl:36
┌ Warning: Incorrect ϵ = NaN; ϵ_previous = 0.07500000000000001 is used instead.
└ @ AdvancedHMC.Adaptation /Users/kcft114/.julia/packages/AdvancedHMC/YWXfk/src/adaptation/stepsize.jl:103
┌ Warning: The current proposal will be rejected due to numerical error(s).
│   isfiniteθ = true
│   isfiniter = true
│   isfiniteℓπ = false
│   isfiniteℓκ = true
└ @ AdvancedHMC /Users/kcft114/.julia/packages/AdvancedHMC/YWXfk/src/hamiltonian.jl:36
┌ Warning: The current proposal will be rejected due to numerical error(s).
│   isfiniteθ = false
│   isfiniter = false
│   isfiniteℓπ = false
│   isfiniteℓκ = false
└ @ AdvancedHMC /Users/kcft114/.julia/packages/AdvancedHMC/YWXfk/src/hamiltonian.jl:36
┌ Warning: Incorrect ϵ = NaN; ϵ_previous = 0.07500000000000001 is used instead.
└ @ AdvancedHMC.Adaptation /Users/kcft114/.julia/packages/AdvancedHMC/YWXfk/src/adaptation/stepsize.jl:103
┌ Warning: The current proposal will be rejected due to numerical error(s).
│   isfiniteθ = true
│   isfiniter = true
│   isfiniteℓπ = false
│   isfiniteℓκ = true
└ @ AdvancedHMC /Users/kcft114/.julia/packages/AdvancedHMC/YWXfk/src/hamiltonian.jl:36
┌ Warning: The current proposal will be rejected due to numerical error(s).
│   isfiniteθ = false
│   isfiniter = false
│   isfiniteℓπ = false
│   isfiniteℓκ = false
└ @ AdvancedHMC /Users/kcft114/.julia/packages/AdvancedHMC/YWXfk/src/hamiltonian.jl:36
┌ Warning: Incorrect ϵ = NaN; ϵ_previous = 0.07500000000000001 is used instead.
└ @ AdvancedHMC.Adaptation /Users/kcft114/.julia/packages/AdvancedHMC/YWXfk/src/adaptation/stepsize.jl:103
┌ Warning: The current proposal will be rejected due to numerical error(s).
│   isfiniteθ = true
│   isfiniter = true
│   isfiniteℓπ = false
│   isfiniteℓκ = true
└ @ AdvancedHMC /Users/kcft114/.julia/packages/AdvancedHMC/YWXfk/src/hamiltonian.jl:36
┌ Warning: The current proposal will be rejected due to numerical error(s).
│   isfiniteθ = false
│   isfiniter = false
│   isfiniteℓπ = false
│   isfiniteℓκ = false
└ @ AdvancedHMC /Users/kcft114/.julia/packages/AdvancedHMC/YWXfk/src/hamiltonian.jl:36
┌ Warning: Incorrect ϵ = NaN; ϵ_previous = 0.07500000000000001 is used instead.
└ @ AdvancedHMC.Adaptation /Users/kcft114/.julia/packages/AdvancedHMC/YWXfk/src/adaptation/stepsize.jl:103
┌ Warning: The current proposal will be rejected due to numerical error(s).
│   isfiniteθ = true
│   isfiniter = true
│   isfiniteℓπ = false
│   isfiniteℓκ = true
└ @ AdvancedHMC /Users/kcft114/.julia/packages/AdvancedHMC/YWXfk/src/hamiltonian.jl:36
┌ Warning: The current proposal will be rejected due to numerical error(s).
│   isfiniteθ = false
│   isfiniter = false
│   isfiniteℓπ = false
│   isfiniteℓκ = false
└ @ AdvancedHMC /Users/kcft114/.julia/packages/AdvancedHMC/YWXfk/src/hamiltonian.jl:36
┌ Warning: Incorrect ϵ = NaN; ϵ_previous = 0.07500000000000001 is used instead.
└ @ AdvancedHMC.Adaptation /Users/kcft114/.julia/packages/AdvancedHMC/YWXfk/src/adaptation/stepsize.jl:103
┌ Warning: The current proposal will be rejected due to numerical error(s).
│   isfiniteθ = true
│   isfiniter = true
│   isfiniteℓπ = false
│   isfiniteℓκ = true
└ @ AdvancedHMC /Users/kcft114/.julia/packages/AdvancedHMC/YWXfk/src/hamiltonian.jl:36
┌ Warning: The current proposal will be rejected due to numerical error(s).
│   isfiniteθ = false
│   isfiniter = false
│   isfiniteℓπ = false
│   isfiniteℓκ = false
└ @ AdvancedHMC /Users/kcft114/.julia/packages/AdvancedHMC/YWXfk/src/hamiltonian.jl:36
┌ Warning: Incorrect ϵ = NaN; ϵ_previous = 0.07500000000000001 is used instead.
└ @ AdvancedHMC.Adaptation /Users/kcft114/.julia/packages/AdvancedHMC/YWXfk/src/adaptation/stepsize.jl:103
┌ Warning: The current proposal will be rejected due to numerical error(s).
│   isfiniteθ = true
│   isfiniter = true
│   isfiniteℓπ = false
│   isfiniteℓκ = true
└ @ AdvancedHMC /Users/kcft114/.julia/packages/AdvancedHMC/YWXfk/src/hamiltonian.jl:36
┌ Warning: The current proposal will be rejected due to numerical error(s).
│   isfiniteθ = false
│   isfiniter = false
│   isfiniteℓπ = false
│   isfiniteℓκ = false
└ @ AdvancedHMC /Users/kcft114/.julia/packages/AdvancedHMC/YWXfk/src/hamiltonian.jl:36
┌ Warning: Incorrect ϵ = NaN; ϵ_previous = 0.07500000000000001 is used instead.
└ @ AdvancedHMC.Adaptation /Users/kcft114/.julia/packages/AdvancedHMC/YWXfk/src/adaptation/stepsize.jl:103
┌ Warning: The current proposal will be rejected due to numerical error(s).
│   isfiniteθ = true
│   isfiniter = true
│   isfiniteℓπ = false
│   isfiniteℓκ = true
└ @ AdvancedHMC /Users/kcft114/.julia/packages/AdvancedHMC/YWXfk/src/hamiltonian.jl:36
┌ Warning: The current proposal will be rejected due to numerical error(s).
│   isfiniteθ = false
│   isfiniter = false
│   isfiniteℓπ = false
│   isfiniteℓκ = false
└ @ AdvancedHMC /Users/kcft114/.julia/packages/AdvancedHMC/YWXfk/src/hamiltonian.jl:36
┌ Warning: Incorrect ϵ = NaN; ϵ_previous = 0.07500000000000001 is used instead.
└ @ AdvancedHMC.Adaptation /Users/kcft114/.julia/packages/AdvancedHMC/YWXfk/src/adaptation/stepsize.jl:103
┌ Warning: The current proposal will be rejected due to numerical error(s).
│   isfiniteθ = true
│   isfiniter = true
│   isfiniteℓπ = false
│   isfiniteℓκ = true
└ @ AdvancedHMC /Users/kcft114/.julia/packages/AdvancedHMC/YWXfk/src/hamiltonian.jl:36
┌ Warning: The current proposal will be rejected due to numerical error(s).
│   isfiniteθ = false
│   isfiniter = false
│   isfiniteℓπ = false
│   isfiniteℓκ = false
└ @ AdvancedHMC /Users/kcft114/.julia/packages/AdvancedHMC/YWXfk/src/hamiltonian.jl:36
┌ Warning: Incorrect ϵ = NaN; ϵ_previous = 0.07500000000000001 is used instead.
└ @ AdvancedHMC.Adaptation /Users/kcft114/.julia/packages/AdvancedHMC/YWXfk/src/adaptation/stepsize.jl:103
┌ Warning: The current proposal will be rejected due to numerical error(s).
│   isfiniteθ = true
│   isfiniter = true
│   isfiniteℓπ = false
│   isfiniteℓκ = true
└ @ AdvancedHMC /Users/kcft114/.julia/packages/AdvancedHMC/YWXfk/src/hamiltonian.jl:36
┌ Warning: The current proposal will be rejected due to numerical error(s).
│   isfiniteθ = false
│   isfiniter = false
│   isfiniteℓπ = false
│   isfiniteℓκ = false
└ @ AdvancedHMC /Users/kcft114/.julia/packages/AdvancedHMC/YWXfk/src/hamiltonian.jl:36
┌ Warning: Incorrect ϵ = NaN; ϵ_previous = 0.07500000000000001 is used instead.
└ @ AdvancedHMC.Adaptation /Users/kcft114/.julia/packages/AdvancedHMC/YWXfk/src/adaptation/stepsize.jl:103
┌ Warning: The current proposal will be rejected due to numerical error(s).
│   isfiniteθ = true
│   isfiniter = true
│   isfiniteℓπ = false
│   isfiniteℓκ = true
└ @ AdvancedHMC /Users/kcft114/.julia/packages/AdvancedHMC/YWXfk/src/hamiltonian.jl:36
┌ Warning: The current proposal will be rejected due to numerical error(s).
│   isfiniteθ = false
│   isfiniter = false
│   isfiniteℓπ = false
│   isfiniteℓκ = false
└ @ AdvancedHMC /Users/kcft114/.julia/packages/AdvancedHMC/YWXfk/src/hamiltonian.jl:36
┌ Warning: Incorrect ϵ = NaN; ϵ_previous = 0.07500000000000001 is used instead.
└ @ AdvancedHMC.Adaptation /Users/kcft114/.julia/packages/AdvancedHMC/YWXfk/src/adaptation/stepsize.jl:103
┌ Warning: The current proposal will be rejected due to numerical error(s).
│   isfiniteθ = true
│   isfiniter = true
│   isfiniteℓπ = false
│   isfiniteℓκ = true
└ @ AdvancedHMC /Users/kcft114/.julia/packages/AdvancedHMC/YWXfk/src/hamiltonian.jl:36
┌ Warning: The current proposal will be rejected due to numerical error(s).
│   isfiniteθ = false
│   isfiniter = false
│   isfiniteℓπ = false
│   isfiniteℓκ = false
└ @ AdvancedHMC /Users/kcft114/.julia/packages/AdvancedHMC/YWXfk/src/hamiltonian.jl:36
┌ Warning: Incorrect ϵ = NaN; ϵ_previous = 0.07500000000000001 is used instead.
└ @ AdvancedHMC.Adaptation /Users/kcft114/.julia/packages/AdvancedHMC/YWXfk/src/adaptation/stepsize.jl:103
┌ Warning: The current proposal will be rejected due to numerical error(s).
│   isfiniteθ = true
│   isfiniter = true
│   isfiniteℓπ = false
│   isfiniteℓκ = true
└ @ AdvancedHMC /Users/kcft114/.julia/packages/AdvancedHMC/YWXfk/src/hamiltonian.jl:36
┌ Warning: The current proposal will be rejected due to numerical error(s).
│   isfiniteθ = false
│   isfiniter = false
│   isfiniteℓπ = false
│   isfiniteℓκ = false
└ @ AdvancedHMC /Users/kcft114/.julia/packages/AdvancedHMC/YWXfk/src/hamiltonian.jl:36
┌ Warning: Incorrect ϵ = NaN; ϵ_previous = 0.07500000000000001 is used instead.
└ @ AdvancedHMC.Adaptation /Users/kcft114/.julia/packages/AdvancedHMC/YWXfk/src/adaptation/stepsize.jl:103
┌ Warning: The current proposal will be rejected due to numerical error(s).
│   isfiniteθ = true
│   isfiniter = true
│   isfiniteℓπ = false
│   isfiniteℓκ = true
└ @ AdvancedHMC /Users/kcft114/.julia/packages/AdvancedHMC/YWXfk/src/hamiltonian.jl:36
┌ Warning: The current proposal will be rejected due to numerical error(s).
│   isfiniteθ = false
│   isfiniter = false
│   isfiniteℓπ = false
│   isfiniteℓκ = false
└ @ AdvancedHMC /Users/kcft114/.julia/packages/AdvancedHMC/YWXfk/src/hamiltonian.jl:36
┌ Warning: Incorrect ϵ = NaN; ϵ_previous = 0.07500000000000001 is used instead.
└ @ AdvancedHMC.Adaptation /Users/kcft114/.julia/packages/AdvancedHMC/YWXfk/src/adaptation/stepsize.jl:103
┌ Warning: The current proposal will be rejected due to numerical error(s).
│   isfiniteθ = true
│   isfiniter = true
│   isfiniteℓπ = false
│   isfiniteℓκ = true
└ @ AdvancedHMC /Users/kcft114/.julia/packages/AdvancedHMC/YWXfk/src/hamiltonian.jl:36
┌ Warning: The current proposal will be rejected due to numerical error(s).
│   isfiniteθ = false
│   isfiniter = false
│   isfiniteℓπ = false
│   isfiniteℓκ = false
└ @ AdvancedHMC /Users/kcft114/.julia/packages/AdvancedHMC/YWXfk/src/hamiltonian.jl:36
┌ Warning: Incorrect ϵ = NaN; ϵ_previous = 0.07500000000000001 is used instead.
└ @ AdvancedHMC.Adaptation /Users/kcft114/.julia/packages/AdvancedHMC/YWXfk/src/adaptation/stepsize.jl:103
┌ Warning: The current proposal will be rejected due to numerical error(s).
│   isfiniteθ = true
│   isfiniter = true
│   isfiniteℓπ = false
│   isfiniteℓκ = true
└ @ AdvancedHMC /Users/kcft114/.julia/packages/AdvancedHMC/YWXfk/src/hamiltonian.jl:36
┌ Warning: The current proposal will be rejected due to numerical error(s).
│   isfiniteθ = false
│   isfiniter = false
│   isfiniteℓπ = false
│   isfiniteℓκ = false
└ @ AdvancedHMC /Users/kcft114/.julia/packages/AdvancedHMC/YWXfk/src/hamiltonian.jl:36
┌ Warning: Incorrect ϵ = NaN; ϵ_previous = 0.07500000000000001 is used instead.
└ @ AdvancedHMC.Adaptation /Users/kcft114/.julia/packages/AdvancedHMC/YWXfk/src/adaptation/stepsize.jl:103
┌ Warning: The current proposal will be rejected due to numerical error(s).
│   isfiniteθ = true
│   isfiniter = true
│   isfiniteℓπ = false
│   isfiniteℓκ = true
└ @ AdvancedHMC /Users/kcft114/.julia/packages/AdvancedHMC/YWXfk/src/hamiltonian.jl:36
┌ Warning: The current proposal will be rejected due to numerical error(s).
│   isfiniteθ = false
│   isfiniter = false
│   isfiniteℓπ = false
│   isfiniteℓκ = false
└ @ AdvancedHMC /Users/kcft114/.julia/packages/AdvancedHMC/YWXfk/src/hamiltonian.jl:36
┌ Warning: Incorrect ϵ = NaN; ϵ_previous = 0.07500000000000001 is used instead.
└ @ AdvancedHMC.Adaptation /Users/kcft114/.julia/packages/AdvancedHMC/YWXfk/src/adaptation/stepsize.jl:103
┌ Warning: The current proposal will be rejected due to numerical error(s).
│   isfiniteθ = true
│   isfiniter = true
│   isfiniteℓπ = false
│   isfiniteℓκ = true
└ @ AdvancedHMC /Users/kcft114/.julia/packages/AdvancedHMC/YWXfk/src/hamiltonian.jl:36
┌ Warning: The current proposal will be rejected due to numerical error(s).
│   isfiniteθ = false
│   isfiniter = false
│   isfiniteℓπ = false
│   isfiniteℓκ = false
└ @ AdvancedHMC /Users/kcft114/.julia/packages/AdvancedHMC/YWXfk/src/hamiltonian.jl:36
┌ Warning: Incorrect ϵ = NaN; ϵ_previous = 0.07500000000000001 is used instead.
└ @ AdvancedHMC.Adaptation /Users/kcft114/.julia/packages/AdvancedHMC/YWXfk/src/adaptation/stepsize.jl:103
┌ Warning: The current proposal will be rejected due to numerical error(s).
│   isfiniteθ = true
│   isfiniter = true
│   isfiniteℓπ = false
│   isfiniteℓκ = true
└ @ AdvancedHMC /Users/kcft114/.julia/packages/AdvancedHMC/YWXfk/src/hamiltonian.jl:36
┌ Warning: The current proposal will be rejected due to numerical error(s).
│   isfiniteθ = false
│   isfiniter = false
│   isfiniteℓπ = false
│   isfiniteℓκ = false
└ @ AdvancedHMC /Users/kcft114/.julia/packages/AdvancedHMC/YWXfk/src/hamiltonian.jl:36
┌ Warning: Incorrect ϵ = NaN; ϵ_previous = 0.07500000000000001 is used instead.
└ @ AdvancedHMC.Adaptation /Users/kcft114/.julia/packages/AdvancedHMC/YWXfk/src/adaptation/stepsize.jl:103
┌ Warning: The current proposal will be rejected due to numerical error(s).
│   isfiniteθ = true
│   isfiniter = true
│   isfiniteℓπ = false
│   isfiniteℓκ = true
└ @ AdvancedHMC /Users/kcft114/.julia/packages/AdvancedHMC/YWXfk/src/hamiltonian.jl:36
┌ Warning: The current proposal will be rejected due to numerical error(s).
│   isfiniteθ = false
│   isfiniter = false
│   isfiniteℓπ = false
│   isfiniteℓκ = false
└ @ AdvancedHMC /Users/kcft114/.julia/packages/AdvancedHMC/YWXfk/src/hamiltonian.jl:36
┌ Warning: Incorrect ϵ = NaN; ϵ_previous = 0.07500000000000001 is used instead.
└ @ AdvancedHMC.Adaptation /Users/kcft114/.julia/packages/AdvancedHMC/YWXfk/src/adaptation/stepsize.jl:103
┌ Warning: The current proposal will be rejected due to numerical error(s).
│   isfiniteθ = true
│   isfiniter = true
│   isfiniteℓπ = false
│   isfiniteℓκ = true
└ @ AdvancedHMC /Users/kcft114/.julia/packages/AdvancedHMC/YWXfk/src/hamiltonian.jl:36
┌ Warning: The current proposal will be rejected due to numerical error(s).
│   isfiniteθ = false
│   isfiniter = false
│   isfiniteℓπ = false
│   isfiniteℓκ = false
└ @ AdvancedHMC /Users/kcft114/.julia/packages/AdvancedHMC/YWXfk/src/hamiltonian.jl:36
┌ Warning: Incorrect ϵ = NaN; ϵ_previous = 0.07500000000000001 is used instead.
└ @ AdvancedHMC.Adaptation /Users/kcft114/.julia/packages/AdvancedHMC/YWXfk/src/adaptation/stepsize.jl:103
┌ Warning: The current proposal will be rejected due to numerical error(s).
│   isfiniteθ = true
│   isfiniter = true
│   isfiniteℓπ = false
│   isfiniteℓκ = true
└ @ AdvancedHMC /Users/kcft114/.julia/packages/AdvancedHMC/YWXfk/src/hamiltonian.jl:36
┌ Warning: The current proposal will be rejected due to numerical error(s).
│   isfiniteθ = false
│   isfiniter = false
│   isfiniteℓπ = false
│   isfiniteℓκ = false
└ @ AdvancedHMC /Users/kcft114/.julia/packages/AdvancedHMC/YWXfk/src/hamiltonian.jl:36
┌ Warning: Incorrect ϵ = NaN; ϵ_previous = 0.07500000000000001 is used instead.
└ @ AdvancedHMC.Adaptation /Users/kcft114/.julia/packages/AdvancedHMC/YWXfk/src/adaptation/stepsize.jl:103
┌ Warning: The current proposal will be rejected due to numerical error(s).
│   isfiniteθ = true
│   isfiniter = true
│   isfiniteℓπ = false
│   isfiniteℓκ = true
└ @ AdvancedHMC /Users/kcft114/.julia/packages/AdvancedHMC/YWXfk/src/hamiltonian.jl:36
┌ Warning: The current proposal will be rejected due to numerical error(s).
│   isfiniteθ = false
│   isfiniter = false
│   isfiniteℓπ = false
│   isfiniteℓκ = false
└ @ AdvancedHMC /Users/kcft114/.julia/packages/AdvancedHMC/YWXfk/src/hamiltonian.jl:36
┌ Warning: Incorrect ϵ = NaN; ϵ_previous = 0.07500000000000001 is used instead.
└ @ AdvancedHMC.Adaptation /Users/kcft114/.julia/packages/AdvancedHMC/YWXfk/src/adaptation/stepsize.jl:103
┌ Warning: The current proposal will be rejected due to numerical error(s).
│   isfiniteθ = true
│   isfiniter = true
│   isfiniteℓπ = false
│   isfiniteℓκ = true
└ @ AdvancedHMC /Users/kcft114/.julia/packages/AdvancedHMC/YWXfk/src/hamiltonian.jl:36
┌ Warning: The current proposal will be rejected due to numerical error(s).
│   isfiniteθ = false
│   isfiniter = false
│   isfiniteℓπ = false
│   isfiniteℓκ = false
└ @ AdvancedHMC /Users/kcft114/.julia/packages/AdvancedHMC/YWXfk/src/hamiltonian.jl:36
┌ Warning: Incorrect ϵ = NaN; ϵ_previous = 0.07500000000000001 is used instead.
└ @ AdvancedHMC.Adaptation /Users/kcft114/.julia/packages/AdvancedHMC/YWXfk/src/adaptation/stepsize.jl:103
┌ Warning: The current proposal will be rejected due to numerical error(s).
│   isfiniteθ = true
│   isfiniter = true
│   isfiniteℓπ = false
│   isfiniteℓκ = true
└ @ AdvancedHMC /Users/kcft114/.julia/packages/AdvancedHMC/YWXfk/src/hamiltonian.jl:36
┌ Warning: The current proposal will be rejected due to numerical error(s).
│   isfiniteθ = false
│   isfiniter = false
│   isfiniteℓπ = false
│   isfiniteℓκ = false
└ @ AdvancedHMC /Users/kcft114/.julia/packages/AdvancedHMC/YWXfk/src/hamiltonian.jl:36
┌ Warning: Incorrect ϵ = NaN; ϵ_previous = 0.07500000000000001 is used instead.
└ @ AdvancedHMC.Adaptation /Users/kcft114/.julia/packages/AdvancedHMC/YWXfk/src/adaptation/stepsize.jl:103
┌ Warning: The current proposal will be rejected due to numerical error(s).
│   isfiniteθ = true
│   isfiniter = true
│   isfiniteℓπ = false
│   isfiniteℓκ = true
└ @ AdvancedHMC /Users/kcft114/.julia/packages/AdvancedHMC/YWXfk/src/hamiltonian.jl:36
┌ Warning: The current proposal will be rejected due to numerical error(s).
│   isfiniteθ = false
│   isfiniter = false
│   isfiniteℓπ = false
│   isfiniteℓκ = false
└ @ AdvancedHMC /Users/kcft114/.julia/packages/AdvancedHMC/YWXfk/src/hamiltonian.jl:36
┌ Warning: Incorrect ϵ = NaN; ϵ_previous = 0.07500000000000001 is used instead.
└ @ AdvancedHMC.Adaptation /Users/kcft114/.julia/packages/AdvancedHMC/YWXfk/src/adaptation/stepsize.jl:103
┌ Warning: The current proposal will be rejected due to numerical error(s).
│   isfiniteθ = true
│   isfiniter = true
│   isfiniteℓπ = false
│   isfiniteℓκ = true
└ @ AdvancedHMC /Users/kcft114/.julia/packages/AdvancedHMC/YWXfk/src/hamiltonian.jl:36
┌ Warning: The current proposal will be rejected due to numerical error(s).
│   isfiniteθ = false
│   isfiniter = false
│   isfiniteℓπ = false
│   isfiniteℓκ = false
└ @ AdvancedHMC /Users/kcft114/.julia/packages/AdvancedHMC/YWXfk/src/hamiltonian.jl:36
┌ Warning: Incorrect ϵ = NaN; ϵ_previous = 0.07500000000000001 is used instead.
└ @ AdvancedHMC.Adaptation /Users/kcft114/.julia/packages/AdvancedHMC/YWXfk/src/adaptation/stepsize.jl:103
┌ Warning: The current proposal will be rejected due to numerical error(s).
│   isfiniteθ = true
│   isfiniter = true
│   isfiniteℓπ = false
│   isfiniteℓκ = true
└ @ AdvancedHMC /Users/kcft114/.julia/packages/AdvancedHMC/YWXfk/src/hamiltonian.jl:36
┌ Warning: The current proposal will be rejected due to numerical error(s).
│   isfiniteθ = false
│   isfiniter = false
│   isfiniteℓπ = false
│   isfiniteℓκ = false
└ @ AdvancedHMC /Users/kcft114/.julia/packages/AdvancedHMC/YWXfk/src/hamiltonian.jl:36
┌ Warning: Incorrect ϵ = NaN; ϵ_previous = 0.07500000000000001 is used instead.
└ @ AdvancedHMC.Adaptation /Users/kcft114/.julia/packages/AdvancedHMC/YWXfk/src/adaptation/stepsize.jl:103
┌ Warning: The current proposal will be rejected due to numerical error(s).
│   isfiniteθ = true
│   isfiniter = true
│   isfiniteℓπ = false
│   isfiniteℓκ = true
└ @ AdvancedHMC /Users/kcft114/.julia/packages/AdvancedHMC/YWXfk/src/hamiltonian.jl:36
┌ Warning: The current proposal will be rejected due to numerical error(s).
│   isfiniteθ = false
│   isfiniter = false
│   isfiniteℓπ = false
│   isfiniteℓκ = false
└ @ AdvancedHMC /Users/kcft114/.julia/packages/AdvancedHMC/YWXfk/src/hamiltonian.jl:36
┌ Warning: Incorrect ϵ = NaN; ϵ_previous = 0.07500000000000001 is used instead.
└ @ AdvancedHMC.Adaptation /Users/kcft114/.julia/packages/AdvancedHMC/YWXfk/src/adaptation/stepsize.jl:103
┌ Warning: The current proposal will be rejected due to numerical error(s).
│   isfiniteθ = true
│   isfiniter = true
│   isfiniteℓπ = false
│   isfiniteℓκ = true
└ @ AdvancedHMC /Users/kcft114/.julia/packages/AdvancedHMC/YWXfk/src/hamiltonian.jl:36
┌ Warning: The current proposal will be rejected due to numerical error(s).
│   isfiniteθ = false
│   isfiniter = false
│   isfiniteℓπ = false
│   isfiniteℓκ = false
└ @ AdvancedHMC /Users/kcft114/.julia/packages/AdvancedHMC/YWXfk/src/hamiltonian.jl:36
┌ Warning: Incorrect ϵ = NaN; ϵ_previous = 0.07500000000000001 is used instead.
└ @ AdvancedHMC.Adaptation /Users/kcft114/.julia/packages/AdvancedHMC/YWXfk/src/adaptation/stepsize.jl:103
┌ Warning: The current proposal will be rejected due to numerical error(s).
│   isfiniteθ = true
│   isfiniter = true
│   isfiniteℓπ = false
│   isfiniteℓκ = true
└ @ AdvancedHMC /Users/kcft114/.julia/packages/AdvancedHMC/YWXfk/src/hamiltonian.jl:36
┌ Warning: The current proposal will be rejected due to numerical error(s).
│   isfiniteθ = false
│   isfiniter = false
│   isfiniteℓπ = false
│   isfiniteℓκ = false
└ @ AdvancedHMC /Users/kcft114/.julia/packages/AdvancedHMC/YWXfk/src/hamiltonian.jl:36
┌ Warning: Incorrect ϵ = NaN; ϵ_previous = 0.07500000000000001 is used instead.
└ @ AdvancedHMC.Adaptation /Users/kcft114/.julia/packages/AdvancedHMC/YWXfk/src/adaptation/stepsize.jl:103
┌ Warning: The current proposal will be rejected due to numerical error(s).
│   isfiniteθ = true
│   isfiniter = true
│   isfiniteℓπ = false
│   isfiniteℓκ = true
└ @ AdvancedHMC /Users/kcft114/.julia/packages/AdvancedHMC/YWXfk/src/hamiltonian.jl:36
┌ Warning: The current proposal will be rejected due to numerical error(s).
│   isfiniteθ = false
│   isfiniter = false
│   isfiniteℓπ = false
│   isfiniteℓκ = false
└ @ AdvancedHMC /Users/kcft114/.julia/packages/AdvancedHMC/YWXfk/src/hamiltonian.jl:36
┌ Warning: Incorrect ϵ = NaN; ϵ_previous = 0.07500000000000001 is used instead.
└ @ AdvancedHMC.Adaptation /Users/kcft114/.julia/packages/AdvancedHMC/YWXfk/src/adaptation/stepsize.jl:103
┌ Warning: The current proposal will be rejected due to numerical error(s).
│   isfiniteθ = true
│   isfiniter = true
│   isfiniteℓπ = false
│   isfiniteℓκ = true
└ @ AdvancedHMC /Users/kcft114/.julia/packages/AdvancedHMC/YWXfk/src/hamiltonian.jl:36
┌ Warning: The current proposal will be rejected due to numerical error(s).
│   isfiniteθ = false
│   isfiniter = false
│   isfiniteℓπ = false
│   isfiniteℓκ = false
└ @ AdvancedHMC /Users/kcft114/.julia/packages/AdvancedHMC/YWXfk/src/hamiltonian.jl:36
┌ Warning: Incorrect ϵ = NaN; ϵ_previous = 0.07500000000000001 is used instead.
└ @ AdvancedHMC.Adaptation /Users/kcft114/.julia/packages/AdvancedHMC/YWXfk/src/adaptation/stepsize.jl:103
┌ Warning: The current proposal will be rejected due to numerical error(s).
│   isfiniteθ = true
│   isfiniter = true
│   isfiniteℓπ = false
│   isfiniteℓκ = true
└ @ AdvancedHMC /Users/kcft114/.julia/packages/AdvancedHMC/YWXfk/src/hamiltonian.jl:36
┌ Warning: The current proposal will be rejected due to numerical error(s).
│   isfiniteθ = false
│   isfiniter = false
│   isfiniteℓπ = false
│   isfiniteℓκ = false
└ @ AdvancedHMC /Users/kcft114/.julia/packages/AdvancedHMC/YWXfk/src/hamiltonian.jl:36
┌ Warning: Incorrect ϵ = NaN; ϵ_previous = 0.07500000000000001 is used instead.
└ @ AdvancedHMC.Adaptation /Users/kcft114/.julia/packages/AdvancedHMC/YWXfk/src/adaptation/stepsize.jl:103
┌ Warning: The current proposal will be rejected due to numerical error(s).
│   isfiniteθ = true
│   isfiniter = true
│   isfiniteℓπ = false
│   isfiniteℓκ = true
└ @ AdvancedHMC /Users/kcft114/.julia/packages/AdvancedHMC/YWXfk/src/hamiltonian.jl:36
┌ Warning: The current proposal will be rejected due to numerical error(s).
│   isfiniteθ = false
│   isfiniter = false
│   isfiniteℓπ = false
│   isfiniteℓκ = false
└ @ AdvancedHMC /Users/kcft114/.julia/packages/AdvancedHMC/YWXfk/src/hamiltonian.jl:36
┌ Warning: Incorrect ϵ = NaN; ϵ_previous = 0.07500000000000001 is used instead.
└ @ AdvancedHMC.Adaptation /Users/kcft114/.julia/packages/AdvancedHMC/YWXfk/src/adaptation/stepsize.jl:103
┌ Warning: The current proposal will be rejected due to numerical error(s).
│   isfiniteθ = true
│   isfiniter = true
│   isfiniteℓπ = false
│   isfiniteℓκ = true
└ @ AdvancedHMC /Users/kcft114/.julia/packages/AdvancedHMC/YWXfk/src/hamiltonian.jl:36
┌ Warning: The current proposal will be rejected due to numerical error(s).
│   isfiniteθ = false
│   isfiniter = false
│   isfiniteℓπ = false
│   isfiniteℓκ = false
└ @ AdvancedHMC /Users/kcft114/.julia/packages/AdvancedHMC/YWXfk/src/hamiltonian.jl:36
┌ Warning: Incorrect ϵ = NaN; ϵ_previous = 0.07500000000000001 is used instead.
└ @ AdvancedHMC.Adaptation /Users/kcft114/.julia/packages/AdvancedHMC/YWXfk/src/adaptation/stepsize.jl:103
┌ Warning: The current proposal will be rejected due to numerical error(s).
│   isfiniteθ = true
│   isfiniter = true
│   isfiniteℓπ = false
│   isfiniteℓκ = true
└ @ AdvancedHMC /Users/kcft114/.julia/packages/AdvancedHMC/YWXfk/src/hamiltonian.jl:36
┌ Warning: The current proposal will be rejected due to numerical error(s).
│   isfiniteθ = false
│   isfiniter = false
│   isfiniteℓπ = false
│   isfiniteℓκ = false
└ @ AdvancedHMC /Users/kcft114/.julia/packages/AdvancedHMC/YWXfk/src/hamiltonian.jl:36
┌ Warning: Incorrect ϵ = NaN; ϵ_previous = 0.07500000000000001 is used instead.
└ @ AdvancedHMC.Adaptation /Users/kcft114/.julia/packages/AdvancedHMC/YWXfk/src/adaptation/stepsize.jl:103
┌ Warning: The current proposal will be rejected due to numerical error(s).
│   isfiniteθ = true
│   isfiniter = true
│   isfiniteℓπ = false
│   isfiniteℓκ = true
└ @ AdvancedHMC /Users/kcft114/.julia/packages/AdvancedHMC/YWXfk/src/hamiltonian.jl:36
┌ Warning: The current proposal will be rejected due to numerical error(s).
│   isfiniteθ = false
│   isfiniter = false
│   isfiniteℓπ = false
│   isfiniteℓκ = false
└ @ AdvancedHMC /Users/kcft114/.julia/packages/AdvancedHMC/YWXfk/src/hamiltonian.jl:36
┌ Warning: Incorrect ϵ = NaN; ϵ_previous = 0.07500000000000001 is used instead.
└ @ AdvancedHMC.Adaptation /Users/kcft114/.julia/packages/AdvancedHMC/YWXfk/src/adaptation/stepsize.jl:103
┌ Warning: The current proposal will be rejected due to numerical error(s).
│   isfiniteθ = true
│   isfiniter = true
│   isfiniteℓπ = false
│   isfiniteℓκ = true
└ @ AdvancedHMC /Users/kcft114/.julia/packages/AdvancedHMC/YWXfk/src/hamiltonian.jl:36
┌ Warning: The current proposal will be rejected due to numerical error(s).
│   isfiniteθ = false
│   isfiniter = false
│   isfiniteℓπ = false
│   isfiniteℓκ = false
└ @ AdvancedHMC /Users/kcft114/.julia/packages/AdvancedHMC/YWXfk/src/hamiltonian.jl:36
┌ Warning: Incorrect ϵ = NaN; ϵ_previous = 0.07500000000000001 is used instead.
└ @ AdvancedHMC.Adaptation /Users/kcft114/.julia/packages/AdvancedHMC/YWXfk/src/adaptation/stepsize.jl:103
┌ Warning: The current proposal will be rejected due to numerical error(s).
│   isfiniteθ = true
│   isfiniter = true
│   isfiniteℓπ = false
│   isfiniteℓκ = true
└ @ AdvancedHMC /Users/kcft114/.julia/packages/AdvancedHMC/YWXfk/src/hamiltonian.jl:36
┌ Warning: The current proposal will be rejected due to numerical error(s).
│   isfiniteθ = false
│   isfiniter = false
│   isfiniteℓπ = false
│   isfiniteℓκ = false
└ @ AdvancedHMC /Users/kcft114/.julia/packages/AdvancedHMC/YWXfk/src/hamiltonian.jl:36
┌ Warning: Incorrect ϵ = NaN; ϵ_previous = 0.07500000000000001 is used instead.
└ @ AdvancedHMC.Adaptation /Users/kcft114/.julia/packages/AdvancedHMC/YWXfk/src/adaptation/stepsize.jl:103
┌ Warning: The current proposal will be rejected due to numerical error(s).
│   isfiniteθ = true
│   isfiniter = true
│   isfiniteℓπ = false
│   isfiniteℓκ = true
└ @ AdvancedHMC /Users/kcft114/.julia/packages/AdvancedHMC/YWXfk/src/hamiltonian.jl:36
┌ Warning: The current proposal will be rejected due to numerical error(s).
│   isfiniteθ = false
│   isfiniter = false
│   isfiniteℓπ = false
│   isfiniteℓκ = false
└ @ AdvancedHMC /Users/kcft114/.julia/packages/AdvancedHMC/YWXfk/src/hamiltonian.jl:36
┌ Warning: Incorrect ϵ = NaN; ϵ_previous = 0.07500000000000001 is used instead.
└ @ AdvancedHMC.Adaptation /Users/kcft114/.julia/packages/AdvancedHMC/YWXfk/src/adaptation/stepsize.jl:103
┌ Warning: The current proposal will be rejected due to numerical error(s).
│   isfiniteθ = true
│   isfiniter = true
│   isfiniteℓπ = false
│   isfiniteℓκ = true
└ @ AdvancedHMC /Users/kcft114/.julia/packages/AdvancedHMC/YWXfk/src/hamiltonian.jl:36
┌ Warning: The current proposal will be rejected due to numerical error(s).
│   isfiniteθ = false
│   isfiniter = false
│   isfiniteℓπ = false
│   isfiniteℓκ = false
└ @ AdvancedHMC /Users/kcft114/.julia/packages/AdvancedHMC/YWXfk/src/hamiltonian.jl:36
┌ Warning: Incorrect ϵ = NaN; ϵ_previous = 0.07500000000000001 is used instead.
└ @ AdvancedHMC.Adaptation /Users/kcft114/.julia/packages/AdvancedHMC/YWXfk/src/adaptation/stepsize.jl:103
┌ Warning: The current proposal will be rejected due to numerical error(s).
│   isfiniteθ = true
│   isfiniter = true
│   isfiniteℓπ = false
│   isfiniteℓκ = true
└ @ AdvancedHMC /Users/kcft114/.julia/packages/AdvancedHMC/YWXfk/src/hamiltonian.jl:36
┌ Warning: The current proposal will be rejected due to numerical error(s).
│   isfiniteθ = false
│   isfiniter = false
│   isfiniteℓπ = false
│   isfiniteℓκ = false
└ @ AdvancedHMC /Users/kcft114/.julia/packages/AdvancedHMC/YWXfk/src/hamiltonian.jl:36
┌ Warning: Incorrect ϵ = NaN; ϵ_previous = 0.07500000000000001 is used instead.
└ @ AdvancedHMC.Adaptation /Users/kcft114/.julia/packages/AdvancedHMC/YWXfk/src/adaptation/stepsize.jl:103
┌ Warning: The current proposal will be rejected due to numerical error(s).
│   isfiniteθ = true
│   isfiniter = true
│   isfiniteℓπ = false
│   isfiniteℓκ = true
└ @ AdvancedHMC /Users/kcft114/.julia/packages/AdvancedHMC/YWXfk/src/hamiltonian.jl:36
┌ Warning: The current proposal will be rejected due to numerical error(s).
│   isfiniteθ = false
│   isfiniter = false
│   isfiniteℓπ = false
│   isfiniteℓκ = false
└ @ AdvancedHMC /Users/kcft114/.julia/packages/AdvancedHMC/YWXfk/src/hamiltonian.jl:36
┌ Warning: Incorrect ϵ = NaN; ϵ_previous = 0.07500000000000001 is used instead.
└ @ AdvancedHMC.Adaptation /Users/kcft114/.julia/packages/AdvancedHMC/YWXfk/src/adaptation/stepsize.jl:103
┌ Warning: The current proposal will be rejected due to numerical error(s).
│   isfiniteθ = true
│   isfiniter = true
│   isfiniteℓπ = false
│   isfiniteℓκ = true
└ @ AdvancedHMC /Users/kcft114/.julia/packages/AdvancedHMC/YWXfk/src/hamiltonian.jl:36
┌ Warning: The current proposal will be rejected due to numerical error(s).
│   isfiniteθ = false
│   isfiniter = false
│   isfiniteℓπ = false
│   isfiniteℓκ = false
└ @ AdvancedHMC /Users/kcft114/.julia/packages/AdvancedHMC/YWXfk/src/hamiltonian.jl:36
┌ Warning: Incorrect ϵ = NaN; ϵ_previous = 0.07500000000000001 is used instead.
└ @ AdvancedHMC.Adaptation /Users/kcft114/.julia/packages/AdvancedHMC/YWXfk/src/adaptation/stepsize.jl:103
┌ Warning: The current proposal will be rejected due to numerical error(s).
│   isfiniteθ = true
│   isfiniter = true
│   isfiniteℓπ = false
│   isfiniteℓκ = true
└ @ AdvancedHMC /Users/kcft114/.julia/packages/AdvancedHMC/YWXfk/src/hamiltonian.jl:36
┌ Warning: The current proposal will be rejected due to numerical error(s).
│   isfiniteθ = false
│   isfiniter = false
│   isfiniteℓπ = false
│   isfiniteℓκ = false
└ @ AdvancedHMC /Users/kcft114/.julia/packages/AdvancedHMC/YWXfk/src/hamiltonian.jl:36
┌ Warning: Incorrect ϵ = NaN; ϵ_previous = 0.07500000000000001 is used instead.
└ @ AdvancedHMC.Adaptation /Users/kcft114/.julia/packages/AdvancedHMC/YWXfk/src/adaptation/stepsize.jl:103
┌ Warning: The current proposal will be rejected due to numerical error(s).
│   isfiniteθ = true
│   isfiniter = true
│   isfiniteℓπ = false
│   isfiniteℓκ = true
└ @ AdvancedHMC /Users/kcft114/.julia/packages/AdvancedHMC/YWXfk/src/hamiltonian.jl:36
┌ Warning: The current proposal will be rejected due to numerical error(s).
│   isfiniteθ = false
│   isfiniter = false
│   isfiniteℓπ = false
│   isfiniteℓκ = false
└ @ AdvancedHMC /Users/kcft114/.julia/packages/AdvancedHMC/YWXfk/src/hamiltonian.jl:36
┌ Warning: Incorrect ϵ = NaN; ϵ_previous = 0.07500000000000001 is used instead.
└ @ AdvancedHMC.Adaptation /Users/kcft114/.julia/packages/AdvancedHMC/YWXfk/src/adaptation/stepsize.jl:103
┌ Warning: The current proposal will be rejected due to numerical error(s).
│   isfiniteθ = true
│   isfiniter = true
│   isfiniteℓπ = false
│   isfiniteℓκ = true
└ @ AdvancedHMC /Users/kcft114/.julia/packages/AdvancedHMC/YWXfk/src/hamiltonian.jl:36
┌ Warning: The current proposal will be rejected due to numerical error(s).
│   isfiniteθ = false
│   isfiniter = false
│   isfiniteℓπ = false
│   isfiniteℓκ = false
└ @ AdvancedHMC /Users/kcft114/.julia/packages/AdvancedHMC/YWXfk/src/hamiltonian.jl:36
┌ Warning: Incorrect ϵ = NaN; ϵ_previous = 0.07500000000000001 is used instead.
└ @ AdvancedHMC.Adaptation /Users/kcft114/.julia/packages/AdvancedHMC/YWXfk/src/adaptation/stepsize.jl:103
┌ Warning: The current proposal will be rejected due to numerical error(s).
│   isfiniteθ = true
│   isfiniter = true
│   isfiniteℓπ = false
│   isfiniteℓκ = true
└ @ AdvancedHMC /Users/kcft114/.julia/packages/AdvancedHMC/YWXfk/src/hamiltonian.jl:36
┌ Warning: The current proposal will be rejected due to numerical error(s).
│   isfiniteθ = false
│   isfiniter = false
│   isfiniteℓπ = false
│   isfiniteℓκ = false
└ @ AdvancedHMC /Users/kcft114/.julia/packages/AdvancedHMC/YWXfk/src/hamiltonian.jl:36
┌ Warning: Incorrect ϵ = NaN; ϵ_previous = 0.07500000000000001 is used instead.
└ @ AdvancedHMC.Adaptation /Users/kcft114/.julia/packages/AdvancedHMC/YWXfk/src/adaptation/stepsize.jl:103
┌ Warning: The current proposal will be rejected due to numerical error(s).
│   isfiniteθ = true
│   isfiniter = true
│   isfiniteℓπ = false
│   isfiniteℓκ = true
└ @ AdvancedHMC /Users/kcft114/.julia/packages/AdvancedHMC/YWXfk/src/hamiltonian.jl:36
┌ Warning: The current proposal will be rejected due to numerical error(s).
│   isfiniteθ = false
│   isfiniter = false
│   isfiniteℓπ = false
│   isfiniteℓκ = false
└ @ AdvancedHMC /Users/kcft114/.julia/packages/AdvancedHMC/YWXfk/src/hamiltonian.jl:36
┌ Warning: Incorrect ϵ = NaN; ϵ_previous = 0.07500000000000001 is used instead.
└ @ AdvancedHMC.Adaptation /Users/kcft114/.julia/packages/AdvancedHMC/YWXfk/src/adaptation/stepsize.jl:103
┌ Warning: The current proposal will be rejected due to numerical error(s).
│   isfiniteθ = true
│   isfiniter = true
│   isfiniteℓπ = false
│   isfiniteℓκ = true
└ @ AdvancedHMC /Users/kcft114/.julia/packages/AdvancedHMC/YWXfk/src/hamiltonian.jl:36
┌ Warning: The current proposal will be rejected due to numerical error(s).
│   isfiniteθ = false
│   isfiniter = false
│   isfiniteℓπ = false
│   isfiniteℓκ = false
└ @ AdvancedHMC /Users/kcft114/.julia/packages/AdvancedHMC/YWXfk/src/hamiltonian.jl:36
┌ Warning: Incorrect ϵ = NaN; ϵ_previous = 0.07500000000000001 is used instead.
└ @ AdvancedHMC.Adaptation /Users/kcft114/.julia/packages/AdvancedHMC/YWXfk/src/adaptation/stepsize.jl:103
┌ Warning: The current proposal will be rejected due to numerical error(s).
│   isfiniteθ = true
│   isfiniter = true
│   isfiniteℓπ = false
│   isfiniteℓκ = true
└ @ AdvancedHMC /Users/kcft114/.julia/packages/AdvancedHMC/YWXfk/src/hamiltonian.jl:36
┌ Warning: The current proposal will be rejected due to numerical error(s).
│   isfiniteθ = false
│   isfiniter = false
│   isfiniteℓπ = false
│   isfiniteℓκ = false
└ @ AdvancedHMC /Users/kcft114/.julia/packages/AdvancedHMC/YWXfk/src/hamiltonian.jl:36
┌ Warning: Incorrect ϵ = NaN; ϵ_previous = 0.07500000000000001 is used instead.
└ @ AdvancedHMC.Adaptation /Users/kcft114/.julia/packages/AdvancedHMC/YWXfk/src/adaptation/stepsize.jl:103
┌ Warning: The current proposal will be rejected due to numerical error(s).
│   isfiniteθ = true
│   isfiniter = true
│   isfiniteℓπ = false
│   isfiniteℓκ = true
└ @ AdvancedHMC /Users/kcft114/.julia/packages/AdvancedHMC/YWXfk/src/hamiltonian.jl:36
┌ Warning: The current proposal will be rejected due to numerical error(s).
│   isfiniteθ = false
│   isfiniter = false
│   isfiniteℓπ = false
│   isfiniteℓκ = false
└ @ AdvancedHMC /Users/kcft114/.julia/packages/AdvancedHMC/YWXfk/src/hamiltonian.jl:36
┌ Warning: Incorrect ϵ = NaN; ϵ_previous = 0.07500000000000001 is used instead.
└ @ AdvancedHMC.Adaptation /Users/kcft114/.julia/packages/AdvancedHMC/YWXfk/src/adaptation/stepsize.jl:103
┌ Warning: The current proposal will be rejected due to numerical error(s).
│   isfiniteθ = true
│   isfiniter = true
│   isfiniteℓπ = false
│   isfiniteℓκ = true
└ @ AdvancedHMC /Users/kcft114/.julia/packages/AdvancedHMC/YWXfk/src/hamiltonian.jl:36
┌ Warning: The current proposal will be rejected due to numerical error(s).
│   isfiniteθ = false
│   isfiniter = false
│   isfiniteℓπ = false
│   isfiniteℓκ = false
└ @ AdvancedHMC /Users/kcft114/.julia/packages/AdvancedHMC/YWXfk/src/hamiltonian.jl:36
┌ Warning: Incorrect ϵ = NaN; ϵ_previous = 0.07500000000000001 is used instead.
└ @ AdvancedHMC.Adaptation /Users/kcft114/.julia/packages/AdvancedHMC/YWXfk/src/adaptation/stepsize.jl:103
┌ Warning: The current proposal will be rejected due to numerical error(s).
│   isfiniteθ = true
│   isfiniter = true
│   isfiniteℓπ = false
│   isfiniteℓκ = true
└ @ AdvancedHMC /Users/kcft114/.julia/packages/AdvancedHMC/YWXfk/src/hamiltonian.jl:36
┌ Warning: The current proposal will be rejected due to numerical error(s).
│   isfiniteθ = false
│   isfiniter = false
│   isfiniteℓπ = false
│   isfiniteℓκ = false
└ @ AdvancedHMC /Users/kcft114/.julia/packages/AdvancedHMC/YWXfk/src/hamiltonian.jl:36
┌ Warning: Incorrect ϵ = NaN; ϵ_previous = 0.07500000000000001 is used instead.
└ @ AdvancedHMC.Adaptation /Users/kcft114/.julia/packages/AdvancedHMC/YWXfk/src/adaptation/stepsize.jl:103
┌ Warning: The current proposal will be rejected due to numerical error(s).
│   isfiniteθ = true
│   isfiniter = true
│   isfiniteℓπ = false
│   isfiniteℓκ = true
└ @ AdvancedHMC /Users/kcft114/.julia/packages/AdvancedHMC/YWXfk/src/hamiltonian.jl:36
┌ Warning: The current proposal will be rejected due to numerical error(s).
│   isfiniteθ = false
│   isfiniter = false
│   isfiniteℓπ = false
│   isfiniteℓκ = false
└ @ AdvancedHMC /Users/kcft114/.julia/packages/AdvancedHMC/YWXfk/src/hamiltonian.jl:36
┌ Warning: Incorrect ϵ = NaN; ϵ_previous = 0.07500000000000001 is used instead.
└ @ AdvancedHMC.Adaptation /Users/kcft114/.julia/packages/AdvancedHMC/YWXfk/src/adaptation/stepsize.jl:103
┌ Warning: The current proposal will be rejected due to numerical error(s).
│   isfiniteθ = true
│   isfiniter = true
│   isfiniteℓπ = false
│   isfiniteℓκ = true
└ @ AdvancedHMC /Users/kcft114/.julia/packages/AdvancedHMC/YWXfk/src/hamiltonian.jl:36
┌ Warning: The current proposal will be rejected due to numerical error(s).
│   isfiniteθ = false
│   isfiniter = false
│   isfiniteℓπ = false
│   isfiniteℓκ = false
└ @ AdvancedHMC /Users/kcft114/.julia/packages/AdvancedHMC/YWXfk/src/hamiltonian.jl:36
┌ Warning: Incorrect ϵ = NaN; ϵ_previous = 0.07500000000000001 is used instead.
└ @ AdvancedHMC.Adaptation /Users/kcft114/.julia/packages/AdvancedHMC/YWXfk/src/adaptation/stepsize.jl:103
┌ Warning: The current proposal will be rejected due to numerical error(s).
│   isfiniteθ = true
│   isfiniter = true
│   isfiniteℓπ = false
│   isfiniteℓκ = true
└ @ AdvancedHMC /Users/kcft114/.julia/packages/AdvancedHMC/YWXfk/src/hamiltonian.jl:36
┌ Warning: The current proposal will be rejected due to numerical error(s).
│   isfiniteθ = false
│   isfiniter = false
│   isfiniteℓπ = false
│   isfiniteℓκ = false
└ @ AdvancedHMC /Users/kcft114/.julia/packages/AdvancedHMC/YWXfk/src/hamiltonian.jl:36
┌ Warning: Incorrect ϵ = NaN; ϵ_previous = 0.07500000000000001 is used instead.
└ @ AdvancedHMC.Adaptation /Users/kcft114/.julia/packages/AdvancedHMC/YWXfk/src/adaptation/stepsize.jl:103
┌ Warning: The current proposal will be rejected due to numerical error(s).
│   isfiniteθ = true
│   isfiniter = true
│   isfiniteℓπ = false
│   isfiniteℓκ = true
└ @ AdvancedHMC /Users/kcft114/.julia/packages/AdvancedHMC/YWXfk/src/hamiltonian.jl:36
┌ Warning: The current proposal will be rejected due to numerical error(s).
│   isfiniteθ = false
│   isfiniter = false
│   isfiniteℓπ = false
│   isfiniteℓκ = false
└ @ AdvancedHMC /Users/kcft114/.julia/packages/AdvancedHMC/YWXfk/src/hamiltonian.jl:36
┌ Warning: Incorrect ϵ = NaN; ϵ_previous = 0.07500000000000001 is used instead.
└ @ AdvancedHMC.Adaptation /Users/kcft114/.julia/packages/AdvancedHMC/YWXfk/src/adaptation/stepsize.jl:103
┌ Warning: The current proposal will be rejected due to numerical error(s).
│   isfiniteθ = true
│   isfiniter = true
│   isfiniteℓπ = false
│   isfiniteℓκ = true
└ @ AdvancedHMC /Users/kcft114/.julia/packages/AdvancedHMC/YWXfk/src/hamiltonian.jl:36
┌ Warning: The current proposal will be rejected due to numerical error(s).
│   isfiniteθ = false
│   isfiniter = false
│   isfiniteℓπ = false
│   isfiniteℓκ = false
└ @ AdvancedHMC /Users/kcft114/.julia/packages/AdvancedHMC/YWXfk/src/hamiltonian.jl:36
┌ Warning: Incorrect ϵ = NaN; ϵ_previous = 0.07500000000000001 is used instead.
└ @ AdvancedHMC.Adaptation /Users/kcft114/.julia/packages/AdvancedHMC/YWXfk/src/adaptation/stepsize.jl:103
┌ Warning: The current proposal will be rejected due to numerical error(s).
│   isfiniteθ = true
│   isfiniter = true
│   isfiniteℓπ = false
│   isfiniteℓκ = true
└ @ AdvancedHMC /Users/kcft114/.julia/packages/AdvancedHMC/YWXfk/src/hamiltonian.jl:36
┌ Warning: The current proposal will be rejected due to numerical error(s).
│   isfiniteθ = false
│   isfiniter = false
│   isfiniteℓπ = false
│   isfiniteℓκ = false
└ @ AdvancedHMC /Users/kcft114/.julia/packages/AdvancedHMC/YWXfk/src/hamiltonian.jl:36
┌ Warning: Incorrect ϵ = NaN; ϵ_previous = 0.07500000000000001 is used instead.
└ @ AdvancedHMC.Adaptation /Users/kcft114/.julia/packages/AdvancedHMC/YWXfk/src/adaptation/stepsize.jl:103
┌ Warning: The current proposal will be rejected due to numerical error(s).
│   isfiniteθ = true
│   isfiniter = true
│   isfiniteℓπ = false
│   isfiniteℓκ = true
└ @ AdvancedHMC /Users/kcft114/.julia/packages/AdvancedHMC/YWXfk/src/hamiltonian.jl:36
┌ Warning: The current proposal will be rejected due to numerical error(s).
│   isfiniteθ = false
│   isfiniter = false
│   isfiniteℓπ = false
│   isfiniteℓκ = false
└ @ AdvancedHMC /Users/kcft114/.julia/packages/AdvancedHMC/YWXfk/src/hamiltonian.jl:36
┌ Warning: Incorrect ϵ = NaN; ϵ_previous = 0.07500000000000001 is used instead.
└ @ AdvancedHMC.Adaptation /Users/kcft114/.julia/packages/AdvancedHMC/YWXfk/src/adaptation/stepsize.jl:103
┌ Warning: The current proposal will be rejected due to numerical error(s).
│   isfiniteθ = true
│   isfiniter = true
│   isfiniteℓπ = false
│   isfiniteℓκ = true
└ @ AdvancedHMC /Users/kcft114/.julia/packages/AdvancedHMC/YWXfk/src/hamiltonian.jl:36
┌ Warning: The current proposal will be rejected due to numerical error(s).
│   isfiniteθ = false
│   isfiniter = false
│   isfiniteℓπ = false
│   isfiniteℓκ = false
└ @ AdvancedHMC /Users/kcft114/.julia/packages/AdvancedHMC/YWXfk/src/hamiltonian.jl:36
┌ Warning: Incorrect ϵ = NaN; ϵ_previous = 0.07500000000000001 is used instead.
└ @ AdvancedHMC.Adaptation /Users/kcft114/.julia/packages/AdvancedHMC/YWXfk/src/adaptation/stepsize.jl:103
┌ Warning: The current proposal will be rejected due to numerical error(s).
│   isfiniteθ = true
│   isfiniter = true
│   isfiniteℓπ = false
│   isfiniteℓκ = true
└ @ AdvancedHMC /Users/kcft114/.julia/packages/AdvancedHMC/YWXfk/src/hamiltonian.jl:36
┌ Warning: The current proposal will be rejected due to numerical error(s).
│   isfiniteθ = false
│   isfiniter = false
│   isfiniteℓπ = false
│   isfiniteℓκ = false
└ @ AdvancedHMC /Users/kcft114/.julia/packages/AdvancedHMC/YWXfk/src/hamiltonian.jl:36
┌ Warning: Incorrect ϵ = NaN; ϵ_previous = 0.07500000000000001 is used instead.
└ @ AdvancedHMC.Adaptation /Users/kcft114/.julia/packages/AdvancedHMC/YWXfk/src/adaptation/stepsize.jl:103
┌ Warning: The current proposal will be rejected due to numerical error(s).
│   isfiniteθ = true
│   isfiniter = true
│   isfiniteℓπ = false
│   isfiniteℓκ = true
└ @ AdvancedHMC /Users/kcft114/.julia/packages/AdvancedHMC/YWXfk/src/hamiltonian.jl:36
┌ Warning: The current proposal will be rejected due to numerical error(s).
│   isfiniteθ = false
│   isfiniter = false
│   isfiniteℓπ = false
│   isfiniteℓκ = false
└ @ AdvancedHMC /Users/kcft114/.julia/packages/AdvancedHMC/YWXfk/src/hamiltonian.jl:36
┌ Warning: Incorrect ϵ = NaN; ϵ_previous = 0.07500000000000001 is used instead.
└ @ AdvancedHMC.Adaptation /Users/kcft114/.julia/packages/AdvancedHMC/YWXfk/src/adaptation/stepsize.jl:103
┌ Warning: The current proposal will be rejected due to numerical error(s).
│   isfiniteθ = true
│   isfiniter = true
│   isfiniteℓπ = false
│   isfiniteℓκ = true
└ @ AdvancedHMC /Users/kcft114/.julia/packages/AdvancedHMC/YWXfk/src/hamiltonian.jl:36
┌ Warning: The current proposal will be rejected due to numerical error(s).
│   isfiniteθ = false
│   isfiniter = false
│   isfiniteℓπ = false
│   isfiniteℓκ = false
└ @ AdvancedHMC /Users/kcft114/.julia/packages/AdvancedHMC/YWXfk/src/hamiltonian.jl:36
┌ Warning: Incorrect ϵ = NaN; ϵ_previous = 0.07500000000000001 is used instead.
└ @ AdvancedHMC.Adaptation /Users/kcft114/.julia/packages/AdvancedHMC/YWXfk/src/adaptation/stepsize.jl:103
┌ Warning: The current proposal will be rejected due to numerical error(s).
│   isfiniteθ = true
│   isfiniter = true
│   isfiniteℓπ = false
│   isfiniteℓκ = true
└ @ AdvancedHMC /Users/kcft114/.julia/packages/AdvancedHMC/YWXfk/src/hamiltonian.jl:36
┌ Warning: The current proposal will be rejected due to numerical error(s).
│   isfiniteθ = false
│   isfiniter = false
│   isfiniteℓπ = false
│   isfiniteℓκ = false
└ @ AdvancedHMC /Users/kcft114/.julia/packages/AdvancedHMC/YWXfk/src/hamiltonian.jl:36
┌ Warning: Incorrect ϵ = NaN; ϵ_previous = 0.07500000000000001 is used instead.
└ @ AdvancedHMC.Adaptation /Users/kcft114/.julia/packages/AdvancedHMC/YWXfk/src/adaptation/stepsize.jl:103
┌ Warning: The current proposal will be rejected due to numerical error(s).
│   isfiniteθ = true
│   isfiniter = true
│   isfiniteℓπ = false
│   isfiniteℓκ = true
└ @ AdvancedHMC /Users/kcft114/.julia/packages/AdvancedHMC/YWXfk/src/hamiltonian.jl:36
┌ Warning: The current proposal will be rejected due to numerical error(s).
│   isfiniteθ = false
│   isfiniter = false
│   isfiniteℓπ = false
│   isfiniteℓκ = false
└ @ AdvancedHMC /Users/kcft114/.julia/packages/AdvancedHMC/YWXfk/src/hamiltonian.jl:36
┌ Warning: Incorrect ϵ = NaN; ϵ_previous = 0.07500000000000001 is used instead.
└ @ AdvancedHMC.Adaptation /Users/kcft114/.julia/packages/AdvancedHMC/YWXfk/src/adaptation/stepsize.jl:103
┌ Warning: The current proposal will be rejected due to numerical error(s).
│   isfiniteθ = true
│   isfiniter = true
│   isfiniteℓπ = false
│   isfiniteℓκ = true
└ @ AdvancedHMC /Users/kcft114/.julia/packages/AdvancedHMC/YWXfk/src/hamiltonian.jl:36
┌ Warning: The current proposal will be rejected due to numerical error(s).
│   isfiniteθ = false
│   isfiniter = false
│   isfiniteℓπ = false
│   isfiniteℓκ = false
└ @ AdvancedHMC /Users/kcft114/.julia/packages/AdvancedHMC/YWXfk/src/hamiltonian.jl:36
┌ Warning: Incorrect ϵ = NaN; ϵ_previous = 0.07500000000000001 is used instead.
└ @ AdvancedHMC.Adaptation /Users/kcft114/.julia/packages/AdvancedHMC/YWXfk/src/adaptation/stepsize.jl:103
┌ Warning: The current proposal will be rejected due to numerical error(s).
│   isfiniteθ = true
│   isfiniter = true
│   isfiniteℓπ = false
│   isfiniteℓκ = true
└ @ AdvancedHMC /Users/kcft114/.julia/packages/AdvancedHMC/YWXfk/src/hamiltonian.jl:36
┌ Warning: The current proposal will be rejected due to numerical error(s).
│   isfiniteθ = false
│   isfiniter = false
│   isfiniteℓπ = false
│   isfiniteℓκ = false
└ @ AdvancedHMC /Users/kcft114/.julia/packages/AdvancedHMC/YWXfk/src/hamiltonian.jl:36
┌ Warning: Incorrect ϵ = NaN; ϵ_previous = 0.07500000000000001 is used instead.
└ @ AdvancedHMC.Adaptation /Users/kcft114/.julia/packages/AdvancedHMC/YWXfk/src/adaptation/stepsize.jl:103
┌ Warning: The current proposal will be rejected due to numerical error(s).
│   isfiniteθ = true
│   isfiniter = true
│   isfiniteℓπ = false
│   isfiniteℓκ = true
└ @ AdvancedHMC /Users/kcft114/.julia/packages/AdvancedHMC/YWXfk/src/hamiltonian.jl:36
┌ Warning: The current proposal will be rejected due to numerical error(s).
│   isfiniteθ = false
│   isfiniter = false
│   isfiniteℓπ = false
│   isfiniteℓκ = false
└ @ AdvancedHMC /Users/kcft114/.julia/packages/AdvancedHMC/YWXfk/src/hamiltonian.jl:36
┌ Warning: Incorrect ϵ = NaN; ϵ_previous = 0.07500000000000001 is used instead.
└ @ AdvancedHMC.Adaptation /Users/kcft114/.julia/packages/AdvancedHMC/YWXfk/src/adaptation/stepsize.jl:103
┌ Warning: The current proposal will be rejected due to numerical error(s).
│   isfiniteθ = true
│   isfiniter = true
│   isfiniteℓπ = false
│   isfiniteℓκ = true
└ @ AdvancedHMC /Users/kcft114/.julia/packages/AdvancedHMC/YWXfk/src/hamiltonian.jl:36
┌ Warning: The current proposal will be rejected due to numerical error(s).
│   isfiniteθ = false
│   isfiniter = false
│   isfiniteℓπ = false
│   isfiniteℓκ = false
└ @ AdvancedHMC /Users/kcft114/.julia/packages/AdvancedHMC/YWXfk/src/hamiltonian.jl:36
┌ Warning: Incorrect ϵ = NaN; ϵ_previous = 0.07500000000000001 is used instead.
└ @ AdvancedHMC.Adaptation /Users/kcft114/.julia/packages/AdvancedHMC/YWXfk/src/adaptation/stepsize.jl:103
┌ Warning: The current proposal will be rejected due to numerical error(s).
│   isfiniteθ = true
│   isfiniter = true
│   isfiniteℓπ = false
│   isfiniteℓκ = true
└ @ AdvancedHMC /Users/kcft114/.julia/packages/AdvancedHMC/YWXfk/src/hamiltonian.jl:36
┌ Warning: The current proposal will be rejected due to numerical error(s).
│   isfiniteθ = false
│   isfiniter = false
│   isfiniteℓπ = false
│   isfiniteℓκ = false
└ @ AdvancedHMC /Users/kcft114/.julia/packages/AdvancedHMC/YWXfk/src/hamiltonian.jl:36
┌ Warning: Incorrect ϵ = NaN; ϵ_previous = 0.07500000000000001 is used instead.
└ @ AdvancedHMC.Adaptation /Users/kcft114/.julia/packages/AdvancedHMC/YWXfk/src/adaptation/stepsize.jl:103
┌ Warning: The current proposal will be rejected due to numerical error(s).
│   isfiniteθ = true
│   isfiniter = true
│   isfiniteℓπ = false
│   isfiniteℓκ = true
└ @ AdvancedHMC /Users/kcft114/.julia/packages/AdvancedHMC/YWXfk/src/hamiltonian.jl:36
┌ Warning: The current proposal will be rejected due to numerical error(s).
│   isfiniteθ = false
│   isfiniter = false
│   isfiniteℓπ = false
│   isfiniteℓκ = false
└ @ AdvancedHMC /Users/kcft114/.julia/packages/AdvancedHMC/YWXfk/src/hamiltonian.jl:36
┌ Warning: Incorrect ϵ = NaN; ϵ_previous = 0.07500000000000001 is used instead.
└ @ AdvancedHMC.Adaptation /Users/kcft114/.julia/packages/AdvancedHMC/YWXfk/src/adaptation/stepsize.jl:103
┌ Warning: The current proposal will be rejected due to numerical error(s).
│   isfiniteθ = true
│   isfiniter = true
│   isfiniteℓπ = false
│   isfiniteℓκ = true
└ @ AdvancedHMC /Users/kcft114/.julia/packages/AdvancedHMC/YWXfk/src/hamiltonian.jl:36
┌ Warning: The current proposal will be rejected due to numerical error(s).
│   isfiniteθ = false
│   isfiniter = false
│   isfiniteℓπ = false
│   isfiniteℓκ = false
└ @ AdvancedHMC /Users/kcft114/.julia/packages/AdvancedHMC/YWXfk/src/hamiltonian.jl:36
┌ Warning: Incorrect ϵ = NaN; ϵ_previous = 0.07500000000000001 is used instead.
└ @ AdvancedHMC.Adaptation /Users/kcft114/.julia/packages/AdvancedHMC/YWXfk/src/adaptation/stepsize.jl:103
┌ Warning: The current proposal will be rejected due to numerical error(s).
│   isfiniteθ = true
│   isfiniter = true
│   isfiniteℓπ = false
│   isfiniteℓκ = true
└ @ AdvancedHMC /Users/kcft114/.julia/packages/AdvancedHMC/YWXfk/src/hamiltonian.jl:36
┌ Warning: The current proposal will be rejected due to numerical error(s).
│   isfiniteθ = false
│   isfiniter = false
│   isfiniteℓπ = false
│   isfiniteℓκ = false
└ @ AdvancedHMC /Users/kcft114/.julia/packages/AdvancedHMC/YWXfk/src/hamiltonian.jl:36
┌ Warning: Incorrect ϵ = NaN; ϵ_previous = 0.07500000000000001 is used instead.
└ @ AdvancedHMC.Adaptation /Users/kcft114/.julia/packages/AdvancedHMC/YWXfk/src/adaptation/stepsize.jl:103
┌ Warning: The current proposal will be rejected due to numerical error(s).
│   isfiniteθ = true
│   isfiniter = true
│   isfiniteℓπ = false
│   isfiniteℓκ = true
└ @ AdvancedHMC /Users/kcft114/.julia/packages/AdvancedHMC/YWXfk/src/hamiltonian.jl:36
┌ Warning: The current proposal will be rejected due to numerical error(s).
│   isfiniteθ = false
│   isfiniter = false
│   isfiniteℓπ = false
│   isfiniteℓκ = false
└ @ AdvancedHMC /Users/kcft114/.julia/packages/AdvancedHMC/YWXfk/src/hamiltonian.jl:36
┌ Warning: Incorrect ϵ = NaN; ϵ_previous = 0.07500000000000001 is used instead.
└ @ AdvancedHMC.Adaptation /Users/kcft114/.julia/packages/AdvancedHMC/YWXfk/src/adaptation/stepsize.jl:103
┌ Warning: The current proposal will be rejected due to numerical error(s).
│   isfiniteθ = true
│   isfiniter = true
│   isfiniteℓπ = false
│   isfiniteℓκ = true
└ @ AdvancedHMC /Users/kcft114/.julia/packages/AdvancedHMC/YWXfk/src/hamiltonian.jl:36
┌ Warning: The current proposal will be rejected due to numerical error(s).
│   isfiniteθ = false
│   isfiniter = false
│   isfiniteℓπ = false
│   isfiniteℓκ = false
└ @ AdvancedHMC /Users/kcft114/.julia/packages/AdvancedHMC/YWXfk/src/hamiltonian.jl:36
┌ Warning: Incorrect ϵ = NaN; ϵ_previous = 0.07500000000000001 is used instead.
└ @ AdvancedHMC.Adaptation /Users/kcft114/.julia/packages/AdvancedHMC/YWXfk/src/adaptation/stepsize.jl:103
┌ Warning: The current proposal will be rejected due to numerical error(s).
│   isfiniteθ = true
│   isfiniter = true
│   isfiniteℓπ = false
│   isfiniteℓκ = true
└ @ AdvancedHMC /Users/kcft114/.julia/packages/AdvancedHMC/YWXfk/src/hamiltonian.jl:36
┌ Warning: The current proposal will be rejected due to numerical error(s).
│   isfiniteθ = false
│   isfiniter = false
│   isfiniteℓπ = false
│   isfiniteℓκ = false
└ @ AdvancedHMC /Users/kcft114/.julia/packages/AdvancedHMC/YWXfk/src/hamiltonian.jl:36
┌ Warning: Incorrect ϵ = NaN; ϵ_previous = 0.07500000000000001 is used instead.
└ @ AdvancedHMC.Adaptation /Users/kcft114/.julia/packages/AdvancedHMC/YWXfk/src/adaptation/stepsize.jl:103
┌ Warning: The current proposal will be rejected due to numerical error(s).
│   isfiniteθ = true
│   isfiniter = true
│   isfiniteℓπ = false
│   isfiniteℓκ = true
└ @ AdvancedHMC /Users/kcft114/.julia/packages/AdvancedHMC/YWXfk/src/hamiltonian.jl:36
┌ Warning: The current proposal will be rejected due to numerical error(s).
│   isfiniteθ = false
│   isfiniter = false
│   isfiniteℓπ = false
│   isfiniteℓκ = false
└ @ AdvancedHMC /Users/kcft114/.julia/packages/AdvancedHMC/YWXfk/src/hamiltonian.jl:36
┌ Warning: Incorrect ϵ = NaN; ϵ_previous = 0.07500000000000001 is used instead.
└ @ AdvancedHMC.Adaptation /Users/kcft114/.julia/packages/AdvancedHMC/YWXfk/src/adaptation/stepsize.jl:103
┌ Warning: The current proposal will be rejected due to numerical error(s).
│   isfiniteθ = true
│   isfiniter = true
│   isfiniteℓπ = false
│   isfiniteℓκ = true
└ @ AdvancedHMC /Users/kcft114/.julia/packages/AdvancedHMC/YWXfk/src/hamiltonian.jl:36
┌ Warning: The current proposal will be rejected due to numerical error(s).
│   isfiniteθ = false
│   isfiniter = false
│   isfiniteℓπ = false
│   isfiniteℓκ = false
└ @ AdvancedHMC /Users/kcft114/.julia/packages/AdvancedHMC/YWXfk/src/hamiltonian.jl:36
┌ Warning: Incorrect ϵ = NaN; ϵ_previous = 0.07500000000000001 is used instead.
└ @ AdvancedHMC.Adaptation /Users/kcft114/.julia/packages/AdvancedHMC/YWXfk/src/adaptation/stepsize.jl:103
┌ Warning: The current proposal will be rejected due to numerical error(s).
│   isfiniteθ = true
│   isfiniter = true
│   isfiniteℓπ = false
│   isfiniteℓκ = true
└ @ AdvancedHMC /Users/kcft114/.julia/packages/AdvancedHMC/YWXfk/src/hamiltonian.jl:36
┌ Warning: The current proposal will be rejected due to numerical error(s).
│   isfiniteθ = false
│   isfiniter = false
│   isfiniteℓπ = false
│   isfiniteℓκ = false
└ @ AdvancedHMC /Users/kcft114/.julia/packages/AdvancedHMC/YWXfk/src/hamiltonian.jl:36
┌ Warning: Incorrect ϵ = NaN; ϵ_previous = 0.07500000000000001 is used instead.
└ @ AdvancedHMC.Adaptation /Users/kcft114/.julia/packages/AdvancedHMC/YWXfk/src/adaptation/stepsize.jl:103
┌ Warning: The current proposal will be rejected due to numerical error(s).
│   isfiniteθ = true
│   isfiniter = true
│   isfiniteℓπ = false
│   isfiniteℓκ = true
└ @ AdvancedHMC /Users/kcft114/.julia/packages/AdvancedHMC/YWXfk/src/hamiltonian.jl:36
┌ Warning: The current proposal will be rejected due to numerical error(s).
│   isfiniteθ = false
│   isfiniter = false
│   isfiniteℓπ = false
│   isfiniteℓκ = false
└ @ AdvancedHMC /Users/kcft114/.julia/packages/AdvancedHMC/YWXfk/src/hamiltonian.jl:36
┌ Warning: Incorrect ϵ = NaN; ϵ_previous = 0.07500000000000001 is used instead.
└ @ AdvancedHMC.Adaptation /Users/kcft114/.julia/packages/AdvancedHMC/YWXfk/src/adaptation/stepsize.jl:103
┌ Warning: The current proposal will be rejected due to numerical error(s).
│   isfiniteθ = true
│   isfiniter = true
│   isfiniteℓπ = false
│   isfiniteℓκ = true
└ @ AdvancedHMC /Users/kcft114/.julia/packages/AdvancedHMC/YWXfk/src/hamiltonian.jl:36
┌ Warning: The current proposal will be rejected due to numerical error(s).
│   isfiniteθ = false
│   isfiniter = false
│   isfiniteℓπ = false
│   isfiniteℓκ = false
└ @ AdvancedHMC /Users/kcft114/.julia/packages/AdvancedHMC/YWXfk/src/hamiltonian.jl:36
┌ Warning: Incorrect ϵ = NaN; ϵ_previous = 0.07500000000000001 is used instead.
└ @ AdvancedHMC.Adaptation /Users/kcft114/.julia/packages/AdvancedHMC/YWXfk/src/adaptation/stepsize.jl:103
┌ Warning: The current proposal will be rejected due to numerical error(s).
│   isfiniteθ = true
│   isfiniter = true
│   isfiniteℓπ = false
│   isfiniteℓκ = true
└ @ AdvancedHMC /Users/kcft114/.julia/packages/AdvancedHMC/YWXfk/src/hamiltonian.jl:36
┌ Warning: The current proposal will be rejected due to numerical error(s).
│   isfiniteθ = false
│   isfiniter = false
│   isfiniteℓπ = false
│   isfiniteℓκ = false
└ @ AdvancedHMC /Users/kcft114/.julia/packages/AdvancedHMC/YWXfk/src/hamiltonian.jl:36
┌ Warning: Incorrect ϵ = NaN; ϵ_previous = 0.07500000000000001 is used instead.
└ @ AdvancedHMC.Adaptation /Users/kcft114/.julia/packages/AdvancedHMC/YWXfk/src/adaptation/stepsize.jl:103
┌ Warning: The current proposal will be rejected due to numerical error(s).
│   isfiniteθ = true
│   isfiniter = true
│   isfiniteℓπ = false
│   isfiniteℓκ = true
└ @ AdvancedHMC /Users/kcft114/.julia/packages/AdvancedHMC/YWXfk/src/hamiltonian.jl:36
┌ Warning: The current proposal will be rejected due to numerical error(s).
│   isfiniteθ = false
│   isfiniter = false
│   isfiniteℓπ = false
│   isfiniteℓκ = false
└ @ AdvancedHMC /Users/kcft114/.julia/packages/AdvancedHMC/YWXfk/src/hamiltonian.jl:36
┌ Warning: Incorrect ϵ = NaN; ϵ_previous = 0.07500000000000001 is used instead.
└ @ AdvancedHMC.Adaptation /Users/kcft114/.julia/packages/AdvancedHMC/YWXfk/src/adaptation/stepsize.jl:103
┌ Warning: The current proposal will be rejected due to numerical error(s).
│   isfiniteθ = true
│   isfiniter = true
│   isfiniteℓπ = false
│   isfiniteℓκ = true
└ @ AdvancedHMC /Users/kcft114/.julia/packages/AdvancedHMC/YWXfk/src/hamiltonian.jl:36
┌ Warning: The current proposal will be rejected due to numerical error(s).
│   isfiniteθ = false
│   isfiniter = false
│   isfiniteℓπ = false
│   isfiniteℓκ = false
└ @ AdvancedHMC /Users/kcft114/.julia/packages/AdvancedHMC/YWXfk/src/hamiltonian.jl:36
┌ Warning: Incorrect ϵ = NaN; ϵ_previous = 0.07500000000000001 is used instead.
└ @ AdvancedHMC.Adaptation /Users/kcft114/.julia/packages/AdvancedHMC/YWXfk/src/adaptation/stepsize.jl:103
┌ Warning: The current proposal will be rejected due to numerical error(s).
│   isfiniteθ = true
│   isfiniter = true
│   isfiniteℓπ = false
│   isfiniteℓκ = true
└ @ AdvancedHMC /Users/kcft114/.julia/packages/AdvancedHMC/YWXfk/src/hamiltonian.jl:36
┌ Warning: The current proposal will be rejected due to numerical error(s).
│   isfiniteθ = false
│   isfiniter = false
│   isfiniteℓπ = false
│   isfiniteℓκ = false
└ @ AdvancedHMC /Users/kcft114/.julia/packages/AdvancedHMC/YWXfk/src/hamiltonian.jl:36
┌ Warning: Incorrect ϵ = NaN; ϵ_previous = 0.07500000000000001 is used instead.
└ @ AdvancedHMC.Adaptation /Users/kcft114/.julia/packages/AdvancedHMC/YWXfk/src/adaptation/stepsize.jl:103
┌ Warning: The current proposal will be rejected due to numerical error(s).
│   isfiniteθ = true
│   isfiniter = true
│   isfiniteℓπ = false
│   isfiniteℓκ = true
└ @ AdvancedHMC /Users/kcft114/.julia/packages/AdvancedHMC/YWXfk/src/hamiltonian.jl:36
┌ Warning: The current proposal will be rejected due to numerical error(s).
│   isfiniteθ = false
│   isfiniter = false
│   isfiniteℓπ = false
│   isfiniteℓκ = false
└ @ AdvancedHMC /Users/kcft114/.julia/packages/AdvancedHMC/YWXfk/src/hamiltonian.jl:36
┌ Warning: Incorrect ϵ = NaN; ϵ_previous = 0.07500000000000001 is used instead.
└ @ AdvancedHMC.Adaptation /Users/kcft114/.julia/packages/AdvancedHMC/YWXfk/src/adaptation/stepsize.jl:103
┌ Warning: The current proposal will be rejected due to numerical error(s).
│   isfiniteθ = true
│   isfiniter = true
│   isfiniteℓπ = false
│   isfiniteℓκ = true
└ @ AdvancedHMC /Users/kcft114/.julia/packages/AdvancedHMC/YWXfk/src/hamiltonian.jl:36
┌ Warning: The current proposal will be rejected due to numerical error(s).
│   isfiniteθ = false
│   isfiniter = false
│   isfiniteℓπ = false
│   isfiniteℓκ = false
└ @ AdvancedHMC /Users/kcft114/.julia/packages/AdvancedHMC/YWXfk/src/hamiltonian.jl:36
┌ Warning: Incorrect ϵ = NaN; ϵ_previous = 0.07500000000000001 is used instead.
└ @ AdvancedHMC.Adaptation /Users/kcft114/.julia/packages/AdvancedHMC/YWXfk/src/adaptation/stepsize.jl:103
┌ Warning: The current proposal will be rejected due to numerical error(s).
│   isfiniteθ = true
│   isfiniter = true
│   isfiniteℓπ = false
│   isfiniteℓκ = true
└ @ AdvancedHMC /Users/kcft114/.julia/packages/AdvancedHMC/YWXfk/src/hamiltonian.jl:36
┌ Warning: The current proposal will be rejected due to numerical error(s).
│   isfiniteθ = false
│   isfiniter = false
│   isfiniteℓπ = false
│   isfiniteℓκ = false
└ @ AdvancedHMC /Users/kcft114/.julia/packages/AdvancedHMC/YWXfk/src/hamiltonian.jl:36
┌ Warning: Incorrect ϵ = NaN; ϵ_previous = 0.07500000000000001 is used instead.
└ @ AdvancedHMC.Adaptation /Users/kcft114/.julia/packages/AdvancedHMC/YWXfk/src/adaptation/stepsize.jl:103
┌ Warning: The current proposal will be rejected due to numerical error(s).
│   isfiniteθ = true
│   isfiniter = true
│   isfiniteℓπ = false
│   isfiniteℓκ = true
└ @ AdvancedHMC /Users/kcft114/.julia/packages/AdvancedHMC/YWXfk/src/hamiltonian.jl:36
┌ Warning: The current proposal will be rejected due to numerical error(s).
│   isfiniteθ = false
│   isfiniter = false
│   isfiniteℓπ = false
│   isfiniteℓκ = false
└ @ AdvancedHMC /Users/kcft114/.julia/packages/AdvancedHMC/YWXfk/src/hamiltonian.jl:36
┌ Warning: Incorrect ϵ = NaN; ϵ_previous = 0.07500000000000001 is used instead.
└ @ AdvancedHMC.Adaptation /Users/kcft114/.julia/packages/AdvancedHMC/YWXfk/src/adaptation/stepsize.jl:103
┌ Warning: The current proposal will be rejected due to numerical error(s).
│   isfiniteθ = true
│   isfiniter = true
│   isfiniteℓπ = false
│   isfiniteℓκ = true
└ @ AdvancedHMC /Users/kcft114/.julia/packages/AdvancedHMC/YWXfk/src/hamiltonian.jl:36
┌ Warning: The current proposal will be rejected due to numerical error(s).
│   isfiniteθ = false
│   isfiniter = false
│   isfiniteℓπ = false
│   isfiniteℓκ = false
└ @ AdvancedHMC /Users/kcft114/.julia/packages/AdvancedHMC/YWXfk/src/hamiltonian.jl:36
┌ Warning: Incorrect ϵ = NaN; ϵ_previous = 0.07500000000000001 is used instead.
└ @ AdvancedHMC.Adaptation /Users/kcft114/.julia/packages/AdvancedHMC/YWXfk/src/adaptation/stepsize.jl:103
┌ Warning: The current proposal will be rejected due to numerical error(s).
│   isfiniteθ = true
│   isfiniter = true
│   isfiniteℓπ = false
│   isfiniteℓκ = true
└ @ AdvancedHMC /Users/kcft114/.julia/packages/AdvancedHMC/YWXfk/src/hamiltonian.jl:36
┌ Warning: The current proposal will be rejected due to numerical error(s).
│   isfiniteθ = false
│   isfiniter = false
│   isfiniteℓπ = false
│   isfiniteℓκ = false
└ @ AdvancedHMC /Users/kcft114/.julia/packages/AdvancedHMC/YWXfk/src/hamiltonian.jl:36
┌ Warning: Incorrect ϵ = NaN; ϵ_previous = 0.07500000000000001 is used instead.
└ @ AdvancedHMC.Adaptation /Users/kcft114/.julia/packages/AdvancedHMC/YWXfk/src/adaptation/stepsize.jl:103
┌ Warning: The current proposal will be rejected due to numerical error(s).
│   isfiniteθ = true
│   isfiniter = true
│   isfiniteℓπ = false
│   isfiniteℓκ = true
└ @ AdvancedHMC /Users/kcft114/.julia/packages/AdvancedHMC/YWXfk/src/hamiltonian.jl:36
┌ Warning: The current proposal will be rejected due to numerical error(s).
│   isfiniteθ = false
│   isfiniter = false
│   isfiniteℓπ = false
│   isfiniteℓκ = false
└ @ AdvancedHMC /Users/kcft114/.julia/packages/AdvancedHMC/YWXfk/src/hamiltonian.jl:36
┌ Warning: Incorrect ϵ = NaN; ϵ_previous = 0.07500000000000001 is used instead.
└ @ AdvancedHMC.Adaptation /Users/kcft114/.julia/packages/AdvancedHMC/YWXfk/src/adaptation/stepsize.jl:103
┌ Warning: The current proposal will be rejected due to numerical error(s).
│   isfiniteθ = true
│   isfiniter = true
│   isfiniteℓπ = false
│   isfiniteℓκ = true
└ @ AdvancedHMC /Users/kcft114/.julia/packages/AdvancedHMC/YWXfk/src/hamiltonian.jl:36
┌ Warning: The current proposal will be rejected due to numerical error(s).
│   isfiniteθ = false
│   isfiniter = false
│   isfiniteℓπ = false
│   isfiniteℓκ = false
└ @ AdvancedHMC /Users/kcft114/.julia/packages/AdvancedHMC/YWXfk/src/hamiltonian.jl:36
┌ Warning: Incorrect ϵ = NaN; ϵ_previous = 0.07500000000000001 is used instead.
└ @ AdvancedHMC.Adaptation /Users/kcft114/.julia/packages/AdvancedHMC/YWXfk/src/adaptation/stepsize.jl:103
┌ Warning: The current proposal will be rejected due to numerical error(s).
│   isfiniteθ = true
│   isfiniter = true
│   isfiniteℓπ = false
│   isfiniteℓκ = true
└ @ AdvancedHMC /Users/kcft114/.julia/packages/AdvancedHMC/YWXfk/src/hamiltonian.jl:36
┌ Warning: The current proposal will be rejected due to numerical error(s).
│   isfiniteθ = false
│   isfiniter = false
│   isfiniteℓπ = false
│   isfiniteℓκ = false
└ @ AdvancedHMC /Users/kcft114/.julia/packages/AdvancedHMC/YWXfk/src/hamiltonian.jl:36
┌ Warning: Incorrect ϵ = NaN; ϵ_previous = 0.07500000000000001 is used instead.
└ @ AdvancedHMC.Adaptation /Users/kcft114/.julia/packages/AdvancedHMC/YWXfk/src/adaptation/stepsize.jl:103
┌ Warning: The current proposal will be rejected due to numerical error(s).
│   isfiniteθ = true
│   isfiniter = true
│   isfiniteℓπ = false
│   isfiniteℓκ = true
└ @ AdvancedHMC /Users/kcft114/.julia/packages/AdvancedHMC/YWXfk/src/hamiltonian.jl:36
┌ Warning: The current proposal will be rejected due to numerical error(s).
│   isfiniteθ = false
│   isfiniter = false
│   isfiniteℓπ = false
│   isfiniteℓκ = false
└ @ AdvancedHMC /Users/kcft114/.julia/packages/AdvancedHMC/YWXfk/src/hamiltonian.jl:36
┌ Warning: Incorrect ϵ = NaN; ϵ_previous = 0.07500000000000001 is used instead.
└ @ AdvancedHMC.Adaptation /Users/kcft114/.julia/packages/AdvancedHMC/YWXfk/src/adaptation/stepsize.jl:103
┌ Warning: The current proposal will be rejected due to numerical error(s).
│   isfiniteθ = true
│   isfiniter = true
│   isfiniteℓπ = false
│   isfiniteℓκ = true
└ @ AdvancedHMC /Users/kcft114/.julia/packages/AdvancedHMC/YWXfk/src/hamiltonian.jl:36
┌ Warning: The current proposal will be rejected due to numerical error(s).
│   isfiniteθ = false
│   isfiniter = false
│   isfiniteℓπ = false
│   isfiniteℓκ = false
└ @ AdvancedHMC /Users/kcft114/.julia/packages/AdvancedHMC/YWXfk/src/hamiltonian.jl:36
┌ Warning: Incorrect ϵ = NaN; ϵ_previous = 0.07500000000000001 is used instead.
└ @ AdvancedHMC.Adaptation /Users/kcft114/.julia/packages/AdvancedHMC/YWXfk/src/adaptation/stepsize.jl:103
┌ Warning: The current proposal will be rejected due to numerical error(s).
│   isfiniteθ = true
│   isfiniter = true
│   isfiniteℓπ = false
│   isfiniteℓκ = true
└ @ AdvancedHMC /Users/kcft114/.julia/packages/AdvancedHMC/YWXfk/src/hamiltonian.jl:36
┌ Warning: The current proposal will be rejected due to numerical error(s).
│   isfiniteθ = false
│   isfiniter = false
│   isfiniteℓπ = false
│   isfiniteℓκ = false
└ @ AdvancedHMC /Users/kcft114/.julia/packages/AdvancedHMC/YWXfk/src/hamiltonian.jl:36
┌ Warning: Incorrect ϵ = NaN; ϵ_previous = 0.07500000000000001 is used instead.
└ @ AdvancedHMC.Adaptation /Users/kcft114/.julia/packages/AdvancedHMC/YWXfk/src/adaptation/stepsize.jl:103
┌ Warning: The current proposal will be rejected due to numerical error(s).
│   isfiniteθ = true
│   isfiniter = true
│   isfiniteℓπ = false
│   isfiniteℓκ = true
└ @ AdvancedHMC /Users/kcft114/.julia/packages/AdvancedHMC/YWXfk/src/hamiltonian.jl:36
┌ Warning: The current proposal will be rejected due to numerical error(s).
│   isfiniteθ = false
│   isfiniter = false
│   isfiniteℓπ = false
│   isfiniteℓκ = false
└ @ AdvancedHMC /Users/kcft114/.julia/packages/AdvancedHMC/YWXfk/src/hamiltonian.jl:36
┌ Warning: Incorrect ϵ = NaN; ϵ_previous = 0.07500000000000001 is used instead.
└ @ AdvancedHMC.Adaptation /Users/kcft114/.julia/packages/AdvancedHMC/YWXfk/src/adaptation/stepsize.jl:103
┌ Warning: The current proposal will be rejected due to numerical error(s).
│   isfiniteθ = true
│   isfiniter = true
│   isfiniteℓπ = false
│   isfiniteℓκ = true
└ @ AdvancedHMC /Users/kcft114/.julia/packages/AdvancedHMC/YWXfk/src/hamiltonian.jl:36
┌ Warning: The current proposal will be rejected due to numerical error(s).
│   isfiniteθ = false
│   isfiniter = false
│   isfiniteℓπ = false
│   isfiniteℓκ = false
└ @ AdvancedHMC /Users/kcft114/.julia/packages/AdvancedHMC/YWXfk/src/hamiltonian.jl:36
┌ Warning: Incorrect ϵ = NaN; ϵ_previous = 0.07500000000000001 is used instead.
└ @ AdvancedHMC.Adaptation /Users/kcft114/.julia/packages/AdvancedHMC/YWXfk/src/adaptation/stepsize.jl:103
┌ Warning: The current proposal will be rejected due to numerical error(s).
│   isfiniteθ = true
│   isfiniter = true
│   isfiniteℓπ = false
│   isfiniteℓκ = true
└ @ AdvancedHMC /Users/kcft114/.julia/packages/AdvancedHMC/YWXfk/src/hamiltonian.jl:36
┌ Warning: The current proposal will be rejected due to numerical error(s).
│   isfiniteθ = false
│   isfiniter = false
│   isfiniteℓπ = false
│   isfiniteℓκ = false
└ @ AdvancedHMC /Users/kcft114/.julia/packages/AdvancedHMC/YWXfk/src/hamiltonian.jl:36
┌ Warning: Incorrect ϵ = NaN; ϵ_previous = 0.07500000000000001 is used instead.
└ @ AdvancedHMC.Adaptation /Users/kcft114/.julia/packages/AdvancedHMC/YWXfk/src/adaptation/stepsize.jl:103
┌ Warning: The current proposal will be rejected due to numerical error(s).
│   isfiniteθ = true
│   isfiniter = true
│   isfiniteℓπ = false
│   isfiniteℓκ = true
└ @ AdvancedHMC /Users/kcft114/.julia/packages/AdvancedHMC/YWXfk/src/hamiltonian.jl:36
┌ Warning: The current proposal will be rejected due to numerical error(s).
│   isfiniteθ = false
│   isfiniter = false
│   isfiniteℓπ = false
│   isfiniteℓκ = false
└ @ AdvancedHMC /Users/kcft114/.julia/packages/AdvancedHMC/YWXfk/src/hamiltonian.jl:36
┌ Warning: Incorrect ϵ = NaN; ϵ_previous = 0.07500000000000001 is used instead.
└ @ AdvancedHMC.Adaptation /Users/kcft114/.julia/packages/AdvancedHMC/YWXfk/src/adaptation/stepsize.jl:103
┌ Warning: The current proposal will be rejected due to numerical error(s).
│   isfiniteθ = true
│   isfiniter = true
│   isfiniteℓπ = false
│   isfiniteℓκ = true
└ @ AdvancedHMC /Users/kcft114/.julia/packages/AdvancedHMC/YWXfk/src/hamiltonian.jl:36
┌ Warning: The current proposal will be rejected due to numerical error(s).
│   isfiniteθ = false
│   isfiniter = false
│   isfiniteℓπ = false
│   isfiniteℓκ = false
└ @ AdvancedHMC /Users/kcft114/.julia/packages/AdvancedHMC/YWXfk/src/hamiltonian.jl:36
┌ Warning: Incorrect ϵ = NaN; ϵ_previous = 0.07500000000000001 is used instead.
└ @ AdvancedHMC.Adaptation /Users/kcft114/.julia/packages/AdvancedHMC/YWXfk/src/adaptation/stepsize.jl:103
┌ Warning: The current proposal will be rejected due to numerical error(s).
│   isfiniteθ = true
│   isfiniter = true
│   isfiniteℓπ = false
│   isfiniteℓκ = true
└ @ AdvancedHMC /Users/kcft114/.julia/packages/AdvancedHMC/YWXfk/src/hamiltonian.jl:36
┌ Warning: The current proposal will be rejected due to numerical error(s).
│   isfiniteθ = false
│   isfiniter = false
│   isfiniteℓπ = false
│   isfiniteℓκ = false
└ @ AdvancedHMC /Users/kcft114/.julia/packages/AdvancedHMC/YWXfk/src/hamiltonian.jl:36
┌ Warning: Incorrect ϵ = NaN; ϵ_previous = 0.07500000000000001 is used instead.
└ @ AdvancedHMC.Adaptation /Users/kcft114/.julia/packages/AdvancedHMC/YWXfk/src/adaptation/stepsize.jl:103
┌ Warning: The current proposal will be rejected due to numerical error(s).
│   isfiniteθ = true
│   isfiniter = true
│   isfiniteℓπ = false
│   isfiniteℓκ = true
└ @ AdvancedHMC /Users/kcft114/.julia/packages/AdvancedHMC/YWXfk/src/hamiltonian.jl:36
┌ Warning: The current proposal will be rejected due to numerical error(s).
│   isfiniteθ = false
│   isfiniter = false
│   isfiniteℓπ = false
│   isfiniteℓκ = false
└ @ AdvancedHMC /Users/kcft114/.julia/packages/AdvancedHMC/YWXfk/src/hamiltonian.jl:36
┌ Warning: Incorrect ϵ = NaN; ϵ_previous = 0.07500000000000001 is used instead.
└ @ AdvancedHMC.Adaptation /Users/kcft114/.julia/packages/AdvancedHMC/YWXfk/src/adaptation/stepsize.jl:103
┌ Warning: The current proposal will be rejected due to numerical error(s).
│   isfiniteθ = true
│   isfiniter = true
│   isfiniteℓπ = false
│   isfiniteℓκ = true
└ @ AdvancedHMC /Users/kcft114/.julia/packages/AdvancedHMC/YWXfk/src/hamiltonian.jl:36
┌ Warning: The current proposal will be rejected due to numerical error(s).
│   isfiniteθ = false
│   isfiniter = false
│   isfiniteℓπ = false
│   isfiniteℓκ = false
└ @ AdvancedHMC /Users/kcft114/.julia/packages/AdvancedHMC/YWXfk/src/hamiltonian.jl:36
┌ Warning: Incorrect ϵ = NaN; ϵ_previous = 0.07500000000000001 is used instead.
└ @ AdvancedHMC.Adaptation /Users/kcft114/.julia/packages/AdvancedHMC/YWXfk/src/adaptation/stepsize.jl:103
┌ Warning: The current proposal will be rejected due to numerical error(s).
│   isfiniteθ = true
│   isfiniter = true
│   isfiniteℓπ = false
│   isfiniteℓκ = true
└ @ AdvancedHMC /Users/kcft114/.julia/packages/AdvancedHMC/YWXfk/src/hamiltonian.jl:36
┌ Warning: The current proposal will be rejected due to numerical error(s).
│   isfiniteθ = false
│   isfiniter = false
│   isfiniteℓπ = false
│   isfiniteℓκ = false
└ @ AdvancedHMC /Users/kcft114/.julia/packages/AdvancedHMC/YWXfk/src/hamiltonian.jl:36
┌ Warning: Incorrect ϵ = NaN; ϵ_previous = 0.07500000000000001 is used instead.
└ @ AdvancedHMC.Adaptation /Users/kcft114/.julia/packages/AdvancedHMC/YWXfk/src/adaptation/stepsize.jl:103
┌ Warning: The current proposal will be rejected due to numerical error(s).
│   isfiniteθ = true
│   isfiniter = true
│   isfiniteℓπ = false
│   isfiniteℓκ = true
└ @ AdvancedHMC /Users/kcft114/.julia/packages/AdvancedHMC/YWXfk/src/hamiltonian.jl:36
┌ Warning: The current proposal will be rejected due to numerical error(s).
│   isfiniteθ = false
│   isfiniter = false
│   isfiniteℓπ = false
│   isfiniteℓκ = false
└ @ AdvancedHMC /Users/kcft114/.julia/packages/AdvancedHMC/YWXfk/src/hamiltonian.jl:36
┌ Warning: Incorrect ϵ = NaN; ϵ_previous = 0.07500000000000001 is used instead.
└ @ AdvancedHMC.Adaptation /Users/kcft114/.julia/packages/AdvancedHMC/YWXfk/src/adaptation/stepsize.jl:103
┌ Warning: The current proposal will be rejected due to numerical error(s).
│   isfiniteθ = true
│   isfiniter = true
│   isfiniteℓπ = false
│   isfiniteℓκ = true
└ @ AdvancedHMC /Users/kcft114/.julia/packages/AdvancedHMC/YWXfk/src/hamiltonian.jl:36
┌ Warning: The current proposal will be rejected due to numerical error(s).
│   isfiniteθ = false
│   isfiniter = false
│   isfiniteℓπ = false
│   isfiniteℓκ = false
└ @ AdvancedHMC /Users/kcft114/.julia/packages/AdvancedHMC/YWXfk/src/hamiltonian.jl:36
┌ Warning: Incorrect ϵ = NaN; ϵ_previous = 0.07500000000000001 is used instead.
└ @ AdvancedHMC.Adaptation /Users/kcft114/.julia/packages/AdvancedHMC/YWXfk/src/adaptation/stepsize.jl:103
┌ Warning: The current proposal will be rejected due to numerical error(s).
│   isfiniteθ = true
│   isfiniter = true
│   isfiniteℓπ = false
│   isfiniteℓκ = true
└ @ AdvancedHMC /Users/kcft114/.julia/packages/AdvancedHMC/YWXfk/src/hamiltonian.jl:36
┌ Warning: The current proposal will be rejected due to numerical error(s).
│   isfiniteθ = false
│   isfiniter = false
│   isfiniteℓπ = false
│   isfiniteℓκ = false
└ @ AdvancedHMC /Users/kcft114/.julia/packages/AdvancedHMC/YWXfk/src/hamiltonian.jl:36
┌ Warning: Incorrect ϵ = NaN; ϵ_previous = 0.07500000000000001 is used instead.
└ @ AdvancedHMC.Adaptation /Users/kcft114/.julia/packages/AdvancedHMC/YWXfk/src/adaptation/stepsize.jl:103
┌ Warning: The current proposal will be rejected due to numerical error(s).
│   isfiniteθ = true
│   isfiniter = true
│   isfiniteℓπ = false
│   isfiniteℓκ = true
└ @ AdvancedHMC /Users/kcft114/.julia/packages/AdvancedHMC/YWXfk/src/hamiltonian.jl:36
┌ Warning: The current proposal will be rejected due to numerical error(s).
│   isfiniteθ = false
│   isfiniter = false
│   isfiniteℓπ = false
│   isfiniteℓκ = false
└ @ AdvancedHMC /Users/kcft114/.julia/packages/AdvancedHMC/YWXfk/src/hamiltonian.jl:36
┌ Warning: Incorrect ϵ = NaN; ϵ_previous = 0.07500000000000001 is used instead.
└ @ AdvancedHMC.Adaptation /Users/kcft114/.julia/packages/AdvancedHMC/YWXfk/src/adaptation/stepsize.jl:103
┌ Warning: The current proposal will be rejected due to numerical error(s).
│   isfiniteθ = true
│   isfiniter = true
│   isfiniteℓπ = false
│   isfiniteℓκ = true
└ @ AdvancedHMC /Users/kcft114/.julia/packages/AdvancedHMC/YWXfk/src/hamiltonian.jl:36
┌ Warning: The current proposal will be rejected due to numerical error(s).
│   isfiniteθ = false
│   isfiniter = false
│   isfiniteℓπ = false
│   isfiniteℓκ = false
└ @ AdvancedHMC /Users/kcft114/.julia/packages/AdvancedHMC/YWXfk/src/hamiltonian.jl:36
┌ Warning: Incorrect ϵ = NaN; ϵ_previous = 0.07500000000000001 is used instead.
└ @ AdvancedHMC.Adaptation /Users/kcft114/.julia/packages/AdvancedHMC/YWXfk/src/adaptation/stepsize.jl:103
┌ Warning: The current proposal will be rejected due to numerical error(s).
│   isfiniteθ = true
│   isfiniter = true
│   isfiniteℓπ = false
│   isfiniteℓκ = true
└ @ AdvancedHMC /Users/kcft114/.julia/packages/AdvancedHMC/YWXfk/src/hamiltonian.jl:36
┌ Warning: The current proposal will be rejected due to numerical error(s).
│   isfiniteθ = false
│   isfiniter = false
│   isfiniteℓπ = false
│   isfiniteℓκ = false
└ @ AdvancedHMC /Users/kcft114/.julia/packages/AdvancedHMC/YWXfk/src/hamiltonian.jl:36
┌ Warning: Incorrect ϵ = NaN; ϵ_previous = 0.07500000000000001 is used instead.
└ @ AdvancedHMC.Adaptation /Users/kcft114/.julia/packages/AdvancedHMC/YWXfk/src/adaptation/stepsize.jl:103
┌ Warning: The current proposal will be rejected due to numerical error(s).
│   isfiniteθ = true
│   isfiniter = true
│   isfiniteℓπ = false
│   isfiniteℓκ = true
└ @ AdvancedHMC /Users/kcft114/.julia/packages/AdvancedHMC/YWXfk/src/hamiltonian.jl:36
┌ Warning: The current proposal will be rejected due to numerical error(s).
│   isfiniteθ = false
│   isfiniter = false
│   isfiniteℓπ = false
│   isfiniteℓκ = false
└ @ AdvancedHMC /Users/kcft114/.julia/packages/AdvancedHMC/YWXfk/src/hamiltonian.jl:36
┌ Warning: Incorrect ϵ = NaN; ϵ_previous = 0.07500000000000001 is used instead.
└ @ AdvancedHMC.Adaptation /Users/kcft114/.julia/packages/AdvancedHMC/YWXfk/src/adaptation/stepsize.jl:103
┌ Warning: The current proposal will be rejected due to numerical error(s).
│   isfiniteθ = true
│   isfiniter = true
│   isfiniteℓπ = false
│   isfiniteℓκ = true
└ @ AdvancedHMC /Users/kcft114/.julia/packages/AdvancedHMC/YWXfk/src/hamiltonian.jl:36
┌ Warning: The current proposal will be rejected due to numerical error(s).
│   isfiniteθ = false
│   isfiniter = false
│   isfiniteℓπ = false
│   isfiniteℓκ = false
└ @ AdvancedHMC /Users/kcft114/.julia/packages/AdvancedHMC/YWXfk/src/hamiltonian.jl:36
┌ Warning: Incorrect ϵ = NaN; ϵ_previous = 0.07500000000000001 is used instead.
└ @ AdvancedHMC.Adaptation /Users/kcft114/.julia/packages/AdvancedHMC/YWXfk/src/adaptation/stepsize.jl:103
┌ Warning: The current proposal will be rejected due to numerical error(s).
│   isfiniteθ = true
│   isfiniter = true
│   isfiniteℓπ = false
│   isfiniteℓκ = true
└ @ AdvancedHMC /Users/kcft114/.julia/packages/AdvancedHMC/YWXfk/src/hamiltonian.jl:36
┌ Warning: The current proposal will be rejected due to numerical error(s).
│   isfiniteθ = false
│   isfiniter = false
│   isfiniteℓπ = false
│   isfiniteℓκ = false
└ @ AdvancedHMC /Users/kcft114/.julia/packages/AdvancedHMC/YWXfk/src/hamiltonian.jl:36
┌ Warning: Incorrect ϵ = NaN; ϵ_previous = 0.07500000000000001 is used instead.
└ @ AdvancedHMC.Adaptation /Users/kcft114/.julia/packages/AdvancedHMC/YWXfk/src/adaptation/stepsize.jl:103
┌ Warning: The current proposal will be rejected due to numerical error(s).
│   isfiniteθ = true
│   isfiniter = true
│   isfiniteℓπ = false
│   isfiniteℓκ = true
└ @ AdvancedHMC /Users/kcft114/.julia/packages/AdvancedHMC/YWXfk/src/hamiltonian.jl:36
┌ Warning: The current proposal will be rejected due to numerical error(s).
│   isfiniteθ = false
│   isfiniter = false
│   isfiniteℓπ = false
│   isfiniteℓκ = false
└ @ AdvancedHMC /Users/kcft114/.julia/packages/AdvancedHMC/YWXfk/src/hamiltonian.jl:36
┌ Warning: Incorrect ϵ = NaN; ϵ_previous = 0.07500000000000001 is used instead.
└ @ AdvancedHMC.Adaptation /Users/kcft114/.julia/packages/AdvancedHMC/YWXfk/src/adaptation/stepsize.jl:103
┌ Warning: The current proposal will be rejected due to numerical error(s).
│   isfiniteθ = true
│   isfiniter = true
│   isfiniteℓπ = false
│   isfiniteℓκ = true
└ @ AdvancedHMC /Users/kcft114/.julia/packages/AdvancedHMC/YWXfk/src/hamiltonian.jl:36
┌ Warning: The current proposal will be rejected due to numerical error(s).
│   isfiniteθ = false
│   isfiniter = false
│   isfiniteℓπ = false
│   isfiniteℓκ = false
└ @ AdvancedHMC /Users/kcft114/.julia/packages/AdvancedHMC/YWXfk/src/hamiltonian.jl:36
┌ Warning: Incorrect ϵ = NaN; ϵ_previous = 0.07500000000000001 is used instead.
└ @ AdvancedHMC.Adaptation /Users/kcft114/.julia/packages/AdvancedHMC/YWXfk/src/adaptation/stepsize.jl:103
┌ Warning: The current proposal will be rejected due to numerical error(s).
│   isfiniteθ = true
│   isfiniter = true
│   isfiniteℓπ = false
│   isfiniteℓκ = true
└ @ AdvancedHMC /Users/kcft114/.julia/packages/AdvancedHMC/YWXfk/src/hamiltonian.jl:36
┌ Warning: The current proposal will be rejected due to numerical error(s).
│   isfiniteθ = false
│   isfiniter = false
│   isfiniteℓπ = false
│   isfiniteℓκ = false
└ @ AdvancedHMC /Users/kcft114/.julia/packages/AdvancedHMC/YWXfk/src/hamiltonian.jl:36
┌ Warning: Incorrect ϵ = NaN; ϵ_previous = 0.07500000000000001 is used instead.
└ @ AdvancedHMC.Adaptation /Users/kcft114/.julia/packages/AdvancedHMC/YWXfk/src/adaptation/stepsize.jl:103
┌ Warning: The current proposal will be rejected due to numerical error(s).
│   isfiniteθ = true
│   isfiniter = true
│   isfiniteℓπ = false
│   isfiniteℓκ = true
└ @ AdvancedHMC /Users/kcft114/.julia/packages/AdvancedHMC/YWXfk/src/hamiltonian.jl:36
┌ Warning: The current proposal will be rejected due to numerical error(s).
│   isfiniteθ = false
│   isfiniter = false
│   isfiniteℓπ = false
│   isfiniteℓκ = false
└ @ AdvancedHMC /Users/kcft114/.julia/packages/AdvancedHMC/YWXfk/src/hamiltonian.jl:36
┌ Warning: Incorrect ϵ = NaN; ϵ_previous = 0.07500000000000001 is used instead.
└ @ AdvancedHMC.Adaptation /Users/kcft114/.julia/packages/AdvancedHMC/YWXfk/src/adaptation/stepsize.jl:103
┌ Warning: The current proposal will be rejected due to numerical error(s).
│   isfiniteθ = true
│   isfiniter = true
│   isfiniteℓπ = false
│   isfiniteℓκ = true
└ @ AdvancedHMC /Users/kcft114/.julia/packages/AdvancedHMC/YWXfk/src/hamiltonian.jl:36
┌ Warning: The current proposal will be rejected due to numerical error(s).
│   isfiniteθ = false
│   isfiniter = false
│   isfiniteℓπ = false
│   isfiniteℓκ = false
└ @ AdvancedHMC /Users/kcft114/.julia/packages/AdvancedHMC/YWXfk/src/hamiltonian.jl:36
┌ Warning: Incorrect ϵ = NaN; ϵ_previous = 0.07500000000000001 is used instead.
└ @ AdvancedHMC.Adaptation /Users/kcft114/.julia/packages/AdvancedHMC/YWXfk/src/adaptation/stepsize.jl:103
┌ Warning: The current proposal will be rejected due to numerical error(s).
│   isfiniteθ = true
│   isfiniter = true
│   isfiniteℓπ = false
│   isfiniteℓκ = true
└ @ AdvancedHMC /Users/kcft114/.julia/packages/AdvancedHMC/YWXfk/src/hamiltonian.jl:36
┌ Warning: The current proposal will be rejected due to numerical error(s).
│   isfiniteθ = false
│   isfiniter = false
│   isfiniteℓπ = false
│   isfiniteℓκ = false
└ @ AdvancedHMC /Users/kcft114/.julia/packages/AdvancedHMC/YWXfk/src/hamiltonian.jl:36
┌ Warning: Incorrect ϵ = NaN; ϵ_previous = 0.07500000000000001 is used instead.
└ @ AdvancedHMC.Adaptation /Users/kcft114/.julia/packages/AdvancedHMC/YWXfk/src/adaptation/stepsize.jl:103
┌ Warning: The current proposal will be rejected due to numerical error(s).
│   isfiniteθ = true
│   isfiniter = true
│   isfiniteℓπ = false
│   isfiniteℓκ = true
└ @ AdvancedHMC /Users/kcft114/.julia/packages/AdvancedHMC/YWXfk/src/hamiltonian.jl:36
┌ Warning: The current proposal will be rejected due to numerical error(s).
│   isfiniteθ = false
│   isfiniter = false
│   isfiniteℓπ = false
│   isfiniteℓκ = false
└ @ AdvancedHMC /Users/kcft114/.julia/packages/AdvancedHMC/YWXfk/src/hamiltonian.jl:36
┌ Warning: Incorrect ϵ = NaN; ϵ_previous = 0.07500000000000001 is used instead.
└ @ AdvancedHMC.Adaptation /Users/kcft114/.julia/packages/AdvancedHMC/YWXfk/src/adaptation/stepsize.jl:103
┌ Warning: The current proposal will be rejected due to numerical error(s).
│   isfiniteθ = true
│   isfiniter = true
│   isfiniteℓπ = false
│   isfiniteℓκ = true
└ @ AdvancedHMC /Users/kcft114/.julia/packages/AdvancedHMC/YWXfk/src/hamiltonian.jl:36
┌ Warning: The current proposal will be rejected due to numerical error(s).
│   isfiniteθ = false
│   isfiniter = false
│   isfiniteℓπ = false
│   isfiniteℓκ = false
└ @ AdvancedHMC /Users/kcft114/.julia/packages/AdvancedHMC/YWXfk/src/hamiltonian.jl:36
┌ Warning: Incorrect ϵ = NaN; ϵ_previous = 0.07500000000000001 is used instead.
└ @ AdvancedHMC.Adaptation /Users/kcft114/.julia/packages/AdvancedHMC/YWXfk/src/adaptation/stepsize.jl:103
┌ Warning: The current proposal will be rejected due to numerical error(s).
│   isfiniteθ = true
│   isfiniter = true
│   isfiniteℓπ = false
│   isfiniteℓκ = true
└ @ AdvancedHMC /Users/kcft114/.julia/packages/AdvancedHMC/YWXfk/src/hamiltonian.jl:36
┌ Warning: The current proposal will be rejected due to numerical error(s).
│   isfiniteθ = false
│   isfiniter = false
│   isfiniteℓπ = false
│   isfiniteℓκ = false
└ @ AdvancedHMC /Users/kcft114/.julia/packages/AdvancedHMC/YWXfk/src/hamiltonian.jl:36
┌ Warning: Incorrect ϵ = NaN; ϵ_previous = 0.07500000000000001 is used instead.
└ @ AdvancedHMC.Adaptation /Users/kcft114/.julia/packages/AdvancedHMC/YWXfk/src/adaptation/stepsize.jl:103
┌ Warning: The current proposal will be rejected due to numerical error(s).
│   isfiniteθ = true
│   isfiniter = true
│   isfiniteℓπ = false
│   isfiniteℓκ = true
└ @ AdvancedHMC /Users/kcft114/.julia/packages/AdvancedHMC/YWXfk/src/hamiltonian.jl:36
┌ Warning: The current proposal will be rejected due to numerical error(s).
│   isfiniteθ = false
│   isfiniter = false
│   isfiniteℓπ = false
│   isfiniteℓκ = false
└ @ AdvancedHMC /Users/kcft114/.julia/packages/AdvancedHMC/YWXfk/src/hamiltonian.jl:36
┌ Warning: Incorrect ϵ = NaN; ϵ_previous = 0.07500000000000001 is used instead.
└ @ AdvancedHMC.Adaptation /Users/kcft114/.julia/packages/AdvancedHMC/YWXfk/src/adaptation/stepsize.jl:103
┌ Warning: The current proposal will be rejected due to numerical error(s).
│   isfiniteθ = true
│   isfiniter = true
│   isfiniteℓπ = false
│   isfiniteℓκ = true
└ @ AdvancedHMC /Users/kcft114/.julia/packages/AdvancedHMC/YWXfk/src/hamiltonian.jl:36
┌ Warning: The current proposal will be rejected due to numerical error(s).
│   isfiniteθ = false
│   isfiniter = false
│   isfiniteℓπ = false
│   isfiniteℓκ = false
└ @ AdvancedHMC /Users/kcft114/.julia/packages/AdvancedHMC/YWXfk/src/hamiltonian.jl:36
┌ Warning: Incorrect ϵ = NaN; ϵ_previous = 0.07500000000000001 is used instead.
└ @ AdvancedHMC.Adaptation /Users/kcft114/.julia/packages/AdvancedHMC/YWXfk/src/adaptation/stepsize.jl:103
┌ Warning: The current proposal will be rejected due to numerical error(s).
│   isfiniteθ = true
│   isfiniter = true
│   isfiniteℓπ = false
│   isfiniteℓκ = true
└ @ AdvancedHMC /Users/kcft114/.julia/packages/AdvancedHMC/YWXfk/src/hamiltonian.jl:36
┌ Warning: The current proposal will be rejected due to numerical error(s).
│   isfiniteθ = false
│   isfiniter = false
│   isfiniteℓπ = false
│   isfiniteℓκ = false
└ @ AdvancedHMC /Users/kcft114/.julia/packages/AdvancedHMC/YWXfk/src/hamiltonian.jl:36
┌ Warning: Incorrect ϵ = NaN; ϵ_previous = 0.07500000000000001 is used instead.
└ @ AdvancedHMC.Adaptation /Users/kcft114/.julia/packages/AdvancedHMC/YWXfk/src/adaptation/stepsize.jl:103
┌ Warning: The current proposal will be rejected due to numerical error(s).
│   isfiniteθ = true
│   isfiniter = true
│   isfiniteℓπ = false
│   isfiniteℓκ = true
└ @ AdvancedHMC /Users/kcft114/.julia/packages/AdvancedHMC/YWXfk/src/hamiltonian.jl:36
┌ Warning: The current proposal will be rejected due to numerical error(s).
│   isfiniteθ = false
│   isfiniter = false
│   isfiniteℓπ = false
│   isfiniteℓκ = false
└ @ AdvancedHMC /Users/kcft114/.julia/packages/AdvancedHMC/YWXfk/src/hamiltonian.jl:36
┌ Warning: Incorrect ϵ = NaN; ϵ_previous = 0.07500000000000001 is used instead.
└ @ AdvancedHMC.Adaptation /Users/kcft114/.julia/packages/AdvancedHMC/YWXfk/src/adaptation/stepsize.jl:103
┌ Warning: The current proposal will be rejected due to numerical error(s).
│   isfiniteθ = true
│   isfiniter = true
│   isfiniteℓπ = false
│   isfiniteℓκ = true
└ @ AdvancedHMC /Users/kcft114/.julia/packages/AdvancedHMC/YWXfk/src/hamiltonian.jl:36
┌ Warning: The current proposal will be rejected due to numerical error(s).
│   isfiniteθ = false
│   isfiniter = false
│   isfiniteℓπ = false
│   isfiniteℓκ = false
└ @ AdvancedHMC /Users/kcft114/.julia/packages/AdvancedHMC/YWXfk/src/hamiltonian.jl:36
┌ Warning: Incorrect ϵ = NaN; ϵ_previous = 0.07500000000000001 is used instead.
└ @ AdvancedHMC.Adaptation /Users/kcft114/.julia/packages/AdvancedHMC/YWXfk/src/adaptation/stepsize.jl:103
┌ Warning: The current proposal will be rejected due to numerical error(s).
│   isfiniteθ = true
│   isfiniter = true
│   isfiniteℓπ = false
│   isfiniteℓκ = true
└ @ AdvancedHMC /Users/kcft114/.julia/packages/AdvancedHMC/YWXfk/src/hamiltonian.jl:36
┌ Warning: The current proposal will be rejected due to numerical error(s).
│   isfiniteθ = false
│   isfiniter = false
│   isfiniteℓπ = false
│   isfiniteℓκ = false
└ @ AdvancedHMC /Users/kcft114/.julia/packages/AdvancedHMC/YWXfk/src/hamiltonian.jl:36
┌ Warning: Incorrect ϵ = NaN; ϵ_previous = 0.07500000000000001 is used instead.
└ @ AdvancedHMC.Adaptation /Users/kcft114/.julia/packages/AdvancedHMC/YWXfk/src/adaptation/stepsize.jl:103
┌ Warning: The current proposal will be rejected due to numerical error(s).
│   isfiniteθ = true
│   isfiniter = true
│   isfiniteℓπ = false
│   isfiniteℓκ = true
└ @ AdvancedHMC /Users/kcft114/.julia/packages/AdvancedHMC/YWXfk/src/hamiltonian.jl:36
┌ Warning: The current proposal will be rejected due to numerical error(s).
│   isfiniteθ = false
│   isfiniter = false
│   isfiniteℓπ = false
│   isfiniteℓκ = false
└ @ AdvancedHMC /Users/kcft114/.julia/packages/AdvancedHMC/YWXfk/src/hamiltonian.jl:36
┌ Warning: Incorrect ϵ = NaN; ϵ_previous = 0.07500000000000001 is used instead.
└ @ AdvancedHMC.Adaptation /Users/kcft114/.julia/packages/AdvancedHMC/YWXfk/src/adaptation/stepsize.jl:103
┌ Warning: The current proposal will be rejected due to numerical error(s).
│   isfiniteθ = true
│   isfiniter = true
│   isfiniteℓπ = false
│   isfiniteℓκ = true
└ @ AdvancedHMC /Users/kcft114/.julia/packages/AdvancedHMC/YWXfk/src/hamiltonian.jl:36
┌ Warning: The current proposal will be rejected due to numerical error(s).
│   isfiniteθ = false
│   isfiniter = false
│   isfiniteℓπ = false
│   isfiniteℓκ = false
└ @ AdvancedHMC /Users/kcft114/.julia/packages/AdvancedHMC/YWXfk/src/hamiltonian.jl:36
┌ Warning: Incorrect ϵ = NaN; ϵ_previous = 0.07500000000000001 is used instead.
└ @ AdvancedHMC.Adaptation /Users/kcft114/.julia/packages/AdvancedHMC/YWXfk/src/adaptation/stepsize.jl:103
┌ Warning: The current proposal will be rejected due to numerical error(s).
│   isfiniteθ = true
│   isfiniter = true
│   isfiniteℓπ = false
│   isfiniteℓκ = true
└ @ AdvancedHMC /Users/kcft114/.julia/packages/AdvancedHMC/YWXfk/src/hamiltonian.jl:36
┌ Warning: The current proposal will be rejected due to numerical error(s).
│   isfiniteθ = false
│   isfiniter = false
│   isfiniteℓπ = false
│   isfiniteℓκ = false
└ @ AdvancedHMC /Users/kcft114/.julia/packages/AdvancedHMC/YWXfk/src/hamiltonian.jl:36
┌ Warning: Incorrect ϵ = NaN; ϵ_previous = 0.07500000000000001 is used instead.
└ @ AdvancedHMC.Adaptation /Users/kcft114/.julia/packages/AdvancedHMC/YWXfk/src/adaptation/stepsize.jl:103
┌ Warning: The current proposal will be rejected due to numerical error(s).
│   isfiniteθ = true
│   isfiniter = true
│   isfiniteℓπ = false
│   isfiniteℓκ = true
└ @ AdvancedHMC /Users/kcft114/.julia/packages/AdvancedHMC/YWXfk/src/hamiltonian.jl:36
┌ Warning: The current proposal will be rejected due to numerical error(s).
│   isfiniteθ = false
│   isfiniter = false
│   isfiniteℓπ = false
│   isfiniteℓκ = false
└ @ AdvancedHMC /Users/kcft114/.julia/packages/AdvancedHMC/YWXfk/src/hamiltonian.jl:36
┌ Warning: Incorrect ϵ = NaN; ϵ_previous = 0.07500000000000001 is used instead.
└ @ AdvancedHMC.Adaptation /Users/kcft114/.julia/packages/AdvancedHMC/YWXfk/src/adaptation/stepsize.jl:103
┌ Warning: The current proposal will be rejected due to numerical error(s).
│   isfiniteθ = true
│   isfiniter = true
│   isfiniteℓπ = false
│   isfiniteℓκ = true
└ @ AdvancedHMC /Users/kcft114/.julia/packages/AdvancedHMC/YWXfk/src/hamiltonian.jl:36
┌ Warning: The current proposal will be rejected due to numerical error(s).
│   isfiniteθ = false
│   isfiniter = false
│   isfiniteℓπ = false
│   isfiniteℓκ = false
└ @ AdvancedHMC /Users/kcft114/.julia/packages/AdvancedHMC/YWXfk/src/hamiltonian.jl:36
┌ Warning: Incorrect ϵ = NaN; ϵ_previous = 0.07500000000000001 is used instead.
└ @ AdvancedHMC.Adaptation /Users/kcft114/.julia/packages/AdvancedHMC/YWXfk/src/adaptation/stepsize.jl:103
┌ Warning: The current proposal will be rejected due to numerical error(s).
│   isfiniteθ = true
│   isfiniter = true
│   isfiniteℓπ = false
│   isfiniteℓκ = true
└ @ AdvancedHMC /Users/kcft114/.julia/packages/AdvancedHMC/YWXfk/src/hamiltonian.jl:36
┌ Warning: The current proposal will be rejected due to numerical error(s).
│   isfiniteθ = false
│   isfiniter = false
│   isfiniteℓπ = false
│   isfiniteℓκ = false
└ @ AdvancedHMC /Users/kcft114/.julia/packages/AdvancedHMC/YWXfk/src/hamiltonian.jl:36
┌ Warning: Incorrect ϵ = NaN; ϵ_previous = 0.07500000000000001 is used instead.
└ @ AdvancedHMC.Adaptation /Users/kcft114/.julia/packages/AdvancedHMC/YWXfk/src/adaptation/stepsize.jl:103
┌ Warning: The current proposal will be rejected due to numerical error(s).
│   isfiniteθ = true
│   isfiniter = true
│   isfiniteℓπ = false
│   isfiniteℓκ = true
└ @ AdvancedHMC /Users/kcft114/.julia/packages/AdvancedHMC/YWXfk/src/hamiltonian.jl:36
┌ Warning: The current proposal will be rejected due to numerical error(s).
│   isfiniteθ = false
│   isfiniter = false
│   isfiniteℓπ = false
│   isfiniteℓκ = false
└ @ AdvancedHMC /Users/kcft114/.julia/packages/AdvancedHMC/YWXfk/src/hamiltonian.jl:36
┌ Warning: Incorrect ϵ = NaN; ϵ_previous = 0.07500000000000001 is used instead.
└ @ AdvancedHMC.Adaptation /Users/kcft114/.julia/packages/AdvancedHMC/YWXfk/src/adaptation/stepsize.jl:103
┌ Warning: The current proposal will be rejected due to numerical error(s).
│   isfiniteθ = true
│   isfiniter = true
│   isfiniteℓπ = false
│   isfiniteℓκ = true
└ @ AdvancedHMC /Users/kcft114/.julia/packages/AdvancedHMC/YWXfk/src/hamiltonian.jl:36
┌ Warning: The current proposal will be rejected due to numerical error(s).
│   isfiniteθ = false
│   isfiniter = false
│   isfiniteℓπ = false
│   isfiniteℓκ = false
└ @ AdvancedHMC /Users/kcft114/.julia/packages/AdvancedHMC/YWXfk/src/hamiltonian.jl:36
┌ Warning: Incorrect ϵ = NaN; ϵ_previous = 0.07500000000000001 is used instead.
└ @ AdvancedHMC.Adaptation /Users/kcft114/.julia/packages/AdvancedHMC/YWXfk/src/adaptation/stepsize.jl:103
┌ Warning: The current proposal will be rejected due to numerical error(s).
│   isfiniteθ = true
│   isfiniter = true
│   isfiniteℓπ = false
│   isfiniteℓκ = true
└ @ AdvancedHMC /Users/kcft114/.julia/packages/AdvancedHMC/YWXfk/src/hamiltonian.jl:36
┌ Warning: The current proposal will be rejected due to numerical error(s).
│   isfiniteθ = false
│   isfiniter = false
│   isfiniteℓπ = false
│   isfiniteℓκ = false
└ @ AdvancedHMC /Users/kcft114/.julia/packages/AdvancedHMC/YWXfk/src/hamiltonian.jl:36
┌ Warning: Incorrect ϵ = NaN; ϵ_previous = 0.07500000000000001 is used instead.
└ @ AdvancedHMC.Adaptation /Users/kcft114/.julia/packages/AdvancedHMC/YWXfk/src/adaptation/stepsize.jl:103
┌ Warning: The current proposal will be rejected due to numerical error(s).
│   isfiniteθ = true
│   isfiniter = true
│   isfiniteℓπ = false
│   isfiniteℓκ = true
└ @ AdvancedHMC /Users/kcft114/.julia/packages/AdvancedHMC/YWXfk/src/hamiltonian.jl:36
┌ Warning: The current proposal will be rejected due to numerical error(s).
│   isfiniteθ = false
│   isfiniter = false
│   isfiniteℓπ = false
│   isfiniteℓκ = false
└ @ AdvancedHMC /Users/kcft114/.julia/packages/AdvancedHMC/YWXfk/src/hamiltonian.jl:36
┌ Warning: Incorrect ϵ = NaN; ϵ_previous = 0.07500000000000001 is used instead.
└ @ AdvancedHMC.Adaptation /Users/kcft114/.julia/packages/AdvancedHMC/YWXfk/src/adaptation/stepsize.jl:103
┌ Warning: The current proposal will be rejected due to numerical error(s).
│   isfiniteθ = true
│   isfiniter = true
│   isfiniteℓπ = false
│   isfiniteℓκ = true
└ @ AdvancedHMC /Users/kcft114/.julia/packages/AdvancedHMC/YWXfk/src/hamiltonian.jl:36
┌ Warning: The current proposal will be rejected due to numerical error(s).
│   isfiniteθ = false
│   isfiniter = false
│   isfiniteℓπ = false
│   isfiniteℓκ = false
└ @ AdvancedHMC /Users/kcft114/.julia/packages/AdvancedHMC/YWXfk/src/hamiltonian.jl:36
┌ Warning: Incorrect ϵ = NaN; ϵ_previous = 0.07500000000000001 is used instead.
└ @ AdvancedHMC.Adaptation /Users/kcft114/.julia/packages/AdvancedHMC/YWXfk/src/adaptation/stepsize.jl:103
┌ Warning: The current proposal will be rejected due to numerical error(s).
│   isfiniteθ = true
│   isfiniter = true
│   isfiniteℓπ = false
│   isfiniteℓκ = true
└ @ AdvancedHMC /Users/kcft114/.julia/packages/AdvancedHMC/YWXfk/src/hamiltonian.jl:36
┌ Warning: The current proposal will be rejected due to numerical error(s).
│   isfiniteθ = false
│   isfiniter = false
│   isfiniteℓπ = false
│   isfiniteℓκ = false
└ @ AdvancedHMC /Users/kcft114/.julia/packages/AdvancedHMC/YWXfk/src/hamiltonian.jl:36
┌ Warning: Incorrect ϵ = NaN; ϵ_previous = 0.07500000000000001 is used instead.
└ @ AdvancedHMC.Adaptation /Users/kcft114/.julia/packages/AdvancedHMC/YWXfk/src/adaptation/stepsize.jl:103
┌ Warning: The current proposal will be rejected due to numerical error(s).
│   isfiniteθ = true
│   isfiniter = true
│   isfiniteℓπ = false
│   isfiniteℓκ = true
└ @ AdvancedHMC /Users/kcft114/.julia/packages/AdvancedHMC/YWXfk/src/hamiltonian.jl:36
┌ Warning: The current proposal will be rejected due to numerical error(s).
│   isfiniteθ = false
│   isfiniter = false
│   isfiniteℓπ = false
│   isfiniteℓκ = false
└ @ AdvancedHMC /Users/kcft114/.julia/packages/AdvancedHMC/YWXfk/src/hamiltonian.jl:36
┌ Warning: Incorrect ϵ = NaN; ϵ_previous = 0.07500000000000001 is used instead.
└ @ AdvancedHMC.Adaptation /Users/kcft114/.julia/packages/AdvancedHMC/YWXfk/src/adaptation/stepsize.jl:103
┌ Warning: The current proposal will be rejected due to numerical error(s).
│   isfiniteθ = true
│   isfiniter = true
│   isfiniteℓπ = false
│   isfiniteℓκ = true
└ @ AdvancedHMC /Users/kcft114/.julia/packages/AdvancedHMC/YWXfk/src/hamiltonian.jl:36
┌ Warning: The current proposal will be rejected due to numerical error(s).
│   isfiniteθ = false
│   isfiniter = false
│   isfiniteℓπ = false
│   isfiniteℓκ = false
└ @ AdvancedHMC /Users/kcft114/.julia/packages/AdvancedHMC/YWXfk/src/hamiltonian.jl:36
┌ Warning: Incorrect ϵ = NaN; ϵ_previous = 0.07500000000000001 is used instead.
└ @ AdvancedHMC.Adaptation /Users/kcft114/.julia/packages/AdvancedHMC/YWXfk/src/adaptation/stepsize.jl:103
┌ Warning: The current proposal will be rejected due to numerical error(s).
│   isfiniteθ = true
│   isfiniter = true
│   isfiniteℓπ = false
│   isfiniteℓκ = true
└ @ AdvancedHMC /Users/kcft114/.julia/packages/AdvancedHMC/YWXfk/src/hamiltonian.jl:36
┌ Warning: The current proposal will be rejected due to numerical error(s).
│   isfiniteθ = false
│   isfiniter = false
│   isfiniteℓπ = false
│   isfiniteℓκ = false
└ @ AdvancedHMC /Users/kcft114/.julia/packages/AdvancedHMC/YWXfk/src/hamiltonian.jl:36
┌ Warning: Incorrect ϵ = NaN; ϵ_previous = 0.07500000000000001 is used instead.
└ @ AdvancedHMC.Adaptation /Users/kcft114/.julia/packages/AdvancedHMC/YWXfk/src/adaptation/stepsize.jl:103
┌ Warning: The current proposal will be rejected due to numerical error(s).
│   isfiniteθ = true
│   isfiniter = true
│   isfiniteℓπ = false
│   isfiniteℓκ = true
└ @ AdvancedHMC /Users/kcft114/.julia/packages/AdvancedHMC/YWXfk/src/hamiltonian.jl:36
┌ Warning: The current proposal will be rejected due to numerical error(s).
│   isfiniteθ = false
│   isfiniter = false
│   isfiniteℓπ = false
│   isfiniteℓκ = false
└ @ AdvancedHMC /Users/kcft114/.julia/packages/AdvancedHMC/YWXfk/src/hamiltonian.jl:36
┌ Warning: Incorrect ϵ = NaN; ϵ_previous = 0.07500000000000001 is used instead.
└ @ AdvancedHMC.Adaptation /Users/kcft114/.julia/packages/AdvancedHMC/YWXfk/src/adaptation/stepsize.jl:103
┌ Warning: The current proposal will be rejected due to numerical error(s).
│   isfiniteθ = true
│   isfiniter = true
│   isfiniteℓπ = false
│   isfiniteℓκ = true
└ @ AdvancedHMC /Users/kcft114/.julia/packages/AdvancedHMC/YWXfk/src/hamiltonian.jl:36
┌ Warning: The current proposal will be rejected due to numerical error(s).
│   isfiniteθ = false
│   isfiniter = false
│   isfiniteℓπ = false
│   isfiniteℓκ = false
└ @ AdvancedHMC /Users/kcft114/.julia/packages/AdvancedHMC/YWXfk/src/hamiltonian.jl:36
┌ Warning: Incorrect ϵ = NaN; ϵ_previous = 0.07500000000000001 is used instead.
└ @ AdvancedHMC.Adaptation /Users/kcft114/.julia/packages/AdvancedHMC/YWXfk/src/adaptation/stepsize.jl:103
┌ Warning: The current proposal will be rejected due to numerical error(s).
│   isfiniteθ = true
│   isfiniter = true
│   isfiniteℓπ = false
│   isfiniteℓκ = true
└ @ AdvancedHMC /Users/kcft114/.julia/packages/AdvancedHMC/YWXfk/src/hamiltonian.jl:36
┌ Warning: The current proposal will be rejected due to numerical error(s).
│   isfiniteθ = false
│   isfiniter = false
│   isfiniteℓπ = false
│   isfiniteℓκ = false
└ @ AdvancedHMC /Users/kcft114/.julia/packages/AdvancedHMC/YWXfk/src/hamiltonian.jl:36
┌ Warning: Incorrect ϵ = NaN; ϵ_previous = 0.07500000000000001 is used instead.
└ @ AdvancedHMC.Adaptation /Users/kcft114/.julia/packages/AdvancedHMC/YWXfk/src/adaptation/stepsize.jl:103
┌ Warning: The current proposal will be rejected due to numerical error(s).
│   isfiniteθ = true
│   isfiniter = true
│   isfiniteℓπ = false
│   isfiniteℓκ = true
└ @ AdvancedHMC /Users/kcft114/.julia/packages/AdvancedHMC/YWXfk/src/hamiltonian.jl:36
┌ Warning: The current proposal will be rejected due to numerical error(s).
│   isfiniteθ = false
│   isfiniter = false
│   isfiniteℓπ = false
│   isfiniteℓκ = false
└ @ AdvancedHMC /Users/kcft114/.julia/packages/AdvancedHMC/YWXfk/src/hamiltonian.jl:36
┌ Warning: Incorrect ϵ = NaN; ϵ_previous = 0.07500000000000001 is used instead.
└ @ AdvancedHMC.Adaptation /Users/kcft114/.julia/packages/AdvancedHMC/YWXfk/src/adaptation/stepsize.jl:103
┌ Warning: The current proposal will be rejected due to numerical error(s).
│   isfiniteθ = true
│   isfiniter = true
│   isfiniteℓπ = false
│   isfiniteℓκ = true
└ @ AdvancedHMC /Users/kcft114/.julia/packages/AdvancedHMC/YWXfk/src/hamiltonian.jl:36
┌ Warning: The current proposal will be rejected due to numerical error(s).
│   isfiniteθ = false
│   isfiniter = false
│   isfiniteℓπ = false
│   isfiniteℓκ = false
└ @ AdvancedHMC /Users/kcft114/.julia/packages/AdvancedHMC/YWXfk/src/hamiltonian.jl:36
┌ Warning: Incorrect ϵ = NaN; ϵ_previous = 0.07500000000000001 is used instead.
└ @ AdvancedHMC.Adaptation /Users/kcft114/.julia/packages/AdvancedHMC/YWXfk/src/adaptation/stepsize.jl:103
┌ Warning: The current proposal will be rejected due to numerical error(s).
│   isfiniteθ = true
│   isfiniter = true
│   isfiniteℓπ = false
│   isfiniteℓκ = true
└ @ AdvancedHMC /Users/kcft114/.julia/packages/AdvancedHMC/YWXfk/src/hamiltonian.jl:36
┌ Warning: The current proposal will be rejected due to numerical error(s).
│   isfiniteθ = false
│   isfiniter = false
│   isfiniteℓπ = false
│   isfiniteℓκ = false
└ @ AdvancedHMC /Users/kcft114/.julia/packages/AdvancedHMC/YWXfk/src/hamiltonian.jl:36
┌ Warning: Incorrect ϵ = NaN; ϵ_previous = 0.07500000000000001 is used instead.
└ @ AdvancedHMC.Adaptation /Users/kcft114/.julia/packages/AdvancedHMC/YWXfk/src/adaptation/stepsize.jl:103
┌ Warning: The current proposal will be rejected due to numerical error(s).
│   isfiniteθ = true
│   isfiniter = true
│   isfiniteℓπ = false
│   isfiniteℓκ = true
└ @ AdvancedHMC /Users/kcft114/.julia/packages/AdvancedHMC/YWXfk/src/hamiltonian.jl:36
┌ Warning: The current proposal will be rejected due to numerical error(s).
│   isfiniteθ = false
│   isfiniter = false
│   isfiniteℓπ = false
│   isfiniteℓκ = false
└ @ AdvancedHMC /Users/kcft114/.julia/packages/AdvancedHMC/YWXfk/src/hamiltonian.jl:36
┌ Warning: Incorrect ϵ = NaN; ϵ_previous = 0.07500000000000001 is used instead.
└ @ AdvancedHMC.Adaptation /Users/kcft114/.julia/packages/AdvancedHMC/YWXfk/src/adaptation/stepsize.jl:103
┌ Warning: The current proposal will be rejected due to numerical error(s).
│   isfiniteθ = true
│   isfiniter = true
│   isfiniteℓπ = false
│   isfiniteℓκ = true
└ @ AdvancedHMC /Users/kcft114/.julia/packages/AdvancedHMC/YWXfk/src/hamiltonian.jl:36
┌ Warning: The current proposal will be rejected due to numerical error(s).
│   isfiniteθ = false
│   isfiniter = false
│   isfiniteℓπ = false
│   isfiniteℓκ = false
└ @ AdvancedHMC /Users/kcft114/.julia/packages/AdvancedHMC/YWXfk/src/hamiltonian.jl:36
┌ Warning: Incorrect ϵ = NaN; ϵ_previous = 0.07500000000000001 is used instead.
└ @ AdvancedHMC.Adaptation /Users/kcft114/.julia/packages/AdvancedHMC/YWXfk/src/adaptation/stepsize.jl:103
┌ Warning: The current proposal will be rejected due to numerical error(s).
│   isfiniteθ = true
│   isfiniter = true
│   isfiniteℓπ = false
│   isfiniteℓκ = true
└ @ AdvancedHMC /Users/kcft114/.julia/packages/AdvancedHMC/YWXfk/src/hamiltonian.jl:36
┌ Warning: The current proposal will be rejected due to numerical error(s).
│   isfiniteθ = false
│   isfiniter = false
│   isfiniteℓπ = false
│   isfiniteℓκ = false
└ @ AdvancedHMC /Users/kcft114/.julia/packages/AdvancedHMC/YWXfk/src/hamiltonian.jl:36
┌ Warning: Incorrect ϵ = NaN; ϵ_previous = 0.07500000000000001 is used instead.
└ @ AdvancedHMC.Adaptation /Users/kcft114/.julia/packages/AdvancedHMC/YWXfk/src/adaptation/stepsize.jl:103
┌ Warning: The current proposal will be rejected due to numerical error(s).
│   isfiniteθ = true
│   isfiniter = true
│   isfiniteℓπ = false
│   isfiniteℓκ = true
└ @ AdvancedHMC /Users/kcft114/.julia/packages/AdvancedHMC/YWXfk/src/hamiltonian.jl:36
┌ Warning: The current proposal will be rejected due to numerical error(s).
│   isfiniteθ = false
│   isfiniter = false
│   isfiniteℓπ = false
│   isfiniteℓκ = false
└ @ AdvancedHMC /Users/kcft114/.julia/packages/AdvancedHMC/YWXfk/src/hamiltonian.jl:36
┌ Warning: Incorrect ϵ = NaN; ϵ_previous = 0.07500000000000001 is used instead.
└ @ AdvancedHMC.Adaptation /Users/kcft114/.julia/packages/AdvancedHMC/YWXfk/src/adaptation/stepsize.jl:103
┌ Warning: The current proposal will be rejected due to numerical error(s).
│   isfiniteθ = true
│   isfiniter = true
│   isfiniteℓπ = false
│   isfiniteℓκ = true
└ @ AdvancedHMC /Users/kcft114/.julia/packages/AdvancedHMC/YWXfk/src/hamiltonian.jl:36
┌ Warning: The current proposal will be rejected due to numerical error(s).
│   isfiniteθ = false
│   isfiniter = false
│   isfiniteℓπ = false
│   isfiniteℓκ = false
└ @ AdvancedHMC /Users/kcft114/.julia/packages/AdvancedHMC/YWXfk/src/hamiltonian.jl:36
┌ Warning: Incorrect ϵ = NaN; ϵ_previous = 0.07500000000000001 is used instead.
└ @ AdvancedHMC.Adaptation /Users/kcft114/.julia/packages/AdvancedHMC/YWXfk/src/adaptation/stepsize.jl:103
┌ Warning: The current proposal will be rejected due to numerical error(s).
│   isfiniteθ = true
│   isfiniter = true
│   isfiniteℓπ = false
│   isfiniteℓκ = true
└ @ AdvancedHMC /Users/kcft114/.julia/packages/AdvancedHMC/YWXfk/src/hamiltonian.jl:36
┌ Warning: The current proposal will be rejected due to numerical error(s).
│   isfiniteθ = false
│   isfiniter = false
│   isfiniteℓπ = false
│   isfiniteℓκ = false
└ @ AdvancedHMC /Users/kcft114/.julia/packages/AdvancedHMC/YWXfk/src/hamiltonian.jl:36
┌ Warning: Incorrect ϵ = NaN; ϵ_previous = 0.07500000000000001 is used instead.
└ @ AdvancedHMC.Adaptation /Users/kcft114/.julia/packages/AdvancedHMC/YWXfk/src/adaptation/stepsize.jl:103
┌ Warning: The current proposal will be rejected due to numerical error(s).
│   isfiniteθ = true
│   isfiniter = true
│   isfiniteℓπ = false
│   isfiniteℓκ = true
└ @ AdvancedHMC /Users/kcft114/.julia/packages/AdvancedHMC/YWXfk/src/hamiltonian.jl:36
┌ Warning: The current proposal will be rejected due to numerical error(s).
│   isfiniteθ = false
│   isfiniter = false
│   isfiniteℓπ = false
│   isfiniteℓκ = false
└ @ AdvancedHMC /Users/kcft114/.julia/packages/AdvancedHMC/YWXfk/src/hamiltonian.jl:36
┌ Warning: Incorrect ϵ = NaN; ϵ_previous = 0.07500000000000001 is used instead.
└ @ AdvancedHMC.Adaptation /Users/kcft114/.julia/packages/AdvancedHMC/YWXfk/src/adaptation/stepsize.jl:103
┌ Warning: The current proposal will be rejected due to numerical error(s).
│   isfiniteθ = true
│   isfiniter = true
│   isfiniteℓπ = false
│   isfiniteℓκ = true
└ @ AdvancedHMC /Users/kcft114/.julia/packages/AdvancedHMC/YWXfk/src/hamiltonian.jl:36
┌ Warning: The current proposal will be rejected due to numerical error(s).
│   isfiniteθ = false
│   isfiniter = false
│   isfiniteℓπ = false
│   isfiniteℓκ = false
└ @ AdvancedHMC /Users/kcft114/.julia/packages/AdvancedHMC/YWXfk/src/hamiltonian.jl:36
┌ Warning: Incorrect ϵ = NaN; ϵ_previous = 0.07500000000000001 is used instead.
└ @ AdvancedHMC.Adaptation /Users/kcft114/.julia/packages/AdvancedHMC/YWXfk/src/adaptation/stepsize.jl:103
┌ Warning: The current proposal will be rejected due to numerical error(s).
│   isfiniteθ = true
│   isfiniter = true
│   isfiniteℓπ = false
│   isfiniteℓκ = true
└ @ AdvancedHMC /Users/kcft114/.julia/packages/AdvancedHMC/YWXfk/src/hamiltonian.jl:36
┌ Warning: The current proposal will be rejected due to numerical error(s).
│   isfiniteθ = false
│   isfiniter = false
│   isfiniteℓπ = false
│   isfiniteℓκ = false
└ @ AdvancedHMC /Users/kcft114/.julia/packages/AdvancedHMC/YWXfk/src/hamiltonian.jl:36
┌ Warning: Incorrect ϵ = NaN; ϵ_previous = 0.07500000000000001 is used instead.
└ @ AdvancedHMC.Adaptation /Users/kcft114/.julia/packages/AdvancedHMC/YWXfk/src/adaptation/stepsize.jl:103
┌ Warning: The current proposal will be rejected due to numerical error(s).
│   isfiniteθ = true
│   isfiniter = true
│   isfiniteℓπ = false
│   isfiniteℓκ = true
└ @ AdvancedHMC /Users/kcft114/.julia/packages/AdvancedHMC/YWXfk/src/hamiltonian.jl:36
┌ Warning: The current proposal will be rejected due to numerical error(s).
│   isfiniteθ = false
│   isfiniter = false
│   isfiniteℓπ = false
│   isfiniteℓκ = false
└ @ AdvancedHMC /Users/kcft114/.julia/packages/AdvancedHMC/YWXfk/src/hamiltonian.jl:36
┌ Warning: Incorrect ϵ = NaN; ϵ_previous = 0.07500000000000001 is used instead.
└ @ AdvancedHMC.Adaptation /Users/kcft114/.julia/packages/AdvancedHMC/YWXfk/src/adaptation/stepsize.jl:103
┌ Warning: The current proposal will be rejected due to numerical error(s).
│   isfiniteθ = true
│   isfiniter = true
│   isfiniteℓπ = false
│   isfiniteℓκ = true
└ @ AdvancedHMC /Users/kcft114/.julia/packages/AdvancedHMC/YWXfk/src/hamiltonian.jl:36
┌ Warning: The current proposal will be rejected due to numerical error(s).
│   isfiniteθ = false
│   isfiniter = false
│   isfiniteℓπ = false
│   isfiniteℓκ = false
└ @ AdvancedHMC /Users/kcft114/.julia/packages/AdvancedHMC/YWXfk/src/hamiltonian.jl:36
┌ Warning: Incorrect ϵ = NaN; ϵ_previous = 0.07500000000000001 is used instead.
└ @ AdvancedHMC.Adaptation /Users/kcft114/.julia/packages/AdvancedHMC/YWXfk/src/adaptation/stepsize.jl:103
┌ Warning: The current proposal will be rejected due to numerical error(s).
│   isfiniteθ = true
│   isfiniter = true
│   isfiniteℓπ = false
│   isfiniteℓκ = true
└ @ AdvancedHMC /Users/kcft114/.julia/packages/AdvancedHMC/YWXfk/src/hamiltonian.jl:36
┌ Warning: The current proposal will be rejected due to numerical error(s).
│   isfiniteθ = false
│   isfiniter = false
│   isfiniteℓπ = false
│   isfiniteℓκ = false
└ @ AdvancedHMC /Users/kcft114/.julia/packages/AdvancedHMC/YWXfk/src/hamiltonian.jl:36
┌ Warning: Incorrect ϵ = NaN; ϵ_previous = 0.07500000000000001 is used instead.
└ @ AdvancedHMC.Adaptation /Users/kcft114/.julia/packages/AdvancedHMC/YWXfk/src/adaptation/stepsize.jl:103
┌ Warning: The current proposal will be rejected due to numerical error(s).
│   isfiniteθ = true
│   isfiniter = true
│   isfiniteℓπ = false
│   isfiniteℓκ = true
└ @ AdvancedHMC /Users/kcft114/.julia/packages/AdvancedHMC/YWXfk/src/hamiltonian.jl:36
┌ Warning: The current proposal will be rejected due to numerical error(s).
│   isfiniteθ = false
│   isfiniter = false
│   isfiniteℓπ = false
│   isfiniteℓκ = false
└ @ AdvancedHMC /Users/kcft114/.julia/packages/AdvancedHMC/YWXfk/src/hamiltonian.jl:36
┌ Warning: Incorrect ϵ = NaN; ϵ_previous = 0.07500000000000001 is used instead.
└ @ AdvancedHMC.Adaptation /Users/kcft114/.julia/packages/AdvancedHMC/YWXfk/src/adaptation/stepsize.jl:103
┌ Warning: The current proposal will be rejected due to numerical error(s).
│   isfiniteθ = true
│   isfiniter = true
│   isfiniteℓπ = false
│   isfiniteℓκ = true
└ @ AdvancedHMC /Users/kcft114/.julia/packages/AdvancedHMC/YWXfk/src/hamiltonian.jl:36
┌ Warning: The current proposal will be rejected due to numerical error(s).
│   isfiniteθ = false
│   isfiniter = false
│   isfiniteℓπ = false
│   isfiniteℓκ = false
└ @ AdvancedHMC /Users/kcft114/.julia/packages/AdvancedHMC/YWXfk/src/hamiltonian.jl:36
┌ Warning: Incorrect ϵ = NaN; ϵ_previous = 0.07500000000000001 is used instead.
└ @ AdvancedHMC.Adaptation /Users/kcft114/.julia/packages/AdvancedHMC/YWXfk/src/adaptation/stepsize.jl:103
┌ Warning: The current proposal will be rejected due to numerical error(s).
│   isfiniteθ = true
│   isfiniter = true
│   isfiniteℓπ = false
│   isfiniteℓκ = true
└ @ AdvancedHMC /Users/kcft114/.julia/packages/AdvancedHMC/YWXfk/src/hamiltonian.jl:36
┌ Warning: The current proposal will be rejected due to numerical error(s).
│   isfiniteθ = false
│   isfiniter = false
│   isfiniteℓπ = false
│   isfiniteℓκ = false
└ @ AdvancedHMC /Users/kcft114/.julia/packages/AdvancedHMC/YWXfk/src/hamiltonian.jl:36
┌ Warning: Incorrect ϵ = NaN; ϵ_previous = 0.07500000000000001 is used instead.
└ @ AdvancedHMC.Adaptation /Users/kcft114/.julia/packages/AdvancedHMC/YWXfk/src/adaptation/stepsize.jl:103
┌ Warning: The current proposal will be rejected due to numerical error(s).
│   isfiniteθ = true
│   isfiniter = true
│   isfiniteℓπ = false
│   isfiniteℓκ = true
└ @ AdvancedHMC /Users/kcft114/.julia/packages/AdvancedHMC/YWXfk/src/hamiltonian.jl:36
┌ Warning: The current proposal will be rejected due to numerical error(s).
│   isfiniteθ = false
│   isfiniter = false
│   isfiniteℓπ = false
│   isfiniteℓκ = false
└ @ AdvancedHMC /Users/kcft114/.julia/packages/AdvancedHMC/YWXfk/src/hamiltonian.jl:36
┌ Warning: Incorrect ϵ = NaN; ϵ_previous = 0.07500000000000001 is used instead.
└ @ AdvancedHMC.Adaptation /Users/kcft114/.julia/packages/AdvancedHMC/YWXfk/src/adaptation/stepsize.jl:103
┌ Warning: The current proposal will be rejected due to numerical error(s).
│   isfiniteθ = true
│   isfiniter = true
│   isfiniteℓπ = false
│   isfiniteℓκ = true
└ @ AdvancedHMC /Users/kcft114/.julia/packages/AdvancedHMC/YWXfk/src/hamiltonian.jl:36
┌ Warning: The current proposal will be rejected due to numerical error(s).
│   isfiniteθ = false
│   isfiniter = false
│   isfiniteℓπ = false
│   isfiniteℓκ = false
└ @ AdvancedHMC /Users/kcft114/.julia/packages/AdvancedHMC/YWXfk/src/hamiltonian.jl:36
┌ Warning: Incorrect ϵ = NaN; ϵ_previous = 0.07500000000000001 is used instead.
└ @ AdvancedHMC.Adaptation /Users/kcft114/.julia/packages/AdvancedHMC/YWXfk/src/adaptation/stepsize.jl:103
┌ Warning: The current proposal will be rejected due to numerical error(s).
│   isfiniteθ = true
│   isfiniter = true
│   isfiniteℓπ = false
│   isfiniteℓκ = true
└ @ AdvancedHMC /Users/kcft114/.julia/packages/AdvancedHMC/YWXfk/src/hamiltonian.jl:36
┌ Warning: The current proposal will be rejected due to numerical error(s).
│   isfiniteθ = false
│   isfiniter = false
│   isfiniteℓπ = false
│   isfiniteℓκ = false
└ @ AdvancedHMC /Users/kcft114/.julia/packages/AdvancedHMC/YWXfk/src/hamiltonian.jl:36
┌ Warning: Incorrect ϵ = NaN; ϵ_previous = 0.07500000000000001 is used instead.
└ @ AdvancedHMC.Adaptation /Users/kcft114/.julia/packages/AdvancedHMC/YWXfk/src/adaptation/stepsize.jl:103
┌ Warning: The current proposal will be rejected due to numerical error(s).
│   isfiniteθ = true
│   isfiniter = true
│   isfiniteℓπ = false
│   isfiniteℓκ = true
└ @ AdvancedHMC /Users/kcft114/.julia/packages/AdvancedHMC/YWXfk/src/hamiltonian.jl:36
┌ Warning: The current proposal will be rejected due to numerical error(s).
│   isfiniteθ = false
│   isfiniter = false
│   isfiniteℓπ = false
│   isfiniteℓκ = false
└ @ AdvancedHMC /Users/kcft114/.julia/packages/AdvancedHMC/YWXfk/src/hamiltonian.jl:36
┌ Warning: Incorrect ϵ = NaN; ϵ_previous = 0.07500000000000001 is used instead.
└ @ AdvancedHMC.Adaptation /Users/kcft114/.julia/packages/AdvancedHMC/YWXfk/src/adaptation/stepsize.jl:103
┌ Warning: The current proposal will be rejected due to numerical error(s).
│   isfiniteθ = true
│   isfiniter = true
│   isfiniteℓπ = false
│   isfiniteℓκ = true
└ @ AdvancedHMC /Users/kcft114/.julia/packages/AdvancedHMC/YWXfk/src/hamiltonian.jl:36
┌ Warning: The current proposal will be rejected due to numerical error(s).
│   isfiniteθ = false
│   isfiniter = false
│   isfiniteℓπ = false
│   isfiniteℓκ = false
└ @ AdvancedHMC /Users/kcft114/.julia/packages/AdvancedHMC/YWXfk/src/hamiltonian.jl:36
┌ Warning: Incorrect ϵ = NaN; ϵ_previous = 0.07500000000000001 is used instead.
└ @ AdvancedHMC.Adaptation /Users/kcft114/.julia/packages/AdvancedHMC/YWXfk/src/adaptation/stepsize.jl:103
┌ Warning: The current proposal will be rejected due to numerical error(s).
│   isfiniteθ = true
│   isfiniter = true
│   isfiniteℓπ = false
│   isfiniteℓκ = true
└ @ AdvancedHMC /Users/kcft114/.julia/packages/AdvancedHMC/YWXfk/src/hamiltonian.jl:36
┌ Warning: The current proposal will be rejected due to numerical error(s).
│   isfiniteθ = false
│   isfiniter = false
│   isfiniteℓπ = false
│   isfiniteℓκ = false
└ @ AdvancedHMC /Users/kcft114/.julia/packages/AdvancedHMC/YWXfk/src/hamiltonian.jl:36
┌ Warning: Incorrect ϵ = NaN; ϵ_previous = 0.07500000000000001 is used instead.
└ @ AdvancedHMC.Adaptation /Users/kcft114/.julia/packages/AdvancedHMC/YWXfk/src/adaptation/stepsize.jl:103
┌ Warning: The current proposal will be rejected due to numerical error(s).
│   isfiniteθ = true
│   isfiniter = true
│   isfiniteℓπ = false
│   isfiniteℓκ = true
└ @ AdvancedHMC /Users/kcft114/.julia/packages/AdvancedHMC/YWXfk/src/hamiltonian.jl:36
┌ Warning: The current proposal will be rejected due to numerical error(s).
│   isfiniteθ = false
│   isfiniter = false
│   isfiniteℓπ = false
│   isfiniteℓκ = false
└ @ AdvancedHMC /Users/kcft114/.julia/packages/AdvancedHMC/YWXfk/src/hamiltonian.jl:36
┌ Warning: Incorrect ϵ = NaN; ϵ_previous = 0.07500000000000001 is used instead.
└ @ AdvancedHMC.Adaptation /Users/kcft114/.julia/packages/AdvancedHMC/YWXfk/src/adaptation/stepsize.jl:103
┌ Warning: The current proposal will be rejected due to numerical error(s).
│   isfiniteθ = true
│   isfiniter = true
│   isfiniteℓπ = false
│   isfiniteℓκ = true
└ @ AdvancedHMC /Users/kcft114/.julia/packages/AdvancedHMC/YWXfk/src/hamiltonian.jl:36
┌ Warning: The current proposal will be rejected due to numerical error(s).
│   isfiniteθ = false
│   isfiniter = false
│   isfiniteℓπ = false
│   isfiniteℓκ = false
└ @ AdvancedHMC /Users/kcft114/.julia/packages/AdvancedHMC/YWXfk/src/hamiltonian.jl:36
┌ Warning: Incorrect ϵ = NaN; ϵ_previous = 0.07500000000000001 is used instead.
└ @ AdvancedHMC.Adaptation /Users/kcft114/.julia/packages/AdvancedHMC/YWXfk/src/adaptation/stepsize.jl:103
┌ Warning: The current proposal will be rejected due to numerical error(s).
│   isfiniteθ = true
│   isfiniter = true
│   isfiniteℓπ = false
│   isfiniteℓκ = true
└ @ AdvancedHMC /Users/kcft114/.julia/packages/AdvancedHMC/YWXfk/src/hamiltonian.jl:36
┌ Warning: The current proposal will be rejected due to numerical error(s).
│   isfiniteθ = false
│   isfiniter = false
│   isfiniteℓπ = false
│   isfiniteℓκ = false
└ @ AdvancedHMC /Users/kcft114/.julia/packages/AdvancedHMC/YWXfk/src/hamiltonian.jl:36
┌ Warning: Incorrect ϵ = NaN; ϵ_previous = 0.07500000000000001 is used instead.
└ @ AdvancedHMC.Adaptation /Users/kcft114/.julia/packages/AdvancedHMC/YWXfk/src/adaptation/stepsize.jl:103
┌ Warning: The current proposal will be rejected due to numerical error(s).
│   isfiniteθ = true
│   isfiniter = true
│   isfiniteℓπ = false
│   isfiniteℓκ = true
└ @ AdvancedHMC /Users/kcft114/.julia/packages/AdvancedHMC/YWXfk/src/hamiltonian.jl:36
┌ Warning: The current proposal will be rejected due to numerical error(s).
│   isfiniteθ = false
│   isfiniter = false
│   isfiniteℓπ = false
│   isfiniteℓκ = false
└ @ AdvancedHMC /Users/kcft114/.julia/packages/AdvancedHMC/YWXfk/src/hamiltonian.jl:36
┌ Warning: Incorrect ϵ = NaN; ϵ_previous = 0.07500000000000001 is used instead.
└ @ AdvancedHMC.Adaptation /Users/kcft114/.julia/packages/AdvancedHMC/YWXfk/src/adaptation/stepsize.jl:103
┌ Warning: The current proposal will be rejected due to numerical error(s).
│   isfiniteθ = true
│   isfiniter = true
│   isfiniteℓπ = false
│   isfiniteℓκ = true
└ @ AdvancedHMC /Users/kcft114/.julia/packages/AdvancedHMC/YWXfk/src/hamiltonian.jl:36
┌ Warning: The current proposal will be rejected due to numerical error(s).
│   isfiniteθ = false
│   isfiniter = false
│   isfiniteℓπ = false
│   isfiniteℓκ = false
└ @ AdvancedHMC /Users/kcft114/.julia/packages/AdvancedHMC/YWXfk/src/hamiltonian.jl:36
┌ Warning: Incorrect ϵ = NaN; ϵ_previous = 0.07500000000000001 is used instead.
└ @ AdvancedHMC.Adaptation /Users/kcft114/.julia/packages/AdvancedHMC/YWXfk/src/adaptation/stepsize.jl:103
┌ Warning: The current proposal will be rejected due to numerical error(s).
│   isfiniteθ = true
│   isfiniter = true
│   isfiniteℓπ = false
│   isfiniteℓκ = true
└ @ AdvancedHMC /Users/kcft114/.julia/packages/AdvancedHMC/YWXfk/src/hamiltonian.jl:36
┌ Warning: The current proposal will be rejected due to numerical error(s).
│   isfiniteθ = false
│   isfiniter = false
│   isfiniteℓπ = false
│   isfiniteℓκ = false
└ @ AdvancedHMC /Users/kcft114/.julia/packages/AdvancedHMC/YWXfk/src/hamiltonian.jl:36
┌ Warning: Incorrect ϵ = NaN; ϵ_previous = 0.07500000000000001 is used instead.
└ @ AdvancedHMC.Adaptation /Users/kcft114/.julia/packages/AdvancedHMC/YWXfk/src/adaptation/stepsize.jl:103
┌ Warning: The current proposal will be rejected due to numerical error(s).
│   isfiniteθ = true
│   isfiniter = true
│   isfiniteℓπ = false
│   isfiniteℓκ = true
└ @ AdvancedHMC /Users/kcft114/.julia/packages/AdvancedHMC/YWXfk/src/hamiltonian.jl:36
┌ Warning: The current proposal will be rejected due to numerical error(s).
│   isfiniteθ = false
│   isfiniter = false
│   isfiniteℓπ = false
│   isfiniteℓκ = false
└ @ AdvancedHMC /Users/kcft114/.julia/packages/AdvancedHMC/YWXfk/src/hamiltonian.jl:36
┌ Warning: Incorrect ϵ = NaN; ϵ_previous = 0.07500000000000001 is used instead.
└ @ AdvancedHMC.Adaptation /Users/kcft114/.julia/packages/AdvancedHMC/YWXfk/src/adaptation/stepsize.jl:103
┌ Warning: The current proposal will be rejected due to numerical error(s).
│   isfiniteθ = true
│   isfiniter = true
│   isfiniteℓπ = false
│   isfiniteℓκ = true
└ @ AdvancedHMC /Users/kcft114/.julia/packages/AdvancedHMC/YWXfk/src/hamiltonian.jl:36
┌ Warning: The current proposal will be rejected due to numerical error(s).
│   isfiniteθ = false
│   isfiniter = false
│   isfiniteℓπ = false
│   isfiniteℓκ = false
└ @ AdvancedHMC /Users/kcft114/.julia/packages/AdvancedHMC/YWXfk/src/hamiltonian.jl:36
┌ Warning: Incorrect ϵ = NaN; ϵ_previous = 0.07500000000000001 is used instead.
└ @ AdvancedHMC.Adaptation /Users/kcft114/.julia/packages/AdvancedHMC/YWXfk/src/adaptation/stepsize.jl:103
┌ Warning: The current proposal will be rejected due to numerical error(s).
│   isfiniteθ = true
│   isfiniter = true
│   isfiniteℓπ = false
│   isfiniteℓκ = true
└ @ AdvancedHMC /Users/kcft114/.julia/packages/AdvancedHMC/YWXfk/src/hamiltonian.jl:36
┌ Warning: The current proposal will be rejected due to numerical error(s).
│   isfiniteθ = false
│   isfiniter = false
│   isfiniteℓπ = false
│   isfiniteℓκ = false
└ @ AdvancedHMC /Users/kcft114/.julia/packages/AdvancedHMC/YWXfk/src/hamiltonian.jl:36
┌ Warning: Incorrect ϵ = NaN; ϵ_previous = 0.07500000000000001 is used instead.
└ @ AdvancedHMC.Adaptation /Users/kcft114/.julia/packages/AdvancedHMC/YWXfk/src/adaptation/stepsize.jl:103
┌ Warning: The current proposal will be rejected due to numerical error(s).
│   isfiniteθ = true
│   isfiniter = true
│   isfiniteℓπ = false
│   isfiniteℓκ = true
└ @ AdvancedHMC /Users/kcft114/.julia/packages/AdvancedHMC/YWXfk/src/hamiltonian.jl:36
┌ Warning: The current proposal will be rejected due to numerical error(s).
│   isfiniteθ = false
│   isfiniter = false
│   isfiniteℓπ = false
│   isfiniteℓκ = false
└ @ AdvancedHMC /Users/kcft114/.julia/packages/AdvancedHMC/YWXfk/src/hamiltonian.jl:36
┌ Warning: Incorrect ϵ = NaN; ϵ_previous = 0.07500000000000001 is used instead.
└ @ AdvancedHMC.Adaptation /Users/kcft114/.julia/packages/AdvancedHMC/YWXfk/src/adaptation/stepsize.jl:103
┌ Warning: The current proposal will be rejected due to numerical error(s).
│   isfiniteθ = true
│   isfiniter = true
│   isfiniteℓπ = false
│   isfiniteℓκ = true
└ @ AdvancedHMC /Users/kcft114/.julia/packages/AdvancedHMC/YWXfk/src/hamiltonian.jl:36
┌ Warning: The current proposal will be rejected due to numerical error(s).
│   isfiniteθ = false
│   isfiniter = false
│   isfiniteℓπ = false
│   isfiniteℓκ = false
└ @ AdvancedHMC /Users/kcft114/.julia/packages/AdvancedHMC/YWXfk/src/hamiltonian.jl:36
┌ Warning: Incorrect ϵ = NaN; ϵ_previous = 0.07500000000000001 is used instead.
└ @ AdvancedHMC.Adaptation /Users/kcft114/.julia/packages/AdvancedHMC/YWXfk/src/adaptation/stepsize.jl:103
┌ Warning: The current proposal will be rejected due to numerical error(s).
│   isfiniteθ = true
│   isfiniter = true
│   isfiniteℓπ = false
│   isfiniteℓκ = true
└ @ AdvancedHMC /Users/kcft114/.julia/packages/AdvancedHMC/YWXfk/src/hamiltonian.jl:36
┌ Warning: The current proposal will be rejected due to numerical error(s).
│   isfiniteθ = false
│   isfiniter = false
│   isfiniteℓπ = false
│   isfiniteℓκ = false
└ @ AdvancedHMC /Users/kcft114/.julia/packages/AdvancedHMC/YWXfk/src/hamiltonian.jl:36
┌ Warning: Incorrect ϵ = NaN; ϵ_previous = 0.07500000000000001 is used instead.
└ @ AdvancedHMC.Adaptation /Users/kcft114/.julia/packages/AdvancedHMC/YWXfk/src/adaptation/stepsize.jl:103
┌ Warning: The current proposal will be rejected due to numerical error(s).
│   isfiniteθ = true
│   isfiniter = true
│   isfiniteℓπ = false
│   isfiniteℓκ = true
└ @ AdvancedHMC /Users/kcft114/.julia/packages/AdvancedHMC/YWXfk/src/hamiltonian.jl:36
┌ Warning: The current proposal will be rejected due to numerical error(s).
│   isfiniteθ = false
│   isfiniter = false
│   isfiniteℓπ = false
│   isfiniteℓκ = false
└ @ AdvancedHMC /Users/kcft114/.julia/packages/AdvancedHMC/YWXfk/src/hamiltonian.jl:36
┌ Warning: Incorrect ϵ = NaN; ϵ_previous = 0.07500000000000001 is used instead.
└ @ AdvancedHMC.Adaptation /Users/kcft114/.julia/packages/AdvancedHMC/YWXfk/src/adaptation/stepsize.jl:103
┌ Warning: The current proposal will be rejected due to numerical error(s).
│   isfiniteθ = true
│   isfiniter = true
│   isfiniteℓπ = false
│   isfiniteℓκ = true
└ @ AdvancedHMC /Users/kcft114/.julia/packages/AdvancedHMC/YWXfk/src/hamiltonian.jl:36
┌ Warning: The current proposal will be rejected due to numerical error(s).
│   isfiniteθ = false
│   isfiniter = false
│   isfiniteℓπ = false
│   isfiniteℓκ = false
└ @ AdvancedHMC /Users/kcft114/.julia/packages/AdvancedHMC/YWXfk/src/hamiltonian.jl:36
┌ Warning: Incorrect ϵ = NaN; ϵ_previous = 0.07500000000000001 is used instead.
└ @ AdvancedHMC.Adaptation /Users/kcft114/.julia/packages/AdvancedHMC/YWXfk/src/adaptation/stepsize.jl:103
┌ Warning: The current proposal will be rejected due to numerical error(s).
│   isfiniteθ = true
│   isfiniter = true
│   isfiniteℓπ = false
│   isfiniteℓκ = true
└ @ AdvancedHMC /Users/kcft114/.julia/packages/AdvancedHMC/YWXfk/src/hamiltonian.jl:36
┌ Warning: The current proposal will be rejected due to numerical error(s).
│   isfiniteθ = false
│   isfiniter = false
│   isfiniteℓπ = false
│   isfiniteℓκ = false
└ @ AdvancedHMC /Users/kcft114/.julia/packages/AdvancedHMC/YWXfk/src/hamiltonian.jl:36
┌ Warning: Incorrect ϵ = NaN; ϵ_previous = 0.07500000000000001 is used instead.
└ @ AdvancedHMC.Adaptation /Users/kcft114/.julia/packages/AdvancedHMC/YWXfk/src/adaptation/stepsize.jl:103
┌ Warning: The current proposal will be rejected due to numerical error(s).
│   isfiniteθ = true
│   isfiniter = true
│   isfiniteℓπ = false
│   isfiniteℓκ = true
└ @ AdvancedHMC /Users/kcft114/.julia/packages/AdvancedHMC/YWXfk/src/hamiltonian.jl:36
┌ Warning: The current proposal will be rejected due to numerical error(s).
│   isfiniteθ = false
│   isfiniter = false
│   isfiniteℓπ = false
│   isfiniteℓκ = false
└ @ AdvancedHMC /Users/kcft114/.julia/packages/AdvancedHMC/YWXfk/src/hamiltonian.jl:36
┌ Warning: Incorrect ϵ = NaN; ϵ_previous = 0.07500000000000001 is used instead.
└ @ AdvancedHMC.Adaptation /Users/kcft114/.julia/packages/AdvancedHMC/YWXfk/src/adaptation/stepsize.jl:103
┌ Warning: The current proposal will be rejected due to numerical error(s).
│   isfiniteθ = true
│   isfiniter = true
│   isfiniteℓπ = false
│   isfiniteℓκ = true
└ @ AdvancedHMC /Users/kcft114/.julia/packages/AdvancedHMC/YWXfk/src/hamiltonian.jl:36
┌ Warning: The current proposal will be rejected due to numerical error(s).
│   isfiniteθ = false
│   isfiniter = false
│   isfiniteℓπ = false
│   isfiniteℓκ = false
└ @ AdvancedHMC /Users/kcft114/.julia/packages/AdvancedHMC/YWXfk/src/hamiltonian.jl:36
┌ Warning: Incorrect ϵ = NaN; ϵ_previous = 0.07500000000000001 is used instead.
└ @ AdvancedHMC.Adaptation /Users/kcft114/.julia/packages/AdvancedHMC/YWXfk/src/adaptation/stepsize.jl:103
┌ Warning: The current proposal will be rejected due to numerical error(s).
│   isfiniteθ = true
│   isfiniter = true
│   isfiniteℓπ = false
│   isfiniteℓκ = true
└ @ AdvancedHMC /Users/kcft114/.julia/packages/AdvancedHMC/YWXfk/src/hamiltonian.jl:36
┌ Warning: The current proposal will be rejected due to numerical error(s).
│   isfiniteθ = false
│   isfiniter = false
│   isfiniteℓπ = false
│   isfiniteℓκ = false
└ @ AdvancedHMC /Users/kcft114/.julia/packages/AdvancedHMC/YWXfk/src/hamiltonian.jl:36
┌ Warning: Incorrect ϵ = NaN; ϵ_previous = 0.07500000000000001 is used instead.
└ @ AdvancedHMC.Adaptation /Users/kcft114/.julia/packages/AdvancedHMC/YWXfk/src/adaptation/stepsize.jl:103
┌ Warning: The current proposal will be rejected due to numerical error(s).
│   isfiniteθ = true
│   isfiniter = true
│   isfiniteℓπ = false
│   isfiniteℓκ = true
└ @ AdvancedHMC /Users/kcft114/.julia/packages/AdvancedHMC/YWXfk/src/hamiltonian.jl:36
┌ Warning: The current proposal will be rejected due to numerical error(s).
│   isfiniteθ = false
│   isfiniter = false
│   isfiniteℓπ = false
│   isfiniteℓκ = false
└ @ AdvancedHMC /Users/kcft114/.julia/packages/AdvancedHMC/YWXfk/src/hamiltonian.jl:36
┌ Warning: Incorrect ϵ = NaN; ϵ_previous = 0.07500000000000001 is used instead.
└ @ AdvancedHMC.Adaptation /Users/kcft114/.julia/packages/AdvancedHMC/YWXfk/src/adaptation/stepsize.jl:103
┌ Warning: The current proposal will be rejected due to numerical error(s).
│   isfiniteθ = true
│   isfiniter = true
│   isfiniteℓπ = false
│   isfiniteℓκ = true
└ @ AdvancedHMC /Users/kcft114/.julia/packages/AdvancedHMC/YWXfk/src/hamiltonian.jl:36
┌ Warning: The current proposal will be rejected due to numerical error(s).
│   isfiniteθ = false
│   isfiniter = false
│   isfiniteℓπ = false
│   isfiniteℓκ = false
└ @ AdvancedHMC /Users/kcft114/.julia/packages/AdvancedHMC/YWXfk/src/hamiltonian.jl:36
┌ Warning: Incorrect ϵ = NaN; ϵ_previous = 0.07500000000000001 is used instead.
└ @ AdvancedHMC.Adaptation /Users/kcft114/.julia/packages/AdvancedHMC/YWXfk/src/adaptation/stepsize.jl:103
┌ Warning: The current proposal will be rejected due to numerical error(s).
│   isfiniteθ = true
│   isfiniter = true
│   isfiniteℓπ = false
│   isfiniteℓκ = true
└ @ AdvancedHMC /Users/kcft114/.julia/packages/AdvancedHMC/YWXfk/src/hamiltonian.jl:36
┌ Warning: The current proposal will be rejected due to numerical error(s).
│   isfiniteθ = false
│   isfiniter = false
│   isfiniteℓπ = false
│   isfiniteℓκ = false
└ @ AdvancedHMC /Users/kcft114/.julia/packages/AdvancedHMC/YWXfk/src/hamiltonian.jl:36
┌ Warning: Incorrect ϵ = NaN; ϵ_previous = 0.07500000000000001 is used instead.
└ @ AdvancedHMC.Adaptation /Users/kcft114/.julia/packages/AdvancedHMC/YWXfk/src/adaptation/stepsize.jl:103
┌ Warning: The current proposal will be rejected due to numerical error(s).
│   isfiniteθ = true
│   isfiniter = true
│   isfiniteℓπ = false
│   isfiniteℓκ = true
└ @ AdvancedHMC /Users/kcft114/.julia/packages/AdvancedHMC/YWXfk/src/hamiltonian.jl:36
┌ Warning: The current proposal will be rejected due to numerical error(s).
│   isfiniteθ = false
│   isfiniter = false
│   isfiniteℓπ = false
│   isfiniteℓκ = false
└ @ AdvancedHMC /Users/kcft114/.julia/packages/AdvancedHMC/YWXfk/src/hamiltonian.jl:36
┌ Warning: Incorrect ϵ = NaN; ϵ_previous = 0.07500000000000001 is used instead.
└ @ AdvancedHMC.Adaptation /Users/kcft114/.julia/packages/AdvancedHMC/YWXfk/src/adaptation/stepsize.jl:103
┌ Warning: The current proposal will be rejected due to numerical error(s).
│   isfiniteθ = true
│   isfiniter = true
│   isfiniteℓπ = false
│   isfiniteℓκ = true
└ @ AdvancedHMC /Users/kcft114/.julia/packages/AdvancedHMC/YWXfk/src/hamiltonian.jl:36
┌ Warning: The current proposal will be rejected due to numerical error(s).
│   isfiniteθ = false
│   isfiniter = false
│   isfiniteℓπ = false
│   isfiniteℓκ = false
└ @ AdvancedHMC /Users/kcft114/.julia/packages/AdvancedHMC/YWXfk/src/hamiltonian.jl:36
┌ Warning: Incorrect ϵ = NaN; ϵ_previous = 0.07500000000000001 is used instead.
└ @ AdvancedHMC.Adaptation /Users/kcft114/.julia/packages/AdvancedHMC/YWXfk/src/adaptation/stepsize.jl:103
┌ Warning: The current proposal will be rejected due to numerical error(s).
│   isfiniteθ = true
│   isfiniter = true
│   isfiniteℓπ = false
│   isfiniteℓκ = true
└ @ AdvancedHMC /Users/kcft114/.julia/packages/AdvancedHMC/YWXfk/src/hamiltonian.jl:36
┌ Warning: The current proposal will be rejected due to numerical error(s).
│   isfiniteθ = false
│   isfiniter = false
│   isfiniteℓπ = false
│   isfiniteℓκ = false
└ @ AdvancedHMC /Users/kcft114/.julia/packages/AdvancedHMC/YWXfk/src/hamiltonian.jl:36
┌ Warning: Incorrect ϵ = NaN; ϵ_previous = 0.07500000000000001 is used instead.
└ @ AdvancedHMC.Adaptation /Users/kcft114/.julia/packages/AdvancedHMC/YWXfk/src/adaptation/stepsize.jl:103
┌ Warning: The current proposal will be rejected due to numerical error(s).
│   isfiniteθ = true
│   isfiniter = true
│   isfiniteℓπ = false
│   isfiniteℓκ = true
└ @ AdvancedHMC /Users/kcft114/.julia/packages/AdvancedHMC/YWXfk/src/hamiltonian.jl:36
┌ Warning: The current proposal will be rejected due to numerical error(s).
│   isfiniteθ = false
│   isfiniter = false
│   isfiniteℓπ = false
│   isfiniteℓκ = false
└ @ AdvancedHMC /Users/kcft114/.julia/packages/AdvancedHMC/YWXfk/src/hamiltonian.jl:36
┌ Warning: Incorrect ϵ = NaN; ϵ_previous = 0.07500000000000001 is used instead.
└ @ AdvancedHMC.Adaptation /Users/kcft114/.julia/packages/AdvancedHMC/YWXfk/src/adaptation/stepsize.jl:103
┌ Warning: The current proposal will be rejected due to numerical error(s).
│   isfiniteθ = true
│   isfiniter = true
│   isfiniteℓπ = false
│   isfiniteℓκ = true
└ @ AdvancedHMC /Users/kcft114/.julia/packages/AdvancedHMC/YWXfk/src/hamiltonian.jl:36
┌ Warning: The current proposal will be rejected due to numerical error(s).
│   isfiniteθ = false
│   isfiniter = false
│   isfiniteℓπ = false
│   isfiniteℓκ = false
└ @ AdvancedHMC /Users/kcft114/.julia/packages/AdvancedHMC/YWXfk/src/hamiltonian.jl:36
┌ Warning: Incorrect ϵ = NaN; ϵ_previous = 0.07500000000000001 is used instead.
└ @ AdvancedHMC.Adaptation /Users/kcft114/.julia/packages/AdvancedHMC/YWXfk/src/adaptation/stepsize.jl:103
┌ Warning: The current proposal will be rejected due to numerical error(s).
│   isfiniteθ = true
│   isfiniter = true
│   isfiniteℓπ = false
│   isfiniteℓκ = true
└ @ AdvancedHMC /Users/kcft114/.julia/packages/AdvancedHMC/YWXfk/src/hamiltonian.jl:36
┌ Warning: The current proposal will be rejected due to numerical error(s).
│   isfiniteθ = false
│   isfiniter = false
│   isfiniteℓπ = false
│   isfiniteℓκ = false
└ @ AdvancedHMC /Users/kcft114/.julia/packages/AdvancedHMC/YWXfk/src/hamiltonian.jl:36
┌ Warning: Incorrect ϵ = NaN; ϵ_previous = 0.07500000000000001 is used instead.
└ @ AdvancedHMC.Adaptation /Users/kcft114/.julia/packages/AdvancedHMC/YWXfk/src/adaptation/stepsize.jl:103
┌ Warning: The current proposal will be rejected due to numerical error(s).
│   isfiniteθ = true
│   isfiniter = true
│   isfiniteℓπ = false
│   isfiniteℓκ = true
└ @ AdvancedHMC /Users/kcft114/.julia/packages/AdvancedHMC/YWXfk/src/hamiltonian.jl:36
┌ Warning: The current proposal will be rejected due to numerical error(s).
│   isfiniteθ = false
│   isfiniter = false
│   isfiniteℓπ = false
│   isfiniteℓκ = false
└ @ AdvancedHMC /Users/kcft114/.julia/packages/AdvancedHMC/YWXfk/src/hamiltonian.jl:36
┌ Warning: Incorrect ϵ = NaN; ϵ_previous = 0.07500000000000001 is used instead.
└ @ AdvancedHMC.Adaptation /Users/kcft114/.julia/packages/AdvancedHMC/YWXfk/src/adaptation/stepsize.jl:103
┌ Warning: The current proposal will be rejected due to numerical error(s).
│   isfiniteθ = true
│   isfiniter = true
│   isfiniteℓπ = false
│   isfiniteℓκ = true
└ @ AdvancedHMC /Users/kcft114/.julia/packages/AdvancedHMC/YWXfk/src/hamiltonian.jl:36
┌ Warning: The current proposal will be rejected due to numerical error(s).
│   isfiniteθ = false
│   isfiniter = false
│   isfiniteℓπ = false
│   isfiniteℓκ = false
└ @ AdvancedHMC /Users/kcft114/.julia/packages/AdvancedHMC/YWXfk/src/hamiltonian.jl:36
┌ Warning: Incorrect ϵ = NaN; ϵ_previous = 0.07500000000000001 is used instead.
└ @ AdvancedHMC.Adaptation /Users/kcft114/.julia/packages/AdvancedHMC/YWXfk/src/adaptation/stepsize.jl:103
┌ Warning: The current proposal will be rejected due to numerical error(s).
│   isfiniteθ = true
│   isfiniter = true
│   isfiniteℓπ = false
│   isfiniteℓκ = true
└ @ AdvancedHMC /Users/kcft114/.julia/packages/AdvancedHMC/YWXfk/src/hamiltonian.jl:36
┌ Warning: The current proposal will be rejected due to numerical error(s).
│   isfiniteθ = false
│   isfiniter = false
│   isfiniteℓπ = false
│   isfiniteℓκ = false
└ @ AdvancedHMC /Users/kcft114/.julia/packages/AdvancedHMC/YWXfk/src/hamiltonian.jl:36
┌ Warning: Incorrect ϵ = NaN; ϵ_previous = 0.07500000000000001 is used instead.
└ @ AdvancedHMC.Adaptation /Users/kcft114/.julia/packages/AdvancedHMC/YWXfk/src/adaptation/stepsize.jl:103
┌ Warning: The current proposal will be rejected due to numerical error(s).
│   isfiniteθ = true
│   isfiniter = true
│   isfiniteℓπ = false
│   isfiniteℓκ = true
└ @ AdvancedHMC /Users/kcft114/.julia/packages/AdvancedHMC/YWXfk/src/hamiltonian.jl:36
┌ Warning: The current proposal will be rejected due to numerical error(s).
│   isfiniteθ = false
│   isfiniter = false
│   isfiniteℓπ = false
│   isfiniteℓκ = false
└ @ AdvancedHMC /Users/kcft114/.julia/packages/AdvancedHMC/YWXfk/src/hamiltonian.jl:36
┌ Warning: Incorrect ϵ = NaN; ϵ_previous = 0.07500000000000001 is used instead.
└ @ AdvancedHMC.Adaptation /Users/kcft114/.julia/packages/AdvancedHMC/YWXfk/src/adaptation/stepsize.jl:103
┌ Warning: The current proposal will be rejected due to numerical error(s).
│   isfiniteθ = true
│   isfiniter = true
│   isfiniteℓπ = false
│   isfiniteℓκ = true
└ @ AdvancedHMC /Users/kcft114/.julia/packages/AdvancedHMC/YWXfk/src/hamiltonian.jl:36
┌ Warning: The current proposal will be rejected due to numerical error(s).
│   isfiniteθ = false
│   isfiniter = false
│   isfiniteℓπ = false
│   isfiniteℓκ = false
└ @ AdvancedHMC /Users/kcft114/.julia/packages/AdvancedHMC/YWXfk/src/hamiltonian.jl:36
┌ Warning: Incorrect ϵ = NaN; ϵ_previous = 0.07500000000000001 is used instead.
└ @ AdvancedHMC.Adaptation /Users/kcft114/.julia/packages/AdvancedHMC/YWXfk/src/adaptation/stepsize.jl:103
┌ Warning: The current proposal will be rejected due to numerical error(s).
│   isfiniteθ = true
│   isfiniter = true
│   isfiniteℓπ = false
│   isfiniteℓκ = true
└ @ AdvancedHMC /Users/kcft114/.julia/packages/AdvancedHMC/YWXfk/src/hamiltonian.jl:36
┌ Warning: The current proposal will be rejected due to numerical error(s).
│   isfiniteθ = false
│   isfiniter = false
│   isfiniteℓπ = false
│   isfiniteℓκ = false
└ @ AdvancedHMC /Users/kcft114/.julia/packages/AdvancedHMC/YWXfk/src/hamiltonian.jl:36
┌ Warning: Incorrect ϵ = NaN; ϵ_previous = 0.07500000000000001 is used instead.
└ @ AdvancedHMC.Adaptation /Users/kcft114/.julia/packages/AdvancedHMC/YWXfk/src/adaptation/stepsize.jl:103
┌ Warning: The current proposal will be rejected due to numerical error(s).
│   isfiniteθ = true
│   isfiniter = true
│   isfiniteℓπ = false
│   isfiniteℓκ = true
└ @ AdvancedHMC /Users/kcft114/.julia/packages/AdvancedHMC/YWXfk/src/hamiltonian.jl:36
┌ Warning: The current proposal will be rejected due to numerical error(s).
│   isfiniteθ = false
│   isfiniter = false
│   isfiniteℓπ = false
│   isfiniteℓκ = false
└ @ AdvancedHMC /Users/kcft114/.julia/packages/AdvancedHMC/YWXfk/src/hamiltonian.jl:36
┌ Warning: Incorrect ϵ = NaN; ϵ_previous = 0.07500000000000001 is used instead.
└ @ AdvancedHMC.Adaptation /Users/kcft114/.julia/packages/AdvancedHMC/YWXfk/src/adaptation/stepsize.jl:103
┌ Warning: The current proposal will be rejected due to numerical error(s).
│   isfiniteθ = true
│   isfiniter = true
│   isfiniteℓπ = false
│   isfiniteℓκ = true
└ @ AdvancedHMC /Users/kcft114/.julia/packages/AdvancedHMC/YWXfk/src/hamiltonian.jl:36
┌ Warning: The current proposal will be rejected due to numerical error(s).
│   isfiniteθ = false
│   isfiniter = false
│   isfiniteℓπ = false
│   isfiniteℓκ = false
└ @ AdvancedHMC /Users/kcft114/.julia/packages/AdvancedHMC/YWXfk/src/hamiltonian.jl:36
┌ Warning: Incorrect ϵ = NaN; ϵ_previous = 0.07500000000000001 is used instead.
└ @ AdvancedHMC.Adaptation /Users/kcft114/.julia/packages/AdvancedHMC/YWXfk/src/adaptation/stepsize.jl:103
┌ Warning: The current proposal will be rejected due to numerical error(s).
│   isfiniteθ = true
│   isfiniter = true
│   isfiniteℓπ = false
│   isfiniteℓκ = true
└ @ AdvancedHMC /Users/kcft114/.julia/packages/AdvancedHMC/YWXfk/src/hamiltonian.jl:36
┌ Warning: The current proposal will be rejected due to numerical error(s).
│   isfiniteθ = false
│   isfiniter = false
│   isfiniteℓπ = false
│   isfiniteℓκ = false
└ @ AdvancedHMC /Users/kcft114/.julia/packages/AdvancedHMC/YWXfk/src/hamiltonian.jl:36
┌ Warning: Incorrect ϵ = NaN; ϵ_previous = 0.07500000000000001 is used instead.
└ @ AdvancedHMC.Adaptation /Users/kcft114/.julia/packages/AdvancedHMC/YWXfk/src/adaptation/stepsize.jl:103
┌ Warning: The current proposal will be rejected due to numerical error(s).
│   isfiniteθ = true
│   isfiniter = true
│   isfiniteℓπ = false
│   isfiniteℓκ = true
└ @ AdvancedHMC /Users/kcft114/.julia/packages/AdvancedHMC/YWXfk/src/hamiltonian.jl:36
┌ Warning: The current proposal will be rejected due to numerical error(s).
│   isfiniteθ = false
│   isfiniter = false
│   isfiniteℓπ = false
│   isfiniteℓκ = false
└ @ AdvancedHMC /Users/kcft114/.julia/packages/AdvancedHMC/YWXfk/src/hamiltonian.jl:36
┌ Warning: Incorrect ϵ = NaN; ϵ_previous = 0.07500000000000001 is used instead.
└ @ AdvancedHMC.Adaptation /Users/kcft114/.julia/packages/AdvancedHMC/YWXfk/src/adaptation/stepsize.jl:103
┌ Warning: The current proposal will be rejected due to numerical error(s).
│   isfiniteθ = true
│   isfiniter = true
│   isfiniteℓπ = false
│   isfiniteℓκ = true
└ @ AdvancedHMC /Users/kcft114/.julia/packages/AdvancedHMC/YWXfk/src/hamiltonian.jl:36
┌ Warning: The current proposal will be rejected due to numerical error(s).
│   isfiniteθ = false
│   isfiniter = false
│   isfiniteℓπ = false
│   isfiniteℓκ = false
└ @ AdvancedHMC /Users/kcft114/.julia/packages/AdvancedHMC/YWXfk/src/hamiltonian.jl:36
┌ Warning: Incorrect ϵ = NaN; ϵ_previous = 0.07500000000000001 is used instead.
└ @ AdvancedHMC.Adaptation /Users/kcft114/.julia/packages/AdvancedHMC/YWXfk/src/adaptation/stepsize.jl:103
┌ Warning: The current proposal will be rejected due to numerical error(s).
│   isfiniteθ = true
│   isfiniter = true
│   isfiniteℓπ = false
│   isfiniteℓκ = true
└ @ AdvancedHMC /Users/kcft114/.julia/packages/AdvancedHMC/YWXfk/src/hamiltonian.jl:36
┌ Warning: The current proposal will be rejected due to numerical error(s).
│   isfiniteθ = false
│   isfiniter = false
│   isfiniteℓπ = false
│   isfiniteℓκ = false
└ @ AdvancedHMC /Users/kcft114/.julia/packages/AdvancedHMC/YWXfk/src/hamiltonian.jl:36
┌ Warning: Incorrect ϵ = NaN; ϵ_previous = 0.07500000000000001 is used instead.
└ @ AdvancedHMC.Adaptation /Users/kcft114/.julia/packages/AdvancedHMC/YWXfk/src/adaptation/stepsize.jl:103
┌ Warning: The current proposal will be rejected due to numerical error(s).
│   isfiniteθ = true
│   isfiniter = true
│   isfiniteℓπ = false
│   isfiniteℓκ = true
└ @ AdvancedHMC /Users/kcft114/.julia/packages/AdvancedHMC/YWXfk/src/hamiltonian.jl:36
┌ Warning: The current proposal will be rejected due to numerical error(s).
│   isfiniteθ = false
│   isfiniter = false
│   isfiniteℓπ = false
│   isfiniteℓκ = false
└ @ AdvancedHMC /Users/kcft114/.julia/packages/AdvancedHMC/YWXfk/src/hamiltonian.jl:36
┌ Warning: Incorrect ϵ = NaN; ϵ_previous = 0.07500000000000001 is used instead.
└ @ AdvancedHMC.Adaptation /Users/kcft114/.julia/packages/AdvancedHMC/YWXfk/src/adaptation/stepsize.jl:103
┌ Warning: The current proposal will be rejected due to numerical error(s).
│   isfiniteθ = true
│   isfiniter = true
│   isfiniteℓπ = false
│   isfiniteℓκ = true
└ @ AdvancedHMC /Users/kcft114/.julia/packages/AdvancedHMC/YWXfk/src/hamiltonian.jl:36
┌ Warning: The current proposal will be rejected due to numerical error(s).
│   isfiniteθ = false
│   isfiniter = false
│   isfiniteℓπ = false
│   isfiniteℓκ = false
└ @ AdvancedHMC /Users/kcft114/.julia/packages/AdvancedHMC/YWXfk/src/hamiltonian.jl:36
┌ Warning: Incorrect ϵ = NaN; ϵ_previous = 0.07500000000000001 is used instead.
└ @ AdvancedHMC.Adaptation /Users/kcft114/.julia/packages/AdvancedHMC/YWXfk/src/adaptation/stepsize.jl:103
┌ Warning: The current proposal will be rejected due to numerical error(s).
│   isfiniteθ = true
│   isfiniter = true
│   isfiniteℓπ = false
│   isfiniteℓκ = true
└ @ AdvancedHMC /Users/kcft114/.julia/packages/AdvancedHMC/YWXfk/src/hamiltonian.jl:36
┌ Warning: The current proposal will be rejected due to numerical error(s).
│   isfiniteθ = false
│   isfiniter = false
│   isfiniteℓπ = false
│   isfiniteℓκ = false
└ @ AdvancedHMC /Users/kcft114/.julia/packages/AdvancedHMC/YWXfk/src/hamiltonian.jl:36
┌ Warning: Incorrect ϵ = NaN; ϵ_previous = 0.07500000000000001 is used instead.
└ @ AdvancedHMC.Adaptation /Users/kcft114/.julia/packages/AdvancedHMC/YWXfk/src/adaptation/stepsize.jl:103
┌ Warning: The current proposal will be rejected due to numerical error(s).
│   isfiniteθ = true
│   isfiniter = true
│   isfiniteℓπ = false
│   isfiniteℓκ = true
└ @ AdvancedHMC /Users/kcft114/.julia/packages/AdvancedHMC/YWXfk/src/hamiltonian.jl:36
┌ Warning: The current proposal will be rejected due to numerical error(s).
│   isfiniteθ = false
│   isfiniter = false
│   isfiniteℓπ = false
│   isfiniteℓκ = false
└ @ AdvancedHMC /Users/kcft114/.julia/packages/AdvancedHMC/YWXfk/src/hamiltonian.jl:36
┌ Warning: Incorrect ϵ = NaN; ϵ_previous = 0.07500000000000001 is used instead.
└ @ AdvancedHMC.Adaptation /Users/kcft114/.julia/packages/AdvancedHMC/YWXfk/src/adaptation/stepsize.jl:103
┌ Warning: The current proposal will be rejected due to numerical error(s).
│   isfiniteθ = true
│   isfiniter = true
│   isfiniteℓπ = false
│   isfiniteℓκ = true
└ @ AdvancedHMC /Users/kcft114/.julia/packages/AdvancedHMC/YWXfk/src/hamiltonian.jl:36
┌ Warning: The current proposal will be rejected due to numerical error(s).
│   isfiniteθ = false
│   isfiniter = false
│   isfiniteℓπ = false
│   isfiniteℓκ = false
└ @ AdvancedHMC /Users/kcft114/.julia/packages/AdvancedHMC/YWXfk/src/hamiltonian.jl:36
┌ Warning: Incorrect ϵ = NaN; ϵ_previous = 0.07500000000000001 is used instead.
└ @ AdvancedHMC.Adaptation /Users/kcft114/.julia/packages/AdvancedHMC/YWXfk/src/adaptation/stepsize.jl:103
┌ Warning: The current proposal will be rejected due to numerical error(s).
│   isfiniteθ = true
│   isfiniter = true
│   isfiniteℓπ = false
│   isfiniteℓκ = true
└ @ AdvancedHMC /Users/kcft114/.julia/packages/AdvancedHMC/YWXfk/src/hamiltonian.jl:36
┌ Warning: The current proposal will be rejected due to numerical error(s).
│   isfiniteθ = false
│   isfiniter = false
│   isfiniteℓπ = false
│   isfiniteℓκ = false
└ @ AdvancedHMC /Users/kcft114/.julia/packages/AdvancedHMC/YWXfk/src/hamiltonian.jl:36
┌ Warning: Incorrect ϵ = NaN; ϵ_previous = 0.07500000000000001 is used instead.
└ @ AdvancedHMC.Adaptation /Users/kcft114/.julia/packages/AdvancedHMC/YWXfk/src/adaptation/stepsize.jl:103
┌ Warning: The current proposal will be rejected due to numerical error(s).
│   isfiniteθ = true
│   isfiniter = true
│   isfiniteℓπ = false
│   isfiniteℓκ = true
└ @ AdvancedHMC /Users/kcft114/.julia/packages/AdvancedHMC/YWXfk/src/hamiltonian.jl:36
┌ Warning: The current proposal will be rejected due to numerical error(s).
│   isfiniteθ = false
│   isfiniter = false
│   isfiniteℓπ = false
│   isfiniteℓκ = false
└ @ AdvancedHMC /Users/kcft114/.julia/packages/AdvancedHMC/YWXfk/src/hamiltonian.jl:36
┌ Warning: Incorrect ϵ = NaN; ϵ_previous = 0.07500000000000001 is used instead.
└ @ AdvancedHMC.Adaptation /Users/kcft114/.julia/packages/AdvancedHMC/YWXfk/src/adaptation/stepsize.jl:103
┌ Warning: The current proposal will be rejected due to numerical error(s).
│   isfiniteθ = true
│   isfiniter = true
│   isfiniteℓπ = false
│   isfiniteℓκ = true
└ @ AdvancedHMC /Users/kcft114/.julia/packages/AdvancedHMC/YWXfk/src/hamiltonian.jl:36
┌ Warning: The current proposal will be rejected due to numerical error(s).
│   isfiniteθ = false
│   isfiniter = false
│   isfiniteℓπ = false
│   isfiniteℓκ = false
└ @ AdvancedHMC /Users/kcft114/.julia/packages/AdvancedHMC/YWXfk/src/hamiltonian.jl:36
┌ Warning: Incorrect ϵ = NaN; ϵ_previous = 0.07500000000000001 is used instead.
└ @ AdvancedHMC.Adaptation /Users/kcft114/.julia/packages/AdvancedHMC/YWXfk/src/adaptation/stepsize.jl:103
┌ Warning: The current proposal will be rejected due to numerical error(s).
│   isfiniteθ = true
│   isfiniter = true
│   isfiniteℓπ = false
│   isfiniteℓκ = true
└ @ AdvancedHMC /Users/kcft114/.julia/packages/AdvancedHMC/YWXfk/src/hamiltonian.jl:36
┌ Warning: The current proposal will be rejected due to numerical error(s).
│   isfiniteθ = false
│   isfiniter = false
│   isfiniteℓπ = false
│   isfiniteℓκ = false
└ @ AdvancedHMC /Users/kcft114/.julia/packages/AdvancedHMC/YWXfk/src/hamiltonian.jl:36
┌ Warning: Incorrect ϵ = NaN; ϵ_previous = 0.07500000000000001 is used instead.
└ @ AdvancedHMC.Adaptation /Users/kcft114/.julia/packages/AdvancedHMC/YWXfk/src/adaptation/stepsize.jl:103
┌ Warning: The current proposal will be rejected due to numerical error(s).
│   isfiniteθ = true
│   isfiniter = true
│   isfiniteℓπ = false
│   isfiniteℓκ = true
└ @ AdvancedHMC /Users/kcft114/.julia/packages/AdvancedHMC/YWXfk/src/hamiltonian.jl:36
┌ Warning: The current proposal will be rejected due to numerical error(s).
│   isfiniteθ = false
│   isfiniter = false
│   isfiniteℓπ = false
│   isfiniteℓκ = false
└ @ AdvancedHMC /Users/kcft114/.julia/packages/AdvancedHMC/YWXfk/src/hamiltonian.jl:36
┌ Warning: Incorrect ϵ = NaN; ϵ_previous = 0.07500000000000001 is used instead.
└ @ AdvancedHMC.Adaptation /Users/kcft114/.julia/packages/AdvancedHMC/YWXfk/src/adaptation/stepsize.jl:103
┌ Warning: The current proposal will be rejected due to numerical error(s).
│   isfiniteθ = true
│   isfiniter = true
│   isfiniteℓπ = false
│   isfiniteℓκ = true
└ @ AdvancedHMC /Users/kcft114/.julia/packages/AdvancedHMC/YWXfk/src/hamiltonian.jl:36
┌ Warning: The current proposal will be rejected due to numerical error(s).
│   isfiniteθ = false
│   isfiniter = false
│   isfiniteℓπ = false
│   isfiniteℓκ = false
└ @ AdvancedHMC /Users/kcft114/.julia/packages/AdvancedHMC/YWXfk/src/hamiltonian.jl:36
┌ Warning: Incorrect ϵ = NaN; ϵ_previous = 0.07500000000000001 is used instead.
└ @ AdvancedHMC.Adaptation /Users/kcft114/.julia/packages/AdvancedHMC/YWXfk/src/adaptation/stepsize.jl:103
┌ Warning: The current proposal will be rejected due to numerical error(s).
│   isfiniteθ = true
│   isfiniter = true
│   isfiniteℓπ = false
│   isfiniteℓκ = true
└ @ AdvancedHMC /Users/kcft114/.julia/packages/AdvancedHMC/YWXfk/src/hamiltonian.jl:36
┌ Warning: The current proposal will be rejected due to numerical error(s).
│   isfiniteθ = false
│   isfiniter = false
│   isfiniteℓπ = false
│   isfiniteℓκ = false
└ @ AdvancedHMC /Users/kcft114/.julia/packages/AdvancedHMC/YWXfk/src/hamiltonian.jl:36
┌ Warning: Incorrect ϵ = NaN; ϵ_previous = 0.07500000000000001 is used instead.
└ @ AdvancedHMC.Adaptation /Users/kcft114/.julia/packages/AdvancedHMC/YWXfk/src/adaptation/stepsize.jl:103
┌ Warning: The current proposal will be rejected due to numerical error(s).
│   isfiniteθ = true
│   isfiniter = true
│   isfiniteℓπ = false
│   isfiniteℓκ = true
└ @ AdvancedHMC /Users/kcft114/.julia/packages/AdvancedHMC/YWXfk/src/hamiltonian.jl:36
┌ Warning: The current proposal will be rejected due to numerical error(s).
│   isfiniteθ = false
│   isfiniter = false
│   isfiniteℓπ = false
│   isfiniteℓκ = false
└ @ AdvancedHMC /Users/kcft114/.julia/packages/AdvancedHMC/YWXfk/src/hamiltonian.jl:36
┌ Warning: Incorrect ϵ = NaN; ϵ_previous = 0.07500000000000001 is used instead.
└ @ AdvancedHMC.Adaptation /Users/kcft114/.julia/packages/AdvancedHMC/YWXfk/src/adaptation/stepsize.jl:103
┌ Warning: The current proposal will be rejected due to numerical error(s).
│   isfiniteθ = true
│   isfiniter = true
│   isfiniteℓπ = false
│   isfiniteℓκ = true
└ @ AdvancedHMC /Users/kcft114/.julia/packages/AdvancedHMC/YWXfk/src/hamiltonian.jl:36
┌ Warning: The current proposal will be rejected due to numerical error(s).
│   isfiniteθ = false
│   isfiniter = false
│   isfiniteℓπ = false
│   isfiniteℓκ = false
└ @ AdvancedHMC /Users/kcft114/.julia/packages/AdvancedHMC/YWXfk/src/hamiltonian.jl:36
┌ Warning: Incorrect ϵ = NaN; ϵ_previous = 0.07500000000000001 is used instead.
└ @ AdvancedHMC.Adaptation /Users/kcft114/.julia/packages/AdvancedHMC/YWXfk/src/adaptation/stepsize.jl:103
┌ Warning: The current proposal will be rejected due to numerical error(s).
│   isfiniteθ = true
│   isfiniter = true
│   isfiniteℓπ = false
│   isfiniteℓκ = true
└ @ AdvancedHMC /Users/kcft114/.julia/packages/AdvancedHMC/YWXfk/src/hamiltonian.jl:36
┌ Warning: The current proposal will be rejected due to numerical error(s).
│   isfiniteθ = false
│   isfiniter = false
│   isfiniteℓπ = false
│   isfiniteℓκ = false
└ @ AdvancedHMC /Users/kcft114/.julia/packages/AdvancedHMC/YWXfk/src/hamiltonian.jl:36
┌ Warning: Incorrect ϵ = NaN; ϵ_previous = 0.07500000000000001 is used instead.
└ @ AdvancedHMC.Adaptation /Users/kcft114/.julia/packages/AdvancedHMC/YWXfk/src/adaptation/stepsize.jl:103
┌ Warning: The current proposal will be rejected due to numerical error(s).
│   isfiniteθ = true
│   isfiniter = true
│   isfiniteℓπ = false
│   isfiniteℓκ = true
└ @ AdvancedHMC /Users/kcft114/.julia/packages/AdvancedHMC/YWXfk/src/hamiltonian.jl:36
┌ Warning: The current proposal will be rejected due to numerical error(s).
│   isfiniteθ = false
│   isfiniter = false
│   isfiniteℓπ = false
│   isfiniteℓκ = false
└ @ AdvancedHMC /Users/kcft114/.julia/packages/AdvancedHMC/YWXfk/src/hamiltonian.jl:36
┌ Warning: Incorrect ϵ = NaN; ϵ_previous = 0.07500000000000001 is used instead.
└ @ AdvancedHMC.Adaptation /Users/kcft114/.julia/packages/AdvancedHMC/YWXfk/src/adaptation/stepsize.jl:103
┌ Warning: The current proposal will be rejected due to numerical error(s).
│   isfiniteθ = true
│   isfiniter = true
│   isfiniteℓπ = false
│   isfiniteℓκ = true
└ @ AdvancedHMC /Users/kcft114/.julia/packages/AdvancedHMC/YWXfk/src/hamiltonian.jl:36
┌ Warning: The current proposal will be rejected due to numerical error(s).
│   isfiniteθ = false
│   isfiniter = false
│   isfiniteℓπ = false
│   isfiniteℓκ = false
└ @ AdvancedHMC /Users/kcft114/.julia/packages/AdvancedHMC/YWXfk/src/hamiltonian.jl:36
┌ Warning: Incorrect ϵ = NaN; ϵ_previous = 0.07500000000000001 is used instead.
└ @ AdvancedHMC.Adaptation /Users/kcft114/.julia/packages/AdvancedHMC/YWXfk/src/adaptation/stepsize.jl:103
┌ Warning: The current proposal will be rejected due to numerical error(s).
│   isfiniteθ = true
│   isfiniter = true
│   isfiniteℓπ = false
│   isfiniteℓκ = true
└ @ AdvancedHMC /Users/kcft114/.julia/packages/AdvancedHMC/YWXfk/src/hamiltonian.jl:36
┌ Warning: The current proposal will be rejected due to numerical error(s).
│   isfiniteθ = false
│   isfiniter = false
│   isfiniteℓπ = false
│   isfiniteℓκ = false
└ @ AdvancedHMC /Users/kcft114/.julia/packages/AdvancedHMC/YWXfk/src/hamiltonian.jl:36
┌ Warning: Incorrect ϵ = NaN; ϵ_previous = 0.07500000000000001 is used instead.
└ @ AdvancedHMC.Adaptation /Users/kcft114/.julia/packages/AdvancedHMC/YWXfk/src/adaptation/stepsize.jl:103
┌ Warning: The current proposal will be rejected due to numerical error(s).
│   isfiniteθ = true
│   isfiniter = true
│   isfiniteℓπ = false
│   isfiniteℓκ = true
└ @ AdvancedHMC /Users/kcft114/.julia/packages/AdvancedHMC/YWXfk/src/hamiltonian.jl:36
┌ Warning: The current proposal will be rejected due to numerical error(s).
│   isfiniteθ = false
│   isfiniter = false
│   isfiniteℓπ = false
│   isfiniteℓκ = false
└ @ AdvancedHMC /Users/kcft114/.julia/packages/AdvancedHMC/YWXfk/src/hamiltonian.jl:36
┌ Warning: Incorrect ϵ = NaN; ϵ_previous = 0.07500000000000001 is used instead.
└ @ AdvancedHMC.Adaptation /Users/kcft114/.julia/packages/AdvancedHMC/YWXfk/src/adaptation/stepsize.jl:103
┌ Warning: The current proposal will be rejected due to numerical error(s).
│   isfiniteθ = true
│   isfiniter = true
│   isfiniteℓπ = false
│   isfiniteℓκ = true
└ @ AdvancedHMC /Users/kcft114/.julia/packages/AdvancedHMC/YWXfk/src/hamiltonian.jl:36
┌ Warning: The current proposal will be rejected due to numerical error(s).
│   isfiniteθ = false
│   isfiniter = false
│   isfiniteℓπ = false
│   isfiniteℓκ = false
└ @ AdvancedHMC /Users/kcft114/.julia/packages/AdvancedHMC/YWXfk/src/hamiltonian.jl:36
┌ Warning: Incorrect ϵ = NaN; ϵ_previous = 0.07500000000000001 is used instead.
└ @ AdvancedHMC.Adaptation /Users/kcft114/.julia/packages/AdvancedHMC/YWXfk/src/adaptation/stepsize.jl:103
┌ Warning: The current proposal will be rejected due to numerical error(s).
│   isfiniteθ = true
│   isfiniter = true
│   isfiniteℓπ = false
│   isfiniteℓκ = true
└ @ AdvancedHMC /Users/kcft114/.julia/packages/AdvancedHMC/YWXfk/src/hamiltonian.jl:36
┌ Warning: The current proposal will be rejected due to numerical error(s).
│   isfiniteθ = false
│   isfiniter = false
│   isfiniteℓπ = false
│   isfiniteℓκ = false
└ @ AdvancedHMC /Users/kcft114/.julia/packages/AdvancedHMC/YWXfk/src/hamiltonian.jl:36
┌ Warning: Incorrect ϵ = NaN; ϵ_previous = 0.07500000000000001 is used instead.
└ @ AdvancedHMC.Adaptation /Users/kcft114/.julia/packages/AdvancedHMC/YWXfk/src/adaptation/stepsize.jl:103
┌ Warning: The current proposal will be rejected due to numerical error(s).
│   isfiniteθ = true
│   isfiniter = true
│   isfiniteℓπ = false
│   isfiniteℓκ = true
└ @ AdvancedHMC /Users/kcft114/.julia/packages/AdvancedHMC/YWXfk/src/hamiltonian.jl:36
┌ Warning: The current proposal will be rejected due to numerical error(s).
│   isfiniteθ = false
│   isfiniter = false
│   isfiniteℓπ = false
│   isfiniteℓκ = false
└ @ AdvancedHMC /Users/kcft114/.julia/packages/AdvancedHMC/YWXfk/src/hamiltonian.jl:36
┌ Warning: Incorrect ϵ = NaN; ϵ_previous = 0.07500000000000001 is used instead.
└ @ AdvancedHMC.Adaptation /Users/kcft114/.julia/packages/AdvancedHMC/YWXfk/src/adaptation/stepsize.jl:103
┌ Warning: The current proposal will be rejected due to numerical error(s).
│   isfiniteθ = true
│   isfiniter = true
│   isfiniteℓπ = false
│   isfiniteℓκ = true
└ @ AdvancedHMC /Users/kcft114/.julia/packages/AdvancedHMC/YWXfk/src/hamiltonian.jl:36
┌ Warning: The current proposal will be rejected due to numerical error(s).
│   isfiniteθ = false
│   isfiniter = false
│   isfiniteℓπ = false
│   isfiniteℓκ = false
└ @ AdvancedHMC /Users/kcft114/.julia/packages/AdvancedHMC/YWXfk/src/hamiltonian.jl:36
┌ Warning: Incorrect ϵ = NaN; ϵ_previous = 0.07500000000000001 is used instead.
└ @ AdvancedHMC.Adaptation /Users/kcft114/.julia/packages/AdvancedHMC/YWXfk/src/adaptation/stepsize.jl:103
┌ Warning: The current proposal will be rejected due to numerical error(s).
│   isfiniteθ = true
│   isfiniter = true
│   isfiniteℓπ = false
│   isfiniteℓκ = true
└ @ AdvancedHMC /Users/kcft114/.julia/packages/AdvancedHMC/YWXfk/src/hamiltonian.jl:36
┌ Warning: The current proposal will be rejected due to numerical error(s).
│   isfiniteθ = false
│   isfiniter = false
│   isfiniteℓπ = false
│   isfiniteℓκ = false
└ @ AdvancedHMC /Users/kcft114/.julia/packages/AdvancedHMC/YWXfk/src/hamiltonian.jl:36
┌ Warning: Incorrect ϵ = NaN; ϵ_previous = 0.07500000000000001 is used instead.
└ @ AdvancedHMC.Adaptation /Users/kcft114/.julia/packages/AdvancedHMC/YWXfk/src/adaptation/stepsize.jl:103
┌ Warning: The current proposal will be rejected due to numerical error(s).
│   isfiniteθ = true
│   isfiniter = true
│   isfiniteℓπ = false
│   isfiniteℓκ = true
└ @ AdvancedHMC /Users/kcft114/.julia/packages/AdvancedHMC/YWXfk/src/hamiltonian.jl:36
┌ Warning: The current proposal will be rejected due to numerical error(s).
│   isfiniteθ = false
│   isfiniter = false
│   isfiniteℓπ = false
│   isfiniteℓκ = false
└ @ AdvancedHMC /Users/kcft114/.julia/packages/AdvancedHMC/YWXfk/src/hamiltonian.jl:36
┌ Warning: Incorrect ϵ = NaN; ϵ_previous = 0.07500000000000001 is used instead.
└ @ AdvancedHMC.Adaptation /Users/kcft114/.julia/packages/AdvancedHMC/YWXfk/src/adaptation/stepsize.jl:103
┌ Warning: The current proposal will be rejected due to numerical error(s).
│   isfiniteθ = true
│   isfiniter = true
│   isfiniteℓπ = false
│   isfiniteℓκ = true
└ @ AdvancedHMC /Users/kcft114/.julia/packages/AdvancedHMC/YWXfk/src/hamiltonian.jl:36
┌ Warning: The current proposal will be rejected due to numerical error(s).
│   isfiniteθ = false
│   isfiniter = false
│   isfiniteℓπ = false
│   isfiniteℓκ = false
└ @ AdvancedHMC /Users/kcft114/.julia/packages/AdvancedHMC/YWXfk/src/hamiltonian.jl:36
┌ Warning: Incorrect ϵ = NaN; ϵ_previous = 0.07500000000000001 is used instead.
└ @ AdvancedHMC.Adaptation /Users/kcft114/.julia/packages/AdvancedHMC/YWXfk/src/adaptation/stepsize.jl:103
┌ Warning: The current proposal will be rejected due to numerical error(s).
│   isfiniteθ = true
│   isfiniter = true
│   isfiniteℓπ = false
│   isfiniteℓκ = true
└ @ AdvancedHMC /Users/kcft114/.julia/packages/AdvancedHMC/YWXfk/src/hamiltonian.jl:36
┌ Warning: The current proposal will be rejected due to numerical error(s).
│   isfiniteθ = false
│   isfiniter = false
│   isfiniteℓπ = false
│   isfiniteℓκ = false
└ @ AdvancedHMC /Users/kcft114/.julia/packages/AdvancedHMC/YWXfk/src/hamiltonian.jl:36
┌ Warning: Incorrect ϵ = NaN; ϵ_previous = 0.07500000000000001 is used instead.
└ @ AdvancedHMC.Adaptation /Users/kcft114/.julia/packages/AdvancedHMC/YWXfk/src/adaptation/stepsize.jl:103
┌ Warning: The current proposal will be rejected due to numerical error(s).
│   isfiniteθ = true
│   isfiniter = true
│   isfiniteℓπ = false
│   isfiniteℓκ = true
└ @ AdvancedHMC /Users/kcft114/.julia/packages/AdvancedHMC/YWXfk/src/hamiltonian.jl:36
┌ Warning: The current proposal will be rejected due to numerical error(s).
│   isfiniteθ = false
│   isfiniter = false
│   isfiniteℓπ = false
│   isfiniteℓκ = false
└ @ AdvancedHMC /Users/kcft114/.julia/packages/AdvancedHMC/YWXfk/src/hamiltonian.jl:36
┌ Warning: Incorrect ϵ = NaN; ϵ_previous = 0.07500000000000001 is used instead.
└ @ AdvancedHMC.Adaptation /Users/kcft114/.julia/packages/AdvancedHMC/YWXfk/src/adaptation/stepsize.jl:103
┌ Warning: The current proposal will be rejected due to numerical error(s).
│   isfiniteθ = true
│   isfiniter = true
│   isfiniteℓπ = false
│   isfiniteℓκ = true
└ @ AdvancedHMC /Users/kcft114/.julia/packages/AdvancedHMC/YWXfk/src/hamiltonian.jl:36
┌ Warning: The current proposal will be rejected due to numerical error(s).
│   isfiniteθ = false
│   isfiniter = false
│   isfiniteℓπ = false
│   isfiniteℓκ = false
└ @ AdvancedHMC /Users/kcft114/.julia/packages/AdvancedHMC/YWXfk/src/hamiltonian.jl:36
┌ Warning: Incorrect ϵ = NaN; ϵ_previous = 0.07500000000000001 is used instead.
└ @ AdvancedHMC.Adaptation /Users/kcft114/.julia/packages/AdvancedHMC/YWXfk/src/adaptation/stepsize.jl:103
┌ Warning: The current proposal will be rejected due to numerical error(s).
│   isfiniteθ = true
│   isfiniter = true
│   isfiniteℓπ = false
│   isfiniteℓκ = true
└ @ AdvancedHMC /Users/kcft114/.julia/packages/AdvancedHMC/YWXfk/src/hamiltonian.jl:36
┌ Warning: The current proposal will be rejected due to numerical error(s).
│   isfiniteθ = false
│   isfiniter = false
│   isfiniteℓπ = false
│   isfiniteℓκ = false
└ @ AdvancedHMC /Users/kcft114/.julia/packages/AdvancedHMC/YWXfk/src/hamiltonian.jl:36
┌ Warning: Incorrect ϵ = NaN; ϵ_previous = 0.07500000000000001 is used instead.
└ @ AdvancedHMC.Adaptation /Users/kcft114/.julia/packages/AdvancedHMC/YWXfk/src/adaptation/stepsize.jl:103
┌ Warning: The current proposal will be rejected due to numerical error(s).
│   isfiniteθ = true
│   isfiniter = true
│   isfiniteℓπ = false
│   isfiniteℓκ = true
└ @ AdvancedHMC /Users/kcft114/.julia/packages/AdvancedHMC/YWXfk/src/hamiltonian.jl:36
┌ Warning: The current proposal will be rejected due to numerical error(s).
│   isfiniteθ = false
│   isfiniter = false
│   isfiniteℓπ = false
│   isfiniteℓκ = false
└ @ AdvancedHMC /Users/kcft114/.julia/packages/AdvancedHMC/YWXfk/src/hamiltonian.jl:36
┌ Warning: Incorrect ϵ = NaN; ϵ_previous = 0.07500000000000001 is used instead.
└ @ AdvancedHMC.Adaptation /Users/kcft114/.julia/packages/AdvancedHMC/YWXfk/src/adaptation/stepsize.jl:103
┌ Warning: The current proposal will be rejected due to numerical error(s).
│   isfiniteθ = true
│   isfiniter = true
│   isfiniteℓπ = false
│   isfiniteℓκ = true
└ @ AdvancedHMC /Users/kcft114/.julia/packages/AdvancedHMC/YWXfk/src/hamiltonian.jl:36
┌ Warning: The current proposal will be rejected due to numerical error(s).
│   isfiniteθ = false
│   isfiniter = false
│   isfiniteℓπ = false
│   isfiniteℓκ = false
└ @ AdvancedHMC /Users/kcft114/.julia/packages/AdvancedHMC/YWXfk/src/hamiltonian.jl:36
┌ Warning: Incorrect ϵ = NaN; ϵ_previous = 0.07500000000000001 is used instead.
└ @ AdvancedHMC.Adaptation /Users/kcft114/.julia/packages/AdvancedHMC/YWXfk/src/adaptation/stepsize.jl:103
┌ Warning: The current proposal will be rejected due to numerical error(s).
│   isfiniteθ = true
│   isfiniter = true
│   isfiniteℓπ = false
│   isfiniteℓκ = true
└ @ AdvancedHMC /Users/kcft114/.julia/packages/AdvancedHMC/YWXfk/src/hamiltonian.jl:36
┌ Warning: The current proposal will be rejected due to numerical error(s).
│   isfiniteθ = false
│   isfiniter = false
│   isfiniteℓπ = false
│   isfiniteℓκ = false
└ @ AdvancedHMC /Users/kcft114/.julia/packages/AdvancedHMC/YWXfk/src/hamiltonian.jl:36
┌ Warning: Incorrect ϵ = NaN; ϵ_previous = 0.07500000000000001 is used instead.
└ @ AdvancedHMC.Adaptation /Users/kcft114/.julia/packages/AdvancedHMC/YWXfk/src/adaptation/stepsize.jl:103
┌ Warning: The current proposal will be rejected due to numerical error(s).
│   isfiniteθ = true
│   isfiniter = true
│   isfiniteℓπ = false
│   isfiniteℓκ = true
└ @ AdvancedHMC /Users/kcft114/.julia/packages/AdvancedHMC/YWXfk/src/hamiltonian.jl:36
┌ Warning: The current proposal will be rejected due to numerical error(s).
│   isfiniteθ = false
│   isfiniter = false
│   isfiniteℓπ = false
│   isfiniteℓκ = false
└ @ AdvancedHMC /Users/kcft114/.julia/packages/AdvancedHMC/YWXfk/src/hamiltonian.jl:36
┌ Warning: Incorrect ϵ = NaN; ϵ_previous = 0.07500000000000001 is used instead.
└ @ AdvancedHMC.Adaptation /Users/kcft114/.julia/packages/AdvancedHMC/YWXfk/src/adaptation/stepsize.jl:103
┌ Warning: The current proposal will be rejected due to numerical error(s).
│   isfiniteθ = true
│   isfiniter = true
│   isfiniteℓπ = false
│   isfiniteℓκ = true
└ @ AdvancedHMC /Users/kcft114/.julia/packages/AdvancedHMC/YWXfk/src/hamiltonian.jl:36
┌ Warning: The current proposal will be rejected due to numerical error(s).
│   isfiniteθ = false
│   isfiniter = false
│   isfiniteℓπ = false
│   isfiniteℓκ = false
└ @ AdvancedHMC /Users/kcft114/.julia/packages/AdvancedHMC/YWXfk/src/hamiltonian.jl:36
┌ Warning: Incorrect ϵ = NaN; ϵ_previous = 0.07500000000000001 is used instead.
└ @ AdvancedHMC.Adaptation /Users/kcft114/.julia/packages/AdvancedHMC/YWXfk/src/adaptation/stepsize.jl:103
┌ Warning: The current proposal will be rejected due to numerical error(s).
│   isfiniteθ = true
│   isfiniter = true
│   isfiniteℓπ = false
│   isfiniteℓκ = true
└ @ AdvancedHMC /Users/kcft114/.julia/packages/AdvancedHMC/YWXfk/src/hamiltonian.jl:36
┌ Warning: The current proposal will be rejected due to numerical error(s).
│   isfiniteθ = false
│   isfiniter = false
│   isfiniteℓπ = false
│   isfiniteℓκ = false
└ @ AdvancedHMC /Users/kcft114/.julia/packages/AdvancedHMC/YWXfk/src/hamiltonian.jl:36
┌ Warning: Incorrect ϵ = NaN; ϵ_previous = 0.07500000000000001 is used instead.
└ @ AdvancedHMC.Adaptation /Users/kcft114/.julia/packages/AdvancedHMC/YWXfk/src/adaptation/stepsize.jl:103
┌ Warning: The current proposal will be rejected due to numerical error(s).
│   isfiniteθ = true
│   isfiniter = true
│   isfiniteℓπ = false
│   isfiniteℓκ = true
└ @ AdvancedHMC /Users/kcft114/.julia/packages/AdvancedHMC/YWXfk/src/hamiltonian.jl:36
┌ Warning: The current proposal will be rejected due to numerical error(s).
│   isfiniteθ = false
│   isfiniter = false
│   isfiniteℓπ = false
│   isfiniteℓκ = false
└ @ AdvancedHMC /Users/kcft114/.julia/packages/AdvancedHMC/YWXfk/src/hamiltonian.jl:36
┌ Warning: Incorrect ϵ = NaN; ϵ_previous = 0.07500000000000001 is used instead.
└ @ AdvancedHMC.Adaptation /Users/kcft114/.julia/packages/AdvancedHMC/YWXfk/src/adaptation/stepsize.jl:103
┌ Warning: The current proposal will be rejected due to numerical error(s).
│   isfiniteθ = true
│   isfiniter = true
│   isfiniteℓπ = false
│   isfiniteℓκ = true
└ @ AdvancedHMC /Users/kcft114/.julia/packages/AdvancedHMC/YWXfk/src/hamiltonian.jl:36
┌ Warning: The current proposal will be rejected due to numerical error(s).
│   isfiniteθ = false
│   isfiniter = false
│   isfiniteℓπ = false
│   isfiniteℓκ = false
└ @ AdvancedHMC /Users/kcft114/.julia/packages/AdvancedHMC/YWXfk/src/hamiltonian.jl:36
┌ Warning: Incorrect ϵ = NaN; ϵ_previous = 0.07500000000000001 is used instead.
└ @ AdvancedHMC.Adaptation /Users/kcft114/.julia/packages/AdvancedHMC/YWXfk/src/adaptation/stepsize.jl:103
┌ Warning: The current proposal will be rejected due to numerical error(s).
│   isfiniteθ = true
│   isfiniter = true
│   isfiniteℓπ = false
│   isfiniteℓκ = true
└ @ AdvancedHMC /Users/kcft114/.julia/packages/AdvancedHMC/YWXfk/src/hamiltonian.jl:36
┌ Warning: The current proposal will be rejected due to numerical error(s).
│   isfiniteθ = false
│   isfiniter = false
│   isfiniteℓπ = false
│   isfiniteℓκ = false
└ @ AdvancedHMC /Users/kcft114/.julia/packages/AdvancedHMC/YWXfk/src/hamiltonian.jl:36
┌ Warning: Incorrect ϵ = NaN; ϵ_previous = 0.07500000000000001 is used instead.
└ @ AdvancedHMC.Adaptation /Users/kcft114/.julia/packages/AdvancedHMC/YWXfk/src/adaptation/stepsize.jl:103
┌ Warning: The current proposal will be rejected due to numerical error(s).
│   isfiniteθ = true
│   isfiniter = true
│   isfiniteℓπ = false
│   isfiniteℓκ = true
└ @ AdvancedHMC /Users/kcft114/.julia/packages/AdvancedHMC/YWXfk/src/hamiltonian.jl:36
┌ Warning: The current proposal will be rejected due to numerical error(s).
│   isfiniteθ = false
│   isfiniter = false
│   isfiniteℓπ = false
│   isfiniteℓκ = false
└ @ AdvancedHMC /Users/kcft114/.julia/packages/AdvancedHMC/YWXfk/src/hamiltonian.jl:36
┌ Warning: Incorrect ϵ = NaN; ϵ_previous = 0.07500000000000001 is used instead.
└ @ AdvancedHMC.Adaptation /Users/kcft114/.julia/packages/AdvancedHMC/YWXfk/src/adaptation/stepsize.jl:103
┌ Warning: The current proposal will be rejected due to numerical error(s).
│   isfiniteθ = true
│   isfiniter = true
│   isfiniteℓπ = false
│   isfiniteℓκ = true
└ @ AdvancedHMC /Users/kcft114/.julia/packages/AdvancedHMC/YWXfk/src/hamiltonian.jl:36
┌ Warning: The current proposal will be rejected due to numerical error(s).
│   isfiniteθ = false
│   isfiniter = false
│   isfiniteℓπ = false
│   isfiniteℓκ = false
└ @ AdvancedHMC /Users/kcft114/.julia/packages/AdvancedHMC/YWXfk/src/hamiltonian.jl:36
┌ Warning: Incorrect ϵ = NaN; ϵ_previous = 0.07500000000000001 is used instead.
└ @ AdvancedHMC.Adaptation /Users/kcft114/.julia/packages/AdvancedHMC/YWXfk/src/adaptation/stepsize.jl:103
┌ Warning: The current proposal will be rejected due to numerical error(s).
│   isfiniteθ = true
│   isfiniter = true
│   isfiniteℓπ = false
│   isfiniteℓκ = true
└ @ AdvancedHMC /Users/kcft114/.julia/packages/AdvancedHMC/YWXfk/src/hamiltonian.jl:36
┌ Warning: The current proposal will be rejected due to numerical error(s).
│   isfiniteθ = false
│   isfiniter = false
│   isfiniteℓπ = false
│   isfiniteℓκ = false
└ @ AdvancedHMC /Users/kcft114/.julia/packages/AdvancedHMC/YWXfk/src/hamiltonian.jl:36
┌ Warning: Incorrect ϵ = NaN; ϵ_previous = 0.07500000000000001 is used instead.
└ @ AdvancedHMC.Adaptation /Users/kcft114/.julia/packages/AdvancedHMC/YWXfk/src/adaptation/stepsize.jl:103
┌ Warning: The current proposal will be rejected due to numerical error(s).
│   isfiniteθ = true
│   isfiniter = true
│   isfiniteℓπ = false
│   isfiniteℓκ = true
└ @ AdvancedHMC /Users/kcft114/.julia/packages/AdvancedHMC/YWXfk/src/hamiltonian.jl:36
┌ Warning: The current proposal will be rejected due to numerical error(s).
│   isfiniteθ = false
│   isfiniter = false
│   isfiniteℓπ = false
│   isfiniteℓκ = false
└ @ AdvancedHMC /Users/kcft114/.julia/packages/AdvancedHMC/YWXfk/src/hamiltonian.jl:36
┌ Warning: Incorrect ϵ = NaN; ϵ_previous = 0.07500000000000001 is used instead.
└ @ AdvancedHMC.Adaptation /Users/kcft114/.julia/packages/AdvancedHMC/YWXfk/src/adaptation/stepsize.jl:103
┌ Warning: The current proposal will be rejected due to numerical error(s).
│   isfiniteθ = true
│   isfiniter = true
│   isfiniteℓπ = false
│   isfiniteℓκ = true
└ @ AdvancedHMC /Users/kcft114/.julia/packages/AdvancedHMC/YWXfk/src/hamiltonian.jl:36
┌ Warning: The current proposal will be rejected due to numerical error(s).
│   isfiniteθ = false
│   isfiniter = false
│   isfiniteℓπ = false
│   isfiniteℓκ = false
└ @ AdvancedHMC /Users/kcft114/.julia/packages/AdvancedHMC/YWXfk/src/hamiltonian.jl:36
┌ Warning: Incorrect ϵ = NaN; ϵ_previous = 0.07500000000000001 is used instead.
└ @ AdvancedHMC.Adaptation /Users/kcft114/.julia/packages/AdvancedHMC/YWXfk/src/adaptation/stepsize.jl:103
┌ Warning: The current proposal will be rejected due to numerical error(s).
│   isfiniteθ = true
│   isfiniter = true
│   isfiniteℓπ = false
│   isfiniteℓκ = true
└ @ AdvancedHMC /Users/kcft114/.julia/packages/AdvancedHMC/YWXfk/src/hamiltonian.jl:36
┌ Warning: The current proposal will be rejected due to numerical error(s).
│   isfiniteθ = false
│   isfiniter = false
│   isfiniteℓπ = false
│   isfiniteℓκ = false
└ @ AdvancedHMC /Users/kcft114/.julia/packages/AdvancedHMC/YWXfk/src/hamiltonian.jl:36
┌ Warning: Incorrect ϵ = NaN; ϵ_previous = 0.07500000000000001 is used instead.
└ @ AdvancedHMC.Adaptation /Users/kcft114/.julia/packages/AdvancedHMC/YWXfk/src/adaptation/stepsize.jl:103
┌ Warning: The current proposal will be rejected due to numerical error(s).
│   isfiniteθ = true
│   isfiniter = true
│   isfiniteℓπ = false
│   isfiniteℓκ = true
└ @ AdvancedHMC /Users/kcft114/.julia/packages/AdvancedHMC/YWXfk/src/hamiltonian.jl:36
┌ Warning: The current proposal will be rejected due to numerical error(s).
│   isfiniteθ = false
│   isfiniter = false
│   isfiniteℓπ = false
│   isfiniteℓκ = false
└ @ AdvancedHMC /Users/kcft114/.julia/packages/AdvancedHMC/YWXfk/src/hamiltonian.jl:36
┌ Warning: Incorrect ϵ = NaN; ϵ_previous = 0.07500000000000001 is used instead.
└ @ AdvancedHMC.Adaptation /Users/kcft114/.julia/packages/AdvancedHMC/YWXfk/src/adaptation/stepsize.jl:103
┌ Warning: The current proposal will be rejected due to numerical error(s).
│   isfiniteθ = true
│   isfiniter = true
│   isfiniteℓπ = false
│   isfiniteℓκ = true
└ @ AdvancedHMC /Users/kcft114/.julia/packages/AdvancedHMC/YWXfk/src/hamiltonian.jl:36
┌ Warning: The current proposal will be rejected due to numerical error(s).
│   isfiniteθ = false
│   isfiniter = false
│   isfiniteℓπ = false
│   isfiniteℓκ = false
└ @ AdvancedHMC /Users/kcft114/.julia/packages/AdvancedHMC/YWXfk/src/hamiltonian.jl:36
┌ Warning: Incorrect ϵ = NaN; ϵ_previous = 0.07500000000000001 is used instead.
└ @ AdvancedHMC.Adaptation /Users/kcft114/.julia/packages/AdvancedHMC/YWXfk/src/adaptation/stepsize.jl:103
┌ Warning: The current proposal will be rejected due to numerical error(s).
│   isfiniteθ = true
│   isfiniter = true
│   isfiniteℓπ = false
│   isfiniteℓκ = true
└ @ AdvancedHMC /Users/kcft114/.julia/packages/AdvancedHMC/YWXfk/src/hamiltonian.jl:36
┌ Warning: The current proposal will be rejected due to numerical error(s).
│   isfiniteθ = false
│   isfiniter = false
│   isfiniteℓπ = false
│   isfiniteℓκ = false
└ @ AdvancedHMC /Users/kcft114/.julia/packages/AdvancedHMC/YWXfk/src/hamiltonian.jl:36
┌ Warning: Incorrect ϵ = NaN; ϵ_previous = 0.07500000000000001 is used instead.
└ @ AdvancedHMC.Adaptation /Users/kcft114/.julia/packages/AdvancedHMC/YWXfk/src/adaptation/stepsize.jl:103
┌ Warning: The current proposal will be rejected due to numerical error(s).
│   isfiniteθ = true
│   isfiniter = true
│   isfiniteℓπ = false
│   isfiniteℓκ = true
└ @ AdvancedHMC /Users/kcft114/.julia/packages/AdvancedHMC/YWXfk/src/hamiltonian.jl:36
┌ Warning: The current proposal will be rejected due to numerical error(s).
│   isfiniteθ = false
│   isfiniter = false
│   isfiniteℓπ = false
│   isfiniteℓκ = false
└ @ AdvancedHMC /Users/kcft114/.julia/packages/AdvancedHMC/YWXfk/src/hamiltonian.jl:36
┌ Warning: Incorrect ϵ = NaN; ϵ_previous = 0.07500000000000001 is used instead.
└ @ AdvancedHMC.Adaptation /Users/kcft114/.julia/packages/AdvancedHMC/YWXfk/src/adaptation/stepsize.jl:103
┌ Warning: The current proposal will be rejected due to numerical error(s).
│   isfiniteθ = true
│   isfiniter = true
│   isfiniteℓπ = false
│   isfiniteℓκ = true
└ @ AdvancedHMC /Users/kcft114/.julia/packages/AdvancedHMC/YWXfk/src/hamiltonian.jl:36
┌ Warning: The current proposal will be rejected due to numerical error(s).
│   isfiniteθ = false
│   isfiniter = false
│   isfiniteℓπ = false
│   isfiniteℓκ = false
└ @ AdvancedHMC /Users/kcft114/.julia/packages/AdvancedHMC/YWXfk/src/hamiltonian.jl:36
┌ Warning: Incorrect ϵ = NaN; ϵ_previous = 0.07500000000000001 is used instead.
└ @ AdvancedHMC.Adaptation /Users/kcft114/.julia/packages/AdvancedHMC/YWXfk/src/adaptation/stepsize.jl:103
┌ Warning: The current proposal will be rejected due to numerical error(s).
│   isfiniteθ = true
│   isfiniter = true
│   isfiniteℓπ = false
│   isfiniteℓκ = true
└ @ AdvancedHMC /Users/kcft114/.julia/packages/AdvancedHMC/YWXfk/src/hamiltonian.jl:36
┌ Warning: The current proposal will be rejected due to numerical error(s).
│   isfiniteθ = false
│   isfiniter = false
│   isfiniteℓπ = false
│   isfiniteℓκ = false
└ @ AdvancedHMC /Users/kcft114/.julia/packages/AdvancedHMC/YWXfk/src/hamiltonian.jl:36
┌ Warning: Incorrect ϵ = NaN; ϵ_previous = 0.07500000000000001 is used instead.
└ @ AdvancedHMC.Adaptation /Users/kcft114/.julia/packages/AdvancedHMC/YWXfk/src/adaptation/stepsize.jl:103
┌ Warning: The current proposal will be rejected due to numerical error(s).
│   isfiniteθ = true
│   isfiniter = true
│   isfiniteℓπ = false
│   isfiniteℓκ = true
└ @ AdvancedHMC /Users/kcft114/.julia/packages/AdvancedHMC/YWXfk/src/hamiltonian.jl:36
┌ Warning: The current proposal will be rejected due to numerical error(s).
│   isfiniteθ = false
│   isfiniter = false
│   isfiniteℓπ = false
│   isfiniteℓκ = false
└ @ AdvancedHMC /Users/kcft114/.julia/packages/AdvancedHMC/YWXfk/src/hamiltonian.jl:36
┌ Warning: Incorrect ϵ = NaN; ϵ_previous = 0.07500000000000001 is used instead.
└ @ AdvancedHMC.Adaptation /Users/kcft114/.julia/packages/AdvancedHMC/YWXfk/src/adaptation/stepsize.jl:103
┌ Warning: The current proposal will be rejected due to numerical error(s).
│   isfiniteθ = true
│   isfiniter = true
│   isfiniteℓπ = false
│   isfiniteℓκ = true
└ @ AdvancedHMC /Users/kcft114/.julia/packages/AdvancedHMC/YWXfk/src/hamiltonian.jl:36
┌ Warning: The current proposal will be rejected due to numerical error(s).
│   isfiniteθ = false
│   isfiniter = false
│   isfiniteℓπ = false
│   isfiniteℓκ = false
└ @ AdvancedHMC /Users/kcft114/.julia/packages/AdvancedHMC/YWXfk/src/hamiltonian.jl:36
┌ Warning: Incorrect ϵ = NaN; ϵ_previous = 0.07500000000000001 is used instead.
└ @ AdvancedHMC.Adaptation /Users/kcft114/.julia/packages/AdvancedHMC/YWXfk/src/adaptation/stepsize.jl:103
┌ Warning: The current proposal will be rejected due to numerical error(s).
│   isfiniteθ = true
│   isfiniter = true
│   isfiniteℓπ = false
│   isfiniteℓκ = true
└ @ AdvancedHMC /Users/kcft114/.julia/packages/AdvancedHMC/YWXfk/src/hamiltonian.jl:36
┌ Warning: The current proposal will be rejected due to numerical error(s).
│   isfiniteθ = false
│   isfiniter = false
│   isfiniteℓπ = false
│   isfiniteℓκ = false
└ @ AdvancedHMC /Users/kcft114/.julia/packages/AdvancedHMC/YWXfk/src/hamiltonian.jl:36
┌ Warning: Incorrect ϵ = NaN; ϵ_previous = 0.07500000000000001 is used instead.
└ @ AdvancedHMC.Adaptation /Users/kcft114/.julia/packages/AdvancedHMC/YWXfk/src/adaptation/stepsize.jl:103
┌ Warning: The current proposal will be rejected due to numerical error(s).
│   isfiniteθ = true
│   isfiniter = true
│   isfiniteℓπ = false
│   isfiniteℓκ = true
└ @ AdvancedHMC /Users/kcft114/.julia/packages/AdvancedHMC/YWXfk/src/hamiltonian.jl:36
┌ Warning: The current proposal will be rejected due to numerical error(s).
│   isfiniteθ = false
│   isfiniter = false
│   isfiniteℓπ = false
│   isfiniteℓκ = false
└ @ AdvancedHMC /Users/kcft114/.julia/packages/AdvancedHMC/YWXfk/src/hamiltonian.jl:36
┌ Warning: Incorrect ϵ = NaN; ϵ_previous = 0.07500000000000001 is used instead.
└ @ AdvancedHMC.Adaptation /Users/kcft114/.julia/packages/AdvancedHMC/YWXfk/src/adaptation/stepsize.jl:103
┌ Warning: The current proposal will be rejected due to numerical error(s).
│   isfiniteθ = true
│   isfiniter = true
│   isfiniteℓπ = false
│   isfiniteℓκ = true
└ @ AdvancedHMC /Users/kcft114/.julia/packages/AdvancedHMC/YWXfk/src/hamiltonian.jl:36
┌ Warning: The current proposal will be rejected due to numerical error(s).
│   isfiniteθ = false
│   isfiniter = false
│   isfiniteℓπ = false
│   isfiniteℓκ = false
└ @ AdvancedHMC /Users/kcft114/.julia/packages/AdvancedHMC/YWXfk/src/hamiltonian.jl:36
┌ Warning: Incorrect ϵ = NaN; ϵ_previous = 0.07500000000000001 is used instead.
└ @ AdvancedHMC.Adaptation /Users/kcft114/.julia/packages/AdvancedHMC/YWXfk/src/adaptation/stepsize.jl:103
┌ Warning: The current proposal will be rejected due to numerical error(s).
│   isfiniteθ = true
│   isfiniter = true
│   isfiniteℓπ = false
│   isfiniteℓκ = true
└ @ AdvancedHMC /Users/kcft114/.julia/packages/AdvancedHMC/YWXfk/src/hamiltonian.jl:36
┌ Warning: The current proposal will be rejected due to numerical error(s).
│   isfiniteθ = false
│   isfiniter = false
│   isfiniteℓπ = false
│   isfiniteℓκ = false
└ @ AdvancedHMC /Users/kcft114/.julia/packages/AdvancedHMC/YWXfk/src/hamiltonian.jl:36
┌ Warning: Incorrect ϵ = NaN; ϵ_previous = 0.07500000000000001 is used instead.
└ @ AdvancedHMC.Adaptation /Users/kcft114/.julia/packages/AdvancedHMC/YWXfk/src/adaptation/stepsize.jl:103
┌ Warning: The current proposal will be rejected due to numerical error(s).
│   isfiniteθ = true
│   isfiniter = true
│   isfiniteℓπ = false
│   isfiniteℓκ = true
└ @ AdvancedHMC /Users/kcft114/.julia/packages/AdvancedHMC/YWXfk/src/hamiltonian.jl:36
┌ Warning: The current proposal will be rejected due to numerical error(s).
│   isfiniteθ = false
│   isfiniter = false
│   isfiniteℓπ = false
│   isfiniteℓκ = false
└ @ AdvancedHMC /Users/kcft114/.julia/packages/AdvancedHMC/YWXfk/src/hamiltonian.jl:36
┌ Warning: Incorrect ϵ = NaN; ϵ_previous = 0.07500000000000001 is used instead.
└ @ AdvancedHMC.Adaptation /Users/kcft114/.julia/packages/AdvancedHMC/YWXfk/src/adaptation/stepsize.jl:103
┌ Warning: The current proposal will be rejected due to numerical error(s).
│   isfiniteθ = true
│   isfiniter = true
│   isfiniteℓπ = false
│   isfiniteℓκ = true
└ @ AdvancedHMC /Users/kcft114/.julia/packages/AdvancedHMC/YWXfk/src/hamiltonian.jl:36
┌ Warning: The current proposal will be rejected due to numerical error(s).
│   isfiniteθ = false
│   isfiniter = false
│   isfiniteℓπ = false
│   isfiniteℓκ = false
└ @ AdvancedHMC /Users/kcft114/.julia/packages/AdvancedHMC/YWXfk/src/hamiltonian.jl:36
┌ Warning: Incorrect ϵ = NaN; ϵ_previous = 0.07500000000000001 is used instead.
└ @ AdvancedHMC.Adaptation /Users/kcft114/.julia/packages/AdvancedHMC/YWXfk/src/adaptation/stepsize.jl:103
┌ Warning: The current proposal will be rejected due to numerical error(s).
│   isfiniteθ = true
│   isfiniter = true
│   isfiniteℓπ = false
│   isfiniteℓκ = true
└ @ AdvancedHMC /Users/kcft114/.julia/packages/AdvancedHMC/YWXfk/src/hamiltonian.jl:36
┌ Warning: The current proposal will be rejected due to numerical error(s).
│   isfiniteθ = false
│   isfiniter = false
│   isfiniteℓπ = false
│   isfiniteℓκ = false
└ @ AdvancedHMC /Users/kcft114/.julia/packages/AdvancedHMC/YWXfk/src/hamiltonian.jl:36
┌ Warning: Incorrect ϵ = NaN; ϵ_previous = 0.07500000000000001 is used instead.
└ @ AdvancedHMC.Adaptation /Users/kcft114/.julia/packages/AdvancedHMC/YWXfk/src/adaptation/stepsize.jl:103
┌ Warning: The current proposal will be rejected due to numerical error(s).
│   isfiniteθ = true
│   isfiniter = true
│   isfiniteℓπ = false
│   isfiniteℓκ = true
└ @ AdvancedHMC /Users/kcft114/.julia/packages/AdvancedHMC/YWXfk/src/hamiltonian.jl:36
┌ Warning: The current proposal will be rejected due to numerical error(s).
│   isfiniteθ = false
│   isfiniter = false
│   isfiniteℓπ = false
│   isfiniteℓκ = false
└ @ AdvancedHMC /Users/kcft114/.julia/packages/AdvancedHMC/YWXfk/src/hamiltonian.jl:36
┌ Warning: Incorrect ϵ = NaN; ϵ_previous = 0.07500000000000001 is used instead.
└ @ AdvancedHMC.Adaptation /Users/kcft114/.julia/packages/AdvancedHMC/YWXfk/src/adaptation/stepsize.jl:103
┌ Warning: The current proposal will be rejected due to numerical error(s).
│   isfiniteθ = true
│   isfiniter = true
│   isfiniteℓπ = false
│   isfiniteℓκ = true
└ @ AdvancedHMC /Users/kcft114/.julia/packages/AdvancedHMC/YWXfk/src/hamiltonian.jl:36
┌ Warning: The current proposal will be rejected due to numerical error(s).
│   isfiniteθ = false
│   isfiniter = false
│   isfiniteℓπ = false
│   isfiniteℓκ = false
└ @ AdvancedHMC /Users/kcft114/.julia/packages/AdvancedHMC/YWXfk/src/hamiltonian.jl:36
┌ Warning: Incorrect ϵ = NaN; ϵ_previous = 0.07500000000000001 is used instead.
└ @ AdvancedHMC.Adaptation /Users/kcft114/.julia/packages/AdvancedHMC/YWXfk/src/adaptation/stepsize.jl:103
┌ Warning: The current proposal will be rejected due to numerical error(s).
│   isfiniteθ = true
│   isfiniter = true
│   isfiniteℓπ = false
│   isfiniteℓκ = true
└ @ AdvancedHMC /Users/kcft114/.julia/packages/AdvancedHMC/YWXfk/src/hamiltonian.jl:36
┌ Warning: The current proposal will be rejected due to numerical error(s).
│   isfiniteθ = false
│   isfiniter = false
│   isfiniteℓπ = false
│   isfiniteℓκ = false
└ @ AdvancedHMC /Users/kcft114/.julia/packages/AdvancedHMC/YWXfk/src/hamiltonian.jl:36
┌ Warning: Incorrect ϵ = NaN; ϵ_previous = 0.07500000000000001 is used instead.
└ @ AdvancedHMC.Adaptation /Users/kcft114/.julia/packages/AdvancedHMC/YWXfk/src/adaptation/stepsize.jl:103
┌ Warning: The current proposal will be rejected due to numerical error(s).
│   isfiniteθ = true
│   isfiniter = true
│   isfiniteℓπ = false
│   isfiniteℓκ = true
└ @ AdvancedHMC /Users/kcft114/.julia/packages/AdvancedHMC/YWXfk/src/hamiltonian.jl:36
┌ Warning: The current proposal will be rejected due to numerical error(s).
│   isfiniteθ = false
│   isfiniter = false
│   isfiniteℓπ = false
│   isfiniteℓκ = false
└ @ AdvancedHMC /Users/kcft114/.julia/packages/AdvancedHMC/YWXfk/src/hamiltonian.jl:36
┌ Warning: Incorrect ϵ = NaN; ϵ_previous = 0.07500000000000001 is used instead.
└ @ AdvancedHMC.Adaptation /Users/kcft114/.julia/packages/AdvancedHMC/YWXfk/src/adaptation/stepsize.jl:103
┌ Warning: The current proposal will be rejected due to numerical error(s).
│   isfiniteθ = true
│   isfiniter = true
│   isfiniteℓπ = false
│   isfiniteℓκ = true
└ @ AdvancedHMC /Users/kcft114/.julia/packages/AdvancedHMC/YWXfk/src/hamiltonian.jl:36
┌ Warning: The current proposal will be rejected due to numerical error(s).
│   isfiniteθ = false
│   isfiniter = false
│   isfiniteℓπ = false
│   isfiniteℓκ = false
└ @ AdvancedHMC /Users/kcft114/.julia/packages/AdvancedHMC/YWXfk/src/hamiltonian.jl:36
┌ Warning: Incorrect ϵ = NaN; ϵ_previous = 0.07500000000000001 is used instead.
└ @ AdvancedHMC.Adaptation /Users/kcft114/.julia/packages/AdvancedHMC/YWXfk/src/adaptation/stepsize.jl:103
┌ Warning: The current proposal will be rejected due to numerical error(s).
│   isfiniteθ = true
│   isfiniter = true
│   isfiniteℓπ = false
│   isfiniteℓκ = true
└ @ AdvancedHMC /Users/kcft114/.julia/packages/AdvancedHMC/YWXfk/src/hamiltonian.jl:36
┌ Warning: The current proposal will be rejected due to numerical error(s).
│   isfiniteθ = false
│   isfiniter = false
│   isfiniteℓπ = false
│   isfiniteℓκ = false
└ @ AdvancedHMC /Users/kcft114/.julia/packages/AdvancedHMC/YWXfk/src/hamiltonian.jl:36
┌ Warning: Incorrect ϵ = NaN; ϵ_previous = 0.07500000000000001 is used instead.
└ @ AdvancedHMC.Adaptation /Users/kcft114/.julia/packages/AdvancedHMC/YWXfk/src/adaptation/stepsize.jl:103
┌ Warning: The current proposal will be rejected due to numerical error(s).
│   isfiniteθ = true
│   isfiniter = true
│   isfiniteℓπ = false
│   isfiniteℓκ = true
└ @ AdvancedHMC /Users/kcft114/.julia/packages/AdvancedHMC/YWXfk/src/hamiltonian.jl:36
┌ Warning: The current proposal will be rejected due to numerical error(s).
│   isfiniteθ = false
│   isfiniter = false
│   isfiniteℓπ = false
│   isfiniteℓκ = false
└ @ AdvancedHMC /Users/kcft114/.julia/packages/AdvancedHMC/YWXfk/src/hamiltonian.jl:36
┌ Warning: Incorrect ϵ = NaN; ϵ_previous = 0.07500000000000001 is used instead.
└ @ AdvancedHMC.Adaptation /Users/kcft114/.julia/packages/AdvancedHMC/YWXfk/src/adaptation/stepsize.jl:103
┌ Warning: The current proposal will be rejected due to numerical error(s).
│   isfiniteθ = true
│   isfiniter = true
│   isfiniteℓπ = false
│   isfiniteℓκ = true
└ @ AdvancedHMC /Users/kcft114/.julia/packages/AdvancedHMC/YWXfk/src/hamiltonian.jl:36
┌ Warning: The current proposal will be rejected due to numerical error(s).
│   isfiniteθ = false
│   isfiniter = false
│   isfiniteℓπ = false
│   isfiniteℓκ = false
└ @ AdvancedHMC /Users/kcft114/.julia/packages/AdvancedHMC/YWXfk/src/hamiltonian.jl:36
┌ Warning: Incorrect ϵ = NaN; ϵ_previous = 0.07500000000000001 is used instead.
└ @ AdvancedHMC.Adaptation /Users/kcft114/.julia/packages/AdvancedHMC/YWXfk/src/adaptation/stepsize.jl:103
┌ Warning: The current proposal will be rejected due to numerical error(s).
│   isfiniteθ = true
│   isfiniter = true
│   isfiniteℓπ = false
│   isfiniteℓκ = true
└ @ AdvancedHMC /Users/kcft114/.julia/packages/AdvancedHMC/YWXfk/src/hamiltonian.jl:36
┌ Warning: The current proposal will be rejected due to numerical error(s).
│   isfiniteθ = false
│   isfiniter = false
│   isfiniteℓπ = false
│   isfiniteℓκ = false
└ @ AdvancedHMC /Users/kcft114/.julia/packages/AdvancedHMC/YWXfk/src/hamiltonian.jl:36
┌ Warning: Incorrect ϵ = NaN; ϵ_previous = 0.07500000000000001 is used instead.
└ @ AdvancedHMC.Adaptation /Users/kcft114/.julia/packages/AdvancedHMC/YWXfk/src/adaptation/stepsize.jl:103
┌ Warning: The current proposal will be rejected due to numerical error(s).
│   isfiniteθ = true
│   isfiniter = true
│   isfiniteℓπ = false
│   isfiniteℓκ = true
└ @ AdvancedHMC /Users/kcft114/.julia/packages/AdvancedHMC/YWXfk/src/hamiltonian.jl:36
Excessive output truncated after 524589 bytes.┌ Warning: The current proposal will be rejected due to numerical error(s).
│   isfiniteθ = false
│   isfiniter = false
│   isfiniteℓπ = false
│   isfiniteℓκ = false
└ @ AdvancedHMC /Users/kcft114/.julia/packages/AdvancedHMC/YWXfk/src/hamiltonian.jl:36
┌ Warning: Incorrect ϵ = NaN; ϵ_previous = 0.07500000000000001 is used instead.
└ @ AdvancedHMC.Adaptation /Users/kcft114/.julia/packages/AdvancedHMC/YWXfk/src/adaptation/stepsize.jl:103
┌ Warning: The current proposal will be rejected due to numerical error(s).
│   isfiniteθ = true
│   isfiniter = true
│   isfiniteℓπ = false
│   isfiniteℓκ = true
└ @ AdvancedHMC /Users/kcft114/.julia/packages/AdvancedHMC/YWXfk/src/hamiltonian.jl:36
┌ Warning: The current proposal will be rejected due to numerical error(s).
│   isfiniteθ = false
│   isfiniter = false
│   isfiniteℓπ = false
│   isfiniteℓκ = false
└ @ AdvancedHMC /Users/kcft114/.julia/packages/AdvancedHMC/YWXfk/src/hamiltonian.jl:36
┌ Warning: Incorrect ϵ = NaN; ϵ_previous = 0.07500000000000001 is used instead.
└ @ AdvancedHMC.Adaptation /Users/kcft114/.julia/packages/AdvancedHMC/YWXfk/src/adaptation/stepsize.jl:103
┌ Warning: The current proposal will be rejected due to numerical error(s).
│   isfiniteθ = true
│   isfiniter = true
│   isfiniteℓπ = false
│   isfiniteℓκ = true
└ @ AdvancedHMC /Users/kcft114/.julia/packages/AdvancedHMC/YWXfk/src/hamiltonian.jl:36
```

In [156]:

```
## ---------------------------------------------------------
## Save/read results
## ---------------------------------------------------------

#write("chains/BNN.jls", ch2)

ch2 = read("chains/BNN.jls", Chains);
```

In [157]:

```
## ---------------------------------------------------------
## Check convergence
## ---------------------------------------------------------
```

In [158]:

```
# should be true
!(sum(isnan.(summarystats(ch2)[:, :r_hat]))>0) | !(sum(abs.(summarystats(ch2)[:, :r_hat] .- 1) .> 0.1) > 0) | !(sum(summarystats(ch2)[:, :std] .< 1e-13) > 0)
```

Out[158]:

```
true
```

In [159]:

```
show(ch2)
```

```
Object of type Chains, with data of type 1000×163×1 Array{Union{Missing, Float64},3}

Log evidence      = 0.0
Iterations        = 1:1000
Thinning interval = 1
Chains            = 1
Samples per chain = 1000
internals         = eval_num, lp, acceptance_rate, hamiltonian_energy, is_accept, log_density, n_steps, numerical_error, step_size, tree_depth
parameters        = θ[142], θ[98], θ[102], θ[105], θ[26], θ[64], θ[106], θ[71], θ[76], θ[74], θ[66], θ[131], θ[83], θ[108], θ[127], θ[125], θ[121], θ[68], θ[135], θ[38], θ[85], θ[67], θ[17], θ[42], θ[69], θ[92], θ[99], θ[100], θ[43], θ[119], θ[72], θ[110], θ[50], θ[90], θ[91], θ[7], θ[44], θ[128], θ[136], θ[73], θ[14], θ[146], θ[148], θ[27], θ[33], θ[134], θ[118], θ[147], θ[82], θ[78], θ[101], c1, θ[104], θ[129], θ[13], θ[138], θ[21], θ[4], θ[81], θ[109], θ[59], θ[112], θ[54], θ[25], θ[1], θ[3], θ[97], θ[87], θ[2], θ[11], θ[88], θ[29], θ[28], θ[114], θ[94], θ[132], θ[57], θ[145], θ[144], θ[20], θ[19], θ[34], θ[58], θ[5], θ[24], θ[12], θ[36], θ[86], θ[124], θ[137], θ[150], θ[96], θ[143], θ[47], θ[62], θ[141], θ[53], θ[55], θ[133], sig, θ[123], θ[79], θ[84], θ[130], θ[117], θ[70], θ[52], θ[60], θ[93], θ[49], θ[80], θ[9], θ[115], θ[140], θ[139], θ[126], θ[103], θ[95], θ[46], θ[6], log_diff_c, θ[61], θ[75], θ[31], θ[48], θ[40], θ[22], θ[111], θ[63], θ[35], θ[30], θ[8], θ[149], θ[89], θ[15], θ[32], θ[16], θ[77], θ[51], θ[113], θ[45], θ[65], θ[23], θ[39], θ[56], θ[122], θ[41], θ[10], θ[116], θ[120], θ[37], θ[18], θ[107]

2-element Array{ChainDataFrame,1}

Summary Statistics

│ Row │ parameters │ mean         │ std      │ naive_se   │ mcse      │ ess     │ r_hat    │
│     │ Symbol     │ Float64      │ Float64  │ Float64    │ Float64   │ Any     │ Any      │
├─────┼────────────┼──────────────┼──────────┼────────────┼───────────┼─────────┼──────────┤
│ 1   │ c1         │ -1.44436     │ 1.47016  │ 0.0464907  │ 0.0977405 │ 300.759 │ 0.999181 │
│ 2   │ log_diff_c │ 0.784309     │ 0.171346 │ 0.00541845 │ 0.0140885 │ 91.669  │ 1.02946  │
│ 3   │ sig        │ 0.686834     │ 0.169961 │ 0.00537464 │ 0.0278532 │ 21.4614 │ 1.12508  │
│ 4   │ θ[1]       │ 0.0422689    │ 0.600846 │ 0.0190004  │ 0.01569   │ 651.117 │ 0.999001 │
│ 5   │ θ[2]       │ -0.0342268   │ 0.677427 │ 0.0214221  │ 0.0228516 │ 660.517 │ 0.999788 │
│ 6   │ θ[3]       │ 0.00167577   │ 0.631282 │ 0.0199629  │ 0.032024  │ 517.662 │ 0.999883 │
│ 7   │ θ[4]       │ 0.0402734    │ 0.664995 │ 0.021029   │ 0.0407456 │ 427.959 │ 1.00032  │
│ 8   │ θ[5]       │ 0.00870489   │ 0.626641 │ 0.0198161  │ 0.0240245 │ 589.892 │ 0.999002 │
│ 9   │ θ[6]       │ 0.019588     │ 0.661977 │ 0.0209335  │ 0.0237971 │ 416.184 │ 0.999019 │
│ 10  │ θ[7]       │ 0.0156781    │ 0.641215 │ 0.020277   │ 0.0174001 │ 745.939 │ 1.00046  │
│ 11  │ θ[8]       │ -0.00545789  │ 0.633486 │ 0.0200326  │ 0.0225871 │ 712.033 │ 1.00196  │
│ 12  │ θ[9]       │ 0.00586565   │ 0.642436 │ 0.0203156  │ 0.0252506 │ 809.086 │ 0.999029 │
│ 13  │ θ[10]      │ 0.00894735   │ 0.588179 │ 0.0185999  │ 0.0255452 │ 658.732 │ 0.999565 │
│ 14  │ θ[11]      │ 0.0251898    │ 0.639672 │ 0.0202282  │ 0.0272448 │ 546.131 │ 1.00069  │
│ 15  │ θ[12]      │ -0.00354834  │ 0.642872 │ 0.0203294  │ 0.0338149 │ 526.421 │ 1.00582  │
│ 16  │ θ[13]      │ 0.0241158    │ 0.688517 │ 0.0217728  │ 0.015765  │ 671.767 │ 0.999224 │
│ 17  │ θ[14]      │ 0.00787196   │ 0.663935 │ 0.0209955  │ 0.0226633 │ 810.544 │ 1.00305  │
│ 18  │ θ[15]      │ 0.00463807   │ 0.640747 │ 0.0202622  │ 0.0191591 │ 613.959 │ 0.999004 │
│ 19  │ θ[16]      │ 0.0246973    │ 0.811446 │ 0.0256602  │ 0.0394912 │ 351.497 │ 0.999822 │
│ 20  │ θ[17]      │ -0.000783169 │ 0.76273  │ 0.0241196  │ 0.0482357 │ 437.792 │ 0.999064 │
│ 21  │ θ[18]      │ -0.079562    │ 0.78084  │ 0.0246923  │ 0.0394498 │ 272.363 │ 1.00107  │
│ 22  │ θ[19]      │ 0.0416398    │ 0.834228 │ 0.0263806  │ 0.0733964 │ 189.269 │ 0.999881 │
│ 23  │ θ[20]      │ -0.0304084   │ 0.720399 │ 0.022781   │ 0.0291334 │ 491.221 │ 1.00207  │
│ 24  │ θ[21]      │ -0.0754394   │ 0.729785 │ 0.0230778  │ 0.0529389 │ 181.535 │ 1.00986  │
│ 25  │ θ[22]      │ 0.0277613    │ 0.760835 │ 0.0240597  │ 0.0460884 │ 536.339 │ 1.00564  │
│ 26  │ θ[23]      │ 0.0591377    │ 0.774903 │ 0.0245046  │ 0.0468759 │ 258.493 │ 1.00708  │
│ 27  │ θ[24]      │ 0.0154541    │ 0.809617 │ 0.0256023  │ 0.0468076 │ 344.816 │ 1.00312  │
│ 28  │ θ[25]      │ 0.0370477    │ 0.788557 │ 0.0249364  │ 0.0405882 │ 490.678 │ 0.999199 │
│ 29  │ θ[26]      │ 0.0481719    │ 0.808325 │ 0.0255615  │ 0.0410734 │ 206.62  │ 0.999094 │
│ 30  │ θ[27]      │ 0.0326515    │ 0.81422  │ 0.0257479  │ 0.0483771 │ 354.242 │ 1.00539  │
│ 31  │ θ[28]      │ 0.0234693    │ 0.730185 │ 0.0230905  │ 0.0460992 │ 211.2   │ 0.99929  │
│ 32  │ θ[29]      │ 0.0617091    │ 0.735885 │ 0.0232707  │ 0.030164  │ 524.463 │ 0.999935 │
│ 33  │ θ[30]      │ 0.0766002    │ 0.813313 │ 0.0257192  │ 0.0511192 │ 212.487 │ 1.00121  │
│ 34  │ θ[31]      │ -0.0282371   │ 0.706069 │ 0.0223279  │ 0.0416743 │ 776.177 │ 1.00721  │
│ 35  │ θ[32]      │ -0.0236984   │ 0.654161 │ 0.0206864  │ 0.0286448 │ 617.77  │ 1.0063   │
│ 36  │ θ[33]      │ -0.0278008   │ 0.703775 │ 0.0222553  │ 0.0323906 │ 496.921 │ 1.00713  │
│ 37  │ θ[34]      │ -0.0712652   │ 0.662199 │ 0.0209406  │ 0.0288917 │ 589.173 │ 1.00149  │
│ 38  │ θ[35]      │ 0.0042509    │ 0.689015 │ 0.0217886  │ 0.0282127 │ 410.904 │ 1.00259  │
│ 39  │ θ[36]      │ -0.0384582   │ 0.684191 │ 0.021636   │ 0.0438625 │ 594.525 │ 1.0006   │
│ 40  │ θ[37]      │ -0.0611018   │ 0.699754 │ 0.0221282  │ 0.0212707 │ 1000.0  │ 0.999619 │
│ 41  │ θ[38]      │ -0.0706976   │ 0.650278 │ 0.0205636  │ 0.0267317 │ 457.055 │ 0.999509 │
│ 42  │ θ[39]      │ -0.0105212   │ 0.657863 │ 0.0208034  │ 0.0243477 │ 668.805 │ 0.999054 │
│ 43  │ θ[40]      │ 0.0123109    │ 0.720139 │ 0.0227728  │ 0.0368649 │ 375.785 │ 1.0111   │
│ 44  │ θ[41]      │ -0.0465102   │ 0.737845 │ 0.0233327  │ 0.0258163 │ 687.854 │ 0.999756 │
│ 45  │ θ[42]      │ -0.0793486   │ 0.670704 │ 0.0212095  │ 0.0240119 │ 804.208 │ 0.999028 │
│ 46  │ θ[43]      │ -0.0395483   │ 0.697921 │ 0.0220702  │ 0.0388775 │ 343.305 │ 0.999    │
│ 47  │ θ[44]      │ -0.00675606  │ 0.678076 │ 0.0214426  │ 0.0405659 │ 293.601 │ 1.00738  │
│ 48  │ θ[45]      │ -0.0220842   │ 0.641296 │ 0.0202796  │ 0.0166522 │ 835.627 │ 0.999823 │
│ 49  │ θ[46]      │ -0.0123402   │ 0.718803 │ 0.0227305  │ 0.029113  │ 555.526 │ 0.999024 │
│ 50  │ θ[47]      │ 0.0474252    │ 0.688874 │ 0.0217841  │ 0.0303535 │ 759.916 │ 1.00322  │
│ 51  │ θ[48]      │ 0.0390722    │ 0.759205 │ 0.0240082  │ 0.0334572 │ 521.338 │ 1.00078  │
│ 52  │ θ[49]      │ 0.0543181    │ 0.71237  │ 0.0225271  │ 0.0298782 │ 689.19  │ 0.999029 │
│ 53  │ θ[50]      │ -0.00387906  │ 0.700704 │ 0.0221582  │ 0.0232028 │ 675.53  │ 1.00041  │
│ 54  │ θ[51]      │ 0.0396675    │ 0.712992 │ 0.0225468  │ 0.0396895 │ 527.342 │ 0.999386 │
│ 55  │ θ[52]      │ 0.067716     │ 0.68361  │ 0.0216176  │ 0.0355076 │ 296.547 │ 1.01061  │
│ 56  │ θ[53]      │ 0.0324577    │ 0.719327 │ 0.0227471  │ 0.0282248 │ 378.703 │ 1.0001   │
│ 57  │ θ[54]      │ -0.0142313   │ 0.700431 │ 0.0221496  │ 0.0229427 │ 591.563 │ 0.999994 │
│ 58  │ θ[55]      │ -0.0154251   │ 0.701044 │ 0.0221689  │ 0.0323623 │ 516.323 │ 1.00183  │
│ 59  │ θ[56]      │ -0.0534623   │ 0.769119 │ 0.0243217  │ 0.0355191 │ 301.154 │ 1.00399  │
│ 60  │ θ[57]      │ -0.0322081   │ 0.700565 │ 0.0221538  │ 0.0294312 │ 768.559 │ 0.999067 │
│ 61  │ θ[58]      │ 0.032069     │ 0.668875 │ 0.0211517  │ 0.0352031 │ 580.497 │ 1.00526  │
│ 62  │ θ[59]      │ 0.0440055    │ 0.745985 │ 0.0235901  │ 0.0426681 │ 440.83  │ 1.00368  │
│ 63  │ θ[60]      │ 0.0412567    │ 0.736958 │ 0.0233047  │ 0.0229848 │ 548.318 │ 0.999616 │
│ 64  │ θ[61]      │ 0.00899984   │ 0.617626 │ 0.019531   │ 0.0300872 │ 587.582 │ 0.999025 │
│ 65  │ θ[62]      │ -0.00679929  │ 0.6407   │ 0.0202607  │ 0.017917  │ 837.683 │ 1.00011  │
│ 66  │ θ[63]      │ -0.0446423   │ 0.666277 │ 0.0210695  │ 0.0253015 │ 537.955 │ 1.00032  │
│ 67  │ θ[64]      │ -0.011716    │ 0.625655 │ 0.019785   │ 0.0128421 │ 836.064 │ 0.999    │
│ 68  │ θ[65]      │ -0.00243486  │ 0.666599 │ 0.0210797  │ 0.0152573 │ 696.948 │ 0.999    │
│ 69  │ θ[66]      │ -0.0470928   │ 0.635999 │ 0.0201121  │ 0.0133674 │ 652.678 │ 0.999661 │
│ 70  │ θ[67]      │ -0.0239345   │ 0.659242 │ 0.0208471  │ 0.0179499 │ 898.26  │ 0.999572 │
│ 71  │ θ[68]      │ -0.0198702   │ 0.671374 │ 0.0212307  │ 0.0278683 │ 700.012 │ 1.00014  │
│ 72  │ θ[69]      │ -0.0756352   │ 0.677782 │ 0.0214333  │ 0.0266876 │ 484.978 │ 1.0022   │
│ 73  │ θ[70]      │ 0.0031994    │ 0.670206 │ 0.0211938  │ 0.0239568 │ 844.888 │ 0.999034 │
│ 74  │ θ[71]      │ 0.00266609   │ 0.676475 │ 0.021392   │ 0.0218369 │ 893.129 │ 0.999712 │
│ 75  │ θ[72]      │ -0.0263279   │ 0.644207 │ 0.0203716  │ 0.0307667 │ 761.061 │ 0.999488 │
│ 76  │ θ[73]      │ -0.0565526   │ 0.662344 │ 0.0209452  │ 0.0309748 │ 705.129 │ 1.00489  │
│ 77  │ θ[74]      │ 0.00397308   │ 0.629948 │ 0.0199207  │ 0.021919  │ 804.165 │ 1.00176  │
│ 78  │ θ[75]      │ 0.00719474   │ 0.653599 │ 0.0206686  │ 0.02038   │ 943.496 │ 0.999009 │
│ 79  │ θ[76]      │ 0.0547359    │ 0.678302 │ 0.0214498  │ 0.0203452 │ 653.594 │ 0.999037 │
│ 80  │ θ[77]      │ 0.0031783    │ 0.665827 │ 0.0210553  │ 0.0270441 │ 625.411 │ 1.0049   │
│ 81  │ θ[78]      │ -0.00817279  │ 0.645199 │ 0.020403   │ 0.019762  │ 909.63  │ 0.999161 │
│ 82  │ θ[79]      │ -0.00706151  │ 0.649715 │ 0.0205458  │ 0.0281078 │ 738.99  │ 1.00538  │
│ 83  │ θ[80]      │ 0.00946464   │ 0.695801 │ 0.0220032  │ 0.0203245 │ 490.636 │ 0.999277 │
│ 84  │ θ[81]      │ -0.00784246  │ 0.691046 │ 0.0218528  │ 0.0181082 │ 782.226 │ 0.999    │
│ 85  │ θ[82]      │ 0.0562211    │ 0.692646 │ 0.0219034  │ 0.0205343 │ 919.721 │ 0.999089 │
│ 86  │ θ[83]      │ 0.0198181    │ 0.684365 │ 0.0216415  │ 0.0330181 │ 759.683 │ 0.99908  │
│ 87  │ θ[84]      │ -0.0167856   │ 0.702297 │ 0.0222086  │ 0.0228027 │ 777.453 │ 0.999066 │
│ 88  │ θ[85]      │ -0.0104739   │ 0.659369 │ 0.0208511  │ 0.021114  │ 1000.0  │ 0.999    │
│ 89  │ θ[86]      │ 0.0282526    │ 0.665026 │ 0.02103    │ 0.0335253 │ 579.283 │ 1.00126  │
│ 90  │ θ[87]      │ 0.0592104    │ 0.705131 │ 0.0222982  │ 0.02501   │ 685.225 │ 1.00036  │
│ 91  │ θ[88]      │ 0.0615105    │ 0.684017 │ 0.0216305  │ 0.0324732 │ 515.509 │ 1.00051  │
│ 92  │ θ[89]      │ -0.0323052   │ 0.656508 │ 0.0207606  │ 0.0135657 │ 724.74  │ 1.00109  │
│ 93  │ θ[90]      │ -0.00531484  │ 0.696068 │ 0.0220116  │ 0.0243403 │ 825.211 │ 0.999034 │
│ 94  │ θ[91]      │ -0.0260274   │ 0.714963 │ 0.0226091  │ 0.0283225 │ 384.544 │ 1.00248  │
│ 95  │ θ[92]      │ -0.0544405   │ 0.697223 │ 0.0220481  │ 0.0316458 │ 547.298 │ 0.999196 │
│ 96  │ θ[93]      │ -0.0649624   │ 0.740742 │ 0.0234243  │ 0.0312782 │ 509.894 │ 0.999778 │
│ 97  │ θ[94]      │ -0.0632387   │ 0.649841 │ 0.0205498  │ 0.0190965 │ 619.471 │ 0.999037 │
│ 98  │ θ[95]      │ -0.103841    │ 0.667539 │ 0.0211094  │ 0.0193038 │ 645.472 │ 0.999979 │
│ 99  │ θ[96]      │ -0.0888712   │ 0.699109 │ 0.0221078  │ 0.0341145 │ 409.258 │ 1.00299  │
│ 100 │ θ[97]      │ -0.077687    │ 0.735863 │ 0.02327    │ 0.039563  │ 246.233 │ 1.00625  │
│ 101 │ θ[98]      │ -0.0564509   │ 0.662083 │ 0.0209369  │ 0.040323  │ 466.841 │ 1.00184  │
│ 102 │ θ[99]      │ -0.0341822   │ 0.671044 │ 0.0212203  │ 0.0306949 │ 362.425 │ 1.00479  │
│ 103 │ θ[100]     │ -0.0390599   │ 0.6919   │ 0.0218798  │ 0.0365071 │ 520.173 │ 1.00118  │
│ 104 │ θ[101]     │ -0.0380851   │ 0.667104 │ 0.0210957  │ 0.0367488 │ 400.031 │ 1.00278  │
│ 105 │ θ[102]     │ -0.0195978   │ 0.669801 │ 0.021181   │ 0.0290515 │ 483.9   │ 0.999925 │
│ 106 │ θ[103]     │ -0.0114919   │ 0.704542 │ 0.0222796  │ 0.0405424 │ 306.744 │ 1.01421  │
│ 107 │ θ[104]     │ -0.108053    │ 0.717275 │ 0.0226822  │ 0.0452797 │ 432.38  │ 0.999013 │
│ 108 │ θ[105]     │ -0.0869239   │ 0.69814  │ 0.0220771  │ 0.0430515 │ 310.805 │ 0.999669 │
│ 109 │ θ[106]     │ 0.0557887    │ 0.743399 │ 0.0235083  │ 0.046391  │ 381.514 │ 0.999398 │
│ 110 │ θ[107]     │ 0.0602319    │ 0.745155 │ 0.0235639  │ 0.0264673 │ 495.325 │ 1.00032  │
│ 111 │ θ[108]     │ 0.0408561    │ 0.754331 │ 0.023854   │ 0.0364359 │ 330.697 │ 1.00279  │
│ 112 │ θ[109]     │ 0.0416592    │ 0.796939 │ 0.0252014  │ 0.0383891 │ 315.946 │ 1.00145  │
│ 113 │ θ[110]     │ 0.0451974    │ 0.748776 │ 0.0236784  │ 0.0220042 │ 479.591 │ 1.00218  │
│ 114 │ θ[111]     │ 0.0758384    │ 0.759247 │ 0.0240095  │ 0.0493337 │ 248.62  │ 0.999089 │
│ 115 │ θ[112]     │ 0.05395      │ 0.733982 │ 0.0232105  │ 0.0437718 │ 290.376 │ 0.999918 │
│ 116 │ θ[113]     │ 0.0786027    │ 0.747083 │ 0.0236249  │ 0.0282164 │ 515.357 │ 1.00018  │
│ 117 │ θ[114]     │ 0.0810854    │ 0.749376 │ 0.0236973  │ 0.0354352 │ 398.882 │ 1.00138  │
│ 118 │ θ[115]     │ 0.0820551    │ 0.724197 │ 0.0229011  │ 0.0371597 │ 589.757 │ 1.00007  │
│ 119 │ θ[116]     │ 0.0756049    │ 0.72511  │ 0.02293    │ 0.0432784 │ 370.918 │ 1.01174  │
│ 120 │ θ[117]     │ 0.0906049    │ 0.754177 │ 0.0238492  │ 0.0604054 │ 309.123 │ 1.01104  │
│ 121 │ θ[118]     │ 0.0686043    │ 0.77714  │ 0.0245753  │ 0.0252362 │ 430.679 │ 1.0016   │
│ 122 │ θ[119]     │ 0.0326688    │ 0.722721 │ 0.0228545  │ 0.0332793 │ 448.862 │ 0.999588 │
│ 123 │ θ[120]     │ 0.101991     │ 0.759777 │ 0.0240263  │ 0.033751  │ 362.058 │ 0.999276 │
│ 124 │ θ[121]     │ -0.033215    │ 0.760709 │ 0.0240557  │ 0.0379605 │ 457.659 │ 1.00163  │
│ 125 │ θ[122]     │ -0.0307778   │ 0.705854 │ 0.0223211  │ 0.0306539 │ 565.652 │ 1.00693  │
│ 126 │ θ[123]     │ 0.0337912    │ 0.712791 │ 0.0225404  │ 0.0296234 │ 639.663 │ 1.00124  │
│ 127 │ θ[124]     │ -0.0272024   │ 0.714434 │ 0.0225924  │ 0.0247458 │ 679.667 │ 0.999411 │
│ 128 │ θ[125]     │ -0.0528841   │ 0.717192 │ 0.0226796  │ 0.0221105 │ 612.635 │ 0.999004 │
│ 129 │ θ[126]     │ 0.0249675    │ 0.709493 │ 0.0224361  │ 0.0227312 │ 661.24  │ 1.00046  │
│ 130 │ θ[127]     │ -0.0274465   │ 0.779926 │ 0.0246634  │ 0.0255844 │ 495.682 │ 0.999011 │
│ 131 │ θ[128]     │ -0.0384062   │ 0.680913 │ 0.0215324  │ 0.0230935 │ 721.214 │ 0.999813 │
│ 132 │ θ[129]     │ 0.0410301    │ 0.705263 │ 0.0223024  │ 0.036584  │ 460.544 │ 1.00565  │
│ 133 │ θ[130]     │ -0.0248907   │ 0.672493 │ 0.0212661  │ 0.0126532 │ 831.86  │ 0.999    │
│ 134 │ θ[131]     │ -0.0153937   │ 0.71342  │ 0.0225603  │ 0.023625  │ 540.109 │ 0.999053 │
│ 135 │ θ[132]     │ -0.0465743   │ 0.741867 │ 0.0234599  │ 0.0410216 │ 414.034 │ 1.00077  │
│ 136 │ θ[133]     │ -0.0343415   │ 0.703653 │ 0.0222515  │ 0.0196545 │ 760.649 │ 0.999457 │
│ 137 │ θ[134]     │ -0.0234433   │ 0.746413 │ 0.0236037  │ 0.0343169 │ 496.867 │ 0.999014 │
│ 138 │ θ[135]     │ -0.0137922   │ 0.694609 │ 0.0219655  │ 0.0277884 │ 567.802 │ 0.999185 │
│ 139 │ θ[136]     │ -0.0339168   │ 0.663431 │ 0.0209795  │ 0.0360375 │ 297.097 │ 0.999241 │
│ 140 │ θ[137]     │ 0.0117195    │ 0.651811 │ 0.0206121  │ 0.0364741 │ 275.267 │ 0.999941 │
│ 141 │ θ[138]     │ 0.0127132    │ 0.667166 │ 0.0210976  │ 0.0478616 │ 276.688 │ 0.999353 │
│ 142 │ θ[139]     │ -0.0908917   │ 0.701265 │ 0.022176   │ 0.0606818 │ 175.539 │ 1.00992  │
│ 143 │ θ[140]     │ 0.0600896    │ 0.653243 │ 0.0206573  │ 0.0496652 │ 217.34  │ 1.00212  │
│ 144 │ θ[141]     │ 0.0255407    │ 0.681931 │ 0.0215645  │ 0.0559289 │ 196.555 │ 0.999014 │
│ 145 │ θ[142]     │ 0.0755346    │ 0.700853 │ 0.0221629  │ 0.0553446 │ 216.984 │ 0.999003 │
│ 146 │ θ[143]     │ -0.0228305   │ 0.678402 │ 0.021453   │ 0.0460689 │ 165.519 │ 0.999364 │
│ 147 │ θ[144]     │ -0.00905815  │ 0.64009  │ 0.0202414  │ 0.0286495 │ 261.892 │ 1.00056  │
│ 148 │ θ[145]     │ 0.0126905    │ 0.673561 │ 0.0212999  │ 0.0444579 │ 305.363 │ 1.00155  │
│ 149 │ θ[146]     │ -0.0317279   │ 0.687796 │ 0.02175    │ 0.0397742 │ 216.54  │ 1.00431  │
│ 150 │ θ[147]     │ -0.00639916  │ 0.672049 │ 0.021252   │ 0.0695574 │ 183.118 │ 1.00012  │
│ 151 │ θ[148]     │ 0.0339678    │ 0.640095 │ 0.0202416  │ 0.0464543 │ 248.521 │ 1.00632  │
│ 152 │ θ[149]     │ 0.00883909   │ 0.670673 │ 0.0212085  │ 0.0422367 │ 264.181 │ 1.00085  │
│ 153 │ θ[150]     │ -0.0207952   │ 0.699408 │ 0.0221172  │ 0.05      │ 194.798 │ 1.00593  │

Quantiles

│ Row │ parameters │ 2.5%     │ 25.0%     │ 50.0%        │ 75.0%     │ 97.5%   │
│     │ Symbol     │ Float64  │ Float64   │ Float64      │ Float64   │ Float64 │
├─────┼────────────┼──────────┼───────────┼──────────────┼───────────┼─────────┤
│ 1   │ c1         │ -4.21364 │ -2.32003  │ -1.43987     │ -0.589687 │ 1.48709 │
│ 2   │ log_diff_c │ 0.445742 │ 0.671916  │ 0.788029     │ 0.900895  │ 1.10933 │
│ 3   │ sig        │ 0.427589 │ 0.563974  │ 0.659366     │ 0.779799  │ 1.09459 │
│ 4   │ θ[1]       │ -1.25416 │ -0.324348 │ 0.0538895    │ 0.401436  │ 1.22416 │
│ 5   │ θ[2]       │ -1.36247 │ -0.455411 │ -0.0238037   │ 0.400606  │ 1.25063 │
│ 6   │ θ[3]       │ -1.21545 │ -0.423453 │ -0.00634231  │ 0.398774  │ 1.22596 │
│ 7   │ θ[4]       │ -1.2319  │ -0.389932 │ 0.041418     │ 0.49485   │ 1.41209 │
│ 8   │ θ[5]       │ -1.12689 │ -0.377898 │ -0.0179172   │ 0.378574  │ 1.30673 │
│ 9   │ θ[6]       │ -1.39716 │ -0.372634 │ 0.0526812    │ 0.406669  │ 1.29357 │
│ 10  │ θ[7]       │ -1.21077 │ -0.36297  │ 0.0056438    │ 0.407002  │ 1.27888 │
│ 11  │ θ[8]       │ -1.42217 │ -0.400678 │ 0.0171975    │ 0.394606  │ 1.16701 │
│ 12  │ θ[9]       │ -1.29808 │ -0.36261  │ 0.0112514    │ 0.3943    │ 1.30887 │
│ 13  │ θ[10]      │ -1.27541 │ -0.331709 │ 0.0120783    │ 0.392133  │ 1.16281 │
│ 14  │ θ[11]      │ -1.26558 │ -0.379235 │ 0.0334503    │ 0.442002  │ 1.31059 │
│ 15  │ θ[12]      │ -1.36783 │ -0.379617 │ -0.0158274   │ 0.385061  │ 1.31941 │
│ 16  │ θ[13]      │ -1.37839 │ -0.419985 │ 0.018096     │ 0.439329  │ 1.5623  │
│ 17  │ θ[14]      │ -1.41721 │ -0.35919  │ 0.0149091    │ 0.392602  │ 1.27922 │
│ 18  │ θ[15]      │ -1.23322 │ -0.422196 │ 0.0108471    │ 0.445735  │ 1.24914 │
│ 19  │ θ[16]      │ -1.44456 │ -0.49668  │ -0.0418525   │ 0.499965  │ 1.7756  │
│ 20  │ θ[17]      │ -1.49047 │ -0.467631 │ -0.0358636   │ 0.462605  │ 1.53146 │
│ 21  │ θ[18]      │ -1.67699 │ -0.547833 │ -0.0964584   │ 0.391646  │ 1.57355 │
│ 22  │ θ[19]      │ -1.7528  │ -0.457881 │ 0.00852779   │ 0.533761  │ 1.86009 │
│ 23  │ θ[20]      │ -1.46166 │ -0.497023 │ -0.0496473   │ 0.414929  │ 1.48604 │
│ 24  │ θ[21]      │ -1.45304 │ -0.542137 │ -0.0745904   │ 0.371745  │ 1.36746 │
│ 25  │ θ[22]      │ -1.40828 │ -0.486317 │ -0.0207834   │ 0.519895  │ 1.6009  │
│ 26  │ θ[23]      │ -1.42833 │ -0.426137 │ 0.0072025    │ 0.512998  │ 1.71383 │
│ 27  │ θ[24]      │ -1.59438 │ -0.484719 │ -0.00792077  │ 0.489556  │ 1.64253 │
│ 28  │ θ[25]      │ -1.47192 │ -0.442635 │ 0.016852     │ 0.502861  │ 1.65945 │
│ 29  │ θ[26]      │ -1.56631 │ -0.428778 │ 0.0197279    │ 0.499965  │ 1.74474 │
│ 30  │ θ[27]      │ -1.50926 │ -0.496323 │ -0.0269343   │ 0.576891  │ 1.57    │
│ 31  │ θ[28]      │ -1.33621 │ -0.416679 │ -0.019913    │ 0.413412  │ 1.71086 │
│ 32  │ θ[29]      │ -1.3976  │ -0.427533 │ 0.0314751    │ 0.524075  │ 1.58438 │
│ 33  │ θ[30]      │ -1.38851 │ -0.487375 │ 0.0296572    │ 0.596397  │ 1.73987 │
│ 34  │ θ[31]      │ -1.39826 │ -0.492174 │ -0.042908    │ 0.40642   │ 1.37578 │
│ 35  │ θ[32]      │ -1.3721  │ -0.451182 │ -0.000873611 │ 0.398027  │ 1.25643 │
│ 36  │ θ[33]      │ -1.35886 │ -0.439169 │ -0.0431794   │ 0.382629  │ 1.53017 │
│ 37  │ θ[34]      │ -1.43406 │ -0.474216 │ -0.0380782   │ 0.364009  │ 1.22332 │
│ 38  │ θ[35]      │ -1.35282 │ -0.434354 │ -0.00125667  │ 0.452544  │ 1.38275 │
│ 39  │ θ[36]      │ -1.28924 │ -0.47434  │ -0.0685111   │ 0.418078  │ 1.35292 │
│ 40  │ θ[37]      │ -1.43596 │ -0.493984 │ -0.0813643   │ 0.381412  │ 1.37586 │
│ 41  │ θ[38]      │ -1.40802 │ -0.470796 │ -0.068352    │ 0.324824  │ 1.1848  │
│ 42  │ θ[39]      │ -1.27464 │ -0.439724 │ -0.0137112   │ 0.361008  │ 1.31437 │
│ 43  │ θ[40]      │ -1.31542 │ -0.442265 │ -0.0240819   │ 0.428622  │ 1.50336 │
│ 44  │ θ[41]      │ -1.5562  │ -0.420535 │ -0.037907    │ 0.36457   │ 1.3671  │
│ 45  │ θ[42]      │ -1.43035 │ -0.497851 │ -0.063536    │ 0.354269  │ 1.1602  │
│ 46  │ θ[43]      │ -1.41299 │ -0.513786 │ -0.0595422   │ 0.408099  │ 1.38084 │
│ 47  │ θ[44]      │ -1.27647 │ -0.437738 │ -0.0433403   │ 0.390484  │ 1.46729 │
│ 48  │ θ[45]      │ -1.30768 │ -0.429345 │ -0.0366154   │ 0.39631   │ 1.27584 │
│ 49  │ θ[46]      │ -1.43443 │ -0.446403 │ 0.0115101    │ 0.435351  │ 1.3473  │
│ 50  │ θ[47]      │ -1.26681 │ -0.369948 │ 0.0532465    │ 0.433483  │ 1.51195 │
│ 51  │ θ[48]      │ -1.5349  │ -0.421898 │ 0.0419424    │ 0.488246  │ 1.62046 │
│ 52  │ θ[49]      │ -1.26394 │ -0.427792 │ 0.0532985    │ 0.513578  │ 1.5607  │
│ 53  │ θ[50]      │ -1.36587 │ -0.462463 │ -0.000147727 │ 0.446508  │ 1.41433 │
│ 54  │ θ[51]      │ -1.31655 │ -0.426419 │ 0.0536954    │ 0.486486  │ 1.41166 │
│ 55  │ θ[52]      │ -1.36168 │ -0.339729 │ 0.0487487    │ 0.504042  │ 1.45539 │
│ 56  │ θ[53]      │ -1.36063 │ -0.405613 │ 0.0362613    │ 0.476177  │ 1.38266 │
│ 57  │ θ[54]      │ -1.49918 │ -0.425988 │ 0.0209319    │ 0.436106  │ 1.22243 │
│ 58  │ θ[55]      │ -1.36074 │ -0.432966 │ -0.0142805   │ 0.435073  │ 1.44817 │
│ 59  │ θ[56]      │ -1.80646 │ -0.435714 │ -0.0151903   │ 0.401264  │ 1.35327 │
│ 60  │ θ[57]      │ -1.39414 │ -0.459353 │ -0.0408775   │ 0.407282  │ 1.37177 │
│ 61  │ θ[58]      │ -1.36199 │ -0.366504 │ 0.0628859    │ 0.453309  │ 1.32909 │
│ 62  │ θ[59]      │ -1.38657 │ -0.403653 │ 0.0153761    │ 0.475938  │ 1.64237 │
│ 63  │ θ[60]      │ -1.37644 │ -0.376139 │ 0.043532     │ 0.484187  │ 1.49435 │
│ 64  │ θ[61]      │ -1.16846 │ -0.389237 │ -0.0044437   │ 0.413147  │ 1.33045 │
│ 65  │ θ[62]      │ -1.25978 │ -0.388525 │ -0.0171203   │ 0.375004  │ 1.29899 │
│ 66  │ θ[63]      │ -1.50828 │ -0.449511 │ -0.0554307   │ 0.367214  │ 1.32266 │
│ 67  │ θ[64]      │ -1.19124 │ -0.418566 │ -0.0131106   │ 0.402886  │ 1.19151 │
│ 68  │ θ[65]      │ -1.22432 │ -0.440608 │ 0.0255283    │ 0.407949  │ 1.32746 │
│ 69  │ θ[66]      │ -1.24968 │ -0.446118 │ -0.0394156   │ 0.363576  │ 1.17229 │
│ 70  │ θ[67]      │ -1.38828 │ -0.419153 │ -0.0123899   │ 0.384713  │ 1.32262 │
│ 71  │ θ[68]      │ -1.44245 │ -0.445027 │ -0.0149068   │ 0.397002  │ 1.32518 │
│ 72  │ θ[69]      │ -1.4278  │ -0.500136 │ -0.06204     │ 0.348355  │ 1.26852 │
│ 73  │ θ[70]      │ -1.30999 │ -0.429199 │ 0.0109176    │ 0.429731  │ 1.30662 │
│ 74  │ θ[71]      │ -1.31854 │ -0.500649 │ 0.0233564    │ 0.434402  │ 1.37161 │
│ 75  │ θ[72]      │ -1.29704 │ -0.452286 │ -0.0269103   │ 0.416172  │ 1.27218 │
│ 76  │ θ[73]      │ -1.40059 │ -0.439499 │ -0.074544    │ 0.313274  │ 1.33752 │
│ 77  │ θ[74]      │ -1.25544 │ -0.396567 │ 0.00941627   │ 0.407257  │ 1.27203 │
│ 78  │ θ[75]      │ -1.26279 │ -0.408757 │ 0.00442037   │ 0.432933  │ 1.26386 │
│ 79  │ θ[76]      │ -1.22183 │ -0.413867 │ 0.0462252    │ 0.498853  │ 1.42741 │
│ 80  │ θ[77]      │ -1.40322 │ -0.390044 │ 0.0150788    │ 0.433529  │ 1.25897 │
│ 81  │ θ[78]      │ -1.28786 │ -0.406944 │ -0.0190515   │ 0.397944  │ 1.27622 │
│ 82  │ θ[79]      │ -1.26381 │ -0.421139 │ -0.00681767  │ 0.408521  │ 1.30408 │
│ 83  │ θ[80]      │ -1.46619 │ -0.402068 │ 0.0183579    │ 0.444363  │ 1.40577 │
│ 84  │ θ[81]      │ -1.4067  │ -0.399517 │ -0.00996047  │ 0.41016   │ 1.41685 │
│ 85  │ θ[82]      │ -1.31966 │ -0.397122 │ 0.0316475    │ 0.50375   │ 1.50074 │
│ 86  │ θ[83]      │ -1.39549 │ -0.412754 │ 0.0227564    │ 0.446502  │ 1.35418 │
│ 87  │ θ[84]      │ -1.47532 │ -0.481888 │ 0.0149138    │ 0.4194    │ 1.36003 │
│ 88  │ θ[85]      │ -1.35885 │ -0.447608 │ 0.00293754   │ 0.405075  │ 1.32195 │
│ 89  │ θ[86]      │ -1.37301 │ -0.351168 │ 0.0256512    │ 0.410591  │ 1.49603 │
│ 90  │ θ[87]      │ -1.32324 │ -0.365562 │ 0.0639581    │ 0.501471  │ 1.43574 │
│ 91  │ θ[88]      │ -1.32991 │ -0.36942  │ 0.0307848    │ 0.499832  │ 1.50343 │
│ 92  │ θ[89]      │ -1.39883 │ -0.42345  │ -0.0251394   │ 0.395259  │ 1.24167 │
│ 93  │ θ[90]      │ -1.46206 │ -0.432951 │ 0.0231079    │ 0.452139  │ 1.35411 │
│ 94  │ θ[91]      │ -1.54341 │ -0.432163 │ -0.00526869  │ 0.389817  │ 1.41775 │
│ 95  │ θ[92]      │ -1.46597 │ -0.470503 │ -0.0552184   │ 0.374649  │ 1.40317 │
│ 96  │ θ[93]      │ -1.45145 │ -0.539605 │ -0.0679053   │ 0.385833  │ 1.4429  │
│ 97  │ θ[94]      │ -1.36085 │ -0.481666 │ -0.0379939   │ 0.348591  │ 1.18677 │
│ 98  │ θ[95]      │ -1.50795 │ -0.521047 │ -0.106486    │ 0.32952   │ 1.36117 │
│ 99  │ θ[96]      │ -1.54254 │ -0.5087   │ -0.0647784   │ 0.345959  │ 1.20161 │
│ 100 │ θ[97]      │ -1.54675 │ -0.550079 │ -0.0660638   │ 0.361072  │ 1.35072 │
│ 101 │ θ[98]      │ -1.48091 │ -0.46252  │ -0.0570559   │ 0.371448  │ 1.23722 │
│ 102 │ θ[99]      │ -1.40707 │ -0.476855 │ -0.0335152   │ 0.40631   │ 1.34284 │
│ 103 │ θ[100]     │ -1.42825 │ -0.436985 │ -0.0521414   │ 0.376361  │ 1.44809 │
│ 104 │ θ[101]     │ -1.40639 │ -0.459675 │ -0.0205963   │ 0.358898  │ 1.35959 │
│ 105 │ θ[102]     │ -1.34182 │ -0.461106 │ 0.0159509    │ 0.406209  │ 1.34012 │
│ 106 │ θ[103]     │ -1.39078 │ -0.434803 │ -0.046909    │ 0.420722  │ 1.48221 │
│ 107 │ θ[104]     │ -1.57108 │ -0.553071 │ -0.101753    │ 0.356449  │ 1.20771 │
│ 108 │ θ[105]     │ -1.49553 │ -0.546387 │ -0.104913    │ 0.387566  │ 1.27723 │
│ 109 │ θ[106]     │ -1.50745 │ -0.398224 │ 0.0570903    │ 0.517026  │ 1.51303 │
│ 110 │ θ[107]     │ -1.43865 │ -0.464008 │ 0.100972     │ 0.572295  │ 1.51399 │
│ 111 │ θ[108]     │ -1.62167 │ -0.403003 │ 0.0731888    │ 0.525827  │ 1.44382 │
│ 112 │ θ[109]     │ -1.51241 │ -0.45751  │ 0.0486727    │ 0.536884  │ 1.64987 │
│ 113 │ θ[110]     │ -1.4133  │ -0.421723 │ 0.0257947    │ 0.508615  │ 1.49423 │
│ 114 │ θ[111]     │ -1.44876 │ -0.430082 │ 0.116955     │ 0.589628  │ 1.53194 │
│ 115 │ θ[112]     │ -1.51398 │ -0.387191 │ 0.0787192    │ 0.519855  │ 1.40561 │
│ 116 │ θ[113]     │ -1.47599 │ -0.395565 │ 0.0987518    │ 0.573555  │ 1.48287 │
│ 117 │ θ[114]     │ -1.39567 │ -0.375855 │ 0.0713862    │ 0.574712  │ 1.51794 │
│ 118 │ θ[115]     │ -1.3982  │ -0.383872 │ 0.0719347    │ 0.55597   │ 1.48001 │
│ 119 │ θ[116]     │ -1.34478 │ -0.40435  │ 0.0948653    │ 0.569317  │ 1.47443 │
│ 120 │ θ[117]     │ -1.30752 │ -0.451254 │ 0.123252     │ 0.578493  │ 1.56601 │
│ 121 │ θ[118]     │ -1.63339 │ -0.368155 │ 0.0938648    │ 0.529361  │ 1.55147 │
│ 122 │ θ[119]     │ -1.43618 │ -0.442946 │ 0.057429     │ 0.497799  │ 1.42088 │
│ 123 │ θ[120]     │ -1.40056 │ -0.421718 │ 0.118576     │ 0.61514   │ 1.55242 │
│ 124 │ θ[121]     │ -1.53841 │ -0.48003  │ -0.0460021   │ 0.482198  │ 1.3591  │
│ 125 │ θ[122]     │ -1.43776 │ -0.459247 │ -0.0542885   │ 0.391443  │ 1.34831 │
│ 126 │ θ[123]     │ -1.37557 │ -0.434857 │ 0.0540424    │ 0.46539   │ 1.53815 │
│ 127 │ θ[124]     │ -1.58577 │ -0.479548 │ -0.00278725  │ 0.449369  │ 1.34597 │
│ 128 │ θ[125]     │ -1.59641 │ -0.471639 │ -0.0186811   │ 0.409118  │ 1.27852 │
│ 129 │ θ[126]     │ -1.35972 │ -0.415552 │ 0.0328366    │ 0.457748  │ 1.38879 │
│ 130 │ θ[127]     │ -1.67586 │ -0.508775 │ 0.0223643    │ 0.445098  │ 1.52992 │
│ 131 │ θ[128]     │ -1.51212 │ -0.488969 │ -0.02526     │ 0.404979  │ 1.24841 │
│ 132 │ θ[129]     │ -1.32946 │ -0.427818 │ 0.0254997    │ 0.486727  │ 1.42918 │
│ 133 │ θ[130]     │ -1.33464 │ -0.455091 │ -0.0445645   │ 0.396423  │ 1.32277 │
│ 134 │ θ[131]     │ -1.3873  │ -0.485731 │ -0.0186891   │ 0.439844  │ 1.42333 │
│ 135 │ θ[132]     │ -1.559   │ -0.525411 │ -0.0286328   │ 0.390707  │ 1.40204 │
│ 136 │ θ[133]     │ -1.43762 │ -0.475699 │ -0.0192159   │ 0.402298  │ 1.33832 │
│ 137 │ θ[134]     │ -1.52055 │ -0.48413  │ -0.0116835   │ 0.408287  │ 1.50888 │
│ 138 │ θ[135]     │ -1.44006 │ -0.465139 │ -0.00630592  │ 0.428209  │ 1.40386 │
│ 139 │ θ[136]     │ -1.33038 │ -0.488569 │ -0.0174244   │ 0.40726   │ 1.24516 │
│ 140 │ θ[137]     │ -1.30491 │ -0.429305 │ 0.0209522    │ 0.466507  │ 1.24567 │
│ 141 │ θ[138]     │ -1.26229 │ -0.390814 │ 0.0340052    │ 0.443904  │ 1.22245 │
│ 142 │ θ[139]     │ -1.54617 │ -0.550395 │ -0.0761326   │ 0.38359   │ 1.27668 │
│ 143 │ θ[140]     │ -1.18642 │ -0.398668 │ 0.0667236    │ 0.518191  │ 1.3328  │
│ 144 │ θ[141]     │ -1.26791 │ -0.435891 │ 0.0207375    │ 0.480737  │ 1.32363 │
│ 145 │ θ[142]     │ -1.24291 │ -0.392791 │ 0.0538593    │ 0.519877  │ 1.5526  │
│ 146 │ θ[143]     │ -1.27978 │ -0.455386 │ -0.0578343   │ 0.428325  │ 1.36699 │
│ 147 │ θ[144]     │ -1.22354 │ -0.429349 │ -0.0299434   │ 0.445095  │ 1.22394 │
│ 148 │ θ[145]     │ -1.32305 │ -0.44079  │ 0.0189598    │ 0.462383  │ 1.30563 │
│ 149 │ θ[146]     │ -1.4327  │ -0.510874 │ -0.0447093   │ 0.467501  │ 1.2445  │
│ 150 │ θ[147]     │ -1.36478 │ -0.47289  │ 0.0170359    │ 0.447094  │ 1.24948 │
│ 151 │ θ[148]     │ -1.17934 │ -0.4258   │ 0.0485161    │ 0.443407  │ 1.2771  │
│ 152 │ θ[149]     │ -1.34658 │ -0.470378 │ -0.0206104   │ 0.478268  │ 1.35794 │
│ 153 │ θ[150]     │ -1.55763 │ -0.477268 │ -0.0269964   │ 0.446997  │ 1.28711 │
```

In [160]:

```
# Extract the θ parameters from the sampled chain.
params2 = ch2[:θ].value.data

# cutpoints
e_log_diff_c = exp.(ch2[:log_diff_c].value.data)[:,1,1]
c1_est = ch2[:c1].value.data[:,1,1]
c2_est = c1_est + e_log_diff_c;

n_samps = size(ch2, 1)
```

Out[160]:

```
1000
```

In [161]:

```
## ---------------------------------------------------------
## predict for training data
## ---------------------------------------------------------

probs_mat_train, y_pred_samps_train, logpdf_mat_train, y_pred_train, eta_post_train = predict(X_train, y_train, n_samps);

## ---------------------------------------------------------
## metrics for training data
## ---------------------------------------------------------

waic = WAIC_logpfd(logpdf_mat_train)
tbl = freqtable(y_train)
ptbl = prop(tbl)
ptbl = convert(Array, ptbl)
v = vcat(fill(ptbl, length(y_train)))
probs_mat_freq = convert(Array{Float64,2}, hcat(v...)')

# model-based Brier score
cBrier_train = cumBrier(probs_mat_train, y_train)

# baseline Brier score
cBrier_BB_train = cumBrier(probs_mat_freq, y_train)

# Brier Skill score
BSS_train = (cBrier_BB_train .- cBrier_train) ./ cBrier_BB_train

y_pred = convert(Array{Int64,1}, y_pred_train);
y_train = convert(Array{Int64,1}, y_train);
C = confusmat(3, y_train, y_pred)
acc = (C[1,1] + C[2,2] + C[3,3]) / sum(C)
bacc = 1/3 *(C[1,1] / sum(C[1, :]) + C[2,2] / sum(C[2, :]) + C[3,3] / sum(C[3, :]))

println("WAIC = ", round(waic, digits=1))
println("mean cumBrier = ", round(mean(cBrier_train), digits=2))
println("median cumBrier = ", round(median(cBrier_train), digits=2))
println("mean BSS = ", round(mean(BSS_train), digits=2))
println("median BSS = ", round(median(BSS_train), digits=2))
println("Balanced accuracy = ", round(bacc, digits=2))
println("====== Confision Matrix:")
C
```

```
WAIC = 252.8
mean cumBrier = 0.12
median cumBrier = 0.08
mean BSS = 0.37
median BSS = 0.47
Balanced accuracy = 0.7
====== Confision Matrix:
```

Out[161]:

```
3×3 Array{Int64,2}:
 28   9   0
  4  24  17
  1  11  53
```

In [ ]:

In [ ]:

In [142]:

```
# there is randomness in predictions, hence need to average over a few of them
n_exper = 10

waic_exp = zeros(n_exper)
cBrier_train_mean = zeros(n_exper)
cBrier_train_median= zeros(n_exper)
BSS_train_mean = zeros(n_exper)
BSS_train_meadian = zeros(n_exper)
bacc_exp = zeros(n_exper)

for exp=1:n_exper
    
    ## ---------------------------------------------------------
    ## predict for training data
    ## ---------------------------------------------------------
    
    probs_mat_train, y_pred_samps_train, logpdf_mat_train, y_pred_train, eta_post_train = predict(X_train, y_train, n_samps);

    ## ---------------------------------------------------------
    ## metrics for training data
    ## ---------------------------------------------------------

    waic = WAIC_logpfd(logpdf_mat_train)
    tbl = freqtable(y_train)
    ptbl = prop(tbl)
    ptbl = convert(Array, ptbl)
    v = vcat(fill(ptbl, length(y_train)))
    probs_mat_freq = convert(Array{Float64,2}, hcat(v...)')

    # model-based Brier score
    cBrier_train = cumBrier(probs_mat_train, y_train)

    # baseline Brier score
    cBrier_BB_train = cumBrier(probs_mat_freq, y_train)

    # Brier Skill score
    BSS_train = (cBrier_BB_train .- cBrier_train) ./ cBrier_BB_train


    y_pred = convert(Array{Int64,1}, y_pred_train);
    y_train = convert(Array{Int64,1}, y_train);
    C = confusmat(3, y_train, y_pred)
    acc = (C[1,1] + C[2,2] + C[3,3]) / sum(C)
    bacc = 1/3 *(C[1,1] / sum(C[1, :]) + C[2,2] / sum(C[2, :]) + C[3,3] / sum(C[3, :]))
    
    waic_exp[exp] = waic
    cBrier_train_mean[exp] = mean(cBrier_train)
    cBrier_train_median[exp] = median(cBrier_train)
    BSS_train_mean[exp] = mean(BSS_train)
    BSS_train_meadian[exp] = median(BSS_train)
    bacc_exp[exp]  = bacc
    
end

println("train WAIC = ", round(mean(waic), digits=2))
println("train mean cumBrier = ", round(mean(cBrier_train_mean), digits = 2))
println("train median cumBrier = ", round(mean(cBrier_train_median), digits = 2))
println("train mean BSS = ", round(mean(BSS_train_mean), digits = 2))
println("train median BSS = ", round(mean(BSS_train_meadian), digits = 2))
println("train Balanced accuracy = ", round(mean(bacc_exp), digits=2))
```

```
train WAIC = 252.77
train mean cumBrier = 0.12
train median cumBrier = 0.08
train mean BSS = 0.37
train median BSS = 0.46
train Balanced accuracy = 0.7
```

In [162]:

```
# Rosiglitazone
probsplot(drug_names_train, y_pred_samps_train, 38, y_train)
```

```
mean cumBrier = 0.14
median cumBrier = 0.1
mean BSS = 0.31
median BSS = 0.39
Balanced accuracy = 0.67
====== Confision Matrix:
```

Out[166]:

```
3×3 Array{Int64,2}:
 8  1   1
 1  5   5
 1  3  12
```

In [167]:

```
sum(C)
```

Out[167]:

```
37
```

In [168]:

```
save("chains/bnn_cBrier.jld", "bnn_cBrier_train", cBrier_train, "bnn_cBrier_test", cBrier_test, 
                              "bnn_BSS_train", BSS_train, "bnn_BSS_test", BSS_test )
```

In [169]:

```
## ---------------------------------------------------------
## Plots
## ---------------------------------------------------------

k1 = kde(cBrier_train)
k2 = kde(cBrier_test)
p1 = plot(k2.x, k2.density,  
          color = :navy,
          fill = (0, 0.6, :navy), 
          label="test data", 
          yticks = true, 
          legend = :topright,
          #legend = false,
          #ylim=(0, 1.05 * maximum(vcat(k1.density, k2.density))),
          framestyle = :box,
          #title="Ordered Brier Score.",
          #titlefont = font(10, "Calibri"),
          size=[350,350],
          xlabel="Ordered Brier Score",
          xguidefontsize=font(7, "Calibri"),
          xlim = (-0.3, 1.2), 
          ylim = (0, 6.5),
          grid = :xy,
          gridopacity = 0.5)
plot!(k1.x, k1.density, color = :purple, fill = (0, 0.6, :purple), label="training data")
```

Out[169]:

xml version="1.0" encoding="utf-8"?


-0.25

0.00

0.25

0.50

0.75

1.00

0

1

2

3

4

5

6


Ordered Brier Score


test data


training data

In [170]:

```
## ---------------------------------------------------------
## Plots
## ---------------------------------------------------------

k1 = kde(cBrier_train)
k2 = kde(cBrier_test)
p1 = plot(k2.x, k2.density,  
          color = :navy,
          fill = (0, 0.6, :navy), 
          label="test data", 
          yticks = true, 
          legend = :topright,
          #legend = false,
          #ylim=(0, 1.05 * maximum(vcat(k1.density, k2.density))),
          framestyle = :box,
          #title="Ordered Brier Score.",
          #titlefont = font(10, "Calibri"),
          size=[350,350],
          xlabel="Ordered Brier Score",
          xguidefontsize=font(7, "Calibri"),
          xlim = (-0.3, 1.2), 
          ylim = (0, 6.5),
          grid = :xy,
          gridopacity = 0.5)
plot!(k1.x, k1.density, color = :purple, fill = (0, 0.6, :purple), label="training data")
```

Out[170]:

xml version="1.0" encoding="utf-8"?


-0.25

0.00

0.25

0.50

0.75

1.00

0

1

2

3

4

5

6


Ordered Brier Score


test data


training data

In [171]:

```
#savefig("figs/OBS_BNN.pdf")
```

In [172]:

```
# Brier Skill score

k1 = kde(BSS_train)
k2 = kde(BSS_test)
p1 = plot(k2.x, k2.density,  
          color = :navy,
          fill = (0, 0.6, :navy), 
          label="test data", 
          yticks = true, 
          legend = :topleft,
          #legend = false,
          #ylim=(0, 1.05 * maximum(vcat(k1.density, k2.density))),
          framestyle = :box,
          #title="Ordered Brier Score.",
          #titlefont = font(10, "Calibri"),
          size=[350,350],
          xlabel="Brier Skill Score",
          xguidefontsize=font(7, "Calibri"),
          xlim = (-2, 2),
          ylim = (0, 1.01),
          grid = :xy,
          gridopacity = 0.5)
plot!(k1.x, k1.density, color = :purple, fill = (0, 0.6, :purple), label="training data")
```

Out[172]:

xml version="1.0" encoding="utf-8"?


-2

-1

0

1

2

0.00

0.25

0.50

0.75

1.00


Brier Skill Score


test data


training data

In [173]:

```
# Brier Skill score

k1 = kde(BSS_train)
k2 = kde(BSS_test)
p1 = plot(k2.x, k2.density,  
          color = :navy,
          fill = (0, 0.6, :navy), 
          label="test data", 
          yticks = true, 
          legend = :topleft,
          #legend = false,
          #ylim=(0, 1.05 * maximum(vcat(k1.density, k2.density))),
          framestyle = :box,
          #title="Ordered Brier Score.",
          #titlefont = font(10, "Calibri"),
          size=[350,350],
          xlabel="Brier Skill Score",
          xguidefontsize=font(7, "Calibri"),
          xlim = (-2, 2),
          ylim = (0, 1.01),
          grid = :xy,
          gridopacity = 0.5)
plot!(k1.x, k1.density, color = :purple, fill = (0, 0.6, :purple), label="training data")
```

Out[173]:

xml version="1.0" encoding="utf-8"?


-2

-1

0

1

2

0.00

0.25

0.50

0.75

1.00


Brier Skill Score


test data


training data

In [174]:

```
#savefig("figs/BSS_BNN.pdf")
```

In [175]:

```
probsplot(drug_names_test, y_pred_samps_test, 6, y_test)
```

xml version="1.0" encoding="utf-8"?


1

2

3

0.0

0.2

0.4

0.6

0.8

1.0


Pioglitazone, true category = 2

DILI class

Predicted probability

In [176]:

```
probsplot(drug_names_test, y_pred_samps_test, 6, y_test)
```

xml version="1.0" encoding="utf-8"?


1

2

3

0.0

0.2

0.4

0.6

0.8

1.0


Pioglitazone, true category = 2

DILI class

Predicted probability

In [177]:

```
## extract predicted values and convert to 0-1 scale
post = logistic.(eta_post_test)
post_test = post

# cutpoints
c1 = logistic(mean(c1_est))
c2 = logistic(mean(c2_est))
```

Out[177]:

```
0.6854278113247833
```

In [178]:

```
postplot(drug_names_test, post, 6, y_test)
```

Out[178]:

xml version="1.0" encoding="utf-8"?


0.00

0.25

0.50

0.75

1.00


Pioglitazone

P(DILI)

In [179]:

```
## ---------------------------------------------------------
## dot plot
## ---------------------------------------------------------

post_med = median(post, dims=1)';

col = []
for i in 1:length(y_test)
    if y_test[i] == 1
        col = vcat(col, :green4)
    elseif y_test[i] == 2
        col = vcat(col, :darkgoldenrod)
    else
        col = vcat(col, :firebrick)
    end
end 

scatter(y_test + randn(length(y_test))/15, post_med, 
        alpha=1.0, 
        xlim=(0.5,3.5), 
        ylim=(0,1), 
        size=(250,250),
        legend = false,
        xlabel="True DILI category" ,
        xguidefontsize=font(11, "Calibri"), 
        xticks = 1:1:3,
        ylabel="Predicted severity", 
        yguidefontsize=font(11, "Calibri"),
        #fillcolor = [:navy, :purple, :magenta],
        color = col,
        framestyle = :box,
        tickfont=font(10))
hline!([c1, c2], color = :black, linestyle = :dash)
```

Out[179]:

xml version="1.0" encoding="utf-8"?


1

2

3

0.0

0.2

0.4

0.6

0.8

1.0


True DILI category

Predicted severity

In [180]:

```
#savefig("figs/preds_dot_BNN.pdf")
```

In [181]:

```
## ---------------------------------------------------------
## increasing human DILI - posteriors
## ---------------------------------------------------------
   
    # cutpoints
    c1 = logistic(mean(c1_est))
    c2 = logistic(mean(c2_est))

#---- Rosiglitazone ---------------------
    post = post_train

    ind = 38

    post_ind = post[:,ind]
    
    post_ind = vcat(0, post_ind, 1)
    
    kde_npoints = 2048
    #dens = kde(post_ind, npoints=kde_npoints, bandwidth=0.1)
    dens = kde(post_ind, npoints=kde_npoints)

    p1 = plot(dens.x, dens.density,  
              fill = (0, 0.2, :darkgoldenrod), 
              color = :darkgoldenrod, 
              #title = drug_names_train[ind], 
              xlabel="P(DILI)", 
              xlims = (0,1.01),
              ylims = (0,6),
              legend=false,
              yticks = false,
              framestyle = :box,
              size=[250,250],
              titlefont = font(14, "Calibri"),
              xguidefontsize=font(14, "Calibri"))
    vline!([c1, c2], color = :black, linestyle = :dash)

    # define a function that returns a Plots.Shape
    rectangle(w, h, x, y) = Shape(x .+ [0,w,w,0], y .+ [0,0,h,h])

    plot!(rectangle(c1+0.05, 0.25, -0.05,0), color = :green4, alpha = 0.9)
    plot!(rectangle(c2-c1,0.25,c1,0), color = :darkgoldenrod, alpha = 0.9)
    plot!(rectangle(1.05-c2,0.25,c2,0), color = :firebrick, alpha = 0.9)

#---- Troglitazone ---------------------
post = post_train

    ind = 107

    post_ind = post[:,ind]
    
    post_ind = vcat(0, post_ind, 1)
    
    kde_npoints = 2048
    #dens = kde(post_ind, npoints=kde_npoints, bandwidth=0.1)
    dens = kde(post_ind, npoints=kde_npoints)

    plot!(dens.x, dens.density,  
              fill = (0, 0.4, :firebrick), 
              color = :firebrick, 
              #title = drug_names_train[ind], 
              xlabel="P(DILI)", 
              xlims = (0,1.01),
              ylims = (0,6),
              legend=false,
              yticks = false,
              framestyle = :box,
              size=[250,250],
              titlefont = font(14, "Calibri"),
              xguidefontsize=font(14, "Calibri"))
    vline!([c1, c2], color = :black, linestyle = :dash)

    # define a function that returns a Plots.Shape
    rectangle(w, h, x, y) = Shape(x .+ [0,w,w,0], y .+ [0,0,h,h])

    plot!(rectangle(c1+0.05, 0.25, -0.05,0), color = :green4, alpha = 0.9)
    plot!(rectangle(c2-c1,0.25,c1,0), color = :darkgoldenrod, alpha = 0.9)
    plot!(rectangle(1.05-c2,0.25,c2,0), color = :firebrick, alpha = 0.9)

#---- Pioglitazone ---------------------
    post = post_test

    ind = 6

    post_ind = post[:,ind]
    
    post_ind = vcat(0, post_ind, 1)
    
    kde_npoints = 2048
    #dens = kde(post_ind, npoints=kde_npoints, bandwidth=0.1)
    dens = kde(post_ind, npoints=kde_npoints)

    plot!(dens.x, dens.density,  
              fill = (0, 0.6, :darkgoldenrod), 
              color = :darkgoldenrod, 
             #title = drug_names_test[ind], 
              xlabel="P(DILI)", 
              xlims = (0,1.01),
              ylims = (0,6),
              legend=false,
              yticks = false,
              framestyle = :box,
              size=[250,250],
              titlefont = font(14, "Calibri"),
              xguidefontsize=font(10, "Calibri"))
    vline!([c1, c2], color = :black, linestyle = :dash)

    # define a function that returns a Plots.Shape
    rectangle(w, h, x, y) = Shape(x .+ [0,w,w,0], y .+ [0,0,h,h])

    plot!(rectangle(c1+0.05, 0.25, -0.05,0), color = :green4, alpha = 0.9)
    plot!(rectangle(c2-c1,0.25,c1,0), color = :darkgoldenrod, alpha = 0.9)
    plot!(rectangle(1.05-c2,0.25,c2,0), color = :firebrick, alpha = 0.9)

#------- annotations ------------
    annotate!(0.87, 5.5, text("Troglitazone", :balck, 9))
    annotate!(0.87, 2.5, text("Pioglitazone", :balck, 9))
    annotate!(0.35, 2, text("Rosiglitazone", :balck, 9))
```

Out[181]:

xml version="1.0" encoding="utf-8"?


0.00

0.25

0.50

0.75

1.00


P(DILI)


Troglitazone

Pioglitazone

Rosiglitazone

In [182]:

```
#savefig("figs/BNN_rosi_pio.pdf")
```

In [183]:

```
## ---------------------------------------------------------
## increasing human DILI - posterior predictive distributions
## ---------------------------------------------------------

#---- Rosiglitazone ---------------------
    y_pred_samps = y_pred_samps_train
    y = y_train
    drug_names = drug_names_train

    j = 38

    tbl = freqtable(y_pred_samps[:,j])
    ptbl = prop(tbl)
    b1 = bar( ptbl, 
             labels="10 sin values", 
             #size=[150,250],
              xlabel=drug_names[j], 
              ylabel="Predicted probability", 
              xticks = 1:1:3,
              legend=false,
              yticks = true,
              framestyle = :grid,
              ylimit = (0,1),
              title = "true category = " * string(y[j]) , 
              titlefont = font(14, "Calibri"),
              xguidefontsize=font(14, "Calibri"),
              yguidefontsize=font(14, "Calibri"),
              #fillcolor = [:navy, :purple, :magenta],
              fillcolor = [:green4, :darkgoldenrod, :firebrick],
              #alpha= [0.9, 0.9,0.6],  
              minorgrid = true,
              #gridcolor = :black,
              gridopacity = 0.5,
              grid = :xy,
              bar_width = 0.9);

#---- Troglitazone ---------------------
    y_pred_samps = y_pred_samps_train
    y = y_train   
    drug_names = drug_names_train

    j = 107

    tbl = freqtable(y_pred_samps[:,j])
    ptbl = prop(tbl)
    b2 = bar( ptbl, 
             labels="10 sin values", 
             #size=[150,250],
              xlabel=drug_names[j], 
              #ylabel="Predicted probability", 
              xticks = 1:1:3,
              legend=false,
              yticks = true,
              framestyle = :grid,
              ylimit = (0,1),
              title = "true category = " * string(y[j]) , 
              titlefont = font(14, "Calibri"),
              xguidefontsize=font(14, "Calibri"),
              yguidefontsize=font(14, "Calibri"),
              #fillcolor = [:navy, :purple, :magenta],
              fillcolor = [:green4, :darkgoldenrod, :firebrick],
              #alpha= [0.9, 0.9,0.6],  
              minorgrid = true,
              #gridcolor = :black,
              gridopacity = 0.5,
              grid = :xy,
              bar_width = 0.9);
#---- Pioglitazone ---------------------
    y_pred_samps = y_pred_samps_test
    y = y_test
    drug_names = drug_names_test

    j = 6

    tbl = freqtable(y_pred_samps[:,j])
    ptbl = prop(tbl)
    b3 = bar( ptbl, 
             labels="10 sin values", 
             #size=[150,250],
              xlabel=drug_names[j], 
              #ylabel="Predicted probability", 
              xticks = 1:1:3,
              legend=false,
              #yticks = true,
              #yticks = false,
              framestyle = :grid,
              ylimit = (0,1),
              title = "true category = " * string(y[j]) , 
              titlefont = font(14, "Calibri"),
              xguidefontsize=font(14, "Calibri"),
              #yguidefontsize=font(14, "Calibri"),
              #fillcolor = [:navy, :purple, :magenta],
              fillcolor = [:green4, :darkgoldenrod, :firebrick],
              #alpha= [0.9, 0.9,0.6],  
              minorgrid = true,
              #gridcolor = :black,
              gridopacity = 0.5,
              grid = :xy,
              bar_width = 0.9);

plot(b1, b3, b2, layout=(1, 3), label="")
```

Out[183]:

xml version="1.0" encoding="utf-8"?


1

2

3

0.0

0.2

0.4

0.6

0.8

1.0

true category = 2

Rosiglitazone

Predicted probability


1

2

3

0.0

0.2

0.4

0.6

0.8

1.0

true category = 2

Pioglitazone


1

2

3

0.0

0.2

0.4

0.6

0.8

1.0

true category = 3

Troglitazone

In [184]:

```
#savefig("figs/BNN_rosi_pio_pred.pdf")
```

In [185]:

```
## ---------------------------------------------------------
## Predict for ambiguous-category data
## ---------------------------------------------------------

probs_mat_ambig, y_pred_samps_ambig, logpdf_mat_ambig, y_pred_ambig, eta_post_ambig = predict(X_ambig, false, n_samps);
```

In [186]:

```
## extract predicted values and convert to 0-1 scale
post = logistic.(eta_post_ambig)
post_ambig = post

# cutpoints
c1 = logistic(mean(c1_est))
c2 = logistic(mean(c2_est))

postplot(drug_names_ambig, post_ambig, 2, false)
```

Out[186]:

xml version="1.0" encoding="utf-8"?


0.00

0.25

0.50

0.75

1.00


Ambrisentan

P(DILI)

In [187]:

```
postplot(drug_names_ambig, post, 3, false)
```

Out[187]:

xml version="1.0" encoding="utf-8"?


0.00

0.25

0.50

0.75

1.00


Buspirone

P(DILI)

In [188]:

```
probsplot(drug_names_ambig, y_pred_samps_ambig, 2, false)
```

Out[192]:

xml version="1.0" encoding="utf-8"?


1

2

3

0.0

0.2

0.4

0.6

0.8

1.0

Buspirone


1

2

3

0.0

0.2

0.4

0.6

0.8

1.0

true category = 3

Nefazodone

Predicted probability

In [193]:

```
drug_names_train[58]
```

Out[193]:

```
"Bosentan"
```

In [194]:

```
drug_names_ambig[3]
```

Out[194]:

```
"Buspirone"
```

In [195]:

```
#---- Bozentan ---------------------
    y_pred_samps = y_pred_samps_train
    y = y_train
    drug_names = drug_names_train

    j = 58

    tbl = freqtable(y_pred_samps[:,j])
    ptbl = prop(tbl)
    b1 = bar( ptbl, 
             labels="10 sin values", 
             #size=[150,250],
              xlabel=drug_names[j], 
              ylabel="Predicted probability", 
              xticks = 1:1:3,
              legend=false,
              yticks = true,
              framestyle = :grid,
              ylimit = (0,1),
              title = "true category = " * string(y[j]) , 
              titlefont = font(7, "Calibri"),
              xguidefontsize=font(7, "Calibri"),
              yguidefontsize=font(7, "Calibri"),
              #fillcolor = [:navy, :purple, :magenta],
              fillcolor = [:green4, :darkgoldenrod, :firebrick],
              #alpha= [0.9, 0.9,0.6],  
              minorgrid = true,
              #gridcolor = :black,
              gridopacity = 0.5,
              grid = :xy,
              bar_width = 0.9);

#---- Bosentan ---------------------
    y_pred_samps = y_pred_samps_ambig
#    y = y_train   
    drug_names = drug_names_ambig

    j = 2

    tbl = freqtable(y_pred_samps[:,j])
    ptbl = prop(tbl)
    b2 = bar( ptbl, 
             labels="10 sin values", 
             #size=[150,250],
              xlabel=drug_names[j], 
              #ylabel="Predicted probability", 
              xticks = 1:1:3,
              legend=false,
              yticks = true,
              framestyle = :grid,
              ylimit = (0,1),
              #title = "true category = " * string(y[j]) , 
              titlefont = font(7, "Calibri"),
              xguidefontsize=font(7, "Calibri"),
              yguidefontsize=font(7, "Calibri"),
              #fillcolor = [:navy, :purple, :magenta],
              fillcolor = [:green4, :darkgoldenrod, :firebrick],
              #alpha= [0.9, 0.9,0.6],  
              minorgrid = true,
              #gridcolor = :black,
              gridopacity = 0.5,
              grid = :xy,
              bar_width = 0.9);

plot(b2, b1, layout=(1, 2), label="")
```

Out[195]:

xml version="1.0" encoding="utf-8"?


1

2

3

0.0

0.2

0.4

0.6

0.8

1.0

Ambrisentan


1

2

3

0.0

0.2

0.4

0.6

0.8

1.0

true category = 3

Bosentan

Predicted probability

In [196]:

```
drug_names_test[30]
```

Out[196]:

```
"Folic Acid"
```

In [197]:

```
#---- Folic acid ---------------------
   # cutpoints
    c1 = logistic(mean(c1_est))
    c2 = logistic(mean(c2_est))

    post = post_test
    drug_names = drug_names_test

    ind = 30

    post_ind = post[:,ind]
  
    post_ind = vcat(0, post_ind, 1)

    kde_npoints = 2048
    dens = kde(post_ind, npoints=kde_npoints)

    p1 = plot(dens.x, dens.density,  
              fill = (0, 0.2, :blue), 
              color = :blue, 
              #title = drug_names[ind], 
              title = "Posterior density distribution", 
              xlabel="P(DILI)", 
              xlims = (0,1.01),
              ylims = (0,6),
              legend=false,
              yticks = false,
              framestyle = :box,
              size=[250,250],
              titlefont = font(14, "Calibri"),
              xguidefontsize=font(14, "Calibri"),
              tickfont=font(9))
   vline!([c1, c2], color = :black, linestyle = :dash)

    # define a function that returns a Plots.Shape
    rectangle(w, h, x, y) = Shape(x .+ [0,w,w,0], y .+ [0,0,h,h])

    plot!(rectangle(c1+0.05, 0.25, -0.05,0), color = :green4, alpha = 0.9)
    plot!(rectangle(c2-c1,0.25,c1,0), color = :darkgoldenrod, alpha = 0.9)
    plot!(rectangle(1.05-c2,0.25,c2,0), color = :firebrick, alpha = 0.9)

#---- Folic acid ---------------------
    y_pred_samps = y_pred_samps_test
    y = y_test

    j = 30

    tbl = freqtable(y_pred_samps[:,j])
    ptbl = prop(tbl)
    p2 = bar( ptbl, 
             labels="10 sin values", 
             #size=[150,250],
              xlabel=drug_names[j], 
              #ylabel="Predicted probability", 
              xticks = 1:1:3,
              legend=false,
              #yticks = true,
            #yticks = false,
              framestyle = :grid,
              ylimit = (0,1),
              title = "true category = " * string(y[j]) , 
              titlefont = font(14, "Calibri"),
              xguidefontsize=font(14, "Calibri"),
              #yguidefontsize=font(7, "Calibri"),
              #fillcolor = [:navy, :purple, :magenta],
              fillcolor = [:green4, :darkgoldenrod, :firebrick],
              #alpha= [0.9, 0.9,0.6],  
              minorgrid = true,
              #gridcolor = :black,
              gridopacity = 0.5,
              grid = :xy,
              bar_width = 0.9,
              tickfont=font(9));

plot(p1, p2, layout=(1, 2), label="")
```

Out[197]:

xml version="1.0" encoding="utf-8"?


0.00

0.25

0.50

0.75

1.00


Posterior density distribution

P(DILI)


1

2

3

0.0

0.2

0.4

0.6

0.8

1.0

true category = 1

Folic Acid

In [48]:

```
#savefig("figs/BNN_FolicAcid.pdf")
```

In [198]:

```
using Pkg
Pkg.status()
```

```
    Status `~/Box Sync/27_BNN_DILI/revision_correct_submit/Project.toml`
  [336ed68f] CSV v0.5.9
  [a93c6f00] DataFrames v0.19.1
  [587475ba] Flux v0.8.3
  [da1fdf0e] FreqTables v0.3.1
  [4138dd39] JLD v0.9.2
  [5ab0869b] KernelDensity v0.5.1
  [f0e99cf1] MLBase v0.8.0
  [91a5bcdd] Plots v0.25.3
  [2913bbd2] StatsBase v0.32.0
  [4c63d2b9] StatsFuns v0.8.0
  [3eaba693] StatsModels v0.6.2
  [fce5fe82] Turing v0.6.23 #master (https://github.com/TuringLang/Turing.jl.git)
  [9a3f8284] Random
```

In [ ]:

```
# Pkg.add(Pkg.PackageSpec(;name="Turing", version="0.6.23"))
```
