## Supplementary material for "A Bayesian neural network for toxicity prediction": 3_BNN_multiclass.html


In [12]:

```
## ---------------------------------------------------------
## Import libraries and functions
## ---------------------------------------------------------

using CSV
using DataFrames
using Turing, Flux, Plots, Random
using StatsBase
using StatsFuns: logistic, logsumexp
using MLBase
using StatsModels
using FreqTables
using KernelDensity
using JLD
using CPUTime
using MLPreprocessing

Turing.turnprogress(true);
Turing.setadbackend(:reverse_diff)

include("functions.jl")
```

```
┌ Info: [Turing]: global PROGRESS is set as true
└ @ Turing /Users/kcft114/.julia/packages/Turing/m05p3/src/Turing.jl:24
```

Out[12]:

```
euc_dist (generic function with 1 method)
```

In [13]:

```
## ---------------------------------------------------------
## Read and prepare data
## ---------------------------------------------------------

# read in data
df_train = CSV.read("data/Aleo_train_match.csv")
df_test = CSV.read("data/Aleo_test_match.csv");
```

In [14]:

```
println(size(df_train))
println(size(df_test))

df_Xtrain = df_train[!, [:ClogP, :BSEP, :Glu, :Glu_Gal, :THLE, :HepG2, :Fsp3, :log10cmax]]
df_Xtest = df_test[!, [:ClogP, :BSEP, :Glu, :Glu_Gal, :THLE, :HepG2, :Fsp3, :log10cmax]]

scaler = fit(StandardScaler, df_Xtrain)
transform!(df_Xtrain, scaler)
transform!(df_Xtest, scaler)

df_y_train = df_train[!, :dili_sev]
df_y_test = df_test[!, :dili_sev];

X_train = convert(Matrix, df_Xtrain)
X_test = convert(Matrix, df_Xtest)

Y_train = Int.(convert(Array, df_y_train)) .- 1
Y_test = Int.(convert(Array, df_y_test)) .- 1

println(sort(countmap(Y_train)))
println(sort(countmap(Y_test)))
```

```
(147, 12)
(37, 12)
OrderedCollections.OrderedDict(0=>37,1=>45,2=>65)
OrderedCollections.OrderedDict(0=>10,1=>11,2=>16)
```

In [15]:

```
n0 = 8
n1 = 15
K = 3
num_params = (n1 * (n0 + 1 + K) + K)
```

Out[15]:

```
183
```

In [16]:

```
## ---------------------------------------------------------
## Define model
## ---------------------------------------------------------

function weights(theta::AbstractVector)
    W0 = reshape(theta[ 1:(n0*n1)], n1, n0); 
    b0 = reshape(theta[(n0*n1 + 1): (n0*n1 + n1)], n1)
    W1 = reshape(theta[(n0*n1 + n1 + 1): (n1 * (n0 + 1 + K))], K, n1); 
    b1 = reshape(theta[(n1 * (n0 + 1 + K) + 1): (n1 * (n0 + 1 + K) + K)], K)
    return W0, b0, W1, b1
end

function feedforward(inp::AbstractArray, theta::AbstractVector)
    W0, b0, W1, b1 = weights(theta)
    model = Chain(
        Dense(W0, b0, relu),
        Dense(W1, b1)
    )
    return model(inp)
end

# Create `CategoricalLogit` to prevent numerical issues
struct CategoricalLogit <: DiscreteUnivariateDistribution
    logitp
end

# `logsumexp` avoids the numerical issue
function Distributions.logpdf(d::CategoricalLogit, k::Int)
    return (d.logitp .- logsumexp(d.logitp))[k+1]    # use k+1 as lab[i] is 0-indexed
end

@model bayesnn(inp, lab) = begin
    
    sig ~ TruncatedNormal(0, 1, 0, Inf)
    theta~ MvNormal(zeros(num_params), sig .* ones(num_params))

    preds = feedforward(inp, theta)
    for i = 1:length(lab)
        lab[i] ~ CategoricalLogit(preds[:,i])
    end
end
```

Out[16]:

```
bayesnn (generic function with 3 methods)
```

In [17]:

```
inputs = X_train'
labels = Y_train;
```

In [18]:

```
## ---------------------------------------------------------
## Inference
## ---------------------------------------------------------

# one chain:
steps = 20_000
chain = sample(bayesnn(Array(inputs), labels), NUTS(steps, 0.65));

# for multiple chains run the following:
#num_chains = 4
#num_samples = 20_000
#chains = mapreduce(c -> sample(bayesnn(Array(inputs), labels), NUTS(steps, 0.65)), chainscat, 1:num_chains)
```

```
┌ Info: Found initial step size
│   init_ϵ = 0.2
└ @ Turing.Inference /Users/kcft114/.julia/packages/Turing/m05p3/src/inference/hmc.jl:365
┌ Warning: The current proposal will be rejected due to numerical error(s).
│   isfiniteθ = true
│   isfiniter = false
│   isfiniteℓπ = false
│   isfiniteℓκ = false
└ @ AdvancedHMC /Users/kcft114/.julia/packages/AdvancedHMC/YWXfk/src/hamiltonian.jl:36
┌ Info: Finished 1000 adapation steps
│   adaptor = StanHMCAdaptor(n_adapts=1000, pc=DiagPreconditioner, ssa=NesterovDualAveraging(γ=0.05, t_0=10.0, κ=0.75, δ=0.65, state.ϵ=0.09908916063682727), init_buffer=75, term_buffer=50)
│   τ.integrator = Leapfrog(ϵ=0.0991)
│   h.metric = DiagEuclideanMetric([0.0250861, 0.189947, 0.178 ...])
└ @ AdvancedHMC /Users/kcft114/.julia/packages/AdvancedHMC/YWXfk/src/sampler.jl:67
┌ Info: Finished 20000 sampling steps in 13062.460452772 (s)
│   h = Hamiltonian(metric=DiagEuclideanMetric([0.0250861, 0.189947, 0.178 ...]))
│   τ = NUTS{Multinomial}(integrator=Leapfrog(ϵ=0.0991), max_depth=5), Δ_max=1000.0)
│   EBFMI(Hs) = 1997.2942857514247
│   mean(αs) = 0.7319993934173448
└ @ AdvancedHMC /Users/kcft114/.julia/packages/AdvancedHMC/YWXfk/src/sampler.jl:77
```

In [33]:

```
## ---------------------------------------------------------
## Save/read results
## ---------------------------------------------------------

#write("chains/BNN_multiclass.jls", chain)

chain = read("chains/BNN_multiclass.jls", Chains);
```

In [20]:

```
!(sum(isnan.(summarystats(chain)[:, :r_hat]))>0) & !(sum(abs.(summarystats(chain)[:, :r_hat] .- 1) .> 0.1) > 0) & !(sum(summarystats(chain)[:, :std] .< 1e-13) > 0)
```

Out[20]:

```
true
```

In [19]:

```
show(chain)
```

```
Object of type Chains, with data of type 19000×194×1 Array{Union{Missing, Float64},3}

Log evidence      = 0.0
Iterations        = 1:19000
Thinning interval = 1
Chains            = 1
Samples per chain = 19000
internals         = eval_num, lp, acceptance_rate, hamiltonian_energy, is_accept, log_density, n_steps, numerical_error, step_size, tree_depth
parameters        = theta[135], theta[145], theta[109], theta[72], theta[96], theta[69], theta[16], theta[87], theta[167], theta[54], theta[121], theta[13], theta[132], theta[113], theta[170], theta[37], theta[115], theta[118], theta[127], theta[142], theta[128], theta[38], theta[175], theta[148], theta[7], theta[77], theta[141], theta[32], theta[101], theta[84], theta[71], theta[164], theta[131], theta[64], theta[17], theta[56], theta[86], theta[4], theta[102], theta[104], theta[49], theta[126], theta[117], theta[14], theta[144], theta[172], theta[28], theta[105], theta[136], theta[183], theta[153], theta[122], theta[33], theta[63], theta[107], theta[161], theta[80], theta[89], theta[151], theta[83], theta[81], theta[157], theta[51], theta[53], theta[78], theta[149], theta[156], theta[11], theta[140], theta[134], theta[12], theta[116], theta[65], theta[173], theta[169], theta[147], theta[120], theta[2], theta[68], theta[139], theta[48], theta[93], theta[166], theta[174], theta[59], theta[30], theta[40], theta[162], theta[57], theta[36], theta[19], theta[177], theta[52], theta[39], theta[70], theta[31], theta[43], theta[125], theta[179], theta[108], theta[23], theta[61], theta[91], theta[150], theta[25], theta[46], theta[92], theta[106], theta[26], theta[155], theta[95], theta[88], theta[21], sig, theta[29], theta[119], theta[79], theta[10], theta[100], theta[178], theta[152], theta[143], theta[176], theta[112], theta[41], theta[1], theta[110], theta[66], theta[76], theta[171], theta[18], theta[180], theta[73], theta[75], theta[3], theta[82], theta[67], theta[9], theta[111], theta[130], theta[35], theta[34], theta[154], theta[114], theta[5], theta[27], theta[74], theta[168], theta[50], theta[138], theta[158], theta[85], theta[22], theta[94], theta[24], theta[99], theta[123], theta[90], theta[62], theta[163], theta[129], theta[44], theta[20], theta[133], theta[165], theta[15], theta[42], theta[60], theta[97], theta[124], theta[146], theta[103], theta[182], theta[98], theta[6], theta[160], theta[8], theta[181], theta[45], theta[159], theta[58], theta[137], theta[47], theta[55]

2-element Array{ChainDataFrame,1}

Summary Statistics

│ Row │ parameters │ mean        │ std       │ naive_se    │ mcse       │ ess     │ r_hat    │
│     │ Symbol     │ Float64     │ Float64   │ Float64     │ Float64    │ Any     │ Any      │
├─────┼────────────┼─────────────┼───────────┼─────────────┼────────────┼─────────┼──────────┤
│ 1   │ sig        │ 0.468703    │ 0.0850081 │ 0.000616714 │ 0.00239869 │ 1266.65 │ 1.00348  │
│ 2   │ theta[1]   │ 0.0208085   │ 0.454165  │ 0.00329486  │ 0.00399293 │ 12677.1 │ 0.999952 │
│ 3   │ theta[2]   │ 0.0295059   │ 0.445342  │ 0.00323085  │ 0.00426133 │ 11900.4 │ 0.999948 │
│ 4   │ theta[3]   │ 0.0205928   │ 0.45007   │ 0.00326515  │ 0.00345674 │ 12470.1 │ 1.00002  │
│ 5   │ theta[4]   │ 0.019907    │ 0.451054  │ 0.00327229  │ 0.00354835 │ 14941.8 │ 0.999949 │
│ 6   │ theta[5]   │ 0.02069     │ 0.450749  │ 0.00327008  │ 0.00396214 │ 13383.9 │ 1.00005  │
│ 7   │ theta[6]   │ 0.0278563   │ 0.4501    │ 0.00326537  │ 0.00368568 │ 12864.2 │ 0.99995  │
│ 8   │ theta[7]   │ 0.0199788   │ 0.457538  │ 0.00331933  │ 0.00411898 │ 13367.9 │ 0.999997 │
│ 9   │ theta[8]   │ 0.0211095   │ 0.447795  │ 0.00324865  │ 0.00417632 │ 12558.7 │ 1.00024  │
│ 10  │ theta[9]   │ 0.0199899   │ 0.452773  │ 0.00328476  │ 0.0039207  │ 13343.8 │ 0.999973 │
│ 11  │ theta[10]  │ 0.0213251   │ 0.454781  │ 0.00329933  │ 0.00373585 │ 13246.0 │ 1.00029  │
│ 12  │ theta[11]  │ 0.0230034   │ 0.451499  │ 0.00327552  │ 0.00449027 │ 10838.9 │ 0.99995  │
│ 13  │ theta[12]  │ 0.0279295   │ 0.453319  │ 0.00328872  │ 0.00396279 │ 12917.2 │ 0.999981 │
│ 14  │ theta[13]  │ 0.0176939   │ 0.444501  │ 0.00322475  │ 0.0039155  │ 13134.7 │ 0.999981 │
│ 15  │ theta[14]  │ 0.019062    │ 0.455973  │ 0.00330798  │ 0.0039751  │ 11485.6 │ 1.00005  │
│ 16  │ theta[15]  │ 0.0186703   │ 0.452705  │ 0.00328426  │ 0.00379193 │ 12958.2 │ 1.00009  │
│ 17  │ theta[16]  │ -0.0112186  │ 0.511631  │ 0.00371176  │ 0.00449733 │ 10046.8 │ 1.00001  │
│ 18  │ theta[17]  │ -0.0173225  │ 0.509902  │ 0.00369922  │ 0.00506699 │ 9640.36 │ 1.0001   │
│ 19  │ theta[18]  │ 0.00723932  │ 0.51571   │ 0.00374135  │ 0.00553253 │ 8131.87 │ 1.00009  │
│ 20  │ theta[19]  │ -0.0155775  │ 0.503437  │ 0.00365232  │ 0.00501206 │ 10313.6 │ 1.00017  │
│ 21  │ theta[20]  │ -0.0090176  │ 0.508276  │ 0.00368742  │ 0.00520838 │ 9012.36 │ 0.999948 │
│ 22  │ theta[21]  │ -0.0105527  │ 0.505225  │ 0.00366528  │ 0.00552423 │ 8486.58 │ 0.999957 │
│ 23  │ theta[22]  │ -0.00958928 │ 0.509247  │ 0.00369447  │ 0.00558309 │ 9410.87 │ 1.00038  │
│ 24  │ theta[23]  │ -0.00532523 │ 0.511479  │ 0.00371066  │ 0.00539408 │ 9922.68 │ 1.00007  │
│ 25  │ theta[24]  │ 0.00128641  │ 0.514122  │ 0.00372983  │ 0.0054625  │ 8305.79 │ 0.99997  │
│ 26  │ theta[25]  │ -0.00527365 │ 0.509293  │ 0.0036948   │ 0.00526803 │ 9017.73 │ 1.00052  │
│ 27  │ theta[26]  │ -0.00616845 │ 0.50926   │ 0.00369456  │ 0.00588819 │ 8700.52 │ 1.00015  │
│ 28  │ theta[27]  │ -0.00915707 │ 0.515744  │ 0.0037416   │ 0.00510121 │ 9220.73 │ 1.00042  │
│ 29  │ theta[28]  │ -0.00488288 │ 0.504038  │ 0.00365668  │ 0.00542514 │ 7673.83 │ 0.999985 │
│ 30  │ theta[29]  │ -0.00207719 │ 0.50956   │ 0.00369674  │ 0.00544785 │ 9123.13 │ 1.00009  │
│ 31  │ theta[30]  │ -0.00366845 │ 0.506649  │ 0.00367562  │ 0.00500212 │ 9724.93 │ 0.999996 │
│ 32  │ theta[31]  │ -0.0316993  │ 0.452503  │ 0.0032828   │ 0.00374055 │ 13856.1 │ 0.999985 │
│ 33  │ theta[32]  │ -0.020453   │ 0.451601  │ 0.00327626  │ 0.00389671 │ 13365.3 │ 0.999964 │
│ 34  │ theta[33]  │ -0.0254607  │ 0.457803  │ 0.00332125  │ 0.00408545 │ 15837.4 │ 1.00001  │
│ 35  │ theta[34]  │ -0.0230883  │ 0.4535    │ 0.00329004  │ 0.0035107  │ 16327.1 │ 0.999955 │
│ 36  │ theta[35]  │ -0.0241311  │ 0.451515  │ 0.00327564  │ 0.00355625 │ 15818.7 │ 0.999954 │
│ 37  │ theta[36]  │ -0.0235982  │ 0.45709   │ 0.00331608  │ 0.00346801 │ 15447.0 │ 0.999984 │
│ 38  │ theta[37]  │ -0.0215983  │ 0.450994  │ 0.00327185  │ 0.00371397 │ 15518.5 │ 0.999992 │
│ 39  │ theta[38]  │ -0.0304967  │ 0.456297  │ 0.00331033  │ 0.00357274 │ 15706.7 │ 0.99999  │
│ 40  │ theta[39]  │ -0.0281294  │ 0.448769  │ 0.00325571  │ 0.00314696 │ 16020.7 │ 0.999963 │
│ 41  │ theta[40]  │ -0.0248234  │ 0.449587  │ 0.00326165  │ 0.00342501 │ 14996.1 │ 0.999948 │
│ 42  │ theta[41]  │ -0.0225861  │ 0.45903   │ 0.00333015  │ 0.00364242 │ 15500.5 │ 1.00004  │
│ 43  │ theta[42]  │ -0.0184193  │ 0.453262  │ 0.00328831  │ 0.00378059 │ 15701.9 │ 0.999993 │
│ 44  │ theta[43]  │ -0.0230133  │ 0.449547  │ 0.00326136  │ 0.00389163 │ 14659.2 │ 0.99997  │
│ 45  │ theta[44]  │ -0.0212217  │ 0.446725  │ 0.00324089  │ 0.00373893 │ 14655.4 │ 1.00005  │
│ 46  │ theta[45]  │ -0.0229562  │ 0.445082  │ 0.00322897  │ 0.0036723  │ 15534.6 │ 1.00001  │
│ 47  │ theta[46]  │ 0.0246008   │ 0.46338   │ 0.00336171  │ 0.00379672 │ 14141.1 │ 0.999972 │
│ 48  │ theta[47]  │ 0.0250476   │ 0.465262  │ 0.00337536  │ 0.00416798 │ 13482.4 │ 0.999948 │
│ 49  │ theta[48]  │ 0.0271944   │ 0.46955   │ 0.00340647  │ 0.0041583  │ 13276.6 │ 0.999955 │
│ 50  │ theta[49]  │ 0.0285089   │ 0.466978  │ 0.00338782  │ 0.00400741 │ 13472.3 │ 0.999969 │
│ 51  │ theta[50]  │ 0.0293338   │ 0.464095  │ 0.0033669   │ 0.00383649 │ 13787.4 │ 1.00007  │
│ 52  │ theta[51]  │ 0.029621    │ 0.465267  │ 0.0033754   │ 0.00437577 │ 13209.1 │ 0.999969 │
│ 53  │ theta[52]  │ 0.0215676   │ 0.469121  │ 0.00340336  │ 0.0040837  │ 13953.7 │ 1.00016  │
│ 54  │ theta[53]  │ 0.0204839   │ 0.467407  │ 0.00339092  │ 0.00396007 │ 14066.6 │ 0.999967 │
│ 55  │ theta[54]  │ 0.0225771   │ 0.47399   │ 0.00343869  │ 0.00410106 │ 13256.5 │ 0.999955 │
│ 56  │ theta[55]  │ 0.0236227   │ 0.46701   │ 0.00338805  │ 0.00351849 │ 15795.8 │ 0.999968 │
│ 57  │ theta[56]  │ 0.0235902   │ 0.469019  │ 0.00340263  │ 0.00383604 │ 13418.5 │ 1.00011  │
│ 58  │ theta[57]  │ 0.0269754   │ 0.463905  │ 0.00336552  │ 0.00365251 │ 15015.7 │ 0.999968 │
│ 59  │ theta[58]  │ 0.0292149   │ 0.469769  │ 0.00340806  │ 0.00410274 │ 13081.6 │ 0.999967 │
│ 60  │ theta[59]  │ 0.0195686   │ 0.464285  │ 0.00336828  │ 0.00368044 │ 14964.3 │ 1.00001  │
│ 61  │ theta[60]  │ 0.0191371   │ 0.471894  │ 0.00342348  │ 0.00363261 │ 15749.9 │ 1.00011  │
│ 62  │ theta[61]  │ -0.0216007  │ 0.449568  │ 0.00326151  │ 0.00373801 │ 13370.4 │ 1.00001  │
│ 63  │ theta[62]  │ -0.0225279  │ 0.446286  │ 0.0032377   │ 0.00378282 │ 14623.6 │ 0.999948 │
│ 64  │ theta[63]  │ -0.0223446  │ 0.444197  │ 0.00322254  │ 0.00370704 │ 15069.5 │ 0.999959 │
│ 65  │ theta[64]  │ -0.0270022  │ 0.446087  │ 0.00323626  │ 0.00398654 │ 13769.2 │ 0.999952 │
│ 66  │ theta[65]  │ -0.0227043  │ 0.45369   │ 0.00329141  │ 0.00416264 │ 13749.6 │ 0.999947 │
│ 67  │ theta[66]  │ -0.0251815  │ 0.446002  │ 0.00323564  │ 0.00348534 │ 15963.1 │ 0.999995 │
│ 68  │ theta[67]  │ -0.0156178  │ 0.448     │ 0.00325013  │ 0.00358072 │ 15805.6 │ 1.00016  │
│ 69  │ theta[68]  │ -0.0214269  │ 0.440947  │ 0.00319897  │ 0.00392215 │ 14080.4 │ 0.999951 │
│ 70  │ theta[69]  │ -0.0238791  │ 0.448281  │ 0.00325217  │ 0.0038014  │ 12767.7 │ 0.999947 │
│ 71  │ theta[70]  │ -0.0229291  │ 0.445931  │ 0.00323513  │ 0.00359525 │ 15295.1 │ 0.999947 │
│ 72  │ theta[71]  │ -0.0317158  │ 0.450913  │ 0.00327126  │ 0.00355988 │ 13560.5 │ 0.999947 │
│ 73  │ theta[72]  │ -0.0251145  │ 0.443873  │ 0.0032202   │ 0.00364288 │ 15499.9 │ 0.999967 │
│ 74  │ theta[73]  │ -0.020272   │ 0.443375  │ 0.00321658  │ 0.0036283  │ 15182.1 │ 0.999961 │
│ 75  │ theta[74]  │ -0.024549   │ 0.441998  │ 0.00320659  │ 0.0035907  │ 14828.1 │ 0.999982 │
│ 76  │ theta[75]  │ -0.0156936  │ 0.446413  │ 0.00323862  │ 0.00381921 │ 14652.2 │ 0.999948 │
│ 77  │ theta[76]  │ 0.0166373   │ 0.457733  │ 0.00332074  │ 0.00399936 │ 13737.1 │ 0.999967 │
│ 78  │ theta[77]  │ 0.0195745   │ 0.451425  │ 0.00327498  │ 0.00399024 │ 14165.8 │ 1.00021  │
│ 79  │ theta[78]  │ 0.0131859   │ 0.456811  │ 0.00331405  │ 0.00394097 │ 13845.2 │ 0.999991 │
│ 80  │ theta[79]  │ 0.0161121   │ 0.447904  │ 0.00324944  │ 0.00371988 │ 14743.2 │ 1.00002  │
│ 81  │ theta[80]  │ 0.0105067   │ 0.45184   │ 0.00327799  │ 0.00387577 │ 13358.6 │ 0.999951 │
│ 82  │ theta[81]  │ 0.0141229   │ 0.456736  │ 0.00331351  │ 0.00388584 │ 13479.5 │ 0.999952 │
│ 83  │ theta[82]  │ 0.0247749   │ 0.456237  │ 0.00330989  │ 0.00390529 │ 13395.8 │ 0.999948 │
│ 84  │ theta[83]  │ 0.00695638  │ 0.456035  │ 0.00330843  │ 0.00363423 │ 14998.6 │ 1.00029  │
│ 85  │ theta[84]  │ 0.0101314   │ 0.4629    │ 0.00335823  │ 0.00384985 │ 14805.5 │ 0.999955 │
│ 86  │ theta[85]  │ 0.02215     │ 0.45495   │ 0.00330055  │ 0.00368327 │ 13894.8 │ 0.999958 │
│ 87  │ theta[86]  │ 0.0146109   │ 0.455304  │ 0.00330313  │ 0.00355616 │ 14019.8 │ 1.00002  │
│ 88  │ theta[87]  │ 0.0174906   │ 0.462474  │ 0.00335514  │ 0.003847   │ 15006.8 │ 0.999948 │
│ 89  │ theta[88]  │ 0.0149032   │ 0.463774  │ 0.00336457  │ 0.00378715 │ 13994.4 │ 0.999949 │
│ 90  │ theta[89]  │ 0.0209952   │ 0.454486  │ 0.00329719  │ 0.00359124 │ 14950.8 │ 0.999956 │
│ 91  │ theta[90]  │ 0.0145156   │ 0.456563  │ 0.00331225  │ 0.00392647 │ 13554.1 │ 1.00004  │
│ 92  │ theta[91]  │ -0.0732574  │ 0.457431  │ 0.00331855  │ 0.00402807 │ 11597.0 │ 0.999965 │
│ 93  │ theta[92]  │ -0.0791093  │ 0.459321  │ 0.00333227  │ 0.00422581 │ 11539.1 │ 0.999968 │
│ 94  │ theta[93]  │ -0.0756725  │ 0.458414  │ 0.00332568  │ 0.00426727 │ 11318.0 │ 0.999952 │
│ 95  │ theta[94]  │ -0.0759509  │ 0.457685  │ 0.00332039  │ 0.00404947 │ 10608.6 │ 0.999955 │
│ 96  │ theta[95]  │ -0.07321    │ 0.462413  │ 0.0033547   │ 0.00462291 │ 10289.1 │ 0.999994 │
│ 97  │ theta[96]  │ -0.0805011  │ 0.456559  │ 0.00331222  │ 0.00440719 │ 10389.5 │ 1.00001  │
│ 98  │ theta[97]  │ -0.0666183  │ 0.465339  │ 0.00337593  │ 0.00419919 │ 11087.8 │ 0.999949 │
│ 99  │ theta[98]  │ -0.067717   │ 0.460809  │ 0.00334306  │ 0.00396906 │ 12004.1 │ 0.999949 │
│ 100 │ theta[99]  │ -0.072749   │ 0.457153  │ 0.00331653  │ 0.00448355 │ 11149.4 │ 0.999969 │
│ 101 │ theta[100] │ -0.0630883  │ 0.46088   │ 0.00334358  │ 0.00410133 │ 11455.7 │ 0.99995  │
│ 102 │ theta[101] │ -0.0676333  │ 0.462388  │ 0.00335452  │ 0.0044021  │ 10465.6 │ 1.00016  │
│ 103 │ theta[102] │ -0.0683296  │ 0.460824  │ 0.00334317  │ 0.00379885 │ 12153.8 │ 0.999982 │
│ 104 │ theta[103] │ -0.0796449  │ 0.467167  │ 0.00338918  │ 0.00443888 │ 10710.8 │ 0.999993 │
│ 105 │ theta[104] │ -0.0689931  │ 0.452807  │ 0.00328501  │ 0.00402074 │ 11895.7 │ 1.00016  │
│ 106 │ theta[105] │ -0.0712451  │ 0.458052  │ 0.00332306  │ 0.00406345 │ 12950.4 │ 0.999947 │
│ 107 │ theta[106] │ 0.0867099   │ 0.532957  │ 0.00386648  │ 0.00645848 │ 7789.55 │ 0.99998  │
│ 108 │ theta[107] │ 0.0734302   │ 0.536014  │ 0.00388866  │ 0.00685544 │ 6726.14 │ 1.00048  │
│ 109 │ theta[108] │ 0.0710462   │ 0.528624  │ 0.00383504  │ 0.00603388 │ 7882.29 │ 1.00014  │
│ 110 │ theta[109] │ 0.0830351   │ 0.529071  │ 0.00383828  │ 0.00604199 │ 8032.25 │ 1.00001  │
│ 111 │ theta[110] │ 0.0831187   │ 0.535456  │ 0.00388461  │ 0.0058882  │ 7722.84 │ 0.999951 │
│ 112 │ theta[111] │ 0.0832261   │ 0.53388   │ 0.00387317  │ 0.00615134 │ 7824.86 │ 1.00011  │
│ 113 │ theta[112] │ 0.0744918   │ 0.53178   │ 0.00385794  │ 0.00526234 │ 8713.39 │ 0.999949 │
│ 114 │ theta[113] │ 0.069373    │ 0.528974  │ 0.00383758  │ 0.00574951 │ 7539.46 │ 1.00049  │
│ 115 │ theta[114] │ 0.084325    │ 0.530442  │ 0.00384823  │ 0.00550193 │ 8287.6  │ 0.999983 │
│ 116 │ theta[115] │ 0.0816117   │ 0.524904  │ 0.00380805  │ 0.00563739 │ 7787.62 │ 0.999959 │
│ 117 │ theta[116] │ 0.0803784   │ 0.526378  │ 0.00381875  │ 0.00636846 │ 7234.12 │ 1.00003  │
│ 118 │ theta[117] │ 0.0744759   │ 0.537811  │ 0.00390169  │ 0.00594636 │ 7769.52 │ 0.999962 │
│ 119 │ theta[118] │ 0.079637    │ 0.528545  │ 0.00383447  │ 0.00551642 │ 8370.65 │ 1.00008  │
│ 120 │ theta[119] │ 0.067142    │ 0.527664  │ 0.00382808  │ 0.00549408 │ 8151.83 │ 0.999961 │
│ 121 │ theta[120] │ 0.0691846   │ 0.527945  │ 0.00383011  │ 0.00590491 │ 8473.89 │ 1.00044  │
│ 122 │ theta[121] │ -0.0172935  │ 0.477983  │ 0.00346765  │ 0.00454783 │ 11800.2 │ 1.00008  │
│ 123 │ theta[122] │ -0.0175839  │ 0.477154  │ 0.00346164  │ 0.00402789 │ 13095.1 │ 1.00004  │
│ 124 │ theta[123] │ -0.0207045  │ 0.476583  │ 0.00345749  │ 0.00420423 │ 13881.7 │ 1.00006  │
│ 125 │ theta[124] │ -0.0171741  │ 0.475413  │ 0.00344901  │ 0.00413846 │ 13590.6 │ 0.999949 │
│ 126 │ theta[125] │ -0.0169448  │ 0.475594  │ 0.00345032  │ 0.00404019 │ 11555.2 │ 0.999977 │
│ 127 │ theta[126] │ -0.0173373  │ 0.474523  │ 0.00344255  │ 0.00420992 │ 14053.5 │ 0.999984 │
│ 128 │ theta[127] │ -0.0262929  │ 0.477547  │ 0.00346449  │ 0.00398828 │ 13924.0 │ 0.999972 │
│ 129 │ theta[128] │ -0.0244488  │ 0.474093  │ 0.00343943  │ 0.0041079  │ 13781.1 │ 0.999959 │
│ 130 │ theta[129] │ -0.0250342  │ 0.47714   │ 0.00346153  │ 0.00430761 │ 12684.9 │ 1.00023  │
│ 131 │ theta[130] │ -0.0243786  │ 0.479034  │ 0.00347528  │ 0.00398821 │ 14182.2 │ 0.999947 │
│ 132 │ theta[131] │ -0.0251655  │ 0.471539  │ 0.0034209   │ 0.00391027 │ 13447.7 │ 1.00001  │
│ 133 │ theta[132] │ -0.020835   │ 0.475911  │ 0.00345262  │ 0.00408661 │ 13135.9 │ 1.00012  │
│ 134 │ theta[133] │ -0.0202957  │ 0.476815  │ 0.00345918  │ 0.00409147 │ 14401.9 │ 1.00018  │
│ 135 │ theta[134] │ -0.0297836  │ 0.476787  │ 0.00345898  │ 0.00453781 │ 12310.0 │ 0.999963 │
│ 136 │ theta[135] │ -0.024864   │ 0.477583  │ 0.00346475  │ 0.00402917 │ 15104.2 │ 0.999963 │
│ 137 │ theta[136] │ -0.0343114  │ 0.500871  │ 0.0036337   │ 0.00551735 │ 8279.92 │ 0.999951 │
│ 138 │ theta[137] │ 0.00856224  │ 0.420185  │ 0.00304834  │ 0.00396315 │ 12830.7 │ 0.999962 │
│ 139 │ theta[138] │ 0.0312448   │ 0.482202  │ 0.00349826  │ 0.00541453 │ 7830.49 │ 1.00018  │
│ 140 │ theta[139] │ -0.0436027  │ 0.504047  │ 0.00365674  │ 0.00594323 │ 7181.83 │ 1.0      │
│ 141 │ theta[140] │ 0.00851702  │ 0.422588  │ 0.00306578  │ 0.00374351 │ 11952.3 │ 0.999992 │
│ 142 │ theta[141] │ 0.0272535   │ 0.489138  │ 0.00354858  │ 0.00574743 │ 6810.47 │ 0.99997  │
│ 143 │ theta[142] │ -0.0207713  │ 0.502988  │ 0.00364906  │ 0.00564577 │ 6974.66 │ 1.0      │
│ 144 │ theta[143] │ 0.00393738  │ 0.419759  │ 0.00304525  │ 0.00369096 │ 12713.1 │ 1.0      │
│ 145 │ theta[144] │ 0.0301668   │ 0.481477  │ 0.003493    │ 0.00535813 │ 7997.73 │ 0.999963 │
│ 146 │ theta[145] │ -0.0393966  │ 0.503412  │ 0.00365214  │ 0.00546696 │ 8391.36 │ 0.999953 │
│ 147 │ theta[146] │ 0.00355615  │ 0.426902  │ 0.00309707  │ 0.00392039 │ 11908.6 │ 0.999954 │
│ 148 │ theta[147] │ 0.0342556   │ 0.482337  │ 0.00349924  │ 0.00537029 │ 8193.77 │ 0.999952 │
│ 149 │ theta[148] │ -0.0441515  │ 0.500971  │ 0.00363443  │ 0.00544754 │ 7496.6  │ 0.999964 │
│ 150 │ theta[149] │ 0.00238136  │ 0.420038  │ 0.00304728  │ 0.00373095 │ 11891.2 │ 0.999948 │
│ 151 │ theta[150] │ 0.0325853   │ 0.482196  │ 0.00349822  │ 0.00557312 │ 7167.62 │ 0.999975 │
│ 152 │ theta[151] │ -0.0442036  │ 0.497634  │ 0.00361022  │ 0.00580689 │ 7178.64 │ 1.00019  │
│ 153 │ theta[152] │ -0.00149792 │ 0.417757  │ 0.00303072  │ 0.00366783 │ 12046.8 │ 1.0001   │
│ 154 │ theta[153] │ 0.0315773   │ 0.487299  │ 0.00353524  │ 0.00526414 │ 7889.71 │ 1.00036  │
│ 155 │ theta[154] │ -0.0265945  │ 0.494366  │ 0.00358651  │ 0.00554699 │ 8388.42 │ 1.00014  │
│ 156 │ theta[155] │ 0.0118999   │ 0.426819  │ 0.00309647  │ 0.00398494 │ 12029.1 │ 1.00017  │
│ 157 │ theta[156] │ 0.022825    │ 0.486221  │ 0.00352742  │ 0.00509448 │ 8760.36 │ 0.999947 │
│ 158 │ theta[157] │ -0.0285127  │ 0.497422  │ 0.00360868  │ 0.00609955 │ 7182.2  │ 1.00011  │
│ 159 │ theta[158] │ 0.0025841   │ 0.422743  │ 0.0030669   │ 0.00409094 │ 11858.6 │ 1.00027  │
│ 160 │ theta[159] │ 0.02182     │ 0.481134  │ 0.00349051  │ 0.00550255 │ 7882.42 │ 1.00003  │
│ 161 │ theta[160] │ -0.0351941  │ 0.504393  │ 0.00365925  │ 0.0053147  │ 7504.2  │ 0.999953 │
│ 162 │ theta[161] │ 0.00262874  │ 0.420388  │ 0.00304981  │ 0.00353241 │ 11843.9 │ 0.999986 │
│ 163 │ theta[162] │ 0.0353971   │ 0.483715  │ 0.00350924  │ 0.00530172 │ 7165.13 │ 0.999959 │
│ 164 │ theta[163] │ -0.0317718  │ 0.507499  │ 0.00368179  │ 0.00548763 │ 7851.64 │ 0.999987 │
│ 165 │ theta[164] │ 0.0023897   │ 0.420045  │ 0.00304733  │ 0.0037908  │ 13103.0 │ 0.99995  │
│ 166 │ theta[165] │ 0.0267763   │ 0.480211  │ 0.00348382  │ 0.00510478 │ 8185.28 │ 1.00022  │
│ 167 │ theta[166] │ -0.036722   │ 0.498034  │ 0.00361312  │ 0.00586579 │ 7717.49 │ 1.00042  │
│ 168 │ theta[167] │ -0.00429574 │ 0.42138   │ 0.00305701  │ 0.00378922 │ 11839.3 │ 1.00009  │
│ 169 │ theta[168] │ 0.0327864   │ 0.479078  │ 0.0034756   │ 0.00596147 │ 7156.3  │ 1.00021  │
│ 170 │ theta[169] │ -0.0318568  │ 0.500773  │ 0.00363299  │ 0.00562137 │ 8045.24 │ 0.99995  │
│ 171 │ theta[170] │ 0.00543743  │ 0.415274  │ 0.00301271  │ 0.00360705 │ 12690.1 │ 0.999949 │
│ 172 │ theta[171] │ 0.0243815   │ 0.479591  │ 0.00347932  │ 0.00514725 │ 7867.13 │ 0.999975 │
│ 173 │ theta[172] │ -0.0347715  │ 0.49913   │ 0.00362107  │ 0.00539593 │ 7782.43 │ 0.999962 │
│ 174 │ theta[173] │ 0.00493459  │ 0.422443  │ 0.00306473  │ 0.00373326 │ 11712.0 │ 1.00002  │
│ 175 │ theta[174] │ 0.0368154   │ 0.482582  │ 0.00350102  │ 0.00518396 │ 7804.34 │ 1.00003  │
│ 176 │ theta[175] │ -0.020204   │ 0.501655  │ 0.00363939  │ 0.00575699 │ 7359.55 │ 0.999947 │
│ 177 │ theta[176] │ 0.00798955  │ 0.423654  │ 0.00307351  │ 0.00386244 │ 12140.1 │ 1.00006  │
│ 178 │ theta[177] │ 0.0218517   │ 0.487727  │ 0.00353835  │ 0.00523926 │ 8418.64 │ 0.999999 │
│ 179 │ theta[178] │ -0.0307315  │ 0.498977  │ 0.00361996  │ 0.00562818 │ 7986.27 │ 1.0002   │
│ 180 │ theta[179] │ 0.00810102  │ 0.424346  │ 0.00307853  │ 0.00372525 │ 12517.2 │ 0.99997  │
│ 181 │ theta[180] │ 0.0211243   │ 0.483527  │ 0.00350788  │ 0.00529627 │ 8480.08 │ 0.99996  │
│ 182 │ theta[181] │ -0.037273   │ 0.436485  │ 0.0031666   │ 0.00365465 │ 13958.9 │ 1.00014  │
│ 183 │ theta[182] │ 0.0196854   │ 0.407581  │ 0.00295691  │ 0.0036384  │ 14785.5 │ 0.999948 │
│ 184 │ theta[183] │ 0.0142924   │ 0.426422  │ 0.00309359  │ 0.00328881 │ 15555.6 │ 0.999949 │

Quantiles

│ Row │ parameters │ 2.5%      │ 25.0%     │ 50.0%        │ 75.0%    │ 97.5%    │
│     │ Symbol     │ Float64   │ Float64   │ Float64      │ Float64  │ Float64  │
├─────┼────────────┼───────────┼───────────┼──────────────┼──────────┼──────────┤
│ 1   │ sig        │ 0.329828  │ 0.409333  │ 0.459263     │ 0.517021 │ 0.66808  │
│ 2   │ theta[1]   │ -0.887287 │ -0.271079 │ 0.0283992    │ 0.314666 │ 0.910063 │
│ 3   │ theta[2]   │ -0.862703 │ -0.259152 │ 0.0344455    │ 0.324244 │ 0.895011 │
│ 4   │ theta[3]   │ -0.878496 │ -0.269019 │ 0.025775     │ 0.315963 │ 0.891611 │
│ 5   │ theta[4]   │ -0.887865 │ -0.272219 │ 0.0271391    │ 0.314449 │ 0.897984 │
│ 6   │ theta[5]   │ -0.877698 │ -0.273513 │ 0.0269631    │ 0.316215 │ 0.91071  │
│ 7   │ theta[6]   │ -0.869793 │ -0.262233 │ 0.0308839    │ 0.323056 │ 0.913506 │
│ 8   │ theta[7]   │ -0.898877 │ -0.273661 │ 0.0243961    │ 0.32415  │ 0.910864 │
│ 9   │ theta[8]   │ -0.864497 │ -0.272101 │ 0.0272869    │ 0.313082 │ 0.894477 │
│ 10  │ theta[9]   │ -0.900888 │ -0.272368 │ 0.0282575    │ 0.319879 │ 0.892476 │
│ 11  │ theta[10]  │ -0.89435  │ -0.271689 │ 0.0280967    │ 0.322986 │ 0.894781 │
│ 12  │ theta[11]  │ -0.894188 │ -0.267874 │ 0.0264728    │ 0.320453 │ 0.900524 │
│ 13  │ theta[12]  │ -0.875292 │ -0.268683 │ 0.0342516    │ 0.32946  │ 0.914442 │
│ 14  │ theta[13]  │ -0.87172  │ -0.272339 │ 0.0223168    │ 0.313698 │ 0.895611 │
│ 15  │ theta[14]  │ -0.900555 │ -0.277909 │ 0.0244373    │ 0.315876 │ 0.900694 │
│ 16  │ theta[15]  │ -0.89296  │ -0.273477 │ 0.0257216    │ 0.315793 │ 0.904023 │
│ 17  │ theta[16]  │ -0.960493 │ -0.348343 │ -0.0450769   │ 0.298436 │ 1.09227  │
│ 18  │ theta[17]  │ -0.961923 │ -0.349926 │ -0.0518437   │ 0.289519 │ 1.08034  │
│ 19  │ theta[18]  │ -0.931968 │ -0.333648 │ -0.0311961   │ 0.318391 │ 1.12549  │
│ 20  │ theta[19]  │ -0.9567   │ -0.346207 │ -0.0461834   │ 0.291103 │ 1.06837  │
│ 21  │ theta[20]  │ -0.96147  │ -0.341409 │ -0.0407663   │ 0.305766 │ 1.05441  │
│ 22  │ theta[21]  │ -0.95381  │ -0.340255 │ -0.0418266   │ 0.29642  │ 1.07892  │
│ 23  │ theta[22]  │ -0.93741  │ -0.349482 │ -0.0422387   │ 0.297073 │ 1.07946  │
│ 24  │ theta[23]  │ -0.957556 │ -0.336646 │ -0.0370671   │ 0.30384  │ 1.08428  │
│ 25  │ theta[24]  │ -0.95446  │ -0.339184 │ -0.0366035   │ 0.315348 │ 1.08997  │
│ 26  │ theta[25]  │ -0.931422 │ -0.347319 │ -0.0440196   │ 0.311562 │ 1.07295  │
│ 27  │ theta[26]  │ -0.934088 │ -0.344541 │ -0.040677    │ 0.303832 │ 1.09524  │
│ 28  │ theta[27]  │ -0.968774 │ -0.349412 │ -0.0446467   │ 0.301931 │ 1.09706  │
│ 29  │ theta[28]  │ -0.93784  │ -0.338582 │ -0.0371636   │ 0.305411 │ 1.06942  │
│ 30  │ theta[29]  │ -0.960921 │ -0.338602 │ -0.0343398   │ 0.309643 │ 1.07039  │
│ 31  │ theta[30]  │ -0.951089 │ -0.338652 │ -0.0331322   │ 0.308854 │ 1.06781  │
│ 32  │ theta[31]  │ -0.923962 │ -0.327857 │ -0.0327188   │ 0.263937 │ 0.870971 │
│ 33  │ theta[32]  │ -0.909393 │ -0.311534 │ -0.0273547   │ 0.268397 │ 0.879447 │
│ 34  │ theta[33]  │ -0.93602  │ -0.316438 │ -0.0269363   │ 0.266847 │ 0.891197 │
│ 35  │ theta[34]  │ -0.921194 │ -0.31855  │ -0.024163    │ 0.267285 │ 0.890502 │
│ 36  │ theta[35]  │ -0.917348 │ -0.319583 │ -0.0256143   │ 0.270635 │ 0.879139 │
│ 37  │ theta[36]  │ -0.930996 │ -0.318896 │ -0.0207848   │ 0.273027 │ 0.87913  │
│ 38  │ theta[37]  │ -0.918904 │ -0.309315 │ -0.0226586   │ 0.272234 │ 0.873971 │
│ 39  │ theta[38]  │ -0.936709 │ -0.332868 │ -0.0289394   │ 0.264268 │ 0.879242 │
│ 40  │ theta[39]  │ -0.925023 │ -0.319174 │ -0.0309289   │ 0.263052 │ 0.869527 │
│ 41  │ theta[40]  │ -0.925765 │ -0.319456 │ -0.0261826   │ 0.272661 │ 0.862295 │
│ 42  │ theta[41]  │ -0.906555 │ -0.321016 │ -0.0274374   │ 0.271472 │ 0.897172 │
│ 43  │ theta[42]  │ -0.928264 │ -0.314305 │ -0.0187171   │ 0.275604 │ 0.892665 │
│ 44  │ theta[43]  │ -0.909861 │ -0.314195 │ -0.0239168   │ 0.267685 │ 0.866548 │
│ 45  │ theta[44]  │ -0.921699 │ -0.308767 │ -0.0223985   │ 0.264824 │ 0.861979 │
│ 46  │ theta[45]  │ -0.897174 │ -0.313283 │ -0.0217144   │ 0.267856 │ 0.851716 │
│ 47  │ theta[46]  │ -0.901061 │ -0.273017 │ 0.0247771    │ 0.327546 │ 0.944123 │
│ 48  │ theta[47]  │ -0.91411  │ -0.276523 │ 0.0278734    │ 0.327007 │ 0.954512 │
│ 49  │ theta[48]  │ -0.907947 │ -0.277279 │ 0.0282806    │ 0.332338 │ 0.956602 │
│ 50  │ theta[49]  │ -0.89731  │ -0.274035 │ 0.029842     │ 0.331414 │ 0.945997 │
│ 51  │ theta[50]  │ -0.885219 │ -0.27305  │ 0.0290999    │ 0.330945 │ 0.955871 │
│ 52  │ theta[51]  │ -0.89368  │ -0.27228  │ 0.0271298    │ 0.333573 │ 0.946404 │
│ 53  │ theta[52]  │ -0.920605 │ -0.277775 │ 0.0183738    │ 0.329355 │ 0.940645 │
│ 54  │ theta[53]  │ -0.897088 │ -0.283176 │ 0.0187122    │ 0.324781 │ 0.956301 │
│ 55  │ theta[54]  │ -0.921261 │ -0.28302  │ 0.0225736    │ 0.330461 │ 0.948422 │
│ 56  │ theta[55]  │ -0.922306 │ -0.279874 │ 0.0278678    │ 0.327695 │ 0.949523 │
│ 57  │ theta[56]  │ -0.888398 │ -0.284217 │ 0.0217309    │ 0.327957 │ 0.938747 │
│ 58  │ theta[57]  │ -0.903685 │ -0.275214 │ 0.0250675    │ 0.336545 │ 0.930868 │
│ 59  │ theta[58]  │ -0.925963 │ -0.275784 │ 0.0285905    │ 0.335018 │ 0.95452  │
│ 60  │ theta[59]  │ -0.90194  │ -0.281693 │ 0.0189313    │ 0.322058 │ 0.942369 │
│ 61  │ theta[60]  │ -0.915733 │ -0.287879 │ 0.0214386    │ 0.323533 │ 0.95911  │
│ 62  │ theta[61]  │ -0.910979 │ -0.311571 │ -0.0221166   │ 0.274845 │ 0.861988 │
│ 63  │ theta[62]  │ -0.905955 │ -0.308969 │ -0.0220363   │ 0.267264 │ 0.871892 │
│ 64  │ theta[63]  │ -0.907734 │ -0.310468 │ -0.0209133   │ 0.265685 │ 0.855778 │
│ 65  │ theta[64]  │ -0.911106 │ -0.317592 │ -0.0248041   │ 0.26286  │ 0.849773 │
│ 66  │ theta[65]  │ -0.90861  │ -0.317488 │ -0.0265589   │ 0.269971 │ 0.874    │
│ 67  │ theta[66]  │ -0.901101 │ -0.313597 │ -0.0228547   │ 0.262211 │ 0.86019  │
│ 68  │ theta[67]  │ -0.910526 │ -0.303108 │ -0.0104162   │ 0.275181 │ 0.878749 │
│ 69  │ theta[68]  │ -0.900187 │ -0.305931 │ -0.0183024   │ 0.266401 │ 0.840471 │
│ 70  │ theta[69]  │ -0.912874 │ -0.309416 │ -0.0257719   │ 0.266863 │ 0.862043 │
│ 71  │ theta[70]  │ -0.916281 │ -0.31749  │ -0.0234316   │ 0.266557 │ 0.881501 │
│ 72  │ theta[71]  │ -0.932294 │ -0.322661 │ -0.0262963   │ 0.262147 │ 0.853884 │
│ 73  │ theta[72]  │ -0.909388 │ -0.317422 │ -0.0241428   │ 0.269594 │ 0.836786 │
│ 74  │ theta[73]  │ -0.90648  │ -0.30819  │ -0.0218921   │ 0.273815 │ 0.85468  │
│ 75  │ theta[74]  │ -0.891846 │ -0.311913 │ -0.0231451   │ 0.263109 │ 0.85212  │
│ 76  │ theta[75]  │ -0.904746 │ -0.30406  │ -0.0109521   │ 0.270872 │ 0.864827 │
│ 77  │ theta[76]  │ -0.886872 │ -0.279336 │ 0.0114971    │ 0.309326 │ 0.933496 │
│ 78  │ theta[77]  │ -0.866478 │ -0.275347 │ 0.0163951    │ 0.310314 │ 0.922362 │
│ 79  │ theta[78]  │ -0.891665 │ -0.28707  │ 0.00949942   │ 0.308538 │ 0.933509 │
│ 80  │ theta[79]  │ -0.858037 │ -0.27663  │ 0.0122929    │ 0.305681 │ 0.912505 │
│ 81  │ theta[80]  │ -0.885343 │ -0.289004 │ 0.00829029   │ 0.306197 │ 0.910139 │
│ 82  │ theta[81]  │ -0.87621  │ -0.287828 │ 0.00970307   │ 0.310744 │ 0.935678 │
│ 83  │ theta[82]  │ -0.864061 │ -0.271203 │ 0.0186707    │ 0.321574 │ 0.94595  │
│ 84  │ theta[83]  │ -0.876798 │ -0.296297 │ -0.000420651 │ 0.304583 │ 0.917683 │
│ 85  │ theta[84]  │ -0.895664 │ -0.292934 │ 0.00671708   │ 0.315574 │ 0.93353  │
│ 86  │ theta[85]  │ -0.884669 │ -0.274102 │ 0.0208328    │ 0.31447  │ 0.937233 │
│ 87  │ theta[86]  │ -0.865154 │ -0.286769 │ 0.00965867   │ 0.31146  │ 0.933503 │
│ 88  │ theta[87]  │ -0.90326  │ -0.282356 │ 0.0133207    │ 0.314608 │ 0.953264 │
│ 89  │ theta[88]  │ -0.894261 │ -0.28729  │ 0.0104721    │ 0.315769 │ 0.939781 │
│ 90  │ theta[89]  │ -0.87969  │ -0.278346 │ 0.0226905    │ 0.317541 │ 0.927837 │
│ 91  │ theta[90]  │ -0.884825 │ -0.285309 │ 0.0109771    │ 0.31185  │ 0.940303 │
│ 92  │ theta[91]  │ -1.00761  │ -0.36679  │ -0.0641131   │ 0.230988 │ 0.810317 │
│ 93  │ theta[92]  │ -1.02103  │ -0.372784 │ -0.0693038   │ 0.220278 │ 0.812505 │
│ 94  │ theta[93]  │ -1.0097   │ -0.369206 │ -0.0628089   │ 0.222148 │ 0.811837 │
│ 95  │ theta[94]  │ -1.00604  │ -0.373131 │ -0.0669905   │ 0.225226 │ 0.810542 │
│ 96  │ theta[95]  │ -1.01209  │ -0.371236 │ -0.0609029   │ 0.233601 │ 0.818539 │
│ 97  │ theta[96]  │ -1.00014  │ -0.380945 │ -0.0688738   │ 0.220635 │ 0.807033 │
│ 98  │ theta[97]  │ -1.01658  │ -0.364422 │ -0.0561036   │ 0.236123 │ 0.839646 │
│ 99  │ theta[98]  │ -0.997997 │ -0.363896 │ -0.0593028   │ 0.234384 │ 0.814997 │
│ 100 │ theta[99]  │ -0.997447 │ -0.366832 │ -0.067957    │ 0.227455 │ 0.815599 │
│ 101 │ theta[100] │ -0.977783 │ -0.359284 │ -0.0572172   │ 0.239143 │ 0.835927 │
│ 102 │ theta[101] │ -1.0131   │ -0.365019 │ -0.0583609   │ 0.239582 │ 0.829551 │
│ 103 │ theta[102] │ -1.00208  │ -0.360868 │ -0.0629239   │ 0.233683 │ 0.822989 │
│ 104 │ theta[103] │ -1.02918  │ -0.379775 │ -0.0712675   │ 0.23143  │ 0.82071  │
│ 105 │ theta[104] │ -0.977448 │ -0.360855 │ -0.0596602   │ 0.232186 │ 0.80152  │
│ 106 │ theta[105] │ -0.989124 │ -0.364276 │ -0.0640509   │ 0.233008 │ 0.806809 │
│ 107 │ theta[106] │ -0.988461 │ -0.262218 │ 0.0941339    │ 0.443905 │ 1.12255  │
│ 108 │ theta[107] │ -1.02446  │ -0.270543 │ 0.0792337    │ 0.433546 │ 1.10328  │
│ 109 │ theta[108] │ -0.995708 │ -0.275061 │ 0.0817714    │ 0.42672  │ 1.08608  │
│ 110 │ theta[109] │ -0.976367 │ -0.261615 │ 0.0875769    │ 0.440779 │ 1.10103  │
│ 111 │ theta[110] │ -1.00781  │ -0.263557 │ 0.0926734    │ 0.445622 │ 1.10557  │
│ 112 │ theta[111] │ -1.01756  │ -0.256466 │ 0.0881154    │ 0.441438 │ 1.09528  │
│ 113 │ theta[112] │ -1.00497  │ -0.271059 │ 0.0819578    │ 0.431294 │ 1.08713  │
│ 114 │ theta[113] │ -0.999124 │ -0.273241 │ 0.075765     │ 0.421797 │ 1.08998  │
│ 115 │ theta[114] │ -0.994608 │ -0.262199 │ 0.0900014    │ 0.439074 │ 1.10458  │
│ 116 │ theta[115] │ -0.965489 │ -0.265884 │ 0.090376     │ 0.439366 │ 1.08035  │
│ 117 │ theta[116] │ -0.988909 │ -0.260284 │ 0.0890234    │ 0.43603  │ 1.08846  │
│ 118 │ theta[117] │ -1.01533  │ -0.273905 │ 0.0840513    │ 0.434632 │ 1.12643  │
│ 119 │ theta[118] │ -0.990528 │ -0.26292  │ 0.0920918    │ 0.434477 │ 1.10185  │
│ 120 │ theta[119] │ -0.998575 │ -0.270288 │ 0.0743675    │ 0.415153 │ 1.09191  │
│ 121 │ theta[120] │ -1.01259  │ -0.272552 │ 0.0817896    │ 0.422338 │ 1.07116  │
│ 122 │ theta[121] │ -0.980052 │ -0.327441 │ -0.00965106  │ 0.290825 │ 0.923437 │
│ 123 │ theta[122] │ -0.974898 │ -0.323732 │ -0.0122773   │ 0.299601 │ 0.903331 │
│ 124 │ theta[123] │ -0.986029 │ -0.327215 │ -0.0125663   │ 0.289688 │ 0.898664 │
│ 125 │ theta[124] │ -0.96702  │ -0.32696  │ -0.0131933   │ 0.299705 │ 0.902232 │
│ 126 │ theta[125] │ -0.978598 │ -0.320516 │ -0.00903537  │ 0.293522 │ 0.906992 │
│ 127 │ theta[126] │ -0.969546 │ -0.329052 │ -0.00911985  │ 0.297926 │ 0.900865 │
│ 128 │ theta[127] │ -0.9976   │ -0.330199 │ -0.0194123   │ 0.289093 │ 0.893966 │
│ 129 │ theta[128] │ -0.967356 │ -0.333779 │ -0.0189808   │ 0.285768 │ 0.908398 │
│ 130 │ theta[129] │ -0.973393 │ -0.330652 │ -0.0150362   │ 0.288048 │ 0.907112 │
│ 131 │ theta[130] │ -0.996919 │ -0.335393 │ -0.0175681   │ 0.29383  │ 0.910152 │
│ 132 │ theta[131] │ -0.973414 │ -0.326468 │ -0.0222863   │ 0.288055 │ 0.890299 │
│ 133 │ theta[132] │ -0.990958 │ -0.324874 │ -0.0097257   │ 0.29645  │ 0.901343 │
│ 134 │ theta[133] │ -0.973508 │ -0.323241 │ -0.0197514   │ 0.29401  │ 0.917731 │
│ 135 │ theta[134] │ -0.988167 │ -0.341099 │ -0.0272434   │ 0.287079 │ 0.894962 │
│ 136 │ theta[135] │ -0.989093 │ -0.334384 │ -0.0188774   │ 0.288963 │ 0.902676 │
│ 137 │ theta[136] │ -1.04883  │ -0.359971 │ -0.0272096   │ 0.29926  │ 0.914927 │
│ 138 │ theta[137] │ -0.823846 │ -0.267764 │ 0.0117356    │ 0.282515 │ 0.833995 │
│ 139 │ theta[138] │ -0.904518 │ -0.289914 │ 0.02719      │ 0.352345 │ 0.970429 │
│ 140 │ theta[139] │ -1.05139  │ -0.37061  │ -0.0314969   │ 0.29538  │ 0.921523 │
│ 141 │ theta[140] │ -0.834123 │ -0.264576 │ 0.00928662   │ 0.286706 │ 0.82939  │
│ 142 │ theta[141] │ -0.945382 │ -0.296065 │ 0.0295432    │ 0.35533  │ 0.979248 │
│ 143 │ theta[142] │ -1.03948  │ -0.347322 │ -0.015915    │ 0.316789 │ 0.94779  │
│ 144 │ theta[143] │ -0.829938 │ -0.268371 │ 0.00346804   │ 0.281615 │ 0.832165 │
│ 145 │ theta[144] │ -0.913143 │ -0.286915 │ 0.0305759    │ 0.350677 │ 0.973133 │
│ 146 │ theta[145] │ -1.05275  │ -0.36825  │ -0.0325153   │ 0.29625  │ 0.937762 │
│ 147 │ theta[146] │ -0.848845 │ -0.277301 │ 0.00694174   │ 0.28412  │ 0.843413 │
│ 148 │ theta[147] │ -0.919377 │ -0.286127 │ 0.0298954    │ 0.352959 │ 0.987375 │
│ 149 │ theta[148] │ -1.05182  │ -0.37452  │ -0.0352453   │ 0.286891 │ 0.918812 │
│ 150 │ theta[149] │ -0.829818 │ -0.271272 │ 0.00322892   │ 0.276317 │ 0.845153 │
│ 151 │ theta[150] │ -0.909346 │ -0.287519 │ 0.030604     │ 0.351053 │ 0.988498 │
│ 152 │ theta[151] │ -1.04669  │ -0.363313 │ -0.0384106   │ 0.292095 │ 0.909435 │
│ 153 │ theta[152] │ -0.829665 │ -0.274808 │ 3.84523e-5   │ 0.272048 │ 0.819743 │
│ 154 │ theta[153] │ -0.938597 │ -0.291398 │ 0.0360848    │ 0.357995 │ 0.973191 │
│ 155 │ theta[154] │ -1.01779  │ -0.350541 │ -0.016909    │ 0.304101 │ 0.929256 │
│ 156 │ theta[155] │ -0.83716  │ -0.264959 │ 0.015483     │ 0.293129 │ 0.851181 │
│ 157 │ theta[156] │ -0.946139 │ -0.300526 │ 0.0249235    │ 0.346865 │ 0.966584 │
│ 158 │ theta[157] │ -1.04392  │ -0.350758 │ -0.0211348   │ 0.302106 │ 0.931763 │
│ 159 │ theta[158] │ -0.818737 │ -0.277329 │ 0.00450001   │ 0.282924 │ 0.829593 │
│ 160 │ theta[159] │ -0.930031 │ -0.294391 │ 0.0150057    │ 0.341408 │ 0.972702 │
│ 161 │ theta[160] │ -1.04522  │ -0.371483 │ -0.0254223   │ 0.298514 │ 0.937034 │
│ 162 │ theta[161] │ -0.824527 │ -0.274019 │ 0.00210998   │ 0.278605 │ 0.83446  │
│ 163 │ theta[162] │ -0.935157 │ -0.281828 │ 0.0416629    │ 0.356517 │ 0.976496 │
│ 164 │ theta[163] │ -1.04301  │ -0.364464 │ -0.0211329   │ 0.310333 │ 0.94681  │
│ 165 │ theta[164] │ -0.844585 │ -0.273175 │ 0.0077837    │ 0.281718 │ 0.821316 │
│ 166 │ theta[165] │ -0.913123 │ -0.294045 │ 0.0252889    │ 0.344876 │ 0.97622  │
│ 167 │ theta[166] │ -1.0349   │ -0.363442 │ -0.0259885   │ 0.298578 │ 0.915965 │
│ 168 │ theta[167] │ -0.848853 │ -0.277633 │ -0.00220983  │ 0.270439 │ 0.819841 │
│ 169 │ theta[168] │ -0.897287 │ -0.287752 │ 0.0302016    │ 0.353143 │ 0.971858 │
│ 170 │ theta[169] │ -1.04485  │ -0.359712 │ -0.023861    │ 0.300655 │ 0.931861 │
│ 171 │ theta[170] │ -0.834215 │ -0.262803 │ 0.0116798    │ 0.275843 │ 0.819847 │
│ 172 │ theta[171] │ -0.934918 │ -0.286729 │ 0.0267653    │ 0.340782 │ 0.966996 │
│ 173 │ theta[172] │ -1.04387  │ -0.353324 │ -0.0274244   │ 0.297881 │ 0.923455 │
│ 174 │ theta[173] │ -0.83731  │ -0.264419 │ 0.00484567   │ 0.280282 │ 0.831996 │
│ 175 │ theta[174] │ -0.902011 │ -0.283399 │ 0.0348523    │ 0.35258  │ 1.00357  │
│ 176 │ theta[175] │ -1.02103  │ -0.347377 │ -0.011938    │ 0.314186 │ 0.94839  │
│ 177 │ theta[176] │ -0.833861 │ -0.265245 │ 0.00829288   │ 0.279918 │ 0.847062 │
│ 178 │ theta[177] │ -0.932983 │ -0.301651 │ 0.0219849    │ 0.341597 │ 0.987273 │
│ 179 │ theta[178] │ -1.03447  │ -0.357271 │ -0.0255201   │ 0.306163 │ 0.928296 │
│ 180 │ theta[179] │ -0.84211  │ -0.264716 │ 0.00836315   │ 0.284879 │ 0.837814 │
│ 181 │ theta[180] │ -0.924213 │ -0.296962 │ 0.0210735    │ 0.347102 │ 0.956842 │
│ 182 │ theta[181] │ -0.891027 │ -0.325476 │ -0.0364037   │ 0.243038 │ 0.837976 │
│ 183 │ theta[182] │ -0.79543  │ -0.242574 │ 0.0235234    │ 0.281035 │ 0.82798  │
│ 184 │ theta[183] │ -0.833494 │ -0.262726 │ 0.0118868    │ 0.290837 │ 0.859885 │
```

In [21]:

```
# Extract all weight and bias parameters.
theta = chain[:theta].value.data[:,:,1];
niter = size(theta,  1)
```

Out[21]:

```
19000
```

In [34]:

```
## ---------------------------------------------------------
## predict for training set
## ---------------------------------------------------------

inputs = X_train'
labels = Y_train;

n_exper = 10
ac_ba_train = zeros(n_exper,2)

for exp=1:n_exper
    
    y_pred_samps = zeros(niter, length(labels))
    y_pred = zeros(length(labels))

    for j in 1:length(labels)
       for i in 1:niter
            preds = softmax(feedforward(inputs[:,j], theta[i,:]))
            dist = Categorical(preds)
            y_pred_samps[i,j] = rand(dist)
        end
        probs = [mean(y_pred_samps[:,j] .== 1), mean(y_pred_samps[:,j] .== 2), mean(y_pred_samps[:,j] .== 3)]
        y_pred[j] = sum((probs .== maximum(probs)) .* [1, 2, 3])
    end

    #println("test accuracy = ", mean(y_pred .- 1 .== labels))

    y_pred = convert(Array{Int64,1}, y_pred);
    C = confusmat(3, labels .+ 1, y_pred)
    bacc = 1/3 *(C[1,1] / sum(C[1, :]) + C[2,2] / sum(C[2, :]) + C[3,3] / sum(C[3, :]))
    #println("train BA = ", round(bacc, digits=2))
    
    ac_ba_train[exp,1] = mean(y_pred .- 1 .== labels)
    ac_ba_train[exp,2] = bacc
    
end

println(round.(mean(ac_ba_train, dims=1), digits=2))
println(round.(std(ac_ba_train, dims=1), digits = 2))
```

```
[0.69 0.68]
[0.0 0.0]
```

In [38]:

```
## ---------------------------------------------------------
## predict for test set
## ---------------------------------------------------------


inputs = X_test'
labels = Y_test

n_exper = 10
ac_ba_test = zeros(n_exper,2)

for exp=1:n_exper
    
    y_pred_samps = zeros(niter, length(labels))
    y_pred = zeros(length(labels))

    for j in 1:length(labels)
       for i in 1:niter
            preds = softmax(feedforward(inputs[:,j], theta[i,:]))
            dist = Categorical(preds)
            y_pred_samps[i,j] = rand(dist)
        end
        probs = [mean(y_pred_samps[:,j] .== 1), mean(y_pred_samps[:,j] .== 2), mean(y_pred_samps[:,j] .== 3)]
        y_pred[j] = sum((probs .== maximum(probs)) .* [1, 2, 3])
    end

    #println("test accuracy = ", mean(y_pred .- 1 .== labels))

    y_pred = convert(Array{Int64,1}, y_pred);
    C = confusmat(3, labels .+ 1, y_pred)
    bacc = 1/3 *(C[1,1] / sum(C[1, :]) + C[2,2] / sum(C[2, :]) + C[3,3] / sum(C[3, :]))
    #println("train BA = ", round(bacc, digits=2))
    
    ac_ba_test[exp,1] = mean(y_pred .- 1 .== labels)
    ac_ba_test[exp,2] = bacc
    
end

println(round.(mean(ac_ba_test, dims=1), digits=2))
println(round.(std(ac_ba_test, dims=1), digits = 2))
```

```
[0.63 0.62]
[0.02 0.02]
```

In [ ]:
