## Supplementary material for "A Bayesian neural network for toxicity prediction": 4_NN_multiclass.html


In [1]:

```
## ---------------------------------------------------------
## Import libraries 
## ---------------------------------------------------------

using CSV
using Statistics
using Flux
using Flux: onehotbatch, throttle, crossentropy, onecold, @epochs
using Base.Iterators: repeated
using MLBase
using LinearAlgebra
using MLPreprocessing

using Random
Random.seed!(1234);
```

In [2]:

```
function res(X, Y, Xt, Yt)
    println("train accuracy = ", accuracy(X, Y))
    println("test accuracy = ", accuracy(Xt, Yt))

    y_pred = onecold(nn(X)) ; y = onecold(Y); C = confusmat(3, y, y_pred)
    bacc_train = 1/3 *(C[1,1] / sum(C[1, :]) + C[2,2] / sum(C[2, :]) + C[3,3] / sum(C[3, :]))
    println("train BA = ", round(bacc_train, digits=2))

    y_pred = onecold(nn(Xt)) ; y = onecold(Yt); C = confusmat(3, y, y_pred)
    bacc_test = 1/3 *(C[1,1] / sum(C[1, :]) + C[2,2] / sum(C[2, :]) + C[3,3] / sum(C[3, :]))
    println("test BA = ", round(bacc_test, digits=2))
    
    return accuracy(X, Y), accuracy(Xt, Yt), bacc_train, bacc_test
end
```

Out[2]:

```
res (generic function with 1 method)
```

In [3]:

```
## ---------------------------------------------------------
## Read and prepare data
## ---------------------------------------------------------

df_train = CSV.read("/Users/kcft114/Box Sync/27_BNN_DILI/data/Aleo_train_match.csv")
df_test = CSV.read("/Users/kcft114/Box Sync/27_BNN_DILI/data/Aleo_test_match.csv")

df_Xtrain = df_train[!, [:ClogP, :BSEP, :Glu, :Glu_Gal, :THLE, :HepG2, :Fsp3, :log10cmax]]
df_Xtest = df_test[!, [:ClogP, :BSEP, :Glu, :Glu_Gal, :THLE, :HepG2, :Fsp3, :log10cmax]]

scaler = fit(StandardScaler, df_Xtrain)
transform!(df_Xtrain, scaler)
transform!(df_Xtest, scaler)

df_y_train = df_train[!, :dili_sev]
df_y_test = df_test[!, :dili_sev];

df_yXtrain = hcat(df_y_train, df_Xtrain)
df_yXtest = hcat(df_y_test, df_Xtest)

X_train = convert(Matrix, df_Xtrain)
X_test = convert(Matrix, df_Xtest)

df_y_train = df_train[!, :dili_sev]
df_y_test = df_test[!, :dili_sev]

Y_train = Int.(convert(Array, df_y_train))
Y_test = Int.(convert(Array, df_y_test))

Y = onehotbatch(Y_train, 1:3)
X = X_train'

Yt = onehotbatch(Y_test, 1:3)
Xt = X_test'

println(size(X))
println(size(Y))
```

```
┌ Warning: `getindex(df::DataFrame, col_ind::ColumnIndex)` is deprecated, use `df[!, col_ind]` instead.
│   caller = valid_columns(::DataFrames.DataFrame) at scaleselection.jl:28
└ @ MLPreprocessing /Users/kcft114/.julia/packages/MLPreprocessing/T8hq3/src/scaleselection.jl:28
┌ Warning: `getindex(df::DataFrame, col_ind::ColumnIndex)` is deprecated, use `df[!, col_ind]` instead.
│   caller = valid_columns(::DataFrames.DataFrame) at scaleselection.jl:28
└ @ MLPreprocessing /Users/kcft114/.julia/packages/MLPreprocessing/T8hq3/src/scaleselection.jl:28
┌ Warning: `getindex(df::DataFrame, col_ind::ColumnIndex)` is deprecated, use `df[!, col_ind]` instead.
│   caller = valid_columns(::DataFrames.DataFrame, ::Array{Symbol,1}) at scaleselection.jl:40
└ @ MLPreprocessing /Users/kcft114/.julia/packages/MLPreprocessing/T8hq3/src/scaleselection.jl:40
┌ Warning: `getindex(df::DataFrame, col_ind::ColumnIndex)` is deprecated, use `df[!, col_ind]` instead.
│   caller = valid_columns(::DataFrames.DataFrame, ::Array{Symbol,1}) at scaleselection.jl:40
└ @ MLPreprocessing /Users/kcft114/.julia/packages/MLPreprocessing/T8hq3/src/scaleselection.jl:40
┌ Warning: `getindex(df::DataFrame, col_ind::ColumnIndex)` is deprecated, use `df[!, col_ind]` instead.
│   caller = StandardScaler(::DataFrames.DataFrame, ::Array{Symbol,1}) at standardize.jl:260
└ @ MLPreprocessing /Users/kcft114/.julia/packages/MLPreprocessing/T8hq3/src/standardize.jl:260
┌ Warning: `getindex(df::DataFrame, col_ind::ColumnIndex)` is deprecated, use `df[!, col_ind]` instead.
│   caller = StandardScaler(::DataFrames.DataFrame, ::Array{Symbol,1}) at standardize.jl:261
└ @ MLPreprocessing /Users/kcft114/.julia/packages/MLPreprocessing/T8hq3/src/standardize.jl:261
┌ Warning: `getindex(df::DataFrame, col_ind::ColumnIndex)` is deprecated, use `df[!, col_ind]` instead.
│   caller = standardize!(::DataFrames.DataFrame, ::Float64, ::Float64, ::Symbol) at standardize.jl:156
└ @ MLPreprocessing /Users/kcft114/.julia/packages/MLPreprocessing/T8hq3/src/standardize.jl:156
┌ Warning: `getindex(df::DataFrame, col_ind::ColumnIndex)` is deprecated, use `df[!, col_ind]` instead.
│   caller = standardize!(::DataFrames.DataFrame, ::Float64, ::Float64, ::Symbol) at standardize.jl:156
└ @ MLPreprocessing /Users/kcft114/.julia/packages/MLPreprocessing/T8hq3/src/standardize.jl:156
┌ Warning: `getindex(df::DataFrame, col_ind::ColumnIndex)` is deprecated, use `df[!, col_ind]` instead.
│   caller = standardize!(::DataFrames.DataFrame, ::Float64, ::Float64, ::Symbol) at standardize.jl:159
└ @ MLPreprocessing /Users/kcft114/.julia/packages/MLPreprocessing/T8hq3/src/standardize.jl:159
┌ Warning: `setindex!(df::DataFrame, v::AbstractVector, col_ind::ColumnIndex)` is deprecated, use `begin
│     df[!, col_ind] = v
│     df
│ end` instead.
│   caller = standardize!(::DataFrames.DataFrame, ::Float64, ::Float64, ::Symbol) at standardize.jl:164
└ @ MLPreprocessing /Users/kcft114/.julia/packages/MLPreprocessing/T8hq3/src/standardize.jl:164
```

```
(8, 147)
(3, 147)
```

In [4]:

```
n_h = 15
```

Out[4]:

```
15
```

In [9]:

```
## ---------------------------------------------------------
## Experiments - no regularization
## ---------------------------------------------------------

n_exper = 100
experiments_none = zeros(n_exper, 4)


for i=1:n_exper
    
    nn = Chain(
          Dense(8, n_h, relu),
          Dense(n_h, 3),
          softmax)
    
    loss(x, y) = crossentropy(nn(x), y)
    accuracy(x, y) = mean(onecold(nn(x)) .== onecold(y))

    dataset = [(X, Y)]
    evalcb = () -> @show(loss(X, Y))
    
    opt = ADAM(0.001)
    
    @epochs 20000 Flux.train!(loss, Flux.params(nn), dataset, opt, cb = throttle(evalcb, 10));
    
    ac_train, ac_test, ba_train, ba_test =  res(X, Y, Xt, Yt);
    global experiments_none[i,:] = [ac_train, ac_test, ba_train, ba_test]
end
```

```
┌ Info: Epoch 1
└ @ Main /Users/kcft114/.julia/packages/Flux/qXNjB/src/optimise/train.jl:105
┌ Info: Epoch 2
└ @ Main /Users/kcft114/.julia/packages/Flux/qXNjB/src/optimise/train.jl:105
┌ Info: Epoch 3
└ @ Main /Users/kcft114/.julia/packages/Flux/qXNjB/src/optimise/train.jl:105
┌ Info: Epoch 4
└ @ Main /Users/kcft114/.julia/packages/Flux/qXNjB/src/optimise/train.jl:105
┌ Info: Epoch 5
└ @ Main /Users/kcft114/.julia/packages/Flux/qXNjB/src/optimise/train.jl:105
┌ Info: Epoch 6
└ @ Main /Users/kcft114/.julia/packages/Flux/qXNjB/src/optimise/train.jl:105
┌ Info: Epoch 7
└ @ Main /Users/kcft114/.julia/packages/Flux/qXNjB/src/optimise/train.jl:105
┌ Info: Epoch 8
└ @ Main /Users/kcft114/.julia/packages/Flux/qXNjB/src/optimise/train.jl:105
┌ Info: Epoch 9
└ @ Main /Users/kcft114/.julia/packages/Flux/qXNjB/src/optimise/train.jl:105
┌ Info: Epoch 10
└ @ Main /Users/kcft114/.julia/packages/Flux/qXNjB/src/optimise/train.jl:105
┌ Info: Epoch 11
└ @ Main /Users/kcft114/.julia/packages/Flux/qXNjB/src/optimise/train.jl:105
┌ Info: Epoch 12
└ @ Main /Users/kcft114/.julia/packages/Flux/qXNjB/src/optimise/train.jl:105
┌ Info: Epoch 13
└ @ Main /Users/kcft114/.julia/packages/Flux/qXNjB/src/optimise/train.jl:105
┌ Info: Epoch 14
└ @ Main /Users/kcft114/.julia/packages/Flux/qXNjB/src/optimise/train.jl:105
┌ Info: Epoch 15
└ @ Main /Users/kcft114/.julia/packages/Flux/qXNjB/src/optimise/train.jl:105
┌ Info: Epoch 16
└ @ Main /Users/kcft114/.julia/packages/Flux/qXNjB/src/optimise/train.jl:105
┌ Info: Epoch 17
└ @ Main /Users/kcft114/.julia/packages/Flux/qXNjB/src/optimise/train.jl:105
┌ Info: Epoch 18
└ @ Main /Users/kcft114/.julia/packages/Flux/qXNjB/src/optimise/train.jl:105
┌ Info: Epoch 19
└ @ Main /Users/kcft114/.julia/packages/Flux/qXNjB/src/optimise/train.jl:105
┌ Info: Epoch 20
└ @ Main /Users/kcft114/.julia/packages/Flux/qXNjB/src/optimise/train.jl:105
┌ Info: Epoch 21
└ @ Main /Users/kcft114/.julia/packages/Flux/qXNjB/src/optimise/train.jl:105
┌ Info: Epoch 22
└ @ Main /Users/kcft114/.julia/packages/Flux/qXNjB/src/optimise/train.jl:105
┌ Info: Epoch 23
└ @ Main /Users/kcft114/.julia/packages/Flux/qXNjB/src/optimise/train.jl:105
┌ Info: Epoch 24
└ @ Main /Users/kcft114/.julia/packages/Flux/qXNjB/src/optimise/train.jl:105
┌ Info: Epoch 25
└ @ Main /Users/kcft114/.julia/packages/Flux/qXNjB/src/optimise/train.jl:105
┌ Info: Epoch 26
└ @ Main /Users/kcft114/.julia/packages/Flux/qXNjB/src/optimise/train.jl:105
┌ Info: Epoch 27
└ @ Main /Users/kcft114/.julia/packages/Flux/qXNjB/src/optimise/train.jl:105
┌ Info: Epoch 28
└ @ Main /Users/kcft114/.julia/packages/Flux/qXNjB/src/optimise/train.jl:105
┌ Info: Epoch 29
└ @ Main /Users/kcft114/.julia/packages/Flux/qXNjB/src/optimise/train.jl:105
┌ Info: Epoch 30
└ @ Main /Users/kcft114/.julia/packages/Flux/qXNjB/src/optimise/train.jl:105
┌ Info: Epoch 31
└ @ Main /Users/kcft114/.julia/packages/Flux/qXNjB/src/optimise/train.jl:105
┌ Info: Epoch 32
└ @ Main /Users/kcft114/.julia/packages/Flux/qXNjB/src/optimise/train.jl:105
┌ Info: Epoch 33
└ @ Main /Users/kcft114/.julia/packages/Flux/qXNjB/src/optimise/train.jl:105
┌ Info: Epoch 34
└ @ Main /Users/kcft114/.julia/packages/Flux/qXNjB/src/optimise/train.jl:105
┌ Info: Epoch 35
└ @ Main /Users/kcft114/.julia/packages/Flux/qXNjB/src/optimise/train.jl:105
┌ Info: Epoch 36
└ @ Main /Users/kcft114/.julia/packages/Flux/qXNjB/src/optimise/train.jl:105
┌ Info: Epoch 37
└ @ Main /Users/kcft114/.julia/packages/Flux/qXNjB/src/optimise/train.jl:105
┌ Info: Epoch 38
└ @ Main /Users/kcft114/.julia/packages/Flux/qXNjB/src/optimise/train.jl:105
┌ Info: Epoch 39
└ @ Main /Users/kcft114/.julia/packages/Flux/qXNjB/src/optimise/train.jl:105
┌ Info: Epoch 40
└ @ Main /Users/kcft114/.julia/packages/Flux/qXNjB/src/optimise/train.jl:105
┌ Info: Epoch 41
└ @ Main /Users/kcft114/.julia/packages/Flux/qXNjB/src/optimise/train.jl:105
┌ Info: Epoch 42
└ @ Main /Users/kcft114/.julia/packages/Flux/qXNjB/src/optimise/train.jl:105
┌ Info: Epoch 43
└ @ Main /Users/kcft114/.julia/packages/Flux/qXNjB/src/optimise/train.jl:105
┌ Info: Epoch 44
└ @ Main /Users/kcft114/.julia/packages/Flux/qXNjB/src/optimise/train.jl:105
┌ Info: Epoch 45
└ @ Main /Users/kcft114/.julia/packages/Flux/qXNjB/src/optimise/train.jl:105
┌ Info: Epoch 46
└ @ Main /Users/kcft114/.julia/packages/Flux/qXNjB/src/optimise/train.jl:105
┌ Info: Epoch 47
└ @ Main /Users/kcft114/.julia/packages/Flux/qXNjB/src/optimise/train.jl:105
┌ Info: Epoch 48
└ @ Main /Users/kcft114/.julia/packages/Flux/qXNjB/src/optimise/train.jl:105
┌ Info: Epoch 49
└ @ Main /Users/kcft114/.julia/packages/Flux/qXNjB/src/optimise/train.jl:105
┌ Info: Epoch 50
└ @ Main /Users/kcft114/.julia/packages/Flux/qXNjB/src/optimise/train.jl:105
┌ Info: Epoch 51
└ @ Main /Users/kcft114/.julia/packages/Flux/qXNjB/src/optimise/train.jl:105
┌ Info: Epoch 52
└ @ Main /Users/kcft114/.julia/packages/Flux/qXNjB/src/optimise/train.jl:105
┌ Info: Epoch 53
└ @ Main /Users/kcft114/.julia/packages/Flux/qXNjB/src/optimise/train.jl:105
┌ Info: Epoch 54
└ @ Main /Users/kcft114/.julia/packages/Flux/qXNjB/src/optimise/train.jl:105
┌ Info: Epoch 55
└ @ Main /Users/kcft114/.julia/packages/Flux/qXNjB/src/optimise/train.jl:105
┌ Info: Epoch 56
└ @ Main /Users/kcft114/.julia/packages/Flux/qXNjB/src/optimise/train.jl:105
┌ Info: Epoch 57
└ @ Main /Users/kcft114/.julia/packages/Flux/qXNjB/src/optimise/train.jl:105
┌ Info: Epoch 58
└ @ Main /Users/kcft114/.julia/packages/Flux/qXNjB/src/optimise/train.jl:105
┌ Info: Epoch 59
└ @ Main /Users/kcft114/.julia/packages/Flux/qXNjB/src/optimise/train.jl:105
┌ Info: Epoch 60
└ @ Main /Users/kcft114/.julia/packages/Flux/qXNjB/src/optimise/train.jl:105
┌ Info: Epoch 61
└ @ Main /Users/kcft114/.julia/packages/Flux/qXNjB/src/optimise/train.jl:105
┌ Info: Epoch 62
└ @ Main /Users/kcft114/.julia/packages/Flux/qXNjB/src/optimise/train.jl:105
┌ Info: Epoch 63
└ @ Main /Users/kcft114/.julia/packages/Flux/qXNjB/src/optimise/train.jl:105
┌ Info: Epoch 64
└ @ Main /Users/kcft114/.julia/packages/Flux/qXNjB/src/optimise/train.jl:105
┌ Info: Epoch 65
└ @ Main /Users/kcft114/.julia/packages/Flux/qXNjB/src/optimise/train.jl:105
┌ Info: Epoch 66
└ @ Main /Users/kcft114/.julia/packages/Flux/qXNjB/src/optimise/train.jl:105
┌ Info: Epoch 67
└ @ Main /Users/kcft114/.julia/packages/Flux/qXNjB/src/optimise/train.jl:105
┌ Info: Epoch 68
└ @ Main /Users/kcft114/.julia/packages/Flux/qXNjB/src/optimise/train.jl:105
┌ Info: Epoch 69
└ @ Main /Users/kcft114/.julia/packages/Flux/qXNjB/src/optimise/train.jl:105
┌ Info: Epoch 70
└ @ Main /Users/kcft114/.julia/packages/Flux/qXNjB/src/optimise/train.jl:105
┌ Info: Epoch 71
└ @ Main /Users/kcft114/.julia/packages/Flux/qXNjB/src/optimise/train.jl:105
┌ Info: Epoch 72
└ @ Main /Users/kcft114/.julia/packages/Flux/qXNjB/src/optimise/train.jl:105
┌ Info: Epoch 73
└ @ Main /Users/kcft114/.julia/packages/Flux/qXNjB/src/optimise/train.jl:105
┌ Info: Epoch 74
└ @ Main /Users/kcft114/.julia/packages/Flux/qXNjB/src/optimise/train.jl:105
┌ Info: Epoch 75
└ @ Main /Users/kcft114/.julia/packages/Flux/qXNjB/src/optimise/train.jl:105
┌ Info: Epoch 76
└ @ Main /Users/kcft114/.julia/packages/Flux/qXNjB/src/optimise/train.jl:105
┌ Info: Epoch 77
└ @ Main /Users/kcft114/.julia/packages/Flux/qXNjB/src/optimise/train.jl:105
┌ Info: Epoch 78
└ @ Main /Users/kcft114/.julia/packages/Flux/qXNjB/src/optimise/train.jl:105
┌ Info: Epoch 79
└ @ Main /Users/kcft114/.julia/packages/Flux/qXNjB/src/optimise/train.jl:105
┌ Info: Epoch 80
└ @ Main /Users/kcft114/.julia/packages/Flux/qXNjB/src/optimise/train.jl:105
┌ Info: Epoch 81
└ @ Main /Users/kcft114/.julia/packages/Flux/qXNjB/src/optimise/train.jl:105
┌ Info: Epoch 82
└ @ Main /Users/kcft114/.julia/packages/Flux/qXNjB/src/optimise/train.jl:105
┌ Info: Epoch 83
└ @ Main /Users/kcft114/.julia/packages/Flux/qXNjB/src/optimise/train.jl:105
┌ Info: Epoch 84
└ @ Main /Users/kcft114/.julia/packages/Flux/qXNjB/src/optimise/train.jl:105
┌ Info: Epoch 85
└ @ Main /Users/kcft114/.julia/packages/Flux/qXNjB/src/optimise/train.jl:105
┌ Info: Epoch 86
└ @ Main /Users/kcft114/.julia/packages/Flux/qXNjB/src/optimise/train.jl:105
┌ Info: Epoch 87
└ @ Main /Users/kcft114/.julia/packages/Flux/qXNjB/src/optimise/train.jl:105
┌ Info: Epoch 88
└ @ Main /Users/kcft114/.julia/packages/Flux/qXNjB/src/optimise/train.jl:105
┌ Info: Epoch 89
└ @ Main /Users/kcft114/.julia/packages/Flux/qXNjB/src/optimise/train.jl:105
┌ Info: Epoch 90
└ @ Main /Users/kcft114/.julia/packages/Flux/qXNjB/src/optimise/train.jl:105
┌ Info: Epoch 91
└ @ Main /Users/kcft114/.julia/packages/Flux/qXNjB/src/optimise/train.jl:105
┌ Info: Epoch 92
└ @ Main /Users/kcft114/.julia/packages/Flux/qXNjB/src/optimise/train.jl:105
┌ Info: Epoch 93
└ @ Main /Users/kcft114/.julia/packages/Flux/qXNjB/src/optimise/train.jl:105
┌ Info: Epoch 94
└ @ Main /Users/kcft114/.julia/packages/Flux/qXNjB/src/optimise/train.jl:105
┌ Info: Epoch 95
└ @ Main /Users/kcft114/.julia/packages/Flux/qXNjB/src/optimise/train.jl:105
┌ Info: Epoch 96
└ @ Main /Users/kcft114/.julia/packages/Flux/qXNjB/src/optimise/train.jl:105
┌ Info: Epoch 97
└ @ Main /Users/kcft114/.julia/packages/Flux/qXNjB/src/optimise/train.jl:105
┌ Info: Epoch 98
└ @ Main /Users/kcft114/.julia/packages/Flux/qXNjB/src/optimise/train.jl:105
┌ Info: Epoch 99
└ @ Main /Users/kcft114/.julia/packages/Flux/qXNjB/src/optimise/train.jl:105
┌ Info: Epoch 100
└ @ Main /Users/kcft114/.julia/packages/Flux/qXNjB/src/optimise/train.jl:105
┌ Info: Epoch 101
└ @ Main /Users/kcft114/.julia/packages/Flux/qXNjB/src/optimise/train.jl:105
┌ Info: Epoch 102
└ @ Main /Users/kcft114/.julia/packages/Flux/qXNjB/src/optimise/train.jl:105
┌ Info: Epoch 103
└ @ Main /Users/kcft114/.julia/packages/Flux/qXNjB/src/optimise/train.jl:105
┌ Info: Epoch 104
└ @ Main /Users/kcft114/.julia/packages/Flux/qXNjB/src/optimise/train.jl:105
┌ Info: Epoch 105
└ @ Main /Users/kcft114/.julia/packages/Flux/qXNjB/src/optimise/train.jl:105
┌ Info: Epoch 106
└ @ Main /Users/kcft114/.julia/packages/Flux/qXNjB/src/optimise/train.jl:105
```

```
loss(X, Y) = 1.3075736f0 (tracked)
loss(X, Y) = 1.3006854f0 (tracked)
loss(X, Y) = 1.2938817f0 (tracked)
loss(X, Y) = 1.2871515f0 (tracked)
loss(X, Y) = 1.2805051f0 (tracked)
loss(X, Y) = 1.2739561f0 (tracked)
loss(X, Y) = 1.267506f0 (tracked)
loss(X, Y) = 1.2611413f0 (tracked)
loss(X, Y) = 1.2548621f0 (tracked)
loss(X, Y) = 1.2486597f0 (tracked)
loss(X, Y) = 1.2425168f0 (tracked)
loss(X, Y) = 1.236432f0 (tracked)
loss(X, Y) = 1.2304165f0 (tracked)
loss(X, Y) = 1.2244939f0 (tracked)
loss(X, Y) = 1.2186538f0 (tracked)
loss(X, Y) = 1.2128806f0 (tracked)
loss(X, Y) = 1.2071487f0 (tracked)
loss(X, Y) = 1.2014692f0 (tracked)
loss(X, Y) = 1.1958678f0 (tracked)
loss(X, Y) = 1.190325f0 (tracked)
loss(X, Y) = 1.1848843f0 (tracked)
loss(X, Y) = 1.1795079f0 (tracked)
loss(X, Y) = 1.174199f0 (tracked)
loss(X, Y) = 1.1689618f0 (tracked)
loss(X, Y) = 1.1637857f0 (tracked)
loss(X, Y) = 1.1586673f0 (tracked)
loss(X, Y) = 1.1536139f0 (tracked)
loss(X, Y) = 1.1486207f0 (tracked)
loss(X, Y) = 1.1436867f0 (tracked)
loss(X, Y) = 1.1388298f0 (tracked)
loss(X, Y) = 1.1340446f0 (tracked)
loss(X, Y) = 1.1293241f0 (tracked)
loss(X, Y) = 1.1246406f0 (tracked)
loss(X, Y) = 1.1200168f0 (tracked)
loss(X, Y) = 1.1154588f0 (tracked)
loss(X, Y) = 1.1109856f0 (tracked)
loss(X, Y) = 1.1065823f0 (tracked)
loss(X, Y) = 1.1022396f0 (tracked)
loss(X, Y) = 1.097951f0 (tracked)
loss(X, Y) = 1.0937216f0 (tracked)
loss(X, Y) = 1.0895444f0 (tracked)
loss(X, Y) = 1.0854226f0 (tracked)
loss(X, Y) = 1.0813596f0 (tracked)
loss(X, Y) = 1.0773551f0 (tracked)
loss(X, Y) = 1.0733947f0 (tracked)
loss(X, Y) = 1.0694888f0 (tracked)
loss(X, Y) = 1.0656428f0 (tracked)
loss(X, Y) = 1.06185f0 (tracked)
loss(X, Y) = 1.0581057f0 (tracked)
loss(X, Y) = 1.0544139f0 (tracked)
loss(X, Y) = 1.0507717f0 (tracked)
loss(X, Y) = 1.0471724f0 (tracked)
loss(X, Y) = 1.0436141f0 (tracked)
loss(X, Y) = 1.0400991f0 (tracked)
loss(X, Y) = 1.0366387f0 (tracked)
loss(X, Y) = 1.0332586f0 (tracked)
loss(X, Y) = 1.029933f0 (tracked)
loss(X, Y) = 1.02667f0 (tracked)
loss(X, Y) = 1.0234591f0 (tracked)
loss(X, Y) = 1.020296f0 (tracked)
loss(X, Y) = 1.0171772f0 (tracked)
loss(X, Y) = 1.0140953f0 (tracked)
loss(X, Y) = 1.0110565f0 (tracked)
loss(X, Y) = 1.0080508f0 (tracked)
loss(X, Y) = 1.0050766f0 (tracked)
loss(X, Y) = 1.0021445f0 (tracked)
loss(X, Y) = 0.99924815f0 (tracked)
loss(X, Y) = 0.99638945f0 (tracked)
loss(X, Y) = 0.9935717f0 (tracked)
loss(X, Y) = 0.99079496f0 (tracked)
loss(X, Y) = 0.9880537f0 (tracked)
loss(X, Y) = 0.98534024f0 (tracked)
loss(X, Y) = 0.98265207f0 (tracked)
loss(X, Y) = 0.97999f0 (tracked)
loss(X, Y) = 0.9773544f0 (tracked)
loss(X, Y) = 0.97474355f0 (tracked)
loss(X, Y) = 0.9721624f0 (tracked)
loss(X, Y) = 0.96961164f0 (tracked)
loss(X, Y) = 0.9670787f0 (tracked)
loss(X, Y) = 0.9645779f0 (tracked)
loss(X, Y) = 0.96211416f0 (tracked)
loss(X, Y) = 0.95968586f0 (tracked)
loss(X, Y) = 0.9572863f0 (tracked)
loss(X, Y) = 0.95489717f0 (tracked)
loss(X, Y) = 0.9525191f0 (tracked)
loss(X, Y) = 0.950149f0 (tracked)
loss(X, Y) = 0.9477952f0 (tracked)
loss(X, Y) = 0.94545543f0 (tracked)
loss(X, Y) = 0.9431391f0 (tracked)
loss(X, Y) = 0.94085056f0 (tracked)
loss(X, Y) = 0.93858594f0 (tracked)
loss(X, Y) = 0.9363461f0 (tracked)
loss(X, Y) = 0.9341285f0 (tracked)
loss(X, Y) = 0.9319338f0 (tracked)
loss(X, Y) = 0.9297729f0 (tracked)
loss(X, Y) = 0.92763054f0 (tracked)
loss(X, Y) = 0.9255115f0 (tracked)
loss(X, Y) = 0.923416f0 (tracked)
loss(X, Y) = 0.92134166f0 (tracked)
loss(X, Y) = 0.9192911f0 (tracked)
loss(X, Y) = 0.9172611f0 (tracked)
loss(X, Y) = 0.9152282f0 (tracked)
loss(X, Y) = 0.913214f0 (tracked)
loss(X, Y) = 0.91122717f0 (tracked)
loss(X, Y) = 0.90926445f0 (tracked)
loss(X, Y) = 0.90731734f0 (tracked)
loss(X, Y) = 0.90539235f0 (tracked)
loss(X, Y) = 0.9034815f0 (tracked)
loss(X, Y) = 0.90159357f0 (tracked)
loss(X, Y) = 0.8997344f0 (tracked)
loss(X, Y) = 0.89790404f0 (tracked)
loss(X, Y) = 0.8960916f0 (tracked)
loss(X, Y) = 0.8942978f0 (tracked)
loss(X, Y) = 0.892524f0 (tracked)
loss(X, Y) = 0.8907813f0 (tracked)
loss(X, Y) = 0.88905823f0 (tracked)
loss(X, Y) = 0.88735485f0 (tracked)
loss(X, Y) = 0.8856953f0 (tracked)
loss(X, Y) = 0.88406545f0 (tracked)
loss(X, Y) = 0.882456f0 (tracked)
loss(X, Y) = 0.880858f0 (tracked)
loss(X, Y) = 0.8792736f0 (tracked)
loss(X, Y) = 0.8777073f0 (tracked)
loss(X, Y) = 0.8761702f0 (tracked)
loss(X, Y) = 0.87465507f0 (tracked)
loss(X, Y) = 0.8731573f0 (tracked)
loss(X, Y) = 0.8716764f0 (tracked)
loss(X, Y) = 0.87021106f0 (tracked)
loss(X, Y) = 0.8687612f0 (tracked)
loss(X, Y) = 0.8673264f0 (tracked)
loss(X, Y) = 0.8659134f0 (tracked)
loss(X, Y) = 0.8645164f0 (tracked)
loss(X, Y) = 0.86313313f0 (tracked)
loss(X, Y) = 0.8617642f0 (tracked)
loss(X, Y) = 0.8604123f0 (tracked)
loss(X, Y) = 0.85908794f0 (tracked)
loss(X, Y) = 0.8577916f0 (tracked)
loss(X, Y) = 0.8565159f0 (tracked)
loss(X, Y) = 0.8552536f0 (tracked)
loss(X, Y) = 0.8540075f0 (tracked)
loss(X, Y) = 0.8527761f0 (tracked)
loss(X, Y) = 0.85155857f0 (tracked)
loss(X, Y) = 0.8503475f0 (tracked)
loss(X, Y) = 0.84913975f0 (tracked)
loss(X, Y) = 0.8479287f0 (tracked)
loss(X, Y) = 0.846725f0 (tracked)
loss(X, Y) = 0.84553f0 (tracked)
loss(X, Y) = 0.84434164f0 (tracked)
loss(X, Y) = 0.8431648f0 (tracked)
loss(X, Y) = 0.8419981f0 (tracked)
loss(X, Y) = 0.84084684f0 (tracked)
loss(X, Y) = 0.8397073f0 (tracked)
loss(X, Y) = 0.83857536f0 (tracked)
loss(X, Y) = 0.8374518f0 (tracked)
loss(X, Y) = 0.8363363f0 (tracked)
loss(X, Y) = 0.8352287f0 (tracked)
loss(X, Y) = 0.83413553f0 (tracked)
loss(X, Y) = 0.83305377f0 (tracked)
loss(X, Y) = 0.8319863f0 (tracked)
loss(X, Y) = 0.83093286f0 (tracked)
```

```
┌ Info: Epoch 107
└ @ Main /Users/kcft114/.julia/packages/Flux/qXNjB/src/optimise/train.jl:105
┌ Info: Epoch 108
└ @ Main /Users/kcft114/.julia/packages/Flux/qXNjB/src/optimise/train.jl:105
┌ Info: Epoch 109
└ @ Main /Users/kcft114/.julia/packages/Flux/qXNjB/src/optimise/train.jl:105
┌ Info: Epoch 110
└ @ Main /Users/kcft114/.julia/packages/Flux/qXNjB/src/optimise/train.jl:105
┌ Info: Epoch 111
└ @ Main /Users/kcft114/.julia/packages/Flux/qXNjB/src/optimise/train.jl:105
┌ Info: Epoch 112
└ @ Main /Users/kcft114/.julia/packages/Flux/qXNjB/src/optimise/train.jl:105
┌ Info: Epoch 113
└ @ Main /Users/kcft114/.julia/packages/Flux/qXNjB/src/optimise/train.jl:105
┌ Info: Epoch 114
└ @ Main /Users/kcft114/.julia/packages/Flux/qXNjB/src/optimise/train.jl:105
┌ Info: Epoch 115
└ @ Main /Users/kcft114/.julia/packages/Flux/qXNjB/src/optimise/train.jl:105
┌ Info: Epoch 116
└ @ Main /Users/kcft114/.julia/packages/Flux/qXNjB/src/optimise/train.jl:105
┌ Info: Epoch 117
└ @ Main /Users/kcft114/.julia/packages/Flux/qXNjB/src/optimise/train.jl:105
┌ Info: Epoch 118
└ @ Main /Users/kcft114/.julia/packages/Flux/qXNjB/src/optimise/train.jl:105
┌ Info: Epoch 119
└ @ Main /Users/kcft114/.julia/packages/Flux/qXNjB/src/optimise/train.jl:105
┌ Info: Epoch 120
└ @ Main /Users/kcft114/.julia/packages/Flux/qXNjB/src/optimise/train.jl:105
┌ Info: Epoch 121
└ @ Main /Users/kcft114/.julia/packages/Flux/qXNjB/src/optimise/train.jl:105
┌ Info: Epoch 122
└ @ Main /Users/kcft114/.julia/packages/Flux/qXNjB/src/optimise/train.jl:105
┌ Info: Epoch 123
└ @ Main /Users/kcft114/.julia/packages/Flux/qXNjB/src/optimise/train.jl:105
┌ Info: Epoch 124
└ @ Main /Users/kcft114/.julia/packages/Flux/qXNjB/src/optimise/train.jl:105
┌ Info: Epoch 125
└ @ Main /Users/kcft114/.julia/packages/Flux/qXNjB/src/optimise/train.jl:105
┌ Info: Epoch 126
└ @ Main /Users/kcft114/.julia/packages/Flux/qXNjB/src/optimise/train.jl:105
┌ Info: Epoch 127
└ @ Main /Users/kcft114/.julia/packages/Flux/qXNjB/src/optimise/train.jl:105
┌ Info: Epoch 128
└ @ Main /Users/kcft114/.julia/packages/Flux/qXNjB/src/optimise/train.jl:105
┌ Info: Epoch 129
└ @ Main /Users/kcft114/.julia/packages/Flux/qXNjB/src/optimise/train.jl:105
┌ Info: Epoch 130
└ @ Main /Users/kcft114/.julia/packages/Flux/qXNjB/src/optimise/train.jl:105
┌ Info: Epoch 131
└ @ Main /Users/kcft114/.julia/packages/Flux/qXNjB/src/optimise/train.jl:105
┌ Info: Epoch 132
└ @ Main /Users/kcft114/.julia/packages/Flux/qXNjB/src/optimise/train.jl:105
┌ Info: Epoch 133
└ @ Main /Users/kcft114/.julia/packages/Flux/qXNjB/src/optimise/train.jl:105
┌ Info: Epoch 134
└ @ Main /Users/kcft114/.julia/packages/Flux/qXNjB/src/optimise/train.jl:105
┌ Info: Epoch 135
└ @ Main /Users/kcft114/.julia/packages/Flux/qXNjB/src/optimise/train.jl:105
┌ Info: Epoch 136
└ @ Main /Users/kcft114/.julia/packages/Flux/qXNjB/src/optimise/train.jl:105
┌ Info: Epoch 137
└ @ Main /Users/kcft114/.julia/packages/Flux/qXNjB/src/optimise/train.jl:105
┌ Info: Epoch 138
└ @ Main /Users/kcft114/.julia/packages/Flux/qXNjB/src/optimise/train.jl:105
┌ Info: Epoch 139
└ @ Main /Users/kcft114/.julia/packages/Flux/qXNjB/src/optimise/train.jl:105
┌ Info: Epoch 140
└ @ Main /Users/kcft114/.julia/packages/Flux/qXNjB/src/optimise/train.jl:105
┌ Info: Epoch 141
└ @ Main /Users/kcft114/.julia/packages/Flux/qXNjB/src/optimise/train.jl:105
┌ Info: Epoch 142
└ @ Main /Users/kcft114/.julia/packages/Flux/qXNjB/src/optimise/train.jl:105
┌ Info: Epoch 143
└ @ Main /Users/kcft114/.julia/packages/Flux/qXNjB/src/optimise/train.jl:105
┌ Info: Epoch 144
└ @ Main /Users/kcft114/.julia/packages/Flux/qXNjB/src/optimise/train.jl:105
┌ Info: Epoch 145
└ @ Main /Users/kcft114/.julia/packages/Flux/qXNjB/src/optimise/train.jl:105
┌ Info: Epoch 146
└ @ Main /Users/kcft114/.julia/packages/Flux/qXNjB/src/optimise/train.jl:105
┌ Info: Epoch 147
└ @ Main /Users/kcft114/.julia/packages/Flux/qXNjB/src/optimise/train.jl:105
┌ Info: Epoch 148
└ @ Main /Users/kcft114/.julia/packages/Flux/qXNjB/src/optimise/train.jl:105
┌ Info: Epoch 149
└ @ Main /Users/kcft114/.julia/packages/Flux/qXNjB/src/optimise/train.jl:105
┌ Info: Epoch 150
└ @ Main /Users/kcft114/.julia/packages/Flux/qXNjB/src/optimise/train.jl:105
┌ Info: Epoch 151
└ @ Main /Users/kcft114/.julia/packages/Flux/qXNjB/src/optimise/train.jl:105
┌ Info: Epoch 152
└ @ Main /Users/kcft114/.julia/packages/Flux/qXNjB/src/optimise/train.jl:105
┌ Info: Epoch 153
└ @ Main /Users/kcft114/.julia/packages/Flux/qXNjB/src/optimise/train.jl:105
┌ Info: Epoch 154
└ @ Main /Users/kcft114/.julia/packages/Flux/qXNjB/src/optimise/train.jl:105
┌ Info: Epoch 155
└ @ Main /Users/kcft114/.julia/packages/Flux/qXNjB/src/optimise/train.jl:105
┌ Info: Epoch 156
└ @ Main /Users/kcft114/.julia/packages/Flux/qXNjB/src/optimise/train.jl:105
┌ Info: Epoch 157
└ @ Main /Users/kcft114/.julia/packages/Flux/qXNjB/src/optimise/train.jl:105
┌ Info: Epoch 158
└ @ Main /Users/kcft114/.julia/packages/Flux/qXNjB/src/optimise/train.jl:105
┌ Info: Epoch 159
└ @ Main /Users/kcft114/.julia/packages/Flux/qXNjB/src/optimise/train.jl:105
┌ Info: Epoch 160
└ @ Main /Users/kcft114/.julia/packages/Flux/qXNjB/src/optimise/train.jl:105
┌ Info: Epoch 161
└ @ Main /Users/kcft114/.julia/packages/Flux/qXNjB/src/optimise/train.jl:105
┌ Info: Epoch 162
└ @ Main /Users/kcft114/.julia/packages/Flux/qXNjB/src/optimise/train.jl:105
┌ Info: Epoch 163
└ @ Main /Users/kcft114/.julia/packages/Flux/qXNjB/src/optimise/train.jl:105
┌ Info: Epoch 164
└ @ Main /Users/kcft114/.julia/packages/Flux/qXNjB/src/optimise/train.jl:105
┌ Info: Epoch 165
└ @ Main /Users/kcft114/.julia/packages/Flux/qXNjB/src/optimise/train.jl:105
┌ Info: Epoch 166
└ @ Main /Users/kcft114/.julia/packages/Flux/qXNjB/src/optimise/train.jl:105
┌ Info: Epoch 167
└ @ Main /Users/kcft114/.julia/packages/Flux/qXNjB/src/optimise/train.jl:105
┌ Info: Epoch 168
└ @ Main /Users/kcft114/.julia/packages/Flux/qXNjB/src/optimise/train.jl:105
┌ Info: Epoch 169
└ @ Main /Users/kcft114/.julia/packages/Flux/qXNjB/src/optimise/train.jl:105
┌ Info: Epoch 170
└ @ Main /Users/kcft114/.julia/packages/Flux/qXNjB/src/optimise/train.jl:105
┌ Info: Epoch 171
└ @ Main /Users/kcft114/.julia/packages/Flux/qXNjB/src/optimise/train.jl:105
┌ Info: Epoch 172
└ @ Main /Users/kcft114/.julia/packages/Flux/qXNjB/src/optimise/train.jl:105
┌ Info: Epoch 173
└ @ Main /Users/kcft114/.julia/packages/Flux/qXNjB/src/optimise/train.jl:105
┌ Info: Epoch 174
└ @ Main /Users/kcft114/.julia/packages/Flux/qXNjB/src/optimise/train.jl:105
┌ Info: Epoch 175
└ @ Main /Users/kcft114/.julia/packages/Flux/qXNjB/src/optimise/train.jl:105
┌ Info: Epoch 176
└ @ Main /Users/kcft114/.julia/packages/Flux/qXNjB/src/optimise/train.jl:105
┌ Info: Epoch 177
└ @ Main /Users/kcft114/.julia/packages/Flux/qXNjB/src/optimise/train.jl:105
┌ Info: Epoch 178
└ @ Main /Users/kcft114/.julia/packages/Flux/qXNjB/src/optimise/train.jl:105
┌ Info: Epoch 179
└ @ Main /Users/kcft114/.julia/packages/Flux/qXNjB/src/optimise/train.jl:105
┌ Info: Epoch 180
└ @ Main /Users/kcft114/.julia/packages/Flux/qXNjB/src/optimise/train.jl:105
┌ Info: Epoch 181
└ @ Main /Users/kcft114/.julia/packages/Flux/qXNjB/src/optimise/train.jl:105
┌ Info: Epoch 182
└ @ Main /Users/kcft114/.julia/packages/Flux/qXNjB/src/optimise/train.jl:105
┌ Info: Epoch 183
└ @ Main /Users/kcft114/.julia/packages/Flux/qXNjB/src/optimise/train.jl:105
┌ Info: Epoch 184
└ @ Main /Users/kcft114/.julia/packages/Flux/qXNjB/src/optimise/train.jl:105
┌ Info: Epoch 185
└ @ Main /Users/kcft114/.julia/packages/Flux/qXNjB/src/optimise/train.jl:105
┌ Info: Epoch 186
└ @ Main /Users/kcft114/.julia/packages/Flux/qXNjB/src/optimise/train.jl:105
┌ Info: Epoch 187
└ @ Main /Users/kcft114/.julia/packages/Flux/qXNjB/src/optimise/train.jl:105
┌ Info: Epoch 188
└ @ Main /Users/kcft114/.julia/packages/Flux/qXNjB/src/optimise/train.jl:105
┌ Info: Epoch 189
└ @ Main /Users/kcft114/.julia/packages/Flux/qXNjB/src/optimise/train.jl:105
┌ Info: Epoch 190
└ @ Main /Users/kcft114/.julia/packages/Flux/qXNjB/src/optimise/train.jl:105
┌ Info: Epoch 191
└ @ Main /Users/kcft114/.julia/packages/Flux/qXNjB/src/optimise/train.jl:105
┌ Info: Epoch 192
└ @ Main /Users/kcft114/.julia/packages/Flux/qXNjB/src/optimise/train.jl:105
┌ Info: Epoch 193
└ @ Main /Users/kcft114/.julia/packages/Flux/qXNjB/src/optimise/train.jl:105
┌ Info: Epoch 194
└ @ Main /Users/kcft114/.julia/packages/Flux/qXNjB/src/optimise/train.jl:105
┌ Info: Epoch 195
└ @ Main /Users/kcft114/.julia/packages/Flux/qXNjB/src/optimise/train.jl:105
┌ Info: Epoch 196
└ @ Main /Users/kcft114/.julia/packages/Flux/qXNjB/src/optimise/train.jl:105
┌ Info: Epoch 197
└ @ Main /Users/kcft114/.julia/packages/Flux/qXNjB/src/optimise/train.jl:105
┌ Info: Epoch 198
└ @ Main /Users/kcft114/.julia/packages/Flux/qXNjB/src/optimise/train.jl:105
┌ Info: Epoch 199
└ @ Main /Users/kcft114/.julia/packages/Flux/qXNjB/src/optimise/train.jl:105
┌ Info: Epoch 200
└ @ Main /Users/kcft114/.julia/packages/Flux/qXNjB/src/optimise/train.jl:105
┌ Info: Epoch 201
└ @ Main /Users/kcft114/.julia/packages/Flux/qXNjB/src/optimise/train.jl:105
┌ Info: Epoch 202
└ @ Main /Users/kcft114/.julia/packages/Flux/qXNjB/src/optimise/train.jl:105
┌ Info: Epoch 203
└ @ Main /Users/kcft114/.julia/packages/Flux/qXNjB/src/optimise/train.jl:105
┌ Info: Epoch 204
└ @ Main /Users/kcft114/.julia/packages/Flux/qXNjB/src/optimise/train.jl:105
┌ Info: Epoch 205
└ @ Main /Users/kcft114/.julia/packages/Flux/qXNjB/src/optimise/train.jl:105
┌ Info: Epoch 206
└ @ Main /Users/kcft114/.julia/packages/Flux/qXNjB/src/optimise/train.jl:105
┌ Info: Epoch 207
└ @ Main /Users/kcft114/.julia/packages/Flux/qXNjB/src/optimise/train.jl:105
┌ Info: Epoch 208
└ @ Main /Users/kcft114/.julia/packages/Flux/qXNjB/src/optimise/train.jl:105
┌ Info: Epoch 209
└ @ Main /Users/kcft114/.julia/packages/Flux/qXNjB/src/optimise/train.jl:105
┌ Info: Epoch 210
└ @ Main /Users/kcft114/.julia/packages/Flux/qXNjB/src/optimise/train.jl:105
┌ Info: Epoch 211
└ @ Main /Users/kcft114/.julia/packages/Flux/qXNjB/src/optimise/train.jl:105
┌ Info: Epoch 212
└ @ Main /Users/kcft114/.julia/packages/Flux/qXNjB/src/optimise/train.jl:105
┌ Info: Epoch 213
└ @ Main /Users/kcft114/.julia/packages/Flux/qXNjB/src/optimise/train.jl:105
┌ Info: Epoch 214
└ @ Main /Users/kcft114/.julia/packages/Flux/qXNjB/src/optimise/train.jl:105
┌ Info: Epoch 215
└ @ Main /Users/kcft114/.julia/packages/Flux/qXNjB/src/optimise/train.jl:105
┌ Info: Epoch 216
└ @ Main /Users/kcft114/.julia/packages/Flux/qXNjB/src/optimise/train.jl:105
┌ Info: Epoch 217
└ @ Main /Users/kcft114/.julia/packages/Flux/qXNjB/src/optimise/train.jl:105
┌ Info: Epoch 218
└ @ Main /Users/kcft114/.julia/packages/Flux/qXNjB/src/optimise/train.jl:105
┌ Info: Epoch 219
└ @ Main /Users/kcft114/.julia/packages/Flux/qXNjB/src/optimise/train.jl:105
┌ Info: Epoch 220
└ @ Main /Users/kcft114/.julia/packages/Flux/qXNjB/src/optimise/train.jl:105
┌ Info: Epoch 221
└ @ Main /Users/kcft114/.julia/packages/Flux/qXNjB/src/optimise/train.jl:105
┌ Info: Epoch 222
└ @ Main /Users/kcft114/.julia/packages/Flux/qXNjB/src/optimise/train.jl:105
┌ Info: Epoch 223
└ @ Main /Users/kcft114/.julia/packages/Flux/qXNjB/src/optimise/train.jl:105
┌ Info: Epoch 224
└ @ Main /Users/kcft114/.julia/packages/Flux/qXNjB/src/optimise/train.jl:105
┌ Info: Epoch 225
└ @ Main /Users/kcft114/.julia/packages/Flux/qXNjB/src/optimise/train.jl:105
┌ Info: Epoch 226
└ @ Main /Users/kcft114/.julia/packages/Flux/qXNjB/src/optimise/train.jl:105
┌ Info: Epoch 227
└ @ Main /Users/kcft114/.julia/packages/Flux/qXNjB/src/optimise/train.jl:105
┌ Info: Epoch 228
└ @ Main /Users/kcft114/.julia/packages/Flux/qXNjB/src/optimise/train.jl:105
┌ Info: Epoch 229
└ @ Main /Users/kcft114/.julia/packages/Flux/qXNjB/src/optimise/train.jl:105
┌ Info: Epoch 230
└ @ Main /Users/kcft114/.julia/packages/Flux/qXNjB/src/optimise/train.jl:105
┌ Info: Epoch 231
└ @ Main /Users/kcft114/.julia/packages/Flux/qXNjB/src/optimise/train.jl:105
┌ Info: Epoch 232
└ @ Main /Users/kcft114/.julia/packages/Flux/qXNjB/src/optimise/train.jl:105
┌ Info: Epoch 233
└ @ Main /Users/kcft114/.julia/packages/Flux/qXNjB/src/optimise/train.jl:105
┌ Info: Epoch 234
└ @ Main /Users/kcft114/.julia/packages/Flux/qXNjB/src/optimise/train.jl:105
┌ Info: Epoch 235
└ @ Main /Users/kcft114/.julia/packages/Flux/qXNjB/src/optimise/train.jl:105
┌ Info: Epoch 236
└ @ Main /Users/kcft114/.julia/packages/Flux/qXNjB/src/optimise/train.jl:105
┌ Info: Epoch 237
└ @ Main /Users/kcft114/.julia/packages/Flux/qXNjB/src/optimise/train.jl:105
┌ Info: Epoch 238
└ @ Main /Users/kcft114/.julia/packages/Flux/qXNjB/src/optimise/train.jl:105
┌ Info: Epoch 239
└ @ Main /Users/kcft114/.julia/packages/Flux/qXNjB/src/optimise/train.jl:105
┌ Info: Epoch 240
└ @ Main /Users/kcft114/.julia/packages/Flux/qXNjB/src/optimise/train.jl:105
┌ Info: Epoch 241
└ @ Main /Users/kcft114/.julia/packages/Flux/qXNjB/src/optimise/train.jl:105
┌ Info: Epoch 242
└ @ Main /Users/kcft114/.julia/packages/Flux/qXNjB/src/optimise/train.jl:105
┌ Info: Epoch 243
└ @ Main /Users/kcft114/.julia/packages/Flux/qXNjB/src/optimise/train.jl:105
┌ Info: Epoch 244
└ @ Main /Users/kcft114/.julia/packages/Flux/qXNjB/src/optimise/train.jl:105
┌ Info: Epoch 245
└ @ Main /Users/kcft114/.julia/packages/Flux/qXNjB/src/optimise/train.jl:105
┌ Info: Epoch 246
└ @ Main /Users/kcft114/.julia/packages/Flux/qXNjB/src/optimise/train.jl:105
┌ Info: Epoch 247
└ @ Main /Users/kcft114/.julia/packages/Flux/qXNjB/src/optimise/train.jl:105
┌ Info: Epoch 248
└ @ Main /Users/kcft114/.julia/packages/Flux/qXNjB/src/optimise/train.jl:105
┌ Info: Epoch 249
└ @ Main /Users/kcft114/.julia/packages/Flux/qXNjB/src/optimise/train.jl:105
┌ Info: Epoch 250
└ @ Main /Users/kcft114/.julia/packages/Flux/qXNjB/src/optimise/train.jl:105
┌ Info: Epoch 251
└ @ Main /Users/kcft114/.julia/packages/Flux/qXNjB/src/optimise/train.jl:105
┌ Info: Epoch 252
└ @ Main /Users/kcft114/.julia/packages/Flux/qXNjB/src/optimise/train.jl:105
┌ Info: Epoch 253
└ @ Main /Users/kcft114/.julia/packages/Flux/qXNjB/src/optimise/train.jl:105
┌ Info: Epoch 254
└ @ Main /Users/kcft114/.julia/packages/Flux/qXNjB/src/optimise/train.jl:105
┌ Info: Epoch 255
└ @ Main /Users/kcft114/.julia/packages/Flux/qXNjB/src/optimise/train.jl:105
┌ Info: Epoch 256
└ @ Main /Users/kcft114/.julia/packages/Flux/qXNjB/src/optimise/train.jl:105
┌ Info: Epoch 257
└ @ Main /Users/kcft114/.julia/packages/Flux/qXNjB/src/optimise/train.jl:105
┌ Info: Epoch 258
└ @ Main /Users/kcft114/.julia/packages/Flux/qXNjB/src/optimise/train.jl:105
┌ Info: Epoch 259
└ @ Main /Users/kcft114/.julia/packages/Flux/qXNjB/src/optimise/train.jl:105
┌ Info: Epoch 260
└ @ Main /Users/kcft114/.julia/packages/Flux/qXNjB/src/optimise/train.jl:105
┌ Info: Epoch 261
└ @ Main /Users/kcft114/.julia/packages/Flux/qXNjB/src/optimise/train.jl:105
┌ Info: Epoch 262
└ @ Main /Users/kcft114/.julia/packages/Flux/qXNjB/src/optimise/train.jl:105
┌ Info: Epoch 263
└ @ Main /Users/kcft114/.julia/packages/Flux/qXNjB/src/optimise/train.jl:105
┌ Info: Epoch 264
└ @ Main /Users/kcft114/.julia/packages/Flux/qXNjB/src/optimise/train.jl:105
┌ Info: Epoch 265
└ @ Main /Users/kcft114/.julia/packages/Flux/qXNjB/src/optimise/train.jl:105
```

```
loss(X, Y) = 0.8298877f0 (tracked)
loss(X, Y) = 0.82885104f0 (tracked)
loss(X, Y) = 0.8278227f0 (tracked)
loss(X, Y) = 0.8268017f0 (tracked)
loss(X, Y) = 0.82579035f0 (tracked)
loss(X, Y) = 0.82478875f0 (tracked)
loss(X, Y) = 0.823805f0 (tracked)
loss(X, Y) = 0.8228271f0 (tracked)
loss(X, Y) = 0.8218509f0 (tracked)
loss(X, Y) = 0.82088083f0 (tracked)
loss(X, Y) = 0.8199253f0 (tracked)
loss(X, Y) = 0.81897616f0 (tracked)
loss(X, Y) = 0.81802624f0 (tracked)
loss(X, Y) = 0.8170875f0 (tracked)
loss(X, Y) = 0.8161659f0 (tracked)
loss(X, Y) = 0.8152571f0 (tracked)
loss(X, Y) = 0.8143609f0 (tracked)
loss(X, Y) = 0.81347317f0 (tracked)
loss(X, Y) = 0.8125944f0 (tracked)
loss(X, Y) = 0.81172615f0 (tracked)
loss(X, Y) = 0.81086475f0 (tracked)
loss(X, Y) = 0.81002337f0 (tracked)
loss(X, Y) = 0.80919963f0 (tracked)
loss(X, Y) = 0.8083839f0 (tracked)
loss(X, Y) = 0.80757576f0 (tracked)
loss(X, Y) = 0.8067733f0 (tracked)
loss(X, Y) = 0.8059766f0 (tracked)
loss(X, Y) = 0.80518544f0 (tracked)
loss(X, Y) = 0.8044002f0 (tracked)
loss(X, Y) = 0.8036246f0 (tracked)
loss(X, Y) = 0.8028559f0 (tracked)
loss(X, Y) = 0.80209607f0 (tracked)
loss(X, Y) = 0.80134445f0 (tracked)
loss(X, Y) = 0.8006026f0 (tracked)
loss(X, Y) = 0.7998675f0 (tracked)
loss(X, Y) = 0.7991402f0 (tracked)
loss(X, Y) = 0.79841864f0 (tracked)
loss(X, Y) = 0.79770195f0 (tracked)
loss(X, Y) = 0.79699165f0 (tracked)
loss(X, Y) = 0.79628986f0 (tracked)
loss(X, Y) = 0.79559875f0 (tracked)
loss(X, Y) = 0.79491466f0 (tracked)
loss(X, Y) = 0.7942377f0 (tracked)
loss(X, Y) = 0.7935652f0 (tracked)
loss(X, Y) = 0.7928995f0 (tracked)
loss(X, Y) = 0.79223907f0 (tracked)
loss(X, Y) = 0.7915844f0 (tracked)
loss(X, Y) = 0.79093677f0 (tracked)
loss(X, Y) = 0.7902961f0 (tracked)
loss(X, Y) = 0.7896591f0 (tracked)
loss(X, Y) = 0.78902847f0 (tracked)
loss(X, Y) = 0.7884025f0 (tracked)
loss(X, Y) = 0.7877792f0 (tracked)
loss(X, Y) = 0.7871612f0 (tracked)
loss(X, Y) = 0.78654814f0 (tracked)
loss(X, Y) = 0.785955f0 (tracked)
loss(X, Y) = 0.78537416f0 (tracked)
loss(X, Y) = 0.7848062f0 (tracked)
loss(X, Y) = 0.78424144f0 (tracked)
loss(X, Y) = 0.78367966f0 (tracked)
loss(X, Y) = 0.78312045f0 (tracked)
loss(X, Y) = 0.782568f0 (tracked)
loss(X, Y) = 0.78201836f0 (tracked)
loss(X, Y) = 0.78147167f0 (tracked)
loss(X, Y) = 0.78093165f0 (tracked)
loss(X, Y) = 0.7803951f0 (tracked)
loss(X, Y) = 0.7798613f0 (tracked)
loss(X, Y) = 0.7793298f0 (tracked)
loss(X, Y) = 0.77880335f0 (tracked)
loss(X, Y) = 0.7782805f0 (tracked)
loss(X, Y) = 0.77775985f0 (tracked)
loss(X, Y) = 0.7772407f0 (tracked)
loss(X, Y) = 0.7767204f0 (tracked)
loss(X, Y) = 0.7761986f0 (tracked)
loss(X, Y) = 0.7756839f0 (tracked)
loss(X, Y) = 0.7751701f0 (tracked)
loss(X, Y) = 0.77465564f0 (tracked)
loss(X, Y) = 0.7741432f0 (tracked)
loss(X, Y) = 0.77363336f0 (tracked)
loss(X, Y) = 0.7731268f0 (tracked)
loss(X, Y) = 0.77262086f0 (tracked)
loss(X, Y) = 0.77211547f0 (tracked)
loss(X, Y) = 0.77161205f0 (tracked)
loss(X, Y) = 0.7711131f0 (tracked)
loss(X, Y) = 0.77061737f0 (tracked)
loss(X, Y) = 0.7701212f0 (tracked)
loss(X, Y) = 0.7696282f0 (tracked)
loss(X, Y) = 0.7691394f0 (tracked)
loss(X, Y) = 0.76865125f0 (tracked)
loss(X, Y) = 0.76816493f0 (tracked)
loss(X, Y) = 0.76767915f0 (tracked)
loss(X, Y) = 0.76719487f0 (tracked)
loss(X, Y) = 0.76671535f0 (tracked)
loss(X, Y) = 0.7662418f0 (tracked)
loss(X, Y) = 0.7657653f0 (tracked)
loss(X, Y) = 0.7652832f0 (tracked)
loss(X, Y) = 0.7648025f0 (tracked)
loss(X, Y) = 0.7643201f0 (tracked)
loss(X, Y) = 0.76383543f0 (tracked)
loss(X, Y) = 0.76334965f0 (tracked)
loss(X, Y) = 0.7628629f0 (tracked)
loss(X, Y) = 0.7623751f0 (tracked)
loss(X, Y) = 0.7618907f0 (tracked)
loss(X, Y) = 0.761407f0 (tracked)
loss(X, Y) = 0.76092255f0 (tracked)
loss(X, Y) = 0.76043934f0 (tracked)
loss(X, Y) = 0.7599604f0 (tracked)
loss(X, Y) = 0.75948375f0 (tracked)
loss(X, Y) = 0.7590112f0 (tracked)
loss(X, Y) = 0.75854003f0 (tracked)
loss(X, Y) = 0.7580677f0 (tracked)
loss(X, Y) = 0.757596f0 (tracked)
loss(X, Y) = 0.75712544f0 (tracked)
loss(X, Y) = 0.7566568f0 (tracked)
loss(X, Y) = 0.75618875f0 (tracked)
loss(X, Y) = 0.7557217f0 (tracked)
loss(X, Y) = 0.75525504f0 (tracked)
loss(X, Y) = 0.7547914f0 (tracked)
loss(X, Y) = 0.75432086f0 (tracked)
loss(X, Y) = 0.7538505f0 (tracked)
loss(X, Y) = 0.7533785f0 (tracked)
loss(X, Y) = 0.75290745f0 (tracked)
loss(X, Y) = 0.7524269f0 (tracked)
loss(X, Y) = 0.7519447f0 (tracked)
loss(X, Y) = 0.7514605f0 (tracked)
loss(X, Y) = 0.7509747f0 (tracked)
loss(X, Y) = 0.750489f0 (tracked)
loss(X, Y) = 0.7500041f0 (tracked)
loss(X, Y) = 0.74951386f0 (tracked)
loss(X, Y) = 0.7490167f0 (tracked)
loss(X, Y) = 0.74851954f0 (tracked)
loss(X, Y) = 0.74802685f0 (tracked)
loss(X, Y) = 0.74753535f0 (tracked)
loss(X, Y) = 0.74705005f0 (tracked)
loss(X, Y) = 0.7465687f0 (tracked)
loss(X, Y) = 0.746087f0 (tracked)
loss(X, Y) = 0.7456052f0 (tracked)
loss(X, Y) = 0.7451237f0 (tracked)
loss(X, Y) = 0.74464625f0 (tracked)
loss(X, Y) = 0.7441687f0 (tracked)
loss(X, Y) = 0.7436917f0 (tracked)
loss(X, Y) = 0.7432193f0 (tracked)
loss(X, Y) = 0.7427413f0 (tracked)
loss(X, Y) = 0.74224746f0 (tracked)
loss(X, Y) = 0.7417535f0 (tracked)
loss(X, Y) = 0.7412582f0 (tracked)
loss(X, Y) = 0.74076355f0 (tracked)
loss(X, Y) = 0.7402669f0 (tracked)
loss(X, Y) = 0.7397699f0 (tracked)
loss(X, Y) = 0.739273f0 (tracked)
loss(X, Y) = 0.7387802f0 (tracked)
loss(X, Y) = 0.7382934f0 (tracked)
loss(X, Y) = 0.73780304f0 (tracked)
loss(X, Y) = 0.7373128f0 (tracked)
loss(X, Y) = 0.73682433f0 (tracked)
loss(X, Y) = 0.7363357f0 (tracked)
loss(X, Y) = 0.73584795f0 (tracked)
loss(X, Y) = 0.73536366f0 (tracked)
loss(X, Y) = 0.73488617f0 (tracked)
loss(X, Y) = 0.7344137f0 (tracked)
loss(X, Y) = 0.7339418f0 (tracked)
loss(X, Y) = 0.7334732f0 (tracked)
loss(X, Y) = 0.7330086f0 (tracked)
loss(X, Y) = 0.7325339f0 (tracked)
loss(X, Y) = 0.7320558f0 (tracked)
loss(X, Y) = 0.7315771f0 (tracked)
```

```
┌ Info: Epoch 266
└ @ Main /Users/kcft114/.julia/packages/Flux/qXNjB/src/optimise/train.jl:105
┌ Info: Epoch 267
└ @ Main /Users/kcft114/.julia/packages/Flux/qXNjB/src/optimise/train.jl:105
┌ Info: Epoch 268
└ @ Main /Users/kcft114/.julia/packages/Flux/qXNjB/src/optimise/train.jl:105
┌ Info: Epoch 269
└ @ Main /Users/kcft114/.julia/packages/Flux/qXNjB/src/optimise/train.jl:105
┌ Info: Epoch 270
└ @ Main /Users/kcft114/.julia/packages/Flux/qXNjB/src/optimise/train.jl:105
┌ Info: Epoch 271
└ @ Main /Users/kcft114/.julia/packages/Flux/qXNjB/src/optimise/train.jl:105
┌ Info: Epoch 272
└ @ Main /Users/kcft114/.julia/packages/Flux/qXNjB/src/optimise/train.jl:105
┌ Info: Epoch 273
└ @ Main /Users/kcft114/.julia/packages/Flux/qXNjB/src/optimise/train.jl:105
┌ Info: Epoch 274
└ @ Main /Users/kcft114/.julia/packages/Flux/qXNjB/src/optimise/train.jl:105
┌ Info: Epoch 275
└ @ Main /Users/kcft114/.julia/packages/Flux/qXNjB/src/optimise/train.jl:105
┌ Info: Epoch 276
└ @ Main /Users/kcft114/.julia/packages/Flux/qXNjB/src/optimise/train.jl:105
┌ Info: Epoch 277
└ @ Main /Users/kcft114/.julia/packages/Flux/qXNjB/src/optimise/train.jl:105
┌ Info: Epoch 278
└ @ Main /Users/kcft114/.julia/packages/Flux/qXNjB/src/optimise/train.jl:105
┌ Info: Epoch 279
└ @ Main /Users/kcft114/.julia/packages/Flux/qXNjB/src/optimise/train.jl:105
┌ Info: Epoch 280
└ @ Main /Users/kcft114/.julia/packages/Flux/qXNjB/src/optimise/train.jl:105
┌ Info: Epoch 281
└ @ Main /Users/kcft114/.julia/packages/Flux/qXNjB/src/optimise/train.jl:105
┌ Info: Epoch 282
└ @ Main /Users/kcft114/.julia/packages/Flux/qXNjB/src/optimise/train.jl:105
┌ Info: Epoch 283
└ @ Main /Users/kcft114/.julia/packages/Flux/qXNjB/src/optimise/train.jl:105
┌ Info: Epoch 284
└ @ Main /Users/kcft114/.julia/packages/Flux/qXNjB/src/optimise/train.jl:105
┌ Info: Epoch 285
└ @ Main /Users/kcft114/.julia/packages/Flux/qXNjB/src/optimise/train.jl:105
┌ Info: Epoch 286
└ @ Main /Users/kcft114/.julia/packages/Flux/qXNjB/src/optimise/train.jl:105
┌ Info: Epoch 287
└ @ Main /Users/kcft114/.julia/packages/Flux/qXNjB/src/optimise/train.jl:105
┌ Info: Epoch 288
└ @ Main /Users/kcft114/.julia/packages/Flux/qXNjB/src/optimise/train.jl:105
┌ Info: Epoch 289
└ @ Main /Users/kcft114/.julia/packages/Flux/qXNjB/src/optimise/train.jl:105
┌ Info: Epoch 290
└ @ Main /Users/kcft114/.julia/packages/Flux/qXNjB/src/optimise/train.jl:105
┌ Info: Epoch 291
└ @ Main /Users/kcft114/.julia/packages/Flux/qXNjB/src/optimise/train.jl:105
┌ Info: Epoch 292
└ @ Main /Users/kcft114/.julia/packages/Flux/qXNjB/src/optimise/train.jl:105
┌ Info: Epoch 293
└ @ Main /Users/kcft114/.julia/packages/Flux/qXNjB/src/optimise/train.jl:105
┌ Info: Epoch 294
└ @ Main /Users/kcft114/.julia/packages/Flux/qXNjB/src/optimise/train.jl:105
┌ Info: Epoch 295
└ @ Main /Users/kcft114/.julia/packages/Flux/qXNjB/src/optimise/train.jl:105
┌ Info: Epoch 296
└ @ Main /Users/kcft114/.julia/packages/Flux/qXNjB/src/optimise/train.jl:105
┌ Info: Epoch 297
└ @ Main /Users/kcft114/.julia/packages/Flux/qXNjB/src/optimise/train.jl:105
┌ Info: Epoch 298
└ @ Main /Users/kcft114/.julia/packages/Flux/qXNjB/src/optimise/train.jl:105
┌ Info: Epoch 299
└ @ Main /Users/kcft114/.julia/packages/Flux/qXNjB/src/optimise/train.jl:105
┌ Info: Epoch 300
└ @ Main /Users/kcft114/.julia/packages/Flux/qXNjB/src/optimise/train.jl:105
┌ Info: Epoch 301
└ @ Main /Users/kcft114/.julia/packages/Flux/qXNjB/src/optimise/train.jl:105
┌ Info: Epoch 302
└ @ Main /Users/kcft114/.julia/packages/Flux/qXNjB/src/optimise/train.jl:105
┌ Info: Epoch 303
└ @ Main /Users/kcft114/.julia/packages/Flux/qXNjB/src/optimise/train.jl:105
┌ Info: Epoch 304
└ @ Main /Users/kcft114/.julia/packages/Flux/qXNjB/src/optimise/train.jl:105
┌ Info: Epoch 305
└ @ Main /Users/kcft114/.julia/packages/Flux/qXNjB/src/optimise/train.jl:105
┌ Info: Epoch 306
└ @ Main /Users/kcft114/.julia/packages/Flux/qXNjB/src/optimise/train.jl:105
┌ Info: Epoch 307
└ @ Main /Users/kcft114/.julia/packages/Flux/qXNjB/src/optimise/train.jl:105
┌ Info: Epoch 308
└ @ Main /Users/kcft114/.julia/packages/Flux/qXNjB/src/optimise/train.jl:105
┌ Info: Epoch 309
└ @ Main /Users/kcft114/.julia/packages/Flux/qXNjB/src/optimise/train.jl:105
┌ Info: Epoch 310
└ @ Main /Users/kcft114/.julia/packages/Flux/qXNjB/src/optimise/train.jl:105
┌ Info: Epoch 311
└ @ Main /Users/kcft114/.julia/packages/Flux/qXNjB/src/optimise/train.jl:105
┌ Info: Epoch 312
└ @ Main /Users/kcft114/.julia/packages/Flux/qXNjB/src/optimise/train.jl:105
┌ Info: Epoch 313
└ @ Main /Users/kcft114/.julia/packages/Flux/qXNjB/src/optimise/train.jl:105
┌ Info: Epoch 314
└ @ Main /Users/kcft114/.julia/packages/Flux/qXNjB/src/optimise/train.jl:105
┌ Info: Epoch 315
└ @ Main /Users/kcft114/.julia/packages/Flux/qXNjB/src/optimise/train.jl:105
┌ Info: Epoch 316
└ @ Main /Users/kcft114/.julia/packages/Flux/qXNjB/src/optimise/train.jl:105
┌ Info: Epoch 317
└ @ Main /Users/kcft114/.julia/packages/Flux/qXNjB/src/optimise/train.jl:105
┌ Info: Epoch 318
└ @ Main /Users/kcft114/.julia/packages/Flux/qXNjB/src/optimise/train.jl:105
┌ Info: Epoch 319
└ @ Main /Users/kcft114/.julia/packages/Flux/qXNjB/src/optimise/train.jl:105
┌ Info: Epoch 320
└ @ Main /Users/kcft114/.julia/packages/Flux/qXNjB/src/optimise/train.jl:105
┌ Info: Epoch 321
└ @ Main /Users/kcft114/.julia/packages/Flux/qXNjB/src/optimise/train.jl:105
┌ Info: Epoch 322
└ @ Main /Users/kcft114/.julia/packages/Flux/qXNjB/src/optimise/train.jl:105
┌ Info: Epoch 323
└ @ Main /Users/kcft114/.julia/packages/Flux/qXNjB/src/optimise/train.jl:105
┌ Info: Epoch 324
└ @ Main /Users/kcft114/.julia/packages/Flux/qXNjB/src/optimise/train.jl:105
┌ Info: Epoch 325
└ @ Main /Users/kcft114/.julia/packages/Flux/qXNjB/src/optimise/train.jl:105
┌ Info: Epoch 326
└ @ Main /Users/kcft114/.julia/packages/Flux/qXNjB/src/optimise/train.jl:105
┌ Info: Epoch 327
└ @ Main /Users/kcft114/.julia/packages/Flux/qXNjB/src/optimise/train.jl:105
┌ Info: Epoch 328
└ @ Main /Users/kcft114/.julia/packages/Flux/qXNjB/src/optimise/train.jl:105
┌ Info: Epoch 329
└ @ Main /Users/kcft114/.julia/packages/Flux/qXNjB/src/optimise/train.jl:105
┌ Info: Epoch 330
└ @ Main /Users/kcft114/.julia/packages/Flux/qXNjB/src/optimise/train.jl:105
┌ Info: Epoch 331
└ @ Main /Users/kcft114/.julia/packages/Flux/qXNjB/src/optimise/train.jl:105
┌ Info: Epoch 332
└ @ Main /Users/kcft114/.julia/packages/Flux/qXNjB/src/optimise/train.jl:105
┌ Info: Epoch 333
└ @ Main /Users/kcft114/.julia/packages/Flux/qXNjB/src/optimise/train.jl:105
┌ Info: Epoch 334
└ @ Main /Users/kcft114/.julia/packages/Flux/qXNjB/src/optimise/train.jl:105
┌ Info: Epoch 335
└ @ Main /Users/kcft114/.julia/packages/Flux/qXNjB/src/optimise/train.jl:105
┌ Info: Epoch 336
└ @ Main /Users/kcft114/.julia/packages/Flux/qXNjB/src/optimise/train.jl:105
┌ Info: Epoch 337
└ @ Main /Users/kcft114/.julia/packages/Flux/qXNjB/src/optimise/train.jl:105
┌ Info: Epoch 338
└ @ Main /Users/kcft114/.julia/packages/Flux/qXNjB/src/optimise/train.jl:105
┌ Info: Epoch 339
└ @ Main /Users/kcft114/.julia/packages/Flux/qXNjB/src/optimise/train.jl:105
┌ Info: Epoch 340
└ @ Main /Users/kcft114/.julia/packages/Flux/qXNjB/src/optimise/train.jl:105
┌ Info: Epoch 341
└ @ Main /Users/kcft114/.julia/packages/Flux/qXNjB/src/optimise/train.jl:105
┌ Info: Epoch 342
└ @ Main /Users/kcft114/.julia/packages/Flux/qXNjB/src/optimise/train.jl:105
┌ Info: Epoch 343
└ @ Main /Users/kcft114/.julia/packages/Flux/qXNjB/src/optimise/train.jl:105
┌ Info: Epoch 344
└ @ Main /Users/kcft114/.julia/packages/Flux/qXNjB/src/optimise/train.jl:105
┌ Info: Epoch 345
└ @ Main /Users/kcft114/.julia/packages/Flux/qXNjB/src/optimise/train.jl:105
┌ Info: Epoch 346
└ @ Main /Users/kcft114/.julia/packages/Flux/qXNjB/src/optimise/train.jl:105
┌ Info: Epoch 347
└ @ Main /Users/kcft114/.julia/packages/Flux/qXNjB/src/optimise/train.jl:105
┌ Info: Epoch 348
└ @ Main /Users/kcft114/.julia/packages/Flux/qXNjB/src/optimise/train.jl:105
┌ Info: Epoch 349
└ @ Main /Users/kcft114/.julia/packages/Flux/qXNjB/src/optimise/train.jl:105
┌ Info: Epoch 350
└ @ Main /Users/kcft114/.julia/packages/Flux/qXNjB/src/optimise/train.jl:105
┌ Info: Epoch 351
└ @ Main /Users/kcft114/.julia/packages/Flux/qXNjB/src/optimise/train.jl:105
┌ Info: Epoch 352
└ @ Main /Users/kcft114/.julia/packages/Flux/qXNjB/src/optimise/train.jl:105
┌ Info: Epoch 353
└ @ Main /Users/kcft114/.julia/packages/Flux/qXNjB/src/optimise/train.jl:105
┌ Info: Epoch 354
└ @ Main /Users/kcft114/.julia/packages/Flux/qXNjB/src/optimise/train.jl:105
┌ Info: Epoch 355
└ @ Main /Users/kcft114/.julia/packages/Flux/qXNjB/src/optimise/train.jl:105
┌ Info: Epoch 356
└ @ Main /Users/kcft114/.julia/packages/Flux/qXNjB/src/optimise/train.jl:105
┌ Info: Epoch 357
└ @ Main /Users/kcft114/.julia/packages/Flux/qXNjB/src/optimise/train.jl:105
┌ Info: Epoch 358
└ @ Main /Users/kcft114/.julia/packages/Flux/qXNjB/src/optimise/train.jl:105
┌ Info: Epoch 359
└ @ Main /Users/kcft114/.julia/packages/Flux/qXNjB/src/optimise/train.jl:105
┌ Info: Epoch 360
└ @ Main /Users/kcft114/.julia/packages/Flux/qXNjB/src/optimise/train.jl:105
┌ Info: Epoch 361
└ @ Main /Users/kcft114/.julia/packages/Flux/qXNjB/src/optimise/train.jl:105
┌ Info: Epoch 362
└ @ Main /Users/kcft114/.julia/packages/Flux/qXNjB/src/optimise/train.jl:105
┌ Info: Epoch 363
└ @ Main /Users/kcft114/.julia/packages/Flux/qXNjB/src/optimise/train.jl:105
┌ Info: Epoch 364
└ @ Main /Users/kcft114/.julia/packages/Flux/qXNjB/src/optimise/train.jl:105
┌ Info: Epoch 365
└ @ Main /Users/kcft114/.julia/packages/Flux/qXNjB/src/optimise/train.jl:105
┌ Info: Epoch 366
└ @ Main /Users/kcft114/.julia/packages/Flux/qXNjB/src/optimise/train.jl:105
┌ Info: Epoch 367
└ @ Main /Users/kcft114/.julia/packages/Flux/qXNjB/src/optimise/train.jl:105
┌ Info: Epoch 368
└ @ Main /Users/kcft114/.julia/packages/Flux/qXNjB/src/optimise/train.jl:105
┌ Info: Epoch 369
└ @ Main /Users/kcft114/.julia/packages/Flux/qXNjB/src/optimise/train.jl:105
┌ Info: Epoch 370
└ @ Main /Users/kcft114/.julia/packages/Flux/qXNjB/src/optimise/train.jl:105
┌ Info: Epoch 371
└ @ Main /Users/kcft114/.julia/packages/Flux/qXNjB/src/optimise/train.jl:105
┌ Info: Epoch 372
└ @ Main /Users/kcft114/.julia/packages/Flux/qXNjB/src/optimise/train.jl:105
┌ Info: Epoch 373
└ @ Main /Users/kcft114/.julia/packages/Flux/qXNjB/src/optimise/train.jl:105
┌ Info: Epoch 374
└ @ Main /Users/kcft114/.julia/packages/Flux/qXNjB/src/optimise/train.jl:105
┌ Info: Epoch 375
└ @ Main /Users/kcft114/.julia/packages/Flux/qXNjB/src/optimise/train.jl:105
┌ Info: Epoch 376
└ @ Main /Users/kcft114/.julia/packages/Flux/qXNjB/src/optimise/train.jl:105
┌ Info: Epoch 377
└ @ Main /Users/kcft114/.julia/packages/Flux/qXNjB/src/optimise/train.jl:105
┌ Info: Epoch 378
└ @ Main /Users/kcft114/.julia/packages/Flux/qXNjB/src/optimise/train.jl:105
┌ Info: Epoch 379
└ @ Main /Users/kcft114/.julia/packages/Flux/qXNjB/src/optimise/train.jl:105
┌ Info: Epoch 380
└ @ Main /Users/kcft114/.julia/packages/Flux/qXNjB/src/optimise/train.jl:105
┌ Info: Epoch 381
└ @ Main /Users/kcft114/.julia/packages/Flux/qXNjB/src/optimise/train.jl:105
┌ Info: Epoch 382
└ @ Main /Users/kcft114/.julia/packages/Flux/qXNjB/src/optimise/train.jl:105
┌ Info: Epoch 383
└ @ Main /Users/kcft114/.julia/packages/Flux/qXNjB/src/optimise/train.jl:105
┌ Info: Epoch 384
└ @ Main /Users/kcft114/.julia/packages/Flux/qXNjB/src/optimise/train.jl:105
┌ Info: Epoch 385
└ @ Main /Users/kcft114/.julia/packages/Flux/qXNjB/src/optimise/train.jl:105
┌ Info: Epoch 386
└ @ Main /Users/kcft114/.julia/packages/Flux/qXNjB/src/optimise/train.jl:105
┌ Info: Epoch 387
└ @ Main /Users/kcft114/.julia/packages/Flux/qXNjB/src/optimise/train.jl:105
┌ Info: Epoch 388
└ @ Main /Users/kcft114/.julia/packages/Flux/qXNjB/src/optimise/train.jl:105
┌ Info: Epoch 389
└ @ Main /Users/kcft114/.julia/packages/Flux/qXNjB/src/optimise/train.jl:105
┌ Info: Epoch 390
└ @ Main /Users/kcft114/.julia/packages/Flux/qXNjB/src/optimise/train.jl:105
┌ Info: Epoch 391
└ @ Main /Users/kcft114/.julia/packages/Flux/qXNjB/src/optimise/train.jl:105
┌ Info: Epoch 392
└ @ Main /Users/kcft114/.julia/packages/Flux/qXNjB/src/optimise/train.jl:105
┌ Info: Epoch 393
└ @ Main /Users/kcft114/.julia/packages/Flux/qXNjB/src/optimise/train.jl:105
┌ Info: Epoch 394
└ @ Main /Users/kcft114/.julia/packages/Flux/qXNjB/src/optimise/train.jl:105
┌ Info: Epoch 395
└ @ Main /Users/kcft114/.julia/packages/Flux/qXNjB/src/optimise/train.jl:105
┌ Info: Epoch 396
└ @ Main /Users/kcft114/.julia/packages/Flux/qXNjB/src/optimise/train.jl:105
┌ Info: Epoch 397
└ @ Main /Users/kcft114/.julia/packages/Flux/qXNjB/src/optimise/train.jl:105
┌ Info: Epoch 398
└ @ Main /Users/kcft114/.julia/packages/Flux/qXNjB/src/optimise/train.jl:105
┌ Info: Epoch 399
└ @ Main /Users/kcft114/.julia/packages/Flux/qXNjB/src/optimise/train.jl:105
┌ Info: Epoch 400
└ @ Main /Users/kcft114/.julia/packages/Flux/qXNjB/src/optimise/train.jl:105
┌ Info: Epoch 401
└ @ Main /Users/kcft114/.julia/packages/Flux/qXNjB/src/optimise/train.jl:105
┌ Info: Epoch 402
└ @ Main /Users/kcft114/.julia/packages/Flux/qXNjB/src/optimise/train.jl:105
┌ Info: Epoch 403
└ @ Main /Users/kcft114/.julia/packages/Flux/qXNjB/src/optimise/train.jl:105
┌ Info: Epoch 404
└ @ Main /Users/kcft114/.julia/packages/Flux/qXNjB/src/optimise/train.jl:105
┌ Info: Epoch 405
└ @ Main /Users/kcft114/.julia/packages/Flux/qXNjB/src/optimise/train.jl:105
┌ Info: Epoch 406
└ @ Main /Users/kcft114/.julia/packages/Flux/qXNjB/src/optimise/train.jl:105
┌ Info: Epoch 407
└ @ Main /Users/kcft114/.julia/packages/Flux/qXNjB/src/optimise/train.jl:105
┌ Info: Epoch 408
└ @ Main /Users/kcft114/.julia/packages/Flux/qXNjB/src/optimise/train.jl:105
┌ Info: Epoch 409
└ @ Main /Users/kcft114/.julia/packages/Flux/qXNjB/src/optimise/train.jl:105
┌ Info: Epoch 410
└ @ Main /Users/kcft114/.julia/packages/Flux/qXNjB/src/optimise/train.jl:105
┌ Info: Epoch 411
└ @ Main /Users/kcft114/.julia/packages/Flux/qXNjB/src/optimise/train.jl:105
┌ Info: Epoch 412
└ @ Main /Users/kcft114/.julia/packages/Flux/qXNjB/src/optimise/train.jl:105
┌ Info: Epoch 413
└ @ Main /Users/kcft114/.julia/packages/Flux/qXNjB/src/optimise/train.jl:105
┌ Info: Epoch 414
└ @ Main /Users/kcft114/.julia/packages/Flux/qXNjB/src/optimise/train.jl:105
┌ Info: Epoch 415
└ @ Main /Users/kcft114/.julia/packages/Flux/qXNjB/src/optimise/train.jl:105
┌ Info: Epoch 416
└ @ Main /Users/kcft114/.julia/packages/Flux/qXNjB/src/optimise/train.jl:105
┌ Info: Epoch 417
└ @ Main /Users/kcft114/.julia/packages/Flux/qXNjB/src/optimise/train.jl:105
┌ Info: Epoch 418
└ @ Main /Users/kcft114/.julia/packages/Flux/qXNjB/src/optimise/train.jl:105
┌ Info: Epoch 419
└ @ Main /Users/kcft114/.julia/packages/Flux/qXNjB/src/optimise/train.jl:105
┌ Info: Epoch 420
└ @ Main /Users/kcft114/.julia/packages/Flux/qXNjB/src/optimise/train.jl:105
┌ Info: Epoch 421
└ @ Main /Users/kcft114/.julia/packages/Flux/qXNjB/src/optimise/train.jl:105
┌ Info: Epoch 422
└ @ Main /Users/kcft114/.julia/packages/Flux/qXNjB/src/optimise/train.jl:105
┌ Info: Epoch 423
└ @ Main /Users/kcft114/.julia/packages/Flux/qXNjB/src/optimise/train.jl:105
┌ Info: Epoch 424
└ @ Main /Users/kcft114/.julia/packages/Flux/qXNjB/src/optimise/train.jl:105
┌ Info: Epoch 425
└ @ Main /Users/kcft114/.julia/packages/Flux/qXNjB/src/optimise/train.jl:105
┌ Info: Epoch 426
└ @ Main /Users/kcft114/.julia/packages/Flux/qXNjB/src/optimise/train.jl:105
┌ Info: Epoch 427
└ @ Main /Users/kcft114/.julia/packages/Flux/qXNjB/src/optimise/train.jl:105
┌ Info: Epoch 428
└ @ Main /Users/kcft114/.julia/packages/Flux/qXNjB/src/optimise/train.jl:105
┌ Info: Epoch 429
└ @ Main /Users/kcft114/.julia/packages/Flux/qXNjB/src/optimise/train.jl:105
┌ Info: Epoch 430
└ @ Main /Users/kcft114/.julia/packages/Flux/qXNjB/src/optimise/train.jl:105
┌ Info: Epoch 431
└ @ Main /Users/kcft114/.julia/packages/Flux/qXNjB/src/optimise/train.jl:105
```

```
loss(X, Y) = 0.7310863f0 (tracked)
loss(X, Y) = 0.7305865f0 (tracked)
loss(X, Y) = 0.73008615f0 (tracked)
loss(X, Y) = 0.7295873f0 (tracked)
loss(X, Y) = 0.7290863f0 (tracked)
loss(X, Y) = 0.72858137f0 (tracked)
loss(X, Y) = 0.7280792f0 (tracked)
loss(X, Y) = 0.7275808f0 (tracked)
loss(X, Y) = 0.7270823f0 (tracked)
loss(X, Y) = 0.7265834f0 (tracked)
loss(X, Y) = 0.72608477f0 (tracked)
loss(X, Y) = 0.7255867f0 (tracked)
loss(X, Y) = 0.72509134f0 (tracked)
loss(X, Y) = 0.7246073f0 (tracked)
loss(X, Y) = 0.724138f0 (tracked)
loss(X, Y) = 0.72366875f0 (tracked)
loss(X, Y) = 0.72319794f0 (tracked)
loss(X, Y) = 0.7227288f0 (tracked)
loss(X, Y) = 0.7222604f0 (tracked)
loss(X, Y) = 0.72179335f0 (tracked)
loss(X, Y) = 0.721332f0 (tracked)
loss(X, Y) = 0.7208738f0 (tracked)
loss(X, Y) = 0.72040063f0 (tracked)
loss(X, Y) = 0.7199027f0 (tracked)
loss(X, Y) = 0.719409f0 (tracked)
loss(X, Y) = 0.7189388f0 (tracked)
loss(X, Y) = 0.71847725f0 (tracked)
loss(X, Y) = 0.7180192f0 (tracked)
loss(X, Y) = 0.7175621f0 (tracked)
loss(X, Y) = 0.71710545f0 (tracked)
loss(X, Y) = 0.71665114f0 (tracked)
loss(X, Y) = 0.7161975f0 (tracked)
loss(X, Y) = 0.7157575f0 (tracked)
loss(X, Y) = 0.7153331f0 (tracked)
loss(X, Y) = 0.7149115f0 (tracked)
loss(X, Y) = 0.7144928f0 (tracked)
loss(X, Y) = 0.71408194f0 (tracked)
loss(X, Y) = 0.71367204f0 (tracked)
loss(X, Y) = 0.71327055f0 (tracked)
loss(X, Y) = 0.7128719f0 (tracked)
loss(X, Y) = 0.7124877f0 (tracked)
loss(X, Y) = 0.7121127f0 (tracked)
loss(X, Y) = 0.71174085f0 (tracked)
loss(X, Y) = 0.7113662f0 (tracked)
loss(X, Y) = 0.7109904f0 (tracked)
loss(X, Y) = 0.7106124f0 (tracked)
loss(X, Y) = 0.7102332f0 (tracked)
loss(X, Y) = 0.7098535f0 (tracked)
loss(X, Y) = 0.7094759f0 (tracked)
loss(X, Y) = 0.7091055f0 (tracked)
loss(X, Y) = 0.7087363f0 (tracked)
loss(X, Y) = 0.7083708f0 (tracked)
loss(X, Y) = 0.7080122f0 (tracked)
loss(X, Y) = 0.707653f0 (tracked)
loss(X, Y) = 0.7072932f0 (tracked)
loss(X, Y) = 0.70693284f0 (tracked)
loss(X, Y) = 0.7065738f0 (tracked)
loss(X, Y) = 0.70621556f0 (tracked)
loss(X, Y) = 0.705861f0 (tracked)
loss(X, Y) = 0.7055107f0 (tracked)
loss(X, Y) = 0.70516014f0 (tracked)
loss(X, Y) = 0.70481384f0 (tracked)
loss(X, Y) = 0.70446825f0 (tracked)
loss(X, Y) = 0.70412093f0 (tracked)
loss(X, Y) = 0.7037714f0 (tracked)
loss(X, Y) = 0.70341885f0 (tracked)
loss(X, Y) = 0.7030671f0 (tracked)
loss(X, Y) = 0.7027161f0 (tracked)
loss(X, Y) = 0.7023624f0 (tracked)
loss(X, Y) = 0.702009f0 (tracked)
loss(X, Y) = 0.7016634f0 (tracked)
loss(X, Y) = 0.70131695f0 (tracked)
loss(X, Y) = 0.70097023f0 (tracked)
loss(X, Y) = 0.7006263f0 (tracked)
loss(X, Y) = 0.7002806f0 (tracked)
loss(X, Y) = 0.6999336f0 (tracked)
loss(X, Y) = 0.6995859f0 (tracked)
loss(X, Y) = 0.6992411f0 (tracked)
loss(X, Y) = 0.698895f0 (tracked)
loss(X, Y) = 0.6985484f0 (tracked)
loss(X, Y) = 0.69820267f0 (tracked)
loss(X, Y) = 0.69785786f0 (tracked)
loss(X, Y) = 0.69751304f0 (tracked)
loss(X, Y) = 0.697171f0 (tracked)
loss(X, Y) = 0.6968302f0 (tracked)
loss(X, Y) = 0.6964874f0 (tracked)
loss(X, Y) = 0.69614255f0 (tracked)
loss(X, Y) = 0.69579685f0 (tracked)
loss(X, Y) = 0.6954501f0 (tracked)
loss(X, Y) = 0.6951083f0 (tracked)
loss(X, Y) = 0.6947682f0 (tracked)
loss(X, Y) = 0.6944282f0 (tracked)
loss(X, Y) = 0.6940906f0 (tracked)
loss(X, Y) = 0.6937542f0 (tracked)
loss(X, Y) = 0.6934166f0 (tracked)
loss(X, Y) = 0.6930792f0 (tracked)
loss(X, Y) = 0.6927446f0 (tracked)
loss(X, Y) = 0.69241095f0 (tracked)
loss(X, Y) = 0.6920743f0 (tracked)
loss(X, Y) = 0.6917366f0 (tracked)
loss(X, Y) = 0.691408f0 (tracked)
loss(X, Y) = 0.6910847f0 (tracked)
loss(X, Y) = 0.6907597f0 (tracked)
loss(X, Y) = 0.6904351f0 (tracked)
loss(X, Y) = 0.69010854f0 (tracked)
loss(X, Y) = 0.68977976f0 (tracked)
loss(X, Y) = 0.6894519f0 (tracked)
loss(X, Y) = 0.68912315f0 (tracked)
loss(X, Y) = 0.68879485f0 (tracked)
loss(X, Y) = 0.68846864f0 (tracked)
loss(X, Y) = 0.68814516f0 (tracked)
loss(X, Y) = 0.6878166f0 (tracked)
loss(X, Y) = 0.68748784f0 (tracked)
loss(X, Y) = 0.6871632f0 (tracked)
loss(X, Y) = 0.68683714f0 (tracked)
loss(X, Y) = 0.68650967f0 (tracked)
loss(X, Y) = 0.6861804f0 (tracked)
loss(X, Y) = 0.6858529f0 (tracked)
loss(X, Y) = 0.6855264f0 (tracked)
loss(X, Y) = 0.68519926f0 (tracked)
loss(X, Y) = 0.68487436f0 (tracked)
loss(X, Y) = 0.68454796f0 (tracked)
loss(X, Y) = 0.68421936f0 (tracked)
loss(X, Y) = 0.68388844f0 (tracked)
loss(X, Y) = 0.6835565f0 (tracked)
loss(X, Y) = 0.68322784f0 (tracked)
loss(X, Y) = 0.6828977f0 (tracked)
loss(X, Y) = 0.6825697f0 (tracked)
loss(X, Y) = 0.68224007f0 (tracked)
loss(X, Y) = 0.68191206f0 (tracked)
loss(X, Y) = 0.6815837f0 (tracked)
loss(X, Y) = 0.6812548f0 (tracked)
loss(X, Y) = 0.6809252f0 (tracked)
loss(X, Y) = 0.680596f0 (tracked)
loss(X, Y) = 0.68027145f0 (tracked)
loss(X, Y) = 0.67994636f0 (tracked)
loss(X, Y) = 0.67961645f0 (tracked)
loss(X, Y) = 0.67928904f0 (tracked)
loss(X, Y) = 0.67896014f0 (tracked)
loss(X, Y) = 0.67863625f0 (tracked)
loss(X, Y) = 0.67831093f0 (tracked)
loss(X, Y) = 0.67798436f0 (tracked)
loss(X, Y) = 0.67765564f0 (tracked)
loss(X, Y) = 0.6773296f0 (tracked)
loss(X, Y) = 0.67700243f0 (tracked)
loss(X, Y) = 0.67667574f0 (tracked)
loss(X, Y) = 0.6763496f0 (tracked)
loss(X, Y) = 0.67602324f0 (tracked)
loss(X, Y) = 0.67569244f0 (tracked)
loss(X, Y) = 0.6753607f0 (tracked)
loss(X, Y) = 0.67503124f0 (tracked)
loss(X, Y) = 0.674701f0 (tracked)
loss(X, Y) = 0.6743719f0 (tracked)
```

```
┌ Info: Epoch 432
└ @ Main /Users/kcft114/.julia/packages/Flux/qXNjB/src/optimise/train.jl:105
┌ Info: Epoch 433
└ @ Main /Users/kcft114/.julia/packages/Flux/qXNjB/src/optimise/train.jl:105
┌ Info: Epoch 434
└ @ Main /Users/kcft114/.julia/packages/Flux/qXNjB/src/optimise/train.jl:105
┌ Info: Epoch 435
└ @ Main /Users/kcft114/.julia/packages/Flux/qXNjB/src/optimise/train.jl:105
┌ Info: Epoch 436
└ @ Main /Users/kcft114/.julia/packages/Flux/qXNjB/src/optimise/train.jl:105
┌ Info: Epoch 437
└ @ Main /Users/kcft114/.julia/packages/Flux/qXNjB/src/optimise/train.jl:105
┌ Info: Epoch 438
└ @ Main /Users/kcft114/.julia/packages/Flux/qXNjB/src/optimise/train.jl:105
┌ Info: Epoch 439
└ @ Main /Users/kcft114/.julia/packages/Flux/qXNjB/src/optimise/train.jl:105
┌ Info: Epoch 440
└ @ Main /Users/kcft114/.julia/packages/Flux/qXNjB/src/optimise/train.jl:105
┌ Info: Epoch 441
└ @ Main /Users/kcft114/.julia/packages/Flux/qXNjB/src/optimise/train.jl:105
┌ Info: Epoch 442
└ @ Main /Users/kcft114/.julia/packages/Flux/qXNjB/src/optimise/train.jl:105
┌ Info: Epoch 443
└ @ Main /Users/kcft114/.julia/packages/Flux/qXNjB/src/optimise/train.jl:105
┌ Info: Epoch 444
└ @ Main /Users/kcft114/.julia/packages/Flux/qXNjB/src/optimise/train.jl:105
┌ Info: Epoch 445
└ @ Main /Users/kcft114/.julia/packages/Flux/qXNjB/src/optimise/train.jl:105
┌ Info: Epoch 446
└ @ Main /Users/kcft114/.julia/packages/Flux/qXNjB/src/optimise/train.jl:105
┌ Info: Epoch 447
└ @ Main /Users/kcft114/.julia/packages/Flux/qXNjB/src/optimise/train.jl:105
┌ Info: Epoch 448
└ @ Main /Users/kcft114/.julia/packages/Flux/qXNjB/src/optimise/train.jl:105
┌ Info: Epoch 449
└ @ Main /Users/kcft114/.julia/packages/Flux/qXNjB/src/optimise/train.jl:105
┌ Info: Epoch 450
└ @ Main /Users/kcft114/.julia/packages/Flux/qXNjB/src/optimise/train.jl:105
┌ Info: Epoch 451
└ @ Main /Users/kcft114/.julia/packages/Flux/qXNjB/src/optimise/train.jl:105
┌ Info: Epoch 452
└ @ Main /Users/kcft114/.julia/packages/Flux/qXNjB/src/optimise/train.jl:105
┌ Info: Epoch 453
└ @ Main /Users/kcft114/.julia/packages/Flux/qXNjB/src/optimise/train.jl:105
┌ Info: Epoch 454
└ @ Main /Users/kcft114/.julia/packages/Flux/qXNjB/src/optimise/train.jl:105
┌ Info: Epoch 455
└ @ Main /Users/kcft114/.julia/packages/Flux/qXNjB/src/optimise/train.jl:105
┌ Info: Epoch 456
└ @ Main /Users/kcft114/.julia/packages/Flux/qXNjB/src/optimise/train.jl:105
┌ Info: Epoch 457
└ @ Main /Users/kcft114/.julia/packages/Flux/qXNjB/src/optimise/train.jl:105
┌ Info: Epoch 458
└ @ Main /Users/kcft114/.julia/packages/Flux/qXNjB/src/optimise/train.jl:105
┌ Info: Epoch 459
└ @ Main /Users/kcft114/.julia/packages/Flux/qXNjB/src/optimise/train.jl:105
┌ Info: Epoch 460
└ @ Main /Users/kcft114/.julia/packages/Flux/qXNjB/src/optimise/train.jl:105
┌ Info: Epoch 461
└ @ Main /Users/kcft114/.julia/packages/Flux/qXNjB/src/optimise/train.jl:105
┌ Info: Epoch 462
└ @ Main /Users/kcft114/.julia/packages/Flux/qXNjB/src/optimise/train.jl:105
┌ Info: Epoch 463
└ @ Main /Users/kcft114/.julia/packages/Flux/qXNjB/src/optimise/train.jl:105
┌ Info: Epoch 464
└ @ Main /Users/kcft114/.julia/packages/Flux/qXNjB/src/optimise/train.jl:105
┌ Info: Epoch 465
└ @ Main /Users/kcft114/.julia/packages/Flux/qXNjB/src/optimise/train.jl:105
┌ Info: Epoch 466
└ @ Main /Users/kcft114/.julia/packages/Flux/qXNjB/src/optimise/train.jl:105
┌ Info: Epoch 467
└ @ Main /Users/kcft114/.julia/packages/Flux/qXNjB/src/optimise/train.jl:105
┌ Info: Epoch 468
└ @ Main /Users/kcft114/.julia/packages/Flux/qXNjB/src/optimise/train.jl:105
┌ Info: Epoch 469
└ @ Main /Users/kcft114/.julia/packages/Flux/qXNjB/src/optimise/train.jl:105
┌ Info: Epoch 470
└ @ Main /Users/kcft114/.julia/packages/Flux/qXNjB/src/optimise/train.jl:105
┌ Info: Epoch 471
└ @ Main /Users/kcft114/.julia/packages/Flux/qXNjB/src/optimise/train.jl:105
┌ Info: Epoch 472
└ @ Main /Users/kcft114/.julia/packages/Flux/qXNjB/src/optimise/train.jl:105
┌ Info: Epoch 473
└ @ Main /Users/kcft114/.julia/packages/Flux/qXNjB/src/optimise/train.jl:105
┌ Info: Epoch 474
└ @ Main /Users/kcft114/.julia/packages/Flux/qXNjB/src/optimise/train.jl:105
┌ Info: Epoch 475
└ @ Main /Users/kcft114/.julia/packages/Flux/qXNjB/src/optimise/train.jl:105
┌ Info: Epoch 476
└ @ Main /Users/kcft114/.julia/packages/Flux/qXNjB/src/optimise/train.jl:105
┌ Info: Epoch 477
└ @ Main /Users/kcft114/.julia/packages/Flux/qXNjB/src/optimise/train.jl:105
┌ Info: Epoch 478
└ @ Main /Users/kcft114/.julia/packages/Flux/qXNjB/src/optimise/train.jl:105
┌ Info: Epoch 479
└ @ Main /Users/kcft114/.julia/packages/Flux/qXNjB/src/optimise/train.jl:105
┌ Info: Epoch 480
└ @ Main /Users/kcft114/.julia/packages/Flux/qXNjB/src/optimise/train.jl:105
┌ Info: Epoch 481
└ @ Main /Users/kcft114/.julia/packages/Flux/qXNjB/src/optimise/train.jl:105
┌ Info: Epoch 482
└ @ Main /Users/kcft114/.julia/packages/Flux/qXNjB/src/optimise/train.jl:105
┌ Info: Epoch 483
└ @ Main /Users/kcft114/.julia/packages/Flux/qXNjB/src/optimise/train.jl:105
┌ Info: Epoch 484
└ @ Main /Users/kcft114/.julia/packages/Flux/qXNjB/src/optimise/train.jl:105
┌ Info: Epoch 485
└ @ Main /Users/kcft114/.julia/packages/Flux/qXNjB/src/optimise/train.jl:105
┌ Info: Epoch 486
└ @ Main /Users/kcft114/.julia/packages/Flux/qXNjB/src/optimise/train.jl:105
┌ Info: Epoch 487
└ @ Main /Users/kcft114/.julia/packages/Flux/qXNjB/src/optimise/train.jl:105
┌ Info: Epoch 488
└ @ Main /Users/kcft114/.julia/packages/Flux/qXNjB/src/optimise/train.jl:105
┌ Info: Epoch 489
└ @ Main /Users/kcft114/.julia/packages/Flux/qXNjB/src/optimise/train.jl:105
┌ Info: Epoch 490
└ @ Main /Users/kcft114/.julia/packages/Flux/qXNjB/src/optimise/train.jl:105
┌ Info: Epoch 491
└ @ Main /Users/kcft114/.julia/packages/Flux/qXNjB/src/optimise/train.jl:105
┌ Info: Epoch 492
└ @ Main /Users/kcft114/.julia/packages/Flux/qXNjB/src/optimise/train.jl:105
┌ Info: Epoch 493
└ @ Main /Users/kcft114/.julia/packages/Flux/qXNjB/src/optimise/train.jl:105
┌ Info: Epoch 494
└ @ Main /Users/kcft114/.julia/packages/Flux/qXNjB/src/optimise/train.jl:105
┌ Info: Epoch 495
└ @ Main /Users/kcft114/.julia/packages/Flux/qXNjB/src/optimise/train.jl:105
┌ Info: Epoch 496
└ @ Main /Users/kcft114/.julia/packages/Flux/qXNjB/src/optimise/train.jl:105
┌ Info: Epoch 497
└ @ Main /Users/kcft114/.julia/packages/Flux/qXNjB/src/optimise/train.jl:105
┌ Info: Epoch 498
└ @ Main /Users/kcft114/.julia/packages/Flux/qXNjB/src/optimise/train.jl:105
┌ Info: Epoch 499
└ @ Main /Users/kcft114/.julia/packages/Flux/qXNjB/src/optimise/train.jl:105
┌ Info: Epoch 500
└ @ Main /Users/kcft114/.julia/packages/Flux/qXNjB/src/optimise/train.jl:105
┌ Info: Epoch 501
└ @ Main /Users/kcft114/.julia/packages/Flux/qXNjB/src/optimise/train.jl:105
┌ Info: Epoch 502
└ @ Main /Users/kcft114/.julia/packages/Flux/qXNjB/src/optimise/train.jl:105
┌ Info: Epoch 503
└ @ Main /Users/kcft114/.julia/packages/Flux/qXNjB/src/optimise/train.jl:105
┌ Info: Epoch 504
└ @ Main /Users/kcft114/.julia/packages/Flux/qXNjB/src/optimise/train.jl:105
┌ Info: Epoch 505
└ @ Main /Users/kcft114/.julia/packages/Flux/qXNjB/src/optimise/train.jl:105
┌ Info: Epoch 506
└ @ Main /Users/kcft114/.julia/packages/Flux/qXNjB/src/optimise/train.jl:105
┌ Info: Epoch 507
└ @ Main /Users/kcft114/.julia/packages/Flux/qXNjB/src/optimise/train.jl:105
┌ Info: Epoch 508
└ @ Main /Users/kcft114/.julia/packages/Flux/qXNjB/src/optimise/train.jl:105
┌ Info: Epoch 509
└ @ Main /Users/kcft114/.julia/packages/Flux/qXNjB/src/optimise/train.jl:105
┌ Info: Epoch 510
└ @ Main /Users/kcft114/.julia/packages/Flux/qXNjB/src/optimise/train.jl:105
┌ Info: Epoch 511
└ @ Main /Users/kcft114/.julia/packages/Flux/qXNjB/src/optimise/train.jl:105
┌ Info: Epoch 512
└ @ Main /Users/kcft114/.julia/packages/Flux/qXNjB/src/optimise/train.jl:105
┌ Info: Epoch 513
└ @ Main /Users/kcft114/.julia/packages/Flux/qXNjB/src/optimise/train.jl:105
┌ Info: Epoch 514
└ @ Main /Users/kcft114/.julia/packages/Flux/qXNjB/src/optimise/train.jl:105
┌ Info: Epoch 515
└ @ Main /Users/kcft114/.julia/packages/Flux/qXNjB/src/optimise/train.jl:105
┌ Info: Epoch 516
└ @ Main /Users/kcft114/.julia/packages/Flux/qXNjB/src/optimise/train.jl:105
┌ Info: Epoch 517
└ @ Main /Users/kcft114/.julia/packages/Flux/qXNjB/src/optimise/train.jl:105
┌ Info: Epoch 518
└ @ Main /Users/kcft114/.julia/packages/Flux/qXNjB/src/optimise/train.jl:105
┌ Info: Epoch 519
└ @ Main /Users/kcft114/.julia/packages/Flux/qXNjB/src/optimise/train.jl:105
┌ Info: Epoch 520
└ @ Main /Users/kcft114/.julia/packages/Flux/qXNjB/src/optimise/train.jl:105
┌ Info: Epoch 521
└ @ Main /Users/kcft114/.julia/packages/Flux/qXNjB/src/optimise/train.jl:105
┌ Info: Epoch 522
└ @ Main /Users/kcft114/.julia/packages/Flux/qXNjB/src/optimise/train.jl:105
┌ Info: Epoch 523
└ @ Main /Users/kcft114/.julia/packages/Flux/qXNjB/src/optimise/train.jl:105
┌ Info: Epoch 524
└ @ Main /Users/kcft114/.julia/packages/Flux/qXNjB/src/optimise/train.jl:105
┌ Info: Epoch 525
└ @ Main /Users/kcft114/.julia/packages/Flux/qXNjB/src/optimise/train.jl:105
┌ Info: Epoch 526
└ @ Main /Users/kcft114/.julia/packages/Flux/qXNjB/src/optimise/train.jl:105
┌ Info: Epoch 527
└ @ Main /Users/kcft114/.julia/packages/Flux/qXNjB/src/optimise/train.jl:105
┌ Info: Epoch 528
└ @ Main /Users/kcft114/.julia/packages/Flux/qXNjB/src/optimise/train.jl:105
┌ Info: Epoch 529
└ @ Main /Users/kcft114/.julia/packages/Flux/qXNjB/src/optimise/train.jl:105
┌ Info: Epoch 530
└ @ Main /Users/kcft114/.julia/packages/Flux/qXNjB/src/optimise/train.jl:105
┌ Info: Epoch 531
└ @ Main /Users/kcft114/.julia/packages/Flux/qXNjB/src/optimise/train.jl:105
┌ Info: Epoch 532
└ @ Main /Users/kcft114/.julia/packages/Flux/qXNjB/src/optimise/train.jl:105
┌ Info: Epoch 533
└ @ Main /Users/kcft114/.julia/packages/Flux/qXNjB/src/optimise/train.jl:105
┌ Info: Epoch 534
└ @ Main /Users/kcft114/.julia/packages/Flux/qXNjB/src/optimise/train.jl:105
┌ Info: Epoch 535
└ @ Main /Users/kcft114/.julia/packages/Flux/qXNjB/src/optimise/train.jl:105
┌ Info: Epoch 536
└ @ Main /Users/kcft114/.julia/packages/Flux/qXNjB/src/optimise/train.jl:105
┌ Info: Epoch 537
└ @ Main /Users/kcft114/.julia/packages/Flux/qXNjB/src/optimise/train.jl:105
┌ Info: Epoch 538
└ @ Main /Users/kcft114/.julia/packages/Flux/qXNjB/src/optimise/train.jl:105
┌ Info: Epoch 539
└ @ Main /Users/kcft114/.julia/packages/Flux/qXNjB/src/optimise/train.jl:105
┌ Info: Epoch 540
└ @ Main /Users/kcft114/.julia/packages/Flux/qXNjB/src/optimise/train.jl:105
┌ Info: Epoch 541
└ @ Main /Users/kcft114/.julia/packages/Flux/qXNjB/src/optimise/train.jl:105
┌ Info: Epoch 542
└ @ Main /Users/kcft114/.julia/packages/Flux/qXNjB/src/optimise/train.jl:105
┌ Info: Epoch 543
└ @ Main /Users/kcft114/.julia/packages/Flux/qXNjB/src/optimise/train.jl:105
┌ Info: Epoch 544
└ @ Main /Users/kcft114/.julia/packages/Flux/qXNjB/src/optimise/train.jl:105
┌ Info: Epoch 545
└ @ Main /Users/kcft114/.julia/packages/Flux/qXNjB/src/optimise/train.jl:105
┌ Info: Epoch 546
└ @ Main /Users/kcft114/.julia/packages/Flux/qXNjB/src/optimise/train.jl:105
┌ Info: Epoch 547
└ @ Main /Users/kcft114/.julia/packages/Flux/qXNjB/src/optimise/train.jl:105
┌ Info: Epoch 548
└ @ Main /Users/kcft114/.julia/packages/Flux/qXNjB/src/optimise/train.jl:105
┌ Info: Epoch 549
└ @ Main /Users/kcft114/.julia/packages/Flux/qXNjB/src/optimise/train.jl:105
┌ Info: Epoch 550
└ @ Main /Users/kcft114/.julia/packages/Flux/qXNjB/src/optimise/train.jl:105
┌ Info: Epoch 551
└ @ Main /Users/kcft114/.julia/packages/Flux/qXNjB/src/optimise/train.jl:105
┌ Info: Epoch 552
└ @ Main /Users/kcft114/.julia/packages/Flux/qXNjB/src/optimise/train.jl:105
┌ Info: Epoch 553
└ @ Main /Users/kcft114/.julia/packages/Flux/qXNjB/src/optimise/train.jl:105
┌ Info: Epoch 554
└ @ Main /Users/kcft114/.julia/packages/Flux/qXNjB/src/optimise/train.jl:105
┌ Info: Epoch 555
└ @ Main /Users/kcft114/.julia/packages/Flux/qXNjB/src/optimise/train.jl:105
┌ Info: Epoch 556
└ @ Main /Users/kcft114/.julia/packages/Flux/qXNjB/src/optimise/train.jl:105
┌ Info: Epoch 557
└ @ Main /Users/kcft114/.julia/packages/Flux/qXNjB/src/optimise/train.jl:105
┌ Info: Epoch 558
└ @ Main /Users/kcft114/.julia/packages/Flux/qXNjB/src/optimise/train.jl:105
┌ Info: Epoch 559
└ @ Main /Users/kcft114/.julia/packages/Flux/qXNjB/src/optimise/train.jl:105
┌ Info: Epoch 560
└ @ Main /Users/kcft114/.julia/packages/Flux/qXNjB/src/optimise/train.jl:105
┌ Info: Epoch 561
└ @ Main /Users/kcft114/.julia/packages/Flux/qXNjB/src/optimise/train.jl:105
┌ Info: Epoch 562
└ @ Main /Users/kcft114/.julia/packages/Flux/qXNjB/src/optimise/train.jl:105
┌ Info: Epoch 563
└ @ Main /Users/kcft114/.julia/packages/Flux/qXNjB/src/optimise/train.jl:105
┌ Info: Epoch 564
└ @ Main /Users/kcft114/.julia/packages/Flux/qXNjB/src/optimise/train.jl:105
┌ Info: Epoch 565
└ @ Main /Users/kcft114/.julia/packages/Flux/qXNjB/src/optimise/train.jl:105
┌ Info: Epoch 566
└ @ Main /Users/kcft114/.julia/packages/Flux/qXNjB/src/optimise/train.jl:105
┌ Info: Epoch 567
└ @ Main /Users/kcft114/.julia/packages/Flux/qXNjB/src/optimise/train.jl:105
┌ Info: Epoch 568
└ @ Main /Users/kcft114/.julia/packages/Flux/qXNjB/src/optimise/train.jl:105
┌ Info: Epoch 569
└ @ Main /Users/kcft114/.julia/packages/Flux/qXNjB/src/optimise/train.jl:105
┌ Info: Epoch 570
└ @ Main /Users/kcft114/.julia/packages/Flux/qXNjB/src/optimise/train.jl:105
┌ Info: Epoch 571
└ @ Main /Users/kcft114/.julia/packages/Flux/qXNjB/src/optimise/train.jl:105
┌ Info: Epoch 572
└ @ Main /Users/kcft114/.julia/packages/Flux/qXNjB/src/optimise/train.jl:105
┌ Info: Epoch 573
└ @ Main /Users/kcft114/.julia/packages/Flux/qXNjB/src/optimise/train.jl:105
┌ Info: Epoch 574
└ @ Main /Users/kcft114/.julia/packages/Flux/qXNjB/src/optimise/train.jl:105
┌ Info: Epoch 575
└ @ Main /Users/kcft114/.julia/packages/Flux/qXNjB/src/optimise/train.jl:105
┌ Info: Epoch 576
└ @ Main /Users/kcft114/.julia/packages/Flux/qXNjB/src/optimise/train.jl:105
┌ Info: Epoch 577
└ @ Main /Users/kcft114/.julia/packages/Flux/qXNjB/src/optimise/train.jl:105
┌ Info: Epoch 578
└ @ Main /Users/kcft114/.julia/packages/Flux/qXNjB/src/optimise/train.jl:105
┌ Info: Epoch 579
└ @ Main /Users/kcft114/.julia/packages/Flux/qXNjB/src/optimise/train.jl:105
┌ Info: Epoch 580
└ @ Main /Users/kcft114/.julia/packages/Flux/qXNjB/src/optimise/train.jl:105
┌ Info: Epoch 581
└ @ Main /Users/kcft114/.julia/packages/Flux/qXNjB/src/optimise/train.jl:105
┌ Info: Epoch 582
└ @ Main /Users/kcft114/.julia/packages/Flux/qXNjB/src/optimise/train.jl:105
┌ Info: Epoch 583
└ @ Main /Users/kcft114/.julia/packages/Flux/qXNjB/src/optimise/train.jl:105
┌ Info: Epoch 584
└ @ Main /Users/kcft114/.julia/packages/Flux/qXNjB/src/optimise/train.jl:105
```

```
loss(X, Y) = 0.67404234f0 (tracked)
loss(X, Y) = 0.673715f0 (tracked)
loss(X, Y) = 0.67338437f0 (tracked)
loss(X, Y) = 0.6730552f0 (tracked)
loss(X, Y) = 0.6727269f0 (tracked)
loss(X, Y) = 0.67239404f0 (tracked)
loss(X, Y) = 0.6720646f0 (tracked)
loss(X, Y) = 0.67173696f0 (tracked)
loss(X, Y) = 0.67139965f0 (tracked)
loss(X, Y) = 0.67105603f0 (tracked)
loss(X, Y) = 0.67070675f0 (tracked)
loss(X, Y) = 0.6703519f0 (tracked)
loss(X, Y) = 0.66999906f0 (tracked)
loss(X, Y) = 0.66964096f0 (tracked)
loss(X, Y) = 0.6692774f0 (tracked)
loss(X, Y) = 0.6689077f0 (tracked)
loss(X, Y) = 0.6685388f0 (tracked)
loss(X, Y) = 0.6681692f0 (tracked)
loss(X, Y) = 0.6677949f0 (tracked)
loss(X, Y) = 0.6674183f0 (tracked)
loss(X, Y) = 0.6670421f0 (tracked)
loss(X, Y) = 0.6666635f0 (tracked)
loss(X, Y) = 0.6662888f0 (tracked)
loss(X, Y) = 0.6659125f0 (tracked)
loss(X, Y) = 0.66552997f0 (tracked)
loss(X, Y) = 0.6651446f0 (tracked)
loss(X, Y) = 0.6647648f0 (tracked)
loss(X, Y) = 0.66438746f0 (tracked)
loss(X, Y) = 0.664005f0 (tracked)
loss(X, Y) = 0.6636191f0 (tracked)
loss(X, Y) = 0.6632366f0 (tracked)
loss(X, Y) = 0.66285247f0 (tracked)
loss(X, Y) = 0.6624646f0 (tracked)
loss(X, Y) = 0.66207874f0 (tracked)
loss(X, Y) = 0.66169095f0 (tracked)
loss(X, Y) = 0.661303f0 (tracked)
loss(X, Y) = 0.660914f0 (tracked)
loss(X, Y) = 0.6605298f0 (tracked)
loss(X, Y) = 0.6601479f0 (tracked)
loss(X, Y) = 0.6597652f0 (tracked)
loss(X, Y) = 0.6593809f0 (tracked)
loss(X, Y) = 0.6589984f0 (tracked)
loss(X, Y) = 0.6586197f0 (tracked)
loss(X, Y) = 0.6582367f0 (tracked)
loss(X, Y) = 0.6578512f0 (tracked)
loss(X, Y) = 0.657468f0 (tracked)
loss(X, Y) = 0.65708625f0 (tracked)
loss(X, Y) = 0.6567032f0 (tracked)
loss(X, Y) = 0.65632504f0 (tracked)
loss(X, Y) = 0.6559497f0 (tracked)
loss(X, Y) = 0.6555723f0 (tracked)
loss(X, Y) = 0.6551911f0 (tracked)
loss(X, Y) = 0.65481144f0 (tracked)
loss(X, Y) = 0.65443164f0 (tracked)
loss(X, Y) = 0.65405387f0 (tracked)
loss(X, Y) = 0.653681f0 (tracked)
loss(X, Y) = 0.65330625f0 (tracked)
loss(X, Y) = 0.6529301f0 (tracked)
loss(X, Y) = 0.65255255f0 (tracked)
loss(X, Y) = 0.6521744f0 (tracked)
loss(X, Y) = 0.65180093f0 (tracked)
loss(X, Y) = 0.6514277f0 (tracked)
loss(X, Y) = 0.6510557f0 (tracked)
loss(X, Y) = 0.6506807f0 (tracked)
loss(X, Y) = 0.6503066f0 (tracked)
loss(X, Y) = 0.6499351f0 (tracked)
loss(X, Y) = 0.6495649f0 (tracked)
loss(X, Y) = 0.649196f0 (tracked)
loss(X, Y) = 0.64882624f0 (tracked)
loss(X, Y) = 0.64845634f0 (tracked)
loss(X, Y) = 0.6480864f0 (tracked)
loss(X, Y) = 0.6477168f0 (tracked)
loss(X, Y) = 0.64734995f0 (tracked)
loss(X, Y) = 0.64698136f0 (tracked)
loss(X, Y) = 0.64661753f0 (tracked)
loss(X, Y) = 0.64625263f0 (tracked)
loss(X, Y) = 0.64588773f0 (tracked)
loss(X, Y) = 0.64552057f0 (tracked)
loss(X, Y) = 0.645156f0 (tracked)
loss(X, Y) = 0.6447833f0 (tracked)
loss(X, Y) = 0.64441067f0 (tracked)
loss(X, Y) = 0.6440382f0 (tracked)
loss(X, Y) = 0.6436642f0 (tracked)
loss(X, Y) = 0.64328897f0 (tracked)
loss(X, Y) = 0.64291793f0 (tracked)
loss(X, Y) = 0.6425413f0 (tracked)
loss(X, Y) = 0.6421641f0 (tracked)
loss(X, Y) = 0.64179f0 (tracked)
loss(X, Y) = 0.64141f0 (tracked)
loss(X, Y) = 0.64104015f0 (tracked)
loss(X, Y) = 0.6406909f0 (tracked)
loss(X, Y) = 0.64033383f0 (tracked)
loss(X, Y) = 0.63997656f0 (tracked)
loss(X, Y) = 0.6396241f0 (tracked)
loss(X, Y) = 0.63927203f0 (tracked)
loss(X, Y) = 0.63892275f0 (tracked)
loss(X, Y) = 0.63857263f0 (tracked)
loss(X, Y) = 0.63822114f0 (tracked)
loss(X, Y) = 0.63786656f0 (tracked)
loss(X, Y) = 0.6375138f0 (tracked)
loss(X, Y) = 0.63716525f0 (tracked)
loss(X, Y) = 0.6368142f0 (tracked)
loss(X, Y) = 0.6364686f0 (tracked)
loss(X, Y) = 0.63612163f0 (tracked)
loss(X, Y) = 0.6357692f0 (tracked)
loss(X, Y) = 0.63541996f0 (tracked)
loss(X, Y) = 0.6350701f0 (tracked)
loss(X, Y) = 0.63472265f0 (tracked)
loss(X, Y) = 0.6343779f0 (tracked)
loss(X, Y) = 0.6340355f0 (tracked)
loss(X, Y) = 0.63368714f0 (tracked)
loss(X, Y) = 0.6333415f0 (tracked)
loss(X, Y) = 0.6329919f0 (tracked)
loss(X, Y) = 0.6326406f0 (tracked)
loss(X, Y) = 0.6322992f0 (tracked)
loss(X, Y) = 0.63195056f0 (tracked)
loss(X, Y) = 0.63160926f0 (tracked)
loss(X, Y) = 0.63126606f0 (tracked)
loss(X, Y) = 0.6309167f0 (tracked)
loss(X, Y) = 0.630564f0 (tracked)
loss(X, Y) = 0.63022166f0 (tracked)
loss(X, Y) = 0.62988394f0 (tracked)
loss(X, Y) = 0.62953484f0 (tracked)
loss(X, Y) = 0.6291874f0 (tracked)
loss(X, Y) = 0.6288522f0 (tracked)
loss(X, Y) = 0.6285125f0 (tracked)
loss(X, Y) = 0.62817353f0 (tracked)
loss(X, Y) = 0.62783766f0 (tracked)
loss(X, Y) = 0.6274967f0 (tracked)
loss(X, Y) = 0.6271644f0 (tracked)
loss(X, Y) = 0.62682796f0 (tracked)
loss(X, Y) = 0.62649167f0 (tracked)
loss(X, Y) = 0.6261518f0 (tracked)
loss(X, Y) = 0.6258111f0 (tracked)
loss(X, Y) = 0.62547344f0 (tracked)
loss(X, Y) = 0.6251317f0 (tracked)
```

```
┌ Info: Epoch 585
└ @ Main /Users/kcft114/.julia/packages/Flux/qXNjB/src/optimise/train.jl:105
┌ Info: Epoch 586
└ @ Main /Users/kcft114/.julia/packages/Flux/qXNjB/src/optimise/train.jl:105
┌ Info: Epoch 587
└ @ Main /Users/kcft114/.julia/packages/Flux/qXNjB/src/optimise/train.jl:105
┌ Info: Epoch 588
└ @ Main /Users/kcft114/.julia/packages/Flux/qXNjB/src/optimise/train.jl:105
┌ Info: Epoch 589
└ @ Main /Users/kcft114/.julia/packages/Flux/qXNjB/src/optimise/train.jl:105
┌ Info: Epoch 590
└ @ Main /Users/kcft114/.julia/packages/Flux/qXNjB/src/optimise/train.jl:105
┌ Info: Epoch 591
└ @ Main /Users/kcft114/.julia/packages/Flux/qXNjB/src/optimise/train.jl:105
┌ Info: Epoch 592
└ @ Main /Users/kcft114/.julia/packages/Flux/qXNjB/src/optimise/train.jl:105
┌ Info: Epoch 593
└ @ Main /Users/kcft114/.julia/packages/Flux/qXNjB/src/optimise/train.jl:105
┌ Info: Epoch 594
└ @ Main /Users/kcft114/.julia/packages/Flux/qXNjB/src/optimise/train.jl:105
┌ Info: Epoch 595
└ @ Main /Users/kcft114/.julia/packages/Flux/qXNjB/src/optimise/train.jl:105
┌ Info: Epoch 596
└ @ Main /Users/kcft114/.julia/packages/Flux/qXNjB/src/optimise/train.jl:105
┌ Info: Epoch 597
└ @ Main /Users/kcft114/.julia/packages/Flux/qXNjB/src/optimise/train.jl:105
┌ Info: Epoch 598
└ @ Main /Users/kcft114/.julia/packages/Flux/qXNjB/src/optimise/train.jl:105
┌ Info: Epoch 599
└ @ Main /Users/kcft114/.julia/packages/Flux/qXNjB/src/optimise/train.jl:105
┌ Info: Epoch 600
└ @ Main /Users/kcft114/.julia/packages/Flux/qXNjB/src/optimise/train.jl:105
┌ Info: Epoch 601
└ @ Main /Users/kcft114/.julia/packages/Flux/qXNjB/src/optimise/train.jl:105
┌ Info: Epoch 602
└ @ Main /Users/kcft114/.julia/packages/Flux/qXNjB/src/optimise/train.jl:105
┌ Info: Epoch 603
└ @ Main /Users/kcft114/.julia/packages/Flux/qXNjB/src/optimise/train.jl:105
┌ Info: Epoch 604
└ @ Main /Users/kcft114/.julia/packages/Flux/qXNjB/src/optimise/train.jl:105
┌ Info: Epoch 605
└ @ Main /Users/kcft114/.julia/packages/Flux/qXNjB/src/optimise/train.jl:105
┌ Info: Epoch 606
└ @ Main /Users/kcft114/.julia/packages/Flux/qXNjB/src/optimise/train.jl:105
┌ Info: Epoch 607
└ @ Main /Users/kcft114/.julia/packages/Flux/qXNjB/src/optimise/train.jl:105
┌ Info: Epoch 608
└ @ Main /Users/kcft114/.julia/packages/Flux/qXNjB/src/optimise/train.jl:105
┌ Info: Epoch 609
└ @ Main /Users/kcft114/.julia/packages/Flux/qXNjB/src/optimise/train.jl:105
┌ Info: Epoch 610
└ @ Main /Users/kcft114/.julia/packages/Flux/qXNjB/src/optimise/train.jl:105
┌ Info: Epoch 611
└ @ Main /Users/kcft114/.julia/packages/Flux/qXNjB/src/optimise/train.jl:105
┌ Info: Epoch 612
└ @ Main /Users/kcft114/.julia/packages/Flux/qXNjB/src/optimise/train.jl:105
┌ Info: Epoch 613
└ @ Main /Users/kcft114/.julia/packages/Flux/qXNjB/src/optimise/train.jl:105
┌ Info: Epoch 614
└ @ Main /Users/kcft114/.julia/packages/Flux/qXNjB/src/optimise/train.jl:105
┌ Info: Epoch 615
└ @ Main /Users/kcft114/.julia/packages/Flux/qXNjB/src/optimise/train.jl:105
┌ Info: Epoch 616
└ @ Main /Users/kcft114/.julia/packages/Flux/qXNjB/src/optimise/train.jl:105
┌ Info: Epoch 617
└ @ Main /Users/kcft114/.julia/packages/Flux/qXNjB/src/optimise/train.jl:105
┌ Info: Epoch 618
└ @ Main /Users/kcft114/.julia/packages/Flux/qXNjB/src/optimise/train.jl:105
┌ Info: Epoch 619
└ @ Main /Users/kcft114/.julia/packages/Flux/qXNjB/src/optimise/train.jl:105
┌ Info: Epoch 620
└ @ Main /Users/kcft114/.julia/packages/Flux/qXNjB/src/optimise/train.jl:105
┌ Info: Epoch 621
└ @ Main /Users/kcft114/.julia/packages/Flux/qXNjB/src/optimise/train.jl:105
┌ Info: Epoch 622
└ @ Main /Users/kcft114/.julia/packages/Flux/qXNjB/src/optimise/train.jl:105
┌ Info: Epoch 623
└ @ Main /Users/kcft114/.julia/packages/Flux/qXNjB/src/optimise/train.jl:105
┌ Info: Epoch 624
└ @ Main /Users/kcft114/.julia/packages/Flux/qXNjB/src/optimise/train.jl:105
┌ Info: Epoch 625
└ @ Main /Users/kcft114/.julia/packages/Flux/qXNjB/src/optimise/train.jl:105
┌ Info: Epoch 626
└ @ Main /Users/kcft114/.julia/packages/Flux/qXNjB/src/optimise/train.jl:105
┌ Info: Epoch 627
└ @ Main /Users/kcft114/.julia/packages/Flux/qXNjB/src/optimise/train.jl:105
┌ Info: Epoch 628
└ @ Main /Users/kcft114/.julia/packages/Flux/qXNjB/src/optimise/train.jl:105
┌ Info: Epoch 629
└ @ Main /Users/kcft114/.julia/packages/Flux/qXNjB/src/optimise/train.jl:105
┌ Info: Epoch 630
└ @ Main /Users/kcft114/.julia/packages/Flux/qXNjB/src/optimise/train.jl:105
┌ Info: Epoch 631
└ @ Main /Users/kcft114/.julia/packages/Flux/qXNjB/src/optimise/train.jl:105
┌ Info: Epoch 632
└ @ Main /Users/kcft114/.julia/packages/Flux/qXNjB/src/optimise/train.jl:105
┌ Info: Epoch 633
└ @ Main /Users/kcft114/.julia/packages/Flux/qXNjB/src/optimise/train.jl:105
┌ Info: Epoch 634
└ @ Main /Users/kcft114/.julia/packages/Flux/qXNjB/src/optimise/train.jl:105
┌ Info: Epoch 635
└ @ Main /Users/kcft114/.julia/packages/Flux/qXNjB/src/optimise/train.jl:105
┌ Info: Epoch 636
└ @ Main /Users/kcft114/.julia/packages/Flux/qXNjB/src/optimise/train.jl:105
┌ Info: Epoch 637
└ @ Main /Users/kcft114/.julia/packages/Flux/qXNjB/src/optimise/train.jl:105
┌ Info: Epoch 638
└ @ Main /Users/kcft114/.julia/packages/Flux/qXNjB/src/optimise/train.jl:105
┌ Info: Epoch 639
└ @ Main /Users/kcft114/.julia/packages/Flux/qXNjB/src/optimise/train.jl:105
┌ Info: Epoch 640
└ @ Main /Users/kcft114/.julia/packages/Flux/qXNjB/src/optimise/train.jl:105
┌ Info: Epoch 641
└ @ Main /Users/kcft114/.julia/packages/Flux/qXNjB/src/optimise/train.jl:105
┌ Info: Epoch 642
└ @ Main /Users/kcft114/.julia/packages/Flux/qXNjB/src/optimise/train.jl:105
┌ Info: Epoch 643
└ @ Main /Users/kcft114/.julia/packages/Flux/qXNjB/src/optimise/train.jl:105
┌ Info: Epoch 644
└ @ Main /Users/kcft114/.julia/packages/Flux/qXNjB/src/optimise/train.jl:105
┌ Info: Epoch 645
└ @ Main /Users/kcft114/.julia/packages/Flux/qXNjB/src/optimise/train.jl:105
┌ Info: Epoch 646
└ @ Main /Users/kcft114/.julia/packages/Flux/qXNjB/src/optimise/train.jl:105
┌ Info: Epoch 647
└ @ Main /Users/kcft114/.julia/packages/Flux/qXNjB/src/optimise/train.jl:105
┌ Info: Epoch 648
└ @ Main /Users/kcft114/.julia/packages/Flux/qXNjB/src/optimise/train.jl:105
┌ Info: Epoch 649
└ @ Main /Users/kcft114/.julia/packages/Flux/qXNjB/src/optimise/train.jl:105
┌ Info: Epoch 650
└ @ Main /Users/kcft114/.julia/packages/Flux/qXNjB/src/optimise/train.jl:105
┌ Info: Epoch 651
└ @ Main /Users/kcft114/.julia/packages/Flux/qXNjB/src/optimise/train.jl:105
┌ Info: Epoch 652
└ @ Main /Users/kcft114/.julia/packages/Flux/qXNjB/src/optimise/train.jl:105
┌ Info: Epoch 653
└ @ Main /Users/kcft114/.julia/packages/Flux/qXNjB/src/optimise/train.jl:105
┌ Info: Epoch 654
└ @ Main /Users/kcft114/.julia/packages/Flux/qXNjB/src/optimise/train.jl:105
┌ Info: Epoch 655
└ @ Main /Users/kcft114/.julia/packages/Flux/qXNjB/src/optimise/train.jl:105
┌ Info: Epoch 656
└ @ Main /Users/kcft114/.julia/packages/Flux/qXNjB/src/optimise/train.jl:105
┌ Info: Epoch 657
└ @ Main /Users/kcft114/.julia/packages/Flux/qXNjB/src/optimise/train.jl:105
┌ Info: Epoch 658
└ @ Main /Users/kcft114/.julia/packages/Flux/qXNjB/src/optimise/train.jl:105
┌ Info: Epoch 659
└ @ Main /Users/kcft114/.julia/packages/Flux/qXNjB/src/optimise/train.jl:105
┌ Info: Epoch 660
└ @ Main /Users/kcft114/.julia/packages/Flux/qXNjB/src/optimise/train.jl:105
┌ Info: Epoch 661
└ @ Main /Users/kcft114/.julia/packages/Flux/qXNjB/src/optimise/train.jl:105
┌ Info: Epoch 662
└ @ Main /Users/kcft114/.julia/packages/Flux/qXNjB/src/optimise/train.jl:105
┌ Info: Epoch 663
└ @ Main /Users/kcft114/.julia/packages/Flux/qXNjB/src/optimise/train.jl:105
┌ Info: Epoch 664
└ @ Main /Users/kcft114/.julia/packages/Flux/qXNjB/src/optimise/train.jl:105
┌ Info: Epoch 665
└ @ Main /Users/kcft114/.julia/packages/Flux/qXNjB/src/optimise/train.jl:105
┌ Info: Epoch 666
└ @ Main /Users/kcft114/.julia/packages/Flux/qXNjB/src/optimise/train.jl:105
┌ Info: Epoch 667
└ @ Main /Users/kcft114/.julia/packages/Flux/qXNjB/src/optimise/train.jl:105
┌ Info: Epoch 668
└ @ Main /Users/kcft114/.julia/packages/Flux/qXNjB/src/optimise/train.jl:105
┌ Info: Epoch 669
└ @ Main /Users/kcft114/.julia/packages/Flux/qXNjB/src/optimise/train.jl:105
┌ Info: Epoch 670
└ @ Main /Users/kcft114/.julia/packages/Flux/qXNjB/src/optimise/train.jl:105
┌ Info: Epoch 671
└ @ Main /Users/kcft114/.julia/packages/Flux/qXNjB/src/optimise/train.jl:105
┌ Info: Epoch 672
└ @ Main /Users/kcft114/.julia/packages/Flux/qXNjB/src/optimise/train.jl:105
┌ Info: Epoch 673
└ @ Main /Users/kcft114/.julia/packages/Flux/qXNjB/src/optimise/train.jl:105
┌ Info: Epoch 674
└ @ Main /Users/kcft114/.julia/packages/Flux/qXNjB/src/optimise/train.jl:105
┌ Info: Epoch 675
└ @ Main /Users/kcft114/.julia/packages/Flux/qXNjB/src/optimise/train.jl:105
┌ Info: Epoch 676
└ @ Main /Users/kcft114/.julia/packages/Flux/qXNjB/src/optimise/train.jl:105
┌ Info: Epoch 677
└ @ Main /Users/kcft114/.julia/packages/Flux/qXNjB/src/optimise/train.jl:105
┌ Info: Epoch 678
└ @ Main /Users/kcft114/.julia/packages/Flux/qXNjB/src/optimise/train.jl:105
┌ Info: Epoch 679
└ @ Main /Users/kcft114/.julia/packages/Flux/qXNjB/src/optimise/train.jl:105
┌ Info: Epoch 680
└ @ Main /Users/kcft114/.julia/packages/Flux/qXNjB/src/optimise/train.jl:105
┌ Info: Epoch 681
└ @ Main /Users/kcft114/.julia/packages/Flux/qXNjB/src/optimise/train.jl:105
┌ Info: Epoch 682
└ @ Main /Users/kcft114/.julia/packages/Flux/qXNjB/src/optimise/train.jl:105
┌ Info: Epoch 683
└ @ Main /Users/kcft114/.julia/packages/Flux/qXNjB/src/optimise/train.jl:105
┌ Info: Epoch 684
└ @ Main /Users/kcft114/.julia/packages/Flux/qXNjB/src/optimise/train.jl:105
┌ Info: Epoch 685
└ @ Main /Users/kcft114/.julia/packages/Flux/qXNjB/src/optimise/train.jl:105
┌ Info: Epoch 686
└ @ Main /Users/kcft114/.julia/packages/Flux/qXNjB/src/optimise/train.jl:105
┌ Info: Epoch 687
└ @ Main /Users/kcft114/.julia/packages/Flux/qXNjB/src/optimise/train.jl:105
┌ Info: Epoch 688
└ @ Main /Users/kcft114/.julia/packages/Flux/qXNjB/src/optimise/train.jl:105
┌ Info: Epoch 689
└ @ Main /Users/kcft114/.julia/packages/Flux/qXNjB/src/optimise/train.jl:105
┌ Info: Epoch 690
└ @ Main /Users/kcft114/.julia/packages/Flux/qXNjB/src/optimise/train.jl:105
┌ Info: Epoch 691
└ @ Main /Users/kcft114/.julia/packages/Flux/qXNjB/src/optimise/train.jl:105
┌ Info: Epoch 692
└ @ Main /Users/kcft114/.julia/packages/Flux/qXNjB/src/optimise/train.jl:105
┌ Info: Epoch 693
└ @ Main /Users/kcft114/.julia/packages/Flux/qXNjB/src/optimise/train.jl:105
┌ Info: Epoch 694
└ @ Main /Users/kcft114/.julia/packages/Flux/qXNjB/src/optimise/train.jl:105
┌ Info: Epoch 695
└ @ Main /Users/kcft114/.julia/packages/Flux/qXNjB/src/optimise/train.jl:105
┌ Info: Epoch 696
└ @ Main /Users/kcft114/.julia/packages/Flux/qXNjB/src/optimise/train.jl:105
┌ Info: Epoch 697
└ @ Main /Users/kcft114/.julia/packages/Flux/qXNjB/src/optimise/train.jl:105
┌ Info: Epoch 698
└ @ Main /Users/kcft114/.julia/packages/Flux/qXNjB/src/optimise/train.jl:105
┌ Info: Epoch 699
└ @ Main /Users/kcft114/.julia/packages/Flux/qXNjB/src/optimise/train.jl:105
┌ Info: Epoch 700
└ @ Main /Users/kcft114/.julia/packages/Flux/qXNjB/src/optimise/train.jl:105
┌ Info: Epoch 701
└ @ Main /Users/kcft114/.julia/packages/Flux/qXNjB/src/optimise/train.jl:105
┌ Info: Epoch 702
└ @ Main /Users/kcft114/.julia/packages/Flux/qXNjB/src/optimise/train.jl:105
┌ Info: Epoch 703
└ @ Main /Users/kcft114/.julia/packages/Flux/qXNjB/src/optimise/train.jl:105
┌ Info: Epoch 704
└ @ Main /Users/kcft114/.julia/packages/Flux/qXNjB/src/optimise/train.jl:105
┌ Info: Epoch 705
└ @ Main /Users/kcft114/.julia/packages/Flux/qXNjB/src/optimise/train.jl:105
┌ Info: Epoch 706
└ @ Main /Users/kcft114/.julia/packages/Flux/qXNjB/src/optimise/train.jl:105
┌ Info: Epoch 707
└ @ Main /Users/kcft114/.julia/packages/Flux/qXNjB/src/optimise/train.jl:105
┌ Info: Epoch 708
└ @ Main /Users/kcft114/.julia/packages/Flux/qXNjB/src/optimise/train.jl:105
┌ Info: Epoch 709
└ @ Main /Users/kcft114/.julia/packages/Flux/qXNjB/src/optimise/train.jl:105
┌ Info: Epoch 710
└ @ Main /Users/kcft114/.julia/packages/Flux/qXNjB/src/optimise/train.jl:105
┌ Info: Epoch 711
└ @ Main /Users/kcft114/.julia/packages/Flux/qXNjB/src/optimise/train.jl:105
┌ Info: Epoch 712
└ @ Main /Users/kcft114/.julia/packages/Flux/qXNjB/src/optimise/train.jl:105
┌ Info: Epoch 713
└ @ Main /Users/kcft114/.julia/packages/Flux/qXNjB/src/optimise/train.jl:105
┌ Info: Epoch 714
└ @ Main /Users/kcft114/.julia/packages/Flux/qXNjB/src/optimise/train.jl:105
┌ Info: Epoch 715
└ @ Main /Users/kcft114/.julia/packages/Flux/qXNjB/src/optimise/train.jl:105
┌ Info: Epoch 716
└ @ Main /Users/kcft114/.julia/packages/Flux/qXNjB/src/optimise/train.jl:105
┌ Info: Epoch 717
└ @ Main /Users/kcft114/.julia/packages/Flux/qXNjB/src/optimise/train.jl:105
┌ Info: Epoch 718
└ @ Main /Users/kcft114/.julia/packages/Flux/qXNjB/src/optimise/train.jl:105
┌ Info: Epoch 719
└ @ Main /Users/kcft114/.julia/packages/Flux/qXNjB/src/optimise/train.jl:105
┌ Info: Epoch 720
└ @ Main /Users/kcft114/.julia/packages/Flux/qXNjB/src/optimise/train.jl:105
```

```
loss(X, Y) = 0.6247904f0 (tracked)
loss(X, Y) = 0.6244503f0 (tracked)
loss(X, Y) = 0.62411374f0 (tracked)
loss(X, Y) = 0.62377083f0 (tracked)
loss(X, Y) = 0.6234291f0 (tracked)
loss(X, Y) = 0.62309235f0 (tracked)
loss(X, Y) = 0.6227445f0 (tracked)
loss(X, Y) = 0.62239164f0 (tracked)
loss(X, Y) = 0.6220399f0 (tracked)
loss(X, Y) = 0.6216875f0 (tracked)
loss(X, Y) = 0.6213336f0 (tracked)
loss(X, Y) = 0.62097627f0 (tracked)
loss(X, Y) = 0.6206128f0 (tracked)
loss(X, Y) = 0.6202542f0 (tracked)
loss(X, Y) = 0.6198971f0 (tracked)
loss(X, Y) = 0.61953926f0 (tracked)
loss(X, Y) = 0.6191721f0 (tracked)
loss(X, Y) = 0.6187992f0 (tracked)
loss(X, Y) = 0.6184326f0 (tracked)
loss(X, Y) = 0.6180606f0 (tracked)
loss(X, Y) = 0.6176859f0 (tracked)
loss(X, Y) = 0.617313f0 (tracked)
loss(X, Y) = 0.6169375f0 (tracked)
loss(X, Y) = 0.61656237f0 (tracked)
loss(X, Y) = 0.6161905f0 (tracked)
loss(X, Y) = 0.61582196f0 (tracked)
loss(X, Y) = 0.61546546f0 (tracked)
loss(X, Y) = 0.6151118f0 (tracked)
loss(X, Y) = 0.61476564f0 (tracked)
loss(X, Y) = 0.61442405f0 (tracked)
loss(X, Y) = 0.6140894f0 (tracked)
loss(X, Y) = 0.6137462f0 (tracked)
loss(X, Y) = 0.61340475f0 (tracked)
loss(X, Y) = 0.61306435f0 (tracked)
loss(X, Y) = 0.61272824f0 (tracked)
loss(X, Y) = 0.61240363f0 (tracked)
loss(X, Y) = 0.6120717f0 (tracked)
loss(X, Y) = 0.61174214f0 (tracked)
loss(X, Y) = 0.61141247f0 (tracked)
loss(X, Y) = 0.6110801f0 (tracked)
loss(X, Y) = 0.61074346f0 (tracked)
loss(X, Y) = 0.61040705f0 (tracked)
loss(X, Y) = 0.61007124f0 (tracked)
loss(X, Y) = 0.609743f0 (tracked)
loss(X, Y) = 0.6094086f0 (tracked)
loss(X, Y) = 0.6090669f0 (tracked)
loss(X, Y) = 0.6087318f0 (tracked)
loss(X, Y) = 0.6083899f0 (tracked)
loss(X, Y) = 0.6080531f0 (tracked)
loss(X, Y) = 0.60772073f0 (tracked)
loss(X, Y) = 0.60738206f0 (tracked)
loss(X, Y) = 0.6070394f0 (tracked)
loss(X, Y) = 0.6067051f0 (tracked)
loss(X, Y) = 0.6063713f0 (tracked)
loss(X, Y) = 0.6060391f0 (tracked)
loss(X, Y) = 0.60570484f0 (tracked)
loss(X, Y) = 0.6053818f0 (tracked)
loss(X, Y) = 0.60506445f0 (tracked)
loss(X, Y) = 0.60474175f0 (tracked)
loss(X, Y) = 0.604418f0 (tracked)
loss(X, Y) = 0.6040954f0 (tracked)
loss(X, Y) = 0.6037655f0 (tracked)
loss(X, Y) = 0.60343355f0 (tracked)
loss(X, Y) = 0.603107f0 (tracked)
loss(X, Y) = 0.6027838f0 (tracked)
loss(X, Y) = 0.6024623f0 (tracked)
loss(X, Y) = 0.60213506f0 (tracked)
loss(X, Y) = 0.6018217f0 (tracked)
loss(X, Y) = 0.60150266f0 (tracked)
loss(X, Y) = 0.60117614f0 (tracked)
loss(X, Y) = 0.60085046f0 (tracked)
loss(X, Y) = 0.60052496f0 (tracked)
loss(X, Y) = 0.60020465f0 (tracked)
loss(X, Y) = 0.59988296f0 (tracked)
loss(X, Y) = 0.59956086f0 (tracked)
loss(X, Y) = 0.5992465f0 (tracked)
loss(X, Y) = 0.5989346f0 (tracked)
loss(X, Y) = 0.5986103f0 (tracked)
loss(X, Y) = 0.5982859f0 (tracked)
loss(X, Y) = 0.59797287f0 (tracked)
loss(X, Y) = 0.59765714f0 (tracked)
loss(X, Y) = 0.59734577f0 (tracked)
loss(X, Y) = 0.59702617f0 (tracked)
loss(X, Y) = 0.5967088f0 (tracked)
loss(X, Y) = 0.5963957f0 (tracked)
loss(X, Y) = 0.59608454f0 (tracked)
loss(X, Y) = 0.5957723f0 (tracked)
loss(X, Y) = 0.595456f0 (tracked)
loss(X, Y) = 0.59513986f0 (tracked)
loss(X, Y) = 0.594825f0 (tracked)
loss(X, Y) = 0.594506f0 (tracked)
loss(X, Y) = 0.5941848f0 (tracked)
loss(X, Y) = 0.59387416f0 (tracked)
loss(X, Y) = 0.5935577f0 (tracked)
loss(X, Y) = 0.5932413f0 (tracked)
loss(X, Y) = 0.5929202f0 (tracked)
loss(X, Y) = 0.5926098f0 (tracked)
loss(X, Y) = 0.5922938f0 (tracked)
loss(X, Y) = 0.5919789f0 (tracked)
loss(X, Y) = 0.5916621f0 (tracked)
loss(X, Y) = 0.5913432f0 (tracked)
loss(X, Y) = 0.5910233f0 (tracked)
loss(X, Y) = 0.59070486f0 (tracked)
loss(X, Y) = 0.5903914f0 (tracked)
loss(X, Y) = 0.5900735f0 (tracked)
loss(X, Y) = 0.5897584f0 (tracked)
loss(X, Y) = 0.5894479f0 (tracked)
loss(X, Y) = 0.58913213f0 (tracked)
loss(X, Y) = 0.5888237f0 (tracked)
loss(X, Y) = 0.58851683f0 (tracked)
loss(X, Y) = 0.5882035f0 (tracked)
loss(X, Y) = 0.5878914f0 (tracked)
loss(X, Y) = 0.5875791f0 (tracked)
loss(X, Y) = 0.587264f0 (tracked)
loss(X, Y) = 0.5869563f0 (tracked)
loss(X, Y) = 0.5866441f0 (tracked)
loss(X, Y) = 0.58632755f0 (tracked)
loss(X, Y) = 0.58601284f0 (tracked)
loss(X, Y) = 0.5856971f0 (tracked)
loss(X, Y) = 0.5853855f0 (tracked)
loss(X, Y) = 0.5850772f0 (tracked)
loss(X, Y) = 0.5847648f0 (tracked)
loss(X, Y) = 0.5844517f0 (tracked)
loss(X, Y) = 0.5841427f0 (tracked)
loss(X, Y) = 0.5838319f0 (tracked)
loss(X, Y) = 0.58351433f0 (tracked)
loss(X, Y) = 0.5831949f0 (tracked)
loss(X, Y) = 0.5828785f0 (tracked)
loss(X, Y) = 0.58256984f0 (tracked)
loss(X, Y) = 0.58225095f0 (tracked)
loss(X, Y) = 0.5819375f0 (tracked)
loss(X, Y) = 0.58162194f0 (tracked)
loss(X, Y) = 0.58130604f0 (tracked)
```

```
┌ Info: Epoch 721
└ @ Main /Users/kcft114/.julia/packages/Flux/qXNjB/src/optimise/train.jl:105
┌ Info: Epoch 722
└ @ Main /Users/kcft114/.julia/packages/Flux/qXNjB/src/optimise/train.jl:105
┌ Info: Epoch 723
└ @ Main /Users/kcft114/.julia/packages/Flux/qXNjB/src/optimise/train.jl:105
┌ Info: Epoch 724
└ @ Main /Users/kcft114/.julia/packages/Flux/qXNjB/src/optimise/train.jl:105
┌ Info: Epoch 725
└ @ Main /Users/kcft114/.julia/packages/Flux/qXNjB/src/optimise/train.jl:105
┌ Info: Epoch 726
└ @ Main /Users/kcft114/.julia/packages/Flux/qXNjB/src/optimise/train.jl:105
┌ Info: Epoch 727
└ @ Main /Users/kcft114/.julia/packages/Flux/qXNjB/src/optimise/train.jl:105
┌ Info: Epoch 728
└ @ Main /Users/kcft114/.julia/packages/Flux/qXNjB/src/optimise/train.jl:105
┌ Info: Epoch 729
└ @ Main /Users/kcft114/.julia/packages/Flux/qXNjB/src/optimise/train.jl:105
┌ Info: Epoch 730
└ @ Main /Users/kcft114/.julia/packages/Flux/qXNjB/src/optimise/train.jl:105
┌ Info: Epoch 731
└ @ Main /Users/kcft114/.julia/packages/Flux/qXNjB/src/optimise/train.jl:105
┌ Info: Epoch 732
└ @ Main /Users/kcft114/.julia/packages/Flux/qXNjB/src/optimise/train.jl:105
┌ Info: Epoch 733
└ @ Main /Users/kcft114/.julia/packages/Flux/qXNjB/src/optimise/train.jl:105
┌ Info: Epoch 734
└ @ Main /Users/kcft114/.julia/packages/Flux/qXNjB/src/optimise/train.jl:105
┌ Info: Epoch 735
└ @ Main /Users/kcft114/.julia/packages/Flux/qXNjB/src/optimise/train.jl:105
┌ Info: Epoch 736
└ @ Main /Users/kcft114/.julia/packages/Flux/qXNjB/src/optimise/train.jl:105
┌ Info: Epoch 737
└ @ Main /Users/kcft114/.julia/packages/Flux/qXNjB/src/optimise/train.jl:105
┌ Info: Epoch 738
└ @ Main /Users/kcft114/.julia/packages/Flux/qXNjB/src/optimise/train.jl:105
┌ Info: Epoch 739
└ @ Main /Users/kcft114/.julia/packages/Flux/qXNjB/src/optimise/train.jl:105
┌ Info: Epoch 740
└ @ Main /Users/kcft114/.julia/packages/Flux/qXNjB/src/optimise/train.jl:105
┌ Info: Epoch 741
└ @ Main /Users/kcft114/.julia/packages/Flux/qXNjB/src/optimise/train.jl:105
┌ Info: Epoch 742
└ @ Main /Users/kcft114/.julia/packages/Flux/qXNjB/src/optimise/train.jl:105
┌ Info: Epoch 743
└ @ Main /Users/kcft114/.julia/packages/Flux/qXNjB/src/optimise/train.jl:105
┌ Info: Epoch 744
└ @ Main /Users/kcft114/.julia/packages/Flux/qXNjB/src/optimise/train.jl:105
┌ Info: Epoch 745
└ @ Main /Users/kcft114/.julia/packages/Flux/qXNjB/src/optimise/train.jl:105
┌ Info: Epoch 746
└ @ Main /Users/kcft114/.julia/packages/Flux/qXNjB/src/optimise/train.jl:105
┌ Info: Epoch 747
└ @ Main /Users/kcft114/.julia/packages/Flux/qXNjB/src/optimise/train.jl:105
┌ Info: Epoch 748
└ @ Main /Users/kcft114/.julia/packages/Flux/qXNjB/src/optimise/train.jl:105
┌ Info: Epoch 749
└ @ Main /Users/kcft114/.julia/packages/Flux/qXNjB/src/optimise/train.jl:105
┌ Info: Epoch 750
└ @ Main /Users/kcft114/.julia/packages/Flux/qXNjB/src/optimise/train.jl:105
┌ Info: Epoch 751
└ @ Main /Users/kcft114/.julia/packages/Flux/qXNjB/src/optimise/train.jl:105
┌ Info: Epoch 752
└ @ Main /Users/kcft114/.julia/packages/Flux/qXNjB/src/optimise/train.jl:105
┌ Info: Epoch 753
└ @ Main /Users/kcft114/.julia/packages/Flux/qXNjB/src/optimise/train.jl:105
┌ Info: Epoch 754
└ @ Main /Users/kcft114/.julia/packages/Flux/qXNjB/src/optimise/train.jl:105
┌ Info: Epoch 755
└ @ Main /Users/kcft114/.julia/packages/Flux/qXNjB/src/optimise/train.jl:105
┌ Info: Epoch 756
└ @ Main /Users/kcft114/.julia/packages/Flux/qXNjB/src/optimise/train.jl:105
┌ Info: Epoch 757
└ @ Main /Users/kcft114/.julia/packages/Flux/qXNjB/src/optimise/train.jl:105
┌ Info: Epoch 758
└ @ Main /Users/kcft114/.julia/packages/Flux/qXNjB/src/optimise/train.jl:105
┌ Info: Epoch 759
└ @ Main /Users/kcft114/.julia/packages/Flux/qXNjB/src/optimise/train.jl:105
┌ Info: Epoch 760
└ @ Main /Users/kcft114/.julia/packages/Flux/qXNjB/src/optimise/train.jl:105
┌ Info: Epoch 761
└ @ Main /Users/kcft114/.julia/packages/Flux/qXNjB/src/optimise/train.jl:105
┌ Info: Epoch 762
└ @ Main /Users/kcft114/.julia/packages/Flux/qXNjB/src/optimise/train.jl:105
┌ Info: Epoch 763
└ @ Main /Users/kcft114/.julia/packages/Flux/qXNjB/src/optimise/train.jl:105
┌ Info: Epoch 764
└ @ Main /Users/kcft114/.julia/packages/Flux/qXNjB/src/optimise/train.jl:105
┌ Info: Epoch 765
└ @ Main /Users/kcft114/.julia/packages/Flux/qXNjB/src/optimise/train.jl:105
┌ Info: Epoch 766
└ @ Main /Users/kcft114/.julia/packages/Flux/qXNjB/src/optimise/train.jl:105
┌ Info: Epoch 767
└ @ Main /Users/kcft114/.julia/packages/Flux/qXNjB/src/optimise/train.jl:105
┌ Info: Epoch 768
└ @ Main /Users/kcft114/.julia/packages/Flux/qXNjB/src/optimise/train.jl:105
┌ Info: Epoch 769
└ @ Main /Users/kcft114/.julia/packages/Flux/qXNjB/src/optimise/train.jl:105
┌ Info: Epoch 770
└ @ Main /Users/kcft114/.julia/packages/Flux/qXNjB/src/optimise/train.jl:105
┌ Info: Epoch 771
└ @ Main /Users/kcft114/.julia/packages/Flux/qXNjB/src/optimise/train.jl:105
┌ Info: Epoch 772
└ @ Main /Users/kcft114/.julia/packages/Flux/qXNjB/src/optimise/train.jl:105
┌ Info: Epoch 773
└ @ Main /Users/kcft114/.julia/packages/Flux/qXNjB/src/optimise/train.jl:105
┌ Info: Epoch 774
└ @ Main /Users/kcft114/.julia/packages/Flux/qXNjB/src/optimise/train.jl:105
┌ Info: Epoch 775
└ @ Main /Users/kcft114/.julia/packages/Flux/qXNjB/src/optimise/train.jl:105
┌ Info: Epoch 776
└ @ Main /Users/kcft114/.julia/packages/Flux/qXNjB/src/optimise/train.jl:105
┌ Info: Epoch 777
└ @ Main /Users/kcft114/.julia/packages/Flux/qXNjB/src/optimise/train.jl:105
┌ Info: Epoch 778
└ @ Main /Users/kcft114/.julia/packages/Flux/qXNjB/src/optimise/train.jl:105
┌ Info: Epoch 779
└ @ Main /Users/kcft114/.julia/packages/Flux/qXNjB/src/optimise/train.jl:105
┌ Info: Epoch 780
└ @ Main /Users/kcft114/.julia/packages/Flux/qXNjB/src/optimise/train.jl:105
┌ Info: Epoch 781
└ @ Main /Users/kcft114/.julia/packages/Flux/qXNjB/src/optimise/train.jl:105
┌ Info: Epoch 782
└ @ Main /Users/kcft114/.julia/packages/Flux/qXNjB/src/optimise/train.jl:105
┌ Info: Epoch 783
└ @ Main /Users/kcft114/.julia/packages/Flux/qXNjB/src/optimise/train.jl:105
┌ Info: Epoch 784
└ @ Main /Users/kcft114/.julia/packages/Flux/qXNjB/src/optimise/train.jl:105
┌ Info: Epoch 785
└ @ Main /Users/kcft114/.julia/packages/Flux/qXNjB/src/optimise/train.jl:105
┌ Info: Epoch 786
└ @ Main /Users/kcft114/.julia/packages/Flux/qXNjB/src/optimise/train.jl:105
┌ Info: Epoch 787
└ @ Main /Users/kcft114/.julia/packages/Flux/qXNjB/src/optimise/train.jl:105
┌ Info: Epoch 788
└ @ Main /Users/kcft114/.julia/packages/Flux/qXNjB/src/optimise/train.jl:105
┌ Info: Epoch 789
└ @ Main /Users/kcft114/.julia/packages/Flux/qXNjB/src/optimise/train.jl:105
┌ Info: Epoch 790
└ @ Main /Users/kcft114/.julia/packages/Flux/qXNjB/src/optimise/train.jl:105
┌ Info: Epoch 791
└ @ Main /Users/kcft114/.julia/packages/Flux/qXNjB/src/optimise/train.jl:105
┌ Info: Epoch 792
└ @ Main /Users/kcft114/.julia/packages/Flux/qXNjB/src/optimise/train.jl:105
┌ Info: Epoch 793
└ @ Main /Users/kcft114/.julia/packages/Flux/qXNjB/src/optimise/train.jl:105
┌ Info: Epoch 794
└ @ Main /Users/kcft114/.julia/packages/Flux/qXNjB/src/optimise/train.jl:105
┌ Info: Epoch 795
└ @ Main /Users/kcft114/.julia/packages/Flux/qXNjB/src/optimise/train.jl:105
┌ Info: Epoch 796
└ @ Main /Users/kcft114/.julia/packages/Flux/qXNjB/src/optimise/train.jl:105
┌ Info: Epoch 797
└ @ Main /Users/kcft114/.julia/packages/Flux/qXNjB/src/optimise/train.jl:105
┌ Info: Epoch 798
└ @ Main /Users/kcft114/.julia/packages/Flux/qXNjB/src/optimise/train.jl:105
┌ Info: Epoch 799
└ @ Main /Users/kcft114/.julia/packages/Flux/qXNjB/src/optimise/train.jl:105
┌ Info: Epoch 800
└ @ Main /Users/kcft114/.julia/packages/Flux/qXNjB/src/optimise/train.jl:105
┌ Info: Epoch 801
└ @ Main /Users/kcft114/.julia/packages/Flux/qXNjB/src/optimise/train.jl:105
┌ Info: Epoch 802
└ @ Main /Users/kcft114/.julia/packages/Flux/qXNjB/src/optimise/train.jl:105
┌ Info: Epoch 803
└ @ Main /Users/kcft114/.julia/packages/Flux/qXNjB/src/optimise/train.jl:105
┌ Info: Epoch 804
└ @ Main /Users/kcft114/.julia/packages/Flux/qXNjB/src/optimise/train.jl:105
┌ Info: Epoch 805
└ @ Main /Users/kcft114/.julia/packages/Flux/qXNjB/src/optimise/train.jl:105
┌ Info: Epoch 806
└ @ Main /Users/kcft114/.julia/packages/Flux/qXNjB/src/optimise/train.jl:105
┌ Info: Epoch 807
└ @ Main /Users/kcft114/.julia/packages/Flux/qXNjB/src/optimise/train.jl:105
┌ Info: Epoch 808
└ @ Main /Users/kcft114/.julia/packages/Flux/qXNjB/src/optimise/train.jl:105
┌ Info: Epoch 809
└ @ Main /Users/kcft114/.julia/packages/Flux/qXNjB/src/optimise/train.jl:105
┌ Info: Epoch 810
└ @ Main /Users/kcft114/.julia/packages/Flux/qXNjB/src/optimise/train.jl:105
┌ Info: Epoch 811
└ @ Main /Users/kcft114/.julia/packages/Flux/qXNjB/src/optimise/train.jl:105
┌ Info: Epoch 812
└ @ Main /Users/kcft114/.julia/packages/Flux/qXNjB/src/optimise/train.jl:105
┌ Info: Epoch 813
└ @ Main /Users/kcft114/.julia/packages/Flux/qXNjB/src/optimise/train.jl:105
┌ Info: Epoch 814
└ @ Main /Users/kcft114/.julia/packages/Flux/qXNjB/src/optimise/train.jl:105
┌ Info: Epoch 815
└ @ Main /Users/kcft114/.julia/packages/Flux/qXNjB/src/optimise/train.jl:105
┌ Info: Epoch 816
└ @ Main /Users/kcft114/.julia/packages/Flux/qXNjB/src/optimise/train.jl:105
┌ Info: Epoch 817
└ @ Main /Users/kcft114/.julia/packages/Flux/qXNjB/src/optimise/train.jl:105
┌ Info: Epoch 818
└ @ Main /Users/kcft114/.julia/packages/Flux/qXNjB/src/optimise/train.jl:105
┌ Info: Epoch 819
└ @ Main /Users/kcft114/.julia/packages/Flux/qXNjB/src/optimise/train.jl:105
┌ Info: Epoch 820
└ @ Main /Users/kcft114/.julia/packages/Flux/qXNjB/src/optimise/train.jl:105
┌ Info: Epoch 821
└ @ Main /Users/kcft114/.julia/packages/Flux/qXNjB/src/optimise/train.jl:105
┌ Info: Epoch 822
└ @ Main /Users/kcft114/.julia/packages/Flux/qXNjB/src/optimise/train.jl:105
┌ Info: Epoch 823
└ @ Main /Users/kcft114/.julia/packages/Flux/qXNjB/src/optimise/train.jl:105
┌ Info: Epoch 824
└ @ Main /Users/kcft114/.julia/packages/Flux/qXNjB/src/optimise/train.jl:105
┌ Info: Epoch 825
└ @ Main /Users/kcft114/.julia/packages/Flux/qXNjB/src/optimise/train.jl:105
┌ Info: Epoch 826
└ @ Main /Users/kcft114/.julia/packages/Flux/qXNjB/src/optimise/train.jl:105
┌ Info: Epoch 827
└ @ Main /Users/kcft114/.julia/packages/Flux/qXNjB/src/optimise/train.jl:105
┌ Info: Epoch 828
└ @ Main /Users/kcft114/.julia/packages/Flux/qXNjB/src/optimise/train.jl:105
┌ Info: Epoch 829
└ @ Main /Users/kcft114/.julia/packages/Flux/qXNjB/src/optimise/train.jl:105
┌ Info: Epoch 830
└ @ Main /Users/kcft114/.julia/packages/Flux/qXNjB/src/optimise/train.jl:105
┌ Info: Epoch 831
└ @ Main /Users/kcft114/.julia/packages/Flux/qXNjB/src/optimise/train.jl:105
┌ Info: Epoch 832
└ @ Main /Users/kcft114/.julia/packages/Flux/qXNjB/src/optimise/train.jl:105
┌ Info: Epoch 833
└ @ Main /Users/kcft114/.julia/packages/Flux/qXNjB/src/optimise/train.jl:105
┌ Info: Epoch 834
└ @ Main /Users/kcft114/.julia/packages/Flux/qXNjB/src/optimise/train.jl:105
┌ Info: Epoch 835
└ @ Main /Users/kcft114/.julia/packages/Flux/qXNjB/src/optimise/train.jl:105
┌ Info: Epoch 836
└ @ Main /Users/kcft114/.julia/packages/Flux/qXNjB/src/optimise/train.jl:105
┌ Info: Epoch 837
└ @ Main /Users/kcft114/.julia/packages/Flux/qXNjB/src/optimise/train.jl:105
┌ Info: Epoch 838
└ @ Main /Users/kcft114/.julia/packages/Flux/qXNjB/src/optimise/train.jl:105
┌ Info: Epoch 839
└ @ Main /Users/kcft114/.julia/packages/Flux/qXNjB/src/optimise/train.jl:105
┌ Info: Epoch 840
└ @ Main /Users/kcft114/.julia/packages/Flux/qXNjB/src/optimise/train.jl:105
┌ Info: Epoch 841
└ @ Main /Users/kcft114/.julia/packages/Flux/qXNjB/src/optimise/train.jl:105
┌ Info: Epoch 842
└ @ Main /Users/kcft114/.julia/packages/Flux/qXNjB/src/optimise/train.jl:105
┌ Info: Epoch 843
└ @ Main /Users/kcft114/.julia/packages/Flux/qXNjB/src/optimise/train.jl:105
┌ Info: Epoch 844
└ @ Main /Users/kcft114/.julia/packages/Flux/qXNjB/src/optimise/train.jl:105
┌ Info: Epoch 845
└ @ Main /Users/kcft114/.julia/packages/Flux/qXNjB/src/optimise/train.jl:105
┌ Info: Epoch 846
└ @ Main /Users/kcft114/.julia/packages/Flux/qXNjB/src/optimise/train.jl:105
┌ Info: Epoch 847
└ @ Main /Users/kcft114/.julia/packages/Flux/qXNjB/src/optimise/train.jl:105
┌ Info: Epoch 848
└ @ Main /Users/kcft114/.julia/packages/Flux/qXNjB/src/optimise/train.jl:105
┌ Info: Epoch 849
└ @ Main /Users/kcft114/.julia/packages/Flux/qXNjB/src/optimise/train.jl:105
┌ Info: Epoch 850
└ @ Main /Users/kcft114/.julia/packages/Flux/qXNjB/src/optimise/train.jl:105
┌ Info: Epoch 851
└ @ Main /Users/kcft114/.julia/packages/Flux/qXNjB/src/optimise/train.jl:105
┌ Info: Epoch 852
└ @ Main /Users/kcft114/.julia/packages/Flux/qXNjB/src/optimise/train.jl:105
┌ Info: Epoch 853
└ @ Main /Users/kcft114/.julia/packages/Flux/qXNjB/src/optimise/train.jl:105
```

```
loss(X, Y) = 0.58099216f0 (tracked)
loss(X, Y) = 0.58068323f0 (tracked)
loss(X, Y) = 0.5803638f0 (tracked)
loss(X, Y) = 0.5800488f0 (tracked)
loss(X, Y) = 0.5797308f0 (tracked)
loss(X, Y) = 0.5794106f0 (tracked)
loss(X, Y) = 0.5790813f0 (tracked)
loss(X, Y) = 0.57875794f0 (tracked)
loss(X, Y) = 0.57843894f0 (tracked)
loss(X, Y) = 0.57810944f0 (tracked)
loss(X, Y) = 0.577783f0 (tracked)
loss(X, Y) = 0.5774548f0 (tracked)
loss(X, Y) = 0.577139f0 (tracked)
loss(X, Y) = 0.57681555f0 (tracked)
loss(X, Y) = 0.57648945f0 (tracked)
loss(X, Y) = 0.57616115f0 (tracked)
loss(X, Y) = 0.5758229f0 (tracked)
loss(X, Y) = 0.57548815f0 (tracked)
loss(X, Y) = 0.5751619f0 (tracked)
loss(X, Y) = 0.5748305f0 (tracked)
loss(X, Y) = 0.57449824f0 (tracked)
loss(X, Y) = 0.5741632f0 (tracked)
loss(X, Y) = 0.5738306f0 (tracked)
loss(X, Y) = 0.573501f0 (tracked)
loss(X, Y) = 0.5731691f0 (tracked)
loss(X, Y) = 0.572838f0 (tracked)
loss(X, Y) = 0.5724992f0 (tracked)
loss(X, Y) = 0.57217f0 (tracked)
loss(X, Y) = 0.5718335f0 (tracked)
loss(X, Y) = 0.5714954f0 (tracked)
loss(X, Y) = 0.57115513f0 (tracked)
loss(X, Y) = 0.5708189f0 (tracked)
loss(X, Y) = 0.5704868f0 (tracked)
loss(X, Y) = 0.57014716f0 (tracked)
loss(X, Y) = 0.5698014f0 (tracked)
loss(X, Y) = 0.56947106f0 (tracked)
loss(X, Y) = 0.5691393f0 (tracked)
loss(X, Y) = 0.5687996f0 (tracked)
loss(X, Y) = 0.56845725f0 (tracked)
loss(X, Y) = 0.5681156f0 (tracked)
loss(X, Y) = 0.5677751f0 (tracked)
loss(X, Y) = 0.56743354f0 (tracked)
loss(X, Y) = 0.56708443f0 (tracked)
loss(X, Y) = 0.5667313f0 (tracked)
loss(X, Y) = 0.5663842f0 (tracked)
loss(X, Y) = 0.5660382f0 (tracked)
loss(X, Y) = 0.5656847f0 (tracked)
loss(X, Y) = 0.56533545f0 (tracked)
loss(X, Y) = 0.5649886f0 (tracked)
loss(X, Y) = 0.56463754f0 (tracked)
loss(X, Y) = 0.5642833f0 (tracked)
loss(X, Y) = 0.5639298f0 (tracked)
loss(X, Y) = 0.5635694f0 (tracked)
loss(X, Y) = 0.5632033f0 (tracked)
loss(X, Y) = 0.5628417f0 (tracked)
loss(X, Y) = 0.56248176f0 (tracked)
loss(X, Y) = 0.56211835f0 (tracked)
loss(X, Y) = 0.56175566f0 (tracked)
loss(X, Y) = 0.561389f0 (tracked)
loss(X, Y) = 0.5610234f0 (tracked)
loss(X, Y) = 0.5606555f0 (tracked)
loss(X, Y) = 0.56028426f0 (tracked)
loss(X, Y) = 0.5599045f0 (tracked)
loss(X, Y) = 0.5595216f0 (tracked)
loss(X, Y) = 0.5591349f0 (tracked)
loss(X, Y) = 0.5587395f0 (tracked)
loss(X, Y) = 0.5583438f0 (tracked)
loss(X, Y) = 0.557953f0 (tracked)
loss(X, Y) = 0.55756676f0 (tracked)
loss(X, Y) = 0.5571828f0 (tracked)
loss(X, Y) = 0.556797f0 (tracked)
loss(X, Y) = 0.5564204f0 (tracked)
loss(X, Y) = 0.556044f0 (tracked)
loss(X, Y) = 0.5556702f0 (tracked)
loss(X, Y) = 0.5552902f0 (tracked)
loss(X, Y) = 0.55490893f0 (tracked)
loss(X, Y) = 0.554542f0 (tracked)
loss(X, Y) = 0.5541831f0 (tracked)
loss(X, Y) = 0.5538249f0 (tracked)
loss(X, Y) = 0.55346256f0 (tracked)
loss(X, Y) = 0.55310094f0 (tracked)
loss(X, Y) = 0.55274093f0 (tracked)
loss(X, Y) = 0.552383f0 (tracked)
loss(X, Y) = 0.55202293f0 (tracked)
loss(X, Y) = 0.5516654f0 (tracked)
loss(X, Y) = 0.55131f0 (tracked)
loss(X, Y) = 0.5509514f0 (tracked)
loss(X, Y) = 0.55059046f0 (tracked)
loss(X, Y) = 0.55023086f0 (tracked)
loss(X, Y) = 0.54986846f0 (tracked)
loss(X, Y) = 0.5495157f0 (tracked)
loss(X, Y) = 0.5491597f0 (tracked)
loss(X, Y) = 0.5488101f0 (tracked)
loss(X, Y) = 0.5484672f0 (tracked)
loss(X, Y) = 0.54812574f0 (tracked)
loss(X, Y) = 0.54778445f0 (tracked)
loss(X, Y) = 0.54744154f0 (tracked)
loss(X, Y) = 0.5471001f0 (tracked)
loss(X, Y) = 0.54675746f0 (tracked)
loss(X, Y) = 0.5464147f0 (tracked)
loss(X, Y) = 0.54606533f0 (tracked)
loss(X, Y) = 0.5457154f0 (tracked)
loss(X, Y) = 0.54537123f0 (tracked)
loss(X, Y) = 0.5450119f0 (tracked)
loss(X, Y) = 0.5446485f0 (tracked)
loss(X, Y) = 0.54429066f0 (tracked)
loss(X, Y) = 0.5439329f0 (tracked)
loss(X, Y) = 0.54357964f0 (tracked)
loss(X, Y) = 0.5432162f0 (tracked)
loss(X, Y) = 0.54285693f0 (tracked)
loss(X, Y) = 0.5424918f0 (tracked)
loss(X, Y) = 0.54212815f0 (tracked)
loss(X, Y) = 0.5417674f0 (tracked)
loss(X, Y) = 0.54140884f0 (tracked)
loss(X, Y) = 0.5410465f0 (tracked)
loss(X, Y) = 0.5406781f0 (tracked)
loss(X, Y) = 0.5403085f0 (tracked)
loss(X, Y) = 0.539946f0 (tracked)
loss(X, Y) = 0.53957915f0 (tracked)
loss(X, Y) = 0.53921235f0 (tracked)
loss(X, Y) = 0.53883827f0 (tracked)
loss(X, Y) = 0.5384782f0 (tracked)
loss(X, Y) = 0.5381223f0 (tracked)
loss(X, Y) = 0.5377669f0 (tracked)
loss(X, Y) = 0.53740835f0 (tracked)
loss(X, Y) = 0.5370472f0 (tracked)
loss(X, Y) = 0.5366843f0 (tracked)
loss(X, Y) = 0.5363152f0 (tracked)
loss(X, Y) = 0.5359467f0 (tracked)
loss(X, Y) = 0.5355804f0 (tracked)
loss(X, Y) = 0.5352151f0 (tracked)
```

```
┌ Info: Epoch 854
└ @ Main /Users/kcft114/.julia/packages/Flux/qXNjB/src/optimise/train.jl:105
┌ Info: Epoch 855
└ @ Main /Users/kcft114/.julia/packages/Flux/qXNjB/src/optimise/train.jl:105
┌ Info: Epoch 856
└ @ Main /Users/kcft114/.julia/packages/Flux/qXNjB/src/optimise/train.jl:105
┌ Info: Epoch 857
└ @ Main /Users/kcft114/.julia/packages/Flux/qXNjB/src/optimise/train.jl:105
┌ Info: Epoch 858
└ @ Main /Users/kcft114/.julia/packages/Flux/qXNjB/src/optimise/train.jl:105
┌ Info: Epoch 859
└ @ Main /Users/kcft114/.julia/packages/Flux/qXNjB/src/optimise/train.jl:105
┌ Info: Epoch 860
└ @ Main /Users/kcft114/.julia/packages/Flux/qXNjB/src/optimise/train.jl:105
┌ Info: Epoch 861
└ @ Main /Users/kcft114/.julia/packages/Flux/qXNjB/src/optimise/train.jl:105
┌ Info: Epoch 862
└ @ Main /Users/kcft114/.julia/packages/Flux/qXNjB/src/optimise/train.jl:105
┌ Info: Epoch 863
└ @ Main /Users/kcft114/.julia/packages/Flux/qXNjB/src/optimise/train.jl:105
┌ Info: Epoch 864
└ @ Main /Users/kcft114/.julia/packages/Flux/qXNjB/src/optimise/train.jl:105
┌ Info: Epoch 865
└ @ Main /Users/kcft114/.julia/packages/Flux/qXNjB/src/optimise/train.jl:105
┌ Info: Epoch 866
└ @ Main /Users/kcft114/.julia/packages/Flux/qXNjB/src/optimise/train.jl:105
┌ Info: Epoch 867
└ @ Main /Users/kcft114/.julia/packages/Flux/qXNjB/src/optimise/train.jl:105
┌ Info: Epoch 868
└ @ Main /Users/kcft114/.julia/packages/Flux/qXNjB/src/optimise/train.jl:105
┌ Info: Epoch 869
└ @ Main /Users/kcft114/.julia/packages/Flux/qXNjB/src/optimise/train.jl:105
┌ Info: Epoch 870
└ @ Main /Users/kcft114/.julia/packages/Flux/qXNjB/src/optimise/train.jl:105
┌ Info: Epoch 871
└ @ Main /Users/kcft114/.julia/packages/Flux/qXNjB/src/optimise/train.jl:105
┌ Info: Epoch 872
└ @ Main /Users/kcft114/.julia/packages/Flux/qXNjB/src/optimise/train.jl:105
┌ Info: Epoch 873
└ @ Main /Users/kcft114/.julia/packages/Flux/qXNjB/src/optimise/train.jl:105
┌ Info: Epoch 874
└ @ Main /Users/kcft114/.julia/packages/Flux/qXNjB/src/optimise/train.jl:105
┌ Info: Epoch 875
└ @ Main /Users/kcft114/.julia/packages/Flux/qXNjB/src/optimise/train.jl:105
┌ Info: Epoch 876
└ @ Main /Users/kcft114/.julia/packages/Flux/qXNjB/src/optimise/train.jl:105
┌ Info: Epoch 877
└ @ Main /Users/kcft114/.julia/packages/Flux/qXNjB/src/optimise/train.jl:105
┌ Info: Epoch 878
└ @ Main /Users/kcft114/.julia/packages/Flux/qXNjB/src/optimise/train.jl:105
┌ Info: Epoch 879
└ @ Main /Users/kcft114/.julia/packages/Flux/qXNjB/src/optimise/train.jl:105
┌ Info: Epoch 880
└ @ Main /Users/kcft114/.julia/packages/Flux/qXNjB/src/optimise/train.jl:105
┌ Info: Epoch 881
└ @ Main /Users/kcft114/.julia/packages/Flux/qXNjB/src/optimise/train.jl:105
┌ Info: Epoch 882
└ @ Main /Users/kcft114/.julia/packages/Flux/qXNjB/src/optimise/train.jl:105
┌ Info: Epoch 883
└ @ Main /Users/kcft114/.julia/packages/Flux/qXNjB/src/optimise/train.jl:105
┌ Info: Epoch 884
└ @ Main /Users/kcft114/.julia/packages/Flux/qXNjB/src/optimise/train.jl:105
┌ Info: Epoch 885
└ @ Main /Users/kcft114/.julia/packages/Flux/qXNjB/src/optimise/train.jl:105
┌ Info: Epoch 886
└ @ Main /Users/kcft114/.julia/packages/Flux/qXNjB/src/optimise/train.jl:105
┌ Info: Epoch 887
└ @ Main /Users/kcft114/.julia/packages/Flux/qXNjB/src/optimise/train.jl:105
┌ Info: Epoch 888
└ @ Main /Users/kcft114/.julia/packages/Flux/qXNjB/src/optimise/train.jl:105
┌ Info: Epoch 889
└ @ Main /Users/kcft114/.julia/packages/Flux/qXNjB/src/optimise/train.jl:105
┌ Info: Epoch 890
└ @ Main /Users/kcft114/.julia/packages/Flux/qXNjB/src/optimise/train.jl:105
┌ Info: Epoch 891
└ @ Main /Users/kcft114/.julia/packages/Flux/qXNjB/src/optimise/train.jl:105
┌ Info: Epoch 892
└ @ Main /Users/kcft114/.julia/packages/Flux/qXNjB/src/optimise/train.jl:105
┌ Info: Epoch 893
└ @ Main /Users/kcft114/.julia/packages/Flux/qXNjB/src/optimise/train.jl:105
┌ Info: Epoch 894
└ @ Main /Users/kcft114/.julia/packages/Flux/qXNjB/src/optimise/train.jl:105
┌ Info: Epoch 895
└ @ Main /Users/kcft114/.julia/packages/Flux/qXNjB/src/optimise/train.jl:105
┌ Info: Epoch 896
└ @ Main /Users/kcft114/.julia/packages/Flux/qXNjB/src/optimise/train.jl:105
┌ Info: Epoch 897
└ @ Main /Users/kcft114/.julia/packages/Flux/qXNjB/src/optimise/train.jl:105
┌ Info: Epoch 898
└ @ Main /Users/kcft114/.julia/packages/Flux/qXNjB/src/optimise/train.jl:105
┌ Info: Epoch 899
└ @ Main /Users/kcft114/.julia/packages/Flux/qXNjB/src/optimise/train.jl:105
┌ Info: Epoch 900
└ @ Main /Users/kcft114/.julia/packages/Flux/qXNjB/src/optimise/train.jl:105
┌ Info: Epoch 901
└ @ Main /Users/kcft114/.julia/packages/Flux/qXNjB/src/optimise/train.jl:105
┌ Info: Epoch 902
└ @ Main /Users/kcft114/.julia/packages/Flux/qXNjB/src/optimise/train.jl:105
┌ Info: Epoch 903
└ @ Main /Users/kcft114/.julia/packages/Flux/qXNjB/src/optimise/train.jl:105
┌ Info: Epoch 904
└ @ Main /Users/kcft114/.julia/packages/Flux/qXNjB/src/optimise/train.jl:105
┌ Info: Epoch 905
└ @ Main /Users/kcft114/.julia/packages/Flux/qXNjB/src/optimise/train.jl:105
┌ Info: Epoch 906
└ @ Main /Users/kcft114/.julia/packages/Flux/qXNjB/src/optimise/train.jl:105
┌ Info: Epoch 907
└ @ Main /Users/kcft114/.julia/packages/Flux/qXNjB/src/optimise/train.jl:105
┌ Info: Epoch 908
└ @ Main /Users/kcft114/.julia/packages/Flux/qXNjB/src/optimise/train.jl:105
┌ Info: Epoch 909
└ @ Main /Users/kcft114/.julia/packages/Flux/qXNjB/src/optimise/train.jl:105
┌ Info: Epoch 910
└ @ Main /Users/kcft114/.julia/packages/Flux/qXNjB/src/optimise/train.jl:105
┌ Info: Epoch 911
└ @ Main /Users/kcft114/.julia/packages/Flux/qXNjB/src/optimise/train.jl:105
┌ Info: Epoch 912
└ @ Main /Users/kcft114/.julia/packages/Flux/qXNjB/src/optimise/train.jl:105
┌ Info: Epoch 913
└ @ Main /Users/kcft114/.julia/packages/Flux/qXNjB/src/optimise/train.jl:105
┌ Info: Epoch 914
└ @ Main /Users/kcft114/.julia/packages/Flux/qXNjB/src/optimise/train.jl:105
┌ Info: Epoch 915
└ @ Main /Users/kcft114/.julia/packages/Flux/qXNjB/src/optimise/train.jl:105
┌ Info: Epoch 916
└ @ Main /Users/kcft114/.julia/packages/Flux/qXNjB/src/optimise/train.jl:105
┌ Info: Epoch 917
└ @ Main /Users/kcft114/.julia/packages/Flux/qXNjB/src/optimise/train.jl:105
┌ Info: Epoch 918
└ @ Main /Users/kcft114/.julia/packages/Flux/qXNjB/src/optimise/train.jl:105
┌ Info: Epoch 919
└ @ Main /Users/kcft114/.julia/packages/Flux/qXNjB/src/optimise/train.jl:105
┌ Info: Epoch 920
└ @ Main /Users/kcft114/.julia/packages/Flux/qXNjB/src/optimise/train.jl:105
┌ Info: Epoch 921
└ @ Main /Users/kcft114/.julia/packages/Flux/qXNjB/src/optimise/train.jl:105
┌ Info: Epoch 922
└ @ Main /Users/kcft114/.julia/packages/Flux/qXNjB/src/optimise/train.jl:105
┌ Info: Epoch 923
└ @ Main /Users/kcft114/.julia/packages/Flux/qXNjB/src/optimise/train.jl:105
┌ Info: Epoch 924
└ @ Main /Users/kcft114/.julia/packages/Flux/qXNjB/src/optimise/train.jl:105
┌ Info: Epoch 925
└ @ Main /Users/kcft114/.julia/packages/Flux/qXNjB/src/optimise/train.jl:105
┌ Info: Epoch 926
└ @ Main /Users/kcft114/.julia/packages/Flux/qXNjB/src/optimise/train.jl:105
┌ Info: Epoch 927
└ @ Main /Users/kcft114/.julia/packages/Flux/qXNjB/src/optimise/train.jl:105
┌ Info: Epoch 928
└ @ Main /Users/kcft114/.julia/packages/Flux/qXNjB/src/optimise/train.jl:105
┌ Info: Epoch 929
└ @ Main /Users/kcft114/.julia/packages/Flux/qXNjB/src/optimise/train.jl:105
┌ Info: Epoch 930
└ @ Main /Users/kcft114/.julia/packages/Flux/qXNjB/src/optimise/train.jl:105
┌ Info: Epoch 931
└ @ Main /Users/kcft114/.julia/packages/Flux/qXNjB/src/optimise/train.jl:105
┌ Info: Epoch 932
└ @ Main /Users/kcft114/.julia/packages/Flux/qXNjB/src/optimise/train.jl:105
┌ Info: Epoch 933
└ @ Main /Users/kcft114/.julia/packages/Flux/qXNjB/src/optimise/train.jl:105
┌ Info: Epoch 934
└ @ Main /Users/kcft114/.julia/packages/Flux/qXNjB/src/optimise/train.jl:105
┌ Info: Epoch 935
└ @ Main /Users/kcft114/.julia/packages/Flux/qXNjB/src/optimise/train.jl:105
┌ Info: Epoch 936
└ @ Main /Users/kcft114/.julia/packages/Flux/qXNjB/src/optimise/train.jl:105
┌ Info: Epoch 937
└ @ Main /Users/kcft114/.julia/packages/Flux/qXNjB/src/optimise/train.jl:105
┌ Info: Epoch 938
└ @ Main /Users/kcft114/.julia/packages/Flux/qXNjB/src/optimise/train.jl:105
┌ Info: Epoch 939
└ @ Main /Users/kcft114/.julia/packages/Flux/qXNjB/src/optimise/train.jl:105
┌ Info: Epoch 940
└ @ Main /Users/kcft114/.julia/packages/Flux/qXNjB/src/optimise/train.jl:105
┌ Info: Epoch 941
└ @ Main /Users/kcft114/.julia/packages/Flux/qXNjB/src/optimise/train.jl:105
┌ Info: Epoch 942
└ @ Main /Users/kcft114/.julia/packages/Flux/qXNjB/src/optimise/train.jl:105
┌ Info: Epoch 943
└ @ Main /Users/kcft114/.julia/packages/Flux/qXNjB/src/optimise/train.jl:105
┌ Info: Epoch 944
└ @ Main /Users/kcft114/.julia/packages/Flux/qXNjB/src/optimise/train.jl:105
┌ Info: Epoch 945
└ @ Main /Users/kcft114/.julia/packages/Flux/qXNjB/src/optimise/train.jl:105
┌ Info: Epoch 946
└ @ Main /Users/kcft114/.julia/packages/Flux/qXNjB/src/optimise/train.jl:105
┌ Info: Epoch 947
└ @ Main /Users/kcft114/.julia/packages/Flux/qXNjB/src/optimise/train.jl:105
┌ Info: Epoch 948
└ @ Main /Users/kcft114/.julia/packages/Flux/qXNjB/src/optimise/train.jl:105
┌ Info: Epoch 949
└ @ Main /Users/kcft114/.julia/packages/Flux/qXNjB/src/optimise/train.jl:105
┌ Info: Epoch 950
└ @ Main /Users/kcft114/.julia/packages/Flux/qXNjB/src/optimise/train.jl:105
┌ Info: Epoch 951
└ @ Main /Users/kcft114/.julia/packages/Flux/qXNjB/src/optimise/train.jl:105
┌ Info: Epoch 952
└ @ Main /Users/kcft114/.julia/packages/Flux/qXNjB/src/optimise/train.jl:105
┌ Info: Epoch 953
└ @ Main /Users/kcft114/.julia/packages/Flux/qXNjB/src/optimise/train.jl:105
┌ Info: Epoch 954
└ @ Main /Users/kcft114/.julia/packages/Flux/qXNjB/src/optimise/train.jl:105
┌ Info: Epoch 955
└ @ Main /Users/kcft114/.julia/packages/Flux/qXNjB/src/optimise/train.jl:105
┌ Info: Epoch 956
└ @ Main /Users/kcft114/.julia/packages/Flux/qXNjB/src/optimise/train.jl:105
┌ Info: Epoch 957
└ @ Main /Users/kcft114/.julia/packages/Flux/qXNjB/src/optimise/train.jl:105
┌ Info: Epoch 958
└ @ Main /Users/kcft114/.julia/packages/Flux/qXNjB/src/optimise/train.jl:105
┌ Info: Epoch 959
└ @ Main /Users/kcft114/.julia/packages/Flux/qXNjB/src/optimise/train.jl:105
┌ Info: Epoch 960
└ @ Main /Users/kcft114/.julia/packages/Flux/qXNjB/src/optimise/train.jl:105
┌ Info: Epoch 961
└ @ Main /Users/kcft114/.julia/packages/Flux/qXNjB/src/optimise/train.jl:105
┌ Info: Epoch 962
└ @ Main /Users/kcft114/.julia/packages/Flux/qXNjB/src/optimise/train.jl:105
┌ Info: Epoch 963
└ @ Main /Users/kcft114/.julia/packages/Flux/qXNjB/src/optimise/train.jl:105
┌ Info: Epoch 964
└ @ Main /Users/kcft114/.julia/packages/Flux/qXNjB/src/optimise/train.jl:105
┌ Info: Epoch 965
└ @ Main /Users/kcft114/.julia/packages/Flux/qXNjB/src/optimise/train.jl:105
┌ Info: Epoch 966
└ @ Main /Users/kcft114/.julia/packages/Flux/qXNjB/src/optimise/train.jl:105
┌ Info: Epoch 967
└ @ Main /Users/kcft114/.julia/packages/Flux/qXNjB/src/optimise/train.jl:105
┌ Info: Epoch 968
└ @ Main /Users/kcft114/.julia/packages/Flux/qXNjB/src/optimise/train.jl:105
┌ Info: Epoch 969
└ @ Main /Users/kcft114/.julia/packages/Flux/qXNjB/src/optimise/train.jl:105
┌ Info: Epoch 970
└ @ Main /Users/kcft114/.julia/packages/Flux/qXNjB/src/optimise/train.jl:105
┌ Info: Epoch 971
└ @ Main /Users/kcft114/.julia/packages/Flux/qXNjB/src/optimise/train.jl:105
┌ Info: Epoch 972
└ @ Main /Users/kcft114/.julia/packages/Flux/qXNjB/src/optimise/train.jl:105
┌ Info: Epoch 973
└ @ Main /Users/kcft114/.julia/packages/Flux/qXNjB/src/optimise/train.jl:105
┌ Info: Epoch 974
└ @ Main /Users/kcft114/.julia/packages/Flux/qXNjB/src/optimise/train.jl:105
┌ Info: Epoch 975
└ @ Main /Users/kcft114/.julia/packages/Flux/qXNjB/src/optimise/train.jl:105
┌ Info: Epoch 976
└ @ Main /Users/kcft114/.julia/packages/Flux/qXNjB/src/optimise/train.jl:105
┌ Info: Epoch 977
└ @ Main /Users/kcft114/.julia/packages/Flux/qXNjB/src/optimise/train.jl:105
┌ Info: Epoch 978
└ @ Main /Users/kcft114/.julia/packages/Flux/qXNjB/src/optimise/train.jl:105
┌ Info: Epoch 979
└ @ Main /Users/kcft114/.julia/packages/Flux/qXNjB/src/optimise/train.jl:105
┌ Info: Epoch 980
└ @ Main /Users/kcft114/.julia/packages/Flux/qXNjB/src/optimise/train.jl:105
┌ Info: Epoch 981
└ @ Main /Users/kcft114/.julia/packages/Flux/qXNjB/src/optimise/train.jl:105
┌ Info: Epoch 982
└ @ Main /Users/kcft114/.julia/packages/Flux/qXNjB/src/optimise/train.jl:105
┌ Info: Epoch 983
└ @ Main /Users/kcft114/.julia/packages/Flux/qXNjB/src/optimise/train.jl:105
┌ Info: Epoch 984
└ @ Main /Users/kcft114/.julia/packages/Flux/qXNjB/src/optimise/train.jl:105
```

```
loss(X, Y) = 0.5348516f0 (tracked)
loss(X, Y) = 0.5344863f0 (tracked)
loss(X, Y) = 0.5341188f0 (tracked)
loss(X, Y) = 0.53375125f0 (tracked)
loss(X, Y) = 0.5333801f0 (tracked)
loss(X, Y) = 0.5330093f0 (tracked)
loss(X, Y) = 0.5326403f0 (tracked)
loss(X, Y) = 0.53227514f0 (tracked)
loss(X, Y) = 0.53190494f0 (tracked)
loss(X, Y) = 0.53154117f0 (tracked)
loss(X, Y) = 0.53117603f0 (tracked)
loss(X, Y) = 0.53080416f0 (tracked)
loss(X, Y) = 0.5304354f0 (tracked)
loss(X, Y) = 0.5300566f0 (tracked)
loss(X, Y) = 0.5296914f0 (tracked)
loss(X, Y) = 0.52932334f0 (tracked)
loss(X, Y) = 0.5289515f0 (tracked)
loss(X, Y) = 0.5285836f0 (tracked)
loss(X, Y) = 0.5282197f0 (tracked)
loss(X, Y) = 0.52785116f0 (tracked)
loss(X, Y) = 0.52748656f0 (tracked)
loss(X, Y) = 0.52712107f0 (tracked)
loss(X, Y) = 0.5267552f0 (tracked)
loss(X, Y) = 0.52639717f0 (tracked)
loss(X, Y) = 0.526035f0 (tracked)
loss(X, Y) = 0.52567345f0 (tracked)
loss(X, Y) = 0.52531123f0 (tracked)
loss(X, Y) = 0.5249506f0 (tracked)
loss(X, Y) = 0.5245986f0 (tracked)
loss(X, Y) = 0.52424455f0 (tracked)
loss(X, Y) = 0.5238915f0 (tracked)
loss(X, Y) = 0.52354324f0 (tracked)
loss(X, Y) = 0.5231947f0 (tracked)
loss(X, Y) = 0.52283955f0 (tracked)
loss(X, Y) = 0.52249485f0 (tracked)
loss(X, Y) = 0.52215064f0 (tracked)
loss(X, Y) = 0.5217951f0 (tracked)
loss(X, Y) = 0.52144766f0 (tracked)
loss(X, Y) = 0.52109486f0 (tracked)
loss(X, Y) = 0.52074873f0 (tracked)
loss(X, Y) = 0.5204017f0 (tracked)
loss(X, Y) = 0.52004814f0 (tracked)
loss(X, Y) = 0.5196962f0 (tracked)
loss(X, Y) = 0.5193481f0 (tracked)
loss(X, Y) = 0.518998f0 (tracked)
loss(X, Y) = 0.518654f0 (tracked)
loss(X, Y) = 0.51830953f0 (tracked)
loss(X, Y) = 0.517957f0 (tracked)
loss(X, Y) = 0.51760846f0 (tracked)
loss(X, Y) = 0.5172606f0 (tracked)
loss(X, Y) = 0.51691073f0 (tracked)
loss(X, Y) = 0.5165614f0 (tracked)
loss(X, Y) = 0.5162063f0 (tracked)
loss(X, Y) = 0.51585156f0 (tracked)
loss(X, Y) = 0.51549196f0 (tracked)
loss(X, Y) = 0.51513565f0 (tracked)
loss(X, Y) = 0.5147841f0 (tracked)
loss(X, Y) = 0.514426f0 (tracked)
loss(X, Y) = 0.5140724f0 (tracked)
loss(X, Y) = 0.5137158f0 (tracked)
loss(X, Y) = 0.51335996f0 (tracked)
loss(X, Y) = 0.513009f0 (tracked)
loss(X, Y) = 0.5126471f0 (tracked)
loss(X, Y) = 0.51229644f0 (tracked)
loss(X, Y) = 0.5119362f0 (tracked)
loss(X, Y) = 0.51158315f0 (tracked)
loss(X, Y) = 0.5112281f0 (tracked)
loss(X, Y) = 0.5108692f0 (tracked)
loss(X, Y) = 0.5105154f0 (tracked)
loss(X, Y) = 0.51015794f0 (tracked)
loss(X, Y) = 0.5098001f0 (tracked)
loss(X, Y) = 0.50944734f0 (tracked)
loss(X, Y) = 0.50908685f0 (tracked)
loss(X, Y) = 0.50873274f0 (tracked)
loss(X, Y) = 0.5083777f0 (tracked)
loss(X, Y) = 0.5080202f0 (tracked)
loss(X, Y) = 0.5076702f0 (tracked)
loss(X, Y) = 0.50731725f0 (tracked)
loss(X, Y) = 0.5069628f0 (tracked)
loss(X, Y) = 0.506608f0 (tracked)
loss(X, Y) = 0.5062507f0 (tracked)
loss(X, Y) = 0.5058948f0 (tracked)
loss(X, Y) = 0.50554496f0 (tracked)
loss(X, Y) = 0.50518954f0 (tracked)
loss(X, Y) = 0.5048311f0 (tracked)
loss(X, Y) = 0.50447875f0 (tracked)
loss(X, Y) = 0.50413f0 (tracked)
loss(X, Y) = 0.5037816f0 (tracked)
loss(X, Y) = 0.50342655f0 (tracked)
loss(X, Y) = 0.50307167f0 (tracked)
loss(X, Y) = 0.5027282f0 (tracked)
loss(X, Y) = 0.5023753f0 (tracked)
loss(X, Y) = 0.5020255f0 (tracked)
loss(X, Y) = 0.5016685f0 (tracked)
loss(X, Y) = 0.50130886f0 (tracked)
loss(X, Y) = 0.5009466f0 (tracked)
loss(X, Y) = 0.5005912f0 (tracked)
loss(X, Y) = 0.50023425f0 (tracked)
loss(X, Y) = 0.4998647f0 (tracked)
loss(X, Y) = 0.49950382f0 (tracked)
loss(X, Y) = 0.49913928f0 (tracked)
loss(X, Y) = 0.49877083f0 (tracked)
loss(X, Y) = 0.4984045f0 (tracked)
loss(X, Y) = 0.49803758f0 (tracked)
loss(X, Y) = 0.49767333f0 (tracked)
loss(X, Y) = 0.4973052f0 (tracked)
loss(X, Y) = 0.4969362f0 (tracked)
loss(X, Y) = 0.4965716f0 (tracked)
loss(X, Y) = 0.49620044f0 (tracked)
loss(X, Y) = 0.49582982f0 (tracked)
loss(X, Y) = 0.49546212f0 (tracked)
loss(X, Y) = 0.49510077f0 (tracked)
loss(X, Y) = 0.49472907f0 (tracked)
loss(X, Y) = 0.49435347f0 (tracked)
loss(X, Y) = 0.49398643f0 (tracked)
loss(X, Y) = 0.49361393f0 (tracked)
loss(X, Y) = 0.4932485f0 (tracked)
loss(X, Y) = 0.49287543f0 (tracked)
loss(X, Y) = 0.4925046f0 (tracked)
loss(X, Y) = 0.49213496f0 (tracked)
loss(X, Y) = 0.49177104f0 (tracked)
loss(X, Y) = 0.49140072f0 (tracked)
loss(X, Y) = 0.49102783f0 (tracked)
loss(X, Y) = 0.49065962f0 (tracked)
loss(X, Y) = 0.49029452f0 (tracked)
loss(X, Y) = 0.4899229f0 (tracked)
loss(X, Y) = 0.48955593f0 (tracked)
loss(X, Y) = 0.48918852f0 (tracked)
loss(X, Y) = 0.48882157f0 (tracked)
```

```
┌ Info: Epoch 985
└ @ Main /Users/kcft114/.julia/packages/Flux/qXNjB/src/optimise/train.jl:105
┌ Info: Epoch 986
└ @ Main /Users/kcft114/.julia/packages/Flux/qXNjB/src/optimise/train.jl:105
┌ Info: Epoch 987
└ @ Main /Users/kcft114/.julia/packages/Flux/qXNjB/src/optimise/train.jl:105
┌ Info: Epoch 988
└ @ Main /Users/kcft114/.julia/packages/Flux/qXNjB/src/optimise/train.jl:105
┌ Info: Epoch 989
└ @ Main /Users/kcft114/.julia/packages/Flux/qXNjB/src/optimise/train.jl:105
┌ Info: Epoch 990
└ @ Main /Users/kcft114/.julia/packages/Flux/qXNjB/src/optimise/train.jl:105
┌ Info: Epoch 991
└ @ Main /Users/kcft114/.julia/packages/Flux/qXNjB/src/optimise/train.jl:105
┌ Info: Epoch 992
└ @ Main /Users/kcft114/.julia/packages/Flux/qXNjB/src/optimise/train.jl:105
┌ Info: Epoch 993
└ @ Main /Users/kcft114/.julia/packages/Flux/qXNjB/src/optimise/train.jl:105
┌ Info: Epoch 994
└ @ Main /Users/kcft114/.julia/packages/Flux/qXNjB/src/optimise/train.jl:105
┌ Info: Epoch 995
└ @ Main /Users/kcft114/.julia/packages/Flux/qXNjB/src/optimise/train.jl:105
┌ Info: Epoch 996
└ @ Main /Users/kcft114/.julia/packages/Flux/qXNjB/src/optimise/train.jl:105
┌ Info: Epoch 997
└ @ Main /Users/kcft114/.julia/packages/Flux/qXNjB/src/optimise/train.jl:105
┌ Info: Epoch 998
└ @ Main /Users/kcft114/.julia/packages/Flux/qXNjB/src/optimise/train.jl:105
┌ Info: Epoch 999
└ @ Main /Users/kcft114/.julia/packages/Flux/qXNjB/src/optimise/train.jl:105
┌ Info: Epoch 1000
└ @ Main /Users/kcft114/.julia/packages/Flux/qXNjB/src/optimise/train.jl:105
┌ Info: Epoch 1001
└ @ Main /Users/kcft114/.julia/packages/Flux/qXNjB/src/optimise/train.jl:105
┌ Info: Epoch 1002
└ @ Main /Users/kcft114/.julia/packages/Flux/qXNjB/src/optimise/train.jl:105
┌ Info: Epoch 1003
└ @ Main /Users/kcft114/.julia/packages/Flux/qXNjB/src/optimise/train.jl:105
┌ Info: Epoch 1004
└ @ Main /Users/kcft114/.julia/packages/Flux/qXNjB/src/optimise/train.jl:105
┌ Info: Epoch 1005
└ @ Main /Users/kcft114/.julia/packages/Flux/qXNjB/src/optimise/train.jl:105
┌ Info: Epoch 1006
└ @ Main /Users/kcft114/.julia/packages/Flux/qXNjB/src/optimise/train.jl:105
┌ Info: Epoch 1007
└ @ Main /Users/kcft114/.julia/packages/Flux/qXNjB/src/optimise/train.jl:105
┌ Info: Epoch 1008
└ @ Main /Users/kcft114/.julia/packages/Flux/qXNjB/src/optimise/train.jl:105
┌ Info: Epoch 1009
└ @ Main /Users/kcft114/.julia/packages/Flux/qXNjB/src/optimise/train.jl:105
┌ Info: Epoch 1010
└ @ Main /Users/kcft114/.julia/packages/Flux/qXNjB/src/optimise/train.jl:105
┌ Info: Epoch 1011
└ @ Main /Users/kcft114/.julia/packages/Flux/qXNjB/src/optimise/train.jl:105
┌ Info: Epoch 1012
└ @ Main /Users/kcft114/.julia/packages/Flux/qXNjB/src/optimise/train.jl:105
┌ Info: Epoch 1013
└ @ Main /Users/kcft114/.julia/packages/Flux/qXNjB/src/optimise/train.jl:105
┌ Info: Epoch 1014
└ @ Main /Users/kcft114/.julia/packages/Flux/qXNjB/src/optimise/train.jl:105
┌ Info: Epoch 1015
└ @ Main /Users/kcft114/.julia/packages/Flux/qXNjB/src/optimise/train.jl:105
┌ Info: Epoch 1016
└ @ Main /Users/kcft114/.julia/packages/Flux/qXNjB/src/optimise/train.jl:105
┌ Info: Epoch 1017
└ @ Main /Users/kcft114/.julia/packages/Flux/qXNjB/src/optimise/train.jl:105
┌ Info: Epoch 1018
└ @ Main /Users/kcft114/.julia/packages/Flux/qXNjB/src/optimise/train.jl:105
┌ Info: Epoch 1019
└ @ Main /Users/kcft114/.julia/packages/Flux/qXNjB/src/optimise/train.jl:105
┌ Info: Epoch 1020
└ @ Main /Users/kcft114/.julia/packages/Flux/qXNjB/src/optimise/train.jl:105
┌ Info: Epoch 1021
└ @ Main /Users/kcft114/.julia/packages/Flux/qXNjB/src/optimise/train.jl:105
┌ Info: Epoch 1022
└ @ Main /Users/kcft114/.julia/packages/Flux/qXNjB/src/optimise/train.jl:105
┌ Info: Epoch 1023
└ @ Main /Users/kcft114/.julia/packages/Flux/qXNjB/src/optimise/train.jl:105
┌ Info: Epoch 1024
└ @ Main /Users/kcft114/.julia/packages/Flux/qXNjB/src/optimise/train.jl:105
┌ Info: Epoch 1025
└ @ Main /Users/kcft114/.julia/packages/Flux/qXNjB/src/optimise/train.jl:105
┌ Info: Epoch 1026
└ @ Main /Users/kcft114/.julia/packages/Flux/qXNjB/src/optimise/train.jl:105
┌ Info: Epoch 1027
└ @ Main /Users/kcft114/.julia/packages/Flux/qXNjB/src/optimise/train.jl:105
┌ Info: Epoch 1028
└ @ Main /Users/kcft114/.julia/packages/Flux/qXNjB/src/optimise/train.jl:105
┌ Info: Epoch 1029
└ @ Main /Users/kcft114/.julia/packages/Flux/qXNjB/src/optimise/train.jl:105
┌ Info: Epoch 1030
└ @ Main /Users/kcft114/.julia/packages/Flux/qXNjB/src/optimise/train.jl:105
┌ Info: Epoch 1031
└ @ Main /Users/kcft114/.julia/packages/Flux/qXNjB/src/optimise/train.jl:105
┌ Info: Epoch 1032
└ @ Main /Users/kcft114/.julia/packages/Flux/qXNjB/src/optimise/train.jl:105
┌ Info: Epoch 1033
└ @ Main /Users/kcft114/.julia/packages/Flux/qXNjB/src/optimise/train.jl:105
┌ Info: Epoch 1034
└ @ Main /Users/kcft114/.julia/packages/Flux/qXNjB/src/optimise/train.jl:105
┌ Info: Epoch 1035
└ @ Main /Users/kcft114/.julia/packages/Flux/qXNjB/src/optimise/train.jl:105
┌ Info: Epoch 1036
└ @ Main /Users/kcft114/.julia/packages/Flux/qXNjB/src/optimise/train.jl:105
┌ Info: Epoch 1037
└ @ Main /Users/kcft114/.julia/packages/Flux/qXNjB/src/optimise/train.jl:105
┌ Info: Epoch 1038
└ @ Main /Users/kcft114/.julia/packages/Flux/qXNjB/src/optimise/train.jl:105
┌ Info: Epoch 1039
└ @ Main /Users/kcft114/.julia/packages/Flux/qXNjB/src/optimise/train.jl:105
┌ Info: Epoch 1040
└ @ Main /Users/kcft114/.julia/packages/Flux/qXNjB/src/optimise/train.jl:105
┌ Info: Epoch 1041
└ @ Main /Users/kcft114/.julia/packages/Flux/qXNjB/src/optimise/train.jl:105
┌ Info: Epoch 1042
└ @ Main /Users/kcft114/.julia/packages/Flux/qXNjB/src/optimise/train.jl:105
┌ Info: Epoch 1043
└ @ Main /Users/kcft114/.julia/packages/Flux/qXNjB/src/optimise/train.jl:105
┌ Info: Epoch 1044
└ @ Main /Users/kcft114/.julia/packages/Flux/qXNjB/src/optimise/train.jl:105
┌ Info: Epoch 1045
└ @ Main /Users/kcft114/.julia/packages/Flux/qXNjB/src/optimise/train.jl:105
┌ Info: Epoch 1046
└ @ Main /Users/kcft114/.julia/packages/Flux/qXNjB/src/optimise/train.jl:105
┌ Info: Epoch 1047
└ @ Main /Users/kcft114/.julia/packages/Flux/qXNjB/src/optimise/train.jl:105
┌ Info: Epoch 1048
└ @ Main /Users/kcft114/.julia/packages/Flux/qXNjB/src/optimise/train.jl:105
┌ Info: Epoch 1049
└ @ Main /Users/kcft114/.julia/packages/Flux/qXNjB/src/optimise/train.jl:105
┌ Info: Epoch 1050
└ @ Main /Users/kcft114/.julia/packages/Flux/qXNjB/src/optimise/train.jl:105
┌ Info: Epoch 1051
└ @ Main /Users/kcft114/.julia/packages/Flux/qXNjB/src/optimise/train.jl:105
┌ Info: Epoch 1052
└ @ Main /Users/kcft114/.julia/packages/Flux/qXNjB/src/optimise/train.jl:105
┌ Info: Epoch 1053
└ @ Main /Users/kcft114/.julia/packages/Flux/qXNjB/src/optimise/train.jl:105
┌ Info: Epoch 1054
└ @ Main /Users/kcft114/.julia/packages/Flux/qXNjB/src/optimise/train.jl:105
┌ Info: Epoch 1055
└ @ Main /Users/kcft114/.julia/packages/Flux/qXNjB/src/optimise/train.jl:105
┌ Info: Epoch 1056
└ @ Main /Users/kcft114/.julia/packages/Flux/qXNjB/src/optimise/train.jl:105
┌ Info: Epoch 1057
└ @ Main /Users/kcft114/.julia/packages/Flux/qXNjB/src/optimise/train.jl:105
┌ Info: Epoch 1058
└ @ Main /Users/kcft114/.julia/packages/Flux/qXNjB/src/optimise/train.jl:105
┌ Info: Epoch 1059
└ @ Main /Users/kcft114/.julia/packages/Flux/qXNjB/src/optimise/train.jl:105
┌ Info: Epoch 1060
└ @ Main /Users/kcft114/.julia/packages/Flux/qXNjB/src/optimise/train.jl:105
┌ Info: Epoch 1061
└ @ Main /Users/kcft114/.julia/packages/Flux/qXNjB/src/optimise/train.jl:105
┌ Info: Epoch 1062
└ @ Main /Users/kcft114/.julia/packages/Flux/qXNjB/src/optimise/train.jl:105
┌ Info: Epoch 1063
└ @ Main /Users/kcft114/.julia/packages/Flux/qXNjB/src/optimise/train.jl:105
┌ Info: Epoch 1064
└ @ Main /Users/kcft114/.julia/packages/Flux/qXNjB/src/optimise/train.jl:105
┌ Info: Epoch 1065
└ @ Main /Users/kcft114/.julia/packages/Flux/qXNjB/src/optimise/train.jl:105
┌ Info: Epoch 1066
└ @ Main /Users/kcft114/.julia/packages/Flux/qXNjB/src/optimise/train.jl:105
┌ Info: Epoch 1067
└ @ Main /Users/kcft114/.julia/packages/Flux/qXNjB/src/optimise/train.jl:105
┌ Info: Epoch 1068
└ @ Main /Users/kcft114/.julia/packages/Flux/qXNjB/src/optimise/train.jl:105
┌ Info: Epoch 1069
└ @ Main /Users/kcft114/.julia/packages/Flux/qXNjB/src/optimise/train.jl:105
┌ Info: Epoch 1070
└ @ Main /Users/kcft114/.julia/packages/Flux/qXNjB/src/optimise/train.jl:105
┌ Info: Epoch 1071
└ @ Main /Users/kcft114/.julia/packages/Flux/qXNjB/src/optimise/train.jl:105
┌ Info: Epoch 1072
└ @ Main /Users/kcft114/.julia/packages/Flux/qXNjB/src/optimise/train.jl:105
┌ Info: Epoch 1073
└ @ Main /Users/kcft114/.julia/packages/Flux/qXNjB/src/optimise/train.jl:105
┌ Info: Epoch 1074
└ @ Main /Users/kcft114/.julia/packages/Flux/qXNjB/src/optimise/train.jl:105
┌ Info: Epoch 1075
└ @ Main /Users/kcft114/.julia/packages/Flux/qXNjB/src/optimise/train.jl:105
┌ Info: Epoch 1076
└ @ Main /Users/kcft114/.julia/packages/Flux/qXNjB/src/optimise/train.jl:105
┌ Info: Epoch 1077
└ @ Main /Users/kcft114/.julia/packages/Flux/qXNjB/src/optimise/train.jl:105
┌ Info: Epoch 1078
└ @ Main /Users/kcft114/.julia/packages/Flux/qXNjB/src/optimise/train.jl:105
┌ Info: Epoch 1079
└ @ Main /Users/kcft114/.julia/packages/Flux/qXNjB/src/optimise/train.jl:105
┌ Info: Epoch 1080
└ @ Main /Users/kcft114/.julia/packages/Flux/qXNjB/src/optimise/train.jl:105
┌ Info: Epoch 1081
└ @ Main /Users/kcft114/.julia/packages/Flux/qXNjB/src/optimise/train.jl:105
┌ Info: Epoch 1082
└ @ Main /Users/kcft114/.julia/packages/Flux/qXNjB/src/optimise/train.jl:105
┌ Info: Epoch 1083
└ @ Main /Users/kcft114/.julia/packages/Flux/qXNjB/src/optimise/train.jl:105
┌ Info: Epoch 1084
└ @ Main /Users/kcft114/.julia/packages/Flux/qXNjB/src/optimise/train.jl:105
┌ Info: Epoch 1085
└ @ Main /Users/kcft114/.julia/packages/Flux/qXNjB/src/optimise/train.jl:105
┌ Info: Epoch 1086
└ @ Main /Users/kcft114/.julia/packages/Flux/qXNjB/src/optimise/train.jl:105
┌ Info: Epoch 1087
└ @ Main /Users/kcft114/.julia/packages/Flux/qXNjB/src/optimise/train.jl:105
┌ Info: Epoch 1088
└ @ Main /Users/kcft114/.julia/packages/Flux/qXNjB/src/optimise/train.jl:105
┌ Info: Epoch 1089
└ @ Main /Users/kcft114/.julia/packages/Flux/qXNjB/src/optimise/train.jl:105
┌ Info: Epoch 1090
└ @ Main /Users/kcft114/.julia/packages/Flux/qXNjB/src/optimise/train.jl:105
┌ Info: Epoch 1091
└ @ Main /Users/kcft114/.julia/packages/Flux/qXNjB/src/optimise/train.jl:105
┌ Info: Epoch 1092
└ @ Main /Users/kcft114/.julia/packages/Flux/qXNjB/src/optimise/train.jl:105
┌ Info: Epoch 1093
└ @ Main /Users/kcft114/.julia/packages/Flux/qXNjB/src/optimise/train.jl:105
┌ Info: Epoch 1094
└ @ Main /Users/kcft114/.julia/packages/Flux/qXNjB/src/optimise/train.jl:105
┌ Info: Epoch 1095
└ @ Main /Users/kcft114/.julia/packages/Flux/qXNjB/src/optimise/train.jl:105
┌ Info: Epoch 1096
└ @ Main /Users/kcft114/.julia/packages/Flux/qXNjB/src/optimise/train.jl:105
┌ Info: Epoch 1097
└ @ Main /Users/kcft114/.julia/packages/Flux/qXNjB/src/optimise/train.jl:105
┌ Info: Epoch 1098
└ @ Main /Users/kcft114/.julia/packages/Flux/qXNjB/src/optimise/train.jl:105
┌ Info: Epoch 1099
└ @ Main /Users/kcft114/.julia/packages/Flux/qXNjB/src/optimise/train.jl:105
┌ Info: Epoch 1100
└ @ Main /Users/kcft114/.julia/packages/Flux/qXNjB/src/optimise/train.jl:105
┌ Info: Epoch 1101
└ @ Main /Users/kcft114/.julia/packages/Flux/qXNjB/src/optimise/train.jl:105
┌ Info: Epoch 1102
└ @ Main /Users/kcft114/.julia/packages/Flux/qXNjB/src/optimise/train.jl:105
┌ Info: Epoch 1103
└ @ Main /Users/kcft114/.julia/packages/Flux/qXNjB/src/optimise/train.jl:105
┌ Info: Epoch 1104
└ @ Main /Users/kcft114/.julia/packages/Flux/qXNjB/src/optimise/train.jl:105
┌ Info: Epoch 1105
└ @ Main /Users/kcft114/.julia/packages/Flux/qXNjB/src/optimise/train.jl:105
┌ Info: Epoch 1106
└ @ Main /Users/kcft114/.julia/packages/Flux/qXNjB/src/optimise/train.jl:105
┌ Info: Epoch 1107
└ @ Main /Users/kcft114/.julia/packages/Flux/qXNjB/src/optimise/train.jl:105
┌ Info: Epoch 1108
└ @ Main /Users/kcft114/.julia/packages/Flux/qXNjB/src/optimise/train.jl:105
┌ Info: Epoch 1109
└ @ Main /Users/kcft114/.julia/packages/Flux/qXNjB/src/optimise/train.jl:105
┌ Info: Epoch 1110
└ @ Main /Users/kcft114/.julia/packages/Flux/qXNjB/src/optimise/train.jl:105
┌ Info: Epoch 1111
└ @ Main /Users/kcft114/.julia/packages/Flux/qXNjB/src/optimise/train.jl:105
┌ Info: Epoch 1112
└ @ Main /Users/kcft114/.julia/packages/Flux/qXNjB/src/optimise/train.jl:105
```

```
loss(X, Y) = 0.48845878f0 (tracked)
loss(X, Y) = 0.48809278f0 (tracked)
loss(X, Y) = 0.48772633f0 (tracked)
loss(X, Y) = 0.48736048f0 (tracked)
loss(X, Y) = 0.48699805f0 (tracked)
loss(X, Y) = 0.48663205f0 (tracked)
loss(X, Y) = 0.48627213f0 (tracked)
loss(X, Y) = 0.48590714f0 (tracked)
loss(X, Y) = 0.48554292f0 (tracked)
loss(X, Y) = 0.48518375f0 (tracked)
loss(X, Y) = 0.48482355f0 (tracked)
loss(X, Y) = 0.48445845f0 (tracked)
loss(X, Y) = 0.48409587f0 (tracked)
loss(X, Y) = 0.4837391f0 (tracked)
loss(X, Y) = 0.48338327f0 (tracked)
loss(X, Y) = 0.48302147f0 (tracked)
loss(X, Y) = 0.48266745f0 (tracked)
loss(X, Y) = 0.4823146f0 (tracked)
loss(X, Y) = 0.48195872f0 (tracked)
loss(X, Y) = 0.48159686f0 (tracked)
loss(X, Y) = 0.48123932f0 (tracked)
loss(X, Y) = 0.48088524f0 (tracked)
loss(X, Y) = 0.4805347f0 (tracked)
loss(X, Y) = 0.48018116f0 (tracked)
loss(X, Y) = 0.4798327f0 (tracked)
loss(X, Y) = 0.4794857f0 (tracked)
loss(X, Y) = 0.4791316f0 (tracked)
loss(X, Y) = 0.47878715f0 (tracked)
loss(X, Y) = 0.4784502f0 (tracked)
loss(X, Y) = 0.47811827f0 (tracked)
loss(X, Y) = 0.47777763f0 (tracked)
loss(X, Y) = 0.47744712f0 (tracked)
loss(X, Y) = 0.47712183f0 (tracked)
loss(X, Y) = 0.47679442f0 (tracked)
loss(X, Y) = 0.47647166f0 (tracked)
loss(X, Y) = 0.4761366f0 (tracked)
loss(X, Y) = 0.4758118f0 (tracked)
loss(X, Y) = 0.4754922f0 (tracked)
loss(X, Y) = 0.47517306f0 (tracked)
loss(X, Y) = 0.4748529f0 (tracked)
loss(X, Y) = 0.47454163f0 (tracked)
loss(X, Y) = 0.47423676f0 (tracked)
loss(X, Y) = 0.4739137f0 (tracked)
loss(X, Y) = 0.47359818f0 (tracked)
loss(X, Y) = 0.4732849f0 (tracked)
loss(X, Y) = 0.47296828f0 (tracked)
loss(X, Y) = 0.47264904f0 (tracked)
loss(X, Y) = 0.47232974f0 (tracked)
loss(X, Y) = 0.47202358f0 (tracked)
loss(X, Y) = 0.47171035f0 (tracked)
loss(X, Y) = 0.47139817f0 (tracked)
loss(X, Y) = 0.4710848f0 (tracked)
loss(X, Y) = 0.4707788f0 (tracked)
loss(X, Y) = 0.4704716f0 (tracked)
loss(X, Y) = 0.47016636f0 (tracked)
loss(X, Y) = 0.4698525f0 (tracked)
loss(X, Y) = 0.46954167f0 (tracked)
loss(X, Y) = 0.46924153f0 (tracked)
loss(X, Y) = 0.46893433f0 (tracked)
loss(X, Y) = 0.46862674f0 (tracked)
loss(X, Y) = 0.46832955f0 (tracked)
loss(X, Y) = 0.4680279f0 (tracked)
loss(X, Y) = 0.4677211f0 (tracked)
loss(X, Y) = 0.46742684f0 (tracked)
loss(X, Y) = 0.46713454f0 (tracked)
loss(X, Y) = 0.46683076f0 (tracked)
loss(X, Y) = 0.4665326f0 (tracked)
loss(X, Y) = 0.46623755f0 (tracked)
loss(X, Y) = 0.46593338f0 (tracked)
loss(X, Y) = 0.4656349f0 (tracked)
loss(X, Y) = 0.46533513f0 (tracked)
loss(X, Y) = 0.46504557f0 (tracked)
loss(X, Y) = 0.46474418f0 (tracked)
loss(X, Y) = 0.46444675f0 (tracked)
loss(X, Y) = 0.46415356f0 (tracked)
loss(X, Y) = 0.4638574f0 (tracked)
loss(X, Y) = 0.46356118f0 (tracked)
loss(X, Y) = 0.46327084f0 (tracked)
loss(X, Y) = 0.4629712f0 (tracked)
loss(X, Y) = 0.46267638f0 (tracked)
loss(X, Y) = 0.46238652f0 (tracked)
loss(X, Y) = 0.46209887f0 (tracked)
loss(X, Y) = 0.46180528f0 (tracked)
loss(X, Y) = 0.46152323f0 (tracked)
loss(X, Y) = 0.4612411f0 (tracked)
loss(X, Y) = 0.4609446f0 (tracked)
loss(X, Y) = 0.46065685f0 (tracked)
loss(X, Y) = 0.46037114f0 (tracked)
loss(X, Y) = 0.46009073f0 (tracked)
loss(X, Y) = 0.4597987f0 (tracked)
loss(X, Y) = 0.4595064f0 (tracked)
loss(X, Y) = 0.45922482f0 (tracked)
loss(X, Y) = 0.45894232f0 (tracked)
loss(X, Y) = 0.45865372f0 (tracked)
loss(X, Y) = 0.45836675f0 (tracked)
loss(X, Y) = 0.45808f0 (tracked)
loss(X, Y) = 0.45779595f0 (tracked)
loss(X, Y) = 0.4575147f0 (tracked)
loss(X, Y) = 0.45722643f0 (tracked)
loss(X, Y) = 0.45693985f0 (tracked)
loss(X, Y) = 0.45666087f0 (tracked)
loss(X, Y) = 0.45638144f0 (tracked)
loss(X, Y) = 0.4560972f0 (tracked)
loss(X, Y) = 0.45581582f0 (tracked)
loss(X, Y) = 0.45553654f0 (tracked)
loss(X, Y) = 0.45525432f0 (tracked)
loss(X, Y) = 0.454971f0 (tracked)
loss(X, Y) = 0.4546858f0 (tracked)
loss(X, Y) = 0.45440745f0 (tracked)
loss(X, Y) = 0.45413664f0 (tracked)
loss(X, Y) = 0.45385918f0 (tracked)
loss(X, Y) = 0.45357174f0 (tracked)
loss(X, Y) = 0.45328438f0 (tracked)
loss(X, Y) = 0.453012f0 (tracked)
loss(X, Y) = 0.45273066f0 (tracked)
loss(X, Y) = 0.45244557f0 (tracked)
loss(X, Y) = 0.45216793f0 (tracked)
loss(X, Y) = 0.45188883f0 (tracked)
loss(X, Y) = 0.45160818f0 (tracked)
loss(X, Y) = 0.45132846f0 (tracked)
loss(X, Y) = 0.45104885f0 (tracked)
loss(X, Y) = 0.45076975f0 (tracked)
loss(X, Y) = 0.45048636f0 (tracked)
loss(X, Y) = 0.45021388f0 (tracked)
loss(X, Y) = 0.4499306f0 (tracked)
loss(X, Y) = 0.44965205f0 (tracked)
loss(X, Y) = 0.44937834f0 (tracked)
loss(X, Y) = 0.4490953f0 (tracked)
loss(X, Y) = 0.44881755f0 (tracked)
loss(X, Y) = 0.44854423f0 (tracked)
loss(X, Y) = 0.44826797f0 (tracked)
loss(X, Y) = 0.44798428f0 (tracked)
loss(X, Y) = 0.44770935f0 (tracked)
loss(X, Y) = 0.44743392f0 (tracked)
loss(X, Y) = 0.44715562f0 (tracked)
loss(X, Y) = 0.4468829f0 (tracked)
loss(X, Y) = 0.44660747f0 (tracked)
loss(X, Y) = 0.44632828f0 (tracked)
loss(X, Y) = 0.44605094f0 (tracked)
loss(X, Y) = 0.44578344f0 (tracked)
loss(X, Y) = 0.4455101f0 (tracked)
loss(X, Y) = 0.44523117f0 (tracked)
loss(X, Y) = 0.44494691f0 (tracked)
loss(X, Y) = 0.44468683f0 (tracked)
loss(X, Y) = 0.44441617f0 (tracked)
loss(X, Y) = 0.44413444f0 (tracked)
loss(X, Y) = 0.4438558f0 (tracked)
loss(X, Y) = 0.44358695f0 (tracked)
loss(X, Y) = 0.44331726f0 (tracked)
loss(X, Y) = 0.44303915f0 (tracked)
loss(X, Y) = 0.44276246f0 (tracked)
```

```
┌ Info: Epoch 1113
└ @ Main /Users/kcft114/.julia/packages/Flux/qXNjB/src/optimise/train.jl:105
┌ Info: Epoch 1114
└ @ Main /Users/kcft114/.julia/packages/Flux/qXNjB/src/optimise/train.jl:105
┌ Info: Epoch 1115
└ @ Main /Users/kcft114/.julia/packages/Flux/qXNjB/src/optimise/train.jl:105
┌ Info: Epoch 1116
└ @ Main /Users/kcft114/.julia/packages/Flux/qXNjB/src/optimise/train.jl:105
┌ Info: Epoch 1117
└ @ Main /Users/kcft114/.julia/packages/Flux/qXNjB/src/optimise/train.jl:105
┌ Info: Epoch 1118
└ @ Main /Users/kcft114/.julia/packages/Flux/qXNjB/src/optimise/train.jl:105
┌ Info: Epoch 1119
└ @ Main /Users/kcft114/.julia/packages/Flux/qXNjB/src/optimise/train.jl:105
┌ Info: Epoch 1120
└ @ Main /Users/kcft114/.julia/packages/Flux/qXNjB/src/optimise/train.jl:105
┌ Info: Epoch 1121
└ @ Main /Users/kcft114/.julia/packages/Flux/qXNjB/src/optimise/train.jl:105
┌ Info: Epoch 1122
└ @ Main /Users/kcft114/.julia/packages/Flux/qXNjB/src/optimise/train.jl:105
┌ Info: Epoch 1123
└ @ Main /Users/kcft114/.julia/packages/Flux/qXNjB/src/optimise/train.jl:105
┌ Info: Epoch 1124
└ @ Main /Users/kcft114/.julia/packages/Flux/qXNjB/src/optimise/train.jl:105
┌ Info: Epoch 1125
└ @ Main /Users/kcft114/.julia/packages/Flux/qXNjB/src/optimise/train.jl:105
┌ Info: Epoch 1126
└ @ Main /Users/kcft114/.julia/packages/Flux/qXNjB/src/optimise/train.jl:105
┌ Info: Epoch 1127
└ @ Main /Users/kcft114/.julia/packages/Flux/qXNjB/src/optimise/train.jl:105
┌ Info: Epoch 1128
└ @ Main /Users/kcft114/.julia/packages/Flux/qXNjB/src/optimise/train.jl:105
┌ Info: Epoch 1129
└ @ Main /Users/kcft114/.julia/packages/Flux/qXNjB/src/optimise/train.jl:105
┌ Info: Epoch 1130
└ @ Main /Users/kcft114/.julia/packages/Flux/qXNjB/src/optimise/train.jl:105
┌ Info: Epoch 1131
└ @ Main /Users/kcft114/.julia/packages/Flux/qXNjB/src/optimise/train.jl:105
┌ Info: Epoch 1132
└ @ Main /Users/kcft114/.julia/packages/Flux/qXNjB/src/optimise/train.jl:105
┌ Info: Epoch 1133
└ @ Main /Users/kcft114/.julia/packages/Flux/qXNjB/src/optimise/train.jl:105
┌ Info: Epoch 1134
└ @ Main /Users/kcft114/.julia/packages/Flux/qXNjB/src/optimise/train.jl:105
┌ Info: Epoch 1135
└ @ Main /Users/kcft114/.julia/packages/Flux/qXNjB/src/optimise/train.jl:105
┌ Info: Epoch 1136
└ @ Main /Users/kcft114/.julia/packages/Flux/qXNjB/src/optimise/train.jl:105
┌ Info: Epoch 1137
└ @ Main /Users/kcft114/.julia/packages/Flux/qXNjB/src/optimise/train.jl:105
┌ Info: Epoch 1138
└ @ Main /Users/kcft114/.julia/packages/Flux/qXNjB/src/optimise/train.jl:105
┌ Info: Epoch 1139
└ @ Main /Users/kcft114/.julia/packages/Flux/qXNjB/src/optimise/train.jl:105
┌ Info: Epoch 1140
└ @ Main /Users/kcft114/.julia/packages/Flux/qXNjB/src/optimise/train.jl:105
┌ Info: Epoch 1141
└ @ Main /Users/kcft114/.julia/packages/Flux/qXNjB/src/optimise/train.jl:105
┌ Info: Epoch 1142
└ @ Main /Users/kcft114/.julia/packages/Flux/qXNjB/src/optimise/train.jl:105
┌ Info: Epoch 1143
└ @ Main /Users/kcft114/.julia/packages/Flux/qXNjB/src/optimise/train.jl:105
┌ Info: Epoch 1144
└ @ Main /Users/kcft114/.julia/packages/Flux/qXNjB/src/optimise/train.jl:105
┌ Info: Epoch 1145
└ @ Main /Users/kcft114/.julia/packages/Flux/qXNjB/src/optimise/train.jl:105
┌ Info: Epoch 1146
└ @ Main /Users/kcft114/.julia/packages/Flux/qXNjB/src/optimise/train.jl:105
┌ Info: Epoch 1147
└ @ Main /Users/kcft114/.julia/packages/Flux/qXNjB/src/optimise/train.jl:105
┌ Info: Epoch 1148
└ @ Main /Users/kcft114/.julia/packages/Flux/qXNjB/src/optimise/train.jl:105
┌ Info: Epoch 1149
└ @ Main /Users/kcft114/.julia/packages/Flux/qXNjB/src/optimise/train.jl:105
┌ Info: Epoch 1150
└ @ Main /Users/kcft114/.julia/packages/Flux/qXNjB/src/optimise/train.jl:105
┌ Info: Epoch 1151
└ @ Main /Users/kcft114/.julia/packages/Flux/qXNjB/src/optimise/train.jl:105
┌ Info: Epoch 1152
└ @ Main /Users/kcft114/.julia/packages/Flux/qXNjB/src/optimise/train.jl:105
┌ Info: Epoch 1153
└ @ Main /Users/kcft114/.julia/packages/Flux/qXNjB/src/optimise/train.jl:105
┌ Info: Epoch 1154
└ @ Main /Users/kcft114/.julia/packages/Flux/qXNjB/src/optimise/train.jl:105
┌ Info: Epoch 1155
└ @ Main /Users/kcft114/.julia/packages/Flux/qXNjB/src/optimise/train.jl:105
┌ Info: Epoch 1156
└ @ Main /Users/kcft114/.julia/packages/Flux/qXNjB/src/optimise/train.jl:105
┌ Info: Epoch 1157
└ @ Main /Users/kcft114/.julia/packages/Flux/qXNjB/src/optimise/train.jl:105
┌ Info: Epoch 1158
└ @ Main /Users/kcft114/.julia/packages/Flux/qXNjB/src/optimise/train.jl:105
┌ Info: Epoch 1159
└ @ Main /Users/kcft114/.julia/packages/Flux/qXNjB/src/optimise/train.jl:105
┌ Info: Epoch 1160
└ @ Main /Users/kcft114/.julia/packages/Flux/qXNjB/src/optimise/train.jl:105
┌ Info: Epoch 1161
└ @ Main /Users/kcft114/.julia/packages/Flux/qXNjB/src/optimise/train.jl:105
┌ Info: Epoch 1162
└ @ Main /Users/kcft114/.julia/packages/Flux/qXNjB/src/optimise/train.jl:105
┌ Info: Epoch 1163
└ @ Main /Users/kcft114/.julia/packages/Flux/qXNjB/src/optimise/train.jl:105
┌ Info: Epoch 1164
└ @ Main /Users/kcft114/.julia/packages/Flux/qXNjB/src/optimise/train.jl:105
┌ Info: Epoch 1165
└ @ Main /Users/kcft114/.julia/packages/Flux/qXNjB/src/optimise/train.jl:105
┌ Info: Epoch 1166
└ @ Main /Users/kcft114/.julia/packages/Flux/qXNjB/src/optimise/train.jl:105
┌ Info: Epoch 1167
└ @ Main /Users/kcft114/.julia/packages/Flux/qXNjB/src/optimise/train.jl:105
┌ Info: Epoch 1168
└ @ Main /Users/kcft114/.julia/packages/Flux/qXNjB/src/optimise/train.jl:105
┌ Info: Epoch 1169
└ @ Main /Users/kcft114/.julia/packages/Flux/qXNjB/src/optimise/train.jl:105
┌ Info: Epoch 1170
└ @ Main /Users/kcft114/.julia/packages/Flux/qXNjB/src/optimise/train.jl:105
┌ Info: Epoch 1171
└ @ Main /Users/kcft114/.julia/packages/Flux/qXNjB/src/optimise/train.jl:105
┌ Info: Epoch 1172
└ @ Main /Users/kcft114/.julia/packages/Flux/qXNjB/src/optimise/train.jl:105
┌ Info: Epoch 1173
└ @ Main /Users/kcft114/.julia/packages/Flux/qXNjB/src/optimise/train.jl:105
┌ Info: Epoch 1174
└ @ Main /Users/kcft114/.julia/packages/Flux/qXNjB/src/optimise/train.jl:105
┌ Info: Epoch 1175
└ @ Main /Users/kcft114/.julia/packages/Flux/qXNjB/src/optimise/train.jl:105
┌ Info: Epoch 1176
└ @ Main /Users/kcft114/.julia/packages/Flux/qXNjB/src/optimise/train.jl:105
┌ Info: Epoch 1177
└ @ Main /Users/kcft114/.julia/packages/Flux/qXNjB/src/optimise/train.jl:105
┌ Info: Epoch 1178
└ @ Main /Users/kcft114/.julia/packages/Flux/qXNjB/src/optimise/train.jl:105
┌ Info: Epoch 1179
└ @ Main /Users/kcft114/.julia/packages/Flux/qXNjB/src/optimise/train.jl:105
┌ Info: Epoch 1180
└ @ Main /Users/kcft114/.julia/packages/Flux/qXNjB/src/optimise/train.jl:105
┌ Info: Epoch 1181
└ @ Main /Users/kcft114/.julia/packages/Flux/qXNjB/src/optimise/train.jl:105
┌ Info: Epoch 1182
└ @ Main /Users/kcft114/.julia/packages/Flux/qXNjB/src/optimise/train.jl:105
┌ Info: Epoch 1183
└ @ Main /Users/kcft114/.julia/packages/Flux/qXNjB/src/optimise/train.jl:105
┌ Info: Epoch 1184
└ @ Main /Users/kcft114/.julia/packages/Flux/qXNjB/src/optimise/train.jl:105
┌ Info: Epoch 1185
└ @ Main /Users/kcft114/.julia/packages/Flux/qXNjB/src/optimise/train.jl:105
┌ Info: Epoch 1186
└ @ Main /Users/kcft114/.julia/packages/Flux/qXNjB/src/optimise/train.jl:105
┌ Info: Epoch 1187
└ @ Main /Users/kcft114/.julia/packages/Flux/qXNjB/src/optimise/train.jl:105
┌ Info: Epoch 1188
└ @ Main /Users/kcft114/.julia/packages/Flux/qXNjB/src/optimise/train.jl:105
┌ Info: Epoch 1189
└ @ Main /Users/kcft114/.julia/packages/Flux/qXNjB/src/optimise/train.jl:105
┌ Info: Epoch 1190
└ @ Main /Users/kcft114/.julia/packages/Flux/qXNjB/src/optimise/train.jl:105
┌ Info: Epoch 1191
└ @ Main /Users/kcft114/.julia/packages/Flux/qXNjB/src/optimise/train.jl:105
┌ Info: Epoch 1192
└ @ Main /Users/kcft114/.julia/packages/Flux/qXNjB/src/optimise/train.jl:105
┌ Info: Epoch 1193
└ @ Main /Users/kcft114/.julia/packages/Flux/qXNjB/src/optimise/train.jl:105
┌ Info: Epoch 1194
└ @ Main /Users/kcft114/.julia/packages/Flux/qXNjB/src/optimise/train.jl:105
┌ Info: Epoch 1195
└ @ Main /Users/kcft114/.julia/packages/Flux/qXNjB/src/optimise/train.jl:105
┌ Info: Epoch 1196
└ @ Main /Users/kcft114/.julia/packages/Flux/qXNjB/src/optimise/train.jl:105
┌ Info: Epoch 1197
└ @ Main /Users/kcft114/.julia/packages/Flux/qXNjB/src/optimise/train.jl:105
┌ Info: Epoch 1198
└ @ Main /Users/kcft114/.julia/packages/Flux/qXNjB/src/optimise/train.jl:105
┌ Info: Epoch 1199
└ @ Main /Users/kcft114/.julia/packages/Flux/qXNjB/src/optimise/train.jl:105
┌ Info: Epoch 1200
└ @ Main /Users/kcft114/.julia/packages/Flux/qXNjB/src/optimise/train.jl:105
┌ Info: Epoch 1201
└ @ Main /Users/kcft114/.julia/packages/Flux/qXNjB/src/optimise/train.jl:105
┌ Info: Epoch 1202
└ @ Main /Users/kcft114/.julia/packages/Flux/qXNjB/src/optimise/train.jl:105
┌ Info: Epoch 1203
└ @ Main /Users/kcft114/.julia/packages/Flux/qXNjB/src/optimise/train.jl:105
┌ Info: Epoch 1204
└ @ Main /Users/kcft114/.julia/packages/Flux/qXNjB/src/optimise/train.jl:105
┌ Info: Epoch 1205
└ @ Main /Users/kcft114/.julia/packages/Flux/qXNjB/src/optimise/train.jl:105
┌ Info: Epoch 1206
└ @ Main /Users/kcft114/.julia/packages/Flux/qXNjB/src/optimise/train.jl:105
┌ Info: Epoch 1207
└ @ Main /Users/kcft114/.julia/packages/Flux/qXNjB/src/optimise/train.jl:105
┌ Info: Epoch 1208
└ @ Main /Users/kcft114/.julia/packages/Flux/qXNjB/src/optimise/train.jl:105
┌ Info: Epoch 1209
└ @ Main /Users/kcft114/.julia/packages/Flux/qXNjB/src/optimise/train.jl:105
┌ Info: Epoch 1210
└ @ Main /Users/kcft114/.julia/packages/Flux/qXNjB/src/optimise/train.jl:105
┌ Info: Epoch 1211
└ @ Main /Users/kcft114/.julia/packages/Flux/qXNjB/src/optimise/train.jl:105
┌ Info: Epoch 1212
└ @ Main /Users/kcft114/.julia/packages/Flux/qXNjB/src/optimise/train.jl:105
┌ Info: Epoch 1213
└ @ Main /Users/kcft114/.julia/packages/Flux/qXNjB/src/optimise/train.jl:105
┌ Info: Epoch 1214
└ @ Main /Users/kcft114/.julia/packages/Flux/qXNjB/src/optimise/train.jl:105
┌ Info: Epoch 1215
└ @ Main /Users/kcft114/.julia/packages/Flux/qXNjB/src/optimise/train.jl:105
┌ Info: Epoch 1216
└ @ Main /Users/kcft114/.julia/packages/Flux/qXNjB/src/optimise/train.jl:105
┌ Info: Epoch 1217
└ @ Main /Users/kcft114/.julia/packages/Flux/qXNjB/src/optimise/train.jl:105
┌ Info: Epoch 1218
└ @ Main /Users/kcft114/.julia/packages/Flux/qXNjB/src/optimise/train.jl:105
┌ Info: Epoch 1219
└ @ Main /Users/kcft114/.julia/packages/Flux/qXNjB/src/optimise/train.jl:105
┌ Info: Epoch 1220
└ @ Main /Users/kcft114/.julia/packages/Flux/qXNjB/src/optimise/train.jl:105
┌ Info: Epoch 1221
└ @ Main /Users/kcft114/.julia/packages/Flux/qXNjB/src/optimise/train.jl:105
┌ Info: Epoch 1222
└ @ Main /Users/kcft114/.julia/packages/Flux/qXNjB/src/optimise/train.jl:105
┌ Info: Epoch 1223
└ @ Main /Users/kcft114/.julia/packages/Flux/qXNjB/src/optimise/train.jl:105
┌ Info: Epoch 1224
└ @ Main /Users/kcft114/.julia/packages/Flux/qXNjB/src/optimise/train.jl:105
┌ Info: Epoch 1225
└ @ Main /Users/kcft114/.julia/packages/Flux/qXNjB/src/optimise/train.jl:105
┌ Info: Epoch 1226
└ @ Main /Users/kcft114/.julia/packages/Flux/qXNjB/src/optimise/train.jl:105
┌ Info: Epoch 1227
└ @ Main /Users/kcft114/.julia/packages/Flux/qXNjB/src/optimise/train.jl:105
┌ Info: Epoch 1228
└ @ Main /Users/kcft114/.julia/packages/Flux/qXNjB/src/optimise/train.jl:105
┌ Info: Epoch 1229
└ @ Main /Users/kcft114/.julia/packages/Flux/qXNjB/src/optimise/train.jl:105
┌ Info: Epoch 1230
└ @ Main /Users/kcft114/.julia/packages/Flux/qXNjB/src/optimise/train.jl:105
┌ Info: Epoch 1231
└ @ Main /Users/kcft114/.julia/packages/Flux/qXNjB/src/optimise/train.jl:105
┌ Info: Epoch 1232
└ @ Main /Users/kcft114/.julia/packages/Flux/qXNjB/src/optimise/train.jl:105
┌ Info: Epoch 1233
└ @ Main /Users/kcft114/.julia/packages/Flux/qXNjB/src/optimise/train.jl:105
┌ Info: Epoch 1234
└ @ Main /Users/kcft114/.julia/packages/Flux/qXNjB/src/optimise/train.jl:105
┌ Info: Epoch 1235
└ @ Main /Users/kcft114/.julia/packages/Flux/qXNjB/src/optimise/train.jl:105
┌ Info: Epoch 1236
└ @ Main /Users/kcft114/.julia/packages/Flux/qXNjB/src/optimise/train.jl:105
┌ Info: Epoch 1237
└ @ Main /Users/kcft114/.julia/packages/Flux/qXNjB/src/optimise/train.jl:105
┌ Info: Epoch 1238
└ @ Main /Users/kcft114/.julia/packages/Flux/qXNjB/src/optimise/train.jl:105
┌ Info: Epoch 1239
└ @ Main /Users/kcft114/.julia/packages/Flux/qXNjB/src/optimise/train.jl:105
┌ Info: Epoch 1240
└ @ Main /Users/kcft114/.julia/packages/Flux/qXNjB/src/optimise/train.jl:105
┌ Info: Epoch 1241
└ @ Main /Users/kcft114/.julia/packages/Flux/qXNjB/src/optimise/train.jl:105
┌ Info: Epoch 1242
└ @ Main /Users/kcft114/.julia/packages/Flux/qXNjB/src/optimise/train.jl:105
┌ Info: Epoch 1243
└ @ Main /Users/kcft114/.julia/packages/Flux/qXNjB/src/optimise/train.jl:105
┌ Info: Epoch 1244
└ @ Main /Users/kcft114/.julia/packages/Flux/qXNjB/src/optimise/train.jl:105
┌ Info: Epoch 1245
└ @ Main /Users/kcft114/.julia/packages/Flux/qXNjB/src/optimise/train.jl:105
┌ Info: Epoch 1246
└ @ Main /Users/kcft114/.julia/packages/Flux/qXNjB/src/optimise/train.jl:105
┌ Info: Epoch 1247
└ @ Main /Users/kcft114/.julia/packages/Flux/qXNjB/src/optimise/train.jl:105
┌ Info: Epoch 1248
└ @ Main /Users/kcft114/.julia/packages/Flux/qXNjB/src/optimise/train.jl:105
┌ Info: Epoch 1249
└ @ Main /Users/kcft114/.julia/packages/Flux/qXNjB/src/optimise/train.jl:105
┌ Info: Epoch 1250
└ @ Main /Users/kcft114/.julia/packages/Flux/qXNjB/src/optimise/train.jl:105
┌ Info: Epoch 1251
└ @ Main /Users/kcft114/.julia/packages/Flux/qXNjB/src/optimise/train.jl:105
┌ Info: Epoch 1252
└ @ Main /Users/kcft114/.julia/packages/Flux/qXNjB/src/optimise/train.jl:105
┌ Info: Epoch 1253
└ @ Main /Users/kcft114/.julia/packages/Flux/qXNjB/src/optimise/train.jl:105
┌ Info: Epoch 1254
└ @ Main /Users/kcft114/.julia/packages/Flux/qXNjB/src/optimise/train.jl:105
┌ Info: Epoch 1255
└ @ Main /Users/kcft114/.julia/packages/Flux/qXNjB/src/optimise/train.jl:105
┌ Info: Epoch 1256
└ @ Main /Users/kcft114/.julia/packages/Flux/qXNjB/src/optimise/train.jl:105
┌ Info: Epoch 1257
└ @ Main /Users/kcft114/.julia/packages/Flux/qXNjB/src/optimise/train.jl:105
┌ Info: Epoch 1258
└ @ Main /Users/kcft114/.julia/packages/Flux/qXNjB/src/optimise/train.jl:105
┌ Info: Epoch 1259
└ @ Main /Users/kcft114/.julia/packages/Flux/qXNjB/src/optimise/train.jl:105
┌ Info: Epoch 1260
└ @ Main /Users/kcft114/.julia/packages/Flux/qXNjB/src/optimise/train.jl:105
┌ Info: Epoch 1261
└ @ Main /Users/kcft114/.julia/packages/Flux/qXNjB/src/optimise/train.jl:105
┌ Info: Epoch 1262
└ @ Main /Users/kcft114/.julia/packages/Flux/qXNjB/src/optimise/train.jl:105
┌ Info: Epoch 1263
└ @ Main /Users/kcft114/.julia/packages/Flux/qXNjB/src/optimise/train.jl:105
```

```
loss(X, Y) = 0.44248858f0 (tracked)
loss(X, Y) = 0.44221377f0 (tracked)
loss(X, Y) = 0.4419388f0 (tracked)
loss(X, Y) = 0.4416741f0 (tracked)
loss(X, Y) = 0.4414095f0 (tracked)
loss(X, Y) = 0.4411302f0 (tracked)
loss(X, Y) = 0.44085297f0 (tracked)
loss(X, Y) = 0.44058648f0 (tracked)
loss(X, Y) = 0.44031456f0 (tracked)
loss(X, Y) = 0.44003955f0 (tracked)
loss(X, Y) = 0.43976435f0 (tracked)
loss(X, Y) = 0.439494f0 (tracked)
loss(X, Y) = 0.43922174f0 (tracked)
loss(X, Y) = 0.43894932f0 (tracked)
loss(X, Y) = 0.4386816f0 (tracked)
loss(X, Y) = 0.43841717f0 (tracked)
loss(X, Y) = 0.43814453f0 (tracked)
loss(X, Y) = 0.43786502f0 (tracked)
loss(X, Y) = 0.43759522f0 (tracked)
loss(X, Y) = 0.43732214f0 (tracked)
loss(X, Y) = 0.43704918f0 (tracked)
loss(X, Y) = 0.43677655f0 (tracked)
loss(X, Y) = 0.43649757f0 (tracked)
loss(X, Y) = 0.43622074f0 (tracked)
loss(X, Y) = 0.4359381f0 (tracked)
loss(X, Y) = 0.4356571f0 (tracked)
loss(X, Y) = 0.43537366f0 (tracked)
loss(X, Y) = 0.4350912f0 (tracked)
loss(X, Y) = 0.4348085f0 (tracked)
loss(X, Y) = 0.434523f0 (tracked)
loss(X, Y) = 0.43423575f0 (tracked)
loss(X, Y) = 0.43395996f0 (tracked)
loss(X, Y) = 0.43369737f0 (tracked)
loss(X, Y) = 0.43343255f0 (tracked)
loss(X, Y) = 0.4331532f0 (tracked)
loss(X, Y) = 0.43286547f0 (tracked)
loss(X, Y) = 0.43257377f0 (tracked)
loss(X, Y) = 0.4322763f0 (tracked)
loss(X, Y) = 0.4319892f0 (tracked)
loss(X, Y) = 0.43171355f0 (tracked)
loss(X, Y) = 0.43143174f0 (tracked)
loss(X, Y) = 0.4311386f0 (tracked)
loss(X, Y) = 0.43084726f0 (tracked)
loss(X, Y) = 0.43055886f0 (tracked)
loss(X, Y) = 0.4302773f0 (tracked)
loss(X, Y) = 0.42998505f0 (tracked)
loss(X, Y) = 0.4296972f0 (tracked)
loss(X, Y) = 0.4294159f0 (tracked)
loss(X, Y) = 0.4291271f0 (tracked)
loss(X, Y) = 0.42883736f0 (tracked)
loss(X, Y) = 0.42855784f0 (tracked)
loss(X, Y) = 0.4282621f0 (tracked)
loss(X, Y) = 0.42796975f0 (tracked)
loss(X, Y) = 0.42768583f0 (tracked)
loss(X, Y) = 0.4274007f0 (tracked)
loss(X, Y) = 0.42711428f0 (tracked)
loss(X, Y) = 0.426828f0 (tracked)
loss(X, Y) = 0.4265434f0 (tracked)
loss(X, Y) = 0.42625588f0 (tracked)
loss(X, Y) = 0.42597142f0 (tracked)
loss(X, Y) = 0.42568818f0 (tracked)
loss(X, Y) = 0.42540088f0 (tracked)
loss(X, Y) = 0.42511237f0 (tracked)
loss(X, Y) = 0.42483187f0 (tracked)
loss(X, Y) = 0.4245465f0 (tracked)
loss(X, Y) = 0.42426202f0 (tracked)
loss(X, Y) = 0.4239749f0 (tracked)
loss(X, Y) = 0.42369384f0 (tracked)
loss(X, Y) = 0.42340702f0 (tracked)
loss(X, Y) = 0.42312962f0 (tracked)
loss(X, Y) = 0.42285213f0 (tracked)
loss(X, Y) = 0.42256525f0 (tracked)
loss(X, Y) = 0.4222913f0 (tracked)
loss(X, Y) = 0.42201048f0 (tracked)
loss(X, Y) = 0.42173052f0 (tracked)
loss(X, Y) = 0.42145097f0 (tracked)
loss(X, Y) = 0.4211619f0 (tracked)
loss(X, Y) = 0.420876f0 (tracked)
loss(X, Y) = 0.42059535f0 (tracked)
loss(X, Y) = 0.42030212f0 (tracked)
loss(X, Y) = 0.42000723f0 (tracked)
loss(X, Y) = 0.41972005f0 (tracked)
loss(X, Y) = 0.41943341f0 (tracked)
loss(X, Y) = 0.4191401f0 (tracked)
loss(X, Y) = 0.4188503f0 (tracked)
loss(X, Y) = 0.4185481f0 (tracked)
loss(X, Y) = 0.41824794f0 (tracked)
loss(X, Y) = 0.41795835f0 (tracked)
loss(X, Y) = 0.41765818f0 (tracked)
loss(X, Y) = 0.41736755f0 (tracked)
loss(X, Y) = 0.4170645f0 (tracked)
loss(X, Y) = 0.41676304f0 (tracked)
loss(X, Y) = 0.41647512f0 (tracked)
loss(X, Y) = 0.4161742f0 (tracked)
loss(X, Y) = 0.41586912f0 (tracked)
loss(X, Y) = 0.41556886f0 (tracked)
loss(X, Y) = 0.41528124f0 (tracked)
loss(X, Y) = 0.41498122f0 (tracked)
loss(X, Y) = 0.4146756f0 (tracked)
loss(X, Y) = 0.41438714f0 (tracked)
loss(X, Y) = 0.41409647f0 (tracked)
loss(X, Y) = 0.41380465f0 (tracked)
loss(X, Y) = 0.41351584f0 (tracked)
loss(X, Y) = 0.4132201f0 (tracked)
loss(X, Y) = 0.41292655f0 (tracked)
loss(X, Y) = 0.41264346f0 (tracked)
loss(X, Y) = 0.4123673f0 (tracked)
loss(X, Y) = 0.41208544f0 (tracked)
loss(X, Y) = 0.41179714f0 (tracked)
loss(X, Y) = 0.41152054f0 (tracked)
loss(X, Y) = 0.41124815f0 (tracked)
loss(X, Y) = 0.41097492f0 (tracked)
loss(X, Y) = 0.41069093f0 (tracked)
loss(X, Y) = 0.41040802f0 (tracked)
loss(X, Y) = 0.41012964f0 (tracked)
loss(X, Y) = 0.40984142f0 (tracked)
loss(X, Y) = 0.40956336f0 (tracked)
loss(X, Y) = 0.40929115f0 (tracked)
loss(X, Y) = 0.4090023f0 (tracked)
loss(X, Y) = 0.4087214f0 (tracked)
loss(X, Y) = 0.40844575f0 (tracked)
loss(X, Y) = 0.40816516f0 (tracked)
loss(X, Y) = 0.40787792f0 (tracked)
loss(X, Y) = 0.40759563f0 (tracked)
loss(X, Y) = 0.40732843f0 (tracked)
loss(X, Y) = 0.40704897f0 (tracked)
loss(X, Y) = 0.40677306f0 (tracked)
loss(X, Y) = 0.4065031f0 (tracked)
loss(X, Y) = 0.40622836f0 (tracked)
loss(X, Y) = 0.40594825f0 (tracked)
loss(X, Y) = 0.4056833f0 (tracked)
loss(X, Y) = 0.4054105f0 (tracked)
loss(X, Y) = 0.40514776f0 (tracked)
loss(X, Y) = 0.40488115f0 (tracked)
loss(X, Y) = 0.40460297f0 (tracked)
loss(X, Y) = 0.40432972f0 (tracked)
loss(X, Y) = 0.4040724f0 (tracked)
loss(X, Y) = 0.4038118f0 (tracked)
loss(X, Y) = 0.40354154f0 (tracked)
loss(X, Y) = 0.4032738f0 (tracked)
loss(X, Y) = 0.40301448f0 (tracked)
loss(X, Y) = 0.4027503f0 (tracked)
loss(X, Y) = 0.40247756f0 (tracked)
loss(X, Y) = 0.40222362f0 (tracked)
loss(X, Y) = 0.40197316f0 (tracked)
loss(X, Y) = 0.40171093f0 (tracked)
```

```
┌ Info: Epoch 1264
└ @ Main /Users/kcft114/.julia/packages/Flux/qXNjB/src/optimise/train.jl:105
┌ Info: Epoch 1265
└ @ Main /Users/kcft114/.julia/packages/Flux/qXNjB/src/optimise/train.jl:105
┌ Info: Epoch 1266
└ @ Main /Users/kcft114/.julia/packages/Flux/qXNjB/src/optimise/train.jl:105
┌ Info: Epoch 1267
└ @ Main /Users/kcft114/.julia/packages/Flux/qXNjB/src/optimise/train.jl:105
┌ Info: Epoch 1268
└ @ Main /Users/kcft114/.julia/packages/Flux/qXNjB/src/optimise/train.jl:105
┌ Info: Epoch 1269
└ @ Main /Users/kcft114/.julia/packages/Flux/qXNjB/src/optimise/train.jl:105
┌ Info: Epoch 1270
└ @ Main /Users/kcft114/.julia/packages/Flux/qXNjB/src/optimise/train.jl:105
┌ Info: Epoch 1271
└ @ Main /Users/kcft114/.julia/packages/Flux/qXNjB/src/optimise/train.jl:105
┌ Info: Epoch 1272
└ @ Main /Users/kcft114/.julia/packages/Flux/qXNjB/src/optimise/train.jl:105
┌ Info: Epoch 1273
└ @ Main /Users/kcft114/.julia/packages/Flux/qXNjB/src/optimise/train.jl:105
┌ Info: Epoch 1274
└ @ Main /Users/kcft114/.julia/packages/Flux/qXNjB/src/optimise/train.jl:105
┌ Info: Epoch 1275
└ @ Main /Users/kcft114/.julia/packages/Flux/qXNjB/src/optimise/train.jl:105
┌ Info: Epoch 1276
└ @ Main /Users/kcft114/.julia/packages/Flux/qXNjB/src/optimise/train.jl:105
┌ Info: Epoch 1277
└ @ Main /Users/kcft114/.julia/packages/Flux/qXNjB/src/optimise/train.jl:105
┌ Info: Epoch 1278
└ @ Main /Users/kcft114/.julia/packages/Flux/qXNjB/src/optimise/train.jl:105
┌ Info: Epoch 1279
└ @ Main /Users/kcft114/.julia/packages/Flux/qXNjB/src/optimise/train.jl:105
┌ Info: Epoch 1280
└ @ Main /Users/kcft114/.julia/packages/Flux/qXNjB/src/optimise/train.jl:105
┌ Info: Epoch 1281
└ @ Main /Users/kcft114/.julia/packages/Flux/qXNjB/src/optimise/train.jl:105
┌ Info: Epoch 1282
└ @ Main /Users/kcft114/.julia/packages/Flux/qXNjB/src/optimise/train.jl:105
┌ Info: Epoch 1283
└ @ Main /Users/kcft114/.julia/packages/Flux/qXNjB/src/optimise/train.jl:105
┌ Info: Epoch 1284
└ @ Main /Users/kcft114/.julia/packages/Flux/qXNjB/src/optimise/train.jl:105
┌ Info: Epoch 1285
└ @ Main /Users/kcft114/.julia/packages/Flux/qXNjB/src/optimise/train.jl:105
┌ Info: Epoch 1286
└ @ Main /Users/kcft114/.julia/packages/Flux/qXNjB/src/optimise/train.jl:105
┌ Info: Epoch 1287
└ @ Main /Users/kcft114/.julia/packages/Flux/qXNjB/src/optimise/train.jl:105
┌ Info: Epoch 1288
└ @ Main /Users/kcft114/.julia/packages/Flux/qXNjB/src/optimise/train.jl:105
┌ Info: Epoch 1289
└ @ Main /Users/kcft114/.julia/packages/Flux/qXNjB/src/optimise/train.jl:105
┌ Info: Epoch 1290
└ @ Main /Users/kcft114/.julia/packages/Flux/qXNjB/src/optimise/train.jl:105
┌ Info: Epoch 1291
└ @ Main /Users/kcft114/.julia/packages/Flux/qXNjB/src/optimise/train.jl:105
┌ Info: Epoch 1292
└ @ Main /Users/kcft114/.julia/packages/Flux/qXNjB/src/optimise/train.jl:105
┌ Info: Epoch 1293
└ @ Main /Users/kcft114/.julia/packages/Flux/qXNjB/src/optimise/train.jl:105
┌ Info: Epoch 1294
└ @ Main /Users/kcft114/.julia/packages/Flux/qXNjB/src/optimise/train.jl:105
┌ Info: Epoch 1295
└ @ Main /Users/kcft114/.julia/packages/Flux/qXNjB/src/optimise/train.jl:105
┌ Info: Epoch 1296
└ @ Main /Users/kcft114/.julia/packages/Flux/qXNjB/src/optimise/train.jl:105
┌ Info: Epoch 1297
└ @ Main /Users/kcft114/.julia/packages/Flux/qXNjB/src/optimise/train.jl:105
┌ Info: Epoch 1298
└ @ Main /Users/kcft114/.julia/packages/Flux/qXNjB/src/optimise/train.jl:105
┌ Info: Epoch 1299
└ @ Main /Users/kcft114/.julia/packages/Flux/qXNjB/src/optimise/train.jl:105
┌ Info: Epoch 1300
└ @ Main /Users/kcft114/.julia/packages/Flux/qXNjB/src/optimise/train.jl:105
┌ Info: Epoch 1301
└ @ Main /Users/kcft114/.julia/packages/Flux/qXNjB/src/optimise/train.jl:105
┌ Info: Epoch 1302
└ @ Main /Users/kcft114/.julia/packages/Flux/qXNjB/src/optimise/train.jl:105
┌ Info: Epoch 1303
└ @ Main /Users/kcft114/.julia/packages/Flux/qXNjB/src/optimise/train.jl:105
┌ Info: Epoch 1304
└ @ Main /Users/kcft114/.julia/packages/Flux/qXNjB/src/optimise/train.jl:105
┌ Info: Epoch 1305
└ @ Main /Users/kcft114/.julia/packages/Flux/qXNjB/src/optimise/train.jl:105
┌ Info: Epoch 1306
└ @ Main /Users/kcft114/.julia/packages/Flux/qXNjB/src/optimise/train.jl:105
┌ Info: Epoch 1307
└ @ Main /Users/kcft114/.julia/packages/Flux/qXNjB/src/optimise/train.jl:105
┌ Info: Epoch 1308
└ @ Main /Users/kcft114/.julia/packages/Flux/qXNjB/src/optimise/train.jl:105
┌ Info: Epoch 1309
└ @ Main /Users/kcft114/.julia/packages/Flux/qXNjB/src/optimise/train.jl:105
┌ Info: Epoch 1310
└ @ Main /Users/kcft114/.julia/packages/Flux/qXNjB/src/optimise/train.jl:105
┌ Info: Epoch 1311
└ @ Main /Users/kcft114/.julia/packages/Flux/qXNjB/src/optimise/train.jl:105
┌ Info: Epoch 1312
└ @ Main /Users/kcft114/.julia/packages/Flux/qXNjB/src/optimise/train.jl:105
┌ Info: Epoch 1313
└ @ Main /Users/kcft114/.julia/packages/Flux/qXNjB/src/optimise/train.jl:105
┌ Info: Epoch 1314
└ @ Main /Users/kcft114/.julia/packages/Flux/qXNjB/src/optimise/train.jl:105
┌ Info: Epoch 1315
└ @ Main /Users/kcft114/.julia/packages/Flux/qXNjB/src/optimise/train.jl:105
┌ Info: Epoch 1316
└ @ Main /Users/kcft114/.julia/packages/Flux/qXNjB/src/optimise/train.jl:105
┌ Info: Epoch 1317
└ @ Main /Users/kcft114/.julia/packages/Flux/qXNjB/src/optimise/train.jl:105
┌ Info: Epoch 1318
└ @ Main /Users/kcft114/.julia/packages/Flux/qXNjB/src/optimise/train.jl:105
┌ Info: Epoch 1319
└ @ Main /Users/kcft114/.julia/packages/Flux/qXNjB/src/optimise/train.jl:105
┌ Info: Epoch 1320
└ @ Main /Users/kcft114/.julia/packages/Flux/qXNjB/src/optimise/train.jl:105
┌ Info: Epoch 1321
└ @ Main /Users/kcft114/.julia/packages/Flux/qXNjB/src/optimise/train.jl:105
┌ Info: Epoch 1322
└ @ Main /Users/kcft114/.julia/packages/Flux/qXNjB/src/optimise/train.jl:105
┌ Info: Epoch 1323
└ @ Main /Users/kcft114/.julia/packages/Flux/qXNjB/src/optimise/train.jl:105
┌ Info: Epoch 1324
└ @ Main /Users/kcft114/.julia/packages/Flux/qXNjB/src/optimise/train.jl:105
┌ Info: Epoch 1325
└ @ Main /Users/kcft114/.julia/packages/Flux/qXNjB/src/optimise/train.jl:105
┌ Info: Epoch 1326
└ @ Main /Users/kcft114/.julia/packages/Flux/qXNjB/src/optimise/train.jl:105
┌ Info: Epoch 1327
└ @ Main /Users/kcft114/.julia/packages/Flux/qXNjB/src/optimise/train.jl:105
┌ Info: Epoch 1328
└ @ Main /Users/kcft114/.julia/packages/Flux/qXNjB/src/optimise/train.jl:105
┌ Info: Epoch 1329
└ @ Main /Users/kcft114/.julia/packages/Flux/qXNjB/src/optimise/train.jl:105
┌ Info: Epoch 1330
└ @ Main /Users/kcft114/.julia/packages/Flux/qXNjB/src/optimise/train.jl:105
┌ Info: Epoch 1331
└ @ Main /Users/kcft114/.julia/packages/Flux/qXNjB/src/optimise/train.jl:105
┌ Info: Epoch 1332
└ @ Main /Users/kcft114/.julia/packages/Flux/qXNjB/src/optimise/train.jl:105
┌ Info: Epoch 1333
└ @ Main /Users/kcft114/.julia/packages/Flux/qXNjB/src/optimise/train.jl:105
┌ Info: Epoch 1334
└ @ Main /Users/kcft114/.julia/packages/Flux/qXNjB/src/optimise/train.jl:105
┌ Info: Epoch 1335
└ @ Main /Users/kcft114/.julia/packages/Flux/qXNjB/src/optimise/train.jl:105
┌ Info: Epoch 1336
└ @ Main /Users/kcft114/.julia/packages/Flux/qXNjB/src/optimise/train.jl:105
┌ Info: Epoch 1337
└ @ Main /Users/kcft114/.julia/packages/Flux/qXNjB/src/optimise/train.jl:105
┌ Info: Epoch 1338
└ @ Main /Users/kcft114/.julia/packages/Flux/qXNjB/src/optimise/train.jl:105
┌ Info: Epoch 1339
└ @ Main /Users/kcft114/.julia/packages/Flux/qXNjB/src/optimise/train.jl:105
┌ Info: Epoch 1340
└ @ Main /Users/kcft114/.julia/packages/Flux/qXNjB/src/optimise/train.jl:105
┌ Info: Epoch 1341
└ @ Main /Users/kcft114/.julia/packages/Flux/qXNjB/src/optimise/train.jl:105
┌ Info: Epoch 1342
└ @ Main /Users/kcft114/.julia/packages/Flux/qXNjB/src/optimise/train.jl:105
┌ Info: Epoch 1343
└ @ Main /Users/kcft114/.julia/packages/Flux/qXNjB/src/optimise/train.jl:105
┌ Info: Epoch 1344
└ @ Main /Users/kcft114/.julia/packages/Flux/qXNjB/src/optimise/train.jl:105
┌ Info: Epoch 1345
└ @ Main /Users/kcft114/.julia/packages/Flux/qXNjB/src/optimise/train.jl:105
┌ Info: Epoch 1346
└ @ Main /Users/kcft114/.julia/packages/Flux/qXNjB/src/optimise/train.jl:105
┌ Info: Epoch 1347
└ @ Main /Users/kcft114/.julia/packages/Flux/qXNjB/src/optimise/train.jl:105
┌ Info: Epoch 1348
└ @ Main /Users/kcft114/.julia/packages/Flux/qXNjB/src/optimise/train.jl:105
┌ Info: Epoch 1349
└ @ Main /Users/kcft114/.julia/packages/Flux/qXNjB/src/optimise/train.jl:105
┌ Info: Epoch 1350
└ @ Main /Users/kcft114/.julia/packages/Flux/qXNjB/src/optimise/train.jl:105
┌ Info: Epoch 1351
└ @ Main /Users/kcft114/.julia/packages/Flux/qXNjB/src/optimise/train.jl:105
┌ Info: Epoch 1352
└ @ Main /Users/kcft114/.julia/packages/Flux/qXNjB/src/optimise/train.jl:105
┌ Info: Epoch 1353
└ @ Main /Users/kcft114/.julia/packages/Flux/qXNjB/src/optimise/train.jl:105
┌ Info: Epoch 1354
└ @ Main /Users/kcft114/.julia/packages/Flux/qXNjB/src/optimise/train.jl:105
┌ Info: Epoch 1355
└ @ Main /Users/kcft114/.julia/packages/Flux/qXNjB/src/optimise/train.jl:105
┌ Info: Epoch 1356
└ @ Main /Users/kcft114/.julia/packages/Flux/qXNjB/src/optimise/train.jl:105
┌ Info: Epoch 1357
└ @ Main /Users/kcft114/.julia/packages/Flux/qXNjB/src/optimise/train.jl:105
┌ Info: Epoch 1358
└ @ Main /Users/kcft114/.julia/packages/Flux/qXNjB/src/optimise/train.jl:105
┌ Info: Epoch 1359
└ @ Main /Users/kcft114/.julia/packages/Flux/qXNjB/src/optimise/train.jl:105
┌ Info: Epoch 1360
└ @ Main /Users/kcft114/.julia/packages/Flux/qXNjB/src/optimise/train.jl:105
┌ Info: Epoch 1361
└ @ Main /Users/kcft114/.julia/packages/Flux/qXNjB/src/optimise/train.jl:105
┌ Info: Epoch 1362
└ @ Main /Users/kcft114/.julia/packages/Flux/qXNjB/src/optimise/train.jl:105
┌ Info: Epoch 1363
└ @ Main /Users/kcft114/.julia/packages/Flux/qXNjB/src/optimise/train.jl:105
┌ Info: Epoch 1364
└ @ Main /Users/kcft114/.julia/packages/Flux/qXNjB/src/optimise/train.jl:105
┌ Info: Epoch 1365
└ @ Main /Users/kcft114/.julia/packages/Flux/qXNjB/src/optimise/train.jl:105
┌ Info: Epoch 1366
└ @ Main /Users/kcft114/.julia/packages/Flux/qXNjB/src/optimise/train.jl:105
┌ Info: Epoch 1367
└ @ Main /Users/kcft114/.julia/packages/Flux/qXNjB/src/optimise/train.jl:105
┌ Info: Epoch 1368
└ @ Main /Users/kcft114/.julia/packages/Flux/qXNjB/src/optimise/train.jl:105
┌ Info: Epoch 1369
└ @ Main /Users/kcft114/.julia/packages/Flux/qXNjB/src/optimise/train.jl:105
┌ Info: Epoch 1370
└ @ Main /Users/kcft114/.julia/packages/Flux/qXNjB/src/optimise/train.jl:105
┌ Info: Epoch 1371
└ @ Main /Users/kcft114/.julia/packages/Flux/qXNjB/src/optimise/train.jl:105
┌ Info: Epoch 1372
└ @ Main /Users/kcft114/.julia/packages/Flux/qXNjB/src/optimise/train.jl:105
┌ Info: Epoch 1373
└ @ Main /Users/kcft114/.julia/packages/Flux/qXNjB/src/optimise/train.jl:105
┌ Info: Epoch 1374
└ @ Main /Users/kcft114/.julia/packages/Flux/qXNjB/src/optimise/train.jl:105
┌ Info: Epoch 1375
└ @ Main /Users/kcft114/.julia/packages/Flux/qXNjB/src/optimise/train.jl:105
┌ Info: Epoch 1376
└ @ Main /Users/kcft114/.julia/packages/Flux/qXNjB/src/optimise/train.jl:105
┌ Info: Epoch 1377
└ @ Main /Users/kcft114/.julia/packages/Flux/qXNjB/src/optimise/train.jl:105
┌ Info: Epoch 1378
└ @ Main /Users/kcft114/.julia/packages/Flux/qXNjB/src/optimise/train.jl:105
┌ Info: Epoch 1379
└ @ Main /Users/kcft114/.julia/packages/Flux/qXNjB/src/optimise/train.jl:105
┌ Info: Epoch 1380
└ @ Main /Users/kcft114/.julia/packages/Flux/qXNjB/src/optimise/train.jl:105
┌ Info: Epoch 1381
└ @ Main /Users/kcft114/.julia/packages/Flux/qXNjB/src/optimise/train.jl:105
┌ Info: Epoch 1382
└ @ Main /Users/kcft114/.julia/packages/Flux/qXNjB/src/optimise/train.jl:105
┌ Info: Epoch 1383
└ @ Main /Users/kcft114/.julia/packages/Flux/qXNjB/src/optimise/train.jl:105
┌ Info: Epoch 1384
└ @ Main /Users/kcft114/.julia/packages/Flux/qXNjB/src/optimise/train.jl:105
┌ Info: Epoch 1385
└ @ Main /Users/kcft114/.julia/packages/Flux/qXNjB/src/optimise/train.jl:105
┌ Info: Epoch 1386
└ @ Main /Users/kcft114/.julia/packages/Flux/qXNjB/src/optimise/train.jl:105
┌ Info: Epoch 1387
└ @ Main /Users/kcft114/.julia/packages/Flux/qXNjB/src/optimise/train.jl:105
┌ Info: Epoch 1388
└ @ Main /Users/kcft114/.julia/packages/Flux/qXNjB/src/optimise/train.jl:105
┌ Info: Epoch 1389
└ @ Main /Users/kcft114/.julia/packages/Flux/qXNjB/src/optimise/train.jl:105
┌ Info: Epoch 1390
└ @ Main /Users/kcft114/.julia/packages/Flux/qXNjB/src/optimise/train.jl:105
┌ Info: Epoch 1391
└ @ Main /Users/kcft114/.julia/packages/Flux/qXNjB/src/optimise/train.jl:105
┌ Info: Epoch 1392
└ @ Main /Users/kcft114/.julia/packages/Flux/qXNjB/src/optimise/train.jl:105
┌ Info: Epoch 1393
└ @ Main /Users/kcft114/.julia/packages/Flux/qXNjB/src/optimise/train.jl:105
┌ Info: Epoch 1394
└ @ Main /Users/kcft114/.julia/packages/Flux/qXNjB/src/optimise/train.jl:105
┌ Info: Epoch 1395
└ @ Main /Users/kcft114/.julia/packages/Flux/qXNjB/src/optimise/train.jl:105
┌ Info: Epoch 1396
└ @ Main /Users/kcft114/.julia/packages/Flux/qXNjB/src/optimise/train.jl:105
┌ Info: Epoch 1397
└ @ Main /Users/kcft114/.julia/packages/Flux/qXNjB/src/optimise/train.jl:105
┌ Info: Epoch 1398
└ @ Main /Users/kcft114/.julia/packages/Flux/qXNjB/src/optimise/train.jl:105
┌ Info: Epoch 1399
└ @ Main /Users/kcft114/.julia/packages/Flux/qXNjB/src/optimise/train.jl:105
┌ Info: Epoch 1400
└ @ Main /Users/kcft114/.julia/packages/Flux/qXNjB/src/optimise/train.jl:105
┌ Info: Epoch 1401
└ @ Main /Users/kcft114/.julia/packages/Flux/qXNjB/src/optimise/train.jl:105
┌ Info: Epoch 1402
└ @ Main /Users/kcft114/.julia/packages/Flux/qXNjB/src/optimise/train.jl:105
┌ Info: Epoch 1403
└ @ Main /Users/kcft114/.julia/packages/Flux/qXNjB/src/optimise/train.jl:105
┌ Info: Epoch 1404
└ @ Main /Users/kcft114/.julia/packages/Flux/qXNjB/src/optimise/train.jl:105
┌ Info: Epoch 1405
└ @ Main /Users/kcft114/.julia/packages/Flux/qXNjB/src/optimise/train.jl:105
┌ Info: Epoch 1406
└ @ Main /Users/kcft114/.julia/packages/Flux/qXNjB/src/optimise/train.jl:105
┌ Info: Epoch 1407
└ @ Main /Users/kcft114/.julia/packages/Flux/qXNjB/src/optimise/train.jl:105
┌ Info: Epoch 1408
└ @ Main /Users/kcft114/.julia/packages/Flux/qXNjB/src/optimise/train.jl:105
┌ Info: Epoch 1409
└ @ Main /Users/kcft114/.julia/packages/Flux/qXNjB/src/optimise/train.jl:105
```

```
loss(X, Y) = 0.40143824f0 (tracked)
loss(X, Y) = 0.40119085f0 (tracked)
loss(X, Y) = 0.4009442f0 (tracked)
loss(X, Y) = 0.40068698f0 (tracked)
loss(X, Y) = 0.40043595f0 (tracked)
loss(X, Y) = 0.400186f0 (tracked)
loss(X, Y) = 0.3999352f0 (tracked)
loss(X, Y) = 0.39968058f0 (tracked)
loss(X, Y) = 0.39942554f0 (tracked)
loss(X, Y) = 0.39917442f0 (tracked)
loss(X, Y) = 0.39892894f0 (tracked)
loss(X, Y) = 0.39867398f0 (tracked)
loss(X, Y) = 0.39842415f0 (tracked)
loss(X, Y) = 0.3981803f0 (tracked)
loss(X, Y) = 0.3979198f0 (tracked)
loss(X, Y) = 0.39766234f0 (tracked)
loss(X, Y) = 0.39739603f0 (tracked)
loss(X, Y) = 0.39713955f0 (tracked)
loss(X, Y) = 0.39688033f0 (tracked)
loss(X, Y) = 0.39661354f0 (tracked)
loss(X, Y) = 0.39636177f0 (tracked)
loss(X, Y) = 0.39611116f0 (tracked)
loss(X, Y) = 0.3958562f0 (tracked)
loss(X, Y) = 0.3956012f0 (tracked)
loss(X, Y) = 0.39535287f0 (tracked)
loss(X, Y) = 0.395098f0 (tracked)
loss(X, Y) = 0.39483306f0 (tracked)
loss(X, Y) = 0.39458278f0 (tracked)
loss(X, Y) = 0.39433485f0 (tracked)
loss(X, Y) = 0.39408833f0 (tracked)
loss(X, Y) = 0.39382544f0 (tracked)
loss(X, Y) = 0.393573f0 (tracked)
loss(X, Y) = 0.39331663f0 (tracked)
loss(X, Y) = 0.39307073f0 (tracked)
loss(X, Y) = 0.3928348f0 (tracked)
loss(X, Y) = 0.3925834f0 (tracked)
loss(X, Y) = 0.39233002f0 (tracked)
loss(X, Y) = 0.39209002f0 (tracked)
loss(X, Y) = 0.39184618f0 (tracked)
loss(X, Y) = 0.39159736f0 (tracked)
loss(X, Y) = 0.39135453f0 (tracked)
loss(X, Y) = 0.39112002f0 (tracked)
loss(X, Y) = 0.39086974f0 (tracked)
loss(X, Y) = 0.39063162f0 (tracked)
loss(X, Y) = 0.3903983f0 (tracked)
loss(X, Y) = 0.39014915f0 (tracked)
loss(X, Y) = 0.38990545f0 (tracked)
loss(X, Y) = 0.38967457f0 (tracked)
loss(X, Y) = 0.38944027f0 (tracked)
loss(X, Y) = 0.3892001f0 (tracked)
loss(X, Y) = 0.38895646f0 (tracked)
loss(X, Y) = 0.38871554f0 (tracked)
loss(X, Y) = 0.3884896f0 (tracked)
loss(X, Y) = 0.3882541f0 (tracked)
loss(X, Y) = 0.38801625f0 (tracked)
loss(X, Y) = 0.38777417f0 (tracked)
loss(X, Y) = 0.3875511f0 (tracked)
loss(X, Y) = 0.38732222f0 (tracked)
loss(X, Y) = 0.38709107f0 (tracked)
loss(X, Y) = 0.38685337f0 (tracked)
loss(X, Y) = 0.38661644f0 (tracked)
loss(X, Y) = 0.38638875f0 (tracked)
loss(X, Y) = 0.3861551f0 (tracked)
loss(X, Y) = 0.38592172f0 (tracked)
loss(X, Y) = 0.38569307f0 (tracked)
loss(X, Y) = 0.38547286f0 (tracked)
loss(X, Y) = 0.38523808f0 (tracked)
loss(X, Y) = 0.38500106f0 (tracked)
loss(X, Y) = 0.3847833f0 (tracked)
loss(X, Y) = 0.38454884f0 (tracked)
loss(X, Y) = 0.38431725f0 (tracked)
loss(X, Y) = 0.38409385f0 (tracked)
loss(X, Y) = 0.38385916f0 (tracked)
loss(X, Y) = 0.38362396f0 (tracked)
loss(X, Y) = 0.38340583f0 (tracked)
loss(X, Y) = 0.38317722f0 (tracked)
loss(X, Y) = 0.38294268f0 (tracked)
loss(X, Y) = 0.38272002f0 (tracked)
loss(X, Y) = 0.38249558f0 (tracked)
loss(X, Y) = 0.38225946f0 (tracked)
loss(X, Y) = 0.38202038f0 (tracked)
loss(X, Y) = 0.3817926f0 (tracked)
loss(X, Y) = 0.38155815f0 (tracked)
loss(X, Y) = 0.3813235f0 (tracked)
loss(X, Y) = 0.38109338f0 (tracked)
loss(X, Y) = 0.38086542f0 (tracked)
loss(X, Y) = 0.3806248f0 (tracked)
loss(X, Y) = 0.3803992f0 (tracked)
loss(X, Y) = 0.3801629f0 (tracked)
loss(X, Y) = 0.37992853f0 (tracked)
loss(X, Y) = 0.37969783f0 (tracked)
loss(X, Y) = 0.3794708f0 (tracked)
loss(X, Y) = 0.37923938f0 (tracked)
loss(X, Y) = 0.3790007f0 (tracked)
loss(X, Y) = 0.3787731f0 (tracked)
loss(X, Y) = 0.37854207f0 (tracked)
loss(X, Y) = 0.37830466f0 (tracked)
loss(X, Y) = 0.3780773f0 (tracked)
loss(X, Y) = 0.37784886f0 (tracked)
loss(X, Y) = 0.37762135f0 (tracked)
loss(X, Y) = 0.3773984f0 (tracked)
loss(X, Y) = 0.37717572f0 (tracked)
loss(X, Y) = 0.37694559f0 (tracked)
loss(X, Y) = 0.37672344f0 (tracked)
loss(X, Y) = 0.3764944f0 (tracked)
loss(X, Y) = 0.376266f0 (tracked)
loss(X, Y) = 0.376046f0 (tracked)
loss(X, Y) = 0.37581822f0 (tracked)
loss(X, Y) = 0.37559268f0 (tracked)
loss(X, Y) = 0.37536263f0 (tracked)
loss(X, Y) = 0.3751424f0 (tracked)
loss(X, Y) = 0.37491357f0 (tracked)
loss(X, Y) = 0.37469277f0 (tracked)
loss(X, Y) = 0.37446353f0 (tracked)
loss(X, Y) = 0.37423447f0 (tracked)
loss(X, Y) = 0.3740119f0 (tracked)
loss(X, Y) = 0.37379438f0 (tracked)
loss(X, Y) = 0.3735712f0 (tracked)
loss(X, Y) = 0.37334573f0 (tracked)
loss(X, Y) = 0.37311992f0 (tracked)
loss(X, Y) = 0.37288746f0 (tracked)
loss(X, Y) = 0.3726668f0 (tracked)
loss(X, Y) = 0.3724484f0 (tracked)
loss(X, Y) = 0.3722239f0 (tracked)
loss(X, Y) = 0.3720016f0 (tracked)
loss(X, Y) = 0.37177354f0 (tracked)
loss(X, Y) = 0.37155694f0 (tracked)
loss(X, Y) = 0.371337f0 (tracked)
loss(X, Y) = 0.3711195f0 (tracked)
loss(X, Y) = 0.3708948f0 (tracked)
loss(X, Y) = 0.37068272f0 (tracked)
loss(X, Y) = 0.37045777f0 (tracked)
loss(X, Y) = 0.37022525f0 (tracked)
loss(X, Y) = 0.37001282f0 (tracked)
loss(X, Y) = 0.36981305f0 (tracked)
loss(X, Y) = 0.36958343f0 (tracked)
loss(X, Y) = 0.36935994f0 (tracked)
loss(X, Y) = 0.36915523f0 (tracked)
loss(X, Y) = 0.36894527f0 (tracked)
loss(X, Y) = 0.36872232f0 (tracked)
loss(X, Y) = 0.36849368f0 (tracked)
loss(X, Y) = 0.36827552f0 (tracked)
loss(X, Y) = 0.36806598f0 (tracked)
loss(X, Y) = 0.3678521f0 (tracked)
loss(X, Y) = 0.36763263f0 (tracked)
loss(X, Y) = 0.36740193f0 (tracked)
loss(X, Y) = 0.3671811f0 (tracked)
```

```
┌ Info: Epoch 1410
└ @ Main /Users/kcft114/.julia/packages/Flux/qXNjB/src/optimise/train.jl:105
┌ Info: Epoch 1411
└ @ Main /Users/kcft114/.julia/packages/Flux/qXNjB/src/optimise/train.jl:105
┌ Info: Epoch 1412
└ @ Main /Users/kcft114/.julia/packages/Flux/qXNjB/src/optimise/train.jl:105
┌ Info: Epoch 1413
└ @ Main /Users/kcft114/.julia/packages/Flux/qXNjB/src/optimise/train.jl:105
┌ Info: Epoch 1414
└ @ Main /Users/kcft114/.julia/packages/Flux/qXNjB/src/optimise/train.jl:105
┌ Info: Epoch 1415
└ @ Main /Users/kcft114/.julia/packages/Flux/qXNjB/src/optimise/train.jl:105
┌ Info: Epoch 1416
└ @ Main /Users/kcft114/.julia/packages/Flux/qXNjB/src/optimise/train.jl:105
┌ Info: Epoch 1417
└ @ Main /Users/kcft114/.julia/packages/Flux/qXNjB/src/optimise/train.jl:105
┌ Info: Epoch 1418
└ @ Main /Users/kcft114/.julia/packages/Flux/qXNjB/src/optimise/train.jl:105
┌ Info: Epoch 1419
└ @ Main /Users/kcft114/.julia/packages/Flux/qXNjB/src/optimise/train.jl:105
┌ Info: Epoch 1420
└ @ Main /Users/kcft114/.julia/packages/Flux/qXNjB/src/optimise/train.jl:105
┌ Info: Epoch 1421
└ @ Main /Users/kcft114/.julia/packages/Flux/qXNjB/src/optimise/train.jl:105
┌ Info: Epoch 1422
└ @ Main /Users/kcft114/.julia/packages/Flux/qXNjB/src/optimise/train.jl:105
┌ Info: Epoch 1423
└ @ Main /Users/kcft114/.julia/packages/Flux/qXNjB/src/optimise/train.jl:105
┌ Info: Epoch 1424
└ @ Main /Users/kcft114/.julia/packages/Flux/qXNjB/src/optimise/train.jl:105
┌ Info: Epoch 1425
└ @ Main /Users/kcft114/.julia/packages/Flux/qXNjB/src/optimise/train.jl:105
┌ Info: Epoch 1426
└ @ Main /Users/kcft114/.julia/packages/Flux/qXNjB/src/optimise/train.jl:105
┌ Info: Epoch 1427
└ @ Main /Users/kcft114/.julia/packages/Flux/qXNjB/src/optimise/train.jl:105
┌ Info: Epoch 1428
└ @ Main /Users/kcft114/.julia/packages/Flux/qXNjB/src/optimise/train.jl:105
┌ Info: Epoch 1429
└ @ Main /Users/kcft114/.julia/packages/Flux/qXNjB/src/optimise/train.jl:105
┌ Info: Epoch 1430
└ @ Main /Users/kcft114/.julia/packages/Flux/qXNjB/src/optimise/train.jl:105
┌ Info: Epoch 1431
└ @ Main /Users/kcft114/.julia/packages/Flux/qXNjB/src/optimise/train.jl:105
┌ Info: Epoch 1432
└ @ Main /Users/kcft114/.julia/packages/Flux/qXNjB/src/optimise/train.jl:105
┌ Info: Epoch 1433
└ @ Main /Users/kcft114/.julia/packages/Flux/qXNjB/src/optimise/train.jl:105
┌ Info: Epoch 1434
└ @ Main /Users/kcft114/.julia/packages/Flux/qXNjB/src/optimise/train.jl:105
┌ Info: Epoch 1435
└ @ Main /Users/kcft114/.julia/packages/Flux/qXNjB/src/optimise/train.jl:105
┌ Info: Epoch 1436
└ @ Main /Users/kcft114/.julia/packages/Flux/qXNjB/src/optimise/train.jl:105
┌ Info: Epoch 1437
└ @ Main /Users/kcft114/.julia/packages/Flux/qXNjB/src/optimise/train.jl:105
┌ Info: Epoch 1438
└ @ Main /Users/kcft114/.julia/packages/Flux/qXNjB/src/optimise/train.jl:105
┌ Info: Epoch 1439
└ @ Main /Users/kcft114/.julia/packages/Flux/qXNjB/src/optimise/train.jl:105
┌ Info: Epoch 1440
└ @ Main /Users/kcft114/.julia/packages/Flux/qXNjB/src/optimise/train.jl:105
┌ Info: Epoch 1441
└ @ Main /Users/kcft114/.julia/packages/Flux/qXNjB/src/optimise/train.jl:105
┌ Info: Epoch 1442
└ @ Main /Users/kcft114/.julia/packages/Flux/qXNjB/src/optimise/train.jl:105
┌ Info: Epoch 1443
└ @ Main /Users/kcft114/.julia/packages/Flux/qXNjB/src/optimise/train.jl:105
┌ Info: Epoch 1444
└ @ Main /Users/kcft114/.julia/packages/Flux/qXNjB/src/optimise/train.jl:105
┌ Info: Epoch 1445
└ @ Main /Users/kcft114/.julia/packages/Flux/qXNjB/src/optimise/train.jl:105
┌ Info: Epoch 1446
└ @ Main /Users/kcft114/.julia/packages/Flux/qXNjB/src/optimise/train.jl:105
┌ Info: Epoch 1447
└ @ Main /Users/kcft114/.julia/packages/Flux/qXNjB/src/optimise/train.jl:105
┌ Info: Epoch 1448
└ @ Main /Users/kcft114/.julia/packages/Flux/qXNjB/src/optimise/train.jl:105
┌ Info: Epoch 1449
└ @ Main /Users/kcft114/.julia/packages/Flux/qXNjB/src/optimise/train.jl:105
┌ Info: Epoch 1450
└ @ Main /Users/kcft114/.julia/packages/Flux/qXNjB/src/optimise/train.jl:105
┌ Info: Epoch 1451
└ @ Main /Users/kcft114/.julia/packages/Flux/qXNjB/src/optimise/train.jl:105
┌ Info: Epoch 1452
└ @ Main /Users/kcft114/.julia/packages/Flux/qXNjB/src/optimise/train.jl:105
┌ Info: Epoch 1453
└ @ Main /Users/kcft114/.julia/packages/Flux/qXNjB/src/optimise/train.jl:105
┌ Info: Epoch 1454
└ @ Main /Users/kcft114/.julia/packages/Flux/qXNjB/src/optimise/train.jl:105
┌ Info: Epoch 1455
└ @ Main /Users/kcft114/.julia/packages/Flux/qXNjB/src/optimise/train.jl:105
┌ Info: Epoch 1456
└ @ Main /Users/kcft114/.julia/packages/Flux/qXNjB/src/optimise/train.jl:105
┌ Info: Epoch 1457
└ @ Main /Users/kcft114/.julia/packages/Flux/qXNjB/src/optimise/train.jl:105
┌ Info: Epoch 1458
└ @ Main /Users/kcft114/.julia/packages/Flux/qXNjB/src/optimise/train.jl:105
┌ Info: Epoch 1459
└ @ Main /Users/kcft114/.julia/packages/Flux/qXNjB/src/optimise/train.jl:105
┌ Info: Epoch 1460
└ @ Main /Users/kcft114/.julia/packages/Flux/qXNjB/src/optimise/train.jl:105
┌ Info: Epoch 1461
└ @ Main /Users/kcft114/.julia/packages/Flux/qXNjB/src/optimise/train.jl:105
┌ Info: Epoch 1462
└ @ Main /Users/kcft114/.julia/packages/Flux/qXNjB/src/optimise/train.jl:105
┌ Info: Epoch 1463
└ @ Main /Users/kcft114/.julia/packages/Flux/qXNjB/src/optimise/train.jl:105
┌ Info: Epoch 1464
└ @ Main /Users/kcft114/.julia/packages/Flux/qXNjB/src/optimise/train.jl:105
┌ Info: Epoch 1465
└ @ Main /Users/kcft114/.julia/packages/Flux/qXNjB/src/optimise/train.jl:105
┌ Info: Epoch 1466
└ @ Main /Users/kcft114/.julia/packages/Flux/qXNjB/src/optimise/train.jl:105
┌ Info: Epoch 1467
└ @ Main /Users/kcft114/.julia/packages/Flux/qXNjB/src/optimise/train.jl:105
┌ Info: Epoch 1468
└ @ Main /Users/kcft114/.julia/packages/Flux/qXNjB/src/optimise/train.jl:105
┌ Info: Epoch 1469
└ @ Main /Users/kcft114/.julia/packages/Flux/qXNjB/src/optimise/train.jl:105
┌ Info: Epoch 1470
└ @ Main /Users/kcft114/.julia/packages/Flux/qXNjB/src/optimise/train.jl:105
┌ Info: Epoch 1471
└ @ Main /Users/kcft114/.julia/packages/Flux/qXNjB/src/optimise/train.jl:105
┌ Info: Epoch 1472
└ @ Main /Users/kcft114/.julia/packages/Flux/qXNjB/src/optimise/train.jl:105
┌ Info: Epoch 1473
└ @ Main /Users/kcft114/.julia/packages/Flux/qXNjB/src/optimise/train.jl:105
┌ Info: Epoch 1474
└ @ Main /Users/kcft114/.julia/packages/Flux/qXNjB/src/optimise/train.jl:105
┌ Info: Epoch 1475
└ @ Main /Users/kcft114/.julia/packages/Flux/qXNjB/src/optimise/train.jl:105
┌ Info: Epoch 1476
└ @ Main /Users/kcft114/.julia/packages/Flux/qXNjB/src/optimise/train.jl:105
┌ Info: Epoch 1477
└ @ Main /Users/kcft114/.julia/packages/Flux/qXNjB/src/optimise/train.jl:105
┌ Info: Epoch 1478
└ @ Main /Users/kcft114/.julia/packages/Flux/qXNjB/src/optimise/train.jl:105
┌ Info: Epoch 1479
└ @ Main /Users/kcft114/.julia/packages/Flux/qXNjB/src/optimise/train.jl:105
┌ Info: Epoch 1480
└ @ Main /Users/kcft114/.julia/packages/Flux/qXNjB/src/optimise/train.jl:105
┌ Info: Epoch 1481
└ @ Main /Users/kcft114/.julia/packages/Flux/qXNjB/src/optimise/train.jl:105
┌ Info: Epoch 1482
└ @ Main /Users/kcft114/.julia/packages/Flux/qXNjB/src/optimise/train.jl:105
┌ Info: Epoch 1483
└ @ Main /Users/kcft114/.julia/packages/Flux/qXNjB/src/optimise/train.jl:105
┌ Info: Epoch 1484
└ @ Main /Users/kcft114/.julia/packages/Flux/qXNjB/src/optimise/train.jl:105
┌ Info: Epoch 1485
└ @ Main /Users/kcft114/.julia/packages/Flux/qXNjB/src/optimise/train.jl:105
┌ Info: Epoch 1486
└ @ Main /Users/kcft114/.julia/packages/Flux/qXNjB/src/optimise/train.jl:105
┌ Info: Epoch 1487
└ @ Main /Users/kcft114/.julia/packages/Flux/qXNjB/src/optimise/train.jl:105
┌ Info: Epoch 1488
└ @ Main /Users/kcft114/.julia/packages/Flux/qXNjB/src/optimise/train.jl:105
┌ Info: Epoch 1489
└ @ Main /Users/kcft114/.julia/packages/Flux/qXNjB/src/optimise/train.jl:105
┌ Info: Epoch 1490
└ @ Main /Users/kcft114/.julia/packages/Flux/qXNjB/src/optimise/train.jl:105
┌ Info: Epoch 1491
└ @ Main /Users/kcft114/.julia/packages/Flux/qXNjB/src/optimise/train.jl:105
┌ Info: Epoch 1492
└ @ Main /Users/kcft114/.julia/packages/Flux/qXNjB/src/optimise/train.jl:105
┌ Info: Epoch 1493
└ @ Main /Users/kcft114/.julia/packages/Flux/qXNjB/src/optimise/train.jl:105
┌ Info: Epoch 1494
└ @ Main /Users/kcft114/.julia/packages/Flux/qXNjB/src/optimise/train.jl:105
┌ Info: Epoch 1495
└ @ Main /Users/kcft114/.julia/packages/Flux/qXNjB/src/optimise/train.jl:105
┌ Info: Epoch 1496
└ @ Main /Users/kcft114/.julia/packages/Flux/qXNjB/src/optimise/train.jl:105
┌ Info: Epoch 1497
└ @ Main /Users/kcft114/.julia/packages/Flux/qXNjB/src/optimise/train.jl:105
┌ Info: Epoch 1498
└ @ Main /Users/kcft114/.julia/packages/Flux/qXNjB/src/optimise/train.jl:105
┌ Info: Epoch 1499
└ @ Main /Users/kcft114/.julia/packages/Flux/qXNjB/src/optimise/train.jl:105
┌ Info: Epoch 1500
└ @ Main /Users/kcft114/.julia/packages/Flux/qXNjB/src/optimise/train.jl:105
┌ Info: Epoch 1501
└ @ Main /Users/kcft114/.julia/packages/Flux/qXNjB/src/optimise/train.jl:105
┌ Info: Epoch 1502
└ @ Main /Users/kcft114/.julia/packages/Flux/qXNjB/src/optimise/train.jl:105
┌ Info: Epoch 1503
└ @ Main /Users/kcft114/.julia/packages/Flux/qXNjB/src/optimise/train.jl:105
┌ Info: Epoch 1504
└ @ Main /Users/kcft114/.julia/packages/Flux/qXNjB/src/optimise/train.jl:105
┌ Info: Epoch 1505
└ @ Main /Users/kcft114/.julia/packages/Flux/qXNjB/src/optimise/train.jl:105
┌ Info: Epoch 1506
└ @ Main /Users/kcft114/.julia/packages/Flux/qXNjB/src/optimise/train.jl:105
┌ Info: Epoch 1507
└ @ Main /Users/kcft114/.julia/packages/Flux/qXNjB/src/optimise/train.jl:105
┌ Info: Epoch 1508
└ @ Main /Users/kcft114/.julia/packages/Flux/qXNjB/src/optimise/train.jl:105
┌ Info: Epoch 1509
└ @ Main /Users/kcft114/.julia/packages/Flux/qXNjB/src/optimise/train.jl:105
┌ Info: Epoch 1510
└ @ Main /Users/kcft114/.julia/packages/Flux/qXNjB/src/optimise/train.jl:105
┌ Info: Epoch 1511
└ @ Main /Users/kcft114/.julia/packages/Flux/qXNjB/src/optimise/train.jl:105
┌ Info: Epoch 1512
└ @ Main /Users/kcft114/.julia/packages/Flux/qXNjB/src/optimise/train.jl:105
┌ Info: Epoch 1513
└ @ Main /Users/kcft114/.julia/packages/Flux/qXNjB/src/optimise/train.jl:105
┌ Info: Epoch 1514
└ @ Main /Users/kcft114/.julia/packages/Flux/qXNjB/src/optimise/train.jl:105
┌ Info: Epoch 1515
└ @ Main /Users/kcft114/.julia/packages/Flux/qXNjB/src/optimise/train.jl:105
┌ Info: Epoch 1516
└ @ Main /Users/kcft114/.julia/packages/Flux/qXNjB/src/optimise/train.jl:105
┌ Info: Epoch 1517
└ @ Main /Users/kcft114/.julia/packages/Flux/qXNjB/src/optimise/train.jl:105
┌ Info: Epoch 1518
└ @ Main /Users/kcft114/.julia/packages/Flux/qXNjB/src/optimise/train.jl:105
┌ Info: Epoch 1519
└ @ Main /Users/kcft114/.julia/packages/Flux/qXNjB/src/optimise/train.jl:105
┌ Info: Epoch 1520
└ @ Main /Users/kcft114/.julia/packages/Flux/qXNjB/src/optimise/train.jl:105
┌ Info: Epoch 1521
└ @ Main /Users/kcft114/.julia/packages/Flux/qXNjB/src/optimise/train.jl:105
┌ Info: Epoch 1522
└ @ Main /Users/kcft114/.julia/packages/Flux/qXNjB/src/optimise/train.jl:105
┌ Info: Epoch 1523
└ @ Main /Users/kcft114/.julia/packages/Flux/qXNjB/src/optimise/train.jl:105
┌ Info: Epoch 1524
└ @ Main /Users/kcft114/.julia/packages/Flux/qXNjB/src/optimise/train.jl:105
┌ Info: Epoch 1525
└ @ Main /Users/kcft114/.julia/packages/Flux/qXNjB/src/optimise/train.jl:105
┌ Info: Epoch 1526
└ @ Main /Users/kcft114/.julia/packages/Flux/qXNjB/src/optimise/train.jl:105
┌ Info: Epoch 1527
└ @ Main /Users/kcft114/.julia/packages/Flux/qXNjB/src/optimise/train.jl:105
┌ Info: Epoch 1528
└ @ Main /Users/kcft114/.julia/packages/Flux/qXNjB/src/optimise/train.jl:105
┌ Info: Epoch 1529
└ @ Main /Users/kcft114/.julia/packages/Flux/qXNjB/src/optimise/train.jl:105
┌ Info: Epoch 1530
└ @ Main /Users/kcft114/.julia/packages/Flux/qXNjB/src/optimise/train.jl:105
┌ Info: Epoch 1531
└ @ Main /Users/kcft114/.julia/packages/Flux/qXNjB/src/optimise/train.jl:105
┌ Info: Epoch 1532
└ @ Main /Users/kcft114/.julia/packages/Flux/qXNjB/src/optimise/train.jl:105
┌ Info: Epoch 1533
└ @ Main /Users/kcft114/.julia/packages/Flux/qXNjB/src/optimise/train.jl:105
┌ Info: Epoch 1534
└ @ Main /Users/kcft114/.julia/packages/Flux/qXNjB/src/optimise/train.jl:105
┌ Info: Epoch 1535
└ @ Main /Users/kcft114/.julia/packages/Flux/qXNjB/src/optimise/train.jl:105
┌ Info: Epoch 1536
└ @ Main /Users/kcft114/.julia/packages/Flux/qXNjB/src/optimise/train.jl:105
┌ Info: Epoch 1537
└ @ Main /Users/kcft114/.julia/packages/Flux/qXNjB/src/optimise/train.jl:105
┌ Info: Epoch 1538
└ @ Main /Users/kcft114/.julia/packages/Flux/qXNjB/src/optimise/train.jl:105
┌ Info: Epoch 1539
└ @ Main /Users/kcft114/.julia/packages/Flux/qXNjB/src/optimise/train.jl:105
┌ Info: Epoch 1540
└ @ Main /Users/kcft114/.julia/packages/Flux/qXNjB/src/optimise/train.jl:105
┌ Info: Epoch 1541
└ @ Main /Users/kcft114/.julia/packages/Flux/qXNjB/src/optimise/train.jl:105
┌ Info: Epoch 1542
└ @ Main /Users/kcft114/.julia/packages/Flux/qXNjB/src/optimise/train.jl:105
┌ Info: Epoch 1543
└ @ Main /Users/kcft114/.julia/packages/Flux/qXNjB/src/optimise/train.jl:105
┌ Info: Epoch 1544
└ @ Main /Users/kcft114/.julia/packages/Flux/qXNjB/src/optimise/train.jl:105
┌ Info: Epoch 1545
└ @ Main /Users/kcft114/.julia/packages/Flux/qXNjB/src/optimise/train.jl:105
┌ Info: Epoch 1546
└ @ Main /Users/kcft114/.julia/packages/Flux/qXNjB/src/optimise/train.jl:105
┌ Info: Epoch 1547
└ @ Main /Users/kcft114/.julia/packages/Flux/qXNjB/src/optimise/train.jl:105
┌ Info: Epoch 1548
└ @ Main /Users/kcft114/.julia/packages/Flux/qXNjB/src/optimise/train.jl:105
┌ Info: Epoch 1549
└ @ Main /Users/kcft114/.julia/packages/Flux/qXNjB/src/optimise/train.jl:105
┌ Info: Epoch 1550
└ @ Main /Users/kcft114/.julia/packages/Flux/qXNjB/src/optimise/train.jl:105
┌ Info: Epoch 1551
└ @ Main /Users/kcft114/.julia/packages/Flux/qXNjB/src/optimise/train.jl:105
┌ Info: Epoch 1552
└ @ Main /Users/kcft114/.julia/packages/Flux/qXNjB/src/optimise/train.jl:105
┌ Info: Epoch 1553
└ @ Main /Users/kcft114/.julia/packages/Flux/qXNjB/src/optimise/train.jl:105
┌ Info: Epoch 1554
└ @ Main /Users/kcft114/.julia/packages/Flux/qXNjB/src/optimise/train.jl:105
┌ Info: Epoch 1555
└ @ Main /Users/kcft114/.julia/packages/Flux/qXNjB/src/optimise/train.jl:105
┌ Info: Epoch 1556
└ @ Main /Users/kcft114/.julia/packages/Flux/qXNjB/src/optimise/train.jl:105
```

```
loss(X, Y) = 0.36696628f0 (tracked)
loss(X, Y) = 0.3667422f0 (tracked)
loss(X, Y) = 0.36651254f0 (tracked)
loss(X, Y) = 0.3662741f0 (tracked)
loss(X, Y) = 0.36604866f0 (tracked)
loss(X, Y) = 0.36581394f0 (tracked)
loss(X, Y) = 0.36558914f0 (tracked)
loss(X, Y) = 0.3653555f0 (tracked)
loss(X, Y) = 0.3651274f0 (tracked)
loss(X, Y) = 0.36489177f0 (tracked)
loss(X, Y) = 0.36467063f0 (tracked)
loss(X, Y) = 0.36444792f0 (tracked)
loss(X, Y) = 0.36422652f0 (tracked)
loss(X, Y) = 0.36399844f0 (tracked)
loss(X, Y) = 0.36379415f0 (tracked)
loss(X, Y) = 0.3635937f0 (tracked)
loss(X, Y) = 0.36337543f0 (tracked)
loss(X, Y) = 0.36317033f0 (tracked)
loss(X, Y) = 0.36296672f0 (tracked)
loss(X, Y) = 0.3627609f0 (tracked)
loss(X, Y) = 0.36255318f0 (tracked)
loss(X, Y) = 0.36235586f0 (tracked)
loss(X, Y) = 0.36213863f0 (tracked)
loss(X, Y) = 0.3619292f0 (tracked)
loss(X, Y) = 0.36173f0 (tracked)
loss(X, Y) = 0.36151797f0 (tracked)
loss(X, Y) = 0.3613103f0 (tracked)
loss(X, Y) = 0.36111516f0 (tracked)
loss(X, Y) = 0.3609052f0 (tracked)
loss(X, Y) = 0.3607154f0 (tracked)
loss(X, Y) = 0.36051852f0 (tracked)
loss(X, Y) = 0.3603108f0 (tracked)
loss(X, Y) = 0.36011598f0 (tracked)
loss(X, Y) = 0.35990918f0 (tracked)
loss(X, Y) = 0.35971177f0 (tracked)
loss(X, Y) = 0.3595162f0 (tracked)
loss(X, Y) = 0.35931748f0 (tracked)
loss(X, Y) = 0.35912865f0 (tracked)
loss(X, Y) = 0.35893327f0 (tracked)
loss(X, Y) = 0.35873327f0 (tracked)
loss(X, Y) = 0.3585397f0 (tracked)
loss(X, Y) = 0.35834914f0 (tracked)
loss(X, Y) = 0.3581565f0 (tracked)
loss(X, Y) = 0.35796335f0 (tracked)
loss(X, Y) = 0.35776532f0 (tracked)
loss(X, Y) = 0.35757226f0 (tracked)
loss(X, Y) = 0.35737675f0 (tracked)
loss(X, Y) = 0.35718715f0 (tracked)
loss(X, Y) = 0.3569955f0 (tracked)
loss(X, Y) = 0.35680294f0 (tracked)
loss(X, Y) = 0.35661295f0 (tracked)
loss(X, Y) = 0.35642952f0 (tracked)
loss(X, Y) = 0.35624143f0 (tracked)
loss(X, Y) = 0.35604134f0 (tracked)
loss(X, Y) = 0.35585228f0 (tracked)
loss(X, Y) = 0.3556616f0 (tracked)
loss(X, Y) = 0.35546938f0 (tracked)
loss(X, Y) = 0.3552882f0 (tracked)
loss(X, Y) = 0.35509443f0 (tracked)
loss(X, Y) = 0.35490403f0 (tracked)
loss(X, Y) = 0.3547161f0 (tracked)
loss(X, Y) = 0.35453796f0 (tracked)
loss(X, Y) = 0.3543464f0 (tracked)
loss(X, Y) = 0.35416237f0 (tracked)
loss(X, Y) = 0.35397848f0 (tracked)
loss(X, Y) = 0.35378632f0 (tracked)
loss(X, Y) = 0.3535984f0 (tracked)
loss(X, Y) = 0.35341284f0 (tracked)
loss(X, Y) = 0.35322374f0 (tracked)
loss(X, Y) = 0.35303685f0 (tracked)
loss(X, Y) = 0.35285363f0 (tracked)
loss(X, Y) = 0.3526676f0 (tracked)
loss(X, Y) = 0.35248148f0 (tracked)
loss(X, Y) = 0.3522976f0 (tracked)
loss(X, Y) = 0.35211623f0 (tracked)
loss(X, Y) = 0.3519238f0 (tracked)
loss(X, Y) = 0.35174412f0 (tracked)
loss(X, Y) = 0.35156456f0 (tracked)
loss(X, Y) = 0.35138333f0 (tracked)
loss(X, Y) = 0.3512004f0 (tracked)
loss(X, Y) = 0.35101512f0 (tracked)
loss(X, Y) = 0.35082248f0 (tracked)
loss(X, Y) = 0.35064968f0 (tracked)
loss(X, Y) = 0.3504661f0 (tracked)
loss(X, Y) = 0.350281f0 (tracked)
loss(X, Y) = 0.35010466f0 (tracked)
loss(X, Y) = 0.3499214f0 (tracked)
loss(X, Y) = 0.3497365f0 (tracked)
loss(X, Y) = 0.34955364f0 (tracked)
loss(X, Y) = 0.3493744f0 (tracked)
loss(X, Y) = 0.34918603f0 (tracked)
loss(X, Y) = 0.3490053f0 (tracked)
loss(X, Y) = 0.3488269f0 (tracked)
loss(X, Y) = 0.3486565f0 (tracked)
loss(X, Y) = 0.34847707f0 (tracked)
loss(X, Y) = 0.34829277f0 (tracked)
loss(X, Y) = 0.34811896f0 (tracked)
loss(X, Y) = 0.34793678f0 (tracked)
loss(X, Y) = 0.34776452f0 (tracked)
loss(X, Y) = 0.3475859f0 (tracked)
loss(X, Y) = 0.34740165f0 (tracked)
loss(X, Y) = 0.3472388f0 (tracked)
loss(X, Y) = 0.3470669f0 (tracked)
loss(X, Y) = 0.34688616f0 (tracked)
loss(X, Y) = 0.3467047f0 (tracked)
loss(X, Y) = 0.34652933f0 (tracked)
loss(X, Y) = 0.34636018f0 (tracked)
loss(X, Y) = 0.34618896f0 (tracked)
loss(X, Y) = 0.34600636f0 (tracked)
loss(X, Y) = 0.34583098f0 (tracked)
loss(X, Y) = 0.34566292f0 (tracked)
loss(X, Y) = 0.34549138f0 (tracked)
loss(X, Y) = 0.34531525f0 (tracked)
loss(X, Y) = 0.34514028f0 (tracked)
loss(X, Y) = 0.34497228f0 (tracked)
loss(X, Y) = 0.34479794f0 (tracked)
loss(X, Y) = 0.34461808f0 (tracked)
loss(X, Y) = 0.34445444f0 (tracked)
loss(X, Y) = 0.344282f0 (tracked)
loss(X, Y) = 0.34409878f0 (tracked)
loss(X, Y) = 0.3439246f0 (tracked)
loss(X, Y) = 0.34377116f0 (tracked)
loss(X, Y) = 0.3435946f0 (tracked)
loss(X, Y) = 0.34340933f0 (tracked)
loss(X, Y) = 0.3432461f0 (tracked)
loss(X, Y) = 0.34308726f0 (tracked)
loss(X, Y) = 0.34291336f0 (tracked)
loss(X, Y) = 0.34273243f0 (tracked)
loss(X, Y) = 0.34256104f0 (tracked)
loss(X, Y) = 0.34240097f0 (tracked)
loss(X, Y) = 0.34223548f0 (tracked)
loss(X, Y) = 0.3420585f0 (tracked)
loss(X, Y) = 0.34189495f0 (tracked)
loss(X, Y) = 0.34172788f0 (tracked)
loss(X, Y) = 0.341565f0 (tracked)
loss(X, Y) = 0.34138814f0 (tracked)
loss(X, Y) = 0.34121126f0 (tracked)
loss(X, Y) = 0.34104156f0 (tracked)
loss(X, Y) = 0.3408774f0 (tracked)
loss(X, Y) = 0.34071234f0 (tracked)
loss(X, Y) = 0.34053627f0 (tracked)
loss(X, Y) = 0.3403606f0 (tracked)
```

```
┌ Info: Epoch 1557
└ @ Main /Users/kcft114/.julia/packages/Flux/qXNjB/src/optimise/train.jl:105
┌ Info: Epoch 1558
└ @ Main /Users/kcft114/.julia/packages/Flux/qXNjB/src/optimise/train.jl:105
┌ Info: Epoch 1559
└ @ Main /Users/kcft114/.julia/packages/Flux/qXNjB/src/optimise/train.jl:105
┌ Info: Epoch 1560
└ @ Main /Users/kcft114/.julia/packages/Flux/qXNjB/src/optimise/train.jl:105
┌ Info: Epoch 1561
└ @ Main /Users/kcft114/.julia/packages/Flux/qXNjB/src/optimise/train.jl:105
┌ Info: Epoch 1562
└ @ Main /Users/kcft114/.julia/packages/Flux/qXNjB/src/optimise/train.jl:105
┌ Info: Epoch 1563
└ @ Main /Users/kcft114/.julia/packages/Flux/qXNjB/src/optimise/train.jl:105
┌ Info: Epoch 1564
└ @ Main /Users/kcft114/.julia/packages/Flux/qXNjB/src/optimise/train.jl:105
┌ Info: Epoch 1565
└ @ Main /Users/kcft114/.julia/packages/Flux/qXNjB/src/optimise/train.jl:105
┌ Info: Epoch 1566
└ @ Main /Users/kcft114/.julia/packages/Flux/qXNjB/src/optimise/train.jl:105
┌ Info: Epoch 1567
└ @ Main /Users/kcft114/.julia/packages/Flux/qXNjB/src/optimise/train.jl:105
┌ Info: Epoch 1568
└ @ Main /Users/kcft114/.julia/packages/Flux/qXNjB/src/optimise/train.jl:105
┌ Info: Epoch 1569
└ @ Main /Users/kcft114/.julia/packages/Flux/qXNjB/src/optimise/train.jl:105
┌ Info: Epoch 1570
└ @ Main /Users/kcft114/.julia/packages/Flux/qXNjB/src/optimise/train.jl:105
┌ Info: Epoch 1571
└ @ Main /Users/kcft114/.julia/packages/Flux/qXNjB/src/optimise/train.jl:105
┌ Info: Epoch 1572
└ @ Main /Users/kcft114/.julia/packages/Flux/qXNjB/src/optimise/train.jl:105
┌ Info: Epoch 1573
└ @ Main /Users/kcft114/.julia/packages/Flux/qXNjB/src/optimise/train.jl:105
┌ Info: Epoch 1574
└ @ Main /Users/kcft114/.julia/packages/Flux/qXNjB/src/optimise/train.jl:105
┌ Info: Epoch 1575
└ @ Main /Users/kcft114/.julia/packages/Flux/qXNjB/src/optimise/train.jl:105
┌ Info: Epoch 1576
└ @ Main /Users/kcft114/.julia/packages/Flux/qXNjB/src/optimise/train.jl:105
┌ Info: Epoch 1577
└ @ Main /Users/kcft114/.julia/packages/Flux/qXNjB/src/optimise/train.jl:105
┌ Info: Epoch 1578
└ @ Main /Users/kcft114/.julia/packages/Flux/qXNjB/src/optimise/train.jl:105
┌ Info: Epoch 1579
└ @ Main /Users/kcft114/.julia/packages/Flux/qXNjB/src/optimise/train.jl:105
┌ Info: Epoch 1580
└ @ Main /Users/kcft114/.julia/packages/Flux/qXNjB/src/optimise/train.jl:105
┌ Info: Epoch 1581
└ @ Main /Users/kcft114/.julia/packages/Flux/qXNjB/src/optimise/train.jl:105
┌ Info: Epoch 1582
└ @ Main /Users/kcft114/.julia/packages/Flux/qXNjB/src/optimise/train.jl:105
┌ Info: Epoch 1583
└ @ Main /Users/kcft114/.julia/packages/Flux/qXNjB/src/optimise/train.jl:105
┌ Info: Epoch 1584
└ @ Main /Users/kcft114/.julia/packages/Flux/qXNjB/src/optimise/train.jl:105
┌ Info: Epoch 1585
└ @ Main /Users/kcft114/.julia/packages/Flux/qXNjB/src/optimise/train.jl:105
┌ Info: Epoch 1586
└ @ Main /Users/kcft114/.julia/packages/Flux/qXNjB/src/optimise/train.jl:105
┌ Info: Epoch 1587
└ @ Main /Users/kcft114/.julia/packages/Flux/qXNjB/src/optimise/train.jl:105
┌ Info: Epoch 1588
└ @ Main /Users/kcft114/.julia/packages/Flux/qXNjB/src/optimise/train.jl:105
┌ Info: Epoch 1589
└ @ Main /Users/kcft114/.julia/packages/Flux/qXNjB/src/optimise/train.jl:105
┌ Info: Epoch 1590
└ @ Main /Users/kcft114/.julia/packages/Flux/qXNjB/src/optimise/train.jl:105
┌ Info: Epoch 1591
└ @ Main /Users/kcft114/.julia/packages/Flux/qXNjB/src/optimise/train.jl:105
┌ Info: Epoch 1592
└ @ Main /Users/kcft114/.julia/packages/Flux/qXNjB/src/optimise/train.jl:105
┌ Info: Epoch 1593
└ @ Main /Users/kcft114/.julia/packages/Flux/qXNjB/src/optimise/train.jl:105
┌ Info: Epoch 1594
└ @ Main /Users/kcft114/.julia/packages/Flux/qXNjB/src/optimise/train.jl:105
┌ Info: Epoch 1595
└ @ Main /Users/kcft114/.julia/packages/Flux/qXNjB/src/optimise/train.jl:105
┌ Info: Epoch 1596
└ @ Main /Users/kcft114/.julia/packages/Flux/qXNjB/src/optimise/train.jl:105
┌ Info: Epoch 1597
└ @ Main /Users/kcft114/.julia/packages/Flux/qXNjB/src/optimise/train.jl:105
┌ Info: Epoch 1598
└ @ Main /Users/kcft114/.julia/packages/Flux/qXNjB/src/optimise/train.jl:105
┌ Info: Epoch 1599
└ @ Main /Users/kcft114/.julia/packages/Flux/qXNjB/src/optimise/train.jl:105
┌ Info: Epoch 1600
└ @ Main /Users/kcft114/.julia/packages/Flux/qXNjB/src/optimise/train.jl:105
┌ Info: Epoch 1601
└ @ Main /Users/kcft114/.julia/packages/Flux/qXNjB/src/optimise/train.jl:105
┌ Info: Epoch 1602
└ @ Main /Users/kcft114/.julia/packages/Flux/qXNjB/src/optimise/train.jl:105
┌ Info: Epoch 1603
└ @ Main /Users/kcft114/.julia/packages/Flux/qXNjB/src/optimise/train.jl:105
┌ Info: Epoch 1604
└ @ Main /Users/kcft114/.julia/packages/Flux/qXNjB/src/optimise/train.jl:105
┌ Info: Epoch 1605
└ @ Main /Users/kcft114/.julia/packages/Flux/qXNjB/src/optimise/train.jl:105
┌ Info: Epoch 1606
└ @ Main /Users/kcft114/.julia/packages/Flux/qXNjB/src/optimise/train.jl:105
┌ Info: Epoch 1607
└ @ Main /Users/kcft114/.julia/packages/Flux/qXNjB/src/optimise/train.jl:105
┌ Info: Epoch 1608
└ @ Main /Users/kcft114/.julia/packages/Flux/qXNjB/src/optimise/train.jl:105
┌ Info: Epoch 1609
└ @ Main /Users/kcft114/.julia/packages/Flux/qXNjB/src/optimise/train.jl:105
┌ Info: Epoch 1610
└ @ Main /Users/kcft114/.julia/packages/Flux/qXNjB/src/optimise/train.jl:105
┌ Info: Epoch 1611
└ @ Main /Users/kcft114/.julia/packages/Flux/qXNjB/src/optimise/train.jl:105
┌ Info: Epoch 1612
└ @ Main /Users/kcft114/.julia/packages/Flux/qXNjB/src/optimise/train.jl:105
┌ Info: Epoch 1613
└ @ Main /Users/kcft114/.julia/packages/Flux/qXNjB/src/optimise/train.jl:105
┌ Info: Epoch 1614
└ @ Main /Users/kcft114/.julia/packages/Flux/qXNjB/src/optimise/train.jl:105
┌ Info: Epoch 1615
└ @ Main /Users/kcft114/.julia/packages/Flux/qXNjB/src/optimise/train.jl:105
┌ Info: Epoch 1616
└ @ Main /Users/kcft114/.julia/packages/Flux/qXNjB/src/optimise/train.jl:105
┌ Info: Epoch 1617
└ @ Main /Users/kcft114/.julia/packages/Flux/qXNjB/src/optimise/train.jl:105
┌ Info: Epoch 1618
└ @ Main /Users/kcft114/.julia/packages/Flux/qXNjB/src/optimise/train.jl:105
┌ Info: Epoch 1619
└ @ Main /Users/kcft114/.julia/packages/Flux/qXNjB/src/optimise/train.jl:105
┌ Info: Epoch 1620
└ @ Main /Users/kcft114/.julia/packages/Flux/qXNjB/src/optimise/train.jl:105
┌ Info: Epoch 1621
└ @ Main /Users/kcft114/.julia/packages/Flux/qXNjB/src/optimise/train.jl:105
┌ Info: Epoch 1622
└ @ Main /Users/kcft114/.julia/packages/Flux/qXNjB/src/optimise/train.jl:105
┌ Info: Epoch 1623
└ @ Main /Users/kcft114/.julia/packages/Flux/qXNjB/src/optimise/train.jl:105
┌ Info: Epoch 1624
└ @ Main /Users/kcft114/.julia/packages/Flux/qXNjB/src/optimise/train.jl:105
┌ Info: Epoch 1625
└ @ Main /Users/kcft114/.julia/packages/Flux/qXNjB/src/optimise/train.jl:105
┌ Info: Epoch 1626
└ @ Main /Users/kcft114/.julia/packages/Flux/qXNjB/src/optimise/train.jl:105
┌ Info: Epoch 1627
└ @ Main /Users/kcft114/.julia/packages/Flux/qXNjB/src/optimise/train.jl:105
┌ Info: Epoch 1628
└ @ Main /Users/kcft114/.julia/packages/Flux/qXNjB/src/optimise/train.jl:105
┌ Info: Epoch 1629
└ @ Main /Users/kcft114/.julia/packages/Flux/qXNjB/src/optimise/train.jl:105
┌ Info: Epoch 1630
└ @ Main /Users/kcft114/.julia/packages/Flux/qXNjB/src/optimise/train.jl:105
┌ Info: Epoch 1631
└ @ Main /Users/kcft114/.julia/packages/Flux/qXNjB/src/optimise/train.jl:105
┌ Info: Epoch 1632
└ @ Main /Users/kcft114/.julia/packages/Flux/qXNjB/src/optimise/train.jl:105
┌ Info: Epoch 1633
└ @ Main /Users/kcft114/.julia/packages/Flux/qXNjB/src/optimise/train.jl:105
┌ Info: Epoch 1634
└ @ Main /Users/kcft114/.julia/packages/Flux/qXNjB/src/optimise/train.jl:105
┌ Info: Epoch 1635
└ @ Main /Users/kcft114/.julia/packages/Flux/qXNjB/src/optimise/train.jl:105
┌ Info: Epoch 1636
└ @ Main /Users/kcft114/.julia/packages/Flux/qXNjB/src/optimise/train.jl:105
┌ Info: Epoch 1637
└ @ Main /Users/kcft114/.julia/packages/Flux/qXNjB/src/optimise/train.jl:105
┌ Info: Epoch 1638
└ @ Main /Users/kcft114/.julia/packages/Flux/qXNjB/src/optimise/train.jl:105
┌ Info: Epoch 1639
└ @ Main /Users/kcft114/.julia/packages/Flux/qXNjB/src/optimise/train.jl:105
┌ Info: Epoch 1640
└ @ Main /Users/kcft114/.julia/packages/Flux/qXNjB/src/optimise/train.jl:105
┌ Info: Epoch 1641
└ @ Main /Users/kcft114/.julia/packages/Flux/qXNjB/src/optimise/train.jl:105
┌ Info: Epoch 1642
└ @ Main /Users/kcft114/.julia/packages/Flux/qXNjB/src/optimise/train.jl:105
┌ Info: Epoch 1643
└ @ Main /Users/kcft114/.julia/packages/Flux/qXNjB/src/optimise/train.jl:105
┌ Info: Epoch 1644
└ @ Main /Users/kcft114/.julia/packages/Flux/qXNjB/src/optimise/train.jl:105
┌ Info: Epoch 1645
└ @ Main /Users/kcft114/.julia/packages/Flux/qXNjB/src/optimise/train.jl:105
┌ Info: Epoch 1646
└ @ Main /Users/kcft114/.julia/packages/Flux/qXNjB/src/optimise/train.jl:105
┌ Info: Epoch 1647
└ @ Main /Users/kcft114/.julia/packages/Flux/qXNjB/src/optimise/train.jl:105
┌ Info: Epoch 1648
└ @ Main /Users/kcft114/.julia/packages/Flux/qXNjB/src/optimise/train.jl:105
┌ Info: Epoch 1649
└ @ Main /Users/kcft114/.julia/packages/Flux/qXNjB/src/optimise/train.jl:105
┌ Info: Epoch 1650
└ @ Main /Users/kcft114/.julia/packages/Flux/qXNjB/src/optimise/train.jl:105
┌ Info: Epoch 1651
└ @ Main /Users/kcft114/.julia/packages/Flux/qXNjB/src/optimise/train.jl:105
┌ Info: Epoch 1652
└ @ Main /Users/kcft114/.julia/packages/Flux/qXNjB/src/optimise/train.jl:105
┌ Info: Epoch 1653
└ @ Main /Users/kcft114/.julia/packages/Flux/qXNjB/src/optimise/train.jl:105
┌ Info: Epoch 1654
└ @ Main /Users/kcft114/.julia/packages/Flux/qXNjB/src/optimise/train.jl:105
┌ Info: Epoch 1655
└ @ Main /Users/kcft114/.julia/packages/Flux/qXNjB/src/optimise/train.jl:105
┌ Info: Epoch 1656
└ @ Main /Users/kcft114/.julia/packages/Flux/qXNjB/src/optimise/train.jl:105
┌ Info: Epoch 1657
└ @ Main /Users/kcft114/.julia/packages/Flux/qXNjB/src/optimise/train.jl:105
┌ Info: Epoch 1658
└ @ Main /Users/kcft114/.julia/packages/Flux/qXNjB/src/optimise/train.jl:105
┌ Info: Epoch 1659
└ @ Main /Users/kcft114/.julia/packages/Flux/qXNjB/src/optimise/train.jl:105
┌ Info: Epoch 1660
└ @ Main /Users/kcft114/.julia/packages/Flux/qXNjB/src/optimise/train.jl:105
┌ Info: Epoch 1661
└ @ Main /Users/kcft114/.julia/packages/Flux/qXNjB/src/optimise/train.jl:105
┌ Info: Epoch 1662
└ @ Main /Users/kcft114/.julia/packages/Flux/qXNjB/src/optimise/train.jl:105
┌ Info: Epoch 1663
└ @ Main /Users/kcft114/.julia/packages/Flux/qXNjB/src/optimise/train.jl:105
┌ Info: Epoch 1664
└ @ Main /Users/kcft114/.julia/packages/Flux/qXNjB/src/optimise/train.jl:105
┌ Info: Epoch 1665
└ @ Main /Users/kcft114/.julia/packages/Flux/qXNjB/src/optimise/train.jl:105
┌ Info: Epoch 1666
└ @ Main /Users/kcft114/.julia/packages/Flux/qXNjB/src/optimise/train.jl:105
┌ Info: Epoch 1667
└ @ Main /Users/kcft114/.julia/packages/Flux/qXNjB/src/optimise/train.jl:105
┌ Info: Epoch 1668
└ @ Main /Users/kcft114/.julia/packages/Flux/qXNjB/src/optimise/train.jl:105
┌ Info: Epoch 1669
└ @ Main /Users/kcft114/.julia/packages/Flux/qXNjB/src/optimise/train.jl:105
┌ Info: Epoch 1670
└ @ Main /Users/kcft114/.julia/packages/Flux/qXNjB/src/optimise/train.jl:105
┌ Info: Epoch 1671
└ @ Main /Users/kcft114/.julia/packages/Flux/qXNjB/src/optimise/train.jl:105
┌ Info: Epoch 1672
└ @ Main /Users/kcft114/.julia/packages/Flux/qXNjB/src/optimise/train.jl:105
┌ Info: Epoch 1673
└ @ Main /Users/kcft114/.julia/packages/Flux/qXNjB/src/optimise/train.jl:105
┌ Info: Epoch 1674
└ @ Main /Users/kcft114/.julia/packages/Flux/qXNjB/src/optimise/train.jl:105
┌ Info: Epoch 1675
└ @ Main /Users/kcft114/.julia/packages/Flux/qXNjB/src/optimise/train.jl:105
┌ Info: Epoch 1676
└ @ Main /Users/kcft114/.julia/packages/Flux/qXNjB/src/optimise/train.jl:105
┌ Info: Epoch 1677
└ @ Main /Users/kcft114/.julia/packages/Flux/qXNjB/src/optimise/train.jl:105
┌ Info: Epoch 1678
└ @ Main /Users/kcft114/.julia/packages/Flux/qXNjB/src/optimise/train.jl:105
┌ Info: Epoch 1679
└ @ Main /Users/kcft114/.julia/packages/Flux/qXNjB/src/optimise/train.jl:105
┌ Info: Epoch 1680
└ @ Main /Users/kcft114/.julia/packages/Flux/qXNjB/src/optimise/train.jl:105
┌ Info: Epoch 1681
└ @ Main /Users/kcft114/.julia/packages/Flux/qXNjB/src/optimise/train.jl:105
┌ Info: Epoch 1682
└ @ Main /Users/kcft114/.julia/packages/Flux/qXNjB/src/optimise/train.jl:105
┌ Info: Epoch 1683
└ @ Main /Users/kcft114/.julia/packages/Flux/qXNjB/src/optimise/train.jl:105
┌ Info: Epoch 1684
└ @ Main /Users/kcft114/.julia/packages/Flux/qXNjB/src/optimise/train.jl:105
┌ Info: Epoch 1685
└ @ Main /Users/kcft114/.julia/packages/Flux/qXNjB/src/optimise/train.jl:105
┌ Info: Epoch 1686
└ @ Main /Users/kcft114/.julia/packages/Flux/qXNjB/src/optimise/train.jl:105
┌ Info: Epoch 1687
└ @ Main /Users/kcft114/.julia/packages/Flux/qXNjB/src/optimise/train.jl:105
┌ Info: Epoch 1688
└ @ Main /Users/kcft114/.julia/packages/Flux/qXNjB/src/optimise/train.jl:105
┌ Info: Epoch 1689
└ @ Main /Users/kcft114/.julia/packages/Flux/qXNjB/src/optimise/train.jl:105
┌ Info: Epoch 1690
└ @ Main /Users/kcft114/.julia/packages/Flux/qXNjB/src/optimise/train.jl:105
┌ Info: Epoch 1691
└ @ Main /Users/kcft114/.julia/packages/Flux/qXNjB/src/optimise/train.jl:105
┌ Info: Epoch 1692
└ @ Main /Users/kcft114/.julia/packages/Flux/qXNjB/src/optimise/train.jl:105
┌ Info: Epoch 1693
└ @ Main /Users/kcft114/.julia/packages/Flux/qXNjB/src/optimise/train.jl:105
┌ Info: Epoch 1694
└ @ Main /Users/kcft114/.julia/packages/Flux/qXNjB/src/optimise/train.jl:105
┌ Info: Epoch 1695
└ @ Main /Users/kcft114/.julia/packages/Flux/qXNjB/src/optimise/train.jl:105
┌ Info: Epoch 1696
└ @ Main /Users/kcft114/.julia/packages/Flux/qXNjB/src/optimise/train.jl:105
┌ Info: Epoch 1697
└ @ Main /Users/kcft114/.julia/packages/Flux/qXNjB/src/optimise/train.jl:105
┌ Info: Epoch 1698
└ @ Main /Users/kcft114/.julia/packages/Flux/qXNjB/src/optimise/train.jl:105
```

```
loss(X, Y) = 0.3402032f0 (tracked)
loss(X, Y) = 0.34004176f0 (tracked)
loss(X, Y) = 0.3398695f0 (tracked)
loss(X, Y) = 0.33970496f0 (tracked)
loss(X, Y) = 0.33954313f0 (tracked)
loss(X, Y) = 0.3393687f0 (tracked)
loss(X, Y) = 0.3392035f0 (tracked)
loss(X, Y) = 0.33903968f0 (tracked)
loss(X, Y) = 0.3388795f0 (tracked)
loss(X, Y) = 0.33870634f0 (tracked)
loss(X, Y) = 0.33853856f0 (tracked)
loss(X, Y) = 0.33837274f0 (tracked)
loss(X, Y) = 0.33821633f0 (tracked)
loss(X, Y) = 0.33805507f0 (tracked)
loss(X, Y) = 0.33788344f0 (tracked)
loss(X, Y) = 0.33771327f0 (tracked)
loss(X, Y) = 0.33756432f0 (tracked)
loss(X, Y) = 0.33739755f0 (tracked)
loss(X, Y) = 0.33723623f0 (tracked)
loss(X, Y) = 0.33707008f0 (tracked)
loss(X, Y) = 0.33691174f0 (tracked)
loss(X, Y) = 0.33674085f0 (tracked)
loss(X, Y) = 0.33657813f0 (tracked)
loss(X, Y) = 0.3364152f0 (tracked)
loss(X, Y) = 0.33625546f0 (tracked)
loss(X, Y) = 0.3360966f0 (tracked)
loss(X, Y) = 0.33592972f0 (tracked)
loss(X, Y) = 0.3357669f0 (tracked)
loss(X, Y) = 0.3356117f0 (tracked)
loss(X, Y) = 0.33544564f0 (tracked)
loss(X, Y) = 0.3352808f0 (tracked)
loss(X, Y) = 0.335111f0 (tracked)
loss(X, Y) = 0.33494687f0 (tracked)
loss(X, Y) = 0.3348031f0 (tracked)
loss(X, Y) = 0.33464298f0 (tracked)
loss(X, Y) = 0.33446494f0 (tracked)
loss(X, Y) = 0.33431336f0 (tracked)
loss(X, Y) = 0.33415234f0 (tracked)
loss(X, Y) = 0.33398688f0 (tracked)
loss(X, Y) = 0.33381945f0 (tracked)
loss(X, Y) = 0.33366126f0 (tracked)
loss(X, Y) = 0.33350188f0 (tracked)
loss(X, Y) = 0.33333683f0 (tracked)
loss(X, Y) = 0.33316794f0 (tracked)
loss(X, Y) = 0.33300385f0 (tracked)
loss(X, Y) = 0.33284798f0 (tracked)
loss(X, Y) = 0.33268523f0 (tracked)
loss(X, Y) = 0.33252287f0 (tracked)
loss(X, Y) = 0.33236003f0 (tracked)
loss(X, Y) = 0.3322034f0 (tracked)
loss(X, Y) = 0.33204535f0 (tracked)
loss(X, Y) = 0.3318852f0 (tracked)
loss(X, Y) = 0.33172005f0 (tracked)
loss(X, Y) = 0.3315629f0 (tracked)
loss(X, Y) = 0.3314071f0 (tracked)
loss(X, Y) = 0.3312425f0 (tracked)
loss(X, Y) = 0.3310733f0 (tracked)
loss(X, Y) = 0.33090833f0 (tracked)
loss(X, Y) = 0.330759f0 (tracked)
loss(X, Y) = 0.33060458f0 (tracked)
loss(X, Y) = 0.3304363f0 (tracked)
loss(X, Y) = 0.3302715f0 (tracked)
loss(X, Y) = 0.33011243f0 (tracked)
loss(X, Y) = 0.32994527f0 (tracked)
loss(X, Y) = 0.3297918f0 (tracked)
loss(X, Y) = 0.3296338f0 (tracked)
loss(X, Y) = 0.3294669f0 (tracked)
loss(X, Y) = 0.32930392f0 (tracked)
loss(X, Y) = 0.329148f0 (tracked)
loss(X, Y) = 0.32897928f0 (tracked)
loss(X, Y) = 0.3288153f0 (tracked)
loss(X, Y) = 0.3286567f0 (tracked)
loss(X, Y) = 0.3284918f0 (tracked)
loss(X, Y) = 0.3283373f0 (tracked)
loss(X, Y) = 0.32817784f0 (tracked)
loss(X, Y) = 0.32801244f0 (tracked)
loss(X, Y) = 0.3278483f0 (tracked)
loss(X, Y) = 0.3276895f0 (tracked)
loss(X, Y) = 0.32753706f0 (tracked)
loss(X, Y) = 0.32737434f0 (tracked)
loss(X, Y) = 0.3272102f0 (tracked)
loss(X, Y) = 0.3270424f0 (tracked)
loss(X, Y) = 0.3268828f0 (tracked)
loss(X, Y) = 0.32672867f0 (tracked)
loss(X, Y) = 0.3265633f0 (tracked)
loss(X, Y) = 0.32640246f0 (tracked)
loss(X, Y) = 0.32624364f0 (tracked)
loss(X, Y) = 0.3260928f0 (tracked)
loss(X, Y) = 0.3259355f0 (tracked)
loss(X, Y) = 0.32577172f0 (tracked)
loss(X, Y) = 0.3256152f0 (tracked)
loss(X, Y) = 0.32546508f0 (tracked)
loss(X, Y) = 0.32530212f0 (tracked)
loss(X, Y) = 0.32513687f0 (tracked)
loss(X, Y) = 0.32498315f0 (tracked)
loss(X, Y) = 0.32483968f0 (tracked)
loss(X, Y) = 0.32467416f0 (tracked)
loss(X, Y) = 0.32450476f0 (tracked)
loss(X, Y) = 0.3243542f0 (tracked)
loss(X, Y) = 0.32420108f0 (tracked)
loss(X, Y) = 0.32403994f0 (tracked)
loss(X, Y) = 0.323871f0 (tracked)
loss(X, Y) = 0.32371792f0 (tracked)
loss(X, Y) = 0.3235736f0 (tracked)
loss(X, Y) = 0.32341978f0 (tracked)
loss(X, Y) = 0.32325083f0 (tracked)
loss(X, Y) = 0.32309094f0 (tracked)
loss(X, Y) = 0.32294884f0 (tracked)
loss(X, Y) = 0.32279018f0 (tracked)
loss(X, Y) = 0.32263216f0 (tracked)
loss(X, Y) = 0.32247308f0 (tracked)
loss(X, Y) = 0.32230675f0 (tracked)
loss(X, Y) = 0.32215405f0 (tracked)
loss(X, Y) = 0.32200265f0 (tracked)
loss(X, Y) = 0.32184523f0 (tracked)
loss(X, Y) = 0.321696f0 (tracked)
loss(X, Y) = 0.32153192f0 (tracked)
loss(X, Y) = 0.32138276f0 (tracked)
loss(X, Y) = 0.3212295f0 (tracked)
loss(X, Y) = 0.3210679f0 (tracked)
loss(X, Y) = 0.32091126f0 (tracked)
loss(X, Y) = 0.32076463f0 (tracked)
loss(X, Y) = 0.32060608f0 (tracked)
loss(X, Y) = 0.32045013f0 (tracked)
loss(X, Y) = 0.3202949f0 (tracked)
loss(X, Y) = 0.32013232f0 (tracked)
loss(X, Y) = 0.3199797f0 (tracked)
loss(X, Y) = 0.31983343f0 (tracked)
loss(X, Y) = 0.31968367f0 (tracked)
loss(X, Y) = 0.31952852f0 (tracked)
loss(X, Y) = 0.31937283f0 (tracked)
loss(X, Y) = 0.31922367f0 (tracked)
loss(X, Y) = 0.3190642f0 (tracked)
loss(X, Y) = 0.31892043f0 (tracked)
loss(X, Y) = 0.31877238f0 (tracked)
loss(X, Y) = 0.3186174f0 (tracked)
loss(X, Y) = 0.31845662f0 (tracked)
loss(X, Y) = 0.31831f0 (tracked)
loss(X, Y) = 0.31816107f0 (tracked)
loss(X, Y) = 0.31800053f0 (tracked)
loss(X, Y) = 0.31785268f0 (tracked)
loss(X, Y) = 0.3177033f0 (tracked)
loss(X, Y) = 0.31754872f0 (tracked)
loss(X, Y) = 0.3173982f0 (tracked)
loss(X, Y) = 0.31725088f0 (tracked)
loss(X, Y) = 0.3171064f0 (tracked)
loss(X, Y) = 0.31695426f0 (tracked)
loss(X, Y) = 0.3167952f0 (tracked)
loss(X, Y) = 0.31664482f0 (tracked)
loss(X, Y) = 0.31650954f0 (tracked)
```

```
┌ Info: Epoch 1699
└ @ Main /Users/kcft114/.julia/packages/Flux/qXNjB/src/optimise/train.jl:105
┌ Info: Epoch 1700
└ @ Main /Users/kcft114/.julia/packages/Flux/qXNjB/src/optimise/train.jl:105
┌ Info: Epoch 1701
└ @ Main /Users/kcft114/.julia/packages/Flux/qXNjB/src/optimise/train.jl:105
┌ Info: Epoch 1702
└ @ Main /Users/kcft114/.julia/packages/Flux/qXNjB/src/optimise/train.jl:105
┌ Info: Epoch 1703
└ @ Main /Users/kcft114/.julia/packages/Flux/qXNjB/src/optimise/train.jl:105
┌ Info: Epoch 1704
└ @ Main /Users/kcft114/.julia/packages/Flux/qXNjB/src/optimise/train.jl:105
┌ Info: Epoch 1705
└ @ Main /Users/kcft114/.julia/packages/Flux/qXNjB/src/optimise/train.jl:105
┌ Info: Epoch 1706
└ @ Main /Users/kcft114/.julia/packages/Flux/qXNjB/src/optimise/train.jl:105
┌ Info: Epoch 1707
└ @ Main /Users/kcft114/.julia/packages/Flux/qXNjB/src/optimise/train.jl:105
┌ Info: Epoch 1708
└ @ Main /Users/kcft114/.julia/packages/Flux/qXNjB/src/optimise/train.jl:105
┌ Info: Epoch 1709
└ @ Main /Users/kcft114/.julia/packages/Flux/qXNjB/src/optimise/train.jl:105
┌ Info: Epoch 1710
└ @ Main /Users/kcft114/.julia/packages/Flux/qXNjB/src/optimise/train.jl:105
┌ Info: Epoch 1711
└ @ Main /Users/kcft114/.julia/packages/Flux/qXNjB/src/optimise/train.jl:105
┌ Info: Epoch 1712
└ @ Main /Users/kcft114/.julia/packages/Flux/qXNjB/src/optimise/train.jl:105
┌ Info: Epoch 1713
└ @ Main /Users/kcft114/.julia/packages/Flux/qXNjB/src/optimise/train.jl:105
┌ Info: Epoch 1714
└ @ Main /Users/kcft114/.julia/packages/Flux/qXNjB/src/optimise/train.jl:105
┌ Info: Epoch 1715
└ @ Main /Users/kcft114/.julia/packages/Flux/qXNjB/src/optimise/train.jl:105
┌ Info: Epoch 1716
└ @ Main /Users/kcft114/.julia/packages/Flux/qXNjB/src/optimise/train.jl:105
┌ Info: Epoch 1717
└ @ Main /Users/kcft114/.julia/packages/Flux/qXNjB/src/optimise/train.jl:105
┌ Info: Epoch 1718
└ @ Main /Users/kcft114/.julia/packages/Flux/qXNjB/src/optimise/train.jl:105
┌ Info: Epoch 1719
└ @ Main /Users/kcft114/.julia/packages/Flux/qXNjB/src/optimise/train.jl:105
┌ Info: Epoch 1720
└ @ Main /Users/kcft114/.julia/packages/Flux/qXNjB/src/optimise/train.jl:105
┌ Info: Epoch 1721
└ @ Main /Users/kcft114/.julia/packages/Flux/qXNjB/src/optimise/train.jl:105
┌ Info: Epoch 1722
└ @ Main /Users/kcft114/.julia/packages/Flux/qXNjB/src/optimise/train.jl:105
┌ Info: Epoch 1723
└ @ Main /Users/kcft114/.julia/packages/Flux/qXNjB/src/optimise/train.jl:105
┌ Info: Epoch 1724
└ @ Main /Users/kcft114/.julia/packages/Flux/qXNjB/src/optimise/train.jl:105
┌ Info: Epoch 1725
└ @ Main /Users/kcft114/.julia/packages/Flux/qXNjB/src/optimise/train.jl:105
┌ Info: Epoch 1726
└ @ Main /Users/kcft114/.julia/packages/Flux/qXNjB/src/optimise/train.jl:105
┌ Info: Epoch 1727
└ @ Main /Users/kcft114/.julia/packages/Flux/qXNjB/src/optimise/train.jl:105
┌ Info: Epoch 1728
└ @ Main /Users/kcft114/.julia/packages/Flux/qXNjB/src/optimise/train.jl:105
┌ Info: Epoch 1729
└ @ Main /Users/kcft114/.julia/packages/Flux/qXNjB/src/optimise/train.jl:105
┌ Info: Epoch 1730
└ @ Main /Users/kcft114/.julia/packages/Flux/qXNjB/src/optimise/train.jl:105
┌ Info: Epoch 1731
└ @ Main /Users/kcft114/.julia/packages/Flux/qXNjB/src/optimise/train.jl:105
┌ Info: Epoch 1732
└ @ Main /Users/kcft114/.julia/packages/Flux/qXNjB/src/optimise/train.jl:105
┌ Info: Epoch 1733
└ @ Main /Users/kcft114/.julia/packages/Flux/qXNjB/src/optimise/train.jl:105
┌ Info: Epoch 1734
└ @ Main /Users/kcft114/.julia/packages/Flux/qXNjB/src/optimise/train.jl:105
┌ Info: Epoch 1735
└ @ Main /Users/kcft114/.julia/packages/Flux/qXNjB/src/optimise/train.jl:105
┌ Info: Epoch 1736
└ @ Main /Users/kcft114/.julia/packages/Flux/qXNjB/src/optimise/train.jl:105
┌ Info: Epoch 1737
└ @ Main /Users/kcft114/.julia/packages/Flux/qXNjB/src/optimise/train.jl:105
┌ Info: Epoch 1738
└ @ Main /Users/kcft114/.julia/packages/Flux/qXNjB/src/optimise/train.jl:105
┌ Info: Epoch 1739
└ @ Main /Users/kcft114/.julia/packages/Flux/qXNjB/src/optimise/train.jl:105
┌ Info: Epoch 1740
└ @ Main /Users/kcft114/.julia/packages/Flux/qXNjB/src/optimise/train.jl:105
┌ Info: Epoch 1741
└ @ Main /Users/kcft114/.julia/packages/Flux/qXNjB/src/optimise/train.jl:105
┌ Info: Epoch 1742
└ @ Main /Users/kcft114/.julia/packages/Flux/qXNjB/src/optimise/train.jl:105
┌ Info: Epoch 1743
└ @ Main /Users/kcft114/.julia/packages/Flux/qXNjB/src/optimise/train.jl:105
┌ Info: Epoch 1744
└ @ Main /Users/kcft114/.julia/packages/Flux/qXNjB/src/optimise/train.jl:105
┌ Info: Epoch 1745
└ @ Main /Users/kcft114/.julia/packages/Flux/qXNjB/src/optimise/train.jl:105
┌ Info: Epoch 1746
└ @ Main /Users/kcft114/.julia/packages/Flux/qXNjB/src/optimise/train.jl:105
┌ Info: Epoch 1747
└ @ Main /Users/kcft114/.julia/packages/Flux/qXNjB/src/optimise/train.jl:105
┌ Info: Epoch 1748
└ @ Main /Users/kcft114/.julia/packages/Flux/qXNjB/src/optimise/train.jl:105
┌ Info: Epoch 1749
└ @ Main /Users/kcft114/.julia/packages/Flux/qXNjB/src/optimise/train.jl:105
┌ Info: Epoch 1750
└ @ Main /Users/kcft114/.julia/packages/Flux/qXNjB/src/optimise/train.jl:105
┌ Info: Epoch 1751
└ @ Main /Users/kcft114/.julia/packages/Flux/qXNjB/src/optimise/train.jl:105
┌ Info: Epoch 1752
└ @ Main /Users/kcft114/.julia/packages/Flux/qXNjB/src/optimise/train.jl:105
┌ Info: Epoch 1753
└ @ Main /Users/kcft114/.julia/packages/Flux/qXNjB/src/optimise/train.jl:105
┌ Info: Epoch 1754
└ @ Main /Users/kcft114/.julia/packages/Flux/qXNjB/src/optimise/train.jl:105
┌ Info: Epoch 1755
└ @ Main /Users/kcft114/.julia/packages/Flux/qXNjB/src/optimise/train.jl:105
┌ Info: Epoch 1756
└ @ Main /Users/kcft114/.julia/packages/Flux/qXNjB/src/optimise/train.jl:105
┌ Info: Epoch 1757
└ @ Main /Users/kcft114/.julia/packages/Flux/qXNjB/src/optimise/train.jl:105
┌ Info: Epoch 1758
└ @ Main /Users/kcft114/.julia/packages/Flux/qXNjB/src/optimise/train.jl:105
┌ Info: Epoch 1759
└ @ Main /Users/kcft114/.julia/packages/Flux/qXNjB/src/optimise/train.jl:105
┌ Info: Epoch 1760
└ @ Main /Users/kcft114/.julia/packages/Flux/qXNjB/src/optimise/train.jl:105
┌ Info: Epoch 1761
└ @ Main /Users/kcft114/.julia/packages/Flux/qXNjB/src/optimise/train.jl:105
┌ Info: Epoch 1762
└ @ Main /Users/kcft114/.julia/packages/Flux/qXNjB/src/optimise/train.jl:105
┌ Info: Epoch 1763
└ @ Main /Users/kcft114/.julia/packages/Flux/qXNjB/src/optimise/train.jl:105
┌ Info: Epoch 1764
└ @ Main /Users/kcft114/.julia/packages/Flux/qXNjB/src/optimise/train.jl:105
┌ Info: Epoch 1765
└ @ Main /Users/kcft114/.julia/packages/Flux/qXNjB/src/optimise/train.jl:105
┌ Info: Epoch 1766
└ @ Main /Users/kcft114/.julia/packages/Flux/qXNjB/src/optimise/train.jl:105
┌ Info: Epoch 1767
└ @ Main /Users/kcft114/.julia/packages/Flux/qXNjB/src/optimise/train.jl:105
┌ Info: Epoch 1768
└ @ Main /Users/kcft114/.julia/packages/Flux/qXNjB/src/optimise/train.jl:105
┌ Info: Epoch 1769
└ @ Main /Users/kcft114/.julia/packages/Flux/qXNjB/src/optimise/train.jl:105
┌ Info: Epoch 1770
└ @ Main /Users/kcft114/.julia/packages/Flux/qXNjB/src/optimise/train.jl:105
┌ Info: Epoch 1771
└ @ Main /Users/kcft114/.julia/packages/Flux/qXNjB/src/optimise/train.jl:105
┌ Info: Epoch 1772
└ @ Main /Users/kcft114/.julia/packages/Flux/qXNjB/src/optimise/train.jl:105
┌ Info: Epoch 1773
└ @ Main /Users/kcft114/.julia/packages/Flux/qXNjB/src/optimise/train.jl:105
┌ Info: Epoch 1774
└ @ Main /Users/kcft114/.julia/packages/Flux/qXNjB/src/optimise/train.jl:105
┌ Info: Epoch 1775
└ @ Main /Users/kcft114/.julia/packages/Flux/qXNjB/src/optimise/train.jl:105
┌ Info: Epoch 1776
└ @ Main /Users/kcft114/.julia/packages/Flux/qXNjB/src/optimise/train.jl:105
┌ Info: Epoch 1777
└ @ Main /Users/kcft114/.julia/packages/Flux/qXNjB/src/optimise/train.jl:105
┌ Info: Epoch 1778
└ @ Main /Users/kcft114/.julia/packages/Flux/qXNjB/src/optimise/train.jl:105
┌ Info: Epoch 1779
└ @ Main /Users/kcft114/.julia/packages/Flux/qXNjB/src/optimise/train.jl:105
┌ Info: Epoch 1780
└ @ Main /Users/kcft114/.julia/packages/Flux/qXNjB/src/optimise/train.jl:105
┌ Info: Epoch 1781
└ @ Main /Users/kcft114/.julia/packages/Flux/qXNjB/src/optimise/train.jl:105
┌ Info: Epoch 1782
└ @ Main /Users/kcft114/.julia/packages/Flux/qXNjB/src/optimise/train.jl:105
┌ Info: Epoch 1783
└ @ Main /Users/kcft114/.julia/packages/Flux/qXNjB/src/optimise/train.jl:105
┌ Info: Epoch 1784
└ @ Main /Users/kcft114/.julia/packages/Flux/qXNjB/src/optimise/train.jl:105
┌ Info: Epoch 1785
└ @ Main /Users/kcft114/.julia/packages/Flux/qXNjB/src/optimise/train.jl:105
┌ Info: Epoch 1786
└ @ Main /Users/kcft114/.julia/packages/Flux/qXNjB/src/optimise/train.jl:105
┌ Info: Epoch 1787
└ @ Main /Users/kcft114/.julia/packages/Flux/qXNjB/src/optimise/train.jl:105
┌ Info: Epoch 1788
└ @ Main /Users/kcft114/.julia/packages/Flux/qXNjB/src/optimise/train.jl:105
┌ Info: Epoch 1789
└ @ Main /Users/kcft114/.julia/packages/Flux/qXNjB/src/optimise/train.jl:105
┌ Info: Epoch 1790
└ @ Main /Users/kcft114/.julia/packages/Flux/qXNjB/src/optimise/train.jl:105
┌ Info: Epoch 1791
└ @ Main /Users/kcft114/.julia/packages/Flux/qXNjB/src/optimise/train.jl:105
┌ Info: Epoch 1792
└ @ Main /Users/kcft114/.julia/packages/Flux/qXNjB/src/optimise/train.jl:105
┌ Info: Epoch 1793
└ @ Main /Users/kcft114/.julia/packages/Flux/qXNjB/src/optimise/train.jl:105
┌ Info: Epoch 1794
└ @ Main /Users/kcft114/.julia/packages/Flux/qXNjB/src/optimise/train.jl:105
┌ Info: Epoch 1795
└ @ Main /Users/kcft114/.julia/packages/Flux/qXNjB/src/optimise/train.jl:105
┌ Info: Epoch 1796
└ @ Main /Users/kcft114/.julia/packages/Flux/qXNjB/src/optimise/train.jl:105
┌ Info: Epoch 1797
└ @ Main /Users/kcft114/.julia/packages/Flux/qXNjB/src/optimise/train.jl:105
┌ Info: Epoch 1798
└ @ Main /Users/kcft114/.julia/packages/Flux/qXNjB/src/optimise/train.jl:105
┌ Info: Epoch 1799
└ @ Main /Users/kcft114/.julia/packages/Flux/qXNjB/src/optimise/train.jl:105
┌ Info: Epoch 1800
└ @ Main /Users/kcft114/.julia/packages/Flux/qXNjB/src/optimise/train.jl:105
┌ Info: Epoch 1801
└ @ Main /Users/kcft114/.julia/packages/Flux/qXNjB/src/optimise/train.jl:105
┌ Info: Epoch 1802
└ @ Main /Users/kcft114/.julia/packages/Flux/qXNjB/src/optimise/train.jl:105
┌ Info: Epoch 1803
└ @ Main /Users/kcft114/.julia/packages/Flux/qXNjB/src/optimise/train.jl:105
┌ Info: Epoch 1804
└ @ Main /Users/kcft114/.julia/packages/Flux/qXNjB/src/optimise/train.jl:105
┌ Info: Epoch 1805
└ @ Main /Users/kcft114/.julia/packages/Flux/qXNjB/src/optimise/train.jl:105
┌ Info: Epoch 1806
└ @ Main /Users/kcft114/.julia/packages/Flux/qXNjB/src/optimise/train.jl:105
┌ Info: Epoch 1807
└ @ Main /Users/kcft114/.julia/packages/Flux/qXNjB/src/optimise/train.jl:105
┌ Info: Epoch 1808
└ @ Main /Users/kcft114/.julia/packages/Flux/qXNjB/src/optimise/train.jl:105
┌ Info: Epoch 1809
└ @ Main /Users/kcft114/.julia/packages/Flux/qXNjB/src/optimise/train.jl:105
┌ Info: Epoch 1810
└ @ Main /Users/kcft114/.julia/packages/Flux/qXNjB/src/optimise/train.jl:105
┌ Info: Epoch 1811
└ @ Main /Users/kcft114/.julia/packages/Flux/qXNjB/src/optimise/train.jl:105
┌ Info: Epoch 1812
└ @ Main /Users/kcft114/.julia/packages/Flux/qXNjB/src/optimise/train.jl:105
┌ Info: Epoch 1813
└ @ Main /Users/kcft114/.julia/packages/Flux/qXNjB/src/optimise/train.jl:105
┌ Info: Epoch 1814
└ @ Main /Users/kcft114/.julia/packages/Flux/qXNjB/src/optimise/train.jl:105
┌ Info: Epoch 1815
└ @ Main /Users/kcft114/.julia/packages/Flux/qXNjB/src/optimise/train.jl:105
┌ Info: Epoch 1816
└ @ Main /Users/kcft114/.julia/packages/Flux/qXNjB/src/optimise/train.jl:105
┌ Info: Epoch 1817
└ @ Main /Users/kcft114/.julia/packages/Flux/qXNjB/src/optimise/train.jl:105
┌ Info: Epoch 1818
└ @ Main /Users/kcft114/.julia/packages/Flux/qXNjB/src/optimise/train.jl:105
┌ Info: Epoch 1819
└ @ Main /Users/kcft114/.julia/packages/Flux/qXNjB/src/optimise/train.jl:105
┌ Info: Epoch 1820
└ @ Main /Users/kcft114/.julia/packages/Flux/qXNjB/src/optimise/train.jl:105
┌ Info: Epoch 1821
└ @ Main /Users/kcft114/.julia/packages/Flux/qXNjB/src/optimise/train.jl:105
┌ Info: Epoch 1822
└ @ Main /Users/kcft114/.julia/packages/Flux/qXNjB/src/optimise/train.jl:105
┌ Info: Epoch 1823
└ @ Main /Users/kcft114/.julia/packages/Flux/qXNjB/src/optimise/train.jl:105
┌ Info: Epoch 1824
└ @ Main /Users/kcft114/.julia/packages/Flux/qXNjB/src/optimise/train.jl:105
┌ Info: Epoch 1825
└ @ Main /Users/kcft114/.julia/packages/Flux/qXNjB/src/optimise/train.jl:105
┌ Info: Epoch 1826
└ @ Main /Users/kcft114/.julia/packages/Flux/qXNjB/src/optimise/train.jl:105
┌ Info: Epoch 1827
└ @ Main /Users/kcft114/.julia/packages/Flux/qXNjB/src/optimise/train.jl:105
┌ Info: Epoch 1828
└ @ Main /Users/kcft114/.julia/packages/Flux/qXNjB/src/optimise/train.jl:105
┌ Info: Epoch 1829
└ @ Main /Users/kcft114/.julia/packages/Flux/qXNjB/src/optimise/train.jl:105
┌ Info: Epoch 1830
└ @ Main /Users/kcft114/.julia/packages/Flux/qXNjB/src/optimise/train.jl:105
┌ Info: Epoch 1831
└ @ Main /Users/kcft114/.julia/packages/Flux/qXNjB/src/optimise/train.jl:105
┌ Info: Epoch 1832
└ @ Main /Users/kcft114/.julia/packages/Flux/qXNjB/src/optimise/train.jl:105
┌ Info: Epoch 1833
└ @ Main /Users/kcft114/.julia/packages/Flux/qXNjB/src/optimise/train.jl:105
┌ Info: Epoch 1834
└ @ Main /Users/kcft114/.julia/packages/Flux/qXNjB/src/optimise/train.jl:105
┌ Info: Epoch 1835
└ @ Main /Users/kcft114/.julia/packages/Flux/qXNjB/src/optimise/train.jl:105
┌ Info: Epoch 1836
└ @ Main /Users/kcft114/.julia/packages/Flux/qXNjB/src/optimise/train.jl:105
┌ Info: Epoch 1837
└ @ Main /Users/kcft114/.julia/packages/Flux/qXNjB/src/optimise/train.jl:105
┌ Info: Epoch 1838
└ @ Main /Users/kcft114/.julia/packages/Flux/qXNjB/src/optimise/train.jl:105
┌ Info: Epoch 1839
└ @ Main /Users/kcft114/.julia/packages/Flux/qXNjB/src/optimise/train.jl:105
┌ Info: Epoch 1840
└ @ Main /Users/kcft114/.julia/packages/Flux/qXNjB/src/optimise/train.jl:105
┌ Info: Epoch 1841
└ @ Main /Users/kcft114/.julia/packages/Flux/qXNjB/src/optimise/train.jl:105
┌ Info: Epoch 1842
└ @ Main /Users/kcft114/.julia/packages/Flux/qXNjB/src/optimise/train.jl:105
┌ Info: Epoch 1843
└ @ Main /Users/kcft114/.julia/packages/Flux/qXNjB/src/optimise/train.jl:105
┌ Info: Epoch 1844
└ @ Main /Users/kcft114/.julia/packages/Flux/qXNjB/src/optimise/train.jl:105
┌ Info: Epoch 1845
└ @ Main /Users/kcft114/.julia/packages/Flux/qXNjB/src/optimise/train.jl:105
┌ Info: Epoch 1846
└ @ Main /Users/kcft114/.julia/packages/Flux/qXNjB/src/optimise/train.jl:105
┌ Info: Epoch 1847
└ @ Main /Users/kcft114/.julia/packages/Flux/qXNjB/src/optimise/train.jl:105
┌ Info: Epoch 1848
└ @ Main /Users/kcft114/.julia/packages/Flux/qXNjB/src/optimise/train.jl:105
```

```
loss(X, Y) = 0.3163586f0 (tracked)
loss(X, Y) = 0.31620374f0 (tracked)
loss(X, Y) = 0.31605658f0 (tracked)
loss(X, Y) = 0.31591237f0 (tracked)
loss(X, Y) = 0.3157579f0 (tracked)
loss(X, Y) = 0.3156105f0 (tracked)
loss(X, Y) = 0.3154619f0 (tracked)
loss(X, Y) = 0.31530735f0 (tracked)
loss(X, Y) = 0.31516138f0 (tracked)
loss(X, Y) = 0.31501454f0 (tracked)
loss(X, Y) = 0.3148668f0 (tracked)
loss(X, Y) = 0.31472665f0 (tracked)
loss(X, Y) = 0.31457856f0 (tracked)
loss(X, Y) = 0.3144259f0 (tracked)
loss(X, Y) = 0.31428316f0 (tracked)
loss(X, Y) = 0.31413886f0 (tracked)
loss(X, Y) = 0.31398657f0 (tracked)
loss(X, Y) = 0.31384102f0 (tracked)
loss(X, Y) = 0.31368724f0 (tracked)
loss(X, Y) = 0.31353593f0 (tracked)
loss(X, Y) = 0.31339458f0 (tracked)
loss(X, Y) = 0.31325114f0 (tracked)
loss(X, Y) = 0.31308815f0 (tracked)
loss(X, Y) = 0.31294388f0 (tracked)
loss(X, Y) = 0.31280562f0 (tracked)
loss(X, Y) = 0.31266147f0 (tracked)
loss(X, Y) = 0.31251186f0 (tracked)
loss(X, Y) = 0.31234765f0 (tracked)
loss(X, Y) = 0.31220022f0 (tracked)
loss(X, Y) = 0.31206524f0 (tracked)
loss(X, Y) = 0.31191352f0 (tracked)
loss(X, Y) = 0.3117662f0 (tracked)
loss(X, Y) = 0.31161264f0 (tracked)
loss(X, Y) = 0.3114666f0 (tracked)
loss(X, Y) = 0.31132284f0 (tracked)
loss(X, Y) = 0.31116977f0 (tracked)
loss(X, Y) = 0.31102148f0 (tracked)
loss(X, Y) = 0.31086898f0 (tracked)
loss(X, Y) = 0.31073084f0 (tracked)
loss(X, Y) = 0.31058142f0 (tracked)
loss(X, Y) = 0.31042874f0 (tracked)
loss(X, Y) = 0.31027845f0 (tracked)
loss(X, Y) = 0.31013635f0 (tracked)
loss(X, Y) = 0.30999112f0 (tracked)
loss(X, Y) = 0.30984414f0 (tracked)
loss(X, Y) = 0.30969313f0 (tracked)
loss(X, Y) = 0.30954885f0 (tracked)
loss(X, Y) = 0.3093984f0 (tracked)
loss(X, Y) = 0.30925173f0 (tracked)
loss(X, Y) = 0.30911434f0 (tracked)
loss(X, Y) = 0.30896696f0 (tracked)
loss(X, Y) = 0.3088231f0 (tracked)
loss(X, Y) = 0.30867535f0 (tracked)
loss(X, Y) = 0.3085311f0 (tracked)
loss(X, Y) = 0.30838376f0 (tracked)
loss(X, Y) = 0.3082416f0 (tracked)
loss(X, Y) = 0.30809548f0 (tracked)
loss(X, Y) = 0.3079564f0 (tracked)
loss(X, Y) = 0.30781263f0 (tracked)
loss(X, Y) = 0.30766502f0 (tracked)
loss(X, Y) = 0.3075168f0 (tracked)
loss(X, Y) = 0.30737463f0 (tracked)
loss(X, Y) = 0.3072379f0 (tracked)
loss(X, Y) = 0.30709887f0 (tracked)
loss(X, Y) = 0.3069654f0 (tracked)
loss(X, Y) = 0.3068133f0 (tracked)
loss(X, Y) = 0.30666503f0 (tracked)
loss(X, Y) = 0.3065183f0 (tracked)
loss(X, Y) = 0.30638072f0 (tracked)
loss(X, Y) = 0.3062454f0 (tracked)
loss(X, Y) = 0.30609962f0 (tracked)
loss(X, Y) = 0.30595642f0 (tracked)
loss(X, Y) = 0.30581725f0 (tracked)
loss(X, Y) = 0.30567715f0 (tracked)
loss(X, Y) = 0.30553794f0 (tracked)
loss(X, Y) = 0.3054023f0 (tracked)
loss(X, Y) = 0.30525935f0 (tracked)
loss(X, Y) = 0.3051082f0 (tracked)
loss(X, Y) = 0.30497077f0 (tracked)
loss(X, Y) = 0.30483294f0 (tracked)
loss(X, Y) = 0.30469424f0 (tracked)
loss(X, Y) = 0.3045565f0 (tracked)
loss(X, Y) = 0.30441168f0 (tracked)
loss(X, Y) = 0.30426607f0 (tracked)
loss(X, Y) = 0.3041277f0 (tracked)
loss(X, Y) = 0.3039927f0 (tracked)
loss(X, Y) = 0.30384195f0 (tracked)
loss(X, Y) = 0.30370036f0 (tracked)
loss(X, Y) = 0.30357528f0 (tracked)
loss(X, Y) = 0.3034319f0 (tracked)
loss(X, Y) = 0.30329716f0 (tracked)
loss(X, Y) = 0.3031575f0 (tracked)
loss(X, Y) = 0.3030085f0 (tracked)
loss(X, Y) = 0.30286548f0 (tracked)
loss(X, Y) = 0.30272844f0 (tracked)
loss(X, Y) = 0.30259126f0 (tracked)
loss(X, Y) = 0.30245632f0 (tracked)
loss(X, Y) = 0.30231172f0 (tracked)
loss(X, Y) = 0.3021744f0 (tracked)
loss(X, Y) = 0.30204794f0 (tracked)
loss(X, Y) = 0.3019072f0 (tracked)
loss(X, Y) = 0.30176926f0 (tracked)
loss(X, Y) = 0.30163014f0 (tracked)
loss(X, Y) = 0.30148724f0 (tracked)
loss(X, Y) = 0.3013521f0 (tracked)
loss(X, Y) = 0.30121732f0 (tracked)
loss(X, Y) = 0.30107236f0 (tracked)
loss(X, Y) = 0.300934f0 (tracked)
loss(X, Y) = 0.3008058f0 (tracked)
loss(X, Y) = 0.3006656f0 (tracked)
loss(X, Y) = 0.3005249f0 (tracked)
loss(X, Y) = 0.3003904f0 (tracked)
loss(X, Y) = 0.3002555f0 (tracked)
loss(X, Y) = 0.3001145f0 (tracked)
loss(X, Y) = 0.29998344f0 (tracked)
loss(X, Y) = 0.29983824f0 (tracked)
loss(X, Y) = 0.29970396f0 (tracked)
loss(X, Y) = 0.29956618f0 (tracked)
loss(X, Y) = 0.29943043f0 (tracked)
loss(X, Y) = 0.29929462f0 (tracked)
loss(X, Y) = 0.29915568f0 (tracked)
loss(X, Y) = 0.29902038f0 (tracked)
loss(X, Y) = 0.29888418f0 (tracked)
loss(X, Y) = 0.29874927f0 (tracked)
loss(X, Y) = 0.298618f0 (tracked)
loss(X, Y) = 0.29848486f0 (tracked)
loss(X, Y) = 0.29834622f0 (tracked)
loss(X, Y) = 0.29821166f0 (tracked)
loss(X, Y) = 0.29807547f0 (tracked)
loss(X, Y) = 0.29794845f0 (tracked)
loss(X, Y) = 0.29781288f0 (tracked)
loss(X, Y) = 0.29767448f0 (tracked)
loss(X, Y) = 0.29753736f0 (tracked)
```

```
┌ Info: Epoch 1849
└ @ Main /Users/kcft114/.julia/packages/Flux/qXNjB/src/optimise/train.jl:105
┌ Info: Epoch 1850
└ @ Main /Users/kcft114/.julia/packages/Flux/qXNjB/src/optimise/train.jl:105
┌ Info: Epoch 1851
└ @ Main /Users/kcft114/.julia/packages/Flux/qXNjB/src/optimise/train.jl:105
┌ Info: Epoch 1852
└ @ Main /Users/kcft114/.julia/packages/Flux/qXNjB/src/optimise/train.jl:105
┌ Info: Epoch 1853
└ @ Main /Users/kcft114/.julia/packages/Flux/qXNjB/src/optimise/train.jl:105
┌ Info: Epoch 1854
└ @ Main /Users/kcft114/.julia/packages/Flux/qXNjB/src/optimise/train.jl:105
┌ Info: Epoch 1855
└ @ Main /Users/kcft114/.julia/packages/Flux/qXNjB/src/optimise/train.jl:105
┌ Info: Epoch 1856
└ @ Main /Users/kcft114/.julia/packages/Flux/qXNjB/src/optimise/train.jl:105
┌ Info: Epoch 1857
└ @ Main /Users/kcft114/.julia/packages/Flux/qXNjB/src/optimise/train.jl:105
┌ Info: Epoch 1858
└ @ Main /Users/kcft114/.julia/packages/Flux/qXNjB/src/optimise/train.jl:105
┌ Info: Epoch 1859
└ @ Main /Users/kcft114/.julia/packages/Flux/qXNjB/src/optimise/train.jl:105
┌ Info: Epoch 1860
└ @ Main /Users/kcft114/.julia/packages/Flux/qXNjB/src/optimise/train.jl:105
┌ Info: Epoch 1861
└ @ Main /Users/kcft114/.julia/packages/Flux/qXNjB/src/optimise/train.jl:105
┌ Info: Epoch 1862
└ @ Main /Users/kcft114/.julia/packages/Flux/qXNjB/src/optimise/train.jl:105
┌ Info: Epoch 1863
└ @ Main /Users/kcft114/.julia/packages/Flux/qXNjB/src/optimise/train.jl:105
┌ Info: Epoch 1864
└ @ Main /Users/kcft114/.julia/packages/Flux/qXNjB/src/optimise/train.jl:105
┌ Info: Epoch 1865
└ @ Main /Users/kcft114/.julia/packages/Flux/qXNjB/src/optimise/train.jl:105
┌ Info: Epoch 1866
└ @ Main /Users/kcft114/.julia/packages/Flux/qXNjB/src/optimise/train.jl:105
┌ Info: Epoch 1867
└ @ Main /Users/kcft114/.julia/packages/Flux/qXNjB/src/optimise/train.jl:105
┌ Info: Epoch 1868
└ @ Main /Users/kcft114/.julia/packages/Flux/qXNjB/src/optimise/train.jl:105
┌ Info: Epoch 1869
└ @ Main /Users/kcft114/.julia/packages/Flux/qXNjB/src/optimise/train.jl:105
┌ Info: Epoch 1870
└ @ Main /Users/kcft114/.julia/packages/Flux/qXNjB/src/optimise/train.jl:105
┌ Info: Epoch 1871
└ @ Main /Users/kcft114/.julia/packages/Flux/qXNjB/src/optimise/train.jl:105
┌ Info: Epoch 1872
└ @ Main /Users/kcft114/.julia/packages/Flux/qXNjB/src/optimise/train.jl:105
┌ Info: Epoch 1873
└ @ Main /Users/kcft114/.julia/packages/Flux/qXNjB/src/optimise/train.jl:105
┌ Info: Epoch 1874
└ @ Main /Users/kcft114/.julia/packages/Flux/qXNjB/src/optimise/train.jl:105
┌ Info: Epoch 1875
└ @ Main /Users/kcft114/.julia/packages/Flux/qXNjB/src/optimise/train.jl:105
┌ Info: Epoch 1876
└ @ Main /Users/kcft114/.julia/packages/Flux/qXNjB/src/optimise/train.jl:105
┌ Info: Epoch 1877
└ @ Main /Users/kcft114/.julia/packages/Flux/qXNjB/src/optimise/train.jl:105
┌ Info: Epoch 1878
└ @ Main /Users/kcft114/.julia/packages/Flux/qXNjB/src/optimise/train.jl:105
┌ Info: Epoch 1879
└ @ Main /Users/kcft114/.julia/packages/Flux/qXNjB/src/optimise/train.jl:105
┌ Info: Epoch 1880
└ @ Main /Users/kcft114/.julia/packages/Flux/qXNjB/src/optimise/train.jl:105
┌ Info: Epoch 1881
└ @ Main /Users/kcft114/.julia/packages/Flux/qXNjB/src/optimise/train.jl:105
┌ Info: Epoch 1882
└ @ Main /Users/kcft114/.julia/packages/Flux/qXNjB/src/optimise/train.jl:105
┌ Info: Epoch 1883
└ @ Main /Users/kcft114/.julia/packages/Flux/qXNjB/src/optimise/train.jl:105
┌ Info: Epoch 1884
└ @ Main /Users/kcft114/.julia/packages/Flux/qXNjB/src/optimise/train.jl:105
┌ Info: Epoch 1885
└ @ Main /Users/kcft114/.julia/packages/Flux/qXNjB/src/optimise/train.jl:105
┌ Info: Epoch 1886
└ @ Main /Users/kcft114/.julia/packages/Flux/qXNjB/src/optimise/train.jl:105
┌ Info: Epoch 1887
└ @ Main /Users/kcft114/.julia/packages/Flux/qXNjB/src/optimise/train.jl:105
┌ Info: Epoch 1888
└ @ Main /Users/kcft114/.julia/packages/Flux/qXNjB/src/optimise/train.jl:105
┌ Info: Epoch 1889
└ @ Main /Users/kcft114/.julia/packages/Flux/qXNjB/src/optimise/train.jl:105
┌ Info: Epoch 1890
└ @ Main /Users/kcft114/.julia/packages/Flux/qXNjB/src/optimise/train.jl:105
┌ Info: Epoch 1891
└ @ Main /Users/kcft114/.julia/packages/Flux/qXNjB/src/optimise/train.jl:105
┌ Info: Epoch 1892
└ @ Main /Users/kcft114/.julia/packages/Flux/qXNjB/src/optimise/train.jl:105
┌ Info: Epoch 1893
└ @ Main /Users/kcft114/.julia/packages/Flux/qXNjB/src/optimise/train.jl:105
┌ Info: Epoch 1894
└ @ Main /Users/kcft114/.julia/packages/Flux/qXNjB/src/optimise/train.jl:105
┌ Info: Epoch 1895
└ @ Main /Users/kcft114/.julia/packages/Flux/qXNjB/src/optimise/train.jl:105
┌ Info: Epoch 1896
└ @ Main /Users/kcft114/.julia/packages/Flux/qXNjB/src/optimise/train.jl:105
┌ Info: Epoch 1897
└ @ Main /Users/kcft114/.julia/packages/Flux/qXNjB/src/optimise/train.jl:105
┌ Info: Epoch 1898
└ @ Main /Users/kcft114/.julia/packages/Flux/qXNjB/src/optimise/train.jl:105
┌ Info: Epoch 1899
└ @ Main /Users/kcft114/.julia/packages/Flux/qXNjB/src/optimise/train.jl:105
┌ Info: Epoch 1900
└ @ Main /Users/kcft114/.julia/packages/Flux/qXNjB/src/optimise/train.jl:105
┌ Info: Epoch 1901
└ @ Main /Users/kcft114/.julia/packages/Flux/qXNjB/src/optimise/train.jl:105
┌ Info: Epoch 1902
└ @ Main /Users/kcft114/.julia/packages/Flux/qXNjB/src/optimise/train.jl:105
┌ Info: Epoch 1903
└ @ Main /Users/kcft114/.julia/packages/Flux/qXNjB/src/optimise/train.jl:105
┌ Info: Epoch 1904
└ @ Main /Users/kcft114/.julia/packages/Flux/qXNjB/src/optimise/train.jl:105
┌ Info: Epoch 1905
└ @ Main /Users/kcft114/.julia/packages/Flux/qXNjB/src/optimise/train.jl:105
┌ Info: Epoch 1906
└ @ Main /Users/kcft114/.julia/packages/Flux/qXNjB/src/optimise/train.jl:105
┌ Info: Epoch 1907
└ @ Main /Users/kcft114/.julia/packages/Flux/qXNjB/src/optimise/train.jl:105
┌ Info: Epoch 1908
└ @ Main /Users/kcft114/.julia/packages/Flux/qXNjB/src/optimise/train.jl:105
┌ Info: Epoch 1909
└ @ Main /Users/kcft114/.julia/packages/Flux/qXNjB/src/optimise/train.jl:105
┌ Info: Epoch 1910
└ @ Main /Users/kcft114/.julia/packages/Flux/qXNjB/src/optimise/train.jl:105
┌ Info: Epoch 1911
└ @ Main /Users/kcft114/.julia/packages/Flux/qXNjB/src/optimise/train.jl:105
┌ Info: Epoch 1912
└ @ Main /Users/kcft114/.julia/packages/Flux/qXNjB/src/optimise/train.jl:105
┌ Info: Epoch 1913
└ @ Main /Users/kcft114/.julia/packages/Flux/qXNjB/src/optimise/train.jl:105
┌ Info: Epoch 1914
└ @ Main /Users/kcft114/.julia/packages/Flux/qXNjB/src/optimise/train.jl:105
┌ Info: Epoch 1915
└ @ Main /Users/kcft114/.julia/packages/Flux/qXNjB/src/optimise/train.jl:105
┌ Info: Epoch 1916
└ @ Main /Users/kcft114/.julia/packages/Flux/qXNjB/src/optimise/train.jl:105
┌ Info: Epoch 1917
└ @ Main /Users/kcft114/.julia/packages/Flux/qXNjB/src/optimise/train.jl:105
┌ Info: Epoch 1918
└ @ Main /Users/kcft114/.julia/packages/Flux/qXNjB/src/optimise/train.jl:105
┌ Info: Epoch 1919
└ @ Main /Users/kcft114/.julia/packages/Flux/qXNjB/src/optimise/train.jl:105
┌ Info: Epoch 1920
└ @ Main /Users/kcft114/.julia/packages/Flux/qXNjB/src/optimise/train.jl:105
┌ Info: Epoch 1921
└ @ Main /Users/kcft114/.julia/packages/Flux/qXNjB/src/optimise/train.jl:105
┌ Info: Epoch 1922
└ @ Main /Users/kcft114/.julia/packages/Flux/qXNjB/src/optimise/train.jl:105
┌ Info: Epoch 1923
└ @ Main /Users/kcft114/.julia/packages/Flux/qXNjB/src/optimise/train.jl:105
┌ Info: Epoch 1924
└ @ Main /Users/kcft114/.julia/packages/Flux/qXNjB/src/optimise/train.jl:105
┌ Info: Epoch 1925
└ @ Main /Users/kcft114/.julia/packages/Flux/qXNjB/src/optimise/train.jl:105
┌ Info: Epoch 1926
└ @ Main /Users/kcft114/.julia/packages/Flux/qXNjB/src/optimise/train.jl:105
┌ Info: Epoch 1927
└ @ Main /Users/kcft114/.julia/packages/Flux/qXNjB/src/optimise/train.jl:105
┌ Info: Epoch 1928
└ @ Main /Users/kcft114/.julia/packages/Flux/qXNjB/src/optimise/train.jl:105
┌ Info: Epoch 1929
└ @ Main /Users/kcft114/.julia/packages/Flux/qXNjB/src/optimise/train.jl:105
┌ Info: Epoch 1930
└ @ Main /Users/kcft114/.julia/packages/Flux/qXNjB/src/optimise/train.jl:105
┌ Info: Epoch 1931
└ @ Main /Users/kcft114/.julia/packages/Flux/qXNjB/src/optimise/train.jl:105
┌ Info: Epoch 1932
└ @ Main /Users/kcft114/.julia/packages/Flux/qXNjB/src/optimise/train.jl:105
┌ Info: Epoch 1933
└ @ Main /Users/kcft114/.julia/packages/Flux/qXNjB/src/optimise/train.jl:105
┌ Info: Epoch 1934
└ @ Main /Users/kcft114/.julia/packages/Flux/qXNjB/src/optimise/train.jl:105
┌ Info: Epoch 1935
└ @ Main /Users/kcft114/.julia/packages/Flux/qXNjB/src/optimise/train.jl:105
┌ Info: Epoch 1936
└ @ Main /Users/kcft114/.julia/packages/Flux/qXNjB/src/optimise/train.jl:105
┌ Info: Epoch 1937
└ @ Main /Users/kcft114/.julia/packages/Flux/qXNjB/src/optimise/train.jl:105
┌ Info: Epoch 1938
└ @ Main /Users/kcft114/.julia/packages/Flux/qXNjB/src/optimise/train.jl:105
┌ Info: Epoch 1939
└ @ Main /Users/kcft114/.julia/packages/Flux/qXNjB/src/optimise/train.jl:105
┌ Info: Epoch 1940
└ @ Main /Users/kcft114/.julia/packages/Flux/qXNjB/src/optimise/train.jl:105
┌ Info: Epoch 1941
└ @ Main /Users/kcft114/.julia/packages/Flux/qXNjB/src/optimise/train.jl:105
┌ Info: Epoch 1942
└ @ Main /Users/kcft114/.julia/packages/Flux/qXNjB/src/optimise/train.jl:105
┌ Info: Epoch 1943
└ @ Main /Users/kcft114/.julia/packages/Flux/qXNjB/src/optimise/train.jl:105
┌ Info: Epoch 1944
└ @ Main /Users/kcft114/.julia/packages/Flux/qXNjB/src/optimise/train.jl:105
┌ Info: Epoch 1945
└ @ Main /Users/kcft114/.julia/packages/Flux/qXNjB/src/optimise/train.jl:105
┌ Info: Epoch 1946
└ @ Main /Users/kcft114/.julia/packages/Flux/qXNjB/src/optimise/train.jl:105
┌ Info: Epoch 1947
└ @ Main /Users/kcft114/.julia/packages/Flux/qXNjB/src/optimise/train.jl:105
┌ Info: Epoch 1948
└ @ Main /Users/kcft114/.julia/packages/Flux/qXNjB/src/optimise/train.jl:105
┌ Info: Epoch 1949
└ @ Main /Users/kcft114/.julia/packages/Flux/qXNjB/src/optimise/train.jl:105
┌ Info: Epoch 1950
└ @ Main /Users/kcft114/.julia/packages/Flux/qXNjB/src/optimise/train.jl:105
┌ Info: Epoch 1951
└ @ Main /Users/kcft114/.julia/packages/Flux/qXNjB/src/optimise/train.jl:105
┌ Info: Epoch 1952
└ @ Main /Users/kcft114/.julia/packages/Flux/qXNjB/src/optimise/train.jl:105
┌ Info: Epoch 1953
└ @ Main /Users/kcft114/.julia/packages/Flux/qXNjB/src/optimise/train.jl:105
┌ Info: Epoch 1954
└ @ Main /Users/kcft114/.julia/packages/Flux/qXNjB/src/optimise/train.jl:105
┌ Info: Epoch 1955
└ @ Main /Users/kcft114/.julia/packages/Flux/qXNjB/src/optimise/train.jl:105
┌ Info: Epoch 1956
└ @ Main /Users/kcft114/.julia/packages/Flux/qXNjB/src/optimise/train.jl:105
┌ Info: Epoch 1957
└ @ Main /Users/kcft114/.julia/packages/Flux/qXNjB/src/optimise/train.jl:105
┌ Info: Epoch 1958
└ @ Main /Users/kcft114/.julia/packages/Flux/qXNjB/src/optimise/train.jl:105
┌ Info: Epoch 1959
└ @ Main /Users/kcft114/.julia/packages/Flux/qXNjB/src/optimise/train.jl:105
┌ Info: Epoch 1960
└ @ Main /Users/kcft114/.julia/packages/Flux/qXNjB/src/optimise/train.jl:105
┌ Info: Epoch 1961
└ @ Main /Users/kcft114/.julia/packages/Flux/qXNjB/src/optimise/train.jl:105
┌ Info: Epoch 1962
└ @ Main /Users/kcft114/.julia/packages/Flux/qXNjB/src/optimise/train.jl:105
┌ Info: Epoch 1963
└ @ Main /Users/kcft114/.julia/packages/Flux/qXNjB/src/optimise/train.jl:105
┌ Info: Epoch 1964
└ @ Main /Users/kcft114/.julia/packages/Flux/qXNjB/src/optimise/train.jl:105
┌ Info: Epoch 1965
└ @ Main /Users/kcft114/.julia/packages/Flux/qXNjB/src/optimise/train.jl:105
┌ Info: Epoch 1966
└ @ Main /Users/kcft114/.julia/packages/Flux/qXNjB/src/optimise/train.jl:105
┌ Info: Epoch 1967
└ @ Main /Users/kcft114/.julia/packages/Flux/qXNjB/src/optimise/train.jl:105
┌ Info: Epoch 1968
└ @ Main /Users/kcft114/.julia/packages/Flux/qXNjB/src/optimise/train.jl:105
┌ Info: Epoch 1969
└ @ Main /Users/kcft114/.julia/packages/Flux/qXNjB/src/optimise/train.jl:105
┌ Info: Epoch 1970
└ @ Main /Users/kcft114/.julia/packages/Flux/qXNjB/src/optimise/train.jl:105
┌ Info: Epoch 1971
└ @ Main /Users/kcft114/.julia/packages/Flux/qXNjB/src/optimise/train.jl:105
┌ Info: Epoch 1972
└ @ Main /Users/kcft114/.julia/packages/Flux/qXNjB/src/optimise/train.jl:105
┌ Info: Epoch 1973
└ @ Main /Users/kcft114/.julia/packages/Flux/qXNjB/src/optimise/train.jl:105
┌ Info: Epoch 1974
└ @ Main /Users/kcft114/.julia/packages/Flux/qXNjB/src/optimise/train.jl:105
┌ Info: Epoch 1975
└ @ Main /Users/kcft114/.julia/packages/Flux/qXNjB/src/optimise/train.jl:105
┌ Info: Epoch 1976
└ @ Main /Users/kcft114/.julia/packages/Flux/qXNjB/src/optimise/train.jl:105
┌ Info: Epoch 1977
└ @ Main /Users/kcft114/.julia/packages/Flux/qXNjB/src/optimise/train.jl:105
┌ Info: Epoch 1978
└ @ Main /Users/kcft114/.julia/packages/Flux/qXNjB/src/optimise/train.jl:105
┌ Info: Epoch 1979
└ @ Main /Users/kcft114/.julia/packages/Flux/qXNjB/src/optimise/train.jl:105
┌ Info: Epoch 1980
└ @ Main /Users/kcft114/.julia/packages/Flux/qXNjB/src/optimise/train.jl:105
┌ Info: Epoch 1981
└ @ Main /Users/kcft114/.julia/packages/Flux/qXNjB/src/optimise/train.jl:105
```

```
loss(X, Y) = 0.29739988f0 (tracked)
loss(X, Y) = 0.29727057f0 (tracked)
loss(X, Y) = 0.2971335f0 (tracked)
loss(X, Y) = 0.29699874f0 (tracked)
loss(X, Y) = 0.29686838f0 (tracked)
loss(X, Y) = 0.2967357f0 (tracked)
loss(X, Y) = 0.29659998f0 (tracked)
loss(X, Y) = 0.29646286f0 (tracked)
loss(X, Y) = 0.29632878f0 (tracked)
loss(X, Y) = 0.29619256f0 (tracked)
loss(X, Y) = 0.2960684f0 (tracked)
loss(X, Y) = 0.29593563f0 (tracked)
loss(X, Y) = 0.2958018f0 (tracked)
loss(X, Y) = 0.29567018f0 (tracked)
loss(X, Y) = 0.2955344f0 (tracked)
loss(X, Y) = 0.29540178f0 (tracked)
loss(X, Y) = 0.29526967f0 (tracked)
loss(X, Y) = 0.29514155f0 (tracked)
loss(X, Y) = 0.29500633f0 (tracked)
loss(X, Y) = 0.29487896f0 (tracked)
loss(X, Y) = 0.29474485f0 (tracked)
loss(X, Y) = 0.29460832f0 (tracked)
loss(X, Y) = 0.29448003f0 (tracked)
loss(X, Y) = 0.29434744f0 (tracked)
loss(X, Y) = 0.2942152f0 (tracked)
loss(X, Y) = 0.2940815f0 (tracked)
loss(X, Y) = 0.29394987f0 (tracked)
loss(X, Y) = 0.2938214f0 (tracked)
loss(X, Y) = 0.29368776f0 (tracked)
loss(X, Y) = 0.29356295f0 (tracked)
loss(X, Y) = 0.29343417f0 (tracked)
loss(X, Y) = 0.29330096f0 (tracked)
loss(X, Y) = 0.29317093f0 (tracked)
loss(X, Y) = 0.29303855f0 (tracked)
loss(X, Y) = 0.2929174f0 (tracked)
loss(X, Y) = 0.29278883f0 (tracked)
loss(X, Y) = 0.2926535f0 (tracked)
loss(X, Y) = 0.29253304f0 (tracked)
loss(X, Y) = 0.29240668f0 (tracked)
loss(X, Y) = 0.29227996f0 (tracked)
loss(X, Y) = 0.29215154f0 (tracked)
loss(X, Y) = 0.29202455f0 (tracked)
loss(X, Y) = 0.2918968f0 (tracked)
loss(X, Y) = 0.29177082f0 (tracked)
loss(X, Y) = 0.29163834f0 (tracked)
loss(X, Y) = 0.2915185f0 (tracked)
loss(X, Y) = 0.29139373f0 (tracked)
loss(X, Y) = 0.29126567f0 (tracked)
loss(X, Y) = 0.29114416f0 (tracked)
loss(X, Y) = 0.29101738f0 (tracked)
loss(X, Y) = 0.29088378f0 (tracked)
loss(X, Y) = 0.29075295f0 (tracked)
loss(X, Y) = 0.2906295f0 (tracked)
loss(X, Y) = 0.29050645f0 (tracked)
loss(X, Y) = 0.2903736f0 (tracked)
loss(X, Y) = 0.2902465f0 (tracked)
loss(X, Y) = 0.29012284f0 (tracked)
loss(X, Y) = 0.28999794f0 (tracked)
loss(X, Y) = 0.28986484f0 (tracked)
loss(X, Y) = 0.2897268f0 (tracked)
loss(X, Y) = 0.28959343f0 (tracked)
loss(X, Y) = 0.2894809f0 (tracked)
loss(X, Y) = 0.2893553f0 (tracked)
loss(X, Y) = 0.2892067f0 (tracked)
loss(X, Y) = 0.28906852f0 (tracked)
loss(X, Y) = 0.28894845f0 (tracked)
loss(X, Y) = 0.28882387f0 (tracked)
loss(X, Y) = 0.28868696f0 (tracked)
loss(X, Y) = 0.2885428f0 (tracked)
loss(X, Y) = 0.28840885f0 (tracked)
loss(X, Y) = 0.28828558f0 (tracked)
loss(X, Y) = 0.28814888f0 (tracked)
loss(X, Y) = 0.28802612f0 (tracked)
loss(X, Y) = 0.28789526f0 (tracked)
loss(X, Y) = 0.287774f0 (tracked)
loss(X, Y) = 0.2876451f0 (tracked)
loss(X, Y) = 0.28751135f0 (tracked)
loss(X, Y) = 0.28737617f0 (tracked)
loss(X, Y) = 0.28725725f0 (tracked)
loss(X, Y) = 0.2871327f0 (tracked)
loss(X, Y) = 0.28699878f0 (tracked)
loss(X, Y) = 0.28687885f0 (tracked)
loss(X, Y) = 0.2867551f0 (tracked)
loss(X, Y) = 0.28662965f0 (tracked)
loss(X, Y) = 0.28650394f0 (tracked)
loss(X, Y) = 0.28636837f0 (tracked)
loss(X, Y) = 0.28623325f0 (tracked)
loss(X, Y) = 0.28611434f0 (tracked)
loss(X, Y) = 0.28599244f0 (tracked)
loss(X, Y) = 0.28586382f0 (tracked)
loss(X, Y) = 0.28572604f0 (tracked)
loss(X, Y) = 0.2856168f0 (tracked)
loss(X, Y) = 0.28550127f0 (tracked)
loss(X, Y) = 0.2853753f0 (tracked)
loss(X, Y) = 0.28524885f0 (tracked)
loss(X, Y) = 0.2851219f0 (tracked)
loss(X, Y) = 0.28499487f0 (tracked)
loss(X, Y) = 0.28486332f0 (tracked)
loss(X, Y) = 0.28474253f0 (tracked)
loss(X, Y) = 0.2846235f0 (tracked)
loss(X, Y) = 0.28449073f0 (tracked)
loss(X, Y) = 0.28435454f0 (tracked)
loss(X, Y) = 0.28423315f0 (tracked)
loss(X, Y) = 0.28411004f0 (tracked)
loss(X, Y) = 0.2839858f0 (tracked)
loss(X, Y) = 0.28385738f0 (tracked)
loss(X, Y) = 0.28373384f0 (tracked)
loss(X, Y) = 0.28361577f0 (tracked)
loss(X, Y) = 0.2834903f0 (tracked)
loss(X, Y) = 0.28336594f0 (tracked)
loss(X, Y) = 0.2832496f0 (tracked)
loss(X, Y) = 0.28312296f0 (tracked)
loss(X, Y) = 0.28300896f0 (tracked)
loss(X, Y) = 0.28288013f0 (tracked)
loss(X, Y) = 0.2827676f0 (tracked)
loss(X, Y) = 0.2826497f0 (tracked)
loss(X, Y) = 0.2825214f0 (tracked)
loss(X, Y) = 0.28239712f0 (tracked)
loss(X, Y) = 0.28228444f0 (tracked)
loss(X, Y) = 0.2821655f0 (tracked)
loss(X, Y) = 0.28203908f0 (tracked)
loss(X, Y) = 0.28191102f0 (tracked)
loss(X, Y) = 0.2817978f0 (tracked)
loss(X, Y) = 0.2816857f0 (tracked)
loss(X, Y) = 0.2815521f0 (tracked)
loss(X, Y) = 0.28143746f0 (tracked)
loss(X, Y) = 0.28132656f0 (tracked)
loss(X, Y) = 0.2812117f0 (tracked)
loss(X, Y) = 0.2810859f0 (tracked)
loss(X, Y) = 0.28096274f0 (tracked)
loss(X, Y) = 0.28083205f0 (tracked)
loss(X, Y) = 0.28072277f0 (tracked)
loss(X, Y) = 0.2806126f0 (tracked)
loss(X, Y) = 0.2804878f0 (tracked)
loss(X, Y) = 0.28036186f0 (tracked)
loss(X, Y) = 0.28024277f0 (tracked)
loss(X, Y) = 0.2801285f0 (tracked)
loss(X, Y) = 0.28002238f0 (tracked)
loss(X, Y) = 0.27990723f0 (tracked)
loss(X, Y) = 0.279794f0 (tracked)
loss(X, Y) = 0.2796698f0 (tracked)
loss(X, Y) = 0.27953845f0 (tracked)
loss(X, Y) = 0.27942637f0 (tracked)
loss(X, Y) = 0.2793155f0 (tracked)
loss(X, Y) = 0.27920312f0 (tracked)
loss(X, Y) = 0.2790783f0 (tracked)
loss(X, Y) = 0.27897316f0 (tracked)
loss(X, Y) = 0.27885f0 (tracked)
loss(X, Y) = 0.27873158f0 (tracked)
loss(X, Y) = 0.27862296f0 (tracked)
loss(X, Y) = 0.27850246f0 (tracked)
loss(X, Y) = 0.27838194f0 (tracked)
loss(X, Y) = 0.2782664f0 (tracked)
loss(X, Y) = 0.27815783f0 (tracked)
loss(X, Y) = 0.27803794f0 (tracked)
```

```
┌ Info: Epoch 1982
└ @ Main /Users/kcft114/.julia/packages/Flux/qXNjB/src/optimise/train.jl:105
┌ Info: Epoch 1983
└ @ Main /Users/kcft114/.julia/packages/Flux/qXNjB/src/optimise/train.jl:105
┌ Info: Epoch 1984
└ @ Main /Users/kcft114/.julia/packages/Flux/qXNjB/src/optimise/train.jl:105
┌ Info: Epoch 1985
└ @ Main /Users/kcft114/.julia/packages/Flux/qXNjB/src/optimise/train.jl:105
┌ Info: Epoch 1986
└ @ Main /Users/kcft114/.julia/packages/Flux/qXNjB/src/optimise/train.jl:105
┌ Info: Epoch 1987
└ @ Main /Users/kcft114/.julia/packages/Flux/qXNjB/src/optimise/train.jl:105
┌ Info: Epoch 1988
└ @ Main /Users/kcft114/.julia/packages/Flux/qXNjB/src/optimise/train.jl:105
┌ Info: Epoch 1989
└ @ Main /Users/kcft114/.julia/packages/Flux/qXNjB/src/optimise/train.jl:105
┌ Info: Epoch 1990
└ @ Main /Users/kcft114/.julia/packages/Flux/qXNjB/src/optimise/train.jl:105
┌ Info: Epoch 1991
└ @ Main /Users/kcft114/.julia/packages/Flux/qXNjB/src/optimise/train.jl:105
┌ Info: Epoch 1992
└ @ Main /Users/kcft114/.julia/packages/Flux/qXNjB/src/optimise/train.jl:105
┌ Info: Epoch 1993
└ @ Main /Users/kcft114/.julia/packages/Flux/qXNjB/src/optimise/train.jl:105
┌ Info: Epoch 1994
└ @ Main /Users/kcft114/.julia/packages/Flux/qXNjB/src/optimise/train.jl:105
┌ Info: Epoch 1995
└ @ Main /Users/kcft114/.julia/packages/Flux/qXNjB/src/optimise/train.jl:105
┌ Info: Epoch 1996
└ @ Main /Users/kcft114/.julia/packages/Flux/qXNjB/src/optimise/train.jl:105
┌ Info: Epoch 1997
└ @ Main /Users/kcft114/.julia/packages/Flux/qXNjB/src/optimise/train.jl:105
┌ Info: Epoch 1998
└ @ Main /Users/kcft114/.julia/packages/Flux/qXNjB/src/optimise/train.jl:105
┌ Info: Epoch 1999
└ @ Main /Users/kcft114/.julia/packages/Flux/qXNjB/src/optimise/train.jl:105
┌ Info: Epoch 2000
└ @ Main /Users/kcft114/.julia/packages/Flux/qXNjB/src/optimise/train.jl:105
┌ Info: Epoch 2001
└ @ Main /Users/kcft114/.julia/packages/Flux/qXNjB/src/optimise/train.jl:105
┌ Info: Epoch 2002
└ @ Main /Users/kcft114/.julia/packages/Flux/qXNjB/src/optimise/train.jl:105
┌ Info: Epoch 2003
└ @ Main /Users/kcft114/.julia/packages/Flux/qXNjB/src/optimise/train.jl:105
┌ Info: Epoch 2004
└ @ Main /Users/kcft114/.julia/packages/Flux/qXNjB/src/optimise/train.jl:105
┌ Info: Epoch 2005
└ @ Main /Users/kcft114/.julia/packages/Flux/qXNjB/src/optimise/train.jl:105
┌ Info: Epoch 2006
└ @ Main /Users/kcft114/.julia/packages/Flux/qXNjB/src/optimise/train.jl:105
┌ Info: Epoch 2007
└ @ Main /Users/kcft114/.julia/packages/Flux/qXNjB/src/optimise/train.jl:105
┌ Info: Epoch 2008
└ @ Main /Users/kcft114/.julia/packages/Flux/qXNjB/src/optimise/train.jl:105
┌ Info: Epoch 2009
└ @ Main /Users/kcft114/.julia/packages/Flux/qXNjB/src/optimise/train.jl:105
┌ Info: Epoch 2010
└ @ Main /Users/kcft114/.julia/packages/Flux/qXNjB/src/optimise/train.jl:105
┌ Info: Epoch 2011
└ @ Main /Users/kcft114/.julia/packages/Flux/qXNjB/src/optimise/train.jl:105
┌ Info: Epoch 2012
└ @ Main /Users/kcft114/.julia/packages/Flux/qXNjB/src/optimise/train.jl:105
┌ Info: Epoch 2013
└ @ Main /Users/kcft114/.julia/packages/Flux/qXNjB/src/optimise/train.jl:105
┌ Info: Epoch 2014
└ @ Main /Users/kcft114/.julia/packages/Flux/qXNjB/src/optimise/train.jl:105
┌ Info: Epoch 2015
└ @ Main /Users/kcft114/.julia/packages/Flux/qXNjB/src/optimise/train.jl:105
┌ Info: Epoch 2016
└ @ Main /Users/kcft114/.julia/packages/Flux/qXNjB/src/optimise/train.jl:105
┌ Info: Epoch 2017
└ @ Main /Users/kcft114/.julia/packages/Flux/qXNjB/src/optimise/train.jl:105
┌ Info: Epoch 2018
└ @ Main /Users/kcft114/.julia/packages/Flux/qXNjB/src/optimise/train.jl:105
┌ Info: Epoch 2019
└ @ Main /Users/kcft114/.julia/packages/Flux/qXNjB/src/optimise/train.jl:105
┌ Info: Epoch 2020
└ @ Main /Users/kcft114/.julia/packages/Flux/qXNjB/src/optimise/train.jl:105
┌ Info: Epoch 2021
└ @ Main /Users/kcft114/.julia/packages/Flux/qXNjB/src/optimise/train.jl:105
┌ Info: Epoch 2022
└ @ Main /Users/kcft114/.julia/packages/Flux/qXNjB/src/optimise/train.jl:105
┌ Info: Epoch 2023
└ @ Main /Users/kcft114/.julia/packages/Flux/qXNjB/src/optimise/train.jl:105
┌ Info: Epoch 2024
└ @ Main /Users/kcft114/.julia/packages/Flux/qXNjB/src/optimise/train.jl:105
┌ Info: Epoch 2025
└ @ Main /Users/kcft114/.julia/packages/Flux/qXNjB/src/optimise/train.jl:105
┌ Info: Epoch 2026
└ @ Main /Users/kcft114/.julia/packages/Flux/qXNjB/src/optimise/train.jl:105
┌ Info: Epoch 2027
└ @ Main /Users/kcft114/.julia/packages/Flux/qXNjB/src/optimise/train.jl:105
┌ Info: Epoch 2028
└ @ Main /Users/kcft114/.julia/packages/Flux/qXNjB/src/optimise/train.jl:105
┌ Info: Epoch 2029
└ @ Main /Users/kcft114/.julia/packages/Flux/qXNjB/src/optimise/train.jl:105
┌ Info: Epoch 2030
└ @ Main /Users/kcft114/.julia/packages/Flux/qXNjB/src/optimise/train.jl:105
┌ Info: Epoch 2031
└ @ Main /Users/kcft114/.julia/packages/Flux/qXNjB/src/optimise/train.jl:105
┌ Info: Epoch 2032
└ @ Main /Users/kcft114/.julia/packages/Flux/qXNjB/src/optimise/train.jl:105
┌ Info: Epoch 2033
└ @ Main /Users/kcft114/.julia/packages/Flux/qXNjB/src/optimise/train.jl:105
┌ Info: Epoch 2034
└ @ Main /Users/kcft114/.julia/packages/Flux/qXNjB/src/optimise/train.jl:105
┌ Info: Epoch 2035
└ @ Main /Users/kcft114/.julia/packages/Flux/qXNjB/src/optimise/train.jl:105
┌ Info: Epoch 2036
└ @ Main /Users/kcft114/.julia/packages/Flux/qXNjB/src/optimise/train.jl:105
┌ Info: Epoch 2037
└ @ Main /Users/kcft114/.julia/packages/Flux/qXNjB/src/optimise/train.jl:105
┌ Info: Epoch 2038
└ @ Main /Users/kcft114/.julia/packages/Flux/qXNjB/src/optimise/train.jl:105
┌ Info: Epoch 2039
└ @ Main /Users/kcft114/.julia/packages/Flux/qXNjB/src/optimise/train.jl:105
┌ Info: Epoch 2040
└ @ Main /Users/kcft114/.julia/packages/Flux/qXNjB/src/optimise/train.jl:105
┌ Info: Epoch 2041
└ @ Main /Users/kcft114/.julia/packages/Flux/qXNjB/src/optimise/train.jl:105
┌ Info: Epoch 2042
└ @ Main /Users/kcft114/.julia/packages/Flux/qXNjB/src/optimise/train.jl:105
┌ Info: Epoch 2043
└ @ Main /Users/kcft114/.julia/packages/Flux/qXNjB/src/optimise/train.jl:105
┌ Info: Epoch 2044
└ @ Main /Users/kcft114/.julia/packages/Flux/qXNjB/src/optimise/train.jl:105
┌ Info: Epoch 2045
└ @ Main /Users/kcft114/.julia/packages/Flux/qXNjB/src/optimise/train.jl:105
┌ Info: Epoch 2046
└ @ Main /Users/kcft114/.julia/packages/Flux/qXNjB/src/optimise/train.jl:105
┌ Info: Epoch 2047
└ @ Main /Users/kcft114/.julia/packages/Flux/qXNjB/src/optimise/train.jl:105
┌ Info: Epoch 2048
└ @ Main /Users/kcft114/.julia/packages/Flux/qXNjB/src/optimise/train.jl:105
┌ Info: Epoch 2049
└ @ Main /Users/kcft114/.julia/packages/Flux/qXNjB/src/optimise/train.jl:105
┌ Info: Epoch 2050
└ @ Main /Users/kcft114/.julia/packages/Flux/qXNjB/src/optimise/train.jl:105
┌ Info: Epoch 2051
└ @ Main /Users/kcft114/.julia/packages/Flux/qXNjB/src/optimise/train.jl:105
┌ Info: Epoch 2052
└ @ Main /Users/kcft114/.julia/packages/Flux/qXNjB/src/optimise/train.jl:105
┌ Info: Epoch 2053
└ @ Main /Users/kcft114/.julia/packages/Flux/qXNjB/src/optimise/train.jl:105
┌ Info: Epoch 2054
└ @ Main /Users/kcft114/.julia/packages/Flux/qXNjB/src/optimise/train.jl:105
┌ Info: Epoch 2055
└ @ Main /Users/kcft114/.julia/packages/Flux/qXNjB/src/optimise/train.jl:105
┌ Info: Epoch 2056
└ @ Main /Users/kcft114/.julia/packages/Flux/qXNjB/src/optimise/train.jl:105
┌ Info: Epoch 2057
└ @ Main /Users/kcft114/.julia/packages/Flux/qXNjB/src/optimise/train.jl:105
┌ Info: Epoch 2058
└ @ Main /Users/kcft114/.julia/packages/Flux/qXNjB/src/optimise/train.jl:105
┌ Info: Epoch 2059
└ @ Main /Users/kcft114/.julia/packages/Flux/qXNjB/src/optimise/train.jl:105
┌ Info: Epoch 2060
└ @ Main /Users/kcft114/.julia/packages/Flux/qXNjB/src/optimise/train.jl:105
┌ Info: Epoch 2061
└ @ Main /Users/kcft114/.julia/packages/Flux/qXNjB/src/optimise/train.jl:105
┌ Info: Epoch 2062
└ @ Main /Users/kcft114/.julia/packages/Flux/qXNjB/src/optimise/train.jl:105
┌ Info: Epoch 2063
└ @ Main /Users/kcft114/.julia/packages/Flux/qXNjB/src/optimise/train.jl:105
┌ Info: Epoch 2064
└ @ Main /Users/kcft114/.julia/packages/Flux/qXNjB/src/optimise/train.jl:105
┌ Info: Epoch 2065
└ @ Main /Users/kcft114/.julia/packages/Flux/qXNjB/src/optimise/train.jl:105
┌ Info: Epoch 2066
└ @ Main /Users/kcft114/.julia/packages/Flux/qXNjB/src/optimise/train.jl:105
┌ Info: Epoch 2067
└ @ Main /Users/kcft114/.julia/packages/Flux/qXNjB/src/optimise/train.jl:105
┌ Info: Epoch 2068
└ @ Main /Users/kcft114/.julia/packages/Flux/qXNjB/src/optimise/train.jl:105
┌ Info: Epoch 2069
└ @ Main /Users/kcft114/.julia/packages/Flux/qXNjB/src/optimise/train.jl:105
┌ Info: Epoch 2070
└ @ Main /Users/kcft114/.julia/packages/Flux/qXNjB/src/optimise/train.jl:105
┌ Info: Epoch 2071
└ @ Main /Users/kcft114/.julia/packages/Flux/qXNjB/src/optimise/train.jl:105
┌ Info: Epoch 2072
└ @ Main /Users/kcft114/.julia/packages/Flux/qXNjB/src/optimise/train.jl:105
┌ Info: Epoch 2073
└ @ Main /Users/kcft114/.julia/packages/Flux/qXNjB/src/optimise/train.jl:105
┌ Info: Epoch 2074
└ @ Main /Users/kcft114/.julia/packages/Flux/qXNjB/src/optimise/train.jl:105
┌ Info: Epoch 2075
└ @ Main /Users/kcft114/.julia/packages/Flux/qXNjB/src/optimise/train.jl:105
┌ Info: Epoch 2076
└ @ Main /Users/kcft114/.julia/packages/Flux/qXNjB/src/optimise/train.jl:105
┌ Info: Epoch 2077
└ @ Main /Users/kcft114/.julia/packages/Flux/qXNjB/src/optimise/train.jl:105
┌ Info: Epoch 2078
└ @ Main /Users/kcft114/.julia/packages/Flux/qXNjB/src/optimise/train.jl:105
┌ Info: Epoch 2079
└ @ Main /Users/kcft114/.julia/packages/Flux/qXNjB/src/optimise/train.jl:105
┌ Info: Epoch 2080
└ @ Main /Users/kcft114/.julia/packages/Flux/qXNjB/src/optimise/train.jl:105
┌ Info: Epoch 2081
└ @ Main /Users/kcft114/.julia/packages/Flux/qXNjB/src/optimise/train.jl:105
┌ Info: Epoch 2082
└ @ Main /Users/kcft114/.julia/packages/Flux/qXNjB/src/optimise/train.jl:105
┌ Info: Epoch 2083
└ @ Main /Users/kcft114/.julia/packages/Flux/qXNjB/src/optimise/train.jl:105
┌ Info: Epoch 2084
└ @ Main /Users/kcft114/.julia/packages/Flux/qXNjB/src/optimise/train.jl:105
┌ Info: Epoch 2085
└ @ Main /Users/kcft114/.julia/packages/Flux/qXNjB/src/optimise/train.jl:105
┌ Info: Epoch 2086
└ @ Main /Users/kcft114/.julia/packages/Flux/qXNjB/src/optimise/train.jl:105
┌ Info: Epoch 2087
└ @ Main /Users/kcft114/.julia/packages/Flux/qXNjB/src/optimise/train.jl:105
┌ Info: Epoch 2088
└ @ Main /Users/kcft114/.julia/packages/Flux/qXNjB/src/optimise/train.jl:105
┌ Info: Epoch 2089
└ @ Main /Users/kcft114/.julia/packages/Flux/qXNjB/src/optimise/train.jl:105
┌ Info: Epoch 2090
└ @ Main /Users/kcft114/.julia/packages/Flux/qXNjB/src/optimise/train.jl:105
┌ Info: Epoch 2091
└ @ Main /Users/kcft114/.julia/packages/Flux/qXNjB/src/optimise/train.jl:105
┌ Info: Epoch 2092
└ @ Main /Users/kcft114/.julia/packages/Flux/qXNjB/src/optimise/train.jl:105
┌ Info: Epoch 2093
└ @ Main /Users/kcft114/.julia/packages/Flux/qXNjB/src/optimise/train.jl:105
┌ Info: Epoch 2094
└ @ Main /Users/kcft114/.julia/packages/Flux/qXNjB/src/optimise/train.jl:105
┌ Info: Epoch 2095
└ @ Main /Users/kcft114/.julia/packages/Flux/qXNjB/src/optimise/train.jl:105
┌ Info: Epoch 2096
└ @ Main /Users/kcft114/.julia/packages/Flux/qXNjB/src/optimise/train.jl:105
┌ Info: Epoch 2097
└ @ Main /Users/kcft114/.julia/packages/Flux/qXNjB/src/optimise/train.jl:105
┌ Info: Epoch 2098
└ @ Main /Users/kcft114/.julia/packages/Flux/qXNjB/src/optimise/train.jl:105
┌ Info: Epoch 2099
└ @ Main /Users/kcft114/.julia/packages/Flux/qXNjB/src/optimise/train.jl:105
┌ Info: Epoch 2100
└ @ Main /Users/kcft114/.julia/packages/Flux/qXNjB/src/optimise/train.jl:105
┌ Info: Epoch 2101
└ @ Main /Users/kcft114/.julia/packages/Flux/qXNjB/src/optimise/train.jl:105
┌ Info: Epoch 2102
└ @ Main /Users/kcft114/.julia/packages/Flux/qXNjB/src/optimise/train.jl:105
┌ Info: Epoch 2103
└ @ Main /Users/kcft114/.julia/packages/Flux/qXNjB/src/optimise/train.jl:105
┌ Info: Epoch 2104
└ @ Main /Users/kcft114/.julia/packages/Flux/qXNjB/src/optimise/train.jl:105
┌ Info: Epoch 2105
└ @ Main /Users/kcft114/.julia/packages/Flux/qXNjB/src/optimise/train.jl:105
┌ Info: Epoch 2106
└ @ Main /Users/kcft114/.julia/packages/Flux/qXNjB/src/optimise/train.jl:105
┌ Info: Epoch 2107
└ @ Main /Users/kcft114/.julia/packages/Flux/qXNjB/src/optimise/train.jl:105
┌ Info: Epoch 2108
└ @ Main /Users/kcft114/.julia/packages/Flux/qXNjB/src/optimise/train.jl:105
┌ Info: Epoch 2109
└ @ Main /Users/kcft114/.julia/packages/Flux/qXNjB/src/optimise/train.jl:105
┌ Info: Epoch 2110
└ @ Main /Users/kcft114/.julia/packages/Flux/qXNjB/src/optimise/train.jl:105
┌ Info: Epoch 2111
└ @ Main /Users/kcft114/.julia/packages/Flux/qXNjB/src/optimise/train.jl:105
┌ Info: Epoch 2112
└ @ Main /Users/kcft114/.julia/packages/Flux/qXNjB/src/optimise/train.jl:105
┌ Info: Epoch 2113
└ @ Main /Users/kcft114/.julia/packages/Flux/qXNjB/src/optimise/train.jl:105
┌ Info: Epoch 2114
└ @ Main /Users/kcft114/.julia/packages/Flux/qXNjB/src/optimise/train.jl:105
┌ Info: Epoch 2115
└ @ Main /Users/kcft114/.julia/packages/Flux/qXNjB/src/optimise/train.jl:105
┌ Info: Epoch 2116
└ @ Main /Users/kcft114/.julia/packages/Flux/qXNjB/src/optimise/train.jl:105
┌ Info: Epoch 2117
└ @ Main /Users/kcft114/.julia/packages/Flux/qXNjB/src/optimise/train.jl:105
┌ Info: Epoch 2118
└ @ Main /Users/kcft114/.julia/packages/Flux/qXNjB/src/optimise/train.jl:105
┌ Info: Epoch 2119
└ @ Main /Users/kcft114/.julia/packages/Flux/qXNjB/src/optimise/train.jl:105
┌ Info: Epoch 2120
└ @ Main /Users/kcft114/.julia/packages/Flux/qXNjB/src/optimise/train.jl:105
┌ Info: Epoch 2121
└ @ Main /Users/kcft114/.julia/packages/Flux/qXNjB/src/optimise/train.jl:105
┌ Info: Epoch 2122
└ @ Main /Users/kcft114/.julia/packages/Flux/qXNjB/src/optimise/train.jl:105
┌ Info: Epoch 2123
└ @ Main /Users/kcft114/.julia/packages/Flux/qXNjB/src/optimise/train.jl:105
┌ Info: Epoch 2124
└ @ Main /Users/kcft114/.julia/packages/Flux/qXNjB/src/optimise/train.jl:105
┌ Info: Epoch 2125
└ @ Main /Users/kcft114/.julia/packages/Flux/qXNjB/src/optimise/train.jl:105
┌ Info: Epoch 2126
└ @ Main /Users/kcft114/.julia/packages/Flux/qXNjB/src/optimise/train.jl:105
┌ Info: Epoch 2127
└ @ Main /Users/kcft114/.julia/packages/Flux/qXNjB/src/optimise/train.jl:105
┌ Info: Epoch 2128
└ @ Main /Users/kcft114/.julia/packages/Flux/qXNjB/src/optimise/train.jl:105
┌ Info: Epoch 2129
└ @ Main /Users/kcft114/.julia/packages/Flux/qXNjB/src/optimise/train.jl:105
┌ Info: Epoch 2130
└ @ Main /Users/kcft114/.julia/packages/Flux/qXNjB/src/optimise/train.jl:105
┌ Info: Epoch 2131
└ @ Main /Users/kcft114/.julia/packages/Flux/qXNjB/src/optimise/train.jl:105
┌ Info: Epoch 2132
└ @ Main /Users/kcft114/.julia/packages/Flux/qXNjB/src/optimise/train.jl:105
┌ Info: Epoch 2133
└ @ Main /Users/kcft114/.julia/packages/Flux/qXNjB/src/optimise/train.jl:105
┌ Info: Epoch 2134
└ @ Main /Users/kcft114/.julia/packages/Flux/qXNjB/src/optimise/train.jl:105
┌ Info: Epoch 2135
└ @ Main /Users/kcft114/.julia/packages/Flux/qXNjB/src/optimise/train.jl:105
┌ Info: Epoch 2136
└ @ Main /Users/kcft114/.julia/packages/Flux/qXNjB/src/optimise/train.jl:105
```

```
loss(X, Y) = 0.277918f0 (tracked)
loss(X, Y) = 0.27780464f0 (tracked)
loss(X, Y) = 0.27770224f0 (tracked)
loss(X, Y) = 0.2775838f0 (tracked)
loss(X, Y) = 0.27745715f0 (tracked)
loss(X, Y) = 0.2773315f0 (tracked)
loss(X, Y) = 0.2772247f0 (tracked)
loss(X, Y) = 0.27711588f0 (tracked)
loss(X, Y) = 0.2769916f0 (tracked)
loss(X, Y) = 0.27687374f0 (tracked)
loss(X, Y) = 0.27676097f0 (tracked)
loss(X, Y) = 0.27665025f0 (tracked)
loss(X, Y) = 0.2765486f0 (tracked)
loss(X, Y) = 0.27643052f0 (tracked)
loss(X, Y) = 0.27629438f0 (tracked)
loss(X, Y) = 0.2761979f0 (tracked)
loss(X, Y) = 0.27609715f0 (tracked)
loss(X, Y) = 0.2759716f0 (tracked)
loss(X, Y) = 0.27584863f0 (tracked)
loss(X, Y) = 0.2757512f0 (tracked)
loss(X, Y) = 0.27563503f0 (tracked)
loss(X, Y) = 0.27551863f0 (tracked)
loss(X, Y) = 0.27541098f0 (tracked)
loss(X, Y) = 0.27528286f0 (tracked)
loss(X, Y) = 0.2751699f0 (tracked)
loss(X, Y) = 0.2750628f0 (tracked)
loss(X, Y) = 0.27494842f0 (tracked)
loss(X, Y) = 0.27483335f0 (tracked)
loss(X, Y) = 0.274712f0 (tracked)
loss(X, Y) = 0.27460027f0 (tracked)
loss(X, Y) = 0.27449813f0 (tracked)
loss(X, Y) = 0.27438357f0 (tracked)
loss(X, Y) = 0.274268f0 (tracked)
loss(X, Y) = 0.27414146f0 (tracked)
loss(X, Y) = 0.2740366f0 (tracked)
loss(X, Y) = 0.27393734f0 (tracked)
loss(X, Y) = 0.2738228f0 (tracked)
loss(X, Y) = 0.2737045f0 (tracked)
loss(X, Y) = 0.27359062f0 (tracked)
loss(X, Y) = 0.273485f0 (tracked)
loss(X, Y) = 0.27337596f0 (tracked)
loss(X, Y) = 0.27325594f0 (tracked)
loss(X, Y) = 0.2731338f0 (tracked)
loss(X, Y) = 0.27303433f0 (tracked)
loss(X, Y) = 0.2729251f0 (tracked)
loss(X, Y) = 0.27280378f0 (tracked)
loss(X, Y) = 0.27268514f0 (tracked)
loss(X, Y) = 0.27258366f0 (tracked)
loss(X, Y) = 0.27247158f0 (tracked)
loss(X, Y) = 0.27236572f0 (tracked)
loss(X, Y) = 0.27224517f0 (tracked)
loss(X, Y) = 0.27213442f0 (tracked)
loss(X, Y) = 0.2720329f0 (tracked)
loss(X, Y) = 0.2719155f0 (tracked)
loss(X, Y) = 0.2718089f0 (tracked)
loss(X, Y) = 0.27169052f0 (tracked)
loss(X, Y) = 0.2715792f0 (tracked)
loss(X, Y) = 0.27147314f0 (tracked)
loss(X, Y) = 0.27136204f0 (tracked)
loss(X, Y) = 0.27125123f0 (tracked)
loss(X, Y) = 0.2711338f0 (tracked)
loss(X, Y) = 0.27102938f0 (tracked)
loss(X, Y) = 0.2709138f0 (tracked)
loss(X, Y) = 0.2707991f0 (tracked)
loss(X, Y) = 0.27069378f0 (tracked)
loss(X, Y) = 0.270586f0 (tracked)
loss(X, Y) = 0.27047667f0 (tracked)
loss(X, Y) = 0.27036119f0 (tracked)
loss(X, Y) = 0.27024376f0 (tracked)
loss(X, Y) = 0.2701212f0 (tracked)
loss(X, Y) = 0.27001184f0 (tracked)
loss(X, Y) = 0.26989508f0 (tracked)
loss(X, Y) = 0.26977658f0 (tracked)
loss(X, Y) = 0.26966023f0 (tracked)
loss(X, Y) = 0.26955616f0 (tracked)
loss(X, Y) = 0.269443f0 (tracked)
loss(X, Y) = 0.26932317f0 (tracked)
loss(X, Y) = 0.2692065f0 (tracked)
loss(X, Y) = 0.26909292f0 (tracked)
loss(X, Y) = 0.2689852f0 (tracked)
loss(X, Y) = 0.26886612f0 (tracked)
loss(X, Y) = 0.26875836f0 (tracked)
loss(X, Y) = 0.2686443f0 (tracked)
loss(X, Y) = 0.26853403f0 (tracked)
loss(X, Y) = 0.2684183f0 (tracked)
loss(X, Y) = 0.26830098f0 (tracked)
loss(X, Y) = 0.2681935f0 (tracked)
loss(X, Y) = 0.26807898f0 (tracked)
loss(X, Y) = 0.26795566f0 (tracked)
loss(X, Y) = 0.2678444f0 (tracked)
loss(X, Y) = 0.2677287f0 (tracked)
loss(X, Y) = 0.26762435f0 (tracked)
loss(X, Y) = 0.2675096f0 (tracked)
loss(X, Y) = 0.26739013f0 (tracked)
loss(X, Y) = 0.26728186f0 (tracked)
loss(X, Y) = 0.2671769f0 (tracked)
loss(X, Y) = 0.2670657f0 (tracked)
loss(X, Y) = 0.2669454f0 (tracked)
loss(X, Y) = 0.26683804f0 (tracked)
loss(X, Y) = 0.26672417f0 (tracked)
loss(X, Y) = 0.2666069f0 (tracked)
loss(X, Y) = 0.26649445f0 (tracked)
loss(X, Y) = 0.2663836f0 (tracked)
loss(X, Y) = 0.26627553f0 (tracked)
loss(X, Y) = 0.26615995f0 (tracked)
loss(X, Y) = 0.2660479f0 (tracked)
loss(X, Y) = 0.26593566f0 (tracked)
loss(X, Y) = 0.26582652f0 (tracked)
loss(X, Y) = 0.26571244f0 (tracked)
loss(X, Y) = 0.26561117f0 (tracked)
loss(X, Y) = 0.26549503f0 (tracked)
loss(X, Y) = 0.2653712f0 (tracked)
loss(X, Y) = 0.26527685f0 (tracked)
loss(X, Y) = 0.26515958f0 (tracked)
loss(X, Y) = 0.26504093f0 (tracked)
loss(X, Y) = 0.26493296f0 (tracked)
loss(X, Y) = 0.26482382f0 (tracked)
loss(X, Y) = 0.26470932f0 (tracked)
loss(X, Y) = 0.26460305f0 (tracked)
loss(X, Y) = 0.2644955f0 (tracked)
loss(X, Y) = 0.26438138f0 (tracked)
loss(X, Y) = 0.26428428f0 (tracked)
loss(X, Y) = 0.26416498f0 (tracked)
loss(X, Y) = 0.26404664f0 (tracked)
loss(X, Y) = 0.26393995f0 (tracked)
loss(X, Y) = 0.2638357f0 (tracked)
loss(X, Y) = 0.26373246f0 (tracked)
loss(X, Y) = 0.26362056f0 (tracked)
loss(X, Y) = 0.2635115f0 (tracked)
loss(X, Y) = 0.26339787f0 (tracked)
loss(X, Y) = 0.26329154f0 (tracked)
loss(X, Y) = 0.2631842f0 (tracked)
loss(X, Y) = 0.2630749f0 (tracked)
loss(X, Y) = 0.26296693f0 (tracked)
loss(X, Y) = 0.26285678f0 (tracked)
loss(X, Y) = 0.26275533f0 (tracked)
loss(X, Y) = 0.26264015f0 (tracked)
loss(X, Y) = 0.26252654f0 (tracked)
loss(X, Y) = 0.2624184f0 (tracked)
loss(X, Y) = 0.26230568f0 (tracked)
loss(X, Y) = 0.26217067f0 (tracked)
loss(X, Y) = 0.2620121f0 (tracked)
loss(X, Y) = 0.2618309f0 (tracked)
loss(X, Y) = 0.26167408f0 (tracked)
loss(X, Y) = 0.26152453f0 (tracked)
loss(X, Y) = 0.2613519f0 (tracked)
loss(X, Y) = 0.2611957f0 (tracked)
loss(X, Y) = 0.2610458f0 (tracked)
loss(X, Y) = 0.26088557f0 (tracked)
loss(X, Y) = 0.2607408f0 (tracked)
loss(X, Y) = 0.2605924f0 (tracked)
loss(X, Y) = 0.26043615f0 (tracked)
loss(X, Y) = 0.2602838f0 (tracked)
loss(X, Y) = 0.26013166f0 (tracked)
loss(X, Y) = 0.26000026f0 (tracked)
loss(X, Y) = 0.25987288f0 (tracked)
loss(X, Y) = 0.2597354f0 (tracked)
loss(X, Y) = 0.2595995f0 (tracked)
loss(X, Y) = 0.25946975f0 (tracked)
loss(X, Y) = 0.25933108f0 (tracked)
loss(X, Y) = 0.25919467f0 (tracked)
loss(X, Y) = 0.25905696f0 (tracked)
loss(X, Y) = 0.25892f0 (tracked)
loss(X, Y) = 0.25877643f0 (tracked)
loss(X, Y) = 0.25864112f0 (tracked)
```

```
┌ Info: Epoch 2137
└ @ Main /Users/kcft114/.julia/packages/Flux/qXNjB/src/optimise/train.jl:105
┌ Info: Epoch 2138
└ @ Main /Users/kcft114/.julia/packages/Flux/qXNjB/src/optimise/train.jl:105
┌ Info: Epoch 2139
└ @ Main /Users/kcft114/.julia/packages/Flux/qXNjB/src/optimise/train.jl:105
┌ Info: Epoch 2140
└ @ Main /Users/kcft114/.julia/packages/Flux/qXNjB/src/optimise/train.jl:105
┌ Info: Epoch 2141
└ @ Main /Users/kcft114/.julia/packages/Flux/qXNjB/src/optimise/train.jl:105
┌ Info: Epoch 2142
└ @ Main /Users/kcft114/.julia/packages/Flux/qXNjB/src/optimise/train.jl:105
┌ Info: Epoch 2143
└ @ Main /Users/kcft114/.julia/packages/Flux/qXNjB/src/optimise/train.jl:105
┌ Info: Epoch 2144
└ @ Main /Users/kcft114/.julia/packages/Flux/qXNjB/src/optimise/train.jl:105
┌ Info: Epoch 2145
└ @ Main /Users/kcft114/.julia/packages/Flux/qXNjB/src/optimise/train.jl:105
┌ Info: Epoch 2146
└ @ Main /Users/kcft114/.julia/packages/Flux/qXNjB/src/optimise/train.jl:105
┌ Info: Epoch 2147
└ @ Main /Users/kcft114/.julia/packages/Flux/qXNjB/src/optimise/train.jl:105
┌ Info: Epoch 2148
└ @ Main /Users/kcft114/.julia/packages/Flux/qXNjB/src/optimise/train.jl:105
┌ Info: Epoch 2149
└ @ Main /Users/kcft114/.julia/packages/Flux/qXNjB/src/optimise/train.jl:105
┌ Info: Epoch 2150
└ @ Main /Users/kcft114/.julia/packages/Flux/qXNjB/src/optimise/train.jl:105
┌ Info: Epoch 2151
└ @ Main /Users/kcft114/.julia/packages/Flux/qXNjB/src/optimise/train.jl:105
┌ Info: Epoch 2152
└ @ Main /Users/kcft114/.julia/packages/Flux/qXNjB/src/optimise/train.jl:105
┌ Info: Epoch 2153
└ @ Main /Users/kcft114/.julia/packages/Flux/qXNjB/src/optimise/train.jl:105
┌ Info: Epoch 2154
└ @ Main /Users/kcft114/.julia/packages/Flux/qXNjB/src/optimise/train.jl:105
┌ Info: Epoch 2155
└ @ Main /Users/kcft114/.julia/packages/Flux/qXNjB/src/optimise/train.jl:105
┌ Info: Epoch 2156
└ @ Main /Users/kcft114/.julia/packages/Flux/qXNjB/src/optimise/train.jl:105
┌ Info: Epoch 2157
└ @ Main /Users/kcft114/.julia/packages/Flux/qXNjB/src/optimise/train.jl:105
┌ Info: Epoch 2158
└ @ Main /Users/kcft114/.julia/packages/Flux/qXNjB/src/optimise/train.jl:105
┌ Info: Epoch 2159
└ @ Main /Users/kcft114/.julia/packages/Flux/qXNjB/src/optimise/train.jl:105
┌ Info: Epoch 2160
└ @ Main /Users/kcft114/.julia/packages/Flux/qXNjB/src/optimise/train.jl:105
┌ Info: Epoch 2161
└ @ Main /Users/kcft114/.julia/packages/Flux/qXNjB/src/optimise/train.jl:105
┌ Info: Epoch 2162
└ @ Main /Users/kcft114/.julia/packages/Flux/qXNjB/src/optimise/train.jl:105
┌ Info: Epoch 2163
└ @ Main /Users/kcft114/.julia/packages/Flux/qXNjB/src/optimise/train.jl:105
┌ Info: Epoch 2164
└ @ Main /Users/kcft114/.julia/packages/Flux/qXNjB/src/optimise/train.jl:105
┌ Info: Epoch 2165
└ @ Main /Users/kcft114/.julia/packages/Flux/qXNjB/src/optimise/train.jl:105
┌ Info: Epoch 2166
└ @ Main /Users/kcft114/.julia/packages/Flux/qXNjB/src/optimise/train.jl:105
┌ Info: Epoch 2167
└ @ Main /Users/kcft114/.julia/packages/Flux/qXNjB/src/optimise/train.jl:105
┌ Info: Epoch 2168
└ @ Main /Users/kcft114/.julia/packages/Flux/qXNjB/src/optimise/train.jl:105
┌ Info: Epoch 2169
└ @ Main /Users/kcft114/.julia/packages/Flux/qXNjB/src/optimise/train.jl:105
┌ Info: Epoch 2170
└ @ Main /Users/kcft114/.julia/packages/Flux/qXNjB/src/optimise/train.jl:105
┌ Info: Epoch 2171
└ @ Main /Users/kcft114/.julia/packages/Flux/qXNjB/src/optimise/train.jl:105
┌ Info: Epoch 2172
└ @ Main /Users/kcft114/.julia/packages/Flux/qXNjB/src/optimise/train.jl:105
┌ Info: Epoch 2173
└ @ Main /Users/kcft114/.julia/packages/Flux/qXNjB/src/optimise/train.jl:105
┌ Info: Epoch 2174
└ @ Main /Users/kcft114/.julia/packages/Flux/qXNjB/src/optimise/train.jl:105
┌ Info: Epoch 2175
└ @ Main /Users/kcft114/.julia/packages/Flux/qXNjB/src/optimise/train.jl:105
┌ Info: Epoch 2176
└ @ Main /Users/kcft114/.julia/packages/Flux/qXNjB/src/optimise/train.jl:105
┌ Info: Epoch 2177
└ @ Main /Users/kcft114/.julia/packages/Flux/qXNjB/src/optimise/train.jl:105
┌ Info: Epoch 2178
└ @ Main /Users/kcft114/.julia/packages/Flux/qXNjB/src/optimise/train.jl:105
┌ Info: Epoch 2179
└ @ Main /Users/kcft114/.julia/packages/Flux/qXNjB/src/optimise/train.jl:105
┌ Info: Epoch 2180
└ @ Main /Users/kcft114/.julia/packages/Flux/qXNjB/src/optimise/train.jl:105
┌ Info: Epoch 2181
└ @ Main /Users/kcft114/.julia/packages/Flux/qXNjB/src/optimise/train.jl:105
┌ Info: Epoch 2182
└ @ Main /Users/kcft114/.julia/packages/Flux/qXNjB/src/optimise/train.jl:105
┌ Info: Epoch 2183
└ @ Main /Users/kcft114/.julia/packages/Flux/qXNjB/src/optimise/train.jl:105
┌ Info: Epoch 2184
└ @ Main /Users/kcft114/.julia/packages/Flux/qXNjB/src/optimise/train.jl:105
┌ Info: Epoch 2185
└ @ Main /Users/kcft114/.julia/packages/Flux/qXNjB/src/optimise/train.jl:105
┌ Info: Epoch 2186
└ @ Main /Users/kcft114/.julia/packages/Flux/qXNjB/src/optimise/train.jl:105
┌ Info: Epoch 2187
└ @ Main /Users/kcft114/.julia/packages/Flux/qXNjB/src/optimise/train.jl:105
┌ Info: Epoch 2188
└ @ Main /Users/kcft114/.julia/packages/Flux/qXNjB/src/optimise/train.jl:105
┌ Info: Epoch 2189
└ @ Main /Users/kcft114/.julia/packages/Flux/qXNjB/src/optimise/train.jl:105
┌ Info: Epoch 2190
└ @ Main /Users/kcft114/.julia/packages/Flux/qXNjB/src/optimise/train.jl:105
┌ Info: Epoch 2191
└ @ Main /Users/kcft114/.julia/packages/Flux/qXNjB/src/optimise/train.jl:105
┌ Info: Epoch 2192
└ @ Main /Users/kcft114/.julia/packages/Flux/qXNjB/src/optimise/train.jl:105
┌ Info: Epoch 2193
└ @ Main /Users/kcft114/.julia/packages/Flux/qXNjB/src/optimise/train.jl:105
┌ Info: Epoch 2194
└ @ Main /Users/kcft114/.julia/packages/Flux/qXNjB/src/optimise/train.jl:105
┌ Info: Epoch 2195
└ @ Main /Users/kcft114/.julia/packages/Flux/qXNjB/src/optimise/train.jl:105
┌ Info: Epoch 2196
└ @ Main /Users/kcft114/.julia/packages/Flux/qXNjB/src/optimise/train.jl:105
┌ Info: Epoch 2197
└ @ Main /Users/kcft114/.julia/packages/Flux/qXNjB/src/optimise/train.jl:105
┌ Info: Epoch 2198
└ @ Main /Users/kcft114/.julia/packages/Flux/qXNjB/src/optimise/train.jl:105
┌ Info: Epoch 2199
└ @ Main /Users/kcft114/.julia/packages/Flux/qXNjB/src/optimise/train.jl:105
┌ Info: Epoch 2200
└ @ Main /Users/kcft114/.julia/packages/Flux/qXNjB/src/optimise/train.jl:105
┌ Info: Epoch 2201
└ @ Main /Users/kcft114/.julia/packages/Flux/qXNjB/src/optimise/train.jl:105
┌ Info: Epoch 2202
└ @ Main /Users/kcft114/.julia/packages/Flux/qXNjB/src/optimise/train.jl:105
┌ Info: Epoch 2203
└ @ Main /Users/kcft114/.julia/packages/Flux/qXNjB/src/optimise/train.jl:105
┌ Info: Epoch 2204
└ @ Main /Users/kcft114/.julia/packages/Flux/qXNjB/src/optimise/train.jl:105
┌ Info: Epoch 2205
└ @ Main /Users/kcft114/.julia/packages/Flux/qXNjB/src/optimise/train.jl:105
┌ Info: Epoch 2206
└ @ Main /Users/kcft114/.julia/packages/Flux/qXNjB/src/optimise/train.jl:105
┌ Info: Epoch 2207
└ @ Main /Users/kcft114/.julia/packages/Flux/qXNjB/src/optimise/train.jl:105
┌ Info: Epoch 2208
└ @ Main /Users/kcft114/.julia/packages/Flux/qXNjB/src/optimise/train.jl:105
┌ Info: Epoch 2209
└ @ Main /Users/kcft114/.julia/packages/Flux/qXNjB/src/optimise/train.jl:105
┌ Info: Epoch 2210
└ @ Main /Users/kcft114/.julia/packages/Flux/qXNjB/src/optimise/train.jl:105
┌ Info: Epoch 2211
└ @ Main /Users/kcft114/.julia/packages/Flux/qXNjB/src/optimise/train.jl:105
┌ Info: Epoch 2212
└ @ Main /Users/kcft114/.julia/packages/Flux/qXNjB/src/optimise/train.jl:105
┌ Info: Epoch 2213
└ @ Main /Users/kcft114/.julia/packages/Flux/qXNjB/src/optimise/train.jl:105
┌ Info: Epoch 2214
└ @ Main /Users/kcft114/.julia/packages/Flux/qXNjB/src/optimise/train.jl:105
┌ Info: Epoch 2215
└ @ Main /Users/kcft114/.julia/packages/Flux/qXNjB/src/optimise/train.jl:105
┌ Info: Epoch 2216
└ @ Main /Users/kcft114/.julia/packages/Flux/qXNjB/src/optimise/train.jl:105
┌ Info: Epoch 2217
└ @ Main /Users/kcft114/.julia/packages/Flux/qXNjB/src/optimise/train.jl:105
┌ Info: Epoch 2218
└ @ Main /Users/kcft114/.julia/packages/Flux/qXNjB/src/optimise/train.jl:105
┌ Info: Epoch 2219
└ @ Main /Users/kcft114/.julia/packages/Flux/qXNjB/src/optimise/train.jl:105
┌ Info: Epoch 2220
└ @ Main /Users/kcft114/.julia/packages/Flux/qXNjB/src/optimise/train.jl:105
┌ Info: Epoch 2221
└ @ Main /Users/kcft114/.julia/packages/Flux/qXNjB/src/optimise/train.jl:105
┌ Info: Epoch 2222
└ @ Main /Users/kcft114/.julia/packages/Flux/qXNjB/src/optimise/train.jl:105
┌ Info: Epoch 2223
└ @ Main /Users/kcft114/.julia/packages/Flux/qXNjB/src/optimise/train.jl:105
┌ Info: Epoch 2224
└ @ Main /Users/kcft114/.julia/packages/Flux/qXNjB/src/optimise/train.jl:105
┌ Info: Epoch 2225
└ @ Main /Users/kcft114/.julia/packages/Flux/qXNjB/src/optimise/train.jl:105
┌ Info: Epoch 2226
└ @ Main /Users/kcft114/.julia/packages/Flux/qXNjB/src/optimise/train.jl:105
┌ Info: Epoch 2227
└ @ Main /Users/kcft114/.julia/packages/Flux/qXNjB/src/optimise/train.jl:105
┌ Info: Epoch 2228
└ @ Main /Users/kcft114/.julia/packages/Flux/qXNjB/src/optimise/train.jl:105
┌ Info: Epoch 2229
└ @ Main /Users/kcft114/.julia/packages/Flux/qXNjB/src/optimise/train.jl:105
┌ Info: Epoch 2230
└ @ Main /Users/kcft114/.julia/packages/Flux/qXNjB/src/optimise/train.jl:105
┌ Info: Epoch 2231
└ @ Main /Users/kcft114/.julia/packages/Flux/qXNjB/src/optimise/train.jl:105
┌ Info: Epoch 2232
└ @ Main /Users/kcft114/.julia/packages/Flux/qXNjB/src/optimise/train.jl:105
┌ Info: Epoch 2233
└ @ Main /Users/kcft114/.julia/packages/Flux/qXNjB/src/optimise/train.jl:105
┌ Info: Epoch 2234
└ @ Main /Users/kcft114/.julia/packages/Flux/qXNjB/src/optimise/train.jl:105
┌ Info: Epoch 2235
└ @ Main /Users/kcft114/.julia/packages/Flux/qXNjB/src/optimise/train.jl:105
┌ Info: Epoch 2236
└ @ Main /Users/kcft114/.julia/packages/Flux/qXNjB/src/optimise/train.jl:105
┌ Info: Epoch 2237
└ @ Main /Users/kcft114/.julia/packages/Flux/qXNjB/src/optimise/train.jl:105
┌ Info: Epoch 2238
└ @ Main /Users/kcft114/.julia/packages/Flux/qXNjB/src/optimise/train.jl:105
┌ Info: Epoch 2239
└ @ Main /Users/kcft114/.julia/packages/Flux/qXNjB/src/optimise/train.jl:105
┌ Info: Epoch 2240
└ @ Main /Users/kcft114/.julia/packages/Flux/qXNjB/src/optimise/train.jl:105
┌ Info: Epoch 2241
└ @ Main /Users/kcft114/.julia/packages/Flux/qXNjB/src/optimise/train.jl:105
┌ Info: Epoch 2242
└ @ Main /Users/kcft114/.julia/packages/Flux/qXNjB/src/optimise/train.jl:105
┌ Info: Epoch 2243
└ @ Main /Users/kcft114/.julia/packages/Flux/qXNjB/src/optimise/train.jl:105
┌ Info: Epoch 2244
└ @ Main /Users/kcft114/.julia/packages/Flux/qXNjB/src/optimise/train.jl:105
┌ Info: Epoch 2245
└ @ Main /Users/kcft114/.julia/packages/Flux/qXNjB/src/optimise/train.jl:105
┌ Info: Epoch 2246
└ @ Main /Users/kcft114/.julia/packages/Flux/qXNjB/src/optimise/train.jl:105
┌ Info: Epoch 2247
└ @ Main /Users/kcft114/.julia/packages/Flux/qXNjB/src/optimise/train.jl:105
┌ Info: Epoch 2248
└ @ Main /Users/kcft114/.julia/packages/Flux/qXNjB/src/optimise/train.jl:105
┌ Info: Epoch 2249
└ @ Main /Users/kcft114/.julia/packages/Flux/qXNjB/src/optimise/train.jl:105
┌ Info: Epoch 2250
└ @ Main /Users/kcft114/.julia/packages/Flux/qXNjB/src/optimise/train.jl:105
┌ Info: Epoch 2251
└ @ Main /Users/kcft114/.julia/packages/Flux/qXNjB/src/optimise/train.jl:105
┌ Info: Epoch 2252
└ @ Main /Users/kcft114/.julia/packages/Flux/qXNjB/src/optimise/train.jl:105
┌ Info: Epoch 2253
└ @ Main /Users/kcft114/.julia/packages/Flux/qXNjB/src/optimise/train.jl:105
┌ Info: Epoch 2254
└ @ Main /Users/kcft114/.julia/packages/Flux/qXNjB/src/optimise/train.jl:105
┌ Info: Epoch 2255
└ @ Main /Users/kcft114/.julia/packages/Flux/qXNjB/src/optimise/train.jl:105
┌ Info: Epoch 2256
└ @ Main /Users/kcft114/.julia/packages/Flux/qXNjB/src/optimise/train.jl:105
┌ Info: Epoch 2257
└ @ Main /Users/kcft114/.julia/packages/Flux/qXNjB/src/optimise/train.jl:105
┌ Info: Epoch 2258
└ @ Main /Users/kcft114/.julia/packages/Flux/qXNjB/src/optimise/train.jl:105
┌ Info: Epoch 2259
└ @ Main /Users/kcft114/.julia/packages/Flux/qXNjB/src/optimise/train.jl:105
┌ Info: Epoch 2260
└ @ Main /Users/kcft114/.julia/packages/Flux/qXNjB/src/optimise/train.jl:105
┌ Info: Epoch 2261
└ @ Main /Users/kcft114/.julia/packages/Flux/qXNjB/src/optimise/train.jl:105
┌ Info: Epoch 2262
└ @ Main /Users/kcft114/.julia/packages/Flux/qXNjB/src/optimise/train.jl:105
┌ Info: Epoch 2263
└ @ Main /Users/kcft114/.julia/packages/Flux/qXNjB/src/optimise/train.jl:105
┌ Info: Epoch 2264
└ @ Main /Users/kcft114/.julia/packages/Flux/qXNjB/src/optimise/train.jl:105
┌ Info: Epoch 2265
└ @ Main /Users/kcft114/.julia/packages/Flux/qXNjB/src/optimise/train.jl:105
┌ Info: Epoch 2266
└ @ Main /Users/kcft114/.julia/packages/Flux/qXNjB/src/optimise/train.jl:105
┌ Info: Epoch 2267
└ @ Main /Users/kcft114/.julia/packages/Flux/qXNjB/src/optimise/train.jl:105
┌ Info: Epoch 2268
└ @ Main /Users/kcft114/.julia/packages/Flux/qXNjB/src/optimise/train.jl:105
┌ Info: Epoch 2269
└ @ Main /Users/kcft114/.julia/packages/Flux/qXNjB/src/optimise/train.jl:105
┌ Info: Epoch 2270
└ @ Main /Users/kcft114/.julia/packages/Flux/qXNjB/src/optimise/train.jl:105
┌ Info: Epoch 2271
└ @ Main /Users/kcft114/.julia/packages/Flux/qXNjB/src/optimise/train.jl:105
┌ Info: Epoch 2272
└ @ Main /Users/kcft114/.julia/packages/Flux/qXNjB/src/optimise/train.jl:105
┌ Info: Epoch 2273
└ @ Main /Users/kcft114/.julia/packages/Flux/qXNjB/src/optimise/train.jl:105
┌ Info: Epoch 2274
└ @ Main /Users/kcft114/.julia/packages/Flux/qXNjB/src/optimise/train.jl:105
┌ Info: Epoch 2275
└ @ Main /Users/kcft114/.julia/packages/Flux/qXNjB/src/optimise/train.jl:105
┌ Info: Epoch 2276
└ @ Main /Users/kcft114/.julia/packages/Flux/qXNjB/src/optimise/train.jl:105
┌ Info: Epoch 2277
└ @ Main /Users/kcft114/.julia/packages/Flux/qXNjB/src/optimise/train.jl:105
┌ Info: Epoch 2278
└ @ Main /Users/kcft114/.julia/packages/Flux/qXNjB/src/optimise/train.jl:105
┌ Info: Epoch 2279
└ @ Main /Users/kcft114/.julia/packages/Flux/qXNjB/src/optimise/train.jl:105
┌ Info: Epoch 2280
└ @ Main /Users/kcft114/.julia/packages/Flux/qXNjB/src/optimise/train.jl:105
┌ Info: Epoch 2281
└ @ Main /Users/kcft114/.julia/packages/Flux/qXNjB/src/optimise/train.jl:105
┌ Info: Epoch 2282
└ @ Main /Users/kcft114/.julia/packages/Flux/qXNjB/src/optimise/train.jl:105
┌ Info: Epoch 2283
└ @ Main /Users/kcft114/.julia/packages/Flux/qXNjB/src/optimise/train.jl:105
┌ Info: Epoch 2284
└ @ Main /Users/kcft114/.julia/packages/Flux/qXNjB/src/optimise/train.jl:105
┌ Info: Epoch 2285
└ @ Main /Users/kcft114/.julia/packages/Flux/qXNjB/src/optimise/train.jl:105
┌ Info: Epoch 2286
└ @ Main /Users/kcft114/.julia/packages/Flux/qXNjB/src/optimise/train.jl:105
┌ Info: Epoch 2287
└ @ Main /Users/kcft114/.julia/packages/Flux/qXNjB/src/optimise/train.jl:105
┌ Info: Epoch 2288
└ @ Main /Users/kcft114/.julia/packages/Flux/qXNjB/src/optimise/train.jl:105
┌ Info: Epoch 2289
└ @ Main /Users/kcft114/.julia/packages/Flux/qXNjB/src/optimise/train.jl:105
┌ Info: Epoch 2290
└ @ Main /Users/kcft114/.julia/packages/Flux/qXNjB/src/optimise/train.jl:105
┌ Info: Epoch 2291
└ @ Main /Users/kcft114/.julia/packages/Flux/qXNjB/src/optimise/train.jl:105
┌ Info: Epoch 2292
└ @ Main /Users/kcft114/.julia/packages/Flux/qXNjB/src/optimise/train.jl:105
┌ Info: Epoch 2293
└ @ Main /Users/kcft114/.julia/packages/Flux/qXNjB/src/optimise/train.jl:105
┌ Info: Epoch 2294
└ @ Main /Users/kcft114/.julia/packages/Flux/qXNjB/src/optimise/train.jl:105
┌ Info: Epoch 2295
└ @ Main /Users/kcft114/.julia/packages/Flux/qXNjB/src/optimise/train.jl:105
┌ Info: Epoch 2296
└ @ Main /Users/kcft114/.julia/packages/Flux/qXNjB/src/optimise/train.jl:105
┌ Info: Epoch 2297
└ @ Main /Users/kcft114/.julia/packages/Flux/qXNjB/src/optimise/train.jl:105
┌ Info: Epoch 2298
└ @ Main /Users/kcft114/.julia/packages/Flux/qXNjB/src/optimise/train.jl:105
┌ Info: Epoch 2299
└ @ Main /Users/kcft114/.julia/packages/Flux/qXNjB/src/optimise/train.jl:105
┌ Info: Epoch 2300
└ @ Main /Users/kcft114/.julia/packages/Flux/qXNjB/src/optimise/train.jl:105
┌ Info: Epoch 2301
└ @ Main /Users/kcft114/.julia/packages/Flux/qXNjB/src/optimise/train.jl:105
```

```
loss(X, Y) = 0.2585141f0 (tracked)
loss(X, Y) = 0.25838467f0 (tracked)
loss(X, Y) = 0.25825378f0 (tracked)
loss(X, Y) = 0.25812638f0 (tracked)
loss(X, Y) = 0.25799426f0 (tracked)
loss(X, Y) = 0.25786835f0 (tracked)
loss(X, Y) = 0.2577317f0 (tracked)
loss(X, Y) = 0.2576043f0 (tracked)
loss(X, Y) = 0.25748372f0 (tracked)
loss(X, Y) = 0.25734434f0 (tracked)
loss(X, Y) = 0.2572202f0 (tracked)
loss(X, Y) = 0.25708663f0 (tracked)
loss(X, Y) = 0.25696078f0 (tracked)
loss(X, Y) = 0.25683206f0 (tracked)
loss(X, Y) = 0.25670013f0 (tracked)
loss(X, Y) = 0.25656527f0 (tracked)
loss(X, Y) = 0.2564473f0 (tracked)
loss(X, Y) = 0.2563282f0 (tracked)
loss(X, Y) = 0.25619924f0 (tracked)
loss(X, Y) = 0.25605774f0 (tracked)
loss(X, Y) = 0.25593227f0 (tracked)
loss(X, Y) = 0.25581205f0 (tracked)
loss(X, Y) = 0.25568488f0 (tracked)
loss(X, Y) = 0.25555107f0 (tracked)
loss(X, Y) = 0.25542235f0 (tracked)
loss(X, Y) = 0.25529015f0 (tracked)
loss(X, Y) = 0.25515518f0 (tracked)
loss(X, Y) = 0.25503024f0 (tracked)
loss(X, Y) = 0.25491473f0 (tracked)
loss(X, Y) = 0.25478727f0 (tracked)
loss(X, Y) = 0.25464824f0 (tracked)
loss(X, Y) = 0.25452548f0 (tracked)
loss(X, Y) = 0.25440866f0 (tracked)
loss(X, Y) = 0.25428417f0 (tracked)
loss(X, Y) = 0.2541453f0 (tracked)
loss(X, Y) = 0.25401247f0 (tracked)
loss(X, Y) = 0.25389722f0 (tracked)
loss(X, Y) = 0.2537714f0 (tracked)
loss(X, Y) = 0.25363967f0 (tracked)
loss(X, Y) = 0.2535114f0 (tracked)
loss(X, Y) = 0.25338864f0 (tracked)
loss(X, Y) = 0.25326192f0 (tracked)
loss(X, Y) = 0.25312385f0 (tracked)
loss(X, Y) = 0.25299433f0 (tracked)
loss(X, Y) = 0.25287607f0 (tracked)
loss(X, Y) = 0.2527396f0 (tracked)
loss(X, Y) = 0.25260413f0 (tracked)
loss(X, Y) = 0.25247648f0 (tracked)
loss(X, Y) = 0.2523667f0 (tracked)
loss(X, Y) = 0.25222784f0 (tracked)
loss(X, Y) = 0.25208187f0 (tracked)
loss(X, Y) = 0.25196287f0 (tracked)
loss(X, Y) = 0.2518306f0 (tracked)
loss(X, Y) = 0.25170046f0 (tracked)
loss(X, Y) = 0.25157434f0 (tracked)
loss(X, Y) = 0.25145024f0 (tracked)
loss(X, Y) = 0.25132126f0 (tracked)
loss(X, Y) = 0.25118503f0 (tracked)
loss(X, Y) = 0.251067f0 (tracked)
loss(X, Y) = 0.25095153f0 (tracked)
loss(X, Y) = 0.25081486f0 (tracked)
loss(X, Y) = 0.25069007f0 (tracked)
loss(X, Y) = 0.2505742f0 (tracked)
loss(X, Y) = 0.25044838f0 (tracked)
loss(X, Y) = 0.2503292f0 (tracked)
loss(X, Y) = 0.2501975f0 (tracked)
loss(X, Y) = 0.25007355f0 (tracked)
loss(X, Y) = 0.24995108f0 (tracked)
loss(X, Y) = 0.24981801f0 (tracked)
loss(X, Y) = 0.24969329f0 (tracked)
loss(X, Y) = 0.2495708f0 (tracked)
loss(X, Y) = 0.24944907f0 (tracked)
loss(X, Y) = 0.24931931f0 (tracked)
loss(X, Y) = 0.24919745f0 (tracked)
loss(X, Y) = 0.24907504f0 (tracked)
loss(X, Y) = 0.24895243f0 (tracked)
loss(X, Y) = 0.24883431f0 (tracked)
loss(X, Y) = 0.24869794f0 (tracked)
loss(X, Y) = 0.24857768f0 (tracked)
loss(X, Y) = 0.24845873f0 (tracked)
loss(X, Y) = 0.24833253f0 (tracked)
loss(X, Y) = 0.24821232f0 (tracked)
loss(X, Y) = 0.24808492f0 (tracked)
loss(X, Y) = 0.24796684f0 (tracked)
loss(X, Y) = 0.24785164f0 (tracked)
loss(X, Y) = 0.24772987f0 (tracked)
loss(X, Y) = 0.24760766f0 (tracked)
loss(X, Y) = 0.2474806f0 (tracked)
loss(X, Y) = 0.24735916f0 (tracked)
loss(X, Y) = 0.24724628f0 (tracked)
loss(X, Y) = 0.24710666f0 (tracked)
loss(X, Y) = 0.24698348f0 (tracked)
loss(X, Y) = 0.24686685f0 (tracked)
loss(X, Y) = 0.24674135f0 (tracked)
loss(X, Y) = 0.24661116f0 (tracked)
loss(X, Y) = 0.24649675f0 (tracked)
loss(X, Y) = 0.24636103f0 (tracked)
loss(X, Y) = 0.24623059f0 (tracked)
loss(X, Y) = 0.246117f0 (tracked)
loss(X, Y) = 0.24599566f0 (tracked)
loss(X, Y) = 0.24587597f0 (tracked)
loss(X, Y) = 0.24575561f0 (tracked)
loss(X, Y) = 0.2456283f0 (tracked)
loss(X, Y) = 0.24550737f0 (tracked)
loss(X, Y) = 0.24538621f0 (tracked)
loss(X, Y) = 0.24526557f0 (tracked)
loss(X, Y) = 0.24513777f0 (tracked)
loss(X, Y) = 0.24500723f0 (tracked)
loss(X, Y) = 0.24488209f0 (tracked)
loss(X, Y) = 0.24477077f0 (tracked)
loss(X, Y) = 0.24464922f0 (tracked)
loss(X, Y) = 0.2445206f0 (tracked)
loss(X, Y) = 0.24438998f0 (tracked)
loss(X, Y) = 0.24426632f0 (tracked)
loss(X, Y) = 0.24414763f0 (tracked)
loss(X, Y) = 0.24401586f0 (tracked)
loss(X, Y) = 0.24388768f0 (tracked)
loss(X, Y) = 0.24378404f0 (tracked)
loss(X, Y) = 0.24365886f0 (tracked)
loss(X, Y) = 0.24352528f0 (tracked)
loss(X, Y) = 0.24339828f0 (tracked)
loss(X, Y) = 0.24327907f0 (tracked)
loss(X, Y) = 0.24315384f0 (tracked)
loss(X, Y) = 0.243024f0 (tracked)
loss(X, Y) = 0.24290621f0 (tracked)
loss(X, Y) = 0.24278474f0 (tracked)
loss(X, Y) = 0.24266575f0 (tracked)
loss(X, Y) = 0.24253455f0 (tracked)
loss(X, Y) = 0.24240595f0 (tracked)
loss(X, Y) = 0.24228017f0 (tracked)
loss(X, Y) = 0.2421481f0 (tracked)
loss(X, Y) = 0.24202074f0 (tracked)
loss(X, Y) = 0.24189898f0 (tracked)
loss(X, Y) = 0.24178101f0 (tracked)
loss(X, Y) = 0.24165744f0 (tracked)
loss(X, Y) = 0.24153827f0 (tracked)
loss(X, Y) = 0.24142604f0 (tracked)
loss(X, Y) = 0.24130632f0 (tracked)
loss(X, Y) = 0.2411826f0 (tracked)
loss(X, Y) = 0.24105601f0 (tracked)
loss(X, Y) = 0.24093306f0 (tracked)
loss(X, Y) = 0.24081005f0 (tracked)
loss(X, Y) = 0.24068312f0 (tracked)
loss(X, Y) = 0.24057022f0 (tracked)
loss(X, Y) = 0.24044584f0 (tracked)
loss(X, Y) = 0.24031396f0 (tracked)
loss(X, Y) = 0.2401983f0 (tracked)
loss(X, Y) = 0.24008428f0 (tracked)
loss(X, Y) = 0.23996376f0 (tracked)
loss(X, Y) = 0.23983756f0 (tracked)
loss(X, Y) = 0.2397178f0 (tracked)
loss(X, Y) = 0.23959693f0 (tracked)
loss(X, Y) = 0.23948127f0 (tracked)
loss(X, Y) = 0.23935801f0 (tracked)
loss(X, Y) = 0.23922817f0 (tracked)
loss(X, Y) = 0.23911142f0 (tracked)
loss(X, Y) = 0.23899584f0 (tracked)
loss(X, Y) = 0.23888189f0 (tracked)
loss(X, Y) = 0.238761f0 (tracked)
loss(X, Y) = 0.23863925f0 (tracked)
loss(X, Y) = 0.23853543f0 (tracked)
loss(X, Y) = 0.23841617f0 (tracked)
loss(X, Y) = 0.23829521f0 (tracked)
loss(X, Y) = 0.23817305f0 (tracked)
loss(X, Y) = 0.2380704f0 (tracked)
loss(X, Y) = 0.23795526f0 (tracked)
loss(X, Y) = 0.23783542f0 (tracked)
loss(X, Y) = 0.23771724f0 (tracked)
loss(X, Y) = 0.23759346f0 (tracked)
loss(X, Y) = 0.23748657f0 (tracked)
```

```
┌ Info: Epoch 2302
└ @ Main /Users/kcft114/.julia/packages/Flux/qXNjB/src/optimise/train.jl:105
┌ Info: Epoch 2303
└ @ Main /Users/kcft114/.julia/packages/Flux/qXNjB/src/optimise/train.jl:105
┌ Info: Epoch 2304
└ @ Main /Users/kcft114/.julia/packages/Flux/qXNjB/src/optimise/train.jl:105
┌ Info: Epoch 2305
└ @ Main /Users/kcft114/.julia/packages/Flux/qXNjB/src/optimise/train.jl:105
┌ Info: Epoch 2306
└ @ Main /Users/kcft114/.julia/packages/Flux/qXNjB/src/optimise/train.jl:105
┌ Info: Epoch 2307
└ @ Main /Users/kcft114/.julia/packages/Flux/qXNjB/src/optimise/train.jl:105
┌ Info: Epoch 2308
└ @ Main /Users/kcft114/.julia/packages/Flux/qXNjB/src/optimise/train.jl:105
┌ Info: Epoch 2309
└ @ Main /Users/kcft114/.julia/packages/Flux/qXNjB/src/optimise/train.jl:105
┌ Info: Epoch 2310
└ @ Main /Users/kcft114/.julia/packages/Flux/qXNjB/src/optimise/train.jl:105
┌ Info: Epoch 2311
└ @ Main /Users/kcft114/.julia/packages/Flux/qXNjB/src/optimise/train.jl:105
┌ Info: Epoch 2312
└ @ Main /Users/kcft114/.julia/packages/Flux/qXNjB/src/optimise/train.jl:105
┌ Info: Epoch 2313
└ @ Main /Users/kcft114/.julia/packages/Flux/qXNjB/src/optimise/train.jl:105
┌ Info: Epoch 2314
└ @ Main /Users/kcft114/.julia/packages/Flux/qXNjB/src/optimise/train.jl:105
┌ Info: Epoch 2315
└ @ Main /Users/kcft114/.julia/packages/Flux/qXNjB/src/optimise/train.jl:105
┌ Info: Epoch 2316
└ @ Main /Users/kcft114/.julia/packages/Flux/qXNjB/src/optimise/train.jl:105
┌ Info: Epoch 2317
└ @ Main /Users/kcft114/.julia/packages/Flux/qXNjB/src/optimise/train.jl:105
┌ Info: Epoch 2318
└ @ Main /Users/kcft114/.julia/packages/Flux/qXNjB/src/optimise/train.jl:105
┌ Info: Epoch 2319
└ @ Main /Users/kcft114/.julia/packages/Flux/qXNjB/src/optimise/train.jl:105
┌ Info: Epoch 2320
└ @ Main /Users/kcft114/.julia/packages/Flux/qXNjB/src/optimise/train.jl:105
┌ Info: Epoch 2321
└ @ Main /Users/kcft114/.julia/packages/Flux/qXNjB/src/optimise/train.jl:105
┌ Info: Epoch 2322
└ @ Main /Users/kcft114/.julia/packages/Flux/qXNjB/src/optimise/train.jl:105
┌ Info: Epoch 2323
└ @ Main /Users/kcft114/.julia/packages/Flux/qXNjB/src/optimise/train.jl:105
┌ Info: Epoch 2324
└ @ Main /Users/kcft114/.julia/packages/Flux/qXNjB/src/optimise/train.jl:105
┌ Info: Epoch 2325
└ @ Main /Users/kcft114/.julia/packages/Flux/qXNjB/src/optimise/train.jl:105
┌ Info: Epoch 2326
└ @ Main /Users/kcft114/.julia/packages/Flux/qXNjB/src/optimise/train.jl:105
┌ Info: Epoch 2327
└ @ Main /Users/kcft114/.julia/packages/Flux/qXNjB/src/optimise/train.jl:105
┌ Info: Epoch 2328
└ @ Main /Users/kcft114/.julia/packages/Flux/qXNjB/src/optimise/train.jl:105
┌ Info: Epoch 2329
└ @ Main /Users/kcft114/.julia/packages/Flux/qXNjB/src/optimise/train.jl:105
┌ Info: Epoch 2330
└ @ Main /Users/kcft114/.julia/packages/Flux/qXNjB/src/optimise/train.jl:105
┌ Info: Epoch 2331
└ @ Main /Users/kcft114/.julia/packages/Flux/qXNjB/src/optimise/train.jl:105
┌ Info: Epoch 2332
└ @ Main /Users/kcft114/.julia/packages/Flux/qXNjB/src/optimise/train.jl:105
┌ Info: Epoch 2333
└ @ Main /Users/kcft114/.julia/packages/Flux/qXNjB/src/optimise/train.jl:105
┌ Info: Epoch 2334
└ @ Main /Users/kcft114/.julia/packages/Flux/qXNjB/src/optimise/train.jl:105
┌ Info: Epoch 2335
└ @ Main /Users/kcft114/.julia/packages/Flux/qXNjB/src/optimise/train.jl:105
┌ Info: Epoch 2336
└ @ Main /Users/kcft114/.julia/packages/Flux/qXNjB/src/optimise/train.jl:105
┌ Info: Epoch 2337
└ @ Main /Users/kcft114/.julia/packages/Flux/qXNjB/src/optimise/train.jl:105
┌ Info: Epoch 2338
└ @ Main /Users/kcft114/.julia/packages/Flux/qXNjB/src/optimise/train.jl:105
┌ Info: Epoch 2339
└ @ Main /Users/kcft114/.julia/packages/Flux/qXNjB/src/optimise/train.jl:105
┌ Info: Epoch 2340
└ @ Main /Users/kcft114/.julia/packages/Flux/qXNjB/src/optimise/train.jl:105
┌ Info: Epoch 2341
└ @ Main /Users/kcft114/.julia/packages/Flux/qXNjB/src/optimise/train.jl:105
┌ Info: Epoch 2342
└ @ Main /Users/kcft114/.julia/packages/Flux/qXNjB/src/optimise/train.jl:105
┌ Info: Epoch 2343
└ @ Main /Users/kcft114/.julia/packages/Flux/qXNjB/src/optimise/train.jl:105
┌ Info: Epoch 2344
└ @ Main /Users/kcft114/.julia/packages/Flux/qXNjB/src/optimise/train.jl:105
┌ Info: Epoch 2345
└ @ Main /Users/kcft114/.julia/packages/Flux/qXNjB/src/optimise/train.jl:105
┌ Info: Epoch 2346
└ @ Main /Users/kcft114/.julia/packages/Flux/qXNjB/src/optimise/train.jl:105
┌ Info: Epoch 2347
└ @ Main /Users/kcft114/.julia/packages/Flux/qXNjB/src/optimise/train.jl:105
┌ Info: Epoch 2348
└ @ Main /Users/kcft114/.julia/packages/Flux/qXNjB/src/optimise/train.jl:105
┌ Info: Epoch 2349
└ @ Main /Users/kcft114/.julia/packages/Flux/qXNjB/src/optimise/train.jl:105
┌ Info: Epoch 2350
└ @ Main /Users/kcft114/.julia/packages/Flux/qXNjB/src/optimise/train.jl:105
┌ Info: Epoch 2351
└ @ Main /Users/kcft114/.julia/packages/Flux/qXNjB/src/optimise/train.jl:105
┌ Info: Epoch 2352
└ @ Main /Users/kcft114/.julia/packages/Flux/qXNjB/src/optimise/train.jl:105
┌ Info: Epoch 2353
└ @ Main /Users/kcft114/.julia/packages/Flux/qXNjB/src/optimise/train.jl:105
┌ Info: Epoch 2354
└ @ Main /Users/kcft114/.julia/packages/Flux/qXNjB/src/optimise/train.jl:105
┌ Info: Epoch 2355
└ @ Main /Users/kcft114/.julia/packages/Flux/qXNjB/src/optimise/train.jl:105
┌ Info: Epoch 2356
└ @ Main /Users/kcft114/.julia/packages/Flux/qXNjB/src/optimise/train.jl:105
┌ Info: Epoch 2357
└ @ Main /Users/kcft114/.julia/packages/Flux/qXNjB/src/optimise/train.jl:105
┌ Info: Epoch 2358
└ @ Main /Users/kcft114/.julia/packages/Flux/qXNjB/src/optimise/train.jl:105
┌ Info: Epoch 2359
└ @ Main /Users/kcft114/.julia/packages/Flux/qXNjB/src/optimise/train.jl:105
┌ Info: Epoch 2360
└ @ Main /Users/kcft114/.julia/packages/Flux/qXNjB/src/optimise/train.jl:105
┌ Info: Epoch 2361
└ @ Main /Users/kcft114/.julia/packages/Flux/qXNjB/src/optimise/train.jl:105
┌ Info: Epoch 2362
└ @ Main /Users/kcft114/.julia/packages/Flux/qXNjB/src/optimise/train.jl:105
┌ Info: Epoch 2363
└ @ Main /Users/kcft114/.julia/packages/Flux/qXNjB/src/optimise/train.jl:105
┌ Info: Epoch 2364
└ @ Main /Users/kcft114/.julia/packages/Flux/qXNjB/src/optimise/train.jl:105
┌ Info: Epoch 2365
└ @ Main /Users/kcft114/.julia/packages/Flux/qXNjB/src/optimise/train.jl:105
┌ Info: Epoch 2366
└ @ Main /Users/kcft114/.julia/packages/Flux/qXNjB/src/optimise/train.jl:105
┌ Info: Epoch 2367
└ @ Main /Users/kcft114/.julia/packages/Flux/qXNjB/src/optimise/train.jl:105
┌ Info: Epoch 2368
└ @ Main /Users/kcft114/.julia/packages/Flux/qXNjB/src/optimise/train.jl:105
┌ Info: Epoch 2369
└ @ Main /Users/kcft114/.julia/packages/Flux/qXNjB/src/optimise/train.jl:105
┌ Info: Epoch 2370
└ @ Main /Users/kcft114/.julia/packages/Flux/qXNjB/src/optimise/train.jl:105
┌ Info: Epoch 2371
└ @ Main /Users/kcft114/.julia/packages/Flux/qXNjB/src/optimise/train.jl:105
┌ Info: Epoch 2372
└ @ Main /Users/kcft114/.julia/packages/Flux/qXNjB/src/optimise/train.jl:105
┌ Info: Epoch 2373
└ @ Main /Users/kcft114/.julia/packages/Flux/qXNjB/src/optimise/train.jl:105
┌ Info: Epoch 2374
└ @ Main /Users/kcft114/.julia/packages/Flux/qXNjB/src/optimise/train.jl:105
┌ Info: Epoch 2375
└ @ Main /Users/kcft114/.julia/packages/Flux/qXNjB/src/optimise/train.jl:105
┌ Info: Epoch 2376
└ @ Main /Users/kcft114/.julia/packages/Flux/qXNjB/src/optimise/train.jl:105
┌ Info: Epoch 2377
└ @ Main /Users/kcft114/.julia/packages/Flux/qXNjB/src/optimise/train.jl:105
┌ Info: Epoch 2378
└ @ Main /Users/kcft114/.julia/packages/Flux/qXNjB/src/optimise/train.jl:105
┌ Info: Epoch 2379
└ @ Main /Users/kcft114/.julia/packages/Flux/qXNjB/src/optimise/train.jl:105
┌ Info: Epoch 2380
└ @ Main /Users/kcft114/.julia/packages/Flux/qXNjB/src/optimise/train.jl:105
┌ Info: Epoch 2381
└ @ Main /Users/kcft114/.julia/packages/Flux/qXNjB/src/optimise/train.jl:105
┌ Info: Epoch 2382
└ @ Main /Users/kcft114/.julia/packages/Flux/qXNjB/src/optimise/train.jl:105
┌ Info: Epoch 2383
└ @ Main /Users/kcft114/.julia/packages/Flux/qXNjB/src/optimise/train.jl:105
┌ Info: Epoch 2384
└ @ Main /Users/kcft114/.julia/packages/Flux/qXNjB/src/optimise/train.jl:105
┌ Info: Epoch 2385
└ @ Main /Users/kcft114/.julia/packages/Flux/qXNjB/src/optimise/train.jl:105
┌ Info: Epoch 2386
└ @ Main /Users/kcft114/.julia/packages/Flux/qXNjB/src/optimise/train.jl:105
┌ Info: Epoch 2387
└ @ Main /Users/kcft114/.julia/packages/Flux/qXNjB/src/optimise/train.jl:105
┌ Info: Epoch 2388
└ @ Main /Users/kcft114/.julia/packages/Flux/qXNjB/src/optimise/train.jl:105
┌ Info: Epoch 2389
└ @ Main /Users/kcft114/.julia/packages/Flux/qXNjB/src/optimise/train.jl:105
┌ Info: Epoch 2390
└ @ Main /Users/kcft114/.julia/packages/Flux/qXNjB/src/optimise/train.jl:105
┌ Info: Epoch 2391
└ @ Main /Users/kcft114/.julia/packages/Flux/qXNjB/src/optimise/train.jl:105
┌ Info: Epoch 2392
└ @ Main /Users/kcft114/.julia/packages/Flux/qXNjB/src/optimise/train.jl:105
┌ Info: Epoch 2393
└ @ Main /Users/kcft114/.julia/packages/Flux/qXNjB/src/optimise/train.jl:105
┌ Info: Epoch 2394
└ @ Main /Users/kcft114/.julia/packages/Flux/qXNjB/src/optimise/train.jl:105
┌ Info: Epoch 2395
└ @ Main /Users/kcft114/.julia/packages/Flux/qXNjB/src/optimise/train.jl:105
┌ Info: Epoch 2396
└ @ Main /Users/kcft114/.julia/packages/Flux/qXNjB/src/optimise/train.jl:105
┌ Info: Epoch 2397
└ @ Main /Users/kcft114/.julia/packages/Flux/qXNjB/src/optimise/train.jl:105
┌ Info: Epoch 2398
└ @ Main /Users/kcft114/.julia/packages/Flux/qXNjB/src/optimise/train.jl:105
┌ Info: Epoch 2399
└ @ Main /Users/kcft114/.julia/packages/Flux/qXNjB/src/optimise/train.jl:105
┌ Info: Epoch 2400
└ @ Main /Users/kcft114/.julia/packages/Flux/qXNjB/src/optimise/train.jl:105
┌ Info: Epoch 2401
└ @ Main /Users/kcft114/.julia/packages/Flux/qXNjB/src/optimise/train.jl:105
┌ Info: Epoch 2402
└ @ Main /Users/kcft114/.julia/packages/Flux/qXNjB/src/optimise/train.jl:105
┌ Info: Epoch 2403
└ @ Main /Users/kcft114/.julia/packages/Flux/qXNjB/src/optimise/train.jl:105
┌ Info: Epoch 2404
└ @ Main /Users/kcft114/.julia/packages/Flux/qXNjB/src/optimise/train.jl:105
┌ Info: Epoch 2405
└ @ Main /Users/kcft114/.julia/packages/Flux/qXNjB/src/optimise/train.jl:105
┌ Info: Epoch 2406
└ @ Main /Users/kcft114/.julia/packages/Flux/qXNjB/src/optimise/train.jl:105
┌ Info: Epoch 2407
└ @ Main /Users/kcft114/.julia/packages/Flux/qXNjB/src/optimise/train.jl:105
┌ Info: Epoch 2408
└ @ Main /Users/kcft114/.julia/packages/Flux/qXNjB/src/optimise/train.jl:105
┌ Info: Epoch 2409
└ @ Main /Users/kcft114/.julia/packages/Flux/qXNjB/src/optimise/train.jl:105
┌ Info: Epoch 2410
└ @ Main /Users/kcft114/.julia/packages/Flux/qXNjB/src/optimise/train.jl:105
┌ Info: Epoch 2411
└ @ Main /Users/kcft114/.julia/packages/Flux/qXNjB/src/optimise/train.jl:105
┌ Info: Epoch 2412
└ @ Main /Users/kcft114/.julia/packages/Flux/qXNjB/src/optimise/train.jl:105
┌ Info: Epoch 2413
└ @ Main /Users/kcft114/.julia/packages/Flux/qXNjB/src/optimise/train.jl:105
┌ Info: Epoch 2414
└ @ Main /Users/kcft114/.julia/packages/Flux/qXNjB/src/optimise/train.jl:105
┌ Info: Epoch 2415
└ @ Main /Users/kcft114/.julia/packages/Flux/qXNjB/src/optimise/train.jl:105
┌ Info: Epoch 2416
└ @ Main /Users/kcft114/.julia/packages/Flux/qXNjB/src/optimise/train.jl:105
┌ Info: Epoch 2417
└ @ Main /Users/kcft114/.julia/packages/Flux/qXNjB/src/optimise/train.jl:105
┌ Info: Epoch 2418
└ @ Main /Users/kcft114/.julia/packages/Flux/qXNjB/src/optimise/train.jl:105
┌ Info: Epoch 2419
└ @ Main /Users/kcft114/.julia/packages/Flux/qXNjB/src/optimise/train.jl:105
┌ Info: Epoch 2420
└ @ Main /Users/kcft114/.julia/packages/Flux/qXNjB/src/optimise/train.jl:105
┌ Info: Epoch 2421
└ @ Main /Users/kcft114/.julia/packages/Flux/qXNjB/src/optimise/train.jl:105
┌ Info: Epoch 2422
└ @ Main /Users/kcft114/.julia/packages/Flux/qXNjB/src/optimise/train.jl:105
┌ Info: Epoch 2423
└ @ Main /Users/kcft114/.julia/packages/Flux/qXNjB/src/optimise/train.jl:105
┌ Info: Epoch 2424
└ @ Main /Users/kcft114/.julia/packages/Flux/qXNjB/src/optimise/train.jl:105
┌ Info: Epoch 2425
└ @ Main /Users/kcft114/.julia/packages/Flux/qXNjB/src/optimise/train.jl:105
┌ Info: Epoch 2426
└ @ Main /Users/kcft114/.julia/packages/Flux/qXNjB/src/optimise/train.jl:105
┌ Info: Epoch 2427
└ @ Main /Users/kcft114/.julia/packages/Flux/qXNjB/src/optimise/train.jl:105
┌ Info: Epoch 2428
└ @ Main /Users/kcft114/.julia/packages/Flux/qXNjB/src/optimise/train.jl:105
┌ Info: Epoch 2429
└ @ Main /Users/kcft114/.julia/packages/Flux/qXNjB/src/optimise/train.jl:105
┌ Info: Epoch 2430
└ @ Main /Users/kcft114/.julia/packages/Flux/qXNjB/src/optimise/train.jl:105
┌ Info: Epoch 2431
└ @ Main /Users/kcft114/.julia/packages/Flux/qXNjB/src/optimise/train.jl:105
┌ Info: Epoch 2432
└ @ Main /Users/kcft114/.julia/packages/Flux/qXNjB/src/optimise/train.jl:105
┌ Info: Epoch 2433
└ @ Main /Users/kcft114/.julia/packages/Flux/qXNjB/src/optimise/train.jl:105
┌ Info: Epoch 2434
└ @ Main /Users/kcft114/.julia/packages/Flux/qXNjB/src/optimise/train.jl:105
┌ Info: Epoch 2435
└ @ Main /Users/kcft114/.julia/packages/Flux/qXNjB/src/optimise/train.jl:105
┌ Info: Epoch 2436
└ @ Main /Users/kcft114/.julia/packages/Flux/qXNjB/src/optimise/train.jl:105
┌ Info: Epoch 2437
└ @ Main /Users/kcft114/.julia/packages/Flux/qXNjB/src/optimise/train.jl:105
┌ Info: Epoch 2438
└ @ Main /Users/kcft114/.julia/packages/Flux/qXNjB/src/optimise/train.jl:105
┌ Info: Epoch 2439
└ @ Main /Users/kcft114/.julia/packages/Flux/qXNjB/src/optimise/train.jl:105
┌ Info: Epoch 2440
└ @ Main /Users/kcft114/.julia/packages/Flux/qXNjB/src/optimise/train.jl:105
┌ Info: Epoch 2441
└ @ Main /Users/kcft114/.julia/packages/Flux/qXNjB/src/optimise/train.jl:105
┌ Info: Epoch 2442
└ @ Main /Users/kcft114/.julia/packages/Flux/qXNjB/src/optimise/train.jl:105
┌ Info: Epoch 2443
└ @ Main /Users/kcft114/.julia/packages/Flux/qXNjB/src/optimise/train.jl:105
┌ Info: Epoch 2444
└ @ Main /Users/kcft114/.julia/packages/Flux/qXNjB/src/optimise/train.jl:105
┌ Info: Epoch 2445
└ @ Main /Users/kcft114/.julia/packages/Flux/qXNjB/src/optimise/train.jl:105
┌ Info: Epoch 2446
└ @ Main /Users/kcft114/.julia/packages/Flux/qXNjB/src/optimise/train.jl:105
┌ Info: Epoch 2447
└ @ Main /Users/kcft114/.julia/packages/Flux/qXNjB/src/optimise/train.jl:105
┌ Info: Epoch 2448
└ @ Main /Users/kcft114/.julia/packages/Flux/qXNjB/src/optimise/train.jl:105
┌ Info: Epoch 2449
└ @ Main /Users/kcft114/.julia/packages/Flux/qXNjB/src/optimise/train.jl:105
┌ Info: Epoch 2450
└ @ Main /Users/kcft114/.julia/packages/Flux/qXNjB/src/optimise/train.jl:105
┌ Info: Epoch 2451
└ @ Main /Users/kcft114/.julia/packages/Flux/qXNjB/src/optimise/train.jl:105
┌ Info: Epoch 2452
└ @ Main /Users/kcft114/.julia/packages/Flux/qXNjB/src/optimise/train.jl:105
┌ Info: Epoch 2453
└ @ Main /Users/kcft114/.julia/packages/Flux/qXNjB/src/optimise/train.jl:105
┌ Info: Epoch 2454
└ @ Main /Users/kcft114/.julia/packages/Flux/qXNjB/src/optimise/train.jl:105
┌ Info: Epoch 2455
└ @ Main /Users/kcft114/.julia/packages/Flux/qXNjB/src/optimise/train.jl:105
┌ Info: Epoch 2456
└ @ Main /Users/kcft114/.julia/packages/Flux/qXNjB/src/optimise/train.jl:105
┌ Info: Epoch 2457
└ @ Main /Users/kcft114/.julia/packages/Flux/qXNjB/src/optimise/train.jl:105
┌ Info: Epoch 2458
└ @ Main /Users/kcft114/.julia/packages/Flux/qXNjB/src/optimise/train.jl:105
┌ Info: Epoch 2459
└ @ Main /Users/kcft114/.julia/packages/Flux/qXNjB/src/optimise/train.jl:105
┌ Info: Epoch 2460
└ @ Main /Users/kcft114/.julia/packages/Flux/qXNjB/src/optimise/train.jl:105
┌ Info: Epoch 2461
└ @ Main /Users/kcft114/.julia/packages/Flux/qXNjB/src/optimise/train.jl:105
┌ Info: Epoch 2462
└ @ Main /Users/kcft114/.julia/packages/Flux/qXNjB/src/optimise/train.jl:105
┌ Info: Epoch 2463
└ @ Main /Users/kcft114/.julia/packages/Flux/qXNjB/src/optimise/train.jl:105
┌ Info: Epoch 2464
└ @ Main /Users/kcft114/.julia/packages/Flux/qXNjB/src/optimise/train.jl:105
┌ Info: Epoch 2465
└ @ Main /Users/kcft114/.julia/packages/Flux/qXNjB/src/optimise/train.jl:105
┌ Info: Epoch 2466
└ @ Main /Users/kcft114/.julia/packages/Flux/qXNjB/src/optimise/train.jl:105
┌ Info: Epoch 2467
└ @ Main /Users/kcft114/.julia/packages/Flux/qXNjB/src/optimise/train.jl:105
┌ Info: Epoch 2468
└ @ Main /Users/kcft114/.julia/packages/Flux/qXNjB/src/optimise/train.jl:105
┌ Info: Epoch 2469
└ @ Main /Users/kcft114/.julia/packages/Flux/qXNjB/src/optimise/train.jl:105
┌ Info: Epoch 2470
└ @ Main /Users/kcft114/.julia/packages/Flux/qXNjB/src/optimise/train.jl:105
┌ Info: Epoch 2471
└ @ Main /Users/kcft114/.julia/packages/Flux/qXNjB/src/optimise/train.jl:105
```

```
loss(X, Y) = 0.23738103f0 (tracked)
loss(X, Y) = 0.23725276f0 (tracked)
loss(X, Y) = 0.23714629f0 (tracked)
loss(X, Y) = 0.2370489f0 (tracked)
loss(X, Y) = 0.23693001f0 (tracked)
loss(X, Y) = 0.23681071f0 (tracked)
loss(X, Y) = 0.23669256f0 (tracked)
loss(X, Y) = 0.23657721f0 (tracked)
loss(X, Y) = 0.23646137f0 (tracked)
loss(X, Y) = 0.23634265f0 (tracked)
loss(X, Y) = 0.23623696f0 (tracked)
loss(X, Y) = 0.23612787f0 (tracked)
loss(X, Y) = 0.23600906f0 (tracked)
loss(X, Y) = 0.23590513f0 (tracked)
loss(X, Y) = 0.23579754f0 (tracked)
loss(X, Y) = 0.2356789f0 (tracked)
loss(X, Y) = 0.2355717f0 (tracked)
loss(X, Y) = 0.23545744f0 (tracked)
loss(X, Y) = 0.2353543f0 (tracked)
loss(X, Y) = 0.23524056f0 (tracked)
loss(X, Y) = 0.2351202f0 (tracked)
loss(X, Y) = 0.23500362f0 (tracked)
loss(X, Y) = 0.23490284f0 (tracked)
loss(X, Y) = 0.23477636f0 (tracked)
loss(X, Y) = 0.23466818f0 (tracked)
loss(X, Y) = 0.23455209f0 (tracked)
loss(X, Y) = 0.23443706f0 (tracked)
loss(X, Y) = 0.23433799f0 (tracked)
loss(X, Y) = 0.23421752f0 (tracked)
loss(X, Y) = 0.23410311f0 (tracked)
loss(X, Y) = 0.23401244f0 (tracked)
loss(X, Y) = 0.23389621f0 (tracked)
loss(X, Y) = 0.233772f0 (tracked)
loss(X, Y) = 0.23366897f0 (tracked)
loss(X, Y) = 0.23357227f0 (tracked)
loss(X, Y) = 0.23345825f0 (tracked)
loss(X, Y) = 0.23334007f0 (tracked)
loss(X, Y) = 0.23321937f0 (tracked)
loss(X, Y) = 0.2331255f0 (tracked)
loss(X, Y) = 0.2330196f0 (tracked)
loss(X, Y) = 0.23288728f0 (tracked)
loss(X, Y) = 0.2327918f0 (tracked)
loss(X, Y) = 0.23268951f0 (tracked)
loss(X, Y) = 0.23259509f0 (tracked)
loss(X, Y) = 0.23247102f0 (tracked)
loss(X, Y) = 0.23235452f0 (tracked)
loss(X, Y) = 0.2322683f0 (tracked)
loss(X, Y) = 0.23216945f0 (tracked)
loss(X, Y) = 0.23204492f0 (tracked)
loss(X, Y) = 0.23194253f0 (tracked)
loss(X, Y) = 0.23184206f0 (tracked)
loss(X, Y) = 0.2317401f0 (tracked)
loss(X, Y) = 0.23162717f0 (tracked)
loss(X, Y) = 0.2315129f0 (tracked)
loss(X, Y) = 0.23140857f0 (tracked)
loss(X, Y) = 0.23130278f0 (tracked)
loss(X, Y) = 0.23119201f0 (tracked)
loss(X, Y) = 0.2310852f0 (tracked)
loss(X, Y) = 0.23098129f0 (tracked)
loss(X, Y) = 0.23087648f0 (tracked)
loss(X, Y) = 0.23076081f0 (tracked)
loss(X, Y) = 0.23065662f0 (tracked)
loss(X, Y) = 0.23056202f0 (tracked)
loss(X, Y) = 0.23044537f0 (tracked)
loss(X, Y) = 0.23033111f0 (tracked)
loss(X, Y) = 0.23023215f0 (tracked)
loss(X, Y) = 0.23012467f0 (tracked)
loss(X, Y) = 0.23002695f0 (tracked)
loss(X, Y) = 0.22991608f0 (tracked)
loss(X, Y) = 0.22981304f0 (tracked)
loss(X, Y) = 0.22971526f0 (tracked)
loss(X, Y) = 0.22959417f0 (tracked)
loss(X, Y) = 0.22949602f0 (tracked)
loss(X, Y) = 0.2293998f0 (tracked)
loss(X, Y) = 0.22929673f0 (tracked)
loss(X, Y) = 0.22919983f0 (tracked)
loss(X, Y) = 0.22909567f0 (tracked)
loss(X, Y) = 0.22897722f0 (tracked)
loss(X, Y) = 0.22886546f0 (tracked)
loss(X, Y) = 0.22879018f0 (tracked)
loss(X, Y) = 0.22869352f0 (tracked)
loss(X, Y) = 0.22857577f0 (tracked)
loss(X, Y) = 0.22846043f0 (tracked)
loss(X, Y) = 0.22835465f0 (tracked)
loss(X, Y) = 0.22826032f0 (tracked)
loss(X, Y) = 0.2281587f0 (tracked)
loss(X, Y) = 0.22803956f0 (tracked)
loss(X, Y) = 0.22792356f0 (tracked)
loss(X, Y) = 0.22781883f0 (tracked)
loss(X, Y) = 0.22772068f0 (tracked)
loss(X, Y) = 0.22761655f0 (tracked)
loss(X, Y) = 0.22750285f0 (tracked)
loss(X, Y) = 0.22740035f0 (tracked)
loss(X, Y) = 0.22732754f0 (tracked)
loss(X, Y) = 0.22721034f0 (tracked)
loss(X, Y) = 0.22709614f0 (tracked)
loss(X, Y) = 0.22699599f0 (tracked)
loss(X, Y) = 0.2268917f0 (tracked)
loss(X, Y) = 0.22678813f0 (tracked)
loss(X, Y) = 0.22667967f0 (tracked)
loss(X, Y) = 0.22657757f0 (tracked)
loss(X, Y) = 0.22648713f0 (tracked)
loss(X, Y) = 0.22638273f0 (tracked)
loss(X, Y) = 0.2262708f0 (tracked)
loss(X, Y) = 0.22617719f0 (tracked)
loss(X, Y) = 0.2260819f0 (tracked)
loss(X, Y) = 0.22597446f0 (tracked)
loss(X, Y) = 0.22586985f0 (tracked)
loss(X, Y) = 0.22576866f0 (tracked)
loss(X, Y) = 0.22566958f0 (tracked)
loss(X, Y) = 0.2255702f0 (tracked)
loss(X, Y) = 0.2254587f0 (tracked)
loss(X, Y) = 0.22534953f0 (tracked)
loss(X, Y) = 0.22526576f0 (tracked)
loss(X, Y) = 0.22517437f0 (tracked)
loss(X, Y) = 0.22506215f0 (tracked)
loss(X, Y) = 0.22496484f0 (tracked)
loss(X, Y) = 0.22486818f0 (tracked)
loss(X, Y) = 0.22475784f0 (tracked)
loss(X, Y) = 0.22463913f0 (tracked)
loss(X, Y) = 0.22455028f0 (tracked)
loss(X, Y) = 0.22445476f0 (tracked)
loss(X, Y) = 0.2243487f0 (tracked)
loss(X, Y) = 0.22424184f0 (tracked)
loss(X, Y) = 0.2241485f0 (tracked)
loss(X, Y) = 0.22404303f0 (tracked)
loss(X, Y) = 0.22394072f0 (tracked)
loss(X, Y) = 0.22384284f0 (tracked)
loss(X, Y) = 0.22374208f0 (tracked)
loss(X, Y) = 0.22364184f0 (tracked)
loss(X, Y) = 0.22353493f0 (tracked)
loss(X, Y) = 0.22344086f0 (tracked)
loss(X, Y) = 0.2233505f0 (tracked)
loss(X, Y) = 0.22324553f0 (tracked)
loss(X, Y) = 0.22315122f0 (tracked)
loss(X, Y) = 0.2230544f0 (tracked)
loss(X, Y) = 0.22294082f0 (tracked)
loss(X, Y) = 0.22285451f0 (tracked)
loss(X, Y) = 0.22275834f0 (tracked)
loss(X, Y) = 0.22265314f0 (tracked)
loss(X, Y) = 0.22256239f0 (tracked)
loss(X, Y) = 0.22246748f0 (tracked)
loss(X, Y) = 0.22236073f0 (tracked)
loss(X, Y) = 0.22225861f0 (tracked)
loss(X, Y) = 0.22216122f0 (tracked)
loss(X, Y) = 0.22206473f0 (tracked)
loss(X, Y) = 0.2219632f0 (tracked)
loss(X, Y) = 0.22186185f0 (tracked)
loss(X, Y) = 0.22177103f0 (tracked)
loss(X, Y) = 0.2216623f0 (tracked)
loss(X, Y) = 0.22156696f0 (tracked)
loss(X, Y) = 0.2214802f0 (tracked)
loss(X, Y) = 0.22137815f0 (tracked)
loss(X, Y) = 0.2212733f0 (tracked)
loss(X, Y) = 0.22117427f0 (tracked)
```

```
┌ Info: Epoch 2472
└ @ Main /Users/kcft114/.julia/packages/Flux/qXNjB/src/optimise/train.jl:105
┌ Info: Epoch 2473
└ @ Main /Users/kcft114/.julia/packages/Flux/qXNjB/src/optimise/train.jl:105
┌ Info: Epoch 2474
└ @ Main /Users/kcft114/.julia/packages/Flux/qXNjB/src/optimise/train.jl:105
┌ Info: Epoch 2475
└ @ Main /Users/kcft114/.julia/packages/Flux/qXNjB/src/optimise/train.jl:105
┌ Info: Epoch 2476
└ @ Main /Users/kcft114/.julia/packages/Flux/qXNjB/src/optimise/train.jl:105
┌ Info: Epoch 2477
└ @ Main /Users/kcft114/.julia/packages/Flux/qXNjB/src/optimise/train.jl:105
┌ Info: Epoch 2478
└ @ Main /Users/kcft114/.julia/packages/Flux/qXNjB/src/optimise/train.jl:105
┌ Info: Epoch 2479
└ @ Main /Users/kcft114/.julia/packages/Flux/qXNjB/src/optimise/train.jl:105
┌ Info: Epoch 2480
└ @ Main /Users/kcft114/.julia/packages/Flux/qXNjB/src/optimise/train.jl:105
┌ Info: Epoch 2481
└ @ Main /Users/kcft114/.julia/packages/Flux/qXNjB/src/optimise/train.jl:105
┌ Info: Epoch 2482
└ @ Main /Users/kcft114/.julia/packages/Flux/qXNjB/src/optimise/train.jl:105
┌ Info: Epoch 2483
└ @ Main /Users/kcft114/.julia/packages/Flux/qXNjB/src/optimise/train.jl:105
┌ Info: Epoch 2484
└ @ Main /Users/kcft114/.julia/packages/Flux/qXNjB/src/optimise/train.jl:105
┌ Info: Epoch 2485
└ @ Main /Users/kcft114/.julia/packages/Flux/qXNjB/src/optimise/train.jl:105
┌ Info: Epoch 2486
└ @ Main /Users/kcft114/.julia/packages/Flux/qXNjB/src/optimise/train.jl:105
┌ Info: Epoch 2487
└ @ Main /Users/kcft114/.julia/packages/Flux/qXNjB/src/optimise/train.jl:105
┌ Info: Epoch 2488
└ @ Main /Users/kcft114/.julia/packages/Flux/qXNjB/src/optimise/train.jl:105
┌ Info: Epoch 2489
└ @ Main /Users/kcft114/.julia/packages/Flux/qXNjB/src/optimise/train.jl:105
┌ Info: Epoch 2490
└ @ Main /Users/kcft114/.julia/packages/Flux/qXNjB/src/optimise/train.jl:105
┌ Info: Epoch 2491
└ @ Main /Users/kcft114/.julia/packages/Flux/qXNjB/src/optimise/train.jl:105
┌ Info: Epoch 2492
└ @ Main /Users/kcft114/.julia/packages/Flux/qXNjB/src/optimise/train.jl:105
┌ Info: Epoch 2493
└ @ Main /Users/kcft114/.julia/packages/Flux/qXNjB/src/optimise/train.jl:105
┌ Info: Epoch 2494
└ @ Main /Users/kcft114/.julia/packages/Flux/qXNjB/src/optimise/train.jl:105
┌ Info: Epoch 2495
└ @ Main /Users/kcft114/.julia/packages/Flux/qXNjB/src/optimise/train.jl:105
┌ Info: Epoch 2496
└ @ Main /Users/kcft114/.julia/packages/Flux/qXNjB/src/optimise/train.jl:105
┌ Info: Epoch 2497
└ @ Main /Users/kcft114/.julia/packages/Flux/qXNjB/src/optimise/train.jl:105
┌ Info: Epoch 2498
└ @ Main /Users/kcft114/.julia/packages/Flux/qXNjB/src/optimise/train.jl:105
┌ Info: Epoch 2499
└ @ Main /Users/kcft114/.julia/packages/Flux/qXNjB/src/optimise/train.jl:105
┌ Info: Epoch 2500
└ @ Main /Users/kcft114/.julia/packages/Flux/qXNjB/src/optimise/train.jl:105
┌ Info: Epoch 2501
└ @ Main /Users/kcft114/.julia/packages/Flux/qXNjB/src/optimise/train.jl:105
┌ Info: Epoch 2502
└ @ Main /Users/kcft114/.julia/packages/Flux/qXNjB/src/optimise/train.jl:105
┌ Info: Epoch 2503
└ @ Main /Users/kcft114/.julia/packages/Flux/qXNjB/src/optimise/train.jl:105
┌ Info: Epoch 2504
└ @ Main /Users/kcft114/.julia/packages/Flux/qXNjB/src/optimise/train.jl:105
┌ Info: Epoch 2505
└ @ Main /Users/kcft114/.julia/packages/Flux/qXNjB/src/optimise/train.jl:105
┌ Info: Epoch 2506
└ @ Main /Users/kcft114/.julia/packages/Flux/qXNjB/src/optimise/train.jl:105
┌ Info: Epoch 2507
└ @ Main /Users/kcft114/.julia/packages/Flux/qXNjB/src/optimise/train.jl:105
┌ Info: Epoch 2508
└ @ Main /Users/kcft114/.julia/packages/Flux/qXNjB/src/optimise/train.jl:105
┌ Info: Epoch 2509
└ @ Main /Users/kcft114/.julia/packages/Flux/qXNjB/src/optimise/train.jl:105
┌ Info: Epoch 2510
└ @ Main /Users/kcft114/.julia/packages/Flux/qXNjB/src/optimise/train.jl:105
┌ Info: Epoch 2511
└ @ Main /Users/kcft114/.julia/packages/Flux/qXNjB/src/optimise/train.jl:105
┌ Info: Epoch 2512
└ @ Main /Users/kcft114/.julia/packages/Flux/qXNjB/src/optimise/train.jl:105
┌ Info: Epoch 2513
└ @ Main /Users/kcft114/.julia/packages/Flux/qXNjB/src/optimise/train.jl:105
┌ Info: Epoch 2514
└ @ Main /Users/kcft114/.julia/packages/Flux/qXNjB/src/optimise/train.jl:105
┌ Info: Epoch 2515
└ @ Main /Users/kcft114/.julia/packages/Flux/qXNjB/src/optimise/train.jl:105
┌ Info: Epoch 2516
└ @ Main /Users/kcft114/.julia/packages/Flux/qXNjB/src/optimise/train.jl:105
┌ Info: Epoch 2517
└ @ Main /Users/kcft114/.julia/packages/Flux/qXNjB/src/optimise/train.jl:105
┌ Info: Epoch 2518
└ @ Main /Users/kcft114/.julia/packages/Flux/qXNjB/src/optimise/train.jl:105
┌ Info: Epoch 2519
└ @ Main /Users/kcft114/.julia/packages/Flux/qXNjB/src/optimise/train.jl:105
┌ Info: Epoch 2520
└ @ Main /Users/kcft114/.julia/packages/Flux/qXNjB/src/optimise/train.jl:105
┌ Info: Epoch 2521
└ @ Main /Users/kcft114/.julia/packages/Flux/qXNjB/src/optimise/train.jl:105
┌ Info: Epoch 2522
└ @ Main /Users/kcft114/.julia/packages/Flux/qXNjB/src/optimise/train.jl:105
┌ Info: Epoch 2523
└ @ Main /Users/kcft114/.julia/packages/Flux/qXNjB/src/optimise/train.jl:105
┌ Info: Epoch 2524
└ @ Main /Users/kcft114/.julia/packages/Flux/qXNjB/src/optimise/train.jl:105
┌ Info: Epoch 2525
└ @ Main /Users/kcft114/.julia/packages/Flux/qXNjB/src/optimise/train.jl:105
┌ Info: Epoch 2526
└ @ Main /Users/kcft114/.julia/packages/Flux/qXNjB/src/optimise/train.jl:105
┌ Info: Epoch 2527
└ @ Main /Users/kcft114/.julia/packages/Flux/qXNjB/src/optimise/train.jl:105
┌ Info: Epoch 2528
└ @ Main /Users/kcft114/.julia/packages/Flux/qXNjB/src/optimise/train.jl:105
┌ Info: Epoch 2529
└ @ Main /Users/kcft114/.julia/packages/Flux/qXNjB/src/optimise/train.jl:105
┌ Info: Epoch 2530
└ @ Main /Users/kcft114/.julia/packages/Flux/qXNjB/src/optimise/train.jl:105
┌ Info: Epoch 2531
└ @ Main /Users/kcft114/.julia/packages/Flux/qXNjB/src/optimise/train.jl:105
┌ Info: Epoch 2532
└ @ Main /Users/kcft114/.julia/packages/Flux/qXNjB/src/optimise/train.jl:105
┌ Info: Epoch 2533
└ @ Main /Users/kcft114/.julia/packages/Flux/qXNjB/src/optimise/train.jl:105
┌ Info: Epoch 2534
└ @ Main /Users/kcft114/.julia/packages/Flux/qXNjB/src/optimise/train.jl:105
┌ Info: Epoch 2535
└ @ Main /Users/kcft114/.julia/packages/Flux/qXNjB/src/optimise/train.jl:105
┌ Info: Epoch 2536
└ @ Main /Users/kcft114/.julia/packages/Flux/qXNjB/src/optimise/train.jl:105
┌ Info: Epoch 2537
└ @ Main /Users/kcft114/.julia/packages/Flux/qXNjB/src/optimise/train.jl:105
┌ Info: Epoch 2538
└ @ Main /Users/kcft114/.julia/packages/Flux/qXNjB/src/optimise/train.jl:105
┌ Info: Epoch 2539
└ @ Main /Users/kcft114/.julia/packages/Flux/qXNjB/src/optimise/train.jl:105
┌ Info: Epoch 2540
└ @ Main /Users/kcft114/.julia/packages/Flux/qXNjB/src/optimise/train.jl:105
┌ Info: Epoch 2541
└ @ Main /Users/kcft114/.julia/packages/Flux/qXNjB/src/optimise/train.jl:105
┌ Info: Epoch 2542
└ @ Main /Users/kcft114/.julia/packages/Flux/qXNjB/src/optimise/train.jl:105
┌ Info: Epoch 2543
└ @ Main /Users/kcft114/.julia/packages/Flux/qXNjB/src/optimise/train.jl:105
┌ Info: Epoch 2544
└ @ Main /Users/kcft114/.julia/packages/Flux/qXNjB/src/optimise/train.jl:105
┌ Info: Epoch 2545
└ @ Main /Users/kcft114/.julia/packages/Flux/qXNjB/src/optimise/train.jl:105
┌ Info: Epoch 2546
└ @ Main /Users/kcft114/.julia/packages/Flux/qXNjB/src/optimise/train.jl:105
┌ Info: Epoch 2547
└ @ Main /Users/kcft114/.julia/packages/Flux/qXNjB/src/optimise/train.jl:105
┌ Info: Epoch 2548
└ @ Main /Users/kcft114/.julia/packages/Flux/qXNjB/src/optimise/train.jl:105
┌ Info: Epoch 2549
└ @ Main /Users/kcft114/.julia/packages/Flux/qXNjB/src/optimise/train.jl:105
┌ Info: Epoch 2550
└ @ Main /Users/kcft114/.julia/packages/Flux/qXNjB/src/optimise/train.jl:105
┌ Info: Epoch 2551
└ @ Main /Users/kcft114/.julia/packages/Flux/qXNjB/src/optimise/train.jl:105
┌ Info: Epoch 2552
└ @ Main /Users/kcft114/.julia/packages/Flux/qXNjB/src/optimise/train.jl:105
┌ Info: Epoch 2553
└ @ Main /Users/kcft114/.julia/packages/Flux/qXNjB/src/optimise/train.jl:105
┌ Info: Epoch 2554
└ @ Main /Users/kcft114/.julia/packages/Flux/qXNjB/src/optimise/train.jl:105
┌ Info: Epoch 2555
└ @ Main /Users/kcft114/.julia/packages/Flux/qXNjB/src/optimise/train.jl:105
┌ Info: Epoch 2556
└ @ Main /Users/kcft114/.julia/packages/Flux/qXNjB/src/optimise/train.jl:105
┌ Info: Epoch 2557
└ @ Main /Users/kcft114/.julia/packages/Flux/qXNjB/src/optimise/train.jl:105
┌ Info: Epoch 2558
└ @ Main /Users/kcft114/.julia/packages/Flux/qXNjB/src/optimise/train.jl:105
┌ Info: Epoch 2559
└ @ Main /Users/kcft114/.julia/packages/Flux/qXNjB/src/optimise/train.jl:105
┌ Info: Epoch 2560
└ @ Main /Users/kcft114/.julia/packages/Flux/qXNjB/src/optimise/train.jl:105
┌ Info: Epoch 2561
└ @ Main /Users/kcft114/.julia/packages/Flux/qXNjB/src/optimise/train.jl:105
┌ Info: Epoch 2562
└ @ Main /Users/kcft114/.julia/packages/Flux/qXNjB/src/optimise/train.jl:105
┌ Info: Epoch 2563
└ @ Main /Users/kcft114/.julia/packages/Flux/qXNjB/src/optimise/train.jl:105
┌ Info: Epoch 2564
└ @ Main /Users/kcft114/.julia/packages/Flux/qXNjB/src/optimise/train.jl:105
┌ Info: Epoch 2565
└ @ Main /Users/kcft114/.julia/packages/Flux/qXNjB/src/optimise/train.jl:105
┌ Info: Epoch 2566
└ @ Main /Users/kcft114/.julia/packages/Flux/qXNjB/src/optimise/train.jl:105
┌ Info: Epoch 2567
└ @ Main /Users/kcft114/.julia/packages/Flux/qXNjB/src/optimise/train.jl:105
┌ Info: Epoch 2568
└ @ Main /Users/kcft114/.julia/packages/Flux/qXNjB/src/optimise/train.jl:105
┌ Info: Epoch 2569
└ @ Main /Users/kcft114/.julia/packages/Flux/qXNjB/src/optimise/train.jl:105
┌ Info: Epoch 2570
└ @ Main /Users/kcft114/.julia/packages/Flux/qXNjB/src/optimise/train.jl:105
┌ Info: Epoch 2571
└ @ Main /Users/kcft114/.julia/packages/Flux/qXNjB/src/optimise/train.jl:105
┌ Info: Epoch 2572
└ @ Main /Users/kcft114/.julia/packages/Flux/qXNjB/src/optimise/train.jl:105
┌ Info: Epoch 2573
└ @ Main /Users/kcft114/.julia/packages/Flux/qXNjB/src/optimise/train.jl:105
┌ Info: Epoch 2574
└ @ Main /Users/kcft114/.julia/packages/Flux/qXNjB/src/optimise/train.jl:105
┌ Info: Epoch 2575
└ @ Main /Users/kcft114/.julia/packages/Flux/qXNjB/src/optimise/train.jl:105
┌ Info: Epoch 2576
└ @ Main /Users/kcft114/.julia/packages/Flux/qXNjB/src/optimise/train.jl:105
┌ Info: Epoch 2577
└ @ Main /Users/kcft114/.julia/packages/Flux/qXNjB/src/optimise/train.jl:105
┌ Info: Epoch 2578
└ @ Main /Users/kcft114/.julia/packages/Flux/qXNjB/src/optimise/train.jl:105
┌ Info: Epoch 2579
└ @ Main /Users/kcft114/.julia/packages/Flux/qXNjB/src/optimise/train.jl:105
┌ Info: Epoch 2580
└ @ Main /Users/kcft114/.julia/packages/Flux/qXNjB/src/optimise/train.jl:105
┌ Info: Epoch 2581
└ @ Main /Users/kcft114/.julia/packages/Flux/qXNjB/src/optimise/train.jl:105
┌ Info: Epoch 2582
└ @ Main /Users/kcft114/.julia/packages/Flux/qXNjB/src/optimise/train.jl:105
┌ Info: Epoch 2583
└ @ Main /Users/kcft114/.julia/packages/Flux/qXNjB/src/optimise/train.jl:105
┌ Info: Epoch 2584
└ @ Main /Users/kcft114/.julia/packages/Flux/qXNjB/src/optimise/train.jl:105
┌ Info: Epoch 2585
└ @ Main /Users/kcft114/.julia/packages/Flux/qXNjB/src/optimise/train.jl:105
┌ Info: Epoch 2586
└ @ Main /Users/kcft114/.julia/packages/Flux/qXNjB/src/optimise/train.jl:105
┌ Info: Epoch 2587
└ @ Main /Users/kcft114/.julia/packages/Flux/qXNjB/src/optimise/train.jl:105
┌ Info: Epoch 2588
└ @ Main /Users/kcft114/.julia/packages/Flux/qXNjB/src/optimise/train.jl:105
┌ Info: Epoch 2589
└ @ Main /Users/kcft114/.julia/packages/Flux/qXNjB/src/optimise/train.jl:105
┌ Info: Epoch 2590
└ @ Main /Users/kcft114/.julia/packages/Flux/qXNjB/src/optimise/train.jl:105
┌ Info: Epoch 2591
└ @ Main /Users/kcft114/.julia/packages/Flux/qXNjB/src/optimise/train.jl:105
┌ Info: Epoch 2592
└ @ Main /Users/kcft114/.julia/packages/Flux/qXNjB/src/optimise/train.jl:105
┌ Info: Epoch 2593
└ @ Main /Users/kcft114/.julia/packages/Flux/qXNjB/src/optimise/train.jl:105
┌ Info: Epoch 2594
└ @ Main /Users/kcft114/.julia/packages/Flux/qXNjB/src/optimise/train.jl:105
┌ Info: Epoch 2595
└ @ Main /Users/kcft114/.julia/packages/Flux/qXNjB/src/optimise/train.jl:105
┌ Info: Epoch 2596
└ @ Main /Users/kcft114/.julia/packages/Flux/qXNjB/src/optimise/train.jl:105
┌ Info: Epoch 2597
└ @ Main /Users/kcft114/.julia/packages/Flux/qXNjB/src/optimise/train.jl:105
┌ Info: Epoch 2598
└ @ Main /Users/kcft114/.julia/packages/Flux/qXNjB/src/optimise/train.jl:105
┌ Info: Epoch 2599
└ @ Main /Users/kcft114/.julia/packages/Flux/qXNjB/src/optimise/train.jl:105
┌ Info: Epoch 2600
└ @ Main /Users/kcft114/.julia/packages/Flux/qXNjB/src/optimise/train.jl:105
┌ Info: Epoch 2601
└ @ Main /Users/kcft114/.julia/packages/Flux/qXNjB/src/optimise/train.jl:105
┌ Info: Epoch 2602
└ @ Main /Users/kcft114/.julia/packages/Flux/qXNjB/src/optimise/train.jl:105
┌ Info: Epoch 2603
└ @ Main /Users/kcft114/.julia/packages/Flux/qXNjB/src/optimise/train.jl:105
┌ Info: Epoch 2604
└ @ Main /Users/kcft114/.julia/packages/Flux/qXNjB/src/optimise/train.jl:105
┌ Info: Epoch 2605
└ @ Main /Users/kcft114/.julia/packages/Flux/qXNjB/src/optimise/train.jl:105
┌ Info: Epoch 2606
└ @ Main /Users/kcft114/.julia/packages/Flux/qXNjB/src/optimise/train.jl:105
┌ Info: Epoch 2607
└ @ Main /Users/kcft114/.julia/packages/Flux/qXNjB/src/optimise/train.jl:105
┌ Info: Epoch 2608
└ @ Main /Users/kcft114/.julia/packages/Flux/qXNjB/src/optimise/train.jl:105
┌ Info: Epoch 2609
└ @ Main /Users/kcft114/.julia/packages/Flux/qXNjB/src/optimise/train.jl:105
┌ Info: Epoch 2610
└ @ Main /Users/kcft114/.julia/packages/Flux/qXNjB/src/optimise/train.jl:105
┌ Info: Epoch 2611
└ @ Main /Users/kcft114/.julia/packages/Flux/qXNjB/src/optimise/train.jl:105
┌ Info: Epoch 2612
└ @ Main /Users/kcft114/.julia/packages/Flux/qXNjB/src/optimise/train.jl:105
┌ Info: Epoch 2613
└ @ Main /Users/kcft114/.julia/packages/Flux/qXNjB/src/optimise/train.jl:105
┌ Info: Epoch 2614
└ @ Main /Users/kcft114/.julia/packages/Flux/qXNjB/src/optimise/train.jl:105
┌ Info: Epoch 2615
└ @ Main /Users/kcft114/.julia/packages/Flux/qXNjB/src/optimise/train.jl:105
┌ Info: Epoch 2616
└ @ Main /Users/kcft114/.julia/packages/Flux/qXNjB/src/optimise/train.jl:105
┌ Info: Epoch 2617
└ @ Main /Users/kcft114/.julia/packages/Flux/qXNjB/src/optimise/train.jl:105
┌ Info: Epoch 2618
└ @ Main /Users/kcft114/.julia/packages/Flux/qXNjB/src/optimise/train.jl:105
┌ Info: Epoch 2619
└ @ Main /Users/kcft114/.julia/packages/Flux/qXNjB/src/optimise/train.jl:105
┌ Info: Epoch 2620
└ @ Main /Users/kcft114/.julia/packages/Flux/qXNjB/src/optimise/train.jl:105
┌ Info: Epoch 2621
└ @ Main /Users/kcft114/.julia/packages/Flux/qXNjB/src/optimise/train.jl:105
┌ Info: Epoch 2622
└ @ Main /Users/kcft114/.julia/packages/Flux/qXNjB/src/optimise/train.jl:105
┌ Info: Epoch 2623
└ @ Main /Users/kcft114/.julia/packages/Flux/qXNjB/src/optimise/train.jl:105
┌ Info: Epoch 2624
└ @ Main /Users/kcft114/.julia/packages/Flux/qXNjB/src/optimise/train.jl:105
┌ Info: Epoch 2625
└ @ Main /Users/kcft114/.julia/packages/Flux/qXNjB/src/optimise/train.jl:105
┌ Info: Epoch 2626
└ @ Main /Users/kcft114/.julia/packages/Flux/qXNjB/src/optimise/train.jl:105
```

```
loss(X, Y) = 0.22108282f0 (tracked)
loss(X, Y) = 0.22099838f0 (tracked)
loss(X, Y) = 0.22088525f0 (tracked)
loss(X, Y) = 0.22079867f0 (tracked)
loss(X, Y) = 0.220706f0 (tracked)
loss(X, Y) = 0.22060215f0 (tracked)
loss(X, Y) = 0.2205081f0 (tracked)
loss(X, Y) = 0.22041087f0 (tracked)
loss(X, Y) = 0.22031415f0 (tracked)
loss(X, Y) = 0.22021717f0 (tracked)
loss(X, Y) = 0.22012588f0 (tracked)
loss(X, Y) = 0.22004038f0 (tracked)
loss(X, Y) = 0.21994446f0 (tracked)
loss(X, Y) = 0.21983996f0 (tracked)
loss(X, Y) = 0.21975054f0 (tracked)
loss(X, Y) = 0.21964817f0 (tracked)
loss(X, Y) = 0.21957308f0 (tracked)
loss(X, Y) = 0.21947813f0 (tracked)
loss(X, Y) = 0.219367f0 (tracked)
loss(X, Y) = 0.21925977f0 (tracked)
loss(X, Y) = 0.2191746f0 (tracked)
loss(X, Y) = 0.2190915f0 (tracked)
loss(X, Y) = 0.21899839f0 (tracked)
loss(X, Y) = 0.21887782f0 (tracked)
loss(X, Y) = 0.21879333f0 (tracked)
loss(X, Y) = 0.2187052f0 (tracked)
loss(X, Y) = 0.21862559f0 (tracked)
loss(X, Y) = 0.21853681f0 (tracked)
loss(X, Y) = 0.2184343f0 (tracked)
loss(X, Y) = 0.21832636f0 (tracked)
loss(X, Y) = 0.2182263f0 (tracked)
loss(X, Y) = 0.21815449f0 (tracked)
loss(X, Y) = 0.21805084f0 (tracked)
loss(X, Y) = 0.21795967f0 (tracked)
loss(X, Y) = 0.21785964f0 (tracked)
loss(X, Y) = 0.21775137f0 (tracked)
loss(X, Y) = 0.21765482f0 (tracked)
loss(X, Y) = 0.21756911f0 (tracked)
loss(X, Y) = 0.21748133f0 (tracked)
loss(X, Y) = 0.21737984f0 (tracked)
loss(X, Y) = 0.21728379f0 (tracked)
loss(X, Y) = 0.21718657f0 (tracked)
loss(X, Y) = 0.21709494f0 (tracked)
loss(X, Y) = 0.2169919f0 (tracked)
loss(X, Y) = 0.216915f0 (tracked)
loss(X, Y) = 0.21681115f0 (tracked)
loss(X, Y) = 0.21672463f0 (tracked)
loss(X, Y) = 0.21664019f0 (tracked)
loss(X, Y) = 0.2165393f0 (tracked)
loss(X, Y) = 0.2164392f0 (tracked)
loss(X, Y) = 0.21634926f0 (tracked)
loss(X, Y) = 0.21626273f0 (tracked)
loss(X, Y) = 0.21616101f0 (tracked)
loss(X, Y) = 0.21606416f0 (tracked)
loss(X, Y) = 0.21597005f0 (tracked)
loss(X, Y) = 0.21587864f0 (tracked)
loss(X, Y) = 0.21578015f0 (tracked)
loss(X, Y) = 0.21569751f0 (tracked)
loss(X, Y) = 0.21559867f0 (tracked)
loss(X, Y) = 0.21550924f0 (tracked)
loss(X, Y) = 0.21542314f0 (tracked)
loss(X, Y) = 0.2153245f0 (tracked)
loss(X, Y) = 0.21522684f0 (tracked)
loss(X, Y) = 0.21514016f0 (tracked)
loss(X, Y) = 0.21505368f0 (tracked)
loss(X, Y) = 0.21495193f0 (tracked)
loss(X, Y) = 0.21485797f0 (tracked)
loss(X, Y) = 0.21476685f0 (tracked)
loss(X, Y) = 0.21468301f0 (tracked)
loss(X, Y) = 0.21458942f0 (tracked)
loss(X, Y) = 0.2144898f0 (tracked)
loss(X, Y) = 0.21441138f0 (tracked)
loss(X, Y) = 0.21432598f0 (tracked)
loss(X, Y) = 0.21422695f0 (tracked)
loss(X, Y) = 0.21412985f0 (tracked)
loss(X, Y) = 0.21405691f0 (tracked)
loss(X, Y) = 0.21396808f0 (tracked)
loss(X, Y) = 0.21387142f0 (tracked)
loss(X, Y) = 0.21376587f0 (tracked)
loss(X, Y) = 0.21367456f0 (tracked)
loss(X, Y) = 0.21357955f0 (tracked)
loss(X, Y) = 0.21349144f0 (tracked)
loss(X, Y) = 0.21340474f0 (tracked)
loss(X, Y) = 0.21330674f0 (tracked)
loss(X, Y) = 0.21322252f0 (tracked)
loss(X, Y) = 0.21313992f0 (tracked)
loss(X, Y) = 0.21303688f0 (tracked)
loss(X, Y) = 0.21296865f0 (tracked)
loss(X, Y) = 0.21285796f0 (tracked)
loss(X, Y) = 0.21275486f0 (tracked)
loss(X, Y) = 0.21268398f0 (tracked)
loss(X, Y) = 0.2126022f0 (tracked)
loss(X, Y) = 0.2124982f0 (tracked)
loss(X, Y) = 0.21241026f0 (tracked)
loss(X, Y) = 0.21233562f0 (tracked)
loss(X, Y) = 0.21224816f0 (tracked)
loss(X, Y) = 0.21214962f0 (tracked)
loss(X, Y) = 0.2120388f0 (tracked)
loss(X, Y) = 0.21195567f0 (tracked)
loss(X, Y) = 0.21187104f0 (tracked)
loss(X, Y) = 0.21176584f0 (tracked)
loss(X, Y) = 0.21167745f0 (tracked)
loss(X, Y) = 0.21158466f0 (tracked)
loss(X, Y) = 0.21149614f0 (tracked)
loss(X, Y) = 0.21140085f0 (tracked)
loss(X, Y) = 0.21130177f0 (tracked)
loss(X, Y) = 0.21122171f0 (tracked)
loss(X, Y) = 0.21112601f0 (tracked)
loss(X, Y) = 0.21103762f0 (tracked)
loss(X, Y) = 0.21094213f0 (tracked)
loss(X, Y) = 0.21086222f0 (tracked)
loss(X, Y) = 0.21077237f0 (tracked)
loss(X, Y) = 0.21068832f0 (tracked)
loss(X, Y) = 0.21058884f0 (tracked)
loss(X, Y) = 0.21049441f0 (tracked)
loss(X, Y) = 0.2104245f0 (tracked)
loss(X, Y) = 0.2103366f0 (tracked)
loss(X, Y) = 0.21022779f0 (tracked)
loss(X, Y) = 0.21013395f0 (tracked)
loss(X, Y) = 0.21003693f0 (tracked)
loss(X, Y) = 0.20995751f0 (tracked)
loss(X, Y) = 0.20988102f0 (tracked)
loss(X, Y) = 0.20977396f0 (tracked)
loss(X, Y) = 0.20965679f0 (tracked)
loss(X, Y) = 0.20957644f0 (tracked)
loss(X, Y) = 0.2095124f0 (tracked)
loss(X, Y) = 0.20942026f0 (tracked)
loss(X, Y) = 0.20933232f0 (tracked)
loss(X, Y) = 0.2092329f0 (tracked)
loss(X, Y) = 0.20913869f0 (tracked)
loss(X, Y) = 0.20905869f0 (tracked)
loss(X, Y) = 0.20896392f0 (tracked)
loss(X, Y) = 0.20887527f0 (tracked)
loss(X, Y) = 0.20877689f0 (tracked)
loss(X, Y) = 0.20868443f0 (tracked)
loss(X, Y) = 0.208597f0 (tracked)
loss(X, Y) = 0.20851277f0 (tracked)
loss(X, Y) = 0.20840973f0 (tracked)
loss(X, Y) = 0.20832306f0 (tracked)
loss(X, Y) = 0.20823859f0 (tracked)
```

```
┌ Info: Epoch 2627
└ @ Main /Users/kcft114/.julia/packages/Flux/qXNjB/src/optimise/train.jl:105
┌ Info: Epoch 2628
└ @ Main /Users/kcft114/.julia/packages/Flux/qXNjB/src/optimise/train.jl:105
┌ Info: Epoch 2629
└ @ Main /Users/kcft114/.julia/packages/Flux/qXNjB/src/optimise/train.jl:105
┌ Info: Epoch 2630
└ @ Main /Users/kcft114/.julia/packages/Flux/qXNjB/src/optimise/train.jl:105
┌ Info: Epoch 2631
└ @ Main /Users/kcft114/.julia/packages/Flux/qXNjB/src/optimise/train.jl:105
┌ Info: Epoch 2632
└ @ Main /Users/kcft114/.julia/packages/Flux/qXNjB/src/optimise/train.jl:105
┌ Info: Epoch 2633
└ @ Main /Users/kcft114/.julia/packages/Flux/qXNjB/src/optimise/train.jl:105
┌ Info: Epoch 2634
└ @ Main /Users/kcft114/.julia/packages/Flux/qXNjB/src/optimise/train.jl:105
┌ Info: Epoch 2635
└ @ Main /Users/kcft114/.julia/packages/Flux/qXNjB/src/optimise/train.jl:105
┌ Info: Epoch 2636
└ @ Main /Users/kcft114/.julia/packages/Flux/qXNjB/src/optimise/train.jl:105
┌ Info: Epoch 2637
└ @ Main /Users/kcft114/.julia/packages/Flux/qXNjB/src/optimise/train.jl:105
┌ Info: Epoch 2638
└ @ Main /Users/kcft114/.julia/packages/Flux/qXNjB/src/optimise/train.jl:105
┌ Info: Epoch 2639
└ @ Main /Users/kcft114/.julia/packages/Flux/qXNjB/src/optimise/train.jl:105
┌ Info: Epoch 2640
└ @ Main /Users/kcft114/.julia/packages/Flux/qXNjB/src/optimise/train.jl:105
┌ Info: Epoch 2641
└ @ Main /Users/kcft114/.julia/packages/Flux/qXNjB/src/optimise/train.jl:105
┌ Info: Epoch 2642
└ @ Main /Users/kcft114/.julia/packages/Flux/qXNjB/src/optimise/train.jl:105
┌ Info: Epoch 2643
└ @ Main /Users/kcft114/.julia/packages/Flux/qXNjB/src/optimise/train.jl:105
┌ Info: Epoch 2644
└ @ Main /Users/kcft114/.julia/packages/Flux/qXNjB/src/optimise/train.jl:105
┌ Info: Epoch 2645
└ @ Main /Users/kcft114/.julia/packages/Flux/qXNjB/src/optimise/train.jl:105
┌ Info: Epoch 2646
└ @ Main /Users/kcft114/.julia/packages/Flux/qXNjB/src/optimise/train.jl:105
┌ Info: Epoch 2647
└ @ Main /Users/kcft114/.julia/packages/Flux/qXNjB/src/optimise/train.jl:105
┌ Info: Epoch 2648
└ @ Main /Users/kcft114/.julia/packages/Flux/qXNjB/src/optimise/train.jl:105
┌ Info: Epoch 2649
└ @ Main /Users/kcft114/.julia/packages/Flux/qXNjB/src/optimise/train.jl:105
┌ Info: Epoch 2650
└ @ Main /Users/kcft114/.julia/packages/Flux/qXNjB/src/optimise/train.jl:105
┌ Info: Epoch 2651
└ @ Main /Users/kcft114/.julia/packages/Flux/qXNjB/src/optimise/train.jl:105
┌ Info: Epoch 2652
└ @ Main /Users/kcft114/.julia/packages/Flux/qXNjB/src/optimise/train.jl:105
┌ Info: Epoch 2653
└ @ Main /Users/kcft114/.julia/packages/Flux/qXNjB/src/optimise/train.jl:105
┌ Info: Epoch 2654
└ @ Main /Users/kcft114/.julia/packages/Flux/qXNjB/src/optimise/train.jl:105
┌ Info: Epoch 2655
└ @ Main /Users/kcft114/.julia/packages/Flux/qXNjB/src/optimise/train.jl:105
┌ Info: Epoch 2656
└ @ Main /Users/kcft114/.julia/packages/Flux/qXNjB/src/optimise/train.jl:105
┌ Info: Epoch 2657
└ @ Main /Users/kcft114/.julia/packages/Flux/qXNjB/src/optimise/train.jl:105
┌ Info: Epoch 2658
└ @ Main /Users/kcft114/.julia/packages/Flux/qXNjB/src/optimise/train.jl:105
┌ Info: Epoch 2659
└ @ Main /Users/kcft114/.julia/packages/Flux/qXNjB/src/optimise/train.jl:105
┌ Info: Epoch 2660
└ @ Main /Users/kcft114/.julia/packages/Flux/qXNjB/src/optimise/train.jl:105
┌ Info: Epoch 2661
└ @ Main /Users/kcft114/.julia/packages/Flux/qXNjB/src/optimise/train.jl:105
┌ Info: Epoch 2662
└ @ Main /Users/kcft114/.julia/packages/Flux/qXNjB/src/optimise/train.jl:105
┌ Info: Epoch 2663
└ @ Main /Users/kcft114/.julia/packages/Flux/qXNjB/src/optimise/train.jl:105
┌ Info: Epoch 2664
└ @ Main /Users/kcft114/.julia/packages/Flux/qXNjB/src/optimise/train.jl:105
┌ Info: Epoch 2665
└ @ Main /Users/kcft114/.julia/packages/Flux/qXNjB/src/optimise/train.jl:105
┌ Info: Epoch 2666
└ @ Main /Users/kcft114/.julia/packages/Flux/qXNjB/src/optimise/train.jl:105
┌ Info: Epoch 2667
└ @ Main /Users/kcft114/.julia/packages/Flux/qXNjB/src/optimise/train.jl:105
┌ Info: Epoch 2668
└ @ Main /Users/kcft114/.julia/packages/Flux/qXNjB/src/optimise/train.jl:105
┌ Info: Epoch 2669
└ @ Main /Users/kcft114/.julia/packages/Flux/qXNjB/src/optimise/train.jl:105
┌ Info: Epoch 2670
└ @ Main /Users/kcft114/.julia/packages/Flux/qXNjB/src/optimise/train.jl:105
┌ Info: Epoch 2671
└ @ Main /Users/kcft114/.julia/packages/Flux/qXNjB/src/optimise/train.jl:105
┌ Info: Epoch 2672
└ @ Main /Users/kcft114/.julia/packages/Flux/qXNjB/src/optimise/train.jl:105
┌ Info: Epoch 2673
└ @ Main /Users/kcft114/.julia/packages/Flux/qXNjB/src/optimise/train.jl:105
┌ Info: Epoch 2674
└ @ Main /Users/kcft114/.julia/packages/Flux/qXNjB/src/optimise/train.jl:105
┌ Info: Epoch 2675
└ @ Main /Users/kcft114/.julia/packages/Flux/qXNjB/src/optimise/train.jl:105
┌ Info: Epoch 2676
└ @ Main /Users/kcft114/.julia/packages/Flux/qXNjB/src/optimise/train.jl:105
┌ Info: Epoch 2677
└ @ Main /Users/kcft114/.julia/packages/Flux/qXNjB/src/optimise/train.jl:105
┌ Info: Epoch 2678
└ @ Main /Users/kcft114/.julia/packages/Flux/qXNjB/src/optimise/train.jl:105
┌ Info: Epoch 2679
└ @ Main /Users/kcft114/.julia/packages/Flux/qXNjB/src/optimise/train.jl:105
┌ Info: Epoch 2680
└ @ Main /Users/kcft114/.julia/packages/Flux/qXNjB/src/optimise/train.jl:105
┌ Info: Epoch 2681
└ @ Main /Users/kcft114/.julia/packages/Flux/qXNjB/src/optimise/train.jl:105
┌ Info: Epoch 2682
└ @ Main /Users/kcft114/.julia/packages/Flux/qXNjB/src/optimise/train.jl:105
┌ Info: Epoch 2683
└ @ Main /Users/kcft114/.julia/packages/Flux/qXNjB/src/optimise/train.jl:105
┌ Info: Epoch 2684
└ @ Main /Users/kcft114/.julia/packages/Flux/qXNjB/src/optimise/train.jl:105
┌ Info: Epoch 2685
└ @ Main /Users/kcft114/.julia/packages/Flux/qXNjB/src/optimise/train.jl:105
┌ Info: Epoch 2686
└ @ Main /Users/kcft114/.julia/packages/Flux/qXNjB/src/optimise/train.jl:105
┌ Info: Epoch 2687
└ @ Main /Users/kcft114/.julia/packages/Flux/qXNjB/src/optimise/train.jl:105
┌ Info: Epoch 2688
└ @ Main /Users/kcft114/.julia/packages/Flux/qXNjB/src/optimise/train.jl:105
┌ Info: Epoch 2689
└ @ Main /Users/kcft114/.julia/packages/Flux/qXNjB/src/optimise/train.jl:105
┌ Info: Epoch 2690
└ @ Main /Users/kcft114/.julia/packages/Flux/qXNjB/src/optimise/train.jl:105
┌ Info: Epoch 2691
└ @ Main /Users/kcft114/.julia/packages/Flux/qXNjB/src/optimise/train.jl:105
┌ Info: Epoch 2692
└ @ Main /Users/kcft114/.julia/packages/Flux/qXNjB/src/optimise/train.jl:105
┌ Info: Epoch 2693
└ @ Main /Users/kcft114/.julia/packages/Flux/qXNjB/src/optimise/train.jl:105
┌ Info: Epoch 2694
└ @ Main /Users/kcft114/.julia/packages/Flux/qXNjB/src/optimise/train.jl:105
┌ Info: Epoch 2695
└ @ Main /Users/kcft114/.julia/packages/Flux/qXNjB/src/optimise/train.jl:105
┌ Info: Epoch 2696
└ @ Main /Users/kcft114/.julia/packages/Flux/qXNjB/src/optimise/train.jl:105
┌ Info: Epoch 2697
└ @ Main /Users/kcft114/.julia/packages/Flux/qXNjB/src/optimise/train.jl:105
┌ Info: Epoch 2698
└ @ Main /Users/kcft114/.julia/packages/Flux/qXNjB/src/optimise/train.jl:105
┌ Info: Epoch 2699
└ @ Main /Users/kcft114/.julia/packages/Flux/qXNjB/src/optimise/train.jl:105
┌ Info: Epoch 2700
└ @ Main /Users/kcft114/.julia/packages/Flux/qXNjB/src/optimise/train.jl:105
┌ Info: Epoch 2701
└ @ Main /Users/kcft114/.julia/packages/Flux/qXNjB/src/optimise/train.jl:105
┌ Info: Epoch 2702
└ @ Main /Users/kcft114/.julia/packages/Flux/qXNjB/src/optimise/train.jl:105
┌ Info: Epoch 2703
└ @ Main /Users/kcft114/.julia/packages/Flux/qXNjB/src/optimise/train.jl:105
┌ Info: Epoch 2704
└ @ Main /Users/kcft114/.julia/packages/Flux/qXNjB/src/optimise/train.jl:105
┌ Info: Epoch 2705
└ @ Main /Users/kcft114/.julia/packages/Flux/qXNjB/src/optimise/train.jl:105
┌ Info: Epoch 2706
└ @ Main /Users/kcft114/.julia/packages/Flux/qXNjB/src/optimise/train.jl:105
┌ Info: Epoch 2707
└ @ Main /Users/kcft114/.julia/packages/Flux/qXNjB/src/optimise/train.jl:105
┌ Info: Epoch 2708
└ @ Main /Users/kcft114/.julia/packages/Flux/qXNjB/src/optimise/train.jl:105
┌ Info: Epoch 2709
└ @ Main /Users/kcft114/.julia/packages/Flux/qXNjB/src/optimise/train.jl:105
┌ Info: Epoch 2710
└ @ Main /Users/kcft114/.julia/packages/Flux/qXNjB/src/optimise/train.jl:105
┌ Info: Epoch 2711
└ @ Main /Users/kcft114/.julia/packages/Flux/qXNjB/src/optimise/train.jl:105
┌ Info: Epoch 2712
└ @ Main /Users/kcft114/.julia/packages/Flux/qXNjB/src/optimise/train.jl:105
┌ Info: Epoch 2713
└ @ Main /Users/kcft114/.julia/packages/Flux/qXNjB/src/optimise/train.jl:105
┌ Info: Epoch 2714
└ @ Main /Users/kcft114/.julia/packages/Flux/qXNjB/src/optimise/train.jl:105
┌ Info: Epoch 2715
└ @ Main /Users/kcft114/.julia/packages/Flux/qXNjB/src/optimise/train.jl:105
┌ Info: Epoch 2716
└ @ Main /Users/kcft114/.julia/packages/Flux/qXNjB/src/optimise/train.jl:105
┌ Info: Epoch 2717
└ @ Main /Users/kcft114/.julia/packages/Flux/qXNjB/src/optimise/train.jl:105
┌ Info: Epoch 2718
└ @ Main /Users/kcft114/.julia/packages/Flux/qXNjB/src/optimise/train.jl:105
┌ Info: Epoch 2719
└ @ Main /Users/kcft114/.julia/packages/Flux/qXNjB/src/optimise/train.jl:105
┌ Info: Epoch 2720
└ @ Main /Users/kcft114/.julia/packages/Flux/qXNjB/src/optimise/train.jl:105
┌ Info: Epoch 2721
└ @ Main /Users/kcft114/.julia/packages/Flux/qXNjB/src/optimise/train.jl:105
┌ Info: Epoch 2722
└ @ Main /Users/kcft114/.julia/packages/Flux/qXNjB/src/optimise/train.jl:105
┌ Info: Epoch 2723
└ @ Main /Users/kcft114/.julia/packages/Flux/qXNjB/src/optimise/train.jl:105
┌ Info: Epoch 2724
└ @ Main /Users/kcft114/.julia/packages/Flux/qXNjB/src/optimise/train.jl:105
┌ Info: Epoch 2725
└ @ Main /Users/kcft114/.julia/packages/Flux/qXNjB/src/optimise/train.jl:105
┌ Info: Epoch 2726
└ @ Main /Users/kcft114/.julia/packages/Flux/qXNjB/src/optimise/train.jl:105
┌ Info: Epoch 2727
└ @ Main /Users/kcft114/.julia/packages/Flux/qXNjB/src/optimise/train.jl:105
┌ Info: Epoch 2728
└ @ Main /Users/kcft114/.julia/packages/Flux/qXNjB/src/optimise/train.jl:105
┌ Info: Epoch 2729
└ @ Main /Users/kcft114/.julia/packages/Flux/qXNjB/src/optimise/train.jl:105
┌ Info: Epoch 2730
└ @ Main /Users/kcft114/.julia/packages/Flux/qXNjB/src/optimise/train.jl:105
┌ Info: Epoch 2731
└ @ Main /Users/kcft114/.julia/packages/Flux/qXNjB/src/optimise/train.jl:105
┌ Info: Epoch 2732
└ @ Main /Users/kcft114/.julia/packages/Flux/qXNjB/src/optimise/train.jl:105
┌ Info: Epoch 2733
└ @ Main /Users/kcft114/.julia/packages/Flux/qXNjB/src/optimise/train.jl:105
┌ Info: Epoch 2734
└ @ Main /Users/kcft114/.julia/packages/Flux/qXNjB/src/optimise/train.jl:105
┌ Info: Epoch 2735
└ @ Main /Users/kcft114/.julia/packages/Flux/qXNjB/src/optimise/train.jl:105
┌ Info: Epoch 2736
└ @ Main /Users/kcft114/.julia/packages/Flux/qXNjB/src/optimise/train.jl:105
┌ Info: Epoch 2737
└ @ Main /Users/kcft114/.julia/packages/Flux/qXNjB/src/optimise/train.jl:105
┌ Info: Epoch 2738
└ @ Main /Users/kcft114/.julia/packages/Flux/qXNjB/src/optimise/train.jl:105
┌ Info: Epoch 2739
└ @ Main /Users/kcft114/.julia/packages/Flux/qXNjB/src/optimise/train.jl:105
┌ Info: Epoch 2740
└ @ Main /Users/kcft114/.julia/packages/Flux/qXNjB/src/optimise/train.jl:105
┌ Info: Epoch 2741
└ @ Main /Users/kcft114/.julia/packages/Flux/qXNjB/src/optimise/train.jl:105
┌ Info: Epoch 2742
└ @ Main /Users/kcft114/.julia/packages/Flux/qXNjB/src/optimise/train.jl:105
┌ Info: Epoch 2743
└ @ Main /Users/kcft114/.julia/packages/Flux/qXNjB/src/optimise/train.jl:105
┌ Info: Epoch 2744
└ @ Main /Users/kcft114/.julia/packages/Flux/qXNjB/src/optimise/train.jl:105
┌ Info: Epoch 2745
└ @ Main /Users/kcft114/.julia/packages/Flux/qXNjB/src/optimise/train.jl:105
┌ Info: Epoch 2746
└ @ Main /Users/kcft114/.julia/packages/Flux/qXNjB/src/optimise/train.jl:105
┌ Info: Epoch 2747
└ @ Main /Users/kcft114/.julia/packages/Flux/qXNjB/src/optimise/train.jl:105
┌ Info: Epoch 2748
└ @ Main /Users/kcft114/.julia/packages/Flux/qXNjB/src/optimise/train.jl:105
┌ Info: Epoch 2749
└ @ Main /Users/kcft114/.julia/packages/Flux/qXNjB/src/optimise/train.jl:105
┌ Info: Epoch 2750
└ @ Main /Users/kcft114/.julia/packages/Flux/qXNjB/src/optimise/train.jl:105
┌ Info: Epoch 2751
└ @ Main /Users/kcft114/.julia/packages/Flux/qXNjB/src/optimise/train.jl:105
┌ Info: Epoch 2752
└ @ Main /Users/kcft114/.julia/packages/Flux/qXNjB/src/optimise/train.jl:105
┌ Info: Epoch 2753
└ @ Main /Users/kcft114/.julia/packages/Flux/qXNjB/src/optimise/train.jl:105
┌ Info: Epoch 2754
└ @ Main /Users/kcft114/.julia/packages/Flux/qXNjB/src/optimise/train.jl:105
┌ Info: Epoch 2755
└ @ Main /Users/kcft114/.julia/packages/Flux/qXNjB/src/optimise/train.jl:105
┌ Info: Epoch 2756
└ @ Main /Users/kcft114/.julia/packages/Flux/qXNjB/src/optimise/train.jl:105
┌ Info: Epoch 2757
└ @ Main /Users/kcft114/.julia/packages/Flux/qXNjB/src/optimise/train.jl:105
┌ Info: Epoch 2758
└ @ Main /Users/kcft114/.julia/packages/Flux/qXNjB/src/optimise/train.jl:105
┌ Info: Epoch 2759
└ @ Main /Users/kcft114/.julia/packages/Flux/qXNjB/src/optimise/train.jl:105
┌ Info: Epoch 2760
└ @ Main /Users/kcft114/.julia/packages/Flux/qXNjB/src/optimise/train.jl:105
┌ Info: Epoch 2761
└ @ Main /Users/kcft114/.julia/packages/Flux/qXNjB/src/optimise/train.jl:105
┌ Info: Epoch 2762
└ @ Main /Users/kcft114/.julia/packages/Flux/qXNjB/src/optimise/train.jl:105
┌ Info: Epoch 2763
└ @ Main /Users/kcft114/.julia/packages/Flux/qXNjB/src/optimise/train.jl:105
┌ Info: Epoch 2764
└ @ Main /Users/kcft114/.julia/packages/Flux/qXNjB/src/optimise/train.jl:105
┌ Info: Epoch 2765
└ @ Main /Users/kcft114/.julia/packages/Flux/qXNjB/src/optimise/train.jl:105
┌ Info: Epoch 2766
└ @ Main /Users/kcft114/.julia/packages/Flux/qXNjB/src/optimise/train.jl:105
```

```
loss(X, Y) = 0.20815276f0 (tracked)
loss(X, Y) = 0.20805517f0 (tracked)
loss(X, Y) = 0.20799251f0 (tracked)
loss(X, Y) = 0.20788747f0 (tracked)
loss(X, Y) = 0.20781274f0 (tracked)
loss(X, Y) = 0.2077228f0 (tracked)
loss(X, Y) = 0.20763238f0 (tracked)
loss(X, Y) = 0.20754628f0 (tracked)
loss(X, Y) = 0.20746917f0 (tracked)
loss(X, Y) = 0.20737642f0 (tracked)
loss(X, Y) = 0.20727663f0 (tracked)
loss(X, Y) = 0.20719759f0 (tracked)
loss(X, Y) = 0.20711248f0 (tracked)
loss(X, Y) = 0.2070092f0 (tracked)
loss(X, Y) = 0.20692548f0 (tracked)
loss(X, Y) = 0.2068526f0 (tracked)
loss(X, Y) = 0.20676304f0 (tracked)
loss(X, Y) = 0.206652f0 (tracked)
loss(X, Y) = 0.20656589f0 (tracked)
loss(X, Y) = 0.20648028f0 (tracked)
loss(X, Y) = 0.20639053f0 (tracked)
loss(X, Y) = 0.20630503f0 (tracked)
loss(X, Y) = 0.20622775f0 (tracked)
loss(X, Y) = 0.20614699f0 (tracked)
loss(X, Y) = 0.20605662f0 (tracked)
loss(X, Y) = 0.20596586f0 (tracked)
loss(X, Y) = 0.20586106f0 (tracked)
loss(X, Y) = 0.20578988f0 (tracked)
loss(X, Y) = 0.20569894f0 (tracked)
loss(X, Y) = 0.20559329f0 (tracked)
loss(X, Y) = 0.20550326f0 (tracked)
loss(X, Y) = 0.20543617f0 (tracked)
loss(X, Y) = 0.20534538f0 (tracked)
loss(X, Y) = 0.20523703f0 (tracked)
loss(X, Y) = 0.20516588f0 (tracked)
loss(X, Y) = 0.20507675f0 (tracked)
loss(X, Y) = 0.20500754f0 (tracked)
loss(X, Y) = 0.20491622f0 (tracked)
loss(X, Y) = 0.20482096f0 (tracked)
loss(X, Y) = 0.20471665f0 (tracked)
loss(X, Y) = 0.20464219f0 (tracked)
loss(X, Y) = 0.20456083f0 (tracked)
loss(X, Y) = 0.20447625f0 (tracked)
loss(X, Y) = 0.20437303f0 (tracked)
loss(X, Y) = 0.2042897f0 (tracked)
loss(X, Y) = 0.20421578f0 (tracked)
loss(X, Y) = 0.20412843f0 (tracked)
loss(X, Y) = 0.20405424f0 (tracked)
loss(X, Y) = 0.20395692f0 (tracked)
loss(X, Y) = 0.20385757f0 (tracked)
loss(X, Y) = 0.20379004f0 (tracked)
loss(X, Y) = 0.20371108f0 (tracked)
loss(X, Y) = 0.20360519f0 (tracked)
loss(X, Y) = 0.20351264f0 (tracked)
loss(X, Y) = 0.20344214f0 (tracked)
loss(X, Y) = 0.20336294f0 (tracked)
loss(X, Y) = 0.20326044f0 (tracked)
loss(X, Y) = 0.20317353f0 (tracked)
loss(X, Y) = 0.20309384f0 (tracked)
loss(X, Y) = 0.20302096f0 (tracked)
loss(X, Y) = 0.20293139f0 (tracked)
loss(X, Y) = 0.20284323f0 (tracked)
loss(X, Y) = 0.20277487f0 (tracked)
loss(X, Y) = 0.20268404f0 (tracked)
loss(X, Y) = 0.20258908f0 (tracked)
loss(X, Y) = 0.20251073f0 (tracked)
loss(X, Y) = 0.20242694f0 (tracked)
loss(X, Y) = 0.20234543f0 (tracked)
loss(X, Y) = 0.20225058f0 (tracked)
loss(X, Y) = 0.20216738f0 (tracked)
loss(X, Y) = 0.20208147f0 (tracked)
loss(X, Y) = 0.20201041f0 (tracked)
loss(X, Y) = 0.20192254f0 (tracked)
loss(X, Y) = 0.20184635f0 (tracked)
loss(X, Y) = 0.20177658f0 (tracked)
loss(X, Y) = 0.20168394f0 (tracked)
loss(X, Y) = 0.20160247f0 (tracked)
loss(X, Y) = 0.20151854f0 (tracked)
loss(X, Y) = 0.2014484f0 (tracked)
loss(X, Y) = 0.20135261f0 (tracked)
loss(X, Y) = 0.20128545f0 (tracked)
loss(X, Y) = 0.2012081f0 (tracked)
loss(X, Y) = 0.20111558f0 (tracked)
loss(X, Y) = 0.20103784f0 (tracked)
loss(X, Y) = 0.20095827f0 (tracked)
loss(X, Y) = 0.20087622f0 (tracked)
loss(X, Y) = 0.20079562f0 (tracked)
loss(X, Y) = 0.20071314f0 (tracked)
loss(X, Y) = 0.20062622f0 (tracked)
loss(X, Y) = 0.20054784f0 (tracked)
loss(X, Y) = 0.20048015f0 (tracked)
loss(X, Y) = 0.2003856f0 (tracked)
loss(X, Y) = 0.20029658f0 (tracked)
loss(X, Y) = 0.20023555f0 (tracked)
loss(X, Y) = 0.20015909f0 (tracked)
loss(X, Y) = 0.20006949f0 (tracked)
loss(X, Y) = 0.19997853f0 (tracked)
loss(X, Y) = 0.19989276f0 (tracked)
loss(X, Y) = 0.19981003f0 (tracked)
loss(X, Y) = 0.19973572f0 (tracked)
loss(X, Y) = 0.19966705f0 (tracked)
loss(X, Y) = 0.1995863f0 (tracked)
loss(X, Y) = 0.19949022f0 (tracked)
loss(X, Y) = 0.19942686f0 (tracked)
loss(X, Y) = 0.19935009f0 (tracked)
loss(X, Y) = 0.19928655f0 (tracked)
loss(X, Y) = 0.19918863f0 (tracked)
loss(X, Y) = 0.19910085f0 (tracked)
loss(X, Y) = 0.19901189f0 (tracked)
loss(X, Y) = 0.19896738f0 (tracked)
loss(X, Y) = 0.19886476f0 (tracked)
loss(X, Y) = 0.19876966f0 (tracked)
loss(X, Y) = 0.19869316f0 (tracked)
loss(X, Y) = 0.19863026f0 (tracked)
loss(X, Y) = 0.19857264f0 (tracked)
loss(X, Y) = 0.1984919f0 (tracked)
loss(X, Y) = 0.19839561f0 (tracked)
loss(X, Y) = 0.19830011f0 (tracked)
loss(X, Y) = 0.19822143f0 (tracked)
loss(X, Y) = 0.19814283f0 (tracked)
loss(X, Y) = 0.19806486f0 (tracked)
loss(X, Y) = 0.19798754f0 (tracked)
loss(X, Y) = 0.19790627f0 (tracked)
loss(X, Y) = 0.19781367f0 (tracked)
loss(X, Y) = 0.19774382f0 (tracked)
loss(X, Y) = 0.19766708f0 (tracked)
loss(X, Y) = 0.19758634f0 (tracked)
loss(X, Y) = 0.19750877f0 (tracked)
```

```
┌ Info: Epoch 2767
└ @ Main /Users/kcft114/.julia/packages/Flux/qXNjB/src/optimise/train.jl:105
┌ Info: Epoch 2768
└ @ Main /Users/kcft114/.julia/packages/Flux/qXNjB/src/optimise/train.jl:105
┌ Info: Epoch 2769
└ @ Main /Users/kcft114/.julia/packages/Flux/qXNjB/src/optimise/train.jl:105
┌ Info: Epoch 2770
└ @ Main /Users/kcft114/.julia/packages/Flux/qXNjB/src/optimise/train.jl:105
┌ Info: Epoch 2771
└ @ Main /Users/kcft114/.julia/packages/Flux/qXNjB/src/optimise/train.jl:105
┌ Info: Epoch 2772
└ @ Main /Users/kcft114/.julia/packages/Flux/qXNjB/src/optimise/train.jl:105
┌ Info: Epoch 2773
└ @ Main /Users/kcft114/.julia/packages/Flux/qXNjB/src/optimise/train.jl:105
┌ Info: Epoch 2774
└ @ Main /Users/kcft114/.julia/packages/Flux/qXNjB/src/optimise/train.jl:105
┌ Info: Epoch 2775
└ @ Main /Users/kcft114/.julia/packages/Flux/qXNjB/src/optimise/train.jl:105
┌ Info: Epoch 2776
└ @ Main /Users/kcft114/.julia/packages/Flux/qXNjB/src/optimise/train.jl:105
┌ Info: Epoch 2777
└ @ Main /Users/kcft114/.julia/packages/Flux/qXNjB/src/optimise/train.jl:105
┌ Info: Epoch 2778
└ @ Main /Users/kcft114/.julia/packages/Flux/qXNjB/src/optimise/train.jl:105
┌ Info: Epoch 2779
└ @ Main /Users/kcft114/.julia/packages/Flux/qXNjB/src/optimise/train.jl:105
┌ Info: Epoch 2780
└ @ Main /Users/kcft114/.julia/packages/Flux/qXNjB/src/optimise/train.jl:105
┌ Info: Epoch 2781
└ @ Main /Users/kcft114/.julia/packages/Flux/qXNjB/src/optimise/train.jl:105
┌ Info: Epoch 2782
└ @ Main /Users/kcft114/.julia/packages/Flux/qXNjB/src/optimise/train.jl:105
┌ Info: Epoch 2783
└ @ Main /Users/kcft114/.julia/packages/Flux/qXNjB/src/optimise/train.jl:105
┌ Info: Epoch 2784
└ @ Main /Users/kcft114/.julia/packages/Flux/qXNjB/src/optimise/train.jl:105
┌ Info: Epoch 2785
└ @ Main /Users/kcft114/.julia/packages/Flux/qXNjB/src/optimise/train.jl:105
┌ Info: Epoch 2786
└ @ Main /Users/kcft114/.julia/packages/Flux/qXNjB/src/optimise/train.jl:105
┌ Info: Epoch 2787
└ @ Main /Users/kcft114/.julia/packages/Flux/qXNjB/src/optimise/train.jl:105
┌ Info: Epoch 2788
└ @ Main /Users/kcft114/.julia/packages/Flux/qXNjB/src/optimise/train.jl:105
┌ Info: Epoch 2789
└ @ Main /Users/kcft114/.julia/packages/Flux/qXNjB/src/optimise/train.jl:105
┌ Info: Epoch 2790
└ @ Main /Users/kcft114/.julia/packages/Flux/qXNjB/src/optimise/train.jl:105
┌ Info: Epoch 2791
└ @ Main /Users/kcft114/.julia/packages/Flux/qXNjB/src/optimise/train.jl:105
┌ Info: Epoch 2792
└ @ Main /Users/kcft114/.julia/packages/Flux/qXNjB/src/optimise/train.jl:105
┌ Info: Epoch 2793
└ @ Main /Users/kcft114/.julia/packages/Flux/qXNjB/src/optimise/train.jl:105
┌ Info: Epoch 2794
└ @ Main /Users/kcft114/.julia/packages/Flux/qXNjB/src/optimise/train.jl:105
┌ Info: Epoch 2795
└ @ Main /Users/kcft114/.julia/packages/Flux/qXNjB/src/optimise/train.jl:105
┌ Info: Epoch 2796
└ @ Main /Users/kcft114/.julia/packages/Flux/qXNjB/src/optimise/train.jl:105
┌ Info: Epoch 2797
└ @ Main /Users/kcft114/.julia/packages/Flux/qXNjB/src/optimise/train.jl:105
┌ Info: Epoch 2798
└ @ Main /Users/kcft114/.julia/packages/Flux/qXNjB/src/optimise/train.jl:105
┌ Info: Epoch 2799
└ @ Main /Users/kcft114/.julia/packages/Flux/qXNjB/src/optimise/train.jl:105
┌ Info: Epoch 2800
└ @ Main /Users/kcft114/.julia/packages/Flux/qXNjB/src/optimise/train.jl:105
┌ Info: Epoch 2801
└ @ Main /Users/kcft114/.julia/packages/Flux/qXNjB/src/optimise/train.jl:105
┌ Info: Epoch 2802
└ @ Main /Users/kcft114/.julia/packages/Flux/qXNjB/src/optimise/train.jl:105
┌ Info: Epoch 2803
└ @ Main /Users/kcft114/.julia/packages/Flux/qXNjB/src/optimise/train.jl:105
┌ Info: Epoch 2804
└ @ Main /Users/kcft114/.julia/packages/Flux/qXNjB/src/optimise/train.jl:105
┌ Info: Epoch 2805
└ @ Main /Users/kcft114/.julia/packages/Flux/qXNjB/src/optimise/train.jl:105
┌ Info: Epoch 2806
└ @ Main /Users/kcft114/.julia/packages/Flux/qXNjB/src/optimise/train.jl:105
┌ Info: Epoch 2807
└ @ Main /Users/kcft114/.julia/packages/Flux/qXNjB/src/optimise/train.jl:105
┌ Info: Epoch 2808
└ @ Main /Users/kcft114/.julia/packages/Flux/qXNjB/src/optimise/train.jl:105
┌ Info: Epoch 2809
└ @ Main /Users/kcft114/.julia/packages/Flux/qXNjB/src/optimise/train.jl:105
┌ Info: Epoch 2810
└ @ Main /Users/kcft114/.julia/packages/Flux/qXNjB/src/optimise/train.jl:105
┌ Info: Epoch 2811
└ @ Main /Users/kcft114/.julia/packages/Flux/qXNjB/src/optimise/train.jl:105
┌ Info: Epoch 2812
└ @ Main /Users/kcft114/.julia/packages/Flux/qXNjB/src/optimise/train.jl:105
┌ Info: Epoch 2813
└ @ Main /Users/kcft114/.julia/packages/Flux/qXNjB/src/optimise/train.jl:105
┌ Info: Epoch 2814
└ @ Main /Users/kcft114/.julia/packages/Flux/qXNjB/src/optimise/train.jl:105
┌ Info: Epoch 2815
└ @ Main /Users/kcft114/.julia/packages/Flux/qXNjB/src/optimise/train.jl:105
┌ Info: Epoch 2816
└ @ Main /Users/kcft114/.julia/packages/Flux/qXNjB/src/optimise/train.jl:105
┌ Info: Epoch 2817
└ @ Main /Users/kcft114/.julia/packages/Flux/qXNjB/src/optimise/train.jl:105
┌ Info: Epoch 2818
└ @ Main /Users/kcft114/.julia/packages/Flux/qXNjB/src/optimise/train.jl:105
┌ Info: Epoch 2819
└ @ Main /Users/kcft114/.julia/packages/Flux/qXNjB/src/optimise/train.jl:105
┌ Info: Epoch 2820
└ @ Main /Users/kcft114/.julia/packages/Flux/qXNjB/src/optimise/train.jl:105
┌ Info: Epoch 2821
└ @ Main /Users/kcft114/.julia/packages/Flux/qXNjB/src/optimise/train.jl:105
┌ Info: Epoch 2822
└ @ Main /Users/kcft114/.julia/packages/Flux/qXNjB/src/optimise/train.jl:105
┌ Info: Epoch 2823
└ @ Main /Users/kcft114/.julia/packages/Flux/qXNjB/src/optimise/train.jl:105
┌ Info: Epoch 2824
└ @ Main /Users/kcft114/.julia/packages/Flux/qXNjB/src/optimise/train.jl:105
┌ Info: Epoch 2825
└ @ Main /Users/kcft114/.julia/packages/Flux/qXNjB/src/optimise/train.jl:105
┌ Info: Epoch 2826
└ @ Main /Users/kcft114/.julia/packages/Flux/qXNjB/src/optimise/train.jl:105
┌ Info: Epoch 2827
└ @ Main /Users/kcft114/.julia/packages/Flux/qXNjB/src/optimise/train.jl:105
┌ Info: Epoch 2828
└ @ Main /Users/kcft114/.julia/packages/Flux/qXNjB/src/optimise/train.jl:105
┌ Info: Epoch 2829
└ @ Main /Users/kcft114/.julia/packages/Flux/qXNjB/src/optimise/train.jl:105
┌ Info: Epoch 2830
└ @ Main /Users/kcft114/.julia/packages/Flux/qXNjB/src/optimise/train.jl:105
┌ Info: Epoch 2831
└ @ Main /Users/kcft114/.julia/packages/Flux/qXNjB/src/optimise/train.jl:105
┌ Info: Epoch 2832
└ @ Main /Users/kcft114/.julia/packages/Flux/qXNjB/src/optimise/train.jl:105
┌ Info: Epoch 2833
└ @ Main /Users/kcft114/.julia/packages/Flux/qXNjB/src/optimise/train.jl:105
┌ Info: Epoch 2834
└ @ Main /Users/kcft114/.julia/packages/Flux/qXNjB/src/optimise/train.jl:105
┌ Info: Epoch 2835
└ @ Main /Users/kcft114/.julia/packages/Flux/qXNjB/src/optimise/train.jl:105
┌ Info: Epoch 2836
└ @ Main /Users/kcft114/.julia/packages/Flux/qXNjB/src/optimise/train.jl:105
┌ Info: Epoch 2837
└ @ Main /Users/kcft114/.julia/packages/Flux/qXNjB/src/optimise/train.jl:105
┌ Info: Epoch 2838
└ @ Main /Users/kcft114/.julia/packages/Flux/qXNjB/src/optimise/train.jl:105
┌ Info: Epoch 2839
└ @ Main /Users/kcft114/.julia/packages/Flux/qXNjB/src/optimise/train.jl:105
┌ Info: Epoch 2840
└ @ Main /Users/kcft114/.julia/packages/Flux/qXNjB/src/optimise/train.jl:105
┌ Info: Epoch 2841
└ @ Main /Users/kcft114/.julia/packages/Flux/qXNjB/src/optimise/train.jl:105
┌ Info: Epoch 2842
└ @ Main /Users/kcft114/.julia/packages/Flux/qXNjB/src/optimise/train.jl:105
┌ Info: Epoch 2843
└ @ Main /Users/kcft114/.julia/packages/Flux/qXNjB/src/optimise/train.jl:105
┌ Info: Epoch 2844
└ @ Main /Users/kcft114/.julia/packages/Flux/qXNjB/src/optimise/train.jl:105
┌ Info: Epoch 2845
└ @ Main /Users/kcft114/.julia/packages/Flux/qXNjB/src/optimise/train.jl:105
┌ Info: Epoch 2846
└ @ Main /Users/kcft114/.julia/packages/Flux/qXNjB/src/optimise/train.jl:105
┌ Info: Epoch 2847
└ @ Main /Users/kcft114/.julia/packages/Flux/qXNjB/src/optimise/train.jl:105
┌ Info: Epoch 2848
└ @ Main /Users/kcft114/.julia/packages/Flux/qXNjB/src/optimise/train.jl:105
┌ Info: Epoch 2849
└ @ Main /Users/kcft114/.julia/packages/Flux/qXNjB/src/optimise/train.jl:105
┌ Info: Epoch 2850
└ @ Main /Users/kcft114/.julia/packages/Flux/qXNjB/src/optimise/train.jl:105
┌ Info: Epoch 2851
└ @ Main /Users/kcft114/.julia/packages/Flux/qXNjB/src/optimise/train.jl:105
┌ Info: Epoch 2852
└ @ Main /Users/kcft114/.julia/packages/Flux/qXNjB/src/optimise/train.jl:105
┌ Info: Epoch 2853
└ @ Main /Users/kcft114/.julia/packages/Flux/qXNjB/src/optimise/train.jl:105
┌ Info: Epoch 2854
└ @ Main /Users/kcft114/.julia/packages/Flux/qXNjB/src/optimise/train.jl:105
┌ Info: Epoch 2855
└ @ Main /Users/kcft114/.julia/packages/Flux/qXNjB/src/optimise/train.jl:105
┌ Info: Epoch 2856
└ @ Main /Users/kcft114/.julia/packages/Flux/qXNjB/src/optimise/train.jl:105
┌ Info: Epoch 2857
└ @ Main /Users/kcft114/.julia/packages/Flux/qXNjB/src/optimise/train.jl:105
┌ Info: Epoch 2858
└ @ Main /Users/kcft114/.julia/packages/Flux/qXNjB/src/optimise/train.jl:105
┌ Info: Epoch 2859
└ @ Main /Users/kcft114/.julia/packages/Flux/qXNjB/src/optimise/train.jl:105
┌ Info: Epoch 2860
└ @ Main /Users/kcft114/.julia/packages/Flux/qXNjB/src/optimise/train.jl:105
┌ Info: Epoch 2861
└ @ Main /Users/kcft114/.julia/packages/Flux/qXNjB/src/optimise/train.jl:105
┌ Info: Epoch 2862
└ @ Main /Users/kcft114/.julia/packages/Flux/qXNjB/src/optimise/train.jl:105
┌ Info: Epoch 2863
└ @ Main /Users/kcft114/.julia/packages/Flux/qXNjB/src/optimise/train.jl:105
┌ Info: Epoch 2864
└ @ Main /Users/kcft114/.julia/packages/Flux/qXNjB/src/optimise/train.jl:105
┌ Info: Epoch 2865
└ @ Main /Users/kcft114/.julia/packages/Flux/qXNjB/src/optimise/train.jl:105
┌ Info: Epoch 2866
└ @ Main /Users/kcft114/.julia/packages/Flux/qXNjB/src/optimise/train.jl:105
┌ Info: Epoch 2867
└ @ Main /Users/kcft114/.julia/packages/Flux/qXNjB/src/optimise/train.jl:105
┌ Info: Epoch 2868
└ @ Main /Users/kcft114/.julia/packages/Flux/qXNjB/src/optimise/train.jl:105
┌ Info: Epoch 2869
└ @ Main /Users/kcft114/.julia/packages/Flux/qXNjB/src/optimise/train.jl:105
┌ Info: Epoch 2870
└ @ Main /Users/kcft114/.julia/packages/Flux/qXNjB/src/optimise/train.jl:105
┌ Info: Epoch 2871
└ @ Main /Users/kcft114/.julia/packages/Flux/qXNjB/src/optimise/train.jl:105
┌ Info: Epoch 2872
└ @ Main /Users/kcft114/.julia/packages/Flux/qXNjB/src/optimise/train.jl:105
┌ Info: Epoch 2873
└ @ Main /Users/kcft114/.julia/packages/Flux/qXNjB/src/optimise/train.jl:105
┌ Info: Epoch 2874
└ @ Main /Users/kcft114/.julia/packages/Flux/qXNjB/src/optimise/train.jl:105
┌ Info: Epoch 2875
└ @ Main /Users/kcft114/.julia/packages/Flux/qXNjB/src/optimise/train.jl:105
┌ Info: Epoch 2876
└ @ Main /Users/kcft114/.julia/packages/Flux/qXNjB/src/optimise/train.jl:105
┌ Info: Epoch 2877
└ @ Main /Users/kcft114/.julia/packages/Flux/qXNjB/src/optimise/train.jl:105
┌ Info: Epoch 2878
└ @ Main /Users/kcft114/.julia/packages/Flux/qXNjB/src/optimise/train.jl:105
┌ Info: Epoch 2879
└ @ Main /Users/kcft114/.julia/packages/Flux/qXNjB/src/optimise/train.jl:105
┌ Info: Epoch 2880
└ @ Main /Users/kcft114/.julia/packages/Flux/qXNjB/src/optimise/train.jl:105
┌ Info: Epoch 2881
└ @ Main /Users/kcft114/.julia/packages/Flux/qXNjB/src/optimise/train.jl:105
┌ Info: Epoch 2882
└ @ Main /Users/kcft114/.julia/packages/Flux/qXNjB/src/optimise/train.jl:105
┌ Info: Epoch 2883
└ @ Main /Users/kcft114/.julia/packages/Flux/qXNjB/src/optimise/train.jl:105
┌ Info: Epoch 2884
└ @ Main /Users/kcft114/.julia/packages/Flux/qXNjB/src/optimise/train.jl:105
┌ Info: Epoch 2885
└ @ Main /Users/kcft114/.julia/packages/Flux/qXNjB/src/optimise/train.jl:105
┌ Info: Epoch 2886
└ @ Main /Users/kcft114/.julia/packages/Flux/qXNjB/src/optimise/train.jl:105
┌ Info: Epoch 2887
└ @ Main /Users/kcft114/.julia/packages/Flux/qXNjB/src/optimise/train.jl:105
┌ Info: Epoch 2888
└ @ Main /Users/kcft114/.julia/packages/Flux/qXNjB/src/optimise/train.jl:105
┌ Info: Epoch 2889
└ @ Main /Users/kcft114/.julia/packages/Flux/qXNjB/src/optimise/train.jl:105
┌ Info: Epoch 2890
└ @ Main /Users/kcft114/.julia/packages/Flux/qXNjB/src/optimise/train.jl:105
┌ Info: Epoch 2891
└ @ Main /Users/kcft114/.julia/packages/Flux/qXNjB/src/optimise/train.jl:105
┌ Info: Epoch 2892
└ @ Main /Users/kcft114/.julia/packages/Flux/qXNjB/src/optimise/train.jl:105
┌ Info: Epoch 2893
└ @ Main /Users/kcft114/.julia/packages/Flux/qXNjB/src/optimise/train.jl:105
┌ Info: Epoch 2894
└ @ Main /Users/kcft114/.julia/packages/Flux/qXNjB/src/optimise/train.jl:105
```

```
loss(X, Y) = 0.19742225f0 (tracked)
loss(X, Y) = 0.1973447f0 (tracked)
loss(X, Y) = 0.19725932f0 (tracked)
loss(X, Y) = 0.1971845f0 (tracked)
loss(X, Y) = 0.19710392f0 (tracked)
loss(X, Y) = 0.19703828f0 (tracked)
loss(X, Y) = 0.19696166f0 (tracked)
loss(X, Y) = 0.19689603f0 (tracked)
loss(X, Y) = 0.19680482f0 (tracked)
loss(X, Y) = 0.19672747f0 (tracked)
loss(X, Y) = 0.19664046f0 (tracked)
loss(X, Y) = 0.19657308f0 (tracked)
loss(X, Y) = 0.19649287f0 (tracked)
loss(X, Y) = 0.19640379f0 (tracked)
loss(X, Y) = 0.1963337f0 (tracked)
loss(X, Y) = 0.196268f0 (tracked)
loss(X, Y) = 0.19618638f0 (tracked)
loss(X, Y) = 0.19608144f0 (tracked)
loss(X, Y) = 0.19603007f0 (tracked)
loss(X, Y) = 0.19596027f0 (tracked)
loss(X, Y) = 0.19588387f0 (tracked)
loss(X, Y) = 0.19577739f0 (tracked)
loss(X, Y) = 0.19571114f0 (tracked)
loss(X, Y) = 0.19565547f0 (tracked)
loss(X, Y) = 0.19557859f0 (tracked)
loss(X, Y) = 0.19548284f0 (tracked)
loss(X, Y) = 0.1954115f0 (tracked)
loss(X, Y) = 0.19534035f0 (tracked)
loss(X, Y) = 0.19526789f0 (tracked)
loss(X, Y) = 0.19518313f0 (tracked)
loss(X, Y) = 0.19508429f0 (tracked)
loss(X, Y) = 0.1950277f0 (tracked)
loss(X, Y) = 0.19495697f0 (tracked)
loss(X, Y) = 0.19485235f0 (tracked)
loss(X, Y) = 0.19478962f0 (tracked)
loss(X, Y) = 0.19470492f0 (tracked)
loss(X, Y) = 0.19461595f0 (tracked)
loss(X, Y) = 0.19454898f0 (tracked)
loss(X, Y) = 0.19447175f0 (tracked)
loss(X, Y) = 0.19438547f0 (tracked)
loss(X, Y) = 0.19429669f0 (tracked)
loss(X, Y) = 0.1942349f0 (tracked)
loss(X, Y) = 0.19414735f0 (tracked)
loss(X, Y) = 0.19408475f0 (tracked)
loss(X, Y) = 0.19399568f0 (tracked)
loss(X, Y) = 0.19391824f0 (tracked)
loss(X, Y) = 0.19385834f0 (tracked)
loss(X, Y) = 0.19378415f0 (tracked)
loss(X, Y) = 0.19371401f0 (tracked)
loss(X, Y) = 0.19363022f0 (tracked)
loss(X, Y) = 0.19353136f0 (tracked)
loss(X, Y) = 0.19348206f0 (tracked)
loss(X, Y) = 0.19340678f0 (tracked)
loss(X, Y) = 0.193321f0 (tracked)
loss(X, Y) = 0.19323534f0 (tracked)
loss(X, Y) = 0.19315621f0 (tracked)
loss(X, Y) = 0.1930828f0 (tracked)
loss(X, Y) = 0.19301479f0 (tracked)
loss(X, Y) = 0.19292502f0 (tracked)
loss(X, Y) = 0.19286117f0 (tracked)
loss(X, Y) = 0.19278386f0 (tracked)
loss(X, Y) = 0.19269915f0 (tracked)
loss(X, Y) = 0.19261436f0 (tracked)
loss(X, Y) = 0.19253612f0 (tracked)
loss(X, Y) = 0.19245902f0 (tracked)
loss(X, Y) = 0.19238658f0 (tracked)
loss(X, Y) = 0.19231015f0 (tracked)
loss(X, Y) = 0.19223697f0 (tracked)
loss(X, Y) = 0.19215313f0 (tracked)
loss(X, Y) = 0.19208993f0 (tracked)
loss(X, Y) = 0.19203378f0 (tracked)
loss(X, Y) = 0.19194174f0 (tracked)
loss(X, Y) = 0.19185674f0 (tracked)
loss(X, Y) = 0.19178513f0 (tracked)
loss(X, Y) = 0.19171453f0 (tracked)
loss(X, Y) = 0.19163768f0 (tracked)
loss(X, Y) = 0.19156317f0 (tracked)
loss(X, Y) = 0.19148274f0 (tracked)
loss(X, Y) = 0.19140796f0 (tracked)
loss(X, Y) = 0.19132815f0 (tracked)
loss(X, Y) = 0.1912558f0 (tracked)
loss(X, Y) = 0.19119325f0 (tracked)
loss(X, Y) = 0.19112477f0 (tracked)
loss(X, Y) = 0.19103992f0 (tracked)
loss(X, Y) = 0.19097483f0 (tracked)
loss(X, Y) = 0.19090226f0 (tracked)
loss(X, Y) = 0.19082835f0 (tracked)
loss(X, Y) = 0.1907599f0 (tracked)
loss(X, Y) = 0.19068117f0 (tracked)
loss(X, Y) = 0.19061372f0 (tracked)
loss(X, Y) = 0.19053158f0 (tracked)
loss(X, Y) = 0.19044402f0 (tracked)
loss(X, Y) = 0.19037056f0 (tracked)
loss(X, Y) = 0.19030473f0 (tracked)
loss(X, Y) = 0.19022076f0 (tracked)
loss(X, Y) = 0.19015306f0 (tracked)
loss(X, Y) = 0.1900891f0 (tracked)
loss(X, Y) = 0.19000885f0 (tracked)
loss(X, Y) = 0.1899085f0 (tracked)
loss(X, Y) = 0.18985806f0 (tracked)
loss(X, Y) = 0.1897877f0 (tracked)
loss(X, Y) = 0.18970874f0 (tracked)
loss(X, Y) = 0.18964912f0 (tracked)
loss(X, Y) = 0.18956596f0 (tracked)
loss(X, Y) = 0.18949963f0 (tracked)
loss(X, Y) = 0.18943122f0 (tracked)
loss(X, Y) = 0.18935882f0 (tracked)
loss(X, Y) = 0.18927255f0 (tracked)
loss(X, Y) = 0.18920709f0 (tracked)
loss(X, Y) = 0.18912588f0 (tracked)
loss(X, Y) = 0.18905437f0 (tracked)
loss(X, Y) = 0.18898092f0 (tracked)
loss(X, Y) = 0.18890102f0 (tracked)
loss(X, Y) = 0.18882471f0 (tracked)
loss(X, Y) = 0.18875457f0 (tracked)
loss(X, Y) = 0.18867593f0 (tracked)
loss(X, Y) = 0.1886071f0 (tracked)
loss(X, Y) = 0.18853146f0 (tracked)
loss(X, Y) = 0.18846099f0 (tracked)
loss(X, Y) = 0.18838231f0 (tracked)
loss(X, Y) = 0.18830983f0 (tracked)
loss(X, Y) = 0.1882388f0 (tracked)
```

```
┌ Info: Epoch 2895
└ @ Main /Users/kcft114/.julia/packages/Flux/qXNjB/src/optimise/train.jl:105
┌ Info: Epoch 2896
└ @ Main /Users/kcft114/.julia/packages/Flux/qXNjB/src/optimise/train.jl:105
┌ Info: Epoch 2897
└ @ Main /Users/kcft114/.julia/packages/Flux/qXNjB/src/optimise/train.jl:105
┌ Info: Epoch 2898
└ @ Main /Users/kcft114/.julia/packages/Flux/qXNjB/src/optimise/train.jl:105
┌ Info: Epoch 2899
└ @ Main /Users/kcft114/.julia/packages/Flux/qXNjB/src/optimise/train.jl:105
┌ Info: Epoch 2900
└ @ Main /Users/kcft114/.julia/packages/Flux/qXNjB/src/optimise/train.jl:105
┌ Info: Epoch 2901
└ @ Main /Users/kcft114/.julia/packages/Flux/qXNjB/src/optimise/train.jl:105
┌ Info: Epoch 2902
└ @ Main /Users/kcft114/.julia/packages/Flux/qXNjB/src/optimise/train.jl:105
┌ Info: Epoch 2903
└ @ Main /Users/kcft114/.julia/packages/Flux/qXNjB/src/optimise/train.jl:105
┌ Info: Epoch 2904
└ @ Main /Users/kcft114/.julia/packages/Flux/qXNjB/src/optimise/train.jl:105
┌ Info: Epoch 2905
└ @ Main /Users/kcft114/.julia/packages/Flux/qXNjB/src/optimise/train.jl:105
┌ Info: Epoch 2906
└ @ Main /Users/kcft114/.julia/packages/Flux/qXNjB/src/optimise/train.jl:105
┌ Info: Epoch 2907
└ @ Main /Users/kcft114/.julia/packages/Flux/qXNjB/src/optimise/train.jl:105
┌ Info: Epoch 2908
└ @ Main /Users/kcft114/.julia/packages/Flux/qXNjB/src/optimise/train.jl:105
┌ Info: Epoch 2909
└ @ Main /Users/kcft114/.julia/packages/Flux/qXNjB/src/optimise/train.jl:105
┌ Info: Epoch 2910
└ @ Main /Users/kcft114/.julia/packages/Flux/qXNjB/src/optimise/train.jl:105
┌ Info: Epoch 2911
└ @ Main /Users/kcft114/.julia/packages/Flux/qXNjB/src/optimise/train.jl:105
┌ Info: Epoch 2912
└ @ Main /Users/kcft114/.julia/packages/Flux/qXNjB/src/optimise/train.jl:105
┌ Info: Epoch 2913
└ @ Main /Users/kcft114/.julia/packages/Flux/qXNjB/src/optimise/train.jl:105
┌ Info: Epoch 2914
└ @ Main /Users/kcft114/.julia/packages/Flux/qXNjB/src/optimise/train.jl:105
┌ Info: Epoch 2915
└ @ Main /Users/kcft114/.julia/packages/Flux/qXNjB/src/optimise/train.jl:105
┌ Info: Epoch 2916
└ @ Main /Users/kcft114/.julia/packages/Flux/qXNjB/src/optimise/train.jl:105
┌ Info: Epoch 2917
└ @ Main /Users/kcft114/.julia/packages/Flux/qXNjB/src/optimise/train.jl:105
┌ Info: Epoch 2918
└ @ Main /Users/kcft114/.julia/packages/Flux/qXNjB/src/optimise/train.jl:105
┌ Info: Epoch 2919
└ @ Main /Users/kcft114/.julia/packages/Flux/qXNjB/src/optimise/train.jl:105
┌ Info: Epoch 2920
└ @ Main /Users/kcft114/.julia/packages/Flux/qXNjB/src/optimise/train.jl:105
┌ Info: Epoch 2921
└ @ Main /Users/kcft114/.julia/packages/Flux/qXNjB/src/optimise/train.jl:105
┌ Info: Epoch 2922
└ @ Main /Users/kcft114/.julia/packages/Flux/qXNjB/src/optimise/train.jl:105
┌ Info: Epoch 2923
└ @ Main /Users/kcft114/.julia/packages/Flux/qXNjB/src/optimise/train.jl:105
┌ Info: Epoch 2924
└ @ Main /Users/kcft114/.julia/packages/Flux/qXNjB/src/optimise/train.jl:105
┌ Info: Epoch 2925
└ @ Main /Users/kcft114/.julia/packages/Flux/qXNjB/src/optimise/train.jl:105
┌ Info: Epoch 2926
└ @ Main /Users/kcft114/.julia/packages/Flux/qXNjB/src/optimise/train.jl:105
┌ Info: Epoch 2927
└ @ Main /Users/kcft114/.julia/packages/Flux/qXNjB/src/optimise/train.jl:105
┌ Info: Epoch 2928
└ @ Main /Users/kcft114/.julia/packages/Flux/qXNjB/src/optimise/train.jl:105
┌ Info: Epoch 2929
└ @ Main /Users/kcft114/.julia/packages/Flux/qXNjB/src/optimise/train.jl:105
┌ Info: Epoch 2930
└ @ Main /Users/kcft114/.julia/packages/Flux/qXNjB/src/optimise/train.jl:105
┌ Info: Epoch 2931
└ @ Main /Users/kcft114/.julia/packages/Flux/qXNjB/src/optimise/train.jl:105
┌ Info: Epoch 2932
└ @ Main /Users/kcft114/.julia/packages/Flux/qXNjB/src/optimise/train.jl:105
┌ Info: Epoch 2933
└ @ Main /Users/kcft114/.julia/packages/Flux/qXNjB/src/optimise/train.jl:105
┌ Info: Epoch 2934
└ @ Main /Users/kcft114/.julia/packages/Flux/qXNjB/src/optimise/train.jl:105
┌ Info: Epoch 2935
└ @ Main /Users/kcft114/.julia/packages/Flux/qXNjB/src/optimise/train.jl:105
┌ Info: Epoch 2936
└ @ Main /Users/kcft114/.julia/packages/Flux/qXNjB/src/optimise/train.jl:105
┌ Info: Epoch 2937
└ @ Main /Users/kcft114/.julia/packages/Flux/qXNjB/src/optimise/train.jl:105
┌ Info: Epoch 2938
└ @ Main /Users/kcft114/.julia/packages/Flux/qXNjB/src/optimise/train.jl:105
┌ Info: Epoch 2939
└ @ Main /Users/kcft114/.julia/packages/Flux/qXNjB/src/optimise/train.jl:105
┌ Info: Epoch 2940
└ @ Main /Users/kcft114/.julia/packages/Flux/qXNjB/src/optimise/train.jl:105
┌ Info: Epoch 2941
└ @ Main /Users/kcft114/.julia/packages/Flux/qXNjB/src/optimise/train.jl:105
┌ Info: Epoch 2942
└ @ Main /Users/kcft114/.julia/packages/Flux/qXNjB/src/optimise/train.jl:105
┌ Info: Epoch 2943
└ @ Main /Users/kcft114/.julia/packages/Flux/qXNjB/src/optimise/train.jl:105
┌ Info: Epoch 2944
└ @ Main /Users/kcft114/.julia/packages/Flux/qXNjB/src/optimise/train.jl:105
┌ Info: Epoch 2945
└ @ Main /Users/kcft114/.julia/packages/Flux/qXNjB/src/optimise/train.jl:105
┌ Info: Epoch 2946
└ @ Main /Users/kcft114/.julia/packages/Flux/qXNjB/src/optimise/train.jl:105
┌ Info: Epoch 2947
└ @ Main /Users/kcft114/.julia/packages/Flux/qXNjB/src/optimise/train.jl:105
┌ Info: Epoch 2948
└ @ Main /Users/kcft114/.julia/packages/Flux/qXNjB/src/optimise/train.jl:105
┌ Info: Epoch 2949
└ @ Main /Users/kcft114/.julia/packages/Flux/qXNjB/src/optimise/train.jl:105
┌ Info: Epoch 2950
└ @ Main /Users/kcft114/.julia/packages/Flux/qXNjB/src/optimise/train.jl:105
┌ Info: Epoch 2951
└ @ Main /Users/kcft114/.julia/packages/Flux/qXNjB/src/optimise/train.jl:105
┌ Info: Epoch 2952
└ @ Main /Users/kcft114/.julia/packages/Flux/qXNjB/src/optimise/train.jl:105
┌ Info: Epoch 2953
└ @ Main /Users/kcft114/.julia/packages/Flux/qXNjB/src/optimise/train.jl:105
┌ Info: Epoch 2954
└ @ Main /Users/kcft114/.julia/packages/Flux/qXNjB/src/optimise/train.jl:105
┌ Info: Epoch 2955
└ @ Main /Users/kcft114/.julia/packages/Flux/qXNjB/src/optimise/train.jl:105
┌ Info: Epoch 2956
└ @ Main /Users/kcft114/.julia/packages/Flux/qXNjB/src/optimise/train.jl:105
┌ Info: Epoch 2957
└ @ Main /Users/kcft114/.julia/packages/Flux/qXNjB/src/optimise/train.jl:105
┌ Info: Epoch 2958
└ @ Main /Users/kcft114/.julia/packages/Flux/qXNjB/src/optimise/train.jl:105
┌ Info: Epoch 2959
└ @ Main /Users/kcft114/.julia/packages/Flux/qXNjB/src/optimise/train.jl:105
┌ Info: Epoch 2960
└ @ Main /Users/kcft114/.julia/packages/Flux/qXNjB/src/optimise/train.jl:105
┌ Info: Epoch 2961
└ @ Main /Users/kcft114/.julia/packages/Flux/qXNjB/src/optimise/train.jl:105
┌ Info: Epoch 2962
└ @ Main /Users/kcft114/.julia/packages/Flux/qXNjB/src/optimise/train.jl:105
┌ Info: Epoch 2963
└ @ Main /Users/kcft114/.julia/packages/Flux/qXNjB/src/optimise/train.jl:105
┌ Info: Epoch 2964
└ @ Main /Users/kcft114/.julia/packages/Flux/qXNjB/src/optimise/train.jl:105
┌ Info: Epoch 2965
└ @ Main /Users/kcft114/.julia/packages/Flux/qXNjB/src/optimise/train.jl:105
┌ Info: Epoch 2966
└ @ Main /Users/kcft114/.julia/packages/Flux/qXNjB/src/optimise/train.jl:105
┌ Info: Epoch 2967
└ @ Main /Users/kcft114/.julia/packages/Flux/qXNjB/src/optimise/train.jl:105
┌ Info: Epoch 2968
└ @ Main /Users/kcft114/.julia/packages/Flux/qXNjB/src/optimise/train.jl:105
┌ Info: Epoch 2969
└ @ Main /Users/kcft114/.julia/packages/Flux/qXNjB/src/optimise/train.jl:105
┌ Info: Epoch 2970
└ @ Main /Users/kcft114/.julia/packages/Flux/qXNjB/src/optimise/train.jl:105
┌ Info: Epoch 2971
└ @ Main /Users/kcft114/.julia/packages/Flux/qXNjB/src/optimise/train.jl:105
┌ Info: Epoch 2972
└ @ Main /Users/kcft114/.julia/packages/Flux/qXNjB/src/optimise/train.jl:105
┌ Info: Epoch 2973
└ @ Main /Users/kcft114/.julia/packages/Flux/qXNjB/src/optimise/train.jl:105
┌ Info: Epoch 2974
└ @ Main /Users/kcft114/.julia/packages/Flux/qXNjB/src/optimise/train.jl:105
┌ Info: Epoch 2975
└ @ Main /Users/kcft114/.julia/packages/Flux/qXNjB/src/optimise/train.jl:105
┌ Info: Epoch 2976
└ @ Main /Users/kcft114/.julia/packages/Flux/qXNjB/src/optimise/train.jl:105
┌ Info: Epoch 2977
└ @ Main /Users/kcft114/.julia/packages/Flux/qXNjB/src/optimise/train.jl:105
┌ Info: Epoch 2978
└ @ Main /Users/kcft114/.julia/packages/Flux/qXNjB/src/optimise/train.jl:105
┌ Info: Epoch 2979
└ @ Main /Users/kcft114/.julia/packages/Flux/qXNjB/src/optimise/train.jl:105
┌ Info: Epoch 2980
└ @ Main /Users/kcft114/.julia/packages/Flux/qXNjB/src/optimise/train.jl:105
┌ Info: Epoch 2981
└ @ Main /Users/kcft114/.julia/packages/Flux/qXNjB/src/optimise/train.jl:105
┌ Info: Epoch 2982
└ @ Main /Users/kcft114/.julia/packages/Flux/qXNjB/src/optimise/train.jl:105
┌ Info: Epoch 2983
└ @ Main /Users/kcft114/.julia/packages/Flux/qXNjB/src/optimise/train.jl:105
┌ Info: Epoch 2984
└ @ Main /Users/kcft114/.julia/packages/Flux/qXNjB/src/optimise/train.jl:105
┌ Info: Epoch 2985
└ @ Main /Users/kcft114/.julia/packages/Flux/qXNjB/src/optimise/train.jl:105
┌ Info: Epoch 2986
└ @ Main /Users/kcft114/.julia/packages/Flux/qXNjB/src/optimise/train.jl:105
┌ Info: Epoch 2987
└ @ Main /Users/kcft114/.julia/packages/Flux/qXNjB/src/optimise/train.jl:105
┌ Info: Epoch 2988
└ @ Main /Users/kcft114/.julia/packages/Flux/qXNjB/src/optimise/train.jl:105
┌ Info: Epoch 2989
└ @ Main /Users/kcft114/.julia/packages/Flux/qXNjB/src/optimise/train.jl:105
┌ Info: Epoch 2990
└ @ Main /Users/kcft114/.julia/packages/Flux/qXNjB/src/optimise/train.jl:105
┌ Info: Epoch 2991
└ @ Main /Users/kcft114/.julia/packages/Flux/qXNjB/src/optimise/train.jl:105
┌ Info: Epoch 2992
└ @ Main /Users/kcft114/.julia/packages/Flux/qXNjB/src/optimise/train.jl:105
┌ Info: Epoch 2993
└ @ Main /Users/kcft114/.julia/packages/Flux/qXNjB/src/optimise/train.jl:105
┌ Info: Epoch 2994
└ @ Main /Users/kcft114/.julia/packages/Flux/qXNjB/src/optimise/train.jl:105
┌ Info: Epoch 2995
└ @ Main /Users/kcft114/.julia/packages/Flux/qXNjB/src/optimise/train.jl:105
┌ Info: Epoch 2996
└ @ Main /Users/kcft114/.julia/packages/Flux/qXNjB/src/optimise/train.jl:105
┌ Info: Epoch 2997
└ @ Main /Users/kcft114/.julia/packages/Flux/qXNjB/src/optimise/train.jl:105
┌ Info: Epoch 2998
└ @ Main /Users/kcft114/.julia/packages/Flux/qXNjB/src/optimise/train.jl:105
┌ Info: Epoch 2999
└ @ Main /Users/kcft114/.julia/packages/Flux/qXNjB/src/optimise/train.jl:105
┌ Info: Epoch 3000
└ @ Main /Users/kcft114/.julia/packages/Flux/qXNjB/src/optimise/train.jl:105
┌ Info: Epoch 3001
└ @ Main /Users/kcft114/.julia/packages/Flux/qXNjB/src/optimise/train.jl:105
┌ Info: Epoch 3002
└ @ Main /Users/kcft114/.julia/packages/Flux/qXNjB/src/optimise/train.jl:105
┌ Info: Epoch 3003
└ @ Main /Users/kcft114/.julia/packages/Flux/qXNjB/src/optimise/train.jl:105
┌ Info: Epoch 3004
└ @ Main /Users/kcft114/.julia/packages/Flux/qXNjB/src/optimise/train.jl:105
┌ Info: Epoch 3005
└ @ Main /Users/kcft114/.julia/packages/Flux/qXNjB/src/optimise/train.jl:105
┌ Info: Epoch 3006
└ @ Main /Users/kcft114/.julia/packages/Flux/qXNjB/src/optimise/train.jl:105
┌ Info: Epoch 3007
└ @ Main /Users/kcft114/.julia/packages/Flux/qXNjB/src/optimise/train.jl:105
┌ Info: Epoch 3008
└ @ Main /Users/kcft114/.julia/packages/Flux/qXNjB/src/optimise/train.jl:105
┌ Info: Epoch 3009
└ @ Main /Users/kcft114/.julia/packages/Flux/qXNjB/src/optimise/train.jl:105
┌ Info: Epoch 3010
└ @ Main /Users/kcft114/.julia/packages/Flux/qXNjB/src/optimise/train.jl:105
┌ Info: Epoch 3011
└ @ Main /Users/kcft114/.julia/packages/Flux/qXNjB/src/optimise/train.jl:105
┌ Info: Epoch 3012
└ @ Main /Users/kcft114/.julia/packages/Flux/qXNjB/src/optimise/train.jl:105
┌ Info: Epoch 3013
└ @ Main /Users/kcft114/.julia/packages/Flux/qXNjB/src/optimise/train.jl:105
┌ Info: Epoch 3014
└ @ Main /Users/kcft114/.julia/packages/Flux/qXNjB/src/optimise/train.jl:105
┌ Info: Epoch 3015
└ @ Main /Users/kcft114/.julia/packages/Flux/qXNjB/src/optimise/train.jl:105
┌ Info: Epoch 3016
└ @ Main /Users/kcft114/.julia/packages/Flux/qXNjB/src/optimise/train.jl:105
```

```
loss(X, Y) = 0.18817365f0 (tracked)
loss(X, Y) = 0.1880945f0 (tracked)
loss(X, Y) = 0.18802798f0 (tracked)
loss(X, Y) = 0.18795423f0 (tracked)
loss(X, Y) = 0.18788135f0 (tracked)
loss(X, Y) = 0.18779938f0 (tracked)
loss(X, Y) = 0.1877278f0 (tracked)
loss(X, Y) = 0.18765813f0 (tracked)
loss(X, Y) = 0.18758635f0 (tracked)
loss(X, Y) = 0.18752883f0 (tracked)
loss(X, Y) = 0.18745151f0 (tracked)
loss(X, Y) = 0.18736668f0 (tracked)
loss(X, Y) = 0.18730475f0 (tracked)
loss(X, Y) = 0.18722466f0 (tracked)
loss(X, Y) = 0.1871573f0 (tracked)
loss(X, Y) = 0.18708621f0 (tracked)
loss(X, Y) = 0.18701772f0 (tracked)
loss(X, Y) = 0.18693846f0 (tracked)
loss(X, Y) = 0.18686254f0 (tracked)
loss(X, Y) = 0.18679872f0 (tracked)
loss(X, Y) = 0.18674317f0 (tracked)
loss(X, Y) = 0.18666379f0 (tracked)
loss(X, Y) = 0.18659048f0 (tracked)
loss(X, Y) = 0.18652016f0 (tracked)
loss(X, Y) = 0.1864385f0 (tracked)
loss(X, Y) = 0.18638025f0 (tracked)
loss(X, Y) = 0.18631108f0 (tracked)
loss(X, Y) = 0.18622953f0 (tracked)
loss(X, Y) = 0.18614419f0 (tracked)
loss(X, Y) = 0.18609609f0 (tracked)
loss(X, Y) = 0.18602438f0 (tracked)
loss(X, Y) = 0.18594375f0 (tracked)
loss(X, Y) = 0.18587898f0 (tracked)
loss(X, Y) = 0.18581852f0 (tracked)
loss(X, Y) = 0.18574806f0 (tracked)
loss(X, Y) = 0.18566366f0 (tracked)
loss(X, Y) = 0.18558678f0 (tracked)
loss(X, Y) = 0.185519f0 (tracked)
loss(X, Y) = 0.18544665f0 (tracked)
loss(X, Y) = 0.18536791f0 (tracked)
loss(X, Y) = 0.18530758f0 (tracked)
loss(X, Y) = 0.18523672f0 (tracked)
loss(X, Y) = 0.18515156f0 (tracked)
loss(X, Y) = 0.18508612f0 (tracked)
loss(X, Y) = 0.1850171f0 (tracked)
loss(X, Y) = 0.18493362f0 (tracked)
loss(X, Y) = 0.18487646f0 (tracked)
loss(X, Y) = 0.18480489f0 (tracked)
loss(X, Y) = 0.18473901f0 (tracked)
loss(X, Y) = 0.1846561f0 (tracked)
loss(X, Y) = 0.1845852f0 (tracked)
loss(X, Y) = 0.18451113f0 (tracked)
loss(X, Y) = 0.18443762f0 (tracked)
loss(X, Y) = 0.18436444f0 (tracked)
loss(X, Y) = 0.18428421f0 (tracked)
loss(X, Y) = 0.18421945f0 (tracked)
loss(X, Y) = 0.18416436f0 (tracked)
loss(X, Y) = 0.18408819f0 (tracked)
loss(X, Y) = 0.18400404f0 (tracked)
loss(X, Y) = 0.18393509f0 (tracked)
loss(X, Y) = 0.18386252f0 (tracked)
loss(X, Y) = 0.18378893f0 (tracked)
loss(X, Y) = 0.18373348f0 (tracked)
loss(X, Y) = 0.18365423f0 (tracked)
loss(X, Y) = 0.18358143f0 (tracked)
loss(X, Y) = 0.1835213f0 (tracked)
loss(X, Y) = 0.18344142f0 (tracked)
loss(X, Y) = 0.18338536f0 (tracked)
loss(X, Y) = 0.18331437f0 (tracked)
loss(X, Y) = 0.18324707f0 (tracked)
loss(X, Y) = 0.1831822f0 (tracked)
loss(X, Y) = 0.1831198f0 (tracked)
loss(X, Y) = 0.1830437f0 (tracked)
loss(X, Y) = 0.18297718f0 (tracked)
loss(X, Y) = 0.18289828f0 (tracked)
loss(X, Y) = 0.1828199f0 (tracked)
loss(X, Y) = 0.18276186f0 (tracked)
loss(X, Y) = 0.1826964f0 (tracked)
loss(X, Y) = 0.18264352f0 (tracked)
loss(X, Y) = 0.18257165f0 (tracked)
loss(X, Y) = 0.18249905f0 (tracked)
loss(X, Y) = 0.18240888f0 (tracked)
loss(X, Y) = 0.18234977f0 (tracked)
loss(X, Y) = 0.18229355f0 (tracked)
loss(X, Y) = 0.18220012f0 (tracked)
loss(X, Y) = 0.18212712f0 (tracked)
loss(X, Y) = 0.1820721f0 (tracked)
loss(X, Y) = 0.18201318f0 (tracked)
loss(X, Y) = 0.18194084f0 (tracked)
loss(X, Y) = 0.18186752f0 (tracked)
loss(X, Y) = 0.18178627f0 (tracked)
loss(X, Y) = 0.18172628f0 (tracked)
loss(X, Y) = 0.18166211f0 (tracked)
loss(X, Y) = 0.18158597f0 (tracked)
loss(X, Y) = 0.18151292f0 (tracked)
loss(X, Y) = 0.18145595f0 (tracked)
loss(X, Y) = 0.1813787f0 (tracked)
loss(X, Y) = 0.18130967f0 (tracked)
loss(X, Y) = 0.1812551f0 (tracked)
loss(X, Y) = 0.18118429f0 (tracked)
loss(X, Y) = 0.18112244f0 (tracked)
loss(X, Y) = 0.18106173f0 (tracked)
loss(X, Y) = 0.18098515f0 (tracked)
loss(X, Y) = 0.18091244f0 (tracked)
loss(X, Y) = 0.1808495f0 (tracked)
loss(X, Y) = 0.18076232f0 (tracked)
loss(X, Y) = 0.18071029f0 (tracked)
loss(X, Y) = 0.18065734f0 (tracked)
loss(X, Y) = 0.18057458f0 (tracked)
loss(X, Y) = 0.18050534f0 (tracked)
loss(X, Y) = 0.18044996f0 (tracked)
loss(X, Y) = 0.1803836f0 (tracked)
loss(X, Y) = 0.18031724f0 (tracked)
loss(X, Y) = 0.18027222f0 (tracked)
loss(X, Y) = 0.18019629f0 (tracked)
```

```
┌ Info: Epoch 3017
└ @ Main /Users/kcft114/.julia/packages/Flux/qXNjB/src/optimise/train.jl:105
┌ Info: Epoch 3018
└ @ Main /Users/kcft114/.julia/packages/Flux/qXNjB/src/optimise/train.jl:105
┌ Info: Epoch 3019
└ @ Main /Users/kcft114/.julia/packages/Flux/qXNjB/src/optimise/train.jl:105
┌ Info: Epoch 3020
└ @ Main /Users/kcft114/.julia/packages/Flux/qXNjB/src/optimise/train.jl:105
┌ Info: Epoch 3021
└ @ Main /Users/kcft114/.julia/packages/Flux/qXNjB/src/optimise/train.jl:105
┌ Info: Epoch 3022
└ @ Main /Users/kcft114/.julia/packages/Flux/qXNjB/src/optimise/train.jl:105
┌ Info: Epoch 3023
└ @ Main /Users/kcft114/.julia/packages/Flux/qXNjB/src/optimise/train.jl:105
┌ Info: Epoch 3024
└ @ Main /Users/kcft114/.julia/packages/Flux/qXNjB/src/optimise/train.jl:105
┌ Info: Epoch 3025
└ @ Main /Users/kcft114/.julia/packages/Flux/qXNjB/src/optimise/train.jl:105
┌ Info: Epoch 3026
└ @ Main /Users/kcft114/.julia/packages/Flux/qXNjB/src/optimise/train.jl:105
┌ Info: Epoch 3027
└ @ Main /Users/kcft114/.julia/packages/Flux/qXNjB/src/optimise/train.jl:105
┌ Info: Epoch 3028
└ @ Main /Users/kcft114/.julia/packages/Flux/qXNjB/src/optimise/train.jl:105
┌ Info: Epoch 3029
└ @ Main /Users/kcft114/.julia/packages/Flux/qXNjB/src/optimise/train.jl:105
┌ Info: Epoch 3030
└ @ Main /Users/kcft114/.julia/packages/Flux/qXNjB/src/optimise/train.jl:105
┌ Info: Epoch 3031
└ @ Main /Users/kcft114/.julia/packages/Flux/qXNjB/src/optimise/train.jl:105
┌ Info: Epoch 3032
└ @ Main /Users/kcft114/.julia/packages/Flux/qXNjB/src/optimise/train.jl:105
┌ Info: Epoch 3033
└ @ Main /Users/kcft114/.julia/packages/Flux/qXNjB/src/optimise/train.jl:105
┌ Info: Epoch 3034
└ @ Main /Users/kcft114/.julia/packages/Flux/qXNjB/src/optimise/train.jl:105
┌ Info: Epoch 3035
└ @ Main /Users/kcft114/.julia/packages/Flux/qXNjB/src/optimise/train.jl:105
┌ Info: Epoch 3036
└ @ Main /Users/kcft114/.julia/packages/Flux/qXNjB/src/optimise/train.jl:105
┌ Info: Epoch 3037
└ @ Main /Users/kcft114/.julia/packages/Flux/qXNjB/src/optimise/train.jl:105
┌ Info: Epoch 3038
└ @ Main /Users/kcft114/.julia/packages/Flux/qXNjB/src/optimise/train.jl:105
┌ Info: Epoch 3039
└ @ Main /Users/kcft114/.julia/packages/Flux/qXNjB/src/optimise/train.jl:105
┌ Info: Epoch 3040
└ @ Main /Users/kcft114/.julia/packages/Flux/qXNjB/src/optimise/train.jl:105
┌ Info: Epoch 3041
└ @ Main /Users/kcft114/.julia/packages/Flux/qXNjB/src/optimise/train.jl:105
┌ Info: Epoch 3042
└ @ Main /Users/kcft114/.julia/packages/Flux/qXNjB/src/optimise/train.jl:105
┌ Info: Epoch 3043
└ @ Main /Users/kcft114/.julia/packages/Flux/qXNjB/src/optimise/train.jl:105
┌ Info: Epoch 3044
└ @ Main /Users/kcft114/.julia/packages/Flux/qXNjB/src/optimise/train.jl:105
┌ Info: Epoch 3045
└ @ Main /Users/kcft114/.julia/packages/Flux/qXNjB/src/optimise/train.jl:105
┌ Info: Epoch 3046
└ @ Main /Users/kcft114/.julia/packages/Flux/qXNjB/src/optimise/train.jl:105
┌ Info: Epoch 3047
└ @ Main /Users/kcft114/.julia/packages/Flux/qXNjB/src/optimise/train.jl:105
┌ Info: Epoch 3048
└ @ Main /Users/kcft114/.julia/packages/Flux/qXNjB/src/optimise/train.jl:105
┌ Info: Epoch 3049
└ @ Main /Users/kcft114/.julia/packages/Flux/qXNjB/src/optimise/train.jl:105
┌ Info: Epoch 3050
└ @ Main /Users/kcft114/.julia/packages/Flux/qXNjB/src/optimise/train.jl:105
┌ Info: Epoch 3051
└ @ Main /Users/kcft114/.julia/packages/Flux/qXNjB/src/optimise/train.jl:105
┌ Info: Epoch 3052
└ @ Main /Users/kcft114/.julia/packages/Flux/qXNjB/src/optimise/train.jl:105
┌ Info: Epoch 3053
└ @ Main /Users/kcft114/.julia/packages/Flux/qXNjB/src/optimise/train.jl:105
┌ Info: Epoch 3054
└ @ Main /Users/kcft114/.julia/packages/Flux/qXNjB/src/optimise/train.jl:105
┌ Info: Epoch 3055
└ @ Main /Users/kcft114/.julia/packages/Flux/qXNjB/src/optimise/train.jl:105
┌ Info: Epoch 3056
└ @ Main /Users/kcft114/.julia/packages/Flux/qXNjB/src/optimise/train.jl:105
┌ Info: Epoch 3057
└ @ Main /Users/kcft114/.julia/packages/Flux/qXNjB/src/optimise/train.jl:105
┌ Info: Epoch 3058
└ @ Main /Users/kcft114/.julia/packages/Flux/qXNjB/src/optimise/train.jl:105
┌ Info: Epoch 3059
└ @ Main /Users/kcft114/.julia/packages/Flux/qXNjB/src/optimise/train.jl:105
┌ Info: Epoch 3060
└ @ Main /Users/kcft114/.julia/packages/Flux/qXNjB/src/optimise/train.jl:105
┌ Info: Epoch 3061
└ @ Main /Users/kcft114/.julia/packages/Flux/qXNjB/src/optimise/train.jl:105
┌ Info: Epoch 3062
└ @ Main /Users/kcft114/.julia/packages/Flux/qXNjB/src/optimise/train.jl:105
┌ Info: Epoch 3063
└ @ Main /Users/kcft114/.julia/packages/Flux/qXNjB/src/optimise/train.jl:105
┌ Info: Epoch 3064
└ @ Main /Users/kcft114/.julia/packages/Flux/qXNjB/src/optimise/train.jl:105
┌ Info: Epoch 3065
└ @ Main /Users/kcft114/.julia/packages/Flux/qXNjB/src/optimise/train.jl:105
┌ Info: Epoch 3066
└ @ Main /Users/kcft114/.julia/packages/Flux/qXNjB/src/optimise/train.jl:105
┌ Info: Epoch 3067
└ @ Main /Users/kcft114/.julia/packages/Flux/qXNjB/src/optimise/train.jl:105
┌ Info: Epoch 3068
└ @ Main /Users/kcft114/.julia/packages/Flux/qXNjB/src/optimise/train.jl:105
┌ Info: Epoch 3069
└ @ Main /Users/kcft114/.julia/packages/Flux/qXNjB/src/optimise/train.jl:105
┌ Info: Epoch 3070
└ @ Main /Users/kcft114/.julia/packages/Flux/qXNjB/src/optimise/train.jl:105
┌ Info: Epoch 3071
└ @ Main /Users/kcft114/.julia/packages/Flux/qXNjB/src/optimise/train.jl:105
┌ Info: Epoch 3072
└ @ Main /Users/kcft114/.julia/packages/Flux/qXNjB/src/optimise/train.jl:105
┌ Info: Epoch 3073
└ @ Main /Users/kcft114/.julia/packages/Flux/qXNjB/src/optimise/train.jl:105
┌ Info: Epoch 3074
└ @ Main /Users/kcft114/.julia/packages/Flux/qXNjB/src/optimise/train.jl:105
┌ Info: Epoch 3075
└ @ Main /Users/kcft114/.julia/packages/Flux/qXNjB/src/optimise/train.jl:105
┌ Info: Epoch 3076
└ @ Main /Users/kcft114/.julia/packages/Flux/qXNjB/src/optimise/train.jl:105
┌ Info: Epoch 3077
└ @ Main /Users/kcft114/.julia/packages/Flux/qXNjB/src/optimise/train.jl:105
┌ Info: Epoch 3078
└ @ Main /Users/kcft114/.julia/packages/Flux/qXNjB/src/optimise/train.jl:105
┌ Info: Epoch 3079
└ @ Main /Users/kcft114/.julia/packages/Flux/qXNjB/src/optimise/train.jl:105
┌ Info: Epoch 3080
└ @ Main /Users/kcft114/.julia/packages/Flux/qXNjB/src/optimise/train.jl:105
┌ Info: Epoch 3081
└ @ Main /Users/kcft114/.julia/packages/Flux/qXNjB/src/optimise/train.jl:105
┌ Info: Epoch 3082
└ @ Main /Users/kcft114/.julia/packages/Flux/qXNjB/src/optimise/train.jl:105
┌ Info: Epoch 3083
└ @ Main /Users/kcft114/.julia/packages/Flux/qXNjB/src/optimise/train.jl:105
┌ Info: Epoch 3084
└ @ Main /Users/kcft114/.julia/packages/Flux/qXNjB/src/optimise/train.jl:105
┌ Info: Epoch 3085
└ @ Main /Users/kcft114/.julia/packages/Flux/qXNjB/src/optimise/train.jl:105
┌ Info: Epoch 3086
└ @ Main /Users/kcft114/.julia/packages/Flux/qXNjB/src/optimise/train.jl:105
┌ Info: Epoch 3087
└ @ Main /Users/kcft114/.julia/packages/Flux/qXNjB/src/optimise/train.jl:105
┌ Info: Epoch 3088
└ @ Main /Users/kcft114/.julia/packages/Flux/qXNjB/src/optimise/train.jl:105
┌ Info: Epoch 3089
└ @ Main /Users/kcft114/.julia/packages/Flux/qXNjB/src/optimise/train.jl:105
┌ Info: Epoch 3090
└ @ Main /Users/kcft114/.julia/packages/Flux/qXNjB/src/optimise/train.jl:105
┌ Info: Epoch 3091
└ @ Main /Users/kcft114/.julia/packages/Flux/qXNjB/src/optimise/train.jl:105
┌ Info: Epoch 3092
└ @ Main /Users/kcft114/.julia/packages/Flux/qXNjB/src/optimise/train.jl:105
┌ Info: Epoch 3093
└ @ Main /Users/kcft114/.julia/packages/Flux/qXNjB/src/optimise/train.jl:105
┌ Info: Epoch 3094
└ @ Main /Users/kcft114/.julia/packages/Flux/qXNjB/src/optimise/train.jl:105
┌ Info: Epoch 3095
└ @ Main /Users/kcft114/.julia/packages/Flux/qXNjB/src/optimise/train.jl:105
┌ Info: Epoch 3096
└ @ Main /Users/kcft114/.julia/packages/Flux/qXNjB/src/optimise/train.jl:105
┌ Info: Epoch 3097
└ @ Main /Users/kcft114/.julia/packages/Flux/qXNjB/src/optimise/train.jl:105
┌ Info: Epoch 3098
└ @ Main /Users/kcft114/.julia/packages/Flux/qXNjB/src/optimise/train.jl:105
┌ Info: Epoch 3099
└ @ Main /Users/kcft114/.julia/packages/Flux/qXNjB/src/optimise/train.jl:105
┌ Info: Epoch 3100
└ @ Main /Users/kcft114/.julia/packages/Flux/qXNjB/src/optimise/train.jl:105
┌ Info: Epoch 3101
└ @ Main /Users/kcft114/.julia/packages/Flux/qXNjB/src/optimise/train.jl:105
┌ Info: Epoch 3102
└ @ Main /Users/kcft114/.julia/packages/Flux/qXNjB/src/optimise/train.jl:105
┌ Info: Epoch 3103
└ @ Main /Users/kcft114/.julia/packages/Flux/qXNjB/src/optimise/train.jl:105
┌ Info: Epoch 3104
└ @ Main /Users/kcft114/.julia/packages/Flux/qXNjB/src/optimise/train.jl:105
┌ Info: Epoch 3105
└ @ Main /Users/kcft114/.julia/packages/Flux/qXNjB/src/optimise/train.jl:105
┌ Info: Epoch 3106
└ @ Main /Users/kcft114/.julia/packages/Flux/qXNjB/src/optimise/train.jl:105
┌ Info: Epoch 3107
└ @ Main /Users/kcft114/.julia/packages/Flux/qXNjB/src/optimise/train.jl:105
┌ Info: Epoch 3108
└ @ Main /Users/kcft114/.julia/packages/Flux/qXNjB/src/optimise/train.jl:105
┌ Info: Epoch 3109
└ @ Main /Users/kcft114/.julia/packages/Flux/qXNjB/src/optimise/train.jl:105
┌ Info: Epoch 3110
└ @ Main /Users/kcft114/.julia/packages/Flux/qXNjB/src/optimise/train.jl:105
┌ Info: Epoch 3111
└ @ Main /Users/kcft114/.julia/packages/Flux/qXNjB/src/optimise/train.jl:105
┌ Info: Epoch 3112
└ @ Main /Users/kcft114/.julia/packages/Flux/qXNjB/src/optimise/train.jl:105
┌ Info: Epoch 3113
└ @ Main /Users/kcft114/.julia/packages/Flux/qXNjB/src/optimise/train.jl:105
┌ Info: Epoch 3114
└ @ Main /Users/kcft114/.julia/packages/Flux/qXNjB/src/optimise/train.jl:105
┌ Info: Epoch 3115
└ @ Main /Users/kcft114/.julia/packages/Flux/qXNjB/src/optimise/train.jl:105
┌ Info: Epoch 3116
└ @ Main /Users/kcft114/.julia/packages/Flux/qXNjB/src/optimise/train.jl:105
┌ Info: Epoch 3117
└ @ Main /Users/kcft114/.julia/packages/Flux/qXNjB/src/optimise/train.jl:105
┌ Info: Epoch 3118
└ @ Main /Users/kcft114/.julia/packages/Flux/qXNjB/src/optimise/train.jl:105
┌ Info: Epoch 3119
└ @ Main /Users/kcft114/.julia/packages/Flux/qXNjB/src/optimise/train.jl:105
┌ Info: Epoch 3120
└ @ Main /Users/kcft114/.julia/packages/Flux/qXNjB/src/optimise/train.jl:105
┌ Info: Epoch 3121
└ @ Main /Users/kcft114/.julia/packages/Flux/qXNjB/src/optimise/train.jl:105
┌ Info: Epoch 3122
└ @ Main /Users/kcft114/.julia/packages/Flux/qXNjB/src/optimise/train.jl:105
┌ Info: Epoch 3123
└ @ Main /Users/kcft114/.julia/packages/Flux/qXNjB/src/optimise/train.jl:105
┌ Info: Epoch 3124
└ @ Main /Users/kcft114/.julia/packages/Flux/qXNjB/src/optimise/train.jl:105
┌ Info: Epoch 3125
└ @ Main /Users/kcft114/.julia/packages/Flux/qXNjB/src/optimise/train.jl:105
┌ Info: Epoch 3126
└ @ Main /Users/kcft114/.julia/packages/Flux/qXNjB/src/optimise/train.jl:105
┌ Info: Epoch 3127
└ @ Main /Users/kcft114/.julia/packages/Flux/qXNjB/src/optimise/train.jl:105
┌ Info: Epoch 3128
└ @ Main /Users/kcft114/.julia/packages/Flux/qXNjB/src/optimise/train.jl:105
┌ Info: Epoch 3129
└ @ Main /Users/kcft114/.julia/packages/Flux/qXNjB/src/optimise/train.jl:105
┌ Info: Epoch 3130
└ @ Main /Users/kcft114/.julia/packages/Flux/qXNjB/src/optimise/train.jl:105
┌ Info: Epoch 3131
└ @ Main /Users/kcft114/.julia/packages/Flux/qXNjB/src/optimise/train.jl:105
```

```
loss(X, Y) = 0.18010952f0 (tracked)
loss(X, Y) = 0.18003224f0 (tracked)
loss(X, Y) = 0.17997262f0 (tracked)
loss(X, Y) = 0.17991255f0 (tracked)
loss(X, Y) = 0.17983884f0 (tracked)
loss(X, Y) = 0.17976269f0 (tracked)
loss(X, Y) = 0.17971586f0 (tracked)
loss(X, Y) = 0.1796483f0 (tracked)
loss(X, Y) = 0.17957774f0 (tracked)
loss(X, Y) = 0.17951289f0 (tracked)
loss(X, Y) = 0.17944258f0 (tracked)
loss(X, Y) = 0.17936821f0 (tracked)
loss(X, Y) = 0.17930827f0 (tracked)
loss(X, Y) = 0.17923582f0 (tracked)
loss(X, Y) = 0.17916822f0 (tracked)
loss(X, Y) = 0.17910086f0 (tracked)
loss(X, Y) = 0.17903817f0 (tracked)
loss(X, Y) = 0.17897677f0 (tracked)
loss(X, Y) = 0.17889929f0 (tracked)
loss(X, Y) = 0.17881931f0 (tracked)
loss(X, Y) = 0.17876151f0 (tracked)
loss(X, Y) = 0.17869785f0 (tracked)
loss(X, Y) = 0.17863154f0 (tracked)
loss(X, Y) = 0.17856802f0 (tracked)
loss(X, Y) = 0.17849693f0 (tracked)
loss(X, Y) = 0.1784357f0 (tracked)
loss(X, Y) = 0.17836308f0 (tracked)
loss(X, Y) = 0.1782869f0 (tracked)
loss(X, Y) = 0.17824496f0 (tracked)
loss(X, Y) = 0.1781804f0 (tracked)
loss(X, Y) = 0.178108f0 (tracked)
loss(X, Y) = 0.178045f0 (tracked)
loss(X, Y) = 0.17798215f0 (tracked)
loss(X, Y) = 0.1779419f0 (tracked)
loss(X, Y) = 0.17785336f0 (tracked)
loss(X, Y) = 0.17778602f0 (tracked)
loss(X, Y) = 0.1777115f0 (tracked)
loss(X, Y) = 0.17765188f0 (tracked)
loss(X, Y) = 0.17759778f0 (tracked)
loss(X, Y) = 0.17752191f0 (tracked)
loss(X, Y) = 0.17746064f0 (tracked)
loss(X, Y) = 0.17740138f0 (tracked)
loss(X, Y) = 0.1773289f0 (tracked)
loss(X, Y) = 0.17726186f0 (tracked)
loss(X, Y) = 0.17719974f0 (tracked)
loss(X, Y) = 0.17714281f0 (tracked)
loss(X, Y) = 0.17706242f0 (tracked)
loss(X, Y) = 0.1769917f0 (tracked)
loss(X, Y) = 0.1769531f0 (tracked)
loss(X, Y) = 0.17689796f0 (tracked)
loss(X, Y) = 0.17681435f0 (tracked)
loss(X, Y) = 0.1767445f0 (tracked)
loss(X, Y) = 0.17669372f0 (tracked)
loss(X, Y) = 0.17663082f0 (tracked)
loss(X, Y) = 0.17656375f0 (tracked)
loss(X, Y) = 0.17649733f0 (tracked)
loss(X, Y) = 0.1764305f0 (tracked)
loss(X, Y) = 0.17636466f0 (tracked)
loss(X, Y) = 0.1763069f0 (tracked)
loss(X, Y) = 0.17622936f0 (tracked)
loss(X, Y) = 0.17616439f0 (tracked)
loss(X, Y) = 0.17610912f0 (tracked)
loss(X, Y) = 0.17604077f0 (tracked)
loss(X, Y) = 0.17596404f0 (tracked)
loss(X, Y) = 0.17590933f0 (tracked)
loss(X, Y) = 0.1758548f0 (tracked)
loss(X, Y) = 0.17579757f0 (tracked)
loss(X, Y) = 0.17572562f0 (tracked)
loss(X, Y) = 0.17566419f0 (tracked)
loss(X, Y) = 0.17559625f0 (tracked)
loss(X, Y) = 0.17553134f0 (tracked)
loss(X, Y) = 0.1754665f0 (tracked)
loss(X, Y) = 0.1753911f0 (tracked)
loss(X, Y) = 0.17532977f0 (tracked)
loss(X, Y) = 0.17526415f0 (tracked)
loss(X, Y) = 0.17520523f0 (tracked)
loss(X, Y) = 0.17516048f0 (tracked)
loss(X, Y) = 0.17507006f0 (tracked)
loss(X, Y) = 0.1750226f0 (tracked)
loss(X, Y) = 0.17496361f0 (tracked)
loss(X, Y) = 0.17488964f0 (tracked)
loss(X, Y) = 0.17482698f0 (tracked)
loss(X, Y) = 0.174778f0 (tracked)
loss(X, Y) = 0.17471498f0 (tracked)
loss(X, Y) = 0.17464626f0 (tracked)
loss(X, Y) = 0.17459227f0 (tracked)
loss(X, Y) = 0.17451805f0 (tracked)
loss(X, Y) = 0.17444561f0 (tracked)
loss(X, Y) = 0.17439125f0 (tracked)
loss(X, Y) = 0.17433725f0 (tracked)
loss(X, Y) = 0.17427716f0 (tracked)
loss(X, Y) = 0.17422137f0 (tracked)
loss(X, Y) = 0.17414849f0 (tracked)
loss(X, Y) = 0.17406516f0 (tracked)
loss(X, Y) = 0.17400402f0 (tracked)
loss(X, Y) = 0.17394443f0 (tracked)
loss(X, Y) = 0.17387187f0 (tracked)
loss(X, Y) = 0.17382112f0 (tracked)
loss(X, Y) = 0.17377675f0 (tracked)
loss(X, Y) = 0.17369233f0 (tracked)
loss(X, Y) = 0.17362916f0 (tracked)
loss(X, Y) = 0.17357036f0 (tracked)
loss(X, Y) = 0.17351092f0 (tracked)
loss(X, Y) = 0.17344253f0 (tracked)
loss(X, Y) = 0.17338502f0 (tracked)
loss(X, Y) = 0.17332365f0 (tracked)
loss(X, Y) = 0.17325503f0 (tracked)
loss(X, Y) = 0.17319001f0 (tracked)
loss(X, Y) = 0.17313938f0 (tracked)
loss(X, Y) = 0.17306669f0 (tracked)
loss(X, Y) = 0.17299272f0 (tracked)
loss(X, Y) = 0.17293642f0 (tracked)
loss(X, Y) = 0.172892f0 (tracked)
loss(X, Y) = 0.17281705f0 (tracked)
loss(X, Y) = 0.17274994f0 (tracked)
loss(X, Y) = 0.17268865f0 (tracked)
loss(X, Y) = 0.17263074f0 (tracked)
loss(X, Y) = 0.1725629f0 (tracked)
loss(X, Y) = 0.17248839f0 (tracked)
loss(X, Y) = 0.17242475f0 (tracked)
```

```
┌ Info: Epoch 3132
└ @ Main /Users/kcft114/.julia/packages/Flux/qXNjB/src/optimise/train.jl:105
┌ Info: Epoch 3133
└ @ Main /Users/kcft114/.julia/packages/Flux/qXNjB/src/optimise/train.jl:105
┌ Info: Epoch 3134
└ @ Main /Users/kcft114/.julia/packages/Flux/qXNjB/src/optimise/train.jl:105
┌ Info: Epoch 3135
└ @ Main /Users/kcft114/.julia/packages/Flux/qXNjB/src/optimise/train.jl:105
┌ Info: Epoch 3136
└ @ Main /Users/kcft114/.julia/packages/Flux/qXNjB/src/optimise/train.jl:105
┌ Info: Epoch 3137
└ @ Main /Users/kcft114/.julia/packages/Flux/qXNjB/src/optimise/train.jl:105
┌ Info: Epoch 3138
└ @ Main /Users/kcft114/.julia/packages/Flux/qXNjB/src/optimise/train.jl:105
┌ Info: Epoch 3139
└ @ Main /Users/kcft114/.julia/packages/Flux/qXNjB/src/optimise/train.jl:105
┌ Info: Epoch 3140
└ @ Main /Users/kcft114/.julia/packages/Flux/qXNjB/src/optimise/train.jl:105
┌ Info: Epoch 3141
└ @ Main /Users/kcft114/.julia/packages/Flux/qXNjB/src/optimise/train.jl:105
┌ Info: Epoch 3142
└ @ Main /Users/kcft114/.julia/packages/Flux/qXNjB/src/optimise/train.jl:105
┌ Info: Epoch 3143
└ @ Main /Users/kcft114/.julia/packages/Flux/qXNjB/src/optimise/train.jl:105
┌ Info: Epoch 3144
└ @ Main /Users/kcft114/.julia/packages/Flux/qXNjB/src/optimise/train.jl:105
┌ Info: Epoch 3145
└ @ Main /Users/kcft114/.julia/packages/Flux/qXNjB/src/optimise/train.jl:105
┌ Info: Epoch 3146
└ @ Main /Users/kcft114/.julia/packages/Flux/qXNjB/src/optimise/train.jl:105
┌ Info: Epoch 3147
└ @ Main /Users/kcft114/.julia/packages/Flux/qXNjB/src/optimise/train.jl:105
┌ Info: Epoch 3148
└ @ Main /Users/kcft114/.julia/packages/Flux/qXNjB/src/optimise/train.jl:105
┌ Info: Epoch 3149
└ @ Main /Users/kcft114/.julia/packages/Flux/qXNjB/src/optimise/train.jl:105
┌ Info: Epoch 3150
└ @ Main /Users/kcft114/.julia/packages/Flux/qXNjB/src/optimise/train.jl:105
┌ Info: Epoch 3151
└ @ Main /Users/kcft114/.julia/packages/Flux/qXNjB/src/optimise/train.jl:105
┌ Info: Epoch 3152
└ @ Main /Users/kcft114/.julia/packages/Flux/qXNjB/src/optimise/train.jl:105
┌ Info: Epoch 3153
└ @ Main /Users/kcft114/.julia/packages/Flux/qXNjB/src/optimise/train.jl:105
┌ Info: Epoch 3154
└ @ Main /Users/kcft114/.julia/packages/Flux/qXNjB/src/optimise/train.jl:105
┌ Info: Epoch 3155
└ @ Main /Users/kcft114/.julia/packages/Flux/qXNjB/src/optimise/train.jl:105
┌ Info: Epoch 3156
└ @ Main /Users/kcft114/.julia/packages/Flux/qXNjB/src/optimise/train.jl:105
┌ Info: Epoch 3157
└ @ Main /Users/kcft114/.julia/packages/Flux/qXNjB/src/optimise/train.jl:105
┌ Info: Epoch 3158
└ @ Main /Users/kcft114/.julia/packages/Flux/qXNjB/src/optimise/train.jl:105
┌ Info: Epoch 3159
└ @ Main /Users/kcft114/.julia/packages/Flux/qXNjB/src/optimise/train.jl:105
┌ Info: Epoch 3160
└ @ Main /Users/kcft114/.julia/packages/Flux/qXNjB/src/optimise/train.jl:105
┌ Info: Epoch 3161
└ @ Main /Users/kcft114/.julia/packages/Flux/qXNjB/src/optimise/train.jl:105
┌ Info: Epoch 3162
└ @ Main /Users/kcft114/.julia/packages/Flux/qXNjB/src/optimise/train.jl:105
┌ Info: Epoch 3163
└ @ Main /Users/kcft114/.julia/packages/Flux/qXNjB/src/optimise/train.jl:105
┌ Info: Epoch 3164
└ @ Main /Users/kcft114/.julia/packages/Flux/qXNjB/src/optimise/train.jl:105
┌ Info: Epoch 3165
└ @ Main /Users/kcft114/.julia/packages/Flux/qXNjB/src/optimise/train.jl:105
┌ Info: Epoch 3166
└ @ Main /Users/kcft114/.julia/packages/Flux/qXNjB/src/optimise/train.jl:105
┌ Info: Epoch 3167
└ @ Main /Users/kcft114/.julia/packages/Flux/qXNjB/src/optimise/train.jl:105
┌ Info: Epoch 3168
└ @ Main /Users/kcft114/.julia/packages/Flux/qXNjB/src/optimise/train.jl:105
┌ Info: Epoch 3169
└ @ Main /Users/kcft114/.julia/packages/Flux/qXNjB/src/optimise/train.jl:105
┌ Info: Epoch 3170
└ @ Main /Users/kcft114/.julia/packages/Flux/qXNjB/src/optimise/train.jl:105
┌ Info: Epoch 3171
└ @ Main /Users/kcft114/.julia/packages/Flux/qXNjB/src/optimise/train.jl:105
┌ Info: Epoch 3172
└ @ Main /Users/kcft114/.julia/packages/Flux/qXNjB/src/optimise/train.jl:105
┌ Info: Epoch 3173
└ @ Main /Users/kcft114/.julia/packages/Flux/qXNjB/src/optimise/train.jl:105
┌ Info: Epoch 3174
└ @ Main /Users/kcft114/.julia/packages/Flux/qXNjB/src/optimise/train.jl:105
┌ Info: Epoch 3175
└ @ Main /Users/kcft114/.julia/packages/Flux/qXNjB/src/optimise/train.jl:105
┌ Info: Epoch 3176
└ @ Main /Users/kcft114/.julia/packages/Flux/qXNjB/src/optimise/train.jl:105
┌ Info: Epoch 3177
└ @ Main /Users/kcft114/.julia/packages/Flux/qXNjB/src/optimise/train.jl:105
┌ Info: Epoch 3178
└ @ Main /Users/kcft114/.julia/packages/Flux/qXNjB/src/optimise/train.jl:105
┌ Info: Epoch 3179
└ @ Main /Users/kcft114/.julia/packages/Flux/qXNjB/src/optimise/train.jl:105
┌ Info: Epoch 3180
└ @ Main /Users/kcft114/.julia/packages/Flux/qXNjB/src/optimise/train.jl:105
┌ Info: Epoch 3181
└ @ Main /Users/kcft114/.julia/packages/Flux/qXNjB/src/optimise/train.jl:105
┌ Info: Epoch 3182
└ @ Main /Users/kcft114/.julia/packages/Flux/qXNjB/src/optimise/train.jl:105
┌ Info: Epoch 3183
└ @ Main /Users/kcft114/.julia/packages/Flux/qXNjB/src/optimise/train.jl:105
┌ Info: Epoch 3184
└ @ Main /Users/kcft114/.julia/packages/Flux/qXNjB/src/optimise/train.jl:105
┌ Info: Epoch 3185
└ @ Main /Users/kcft114/.julia/packages/Flux/qXNjB/src/optimise/train.jl:105
┌ Info: Epoch 3186
└ @ Main /Users/kcft114/.julia/packages/Flux/qXNjB/src/optimise/train.jl:105
┌ Info: Epoch 3187
└ @ Main /Users/kcft114/.julia/packages/Flux/qXNjB/src/optimise/train.jl:105
┌ Info: Epoch 3188
└ @ Main /Users/kcft114/.julia/packages/Flux/qXNjB/src/optimise/train.jl:105
┌ Info: Epoch 3189
└ @ Main /Users/kcft114/.julia/packages/Flux/qXNjB/src/optimise/train.jl:105
┌ Info: Epoch 3190
└ @ Main /Users/kcft114/.julia/packages/Flux/qXNjB/src/optimise/train.jl:105
┌ Info: Epoch 3191
└ @ Main /Users/kcft114/.julia/packages/Flux/qXNjB/src/optimise/train.jl:105
┌ Info: Epoch 3192
└ @ Main /Users/kcft114/.julia/packages/Flux/qXNjB/src/optimise/train.jl:105
┌ Info: Epoch 3193
└ @ Main /Users/kcft114/.julia/packages/Flux/qXNjB/src/optimise/train.jl:105
┌ Info: Epoch 3194
└ @ Main /Users/kcft114/.julia/packages/Flux/qXNjB/src/optimise/train.jl:105
┌ Info: Epoch 3195
└ @ Main /Users/kcft114/.julia/packages/Flux/qXNjB/src/optimise/train.jl:105
┌ Info: Epoch 3196
└ @ Main /Users/kcft114/.julia/packages/Flux/qXNjB/src/optimise/train.jl:105
┌ Info: Epoch 3197
└ @ Main /Users/kcft114/.julia/packages/Flux/qXNjB/src/optimise/train.jl:105
┌ Info: Epoch 3198
└ @ Main /Users/kcft114/.julia/packages/Flux/qXNjB/src/optimise/train.jl:105
┌ Info: Epoch 3199
└ @ Main /Users/kcft114/.julia/packages/Flux/qXNjB/src/optimise/train.jl:105
┌ Info: Epoch 3200
└ @ Main /Users/kcft114/.julia/packages/Flux/qXNjB/src/optimise/train.jl:105
┌ Info: Epoch 3201
└ @ Main /Users/kcft114/.julia/packages/Flux/qXNjB/src/optimise/train.jl:105
┌ Info: Epoch 3202
└ @ Main /Users/kcft114/.julia/packages/Flux/qXNjB/src/optimise/train.jl:105
┌ Info: Epoch 3203
└ @ Main /Users/kcft114/.julia/packages/Flux/qXNjB/src/optimise/train.jl:105
┌ Info: Epoch 3204
└ @ Main /Users/kcft114/.julia/packages/Flux/qXNjB/src/optimise/train.jl:105
┌ Info: Epoch 3205
└ @ Main /Users/kcft114/.julia/packages/Flux/qXNjB/src/optimise/train.jl:105
┌ Info: Epoch 3206
└ @ Main /Users/kcft114/.julia/packages/Flux/qXNjB/src/optimise/train.jl:105
┌ Info: Epoch 3207
└ @ Main /Users/kcft114/.julia/packages/Flux/qXNjB/src/optimise/train.jl:105
┌ Info: Epoch 3208
└ @ Main /Users/kcft114/.julia/packages/Flux/qXNjB/src/optimise/train.jl:105
┌ Info: Epoch 3209
└ @ Main /Users/kcft114/.julia/packages/Flux/qXNjB/src/optimise/train.jl:105
┌ Info: Epoch 3210
└ @ Main /Users/kcft114/.julia/packages/Flux/qXNjB/src/optimise/train.jl:105
┌ Info: Epoch 3211
└ @ Main /Users/kcft114/.julia/packages/Flux/qXNjB/src/optimise/train.jl:105
┌ Info: Epoch 3212
└ @ Main /Users/kcft114/.julia/packages/Flux/qXNjB/src/optimise/train.jl:105
┌ Info: Epoch 3213
└ @ Main /Users/kcft114/.julia/packages/Flux/qXNjB/src/optimise/train.jl:105
┌ Info: Epoch 3214
└ @ Main /Users/kcft114/.julia/packages/Flux/qXNjB/src/optimise/train.jl:105
┌ Info: Epoch 3215
└ @ Main /Users/kcft114/.julia/packages/Flux/qXNjB/src/optimise/train.jl:105
┌ Info: Epoch 3216
└ @ Main /Users/kcft114/.julia/packages/Flux/qXNjB/src/optimise/train.jl:105
┌ Info: Epoch 3217
└ @ Main /Users/kcft114/.julia/packages/Flux/qXNjB/src/optimise/train.jl:105
┌ Info: Epoch 3218
└ @ Main /Users/kcft114/.julia/packages/Flux/qXNjB/src/optimise/train.jl:105
┌ Info: Epoch 3219
└ @ Main /Users/kcft114/.julia/packages/Flux/qXNjB/src/optimise/train.jl:105
┌ Info: Epoch 3220
└ @ Main /Users/kcft114/.julia/packages/Flux/qXNjB/src/optimise/train.jl:105
┌ Info: Epoch 3221
└ @ Main /Users/kcft114/.julia/packages/Flux/qXNjB/src/optimise/train.jl:105
┌ Info: Epoch 3222
└ @ Main /Users/kcft114/.julia/packages/Flux/qXNjB/src/optimise/train.jl:105
┌ Info: Epoch 3223
└ @ Main /Users/kcft114/.julia/packages/Flux/qXNjB/src/optimise/train.jl:105
┌ Info: Epoch 3224
└ @ Main /Users/kcft114/.julia/packages/Flux/qXNjB/src/optimise/train.jl:105
┌ Info: Epoch 3225
└ @ Main /Users/kcft114/.julia/packages/Flux/qXNjB/src/optimise/train.jl:105
┌ Info: Epoch 3226
└ @ Main /Users/kcft114/.julia/packages/Flux/qXNjB/src/optimise/train.jl:105
┌ Info: Epoch 3227
└ @ Main /Users/kcft114/.julia/packages/Flux/qXNjB/src/optimise/train.jl:105
┌ Info: Epoch 3228
└ @ Main /Users/kcft114/.julia/packages/Flux/qXNjB/src/optimise/train.jl:105
┌ Info: Epoch 3229
└ @ Main /Users/kcft114/.julia/packages/Flux/qXNjB/src/optimise/train.jl:105
┌ Info: Epoch 3230
└ @ Main /Users/kcft114/.julia/packages/Flux/qXNjB/src/optimise/train.jl:105
┌ Info: Epoch 3231
└ @ Main /Users/kcft114/.julia/packages/Flux/qXNjB/src/optimise/train.jl:105
┌ Info: Epoch 3232
└ @ Main /Users/kcft114/.julia/packages/Flux/qXNjB/src/optimise/train.jl:105
┌ Info: Epoch 3233
└ @ Main /Users/kcft114/.julia/packages/Flux/qXNjB/src/optimise/train.jl:105
┌ Info: Epoch 3234
└ @ Main /Users/kcft114/.julia/packages/Flux/qXNjB/src/optimise/train.jl:105
┌ Info: Epoch 3235
└ @ Main /Users/kcft114/.julia/packages/Flux/qXNjB/src/optimise/train.jl:105
┌ Info: Epoch 3236
└ @ Main /Users/kcft114/.julia/packages/Flux/qXNjB/src/optimise/train.jl:105
┌ Info: Epoch 3237
└ @ Main /Users/kcft114/.julia/packages/Flux/qXNjB/src/optimise/train.jl:105
┌ Info: Epoch 3238
└ @ Main /Users/kcft114/.julia/packages/Flux/qXNjB/src/optimise/train.jl:105
┌ Info: Epoch 3239
└ @ Main /Users/kcft114/.julia/packages/Flux/qXNjB/src/optimise/train.jl:105
┌ Info: Epoch 3240
└ @ Main /Users/kcft114/.julia/packages/Flux/qXNjB/src/optimise/train.jl:105
┌ Info: Epoch 3241
└ @ Main /Users/kcft114/.julia/packages/Flux/qXNjB/src/optimise/train.jl:105
┌ Info: Epoch 3242
└ @ Main /Users/kcft114/.julia/packages/Flux/qXNjB/src/optimise/train.jl:105
┌ Info: Epoch 3243
└ @ Main /Users/kcft114/.julia/packages/Flux/qXNjB/src/optimise/train.jl:105
┌ Info: Epoch 3244
└ @ Main /Users/kcft114/.julia/packages/Flux/qXNjB/src/optimise/train.jl:105
┌ Info: Epoch 3245
└ @ Main /Users/kcft114/.julia/packages/Flux/qXNjB/src/optimise/train.jl:105
┌ Info: Epoch 3246
└ @ Main /Users/kcft114/.julia/packages/Flux/qXNjB/src/optimise/train.jl:105
┌ Info: Epoch 3247
└ @ Main /Users/kcft114/.julia/packages/Flux/qXNjB/src/optimise/train.jl:105
┌ Info: Epoch 3248
└ @ Main /Users/kcft114/.julia/packages/Flux/qXNjB/src/optimise/train.jl:105
┌ Info: Epoch 3249
└ @ Main /Users/kcft114/.julia/packages/Flux/qXNjB/src/optimise/train.jl:105
┌ Info: Epoch 3250
└ @ Main /Users/kcft114/.julia/packages/Flux/qXNjB/src/optimise/train.jl:105
```

```
loss(X, Y) = 0.1723905f0 (tracked)
loss(X, Y) = 0.17232777f0 (tracked)
loss(X, Y) = 0.172244f0 (tracked)
loss(X, Y) = 0.17218514f0 (tracked)
loss(X, Y) = 0.17212586f0 (tracked)
loss(X, Y) = 0.17205818f0 (tracked)
loss(X, Y) = 0.17201398f0 (tracked)
loss(X, Y) = 0.17194943f0 (tracked)
loss(X, Y) = 0.17187089f0 (tracked)
loss(X, Y) = 0.17182949f0 (tracked)
loss(X, Y) = 0.17176068f0 (tracked)
loss(X, Y) = 0.17169383f0 (tracked)
loss(X, Y) = 0.17163394f0 (tracked)
loss(X, Y) = 0.17157513f0 (tracked)
loss(X, Y) = 0.17151654f0 (tracked)
loss(X, Y) = 0.17144924f0 (tracked)
loss(X, Y) = 0.17137814f0 (tracked)
loss(X, Y) = 0.17131601f0 (tracked)
loss(X, Y) = 0.17127077f0 (tracked)
loss(X, Y) = 0.1712078f0 (tracked)
loss(X, Y) = 0.17114338f0 (tracked)
loss(X, Y) = 0.17108914f0 (tracked)
loss(X, Y) = 0.17101541f0 (tracked)
loss(X, Y) = 0.1709865f0 (tracked)
loss(X, Y) = 0.17090538f0 (tracked)
loss(X, Y) = 0.17081688f0 (tracked)
loss(X, Y) = 0.17077254f0 (tracked)
loss(X, Y) = 0.17072396f0 (tracked)
loss(X, Y) = 0.17067277f0 (tracked)
loss(X, Y) = 0.17060305f0 (tracked)
loss(X, Y) = 0.1705483f0 (tracked)
loss(X, Y) = 0.17048788f0 (tracked)
loss(X, Y) = 0.17042209f0 (tracked)
loss(X, Y) = 0.17036025f0 (tracked)
loss(X, Y) = 0.17028774f0 (tracked)
loss(X, Y) = 0.17022303f0 (tracked)
loss(X, Y) = 0.17015389f0 (tracked)
loss(X, Y) = 0.17008752f0 (tracked)
loss(X, Y) = 0.17003267f0 (tracked)
loss(X, Y) = 0.1699681f0 (tracked)
loss(X, Y) = 0.16990504f0 (tracked)
loss(X, Y) = 0.1698487f0 (tracked)
loss(X, Y) = 0.16978516f0 (tracked)
loss(X, Y) = 0.1697217f0 (tracked)
loss(X, Y) = 0.16966517f0 (tracked)
loss(X, Y) = 0.16959786f0 (tracked)
loss(X, Y) = 0.16954117f0 (tracked)
loss(X, Y) = 0.16948095f0 (tracked)
loss(X, Y) = 0.16940229f0 (tracked)
loss(X, Y) = 0.16932988f0 (tracked)
loss(X, Y) = 0.16927983f0 (tracked)
loss(X, Y) = 0.16921048f0 (tracked)
loss(X, Y) = 0.16915374f0 (tracked)
loss(X, Y) = 0.16910517f0 (tracked)
loss(X, Y) = 0.16904412f0 (tracked)
loss(X, Y) = 0.16898152f0 (tracked)
loss(X, Y) = 0.16890271f0 (tracked)
loss(X, Y) = 0.16883227f0 (tracked)
loss(X, Y) = 0.16878487f0 (tracked)
loss(X, Y) = 0.16871057f0 (tracked)
loss(X, Y) = 0.1686265f0 (tracked)
loss(X, Y) = 0.16857749f0 (tracked)
loss(X, Y) = 0.16852f0 (tracked)
loss(X, Y) = 0.16845132f0 (tracked)
loss(X, Y) = 0.16838756f0 (tracked)
loss(X, Y) = 0.16830687f0 (tracked)
loss(X, Y) = 0.1682524f0 (tracked)
loss(X, Y) = 0.1681903f0 (tracked)
loss(X, Y) = 0.16813323f0 (tracked)
loss(X, Y) = 0.16804948f0 (tracked)
loss(X, Y) = 0.16797988f0 (tracked)
loss(X, Y) = 0.16792151f0 (tracked)
loss(X, Y) = 0.16785581f0 (tracked)
loss(X, Y) = 0.1677937f0 (tracked)
loss(X, Y) = 0.16772553f0 (tracked)
loss(X, Y) = 0.16766143f0 (tracked)
loss(X, Y) = 0.16760187f0 (tracked)
loss(X, Y) = 0.16753611f0 (tracked)
loss(X, Y) = 0.1674565f0 (tracked)
loss(X, Y) = 0.1674101f0 (tracked)
loss(X, Y) = 0.16734318f0 (tracked)
loss(X, Y) = 0.16725956f0 (tracked)
loss(X, Y) = 0.16719888f0 (tracked)
loss(X, Y) = 0.16712996f0 (tracked)
loss(X, Y) = 0.16706128f0 (tracked)
loss(X, Y) = 0.16699678f0 (tracked)
loss(X, Y) = 0.16693282f0 (tracked)
loss(X, Y) = 0.16686718f0 (tracked)
loss(X, Y) = 0.16679464f0 (tracked)
loss(X, Y) = 0.1667278f0 (tracked)
loss(X, Y) = 0.1666634f0 (tracked)
loss(X, Y) = 0.1666092f0 (tracked)
loss(X, Y) = 0.1665411f0 (tracked)
loss(X, Y) = 0.16647096f0 (tracked)
loss(X, Y) = 0.16641596f0 (tracked)
loss(X, Y) = 0.16636439f0 (tracked)
loss(X, Y) = 0.1663042f0 (tracked)
loss(X, Y) = 0.16622776f0 (tracked)
loss(X, Y) = 0.16616206f0 (tracked)
loss(X, Y) = 0.1660965f0 (tracked)
loss(X, Y) = 0.1660311f0 (tracked)
loss(X, Y) = 0.16597083f0 (tracked)
loss(X, Y) = 0.16589542f0 (tracked)
loss(X, Y) = 0.16583897f0 (tracked)
loss(X, Y) = 0.16576925f0 (tracked)
loss(X, Y) = 0.16570354f0 (tracked)
loss(X, Y) = 0.16565448f0 (tracked)
loss(X, Y) = 0.16557471f0 (tracked)
loss(X, Y) = 0.16550368f0 (tracked)
loss(X, Y) = 0.16545871f0 (tracked)
loss(X, Y) = 0.16539416f0 (tracked)
loss(X, Y) = 0.16534206f0 (tracked)
loss(X, Y) = 0.16525538f0 (tracked)
loss(X, Y) = 0.16519584f0 (tracked)
loss(X, Y) = 0.16512059f0 (tracked)
loss(X, Y) = 0.1650535f0 (tracked)
loss(X, Y) = 0.16499947f0 (tracked)
loss(X, Y) = 0.16493237f0 (tracked)
loss(X, Y) = 0.16486524f0 (tracked)
loss(X, Y) = 0.16480054f0 (tracked)
loss(X, Y) = 0.16473249f0 (tracked)
loss(X, Y) = 0.16465698f0 (tracked)
loss(X, Y) = 0.16460627f0 (tracked)
loss(X, Y) = 0.16455108f0 (tracked)
loss(X, Y) = 0.1644738f0 (tracked)
loss(X, Y) = 0.1643985f0 (tracked)
loss(X, Y) = 0.1643411f0 (tracked)
loss(X, Y) = 0.16427934f0 (tracked)
loss(X, Y) = 0.16421305f0 (tracked)
loss(X, Y) = 0.16414285f0 (tracked)
loss(X, Y) = 0.16407287f0 (tracked)
loss(X, Y) = 0.16400945f0 (tracked)
loss(X, Y) = 0.16395678f0 (tracked)
loss(X, Y) = 0.16387902f0 (tracked)
loss(X, Y) = 0.16383104f0 (tracked)
loss(X, Y) = 0.16377321f0 (tracked)
loss(X, Y) = 0.16371015f0 (tracked)
loss(X, Y) = 0.16363809f0 (tracked)
loss(X, Y) = 0.16358283f0 (tracked)
loss(X, Y) = 0.16351591f0 (tracked)
loss(X, Y) = 0.16344543f0 (tracked)
loss(X, Y) = 0.16338629f0 (tracked)
loss(X, Y) = 0.16330855f0 (tracked)
loss(X, Y) = 0.16324934f0 (tracked)
loss(X, Y) = 0.16318448f0 (tracked)
```

```
┌ Info: Epoch 3251
└ @ Main /Users/kcft114/.julia/packages/Flux/qXNjB/src/optimise/train.jl:105
┌ Info: Epoch 3252
└ @ Main /Users/kcft114/.julia/packages/Flux/qXNjB/src/optimise/train.jl:105
┌ Info: Epoch 3253
└ @ Main /Users/kcft114/.julia/packages/Flux/qXNjB/src/optimise/train.jl:105
┌ Info: Epoch 3254
└ @ Main /Users/kcft114/.julia/packages/Flux/qXNjB/src/optimise/train.jl:105
┌ Info: Epoch 3255
└ @ Main /Users/kcft114/.julia/packages/Flux/qXNjB/src/optimise/train.jl:105
┌ Info: Epoch 3256
└ @ Main /Users/kcft114/.julia/packages/Flux/qXNjB/src/optimise/train.jl:105
┌ Info: Epoch 3257
└ @ Main /Users/kcft114/.julia/packages/Flux/qXNjB/src/optimise/train.jl:105
┌ Info: Epoch 3258
└ @ Main /Users/kcft114/.julia/packages/Flux/qXNjB/src/optimise/train.jl:105
┌ Info: Epoch 3259
└ @ Main /Users/kcft114/.julia/packages/Flux/qXNjB/src/optimise/train.jl:105
┌ Info: Epoch 3260
└ @ Main /Users/kcft114/.julia/packages/Flux/qXNjB/src/optimise/train.jl:105
┌ Info: Epoch 3261
└ @ Main /Users/kcft114/.julia/packages/Flux/qXNjB/src/optimise/train.jl:105
┌ Info: Epoch 3262
└ @ Main /Users/kcft114/.julia/packages/Flux/qXNjB/src/optimise/train.jl:105
┌ Info: Epoch 3263
└ @ Main /Users/kcft114/.julia/packages/Flux/qXNjB/src/optimise/train.jl:105
┌ Info: Epoch 3264
└ @ Main /Users/kcft114/.julia/packages/Flux/qXNjB/src/optimise/train.jl:105
┌ Info: Epoch 3265
└ @ Main /Users/kcft114/.julia/packages/Flux/qXNjB/src/optimise/train.jl:105
┌ Info: Epoch 3266
└ @ Main /Users/kcft114/.julia/packages/Flux/qXNjB/src/optimise/train.jl:105
┌ Info: Epoch 3267
└ @ Main /Users/kcft114/.julia/packages/Flux/qXNjB/src/optimise/train.jl:105
┌ Info: Epoch 3268
└ @ Main /Users/kcft114/.julia/packages/Flux/qXNjB/src/optimise/train.jl:105
┌ Info: Epoch 3269
└ @ Main /Users/kcft114/.julia/packages/Flux/qXNjB/src/optimise/train.jl:105
┌ Info: Epoch 3270
└ @ Main /Users/kcft114/.julia/packages/Flux/qXNjB/src/optimise/train.jl:105
┌ Info: Epoch 3271
└ @ Main /Users/kcft114/.julia/packages/Flux/qXNjB/src/optimise/train.jl:105
┌ Info: Epoch 3272
└ @ Main /Users/kcft114/.julia/packages/Flux/qXNjB/src/optimise/train.jl:105
┌ Info: Epoch 3273
└ @ Main /Users/kcft114/.julia/packages/Flux/qXNjB/src/optimise/train.jl:105
┌ Info: Epoch 3274
└ @ Main /Users/kcft114/.julia/packages/Flux/qXNjB/src/optimise/train.jl:105
┌ Info: Epoch 3275
└ @ Main /Users/kcft114/.julia/packages/Flux/qXNjB/src/optimise/train.jl:105
┌ Info: Epoch 3276
└ @ Main /Users/kcft114/.julia/packages/Flux/qXNjB/src/optimise/train.jl:105
┌ Info: Epoch 3277
└ @ Main /Users/kcft114/.julia/packages/Flux/qXNjB/src/optimise/train.jl:105
┌ Info: Epoch 3278
└ @ Main /Users/kcft114/.julia/packages/Flux/qXNjB/src/optimise/train.jl:105
┌ Info: Epoch 3279
└ @ Main /Users/kcft114/.julia/packages/Flux/qXNjB/src/optimise/train.jl:105
┌ Info: Epoch 3280
└ @ Main /Users/kcft114/.julia/packages/Flux/qXNjB/src/optimise/train.jl:105
┌ Info: Epoch 3281
└ @ Main /Users/kcft114/.julia/packages/Flux/qXNjB/src/optimise/train.jl:105
┌ Info: Epoch 3282
└ @ Main /Users/kcft114/.julia/packages/Flux/qXNjB/src/optimise/train.jl:105
┌ Info: Epoch 3283
└ @ Main /Users/kcft114/.julia/packages/Flux/qXNjB/src/optimise/train.jl:105
┌ Info: Epoch 3284
└ @ Main /Users/kcft114/.julia/packages/Flux/qXNjB/src/optimise/train.jl:105
┌ Info: Epoch 3285
└ @ Main /Users/kcft114/.julia/packages/Flux/qXNjB/src/optimise/train.jl:105
┌ Info: Epoch 3286
└ @ Main /Users/kcft114/.julia/packages/Flux/qXNjB/src/optimise/train.jl:105
┌ Info: Epoch 3287
└ @ Main /Users/kcft114/.julia/packages/Flux/qXNjB/src/optimise/train.jl:105
┌ Info: Epoch 3288
└ @ Main /Users/kcft114/.julia/packages/Flux/qXNjB/src/optimise/train.jl:105
┌ Info: Epoch 3289
└ @ Main /Users/kcft114/.julia/packages/Flux/qXNjB/src/optimise/train.jl:105
┌ Info: Epoch 3290
└ @ Main /Users/kcft114/.julia/packages/Flux/qXNjB/src/optimise/train.jl:105
┌ Info: Epoch 3291
└ @ Main /Users/kcft114/.julia/packages/Flux/qXNjB/src/optimise/train.jl:105
┌ Info: Epoch 3292
└ @ Main /Users/kcft114/.julia/packages/Flux/qXNjB/src/optimise/train.jl:105
┌ Info: Epoch 3293
└ @ Main /Users/kcft114/.julia/packages/Flux/qXNjB/src/optimise/train.jl:105
┌ Info: Epoch 3294
└ @ Main /Users/kcft114/.julia/packages/Flux/qXNjB/src/optimise/train.jl:105
┌ Info: Epoch 3295
└ @ Main /Users/kcft114/.julia/packages/Flux/qXNjB/src/optimise/train.jl:105
┌ Info: Epoch 3296
└ @ Main /Users/kcft114/.julia/packages/Flux/qXNjB/src/optimise/train.jl:105
┌ Info: Epoch 3297
└ @ Main /Users/kcft114/.julia/packages/Flux/qXNjB/src/optimise/train.jl:105
┌ Info: Epoch 3298
└ @ Main /Users/kcft114/.julia/packages/Flux/qXNjB/src/optimise/train.jl:105
┌ Info: Epoch 3299
└ @ Main /Users/kcft114/.julia/packages/Flux/qXNjB/src/optimise/train.jl:105
┌ Info: Epoch 3300
└ @ Main /Users/kcft114/.julia/packages/Flux/qXNjB/src/optimise/train.jl:105
┌ Info: Epoch 3301
└ @ Main /Users/kcft114/.julia/packages/Flux/qXNjB/src/optimise/train.jl:105
┌ Info: Epoch 3302
└ @ Main /Users/kcft114/.julia/packages/Flux/qXNjB/src/optimise/train.jl:105
┌ Info: Epoch 3303
└ @ Main /Users/kcft114/.julia/packages/Flux/qXNjB/src/optimise/train.jl:105
┌ Info: Epoch 3304
└ @ Main /Users/kcft114/.julia/packages/Flux/qXNjB/src/optimise/train.jl:105
┌ Info: Epoch 3305
└ @ Main /Users/kcft114/.julia/packages/Flux/qXNjB/src/optimise/train.jl:105
┌ Info: Epoch 3306
└ @ Main /Users/kcft114/.julia/packages/Flux/qXNjB/src/optimise/train.jl:105
┌ Info: Epoch 3307
└ @ Main /Users/kcft114/.julia/packages/Flux/qXNjB/src/optimise/train.jl:105
┌ Info: Epoch 3308
└ @ Main /Users/kcft114/.julia/packages/Flux/qXNjB/src/optimise/train.jl:105
┌ Info: Epoch 3309
└ @ Main /Users/kcft114/.julia/packages/Flux/qXNjB/src/optimise/train.jl:105
┌ Info: Epoch 3310
└ @ Main /Users/kcft114/.julia/packages/Flux/qXNjB/src/optimise/train.jl:105
┌ Info: Epoch 3311
└ @ Main /Users/kcft114/.julia/packages/Flux/qXNjB/src/optimise/train.jl:105
┌ Info: Epoch 3312
└ @ Main /Users/kcft114/.julia/packages/Flux/qXNjB/src/optimise/train.jl:105
┌ Info: Epoch 3313
└ @ Main /Users/kcft114/.julia/packages/Flux/qXNjB/src/optimise/train.jl:105
┌ Info: Epoch 3314
└ @ Main /Users/kcft114/.julia/packages/Flux/qXNjB/src/optimise/train.jl:105
┌ Info: Epoch 3315
└ @ Main /Users/kcft114/.julia/packages/Flux/qXNjB/src/optimise/train.jl:105
┌ Info: Epoch 3316
└ @ Main /Users/kcft114/.julia/packages/Flux/qXNjB/src/optimise/train.jl:105
┌ Info: Epoch 3317
└ @ Main /Users/kcft114/.julia/packages/Flux/qXNjB/src/optimise/train.jl:105
┌ Info: Epoch 3318
└ @ Main /Users/kcft114/.julia/packages/Flux/qXNjB/src/optimise/train.jl:105
┌ Info: Epoch 3319
└ @ Main /Users/kcft114/.julia/packages/Flux/qXNjB/src/optimise/train.jl:105
┌ Info: Epoch 3320
└ @ Main /Users/kcft114/.julia/packages/Flux/qXNjB/src/optimise/train.jl:105
┌ Info: Epoch 3321
└ @ Main /Users/kcft114/.julia/packages/Flux/qXNjB/src/optimise/train.jl:105
┌ Info: Epoch 3322
└ @ Main /Users/kcft114/.julia/packages/Flux/qXNjB/src/optimise/train.jl:105
┌ Info: Epoch 3323
└ @ Main /Users/kcft114/.julia/packages/Flux/qXNjB/src/optimise/train.jl:105
┌ Info: Epoch 3324
└ @ Main /Users/kcft114/.julia/packages/Flux/qXNjB/src/optimise/train.jl:105
┌ Info: Epoch 3325
└ @ Main /Users/kcft114/.julia/packages/Flux/qXNjB/src/optimise/train.jl:105
┌ Info: Epoch 3326
└ @ Main /Users/kcft114/.julia/packages/Flux/qXNjB/src/optimise/train.jl:105
┌ Info: Epoch 3327
└ @ Main /Users/kcft114/.julia/packages/Flux/qXNjB/src/optimise/train.jl:105
┌ Info: Epoch 3328
└ @ Main /Users/kcft114/.julia/packages/Flux/qXNjB/src/optimise/train.jl:105
┌ Info: Epoch 3329
└ @ Main /Users/kcft114/.julia/packages/Flux/qXNjB/src/optimise/train.jl:105
┌ Info: Epoch 3330
└ @ Main /Users/kcft114/.julia/packages/Flux/qXNjB/src/optimise/train.jl:105
┌ Info: Epoch 3331
└ @ Main /Users/kcft114/.julia/packages/Flux/qXNjB/src/optimise/train.jl:105
┌ Info: Epoch 3332
└ @ Main /Users/kcft114/.julia/packages/Flux/qXNjB/src/optimise/train.jl:105
┌ Info: Epoch 3333
└ @ Main /Users/kcft114/.julia/packages/Flux/qXNjB/src/optimise/train.jl:105
┌ Info: Epoch 3334
└ @ Main /Users/kcft114/.julia/packages/Flux/qXNjB/src/optimise/train.jl:105
┌ Info: Epoch 3335
└ @ Main /Users/kcft114/.julia/packages/Flux/qXNjB/src/optimise/train.jl:105
┌ Info: Epoch 3336
└ @ Main /Users/kcft114/.julia/packages/Flux/qXNjB/src/optimise/train.jl:105
┌ Info: Epoch 3337
└ @ Main /Users/kcft114/.julia/packages/Flux/qXNjB/src/optimise/train.jl:105
┌ Info: Epoch 3338
└ @ Main /Users/kcft114/.julia/packages/Flux/qXNjB/src/optimise/train.jl:105
┌ Info: Epoch 3339
└ @ Main /Users/kcft114/.julia/packages/Flux/qXNjB/src/optimise/train.jl:105
┌ Info: Epoch 3340
└ @ Main /Users/kcft114/.julia/packages/Flux/qXNjB/src/optimise/train.jl:105
┌ Info: Epoch 3341
└ @ Main /Users/kcft114/.julia/packages/Flux/qXNjB/src/optimise/train.jl:105
┌ Info: Epoch 3342
└ @ Main /Users/kcft114/.julia/packages/Flux/qXNjB/src/optimise/train.jl:105
┌ Info: Epoch 3343
└ @ Main /Users/kcft114/.julia/packages/Flux/qXNjB/src/optimise/train.jl:105
┌ Info: Epoch 3344
└ @ Main /Users/kcft114/.julia/packages/Flux/qXNjB/src/optimise/train.jl:105
┌ Info: Epoch 3345
└ @ Main /Users/kcft114/.julia/packages/Flux/qXNjB/src/optimise/train.jl:105
┌ Info: Epoch 3346
└ @ Main /Users/kcft114/.julia/packages/Flux/qXNjB/src/optimise/train.jl:105
┌ Info: Epoch 3347
└ @ Main /Users/kcft114/.julia/packages/Flux/qXNjB/src/optimise/train.jl:105
┌ Info: Epoch 3348
└ @ Main /Users/kcft114/.julia/packages/Flux/qXNjB/src/optimise/train.jl:105
┌ Info: Epoch 3349
└ @ Main /Users/kcft114/.julia/packages/Flux/qXNjB/src/optimise/train.jl:105
┌ Info: Epoch 3350
└ @ Main /Users/kcft114/.julia/packages/Flux/qXNjB/src/optimise/train.jl:105
┌ Info: Epoch 3351
└ @ Main /Users/kcft114/.julia/packages/Flux/qXNjB/src/optimise/train.jl:105
┌ Info: Epoch 3352
└ @ Main /Users/kcft114/.julia/packages/Flux/qXNjB/src/optimise/train.jl:105
┌ Info: Epoch 3353
└ @ Main /Users/kcft114/.julia/packages/Flux/qXNjB/src/optimise/train.jl:105
┌ Info: Epoch 3354
└ @ Main /Users/kcft114/.julia/packages/Flux/qXNjB/src/optimise/train.jl:105
┌ Info: Epoch 3355
└ @ Main /Users/kcft114/.julia/packages/Flux/qXNjB/src/optimise/train.jl:105
┌ Info: Epoch 3356
└ @ Main /Users/kcft114/.julia/packages/Flux/qXNjB/src/optimise/train.jl:105
┌ Info: Epoch 3357
└ @ Main /Users/kcft114/.julia/packages/Flux/qXNjB/src/optimise/train.jl:105
┌ Info: Epoch 3358
└ @ Main /Users/kcft114/.julia/packages/Flux/qXNjB/src/optimise/train.jl:105
┌ Info: Epoch 3359
└ @ Main /Users/kcft114/.julia/packages/Flux/qXNjB/src/optimise/train.jl:105
┌ Info: Epoch 3360
└ @ Main /Users/kcft114/.julia/packages/Flux/qXNjB/src/optimise/train.jl:105
┌ Info: Epoch 3361
└ @ Main /Users/kcft114/.julia/packages/Flux/qXNjB/src/optimise/train.jl:105
┌ Info: Epoch 3362
└ @ Main /Users/kcft114/.julia/packages/Flux/qXNjB/src/optimise/train.jl:105
┌ Info: Epoch 3363
└ @ Main /Users/kcft114/.julia/packages/Flux/qXNjB/src/optimise/train.jl:105
┌ Info: Epoch 3364
└ @ Main /Users/kcft114/.julia/packages/Flux/qXNjB/src/optimise/train.jl:105
┌ Info: Epoch 3365
└ @ Main /Users/kcft114/.julia/packages/Flux/qXNjB/src/optimise/train.jl:105
┌ Info: Epoch 3366
└ @ Main /Users/kcft114/.julia/packages/Flux/qXNjB/src/optimise/train.jl:105
┌ Info: Epoch 3367
└ @ Main /Users/kcft114/.julia/packages/Flux/qXNjB/src/optimise/train.jl:105
┌ Info: Epoch 3368
└ @ Main /Users/kcft114/.julia/packages/Flux/qXNjB/src/optimise/train.jl:105
┌ Info: Epoch 3369
└ @ Main /Users/kcft114/.julia/packages/Flux/qXNjB/src/optimise/train.jl:105
┌ Info: Epoch 3370
└ @ Main /Users/kcft114/.julia/packages/Flux/qXNjB/src/optimise/train.jl:105
┌ Info: Epoch 3371
└ @ Main /Users/kcft114/.julia/packages/Flux/qXNjB/src/optimise/train.jl:105
┌ Info: Epoch 3372
└ @ Main /Users/kcft114/.julia/packages/Flux/qXNjB/src/optimise/train.jl:105
┌ Info: Epoch 3373
└ @ Main /Users/kcft114/.julia/packages/Flux/qXNjB/src/optimise/train.jl:105
┌ Info: Epoch 3374
└ @ Main /Users/kcft114/.julia/packages/Flux/qXNjB/src/optimise/train.jl:105
┌ Info: Epoch 3375
└ @ Main /Users/kcft114/.julia/packages/Flux/qXNjB/src/optimise/train.jl:105
┌ Info: Epoch 3376
└ @ Main /Users/kcft114/.julia/packages/Flux/qXNjB/src/optimise/train.jl:105
┌ Info: Epoch 3377
└ @ Main /Users/kcft114/.julia/packages/Flux/qXNjB/src/optimise/train.jl:105
┌ Info: Epoch 3378
└ @ Main /Users/kcft114/.julia/packages/Flux/qXNjB/src/optimise/train.jl:105
┌ Info: Epoch 3379
└ @ Main /Users/kcft114/.julia/packages/Flux/qXNjB/src/optimise/train.jl:105
┌ Info: Epoch 3380
└ @ Main /Users/kcft114/.julia/packages/Flux/qXNjB/src/optimise/train.jl:105
┌ Info: Epoch 3381
└ @ Main /Users/kcft114/.julia/packages/Flux/qXNjB/src/optimise/train.jl:105
┌ Info: Epoch 3382
└ @ Main /Users/kcft114/.julia/packages/Flux/qXNjB/src/optimise/train.jl:105
┌ Info: Epoch 3383
└ @ Main /Users/kcft114/.julia/packages/Flux/qXNjB/src/optimise/train.jl:105
┌ Info: Epoch 3384
└ @ Main /Users/kcft114/.julia/packages/Flux/qXNjB/src/optimise/train.jl:105
┌ Info: Epoch 3385
└ @ Main /Users/kcft114/.julia/packages/Flux/qXNjB/src/optimise/train.jl:105
┌ Info: Epoch 3386
└ @ Main /Users/kcft114/.julia/packages/Flux/qXNjB/src/optimise/train.jl:105
┌ Info: Epoch 3387
└ @ Main /Users/kcft114/.julia/packages/Flux/qXNjB/src/optimise/train.jl:105
┌ Info: Epoch 3388
└ @ Main /Users/kcft114/.julia/packages/Flux/qXNjB/src/optimise/train.jl:105
┌ Info: Epoch 3389
└ @ Main /Users/kcft114/.julia/packages/Flux/qXNjB/src/optimise/train.jl:105
┌ Info: Epoch 3390
└ @ Main /Users/kcft114/.julia/packages/Flux/qXNjB/src/optimise/train.jl:105
┌ Info: Epoch 3391
└ @ Main /Users/kcft114/.julia/packages/Flux/qXNjB/src/optimise/train.jl:105
┌ Info: Epoch 3392
└ @ Main /Users/kcft114/.julia/packages/Flux/qXNjB/src/optimise/train.jl:105
┌ Info: Epoch 3393
└ @ Main /Users/kcft114/.julia/packages/Flux/qXNjB/src/optimise/train.jl:105
┌ Info: Epoch 3394
└ @ Main /Users/kcft114/.julia/packages/Flux/qXNjB/src/optimise/train.jl:105
┌ Info: Epoch 3395
└ @ Main /Users/kcft114/.julia/packages/Flux/qXNjB/src/optimise/train.jl:105
┌ Info: Epoch 3396
└ @ Main /Users/kcft114/.julia/packages/Flux/qXNjB/src/optimise/train.jl:105
```

```
loss(X, Y) = 0.16312863f0 (tracked)
loss(X, Y) = 0.16306485f0 (tracked)
loss(X, Y) = 0.1629901f0 (tracked)
loss(X, Y) = 0.16291995f0 (tracked)
loss(X, Y) = 0.16285889f0 (tracked)
loss(X, Y) = 0.16281392f0 (tracked)
loss(X, Y) = 0.1627307f0 (tracked)
loss(X, Y) = 0.16267256f0 (tracked)
loss(X, Y) = 0.16261454f0 (tracked)
loss(X, Y) = 0.1625438f0 (tracked)
loss(X, Y) = 0.16247547f0 (tracked)
loss(X, Y) = 0.16241996f0 (tracked)
loss(X, Y) = 0.16235244f0 (tracked)
loss(X, Y) = 0.1622788f0 (tracked)
loss(X, Y) = 0.1622167f0 (tracked)
loss(X, Y) = 0.16215605f0 (tracked)
loss(X, Y) = 0.16207701f0 (tracked)
loss(X, Y) = 0.16202964f0 (tracked)
loss(X, Y) = 0.16196938f0 (tracked)
loss(X, Y) = 0.16189198f0 (tracked)
loss(X, Y) = 0.16182663f0 (tracked)
loss(X, Y) = 0.16176985f0 (tracked)
loss(X, Y) = 0.16171181f0 (tracked)
loss(X, Y) = 0.16167207f0 (tracked)
loss(X, Y) = 0.16159241f0 (tracked)
loss(X, Y) = 0.1615316f0 (tracked)
loss(X, Y) = 0.16147614f0 (tracked)
loss(X, Y) = 0.16142507f0 (tracked)
loss(X, Y) = 0.1613515f0 (tracked)
loss(X, Y) = 0.16128537f0 (tracked)
loss(X, Y) = 0.16121897f0 (tracked)
loss(X, Y) = 0.16116263f0 (tracked)
loss(X, Y) = 0.16110237f0 (tracked)
loss(X, Y) = 0.16102894f0 (tracked)
loss(X, Y) = 0.16098383f0 (tracked)
loss(X, Y) = 0.16092281f0 (tracked)
loss(X, Y) = 0.16084826f0 (tracked)
loss(X, Y) = 0.16078666f0 (tracked)
loss(X, Y) = 0.16073579f0 (tracked)
loss(X, Y) = 0.16068384f0 (tracked)
loss(X, Y) = 0.16060525f0 (tracked)
loss(X, Y) = 0.16056281f0 (tracked)
loss(X, Y) = 0.16049483f0 (tracked)
loss(X, Y) = 0.16042803f0 (tracked)
loss(X, Y) = 0.1603615f0 (tracked)
loss(X, Y) = 0.16029541f0 (tracked)
loss(X, Y) = 0.16023578f0 (tracked)
loss(X, Y) = 0.16018534f0 (tracked)
loss(X, Y) = 0.16012418f0 (tracked)
loss(X, Y) = 0.16005391f0 (tracked)
loss(X, Y) = 0.16000588f0 (tracked)
loss(X, Y) = 0.15995361f0 (tracked)
loss(X, Y) = 0.1598825f0 (tracked)
loss(X, Y) = 0.15981318f0 (tracked)
loss(X, Y) = 0.15976606f0 (tracked)
loss(X, Y) = 0.15971376f0 (tracked)
loss(X, Y) = 0.15964368f0 (tracked)
loss(X, Y) = 0.15958957f0 (tracked)
loss(X, Y) = 0.15952525f0 (tracked)
loss(X, Y) = 0.15946777f0 (tracked)
loss(X, Y) = 0.15940863f0 (tracked)
loss(X, Y) = 0.15933006f0 (tracked)
loss(X, Y) = 0.15928175f0 (tracked)
loss(X, Y) = 0.15923446f0 (tracked)
loss(X, Y) = 0.1591786f0 (tracked)
loss(X, Y) = 0.15911332f0 (tracked)
loss(X, Y) = 0.15905721f0 (tracked)
loss(X, Y) = 0.15898316f0 (tracked)
loss(X, Y) = 0.15892987f0 (tracked)
loss(X, Y) = 0.15887265f0 (tracked)
loss(X, Y) = 0.15879233f0 (tracked)
loss(X, Y) = 0.15874037f0 (tracked)
loss(X, Y) = 0.15868472f0 (tracked)
loss(X, Y) = 0.15861039f0 (tracked)
loss(X, Y) = 0.15853578f0 (tracked)
loss(X, Y) = 0.15849961f0 (tracked)
loss(X, Y) = 0.15842961f0 (tracked)
loss(X, Y) = 0.1583462f0 (tracked)
loss(X, Y) = 0.15831198f0 (tracked)
loss(X, Y) = 0.15824641f0 (tracked)
loss(X, Y) = 0.15817867f0 (tracked)
loss(X, Y) = 0.15811737f0 (tracked)
loss(X, Y) = 0.15806614f0 (tracked)
loss(X, Y) = 0.15797544f0 (tracked)
loss(X, Y) = 0.157905f0 (tracked)
loss(X, Y) = 0.15786485f0 (tracked)
loss(X, Y) = 0.1578042f0 (tracked)
loss(X, Y) = 0.15772496f0 (tracked)
loss(X, Y) = 0.15765995f0 (tracked)
loss(X, Y) = 0.15758717f0 (tracked)
loss(X, Y) = 0.15751377f0 (tracked)
loss(X, Y) = 0.15743433f0 (tracked)
loss(X, Y) = 0.15737355f0 (tracked)
loss(X, Y) = 0.15730715f0 (tracked)
loss(X, Y) = 0.15723503f0 (tracked)
loss(X, Y) = 0.15715522f0 (tracked)
loss(X, Y) = 0.1570887f0 (tracked)
loss(X, Y) = 0.15703282f0 (tracked)
loss(X, Y) = 0.15694739f0 (tracked)
loss(X, Y) = 0.1568811f0 (tracked)
loss(X, Y) = 0.15683332f0 (tracked)
loss(X, Y) = 0.15676014f0 (tracked)
loss(X, Y) = 0.15668269f0 (tracked)
loss(X, Y) = 0.15660259f0 (tracked)
loss(X, Y) = 0.15652946f0 (tracked)
loss(X, Y) = 0.15647462f0 (tracked)
loss(X, Y) = 0.15641484f0 (tracked)
loss(X, Y) = 0.15632747f0 (tracked)
loss(X, Y) = 0.15626854f0 (tracked)
loss(X, Y) = 0.15620616f0 (tracked)
loss(X, Y) = 0.1561384f0 (tracked)
loss(X, Y) = 0.15606192f0 (tracked)
loss(X, Y) = 0.15597695f0 (tracked)
loss(X, Y) = 0.15591542f0 (tracked)
loss(X, Y) = 0.15585741f0 (tracked)
loss(X, Y) = 0.15578495f0 (tracked)
loss(X, Y) = 0.1557015f0 (tracked)
loss(X, Y) = 0.15561405f0 (tracked)
loss(X, Y) = 0.15555634f0 (tracked)
loss(X, Y) = 0.15548497f0 (tracked)
loss(X, Y) = 0.1554044f0 (tracked)
loss(X, Y) = 0.15534511f0 (tracked)
loss(X, Y) = 0.15528633f0 (tracked)
loss(X, Y) = 0.15521377f0 (tracked)
loss(X, Y) = 0.15514101f0 (tracked)
loss(X, Y) = 0.15507732f0 (tracked)
loss(X, Y) = 0.15502344f0 (tracked)
loss(X, Y) = 0.15494142f0 (tracked)
loss(X, Y) = 0.1548726f0 (tracked)
loss(X, Y) = 0.15481466f0 (tracked)
loss(X, Y) = 0.15474167f0 (tracked)
loss(X, Y) = 0.15466572f0 (tracked)
loss(X, Y) = 0.15461168f0 (tracked)
loss(X, Y) = 0.1545409f0 (tracked)
loss(X, Y) = 0.15446258f0 (tracked)
loss(X, Y) = 0.1544006f0 (tracked)
loss(X, Y) = 0.15432946f0 (tracked)
loss(X, Y) = 0.15427086f0 (tracked)
loss(X, Y) = 0.15422434f0 (tracked)
loss(X, Y) = 0.15415013f0 (tracked)
loss(X, Y) = 0.1540695f0 (tracked)
loss(X, Y) = 0.1540064f0 (tracked)
loss(X, Y) = 0.1539386f0 (tracked)
loss(X, Y) = 0.15389791f0 (tracked)
loss(X, Y) = 0.15382919f0 (tracked)
loss(X, Y) = 0.153737f0 (tracked)
loss(X, Y) = 0.15368111f0 (tracked)
loss(X, Y) = 0.15362652f0 (tracked)
loss(X, Y) = 0.15356846f0 (tracked)
loss(X, Y) = 0.15350337f0 (tracked)
```

```
┌ Info: Epoch 3397
└ @ Main /Users/kcft114/.julia/packages/Flux/qXNjB/src/optimise/train.jl:105
┌ Info: Epoch 3398
└ @ Main /Users/kcft114/.julia/packages/Flux/qXNjB/src/optimise/train.jl:105
┌ Info: Epoch 3399
└ @ Main /Users/kcft114/.julia/packages/Flux/qXNjB/src/optimise/train.jl:105
┌ Info: Epoch 3400
└ @ Main /Users/kcft114/.julia/packages/Flux/qXNjB/src/optimise/train.jl:105
┌ Info: Epoch 3401
└ @ Main /Users/kcft114/.julia/packages/Flux/qXNjB/src/optimise/train.jl:105
┌ Info: Epoch 3402
└ @ Main /Users/kcft114/.julia/packages/Flux/qXNjB/src/optimise/train.jl:105
┌ Info: Epoch 3403
└ @ Main /Users/kcft114/.julia/packages/Flux/qXNjB/src/optimise/train.jl:105
┌ Info: Epoch 3404
└ @ Main /Users/kcft114/.julia/packages/Flux/qXNjB/src/optimise/train.jl:105
┌ Info: Epoch 3405
└ @ Main /Users/kcft114/.julia/packages/Flux/qXNjB/src/optimise/train.jl:105
┌ Info: Epoch 3406
└ @ Main /Users/kcft114/.julia/packages/Flux/qXNjB/src/optimise/train.jl:105
┌ Info: Epoch 3407
└ @ Main /Users/kcft114/.julia/packages/Flux/qXNjB/src/optimise/train.jl:105
┌ Info: Epoch 3408
└ @ Main /Users/kcft114/.julia/packages/Flux/qXNjB/src/optimise/train.jl:105
┌ Info: Epoch 3409
└ @ Main /Users/kcft114/.julia/packages/Flux/qXNjB/src/optimise/train.jl:105
┌ Info: Epoch 3410
└ @ Main /Users/kcft114/.julia/packages/Flux/qXNjB/src/optimise/train.jl:105
┌ Info: Epoch 3411
└ @ Main /Users/kcft114/.julia/packages/Flux/qXNjB/src/optimise/train.jl:105
┌ Info: Epoch 3412
└ @ Main /Users/kcft114/.julia/packages/Flux/qXNjB/src/optimise/train.jl:105
┌ Info: Epoch 3413
└ @ Main /Users/kcft114/.julia/packages/Flux/qXNjB/src/optimise/train.jl:105
┌ Info: Epoch 3414
└ @ Main /Users/kcft114/.julia/packages/Flux/qXNjB/src/optimise/train.jl:105
┌ Info: Epoch 3415
└ @ Main /Users/kcft114/.julia/packages/Flux/qXNjB/src/optimise/train.jl:105
┌ Info: Epoch 3416
└ @ Main /Users/kcft114/.julia/packages/Flux/qXNjB/src/optimise/train.jl:105
┌ Info: Epoch 3417
└ @ Main /Users/kcft114/.julia/packages/Flux/qXNjB/src/optimise/train.jl:105
┌ Info: Epoch 3418
└ @ Main /Users/kcft114/.julia/packages/Flux/qXNjB/src/optimise/train.jl:105
┌ Info: Epoch 3419
└ @ Main /Users/kcft114/.julia/packages/Flux/qXNjB/src/optimise/train.jl:105
┌ Info: Epoch 3420
└ @ Main /Users/kcft114/.julia/packages/Flux/qXNjB/src/optimise/train.jl:105
┌ Info: Epoch 3421
└ @ Main /Users/kcft114/.julia/packages/Flux/qXNjB/src/optimise/train.jl:105
┌ Info: Epoch 3422
└ @ Main /Users/kcft114/.julia/packages/Flux/qXNjB/src/optimise/train.jl:105
┌ Info: Epoch 3423
└ @ Main /Users/kcft114/.julia/packages/Flux/qXNjB/src/optimise/train.jl:105
┌ Info: Epoch 3424
└ @ Main /Users/kcft114/.julia/packages/Flux/qXNjB/src/optimise/train.jl:105
┌ Info: Epoch 3425
└ @ Main /Users/kcft114/.julia/packages/Flux/qXNjB/src/optimise/train.jl:105
┌ Info: Epoch 3426
└ @ Main /Users/kcft114/.julia/packages/Flux/qXNjB/src/optimise/train.jl:105
┌ Info: Epoch 3427
└ @ Main /Users/kcft114/.julia/packages/Flux/qXNjB/src/optimise/train.jl:105
┌ Info: Epoch 3428
└ @ Main /Users/kcft114/.julia/packages/Flux/qXNjB/src/optimise/train.jl:105
┌ Info: Epoch 3429
└ @ Main /Users/kcft114/.julia/packages/Flux/qXNjB/src/optimise/train.jl:105
┌ Info: Epoch 3430
└ @ Main /Users/kcft114/.julia/packages/Flux/qXNjB/src/optimise/train.jl:105
┌ Info: Epoch 3431
└ @ Main /Users/kcft114/.julia/packages/Flux/qXNjB/src/optimise/train.jl:105
┌ Info: Epoch 3432
└ @ Main /Users/kcft114/.julia/packages/Flux/qXNjB/src/optimise/train.jl:105
┌ Info: Epoch 3433
└ @ Main /Users/kcft114/.julia/packages/Flux/qXNjB/src/optimise/train.jl:105
┌ Info: Epoch 3434
└ @ Main /Users/kcft114/.julia/packages/Flux/qXNjB/src/optimise/train.jl:105
┌ Info: Epoch 3435
└ @ Main /Users/kcft114/.julia/packages/Flux/qXNjB/src/optimise/train.jl:105
┌ Info: Epoch 3436
└ @ Main /Users/kcft114/.julia/packages/Flux/qXNjB/src/optimise/train.jl:105
┌ Info: Epoch 3437
└ @ Main /Users/kcft114/.julia/packages/Flux/qXNjB/src/optimise/train.jl:105
┌ Info: Epoch 3438
└ @ Main /Users/kcft114/.julia/packages/Flux/qXNjB/src/optimise/train.jl:105
┌ Info: Epoch 3439
└ @ Main /Users/kcft114/.julia/packages/Flux/qXNjB/src/optimise/train.jl:105
┌ Info: Epoch 3440
└ @ Main /Users/kcft114/.julia/packages/Flux/qXNjB/src/optimise/train.jl:105
┌ Info: Epoch 3441
└ @ Main /Users/kcft114/.julia/packages/Flux/qXNjB/src/optimise/train.jl:105
┌ Info: Epoch 3442
└ @ Main /Users/kcft114/.julia/packages/Flux/qXNjB/src/optimise/train.jl:105
┌ Info: Epoch 3443
└ @ Main /Users/kcft114/.julia/packages/Flux/qXNjB/src/optimise/train.jl:105
┌ Info: Epoch 3444
└ @ Main /Users/kcft114/.julia/packages/Flux/qXNjB/src/optimise/train.jl:105
┌ Info: Epoch 3445
└ @ Main /Users/kcft114/.julia/packages/Flux/qXNjB/src/optimise/train.jl:105
┌ Info: Epoch 3446
└ @ Main /Users/kcft114/.julia/packages/Flux/qXNjB/src/optimise/train.jl:105
┌ Info: Epoch 3447
└ @ Main /Users/kcft114/.julia/packages/Flux/qXNjB/src/optimise/train.jl:105
┌ Info: Epoch 3448
└ @ Main /Users/kcft114/.julia/packages/Flux/qXNjB/src/optimise/train.jl:105
┌ Info: Epoch 3449
└ @ Main /Users/kcft114/.julia/packages/Flux/qXNjB/src/optimise/train.jl:105
┌ Info: Epoch 3450
└ @ Main /Users/kcft114/.julia/packages/Flux/qXNjB/src/optimise/train.jl:105
┌ Info: Epoch 3451
└ @ Main /Users/kcft114/.julia/packages/Flux/qXNjB/src/optimise/train.jl:105
┌ Info: Epoch 3452
└ @ Main /Users/kcft114/.julia/packages/Flux/qXNjB/src/optimise/train.jl:105
┌ Info: Epoch 3453
└ @ Main /Users/kcft114/.julia/packages/Flux/qXNjB/src/optimise/train.jl:105
┌ Info: Epoch 3454
└ @ Main /Users/kcft114/.julia/packages/Flux/qXNjB/src/optimise/train.jl:105
┌ Info: Epoch 3455
└ @ Main /Users/kcft114/.julia/packages/Flux/qXNjB/src/optimise/train.jl:105
┌ Info: Epoch 3456
└ @ Main /Users/kcft114/.julia/packages/Flux/qXNjB/src/optimise/train.jl:105
┌ Info: Epoch 3457
└ @ Main /Users/kcft114/.julia/packages/Flux/qXNjB/src/optimise/train.jl:105
┌ Info: Epoch 3458
└ @ Main /Users/kcft114/.julia/packages/Flux/qXNjB/src/optimise/train.jl:105
┌ Info: Epoch 3459
└ @ Main /Users/kcft114/.julia/packages/Flux/qXNjB/src/optimise/train.jl:105
┌ Info: Epoch 3460
└ @ Main /Users/kcft114/.julia/packages/Flux/qXNjB/src/optimise/train.jl:105
┌ Info: Epoch 3461
└ @ Main /Users/kcft114/.julia/packages/Flux/qXNjB/src/optimise/train.jl:105
┌ Info: Epoch 3462
└ @ Main /Users/kcft114/.julia/packages/Flux/qXNjB/src/optimise/train.jl:105
┌ Info: Epoch 3463
└ @ Main /Users/kcft114/.julia/packages/Flux/qXNjB/src/optimise/train.jl:105
┌ Info: Epoch 3464
└ @ Main /Users/kcft114/.julia/packages/Flux/qXNjB/src/optimise/train.jl:105
┌ Info: Epoch 3465
└ @ Main /Users/kcft114/.julia/packages/Flux/qXNjB/src/optimise/train.jl:105
┌ Info: Epoch 3466
└ @ Main /Users/kcft114/.julia/packages/Flux/qXNjB/src/optimise/train.jl:105
┌ Info: Epoch 3467
└ @ Main /Users/kcft114/.julia/packages/Flux/qXNjB/src/optimise/train.jl:105
┌ Info: Epoch 3468
└ @ Main /Users/kcft114/.julia/packages/Flux/qXNjB/src/optimise/train.jl:105
┌ Info: Epoch 3469
└ @ Main /Users/kcft114/.julia/packages/Flux/qXNjB/src/optimise/train.jl:105
┌ Info: Epoch 3470
└ @ Main /Users/kcft114/.julia/packages/Flux/qXNjB/src/optimise/train.jl:105
┌ Info: Epoch 3471
└ @ Main /Users/kcft114/.julia/packages/Flux/qXNjB/src/optimise/train.jl:105
┌ Info: Epoch 3472
└ @ Main /Users/kcft114/.julia/packages/Flux/qXNjB/src/optimise/train.jl:105
┌ Info: Epoch 3473
└ @ Main /Users/kcft114/.julia/packages/Flux/qXNjB/src/optimise/train.jl:105
┌ Info: Epoch 3474
└ @ Main /Users/kcft114/.julia/packages/Flux/qXNjB/src/optimise/train.jl:105
┌ Info: Epoch 3475
└ @ Main /Users/kcft114/.julia/packages/Flux/qXNjB/src/optimise/train.jl:105
┌ Info: Epoch 3476
└ @ Main /Users/kcft114/.julia/packages/Flux/qXNjB/src/optimise/train.jl:105
┌ Info: Epoch 3477
└ @ Main /Users/kcft114/.julia/packages/Flux/qXNjB/src/optimise/train.jl:105
┌ Info: Epoch 3478
└ @ Main /Users/kcft114/.julia/packages/Flux/qXNjB/src/optimise/train.jl:105
┌ Info: Epoch 3479
└ @ Main /Users/kcft114/.julia/packages/Flux/qXNjB/src/optimise/train.jl:105
┌ Info: Epoch 3480
└ @ Main /Users/kcft114/.julia/packages/Flux/qXNjB/src/optimise/train.jl:105
┌ Info: Epoch 3481
└ @ Main /Users/kcft114/.julia/packages/Flux/qXNjB/src/optimise/train.jl:105
┌ Info: Epoch 3482
└ @ Main /Users/kcft114/.julia/packages/Flux/qXNjB/src/optimise/train.jl:105
┌ Info: Epoch 3483
└ @ Main /Users/kcft114/.julia/packages/Flux/qXNjB/src/optimise/train.jl:105
┌ Info: Epoch 3484
└ @ Main /Users/kcft114/.julia/packages/Flux/qXNjB/src/optimise/train.jl:105
┌ Info: Epoch 3485
└ @ Main /Users/kcft114/.julia/packages/Flux/qXNjB/src/optimise/train.jl:105
┌ Info: Epoch 3486
└ @ Main /Users/kcft114/.julia/packages/Flux/qXNjB/src/optimise/train.jl:105
┌ Info: Epoch 3487
└ @ Main /Users/kcft114/.julia/packages/Flux/qXNjB/src/optimise/train.jl:105
┌ Info: Epoch 3488
└ @ Main /Users/kcft114/.julia/packages/Flux/qXNjB/src/optimise/train.jl:105
┌ Info: Epoch 3489
└ @ Main /Users/kcft114/.julia/packages/Flux/qXNjB/src/optimise/train.jl:105
┌ Info: Epoch 3490
└ @ Main /Users/kcft114/.julia/packages/Flux/qXNjB/src/optimise/train.jl:105
┌ Info: Epoch 3491
└ @ Main /Users/kcft114/.julia/packages/Flux/qXNjB/src/optimise/train.jl:105
┌ Info: Epoch 3492
└ @ Main /Users/kcft114/.julia/packages/Flux/qXNjB/src/optimise/train.jl:105
┌ Info: Epoch 3493
└ @ Main /Users/kcft114/.julia/packages/Flux/qXNjB/src/optimise/train.jl:105
┌ Info: Epoch 3494
└ @ Main /Users/kcft114/.julia/packages/Flux/qXNjB/src/optimise/train.jl:105
┌ Info: Epoch 3495
└ @ Main /Users/kcft114/.julia/packages/Flux/qXNjB/src/optimise/train.jl:105
┌ Info: Epoch 3496
└ @ Main /Users/kcft114/.julia/packages/Flux/qXNjB/src/optimise/train.jl:105
┌ Info: Epoch 3497
└ @ Main /Users/kcft114/.julia/packages/Flux/qXNjB/src/optimise/train.jl:105
┌ Info: Epoch 3498
└ @ Main /Users/kcft114/.julia/packages/Flux/qXNjB/src/optimise/train.jl:105
┌ Info: Epoch 3499
└ @ Main /Users/kcft114/.julia/packages/Flux/qXNjB/src/optimise/train.jl:105
┌ Info: Epoch 3500
└ @ Main /Users/kcft114/.julia/packages/Flux/qXNjB/src/optimise/train.jl:105
┌ Info: Epoch 3501
└ @ Main /Users/kcft114/.julia/packages/Flux/qXNjB/src/optimise/train.jl:105
┌ Info: Epoch 3502
└ @ Main /Users/kcft114/.julia/packages/Flux/qXNjB/src/optimise/train.jl:105
┌ Info: Epoch 3503
└ @ Main /Users/kcft114/.julia/packages/Flux/qXNjB/src/optimise/train.jl:105
┌ Info: Epoch 3504
└ @ Main /Users/kcft114/.julia/packages/Flux/qXNjB/src/optimise/train.jl:105
┌ Info: Epoch 3505
└ @ Main /Users/kcft114/.julia/packages/Flux/qXNjB/src/optimise/train.jl:105
┌ Info: Epoch 3506
└ @ Main /Users/kcft114/.julia/packages/Flux/qXNjB/src/optimise/train.jl:105
┌ Info: Epoch 3507
└ @ Main /Users/kcft114/.julia/packages/Flux/qXNjB/src/optimise/train.jl:105
┌ Info: Epoch 3508
└ @ Main /Users/kcft114/.julia/packages/Flux/qXNjB/src/optimise/train.jl:105
┌ Info: Epoch 3509
└ @ Main /Users/kcft114/.julia/packages/Flux/qXNjB/src/optimise/train.jl:105
┌ Info: Epoch 3510
└ @ Main /Users/kcft114/.julia/packages/Flux/qXNjB/src/optimise/train.jl:105
┌ Info: Epoch 3511
└ @ Main /Users/kcft114/.julia/packages/Flux/qXNjB/src/optimise/train.jl:105
┌ Info: Epoch 3512
└ @ Main /Users/kcft114/.julia/packages/Flux/qXNjB/src/optimise/train.jl:105
┌ Info: Epoch 3513
└ @ Main /Users/kcft114/.julia/packages/Flux/qXNjB/src/optimise/train.jl:105
┌ Info: Epoch 3514
└ @ Main /Users/kcft114/.julia/packages/Flux/qXNjB/src/optimise/train.jl:105
┌ Info: Epoch 3515
└ @ Main /Users/kcft114/.julia/packages/Flux/qXNjB/src/optimise/train.jl:105
┌ Info: Epoch 3516
└ @ Main /Users/kcft114/.julia/packages/Flux/qXNjB/src/optimise/train.jl:105
┌ Info: Epoch 3517
└ @ Main /Users/kcft114/.julia/packages/Flux/qXNjB/src/optimise/train.jl:105
┌ Info: Epoch 3518
└ @ Main /Users/kcft114/.julia/packages/Flux/qXNjB/src/optimise/train.jl:105
┌ Info: Epoch 3519
└ @ Main /Users/kcft114/.julia/packages/Flux/qXNjB/src/optimise/train.jl:105
┌ Info: Epoch 3520
└ @ Main /Users/kcft114/.julia/packages/Flux/qXNjB/src/optimise/train.jl:105
┌ Info: Epoch 3521
└ @ Main /Users/kcft114/.julia/packages/Flux/qXNjB/src/optimise/train.jl:105
┌ Info: Epoch 3522
└ @ Main /Users/kcft114/.julia/packages/Flux/qXNjB/src/optimise/train.jl:105
┌ Info: Epoch 3523
└ @ Main /Users/kcft114/.julia/packages/Flux/qXNjB/src/optimise/train.jl:105
┌ Info: Epoch 3524
└ @ Main /Users/kcft114/.julia/packages/Flux/qXNjB/src/optimise/train.jl:105
┌ Info: Epoch 3525
└ @ Main /Users/kcft114/.julia/packages/Flux/qXNjB/src/optimise/train.jl:105
┌ Info: Epoch 3526
└ @ Main /Users/kcft114/.julia/packages/Flux/qXNjB/src/optimise/train.jl:105
┌ Info: Epoch 3527
└ @ Main /Users/kcft114/.julia/packages/Flux/qXNjB/src/optimise/train.jl:105
┌ Info: Epoch 3528
└ @ Main /Users/kcft114/.julia/packages/Flux/qXNjB/src/optimise/train.jl:105
┌ Info: Epoch 3529
└ @ Main /Users/kcft114/.julia/packages/Flux/qXNjB/src/optimise/train.jl:105
┌ Info: Epoch 3530
└ @ Main /Users/kcft114/.julia/packages/Flux/qXNjB/src/optimise/train.jl:105
┌ Info: Epoch 3531
└ @ Main /Users/kcft114/.julia/packages/Flux/qXNjB/src/optimise/train.jl:105
┌ Info: Epoch 3532
└ @ Main /Users/kcft114/.julia/packages/Flux/qXNjB/src/optimise/train.jl:105
┌ Info: Epoch 3533
└ @ Main /Users/kcft114/.julia/packages/Flux/qXNjB/src/optimise/train.jl:105
┌ Info: Epoch 3534
└ @ Main /Users/kcft114/.julia/packages/Flux/qXNjB/src/optimise/train.jl:105
┌ Info: Epoch 3535
└ @ Main /Users/kcft114/.julia/packages/Flux/qXNjB/src/optimise/train.jl:105
┌ Info: Epoch 3536
└ @ Main /Users/kcft114/.julia/packages/Flux/qXNjB/src/optimise/train.jl:105
┌ Info: Epoch 3537
└ @ Main /Users/kcft114/.julia/packages/Flux/qXNjB/src/optimise/train.jl:105
┌ Info: Epoch 3538
└ @ Main /Users/kcft114/.julia/packages/Flux/qXNjB/src/optimise/train.jl:105
┌ Info: Epoch 3539
└ @ Main /Users/kcft114/.julia/packages/Flux/qXNjB/src/optimise/train.jl:105
┌ Info: Epoch 3540
└ @ Main /Users/kcft114/.julia/packages/Flux/qXNjB/src/optimise/train.jl:105
┌ Info: Epoch 3541
└ @ Main /Users/kcft114/.julia/packages/Flux/qXNjB/src/optimise/train.jl:105
┌ Info: Epoch 3542
└ @ Main /Users/kcft114/.julia/packages/Flux/qXNjB/src/optimise/train.jl:105
┌ Info: Epoch 3543
└ @ Main /Users/kcft114/.julia/packages/Flux/qXNjB/src/optimise/train.jl:105
┌ Info: Epoch 3544
└ @ Main /Users/kcft114/.julia/packages/Flux/qXNjB/src/optimise/train.jl:105
┌ Info: Epoch 3545
└ @ Main /Users/kcft114/.julia/packages/Flux/qXNjB/src/optimise/train.jl:105
┌ Info: Epoch 3546
└ @ Main /Users/kcft114/.julia/packages/Flux/qXNjB/src/optimise/train.jl:105
```

```
loss(X, Y) = 0.15342636f0 (tracked)
loss(X, Y) = 0.15335949f0 (tracked)
loss(X, Y) = 0.15328965f0 (tracked)
loss(X, Y) = 0.1532331f0 (tracked)
loss(X, Y) = 0.15316097f0 (tracked)
loss(X, Y) = 0.15309678f0 (tracked)
loss(X, Y) = 0.15303755f0 (tracked)
loss(X, Y) = 0.15296488f0 (tracked)
loss(X, Y) = 0.1529112f0 (tracked)
loss(X, Y) = 0.1528333f0 (tracked)
loss(X, Y) = 0.15276964f0 (tracked)
loss(X, Y) = 0.15272172f0 (tracked)
loss(X, Y) = 0.15267073f0 (tracked)
loss(X, Y) = 0.15258588f0 (tracked)
loss(X, Y) = 0.15252702f0 (tracked)
loss(X, Y) = 0.15247247f0 (tracked)
loss(X, Y) = 0.15240914f0 (tracked)
loss(X, Y) = 0.1523449f0 (tracked)
loss(X, Y) = 0.15228145f0 (tracked)
loss(X, Y) = 0.15221183f0 (tracked)
loss(X, Y) = 0.1521411f0 (tracked)
loss(X, Y) = 0.15210766f0 (tracked)
loss(X, Y) = 0.15204635f0 (tracked)
loss(X, Y) = 0.15196593f0 (tracked)
loss(X, Y) = 0.1518944f0 (tracked)
loss(X, Y) = 0.15183926f0 (tracked)
loss(X, Y) = 0.15177087f0 (tracked)
loss(X, Y) = 0.15174562f0 (tracked)
loss(X, Y) = 0.15166123f0 (tracked)
loss(X, Y) = 0.15157905f0 (tracked)
loss(X, Y) = 0.15152201f0 (tracked)
loss(X, Y) = 0.15147947f0 (tracked)
loss(X, Y) = 0.1514054f0 (tracked)
loss(X, Y) = 0.1513356f0 (tracked)
loss(X, Y) = 0.15129408f0 (tracked)
loss(X, Y) = 0.15122974f0 (tracked)
loss(X, Y) = 0.15115835f0 (tracked)
loss(X, Y) = 0.15109597f0 (tracked)
loss(X, Y) = 0.15103032f0 (tracked)
loss(X, Y) = 0.15097696f0 (tracked)
loss(X, Y) = 0.1509176f0 (tracked)
loss(X, Y) = 0.15084736f0 (tracked)
loss(X, Y) = 0.15077172f0 (tracked)
loss(X, Y) = 0.15072374f0 (tracked)
loss(X, Y) = 0.15066151f0 (tracked)
loss(X, Y) = 0.15059008f0 (tracked)
loss(X, Y) = 0.1505326f0 (tracked)
loss(X, Y) = 0.15046483f0 (tracked)
loss(X, Y) = 0.15039226f0 (tracked)
loss(X, Y) = 0.15033683f0 (tracked)
loss(X, Y) = 0.15028012f0 (tracked)
loss(X, Y) = 0.15022139f0 (tracked)
loss(X, Y) = 0.15016215f0 (tracked)
loss(X, Y) = 0.15010345f0 (tracked)
loss(X, Y) = 0.15005486f0 (tracked)
loss(X, Y) = 0.150005f0 (tracked)
loss(X, Y) = 0.1499247f0 (tracked)
loss(X, Y) = 0.14986911f0 (tracked)
loss(X, Y) = 0.14981534f0 (tracked)
loss(X, Y) = 0.14975585f0 (tracked)
loss(X, Y) = 0.14969364f0 (tracked)
loss(X, Y) = 0.14963065f0 (tracked)
loss(X, Y) = 0.14956276f0 (tracked)
loss(X, Y) = 0.1495131f0 (tracked)
loss(X, Y) = 0.14945523f0 (tracked)
loss(X, Y) = 0.1493851f0 (tracked)
loss(X, Y) = 0.14932314f0 (tracked)
loss(X, Y) = 0.14926097f0 (tracked)
loss(X, Y) = 0.14921746f0 (tracked)
loss(X, Y) = 0.14914976f0 (tracked)
loss(X, Y) = 0.14909568f0 (tracked)
loss(X, Y) = 0.14902037f0 (tracked)
loss(X, Y) = 0.14897768f0 (tracked)
loss(X, Y) = 0.14892076f0 (tracked)
loss(X, Y) = 0.14884844f0 (tracked)
loss(X, Y) = 0.14880084f0 (tracked)
loss(X, Y) = 0.1487483f0 (tracked)
loss(X, Y) = 0.14869782f0 (tracked)
loss(X, Y) = 0.14864199f0 (tracked)
loss(X, Y) = 0.14856318f0 (tracked)
loss(X, Y) = 0.14849547f0 (tracked)
loss(X, Y) = 0.14845777f0 (tracked)
loss(X, Y) = 0.14838728f0 (tracked)
loss(X, Y) = 0.14831154f0 (tracked)
loss(X, Y) = 0.14825533f0 (tracked)
loss(X, Y) = 0.14819126f0 (tracked)
loss(X, Y) = 0.14813381f0 (tracked)
loss(X, Y) = 0.14806837f0 (tracked)
loss(X, Y) = 0.14799932f0 (tracked)
loss(X, Y) = 0.147951f0 (tracked)
loss(X, Y) = 0.14788476f0 (tracked)
loss(X, Y) = 0.14782774f0 (tracked)
loss(X, Y) = 0.14777502f0 (tracked)
loss(X, Y) = 0.14772473f0 (tracked)
loss(X, Y) = 0.14765385f0 (tracked)
loss(X, Y) = 0.14760563f0 (tracked)
loss(X, Y) = 0.1475454f0 (tracked)
loss(X, Y) = 0.14749047f0 (tracked)
loss(X, Y) = 0.14742588f0 (tracked)
loss(X, Y) = 0.14737311f0 (tracked)
loss(X, Y) = 0.1473128f0 (tracked)
loss(X, Y) = 0.14724778f0 (tracked)
loss(X, Y) = 0.14721474f0 (tracked)
loss(X, Y) = 0.14716238f0 (tracked)
loss(X, Y) = 0.14709295f0 (tracked)
loss(X, Y) = 0.1470252f0 (tracked)
loss(X, Y) = 0.14696234f0 (tracked)
loss(X, Y) = 0.14690918f0 (tracked)
loss(X, Y) = 0.14683099f0 (tracked)
loss(X, Y) = 0.14679296f0 (tracked)
loss(X, Y) = 0.1467433f0 (tracked)
loss(X, Y) = 0.14666304f0 (tracked)
loss(X, Y) = 0.14661248f0 (tracked)
loss(X, Y) = 0.14656259f0 (tracked)
loss(X, Y) = 0.1464844f0 (tracked)
loss(X, Y) = 0.14642487f0 (tracked)
loss(X, Y) = 0.14639825f0 (tracked)
loss(X, Y) = 0.14634874f0 (tracked)
loss(X, Y) = 0.14625517f0 (tracked)
loss(X, Y) = 0.14620605f0 (tracked)
loss(X, Y) = 0.14616892f0 (tracked)
loss(X, Y) = 0.1461072f0 (tracked)
loss(X, Y) = 0.14604056f0 (tracked)
loss(X, Y) = 0.14598483f0 (tracked)
loss(X, Y) = 0.14593178f0 (tracked)
loss(X, Y) = 0.14586866f0 (tracked)
loss(X, Y) = 0.14581263f0 (tracked)
loss(X, Y) = 0.14574447f0 (tracked)
loss(X, Y) = 0.14569753f0 (tracked)
loss(X, Y) = 0.14564477f0 (tracked)
loss(X, Y) = 0.14558542f0 (tracked)
loss(X, Y) = 0.14553402f0 (tracked)
loss(X, Y) = 0.14548016f0 (tracked)
loss(X, Y) = 0.14541657f0 (tracked)
loss(X, Y) = 0.14535876f0 (tracked)
loss(X, Y) = 0.14530872f0 (tracked)
loss(X, Y) = 0.14525245f0 (tracked)
loss(X, Y) = 0.14517939f0 (tracked)
loss(X, Y) = 0.14511451f0 (tracked)
loss(X, Y) = 0.14507358f0 (tracked)
loss(X, Y) = 0.1450152f0 (tracked)
loss(X, Y) = 0.14496267f0 (tracked)
loss(X, Y) = 0.14488898f0 (tracked)
loss(X, Y) = 0.14484008f0 (tracked)
loss(X, Y) = 0.1447926f0 (tracked)
loss(X, Y) = 0.14471558f0 (tracked)
loss(X, Y) = 0.14465523f0 (tracked)
loss(X, Y) = 0.14460625f0 (tracked)
loss(X, Y) = 0.14454167f0 (tracked)
loss(X, Y) = 0.14449269f0 (tracked)
loss(X, Y) = 0.1444443f0 (tracked)
loss(X, Y) = 0.14439529f0 (tracked)
loss(X, Y) = 0.14432819f0 (tracked)
loss(X, Y) = 0.14425014f0 (tracked)
loss(X, Y) = 0.14421108f0 (tracked)
loss(X, Y) = 0.1441625f0 (tracked)
loss(X, Y) = 0.14409655f0 (tracked)
loss(X, Y) = 0.14404044f0 (tracked)
loss(X, Y) = 0.14398406f0 (tracked)
```

```
┌ Info: Epoch 3547
└ @ Main /Users/kcft114/.julia/packages/Flux/qXNjB/src/optimise/train.jl:105
┌ Info: Epoch 3548
└ @ Main /Users/kcft114/.julia/packages/Flux/qXNjB/src/optimise/train.jl:105
┌ Info: Epoch 3549
└ @ Main /Users/kcft114/.julia/packages/Flux/qXNjB/src/optimise/train.jl:105
┌ Info: Epoch 3550
└ @ Main /Users/kcft114/.julia/packages/Flux/qXNjB/src/optimise/train.jl:105
┌ Info: Epoch 3551
└ @ Main /Users/kcft114/.julia/packages/Flux/qXNjB/src/optimise/train.jl:105
┌ Info: Epoch 3552
└ @ Main /Users/kcft114/.julia/packages/Flux/qXNjB/src/optimise/train.jl:105
┌ Info: Epoch 3553
└ @ Main /Users/kcft114/.julia/packages/Flux/qXNjB/src/optimise/train.jl:105
┌ Info: Epoch 3554
└ @ Main /Users/kcft114/.julia/packages/Flux/qXNjB/src/optimise/train.jl:105
┌ Info: Epoch 3555
└ @ Main /Users/kcft114/.julia/packages/Flux/qXNjB/src/optimise/train.jl:105
┌ Info: Epoch 3556
└ @ Main /Users/kcft114/.julia/packages/Flux/qXNjB/src/optimise/train.jl:105
┌ Info: Epoch 3557
└ @ Main /Users/kcft114/.julia/packages/Flux/qXNjB/src/optimise/train.jl:105
┌ Info: Epoch 3558
└ @ Main /Users/kcft114/.julia/packages/Flux/qXNjB/src/optimise/train.jl:105
┌ Info: Epoch 3559
└ @ Main /Users/kcft114/.julia/packages/Flux/qXNjB/src/optimise/train.jl:105
┌ Info: Epoch 3560
└ @ Main /Users/kcft114/.julia/packages/Flux/qXNjB/src/optimise/train.jl:105
┌ Info: Epoch 3561
└ @ Main /Users/kcft114/.julia/packages/Flux/qXNjB/src/optimise/train.jl:105
┌ Info: Epoch 3562
└ @ Main /Users/kcft114/.julia/packages/Flux/qXNjB/src/optimise/train.jl:105
┌ Info: Epoch 3563
└ @ Main /Users/kcft114/.julia/packages/Flux/qXNjB/src/optimise/train.jl:105
┌ Info: Epoch 3564
└ @ Main /Users/kcft114/.julia/packages/Flux/qXNjB/src/optimise/train.jl:105
┌ Info: Epoch 3565
└ @ Main /Users/kcft114/.julia/packages/Flux/qXNjB/src/optimise/train.jl:105
┌ Info: Epoch 3566
└ @ Main /Users/kcft114/.julia/packages/Flux/qXNjB/src/optimise/train.jl:105
┌ Info: Epoch 3567
└ @ Main /Users/kcft114/.julia/packages/Flux/qXNjB/src/optimise/train.jl:105
┌ Info: Epoch 3568
└ @ Main /Users/kcft114/.julia/packages/Flux/qXNjB/src/optimise/train.jl:105
┌ Info: Epoch 3569
└ @ Main /Users/kcft114/.julia/packages/Flux/qXNjB/src/optimise/train.jl:105
┌ Info: Epoch 3570
└ @ Main /Users/kcft114/.julia/packages/Flux/qXNjB/src/optimise/train.jl:105
┌ Info: Epoch 3571
└ @ Main /Users/kcft114/.julia/packages/Flux/qXNjB/src/optimise/train.jl:105
┌ Info: Epoch 3572
└ @ Main /Users/kcft114/.julia/packages/Flux/qXNjB/src/optimise/train.jl:105
┌ Info: Epoch 3573
└ @ Main /Users/kcft114/.julia/packages/Flux/qXNjB/src/optimise/train.jl:105
┌ Info: Epoch 3574
└ @ Main /Users/kcft114/.julia/packages/Flux/qXNjB/src/optimise/train.jl:105
┌ Info: Epoch 3575
└ @ Main /Users/kcft114/.julia/packages/Flux/qXNjB/src/optimise/train.jl:105
┌ Info: Epoch 3576
└ @ Main /Users/kcft114/.julia/packages/Flux/qXNjB/src/optimise/train.jl:105
┌ Info: Epoch 3577
└ @ Main /Users/kcft114/.julia/packages/Flux/qXNjB/src/optimise/train.jl:105
┌ Info: Epoch 3578
└ @ Main /Users/kcft114/.julia/packages/Flux/qXNjB/src/optimise/train.jl:105
┌ Info: Epoch 3579
└ @ Main /Users/kcft114/.julia/packages/Flux/qXNjB/src/optimise/train.jl:105
┌ Info: Epoch 3580
└ @ Main /Users/kcft114/.julia/packages/Flux/qXNjB/src/optimise/train.jl:105
┌ Info: Epoch 3581
└ @ Main /Users/kcft114/.julia/packages/Flux/qXNjB/src/optimise/train.jl:105
┌ Info: Epoch 3582
└ @ Main /Users/kcft114/.julia/packages/Flux/qXNjB/src/optimise/train.jl:105
┌ Info: Epoch 3583
└ @ Main /Users/kcft114/.julia/packages/Flux/qXNjB/src/optimise/train.jl:105
┌ Info: Epoch 3584
└ @ Main /Users/kcft114/.julia/packages/Flux/qXNjB/src/optimise/train.jl:105
┌ Info: Epoch 3585
└ @ Main /Users/kcft114/.julia/packages/Flux/qXNjB/src/optimise/train.jl:105
┌ Info: Epoch 3586
└ @ Main /Users/kcft114/.julia/packages/Flux/qXNjB/src/optimise/train.jl:105
┌ Info: Epoch 3587
└ @ Main /Users/kcft114/.julia/packages/Flux/qXNjB/src/optimise/train.jl:105
┌ Info: Epoch 3588
└ @ Main /Users/kcft114/.julia/packages/Flux/qXNjB/src/optimise/train.jl:105
┌ Info: Epoch 3589
└ @ Main /Users/kcft114/.julia/packages/Flux/qXNjB/src/optimise/train.jl:105
┌ Info: Epoch 3590
└ @ Main /Users/kcft114/.julia/packages/Flux/qXNjB/src/optimise/train.jl:105
┌ Info: Epoch 3591
└ @ Main /Users/kcft114/.julia/packages/Flux/qXNjB/src/optimise/train.jl:105
┌ Info: Epoch 3592
└ @ Main /Users/kcft114/.julia/packages/Flux/qXNjB/src/optimise/train.jl:105
┌ Info: Epoch 3593
└ @ Main /Users/kcft114/.julia/packages/Flux/qXNjB/src/optimise/train.jl:105
┌ Info: Epoch 3594
└ @ Main /Users/kcft114/.julia/packages/Flux/qXNjB/src/optimise/train.jl:105
┌ Info: Epoch 3595
└ @ Main /Users/kcft114/.julia/packages/Flux/qXNjB/src/optimise/train.jl:105
┌ Info: Epoch 3596
└ @ Main /Users/kcft114/.julia/packages/Flux/qXNjB/src/optimise/train.jl:105
┌ Info: Epoch 3597
└ @ Main /Users/kcft114/.julia/packages/Flux/qXNjB/src/optimise/train.jl:105
┌ Info: Epoch 3598
└ @ Main /Users/kcft114/.julia/packages/Flux/qXNjB/src/optimise/train.jl:105
┌ Info: Epoch 3599
└ @ Main /Users/kcft114/.julia/packages/Flux/qXNjB/src/optimise/train.jl:105
┌ Info: Epoch 3600
└ @ Main /Users/kcft114/.julia/packages/Flux/qXNjB/src/optimise/train.jl:105
┌ Info: Epoch 3601
└ @ Main /Users/kcft114/.julia/packages/Flux/qXNjB/src/optimise/train.jl:105
┌ Info: Epoch 3602
└ @ Main /Users/kcft114/.julia/packages/Flux/qXNjB/src/optimise/train.jl:105
┌ Info: Epoch 3603
└ @ Main /Users/kcft114/.julia/packages/Flux/qXNjB/src/optimise/train.jl:105
┌ Info: Epoch 3604
└ @ Main /Users/kcft114/.julia/packages/Flux/qXNjB/src/optimise/train.jl:105
┌ Info: Epoch 3605
└ @ Main /Users/kcft114/.julia/packages/Flux/qXNjB/src/optimise/train.jl:105
┌ Info: Epoch 3606
└ @ Main /Users/kcft114/.julia/packages/Flux/qXNjB/src/optimise/train.jl:105
┌ Info: Epoch 3607
└ @ Main /Users/kcft114/.julia/packages/Flux/qXNjB/src/optimise/train.jl:105
┌ Info: Epoch 3608
└ @ Main /Users/kcft114/.julia/packages/Flux/qXNjB/src/optimise/train.jl:105
┌ Info: Epoch 3609
└ @ Main /Users/kcft114/.julia/packages/Flux/qXNjB/src/optimise/train.jl:105
┌ Info: Epoch 3610
└ @ Main /Users/kcft114/.julia/packages/Flux/qXNjB/src/optimise/train.jl:105
┌ Info: Epoch 3611
└ @ Main /Users/kcft114/.julia/packages/Flux/qXNjB/src/optimise/train.jl:105
┌ Info: Epoch 3612
└ @ Main /Users/kcft114/.julia/packages/Flux/qXNjB/src/optimise/train.jl:105
┌ Info: Epoch 3613
└ @ Main /Users/kcft114/.julia/packages/Flux/qXNjB/src/optimise/train.jl:105
┌ Info: Epoch 3614
└ @ Main /Users/kcft114/.julia/packages/Flux/qXNjB/src/optimise/train.jl:105
┌ Info: Epoch 3615
└ @ Main /Users/kcft114/.julia/packages/Flux/qXNjB/src/optimise/train.jl:105
┌ Info: Epoch 3616
└ @ Main /Users/kcft114/.julia/packages/Flux/qXNjB/src/optimise/train.jl:105
┌ Info: Epoch 3617
└ @ Main /Users/kcft114/.julia/packages/Flux/qXNjB/src/optimise/train.jl:105
┌ Info: Epoch 3618
└ @ Main /Users/kcft114/.julia/packages/Flux/qXNjB/src/optimise/train.jl:105
┌ Info: Epoch 3619
└ @ Main /Users/kcft114/.julia/packages/Flux/qXNjB/src/optimise/train.jl:105
┌ Info: Epoch 3620
└ @ Main /Users/kcft114/.julia/packages/Flux/qXNjB/src/optimise/train.jl:105
┌ Info: Epoch 3621
└ @ Main /Users/kcft114/.julia/packages/Flux/qXNjB/src/optimise/train.jl:105
┌ Info: Epoch 3622
└ @ Main /Users/kcft114/.julia/packages/Flux/qXNjB/src/optimise/train.jl:105
┌ Info: Epoch 3623
└ @ Main /Users/kcft114/.julia/packages/Flux/qXNjB/src/optimise/train.jl:105
┌ Info: Epoch 3624
└ @ Main /Users/kcft114/.julia/packages/Flux/qXNjB/src/optimise/train.jl:105
┌ Info: Epoch 3625
└ @ Main /Users/kcft114/.julia/packages/Flux/qXNjB/src/optimise/train.jl:105
┌ Info: Epoch 3626
└ @ Main /Users/kcft114/.julia/packages/Flux/qXNjB/src/optimise/train.jl:105
┌ Info: Epoch 3627
└ @ Main /Users/kcft114/.julia/packages/Flux/qXNjB/src/optimise/train.jl:105
┌ Info: Epoch 3628
└ @ Main /Users/kcft114/.julia/packages/Flux/qXNjB/src/optimise/train.jl:105
┌ Info: Epoch 3629
└ @ Main /Users/kcft114/.julia/packages/Flux/qXNjB/src/optimise/train.jl:105
┌ Info: Epoch 3630
└ @ Main /Users/kcft114/.julia/packages/Flux/qXNjB/src/optimise/train.jl:105
┌ Info: Epoch 3631
└ @ Main /Users/kcft114/.julia/packages/Flux/qXNjB/src/optimise/train.jl:105
┌ Info: Epoch 3632
└ @ Main /Users/kcft114/.julia/packages/Flux/qXNjB/src/optimise/train.jl:105
┌ Info: Epoch 3633
└ @ Main /Users/kcft114/.julia/packages/Flux/qXNjB/src/optimise/train.jl:105
┌ Info: Epoch 3634
└ @ Main /Users/kcft114/.julia/packages/Flux/qXNjB/src/optimise/train.jl:105
┌ Info: Epoch 3635
└ @ Main /Users/kcft114/.julia/packages/Flux/qXNjB/src/optimise/train.jl:105
┌ Info: Epoch 3636
└ @ Main /Users/kcft114/.julia/packages/Flux/qXNjB/src/optimise/train.jl:105
┌ Info: Epoch 3637
└ @ Main /Users/kcft114/.julia/packages/Flux/qXNjB/src/optimise/train.jl:105
┌ Info: Epoch 3638
└ @ Main /Users/kcft114/.julia/packages/Flux/qXNjB/src/optimise/train.jl:105
┌ Info: Epoch 3639
└ @ Main /Users/kcft114/.julia/packages/Flux/qXNjB/src/optimise/train.jl:105
┌ Info: Epoch 3640
└ @ Main /Users/kcft114/.julia/packages/Flux/qXNjB/src/optimise/train.jl:105
┌ Info: Epoch 3641
└ @ Main /Users/kcft114/.julia/packages/Flux/qXNjB/src/optimise/train.jl:105
┌ Info: Epoch 3642
└ @ Main /Users/kcft114/.julia/packages/Flux/qXNjB/src/optimise/train.jl:105
┌ Info: Epoch 3643
└ @ Main /Users/kcft114/.julia/packages/Flux/qXNjB/src/optimise/train.jl:105
┌ Info: Epoch 3644
└ @ Main /Users/kcft114/.julia/packages/Flux/qXNjB/src/optimise/train.jl:105
┌ Info: Epoch 3645
└ @ Main /Users/kcft114/.julia/packages/Flux/qXNjB/src/optimise/train.jl:105
┌ Info: Epoch 3646
└ @ Main /Users/kcft114/.julia/packages/Flux/qXNjB/src/optimise/train.jl:105
┌ Info: Epoch 3647
└ @ Main /Users/kcft114/.julia/packages/Flux/qXNjB/src/optimise/train.jl:105
┌ Info: Epoch 3648
└ @ Main /Users/kcft114/.julia/packages/Flux/qXNjB/src/optimise/train.jl:105
┌ Info: Epoch 3649
└ @ Main /Users/kcft114/.julia/packages/Flux/qXNjB/src/optimise/train.jl:105
┌ Info: Epoch 3650
└ @ Main /Users/kcft114/.julia/packages/Flux/qXNjB/src/optimise/train.jl:105
┌ Info: Epoch 3651
└ @ Main /Users/kcft114/.julia/packages/Flux/qXNjB/src/optimise/train.jl:105
┌ Info: Epoch 3652
└ @ Main /Users/kcft114/.julia/packages/Flux/qXNjB/src/optimise/train.jl:105
┌ Info: Epoch 3653
└ @ Main /Users/kcft114/.julia/packages/Flux/qXNjB/src/optimise/train.jl:105
┌ Info: Epoch 3654
└ @ Main /Users/kcft114/.julia/packages/Flux/qXNjB/src/optimise/train.jl:105
┌ Info: Epoch 3655
└ @ Main /Users/kcft114/.julia/packages/Flux/qXNjB/src/optimise/train.jl:105
┌ Info: Epoch 3656
└ @ Main /Users/kcft114/.julia/packages/Flux/qXNjB/src/optimise/train.jl:105
┌ Info: Epoch 3657
└ @ Main /Users/kcft114/.julia/packages/Flux/qXNjB/src/optimise/train.jl:105
┌ Info: Epoch 3658
└ @ Main /Users/kcft114/.julia/packages/Flux/qXNjB/src/optimise/train.jl:105
┌ Info: Epoch 3659
└ @ Main /Users/kcft114/.julia/packages/Flux/qXNjB/src/optimise/train.jl:105
┌ Info: Epoch 3660
└ @ Main /Users/kcft114/.julia/packages/Flux/qXNjB/src/optimise/train.jl:105
┌ Info: Epoch 3661
└ @ Main /Users/kcft114/.julia/packages/Flux/qXNjB/src/optimise/train.jl:105
┌ Info: Epoch 3662
└ @ Main /Users/kcft114/.julia/packages/Flux/qXNjB/src/optimise/train.jl:105
┌ Info: Epoch 3663
└ @ Main /Users/kcft114/.julia/packages/Flux/qXNjB/src/optimise/train.jl:105
┌ Info: Epoch 3664
└ @ Main /Users/kcft114/.julia/packages/Flux/qXNjB/src/optimise/train.jl:105
┌ Info: Epoch 3665
└ @ Main /Users/kcft114/.julia/packages/Flux/qXNjB/src/optimise/train.jl:105
┌ Info: Epoch 3666
└ @ Main /Users/kcft114/.julia/packages/Flux/qXNjB/src/optimise/train.jl:105
┌ Info: Epoch 3667
└ @ Main /Users/kcft114/.julia/packages/Flux/qXNjB/src/optimise/train.jl:105
┌ Info: Epoch 3668
└ @ Main /Users/kcft114/.julia/packages/Flux/qXNjB/src/optimise/train.jl:105
┌ Info: Epoch 3669
└ @ Main /Users/kcft114/.julia/packages/Flux/qXNjB/src/optimise/train.jl:105
┌ Info: Epoch 3670
└ @ Main /Users/kcft114/.julia/packages/Flux/qXNjB/src/optimise/train.jl:105
┌ Info: Epoch 3671
└ @ Main /Users/kcft114/.julia/packages/Flux/qXNjB/src/optimise/train.jl:105
┌ Info: Epoch 3672
└ @ Main /Users/kcft114/.julia/packages/Flux/qXNjB/src/optimise/train.jl:105
┌ Info: Epoch 3673
└ @ Main /Users/kcft114/.julia/packages/Flux/qXNjB/src/optimise/train.jl:105
┌ Info: Epoch 3674
└ @ Main /Users/kcft114/.julia/packages/Flux/qXNjB/src/optimise/train.jl:105
┌ Info: Epoch 3675
└ @ Main /Users/kcft114/.julia/packages/Flux/qXNjB/src/optimise/train.jl:105
┌ Info: Epoch 3676
└ @ Main /Users/kcft114/.julia/packages/Flux/qXNjB/src/optimise/train.jl:105
┌ Info: Epoch 3677
└ @ Main /Users/kcft114/.julia/packages/Flux/qXNjB/src/optimise/train.jl:105
┌ Info: Epoch 3678
└ @ Main /Users/kcft114/.julia/packages/Flux/qXNjB/src/optimise/train.jl:105
┌ Info: Epoch 3679
└ @ Main /Users/kcft114/.julia/packages/Flux/qXNjB/src/optimise/train.jl:105
┌ Info: Epoch 3680
└ @ Main /Users/kcft114/.julia/packages/Flux/qXNjB/src/optimise/train.jl:105
┌ Info: Epoch 3681
└ @ Main /Users/kcft114/.julia/packages/Flux/qXNjB/src/optimise/train.jl:105
┌ Info: Epoch 3682
└ @ Main /Users/kcft114/.julia/packages/Flux/qXNjB/src/optimise/train.jl:105
┌ Info: Epoch 3683
└ @ Main /Users/kcft114/.julia/packages/Flux/qXNjB/src/optimise/train.jl:105
┌ Info: Epoch 3684
└ @ Main /Users/kcft114/.julia/packages/Flux/qXNjB/src/optimise/train.jl:105
┌ Info: Epoch 3685
└ @ Main /Users/kcft114/.julia/packages/Flux/qXNjB/src/optimise/train.jl:105
┌ Info: Epoch 3686
└ @ Main /Users/kcft114/.julia/packages/Flux/qXNjB/src/optimise/train.jl:105
┌ Info: Epoch 3687
└ @ Main /Users/kcft114/.julia/packages/Flux/qXNjB/src/optimise/train.jl:105
┌ Info: Epoch 3688
└ @ Main /Users/kcft114/.julia/packages/Flux/qXNjB/src/optimise/train.jl:105
┌ Info: Epoch 3689
└ @ Main /Users/kcft114/.julia/packages/Flux/qXNjB/src/optimise/train.jl:105
┌ Info: Epoch 3690
└ @ Main /Users/kcft114/.julia/packages/Flux/qXNjB/src/optimise/train.jl:105
┌ Info: Epoch 3691
└ @ Main /Users/kcft114/.julia/packages/Flux/qXNjB/src/optimise/train.jl:105
┌ Info: Epoch 3692
└ @ Main /Users/kcft114/.julia/packages/Flux/qXNjB/src/optimise/train.jl:105
┌ Info: Epoch 3693
└ @ Main /Users/kcft114/.julia/packages/Flux/qXNjB/src/optimise/train.jl:105
┌ Info: Epoch 3694
└ @ Main /Users/kcft114/.julia/packages/Flux/qXNjB/src/optimise/train.jl:105
┌ Info: Epoch 3695
└ @ Main /Users/kcft114/.julia/packages/Flux/qXNjB/src/optimise/train.jl:105
┌ Info: Epoch 3696
└ @ Main /Users/kcft114/.julia/packages/Flux/qXNjB/src/optimise/train.jl:105
┌ Info: Epoch 3697
└ @ Main /Users/kcft114/.julia/packages/Flux/qXNjB/src/optimise/train.jl:105
┌ Info: Epoch 3698
└ @ Main /Users/kcft114/.julia/packages/Flux/qXNjB/src/optimise/train.jl:105
┌ Info: Epoch 3699
└ @ Main /Users/kcft114/.julia/packages/Flux/qXNjB/src/optimise/train.jl:105
┌ Info: Epoch 3700
└ @ Main /Users/kcft114/.julia/packages/Flux/qXNjB/src/optimise/train.jl:105
┌ Info: Epoch 3701
└ @ Main /Users/kcft114/.julia/packages/Flux/qXNjB/src/optimise/train.jl:105
┌ Info: Epoch 3702
└ @ Main /Users/kcft114/.julia/packages/Flux/qXNjB/src/optimise/train.jl:105
┌ Info: Epoch 3703
└ @ Main /Users/kcft114/.julia/packages/Flux/qXNjB/src/optimise/train.jl:105
┌ Info: Epoch 3704
└ @ Main /Users/kcft114/.julia/packages/Flux/qXNjB/src/optimise/train.jl:105
┌ Info: Epoch 3705
└ @ Main /Users/kcft114/.julia/packages/Flux/qXNjB/src/optimise/train.jl:105
```

```
loss(X, Y) = 0.14390887f0 (tracked)
loss(X, Y) = 0.14385451f0 (tracked)
loss(X, Y) = 0.14383999f0 (tracked)
loss(X, Y) = 0.14378345f0 (tracked)
loss(X, Y) = 0.1436917f0 (tracked)
loss(X, Y) = 0.14365494f0 (tracked)
loss(X, Y) = 0.14360519f0 (tracked)
loss(X, Y) = 0.1435305f0 (tracked)
loss(X, Y) = 0.14346828f0 (tracked)
loss(X, Y) = 0.14340195f0 (tracked)
loss(X, Y) = 0.14335637f0 (tracked)
loss(X, Y) = 0.14331359f0 (tracked)
loss(X, Y) = 0.14325914f0 (tracked)
loss(X, Y) = 0.14319934f0 (tracked)
loss(X, Y) = 0.14312495f0 (tracked)
loss(X, Y) = 0.14307426f0 (tracked)
loss(X, Y) = 0.14303218f0 (tracked)
loss(X, Y) = 0.14295745f0 (tracked)
loss(X, Y) = 0.14288935f0 (tracked)
loss(X, Y) = 0.14284703f0 (tracked)
loss(X, Y) = 0.14279445f0 (tracked)
loss(X, Y) = 0.14271812f0 (tracked)
loss(X, Y) = 0.14266166f0 (tracked)
loss(X, Y) = 0.14261763f0 (tracked)
loss(X, Y) = 0.14255579f0 (tracked)
loss(X, Y) = 0.14249407f0 (tracked)
loss(X, Y) = 0.14243962f0 (tracked)
loss(X, Y) = 0.14240418f0 (tracked)
loss(X, Y) = 0.14234026f0 (tracked)
loss(X, Y) = 0.14228855f0 (tracked)
loss(X, Y) = 0.14223251f0 (tracked)
loss(X, Y) = 0.14217103f0 (tracked)
loss(X, Y) = 0.14210105f0 (tracked)
loss(X, Y) = 0.14205717f0 (tracked)
loss(X, Y) = 0.14200434f0 (tracked)
loss(X, Y) = 0.14194301f0 (tracked)
loss(X, Y) = 0.1418988f0 (tracked)
loss(X, Y) = 0.14185648f0 (tracked)
loss(X, Y) = 0.14179695f0 (tracked)
loss(X, Y) = 0.14174253f0 (tracked)
loss(X, Y) = 0.14169107f0 (tracked)
loss(X, Y) = 0.1416297f0 (tracked)
loss(X, Y) = 0.14158072f0 (tracked)
loss(X, Y) = 0.14152315f0 (tracked)
loss(X, Y) = 0.14146292f0 (tracked)
loss(X, Y) = 0.14140627f0 (tracked)
loss(X, Y) = 0.14134558f0 (tracked)
loss(X, Y) = 0.14128406f0 (tracked)
loss(X, Y) = 0.14122218f0 (tracked)
loss(X, Y) = 0.14117499f0 (tracked)
loss(X, Y) = 0.14110844f0 (tracked)
loss(X, Y) = 0.14106151f0 (tracked)
loss(X, Y) = 0.14100167f0 (tracked)
loss(X, Y) = 0.14094779f0 (tracked)
loss(X, Y) = 0.14090551f0 (tracked)
loss(X, Y) = 0.1408335f0 (tracked)
loss(X, Y) = 0.14079233f0 (tracked)
loss(X, Y) = 0.14074077f0 (tracked)
loss(X, Y) = 0.14067398f0 (tracked)
loss(X, Y) = 0.14061312f0 (tracked)
loss(X, Y) = 0.14057514f0 (tracked)
loss(X, Y) = 0.14052404f0 (tracked)
loss(X, Y) = 0.14045744f0 (tracked)
loss(X, Y) = 0.14039755f0 (tracked)
loss(X, Y) = 0.14035037f0 (tracked)
loss(X, Y) = 0.1403006f0 (tracked)
loss(X, Y) = 0.14022599f0 (tracked)
loss(X, Y) = 0.14017461f0 (tracked)
loss(X, Y) = 0.14012639f0 (tracked)
loss(X, Y) = 0.14007974f0 (tracked)
loss(X, Y) = 0.14000936f0 (tracked)
loss(X, Y) = 0.13994056f0 (tracked)
loss(X, Y) = 0.13989224f0 (tracked)
loss(X, Y) = 0.13982525f0 (tracked)
loss(X, Y) = 0.13978545f0 (tracked)
loss(X, Y) = 0.13972178f0 (tracked)
loss(X, Y) = 0.13966091f0 (tracked)
loss(X, Y) = 0.13961385f0 (tracked)
loss(X, Y) = 0.13954233f0 (tracked)
loss(X, Y) = 0.13947421f0 (tracked)
loss(X, Y) = 0.13943876f0 (tracked)
loss(X, Y) = 0.13937967f0 (tracked)
loss(X, Y) = 0.13931352f0 (tracked)
loss(X, Y) = 0.1392396f0 (tracked)
loss(X, Y) = 0.13919914f0 (tracked)
loss(X, Y) = 0.13914444f0 (tracked)
loss(X, Y) = 0.13907947f0 (tracked)
loss(X, Y) = 0.13902129f0 (tracked)
loss(X, Y) = 0.13895892f0 (tracked)
loss(X, Y) = 0.13891813f0 (tracked)
loss(X, Y) = 0.13885686f0 (tracked)
loss(X, Y) = 0.138801f0 (tracked)
loss(X, Y) = 0.13873418f0 (tracked)
loss(X, Y) = 0.13868812f0 (tracked)
loss(X, Y) = 0.13864441f0 (tracked)
loss(X, Y) = 0.13857634f0 (tracked)
loss(X, Y) = 0.13852535f0 (tracked)
loss(X, Y) = 0.13846059f0 (tracked)
loss(X, Y) = 0.13839313f0 (tracked)
loss(X, Y) = 0.13834901f0 (tracked)
loss(X, Y) = 0.13829492f0 (tracked)
loss(X, Y) = 0.13821688f0 (tracked)
loss(X, Y) = 0.13815902f0 (tracked)
loss(X, Y) = 0.1381278f0 (tracked)
loss(X, Y) = 0.13805771f0 (tracked)
loss(X, Y) = 0.13800882f0 (tracked)
loss(X, Y) = 0.13796194f0 (tracked)
loss(X, Y) = 0.13790867f0 (tracked)
loss(X, Y) = 0.13785928f0 (tracked)
loss(X, Y) = 0.13779573f0 (tracked)
loss(X, Y) = 0.13775088f0 (tracked)
loss(X, Y) = 0.13769072f0 (tracked)
```

```
┌ Info: Epoch 3706
└ @ Main /Users/kcft114/.julia/packages/Flux/qXNjB/src/optimise/train.jl:105
┌ Info: Epoch 3707
└ @ Main /Users/kcft114/.julia/packages/Flux/qXNjB/src/optimise/train.jl:105
┌ Info: Epoch 3708
└ @ Main /Users/kcft114/.julia/packages/Flux/qXNjB/src/optimise/train.jl:105
┌ Info: Epoch 3709
└ @ Main /Users/kcft114/.julia/packages/Flux/qXNjB/src/optimise/train.jl:105
┌ Info: Epoch 3710
└ @ Main /Users/kcft114/.julia/packages/Flux/qXNjB/src/optimise/train.jl:105
┌ Info: Epoch 3711
└ @ Main /Users/kcft114/.julia/packages/Flux/qXNjB/src/optimise/train.jl:105
┌ Info: Epoch 3712
└ @ Main /Users/kcft114/.julia/packages/Flux/qXNjB/src/optimise/train.jl:105
┌ Info: Epoch 3713
└ @ Main /Users/kcft114/.julia/packages/Flux/qXNjB/src/optimise/train.jl:105
┌ Info: Epoch 3714
└ @ Main /Users/kcft114/.julia/packages/Flux/qXNjB/src/optimise/train.jl:105
┌ Info: Epoch 3715
└ @ Main /Users/kcft114/.julia/packages/Flux/qXNjB/src/optimise/train.jl:105
┌ Info: Epoch 3716
└ @ Main /Users/kcft114/.julia/packages/Flux/qXNjB/src/optimise/train.jl:105
┌ Info: Epoch 3717
└ @ Main /Users/kcft114/.julia/packages/Flux/qXNjB/src/optimise/train.jl:105
┌ Info: Epoch 3718
└ @ Main /Users/kcft114/.julia/packages/Flux/qXNjB/src/optimise/train.jl:105
┌ Info: Epoch 3719
└ @ Main /Users/kcft114/.julia/packages/Flux/qXNjB/src/optimise/train.jl:105
┌ Info: Epoch 3720
└ @ Main /Users/kcft114/.julia/packages/Flux/qXNjB/src/optimise/train.jl:105
┌ Info: Epoch 3721
└ @ Main /Users/kcft114/.julia/packages/Flux/qXNjB/src/optimise/train.jl:105
┌ Info: Epoch 3722
└ @ Main /Users/kcft114/.julia/packages/Flux/qXNjB/src/optimise/train.jl:105
┌ Info: Epoch 3723
└ @ Main /Users/kcft114/.julia/packages/Flux/qXNjB/src/optimise/train.jl:105
┌ Info: Epoch 3724
└ @ Main /Users/kcft114/.julia/packages/Flux/qXNjB/src/optimise/train.jl:105
┌ Info: Epoch 3725
└ @ Main /Users/kcft114/.julia/packages/Flux/qXNjB/src/optimise/train.jl:105
┌ Info: Epoch 3726
└ @ Main /Users/kcft114/.julia/packages/Flux/qXNjB/src/optimise/train.jl:105
┌ Info: Epoch 3727
└ @ Main /Users/kcft114/.julia/packages/Flux/qXNjB/src/optimise/train.jl:105
┌ Info: Epoch 3728
└ @ Main /Users/kcft114/.julia/packages/Flux/qXNjB/src/optimise/train.jl:105
┌ Info: Epoch 3729
└ @ Main /Users/kcft114/.julia/packages/Flux/qXNjB/src/optimise/train.jl:105
┌ Info: Epoch 3730
└ @ Main /Users/kcft114/.julia/packages/Flux/qXNjB/src/optimise/train.jl:105
┌ Info: Epoch 3731
└ @ Main /Users/kcft114/.julia/packages/Flux/qXNjB/src/optimise/train.jl:105
┌ Info: Epoch 3732
└ @ Main /Users/kcft114/.julia/packages/Flux/qXNjB/src/optimise/train.jl:105
┌ Info: Epoch 3733
└ @ Main /Users/kcft114/.julia/packages/Flux/qXNjB/src/optimise/train.jl:105
┌ Info: Epoch 3734
└ @ Main /Users/kcft114/.julia/packages/Flux/qXNjB/src/optimise/train.jl:105
┌ Info: Epoch 3735
└ @ Main /Users/kcft114/.julia/packages/Flux/qXNjB/src/optimise/train.jl:105
┌ Info: Epoch 3736
└ @ Main /Users/kcft114/.julia/packages/Flux/qXNjB/src/optimise/train.jl:105
┌ Info: Epoch 3737
└ @ Main /Users/kcft114/.julia/packages/Flux/qXNjB/src/optimise/train.jl:105
┌ Info: Epoch 3738
└ @ Main /Users/kcft114/.julia/packages/Flux/qXNjB/src/optimise/train.jl:105
┌ Info: Epoch 3739
└ @ Main /Users/kcft114/.julia/packages/Flux/qXNjB/src/optimise/train.jl:105
┌ Info: Epoch 3740
└ @ Main /Users/kcft114/.julia/packages/Flux/qXNjB/src/optimise/train.jl:105
┌ Info: Epoch 3741
└ @ Main /Users/kcft114/.julia/packages/Flux/qXNjB/src/optimise/train.jl:105
┌ Info: Epoch 3742
└ @ Main /Users/kcft114/.julia/packages/Flux/qXNjB/src/optimise/train.jl:105
┌ Info: Epoch 3743
└ @ Main /Users/kcft114/.julia/packages/Flux/qXNjB/src/optimise/train.jl:105
┌ Info: Epoch 3744
└ @ Main /Users/kcft114/.julia/packages/Flux/qXNjB/src/optimise/train.jl:105
┌ Info: Epoch 3745
└ @ Main /Users/kcft114/.julia/packages/Flux/qXNjB/src/optimise/train.jl:105
┌ Info: Epoch 3746
└ @ Main /Users/kcft114/.julia/packages/Flux/qXNjB/src/optimise/train.jl:105
┌ Info: Epoch 3747
└ @ Main /Users/kcft114/.julia/packages/Flux/qXNjB/src/optimise/train.jl:105
┌ Info: Epoch 3748
└ @ Main /Users/kcft114/.julia/packages/Flux/qXNjB/src/optimise/train.jl:105
┌ Info: Epoch 3749
└ @ Main /Users/kcft114/.julia/packages/Flux/qXNjB/src/optimise/train.jl:105
┌ Info: Epoch 3750
└ @ Main /Users/kcft114/.julia/packages/Flux/qXNjB/src/optimise/train.jl:105
┌ Info: Epoch 3751
└ @ Main /Users/kcft114/.julia/packages/Flux/qXNjB/src/optimise/train.jl:105
┌ Info: Epoch 3752
└ @ Main /Users/kcft114/.julia/packages/Flux/qXNjB/src/optimise/train.jl:105
┌ Info: Epoch 3753
└ @ Main /Users/kcft114/.julia/packages/Flux/qXNjB/src/optimise/train.jl:105
┌ Info: Epoch 3754
└ @ Main /Users/kcft114/.julia/packages/Flux/qXNjB/src/optimise/train.jl:105
┌ Info: Epoch 3755
└ @ Main /Users/kcft114/.julia/packages/Flux/qXNjB/src/optimise/train.jl:105
┌ Info: Epoch 3756
└ @ Main /Users/kcft114/.julia/packages/Flux/qXNjB/src/optimise/train.jl:105
┌ Info: Epoch 3757
└ @ Main /Users/kcft114/.julia/packages/Flux/qXNjB/src/optimise/train.jl:105
┌ Info: Epoch 3758
└ @ Main /Users/kcft114/.julia/packages/Flux/qXNjB/src/optimise/train.jl:105
┌ Info: Epoch 3759
└ @ Main /Users/kcft114/.julia/packages/Flux/qXNjB/src/optimise/train.jl:105
┌ Info: Epoch 3760
└ @ Main /Users/kcft114/.julia/packages/Flux/qXNjB/src/optimise/train.jl:105
┌ Info: Epoch 3761
└ @ Main /Users/kcft114/.julia/packages/Flux/qXNjB/src/optimise/train.jl:105
┌ Info: Epoch 3762
└ @ Main /Users/kcft114/.julia/packages/Flux/qXNjB/src/optimise/train.jl:105
┌ Info: Epoch 3763
└ @ Main /Users/kcft114/.julia/packages/Flux/qXNjB/src/optimise/train.jl:105
┌ Info: Epoch 3764
└ @ Main /Users/kcft114/.julia/packages/Flux/qXNjB/src/optimise/train.jl:105
┌ Info: Epoch 3765
└ @ Main /Users/kcft114/.julia/packages/Flux/qXNjB/src/optimise/train.jl:105
┌ Info: Epoch 3766
└ @ Main /Users/kcft114/.julia/packages/Flux/qXNjB/src/optimise/train.jl:105
┌ Info: Epoch 3767
└ @ Main /Users/kcft114/.julia/packages/Flux/qXNjB/src/optimise/train.jl:105
┌ Info: Epoch 3768
└ @ Main /Users/kcft114/.julia/packages/Flux/qXNjB/src/optimise/train.jl:105
┌ Info: Epoch 3769
└ @ Main /Users/kcft114/.julia/packages/Flux/qXNjB/src/optimise/train.jl:105
┌ Info: Epoch 3770
└ @ Main /Users/kcft114/.julia/packages/Flux/qXNjB/src/optimise/train.jl:105
┌ Info: Epoch 3771
└ @ Main /Users/kcft114/.julia/packages/Flux/qXNjB/src/optimise/train.jl:105
┌ Info: Epoch 3772
└ @ Main /Users/kcft114/.julia/packages/Flux/qXNjB/src/optimise/train.jl:105
┌ Info: Epoch 3773
└ @ Main /Users/kcft114/.julia/packages/Flux/qXNjB/src/optimise/train.jl:105
┌ Info: Epoch 3774
└ @ Main /Users/kcft114/.julia/packages/Flux/qXNjB/src/optimise/train.jl:105
┌ Info: Epoch 3775
└ @ Main /Users/kcft114/.julia/packages/Flux/qXNjB/src/optimise/train.jl:105
┌ Info: Epoch 3776
└ @ Main /Users/kcft114/.julia/packages/Flux/qXNjB/src/optimise/train.jl:105
┌ Info: Epoch 3777
└ @ Main /Users/kcft114/.julia/packages/Flux/qXNjB/src/optimise/train.jl:105
┌ Info: Epoch 3778
└ @ Main /Users/kcft114/.julia/packages/Flux/qXNjB/src/optimise/train.jl:105
┌ Info: Epoch 3779
└ @ Main /Users/kcft114/.julia/packages/Flux/qXNjB/src/optimise/train.jl:105
┌ Info: Epoch 3780
└ @ Main /Users/kcft114/.julia/packages/Flux/qXNjB/src/optimise/train.jl:105
┌ Info: Epoch 3781
└ @ Main /Users/kcft114/.julia/packages/Flux/qXNjB/src/optimise/train.jl:105
┌ Info: Epoch 3782
└ @ Main /Users/kcft114/.julia/packages/Flux/qXNjB/src/optimise/train.jl:105
┌ Info: Epoch 3783
└ @ Main /Users/kcft114/.julia/packages/Flux/qXNjB/src/optimise/train.jl:105
┌ Info: Epoch 3784
└ @ Main /Users/kcft114/.julia/packages/Flux/qXNjB/src/optimise/train.jl:105
┌ Info: Epoch 3785
└ @ Main /Users/kcft114/.julia/packages/Flux/qXNjB/src/optimise/train.jl:105
┌ Info: Epoch 3786
└ @ Main /Users/kcft114/.julia/packages/Flux/qXNjB/src/optimise/train.jl:105
┌ Info: Epoch 3787
└ @ Main /Users/kcft114/.julia/packages/Flux/qXNjB/src/optimise/train.jl:105
┌ Info: Epoch 3788
└ @ Main /Users/kcft114/.julia/packages/Flux/qXNjB/src/optimise/train.jl:105
┌ Info: Epoch 3789
└ @ Main /Users/kcft114/.julia/packages/Flux/qXNjB/src/optimise/train.jl:105
┌ Info: Epoch 3790
└ @ Main /Users/kcft114/.julia/packages/Flux/qXNjB/src/optimise/train.jl:105
┌ Info: Epoch 3791
└ @ Main /Users/kcft114/.julia/packages/Flux/qXNjB/src/optimise/train.jl:105
┌ Info: Epoch 3792
└ @ Main /Users/kcft114/.julia/packages/Flux/qXNjB/src/optimise/train.jl:105
┌ Info: Epoch 3793
└ @ Main /Users/kcft114/.julia/packages/Flux/qXNjB/src/optimise/train.jl:105
┌ Info: Epoch 3794
└ @ Main /Users/kcft114/.julia/packages/Flux/qXNjB/src/optimise/train.jl:105
┌ Info: Epoch 3795
└ @ Main /Users/kcft114/.julia/packages/Flux/qXNjB/src/optimise/train.jl:105
┌ Info: Epoch 3796
└ @ Main /Users/kcft114/.julia/packages/Flux/qXNjB/src/optimise/train.jl:105
┌ Info: Epoch 3797
└ @ Main /Users/kcft114/.julia/packages/Flux/qXNjB/src/optimise/train.jl:105
┌ Info: Epoch 3798
└ @ Main /Users/kcft114/.julia/packages/Flux/qXNjB/src/optimise/train.jl:105
┌ Info: Epoch 3799
└ @ Main /Users/kcft114/.julia/packages/Flux/qXNjB/src/optimise/train.jl:105
┌ Info: Epoch 3800
└ @ Main /Users/kcft114/.julia/packages/Flux/qXNjB/src/optimise/train.jl:105
┌ Info: Epoch 3801
└ @ Main /Users/kcft114/.julia/packages/Flux/qXNjB/src/optimise/train.jl:105
┌ Info: Epoch 3802
└ @ Main /Users/kcft114/.julia/packages/Flux/qXNjB/src/optimise/train.jl:105
┌ Info: Epoch 3803
└ @ Main /Users/kcft114/.julia/packages/Flux/qXNjB/src/optimise/train.jl:105
┌ Info: Epoch 3804
└ @ Main /Users/kcft114/.julia/packages/Flux/qXNjB/src/optimise/train.jl:105
┌ Info: Epoch 3805
└ @ Main /Users/kcft114/.julia/packages/Flux/qXNjB/src/optimise/train.jl:105
┌ Info: Epoch 3806
└ @ Main /Users/kcft114/.julia/packages/Flux/qXNjB/src/optimise/train.jl:105
┌ Info: Epoch 3807
└ @ Main /Users/kcft114/.julia/packages/Flux/qXNjB/src/optimise/train.jl:105
┌ Info: Epoch 3808
└ @ Main /Users/kcft114/.julia/packages/Flux/qXNjB/src/optimise/train.jl:105
┌ Info: Epoch 3809
└ @ Main /Users/kcft114/.julia/packages/Flux/qXNjB/src/optimise/train.jl:105
┌ Info: Epoch 3810
└ @ Main /Users/kcft114/.julia/packages/Flux/qXNjB/src/optimise/train.jl:105
┌ Info: Epoch 3811
└ @ Main /Users/kcft114/.julia/packages/Flux/qXNjB/src/optimise/train.jl:105
┌ Info: Epoch 3812
└ @ Main /Users/kcft114/.julia/packages/Flux/qXNjB/src/optimise/train.jl:105
┌ Info: Epoch 3813
└ @ Main /Users/kcft114/.julia/packages/Flux/qXNjB/src/optimise/train.jl:105
┌ Info: Epoch 3814
└ @ Main /Users/kcft114/.julia/packages/Flux/qXNjB/src/optimise/train.jl:105
┌ Info: Epoch 3815
└ @ Main /Users/kcft114/.julia/packages/Flux/qXNjB/src/optimise/train.jl:105
┌ Info: Epoch 3816
└ @ Main /Users/kcft114/.julia/packages/Flux/qXNjB/src/optimise/train.jl:105
┌ Info: Epoch 3817
└ @ Main /Users/kcft114/.julia/packages/Flux/qXNjB/src/optimise/train.jl:105
```

```
loss(X, Y) = 0.13763331f0 (tracked)
loss(X, Y) = 0.13758999f0 (tracked)
loss(X, Y) = 0.1375154f0 (tracked)
loss(X, Y) = 0.13746034f0 (tracked)
loss(X, Y) = 0.13742681f0 (tracked)
loss(X, Y) = 0.13736257f0 (tracked)
loss(X, Y) = 0.13728492f0 (tracked)
loss(X, Y) = 0.1372229f0 (tracked)
loss(X, Y) = 0.1371601f0 (tracked)
loss(X, Y) = 0.13709845f0 (tracked)
loss(X, Y) = 0.13703564f0 (tracked)
loss(X, Y) = 0.13699545f0 (tracked)
loss(X, Y) = 0.13693607f0 (tracked)
loss(X, Y) = 0.13687757f0 (tracked)
loss(X, Y) = 0.13683504f0 (tracked)
loss(X, Y) = 0.13677149f0 (tracked)
loss(X, Y) = 0.13670869f0 (tracked)
loss(X, Y) = 0.13666287f0 (tracked)
loss(X, Y) = 0.13661799f0 (tracked)
loss(X, Y) = 0.1365472f0 (tracked)
loss(X, Y) = 0.13649118f0 (tracked)
loss(X, Y) = 0.13642451f0 (tracked)
loss(X, Y) = 0.13637696f0 (tracked)
loss(X, Y) = 0.13635787f0 (tracked)
loss(X, Y) = 0.13627122f0 (tracked)
loss(X, Y) = 0.1362103f0 (tracked)
loss(X, Y) = 0.13616356f0 (tracked)
loss(X, Y) = 0.13612115f0 (tracked)
loss(X, Y) = 0.13606705f0 (tracked)
loss(X, Y) = 0.13600492f0 (tracked)
loss(X, Y) = 0.1359469f0 (tracked)
loss(X, Y) = 0.1358857f0 (tracked)
loss(X, Y) = 0.13584036f0 (tracked)
loss(X, Y) = 0.13579623f0 (tracked)
loss(X, Y) = 0.13573302f0 (tracked)
loss(X, Y) = 0.13567704f0 (tracked)
loss(X, Y) = 0.13561822f0 (tracked)
loss(X, Y) = 0.13555616f0 (tracked)
loss(X, Y) = 0.13550082f0 (tracked)
loss(X, Y) = 0.13544773f0 (tracked)
loss(X, Y) = 0.13539238f0 (tracked)
loss(X, Y) = 0.13532652f0 (tracked)
loss(X, Y) = 0.13528483f0 (tracked)
loss(X, Y) = 0.13523155f0 (tracked)
loss(X, Y) = 0.13515548f0 (tracked)
loss(X, Y) = 0.13510226f0 (tracked)
loss(X, Y) = 0.13505293f0 (tracked)
loss(X, Y) = 0.1350006f0 (tracked)
loss(X, Y) = 0.13494997f0 (tracked)
loss(X, Y) = 0.13489339f0 (tracked)
loss(X, Y) = 0.13483253f0 (tracked)
loss(X, Y) = 0.13477951f0 (tracked)
loss(X, Y) = 0.1347358f0 (tracked)
loss(X, Y) = 0.13468093f0 (tracked)
loss(X, Y) = 0.13461532f0 (tracked)
loss(X, Y) = 0.13456525f0 (tracked)
loss(X, Y) = 0.13452262f0 (tracked)
loss(X, Y) = 0.13446377f0 (tracked)
loss(X, Y) = 0.13440949f0 (tracked)
loss(X, Y) = 0.13434839f0 (tracked)
loss(X, Y) = 0.13427643f0 (tracked)
loss(X, Y) = 0.1342293f0 (tracked)
loss(X, Y) = 0.13418627f0 (tracked)
loss(X, Y) = 0.134128f0 (tracked)
loss(X, Y) = 0.13406646f0 (tracked)
loss(X, Y) = 0.13401456f0 (tracked)
loss(X, Y) = 0.13395406f0 (tracked)
loss(X, Y) = 0.13391598f0 (tracked)
loss(X, Y) = 0.13385701f0 (tracked)
loss(X, Y) = 0.13380314f0 (tracked)
loss(X, Y) = 0.13374764f0 (tracked)
loss(X, Y) = 0.13366869f0 (tracked)
loss(X, Y) = 0.13363351f0 (tracked)
loss(X, Y) = 0.13359731f0 (tracked)
loss(X, Y) = 0.13353938f0 (tracked)
loss(X, Y) = 0.13348041f0 (tracked)
loss(X, Y) = 0.13343377f0 (tracked)
loss(X, Y) = 0.13337612f0 (tracked)
loss(X, Y) = 0.13331497f0 (tracked)
loss(X, Y) = 0.13327262f0 (tracked)
loss(X, Y) = 0.13320784f0 (tracked)
loss(X, Y) = 0.13314626f0 (tracked)
loss(X, Y) = 0.13311312f0 (tracked)
loss(X, Y) = 0.13306084f0 (tracked)
loss(X, Y) = 0.13299935f0 (tracked)
loss(X, Y) = 0.13294905f0 (tracked)
loss(X, Y) = 0.13288845f0 (tracked)
loss(X, Y) = 0.13284595f0 (tracked)
loss(X, Y) = 0.13280186f0 (tracked)
loss(X, Y) = 0.13272822f0 (tracked)
loss(X, Y) = 0.13265572f0 (tracked)
loss(X, Y) = 0.13260111f0 (tracked)
loss(X, Y) = 0.13256548f0 (tracked)
loss(X, Y) = 0.13251895f0 (tracked)
loss(X, Y) = 0.13244079f0 (tracked)
loss(X, Y) = 0.13240287f0 (tracked)
loss(X, Y) = 0.13235682f0 (tracked)
loss(X, Y) = 0.13230437f0 (tracked)
loss(X, Y) = 0.1322424f0 (tracked)
loss(X, Y) = 0.1321845f0 (tracked)
loss(X, Y) = 0.13213673f0 (tracked)
loss(X, Y) = 0.13209215f0 (tracked)
loss(X, Y) = 0.13204287f0 (tracked)
loss(X, Y) = 0.13195503f0 (tracked)
loss(X, Y) = 0.13192411f0 (tracked)
loss(X, Y) = 0.13188621f0 (tracked)
loss(X, Y) = 0.13183025f0 (tracked)
loss(X, Y) = 0.13176891f0 (tracked)
loss(X, Y) = 0.13171282f0 (tracked)
loss(X, Y) = 0.13165087f0 (tracked)
loss(X, Y) = 0.13159996f0 (tracked)
loss(X, Y) = 0.13155048f0 (tracked)
loss(X, Y) = 0.13148515f0 (tracked)
loss(X, Y) = 0.13142717f0 (tracked)
loss(X, Y) = 0.13138549f0 (tracked)
loss(X, Y) = 0.13133803f0 (tracked)
loss(X, Y) = 0.13127021f0 (tracked)
loss(X, Y) = 0.13122997f0 (tracked)
loss(X, Y) = 0.13116641f0 (tracked)
loss(X, Y) = 0.13112055f0 (tracked)
loss(X, Y) = 0.13106109f0 (tracked)
loss(X, Y) = 0.13099138f0 (tracked)
loss(X, Y) = 0.13097185f0 (tracked)
loss(X, Y) = 0.13091703f0 (tracked)
loss(X, Y) = 0.13084035f0 (tracked)
loss(X, Y) = 0.13079052f0 (tracked)
loss(X, Y) = 0.1307586f0 (tracked)
loss(X, Y) = 0.13069035f0 (tracked)
```

```
┌ Info: Epoch 3818
└ @ Main /Users/kcft114/.julia/packages/Flux/qXNjB/src/optimise/train.jl:105
┌ Info: Epoch 3819
└ @ Main /Users/kcft114/.julia/packages/Flux/qXNjB/src/optimise/train.jl:105
┌ Info: Epoch 3820
└ @ Main /Users/kcft114/.julia/packages/Flux/qXNjB/src/optimise/train.jl:105
┌ Info: Epoch 3821
└ @ Main /Users/kcft114/.julia/packages/Flux/qXNjB/src/optimise/train.jl:105
┌ Info: Epoch 3822
└ @ Main /Users/kcft114/.julia/packages/Flux/qXNjB/src/optimise/train.jl:105
┌ Info: Epoch 3823
└ @ Main /Users/kcft114/.julia/packages/Flux/qXNjB/src/optimise/train.jl:105
┌ Info: Epoch 3824
└ @ Main /Users/kcft114/.julia/packages/Flux/qXNjB/src/optimise/train.jl:105
┌ Info: Epoch 3825
└ @ Main /Users/kcft114/.julia/packages/Flux/qXNjB/src/optimise/train.jl:105
┌ Info: Epoch 3826
└ @ Main /Users/kcft114/.julia/packages/Flux/qXNjB/src/optimise/train.jl:105
┌ Info: Epoch 3827
└ @ Main /Users/kcft114/.julia/packages/Flux/qXNjB/src/optimise/train.jl:105
┌ Info: Epoch 3828
└ @ Main /Users/kcft114/.julia/packages/Flux/qXNjB/src/optimise/train.jl:105
┌ Info: Epoch 3829
└ @ Main /Users/kcft114/.julia/packages/Flux/qXNjB/src/optimise/train.jl:105
┌ Info: Epoch 3830
└ @ Main /Users/kcft114/.julia/packages/Flux/qXNjB/src/optimise/train.jl:105
┌ Info: Epoch 3831
└ @ Main /Users/kcft114/.julia/packages/Flux/qXNjB/src/optimise/train.jl:105
┌ Info: Epoch 3832
└ @ Main /Users/kcft114/.julia/packages/Flux/qXNjB/src/optimise/train.jl:105
┌ Info: Epoch 3833
└ @ Main /Users/kcft114/.julia/packages/Flux/qXNjB/src/optimise/train.jl:105
┌ Info: Epoch 3834
└ @ Main /Users/kcft114/.julia/packages/Flux/qXNjB/src/optimise/train.jl:105
┌ Info: Epoch 3835
└ @ Main /Users/kcft114/.julia/packages/Flux/qXNjB/src/optimise/train.jl:105
┌ Info: Epoch 3836
└ @ Main /Users/kcft114/.julia/packages/Flux/qXNjB/src/optimise/train.jl:105
┌ Info: Epoch 3837
└ @ Main /Users/kcft114/.julia/packages/Flux/qXNjB/src/optimise/train.jl:105
┌ Info: Epoch 3838
└ @ Main /Users/kcft114/.julia/packages/Flux/qXNjB/src/optimise/train.jl:105
┌ Info: Epoch 3839
└ @ Main /Users/kcft114/.julia/packages/Flux/qXNjB/src/optimise/train.jl:105
┌ Info: Epoch 3840
└ @ Main /Users/kcft114/.julia/packages/Flux/qXNjB/src/optimise/train.jl:105
┌ Info: Epoch 3841
└ @ Main /Users/kcft114/.julia/packages/Flux/qXNjB/src/optimise/train.jl:105
┌ Info: Epoch 3842
└ @ Main /Users/kcft114/.julia/packages/Flux/qXNjB/src/optimise/train.jl:105
Excessive output truncated after 524324 bytes.
```

```
loss(X, Y) = 0.13062036f0 (tracked)
loss(X, Y) = 0.13058124f0 (tracked)
loss(X, Y) = 0.13055201f0 (tracked)
loss(X, Y) = 0.1304863f0 (tracked)
loss(X, Y) = 0.13042378f0 (tracked)
loss(X, Y) = 0.13037722f0 (tracked)
loss(X, Y) = 0.13033068f0 (tracked)
loss(X, Y) = 0.13026457f0 (tracked)
loss(X, Y) = 0.13021697f0 (tracked)
loss(X, Y) = 0.13015981f0 (tracked)
loss(X, Y) = 0.13012286f0 (tracked)
loss(X, Y) = 0.1300623f0 (tracked)
loss(X, Y) = 0.12999804f0 (tracked)
loss(X, Y) = 0.12996343f0 (tracked)
loss(X, Y) = 0.12992807f0 (tracked)
loss(X, Y) = 0.12985618f0 (tracked)
loss(X, Y) = 0.12979311f0 (tracked)
loss(X, Y) = 0.129762f0 (tracked)
loss(X, Y) = 0.1297032f0 (tracked)
loss(X, Y) = 0.12964013f0 (tracked)
loss(X, Y) = 0.1295741f0 (tracked)
loss(X, Y) = 0.1295424f0 (tracked)
loss(X, Y) = 0.12949319f0 (tracked)
loss(X, Y) = 0.12942249f0 (tracked)
loss(X, Y) = 0.12936777f0 (tracked)
loss(X, Y) = 0.12931907f0 (tracked)
loss(X, Y) = 0.12926118f0 (tracked)
loss(X, Y) = 0.12921882f0 (tracked)
loss(X, Y) = 0.12915693f0 (tracked)
loss(X, Y) = 0.12911302f0 (tracked)
loss(X, Y) = 0.12907529f0 (tracked)
loss(X, Y) = 0.12902491f0 (tracked)
```

```
┌ Info: Epoch 3843
└ @ Main /Users/kcft114/.julia/packages/Flux/qXNjB/src/optimise/train.jl:105
┌ Info: Epoch 3844
└ @ Main /Users/kcft114/.julia/packages/Flux/qXNjB/src/optimise/train.jl:105
┌ Info: Epoch 3845
└ @ Main /Users/kcft114/.julia/packages/Flux/qXNjB/src/optimise/train.jl:105
┌ Info: Epoch 3846
└ @ Main /Users/kcft114/.julia/packages/Flux/qXNjB/src/optimise/train.jl:105
┌ Info: Epoch 3847
└ @ Main /Users/kcft114/.julia/packages/Flux/qXNjB/src/optimise/train.jl:105
┌ Info: Epoch 3848
└ @ Main /Users/kcft114/.julia/packages/Flux/qXNjB/src/optimise/train.jl:105
┌ Info: Epoch 3849
└ @ Main /Users/kcft114/.julia/packages/Flux/qXNjB/src/optimise/train.jl:105
┌ Info: Epoch 3850
└ @ Main /Users/kcft114/.julia/packages/Flux/qXNjB/src/optimise/train.jl:105
┌ Info: Epoch 3851
└ @ Main /Users/kcft114/.julia/packages/Flux/qXNjB/src/optimise/train.jl:105
┌ Info: Epoch 3852
└ @ Main /Users/kcft114/.julia/packages/Flux/qXNjB/src/optimise/train.jl:105
┌ Info: Epoch 3853
└ @ Main /Users/kcft114/.julia/packages/Flux/qXNjB/src/optimise/train.jl:105
┌ Info: Epoch 3854
└ @ Main /Users/kcft114/.julia/packages/Flux/qXNjB/src/optimise/train.jl:105
┌ Info: Epoch 3855
└ @ Main /Users/kcft114/.julia/packages/Flux/qXNjB/src/optimise/train.jl:105
┌ Info: Epoch 3856
└ @ Main /Users/kcft114/.julia/packages/Flux/qXNjB/src/optimise/train.jl:105
┌ Info: Epoch 3857
└ @ Main /Users/kcft114/.julia/packages/Flux/qXNjB/src/optimise/train.jl:105
┌ Info: Epoch 3858
└ @ Main /Users/kcft114/.julia/packages/Flux/qXNjB/src/optimise/train.jl:105
┌ Info: Epoch 3859
└ @ Main /Users/kcft114/.julia/packages/Flux/qXNjB/src/optimise/train.jl:105
┌ Info: Epoch 3860
└ @ Main /Users/kcft114/.julia/packages/Flux/qXNjB/src/optimise/train.jl:105
┌ Info: Epoch 3861
└ @ Main /Users/kcft114/.julia/packages/Flux/qXNjB/src/optimise/train.jl:105
┌ Info: Epoch 3862
└ @ Main /Users/kcft114/.julia/packages/Flux/qXNjB/src/optimise/train.jl:105
┌ Info: Epoch 3863
└ @ Main /Users/kcft114/.julia/packages/Flux/qXNjB/src/optimise/train.jl:105
┌ Info: Epoch 3864
└ @ Main /Users/kcft114/.julia/packages/Flux/qXNjB/src/optimise/train.jl:105
┌ Info: Epoch 3865
└ @ Main /Users/kcft114/.julia/packages/Flux/qXNjB/src/optimise/train.jl:105
┌ Info: Epoch 3866
└ @ Main /Users/kcft114/.julia/packages/Flux/qXNjB/src/optimise/train.jl:105
┌ Info: Epoch 3867
└ @ Main /Users/kcft114/.julia/packages/Flux/qXNjB/src/optimise/train.jl:105
┌ Info: Epoch 3868
└ @ Main /Users/kcft114/.julia/packages/Flux/qXNjB/src/optimise/train.jl:105
┌ Info: Epoch 3869
└ @ Main /Users/kcft114/.julia/packages/Flux/qXNjB/src/optimise/train.jl:105
┌ Info: Epoch 3870
└ @ Main /Users/kcft114/.julia/packages/Flux/qXNjB/src/optimise/train.jl:105
┌ Info: Epoch 3871
└ @ Main /Users/kcft114/.julia/packages/Flux/qXNjB/src/optimise/train.jl:105
┌ Info: Epoch 3872
└ @ Main /Users/kcft114/.julia/packages/Flux/qXNjB/src/optimise/train.jl:105
┌ Info: Epoch 3873
└ @ Main /Users/kcft114/.julia/packages/Flux/qXNjB/src/optimise/train.jl:105
```

In [11]:

```
## ---------------------------------------------------------
## Experiments - Regularization via Dropout
## ---------------------------------------------------------
```

In [11]:

```
p_dropout=0.1

n_exper = 100
experiments_01 = zeros(n_exper, 4)


for i=1:n_exper
    
    nn = Chain(
              Dense(8, n_h, relu),
              Dropout(p_dropout),
              Dense(n_h, 3),
              softmax)
    
    loss(x, y) = crossentropy(nn(x), y)
    accuracy(x, y) = mean(
    onecold(nn(x)) .== onecold(y))

    dataset = [(X, Y)]

    evalcb = () -> @show(loss(X, Y))
    
    opt = ADAM(0.001)
    
    @epochs 2000 Flux.train!(loss, Flux.params(nn), dataset, opt, cb = throttle(evalcb, 10));
    
    ac_train, ac_test, ba_train, ba_test =  res(X, Y, Xt, Yt);
    experiments_01[i,:] = [ac_train, ac_test, ba_train, ba_test]
end
```

```
┌ Info: Epoch 1
└ @ Main /Users/kcft114/.julia/packages/Flux/qXNjB/src/optimise/train.jl:105
```

```
loss(X, Y) = 1.1637123f0 (tracked)
loss(X, Y) = 1.1320153f0 (tracked)
loss(X, Y) = 1.1606095f0 (tracked)
loss(X, Y) = 1.1303873f0 (tracked)
loss(X, Y) = 1.1283019f0 (tracked)
loss(X, Y) = 1.1342638f0 (tracked)
loss(X, Y) = 1.1209987f0 (tracked)
loss(X, Y) = 1.1340778f0 (tracked)
loss(X, Y) = 1.1120175f0 (tracked)
loss(X, Y) = 1.0951488f0 (tracked)
loss(X, Y) = 1.097836f0 (tracked)
loss(X, Y) = 1.0770315f0 (tracked)
loss(X, Y) = 1.1100444f0 (tracked)
loss(X, Y) = 1.0889207f0 (tracked)
loss(X, Y) = 1.0569634f0 (tracked)
loss(X, Y) = 1.0720273f0 (tracked)
loss(X, Y) = 1.070278f0 (tracked)
loss(X, Y) = 1.0915045f0 (tracked)
loss(X, Y) = 1.0589314f0 (tracked)
loss(X, Y) = 1.0712459f0 (tracked)
loss(X, Y) = 1.0735706f0 (tracked)
loss(X, Y) = 1.0671486f0 (tracked)
loss(X, Y) = 1.0955693f0 (tracked)
loss(X, Y) = 1.0619891f0 (tracked)
loss(X, Y) = 1.0419513f0 (tracked)
loss(X, Y) = 1.0475029f0 (tracked)
loss(X, Y) = 1.0624322f0 (tracked)
loss(X, Y) = 1.02566f0 (tracked)
loss(X, Y) = 1.0613681f0 (tracked)
loss(X, Y) = 1.0605611f0 (tracked)
loss(X, Y) = 1.0137303f0 (tracked)
loss(X, Y) = 1.0488304f0 (tracked)
loss(X, Y) = 1.050647f0 (tracked)
loss(X, Y) = 1.0286999f0 (tracked)
loss(X, Y) = 1.0410885f0 (tracked)
loss(X, Y) = 1.0138589f0 (tracked)
loss(X, Y) = 1.0080714f0 (tracked)
loss(X, Y) = 0.9994997f0 (tracked)
loss(X, Y) = 1.0152645f0 (tracked)
loss(X, Y) = 1.025045f0 (tracked)
loss(X, Y) = 1.0098901f0 (tracked)
loss(X, Y) = 1.031713f0 (tracked)
loss(X, Y) = 1.0065614f0 (tracked)
loss(X, Y) = 1.0265001f0 (tracked)
loss(X, Y) = 1.0177029f0 (tracked)
loss(X, Y) = 1.0078683f0 (tracked)
loss(X, Y) = 0.99177873f0 (tracked)
loss(X, Y) = 0.9788679f0 (tracked)
loss(X, Y) = 0.99554086f0 (tracked)
loss(X, Y) = 1.009312f0 (tracked)
loss(X, Y) = 1.0284593f0 (tracked)
loss(X, Y) = 0.97713846f0 (tracked)
loss(X, Y) = 1.0028137f0 (tracked)
loss(X, Y) = 0.95823926f0 (tracked)
loss(X, Y) = 0.9730082f0 (tracked)
loss(X, Y) = 0.9804186f0 (tracked)
loss(X, Y) = 0.97615373f0 (tracked)
loss(X, Y) = 0.97260165f0 (tracked)
loss(X, Y) = 0.99974805f0 (tracked)
loss(X, Y) = 0.98085386f0 (tracked)
loss(X, Y) = 0.9688364f0 (tracked)
loss(X, Y) = 0.9631932f0 (tracked)
loss(X, Y) = 0.9650679f0 (tracked)
loss(X, Y) = 0.97077316f0 (tracked)
loss(X, Y) = 0.9719453f0 (tracked)
loss(X, Y) = 0.94464505f0 (tracked)
loss(X, Y) = 0.94778126f0 (tracked)
loss(X, Y) = 0.9655678f0 (tracked)
loss(X, Y) = 0.94828004f0 (tracked)
loss(X, Y) = 0.9376315f0 (tracked)
loss(X, Y) = 0.95652497f0 (tracked)
loss(X, Y) = 0.9548931f0 (tracked)
loss(X, Y) = 0.93975914f0 (tracked)
loss(X, Y) = 0.9346455f0 (tracked)
loss(X, Y) = 0.93831855f0 (tracked)
loss(X, Y) = 0.95290595f0 (tracked)
loss(X, Y) = 0.9288217f0 (tracked)
loss(X, Y) = 0.91942537f0 (tracked)
loss(X, Y) = 0.9213048f0 (tracked)
```

```
┌ Info: Epoch 2
└ @ Main /Users/kcft114/.julia/packages/Flux/qXNjB/src/optimise/train.jl:105
┌ Info: Epoch 3
└ @ Main /Users/kcft114/.julia/packages/Flux/qXNjB/src/optimise/train.jl:105
┌ Info: Epoch 4
└ @ Main /Users/kcft114/.julia/packages/Flux/qXNjB/src/optimise/train.jl:105
┌ Info: Epoch 5
└ @ Main /Users/kcft114/.julia/packages/Flux/qXNjB/src/optimise/train.jl:105
┌ Info: Epoch 6
└ @ Main /Users/kcft114/.julia/packages/Flux/qXNjB/src/optimise/train.jl:105
┌ Info: Epoch 7
└ @ Main /Users/kcft114/.julia/packages/Flux/qXNjB/src/optimise/train.jl:105
┌ Info: Epoch 8
└ @ Main /Users/kcft114/.julia/packages/Flux/qXNjB/src/optimise/train.jl:105
┌ Info: Epoch 9
└ @ Main /Users/kcft114/.julia/packages/Flux/qXNjB/src/optimise/train.jl:105
┌ Info: Epoch 10
└ @ Main /Users/kcft114/.julia/packages/Flux/qXNjB/src/optimise/train.jl:105
┌ Info: Epoch 11
└ @ Main /Users/kcft114/.julia/packages/Flux/qXNjB/src/optimise/train.jl:105
┌ Info: Epoch 12
└ @ Main /Users/kcft114/.julia/packages/Flux/qXNjB/src/optimise/train.jl:105
┌ Info: Epoch 13
└ @ Main /Users/kcft114/.julia/packages/Flux/qXNjB/src/optimise/train.jl:105
┌ Info: Epoch 14
└ @ Main /Users/kcft114/.julia/packages/Flux/qXNjB/src/optimise/train.jl:105
┌ Info: Epoch 15
└ @ Main /Users/kcft114/.julia/packages/Flux/qXNjB/src/optimise/train.jl:105
┌ Info: Epoch 16
└ @ Main /Users/kcft114/.julia/packages/Flux/qXNjB/src/optimise/train.jl:105
┌ Info: Epoch 17
└ @ Main /Users/kcft114/.julia/packages/Flux/qXNjB/src/optimise/train.jl:105
┌ Info: Epoch 18
└ @ Main /Users/kcft114/.julia/packages/Flux/qXNjB/src/optimise/train.jl:105
┌ Info: Epoch 19
└ @ Main /Users/kcft114/.julia/packages/Flux/qXNjB/src/optimise/train.jl:105
┌ Info: Epoch 20
└ @ Main /Users/kcft114/.julia/packages/Flux/qXNjB/src/optimise/train.jl:105
┌ Info: Epoch 21
└ @ Main /Users/kcft114/.julia/packages/Flux/qXNjB/src/optimise/train.jl:105
┌ Info: Epoch 22
└ @ Main /Users/kcft114/.julia/packages/Flux/qXNjB/src/optimise/train.jl:105
┌ Info: Epoch 23
└ @ Main /Users/kcft114/.julia/packages/Flux/qXNjB/src/optimise/train.jl:105
┌ Info: Epoch 24
└ @ Main /Users/kcft114/.julia/packages/Flux/qXNjB/src/optimise/train.jl:105
┌ Info: Epoch 25
└ @ Main /Users/kcft114/.julia/packages/Flux/qXNjB/src/optimise/train.jl:105
┌ Info: Epoch 26
└ @ Main /Users/kcft114/.julia/packages/Flux/qXNjB/src/optimise/train.jl:105
┌ Info: Epoch 27
└ @ Main /Users/kcft114/.julia/packages/Flux/qXNjB/src/optimise/train.jl:105
┌ Info: Epoch 28
└ @ Main /Users/kcft114/.julia/packages/Flux/qXNjB/src/optimise/train.jl:105
┌ Info: Epoch 29
└ @ Main /Users/kcft114/.julia/packages/Flux/qXNjB/src/optimise/train.jl:105
┌ Info: Epoch 30
└ @ Main /Users/kcft114/.julia/packages/Flux/qXNjB/src/optimise/train.jl:105
┌ Info: Epoch 31
└ @ Main /Users/kcft114/.julia/packages/Flux/qXNjB/src/optimise/train.jl:105
┌ Info: Epoch 32
└ @ Main /Users/kcft114/.julia/packages/Flux/qXNjB/src/optimise/train.jl:105
┌ Info: Epoch 33
└ @ Main /Users/kcft114/.julia/packages/Flux/qXNjB/src/optimise/train.jl:105
┌ Info: Epoch 34
└ @ Main /Users/kcft114/.julia/packages/Flux/qXNjB/src/optimise/train.jl:105
┌ Info: Epoch 35
└ @ Main /Users/kcft114/.julia/packages/Flux/qXNjB/src/optimise/train.jl:105
┌ Info: Epoch 36
└ @ Main /Users/kcft114/.julia/packages/Flux/qXNjB/src/optimise/train.jl:105
┌ Info: Epoch 37
└ @ Main /Users/kcft114/.julia/packages/Flux/qXNjB/src/optimise/train.jl:105
┌ Info: Epoch 38
└ @ Main /Users/kcft114/.julia/packages/Flux/qXNjB/src/optimise/train.jl:105
┌ Info: Epoch 39
└ @ Main /Users/kcft114/.julia/packages/Flux/qXNjB/src/optimise/train.jl:105
┌ Info: Epoch 40
└ @ Main /Users/kcft114/.julia/packages/Flux/qXNjB/src/optimise/train.jl:105
┌ Info: Epoch 41
└ @ Main /Users/kcft114/.julia/packages/Flux/qXNjB/src/optimise/train.jl:105
┌ Info: Epoch 42
└ @ Main /Users/kcft114/.julia/packages/Flux/qXNjB/src/optimise/train.jl:105
┌ Info: Epoch 43
└ @ Main /Users/kcft114/.julia/packages/Flux/qXNjB/src/optimise/train.jl:105
┌ Info: Epoch 44
└ @ Main /Users/kcft114/.julia/packages/Flux/qXNjB/src/optimise/train.jl:105
┌ Info: Epoch 45
└ @ Main /Users/kcft114/.julia/packages/Flux/qXNjB/src/optimise/train.jl:105
┌ Info: Epoch 46
└ @ Main /Users/kcft114/.julia/packages/Flux/qXNjB/src/optimise/train.jl:105
┌ Info: Epoch 47
└ @ Main /Users/kcft114/.julia/packages/Flux/qXNjB/src/optimise/train.jl:105
┌ Info: Epoch 48
└ @ Main /Users/kcft114/.julia/packages/Flux/qXNjB/src/optimise/train.jl:105
┌ Info: Epoch 49
└ @ Main /Users/kcft114/.julia/packages/Flux/qXNjB/src/optimise/train.jl:105
┌ Info: Epoch 50
└ @ Main /Users/kcft114/.julia/packages/Flux/qXNjB/src/optimise/train.jl:105
┌ Info: Epoch 51
└ @ Main /Users/kcft114/.julia/packages/Flux/qXNjB/src/optimise/train.jl:105
┌ Info: Epoch 52
└ @ Main /Users/kcft114/.julia/packages/Flux/qXNjB/src/optimise/train.jl:105
┌ Info: Epoch 53
└ @ Main /Users/kcft114/.julia/packages/Flux/qXNjB/src/optimise/train.jl:105
┌ Info: Epoch 54
└ @ Main /Users/kcft114/.julia/packages/Flux/qXNjB/src/optimise/train.jl:105
┌ Info: Epoch 55
└ @ Main /Users/kcft114/.julia/packages/Flux/qXNjB/src/optimise/train.jl:105
┌ Info: Epoch 56
└ @ Main /Users/kcft114/.julia/packages/Flux/qXNjB/src/optimise/train.jl:105
┌ Info: Epoch 57
└ @ Main /Users/kcft114/.julia/packages/Flux/qXNjB/src/optimise/train.jl:105
┌ Info: Epoch 58
└ @ Main /Users/kcft114/.julia/packages/Flux/qXNjB/src/optimise/train.jl:105
┌ Info: Epoch 59
└ @ Main /Users/kcft114/.julia/packages/Flux/qXNjB/src/optimise/train.jl:105
┌ Info: Epoch 60
└ @ Main /Users/kcft114/.julia/packages/Flux/qXNjB/src/optimise/train.jl:105
┌ Info: Epoch 61
└ @ Main /Users/kcft114/.julia/packages/Flux/qXNjB/src/optimise/train.jl:105
┌ Info: Epoch 62
└ @ Main /Users/kcft114/.julia/packages/Flux/qXNjB/src/optimise/train.jl:105
┌ Info: Epoch 63
└ @ Main /Users/kcft114/.julia/packages/Flux/qXNjB/src/optimise/train.jl:105
┌ Info: Epoch 64
└ @ Main /Users/kcft114/.julia/packages/Flux/qXNjB/src/optimise/train.jl:105
┌ Info: Epoch 65
└ @ Main /Users/kcft114/.julia/packages/Flux/qXNjB/src/optimise/train.jl:105
┌ Info: Epoch 66
└ @ Main /Users/kcft114/.julia/packages/Flux/qXNjB/src/optimise/train.jl:105
┌ Info: Epoch 67
└ @ Main /Users/kcft114/.julia/packages/Flux/qXNjB/src/optimise/train.jl:105
┌ Info: Epoch 68
└ @ Main /Users/kcft114/.julia/packages/Flux/qXNjB/src/optimise/train.jl:105
┌ Info: Epoch 69
└ @ Main /Users/kcft114/.julia/packages/Flux/qXNjB/src/optimise/train.jl:105
┌ Info: Epoch 70
└ @ Main /Users/kcft114/.julia/packages/Flux/qXNjB/src/optimise/train.jl:105
┌ Info: Epoch 71
└ @ Main /Users/kcft114/.julia/packages/Flux/qXNjB/src/optimise/train.jl:105
┌ Info: Epoch 72
└ @ Main /Users/kcft114/.julia/packages/Flux/qXNjB/src/optimise/train.jl:105
┌ Info: Epoch 73
└ @ Main /Users/kcft114/.julia/packages/Flux/qXNjB/src/optimise/train.jl:105
┌ Info: Epoch 74
└ @ Main /Users/kcft114/.julia/packages/Flux/qXNjB/src/optimise/train.jl:105
┌ Info: Epoch 75
└ @ Main /Users/kcft114/.julia/packages/Flux/qXNjB/src/optimise/train.jl:105
┌ Info: Epoch 76
└ @ Main /Users/kcft114/.julia/packages/Flux/qXNjB/src/optimise/train.jl:105
┌ Info: Epoch 77
└ @ Main /Users/kcft114/.julia/packages/Flux/qXNjB/src/optimise/train.jl:105
┌ Info: Epoch 78
└ @ Main /Users/kcft114/.julia/packages/Flux/qXNjB/src/optimise/train.jl:105
┌ Info: Epoch 79
└ @ Main /Users/kcft114/.julia/packages/Flux/qXNjB/src/optimise/train.jl:105
┌ Info: Epoch 80
└ @ Main /Users/kcft114/.julia/packages/Flux/qXNjB/src/optimise/train.jl:105
```

```
loss(X, Y) = 0.91311973f0 (tracked)
loss(X, Y) = 0.92193997f0 (tracked)
loss(X, Y) = 0.92630976f0 (tracked)
loss(X, Y) = 0.92606264f0 (tracked)
loss(X, Y) = 0.914451f0 (tracked)
loss(X, Y) = 0.96589607f0 (tracked)
loss(X, Y) = 0.90077454f0 (tracked)
loss(X, Y) = 0.92771465f0 (tracked)
loss(X, Y) = 0.92039365f0 (tracked)
loss(X, Y) = 0.9238458f0 (tracked)
loss(X, Y) = 0.9119355f0 (tracked)
loss(X, Y) = 0.9181061f0 (tracked)
loss(X, Y) = 0.9197597f0 (tracked)
loss(X, Y) = 0.9216824f0 (tracked)
loss(X, Y) = 0.90302354f0 (tracked)
loss(X, Y) = 0.8895437f0 (tracked)
loss(X, Y) = 0.8909751f0 (tracked)
loss(X, Y) = 0.9194037f0 (tracked)
loss(X, Y) = 0.91344786f0 (tracked)
loss(X, Y) = 0.9344656f0 (tracked)
loss(X, Y) = 0.88012093f0 (tracked)
loss(X, Y) = 0.8991274f0 (tracked)
loss(X, Y) = 0.91990983f0 (tracked)
loss(X, Y) = 0.90353984f0 (tracked)
loss(X, Y) = 0.89850247f0 (tracked)
loss(X, Y) = 0.92583156f0 (tracked)
loss(X, Y) = 0.8973839f0 (tracked)
loss(X, Y) = 0.8944214f0 (tracked)
loss(X, Y) = 0.8588153f0 (tracked)
loss(X, Y) = 0.87703854f0 (tracked)
loss(X, Y) = 0.9022595f0 (tracked)
loss(X, Y) = 0.9140767f0 (tracked)
loss(X, Y) = 0.87372303f0 (tracked)
loss(X, Y) = 0.9057865f0 (tracked)
loss(X, Y) = 0.9011543f0 (tracked)
loss(X, Y) = 0.87943834f0 (tracked)
loss(X, Y) = 0.8761911f0 (tracked)
loss(X, Y) = 0.86436194f0 (tracked)
loss(X, Y) = 0.88800335f0 (tracked)
loss(X, Y) = 0.8709889f0 (tracked)
loss(X, Y) = 0.88066775f0 (tracked)
loss(X, Y) = 0.8581471f0 (tracked)
loss(X, Y) = 0.89150876f0 (tracked)
loss(X, Y) = 0.89514893f0 (tracked)
loss(X, Y) = 0.86375874f0 (tracked)
loss(X, Y) = 0.84778506f0 (tracked)
loss(X, Y) = 0.8713333f0 (tracked)
loss(X, Y) = 0.86913514f0 (tracked)
loss(X, Y) = 0.86461204f0 (tracked)
loss(X, Y) = 0.8597678f0 (tracked)
loss(X, Y) = 0.88930005f0 (tracked)
loss(X, Y) = 0.8627245f0 (tracked)
loss(X, Y) = 0.9014527f0 (tracked)
loss(X, Y) = 0.87099504f0 (tracked)
loss(X, Y) = 0.83584243f0 (tracked)
loss(X, Y) = 0.88197494f0 (tracked)
loss(X, Y) = 0.8575107f0 (tracked)
loss(X, Y) = 0.874681f0 (tracked)
loss(X, Y) = 0.8531673f0 (tracked)
loss(X, Y) = 0.8648417f0 (tracked)
loss(X, Y) = 0.83393776f0 (tracked)
loss(X, Y) = 0.83480775f0 (tracked)
loss(X, Y) = 0.8414547f0 (tracked)
loss(X, Y) = 0.8457453f0 (tracked)
loss(X, Y) = 0.83006364f0 (tracked)
loss(X, Y) = 0.8534563f0 (tracked)
loss(X, Y) = 0.8645224f0 (tracked)
loss(X, Y) = 0.82161707f0 (tracked)
loss(X, Y) = 0.8475952f0 (tracked)
loss(X, Y) = 0.8261213f0 (tracked)
loss(X, Y) = 0.8316294f0 (tracked)
```

```
┌ Info: Epoch 81
└ @ Main /Users/kcft114/.julia/packages/Flux/qXNjB/src/optimise/train.jl:105
┌ Info: Epoch 82
└ @ Main /Users/kcft114/.julia/packages/Flux/qXNjB/src/optimise/train.jl:105
┌ Info: Epoch 83
└ @ Main /Users/kcft114/.julia/packages/Flux/qXNjB/src/optimise/train.jl:105
┌ Info: Epoch 84
└ @ Main /Users/kcft114/.julia/packages/Flux/qXNjB/src/optimise/train.jl:105
┌ Info: Epoch 85
└ @ Main /Users/kcft114/.julia/packages/Flux/qXNjB/src/optimise/train.jl:105
┌ Info: Epoch 86
└ @ Main /Users/kcft114/.julia/packages/Flux/qXNjB/src/optimise/train.jl:105
┌ Info: Epoch 87
└ @ Main /Users/kcft114/.julia/packages/Flux/qXNjB/src/optimise/train.jl:105
┌ Info: Epoch 88
└ @ Main /Users/kcft114/.julia/packages/Flux/qXNjB/src/optimise/train.jl:105
┌ Info: Epoch 89
└ @ Main /Users/kcft114/.julia/packages/Flux/qXNjB/src/optimise/train.jl:105
┌ Info: Epoch 90
└ @ Main /Users/kcft114/.julia/packages/Flux/qXNjB/src/optimise/train.jl:105
┌ Info: Epoch 91
└ @ Main /Users/kcft114/.julia/packages/Flux/qXNjB/src/optimise/train.jl:105
┌ Info: Epoch 92
└ @ Main /Users/kcft114/.julia/packages/Flux/qXNjB/src/optimise/train.jl:105
┌ Info: Epoch 93
└ @ Main /Users/kcft114/.julia/packages/Flux/qXNjB/src/optimise/train.jl:105
┌ Info: Epoch 94
└ @ Main /Users/kcft114/.julia/packages/Flux/qXNjB/src/optimise/train.jl:105
┌ Info: Epoch 95
└ @ Main /Users/kcft114/.julia/packages/Flux/qXNjB/src/optimise/train.jl:105
┌ Info: Epoch 96
└ @ Main /Users/kcft114/.julia/packages/Flux/qXNjB/src/optimise/train.jl:105
┌ Info: Epoch 97
└ @ Main /Users/kcft114/.julia/packages/Flux/qXNjB/src/optimise/train.jl:105
┌ Info: Epoch 98
└ @ Main /Users/kcft114/.julia/packages/Flux/qXNjB/src/optimise/train.jl:105
┌ Info: Epoch 99
└ @ Main /Users/kcft114/.julia/packages/Flux/qXNjB/src/optimise/train.jl:105
┌ Info: Epoch 100
└ @ Main /Users/kcft114/.julia/packages/Flux/qXNjB/src/optimise/train.jl:105
┌ Info: Epoch 101
└ @ Main /Users/kcft114/.julia/packages/Flux/qXNjB/src/optimise/train.jl:105
┌ Info: Epoch 102
└ @ Main /Users/kcft114/.julia/packages/Flux/qXNjB/src/optimise/train.jl:105
┌ Info: Epoch 103
└ @ Main /Users/kcft114/.julia/packages/Flux/qXNjB/src/optimise/train.jl:105
┌ Info: Epoch 104
└ @ Main /Users/kcft114/.julia/packages/Flux/qXNjB/src/optimise/train.jl:105
┌ Info: Epoch 105
└ @ Main /Users/kcft114/.julia/packages/Flux/qXNjB/src/optimise/train.jl:105
┌ Info: Epoch 106
└ @ Main /Users/kcft114/.julia/packages/Flux/qXNjB/src/optimise/train.jl:105
┌ Info: Epoch 107
└ @ Main /Users/kcft114/.julia/packages/Flux/qXNjB/src/optimise/train.jl:105
┌ Info: Epoch 108
└ @ Main /Users/kcft114/.julia/packages/Flux/qXNjB/src/optimise/train.jl:105
┌ Info: Epoch 109
└ @ Main /Users/kcft114/.julia/packages/Flux/qXNjB/src/optimise/train.jl:105
┌ Info: Epoch 110
└ @ Main /Users/kcft114/.julia/packages/Flux/qXNjB/src/optimise/train.jl:105
┌ Info: Epoch 111
└ @ Main /Users/kcft114/.julia/packages/Flux/qXNjB/src/optimise/train.jl:105
┌ Info: Epoch 112
└ @ Main /Users/kcft114/.julia/packages/Flux/qXNjB/src/optimise/train.jl:105
┌ Info: Epoch 113
└ @ Main /Users/kcft114/.julia/packages/Flux/qXNjB/src/optimise/train.jl:105
┌ Info: Epoch 114
└ @ Main /Users/kcft114/.julia/packages/Flux/qXNjB/src/optimise/train.jl:105
┌ Info: Epoch 115
└ @ Main /Users/kcft114/.julia/packages/Flux/qXNjB/src/optimise/train.jl:105
┌ Info: Epoch 116
└ @ Main /Users/kcft114/.julia/packages/Flux/qXNjB/src/optimise/train.jl:105
┌ Info: Epoch 117
└ @ Main /Users/kcft114/.julia/packages/Flux/qXNjB/src/optimise/train.jl:105
┌ Info: Epoch 118
└ @ Main /Users/kcft114/.julia/packages/Flux/qXNjB/src/optimise/train.jl:105
┌ Info: Epoch 119
└ @ Main /Users/kcft114/.julia/packages/Flux/qXNjB/src/optimise/train.jl:105
┌ Info: Epoch 120
└ @ Main /Users/kcft114/.julia/packages/Flux/qXNjB/src/optimise/train.jl:105
┌ Info: Epoch 121
└ @ Main /Users/kcft114/.julia/packages/Flux/qXNjB/src/optimise/train.jl:105
┌ Info: Epoch 122
└ @ Main /Users/kcft114/.julia/packages/Flux/qXNjB/src/optimise/train.jl:105
┌ Info: Epoch 123
└ @ Main /Users/kcft114/.julia/packages/Flux/qXNjB/src/optimise/train.jl:105
┌ Info: Epoch 124
└ @ Main /Users/kcft114/.julia/packages/Flux/qXNjB/src/optimise/train.jl:105
┌ Info: Epoch 125
└ @ Main /Users/kcft114/.julia/packages/Flux/qXNjB/src/optimise/train.jl:105
┌ Info: Epoch 126
└ @ Main /Users/kcft114/.julia/packages/Flux/qXNjB/src/optimise/train.jl:105
┌ Info: Epoch 127
└ @ Main /Users/kcft114/.julia/packages/Flux/qXNjB/src/optimise/train.jl:105
┌ Info: Epoch 128
└ @ Main /Users/kcft114/.julia/packages/Flux/qXNjB/src/optimise/train.jl:105
┌ Info: Epoch 129
└ @ Main /Users/kcft114/.julia/packages/Flux/qXNjB/src/optimise/train.jl:105
┌ Info: Epoch 130
└ @ Main /Users/kcft114/.julia/packages/Flux/qXNjB/src/optimise/train.jl:105
┌ Info: Epoch 131
└ @ Main /Users/kcft114/.julia/packages/Flux/qXNjB/src/optimise/train.jl:105
┌ Info: Epoch 132
└ @ Main /Users/kcft114/.julia/packages/Flux/qXNjB/src/optimise/train.jl:105
┌ Info: Epoch 133
└ @ Main /Users/kcft114/.julia/packages/Flux/qXNjB/src/optimise/train.jl:105
┌ Info: Epoch 134
└ @ Main /Users/kcft114/.julia/packages/Flux/qXNjB/src/optimise/train.jl:105
┌ Info: Epoch 135
└ @ Main /Users/kcft114/.julia/packages/Flux/qXNjB/src/optimise/train.jl:105
┌ Info: Epoch 136
└ @ Main /Users/kcft114/.julia/packages/Flux/qXNjB/src/optimise/train.jl:105
┌ Info: Epoch 137
└ @ Main /Users/kcft114/.julia/packages/Flux/qXNjB/src/optimise/train.jl:105
┌ Info: Epoch 138
└ @ Main /Users/kcft114/.julia/packages/Flux/qXNjB/src/optimise/train.jl:105
┌ Info: Epoch 139
└ @ Main /Users/kcft114/.julia/packages/Flux/qXNjB/src/optimise/train.jl:105
┌ Info: Epoch 140
└ @ Main /Users/kcft114/.julia/packages/Flux/qXNjB/src/optimise/train.jl:105
┌ Info: Epoch 141
└ @ Main /Users/kcft114/.julia/packages/Flux/qXNjB/src/optimise/train.jl:105
┌ Info: Epoch 142
└ @ Main /Users/kcft114/.julia/packages/Flux/qXNjB/src/optimise/train.jl:105
┌ Info: Epoch 143
└ @ Main /Users/kcft114/.julia/packages/Flux/qXNjB/src/optimise/train.jl:105
┌ Info: Epoch 144
└ @ Main /Users/kcft114/.julia/packages/Flux/qXNjB/src/optimise/train.jl:105
┌ Info: Epoch 145
└ @ Main /Users/kcft114/.julia/packages/Flux/qXNjB/src/optimise/train.jl:105
┌ Info: Epoch 146
└ @ Main /Users/kcft114/.julia/packages/Flux/qXNjB/src/optimise/train.jl:105
┌ Info: Epoch 147
└ @ Main /Users/kcft114/.julia/packages/Flux/qXNjB/src/optimise/train.jl:105
┌ Info: Epoch 148
└ @ Main /Users/kcft114/.julia/packages/Flux/qXNjB/src/optimise/train.jl:105
┌ Info: Epoch 149
└ @ Main /Users/kcft114/.julia/packages/Flux/qXNjB/src/optimise/train.jl:105
┌ Info: Epoch 150
└ @ Main /Users/kcft114/.julia/packages/Flux/qXNjB/src/optimise/train.jl:105
┌ Info: Epoch 151
└ @ Main /Users/kcft114/.julia/packages/Flux/qXNjB/src/optimise/train.jl:105
```

```
loss(X, Y) = 0.8263668f0 (tracked)
loss(X, Y) = 0.8502417f0 (tracked)
loss(X, Y) = 0.8371888f0 (tracked)
loss(X, Y) = 0.83555883f0 (tracked)
loss(X, Y) = 0.85133976f0 (tracked)
loss(X, Y) = 0.841604f0 (tracked)
loss(X, Y) = 0.8428897f0 (tracked)
loss(X, Y) = 0.84401655f0 (tracked)
loss(X, Y) = 0.82276565f0 (tracked)
loss(X, Y) = 0.8224795f0 (tracked)
loss(X, Y) = 0.8423564f0 (tracked)
loss(X, Y) = 0.8372273f0 (tracked)
loss(X, Y) = 0.8538842f0 (tracked)
loss(X, Y) = 0.83115727f0 (tracked)
loss(X, Y) = 0.85735667f0 (tracked)
loss(X, Y) = 0.83697605f0 (tracked)
loss(X, Y) = 0.8263928f0 (tracked)
loss(X, Y) = 0.82154346f0 (tracked)
loss(X, Y) = 0.84755194f0 (tracked)
loss(X, Y) = 0.8021015f0 (tracked)
loss(X, Y) = 0.81158054f0 (tracked)
loss(X, Y) = 0.8317987f0 (tracked)
loss(X, Y) = 0.8357584f0 (tracked)
loss(X, Y) = 0.8292734f0 (tracked)
loss(X, Y) = 0.8345401f0 (tracked)
loss(X, Y) = 0.83264446f0 (tracked)
loss(X, Y) = 0.79629374f0 (tracked)
loss(X, Y) = 0.80903256f0 (tracked)
loss(X, Y) = 0.82410216f0 (tracked)
loss(X, Y) = 0.8249903f0 (tracked)
loss(X, Y) = 0.8138525f0 (tracked)
loss(X, Y) = 0.8305749f0 (tracked)
loss(X, Y) = 0.82025164f0 (tracked)
loss(X, Y) = 0.8100515f0 (tracked)
loss(X, Y) = 0.81342256f0 (tracked)
loss(X, Y) = 0.8264798f0 (tracked)
loss(X, Y) = 0.8003007f0 (tracked)
loss(X, Y) = 0.8370451f0 (tracked)
loss(X, Y) = 0.8131683f0 (tracked)
loss(X, Y) = 0.8022948f0 (tracked)
loss(X, Y) = 0.8164993f0 (tracked)
loss(X, Y) = 0.81504244f0 (tracked)
loss(X, Y) = 0.7945789f0 (tracked)
loss(X, Y) = 0.820239f0 (tracked)
loss(X, Y) = 0.815632f0 (tracked)
loss(X, Y) = 0.80094993f0 (tracked)
loss(X, Y) = 0.7979246f0 (tracked)
loss(X, Y) = 0.82412755f0 (tracked)
loss(X, Y) = 0.81077224f0 (tracked)
loss(X, Y) = 0.82411164f0 (tracked)
loss(X, Y) = 0.8200816f0 (tracked)
loss(X, Y) = 0.8208534f0 (tracked)
loss(X, Y) = 0.8458522f0 (tracked)
loss(X, Y) = 0.80540866f0 (tracked)
loss(X, Y) = 0.8058943f0 (tracked)
loss(X, Y) = 0.8225757f0 (tracked)
loss(X, Y) = 0.7814904f0 (tracked)
loss(X, Y) = 0.78358424f0 (tracked)
loss(X, Y) = 0.82354957f0 (tracked)
loss(X, Y) = 0.7998036f0 (tracked)
loss(X, Y) = 0.8165597f0 (tracked)
loss(X, Y) = 0.7806884f0 (tracked)
loss(X, Y) = 0.80575126f0 (tracked)
loss(X, Y) = 0.81468195f0 (tracked)
loss(X, Y) = 0.80316746f0 (tracked)
loss(X, Y) = 0.7888205f0 (tracked)
loss(X, Y) = 0.7747273f0 (tracked)
loss(X, Y) = 0.81316483f0 (tracked)
loss(X, Y) = 0.79712605f0 (tracked)
loss(X, Y) = 0.8188373f0 (tracked)
loss(X, Y) = 0.78068465f0 (tracked)
loss(X, Y) = 0.78821534f0 (tracked)
loss(X, Y) = 0.78898865f0 (tracked)
loss(X, Y) = 0.79988647f0 (tracked)
loss(X, Y) = 0.7939209f0 (tracked)
loss(X, Y) = 0.7717598f0 (tracked)
loss(X, Y) = 0.79607093f0 (tracked)
loss(X, Y) = 0.8119316f0 (tracked)
loss(X, Y) = 0.7982038f0 (tracked)
loss(X, Y) = 0.7873009f0 (tracked)
loss(X, Y) = 0.79300374f0 (tracked)
loss(X, Y) = 0.78813475f0 (tracked)
loss(X, Y) = 0.7807382f0 (tracked)
loss(X, Y) = 0.7932986f0 (tracked)
loss(X, Y) = 0.7817526f0 (tracked)
loss(X, Y) = 0.7941582f0 (tracked)
loss(X, Y) = 0.7942401f0 (tracked)
loss(X, Y) = 0.7655133f0 (tracked)
loss(X, Y) = 0.77095544f0 (tracked)
loss(X, Y) = 0.78624636f0 (tracked)
loss(X, Y) = 0.7907049f0 (tracked)
loss(X, Y) = 0.7749114f0 (tracked)
loss(X, Y) = 0.7555177f0 (tracked)
loss(X, Y) = 0.77263105f0 (tracked)
loss(X, Y) = 0.80681914f0 (tracked)
loss(X, Y) = 0.7792549f0 (tracked)
loss(X, Y) = 0.79521704f0 (tracked)
```

```
┌ Info: Epoch 152
└ @ Main /Users/kcft114/.julia/packages/Flux/qXNjB/src/optimise/train.jl:105
┌ Info: Epoch 153
└ @ Main /Users/kcft114/.julia/packages/Flux/qXNjB/src/optimise/train.jl:105
┌ Info: Epoch 154
└ @ Main /Users/kcft114/.julia/packages/Flux/qXNjB/src/optimise/train.jl:105
┌ Info: Epoch 155
└ @ Main /Users/kcft114/.julia/packages/Flux/qXNjB/src/optimise/train.jl:105
┌ Info: Epoch 156
└ @ Main /Users/kcft114/.julia/packages/Flux/qXNjB/src/optimise/train.jl:105
┌ Info: Epoch 157
└ @ Main /Users/kcft114/.julia/packages/Flux/qXNjB/src/optimise/train.jl:105
┌ Info: Epoch 158
└ @ Main /Users/kcft114/.julia/packages/Flux/qXNjB/src/optimise/train.jl:105
┌ Info: Epoch 159
└ @ Main /Users/kcft114/.julia/packages/Flux/qXNjB/src/optimise/train.jl:105
┌ Info: Epoch 160
└ @ Main /Users/kcft114/.julia/packages/Flux/qXNjB/src/optimise/train.jl:105
┌ Info: Epoch 161
└ @ Main /Users/kcft114/.julia/packages/Flux/qXNjB/src/optimise/train.jl:105
┌ Info: Epoch 162
└ @ Main /Users/kcft114/.julia/packages/Flux/qXNjB/src/optimise/train.jl:105
┌ Info: Epoch 163
└ @ Main /Users/kcft114/.julia/packages/Flux/qXNjB/src/optimise/train.jl:105
┌ Info: Epoch 164
└ @ Main /Users/kcft114/.julia/packages/Flux/qXNjB/src/optimise/train.jl:105
┌ Info: Epoch 165
└ @ Main /Users/kcft114/.julia/packages/Flux/qXNjB/src/optimise/train.jl:105
┌ Info: Epoch 166
└ @ Main /Users/kcft114/.julia/packages/Flux/qXNjB/src/optimise/train.jl:105
┌ Info: Epoch 167
└ @ Main /Users/kcft114/.julia/packages/Flux/qXNjB/src/optimise/train.jl:105
┌ Info: Epoch 168
└ @ Main /Users/kcft114/.julia/packages/Flux/qXNjB/src/optimise/train.jl:105
┌ Info: Epoch 169
└ @ Main /Users/kcft114/.julia/packages/Flux/qXNjB/src/optimise/train.jl:105
┌ Info: Epoch 170
└ @ Main /Users/kcft114/.julia/packages/Flux/qXNjB/src/optimise/train.jl:105
┌ Info: Epoch 171
└ @ Main /Users/kcft114/.julia/packages/Flux/qXNjB/src/optimise/train.jl:105
┌ Info: Epoch 172
└ @ Main /Users/kcft114/.julia/packages/Flux/qXNjB/src/optimise/train.jl:105
┌ Info: Epoch 173
└ @ Main /Users/kcft114/.julia/packages/Flux/qXNjB/src/optimise/train.jl:105
┌ Info: Epoch 174
└ @ Main /Users/kcft114/.julia/packages/Flux/qXNjB/src/optimise/train.jl:105
┌ Info: Epoch 175
└ @ Main /Users/kcft114/.julia/packages/Flux/qXNjB/src/optimise/train.jl:105
┌ Info: Epoch 176
└ @ Main /Users/kcft114/.julia/packages/Flux/qXNjB/src/optimise/train.jl:105
┌ Info: Epoch 177
└ @ Main /Users/kcft114/.julia/packages/Flux/qXNjB/src/optimise/train.jl:105
┌ Info: Epoch 178
└ @ Main /Users/kcft114/.julia/packages/Flux/qXNjB/src/optimise/train.jl:105
┌ Info: Epoch 179
└ @ Main /Users/kcft114/.julia/packages/Flux/qXNjB/src/optimise/train.jl:105
┌ Info: Epoch 180
└ @ Main /Users/kcft114/.julia/packages/Flux/qXNjB/src/optimise/train.jl:105
┌ Info: Epoch 181
└ @ Main /Users/kcft114/.julia/packages/Flux/qXNjB/src/optimise/train.jl:105
┌ Info: Epoch 182
└ @ Main /Users/kcft114/.julia/packages/Flux/qXNjB/src/optimise/train.jl:105
┌ Info: Epoch 183
└ @ Main /Users/kcft114/.julia/packages/Flux/qXNjB/src/optimise/train.jl:105
┌ Info: Epoch 184
└ @ Main /Users/kcft114/.julia/packages/Flux/qXNjB/src/optimise/train.jl:105
┌ Info: Epoch 185
└ @ Main /Users/kcft114/.julia/packages/Flux/qXNjB/src/optimise/train.jl:105
┌ Info: Epoch 186
└ @ Main /Users/kcft114/.julia/packages/Flux/qXNjB/src/optimise/train.jl:105
┌ Info: Epoch 187
└ @ Main /Users/kcft114/.julia/packages/Flux/qXNjB/src/optimise/train.jl:105
┌ Info: Epoch 188
└ @ Main /Users/kcft114/.julia/packages/Flux/qXNjB/src/optimise/train.jl:105
┌ Info: Epoch 189
└ @ Main /Users/kcft114/.julia/packages/Flux/qXNjB/src/optimise/train.jl:105
┌ Info: Epoch 190
└ @ Main /Users/kcft114/.julia/packages/Flux/qXNjB/src/optimise/train.jl:105
┌ Info: Epoch 191
└ @ Main /Users/kcft114/.julia/packages/Flux/qXNjB/src/optimise/train.jl:105
┌ Info: Epoch 192
└ @ Main /Users/kcft114/.julia/packages/Flux/qXNjB/src/optimise/train.jl:105
┌ Info: Epoch 193
└ @ Main /Users/kcft114/.julia/packages/Flux/qXNjB/src/optimise/train.jl:105
┌ Info: Epoch 194
└ @ Main /Users/kcft114/.julia/packages/Flux/qXNjB/src/optimise/train.jl:105
┌ Info: Epoch 195
└ @ Main /Users/kcft114/.julia/packages/Flux/qXNjB/src/optimise/train.jl:105
┌ Info: Epoch 196
└ @ Main /Users/kcft114/.julia/packages/Flux/qXNjB/src/optimise/train.jl:105
┌ Info: Epoch 197
└ @ Main /Users/kcft114/.julia/packages/Flux/qXNjB/src/optimise/train.jl:105
┌ Info: Epoch 198
└ @ Main /Users/kcft114/.julia/packages/Flux/qXNjB/src/optimise/train.jl:105
┌ Info: Epoch 199
└ @ Main /Users/kcft114/.julia/packages/Flux/qXNjB/src/optimise/train.jl:105
┌ Info: Epoch 200
└ @ Main /Users/kcft114/.julia/packages/Flux/qXNjB/src/optimise/train.jl:105
┌ Info: Epoch 201
└ @ Main /Users/kcft114/.julia/packages/Flux/qXNjB/src/optimise/train.jl:105
┌ Info: Epoch 202
└ @ Main /Users/kcft114/.julia/packages/Flux/qXNjB/src/optimise/train.jl:105
┌ Info: Epoch 203
└ @ Main /Users/kcft114/.julia/packages/Flux/qXNjB/src/optimise/train.jl:105
┌ Info: Epoch 204
└ @ Main /Users/kcft114/.julia/packages/Flux/qXNjB/src/optimise/train.jl:105
┌ Info: Epoch 205
└ @ Main /Users/kcft114/.julia/packages/Flux/qXNjB/src/optimise/train.jl:105
┌ Info: Epoch 206
└ @ Main /Users/kcft114/.julia/packages/Flux/qXNjB/src/optimise/train.jl:105
┌ Info: Epoch 207
└ @ Main /Users/kcft114/.julia/packages/Flux/qXNjB/src/optimise/train.jl:105
┌ Info: Epoch 208
└ @ Main /Users/kcft114/.julia/packages/Flux/qXNjB/src/optimise/train.jl:105
┌ Info: Epoch 209
└ @ Main /Users/kcft114/.julia/packages/Flux/qXNjB/src/optimise/train.jl:105
┌ Info: Epoch 210
└ @ Main /Users/kcft114/.julia/packages/Flux/qXNjB/src/optimise/train.jl:105
┌ Info: Epoch 211
└ @ Main /Users/kcft114/.julia/packages/Flux/qXNjB/src/optimise/train.jl:105
┌ Info: Epoch 212
└ @ Main /Users/kcft114/.julia/packages/Flux/qXNjB/src/optimise/train.jl:105
┌ Info: Epoch 213
└ @ Main /Users/kcft114/.julia/packages/Flux/qXNjB/src/optimise/train.jl:105
┌ Info: Epoch 214
└ @ Main /Users/kcft114/.julia/packages/Flux/qXNjB/src/optimise/train.jl:105
┌ Info: Epoch 215
└ @ Main /Users/kcft114/.julia/packages/Flux/qXNjB/src/optimise/train.jl:105
┌ Info: Epoch 216
└ @ Main /Users/kcft114/.julia/packages/Flux/qXNjB/src/optimise/train.jl:105
┌ Info: Epoch 217
└ @ Main /Users/kcft114/.julia/packages/Flux/qXNjB/src/optimise/train.jl:105
┌ Info: Epoch 218
└ @ Main /Users/kcft114/.julia/packages/Flux/qXNjB/src/optimise/train.jl:105
┌ Info: Epoch 219
└ @ Main /Users/kcft114/.julia/packages/Flux/qXNjB/src/optimise/train.jl:105
┌ Info: Epoch 220
└ @ Main /Users/kcft114/.julia/packages/Flux/qXNjB/src/optimise/train.jl:105
┌ Info: Epoch 221
└ @ Main /Users/kcft114/.julia/packages/Flux/qXNjB/src/optimise/train.jl:105
┌ Info: Epoch 222
└ @ Main /Users/kcft114/.julia/packages/Flux/qXNjB/src/optimise/train.jl:105
┌ Info: Epoch 223
└ @ Main /Users/kcft114/.julia/packages/Flux/qXNjB/src/optimise/train.jl:105
┌ Info: Epoch 224
└ @ Main /Users/kcft114/.julia/packages/Flux/qXNjB/src/optimise/train.jl:105
┌ Info: Epoch 225
└ @ Main /Users/kcft114/.julia/packages/Flux/qXNjB/src/optimise/train.jl:105
┌ Info: Epoch 226
└ @ Main /Users/kcft114/.julia/packages/Flux/qXNjB/src/optimise/train.jl:105
┌ Info: Epoch 227
└ @ Main /Users/kcft114/.julia/packages/Flux/qXNjB/src/optimise/train.jl:105
┌ Info: Epoch 228
└ @ Main /Users/kcft114/.julia/packages/Flux/qXNjB/src/optimise/train.jl:105
┌ Info: Epoch 229
└ @ Main /Users/kcft114/.julia/packages/Flux/qXNjB/src/optimise/train.jl:105
┌ Info: Epoch 230
└ @ Main /Users/kcft114/.julia/packages/Flux/qXNjB/src/optimise/train.jl:105
┌ Info: Epoch 231
└ @ Main /Users/kcft114/.julia/packages/Flux/qXNjB/src/optimise/train.jl:105
┌ Info: Epoch 232
└ @ Main /Users/kcft114/.julia/packages/Flux/qXNjB/src/optimise/train.jl:105
┌ Info: Epoch 233
└ @ Main /Users/kcft114/.julia/packages/Flux/qXNjB/src/optimise/train.jl:105
┌ Info: Epoch 234
└ @ Main /Users/kcft114/.julia/packages/Flux/qXNjB/src/optimise/train.jl:105
┌ Info: Epoch 235
└ @ Main /Users/kcft114/.julia/packages/Flux/qXNjB/src/optimise/train.jl:105
┌ Info: Epoch 236
└ @ Main /Users/kcft114/.julia/packages/Flux/qXNjB/src/optimise/train.jl:105
┌ Info: Epoch 237
└ @ Main /Users/kcft114/.julia/packages/Flux/qXNjB/src/optimise/train.jl:105
┌ Info: Epoch 238
└ @ Main /Users/kcft114/.julia/packages/Flux/qXNjB/src/optimise/train.jl:105
┌ Info: Epoch 239
└ @ Main /Users/kcft114/.julia/packages/Flux/qXNjB/src/optimise/train.jl:105
┌ Info: Epoch 240
└ @ Main /Users/kcft114/.julia/packages/Flux/qXNjB/src/optimise/train.jl:105
┌ Info: Epoch 241
└ @ Main /Users/kcft114/.julia/packages/Flux/qXNjB/src/optimise/train.jl:105
┌ Info: Epoch 242
└ @ Main /Users/kcft114/.julia/packages/Flux/qXNjB/src/optimise/train.jl:105
┌ Info: Epoch 243
└ @ Main /Users/kcft114/.julia/packages/Flux/qXNjB/src/optimise/train.jl:105
┌ Info: Epoch 244
└ @ Main /Users/kcft114/.julia/packages/Flux/qXNjB/src/optimise/train.jl:105
┌ Info: Epoch 245
└ @ Main /Users/kcft114/.julia/packages/Flux/qXNjB/src/optimise/train.jl:105
┌ Info: Epoch 246
└ @ Main /Users/kcft114/.julia/packages/Flux/qXNjB/src/optimise/train.jl:105
┌ Info: Epoch 247
└ @ Main /Users/kcft114/.julia/packages/Flux/qXNjB/src/optimise/train.jl:105
┌ Info: Epoch 248
└ @ Main /Users/kcft114/.julia/packages/Flux/qXNjB/src/optimise/train.jl:105
┌ Info: Epoch 249
└ @ Main /Users/kcft114/.julia/packages/Flux/qXNjB/src/optimise/train.jl:105
┌ Info: Epoch 250
└ @ Main /Users/kcft114/.julia/packages/Flux/qXNjB/src/optimise/train.jl:105
┌ Info: Epoch 251
└ @ Main /Users/kcft114/.julia/packages/Flux/qXNjB/src/optimise/train.jl:105
┌ Info: Epoch 252
└ @ Main /Users/kcft114/.julia/packages/Flux/qXNjB/src/optimise/train.jl:105
┌ Info: Epoch 253
└ @ Main /Users/kcft114/.julia/packages/Flux/qXNjB/src/optimise/train.jl:105
┌ Info: Epoch 254
└ @ Main /Users/kcft114/.julia/packages/Flux/qXNjB/src/optimise/train.jl:105
┌ Info: Epoch 255
└ @ Main /Users/kcft114/.julia/packages/Flux/qXNjB/src/optimise/train.jl:105
┌ Info: Epoch 256
└ @ Main /Users/kcft114/.julia/packages/Flux/qXNjB/src/optimise/train.jl:105
┌ Info: Epoch 257
└ @ Main /Users/kcft114/.julia/packages/Flux/qXNjB/src/optimise/train.jl:105
┌ Info: Epoch 258
└ @ Main /Users/kcft114/.julia/packages/Flux/qXNjB/src/optimise/train.jl:105
┌ Info: Epoch 259
└ @ Main /Users/kcft114/.julia/packages/Flux/qXNjB/src/optimise/train.jl:105
┌ Info: Epoch 260
└ @ Main /Users/kcft114/.julia/packages/Flux/qXNjB/src/optimise/train.jl:105
┌ Info: Epoch 261
└ @ Main /Users/kcft114/.julia/packages/Flux/qXNjB/src/optimise/train.jl:105
┌ Info: Epoch 262
└ @ Main /Users/kcft114/.julia/packages/Flux/qXNjB/src/optimise/train.jl:105
┌ Info: Epoch 263
└ @ Main /Users/kcft114/.julia/packages/Flux/qXNjB/src/optimise/train.jl:105
┌ Info: Epoch 264
└ @ Main /Users/kcft114/.julia/packages/Flux/qXNjB/src/optimise/train.jl:105
┌ Info: Epoch 265
└ @ Main /Users/kcft114/.julia/packages/Flux/qXNjB/src/optimise/train.jl:105
┌ Info: Epoch 266
└ @ Main /Users/kcft114/.julia/packages/Flux/qXNjB/src/optimise/train.jl:105
┌ Info: Epoch 267
└ @ Main /Users/kcft114/.julia/packages/Flux/qXNjB/src/optimise/train.jl:105
┌ Info: Epoch 268
└ @ Main /Users/kcft114/.julia/packages/Flux/qXNjB/src/optimise/train.jl:105
┌ Info: Epoch 269
└ @ Main /Users/kcft114/.julia/packages/Flux/qXNjB/src/optimise/train.jl:105
┌ Info: Epoch 270
└ @ Main /Users/kcft114/.julia/packages/Flux/qXNjB/src/optimise/train.jl:105
┌ Info: Epoch 271
└ @ Main /Users/kcft114/.julia/packages/Flux/qXNjB/src/optimise/train.jl:105
┌ Info: Epoch 272
└ @ Main /Users/kcft114/.julia/packages/Flux/qXNjB/src/optimise/train.jl:105
┌ Info: Epoch 273
└ @ Main /Users/kcft114/.julia/packages/Flux/qXNjB/src/optimise/train.jl:105
┌ Info: Epoch 274
└ @ Main /Users/kcft114/.julia/packages/Flux/qXNjB/src/optimise/train.jl:105
┌ Info: Epoch 275
└ @ Main /Users/kcft114/.julia/packages/Flux/qXNjB/src/optimise/train.jl:105
┌ Info: Epoch 276
└ @ Main /Users/kcft114/.julia/packages/Flux/qXNjB/src/optimise/train.jl:105
┌ Info: Epoch 277
└ @ Main /Users/kcft114/.julia/packages/Flux/qXNjB/src/optimise/train.jl:105
┌ Info: Epoch 278
└ @ Main /Users/kcft114/.julia/packages/Flux/qXNjB/src/optimise/train.jl:105
┌ Info: Epoch 279
└ @ Main /Users/kcft114/.julia/packages/Flux/qXNjB/src/optimise/train.jl:105
┌ Info: Epoch 280
└ @ Main /Users/kcft114/.julia/packages/Flux/qXNjB/src/optimise/train.jl:105
┌ Info: Epoch 281
└ @ Main /Users/kcft114/.julia/packages/Flux/qXNjB/src/optimise/train.jl:105
┌ Info: Epoch 282
└ @ Main /Users/kcft114/.julia/packages/Flux/qXNjB/src/optimise/train.jl:105
┌ Info: Epoch 283
└ @ Main /Users/kcft114/.julia/packages/Flux/qXNjB/src/optimise/train.jl:105
┌ Info: Epoch 284
└ @ Main /Users/kcft114/.julia/packages/Flux/qXNjB/src/optimise/train.jl:105
┌ Info: Epoch 285
└ @ Main /Users/kcft114/.julia/packages/Flux/qXNjB/src/optimise/train.jl:105
┌ Info: Epoch 286
└ @ Main /Users/kcft114/.julia/packages/Flux/qXNjB/src/optimise/train.jl:105
┌ Info: Epoch 287
└ @ Main /Users/kcft114/.julia/packages/Flux/qXNjB/src/optimise/train.jl:105
┌ Info: Epoch 288
└ @ Main /Users/kcft114/.julia/packages/Flux/qXNjB/src/optimise/train.jl:105
┌ Info: Epoch 289
└ @ Main /Users/kcft114/.julia/packages/Flux/qXNjB/src/optimise/train.jl:105
┌ Info: Epoch 290
└ @ Main /Users/kcft114/.julia/packages/Flux/qXNjB/src/optimise/train.jl:105
┌ Info: Epoch 291
└ @ Main /Users/kcft114/.julia/packages/Flux/qXNjB/src/optimise/train.jl:105
┌ Info: Epoch 292
└ @ Main /Users/kcft114/.julia/packages/Flux/qXNjB/src/optimise/train.jl:105
┌ Info: Epoch 293
└ @ Main /Users/kcft114/.julia/packages/Flux/qXNjB/src/optimise/train.jl:105
┌ Info: Epoch 294
└ @ Main /Users/kcft114/.julia/packages/Flux/qXNjB/src/optimise/train.jl:105
┌ Info: Epoch 295
└ @ Main /Users/kcft114/.julia/packages/Flux/qXNjB/src/optimise/train.jl:105
┌ Info: Epoch 296
└ @ Main /Users/kcft114/.julia/packages/Flux/qXNjB/src/optimise/train.jl:105
┌ Info: Epoch 297
└ @ Main /Users/kcft114/.julia/packages/Flux/qXNjB/src/optimise/train.jl:105
┌ Info: Epoch 298
└ @ Main /Users/kcft114/.julia/packages/Flux/qXNjB/src/optimise/train.jl:105
┌ Info: Epoch 299
└ @ Main /Users/kcft114/.julia/packages/Flux/qXNjB/src/optimise/train.jl:105
┌ Info: Epoch 300
└ @ Main /Users/kcft114/.julia/packages/Flux/qXNjB/src/optimise/train.jl:105
┌ Info: Epoch 301
└ @ Main /Users/kcft114/.julia/packages/Flux/qXNjB/src/optimise/train.jl:105
┌ Info: Epoch 302
└ @ Main /Users/kcft114/.julia/packages/Flux/qXNjB/src/optimise/train.jl:105
┌ Info: Epoch 303
└ @ Main /Users/kcft114/.julia/packages/Flux/qXNjB/src/optimise/train.jl:105
┌ Info: Epoch 304
└ @ Main /Users/kcft114/.julia/packages/Flux/qXNjB/src/optimise/train.jl:105
┌ Info: Epoch 305
└ @ Main /Users/kcft114/.julia/packages/Flux/qXNjB/src/optimise/train.jl:105
┌ Info: Epoch 306
└ @ Main /Users/kcft114/.julia/packages/Flux/qXNjB/src/optimise/train.jl:105
┌ Info: Epoch 307
└ @ Main /Users/kcft114/.julia/packages/Flux/qXNjB/src/optimise/train.jl:105
┌ Info: Epoch 308
└ @ Main /Users/kcft114/.julia/packages/Flux/qXNjB/src/optimise/train.jl:105
┌ Info: Epoch 309
└ @ Main /Users/kcft114/.julia/packages/Flux/qXNjB/src/optimise/train.jl:105
┌ Info: Epoch 310
└ @ Main /Users/kcft114/.julia/packages/Flux/qXNjB/src/optimise/train.jl:105
┌ Info: Epoch 311
└ @ Main /Users/kcft114/.julia/packages/Flux/qXNjB/src/optimise/train.jl:105
┌ Info: Epoch 312
└ @ Main /Users/kcft114/.julia/packages/Flux/qXNjB/src/optimise/train.jl:105
┌ Info: Epoch 313
└ @ Main /Users/kcft114/.julia/packages/Flux/qXNjB/src/optimise/train.jl:105
┌ Info: Epoch 314
└ @ Main /Users/kcft114/.julia/packages/Flux/qXNjB/src/optimise/train.jl:105
┌ Info: Epoch 315
└ @ Main /Users/kcft114/.julia/packages/Flux/qXNjB/src/optimise/train.jl:105
┌ Info: Epoch 316
└ @ Main /Users/kcft114/.julia/packages/Flux/qXNjB/src/optimise/train.jl:105
┌ Info: Epoch 317
└ @ Main /Users/kcft114/.julia/packages/Flux/qXNjB/src/optimise/train.jl:105
┌ Info: Epoch 318
└ @ Main /Users/kcft114/.julia/packages/Flux/qXNjB/src/optimise/train.jl:105
┌ Info: Epoch 319
└ @ Main /Users/kcft114/.julia/packages/Flux/qXNjB/src/optimise/train.jl:105
┌ Info: Epoch 320
└ @ Main /Users/kcft114/.julia/packages/Flux/qXNjB/src/optimise/train.jl:105
┌ Info: Epoch 321
└ @ Main /Users/kcft114/.julia/packages/Flux/qXNjB/src/optimise/train.jl:105
┌ Info: Epoch 322
└ @ Main /Users/kcft114/.julia/packages/Flux/qXNjB/src/optimise/train.jl:105
┌ Info: Epoch 323
└ @ Main /Users/kcft114/.julia/packages/Flux/qXNjB/src/optimise/train.jl:105
┌ Info: Epoch 324
└ @ Main /Users/kcft114/.julia/packages/Flux/qXNjB/src/optimise/train.jl:105
┌ Info: Epoch 325
└ @ Main /Users/kcft114/.julia/packages/Flux/qXNjB/src/optimise/train.jl:105
┌ Info: Epoch 326
└ @ Main /Users/kcft114/.julia/packages/Flux/qXNjB/src/optimise/train.jl:105
┌ Info: Epoch 327
└ @ Main /Users/kcft114/.julia/packages/Flux/qXNjB/src/optimise/train.jl:105
┌ Info: Epoch 328
└ @ Main /Users/kcft114/.julia/packages/Flux/qXNjB/src/optimise/train.jl:105
┌ Info: Epoch 329
└ @ Main /Users/kcft114/.julia/packages/Flux/qXNjB/src/optimise/train.jl:105
┌ Info: Epoch 330
└ @ Main /Users/kcft114/.julia/packages/Flux/qXNjB/src/optimise/train.jl:105
┌ Info: Epoch 331
└ @ Main /Users/kcft114/.julia/packages/Flux/qXNjB/src/optimise/train.jl:105
┌ Info: Epoch 332
└ @ Main /Users/kcft114/.julia/packages/Flux/qXNjB/src/optimise/train.jl:105
┌ Info: Epoch 333
└ @ Main /Users/kcft114/.julia/packages/Flux/qXNjB/src/optimise/train.jl:105
┌ Info: Epoch 334
└ @ Main /Users/kcft114/.julia/packages/Flux/qXNjB/src/optimise/train.jl:105
┌ Info: Epoch 335
└ @ Main /Users/kcft114/.julia/packages/Flux/qXNjB/src/optimise/train.jl:105
┌ Info: Epoch 336
└ @ Main /Users/kcft114/.julia/packages/Flux/qXNjB/src/optimise/train.jl:105
┌ Info: Epoch 337
└ @ Main /Users/kcft114/.julia/packages/Flux/qXNjB/src/optimise/train.jl:105
┌ Info: Epoch 338
└ @ Main /Users/kcft114/.julia/packages/Flux/qXNjB/src/optimise/train.jl:105
┌ Info: Epoch 339
└ @ Main /Users/kcft114/.julia/packages/Flux/qXNjB/src/optimise/train.jl:105
┌ Info: Epoch 340
└ @ Main /Users/kcft114/.julia/packages/Flux/qXNjB/src/optimise/train.jl:105
┌ Info: Epoch 341
└ @ Main /Users/kcft114/.julia/packages/Flux/qXNjB/src/optimise/train.jl:105
┌ Info: Epoch 342
└ @ Main /Users/kcft114/.julia/packages/Flux/qXNjB/src/optimise/train.jl:105
┌ Info: Epoch 343
└ @ Main /Users/kcft114/.julia/packages/Flux/qXNjB/src/optimise/train.jl:105
┌ Info: Epoch 344
└ @ Main /Users/kcft114/.julia/packages/Flux/qXNjB/src/optimise/train.jl:105
┌ Info: Epoch 345
└ @ Main /Users/kcft114/.julia/packages/Flux/qXNjB/src/optimise/train.jl:105
┌ Info: Epoch 346
└ @ Main /Users/kcft114/.julia/packages/Flux/qXNjB/src/optimise/train.jl:105
┌ Info: Epoch 347
└ @ Main /Users/kcft114/.julia/packages/Flux/qXNjB/src/optimise/train.jl:105
┌ Info: Epoch 348
└ @ Main /Users/kcft114/.julia/packages/Flux/qXNjB/src/optimise/train.jl:105
┌ Info: Epoch 349
└ @ Main /Users/kcft114/.julia/packages/Flux/qXNjB/src/optimise/train.jl:105
┌ Info: Epoch 350
└ @ Main /Users/kcft114/.julia/packages/Flux/qXNjB/src/optimise/train.jl:105
┌ Info: Epoch 351
└ @ Main /Users/kcft114/.julia/packages/Flux/qXNjB/src/optimise/train.jl:105
┌ Info: Epoch 352
└ @ Main /Users/kcft114/.julia/packages/Flux/qXNjB/src/optimise/train.jl:105
```

```
loss(X, Y) = 0.77869004f0 (tracked)
loss(X, Y) = 0.7915227f0 (tracked)
loss(X, Y) = 0.7675314f0 (tracked)
loss(X, Y) = 0.77727157f0 (tracked)
loss(X, Y) = 0.7632062f0 (tracked)
loss(X, Y) = 0.78181165f0 (tracked)
loss(X, Y) = 0.7401672f0 (tracked)
loss(X, Y) = 0.77884513f0 (tracked)
loss(X, Y) = 0.7947815f0 (tracked)
loss(X, Y) = 0.7768493f0 (tracked)
loss(X, Y) = 0.7846086f0 (tracked)
loss(X, Y) = 0.7785148f0 (tracked)
loss(X, Y) = 0.768501f0 (tracked)
loss(X, Y) = 0.77829033f0 (tracked)
loss(X, Y) = 0.80172163f0 (tracked)
loss(X, Y) = 0.7619839f0 (tracked)
loss(X, Y) = 0.77321476f0 (tracked)
loss(X, Y) = 0.7805431f0 (tracked)
loss(X, Y) = 0.76038784f0 (tracked)
loss(X, Y) = 0.7839439f0 (tracked)
loss(X, Y) = 0.76866794f0 (tracked)
loss(X, Y) = 0.77705973f0 (tracked)
loss(X, Y) = 0.7623481f0 (tracked)
loss(X, Y) = 0.78744215f0 (tracked)
loss(X, Y) = 0.79068637f0 (tracked)
loss(X, Y) = 0.76900005f0 (tracked)
loss(X, Y) = 0.7997253f0 (tracked)
loss(X, Y) = 0.79271126f0 (tracked)
loss(X, Y) = 0.7689978f0 (tracked)
loss(X, Y) = 0.7841753f0 (tracked)
loss(X, Y) = 0.7718754f0 (tracked)
loss(X, Y) = 0.76237094f0 (tracked)
loss(X, Y) = 0.7546148f0 (tracked)
loss(X, Y) = 0.76062f0 (tracked)
loss(X, Y) = 0.76815206f0 (tracked)
loss(X, Y) = 0.7328653f0 (tracked)
loss(X, Y) = 0.78078413f0 (tracked)
loss(X, Y) = 0.76428527f0 (tracked)
loss(X, Y) = 0.7578797f0 (tracked)
loss(X, Y) = 0.74161565f0 (tracked)
loss(X, Y) = 0.768747f0 (tracked)
loss(X, Y) = 0.7586273f0 (tracked)
loss(X, Y) = 0.7628258f0 (tracked)
loss(X, Y) = 0.7399581f0 (tracked)
loss(X, Y) = 0.7656585f0 (tracked)
loss(X, Y) = 0.72927064f0 (tracked)
loss(X, Y) = 0.7796014f0 (tracked)
loss(X, Y) = 0.7678597f0 (tracked)
loss(X, Y) = 0.76461315f0 (tracked)
loss(X, Y) = 0.7714609f0 (tracked)
loss(X, Y) = 0.7521853f0 (tracked)
loss(X, Y) = 0.7668909f0 (tracked)
loss(X, Y) = 0.77827454f0 (tracked)
loss(X, Y) = 0.7620359f0 (tracked)
loss(X, Y) = 0.7446181f0 (tracked)
loss(X, Y) = 0.7562145f0 (tracked)
loss(X, Y) = 0.76644343f0 (tracked)
loss(X, Y) = 0.78290164f0 (tracked)
loss(X, Y) = 0.75771856f0 (tracked)
loss(X, Y) = 0.75999546f0 (tracked)
loss(X, Y) = 0.76509064f0 (tracked)
loss(X, Y) = 0.744753f0 (tracked)
loss(X, Y) = 0.75166726f0 (tracked)
loss(X, Y) = 0.73736894f0 (tracked)
loss(X, Y) = 0.74407357f0 (tracked)
loss(X, Y) = 0.7407831f0 (tracked)
loss(X, Y) = 0.7919056f0 (tracked)
loss(X, Y) = 0.7727997f0 (tracked)
loss(X, Y) = 0.7637378f0 (tracked)
loss(X, Y) = 0.7572772f0 (tracked)
loss(X, Y) = 0.75644225f0 (tracked)
loss(X, Y) = 0.75773066f0 (tracked)
loss(X, Y) = 0.76130134f0 (tracked)
loss(X, Y) = 0.7610442f0 (tracked)
loss(X, Y) = 0.7695991f0 (tracked)
loss(X, Y) = 0.7437436f0 (tracked)
loss(X, Y) = 0.7384328f0 (tracked)
loss(X, Y) = 0.7934174f0 (tracked)
loss(X, Y) = 0.7678597f0 (tracked)
loss(X, Y) = 0.75481164f0 (tracked)
loss(X, Y) = 0.76311606f0 (tracked)
loss(X, Y) = 0.75046086f0 (tracked)
loss(X, Y) = 0.75331235f0 (tracked)
loss(X, Y) = 0.73856926f0 (tracked)
loss(X, Y) = 0.7763051f0 (tracked)
loss(X, Y) = 0.7302096f0 (tracked)
loss(X, Y) = 0.76063997f0 (tracked)
loss(X, Y) = 0.756287f0 (tracked)
loss(X, Y) = 0.74924874f0 (tracked)
loss(X, Y) = 0.73626685f0 (tracked)
loss(X, Y) = 0.74302864f0 (tracked)
loss(X, Y) = 0.7664034f0 (tracked)
loss(X, Y) = 0.739285f0 (tracked)
loss(X, Y) = 0.7209301f0 (tracked)
loss(X, Y) = 0.7135487f0 (tracked)
loss(X, Y) = 0.72816795f0 (tracked)
loss(X, Y) = 0.7434498f0 (tracked)
loss(X, Y) = 0.75337225f0 (tracked)
loss(X, Y) = 0.7395696f0 (tracked)
loss(X, Y) = 0.7493833f0 (tracked)
loss(X, Y) = 0.7356247f0 (tracked)
loss(X, Y) = 0.7089574f0 (tracked)
loss(X, Y) = 0.73362935f0 (tracked)
loss(X, Y) = 0.750315f0 (tracked)
loss(X, Y) = 0.7652483f0 (tracked)
loss(X, Y) = 0.7714645f0 (tracked)
loss(X, Y) = 0.7445283f0 (tracked)
loss(X, Y) = 0.7727755f0 (tracked)
loss(X, Y) = 0.7403745f0 (tracked)
loss(X, Y) = 0.7413397f0 (tracked)
loss(X, Y) = 0.72359926f0 (tracked)
loss(X, Y) = 0.7564243f0 (tracked)
loss(X, Y) = 0.7378303f0 (tracked)
loss(X, Y) = 0.724175f0 (tracked)
loss(X, Y) = 0.70977676f0 (tracked)
loss(X, Y) = 0.7235635f0 (tracked)
loss(X, Y) = 0.7460617f0 (tracked)
loss(X, Y) = 0.7337501f0 (tracked)
loss(X, Y) = 0.74088115f0 (tracked)
loss(X, Y) = 0.7103338f0 (tracked)
loss(X, Y) = 0.751793f0 (tracked)
```

```
┌ Info: Epoch 353
└ @ Main /Users/kcft114/.julia/packages/Flux/qXNjB/src/optimise/train.jl:105
┌ Info: Epoch 354
└ @ Main /Users/kcft114/.julia/packages/Flux/qXNjB/src/optimise/train.jl:105
┌ Info: Epoch 355
└ @ Main /Users/kcft114/.julia/packages/Flux/qXNjB/src/optimise/train.jl:105
┌ Info: Epoch 356
└ @ Main /Users/kcft114/.julia/packages/Flux/qXNjB/src/optimise/train.jl:105
┌ Info: Epoch 357
└ @ Main /Users/kcft114/.julia/packages/Flux/qXNjB/src/optimise/train.jl:105
┌ Info: Epoch 358
└ @ Main /Users/kcft114/.julia/packages/Flux/qXNjB/src/optimise/train.jl:105
┌ Info: Epoch 359
└ @ Main /Users/kcft114/.julia/packages/Flux/qXNjB/src/optimise/train.jl:105
┌ Info: Epoch 360
└ @ Main /Users/kcft114/.julia/packages/Flux/qXNjB/src/optimise/train.jl:105
┌ Info: Epoch 361
└ @ Main /Users/kcft114/.julia/packages/Flux/qXNjB/src/optimise/train.jl:105
┌ Info: Epoch 362
└ @ Main /Users/kcft114/.julia/packages/Flux/qXNjB/src/optimise/train.jl:105
┌ Info: Epoch 363
└ @ Main /Users/kcft114/.julia/packages/Flux/qXNjB/src/optimise/train.jl:105
┌ Info: Epoch 364
└ @ Main /Users/kcft114/.julia/packages/Flux/qXNjB/src/optimise/train.jl:105
┌ Info: Epoch 365
└ @ Main /Users/kcft114/.julia/packages/Flux/qXNjB/src/optimise/train.jl:105
┌ Info: Epoch 366
└ @ Main /Users/kcft114/.julia/packages/Flux/qXNjB/src/optimise/train.jl:105
┌ Info: Epoch 367
└ @ Main /Users/kcft114/.julia/packages/Flux/qXNjB/src/optimise/train.jl:105
┌ Info: Epoch 368
└ @ Main /Users/kcft114/.julia/packages/Flux/qXNjB/src/optimise/train.jl:105
┌ Info: Epoch 369
└ @ Main /Users/kcft114/.julia/packages/Flux/qXNjB/src/optimise/train.jl:105
┌ Info: Epoch 370
└ @ Main /Users/kcft114/.julia/packages/Flux/qXNjB/src/optimise/train.jl:105
┌ Info: Epoch 371
└ @ Main /Users/kcft114/.julia/packages/Flux/qXNjB/src/optimise/train.jl:105
┌ Info: Epoch 372
└ @ Main /Users/kcft114/.julia/packages/Flux/qXNjB/src/optimise/train.jl:105
┌ Info: Epoch 373
└ @ Main /Users/kcft114/.julia/packages/Flux/qXNjB/src/optimise/train.jl:105
┌ Info: Epoch 374
└ @ Main /Users/kcft114/.julia/packages/Flux/qXNjB/src/optimise/train.jl:105
┌ Info: Epoch 375
└ @ Main /Users/kcft114/.julia/packages/Flux/qXNjB/src/optimise/train.jl:105
┌ Info: Epoch 376
└ @ Main /Users/kcft114/.julia/packages/Flux/qXNjB/src/optimise/train.jl:105
┌ Info: Epoch 377
└ @ Main /Users/kcft114/.julia/packages/Flux/qXNjB/src/optimise/train.jl:105
┌ Info: Epoch 378
└ @ Main /Users/kcft114/.julia/packages/Flux/qXNjB/src/optimise/train.jl:105
┌ Info: Epoch 379
└ @ Main /Users/kcft114/.julia/packages/Flux/qXNjB/src/optimise/train.jl:105
┌ Info: Epoch 380
└ @ Main /Users/kcft114/.julia/packages/Flux/qXNjB/src/optimise/train.jl:105
┌ Info: Epoch 381
└ @ Main /Users/kcft114/.julia/packages/Flux/qXNjB/src/optimise/train.jl:105
┌ Info: Epoch 382
└ @ Main /Users/kcft114/.julia/packages/Flux/qXNjB/src/optimise/train.jl:105
┌ Info: Epoch 383
└ @ Main /Users/kcft114/.julia/packages/Flux/qXNjB/src/optimise/train.jl:105
┌ Info: Epoch 384
└ @ Main /Users/kcft114/.julia/packages/Flux/qXNjB/src/optimise/train.jl:105
┌ Info: Epoch 385
└ @ Main /Users/kcft114/.julia/packages/Flux/qXNjB/src/optimise/train.jl:105
┌ Info: Epoch 386
└ @ Main /Users/kcft114/.julia/packages/Flux/qXNjB/src/optimise/train.jl:105
┌ Info: Epoch 387
└ @ Main /Users/kcft114/.julia/packages/Flux/qXNjB/src/optimise/train.jl:105
┌ Info: Epoch 388
└ @ Main /Users/kcft114/.julia/packages/Flux/qXNjB/src/optimise/train.jl:105
┌ Info: Epoch 389
└ @ Main /Users/kcft114/.julia/packages/Flux/qXNjB/src/optimise/train.jl:105
┌ Info: Epoch 390
└ @ Main /Users/kcft114/.julia/packages/Flux/qXNjB/src/optimise/train.jl:105
┌ Info: Epoch 391
└ @ Main /Users/kcft114/.julia/packages/Flux/qXNjB/src/optimise/train.jl:105
┌ Info: Epoch 392
└ @ Main /Users/kcft114/.julia/packages/Flux/qXNjB/src/optimise/train.jl:105
┌ Info: Epoch 393
└ @ Main /Users/kcft114/.julia/packages/Flux/qXNjB/src/optimise/train.jl:105
┌ Info: Epoch 394
└ @ Main /Users/kcft114/.julia/packages/Flux/qXNjB/src/optimise/train.jl:105
┌ Info: Epoch 395
└ @ Main /Users/kcft114/.julia/packages/Flux/qXNjB/src/optimise/train.jl:105
┌ Info: Epoch 396
└ @ Main /Users/kcft114/.julia/packages/Flux/qXNjB/src/optimise/train.jl:105
┌ Info: Epoch 397
└ @ Main /Users/kcft114/.julia/packages/Flux/qXNjB/src/optimise/train.jl:105
┌ Info: Epoch 398
└ @ Main /Users/kcft114/.julia/packages/Flux/qXNjB/src/optimise/train.jl:105
┌ Info: Epoch 399
└ @ Main /Users/kcft114/.julia/packages/Flux/qXNjB/src/optimise/train.jl:105
┌ Info: Epoch 400
└ @ Main /Users/kcft114/.julia/packages/Flux/qXNjB/src/optimise/train.jl:105
┌ Info: Epoch 401
└ @ Main /Users/kcft114/.julia/packages/Flux/qXNjB/src/optimise/train.jl:105
┌ Info: Epoch 402
└ @ Main /Users/kcft114/.julia/packages/Flux/qXNjB/src/optimise/train.jl:105
┌ Info: Epoch 403
└ @ Main /Users/kcft114/.julia/packages/Flux/qXNjB/src/optimise/train.jl:105
┌ Info: Epoch 404
└ @ Main /Users/kcft114/.julia/packages/Flux/qXNjB/src/optimise/train.jl:105
┌ Info: Epoch 405
└ @ Main /Users/kcft114/.julia/packages/Flux/qXNjB/src/optimise/train.jl:105
┌ Info: Epoch 406
└ @ Main /Users/kcft114/.julia/packages/Flux/qXNjB/src/optimise/train.jl:105
┌ Info: Epoch 407
└ @ Main /Users/kcft114/.julia/packages/Flux/qXNjB/src/optimise/train.jl:105
┌ Info: Epoch 408
└ @ Main /Users/kcft114/.julia/packages/Flux/qXNjB/src/optimise/train.jl:105
┌ Info: Epoch 409
└ @ Main /Users/kcft114/.julia/packages/Flux/qXNjB/src/optimise/train.jl:105
┌ Info: Epoch 410
└ @ Main /Users/kcft114/.julia/packages/Flux/qXNjB/src/optimise/train.jl:105
┌ Info: Epoch 411
└ @ Main /Users/kcft114/.julia/packages/Flux/qXNjB/src/optimise/train.jl:105
┌ Info: Epoch 412
└ @ Main /Users/kcft114/.julia/packages/Flux/qXNjB/src/optimise/train.jl:105
┌ Info: Epoch 413
└ @ Main /Users/kcft114/.julia/packages/Flux/qXNjB/src/optimise/train.jl:105
┌ Info: Epoch 414
└ @ Main /Users/kcft114/.julia/packages/Flux/qXNjB/src/optimise/train.jl:105
┌ Info: Epoch 415
└ @ Main /Users/kcft114/.julia/packages/Flux/qXNjB/src/optimise/train.jl:105
┌ Info: Epoch 416
└ @ Main /Users/kcft114/.julia/packages/Flux/qXNjB/src/optimise/train.jl:105
┌ Info: Epoch 417
└ @ Main /Users/kcft114/.julia/packages/Flux/qXNjB/src/optimise/train.jl:105
┌ Info: Epoch 418
└ @ Main /Users/kcft114/.julia/packages/Flux/qXNjB/src/optimise/train.jl:105
┌ Info: Epoch 419
└ @ Main /Users/kcft114/.julia/packages/Flux/qXNjB/src/optimise/train.jl:105
┌ Info: Epoch 420
└ @ Main /Users/kcft114/.julia/packages/Flux/qXNjB/src/optimise/train.jl:105
┌ Info: Epoch 421
└ @ Main /Users/kcft114/.julia/packages/Flux/qXNjB/src/optimise/train.jl:105
┌ Info: Epoch 422
└ @ Main /Users/kcft114/.julia/packages/Flux/qXNjB/src/optimise/train.jl:105
┌ Info: Epoch 423
└ @ Main /Users/kcft114/.julia/packages/Flux/qXNjB/src/optimise/train.jl:105
┌ Info: Epoch 424
└ @ Main /Users/kcft114/.julia/packages/Flux/qXNjB/src/optimise/train.jl:105
┌ Info: Epoch 425
└ @ Main /Users/kcft114/.julia/packages/Flux/qXNjB/src/optimise/train.jl:105
┌ Info: Epoch 426
└ @ Main /Users/kcft114/.julia/packages/Flux/qXNjB/src/optimise/train.jl:105
┌ Info: Epoch 427
└ @ Main /Users/kcft114/.julia/packages/Flux/qXNjB/src/optimise/train.jl:105
┌ Info: Epoch 428
└ @ Main /Users/kcft114/.julia/packages/Flux/qXNjB/src/optimise/train.jl:105
┌ Info: Epoch 429
└ @ Main /Users/kcft114/.julia/packages/Flux/qXNjB/src/optimise/train.jl:105
┌ Info: Epoch 430
└ @ Main /Users/kcft114/.julia/packages/Flux/qXNjB/src/optimise/train.jl:105
┌ Info: Epoch 431
└ @ Main /Users/kcft114/.julia/packages/Flux/qXNjB/src/optimise/train.jl:105
┌ Info: Epoch 432
└ @ Main /Users/kcft114/.julia/packages/Flux/qXNjB/src/optimise/train.jl:105
┌ Info: Epoch 433
└ @ Main /Users/kcft114/.julia/packages/Flux/qXNjB/src/optimise/train.jl:105
┌ Info: Epoch 434
└ @ Main /Users/kcft114/.julia/packages/Flux/qXNjB/src/optimise/train.jl:105
┌ Info: Epoch 435
└ @ Main /Users/kcft114/.julia/packages/Flux/qXNjB/src/optimise/train.jl:105
┌ Info: Epoch 436
└ @ Main /Users/kcft114/.julia/packages/Flux/qXNjB/src/optimise/train.jl:105
┌ Info: Epoch 437
└ @ Main /Users/kcft114/.julia/packages/Flux/qXNjB/src/optimise/train.jl:105
┌ Info: Epoch 438
└ @ Main /Users/kcft114/.julia/packages/Flux/qXNjB/src/optimise/train.jl:105
┌ Info: Epoch 439
└ @ Main /Users/kcft114/.julia/packages/Flux/qXNjB/src/optimise/train.jl:105
┌ Info: Epoch 440
└ @ Main /Users/kcft114/.julia/packages/Flux/qXNjB/src/optimise/train.jl:105
┌ Info: Epoch 441
└ @ Main /Users/kcft114/.julia/packages/Flux/qXNjB/src/optimise/train.jl:105
┌ Info: Epoch 442
└ @ Main /Users/kcft114/.julia/packages/Flux/qXNjB/src/optimise/train.jl:105
┌ Info: Epoch 443
└ @ Main /Users/kcft114/.julia/packages/Flux/qXNjB/src/optimise/train.jl:105
┌ Info: Epoch 444
└ @ Main /Users/kcft114/.julia/packages/Flux/qXNjB/src/optimise/train.jl:105
┌ Info: Epoch 445
└ @ Main /Users/kcft114/.julia/packages/Flux/qXNjB/src/optimise/train.jl:105
┌ Info: Epoch 446
└ @ Main /Users/kcft114/.julia/packages/Flux/qXNjB/src/optimise/train.jl:105
┌ Info: Epoch 447
└ @ Main /Users/kcft114/.julia/packages/Flux/qXNjB/src/optimise/train.jl:105
┌ Info: Epoch 448
└ @ Main /Users/kcft114/.julia/packages/Flux/qXNjB/src/optimise/train.jl:105
┌ Info: Epoch 449
└ @ Main /Users/kcft114/.julia/packages/Flux/qXNjB/src/optimise/train.jl:105
┌ Info: Epoch 450
└ @ Main /Users/kcft114/.julia/packages/Flux/qXNjB/src/optimise/train.jl:105
┌ Info: Epoch 451
└ @ Main /Users/kcft114/.julia/packages/Flux/qXNjB/src/optimise/train.jl:105
┌ Info: Epoch 452
└ @ Main /Users/kcft114/.julia/packages/Flux/qXNjB/src/optimise/train.jl:105
┌ Info: Epoch 453
└ @ Main /Users/kcft114/.julia/packages/Flux/qXNjB/src/optimise/train.jl:105
┌ Info: Epoch 454
└ @ Main /Users/kcft114/.julia/packages/Flux/qXNjB/src/optimise/train.jl:105
┌ Info: Epoch 455
└ @ Main /Users/kcft114/.julia/packages/Flux/qXNjB/src/optimise/train.jl:105
┌ Info: Epoch 456
└ @ Main /Users/kcft114/.julia/packages/Flux/qXNjB/src/optimise/train.jl:105
┌ Info: Epoch 457
└ @ Main /Users/kcft114/.julia/packages/Flux/qXNjB/src/optimise/train.jl:105
┌ Info: Epoch 458
└ @ Main /Users/kcft114/.julia/packages/Flux/qXNjB/src/optimise/train.jl:105
┌ Info: Epoch 459
└ @ Main /Users/kcft114/.julia/packages/Flux/qXNjB/src/optimise/train.jl:105
┌ Info: Epoch 460
└ @ Main /Users/kcft114/.julia/packages/Flux/qXNjB/src/optimise/train.jl:105
┌ Info: Epoch 461
└ @ Main /Users/kcft114/.julia/packages/Flux/qXNjB/src/optimise/train.jl:105
┌ Info: Epoch 462
└ @ Main /Users/kcft114/.julia/packages/Flux/qXNjB/src/optimise/train.jl:105
┌ Info: Epoch 463
└ @ Main /Users/kcft114/.julia/packages/Flux/qXNjB/src/optimise/train.jl:105
┌ Info: Epoch 464
└ @ Main /Users/kcft114/.julia/packages/Flux/qXNjB/src/optimise/train.jl:105
┌ Info: Epoch 465
└ @ Main /Users/kcft114/.julia/packages/Flux/qXNjB/src/optimise/train.jl:105
┌ Info: Epoch 466
└ @ Main /Users/kcft114/.julia/packages/Flux/qXNjB/src/optimise/train.jl:105
┌ Info: Epoch 467
└ @ Main /Users/kcft114/.julia/packages/Flux/qXNjB/src/optimise/train.jl:105
┌ Info: Epoch 468
└ @ Main /Users/kcft114/.julia/packages/Flux/qXNjB/src/optimise/train.jl:105
┌ Info: Epoch 469
└ @ Main /Users/kcft114/.julia/packages/Flux/qXNjB/src/optimise/train.jl:105
┌ Info: Epoch 470
└ @ Main /Users/kcft114/.julia/packages/Flux/qXNjB/src/optimise/train.jl:105
┌ Info: Epoch 471
└ @ Main /Users/kcft114/.julia/packages/Flux/qXNjB/src/optimise/train.jl:105
┌ Info: Epoch 472
└ @ Main /Users/kcft114/.julia/packages/Flux/qXNjB/src/optimise/train.jl:105
┌ Info: Epoch 473
└ @ Main /Users/kcft114/.julia/packages/Flux/qXNjB/src/optimise/train.jl:105
```

```
loss(X, Y) = 0.7574879f0 (tracked)
loss(X, Y) = 0.73260456f0 (tracked)
loss(X, Y) = 0.73047584f0 (tracked)
loss(X, Y) = 0.73340523f0 (tracked)
loss(X, Y) = 0.7379578f0 (tracked)
loss(X, Y) = 0.7205872f0 (tracked)
loss(X, Y) = 0.74329346f0 (tracked)
loss(X, Y) = 0.74754655f0 (tracked)
loss(X, Y) = 0.7641802f0 (tracked)
loss(X, Y) = 0.7378626f0 (tracked)
loss(X, Y) = 0.72483766f0 (tracked)
loss(X, Y) = 0.7543722f0 (tracked)
loss(X, Y) = 0.73164487f0 (tracked)
loss(X, Y) = 0.7285433f0 (tracked)
loss(X, Y) = 0.726643f0 (tracked)
loss(X, Y) = 0.7014931f0 (tracked)
loss(X, Y) = 0.7242042f0 (tracked)
loss(X, Y) = 0.7414744f0 (tracked)
loss(X, Y) = 0.7332157f0 (tracked)
loss(X, Y) = 0.71378773f0 (tracked)
loss(X, Y) = 0.70179987f0 (tracked)
loss(X, Y) = 0.73974013f0 (tracked)
loss(X, Y) = 0.70660824f0 (tracked)
loss(X, Y) = 0.7069557f0 (tracked)
loss(X, Y) = 0.7386596f0 (tracked)
loss(X, Y) = 0.7068744f0 (tracked)
loss(X, Y) = 0.73381406f0 (tracked)
loss(X, Y) = 0.7321629f0 (tracked)
loss(X, Y) = 0.7660294f0 (tracked)
loss(X, Y) = 0.7307133f0 (tracked)
loss(X, Y) = 0.7152128f0 (tracked)
loss(X, Y) = 0.72460634f0 (tracked)
loss(X, Y) = 0.7204339f0 (tracked)
loss(X, Y) = 0.7294541f0 (tracked)
loss(X, Y) = 0.71571785f0 (tracked)
loss(X, Y) = 0.7276245f0 (tracked)
loss(X, Y) = 0.7072393f0 (tracked)
loss(X, Y) = 0.73197955f0 (tracked)
loss(X, Y) = 0.7132338f0 (tracked)
loss(X, Y) = 0.7339389f0 (tracked)
loss(X, Y) = 0.7298479f0 (tracked)
loss(X, Y) = 0.6899608f0 (tracked)
loss(X, Y) = 0.70783126f0 (tracked)
loss(X, Y) = 0.718f0 (tracked)
loss(X, Y) = 0.7090443f0 (tracked)
loss(X, Y) = 0.7159651f0 (tracked)
loss(X, Y) = 0.7094153f0 (tracked)
loss(X, Y) = 0.7410815f0 (tracked)
loss(X, Y) = 0.71233225f0 (tracked)
loss(X, Y) = 0.73807657f0 (tracked)
loss(X, Y) = 0.7042805f0 (tracked)
loss(X, Y) = 0.73283106f0 (tracked)
loss(X, Y) = 0.70156854f0 (tracked)
loss(X, Y) = 0.7285316f0 (tracked)
loss(X, Y) = 0.72505444f0 (tracked)
loss(X, Y) = 0.7349508f0 (tracked)
loss(X, Y) = 0.73051393f0 (tracked)
loss(X, Y) = 0.7206862f0 (tracked)
loss(X, Y) = 0.7103234f0 (tracked)
loss(X, Y) = 0.7157333f0 (tracked)
loss(X, Y) = 0.7102862f0 (tracked)
loss(X, Y) = 0.7299276f0 (tracked)
loss(X, Y) = 0.69259644f0 (tracked)
loss(X, Y) = 0.70473975f0 (tracked)
loss(X, Y) = 0.71285164f0 (tracked)
loss(X, Y) = 0.7101116f0 (tracked)
loss(X, Y) = 0.70295924f0 (tracked)
loss(X, Y) = 0.7201127f0 (tracked)
loss(X, Y) = 0.70669633f0 (tracked)
loss(X, Y) = 0.70460325f0 (tracked)
loss(X, Y) = 0.71847063f0 (tracked)
loss(X, Y) = 0.6932574f0 (tracked)
loss(X, Y) = 0.7131707f0 (tracked)
loss(X, Y) = 0.7196512f0 (tracked)
loss(X, Y) = 0.7176028f0 (tracked)
loss(X, Y) = 0.7243696f0 (tracked)
loss(X, Y) = 0.7219331f0 (tracked)
loss(X, Y) = 0.69988704f0 (tracked)
loss(X, Y) = 0.70062506f0 (tracked)
loss(X, Y) = 0.7112386f0 (tracked)
loss(X, Y) = 0.72558165f0 (tracked)
loss(X, Y) = 0.7151486f0 (tracked)
loss(X, Y) = 0.7279414f0 (tracked)
loss(X, Y) = 0.71482354f0 (tracked)
loss(X, Y) = 0.70208836f0 (tracked)
loss(X, Y) = 0.7032086f0 (tracked)
loss(X, Y) = 0.709001f0 (tracked)
loss(X, Y) = 0.6985993f0 (tracked)
loss(X, Y) = 0.7097909f0 (tracked)
loss(X, Y) = 0.7461868f0 (tracked)
loss(X, Y) = 0.6848196f0 (tracked)
loss(X, Y) = 0.7184228f0 (tracked)
loss(X, Y) = 0.7209412f0 (tracked)
loss(X, Y) = 0.70139575f0 (tracked)
loss(X, Y) = 0.70985436f0 (tracked)
loss(X, Y) = 0.7010731f0 (tracked)
loss(X, Y) = 0.7046641f0 (tracked)
loss(X, Y) = 0.7063029f0 (tracked)
loss(X, Y) = 0.6920866f0 (tracked)
loss(X, Y) = 0.6877041f0 (tracked)
loss(X, Y) = 0.6901439f0 (tracked)
loss(X, Y) = 0.7117483f0 (tracked)
loss(X, Y) = 0.72249836f0 (tracked)
loss(X, Y) = 0.71520627f0 (tracked)
loss(X, Y) = 0.7176721f0 (tracked)
loss(X, Y) = 0.68677795f0 (tracked)
loss(X, Y) = 0.71221185f0 (tracked)
loss(X, Y) = 0.71274865f0 (tracked)
loss(X, Y) = 0.692876f0 (tracked)
loss(X, Y) = 0.72226584f0 (tracked)
loss(X, Y) = 0.70716405f0 (tracked)
loss(X, Y) = 0.69179577f0 (tracked)
loss(X, Y) = 0.7132075f0 (tracked)
loss(X, Y) = 0.6894981f0 (tracked)
loss(X, Y) = 0.6945404f0 (tracked)
loss(X, Y) = 0.68506545f0 (tracked)
loss(X, Y) = 0.71453017f0 (tracked)
loss(X, Y) = 0.7066427f0 (tracked)
loss(X, Y) = 0.69587183f0 (tracked)
loss(X, Y) = 0.71575534f0 (tracked)
loss(X, Y) = 0.69122213f0 (tracked)
loss(X, Y) = 0.7078901f0 (tracked)
loss(X, Y) = 0.71428007f0 (tracked)
loss(X, Y) = 0.72186756f0 (tracked)
loss(X, Y) = 0.6887859f0 (tracked)
loss(X, Y) = 0.67774445f0 (tracked)
loss(X, Y) = 0.68917495f0 (tracked)
loss(X, Y) = 0.73025006f0 (tracked)
loss(X, Y) = 0.7064749f0 (tracked)
loss(X, Y) = 0.66513044f0 (tracked)
loss(X, Y) = 0.6965196f0 (tracked)
```

```
┌ Info: Epoch 474
└ @ Main /Users/kcft114/.julia/packages/Flux/qXNjB/src/optimise/train.jl:105
┌ Info: Epoch 475
└ @ Main /Users/kcft114/.julia/packages/Flux/qXNjB/src/optimise/train.jl:105
┌ Info: Epoch 476
└ @ Main /Users/kcft114/.julia/packages/Flux/qXNjB/src/optimise/train.jl:105
┌ Info: Epoch 477
└ @ Main /Users/kcft114/.julia/packages/Flux/qXNjB/src/optimise/train.jl:105
┌ Info: Epoch 478
└ @ Main /Users/kcft114/.julia/packages/Flux/qXNjB/src/optimise/train.jl:105
┌ Info: Epoch 479
└ @ Main /Users/kcft114/.julia/packages/Flux/qXNjB/src/optimise/train.jl:105
┌ Info: Epoch 480
└ @ Main /Users/kcft114/.julia/packages/Flux/qXNjB/src/optimise/train.jl:105
┌ Info: Epoch 481
└ @ Main /Users/kcft114/.julia/packages/Flux/qXNjB/src/optimise/train.jl:105
┌ Info: Epoch 482
└ @ Main /Users/kcft114/.julia/packages/Flux/qXNjB/src/optimise/train.jl:105
┌ Info: Epoch 483
└ @ Main /Users/kcft114/.julia/packages/Flux/qXNjB/src/optimise/train.jl:105
┌ Info: Epoch 484
└ @ Main /Users/kcft114/.julia/packages/Flux/qXNjB/src/optimise/train.jl:105
┌ Info: Epoch 485
└ @ Main /Users/kcft114/.julia/packages/Flux/qXNjB/src/optimise/train.jl:105
┌ Info: Epoch 486
└ @ Main /Users/kcft114/.julia/packages/Flux/qXNjB/src/optimise/train.jl:105
┌ Info: Epoch 487
└ @ Main /Users/kcft114/.julia/packages/Flux/qXNjB/src/optimise/train.jl:105
┌ Info: Epoch 488
└ @ Main /Users/kcft114/.julia/packages/Flux/qXNjB/src/optimise/train.jl:105
┌ Info: Epoch 489
└ @ Main /Users/kcft114/.julia/packages/Flux/qXNjB/src/optimise/train.jl:105
┌ Info: Epoch 490
└ @ Main /Users/kcft114/.julia/packages/Flux/qXNjB/src/optimise/train.jl:105
┌ Info: Epoch 491
└ @ Main /Users/kcft114/.julia/packages/Flux/qXNjB/src/optimise/train.jl:105
┌ Info: Epoch 492
└ @ Main /Users/kcft114/.julia/packages/Flux/qXNjB/src/optimise/train.jl:105
┌ Info: Epoch 493
└ @ Main /Users/kcft114/.julia/packages/Flux/qXNjB/src/optimise/train.jl:105
┌ Info: Epoch 494
└ @ Main /Users/kcft114/.julia/packages/Flux/qXNjB/src/optimise/train.jl:105
┌ Info: Epoch 495
└ @ Main /Users/kcft114/.julia/packages/Flux/qXNjB/src/optimise/train.jl:105
┌ Info: Epoch 496
└ @ Main /Users/kcft114/.julia/packages/Flux/qXNjB/src/optimise/train.jl:105
┌ Info: Epoch 497
└ @ Main /Users/kcft114/.julia/packages/Flux/qXNjB/src/optimise/train.jl:105
┌ Info: Epoch 498
└ @ Main /Users/kcft114/.julia/packages/Flux/qXNjB/src/optimise/train.jl:105
┌ Info: Epoch 499
└ @ Main /Users/kcft114/.julia/packages/Flux/qXNjB/src/optimise/train.jl:105
┌ Info: Epoch 500
└ @ Main /Users/kcft114/.julia/packages/Flux/qXNjB/src/optimise/train.jl:105
┌ Info: Epoch 501
└ @ Main /Users/kcft114/.julia/packages/Flux/qXNjB/src/optimise/train.jl:105
┌ Info: Epoch 502
└ @ Main /Users/kcft114/.julia/packages/Flux/qXNjB/src/optimise/train.jl:105
┌ Info: Epoch 503
└ @ Main /Users/kcft114/.julia/packages/Flux/qXNjB/src/optimise/train.jl:105
┌ Info: Epoch 504
└ @ Main /Users/kcft114/.julia/packages/Flux/qXNjB/src/optimise/train.jl:105
┌ Info: Epoch 505
└ @ Main /Users/kcft114/.julia/packages/Flux/qXNjB/src/optimise/train.jl:105
┌ Info: Epoch 506
└ @ Main /Users/kcft114/.julia/packages/Flux/qXNjB/src/optimise/train.jl:105
┌ Info: Epoch 507
└ @ Main /Users/kcft114/.julia/packages/Flux/qXNjB/src/optimise/train.jl:105
┌ Info: Epoch 508
└ @ Main /Users/kcft114/.julia/packages/Flux/qXNjB/src/optimise/train.jl:105
┌ Info: Epoch 509
└ @ Main /Users/kcft114/.julia/packages/Flux/qXNjB/src/optimise/train.jl:105
┌ Info: Epoch 510
└ @ Main /Users/kcft114/.julia/packages/Flux/qXNjB/src/optimise/train.jl:105
┌ Info: Epoch 511
└ @ Main /Users/kcft114/.julia/packages/Flux/qXNjB/src/optimise/train.jl:105
┌ Info: Epoch 512
└ @ Main /Users/kcft114/.julia/packages/Flux/qXNjB/src/optimise/train.jl:105
┌ Info: Epoch 513
└ @ Main /Users/kcft114/.julia/packages/Flux/qXNjB/src/optimise/train.jl:105
┌ Info: Epoch 514
└ @ Main /Users/kcft114/.julia/packages/Flux/qXNjB/src/optimise/train.jl:105
┌ Info: Epoch 515
└ @ Main /Users/kcft114/.julia/packages/Flux/qXNjB/src/optimise/train.jl:105
┌ Info: Epoch 516
└ @ Main /Users/kcft114/.julia/packages/Flux/qXNjB/src/optimise/train.jl:105
┌ Info: Epoch 517
└ @ Main /Users/kcft114/.julia/packages/Flux/qXNjB/src/optimise/train.jl:105
┌ Info: Epoch 518
└ @ Main /Users/kcft114/.julia/packages/Flux/qXNjB/src/optimise/train.jl:105
┌ Info: Epoch 519
└ @ Main /Users/kcft114/.julia/packages/Flux/qXNjB/src/optimise/train.jl:105
┌ Info: Epoch 520
└ @ Main /Users/kcft114/.julia/packages/Flux/qXNjB/src/optimise/train.jl:105
┌ Info: Epoch 521
└ @ Main /Users/kcft114/.julia/packages/Flux/qXNjB/src/optimise/train.jl:105
┌ Info: Epoch 522
└ @ Main /Users/kcft114/.julia/packages/Flux/qXNjB/src/optimise/train.jl:105
┌ Info: Epoch 523
└ @ Main /Users/kcft114/.julia/packages/Flux/qXNjB/src/optimise/train.jl:105
┌ Info: Epoch 524
└ @ Main /Users/kcft114/.julia/packages/Flux/qXNjB/src/optimise/train.jl:105
┌ Info: Epoch 525
└ @ Main /Users/kcft114/.julia/packages/Flux/qXNjB/src/optimise/train.jl:105
┌ Info: Epoch 526
└ @ Main /Users/kcft114/.julia/packages/Flux/qXNjB/src/optimise/train.jl:105
┌ Info: Epoch 527
└ @ Main /Users/kcft114/.julia/packages/Flux/qXNjB/src/optimise/train.jl:105
┌ Info: Epoch 528
└ @ Main /Users/kcft114/.julia/packages/Flux/qXNjB/src/optimise/train.jl:105
┌ Info: Epoch 529
└ @ Main /Users/kcft114/.julia/packages/Flux/qXNjB/src/optimise/train.jl:105
┌ Info: Epoch 530
└ @ Main /Users/kcft114/.julia/packages/Flux/qXNjB/src/optimise/train.jl:105
┌ Info: Epoch 531
└ @ Main /Users/kcft114/.julia/packages/Flux/qXNjB/src/optimise/train.jl:105
┌ Info: Epoch 532
└ @ Main /Users/kcft114/.julia/packages/Flux/qXNjB/src/optimise/train.jl:105
┌ Info: Epoch 533
└ @ Main /Users/kcft114/.julia/packages/Flux/qXNjB/src/optimise/train.jl:105
┌ Info: Epoch 534
└ @ Main /Users/kcft114/.julia/packages/Flux/qXNjB/src/optimise/train.jl:105
┌ Info: Epoch 535
└ @ Main /Users/kcft114/.julia/packages/Flux/qXNjB/src/optimise/train.jl:105
┌ Info: Epoch 536
└ @ Main /Users/kcft114/.julia/packages/Flux/qXNjB/src/optimise/train.jl:105
┌ Info: Epoch 537
└ @ Main /Users/kcft114/.julia/packages/Flux/qXNjB/src/optimise/train.jl:105
┌ Info: Epoch 538
└ @ Main /Users/kcft114/.julia/packages/Flux/qXNjB/src/optimise/train.jl:105
┌ Info: Epoch 539
└ @ Main /Users/kcft114/.julia/packages/Flux/qXNjB/src/optimise/train.jl:105
┌ Info: Epoch 540
└ @ Main /Users/kcft114/.julia/packages/Flux/qXNjB/src/optimise/train.jl:105
┌ Info: Epoch 541
└ @ Main /Users/kcft114/.julia/packages/Flux/qXNjB/src/optimise/train.jl:105
┌ Info: Epoch 542
└ @ Main /Users/kcft114/.julia/packages/Flux/qXNjB/src/optimise/train.jl:105
┌ Info: Epoch 543
└ @ Main /Users/kcft114/.julia/packages/Flux/qXNjB/src/optimise/train.jl:105
┌ Info: Epoch 544
└ @ Main /Users/kcft114/.julia/packages/Flux/qXNjB/src/optimise/train.jl:105
┌ Info: Epoch 545
└ @ Main /Users/kcft114/.julia/packages/Flux/qXNjB/src/optimise/train.jl:105
┌ Info: Epoch 546
└ @ Main /Users/kcft114/.julia/packages/Flux/qXNjB/src/optimise/train.jl:105
┌ Info: Epoch 547
└ @ Main /Users/kcft114/.julia/packages/Flux/qXNjB/src/optimise/train.jl:105
┌ Info: Epoch 548
└ @ Main /Users/kcft114/.julia/packages/Flux/qXNjB/src/optimise/train.jl:105
┌ Info: Epoch 549
└ @ Main /Users/kcft114/.julia/packages/Flux/qXNjB/src/optimise/train.jl:105
┌ Info: Epoch 550
└ @ Main /Users/kcft114/.julia/packages/Flux/qXNjB/src/optimise/train.jl:105
┌ Info: Epoch 551
└ @ Main /Users/kcft114/.julia/packages/Flux/qXNjB/src/optimise/train.jl:105
┌ Info: Epoch 552
└ @ Main /Users/kcft114/.julia/packages/Flux/qXNjB/src/optimise/train.jl:105
┌ Info: Epoch 553
└ @ Main /Users/kcft114/.julia/packages/Flux/qXNjB/src/optimise/train.jl:105
┌ Info: Epoch 554
└ @ Main /Users/kcft114/.julia/packages/Flux/qXNjB/src/optimise/train.jl:105
┌ Info: Epoch 555
└ @ Main /Users/kcft114/.julia/packages/Flux/qXNjB/src/optimise/train.jl:105
┌ Info: Epoch 556
└ @ Main /Users/kcft114/.julia/packages/Flux/qXNjB/src/optimise/train.jl:105
┌ Info: Epoch 557
└ @ Main /Users/kcft114/.julia/packages/Flux/qXNjB/src/optimise/train.jl:105
┌ Info: Epoch 558
└ @ Main /Users/kcft114/.julia/packages/Flux/qXNjB/src/optimise/train.jl:105
┌ Info: Epoch 559
└ @ Main /Users/kcft114/.julia/packages/Flux/qXNjB/src/optimise/train.jl:105
┌ Info: Epoch 560
└ @ Main /Users/kcft114/.julia/packages/Flux/qXNjB/src/optimise/train.jl:105
┌ Info: Epoch 561
└ @ Main /Users/kcft114/.julia/packages/Flux/qXNjB/src/optimise/train.jl:105
┌ Info: Epoch 562
└ @ Main /Users/kcft114/.julia/packages/Flux/qXNjB/src/optimise/train.jl:105
┌ Info: Epoch 563
└ @ Main /Users/kcft114/.julia/packages/Flux/qXNjB/src/optimise/train.jl:105
┌ Info: Epoch 564
└ @ Main /Users/kcft114/.julia/packages/Flux/qXNjB/src/optimise/train.jl:105
┌ Info: Epoch 565
└ @ Main /Users/kcft114/.julia/packages/Flux/qXNjB/src/optimise/train.jl:105
┌ Info: Epoch 566
└ @ Main /Users/kcft114/.julia/packages/Flux/qXNjB/src/optimise/train.jl:105
┌ Info: Epoch 567
└ @ Main /Users/kcft114/.julia/packages/Flux/qXNjB/src/optimise/train.jl:105
┌ Info: Epoch 568
└ @ Main /Users/kcft114/.julia/packages/Flux/qXNjB/src/optimise/train.jl:105
┌ Info: Epoch 569
└ @ Main /Users/kcft114/.julia/packages/Flux/qXNjB/src/optimise/train.jl:105
┌ Info: Epoch 570
└ @ Main /Users/kcft114/.julia/packages/Flux/qXNjB/src/optimise/train.jl:105
┌ Info: Epoch 571
└ @ Main /Users/kcft114/.julia/packages/Flux/qXNjB/src/optimise/train.jl:105
┌ Info: Epoch 572
└ @ Main /Users/kcft114/.julia/packages/Flux/qXNjB/src/optimise/train.jl:105
┌ Info: Epoch 573
└ @ Main /Users/kcft114/.julia/packages/Flux/qXNjB/src/optimise/train.jl:105
┌ Info: Epoch 574
└ @ Main /Users/kcft114/.julia/packages/Flux/qXNjB/src/optimise/train.jl:105
┌ Info: Epoch 575
└ @ Main /Users/kcft114/.julia/packages/Flux/qXNjB/src/optimise/train.jl:105
┌ Info: Epoch 576
└ @ Main /Users/kcft114/.julia/packages/Flux/qXNjB/src/optimise/train.jl:105
┌ Info: Epoch 577
└ @ Main /Users/kcft114/.julia/packages/Flux/qXNjB/src/optimise/train.jl:105
┌ Info: Epoch 578
└ @ Main /Users/kcft114/.julia/packages/Flux/qXNjB/src/optimise/train.jl:105
┌ Info: Epoch 579
└ @ Main /Users/kcft114/.julia/packages/Flux/qXNjB/src/optimise/train.jl:105
┌ Info: Epoch 580
└ @ Main /Users/kcft114/.julia/packages/Flux/qXNjB/src/optimise/train.jl:105
┌ Info: Epoch 581
└ @ Main /Users/kcft114/.julia/packages/Flux/qXNjB/src/optimise/train.jl:105
┌ Info: Epoch 582
└ @ Main /Users/kcft114/.julia/packages/Flux/qXNjB/src/optimise/train.jl:105
┌ Info: Epoch 583
└ @ Main /Users/kcft114/.julia/packages/Flux/qXNjB/src/optimise/train.jl:105
┌ Info: Epoch 584
└ @ Main /Users/kcft114/.julia/packages/Flux/qXNjB/src/optimise/train.jl:105
┌ Info: Epoch 585
└ @ Main /Users/kcft114/.julia/packages/Flux/qXNjB/src/optimise/train.jl:105
┌ Info: Epoch 586
└ @ Main /Users/kcft114/.julia/packages/Flux/qXNjB/src/optimise/train.jl:105
┌ Info: Epoch 587
└ @ Main /Users/kcft114/.julia/packages/Flux/qXNjB/src/optimise/train.jl:105
┌ Info: Epoch 588
└ @ Main /Users/kcft114/.julia/packages/Flux/qXNjB/src/optimise/train.jl:105
┌ Info: Epoch 589
└ @ Main /Users/kcft114/.julia/packages/Flux/qXNjB/src/optimise/train.jl:105
┌ Info: Epoch 590
└ @ Main /Users/kcft114/.julia/packages/Flux/qXNjB/src/optimise/train.jl:105
┌ Info: Epoch 591
└ @ Main /Users/kcft114/.julia/packages/Flux/qXNjB/src/optimise/train.jl:105
┌ Info: Epoch 592
└ @ Main /Users/kcft114/.julia/packages/Flux/qXNjB/src/optimise/train.jl:105
┌ Info: Epoch 593
└ @ Main /Users/kcft114/.julia/packages/Flux/qXNjB/src/optimise/train.jl:105
┌ Info: Epoch 594
└ @ Main /Users/kcft114/.julia/packages/Flux/qXNjB/src/optimise/train.jl:105
┌ Info: Epoch 595
└ @ Main /Users/kcft114/.julia/packages/Flux/qXNjB/src/optimise/train.jl:105
┌ Info: Epoch 596
└ @ Main /Users/kcft114/.julia/packages/Flux/qXNjB/src/optimise/train.jl:105
┌ Info: Epoch 597
└ @ Main /Users/kcft114/.julia/packages/Flux/qXNjB/src/optimise/train.jl:105
┌ Info: Epoch 598
└ @ Main /Users/kcft114/.julia/packages/Flux/qXNjB/src/optimise/train.jl:105
┌ Info: Epoch 599
└ @ Main /Users/kcft114/.julia/packages/Flux/qXNjB/src/optimise/train.jl:105
┌ Info: Epoch 600
└ @ Main /Users/kcft114/.julia/packages/Flux/qXNjB/src/optimise/train.jl:105
┌ Info: Epoch 601
└ @ Main /Users/kcft114/.julia/packages/Flux/qXNjB/src/optimise/train.jl:105
┌ Info: Epoch 602
└ @ Main /Users/kcft114/.julia/packages/Flux/qXNjB/src/optimise/train.jl:105
┌ Info: Epoch 603
└ @ Main /Users/kcft114/.julia/packages/Flux/qXNjB/src/optimise/train.jl:105
┌ Info: Epoch 604
└ @ Main /Users/kcft114/.julia/packages/Flux/qXNjB/src/optimise/train.jl:105
```

```
loss(X, Y) = 0.7051413f0 (tracked)
loss(X, Y) = 0.70464015f0 (tracked)
loss(X, Y) = 0.7085544f0 (tracked)
loss(X, Y) = 0.692609f0 (tracked)
loss(X, Y) = 0.69186974f0 (tracked)
loss(X, Y) = 0.70328695f0 (tracked)
loss(X, Y) = 0.6905081f0 (tracked)
loss(X, Y) = 0.71175814f0 (tracked)
loss(X, Y) = 0.690962f0 (tracked)
loss(X, Y) = 0.6938967f0 (tracked)
loss(X, Y) = 0.7027793f0 (tracked)
loss(X, Y) = 0.7102737f0 (tracked)
loss(X, Y) = 0.68320644f0 (tracked)
loss(X, Y) = 0.7072818f0 (tracked)
loss(X, Y) = 0.7012206f0 (tracked)
loss(X, Y) = 0.69777775f0 (tracked)
loss(X, Y) = 0.69744426f0 (tracked)
loss(X, Y) = 0.68851566f0 (tracked)
loss(X, Y) = 0.68481976f0 (tracked)
loss(X, Y) = 0.6617759f0 (tracked)
loss(X, Y) = 0.69396204f0 (tracked)
loss(X, Y) = 0.6786191f0 (tracked)
loss(X, Y) = 0.6953562f0 (tracked)
loss(X, Y) = 0.68409044f0 (tracked)
loss(X, Y) = 0.6926668f0 (tracked)
loss(X, Y) = 0.684856f0 (tracked)
loss(X, Y) = 0.6791748f0 (tracked)
loss(X, Y) = 0.69967574f0 (tracked)
loss(X, Y) = 0.6859555f0 (tracked)
loss(X, Y) = 0.70985824f0 (tracked)
loss(X, Y) = 0.6804431f0 (tracked)
loss(X, Y) = 0.6946797f0 (tracked)
loss(X, Y) = 0.6754512f0 (tracked)
loss(X, Y) = 0.68065196f0 (tracked)
loss(X, Y) = 0.69426095f0 (tracked)
loss(X, Y) = 0.6874089f0 (tracked)
loss(X, Y) = 0.6802104f0 (tracked)
loss(X, Y) = 0.6784664f0 (tracked)
loss(X, Y) = 0.68094486f0 (tracked)
loss(X, Y) = 0.6647741f0 (tracked)
loss(X, Y) = 0.7008892f0 (tracked)
loss(X, Y) = 0.70499545f0 (tracked)
loss(X, Y) = 0.71546924f0 (tracked)
loss(X, Y) = 0.66720873f0 (tracked)
loss(X, Y) = 0.66839135f0 (tracked)
loss(X, Y) = 0.70800185f0 (tracked)
loss(X, Y) = 0.6822145f0 (tracked)
loss(X, Y) = 0.6796431f0 (tracked)
loss(X, Y) = 0.66476434f0 (tracked)
loss(X, Y) = 0.6871859f0 (tracked)
loss(X, Y) = 0.6845309f0 (tracked)
loss(X, Y) = 0.6652451f0 (tracked)
loss(X, Y) = 0.69257957f0 (tracked)
loss(X, Y) = 0.6786778f0 (tracked)
loss(X, Y) = 0.69411576f0 (tracked)
loss(X, Y) = 0.6817103f0 (tracked)
loss(X, Y) = 0.6670808f0 (tracked)
loss(X, Y) = 0.67382985f0 (tracked)
loss(X, Y) = 0.68954057f0 (tracked)
loss(X, Y) = 0.6990203f0 (tracked)
loss(X, Y) = 0.6863619f0 (tracked)
loss(X, Y) = 0.6731002f0 (tracked)
loss(X, Y) = 0.658472f0 (tracked)
loss(X, Y) = 0.67850363f0 (tracked)
loss(X, Y) = 0.6798423f0 (tracked)
loss(X, Y) = 0.6901927f0 (tracked)
loss(X, Y) = 0.6986719f0 (tracked)
loss(X, Y) = 0.65987825f0 (tracked)
loss(X, Y) = 0.6938429f0 (tracked)
loss(X, Y) = 0.6816877f0 (tracked)
loss(X, Y) = 0.6628233f0 (tracked)
loss(X, Y) = 0.6969991f0 (tracked)
loss(X, Y) = 0.68842256f0 (tracked)
loss(X, Y) = 0.69381857f0 (tracked)
loss(X, Y) = 0.67651945f0 (tracked)
loss(X, Y) = 0.68604136f0 (tracked)
loss(X, Y) = 0.6292271f0 (tracked)
loss(X, Y) = 0.67718714f0 (tracked)
loss(X, Y) = 0.68489593f0 (tracked)
loss(X, Y) = 0.703885f0 (tracked)
loss(X, Y) = 0.6816541f0 (tracked)
loss(X, Y) = 0.67201036f0 (tracked)
loss(X, Y) = 0.67329675f0 (tracked)
loss(X, Y) = 0.65994537f0 (tracked)
loss(X, Y) = 0.6905435f0 (tracked)
loss(X, Y) = 0.68154734f0 (tracked)
loss(X, Y) = 0.6935396f0 (tracked)
loss(X, Y) = 0.70283824f0 (tracked)
loss(X, Y) = 0.6772171f0 (tracked)
loss(X, Y) = 0.70362234f0 (tracked)
loss(X, Y) = 0.6847887f0 (tracked)
loss(X, Y) = 0.69185567f0 (tracked)
loss(X, Y) = 0.6624953f0 (tracked)
loss(X, Y) = 0.6559354f0 (tracked)
loss(X, Y) = 0.6572151f0 (tracked)
loss(X, Y) = 0.70060325f0 (tracked)
loss(X, Y) = 0.66793334f0 (tracked)
loss(X, Y) = 0.6744623f0 (tracked)
loss(X, Y) = 0.66879135f0 (tracked)
loss(X, Y) = 0.6811983f0 (tracked)
loss(X, Y) = 0.6604619f0 (tracked)
loss(X, Y) = 0.6781281f0 (tracked)
loss(X, Y) = 0.63495857f0 (tracked)
loss(X, Y) = 0.6573695f0 (tracked)
loss(X, Y) = 0.6743908f0 (tracked)
loss(X, Y) = 0.6894834f0 (tracked)
loss(X, Y) = 0.67535406f0 (tracked)
loss(X, Y) = 0.6468551f0 (tracked)
loss(X, Y) = 0.67622197f0 (tracked)
loss(X, Y) = 0.6566234f0 (tracked)
loss(X, Y) = 0.6916817f0 (tracked)
loss(X, Y) = 0.6775579f0 (tracked)
loss(X, Y) = 0.6716293f0 (tracked)
loss(X, Y) = 0.67405427f0 (tracked)
loss(X, Y) = 0.6545109f0 (tracked)
loss(X, Y) = 0.6911803f0 (tracked)
loss(X, Y) = 0.6556635f0 (tracked)
loss(X, Y) = 0.683747f0 (tracked)
loss(X, Y) = 0.6800154f0 (tracked)
loss(X, Y) = 0.6394303f0 (tracked)
loss(X, Y) = 0.6882562f0 (tracked)
loss(X, Y) = 0.6767365f0 (tracked)
loss(X, Y) = 0.68865234f0 (tracked)
loss(X, Y) = 0.65968066f0 (tracked)
loss(X, Y) = 0.68292886f0 (tracked)
loss(X, Y) = 0.6583662f0 (tracked)
loss(X, Y) = 0.68913203f0 (tracked)
loss(X, Y) = 0.6461812f0 (tracked)
loss(X, Y) = 0.68533987f0 (tracked)
loss(X, Y) = 0.651528f0 (tracked)
loss(X, Y) = 0.6909276f0 (tracked)
loss(X, Y) = 0.66868997f0 (tracked)
loss(X, Y) = 0.6990647f0 (tracked)
loss(X, Y) = 0.64997715f0 (tracked)
loss(X, Y) = 0.6853781f0 (tracked)
loss(X, Y) = 0.6715952f0 (tracked)
loss(X, Y) = 0.6710118f0 (tracked)
loss(X, Y) = 0.6831397f0 (tracked)
loss(X, Y) = 0.67646694f0 (tracked)
loss(X, Y) = 0.64704293f0 (tracked)
loss(X, Y) = 0.6499549f0 (tracked)
loss(X, Y) = 0.64002955f0 (tracked)
loss(X, Y) = 0.6609028f0 (tracked)
loss(X, Y) = 0.66939014f0 (tracked)
```

```
┌ Info: Epoch 605
└ @ Main /Users/kcft114/.julia/packages/Flux/qXNjB/src/optimise/train.jl:105
┌ Info: Epoch 606
└ @ Main /Users/kcft114/.julia/packages/Flux/qXNjB/src/optimise/train.jl:105
┌ Info: Epoch 607
└ @ Main /Users/kcft114/.julia/packages/Flux/qXNjB/src/optimise/train.jl:105
┌ Info: Epoch 608
└ @ Main /Users/kcft114/.julia/packages/Flux/qXNjB/src/optimise/train.jl:105
┌ Info: Epoch 609
└ @ Main /Users/kcft114/.julia/packages/Flux/qXNjB/src/optimise/train.jl:105
┌ Info: Epoch 610
└ @ Main /Users/kcft114/.julia/packages/Flux/qXNjB/src/optimise/train.jl:105
┌ Info: Epoch 611
└ @ Main /Users/kcft114/.julia/packages/Flux/qXNjB/src/optimise/train.jl:105
┌ Info: Epoch 612
└ @ Main /Users/kcft114/.julia/packages/Flux/qXNjB/src/optimise/train.jl:105
┌ Info: Epoch 613
└ @ Main /Users/kcft114/.julia/packages/Flux/qXNjB/src/optimise/train.jl:105
┌ Info: Epoch 614
└ @ Main /Users/kcft114/.julia/packages/Flux/qXNjB/src/optimise/train.jl:105
┌ Info: Epoch 615
└ @ Main /Users/kcft114/.julia/packages/Flux/qXNjB/src/optimise/train.jl:105
┌ Info: Epoch 616
└ @ Main /Users/kcft114/.julia/packages/Flux/qXNjB/src/optimise/train.jl:105
┌ Info: Epoch 617
└ @ Main /Users/kcft114/.julia/packages/Flux/qXNjB/src/optimise/train.jl:105
┌ Info: Epoch 618
└ @ Main /Users/kcft114/.julia/packages/Flux/qXNjB/src/optimise/train.jl:105
┌ Info: Epoch 619
└ @ Main /Users/kcft114/.julia/packages/Flux/qXNjB/src/optimise/train.jl:105
┌ Info: Epoch 620
└ @ Main /Users/kcft114/.julia/packages/Flux/qXNjB/src/optimise/train.jl:105
┌ Info: Epoch 621
└ @ Main /Users/kcft114/.julia/packages/Flux/qXNjB/src/optimise/train.jl:105
┌ Info: Epoch 622
└ @ Main /Users/kcft114/.julia/packages/Flux/qXNjB/src/optimise/train.jl:105
┌ Info: Epoch 623
└ @ Main /Users/kcft114/.julia/packages/Flux/qXNjB/src/optimise/train.jl:105
┌ Info: Epoch 624
└ @ Main /Users/kcft114/.julia/packages/Flux/qXNjB/src/optimise/train.jl:105
┌ Info: Epoch 625
└ @ Main /Users/kcft114/.julia/packages/Flux/qXNjB/src/optimise/train.jl:105
┌ Info: Epoch 626
└ @ Main /Users/kcft114/.julia/packages/Flux/qXNjB/src/optimise/train.jl:105
┌ Info: Epoch 627
└ @ Main /Users/kcft114/.julia/packages/Flux/qXNjB/src/optimise/train.jl:105
┌ Info: Epoch 628
└ @ Main /Users/kcft114/.julia/packages/Flux/qXNjB/src/optimise/train.jl:105
┌ Info: Epoch 629
└ @ Main /Users/kcft114/.julia/packages/Flux/qXNjB/src/optimise/train.jl:105
┌ Info: Epoch 630
└ @ Main /Users/kcft114/.julia/packages/Flux/qXNjB/src/optimise/train.jl:105
┌ Info: Epoch 631
└ @ Main /Users/kcft114/.julia/packages/Flux/qXNjB/src/optimise/train.jl:105
┌ Info: Epoch 632
└ @ Main /Users/kcft114/.julia/packages/Flux/qXNjB/src/optimise/train.jl:105
┌ Info: Epoch 633
└ @ Main /Users/kcft114/.julia/packages/Flux/qXNjB/src/optimise/train.jl:105
┌ Info: Epoch 634
└ @ Main /Users/kcft114/.julia/packages/Flux/qXNjB/src/optimise/train.jl:105
┌ Info: Epoch 635
└ @ Main /Users/kcft114/.julia/packages/Flux/qXNjB/src/optimise/train.jl:105
┌ Info: Epoch 636
└ @ Main /Users/kcft114/.julia/packages/Flux/qXNjB/src/optimise/train.jl:105
┌ Info: Epoch 637
└ @ Main /Users/kcft114/.julia/packages/Flux/qXNjB/src/optimise/train.jl:105
┌ Info: Epoch 638
└ @ Main /Users/kcft114/.julia/packages/Flux/qXNjB/src/optimise/train.jl:105
┌ Info: Epoch 639
└ @ Main /Users/kcft114/.julia/packages/Flux/qXNjB/src/optimise/train.jl:105
┌ Info: Epoch 640
└ @ Main /Users/kcft114/.julia/packages/Flux/qXNjB/src/optimise/train.jl:105
┌ Info: Epoch 641
└ @ Main /Users/kcft114/.julia/packages/Flux/qXNjB/src/optimise/train.jl:105
┌ Info: Epoch 642
└ @ Main /Users/kcft114/.julia/packages/Flux/qXNjB/src/optimise/train.jl:105
┌ Info: Epoch 643
└ @ Main /Users/kcft114/.julia/packages/Flux/qXNjB/src/optimise/train.jl:105
┌ Info: Epoch 644
└ @ Main /Users/kcft114/.julia/packages/Flux/qXNjB/src/optimise/train.jl:105
┌ Info: Epoch 645
└ @ Main /Users/kcft114/.julia/packages/Flux/qXNjB/src/optimise/train.jl:105
┌ Info: Epoch 646
└ @ Main /Users/kcft114/.julia/packages/Flux/qXNjB/src/optimise/train.jl:105
┌ Info: Epoch 647
└ @ Main /Users/kcft114/.julia/packages/Flux/qXNjB/src/optimise/train.jl:105
┌ Info: Epoch 648
└ @ Main /Users/kcft114/.julia/packages/Flux/qXNjB/src/optimise/train.jl:105
┌ Info: Epoch 649
└ @ Main /Users/kcft114/.julia/packages/Flux/qXNjB/src/optimise/train.jl:105
┌ Info: Epoch 650
└ @ Main /Users/kcft114/.julia/packages/Flux/qXNjB/src/optimise/train.jl:105
┌ Info: Epoch 651
└ @ Main /Users/kcft114/.julia/packages/Flux/qXNjB/src/optimise/train.jl:105
┌ Info: Epoch 652
└ @ Main /Users/kcft114/.julia/packages/Flux/qXNjB/src/optimise/train.jl:105
┌ Info: Epoch 653
└ @ Main /Users/kcft114/.julia/packages/Flux/qXNjB/src/optimise/train.jl:105
┌ Info: Epoch 654
└ @ Main /Users/kcft114/.julia/packages/Flux/qXNjB/src/optimise/train.jl:105
┌ Info: Epoch 655
└ @ Main /Users/kcft114/.julia/packages/Flux/qXNjB/src/optimise/train.jl:105
┌ Info: Epoch 656
└ @ Main /Users/kcft114/.julia/packages/Flux/qXNjB/src/optimise/train.jl:105
┌ Info: Epoch 657
└ @ Main /Users/kcft114/.julia/packages/Flux/qXNjB/src/optimise/train.jl:105
┌ Info: Epoch 658
└ @ Main /Users/kcft114/.julia/packages/Flux/qXNjB/src/optimise/train.jl:105
┌ Info: Epoch 659
└ @ Main /Users/kcft114/.julia/packages/Flux/qXNjB/src/optimise/train.jl:105
┌ Info: Epoch 660
└ @ Main /Users/kcft114/.julia/packages/Flux/qXNjB/src/optimise/train.jl:105
┌ Info: Epoch 661
└ @ Main /Users/kcft114/.julia/packages/Flux/qXNjB/src/optimise/train.jl:105
┌ Info: Epoch 662
└ @ Main /Users/kcft114/.julia/packages/Flux/qXNjB/src/optimise/train.jl:105
┌ Info: Epoch 663
└ @ Main /Users/kcft114/.julia/packages/Flux/qXNjB/src/optimise/train.jl:105
┌ Info: Epoch 664
└ @ Main /Users/kcft114/.julia/packages/Flux/qXNjB/src/optimise/train.jl:105
┌ Info: Epoch 665
└ @ Main /Users/kcft114/.julia/packages/Flux/qXNjB/src/optimise/train.jl:105
┌ Info: Epoch 666
└ @ Main /Users/kcft114/.julia/packages/Flux/qXNjB/src/optimise/train.jl:105
┌ Info: Epoch 667
└ @ Main /Users/kcft114/.julia/packages/Flux/qXNjB/src/optimise/train.jl:105
┌ Info: Epoch 668
└ @ Main /Users/kcft114/.julia/packages/Flux/qXNjB/src/optimise/train.jl:105
┌ Info: Epoch 669
└ @ Main /Users/kcft114/.julia/packages/Flux/qXNjB/src/optimise/train.jl:105
┌ Info: Epoch 670
└ @ Main /Users/kcft114/.julia/packages/Flux/qXNjB/src/optimise/train.jl:105
┌ Info: Epoch 671
└ @ Main /Users/kcft114/.julia/packages/Flux/qXNjB/src/optimise/train.jl:105
┌ Info: Epoch 672
└ @ Main /Users/kcft114/.julia/packages/Flux/qXNjB/src/optimise/train.jl:105
┌ Info: Epoch 673
└ @ Main /Users/kcft114/.julia/packages/Flux/qXNjB/src/optimise/train.jl:105
┌ Info: Epoch 674
└ @ Main /Users/kcft114/.julia/packages/Flux/qXNjB/src/optimise/train.jl:105
┌ Info: Epoch 675
└ @ Main /Users/kcft114/.julia/packages/Flux/qXNjB/src/optimise/train.jl:105
┌ Info: Epoch 676
└ @ Main /Users/kcft114/.julia/packages/Flux/qXNjB/src/optimise/train.jl:105
┌ Info: Epoch 677
└ @ Main /Users/kcft114/.julia/packages/Flux/qXNjB/src/optimise/train.jl:105
┌ Info: Epoch 678
└ @ Main /Users/kcft114/.julia/packages/Flux/qXNjB/src/optimise/train.jl:105
┌ Info: Epoch 679
└ @ Main /Users/kcft114/.julia/packages/Flux/qXNjB/src/optimise/train.jl:105
┌ Info: Epoch 680
└ @ Main /Users/kcft114/.julia/packages/Flux/qXNjB/src/optimise/train.jl:105
┌ Info: Epoch 681
└ @ Main /Users/kcft114/.julia/packages/Flux/qXNjB/src/optimise/train.jl:105
┌ Info: Epoch 682
└ @ Main /Users/kcft114/.julia/packages/Flux/qXNjB/src/optimise/train.jl:105
┌ Info: Epoch 683
└ @ Main /Users/kcft114/.julia/packages/Flux/qXNjB/src/optimise/train.jl:105
┌ Info: Epoch 684
└ @ Main /Users/kcft114/.julia/packages/Flux/qXNjB/src/optimise/train.jl:105
┌ Info: Epoch 685
└ @ Main /Users/kcft114/.julia/packages/Flux/qXNjB/src/optimise/train.jl:105
┌ Info: Epoch 686
└ @ Main /Users/kcft114/.julia/packages/Flux/qXNjB/src/optimise/train.jl:105
┌ Info: Epoch 687
└ @ Main /Users/kcft114/.julia/packages/Flux/qXNjB/src/optimise/train.jl:105
┌ Info: Epoch 688
└ @ Main /Users/kcft114/.julia/packages/Flux/qXNjB/src/optimise/train.jl:105
┌ Info: Epoch 689
└ @ Main /Users/kcft114/.julia/packages/Flux/qXNjB/src/optimise/train.jl:105
┌ Info: Epoch 690
└ @ Main /Users/kcft114/.julia/packages/Flux/qXNjB/src/optimise/train.jl:105
┌ Info: Epoch 691
└ @ Main /Users/kcft114/.julia/packages/Flux/qXNjB/src/optimise/train.jl:105
┌ Info: Epoch 692
└ @ Main /Users/kcft114/.julia/packages/Flux/qXNjB/src/optimise/train.jl:105
┌ Info: Epoch 693
└ @ Main /Users/kcft114/.julia/packages/Flux/qXNjB/src/optimise/train.jl:105
┌ Info: Epoch 694
└ @ Main /Users/kcft114/.julia/packages/Flux/qXNjB/src/optimise/train.jl:105
┌ Info: Epoch 695
└ @ Main /Users/kcft114/.julia/packages/Flux/qXNjB/src/optimise/train.jl:105
┌ Info: Epoch 696
└ @ Main /Users/kcft114/.julia/packages/Flux/qXNjB/src/optimise/train.jl:105
┌ Info: Epoch 697
└ @ Main /Users/kcft114/.julia/packages/Flux/qXNjB/src/optimise/train.jl:105
┌ Info: Epoch 698
└ @ Main /Users/kcft114/.julia/packages/Flux/qXNjB/src/optimise/train.jl:105
┌ Info: Epoch 699
└ @ Main /Users/kcft114/.julia/packages/Flux/qXNjB/src/optimise/train.jl:105
┌ Info: Epoch 700
└ @ Main /Users/kcft114/.julia/packages/Flux/qXNjB/src/optimise/train.jl:105
┌ Info: Epoch 701
└ @ Main /Users/kcft114/.julia/packages/Flux/qXNjB/src/optimise/train.jl:105
┌ Info: Epoch 702
└ @ Main /Users/kcft114/.julia/packages/Flux/qXNjB/src/optimise/train.jl:105
┌ Info: Epoch 703
└ @ Main /Users/kcft114/.julia/packages/Flux/qXNjB/src/optimise/train.jl:105
┌ Info: Epoch 704
└ @ Main /Users/kcft114/.julia/packages/Flux/qXNjB/src/optimise/train.jl:105
┌ Info: Epoch 705
└ @ Main /Users/kcft114/.julia/packages/Flux/qXNjB/src/optimise/train.jl:105
┌ Info: Epoch 706
└ @ Main /Users/kcft114/.julia/packages/Flux/qXNjB/src/optimise/train.jl:105
┌ Info: Epoch 707
└ @ Main /Users/kcft114/.julia/packages/Flux/qXNjB/src/optimise/train.jl:105
┌ Info: Epoch 708
└ @ Main /Users/kcft114/.julia/packages/Flux/qXNjB/src/optimise/train.jl:105
┌ Info: Epoch 709
└ @ Main /Users/kcft114/.julia/packages/Flux/qXNjB/src/optimise/train.jl:105
┌ Info: Epoch 710
└ @ Main /Users/kcft114/.julia/packages/Flux/qXNjB/src/optimise/train.jl:105
┌ Info: Epoch 711
└ @ Main /Users/kcft114/.julia/packages/Flux/qXNjB/src/optimise/train.jl:105
┌ Info: Epoch 712
└ @ Main /Users/kcft114/.julia/packages/Flux/qXNjB/src/optimise/train.jl:105
┌ Info: Epoch 713
└ @ Main /Users/kcft114/.julia/packages/Flux/qXNjB/src/optimise/train.jl:105
┌ Info: Epoch 714
└ @ Main /Users/kcft114/.julia/packages/Flux/qXNjB/src/optimise/train.jl:105
┌ Info: Epoch 715
└ @ Main /Users/kcft114/.julia/packages/Flux/qXNjB/src/optimise/train.jl:105
┌ Info: Epoch 716
└ @ Main /Users/kcft114/.julia/packages/Flux/qXNjB/src/optimise/train.jl:105
┌ Info: Epoch 717
└ @ Main /Users/kcft114/.julia/packages/Flux/qXNjB/src/optimise/train.jl:105
┌ Info: Epoch 718
└ @ Main /Users/kcft114/.julia/packages/Flux/qXNjB/src/optimise/train.jl:105
┌ Info: Epoch 719
└ @ Main /Users/kcft114/.julia/packages/Flux/qXNjB/src/optimise/train.jl:105
┌ Info: Epoch 720
└ @ Main /Users/kcft114/.julia/packages/Flux/qXNjB/src/optimise/train.jl:105
┌ Info: Epoch 721
└ @ Main /Users/kcft114/.julia/packages/Flux/qXNjB/src/optimise/train.jl:105
┌ Info: Epoch 722
└ @ Main /Users/kcft114/.julia/packages/Flux/qXNjB/src/optimise/train.jl:105
┌ Info: Epoch 723
└ @ Main /Users/kcft114/.julia/packages/Flux/qXNjB/src/optimise/train.jl:105
┌ Info: Epoch 724
└ @ Main /Users/kcft114/.julia/packages/Flux/qXNjB/src/optimise/train.jl:105
┌ Info: Epoch 725
└ @ Main /Users/kcft114/.julia/packages/Flux/qXNjB/src/optimise/train.jl:105
┌ Info: Epoch 726
└ @ Main /Users/kcft114/.julia/packages/Flux/qXNjB/src/optimise/train.jl:105
┌ Info: Epoch 727
└ @ Main /Users/kcft114/.julia/packages/Flux/qXNjB/src/optimise/train.jl:105
┌ Info: Epoch 728
└ @ Main /Users/kcft114/.julia/packages/Flux/qXNjB/src/optimise/train.jl:105
┌ Info: Epoch 729
└ @ Main /Users/kcft114/.julia/packages/Flux/qXNjB/src/optimise/train.jl:105
┌ Info: Epoch 730
└ @ Main /Users/kcft114/.julia/packages/Flux/qXNjB/src/optimise/train.jl:105
┌ Info: Epoch 731
└ @ Main /Users/kcft114/.julia/packages/Flux/qXNjB/src/optimise/train.jl:105
┌ Info: Epoch 732
└ @ Main /Users/kcft114/.julia/packages/Flux/qXNjB/src/optimise/train.jl:105
┌ Info: Epoch 733
└ @ Main /Users/kcft114/.julia/packages/Flux/qXNjB/src/optimise/train.jl:105
┌ Info: Epoch 734
└ @ Main /Users/kcft114/.julia/packages/Flux/qXNjB/src/optimise/train.jl:105
┌ Info: Epoch 735
└ @ Main /Users/kcft114/.julia/packages/Flux/qXNjB/src/optimise/train.jl:105
┌ Info: Epoch 736
└ @ Main /Users/kcft114/.julia/packages/Flux/qXNjB/src/optimise/train.jl:105
┌ Info: Epoch 737
└ @ Main /Users/kcft114/.julia/packages/Flux/qXNjB/src/optimise/train.jl:105
┌ Info: Epoch 738
└ @ Main /Users/kcft114/.julia/packages/Flux/qXNjB/src/optimise/train.jl:105
┌ Info: Epoch 739
└ @ Main /Users/kcft114/.julia/packages/Flux/qXNjB/src/optimise/train.jl:105
┌ Info: Epoch 740
└ @ Main /Users/kcft114/.julia/packages/Flux/qXNjB/src/optimise/train.jl:105
┌ Info: Epoch 741
└ @ Main /Users/kcft114/.julia/packages/Flux/qXNjB/src/optimise/train.jl:105
┌ Info: Epoch 742
└ @ Main /Users/kcft114/.julia/packages/Flux/qXNjB/src/optimise/train.jl:105
┌ Info: Epoch 743
└ @ Main /Users/kcft114/.julia/packages/Flux/qXNjB/src/optimise/train.jl:105
┌ Info: Epoch 744
└ @ Main /Users/kcft114/.julia/packages/Flux/qXNjB/src/optimise/train.jl:105
┌ Info: Epoch 745
└ @ Main /Users/kcft114/.julia/packages/Flux/qXNjB/src/optimise/train.jl:105
┌ Info: Epoch 746
└ @ Main /Users/kcft114/.julia/packages/Flux/qXNjB/src/optimise/train.jl:105
┌ Info: Epoch 747
└ @ Main /Users/kcft114/.julia/packages/Flux/qXNjB/src/optimise/train.jl:105
┌ Info: Epoch 748
└ @ Main /Users/kcft114/.julia/packages/Flux/qXNjB/src/optimise/train.jl:105
```

```
loss(X, Y) = 0.6479291f0 (tracked)
loss(X, Y) = 0.6970686f0 (tracked)
loss(X, Y) = 0.66425604f0 (tracked)
loss(X, Y) = 0.68926835f0 (tracked)
loss(X, Y) = 0.65089613f0 (tracked)
loss(X, Y) = 0.66994345f0 (tracked)
loss(X, Y) = 0.65575093f0 (tracked)
loss(X, Y) = 0.64146626f0 (tracked)
loss(X, Y) = 0.67580026f0 (tracked)
loss(X, Y) = 0.6599303f0 (tracked)
loss(X, Y) = 0.66353387f0 (tracked)
loss(X, Y) = 0.6229683f0 (tracked)
loss(X, Y) = 0.6809167f0 (tracked)
loss(X, Y) = 0.67071176f0 (tracked)
loss(X, Y) = 0.66519237f0 (tracked)
loss(X, Y) = 0.6659176f0 (tracked)
loss(X, Y) = 0.6182294f0 (tracked)
loss(X, Y) = 0.66354096f0 (tracked)
loss(X, Y) = 0.61157167f0 (tracked)
loss(X, Y) = 0.6503172f0 (tracked)
loss(X, Y) = 0.68001837f0 (tracked)
loss(X, Y) = 0.65735704f0 (tracked)
loss(X, Y) = 0.67964524f0 (tracked)
loss(X, Y) = 0.68977094f0 (tracked)
loss(X, Y) = 0.6557984f0 (tracked)
loss(X, Y) = 0.665565f0 (tracked)
loss(X, Y) = 0.65433073f0 (tracked)
loss(X, Y) = 0.6671948f0 (tracked)
loss(X, Y) = 0.6407065f0 (tracked)
loss(X, Y) = 0.6333424f0 (tracked)
loss(X, Y) = 0.67591035f0 (tracked)
loss(X, Y) = 0.67455155f0 (tracked)
loss(X, Y) = 0.6539016f0 (tracked)
loss(X, Y) = 0.67386955f0 (tracked)
loss(X, Y) = 0.6563488f0 (tracked)
loss(X, Y) = 0.651363f0 (tracked)
loss(X, Y) = 0.65053576f0 (tracked)
loss(X, Y) = 0.6554686f0 (tracked)
loss(X, Y) = 0.67643684f0 (tracked)
loss(X, Y) = 0.65800077f0 (tracked)
loss(X, Y) = 0.66421837f0 (tracked)
loss(X, Y) = 0.6658221f0 (tracked)
loss(X, Y) = 0.6583657f0 (tracked)
loss(X, Y) = 0.636251f0 (tracked)
loss(X, Y) = 0.65544784f0 (tracked)
loss(X, Y) = 0.6551903f0 (tracked)
loss(X, Y) = 0.657445f0 (tracked)
loss(X, Y) = 0.65052044f0 (tracked)
loss(X, Y) = 0.65774596f0 (tracked)
loss(X, Y) = 0.64950067f0 (tracked)
loss(X, Y) = 0.63905287f0 (tracked)
loss(X, Y) = 0.66983867f0 (tracked)
loss(X, Y) = 0.6607669f0 (tracked)
loss(X, Y) = 0.6307671f0 (tracked)
loss(X, Y) = 0.6389094f0 (tracked)
loss(X, Y) = 0.61614114f0 (tracked)
loss(X, Y) = 0.64453185f0 (tracked)
loss(X, Y) = 0.6548576f0 (tracked)
loss(X, Y) = 0.6575469f0 (tracked)
loss(X, Y) = 0.654783f0 (tracked)
loss(X, Y) = 0.6783067f0 (tracked)
loss(X, Y) = 0.6678801f0 (tracked)
loss(X, Y) = 0.65683955f0 (tracked)
loss(X, Y) = 0.67504185f0 (tracked)
loss(X, Y) = 0.6494264f0 (tracked)
loss(X, Y) = 0.63259554f0 (tracked)
loss(X, Y) = 0.6554976f0 (tracked)
loss(X, Y) = 0.65132636f0 (tracked)
loss(X, Y) = 0.68055224f0 (tracked)
loss(X, Y) = 0.6445431f0 (tracked)
loss(X, Y) = 0.64603376f0 (tracked)
loss(X, Y) = 0.6396781f0 (tracked)
loss(X, Y) = 0.63539696f0 (tracked)
loss(X, Y) = 0.6636959f0 (tracked)
loss(X, Y) = 0.65485275f0 (tracked)
loss(X, Y) = 0.6606256f0 (tracked)
loss(X, Y) = 0.64473265f0 (tracked)
loss(X, Y) = 0.6534401f0 (tracked)
loss(X, Y) = 0.67682636f0 (tracked)
loss(X, Y) = 0.65873694f0 (tracked)
loss(X, Y) = 0.62088543f0 (tracked)
loss(X, Y) = 0.66662085f0 (tracked)
loss(X, Y) = 0.62060094f0 (tracked)
loss(X, Y) = 0.63359267f0 (tracked)
loss(X, Y) = 0.66695994f0 (tracked)
loss(X, Y) = 0.64939386f0 (tracked)
loss(X, Y) = 0.6315095f0 (tracked)
loss(X, Y) = 0.6305164f0 (tracked)
loss(X, Y) = 0.64560163f0 (tracked)
loss(X, Y) = 0.6614945f0 (tracked)
loss(X, Y) = 0.659042f0 (tracked)
loss(X, Y) = 0.63973075f0 (tracked)
loss(X, Y) = 0.6153837f0 (tracked)
loss(X, Y) = 0.64382464f0 (tracked)
loss(X, Y) = 0.6679204f0 (tracked)
loss(X, Y) = 0.6406295f0 (tracked)
loss(X, Y) = 0.65709275f0 (tracked)
loss(X, Y) = 0.66612655f0 (tracked)
loss(X, Y) = 0.6812777f0 (tracked)
loss(X, Y) = 0.65896803f0 (tracked)
loss(X, Y) = 0.6175118f0 (tracked)
loss(X, Y) = 0.6253825f0 (tracked)
loss(X, Y) = 0.67369777f0 (tracked)
loss(X, Y) = 0.65106577f0 (tracked)
loss(X, Y) = 0.6493241f0 (tracked)
loss(X, Y) = 0.64202696f0 (tracked)
loss(X, Y) = 0.6288426f0 (tracked)
loss(X, Y) = 0.64930767f0 (tracked)
loss(X, Y) = 0.66575974f0 (tracked)
loss(X, Y) = 0.62914383f0 (tracked)
loss(X, Y) = 0.6470952f0 (tracked)
loss(X, Y) = 0.6732793f0 (tracked)
loss(X, Y) = 0.65891397f0 (tracked)
loss(X, Y) = 0.64991f0 (tracked)
loss(X, Y) = 0.65157616f0 (tracked)
loss(X, Y) = 0.6321344f0 (tracked)
loss(X, Y) = 0.68355715f0 (tracked)
loss(X, Y) = 0.64359236f0 (tracked)
loss(X, Y) = 0.6432501f0 (tracked)
loss(X, Y) = 0.63280165f0 (tracked)
loss(X, Y) = 0.63566345f0 (tracked)
loss(X, Y) = 0.6328143f0 (tracked)
loss(X, Y) = 0.62328345f0 (tracked)
loss(X, Y) = 0.64853114f0 (tracked)
loss(X, Y) = 0.64305407f0 (tracked)
loss(X, Y) = 0.5983721f0 (tracked)
loss(X, Y) = 0.6526896f0 (tracked)
```

```
┌ Info: Epoch 749
└ @ Main /Users/kcft114/.julia/packages/Flux/qXNjB/src/optimise/train.jl:105
┌ Info: Epoch 750
└ @ Main /Users/kcft114/.julia/packages/Flux/qXNjB/src/optimise/train.jl:105
┌ Info: Epoch 751
└ @ Main /Users/kcft114/.julia/packages/Flux/qXNjB/src/optimise/train.jl:105
┌ Info: Epoch 752
└ @ Main /Users/kcft114/.julia/packages/Flux/qXNjB/src/optimise/train.jl:105
┌ Info: Epoch 753
└ @ Main /Users/kcft114/.julia/packages/Flux/qXNjB/src/optimise/train.jl:105
┌ Info: Epoch 754
└ @ Main /Users/kcft114/.julia/packages/Flux/qXNjB/src/optimise/train.jl:105
┌ Info: Epoch 755
└ @ Main /Users/kcft114/.julia/packages/Flux/qXNjB/src/optimise/train.jl:105
┌ Info: Epoch 756
└ @ Main /Users/kcft114/.julia/packages/Flux/qXNjB/src/optimise/train.jl:105
┌ Info: Epoch 757
└ @ Main /Users/kcft114/.julia/packages/Flux/qXNjB/src/optimise/train.jl:105
┌ Info: Epoch 758
└ @ Main /Users/kcft114/.julia/packages/Flux/qXNjB/src/optimise/train.jl:105
┌ Info: Epoch 759
└ @ Main /Users/kcft114/.julia/packages/Flux/qXNjB/src/optimise/train.jl:105
┌ Info: Epoch 760
└ @ Main /Users/kcft114/.julia/packages/Flux/qXNjB/src/optimise/train.jl:105
┌ Info: Epoch 761
└ @ Main /Users/kcft114/.julia/packages/Flux/qXNjB/src/optimise/train.jl:105
┌ Info: Epoch 762
└ @ Main /Users/kcft114/.julia/packages/Flux/qXNjB/src/optimise/train.jl:105
┌ Info: Epoch 763
└ @ Main /Users/kcft114/.julia/packages/Flux/qXNjB/src/optimise/train.jl:105
┌ Info: Epoch 764
└ @ Main /Users/kcft114/.julia/packages/Flux/qXNjB/src/optimise/train.jl:105
┌ Info: Epoch 765
└ @ Main /Users/kcft114/.julia/packages/Flux/qXNjB/src/optimise/train.jl:105
┌ Info: Epoch 766
└ @ Main /Users/kcft114/.julia/packages/Flux/qXNjB/src/optimise/train.jl:105
┌ Info: Epoch 767
└ @ Main /Users/kcft114/.julia/packages/Flux/qXNjB/src/optimise/train.jl:105
┌ Info: Epoch 768
└ @ Main /Users/kcft114/.julia/packages/Flux/qXNjB/src/optimise/train.jl:105
┌ Info: Epoch 769
└ @ Main /Users/kcft114/.julia/packages/Flux/qXNjB/src/optimise/train.jl:105
┌ Info: Epoch 770
└ @ Main /Users/kcft114/.julia/packages/Flux/qXNjB/src/optimise/train.jl:105
┌ Info: Epoch 771
└ @ Main /Users/kcft114/.julia/packages/Flux/qXNjB/src/optimise/train.jl:105
┌ Info: Epoch 772
└ @ Main /Users/kcft114/.julia/packages/Flux/qXNjB/src/optimise/train.jl:105
┌ Info: Epoch 773
└ @ Main /Users/kcft114/.julia/packages/Flux/qXNjB/src/optimise/train.jl:105
┌ Info: Epoch 774
└ @ Main /Users/kcft114/.julia/packages/Flux/qXNjB/src/optimise/train.jl:105
┌ Info: Epoch 775
└ @ Main /Users/kcft114/.julia/packages/Flux/qXNjB/src/optimise/train.jl:105
┌ Info: Epoch 776
└ @ Main /Users/kcft114/.julia/packages/Flux/qXNjB/src/optimise/train.jl:105
┌ Info: Epoch 777
└ @ Main /Users/kcft114/.julia/packages/Flux/qXNjB/src/optimise/train.jl:105
┌ Info: Epoch 778
└ @ Main /Users/kcft114/.julia/packages/Flux/qXNjB/src/optimise/train.jl:105
┌ Info: Epoch 779
└ @ Main /Users/kcft114/.julia/packages/Flux/qXNjB/src/optimise/train.jl:105
┌ Info: Epoch 780
└ @ Main /Users/kcft114/.julia/packages/Flux/qXNjB/src/optimise/train.jl:105
┌ Info: Epoch 781
└ @ Main /Users/kcft114/.julia/packages/Flux/qXNjB/src/optimise/train.jl:105
┌ Info: Epoch 782
└ @ Main /Users/kcft114/.julia/packages/Flux/qXNjB/src/optimise/train.jl:105
┌ Info: Epoch 783
└ @ Main /Users/kcft114/.julia/packages/Flux/qXNjB/src/optimise/train.jl:105
┌ Info: Epoch 784
└ @ Main /Users/kcft114/.julia/packages/Flux/qXNjB/src/optimise/train.jl:105
┌ Info: Epoch 785
└ @ Main /Users/kcft114/.julia/packages/Flux/qXNjB/src/optimise/train.jl:105
┌ Info: Epoch 786
└ @ Main /Users/kcft114/.julia/packages/Flux/qXNjB/src/optimise/train.jl:105
┌ Info: Epoch 787
└ @ Main /Users/kcft114/.julia/packages/Flux/qXNjB/src/optimise/train.jl:105
┌ Info: Epoch 788
└ @ Main /Users/kcft114/.julia/packages/Flux/qXNjB/src/optimise/train.jl:105
┌ Info: Epoch 789
└ @ Main /Users/kcft114/.julia/packages/Flux/qXNjB/src/optimise/train.jl:105
┌ Info: Epoch 790
└ @ Main /Users/kcft114/.julia/packages/Flux/qXNjB/src/optimise/train.jl:105
┌ Info: Epoch 791
└ @ Main /Users/kcft114/.julia/packages/Flux/qXNjB/src/optimise/train.jl:105
┌ Info: Epoch 792
└ @ Main /Users/kcft114/.julia/packages/Flux/qXNjB/src/optimise/train.jl:105
┌ Info: Epoch 793
└ @ Main /Users/kcft114/.julia/packages/Flux/qXNjB/src/optimise/train.jl:105
┌ Info: Epoch 794
└ @ Main /Users/kcft114/.julia/packages/Flux/qXNjB/src/optimise/train.jl:105
┌ Info: Epoch 795
└ @ Main /Users/kcft114/.julia/packages/Flux/qXNjB/src/optimise/train.jl:105
┌ Info: Epoch 796
└ @ Main /Users/kcft114/.julia/packages/Flux/qXNjB/src/optimise/train.jl:105
┌ Info: Epoch 797
└ @ Main /Users/kcft114/.julia/packages/Flux/qXNjB/src/optimise/train.jl:105
┌ Info: Epoch 798
└ @ Main /Users/kcft114/.julia/packages/Flux/qXNjB/src/optimise/train.jl:105
┌ Info: Epoch 799
└ @ Main /Users/kcft114/.julia/packages/Flux/qXNjB/src/optimise/train.jl:105
┌ Info: Epoch 800
└ @ Main /Users/kcft114/.julia/packages/Flux/qXNjB/src/optimise/train.jl:105
┌ Info: Epoch 801
└ @ Main /Users/kcft114/.julia/packages/Flux/qXNjB/src/optimise/train.jl:105
┌ Info: Epoch 802
└ @ Main /Users/kcft114/.julia/packages/Flux/qXNjB/src/optimise/train.jl:105
┌ Info: Epoch 803
└ @ Main /Users/kcft114/.julia/packages/Flux/qXNjB/src/optimise/train.jl:105
┌ Info: Epoch 804
└ @ Main /Users/kcft114/.julia/packages/Flux/qXNjB/src/optimise/train.jl:105
┌ Info: Epoch 805
└ @ Main /Users/kcft114/.julia/packages/Flux/qXNjB/src/optimise/train.jl:105
┌ Info: Epoch 806
└ @ Main /Users/kcft114/.julia/packages/Flux/qXNjB/src/optimise/train.jl:105
┌ Info: Epoch 807
└ @ Main /Users/kcft114/.julia/packages/Flux/qXNjB/src/optimise/train.jl:105
┌ Info: Epoch 808
└ @ Main /Users/kcft114/.julia/packages/Flux/qXNjB/src/optimise/train.jl:105
┌ Info: Epoch 809
└ @ Main /Users/kcft114/.julia/packages/Flux/qXNjB/src/optimise/train.jl:105
┌ Info: Epoch 810
└ @ Main /Users/kcft114/.julia/packages/Flux/qXNjB/src/optimise/train.jl:105
┌ Info: Epoch 811
└ @ Main /Users/kcft114/.julia/packages/Flux/qXNjB/src/optimise/train.jl:105
┌ Info: Epoch 812
└ @ Main /Users/kcft114/.julia/packages/Flux/qXNjB/src/optimise/train.jl:105
┌ Info: Epoch 813
└ @ Main /Users/kcft114/.julia/packages/Flux/qXNjB/src/optimise/train.jl:105
┌ Info: Epoch 814
└ @ Main /Users/kcft114/.julia/packages/Flux/qXNjB/src/optimise/train.jl:105
┌ Info: Epoch 815
└ @ Main /Users/kcft114/.julia/packages/Flux/qXNjB/src/optimise/train.jl:105
┌ Info: Epoch 816
└ @ Main /Users/kcft114/.julia/packages/Flux/qXNjB/src/optimise/train.jl:105
┌ Info: Epoch 817
└ @ Main /Users/kcft114/.julia/packages/Flux/qXNjB/src/optimise/train.jl:105
┌ Info: Epoch 818
└ @ Main /Users/kcft114/.julia/packages/Flux/qXNjB/src/optimise/train.jl:105
┌ Info: Epoch 819
└ @ Main /Users/kcft114/.julia/packages/Flux/qXNjB/src/optimise/train.jl:105
┌ Info: Epoch 820
└ @ Main /Users/kcft114/.julia/packages/Flux/qXNjB/src/optimise/train.jl:105
┌ Info: Epoch 821
└ @ Main /Users/kcft114/.julia/packages/Flux/qXNjB/src/optimise/train.jl:105
┌ Info: Epoch 822
└ @ Main /Users/kcft114/.julia/packages/Flux/qXNjB/src/optimise/train.jl:105
┌ Info: Epoch 823
└ @ Main /Users/kcft114/.julia/packages/Flux/qXNjB/src/optimise/train.jl:105
┌ Info: Epoch 824
└ @ Main /Users/kcft114/.julia/packages/Flux/qXNjB/src/optimise/train.jl:105
┌ Info: Epoch 825
└ @ Main /Users/kcft114/.julia/packages/Flux/qXNjB/src/optimise/train.jl:105
┌ Info: Epoch 826
└ @ Main /Users/kcft114/.julia/packages/Flux/qXNjB/src/optimise/train.jl:105
┌ Info: Epoch 827
└ @ Main /Users/kcft114/.julia/packages/Flux/qXNjB/src/optimise/train.jl:105
┌ Info: Epoch 828
└ @ Main /Users/kcft114/.julia/packages/Flux/qXNjB/src/optimise/train.jl:105
┌ Info: Epoch 829
└ @ Main /Users/kcft114/.julia/packages/Flux/qXNjB/src/optimise/train.jl:105
┌ Info: Epoch 830
└ @ Main /Users/kcft114/.julia/packages/Flux/qXNjB/src/optimise/train.jl:105
┌ Info: Epoch 831
└ @ Main /Users/kcft114/.julia/packages/Flux/qXNjB/src/optimise/train.jl:105
┌ Info: Epoch 832
└ @ Main /Users/kcft114/.julia/packages/Flux/qXNjB/src/optimise/train.jl:105
┌ Info: Epoch 833
└ @ Main /Users/kcft114/.julia/packages/Flux/qXNjB/src/optimise/train.jl:105
┌ Info: Epoch 834
└ @ Main /Users/kcft114/.julia/packages/Flux/qXNjB/src/optimise/train.jl:105
┌ Info: Epoch 835
└ @ Main /Users/kcft114/.julia/packages/Flux/qXNjB/src/optimise/train.jl:105
┌ Info: Epoch 836
└ @ Main /Users/kcft114/.julia/packages/Flux/qXNjB/src/optimise/train.jl:105
┌ Info: Epoch 837
└ @ Main /Users/kcft114/.julia/packages/Flux/qXNjB/src/optimise/train.jl:105
┌ Info: Epoch 838
└ @ Main /Users/kcft114/.julia/packages/Flux/qXNjB/src/optimise/train.jl:105
┌ Info: Epoch 839
└ @ Main /Users/kcft114/.julia/packages/Flux/qXNjB/src/optimise/train.jl:105
┌ Info: Epoch 840
└ @ Main /Users/kcft114/.julia/packages/Flux/qXNjB/src/optimise/train.jl:105
┌ Info: Epoch 841
└ @ Main /Users/kcft114/.julia/packages/Flux/qXNjB/src/optimise/train.jl:105
┌ Info: Epoch 842
└ @ Main /Users/kcft114/.julia/packages/Flux/qXNjB/src/optimise/train.jl:105
┌ Info: Epoch 843
└ @ Main /Users/kcft114/.julia/packages/Flux/qXNjB/src/optimise/train.jl:105
┌ Info: Epoch 844
└ @ Main /Users/kcft114/.julia/packages/Flux/qXNjB/src/optimise/train.jl:105
┌ Info: Epoch 845
└ @ Main /Users/kcft114/.julia/packages/Flux/qXNjB/src/optimise/train.jl:105
┌ Info: Epoch 846
└ @ Main /Users/kcft114/.julia/packages/Flux/qXNjB/src/optimise/train.jl:105
┌ Info: Epoch 847
└ @ Main /Users/kcft114/.julia/packages/Flux/qXNjB/src/optimise/train.jl:105
┌ Info: Epoch 848
└ @ Main /Users/kcft114/.julia/packages/Flux/qXNjB/src/optimise/train.jl:105
┌ Info: Epoch 849
└ @ Main /Users/kcft114/.julia/packages/Flux/qXNjB/src/optimise/train.jl:105
┌ Info: Epoch 850
└ @ Main /Users/kcft114/.julia/packages/Flux/qXNjB/src/optimise/train.jl:105
┌ Info: Epoch 851
└ @ Main /Users/kcft114/.julia/packages/Flux/qXNjB/src/optimise/train.jl:105
┌ Info: Epoch 852
└ @ Main /Users/kcft114/.julia/packages/Flux/qXNjB/src/optimise/train.jl:105
┌ Info: Epoch 853
└ @ Main /Users/kcft114/.julia/packages/Flux/qXNjB/src/optimise/train.jl:105
┌ Info: Epoch 854
└ @ Main /Users/kcft114/.julia/packages/Flux/qXNjB/src/optimise/train.jl:105
┌ Info: Epoch 855
└ @ Main /Users/kcft114/.julia/packages/Flux/qXNjB/src/optimise/train.jl:105
┌ Info: Epoch 856
└ @ Main /Users/kcft114/.julia/packages/Flux/qXNjB/src/optimise/train.jl:105
┌ Info: Epoch 857
└ @ Main /Users/kcft114/.julia/packages/Flux/qXNjB/src/optimise/train.jl:105
┌ Info: Epoch 858
└ @ Main /Users/kcft114/.julia/packages/Flux/qXNjB/src/optimise/train.jl:105
┌ Info: Epoch 859
└ @ Main /Users/kcft114/.julia/packages/Flux/qXNjB/src/optimise/train.jl:105
┌ Info: Epoch 860
└ @ Main /Users/kcft114/.julia/packages/Flux/qXNjB/src/optimise/train.jl:105
┌ Info: Epoch 861
└ @ Main /Users/kcft114/.julia/packages/Flux/qXNjB/src/optimise/train.jl:105
┌ Info: Epoch 862
└ @ Main /Users/kcft114/.julia/packages/Flux/qXNjB/src/optimise/train.jl:105
┌ Info: Epoch 863
└ @ Main /Users/kcft114/.julia/packages/Flux/qXNjB/src/optimise/train.jl:105
┌ Info: Epoch 864
└ @ Main /Users/kcft114/.julia/packages/Flux/qXNjB/src/optimise/train.jl:105
┌ Info: Epoch 865
└ @ Main /Users/kcft114/.julia/packages/Flux/qXNjB/src/optimise/train.jl:105
┌ Info: Epoch 866
└ @ Main /Users/kcft114/.julia/packages/Flux/qXNjB/src/optimise/train.jl:105
┌ Info: Epoch 867
└ @ Main /Users/kcft114/.julia/packages/Flux/qXNjB/src/optimise/train.jl:105
┌ Info: Epoch 868
└ @ Main /Users/kcft114/.julia/packages/Flux/qXNjB/src/optimise/train.jl:105
┌ Info: Epoch 869
└ @ Main /Users/kcft114/.julia/packages/Flux/qXNjB/src/optimise/train.jl:105
┌ Info: Epoch 870
└ @ Main /Users/kcft114/.julia/packages/Flux/qXNjB/src/optimise/train.jl:105
┌ Info: Epoch 871
└ @ Main /Users/kcft114/.julia/packages/Flux/qXNjB/src/optimise/train.jl:105
┌ Info: Epoch 872
└ @ Main /Users/kcft114/.julia/packages/Flux/qXNjB/src/optimise/train.jl:105
┌ Info: Epoch 873
└ @ Main /Users/kcft114/.julia/packages/Flux/qXNjB/src/optimise/train.jl:105
┌ Info: Epoch 874
└ @ Main /Users/kcft114/.julia/packages/Flux/qXNjB/src/optimise/train.jl:105
┌ Info: Epoch 875
└ @ Main /Users/kcft114/.julia/packages/Flux/qXNjB/src/optimise/train.jl:105
```

```
loss(X, Y) = 0.6462668f0 (tracked)
loss(X, Y) = 0.6177828f0 (tracked)
loss(X, Y) = 0.63507825f0 (tracked)
loss(X, Y) = 0.6305802f0 (tracked)
loss(X, Y) = 0.66246605f0 (tracked)
loss(X, Y) = 0.6399439f0 (tracked)
loss(X, Y) = 0.6358686f0 (tracked)
loss(X, Y) = 0.60884225f0 (tracked)
loss(X, Y) = 0.62649107f0 (tracked)
loss(X, Y) = 0.65117574f0 (tracked)
loss(X, Y) = 0.6377126f0 (tracked)
loss(X, Y) = 0.63067466f0 (tracked)
loss(X, Y) = 0.6803328f0 (tracked)
loss(X, Y) = 0.6393069f0 (tracked)
loss(X, Y) = 0.6507473f0 (tracked)
loss(X, Y) = 0.6534808f0 (tracked)
loss(X, Y) = 0.63207626f0 (tracked)
loss(X, Y) = 0.6230227f0 (tracked)
loss(X, Y) = 0.6236069f0 (tracked)
loss(X, Y) = 0.6098619f0 (tracked)
loss(X, Y) = 0.62856853f0 (tracked)
loss(X, Y) = 0.64357954f0 (tracked)
loss(X, Y) = 0.6413473f0 (tracked)
loss(X, Y) = 0.6296031f0 (tracked)
loss(X, Y) = 0.6433191f0 (tracked)
loss(X, Y) = 0.6154864f0 (tracked)
loss(X, Y) = 0.6492241f0 (tracked)
loss(X, Y) = 0.64780843f0 (tracked)
loss(X, Y) = 0.6301678f0 (tracked)
loss(X, Y) = 0.60612184f0 (tracked)
loss(X, Y) = 0.64410025f0 (tracked)
loss(X, Y) = 0.63656425f0 (tracked)
loss(X, Y) = 0.6217725f0 (tracked)
loss(X, Y) = 0.61255765f0 (tracked)
loss(X, Y) = 0.634813f0 (tracked)
loss(X, Y) = 0.6356245f0 (tracked)
loss(X, Y) = 0.6613065f0 (tracked)
loss(X, Y) = 0.63078743f0 (tracked)
loss(X, Y) = 0.6314635f0 (tracked)
loss(X, Y) = 0.63880444f0 (tracked)
loss(X, Y) = 0.6226299f0 (tracked)
loss(X, Y) = 0.6567125f0 (tracked)
loss(X, Y) = 0.6378731f0 (tracked)
loss(X, Y) = 0.63503116f0 (tracked)
loss(X, Y) = 0.64955527f0 (tracked)
loss(X, Y) = 0.63282406f0 (tracked)
loss(X, Y) = 0.67964137f0 (tracked)
loss(X, Y) = 0.64050645f0 (tracked)
loss(X, Y) = 0.6548587f0 (tracked)
loss(X, Y) = 0.6466519f0 (tracked)
loss(X, Y) = 0.6484073f0 (tracked)
loss(X, Y) = 0.65260637f0 (tracked)
loss(X, Y) = 0.6124423f0 (tracked)
loss(X, Y) = 0.6283012f0 (tracked)
loss(X, Y) = 0.6471414f0 (tracked)
loss(X, Y) = 0.6090675f0 (tracked)
loss(X, Y) = 0.6176067f0 (tracked)
loss(X, Y) = 0.65067303f0 (tracked)
loss(X, Y) = 0.64359796f0 (tracked)
loss(X, Y) = 0.6336236f0 (tracked)
loss(X, Y) = 0.6096135f0 (tracked)
loss(X, Y) = 0.6658887f0 (tracked)
loss(X, Y) = 0.6159546f0 (tracked)
loss(X, Y) = 0.6512206f0 (tracked)
loss(X, Y) = 0.62804264f0 (tracked)
loss(X, Y) = 0.611245f0 (tracked)
loss(X, Y) = 0.62255096f0 (tracked)
loss(X, Y) = 0.6094234f0 (tracked)
loss(X, Y) = 0.60039085f0 (tracked)
loss(X, Y) = 0.6367468f0 (tracked)
loss(X, Y) = 0.6048876f0 (tracked)
loss(X, Y) = 0.610935f0 (tracked)
loss(X, Y) = 0.58771265f0 (tracked)
loss(X, Y) = 0.62570196f0 (tracked)
loss(X, Y) = 0.6619876f0 (tracked)
loss(X, Y) = 0.6451543f0 (tracked)
loss(X, Y) = 0.6317955f0 (tracked)
loss(X, Y) = 0.625559f0 (tracked)
loss(X, Y) = 0.64069575f0 (tracked)
loss(X, Y) = 0.63578445f0 (tracked)
loss(X, Y) = 0.5847988f0 (tracked)
loss(X, Y) = 0.64886445f0 (tracked)
loss(X, Y) = 0.63139105f0 (tracked)
loss(X, Y) = 0.63673884f0 (tracked)
loss(X, Y) = 0.6497269f0 (tracked)
loss(X, Y) = 0.61144507f0 (tracked)
loss(X, Y) = 0.6090248f0 (tracked)
loss(X, Y) = 0.6085209f0 (tracked)
loss(X, Y) = 0.61735344f0 (tracked)
loss(X, Y) = 0.65501666f0 (tracked)
loss(X, Y) = 0.632443f0 (tracked)
loss(X, Y) = 0.60545355f0 (tracked)
loss(X, Y) = 0.6336497f0 (tracked)
loss(X, Y) = 0.62805736f0 (tracked)
loss(X, Y) = 0.6380783f0 (tracked)
loss(X, Y) = 0.6207579f0 (tracked)
loss(X, Y) = 0.6237862f0 (tracked)
loss(X, Y) = 0.62236315f0 (tracked)
loss(X, Y) = 0.61393154f0 (tracked)
loss(X, Y) = 0.6236832f0 (tracked)
loss(X, Y) = 0.6615897f0 (tracked)
loss(X, Y) = 0.62622654f0 (tracked)
loss(X, Y) = 0.5943105f0 (tracked)
loss(X, Y) = 0.6187191f0 (tracked)
loss(X, Y) = 0.63519984f0 (tracked)
loss(X, Y) = 0.61419266f0 (tracked)
loss(X, Y) = 0.62151486f0 (tracked)
loss(X, Y) = 0.636115f0 (tracked)
loss(X, Y) = 0.64266485f0 (tracked)
loss(X, Y) = 0.65913177f0 (tracked)
loss(X, Y) = 0.66267014f0 (tracked)
loss(X, Y) = 0.6899247f0 (tracked)
loss(X, Y) = 0.6476216f0 (tracked)
loss(X, Y) = 0.6415571f0 (tracked)
loss(X, Y) = 0.58970386f0 (tracked)
loss(X, Y) = 0.62416494f0 (tracked)
loss(X, Y) = 0.623761f0 (tracked)
loss(X, Y) = 0.6159553f0 (tracked)
loss(X, Y) = 0.6417036f0 (tracked)
```

```
┌ Info: Epoch 876
└ @ Main /Users/kcft114/.julia/packages/Flux/qXNjB/src/optimise/train.jl:105
┌ Info: Epoch 877
└ @ Main /Users/kcft114/.julia/packages/Flux/qXNjB/src/optimise/train.jl:105
┌ Info: Epoch 878
└ @ Main /Users/kcft114/.julia/packages/Flux/qXNjB/src/optimise/train.jl:105
┌ Info: Epoch 879
└ @ Main /Users/kcft114/.julia/packages/Flux/qXNjB/src/optimise/train.jl:105
┌ Info: Epoch 880
└ @ Main /Users/kcft114/.julia/packages/Flux/qXNjB/src/optimise/train.jl:105
┌ Info: Epoch 881
└ @ Main /Users/kcft114/.julia/packages/Flux/qXNjB/src/optimise/train.jl:105
┌ Info: Epoch 882
└ @ Main /Users/kcft114/.julia/packages/Flux/qXNjB/src/optimise/train.jl:105
┌ Info: Epoch 883
└ @ Main /Users/kcft114/.julia/packages/Flux/qXNjB/src/optimise/train.jl:105
┌ Info: Epoch 884
└ @ Main /Users/kcft114/.julia/packages/Flux/qXNjB/src/optimise/train.jl:105
┌ Info: Epoch 885
└ @ Main /Users/kcft114/.julia/packages/Flux/qXNjB/src/optimise/train.jl:105
┌ Info: Epoch 886
└ @ Main /Users/kcft114/.julia/packages/Flux/qXNjB/src/optimise/train.jl:105
┌ Info: Epoch 887
└ @ Main /Users/kcft114/.julia/packages/Flux/qXNjB/src/optimise/train.jl:105
┌ Info: Epoch 888
└ @ Main /Users/kcft114/.julia/packages/Flux/qXNjB/src/optimise/train.jl:105
┌ Info: Epoch 889
└ @ Main /Users/kcft114/.julia/packages/Flux/qXNjB/src/optimise/train.jl:105
┌ Info: Epoch 890
└ @ Main /Users/kcft114/.julia/packages/Flux/qXNjB/src/optimise/train.jl:105
┌ Info: Epoch 891
└ @ Main /Users/kcft114/.julia/packages/Flux/qXNjB/src/optimise/train.jl:105
┌ Info: Epoch 892
└ @ Main /Users/kcft114/.julia/packages/Flux/qXNjB/src/optimise/train.jl:105
┌ Info: Epoch 893
└ @ Main /Users/kcft114/.julia/packages/Flux/qXNjB/src/optimise/train.jl:105
┌ Info: Epoch 894
└ @ Main /Users/kcft114/.julia/packages/Flux/qXNjB/src/optimise/train.jl:105
┌ Info: Epoch 895
└ @ Main /Users/kcft114/.julia/packages/Flux/qXNjB/src/optimise/train.jl:105
┌ Info: Epoch 896
└ @ Main /Users/kcft114/.julia/packages/Flux/qXNjB/src/optimise/train.jl:105
┌ Info: Epoch 897
└ @ Main /Users/kcft114/.julia/packages/Flux/qXNjB/src/optimise/train.jl:105
┌ Info: Epoch 898
└ @ Main /Users/kcft114/.julia/packages/Flux/qXNjB/src/optimise/train.jl:105
┌ Info: Epoch 899
└ @ Main /Users/kcft114/.julia/packages/Flux/qXNjB/src/optimise/train.jl:105
┌ Info: Epoch 900
└ @ Main /Users/kcft114/.julia/packages/Flux/qXNjB/src/optimise/train.jl:105
┌ Info: Epoch 901
└ @ Main /Users/kcft114/.julia/packages/Flux/qXNjB/src/optimise/train.jl:105
┌ Info: Epoch 902
└ @ Main /Users/kcft114/.julia/packages/Flux/qXNjB/src/optimise/train.jl:105
┌ Info: Epoch 903
└ @ Main /Users/kcft114/.julia/packages/Flux/qXNjB/src/optimise/train.jl:105
┌ Info: Epoch 904
└ @ Main /Users/kcft114/.julia/packages/Flux/qXNjB/src/optimise/train.jl:105
┌ Info: Epoch 905
└ @ Main /Users/kcft114/.julia/packages/Flux/qXNjB/src/optimise/train.jl:105
┌ Info: Epoch 906
└ @ Main /Users/kcft114/.julia/packages/Flux/qXNjB/src/optimise/train.jl:105
┌ Info: Epoch 907
└ @ Main /Users/kcft114/.julia/packages/Flux/qXNjB/src/optimise/train.jl:105
┌ Info: Epoch 908
└ @ Main /Users/kcft114/.julia/packages/Flux/qXNjB/src/optimise/train.jl:105
┌ Info: Epoch 909
└ @ Main /Users/kcft114/.julia/packages/Flux/qXNjB/src/optimise/train.jl:105
┌ Info: Epoch 910
└ @ Main /Users/kcft114/.julia/packages/Flux/qXNjB/src/optimise/train.jl:105
┌ Info: Epoch 911
└ @ Main /Users/kcft114/.julia/packages/Flux/qXNjB/src/optimise/train.jl:105
┌ Info: Epoch 912
└ @ Main /Users/kcft114/.julia/packages/Flux/qXNjB/src/optimise/train.jl:105
┌ Info: Epoch 913
└ @ Main /Users/kcft114/.julia/packages/Flux/qXNjB/src/optimise/train.jl:105
┌ Info: Epoch 914
└ @ Main /Users/kcft114/.julia/packages/Flux/qXNjB/src/optimise/train.jl:105
┌ Info: Epoch 915
└ @ Main /Users/kcft114/.julia/packages/Flux/qXNjB/src/optimise/train.jl:105
┌ Info: Epoch 916
└ @ Main /Users/kcft114/.julia/packages/Flux/qXNjB/src/optimise/train.jl:105
┌ Info: Epoch 917
└ @ Main /Users/kcft114/.julia/packages/Flux/qXNjB/src/optimise/train.jl:105
┌ Info: Epoch 918
└ @ Main /Users/kcft114/.julia/packages/Flux/qXNjB/src/optimise/train.jl:105
┌ Info: Epoch 919
└ @ Main /Users/kcft114/.julia/packages/Flux/qXNjB/src/optimise/train.jl:105
┌ Info: Epoch 920
└ @ Main /Users/kcft114/.julia/packages/Flux/qXNjB/src/optimise/train.jl:105
┌ Info: Epoch 921
└ @ Main /Users/kcft114/.julia/packages/Flux/qXNjB/src/optimise/train.jl:105
┌ Info: Epoch 922
└ @ Main /Users/kcft114/.julia/packages/Flux/qXNjB/src/optimise/train.jl:105
┌ Info: Epoch 923
└ @ Main /Users/kcft114/.julia/packages/Flux/qXNjB/src/optimise/train.jl:105
┌ Info: Epoch 924
└ @ Main /Users/kcft114/.julia/packages/Flux/qXNjB/src/optimise/train.jl:105
┌ Info: Epoch 925
└ @ Main /Users/kcft114/.julia/packages/Flux/qXNjB/src/optimise/train.jl:105
┌ Info: Epoch 926
└ @ Main /Users/kcft114/.julia/packages/Flux/qXNjB/src/optimise/train.jl:105
┌ Info: Epoch 927
└ @ Main /Users/kcft114/.julia/packages/Flux/qXNjB/src/optimise/train.jl:105
┌ Info: Epoch 928
└ @ Main /Users/kcft114/.julia/packages/Flux/qXNjB/src/optimise/train.jl:105
┌ Info: Epoch 929
└ @ Main /Users/kcft114/.julia/packages/Flux/qXNjB/src/optimise/train.jl:105
┌ Info: Epoch 930
└ @ Main /Users/kcft114/.julia/packages/Flux/qXNjB/src/optimise/train.jl:105
┌ Info: Epoch 931
└ @ Main /Users/kcft114/.julia/packages/Flux/qXNjB/src/optimise/train.jl:105
┌ Info: Epoch 932
└ @ Main /Users/kcft114/.julia/packages/Flux/qXNjB/src/optimise/train.jl:105
┌ Info: Epoch 933
└ @ Main /Users/kcft114/.julia/packages/Flux/qXNjB/src/optimise/train.jl:105
┌ Info: Epoch 934
└ @ Main /Users/kcft114/.julia/packages/Flux/qXNjB/src/optimise/train.jl:105
┌ Info: Epoch 935
└ @ Main /Users/kcft114/.julia/packages/Flux/qXNjB/src/optimise/train.jl:105
┌ Info: Epoch 936
└ @ Main /Users/kcft114/.julia/packages/Flux/qXNjB/src/optimise/train.jl:105
┌ Info: Epoch 937
└ @ Main /Users/kcft114/.julia/packages/Flux/qXNjB/src/optimise/train.jl:105
┌ Info: Epoch 938
└ @ Main /Users/kcft114/.julia/packages/Flux/qXNjB/src/optimise/train.jl:105
┌ Info: Epoch 939
└ @ Main /Users/kcft114/.julia/packages/Flux/qXNjB/src/optimise/train.jl:105
┌ Info: Epoch 940
└ @ Main /Users/kcft114/.julia/packages/Flux/qXNjB/src/optimise/train.jl:105
┌ Info: Epoch 941
└ @ Main /Users/kcft114/.julia/packages/Flux/qXNjB/src/optimise/train.jl:105
┌ Info: Epoch 942
└ @ Main /Users/kcft114/.julia/packages/Flux/qXNjB/src/optimise/train.jl:105
┌ Info: Epoch 943
└ @ Main /Users/kcft114/.julia/packages/Flux/qXNjB/src/optimise/train.jl:105
┌ Info: Epoch 944
└ @ Main /Users/kcft114/.julia/packages/Flux/qXNjB/src/optimise/train.jl:105
┌ Info: Epoch 945
└ @ Main /Users/kcft114/.julia/packages/Flux/qXNjB/src/optimise/train.jl:105
┌ Info: Epoch 946
└ @ Main /Users/kcft114/.julia/packages/Flux/qXNjB/src/optimise/train.jl:105
┌ Info: Epoch 947
└ @ Main /Users/kcft114/.julia/packages/Flux/qXNjB/src/optimise/train.jl:105
┌ Info: Epoch 948
└ @ Main /Users/kcft114/.julia/packages/Flux/qXNjB/src/optimise/train.jl:105
┌ Info: Epoch 949
└ @ Main /Users/kcft114/.julia/packages/Flux/qXNjB/src/optimise/train.jl:105
┌ Info: Epoch 950
└ @ Main /Users/kcft114/.julia/packages/Flux/qXNjB/src/optimise/train.jl:105
┌ Info: Epoch 951
└ @ Main /Users/kcft114/.julia/packages/Flux/qXNjB/src/optimise/train.jl:105
┌ Info: Epoch 952
└ @ Main /Users/kcft114/.julia/packages/Flux/qXNjB/src/optimise/train.jl:105
┌ Info: Epoch 953
└ @ Main /Users/kcft114/.julia/packages/Flux/qXNjB/src/optimise/train.jl:105
┌ Info: Epoch 954
└ @ Main /Users/kcft114/.julia/packages/Flux/qXNjB/src/optimise/train.jl:105
┌ Info: Epoch 955
└ @ Main /Users/kcft114/.julia/packages/Flux/qXNjB/src/optimise/train.jl:105
┌ Info: Epoch 956
└ @ Main /Users/kcft114/.julia/packages/Flux/qXNjB/src/optimise/train.jl:105
┌ Info: Epoch 957
└ @ Main /Users/kcft114/.julia/packages/Flux/qXNjB/src/optimise/train.jl:105
┌ Info: Epoch 958
└ @ Main /Users/kcft114/.julia/packages/Flux/qXNjB/src/optimise/train.jl:105
┌ Info: Epoch 959
└ @ Main /Users/kcft114/.julia/packages/Flux/qXNjB/src/optimise/train.jl:105
┌ Info: Epoch 960
└ @ Main /Users/kcft114/.julia/packages/Flux/qXNjB/src/optimise/train.jl:105
┌ Info: Epoch 961
└ @ Main /Users/kcft114/.julia/packages/Flux/qXNjB/src/optimise/train.jl:105
┌ Info: Epoch 962
└ @ Main /Users/kcft114/.julia/packages/Flux/qXNjB/src/optimise/train.jl:105
┌ Info: Epoch 963
└ @ Main /Users/kcft114/.julia/packages/Flux/qXNjB/src/optimise/train.jl:105
┌ Info: Epoch 964
└ @ Main /Users/kcft114/.julia/packages/Flux/qXNjB/src/optimise/train.jl:105
┌ Info: Epoch 965
└ @ Main /Users/kcft114/.julia/packages/Flux/qXNjB/src/optimise/train.jl:105
┌ Info: Epoch 966
└ @ Main /Users/kcft114/.julia/packages/Flux/qXNjB/src/optimise/train.jl:105
┌ Info: Epoch 967
└ @ Main /Users/kcft114/.julia/packages/Flux/qXNjB/src/optimise/train.jl:105
┌ Info: Epoch 968
└ @ Main /Users/kcft114/.julia/packages/Flux/qXNjB/src/optimise/train.jl:105
┌ Info: Epoch 969
└ @ Main /Users/kcft114/.julia/packages/Flux/qXNjB/src/optimise/train.jl:105
┌ Info: Epoch 970
└ @ Main /Users/kcft114/.julia/packages/Flux/qXNjB/src/optimise/train.jl:105
┌ Info: Epoch 971
└ @ Main /Users/kcft114/.julia/packages/Flux/qXNjB/src/optimise/train.jl:105
┌ Info: Epoch 972
└ @ Main /Users/kcft114/.julia/packages/Flux/qXNjB/src/optimise/train.jl:105
┌ Info: Epoch 973
└ @ Main /Users/kcft114/.julia/packages/Flux/qXNjB/src/optimise/train.jl:105
┌ Info: Epoch 974
└ @ Main /Users/kcft114/.julia/packages/Flux/qXNjB/src/optimise/train.jl:105
┌ Info: Epoch 975
└ @ Main /Users/kcft114/.julia/packages/Flux/qXNjB/src/optimise/train.jl:105
┌ Info: Epoch 976
└ @ Main /Users/kcft114/.julia/packages/Flux/qXNjB/src/optimise/train.jl:105
┌ Info: Epoch 977
└ @ Main /Users/kcft114/.julia/packages/Flux/qXNjB/src/optimise/train.jl:105
┌ Info: Epoch 978
└ @ Main /Users/kcft114/.julia/packages/Flux/qXNjB/src/optimise/train.jl:105
┌ Info: Epoch 979
└ @ Main /Users/kcft114/.julia/packages/Flux/qXNjB/src/optimise/train.jl:105
┌ Info: Epoch 980
└ @ Main /Users/kcft114/.julia/packages/Flux/qXNjB/src/optimise/train.jl:105
┌ Info: Epoch 981
└ @ Main /Users/kcft114/.julia/packages/Flux/qXNjB/src/optimise/train.jl:105
┌ Info: Epoch 982
└ @ Main /Users/kcft114/.julia/packages/Flux/qXNjB/src/optimise/train.jl:105
┌ Info: Epoch 983
└ @ Main /Users/kcft114/.julia/packages/Flux/qXNjB/src/optimise/train.jl:105
┌ Info: Epoch 984
└ @ Main /Users/kcft114/.julia/packages/Flux/qXNjB/src/optimise/train.jl:105
┌ Info: Epoch 985
└ @ Main /Users/kcft114/.julia/packages/Flux/qXNjB/src/optimise/train.jl:105
┌ Info: Epoch 986
└ @ Main /Users/kcft114/.julia/packages/Flux/qXNjB/src/optimise/train.jl:105
┌ Info: Epoch 987
└ @ Main /Users/kcft114/.julia/packages/Flux/qXNjB/src/optimise/train.jl:105
┌ Info: Epoch 988
└ @ Main /Users/kcft114/.julia/packages/Flux/qXNjB/src/optimise/train.jl:105
┌ Info: Epoch 989
└ @ Main /Users/kcft114/.julia/packages/Flux/qXNjB/src/optimise/train.jl:105
┌ Info: Epoch 990
└ @ Main /Users/kcft114/.julia/packages/Flux/qXNjB/src/optimise/train.jl:105
┌ Info: Epoch 991
└ @ Main /Users/kcft114/.julia/packages/Flux/qXNjB/src/optimise/train.jl:105
┌ Info: Epoch 992
└ @ Main /Users/kcft114/.julia/packages/Flux/qXNjB/src/optimise/train.jl:105
┌ Info: Epoch 993
└ @ Main /Users/kcft114/.julia/packages/Flux/qXNjB/src/optimise/train.jl:105
┌ Info: Epoch 994
└ @ Main /Users/kcft114/.julia/packages/Flux/qXNjB/src/optimise/train.jl:105
```

```
loss(X, Y) = 0.647674f0 (tracked)
loss(X, Y) = 0.6290396f0 (tracked)
loss(X, Y) = 0.6159215f0 (tracked)
loss(X, Y) = 0.5972236f0 (tracked)
loss(X, Y) = 0.62126887f0 (tracked)
loss(X, Y) = 0.63359964f0 (tracked)
loss(X, Y) = 0.6047247f0 (tracked)
loss(X, Y) = 0.6017758f0 (tracked)
loss(X, Y) = 0.64228874f0 (tracked)
loss(X, Y) = 0.58158684f0 (tracked)
loss(X, Y) = 0.6101485f0 (tracked)
loss(X, Y) = 0.62580943f0 (tracked)
loss(X, Y) = 0.61855567f0 (tracked)
loss(X, Y) = 0.6297737f0 (tracked)
loss(X, Y) = 0.6179066f0 (tracked)
loss(X, Y) = 0.6098212f0 (tracked)
loss(X, Y) = 0.6038305f0 (tracked)
loss(X, Y) = 0.5853289f0 (tracked)
loss(X, Y) = 0.63628304f0 (tracked)
loss(X, Y) = 0.582633f0 (tracked)
loss(X, Y) = 0.6259253f0 (tracked)
loss(X, Y) = 0.6358006f0 (tracked)
loss(X, Y) = 0.58430135f0 (tracked)
loss(X, Y) = 0.5888205f0 (tracked)
loss(X, Y) = 0.6057889f0 (tracked)
loss(X, Y) = 0.6363069f0 (tracked)
loss(X, Y) = 0.6121743f0 (tracked)
loss(X, Y) = 0.6111897f0 (tracked)
loss(X, Y) = 0.63385797f0 (tracked)
loss(X, Y) = 0.59473634f0 (tracked)
loss(X, Y) = 0.6133027f0 (tracked)
loss(X, Y) = 0.6383492f0 (tracked)
loss(X, Y) = 0.6060695f0 (tracked)
loss(X, Y) = 0.62957394f0 (tracked)
loss(X, Y) = 0.623702f0 (tracked)
loss(X, Y) = 0.636762f0 (tracked)
loss(X, Y) = 0.6159699f0 (tracked)
loss(X, Y) = 0.62072736f0 (tracked)
loss(X, Y) = 0.6243266f0 (tracked)
loss(X, Y) = 0.6137454f0 (tracked)
loss(X, Y) = 0.63695115f0 (tracked)
loss(X, Y) = 0.6308365f0 (tracked)
loss(X, Y) = 0.6128749f0 (tracked)
loss(X, Y) = 0.59734565f0 (tracked)
loss(X, Y) = 0.60221094f0 (tracked)
loss(X, Y) = 0.6301968f0 (tracked)
loss(X, Y) = 0.57591087f0 (tracked)
loss(X, Y) = 0.62670046f0 (tracked)
loss(X, Y) = 0.60257626f0 (tracked)
loss(X, Y) = 0.5602645f0 (tracked)
loss(X, Y) = 0.6056905f0 (tracked)
loss(X, Y) = 0.6146852f0 (tracked)
loss(X, Y) = 0.5900425f0 (tracked)
loss(X, Y) = 0.63375604f0 (tracked)
loss(X, Y) = 0.59810877f0 (tracked)
loss(X, Y) = 0.6237943f0 (tracked)
loss(X, Y) = 0.61329633f0 (tracked)
loss(X, Y) = 0.63471746f0 (tracked)
loss(X, Y) = 0.6237792f0 (tracked)
loss(X, Y) = 0.6287716f0 (tracked)
loss(X, Y) = 0.6181585f0 (tracked)
loss(X, Y) = 0.592493f0 (tracked)
loss(X, Y) = 0.6277765f0 (tracked)
loss(X, Y) = 0.6107573f0 (tracked)
loss(X, Y) = 0.6426138f0 (tracked)
loss(X, Y) = 0.6030016f0 (tracked)
loss(X, Y) = 0.619277f0 (tracked)
loss(X, Y) = 0.63667095f0 (tracked)
loss(X, Y) = 0.62854403f0 (tracked)
loss(X, Y) = 0.61803424f0 (tracked)
loss(X, Y) = 0.6181934f0 (tracked)
loss(X, Y) = 0.60783947f0 (tracked)
loss(X, Y) = 0.6052207f0 (tracked)
loss(X, Y) = 0.60188264f0 (tracked)
loss(X, Y) = 0.5897451f0 (tracked)
loss(X, Y) = 0.634391f0 (tracked)
loss(X, Y) = 0.5771255f0 (tracked)
loss(X, Y) = 0.62968653f0 (tracked)
loss(X, Y) = 0.6182665f0 (tracked)
loss(X, Y) = 0.6187412f0 (tracked)
loss(X, Y) = 0.6127198f0 (tracked)
loss(X, Y) = 0.62214696f0 (tracked)
loss(X, Y) = 0.6181289f0 (tracked)
loss(X, Y) = 0.5961185f0 (tracked)
loss(X, Y) = 0.6111024f0 (tracked)
loss(X, Y) = 0.57441294f0 (tracked)
loss(X, Y) = 0.5720224f0 (tracked)
loss(X, Y) = 0.6239618f0 (tracked)
loss(X, Y) = 0.62134653f0 (tracked)
loss(X, Y) = 0.61463475f0 (tracked)
loss(X, Y) = 0.601816f0 (tracked)
loss(X, Y) = 0.5999829f0 (tracked)
loss(X, Y) = 0.6166287f0 (tracked)
loss(X, Y) = 0.60437506f0 (tracked)
loss(X, Y) = 0.6044925f0 (tracked)
loss(X, Y) = 0.6328352f0 (tracked)
loss(X, Y) = 0.6375157f0 (tracked)
loss(X, Y) = 0.5999544f0 (tracked)
loss(X, Y) = 0.6180188f0 (tracked)
loss(X, Y) = 0.59197986f0 (tracked)
loss(X, Y) = 0.63641274f0 (tracked)
loss(X, Y) = 0.5834672f0 (tracked)
loss(X, Y) = 0.6089888f0 (tracked)
loss(X, Y) = 0.6097108f0 (tracked)
loss(X, Y) = 0.6166704f0 (tracked)
loss(X, Y) = 0.6045108f0 (tracked)
loss(X, Y) = 0.6281979f0 (tracked)
loss(X, Y) = 0.6033306f0 (tracked)
loss(X, Y) = 0.6269878f0 (tracked)
loss(X, Y) = 0.63040286f0 (tracked)
loss(X, Y) = 0.5891999f0 (tracked)
loss(X, Y) = 0.6256449f0 (tracked)
loss(X, Y) = 0.6354899f0 (tracked)
loss(X, Y) = 0.61207753f0 (tracked)
loss(X, Y) = 0.57676864f0 (tracked)
loss(X, Y) = 0.59020656f0 (tracked)
loss(X, Y) = 0.63612574f0 (tracked)
loss(X, Y) = 0.56057626f0 (tracked)
loss(X, Y) = 0.63343304f0 (tracked)
loss(X, Y) = 0.6147921f0 (tracked)
loss(X, Y) = 0.6090546f0 (tracked)
loss(X, Y) = 0.6477091f0 (tracked)
loss(X, Y) = 0.6547448f0 (tracked)
loss(X, Y) = 0.6068327f0 (tracked)
loss(X, Y) = 0.6030481f0 (tracked)
loss(X, Y) = 0.6221286f0 (tracked)
loss(X, Y) = 0.6110851f0 (tracked)
loss(X, Y) = 0.6253525f0 (tracked)
loss(X, Y) = 0.59378517f0 (tracked)
loss(X, Y) = 0.5998568f0 (tracked)
loss(X, Y) = 0.55253786f0 (tracked)
loss(X, Y) = 0.6232992f0 (tracked)
loss(X, Y) = 0.6076025f0 (tracked)
loss(X, Y) = 0.62357366f0 (tracked)
```

```
┌ Info: Epoch 995
└ @ Main /Users/kcft114/.julia/packages/Flux/qXNjB/src/optimise/train.jl:105
┌ Info: Epoch 996
└ @ Main /Users/kcft114/.julia/packages/Flux/qXNjB/src/optimise/train.jl:105
┌ Info: Epoch 997
└ @ Main /Users/kcft114/.julia/packages/Flux/qXNjB/src/optimise/train.jl:105
┌ Info: Epoch 998
└ @ Main /Users/kcft114/.julia/packages/Flux/qXNjB/src/optimise/train.jl:105
┌ Info: Epoch 999
└ @ Main /Users/kcft114/.julia/packages/Flux/qXNjB/src/optimise/train.jl:105
┌ Info: Epoch 1000
└ @ Main /Users/kcft114/.julia/packages/Flux/qXNjB/src/optimise/train.jl:105
┌ Info: Epoch 1001
└ @ Main /Users/kcft114/.julia/packages/Flux/qXNjB/src/optimise/train.jl:105
┌ Info: Epoch 1002
└ @ Main /Users/kcft114/.julia/packages/Flux/qXNjB/src/optimise/train.jl:105
┌ Info: Epoch 1003
└ @ Main /Users/kcft114/.julia/packages/Flux/qXNjB/src/optimise/train.jl:105
┌ Info: Epoch 1004
└ @ Main /Users/kcft114/.julia/packages/Flux/qXNjB/src/optimise/train.jl:105
┌ Info: Epoch 1005
└ @ Main /Users/kcft114/.julia/packages/Flux/qXNjB/src/optimise/train.jl:105
┌ Info: Epoch 1006
└ @ Main /Users/kcft114/.julia/packages/Flux/qXNjB/src/optimise/train.jl:105
┌ Info: Epoch 1007
└ @ Main /Users/kcft114/.julia/packages/Flux/qXNjB/src/optimise/train.jl:105
┌ Info: Epoch 1008
└ @ Main /Users/kcft114/.julia/packages/Flux/qXNjB/src/optimise/train.jl:105
┌ Info: Epoch 1009
└ @ Main /Users/kcft114/.julia/packages/Flux/qXNjB/src/optimise/train.jl:105
┌ Info: Epoch 1010
└ @ Main /Users/kcft114/.julia/packages/Flux/qXNjB/src/optimise/train.jl:105
┌ Info: Epoch 1011
└ @ Main /Users/kcft114/.julia/packages/Flux/qXNjB/src/optimise/train.jl:105
┌ Info: Epoch 1012
└ @ Main /Users/kcft114/.julia/packages/Flux/qXNjB/src/optimise/train.jl:105
┌ Info: Epoch 1013
└ @ Main /Users/kcft114/.julia/packages/Flux/qXNjB/src/optimise/train.jl:105
┌ Info: Epoch 1014
└ @ Main /Users/kcft114/.julia/packages/Flux/qXNjB/src/optimise/train.jl:105
┌ Info: Epoch 1015
└ @ Main /Users/kcft114/.julia/packages/Flux/qXNjB/src/optimise/train.jl:105
┌ Info: Epoch 1016
└ @ Main /Users/kcft114/.julia/packages/Flux/qXNjB/src/optimise/train.jl:105
┌ Info: Epoch 1017
└ @ Main /Users/kcft114/.julia/packages/Flux/qXNjB/src/optimise/train.jl:105
┌ Info: Epoch 1018
└ @ Main /Users/kcft114/.julia/packages/Flux/qXNjB/src/optimise/train.jl:105
┌ Info: Epoch 1019
└ @ Main /Users/kcft114/.julia/packages/Flux/qXNjB/src/optimise/train.jl:105
┌ Info: Epoch 1020
└ @ Main /Users/kcft114/.julia/packages/Flux/qXNjB/src/optimise/train.jl:105
┌ Info: Epoch 1021
└ @ Main /Users/kcft114/.julia/packages/Flux/qXNjB/src/optimise/train.jl:105
┌ Info: Epoch 1022
└ @ Main /Users/kcft114/.julia/packages/Flux/qXNjB/src/optimise/train.jl:105
┌ Info: Epoch 1023
└ @ Main /Users/kcft114/.julia/packages/Flux/qXNjB/src/optimise/train.jl:105
┌ Info: Epoch 1024
└ @ Main /Users/kcft114/.julia/packages/Flux/qXNjB/src/optimise/train.jl:105
┌ Info: Epoch 1025
└ @ Main /Users/kcft114/.julia/packages/Flux/qXNjB/src/optimise/train.jl:105
┌ Info: Epoch 1026
└ @ Main /Users/kcft114/.julia/packages/Flux/qXNjB/src/optimise/train.jl:105
┌ Info: Epoch 1027
└ @ Main /Users/kcft114/.julia/packages/Flux/qXNjB/src/optimise/train.jl:105
┌ Info: Epoch 1028
└ @ Main /Users/kcft114/.julia/packages/Flux/qXNjB/src/optimise/train.jl:105
┌ Info: Epoch 1029
└ @ Main /Users/kcft114/.julia/packages/Flux/qXNjB/src/optimise/train.jl:105
┌ Info: Epoch 1030
└ @ Main /Users/kcft114/.julia/packages/Flux/qXNjB/src/optimise/train.jl:105
┌ Info: Epoch 1031
└ @ Main /Users/kcft114/.julia/packages/Flux/qXNjB/src/optimise/train.jl:105
┌ Info: Epoch 1032
└ @ Main /Users/kcft114/.julia/packages/Flux/qXNjB/src/optimise/train.jl:105
┌ Info: Epoch 1033
└ @ Main /Users/kcft114/.julia/packages/Flux/qXNjB/src/optimise/train.jl:105
┌ Info: Epoch 1034
└ @ Main /Users/kcft114/.julia/packages/Flux/qXNjB/src/optimise/train.jl:105
┌ Info: Epoch 1035
└ @ Main /Users/kcft114/.julia/packages/Flux/qXNjB/src/optimise/train.jl:105
┌ Info: Epoch 1036
└ @ Main /Users/kcft114/.julia/packages/Flux/qXNjB/src/optimise/train.jl:105
┌ Info: Epoch 1037
└ @ Main /Users/kcft114/.julia/packages/Flux/qXNjB/src/optimise/train.jl:105
┌ Info: Epoch 1038
└ @ Main /Users/kcft114/.julia/packages/Flux/qXNjB/src/optimise/train.jl:105
┌ Info: Epoch 1039
└ @ Main /Users/kcft114/.julia/packages/Flux/qXNjB/src/optimise/train.jl:105
┌ Info: Epoch 1040
└ @ Main /Users/kcft114/.julia/packages/Flux/qXNjB/src/optimise/train.jl:105
┌ Info: Epoch 1041
└ @ Main /Users/kcft114/.julia/packages/Flux/qXNjB/src/optimise/train.jl:105
┌ Info: Epoch 1042
└ @ Main /Users/kcft114/.julia/packages/Flux/qXNjB/src/optimise/train.jl:105
┌ Info: Epoch 1043
└ @ Main /Users/kcft114/.julia/packages/Flux/qXNjB/src/optimise/train.jl:105
┌ Info: Epoch 1044
└ @ Main /Users/kcft114/.julia/packages/Flux/qXNjB/src/optimise/train.jl:105
┌ Info: Epoch 1045
└ @ Main /Users/kcft114/.julia/packages/Flux/qXNjB/src/optimise/train.jl:105
┌ Info: Epoch 1046
└ @ Main /Users/kcft114/.julia/packages/Flux/qXNjB/src/optimise/train.jl:105
┌ Info: Epoch 1047
└ @ Main /Users/kcft114/.julia/packages/Flux/qXNjB/src/optimise/train.jl:105
┌ Info: Epoch 1048
└ @ Main /Users/kcft114/.julia/packages/Flux/qXNjB/src/optimise/train.jl:105
┌ Info: Epoch 1049
└ @ Main /Users/kcft114/.julia/packages/Flux/qXNjB/src/optimise/train.jl:105
┌ Info: Epoch 1050
└ @ Main /Users/kcft114/.julia/packages/Flux/qXNjB/src/optimise/train.jl:105
┌ Info: Epoch 1051
└ @ Main /Users/kcft114/.julia/packages/Flux/qXNjB/src/optimise/train.jl:105
┌ Info: Epoch 1052
└ @ Main /Users/kcft114/.julia/packages/Flux/qXNjB/src/optimise/train.jl:105
┌ Info: Epoch 1053
└ @ Main /Users/kcft114/.julia/packages/Flux/qXNjB/src/optimise/train.jl:105
┌ Info: Epoch 1054
└ @ Main /Users/kcft114/.julia/packages/Flux/qXNjB/src/optimise/train.jl:105
┌ Info: Epoch 1055
└ @ Main /Users/kcft114/.julia/packages/Flux/qXNjB/src/optimise/train.jl:105
┌ Info: Epoch 1056
└ @ Main /Users/kcft114/.julia/packages/Flux/qXNjB/src/optimise/train.jl:105
┌ Info: Epoch 1057
└ @ Main /Users/kcft114/.julia/packages/Flux/qXNjB/src/optimise/train.jl:105
┌ Info: Epoch 1058
└ @ Main /Users/kcft114/.julia/packages/Flux/qXNjB/src/optimise/train.jl:105
┌ Info: Epoch 1059
└ @ Main /Users/kcft114/.julia/packages/Flux/qXNjB/src/optimise/train.jl:105
┌ Info: Epoch 1060
└ @ Main /Users/kcft114/.julia/packages/Flux/qXNjB/src/optimise/train.jl:105
┌ Info: Epoch 1061
└ @ Main /Users/kcft114/.julia/packages/Flux/qXNjB/src/optimise/train.jl:105
┌ Info: Epoch 1062
└ @ Main /Users/kcft114/.julia/packages/Flux/qXNjB/src/optimise/train.jl:105
┌ Info: Epoch 1063
└ @ Main /Users/kcft114/.julia/packages/Flux/qXNjB/src/optimise/train.jl:105
┌ Info: Epoch 1064
└ @ Main /Users/kcft114/.julia/packages/Flux/qXNjB/src/optimise/train.jl:105
┌ Info: Epoch 1065
└ @ Main /Users/kcft114/.julia/packages/Flux/qXNjB/src/optimise/train.jl:105
┌ Info: Epoch 1066
└ @ Main /Users/kcft114/.julia/packages/Flux/qXNjB/src/optimise/train.jl:105
┌ Info: Epoch 1067
└ @ Main /Users/kcft114/.julia/packages/Flux/qXNjB/src/optimise/train.jl:105
┌ Info: Epoch 1068
└ @ Main /Users/kcft114/.julia/packages/Flux/qXNjB/src/optimise/train.jl:105
┌ Info: Epoch 1069
└ @ Main /Users/kcft114/.julia/packages/Flux/qXNjB/src/optimise/train.jl:105
┌ Info: Epoch 1070
└ @ Main /Users/kcft114/.julia/packages/Flux/qXNjB/src/optimise/train.jl:105
┌ Info: Epoch 1071
└ @ Main /Users/kcft114/.julia/packages/Flux/qXNjB/src/optimise/train.jl:105
┌ Info: Epoch 1072
└ @ Main /Users/kcft114/.julia/packages/Flux/qXNjB/src/optimise/train.jl:105
┌ Info: Epoch 1073
└ @ Main /Users/kcft114/.julia/packages/Flux/qXNjB/src/optimise/train.jl:105
┌ Info: Epoch 1074
└ @ Main /Users/kcft114/.julia/packages/Flux/qXNjB/src/optimise/train.jl:105
┌ Info: Epoch 1075
└ @ Main /Users/kcft114/.julia/packages/Flux/qXNjB/src/optimise/train.jl:105
┌ Info: Epoch 1076
└ @ Main /Users/kcft114/.julia/packages/Flux/qXNjB/src/optimise/train.jl:105
┌ Info: Epoch 1077
└ @ Main /Users/kcft114/.julia/packages/Flux/qXNjB/src/optimise/train.jl:105
┌ Info: Epoch 1078
└ @ Main /Users/kcft114/.julia/packages/Flux/qXNjB/src/optimise/train.jl:105
┌ Info: Epoch 1079
└ @ Main /Users/kcft114/.julia/packages/Flux/qXNjB/src/optimise/train.jl:105
┌ Info: Epoch 1080
└ @ Main /Users/kcft114/.julia/packages/Flux/qXNjB/src/optimise/train.jl:105
┌ Info: Epoch 1081
└ @ Main /Users/kcft114/.julia/packages/Flux/qXNjB/src/optimise/train.jl:105
┌ Info: Epoch 1082
└ @ Main /Users/kcft114/.julia/packages/Flux/qXNjB/src/optimise/train.jl:105
┌ Info: Epoch 1083
└ @ Main /Users/kcft114/.julia/packages/Flux/qXNjB/src/optimise/train.jl:105
┌ Info: Epoch 1084
└ @ Main /Users/kcft114/.julia/packages/Flux/qXNjB/src/optimise/train.jl:105
┌ Info: Epoch 1085
└ @ Main /Users/kcft114/.julia/packages/Flux/qXNjB/src/optimise/train.jl:105
┌ Info: Epoch 1086
└ @ Main /Users/kcft114/.julia/packages/Flux/qXNjB/src/optimise/train.jl:105
┌ Info: Epoch 1087
└ @ Main /Users/kcft114/.julia/packages/Flux/qXNjB/src/optimise/train.jl:105
┌ Info: Epoch 1088
└ @ Main /Users/kcft114/.julia/packages/Flux/qXNjB/src/optimise/train.jl:105
┌ Info: Epoch 1089
└ @ Main /Users/kcft114/.julia/packages/Flux/qXNjB/src/optimise/train.jl:105
┌ Info: Epoch 1090
└ @ Main /Users/kcft114/.julia/packages/Flux/qXNjB/src/optimise/train.jl:105
┌ Info: Epoch 1091
└ @ Main /Users/kcft114/.julia/packages/Flux/qXNjB/src/optimise/train.jl:105
┌ Info: Epoch 1092
└ @ Main /Users/kcft114/.julia/packages/Flux/qXNjB/src/optimise/train.jl:105
┌ Info: Epoch 1093
└ @ Main /Users/kcft114/.julia/packages/Flux/qXNjB/src/optimise/train.jl:105
┌ Info: Epoch 1094
└ @ Main /Users/kcft114/.julia/packages/Flux/qXNjB/src/optimise/train.jl:105
┌ Info: Epoch 1095
└ @ Main /Users/kcft114/.julia/packages/Flux/qXNjB/src/optimise/train.jl:105
┌ Info: Epoch 1096
└ @ Main /Users/kcft114/.julia/packages/Flux/qXNjB/src/optimise/train.jl:105
┌ Info: Epoch 1097
└ @ Main /Users/kcft114/.julia/packages/Flux/qXNjB/src/optimise/train.jl:105
┌ Info: Epoch 1098
└ @ Main /Users/kcft114/.julia/packages/Flux/qXNjB/src/optimise/train.jl:105
┌ Info: Epoch 1099
└ @ Main /Users/kcft114/.julia/packages/Flux/qXNjB/src/optimise/train.jl:105
┌ Info: Epoch 1100
└ @ Main /Users/kcft114/.julia/packages/Flux/qXNjB/src/optimise/train.jl:105
┌ Info: Epoch 1101
└ @ Main /Users/kcft114/.julia/packages/Flux/qXNjB/src/optimise/train.jl:105
┌ Info: Epoch 1102
└ @ Main /Users/kcft114/.julia/packages/Flux/qXNjB/src/optimise/train.jl:105
┌ Info: Epoch 1103
└ @ Main /Users/kcft114/.julia/packages/Flux/qXNjB/src/optimise/train.jl:105
┌ Info: Epoch 1104
└ @ Main /Users/kcft114/.julia/packages/Flux/qXNjB/src/optimise/train.jl:105
┌ Info: Epoch 1105
└ @ Main /Users/kcft114/.julia/packages/Flux/qXNjB/src/optimise/train.jl:105
┌ Info: Epoch 1106
└ @ Main /Users/kcft114/.julia/packages/Flux/qXNjB/src/optimise/train.jl:105
┌ Info: Epoch 1107
└ @ Main /Users/kcft114/.julia/packages/Flux/qXNjB/src/optimise/train.jl:105
┌ Info: Epoch 1108
└ @ Main /Users/kcft114/.julia/packages/Flux/qXNjB/src/optimise/train.jl:105
┌ Info: Epoch 1109
└ @ Main /Users/kcft114/.julia/packages/Flux/qXNjB/src/optimise/train.jl:105
┌ Info: Epoch 1110
└ @ Main /Users/kcft114/.julia/packages/Flux/qXNjB/src/optimise/train.jl:105
┌ Info: Epoch 1111
└ @ Main /Users/kcft114/.julia/packages/Flux/qXNjB/src/optimise/train.jl:105
┌ Info: Epoch 1112
└ @ Main /Users/kcft114/.julia/packages/Flux/qXNjB/src/optimise/train.jl:105
┌ Info: Epoch 1113
└ @ Main /Users/kcft114/.julia/packages/Flux/qXNjB/src/optimise/train.jl:105
┌ Info: Epoch 1114
└ @ Main /Users/kcft114/.julia/packages/Flux/qXNjB/src/optimise/train.jl:105
┌ Info: Epoch 1115
└ @ Main /Users/kcft114/.julia/packages/Flux/qXNjB/src/optimise/train.jl:105
┌ Info: Epoch 1116
└ @ Main /Users/kcft114/.julia/packages/Flux/qXNjB/src/optimise/train.jl:105
┌ Info: Epoch 1117
└ @ Main /Users/kcft114/.julia/packages/Flux/qXNjB/src/optimise/train.jl:105
┌ Info: Epoch 1118
└ @ Main /Users/kcft114/.julia/packages/Flux/qXNjB/src/optimise/train.jl:105
┌ Info: Epoch 1119
└ @ Main /Users/kcft114/.julia/packages/Flux/qXNjB/src/optimise/train.jl:105
┌ Info: Epoch 1120
└ @ Main /Users/kcft114/.julia/packages/Flux/qXNjB/src/optimise/train.jl:105
┌ Info: Epoch 1121
└ @ Main /Users/kcft114/.julia/packages/Flux/qXNjB/src/optimise/train.jl:105
┌ Info: Epoch 1122
└ @ Main /Users/kcft114/.julia/packages/Flux/qXNjB/src/optimise/train.jl:105
┌ Info: Epoch 1123
└ @ Main /Users/kcft114/.julia/packages/Flux/qXNjB/src/optimise/train.jl:105
┌ Info: Epoch 1124
└ @ Main /Users/kcft114/.julia/packages/Flux/qXNjB/src/optimise/train.jl:105
┌ Info: Epoch 1125
└ @ Main /Users/kcft114/.julia/packages/Flux/qXNjB/src/optimise/train.jl:105
┌ Info: Epoch 1126
└ @ Main /Users/kcft114/.julia/packages/Flux/qXNjB/src/optimise/train.jl:105
```

```
loss(X, Y) = 0.6342584f0 (tracked)
loss(X, Y) = 0.6100471f0 (tracked)
loss(X, Y) = 0.5743927f0 (tracked)
loss(X, Y) = 0.60338724f0 (tracked)
loss(X, Y) = 0.66103387f0 (tracked)
loss(X, Y) = 0.6332143f0 (tracked)
loss(X, Y) = 0.5836014f0 (tracked)
loss(X, Y) = 0.627687f0 (tracked)
loss(X, Y) = 0.59136176f0 (tracked)
loss(X, Y) = 0.59577775f0 (tracked)
loss(X, Y) = 0.6291687f0 (tracked)
loss(X, Y) = 0.571512f0 (tracked)
loss(X, Y) = 0.6088274f0 (tracked)
loss(X, Y) = 0.6086012f0 (tracked)
loss(X, Y) = 0.62607485f0 (tracked)
loss(X, Y) = 0.6059503f0 (tracked)
loss(X, Y) = 0.5492709f0 (tracked)
loss(X, Y) = 0.6063944f0 (tracked)
loss(X, Y) = 0.5963595f0 (tracked)
loss(X, Y) = 0.61235654f0 (tracked)
loss(X, Y) = 0.6623973f0 (tracked)
loss(X, Y) = 0.6065814f0 (tracked)
loss(X, Y) = 0.62160856f0 (tracked)
loss(X, Y) = 0.5958918f0 (tracked)
loss(X, Y) = 0.62731594f0 (tracked)
loss(X, Y) = 0.5778279f0 (tracked)
loss(X, Y) = 0.61717194f0 (tracked)
loss(X, Y) = 0.61013323f0 (tracked)
loss(X, Y) = 0.59664375f0 (tracked)
loss(X, Y) = 0.6035633f0 (tracked)
loss(X, Y) = 0.5869877f0 (tracked)
loss(X, Y) = 0.6253967f0 (tracked)
loss(X, Y) = 0.60060114f0 (tracked)
loss(X, Y) = 0.585991f0 (tracked)
loss(X, Y) = 0.6145529f0 (tracked)
loss(X, Y) = 0.58903414f0 (tracked)
loss(X, Y) = 0.59844184f0 (tracked)
loss(X, Y) = 0.5792117f0 (tracked)
loss(X, Y) = 0.62106085f0 (tracked)
loss(X, Y) = 0.62147385f0 (tracked)
loss(X, Y) = 0.59311604f0 (tracked)
loss(X, Y) = 0.5863258f0 (tracked)
loss(X, Y) = 0.5947829f0 (tracked)
loss(X, Y) = 0.6000407f0 (tracked)
loss(X, Y) = 0.6005429f0 (tracked)
loss(X, Y) = 0.62032396f0 (tracked)
loss(X, Y) = 0.59938455f0 (tracked)
loss(X, Y) = 0.5961022f0 (tracked)
loss(X, Y) = 0.6244624f0 (tracked)
loss(X, Y) = 0.6149635f0 (tracked)
loss(X, Y) = 0.62472373f0 (tracked)
loss(X, Y) = 0.5846308f0 (tracked)
loss(X, Y) = 0.6147474f0 (tracked)
loss(X, Y) = 0.57550585f0 (tracked)
loss(X, Y) = 0.6069058f0 (tracked)
loss(X, Y) = 0.61042464f0 (tracked)
loss(X, Y) = 0.58896524f0 (tracked)
loss(X, Y) = 0.63808244f0 (tracked)
loss(X, Y) = 0.58152395f0 (tracked)
loss(X, Y) = 0.58154225f0 (tracked)
loss(X, Y) = 0.63306373f0 (tracked)
loss(X, Y) = 0.6075155f0 (tracked)
loss(X, Y) = 0.62971085f0 (tracked)
loss(X, Y) = 0.62086225f0 (tracked)
loss(X, Y) = 0.5976559f0 (tracked)
loss(X, Y) = 0.5551305f0 (tracked)
loss(X, Y) = 0.57361573f0 (tracked)
loss(X, Y) = 0.57305986f0 (tracked)
loss(X, Y) = 0.60185045f0 (tracked)
loss(X, Y) = 0.5981744f0 (tracked)
loss(X, Y) = 0.59820706f0 (tracked)
loss(X, Y) = 0.5810761f0 (tracked)
loss(X, Y) = 0.6032802f0 (tracked)
loss(X, Y) = 0.56740534f0 (tracked)
loss(X, Y) = 0.6120753f0 (tracked)
loss(X, Y) = 0.5433173f0 (tracked)
loss(X, Y) = 0.6423448f0 (tracked)
loss(X, Y) = 0.66343915f0 (tracked)
loss(X, Y) = 0.59221876f0 (tracked)
loss(X, Y) = 0.5800054f0 (tracked)
loss(X, Y) = 0.5969097f0 (tracked)
loss(X, Y) = 0.6274296f0 (tracked)
loss(X, Y) = 0.5639845f0 (tracked)
loss(X, Y) = 0.5740802f0 (tracked)
loss(X, Y) = 0.5681425f0 (tracked)
loss(X, Y) = 0.56226784f0 (tracked)
loss(X, Y) = 0.61768526f0 (tracked)
loss(X, Y) = 0.5841321f0 (tracked)
loss(X, Y) = 0.57400024f0 (tracked)
loss(X, Y) = 0.58986735f0 (tracked)
loss(X, Y) = 0.5750614f0 (tracked)
loss(X, Y) = 0.5902486f0 (tracked)
loss(X, Y) = 0.59133214f0 (tracked)
loss(X, Y) = 0.6061598f0 (tracked)
loss(X, Y) = 0.62975824f0 (tracked)
loss(X, Y) = 0.62122166f0 (tracked)
loss(X, Y) = 0.6304015f0 (tracked)
loss(X, Y) = 0.5886102f0 (tracked)
loss(X, Y) = 0.62842256f0 (tracked)
loss(X, Y) = 0.61559284f0 (tracked)
loss(X, Y) = 0.62092745f0 (tracked)
loss(X, Y) = 0.59126455f0 (tracked)
loss(X, Y) = 0.59060466f0 (tracked)
loss(X, Y) = 0.6230447f0 (tracked)
loss(X, Y) = 0.5982687f0 (tracked)
loss(X, Y) = 0.5848652f0 (tracked)
loss(X, Y) = 0.6022331f0 (tracked)
loss(X, Y) = 0.57191604f0 (tracked)
loss(X, Y) = 0.60250294f0 (tracked)
loss(X, Y) = 0.5655977f0 (tracked)
loss(X, Y) = 0.56180197f0 (tracked)
loss(X, Y) = 0.55513096f0 (tracked)
loss(X, Y) = 0.60521305f0 (tracked)
loss(X, Y) = 0.5719864f0 (tracked)
loss(X, Y) = 0.6095279f0 (tracked)
loss(X, Y) = 0.6154267f0 (tracked)
loss(X, Y) = 0.5878152f0 (tracked)
loss(X, Y) = 0.5764073f0 (tracked)
loss(X, Y) = 0.58760756f0 (tracked)
loss(X, Y) = 0.5651716f0 (tracked)
loss(X, Y) = 0.5426626f0 (tracked)
```

```
┌ Info: Epoch 1127
└ @ Main /Users/kcft114/.julia/packages/Flux/qXNjB/src/optimise/train.jl:105
┌ Info: Epoch 1128
└ @ Main /Users/kcft114/.julia/packages/Flux/qXNjB/src/optimise/train.jl:105
┌ Info: Epoch 1129
└ @ Main /Users/kcft114/.julia/packages/Flux/qXNjB/src/optimise/train.jl:105
┌ Info: Epoch 1130
└ @ Main /Users/kcft114/.julia/packages/Flux/qXNjB/src/optimise/train.jl:105
┌ Info: Epoch 1131
└ @ Main /Users/kcft114/.julia/packages/Flux/qXNjB/src/optimise/train.jl:105
┌ Info: Epoch 1132
└ @ Main /Users/kcft114/.julia/packages/Flux/qXNjB/src/optimise/train.jl:105
┌ Info: Epoch 1133
└ @ Main /Users/kcft114/.julia/packages/Flux/qXNjB/src/optimise/train.jl:105
┌ Info: Epoch 1134
└ @ Main /Users/kcft114/.julia/packages/Flux/qXNjB/src/optimise/train.jl:105
┌ Info: Epoch 1135
└ @ Main /Users/kcft114/.julia/packages/Flux/qXNjB/src/optimise/train.jl:105
┌ Info: Epoch 1136
└ @ Main /Users/kcft114/.julia/packages/Flux/qXNjB/src/optimise/train.jl:105
┌ Info: Epoch 1137
└ @ Main /Users/kcft114/.julia/packages/Flux/qXNjB/src/optimise/train.jl:105
┌ Info: Epoch 1138
└ @ Main /Users/kcft114/.julia/packages/Flux/qXNjB/src/optimise/train.jl:105
┌ Info: Epoch 1139
└ @ Main /Users/kcft114/.julia/packages/Flux/qXNjB/src/optimise/train.jl:105
┌ Info: Epoch 1140
└ @ Main /Users/kcft114/.julia/packages/Flux/qXNjB/src/optimise/train.jl:105
┌ Info: Epoch 1141
└ @ Main /Users/kcft114/.julia/packages/Flux/qXNjB/src/optimise/train.jl:105
┌ Info: Epoch 1142
└ @ Main /Users/kcft114/.julia/packages/Flux/qXNjB/src/optimise/train.jl:105
┌ Info: Epoch 1143
└ @ Main /Users/kcft114/.julia/packages/Flux/qXNjB/src/optimise/train.jl:105
┌ Info: Epoch 1144
└ @ Main /Users/kcft114/.julia/packages/Flux/qXNjB/src/optimise/train.jl:105
┌ Info: Epoch 1145
└ @ Main /Users/kcft114/.julia/packages/Flux/qXNjB/src/optimise/train.jl:105
┌ Info: Epoch 1146
└ @ Main /Users/kcft114/.julia/packages/Flux/qXNjB/src/optimise/train.jl:105
┌ Info: Epoch 1147
└ @ Main /Users/kcft114/.julia/packages/Flux/qXNjB/src/optimise/train.jl:105
┌ Info: Epoch 1148
└ @ Main /Users/kcft114/.julia/packages/Flux/qXNjB/src/optimise/train.jl:105
┌ Info: Epoch 1149
└ @ Main /Users/kcft114/.julia/packages/Flux/qXNjB/src/optimise/train.jl:105
┌ Info: Epoch 1150
└ @ Main /Users/kcft114/.julia/packages/Flux/qXNjB/src/optimise/train.jl:105
┌ Info: Epoch 1151
└ @ Main /Users/kcft114/.julia/packages/Flux/qXNjB/src/optimise/train.jl:105
┌ Info: Epoch 1152
└ @ Main /Users/kcft114/.julia/packages/Flux/qXNjB/src/optimise/train.jl:105
┌ Info: Epoch 1153
└ @ Main /Users/kcft114/.julia/packages/Flux/qXNjB/src/optimise/train.jl:105
┌ Info: Epoch 1154
└ @ Main /Users/kcft114/.julia/packages/Flux/qXNjB/src/optimise/train.jl:105
┌ Info: Epoch 1155
└ @ Main /Users/kcft114/.julia/packages/Flux/qXNjB/src/optimise/train.jl:105
┌ Info: Epoch 1156
└ @ Main /Users/kcft114/.julia/packages/Flux/qXNjB/src/optimise/train.jl:105
┌ Info: Epoch 1157
└ @ Main /Users/kcft114/.julia/packages/Flux/qXNjB/src/optimise/train.jl:105
┌ Info: Epoch 1158
└ @ Main /Users/kcft114/.julia/packages/Flux/qXNjB/src/optimise/train.jl:105
┌ Info: Epoch 1159
└ @ Main /Users/kcft114/.julia/packages/Flux/qXNjB/src/optimise/train.jl:105
┌ Info: Epoch 1160
└ @ Main /Users/kcft114/.julia/packages/Flux/qXNjB/src/optimise/train.jl:105
┌ Info: Epoch 1161
└ @ Main /Users/kcft114/.julia/packages/Flux/qXNjB/src/optimise/train.jl:105
┌ Info: Epoch 1162
└ @ Main /Users/kcft114/.julia/packages/Flux/qXNjB/src/optimise/train.jl:105
┌ Info: Epoch 1163
└ @ Main /Users/kcft114/.julia/packages/Flux/qXNjB/src/optimise/train.jl:105
┌ Info: Epoch 1164
└ @ Main /Users/kcft114/.julia/packages/Flux/qXNjB/src/optimise/train.jl:105
┌ Info: Epoch 1165
└ @ Main /Users/kcft114/.julia/packages/Flux/qXNjB/src/optimise/train.jl:105
┌ Info: Epoch 1166
└ @ Main /Users/kcft114/.julia/packages/Flux/qXNjB/src/optimise/train.jl:105
┌ Info: Epoch 1167
└ @ Main /Users/kcft114/.julia/packages/Flux/qXNjB/src/optimise/train.jl:105
┌ Info: Epoch 1168
└ @ Main /Users/kcft114/.julia/packages/Flux/qXNjB/src/optimise/train.jl:105
┌ Info: Epoch 1169
└ @ Main /Users/kcft114/.julia/packages/Flux/qXNjB/src/optimise/train.jl:105
┌ Info: Epoch 1170
└ @ Main /Users/kcft114/.julia/packages/Flux/qXNjB/src/optimise/train.jl:105
┌ Info: Epoch 1171
└ @ Main /Users/kcft114/.julia/packages/Flux/qXNjB/src/optimise/train.jl:105
┌ Info: Epoch 1172
└ @ Main /Users/kcft114/.julia/packages/Flux/qXNjB/src/optimise/train.jl:105
┌ Info: Epoch 1173
└ @ Main /Users/kcft114/.julia/packages/Flux/qXNjB/src/optimise/train.jl:105
┌ Info: Epoch 1174
└ @ Main /Users/kcft114/.julia/packages/Flux/qXNjB/src/optimise/train.jl:105
┌ Info: Epoch 1175
└ @ Main /Users/kcft114/.julia/packages/Flux/qXNjB/src/optimise/train.jl:105
┌ Info: Epoch 1176
└ @ Main /Users/kcft114/.julia/packages/Flux/qXNjB/src/optimise/train.jl:105
┌ Info: Epoch 1177
└ @ Main /Users/kcft114/.julia/packages/Flux/qXNjB/src/optimise/train.jl:105
┌ Info: Epoch 1178
└ @ Main /Users/kcft114/.julia/packages/Flux/qXNjB/src/optimise/train.jl:105
┌ Info: Epoch 1179
└ @ Main /Users/kcft114/.julia/packages/Flux/qXNjB/src/optimise/train.jl:105
┌ Info: Epoch 1180
└ @ Main /Users/kcft114/.julia/packages/Flux/qXNjB/src/optimise/train.jl:105
┌ Info: Epoch 1181
└ @ Main /Users/kcft114/.julia/packages/Flux/qXNjB/src/optimise/train.jl:105
┌ Info: Epoch 1182
└ @ Main /Users/kcft114/.julia/packages/Flux/qXNjB/src/optimise/train.jl:105
┌ Info: Epoch 1183
└ @ Main /Users/kcft114/.julia/packages/Flux/qXNjB/src/optimise/train.jl:105
┌ Info: Epoch 1184
└ @ Main /Users/kcft114/.julia/packages/Flux/qXNjB/src/optimise/train.jl:105
┌ Info: Epoch 1185
└ @ Main /Users/kcft114/.julia/packages/Flux/qXNjB/src/optimise/train.jl:105
┌ Info: Epoch 1186
└ @ Main /Users/kcft114/.julia/packages/Flux/qXNjB/src/optimise/train.jl:105
┌ Info: Epoch 1187
└ @ Main /Users/kcft114/.julia/packages/Flux/qXNjB/src/optimise/train.jl:105
┌ Info: Epoch 1188
└ @ Main /Users/kcft114/.julia/packages/Flux/qXNjB/src/optimise/train.jl:105
┌ Info: Epoch 1189
└ @ Main /Users/kcft114/.julia/packages/Flux/qXNjB/src/optimise/train.jl:105
┌ Info: Epoch 1190
└ @ Main /Users/kcft114/.julia/packages/Flux/qXNjB/src/optimise/train.jl:105
┌ Info: Epoch 1191
└ @ Main /Users/kcft114/.julia/packages/Flux/qXNjB/src/optimise/train.jl:105
┌ Info: Epoch 1192
└ @ Main /Users/kcft114/.julia/packages/Flux/qXNjB/src/optimise/train.jl:105
┌ Info: Epoch 1193
└ @ Main /Users/kcft114/.julia/packages/Flux/qXNjB/src/optimise/train.jl:105
┌ Info: Epoch 1194
└ @ Main /Users/kcft114/.julia/packages/Flux/qXNjB/src/optimise/train.jl:105
┌ Info: Epoch 1195
└ @ Main /Users/kcft114/.julia/packages/Flux/qXNjB/src/optimise/train.jl:105
┌ Info: Epoch 1196
└ @ Main /Users/kcft114/.julia/packages/Flux/qXNjB/src/optimise/train.jl:105
┌ Info: Epoch 1197
└ @ Main /Users/kcft114/.julia/packages/Flux/qXNjB/src/optimise/train.jl:105
┌ Info: Epoch 1198
└ @ Main /Users/kcft114/.julia/packages/Flux/qXNjB/src/optimise/train.jl:105
┌ Info: Epoch 1199
└ @ Main /Users/kcft114/.julia/packages/Flux/qXNjB/src/optimise/train.jl:105
┌ Info: Epoch 1200
└ @ Main /Users/kcft114/.julia/packages/Flux/qXNjB/src/optimise/train.jl:105
┌ Info: Epoch 1201
└ @ Main /Users/kcft114/.julia/packages/Flux/qXNjB/src/optimise/train.jl:105
┌ Info: Epoch 1202
└ @ Main /Users/kcft114/.julia/packages/Flux/qXNjB/src/optimise/train.jl:105
┌ Info: Epoch 1203
└ @ Main /Users/kcft114/.julia/packages/Flux/qXNjB/src/optimise/train.jl:105
┌ Info: Epoch 1204
└ @ Main /Users/kcft114/.julia/packages/Flux/qXNjB/src/optimise/train.jl:105
┌ Info: Epoch 1205
└ @ Main /Users/kcft114/.julia/packages/Flux/qXNjB/src/optimise/train.jl:105
┌ Info: Epoch 1206
└ @ Main /Users/kcft114/.julia/packages/Flux/qXNjB/src/optimise/train.jl:105
┌ Info: Epoch 1207
└ @ Main /Users/kcft114/.julia/packages/Flux/qXNjB/src/optimise/train.jl:105
┌ Info: Epoch 1208
└ @ Main /Users/kcft114/.julia/packages/Flux/qXNjB/src/optimise/train.jl:105
┌ Info: Epoch 1209
└ @ Main /Users/kcft114/.julia/packages/Flux/qXNjB/src/optimise/train.jl:105
┌ Info: Epoch 1210
└ @ Main /Users/kcft114/.julia/packages/Flux/qXNjB/src/optimise/train.jl:105
┌ Info: Epoch 1211
└ @ Main /Users/kcft114/.julia/packages/Flux/qXNjB/src/optimise/train.jl:105
┌ Info: Epoch 1212
└ @ Main /Users/kcft114/.julia/packages/Flux/qXNjB/src/optimise/train.jl:105
┌ Info: Epoch 1213
└ @ Main /Users/kcft114/.julia/packages/Flux/qXNjB/src/optimise/train.jl:105
┌ Info: Epoch 1214
└ @ Main /Users/kcft114/.julia/packages/Flux/qXNjB/src/optimise/train.jl:105
┌ Info: Epoch 1215
└ @ Main /Users/kcft114/.julia/packages/Flux/qXNjB/src/optimise/train.jl:105
┌ Info: Epoch 1216
└ @ Main /Users/kcft114/.julia/packages/Flux/qXNjB/src/optimise/train.jl:105
┌ Info: Epoch 1217
└ @ Main /Users/kcft114/.julia/packages/Flux/qXNjB/src/optimise/train.jl:105
┌ Info: Epoch 1218
└ @ Main /Users/kcft114/.julia/packages/Flux/qXNjB/src/optimise/train.jl:105
┌ Info: Epoch 1219
└ @ Main /Users/kcft114/.julia/packages/Flux/qXNjB/src/optimise/train.jl:105
┌ Info: Epoch 1220
└ @ Main /Users/kcft114/.julia/packages/Flux/qXNjB/src/optimise/train.jl:105
┌ Info: Epoch 1221
└ @ Main /Users/kcft114/.julia/packages/Flux/qXNjB/src/optimise/train.jl:105
┌ Info: Epoch 1222
└ @ Main /Users/kcft114/.julia/packages/Flux/qXNjB/src/optimise/train.jl:105
┌ Info: Epoch 1223
└ @ Main /Users/kcft114/.julia/packages/Flux/qXNjB/src/optimise/train.jl:105
┌ Info: Epoch 1224
└ @ Main /Users/kcft114/.julia/packages/Flux/qXNjB/src/optimise/train.jl:105
┌ Info: Epoch 1225
└ @ Main /Users/kcft114/.julia/packages/Flux/qXNjB/src/optimise/train.jl:105
┌ Info: Epoch 1226
└ @ Main /Users/kcft114/.julia/packages/Flux/qXNjB/src/optimise/train.jl:105
┌ Info: Epoch 1227
└ @ Main /Users/kcft114/.julia/packages/Flux/qXNjB/src/optimise/train.jl:105
┌ Info: Epoch 1228
└ @ Main /Users/kcft114/.julia/packages/Flux/qXNjB/src/optimise/train.jl:105
┌ Info: Epoch 1229
└ @ Main /Users/kcft114/.julia/packages/Flux/qXNjB/src/optimise/train.jl:105
┌ Info: Epoch 1230
└ @ Main /Users/kcft114/.julia/packages/Flux/qXNjB/src/optimise/train.jl:105
┌ Info: Epoch 1231
└ @ Main /Users/kcft114/.julia/packages/Flux/qXNjB/src/optimise/train.jl:105
┌ Info: Epoch 1232
└ @ Main /Users/kcft114/.julia/packages/Flux/qXNjB/src/optimise/train.jl:105
┌ Info: Epoch 1233
└ @ Main /Users/kcft114/.julia/packages/Flux/qXNjB/src/optimise/train.jl:105
┌ Info: Epoch 1234
└ @ Main /Users/kcft114/.julia/packages/Flux/qXNjB/src/optimise/train.jl:105
┌ Info: Epoch 1235
└ @ Main /Users/kcft114/.julia/packages/Flux/qXNjB/src/optimise/train.jl:105
┌ Info: Epoch 1236
└ @ Main /Users/kcft114/.julia/packages/Flux/qXNjB/src/optimise/train.jl:105
┌ Info: Epoch 1237
└ @ Main /Users/kcft114/.julia/packages/Flux/qXNjB/src/optimise/train.jl:105
┌ Info: Epoch 1238
└ @ Main /Users/kcft114/.julia/packages/Flux/qXNjB/src/optimise/train.jl:105
┌ Info: Epoch 1239
└ @ Main /Users/kcft114/.julia/packages/Flux/qXNjB/src/optimise/train.jl:105
┌ Info: Epoch 1240
└ @ Main /Users/kcft114/.julia/packages/Flux/qXNjB/src/optimise/train.jl:105
┌ Info: Epoch 1241
└ @ Main /Users/kcft114/.julia/packages/Flux/qXNjB/src/optimise/train.jl:105
┌ Info: Epoch 1242
└ @ Main /Users/kcft114/.julia/packages/Flux/qXNjB/src/optimise/train.jl:105
┌ Info: Epoch 1243
└ @ Main /Users/kcft114/.julia/packages/Flux/qXNjB/src/optimise/train.jl:105
┌ Info: Epoch 1244
└ @ Main /Users/kcft114/.julia/packages/Flux/qXNjB/src/optimise/train.jl:105
┌ Info: Epoch 1245
└ @ Main /Users/kcft114/.julia/packages/Flux/qXNjB/src/optimise/train.jl:105
┌ Info: Epoch 1246
└ @ Main /Users/kcft114/.julia/packages/Flux/qXNjB/src/optimise/train.jl:105
┌ Info: Epoch 1247
└ @ Main /Users/kcft114/.julia/packages/Flux/qXNjB/src/optimise/train.jl:105
```

```
loss(X, Y) = 0.56541705f0 (tracked)
loss(X, Y) = 0.5724768f0 (tracked)
loss(X, Y) = 0.5808937f0 (tracked)
loss(X, Y) = 0.59601957f0 (tracked)
loss(X, Y) = 0.5806582f0 (tracked)
loss(X, Y) = 0.6368052f0 (tracked)
loss(X, Y) = 0.61427444f0 (tracked)
loss(X, Y) = 0.60330397f0 (tracked)
loss(X, Y) = 0.5854293f0 (tracked)
loss(X, Y) = 0.6156212f0 (tracked)
loss(X, Y) = 0.54404f0 (tracked)
loss(X, Y) = 0.59334713f0 (tracked)
loss(X, Y) = 0.59659314f0 (tracked)
loss(X, Y) = 0.6046323f0 (tracked)
loss(X, Y) = 0.59667903f0 (tracked)
loss(X, Y) = 0.59289646f0 (tracked)
loss(X, Y) = 0.5352826f0 (tracked)
loss(X, Y) = 0.58465534f0 (tracked)
loss(X, Y) = 0.5993071f0 (tracked)
loss(X, Y) = 0.5947255f0 (tracked)
loss(X, Y) = 0.59006536f0 (tracked)
loss(X, Y) = 0.58037657f0 (tracked)
loss(X, Y) = 0.586942f0 (tracked)
loss(X, Y) = 0.57097614f0 (tracked)
loss(X, Y) = 0.6097242f0 (tracked)
loss(X, Y) = 0.6082016f0 (tracked)
loss(X, Y) = 0.6074071f0 (tracked)
loss(X, Y) = 0.5628141f0 (tracked)
loss(X, Y) = 0.6000342f0 (tracked)
loss(X, Y) = 0.5725687f0 (tracked)
loss(X, Y) = 0.6284488f0 (tracked)
loss(X, Y) = 0.5816038f0 (tracked)
loss(X, Y) = 0.5933464f0 (tracked)
loss(X, Y) = 0.60219747f0 (tracked)
loss(X, Y) = 0.58219194f0 (tracked)
loss(X, Y) = 0.57511765f0 (tracked)
loss(X, Y) = 0.5843098f0 (tracked)
loss(X, Y) = 0.5951559f0 (tracked)
loss(X, Y) = 0.5793516f0 (tracked)
loss(X, Y) = 0.5824401f0 (tracked)
loss(X, Y) = 0.5993062f0 (tracked)
loss(X, Y) = 0.5781801f0 (tracked)
loss(X, Y) = 0.5795268f0 (tracked)
loss(X, Y) = 0.593411f0 (tracked)
loss(X, Y) = 0.5951116f0 (tracked)
loss(X, Y) = 0.59530073f0 (tracked)
loss(X, Y) = 0.60652465f0 (tracked)
loss(X, Y) = 0.5881605f0 (tracked)
loss(X, Y) = 0.5859937f0 (tracked)
loss(X, Y) = 0.6216711f0 (tracked)
loss(X, Y) = 0.55040276f0 (tracked)
loss(X, Y) = 0.60249925f0 (tracked)
loss(X, Y) = 0.5981664f0 (tracked)
loss(X, Y) = 0.584434f0 (tracked)
loss(X, Y) = 0.57729536f0 (tracked)
loss(X, Y) = 0.5770302f0 (tracked)
loss(X, Y) = 0.5926836f0 (tracked)
loss(X, Y) = 0.6119087f0 (tracked)
loss(X, Y) = 0.5665427f0 (tracked)
loss(X, Y) = 0.60629404f0 (tracked)
loss(X, Y) = 0.59033835f0 (tracked)
loss(X, Y) = 0.6196918f0 (tracked)
loss(X, Y) = 0.5791689f0 (tracked)
loss(X, Y) = 0.59988075f0 (tracked)
loss(X, Y) = 0.5652021f0 (tracked)
loss(X, Y) = 0.5725986f0 (tracked)
loss(X, Y) = 0.5985429f0 (tracked)
loss(X, Y) = 0.542277f0 (tracked)
loss(X, Y) = 0.59417456f0 (tracked)
loss(X, Y) = 0.5733094f0 (tracked)
loss(X, Y) = 0.54991555f0 (tracked)
loss(X, Y) = 0.58059f0 (tracked)
loss(X, Y) = 0.5616037f0 (tracked)
loss(X, Y) = 0.5600451f0 (tracked)
loss(X, Y) = 0.59404397f0 (tracked)
loss(X, Y) = 0.5720818f0 (tracked)
loss(X, Y) = 0.61510414f0 (tracked)
loss(X, Y) = 0.5766286f0 (tracked)
loss(X, Y) = 0.59967244f0 (tracked)
loss(X, Y) = 0.5648584f0 (tracked)
loss(X, Y) = 0.5615878f0 (tracked)
loss(X, Y) = 0.598879f0 (tracked)
loss(X, Y) = 0.5780973f0 (tracked)
loss(X, Y) = 0.61021614f0 (tracked)
loss(X, Y) = 0.54497504f0 (tracked)
loss(X, Y) = 0.58620983f0 (tracked)
loss(X, Y) = 0.58695686f0 (tracked)
loss(X, Y) = 0.5885984f0 (tracked)
loss(X, Y) = 0.5853321f0 (tracked)
loss(X, Y) = 0.55765104f0 (tracked)
loss(X, Y) = 0.5579425f0 (tracked)
loss(X, Y) = 0.5843567f0 (tracked)
loss(X, Y) = 0.56015f0 (tracked)
loss(X, Y) = 0.54815274f0 (tracked)
loss(X, Y) = 0.62862647f0 (tracked)
loss(X, Y) = 0.56059515f0 (tracked)
loss(X, Y) = 0.62263167f0 (tracked)
loss(X, Y) = 0.5754779f0 (tracked)
loss(X, Y) = 0.59117466f0 (tracked)
loss(X, Y) = 0.55342877f0 (tracked)
loss(X, Y) = 0.56497955f0 (tracked)
loss(X, Y) = 0.5942473f0 (tracked)
loss(X, Y) = 0.6081458f0 (tracked)
loss(X, Y) = 0.5552867f0 (tracked)
loss(X, Y) = 0.5879219f0 (tracked)
loss(X, Y) = 0.59876287f0 (tracked)
loss(X, Y) = 0.6016442f0 (tracked)
loss(X, Y) = 0.60264355f0 (tracked)
loss(X, Y) = 0.59362996f0 (tracked)
```

```
┌ Info: Epoch 1248
└ @ Main /Users/kcft114/.julia/packages/Flux/qXNjB/src/optimise/train.jl:105
┌ Info: Epoch 1249
└ @ Main /Users/kcft114/.julia/packages/Flux/qXNjB/src/optimise/train.jl:105
┌ Info: Epoch 1250
└ @ Main /Users/kcft114/.julia/packages/Flux/qXNjB/src/optimise/train.jl:105
┌ Info: Epoch 1251
└ @ Main /Users/kcft114/.julia/packages/Flux/qXNjB/src/optimise/train.jl:105
┌ Info: Epoch 1252
└ @ Main /Users/kcft114/.julia/packages/Flux/qXNjB/src/optimise/train.jl:105
┌ Info: Epoch 1253
└ @ Main /Users/kcft114/.julia/packages/Flux/qXNjB/src/optimise/train.jl:105
┌ Info: Epoch 1254
└ @ Main /Users/kcft114/.julia/packages/Flux/qXNjB/src/optimise/train.jl:105
┌ Info: Epoch 1255
└ @ Main /Users/kcft114/.julia/packages/Flux/qXNjB/src/optimise/train.jl:105
┌ Info: Epoch 1256
└ @ Main /Users/kcft114/.julia/packages/Flux/qXNjB/src/optimise/train.jl:105
┌ Info: Epoch 1257
└ @ Main /Users/kcft114/.julia/packages/Flux/qXNjB/src/optimise/train.jl:105
┌ Info: Epoch 1258
└ @ Main /Users/kcft114/.julia/packages/Flux/qXNjB/src/optimise/train.jl:105
┌ Info: Epoch 1259
└ @ Main /Users/kcft114/.julia/packages/Flux/qXNjB/src/optimise/train.jl:105
┌ Info: Epoch 1260
└ @ Main /Users/kcft114/.julia/packages/Flux/qXNjB/src/optimise/train.jl:105
┌ Info: Epoch 1261
└ @ Main /Users/kcft114/.julia/packages/Flux/qXNjB/src/optimise/train.jl:105
┌ Info: Epoch 1262
└ @ Main /Users/kcft114/.julia/packages/Flux/qXNjB/src/optimise/train.jl:105
┌ Info: Epoch 1263
└ @ Main /Users/kcft114/.julia/packages/Flux/qXNjB/src/optimise/train.jl:105
┌ Info: Epoch 1264
└ @ Main /Users/kcft114/.julia/packages/Flux/qXNjB/src/optimise/train.jl:105
┌ Info: Epoch 1265
└ @ Main /Users/kcft114/.julia/packages/Flux/qXNjB/src/optimise/train.jl:105
┌ Info: Epoch 1266
└ @ Main /Users/kcft114/.julia/packages/Flux/qXNjB/src/optimise/train.jl:105
┌ Info: Epoch 1267
└ @ Main /Users/kcft114/.julia/packages/Flux/qXNjB/src/optimise/train.jl:105
┌ Info: Epoch 1268
└ @ Main /Users/kcft114/.julia/packages/Flux/qXNjB/src/optimise/train.jl:105
┌ Info: Epoch 1269
└ @ Main /Users/kcft114/.julia/packages/Flux/qXNjB/src/optimise/train.jl:105
┌ Info: Epoch 1270
└ @ Main /Users/kcft114/.julia/packages/Flux/qXNjB/src/optimise/train.jl:105
┌ Info: Epoch 1271
└ @ Main /Users/kcft114/.julia/packages/Flux/qXNjB/src/optimise/train.jl:105
┌ Info: Epoch 1272
└ @ Main /Users/kcft114/.julia/packages/Flux/qXNjB/src/optimise/train.jl:105
┌ Info: Epoch 1273
└ @ Main /Users/kcft114/.julia/packages/Flux/qXNjB/src/optimise/train.jl:105
┌ Info: Epoch 1274
└ @ Main /Users/kcft114/.julia/packages/Flux/qXNjB/src/optimise/train.jl:105
┌ Info: Epoch 1275
└ @ Main /Users/kcft114/.julia/packages/Flux/qXNjB/src/optimise/train.jl:105
┌ Info: Epoch 1276
└ @ Main /Users/kcft114/.julia/packages/Flux/qXNjB/src/optimise/train.jl:105
┌ Info: Epoch 1277
└ @ Main /Users/kcft114/.julia/packages/Flux/qXNjB/src/optimise/train.jl:105
┌ Info: Epoch 1278
└ @ Main /Users/kcft114/.julia/packages/Flux/qXNjB/src/optimise/train.jl:105
┌ Info: Epoch 1279
└ @ Main /Users/kcft114/.julia/packages/Flux/qXNjB/src/optimise/train.jl:105
┌ Info: Epoch 1280
└ @ Main /Users/kcft114/.julia/packages/Flux/qXNjB/src/optimise/train.jl:105
┌ Info: Epoch 1281
└ @ Main /Users/kcft114/.julia/packages/Flux/qXNjB/src/optimise/train.jl:105
┌ Info: Epoch 1282
└ @ Main /Users/kcft114/.julia/packages/Flux/qXNjB/src/optimise/train.jl:105
┌ Info: Epoch 1283
└ @ Main /Users/kcft114/.julia/packages/Flux/qXNjB/src/optimise/train.jl:105
┌ Info: Epoch 1284
└ @ Main /Users/kcft114/.julia/packages/Flux/qXNjB/src/optimise/train.jl:105
┌ Info: Epoch 1285
└ @ Main /Users/kcft114/.julia/packages/Flux/qXNjB/src/optimise/train.jl:105
┌ Info: Epoch 1286
└ @ Main /Users/kcft114/.julia/packages/Flux/qXNjB/src/optimise/train.jl:105
┌ Info: Epoch 1287
└ @ Main /Users/kcft114/.julia/packages/Flux/qXNjB/src/optimise/train.jl:105
┌ Info: Epoch 1288
└ @ Main /Users/kcft114/.julia/packages/Flux/qXNjB/src/optimise/train.jl:105
┌ Info: Epoch 1289
└ @ Main /Users/kcft114/.julia/packages/Flux/qXNjB/src/optimise/train.jl:105
┌ Info: Epoch 1290
└ @ Main /Users/kcft114/.julia/packages/Flux/qXNjB/src/optimise/train.jl:105
┌ Info: Epoch 1291
└ @ Main /Users/kcft114/.julia/packages/Flux/qXNjB/src/optimise/train.jl:105
┌ Info: Epoch 1292
└ @ Main /Users/kcft114/.julia/packages/Flux/qXNjB/src/optimise/train.jl:105
┌ Info: Epoch 1293
└ @ Main /Users/kcft114/.julia/packages/Flux/qXNjB/src/optimise/train.jl:105
┌ Info: Epoch 1294
└ @ Main /Users/kcft114/.julia/packages/Flux/qXNjB/src/optimise/train.jl:105
┌ Info: Epoch 1295
└ @ Main /Users/kcft114/.julia/packages/Flux/qXNjB/src/optimise/train.jl:105
┌ Info: Epoch 1296
└ @ Main /Users/kcft114/.julia/packages/Flux/qXNjB/src/optimise/train.jl:105
┌ Info: Epoch 1297
└ @ Main /Users/kcft114/.julia/packages/Flux/qXNjB/src/optimise/train.jl:105
┌ Info: Epoch 1298
└ @ Main /Users/kcft114/.julia/packages/Flux/qXNjB/src/optimise/train.jl:105
┌ Info: Epoch 1299
└ @ Main /Users/kcft114/.julia/packages/Flux/qXNjB/src/optimise/train.jl:105
┌ Info: Epoch 1300
└ @ Main /Users/kcft114/.julia/packages/Flux/qXNjB/src/optimise/train.jl:105
┌ Info: Epoch 1301
└ @ Main /Users/kcft114/.julia/packages/Flux/qXNjB/src/optimise/train.jl:105
┌ Info: Epoch 1302
└ @ Main /Users/kcft114/.julia/packages/Flux/qXNjB/src/optimise/train.jl:105
┌ Info: Epoch 1303
└ @ Main /Users/kcft114/.julia/packages/Flux/qXNjB/src/optimise/train.jl:105
┌ Info: Epoch 1304
└ @ Main /Users/kcft114/.julia/packages/Flux/qXNjB/src/optimise/train.jl:105
┌ Info: Epoch 1305
└ @ Main /Users/kcft114/.julia/packages/Flux/qXNjB/src/optimise/train.jl:105
┌ Info: Epoch 1306
└ @ Main /Users/kcft114/.julia/packages/Flux/qXNjB/src/optimise/train.jl:105
┌ Info: Epoch 1307
└ @ Main /Users/kcft114/.julia/packages/Flux/qXNjB/src/optimise/train.jl:105
┌ Info: Epoch 1308
└ @ Main /Users/kcft114/.julia/packages/Flux/qXNjB/src/optimise/train.jl:105
┌ Info: Epoch 1309
└ @ Main /Users/kcft114/.julia/packages/Flux/qXNjB/src/optimise/train.jl:105
┌ Info: Epoch 1310
└ @ Main /Users/kcft114/.julia/packages/Flux/qXNjB/src/optimise/train.jl:105
┌ Info: Epoch 1311
└ @ Main /Users/kcft114/.julia/packages/Flux/qXNjB/src/optimise/train.jl:105
┌ Info: Epoch 1312
└ @ Main /Users/kcft114/.julia/packages/Flux/qXNjB/src/optimise/train.jl:105
┌ Info: Epoch 1313
└ @ Main /Users/kcft114/.julia/packages/Flux/qXNjB/src/optimise/train.jl:105
┌ Info: Epoch 1314
└ @ Main /Users/kcft114/.julia/packages/Flux/qXNjB/src/optimise/train.jl:105
┌ Info: Epoch 1315
└ @ Main /Users/kcft114/.julia/packages/Flux/qXNjB/src/optimise/train.jl:105
┌ Info: Epoch 1316
└ @ Main /Users/kcft114/.julia/packages/Flux/qXNjB/src/optimise/train.jl:105
┌ Info: Epoch 1317
└ @ Main /Users/kcft114/.julia/packages/Flux/qXNjB/src/optimise/train.jl:105
┌ Info: Epoch 1318
└ @ Main /Users/kcft114/.julia/packages/Flux/qXNjB/src/optimise/train.jl:105
┌ Info: Epoch 1319
└ @ Main /Users/kcft114/.julia/packages/Flux/qXNjB/src/optimise/train.jl:105
┌ Info: Epoch 1320
└ @ Main /Users/kcft114/.julia/packages/Flux/qXNjB/src/optimise/train.jl:105
┌ Info: Epoch 1321
└ @ Main /Users/kcft114/.julia/packages/Flux/qXNjB/src/optimise/train.jl:105
┌ Info: Epoch 1322
└ @ Main /Users/kcft114/.julia/packages/Flux/qXNjB/src/optimise/train.jl:105
┌ Info: Epoch 1323
└ @ Main /Users/kcft114/.julia/packages/Flux/qXNjB/src/optimise/train.jl:105
┌ Info: Epoch 1324
└ @ Main /Users/kcft114/.julia/packages/Flux/qXNjB/src/optimise/train.jl:105
┌ Info: Epoch 1325
└ @ Main /Users/kcft114/.julia/packages/Flux/qXNjB/src/optimise/train.jl:105
┌ Info: Epoch 1326
└ @ Main /Users/kcft114/.julia/packages/Flux/qXNjB/src/optimise/train.jl:105
┌ Info: Epoch 1327
└ @ Main /Users/kcft114/.julia/packages/Flux/qXNjB/src/optimise/train.jl:105
┌ Info: Epoch 1328
└ @ Main /Users/kcft114/.julia/packages/Flux/qXNjB/src/optimise/train.jl:105
┌ Info: Epoch 1329
└ @ Main /Users/kcft114/.julia/packages/Flux/qXNjB/src/optimise/train.jl:105
┌ Info: Epoch 1330
└ @ Main /Users/kcft114/.julia/packages/Flux/qXNjB/src/optimise/train.jl:105
┌ Info: Epoch 1331
└ @ Main /Users/kcft114/.julia/packages/Flux/qXNjB/src/optimise/train.jl:105
┌ Info: Epoch 1332
└ @ Main /Users/kcft114/.julia/packages/Flux/qXNjB/src/optimise/train.jl:105
┌ Info: Epoch 1333
└ @ Main /Users/kcft114/.julia/packages/Flux/qXNjB/src/optimise/train.jl:105
┌ Info: Epoch 1334
└ @ Main /Users/kcft114/.julia/packages/Flux/qXNjB/src/optimise/train.jl:105
┌ Info: Epoch 1335
└ @ Main /Users/kcft114/.julia/packages/Flux/qXNjB/src/optimise/train.jl:105
┌ Info: Epoch 1336
└ @ Main /Users/kcft114/.julia/packages/Flux/qXNjB/src/optimise/train.jl:105
┌ Info: Epoch 1337
└ @ Main /Users/kcft114/.julia/packages/Flux/qXNjB/src/optimise/train.jl:105
┌ Info: Epoch 1338
└ @ Main /Users/kcft114/.julia/packages/Flux/qXNjB/src/optimise/train.jl:105
┌ Info: Epoch 1339
└ @ Main /Users/kcft114/.julia/packages/Flux/qXNjB/src/optimise/train.jl:105
┌ Info: Epoch 1340
└ @ Main /Users/kcft114/.julia/packages/Flux/qXNjB/src/optimise/train.jl:105
┌ Info: Epoch 1341
└ @ Main /Users/kcft114/.julia/packages/Flux/qXNjB/src/optimise/train.jl:105
┌ Info: Epoch 1342
└ @ Main /Users/kcft114/.julia/packages/Flux/qXNjB/src/optimise/train.jl:105
┌ Info: Epoch 1343
└ @ Main /Users/kcft114/.julia/packages/Flux/qXNjB/src/optimise/train.jl:105
┌ Info: Epoch 1344
└ @ Main /Users/kcft114/.julia/packages/Flux/qXNjB/src/optimise/train.jl:105
┌ Info: Epoch 1345
└ @ Main /Users/kcft114/.julia/packages/Flux/qXNjB/src/optimise/train.jl:105
┌ Info: Epoch 1346
└ @ Main /Users/kcft114/.julia/packages/Flux/qXNjB/src/optimise/train.jl:105
┌ Info: Epoch 1347
└ @ Main /Users/kcft114/.julia/packages/Flux/qXNjB/src/optimise/train.jl:105
┌ Info: Epoch 1348
└ @ Main /Users/kcft114/.julia/packages/Flux/qXNjB/src/optimise/train.jl:105
┌ Info: Epoch 1349
└ @ Main /Users/kcft114/.julia/packages/Flux/qXNjB/src/optimise/train.jl:105
┌ Info: Epoch 1350
└ @ Main /Users/kcft114/.julia/packages/Flux/qXNjB/src/optimise/train.jl:105
┌ Info: Epoch 1351
└ @ Main /Users/kcft114/.julia/packages/Flux/qXNjB/src/optimise/train.jl:105
┌ Info: Epoch 1352
└ @ Main /Users/kcft114/.julia/packages/Flux/qXNjB/src/optimise/train.jl:105
┌ Info: Epoch 1353
└ @ Main /Users/kcft114/.julia/packages/Flux/qXNjB/src/optimise/train.jl:105
┌ Info: Epoch 1354
└ @ Main /Users/kcft114/.julia/packages/Flux/qXNjB/src/optimise/train.jl:105
┌ Info: Epoch 1355
└ @ Main /Users/kcft114/.julia/packages/Flux/qXNjB/src/optimise/train.jl:105
┌ Info: Epoch 1356
└ @ Main /Users/kcft114/.julia/packages/Flux/qXNjB/src/optimise/train.jl:105
┌ Info: Epoch 1357
└ @ Main /Users/kcft114/.julia/packages/Flux/qXNjB/src/optimise/train.jl:105
```

```
loss(X, Y) = 0.5571126f0 (tracked)
loss(X, Y) = 0.60126173f0 (tracked)
loss(X, Y) = 0.56859326f0 (tracked)
loss(X, Y) = 0.5709616f0 (tracked)
loss(X, Y) = 0.57994264f0 (tracked)
loss(X, Y) = 0.5693078f0 (tracked)
loss(X, Y) = 0.54849404f0 (tracked)
loss(X, Y) = 0.5599943f0 (tracked)
loss(X, Y) = 0.5980838f0 (tracked)
loss(X, Y) = 0.59045905f0 (tracked)
loss(X, Y) = 0.5788911f0 (tracked)
loss(X, Y) = 0.5434814f0 (tracked)
loss(X, Y) = 0.5471271f0 (tracked)
loss(X, Y) = 0.59210634f0 (tracked)
loss(X, Y) = 0.5407355f0 (tracked)
loss(X, Y) = 0.5600149f0 (tracked)
loss(X, Y) = 0.56589913f0 (tracked)
loss(X, Y) = 0.5616453f0 (tracked)
loss(X, Y) = 0.57202274f0 (tracked)
loss(X, Y) = 0.5910732f0 (tracked)
loss(X, Y) = 0.56220984f0 (tracked)
loss(X, Y) = 0.59667194f0 (tracked)
loss(X, Y) = 0.60099643f0 (tracked)
loss(X, Y) = 0.561116f0 (tracked)
loss(X, Y) = 0.530143f0 (tracked)
loss(X, Y) = 0.5797102f0 (tracked)
loss(X, Y) = 0.62658334f0 (tracked)
loss(X, Y) = 0.5936493f0 (tracked)
loss(X, Y) = 0.5752187f0 (tracked)
loss(X, Y) = 0.5685297f0 (tracked)
loss(X, Y) = 0.562441f0 (tracked)
loss(X, Y) = 0.5551787f0 (tracked)
loss(X, Y) = 0.5689754f0 (tracked)
loss(X, Y) = 0.5728394f0 (tracked)
loss(X, Y) = 0.583244f0 (tracked)
loss(X, Y) = 0.5734752f0 (tracked)
loss(X, Y) = 0.6011585f0 (tracked)
loss(X, Y) = 0.5851446f0 (tracked)
loss(X, Y) = 0.58940023f0 (tracked)
loss(X, Y) = 0.5666196f0 (tracked)
loss(X, Y) = 0.55992466f0 (tracked)
loss(X, Y) = 0.5761431f0 (tracked)
loss(X, Y) = 0.615518f0 (tracked)
loss(X, Y) = 0.56503004f0 (tracked)
loss(X, Y) = 0.5772511f0 (tracked)
loss(X, Y) = 0.5767643f0 (tracked)
loss(X, Y) = 0.59658706f0 (tracked)
loss(X, Y) = 0.602058f0 (tracked)
loss(X, Y) = 0.5719899f0 (tracked)
loss(X, Y) = 0.5428865f0 (tracked)
loss(X, Y) = 0.572569f0 (tracked)
loss(X, Y) = 0.5990351f0 (tracked)
loss(X, Y) = 0.60918933f0 (tracked)
loss(X, Y) = 0.60273916f0 (tracked)
loss(X, Y) = 0.5788836f0 (tracked)
loss(X, Y) = 0.5561518f0 (tracked)
loss(X, Y) = 0.5707425f0 (tracked)
loss(X, Y) = 0.5906954f0 (tracked)
loss(X, Y) = 0.5660107f0 (tracked)
loss(X, Y) = 0.61152154f0 (tracked)
loss(X, Y) = 0.58309346f0 (tracked)
loss(X, Y) = 0.57502943f0 (tracked)
loss(X, Y) = 0.60292f0 (tracked)
loss(X, Y) = 0.58096075f0 (tracked)
loss(X, Y) = 0.5591439f0 (tracked)
loss(X, Y) = 0.5861309f0 (tracked)
loss(X, Y) = 0.55824316f0 (tracked)
loss(X, Y) = 0.5871869f0 (tracked)
loss(X, Y) = 0.5650655f0 (tracked)
loss(X, Y) = 0.55177665f0 (tracked)
loss(X, Y) = 0.5875009f0 (tracked)
loss(X, Y) = 0.5671796f0 (tracked)
loss(X, Y) = 0.5843062f0 (tracked)
loss(X, Y) = 0.5322078f0 (tracked)
loss(X, Y) = 0.5501069f0 (tracked)
loss(X, Y) = 0.57099324f0 (tracked)
loss(X, Y) = 0.5620876f0 (tracked)
loss(X, Y) = 0.55908394f0 (tracked)
loss(X, Y) = 0.52647257f0 (tracked)
loss(X, Y) = 0.6296977f0 (tracked)
loss(X, Y) = 0.56085336f0 (tracked)
loss(X, Y) = 0.604435f0 (tracked)
loss(X, Y) = 0.53000236f0 (tracked)
loss(X, Y) = 0.57377005f0 (tracked)
loss(X, Y) = 0.5850652f0 (tracked)
loss(X, Y) = 0.6083361f0 (tracked)
loss(X, Y) = 0.5523299f0 (tracked)
loss(X, Y) = 0.5841572f0 (tracked)
loss(X, Y) = 0.58865607f0 (tracked)
loss(X, Y) = 0.57274157f0 (tracked)
loss(X, Y) = 0.5563275f0 (tracked)
loss(X, Y) = 0.59102076f0 (tracked)
loss(X, Y) = 0.5741176f0 (tracked)
loss(X, Y) = 0.57760346f0 (tracked)
loss(X, Y) = 0.58425707f0 (tracked)
loss(X, Y) = 0.53366476f0 (tracked)
loss(X, Y) = 0.60491645f0 (tracked)
loss(X, Y) = 0.58357364f0 (tracked)
loss(X, Y) = 0.57195693f0 (tracked)
loss(X, Y) = 0.5652915f0 (tracked)
loss(X, Y) = 0.5667975f0 (tracked)
loss(X, Y) = 0.55971724f0 (tracked)
loss(X, Y) = 0.55679214f0 (tracked)
loss(X, Y) = 0.59409183f0 (tracked)
loss(X, Y) = 0.59243554f0 (tracked)
loss(X, Y) = 0.5851747f0 (tracked)
loss(X, Y) = 0.55817455f0 (tracked)
loss(X, Y) = 0.5716357f0 (tracked)
loss(X, Y) = 0.54397124f0 (tracked)
loss(X, Y) = 0.56245285f0 (tracked)
loss(X, Y) = 0.52189684f0 (tracked)
loss(X, Y) = 0.56143546f0 (tracked)
loss(X, Y) = 0.615014f0 (tracked)
loss(X, Y) = 0.5325562f0 (tracked)
loss(X, Y) = 0.525386f0 (tracked)
loss(X, Y) = 0.5536955f0 (tracked)
loss(X, Y) = 0.56217927f0 (tracked)
loss(X, Y) = 0.5968706f0 (tracked)
loss(X, Y) = 0.5804641f0 (tracked)
loss(X, Y) = 0.5723797f0 (tracked)
```

```
┌ Info: Epoch 1358
└ @ Main /Users/kcft114/.julia/packages/Flux/qXNjB/src/optimise/train.jl:105
┌ Info: Epoch 1359
└ @ Main /Users/kcft114/.julia/packages/Flux/qXNjB/src/optimise/train.jl:105
┌ Info: Epoch 1360
└ @ Main /Users/kcft114/.julia/packages/Flux/qXNjB/src/optimise/train.jl:105
┌ Info: Epoch 1361
└ @ Main /Users/kcft114/.julia/packages/Flux/qXNjB/src/optimise/train.jl:105
┌ Info: Epoch 1362
└ @ Main /Users/kcft114/.julia/packages/Flux/qXNjB/src/optimise/train.jl:105
┌ Info: Epoch 1363
└ @ Main /Users/kcft114/.julia/packages/Flux/qXNjB/src/optimise/train.jl:105
┌ Info: Epoch 1364
└ @ Main /Users/kcft114/.julia/packages/Flux/qXNjB/src/optimise/train.jl:105
┌ Info: Epoch 1365
└ @ Main /Users/kcft114/.julia/packages/Flux/qXNjB/src/optimise/train.jl:105
┌ Info: Epoch 1366
└ @ Main /Users/kcft114/.julia/packages/Flux/qXNjB/src/optimise/train.jl:105
┌ Info: Epoch 1367
└ @ Main /Users/kcft114/.julia/packages/Flux/qXNjB/src/optimise/train.jl:105
┌ Info: Epoch 1368
└ @ Main /Users/kcft114/.julia/packages/Flux/qXNjB/src/optimise/train.jl:105
┌ Info: Epoch 1369
└ @ Main /Users/kcft114/.julia/packages/Flux/qXNjB/src/optimise/train.jl:105
┌ Info: Epoch 1370
└ @ Main /Users/kcft114/.julia/packages/Flux/qXNjB/src/optimise/train.jl:105
┌ Info: Epoch 1371
└ @ Main /Users/kcft114/.julia/packages/Flux/qXNjB/src/optimise/train.jl:105
┌ Info: Epoch 1372
└ @ Main /Users/kcft114/.julia/packages/Flux/qXNjB/src/optimise/train.jl:105
┌ Info: Epoch 1373
└ @ Main /Users/kcft114/.julia/packages/Flux/qXNjB/src/optimise/train.jl:105
┌ Info: Epoch 1374
└ @ Main /Users/kcft114/.julia/packages/Flux/qXNjB/src/optimise/train.jl:105
┌ Info: Epoch 1375
└ @ Main /Users/kcft114/.julia/packages/Flux/qXNjB/src/optimise/train.jl:105
┌ Info: Epoch 1376
└ @ Main /Users/kcft114/.julia/packages/Flux/qXNjB/src/optimise/train.jl:105
┌ Info: Epoch 1377
└ @ Main /Users/kcft114/.julia/packages/Flux/qXNjB/src/optimise/train.jl:105
┌ Info: Epoch 1378
└ @ Main /Users/kcft114/.julia/packages/Flux/qXNjB/src/optimise/train.jl:105
┌ Info: Epoch 1379
└ @ Main /Users/kcft114/.julia/packages/Flux/qXNjB/src/optimise/train.jl:105
┌ Info: Epoch 1380
└ @ Main /Users/kcft114/.julia/packages/Flux/qXNjB/src/optimise/train.jl:105
┌ Info: Epoch 1381
└ @ Main /Users/kcft114/.julia/packages/Flux/qXNjB/src/optimise/train.jl:105
┌ Info: Epoch 1382
└ @ Main /Users/kcft114/.julia/packages/Flux/qXNjB/src/optimise/train.jl:105
┌ Info: Epoch 1383
└ @ Main /Users/kcft114/.julia/packages/Flux/qXNjB/src/optimise/train.jl:105
┌ Info: Epoch 1384
└ @ Main /Users/kcft114/.julia/packages/Flux/qXNjB/src/optimise/train.jl:105
┌ Info: Epoch 1385
└ @ Main /Users/kcft114/.julia/packages/Flux/qXNjB/src/optimise/train.jl:105
┌ Info: Epoch 1386
└ @ Main /Users/kcft114/.julia/packages/Flux/qXNjB/src/optimise/train.jl:105
┌ Info: Epoch 1387
└ @ Main /Users/kcft114/.julia/packages/Flux/qXNjB/src/optimise/train.jl:105
┌ Info: Epoch 1388
└ @ Main /Users/kcft114/.julia/packages/Flux/qXNjB/src/optimise/train.jl:105
┌ Info: Epoch 1389
└ @ Main /Users/kcft114/.julia/packages/Flux/qXNjB/src/optimise/train.jl:105
┌ Info: Epoch 1390
└ @ Main /Users/kcft114/.julia/packages/Flux/qXNjB/src/optimise/train.jl:105
┌ Info: Epoch 1391
└ @ Main /Users/kcft114/.julia/packages/Flux/qXNjB/src/optimise/train.jl:105
┌ Info: Epoch 1392
└ @ Main /Users/kcft114/.julia/packages/Flux/qXNjB/src/optimise/train.jl:105
┌ Info: Epoch 1393
└ @ Main /Users/kcft114/.julia/packages/Flux/qXNjB/src/optimise/train.jl:105
┌ Info: Epoch 1394
└ @ Main /Users/kcft114/.julia/packages/Flux/qXNjB/src/optimise/train.jl:105
┌ Info: Epoch 1395
└ @ Main /Users/kcft114/.julia/packages/Flux/qXNjB/src/optimise/train.jl:105
┌ Info: Epoch 1396
└ @ Main /Users/kcft114/.julia/packages/Flux/qXNjB/src/optimise/train.jl:105
┌ Info: Epoch 1397
└ @ Main /Users/kcft114/.julia/packages/Flux/qXNjB/src/optimise/train.jl:105
┌ Info: Epoch 1398
└ @ Main /Users/kcft114/.julia/packages/Flux/qXNjB/src/optimise/train.jl:105
┌ Info: Epoch 1399
└ @ Main /Users/kcft114/.julia/packages/Flux/qXNjB/src/optimise/train.jl:105
┌ Info: Epoch 1400
└ @ Main /Users/kcft114/.julia/packages/Flux/qXNjB/src/optimise/train.jl:105
┌ Info: Epoch 1401
└ @ Main /Users/kcft114/.julia/packages/Flux/qXNjB/src/optimise/train.jl:105
┌ Info: Epoch 1402
└ @ Main /Users/kcft114/.julia/packages/Flux/qXNjB/src/optimise/train.jl:105
┌ Info: Epoch 1403
└ @ Main /Users/kcft114/.julia/packages/Flux/qXNjB/src/optimise/train.jl:105
┌ Info: Epoch 1404
└ @ Main /Users/kcft114/.julia/packages/Flux/qXNjB/src/optimise/train.jl:105
┌ Info: Epoch 1405
└ @ Main /Users/kcft114/.julia/packages/Flux/qXNjB/src/optimise/train.jl:105
┌ Info: Epoch 1406
└ @ Main /Users/kcft114/.julia/packages/Flux/qXNjB/src/optimise/train.jl:105
┌ Info: Epoch 1407
└ @ Main /Users/kcft114/.julia/packages/Flux/qXNjB/src/optimise/train.jl:105
┌ Info: Epoch 1408
└ @ Main /Users/kcft114/.julia/packages/Flux/qXNjB/src/optimise/train.jl:105
┌ Info: Epoch 1409
└ @ Main /Users/kcft114/.julia/packages/Flux/qXNjB/src/optimise/train.jl:105
┌ Info: Epoch 1410
└ @ Main /Users/kcft114/.julia/packages/Flux/qXNjB/src/optimise/train.jl:105
┌ Info: Epoch 1411
└ @ Main /Users/kcft114/.julia/packages/Flux/qXNjB/src/optimise/train.jl:105
┌ Info: Epoch 1412
└ @ Main /Users/kcft114/.julia/packages/Flux/qXNjB/src/optimise/train.jl:105
┌ Info: Epoch 1413
└ @ Main /Users/kcft114/.julia/packages/Flux/qXNjB/src/optimise/train.jl:105
┌ Info: Epoch 1414
└ @ Main /Users/kcft114/.julia/packages/Flux/qXNjB/src/optimise/train.jl:105
┌ Info: Epoch 1415
└ @ Main /Users/kcft114/.julia/packages/Flux/qXNjB/src/optimise/train.jl:105
┌ Info: Epoch 1416
└ @ Main /Users/kcft114/.julia/packages/Flux/qXNjB/src/optimise/train.jl:105
┌ Info: Epoch 1417
└ @ Main /Users/kcft114/.julia/packages/Flux/qXNjB/src/optimise/train.jl:105
┌ Info: Epoch 1418
└ @ Main /Users/kcft114/.julia/packages/Flux/qXNjB/src/optimise/train.jl:105
┌ Info: Epoch 1419
└ @ Main /Users/kcft114/.julia/packages/Flux/qXNjB/src/optimise/train.jl:105
┌ Info: Epoch 1420
└ @ Main /Users/kcft114/.julia/packages/Flux/qXNjB/src/optimise/train.jl:105
┌ Info: Epoch 1421
└ @ Main /Users/kcft114/.julia/packages/Flux/qXNjB/src/optimise/train.jl:105
┌ Info: Epoch 1422
└ @ Main /Users/kcft114/.julia/packages/Flux/qXNjB/src/optimise/train.jl:105
┌ Info: Epoch 1423
└ @ Main /Users/kcft114/.julia/packages/Flux/qXNjB/src/optimise/train.jl:105
┌ Info: Epoch 1424
└ @ Main /Users/kcft114/.julia/packages/Flux/qXNjB/src/optimise/train.jl:105
┌ Info: Epoch 1425
└ @ Main /Users/kcft114/.julia/packages/Flux/qXNjB/src/optimise/train.jl:105
┌ Info: Epoch 1426
└ @ Main /Users/kcft114/.julia/packages/Flux/qXNjB/src/optimise/train.jl:105
┌ Info: Epoch 1427
└ @ Main /Users/kcft114/.julia/packages/Flux/qXNjB/src/optimise/train.jl:105
┌ Info: Epoch 1428
└ @ Main /Users/kcft114/.julia/packages/Flux/qXNjB/src/optimise/train.jl:105
┌ Info: Epoch 1429
└ @ Main /Users/kcft114/.julia/packages/Flux/qXNjB/src/optimise/train.jl:105
┌ Info: Epoch 1430
└ @ Main /Users/kcft114/.julia/packages/Flux/qXNjB/src/optimise/train.jl:105
┌ Info: Epoch 1431
└ @ Main /Users/kcft114/.julia/packages/Flux/qXNjB/src/optimise/train.jl:105
┌ Info: Epoch 1432
└ @ Main /Users/kcft114/.julia/packages/Flux/qXNjB/src/optimise/train.jl:105
┌ Info: Epoch 1433
└ @ Main /Users/kcft114/.julia/packages/Flux/qXNjB/src/optimise/train.jl:105
┌ Info: Epoch 1434
└ @ Main /Users/kcft114/.julia/packages/Flux/qXNjB/src/optimise/train.jl:105
┌ Info: Epoch 1435
└ @ Main /Users/kcft114/.julia/packages/Flux/qXNjB/src/optimise/train.jl:105
┌ Info: Epoch 1436
└ @ Main /Users/kcft114/.julia/packages/Flux/qXNjB/src/optimise/train.jl:105
┌ Info: Epoch 1437
└ @ Main /Users/kcft114/.julia/packages/Flux/qXNjB/src/optimise/train.jl:105
┌ Info: Epoch 1438
└ @ Main /Users/kcft114/.julia/packages/Flux/qXNjB/src/optimise/train.jl:105
┌ Info: Epoch 1439
└ @ Main /Users/kcft114/.julia/packages/Flux/qXNjB/src/optimise/train.jl:105
┌ Info: Epoch 1440
└ @ Main /Users/kcft114/.julia/packages/Flux/qXNjB/src/optimise/train.jl:105
┌ Info: Epoch 1441
└ @ Main /Users/kcft114/.julia/packages/Flux/qXNjB/src/optimise/train.jl:105
┌ Info: Epoch 1442
└ @ Main /Users/kcft114/.julia/packages/Flux/qXNjB/src/optimise/train.jl:105
┌ Info: Epoch 1443
└ @ Main /Users/kcft114/.julia/packages/Flux/qXNjB/src/optimise/train.jl:105
┌ Info: Epoch 1444
└ @ Main /Users/kcft114/.julia/packages/Flux/qXNjB/src/optimise/train.jl:105
┌ Info: Epoch 1445
└ @ Main /Users/kcft114/.julia/packages/Flux/qXNjB/src/optimise/train.jl:105
┌ Info: Epoch 1446
└ @ Main /Users/kcft114/.julia/packages/Flux/qXNjB/src/optimise/train.jl:105
┌ Info: Epoch 1447
└ @ Main /Users/kcft114/.julia/packages/Flux/qXNjB/src/optimise/train.jl:105
┌ Info: Epoch 1448
└ @ Main /Users/kcft114/.julia/packages/Flux/qXNjB/src/optimise/train.jl:105
┌ Info: Epoch 1449
└ @ Main /Users/kcft114/.julia/packages/Flux/qXNjB/src/optimise/train.jl:105
┌ Info: Epoch 1450
└ @ Main /Users/kcft114/.julia/packages/Flux/qXNjB/src/optimise/train.jl:105
┌ Info: Epoch 1451
└ @ Main /Users/kcft114/.julia/packages/Flux/qXNjB/src/optimise/train.jl:105
┌ Info: Epoch 1452
└ @ Main /Users/kcft114/.julia/packages/Flux/qXNjB/src/optimise/train.jl:105
┌ Info: Epoch 1453
└ @ Main /Users/kcft114/.julia/packages/Flux/qXNjB/src/optimise/train.jl:105
┌ Info: Epoch 1454
└ @ Main /Users/kcft114/.julia/packages/Flux/qXNjB/src/optimise/train.jl:105
┌ Info: Epoch 1455
└ @ Main /Users/kcft114/.julia/packages/Flux/qXNjB/src/optimise/train.jl:105
┌ Info: Epoch 1456
└ @ Main /Users/kcft114/.julia/packages/Flux/qXNjB/src/optimise/train.jl:105
┌ Info: Epoch 1457
└ @ Main /Users/kcft114/.julia/packages/Flux/qXNjB/src/optimise/train.jl:105
┌ Info: Epoch 1458
└ @ Main /Users/kcft114/.julia/packages/Flux/qXNjB/src/optimise/train.jl:105
┌ Info: Epoch 1459
└ @ Main /Users/kcft114/.julia/packages/Flux/qXNjB/src/optimise/train.jl:105
┌ Info: Epoch 1460
└ @ Main /Users/kcft114/.julia/packages/Flux/qXNjB/src/optimise/train.jl:105
┌ Info: Epoch 1461
└ @ Main /Users/kcft114/.julia/packages/Flux/qXNjB/src/optimise/train.jl:105
┌ Info: Epoch 1462
└ @ Main /Users/kcft114/.julia/packages/Flux/qXNjB/src/optimise/train.jl:105
┌ Info: Epoch 1463
└ @ Main /Users/kcft114/.julia/packages/Flux/qXNjB/src/optimise/train.jl:105
┌ Info: Epoch 1464
└ @ Main /Users/kcft114/.julia/packages/Flux/qXNjB/src/optimise/train.jl:105
┌ Info: Epoch 1465
└ @ Main /Users/kcft114/.julia/packages/Flux/qXNjB/src/optimise/train.jl:105
┌ Info: Epoch 1466
└ @ Main /Users/kcft114/.julia/packages/Flux/qXNjB/src/optimise/train.jl:105
┌ Info: Epoch 1467
└ @ Main /Users/kcft114/.julia/packages/Flux/qXNjB/src/optimise/train.jl:105
┌ Info: Epoch 1468
└ @ Main /Users/kcft114/.julia/packages/Flux/qXNjB/src/optimise/train.jl:105
┌ Info: Epoch 1469
└ @ Main /Users/kcft114/.julia/packages/Flux/qXNjB/src/optimise/train.jl:105
┌ Info: Epoch 1470
└ @ Main /Users/kcft114/.julia/packages/Flux/qXNjB/src/optimise/train.jl:105
┌ Info: Epoch 1471
└ @ Main /Users/kcft114/.julia/packages/Flux/qXNjB/src/optimise/train.jl:105
┌ Info: Epoch 1472
└ @ Main /Users/kcft114/.julia/packages/Flux/qXNjB/src/optimise/train.jl:105
┌ Info: Epoch 1473
└ @ Main /Users/kcft114/.julia/packages/Flux/qXNjB/src/optimise/train.jl:105
┌ Info: Epoch 1474
└ @ Main /Users/kcft114/.julia/packages/Flux/qXNjB/src/optimise/train.jl:105
┌ Info: Epoch 1475
└ @ Main /Users/kcft114/.julia/packages/Flux/qXNjB/src/optimise/train.jl:105
┌ Info: Epoch 1476
└ @ Main /Users/kcft114/.julia/packages/Flux/qXNjB/src/optimise/train.jl:105
┌ Info: Epoch 1477
└ @ Main /Users/kcft114/.julia/packages/Flux/qXNjB/src/optimise/train.jl:105
```

```
loss(X, Y) = 0.5514951f0 (tracked)
loss(X, Y) = 0.55080897f0 (tracked)
loss(X, Y) = 0.56900835f0 (tracked)
loss(X, Y) = 0.5533264f0 (tracked)
loss(X, Y) = 0.52314055f0 (tracked)
loss(X, Y) = 0.5495971f0 (tracked)
loss(X, Y) = 0.52240825f0 (tracked)
loss(X, Y) = 0.5511388f0 (tracked)
loss(X, Y) = 0.614064f0 (tracked)
loss(X, Y) = 0.5609705f0 (tracked)
loss(X, Y) = 0.54433876f0 (tracked)
loss(X, Y) = 0.53852856f0 (tracked)
loss(X, Y) = 0.5566279f0 (tracked)
loss(X, Y) = 0.59705526f0 (tracked)
loss(X, Y) = 0.5617155f0 (tracked)
loss(X, Y) = 0.5790571f0 (tracked)
loss(X, Y) = 0.5198644f0 (tracked)
loss(X, Y) = 0.55153793f0 (tracked)
loss(X, Y) = 0.5564333f0 (tracked)
loss(X, Y) = 0.57253027f0 (tracked)
loss(X, Y) = 0.5917568f0 (tracked)
loss(X, Y) = 0.5738376f0 (tracked)
loss(X, Y) = 0.52571017f0 (tracked)
loss(X, Y) = 0.5414072f0 (tracked)
loss(X, Y) = 0.558963f0 (tracked)
loss(X, Y) = 0.54600304f0 (tracked)
loss(X, Y) = 0.5470759f0 (tracked)
loss(X, Y) = 0.5539421f0 (tracked)
loss(X, Y) = 0.54427856f0 (tracked)
loss(X, Y) = 0.5672531f0 (tracked)
loss(X, Y) = 0.5762415f0 (tracked)
loss(X, Y) = 0.58018357f0 (tracked)
loss(X, Y) = 0.57948625f0 (tracked)
loss(X, Y) = 0.5893142f0 (tracked)
loss(X, Y) = 0.5574788f0 (tracked)
loss(X, Y) = 0.56110215f0 (tracked)
loss(X, Y) = 0.5609954f0 (tracked)
loss(X, Y) = 0.5583697f0 (tracked)
loss(X, Y) = 0.61797464f0 (tracked)
loss(X, Y) = 0.5634282f0 (tracked)
loss(X, Y) = 0.59012413f0 (tracked)
loss(X, Y) = 0.6075965f0 (tracked)
loss(X, Y) = 0.6067553f0 (tracked)
loss(X, Y) = 0.5395504f0 (tracked)
loss(X, Y) = 0.5716582f0 (tracked)
loss(X, Y) = 0.5354849f0 (tracked)
loss(X, Y) = 0.5501178f0 (tracked)
loss(X, Y) = 0.5479272f0 (tracked)
loss(X, Y) = 0.5174204f0 (tracked)
loss(X, Y) = 0.5570159f0 (tracked)
loss(X, Y) = 0.54282403f0 (tracked)
loss(X, Y) = 0.5640159f0 (tracked)
loss(X, Y) = 0.5472239f0 (tracked)
loss(X, Y) = 0.56223327f0 (tracked)
loss(X, Y) = 0.5385872f0 (tracked)
loss(X, Y) = 0.5234544f0 (tracked)
loss(X, Y) = 0.58306795f0 (tracked)
loss(X, Y) = 0.5312479f0 (tracked)
loss(X, Y) = 0.5671994f0 (tracked)
loss(X, Y) = 0.5623636f0 (tracked)
loss(X, Y) = 0.5929025f0 (tracked)
loss(X, Y) = 0.54919136f0 (tracked)
loss(X, Y) = 0.55539995f0 (tracked)
loss(X, Y) = 0.5988652f0 (tracked)
loss(X, Y) = 0.5769618f0 (tracked)
loss(X, Y) = 0.5803049f0 (tracked)
loss(X, Y) = 0.5188106f0 (tracked)
loss(X, Y) = 0.5571085f0 (tracked)
loss(X, Y) = 0.5671391f0 (tracked)
loss(X, Y) = 0.5445621f0 (tracked)
loss(X, Y) = 0.5859621f0 (tracked)
loss(X, Y) = 0.58193636f0 (tracked)
loss(X, Y) = 0.60218686f0 (tracked)
loss(X, Y) = 0.53489274f0 (tracked)
loss(X, Y) = 0.55906713f0 (tracked)
loss(X, Y) = 0.51374185f0 (tracked)
loss(X, Y) = 0.5825858f0 (tracked)
loss(X, Y) = 0.5581217f0 (tracked)
loss(X, Y) = 0.55401254f0 (tracked)
loss(X, Y) = 0.5621689f0 (tracked)
loss(X, Y) = 0.565856f0 (tracked)
loss(X, Y) = 0.55490714f0 (tracked)
loss(X, Y) = 0.53405535f0 (tracked)
loss(X, Y) = 0.585103f0 (tracked)
loss(X, Y) = 0.51783925f0 (tracked)
loss(X, Y) = 0.5458106f0 (tracked)
loss(X, Y) = 0.58238864f0 (tracked)
loss(X, Y) = 0.5490778f0 (tracked)
loss(X, Y) = 0.54604405f0 (tracked)
loss(X, Y) = 0.5325968f0 (tracked)
loss(X, Y) = 0.5524759f0 (tracked)
loss(X, Y) = 0.57645184f0 (tracked)
loss(X, Y) = 0.56708455f0 (tracked)
loss(X, Y) = 0.5526721f0 (tracked)
loss(X, Y) = 0.50923955f0 (tracked)
loss(X, Y) = 0.5318709f0 (tracked)
loss(X, Y) = 0.5908952f0 (tracked)
loss(X, Y) = 0.5549035f0 (tracked)
loss(X, Y) = 0.5874425f0 (tracked)
loss(X, Y) = 0.51991117f0 (tracked)
loss(X, Y) = 0.5729532f0 (tracked)
loss(X, Y) = 0.5215764f0 (tracked)
loss(X, Y) = 0.6038299f0 (tracked)
loss(X, Y) = 0.57721615f0 (tracked)
loss(X, Y) = 0.6213169f0 (tracked)
loss(X, Y) = 0.5449814f0 (tracked)
loss(X, Y) = 0.5243829f0 (tracked)
loss(X, Y) = 0.55772084f0 (tracked)
loss(X, Y) = 0.5161825f0 (tracked)
loss(X, Y) = 0.55183524f0 (tracked)
loss(X, Y) = 0.57215506f0 (tracked)
loss(X, Y) = 0.60077196f0 (tracked)
loss(X, Y) = 0.59158975f0 (tracked)
loss(X, Y) = 0.5572201f0 (tracked)
loss(X, Y) = 0.54323214f0 (tracked)
loss(X, Y) = 0.5911378f0 (tracked)
loss(X, Y) = 0.5592029f0 (tracked)
loss(X, Y) = 0.5477031f0 (tracked)
loss(X, Y) = 0.52076733f0 (tracked)
loss(X, Y) = 0.56321484f0 (tracked)
loss(X, Y) = 0.58599144f0 (tracked)
loss(X, Y) = 0.57141775f0 (tracked)
loss(X, Y) = 0.5662069f0 (tracked)
loss(X, Y) = 0.52620625f0 (tracked)
loss(X, Y) = 0.5588087f0 (tracked)
loss(X, Y) = 0.57544196f0 (tracked)
loss(X, Y) = 0.5689148f0 (tracked)
loss(X, Y) = 0.5308664f0 (tracked)
loss(X, Y) = 0.5204285f0 (tracked)
loss(X, Y) = 0.5576474f0 (tracked)
loss(X, Y) = 0.53212f0 (tracked)
loss(X, Y) = 0.5397812f0 (tracked)
```

```
┌ Info: Epoch 1478
└ @ Main /Users/kcft114/.julia/packages/Flux/qXNjB/src/optimise/train.jl:105
┌ Info: Epoch 1479
└ @ Main /Users/kcft114/.julia/packages/Flux/qXNjB/src/optimise/train.jl:105
┌ Info: Epoch 1480
└ @ Main /Users/kcft114/.julia/packages/Flux/qXNjB/src/optimise/train.jl:105
┌ Info: Epoch 1481
└ @ Main /Users/kcft114/.julia/packages/Flux/qXNjB/src/optimise/train.jl:105
┌ Info: Epoch 1482
└ @ Main /Users/kcft114/.julia/packages/Flux/qXNjB/src/optimise/train.jl:105
┌ Info: Epoch 1483
└ @ Main /Users/kcft114/.julia/packages/Flux/qXNjB/src/optimise/train.jl:105
┌ Info: Epoch 1484
└ @ Main /Users/kcft114/.julia/packages/Flux/qXNjB/src/optimise/train.jl:105
┌ Info: Epoch 1485
└ @ Main /Users/kcft114/.julia/packages/Flux/qXNjB/src/optimise/train.jl:105
┌ Info: Epoch 1486
└ @ Main /Users/kcft114/.julia/packages/Flux/qXNjB/src/optimise/train.jl:105
┌ Info: Epoch 1487
└ @ Main /Users/kcft114/.julia/packages/Flux/qXNjB/src/optimise/train.jl:105
┌ Info: Epoch 1488
└ @ Main /Users/kcft114/.julia/packages/Flux/qXNjB/src/optimise/train.jl:105
┌ Info: Epoch 1489
└ @ Main /Users/kcft114/.julia/packages/Flux/qXNjB/src/optimise/train.jl:105
┌ Info: Epoch 1490
└ @ Main /Users/kcft114/.julia/packages/Flux/qXNjB/src/optimise/train.jl:105
┌ Info: Epoch 1491
└ @ Main /Users/kcft114/.julia/packages/Flux/qXNjB/src/optimise/train.jl:105
┌ Info: Epoch 1492
└ @ Main /Users/kcft114/.julia/packages/Flux/qXNjB/src/optimise/train.jl:105
┌ Info: Epoch 1493
└ @ Main /Users/kcft114/.julia/packages/Flux/qXNjB/src/optimise/train.jl:105
┌ Info: Epoch 1494
└ @ Main /Users/kcft114/.julia/packages/Flux/qXNjB/src/optimise/train.jl:105
┌ Info: Epoch 1495
└ @ Main /Users/kcft114/.julia/packages/Flux/qXNjB/src/optimise/train.jl:105
┌ Info: Epoch 1496
└ @ Main /Users/kcft114/.julia/packages/Flux/qXNjB/src/optimise/train.jl:105
┌ Info: Epoch 1497
└ @ Main /Users/kcft114/.julia/packages/Flux/qXNjB/src/optimise/train.jl:105
┌ Info: Epoch 1498
└ @ Main /Users/kcft114/.julia/packages/Flux/qXNjB/src/optimise/train.jl:105
┌ Info: Epoch 1499
└ @ Main /Users/kcft114/.julia/packages/Flux/qXNjB/src/optimise/train.jl:105
┌ Info: Epoch 1500
└ @ Main /Users/kcft114/.julia/packages/Flux/qXNjB/src/optimise/train.jl:105
┌ Info: Epoch 1501
└ @ Main /Users/kcft114/.julia/packages/Flux/qXNjB/src/optimise/train.jl:105
┌ Info: Epoch 1502
└ @ Main /Users/kcft114/.julia/packages/Flux/qXNjB/src/optimise/train.jl:105
┌ Info: Epoch 1503
└ @ Main /Users/kcft114/.julia/packages/Flux/qXNjB/src/optimise/train.jl:105
┌ Info: Epoch 1504
└ @ Main /Users/kcft114/.julia/packages/Flux/qXNjB/src/optimise/train.jl:105
┌ Info: Epoch 1505
└ @ Main /Users/kcft114/.julia/packages/Flux/qXNjB/src/optimise/train.jl:105
┌ Info: Epoch 1506
└ @ Main /Users/kcft114/.julia/packages/Flux/qXNjB/src/optimise/train.jl:105
┌ Info: Epoch 1507
└ @ Main /Users/kcft114/.julia/packages/Flux/qXNjB/src/optimise/train.jl:105
┌ Info: Epoch 1508
└ @ Main /Users/kcft114/.julia/packages/Flux/qXNjB/src/optimise/train.jl:105
┌ Info: Epoch 1509
└ @ Main /Users/kcft114/.julia/packages/Flux/qXNjB/src/optimise/train.jl:105
┌ Info: Epoch 1510
└ @ Main /Users/kcft114/.julia/packages/Flux/qXNjB/src/optimise/train.jl:105
┌ Info: Epoch 1511
└ @ Main /Users/kcft114/.julia/packages/Flux/qXNjB/src/optimise/train.jl:105
┌ Info: Epoch 1512
└ @ Main /Users/kcft114/.julia/packages/Flux/qXNjB/src/optimise/train.jl:105
┌ Info: Epoch 1513
└ @ Main /Users/kcft114/.julia/packages/Flux/qXNjB/src/optimise/train.jl:105
┌ Info: Epoch 1514
└ @ Main /Users/kcft114/.julia/packages/Flux/qXNjB/src/optimise/train.jl:105
┌ Info: Epoch 1515
└ @ Main /Users/kcft114/.julia/packages/Flux/qXNjB/src/optimise/train.jl:105
┌ Info: Epoch 1516
└ @ Main /Users/kcft114/.julia/packages/Flux/qXNjB/src/optimise/train.jl:105
┌ Info: Epoch 1517
└ @ Main /Users/kcft114/.julia/packages/Flux/qXNjB/src/optimise/train.jl:105
┌ Info: Epoch 1518
└ @ Main /Users/kcft114/.julia/packages/Flux/qXNjB/src/optimise/train.jl:105
┌ Info: Epoch 1519
└ @ Main /Users/kcft114/.julia/packages/Flux/qXNjB/src/optimise/train.jl:105
┌ Info: Epoch 1520
└ @ Main /Users/kcft114/.julia/packages/Flux/qXNjB/src/optimise/train.jl:105
┌ Info: Epoch 1521
└ @ Main /Users/kcft114/.julia/packages/Flux/qXNjB/src/optimise/train.jl:105
┌ Info: Epoch 1522
└ @ Main /Users/kcft114/.julia/packages/Flux/qXNjB/src/optimise/train.jl:105
┌ Info: Epoch 1523
└ @ Main /Users/kcft114/.julia/packages/Flux/qXNjB/src/optimise/train.jl:105
┌ Info: Epoch 1524
└ @ Main /Users/kcft114/.julia/packages/Flux/qXNjB/src/optimise/train.jl:105
┌ Info: Epoch 1525
└ @ Main /Users/kcft114/.julia/packages/Flux/qXNjB/src/optimise/train.jl:105
┌ Info: Epoch 1526
└ @ Main /Users/kcft114/.julia/packages/Flux/qXNjB/src/optimise/train.jl:105
┌ Info: Epoch 1527
└ @ Main /Users/kcft114/.julia/packages/Flux/qXNjB/src/optimise/train.jl:105
┌ Info: Epoch 1528
└ @ Main /Users/kcft114/.julia/packages/Flux/qXNjB/src/optimise/train.jl:105
┌ Info: Epoch 1529
└ @ Main /Users/kcft114/.julia/packages/Flux/qXNjB/src/optimise/train.jl:105
┌ Info: Epoch 1530
└ @ Main /Users/kcft114/.julia/packages/Flux/qXNjB/src/optimise/train.jl:105
┌ Info: Epoch 1531
└ @ Main /Users/kcft114/.julia/packages/Flux/qXNjB/src/optimise/train.jl:105
┌ Info: Epoch 1532
└ @ Main /Users/kcft114/.julia/packages/Flux/qXNjB/src/optimise/train.jl:105
┌ Info: Epoch 1533
└ @ Main /Users/kcft114/.julia/packages/Flux/qXNjB/src/optimise/train.jl:105
┌ Info: Epoch 1534
└ @ Main /Users/kcft114/.julia/packages/Flux/qXNjB/src/optimise/train.jl:105
┌ Info: Epoch 1535
└ @ Main /Users/kcft114/.julia/packages/Flux/qXNjB/src/optimise/train.jl:105
┌ Info: Epoch 1536
└ @ Main /Users/kcft114/.julia/packages/Flux/qXNjB/src/optimise/train.jl:105
┌ Info: Epoch 1537
└ @ Main /Users/kcft114/.julia/packages/Flux/qXNjB/src/optimise/train.jl:105
┌ Info: Epoch 1538
└ @ Main /Users/kcft114/.julia/packages/Flux/qXNjB/src/optimise/train.jl:105
┌ Info: Epoch 1539
└ @ Main /Users/kcft114/.julia/packages/Flux/qXNjB/src/optimise/train.jl:105
┌ Info: Epoch 1540
└ @ Main /Users/kcft114/.julia/packages/Flux/qXNjB/src/optimise/train.jl:105
┌ Info: Epoch 1541
└ @ Main /Users/kcft114/.julia/packages/Flux/qXNjB/src/optimise/train.jl:105
┌ Info: Epoch 1542
└ @ Main /Users/kcft114/.julia/packages/Flux/qXNjB/src/optimise/train.jl:105
┌ Info: Epoch 1543
└ @ Main /Users/kcft114/.julia/packages/Flux/qXNjB/src/optimise/train.jl:105
┌ Info: Epoch 1544
└ @ Main /Users/kcft114/.julia/packages/Flux/qXNjB/src/optimise/train.jl:105
┌ Info: Epoch 1545
└ @ Main /Users/kcft114/.julia/packages/Flux/qXNjB/src/optimise/train.jl:105
┌ Info: Epoch 1546
└ @ Main /Users/kcft114/.julia/packages/Flux/qXNjB/src/optimise/train.jl:105
┌ Info: Epoch 1547
└ @ Main /Users/kcft114/.julia/packages/Flux/qXNjB/src/optimise/train.jl:105
┌ Info: Epoch 1548
└ @ Main /Users/kcft114/.julia/packages/Flux/qXNjB/src/optimise/train.jl:105
┌ Info: Epoch 1549
└ @ Main /Users/kcft114/.julia/packages/Flux/qXNjB/src/optimise/train.jl:105
┌ Info: Epoch 1550
└ @ Main /Users/kcft114/.julia/packages/Flux/qXNjB/src/optimise/train.jl:105
┌ Info: Epoch 1551
└ @ Main /Users/kcft114/.julia/packages/Flux/qXNjB/src/optimise/train.jl:105
┌ Info: Epoch 1552
└ @ Main /Users/kcft114/.julia/packages/Flux/qXNjB/src/optimise/train.jl:105
┌ Info: Epoch 1553
└ @ Main /Users/kcft114/.julia/packages/Flux/qXNjB/src/optimise/train.jl:105
┌ Info: Epoch 1554
└ @ Main /Users/kcft114/.julia/packages/Flux/qXNjB/src/optimise/train.jl:105
┌ Info: Epoch 1555
└ @ Main /Users/kcft114/.julia/packages/Flux/qXNjB/src/optimise/train.jl:105
┌ Info: Epoch 1556
└ @ Main /Users/kcft114/.julia/packages/Flux/qXNjB/src/optimise/train.jl:105
┌ Info: Epoch 1557
└ @ Main /Users/kcft114/.julia/packages/Flux/qXNjB/src/optimise/train.jl:105
┌ Info: Epoch 1558
└ @ Main /Users/kcft114/.julia/packages/Flux/qXNjB/src/optimise/train.jl:105
┌ Info: Epoch 1559
└ @ Main /Users/kcft114/.julia/packages/Flux/qXNjB/src/optimise/train.jl:105
┌ Info: Epoch 1560
└ @ Main /Users/kcft114/.julia/packages/Flux/qXNjB/src/optimise/train.jl:105
┌ Info: Epoch 1561
└ @ Main /Users/kcft114/.julia/packages/Flux/qXNjB/src/optimise/train.jl:105
┌ Info: Epoch 1562
└ @ Main /Users/kcft114/.julia/packages/Flux/qXNjB/src/optimise/train.jl:105
┌ Info: Epoch 1563
└ @ Main /Users/kcft114/.julia/packages/Flux/qXNjB/src/optimise/train.jl:105
┌ Info: Epoch 1564
└ @ Main /Users/kcft114/.julia/packages/Flux/qXNjB/src/optimise/train.jl:105
┌ Info: Epoch 1565
└ @ Main /Users/kcft114/.julia/packages/Flux/qXNjB/src/optimise/train.jl:105
┌ Info: Epoch 1566
└ @ Main /Users/kcft114/.julia/packages/Flux/qXNjB/src/optimise/train.jl:105
┌ Info: Epoch 1567
└ @ Main /Users/kcft114/.julia/packages/Flux/qXNjB/src/optimise/train.jl:105
┌ Info: Epoch 1568
└ @ Main /Users/kcft114/.julia/packages/Flux/qXNjB/src/optimise/train.jl:105
┌ Info: Epoch 1569
└ @ Main /Users/kcft114/.julia/packages/Flux/qXNjB/src/optimise/train.jl:105
┌ Info: Epoch 1570
└ @ Main /Users/kcft114/.julia/packages/Flux/qXNjB/src/optimise/train.jl:105
┌ Info: Epoch 1571
└ @ Main /Users/kcft114/.julia/packages/Flux/qXNjB/src/optimise/train.jl:105
┌ Info: Epoch 1572
└ @ Main /Users/kcft114/.julia/packages/Flux/qXNjB/src/optimise/train.jl:105
┌ Info: Epoch 1573
└ @ Main /Users/kcft114/.julia/packages/Flux/qXNjB/src/optimise/train.jl:105
┌ Info: Epoch 1574
└ @ Main /Users/kcft114/.julia/packages/Flux/qXNjB/src/optimise/train.jl:105
┌ Info: Epoch 1575
└ @ Main /Users/kcft114/.julia/packages/Flux/qXNjB/src/optimise/train.jl:105
┌ Info: Epoch 1576
└ @ Main /Users/kcft114/.julia/packages/Flux/qXNjB/src/optimise/train.jl:105
┌ Info: Epoch 1577
└ @ Main /Users/kcft114/.julia/packages/Flux/qXNjB/src/optimise/train.jl:105
┌ Info: Epoch 1578
└ @ Main /Users/kcft114/.julia/packages/Flux/qXNjB/src/optimise/train.jl:105
┌ Info: Epoch 1579
└ @ Main /Users/kcft114/.julia/packages/Flux/qXNjB/src/optimise/train.jl:105
┌ Info: Epoch 1580
└ @ Main /Users/kcft114/.julia/packages/Flux/qXNjB/src/optimise/train.jl:105
┌ Info: Epoch 1581
└ @ Main /Users/kcft114/.julia/packages/Flux/qXNjB/src/optimise/train.jl:105
┌ Info: Epoch 1582
└ @ Main /Users/kcft114/.julia/packages/Flux/qXNjB/src/optimise/train.jl:105
┌ Info: Epoch 1583
└ @ Main /Users/kcft114/.julia/packages/Flux/qXNjB/src/optimise/train.jl:105
┌ Info: Epoch 1584
└ @ Main /Users/kcft114/.julia/packages/Flux/qXNjB/src/optimise/train.jl:105
┌ Info: Epoch 1585
└ @ Main /Users/kcft114/.julia/packages/Flux/qXNjB/src/optimise/train.jl:105
┌ Info: Epoch 1586
└ @ Main /Users/kcft114/.julia/packages/Flux/qXNjB/src/optimise/train.jl:105
┌ Info: Epoch 1587
└ @ Main /Users/kcft114/.julia/packages/Flux/qXNjB/src/optimise/train.jl:105
┌ Info: Epoch 1588
└ @ Main /Users/kcft114/.julia/packages/Flux/qXNjB/src/optimise/train.jl:105
┌ Info: Epoch 1589
└ @ Main /Users/kcft114/.julia/packages/Flux/qXNjB/src/optimise/train.jl:105
┌ Info: Epoch 1590
└ @ Main /Users/kcft114/.julia/packages/Flux/qXNjB/src/optimise/train.jl:105
┌ Info: Epoch 1591
└ @ Main /Users/kcft114/.julia/packages/Flux/qXNjB/src/optimise/train.jl:105
┌ Info: Epoch 1592
└ @ Main /Users/kcft114/.julia/packages/Flux/qXNjB/src/optimise/train.jl:105
┌ Info: Epoch 1593
└ @ Main /Users/kcft114/.julia/packages/Flux/qXNjB/src/optimise/train.jl:105
┌ Info: Epoch 1594
└ @ Main /Users/kcft114/.julia/packages/Flux/qXNjB/src/optimise/train.jl:105
┌ Info: Epoch 1595
└ @ Main /Users/kcft114/.julia/packages/Flux/qXNjB/src/optimise/train.jl:105
┌ Info: Epoch 1596
└ @ Main /Users/kcft114/.julia/packages/Flux/qXNjB/src/optimise/train.jl:105
┌ Info: Epoch 1597
└ @ Main /Users/kcft114/.julia/packages/Flux/qXNjB/src/optimise/train.jl:105
┌ Info: Epoch 1598
└ @ Main /Users/kcft114/.julia/packages/Flux/qXNjB/src/optimise/train.jl:105
┌ Info: Epoch 1599
└ @ Main /Users/kcft114/.julia/packages/Flux/qXNjB/src/optimise/train.jl:105
┌ Info: Epoch 1600
└ @ Main /Users/kcft114/.julia/packages/Flux/qXNjB/src/optimise/train.jl:105
┌ Info: Epoch 1601
└ @ Main /Users/kcft114/.julia/packages/Flux/qXNjB/src/optimise/train.jl:105
┌ Info: Epoch 1602
└ @ Main /Users/kcft114/.julia/packages/Flux/qXNjB/src/optimise/train.jl:105
┌ Info: Epoch 1603
└ @ Main /Users/kcft114/.julia/packages/Flux/qXNjB/src/optimise/train.jl:105
┌ Info: Epoch 1604
└ @ Main /Users/kcft114/.julia/packages/Flux/qXNjB/src/optimise/train.jl:105
┌ Info: Epoch 1605
└ @ Main /Users/kcft114/.julia/packages/Flux/qXNjB/src/optimise/train.jl:105
┌ Info: Epoch 1606
└ @ Main /Users/kcft114/.julia/packages/Flux/qXNjB/src/optimise/train.jl:105
┌ Info: Epoch 1607
└ @ Main /Users/kcft114/.julia/packages/Flux/qXNjB/src/optimise/train.jl:105
┌ Info: Epoch 1608
└ @ Main /Users/kcft114/.julia/packages/Flux/qXNjB/src/optimise/train.jl:105
┌ Info: Epoch 1609
└ @ Main /Users/kcft114/.julia/packages/Flux/qXNjB/src/optimise/train.jl:105
```

```
loss(X, Y) = 0.50999403f0 (tracked)
loss(X, Y) = 0.516331f0 (tracked)
loss(X, Y) = 0.564224f0 (tracked)
loss(X, Y) = 0.5118413f0 (tracked)
loss(X, Y) = 0.5432744f0 (tracked)
loss(X, Y) = 0.5142909f0 (tracked)
loss(X, Y) = 0.5750653f0 (tracked)
loss(X, Y) = 0.5475754f0 (tracked)
loss(X, Y) = 0.53091305f0 (tracked)
loss(X, Y) = 0.5576699f0 (tracked)
loss(X, Y) = 0.5497538f0 (tracked)
loss(X, Y) = 0.54553f0 (tracked)
loss(X, Y) = 0.55392396f0 (tracked)
loss(X, Y) = 0.6083039f0 (tracked)
loss(X, Y) = 0.51239824f0 (tracked)
loss(X, Y) = 0.5272981f0 (tracked)
loss(X, Y) = 0.54044634f0 (tracked)
loss(X, Y) = 0.5553087f0 (tracked)
loss(X, Y) = 0.5516722f0 (tracked)
loss(X, Y) = 0.5283976f0 (tracked)
loss(X, Y) = 0.5364767f0 (tracked)
loss(X, Y) = 0.5339157f0 (tracked)
loss(X, Y) = 0.544301f0 (tracked)
loss(X, Y) = 0.5705886f0 (tracked)
loss(X, Y) = 0.5826371f0 (tracked)
loss(X, Y) = 0.5806066f0 (tracked)
loss(X, Y) = 0.56215405f0 (tracked)
loss(X, Y) = 0.57166505f0 (tracked)
loss(X, Y) = 0.55828655f0 (tracked)
loss(X, Y) = 0.50780493f0 (tracked)
loss(X, Y) = 0.546381f0 (tracked)
loss(X, Y) = 0.5390115f0 (tracked)
loss(X, Y) = 0.52788186f0 (tracked)
loss(X, Y) = 0.52971613f0 (tracked)
loss(X, Y) = 0.5674003f0 (tracked)
loss(X, Y) = 0.55114573f0 (tracked)
loss(X, Y) = 0.52724135f0 (tracked)
loss(X, Y) = 0.5847302f0 (tracked)
loss(X, Y) = 0.57245356f0 (tracked)
loss(X, Y) = 0.5494594f0 (tracked)
loss(X, Y) = 0.5478447f0 (tracked)
loss(X, Y) = 0.547581f0 (tracked)
loss(X, Y) = 0.54588354f0 (tracked)
loss(X, Y) = 0.53450906f0 (tracked)
loss(X, Y) = 0.52745384f0 (tracked)
loss(X, Y) = 0.54980916f0 (tracked)
loss(X, Y) = 0.5376272f0 (tracked)
loss(X, Y) = 0.55464154f0 (tracked)
loss(X, Y) = 0.5706599f0 (tracked)
loss(X, Y) = 0.55358464f0 (tracked)
loss(X, Y) = 0.57059294f0 (tracked)
loss(X, Y) = 0.57810587f0 (tracked)
loss(X, Y) = 0.5763968f0 (tracked)
loss(X, Y) = 0.56791764f0 (tracked)
loss(X, Y) = 0.54745984f0 (tracked)
loss(X, Y) = 0.59156454f0 (tracked)
loss(X, Y) = 0.5396658f0 (tracked)
loss(X, Y) = 0.5160373f0 (tracked)
loss(X, Y) = 0.58200103f0 (tracked)
loss(X, Y) = 0.52761036f0 (tracked)
loss(X, Y) = 0.5852228f0 (tracked)
loss(X, Y) = 0.5629475f0 (tracked)
loss(X, Y) = 0.5487813f0 (tracked)
loss(X, Y) = 0.54638296f0 (tracked)
loss(X, Y) = 0.54615515f0 (tracked)
loss(X, Y) = 0.5651086f0 (tracked)
loss(X, Y) = 0.54056126f0 (tracked)
loss(X, Y) = 0.5824896f0 (tracked)
loss(X, Y) = 0.51423657f0 (tracked)
loss(X, Y) = 0.5575434f0 (tracked)
loss(X, Y) = 0.55768436f0 (tracked)
loss(X, Y) = 0.53842044f0 (tracked)
loss(X, Y) = 0.511406f0 (tracked)
loss(X, Y) = 0.5472526f0 (tracked)
loss(X, Y) = 0.55916214f0 (tracked)
loss(X, Y) = 0.54355955f0 (tracked)
loss(X, Y) = 0.5777884f0 (tracked)
loss(X, Y) = 0.4892098f0 (tracked)
loss(X, Y) = 0.5388048f0 (tracked)
loss(X, Y) = 0.56443274f0 (tracked)
loss(X, Y) = 0.5303576f0 (tracked)
loss(X, Y) = 0.5352421f0 (tracked)
loss(X, Y) = 0.52466965f0 (tracked)
loss(X, Y) = 0.5322396f0 (tracked)
loss(X, Y) = 0.5307759f0 (tracked)
loss(X, Y) = 0.55873f0 (tracked)
loss(X, Y) = 0.516089f0 (tracked)
loss(X, Y) = 0.5473344f0 (tracked)
loss(X, Y) = 0.5772023f0 (tracked)
loss(X, Y) = 0.5406557f0 (tracked)
loss(X, Y) = 0.56022424f0 (tracked)
loss(X, Y) = 0.5115182f0 (tracked)
loss(X, Y) = 0.5199169f0 (tracked)
loss(X, Y) = 0.5469101f0 (tracked)
loss(X, Y) = 0.5312818f0 (tracked)
loss(X, Y) = 0.5473263f0 (tracked)
loss(X, Y) = 0.5759889f0 (tracked)
loss(X, Y) = 0.5662354f0 (tracked)
loss(X, Y) = 0.5453585f0 (tracked)
loss(X, Y) = 0.5440429f0 (tracked)
loss(X, Y) = 0.5434383f0 (tracked)
loss(X, Y) = 0.53839993f0 (tracked)
loss(X, Y) = 0.5338259f0 (tracked)
loss(X, Y) = 0.5341477f0 (tracked)
loss(X, Y) = 0.56283164f0 (tracked)
loss(X, Y) = 0.5817171f0 (tracked)
loss(X, Y) = 0.5297638f0 (tracked)
loss(X, Y) = 0.5779065f0 (tracked)
loss(X, Y) = 0.5268744f0 (tracked)
loss(X, Y) = 0.54534215f0 (tracked)
loss(X, Y) = 0.53896016f0 (tracked)
loss(X, Y) = 0.53658104f0 (tracked)
loss(X, Y) = 0.52622396f0 (tracked)
loss(X, Y) = 0.54453397f0 (tracked)
loss(X, Y) = 0.59775937f0 (tracked)
loss(X, Y) = 0.5605956f0 (tracked)
loss(X, Y) = 0.5543707f0 (tracked)
loss(X, Y) = 0.53156865f0 (tracked)
loss(X, Y) = 0.5330781f0 (tracked)
loss(X, Y) = 0.53389627f0 (tracked)
loss(X, Y) = 0.49769774f0 (tracked)
loss(X, Y) = 0.5447226f0 (tracked)
loss(X, Y) = 0.521821f0 (tracked)
```

```
┌ Info: Epoch 1610
└ @ Main /Users/kcft114/.julia/packages/Flux/qXNjB/src/optimise/train.jl:105
┌ Info: Epoch 1611
└ @ Main /Users/kcft114/.julia/packages/Flux/qXNjB/src/optimise/train.jl:105
┌ Info: Epoch 1612
└ @ Main /Users/kcft114/.julia/packages/Flux/qXNjB/src/optimise/train.jl:105
┌ Info: Epoch 1613
└ @ Main /Users/kcft114/.julia/packages/Flux/qXNjB/src/optimise/train.jl:105
┌ Info: Epoch 1614
└ @ Main /Users/kcft114/.julia/packages/Flux/qXNjB/src/optimise/train.jl:105
┌ Info: Epoch 1615
└ @ Main /Users/kcft114/.julia/packages/Flux/qXNjB/src/optimise/train.jl:105
┌ Info: Epoch 1616
└ @ Main /Users/kcft114/.julia/packages/Flux/qXNjB/src/optimise/train.jl:105
┌ Info: Epoch 1617
└ @ Main /Users/kcft114/.julia/packages/Flux/qXNjB/src/optimise/train.jl:105
┌ Info: Epoch 1618
└ @ Main /Users/kcft114/.julia/packages/Flux/qXNjB/src/optimise/train.jl:105
┌ Info: Epoch 1619
└ @ Main /Users/kcft114/.julia/packages/Flux/qXNjB/src/optimise/train.jl:105
┌ Info: Epoch 1620
└ @ Main /Users/kcft114/.julia/packages/Flux/qXNjB/src/optimise/train.jl:105
┌ Info: Epoch 1621
└ @ Main /Users/kcft114/.julia/packages/Flux/qXNjB/src/optimise/train.jl:105
┌ Info: Epoch 1622
└ @ Main /Users/kcft114/.julia/packages/Flux/qXNjB/src/optimise/train.jl:105
┌ Info: Epoch 1623
└ @ Main /Users/kcft114/.julia/packages/Flux/qXNjB/src/optimise/train.jl:105
┌ Info: Epoch 1624
└ @ Main /Users/kcft114/.julia/packages/Flux/qXNjB/src/optimise/train.jl:105
┌ Info: Epoch 1625
└ @ Main /Users/kcft114/.julia/packages/Flux/qXNjB/src/optimise/train.jl:105
┌ Info: Epoch 1626
└ @ Main /Users/kcft114/.julia/packages/Flux/qXNjB/src/optimise/train.jl:105
┌ Info: Epoch 1627
└ @ Main /Users/kcft114/.julia/packages/Flux/qXNjB/src/optimise/train.jl:105
┌ Info: Epoch 1628
└ @ Main /Users/kcft114/.julia/packages/Flux/qXNjB/src/optimise/train.jl:105
┌ Info: Epoch 1629
└ @ Main /Users/kcft114/.julia/packages/Flux/qXNjB/src/optimise/train.jl:105
┌ Info: Epoch 1630
└ @ Main /Users/kcft114/.julia/packages/Flux/qXNjB/src/optimise/train.jl:105
┌ Info: Epoch 1631
└ @ Main /Users/kcft114/.julia/packages/Flux/qXNjB/src/optimise/train.jl:105
┌ Info: Epoch 1632
└ @ Main /Users/kcft114/.julia/packages/Flux/qXNjB/src/optimise/train.jl:105
┌ Info: Epoch 1633
└ @ Main /Users/kcft114/.julia/packages/Flux/qXNjB/src/optimise/train.jl:105
┌ Info: Epoch 1634
└ @ Main /Users/kcft114/.julia/packages/Flux/qXNjB/src/optimise/train.jl:105
┌ Info: Epoch 1635
└ @ Main /Users/kcft114/.julia/packages/Flux/qXNjB/src/optimise/train.jl:105
┌ Info: Epoch 1636
└ @ Main /Users/kcft114/.julia/packages/Flux/qXNjB/src/optimise/train.jl:105
┌ Info: Epoch 1637
└ @ Main /Users/kcft114/.julia/packages/Flux/qXNjB/src/optimise/train.jl:105
┌ Info: Epoch 1638
└ @ Main /Users/kcft114/.julia/packages/Flux/qXNjB/src/optimise/train.jl:105
┌ Info: Epoch 1639
└ @ Main /Users/kcft114/.julia/packages/Flux/qXNjB/src/optimise/train.jl:105
┌ Info: Epoch 1640
└ @ Main /Users/kcft114/.julia/packages/Flux/qXNjB/src/optimise/train.jl:105
┌ Info: Epoch 1641
└ @ Main /Users/kcft114/.julia/packages/Flux/qXNjB/src/optimise/train.jl:105
┌ Info: Epoch 1642
└ @ Main /Users/kcft114/.julia/packages/Flux/qXNjB/src/optimise/train.jl:105
┌ Info: Epoch 1643
└ @ Main /Users/kcft114/.julia/packages/Flux/qXNjB/src/optimise/train.jl:105
┌ Info: Epoch 1644
└ @ Main /Users/kcft114/.julia/packages/Flux/qXNjB/src/optimise/train.jl:105
┌ Info: Epoch 1645
└ @ Main /Users/kcft114/.julia/packages/Flux/qXNjB/src/optimise/train.jl:105
┌ Info: Epoch 1646
└ @ Main /Users/kcft114/.julia/packages/Flux/qXNjB/src/optimise/train.jl:105
┌ Info: Epoch 1647
└ @ Main /Users/kcft114/.julia/packages/Flux/qXNjB/src/optimise/train.jl:105
┌ Info: Epoch 1648
└ @ Main /Users/kcft114/.julia/packages/Flux/qXNjB/src/optimise/train.jl:105
┌ Info: Epoch 1649
└ @ Main /Users/kcft114/.julia/packages/Flux/qXNjB/src/optimise/train.jl:105
┌ Info: Epoch 1650
└ @ Main /Users/kcft114/.julia/packages/Flux/qXNjB/src/optimise/train.jl:105
┌ Info: Epoch 1651
└ @ Main /Users/kcft114/.julia/packages/Flux/qXNjB/src/optimise/train.jl:105
┌ Info: Epoch 1652
└ @ Main /Users/kcft114/.julia/packages/Flux/qXNjB/src/optimise/train.jl:105
┌ Info: Epoch 1653
└ @ Main /Users/kcft114/.julia/packages/Flux/qXNjB/src/optimise/train.jl:105
┌ Info: Epoch 1654
└ @ Main /Users/kcft114/.julia/packages/Flux/qXNjB/src/optimise/train.jl:105
┌ Info: Epoch 1655
└ @ Main /Users/kcft114/.julia/packages/Flux/qXNjB/src/optimise/train.jl:105
┌ Info: Epoch 1656
└ @ Main /Users/kcft114/.julia/packages/Flux/qXNjB/src/optimise/train.jl:105
┌ Info: Epoch 1657
└ @ Main /Users/kcft114/.julia/packages/Flux/qXNjB/src/optimise/train.jl:105
┌ Info: Epoch 1658
└ @ Main /Users/kcft114/.julia/packages/Flux/qXNjB/src/optimise/train.jl:105
┌ Info: Epoch 1659
└ @ Main /Users/kcft114/.julia/packages/Flux/qXNjB/src/optimise/train.jl:105
┌ Info: Epoch 1660
└ @ Main /Users/kcft114/.julia/packages/Flux/qXNjB/src/optimise/train.jl:105
┌ Info: Epoch 1661
└ @ Main /Users/kcft114/.julia/packages/Flux/qXNjB/src/optimise/train.jl:105
┌ Info: Epoch 1662
└ @ Main /Users/kcft114/.julia/packages/Flux/qXNjB/src/optimise/train.jl:105
┌ Info: Epoch 1663
└ @ Main /Users/kcft114/.julia/packages/Flux/qXNjB/src/optimise/train.jl:105
┌ Info: Epoch 1664
└ @ Main /Users/kcft114/.julia/packages/Flux/qXNjB/src/optimise/train.jl:105
┌ Info: Epoch 1665
└ @ Main /Users/kcft114/.julia/packages/Flux/qXNjB/src/optimise/train.jl:105
┌ Info: Epoch 1666
└ @ Main /Users/kcft114/.julia/packages/Flux/qXNjB/src/optimise/train.jl:105
┌ Info: Epoch 1667
└ @ Main /Users/kcft114/.julia/packages/Flux/qXNjB/src/optimise/train.jl:105
┌ Info: Epoch 1668
└ @ Main /Users/kcft114/.julia/packages/Flux/qXNjB/src/optimise/train.jl:105
┌ Info: Epoch 1669
└ @ Main /Users/kcft114/.julia/packages/Flux/qXNjB/src/optimise/train.jl:105
┌ Info: Epoch 1670
└ @ Main /Users/kcft114/.julia/packages/Flux/qXNjB/src/optimise/train.jl:105
┌ Info: Epoch 1671
└ @ Main /Users/kcft114/.julia/packages/Flux/qXNjB/src/optimise/train.jl:105
┌ Info: Epoch 1672
└ @ Main /Users/kcft114/.julia/packages/Flux/qXNjB/src/optimise/train.jl:105
┌ Info: Epoch 1673
└ @ Main /Users/kcft114/.julia/packages/Flux/qXNjB/src/optimise/train.jl:105
┌ Info: Epoch 1674
└ @ Main /Users/kcft114/.julia/packages/Flux/qXNjB/src/optimise/train.jl:105
┌ Info: Epoch 1675
└ @ Main /Users/kcft114/.julia/packages/Flux/qXNjB/src/optimise/train.jl:105
┌ Info: Epoch 1676
└ @ Main /Users/kcft114/.julia/packages/Flux/qXNjB/src/optimise/train.jl:105
┌ Info: Epoch 1677
└ @ Main /Users/kcft114/.julia/packages/Flux/qXNjB/src/optimise/train.jl:105
┌ Info: Epoch 1678
└ @ Main /Users/kcft114/.julia/packages/Flux/qXNjB/src/optimise/train.jl:105
┌ Info: Epoch 1679
└ @ Main /Users/kcft114/.julia/packages/Flux/qXNjB/src/optimise/train.jl:105
┌ Info: Epoch 1680
└ @ Main /Users/kcft114/.julia/packages/Flux/qXNjB/src/optimise/train.jl:105
┌ Info: Epoch 1681
└ @ Main /Users/kcft114/.julia/packages/Flux/qXNjB/src/optimise/train.jl:105
┌ Info: Epoch 1682
└ @ Main /Users/kcft114/.julia/packages/Flux/qXNjB/src/optimise/train.jl:105
┌ Info: Epoch 1683
└ @ Main /Users/kcft114/.julia/packages/Flux/qXNjB/src/optimise/train.jl:105
┌ Info: Epoch 1684
└ @ Main /Users/kcft114/.julia/packages/Flux/qXNjB/src/optimise/train.jl:105
┌ Info: Epoch 1685
└ @ Main /Users/kcft114/.julia/packages/Flux/qXNjB/src/optimise/train.jl:105
┌ Info: Epoch 1686
└ @ Main /Users/kcft114/.julia/packages/Flux/qXNjB/src/optimise/train.jl:105
┌ Info: Epoch 1687
└ @ Main /Users/kcft114/.julia/packages/Flux/qXNjB/src/optimise/train.jl:105
┌ Info: Epoch 1688
└ @ Main /Users/kcft114/.julia/packages/Flux/qXNjB/src/optimise/train.jl:105
┌ Info: Epoch 1689
└ @ Main /Users/kcft114/.julia/packages/Flux/qXNjB/src/optimise/train.jl:105
┌ Info: Epoch 1690
└ @ Main /Users/kcft114/.julia/packages/Flux/qXNjB/src/optimise/train.jl:105
┌ Info: Epoch 1691
└ @ Main /Users/kcft114/.julia/packages/Flux/qXNjB/src/optimise/train.jl:105
┌ Info: Epoch 1692
└ @ Main /Users/kcft114/.julia/packages/Flux/qXNjB/src/optimise/train.jl:105
┌ Info: Epoch 1693
└ @ Main /Users/kcft114/.julia/packages/Flux/qXNjB/src/optimise/train.jl:105
┌ Info: Epoch 1694
└ @ Main /Users/kcft114/.julia/packages/Flux/qXNjB/src/optimise/train.jl:105
┌ Info: Epoch 1695
└ @ Main /Users/kcft114/.julia/packages/Flux/qXNjB/src/optimise/train.jl:105
┌ Info: Epoch 1696
└ @ Main /Users/kcft114/.julia/packages/Flux/qXNjB/src/optimise/train.jl:105
┌ Info: Epoch 1697
└ @ Main /Users/kcft114/.julia/packages/Flux/qXNjB/src/optimise/train.jl:105
┌ Info: Epoch 1698
└ @ Main /Users/kcft114/.julia/packages/Flux/qXNjB/src/optimise/train.jl:105
┌ Info: Epoch 1699
└ @ Main /Users/kcft114/.julia/packages/Flux/qXNjB/src/optimise/train.jl:105
┌ Info: Epoch 1700
└ @ Main /Users/kcft114/.julia/packages/Flux/qXNjB/src/optimise/train.jl:105
┌ Info: Epoch 1701
└ @ Main /Users/kcft114/.julia/packages/Flux/qXNjB/src/optimise/train.jl:105
┌ Info: Epoch 1702
└ @ Main /Users/kcft114/.julia/packages/Flux/qXNjB/src/optimise/train.jl:105
┌ Info: Epoch 1703
└ @ Main /Users/kcft114/.julia/packages/Flux/qXNjB/src/optimise/train.jl:105
┌ Info: Epoch 1704
└ @ Main /Users/kcft114/.julia/packages/Flux/qXNjB/src/optimise/train.jl:105
┌ Info: Epoch 1705
└ @ Main /Users/kcft114/.julia/packages/Flux/qXNjB/src/optimise/train.jl:105
┌ Info: Epoch 1706
└ @ Main /Users/kcft114/.julia/packages/Flux/qXNjB/src/optimise/train.jl:105
┌ Info: Epoch 1707
└ @ Main /Users/kcft114/.julia/packages/Flux/qXNjB/src/optimise/train.jl:105
┌ Info: Epoch 1708
└ @ Main /Users/kcft114/.julia/packages/Flux/qXNjB/src/optimise/train.jl:105
┌ Info: Epoch 1709
└ @ Main /Users/kcft114/.julia/packages/Flux/qXNjB/src/optimise/train.jl:105
┌ Info: Epoch 1710
└ @ Main /Users/kcft114/.julia/packages/Flux/qXNjB/src/optimise/train.jl:105
┌ Info: Epoch 1711
└ @ Main /Users/kcft114/.julia/packages/Flux/qXNjB/src/optimise/train.jl:105
┌ Info: Epoch 1712
└ @ Main /Users/kcft114/.julia/packages/Flux/qXNjB/src/optimise/train.jl:105
┌ Info: Epoch 1713
└ @ Main /Users/kcft114/.julia/packages/Flux/qXNjB/src/optimise/train.jl:105
┌ Info: Epoch 1714
└ @ Main /Users/kcft114/.julia/packages/Flux/qXNjB/src/optimise/train.jl:105
┌ Info: Epoch 1715
└ @ Main /Users/kcft114/.julia/packages/Flux/qXNjB/src/optimise/train.jl:105
┌ Info: Epoch 1716
└ @ Main /Users/kcft114/.julia/packages/Flux/qXNjB/src/optimise/train.jl:105
┌ Info: Epoch 1717
└ @ Main /Users/kcft114/.julia/packages/Flux/qXNjB/src/optimise/train.jl:105
┌ Info: Epoch 1718
└ @ Main /Users/kcft114/.julia/packages/Flux/qXNjB/src/optimise/train.jl:105
┌ Info: Epoch 1719
└ @ Main /Users/kcft114/.julia/packages/Flux/qXNjB/src/optimise/train.jl:105
┌ Info: Epoch 1720
└ @ Main /Users/kcft114/.julia/packages/Flux/qXNjB/src/optimise/train.jl:105
┌ Info: Epoch 1721
└ @ Main /Users/kcft114/.julia/packages/Flux/qXNjB/src/optimise/train.jl:105
┌ Info: Epoch 1722
└ @ Main /Users/kcft114/.julia/packages/Flux/qXNjB/src/optimise/train.jl:105
┌ Info: Epoch 1723
└ @ Main /Users/kcft114/.julia/packages/Flux/qXNjB/src/optimise/train.jl:105
┌ Info: Epoch 1724
└ @ Main /Users/kcft114/.julia/packages/Flux/qXNjB/src/optimise/train.jl:105
┌ Info: Epoch 1725
└ @ Main /Users/kcft114/.julia/packages/Flux/qXNjB/src/optimise/train.jl:105
┌ Info: Epoch 1726
└ @ Main /Users/kcft114/.julia/packages/Flux/qXNjB/src/optimise/train.jl:105
┌ Info: Epoch 1727
└ @ Main /Users/kcft114/.julia/packages/Flux/qXNjB/src/optimise/train.jl:105
┌ Info: Epoch 1728
└ @ Main /Users/kcft114/.julia/packages/Flux/qXNjB/src/optimise/train.jl:105
┌ Info: Epoch 1729
└ @ Main /Users/kcft114/.julia/packages/Flux/qXNjB/src/optimise/train.jl:105
┌ Info: Epoch 1730
└ @ Main /Users/kcft114/.julia/packages/Flux/qXNjB/src/optimise/train.jl:105
┌ Info: Epoch 1731
└ @ Main /Users/kcft114/.julia/packages/Flux/qXNjB/src/optimise/train.jl:105
┌ Info: Epoch 1732
└ @ Main /Users/kcft114/.julia/packages/Flux/qXNjB/src/optimise/train.jl:105
```

```
loss(X, Y) = 0.5256867f0 (tracked)
loss(X, Y) = 0.5550315f0 (tracked)
loss(X, Y) = 0.5486861f0 (tracked)
loss(X, Y) = 0.65165067f0 (tracked)
loss(X, Y) = 0.48767796f0 (tracked)
loss(X, Y) = 0.54699457f0 (tracked)
loss(X, Y) = 0.5270414f0 (tracked)
loss(X, Y) = 0.56189436f0 (tracked)
loss(X, Y) = 0.5517149f0 (tracked)
loss(X, Y) = 0.50744617f0 (tracked)
loss(X, Y) = 0.5268815f0 (tracked)
loss(X, Y) = 0.5547465f0 (tracked)
loss(X, Y) = 0.525171f0 (tracked)
loss(X, Y) = 0.5293804f0 (tracked)
loss(X, Y) = 0.5554076f0 (tracked)
loss(X, Y) = 0.5334667f0 (tracked)
loss(X, Y) = 0.56127125f0 (tracked)
loss(X, Y) = 0.5365663f0 (tracked)
loss(X, Y) = 0.5257443f0 (tracked)
loss(X, Y) = 0.5639509f0 (tracked)
loss(X, Y) = 0.51622957f0 (tracked)
loss(X, Y) = 0.5397695f0 (tracked)
loss(X, Y) = 0.57891446f0 (tracked)
loss(X, Y) = 0.51946574f0 (tracked)
loss(X, Y) = 0.5312483f0 (tracked)
loss(X, Y) = 0.54985267f0 (tracked)
loss(X, Y) = 0.57063544f0 (tracked)
loss(X, Y) = 0.5465898f0 (tracked)
loss(X, Y) = 0.5291641f0 (tracked)
loss(X, Y) = 0.5730849f0 (tracked)
loss(X, Y) = 0.57591f0 (tracked)
loss(X, Y) = 0.5227734f0 (tracked)
loss(X, Y) = 0.5454228f0 (tracked)
loss(X, Y) = 0.55161494f0 (tracked)
loss(X, Y) = 0.53146493f0 (tracked)
loss(X, Y) = 0.5351157f0 (tracked)
loss(X, Y) = 0.53703684f0 (tracked)
loss(X, Y) = 0.52257806f0 (tracked)
loss(X, Y) = 0.53547573f0 (tracked)
loss(X, Y) = 0.59773755f0 (tracked)
loss(X, Y) = 0.54294443f0 (tracked)
loss(X, Y) = 0.55127615f0 (tracked)
loss(X, Y) = 0.55646026f0 (tracked)
loss(X, Y) = 0.5290107f0 (tracked)
loss(X, Y) = 0.5147555f0 (tracked)
loss(X, Y) = 0.5201734f0 (tracked)
loss(X, Y) = 0.5822257f0 (tracked)
loss(X, Y) = 0.5528267f0 (tracked)
loss(X, Y) = 0.56164336f0 (tracked)
loss(X, Y) = 0.561497f0 (tracked)
loss(X, Y) = 0.5921182f0 (tracked)
loss(X, Y) = 0.5564131f0 (tracked)
loss(X, Y) = 0.5571427f0 (tracked)
loss(X, Y) = 0.5575695f0 (tracked)
loss(X, Y) = 0.53164124f0 (tracked)
loss(X, Y) = 0.5085065f0 (tracked)
loss(X, Y) = 0.5796438f0 (tracked)
loss(X, Y) = 0.57062817f0 (tracked)
loss(X, Y) = 0.5452965f0 (tracked)
loss(X, Y) = 0.5433279f0 (tracked)
loss(X, Y) = 0.5601152f0 (tracked)
loss(X, Y) = 0.54647666f0 (tracked)
loss(X, Y) = 0.4965676f0 (tracked)
loss(X, Y) = 0.49027827f0 (tracked)
loss(X, Y) = 0.5444282f0 (tracked)
loss(X, Y) = 0.56582665f0 (tracked)
loss(X, Y) = 0.54074365f0 (tracked)
loss(X, Y) = 0.5167932f0 (tracked)
loss(X, Y) = 0.5589141f0 (tracked)
loss(X, Y) = 0.5443369f0 (tracked)
loss(X, Y) = 0.56782943f0 (tracked)
loss(X, Y) = 0.55991566f0 (tracked)
loss(X, Y) = 0.5422488f0 (tracked)
loss(X, Y) = 0.52864516f0 (tracked)
loss(X, Y) = 0.5116564f0 (tracked)
loss(X, Y) = 0.53611505f0 (tracked)
loss(X, Y) = 0.51738876f0 (tracked)
loss(X, Y) = 0.4990844f0 (tracked)
loss(X, Y) = 0.5751619f0 (tracked)
loss(X, Y) = 0.5157038f0 (tracked)
loss(X, Y) = 0.5364676f0 (tracked)
loss(X, Y) = 0.5428448f0 (tracked)
loss(X, Y) = 0.5551473f0 (tracked)
loss(X, Y) = 0.53392404f0 (tracked)
loss(X, Y) = 0.48570618f0 (tracked)
loss(X, Y) = 0.5570928f0 (tracked)
loss(X, Y) = 0.50332946f0 (tracked)
loss(X, Y) = 0.53351104f0 (tracked)
loss(X, Y) = 0.5397679f0 (tracked)
loss(X, Y) = 0.54660094f0 (tracked)
loss(X, Y) = 0.5674879f0 (tracked)
loss(X, Y) = 0.53713685f0 (tracked)
loss(X, Y) = 0.55508995f0 (tracked)
loss(X, Y) = 0.5465174f0 (tracked)
loss(X, Y) = 0.51829773f0 (tracked)
loss(X, Y) = 0.5524954f0 (tracked)
loss(X, Y) = 0.5342299f0 (tracked)
loss(X, Y) = 0.51283115f0 (tracked)
loss(X, Y) = 0.5347894f0 (tracked)
loss(X, Y) = 0.53303957f0 (tracked)
loss(X, Y) = 0.5040496f0 (tracked)
loss(X, Y) = 0.53159547f0 (tracked)
loss(X, Y) = 0.5254935f0 (tracked)
loss(X, Y) = 0.60559225f0 (tracked)
loss(X, Y) = 0.5066007f0 (tracked)
loss(X, Y) = 0.5088214f0 (tracked)
loss(X, Y) = 0.57159966f0 (tracked)
loss(X, Y) = 0.55441594f0 (tracked)
loss(X, Y) = 0.5119454f0 (tracked)
loss(X, Y) = 0.520976f0 (tracked)
loss(X, Y) = 0.51722425f0 (tracked)
loss(X, Y) = 0.50634325f0 (tracked)
loss(X, Y) = 0.5176011f0 (tracked)
loss(X, Y) = 0.5392437f0 (tracked)
loss(X, Y) = 0.5224314f0 (tracked)
loss(X, Y) = 0.500653f0 (tracked)
loss(X, Y) = 0.53479326f0 (tracked)
loss(X, Y) = 0.52166736f0 (tracked)
loss(X, Y) = 0.566205f0 (tracked)
loss(X, Y) = 0.5492542f0 (tracked)
loss(X, Y) = 0.5169437f0 (tracked)
loss(X, Y) = 0.5473682f0 (tracked)
loss(X, Y) = 0.56342566f0 (tracked)
loss(X, Y) = 0.5296132f0 (tracked)
loss(X, Y) = 0.6316408f0 (tracked)
loss(X, Y) = 0.5696926f0 (tracked)
loss(X, Y) = 0.5083523f0 (tracked)
loss(X, Y) = 0.5093929f0 (tracked)
loss(X, Y) = 0.61532205f0 (tracked)
loss(X, Y) = 0.53802603f0 (tracked)
loss(X, Y) = 0.5331626f0 (tracked)
loss(X, Y) = 0.557847f0 (tracked)
loss(X, Y) = 0.50215274f0 (tracked)
loss(X, Y) = 0.5578422f0 (tracked)
loss(X, Y) = 0.56380635f0 (tracked)
loss(X, Y) = 0.5137854f0 (tracked)
```

```
┌ Info: Epoch 1733
└ @ Main /Users/kcft114/.julia/packages/Flux/qXNjB/src/optimise/train.jl:105
┌ Info: Epoch 1734
└ @ Main /Users/kcft114/.julia/packages/Flux/qXNjB/src/optimise/train.jl:105
┌ Info: Epoch 1735
└ @ Main /Users/kcft114/.julia/packages/Flux/qXNjB/src/optimise/train.jl:105
┌ Info: Epoch 1736
└ @ Main /Users/kcft114/.julia/packages/Flux/qXNjB/src/optimise/train.jl:105
┌ Info: Epoch 1737
└ @ Main /Users/kcft114/.julia/packages/Flux/qXNjB/src/optimise/train.jl:105
┌ Info: Epoch 1738
└ @ Main /Users/kcft114/.julia/packages/Flux/qXNjB/src/optimise/train.jl:105
┌ Info: Epoch 1739
└ @ Main /Users/kcft114/.julia/packages/Flux/qXNjB/src/optimise/train.jl:105
┌ Info: Epoch 1740
└ @ Main /Users/kcft114/.julia/packages/Flux/qXNjB/src/optimise/train.jl:105
┌ Info: Epoch 1741
└ @ Main /Users/kcft114/.julia/packages/Flux/qXNjB/src/optimise/train.jl:105
┌ Info: Epoch 1742
└ @ Main /Users/kcft114/.julia/packages/Flux/qXNjB/src/optimise/train.jl:105
┌ Info: Epoch 1743
└ @ Main /Users/kcft114/.julia/packages/Flux/qXNjB/src/optimise/train.jl:105
┌ Info: Epoch 1744
└ @ Main /Users/kcft114/.julia/packages/Flux/qXNjB/src/optimise/train.jl:105
┌ Info: Epoch 1745
└ @ Main /Users/kcft114/.julia/packages/Flux/qXNjB/src/optimise/train.jl:105
┌ Info: Epoch 1746
└ @ Main /Users/kcft114/.julia/packages/Flux/qXNjB/src/optimise/train.jl:105
┌ Info: Epoch 1747
└ @ Main /Users/kcft114/.julia/packages/Flux/qXNjB/src/optimise/train.jl:105
┌ Info: Epoch 1748
└ @ Main /Users/kcft114/.julia/packages/Flux/qXNjB/src/optimise/train.jl:105
┌ Info: Epoch 1749
└ @ Main /Users/kcft114/.julia/packages/Flux/qXNjB/src/optimise/train.jl:105
┌ Info: Epoch 1750
└ @ Main /Users/kcft114/.julia/packages/Flux/qXNjB/src/optimise/train.jl:105
┌ Info: Epoch 1751
└ @ Main /Users/kcft114/.julia/packages/Flux/qXNjB/src/optimise/train.jl:105
┌ Info: Epoch 1752
└ @ Main /Users/kcft114/.julia/packages/Flux/qXNjB/src/optimise/train.jl:105
┌ Info: Epoch 1753
└ @ Main /Users/kcft114/.julia/packages/Flux/qXNjB/src/optimise/train.jl:105
┌ Info: Epoch 1754
└ @ Main /Users/kcft114/.julia/packages/Flux/qXNjB/src/optimise/train.jl:105
┌ Info: Epoch 1755
└ @ Main /Users/kcft114/.julia/packages/Flux/qXNjB/src/optimise/train.jl:105
┌ Info: Epoch 1756
└ @ Main /Users/kcft114/.julia/packages/Flux/qXNjB/src/optimise/train.jl:105
┌ Info: Epoch 1757
└ @ Main /Users/kcft114/.julia/packages/Flux/qXNjB/src/optimise/train.jl:105
┌ Info: Epoch 1758
└ @ Main /Users/kcft114/.julia/packages/Flux/qXNjB/src/optimise/train.jl:105
┌ Info: Epoch 1759
└ @ Main /Users/kcft114/.julia/packages/Flux/qXNjB/src/optimise/train.jl:105
┌ Info: Epoch 1760
└ @ Main /Users/kcft114/.julia/packages/Flux/qXNjB/src/optimise/train.jl:105
┌ Info: Epoch 1761
└ @ Main /Users/kcft114/.julia/packages/Flux/qXNjB/src/optimise/train.jl:105
┌ Info: Epoch 1762
└ @ Main /Users/kcft114/.julia/packages/Flux/qXNjB/src/optimise/train.jl:105
┌ Info: Epoch 1763
└ @ Main /Users/kcft114/.julia/packages/Flux/qXNjB/src/optimise/train.jl:105
┌ Info: Epoch 1764
└ @ Main /Users/kcft114/.julia/packages/Flux/qXNjB/src/optimise/train.jl:105
┌ Info: Epoch 1765
└ @ Main /Users/kcft114/.julia/packages/Flux/qXNjB/src/optimise/train.jl:105
┌ Info: Epoch 1766
└ @ Main /Users/kcft114/.julia/packages/Flux/qXNjB/src/optimise/train.jl:105
┌ Info: Epoch 1767
└ @ Main /Users/kcft114/.julia/packages/Flux/qXNjB/src/optimise/train.jl:105
┌ Info: Epoch 1768
└ @ Main /Users/kcft114/.julia/packages/Flux/qXNjB/src/optimise/train.jl:105
┌ Info: Epoch 1769
└ @ Main /Users/kcft114/.julia/packages/Flux/qXNjB/src/optimise/train.jl:105
┌ Info: Epoch 1770
└ @ Main /Users/kcft114/.julia/packages/Flux/qXNjB/src/optimise/train.jl:105
┌ Info: Epoch 1771
└ @ Main /Users/kcft114/.julia/packages/Flux/qXNjB/src/optimise/train.jl:105
┌ Info: Epoch 1772
└ @ Main /Users/kcft114/.julia/packages/Flux/qXNjB/src/optimise/train.jl:105
┌ Info: Epoch 1773
└ @ Main /Users/kcft114/.julia/packages/Flux/qXNjB/src/optimise/train.jl:105
┌ Info: Epoch 1774
└ @ Main /Users/kcft114/.julia/packages/Flux/qXNjB/src/optimise/train.jl:105
┌ Info: Epoch 1775
└ @ Main /Users/kcft114/.julia/packages/Flux/qXNjB/src/optimise/train.jl:105
┌ Info: Epoch 1776
└ @ Main /Users/kcft114/.julia/packages/Flux/qXNjB/src/optimise/train.jl:105
┌ Info: Epoch 1777
└ @ Main /Users/kcft114/.julia/packages/Flux/qXNjB/src/optimise/train.jl:105
┌ Info: Epoch 1778
└ @ Main /Users/kcft114/.julia/packages/Flux/qXNjB/src/optimise/train.jl:105
┌ Info: Epoch 1779
└ @ Main /Users/kcft114/.julia/packages/Flux/qXNjB/src/optimise/train.jl:105
┌ Info: Epoch 1780
└ @ Main /Users/kcft114/.julia/packages/Flux/qXNjB/src/optimise/train.jl:105
┌ Info: Epoch 1781
└ @ Main /Users/kcft114/.julia/packages/Flux/qXNjB/src/optimise/train.jl:105
┌ Info: Epoch 1782
└ @ Main /Users/kcft114/.julia/packages/Flux/qXNjB/src/optimise/train.jl:105
┌ Info: Epoch 1783
└ @ Main /Users/kcft114/.julia/packages/Flux/qXNjB/src/optimise/train.jl:105
┌ Info: Epoch 1784
└ @ Main /Users/kcft114/.julia/packages/Flux/qXNjB/src/optimise/train.jl:105
┌ Info: Epoch 1785
└ @ Main /Users/kcft114/.julia/packages/Flux/qXNjB/src/optimise/train.jl:105
┌ Info: Epoch 1786
└ @ Main /Users/kcft114/.julia/packages/Flux/qXNjB/src/optimise/train.jl:105
┌ Info: Epoch 1787
└ @ Main /Users/kcft114/.julia/packages/Flux/qXNjB/src/optimise/train.jl:105
┌ Info: Epoch 1788
└ @ Main /Users/kcft114/.julia/packages/Flux/qXNjB/src/optimise/train.jl:105
┌ Info: Epoch 1789
└ @ Main /Users/kcft114/.julia/packages/Flux/qXNjB/src/optimise/train.jl:105
┌ Info: Epoch 1790
└ @ Main /Users/kcft114/.julia/packages/Flux/qXNjB/src/optimise/train.jl:105
┌ Info: Epoch 1791
└ @ Main /Users/kcft114/.julia/packages/Flux/qXNjB/src/optimise/train.jl:105
┌ Info: Epoch 1792
└ @ Main /Users/kcft114/.julia/packages/Flux/qXNjB/src/optimise/train.jl:105
┌ Info: Epoch 1793
└ @ Main /Users/kcft114/.julia/packages/Flux/qXNjB/src/optimise/train.jl:105
┌ Info: Epoch 1794
└ @ Main /Users/kcft114/.julia/packages/Flux/qXNjB/src/optimise/train.jl:105
┌ Info: Epoch 1795
└ @ Main /Users/kcft114/.julia/packages/Flux/qXNjB/src/optimise/train.jl:105
┌ Info: Epoch 1796
└ @ Main /Users/kcft114/.julia/packages/Flux/qXNjB/src/optimise/train.jl:105
┌ Info: Epoch 1797
└ @ Main /Users/kcft114/.julia/packages/Flux/qXNjB/src/optimise/train.jl:105
┌ Info: Epoch 1798
└ @ Main /Users/kcft114/.julia/packages/Flux/qXNjB/src/optimise/train.jl:105
┌ Info: Epoch 1799
└ @ Main /Users/kcft114/.julia/packages/Flux/qXNjB/src/optimise/train.jl:105
┌ Info: Epoch 1800
└ @ Main /Users/kcft114/.julia/packages/Flux/qXNjB/src/optimise/train.jl:105
┌ Info: Epoch 1801
└ @ Main /Users/kcft114/.julia/packages/Flux/qXNjB/src/optimise/train.jl:105
┌ Info: Epoch 1802
└ @ Main /Users/kcft114/.julia/packages/Flux/qXNjB/src/optimise/train.jl:105
┌ Info: Epoch 1803
└ @ Main /Users/kcft114/.julia/packages/Flux/qXNjB/src/optimise/train.jl:105
┌ Info: Epoch 1804
└ @ Main /Users/kcft114/.julia/packages/Flux/qXNjB/src/optimise/train.jl:105
┌ Info: Epoch 1805
└ @ Main /Users/kcft114/.julia/packages/Flux/qXNjB/src/optimise/train.jl:105
┌ Info: Epoch 1806
└ @ Main /Users/kcft114/.julia/packages/Flux/qXNjB/src/optimise/train.jl:105
┌ Info: Epoch 1807
└ @ Main /Users/kcft114/.julia/packages/Flux/qXNjB/src/optimise/train.jl:105
┌ Info: Epoch 1808
└ @ Main /Users/kcft114/.julia/packages/Flux/qXNjB/src/optimise/train.jl:105
┌ Info: Epoch 1809
└ @ Main /Users/kcft114/.julia/packages/Flux/qXNjB/src/optimise/train.jl:105
┌ Info: Epoch 1810
└ @ Main /Users/kcft114/.julia/packages/Flux/qXNjB/src/optimise/train.jl:105
┌ Info: Epoch 1811
└ @ Main /Users/kcft114/.julia/packages/Flux/qXNjB/src/optimise/train.jl:105
┌ Info: Epoch 1812
└ @ Main /Users/kcft114/.julia/packages/Flux/qXNjB/src/optimise/train.jl:105
┌ Info: Epoch 1813
└ @ Main /Users/kcft114/.julia/packages/Flux/qXNjB/src/optimise/train.jl:105
┌ Info: Epoch 1814
└ @ Main /Users/kcft114/.julia/packages/Flux/qXNjB/src/optimise/train.jl:105
┌ Info: Epoch 1815
└ @ Main /Users/kcft114/.julia/packages/Flux/qXNjB/src/optimise/train.jl:105
┌ Info: Epoch 1816
└ @ Main /Users/kcft114/.julia/packages/Flux/qXNjB/src/optimise/train.jl:105
┌ Info: Epoch 1817
└ @ Main /Users/kcft114/.julia/packages/Flux/qXNjB/src/optimise/train.jl:105
┌ Info: Epoch 1818
└ @ Main /Users/kcft114/.julia/packages/Flux/qXNjB/src/optimise/train.jl:105
┌ Info: Epoch 1819
└ @ Main /Users/kcft114/.julia/packages/Flux/qXNjB/src/optimise/train.jl:105
┌ Info: Epoch 1820
└ @ Main /Users/kcft114/.julia/packages/Flux/qXNjB/src/optimise/train.jl:105
┌ Info: Epoch 1821
└ @ Main /Users/kcft114/.julia/packages/Flux/qXNjB/src/optimise/train.jl:105
┌ Info: Epoch 1822
└ @ Main /Users/kcft114/.julia/packages/Flux/qXNjB/src/optimise/train.jl:105
┌ Info: Epoch 1823
└ @ Main /Users/kcft114/.julia/packages/Flux/qXNjB/src/optimise/train.jl:105
┌ Info: Epoch 1824
└ @ Main /Users/kcft114/.julia/packages/Flux/qXNjB/src/optimise/train.jl:105
┌ Info: Epoch 1825
└ @ Main /Users/kcft114/.julia/packages/Flux/qXNjB/src/optimise/train.jl:105
┌ Info: Epoch 1826
└ @ Main /Users/kcft114/.julia/packages/Flux/qXNjB/src/optimise/train.jl:105
┌ Info: Epoch 1827
└ @ Main /Users/kcft114/.julia/packages/Flux/qXNjB/src/optimise/train.jl:105
┌ Info: Epoch 1828
└ @ Main /Users/kcft114/.julia/packages/Flux/qXNjB/src/optimise/train.jl:105
┌ Info: Epoch 1829
└ @ Main /Users/kcft114/.julia/packages/Flux/qXNjB/src/optimise/train.jl:105
┌ Info: Epoch 1830
└ @ Main /Users/kcft114/.julia/packages/Flux/qXNjB/src/optimise/train.jl:105
┌ Info: Epoch 1831
└ @ Main /Users/kcft114/.julia/packages/Flux/qXNjB/src/optimise/train.jl:105
┌ Info: Epoch 1832
└ @ Main /Users/kcft114/.julia/packages/Flux/qXNjB/src/optimise/train.jl:105
┌ Info: Epoch 1833
└ @ Main /Users/kcft114/.julia/packages/Flux/qXNjB/src/optimise/train.jl:105
┌ Info: Epoch 1834
└ @ Main /Users/kcft114/.julia/packages/Flux/qXNjB/src/optimise/train.jl:105
┌ Info: Epoch 1835
└ @ Main /Users/kcft114/.julia/packages/Flux/qXNjB/src/optimise/train.jl:105
┌ Info: Epoch 1836
└ @ Main /Users/kcft114/.julia/packages/Flux/qXNjB/src/optimise/train.jl:105
┌ Info: Epoch 1837
└ @ Main /Users/kcft114/.julia/packages/Flux/qXNjB/src/optimise/train.jl:105
┌ Info: Epoch 1838
└ @ Main /Users/kcft114/.julia/packages/Flux/qXNjB/src/optimise/train.jl:105
┌ Info: Epoch 1839
└ @ Main /Users/kcft114/.julia/packages/Flux/qXNjB/src/optimise/train.jl:105
┌ Info: Epoch 1840
└ @ Main /Users/kcft114/.julia/packages/Flux/qXNjB/src/optimise/train.jl:105
┌ Info: Epoch 1841
└ @ Main /Users/kcft114/.julia/packages/Flux/qXNjB/src/optimise/train.jl:105
┌ Info: Epoch 1842
└ @ Main /Users/kcft114/.julia/packages/Flux/qXNjB/src/optimise/train.jl:105
┌ Info: Epoch 1843
└ @ Main /Users/kcft114/.julia/packages/Flux/qXNjB/src/optimise/train.jl:105
┌ Info: Epoch 1844
└ @ Main /Users/kcft114/.julia/packages/Flux/qXNjB/src/optimise/train.jl:105
┌ Info: Epoch 1845
└ @ Main /Users/kcft114/.julia/packages/Flux/qXNjB/src/optimise/train.jl:105
┌ Info: Epoch 1846
└ @ Main /Users/kcft114/.julia/packages/Flux/qXNjB/src/optimise/train.jl:105
┌ Info: Epoch 1847
└ @ Main /Users/kcft114/.julia/packages/Flux/qXNjB/src/optimise/train.jl:105
┌ Info: Epoch 1848
└ @ Main /Users/kcft114/.julia/packages/Flux/qXNjB/src/optimise/train.jl:105
┌ Info: Epoch 1849
└ @ Main /Users/kcft114/.julia/packages/Flux/qXNjB/src/optimise/train.jl:105
┌ Info: Epoch 1850
└ @ Main /Users/kcft114/.julia/packages/Flux/qXNjB/src/optimise/train.jl:105
┌ Info: Epoch 1851
└ @ Main /Users/kcft114/.julia/packages/Flux/qXNjB/src/optimise/train.jl:105
┌ Info: Epoch 1852
└ @ Main /Users/kcft114/.julia/packages/Flux/qXNjB/src/optimise/train.jl:105
┌ Info: Epoch 1853
└ @ Main /Users/kcft114/.julia/packages/Flux/qXNjB/src/optimise/train.jl:105
┌ Info: Epoch 1854
└ @ Main /Users/kcft114/.julia/packages/Flux/qXNjB/src/optimise/train.jl:105
┌ Info: Epoch 1855
└ @ Main /Users/kcft114/.julia/packages/Flux/qXNjB/src/optimise/train.jl:105
┌ Info: Epoch 1856
└ @ Main /Users/kcft114/.julia/packages/Flux/qXNjB/src/optimise/train.jl:105
┌ Info: Epoch 1857
└ @ Main /Users/kcft114/.julia/packages/Flux/qXNjB/src/optimise/train.jl:105
┌ Info: Epoch 1858
└ @ Main /Users/kcft114/.julia/packages/Flux/qXNjB/src/optimise/train.jl:105
┌ Info: Epoch 1859
└ @ Main /Users/kcft114/.julia/packages/Flux/qXNjB/src/optimise/train.jl:105
┌ Info: Epoch 1860
└ @ Main /Users/kcft114/.julia/packages/Flux/qXNjB/src/optimise/train.jl:105
┌ Info: Epoch 1861
└ @ Main /Users/kcft114/.julia/packages/Flux/qXNjB/src/optimise/train.jl:105
┌ Info: Epoch 1862
└ @ Main /Users/kcft114/.julia/packages/Flux/qXNjB/src/optimise/train.jl:105
┌ Info: Epoch 1863
└ @ Main /Users/kcft114/.julia/packages/Flux/qXNjB/src/optimise/train.jl:105
┌ Info: Epoch 1864
└ @ Main /Users/kcft114/.julia/packages/Flux/qXNjB/src/optimise/train.jl:105
┌ Info: Epoch 1865
└ @ Main /Users/kcft114/.julia/packages/Flux/qXNjB/src/optimise/train.jl:105
┌ Info: Epoch 1866
└ @ Main /Users/kcft114/.julia/packages/Flux/qXNjB/src/optimise/train.jl:105
┌ Info: Epoch 1867
└ @ Main /Users/kcft114/.julia/packages/Flux/qXNjB/src/optimise/train.jl:105
┌ Info: Epoch 1868
└ @ Main /Users/kcft114/.julia/packages/Flux/qXNjB/src/optimise/train.jl:105
```

```
loss(X, Y) = 0.5527046f0 (tracked)
loss(X, Y) = 0.53079927f0 (tracked)
loss(X, Y) = 0.53976053f0 (tracked)
loss(X, Y) = 0.5176927f0 (tracked)
loss(X, Y) = 0.536302f0 (tracked)
loss(X, Y) = 0.51481336f0 (tracked)
loss(X, Y) = 0.5147977f0 (tracked)
loss(X, Y) = 0.52204424f0 (tracked)
loss(X, Y) = 0.5602457f0 (tracked)
loss(X, Y) = 0.53994876f0 (tracked)
loss(X, Y) = 0.49292713f0 (tracked)
loss(X, Y) = 0.56032234f0 (tracked)
loss(X, Y) = 0.5389887f0 (tracked)
loss(X, Y) = 0.5471452f0 (tracked)
loss(X, Y) = 0.52710736f0 (tracked)
loss(X, Y) = 0.51401275f0 (tracked)
loss(X, Y) = 0.56959635f0 (tracked)
loss(X, Y) = 0.5530013f0 (tracked)
loss(X, Y) = 0.5465302f0 (tracked)
loss(X, Y) = 0.50528854f0 (tracked)
loss(X, Y) = 0.5211467f0 (tracked)
loss(X, Y) = 0.52168155f0 (tracked)
loss(X, Y) = 0.55918115f0 (tracked)
loss(X, Y) = 0.5762688f0 (tracked)
loss(X, Y) = 0.5388483f0 (tracked)
loss(X, Y) = 0.576783f0 (tracked)
loss(X, Y) = 0.53883237f0 (tracked)
loss(X, Y) = 0.5216762f0 (tracked)
loss(X, Y) = 0.5082278f0 (tracked)
loss(X, Y) = 0.5198659f0 (tracked)
loss(X, Y) = 0.52283037f0 (tracked)
loss(X, Y) = 0.5261408f0 (tracked)
loss(X, Y) = 0.5711594f0 (tracked)
loss(X, Y) = 0.55239725f0 (tracked)
loss(X, Y) = 0.55093944f0 (tracked)
loss(X, Y) = 0.5330059f0 (tracked)
loss(X, Y) = 0.5216743f0 (tracked)
loss(X, Y) = 0.5488026f0 (tracked)
loss(X, Y) = 0.5097821f0 (tracked)
loss(X, Y) = 0.49980888f0 (tracked)
loss(X, Y) = 0.5285417f0 (tracked)
loss(X, Y) = 0.5154873f0 (tracked)
loss(X, Y) = 0.5340903f0 (tracked)
loss(X, Y) = 0.4673169f0 (tracked)
loss(X, Y) = 0.5203717f0 (tracked)
loss(X, Y) = 0.54769564f0 (tracked)
loss(X, Y) = 0.5402453f0 (tracked)
loss(X, Y) = 0.5117099f0 (tracked)
loss(X, Y) = 0.47691172f0 (tracked)
loss(X, Y) = 0.5179865f0 (tracked)
loss(X, Y) = 0.5314163f0 (tracked)
loss(X, Y) = 0.55212474f0 (tracked)
loss(X, Y) = 0.537702f0 (tracked)
loss(X, Y) = 0.50672317f0 (tracked)
loss(X, Y) = 0.55256385f0 (tracked)
loss(X, Y) = 0.55960935f0 (tracked)
loss(X, Y) = 0.52196157f0 (tracked)
loss(X, Y) = 0.55318624f0 (tracked)
loss(X, Y) = 0.5049461f0 (tracked)
loss(X, Y) = 0.5228149f0 (tracked)
loss(X, Y) = 0.5089426f0 (tracked)
loss(X, Y) = 0.5241344f0 (tracked)
loss(X, Y) = 0.5346408f0 (tracked)
loss(X, Y) = 0.55722314f0 (tracked)
loss(X, Y) = 0.50860447f0 (tracked)
loss(X, Y) = 0.47952244f0 (tracked)
loss(X, Y) = 0.5837585f0 (tracked)
loss(X, Y) = 0.56389546f0 (tracked)
loss(X, Y) = 0.54428375f0 (tracked)
loss(X, Y) = 0.5030771f0 (tracked)
loss(X, Y) = 0.55250716f0 (tracked)
loss(X, Y) = 0.527684f0 (tracked)
loss(X, Y) = 0.5296466f0 (tracked)
loss(X, Y) = 0.51640034f0 (tracked)
loss(X, Y) = 0.5163345f0 (tracked)
loss(X, Y) = 0.49079275f0 (tracked)
loss(X, Y) = 0.52639973f0 (tracked)
loss(X, Y) = 0.5124959f0 (tracked)
loss(X, Y) = 0.49647048f0 (tracked)
loss(X, Y) = 0.5200247f0 (tracked)
loss(X, Y) = 0.52985674f0 (tracked)
loss(X, Y) = 0.53252184f0 (tracked)
loss(X, Y) = 0.51669616f0 (tracked)
loss(X, Y) = 0.54790324f0 (tracked)
loss(X, Y) = 0.5046046f0 (tracked)
loss(X, Y) = 0.51024306f0 (tracked)
loss(X, Y) = 0.6122921f0 (tracked)
loss(X, Y) = 0.54999393f0 (tracked)
loss(X, Y) = 0.5395976f0 (tracked)
loss(X, Y) = 0.49882713f0 (tracked)
loss(X, Y) = 0.5234639f0 (tracked)
loss(X, Y) = 0.51707566f0 (tracked)
loss(X, Y) = 0.49231595f0 (tracked)
loss(X, Y) = 0.50825286f0 (tracked)
loss(X, Y) = 0.514139f0 (tracked)
loss(X, Y) = 0.5245405f0 (tracked)
loss(X, Y) = 0.4985465f0 (tracked)
loss(X, Y) = 0.5069226f0 (tracked)
loss(X, Y) = 0.508448f0 (tracked)
loss(X, Y) = 0.56558985f0 (tracked)
loss(X, Y) = 0.4590635f0 (tracked)
loss(X, Y) = 0.4926455f0 (tracked)
loss(X, Y) = 0.5390785f0 (tracked)
loss(X, Y) = 0.5070784f0 (tracked)
loss(X, Y) = 0.51523876f0 (tracked)
loss(X, Y) = 0.52011234f0 (tracked)
loss(X, Y) = 0.5161337f0 (tracked)
loss(X, Y) = 0.5377941f0 (tracked)
loss(X, Y) = 0.5199068f0 (tracked)
loss(X, Y) = 0.52663493f0 (tracked)
loss(X, Y) = 0.5395999f0 (tracked)
loss(X, Y) = 0.5452992f0 (tracked)
loss(X, Y) = 0.49203905f0 (tracked)
loss(X, Y) = 0.48487163f0 (tracked)
loss(X, Y) = 0.508304f0 (tracked)
loss(X, Y) = 0.53854346f0 (tracked)
loss(X, Y) = 0.515481f0 (tracked)
loss(X, Y) = 0.5499518f0 (tracked)
loss(X, Y) = 0.5310388f0 (tracked)
loss(X, Y) = 0.5312975f0 (tracked)
loss(X, Y) = 0.4993999f0 (tracked)
```

```
┌ Info: Epoch 1869
└ @ Main /Users/kcft114/.julia/packages/Flux/qXNjB/src/optimise/train.jl:105
┌ Info: Epoch 1870
└ @ Main /Users/kcft114/.julia/packages/Flux/qXNjB/src/optimise/train.jl:105
┌ Info: Epoch 1871
└ @ Main /Users/kcft114/.julia/packages/Flux/qXNjB/src/optimise/train.jl:105
┌ Info: Epoch 1872
└ @ Main /Users/kcft114/.julia/packages/Flux/qXNjB/src/optimise/train.jl:105
┌ Info: Epoch 1873
└ @ Main /Users/kcft114/.julia/packages/Flux/qXNjB/src/optimise/train.jl:105
┌ Info: Epoch 1874
└ @ Main /Users/kcft114/.julia/packages/Flux/qXNjB/src/optimise/train.jl:105
┌ Info: Epoch 1875
└ @ Main /Users/kcft114/.julia/packages/Flux/qXNjB/src/optimise/train.jl:105
┌ Info: Epoch 1876
└ @ Main /Users/kcft114/.julia/packages/Flux/qXNjB/src/optimise/train.jl:105
┌ Info: Epoch 1877
└ @ Main /Users/kcft114/.julia/packages/Flux/qXNjB/src/optimise/train.jl:105
┌ Info: Epoch 1878
└ @ Main /Users/kcft114/.julia/packages/Flux/qXNjB/src/optimise/train.jl:105
┌ Info: Epoch 1879
└ @ Main /Users/kcft114/.julia/packages/Flux/qXNjB/src/optimise/train.jl:105
┌ Info: Epoch 1880
└ @ Main /Users/kcft114/.julia/packages/Flux/qXNjB/src/optimise/train.jl:105
┌ Info: Epoch 1881
└ @ Main /Users/kcft114/.julia/packages/Flux/qXNjB/src/optimise/train.jl:105
┌ Info: Epoch 1882
└ @ Main /Users/kcft114/.julia/packages/Flux/qXNjB/src/optimise/train.jl:105
┌ Info: Epoch 1883
└ @ Main /Users/kcft114/.julia/packages/Flux/qXNjB/src/optimise/train.jl:105
┌ Info: Epoch 1884
└ @ Main /Users/kcft114/.julia/packages/Flux/qXNjB/src/optimise/train.jl:105
┌ Info: Epoch 1885
└ @ Main /Users/kcft114/.julia/packages/Flux/qXNjB/src/optimise/train.jl:105
┌ Info: Epoch 1886
└ @ Main /Users/kcft114/.julia/packages/Flux/qXNjB/src/optimise/train.jl:105
┌ Info: Epoch 1887
└ @ Main /Users/kcft114/.julia/packages/Flux/qXNjB/src/optimise/train.jl:105
┌ Info: Epoch 1888
└ @ Main /Users/kcft114/.julia/packages/Flux/qXNjB/src/optimise/train.jl:105
┌ Info: Epoch 1889
└ @ Main /Users/kcft114/.julia/packages/Flux/qXNjB/src/optimise/train.jl:105
┌ Info: Epoch 1890
└ @ Main /Users/kcft114/.julia/packages/Flux/qXNjB/src/optimise/train.jl:105
┌ Info: Epoch 1891
└ @ Main /Users/kcft114/.julia/packages/Flux/qXNjB/src/optimise/train.jl:105
┌ Info: Epoch 1892
└ @ Main /Users/kcft114/.julia/packages/Flux/qXNjB/src/optimise/train.jl:105
┌ Info: Epoch 1893
└ @ Main /Users/kcft114/.julia/packages/Flux/qXNjB/src/optimise/train.jl:105
┌ Info: Epoch 1894
└ @ Main /Users/kcft114/.julia/packages/Flux/qXNjB/src/optimise/train.jl:105
┌ Info: Epoch 1895
└ @ Main /Users/kcft114/.julia/packages/Flux/qXNjB/src/optimise/train.jl:105
┌ Info: Epoch 1896
└ @ Main /Users/kcft114/.julia/packages/Flux/qXNjB/src/optimise/train.jl:105
┌ Info: Epoch 1897
└ @ Main /Users/kcft114/.julia/packages/Flux/qXNjB/src/optimise/train.jl:105
┌ Info: Epoch 1898
└ @ Main /Users/kcft114/.julia/packages/Flux/qXNjB/src/optimise/train.jl:105
┌ Info: Epoch 1899
└ @ Main /Users/kcft114/.julia/packages/Flux/qXNjB/src/optimise/train.jl:105
┌ Info: Epoch 1900
└ @ Main /Users/kcft114/.julia/packages/Flux/qXNjB/src/optimise/train.jl:105
┌ Info: Epoch 1901
└ @ Main /Users/kcft114/.julia/packages/Flux/qXNjB/src/optimise/train.jl:105
┌ Info: Epoch 1902
└ @ Main /Users/kcft114/.julia/packages/Flux/qXNjB/src/optimise/train.jl:105
┌ Info: Epoch 1903
└ @ Main /Users/kcft114/.julia/packages/Flux/qXNjB/src/optimise/train.jl:105
┌ Info: Epoch 1904
└ @ Main /Users/kcft114/.julia/packages/Flux/qXNjB/src/optimise/train.jl:105
┌ Info: Epoch 1905
└ @ Main /Users/kcft114/.julia/packages/Flux/qXNjB/src/optimise/train.jl:105
┌ Info: Epoch 1906
└ @ Main /Users/kcft114/.julia/packages/Flux/qXNjB/src/optimise/train.jl:105
┌ Info: Epoch 1907
└ @ Main /Users/kcft114/.julia/packages/Flux/qXNjB/src/optimise/train.jl:105
┌ Info: Epoch 1908
└ @ Main /Users/kcft114/.julia/packages/Flux/qXNjB/src/optimise/train.jl:105
┌ Info: Epoch 1909
└ @ Main /Users/kcft114/.julia/packages/Flux/qXNjB/src/optimise/train.jl:105
┌ Info: Epoch 1910
└ @ Main /Users/kcft114/.julia/packages/Flux/qXNjB/src/optimise/train.jl:105
┌ Info: Epoch 1911
└ @ Main /Users/kcft114/.julia/packages/Flux/qXNjB/src/optimise/train.jl:105
┌ Info: Epoch 1912
└ @ Main /Users/kcft114/.julia/packages/Flux/qXNjB/src/optimise/train.jl:105
┌ Info: Epoch 1913
└ @ Main /Users/kcft114/.julia/packages/Flux/qXNjB/src/optimise/train.jl:105
┌ Info: Epoch 1914
└ @ Main /Users/kcft114/.julia/packages/Flux/qXNjB/src/optimise/train.jl:105
┌ Info: Epoch 1915
└ @ Main /Users/kcft114/.julia/packages/Flux/qXNjB/src/optimise/train.jl:105
┌ Info: Epoch 1916
└ @ Main /Users/kcft114/.julia/packages/Flux/qXNjB/src/optimise/train.jl:105
┌ Info: Epoch 1917
└ @ Main /Users/kcft114/.julia/packages/Flux/qXNjB/src/optimise/train.jl:105
┌ Info: Epoch 1918
└ @ Main /Users/kcft114/.julia/packages/Flux/qXNjB/src/optimise/train.jl:105
┌ Info: Epoch 1919
└ @ Main /Users/kcft114/.julia/packages/Flux/qXNjB/src/optimise/train.jl:105
┌ Info: Epoch 1920
└ @ Main /Users/kcft114/.julia/packages/Flux/qXNjB/src/optimise/train.jl:105
┌ Info: Epoch 1921
└ @ Main /Users/kcft114/.julia/packages/Flux/qXNjB/src/optimise/train.jl:105
┌ Info: Epoch 1922
└ @ Main /Users/kcft114/.julia/packages/Flux/qXNjB/src/optimise/train.jl:105
┌ Info: Epoch 1923
└ @ Main /Users/kcft114/.julia/packages/Flux/qXNjB/src/optimise/train.jl:105
┌ Info: Epoch 1924
└ @ Main /Users/kcft114/.julia/packages/Flux/qXNjB/src/optimise/train.jl:105
┌ Info: Epoch 1925
└ @ Main /Users/kcft114/.julia/packages/Flux/qXNjB/src/optimise/train.jl:105
┌ Info: Epoch 1926
└ @ Main /Users/kcft114/.julia/packages/Flux/qXNjB/src/optimise/train.jl:105
┌ Info: Epoch 1927
└ @ Main /Users/kcft114/.julia/packages/Flux/qXNjB/src/optimise/train.jl:105
┌ Info: Epoch 1928
└ @ Main /Users/kcft114/.julia/packages/Flux/qXNjB/src/optimise/train.jl:105
┌ Info: Epoch 1929
└ @ Main /Users/kcft114/.julia/packages/Flux/qXNjB/src/optimise/train.jl:105
┌ Info: Epoch 1930
└ @ Main /Users/kcft114/.julia/packages/Flux/qXNjB/src/optimise/train.jl:105
┌ Info: Epoch 1931
└ @ Main /Users/kcft114/.julia/packages/Flux/qXNjB/src/optimise/train.jl:105
┌ Info: Epoch 1932
└ @ Main /Users/kcft114/.julia/packages/Flux/qXNjB/src/optimise/train.jl:105
┌ Info: Epoch 1933
└ @ Main /Users/kcft114/.julia/packages/Flux/qXNjB/src/optimise/train.jl:105
┌ Info: Epoch 1934
└ @ Main /Users/kcft114/.julia/packages/Flux/qXNjB/src/optimise/train.jl:105
┌ Info: Epoch 1935
└ @ Main /Users/kcft114/.julia/packages/Flux/qXNjB/src/optimise/train.jl:105
┌ Info: Epoch 1936
└ @ Main /Users/kcft114/.julia/packages/Flux/qXNjB/src/optimise/train.jl:105
┌ Info: Epoch 1937
└ @ Main /Users/kcft114/.julia/packages/Flux/qXNjB/src/optimise/train.jl:105
┌ Info: Epoch 1938
└ @ Main /Users/kcft114/.julia/packages/Flux/qXNjB/src/optimise/train.jl:105
┌ Info: Epoch 1939
└ @ Main /Users/kcft114/.julia/packages/Flux/qXNjB/src/optimise/train.jl:105
┌ Info: Epoch 1940
└ @ Main /Users/kcft114/.julia/packages/Flux/qXNjB/src/optimise/train.jl:105
┌ Info: Epoch 1941
└ @ Main /Users/kcft114/.julia/packages/Flux/qXNjB/src/optimise/train.jl:105
┌ Info: Epoch 1942
└ @ Main /Users/kcft114/.julia/packages/Flux/qXNjB/src/optimise/train.jl:105
┌ Info: Epoch 1943
└ @ Main /Users/kcft114/.julia/packages/Flux/qXNjB/src/optimise/train.jl:105
┌ Info: Epoch 1944
└ @ Main /Users/kcft114/.julia/packages/Flux/qXNjB/src/optimise/train.jl:105
┌ Info: Epoch 1945
└ @ Main /Users/kcft114/.julia/packages/Flux/qXNjB/src/optimise/train.jl:105
┌ Info: Epoch 1946
└ @ Main /Users/kcft114/.julia/packages/Flux/qXNjB/src/optimise/train.jl:105
┌ Info: Epoch 1947
└ @ Main /Users/kcft114/.julia/packages/Flux/qXNjB/src/optimise/train.jl:105
┌ Info: Epoch 1948
└ @ Main /Users/kcft114/.julia/packages/Flux/qXNjB/src/optimise/train.jl:105
┌ Info: Epoch 1949
└ @ Main /Users/kcft114/.julia/packages/Flux/qXNjB/src/optimise/train.jl:105
┌ Info: Epoch 1950
└ @ Main /Users/kcft114/.julia/packages/Flux/qXNjB/src/optimise/train.jl:105
┌ Info: Epoch 1951
└ @ Main /Users/kcft114/.julia/packages/Flux/qXNjB/src/optimise/train.jl:105
┌ Info: Epoch 1952
└ @ Main /Users/kcft114/.julia/packages/Flux/qXNjB/src/optimise/train.jl:105
┌ Info: Epoch 1953
└ @ Main /Users/kcft114/.julia/packages/Flux/qXNjB/src/optimise/train.jl:105
┌ Info: Epoch 1954
└ @ Main /Users/kcft114/.julia/packages/Flux/qXNjB/src/optimise/train.jl:105
┌ Info: Epoch 1955
└ @ Main /Users/kcft114/.julia/packages/Flux/qXNjB/src/optimise/train.jl:105
┌ Info: Epoch 1956
└ @ Main /Users/kcft114/.julia/packages/Flux/qXNjB/src/optimise/train.jl:105
┌ Info: Epoch 1957
└ @ Main /Users/kcft114/.julia/packages/Flux/qXNjB/src/optimise/train.jl:105
┌ Info: Epoch 1958
└ @ Main /Users/kcft114/.julia/packages/Flux/qXNjB/src/optimise/train.jl:105
┌ Info: Epoch 1959
└ @ Main /Users/kcft114/.julia/packages/Flux/qXNjB/src/optimise/train.jl:105
┌ Info: Epoch 1960
└ @ Main /Users/kcft114/.julia/packages/Flux/qXNjB/src/optimise/train.jl:105
┌ Info: Epoch 1961
└ @ Main /Users/kcft114/.julia/packages/Flux/qXNjB/src/optimise/train.jl:105
┌ Info: Epoch 1962
└ @ Main /Users/kcft114/.julia/packages/Flux/qXNjB/src/optimise/train.jl:105
┌ Info: Epoch 1963
└ @ Main /Users/kcft114/.julia/packages/Flux/qXNjB/src/optimise/train.jl:105
┌ Info: Epoch 1964
└ @ Main /Users/kcft114/.julia/packages/Flux/qXNjB/src/optimise/train.jl:105
┌ Info: Epoch 1965
└ @ Main /Users/kcft114/.julia/packages/Flux/qXNjB/src/optimise/train.jl:105
┌ Info: Epoch 1966
└ @ Main /Users/kcft114/.julia/packages/Flux/qXNjB/src/optimise/train.jl:105
┌ Info: Epoch 1967
└ @ Main /Users/kcft114/.julia/packages/Flux/qXNjB/src/optimise/train.jl:105
┌ Info: Epoch 1968
└ @ Main /Users/kcft114/.julia/packages/Flux/qXNjB/src/optimise/train.jl:105
┌ Info: Epoch 1969
└ @ Main /Users/kcft114/.julia/packages/Flux/qXNjB/src/optimise/train.jl:105
┌ Info: Epoch 1970
└ @ Main /Users/kcft114/.julia/packages/Flux/qXNjB/src/optimise/train.jl:105
┌ Info: Epoch 1971
└ @ Main /Users/kcft114/.julia/packages/Flux/qXNjB/src/optimise/train.jl:105
┌ Info: Epoch 1972
└ @ Main /Users/kcft114/.julia/packages/Flux/qXNjB/src/optimise/train.jl:105
┌ Info: Epoch 1973
└ @ Main /Users/kcft114/.julia/packages/Flux/qXNjB/src/optimise/train.jl:105
┌ Info: Epoch 1974
└ @ Main /Users/kcft114/.julia/packages/Flux/qXNjB/src/optimise/train.jl:105
┌ Info: Epoch 1975
└ @ Main /Users/kcft114/.julia/packages/Flux/qXNjB/src/optimise/train.jl:105
┌ Info: Epoch 1976
└ @ Main /Users/kcft114/.julia/packages/Flux/qXNjB/src/optimise/train.jl:105
┌ Info: Epoch 1977
└ @ Main /Users/kcft114/.julia/packages/Flux/qXNjB/src/optimise/train.jl:105
┌ Info: Epoch 1978
└ @ Main /Users/kcft114/.julia/packages/Flux/qXNjB/src/optimise/train.jl:105
┌ Info: Epoch 1979
└ @ Main /Users/kcft114/.julia/packages/Flux/qXNjB/src/optimise/train.jl:105
┌ Info: Epoch 1980
└ @ Main /Users/kcft114/.julia/packages/Flux/qXNjB/src/optimise/train.jl:105
┌ Info: Epoch 1981
└ @ Main /Users/kcft114/.julia/packages/Flux/qXNjB/src/optimise/train.jl:105
┌ Info: Epoch 1982
└ @ Main /Users/kcft114/.julia/packages/Flux/qXNjB/src/optimise/train.jl:105
┌ Info: Epoch 1983
└ @ Main /Users/kcft114/.julia/packages/Flux/qXNjB/src/optimise/train.jl:105
┌ Info: Epoch 1984
└ @ Main /Users/kcft114/.julia/packages/Flux/qXNjB/src/optimise/train.jl:105
┌ Info: Epoch 1985
└ @ Main /Users/kcft114/.julia/packages/Flux/qXNjB/src/optimise/train.jl:105
┌ Info: Epoch 1986
└ @ Main /Users/kcft114/.julia/packages/Flux/qXNjB/src/optimise/train.jl:105
┌ Info: Epoch 1987
└ @ Main /Users/kcft114/.julia/packages/Flux/qXNjB/src/optimise/train.jl:105
┌ Info: Epoch 1988
└ @ Main /Users/kcft114/.julia/packages/Flux/qXNjB/src/optimise/train.jl:105
```

```
loss(X, Y) = 0.541009f0 (tracked)
loss(X, Y) = 0.5066454f0 (tracked)
loss(X, Y) = 0.5081811f0 (tracked)
loss(X, Y) = 0.5933181f0 (tracked)
loss(X, Y) = 0.50254714f0 (tracked)
loss(X, Y) = 0.5130168f0 (tracked)
loss(X, Y) = 0.5203402f0 (tracked)
loss(X, Y) = 0.47619534f0 (tracked)
loss(X, Y) = 0.5099831f0 (tracked)
loss(X, Y) = 0.528731f0 (tracked)
loss(X, Y) = 0.526762f0 (tracked)
loss(X, Y) = 0.50223756f0 (tracked)
loss(X, Y) = 0.52280354f0 (tracked)
loss(X, Y) = 0.58256996f0 (tracked)
loss(X, Y) = 0.5480328f0 (tracked)
loss(X, Y) = 0.5168136f0 (tracked)
loss(X, Y) = 0.5734134f0 (tracked)
loss(X, Y) = 0.54443604f0 (tracked)
loss(X, Y) = 0.52863026f0 (tracked)
loss(X, Y) = 0.4973974f0 (tracked)
loss(X, Y) = 0.4922971f0 (tracked)
loss(X, Y) = 0.52297807f0 (tracked)
loss(X, Y) = 0.5455332f0 (tracked)
loss(X, Y) = 0.4813307f0 (tracked)
loss(X, Y) = 0.54400605f0 (tracked)
loss(X, Y) = 0.55525655f0 (tracked)
loss(X, Y) = 0.49923503f0 (tracked)
loss(X, Y) = 0.5323656f0 (tracked)
loss(X, Y) = 0.514364f0 (tracked)
loss(X, Y) = 0.50269836f0 (tracked)
loss(X, Y) = 0.51538646f0 (tracked)
loss(X, Y) = 0.5135917f0 (tracked)
loss(X, Y) = 0.536411f0 (tracked)
loss(X, Y) = 0.54547894f0 (tracked)
loss(X, Y) = 0.54458743f0 (tracked)
loss(X, Y) = 0.50964266f0 (tracked)
loss(X, Y) = 0.52241015f0 (tracked)
loss(X, Y) = 0.5027888f0 (tracked)
loss(X, Y) = 0.54779905f0 (tracked)
loss(X, Y) = 0.5139568f0 (tracked)
loss(X, Y) = 0.5153744f0 (tracked)
loss(X, Y) = 0.52315557f0 (tracked)
loss(X, Y) = 0.48168024f0 (tracked)
loss(X, Y) = 0.5198468f0 (tracked)
loss(X, Y) = 0.54086274f0 (tracked)
loss(X, Y) = 0.51378644f0 (tracked)
loss(X, Y) = 0.530546f0 (tracked)
loss(X, Y) = 0.48621082f0 (tracked)
loss(X, Y) = 0.5063923f0 (tracked)
loss(X, Y) = 0.5073285f0 (tracked)
loss(X, Y) = 0.53519005f0 (tracked)
loss(X, Y) = 0.56318605f0 (tracked)
loss(X, Y) = 0.5047015f0 (tracked)
loss(X, Y) = 0.533651f0 (tracked)
loss(X, Y) = 0.52099985f0 (tracked)
loss(X, Y) = 0.512014f0 (tracked)
loss(X, Y) = 0.49784446f0 (tracked)
loss(X, Y) = 0.5351987f0 (tracked)
loss(X, Y) = 0.5304538f0 (tracked)
loss(X, Y) = 0.50752175f0 (tracked)
loss(X, Y) = 0.52293646f0 (tracked)
loss(X, Y) = 0.52854264f0 (tracked)
loss(X, Y) = 0.53000927f0 (tracked)
loss(X, Y) = 0.5268481f0 (tracked)
loss(X, Y) = 0.50715935f0 (tracked)
loss(X, Y) = 0.52357405f0 (tracked)
loss(X, Y) = 0.54983705f0 (tracked)
loss(X, Y) = 0.49201518f0 (tracked)
loss(X, Y) = 0.53216445f0 (tracked)
loss(X, Y) = 0.4768236f0 (tracked)
loss(X, Y) = 0.51508313f0 (tracked)
loss(X, Y) = 0.5644928f0 (tracked)
loss(X, Y) = 0.52868325f0 (tracked)
loss(X, Y) = 0.5274844f0 (tracked)
loss(X, Y) = 0.49030754f0 (tracked)
loss(X, Y) = 0.5564707f0 (tracked)
loss(X, Y) = 0.565618f0 (tracked)
loss(X, Y) = 0.48697257f0 (tracked)
loss(X, Y) = 0.50964046f0 (tracked)
loss(X, Y) = 0.51356006f0 (tracked)
loss(X, Y) = 0.5146231f0 (tracked)
loss(X, Y) = 0.49732113f0 (tracked)
loss(X, Y) = 0.51245356f0 (tracked)
loss(X, Y) = 0.50658923f0 (tracked)
loss(X, Y) = 0.49039832f0 (tracked)
loss(X, Y) = 0.50087833f0 (tracked)
loss(X, Y) = 0.4762718f0 (tracked)
loss(X, Y) = 0.5139801f0 (tracked)
loss(X, Y) = 0.5249716f0 (tracked)
loss(X, Y) = 0.4777114f0 (tracked)
loss(X, Y) = 0.5013846f0 (tracked)
loss(X, Y) = 0.5514098f0 (tracked)
loss(X, Y) = 0.52302516f0 (tracked)
loss(X, Y) = 0.49374658f0 (tracked)
loss(X, Y) = 0.5129288f0 (tracked)
loss(X, Y) = 0.5215756f0 (tracked)
loss(X, Y) = 0.51183933f0 (tracked)
loss(X, Y) = 0.4822824f0 (tracked)
loss(X, Y) = 0.47952917f0 (tracked)
loss(X, Y) = 0.53521466f0 (tracked)
loss(X, Y) = 0.51441723f0 (tracked)
loss(X, Y) = 0.474f0 (tracked)
loss(X, Y) = 0.49711758f0 (tracked)
loss(X, Y) = 0.49337876f0 (tracked)
loss(X, Y) = 0.49387684f0 (tracked)
```

```
┌ Info: Epoch 1989
└ @ Main /Users/kcft114/.julia/packages/Flux/qXNjB/src/optimise/train.jl:105
┌ Info: Epoch 1990
└ @ Main /Users/kcft114/.julia/packages/Flux/qXNjB/src/optimise/train.jl:105
┌ Info: Epoch 1991
└ @ Main /Users/kcft114/.julia/packages/Flux/qXNjB/src/optimise/train.jl:105
┌ Info: Epoch 1992
└ @ Main /Users/kcft114/.julia/packages/Flux/qXNjB/src/optimise/train.jl:105
┌ Info: Epoch 1993
└ @ Main /Users/kcft114/.julia/packages/Flux/qXNjB/src/optimise/train.jl:105
┌ Info: Epoch 1994
└ @ Main /Users/kcft114/.julia/packages/Flux/qXNjB/src/optimise/train.jl:105
┌ Info: Epoch 1995
└ @ Main /Users/kcft114/.julia/packages/Flux/qXNjB/src/optimise/train.jl:105
┌ Info: Epoch 1996
└ @ Main /Users/kcft114/.julia/packages/Flux/qXNjB/src/optimise/train.jl:105
┌ Info: Epoch 1997
└ @ Main /Users/kcft114/.julia/packages/Flux/qXNjB/src/optimise/train.jl:105
┌ Info: Epoch 1998
└ @ Main /Users/kcft114/.julia/packages/Flux/qXNjB/src/optimise/train.jl:105
┌ Info: Epoch 1999
└ @ Main /Users/kcft114/.julia/packages/Flux/qXNjB/src/optimise/train.jl:105
┌ Info: Epoch 2000
└ @ Main /Users/kcft114/.julia/packages/Flux/qXNjB/src/optimise/train.jl:105
┌ Info: Epoch 1
└ @ Main /Users/kcft114/.julia/packages/Flux/qXNjB/src/optimise/train.jl:105
┌ Info: Epoch 2
└ @ Main /Users/kcft114/.julia/packages/Flux/qXNjB/src/optimise/train.jl:105
┌ Info: Epoch 3
└ @ Main /Users/kcft114/.julia/packages/Flux/qXNjB/src/optimise/train.jl:105
┌ Info: Epoch 4
└ @ Main /Users/kcft114/.julia/packages/Flux/qXNjB/src/optimise/train.jl:105
┌ Info: Epoch 5
└ @ Main /Users/kcft114/.julia/packages/Flux/qXNjB/src/optimise/train.jl:105
┌ Info: Epoch 6
└ @ Main /Users/kcft114/.julia/packages/Flux/qXNjB/src/optimise/train.jl:105
┌ Info: Epoch 7
└ @ Main /Users/kcft114/.julia/packages/Flux/qXNjB/src/optimise/train.jl:105
┌ Info: Epoch 8
└ @ Main /Users/kcft114/.julia/packages/Flux/qXNjB/src/optimise/train.jl:105
┌ Info: Epoch 9
└ @ Main /Users/kcft114/.julia/packages/Flux/qXNjB/src/optimise/train.jl:105
┌ Info: Epoch 10
└ @ Main /Users/kcft114/.julia/packages/Flux/qXNjB/src/optimise/train.jl:105
┌ Info: Epoch 11
└ @ Main /Users/kcft114/.julia/packages/Flux/qXNjB/src/optimise/train.jl:105
┌ Info: Epoch 12
└ @ Main /Users/kcft114/.julia/packages/Flux/qXNjB/src/optimise/train.jl:105
┌ Info: Epoch 13
└ @ Main /Users/kcft114/.julia/packages/Flux/qXNjB/src/optimise/train.jl:105
┌ Info: Epoch 14
└ @ Main /Users/kcft114/.julia/packages/Flux/qXNjB/src/optimise/train.jl:105
┌ Info: Epoch 15
└ @ Main /Users/kcft114/.julia/packages/Flux/qXNjB/src/optimise/train.jl:105
┌ Info: Epoch 16
└ @ Main /Users/kcft114/.julia/packages/Flux/qXNjB/src/optimise/train.jl:105
┌ Info: Epoch 17
└ @ Main /Users/kcft114/.julia/packages/Flux/qXNjB/src/optimise/train.jl:105
┌ Info: Epoch 18
└ @ Main /Users/kcft114/.julia/packages/Flux/qXNjB/src/optimise/train.jl:105
┌ Info: Epoch 19
└ @ Main /Users/kcft114/.julia/packages/Flux/qXNjB/src/optimise/train.jl:105
┌ Info: Epoch 20
└ @ Main /Users/kcft114/.julia/packages/Flux/qXNjB/src/optimise/train.jl:105
┌ Info: Epoch 21
└ @ Main /Users/kcft114/.julia/packages/Flux/qXNjB/src/optimise/train.jl:105
┌ Info: Epoch 22
└ @ Main /Users/kcft114/.julia/packages/Flux/qXNjB/src/optimise/train.jl:105
┌ Info: Epoch 23
└ @ Main /Users/kcft114/.julia/packages/Flux/qXNjB/src/optimise/train.jl:105
┌ Info: Epoch 24
└ @ Main /Users/kcft114/.julia/packages/Flux/qXNjB/src/optimise/train.jl:105
┌ Info: Epoch 25
└ @ Main /Users/kcft114/.julia/packages/Flux/qXNjB/src/optimise/train.jl:105
┌ Info: Epoch 26
└ @ Main /Users/kcft114/.julia/packages/Flux/qXNjB/src/optimise/train.jl:105
┌ Info: Epoch 27
└ @ Main /Users/kcft114/.julia/packages/Flux/qXNjB/src/optimise/train.jl:105
┌ Info: Epoch 28
└ @ Main /Users/kcft114/.julia/packages/Flux/qXNjB/src/optimise/train.jl:105
┌ Info: Epoch 29
└ @ Main /Users/kcft114/.julia/packages/Flux/qXNjB/src/optimise/train.jl:105
┌ Info: Epoch 30
└ @ Main /Users/kcft114/.julia/packages/Flux/qXNjB/src/optimise/train.jl:105
┌ Info: Epoch 31
└ @ Main /Users/kcft114/.julia/packages/Flux/qXNjB/src/optimise/train.jl:105
┌ Info: Epoch 32
└ @ Main /Users/kcft114/.julia/packages/Flux/qXNjB/src/optimise/train.jl:105
┌ Info: Epoch 33
└ @ Main /Users/kcft114/.julia/packages/Flux/qXNjB/src/optimise/train.jl:105
┌ Info: Epoch 34
└ @ Main /Users/kcft114/.julia/packages/Flux/qXNjB/src/optimise/train.jl:105
┌ Info: Epoch 35
└ @ Main /Users/kcft114/.julia/packages/Flux/qXNjB/src/optimise/train.jl:105
┌ Info: Epoch 36
└ @ Main /Users/kcft114/.julia/packages/Flux/qXNjB/src/optimise/train.jl:105
┌ Info: Epoch 37
└ @ Main /Users/kcft114/.julia/packages/Flux/qXNjB/src/optimise/train.jl:105
┌ Info: Epoch 38
└ @ Main /Users/kcft114/.julia/packages/Flux/qXNjB/src/optimise/train.jl:105
┌ Info: Epoch 39
└ @ Main /Users/kcft114/.julia/packages/Flux/qXNjB/src/optimise/train.jl:105
┌ Info: Epoch 40
└ @ Main /Users/kcft114/.julia/packages/Flux/qXNjB/src/optimise/train.jl:105
┌ Info: Epoch 41
└ @ Main /Users/kcft114/.julia/packages/Flux/qXNjB/src/optimise/train.jl:105
┌ Info: Epoch 42
└ @ Main /Users/kcft114/.julia/packages/Flux/qXNjB/src/optimise/train.jl:105
┌ Info: Epoch 43
└ @ Main /Users/kcft114/.julia/packages/Flux/qXNjB/src/optimise/train.jl:105
┌ Info: Epoch 44
└ @ Main /Users/kcft114/.julia/packages/Flux/qXNjB/src/optimise/train.jl:105
┌ Info: Epoch 45
└ @ Main /Users/kcft114/.julia/packages/Flux/qXNjB/src/optimise/train.jl:105
┌ Info: Epoch 46
└ @ Main /Users/kcft114/.julia/packages/Flux/qXNjB/src/optimise/train.jl:105
┌ Info: Epoch 47
└ @ Main /Users/kcft114/.julia/packages/Flux/qXNjB/src/optimise/train.jl:105
┌ Info: Epoch 48
└ @ Main /Users/kcft114/.julia/packages/Flux/qXNjB/src/optimise/train.jl:105
┌ Info: Epoch 49
└ @ Main /Users/kcft114/.julia/packages/Flux/qXNjB/src/optimise/train.jl:105
┌ Info: Epoch 50
└ @ Main /Users/kcft114/.julia/packages/Flux/qXNjB/src/optimise/train.jl:105
┌ Info: Epoch 51
└ @ Main /Users/kcft114/.julia/packages/Flux/qXNjB/src/optimise/train.jl:105
┌ Info: Epoch 52
└ @ Main /Users/kcft114/.julia/packages/Flux/qXNjB/src/optimise/train.jl:105
┌ Info: Epoch 53
└ @ Main /Users/kcft114/.julia/packages/Flux/qXNjB/src/optimise/train.jl:105
┌ Info: Epoch 54
└ @ Main /Users/kcft114/.julia/packages/Flux/qXNjB/src/optimise/train.jl:105
┌ Info: Epoch 55
└ @ Main /Users/kcft114/.julia/packages/Flux/qXNjB/src/optimise/train.jl:105
┌ Info: Epoch 56
└ @ Main /Users/kcft114/.julia/packages/Flux/qXNjB/src/optimise/train.jl:105
┌ Info: Epoch 57
└ @ Main /Users/kcft114/.julia/packages/Flux/qXNjB/src/optimise/train.jl:105
┌ Info: Epoch 58
└ @ Main /Users/kcft114/.julia/packages/Flux/qXNjB/src/optimise/train.jl:105
┌ Info: Epoch 59
└ @ Main /Users/kcft114/.julia/packages/Flux/qXNjB/src/optimise/train.jl:105
┌ Info: Epoch 60
└ @ Main /Users/kcft114/.julia/packages/Flux/qXNjB/src/optimise/train.jl:105
┌ Info: Epoch 61
└ @ Main /Users/kcft114/.julia/packages/Flux/qXNjB/src/optimise/train.jl:105
┌ Info: Epoch 62
└ @ Main /Users/kcft114/.julia/packages/Flux/qXNjB/src/optimise/train.jl:105
┌ Info: Epoch 63
└ @ Main /Users/kcft114/.julia/packages/Flux/qXNjB/src/optimise/train.jl:105
┌ Info: Epoch 64
└ @ Main /Users/kcft114/.julia/packages/Flux/qXNjB/src/optimise/train.jl:105
┌ Info: Epoch 65
└ @ Main /Users/kcft114/.julia/packages/Flux/qXNjB/src/optimise/train.jl:105
┌ Info: Epoch 66
└ @ Main /Users/kcft114/.julia/packages/Flux/qXNjB/src/optimise/train.jl:105
┌ Info: Epoch 67
└ @ Main /Users/kcft114/.julia/packages/Flux/qXNjB/src/optimise/train.jl:105
┌ Info: Epoch 68
└ @ Main /Users/kcft114/.julia/packages/Flux/qXNjB/src/optimise/train.jl:105
┌ Info: Epoch 69
└ @ Main /Users/kcft114/.julia/packages/Flux/qXNjB/src/optimise/train.jl:105
┌ Info: Epoch 70
└ @ Main /Users/kcft114/.julia/packages/Flux/qXNjB/src/optimise/train.jl:105
┌ Info: Epoch 71
└ @ Main /Users/kcft114/.julia/packages/Flux/qXNjB/src/optimise/train.jl:105
┌ Info: Epoch 72
└ @ Main /Users/kcft114/.julia/packages/Flux/qXNjB/src/optimise/train.jl:105
┌ Info: Epoch 73
└ @ Main /Users/kcft114/.julia/packages/Flux/qXNjB/src/optimise/train.jl:105
┌ Info: Epoch 74
└ @ Main /Users/kcft114/.julia/packages/Flux/qXNjB/src/optimise/train.jl:105
┌ Info: Epoch 75
└ @ Main /Users/kcft114/.julia/packages/Flux/qXNjB/src/optimise/train.jl:105
┌ Info: Epoch 76
└ @ Main /Users/kcft114/.julia/packages/Flux/qXNjB/src/optimise/train.jl:105
┌ Info: Epoch 77
└ @ Main /Users/kcft114/.julia/packages/Flux/qXNjB/src/optimise/train.jl:105
┌ Info: Epoch 78
└ @ Main /Users/kcft114/.julia/packages/Flux/qXNjB/src/optimise/train.jl:105
┌ Info: Epoch 79
└ @ Main /Users/kcft114/.julia/packages/Flux/qXNjB/src/optimise/train.jl:105
┌ Info: Epoch 80
└ @ Main /Users/kcft114/.julia/packages/Flux/qXNjB/src/optimise/train.jl:105
┌ Info: Epoch 81
└ @ Main /Users/kcft114/.julia/packages/Flux/qXNjB/src/optimise/train.jl:105
┌ Info: Epoch 82
└ @ Main /Users/kcft114/.julia/packages/Flux/qXNjB/src/optimise/train.jl:105
┌ Info: Epoch 83
└ @ Main /Users/kcft114/.julia/packages/Flux/qXNjB/src/optimise/train.jl:105
┌ Info: Epoch 84
└ @ Main /Users/kcft114/.julia/packages/Flux/qXNjB/src/optimise/train.jl:105
┌ Info: Epoch 85
└ @ Main /Users/kcft114/.julia/packages/Flux/qXNjB/src/optimise/train.jl:105
┌ Info: Epoch 86
└ @ Main /Users/kcft114/.julia/packages/Flux/qXNjB/src/optimise/train.jl:105
┌ Info: Epoch 87
└ @ Main /Users/kcft114/.julia/packages/Flux/qXNjB/src/optimise/train.jl:105
┌ Info: Epoch 88
└ @ Main /Users/kcft114/.julia/packages/Flux/qXNjB/src/optimise/train.jl:105
┌ Info: Epoch 89
└ @ Main /Users/kcft114/.julia/packages/Flux/qXNjB/src/optimise/train.jl:105
┌ Info: Epoch 90
└ @ Main /Users/kcft114/.julia/packages/Flux/qXNjB/src/optimise/train.jl:105
┌ Info: Epoch 91
└ @ Main /Users/kcft114/.julia/packages/Flux/qXNjB/src/optimise/train.jl:105
┌ Info: Epoch 92
└ @ Main /Users/kcft114/.julia/packages/Flux/qXNjB/src/optimise/train.jl:105
┌ Info: Epoch 93
└ @ Main /Users/kcft114/.julia/packages/Flux/qXNjB/src/optimise/train.jl:105
┌ Info: Epoch 94
└ @ Main /Users/kcft114/.julia/packages/Flux/qXNjB/src/optimise/train.jl:105
┌ Info: Epoch 95
└ @ Main /Users/kcft114/.julia/packages/Flux/qXNjB/src/optimise/train.jl:105
```

```
loss(X, Y) = 0.47449455f0 (tracked)
loss(X, Y) = 0.540901f0 (tracked)
loss(X, Y) = 0.49212313f0 (tracked)
loss(X, Y) = 0.52217996f0 (tracked)
loss(X, Y) = 0.5634943f0 (tracked)
loss(X, Y) = 0.5377652f0 (tracked)
loss(X, Y) = 0.4763579f0 (tracked)
loss(X, Y) = 0.5420434f0 (tracked)
loss(X, Y) = 0.52346736f0 (tracked)
loss(X, Y) = 0.5765263f0 (tracked)
train accuracy = 0.782312925170068
test accuracy = 0.4594594594594595
train BA = 0.8
test BA = 0.52
loss(X, Y) = 1.1924303f0 (tracked)
loss(X, Y) = 1.1670403f0 (tracked)
loss(X, Y) = 1.1839769f0 (tracked)
loss(X, Y) = 1.1551604f0 (tracked)
loss(X, Y) = 1.1754657f0 (tracked)
loss(X, Y) = 1.1729093f0 (tracked)
loss(X, Y) = 1.1539807f0 (tracked)
loss(X, Y) = 1.1697649f0 (tracked)
loss(X, Y) = 1.1596665f0 (tracked)
loss(X, Y) = 1.1446366f0 (tracked)
loss(X, Y) = 1.1513225f0 (tracked)
loss(X, Y) = 1.1365806f0 (tracked)
loss(X, Y) = 1.1131407f0 (tracked)
loss(X, Y) = 1.1310139f0 (tracked)
loss(X, Y) = 1.1053243f0 (tracked)
loss(X, Y) = 1.141161f0 (tracked)
loss(X, Y) = 1.1274886f0 (tracked)
loss(X, Y) = 1.1035318f0 (tracked)
loss(X, Y) = 1.1398599f0 (tracked)
loss(X, Y) = 1.1167977f0 (tracked)
loss(X, Y) = 1.0974493f0 (tracked)
loss(X, Y) = 1.1102148f0 (tracked)
loss(X, Y) = 1.1224184f0 (tracked)
loss(X, Y) = 1.1009597f0 (tracked)
loss(X, Y) = 1.0990167f0 (tracked)
loss(X, Y) = 1.0989705f0 (tracked)
loss(X, Y) = 1.0684087f0 (tracked)
loss(X, Y) = 1.1032207f0 (tracked)
loss(X, Y) = 1.0755585f0 (tracked)
loss(X, Y) = 1.0706838f0 (tracked)
loss(X, Y) = 1.0732312f0 (tracked)
loss(X, Y) = 1.0689998f0 (tracked)
loss(X, Y) = 1.044254f0 (tracked)
loss(X, Y) = 1.0671476f0 (tracked)
loss(X, Y) = 1.0576847f0 (tracked)
loss(X, Y) = 1.0593337f0 (tracked)
loss(X, Y) = 1.0523466f0 (tracked)
loss(X, Y) = 1.0496161f0 (tracked)
loss(X, Y) = 1.0694455f0 (tracked)
loss(X, Y) = 1.0499622f0 (tracked)
loss(X, Y) = 1.0525339f0 (tracked)
loss(X, Y) = 1.0581651f0 (tracked)
loss(X, Y) = 1.0670453f0 (tracked)
loss(X, Y) = 1.0430114f0 (tracked)
loss(X, Y) = 1.024181f0 (tracked)
loss(X, Y) = 1.0247699f0 (tracked)
loss(X, Y) = 1.0248165f0 (tracked)
loss(X, Y) = 1.0310365f0 (tracked)
loss(X, Y) = 1.0312792f0 (tracked)
loss(X, Y) = 1.013583f0 (tracked)
loss(X, Y) = 1.0239927f0 (tracked)
loss(X, Y) = 1.0228463f0 (tracked)
loss(X, Y) = 1.0128883f0 (tracked)
loss(X, Y) = 1.0128149f0 (tracked)
loss(X, Y) = 1.021116f0 (tracked)
loss(X, Y) = 1.0246103f0 (tracked)
loss(X, Y) = 1.0309224f0 (tracked)
loss(X, Y) = 1.0177629f0 (tracked)
loss(X, Y) = 0.9919718f0 (tracked)
loss(X, Y) = 1.011944f0 (tracked)
loss(X, Y) = 1.0103476f0 (tracked)
loss(X, Y) = 0.9999282f0 (tracked)
loss(X, Y) = 0.9983007f0 (tracked)
loss(X, Y) = 0.98852164f0 (tracked)
loss(X, Y) = 0.9978602f0 (tracked)
loss(X, Y) = 0.99554896f0 (tracked)
loss(X, Y) = 0.9942924f0 (tracked)
loss(X, Y) = 1.0019269f0 (tracked)
loss(X, Y) = 1.001684f0 (tracked)
loss(X, Y) = 0.97977656f0 (tracked)
loss(X, Y) = 0.9795937f0 (tracked)
loss(X, Y) = 0.9934992f0 (tracked)
loss(X, Y) = 0.999473f0 (tracked)
loss(X, Y) = 0.9773179f0 (tracked)
loss(X, Y) = 0.9681373f0 (tracked)
loss(X, Y) = 0.99434954f0 (tracked)
loss(X, Y) = 0.98679596f0 (tracked)
loss(X, Y) = 0.9708449f0 (tracked)
loss(X, Y) = 0.9805126f0 (tracked)
loss(X, Y) = 0.95539397f0 (tracked)
loss(X, Y) = 0.96548027f0 (tracked)
loss(X, Y) = 0.969203f0 (tracked)
loss(X, Y) = 0.96261823f0 (tracked)
loss(X, Y) = 0.9581603f0 (tracked)
loss(X, Y) = 0.96451044f0 (tracked)
loss(X, Y) = 0.97139764f0 (tracked)
loss(X, Y) = 0.97843695f0 (tracked)
loss(X, Y) = 0.9495339f0 (tracked)
loss(X, Y) = 0.9620201f0 (tracked)
loss(X, Y) = 0.950853f0 (tracked)
loss(X, Y) = 0.95901465f0 (tracked)
loss(X, Y) = 0.97490895f0 (tracked)
loss(X, Y) = 0.9430436f0 (tracked)
loss(X, Y) = 0.94463557f0 (tracked)
loss(X, Y) = 0.9479321f0 (tracked)
loss(X, Y) = 0.9360145f0 (tracked)
loss(X, Y) = 0.9484025f0 (tracked)
loss(X, Y) = 0.9409691f0 (tracked)
loss(X, Y) = 0.92372876f0 (tracked)
loss(X, Y) = 0.9279392f0 (tracked)
loss(X, Y) = 0.933584f0 (tracked)
loss(X, Y) = 0.93409103f0 (tracked)
loss(X, Y) = 0.94152766f0 (tracked)
loss(X, Y) = 0.9188255f0 (tracked)
loss(X, Y) = 0.94296575f0 (tracked)
loss(X, Y) = 0.9293013f0 (tracked)
loss(X, Y) = 0.9223118f0 (tracked)
loss(X, Y) = 0.9290828f0 (tracked)
loss(X, Y) = 0.94989705f0 (tracked)
loss(X, Y) = 0.93284875f0 (tracked)
loss(X, Y) = 0.9278125f0 (tracked)
```

```
┌ Info: Epoch 96
└ @ Main /Users/kcft114/.julia/packages/Flux/qXNjB/src/optimise/train.jl:105
┌ Info: Epoch 97
└ @ Main /Users/kcft114/.julia/packages/Flux/qXNjB/src/optimise/train.jl:105
┌ Info: Epoch 98
└ @ Main /Users/kcft114/.julia/packages/Flux/qXNjB/src/optimise/train.jl:105
┌ Info: Epoch 99
└ @ Main /Users/kcft114/.julia/packages/Flux/qXNjB/src/optimise/train.jl:105
┌ Info: Epoch 100
└ @ Main /Users/kcft114/.julia/packages/Flux/qXNjB/src/optimise/train.jl:105
┌ Info: Epoch 101
└ @ Main /Users/kcft114/.julia/packages/Flux/qXNjB/src/optimise/train.jl:105
┌ Info: Epoch 102
└ @ Main /Users/kcft114/.julia/packages/Flux/qXNjB/src/optimise/train.jl:105
┌ Info: Epoch 103
└ @ Main /Users/kcft114/.julia/packages/Flux/qXNjB/src/optimise/train.jl:105
┌ Info: Epoch 104
└ @ Main /Users/kcft114/.julia/packages/Flux/qXNjB/src/optimise/train.jl:105
┌ Info: Epoch 105
└ @ Main /Users/kcft114/.julia/packages/Flux/qXNjB/src/optimise/train.jl:105
┌ Info: Epoch 106
└ @ Main /Users/kcft114/.julia/packages/Flux/qXNjB/src/optimise/train.jl:105
┌ Info: Epoch 107
└ @ Main /Users/kcft114/.julia/packages/Flux/qXNjB/src/optimise/train.jl:105
┌ Info: Epoch 108
└ @ Main /Users/kcft114/.julia/packages/Flux/qXNjB/src/optimise/train.jl:105
┌ Info: Epoch 109
└ @ Main /Users/kcft114/.julia/packages/Flux/qXNjB/src/optimise/train.jl:105
┌ Info: Epoch 110
└ @ Main /Users/kcft114/.julia/packages/Flux/qXNjB/src/optimise/train.jl:105
┌ Info: Epoch 111
└ @ Main /Users/kcft114/.julia/packages/Flux/qXNjB/src/optimise/train.jl:105
┌ Info: Epoch 112
└ @ Main /Users/kcft114/.julia/packages/Flux/qXNjB/src/optimise/train.jl:105
┌ Info: Epoch 113
└ @ Main /Users/kcft114/.julia/packages/Flux/qXNjB/src/optimise/train.jl:105
┌ Info: Epoch 114
└ @ Main /Users/kcft114/.julia/packages/Flux/qXNjB/src/optimise/train.jl:105
┌ Info: Epoch 115
└ @ Main /Users/kcft114/.julia/packages/Flux/qXNjB/src/optimise/train.jl:105
┌ Info: Epoch 116
└ @ Main /Users/kcft114/.julia/packages/Flux/qXNjB/src/optimise/train.jl:105
┌ Info: Epoch 117
└ @ Main /Users/kcft114/.julia/packages/Flux/qXNjB/src/optimise/train.jl:105
┌ Info: Epoch 118
└ @ Main /Users/kcft114/.julia/packages/Flux/qXNjB/src/optimise/train.jl:105
┌ Info: Epoch 119
└ @ Main /Users/kcft114/.julia/packages/Flux/qXNjB/src/optimise/train.jl:105
┌ Info: Epoch 120
└ @ Main /Users/kcft114/.julia/packages/Flux/qXNjB/src/optimise/train.jl:105
┌ Info: Epoch 121
└ @ Main /Users/kcft114/.julia/packages/Flux/qXNjB/src/optimise/train.jl:105
┌ Info: Epoch 122
└ @ Main /Users/kcft114/.julia/packages/Flux/qXNjB/src/optimise/train.jl:105
┌ Info: Epoch 123
└ @ Main /Users/kcft114/.julia/packages/Flux/qXNjB/src/optimise/train.jl:105
┌ Info: Epoch 124
└ @ Main /Users/kcft114/.julia/packages/Flux/qXNjB/src/optimise/train.jl:105
┌ Info: Epoch 125
└ @ Main /Users/kcft114/.julia/packages/Flux/qXNjB/src/optimise/train.jl:105
┌ Info: Epoch 126
└ @ Main /Users/kcft114/.julia/packages/Flux/qXNjB/src/optimise/train.jl:105
┌ Info: Epoch 127
└ @ Main /Users/kcft114/.julia/packages/Flux/qXNjB/src/optimise/train.jl:105
┌ Info: Epoch 128
└ @ Main /Users/kcft114/.julia/packages/Flux/qXNjB/src/optimise/train.jl:105
┌ Info: Epoch 129
└ @ Main /Users/kcft114/.julia/packages/Flux/qXNjB/src/optimise/train.jl:105
┌ Info: Epoch 130
└ @ Main /Users/kcft114/.julia/packages/Flux/qXNjB/src/optimise/train.jl:105
┌ Info: Epoch 131
└ @ Main /Users/kcft114/.julia/packages/Flux/qXNjB/src/optimise/train.jl:105
┌ Info: Epoch 132
└ @ Main /Users/kcft114/.julia/packages/Flux/qXNjB/src/optimise/train.jl:105
┌ Info: Epoch 133
└ @ Main /Users/kcft114/.julia/packages/Flux/qXNjB/src/optimise/train.jl:105
┌ Info: Epoch 134
└ @ Main /Users/kcft114/.julia/packages/Flux/qXNjB/src/optimise/train.jl:105
┌ Info: Epoch 135
└ @ Main /Users/kcft114/.julia/packages/Flux/qXNjB/src/optimise/train.jl:105
┌ Info: Epoch 136
└ @ Main /Users/kcft114/.julia/packages/Flux/qXNjB/src/optimise/train.jl:105
┌ Info: Epoch 137
└ @ Main /Users/kcft114/.julia/packages/Flux/qXNjB/src/optimise/train.jl:105
┌ Info: Epoch 138
└ @ Main /Users/kcft114/.julia/packages/Flux/qXNjB/src/optimise/train.jl:105
┌ Info: Epoch 139
└ @ Main /Users/kcft114/.julia/packages/Flux/qXNjB/src/optimise/train.jl:105
┌ Info: Epoch 140
└ @ Main /Users/kcft114/.julia/packages/Flux/qXNjB/src/optimise/train.jl:105
┌ Info: Epoch 141
└ @ Main /Users/kcft114/.julia/packages/Flux/qXNjB/src/optimise/train.jl:105
┌ Info: Epoch 142
└ @ Main /Users/kcft114/.julia/packages/Flux/qXNjB/src/optimise/train.jl:105
┌ Info: Epoch 143
└ @ Main /Users/kcft114/.julia/packages/Flux/qXNjB/src/optimise/train.jl:105
┌ Info: Epoch 144
└ @ Main /Users/kcft114/.julia/packages/Flux/qXNjB/src/optimise/train.jl:105
┌ Info: Epoch 145
└ @ Main /Users/kcft114/.julia/packages/Flux/qXNjB/src/optimise/train.jl:105
┌ Info: Epoch 146
└ @ Main /Users/kcft114/.julia/packages/Flux/qXNjB/src/optimise/train.jl:105
┌ Info: Epoch 147
└ @ Main /Users/kcft114/.julia/packages/Flux/qXNjB/src/optimise/train.jl:105
┌ Info: Epoch 148
└ @ Main /Users/kcft114/.julia/packages/Flux/qXNjB/src/optimise/train.jl:105
┌ Info: Epoch 149
└ @ Main /Users/kcft114/.julia/packages/Flux/qXNjB/src/optimise/train.jl:105
┌ Info: Epoch 150
└ @ Main /Users/kcft114/.julia/packages/Flux/qXNjB/src/optimise/train.jl:105
┌ Info: Epoch 151
└ @ Main /Users/kcft114/.julia/packages/Flux/qXNjB/src/optimise/train.jl:105
┌ Info: Epoch 152
└ @ Main /Users/kcft114/.julia/packages/Flux/qXNjB/src/optimise/train.jl:105
┌ Info: Epoch 153
└ @ Main /Users/kcft114/.julia/packages/Flux/qXNjB/src/optimise/train.jl:105
┌ Info: Epoch 154
└ @ Main /Users/kcft114/.julia/packages/Flux/qXNjB/src/optimise/train.jl:105
┌ Info: Epoch 155
└ @ Main /Users/kcft114/.julia/packages/Flux/qXNjB/src/optimise/train.jl:105
┌ Info: Epoch 156
└ @ Main /Users/kcft114/.julia/packages/Flux/qXNjB/src/optimise/train.jl:105
┌ Info: Epoch 157
└ @ Main /Users/kcft114/.julia/packages/Flux/qXNjB/src/optimise/train.jl:105
┌ Info: Epoch 158
└ @ Main /Users/kcft114/.julia/packages/Flux/qXNjB/src/optimise/train.jl:105
┌ Info: Epoch 159
└ @ Main /Users/kcft114/.julia/packages/Flux/qXNjB/src/optimise/train.jl:105
┌ Info: Epoch 160
└ @ Main /Users/kcft114/.julia/packages/Flux/qXNjB/src/optimise/train.jl:105
┌ Info: Epoch 161
└ @ Main /Users/kcft114/.julia/packages/Flux/qXNjB/src/optimise/train.jl:105
┌ Info: Epoch 162
└ @ Main /Users/kcft114/.julia/packages/Flux/qXNjB/src/optimise/train.jl:105
┌ Info: Epoch 163
└ @ Main /Users/kcft114/.julia/packages/Flux/qXNjB/src/optimise/train.jl:105
┌ Info: Epoch 164
└ @ Main /Users/kcft114/.julia/packages/Flux/qXNjB/src/optimise/train.jl:105
┌ Info: Epoch 165
└ @ Main /Users/kcft114/.julia/packages/Flux/qXNjB/src/optimise/train.jl:105
┌ Info: Epoch 166
└ @ Main /Users/kcft114/.julia/packages/Flux/qXNjB/src/optimise/train.jl:105
┌ Info: Epoch 167
└ @ Main /Users/kcft114/.julia/packages/Flux/qXNjB/src/optimise/train.jl:105
┌ Info: Epoch 168
└ @ Main /Users/kcft114/.julia/packages/Flux/qXNjB/src/optimise/train.jl:105
┌ Info: Epoch 169
└ @ Main /Users/kcft114/.julia/packages/Flux/qXNjB/src/optimise/train.jl:105
┌ Info: Epoch 170
└ @ Main /Users/kcft114/.julia/packages/Flux/qXNjB/src/optimise/train.jl:105
┌ Info: Epoch 171
└ @ Main /Users/kcft114/.julia/packages/Flux/qXNjB/src/optimise/train.jl:105
┌ Info: Epoch 172
└ @ Main /Users/kcft114/.julia/packages/Flux/qXNjB/src/optimise/train.jl:105
┌ Info: Epoch 173
└ @ Main /Users/kcft114/.julia/packages/Flux/qXNjB/src/optimise/train.jl:105
┌ Info: Epoch 174
└ @ Main /Users/kcft114/.julia/packages/Flux/qXNjB/src/optimise/train.jl:105
┌ Info: Epoch 175
└ @ Main /Users/kcft114/.julia/packages/Flux/qXNjB/src/optimise/train.jl:105
┌ Info: Epoch 176
└ @ Main /Users/kcft114/.julia/packages/Flux/qXNjB/src/optimise/train.jl:105
┌ Info: Epoch 177
└ @ Main /Users/kcft114/.julia/packages/Flux/qXNjB/src/optimise/train.jl:105
┌ Info: Epoch 178
└ @ Main /Users/kcft114/.julia/packages/Flux/qXNjB/src/optimise/train.jl:105
┌ Info: Epoch 179
└ @ Main /Users/kcft114/.julia/packages/Flux/qXNjB/src/optimise/train.jl:105
┌ Info: Epoch 180
└ @ Main /Users/kcft114/.julia/packages/Flux/qXNjB/src/optimise/train.jl:105
┌ Info: Epoch 181
└ @ Main /Users/kcft114/.julia/packages/Flux/qXNjB/src/optimise/train.jl:105
┌ Info: Epoch 182
└ @ Main /Users/kcft114/.julia/packages/Flux/qXNjB/src/optimise/train.jl:105
┌ Info: Epoch 183
└ @ Main /Users/kcft114/.julia/packages/Flux/qXNjB/src/optimise/train.jl:105
┌ Info: Epoch 184
└ @ Main /Users/kcft114/.julia/packages/Flux/qXNjB/src/optimise/train.jl:105
┌ Info: Epoch 185
└ @ Main /Users/kcft114/.julia/packages/Flux/qXNjB/src/optimise/train.jl:105
┌ Info: Epoch 186
└ @ Main /Users/kcft114/.julia/packages/Flux/qXNjB/src/optimise/train.jl:105
┌ Info: Epoch 187
└ @ Main /Users/kcft114/.julia/packages/Flux/qXNjB/src/optimise/train.jl:105
┌ Info: Epoch 188
└ @ Main /Users/kcft114/.julia/packages/Flux/qXNjB/src/optimise/train.jl:105
┌ Info: Epoch 189
└ @ Main /Users/kcft114/.julia/packages/Flux/qXNjB/src/optimise/train.jl:105
┌ Info: Epoch 190
└ @ Main /Users/kcft114/.julia/packages/Flux/qXNjB/src/optimise/train.jl:105
┌ Info: Epoch 191
└ @ Main /Users/kcft114/.julia/packages/Flux/qXNjB/src/optimise/train.jl:105
┌ Info: Epoch 192
└ @ Main /Users/kcft114/.julia/packages/Flux/qXNjB/src/optimise/train.jl:105
┌ Info: Epoch 193
└ @ Main /Users/kcft114/.julia/packages/Flux/qXNjB/src/optimise/train.jl:105
┌ Info: Epoch 194
└ @ Main /Users/kcft114/.julia/packages/Flux/qXNjB/src/optimise/train.jl:105
┌ Info: Epoch 195
└ @ Main /Users/kcft114/.julia/packages/Flux/qXNjB/src/optimise/train.jl:105
┌ Info: Epoch 196
└ @ Main /Users/kcft114/.julia/packages/Flux/qXNjB/src/optimise/train.jl:105
┌ Info: Epoch 197
└ @ Main /Users/kcft114/.julia/packages/Flux/qXNjB/src/optimise/train.jl:105
┌ Info: Epoch 198
└ @ Main /Users/kcft114/.julia/packages/Flux/qXNjB/src/optimise/train.jl:105
┌ Info: Epoch 199
└ @ Main /Users/kcft114/.julia/packages/Flux/qXNjB/src/optimise/train.jl:105
┌ Info: Epoch 200
└ @ Main /Users/kcft114/.julia/packages/Flux/qXNjB/src/optimise/train.jl:105
┌ Info: Epoch 201
└ @ Main /Users/kcft114/.julia/packages/Flux/qXNjB/src/optimise/train.jl:105
┌ Info: Epoch 202
└ @ Main /Users/kcft114/.julia/packages/Flux/qXNjB/src/optimise/train.jl:105
┌ Info: Epoch 203
└ @ Main /Users/kcft114/.julia/packages/Flux/qXNjB/src/optimise/train.jl:105
┌ Info: Epoch 204
└ @ Main /Users/kcft114/.julia/packages/Flux/qXNjB/src/optimise/train.jl:105
┌ Info: Epoch 205
└ @ Main /Users/kcft114/.julia/packages/Flux/qXNjB/src/optimise/train.jl:105
┌ Info: Epoch 206
└ @ Main /Users/kcft114/.julia/packages/Flux/qXNjB/src/optimise/train.jl:105
┌ Info: Epoch 207
└ @ Main /Users/kcft114/.julia/packages/Flux/qXNjB/src/optimise/train.jl:105
┌ Info: Epoch 208
└ @ Main /Users/kcft114/.julia/packages/Flux/qXNjB/src/optimise/train.jl:105
┌ Info: Epoch 209
└ @ Main /Users/kcft114/.julia/packages/Flux/qXNjB/src/optimise/train.jl:105
┌ Info: Epoch 210
└ @ Main /Users/kcft114/.julia/packages/Flux/qXNjB/src/optimise/train.jl:105
┌ Info: Epoch 211
└ @ Main /Users/kcft114/.julia/packages/Flux/qXNjB/src/optimise/train.jl:105
┌ Info: Epoch 212
└ @ Main /Users/kcft114/.julia/packages/Flux/qXNjB/src/optimise/train.jl:105
┌ Info: Epoch 213
└ @ Main /Users/kcft114/.julia/packages/Flux/qXNjB/src/optimise/train.jl:105
┌ Info: Epoch 214
└ @ Main /Users/kcft114/.julia/packages/Flux/qXNjB/src/optimise/train.jl:105
┌ Info: Epoch 215
└ @ Main /Users/kcft114/.julia/packages/Flux/qXNjB/src/optimise/train.jl:105
┌ Info: Epoch 216
└ @ Main /Users/kcft114/.julia/packages/Flux/qXNjB/src/optimise/train.jl:105
```

```
loss(X, Y) = 0.9137103f0 (tracked)
loss(X, Y) = 0.91485655f0 (tracked)
loss(X, Y) = 0.917979f0 (tracked)
loss(X, Y) = 0.90084577f0 (tracked)
loss(X, Y) = 0.9064106f0 (tracked)
loss(X, Y) = 0.8996615f0 (tracked)
loss(X, Y) = 0.9080731f0 (tracked)
loss(X, Y) = 0.9006961f0 (tracked)
loss(X, Y) = 0.91146135f0 (tracked)
loss(X, Y) = 0.91477853f0 (tracked)
loss(X, Y) = 0.8981624f0 (tracked)
loss(X, Y) = 0.8900697f0 (tracked)
loss(X, Y) = 0.890227f0 (tracked)
loss(X, Y) = 0.9093781f0 (tracked)
loss(X, Y) = 0.90225583f0 (tracked)
loss(X, Y) = 0.90508795f0 (tracked)
loss(X, Y) = 0.89381087f0 (tracked)
loss(X, Y) = 0.9082359f0 (tracked)
loss(X, Y) = 0.90074384f0 (tracked)
loss(X, Y) = 0.88047475f0 (tracked)
loss(X, Y) = 0.91694754f0 (tracked)
loss(X, Y) = 0.8862893f0 (tracked)
loss(X, Y) = 0.8851692f0 (tracked)
loss(X, Y) = 0.88853836f0 (tracked)
loss(X, Y) = 0.8726931f0 (tracked)
loss(X, Y) = 0.8898573f0 (tracked)
loss(X, Y) = 0.8792392f0 (tracked)
loss(X, Y) = 0.8941343f0 (tracked)
loss(X, Y) = 0.87214804f0 (tracked)
loss(X, Y) = 0.87179875f0 (tracked)
loss(X, Y) = 0.8781468f0 (tracked)
loss(X, Y) = 0.8898169f0 (tracked)
loss(X, Y) = 0.89315057f0 (tracked)
loss(X, Y) = 0.87765867f0 (tracked)
loss(X, Y) = 0.89213324f0 (tracked)
loss(X, Y) = 0.8680729f0 (tracked)
loss(X, Y) = 0.87760043f0 (tracked)
loss(X, Y) = 0.881171f0 (tracked)
loss(X, Y) = 0.88390726f0 (tracked)
loss(X, Y) = 0.86993665f0 (tracked)
loss(X, Y) = 0.86276793f0 (tracked)
loss(X, Y) = 0.8693929f0 (tracked)
loss(X, Y) = 0.88138574f0 (tracked)
loss(X, Y) = 0.863861f0 (tracked)
loss(X, Y) = 0.8638627f0 (tracked)
loss(X, Y) = 0.861278f0 (tracked)
loss(X, Y) = 0.8704945f0 (tracked)
loss(X, Y) = 0.8454314f0 (tracked)
loss(X, Y) = 0.8595106f0 (tracked)
loss(X, Y) = 0.8432274f0 (tracked)
loss(X, Y) = 0.85090566f0 (tracked)
loss(X, Y) = 0.853925f0 (tracked)
loss(X, Y) = 0.8490393f0 (tracked)
loss(X, Y) = 0.8746863f0 (tracked)
loss(X, Y) = 0.8532852f0 (tracked)
loss(X, Y) = 0.8349711f0 (tracked)
loss(X, Y) = 0.82673174f0 (tracked)
loss(X, Y) = 0.83797127f0 (tracked)
loss(X, Y) = 0.85635555f0 (tracked)
loss(X, Y) = 0.8357984f0 (tracked)
loss(X, Y) = 0.8334588f0 (tracked)
loss(X, Y) = 0.86587375f0 (tracked)
loss(X, Y) = 0.83501107f0 (tracked)
loss(X, Y) = 0.8319402f0 (tracked)
loss(X, Y) = 0.83191806f0 (tracked)
loss(X, Y) = 0.83261275f0 (tracked)
loss(X, Y) = 0.8480857f0 (tracked)
loss(X, Y) = 0.84143156f0 (tracked)
loss(X, Y) = 0.8258314f0 (tracked)
loss(X, Y) = 0.84514207f0 (tracked)
loss(X, Y) = 0.82795966f0 (tracked)
loss(X, Y) = 0.8250829f0 (tracked)
loss(X, Y) = 0.82581675f0 (tracked)
loss(X, Y) = 0.84869057f0 (tracked)
loss(X, Y) = 0.8279628f0 (tracked)
loss(X, Y) = 0.8510843f0 (tracked)
loss(X, Y) = 0.8348006f0 (tracked)
loss(X, Y) = 0.8259056f0 (tracked)
loss(X, Y) = 0.83752453f0 (tracked)
loss(X, Y) = 0.8324687f0 (tracked)
loss(X, Y) = 0.8122195f0 (tracked)
loss(X, Y) = 0.82270306f0 (tracked)
loss(X, Y) = 0.8330369f0 (tracked)
loss(X, Y) = 0.8311548f0 (tracked)
loss(X, Y) = 0.8144833f0 (tracked)
loss(X, Y) = 0.8288756f0 (tracked)
loss(X, Y) = 0.80973023f0 (tracked)
loss(X, Y) = 0.8339986f0 (tracked)
loss(X, Y) = 0.82896817f0 (tracked)
loss(X, Y) = 0.82723004f0 (tracked)
loss(X, Y) = 0.81791f0 (tracked)
loss(X, Y) = 0.8174771f0 (tracked)
loss(X, Y) = 0.8010329f0 (tracked)
loss(X, Y) = 0.8207004f0 (tracked)
loss(X, Y) = 0.8424663f0 (tracked)
loss(X, Y) = 0.8121807f0 (tracked)
loss(X, Y) = 0.80566305f0 (tracked)
loss(X, Y) = 0.8157831f0 (tracked)
loss(X, Y) = 0.81781256f0 (tracked)
loss(X, Y) = 0.8075572f0 (tracked)
loss(X, Y) = 0.8224301f0 (tracked)
loss(X, Y) = 0.80418694f0 (tracked)
loss(X, Y) = 0.7942427f0 (tracked)
loss(X, Y) = 0.7936818f0 (tracked)
loss(X, Y) = 0.80042356f0 (tracked)
loss(X, Y) = 0.8137931f0 (tracked)
loss(X, Y) = 0.8050827f0 (tracked)
loss(X, Y) = 0.8001637f0 (tracked)
loss(X, Y) = 0.80107635f0 (tracked)
loss(X, Y) = 0.81809115f0 (tracked)
loss(X, Y) = 0.8052836f0 (tracked)
loss(X, Y) = 0.8218842f0 (tracked)
loss(X, Y) = 0.79566824f0 (tracked)
loss(X, Y) = 0.7895448f0 (tracked)
loss(X, Y) = 0.8037066f0 (tracked)
loss(X, Y) = 0.8040405f0 (tracked)
loss(X, Y) = 0.78778124f0 (tracked)
loss(X, Y) = 0.79012644f0 (tracked)
loss(X, Y) = 0.80355257f0 (tracked)
loss(X, Y) = 0.77568996f0 (tracked)
loss(X, Y) = 0.80087435f0 (tracked)
loss(X, Y) = 0.79570407f0 (tracked)
loss(X, Y) = 0.7864337f0 (tracked)
loss(X, Y) = 0.80021113f0 (tracked)
```

```
┌ Info: Epoch 217
└ @ Main /Users/kcft114/.julia/packages/Flux/qXNjB/src/optimise/train.jl:105
┌ Info: Epoch 218
└ @ Main /Users/kcft114/.julia/packages/Flux/qXNjB/src/optimise/train.jl:105
┌ Info: Epoch 219
└ @ Main /Users/kcft114/.julia/packages/Flux/qXNjB/src/optimise/train.jl:105
┌ Info: Epoch 220
└ @ Main /Users/kcft114/.julia/packages/Flux/qXNjB/src/optimise/train.jl:105
┌ Info: Epoch 221
└ @ Main /Users/kcft114/.julia/packages/Flux/qXNjB/src/optimise/train.jl:105
┌ Info: Epoch 222
└ @ Main /Users/kcft114/.julia/packages/Flux/qXNjB/src/optimise/train.jl:105
┌ Info: Epoch 223
└ @ Main /Users/kcft114/.julia/packages/Flux/qXNjB/src/optimise/train.jl:105
┌ Info: Epoch 224
└ @ Main /Users/kcft114/.julia/packages/Flux/qXNjB/src/optimise/train.jl:105
┌ Info: Epoch 225
└ @ Main /Users/kcft114/.julia/packages/Flux/qXNjB/src/optimise/train.jl:105
┌ Info: Epoch 226
└ @ Main /Users/kcft114/.julia/packages/Flux/qXNjB/src/optimise/train.jl:105
┌ Info: Epoch 227
└ @ Main /Users/kcft114/.julia/packages/Flux/qXNjB/src/optimise/train.jl:105
┌ Info: Epoch 228
└ @ Main /Users/kcft114/.julia/packages/Flux/qXNjB/src/optimise/train.jl:105
┌ Info: Epoch 229
└ @ Main /Users/kcft114/.julia/packages/Flux/qXNjB/src/optimise/train.jl:105
┌ Info: Epoch 230
└ @ Main /Users/kcft114/.julia/packages/Flux/qXNjB/src/optimise/train.jl:105
┌ Info: Epoch 231
└ @ Main /Users/kcft114/.julia/packages/Flux/qXNjB/src/optimise/train.jl:105
┌ Info: Epoch 232
└ @ Main /Users/kcft114/.julia/packages/Flux/qXNjB/src/optimise/train.jl:105
┌ Info: Epoch 233
└ @ Main /Users/kcft114/.julia/packages/Flux/qXNjB/src/optimise/train.jl:105
┌ Info: Epoch 234
└ @ Main /Users/kcft114/.julia/packages/Flux/qXNjB/src/optimise/train.jl:105
┌ Info: Epoch 235
└ @ Main /Users/kcft114/.julia/packages/Flux/qXNjB/src/optimise/train.jl:105
┌ Info: Epoch 236
└ @ Main /Users/kcft114/.julia/packages/Flux/qXNjB/src/optimise/train.jl:105
```

```
loss(X, Y) = 0.79357636f0 (tracked)
loss(X, Y) = 0.784876f0 (tracked)
loss(X, Y) = 0.78023255f0 (tracked)
loss(X, Y) = 0.7870772f0 (tracked)
loss(X, Y) = 0.8065367f0 (tracked)
loss(X, Y) = 0.79432386f0 (tracked)
loss(X, Y) = 0.7953339f0 (tracked)
loss(X, Y) = 0.79584557f0 (tracked)
loss(X, Y) = 0.79065996f0 (tracked)
loss(X, Y) = 0.7975561f0 (tracked)
loss(X, Y) = 0.7748279f0 (tracked)
loss(X, Y) = 0.7948928f0 (tracked)
loss(X, Y) = 0.7943426f0 (tracked)
loss(X, Y) = 0.78609407f0 (tracked)
loss(X, Y) = 0.7900283f0 (tracked)
loss(X, Y) = 0.7818901f0 (tracked)
loss(X, Y) = 0.79338324f0 (tracked)
loss(X, Y) = 0.7637625f0 (tracked)
loss(X, Y) = 0.75994426f0 (tracked)
loss(X, Y) = 0.77244467f0 (tracked)
loss(X, Y) = 0.77941024f0 (tracked)
loss(X, Y) = 0.7763973f0 (tracked)
loss(X, Y) = 0.7798388f0 (tracked)
loss(X, Y) = 0.7839393f0 (tracked)
loss(X, Y) = 0.76528853f0 (tracked)
loss(X, Y) = 0.7666518f0 (tracked)
loss(X, Y) = 0.7836718f0 (tracked)
loss(X, Y) = 0.7966912f0 (tracked)
loss(X, Y) = 0.7706886f0 (tracked)
loss(X, Y) = 0.77618724f0 (tracked)
loss(X, Y) = 0.77559614f0 (tracked)
loss(X, Y) = 0.7687114f0 (tracked)
loss(X, Y) = 0.76352584f0 (tracked)
loss(X, Y) = 0.7611739f0 (tracked)
loss(X, Y) = 0.7655485f0 (tracked)
loss(X, Y) = 0.76146877f0 (tracked)
loss(X, Y) = 0.7760205f0 (tracked)
loss(X, Y) = 0.75214094f0 (tracked)
loss(X, Y) = 0.7915409f0 (tracked)
loss(X, Y) = 0.7586422f0 (tracked)
loss(X, Y) = 0.76161766f0 (tracked)
loss(X, Y) = 0.74973625f0 (tracked)
loss(X, Y) = 0.741476f0 (tracked)
loss(X, Y) = 0.7595565f0 (tracked)
loss(X, Y) = 0.759938f0 (tracked)
loss(X, Y) = 0.7600793f0 (tracked)
loss(X, Y) = 0.7702259f0 (tracked)
loss(X, Y) = 0.74684477f0 (tracked)
loss(X, Y) = 0.7576364f0 (tracked)
loss(X, Y) = 0.75161386f0 (tracked)
loss(X, Y) = 0.7649641f0 (tracked)
loss(X, Y) = 0.7644843f0 (tracked)
loss(X, Y) = 0.7397854f0 (tracked)
loss(X, Y) = 0.76116914f0 (tracked)
loss(X, Y) = 0.776892f0 (tracked)
loss(X, Y) = 0.75689775f0 (tracked)
loss(X, Y) = 0.7533236f0 (tracked)
loss(X, Y) = 0.77433825f0 (tracked)
loss(X, Y) = 0.77225804f0 (tracked)
loss(X, Y) = 0.76523274f0 (tracked)
loss(X, Y) = 0.72612447f0 (tracked)
loss(X, Y) = 0.7506316f0 (tracked)
loss(X, Y) = 0.75949234f0 (tracked)
loss(X, Y) = 0.766301f0 (tracked)
loss(X, Y) = 0.7187339f0 (tracked)
loss(X, Y) = 0.7497289f0 (tracked)
loss(X, Y) = 0.7396994f0 (tracked)
loss(X, Y) = 0.7661951f0 (tracked)
loss(X, Y) = 0.7653021f0 (tracked)
loss(X, Y) = 0.77491117f0 (tracked)
loss(X, Y) = 0.718027f0 (tracked)
loss(X, Y) = 0.729895f0 (tracked)
loss(X, Y) = 0.75503236f0 (tracked)
loss(X, Y) = 0.73056275f0 (tracked)
loss(X, Y) = 0.7636205f0 (tracked)
loss(X, Y) = 0.73591906f0 (tracked)
loss(X, Y) = 0.7458878f0 (tracked)
loss(X, Y) = 0.73797035f0 (tracked)
loss(X, Y) = 0.7444333f0 (tracked)
loss(X, Y) = 0.74915606f0 (tracked)
loss(X, Y) = 0.744847f0 (tracked)
loss(X, Y) = 0.7532718f0 (tracked)
loss(X, Y) = 0.7597923f0 (tracked)
loss(X, Y) = 0.7425549f0 (tracked)
loss(X, Y) = 0.74450743f0 (tracked)
loss(X, Y) = 0.7346986f0 (tracked)
loss(X, Y) = 0.74289966f0 (tracked)
loss(X, Y) = 0.7451025f0 (tracked)
loss(X, Y) = 0.73514444f0 (tracked)
loss(X, Y) = 0.72258997f0 (tracked)
loss(X, Y) = 0.73615354f0 (tracked)
loss(X, Y) = 0.74763715f0 (tracked)
loss(X, Y) = 0.73004305f0 (tracked)
loss(X, Y) = 0.71867126f0 (tracked)
loss(X, Y) = 0.73386514f0 (tracked)
loss(X, Y) = 0.7429601f0 (tracked)
loss(X, Y) = 0.7243595f0 (tracked)
loss(X, Y) = 0.74723184f0 (tracked)
loss(X, Y) = 0.75421566f0 (tracked)
loss(X, Y) = 0.7222218f0 (tracked)
loss(X, Y) = 0.72952026f0 (tracked)
loss(X, Y) = 0.7266088f0 (tracked)
loss(X, Y) = 0.7247975f0 (tracked)
```

```
┌ Info: Epoch 237
└ @ Main /Users/kcft114/.julia/packages/Flux/qXNjB/src/optimise/train.jl:105
┌ Info: Epoch 238
└ @ Main /Users/kcft114/.julia/packages/Flux/qXNjB/src/optimise/train.jl:105
┌ Info: Epoch 239
└ @ Main /Users/kcft114/.julia/packages/Flux/qXNjB/src/optimise/train.jl:105
┌ Info: Epoch 240
└ @ Main /Users/kcft114/.julia/packages/Flux/qXNjB/src/optimise/train.jl:105
┌ Info: Epoch 241
└ @ Main /Users/kcft114/.julia/packages/Flux/qXNjB/src/optimise/train.jl:105
┌ Info: Epoch 242
└ @ Main /Users/kcft114/.julia/packages/Flux/qXNjB/src/optimise/train.jl:105
┌ Info: Epoch 243
└ @ Main /Users/kcft114/.julia/packages/Flux/qXNjB/src/optimise/train.jl:105
┌ Info: Epoch 244
└ @ Main /Users/kcft114/.julia/packages/Flux/qXNjB/src/optimise/train.jl:105
┌ Info: Epoch 245
└ @ Main /Users/kcft114/.julia/packages/Flux/qXNjB/src/optimise/train.jl:105
┌ Info: Epoch 246
└ @ Main /Users/kcft114/.julia/packages/Flux/qXNjB/src/optimise/train.jl:105
┌ Info: Epoch 247
└ @ Main /Users/kcft114/.julia/packages/Flux/qXNjB/src/optimise/train.jl:105
┌ Info: Epoch 248
└ @ Main /Users/kcft114/.julia/packages/Flux/qXNjB/src/optimise/train.jl:105
┌ Info: Epoch 249
└ @ Main /Users/kcft114/.julia/packages/Flux/qXNjB/src/optimise/train.jl:105
┌ Info: Epoch 250
└ @ Main /Users/kcft114/.julia/packages/Flux/qXNjB/src/optimise/train.jl:105
┌ Info: Epoch 251
└ @ Main /Users/kcft114/.julia/packages/Flux/qXNjB/src/optimise/train.jl:105
┌ Info: Epoch 252
└ @ Main /Users/kcft114/.julia/packages/Flux/qXNjB/src/optimise/train.jl:105
┌ Info: Epoch 253
└ @ Main /Users/kcft114/.julia/packages/Flux/qXNjB/src/optimise/train.jl:105
┌ Info: Epoch 254
└ @ Main /Users/kcft114/.julia/packages/Flux/qXNjB/src/optimise/train.jl:105
┌ Info: Epoch 255
└ @ Main /Users/kcft114/.julia/packages/Flux/qXNjB/src/optimise/train.jl:105
┌ Info: Epoch 256
└ @ Main /Users/kcft114/.julia/packages/Flux/qXNjB/src/optimise/train.jl:105
┌ Info: Epoch 257
└ @ Main /Users/kcft114/.julia/packages/Flux/qXNjB/src/optimise/train.jl:105
┌ Info: Epoch 258
└ @ Main /Users/kcft114/.julia/packages/Flux/qXNjB/src/optimise/train.jl:105
┌ Info: Epoch 259
└ @ Main /Users/kcft114/.julia/packages/Flux/qXNjB/src/optimise/train.jl:105
┌ Info: Epoch 260
└ @ Main /Users/kcft114/.julia/packages/Flux/qXNjB/src/optimise/train.jl:105
┌ Info: Epoch 261
└ @ Main /Users/kcft114/.julia/packages/Flux/qXNjB/src/optimise/train.jl:105
┌ Info: Epoch 262
└ @ Main /Users/kcft114/.julia/packages/Flux/qXNjB/src/optimise/train.jl:105
┌ Info: Epoch 263
└ @ Main /Users/kcft114/.julia/packages/Flux/qXNjB/src/optimise/train.jl:105
┌ Info: Epoch 264
└ @ Main /Users/kcft114/.julia/packages/Flux/qXNjB/src/optimise/train.jl:105
┌ Info: Epoch 265
└ @ Main /Users/kcft114/.julia/packages/Flux/qXNjB/src/optimise/train.jl:105
┌ Info: Epoch 266
└ @ Main /Users/kcft114/.julia/packages/Flux/qXNjB/src/optimise/train.jl:105
┌ Info: Epoch 267
└ @ Main /Users/kcft114/.julia/packages/Flux/qXNjB/src/optimise/train.jl:105
┌ Info: Epoch 268
└ @ Main /Users/kcft114/.julia/packages/Flux/qXNjB/src/optimise/train.jl:105
┌ Info: Epoch 269
└ @ Main /Users/kcft114/.julia/packages/Flux/qXNjB/src/optimise/train.jl:105
┌ Info: Epoch 270
└ @ Main /Users/kcft114/.julia/packages/Flux/qXNjB/src/optimise/train.jl:105
┌ Info: Epoch 271
└ @ Main /Users/kcft114/.julia/packages/Flux/qXNjB/src/optimise/train.jl:105
┌ Info: Epoch 272
└ @ Main /Users/kcft114/.julia/packages/Flux/qXNjB/src/optimise/train.jl:105
┌ Info: Epoch 273
└ @ Main /Users/kcft114/.julia/packages/Flux/qXNjB/src/optimise/train.jl:105
┌ Info: Epoch 274
└ @ Main /Users/kcft114/.julia/packages/Flux/qXNjB/src/optimise/train.jl:105
┌ Info: Epoch 275
└ @ Main /Users/kcft114/.julia/packages/Flux/qXNjB/src/optimise/train.jl:105
┌ Info: Epoch 276
└ @ Main /Users/kcft114/.julia/packages/Flux/qXNjB/src/optimise/train.jl:105
┌ Info: Epoch 277
└ @ Main /Users/kcft114/.julia/packages/Flux/qXNjB/src/optimise/train.jl:105
┌ Info: Epoch 278
└ @ Main /Users/kcft114/.julia/packages/Flux/qXNjB/src/optimise/train.jl:105
┌ Info: Epoch 279
└ @ Main /Users/kcft114/.julia/packages/Flux/qXNjB/src/optimise/train.jl:105
┌ Info: Epoch 280
└ @ Main /Users/kcft114/.julia/packages/Flux/qXNjB/src/optimise/train.jl:105
┌ Info: Epoch 281
└ @ Main /Users/kcft114/.julia/packages/Flux/qXNjB/src/optimise/train.jl:105
┌ Info: Epoch 282
└ @ Main /Users/kcft114/.julia/packages/Flux/qXNjB/src/optimise/train.jl:105
┌ Info: Epoch 283
└ @ Main /Users/kcft114/.julia/packages/Flux/qXNjB/src/optimise/train.jl:105
┌ Info: Epoch 284
└ @ Main /Users/kcft114/.julia/packages/Flux/qXNjB/src/optimise/train.jl:105
┌ Info: Epoch 285
└ @ Main /Users/kcft114/.julia/packages/Flux/qXNjB/src/optimise/train.jl:105
┌ Info: Epoch 286
└ @ Main /Users/kcft114/.julia/packages/Flux/qXNjB/src/optimise/train.jl:105
┌ Info: Epoch 287
└ @ Main /Users/kcft114/.julia/packages/Flux/qXNjB/src/optimise/train.jl:105
┌ Info: Epoch 288
└ @ Main /Users/kcft114/.julia/packages/Flux/qXNjB/src/optimise/train.jl:105
┌ Info: Epoch 289
└ @ Main /Users/kcft114/.julia/packages/Flux/qXNjB/src/optimise/train.jl:105
┌ Info: Epoch 290
└ @ Main /Users/kcft114/.julia/packages/Flux/qXNjB/src/optimise/train.jl:105
┌ Info: Epoch 291
└ @ Main /Users/kcft114/.julia/packages/Flux/qXNjB/src/optimise/train.jl:105
┌ Info: Epoch 292
└ @ Main /Users/kcft114/.julia/packages/Flux/qXNjB/src/optimise/train.jl:105
┌ Info: Epoch 293
└ @ Main /Users/kcft114/.julia/packages/Flux/qXNjB/src/optimise/train.jl:105
┌ Info: Epoch 294
└ @ Main /Users/kcft114/.julia/packages/Flux/qXNjB/src/optimise/train.jl:105
┌ Info: Epoch 295
└ @ Main /Users/kcft114/.julia/packages/Flux/qXNjB/src/optimise/train.jl:105
┌ Info: Epoch 296
└ @ Main /Users/kcft114/.julia/packages/Flux/qXNjB/src/optimise/train.jl:105
┌ Info: Epoch 297
└ @ Main /Users/kcft114/.julia/packages/Flux/qXNjB/src/optimise/train.jl:105
┌ Info: Epoch 298
└ @ Main /Users/kcft114/.julia/packages/Flux/qXNjB/src/optimise/train.jl:105
┌ Info: Epoch 299
└ @ Main /Users/kcft114/.julia/packages/Flux/qXNjB/src/optimise/train.jl:105
┌ Info: Epoch 300
└ @ Main /Users/kcft114/.julia/packages/Flux/qXNjB/src/optimise/train.jl:105
┌ Info: Epoch 301
└ @ Main /Users/kcft114/.julia/packages/Flux/qXNjB/src/optimise/train.jl:105
┌ Info: Epoch 302
└ @ Main /Users/kcft114/.julia/packages/Flux/qXNjB/src/optimise/train.jl:105
┌ Info: Epoch 303
└ @ Main /Users/kcft114/.julia/packages/Flux/qXNjB/src/optimise/train.jl:105
┌ Info: Epoch 304
└ @ Main /Users/kcft114/.julia/packages/Flux/qXNjB/src/optimise/train.jl:105
┌ Info: Epoch 305
└ @ Main /Users/kcft114/.julia/packages/Flux/qXNjB/src/optimise/train.jl:105
┌ Info: Epoch 306
└ @ Main /Users/kcft114/.julia/packages/Flux/qXNjB/src/optimise/train.jl:105
┌ Info: Epoch 307
└ @ Main /Users/kcft114/.julia/packages/Flux/qXNjB/src/optimise/train.jl:105
┌ Info: Epoch 308
└ @ Main /Users/kcft114/.julia/packages/Flux/qXNjB/src/optimise/train.jl:105
┌ Info: Epoch 309
└ @ Main /Users/kcft114/.julia/packages/Flux/qXNjB/src/optimise/train.jl:105
┌ Info: Epoch 310
└ @ Main /Users/kcft114/.julia/packages/Flux/qXNjB/src/optimise/train.jl:105
┌ Info: Epoch 311
└ @ Main /Users/kcft114/.julia/packages/Flux/qXNjB/src/optimise/train.jl:105
┌ Info: Epoch 312
└ @ Main /Users/kcft114/.julia/packages/Flux/qXNjB/src/optimise/train.jl:105
┌ Info: Epoch 313
└ @ Main /Users/kcft114/.julia/packages/Flux/qXNjB/src/optimise/train.jl:105
┌ Info: Epoch 314
└ @ Main /Users/kcft114/.julia/packages/Flux/qXNjB/src/optimise/train.jl:105
┌ Info: Epoch 315
└ @ Main /Users/kcft114/.julia/packages/Flux/qXNjB/src/optimise/train.jl:105
┌ Info: Epoch 316
└ @ Main /Users/kcft114/.julia/packages/Flux/qXNjB/src/optimise/train.jl:105
┌ Info: Epoch 317
└ @ Main /Users/kcft114/.julia/packages/Flux/qXNjB/src/optimise/train.jl:105
┌ Info: Epoch 318
└ @ Main /Users/kcft114/.julia/packages/Flux/qXNjB/src/optimise/train.jl:105
┌ Info: Epoch 319
└ @ Main /Users/kcft114/.julia/packages/Flux/qXNjB/src/optimise/train.jl:105
┌ Info: Epoch 320
└ @ Main /Users/kcft114/.julia/packages/Flux/qXNjB/src/optimise/train.jl:105
┌ Info: Epoch 321
└ @ Main /Users/kcft114/.julia/packages/Flux/qXNjB/src/optimise/train.jl:105
┌ Info: Epoch 322
└ @ Main /Users/kcft114/.julia/packages/Flux/qXNjB/src/optimise/train.jl:105
┌ Info: Epoch 323
└ @ Main /Users/kcft114/.julia/packages/Flux/qXNjB/src/optimise/train.jl:105
┌ Info: Epoch 324
└ @ Main /Users/kcft114/.julia/packages/Flux/qXNjB/src/optimise/train.jl:105
┌ Info: Epoch 325
└ @ Main /Users/kcft114/.julia/packages/Flux/qXNjB/src/optimise/train.jl:105
┌ Info: Epoch 326
└ @ Main /Users/kcft114/.julia/packages/Flux/qXNjB/src/optimise/train.jl:105
┌ Info: Epoch 327
└ @ Main /Users/kcft114/.julia/packages/Flux/qXNjB/src/optimise/train.jl:105
┌ Info: Epoch 328
└ @ Main /Users/kcft114/.julia/packages/Flux/qXNjB/src/optimise/train.jl:105
┌ Info: Epoch 329
└ @ Main /Users/kcft114/.julia/packages/Flux/qXNjB/src/optimise/train.jl:105
┌ Info: Epoch 330
└ @ Main /Users/kcft114/.julia/packages/Flux/qXNjB/src/optimise/train.jl:105
┌ Info: Epoch 331
└ @ Main /Users/kcft114/.julia/packages/Flux/qXNjB/src/optimise/train.jl:105
┌ Info: Epoch 332
└ @ Main /Users/kcft114/.julia/packages/Flux/qXNjB/src/optimise/train.jl:105
┌ Info: Epoch 333
└ @ Main /Users/kcft114/.julia/packages/Flux/qXNjB/src/optimise/train.jl:105
┌ Info: Epoch 334
└ @ Main /Users/kcft114/.julia/packages/Flux/qXNjB/src/optimise/train.jl:105
┌ Info: Epoch 335
└ @ Main /Users/kcft114/.julia/packages/Flux/qXNjB/src/optimise/train.jl:105
┌ Info: Epoch 336
└ @ Main /Users/kcft114/.julia/packages/Flux/qXNjB/src/optimise/train.jl:105
┌ Info: Epoch 337
└ @ Main /Users/kcft114/.julia/packages/Flux/qXNjB/src/optimise/train.jl:105
┌ Info: Epoch 338
└ @ Main /Users/kcft114/.julia/packages/Flux/qXNjB/src/optimise/train.jl:105
┌ Info: Epoch 339
└ @ Main /Users/kcft114/.julia/packages/Flux/qXNjB/src/optimise/train.jl:105
┌ Info: Epoch 340
└ @ Main /Users/kcft114/.julia/packages/Flux/qXNjB/src/optimise/train.jl:105
┌ Info: Epoch 341
└ @ Main /Users/kcft114/.julia/packages/Flux/qXNjB/src/optimise/train.jl:105
┌ Info: Epoch 342
└ @ Main /Users/kcft114/.julia/packages/Flux/qXNjB/src/optimise/train.jl:105
┌ Info: Epoch 343
└ @ Main /Users/kcft114/.julia/packages/Flux/qXNjB/src/optimise/train.jl:105
┌ Info: Epoch 344
└ @ Main /Users/kcft114/.julia/packages/Flux/qXNjB/src/optimise/train.jl:105
┌ Info: Epoch 345
└ @ Main /Users/kcft114/.julia/packages/Flux/qXNjB/src/optimise/train.jl:105
┌ Info: Epoch 346
└ @ Main /Users/kcft114/.julia/packages/Flux/qXNjB/src/optimise/train.jl:105
┌ Info: Epoch 347
└ @ Main /Users/kcft114/.julia/packages/Flux/qXNjB/src/optimise/train.jl:105
┌ Info: Epoch 348
└ @ Main /Users/kcft114/.julia/packages/Flux/qXNjB/src/optimise/train.jl:105
┌ Info: Epoch 349
└ @ Main /Users/kcft114/.julia/packages/Flux/qXNjB/src/optimise/train.jl:105
┌ Info: Epoch 350
└ @ Main /Users/kcft114/.julia/packages/Flux/qXNjB/src/optimise/train.jl:105
┌ Info: Epoch 351
└ @ Main /Users/kcft114/.julia/packages/Flux/qXNjB/src/optimise/train.jl:105
┌ Info: Epoch 352
└ @ Main /Users/kcft114/.julia/packages/Flux/qXNjB/src/optimise/train.jl:105
┌ Info: Epoch 353
└ @ Main /Users/kcft114/.julia/packages/Flux/qXNjB/src/optimise/train.jl:105
┌ Info: Epoch 354
└ @ Main /Users/kcft114/.julia/packages/Flux/qXNjB/src/optimise/train.jl:105
┌ Info: Epoch 355
└ @ Main /Users/kcft114/.julia/packages/Flux/qXNjB/src/optimise/train.jl:105
┌ Info: Epoch 356
└ @ Main /Users/kcft114/.julia/packages/Flux/qXNjB/src/optimise/train.jl:105
┌ Info: Epoch 357
└ @ Main /Users/kcft114/.julia/packages/Flux/qXNjB/src/optimise/train.jl:105
┌ Info: Epoch 358
└ @ Main /Users/kcft114/.julia/packages/Flux/qXNjB/src/optimise/train.jl:105
┌ Info: Epoch 359
└ @ Main /Users/kcft114/.julia/packages/Flux/qXNjB/src/optimise/train.jl:105
┌ Info: Epoch 360
└ @ Main /Users/kcft114/.julia/packages/Flux/qXNjB/src/optimise/train.jl:105
┌ Info: Epoch 361
└ @ Main /Users/kcft114/.julia/packages/Flux/qXNjB/src/optimise/train.jl:105
┌ Info: Epoch 362
└ @ Main /Users/kcft114/.julia/packages/Flux/qXNjB/src/optimise/train.jl:105
┌ Info: Epoch 363
└ @ Main /Users/kcft114/.julia/packages/Flux/qXNjB/src/optimise/train.jl:105
┌ Info: Epoch 364
└ @ Main /Users/kcft114/.julia/packages/Flux/qXNjB/src/optimise/train.jl:105
┌ Info: Epoch 365
└ @ Main /Users/kcft114/.julia/packages/Flux/qXNjB/src/optimise/train.jl:105
┌ Info: Epoch 366
└ @ Main /Users/kcft114/.julia/packages/Flux/qXNjB/src/optimise/train.jl:105
┌ Info: Epoch 367
└ @ Main /Users/kcft114/.julia/packages/Flux/qXNjB/src/optimise/train.jl:105
┌ Info: Epoch 368
└ @ Main /Users/kcft114/.julia/packages/Flux/qXNjB/src/optimise/train.jl:105
┌ Info: Epoch 369
└ @ Main /Users/kcft114/.julia/packages/Flux/qXNjB/src/optimise/train.jl:105
┌ Info: Epoch 370
└ @ Main /Users/kcft114/.julia/packages/Flux/qXNjB/src/optimise/train.jl:105
┌ Info: Epoch 371
└ @ Main /Users/kcft114/.julia/packages/Flux/qXNjB/src/optimise/train.jl:105
┌ Info: Epoch 372
└ @ Main /Users/kcft114/.julia/packages/Flux/qXNjB/src/optimise/train.jl:105
┌ Info: Epoch 373
└ @ Main /Users/kcft114/.julia/packages/Flux/qXNjB/src/optimise/train.jl:105
┌ Info: Epoch 374
└ @ Main /Users/kcft114/.julia/packages/Flux/qXNjB/src/optimise/train.jl:105
┌ Info: Epoch 375
└ @ Main /Users/kcft114/.julia/packages/Flux/qXNjB/src/optimise/train.jl:105
┌ Info: Epoch 376
└ @ Main /Users/kcft114/.julia/packages/Flux/qXNjB/src/optimise/train.jl:105
┌ Info: Epoch 377
└ @ Main /Users/kcft114/.julia/packages/Flux/qXNjB/src/optimise/train.jl:105
┌ Info: Epoch 378
└ @ Main /Users/kcft114/.julia/packages/Flux/qXNjB/src/optimise/train.jl:105
┌ Info: Epoch 379
└ @ Main /Users/kcft114/.julia/packages/Flux/qXNjB/src/optimise/train.jl:105
┌ Info: Epoch 380
└ @ Main /Users/kcft114/.julia/packages/Flux/qXNjB/src/optimise/train.jl:105
┌ Info: Epoch 381
└ @ Main /Users/kcft114/.julia/packages/Flux/qXNjB/src/optimise/train.jl:105
┌ Info: Epoch 382
└ @ Main /Users/kcft114/.julia/packages/Flux/qXNjB/src/optimise/train.jl:105
┌ Info: Epoch 383
└ @ Main /Users/kcft114/.julia/packages/Flux/qXNjB/src/optimise/train.jl:105
┌ Info: Epoch 384
└ @ Main /Users/kcft114/.julia/packages/Flux/qXNjB/src/optimise/train.jl:105
┌ Info: Epoch 385
└ @ Main /Users/kcft114/.julia/packages/Flux/qXNjB/src/optimise/train.jl:105
┌ Info: Epoch 386
└ @ Main /Users/kcft114/.julia/packages/Flux/qXNjB/src/optimise/train.jl:105
┌ Info: Epoch 387
└ @ Main /Users/kcft114/.julia/packages/Flux/qXNjB/src/optimise/train.jl:105
┌ Info: Epoch 388
└ @ Main /Users/kcft114/.julia/packages/Flux/qXNjB/src/optimise/train.jl:105
┌ Info: Epoch 389
└ @ Main /Users/kcft114/.julia/packages/Flux/qXNjB/src/optimise/train.jl:105
┌ Info: Epoch 390
└ @ Main /Users/kcft114/.julia/packages/Flux/qXNjB/src/optimise/train.jl:105
┌ Info: Epoch 391
└ @ Main /Users/kcft114/.julia/packages/Flux/qXNjB/src/optimise/train.jl:105
┌ Info: Epoch 392
└ @ Main /Users/kcft114/.julia/packages/Flux/qXNjB/src/optimise/train.jl:105
┌ Info: Epoch 393
└ @ Main /Users/kcft114/.julia/packages/Flux/qXNjB/src/optimise/train.jl:105
┌ Info: Epoch 394
└ @ Main /Users/kcft114/.julia/packages/Flux/qXNjB/src/optimise/train.jl:105
┌ Info: Epoch 395
└ @ Main /Users/kcft114/.julia/packages/Flux/qXNjB/src/optimise/train.jl:105
┌ Info: Epoch 396
└ @ Main /Users/kcft114/.julia/packages/Flux/qXNjB/src/optimise/train.jl:105
┌ Info: Epoch 397
└ @ Main /Users/kcft114/.julia/packages/Flux/qXNjB/src/optimise/train.jl:105
┌ Info: Epoch 398
└ @ Main /Users/kcft114/.julia/packages/Flux/qXNjB/src/optimise/train.jl:105
┌ Info: Epoch 399
└ @ Main /Users/kcft114/.julia/packages/Flux/qXNjB/src/optimise/train.jl:105
┌ Info: Epoch 400
└ @ Main /Users/kcft114/.julia/packages/Flux/qXNjB/src/optimise/train.jl:105
┌ Info: Epoch 401
└ @ Main /Users/kcft114/.julia/packages/Flux/qXNjB/src/optimise/train.jl:105
┌ Info: Epoch 402
└ @ Main /Users/kcft114/.julia/packages/Flux/qXNjB/src/optimise/train.jl:105
┌ Info: Epoch 403
└ @ Main /Users/kcft114/.julia/packages/Flux/qXNjB/src/optimise/train.jl:105
┌ Info: Epoch 404
└ @ Main /Users/kcft114/.julia/packages/Flux/qXNjB/src/optimise/train.jl:105
┌ Info: Epoch 405
└ @ Main /Users/kcft114/.julia/packages/Flux/qXNjB/src/optimise/train.jl:105
┌ Info: Epoch 406
└ @ Main /Users/kcft114/.julia/packages/Flux/qXNjB/src/optimise/train.jl:105
┌ Info: Epoch 407
└ @ Main /Users/kcft114/.julia/packages/Flux/qXNjB/src/optimise/train.jl:105
┌ Info: Epoch 408
└ @ Main /Users/kcft114/.julia/packages/Flux/qXNjB/src/optimise/train.jl:105
┌ Info: Epoch 409
└ @ Main /Users/kcft114/.julia/packages/Flux/qXNjB/src/optimise/train.jl:105
┌ Info: Epoch 410
└ @ Main /Users/kcft114/.julia/packages/Flux/qXNjB/src/optimise/train.jl:105
┌ Info: Epoch 411
└ @ Main /Users/kcft114/.julia/packages/Flux/qXNjB/src/optimise/train.jl:105
┌ Info: Epoch 412
└ @ Main /Users/kcft114/.julia/packages/Flux/qXNjB/src/optimise/train.jl:105
┌ Info: Epoch 413
└ @ Main /Users/kcft114/.julia/packages/Flux/qXNjB/src/optimise/train.jl:105
┌ Info: Epoch 414
└ @ Main /Users/kcft114/.julia/packages/Flux/qXNjB/src/optimise/train.jl:105
┌ Info: Epoch 415
└ @ Main /Users/kcft114/.julia/packages/Flux/qXNjB/src/optimise/train.jl:105
┌ Info: Epoch 416
└ @ Main /Users/kcft114/.julia/packages/Flux/qXNjB/src/optimise/train.jl:105
┌ Info: Epoch 417
└ @ Main /Users/kcft114/.julia/packages/Flux/qXNjB/src/optimise/train.jl:105
┌ Info: Epoch 418
└ @ Main /Users/kcft114/.julia/packages/Flux/qXNjB/src/optimise/train.jl:105
┌ Info: Epoch 419
└ @ Main /Users/kcft114/.julia/packages/Flux/qXNjB/src/optimise/train.jl:105
┌ Info: Epoch 420
└ @ Main /Users/kcft114/.julia/packages/Flux/qXNjB/src/optimise/train.jl:105
┌ Info: Epoch 421
└ @ Main /Users/kcft114/.julia/packages/Flux/qXNjB/src/optimise/train.jl:105
┌ Info: Epoch 422
└ @ Main /Users/kcft114/.julia/packages/Flux/qXNjB/src/optimise/train.jl:105
┌ Info: Epoch 423
└ @ Main /Users/kcft114/.julia/packages/Flux/qXNjB/src/optimise/train.jl:105
┌ Info: Epoch 424
└ @ Main /Users/kcft114/.julia/packages/Flux/qXNjB/src/optimise/train.jl:105
┌ Info: Epoch 425
└ @ Main /Users/kcft114/.julia/packages/Flux/qXNjB/src/optimise/train.jl:105
┌ Info: Epoch 426
└ @ Main /Users/kcft114/.julia/packages/Flux/qXNjB/src/optimise/train.jl:105
┌ Info: Epoch 427
└ @ Main /Users/kcft114/.julia/packages/Flux/qXNjB/src/optimise/train.jl:105
┌ Info: Epoch 428
└ @ Main /Users/kcft114/.julia/packages/Flux/qXNjB/src/optimise/train.jl:105
┌ Info: Epoch 429
└ @ Main /Users/kcft114/.julia/packages/Flux/qXNjB/src/optimise/train.jl:105
┌ Info: Epoch 430
└ @ Main /Users/kcft114/.julia/packages/Flux/qXNjB/src/optimise/train.jl:105
┌ Info: Epoch 431
└ @ Main /Users/kcft114/.julia/packages/Flux/qXNjB/src/optimise/train.jl:105
┌ Info: Epoch 432
└ @ Main /Users/kcft114/.julia/packages/Flux/qXNjB/src/optimise/train.jl:105
┌ Info: Epoch 433
└ @ Main /Users/kcft114/.julia/packages/Flux/qXNjB/src/optimise/train.jl:105
┌ Info: Epoch 434
└ @ Main /Users/kcft114/.julia/packages/Flux/qXNjB/src/optimise/train.jl:105
┌ Info: Epoch 435
└ @ Main /Users/kcft114/.julia/packages/Flux/qXNjB/src/optimise/train.jl:105
┌ Info: Epoch 436
└ @ Main /Users/kcft114/.julia/packages/Flux/qXNjB/src/optimise/train.jl:105
┌ Info: Epoch 437
└ @ Main /Users/kcft114/.julia/packages/Flux/qXNjB/src/optimise/train.jl:105
┌ Info: Epoch 438
└ @ Main /Users/kcft114/.julia/packages/Flux/qXNjB/src/optimise/train.jl:105
┌ Info: Epoch 439
└ @ Main /Users/kcft114/.julia/packages/Flux/qXNjB/src/optimise/train.jl:105
┌ Info: Epoch 440
└ @ Main /Users/kcft114/.julia/packages/Flux/qXNjB/src/optimise/train.jl:105
┌ Info: Epoch 441
└ @ Main /Users/kcft114/.julia/packages/Flux/qXNjB/src/optimise/train.jl:105
┌ Info: Epoch 442
└ @ Main /Users/kcft114/.julia/packages/Flux/qXNjB/src/optimise/train.jl:105
┌ Info: Epoch 443
└ @ Main /Users/kcft114/.julia/packages/Flux/qXNjB/src/optimise/train.jl:105
```

```
loss(X, Y) = 0.72674257f0 (tracked)
loss(X, Y) = 0.71044874f0 (tracked)
loss(X, Y) = 0.7490849f0 (tracked)
loss(X, Y) = 0.71549666f0 (tracked)
loss(X, Y) = 0.72951555f0 (tracked)
loss(X, Y) = 0.712154f0 (tracked)
loss(X, Y) = 0.7176719f0 (tracked)
loss(X, Y) = 0.720596f0 (tracked)
loss(X, Y) = 0.71087515f0 (tracked)
loss(X, Y) = 0.74394757f0 (tracked)
loss(X, Y) = 0.7320468f0 (tracked)
loss(X, Y) = 0.7536637f0 (tracked)
loss(X, Y) = 0.73274267f0 (tracked)
loss(X, Y) = 0.7547554f0 (tracked)
loss(X, Y) = 0.7311066f0 (tracked)
loss(X, Y) = 0.7167266f0 (tracked)
loss(X, Y) = 0.7063448f0 (tracked)
loss(X, Y) = 0.7182323f0 (tracked)
loss(X, Y) = 0.7329014f0 (tracked)
loss(X, Y) = 0.73999965f0 (tracked)
loss(X, Y) = 0.7266256f0 (tracked)
loss(X, Y) = 0.7204033f0 (tracked)
loss(X, Y) = 0.727861f0 (tracked)
loss(X, Y) = 0.70843464f0 (tracked)
loss(X, Y) = 0.72189766f0 (tracked)
loss(X, Y) = 0.72449255f0 (tracked)
loss(X, Y) = 0.7329182f0 (tracked)
loss(X, Y) = 0.71775717f0 (tracked)
loss(X, Y) = 0.70909685f0 (tracked)
loss(X, Y) = 0.6903691f0 (tracked)
loss(X, Y) = 0.7381665f0 (tracked)
loss(X, Y) = 0.7154842f0 (tracked)
loss(X, Y) = 0.70702666f0 (tracked)
loss(X, Y) = 0.72303396f0 (tracked)
loss(X, Y) = 0.70954424f0 (tracked)
loss(X, Y) = 0.729514f0 (tracked)
loss(X, Y) = 0.71172863f0 (tracked)
loss(X, Y) = 0.71145517f0 (tracked)
loss(X, Y) = 0.75132287f0 (tracked)
loss(X, Y) = 0.713619f0 (tracked)
loss(X, Y) = 0.70156336f0 (tracked)
loss(X, Y) = 0.71859473f0 (tracked)
loss(X, Y) = 0.7064498f0 (tracked)
loss(X, Y) = 0.6916704f0 (tracked)
loss(X, Y) = 0.700617f0 (tracked)
loss(X, Y) = 0.7132485f0 (tracked)
loss(X, Y) = 0.733615f0 (tracked)
loss(X, Y) = 0.70378727f0 (tracked)
loss(X, Y) = 0.70530677f0 (tracked)
loss(X, Y) = 0.70050114f0 (tracked)
loss(X, Y) = 0.7249211f0 (tracked)
loss(X, Y) = 0.69962525f0 (tracked)
loss(X, Y) = 0.7323017f0 (tracked)
loss(X, Y) = 0.6983791f0 (tracked)
loss(X, Y) = 0.70679945f0 (tracked)
loss(X, Y) = 0.70989424f0 (tracked)
loss(X, Y) = 0.69363993f0 (tracked)
loss(X, Y) = 0.6968691f0 (tracked)
loss(X, Y) = 0.6840073f0 (tracked)
loss(X, Y) = 0.704226f0 (tracked)
loss(X, Y) = 0.7185279f0 (tracked)
loss(X, Y) = 0.69725436f0 (tracked)
loss(X, Y) = 0.7380604f0 (tracked)
loss(X, Y) = 0.7120918f0 (tracked)
loss(X, Y) = 0.6920376f0 (tracked)
loss(X, Y) = 0.71025914f0 (tracked)
loss(X, Y) = 0.6889067f0 (tracked)
loss(X, Y) = 0.6909675f0 (tracked)
loss(X, Y) = 0.71509737f0 (tracked)
loss(X, Y) = 0.70622176f0 (tracked)
loss(X, Y) = 0.7448438f0 (tracked)
loss(X, Y) = 0.71169955f0 (tracked)
loss(X, Y) = 0.68188286f0 (tracked)
loss(X, Y) = 0.7176523f0 (tracked)
loss(X, Y) = 0.6902553f0 (tracked)
loss(X, Y) = 0.704122f0 (tracked)
loss(X, Y) = 0.70958f0 (tracked)
loss(X, Y) = 0.6956539f0 (tracked)
loss(X, Y) = 0.67150366f0 (tracked)
loss(X, Y) = 0.6801504f0 (tracked)
loss(X, Y) = 0.6925711f0 (tracked)
loss(X, Y) = 0.70093286f0 (tracked)
loss(X, Y) = 0.6783996f0 (tracked)
loss(X, Y) = 0.7157417f0 (tracked)
loss(X, Y) = 0.68610454f0 (tracked)
loss(X, Y) = 0.69659954f0 (tracked)
loss(X, Y) = 0.69384056f0 (tracked)
loss(X, Y) = 0.68125594f0 (tracked)
loss(X, Y) = 0.6844355f0 (tracked)
loss(X, Y) = 0.6736165f0 (tracked)
loss(X, Y) = 0.70271677f0 (tracked)
loss(X, Y) = 0.7096184f0 (tracked)
loss(X, Y) = 0.6978997f0 (tracked)
loss(X, Y) = 0.6996121f0 (tracked)
loss(X, Y) = 0.7079449f0 (tracked)
loss(X, Y) = 0.71303004f0 (tracked)
loss(X, Y) = 0.68912184f0 (tracked)
loss(X, Y) = 0.6868074f0 (tracked)
loss(X, Y) = 0.6902582f0 (tracked)
loss(X, Y) = 0.6732941f0 (tracked)
loss(X, Y) = 0.6644912f0 (tracked)
loss(X, Y) = 0.70155305f0 (tracked)
loss(X, Y) = 0.7389185f0 (tracked)
loss(X, Y) = 0.7216031f0 (tracked)
loss(X, Y) = 0.705387f0 (tracked)
loss(X, Y) = 0.6774981f0 (tracked)
loss(X, Y) = 0.68906456f0 (tracked)
loss(X, Y) = 0.6881328f0 (tracked)
loss(X, Y) = 0.69433135f0 (tracked)
loss(X, Y) = 0.68736756f0 (tracked)
loss(X, Y) = 0.6745681f0 (tracked)
```

```
┌ Info: Epoch 444
└ @ Main /Users/kcft114/.julia/packages/Flux/qXNjB/src/optimise/train.jl:105
┌ Info: Epoch 445
└ @ Main /Users/kcft114/.julia/packages/Flux/qXNjB/src/optimise/train.jl:105
┌ Info: Epoch 446
└ @ Main /Users/kcft114/.julia/packages/Flux/qXNjB/src/optimise/train.jl:105
┌ Info: Epoch 447
└ @ Main /Users/kcft114/.julia/packages/Flux/qXNjB/src/optimise/train.jl:105
┌ Info: Epoch 448
└ @ Main /Users/kcft114/.julia/packages/Flux/qXNjB/src/optimise/train.jl:105
┌ Info: Epoch 449
└ @ Main /Users/kcft114/.julia/packages/Flux/qXNjB/src/optimise/train.jl:105
┌ Info: Epoch 450
└ @ Main /Users/kcft114/.julia/packages/Flux/qXNjB/src/optimise/train.jl:105
┌ Info: Epoch 451
└ @ Main /Users/kcft114/.julia/packages/Flux/qXNjB/src/optimise/train.jl:105
┌ Info: Epoch 452
└ @ Main /Users/kcft114/.julia/packages/Flux/qXNjB/src/optimise/train.jl:105
┌ Info: Epoch 453
└ @ Main /Users/kcft114/.julia/packages/Flux/qXNjB/src/optimise/train.jl:105
┌ Info: Epoch 454
└ @ Main /Users/kcft114/.julia/packages/Flux/qXNjB/src/optimise/train.jl:105
┌ Info: Epoch 455
└ @ Main /Users/kcft114/.julia/packages/Flux/qXNjB/src/optimise/train.jl:105
┌ Info: Epoch 456
└ @ Main /Users/kcft114/.julia/packages/Flux/qXNjB/src/optimise/train.jl:105
┌ Info: Epoch 457
└ @ Main /Users/kcft114/.julia/packages/Flux/qXNjB/src/optimise/train.jl:105
┌ Info: Epoch 458
└ @ Main /Users/kcft114/.julia/packages/Flux/qXNjB/src/optimise/train.jl:105
┌ Info: Epoch 459
└ @ Main /Users/kcft114/.julia/packages/Flux/qXNjB/src/optimise/train.jl:105
┌ Info: Epoch 460
└ @ Main /Users/kcft114/.julia/packages/Flux/qXNjB/src/optimise/train.jl:105
┌ Info: Epoch 461
└ @ Main /Users/kcft114/.julia/packages/Flux/qXNjB/src/optimise/train.jl:105
┌ Info: Epoch 462
└ @ Main /Users/kcft114/.julia/packages/Flux/qXNjB/src/optimise/train.jl:105
┌ Info: Epoch 463
└ @ Main /Users/kcft114/.julia/packages/Flux/qXNjB/src/optimise/train.jl:105
┌ Info: Epoch 464
└ @ Main /Users/kcft114/.julia/packages/Flux/qXNjB/src/optimise/train.jl:105
┌ Info: Epoch 465
└ @ Main /Users/kcft114/.julia/packages/Flux/qXNjB/src/optimise/train.jl:105
┌ Info: Epoch 466
└ @ Main /Users/kcft114/.julia/packages/Flux/qXNjB/src/optimise/train.jl:105
┌ Info: Epoch 467
└ @ Main /Users/kcft114/.julia/packages/Flux/qXNjB/src/optimise/train.jl:105
┌ Info: Epoch 468
└ @ Main /Users/kcft114/.julia/packages/Flux/qXNjB/src/optimise/train.jl:105
┌ Info: Epoch 469
└ @ Main /Users/kcft114/.julia/packages/Flux/qXNjB/src/optimise/train.jl:105
┌ Info: Epoch 470
└ @ Main /Users/kcft114/.julia/packages/Flux/qXNjB/src/optimise/train.jl:105
┌ Info: Epoch 471
└ @ Main /Users/kcft114/.julia/packages/Flux/qXNjB/src/optimise/train.jl:105
┌ Info: Epoch 472
└ @ Main /Users/kcft114/.julia/packages/Flux/qXNjB/src/optimise/train.jl:105
┌ Info: Epoch 473
└ @ Main /Users/kcft114/.julia/packages/Flux/qXNjB/src/optimise/train.jl:105
┌ Info: Epoch 474
└ @ Main /Users/kcft114/.julia/packages/Flux/qXNjB/src/optimise/train.jl:105
┌ Info: Epoch 475
└ @ Main /Users/kcft114/.julia/packages/Flux/qXNjB/src/optimise/train.jl:105
┌ Info: Epoch 476
└ @ Main /Users/kcft114/.julia/packages/Flux/qXNjB/src/optimise/train.jl:105
┌ Info: Epoch 477
└ @ Main /Users/kcft114/.julia/packages/Flux/qXNjB/src/optimise/train.jl:105
┌ Info: Epoch 478
└ @ Main /Users/kcft114/.julia/packages/Flux/qXNjB/src/optimise/train.jl:105
┌ Info: Epoch 479
└ @ Main /Users/kcft114/.julia/packages/Flux/qXNjB/src/optimise/train.jl:105
┌ Info: Epoch 480
└ @ Main /Users/kcft114/.julia/packages/Flux/qXNjB/src/optimise/train.jl:105
┌ Info: Epoch 481
└ @ Main /Users/kcft114/.julia/packages/Flux/qXNjB/src/optimise/train.jl:105
┌ Info: Epoch 482
└ @ Main /Users/kcft114/.julia/packages/Flux/qXNjB/src/optimise/train.jl:105
┌ Info: Epoch 483
└ @ Main /Users/kcft114/.julia/packages/Flux/qXNjB/src/optimise/train.jl:105
┌ Info: Epoch 484
└ @ Main /Users/kcft114/.julia/packages/Flux/qXNjB/src/optimise/train.jl:105
┌ Info: Epoch 485
└ @ Main /Users/kcft114/.julia/packages/Flux/qXNjB/src/optimise/train.jl:105
┌ Info: Epoch 486
└ @ Main /Users/kcft114/.julia/packages/Flux/qXNjB/src/optimise/train.jl:105
┌ Info: Epoch 487
└ @ Main /Users/kcft114/.julia/packages/Flux/qXNjB/src/optimise/train.jl:105
┌ Info: Epoch 488
└ @ Main /Users/kcft114/.julia/packages/Flux/qXNjB/src/optimise/train.jl:105
┌ Info: Epoch 489
└ @ Main /Users/kcft114/.julia/packages/Flux/qXNjB/src/optimise/train.jl:105
┌ Info: Epoch 490
└ @ Main /Users/kcft114/.julia/packages/Flux/qXNjB/src/optimise/train.jl:105
┌ Info: Epoch 491
└ @ Main /Users/kcft114/.julia/packages/Flux/qXNjB/src/optimise/train.jl:105
┌ Info: Epoch 492
└ @ Main /Users/kcft114/.julia/packages/Flux/qXNjB/src/optimise/train.jl:105
┌ Info: Epoch 493
└ @ Main /Users/kcft114/.julia/packages/Flux/qXNjB/src/optimise/train.jl:105
┌ Info: Epoch 494
└ @ Main /Users/kcft114/.julia/packages/Flux/qXNjB/src/optimise/train.jl:105
┌ Info: Epoch 495
└ @ Main /Users/kcft114/.julia/packages/Flux/qXNjB/src/optimise/train.jl:105
┌ Info: Epoch 496
└ @ Main /Users/kcft114/.julia/packages/Flux/qXNjB/src/optimise/train.jl:105
┌ Info: Epoch 497
└ @ Main /Users/kcft114/.julia/packages/Flux/qXNjB/src/optimise/train.jl:105
┌ Info: Epoch 498
└ @ Main /Users/kcft114/.julia/packages/Flux/qXNjB/src/optimise/train.jl:105
┌ Info: Epoch 499
└ @ Main /Users/kcft114/.julia/packages/Flux/qXNjB/src/optimise/train.jl:105
┌ Info: Epoch 500
└ @ Main /Users/kcft114/.julia/packages/Flux/qXNjB/src/optimise/train.jl:105
┌ Info: Epoch 501
└ @ Main /Users/kcft114/.julia/packages/Flux/qXNjB/src/optimise/train.jl:105
┌ Info: Epoch 502
└ @ Main /Users/kcft114/.julia/packages/Flux/qXNjB/src/optimise/train.jl:105
┌ Info: Epoch 503
└ @ Main /Users/kcft114/.julia/packages/Flux/qXNjB/src/optimise/train.jl:105
┌ Info: Epoch 504
└ @ Main /Users/kcft114/.julia/packages/Flux/qXNjB/src/optimise/train.jl:105
┌ Info: Epoch 505
└ @ Main /Users/kcft114/.julia/packages/Flux/qXNjB/src/optimise/train.jl:105
┌ Info: Epoch 506
└ @ Main /Users/kcft114/.julia/packages/Flux/qXNjB/src/optimise/train.jl:105
┌ Info: Epoch 507
└ @ Main /Users/kcft114/.julia/packages/Flux/qXNjB/src/optimise/train.jl:105
┌ Info: Epoch 508
└ @ Main /Users/kcft114/.julia/packages/Flux/qXNjB/src/optimise/train.jl:105
┌ Info: Epoch 509
└ @ Main /Users/kcft114/.julia/packages/Flux/qXNjB/src/optimise/train.jl:105
┌ Info: Epoch 510
└ @ Main /Users/kcft114/.julia/packages/Flux/qXNjB/src/optimise/train.jl:105
┌ Info: Epoch 511
└ @ Main /Users/kcft114/.julia/packages/Flux/qXNjB/src/optimise/train.jl:105
┌ Info: Epoch 512
└ @ Main /Users/kcft114/.julia/packages/Flux/qXNjB/src/optimise/train.jl:105
┌ Info: Epoch 513
└ @ Main /Users/kcft114/.julia/packages/Flux/qXNjB/src/optimise/train.jl:105
┌ Info: Epoch 514
└ @ Main /Users/kcft114/.julia/packages/Flux/qXNjB/src/optimise/train.jl:105
┌ Info: Epoch 515
└ @ Main /Users/kcft114/.julia/packages/Flux/qXNjB/src/optimise/train.jl:105
┌ Info: Epoch 516
└ @ Main /Users/kcft114/.julia/packages/Flux/qXNjB/src/optimise/train.jl:105
┌ Info: Epoch 517
└ @ Main /Users/kcft114/.julia/packages/Flux/qXNjB/src/optimise/train.jl:105
┌ Info: Epoch 518
└ @ Main /Users/kcft114/.julia/packages/Flux/qXNjB/src/optimise/train.jl:105
┌ Info: Epoch 519
└ @ Main /Users/kcft114/.julia/packages/Flux/qXNjB/src/optimise/train.jl:105
┌ Info: Epoch 520
└ @ Main /Users/kcft114/.julia/packages/Flux/qXNjB/src/optimise/train.jl:105
┌ Info: Epoch 521
└ @ Main /Users/kcft114/.julia/packages/Flux/qXNjB/src/optimise/train.jl:105
┌ Info: Epoch 522
└ @ Main /Users/kcft114/.julia/packages/Flux/qXNjB/src/optimise/train.jl:105
┌ Info: Epoch 523
└ @ Main /Users/kcft114/.julia/packages/Flux/qXNjB/src/optimise/train.jl:105
┌ Info: Epoch 524
└ @ Main /Users/kcft114/.julia/packages/Flux/qXNjB/src/optimise/train.jl:105
┌ Info: Epoch 525
└ @ Main /Users/kcft114/.julia/packages/Flux/qXNjB/src/optimise/train.jl:105
┌ Info: Epoch 526
└ @ Main /Users/kcft114/.julia/packages/Flux/qXNjB/src/optimise/train.jl:105
┌ Info: Epoch 527
└ @ Main /Users/kcft114/.julia/packages/Flux/qXNjB/src/optimise/train.jl:105
┌ Info: Epoch 528
└ @ Main /Users/kcft114/.julia/packages/Flux/qXNjB/src/optimise/train.jl:105
┌ Info: Epoch 529
└ @ Main /Users/kcft114/.julia/packages/Flux/qXNjB/src/optimise/train.jl:105
┌ Info: Epoch 530
└ @ Main /Users/kcft114/.julia/packages/Flux/qXNjB/src/optimise/train.jl:105
┌ Info: Epoch 531
└ @ Main /Users/kcft114/.julia/packages/Flux/qXNjB/src/optimise/train.jl:105
┌ Info: Epoch 532
└ @ Main /Users/kcft114/.julia/packages/Flux/qXNjB/src/optimise/train.jl:105
┌ Info: Epoch 533
└ @ Main /Users/kcft114/.julia/packages/Flux/qXNjB/src/optimise/train.jl:105
┌ Info: Epoch 534
└ @ Main /Users/kcft114/.julia/packages/Flux/qXNjB/src/optimise/train.jl:105
┌ Info: Epoch 535
└ @ Main /Users/kcft114/.julia/packages/Flux/qXNjB/src/optimise/train.jl:105
┌ Info: Epoch 536
└ @ Main /Users/kcft114/.julia/packages/Flux/qXNjB/src/optimise/train.jl:105
┌ Info: Epoch 537
└ @ Main /Users/kcft114/.julia/packages/Flux/qXNjB/src/optimise/train.jl:105
┌ Info: Epoch 538
└ @ Main /Users/kcft114/.julia/packages/Flux/qXNjB/src/optimise/train.jl:105
┌ Info: Epoch 539
└ @ Main /Users/kcft114/.julia/packages/Flux/qXNjB/src/optimise/train.jl:105
┌ Info: Epoch 540
└ @ Main /Users/kcft114/.julia/packages/Flux/qXNjB/src/optimise/train.jl:105
┌ Info: Epoch 541
└ @ Main /Users/kcft114/.julia/packages/Flux/qXNjB/src/optimise/train.jl:105
┌ Info: Epoch 542
└ @ Main /Users/kcft114/.julia/packages/Flux/qXNjB/src/optimise/train.jl:105
┌ Info: Epoch 543
└ @ Main /Users/kcft114/.julia/packages/Flux/qXNjB/src/optimise/train.jl:105
┌ Info: Epoch 544
└ @ Main /Users/kcft114/.julia/packages/Flux/qXNjB/src/optimise/train.jl:105
┌ Info: Epoch 545
└ @ Main /Users/kcft114/.julia/packages/Flux/qXNjB/src/optimise/train.jl:105
┌ Info: Epoch 546
└ @ Main /Users/kcft114/.julia/packages/Flux/qXNjB/src/optimise/train.jl:105
┌ Info: Epoch 547
└ @ Main /Users/kcft114/.julia/packages/Flux/qXNjB/src/optimise/train.jl:105
┌ Info: Epoch 548
└ @ Main /Users/kcft114/.julia/packages/Flux/qXNjB/src/optimise/train.jl:105
┌ Info: Epoch 549
└ @ Main /Users/kcft114/.julia/packages/Flux/qXNjB/src/optimise/train.jl:105
┌ Info: Epoch 550
└ @ Main /Users/kcft114/.julia/packages/Flux/qXNjB/src/optimise/train.jl:105
┌ Info: Epoch 551
└ @ Main /Users/kcft114/.julia/packages/Flux/qXNjB/src/optimise/train.jl:105
┌ Info: Epoch 552
└ @ Main /Users/kcft114/.julia/packages/Flux/qXNjB/src/optimise/train.jl:105
┌ Info: Epoch 553
└ @ Main /Users/kcft114/.julia/packages/Flux/qXNjB/src/optimise/train.jl:105
┌ Info: Epoch 554
└ @ Main /Users/kcft114/.julia/packages/Flux/qXNjB/src/optimise/train.jl:105
```

```
loss(X, Y) = 0.7097386f0 (tracked)
loss(X, Y) = 0.64466083f0 (tracked)
loss(X, Y) = 0.66504276f0 (tracked)
loss(X, Y) = 0.6786265f0 (tracked)
loss(X, Y) = 0.70325416f0 (tracked)
loss(X, Y) = 0.6912435f0 (tracked)
loss(X, Y) = 0.71588653f0 (tracked)
loss(X, Y) = 0.68113357f0 (tracked)
loss(X, Y) = 0.6864241f0 (tracked)
loss(X, Y) = 0.70627797f0 (tracked)
loss(X, Y) = 0.66891986f0 (tracked)
loss(X, Y) = 0.6737836f0 (tracked)
loss(X, Y) = 0.6973656f0 (tracked)
loss(X, Y) = 0.6491162f0 (tracked)
loss(X, Y) = 0.6719166f0 (tracked)
loss(X, Y) = 0.6487817f0 (tracked)
loss(X, Y) = 0.69327015f0 (tracked)
loss(X, Y) = 0.6792606f0 (tracked)
loss(X, Y) = 0.6743198f0 (tracked)
loss(X, Y) = 0.6896071f0 (tracked)
loss(X, Y) = 0.68310696f0 (tracked)
loss(X, Y) = 0.67011434f0 (tracked)
loss(X, Y) = 0.6672509f0 (tracked)
loss(X, Y) = 0.670822f0 (tracked)
loss(X, Y) = 0.70403624f0 (tracked)
loss(X, Y) = 0.6857355f0 (tracked)
loss(X, Y) = 0.67564267f0 (tracked)
loss(X, Y) = 0.6699415f0 (tracked)
loss(X, Y) = 0.66754913f0 (tracked)
loss(X, Y) = 0.67444444f0 (tracked)
loss(X, Y) = 0.67785335f0 (tracked)
loss(X, Y) = 0.6831029f0 (tracked)
loss(X, Y) = 0.6818736f0 (tracked)
loss(X, Y) = 0.7089647f0 (tracked)
loss(X, Y) = 0.6781293f0 (tracked)
loss(X, Y) = 0.67037773f0 (tracked)
loss(X, Y) = 0.6724777f0 (tracked)
loss(X, Y) = 0.6733937f0 (tracked)
loss(X, Y) = 0.68776304f0 (tracked)
loss(X, Y) = 0.6868225f0 (tracked)
loss(X, Y) = 0.6793095f0 (tracked)
loss(X, Y) = 0.66306597f0 (tracked)
loss(X, Y) = 0.66104937f0 (tracked)
loss(X, Y) = 0.6465855f0 (tracked)
loss(X, Y) = 0.70911175f0 (tracked)
loss(X, Y) = 0.69974524f0 (tracked)
loss(X, Y) = 0.6439581f0 (tracked)
loss(X, Y) = 0.68910587f0 (tracked)
loss(X, Y) = 0.68025196f0 (tracked)
loss(X, Y) = 0.6985691f0 (tracked)
loss(X, Y) = 0.6952274f0 (tracked)
loss(X, Y) = 0.6757511f0 (tracked)
loss(X, Y) = 0.67019844f0 (tracked)
loss(X, Y) = 0.6892013f0 (tracked)
loss(X, Y) = 0.67345214f0 (tracked)
loss(X, Y) = 0.68572086f0 (tracked)
loss(X, Y) = 0.6788912f0 (tracked)
loss(X, Y) = 0.68513674f0 (tracked)
loss(X, Y) = 0.66016746f0 (tracked)
loss(X, Y) = 0.65040153f0 (tracked)
loss(X, Y) = 0.68429494f0 (tracked)
loss(X, Y) = 0.67678845f0 (tracked)
loss(X, Y) = 0.67126065f0 (tracked)
loss(X, Y) = 0.68123084f0 (tracked)
loss(X, Y) = 0.6550959f0 (tracked)
loss(X, Y) = 0.67046714f0 (tracked)
loss(X, Y) = 0.6498539f0 (tracked)
loss(X, Y) = 0.6869533f0 (tracked)
loss(X, Y) = 0.65767014f0 (tracked)
loss(X, Y) = 0.65837497f0 (tracked)
loss(X, Y) = 0.6569271f0 (tracked)
loss(X, Y) = 0.66062784f0 (tracked)
loss(X, Y) = 0.6691582f0 (tracked)
loss(X, Y) = 0.6709423f0 (tracked)
loss(X, Y) = 0.64732945f0 (tracked)
loss(X, Y) = 0.66528434f0 (tracked)
loss(X, Y) = 0.65938413f0 (tracked)
loss(X, Y) = 0.6566404f0 (tracked)
loss(X, Y) = 0.68069583f0 (tracked)
loss(X, Y) = 0.68123233f0 (tracked)
loss(X, Y) = 0.6814058f0 (tracked)
loss(X, Y) = 0.6785302f0 (tracked)
loss(X, Y) = 0.6317856f0 (tracked)
loss(X, Y) = 0.64179623f0 (tracked)
loss(X, Y) = 0.67779565f0 (tracked)
loss(X, Y) = 0.67659277f0 (tracked)
loss(X, Y) = 0.6704649f0 (tracked)
loss(X, Y) = 0.66838f0 (tracked)
loss(X, Y) = 0.6721494f0 (tracked)
loss(X, Y) = 0.66935426f0 (tracked)
loss(X, Y) = 0.640466f0 (tracked)
loss(X, Y) = 0.64885247f0 (tracked)
loss(X, Y) = 0.6864165f0 (tracked)
loss(X, Y) = 0.6714123f0 (tracked)
loss(X, Y) = 0.67274314f0 (tracked)
loss(X, Y) = 0.63494486f0 (tracked)
loss(X, Y) = 0.6389519f0 (tracked)
loss(X, Y) = 0.68471736f0 (tracked)
loss(X, Y) = 0.6391946f0 (tracked)
loss(X, Y) = 0.6578228f0 (tracked)
loss(X, Y) = 0.6642234f0 (tracked)
loss(X, Y) = 0.64428353f0 (tracked)
loss(X, Y) = 0.68978596f0 (tracked)
loss(X, Y) = 0.6663936f0 (tracked)
loss(X, Y) = 0.6451119f0 (tracked)
loss(X, Y) = 0.6625085f0 (tracked)
loss(X, Y) = 0.6710424f0 (tracked)
loss(X, Y) = 0.6670204f0 (tracked)
loss(X, Y) = 0.6717111f0 (tracked)
loss(X, Y) = 0.6539234f0 (tracked)
loss(X, Y) = 0.6540144f0 (tracked)
loss(X, Y) = 0.638165f0 (tracked)
loss(X, Y) = 0.6556502f0 (tracked)
loss(X, Y) = 0.63306296f0 (tracked)
loss(X, Y) = 0.6803825f0 (tracked)
loss(X, Y) = 0.63757455f0 (tracked)
loss(X, Y) = 0.64590734f0 (tracked)
loss(X, Y) = 0.6670532f0 (tracked)
loss(X, Y) = 0.6553714f0 (tracked)
loss(X, Y) = 0.6709536f0 (tracked)
loss(X, Y) = 0.6601443f0 (tracked)
loss(X, Y) = 0.663845f0 (tracked)
loss(X, Y) = 0.67097664f0 (tracked)
loss(X, Y) = 0.65467244f0 (tracked)
loss(X, Y) = 0.6858397f0 (tracked)
loss(X, Y) = 0.6323507f0 (tracked)
loss(X, Y) = 0.671834f0 (tracked)
```

```
┌ Info: Epoch 555
└ @ Main /Users/kcft114/.julia/packages/Flux/qXNjB/src/optimise/train.jl:105
┌ Info: Epoch 556
└ @ Main /Users/kcft114/.julia/packages/Flux/qXNjB/src/optimise/train.jl:105
┌ Info: Epoch 557
└ @ Main /Users/kcft114/.julia/packages/Flux/qXNjB/src/optimise/train.jl:105
┌ Info: Epoch 558
└ @ Main /Users/kcft114/.julia/packages/Flux/qXNjB/src/optimise/train.jl:105
┌ Info: Epoch 559
└ @ Main /Users/kcft114/.julia/packages/Flux/qXNjB/src/optimise/train.jl:105
┌ Info: Epoch 560
└ @ Main /Users/kcft114/.julia/packages/Flux/qXNjB/src/optimise/train.jl:105
┌ Info: Epoch 561
└ @ Main /Users/kcft114/.julia/packages/Flux/qXNjB/src/optimise/train.jl:105
┌ Info: Epoch 562
└ @ Main /Users/kcft114/.julia/packages/Flux/qXNjB/src/optimise/train.jl:105
┌ Info: Epoch 563
└ @ Main /Users/kcft114/.julia/packages/Flux/qXNjB/src/optimise/train.jl:105
┌ Info: Epoch 564
└ @ Main /Users/kcft114/.julia/packages/Flux/qXNjB/src/optimise/train.jl:105
┌ Info: Epoch 565
└ @ Main /Users/kcft114/.julia/packages/Flux/qXNjB/src/optimise/train.jl:105
┌ Info: Epoch 566
└ @ Main /Users/kcft114/.julia/packages/Flux/qXNjB/src/optimise/train.jl:105
┌ Info: Epoch 567
└ @ Main /Users/kcft114/.julia/packages/Flux/qXNjB/src/optimise/train.jl:105
┌ Info: Epoch 568
└ @ Main /Users/kcft114/.julia/packages/Flux/qXNjB/src/optimise/train.jl:105
┌ Info: Epoch 569
└ @ Main /Users/kcft114/.julia/packages/Flux/qXNjB/src/optimise/train.jl:105
┌ Info: Epoch 570
└ @ Main /Users/kcft114/.julia/packages/Flux/qXNjB/src/optimise/train.jl:105
┌ Info: Epoch 571
└ @ Main /Users/kcft114/.julia/packages/Flux/qXNjB/src/optimise/train.jl:105
┌ Info: Epoch 572
└ @ Main /Users/kcft114/.julia/packages/Flux/qXNjB/src/optimise/train.jl:105
┌ Info: Epoch 573
└ @ Main /Users/kcft114/.julia/packages/Flux/qXNjB/src/optimise/train.jl:105
┌ Info: Epoch 574
└ @ Main /Users/kcft114/.julia/packages/Flux/qXNjB/src/optimise/train.jl:105
┌ Info: Epoch 575
└ @ Main /Users/kcft114/.julia/packages/Flux/qXNjB/src/optimise/train.jl:105
┌ Info: Epoch 576
└ @ Main /Users/kcft114/.julia/packages/Flux/qXNjB/src/optimise/train.jl:105
┌ Info: Epoch 577
└ @ Main /Users/kcft114/.julia/packages/Flux/qXNjB/src/optimise/train.jl:105
┌ Info: Epoch 578
└ @ Main /Users/kcft114/.julia/packages/Flux/qXNjB/src/optimise/train.jl:105
┌ Info: Epoch 579
└ @ Main /Users/kcft114/.julia/packages/Flux/qXNjB/src/optimise/train.jl:105
┌ Info: Epoch 580
└ @ Main /Users/kcft114/.julia/packages/Flux/qXNjB/src/optimise/train.jl:105
┌ Info: Epoch 581
└ @ Main /Users/kcft114/.julia/packages/Flux/qXNjB/src/optimise/train.jl:105
┌ Info: Epoch 582
└ @ Main /Users/kcft114/.julia/packages/Flux/qXNjB/src/optimise/train.jl:105
┌ Info: Epoch 583
└ @ Main /Users/kcft114/.julia/packages/Flux/qXNjB/src/optimise/train.jl:105
┌ Info: Epoch 584
└ @ Main /Users/kcft114/.julia/packages/Flux/qXNjB/src/optimise/train.jl:105
┌ Info: Epoch 585
└ @ Main /Users/kcft114/.julia/packages/Flux/qXNjB/src/optimise/train.jl:105
┌ Info: Epoch 586
└ @ Main /Users/kcft114/.julia/packages/Flux/qXNjB/src/optimise/train.jl:105
┌ Info: Epoch 587
└ @ Main /Users/kcft114/.julia/packages/Flux/qXNjB/src/optimise/train.jl:105
┌ Info: Epoch 588
└ @ Main /Users/kcft114/.julia/packages/Flux/qXNjB/src/optimise/train.jl:105
┌ Info: Epoch 589
└ @ Main /Users/kcft114/.julia/packages/Flux/qXNjB/src/optimise/train.jl:105
┌ Info: Epoch 590
└ @ Main /Users/kcft114/.julia/packages/Flux/qXNjB/src/optimise/train.jl:105
┌ Info: Epoch 591
└ @ Main /Users/kcft114/.julia/packages/Flux/qXNjB/src/optimise/train.jl:105
┌ Info: Epoch 592
└ @ Main /Users/kcft114/.julia/packages/Flux/qXNjB/src/optimise/train.jl:105
┌ Info: Epoch 593
└ @ Main /Users/kcft114/.julia/packages/Flux/qXNjB/src/optimise/train.jl:105
┌ Info: Epoch 594
└ @ Main /Users/kcft114/.julia/packages/Flux/qXNjB/src/optimise/train.jl:105
┌ Info: Epoch 595
└ @ Main /Users/kcft114/.julia/packages/Flux/qXNjB/src/optimise/train.jl:105
┌ Info: Epoch 596
└ @ Main /Users/kcft114/.julia/packages/Flux/qXNjB/src/optimise/train.jl:105
┌ Info: Epoch 597
└ @ Main /Users/kcft114/.julia/packages/Flux/qXNjB/src/optimise/train.jl:105
┌ Info: Epoch 598
└ @ Main /Users/kcft114/.julia/packages/Flux/qXNjB/src/optimise/train.jl:105
┌ Info: Epoch 599
└ @ Main /Users/kcft114/.julia/packages/Flux/qXNjB/src/optimise/train.jl:105
┌ Info: Epoch 600
└ @ Main /Users/kcft114/.julia/packages/Flux/qXNjB/src/optimise/train.jl:105
┌ Info: Epoch 601
└ @ Main /Users/kcft114/.julia/packages/Flux/qXNjB/src/optimise/train.jl:105
┌ Info: Epoch 602
└ @ Main /Users/kcft114/.julia/packages/Flux/qXNjB/src/optimise/train.jl:105
┌ Info: Epoch 603
└ @ Main /Users/kcft114/.julia/packages/Flux/qXNjB/src/optimise/train.jl:105
┌ Info: Epoch 604
└ @ Main /Users/kcft114/.julia/packages/Flux/qXNjB/src/optimise/train.jl:105
┌ Info: Epoch 605
└ @ Main /Users/kcft114/.julia/packages/Flux/qXNjB/src/optimise/train.jl:105
┌ Info: Epoch 606
└ @ Main /Users/kcft114/.julia/packages/Flux/qXNjB/src/optimise/train.jl:105
┌ Info: Epoch 607
└ @ Main /Users/kcft114/.julia/packages/Flux/qXNjB/src/optimise/train.jl:105
┌ Info: Epoch 608
└ @ Main /Users/kcft114/.julia/packages/Flux/qXNjB/src/optimise/train.jl:105
┌ Info: Epoch 609
└ @ Main /Users/kcft114/.julia/packages/Flux/qXNjB/src/optimise/train.jl:105
┌ Info: Epoch 610
└ @ Main /Users/kcft114/.julia/packages/Flux/qXNjB/src/optimise/train.jl:105
┌ Info: Epoch 611
└ @ Main /Users/kcft114/.julia/packages/Flux/qXNjB/src/optimise/train.jl:105
┌ Info: Epoch 612
└ @ Main /Users/kcft114/.julia/packages/Flux/qXNjB/src/optimise/train.jl:105
┌ Info: Epoch 613
└ @ Main /Users/kcft114/.julia/packages/Flux/qXNjB/src/optimise/train.jl:105
┌ Info: Epoch 614
└ @ Main /Users/kcft114/.julia/packages/Flux/qXNjB/src/optimise/train.jl:105
┌ Info: Epoch 615
└ @ Main /Users/kcft114/.julia/packages/Flux/qXNjB/src/optimise/train.jl:105
┌ Info: Epoch 616
└ @ Main /Users/kcft114/.julia/packages/Flux/qXNjB/src/optimise/train.jl:105
┌ Info: Epoch 617
└ @ Main /Users/kcft114/.julia/packages/Flux/qXNjB/src/optimise/train.jl:105
┌ Info: Epoch 618
└ @ Main /Users/kcft114/.julia/packages/Flux/qXNjB/src/optimise/train.jl:105
┌ Info: Epoch 619
└ @ Main /Users/kcft114/.julia/packages/Flux/qXNjB/src/optimise/train.jl:105
┌ Info: Epoch 620
└ @ Main /Users/kcft114/.julia/packages/Flux/qXNjB/src/optimise/train.jl:105
┌ Info: Epoch 621
└ @ Main /Users/kcft114/.julia/packages/Flux/qXNjB/src/optimise/train.jl:105
┌ Info: Epoch 622
└ @ Main /Users/kcft114/.julia/packages/Flux/qXNjB/src/optimise/train.jl:105
┌ Info: Epoch 623
└ @ Main /Users/kcft114/.julia/packages/Flux/qXNjB/src/optimise/train.jl:105
┌ Info: Epoch 624
└ @ Main /Users/kcft114/.julia/packages/Flux/qXNjB/src/optimise/train.jl:105
┌ Info: Epoch 625
└ @ Main /Users/kcft114/.julia/packages/Flux/qXNjB/src/optimise/train.jl:105
┌ Info: Epoch 626
└ @ Main /Users/kcft114/.julia/packages/Flux/qXNjB/src/optimise/train.jl:105
┌ Info: Epoch 627
└ @ Main /Users/kcft114/.julia/packages/Flux/qXNjB/src/optimise/train.jl:105
┌ Info: Epoch 628
└ @ Main /Users/kcft114/.julia/packages/Flux/qXNjB/src/optimise/train.jl:105
┌ Info: Epoch 629
└ @ Main /Users/kcft114/.julia/packages/Flux/qXNjB/src/optimise/train.jl:105
┌ Info: Epoch 630
└ @ Main /Users/kcft114/.julia/packages/Flux/qXNjB/src/optimise/train.jl:105
┌ Info: Epoch 631
└ @ Main /Users/kcft114/.julia/packages/Flux/qXNjB/src/optimise/train.jl:105
┌ Info: Epoch 632
└ @ Main /Users/kcft114/.julia/packages/Flux/qXNjB/src/optimise/train.jl:105
┌ Info: Epoch 633
└ @ Main /Users/kcft114/.julia/packages/Flux/qXNjB/src/optimise/train.jl:105
┌ Info: Epoch 634
└ @ Main /Users/kcft114/.julia/packages/Flux/qXNjB/src/optimise/train.jl:105
┌ Info: Epoch 635
└ @ Main /Users/kcft114/.julia/packages/Flux/qXNjB/src/optimise/train.jl:105
┌ Info: Epoch 636
└ @ Main /Users/kcft114/.julia/packages/Flux/qXNjB/src/optimise/train.jl:105
┌ Info: Epoch 637
└ @ Main /Users/kcft114/.julia/packages/Flux/qXNjB/src/optimise/train.jl:105
┌ Info: Epoch 638
└ @ Main /Users/kcft114/.julia/packages/Flux/qXNjB/src/optimise/train.jl:105
┌ Info: Epoch 639
└ @ Main /Users/kcft114/.julia/packages/Flux/qXNjB/src/optimise/train.jl:105
┌ Info: Epoch 640
└ @ Main /Users/kcft114/.julia/packages/Flux/qXNjB/src/optimise/train.jl:105
┌ Info: Epoch 641
└ @ Main /Users/kcft114/.julia/packages/Flux/qXNjB/src/optimise/train.jl:105
┌ Info: Epoch 642
└ @ Main /Users/kcft114/.julia/packages/Flux/qXNjB/src/optimise/train.jl:105
┌ Info: Epoch 643
└ @ Main /Users/kcft114/.julia/packages/Flux/qXNjB/src/optimise/train.jl:105
┌ Info: Epoch 644
└ @ Main /Users/kcft114/.julia/packages/Flux/qXNjB/src/optimise/train.jl:105
┌ Info: Epoch 645
└ @ Main /Users/kcft114/.julia/packages/Flux/qXNjB/src/optimise/train.jl:105
┌ Info: Epoch 646
└ @ Main /Users/kcft114/.julia/packages/Flux/qXNjB/src/optimise/train.jl:105
┌ Info: Epoch 647
└ @ Main /Users/kcft114/.julia/packages/Flux/qXNjB/src/optimise/train.jl:105
┌ Info: Epoch 648
└ @ Main /Users/kcft114/.julia/packages/Flux/qXNjB/src/optimise/train.jl:105
┌ Info: Epoch 649
└ @ Main /Users/kcft114/.julia/packages/Flux/qXNjB/src/optimise/train.jl:105
┌ Info: Epoch 650
└ @ Main /Users/kcft114/.julia/packages/Flux/qXNjB/src/optimise/train.jl:105
┌ Info: Epoch 651
└ @ Main /Users/kcft114/.julia/packages/Flux/qXNjB/src/optimise/train.jl:105
┌ Info: Epoch 652
└ @ Main /Users/kcft114/.julia/packages/Flux/qXNjB/src/optimise/train.jl:105
┌ Info: Epoch 653
└ @ Main /Users/kcft114/.julia/packages/Flux/qXNjB/src/optimise/train.jl:105
┌ Info: Epoch 654
└ @ Main /Users/kcft114/.julia/packages/Flux/qXNjB/src/optimise/train.jl:105
┌ Info: Epoch 655
└ @ Main /Users/kcft114/.julia/packages/Flux/qXNjB/src/optimise/train.jl:105
┌ Info: Epoch 656
└ @ Main /Users/kcft114/.julia/packages/Flux/qXNjB/src/optimise/train.jl:105
┌ Info: Epoch 657
└ @ Main /Users/kcft114/.julia/packages/Flux/qXNjB/src/optimise/train.jl:105
┌ Info: Epoch 658
└ @ Main /Users/kcft114/.julia/packages/Flux/qXNjB/src/optimise/train.jl:105
┌ Info: Epoch 659
└ @ Main /Users/kcft114/.julia/packages/Flux/qXNjB/src/optimise/train.jl:105
┌ Info: Epoch 660
└ @ Main /Users/kcft114/.julia/packages/Flux/qXNjB/src/optimise/train.jl:105
┌ Info: Epoch 661
└ @ Main /Users/kcft114/.julia/packages/Flux/qXNjB/src/optimise/train.jl:105
┌ Info: Epoch 662
└ @ Main /Users/kcft114/.julia/packages/Flux/qXNjB/src/optimise/train.jl:105
┌ Info: Epoch 663
└ @ Main /Users/kcft114/.julia/packages/Flux/qXNjB/src/optimise/train.jl:105
┌ Info: Epoch 664
└ @ Main /Users/kcft114/.julia/packages/Flux/qXNjB/src/optimise/train.jl:105
┌ Info: Epoch 665
└ @ Main /Users/kcft114/.julia/packages/Flux/qXNjB/src/optimise/train.jl:105
┌ Info: Epoch 666
└ @ Main /Users/kcft114/.julia/packages/Flux/qXNjB/src/optimise/train.jl:105
┌ Info: Epoch 667
└ @ Main /Users/kcft114/.julia/packages/Flux/qXNjB/src/optimise/train.jl:105
┌ Info: Epoch 668
└ @ Main /Users/kcft114/.julia/packages/Flux/qXNjB/src/optimise/train.jl:105
┌ Info: Epoch 669
└ @ Main /Users/kcft114/.julia/packages/Flux/qXNjB/src/optimise/train.jl:105
┌ Info: Epoch 670
└ @ Main /Users/kcft114/.julia/packages/Flux/qXNjB/src/optimise/train.jl:105
┌ Info: Epoch 671
└ @ Main /Users/kcft114/.julia/packages/Flux/qXNjB/src/optimise/train.jl:105
┌ Info: Epoch 672
└ @ Main /Users/kcft114/.julia/packages/Flux/qXNjB/src/optimise/train.jl:105
┌ Info: Epoch 673
└ @ Main /Users/kcft114/.julia/packages/Flux/qXNjB/src/optimise/train.jl:105
┌ Info: Epoch 674
└ @ Main /Users/kcft114/.julia/packages/Flux/qXNjB/src/optimise/train.jl:105
┌ Info: Epoch 675
└ @ Main /Users/kcft114/.julia/packages/Flux/qXNjB/src/optimise/train.jl:105
┌ Info: Epoch 676
└ @ Main /Users/kcft114/.julia/packages/Flux/qXNjB/src/optimise/train.jl:105
┌ Info: Epoch 677
└ @ Main /Users/kcft114/.julia/packages/Flux/qXNjB/src/optimise/train.jl:105
┌ Info: Epoch 678
└ @ Main /Users/kcft114/.julia/packages/Flux/qXNjB/src/optimise/train.jl:105
┌ Info: Epoch 679
└ @ Main /Users/kcft114/.julia/packages/Flux/qXNjB/src/optimise/train.jl:105
┌ Info: Epoch 680
└ @ Main /Users/kcft114/.julia/packages/Flux/qXNjB/src/optimise/train.jl:105
┌ Info: Epoch 681
└ @ Main /Users/kcft114/.julia/packages/Flux/qXNjB/src/optimise/train.jl:105
```

```
loss(X, Y) = 0.63432634f0 (tracked)
loss(X, Y) = 0.64482445f0 (tracked)
loss(X, Y) = 0.6614019f0 (tracked)
loss(X, Y) = 0.6662789f0 (tracked)
loss(X, Y) = 0.65417653f0 (tracked)
loss(X, Y) = 0.6807343f0 (tracked)
loss(X, Y) = 0.6573998f0 (tracked)
loss(X, Y) = 0.6352472f0 (tracked)
loss(X, Y) = 0.68533534f0 (tracked)
loss(X, Y) = 0.66670716f0 (tracked)
loss(X, Y) = 0.6604453f0 (tracked)
loss(X, Y) = 0.6584827f0 (tracked)
loss(X, Y) = 0.6690023f0 (tracked)
loss(X, Y) = 0.6621528f0 (tracked)
loss(X, Y) = 0.6652444f0 (tracked)
loss(X, Y) = 0.64112765f0 (tracked)
loss(X, Y) = 0.6393881f0 (tracked)
loss(X, Y) = 0.66528344f0 (tracked)
loss(X, Y) = 0.6432315f0 (tracked)
loss(X, Y) = 0.62853265f0 (tracked)
loss(X, Y) = 0.63287807f0 (tracked)
loss(X, Y) = 0.66073155f0 (tracked)
loss(X, Y) = 0.6321413f0 (tracked)
loss(X, Y) = 0.66367024f0 (tracked)
loss(X, Y) = 0.65817875f0 (tracked)
loss(X, Y) = 0.6618089f0 (tracked)
loss(X, Y) = 0.63717985f0 (tracked)
loss(X, Y) = 0.64318556f0 (tracked)
loss(X, Y) = 0.64939165f0 (tracked)
loss(X, Y) = 0.65592843f0 (tracked)
loss(X, Y) = 0.6236179f0 (tracked)
loss(X, Y) = 0.6262242f0 (tracked)
loss(X, Y) = 0.6464139f0 (tracked)
loss(X, Y) = 0.6586121f0 (tracked)
loss(X, Y) = 0.6504676f0 (tracked)
loss(X, Y) = 0.65518236f0 (tracked)
loss(X, Y) = 0.67528844f0 (tracked)
loss(X, Y) = 0.6517382f0 (tracked)
loss(X, Y) = 0.63030577f0 (tracked)
loss(X, Y) = 0.64338243f0 (tracked)
loss(X, Y) = 0.630092f0 (tracked)
loss(X, Y) = 0.63863873f0 (tracked)
loss(X, Y) = 0.64991057f0 (tracked)
loss(X, Y) = 0.63184536f0 (tracked)
loss(X, Y) = 0.6267675f0 (tracked)
loss(X, Y) = 0.63316584f0 (tracked)
loss(X, Y) = 0.62388253f0 (tracked)
loss(X, Y) = 0.6593773f0 (tracked)
loss(X, Y) = 0.63774353f0 (tracked)
loss(X, Y) = 0.64086956f0 (tracked)
loss(X, Y) = 0.6410664f0 (tracked)
loss(X, Y) = 0.64134836f0 (tracked)
loss(X, Y) = 0.64192766f0 (tracked)
loss(X, Y) = 0.62877905f0 (tracked)
loss(X, Y) = 0.6402259f0 (tracked)
loss(X, Y) = 0.6445795f0 (tracked)
loss(X, Y) = 0.6689606f0 (tracked)
loss(X, Y) = 0.6485689f0 (tracked)
loss(X, Y) = 0.62158453f0 (tracked)
loss(X, Y) = 0.6392571f0 (tracked)
loss(X, Y) = 0.664144f0 (tracked)
loss(X, Y) = 0.6378508f0 (tracked)
loss(X, Y) = 0.6605162f0 (tracked)
loss(X, Y) = 0.6597331f0 (tracked)
loss(X, Y) = 0.6590527f0 (tracked)
loss(X, Y) = 0.594485f0 (tracked)
loss(X, Y) = 0.6359112f0 (tracked)
loss(X, Y) = 0.63283575f0 (tracked)
loss(X, Y) = 0.65013057f0 (tracked)
loss(X, Y) = 0.6390733f0 (tracked)
loss(X, Y) = 0.65452737f0 (tracked)
loss(X, Y) = 0.63425446f0 (tracked)
loss(X, Y) = 0.6318753f0 (tracked)
loss(X, Y) = 0.6401783f0 (tracked)
loss(X, Y) = 0.6711306f0 (tracked)
loss(X, Y) = 0.6445382f0 (tracked)
loss(X, Y) = 0.6214775f0 (tracked)
loss(X, Y) = 0.6445646f0 (tracked)
loss(X, Y) = 0.6492815f0 (tracked)
loss(X, Y) = 0.6224153f0 (tracked)
loss(X, Y) = 0.6416607f0 (tracked)
loss(X, Y) = 0.63689464f0 (tracked)
loss(X, Y) = 0.64410233f0 (tracked)
loss(X, Y) = 0.62792087f0 (tracked)
loss(X, Y) = 0.63469183f0 (tracked)
loss(X, Y) = 0.6420783f0 (tracked)
loss(X, Y) = 0.6384941f0 (tracked)
loss(X, Y) = 0.62793124f0 (tracked)
loss(X, Y) = 0.6390582f0 (tracked)
loss(X, Y) = 0.61981624f0 (tracked)
loss(X, Y) = 0.64760804f0 (tracked)
loss(X, Y) = 0.6360059f0 (tracked)
loss(X, Y) = 0.6441485f0 (tracked)
loss(X, Y) = 0.6298366f0 (tracked)
loss(X, Y) = 0.6404661f0 (tracked)
loss(X, Y) = 0.63148695f0 (tracked)
loss(X, Y) = 0.6381342f0 (tracked)
loss(X, Y) = 0.6544415f0 (tracked)
loss(X, Y) = 0.6209946f0 (tracked)
loss(X, Y) = 0.6502823f0 (tracked)
loss(X, Y) = 0.60323054f0 (tracked)
loss(X, Y) = 0.61882544f0 (tracked)
loss(X, Y) = 0.6154863f0 (tracked)
loss(X, Y) = 0.63944423f0 (tracked)
loss(X, Y) = 0.61521804f0 (tracked)
loss(X, Y) = 0.6355322f0 (tracked)
loss(X, Y) = 0.64863133f0 (tracked)
loss(X, Y) = 0.62841946f0 (tracked)
loss(X, Y) = 0.64882916f0 (tracked)
loss(X, Y) = 0.6118652f0 (tracked)
loss(X, Y) = 0.63950557f0 (tracked)
loss(X, Y) = 0.6349218f0 (tracked)
loss(X, Y) = 0.6451394f0 (tracked)
loss(X, Y) = 0.6355131f0 (tracked)
```

```
┌ Info: Epoch 682
└ @ Main /Users/kcft114/.julia/packages/Flux/qXNjB/src/optimise/train.jl:105
┌ Info: Epoch 683
└ @ Main /Users/kcft114/.julia/packages/Flux/qXNjB/src/optimise/train.jl:105
┌ Info: Epoch 684
└ @ Main /Users/kcft114/.julia/packages/Flux/qXNjB/src/optimise/train.jl:105
┌ Info: Epoch 685
└ @ Main /Users/kcft114/.julia/packages/Flux/qXNjB/src/optimise/train.jl:105
┌ Info: Epoch 686
└ @ Main /Users/kcft114/.julia/packages/Flux/qXNjB/src/optimise/train.jl:105
┌ Info: Epoch 687
└ @ Main /Users/kcft114/.julia/packages/Flux/qXNjB/src/optimise/train.jl:105
┌ Info: Epoch 688
└ @ Main /Users/kcft114/.julia/packages/Flux/qXNjB/src/optimise/train.jl:105
┌ Info: Epoch 689
└ @ Main /Users/kcft114/.julia/packages/Flux/qXNjB/src/optimise/train.jl:105
┌ Info: Epoch 690
└ @ Main /Users/kcft114/.julia/packages/Flux/qXNjB/src/optimise/train.jl:105
┌ Info: Epoch 691
└ @ Main /Users/kcft114/.julia/packages/Flux/qXNjB/src/optimise/train.jl:105
┌ Info: Epoch 692
└ @ Main /Users/kcft114/.julia/packages/Flux/qXNjB/src/optimise/train.jl:105
┌ Info: Epoch 693
└ @ Main /Users/kcft114/.julia/packages/Flux/qXNjB/src/optimise/train.jl:105
┌ Info: Epoch 694
└ @ Main /Users/kcft114/.julia/packages/Flux/qXNjB/src/optimise/train.jl:105
┌ Info: Epoch 695
└ @ Main /Users/kcft114/.julia/packages/Flux/qXNjB/src/optimise/train.jl:105
┌ Info: Epoch 696
└ @ Main /Users/kcft114/.julia/packages/Flux/qXNjB/src/optimise/train.jl:105
┌ Info: Epoch 697
└ @ Main /Users/kcft114/.julia/packages/Flux/qXNjB/src/optimise/train.jl:105
┌ Info: Epoch 698
└ @ Main /Users/kcft114/.julia/packages/Flux/qXNjB/src/optimise/train.jl:105
┌ Info: Epoch 699
└ @ Main /Users/kcft114/.julia/packages/Flux/qXNjB/src/optimise/train.jl:105
┌ Info: Epoch 700
└ @ Main /Users/kcft114/.julia/packages/Flux/qXNjB/src/optimise/train.jl:105
┌ Info: Epoch 701
└ @ Main /Users/kcft114/.julia/packages/Flux/qXNjB/src/optimise/train.jl:105
┌ Info: Epoch 702
└ @ Main /Users/kcft114/.julia/packages/Flux/qXNjB/src/optimise/train.jl:105
┌ Info: Epoch 703
└ @ Main /Users/kcft114/.julia/packages/Flux/qXNjB/src/optimise/train.jl:105
┌ Info: Epoch 704
└ @ Main /Users/kcft114/.julia/packages/Flux/qXNjB/src/optimise/train.jl:105
┌ Info: Epoch 705
└ @ Main /Users/kcft114/.julia/packages/Flux/qXNjB/src/optimise/train.jl:105
┌ Info: Epoch 706
└ @ Main /Users/kcft114/.julia/packages/Flux/qXNjB/src/optimise/train.jl:105
┌ Info: Epoch 707
└ @ Main /Users/kcft114/.julia/packages/Flux/qXNjB/src/optimise/train.jl:105
┌ Info: Epoch 708
└ @ Main /Users/kcft114/.julia/packages/Flux/qXNjB/src/optimise/train.jl:105
┌ Info: Epoch 709
└ @ Main /Users/kcft114/.julia/packages/Flux/qXNjB/src/optimise/train.jl:105
┌ Info: Epoch 710
└ @ Main /Users/kcft114/.julia/packages/Flux/qXNjB/src/optimise/train.jl:105
┌ Info: Epoch 711
└ @ Main /Users/kcft114/.julia/packages/Flux/qXNjB/src/optimise/train.jl:105
┌ Info: Epoch 712
└ @ Main /Users/kcft114/.julia/packages/Flux/qXNjB/src/optimise/train.jl:105
┌ Info: Epoch 713
└ @ Main /Users/kcft114/.julia/packages/Flux/qXNjB/src/optimise/train.jl:105
┌ Info: Epoch 714
└ @ Main /Users/kcft114/.julia/packages/Flux/qXNjB/src/optimise/train.jl:105
┌ Info: Epoch 715
└ @ Main /Users/kcft114/.julia/packages/Flux/qXNjB/src/optimise/train.jl:105
┌ Info: Epoch 716
└ @ Main /Users/kcft114/.julia/packages/Flux/qXNjB/src/optimise/train.jl:105
┌ Info: Epoch 717
└ @ Main /Users/kcft114/.julia/packages/Flux/qXNjB/src/optimise/train.jl:105
┌ Info: Epoch 718
└ @ Main /Users/kcft114/.julia/packages/Flux/qXNjB/src/optimise/train.jl:105
┌ Info: Epoch 719
└ @ Main /Users/kcft114/.julia/packages/Flux/qXNjB/src/optimise/train.jl:105
┌ Info: Epoch 720
└ @ Main /Users/kcft114/.julia/packages/Flux/qXNjB/src/optimise/train.jl:105
┌ Info: Epoch 721
└ @ Main /Users/kcft114/.julia/packages/Flux/qXNjB/src/optimise/train.jl:105
┌ Info: Epoch 722
└ @ Main /Users/kcft114/.julia/packages/Flux/qXNjB/src/optimise/train.jl:105
┌ Info: Epoch 723
└ @ Main /Users/kcft114/.julia/packages/Flux/qXNjB/src/optimise/train.jl:105
┌ Info: Epoch 724
└ @ Main /Users/kcft114/.julia/packages/Flux/qXNjB/src/optimise/train.jl:105
┌ Info: Epoch 725
└ @ Main /Users/kcft114/.julia/packages/Flux/qXNjB/src/optimise/train.jl:105
┌ Info: Epoch 726
└ @ Main /Users/kcft114/.julia/packages/Flux/qXNjB/src/optimise/train.jl:105
┌ Info: Epoch 727
└ @ Main /Users/kcft114/.julia/packages/Flux/qXNjB/src/optimise/train.jl:105
┌ Info: Epoch 728
└ @ Main /Users/kcft114/.julia/packages/Flux/qXNjB/src/optimise/train.jl:105
┌ Info: Epoch 729
└ @ Main /Users/kcft114/.julia/packages/Flux/qXNjB/src/optimise/train.jl:105
┌ Info: Epoch 730
└ @ Main /Users/kcft114/.julia/packages/Flux/qXNjB/src/optimise/train.jl:105
┌ Info: Epoch 731
└ @ Main /Users/kcft114/.julia/packages/Flux/qXNjB/src/optimise/train.jl:105
┌ Info: Epoch 732
└ @ Main /Users/kcft114/.julia/packages/Flux/qXNjB/src/optimise/train.jl:105
┌ Info: Epoch 733
└ @ Main /Users/kcft114/.julia/packages/Flux/qXNjB/src/optimise/train.jl:105
┌ Info: Epoch 734
└ @ Main /Users/kcft114/.julia/packages/Flux/qXNjB/src/optimise/train.jl:105
┌ Info: Epoch 735
└ @ Main /Users/kcft114/.julia/packages/Flux/qXNjB/src/optimise/train.jl:105
┌ Info: Epoch 736
└ @ Main /Users/kcft114/.julia/packages/Flux/qXNjB/src/optimise/train.jl:105
┌ Info: Epoch 737
└ @ Main /Users/kcft114/.julia/packages/Flux/qXNjB/src/optimise/train.jl:105
┌ Info: Epoch 738
└ @ Main /Users/kcft114/.julia/packages/Flux/qXNjB/src/optimise/train.jl:105
┌ Info: Epoch 739
└ @ Main /Users/kcft114/.julia/packages/Flux/qXNjB/src/optimise/train.jl:105
┌ Info: Epoch 740
└ @ Main /Users/kcft114/.julia/packages/Flux/qXNjB/src/optimise/train.jl:105
┌ Info: Epoch 741
└ @ Main /Users/kcft114/.julia/packages/Flux/qXNjB/src/optimise/train.jl:105
┌ Info: Epoch 742
└ @ Main /Users/kcft114/.julia/packages/Flux/qXNjB/src/optimise/train.jl:105
┌ Info: Epoch 743
└ @ Main /Users/kcft114/.julia/packages/Flux/qXNjB/src/optimise/train.jl:105
┌ Info: Epoch 744
└ @ Main /Users/kcft114/.julia/packages/Flux/qXNjB/src/optimise/train.jl:105
┌ Info: Epoch 745
└ @ Main /Users/kcft114/.julia/packages/Flux/qXNjB/src/optimise/train.jl:105
┌ Info: Epoch 746
└ @ Main /Users/kcft114/.julia/packages/Flux/qXNjB/src/optimise/train.jl:105
┌ Info: Epoch 747
└ @ Main /Users/kcft114/.julia/packages/Flux/qXNjB/src/optimise/train.jl:105
┌ Info: Epoch 748
└ @ Main /Users/kcft114/.julia/packages/Flux/qXNjB/src/optimise/train.jl:105
┌ Info: Epoch 749
└ @ Main /Users/kcft114/.julia/packages/Flux/qXNjB/src/optimise/train.jl:105
┌ Info: Epoch 750
└ @ Main /Users/kcft114/.julia/packages/Flux/qXNjB/src/optimise/train.jl:105
┌ Info: Epoch 751
└ @ Main /Users/kcft114/.julia/packages/Flux/qXNjB/src/optimise/train.jl:105
┌ Info: Epoch 752
└ @ Main /Users/kcft114/.julia/packages/Flux/qXNjB/src/optimise/train.jl:105
┌ Info: Epoch 753
└ @ Main /Users/kcft114/.julia/packages/Flux/qXNjB/src/optimise/train.jl:105
┌ Info: Epoch 754
└ @ Main /Users/kcft114/.julia/packages/Flux/qXNjB/src/optimise/train.jl:105
┌ Info: Epoch 755
└ @ Main /Users/kcft114/.julia/packages/Flux/qXNjB/src/optimise/train.jl:105
┌ Info: Epoch 756
└ @ Main /Users/kcft114/.julia/packages/Flux/qXNjB/src/optimise/train.jl:105
┌ Info: Epoch 757
└ @ Main /Users/kcft114/.julia/packages/Flux/qXNjB/src/optimise/train.jl:105
┌ Info: Epoch 758
└ @ Main /Users/kcft114/.julia/packages/Flux/qXNjB/src/optimise/train.jl:105
┌ Info: Epoch 759
└ @ Main /Users/kcft114/.julia/packages/Flux/qXNjB/src/optimise/train.jl:105
┌ Info: Epoch 760
└ @ Main /Users/kcft114/.julia/packages/Flux/qXNjB/src/optimise/train.jl:105
┌ Info: Epoch 761
└ @ Main /Users/kcft114/.julia/packages/Flux/qXNjB/src/optimise/train.jl:105
┌ Info: Epoch 762
└ @ Main /Users/kcft114/.julia/packages/Flux/qXNjB/src/optimise/train.jl:105
┌ Info: Epoch 763
└ @ Main /Users/kcft114/.julia/packages/Flux/qXNjB/src/optimise/train.jl:105
┌ Info: Epoch 764
└ @ Main /Users/kcft114/.julia/packages/Flux/qXNjB/src/optimise/train.jl:105
┌ Info: Epoch 765
└ @ Main /Users/kcft114/.julia/packages/Flux/qXNjB/src/optimise/train.jl:105
┌ Info: Epoch 766
└ @ Main /Users/kcft114/.julia/packages/Flux/qXNjB/src/optimise/train.jl:105
┌ Info: Epoch 767
└ @ Main /Users/kcft114/.julia/packages/Flux/qXNjB/src/optimise/train.jl:105
┌ Info: Epoch 768
└ @ Main /Users/kcft114/.julia/packages/Flux/qXNjB/src/optimise/train.jl:105
┌ Info: Epoch 769
└ @ Main /Users/kcft114/.julia/packages/Flux/qXNjB/src/optimise/train.jl:105
┌ Info: Epoch 770
└ @ Main /Users/kcft114/.julia/packages/Flux/qXNjB/src/optimise/train.jl:105
┌ Info: Epoch 771
└ @ Main /Users/kcft114/.julia/packages/Flux/qXNjB/src/optimise/train.jl:105
┌ Info: Epoch 772
└ @ Main /Users/kcft114/.julia/packages/Flux/qXNjB/src/optimise/train.jl:105
┌ Info: Epoch 773
└ @ Main /Users/kcft114/.julia/packages/Flux/qXNjB/src/optimise/train.jl:105
┌ Info: Epoch 774
└ @ Main /Users/kcft114/.julia/packages/Flux/qXNjB/src/optimise/train.jl:105
┌ Info: Epoch 775
└ @ Main /Users/kcft114/.julia/packages/Flux/qXNjB/src/optimise/train.jl:105
┌ Info: Epoch 776
└ @ Main /Users/kcft114/.julia/packages/Flux/qXNjB/src/optimise/train.jl:105
┌ Info: Epoch 777
└ @ Main /Users/kcft114/.julia/packages/Flux/qXNjB/src/optimise/train.jl:105
┌ Info: Epoch 778
└ @ Main /Users/kcft114/.julia/packages/Flux/qXNjB/src/optimise/train.jl:105
┌ Info: Epoch 779
└ @ Main /Users/kcft114/.julia/packages/Flux/qXNjB/src/optimise/train.jl:105
┌ Info: Epoch 780
└ @ Main /Users/kcft114/.julia/packages/Flux/qXNjB/src/optimise/train.jl:105
┌ Info: Epoch 781
└ @ Main /Users/kcft114/.julia/packages/Flux/qXNjB/src/optimise/train.jl:105
┌ Info: Epoch 782
└ @ Main /Users/kcft114/.julia/packages/Flux/qXNjB/src/optimise/train.jl:105
┌ Info: Epoch 783
└ @ Main /Users/kcft114/.julia/packages/Flux/qXNjB/src/optimise/train.jl:105
┌ Info: Epoch 784
└ @ Main /Users/kcft114/.julia/packages/Flux/qXNjB/src/optimise/train.jl:105
┌ Info: Epoch 785
└ @ Main /Users/kcft114/.julia/packages/Flux/qXNjB/src/optimise/train.jl:105
┌ Info: Epoch 786
└ @ Main /Users/kcft114/.julia/packages/Flux/qXNjB/src/optimise/train.jl:105
┌ Info: Epoch 787
└ @ Main /Users/kcft114/.julia/packages/Flux/qXNjB/src/optimise/train.jl:105
┌ Info: Epoch 788
└ @ Main /Users/kcft114/.julia/packages/Flux/qXNjB/src/optimise/train.jl:105
┌ Info: Epoch 789
└ @ Main /Users/kcft114/.julia/packages/Flux/qXNjB/src/optimise/train.jl:105
┌ Info: Epoch 790
└ @ Main /Users/kcft114/.julia/packages/Flux/qXNjB/src/optimise/train.jl:105
┌ Info: Epoch 791
└ @ Main /Users/kcft114/.julia/packages/Flux/qXNjB/src/optimise/train.jl:105
┌ Info: Epoch 792
└ @ Main /Users/kcft114/.julia/packages/Flux/qXNjB/src/optimise/train.jl:105
┌ Info: Epoch 793
└ @ Main /Users/kcft114/.julia/packages/Flux/qXNjB/src/optimise/train.jl:105
┌ Info: Epoch 794
└ @ Main /Users/kcft114/.julia/packages/Flux/qXNjB/src/optimise/train.jl:105
┌ Info: Epoch 795
└ @ Main /Users/kcft114/.julia/packages/Flux/qXNjB/src/optimise/train.jl:105
```

```
loss(X, Y) = 0.62195015f0 (tracked)
loss(X, Y) = 0.63119596f0 (tracked)
loss(X, Y) = 0.64906013f0 (tracked)
loss(X, Y) = 0.64635926f0 (tracked)
loss(X, Y) = 0.6305257f0 (tracked)
loss(X, Y) = 0.6275637f0 (tracked)
loss(X, Y) = 0.60977983f0 (tracked)
loss(X, Y) = 0.6276244f0 (tracked)
loss(X, Y) = 0.6139429f0 (tracked)
loss(X, Y) = 0.59867793f0 (tracked)
loss(X, Y) = 0.6478198f0 (tracked)
loss(X, Y) = 0.63517636f0 (tracked)
loss(X, Y) = 0.6066221f0 (tracked)
loss(X, Y) = 0.60553783f0 (tracked)
loss(X, Y) = 0.6325984f0 (tracked)
loss(X, Y) = 0.6420202f0 (tracked)
loss(X, Y) = 0.6412153f0 (tracked)
loss(X, Y) = 0.586027f0 (tracked)
loss(X, Y) = 0.6028066f0 (tracked)
loss(X, Y) = 0.61498517f0 (tracked)
loss(X, Y) = 0.600845f0 (tracked)
loss(X, Y) = 0.62603f0 (tracked)
loss(X, Y) = 0.6328335f0 (tracked)
loss(X, Y) = 0.6130504f0 (tracked)
loss(X, Y) = 0.6001533f0 (tracked)
loss(X, Y) = 0.63757604f0 (tracked)
loss(X, Y) = 0.6595435f0 (tracked)
loss(X, Y) = 0.6324739f0 (tracked)
loss(X, Y) = 0.61698294f0 (tracked)
loss(X, Y) = 0.6188863f0 (tracked)
loss(X, Y) = 0.62701523f0 (tracked)
loss(X, Y) = 0.6315136f0 (tracked)
loss(X, Y) = 0.6146674f0 (tracked)
loss(X, Y) = 0.6181551f0 (tracked)
loss(X, Y) = 0.61919796f0 (tracked)
loss(X, Y) = 0.6369463f0 (tracked)
loss(X, Y) = 0.605776f0 (tracked)
loss(X, Y) = 0.58927953f0 (tracked)
loss(X, Y) = 0.6174287f0 (tracked)
loss(X, Y) = 0.62829643f0 (tracked)
loss(X, Y) = 0.62138534f0 (tracked)
loss(X, Y) = 0.6349811f0 (tracked)
loss(X, Y) = 0.62801486f0 (tracked)
loss(X, Y) = 0.5927878f0 (tracked)
loss(X, Y) = 0.61372197f0 (tracked)
loss(X, Y) = 0.6206623f0 (tracked)
loss(X, Y) = 0.599279f0 (tracked)
loss(X, Y) = 0.59939307f0 (tracked)
loss(X, Y) = 0.5984442f0 (tracked)
loss(X, Y) = 0.63456017f0 (tracked)
loss(X, Y) = 0.6219612f0 (tracked)
loss(X, Y) = 0.6196575f0 (tracked)
loss(X, Y) = 0.61088294f0 (tracked)
loss(X, Y) = 0.6233094f0 (tracked)
loss(X, Y) = 0.6221798f0 (tracked)
loss(X, Y) = 0.61485183f0 (tracked)
loss(X, Y) = 0.58110934f0 (tracked)
loss(X, Y) = 0.62799525f0 (tracked)
loss(X, Y) = 0.6242856f0 (tracked)
loss(X, Y) = 0.6194446f0 (tracked)
loss(X, Y) = 0.60191214f0 (tracked)
loss(X, Y) = 0.634122f0 (tracked)
loss(X, Y) = 0.6078242f0 (tracked)
loss(X, Y) = 0.60262316f0 (tracked)
loss(X, Y) = 0.6060326f0 (tracked)
loss(X, Y) = 0.5973078f0 (tracked)
loss(X, Y) = 0.61521417f0 (tracked)
loss(X, Y) = 0.61162376f0 (tracked)
loss(X, Y) = 0.6181073f0 (tracked)
loss(X, Y) = 0.59661543f0 (tracked)
loss(X, Y) = 0.61583704f0 (tracked)
loss(X, Y) = 0.6126923f0 (tracked)
loss(X, Y) = 0.6302519f0 (tracked)
loss(X, Y) = 0.6086996f0 (tracked)
loss(X, Y) = 0.6066852f0 (tracked)
loss(X, Y) = 0.6379923f0 (tracked)
loss(X, Y) = 0.57191175f0 (tracked)
loss(X, Y) = 0.5875999f0 (tracked)
loss(X, Y) = 0.60848707f0 (tracked)
loss(X, Y) = 0.58626366f0 (tracked)
loss(X, Y) = 0.58253723f0 (tracked)
loss(X, Y) = 0.6509119f0 (tracked)
loss(X, Y) = 0.60902935f0 (tracked)
loss(X, Y) = 0.62235075f0 (tracked)
loss(X, Y) = 0.62245834f0 (tracked)
loss(X, Y) = 0.6331079f0 (tracked)
loss(X, Y) = 0.60109156f0 (tracked)
loss(X, Y) = 0.6169783f0 (tracked)
loss(X, Y) = 0.60723084f0 (tracked)
loss(X, Y) = 0.6150008f0 (tracked)
loss(X, Y) = 0.62062556f0 (tracked)
loss(X, Y) = 0.607059f0 (tracked)
loss(X, Y) = 0.6135237f0 (tracked)
loss(X, Y) = 0.59212047f0 (tracked)
loss(X, Y) = 0.60399646f0 (tracked)
loss(X, Y) = 0.6184469f0 (tracked)
loss(X, Y) = 0.60483986f0 (tracked)
loss(X, Y) = 0.61742246f0 (tracked)
loss(X, Y) = 0.6324121f0 (tracked)
loss(X, Y) = 0.60078686f0 (tracked)
loss(X, Y) = 0.6007616f0 (tracked)
loss(X, Y) = 0.6110016f0 (tracked)
loss(X, Y) = 0.6196802f0 (tracked)
loss(X, Y) = 0.62830067f0 (tracked)
loss(X, Y) = 0.601962f0 (tracked)
loss(X, Y) = 0.6150685f0 (tracked)
loss(X, Y) = 0.5754709f0 (tracked)
loss(X, Y) = 0.62214357f0 (tracked)
loss(X, Y) = 0.62126654f0 (tracked)
loss(X, Y) = 0.6015092f0 (tracked)
loss(X, Y) = 0.60983926f0 (tracked)
loss(X, Y) = 0.6219482f0 (tracked)
loss(X, Y) = 0.60158694f0 (tracked)
loss(X, Y) = 0.60257596f0 (tracked)
loss(X, Y) = 0.612937f0 (tracked)
loss(X, Y) = 0.6158083f0 (tracked)
loss(X, Y) = 0.6122574f0 (tracked)
loss(X, Y) = 0.64344776f0 (tracked)
```

```
┌ Info: Epoch 796
└ @ Main /Users/kcft114/.julia/packages/Flux/qXNjB/src/optimise/train.jl:105
┌ Info: Epoch 797
└ @ Main /Users/kcft114/.julia/packages/Flux/qXNjB/src/optimise/train.jl:105
┌ Info: Epoch 798
└ @ Main /Users/kcft114/.julia/packages/Flux/qXNjB/src/optimise/train.jl:105
┌ Info: Epoch 799
└ @ Main /Users/kcft114/.julia/packages/Flux/qXNjB/src/optimise/train.jl:105
┌ Info: Epoch 800
└ @ Main /Users/kcft114/.julia/packages/Flux/qXNjB/src/optimise/train.jl:105
┌ Info: Epoch 801
└ @ Main /Users/kcft114/.julia/packages/Flux/qXNjB/src/optimise/train.jl:105
┌ Info: Epoch 802
└ @ Main /Users/kcft114/.julia/packages/Flux/qXNjB/src/optimise/train.jl:105
┌ Info: Epoch 803
└ @ Main /Users/kcft114/.julia/packages/Flux/qXNjB/src/optimise/train.jl:105
┌ Info: Epoch 804
└ @ Main /Users/kcft114/.julia/packages/Flux/qXNjB/src/optimise/train.jl:105
┌ Info: Epoch 805
└ @ Main /Users/kcft114/.julia/packages/Flux/qXNjB/src/optimise/train.jl:105
┌ Info: Epoch 806
└ @ Main /Users/kcft114/.julia/packages/Flux/qXNjB/src/optimise/train.jl:105
┌ Info: Epoch 807
└ @ Main /Users/kcft114/.julia/packages/Flux/qXNjB/src/optimise/train.jl:105
┌ Info: Epoch 808
└ @ Main /Users/kcft114/.julia/packages/Flux/qXNjB/src/optimise/train.jl:105
┌ Info: Epoch 809
└ @ Main /Users/kcft114/.julia/packages/Flux/qXNjB/src/optimise/train.jl:105
┌ Info: Epoch 810
└ @ Main /Users/kcft114/.julia/packages/Flux/qXNjB/src/optimise/train.jl:105
┌ Info: Epoch 811
└ @ Main /Users/kcft114/.julia/packages/Flux/qXNjB/src/optimise/train.jl:105
┌ Info: Epoch 812
└ @ Main /Users/kcft114/.julia/packages/Flux/qXNjB/src/optimise/train.jl:105
┌ Info: Epoch 813
└ @ Main /Users/kcft114/.julia/packages/Flux/qXNjB/src/optimise/train.jl:105
┌ Info: Epoch 814
└ @ Main /Users/kcft114/.julia/packages/Flux/qXNjB/src/optimise/train.jl:105
┌ Info: Epoch 815
└ @ Main /Users/kcft114/.julia/packages/Flux/qXNjB/src/optimise/train.jl:105
┌ Info: Epoch 816
└ @ Main /Users/kcft114/.julia/packages/Flux/qXNjB/src/optimise/train.jl:105
┌ Info: Epoch 817
└ @ Main /Users/kcft114/.julia/packages/Flux/qXNjB/src/optimise/train.jl:105
┌ Info: Epoch 818
└ @ Main /Users/kcft114/.julia/packages/Flux/qXNjB/src/optimise/train.jl:105
┌ Info: Epoch 819
└ @ Main /Users/kcft114/.julia/packages/Flux/qXNjB/src/optimise/train.jl:105
┌ Info: Epoch 820
└ @ Main /Users/kcft114/.julia/packages/Flux/qXNjB/src/optimise/train.jl:105
┌ Info: Epoch 821
└ @ Main /Users/kcft114/.julia/packages/Flux/qXNjB/src/optimise/train.jl:105
┌ Info: Epoch 822
└ @ Main /Users/kcft114/.julia/packages/Flux/qXNjB/src/optimise/train.jl:105
┌ Info: Epoch 823
└ @ Main /Users/kcft114/.julia/packages/Flux/qXNjB/src/optimise/train.jl:105
┌ Info: Epoch 824
└ @ Main /Users/kcft114/.julia/packages/Flux/qXNjB/src/optimise/train.jl:105
┌ Info: Epoch 825
└ @ Main /Users/kcft114/.julia/packages/Flux/qXNjB/src/optimise/train.jl:105
┌ Info: Epoch 826
└ @ Main /Users/kcft114/.julia/packages/Flux/qXNjB/src/optimise/train.jl:105
┌ Info: Epoch 827
└ @ Main /Users/kcft114/.julia/packages/Flux/qXNjB/src/optimise/train.jl:105
┌ Info: Epoch 828
└ @ Main /Users/kcft114/.julia/packages/Flux/qXNjB/src/optimise/train.jl:105
┌ Info: Epoch 829
└ @ Main /Users/kcft114/.julia/packages/Flux/qXNjB/src/optimise/train.jl:105
┌ Info: Epoch 830
└ @ Main /Users/kcft114/.julia/packages/Flux/qXNjB/src/optimise/train.jl:105
┌ Info: Epoch 831
└ @ Main /Users/kcft114/.julia/packages/Flux/qXNjB/src/optimise/train.jl:105
┌ Info: Epoch 832
└ @ Main /Users/kcft114/.julia/packages/Flux/qXNjB/src/optimise/train.jl:105
┌ Info: Epoch 833
└ @ Main /Users/kcft114/.julia/packages/Flux/qXNjB/src/optimise/train.jl:105
┌ Info: Epoch 834
└ @ Main /Users/kcft114/.julia/packages/Flux/qXNjB/src/optimise/train.jl:105
┌ Info: Epoch 835
└ @ Main /Users/kcft114/.julia/packages/Flux/qXNjB/src/optimise/train.jl:105
┌ Info: Epoch 836
└ @ Main /Users/kcft114/.julia/packages/Flux/qXNjB/src/optimise/train.jl:105
┌ Info: Epoch 837
└ @ Main /Users/kcft114/.julia/packages/Flux/qXNjB/src/optimise/train.jl:105
┌ Info: Epoch 838
└ @ Main /Users/kcft114/.julia/packages/Flux/qXNjB/src/optimise/train.jl:105
┌ Info: Epoch 839
└ @ Main /Users/kcft114/.julia/packages/Flux/qXNjB/src/optimise/train.jl:105
┌ Info: Epoch 840
└ @ Main /Users/kcft114/.julia/packages/Flux/qXNjB/src/optimise/train.jl:105
┌ Info: Epoch 841
└ @ Main /Users/kcft114/.julia/packages/Flux/qXNjB/src/optimise/train.jl:105
┌ Info: Epoch 842
└ @ Main /Users/kcft114/.julia/packages/Flux/qXNjB/src/optimise/train.jl:105
┌ Info: Epoch 843
└ @ Main /Users/kcft114/.julia/packages/Flux/qXNjB/src/optimise/train.jl:105
┌ Info: Epoch 844
└ @ Main /Users/kcft114/.julia/packages/Flux/qXNjB/src/optimise/train.jl:105
┌ Info: Epoch 845
└ @ Main /Users/kcft114/.julia/packages/Flux/qXNjB/src/optimise/train.jl:105
┌ Info: Epoch 846
└ @ Main /Users/kcft114/.julia/packages/Flux/qXNjB/src/optimise/train.jl:105
┌ Info: Epoch 847
└ @ Main /Users/kcft114/.julia/packages/Flux/qXNjB/src/optimise/train.jl:105
┌ Info: Epoch 848
└ @ Main /Users/kcft114/.julia/packages/Flux/qXNjB/src/optimise/train.jl:105
┌ Info: Epoch 849
└ @ Main /Users/kcft114/.julia/packages/Flux/qXNjB/src/optimise/train.jl:105
┌ Info: Epoch 850
└ @ Main /Users/kcft114/.julia/packages/Flux/qXNjB/src/optimise/train.jl:105
┌ Info: Epoch 851
└ @ Main /Users/kcft114/.julia/packages/Flux/qXNjB/src/optimise/train.jl:105
┌ Info: Epoch 852
└ @ Main /Users/kcft114/.julia/packages/Flux/qXNjB/src/optimise/train.jl:105
┌ Info: Epoch 853
└ @ Main /Users/kcft114/.julia/packages/Flux/qXNjB/src/optimise/train.jl:105
┌ Info: Epoch 854
└ @ Main /Users/kcft114/.julia/packages/Flux/qXNjB/src/optimise/train.jl:105
┌ Info: Epoch 855
└ @ Main /Users/kcft114/.julia/packages/Flux/qXNjB/src/optimise/train.jl:105
┌ Info: Epoch 856
└ @ Main /Users/kcft114/.julia/packages/Flux/qXNjB/src/optimise/train.jl:105
┌ Info: Epoch 857
└ @ Main /Users/kcft114/.julia/packages/Flux/qXNjB/src/optimise/train.jl:105
┌ Info: Epoch 858
└ @ Main /Users/kcft114/.julia/packages/Flux/qXNjB/src/optimise/train.jl:105
┌ Info: Epoch 859
└ @ Main /Users/kcft114/.julia/packages/Flux/qXNjB/src/optimise/train.jl:105
┌ Info: Epoch 860
└ @ Main /Users/kcft114/.julia/packages/Flux/qXNjB/src/optimise/train.jl:105
┌ Info: Epoch 861
└ @ Main /Users/kcft114/.julia/packages/Flux/qXNjB/src/optimise/train.jl:105
┌ Info: Epoch 862
└ @ Main /Users/kcft114/.julia/packages/Flux/qXNjB/src/optimise/train.jl:105
┌ Info: Epoch 863
└ @ Main /Users/kcft114/.julia/packages/Flux/qXNjB/src/optimise/train.jl:105
┌ Info: Epoch 864
└ @ Main /Users/kcft114/.julia/packages/Flux/qXNjB/src/optimise/train.jl:105
┌ Info: Epoch 865
└ @ Main /Users/kcft114/.julia/packages/Flux/qXNjB/src/optimise/train.jl:105
┌ Info: Epoch 866
└ @ Main /Users/kcft114/.julia/packages/Flux/qXNjB/src/optimise/train.jl:105
┌ Info: Epoch 867
└ @ Main /Users/kcft114/.julia/packages/Flux/qXNjB/src/optimise/train.jl:105
┌ Info: Epoch 868
└ @ Main /Users/kcft114/.julia/packages/Flux/qXNjB/src/optimise/train.jl:105
┌ Info: Epoch 869
└ @ Main /Users/kcft114/.julia/packages/Flux/qXNjB/src/optimise/train.jl:105
┌ Info: Epoch 870
└ @ Main /Users/kcft114/.julia/packages/Flux/qXNjB/src/optimise/train.jl:105
┌ Info: Epoch 871
└ @ Main /Users/kcft114/.julia/packages/Flux/qXNjB/src/optimise/train.jl:105
┌ Info: Epoch 872
└ @ Main /Users/kcft114/.julia/packages/Flux/qXNjB/src/optimise/train.jl:105
┌ Info: Epoch 873
└ @ Main /Users/kcft114/.julia/packages/Flux/qXNjB/src/optimise/train.jl:105
┌ Info: Epoch 874
└ @ Main /Users/kcft114/.julia/packages/Flux/qXNjB/src/optimise/train.jl:105
┌ Info: Epoch 875
└ @ Main /Users/kcft114/.julia/packages/Flux/qXNjB/src/optimise/train.jl:105
┌ Info: Epoch 876
└ @ Main /Users/kcft114/.julia/packages/Flux/qXNjB/src/optimise/train.jl:105
┌ Info: Epoch 877
└ @ Main /Users/kcft114/.julia/packages/Flux/qXNjB/src/optimise/train.jl:105
┌ Info: Epoch 878
└ @ Main /Users/kcft114/.julia/packages/Flux/qXNjB/src/optimise/train.jl:105
┌ Info: Epoch 879
└ @ Main /Users/kcft114/.julia/packages/Flux/qXNjB/src/optimise/train.jl:105
┌ Info: Epoch 880
└ @ Main /Users/kcft114/.julia/packages/Flux/qXNjB/src/optimise/train.jl:105
┌ Info: Epoch 881
└ @ Main /Users/kcft114/.julia/packages/Flux/qXNjB/src/optimise/train.jl:105
┌ Info: Epoch 882
└ @ Main /Users/kcft114/.julia/packages/Flux/qXNjB/src/optimise/train.jl:105
┌ Info: Epoch 883
└ @ Main /Users/kcft114/.julia/packages/Flux/qXNjB/src/optimise/train.jl:105
┌ Info: Epoch 884
└ @ Main /Users/kcft114/.julia/packages/Flux/qXNjB/src/optimise/train.jl:105
┌ Info: Epoch 885
└ @ Main /Users/kcft114/.julia/packages/Flux/qXNjB/src/optimise/train.jl:105
┌ Info: Epoch 886
└ @ Main /Users/kcft114/.julia/packages/Flux/qXNjB/src/optimise/train.jl:105
┌ Info: Epoch 887
└ @ Main /Users/kcft114/.julia/packages/Flux/qXNjB/src/optimise/train.jl:105
┌ Info: Epoch 888
└ @ Main /Users/kcft114/.julia/packages/Flux/qXNjB/src/optimise/train.jl:105
┌ Info: Epoch 889
└ @ Main /Users/kcft114/.julia/packages/Flux/qXNjB/src/optimise/train.jl:105
┌ Info: Epoch 890
└ @ Main /Users/kcft114/.julia/packages/Flux/qXNjB/src/optimise/train.jl:105
┌ Info: Epoch 891
└ @ Main /Users/kcft114/.julia/packages/Flux/qXNjB/src/optimise/train.jl:105
┌ Info: Epoch 892
└ @ Main /Users/kcft114/.julia/packages/Flux/qXNjB/src/optimise/train.jl:105
┌ Info: Epoch 893
└ @ Main /Users/kcft114/.julia/packages/Flux/qXNjB/src/optimise/train.jl:105
┌ Info: Epoch 894
└ @ Main /Users/kcft114/.julia/packages/Flux/qXNjB/src/optimise/train.jl:105
┌ Info: Epoch 895
└ @ Main /Users/kcft114/.julia/packages/Flux/qXNjB/src/optimise/train.jl:105
┌ Info: Epoch 896
└ @ Main /Users/kcft114/.julia/packages/Flux/qXNjB/src/optimise/train.jl:105
┌ Info: Epoch 897
└ @ Main /Users/kcft114/.julia/packages/Flux/qXNjB/src/optimise/train.jl:105
┌ Info: Epoch 898
└ @ Main /Users/kcft114/.julia/packages/Flux/qXNjB/src/optimise/train.jl:105
┌ Info: Epoch 899
└ @ Main /Users/kcft114/.julia/packages/Flux/qXNjB/src/optimise/train.jl:105
┌ Info: Epoch 900
└ @ Main /Users/kcft114/.julia/packages/Flux/qXNjB/src/optimise/train.jl:105
┌ Info: Epoch 901
└ @ Main /Users/kcft114/.julia/packages/Flux/qXNjB/src/optimise/train.jl:105
┌ Info: Epoch 902
└ @ Main /Users/kcft114/.julia/packages/Flux/qXNjB/src/optimise/train.jl:105
┌ Info: Epoch 903
└ @ Main /Users/kcft114/.julia/packages/Flux/qXNjB/src/optimise/train.jl:105
┌ Info: Epoch 904
└ @ Main /Users/kcft114/.julia/packages/Flux/qXNjB/src/optimise/train.jl:105
┌ Info: Epoch 905
└ @ Main /Users/kcft114/.julia/packages/Flux/qXNjB/src/optimise/train.jl:105
┌ Info: Epoch 906
└ @ Main /Users/kcft114/.julia/packages/Flux/qXNjB/src/optimise/train.jl:105
┌ Info: Epoch 907
└ @ Main /Users/kcft114/.julia/packages/Flux/qXNjB/src/optimise/train.jl:105
┌ Info: Epoch 908
└ @ Main /Users/kcft114/.julia/packages/Flux/qXNjB/src/optimise/train.jl:105
┌ Info: Epoch 909
└ @ Main /Users/kcft114/.julia/packages/Flux/qXNjB/src/optimise/train.jl:105
┌ Info: Epoch 910
└ @ Main /Users/kcft114/.julia/packages/Flux/qXNjB/src/optimise/train.jl:105
┌ Info: Epoch 911
└ @ Main /Users/kcft114/.julia/packages/Flux/qXNjB/src/optimise/train.jl:105
┌ Info: Epoch 912
└ @ Main /Users/kcft114/.julia/packages/Flux/qXNjB/src/optimise/train.jl:105
┌ Info: Epoch 913
└ @ Main /Users/kcft114/.julia/packages/Flux/qXNjB/src/optimise/train.jl:105
```

```
loss(X, Y) = 0.59431356f0 (tracked)
loss(X, Y) = 0.6024759f0 (tracked)
loss(X, Y) = 0.6014276f0 (tracked)
loss(X, Y) = 0.6194006f0 (tracked)
loss(X, Y) = 0.5989219f0 (tracked)
loss(X, Y) = 0.61610216f0 (tracked)
loss(X, Y) = 0.6012945f0 (tracked)
loss(X, Y) = 0.5877109f0 (tracked)
loss(X, Y) = 0.60402596f0 (tracked)
loss(X, Y) = 0.5877318f0 (tracked)
loss(X, Y) = 0.62656355f0 (tracked)
loss(X, Y) = 0.62031025f0 (tracked)
loss(X, Y) = 0.6036674f0 (tracked)
loss(X, Y) = 0.6331421f0 (tracked)
loss(X, Y) = 0.5949579f0 (tracked)
loss(X, Y) = 0.6058395f0 (tracked)
loss(X, Y) = 0.63211936f0 (tracked)
loss(X, Y) = 0.5769044f0 (tracked)
loss(X, Y) = 0.58159214f0 (tracked)
loss(X, Y) = 0.61277413f0 (tracked)
loss(X, Y) = 0.5968998f0 (tracked)
loss(X, Y) = 0.5943237f0 (tracked)
loss(X, Y) = 0.6406587f0 (tracked)
loss(X, Y) = 0.6025805f0 (tracked)
loss(X, Y) = 0.6462329f0 (tracked)
loss(X, Y) = 0.6075124f0 (tracked)
loss(X, Y) = 0.5899669f0 (tracked)
loss(X, Y) = 0.61466736f0 (tracked)
loss(X, Y) = 0.60084516f0 (tracked)
loss(X, Y) = 0.5966145f0 (tracked)
loss(X, Y) = 0.59919345f0 (tracked)
loss(X, Y) = 0.5875514f0 (tracked)
loss(X, Y) = 0.57794255f0 (tracked)
loss(X, Y) = 0.5870667f0 (tracked)
loss(X, Y) = 0.5632932f0 (tracked)
loss(X, Y) = 0.58291787f0 (tracked)
loss(X, Y) = 0.6119592f0 (tracked)
loss(X, Y) = 0.62077427f0 (tracked)
loss(X, Y) = 0.60733247f0 (tracked)
loss(X, Y) = 0.5885618f0 (tracked)
loss(X, Y) = 0.60027426f0 (tracked)
loss(X, Y) = 0.63959795f0 (tracked)
loss(X, Y) = 0.6170187f0 (tracked)
loss(X, Y) = 0.6332229f0 (tracked)
loss(X, Y) = 0.5924904f0 (tracked)
loss(X, Y) = 0.6380379f0 (tracked)
loss(X, Y) = 0.5744294f0 (tracked)
loss(X, Y) = 0.58582354f0 (tracked)
loss(X, Y) = 0.6255565f0 (tracked)
loss(X, Y) = 0.57476944f0 (tracked)
loss(X, Y) = 0.5877393f0 (tracked)
loss(X, Y) = 0.59874564f0 (tracked)
loss(X, Y) = 0.5898856f0 (tracked)
loss(X, Y) = 0.5672936f0 (tracked)
loss(X, Y) = 0.62455237f0 (tracked)
loss(X, Y) = 0.5592279f0 (tracked)
loss(X, Y) = 0.60324115f0 (tracked)
loss(X, Y) = 0.57302505f0 (tracked)
loss(X, Y) = 0.5937042f0 (tracked)
loss(X, Y) = 0.59288806f0 (tracked)
loss(X, Y) = 0.6179993f0 (tracked)
loss(X, Y) = 0.5942689f0 (tracked)
loss(X, Y) = 0.60090435f0 (tracked)
loss(X, Y) = 0.61772066f0 (tracked)
loss(X, Y) = 0.61054146f0 (tracked)
loss(X, Y) = 0.63296086f0 (tracked)
loss(X, Y) = 0.59968084f0 (tracked)
loss(X, Y) = 0.57369936f0 (tracked)
loss(X, Y) = 0.59959877f0 (tracked)
loss(X, Y) = 0.5737447f0 (tracked)
loss(X, Y) = 0.5557761f0 (tracked)
loss(X, Y) = 0.61551f0 (tracked)
loss(X, Y) = 0.5849341f0 (tracked)
loss(X, Y) = 0.63049704f0 (tracked)
loss(X, Y) = 0.60474443f0 (tracked)
loss(X, Y) = 0.60254884f0 (tracked)
loss(X, Y) = 0.55759835f0 (tracked)
loss(X, Y) = 0.58716017f0 (tracked)
loss(X, Y) = 0.6134113f0 (tracked)
loss(X, Y) = 0.60354203f0 (tracked)
loss(X, Y) = 0.57857925f0 (tracked)
loss(X, Y) = 0.58837944f0 (tracked)
loss(X, Y) = 0.59960455f0 (tracked)
loss(X, Y) = 0.5869877f0 (tracked)
loss(X, Y) = 0.57089806f0 (tracked)
loss(X, Y) = 0.60035586f0 (tracked)
loss(X, Y) = 0.5941015f0 (tracked)
loss(X, Y) = 0.5947211f0 (tracked)
loss(X, Y) = 0.5778298f0 (tracked)
loss(X, Y) = 0.60751295f0 (tracked)
loss(X, Y) = 0.57773924f0 (tracked)
loss(X, Y) = 0.5798222f0 (tracked)
loss(X, Y) = 0.605981f0 (tracked)
loss(X, Y) = 0.59089375f0 (tracked)
loss(X, Y) = 0.54771644f0 (tracked)
loss(X, Y) = 0.63679814f0 (tracked)
loss(X, Y) = 0.5863811f0 (tracked)
loss(X, Y) = 0.59014356f0 (tracked)
loss(X, Y) = 0.6042798f0 (tracked)
loss(X, Y) = 0.59876704f0 (tracked)
loss(X, Y) = 0.5775376f0 (tracked)
loss(X, Y) = 0.5720533f0 (tracked)
loss(X, Y) = 0.58937734f0 (tracked)
loss(X, Y) = 0.5880883f0 (tracked)
loss(X, Y) = 0.574393f0 (tracked)
loss(X, Y) = 0.5834947f0 (tracked)
loss(X, Y) = 0.6281153f0 (tracked)
loss(X, Y) = 0.6182996f0 (tracked)
loss(X, Y) = 0.5795159f0 (tracked)
```

```
┌ Info: Epoch 914
└ @ Main /Users/kcft114/.julia/packages/Flux/qXNjB/src/optimise/train.jl:105
┌ Info: Epoch 915
└ @ Main /Users/kcft114/.julia/packages/Flux/qXNjB/src/optimise/train.jl:105
┌ Info: Epoch 916
└ @ Main /Users/kcft114/.julia/packages/Flux/qXNjB/src/optimise/train.jl:105
┌ Info: Epoch 917
└ @ Main /Users/kcft114/.julia/packages/Flux/qXNjB/src/optimise/train.jl:105
┌ Info: Epoch 918
└ @ Main /Users/kcft114/.julia/packages/Flux/qXNjB/src/optimise/train.jl:105
```

```
loss(X, Y) = 0.60278815f0 (tracked)
loss(X, Y) = 0.56703603f0 (tracked)
loss(X, Y) = 0.58243674f0 (tracked)
loss(X, Y) = 0.60624546f0 (tracked)
loss(X, Y) = 0.56559455f0 (tracked)
loss(X, Y) = 0.6112969f0 (tracked)
loss(X, Y) = 0.61470187f0 (tracked)
loss(X, Y) = 0.5818308f0 (tracked)
loss(X, Y) = 0.5634671f0 (tracked)
loss(X, Y) = 0.5919681f0 (tracked)
loss(X, Y) = 0.5879863f0 (tracked)
loss(X, Y) = 0.5921957f0 (tracked)
loss(X, Y) = 0.60297877f0 (tracked)
loss(X, Y) = 0.61267465f0 (tracked)
loss(X, Y) = 0.62169474f0 (tracked)
loss(X, Y) = 0.5619561f0 (tracked)
loss(X, Y) = 0.5978977f0 (tracked)
loss(X, Y) = 0.5842493f0 (tracked)
loss(X, Y) = 0.5725514f0 (tracked)
loss(X, Y) = 0.59287333f0 (tracked)
loss(X, Y) = 0.61329967f0 (tracked)
loss(X, Y) = 0.5873978f0 (tracked)
loss(X, Y) = 0.58950114f0 (tracked)
loss(X, Y) = 0.58275884f0 (tracked)
loss(X, Y) = 0.54673487f0 (tracked)
loss(X, Y) = 0.5948154f0 (tracked)
loss(X, Y) = 0.58603144f0 (tracked)
loss(X, Y) = 0.59687424f0 (tracked)
loss(X, Y) = 0.5680833f0 (tracked)
loss(X, Y) = 0.5817865f0 (tracked)
loss(X, Y) = 0.5822812f0 (tracked)
loss(X, Y) = 0.6087545f0 (tracked)
loss(X, Y) = 0.58497036f0 (tracked)
loss(X, Y) = 0.59104687f0 (tracked)
loss(X, Y) = 0.583749f0 (tracked)
loss(X, Y) = 0.5644699f0 (tracked)
loss(X, Y) = 0.5748415f0 (tracked)
loss(X, Y) = 0.6389044f0 (tracked)
loss(X, Y) = 0.5426653f0 (tracked)
loss(X, Y) = 0.53890294f0 (tracked)
loss(X, Y) = 0.613075f0 (tracked)
loss(X, Y) = 0.56824136f0 (tracked)
loss(X, Y) = 0.6229677f0 (tracked)
loss(X, Y) = 0.6502636f0 (tracked)
loss(X, Y) = 0.61916107f0 (tracked)
loss(X, Y) = 0.5724212f0 (tracked)
loss(X, Y) = 0.59602386f0 (tracked)
loss(X, Y) = 0.56946456f0 (tracked)
loss(X, Y) = 0.57252616f0 (tracked)
loss(X, Y) = 0.6026097f0 (tracked)
loss(X, Y) = 0.6041095f0 (tracked)
loss(X, Y) = 0.5403869f0 (tracked)
loss(X, Y) = 0.58777535f0 (tracked)
loss(X, Y) = 0.57354623f0 (tracked)
loss(X, Y) = 0.56439465f0 (tracked)
loss(X, Y) = 0.57034326f0 (tracked)
loss(X, Y) = 0.6010414f0 (tracked)
loss(X, Y) = 0.593985f0 (tracked)
loss(X, Y) = 0.6155936f0 (tracked)
loss(X, Y) = 0.59742826f0 (tracked)
loss(X, Y) = 0.5666267f0 (tracked)
loss(X, Y) = 0.55406487f0 (tracked)
loss(X, Y) = 0.6549463f0 (tracked)
loss(X, Y) = 0.5844956f0 (tracked)
loss(X, Y) = 0.56577337f0 (tracked)
loss(X, Y) = 0.5701935f0 (tracked)
loss(X, Y) = 0.5711354f0 (tracked)
loss(X, Y) = 0.561622f0 (tracked)
loss(X, Y) = 0.5691259f0 (tracked)
loss(X, Y) = 0.58696383f0 (tracked)
loss(X, Y) = 0.5711695f0 (tracked)
loss(X, Y) = 0.5836207f0 (tracked)
loss(X, Y) = 0.582188f0 (tracked)
```

```
┌ Info: Epoch 919
└ @ Main /Users/kcft114/.julia/packages/Flux/qXNjB/src/optimise/train.jl:105
┌ Info: Epoch 920
└ @ Main /Users/kcft114/.julia/packages/Flux/qXNjB/src/optimise/train.jl:105
┌ Info: Epoch 921
└ @ Main /Users/kcft114/.julia/packages/Flux/qXNjB/src/optimise/train.jl:105
┌ Info: Epoch 922
└ @ Main /Users/kcft114/.julia/packages/Flux/qXNjB/src/optimise/train.jl:105
┌ Info: Epoch 923
└ @ Main /Users/kcft114/.julia/packages/Flux/qXNjB/src/optimise/train.jl:105
┌ Info: Epoch 924
└ @ Main /Users/kcft114/.julia/packages/Flux/qXNjB/src/optimise/train.jl:105
┌ Info: Epoch 925
└ @ Main /Users/kcft114/.julia/packages/Flux/qXNjB/src/optimise/train.jl:105
┌ Info: Epoch 926
└ @ Main /Users/kcft114/.julia/packages/Flux/qXNjB/src/optimise/train.jl:105
┌ Info: Epoch 927
└ @ Main /Users/kcft114/.julia/packages/Flux/qXNjB/src/optimise/train.jl:105
┌ Info: Epoch 928
└ @ Main /Users/kcft114/.julia/packages/Flux/qXNjB/src/optimise/train.jl:105
┌ Info: Epoch 929
└ @ Main /Users/kcft114/.julia/packages/Flux/qXNjB/src/optimise/train.jl:105
┌ Info: Epoch 930
└ @ Main /Users/kcft114/.julia/packages/Flux/qXNjB/src/optimise/train.jl:105
┌ Info: Epoch 931
└ @ Main /Users/kcft114/.julia/packages/Flux/qXNjB/src/optimise/train.jl:105
┌ Info: Epoch 932
└ @ Main /Users/kcft114/.julia/packages/Flux/qXNjB/src/optimise/train.jl:105
┌ Info: Epoch 933
└ @ Main /Users/kcft114/.julia/packages/Flux/qXNjB/src/optimise/train.jl:105
┌ Info: Epoch 934
└ @ Main /Users/kcft114/.julia/packages/Flux/qXNjB/src/optimise/train.jl:105
┌ Info: Epoch 935
└ @ Main /Users/kcft114/.julia/packages/Flux/qXNjB/src/optimise/train.jl:105
┌ Info: Epoch 936
└ @ Main /Users/kcft114/.julia/packages/Flux/qXNjB/src/optimise/train.jl:105
┌ Info: Epoch 937
└ @ Main /Users/kcft114/.julia/packages/Flux/qXNjB/src/optimise/train.jl:105
┌ Info: Epoch 938
└ @ Main /Users/kcft114/.julia/packages/Flux/qXNjB/src/optimise/train.jl:105
┌ Info: Epoch 939
└ @ Main /Users/kcft114/.julia/packages/Flux/qXNjB/src/optimise/train.jl:105
┌ Info: Epoch 940
└ @ Main /Users/kcft114/.julia/packages/Flux/qXNjB/src/optimise/train.jl:105
┌ Info: Epoch 941
└ @ Main /Users/kcft114/.julia/packages/Flux/qXNjB/src/optimise/train.jl:105
┌ Info: Epoch 942
└ @ Main /Users/kcft114/.julia/packages/Flux/qXNjB/src/optimise/train.jl:105
┌ Info: Epoch 943
└ @ Main /Users/kcft114/.julia/packages/Flux/qXNjB/src/optimise/train.jl:105
┌ Info: Epoch 944
└ @ Main /Users/kcft114/.julia/packages/Flux/qXNjB/src/optimise/train.jl:105
┌ Info: Epoch 945
└ @ Main /Users/kcft114/.julia/packages/Flux/qXNjB/src/optimise/train.jl:105
┌ Info: Epoch 946
└ @ Main /Users/kcft114/.julia/packages/Flux/qXNjB/src/optimise/train.jl:105
┌ Info: Epoch 947
└ @ Main /Users/kcft114/.julia/packages/Flux/qXNjB/src/optimise/train.jl:105
┌ Info: Epoch 948
└ @ Main /Users/kcft114/.julia/packages/Flux/qXNjB/src/optimise/train.jl:105
┌ Info: Epoch 949
└ @ Main /Users/kcft114/.julia/packages/Flux/qXNjB/src/optimise/train.jl:105
┌ Info: Epoch 950
└ @ Main /Users/kcft114/.julia/packages/Flux/qXNjB/src/optimise/train.jl:105
┌ Info: Epoch 951
└ @ Main /Users/kcft114/.julia/packages/Flux/qXNjB/src/optimise/train.jl:105
┌ Info: Epoch 952
└ @ Main /Users/kcft114/.julia/packages/Flux/qXNjB/src/optimise/train.jl:105
┌ Info: Epoch 953
└ @ Main /Users/kcft114/.julia/packages/Flux/qXNjB/src/optimise/train.jl:105
┌ Info: Epoch 954
└ @ Main /Users/kcft114/.julia/packages/Flux/qXNjB/src/optimise/train.jl:105
┌ Info: Epoch 955
└ @ Main /Users/kcft114/.julia/packages/Flux/qXNjB/src/optimise/train.jl:105
┌ Info: Epoch 956
└ @ Main /Users/kcft114/.julia/packages/Flux/qXNjB/src/optimise/train.jl:105
┌ Info: Epoch 957
└ @ Main /Users/kcft114/.julia/packages/Flux/qXNjB/src/optimise/train.jl:105
┌ Info: Epoch 958
└ @ Main /Users/kcft114/.julia/packages/Flux/qXNjB/src/optimise/train.jl:105
┌ Info: Epoch 959
└ @ Main /Users/kcft114/.julia/packages/Flux/qXNjB/src/optimise/train.jl:105
┌ Info: Epoch 960
└ @ Main /Users/kcft114/.julia/packages/Flux/qXNjB/src/optimise/train.jl:105
┌ Info: Epoch 961
└ @ Main /Users/kcft114/.julia/packages/Flux/qXNjB/src/optimise/train.jl:105
┌ Info: Epoch 962
└ @ Main /Users/kcft114/.julia/packages/Flux/qXNjB/src/optimise/train.jl:105
┌ Info: Epoch 963
└ @ Main /Users/kcft114/.julia/packages/Flux/qXNjB/src/optimise/train.jl:105
┌ Info: Epoch 964
└ @ Main /Users/kcft114/.julia/packages/Flux/qXNjB/src/optimise/train.jl:105
┌ Info: Epoch 965
└ @ Main /Users/kcft114/.julia/packages/Flux/qXNjB/src/optimise/train.jl:105
┌ Info: Epoch 966
└ @ Main /Users/kcft114/.julia/packages/Flux/qXNjB/src/optimise/train.jl:105
┌ Info: Epoch 967
└ @ Main /Users/kcft114/.julia/packages/Flux/qXNjB/src/optimise/train.jl:105
┌ Info: Epoch 968
└ @ Main /Users/kcft114/.julia/packages/Flux/qXNjB/src/optimise/train.jl:105
┌ Info: Epoch 969
└ @ Main /Users/kcft114/.julia/packages/Flux/qXNjB/src/optimise/train.jl:105
┌ Info: Epoch 970
└ @ Main /Users/kcft114/.julia/packages/Flux/qXNjB/src/optimise/train.jl:105
┌ Info: Epoch 971
└ @ Main /Users/kcft114/.julia/packages/Flux/qXNjB/src/optimise/train.jl:105
┌ Info: Epoch 972
└ @ Main /Users/kcft114/.julia/packages/Flux/qXNjB/src/optimise/train.jl:105
┌ Info: Epoch 973
└ @ Main /Users/kcft114/.julia/packages/Flux/qXNjB/src/optimise/train.jl:105
┌ Info: Epoch 974
└ @ Main /Users/kcft114/.julia/packages/Flux/qXNjB/src/optimise/train.jl:105
┌ Info: Epoch 975
└ @ Main /Users/kcft114/.julia/packages/Flux/qXNjB/src/optimise/train.jl:105
┌ Info: Epoch 976
└ @ Main /Users/kcft114/.julia/packages/Flux/qXNjB/src/optimise/train.jl:105
┌ Info: Epoch 977
└ @ Main /Users/kcft114/.julia/packages/Flux/qXNjB/src/optimise/train.jl:105
┌ Info: Epoch 978
└ @ Main /Users/kcft114/.julia/packages/Flux/qXNjB/src/optimise/train.jl:105
┌ Info: Epoch 979
└ @ Main /Users/kcft114/.julia/packages/Flux/qXNjB/src/optimise/train.jl:105
┌ Info: Epoch 980
└ @ Main /Users/kcft114/.julia/packages/Flux/qXNjB/src/optimise/train.jl:105
┌ Info: Epoch 981
└ @ Main /Users/kcft114/.julia/packages/Flux/qXNjB/src/optimise/train.jl:105
┌ Info: Epoch 982
└ @ Main /Users/kcft114/.julia/packages/Flux/qXNjB/src/optimise/train.jl:105
┌ Info: Epoch 983
└ @ Main /Users/kcft114/.julia/packages/Flux/qXNjB/src/optimise/train.jl:105
┌ Info: Epoch 984
└ @ Main /Users/kcft114/.julia/packages/Flux/qXNjB/src/optimise/train.jl:105
┌ Info: Epoch 985
└ @ Main /Users/kcft114/.julia/packages/Flux/qXNjB/src/optimise/train.jl:105
┌ Info: Epoch 986
└ @ Main /Users/kcft114/.julia/packages/Flux/qXNjB/src/optimise/train.jl:105
┌ Info: Epoch 987
└ @ Main /Users/kcft114/.julia/packages/Flux/qXNjB/src/optimise/train.jl:105
┌ Info: Epoch 988
└ @ Main /Users/kcft114/.julia/packages/Flux/qXNjB/src/optimise/train.jl:105
┌ Info: Epoch 989
└ @ Main /Users/kcft114/.julia/packages/Flux/qXNjB/src/optimise/train.jl:105
┌ Info: Epoch 990
└ @ Main /Users/kcft114/.julia/packages/Flux/qXNjB/src/optimise/train.jl:105
┌ Info: Epoch 991
└ @ Main /Users/kcft114/.julia/packages/Flux/qXNjB/src/optimise/train.jl:105
```

```
loss(X, Y) = 0.5639461f0 (tracked)
loss(X, Y) = 0.5724914f0 (tracked)
loss(X, Y) = 0.56087464f0 (tracked)
loss(X, Y) = 0.58062106f0 (tracked)
loss(X, Y) = 0.5748894f0 (tracked)
loss(X, Y) = 0.5685221f0 (tracked)
loss(X, Y) = 0.56916726f0 (tracked)
loss(X, Y) = 0.57077885f0 (tracked)
loss(X, Y) = 0.5675151f0 (tracked)
loss(X, Y) = 0.57000154f0 (tracked)
loss(X, Y) = 0.5475717f0 (tracked)
loss(X, Y) = 0.58084035f0 (tracked)
loss(X, Y) = 0.6076143f0 (tracked)
loss(X, Y) = 0.56032974f0 (tracked)
loss(X, Y) = 0.57389045f0 (tracked)
loss(X, Y) = 0.56889623f0 (tracked)
loss(X, Y) = 0.5602689f0 (tracked)
loss(X, Y) = 0.5590999f0 (tracked)
loss(X, Y) = 0.58723146f0 (tracked)
loss(X, Y) = 0.5726879f0 (tracked)
loss(X, Y) = 0.6057137f0 (tracked)
loss(X, Y) = 0.5517181f0 (tracked)
loss(X, Y) = 0.5635589f0 (tracked)
loss(X, Y) = 0.6443458f0 (tracked)
loss(X, Y) = 0.5844961f0 (tracked)
loss(X, Y) = 0.61167085f0 (tracked)
loss(X, Y) = 0.5577614f0 (tracked)
loss(X, Y) = 0.6186533f0 (tracked)
loss(X, Y) = 0.5843488f0 (tracked)
loss(X, Y) = 0.5365396f0 (tracked)
loss(X, Y) = 0.59263575f0 (tracked)
loss(X, Y) = 0.5854242f0 (tracked)
loss(X, Y) = 0.57086986f0 (tracked)
loss(X, Y) = 0.59069663f0 (tracked)
loss(X, Y) = 0.5577143f0 (tracked)
loss(X, Y) = 0.5529705f0 (tracked)
loss(X, Y) = 0.5737455f0 (tracked)
loss(X, Y) = 0.5938488f0 (tracked)
loss(X, Y) = 0.5980905f0 (tracked)
loss(X, Y) = 0.56227714f0 (tracked)
loss(X, Y) = 0.53273433f0 (tracked)
loss(X, Y) = 0.5726742f0 (tracked)
loss(X, Y) = 0.58177817f0 (tracked)
loss(X, Y) = 0.5790654f0 (tracked)
loss(X, Y) = 0.5549213f0 (tracked)
loss(X, Y) = 0.57486457f0 (tracked)
loss(X, Y) = 0.57928395f0 (tracked)
loss(X, Y) = 0.58166397f0 (tracked)
loss(X, Y) = 0.6166594f0 (tracked)
loss(X, Y) = 0.5817302f0 (tracked)
loss(X, Y) = 0.6150512f0 (tracked)
loss(X, Y) = 0.5723804f0 (tracked)
loss(X, Y) = 0.5972594f0 (tracked)
loss(X, Y) = 0.57071203f0 (tracked)
loss(X, Y) = 0.5858236f0 (tracked)
loss(X, Y) = 0.5806153f0 (tracked)
loss(X, Y) = 0.6054768f0 (tracked)
loss(X, Y) = 0.5553248f0 (tracked)
loss(X, Y) = 0.55567944f0 (tracked)
loss(X, Y) = 0.55708057f0 (tracked)
loss(X, Y) = 0.58740914f0 (tracked)
loss(X, Y) = 0.574707f0 (tracked)
loss(X, Y) = 0.56762433f0 (tracked)
loss(X, Y) = 0.53038216f0 (tracked)
loss(X, Y) = 0.5656715f0 (tracked)
loss(X, Y) = 0.55177534f0 (tracked)
loss(X, Y) = 0.60219675f0 (tracked)
loss(X, Y) = 0.5813153f0 (tracked)
loss(X, Y) = 0.5818536f0 (tracked)
loss(X, Y) = 0.58198214f0 (tracked)
loss(X, Y) = 0.6039542f0 (tracked)
loss(X, Y) = 0.5591231f0 (tracked)
loss(X, Y) = 0.55409855f0 (tracked)
loss(X, Y) = 0.52098066f0 (tracked)
loss(X, Y) = 0.57829076f0 (tracked)
loss(X, Y) = 0.5355115f0 (tracked)
loss(X, Y) = 0.5411448f0 (tracked)
loss(X, Y) = 0.5468677f0 (tracked)
loss(X, Y) = 0.54980093f0 (tracked)
loss(X, Y) = 0.5890677f0 (tracked)
loss(X, Y) = 0.587056f0 (tracked)
loss(X, Y) = 0.58873314f0 (tracked)
loss(X, Y) = 0.5196281f0 (tracked)
loss(X, Y) = 0.5619045f0 (tracked)
loss(X, Y) = 0.58627945f0 (tracked)
loss(X, Y) = 0.56784666f0 (tracked)
loss(X, Y) = 0.5670671f0 (tracked)
loss(X, Y) = 0.6047864f0 (tracked)
```

```
┌ Info: Epoch 992
└ @ Main /Users/kcft114/.julia/packages/Flux/qXNjB/src/optimise/train.jl:105
┌ Info: Epoch 993
└ @ Main /Users/kcft114/.julia/packages/Flux/qXNjB/src/optimise/train.jl:105
┌ Info: Epoch 994
└ @ Main /Users/kcft114/.julia/packages/Flux/qXNjB/src/optimise/train.jl:105
┌ Info: Epoch 995
└ @ Main /Users/kcft114/.julia/packages/Flux/qXNjB/src/optimise/train.jl:105
┌ Info: Epoch 996
└ @ Main /Users/kcft114/.julia/packages/Flux/qXNjB/src/optimise/train.jl:105
┌ Info: Epoch 997
└ @ Main /Users/kcft114/.julia/packages/Flux/qXNjB/src/optimise/train.jl:105
┌ Info: Epoch 998
└ @ Main /Users/kcft114/.julia/packages/Flux/qXNjB/src/optimise/train.jl:105
┌ Info: Epoch 999
└ @ Main /Users/kcft114/.julia/packages/Flux/qXNjB/src/optimise/train.jl:105
┌ Info: Epoch 1000
└ @ Main /Users/kcft114/.julia/packages/Flux/qXNjB/src/optimise/train.jl:105
┌ Info: Epoch 1001
└ @ Main /Users/kcft114/.julia/packages/Flux/qXNjB/src/optimise/train.jl:105
┌ Info: Epoch 1002
└ @ Main /Users/kcft114/.julia/packages/Flux/qXNjB/src/optimise/train.jl:105
┌ Info: Epoch 1003
└ @ Main /Users/kcft114/.julia/packages/Flux/qXNjB/src/optimise/train.jl:105
┌ Info: Epoch 1004
└ @ Main /Users/kcft114/.julia/packages/Flux/qXNjB/src/optimise/train.jl:105
┌ Info: Epoch 1005
└ @ Main /Users/kcft114/.julia/packages/Flux/qXNjB/src/optimise/train.jl:105
┌ Info: Epoch 1006
└ @ Main /Users/kcft114/.julia/packages/Flux/qXNjB/src/optimise/train.jl:105
┌ Info: Epoch 1007
└ @ Main /Users/kcft114/.julia/packages/Flux/qXNjB/src/optimise/train.jl:105
┌ Info: Epoch 1008
└ @ Main /Users/kcft114/.julia/packages/Flux/qXNjB/src/optimise/train.jl:105
┌ Info: Epoch 1009
└ @ Main /Users/kcft114/.julia/packages/Flux/qXNjB/src/optimise/train.jl:105
┌ Info: Epoch 1010
└ @ Main /Users/kcft114/.julia/packages/Flux/qXNjB/src/optimise/train.jl:105
┌ Info: Epoch 1011
└ @ Main /Users/kcft114/.julia/packages/Flux/qXNjB/src/optimise/train.jl:105
┌ Info: Epoch 1012
└ @ Main /Users/kcft114/.julia/packages/Flux/qXNjB/src/optimise/train.jl:105
┌ Info: Epoch 1013
└ @ Main /Users/kcft114/.julia/packages/Flux/qXNjB/src/optimise/train.jl:105
┌ Info: Epoch 1014
└ @ Main /Users/kcft114/.julia/packages/Flux/qXNjB/src/optimise/train.jl:105
┌ Info: Epoch 1015
└ @ Main /Users/kcft114/.julia/packages/Flux/qXNjB/src/optimise/train.jl:105
┌ Info: Epoch 1016
└ @ Main /Users/kcft114/.julia/packages/Flux/qXNjB/src/optimise/train.jl:105
┌ Info: Epoch 1017
└ @ Main /Users/kcft114/.julia/packages/Flux/qXNjB/src/optimise/train.jl:105
┌ Info: Epoch 1018
└ @ Main /Users/kcft114/.julia/packages/Flux/qXNjB/src/optimise/train.jl:105
┌ Info: Epoch 1019
└ @ Main /Users/kcft114/.julia/packages/Flux/qXNjB/src/optimise/train.jl:105
┌ Info: Epoch 1020
└ @ Main /Users/kcft114/.julia/packages/Flux/qXNjB/src/optimise/train.jl:105
┌ Info: Epoch 1021
└ @ Main /Users/kcft114/.julia/packages/Flux/qXNjB/src/optimise/train.jl:105
┌ Info: Epoch 1022
└ @ Main /Users/kcft114/.julia/packages/Flux/qXNjB/src/optimise/train.jl:105
┌ Info: Epoch 1023
└ @ Main /Users/kcft114/.julia/packages/Flux/qXNjB/src/optimise/train.jl:105
┌ Info: Epoch 1024
└ @ Main /Users/kcft114/.julia/packages/Flux/qXNjB/src/optimise/train.jl:105
┌ Info: Epoch 1025
└ @ Main /Users/kcft114/.julia/packages/Flux/qXNjB/src/optimise/train.jl:105
┌ Info: Epoch 1026
└ @ Main /Users/kcft114/.julia/packages/Flux/qXNjB/src/optimise/train.jl:105
┌ Info: Epoch 1027
└ @ Main /Users/kcft114/.julia/packages/Flux/qXNjB/src/optimise/train.jl:105
┌ Info: Epoch 1028
└ @ Main /Users/kcft114/.julia/packages/Flux/qXNjB/src/optimise/train.jl:105
┌ Info: Epoch 1029
└ @ Main /Users/kcft114/.julia/packages/Flux/qXNjB/src/optimise/train.jl:105
┌ Info: Epoch 1030
└ @ Main /Users/kcft114/.julia/packages/Flux/qXNjB/src/optimise/train.jl:105
┌ Info: Epoch 1031
└ @ Main /Users/kcft114/.julia/packages/Flux/qXNjB/src/optimise/train.jl:105
┌ Info: Epoch 1032
└ @ Main /Users/kcft114/.julia/packages/Flux/qXNjB/src/optimise/train.jl:105
┌ Info: Epoch 1033
└ @ Main /Users/kcft114/.julia/packages/Flux/qXNjB/src/optimise/train.jl:105
┌ Info: Epoch 1034
└ @ Main /Users/kcft114/.julia/packages/Flux/qXNjB/src/optimise/train.jl:105
┌ Info: Epoch 1035
└ @ Main /Users/kcft114/.julia/packages/Flux/qXNjB/src/optimise/train.jl:105
┌ Info: Epoch 1036
└ @ Main /Users/kcft114/.julia/packages/Flux/qXNjB/src/optimise/train.jl:105
┌ Info: Epoch 1037
└ @ Main /Users/kcft114/.julia/packages/Flux/qXNjB/src/optimise/train.jl:105
┌ Info: Epoch 1038
└ @ Main /Users/kcft114/.julia/packages/Flux/qXNjB/src/optimise/train.jl:105
┌ Info: Epoch 1039
└ @ Main /Users/kcft114/.julia/packages/Flux/qXNjB/src/optimise/train.jl:105
┌ Info: Epoch 1040
└ @ Main /Users/kcft114/.julia/packages/Flux/qXNjB/src/optimise/train.jl:105
┌ Info: Epoch 1041
└ @ Main /Users/kcft114/.julia/packages/Flux/qXNjB/src/optimise/train.jl:105
┌ Info: Epoch 1042
└ @ Main /Users/kcft114/.julia/packages/Flux/qXNjB/src/optimise/train.jl:105
┌ Info: Epoch 1043
└ @ Main /Users/kcft114/.julia/packages/Flux/qXNjB/src/optimise/train.jl:105
┌ Info: Epoch 1044
└ @ Main /Users/kcft114/.julia/packages/Flux/qXNjB/src/optimise/train.jl:105
┌ Info: Epoch 1045
└ @ Main /Users/kcft114/.julia/packages/Flux/qXNjB/src/optimise/train.jl:105
┌ Info: Epoch 1046
└ @ Main /Users/kcft114/.julia/packages/Flux/qXNjB/src/optimise/train.jl:105
┌ Info: Epoch 1047
└ @ Main /Users/kcft114/.julia/packages/Flux/qXNjB/src/optimise/train.jl:105
┌ Info: Epoch 1048
└ @ Main /Users/kcft114/.julia/packages/Flux/qXNjB/src/optimise/train.jl:105
┌ Info: Epoch 1049
└ @ Main /Users/kcft114/.julia/packages/Flux/qXNjB/src/optimise/train.jl:105
┌ Info: Epoch 1050
└ @ Main /Users/kcft114/.julia/packages/Flux/qXNjB/src/optimise/train.jl:105
┌ Info: Epoch 1051
└ @ Main /Users/kcft114/.julia/packages/Flux/qXNjB/src/optimise/train.jl:105
┌ Info: Epoch 1052
└ @ Main /Users/kcft114/.julia/packages/Flux/qXNjB/src/optimise/train.jl:105
┌ Info: Epoch 1053
└ @ Main /Users/kcft114/.julia/packages/Flux/qXNjB/src/optimise/train.jl:105
┌ Info: Epoch 1054
└ @ Main /Users/kcft114/.julia/packages/Flux/qXNjB/src/optimise/train.jl:105
┌ Info: Epoch 1055
└ @ Main /Users/kcft114/.julia/packages/Flux/qXNjB/src/optimise/train.jl:105
┌ Info: Epoch 1056
└ @ Main /Users/kcft114/.julia/packages/Flux/qXNjB/src/optimise/train.jl:105
┌ Info: Epoch 1057
└ @ Main /Users/kcft114/.julia/packages/Flux/qXNjB/src/optimise/train.jl:105
┌ Info: Epoch 1058
└ @ Main /Users/kcft114/.julia/packages/Flux/qXNjB/src/optimise/train.jl:105
┌ Info: Epoch 1059
└ @ Main /Users/kcft114/.julia/packages/Flux/qXNjB/src/optimise/train.jl:105
┌ Info: Epoch 1060
└ @ Main /Users/kcft114/.julia/packages/Flux/qXNjB/src/optimise/train.jl:105
┌ Info: Epoch 1061
└ @ Main /Users/kcft114/.julia/packages/Flux/qXNjB/src/optimise/train.jl:105
┌ Info: Epoch 1062
└ @ Main /Users/kcft114/.julia/packages/Flux/qXNjB/src/optimise/train.jl:105
┌ Info: Epoch 1063
└ @ Main /Users/kcft114/.julia/packages/Flux/qXNjB/src/optimise/train.jl:105
┌ Info: Epoch 1064
└ @ Main /Users/kcft114/.julia/packages/Flux/qXNjB/src/optimise/train.jl:105
┌ Info: Epoch 1065
└ @ Main /Users/kcft114/.julia/packages/Flux/qXNjB/src/optimise/train.jl:105
┌ Info: Epoch 1066
└ @ Main /Users/kcft114/.julia/packages/Flux/qXNjB/src/optimise/train.jl:105
┌ Info: Epoch 1067
└ @ Main /Users/kcft114/.julia/packages/Flux/qXNjB/src/optimise/train.jl:105
┌ Info: Epoch 1068
└ @ Main /Users/kcft114/.julia/packages/Flux/qXNjB/src/optimise/train.jl:105
┌ Info: Epoch 1069
└ @ Main /Users/kcft114/.julia/packages/Flux/qXNjB/src/optimise/train.jl:105
┌ Info: Epoch 1070
└ @ Main /Users/kcft114/.julia/packages/Flux/qXNjB/src/optimise/train.jl:105
┌ Info: Epoch 1071
└ @ Main /Users/kcft114/.julia/packages/Flux/qXNjB/src/optimise/train.jl:105
┌ Info: Epoch 1072
└ @ Main /Users/kcft114/.julia/packages/Flux/qXNjB/src/optimise/train.jl:105
┌ Info: Epoch 1073
└ @ Main /Users/kcft114/.julia/packages/Flux/qXNjB/src/optimise/train.jl:105
┌ Info: Epoch 1074
└ @ Main /Users/kcft114/.julia/packages/Flux/qXNjB/src/optimise/train.jl:105
┌ Info: Epoch 1075
└ @ Main /Users/kcft114/.julia/packages/Flux/qXNjB/src/optimise/train.jl:105
┌ Info: Epoch 1076
└ @ Main /Users/kcft114/.julia/packages/Flux/qXNjB/src/optimise/train.jl:105
┌ Info: Epoch 1077
└ @ Main /Users/kcft114/.julia/packages/Flux/qXNjB/src/optimise/train.jl:105
┌ Info: Epoch 1078
└ @ Main /Users/kcft114/.julia/packages/Flux/qXNjB/src/optimise/train.jl:105
┌ Info: Epoch 1079
└ @ Main /Users/kcft114/.julia/packages/Flux/qXNjB/src/optimise/train.jl:105
```

```
loss(X, Y) = 0.5526562f0 (tracked)
loss(X, Y) = 0.52512187f0 (tracked)
loss(X, Y) = 0.57977307f0 (tracked)
loss(X, Y) = 0.5919703f0 (tracked)
loss(X, Y) = 0.570269f0 (tracked)
loss(X, Y) = 0.56401294f0 (tracked)
loss(X, Y) = 0.5653641f0 (tracked)
loss(X, Y) = 0.57132477f0 (tracked)
loss(X, Y) = 0.55128014f0 (tracked)
loss(X, Y) = 0.5408676f0 (tracked)
loss(X, Y) = 0.58449036f0 (tracked)
loss(X, Y) = 0.58163863f0 (tracked)
loss(X, Y) = 0.5581209f0 (tracked)
loss(X, Y) = 0.5780012f0 (tracked)
loss(X, Y) = 0.5842691f0 (tracked)
loss(X, Y) = 0.5650284f0 (tracked)
loss(X, Y) = 0.56147724f0 (tracked)
loss(X, Y) = 0.5754111f0 (tracked)
loss(X, Y) = 0.5778693f0 (tracked)
loss(X, Y) = 0.52825546f0 (tracked)
loss(X, Y) = 0.5681066f0 (tracked)
loss(X, Y) = 0.5413892f0 (tracked)
loss(X, Y) = 0.57117045f0 (tracked)
loss(X, Y) = 0.5811172f0 (tracked)
loss(X, Y) = 0.534348f0 (tracked)
loss(X, Y) = 0.5768646f0 (tracked)
loss(X, Y) = 0.55636925f0 (tracked)
loss(X, Y) = 0.5353832f0 (tracked)
loss(X, Y) = 0.5144803f0 (tracked)
loss(X, Y) = 0.57047004f0 (tracked)
loss(X, Y) = 0.5657764f0 (tracked)
loss(X, Y) = 0.5685336f0 (tracked)
loss(X, Y) = 0.5861222f0 (tracked)
loss(X, Y) = 0.60373646f0 (tracked)
loss(X, Y) = 0.53780997f0 (tracked)
loss(X, Y) = 0.56351775f0 (tracked)
loss(X, Y) = 0.55777127f0 (tracked)
loss(X, Y) = 0.59060675f0 (tracked)
loss(X, Y) = 0.5598297f0 (tracked)
loss(X, Y) = 0.57602847f0 (tracked)
loss(X, Y) = 0.5815844f0 (tracked)
loss(X, Y) = 0.5409806f0 (tracked)
loss(X, Y) = 0.57946956f0 (tracked)
loss(X, Y) = 0.5643566f0 (tracked)
loss(X, Y) = 0.60105914f0 (tracked)
loss(X, Y) = 0.55935544f0 (tracked)
loss(X, Y) = 0.53901005f0 (tracked)
loss(X, Y) = 0.53951436f0 (tracked)
loss(X, Y) = 0.57457584f0 (tracked)
loss(X, Y) = 0.59258837f0 (tracked)
loss(X, Y) = 0.6049273f0 (tracked)
loss(X, Y) = 0.5910025f0 (tracked)
loss(X, Y) = 0.5478019f0 (tracked)
loss(X, Y) = 0.5872019f0 (tracked)
loss(X, Y) = 0.5736275f0 (tracked)
loss(X, Y) = 0.5749778f0 (tracked)
loss(X, Y) = 0.5985416f0 (tracked)
loss(X, Y) = 0.55330503f0 (tracked)
loss(X, Y) = 0.5685457f0 (tracked)
loss(X, Y) = 0.54712766f0 (tracked)
loss(X, Y) = 0.56171054f0 (tracked)
loss(X, Y) = 0.63341254f0 (tracked)
loss(X, Y) = 0.57457864f0 (tracked)
loss(X, Y) = 0.5580346f0 (tracked)
loss(X, Y) = 0.5778533f0 (tracked)
loss(X, Y) = 0.5605976f0 (tracked)
loss(X, Y) = 0.5652007f0 (tracked)
loss(X, Y) = 0.6102457f0 (tracked)
loss(X, Y) = 0.5408121f0 (tracked)
loss(X, Y) = 0.56160665f0 (tracked)
loss(X, Y) = 0.53822f0 (tracked)
loss(X, Y) = 0.5762731f0 (tracked)
loss(X, Y) = 0.57543796f0 (tracked)
loss(X, Y) = 0.5742852f0 (tracked)
loss(X, Y) = 0.5524543f0 (tracked)
loss(X, Y) = 0.5400472f0 (tracked)
loss(X, Y) = 0.55271643f0 (tracked)
loss(X, Y) = 0.5570395f0 (tracked)
loss(X, Y) = 0.5806083f0 (tracked)
loss(X, Y) = 0.6134938f0 (tracked)
loss(X, Y) = 0.52420664f0 (tracked)
loss(X, Y) = 0.57239825f0 (tracked)
loss(X, Y) = 0.52076906f0 (tracked)
loss(X, Y) = 0.5726526f0 (tracked)
loss(X, Y) = 0.5934797f0 (tracked)
loss(X, Y) = 0.58571756f0 (tracked)
loss(X, Y) = 0.5721652f0 (tracked)
loss(X, Y) = 0.5365843f0 (tracked)
loss(X, Y) = 0.58405864f0 (tracked)
loss(X, Y) = 0.5810503f0 (tracked)
loss(X, Y) = 0.551143f0 (tracked)
loss(X, Y) = 0.5635574f0 (tracked)
loss(X, Y) = 0.59538805f0 (tracked)
```

```
┌ Info: Epoch 1080
└ @ Main /Users/kcft114/.julia/packages/Flux/qXNjB/src/optimise/train.jl:105
┌ Info: Epoch 1081
└ @ Main /Users/kcft114/.julia/packages/Flux/qXNjB/src/optimise/train.jl:105
┌ Info: Epoch 1082
└ @ Main /Users/kcft114/.julia/packages/Flux/qXNjB/src/optimise/train.jl:105
┌ Info: Epoch 1083
└ @ Main /Users/kcft114/.julia/packages/Flux/qXNjB/src/optimise/train.jl:105
┌ Info: Epoch 1084
└ @ Main /Users/kcft114/.julia/packages/Flux/qXNjB/src/optimise/train.jl:105
┌ Info: Epoch 1085
└ @ Main /Users/kcft114/.julia/packages/Flux/qXNjB/src/optimise/train.jl:105
┌ Info: Epoch 1086
└ @ Main /Users/kcft114/.julia/packages/Flux/qXNjB/src/optimise/train.jl:105
┌ Info: Epoch 1087
└ @ Main /Users/kcft114/.julia/packages/Flux/qXNjB/src/optimise/train.jl:105
┌ Info: Epoch 1088
└ @ Main /Users/kcft114/.julia/packages/Flux/qXNjB/src/optimise/train.jl:105
┌ Info: Epoch 1089
└ @ Main /Users/kcft114/.julia/packages/Flux/qXNjB/src/optimise/train.jl:105
┌ Info: Epoch 1090
└ @ Main /Users/kcft114/.julia/packages/Flux/qXNjB/src/optimise/train.jl:105
┌ Info: Epoch 1091
└ @ Main /Users/kcft114/.julia/packages/Flux/qXNjB/src/optimise/train.jl:105
┌ Info: Epoch 1092
└ @ Main /Users/kcft114/.julia/packages/Flux/qXNjB/src/optimise/train.jl:105
┌ Info: Epoch 1093
└ @ Main /Users/kcft114/.julia/packages/Flux/qXNjB/src/optimise/train.jl:105
┌ Info: Epoch 1094
└ @ Main /Users/kcft114/.julia/packages/Flux/qXNjB/src/optimise/train.jl:105
┌ Info: Epoch 1095
└ @ Main /Users/kcft114/.julia/packages/Flux/qXNjB/src/optimise/train.jl:105
┌ Info: Epoch 1096
└ @ Main /Users/kcft114/.julia/packages/Flux/qXNjB/src/optimise/train.jl:105
┌ Info: Epoch 1097
└ @ Main /Users/kcft114/.julia/packages/Flux/qXNjB/src/optimise/train.jl:105
┌ Info: Epoch 1098
└ @ Main /Users/kcft114/.julia/packages/Flux/qXNjB/src/optimise/train.jl:105
┌ Info: Epoch 1099
└ @ Main /Users/kcft114/.julia/packages/Flux/qXNjB/src/optimise/train.jl:105
┌ Info: Epoch 1100
└ @ Main /Users/kcft114/.julia/packages/Flux/qXNjB/src/optimise/train.jl:105
┌ Info: Epoch 1101
└ @ Main /Users/kcft114/.julia/packages/Flux/qXNjB/src/optimise/train.jl:105
┌ Info: Epoch 1102
└ @ Main /Users/kcft114/.julia/packages/Flux/qXNjB/src/optimise/train.jl:105
┌ Info: Epoch 1103
└ @ Main /Users/kcft114/.julia/packages/Flux/qXNjB/src/optimise/train.jl:105
┌ Info: Epoch 1104
└ @ Main /Users/kcft114/.julia/packages/Flux/qXNjB/src/optimise/train.jl:105
┌ Info: Epoch 1105
└ @ Main /Users/kcft114/.julia/packages/Flux/qXNjB/src/optimise/train.jl:105
┌ Info: Epoch 1106
└ @ Main /Users/kcft114/.julia/packages/Flux/qXNjB/src/optimise/train.jl:105
┌ Info: Epoch 1107
└ @ Main /Users/kcft114/.julia/packages/Flux/qXNjB/src/optimise/train.jl:105
┌ Info: Epoch 1108
└ @ Main /Users/kcft114/.julia/packages/Flux/qXNjB/src/optimise/train.jl:105
┌ Info: Epoch 1109
└ @ Main /Users/kcft114/.julia/packages/Flux/qXNjB/src/optimise/train.jl:105
┌ Info: Epoch 1110
└ @ Main /Users/kcft114/.julia/packages/Flux/qXNjB/src/optimise/train.jl:105
┌ Info: Epoch 1111
└ @ Main /Users/kcft114/.julia/packages/Flux/qXNjB/src/optimise/train.jl:105
┌ Info: Epoch 1112
└ @ Main /Users/kcft114/.julia/packages/Flux/qXNjB/src/optimise/train.jl:105
┌ Info: Epoch 1113
└ @ Main /Users/kcft114/.julia/packages/Flux/qXNjB/src/optimise/train.jl:105
┌ Info: Epoch 1114
└ @ Main /Users/kcft114/.julia/packages/Flux/qXNjB/src/optimise/train.jl:105
┌ Info: Epoch 1115
└ @ Main /Users/kcft114/.julia/packages/Flux/qXNjB/src/optimise/train.jl:105
┌ Info: Epoch 1116
└ @ Main /Users/kcft114/.julia/packages/Flux/qXNjB/src/optimise/train.jl:105
┌ Info: Epoch 1117
└ @ Main /Users/kcft114/.julia/packages/Flux/qXNjB/src/optimise/train.jl:105
┌ Info: Epoch 1118
└ @ Main /Users/kcft114/.julia/packages/Flux/qXNjB/src/optimise/train.jl:105
┌ Info: Epoch 1119
└ @ Main /Users/kcft114/.julia/packages/Flux/qXNjB/src/optimise/train.jl:105
┌ Info: Epoch 1120
└ @ Main /Users/kcft114/.julia/packages/Flux/qXNjB/src/optimise/train.jl:105
┌ Info: Epoch 1121
└ @ Main /Users/kcft114/.julia/packages/Flux/qXNjB/src/optimise/train.jl:105
┌ Info: Epoch 1122
└ @ Main /Users/kcft114/.julia/packages/Flux/qXNjB/src/optimise/train.jl:105
┌ Info: Epoch 1123
└ @ Main /Users/kcft114/.julia/packages/Flux/qXNjB/src/optimise/train.jl:105
┌ Info: Epoch 1124
└ @ Main /Users/kcft114/.julia/packages/Flux/qXNjB/src/optimise/train.jl:105
┌ Info: Epoch 1125
└ @ Main /Users/kcft114/.julia/packages/Flux/qXNjB/src/optimise/train.jl:105
┌ Info: Epoch 1126
└ @ Main /Users/kcft114/.julia/packages/Flux/qXNjB/src/optimise/train.jl:105
┌ Info: Epoch 1127
└ @ Main /Users/kcft114/.julia/packages/Flux/qXNjB/src/optimise/train.jl:105
┌ Info: Epoch 1128
└ @ Main /Users/kcft114/.julia/packages/Flux/qXNjB/src/optimise/train.jl:105
┌ Info: Epoch 1129
└ @ Main /Users/kcft114/.julia/packages/Flux/qXNjB/src/optimise/train.jl:105
┌ Info: Epoch 1130
└ @ Main /Users/kcft114/.julia/packages/Flux/qXNjB/src/optimise/train.jl:105
┌ Info: Epoch 1131
└ @ Main /Users/kcft114/.julia/packages/Flux/qXNjB/src/optimise/train.jl:105
┌ Info: Epoch 1132
└ @ Main /Users/kcft114/.julia/packages/Flux/qXNjB/src/optimise/train.jl:105
┌ Info: Epoch 1133
└ @ Main /Users/kcft114/.julia/packages/Flux/qXNjB/src/optimise/train.jl:105
┌ Info: Epoch 1134
└ @ Main /Users/kcft114/.julia/packages/Flux/qXNjB/src/optimise/train.jl:105
┌ Info: Epoch 1135
└ @ Main /Users/kcft114/.julia/packages/Flux/qXNjB/src/optimise/train.jl:105
┌ Info: Epoch 1136
└ @ Main /Users/kcft114/.julia/packages/Flux/qXNjB/src/optimise/train.jl:105
┌ Info: Epoch 1137
└ @ Main /Users/kcft114/.julia/packages/Flux/qXNjB/src/optimise/train.jl:105
┌ Info: Epoch 1138
└ @ Main /Users/kcft114/.julia/packages/Flux/qXNjB/src/optimise/train.jl:105
┌ Info: Epoch 1139
└ @ Main /Users/kcft114/.julia/packages/Flux/qXNjB/src/optimise/train.jl:105
┌ Info: Epoch 1140
└ @ Main /Users/kcft114/.julia/packages/Flux/qXNjB/src/optimise/train.jl:105
┌ Info: Epoch 1141
└ @ Main /Users/kcft114/.julia/packages/Flux/qXNjB/src/optimise/train.jl:105
┌ Info: Epoch 1142
└ @ Main /Users/kcft114/.julia/packages/Flux/qXNjB/src/optimise/train.jl:105
┌ Info: Epoch 1143
└ @ Main /Users/kcft114/.julia/packages/Flux/qXNjB/src/optimise/train.jl:105
┌ Info: Epoch 1144
└ @ Main /Users/kcft114/.julia/packages/Flux/qXNjB/src/optimise/train.jl:105
┌ Info: Epoch 1145
└ @ Main /Users/kcft114/.julia/packages/Flux/qXNjB/src/optimise/train.jl:105
┌ Info: Epoch 1146
└ @ Main /Users/kcft114/.julia/packages/Flux/qXNjB/src/optimise/train.jl:105
┌ Info: Epoch 1147
└ @ Main /Users/kcft114/.julia/packages/Flux/qXNjB/src/optimise/train.jl:105
┌ Info: Epoch 1148
└ @ Main /Users/kcft114/.julia/packages/Flux/qXNjB/src/optimise/train.jl:105
┌ Info: Epoch 1149
└ @ Main /Users/kcft114/.julia/packages/Flux/qXNjB/src/optimise/train.jl:105
┌ Info: Epoch 1150
└ @ Main /Users/kcft114/.julia/packages/Flux/qXNjB/src/optimise/train.jl:105
┌ Info: Epoch 1151
└ @ Main /Users/kcft114/.julia/packages/Flux/qXNjB/src/optimise/train.jl:105
┌ Info: Epoch 1152
└ @ Main /Users/kcft114/.julia/packages/Flux/qXNjB/src/optimise/train.jl:105
┌ Info: Epoch 1153
└ @ Main /Users/kcft114/.julia/packages/Flux/qXNjB/src/optimise/train.jl:105
┌ Info: Epoch 1154
└ @ Main /Users/kcft114/.julia/packages/Flux/qXNjB/src/optimise/train.jl:105
┌ Info: Epoch 1155
└ @ Main /Users/kcft114/.julia/packages/Flux/qXNjB/src/optimise/train.jl:105
┌ Info: Epoch 1156
└ @ Main /Users/kcft114/.julia/packages/Flux/qXNjB/src/optimise/train.jl:105
┌ Info: Epoch 1157
└ @ Main /Users/kcft114/.julia/packages/Flux/qXNjB/src/optimise/train.jl:105
┌ Info: Epoch 1158
└ @ Main /Users/kcft114/.julia/packages/Flux/qXNjB/src/optimise/train.jl:105
┌ Info: Epoch 1159
└ @ Main /Users/kcft114/.julia/packages/Flux/qXNjB/src/optimise/train.jl:105
┌ Info: Epoch 1160
└ @ Main /Users/kcft114/.julia/packages/Flux/qXNjB/src/optimise/train.jl:105
┌ Info: Epoch 1161
└ @ Main /Users/kcft114/.julia/packages/Flux/qXNjB/src/optimise/train.jl:105
┌ Info: Epoch 1162
└ @ Main /Users/kcft114/.julia/packages/Flux/qXNjB/src/optimise/train.jl:105
┌ Info: Epoch 1163
└ @ Main /Users/kcft114/.julia/packages/Flux/qXNjB/src/optimise/train.jl:105
┌ Info: Epoch 1164
└ @ Main /Users/kcft114/.julia/packages/Flux/qXNjB/src/optimise/train.jl:105
┌ Info: Epoch 1165
└ @ Main /Users/kcft114/.julia/packages/Flux/qXNjB/src/optimise/train.jl:105
┌ Info: Epoch 1166
└ @ Main /Users/kcft114/.julia/packages/Flux/qXNjB/src/optimise/train.jl:105
┌ Info: Epoch 1167
└ @ Main /Users/kcft114/.julia/packages/Flux/qXNjB/src/optimise/train.jl:105
┌ Info: Epoch 1168
└ @ Main /Users/kcft114/.julia/packages/Flux/qXNjB/src/optimise/train.jl:105
┌ Info: Epoch 1169
└ @ Main /Users/kcft114/.julia/packages/Flux/qXNjB/src/optimise/train.jl:105
┌ Info: Epoch 1170
└ @ Main /Users/kcft114/.julia/packages/Flux/qXNjB/src/optimise/train.jl:105
┌ Info: Epoch 1171
└ @ Main /Users/kcft114/.julia/packages/Flux/qXNjB/src/optimise/train.jl:105
┌ Info: Epoch 1172
└ @ Main /Users/kcft114/.julia/packages/Flux/qXNjB/src/optimise/train.jl:105
```

```
loss(X, Y) = 0.561482f0 (tracked)
loss(X, Y) = 0.6006574f0 (tracked)
loss(X, Y) = 0.5366088f0 (tracked)
loss(X, Y) = 0.58496255f0 (tracked)
loss(X, Y) = 0.58693224f0 (tracked)
loss(X, Y) = 0.6052325f0 (tracked)
loss(X, Y) = 0.5757564f0 (tracked)
loss(X, Y) = 0.58995354f0 (tracked)
loss(X, Y) = 0.5303965f0 (tracked)
loss(X, Y) = 0.56046706f0 (tracked)
loss(X, Y) = 0.58445656f0 (tracked)
loss(X, Y) = 0.5804919f0 (tracked)
loss(X, Y) = 0.5340974f0 (tracked)
loss(X, Y) = 0.5472995f0 (tracked)
loss(X, Y) = 0.56067634f0 (tracked)
loss(X, Y) = 0.57025236f0 (tracked)
loss(X, Y) = 0.5340312f0 (tracked)
loss(X, Y) = 0.5724143f0 (tracked)
loss(X, Y) = 0.5595484f0 (tracked)
loss(X, Y) = 0.58985585f0 (tracked)
loss(X, Y) = 0.5955379f0 (tracked)
loss(X, Y) = 0.50244725f0 (tracked)
loss(X, Y) = 0.5903054f0 (tracked)
loss(X, Y) = 0.58100706f0 (tracked)
loss(X, Y) = 0.5315344f0 (tracked)
loss(X, Y) = 0.54494435f0 (tracked)
loss(X, Y) = 0.5788816f0 (tracked)
loss(X, Y) = 0.58194035f0 (tracked)
loss(X, Y) = 0.59480757f0 (tracked)
loss(X, Y) = 0.5611743f0 (tracked)
loss(X, Y) = 0.5777747f0 (tracked)
loss(X, Y) = 0.55785984f0 (tracked)
loss(X, Y) = 0.5149618f0 (tracked)
loss(X, Y) = 0.5548784f0 (tracked)
loss(X, Y) = 0.556571f0 (tracked)
loss(X, Y) = 0.5575256f0 (tracked)
loss(X, Y) = 0.54698527f0 (tracked)
loss(X, Y) = 0.4984261f0 (tracked)
loss(X, Y) = 0.57991916f0 (tracked)
loss(X, Y) = 0.6028973f0 (tracked)
loss(X, Y) = 0.5363197f0 (tracked)
loss(X, Y) = 0.54004616f0 (tracked)
loss(X, Y) = 0.55705833f0 (tracked)
loss(X, Y) = 0.61041856f0 (tracked)
loss(X, Y) = 0.5692531f0 (tracked)
loss(X, Y) = 0.5369849f0 (tracked)
loss(X, Y) = 0.5982506f0 (tracked)
loss(X, Y) = 0.54903543f0 (tracked)
loss(X, Y) = 0.54136306f0 (tracked)
loss(X, Y) = 0.54285884f0 (tracked)
loss(X, Y) = 0.58151376f0 (tracked)
loss(X, Y) = 0.58297163f0 (tracked)
loss(X, Y) = 0.5559161f0 (tracked)
loss(X, Y) = 0.56354415f0 (tracked)
loss(X, Y) = 0.6003874f0 (tracked)
loss(X, Y) = 0.5957664f0 (tracked)
loss(X, Y) = 0.577852f0 (tracked)
loss(X, Y) = 0.57021266f0 (tracked)
loss(X, Y) = 0.5389068f0 (tracked)
loss(X, Y) = 0.5525748f0 (tracked)
loss(X, Y) = 0.56936413f0 (tracked)
loss(X, Y) = 0.550165f0 (tracked)
loss(X, Y) = 0.5232059f0 (tracked)
loss(X, Y) = 0.5647871f0 (tracked)
loss(X, Y) = 0.55045825f0 (tracked)
loss(X, Y) = 0.53774166f0 (tracked)
loss(X, Y) = 0.56129104f0 (tracked)
loss(X, Y) = 0.56889194f0 (tracked)
loss(X, Y) = 0.53621566f0 (tracked)
loss(X, Y) = 0.52470356f0 (tracked)
loss(X, Y) = 0.6069544f0 (tracked)
loss(X, Y) = 0.5306784f0 (tracked)
loss(X, Y) = 0.5282018f0 (tracked)
loss(X, Y) = 0.5513206f0 (tracked)
loss(X, Y) = 0.56660247f0 (tracked)
loss(X, Y) = 0.5098078f0 (tracked)
loss(X, Y) = 0.59208184f0 (tracked)
loss(X, Y) = 0.5520917f0 (tracked)
loss(X, Y) = 0.59577763f0 (tracked)
loss(X, Y) = 0.5298547f0 (tracked)
loss(X, Y) = 0.5671906f0 (tracked)
loss(X, Y) = 0.55788755f0 (tracked)
loss(X, Y) = 0.5566145f0 (tracked)
loss(X, Y) = 0.53549135f0 (tracked)
loss(X, Y) = 0.53202194f0 (tracked)
loss(X, Y) = 0.5602137f0 (tracked)
loss(X, Y) = 0.548122f0 (tracked)
loss(X, Y) = 0.5403954f0 (tracked)
loss(X, Y) = 0.57059216f0 (tracked)
loss(X, Y) = 0.5387644f0 (tracked)
loss(X, Y) = 0.56075454f0 (tracked)
loss(X, Y) = 0.547882f0 (tracked)
```

```
┌ Info: Epoch 1173
└ @ Main /Users/kcft114/.julia/packages/Flux/qXNjB/src/optimise/train.jl:105
┌ Info: Epoch 1174
└ @ Main /Users/kcft114/.julia/packages/Flux/qXNjB/src/optimise/train.jl:105
┌ Info: Epoch 1175
└ @ Main /Users/kcft114/.julia/packages/Flux/qXNjB/src/optimise/train.jl:105
┌ Info: Epoch 1176
└ @ Main /Users/kcft114/.julia/packages/Flux/qXNjB/src/optimise/train.jl:105
┌ Info: Epoch 1177
└ @ Main /Users/kcft114/.julia/packages/Flux/qXNjB/src/optimise/train.jl:105
┌ Info: Epoch 1178
└ @ Main /Users/kcft114/.julia/packages/Flux/qXNjB/src/optimise/train.jl:105
┌ Info: Epoch 1179
└ @ Main /Users/kcft114/.julia/packages/Flux/qXNjB/src/optimise/train.jl:105
┌ Info: Epoch 1180
└ @ Main /Users/kcft114/.julia/packages/Flux/qXNjB/src/optimise/train.jl:105
┌ Info: Epoch 1181
└ @ Main /Users/kcft114/.julia/packages/Flux/qXNjB/src/optimise/train.jl:105
┌ Info: Epoch 1182
└ @ Main /Users/kcft114/.julia/packages/Flux/qXNjB/src/optimise/train.jl:105
┌ Info: Epoch 1183
└ @ Main /Users/kcft114/.julia/packages/Flux/qXNjB/src/optimise/train.jl:105
┌ Info: Epoch 1184
└ @ Main /Users/kcft114/.julia/packages/Flux/qXNjB/src/optimise/train.jl:105
┌ Info: Epoch 1185
└ @ Main /Users/kcft114/.julia/packages/Flux/qXNjB/src/optimise/train.jl:105
┌ Info: Epoch 1186
└ @ Main /Users/kcft114/.julia/packages/Flux/qXNjB/src/optimise/train.jl:105
┌ Info: Epoch 1187
└ @ Main /Users/kcft114/.julia/packages/Flux/qXNjB/src/optimise/train.jl:105
┌ Info: Epoch 1188
└ @ Main /Users/kcft114/.julia/packages/Flux/qXNjB/src/optimise/train.jl:105
┌ Info: Epoch 1189
└ @ Main /Users/kcft114/.julia/packages/Flux/qXNjB/src/optimise/train.jl:105
┌ Info: Epoch 1190
└ @ Main /Users/kcft114/.julia/packages/Flux/qXNjB/src/optimise/train.jl:105
┌ Info: Epoch 1191
└ @ Main /Users/kcft114/.julia/packages/Flux/qXNjB/src/optimise/train.jl:105
┌ Info: Epoch 1192
└ @ Main /Users/kcft114/.julia/packages/Flux/qXNjB/src/optimise/train.jl:105
┌ Info: Epoch 1193
└ @ Main /Users/kcft114/.julia/packages/Flux/qXNjB/src/optimise/train.jl:105
┌ Info: Epoch 1194
└ @ Main /Users/kcft114/.julia/packages/Flux/qXNjB/src/optimise/train.jl:105
┌ Info: Epoch 1195
└ @ Main /Users/kcft114/.julia/packages/Flux/qXNjB/src/optimise/train.jl:105
┌ Info: Epoch 1196
└ @ Main /Users/kcft114/.julia/packages/Flux/qXNjB/src/optimise/train.jl:105
┌ Info: Epoch 1197
└ @ Main /Users/kcft114/.julia/packages/Flux/qXNjB/src/optimise/train.jl:105
┌ Info: Epoch 1198
└ @ Main /Users/kcft114/.julia/packages/Flux/qXNjB/src/optimise/train.jl:105
┌ Info: Epoch 1199
└ @ Main /Users/kcft114/.julia/packages/Flux/qXNjB/src/optimise/train.jl:105
┌ Info: Epoch 1200
└ @ Main /Users/kcft114/.julia/packages/Flux/qXNjB/src/optimise/train.jl:105
┌ Info: Epoch 1201
└ @ Main /Users/kcft114/.julia/packages/Flux/qXNjB/src/optimise/train.jl:105
┌ Info: Epoch 1202
└ @ Main /Users/kcft114/.julia/packages/Flux/qXNjB/src/optimise/train.jl:105
┌ Info: Epoch 1203
└ @ Main /Users/kcft114/.julia/packages/Flux/qXNjB/src/optimise/train.jl:105
┌ Info: Epoch 1204
└ @ Main /Users/kcft114/.julia/packages/Flux/qXNjB/src/optimise/train.jl:105
┌ Info: Epoch 1205
└ @ Main /Users/kcft114/.julia/packages/Flux/qXNjB/src/optimise/train.jl:105
┌ Info: Epoch 1206
└ @ Main /Users/kcft114/.julia/packages/Flux/qXNjB/src/optimise/train.jl:105
┌ Info: Epoch 1207
└ @ Main /Users/kcft114/.julia/packages/Flux/qXNjB/src/optimise/train.jl:105
┌ Info: Epoch 1208
└ @ Main /Users/kcft114/.julia/packages/Flux/qXNjB/src/optimise/train.jl:105
┌ Info: Epoch 1209
└ @ Main /Users/kcft114/.julia/packages/Flux/qXNjB/src/optimise/train.jl:105
┌ Info: Epoch 1210
└ @ Main /Users/kcft114/.julia/packages/Flux/qXNjB/src/optimise/train.jl:105
┌ Info: Epoch 1211
└ @ Main /Users/kcft114/.julia/packages/Flux/qXNjB/src/optimise/train.jl:105
┌ Info: Epoch 1212
└ @ Main /Users/kcft114/.julia/packages/Flux/qXNjB/src/optimise/train.jl:105
┌ Info: Epoch 1213
└ @ Main /Users/kcft114/.julia/packages/Flux/qXNjB/src/optimise/train.jl:105
┌ Info: Epoch 1214
└ @ Main /Users/kcft114/.julia/packages/Flux/qXNjB/src/optimise/train.jl:105
┌ Info: Epoch 1215
└ @ Main /Users/kcft114/.julia/packages/Flux/qXNjB/src/optimise/train.jl:105
┌ Info: Epoch 1216
└ @ Main /Users/kcft114/.julia/packages/Flux/qXNjB/src/optimise/train.jl:105
┌ Info: Epoch 1217
└ @ Main /Users/kcft114/.julia/packages/Flux/qXNjB/src/optimise/train.jl:105
┌ Info: Epoch 1218
└ @ Main /Users/kcft114/.julia/packages/Flux/qXNjB/src/optimise/train.jl:105
┌ Info: Epoch 1219
└ @ Main /Users/kcft114/.julia/packages/Flux/qXNjB/src/optimise/train.jl:105
┌ Info: Epoch 1220
└ @ Main /Users/kcft114/.julia/packages/Flux/qXNjB/src/optimise/train.jl:105
┌ Info: Epoch 1221
└ @ Main /Users/kcft114/.julia/packages/Flux/qXNjB/src/optimise/train.jl:105
┌ Info: Epoch 1222
└ @ Main /Users/kcft114/.julia/packages/Flux/qXNjB/src/optimise/train.jl:105
┌ Info: Epoch 1223
└ @ Main /Users/kcft114/.julia/packages/Flux/qXNjB/src/optimise/train.jl:105
┌ Info: Epoch 1224
└ @ Main /Users/kcft114/.julia/packages/Flux/qXNjB/src/optimise/train.jl:105
┌ Info: Epoch 1225
└ @ Main /Users/kcft114/.julia/packages/Flux/qXNjB/src/optimise/train.jl:105
┌ Info: Epoch 1226
└ @ Main /Users/kcft114/.julia/packages/Flux/qXNjB/src/optimise/train.jl:105
┌ Info: Epoch 1227
└ @ Main /Users/kcft114/.julia/packages/Flux/qXNjB/src/optimise/train.jl:105
┌ Info: Epoch 1228
└ @ Main /Users/kcft114/.julia/packages/Flux/qXNjB/src/optimise/train.jl:105
┌ Info: Epoch 1229
└ @ Main /Users/kcft114/.julia/packages/Flux/qXNjB/src/optimise/train.jl:105
┌ Info: Epoch 1230
└ @ Main /Users/kcft114/.julia/packages/Flux/qXNjB/src/optimise/train.jl:105
┌ Info: Epoch 1231
└ @ Main /Users/kcft114/.julia/packages/Flux/qXNjB/src/optimise/train.jl:105
┌ Info: Epoch 1232
└ @ Main /Users/kcft114/.julia/packages/Flux/qXNjB/src/optimise/train.jl:105
┌ Info: Epoch 1233
└ @ Main /Users/kcft114/.julia/packages/Flux/qXNjB/src/optimise/train.jl:105
┌ Info: Epoch 1234
└ @ Main /Users/kcft114/.julia/packages/Flux/qXNjB/src/optimise/train.jl:105
┌ Info: Epoch 1235
└ @ Main /Users/kcft114/.julia/packages/Flux/qXNjB/src/optimise/train.jl:105
┌ Info: Epoch 1236
└ @ Main /Users/kcft114/.julia/packages/Flux/qXNjB/src/optimise/train.jl:105
┌ Info: Epoch 1237
└ @ Main /Users/kcft114/.julia/packages/Flux/qXNjB/src/optimise/train.jl:105
┌ Info: Epoch 1238
└ @ Main /Users/kcft114/.julia/packages/Flux/qXNjB/src/optimise/train.jl:105
┌ Info: Epoch 1239
└ @ Main /Users/kcft114/.julia/packages/Flux/qXNjB/src/optimise/train.jl:105
┌ Info: Epoch 1240
└ @ Main /Users/kcft114/.julia/packages/Flux/qXNjB/src/optimise/train.jl:105
┌ Info: Epoch 1241
└ @ Main /Users/kcft114/.julia/packages/Flux/qXNjB/src/optimise/train.jl:105
┌ Info: Epoch 1242
└ @ Main /Users/kcft114/.julia/packages/Flux/qXNjB/src/optimise/train.jl:105
┌ Info: Epoch 1243
└ @ Main /Users/kcft114/.julia/packages/Flux/qXNjB/src/optimise/train.jl:105
┌ Info: Epoch 1244
└ @ Main /Users/kcft114/.julia/packages/Flux/qXNjB/src/optimise/train.jl:105
┌ Info: Epoch 1245
└ @ Main /Users/kcft114/.julia/packages/Flux/qXNjB/src/optimise/train.jl:105
┌ Info: Epoch 1246
└ @ Main /Users/kcft114/.julia/packages/Flux/qXNjB/src/optimise/train.jl:105
┌ Info: Epoch 1247
└ @ Main /Users/kcft114/.julia/packages/Flux/qXNjB/src/optimise/train.jl:105
┌ Info: Epoch 1248
└ @ Main /Users/kcft114/.julia/packages/Flux/qXNjB/src/optimise/train.jl:105
┌ Info: Epoch 1249
└ @ Main /Users/kcft114/.julia/packages/Flux/qXNjB/src/optimise/train.jl:105
┌ Info: Epoch 1250
└ @ Main /Users/kcft114/.julia/packages/Flux/qXNjB/src/optimise/train.jl:105
┌ Info: Epoch 1251
└ @ Main /Users/kcft114/.julia/packages/Flux/qXNjB/src/optimise/train.jl:105
┌ Info: Epoch 1252
└ @ Main /Users/kcft114/.julia/packages/Flux/qXNjB/src/optimise/train.jl:105
┌ Info: Epoch 1253
└ @ Main /Users/kcft114/.julia/packages/Flux/qXNjB/src/optimise/train.jl:105
┌ Info: Epoch 1254
└ @ Main /Users/kcft114/.julia/packages/Flux/qXNjB/src/optimise/train.jl:105
┌ Info: Epoch 1255
└ @ Main /Users/kcft114/.julia/packages/Flux/qXNjB/src/optimise/train.jl:105
┌ Info: Epoch 1256
└ @ Main /Users/kcft114/.julia/packages/Flux/qXNjB/src/optimise/train.jl:105
┌ Info: Epoch 1257
└ @ Main /Users/kcft114/.julia/packages/Flux/qXNjB/src/optimise/train.jl:105
┌ Info: Epoch 1258
└ @ Main /Users/kcft114/.julia/packages/Flux/qXNjB/src/optimise/train.jl:105
┌ Info: Epoch 1259
└ @ Main /Users/kcft114/.julia/packages/Flux/qXNjB/src/optimise/train.jl:105
┌ Info: Epoch 1260
└ @ Main /Users/kcft114/.julia/packages/Flux/qXNjB/src/optimise/train.jl:105
┌ Info: Epoch 1261
└ @ Main /Users/kcft114/.julia/packages/Flux/qXNjB/src/optimise/train.jl:105
┌ Info: Epoch 1262
└ @ Main /Users/kcft114/.julia/packages/Flux/qXNjB/src/optimise/train.jl:105
┌ Info: Epoch 1263
└ @ Main /Users/kcft114/.julia/packages/Flux/qXNjB/src/optimise/train.jl:105
┌ Info: Epoch 1264
└ @ Main /Users/kcft114/.julia/packages/Flux/qXNjB/src/optimise/train.jl:105
```

```
loss(X, Y) = 0.6246625f0 (tracked)
loss(X, Y) = 0.54926074f0 (tracked)
loss(X, Y) = 0.58727753f0 (tracked)
loss(X, Y) = 0.5602987f0 (tracked)
loss(X, Y) = 0.56401324f0 (tracked)
loss(X, Y) = 0.5639232f0 (tracked)
loss(X, Y) = 0.5852392f0 (tracked)
loss(X, Y) = 0.54414064f0 (tracked)
loss(X, Y) = 0.55014414f0 (tracked)
loss(X, Y) = 0.57165056f0 (tracked)
loss(X, Y) = 0.50548327f0 (tracked)
loss(X, Y) = 0.5779065f0 (tracked)
loss(X, Y) = 0.548062f0 (tracked)
loss(X, Y) = 0.5241015f0 (tracked)
loss(X, Y) = 0.5189106f0 (tracked)
loss(X, Y) = 0.57346517f0 (tracked)
loss(X, Y) = 0.54880625f0 (tracked)
loss(X, Y) = 0.52304137f0 (tracked)
loss(X, Y) = 0.5433782f0 (tracked)
loss(X, Y) = 0.5300316f0 (tracked)
loss(X, Y) = 0.56485784f0 (tracked)
loss(X, Y) = 0.5919087f0 (tracked)
loss(X, Y) = 0.5278232f0 (tracked)
loss(X, Y) = 0.53051007f0 (tracked)
loss(X, Y) = 0.55493134f0 (tracked)
loss(X, Y) = 0.5500862f0 (tracked)
loss(X, Y) = 0.56399924f0 (tracked)
loss(X, Y) = 0.5206266f0 (tracked)
loss(X, Y) = 0.58619064f0 (tracked)
loss(X, Y) = 0.58376545f0 (tracked)
loss(X, Y) = 0.50483894f0 (tracked)
loss(X, Y) = 0.51064736f0 (tracked)
loss(X, Y) = 0.5448272f0 (tracked)
loss(X, Y) = 0.58331007f0 (tracked)
loss(X, Y) = 0.5008075f0 (tracked)
loss(X, Y) = 0.5461628f0 (tracked)
loss(X, Y) = 0.5501454f0 (tracked)
loss(X, Y) = 0.5517621f0 (tracked)
loss(X, Y) = 0.5818008f0 (tracked)
loss(X, Y) = 0.5715256f0 (tracked)
loss(X, Y) = 0.5243068f0 (tracked)
loss(X, Y) = 0.60330164f0 (tracked)
loss(X, Y) = 0.5957588f0 (tracked)
loss(X, Y) = 0.614041f0 (tracked)
loss(X, Y) = 0.55904555f0 (tracked)
loss(X, Y) = 0.5390633f0 (tracked)
loss(X, Y) = 0.5439117f0 (tracked)
loss(X, Y) = 0.52736723f0 (tracked)
loss(X, Y) = 0.60059977f0 (tracked)
loss(X, Y) = 0.552553f0 (tracked)
loss(X, Y) = 0.5451206f0 (tracked)
loss(X, Y) = 0.5593994f0 (tracked)
loss(X, Y) = 0.5338223f0 (tracked)
loss(X, Y) = 0.6097455f0 (tracked)
loss(X, Y) = 0.5295635f0 (tracked)
loss(X, Y) = 0.5733551f0 (tracked)
loss(X, Y) = 0.58040184f0 (tracked)
loss(X, Y) = 0.51717f0 (tracked)
loss(X, Y) = 0.5480338f0 (tracked)
loss(X, Y) = 0.53020614f0 (tracked)
loss(X, Y) = 0.5532515f0 (tracked)
loss(X, Y) = 0.5801937f0 (tracked)
loss(X, Y) = 0.5609465f0 (tracked)
loss(X, Y) = 0.5791602f0 (tracked)
loss(X, Y) = 0.5124601f0 (tracked)
loss(X, Y) = 0.5322447f0 (tracked)
loss(X, Y) = 0.55198413f0 (tracked)
loss(X, Y) = 0.5639404f0 (tracked)
loss(X, Y) = 0.5010725f0 (tracked)
loss(X, Y) = 0.5708453f0 (tracked)
loss(X, Y) = 0.5706305f0 (tracked)
loss(X, Y) = 0.56805366f0 (tracked)
loss(X, Y) = 0.4997996f0 (tracked)
loss(X, Y) = 0.5587921f0 (tracked)
loss(X, Y) = 0.535182f0 (tracked)
loss(X, Y) = 0.5485982f0 (tracked)
loss(X, Y) = 0.5267511f0 (tracked)
loss(X, Y) = 0.52276826f0 (tracked)
loss(X, Y) = 0.50721544f0 (tracked)
loss(X, Y) = 0.6134742f0 (tracked)
loss(X, Y) = 0.5213879f0 (tracked)
loss(X, Y) = 0.5628739f0 (tracked)
loss(X, Y) = 0.5477957f0 (tracked)
loss(X, Y) = 0.5599267f0 (tracked)
loss(X, Y) = 0.555201f0 (tracked)
loss(X, Y) = 0.57039344f0 (tracked)
loss(X, Y) = 0.53174067f0 (tracked)
loss(X, Y) = 0.58285826f0 (tracked)
loss(X, Y) = 0.5701324f0 (tracked)
```

```
┌ Info: Epoch 1265
└ @ Main /Users/kcft114/.julia/packages/Flux/qXNjB/src/optimise/train.jl:105
┌ Info: Epoch 1266
└ @ Main /Users/kcft114/.julia/packages/Flux/qXNjB/src/optimise/train.jl:105
┌ Info: Epoch 1267
└ @ Main /Users/kcft114/.julia/packages/Flux/qXNjB/src/optimise/train.jl:105
┌ Info: Epoch 1268
└ @ Main /Users/kcft114/.julia/packages/Flux/qXNjB/src/optimise/train.jl:105
┌ Info: Epoch 1269
└ @ Main /Users/kcft114/.julia/packages/Flux/qXNjB/src/optimise/train.jl:105
┌ Info: Epoch 1270
└ @ Main /Users/kcft114/.julia/packages/Flux/qXNjB/src/optimise/train.jl:105
┌ Info: Epoch 1271
└ @ Main /Users/kcft114/.julia/packages/Flux/qXNjB/src/optimise/train.jl:105
┌ Info: Epoch 1272
└ @ Main /Users/kcft114/.julia/packages/Flux/qXNjB/src/optimise/train.jl:105
┌ Info: Epoch 1273
└ @ Main /Users/kcft114/.julia/packages/Flux/qXNjB/src/optimise/train.jl:105
┌ Info: Epoch 1274
└ @ Main /Users/kcft114/.julia/packages/Flux/qXNjB/src/optimise/train.jl:105
┌ Info: Epoch 1275
└ @ Main /Users/kcft114/.julia/packages/Flux/qXNjB/src/optimise/train.jl:105
┌ Info: Epoch 1276
└ @ Main /Users/kcft114/.julia/packages/Flux/qXNjB/src/optimise/train.jl:105
┌ Info: Epoch 1277
└ @ Main /Users/kcft114/.julia/packages/Flux/qXNjB/src/optimise/train.jl:105
┌ Info: Epoch 1278
└ @ Main /Users/kcft114/.julia/packages/Flux/qXNjB/src/optimise/train.jl:105
┌ Info: Epoch 1279
└ @ Main /Users/kcft114/.julia/packages/Flux/qXNjB/src/optimise/train.jl:105
┌ Info: Epoch 1280
└ @ Main /Users/kcft114/.julia/packages/Flux/qXNjB/src/optimise/train.jl:105
┌ Info: Epoch 1281
└ @ Main /Users/kcft114/.julia/packages/Flux/qXNjB/src/optimise/train.jl:105
┌ Info: Epoch 1282
└ @ Main /Users/kcft114/.julia/packages/Flux/qXNjB/src/optimise/train.jl:105
┌ Info: Epoch 1283
└ @ Main /Users/kcft114/.julia/packages/Flux/qXNjB/src/optimise/train.jl:105
┌ Info: Epoch 1284
└ @ Main /Users/kcft114/.julia/packages/Flux/qXNjB/src/optimise/train.jl:105
┌ Info: Epoch 1285
└ @ Main /Users/kcft114/.julia/packages/Flux/qXNjB/src/optimise/train.jl:105
┌ Info: Epoch 1286
└ @ Main /Users/kcft114/.julia/packages/Flux/qXNjB/src/optimise/train.jl:105
┌ Info: Epoch 1287
└ @ Main /Users/kcft114/.julia/packages/Flux/qXNjB/src/optimise/train.jl:105
┌ Info: Epoch 1288
└ @ Main /Users/kcft114/.julia/packages/Flux/qXNjB/src/optimise/train.jl:105
┌ Info: Epoch 1289
└ @ Main /Users/kcft114/.julia/packages/Flux/qXNjB/src/optimise/train.jl:105
┌ Info: Epoch 1290
└ @ Main /Users/kcft114/.julia/packages/Flux/qXNjB/src/optimise/train.jl:105
┌ Info: Epoch 1291
└ @ Main /Users/kcft114/.julia/packages/Flux/qXNjB/src/optimise/train.jl:105
┌ Info: Epoch 1292
└ @ Main /Users/kcft114/.julia/packages/Flux/qXNjB/src/optimise/train.jl:105
┌ Info: Epoch 1293
└ @ Main /Users/kcft114/.julia/packages/Flux/qXNjB/src/optimise/train.jl:105
┌ Info: Epoch 1294
└ @ Main /Users/kcft114/.julia/packages/Flux/qXNjB/src/optimise/train.jl:105
┌ Info: Epoch 1295
└ @ Main /Users/kcft114/.julia/packages/Flux/qXNjB/src/optimise/train.jl:105
┌ Info: Epoch 1296
└ @ Main /Users/kcft114/.julia/packages/Flux/qXNjB/src/optimise/train.jl:105
┌ Info: Epoch 1297
└ @ Main /Users/kcft114/.julia/packages/Flux/qXNjB/src/optimise/train.jl:105
┌ Info: Epoch 1298
└ @ Main /Users/kcft114/.julia/packages/Flux/qXNjB/src/optimise/train.jl:105
┌ Info: Epoch 1299
└ @ Main /Users/kcft114/.julia/packages/Flux/qXNjB/src/optimise/train.jl:105
┌ Info: Epoch 1300
└ @ Main /Users/kcft114/.julia/packages/Flux/qXNjB/src/optimise/train.jl:105
┌ Info: Epoch 1301
└ @ Main /Users/kcft114/.julia/packages/Flux/qXNjB/src/optimise/train.jl:105
┌ Info: Epoch 1302
└ @ Main /Users/kcft114/.julia/packages/Flux/qXNjB/src/optimise/train.jl:105
┌ Info: Epoch 1303
└ @ Main /Users/kcft114/.julia/packages/Flux/qXNjB/src/optimise/train.jl:105
┌ Info: Epoch 1304
└ @ Main /Users/kcft114/.julia/packages/Flux/qXNjB/src/optimise/train.jl:105
┌ Info: Epoch 1305
└ @ Main /Users/kcft114/.julia/packages/Flux/qXNjB/src/optimise/train.jl:105
┌ Info: Epoch 1306
└ @ Main /Users/kcft114/.julia/packages/Flux/qXNjB/src/optimise/train.jl:105
┌ Info: Epoch 1307
└ @ Main /Users/kcft114/.julia/packages/Flux/qXNjB/src/optimise/train.jl:105
┌ Info: Epoch 1308
└ @ Main /Users/kcft114/.julia/packages/Flux/qXNjB/src/optimise/train.jl:105
┌ Info: Epoch 1309
└ @ Main /Users/kcft114/.julia/packages/Flux/qXNjB/src/optimise/train.jl:105
┌ Info: Epoch 1310
└ @ Main /Users/kcft114/.julia/packages/Flux/qXNjB/src/optimise/train.jl:105
┌ Info: Epoch 1311
└ @ Main /Users/kcft114/.julia/packages/Flux/qXNjB/src/optimise/train.jl:105
┌ Info: Epoch 1312
└ @ Main /Users/kcft114/.julia/packages/Flux/qXNjB/src/optimise/train.jl:105
┌ Info: Epoch 1313
└ @ Main /Users/kcft114/.julia/packages/Flux/qXNjB/src/optimise/train.jl:105
┌ Info: Epoch 1314
└ @ Main /Users/kcft114/.julia/packages/Flux/qXNjB/src/optimise/train.jl:105
┌ Info: Epoch 1315
└ @ Main /Users/kcft114/.julia/packages/Flux/qXNjB/src/optimise/train.jl:105
┌ Info: Epoch 1316
└ @ Main /Users/kcft114/.julia/packages/Flux/qXNjB/src/optimise/train.jl:105
┌ Info: Epoch 1317
└ @ Main /Users/kcft114/.julia/packages/Flux/qXNjB/src/optimise/train.jl:105
┌ Info: Epoch 1318
└ @ Main /Users/kcft114/.julia/packages/Flux/qXNjB/src/optimise/train.jl:105
┌ Info: Epoch 1319
└ @ Main /Users/kcft114/.julia/packages/Flux/qXNjB/src/optimise/train.jl:105
┌ Info: Epoch 1320
└ @ Main /Users/kcft114/.julia/packages/Flux/qXNjB/src/optimise/train.jl:105
┌ Info: Epoch 1321
└ @ Main /Users/kcft114/.julia/packages/Flux/qXNjB/src/optimise/train.jl:105
┌ Info: Epoch 1322
└ @ Main /Users/kcft114/.julia/packages/Flux/qXNjB/src/optimise/train.jl:105
┌ Info: Epoch 1323
└ @ Main /Users/kcft114/.julia/packages/Flux/qXNjB/src/optimise/train.jl:105
┌ Info: Epoch 1324
└ @ Main /Users/kcft114/.julia/packages/Flux/qXNjB/src/optimise/train.jl:105
┌ Info: Epoch 1325
└ @ Main /Users/kcft114/.julia/packages/Flux/qXNjB/src/optimise/train.jl:105
┌ Info: Epoch 1326
└ @ Main /Users/kcft114/.julia/packages/Flux/qXNjB/src/optimise/train.jl:105
┌ Info: Epoch 1327
└ @ Main /Users/kcft114/.julia/packages/Flux/qXNjB/src/optimise/train.jl:105
┌ Info: Epoch 1328
└ @ Main /Users/kcft114/.julia/packages/Flux/qXNjB/src/optimise/train.jl:105
┌ Info: Epoch 1329
└ @ Main /Users/kcft114/.julia/packages/Flux/qXNjB/src/optimise/train.jl:105
┌ Info: Epoch 1330
└ @ Main /Users/kcft114/.julia/packages/Flux/qXNjB/src/optimise/train.jl:105
┌ Info: Epoch 1331
└ @ Main /Users/kcft114/.julia/packages/Flux/qXNjB/src/optimise/train.jl:105
┌ Info: Epoch 1332
└ @ Main /Users/kcft114/.julia/packages/Flux/qXNjB/src/optimise/train.jl:105
┌ Info: Epoch 1333
└ @ Main /Users/kcft114/.julia/packages/Flux/qXNjB/src/optimise/train.jl:105
┌ Info: Epoch 1334
└ @ Main /Users/kcft114/.julia/packages/Flux/qXNjB/src/optimise/train.jl:105
┌ Info: Epoch 1335
└ @ Main /Users/kcft114/.julia/packages/Flux/qXNjB/src/optimise/train.jl:105
┌ Info: Epoch 1336
└ @ Main /Users/kcft114/.julia/packages/Flux/qXNjB/src/optimise/train.jl:105
┌ Info: Epoch 1337
└ @ Main /Users/kcft114/.julia/packages/Flux/qXNjB/src/optimise/train.jl:105
┌ Info: Epoch 1338
└ @ Main /Users/kcft114/.julia/packages/Flux/qXNjB/src/optimise/train.jl:105
┌ Info: Epoch 1339
└ @ Main /Users/kcft114/.julia/packages/Flux/qXNjB/src/optimise/train.jl:105
┌ Info: Epoch 1340
└ @ Main /Users/kcft114/.julia/packages/Flux/qXNjB/src/optimise/train.jl:105
┌ Info: Epoch 1341
└ @ Main /Users/kcft114/.julia/packages/Flux/qXNjB/src/optimise/train.jl:105
┌ Info: Epoch 1342
└ @ Main /Users/kcft114/.julia/packages/Flux/qXNjB/src/optimise/train.jl:105
┌ Info: Epoch 1343
└ @ Main /Users/kcft114/.julia/packages/Flux/qXNjB/src/optimise/train.jl:105
┌ Info: Epoch 1344
└ @ Main /Users/kcft114/.julia/packages/Flux/qXNjB/src/optimise/train.jl:105
┌ Info: Epoch 1345
└ @ Main /Users/kcft114/.julia/packages/Flux/qXNjB/src/optimise/train.jl:105
┌ Info: Epoch 1346
└ @ Main /Users/kcft114/.julia/packages/Flux/qXNjB/src/optimise/train.jl:105
┌ Info: Epoch 1347
└ @ Main /Users/kcft114/.julia/packages/Flux/qXNjB/src/optimise/train.jl:105
┌ Info: Epoch 1348
└ @ Main /Users/kcft114/.julia/packages/Flux/qXNjB/src/optimise/train.jl:105
┌ Info: Epoch 1349
└ @ Main /Users/kcft114/.julia/packages/Flux/qXNjB/src/optimise/train.jl:105
┌ Info: Epoch 1350
└ @ Main /Users/kcft114/.julia/packages/Flux/qXNjB/src/optimise/train.jl:105
┌ Info: Epoch 1351
└ @ Main /Users/kcft114/.julia/packages/Flux/qXNjB/src/optimise/train.jl:105
┌ Info: Epoch 1352
└ @ Main /Users/kcft114/.julia/packages/Flux/qXNjB/src/optimise/train.jl:105
┌ Info: Epoch 1353
└ @ Main /Users/kcft114/.julia/packages/Flux/qXNjB/src/optimise/train.jl:105
```

```
loss(X, Y) = 0.5071108f0 (tracked)
loss(X, Y) = 0.53513414f0 (tracked)
loss(X, Y) = 0.5120013f0 (tracked)
loss(X, Y) = 0.54960537f0 (tracked)
loss(X, Y) = 0.57380646f0 (tracked)
loss(X, Y) = 0.55612546f0 (tracked)
loss(X, Y) = 0.5711757f0 (tracked)
loss(X, Y) = 0.53503627f0 (tracked)
loss(X, Y) = 0.5823492f0 (tracked)
loss(X, Y) = 0.5335161f0 (tracked)
loss(X, Y) = 0.5956653f0 (tracked)
loss(X, Y) = 0.48495576f0 (tracked)
loss(X, Y) = 0.5394709f0 (tracked)
loss(X, Y) = 0.55313575f0 (tracked)
loss(X, Y) = 0.5464409f0 (tracked)
loss(X, Y) = 0.54237324f0 (tracked)
loss(X, Y) = 0.5461546f0 (tracked)
loss(X, Y) = 0.57320356f0 (tracked)
loss(X, Y) = 0.5336814f0 (tracked)
loss(X, Y) = 0.52277106f0 (tracked)
loss(X, Y) = 0.5274322f0 (tracked)
loss(X, Y) = 0.5252136f0 (tracked)
loss(X, Y) = 0.4905589f0 (tracked)
loss(X, Y) = 0.55869263f0 (tracked)
loss(X, Y) = 0.51901984f0 (tracked)
loss(X, Y) = 0.536699f0 (tracked)
loss(X, Y) = 0.5108556f0 (tracked)
loss(X, Y) = 0.55109817f0 (tracked)
loss(X, Y) = 0.5748403f0 (tracked)
loss(X, Y) = 0.5757089f0 (tracked)
loss(X, Y) = 0.5872001f0 (tracked)
loss(X, Y) = 0.56122494f0 (tracked)
loss(X, Y) = 0.55382514f0 (tracked)
loss(X, Y) = 0.6023907f0 (tracked)
loss(X, Y) = 0.58412826f0 (tracked)
loss(X, Y) = 0.58538055f0 (tracked)
loss(X, Y) = 0.51892644f0 (tracked)
loss(X, Y) = 0.51450837f0 (tracked)
loss(X, Y) = 0.5596769f0 (tracked)
loss(X, Y) = 0.57302964f0 (tracked)
loss(X, Y) = 0.5542907f0 (tracked)
loss(X, Y) = 0.5808295f0 (tracked)
loss(X, Y) = 0.63025415f0 (tracked)
loss(X, Y) = 0.53202f0 (tracked)
loss(X, Y) = 0.53964245f0 (tracked)
loss(X, Y) = 0.5329038f0 (tracked)
loss(X, Y) = 0.51048964f0 (tracked)
loss(X, Y) = 0.5404319f0 (tracked)
loss(X, Y) = 0.62328464f0 (tracked)
loss(X, Y) = 0.5549265f0 (tracked)
loss(X, Y) = 0.5695018f0 (tracked)
loss(X, Y) = 0.55889344f0 (tracked)
loss(X, Y) = 0.5778562f0 (tracked)
loss(X, Y) = 0.53110623f0 (tracked)
loss(X, Y) = 0.5393358f0 (tracked)
loss(X, Y) = 0.55180985f0 (tracked)
loss(X, Y) = 0.49278107f0 (tracked)
loss(X, Y) = 0.53500646f0 (tracked)
loss(X, Y) = 0.53779453f0 (tracked)
loss(X, Y) = 0.51418495f0 (tracked)
loss(X, Y) = 0.5362015f0 (tracked)
loss(X, Y) = 0.5710364f0 (tracked)
loss(X, Y) = 0.53873914f0 (tracked)
loss(X, Y) = 0.5659302f0 (tracked)
loss(X, Y) = 0.5515142f0 (tracked)
loss(X, Y) = 0.5564014f0 (tracked)
loss(X, Y) = 0.5411782f0 (tracked)
loss(X, Y) = 0.5793301f0 (tracked)
loss(X, Y) = 0.55651265f0 (tracked)
loss(X, Y) = 0.5296885f0 (tracked)
loss(X, Y) = 0.53873444f0 (tracked)
loss(X, Y) = 0.53979707f0 (tracked)
loss(X, Y) = 0.5563566f0 (tracked)
loss(X, Y) = 0.4597606f0 (tracked)
loss(X, Y) = 0.53741807f0 (tracked)
loss(X, Y) = 0.55263406f0 (tracked)
loss(X, Y) = 0.5350498f0 (tracked)
loss(X, Y) = 0.5461972f0 (tracked)
loss(X, Y) = 0.57490563f0 (tracked)
loss(X, Y) = 0.57771146f0 (tracked)
loss(X, Y) = 0.49549708f0 (tracked)
```

```
┌ Info: Epoch 1354
└ @ Main /Users/kcft114/.julia/packages/Flux/qXNjB/src/optimise/train.jl:105
┌ Info: Epoch 1355
└ @ Main /Users/kcft114/.julia/packages/Flux/qXNjB/src/optimise/train.jl:105
┌ Info: Epoch 1356
└ @ Main /Users/kcft114/.julia/packages/Flux/qXNjB/src/optimise/train.jl:105
┌ Info: Epoch 1357
└ @ Main /Users/kcft114/.julia/packages/Flux/qXNjB/src/optimise/train.jl:105
┌ Info: Epoch 1358
└ @ Main /Users/kcft114/.julia/packages/Flux/qXNjB/src/optimise/train.jl:105
┌ Info: Epoch 1359
└ @ Main /Users/kcft114/.julia/packages/Flux/qXNjB/src/optimise/train.jl:105
┌ Info: Epoch 1360
└ @ Main /Users/kcft114/.julia/packages/Flux/qXNjB/src/optimise/train.jl:105
┌ Info: Epoch 1361
└ @ Main /Users/kcft114/.julia/packages/Flux/qXNjB/src/optimise/train.jl:105
┌ Info: Epoch 1362
└ @ Main /Users/kcft114/.julia/packages/Flux/qXNjB/src/optimise/train.jl:105
┌ Info: Epoch 1363
└ @ Main /Users/kcft114/.julia/packages/Flux/qXNjB/src/optimise/train.jl:105
┌ Info: Epoch 1364
└ @ Main /Users/kcft114/.julia/packages/Flux/qXNjB/src/optimise/train.jl:105
┌ Info: Epoch 1365
└ @ Main /Users/kcft114/.julia/packages/Flux/qXNjB/src/optimise/train.jl:105
┌ Info: Epoch 1366
└ @ Main /Users/kcft114/.julia/packages/Flux/qXNjB/src/optimise/train.jl:105
┌ Info: Epoch 1367
└ @ Main /Users/kcft114/.julia/packages/Flux/qXNjB/src/optimise/train.jl:105
┌ Info: Epoch 1368
└ @ Main /Users/kcft114/.julia/packages/Flux/qXNjB/src/optimise/train.jl:105
┌ Info: Epoch 1369
└ @ Main /Users/kcft114/.julia/packages/Flux/qXNjB/src/optimise/train.jl:105
┌ Info: Epoch 1370
└ @ Main /Users/kcft114/.julia/packages/Flux/qXNjB/src/optimise/train.jl:105
┌ Info: Epoch 1371
└ @ Main /Users/kcft114/.julia/packages/Flux/qXNjB/src/optimise/train.jl:105
┌ Info: Epoch 1372
└ @ Main /Users/kcft114/.julia/packages/Flux/qXNjB/src/optimise/train.jl:105
┌ Info: Epoch 1373
└ @ Main /Users/kcft114/.julia/packages/Flux/qXNjB/src/optimise/train.jl:105
┌ Info: Epoch 1374
└ @ Main /Users/kcft114/.julia/packages/Flux/qXNjB/src/optimise/train.jl:105
┌ Info: Epoch 1375
└ @ Main /Users/kcft114/.julia/packages/Flux/qXNjB/src/optimise/train.jl:105
┌ Info: Epoch 1376
└ @ Main /Users/kcft114/.julia/packages/Flux/qXNjB/src/optimise/train.jl:105
┌ Info: Epoch 1377
└ @ Main /Users/kcft114/.julia/packages/Flux/qXNjB/src/optimise/train.jl:105
┌ Info: Epoch 1378
└ @ Main /Users/kcft114/.julia/packages/Flux/qXNjB/src/optimise/train.jl:105
┌ Info: Epoch 1379
└ @ Main /Users/kcft114/.julia/packages/Flux/qXNjB/src/optimise/train.jl:105
┌ Info: Epoch 1380
└ @ Main /Users/kcft114/.julia/packages/Flux/qXNjB/src/optimise/train.jl:105
┌ Info: Epoch 1381
└ @ Main /Users/kcft114/.julia/packages/Flux/qXNjB/src/optimise/train.jl:105
┌ Info: Epoch 1382
└ @ Main /Users/kcft114/.julia/packages/Flux/qXNjB/src/optimise/train.jl:105
┌ Info: Epoch 1383
└ @ Main /Users/kcft114/.julia/packages/Flux/qXNjB/src/optimise/train.jl:105
┌ Info: Epoch 1384
└ @ Main /Users/kcft114/.julia/packages/Flux/qXNjB/src/optimise/train.jl:105
┌ Info: Epoch 1385
└ @ Main /Users/kcft114/.julia/packages/Flux/qXNjB/src/optimise/train.jl:105
┌ Info: Epoch 1386
└ @ Main /Users/kcft114/.julia/packages/Flux/qXNjB/src/optimise/train.jl:105
┌ Info: Epoch 1387
└ @ Main /Users/kcft114/.julia/packages/Flux/qXNjB/src/optimise/train.jl:105
┌ Info: Epoch 1388
└ @ Main /Users/kcft114/.julia/packages/Flux/qXNjB/src/optimise/train.jl:105
┌ Info: Epoch 1389
└ @ Main /Users/kcft114/.julia/packages/Flux/qXNjB/src/optimise/train.jl:105
┌ Info: Epoch 1390
└ @ Main /Users/kcft114/.julia/packages/Flux/qXNjB/src/optimise/train.jl:105
┌ Info: Epoch 1391
└ @ Main /Users/kcft114/.julia/packages/Flux/qXNjB/src/optimise/train.jl:105
┌ Info: Epoch 1392
└ @ Main /Users/kcft114/.julia/packages/Flux/qXNjB/src/optimise/train.jl:105
┌ Info: Epoch 1393
└ @ Main /Users/kcft114/.julia/packages/Flux/qXNjB/src/optimise/train.jl:105
┌ Info: Epoch 1394
└ @ Main /Users/kcft114/.julia/packages/Flux/qXNjB/src/optimise/train.jl:105
┌ Info: Epoch 1395
└ @ Main /Users/kcft114/.julia/packages/Flux/qXNjB/src/optimise/train.jl:105
┌ Info: Epoch 1396
└ @ Main /Users/kcft114/.julia/packages/Flux/qXNjB/src/optimise/train.jl:105
┌ Info: Epoch 1397
└ @ Main /Users/kcft114/.julia/packages/Flux/qXNjB/src/optimise/train.jl:105
┌ Info: Epoch 1398
└ @ Main /Users/kcft114/.julia/packages/Flux/qXNjB/src/optimise/train.jl:105
┌ Info: Epoch 1399
└ @ Main /Users/kcft114/.julia/packages/Flux/qXNjB/src/optimise/train.jl:105
┌ Info: Epoch 1400
└ @ Main /Users/kcft114/.julia/packages/Flux/qXNjB/src/optimise/train.jl:105
┌ Info: Epoch 1401
└ @ Main /Users/kcft114/.julia/packages/Flux/qXNjB/src/optimise/train.jl:105
┌ Info: Epoch 1402
└ @ Main /Users/kcft114/.julia/packages/Flux/qXNjB/src/optimise/train.jl:105
┌ Info: Epoch 1403
└ @ Main /Users/kcft114/.julia/packages/Flux/qXNjB/src/optimise/train.jl:105
┌ Info: Epoch 1404
└ @ Main /Users/kcft114/.julia/packages/Flux/qXNjB/src/optimise/train.jl:105
┌ Info: Epoch 1405
└ @ Main /Users/kcft114/.julia/packages/Flux/qXNjB/src/optimise/train.jl:105
┌ Info: Epoch 1406
└ @ Main /Users/kcft114/.julia/packages/Flux/qXNjB/src/optimise/train.jl:105
┌ Info: Epoch 1407
└ @ Main /Users/kcft114/.julia/packages/Flux/qXNjB/src/optimise/train.jl:105
┌ Info: Epoch 1408
└ @ Main /Users/kcft114/.julia/packages/Flux/qXNjB/src/optimise/train.jl:105
┌ Info: Epoch 1409
└ @ Main /Users/kcft114/.julia/packages/Flux/qXNjB/src/optimise/train.jl:105
┌ Info: Epoch 1410
└ @ Main /Users/kcft114/.julia/packages/Flux/qXNjB/src/optimise/train.jl:105
┌ Info: Epoch 1411
└ @ Main /Users/kcft114/.julia/packages/Flux/qXNjB/src/optimise/train.jl:105
┌ Info: Epoch 1412
└ @ Main /Users/kcft114/.julia/packages/Flux/qXNjB/src/optimise/train.jl:105
┌ Info: Epoch 1413
└ @ Main /Users/kcft114/.julia/packages/Flux/qXNjB/src/optimise/train.jl:105
┌ Info: Epoch 1414
└ @ Main /Users/kcft114/.julia/packages/Flux/qXNjB/src/optimise/train.jl:105
┌ Info: Epoch 1415
└ @ Main /Users/kcft114/.julia/packages/Flux/qXNjB/src/optimise/train.jl:105
┌ Info: Epoch 1416
└ @ Main /Users/kcft114/.julia/packages/Flux/qXNjB/src/optimise/train.jl:105
┌ Info: Epoch 1417
└ @ Main /Users/kcft114/.julia/packages/Flux/qXNjB/src/optimise/train.jl:105
┌ Info: Epoch 1418
└ @ Main /Users/kcft114/.julia/packages/Flux/qXNjB/src/optimise/train.jl:105
┌ Info: Epoch 1419
└ @ Main /Users/kcft114/.julia/packages/Flux/qXNjB/src/optimise/train.jl:105
┌ Info: Epoch 1420
└ @ Main /Users/kcft114/.julia/packages/Flux/qXNjB/src/optimise/train.jl:105
┌ Info: Epoch 1421
└ @ Main /Users/kcft114/.julia/packages/Flux/qXNjB/src/optimise/train.jl:105
┌ Info: Epoch 1422
└ @ Main /Users/kcft114/.julia/packages/Flux/qXNjB/src/optimise/train.jl:105
┌ Info: Epoch 1423
└ @ Main /Users/kcft114/.julia/packages/Flux/qXNjB/src/optimise/train.jl:105
┌ Info: Epoch 1424
└ @ Main /Users/kcft114/.julia/packages/Flux/qXNjB/src/optimise/train.jl:105
┌ Info: Epoch 1425
└ @ Main /Users/kcft114/.julia/packages/Flux/qXNjB/src/optimise/train.jl:105
┌ Info: Epoch 1426
└ @ Main /Users/kcft114/.julia/packages/Flux/qXNjB/src/optimise/train.jl:105
┌ Info: Epoch 1427
└ @ Main /Users/kcft114/.julia/packages/Flux/qXNjB/src/optimise/train.jl:105
┌ Info: Epoch 1428
└ @ Main /Users/kcft114/.julia/packages/Flux/qXNjB/src/optimise/train.jl:105
┌ Info: Epoch 1429
└ @ Main /Users/kcft114/.julia/packages/Flux/qXNjB/src/optimise/train.jl:105
┌ Info: Epoch 1430
└ @ Main /Users/kcft114/.julia/packages/Flux/qXNjB/src/optimise/train.jl:105
┌ Info: Epoch 1431
└ @ Main /Users/kcft114/.julia/packages/Flux/qXNjB/src/optimise/train.jl:105
┌ Info: Epoch 1432
└ @ Main /Users/kcft114/.julia/packages/Flux/qXNjB/src/optimise/train.jl:105
┌ Info: Epoch 1433
└ @ Main /Users/kcft114/.julia/packages/Flux/qXNjB/src/optimise/train.jl:105
┌ Info: Epoch 1434
└ @ Main /Users/kcft114/.julia/packages/Flux/qXNjB/src/optimise/train.jl:105
```

```
loss(X, Y) = 0.5380105f0 (tracked)
loss(X, Y) = 0.6314107f0 (tracked)
loss(X, Y) = 0.57572037f0 (tracked)
loss(X, Y) = 0.5504293f0 (tracked)
loss(X, Y) = 0.5567913f0 (tracked)
loss(X, Y) = 0.52051216f0 (tracked)
loss(X, Y) = 0.56858927f0 (tracked)
loss(X, Y) = 0.5368006f0 (tracked)
loss(X, Y) = 0.5504528f0 (tracked)
loss(X, Y) = 0.5286815f0 (tracked)
loss(X, Y) = 0.56199014f0 (tracked)
loss(X, Y) = 0.5438944f0 (tracked)
loss(X, Y) = 0.53630155f0 (tracked)
loss(X, Y) = 0.5682803f0 (tracked)
loss(X, Y) = 0.534137f0 (tracked)
loss(X, Y) = 0.52725285f0 (tracked)
loss(X, Y) = 0.549938f0 (tracked)
loss(X, Y) = 0.5235332f0 (tracked)
loss(X, Y) = 0.5377496f0 (tracked)
loss(X, Y) = 0.52816886f0 (tracked)
loss(X, Y) = 0.48857433f0 (tracked)
loss(X, Y) = 0.5574283f0 (tracked)
loss(X, Y) = 0.51514524f0 (tracked)
loss(X, Y) = 0.5332137f0 (tracked)
loss(X, Y) = 0.54805106f0 (tracked)
loss(X, Y) = 0.5455563f0 (tracked)
loss(X, Y) = 0.53372645f0 (tracked)
loss(X, Y) = 0.6345957f0 (tracked)
loss(X, Y) = 0.5626086f0 (tracked)
loss(X, Y) = 0.5035579f0 (tracked)
loss(X, Y) = 0.5286125f0 (tracked)
loss(X, Y) = 0.5181349f0 (tracked)
loss(X, Y) = 0.54330295f0 (tracked)
loss(X, Y) = 0.51897f0 (tracked)
loss(X, Y) = 0.52076536f0 (tracked)
loss(X, Y) = 0.55760103f0 (tracked)
loss(X, Y) = 0.5706565f0 (tracked)
loss(X, Y) = 0.5718991f0 (tracked)
loss(X, Y) = 0.55608636f0 (tracked)
loss(X, Y) = 0.53635055f0 (tracked)
loss(X, Y) = 0.50766367f0 (tracked)
loss(X, Y) = 0.5040716f0 (tracked)
loss(X, Y) = 0.57823646f0 (tracked)
loss(X, Y) = 0.51362133f0 (tracked)
loss(X, Y) = 0.52766335f0 (tracked)
loss(X, Y) = 0.5446242f0 (tracked)
loss(X, Y) = 0.5185816f0 (tracked)
loss(X, Y) = 0.55594987f0 (tracked)
loss(X, Y) = 0.5373677f0 (tracked)
loss(X, Y) = 0.5338954f0 (tracked)
loss(X, Y) = 0.52537805f0 (tracked)
loss(X, Y) = 0.5377366f0 (tracked)
loss(X, Y) = 0.5449623f0 (tracked)
loss(X, Y) = 0.51434016f0 (tracked)
loss(X, Y) = 0.5048291f0 (tracked)
loss(X, Y) = 0.52851844f0 (tracked)
loss(X, Y) = 0.57154953f0 (tracked)
loss(X, Y) = 0.5497488f0 (tracked)
loss(X, Y) = 0.54554105f0 (tracked)
loss(X, Y) = 0.5140789f0 (tracked)
loss(X, Y) = 0.511926f0 (tracked)
loss(X, Y) = 0.5332309f0 (tracked)
loss(X, Y) = 0.5438466f0 (tracked)
loss(X, Y) = 0.54411805f0 (tracked)
loss(X, Y) = 0.5431729f0 (tracked)
loss(X, Y) = 0.54925764f0 (tracked)
loss(X, Y) = 0.49463192f0 (tracked)
loss(X, Y) = 0.5313759f0 (tracked)
loss(X, Y) = 0.54966754f0 (tracked)
loss(X, Y) = 0.5273304f0 (tracked)
loss(X, Y) = 0.5795621f0 (tracked)
loss(X, Y) = 0.5623487f0 (tracked)
loss(X, Y) = 0.5332595f0 (tracked)
loss(X, Y) = 0.53480333f0 (tracked)
loss(X, Y) = 0.52887684f0 (tracked)
loss(X, Y) = 0.50378394f0 (tracked)
loss(X, Y) = 0.5321934f0 (tracked)
loss(X, Y) = 0.6150602f0 (tracked)
loss(X, Y) = 0.5705099f0 (tracked)
loss(X, Y) = 0.51797634f0 (tracked)
loss(X, Y) = 0.53419936f0 (tracked)
loss(X, Y) = 0.5325904f0 (tracked)
loss(X, Y) = 0.55569977f0 (tracked)
loss(X, Y) = 0.48352954f0 (tracked)
loss(X, Y) = 0.5289133f0 (tracked)
loss(X, Y) = 0.56774944f0 (tracked)
loss(X, Y) = 0.6065749f0 (tracked)
loss(X, Y) = 0.5192651f0 (tracked)
loss(X, Y) = 0.55745786f0 (tracked)
loss(X, Y) = 0.5504351f0 (tracked)
loss(X, Y) = 0.4849587f0 (tracked)
loss(X, Y) = 0.51211137f0 (tracked)
```

```
┌ Info: Epoch 1435
└ @ Main /Users/kcft114/.julia/packages/Flux/qXNjB/src/optimise/train.jl:105
┌ Info: Epoch 1436
└ @ Main /Users/kcft114/.julia/packages/Flux/qXNjB/src/optimise/train.jl:105
┌ Info: Epoch 1437
└ @ Main /Users/kcft114/.julia/packages/Flux/qXNjB/src/optimise/train.jl:105
┌ Info: Epoch 1438
└ @ Main /Users/kcft114/.julia/packages/Flux/qXNjB/src/optimise/train.jl:105
┌ Info: Epoch 1439
└ @ Main /Users/kcft114/.julia/packages/Flux/qXNjB/src/optimise/train.jl:105
┌ Info: Epoch 1440
└ @ Main /Users/kcft114/.julia/packages/Flux/qXNjB/src/optimise/train.jl:105
┌ Info: Epoch 1441
└ @ Main /Users/kcft114/.julia/packages/Flux/qXNjB/src/optimise/train.jl:105
┌ Info: Epoch 1442
└ @ Main /Users/kcft114/.julia/packages/Flux/qXNjB/src/optimise/train.jl:105
┌ Info: Epoch 1443
└ @ Main /Users/kcft114/.julia/packages/Flux/qXNjB/src/optimise/train.jl:105
┌ Info: Epoch 1444
└ @ Main /Users/kcft114/.julia/packages/Flux/qXNjB/src/optimise/train.jl:105
┌ Info: Epoch 1445
└ @ Main /Users/kcft114/.julia/packages/Flux/qXNjB/src/optimise/train.jl:105
┌ Info: Epoch 1446
└ @ Main /Users/kcft114/.julia/packages/Flux/qXNjB/src/optimise/train.jl:105
┌ Info: Epoch 1447
└ @ Main /Users/kcft114/.julia/packages/Flux/qXNjB/src/optimise/train.jl:105
┌ Info: Epoch 1448
└ @ Main /Users/kcft114/.julia/packages/Flux/qXNjB/src/optimise/train.jl:105
┌ Info: Epoch 1449
└ @ Main /Users/kcft114/.julia/packages/Flux/qXNjB/src/optimise/train.jl:105
┌ Info: Epoch 1450
└ @ Main /Users/kcft114/.julia/packages/Flux/qXNjB/src/optimise/train.jl:105
┌ Info: Epoch 1451
└ @ Main /Users/kcft114/.julia/packages/Flux/qXNjB/src/optimise/train.jl:105
┌ Info: Epoch 1452
└ @ Main /Users/kcft114/.julia/packages/Flux/qXNjB/src/optimise/train.jl:105
┌ Info: Epoch 1453
└ @ Main /Users/kcft114/.julia/packages/Flux/qXNjB/src/optimise/train.jl:105
┌ Info: Epoch 1454
└ @ Main /Users/kcft114/.julia/packages/Flux/qXNjB/src/optimise/train.jl:105
┌ Info: Epoch 1455
└ @ Main /Users/kcft114/.julia/packages/Flux/qXNjB/src/optimise/train.jl:105
┌ Info: Epoch 1456
└ @ Main /Users/kcft114/.julia/packages/Flux/qXNjB/src/optimise/train.jl:105
┌ Info: Epoch 1457
└ @ Main /Users/kcft114/.julia/packages/Flux/qXNjB/src/optimise/train.jl:105
┌ Info: Epoch 1458
└ @ Main /Users/kcft114/.julia/packages/Flux/qXNjB/src/optimise/train.jl:105
┌ Info: Epoch 1459
└ @ Main /Users/kcft114/.julia/packages/Flux/qXNjB/src/optimise/train.jl:105
┌ Info: Epoch 1460
└ @ Main /Users/kcft114/.julia/packages/Flux/qXNjB/src/optimise/train.jl:105
┌ Info: Epoch 1461
└ @ Main /Users/kcft114/.julia/packages/Flux/qXNjB/src/optimise/train.jl:105
┌ Info: Epoch 1462
└ @ Main /Users/kcft114/.julia/packages/Flux/qXNjB/src/optimise/train.jl:105
┌ Info: Epoch 1463
└ @ Main /Users/kcft114/.julia/packages/Flux/qXNjB/src/optimise/train.jl:105
┌ Info: Epoch 1464
└ @ Main /Users/kcft114/.julia/packages/Flux/qXNjB/src/optimise/train.jl:105
┌ Info: Epoch 1465
└ @ Main /Users/kcft114/.julia/packages/Flux/qXNjB/src/optimise/train.jl:105
┌ Info: Epoch 1466
└ @ Main /Users/kcft114/.julia/packages/Flux/qXNjB/src/optimise/train.jl:105
┌ Info: Epoch 1467
└ @ Main /Users/kcft114/.julia/packages/Flux/qXNjB/src/optimise/train.jl:105
┌ Info: Epoch 1468
└ @ Main /Users/kcft114/.julia/packages/Flux/qXNjB/src/optimise/train.jl:105
┌ Info: Epoch 1469
└ @ Main /Users/kcft114/.julia/packages/Flux/qXNjB/src/optimise/train.jl:105
┌ Info: Epoch 1470
└ @ Main /Users/kcft114/.julia/packages/Flux/qXNjB/src/optimise/train.jl:105
┌ Info: Epoch 1471
└ @ Main /Users/kcft114/.julia/packages/Flux/qXNjB/src/optimise/train.jl:105
┌ Info: Epoch 1472
└ @ Main /Users/kcft114/.julia/packages/Flux/qXNjB/src/optimise/train.jl:105
┌ Info: Epoch 1473
└ @ Main /Users/kcft114/.julia/packages/Flux/qXNjB/src/optimise/train.jl:105
┌ Info: Epoch 1474
└ @ Main /Users/kcft114/.julia/packages/Flux/qXNjB/src/optimise/train.jl:105
┌ Info: Epoch 1475
└ @ Main /Users/kcft114/.julia/packages/Flux/qXNjB/src/optimise/train.jl:105
┌ Info: Epoch 1476
└ @ Main /Users/kcft114/.julia/packages/Flux/qXNjB/src/optimise/train.jl:105
┌ Info: Epoch 1477
└ @ Main /Users/kcft114/.julia/packages/Flux/qXNjB/src/optimise/train.jl:105
┌ Info: Epoch 1478
└ @ Main /Users/kcft114/.julia/packages/Flux/qXNjB/src/optimise/train.jl:105
┌ Info: Epoch 1479
└ @ Main /Users/kcft114/.julia/packages/Flux/qXNjB/src/optimise/train.jl:105
┌ Info: Epoch 1480
└ @ Main /Users/kcft114/.julia/packages/Flux/qXNjB/src/optimise/train.jl:105
┌ Info: Epoch 1481
└ @ Main /Users/kcft114/.julia/packages/Flux/qXNjB/src/optimise/train.jl:105
┌ Info: Epoch 1482
└ @ Main /Users/kcft114/.julia/packages/Flux/qXNjB/src/optimise/train.jl:105
┌ Info: Epoch 1483
└ @ Main /Users/kcft114/.julia/packages/Flux/qXNjB/src/optimise/train.jl:105
┌ Info: Epoch 1484
└ @ Main /Users/kcft114/.julia/packages/Flux/qXNjB/src/optimise/train.jl:105
┌ Info: Epoch 1485
└ @ Main /Users/kcft114/.julia/packages/Flux/qXNjB/src/optimise/train.jl:105
┌ Info: Epoch 1486
└ @ Main /Users/kcft114/.julia/packages/Flux/qXNjB/src/optimise/train.jl:105
┌ Info: Epoch 1487
└ @ Main /Users/kcft114/.julia/packages/Flux/qXNjB/src/optimise/train.jl:105
┌ Info: Epoch 1488
└ @ Main /Users/kcft114/.julia/packages/Flux/qXNjB/src/optimise/train.jl:105
┌ Info: Epoch 1489
└ @ Main /Users/kcft114/.julia/packages/Flux/qXNjB/src/optimise/train.jl:105
┌ Info: Epoch 1490
└ @ Main /Users/kcft114/.julia/packages/Flux/qXNjB/src/optimise/train.jl:105
┌ Info: Epoch 1491
└ @ Main /Users/kcft114/.julia/packages/Flux/qXNjB/src/optimise/train.jl:105
┌ Info: Epoch 1492
└ @ Main /Users/kcft114/.julia/packages/Flux/qXNjB/src/optimise/train.jl:105
┌ Info: Epoch 1493
└ @ Main /Users/kcft114/.julia/packages/Flux/qXNjB/src/optimise/train.jl:105
┌ Info: Epoch 1494
└ @ Main /Users/kcft114/.julia/packages/Flux/qXNjB/src/optimise/train.jl:105
┌ Info: Epoch 1495
└ @ Main /Users/kcft114/.julia/packages/Flux/qXNjB/src/optimise/train.jl:105
┌ Info: Epoch 1496
└ @ Main /Users/kcft114/.julia/packages/Flux/qXNjB/src/optimise/train.jl:105
┌ Info: Epoch 1497
└ @ Main /Users/kcft114/.julia/packages/Flux/qXNjB/src/optimise/train.jl:105
┌ Info: Epoch 1498
└ @ Main /Users/kcft114/.julia/packages/Flux/qXNjB/src/optimise/train.jl:105
┌ Info: Epoch 1499
└ @ Main /Users/kcft114/.julia/packages/Flux/qXNjB/src/optimise/train.jl:105
┌ Info: Epoch 1500
└ @ Main /Users/kcft114/.julia/packages/Flux/qXNjB/src/optimise/train.jl:105
┌ Info: Epoch 1501
└ @ Main /Users/kcft114/.julia/packages/Flux/qXNjB/src/optimise/train.jl:105
┌ Info: Epoch 1502
└ @ Main /Users/kcft114/.julia/packages/Flux/qXNjB/src/optimise/train.jl:105
┌ Info: Epoch 1503
└ @ Main /Users/kcft114/.julia/packages/Flux/qXNjB/src/optimise/train.jl:105
┌ Info: Epoch 1504
└ @ Main /Users/kcft114/.julia/packages/Flux/qXNjB/src/optimise/train.jl:105
┌ Info: Epoch 1505
└ @ Main /Users/kcft114/.julia/packages/Flux/qXNjB/src/optimise/train.jl:105
┌ Info: Epoch 1506
└ @ Main /Users/kcft114/.julia/packages/Flux/qXNjB/src/optimise/train.jl:105
┌ Info: Epoch 1507
└ @ Main /Users/kcft114/.julia/packages/Flux/qXNjB/src/optimise/train.jl:105
┌ Info: Epoch 1508
└ @ Main /Users/kcft114/.julia/packages/Flux/qXNjB/src/optimise/train.jl:105
┌ Info: Epoch 1509
└ @ Main /Users/kcft114/.julia/packages/Flux/qXNjB/src/optimise/train.jl:105
┌ Info: Epoch 1510
└ @ Main /Users/kcft114/.julia/packages/Flux/qXNjB/src/optimise/train.jl:105
┌ Info: Epoch 1511
└ @ Main /Users/kcft114/.julia/packages/Flux/qXNjB/src/optimise/train.jl:105
┌ Info: Epoch 1512
└ @ Main /Users/kcft114/.julia/packages/Flux/qXNjB/src/optimise/train.jl:105
┌ Info: Epoch 1513
└ @ Main /Users/kcft114/.julia/packages/Flux/qXNjB/src/optimise/train.jl:105
┌ Info: Epoch 1514
└ @ Main /Users/kcft114/.julia/packages/Flux/qXNjB/src/optimise/train.jl:105
┌ Info: Epoch 1515
└ @ Main /Users/kcft114/.julia/packages/Flux/qXNjB/src/optimise/train.jl:105
┌ Info: Epoch 1516
└ @ Main /Users/kcft114/.julia/packages/Flux/qXNjB/src/optimise/train.jl:105
┌ Info: Epoch 1517
└ @ Main /Users/kcft114/.julia/packages/Flux/qXNjB/src/optimise/train.jl:105
┌ Info: Epoch 1518
└ @ Main /Users/kcft114/.julia/packages/Flux/qXNjB/src/optimise/train.jl:105
┌ Info: Epoch 1519
└ @ Main /Users/kcft114/.julia/packages/Flux/qXNjB/src/optimise/train.jl:105
┌ Info: Epoch 1520
└ @ Main /Users/kcft114/.julia/packages/Flux/qXNjB/src/optimise/train.jl:105
┌ Info: Epoch 1521
└ @ Main /Users/kcft114/.julia/packages/Flux/qXNjB/src/optimise/train.jl:105
┌ Info: Epoch 1522
└ @ Main /Users/kcft114/.julia/packages/Flux/qXNjB/src/optimise/train.jl:105
┌ Info: Epoch 1523
└ @ Main /Users/kcft114/.julia/packages/Flux/qXNjB/src/optimise/train.jl:105
┌ Info: Epoch 1524
└ @ Main /Users/kcft114/.julia/packages/Flux/qXNjB/src/optimise/train.jl:105
┌ Info: Epoch 1525
└ @ Main /Users/kcft114/.julia/packages/Flux/qXNjB/src/optimise/train.jl:105
┌ Info: Epoch 1526
└ @ Main /Users/kcft114/.julia/packages/Flux/qXNjB/src/optimise/train.jl:105
```

```
loss(X, Y) = 0.57188565f0 (tracked)
loss(X, Y) = 0.5627993f0 (tracked)
loss(X, Y) = 0.5632474f0 (tracked)
loss(X, Y) = 0.5777824f0 (tracked)
loss(X, Y) = 0.55181295f0 (tracked)
loss(X, Y) = 0.5307853f0 (tracked)
loss(X, Y) = 0.5306164f0 (tracked)
loss(X, Y) = 0.5417793f0 (tracked)
loss(X, Y) = 0.5580903f0 (tracked)
loss(X, Y) = 0.5075636f0 (tracked)
loss(X, Y) = 0.6101051f0 (tracked)
loss(X, Y) = 0.5477189f0 (tracked)
loss(X, Y) = 0.49464887f0 (tracked)
loss(X, Y) = 0.56202465f0 (tracked)
loss(X, Y) = 0.52455175f0 (tracked)
loss(X, Y) = 0.54025f0 (tracked)
loss(X, Y) = 0.5217644f0 (tracked)
loss(X, Y) = 0.53092027f0 (tracked)
loss(X, Y) = 0.50512815f0 (tracked)
loss(X, Y) = 0.5684621f0 (tracked)
loss(X, Y) = 0.5368554f0 (tracked)
loss(X, Y) = 0.4971247f0 (tracked)
loss(X, Y) = 0.52564514f0 (tracked)
loss(X, Y) = 0.48273125f0 (tracked)
loss(X, Y) = 0.5068711f0 (tracked)
loss(X, Y) = 0.53662765f0 (tracked)
loss(X, Y) = 0.5892467f0 (tracked)
loss(X, Y) = 0.55336773f0 (tracked)
loss(X, Y) = 0.49778917f0 (tracked)
loss(X, Y) = 0.5259295f0 (tracked)
loss(X, Y) = 0.54024863f0 (tracked)
loss(X, Y) = 0.5186846f0 (tracked)
loss(X, Y) = 0.5435552f0 (tracked)
loss(X, Y) = 0.52307224f0 (tracked)
loss(X, Y) = 0.52501893f0 (tracked)
loss(X, Y) = 0.55664015f0 (tracked)
loss(X, Y) = 0.53821087f0 (tracked)
loss(X, Y) = 0.5015536f0 (tracked)
loss(X, Y) = 0.5410772f0 (tracked)
loss(X, Y) = 0.5771706f0 (tracked)
loss(X, Y) = 0.54060227f0 (tracked)
loss(X, Y) = 0.5128765f0 (tracked)
loss(X, Y) = 0.5559201f0 (tracked)
loss(X, Y) = 0.53454906f0 (tracked)
loss(X, Y) = 0.5791253f0 (tracked)
loss(X, Y) = 0.511367f0 (tracked)
loss(X, Y) = 0.5085699f0 (tracked)
loss(X, Y) = 0.54676604f0 (tracked)
loss(X, Y) = 0.5441634f0 (tracked)
loss(X, Y) = 0.49928272f0 (tracked)
loss(X, Y) = 0.51287705f0 (tracked)
loss(X, Y) = 0.57380706f0 (tracked)
loss(X, Y) = 0.54418874f0 (tracked)
loss(X, Y) = 0.49628583f0 (tracked)
loss(X, Y) = 0.5309781f0 (tracked)
loss(X, Y) = 0.5531082f0 (tracked)
loss(X, Y) = 0.5665225f0 (tracked)
loss(X, Y) = 0.5047099f0 (tracked)
loss(X, Y) = 0.53752106f0 (tracked)
loss(X, Y) = 0.55475515f0 (tracked)
loss(X, Y) = 0.58147013f0 (tracked)
loss(X, Y) = 0.573221f0 (tracked)
loss(X, Y) = 0.5575017f0 (tracked)
loss(X, Y) = 0.53867775f0 (tracked)
loss(X, Y) = 0.57304543f0 (tracked)
loss(X, Y) = 0.5563476f0 (tracked)
loss(X, Y) = 0.53232104f0 (tracked)
loss(X, Y) = 0.57081294f0 (tracked)
loss(X, Y) = 0.49772668f0 (tracked)
loss(X, Y) = 0.49328354f0 (tracked)
loss(X, Y) = 0.52071816f0 (tracked)
loss(X, Y) = 0.48064774f0 (tracked)
loss(X, Y) = 0.510426f0 (tracked)
loss(X, Y) = 0.5296913f0 (tracked)
loss(X, Y) = 0.5489964f0 (tracked)
loss(X, Y) = 0.49926108f0 (tracked)
loss(X, Y) = 0.555664f0 (tracked)
loss(X, Y) = 0.5343846f0 (tracked)
loss(X, Y) = 0.49615026f0 (tracked)
loss(X, Y) = 0.54347813f0 (tracked)
loss(X, Y) = 0.53057694f0 (tracked)
loss(X, Y) = 0.54910785f0 (tracked)
loss(X, Y) = 0.4848283f0 (tracked)
loss(X, Y) = 0.5035509f0 (tracked)
loss(X, Y) = 0.52994156f0 (tracked)
loss(X, Y) = 0.5360849f0 (tracked)
loss(X, Y) = 0.5786289f0 (tracked)
loss(X, Y) = 0.48936424f0 (tracked)
loss(X, Y) = 0.50473875f0 (tracked)
loss(X, Y) = 0.5135057f0 (tracked)
loss(X, Y) = 0.5397783f0 (tracked)
loss(X, Y) = 0.5164984f0 (tracked)
loss(X, Y) = 0.52113146f0 (tracked)
loss(X, Y) = 0.5336631f0 (tracked)
```

```
┌ Info: Epoch 1527
└ @ Main /Users/kcft114/.julia/packages/Flux/qXNjB/src/optimise/train.jl:105
┌ Info: Epoch 1528
└ @ Main /Users/kcft114/.julia/packages/Flux/qXNjB/src/optimise/train.jl:105
┌ Info: Epoch 1529
└ @ Main /Users/kcft114/.julia/packages/Flux/qXNjB/src/optimise/train.jl:105
┌ Info: Epoch 1530
└ @ Main /Users/kcft114/.julia/packages/Flux/qXNjB/src/optimise/train.jl:105
┌ Info: Epoch 1531
└ @ Main /Users/kcft114/.julia/packages/Flux/qXNjB/src/optimise/train.jl:105
┌ Info: Epoch 1532
└ @ Main /Users/kcft114/.julia/packages/Flux/qXNjB/src/optimise/train.jl:105
┌ Info: Epoch 1533
└ @ Main /Users/kcft114/.julia/packages/Flux/qXNjB/src/optimise/train.jl:105
┌ Info: Epoch 1534
└ @ Main /Users/kcft114/.julia/packages/Flux/qXNjB/src/optimise/train.jl:105
┌ Info: Epoch 1535
└ @ Main /Users/kcft114/.julia/packages/Flux/qXNjB/src/optimise/train.jl:105
┌ Info: Epoch 1536
└ @ Main /Users/kcft114/.julia/packages/Flux/qXNjB/src/optimise/train.jl:105
┌ Info: Epoch 1537
└ @ Main /Users/kcft114/.julia/packages/Flux/qXNjB/src/optimise/train.jl:105
┌ Info: Epoch 1538
└ @ Main /Users/kcft114/.julia/packages/Flux/qXNjB/src/optimise/train.jl:105
┌ Info: Epoch 1539
└ @ Main /Users/kcft114/.julia/packages/Flux/qXNjB/src/optimise/train.jl:105
┌ Info: Epoch 1540
└ @ Main /Users/kcft114/.julia/packages/Flux/qXNjB/src/optimise/train.jl:105
┌ Info: Epoch 1541
└ @ Main /Users/kcft114/.julia/packages/Flux/qXNjB/src/optimise/train.jl:105
┌ Info: Epoch 1542
└ @ Main /Users/kcft114/.julia/packages/Flux/qXNjB/src/optimise/train.jl:105
┌ Info: Epoch 1543
└ @ Main /Users/kcft114/.julia/packages/Flux/qXNjB/src/optimise/train.jl:105
┌ Info: Epoch 1544
└ @ Main /Users/kcft114/.julia/packages/Flux/qXNjB/src/optimise/train.jl:105
┌ Info: Epoch 1545
└ @ Main /Users/kcft114/.julia/packages/Flux/qXNjB/src/optimise/train.jl:105
┌ Info: Epoch 1546
└ @ Main /Users/kcft114/.julia/packages/Flux/qXNjB/src/optimise/train.jl:105
┌ Info: Epoch 1547
└ @ Main /Users/kcft114/.julia/packages/Flux/qXNjB/src/optimise/train.jl:105
┌ Info: Epoch 1548
└ @ Main /Users/kcft114/.julia/packages/Flux/qXNjB/src/optimise/train.jl:105
┌ Info: Epoch 1549
└ @ Main /Users/kcft114/.julia/packages/Flux/qXNjB/src/optimise/train.jl:105
┌ Info: Epoch 1550
└ @ Main /Users/kcft114/.julia/packages/Flux/qXNjB/src/optimise/train.jl:105
┌ Info: Epoch 1551
└ @ Main /Users/kcft114/.julia/packages/Flux/qXNjB/src/optimise/train.jl:105
┌ Info: Epoch 1552
└ @ Main /Users/kcft114/.julia/packages/Flux/qXNjB/src/optimise/train.jl:105
┌ Info: Epoch 1553
└ @ Main /Users/kcft114/.julia/packages/Flux/qXNjB/src/optimise/train.jl:105
┌ Info: Epoch 1554
└ @ Main /Users/kcft114/.julia/packages/Flux/qXNjB/src/optimise/train.jl:105
┌ Info: Epoch 1555
└ @ Main /Users/kcft114/.julia/packages/Flux/qXNjB/src/optimise/train.jl:105
┌ Info: Epoch 1556
└ @ Main /Users/kcft114/.julia/packages/Flux/qXNjB/src/optimise/train.jl:105
┌ Info: Epoch 1557
└ @ Main /Users/kcft114/.julia/packages/Flux/qXNjB/src/optimise/train.jl:105
┌ Info: Epoch 1558
└ @ Main /Users/kcft114/.julia/packages/Flux/qXNjB/src/optimise/train.jl:105
┌ Info: Epoch 1559
└ @ Main /Users/kcft114/.julia/packages/Flux/qXNjB/src/optimise/train.jl:105
┌ Info: Epoch 1560
└ @ Main /Users/kcft114/.julia/packages/Flux/qXNjB/src/optimise/train.jl:105
┌ Info: Epoch 1561
└ @ Main /Users/kcft114/.julia/packages/Flux/qXNjB/src/optimise/train.jl:105
┌ Info: Epoch 1562
└ @ Main /Users/kcft114/.julia/packages/Flux/qXNjB/src/optimise/train.jl:105
┌ Info: Epoch 1563
└ @ Main /Users/kcft114/.julia/packages/Flux/qXNjB/src/optimise/train.jl:105
┌ Info: Epoch 1564
└ @ Main /Users/kcft114/.julia/packages/Flux/qXNjB/src/optimise/train.jl:105
┌ Info: Epoch 1565
└ @ Main /Users/kcft114/.julia/packages/Flux/qXNjB/src/optimise/train.jl:105
┌ Info: Epoch 1566
└ @ Main /Users/kcft114/.julia/packages/Flux/qXNjB/src/optimise/train.jl:105
┌ Info: Epoch 1567
└ @ Main /Users/kcft114/.julia/packages/Flux/qXNjB/src/optimise/train.jl:105
┌ Info: Epoch 1568
└ @ Main /Users/kcft114/.julia/packages/Flux/qXNjB/src/optimise/train.jl:105
┌ Info: Epoch 1569
└ @ Main /Users/kcft114/.julia/packages/Flux/qXNjB/src/optimise/train.jl:105
┌ Info: Epoch 1570
└ @ Main /Users/kcft114/.julia/packages/Flux/qXNjB/src/optimise/train.jl:105
┌ Info: Epoch 1571
└ @ Main /Users/kcft114/.julia/packages/Flux/qXNjB/src/optimise/train.jl:105
┌ Info: Epoch 1572
└ @ Main /Users/kcft114/.julia/packages/Flux/qXNjB/src/optimise/train.jl:105
┌ Info: Epoch 1573
└ @ Main /Users/kcft114/.julia/packages/Flux/qXNjB/src/optimise/train.jl:105
┌ Info: Epoch 1574
└ @ Main /Users/kcft114/.julia/packages/Flux/qXNjB/src/optimise/train.jl:105
┌ Info: Epoch 1575
└ @ Main /Users/kcft114/.julia/packages/Flux/qXNjB/src/optimise/train.jl:105
┌ Info: Epoch 1576
└ @ Main /Users/kcft114/.julia/packages/Flux/qXNjB/src/optimise/train.jl:105
┌ Info: Epoch 1577
└ @ Main /Users/kcft114/.julia/packages/Flux/qXNjB/src/optimise/train.jl:105
┌ Info: Epoch 1578
└ @ Main /Users/kcft114/.julia/packages/Flux/qXNjB/src/optimise/train.jl:105
┌ Info: Epoch 1579
└ @ Main /Users/kcft114/.julia/packages/Flux/qXNjB/src/optimise/train.jl:105
┌ Info: Epoch 1580
└ @ Main /Users/kcft114/.julia/packages/Flux/qXNjB/src/optimise/train.jl:105
┌ Info: Epoch 1581
└ @ Main /Users/kcft114/.julia/packages/Flux/qXNjB/src/optimise/train.jl:105
┌ Info: Epoch 1582
└ @ Main /Users/kcft114/.julia/packages/Flux/qXNjB/src/optimise/train.jl:105
┌ Info: Epoch 1583
└ @ Main /Users/kcft114/.julia/packages/Flux/qXNjB/src/optimise/train.jl:105
┌ Info: Epoch 1584
└ @ Main /Users/kcft114/.julia/packages/Flux/qXNjB/src/optimise/train.jl:105
┌ Info: Epoch 1585
└ @ Main /Users/kcft114/.julia/packages/Flux/qXNjB/src/optimise/train.jl:105
┌ Info: Epoch 1586
└ @ Main /Users/kcft114/.julia/packages/Flux/qXNjB/src/optimise/train.jl:105
┌ Info: Epoch 1587
└ @ Main /Users/kcft114/.julia/packages/Flux/qXNjB/src/optimise/train.jl:105
┌ Info: Epoch 1588
└ @ Main /Users/kcft114/.julia/packages/Flux/qXNjB/src/optimise/train.jl:105
┌ Info: Epoch 1589
└ @ Main /Users/kcft114/.julia/packages/Flux/qXNjB/src/optimise/train.jl:105
┌ Info: Epoch 1590
└ @ Main /Users/kcft114/.julia/packages/Flux/qXNjB/src/optimise/train.jl:105
┌ Info: Epoch 1591
└ @ Main /Users/kcft114/.julia/packages/Flux/qXNjB/src/optimise/train.jl:105
┌ Info: Epoch 1592
└ @ Main /Users/kcft114/.julia/packages/Flux/qXNjB/src/optimise/train.jl:105
┌ Info: Epoch 1593
└ @ Main /Users/kcft114/.julia/packages/Flux/qXNjB/src/optimise/train.jl:105
┌ Info: Epoch 1594
└ @ Main /Users/kcft114/.julia/packages/Flux/qXNjB/src/optimise/train.jl:105
┌ Info: Epoch 1595
└ @ Main /Users/kcft114/.julia/packages/Flux/qXNjB/src/optimise/train.jl:105
┌ Info: Epoch 1596
└ @ Main /Users/kcft114/.julia/packages/Flux/qXNjB/src/optimise/train.jl:105
┌ Info: Epoch 1597
└ @ Main /Users/kcft114/.julia/packages/Flux/qXNjB/src/optimise/train.jl:105
┌ Info: Epoch 1598
└ @ Main /Users/kcft114/.julia/packages/Flux/qXNjB/src/optimise/train.jl:105
┌ Info: Epoch 1599
└ @ Main /Users/kcft114/.julia/packages/Flux/qXNjB/src/optimise/train.jl:105
┌ Info: Epoch 1600
└ @ Main /Users/kcft114/.julia/packages/Flux/qXNjB/src/optimise/train.jl:105
┌ Info: Epoch 1601
└ @ Main /Users/kcft114/.julia/packages/Flux/qXNjB/src/optimise/train.jl:105
┌ Info: Epoch 1602
└ @ Main /Users/kcft114/.julia/packages/Flux/qXNjB/src/optimise/train.jl:105
┌ Info: Epoch 1603
└ @ Main /Users/kcft114/.julia/packages/Flux/qXNjB/src/optimise/train.jl:105
┌ Info: Epoch 1604
└ @ Main /Users/kcft114/.julia/packages/Flux/qXNjB/src/optimise/train.jl:105
┌ Info: Epoch 1605
└ @ Main /Users/kcft114/.julia/packages/Flux/qXNjB/src/optimise/train.jl:105
┌ Info: Epoch 1606
└ @ Main /Users/kcft114/.julia/packages/Flux/qXNjB/src/optimise/train.jl:105
┌ Info: Epoch 1607
└ @ Main /Users/kcft114/.julia/packages/Flux/qXNjB/src/optimise/train.jl:105
┌ Info: Epoch 1608
└ @ Main /Users/kcft114/.julia/packages/Flux/qXNjB/src/optimise/train.jl:105
┌ Info: Epoch 1609
└ @ Main /Users/kcft114/.julia/packages/Flux/qXNjB/src/optimise/train.jl:105
┌ Info: Epoch 1610
└ @ Main /Users/kcft114/.julia/packages/Flux/qXNjB/src/optimise/train.jl:105
┌ Info: Epoch 1611
└ @ Main /Users/kcft114/.julia/packages/Flux/qXNjB/src/optimise/train.jl:105
┌ Info: Epoch 1612
└ @ Main /Users/kcft114/.julia/packages/Flux/qXNjB/src/optimise/train.jl:105
┌ Info: Epoch 1613
└ @ Main /Users/kcft114/.julia/packages/Flux/qXNjB/src/optimise/train.jl:105
┌ Info: Epoch 1614
└ @ Main /Users/kcft114/.julia/packages/Flux/qXNjB/src/optimise/train.jl:105
┌ Info: Epoch 1615
└ @ Main /Users/kcft114/.julia/packages/Flux/qXNjB/src/optimise/train.jl:105
┌ Info: Epoch 1616
└ @ Main /Users/kcft114/.julia/packages/Flux/qXNjB/src/optimise/train.jl:105
┌ Info: Epoch 1617
└ @ Main /Users/kcft114/.julia/packages/Flux/qXNjB/src/optimise/train.jl:105
┌ Info: Epoch 1618
└ @ Main /Users/kcft114/.julia/packages/Flux/qXNjB/src/optimise/train.jl:105
┌ Info: Epoch 1619
└ @ Main /Users/kcft114/.julia/packages/Flux/qXNjB/src/optimise/train.jl:105
┌ Info: Epoch 1620
└ @ Main /Users/kcft114/.julia/packages/Flux/qXNjB/src/optimise/train.jl:105
┌ Info: Epoch 1621
└ @ Main /Users/kcft114/.julia/packages/Flux/qXNjB/src/optimise/train.jl:105
┌ Info: Epoch 1622
└ @ Main /Users/kcft114/.julia/packages/Flux/qXNjB/src/optimise/train.jl:105
┌ Info: Epoch 1623
└ @ Main /Users/kcft114/.julia/packages/Flux/qXNjB/src/optimise/train.jl:105
┌ Info: Epoch 1624
└ @ Main /Users/kcft114/.julia/packages/Flux/qXNjB/src/optimise/train.jl:105
┌ Info: Epoch 1625
└ @ Main /Users/kcft114/.julia/packages/Flux/qXNjB/src/optimise/train.jl:105
┌ Info: Epoch 1626
└ @ Main /Users/kcft114/.julia/packages/Flux/qXNjB/src/optimise/train.jl:105
┌ Info: Epoch 1627
└ @ Main /Users/kcft114/.julia/packages/Flux/qXNjB/src/optimise/train.jl:105
┌ Info: Epoch 1628
└ @ Main /Users/kcft114/.julia/packages/Flux/qXNjB/src/optimise/train.jl:105
┌ Info: Epoch 1629
└ @ Main /Users/kcft114/.julia/packages/Flux/qXNjB/src/optimise/train.jl:105
┌ Info: Epoch 1630
└ @ Main /Users/kcft114/.julia/packages/Flux/qXNjB/src/optimise/train.jl:105
┌ Info: Epoch 1631
└ @ Main /Users/kcft114/.julia/packages/Flux/qXNjB/src/optimise/train.jl:105
┌ Info: Epoch 1632
└ @ Main /Users/kcft114/.julia/packages/Flux/qXNjB/src/optimise/train.jl:105
┌ Info: Epoch 1633
└ @ Main /Users/kcft114/.julia/packages/Flux/qXNjB/src/optimise/train.jl:105
┌ Info: Epoch 1634
└ @ Main /Users/kcft114/.julia/packages/Flux/qXNjB/src/optimise/train.jl:105
┌ Info: Epoch 1635
└ @ Main /Users/kcft114/.julia/packages/Flux/qXNjB/src/optimise/train.jl:105
┌ Info: Epoch 1636
└ @ Main /Users/kcft114/.julia/packages/Flux/qXNjB/src/optimise/train.jl:105
┌ Info: Epoch 1637
└ @ Main /Users/kcft114/.julia/packages/Flux/qXNjB/src/optimise/train.jl:105
┌ Info: Epoch 1638
└ @ Main /Users/kcft114/.julia/packages/Flux/qXNjB/src/optimise/train.jl:105
┌ Info: Epoch 1639
└ @ Main /Users/kcft114/.julia/packages/Flux/qXNjB/src/optimise/train.jl:105
┌ Info: Epoch 1640
└ @ Main /Users/kcft114/.julia/packages/Flux/qXNjB/src/optimise/train.jl:105
┌ Info: Epoch 1641
└ @ Main /Users/kcft114/.julia/packages/Flux/qXNjB/src/optimise/train.jl:105
┌ Info: Epoch 1642
└ @ Main /Users/kcft114/.julia/packages/Flux/qXNjB/src/optimise/train.jl:105
┌ Info: Epoch 1643
└ @ Main /Users/kcft114/.julia/packages/Flux/qXNjB/src/optimise/train.jl:105
┌ Info: Epoch 1644
└ @ Main /Users/kcft114/.julia/packages/Flux/qXNjB/src/optimise/train.jl:105
┌ Info: Epoch 1645
└ @ Main /Users/kcft114/.julia/packages/Flux/qXNjB/src/optimise/train.jl:105
┌ Info: Epoch 1646
└ @ Main /Users/kcft114/.julia/packages/Flux/qXNjB/src/optimise/train.jl:105
┌ Info: Epoch 1647
└ @ Main /Users/kcft114/.julia/packages/Flux/qXNjB/src/optimise/train.jl:105
┌ Info: Epoch 1648
└ @ Main /Users/kcft114/.julia/packages/Flux/qXNjB/src/optimise/train.jl:105
┌ Info: Epoch 1649
└ @ Main /Users/kcft114/.julia/packages/Flux/qXNjB/src/optimise/train.jl:105
┌ Info: Epoch 1650
└ @ Main /Users/kcft114/.julia/packages/Flux/qXNjB/src/optimise/train.jl:105
┌ Info: Epoch 1651
└ @ Main /Users/kcft114/.julia/packages/Flux/qXNjB/src/optimise/train.jl:105
┌ Info: Epoch 1652
└ @ Main /Users/kcft114/.julia/packages/Flux/qXNjB/src/optimise/train.jl:105
┌ Info: Epoch 1653
└ @ Main /Users/kcft114/.julia/packages/Flux/qXNjB/src/optimise/train.jl:105
┌ Info: Epoch 1654
└ @ Main /Users/kcft114/.julia/packages/Flux/qXNjB/src/optimise/train.jl:105
┌ Info: Epoch 1655
└ @ Main /Users/kcft114/.julia/packages/Flux/qXNjB/src/optimise/train.jl:105
┌ Info: Epoch 1656
└ @ Main /Users/kcft114/.julia/packages/Flux/qXNjB/src/optimise/train.jl:105
┌ Info: Epoch 1657
└ @ Main /Users/kcft114/.julia/packages/Flux/qXNjB/src/optimise/train.jl:105
┌ Info: Epoch 1658
└ @ Main /Users/kcft114/.julia/packages/Flux/qXNjB/src/optimise/train.jl:105
┌ Info: Epoch 1659
└ @ Main /Users/kcft114/.julia/packages/Flux/qXNjB/src/optimise/train.jl:105
┌ Info: Epoch 1660
└ @ Main /Users/kcft114/.julia/packages/Flux/qXNjB/src/optimise/train.jl:105
┌ Info: Epoch 1661
└ @ Main /Users/kcft114/.julia/packages/Flux/qXNjB/src/optimise/train.jl:105
┌ Info: Epoch 1662
└ @ Main /Users/kcft114/.julia/packages/Flux/qXNjB/src/optimise/train.jl:105
┌ Info: Epoch 1663
└ @ Main /Users/kcft114/.julia/packages/Flux/qXNjB/src/optimise/train.jl:105
┌ Info: Epoch 1664
└ @ Main /Users/kcft114/.julia/packages/Flux/qXNjB/src/optimise/train.jl:105
┌ Info: Epoch 1665
└ @ Main /Users/kcft114/.julia/packages/Flux/qXNjB/src/optimise/train.jl:105
┌ Info: Epoch 1666
└ @ Main /Users/kcft114/.julia/packages/Flux/qXNjB/src/optimise/train.jl:105
┌ Info: Epoch 1667
└ @ Main /Users/kcft114/.julia/packages/Flux/qXNjB/src/optimise/train.jl:105
┌ Info: Epoch 1668
└ @ Main /Users/kcft114/.julia/packages/Flux/qXNjB/src/optimise/train.jl:105
┌ Info: Epoch 1669
└ @ Main /Users/kcft114/.julia/packages/Flux/qXNjB/src/optimise/train.jl:105
┌ Info: Epoch 1670
└ @ Main /Users/kcft114/.julia/packages/Flux/qXNjB/src/optimise/train.jl:105
┌ Info: Epoch 1671
└ @ Main /Users/kcft114/.julia/packages/Flux/qXNjB/src/optimise/train.jl:105
┌ Info: Epoch 1672
└ @ Main /Users/kcft114/.julia/packages/Flux/qXNjB/src/optimise/train.jl:105
┌ Info: Epoch 1673
└ @ Main /Users/kcft114/.julia/packages/Flux/qXNjB/src/optimise/train.jl:105
┌ Info: Epoch 1674
└ @ Main /Users/kcft114/.julia/packages/Flux/qXNjB/src/optimise/train.jl:105
┌ Info: Epoch 1675
└ @ Main /Users/kcft114/.julia/packages/Flux/qXNjB/src/optimise/train.jl:105
┌ Info: Epoch 1676
└ @ Main /Users/kcft114/.julia/packages/Flux/qXNjB/src/optimise/train.jl:105
┌ Info: Epoch 1677
└ @ Main /Users/kcft114/.julia/packages/Flux/qXNjB/src/optimise/train.jl:105
┌ Info: Epoch 1678
└ @ Main /Users/kcft114/.julia/packages/Flux/qXNjB/src/optimise/train.jl:105
┌ Info: Epoch 1679
└ @ Main /Users/kcft114/.julia/packages/Flux/qXNjB/src/optimise/train.jl:105
┌ Info: Epoch 1680
└ @ Main /Users/kcft114/.julia/packages/Flux/qXNjB/src/optimise/train.jl:105
┌ Info: Epoch 1681
└ @ Main /Users/kcft114/.julia/packages/Flux/qXNjB/src/optimise/train.jl:105
┌ Info: Epoch 1682
└ @ Main /Users/kcft114/.julia/packages/Flux/qXNjB/src/optimise/train.jl:105
┌ Info: Epoch 1683
└ @ Main /Users/kcft114/.julia/packages/Flux/qXNjB/src/optimise/train.jl:105
┌ Info: Epoch 1684
└ @ Main /Users/kcft114/.julia/packages/Flux/qXNjB/src/optimise/train.jl:105
┌ Info: Epoch 1685
└ @ Main /Users/kcft114/.julia/packages/Flux/qXNjB/src/optimise/train.jl:105
┌ Info: Epoch 1686
└ @ Main /Users/kcft114/.julia/packages/Flux/qXNjB/src/optimise/train.jl:105
┌ Info: Epoch 1687
└ @ Main /Users/kcft114/.julia/packages/Flux/qXNjB/src/optimise/train.jl:105
┌ Info: Epoch 1688
└ @ Main /Users/kcft114/.julia/packages/Flux/qXNjB/src/optimise/train.jl:105
┌ Info: Epoch 1689
└ @ Main /Users/kcft114/.julia/packages/Flux/qXNjB/src/optimise/train.jl:105
┌ Info: Epoch 1690
└ @ Main /Users/kcft114/.julia/packages/Flux/qXNjB/src/optimise/train.jl:105
┌ Info: Epoch 1691
└ @ Main /Users/kcft114/.julia/packages/Flux/qXNjB/src/optimise/train.jl:105
┌ Info: Epoch 1692
└ @ Main /Users/kcft114/.julia/packages/Flux/qXNjB/src/optimise/train.jl:105
┌ Info: Epoch 1693
└ @ Main /Users/kcft114/.julia/packages/Flux/qXNjB/src/optimise/train.jl:105
┌ Info: Epoch 1694
└ @ Main /Users/kcft114/.julia/packages/Flux/qXNjB/src/optimise/train.jl:105
┌ Info: Epoch 1695
└ @ Main /Users/kcft114/.julia/packages/Flux/qXNjB/src/optimise/train.jl:105
┌ Info: Epoch 1696
└ @ Main /Users/kcft114/.julia/packages/Flux/qXNjB/src/optimise/train.jl:105
┌ Info: Epoch 1697
└ @ Main /Users/kcft114/.julia/packages/Flux/qXNjB/src/optimise/train.jl:105
┌ Info: Epoch 1698
└ @ Main /Users/kcft114/.julia/packages/Flux/qXNjB/src/optimise/train.jl:105
┌ Info: Epoch 1699
└ @ Main /Users/kcft114/.julia/packages/Flux/qXNjB/src/optimise/train.jl:105
┌ Info: Epoch 1700
└ @ Main /Users/kcft114/.julia/packages/Flux/qXNjB/src/optimise/train.jl:105
┌ Info: Epoch 1701
└ @ Main /Users/kcft114/.julia/packages/Flux/qXNjB/src/optimise/train.jl:105
┌ Info: Epoch 1702
└ @ Main /Users/kcft114/.julia/packages/Flux/qXNjB/src/optimise/train.jl:105
┌ Info: Epoch 1703
└ @ Main /Users/kcft114/.julia/packages/Flux/qXNjB/src/optimise/train.jl:105
┌ Info: Epoch 1704
└ @ Main /Users/kcft114/.julia/packages/Flux/qXNjB/src/optimise/train.jl:105
┌ Info: Epoch 1705
└ @ Main /Users/kcft114/.julia/packages/Flux/qXNjB/src/optimise/train.jl:105
┌ Info: Epoch 1706
└ @ Main /Users/kcft114/.julia/packages/Flux/qXNjB/src/optimise/train.jl:105
┌ Info: Epoch 1707
└ @ Main /Users/kcft114/.julia/packages/Flux/qXNjB/src/optimise/train.jl:105
┌ Info: Epoch 1708
└ @ Main /Users/kcft114/.julia/packages/Flux/qXNjB/src/optimise/train.jl:105
┌ Info: Epoch 1709
└ @ Main /Users/kcft114/.julia/packages/Flux/qXNjB/src/optimise/train.jl:105
┌ Info: Epoch 1710
└ @ Main /Users/kcft114/.julia/packages/Flux/qXNjB/src/optimise/train.jl:105
┌ Info: Epoch 1711
└ @ Main /Users/kcft114/.julia/packages/Flux/qXNjB/src/optimise/train.jl:105
┌ Info: Epoch 1712
└ @ Main /Users/kcft114/.julia/packages/Flux/qXNjB/src/optimise/train.jl:105
┌ Info: Epoch 1713
└ @ Main /Users/kcft114/.julia/packages/Flux/qXNjB/src/optimise/train.jl:105
┌ Info: Epoch 1714
└ @ Main /Users/kcft114/.julia/packages/Flux/qXNjB/src/optimise/train.jl:105
┌ Info: Epoch 1715
└ @ Main /Users/kcft114/.julia/packages/Flux/qXNjB/src/optimise/train.jl:105
┌ Info: Epoch 1716
└ @ Main /Users/kcft114/.julia/packages/Flux/qXNjB/src/optimise/train.jl:105
┌ Info: Epoch 1717
└ @ Main /Users/kcft114/.julia/packages/Flux/qXNjB/src/optimise/train.jl:105
┌ Info: Epoch 1718
└ @ Main /Users/kcft114/.julia/packages/Flux/qXNjB/src/optimise/train.jl:105
┌ Info: Epoch 1719
└ @ Main /Users/kcft114/.julia/packages/Flux/qXNjB/src/optimise/train.jl:105
┌ Info: Epoch 1720
└ @ Main /Users/kcft114/.julia/packages/Flux/qXNjB/src/optimise/train.jl:105
┌ Info: Epoch 1721
└ @ Main /Users/kcft114/.julia/packages/Flux/qXNjB/src/optimise/train.jl:105
┌ Info: Epoch 1722
└ @ Main /Users/kcft114/.julia/packages/Flux/qXNjB/src/optimise/train.jl:105
┌ Info: Epoch 1723
└ @ Main /Users/kcft114/.julia/packages/Flux/qXNjB/src/optimise/train.jl:105
```

```
loss(X, Y) = 0.49766177f0 (tracked)
loss(X, Y) = 0.5767595f0 (tracked)
loss(X, Y) = 0.5098359f0 (tracked)
loss(X, Y) = 0.5292928f0 (tracked)
loss(X, Y) = 0.5007277f0 (tracked)
loss(X, Y) = 0.5540479f0 (tracked)
loss(X, Y) = 0.49652317f0 (tracked)
loss(X, Y) = 0.56017673f0 (tracked)
loss(X, Y) = 0.5356362f0 (tracked)
loss(X, Y) = 0.49333438f0 (tracked)
loss(X, Y) = 0.5121678f0 (tracked)
loss(X, Y) = 0.5029285f0 (tracked)
loss(X, Y) = 0.5108942f0 (tracked)
loss(X, Y) = 0.5282889f0 (tracked)
loss(X, Y) = 0.5672428f0 (tracked)
loss(X, Y) = 0.4922995f0 (tracked)
loss(X, Y) = 0.53289247f0 (tracked)
loss(X, Y) = 0.55068755f0 (tracked)
loss(X, Y) = 0.5145313f0 (tracked)
loss(X, Y) = 0.49653095f0 (tracked)
loss(X, Y) = 0.52194124f0 (tracked)
loss(X, Y) = 0.55566216f0 (tracked)
loss(X, Y) = 0.5690028f0 (tracked)
loss(X, Y) = 0.55723923f0 (tracked)
loss(X, Y) = 0.5353102f0 (tracked)
loss(X, Y) = 0.5111555f0 (tracked)
loss(X, Y) = 0.5627494f0 (tracked)
loss(X, Y) = 0.485456f0 (tracked)
loss(X, Y) = 0.5134825f0 (tracked)
loss(X, Y) = 0.4642447f0 (tracked)
loss(X, Y) = 0.5413218f0 (tracked)
loss(X, Y) = 0.5483948f0 (tracked)
loss(X, Y) = 0.55401343f0 (tracked)
loss(X, Y) = 0.4673285f0 (tracked)
loss(X, Y) = 0.5311903f0 (tracked)
loss(X, Y) = 0.5571619f0 (tracked)
loss(X, Y) = 0.49877498f0 (tracked)
loss(X, Y) = 0.5236636f0 (tracked)
loss(X, Y) = 0.5844708f0 (tracked)
loss(X, Y) = 0.50675696f0 (tracked)
loss(X, Y) = 0.482967f0 (tracked)
loss(X, Y) = 0.55544794f0 (tracked)
loss(X, Y) = 0.54865885f0 (tracked)
loss(X, Y) = 0.5684682f0 (tracked)
loss(X, Y) = 0.50583756f0 (tracked)
loss(X, Y) = 0.53058213f0 (tracked)
loss(X, Y) = 0.4842553f0 (tracked)
loss(X, Y) = 0.49224105f0 (tracked)
loss(X, Y) = 0.5106798f0 (tracked)
loss(X, Y) = 0.5403337f0 (tracked)
loss(X, Y) = 0.5576522f0 (tracked)
loss(X, Y) = 0.56235504f0 (tracked)
loss(X, Y) = 0.5227743f0 (tracked)
loss(X, Y) = 0.59852344f0 (tracked)
loss(X, Y) = 0.52286863f0 (tracked)
loss(X, Y) = 0.4866886f0 (tracked)
loss(X, Y) = 0.55312157f0 (tracked)
loss(X, Y) = 0.48263466f0 (tracked)
loss(X, Y) = 0.5271701f0 (tracked)
loss(X, Y) = 0.55418175f0 (tracked)
loss(X, Y) = 0.5439401f0 (tracked)
loss(X, Y) = 0.5517573f0 (tracked)
loss(X, Y) = 0.53064466f0 (tracked)
loss(X, Y) = 0.5438367f0 (tracked)
loss(X, Y) = 0.57906616f0 (tracked)
loss(X, Y) = 0.5164556f0 (tracked)
loss(X, Y) = 0.5165103f0 (tracked)
loss(X, Y) = 0.5359695f0 (tracked)
loss(X, Y) = 0.5070101f0 (tracked)
loss(X, Y) = 0.5404482f0 (tracked)
loss(X, Y) = 0.5225846f0 (tracked)
loss(X, Y) = 0.53777343f0 (tracked)
loss(X, Y) = 0.5533723f0 (tracked)
loss(X, Y) = 0.5627687f0 (tracked)
loss(X, Y) = 0.4928279f0 (tracked)
loss(X, Y) = 0.5268646f0 (tracked)
loss(X, Y) = 0.5289431f0 (tracked)
loss(X, Y) = 0.50748813f0 (tracked)
loss(X, Y) = 0.51675665f0 (tracked)
loss(X, Y) = 0.50798285f0 (tracked)
loss(X, Y) = 0.5120587f0 (tracked)
loss(X, Y) = 0.5075823f0 (tracked)
loss(X, Y) = 0.4944693f0 (tracked)
loss(X, Y) = 0.5686252f0 (tracked)
loss(X, Y) = 0.49660435f0 (tracked)
loss(X, Y) = 0.5724112f0 (tracked)
loss(X, Y) = 0.47958523f0 (tracked)
loss(X, Y) = 0.54508704f0 (tracked)
loss(X, Y) = 0.5629774f0 (tracked)
loss(X, Y) = 0.5705071f0 (tracked)
loss(X, Y) = 0.5355767f0 (tracked)
loss(X, Y) = 0.54666954f0 (tracked)
loss(X, Y) = 0.50668263f0 (tracked)
loss(X, Y) = 0.51713544f0 (tracked)
loss(X, Y) = 0.5162842f0 (tracked)
loss(X, Y) = 0.49152634f0 (tracked)
loss(X, Y) = 0.51422614f0 (tracked)
loss(X, Y) = 0.54291964f0 (tracked)
loss(X, Y) = 0.5000578f0 (tracked)
loss(X, Y) = 0.56842995f0 (tracked)
loss(X, Y) = 0.5240021f0 (tracked)
loss(X, Y) = 0.5086908f0 (tracked)
loss(X, Y) = 0.5563064f0 (tracked)
loss(X, Y) = 0.5267874f0 (tracked)
loss(X, Y) = 0.50876635f0 (tracked)
loss(X, Y) = 0.52136856f0 (tracked)
loss(X, Y) = 0.50415313f0 (tracked)
loss(X, Y) = 0.54555976f0 (tracked)
loss(X, Y) = 0.5373685f0 (tracked)
loss(X, Y) = 0.53852165f0 (tracked)
loss(X, Y) = 0.5537315f0 (tracked)
loss(X, Y) = 0.5224941f0 (tracked)
loss(X, Y) = 0.5478623f0 (tracked)
loss(X, Y) = 0.49456733f0 (tracked)
loss(X, Y) = 0.49914026f0 (tracked)
loss(X, Y) = 0.5389948f0 (tracked)
loss(X, Y) = 0.5649389f0 (tracked)
loss(X, Y) = 0.52615f0 (tracked)
loss(X, Y) = 0.52315277f0 (tracked)
loss(X, Y) = 0.5196996f0 (tracked)
loss(X, Y) = 0.51329774f0 (tracked)
loss(X, Y) = 0.51005524f0 (tracked)
loss(X, Y) = 0.5464948f0 (tracked)
loss(X, Y) = 0.51906025f0 (tracked)
loss(X, Y) = 0.46680647f0 (tracked)
loss(X, Y) = 0.52286613f0 (tracked)
loss(X, Y) = 0.5347015f0 (tracked)
loss(X, Y) = 0.5066642f0 (tracked)
loss(X, Y) = 0.57527524f0 (tracked)
loss(X, Y) = 0.5288455f0 (tracked)
loss(X, Y) = 0.53907615f0 (tracked)
loss(X, Y) = 0.48580563f0 (tracked)
loss(X, Y) = 0.53408927f0 (tracked)
loss(X, Y) = 0.51393765f0 (tracked)
loss(X, Y) = 0.5330081f0 (tracked)
loss(X, Y) = 0.5593658f0 (tracked)
loss(X, Y) = 0.5487383f0 (tracked)
loss(X, Y) = 0.4872375f0 (tracked)
loss(X, Y) = 0.50214505f0 (tracked)
loss(X, Y) = 0.5492593f0 (tracked)
loss(X, Y) = 0.5074822f0 (tracked)
loss(X, Y) = 0.5117682f0 (tracked)
loss(X, Y) = 0.48125803f0 (tracked)
loss(X, Y) = 0.5096789f0 (tracked)
loss(X, Y) = 0.4519543f0 (tracked)
loss(X, Y) = 0.5488396f0 (tracked)
loss(X, Y) = 0.54392105f0 (tracked)
loss(X, Y) = 0.49692523f0 (tracked)
loss(X, Y) = 0.5577598f0 (tracked)
loss(X, Y) = 0.51384693f0 (tracked)
loss(X, Y) = 0.5117915f0 (tracked)
loss(X, Y) = 0.4826743f0 (tracked)
loss(X, Y) = 0.5320066f0 (tracked)
loss(X, Y) = 0.52273613f0 (tracked)
loss(X, Y) = 0.5253358f0 (tracked)
loss(X, Y) = 0.53050256f0 (tracked)
loss(X, Y) = 0.5570835f0 (tracked)
loss(X, Y) = 0.5244174f0 (tracked)
loss(X, Y) = 0.5195885f0 (tracked)
loss(X, Y) = 0.4807391f0 (tracked)
loss(X, Y) = 0.52967083f0 (tracked)
loss(X, Y) = 0.55707645f0 (tracked)
loss(X, Y) = 0.54728884f0 (tracked)
loss(X, Y) = 0.473604f0 (tracked)
loss(X, Y) = 0.54135424f0 (tracked)
loss(X, Y) = 0.47731623f0 (tracked)
loss(X, Y) = 0.5355054f0 (tracked)
loss(X, Y) = 0.54456705f0 (tracked)
loss(X, Y) = 0.5141058f0 (tracked)
loss(X, Y) = 0.51049274f0 (tracked)
loss(X, Y) = 0.5580953f0 (tracked)
loss(X, Y) = 0.49939877f0 (tracked)
loss(X, Y) = 0.48867574f0 (tracked)
loss(X, Y) = 0.56006914f0 (tracked)
loss(X, Y) = 0.5863282f0 (tracked)
loss(X, Y) = 0.52968895f0 (tracked)
loss(X, Y) = 0.49587092f0 (tracked)
loss(X, Y) = 0.5473305f0 (tracked)
loss(X, Y) = 0.5763673f0 (tracked)
loss(X, Y) = 0.5645651f0 (tracked)
loss(X, Y) = 0.5286802f0 (tracked)
loss(X, Y) = 0.48996785f0 (tracked)
loss(X, Y) = 0.5309934f0 (tracked)
loss(X, Y) = 0.48789784f0 (tracked)
loss(X, Y) = 0.4763648f0 (tracked)
loss(X, Y) = 0.5182404f0 (tracked)
loss(X, Y) = 0.5162475f0 (tracked)
loss(X, Y) = 0.48241627f0 (tracked)
loss(X, Y) = 0.5484774f0 (tracked)
loss(X, Y) = 0.5233044f0 (tracked)
loss(X, Y) = 0.48012465f0 (tracked)
loss(X, Y) = 0.509008f0 (tracked)
loss(X, Y) = 0.5398838f0 (tracked)
loss(X, Y) = 0.5989769f0 (tracked)
loss(X, Y) = 0.50650775f0 (tracked)
loss(X, Y) = 0.5065877f0 (tracked)
loss(X, Y) = 0.5079106f0 (tracked)
loss(X, Y) = 0.5191385f0 (tracked)
```

```
┌ Info: Epoch 1724
└ @ Main /Users/kcft114/.julia/packages/Flux/qXNjB/src/optimise/train.jl:105
┌ Info: Epoch 1725
└ @ Main /Users/kcft114/.julia/packages/Flux/qXNjB/src/optimise/train.jl:105
┌ Info: Epoch 1726
└ @ Main /Users/kcft114/.julia/packages/Flux/qXNjB/src/optimise/train.jl:105
┌ Info: Epoch 1727
└ @ Main /Users/kcft114/.julia/packages/Flux/qXNjB/src/optimise/train.jl:105
┌ Info: Epoch 1728
└ @ Main /Users/kcft114/.julia/packages/Flux/qXNjB/src/optimise/train.jl:105
┌ Info: Epoch 1729
└ @ Main /Users/kcft114/.julia/packages/Flux/qXNjB/src/optimise/train.jl:105
┌ Info: Epoch 1730
└ @ Main /Users/kcft114/.julia/packages/Flux/qXNjB/src/optimise/train.jl:105
┌ Info: Epoch 1731
└ @ Main /Users/kcft114/.julia/packages/Flux/qXNjB/src/optimise/train.jl:105
┌ Info: Epoch 1732
└ @ Main /Users/kcft114/.julia/packages/Flux/qXNjB/src/optimise/train.jl:105
┌ Info: Epoch 1733
└ @ Main /Users/kcft114/.julia/packages/Flux/qXNjB/src/optimise/train.jl:105
┌ Info: Epoch 1734
└ @ Main /Users/kcft114/.julia/packages/Flux/qXNjB/src/optimise/train.jl:105
┌ Info: Epoch 1735
└ @ Main /Users/kcft114/.julia/packages/Flux/qXNjB/src/optimise/train.jl:105
┌ Info: Epoch 1736
└ @ Main /Users/kcft114/.julia/packages/Flux/qXNjB/src/optimise/train.jl:105
┌ Info: Epoch 1737
└ @ Main /Users/kcft114/.julia/packages/Flux/qXNjB/src/optimise/train.jl:105
┌ Info: Epoch 1738
└ @ Main /Users/kcft114/.julia/packages/Flux/qXNjB/src/optimise/train.jl:105
┌ Info: Epoch 1739
└ @ Main /Users/kcft114/.julia/packages/Flux/qXNjB/src/optimise/train.jl:105
┌ Info: Epoch 1740
└ @ Main /Users/kcft114/.julia/packages/Flux/qXNjB/src/optimise/train.jl:105
┌ Info: Epoch 1741
└ @ Main /Users/kcft114/.julia/packages/Flux/qXNjB/src/optimise/train.jl:105
┌ Info: Epoch 1742
└ @ Main /Users/kcft114/.julia/packages/Flux/qXNjB/src/optimise/train.jl:105
┌ Info: Epoch 1743
└ @ Main /Users/kcft114/.julia/packages/Flux/qXNjB/src/optimise/train.jl:105
┌ Info: Epoch 1744
└ @ Main /Users/kcft114/.julia/packages/Flux/qXNjB/src/optimise/train.jl:105
┌ Info: Epoch 1745
└ @ Main /Users/kcft114/.julia/packages/Flux/qXNjB/src/optimise/train.jl:105
┌ Info: Epoch 1746
└ @ Main /Users/kcft114/.julia/packages/Flux/qXNjB/src/optimise/train.jl:105
┌ Info: Epoch 1747
└ @ Main /Users/kcft114/.julia/packages/Flux/qXNjB/src/optimise/train.jl:105
┌ Info: Epoch 1748
└ @ Main /Users/kcft114/.julia/packages/Flux/qXNjB/src/optimise/train.jl:105
┌ Info: Epoch 1749
└ @ Main /Users/kcft114/.julia/packages/Flux/qXNjB/src/optimise/train.jl:105
┌ Info: Epoch 1750
└ @ Main /Users/kcft114/.julia/packages/Flux/qXNjB/src/optimise/train.jl:105
┌ Info: Epoch 1751
└ @ Main /Users/kcft114/.julia/packages/Flux/qXNjB/src/optimise/train.jl:105
┌ Info: Epoch 1752
└ @ Main /Users/kcft114/.julia/packages/Flux/qXNjB/src/optimise/train.jl:105
┌ Info: Epoch 1753
└ @ Main /Users/kcft114/.julia/packages/Flux/qXNjB/src/optimise/train.jl:105
┌ Info: Epoch 1754
└ @ Main /Users/kcft114/.julia/packages/Flux/qXNjB/src/optimise/train.jl:105
┌ Info: Epoch 1755
└ @ Main /Users/kcft114/.julia/packages/Flux/qXNjB/src/optimise/train.jl:105
┌ Info: Epoch 1756
└ @ Main /Users/kcft114/.julia/packages/Flux/qXNjB/src/optimise/train.jl:105
┌ Info: Epoch 1757
└ @ Main /Users/kcft114/.julia/packages/Flux/qXNjB/src/optimise/train.jl:105
┌ Info: Epoch 1758
└ @ Main /Users/kcft114/.julia/packages/Flux/qXNjB/src/optimise/train.jl:105
┌ Info: Epoch 1759
└ @ Main /Users/kcft114/.julia/packages/Flux/qXNjB/src/optimise/train.jl:105
┌ Info: Epoch 1760
└ @ Main /Users/kcft114/.julia/packages/Flux/qXNjB/src/optimise/train.jl:105
┌ Info: Epoch 1761
└ @ Main /Users/kcft114/.julia/packages/Flux/qXNjB/src/optimise/train.jl:105
┌ Info: Epoch 1762
└ @ Main /Users/kcft114/.julia/packages/Flux/qXNjB/src/optimise/train.jl:105
┌ Info: Epoch 1763
└ @ Main /Users/kcft114/.julia/packages/Flux/qXNjB/src/optimise/train.jl:105
┌ Info: Epoch 1764
└ @ Main /Users/kcft114/.julia/packages/Flux/qXNjB/src/optimise/train.jl:105
┌ Info: Epoch 1765
└ @ Main /Users/kcft114/.julia/packages/Flux/qXNjB/src/optimise/train.jl:105
┌ Info: Epoch 1766
└ @ Main /Users/kcft114/.julia/packages/Flux/qXNjB/src/optimise/train.jl:105
┌ Info: Epoch 1767
└ @ Main /Users/kcft114/.julia/packages/Flux/qXNjB/src/optimise/train.jl:105
┌ Info: Epoch 1768
└ @ Main /Users/kcft114/.julia/packages/Flux/qXNjB/src/optimise/train.jl:105
┌ Info: Epoch 1769
└ @ Main /Users/kcft114/.julia/packages/Flux/qXNjB/src/optimise/train.jl:105
┌ Info: Epoch 1770
└ @ Main /Users/kcft114/.julia/packages/Flux/qXNjB/src/optimise/train.jl:105
┌ Info: Epoch 1771
└ @ Main /Users/kcft114/.julia/packages/Flux/qXNjB/src/optimise/train.jl:105
┌ Info: Epoch 1772
└ @ Main /Users/kcft114/.julia/packages/Flux/qXNjB/src/optimise/train.jl:105
┌ Info: Epoch 1773
└ @ Main /Users/kcft114/.julia/packages/Flux/qXNjB/src/optimise/train.jl:105
┌ Info: Epoch 1774
└ @ Main /Users/kcft114/.julia/packages/Flux/qXNjB/src/optimise/train.jl:105
┌ Info: Epoch 1775
└ @ Main /Users/kcft114/.julia/packages/Flux/qXNjB/src/optimise/train.jl:105
┌ Info: Epoch 1776
└ @ Main /Users/kcft114/.julia/packages/Flux/qXNjB/src/optimise/train.jl:105
┌ Info: Epoch 1777
└ @ Main /Users/kcft114/.julia/packages/Flux/qXNjB/src/optimise/train.jl:105
┌ Info: Epoch 1778
└ @ Main /Users/kcft114/.julia/packages/Flux/qXNjB/src/optimise/train.jl:105
┌ Info: Epoch 1779
└ @ Main /Users/kcft114/.julia/packages/Flux/qXNjB/src/optimise/train.jl:105
┌ Info: Epoch 1780
└ @ Main /Users/kcft114/.julia/packages/Flux/qXNjB/src/optimise/train.jl:105
┌ Info: Epoch 1781
└ @ Main /Users/kcft114/.julia/packages/Flux/qXNjB/src/optimise/train.jl:105
┌ Info: Epoch 1782
└ @ Main /Users/kcft114/.julia/packages/Flux/qXNjB/src/optimise/train.jl:105
┌ Info: Epoch 1783
└ @ Main /Users/kcft114/.julia/packages/Flux/qXNjB/src/optimise/train.jl:105
┌ Info: Epoch 1784
└ @ Main /Users/kcft114/.julia/packages/Flux/qXNjB/src/optimise/train.jl:105
┌ Info: Epoch 1785
└ @ Main /Users/kcft114/.julia/packages/Flux/qXNjB/src/optimise/train.jl:105
┌ Info: Epoch 1786
└ @ Main /Users/kcft114/.julia/packages/Flux/qXNjB/src/optimise/train.jl:105
┌ Info: Epoch 1787
└ @ Main /Users/kcft114/.julia/packages/Flux/qXNjB/src/optimise/train.jl:105
┌ Info: Epoch 1788
└ @ Main /Users/kcft114/.julia/packages/Flux/qXNjB/src/optimise/train.jl:105
┌ Info: Epoch 1789
└ @ Main /Users/kcft114/.julia/packages/Flux/qXNjB/src/optimise/train.jl:105
┌ Info: Epoch 1790
└ @ Main /Users/kcft114/.julia/packages/Flux/qXNjB/src/optimise/train.jl:105
┌ Info: Epoch 1791
└ @ Main /Users/kcft114/.julia/packages/Flux/qXNjB/src/optimise/train.jl:105
┌ Info: Epoch 1792
└ @ Main /Users/kcft114/.julia/packages/Flux/qXNjB/src/optimise/train.jl:105
┌ Info: Epoch 1793
└ @ Main /Users/kcft114/.julia/packages/Flux/qXNjB/src/optimise/train.jl:105
┌ Info: Epoch 1794
└ @ Main /Users/kcft114/.julia/packages/Flux/qXNjB/src/optimise/train.jl:105
┌ Info: Epoch 1795
└ @ Main /Users/kcft114/.julia/packages/Flux/qXNjB/src/optimise/train.jl:105
┌ Info: Epoch 1796
└ @ Main /Users/kcft114/.julia/packages/Flux/qXNjB/src/optimise/train.jl:105
┌ Info: Epoch 1797
└ @ Main /Users/kcft114/.julia/packages/Flux/qXNjB/src/optimise/train.jl:105
┌ Info: Epoch 1798
└ @ Main /Users/kcft114/.julia/packages/Flux/qXNjB/src/optimise/train.jl:105
┌ Info: Epoch 1799
└ @ Main /Users/kcft114/.julia/packages/Flux/qXNjB/src/optimise/train.jl:105
┌ Info: Epoch 1800
└ @ Main /Users/kcft114/.julia/packages/Flux/qXNjB/src/optimise/train.jl:105
┌ Info: Epoch 1801
└ @ Main /Users/kcft114/.julia/packages/Flux/qXNjB/src/optimise/train.jl:105
┌ Info: Epoch 1802
└ @ Main /Users/kcft114/.julia/packages/Flux/qXNjB/src/optimise/train.jl:105
┌ Info: Epoch 1803
└ @ Main /Users/kcft114/.julia/packages/Flux/qXNjB/src/optimise/train.jl:105
┌ Info: Epoch 1804
└ @ Main /Users/kcft114/.julia/packages/Flux/qXNjB/src/optimise/train.jl:105
┌ Info: Epoch 1805
└ @ Main /Users/kcft114/.julia/packages/Flux/qXNjB/src/optimise/train.jl:105
┌ Info: Epoch 1806
└ @ Main /Users/kcft114/.julia/packages/Flux/qXNjB/src/optimise/train.jl:105
┌ Info: Epoch 1807
└ @ Main /Users/kcft114/.julia/packages/Flux/qXNjB/src/optimise/train.jl:105
┌ Info: Epoch 1808
└ @ Main /Users/kcft114/.julia/packages/Flux/qXNjB/src/optimise/train.jl:105
┌ Info: Epoch 1809
└ @ Main /Users/kcft114/.julia/packages/Flux/qXNjB/src/optimise/train.jl:105
┌ Info: Epoch 1810
└ @ Main /Users/kcft114/.julia/packages/Flux/qXNjB/src/optimise/train.jl:105
┌ Info: Epoch 1811
└ @ Main /Users/kcft114/.julia/packages/Flux/qXNjB/src/optimise/train.jl:105
┌ Info: Epoch 1812
└ @ Main /Users/kcft114/.julia/packages/Flux/qXNjB/src/optimise/train.jl:105
┌ Info: Epoch 1813
└ @ Main /Users/kcft114/.julia/packages/Flux/qXNjB/src/optimise/train.jl:105
┌ Info: Epoch 1814
└ @ Main /Users/kcft114/.julia/packages/Flux/qXNjB/src/optimise/train.jl:105
┌ Info: Epoch 1815
└ @ Main /Users/kcft114/.julia/packages/Flux/qXNjB/src/optimise/train.jl:105
┌ Info: Epoch 1816
└ @ Main /Users/kcft114/.julia/packages/Flux/qXNjB/src/optimise/train.jl:105
┌ Info: Epoch 1817
└ @ Main /Users/kcft114/.julia/packages/Flux/qXNjB/src/optimise/train.jl:105
┌ Info: Epoch 1818
└ @ Main /Users/kcft114/.julia/packages/Flux/qXNjB/src/optimise/train.jl:105
Excessive output truncated after 524381 bytes.
```

```
loss(X, Y) = 0.5199574f0 (tracked)
loss(X, Y) = 0.56694067f0 (tracked)
loss(X, Y) = 0.5254522f0 (tracked)
loss(X, Y) = 0.5380213f0 (tracked)
loss(X, Y) = 0.55485004f0 (tracked)
loss(X, Y) = 0.4828158f0 (tracked)
loss(X, Y) = 0.54848707f0 (tracked)
loss(X, Y) = 0.5389753f0 (tracked)
loss(X, Y) = 0.49384308f0 (tracked)
loss(X, Y) = 0.51260203f0 (tracked)
loss(X, Y) = 0.5330589f0 (tracked)
loss(X, Y) = 0.4865585f0 (tracked)
loss(X, Y) = 0.50366414f0 (tracked)
loss(X, Y) = 0.5697868f0 (tracked)
loss(X, Y) = 0.5893416f0 (tracked)
loss(X, Y) = 0.5271586f0 (tracked)
loss(X, Y) = 0.50030005f0 (tracked)
loss(X, Y) = 0.52616364f0 (tracked)
loss(X, Y) = 0.4990564f0 (tracked)
loss(X, Y) = 0.50080943f0 (tracked)
loss(X, Y) = 0.515071f0 (tracked)
loss(X, Y) = 0.4720558f0 (tracked)
loss(X, Y) = 0.5150554f0 (tracked)
loss(X, Y) = 0.53934073f0 (tracked)
loss(X, Y) = 0.5109552f0 (tracked)
loss(X, Y) = 0.4808869f0 (tracked)
loss(X, Y) = 0.5143465f0 (tracked)
loss(X, Y) = 0.5554795f0 (tracked)
loss(X, Y) = 0.49119925f0 (tracked)
loss(X, Y) = 0.53052306f0 (tracked)
loss(X, Y) = 0.5309513f0 (tracked)
loss(X, Y) = 0.6024628f0 (tracked)
loss(X, Y) = 0.49941766f0 (tracked)
loss(X, Y) = 0.4752831f0 (tracked)
loss(X, Y) = 0.49182522f0 (tracked)
loss(X, Y) = 0.5368246f0 (tracked)
loss(X, Y) = 0.51331085f0 (tracked)
loss(X, Y) = 0.48300496f0 (tracked)
loss(X, Y) = 0.49627045f0 (tracked)
loss(X, Y) = 0.52177644f0 (tracked)
loss(X, Y) = 0.5338519f0 (tracked)
loss(X, Y) = 0.5429963f0 (tracked)
loss(X, Y) = 0.519486f0 (tracked)
loss(X, Y) = 0.57000375f0 (tracked)
loss(X, Y) = 0.4871512f0 (tracked)
loss(X, Y) = 0.52734023f0 (tracked)
loss(X, Y) = 0.5270099f0 (tracked)
loss(X, Y) = 0.5286549f0 (tracked)
loss(X, Y) = 0.47789446f0 (tracked)
loss(X, Y) = 0.48371315f0 (tracked)
loss(X, Y) = 0.51814646f0 (tracked)
loss(X, Y) = 0.51444054f0 (tracked)
loss(X, Y) = 0.50976545f0 (tracked)
loss(X, Y) = 0.5159013f0 (tracked)
loss(X, Y) = 0.49816614f0 (tracked)
loss(X, Y) = 0.48078337f0 (tracked)
loss(X, Y) = 0.502165f0 (tracked)
loss(X, Y) = 0.5024865f0 (tracked)
loss(X, Y) = 0.512472f0 (tracked)
loss(X, Y) = 0.5349612f0 (tracked)
loss(X, Y) = 0.4515831f0 (tracked)
loss(X, Y) = 0.5352235f0 (tracked)
loss(X, Y) = 0.49660444f0 (tracked)
loss(X, Y) = 0.46703845f0 (tracked)
loss(X, Y) = 0.5349447f0 (tracked)
loss(X, Y) = 0.5405726f0 (tracked)
loss(X, Y) = 0.5171532f0 (tracked)
loss(X, Y) = 0.5590195f0 (tracked)
loss(X, Y) = 0.50008065f0 (tracked)
loss(X, Y) = 0.5414509f0 (tracked)
loss(X, Y) = 0.5476699f0 (tracked)
loss(X, Y) = 0.46738216f0 (tracked)
```

```
┌ Info: Epoch 1819
└ @ Main /Users/kcft114/.julia/packages/Flux/qXNjB/src/optimise/train.jl:105
┌ Info: Epoch 1820
└ @ Main /Users/kcft114/.julia/packages/Flux/qXNjB/src/optimise/train.jl:105
┌ Info: Epoch 1821
└ @ Main /Users/kcft114/.julia/packages/Flux/qXNjB/src/optimise/train.jl:105
┌ Info: Epoch 1822
└ @ Main /Users/kcft114/.julia/packages/Flux/qXNjB/src/optimise/train.jl:105
┌ Info: Epoch 1823
└ @ Main /Users/kcft114/.julia/packages/Flux/qXNjB/src/optimise/train.jl:105
┌ Info: Epoch 1824
└ @ Main /Users/kcft114/.julia/packages/Flux/qXNjB/src/optimise/train.jl:105
┌ Info: Epoch 1825
└ @ Main /Users/kcft114/.julia/packages/Flux/qXNjB/src/optimise/train.jl:105
┌ Info: Epoch 1826
└ @ Main /Users/kcft114/.julia/packages/Flux/qXNjB/src/optimise/train.jl:105
┌ Info: Epoch 1827
└ @ Main /Users/kcft114/.julia/packages/Flux/qXNjB/src/optimise/train.jl:105
┌ Info: Epoch 1828
└ @ Main /Users/kcft114/.julia/packages/Flux/qXNjB/src/optimise/train.jl:105
┌ Info: Epoch 1829
└ @ Main /Users/kcft114/.julia/packages/Flux/qXNjB/src/optimise/train.jl:105
┌ Info: Epoch 1830
└ @ Main /Users/kcft114/.julia/packages/Flux/qXNjB/src/optimise/train.jl:105
┌ Info: Epoch 1831
└ @ Main /Users/kcft114/.julia/packages/Flux/qXNjB/src/optimise/train.jl:105
┌ Info: Epoch 1832
└ @ Main /Users/kcft114/.julia/packages/Flux/qXNjB/src/optimise/train.jl:105
┌ Info: Epoch 1833
└ @ Main /Users/kcft114/.julia/packages/Flux/qXNjB/src/optimise/train.jl:105
┌ Info: Epoch 1834
└ @ Main /Users/kcft114/.julia/packages/Flux/qXNjB/src/optimise/train.jl:105
┌ Info: Epoch 1835
└ @ Main /Users/kcft114/.julia/packages/Flux/qXNjB/src/optimise/train.jl:105
┌ Info: Epoch 1836
└ @ Main /Users/kcft114/.julia/packages/Flux/qXNjB/src/optimise/train.jl:105
┌ Info: Epoch 1837
└ @ Main /Users/kcft114/.julia/packages/Flux/qXNjB/src/optimise/train.jl:105
┌ Info: Epoch 1838
└ @ Main /Users/kcft114/.julia/packages/Flux/qXNjB/src/optimise/train.jl:105
┌ Info: Epoch 1839
└ @ Main /Users/kcft114/.julia/packages/Flux/qXNjB/src/optimise/train.jl:105
┌ Info: Epoch 1840
└ @ Main /Users/kcft114/.julia/packages/Flux/qXNjB/src/optimise/train.jl:105
┌ Info: Epoch 1841
└ @ Main /Users/kcft114/.julia/packages/Flux/qXNjB/src/optimise/train.jl:105
┌ Info: Epoch 1842
└ @ Main /Users/kcft114/.julia/packages/Flux/qXNjB/src/optimise/train.jl:105
┌ Info: Epoch 1843
└ @ Main /Users/kcft114/.julia/packages/Flux/qXNjB/src/optimise/train.jl:105
┌ Info: Epoch 1844
└ @ Main /Users/kcft114/.julia/packages/Flux/qXNjB/src/optimise/train.jl:105
┌ Info: Epoch 1845
└ @ Main /Users/kcft114/.julia/packages/Flux/qXNjB/src/optimise/train.jl:105
┌ Info: Epoch 1846
└ @ Main /Users/kcft114/.julia/packages/Flux/qXNjB/src/optimise/train.jl:105
┌ Info: Epoch 1847
└ @ Main /Users/kcft114/.julia/packages/Flux/qXNjB/src/optimise/train.jl:105
┌ Info: Epoch 1848
└ @ Main /Users/kcft114/.julia/packages/Flux/qXNjB/src/optimise/train.jl:105
┌ Info: Epoch 1849
└ @ Main /Users/kcft114/.julia/packages/Flux/qXNjB/src/optimise/train.jl:105
┌ Info: Epoch 1850
└ @ Main /Users/kcft114/.julia/packages/Flux/qXNjB/src/optimise/train.jl:105
┌ Info: Epoch 1851
└ @ Main /Users/kcft114/.julia/packages/Flux/qXNjB/src/optimise/train.jl:105
┌ Info: Epoch 1852
└ @ Main /Users/kcft114/.julia/packages/Flux/qXNjB/src/optimise/train.jl:105
┌ Info: Epoch 1853
└ @ Main /Users/kcft114/.julia/packages/Flux/qXNjB/src/optimise/train.jl:105
┌ Info: Epoch 1854
└ @ Main /Users/kcft114/.julia/packages/Flux/qXNjB/src/optimise/train.jl:105
┌ Info: Epoch 1855
└ @ Main /Users/kcft114/.julia/packages/Flux/qXNjB/src/optimise/train.jl:105
┌ Info: Epoch 1856
└ @ Main /Users/kcft114/.julia/packages/Flux/qXNjB/src/optimise/train.jl:105
┌ Info: Epoch 1857
└ @ Main /Users/kcft114/.julia/packages/Flux/qXNjB/src/optimise/train.jl:105
┌ Info: Epoch 1858
└ @ Main /Users/kcft114/.julia/packages/Flux/qXNjB/src/optimise/train.jl:105
┌ Info: Epoch 1859
└ @ Main /Users/kcft114/.julia/packages/Flux/qXNjB/src/optimise/train.jl:105
┌ Info: Epoch 1860
└ @ Main /Users/kcft114/.julia/packages/Flux/qXNjB/src/optimise/train.jl:105
┌ Info: Epoch 1861
└ @ Main /Users/kcft114/.julia/packages/Flux/qXNjB/src/optimise/train.jl:105
┌ Info: Epoch 1862
└ @ Main /Users/kcft114/.julia/packages/Flux/qXNjB/src/optimise/train.jl:105
┌ Info: Epoch 1863
└ @ Main /Users/kcft114/.julia/packages/Flux/qXNjB/src/optimise/train.jl:105
┌ Info: Epoch 1864
└ @ Main /Users/kcft114/.julia/packages/Flux/qXNjB/src/optimise/train.jl:105
┌ Info: Epoch 1865
└ @ Main /Users/kcft114/.julia/packages/Flux/qXNjB/src/optimise/train.jl:105
┌ Info: Epoch 1866
└ @ Main /Users/kcft114/.julia/packages/Flux/qXNjB/src/optimise/train.jl:105
┌ Info: Epoch 1867
└ @ Main /Users/kcft114/.julia/packages/Flux/qXNjB/src/optimise/train.jl:105
┌ Info: Epoch 1868
└ @ Main /Users/kcft114/.julia/packages/Flux/qXNjB/src/optimise/train.jl:105
┌ Info: Epoch 1869
└ @ Main /Users/kcft114/.julia/packages/Flux/qXNjB/src/optimise/train.jl:105
┌ Info: Epoch 1870
└ @ Main /Users/kcft114/.julia/packages/Flux/qXNjB/src/optimise/train.jl:105
┌ Info: Epoch 1871
└ @ Main /Users/kcft114/.julia/packages/Flux/qXNjB/src/optimise/train.jl:105
┌ Info: Epoch 1872
└ @ Main /Users/kcft114/.julia/packages/Flux/qXNjB/src/optimise/train.jl:105
┌ Info: Epoch 1873
└ @ Main /Users/kcft114/.julia/packages/Flux/qXNjB/src/optimise/train.jl:105
┌ Info: Epoch 1874
└ @ Main /Users/kcft114/.julia/packages/Flux/qXNjB/src/optimise/train.jl:105
┌ Info: Epoch 1875
└ @ Main /Users/kcft114/.julia/packages/Flux/qXNjB/src/optimise/train.jl:105
┌ Info: Epoch 1876
└ @ Main /Users/kcft114/.julia/packages/Flux/qXNjB/src/optimise/train.jl:105
┌ Info: Epoch 1877
└ @ Main /Users/kcft114/.julia/packages/Flux/qXNjB/src/optimise/train.jl:105
┌ Info: Epoch 1878
└ @ Main /Users/kcft114/.julia/packages/Flux/qXNjB/src/optimise/train.jl:105
┌ Info: Epoch 1879
└ @ Main /Users/kcft114/.julia/packages/Flux/qXNjB/src/optimise/train.jl:105
┌ Info: Epoch 1880
└ @ Main /Users/kcft114/.julia/packages/Flux/qXNjB/src/optimise/train.jl:105
┌ Info: Epoch 1881
└ @ Main /Users/kcft114/.julia/packages/Flux/qXNjB/src/optimise/train.jl:105
┌ Info: Epoch 1882
└ @ Main /Users/kcft114/.julia/packages/Flux/qXNjB/src/optimise/train.jl:105
┌ Info: Epoch 1883
└ @ Main /Users/kcft114/.julia/packages/Flux/qXNjB/src/optimise/train.jl:105
┌ Info: Epoch 1884
└ @ Main /Users/kcft114/.julia/packages/Flux/qXNjB/src/optimise/train.jl:105
┌ Info: Epoch 1885
└ @ Main /Users/kcft114/.julia/packages/Flux/qXNjB/src/optimise/train.jl:105
┌ Info: Epoch 1886
└ @ Main /Users/kcft114/.julia/packages/Flux/qXNjB/src/optimise/train.jl:105
┌ Info: Epoch 1887
└ @ Main /Users/kcft114/.julia/packages/Flux/qXNjB/src/optimise/train.jl:105
┌ Info: Epoch 1888
└ @ Main /Users/kcft114/.julia/packages/Flux/qXNjB/src/optimise/train.jl:105
┌ Info: Epoch 1889
└ @ Main /Users/kcft114/.julia/packages/Flux/qXNjB/src/optimise/train.jl:105
```

In [13]:

```
p_dropout=0.2

n_exper = 100
experiments_02 = zeros(n_exper, 4)

for i=1:n_exper
    
    nn = Chain(
              Dense(8, n_h, relu),
              Dropout(p_dropout),
              Dense(n_h, 3),
              softmax)
    
    loss(x, y) = crossentropy(nn(x), y)
    accuracy(x, y) = mean(onecold(nn(x)) .== onecold(y))

    dataset = [(X, Y)]
    evalcb = () -> @show(loss(X, Y))
    
    opt = ADAM(0.001)
    
    @epochs 2000 Flux.train!(loss, Flux.params(nn), dataset, opt, cb = throttle(evalcb, 10));
    
    ac_train, ac_test, ba_train, ba_test =  res(X, Y, Xt, Yt);
    experiments_02[i,:] = [ac_train, ac_test, ba_train, ba_test]
end
```

```
┌ Info: Epoch 1
└ @ Main /Users/kcft114/.julia/packages/Flux/qXNjB/src/optimise/train.jl:105
┌ Info: Epoch 2
└ @ Main /Users/kcft114/.julia/packages/Flux/qXNjB/src/optimise/train.jl:105
┌ Info: Epoch 3
└ @ Main /Users/kcft114/.julia/packages/Flux/qXNjB/src/optimise/train.jl:105
┌ Info: Epoch 4
└ @ Main /Users/kcft114/.julia/packages/Flux/qXNjB/src/optimise/train.jl:105
┌ Info: Epoch 5
└ @ Main /Users/kcft114/.julia/packages/Flux/qXNjB/src/optimise/train.jl:105
┌ Info: Epoch 6
└ @ Main /Users/kcft114/.julia/packages/Flux/qXNjB/src/optimise/train.jl:105
┌ Info: Epoch 7
└ @ Main /Users/kcft114/.julia/packages/Flux/qXNjB/src/optimise/train.jl:105
┌ Info: Epoch 8
└ @ Main /Users/kcft114/.julia/packages/Flux/qXNjB/src/optimise/train.jl:105
┌ Info: Epoch 9
└ @ Main /Users/kcft114/.julia/packages/Flux/qXNjB/src/optimise/train.jl:105
┌ Info: Epoch 10
└ @ Main /Users/kcft114/.julia/packages/Flux/qXNjB/src/optimise/train.jl:105
┌ Info: Epoch 11
└ @ Main /Users/kcft114/.julia/packages/Flux/qXNjB/src/optimise/train.jl:105
┌ Info: Epoch 12
└ @ Main /Users/kcft114/.julia/packages/Flux/qXNjB/src/optimise/train.jl:105
┌ Info: Epoch 13
└ @ Main /Users/kcft114/.julia/packages/Flux/qXNjB/src/optimise/train.jl:105
┌ Info: Epoch 14
└ @ Main /Users/kcft114/.julia/packages/Flux/qXNjB/src/optimise/train.jl:105
┌ Info: Epoch 15
└ @ Main /Users/kcft114/.julia/packages/Flux/qXNjB/src/optimise/train.jl:105
┌ Info: Epoch 16
└ @ Main /Users/kcft114/.julia/packages/Flux/qXNjB/src/optimise/train.jl:105
┌ Info: Epoch 17
└ @ Main /Users/kcft114/.julia/packages/Flux/qXNjB/src/optimise/train.jl:105
┌ Info: Epoch 18
└ @ Main /Users/kcft114/.julia/packages/Flux/qXNjB/src/optimise/train.jl:105
┌ Info: Epoch 19
└ @ Main /Users/kcft114/.julia/packages/Flux/qXNjB/src/optimise/train.jl:105
┌ Info: Epoch 20
└ @ Main /Users/kcft114/.julia/packages/Flux/qXNjB/src/optimise/train.jl:105
┌ Info: Epoch 21
└ @ Main /Users/kcft114/.julia/packages/Flux/qXNjB/src/optimise/train.jl:105
┌ Info: Epoch 22
└ @ Main /Users/kcft114/.julia/packages/Flux/qXNjB/src/optimise/train.jl:105
┌ Info: Epoch 23
└ @ Main /Users/kcft114/.julia/packages/Flux/qXNjB/src/optimise/train.jl:105
┌ Info: Epoch 24
└ @ Main /Users/kcft114/.julia/packages/Flux/qXNjB/src/optimise/train.jl:105
┌ Info: Epoch 25
└ @ Main /Users/kcft114/.julia/packages/Flux/qXNjB/src/optimise/train.jl:105
┌ Info: Epoch 26
└ @ Main /Users/kcft114/.julia/packages/Flux/qXNjB/src/optimise/train.jl:105
┌ Info: Epoch 27
└ @ Main /Users/kcft114/.julia/packages/Flux/qXNjB/src/optimise/train.jl:105
┌ Info: Epoch 28
└ @ Main /Users/kcft114/.julia/packages/Flux/qXNjB/src/optimise/train.jl:105
┌ Info: Epoch 29
└ @ Main /Users/kcft114/.julia/packages/Flux/qXNjB/src/optimise/train.jl:105
┌ Info: Epoch 30
└ @ Main /Users/kcft114/.julia/packages/Flux/qXNjB/src/optimise/train.jl:105
┌ Info: Epoch 31
└ @ Main /Users/kcft114/.julia/packages/Flux/qXNjB/src/optimise/train.jl:105
┌ Info: Epoch 32
└ @ Main /Users/kcft114/.julia/packages/Flux/qXNjB/src/optimise/train.jl:105
┌ Info: Epoch 33
└ @ Main /Users/kcft114/.julia/packages/Flux/qXNjB/src/optimise/train.jl:105
┌ Info: Epoch 34
└ @ Main /Users/kcft114/.julia/packages/Flux/qXNjB/src/optimise/train.jl:105
┌ Info: Epoch 35
└ @ Main /Users/kcft114/.julia/packages/Flux/qXNjB/src/optimise/train.jl:105
┌ Info: Epoch 36
└ @ Main /Users/kcft114/.julia/packages/Flux/qXNjB/src/optimise/train.jl:105
┌ Info: Epoch 37
└ @ Main /Users/kcft114/.julia/packages/Flux/qXNjB/src/optimise/train.jl:105
┌ Info: Epoch 38
└ @ Main /Users/kcft114/.julia/packages/Flux/qXNjB/src/optimise/train.jl:105
┌ Info: Epoch 39
└ @ Main /Users/kcft114/.julia/packages/Flux/qXNjB/src/optimise/train.jl:105
┌ Info: Epoch 40
└ @ Main /Users/kcft114/.julia/packages/Flux/qXNjB/src/optimise/train.jl:105
┌ Info: Epoch 41
└ @ Main /Users/kcft114/.julia/packages/Flux/qXNjB/src/optimise/train.jl:105
┌ Info: Epoch 42
└ @ Main /Users/kcft114/.julia/packages/Flux/qXNjB/src/optimise/train.jl:105
┌ Info: Epoch 43
└ @ Main /Users/kcft114/.julia/packages/Flux/qXNjB/src/optimise/train.jl:105
┌ Info: Epoch 44
└ @ Main /Users/kcft114/.julia/packages/Flux/qXNjB/src/optimise/train.jl:105
┌ Info: Epoch 45
└ @ Main /Users/kcft114/.julia/packages/Flux/qXNjB/src/optimise/train.jl:105
┌ Info: Epoch 46
└ @ Main /Users/kcft114/.julia/packages/Flux/qXNjB/src/optimise/train.jl:105
┌ Info: Epoch 47
└ @ Main /Users/kcft114/.julia/packages/Flux/qXNjB/src/optimise/train.jl:105
┌ Info: Epoch 48
└ @ Main /Users/kcft114/.julia/packages/Flux/qXNjB/src/optimise/train.jl:105
┌ Info: Epoch 49
└ @ Main /Users/kcft114/.julia/packages/Flux/qXNjB/src/optimise/train.jl:105
┌ Info: Epoch 50
└ @ Main /Users/kcft114/.julia/packages/Flux/qXNjB/src/optimise/train.jl:105
┌ Info: Epoch 51
└ @ Main /Users/kcft114/.julia/packages/Flux/qXNjB/src/optimise/train.jl:105
┌ Info: Epoch 52
└ @ Main /Users/kcft114/.julia/packages/Flux/qXNjB/src/optimise/train.jl:105
┌ Info: Epoch 53
└ @ Main /Users/kcft114/.julia/packages/Flux/qXNjB/src/optimise/train.jl:105
┌ Info: Epoch 54
└ @ Main /Users/kcft114/.julia/packages/Flux/qXNjB/src/optimise/train.jl:105
┌ Info: Epoch 55
└ @ Main /Users/kcft114/.julia/packages/Flux/qXNjB/src/optimise/train.jl:105
┌ Info: Epoch 56
└ @ Main /Users/kcft114/.julia/packages/Flux/qXNjB/src/optimise/train.jl:105
┌ Info: Epoch 57
└ @ Main /Users/kcft114/.julia/packages/Flux/qXNjB/src/optimise/train.jl:105
┌ Info: Epoch 58
└ @ Main /Users/kcft114/.julia/packages/Flux/qXNjB/src/optimise/train.jl:105
┌ Info: Epoch 59
└ @ Main /Users/kcft114/.julia/packages/Flux/qXNjB/src/optimise/train.jl:105
┌ Info: Epoch 60
└ @ Main /Users/kcft114/.julia/packages/Flux/qXNjB/src/optimise/train.jl:105
┌ Info: Epoch 61
└ @ Main /Users/kcft114/.julia/packages/Flux/qXNjB/src/optimise/train.jl:105
┌ Info: Epoch 62
└ @ Main /Users/kcft114/.julia/packages/Flux/qXNjB/src/optimise/train.jl:105
┌ Info: Epoch 63
└ @ Main /Users/kcft114/.julia/packages/Flux/qXNjB/src/optimise/train.jl:105
┌ Info: Epoch 64
└ @ Main /Users/kcft114/.julia/packages/Flux/qXNjB/src/optimise/train.jl:105
┌ Info: Epoch 65
└ @ Main /Users/kcft114/.julia/packages/Flux/qXNjB/src/optimise/train.jl:105
┌ Info: Epoch 66
└ @ Main /Users/kcft114/.julia/packages/Flux/qXNjB/src/optimise/train.jl:105
┌ Info: Epoch 67
└ @ Main /Users/kcft114/.julia/packages/Flux/qXNjB/src/optimise/train.jl:105
┌ Info: Epoch 68
└ @ Main /Users/kcft114/.julia/packages/Flux/qXNjB/src/optimise/train.jl:105
┌ Info: Epoch 69
└ @ Main /Users/kcft114/.julia/packages/Flux/qXNjB/src/optimise/train.jl:105
┌ Info: Epoch 70
└ @ Main /Users/kcft114/.julia/packages/Flux/qXNjB/src/optimise/train.jl:105
┌ Info: Epoch 71
└ @ Main /Users/kcft114/.julia/packages/Flux/qXNjB/src/optimise/train.jl:105
┌ Info: Epoch 72
└ @ Main /Users/kcft114/.julia/packages/Flux/qXNjB/src/optimise/train.jl:105
┌ Info: Epoch 73
└ @ Main /Users/kcft114/.julia/packages/Flux/qXNjB/src/optimise/train.jl:105
┌ Info: Epoch 74
└ @ Main /Users/kcft114/.julia/packages/Flux/qXNjB/src/optimise/train.jl:105
┌ Info: Epoch 75
└ @ Main /Users/kcft114/.julia/packages/Flux/qXNjB/src/optimise/train.jl:105
┌ Info: Epoch 76
└ @ Main /Users/kcft114/.julia/packages/Flux/qXNjB/src/optimise/train.jl:105
┌ Info: Epoch 77
└ @ Main /Users/kcft114/.julia/packages/Flux/qXNjB/src/optimise/train.jl:105
┌ Info: Epoch 78
└ @ Main /Users/kcft114/.julia/packages/Flux/qXNjB/src/optimise/train.jl:105
┌ Info: Epoch 79
└ @ Main /Users/kcft114/.julia/packages/Flux/qXNjB/src/optimise/train.jl:105
┌ Info: Epoch 80
└ @ Main /Users/kcft114/.julia/packages/Flux/qXNjB/src/optimise/train.jl:105
┌ Info: Epoch 81
└ @ Main /Users/kcft114/.julia/packages/Flux/qXNjB/src/optimise/train.jl:105
┌ Info: Epoch 82
└ @ Main /Users/kcft114/.julia/packages/Flux/qXNjB/src/optimise/train.jl:105
┌ Info: Epoch 83
└ @ Main /Users/kcft114/.julia/packages/Flux/qXNjB/src/optimise/train.jl:105
┌ Info: Epoch 84
└ @ Main /Users/kcft114/.julia/packages/Flux/qXNjB/src/optimise/train.jl:105
┌ Info: Epoch 85
└ @ Main /Users/kcft114/.julia/packages/Flux/qXNjB/src/optimise/train.jl:105
┌ Info: Epoch 86
└ @ Main /Users/kcft114/.julia/packages/Flux/qXNjB/src/optimise/train.jl:105
┌ Info: Epoch 87
└ @ Main /Users/kcft114/.julia/packages/Flux/qXNjB/src/optimise/train.jl:105
┌ Info: Epoch 88
└ @ Main /Users/kcft114/.julia/packages/Flux/qXNjB/src/optimise/train.jl:105
┌ Info: Epoch 89
└ @ Main /Users/kcft114/.julia/packages/Flux/qXNjB/src/optimise/train.jl:105
┌ Info: Epoch 90
└ @ Main /Users/kcft114/.julia/packages/Flux/qXNjB/src/optimise/train.jl:105
┌ Info: Epoch 91
└ @ Main /Users/kcft114/.julia/packages/Flux/qXNjB/src/optimise/train.jl:105
┌ Info: Epoch 92
└ @ Main /Users/kcft114/.julia/packages/Flux/qXNjB/src/optimise/train.jl:105
┌ Info: Epoch 93
└ @ Main /Users/kcft114/.julia/packages/Flux/qXNjB/src/optimise/train.jl:105
┌ Info: Epoch 94
└ @ Main /Users/kcft114/.julia/packages/Flux/qXNjB/src/optimise/train.jl:105
┌ Info: Epoch 95
└ @ Main /Users/kcft114/.julia/packages/Flux/qXNjB/src/optimise/train.jl:105
┌ Info: Epoch 96
└ @ Main /Users/kcft114/.julia/packages/Flux/qXNjB/src/optimise/train.jl:105
┌ Info: Epoch 97
└ @ Main /Users/kcft114/.julia/packages/Flux/qXNjB/src/optimise/train.jl:105
┌ Info: Epoch 98
└ @ Main /Users/kcft114/.julia/packages/Flux/qXNjB/src/optimise/train.jl:105
┌ Info: Epoch 99
└ @ Main /Users/kcft114/.julia/packages/Flux/qXNjB/src/optimise/train.jl:105
┌ Info: Epoch 100
└ @ Main /Users/kcft114/.julia/packages/Flux/qXNjB/src/optimise/train.jl:105
┌ Info: Epoch 101
└ @ Main /Users/kcft114/.julia/packages/Flux/qXNjB/src/optimise/train.jl:105
┌ Info: Epoch 102
└ @ Main /Users/kcft114/.julia/packages/Flux/qXNjB/src/optimise/train.jl:105
┌ Info: Epoch 103
└ @ Main /Users/kcft114/.julia/packages/Flux/qXNjB/src/optimise/train.jl:105
┌ Info: Epoch 104
└ @ Main /Users/kcft114/.julia/packages/Flux/qXNjB/src/optimise/train.jl:105
┌ Info: Epoch 105
└ @ Main /Users/kcft114/.julia/packages/Flux/qXNjB/src/optimise/train.jl:105
┌ Info: Epoch 106
└ @ Main /Users/kcft114/.julia/packages/Flux/qXNjB/src/optimise/train.jl:105
```

```
loss(X, Y) = 1.1113478f0 (tracked)
loss(X, Y) = 1.1492695f0 (tracked)
loss(X, Y) = 1.1534144f0 (tracked)
loss(X, Y) = 1.1448128f0 (tracked)
loss(X, Y) = 1.1126723f0 (tracked)
loss(X, Y) = 1.0870438f0 (tracked)
loss(X, Y) = 1.1250849f0 (tracked)
loss(X, Y) = 1.0969273f0 (tracked)
loss(X, Y) = 1.103306f0 (tracked)
loss(X, Y) = 1.1031895f0 (tracked)
loss(X, Y) = 1.1237857f0 (tracked)
loss(X, Y) = 1.0653992f0 (tracked)
loss(X, Y) = 1.1205912f0 (tracked)
loss(X, Y) = 1.0894563f0 (tracked)
loss(X, Y) = 1.0807567f0 (tracked)
loss(X, Y) = 1.0694877f0 (tracked)
loss(X, Y) = 1.0575188f0 (tracked)
loss(X, Y) = 1.079952f0 (tracked)
loss(X, Y) = 1.0508941f0 (tracked)
loss(X, Y) = 1.0704479f0 (tracked)
loss(X, Y) = 1.0253327f0 (tracked)
loss(X, Y) = 1.0230862f0 (tracked)
loss(X, Y) = 1.0566608f0 (tracked)
loss(X, Y) = 1.0337061f0 (tracked)
loss(X, Y) = 1.0397117f0 (tracked)
loss(X, Y) = 1.0104371f0 (tracked)
loss(X, Y) = 1.0672498f0 (tracked)
loss(X, Y) = 1.0583181f0 (tracked)
loss(X, Y) = 1.0228127f0 (tracked)
loss(X, Y) = 1.0341092f0 (tracked)
loss(X, Y) = 1.0012662f0 (tracked)
loss(X, Y) = 1.0427389f0 (tracked)
loss(X, Y) = 1.0347512f0 (tracked)
loss(X, Y) = 1.0152639f0 (tracked)
loss(X, Y) = 1.010108f0 (tracked)
loss(X, Y) = 1.0280327f0 (tracked)
loss(X, Y) = 0.9674719f0 (tracked)
loss(X, Y) = 1.0283387f0 (tracked)
loss(X, Y) = 0.9795283f0 (tracked)
loss(X, Y) = 1.0031571f0 (tracked)
loss(X, Y) = 1.0357429f0 (tracked)
loss(X, Y) = 0.9909611f0 (tracked)
loss(X, Y) = 1.0453726f0 (tracked)
loss(X, Y) = 1.0180087f0 (tracked)
loss(X, Y) = 0.9794525f0 (tracked)
loss(X, Y) = 0.99696547f0 (tracked)
loss(X, Y) = 0.9776134f0 (tracked)
loss(X, Y) = 0.9938584f0 (tracked)
loss(X, Y) = 1.0062002f0 (tracked)
loss(X, Y) = 1.0326635f0 (tracked)
loss(X, Y) = 0.9918023f0 (tracked)
loss(X, Y) = 0.9759951f0 (tracked)
loss(X, Y) = 0.97690773f0 (tracked)
loss(X, Y) = 0.9574834f0 (tracked)
loss(X, Y) = 0.9930047f0 (tracked)
loss(X, Y) = 0.9706863f0 (tracked)
loss(X, Y) = 0.986235f0 (tracked)
loss(X, Y) = 0.96883833f0 (tracked)
loss(X, Y) = 0.9532915f0 (tracked)
loss(X, Y) = 0.9568286f0 (tracked)
loss(X, Y) = 0.9513425f0 (tracked)
loss(X, Y) = 0.97284985f0 (tracked)
loss(X, Y) = 0.95871043f0 (tracked)
loss(X, Y) = 0.9517722f0 (tracked)
loss(X, Y) = 0.923348f0 (tracked)
loss(X, Y) = 0.9464964f0 (tracked)
loss(X, Y) = 0.952167f0 (tracked)
loss(X, Y) = 0.967156f0 (tracked)
loss(X, Y) = 0.9556727f0 (tracked)
loss(X, Y) = 0.9483724f0 (tracked)
loss(X, Y) = 0.92465353f0 (tracked)
loss(X, Y) = 0.98458433f0 (tracked)
loss(X, Y) = 0.93487275f0 (tracked)
loss(X, Y) = 0.9215784f0 (tracked)
loss(X, Y) = 0.91227126f0 (tracked)
loss(X, Y) = 0.9294658f0 (tracked)
loss(X, Y) = 0.92100805f0 (tracked)
loss(X, Y) = 0.9435561f0 (tracked)
loss(X, Y) = 0.92237216f0 (tracked)
loss(X, Y) = 0.91704965f0 (tracked)
loss(X, Y) = 0.94301337f0 (tracked)
loss(X, Y) = 0.9431028f0 (tracked)
loss(X, Y) = 0.92744297f0 (tracked)
loss(X, Y) = 0.9157876f0 (tracked)
loss(X, Y) = 0.9064448f0 (tracked)
loss(X, Y) = 0.9524017f0 (tracked)
loss(X, Y) = 0.9292033f0 (tracked)
loss(X, Y) = 0.930175f0 (tracked)
loss(X, Y) = 0.93190247f0 (tracked)
loss(X, Y) = 0.88748145f0 (tracked)
loss(X, Y) = 0.9264088f0 (tracked)
loss(X, Y) = 0.90818983f0 (tracked)
loss(X, Y) = 0.93878764f0 (tracked)
loss(X, Y) = 0.918321f0 (tracked)
loss(X, Y) = 0.9010954f0 (tracked)
loss(X, Y) = 0.912588f0 (tracked)
loss(X, Y) = 0.9161818f0 (tracked)
loss(X, Y) = 0.9255636f0 (tracked)
loss(X, Y) = 0.91858315f0 (tracked)
loss(X, Y) = 0.8940776f0 (tracked)
loss(X, Y) = 0.8995177f0 (tracked)
loss(X, Y) = 0.9190555f0 (tracked)
loss(X, Y) = 0.91020626f0 (tracked)
loss(X, Y) = 0.8981625f0 (tracked)
loss(X, Y) = 0.9151942f0 (tracked)
loss(X, Y) = 0.9389174f0 (tracked)
loss(X, Y) = 0.8929245f0 (tracked)
loss(X, Y) = 0.8739648f0 (tracked)
loss(X, Y) = 0.8879576f0 (tracked)
loss(X, Y) = 0.8900267f0 (tracked)
loss(X, Y) = 0.9256084f0 (tracked)
loss(X, Y) = 0.8807229f0 (tracked)
loss(X, Y) = 0.87565154f0 (tracked)
loss(X, Y) = 0.8905283f0 (tracked)
loss(X, Y) = 0.8878864f0 (tracked)
loss(X, Y) = 0.88600385f0 (tracked)
loss(X, Y) = 0.88487387f0 (tracked)
loss(X, Y) = 0.89618677f0 (tracked)
loss(X, Y) = 0.8787115f0 (tracked)
loss(X, Y) = 0.8722152f0 (tracked)
loss(X, Y) = 0.8871686f0 (tracked)
loss(X, Y) = 0.8883888f0 (tracked)
loss(X, Y) = 0.85588324f0 (tracked)
loss(X, Y) = 0.8694663f0 (tracked)
loss(X, Y) = 0.84034866f0 (tracked)
loss(X, Y) = 0.89918214f0 (tracked)
loss(X, Y) = 0.88909787f0 (tracked)
loss(X, Y) = 0.86690557f0 (tracked)
loss(X, Y) = 0.8600706f0 (tracked)
loss(X, Y) = 0.8613163f0 (tracked)
```

```
┌ Info: Epoch 107
└ @ Main /Users/kcft114/.julia/packages/Flux/qXNjB/src/optimise/train.jl:105
┌ Info: Epoch 108
└ @ Main /Users/kcft114/.julia/packages/Flux/qXNjB/src/optimise/train.jl:105
┌ Info: Epoch 109
└ @ Main /Users/kcft114/.julia/packages/Flux/qXNjB/src/optimise/train.jl:105
┌ Info: Epoch 110
└ @ Main /Users/kcft114/.julia/packages/Flux/qXNjB/src/optimise/train.jl:105
┌ Info: Epoch 111
└ @ Main /Users/kcft114/.julia/packages/Flux/qXNjB/src/optimise/train.jl:105
┌ Info: Epoch 112
└ @ Main /Users/kcft114/.julia/packages/Flux/qXNjB/src/optimise/train.jl:105
┌ Info: Epoch 113
└ @ Main /Users/kcft114/.julia/packages/Flux/qXNjB/src/optimise/train.jl:105
┌ Info: Epoch 114
└ @ Main /Users/kcft114/.julia/packages/Flux/qXNjB/src/optimise/train.jl:105
┌ Info: Epoch 115
└ @ Main /Users/kcft114/.julia/packages/Flux/qXNjB/src/optimise/train.jl:105
┌ Info: Epoch 116
└ @ Main /Users/kcft114/.julia/packages/Flux/qXNjB/src/optimise/train.jl:105
┌ Info: Epoch 117
└ @ Main /Users/kcft114/.julia/packages/Flux/qXNjB/src/optimise/train.jl:105
┌ Info: Epoch 118
└ @ Main /Users/kcft114/.julia/packages/Flux/qXNjB/src/optimise/train.jl:105
┌ Info: Epoch 119
└ @ Main /Users/kcft114/.julia/packages/Flux/qXNjB/src/optimise/train.jl:105
┌ Info: Epoch 120
└ @ Main /Users/kcft114/.julia/packages/Flux/qXNjB/src/optimise/train.jl:105
┌ Info: Epoch 121
└ @ Main /Users/kcft114/.julia/packages/Flux/qXNjB/src/optimise/train.jl:105
┌ Info: Epoch 122
└ @ Main /Users/kcft114/.julia/packages/Flux/qXNjB/src/optimise/train.jl:105
┌ Info: Epoch 123
└ @ Main /Users/kcft114/.julia/packages/Flux/qXNjB/src/optimise/train.jl:105
┌ Info: Epoch 124
└ @ Main /Users/kcft114/.julia/packages/Flux/qXNjB/src/optimise/train.jl:105
┌ Info: Epoch 125
└ @ Main /Users/kcft114/.julia/packages/Flux/qXNjB/src/optimise/train.jl:105
┌ Info: Epoch 126
└ @ Main /Users/kcft114/.julia/packages/Flux/qXNjB/src/optimise/train.jl:105
┌ Info: Epoch 127
└ @ Main /Users/kcft114/.julia/packages/Flux/qXNjB/src/optimise/train.jl:105
┌ Info: Epoch 128
└ @ Main /Users/kcft114/.julia/packages/Flux/qXNjB/src/optimise/train.jl:105
┌ Info: Epoch 129
└ @ Main /Users/kcft114/.julia/packages/Flux/qXNjB/src/optimise/train.jl:105
┌ Info: Epoch 130
└ @ Main /Users/kcft114/.julia/packages/Flux/qXNjB/src/optimise/train.jl:105
┌ Info: Epoch 131
└ @ Main /Users/kcft114/.julia/packages/Flux/qXNjB/src/optimise/train.jl:105
┌ Info: Epoch 132
└ @ Main /Users/kcft114/.julia/packages/Flux/qXNjB/src/optimise/train.jl:105
┌ Info: Epoch 133
└ @ Main /Users/kcft114/.julia/packages/Flux/qXNjB/src/optimise/train.jl:105
┌ Info: Epoch 134
└ @ Main /Users/kcft114/.julia/packages/Flux/qXNjB/src/optimise/train.jl:105
┌ Info: Epoch 135
└ @ Main /Users/kcft114/.julia/packages/Flux/qXNjB/src/optimise/train.jl:105
┌ Info: Epoch 136
└ @ Main /Users/kcft114/.julia/packages/Flux/qXNjB/src/optimise/train.jl:105
┌ Info: Epoch 137
└ @ Main /Users/kcft114/.julia/packages/Flux/qXNjB/src/optimise/train.jl:105
┌ Info: Epoch 138
└ @ Main /Users/kcft114/.julia/packages/Flux/qXNjB/src/optimise/train.jl:105
┌ Info: Epoch 139
└ @ Main /Users/kcft114/.julia/packages/Flux/qXNjB/src/optimise/train.jl:105
┌ Info: Epoch 140
└ @ Main /Users/kcft114/.julia/packages/Flux/qXNjB/src/optimise/train.jl:105
┌ Info: Epoch 141
└ @ Main /Users/kcft114/.julia/packages/Flux/qXNjB/src/optimise/train.jl:105
┌ Info: Epoch 142
└ @ Main /Users/kcft114/.julia/packages/Flux/qXNjB/src/optimise/train.jl:105
┌ Info: Epoch 143
└ @ Main /Users/kcft114/.julia/packages/Flux/qXNjB/src/optimise/train.jl:105
┌ Info: Epoch 144
└ @ Main /Users/kcft114/.julia/packages/Flux/qXNjB/src/optimise/train.jl:105
┌ Info: Epoch 145
└ @ Main /Users/kcft114/.julia/packages/Flux/qXNjB/src/optimise/train.jl:105
┌ Info: Epoch 146
└ @ Main /Users/kcft114/.julia/packages/Flux/qXNjB/src/optimise/train.jl:105
┌ Info: Epoch 147
└ @ Main /Users/kcft114/.julia/packages/Flux/qXNjB/src/optimise/train.jl:105
┌ Info: Epoch 148
└ @ Main /Users/kcft114/.julia/packages/Flux/qXNjB/src/optimise/train.jl:105
┌ Info: Epoch 149
└ @ Main /Users/kcft114/.julia/packages/Flux/qXNjB/src/optimise/train.jl:105
┌ Info: Epoch 150
└ @ Main /Users/kcft114/.julia/packages/Flux/qXNjB/src/optimise/train.jl:105
┌ Info: Epoch 151
└ @ Main /Users/kcft114/.julia/packages/Flux/qXNjB/src/optimise/train.jl:105
┌ Info: Epoch 152
└ @ Main /Users/kcft114/.julia/packages/Flux/qXNjB/src/optimise/train.jl:105
┌ Info: Epoch 153
└ @ Main /Users/kcft114/.julia/packages/Flux/qXNjB/src/optimise/train.jl:105
┌ Info: Epoch 154
└ @ Main /Users/kcft114/.julia/packages/Flux/qXNjB/src/optimise/train.jl:105
┌ Info: Epoch 155
└ @ Main /Users/kcft114/.julia/packages/Flux/qXNjB/src/optimise/train.jl:105
┌ Info: Epoch 156
└ @ Main /Users/kcft114/.julia/packages/Flux/qXNjB/src/optimise/train.jl:105
┌ Info: Epoch 157
└ @ Main /Users/kcft114/.julia/packages/Flux/qXNjB/src/optimise/train.jl:105
┌ Info: Epoch 158
└ @ Main /Users/kcft114/.julia/packages/Flux/qXNjB/src/optimise/train.jl:105
┌ Info: Epoch 159
└ @ Main /Users/kcft114/.julia/packages/Flux/qXNjB/src/optimise/train.jl:105
┌ Info: Epoch 160
└ @ Main /Users/kcft114/.julia/packages/Flux/qXNjB/src/optimise/train.jl:105
┌ Info: Epoch 161
└ @ Main /Users/kcft114/.julia/packages/Flux/qXNjB/src/optimise/train.jl:105
┌ Info: Epoch 162
└ @ Main /Users/kcft114/.julia/packages/Flux/qXNjB/src/optimise/train.jl:105
┌ Info: Epoch 163
└ @ Main /Users/kcft114/.julia/packages/Flux/qXNjB/src/optimise/train.jl:105
┌ Info: Epoch 164
└ @ Main /Users/kcft114/.julia/packages/Flux/qXNjB/src/optimise/train.jl:105
┌ Info: Epoch 165
└ @ Main /Users/kcft114/.julia/packages/Flux/qXNjB/src/optimise/train.jl:105
┌ Info: Epoch 166
└ @ Main /Users/kcft114/.julia/packages/Flux/qXNjB/src/optimise/train.jl:105
┌ Info: Epoch 167
└ @ Main /Users/kcft114/.julia/packages/Flux/qXNjB/src/optimise/train.jl:105
┌ Info: Epoch 168
└ @ Main /Users/kcft114/.julia/packages/Flux/qXNjB/src/optimise/train.jl:105
┌ Info: Epoch 169
└ @ Main /Users/kcft114/.julia/packages/Flux/qXNjB/src/optimise/train.jl:105
┌ Info: Epoch 170
└ @ Main /Users/kcft114/.julia/packages/Flux/qXNjB/src/optimise/train.jl:105
┌ Info: Epoch 171
└ @ Main /Users/kcft114/.julia/packages/Flux/qXNjB/src/optimise/train.jl:105
┌ Info: Epoch 172
└ @ Main /Users/kcft114/.julia/packages/Flux/qXNjB/src/optimise/train.jl:105
┌ Info: Epoch 173
└ @ Main /Users/kcft114/.julia/packages/Flux/qXNjB/src/optimise/train.jl:105
┌ Info: Epoch 174
└ @ Main /Users/kcft114/.julia/packages/Flux/qXNjB/src/optimise/train.jl:105
┌ Info: Epoch 175
└ @ Main /Users/kcft114/.julia/packages/Flux/qXNjB/src/optimise/train.jl:105
┌ Info: Epoch 176
└ @ Main /Users/kcft114/.julia/packages/Flux/qXNjB/src/optimise/train.jl:105
┌ Info: Epoch 177
└ @ Main /Users/kcft114/.julia/packages/Flux/qXNjB/src/optimise/train.jl:105
┌ Info: Epoch 178
└ @ Main /Users/kcft114/.julia/packages/Flux/qXNjB/src/optimise/train.jl:105
┌ Info: Epoch 179
└ @ Main /Users/kcft114/.julia/packages/Flux/qXNjB/src/optimise/train.jl:105
┌ Info: Epoch 180
└ @ Main /Users/kcft114/.julia/packages/Flux/qXNjB/src/optimise/train.jl:105
┌ Info: Epoch 181
└ @ Main /Users/kcft114/.julia/packages/Flux/qXNjB/src/optimise/train.jl:105
┌ Info: Epoch 182
└ @ Main /Users/kcft114/.julia/packages/Flux/qXNjB/src/optimise/train.jl:105
┌ Info: Epoch 183
└ @ Main /Users/kcft114/.julia/packages/Flux/qXNjB/src/optimise/train.jl:105
┌ Info: Epoch 184
└ @ Main /Users/kcft114/.julia/packages/Flux/qXNjB/src/optimise/train.jl:105
┌ Info: Epoch 185
└ @ Main /Users/kcft114/.julia/packages/Flux/qXNjB/src/optimise/train.jl:105
┌ Info: Epoch 186
└ @ Main /Users/kcft114/.julia/packages/Flux/qXNjB/src/optimise/train.jl:105
┌ Info: Epoch 187
└ @ Main /Users/kcft114/.julia/packages/Flux/qXNjB/src/optimise/train.jl:105
┌ Info: Epoch 188
└ @ Main /Users/kcft114/.julia/packages/Flux/qXNjB/src/optimise/train.jl:105
┌ Info: Epoch 189
└ @ Main /Users/kcft114/.julia/packages/Flux/qXNjB/src/optimise/train.jl:105
┌ Info: Epoch 190
└ @ Main /Users/kcft114/.julia/packages/Flux/qXNjB/src/optimise/train.jl:105
┌ Info: Epoch 191
└ @ Main /Users/kcft114/.julia/packages/Flux/qXNjB/src/optimise/train.jl:105
┌ Info: Epoch 192
└ @ Main /Users/kcft114/.julia/packages/Flux/qXNjB/src/optimise/train.jl:105
┌ Info: Epoch 193
└ @ Main /Users/kcft114/.julia/packages/Flux/qXNjB/src/optimise/train.jl:105
┌ Info: Epoch 194
└ @ Main /Users/kcft114/.julia/packages/Flux/qXNjB/src/optimise/train.jl:105
┌ Info: Epoch 195
└ @ Main /Users/kcft114/.julia/packages/Flux/qXNjB/src/optimise/train.jl:105
┌ Info: Epoch 196
└ @ Main /Users/kcft114/.julia/packages/Flux/qXNjB/src/optimise/train.jl:105
┌ Info: Epoch 197
└ @ Main /Users/kcft114/.julia/packages/Flux/qXNjB/src/optimise/train.jl:105
┌ Info: Epoch 198
└ @ Main /Users/kcft114/.julia/packages/Flux/qXNjB/src/optimise/train.jl:105
┌ Info: Epoch 199
└ @ Main /Users/kcft114/.julia/packages/Flux/qXNjB/src/optimise/train.jl:105
┌ Info: Epoch 200
└ @ Main /Users/kcft114/.julia/packages/Flux/qXNjB/src/optimise/train.jl:105
┌ Info: Epoch 201
└ @ Main /Users/kcft114/.julia/packages/Flux/qXNjB/src/optimise/train.jl:105
┌ Info: Epoch 202
└ @ Main /Users/kcft114/.julia/packages/Flux/qXNjB/src/optimise/train.jl:105
┌ Info: Epoch 203
└ @ Main /Users/kcft114/.julia/packages/Flux/qXNjB/src/optimise/train.jl:105
┌ Info: Epoch 204
└ @ Main /Users/kcft114/.julia/packages/Flux/qXNjB/src/optimise/train.jl:105
┌ Info: Epoch 205
└ @ Main /Users/kcft114/.julia/packages/Flux/qXNjB/src/optimise/train.jl:105
┌ Info: Epoch 206
└ @ Main /Users/kcft114/.julia/packages/Flux/qXNjB/src/optimise/train.jl:105
┌ Info: Epoch 207
└ @ Main /Users/kcft114/.julia/packages/Flux/qXNjB/src/optimise/train.jl:105
┌ Info: Epoch 208
└ @ Main /Users/kcft114/.julia/packages/Flux/qXNjB/src/optimise/train.jl:105
┌ Info: Epoch 209
└ @ Main /Users/kcft114/.julia/packages/Flux/qXNjB/src/optimise/train.jl:105
┌ Info: Epoch 210
└ @ Main /Users/kcft114/.julia/packages/Flux/qXNjB/src/optimise/train.jl:105
┌ Info: Epoch 211
└ @ Main /Users/kcft114/.julia/packages/Flux/qXNjB/src/optimise/train.jl:105
┌ Info: Epoch 212
└ @ Main /Users/kcft114/.julia/packages/Flux/qXNjB/src/optimise/train.jl:105
┌ Info: Epoch 213
└ @ Main /Users/kcft114/.julia/packages/Flux/qXNjB/src/optimise/train.jl:105
┌ Info: Epoch 214
└ @ Main /Users/kcft114/.julia/packages/Flux/qXNjB/src/optimise/train.jl:105
┌ Info: Epoch 215
└ @ Main /Users/kcft114/.julia/packages/Flux/qXNjB/src/optimise/train.jl:105
┌ Info: Epoch 216
└ @ Main /Users/kcft114/.julia/packages/Flux/qXNjB/src/optimise/train.jl:105
┌ Info: Epoch 217
└ @ Main /Users/kcft114/.julia/packages/Flux/qXNjB/src/optimise/train.jl:105
┌ Info: Epoch 218
└ @ Main /Users/kcft114/.julia/packages/Flux/qXNjB/src/optimise/train.jl:105
┌ Info: Epoch 219
└ @ Main /Users/kcft114/.julia/packages/Flux/qXNjB/src/optimise/train.jl:105
┌ Info: Epoch 220
└ @ Main /Users/kcft114/.julia/packages/Flux/qXNjB/src/optimise/train.jl:105
┌ Info: Epoch 221
└ @ Main /Users/kcft114/.julia/packages/Flux/qXNjB/src/optimise/train.jl:105
┌ Info: Epoch 222
└ @ Main /Users/kcft114/.julia/packages/Flux/qXNjB/src/optimise/train.jl:105
┌ Info: Epoch 223
└ @ Main /Users/kcft114/.julia/packages/Flux/qXNjB/src/optimise/train.jl:105
┌ Info: Epoch 224
└ @ Main /Users/kcft114/.julia/packages/Flux/qXNjB/src/optimise/train.jl:105
┌ Info: Epoch 225
└ @ Main /Users/kcft114/.julia/packages/Flux/qXNjB/src/optimise/train.jl:105
┌ Info: Epoch 226
└ @ Main /Users/kcft114/.julia/packages/Flux/qXNjB/src/optimise/train.jl:105
┌ Info: Epoch 227
└ @ Main /Users/kcft114/.julia/packages/Flux/qXNjB/src/optimise/train.jl:105
┌ Info: Epoch 228
└ @ Main /Users/kcft114/.julia/packages/Flux/qXNjB/src/optimise/train.jl:105
┌ Info: Epoch 229
└ @ Main /Users/kcft114/.julia/packages/Flux/qXNjB/src/optimise/train.jl:105
┌ Info: Epoch 230
└ @ Main /Users/kcft114/.julia/packages/Flux/qXNjB/src/optimise/train.jl:105
┌ Info: Epoch 231
└ @ Main /Users/kcft114/.julia/packages/Flux/qXNjB/src/optimise/train.jl:105
┌ Info: Epoch 232
└ @ Main /Users/kcft114/.julia/packages/Flux/qXNjB/src/optimise/train.jl:105
┌ Info: Epoch 233
└ @ Main /Users/kcft114/.julia/packages/Flux/qXNjB/src/optimise/train.jl:105
┌ Info: Epoch 234
└ @ Main /Users/kcft114/.julia/packages/Flux/qXNjB/src/optimise/train.jl:105
┌ Info: Epoch 235
└ @ Main /Users/kcft114/.julia/packages/Flux/qXNjB/src/optimise/train.jl:105
```

```
loss(X, Y) = 0.88257635f0 (tracked)
loss(X, Y) = 0.8941046f0 (tracked)
loss(X, Y) = 0.8432511f0 (tracked)
loss(X, Y) = 0.9201223f0 (tracked)
loss(X, Y) = 0.83824193f0 (tracked)
loss(X, Y) = 0.8584487f0 (tracked)
loss(X, Y) = 0.8547475f0 (tracked)
loss(X, Y) = 0.868571f0 (tracked)
loss(X, Y) = 0.8592812f0 (tracked)
loss(X, Y) = 0.8651839f0 (tracked)
loss(X, Y) = 0.8522899f0 (tracked)
loss(X, Y) = 0.85400885f0 (tracked)
loss(X, Y) = 0.8667608f0 (tracked)
loss(X, Y) = 0.8206279f0 (tracked)
loss(X, Y) = 0.85266435f0 (tracked)
loss(X, Y) = 0.85203224f0 (tracked)
loss(X, Y) = 0.8485689f0 (tracked)
loss(X, Y) = 0.84195507f0 (tracked)
loss(X, Y) = 0.83808696f0 (tracked)
loss(X, Y) = 0.8522092f0 (tracked)
loss(X, Y) = 0.8351238f0 (tracked)
loss(X, Y) = 0.8503944f0 (tracked)
loss(X, Y) = 0.87995523f0 (tracked)
loss(X, Y) = 0.853232f0 (tracked)
loss(X, Y) = 0.8491866f0 (tracked)
loss(X, Y) = 0.82714784f0 (tracked)
loss(X, Y) = 0.8262466f0 (tracked)
loss(X, Y) = 0.84319687f0 (tracked)
loss(X, Y) = 0.8213619f0 (tracked)
loss(X, Y) = 0.8206196f0 (tracked)
loss(X, Y) = 0.84258556f0 (tracked)
loss(X, Y) = 0.83452857f0 (tracked)
loss(X, Y) = 0.832275f0 (tracked)
loss(X, Y) = 0.84268165f0 (tracked)
loss(X, Y) = 0.85539246f0 (tracked)
loss(X, Y) = 0.845649f0 (tracked)
loss(X, Y) = 0.82629097f0 (tracked)
loss(X, Y) = 0.8222732f0 (tracked)
loss(X, Y) = 0.8235775f0 (tracked)
loss(X, Y) = 0.84951705f0 (tracked)
loss(X, Y) = 0.82111645f0 (tracked)
loss(X, Y) = 0.80833846f0 (tracked)
loss(X, Y) = 0.8104602f0 (tracked)
loss(X, Y) = 0.8597041f0 (tracked)
loss(X, Y) = 0.8016814f0 (tracked)
loss(X, Y) = 0.83050054f0 (tracked)
loss(X, Y) = 0.85007966f0 (tracked)
loss(X, Y) = 0.8320186f0 (tracked)
loss(X, Y) = 0.82904303f0 (tracked)
loss(X, Y) = 0.8253781f0 (tracked)
loss(X, Y) = 0.8194006f0 (tracked)
loss(X, Y) = 0.82544905f0 (tracked)
loss(X, Y) = 0.8206333f0 (tracked)
loss(X, Y) = 0.82963127f0 (tracked)
loss(X, Y) = 0.8411216f0 (tracked)
loss(X, Y) = 0.8308805f0 (tracked)
loss(X, Y) = 0.8042793f0 (tracked)
loss(X, Y) = 0.8251027f0 (tracked)
loss(X, Y) = 0.82732695f0 (tracked)
loss(X, Y) = 0.8317874f0 (tracked)
loss(X, Y) = 0.82402545f0 (tracked)
loss(X, Y) = 0.81669f0 (tracked)
loss(X, Y) = 0.83776027f0 (tracked)
loss(X, Y) = 0.80094445f0 (tracked)
loss(X, Y) = 0.78075004f0 (tracked)
loss(X, Y) = 0.8007416f0 (tracked)
loss(X, Y) = 0.8290643f0 (tracked)
loss(X, Y) = 0.80640244f0 (tracked)
loss(X, Y) = 0.8491944f0 (tracked)
loss(X, Y) = 0.79862136f0 (tracked)
loss(X, Y) = 0.80656767f0 (tracked)
loss(X, Y) = 0.7975205f0 (tracked)
loss(X, Y) = 0.7865899f0 (tracked)
loss(X, Y) = 0.7783908f0 (tracked)
loss(X, Y) = 0.78723794f0 (tracked)
loss(X, Y) = 0.8046142f0 (tracked)
loss(X, Y) = 0.7955858f0 (tracked)
loss(X, Y) = 0.7708917f0 (tracked)
loss(X, Y) = 0.8124183f0 (tracked)
loss(X, Y) = 0.83587545f0 (tracked)
loss(X, Y) = 0.7832502f0 (tracked)
loss(X, Y) = 0.81168085f0 (tracked)
loss(X, Y) = 0.7909828f0 (tracked)
loss(X, Y) = 0.7832199f0 (tracked)
loss(X, Y) = 0.8118319f0 (tracked)
loss(X, Y) = 0.78808457f0 (tracked)
loss(X, Y) = 0.8073761f0 (tracked)
loss(X, Y) = 0.7801225f0 (tracked)
loss(X, Y) = 0.8196234f0 (tracked)
loss(X, Y) = 0.8132583f0 (tracked)
loss(X, Y) = 0.80340743f0 (tracked)
loss(X, Y) = 0.80184f0 (tracked)
loss(X, Y) = 0.8189802f0 (tracked)
loss(X, Y) = 0.7652894f0 (tracked)
loss(X, Y) = 0.83623505f0 (tracked)
loss(X, Y) = 0.7802271f0 (tracked)
loss(X, Y) = 0.78442764f0 (tracked)
loss(X, Y) = 0.776407f0 (tracked)
loss(X, Y) = 0.8238625f0 (tracked)
loss(X, Y) = 0.7937146f0 (tracked)
loss(X, Y) = 0.7999393f0 (tracked)
loss(X, Y) = 0.780249f0 (tracked)
loss(X, Y) = 0.78960717f0 (tracked)
loss(X, Y) = 0.83347887f0 (tracked)
loss(X, Y) = 0.81075126f0 (tracked)
loss(X, Y) = 0.8042198f0 (tracked)
loss(X, Y) = 0.79091066f0 (tracked)
loss(X, Y) = 0.7914785f0 (tracked)
loss(X, Y) = 0.7765451f0 (tracked)
loss(X, Y) = 0.790272f0 (tracked)
loss(X, Y) = 0.79811025f0 (tracked)
loss(X, Y) = 0.7793105f0 (tracked)
loss(X, Y) = 0.7880194f0 (tracked)
loss(X, Y) = 0.7935223f0 (tracked)
loss(X, Y) = 0.78746647f0 (tracked)
loss(X, Y) = 0.76882833f0 (tracked)
loss(X, Y) = 0.7941324f0 (tracked)
loss(X, Y) = 0.77783376f0 (tracked)
loss(X, Y) = 0.81065255f0 (tracked)
loss(X, Y) = 0.7689309f0 (tracked)
loss(X, Y) = 0.76941866f0 (tracked)
```

```
┌ Info: Epoch 236
└ @ Main /Users/kcft114/.julia/packages/Flux/qXNjB/src/optimise/train.jl:105
┌ Info: Epoch 237
└ @ Main /Users/kcft114/.julia/packages/Flux/qXNjB/src/optimise/train.jl:105
┌ Info: Epoch 238
└ @ Main /Users/kcft114/.julia/packages/Flux/qXNjB/src/optimise/train.jl:105
┌ Info: Epoch 239
└ @ Main /Users/kcft114/.julia/packages/Flux/qXNjB/src/optimise/train.jl:105
┌ Info: Epoch 240
└ @ Main /Users/kcft114/.julia/packages/Flux/qXNjB/src/optimise/train.jl:105
┌ Info: Epoch 241
└ @ Main /Users/kcft114/.julia/packages/Flux/qXNjB/src/optimise/train.jl:105
┌ Info: Epoch 242
└ @ Main /Users/kcft114/.julia/packages/Flux/qXNjB/src/optimise/train.jl:105
┌ Info: Epoch 243
└ @ Main /Users/kcft114/.julia/packages/Flux/qXNjB/src/optimise/train.jl:105
┌ Info: Epoch 244
└ @ Main /Users/kcft114/.julia/packages/Flux/qXNjB/src/optimise/train.jl:105
┌ Info: Epoch 245
└ @ Main /Users/kcft114/.julia/packages/Flux/qXNjB/src/optimise/train.jl:105
┌ Info: Epoch 246
└ @ Main /Users/kcft114/.julia/packages/Flux/qXNjB/src/optimise/train.jl:105
┌ Info: Epoch 247
└ @ Main /Users/kcft114/.julia/packages/Flux/qXNjB/src/optimise/train.jl:105
┌ Info: Epoch 248
└ @ Main /Users/kcft114/.julia/packages/Flux/qXNjB/src/optimise/train.jl:105
┌ Info: Epoch 249
└ @ Main /Users/kcft114/.julia/packages/Flux/qXNjB/src/optimise/train.jl:105
┌ Info: Epoch 250
└ @ Main /Users/kcft114/.julia/packages/Flux/qXNjB/src/optimise/train.jl:105
┌ Info: Epoch 251
└ @ Main /Users/kcft114/.julia/packages/Flux/qXNjB/src/optimise/train.jl:105
┌ Info: Epoch 252
└ @ Main /Users/kcft114/.julia/packages/Flux/qXNjB/src/optimise/train.jl:105
┌ Info: Epoch 253
└ @ Main /Users/kcft114/.julia/packages/Flux/qXNjB/src/optimise/train.jl:105
┌ Info: Epoch 254
└ @ Main /Users/kcft114/.julia/packages/Flux/qXNjB/src/optimise/train.jl:105
┌ Info: Epoch 255
└ @ Main /Users/kcft114/.julia/packages/Flux/qXNjB/src/optimise/train.jl:105
┌ Info: Epoch 256
└ @ Main /Users/kcft114/.julia/packages/Flux/qXNjB/src/optimise/train.jl:105
┌ Info: Epoch 257
└ @ Main /Users/kcft114/.julia/packages/Flux/qXNjB/src/optimise/train.jl:105
┌ Info: Epoch 258
└ @ Main /Users/kcft114/.julia/packages/Flux/qXNjB/src/optimise/train.jl:105
┌ Info: Epoch 259
└ @ Main /Users/kcft114/.julia/packages/Flux/qXNjB/src/optimise/train.jl:105
┌ Info: Epoch 260
└ @ Main /Users/kcft114/.julia/packages/Flux/qXNjB/src/optimise/train.jl:105
┌ Info: Epoch 261
└ @ Main /Users/kcft114/.julia/packages/Flux/qXNjB/src/optimise/train.jl:105
┌ Info: Epoch 262
└ @ Main /Users/kcft114/.julia/packages/Flux/qXNjB/src/optimise/train.jl:105
┌ Info: Epoch 263
└ @ Main /Users/kcft114/.julia/packages/Flux/qXNjB/src/optimise/train.jl:105
┌ Info: Epoch 264
└ @ Main /Users/kcft114/.julia/packages/Flux/qXNjB/src/optimise/train.jl:105
┌ Info: Epoch 265
└ @ Main /Users/kcft114/.julia/packages/Flux/qXNjB/src/optimise/train.jl:105
┌ Info: Epoch 266
└ @ Main /Users/kcft114/.julia/packages/Flux/qXNjB/src/optimise/train.jl:105
┌ Info: Epoch 267
└ @ Main /Users/kcft114/.julia/packages/Flux/qXNjB/src/optimise/train.jl:105
┌ Info: Epoch 268
└ @ Main /Users/kcft114/.julia/packages/Flux/qXNjB/src/optimise/train.jl:105
┌ Info: Epoch 269
└ @ Main /Users/kcft114/.julia/packages/Flux/qXNjB/src/optimise/train.jl:105
┌ Info: Epoch 270
└ @ Main /Users/kcft114/.julia/packages/Flux/qXNjB/src/optimise/train.jl:105
┌ Info: Epoch 271
└ @ Main /Users/kcft114/.julia/packages/Flux/qXNjB/src/optimise/train.jl:105
┌ Info: Epoch 272
└ @ Main /Users/kcft114/.julia/packages/Flux/qXNjB/src/optimise/train.jl:105
┌ Info: Epoch 273
└ @ Main /Users/kcft114/.julia/packages/Flux/qXNjB/src/optimise/train.jl:105
┌ Info: Epoch 274
└ @ Main /Users/kcft114/.julia/packages/Flux/qXNjB/src/optimise/train.jl:105
┌ Info: Epoch 275
└ @ Main /Users/kcft114/.julia/packages/Flux/qXNjB/src/optimise/train.jl:105
┌ Info: Epoch 276
└ @ Main /Users/kcft114/.julia/packages/Flux/qXNjB/src/optimise/train.jl:105
┌ Info: Epoch 277
└ @ Main /Users/kcft114/.julia/packages/Flux/qXNjB/src/optimise/train.jl:105
┌ Info: Epoch 278
└ @ Main /Users/kcft114/.julia/packages/Flux/qXNjB/src/optimise/train.jl:105
┌ Info: Epoch 279
└ @ Main /Users/kcft114/.julia/packages/Flux/qXNjB/src/optimise/train.jl:105
┌ Info: Epoch 280
└ @ Main /Users/kcft114/.julia/packages/Flux/qXNjB/src/optimise/train.jl:105
┌ Info: Epoch 281
└ @ Main /Users/kcft114/.julia/packages/Flux/qXNjB/src/optimise/train.jl:105
┌ Info: Epoch 282
└ @ Main /Users/kcft114/.julia/packages/Flux/qXNjB/src/optimise/train.jl:105
┌ Info: Epoch 283
└ @ Main /Users/kcft114/.julia/packages/Flux/qXNjB/src/optimise/train.jl:105
┌ Info: Epoch 284
└ @ Main /Users/kcft114/.julia/packages/Flux/qXNjB/src/optimise/train.jl:105
┌ Info: Epoch 285
└ @ Main /Users/kcft114/.julia/packages/Flux/qXNjB/src/optimise/train.jl:105
┌ Info: Epoch 286
└ @ Main /Users/kcft114/.julia/packages/Flux/qXNjB/src/optimise/train.jl:105
┌ Info: Epoch 287
└ @ Main /Users/kcft114/.julia/packages/Flux/qXNjB/src/optimise/train.jl:105
┌ Info: Epoch 288
└ @ Main /Users/kcft114/.julia/packages/Flux/qXNjB/src/optimise/train.jl:105
┌ Info: Epoch 289
└ @ Main /Users/kcft114/.julia/packages/Flux/qXNjB/src/optimise/train.jl:105
┌ Info: Epoch 290
└ @ Main /Users/kcft114/.julia/packages/Flux/qXNjB/src/optimise/train.jl:105
┌ Info: Epoch 291
└ @ Main /Users/kcft114/.julia/packages/Flux/qXNjB/src/optimise/train.jl:105
┌ Info: Epoch 292
└ @ Main /Users/kcft114/.julia/packages/Flux/qXNjB/src/optimise/train.jl:105
┌ Info: Epoch 293
└ @ Main /Users/kcft114/.julia/packages/Flux/qXNjB/src/optimise/train.jl:105
┌ Info: Epoch 294
└ @ Main /Users/kcft114/.julia/packages/Flux/qXNjB/src/optimise/train.jl:105
┌ Info: Epoch 295
└ @ Main /Users/kcft114/.julia/packages/Flux/qXNjB/src/optimise/train.jl:105
┌ Info: Epoch 296
└ @ Main /Users/kcft114/.julia/packages/Flux/qXNjB/src/optimise/train.jl:105
┌ Info: Epoch 297
└ @ Main /Users/kcft114/.julia/packages/Flux/qXNjB/src/optimise/train.jl:105
┌ Info: Epoch 298
└ @ Main /Users/kcft114/.julia/packages/Flux/qXNjB/src/optimise/train.jl:105
┌ Info: Epoch 299
└ @ Main /Users/kcft114/.julia/packages/Flux/qXNjB/src/optimise/train.jl:105
┌ Info: Epoch 300
└ @ Main /Users/kcft114/.julia/packages/Flux/qXNjB/src/optimise/train.jl:105
┌ Info: Epoch 301
└ @ Main /Users/kcft114/.julia/packages/Flux/qXNjB/src/optimise/train.jl:105
┌ Info: Epoch 302
└ @ Main /Users/kcft114/.julia/packages/Flux/qXNjB/src/optimise/train.jl:105
┌ Info: Epoch 303
└ @ Main /Users/kcft114/.julia/packages/Flux/qXNjB/src/optimise/train.jl:105
┌ Info: Epoch 304
└ @ Main /Users/kcft114/.julia/packages/Flux/qXNjB/src/optimise/train.jl:105
┌ Info: Epoch 305
└ @ Main /Users/kcft114/.julia/packages/Flux/qXNjB/src/optimise/train.jl:105
┌ Info: Epoch 306
└ @ Main /Users/kcft114/.julia/packages/Flux/qXNjB/src/optimise/train.jl:105
┌ Info: Epoch 307
└ @ Main /Users/kcft114/.julia/packages/Flux/qXNjB/src/optimise/train.jl:105
┌ Info: Epoch 308
└ @ Main /Users/kcft114/.julia/packages/Flux/qXNjB/src/optimise/train.jl:105
┌ Info: Epoch 309
└ @ Main /Users/kcft114/.julia/packages/Flux/qXNjB/src/optimise/train.jl:105
┌ Info: Epoch 310
└ @ Main /Users/kcft114/.julia/packages/Flux/qXNjB/src/optimise/train.jl:105
┌ Info: Epoch 311
└ @ Main /Users/kcft114/.julia/packages/Flux/qXNjB/src/optimise/train.jl:105
┌ Info: Epoch 312
└ @ Main /Users/kcft114/.julia/packages/Flux/qXNjB/src/optimise/train.jl:105
┌ Info: Epoch 313
└ @ Main /Users/kcft114/.julia/packages/Flux/qXNjB/src/optimise/train.jl:105
┌ Info: Epoch 314
└ @ Main /Users/kcft114/.julia/packages/Flux/qXNjB/src/optimise/train.jl:105
┌ Info: Epoch 315
└ @ Main /Users/kcft114/.julia/packages/Flux/qXNjB/src/optimise/train.jl:105
┌ Info: Epoch 316
└ @ Main /Users/kcft114/.julia/packages/Flux/qXNjB/src/optimise/train.jl:105
┌ Info: Epoch 317
└ @ Main /Users/kcft114/.julia/packages/Flux/qXNjB/src/optimise/train.jl:105
┌ Info: Epoch 318
└ @ Main /Users/kcft114/.julia/packages/Flux/qXNjB/src/optimise/train.jl:105
┌ Info: Epoch 319
└ @ Main /Users/kcft114/.julia/packages/Flux/qXNjB/src/optimise/train.jl:105
┌ Info: Epoch 320
└ @ Main /Users/kcft114/.julia/packages/Flux/qXNjB/src/optimise/train.jl:105
┌ Info: Epoch 321
└ @ Main /Users/kcft114/.julia/packages/Flux/qXNjB/src/optimise/train.jl:105
┌ Info: Epoch 322
└ @ Main /Users/kcft114/.julia/packages/Flux/qXNjB/src/optimise/train.jl:105
┌ Info: Epoch 323
└ @ Main /Users/kcft114/.julia/packages/Flux/qXNjB/src/optimise/train.jl:105
┌ Info: Epoch 324
└ @ Main /Users/kcft114/.julia/packages/Flux/qXNjB/src/optimise/train.jl:105
┌ Info: Epoch 325
└ @ Main /Users/kcft114/.julia/packages/Flux/qXNjB/src/optimise/train.jl:105
┌ Info: Epoch 326
└ @ Main /Users/kcft114/.julia/packages/Flux/qXNjB/src/optimise/train.jl:105
┌ Info: Epoch 327
└ @ Main /Users/kcft114/.julia/packages/Flux/qXNjB/src/optimise/train.jl:105
┌ Info: Epoch 328
└ @ Main /Users/kcft114/.julia/packages/Flux/qXNjB/src/optimise/train.jl:105
┌ Info: Epoch 329
└ @ Main /Users/kcft114/.julia/packages/Flux/qXNjB/src/optimise/train.jl:105
┌ Info: Epoch 330
└ @ Main /Users/kcft114/.julia/packages/Flux/qXNjB/src/optimise/train.jl:105
┌ Info: Epoch 331
└ @ Main /Users/kcft114/.julia/packages/Flux/qXNjB/src/optimise/train.jl:105
┌ Info: Epoch 332
└ @ Main /Users/kcft114/.julia/packages/Flux/qXNjB/src/optimise/train.jl:105
┌ Info: Epoch 333
└ @ Main /Users/kcft114/.julia/packages/Flux/qXNjB/src/optimise/train.jl:105
┌ Info: Epoch 334
└ @ Main /Users/kcft114/.julia/packages/Flux/qXNjB/src/optimise/train.jl:105
┌ Info: Epoch 335
└ @ Main /Users/kcft114/.julia/packages/Flux/qXNjB/src/optimise/train.jl:105
┌ Info: Epoch 336
└ @ Main /Users/kcft114/.julia/packages/Flux/qXNjB/src/optimise/train.jl:105
┌ Info: Epoch 337
└ @ Main /Users/kcft114/.julia/packages/Flux/qXNjB/src/optimise/train.jl:105
┌ Info: Epoch 338
└ @ Main /Users/kcft114/.julia/packages/Flux/qXNjB/src/optimise/train.jl:105
┌ Info: Epoch 339
└ @ Main /Users/kcft114/.julia/packages/Flux/qXNjB/src/optimise/train.jl:105
┌ Info: Epoch 340
└ @ Main /Users/kcft114/.julia/packages/Flux/qXNjB/src/optimise/train.jl:105
┌ Info: Epoch 341
└ @ Main /Users/kcft114/.julia/packages/Flux/qXNjB/src/optimise/train.jl:105
┌ Info: Epoch 342
└ @ Main /Users/kcft114/.julia/packages/Flux/qXNjB/src/optimise/train.jl:105
┌ Info: Epoch 343
└ @ Main /Users/kcft114/.julia/packages/Flux/qXNjB/src/optimise/train.jl:105
┌ Info: Epoch 344
└ @ Main /Users/kcft114/.julia/packages/Flux/qXNjB/src/optimise/train.jl:105
┌ Info: Epoch 345
└ @ Main /Users/kcft114/.julia/packages/Flux/qXNjB/src/optimise/train.jl:105
┌ Info: Epoch 346
└ @ Main /Users/kcft114/.julia/packages/Flux/qXNjB/src/optimise/train.jl:105
┌ Info: Epoch 347
└ @ Main /Users/kcft114/.julia/packages/Flux/qXNjB/src/optimise/train.jl:105
┌ Info: Epoch 348
└ @ Main /Users/kcft114/.julia/packages/Flux/qXNjB/src/optimise/train.jl:105
┌ Info: Epoch 349
└ @ Main /Users/kcft114/.julia/packages/Flux/qXNjB/src/optimise/train.jl:105
┌ Info: Epoch 350
└ @ Main /Users/kcft114/.julia/packages/Flux/qXNjB/src/optimise/train.jl:105
┌ Info: Epoch 351
└ @ Main /Users/kcft114/.julia/packages/Flux/qXNjB/src/optimise/train.jl:105
┌ Info: Epoch 352
└ @ Main /Users/kcft114/.julia/packages/Flux/qXNjB/src/optimise/train.jl:105
┌ Info: Epoch 353
└ @ Main /Users/kcft114/.julia/packages/Flux/qXNjB/src/optimise/train.jl:105
┌ Info: Epoch 354
└ @ Main /Users/kcft114/.julia/packages/Flux/qXNjB/src/optimise/train.jl:105
┌ Info: Epoch 355
└ @ Main /Users/kcft114/.julia/packages/Flux/qXNjB/src/optimise/train.jl:105
┌ Info: Epoch 356
└ @ Main /Users/kcft114/.julia/packages/Flux/qXNjB/src/optimise/train.jl:105
```

```
loss(X, Y) = 0.7921099f0 (tracked)
loss(X, Y) = 0.7837072f0 (tracked)
loss(X, Y) = 0.80166554f0 (tracked)
loss(X, Y) = 0.78587085f0 (tracked)
loss(X, Y) = 0.79118085f0 (tracked)
loss(X, Y) = 0.79419094f0 (tracked)
loss(X, Y) = 0.7835252f0 (tracked)
loss(X, Y) = 0.7849315f0 (tracked)
loss(X, Y) = 0.78208435f0 (tracked)
loss(X, Y) = 0.8017802f0 (tracked)
loss(X, Y) = 0.75471264f0 (tracked)
loss(X, Y) = 0.7531905f0 (tracked)
loss(X, Y) = 0.80289143f0 (tracked)
loss(X, Y) = 0.7733814f0 (tracked)
loss(X, Y) = 0.7574645f0 (tracked)
loss(X, Y) = 0.78756136f0 (tracked)
loss(X, Y) = 0.7629026f0 (tracked)
loss(X, Y) = 0.7696289f0 (tracked)
loss(X, Y) = 0.7901974f0 (tracked)
loss(X, Y) = 0.77070826f0 (tracked)
loss(X, Y) = 0.7646903f0 (tracked)
loss(X, Y) = 0.7707862f0 (tracked)
loss(X, Y) = 0.7484944f0 (tracked)
loss(X, Y) = 0.78835255f0 (tracked)
loss(X, Y) = 0.78910154f0 (tracked)
loss(X, Y) = 0.77336466f0 (tracked)
loss(X, Y) = 0.7844243f0 (tracked)
loss(X, Y) = 0.77026695f0 (tracked)
loss(X, Y) = 0.749443f0 (tracked)
loss(X, Y) = 0.764494f0 (tracked)
loss(X, Y) = 0.7831767f0 (tracked)
loss(X, Y) = 0.76344675f0 (tracked)
loss(X, Y) = 0.7604459f0 (tracked)
loss(X, Y) = 0.76928174f0 (tracked)
loss(X, Y) = 0.7638524f0 (tracked)
loss(X, Y) = 0.77187365f0 (tracked)
loss(X, Y) = 0.7564281f0 (tracked)
loss(X, Y) = 0.75255805f0 (tracked)
loss(X, Y) = 0.75905323f0 (tracked)
loss(X, Y) = 0.73888373f0 (tracked)
loss(X, Y) = 0.78246075f0 (tracked)
loss(X, Y) = 0.7587052f0 (tracked)
loss(X, Y) = 0.7596579f0 (tracked)
loss(X, Y) = 0.73375493f0 (tracked)
loss(X, Y) = 0.7652463f0 (tracked)
loss(X, Y) = 0.7533731f0 (tracked)
loss(X, Y) = 0.74677676f0 (tracked)
loss(X, Y) = 0.7532315f0 (tracked)
loss(X, Y) = 0.74197596f0 (tracked)
loss(X, Y) = 0.76300776f0 (tracked)
loss(X, Y) = 0.765548f0 (tracked)
loss(X, Y) = 0.76515967f0 (tracked)
loss(X, Y) = 0.76897085f0 (tracked)
loss(X, Y) = 0.77401865f0 (tracked)
loss(X, Y) = 0.72058827f0 (tracked)
loss(X, Y) = 0.7480129f0 (tracked)
loss(X, Y) = 0.771096f0 (tracked)
loss(X, Y) = 0.8003042f0 (tracked)
loss(X, Y) = 0.74623847f0 (tracked)
loss(X, Y) = 0.7588491f0 (tracked)
loss(X, Y) = 0.7721027f0 (tracked)
loss(X, Y) = 0.7701784f0 (tracked)
loss(X, Y) = 0.76849014f0 (tracked)
loss(X, Y) = 0.75399745f0 (tracked)
loss(X, Y) = 0.7292208f0 (tracked)
loss(X, Y) = 0.7546197f0 (tracked)
loss(X, Y) = 0.77720106f0 (tracked)
loss(X, Y) = 0.75062543f0 (tracked)
loss(X, Y) = 0.75790167f0 (tracked)
loss(X, Y) = 0.7415742f0 (tracked)
loss(X, Y) = 0.7627325f0 (tracked)
loss(X, Y) = 0.73586583f0 (tracked)
loss(X, Y) = 0.77450967f0 (tracked)
loss(X, Y) = 0.75616485f0 (tracked)
loss(X, Y) = 0.75246435f0 (tracked)
loss(X, Y) = 0.7520823f0 (tracked)
loss(X, Y) = 0.74945265f0 (tracked)
loss(X, Y) = 0.74672306f0 (tracked)
loss(X, Y) = 0.7418408f0 (tracked)
loss(X, Y) = 0.7888396f0 (tracked)
loss(X, Y) = 0.7668653f0 (tracked)
loss(X, Y) = 0.74515843f0 (tracked)
loss(X, Y) = 0.7555436f0 (tracked)
loss(X, Y) = 0.7563837f0 (tracked)
loss(X, Y) = 0.76755977f0 (tracked)
loss(X, Y) = 0.7646646f0 (tracked)
loss(X, Y) = 0.7406469f0 (tracked)
loss(X, Y) = 0.7632015f0 (tracked)
loss(X, Y) = 0.7327693f0 (tracked)
loss(X, Y) = 0.7321269f0 (tracked)
loss(X, Y) = 0.7403862f0 (tracked)
loss(X, Y) = 0.74030954f0 (tracked)
loss(X, Y) = 0.7513976f0 (tracked)
loss(X, Y) = 0.7870315f0 (tracked)
loss(X, Y) = 0.73634577f0 (tracked)
loss(X, Y) = 0.72851765f0 (tracked)
loss(X, Y) = 0.7472997f0 (tracked)
loss(X, Y) = 0.74309576f0 (tracked)
loss(X, Y) = 0.7515338f0 (tracked)
loss(X, Y) = 0.7218366f0 (tracked)
loss(X, Y) = 0.74938524f0 (tracked)
loss(X, Y) = 0.7471119f0 (tracked)
loss(X, Y) = 0.73706585f0 (tracked)
loss(X, Y) = 0.7319288f0 (tracked)
loss(X, Y) = 0.73041874f0 (tracked)
loss(X, Y) = 0.7521726f0 (tracked)
loss(X, Y) = 0.74093574f0 (tracked)
loss(X, Y) = 0.7077182f0 (tracked)
loss(X, Y) = 0.74351925f0 (tracked)
loss(X, Y) = 0.7259629f0 (tracked)
loss(X, Y) = 0.74183685f0 (tracked)
```

```
┌ Info: Epoch 357
└ @ Main /Users/kcft114/.julia/packages/Flux/qXNjB/src/optimise/train.jl:105
┌ Info: Epoch 358
└ @ Main /Users/kcft114/.julia/packages/Flux/qXNjB/src/optimise/train.jl:105
┌ Info: Epoch 359
└ @ Main /Users/kcft114/.julia/packages/Flux/qXNjB/src/optimise/train.jl:105
┌ Info: Epoch 360
└ @ Main /Users/kcft114/.julia/packages/Flux/qXNjB/src/optimise/train.jl:105
┌ Info: Epoch 361
└ @ Main /Users/kcft114/.julia/packages/Flux/qXNjB/src/optimise/train.jl:105
┌ Info: Epoch 362
└ @ Main /Users/kcft114/.julia/packages/Flux/qXNjB/src/optimise/train.jl:105
┌ Info: Epoch 363
└ @ Main /Users/kcft114/.julia/packages/Flux/qXNjB/src/optimise/train.jl:105
```

```
loss(X, Y) = 0.7182704f0 (tracked)
loss(X, Y) = 0.7292726f0 (tracked)
loss(X, Y) = 0.7656302f0 (tracked)
loss(X, Y) = 0.7130085f0 (tracked)
loss(X, Y) = 0.7441958f0 (tracked)
loss(X, Y) = 0.7351872f0 (tracked)
loss(X, Y) = 0.7277545f0 (tracked)
loss(X, Y) = 0.74570125f0 (tracked)
loss(X, Y) = 0.73636013f0 (tracked)
loss(X, Y) = 0.7391541f0 (tracked)
loss(X, Y) = 0.72211945f0 (tracked)
loss(X, Y) = 0.7507459f0 (tracked)
loss(X, Y) = 0.708476f0 (tracked)
loss(X, Y) = 0.70451045f0 (tracked)
loss(X, Y) = 0.7440689f0 (tracked)
loss(X, Y) = 0.7514486f0 (tracked)
loss(X, Y) = 0.73894346f0 (tracked)
loss(X, Y) = 0.75489706f0 (tracked)
loss(X, Y) = 0.7523526f0 (tracked)
loss(X, Y) = 0.7661804f0 (tracked)
loss(X, Y) = 0.72895664f0 (tracked)
loss(X, Y) = 0.7322571f0 (tracked)
loss(X, Y) = 0.7138413f0 (tracked)
loss(X, Y) = 0.7414525f0 (tracked)
loss(X, Y) = 0.7131963f0 (tracked)
loss(X, Y) = 0.7418677f0 (tracked)
loss(X, Y) = 0.7216176f0 (tracked)
loss(X, Y) = 0.73283947f0 (tracked)
loss(X, Y) = 0.7259317f0 (tracked)
loss(X, Y) = 0.71372414f0 (tracked)
loss(X, Y) = 0.72312456f0 (tracked)
loss(X, Y) = 0.73530906f0 (tracked)
loss(X, Y) = 0.7409443f0 (tracked)
loss(X, Y) = 0.75059134f0 (tracked)
loss(X, Y) = 0.7071032f0 (tracked)
loss(X, Y) = 0.7217097f0 (tracked)
loss(X, Y) = 0.7217257f0 (tracked)
loss(X, Y) = 0.71434593f0 (tracked)
loss(X, Y) = 0.7257962f0 (tracked)
loss(X, Y) = 0.71572214f0 (tracked)
loss(X, Y) = 0.7649252f0 (tracked)
loss(X, Y) = 0.72361404f0 (tracked)
loss(X, Y) = 0.7318265f0 (tracked)
loss(X, Y) = 0.7282835f0 (tracked)
loss(X, Y) = 0.7413142f0 (tracked)
loss(X, Y) = 0.73176765f0 (tracked)
loss(X, Y) = 0.73666406f0 (tracked)
loss(X, Y) = 0.7375829f0 (tracked)
loss(X, Y) = 0.72684616f0 (tracked)
loss(X, Y) = 0.70843756f0 (tracked)
loss(X, Y) = 0.7267443f0 (tracked)
loss(X, Y) = 0.73352265f0 (tracked)
loss(X, Y) = 0.73330516f0 (tracked)
loss(X, Y) = 0.7174133f0 (tracked)
loss(X, Y) = 0.7069299f0 (tracked)
loss(X, Y) = 0.77329457f0 (tracked)
loss(X, Y) = 0.6988699f0 (tracked)
loss(X, Y) = 0.7229214f0 (tracked)
loss(X, Y) = 0.70889837f0 (tracked)
loss(X, Y) = 0.73382336f0 (tracked)
loss(X, Y) = 0.73708194f0 (tracked)
loss(X, Y) = 0.68362117f0 (tracked)
loss(X, Y) = 0.72952515f0 (tracked)
loss(X, Y) = 0.70500255f0 (tracked)
loss(X, Y) = 0.71366507f0 (tracked)
loss(X, Y) = 0.7326777f0 (tracked)
loss(X, Y) = 0.7068612f0 (tracked)
loss(X, Y) = 0.71383363f0 (tracked)
loss(X, Y) = 0.7110864f0 (tracked)
loss(X, Y) = 0.7059664f0 (tracked)
loss(X, Y) = 0.7000369f0 (tracked)
loss(X, Y) = 0.7081352f0 (tracked)
loss(X, Y) = 0.7240244f0 (tracked)
loss(X, Y) = 0.693644f0 (tracked)
loss(X, Y) = 0.69927263f0 (tracked)
loss(X, Y) = 0.7107599f0 (tracked)
loss(X, Y) = 0.71619755f0 (tracked)
loss(X, Y) = 0.69693124f0 (tracked)
loss(X, Y) = 0.72171676f0 (tracked)
loss(X, Y) = 0.71050966f0 (tracked)
loss(X, Y) = 0.6999358f0 (tracked)
loss(X, Y) = 0.70798546f0 (tracked)
loss(X, Y) = 0.7166876f0 (tracked)
loss(X, Y) = 0.72071207f0 (tracked)
loss(X, Y) = 0.7208266f0 (tracked)
loss(X, Y) = 0.7143612f0 (tracked)
loss(X, Y) = 0.7191428f0 (tracked)
loss(X, Y) = 0.70445615f0 (tracked)
loss(X, Y) = 0.7032702f0 (tracked)
loss(X, Y) = 0.7040572f0 (tracked)
loss(X, Y) = 0.7452208f0 (tracked)
loss(X, Y) = 0.745844f0 (tracked)
loss(X, Y) = 0.7366503f0 (tracked)
loss(X, Y) = 0.669532f0 (tracked)
loss(X, Y) = 0.697757f0 (tracked)
loss(X, Y) = 0.6839629f0 (tracked)
loss(X, Y) = 0.7057353f0 (tracked)
```

```
┌ Info: Epoch 364
└ @ Main /Users/kcft114/.julia/packages/Flux/qXNjB/src/optimise/train.jl:105
┌ Info: Epoch 365
└ @ Main /Users/kcft114/.julia/packages/Flux/qXNjB/src/optimise/train.jl:105
┌ Info: Epoch 366
└ @ Main /Users/kcft114/.julia/packages/Flux/qXNjB/src/optimise/train.jl:105
┌ Info: Epoch 367
└ @ Main /Users/kcft114/.julia/packages/Flux/qXNjB/src/optimise/train.jl:105
┌ Info: Epoch 368
└ @ Main /Users/kcft114/.julia/packages/Flux/qXNjB/src/optimise/train.jl:105
┌ Info: Epoch 369
└ @ Main /Users/kcft114/.julia/packages/Flux/qXNjB/src/optimise/train.jl:105
┌ Info: Epoch 370
└ @ Main /Users/kcft114/.julia/packages/Flux/qXNjB/src/optimise/train.jl:105
┌ Info: Epoch 371
└ @ Main /Users/kcft114/.julia/packages/Flux/qXNjB/src/optimise/train.jl:105
┌ Info: Epoch 372
└ @ Main /Users/kcft114/.julia/packages/Flux/qXNjB/src/optimise/train.jl:105
┌ Info: Epoch 373
└ @ Main /Users/kcft114/.julia/packages/Flux/qXNjB/src/optimise/train.jl:105
┌ Info: Epoch 374
└ @ Main /Users/kcft114/.julia/packages/Flux/qXNjB/src/optimise/train.jl:105
┌ Info: Epoch 375
└ @ Main /Users/kcft114/.julia/packages/Flux/qXNjB/src/optimise/train.jl:105
┌ Info: Epoch 376
└ @ Main /Users/kcft114/.julia/packages/Flux/qXNjB/src/optimise/train.jl:105
┌ Info: Epoch 377
└ @ Main /Users/kcft114/.julia/packages/Flux/qXNjB/src/optimise/train.jl:105
┌ Info: Epoch 378
└ @ Main /Users/kcft114/.julia/packages/Flux/qXNjB/src/optimise/train.jl:105
┌ Info: Epoch 379
└ @ Main /Users/kcft114/.julia/packages/Flux/qXNjB/src/optimise/train.jl:105
┌ Info: Epoch 380
└ @ Main /Users/kcft114/.julia/packages/Flux/qXNjB/src/optimise/train.jl:105
┌ Info: Epoch 381
└ @ Main /Users/kcft114/.julia/packages/Flux/qXNjB/src/optimise/train.jl:105
┌ Info: Epoch 382
└ @ Main /Users/kcft114/.julia/packages/Flux/qXNjB/src/optimise/train.jl:105
┌ Info: Epoch 383
└ @ Main /Users/kcft114/.julia/packages/Flux/qXNjB/src/optimise/train.jl:105
┌ Info: Epoch 384
└ @ Main /Users/kcft114/.julia/packages/Flux/qXNjB/src/optimise/train.jl:105
┌ Info: Epoch 385
└ @ Main /Users/kcft114/.julia/packages/Flux/qXNjB/src/optimise/train.jl:105
┌ Info: Epoch 386
└ @ Main /Users/kcft114/.julia/packages/Flux/qXNjB/src/optimise/train.jl:105
┌ Info: Epoch 387
└ @ Main /Users/kcft114/.julia/packages/Flux/qXNjB/src/optimise/train.jl:105
┌ Info: Epoch 388
└ @ Main /Users/kcft114/.julia/packages/Flux/qXNjB/src/optimise/train.jl:105
┌ Info: Epoch 389
└ @ Main /Users/kcft114/.julia/packages/Flux/qXNjB/src/optimise/train.jl:105
┌ Info: Epoch 390
└ @ Main /Users/kcft114/.julia/packages/Flux/qXNjB/src/optimise/train.jl:105
┌ Info: Epoch 391
└ @ Main /Users/kcft114/.julia/packages/Flux/qXNjB/src/optimise/train.jl:105
┌ Info: Epoch 392
└ @ Main /Users/kcft114/.julia/packages/Flux/qXNjB/src/optimise/train.jl:105
┌ Info: Epoch 393
└ @ Main /Users/kcft114/.julia/packages/Flux/qXNjB/src/optimise/train.jl:105
┌ Info: Epoch 394
└ @ Main /Users/kcft114/.julia/packages/Flux/qXNjB/src/optimise/train.jl:105
┌ Info: Epoch 395
└ @ Main /Users/kcft114/.julia/packages/Flux/qXNjB/src/optimise/train.jl:105
┌ Info: Epoch 396
└ @ Main /Users/kcft114/.julia/packages/Flux/qXNjB/src/optimise/train.jl:105
┌ Info: Epoch 397
└ @ Main /Users/kcft114/.julia/packages/Flux/qXNjB/src/optimise/train.jl:105
┌ Info: Epoch 398
└ @ Main /Users/kcft114/.julia/packages/Flux/qXNjB/src/optimise/train.jl:105
┌ Info: Epoch 399
└ @ Main /Users/kcft114/.julia/packages/Flux/qXNjB/src/optimise/train.jl:105
┌ Info: Epoch 400
└ @ Main /Users/kcft114/.julia/packages/Flux/qXNjB/src/optimise/train.jl:105
┌ Info: Epoch 401
└ @ Main /Users/kcft114/.julia/packages/Flux/qXNjB/src/optimise/train.jl:105
┌ Info: Epoch 402
└ @ Main /Users/kcft114/.julia/packages/Flux/qXNjB/src/optimise/train.jl:105
┌ Info: Epoch 403
└ @ Main /Users/kcft114/.julia/packages/Flux/qXNjB/src/optimise/train.jl:105
┌ Info: Epoch 404
└ @ Main /Users/kcft114/.julia/packages/Flux/qXNjB/src/optimise/train.jl:105
┌ Info: Epoch 405
└ @ Main /Users/kcft114/.julia/packages/Flux/qXNjB/src/optimise/train.jl:105
┌ Info: Epoch 406
└ @ Main /Users/kcft114/.julia/packages/Flux/qXNjB/src/optimise/train.jl:105
┌ Info: Epoch 407
└ @ Main /Users/kcft114/.julia/packages/Flux/qXNjB/src/optimise/train.jl:105
┌ Info: Epoch 408
└ @ Main /Users/kcft114/.julia/packages/Flux/qXNjB/src/optimise/train.jl:105
┌ Info: Epoch 409
└ @ Main /Users/kcft114/.julia/packages/Flux/qXNjB/src/optimise/train.jl:105
┌ Info: Epoch 410
└ @ Main /Users/kcft114/.julia/packages/Flux/qXNjB/src/optimise/train.jl:105
┌ Info: Epoch 411
└ @ Main /Users/kcft114/.julia/packages/Flux/qXNjB/src/optimise/train.jl:105
┌ Info: Epoch 412
└ @ Main /Users/kcft114/.julia/packages/Flux/qXNjB/src/optimise/train.jl:105
┌ Info: Epoch 413
└ @ Main /Users/kcft114/.julia/packages/Flux/qXNjB/src/optimise/train.jl:105
┌ Info: Epoch 414
└ @ Main /Users/kcft114/.julia/packages/Flux/qXNjB/src/optimise/train.jl:105
┌ Info: Epoch 415
└ @ Main /Users/kcft114/.julia/packages/Flux/qXNjB/src/optimise/train.jl:105
┌ Info: Epoch 416
└ @ Main /Users/kcft114/.julia/packages/Flux/qXNjB/src/optimise/train.jl:105
┌ Info: Epoch 417
└ @ Main /Users/kcft114/.julia/packages/Flux/qXNjB/src/optimise/train.jl:105
┌ Info: Epoch 418
└ @ Main /Users/kcft114/.julia/packages/Flux/qXNjB/src/optimise/train.jl:105
┌ Info: Epoch 419
└ @ Main /Users/kcft114/.julia/packages/Flux/qXNjB/src/optimise/train.jl:105
┌ Info: Epoch 420
└ @ Main /Users/kcft114/.julia/packages/Flux/qXNjB/src/optimise/train.jl:105
┌ Info: Epoch 421
└ @ Main /Users/kcft114/.julia/packages/Flux/qXNjB/src/optimise/train.jl:105
┌ Info: Epoch 422
└ @ Main /Users/kcft114/.julia/packages/Flux/qXNjB/src/optimise/train.jl:105
┌ Info: Epoch 423
└ @ Main /Users/kcft114/.julia/packages/Flux/qXNjB/src/optimise/train.jl:105
┌ Info: Epoch 424
└ @ Main /Users/kcft114/.julia/packages/Flux/qXNjB/src/optimise/train.jl:105
┌ Info: Epoch 425
└ @ Main /Users/kcft114/.julia/packages/Flux/qXNjB/src/optimise/train.jl:105
┌ Info: Epoch 426
└ @ Main /Users/kcft114/.julia/packages/Flux/qXNjB/src/optimise/train.jl:105
┌ Info: Epoch 427
└ @ Main /Users/kcft114/.julia/packages/Flux/qXNjB/src/optimise/train.jl:105
┌ Info: Epoch 428
└ @ Main /Users/kcft114/.julia/packages/Flux/qXNjB/src/optimise/train.jl:105
┌ Info: Epoch 429
└ @ Main /Users/kcft114/.julia/packages/Flux/qXNjB/src/optimise/train.jl:105
┌ Info: Epoch 430
└ @ Main /Users/kcft114/.julia/packages/Flux/qXNjB/src/optimise/train.jl:105
┌ Info: Epoch 431
└ @ Main /Users/kcft114/.julia/packages/Flux/qXNjB/src/optimise/train.jl:105
┌ Info: Epoch 432
└ @ Main /Users/kcft114/.julia/packages/Flux/qXNjB/src/optimise/train.jl:105
┌ Info: Epoch 433
└ @ Main /Users/kcft114/.julia/packages/Flux/qXNjB/src/optimise/train.jl:105
┌ Info: Epoch 434
└ @ Main /Users/kcft114/.julia/packages/Flux/qXNjB/src/optimise/train.jl:105
┌ Info: Epoch 435
└ @ Main /Users/kcft114/.julia/packages/Flux/qXNjB/src/optimise/train.jl:105
┌ Info: Epoch 436
└ @ Main /Users/kcft114/.julia/packages/Flux/qXNjB/src/optimise/train.jl:105
┌ Info: Epoch 437
└ @ Main /Users/kcft114/.julia/packages/Flux/qXNjB/src/optimise/train.jl:105
┌ Info: Epoch 438
└ @ Main /Users/kcft114/.julia/packages/Flux/qXNjB/src/optimise/train.jl:105
┌ Info: Epoch 439
└ @ Main /Users/kcft114/.julia/packages/Flux/qXNjB/src/optimise/train.jl:105
┌ Info: Epoch 440
└ @ Main /Users/kcft114/.julia/packages/Flux/qXNjB/src/optimise/train.jl:105
┌ Info: Epoch 441
└ @ Main /Users/kcft114/.julia/packages/Flux/qXNjB/src/optimise/train.jl:105
┌ Info: Epoch 442
└ @ Main /Users/kcft114/.julia/packages/Flux/qXNjB/src/optimise/train.jl:105
┌ Info: Epoch 443
└ @ Main /Users/kcft114/.julia/packages/Flux/qXNjB/src/optimise/train.jl:105
┌ Info: Epoch 444
└ @ Main /Users/kcft114/.julia/packages/Flux/qXNjB/src/optimise/train.jl:105
┌ Info: Epoch 445
└ @ Main /Users/kcft114/.julia/packages/Flux/qXNjB/src/optimise/train.jl:105
┌ Info: Epoch 446
└ @ Main /Users/kcft114/.julia/packages/Flux/qXNjB/src/optimise/train.jl:105
┌ Info: Epoch 447
└ @ Main /Users/kcft114/.julia/packages/Flux/qXNjB/src/optimise/train.jl:105
┌ Info: Epoch 448
└ @ Main /Users/kcft114/.julia/packages/Flux/qXNjB/src/optimise/train.jl:105
┌ Info: Epoch 449
└ @ Main /Users/kcft114/.julia/packages/Flux/qXNjB/src/optimise/train.jl:105
┌ Info: Epoch 450
└ @ Main /Users/kcft114/.julia/packages/Flux/qXNjB/src/optimise/train.jl:105
┌ Info: Epoch 451
└ @ Main /Users/kcft114/.julia/packages/Flux/qXNjB/src/optimise/train.jl:105
┌ Info: Epoch 452
└ @ Main /Users/kcft114/.julia/packages/Flux/qXNjB/src/optimise/train.jl:105
┌ Info: Epoch 453
└ @ Main /Users/kcft114/.julia/packages/Flux/qXNjB/src/optimise/train.jl:105
┌ Info: Epoch 454
└ @ Main /Users/kcft114/.julia/packages/Flux/qXNjB/src/optimise/train.jl:105
┌ Info: Epoch 455
└ @ Main /Users/kcft114/.julia/packages/Flux/qXNjB/src/optimise/train.jl:105
┌ Info: Epoch 456
└ @ Main /Users/kcft114/.julia/packages/Flux/qXNjB/src/optimise/train.jl:105
┌ Info: Epoch 457
└ @ Main /Users/kcft114/.julia/packages/Flux/qXNjB/src/optimise/train.jl:105
┌ Info: Epoch 458
└ @ Main /Users/kcft114/.julia/packages/Flux/qXNjB/src/optimise/train.jl:105
┌ Info: Epoch 459
└ @ Main /Users/kcft114/.julia/packages/Flux/qXNjB/src/optimise/train.jl:105
┌ Info: Epoch 460
└ @ Main /Users/kcft114/.julia/packages/Flux/qXNjB/src/optimise/train.jl:105
```

```
loss(X, Y) = 0.6884932f0 (tracked)
loss(X, Y) = 0.70809615f0 (tracked)
loss(X, Y) = 0.6851491f0 (tracked)
loss(X, Y) = 0.716466f0 (tracked)
loss(X, Y) = 0.6917794f0 (tracked)
loss(X, Y) = 0.7073826f0 (tracked)
loss(X, Y) = 0.71886486f0 (tracked)
loss(X, Y) = 0.6920383f0 (tracked)
loss(X, Y) = 0.6880775f0 (tracked)
loss(X, Y) = 0.7150735f0 (tracked)
loss(X, Y) = 0.7296476f0 (tracked)
loss(X, Y) = 0.7237244f0 (tracked)
loss(X, Y) = 0.72541666f0 (tracked)
loss(X, Y) = 0.71104884f0 (tracked)
loss(X, Y) = 0.7154587f0 (tracked)
loss(X, Y) = 0.6877275f0 (tracked)
loss(X, Y) = 0.7353712f0 (tracked)
loss(X, Y) = 0.70469964f0 (tracked)
loss(X, Y) = 0.714882f0 (tracked)
loss(X, Y) = 0.7376424f0 (tracked)
loss(X, Y) = 0.7160187f0 (tracked)
loss(X, Y) = 0.70995706f0 (tracked)
loss(X, Y) = 0.7098313f0 (tracked)
loss(X, Y) = 0.7321763f0 (tracked)
loss(X, Y) = 0.697226f0 (tracked)
loss(X, Y) = 0.69364816f0 (tracked)
loss(X, Y) = 0.713038f0 (tracked)
loss(X, Y) = 0.6977931f0 (tracked)
loss(X, Y) = 0.7024135f0 (tracked)
loss(X, Y) = 0.6878433f0 (tracked)
loss(X, Y) = 0.69714767f0 (tracked)
loss(X, Y) = 0.6639751f0 (tracked)
loss(X, Y) = 0.7204876f0 (tracked)
loss(X, Y) = 0.74077576f0 (tracked)
loss(X, Y) = 0.6924138f0 (tracked)
loss(X, Y) = 0.7032279f0 (tracked)
loss(X, Y) = 0.7032713f0 (tracked)
loss(X, Y) = 0.68268985f0 (tracked)
loss(X, Y) = 0.67920685f0 (tracked)
loss(X, Y) = 0.7298005f0 (tracked)
loss(X, Y) = 0.6774819f0 (tracked)
loss(X, Y) = 0.7292235f0 (tracked)
loss(X, Y) = 0.69823426f0 (tracked)
loss(X, Y) = 0.7168259f0 (tracked)
loss(X, Y) = 0.68708485f0 (tracked)
loss(X, Y) = 0.714912f0 (tracked)
loss(X, Y) = 0.7082247f0 (tracked)
loss(X, Y) = 0.6935947f0 (tracked)
loss(X, Y) = 0.67753565f0 (tracked)
loss(X, Y) = 0.7178832f0 (tracked)
loss(X, Y) = 0.6955384f0 (tracked)
loss(X, Y) = 0.6960391f0 (tracked)
loss(X, Y) = 0.6985563f0 (tracked)
loss(X, Y) = 0.67897004f0 (tracked)
loss(X, Y) = 0.6929695f0 (tracked)
loss(X, Y) = 0.68199f0 (tracked)
loss(X, Y) = 0.68078136f0 (tracked)
loss(X, Y) = 0.7492181f0 (tracked)
loss(X, Y) = 0.7163653f0 (tracked)
loss(X, Y) = 0.6775329f0 (tracked)
loss(X, Y) = 0.7104631f0 (tracked)
loss(X, Y) = 0.7121f0 (tracked)
loss(X, Y) = 0.7163881f0 (tracked)
loss(X, Y) = 0.7121201f0 (tracked)
loss(X, Y) = 0.7029744f0 (tracked)
loss(X, Y) = 0.7104666f0 (tracked)
loss(X, Y) = 0.72073275f0 (tracked)
loss(X, Y) = 0.6935743f0 (tracked)
loss(X, Y) = 0.6806164f0 (tracked)
loss(X, Y) = 0.67376477f0 (tracked)
loss(X, Y) = 0.72277415f0 (tracked)
loss(X, Y) = 0.693914f0 (tracked)
loss(X, Y) = 0.6830842f0 (tracked)
loss(X, Y) = 0.7017613f0 (tracked)
loss(X, Y) = 0.7034446f0 (tracked)
loss(X, Y) = 0.6929571f0 (tracked)
loss(X, Y) = 0.70291007f0 (tracked)
loss(X, Y) = 0.6658184f0 (tracked)
loss(X, Y) = 0.72725004f0 (tracked)
loss(X, Y) = 0.6759496f0 (tracked)
loss(X, Y) = 0.7390278f0 (tracked)
loss(X, Y) = 0.71687645f0 (tracked)
loss(X, Y) = 0.6801093f0 (tracked)
loss(X, Y) = 0.68873006f0 (tracked)
loss(X, Y) = 0.6567411f0 (tracked)
loss(X, Y) = 0.6630326f0 (tracked)
loss(X, Y) = 0.69965035f0 (tracked)
loss(X, Y) = 0.69858176f0 (tracked)
loss(X, Y) = 0.7007929f0 (tracked)
loss(X, Y) = 0.69072616f0 (tracked)
loss(X, Y) = 0.6814263f0 (tracked)
loss(X, Y) = 0.7232098f0 (tracked)
loss(X, Y) = 0.70467585f0 (tracked)
loss(X, Y) = 0.6759303f0 (tracked)
loss(X, Y) = 0.67970854f0 (tracked)
loss(X, Y) = 0.66709644f0 (tracked)
loss(X, Y) = 0.69833744f0 (tracked)
loss(X, Y) = 0.6770774f0 (tracked)
loss(X, Y) = 0.68364054f0 (tracked)
loss(X, Y) = 0.7056117f0 (tracked)
loss(X, Y) = 0.70698476f0 (tracked)
loss(X, Y) = 0.6653902f0 (tracked)
loss(X, Y) = 0.7205131f0 (tracked)
```

```
┌ Info: Epoch 461
└ @ Main /Users/kcft114/.julia/packages/Flux/qXNjB/src/optimise/train.jl:105
┌ Info: Epoch 462
└ @ Main /Users/kcft114/.julia/packages/Flux/qXNjB/src/optimise/train.jl:105
┌ Info: Epoch 463
└ @ Main /Users/kcft114/.julia/packages/Flux/qXNjB/src/optimise/train.jl:105
┌ Info: Epoch 464
└ @ Main /Users/kcft114/.julia/packages/Flux/qXNjB/src/optimise/train.jl:105
┌ Info: Epoch 465
└ @ Main /Users/kcft114/.julia/packages/Flux/qXNjB/src/optimise/train.jl:105
┌ Info: Epoch 466
└ @ Main /Users/kcft114/.julia/packages/Flux/qXNjB/src/optimise/train.jl:105
┌ Info: Epoch 467
└ @ Main /Users/kcft114/.julia/packages/Flux/qXNjB/src/optimise/train.jl:105
┌ Info: Epoch 468
└ @ Main /Users/kcft114/.julia/packages/Flux/qXNjB/src/optimise/train.jl:105
┌ Info: Epoch 469
└ @ Main /Users/kcft114/.julia/packages/Flux/qXNjB/src/optimise/train.jl:105
┌ Info: Epoch 470
└ @ Main /Users/kcft114/.julia/packages/Flux/qXNjB/src/optimise/train.jl:105
┌ Info: Epoch 471
└ @ Main /Users/kcft114/.julia/packages/Flux/qXNjB/src/optimise/train.jl:105
┌ Info: Epoch 472
└ @ Main /Users/kcft114/.julia/packages/Flux/qXNjB/src/optimise/train.jl:105
┌ Info: Epoch 473
└ @ Main /Users/kcft114/.julia/packages/Flux/qXNjB/src/optimise/train.jl:105
┌ Info: Epoch 474
└ @ Main /Users/kcft114/.julia/packages/Flux/qXNjB/src/optimise/train.jl:105
┌ Info: Epoch 475
└ @ Main /Users/kcft114/.julia/packages/Flux/qXNjB/src/optimise/train.jl:105
┌ Info: Epoch 476
└ @ Main /Users/kcft114/.julia/packages/Flux/qXNjB/src/optimise/train.jl:105
┌ Info: Epoch 477
└ @ Main /Users/kcft114/.julia/packages/Flux/qXNjB/src/optimise/train.jl:105
┌ Info: Epoch 478
└ @ Main /Users/kcft114/.julia/packages/Flux/qXNjB/src/optimise/train.jl:105
┌ Info: Epoch 479
└ @ Main /Users/kcft114/.julia/packages/Flux/qXNjB/src/optimise/train.jl:105
┌ Info: Epoch 480
└ @ Main /Users/kcft114/.julia/packages/Flux/qXNjB/src/optimise/train.jl:105
┌ Info: Epoch 481
└ @ Main /Users/kcft114/.julia/packages/Flux/qXNjB/src/optimise/train.jl:105
┌ Info: Epoch 482
└ @ Main /Users/kcft114/.julia/packages/Flux/qXNjB/src/optimise/train.jl:105
┌ Info: Epoch 483
└ @ Main /Users/kcft114/.julia/packages/Flux/qXNjB/src/optimise/train.jl:105
┌ Info: Epoch 484
└ @ Main /Users/kcft114/.julia/packages/Flux/qXNjB/src/optimise/train.jl:105
┌ Info: Epoch 485
└ @ Main /Users/kcft114/.julia/packages/Flux/qXNjB/src/optimise/train.jl:105
┌ Info: Epoch 486
└ @ Main /Users/kcft114/.julia/packages/Flux/qXNjB/src/optimise/train.jl:105
┌ Info: Epoch 487
└ @ Main /Users/kcft114/.julia/packages/Flux/qXNjB/src/optimise/train.jl:105
┌ Info: Epoch 488
└ @ Main /Users/kcft114/.julia/packages/Flux/qXNjB/src/optimise/train.jl:105
┌ Info: Epoch 489
└ @ Main /Users/kcft114/.julia/packages/Flux/qXNjB/src/optimise/train.jl:105
┌ Info: Epoch 490
└ @ Main /Users/kcft114/.julia/packages/Flux/qXNjB/src/optimise/train.jl:105
┌ Info: Epoch 491
└ @ Main /Users/kcft114/.julia/packages/Flux/qXNjB/src/optimise/train.jl:105
┌ Info: Epoch 492
└ @ Main /Users/kcft114/.julia/packages/Flux/qXNjB/src/optimise/train.jl:105
┌ Info: Epoch 493
└ @ Main /Users/kcft114/.julia/packages/Flux/qXNjB/src/optimise/train.jl:105
┌ Info: Epoch 494
└ @ Main /Users/kcft114/.julia/packages/Flux/qXNjB/src/optimise/train.jl:105
┌ Info: Epoch 495
└ @ Main /Users/kcft114/.julia/packages/Flux/qXNjB/src/optimise/train.jl:105
┌ Info: Epoch 496
└ @ Main /Users/kcft114/.julia/packages/Flux/qXNjB/src/optimise/train.jl:105
┌ Info: Epoch 497
└ @ Main /Users/kcft114/.julia/packages/Flux/qXNjB/src/optimise/train.jl:105
┌ Info: Epoch 498
└ @ Main /Users/kcft114/.julia/packages/Flux/qXNjB/src/optimise/train.jl:105
┌ Info: Epoch 499
└ @ Main /Users/kcft114/.julia/packages/Flux/qXNjB/src/optimise/train.jl:105
┌ Info: Epoch 500
└ @ Main /Users/kcft114/.julia/packages/Flux/qXNjB/src/optimise/train.jl:105
┌ Info: Epoch 501
└ @ Main /Users/kcft114/.julia/packages/Flux/qXNjB/src/optimise/train.jl:105
┌ Info: Epoch 502
└ @ Main /Users/kcft114/.julia/packages/Flux/qXNjB/src/optimise/train.jl:105
┌ Info: Epoch 503
└ @ Main /Users/kcft114/.julia/packages/Flux/qXNjB/src/optimise/train.jl:105
┌ Info: Epoch 504
└ @ Main /Users/kcft114/.julia/packages/Flux/qXNjB/src/optimise/train.jl:105
┌ Info: Epoch 505
└ @ Main /Users/kcft114/.julia/packages/Flux/qXNjB/src/optimise/train.jl:105
┌ Info: Epoch 506
└ @ Main /Users/kcft114/.julia/packages/Flux/qXNjB/src/optimise/train.jl:105
┌ Info: Epoch 507
└ @ Main /Users/kcft114/.julia/packages/Flux/qXNjB/src/optimise/train.jl:105
┌ Info: Epoch 508
└ @ Main /Users/kcft114/.julia/packages/Flux/qXNjB/src/optimise/train.jl:105
┌ Info: Epoch 509
└ @ Main /Users/kcft114/.julia/packages/Flux/qXNjB/src/optimise/train.jl:105
┌ Info: Epoch 510
└ @ Main /Users/kcft114/.julia/packages/Flux/qXNjB/src/optimise/train.jl:105
┌ Info: Epoch 511
└ @ Main /Users/kcft114/.julia/packages/Flux/qXNjB/src/optimise/train.jl:105
┌ Info: Epoch 512
└ @ Main /Users/kcft114/.julia/packages/Flux/qXNjB/src/optimise/train.jl:105
┌ Info: Epoch 513
└ @ Main /Users/kcft114/.julia/packages/Flux/qXNjB/src/optimise/train.jl:105
┌ Info: Epoch 514
└ @ Main /Users/kcft114/.julia/packages/Flux/qXNjB/src/optimise/train.jl:105
┌ Info: Epoch 515
└ @ Main /Users/kcft114/.julia/packages/Flux/qXNjB/src/optimise/train.jl:105
┌ Info: Epoch 516
└ @ Main /Users/kcft114/.julia/packages/Flux/qXNjB/src/optimise/train.jl:105
┌ Info: Epoch 517
└ @ Main /Users/kcft114/.julia/packages/Flux/qXNjB/src/optimise/train.jl:105
┌ Info: Epoch 518
└ @ Main /Users/kcft114/.julia/packages/Flux/qXNjB/src/optimise/train.jl:105
┌ Info: Epoch 519
└ @ Main /Users/kcft114/.julia/packages/Flux/qXNjB/src/optimise/train.jl:105
┌ Info: Epoch 520
└ @ Main /Users/kcft114/.julia/packages/Flux/qXNjB/src/optimise/train.jl:105
┌ Info: Epoch 521
└ @ Main /Users/kcft114/.julia/packages/Flux/qXNjB/src/optimise/train.jl:105
┌ Info: Epoch 522
└ @ Main /Users/kcft114/.julia/packages/Flux/qXNjB/src/optimise/train.jl:105
┌ Info: Epoch 523
└ @ Main /Users/kcft114/.julia/packages/Flux/qXNjB/src/optimise/train.jl:105
┌ Info: Epoch 524
└ @ Main /Users/kcft114/.julia/packages/Flux/qXNjB/src/optimise/train.jl:105
┌ Info: Epoch 525
└ @ Main /Users/kcft114/.julia/packages/Flux/qXNjB/src/optimise/train.jl:105
┌ Info: Epoch 526
└ @ Main /Users/kcft114/.julia/packages/Flux/qXNjB/src/optimise/train.jl:105
┌ Info: Epoch 527
└ @ Main /Users/kcft114/.julia/packages/Flux/qXNjB/src/optimise/train.jl:105
┌ Info: Epoch 528
└ @ Main /Users/kcft114/.julia/packages/Flux/qXNjB/src/optimise/train.jl:105
┌ Info: Epoch 529
└ @ Main /Users/kcft114/.julia/packages/Flux/qXNjB/src/optimise/train.jl:105
┌ Info: Epoch 530
└ @ Main /Users/kcft114/.julia/packages/Flux/qXNjB/src/optimise/train.jl:105
┌ Info: Epoch 531
└ @ Main /Users/kcft114/.julia/packages/Flux/qXNjB/src/optimise/train.jl:105
┌ Info: Epoch 532
└ @ Main /Users/kcft114/.julia/packages/Flux/qXNjB/src/optimise/train.jl:105
┌ Info: Epoch 533
└ @ Main /Users/kcft114/.julia/packages/Flux/qXNjB/src/optimise/train.jl:105
┌ Info: Epoch 534
└ @ Main /Users/kcft114/.julia/packages/Flux/qXNjB/src/optimise/train.jl:105
┌ Info: Epoch 535
└ @ Main /Users/kcft114/.julia/packages/Flux/qXNjB/src/optimise/train.jl:105
┌ Info: Epoch 536
└ @ Main /Users/kcft114/.julia/packages/Flux/qXNjB/src/optimise/train.jl:105
┌ Info: Epoch 537
└ @ Main /Users/kcft114/.julia/packages/Flux/qXNjB/src/optimise/train.jl:105
┌ Info: Epoch 538
└ @ Main /Users/kcft114/.julia/packages/Flux/qXNjB/src/optimise/train.jl:105
┌ Info: Epoch 539
└ @ Main /Users/kcft114/.julia/packages/Flux/qXNjB/src/optimise/train.jl:105
┌ Info: Epoch 540
└ @ Main /Users/kcft114/.julia/packages/Flux/qXNjB/src/optimise/train.jl:105
┌ Info: Epoch 541
└ @ Main /Users/kcft114/.julia/packages/Flux/qXNjB/src/optimise/train.jl:105
┌ Info: Epoch 542
└ @ Main /Users/kcft114/.julia/packages/Flux/qXNjB/src/optimise/train.jl:105
┌ Info: Epoch 543
└ @ Main /Users/kcft114/.julia/packages/Flux/qXNjB/src/optimise/train.jl:105
┌ Info: Epoch 544
└ @ Main /Users/kcft114/.julia/packages/Flux/qXNjB/src/optimise/train.jl:105
┌ Info: Epoch 545
└ @ Main /Users/kcft114/.julia/packages/Flux/qXNjB/src/optimise/train.jl:105
┌ Info: Epoch 546
└ @ Main /Users/kcft114/.julia/packages/Flux/qXNjB/src/optimise/train.jl:105
┌ Info: Epoch 547
└ @ Main /Users/kcft114/.julia/packages/Flux/qXNjB/src/optimise/train.jl:105
┌ Info: Epoch 548
└ @ Main /Users/kcft114/.julia/packages/Flux/qXNjB/src/optimise/train.jl:105
┌ Info: Epoch 549
└ @ Main /Users/kcft114/.julia/packages/Flux/qXNjB/src/optimise/train.jl:105
┌ Info: Epoch 550
└ @ Main /Users/kcft114/.julia/packages/Flux/qXNjB/src/optimise/train.jl:105
┌ Info: Epoch 551
└ @ Main /Users/kcft114/.julia/packages/Flux/qXNjB/src/optimise/train.jl:105
┌ Info: Epoch 552
└ @ Main /Users/kcft114/.julia/packages/Flux/qXNjB/src/optimise/train.jl:105
┌ Info: Epoch 553
└ @ Main /Users/kcft114/.julia/packages/Flux/qXNjB/src/optimise/train.jl:105
┌ Info: Epoch 554
└ @ Main /Users/kcft114/.julia/packages/Flux/qXNjB/src/optimise/train.jl:105
┌ Info: Epoch 555
└ @ Main /Users/kcft114/.julia/packages/Flux/qXNjB/src/optimise/train.jl:105
┌ Info: Epoch 556
└ @ Main /Users/kcft114/.julia/packages/Flux/qXNjB/src/optimise/train.jl:105
┌ Info: Epoch 557
└ @ Main /Users/kcft114/.julia/packages/Flux/qXNjB/src/optimise/train.jl:105
┌ Info: Epoch 558
└ @ Main /Users/kcft114/.julia/packages/Flux/qXNjB/src/optimise/train.jl:105
┌ Info: Epoch 559
└ @ Main /Users/kcft114/.julia/packages/Flux/qXNjB/src/optimise/train.jl:105
┌ Info: Epoch 560
└ @ Main /Users/kcft114/.julia/packages/Flux/qXNjB/src/optimise/train.jl:105
┌ Info: Epoch 561
└ @ Main /Users/kcft114/.julia/packages/Flux/qXNjB/src/optimise/train.jl:105
┌ Info: Epoch 562
└ @ Main /Users/kcft114/.julia/packages/Flux/qXNjB/src/optimise/train.jl:105
┌ Info: Epoch 563
└ @ Main /Users/kcft114/.julia/packages/Flux/qXNjB/src/optimise/train.jl:105
┌ Info: Epoch 564
└ @ Main /Users/kcft114/.julia/packages/Flux/qXNjB/src/optimise/train.jl:105
┌ Info: Epoch 565
└ @ Main /Users/kcft114/.julia/packages/Flux/qXNjB/src/optimise/train.jl:105
┌ Info: Epoch 566
└ @ Main /Users/kcft114/.julia/packages/Flux/qXNjB/src/optimise/train.jl:105
┌ Info: Epoch 567
└ @ Main /Users/kcft114/.julia/packages/Flux/qXNjB/src/optimise/train.jl:105
┌ Info: Epoch 568
└ @ Main /Users/kcft114/.julia/packages/Flux/qXNjB/src/optimise/train.jl:105
┌ Info: Epoch 569
└ @ Main /Users/kcft114/.julia/packages/Flux/qXNjB/src/optimise/train.jl:105
┌ Info: Epoch 570
└ @ Main /Users/kcft114/.julia/packages/Flux/qXNjB/src/optimise/train.jl:105
┌ Info: Epoch 571
└ @ Main /Users/kcft114/.julia/packages/Flux/qXNjB/src/optimise/train.jl:105
┌ Info: Epoch 572
└ @ Main /Users/kcft114/.julia/packages/Flux/qXNjB/src/optimise/train.jl:105
┌ Info: Epoch 573
└ @ Main /Users/kcft114/.julia/packages/Flux/qXNjB/src/optimise/train.jl:105
┌ Info: Epoch 574
└ @ Main /Users/kcft114/.julia/packages/Flux/qXNjB/src/optimise/train.jl:105
┌ Info: Epoch 575
└ @ Main /Users/kcft114/.julia/packages/Flux/qXNjB/src/optimise/train.jl:105
┌ Info: Epoch 576
└ @ Main /Users/kcft114/.julia/packages/Flux/qXNjB/src/optimise/train.jl:105
┌ Info: Epoch 577
└ @ Main /Users/kcft114/.julia/packages/Flux/qXNjB/src/optimise/train.jl:105
┌ Info: Epoch 578
└ @ Main /Users/kcft114/.julia/packages/Flux/qXNjB/src/optimise/train.jl:105
┌ Info: Epoch 579
└ @ Main /Users/kcft114/.julia/packages/Flux/qXNjB/src/optimise/train.jl:105
┌ Info: Epoch 580
└ @ Main /Users/kcft114/.julia/packages/Flux/qXNjB/src/optimise/train.jl:105
┌ Info: Epoch 581
└ @ Main /Users/kcft114/.julia/packages/Flux/qXNjB/src/optimise/train.jl:105
┌ Info: Epoch 582
└ @ Main /Users/kcft114/.julia/packages/Flux/qXNjB/src/optimise/train.jl:105
┌ Info: Epoch 583
└ @ Main /Users/kcft114/.julia/packages/Flux/qXNjB/src/optimise/train.jl:105
┌ Info: Epoch 584
└ @ Main /Users/kcft114/.julia/packages/Flux/qXNjB/src/optimise/train.jl:105
┌ Info: Epoch 585
└ @ Main /Users/kcft114/.julia/packages/Flux/qXNjB/src/optimise/train.jl:105
┌ Info: Epoch 586
└ @ Main /Users/kcft114/.julia/packages/Flux/qXNjB/src/optimise/train.jl:105
┌ Info: Epoch 587
└ @ Main /Users/kcft114/.julia/packages/Flux/qXNjB/src/optimise/train.jl:105
┌ Info: Epoch 588
└ @ Main /Users/kcft114/.julia/packages/Flux/qXNjB/src/optimise/train.jl:105
┌ Info: Epoch 589
└ @ Main /Users/kcft114/.julia/packages/Flux/qXNjB/src/optimise/train.jl:105
┌ Info: Epoch 590
└ @ Main /Users/kcft114/.julia/packages/Flux/qXNjB/src/optimise/train.jl:105
┌ Info: Epoch 591
└ @ Main /Users/kcft114/.julia/packages/Flux/qXNjB/src/optimise/train.jl:105
┌ Info: Epoch 592
└ @ Main /Users/kcft114/.julia/packages/Flux/qXNjB/src/optimise/train.jl:105
┌ Info: Epoch 593
└ @ Main /Users/kcft114/.julia/packages/Flux/qXNjB/src/optimise/train.jl:105
┌ Info: Epoch 594
└ @ Main /Users/kcft114/.julia/packages/Flux/qXNjB/src/optimise/train.jl:105
┌ Info: Epoch 595
└ @ Main /Users/kcft114/.julia/packages/Flux/qXNjB/src/optimise/train.jl:105
┌ Info: Epoch 596
└ @ Main /Users/kcft114/.julia/packages/Flux/qXNjB/src/optimise/train.jl:105
┌ Info: Epoch 597
└ @ Main /Users/kcft114/.julia/packages/Flux/qXNjB/src/optimise/train.jl:105
┌ Info: Epoch 598
└ @ Main /Users/kcft114/.julia/packages/Flux/qXNjB/src/optimise/train.jl:105
┌ Info: Epoch 599
└ @ Main /Users/kcft114/.julia/packages/Flux/qXNjB/src/optimise/train.jl:105
┌ Info: Epoch 600
└ @ Main /Users/kcft114/.julia/packages/Flux/qXNjB/src/optimise/train.jl:105
┌ Info: Epoch 601
└ @ Main /Users/kcft114/.julia/packages/Flux/qXNjB/src/optimise/train.jl:105
┌ Info: Epoch 602
└ @ Main /Users/kcft114/.julia/packages/Flux/qXNjB/src/optimise/train.jl:105
┌ Info: Epoch 603
└ @ Main /Users/kcft114/.julia/packages/Flux/qXNjB/src/optimise/train.jl:105
┌ Info: Epoch 604
└ @ Main /Users/kcft114/.julia/packages/Flux/qXNjB/src/optimise/train.jl:105
┌ Info: Epoch 605
└ @ Main /Users/kcft114/.julia/packages/Flux/qXNjB/src/optimise/train.jl:105
┌ Info: Epoch 606
└ @ Main /Users/kcft114/.julia/packages/Flux/qXNjB/src/optimise/train.jl:105
┌ Info: Epoch 607
└ @ Main /Users/kcft114/.julia/packages/Flux/qXNjB/src/optimise/train.jl:105
┌ Info: Epoch 608
└ @ Main /Users/kcft114/.julia/packages/Flux/qXNjB/src/optimise/train.jl:105
┌ Info: Epoch 609
└ @ Main /Users/kcft114/.julia/packages/Flux/qXNjB/src/optimise/train.jl:105
┌ Info: Epoch 610
└ @ Main /Users/kcft114/.julia/packages/Flux/qXNjB/src/optimise/train.jl:105
┌ Info: Epoch 611
└ @ Main /Users/kcft114/.julia/packages/Flux/qXNjB/src/optimise/train.jl:105
┌ Info: Epoch 612
└ @ Main /Users/kcft114/.julia/packages/Flux/qXNjB/src/optimise/train.jl:105
┌ Info: Epoch 613
└ @ Main /Users/kcft114/.julia/packages/Flux/qXNjB/src/optimise/train.jl:105
┌ Info: Epoch 614
└ @ Main /Users/kcft114/.julia/packages/Flux/qXNjB/src/optimise/train.jl:105
┌ Info: Epoch 615
└ @ Main /Users/kcft114/.julia/packages/Flux/qXNjB/src/optimise/train.jl:105
┌ Info: Epoch 616
└ @ Main /Users/kcft114/.julia/packages/Flux/qXNjB/src/optimise/train.jl:105
┌ Info: Epoch 617
└ @ Main /Users/kcft114/.julia/packages/Flux/qXNjB/src/optimise/train.jl:105
┌ Info: Epoch 618
└ @ Main /Users/kcft114/.julia/packages/Flux/qXNjB/src/optimise/train.jl:105
┌ Info: Epoch 619
└ @ Main /Users/kcft114/.julia/packages/Flux/qXNjB/src/optimise/train.jl:105
┌ Info: Epoch 620
└ @ Main /Users/kcft114/.julia/packages/Flux/qXNjB/src/optimise/train.jl:105
┌ Info: Epoch 621
└ @ Main /Users/kcft114/.julia/packages/Flux/qXNjB/src/optimise/train.jl:105
┌ Info: Epoch 622
└ @ Main /Users/kcft114/.julia/packages/Flux/qXNjB/src/optimise/train.jl:105
┌ Info: Epoch 623
└ @ Main /Users/kcft114/.julia/packages/Flux/qXNjB/src/optimise/train.jl:105
┌ Info: Epoch 624
└ @ Main /Users/kcft114/.julia/packages/Flux/qXNjB/src/optimise/train.jl:105
┌ Info: Epoch 625
└ @ Main /Users/kcft114/.julia/packages/Flux/qXNjB/src/optimise/train.jl:105
┌ Info: Epoch 626
└ @ Main /Users/kcft114/.julia/packages/Flux/qXNjB/src/optimise/train.jl:105
┌ Info: Epoch 627
└ @ Main /Users/kcft114/.julia/packages/Flux/qXNjB/src/optimise/train.jl:105
┌ Info: Epoch 628
└ @ Main /Users/kcft114/.julia/packages/Flux/qXNjB/src/optimise/train.jl:105
┌ Info: Epoch 629
└ @ Main /Users/kcft114/.julia/packages/Flux/qXNjB/src/optimise/train.jl:105
┌ Info: Epoch 630
└ @ Main /Users/kcft114/.julia/packages/Flux/qXNjB/src/optimise/train.jl:105
┌ Info: Epoch 631
└ @ Main /Users/kcft114/.julia/packages/Flux/qXNjB/src/optimise/train.jl:105
┌ Info: Epoch 632
└ @ Main /Users/kcft114/.julia/packages/Flux/qXNjB/src/optimise/train.jl:105
┌ Info: Epoch 633
└ @ Main /Users/kcft114/.julia/packages/Flux/qXNjB/src/optimise/train.jl:105
┌ Info: Epoch 634
└ @ Main /Users/kcft114/.julia/packages/Flux/qXNjB/src/optimise/train.jl:105
┌ Info: Epoch 635
└ @ Main /Users/kcft114/.julia/packages/Flux/qXNjB/src/optimise/train.jl:105
┌ Info: Epoch 636
└ @ Main /Users/kcft114/.julia/packages/Flux/qXNjB/src/optimise/train.jl:105
┌ Info: Epoch 637
└ @ Main /Users/kcft114/.julia/packages/Flux/qXNjB/src/optimise/train.jl:105
┌ Info: Epoch 638
└ @ Main /Users/kcft114/.julia/packages/Flux/qXNjB/src/optimise/train.jl:105
┌ Info: Epoch 639
└ @ Main /Users/kcft114/.julia/packages/Flux/qXNjB/src/optimise/train.jl:105
┌ Info: Epoch 640
└ @ Main /Users/kcft114/.julia/packages/Flux/qXNjB/src/optimise/train.jl:105
┌ Info: Epoch 641
└ @ Main /Users/kcft114/.julia/packages/Flux/qXNjB/src/optimise/train.jl:105
┌ Info: Epoch 642
└ @ Main /Users/kcft114/.julia/packages/Flux/qXNjB/src/optimise/train.jl:105
┌ Info: Epoch 643
└ @ Main /Users/kcft114/.julia/packages/Flux/qXNjB/src/optimise/train.jl:105
┌ Info: Epoch 644
└ @ Main /Users/kcft114/.julia/packages/Flux/qXNjB/src/optimise/train.jl:105
┌ Info: Epoch 645
└ @ Main /Users/kcft114/.julia/packages/Flux/qXNjB/src/optimise/train.jl:105
┌ Info: Epoch 646
└ @ Main /Users/kcft114/.julia/packages/Flux/qXNjB/src/optimise/train.jl:105
┌ Info: Epoch 647
└ @ Main /Users/kcft114/.julia/packages/Flux/qXNjB/src/optimise/train.jl:105
┌ Info: Epoch 648
└ @ Main /Users/kcft114/.julia/packages/Flux/qXNjB/src/optimise/train.jl:105
┌ Info: Epoch 649
└ @ Main /Users/kcft114/.julia/packages/Flux/qXNjB/src/optimise/train.jl:105
┌ Info: Epoch 650
└ @ Main /Users/kcft114/.julia/packages/Flux/qXNjB/src/optimise/train.jl:105
┌ Info: Epoch 651
└ @ Main /Users/kcft114/.julia/packages/Flux/qXNjB/src/optimise/train.jl:105
┌ Info: Epoch 652
└ @ Main /Users/kcft114/.julia/packages/Flux/qXNjB/src/optimise/train.jl:105
┌ Info: Epoch 653
└ @ Main /Users/kcft114/.julia/packages/Flux/qXNjB/src/optimise/train.jl:105
┌ Info: Epoch 654
└ @ Main /Users/kcft114/.julia/packages/Flux/qXNjB/src/optimise/train.jl:105
┌ Info: Epoch 655
└ @ Main /Users/kcft114/.julia/packages/Flux/qXNjB/src/optimise/train.jl:105
┌ Info: Epoch 656
└ @ Main /Users/kcft114/.julia/packages/Flux/qXNjB/src/optimise/train.jl:105
┌ Info: Epoch 657
└ @ Main /Users/kcft114/.julia/packages/Flux/qXNjB/src/optimise/train.jl:105
┌ Info: Epoch 658
└ @ Main /Users/kcft114/.julia/packages/Flux/qXNjB/src/optimise/train.jl:105
┌ Info: Epoch 659
└ @ Main /Users/kcft114/.julia/packages/Flux/qXNjB/src/optimise/train.jl:105
┌ Info: Epoch 660
└ @ Main /Users/kcft114/.julia/packages/Flux/qXNjB/src/optimise/train.jl:105
┌ Info: Epoch 661
└ @ Main /Users/kcft114/.julia/packages/Flux/qXNjB/src/optimise/train.jl:105
┌ Info: Epoch 662
└ @ Main /Users/kcft114/.julia/packages/Flux/qXNjB/src/optimise/train.jl:105
┌ Info: Epoch 663
└ @ Main /Users/kcft114/.julia/packages/Flux/qXNjB/src/optimise/train.jl:105
┌ Info: Epoch 664
└ @ Main /Users/kcft114/.julia/packages/Flux/qXNjB/src/optimise/train.jl:105
┌ Info: Epoch 665
└ @ Main /Users/kcft114/.julia/packages/Flux/qXNjB/src/optimise/train.jl:105
┌ Info: Epoch 666
└ @ Main /Users/kcft114/.julia/packages/Flux/qXNjB/src/optimise/train.jl:105
┌ Info: Epoch 667
└ @ Main /Users/kcft114/.julia/packages/Flux/qXNjB/src/optimise/train.jl:105
```

```
loss(X, Y) = 0.70195127f0 (tracked)
loss(X, Y) = 0.6498978f0 (tracked)
loss(X, Y) = 0.69707114f0 (tracked)
loss(X, Y) = 0.70271695f0 (tracked)
loss(X, Y) = 0.682165f0 (tracked)
loss(X, Y) = 0.7079188f0 (tracked)
loss(X, Y) = 0.709219f0 (tracked)
loss(X, Y) = 0.6761581f0 (tracked)
loss(X, Y) = 0.7101185f0 (tracked)
loss(X, Y) = 0.6749173f0 (tracked)
loss(X, Y) = 0.6650719f0 (tracked)
loss(X, Y) = 0.68010044f0 (tracked)
loss(X, Y) = 0.713835f0 (tracked)
loss(X, Y) = 0.68635327f0 (tracked)
loss(X, Y) = 0.6826041f0 (tracked)
loss(X, Y) = 0.6515405f0 (tracked)
loss(X, Y) = 0.72708905f0 (tracked)
loss(X, Y) = 0.6760807f0 (tracked)
loss(X, Y) = 0.70265204f0 (tracked)
loss(X, Y) = 0.67993325f0 (tracked)
loss(X, Y) = 0.7172888f0 (tracked)
loss(X, Y) = 0.6581323f0 (tracked)
loss(X, Y) = 0.6690989f0 (tracked)
loss(X, Y) = 0.6955575f0 (tracked)
loss(X, Y) = 0.668928f0 (tracked)
loss(X, Y) = 0.6867788f0 (tracked)
loss(X, Y) = 0.64543283f0 (tracked)
loss(X, Y) = 0.68211263f0 (tracked)
loss(X, Y) = 0.7205759f0 (tracked)
loss(X, Y) = 0.70064914f0 (tracked)
loss(X, Y) = 0.6753361f0 (tracked)
loss(X, Y) = 0.67617965f0 (tracked)
loss(X, Y) = 0.6988686f0 (tracked)
loss(X, Y) = 0.68952584f0 (tracked)
loss(X, Y) = 0.7191972f0 (tracked)
loss(X, Y) = 0.71045935f0 (tracked)
loss(X, Y) = 0.6772498f0 (tracked)
loss(X, Y) = 0.6797468f0 (tracked)
loss(X, Y) = 0.68462783f0 (tracked)
loss(X, Y) = 0.6491883f0 (tracked)
loss(X, Y) = 0.6636111f0 (tracked)
loss(X, Y) = 0.6656233f0 (tracked)
loss(X, Y) = 0.673833f0 (tracked)
loss(X, Y) = 0.6808934f0 (tracked)
loss(X, Y) = 0.69831854f0 (tracked)
loss(X, Y) = 0.6668276f0 (tracked)
loss(X, Y) = 0.6427355f0 (tracked)
loss(X, Y) = 0.63552225f0 (tracked)
loss(X, Y) = 0.7016964f0 (tracked)
loss(X, Y) = 0.672654f0 (tracked)
loss(X, Y) = 0.7032568f0 (tracked)
loss(X, Y) = 0.7097661f0 (tracked)
loss(X, Y) = 0.6750288f0 (tracked)
loss(X, Y) = 0.6931418f0 (tracked)
loss(X, Y) = 0.6502554f0 (tracked)
loss(X, Y) = 0.69192886f0 (tracked)
loss(X, Y) = 0.70730597f0 (tracked)
loss(X, Y) = 0.6573797f0 (tracked)
loss(X, Y) = 0.7111882f0 (tracked)
loss(X, Y) = 0.6492298f0 (tracked)
loss(X, Y) = 0.6611758f0 (tracked)
loss(X, Y) = 0.69942355f0 (tracked)
loss(X, Y) = 0.674386f0 (tracked)
loss(X, Y) = 0.70906734f0 (tracked)
loss(X, Y) = 0.66584164f0 (tracked)
loss(X, Y) = 0.6757075f0 (tracked)
loss(X, Y) = 0.6718723f0 (tracked)
loss(X, Y) = 0.64071876f0 (tracked)
loss(X, Y) = 0.64223355f0 (tracked)
loss(X, Y) = 0.6726123f0 (tracked)
loss(X, Y) = 0.6875004f0 (tracked)
loss(X, Y) = 0.68400544f0 (tracked)
loss(X, Y) = 0.6217009f0 (tracked)
loss(X, Y) = 0.66110283f0 (tracked)
loss(X, Y) = 0.6767547f0 (tracked)
loss(X, Y) = 0.70041335f0 (tracked)
loss(X, Y) = 0.632219f0 (tracked)
loss(X, Y) = 0.6543658f0 (tracked)
loss(X, Y) = 0.65267724f0 (tracked)
loss(X, Y) = 0.6743837f0 (tracked)
loss(X, Y) = 0.6733594f0 (tracked)
loss(X, Y) = 0.68345565f0 (tracked)
loss(X, Y) = 0.6413137f0 (tracked)
loss(X, Y) = 0.67078257f0 (tracked)
loss(X, Y) = 0.6680917f0 (tracked)
loss(X, Y) = 0.66659755f0 (tracked)
loss(X, Y) = 0.67270035f0 (tracked)
loss(X, Y) = 0.71072173f0 (tracked)
loss(X, Y) = 0.6355985f0 (tracked)
loss(X, Y) = 0.6711714f0 (tracked)
loss(X, Y) = 0.67210764f0 (tracked)
loss(X, Y) = 0.65602565f0 (tracked)
loss(X, Y) = 0.6600971f0 (tracked)
loss(X, Y) = 0.6844343f0 (tracked)
loss(X, Y) = 0.6871647f0 (tracked)
loss(X, Y) = 0.6796711f0 (tracked)
loss(X, Y) = 0.6762042f0 (tracked)
loss(X, Y) = 0.6794191f0 (tracked)
loss(X, Y) = 0.69131804f0 (tracked)
loss(X, Y) = 0.67215276f0 (tracked)
loss(X, Y) = 0.6356944f0 (tracked)
loss(X, Y) = 0.6517763f0 (tracked)
loss(X, Y) = 0.66532415f0 (tracked)
loss(X, Y) = 0.66933995f0 (tracked)
loss(X, Y) = 0.6180109f0 (tracked)
loss(X, Y) = 0.68386686f0 (tracked)
loss(X, Y) = 0.7238665f0 (tracked)
loss(X, Y) = 0.64909565f0 (tracked)
loss(X, Y) = 0.67426586f0 (tracked)
loss(X, Y) = 0.67255574f0 (tracked)
loss(X, Y) = 0.7189802f0 (tracked)
loss(X, Y) = 0.6756533f0 (tracked)
```

```
┌ Info: Epoch 668
└ @ Main /Users/kcft114/.julia/packages/Flux/qXNjB/src/optimise/train.jl:105
┌ Info: Epoch 669
└ @ Main /Users/kcft114/.julia/packages/Flux/qXNjB/src/optimise/train.jl:105
┌ Info: Epoch 670
└ @ Main /Users/kcft114/.julia/packages/Flux/qXNjB/src/optimise/train.jl:105
┌ Info: Epoch 671
└ @ Main /Users/kcft114/.julia/packages/Flux/qXNjB/src/optimise/train.jl:105
┌ Info: Epoch 672
└ @ Main /Users/kcft114/.julia/packages/Flux/qXNjB/src/optimise/train.jl:105
┌ Info: Epoch 673
└ @ Main /Users/kcft114/.julia/packages/Flux/qXNjB/src/optimise/train.jl:105
┌ Info: Epoch 674
└ @ Main /Users/kcft114/.julia/packages/Flux/qXNjB/src/optimise/train.jl:105
┌ Info: Epoch 675
└ @ Main /Users/kcft114/.julia/packages/Flux/qXNjB/src/optimise/train.jl:105
┌ Info: Epoch 676
└ @ Main /Users/kcft114/.julia/packages/Flux/qXNjB/src/optimise/train.jl:105
┌ Info: Epoch 677
└ @ Main /Users/kcft114/.julia/packages/Flux/qXNjB/src/optimise/train.jl:105
┌ Info: Epoch 678
└ @ Main /Users/kcft114/.julia/packages/Flux/qXNjB/src/optimise/train.jl:105
┌ Info: Epoch 679
└ @ Main /Users/kcft114/.julia/packages/Flux/qXNjB/src/optimise/train.jl:105
┌ Info: Epoch 680
└ @ Main /Users/kcft114/.julia/packages/Flux/qXNjB/src/optimise/train.jl:105
┌ Info: Epoch 681
└ @ Main /Users/kcft114/.julia/packages/Flux/qXNjB/src/optimise/train.jl:105
┌ Info: Epoch 682
└ @ Main /Users/kcft114/.julia/packages/Flux/qXNjB/src/optimise/train.jl:105
┌ Info: Epoch 683
└ @ Main /Users/kcft114/.julia/packages/Flux/qXNjB/src/optimise/train.jl:105
┌ Info: Epoch 684
└ @ Main /Users/kcft114/.julia/packages/Flux/qXNjB/src/optimise/train.jl:105
┌ Info: Epoch 685
└ @ Main /Users/kcft114/.julia/packages/Flux/qXNjB/src/optimise/train.jl:105
┌ Info: Epoch 686
└ @ Main /Users/kcft114/.julia/packages/Flux/qXNjB/src/optimise/train.jl:105
┌ Info: Epoch 687
└ @ Main /Users/kcft114/.julia/packages/Flux/qXNjB/src/optimise/train.jl:105
┌ Info: Epoch 688
└ @ Main /Users/kcft114/.julia/packages/Flux/qXNjB/src/optimise/train.jl:105
┌ Info: Epoch 689
└ @ Main /Users/kcft114/.julia/packages/Flux/qXNjB/src/optimise/train.jl:105
┌ Info: Epoch 690
└ @ Main /Users/kcft114/.julia/packages/Flux/qXNjB/src/optimise/train.jl:105
┌ Info: Epoch 691
└ @ Main /Users/kcft114/.julia/packages/Flux/qXNjB/src/optimise/train.jl:105
┌ Info: Epoch 692
└ @ Main /Users/kcft114/.julia/packages/Flux/qXNjB/src/optimise/train.jl:105
┌ Info: Epoch 693
└ @ Main /Users/kcft114/.julia/packages/Flux/qXNjB/src/optimise/train.jl:105
┌ Info: Epoch 694
└ @ Main /Users/kcft114/.julia/packages/Flux/qXNjB/src/optimise/train.jl:105
┌ Info: Epoch 695
└ @ Main /Users/kcft114/.julia/packages/Flux/qXNjB/src/optimise/train.jl:105
┌ Info: Epoch 696
└ @ Main /Users/kcft114/.julia/packages/Flux/qXNjB/src/optimise/train.jl:105
┌ Info: Epoch 697
└ @ Main /Users/kcft114/.julia/packages/Flux/qXNjB/src/optimise/train.jl:105
┌ Info: Epoch 698
└ @ Main /Users/kcft114/.julia/packages/Flux/qXNjB/src/optimise/train.jl:105
┌ Info: Epoch 699
└ @ Main /Users/kcft114/.julia/packages/Flux/qXNjB/src/optimise/train.jl:105
┌ Info: Epoch 700
└ @ Main /Users/kcft114/.julia/packages/Flux/qXNjB/src/optimise/train.jl:105
┌ Info: Epoch 701
└ @ Main /Users/kcft114/.julia/packages/Flux/qXNjB/src/optimise/train.jl:105
┌ Info: Epoch 702
└ @ Main /Users/kcft114/.julia/packages/Flux/qXNjB/src/optimise/train.jl:105
┌ Info: Epoch 703
└ @ Main /Users/kcft114/.julia/packages/Flux/qXNjB/src/optimise/train.jl:105
┌ Info: Epoch 704
└ @ Main /Users/kcft114/.julia/packages/Flux/qXNjB/src/optimise/train.jl:105
┌ Info: Epoch 705
└ @ Main /Users/kcft114/.julia/packages/Flux/qXNjB/src/optimise/train.jl:105
┌ Info: Epoch 706
└ @ Main /Users/kcft114/.julia/packages/Flux/qXNjB/src/optimise/train.jl:105
┌ Info: Epoch 707
└ @ Main /Users/kcft114/.julia/packages/Flux/qXNjB/src/optimise/train.jl:105
┌ Info: Epoch 708
└ @ Main /Users/kcft114/.julia/packages/Flux/qXNjB/src/optimise/train.jl:105
┌ Info: Epoch 709
└ @ Main /Users/kcft114/.julia/packages/Flux/qXNjB/src/optimise/train.jl:105
┌ Info: Epoch 710
└ @ Main /Users/kcft114/.julia/packages/Flux/qXNjB/src/optimise/train.jl:105
┌ Info: Epoch 711
└ @ Main /Users/kcft114/.julia/packages/Flux/qXNjB/src/optimise/train.jl:105
┌ Info: Epoch 712
└ @ Main /Users/kcft114/.julia/packages/Flux/qXNjB/src/optimise/train.jl:105
┌ Info: Epoch 713
└ @ Main /Users/kcft114/.julia/packages/Flux/qXNjB/src/optimise/train.jl:105
┌ Info: Epoch 714
└ @ Main /Users/kcft114/.julia/packages/Flux/qXNjB/src/optimise/train.jl:105
┌ Info: Epoch 715
└ @ Main /Users/kcft114/.julia/packages/Flux/qXNjB/src/optimise/train.jl:105
┌ Info: Epoch 716
└ @ Main /Users/kcft114/.julia/packages/Flux/qXNjB/src/optimise/train.jl:105
┌ Info: Epoch 717
└ @ Main /Users/kcft114/.julia/packages/Flux/qXNjB/src/optimise/train.jl:105
┌ Info: Epoch 718
└ @ Main /Users/kcft114/.julia/packages/Flux/qXNjB/src/optimise/train.jl:105
┌ Info: Epoch 719
└ @ Main /Users/kcft114/.julia/packages/Flux/qXNjB/src/optimise/train.jl:105
┌ Info: Epoch 720
└ @ Main /Users/kcft114/.julia/packages/Flux/qXNjB/src/optimise/train.jl:105
┌ Info: Epoch 721
└ @ Main /Users/kcft114/.julia/packages/Flux/qXNjB/src/optimise/train.jl:105
┌ Info: Epoch 722
└ @ Main /Users/kcft114/.julia/packages/Flux/qXNjB/src/optimise/train.jl:105
┌ Info: Epoch 723
└ @ Main /Users/kcft114/.julia/packages/Flux/qXNjB/src/optimise/train.jl:105
┌ Info: Epoch 724
└ @ Main /Users/kcft114/.julia/packages/Flux/qXNjB/src/optimise/train.jl:105
┌ Info: Epoch 725
└ @ Main /Users/kcft114/.julia/packages/Flux/qXNjB/src/optimise/train.jl:105
┌ Info: Epoch 726
└ @ Main /Users/kcft114/.julia/packages/Flux/qXNjB/src/optimise/train.jl:105
┌ Info: Epoch 727
└ @ Main /Users/kcft114/.julia/packages/Flux/qXNjB/src/optimise/train.jl:105
┌ Info: Epoch 728
└ @ Main /Users/kcft114/.julia/packages/Flux/qXNjB/src/optimise/train.jl:105
┌ Info: Epoch 729
└ @ Main /Users/kcft114/.julia/packages/Flux/qXNjB/src/optimise/train.jl:105
┌ Info: Epoch 730
└ @ Main /Users/kcft114/.julia/packages/Flux/qXNjB/src/optimise/train.jl:105
┌ Info: Epoch 731
└ @ Main /Users/kcft114/.julia/packages/Flux/qXNjB/src/optimise/train.jl:105
┌ Info: Epoch 732
└ @ Main /Users/kcft114/.julia/packages/Flux/qXNjB/src/optimise/train.jl:105
┌ Info: Epoch 733
└ @ Main /Users/kcft114/.julia/packages/Flux/qXNjB/src/optimise/train.jl:105
┌ Info: Epoch 734
└ @ Main /Users/kcft114/.julia/packages/Flux/qXNjB/src/optimise/train.jl:105
┌ Info: Epoch 735
└ @ Main /Users/kcft114/.julia/packages/Flux/qXNjB/src/optimise/train.jl:105
┌ Info: Epoch 736
└ @ Main /Users/kcft114/.julia/packages/Flux/qXNjB/src/optimise/train.jl:105
┌ Info: Epoch 737
└ @ Main /Users/kcft114/.julia/packages/Flux/qXNjB/src/optimise/train.jl:105
┌ Info: Epoch 738
└ @ Main /Users/kcft114/.julia/packages/Flux/qXNjB/src/optimise/train.jl:105
┌ Info: Epoch 739
└ @ Main /Users/kcft114/.julia/packages/Flux/qXNjB/src/optimise/train.jl:105
┌ Info: Epoch 740
└ @ Main /Users/kcft114/.julia/packages/Flux/qXNjB/src/optimise/train.jl:105
┌ Info: Epoch 741
└ @ Main /Users/kcft114/.julia/packages/Flux/qXNjB/src/optimise/train.jl:105
┌ Info: Epoch 742
└ @ Main /Users/kcft114/.julia/packages/Flux/qXNjB/src/optimise/train.jl:105
┌ Info: Epoch 743
└ @ Main /Users/kcft114/.julia/packages/Flux/qXNjB/src/optimise/train.jl:105
┌ Info: Epoch 744
└ @ Main /Users/kcft114/.julia/packages/Flux/qXNjB/src/optimise/train.jl:105
┌ Info: Epoch 745
└ @ Main /Users/kcft114/.julia/packages/Flux/qXNjB/src/optimise/train.jl:105
┌ Info: Epoch 746
└ @ Main /Users/kcft114/.julia/packages/Flux/qXNjB/src/optimise/train.jl:105
┌ Info: Epoch 747
└ @ Main /Users/kcft114/.julia/packages/Flux/qXNjB/src/optimise/train.jl:105
┌ Info: Epoch 748
└ @ Main /Users/kcft114/.julia/packages/Flux/qXNjB/src/optimise/train.jl:105
┌ Info: Epoch 749
└ @ Main /Users/kcft114/.julia/packages/Flux/qXNjB/src/optimise/train.jl:105
┌ Info: Epoch 750
└ @ Main /Users/kcft114/.julia/packages/Flux/qXNjB/src/optimise/train.jl:105
┌ Info: Epoch 751
└ @ Main /Users/kcft114/.julia/packages/Flux/qXNjB/src/optimise/train.jl:105
┌ Info: Epoch 752
└ @ Main /Users/kcft114/.julia/packages/Flux/qXNjB/src/optimise/train.jl:105
┌ Info: Epoch 753
└ @ Main /Users/kcft114/.julia/packages/Flux/qXNjB/src/optimise/train.jl:105
┌ Info: Epoch 754
└ @ Main /Users/kcft114/.julia/packages/Flux/qXNjB/src/optimise/train.jl:105
┌ Info: Epoch 755
└ @ Main /Users/kcft114/.julia/packages/Flux/qXNjB/src/optimise/train.jl:105
┌ Info: Epoch 756
└ @ Main /Users/kcft114/.julia/packages/Flux/qXNjB/src/optimise/train.jl:105
┌ Info: Epoch 757
└ @ Main /Users/kcft114/.julia/packages/Flux/qXNjB/src/optimise/train.jl:105
┌ Info: Epoch 758
└ @ Main /Users/kcft114/.julia/packages/Flux/qXNjB/src/optimise/train.jl:105
┌ Info: Epoch 759
└ @ Main /Users/kcft114/.julia/packages/Flux/qXNjB/src/optimise/train.jl:105
┌ Info: Epoch 760
└ @ Main /Users/kcft114/.julia/packages/Flux/qXNjB/src/optimise/train.jl:105
┌ Info: Epoch 761
└ @ Main /Users/kcft114/.julia/packages/Flux/qXNjB/src/optimise/train.jl:105
┌ Info: Epoch 762
└ @ Main /Users/kcft114/.julia/packages/Flux/qXNjB/src/optimise/train.jl:105
┌ Info: Epoch 763
└ @ Main /Users/kcft114/.julia/packages/Flux/qXNjB/src/optimise/train.jl:105
┌ Info: Epoch 764
└ @ Main /Users/kcft114/.julia/packages/Flux/qXNjB/src/optimise/train.jl:105
┌ Info: Epoch 765
└ @ Main /Users/kcft114/.julia/packages/Flux/qXNjB/src/optimise/train.jl:105
┌ Info: Epoch 766
└ @ Main /Users/kcft114/.julia/packages/Flux/qXNjB/src/optimise/train.jl:105
┌ Info: Epoch 767
└ @ Main /Users/kcft114/.julia/packages/Flux/qXNjB/src/optimise/train.jl:105
┌ Info: Epoch 768
└ @ Main /Users/kcft114/.julia/packages/Flux/qXNjB/src/optimise/train.jl:105
┌ Info: Epoch 769
└ @ Main /Users/kcft114/.julia/packages/Flux/qXNjB/src/optimise/train.jl:105
┌ Info: Epoch 770
└ @ Main /Users/kcft114/.julia/packages/Flux/qXNjB/src/optimise/train.jl:105
┌ Info: Epoch 771
└ @ Main /Users/kcft114/.julia/packages/Flux/qXNjB/src/optimise/train.jl:105
┌ Info: Epoch 772
└ @ Main /Users/kcft114/.julia/packages/Flux/qXNjB/src/optimise/train.jl:105
┌ Info: Epoch 773
└ @ Main /Users/kcft114/.julia/packages/Flux/qXNjB/src/optimise/train.jl:105
┌ Info: Epoch 774
└ @ Main /Users/kcft114/.julia/packages/Flux/qXNjB/src/optimise/train.jl:105
┌ Info: Epoch 775
└ @ Main /Users/kcft114/.julia/packages/Flux/qXNjB/src/optimise/train.jl:105
┌ Info: Epoch 776
└ @ Main /Users/kcft114/.julia/packages/Flux/qXNjB/src/optimise/train.jl:105
┌ Info: Epoch 777
└ @ Main /Users/kcft114/.julia/packages/Flux/qXNjB/src/optimise/train.jl:105
┌ Info: Epoch 778
└ @ Main /Users/kcft114/.julia/packages/Flux/qXNjB/src/optimise/train.jl:105
┌ Info: Epoch 779
└ @ Main /Users/kcft114/.julia/packages/Flux/qXNjB/src/optimise/train.jl:105
```

```
loss(X, Y) = 0.6743744f0 (tracked)
loss(X, Y) = 0.6273399f0 (tracked)
loss(X, Y) = 0.7085195f0 (tracked)
loss(X, Y) = 0.65922827f0 (tracked)
loss(X, Y) = 0.6304504f0 (tracked)
loss(X, Y) = 0.6841645f0 (tracked)
loss(X, Y) = 0.71427846f0 (tracked)
loss(X, Y) = 0.6412996f0 (tracked)
loss(X, Y) = 0.6434337f0 (tracked)
loss(X, Y) = 0.6773304f0 (tracked)
loss(X, Y) = 0.66610104f0 (tracked)
loss(X, Y) = 0.621158f0 (tracked)
loss(X, Y) = 0.65934426f0 (tracked)
loss(X, Y) = 0.6463872f0 (tracked)
loss(X, Y) = 0.6411946f0 (tracked)
loss(X, Y) = 0.64963996f0 (tracked)
loss(X, Y) = 0.65296626f0 (tracked)
loss(X, Y) = 0.6391077f0 (tracked)
loss(X, Y) = 0.6842801f0 (tracked)
loss(X, Y) = 0.6843457f0 (tracked)
loss(X, Y) = 0.6543569f0 (tracked)
loss(X, Y) = 0.63375366f0 (tracked)
loss(X, Y) = 0.6991514f0 (tracked)
loss(X, Y) = 0.68383735f0 (tracked)
loss(X, Y) = 0.6806277f0 (tracked)
loss(X, Y) = 0.66489184f0 (tracked)
loss(X, Y) = 0.69408184f0 (tracked)
loss(X, Y) = 0.661078f0 (tracked)
loss(X, Y) = 0.66743314f0 (tracked)
loss(X, Y) = 0.66577417f0 (tracked)
loss(X, Y) = 0.66731095f0 (tracked)
loss(X, Y) = 0.6939401f0 (tracked)
loss(X, Y) = 0.6806882f0 (tracked)
loss(X, Y) = 0.6392265f0 (tracked)
loss(X, Y) = 0.65907353f0 (tracked)
loss(X, Y) = 0.6409842f0 (tracked)
loss(X, Y) = 0.65861994f0 (tracked)
loss(X, Y) = 0.6378046f0 (tracked)
loss(X, Y) = 0.66031975f0 (tracked)
loss(X, Y) = 0.6568578f0 (tracked)
loss(X, Y) = 0.700591f0 (tracked)
loss(X, Y) = 0.6602131f0 (tracked)
loss(X, Y) = 0.6702538f0 (tracked)
loss(X, Y) = 0.6730199f0 (tracked)
loss(X, Y) = 0.64937764f0 (tracked)
loss(X, Y) = 0.6423036f0 (tracked)
loss(X, Y) = 0.69661653f0 (tracked)
loss(X, Y) = 0.63788f0 (tracked)
loss(X, Y) = 0.6631819f0 (tracked)
loss(X, Y) = 0.64773244f0 (tracked)
loss(X, Y) = 0.6482763f0 (tracked)
loss(X, Y) = 0.6749493f0 (tracked)
loss(X, Y) = 0.64275074f0 (tracked)
loss(X, Y) = 0.6490342f0 (tracked)
loss(X, Y) = 0.6479355f0 (tracked)
loss(X, Y) = 0.68493617f0 (tracked)
loss(X, Y) = 0.6614562f0 (tracked)
loss(X, Y) = 0.644813f0 (tracked)
loss(X, Y) = 0.6660304f0 (tracked)
loss(X, Y) = 0.6501245f0 (tracked)
loss(X, Y) = 0.6900486f0 (tracked)
loss(X, Y) = 0.6966705f0 (tracked)
loss(X, Y) = 0.64285165f0 (tracked)
loss(X, Y) = 0.6255735f0 (tracked)
loss(X, Y) = 0.66375554f0 (tracked)
loss(X, Y) = 0.6454525f0 (tracked)
loss(X, Y) = 0.6877579f0 (tracked)
loss(X, Y) = 0.66496134f0 (tracked)
loss(X, Y) = 0.68927985f0 (tracked)
loss(X, Y) = 0.652049f0 (tracked)
loss(X, Y) = 0.6760179f0 (tracked)
loss(X, Y) = 0.64879537f0 (tracked)
loss(X, Y) = 0.6421911f0 (tracked)
loss(X, Y) = 0.6917754f0 (tracked)
loss(X, Y) = 0.6515363f0 (tracked)
loss(X, Y) = 0.65015644f0 (tracked)
loss(X, Y) = 0.6788191f0 (tracked)
loss(X, Y) = 0.6739272f0 (tracked)
loss(X, Y) = 0.65109026f0 (tracked)
loss(X, Y) = 0.682486f0 (tracked)
loss(X, Y) = 0.64442766f0 (tracked)
loss(X, Y) = 0.65008754f0 (tracked)
loss(X, Y) = 0.63106084f0 (tracked)
loss(X, Y) = 0.6660221f0 (tracked)
loss(X, Y) = 0.6702277f0 (tracked)
loss(X, Y) = 0.6640899f0 (tracked)
loss(X, Y) = 0.6686103f0 (tracked)
loss(X, Y) = 0.6580889f0 (tracked)
loss(X, Y) = 0.6767527f0 (tracked)
loss(X, Y) = 0.6612422f0 (tracked)
loss(X, Y) = 0.63008595f0 (tracked)
loss(X, Y) = 0.6217894f0 (tracked)
loss(X, Y) = 0.6717378f0 (tracked)
loss(X, Y) = 0.66351527f0 (tracked)
loss(X, Y) = 0.6630527f0 (tracked)
loss(X, Y) = 0.6320064f0 (tracked)
loss(X, Y) = 0.6513382f0 (tracked)
loss(X, Y) = 0.63878965f0 (tracked)
loss(X, Y) = 0.6483192f0 (tracked)
loss(X, Y) = 0.6827972f0 (tracked)
loss(X, Y) = 0.67229015f0 (tracked)
loss(X, Y) = 0.66841346f0 (tracked)
loss(X, Y) = 0.6892826f0 (tracked)
loss(X, Y) = 0.64388424f0 (tracked)
loss(X, Y) = 0.66374147f0 (tracked)
loss(X, Y) = 0.6539732f0 (tracked)
loss(X, Y) = 0.6443856f0 (tracked)
loss(X, Y) = 0.64016604f0 (tracked)
loss(X, Y) = 0.6247715f0 (tracked)
loss(X, Y) = 0.63742197f0 (tracked)
loss(X, Y) = 0.65154064f0 (tracked)
loss(X, Y) = 0.6141447f0 (tracked)
loss(X, Y) = 0.6835553f0 (tracked)
loss(X, Y) = 0.6528933f0 (tracked)
loss(X, Y) = 0.6579794f0 (tracked)
loss(X, Y) = 0.69948816f0 (tracked)
loss(X, Y) = 0.62051314f0 (tracked)
loss(X, Y) = 0.6554496f0 (tracked)
loss(X, Y) = 0.64396954f0 (tracked)
loss(X, Y) = 0.6945884f0 (tracked)
```

```
┌ Info: Epoch 780
└ @ Main /Users/kcft114/.julia/packages/Flux/qXNjB/src/optimise/train.jl:105
┌ Info: Epoch 781
└ @ Main /Users/kcft114/.julia/packages/Flux/qXNjB/src/optimise/train.jl:105
┌ Info: Epoch 782
└ @ Main /Users/kcft114/.julia/packages/Flux/qXNjB/src/optimise/train.jl:105
┌ Info: Epoch 783
└ @ Main /Users/kcft114/.julia/packages/Flux/qXNjB/src/optimise/train.jl:105
┌ Info: Epoch 784
└ @ Main /Users/kcft114/.julia/packages/Flux/qXNjB/src/optimise/train.jl:105
┌ Info: Epoch 785
└ @ Main /Users/kcft114/.julia/packages/Flux/qXNjB/src/optimise/train.jl:105
┌ Info: Epoch 786
└ @ Main /Users/kcft114/.julia/packages/Flux/qXNjB/src/optimise/train.jl:105
┌ Info: Epoch 787
└ @ Main /Users/kcft114/.julia/packages/Flux/qXNjB/src/optimise/train.jl:105
┌ Info: Epoch 788
└ @ Main /Users/kcft114/.julia/packages/Flux/qXNjB/src/optimise/train.jl:105
┌ Info: Epoch 789
└ @ Main /Users/kcft114/.julia/packages/Flux/qXNjB/src/optimise/train.jl:105
┌ Info: Epoch 790
└ @ Main /Users/kcft114/.julia/packages/Flux/qXNjB/src/optimise/train.jl:105
┌ Info: Epoch 791
└ @ Main /Users/kcft114/.julia/packages/Flux/qXNjB/src/optimise/train.jl:105
┌ Info: Epoch 792
└ @ Main /Users/kcft114/.julia/packages/Flux/qXNjB/src/optimise/train.jl:105
┌ Info: Epoch 793
└ @ Main /Users/kcft114/.julia/packages/Flux/qXNjB/src/optimise/train.jl:105
┌ Info: Epoch 794
└ @ Main /Users/kcft114/.julia/packages/Flux/qXNjB/src/optimise/train.jl:105
┌ Info: Epoch 795
└ @ Main /Users/kcft114/.julia/packages/Flux/qXNjB/src/optimise/train.jl:105
┌ Info: Epoch 796
└ @ Main /Users/kcft114/.julia/packages/Flux/qXNjB/src/optimise/train.jl:105
┌ Info: Epoch 797
└ @ Main /Users/kcft114/.julia/packages/Flux/qXNjB/src/optimise/train.jl:105
┌ Info: Epoch 798
└ @ Main /Users/kcft114/.julia/packages/Flux/qXNjB/src/optimise/train.jl:105
┌ Info: Epoch 799
└ @ Main /Users/kcft114/.julia/packages/Flux/qXNjB/src/optimise/train.jl:105
┌ Info: Epoch 800
└ @ Main /Users/kcft114/.julia/packages/Flux/qXNjB/src/optimise/train.jl:105
┌ Info: Epoch 801
└ @ Main /Users/kcft114/.julia/packages/Flux/qXNjB/src/optimise/train.jl:105
┌ Info: Epoch 802
└ @ Main /Users/kcft114/.julia/packages/Flux/qXNjB/src/optimise/train.jl:105
┌ Info: Epoch 803
└ @ Main /Users/kcft114/.julia/packages/Flux/qXNjB/src/optimise/train.jl:105
┌ Info: Epoch 804
└ @ Main /Users/kcft114/.julia/packages/Flux/qXNjB/src/optimise/train.jl:105
┌ Info: Epoch 805
└ @ Main /Users/kcft114/.julia/packages/Flux/qXNjB/src/optimise/train.jl:105
┌ Info: Epoch 806
└ @ Main /Users/kcft114/.julia/packages/Flux/qXNjB/src/optimise/train.jl:105
┌ Info: Epoch 807
└ @ Main /Users/kcft114/.julia/packages/Flux/qXNjB/src/optimise/train.jl:105
┌ Info: Epoch 808
└ @ Main /Users/kcft114/.julia/packages/Flux/qXNjB/src/optimise/train.jl:105
┌ Info: Epoch 809
└ @ Main /Users/kcft114/.julia/packages/Flux/qXNjB/src/optimise/train.jl:105
┌ Info: Epoch 810
└ @ Main /Users/kcft114/.julia/packages/Flux/qXNjB/src/optimise/train.jl:105
┌ Info: Epoch 811
└ @ Main /Users/kcft114/.julia/packages/Flux/qXNjB/src/optimise/train.jl:105
┌ Info: Epoch 812
└ @ Main /Users/kcft114/.julia/packages/Flux/qXNjB/src/optimise/train.jl:105
┌ Info: Epoch 813
└ @ Main /Users/kcft114/.julia/packages/Flux/qXNjB/src/optimise/train.jl:105
┌ Info: Epoch 814
└ @ Main /Users/kcft114/.julia/packages/Flux/qXNjB/src/optimise/train.jl:105
┌ Info: Epoch 815
└ @ Main /Users/kcft114/.julia/packages/Flux/qXNjB/src/optimise/train.jl:105
┌ Info: Epoch 816
└ @ Main /Users/kcft114/.julia/packages/Flux/qXNjB/src/optimise/train.jl:105
┌ Info: Epoch 817
└ @ Main /Users/kcft114/.julia/packages/Flux/qXNjB/src/optimise/train.jl:105
┌ Info: Epoch 818
└ @ Main /Users/kcft114/.julia/packages/Flux/qXNjB/src/optimise/train.jl:105
┌ Info: Epoch 819
└ @ Main /Users/kcft114/.julia/packages/Flux/qXNjB/src/optimise/train.jl:105
┌ Info: Epoch 820
└ @ Main /Users/kcft114/.julia/packages/Flux/qXNjB/src/optimise/train.jl:105
┌ Info: Epoch 821
└ @ Main /Users/kcft114/.julia/packages/Flux/qXNjB/src/optimise/train.jl:105
┌ Info: Epoch 822
└ @ Main /Users/kcft114/.julia/packages/Flux/qXNjB/src/optimise/train.jl:105
┌ Info: Epoch 823
└ @ Main /Users/kcft114/.julia/packages/Flux/qXNjB/src/optimise/train.jl:105
┌ Info: Epoch 824
└ @ Main /Users/kcft114/.julia/packages/Flux/qXNjB/src/optimise/train.jl:105
┌ Info: Epoch 825
└ @ Main /Users/kcft114/.julia/packages/Flux/qXNjB/src/optimise/train.jl:105
┌ Info: Epoch 826
└ @ Main /Users/kcft114/.julia/packages/Flux/qXNjB/src/optimise/train.jl:105
┌ Info: Epoch 827
└ @ Main /Users/kcft114/.julia/packages/Flux/qXNjB/src/optimise/train.jl:105
┌ Info: Epoch 828
└ @ Main /Users/kcft114/.julia/packages/Flux/qXNjB/src/optimise/train.jl:105
┌ Info: Epoch 829
└ @ Main /Users/kcft114/.julia/packages/Flux/qXNjB/src/optimise/train.jl:105
┌ Info: Epoch 830
└ @ Main /Users/kcft114/.julia/packages/Flux/qXNjB/src/optimise/train.jl:105
┌ Info: Epoch 831
└ @ Main /Users/kcft114/.julia/packages/Flux/qXNjB/src/optimise/train.jl:105
┌ Info: Epoch 832
└ @ Main /Users/kcft114/.julia/packages/Flux/qXNjB/src/optimise/train.jl:105
┌ Info: Epoch 833
└ @ Main /Users/kcft114/.julia/packages/Flux/qXNjB/src/optimise/train.jl:105
┌ Info: Epoch 834
└ @ Main /Users/kcft114/.julia/packages/Flux/qXNjB/src/optimise/train.jl:105
┌ Info: Epoch 835
└ @ Main /Users/kcft114/.julia/packages/Flux/qXNjB/src/optimise/train.jl:105
┌ Info: Epoch 836
└ @ Main /Users/kcft114/.julia/packages/Flux/qXNjB/src/optimise/train.jl:105
┌ Info: Epoch 837
└ @ Main /Users/kcft114/.julia/packages/Flux/qXNjB/src/optimise/train.jl:105
┌ Info: Epoch 838
└ @ Main /Users/kcft114/.julia/packages/Flux/qXNjB/src/optimise/train.jl:105
┌ Info: Epoch 839
└ @ Main /Users/kcft114/.julia/packages/Flux/qXNjB/src/optimise/train.jl:105
┌ Info: Epoch 840
└ @ Main /Users/kcft114/.julia/packages/Flux/qXNjB/src/optimise/train.jl:105
┌ Info: Epoch 841
└ @ Main /Users/kcft114/.julia/packages/Flux/qXNjB/src/optimise/train.jl:105
┌ Info: Epoch 842
└ @ Main /Users/kcft114/.julia/packages/Flux/qXNjB/src/optimise/train.jl:105
┌ Info: Epoch 843
└ @ Main /Users/kcft114/.julia/packages/Flux/qXNjB/src/optimise/train.jl:105
┌ Info: Epoch 844
└ @ Main /Users/kcft114/.julia/packages/Flux/qXNjB/src/optimise/train.jl:105
┌ Info: Epoch 845
└ @ Main /Users/kcft114/.julia/packages/Flux/qXNjB/src/optimise/train.jl:105
┌ Info: Epoch 846
└ @ Main /Users/kcft114/.julia/packages/Flux/qXNjB/src/optimise/train.jl:105
┌ Info: Epoch 847
└ @ Main /Users/kcft114/.julia/packages/Flux/qXNjB/src/optimise/train.jl:105
┌ Info: Epoch 848
└ @ Main /Users/kcft114/.julia/packages/Flux/qXNjB/src/optimise/train.jl:105
┌ Info: Epoch 849
└ @ Main /Users/kcft114/.julia/packages/Flux/qXNjB/src/optimise/train.jl:105
┌ Info: Epoch 850
└ @ Main /Users/kcft114/.julia/packages/Flux/qXNjB/src/optimise/train.jl:105
┌ Info: Epoch 851
└ @ Main /Users/kcft114/.julia/packages/Flux/qXNjB/src/optimise/train.jl:105
┌ Info: Epoch 852
└ @ Main /Users/kcft114/.julia/packages/Flux/qXNjB/src/optimise/train.jl:105
┌ Info: Epoch 853
└ @ Main /Users/kcft114/.julia/packages/Flux/qXNjB/src/optimise/train.jl:105
┌ Info: Epoch 854
└ @ Main /Users/kcft114/.julia/packages/Flux/qXNjB/src/optimise/train.jl:105
┌ Info: Epoch 855
└ @ Main /Users/kcft114/.julia/packages/Flux/qXNjB/src/optimise/train.jl:105
┌ Info: Epoch 856
└ @ Main /Users/kcft114/.julia/packages/Flux/qXNjB/src/optimise/train.jl:105
┌ Info: Epoch 857
└ @ Main /Users/kcft114/.julia/packages/Flux/qXNjB/src/optimise/train.jl:105
┌ Info: Epoch 858
└ @ Main /Users/kcft114/.julia/packages/Flux/qXNjB/src/optimise/train.jl:105
┌ Info: Epoch 859
└ @ Main /Users/kcft114/.julia/packages/Flux/qXNjB/src/optimise/train.jl:105
┌ Info: Epoch 860
└ @ Main /Users/kcft114/.julia/packages/Flux/qXNjB/src/optimise/train.jl:105
┌ Info: Epoch 861
└ @ Main /Users/kcft114/.julia/packages/Flux/qXNjB/src/optimise/train.jl:105
┌ Info: Epoch 862
└ @ Main /Users/kcft114/.julia/packages/Flux/qXNjB/src/optimise/train.jl:105
┌ Info: Epoch 863
└ @ Main /Users/kcft114/.julia/packages/Flux/qXNjB/src/optimise/train.jl:105
┌ Info: Epoch 864
└ @ Main /Users/kcft114/.julia/packages/Flux/qXNjB/src/optimise/train.jl:105
┌ Info: Epoch 865
└ @ Main /Users/kcft114/.julia/packages/Flux/qXNjB/src/optimise/train.jl:105
┌ Info: Epoch 866
└ @ Main /Users/kcft114/.julia/packages/Flux/qXNjB/src/optimise/train.jl:105
┌ Info: Epoch 867
└ @ Main /Users/kcft114/.julia/packages/Flux/qXNjB/src/optimise/train.jl:105
┌ Info: Epoch 868
└ @ Main /Users/kcft114/.julia/packages/Flux/qXNjB/src/optimise/train.jl:105
┌ Info: Epoch 869
└ @ Main /Users/kcft114/.julia/packages/Flux/qXNjB/src/optimise/train.jl:105
┌ Info: Epoch 870
└ @ Main /Users/kcft114/.julia/packages/Flux/qXNjB/src/optimise/train.jl:105
┌ Info: Epoch 871
└ @ Main /Users/kcft114/.julia/packages/Flux/qXNjB/src/optimise/train.jl:105
┌ Info: Epoch 872
└ @ Main /Users/kcft114/.julia/packages/Flux/qXNjB/src/optimise/train.jl:105
┌ Info: Epoch 873
└ @ Main /Users/kcft114/.julia/packages/Flux/qXNjB/src/optimise/train.jl:105
┌ Info: Epoch 874
└ @ Main /Users/kcft114/.julia/packages/Flux/qXNjB/src/optimise/train.jl:105
┌ Info: Epoch 875
└ @ Main /Users/kcft114/.julia/packages/Flux/qXNjB/src/optimise/train.jl:105
┌ Info: Epoch 876
└ @ Main /Users/kcft114/.julia/packages/Flux/qXNjB/src/optimise/train.jl:105
┌ Info: Epoch 877
└ @ Main /Users/kcft114/.julia/packages/Flux/qXNjB/src/optimise/train.jl:105
┌ Info: Epoch 878
└ @ Main /Users/kcft114/.julia/packages/Flux/qXNjB/src/optimise/train.jl:105
┌ Info: Epoch 879
└ @ Main /Users/kcft114/.julia/packages/Flux/qXNjB/src/optimise/train.jl:105
┌ Info: Epoch 880
└ @ Main /Users/kcft114/.julia/packages/Flux/qXNjB/src/optimise/train.jl:105
┌ Info: Epoch 881
└ @ Main /Users/kcft114/.julia/packages/Flux/qXNjB/src/optimise/train.jl:105
┌ Info: Epoch 882
└ @ Main /Users/kcft114/.julia/packages/Flux/qXNjB/src/optimise/train.jl:105
┌ Info: Epoch 883
└ @ Main /Users/kcft114/.julia/packages/Flux/qXNjB/src/optimise/train.jl:105
┌ Info: Epoch 884
└ @ Main /Users/kcft114/.julia/packages/Flux/qXNjB/src/optimise/train.jl:105
┌ Info: Epoch 885
└ @ Main /Users/kcft114/.julia/packages/Flux/qXNjB/src/optimise/train.jl:105
┌ Info: Epoch 886
└ @ Main /Users/kcft114/.julia/packages/Flux/qXNjB/src/optimise/train.jl:105
┌ Info: Epoch 887
└ @ Main /Users/kcft114/.julia/packages/Flux/qXNjB/src/optimise/train.jl:105
┌ Info: Epoch 888
└ @ Main /Users/kcft114/.julia/packages/Flux/qXNjB/src/optimise/train.jl:105
┌ Info: Epoch 889
└ @ Main /Users/kcft114/.julia/packages/Flux/qXNjB/src/optimise/train.jl:105
┌ Info: Epoch 890
└ @ Main /Users/kcft114/.julia/packages/Flux/qXNjB/src/optimise/train.jl:105
┌ Info: Epoch 891
└ @ Main /Users/kcft114/.julia/packages/Flux/qXNjB/src/optimise/train.jl:105
┌ Info: Epoch 892
└ @ Main /Users/kcft114/.julia/packages/Flux/qXNjB/src/optimise/train.jl:105
┌ Info: Epoch 893
└ @ Main /Users/kcft114/.julia/packages/Flux/qXNjB/src/optimise/train.jl:105
┌ Info: Epoch 894
└ @ Main /Users/kcft114/.julia/packages/Flux/qXNjB/src/optimise/train.jl:105
┌ Info: Epoch 895
└ @ Main /Users/kcft114/.julia/packages/Flux/qXNjB/src/optimise/train.jl:105
┌ Info: Epoch 896
└ @ Main /Users/kcft114/.julia/packages/Flux/qXNjB/src/optimise/train.jl:105
┌ Info: Epoch 897
└ @ Main /Users/kcft114/.julia/packages/Flux/qXNjB/src/optimise/train.jl:105
┌ Info: Epoch 898
└ @ Main /Users/kcft114/.julia/packages/Flux/qXNjB/src/optimise/train.jl:105
┌ Info: Epoch 899
└ @ Main /Users/kcft114/.julia/packages/Flux/qXNjB/src/optimise/train.jl:105
```

```
loss(X, Y) = 0.6633836f0 (tracked)
loss(X, Y) = 0.6426086f0 (tracked)
loss(X, Y) = 0.66600317f0 (tracked)
loss(X, Y) = 0.6738426f0 (tracked)
loss(X, Y) = 0.62881064f0 (tracked)
loss(X, Y) = 0.64669126f0 (tracked)
loss(X, Y) = 0.62426215f0 (tracked)
loss(X, Y) = 0.65573144f0 (tracked)
loss(X, Y) = 0.6436581f0 (tracked)
loss(X, Y) = 0.6533072f0 (tracked)
loss(X, Y) = 0.67070836f0 (tracked)
loss(X, Y) = 0.6299786f0 (tracked)
loss(X, Y) = 0.6261297f0 (tracked)
loss(X, Y) = 0.63437504f0 (tracked)
loss(X, Y) = 0.6923404f0 (tracked)
loss(X, Y) = 0.6513335f0 (tracked)
loss(X, Y) = 0.62046754f0 (tracked)
loss(X, Y) = 0.6383459f0 (tracked)
loss(X, Y) = 0.6626461f0 (tracked)
loss(X, Y) = 0.6545994f0 (tracked)
loss(X, Y) = 0.6312809f0 (tracked)
loss(X, Y) = 0.6196001f0 (tracked)
loss(X, Y) = 0.64214295f0 (tracked)
loss(X, Y) = 0.66104627f0 (tracked)
loss(X, Y) = 0.6785733f0 (tracked)
loss(X, Y) = 0.6615233f0 (tracked)
loss(X, Y) = 0.6540682f0 (tracked)
loss(X, Y) = 0.582956f0 (tracked)
loss(X, Y) = 0.6840165f0 (tracked)
loss(X, Y) = 0.6201558f0 (tracked)
loss(X, Y) = 0.65479034f0 (tracked)
loss(X, Y) = 0.66190386f0 (tracked)
loss(X, Y) = 0.6353485f0 (tracked)
loss(X, Y) = 0.6258757f0 (tracked)
loss(X, Y) = 0.6504409f0 (tracked)
loss(X, Y) = 0.6107317f0 (tracked)
loss(X, Y) = 0.6549172f0 (tracked)
loss(X, Y) = 0.6420727f0 (tracked)
loss(X, Y) = 0.6062559f0 (tracked)
loss(X, Y) = 0.65609246f0 (tracked)
loss(X, Y) = 0.65042466f0 (tracked)
loss(X, Y) = 0.665911f0 (tracked)
loss(X, Y) = 0.61868936f0 (tracked)
loss(X, Y) = 0.6450105f0 (tracked)
loss(X, Y) = 0.62713057f0 (tracked)
loss(X, Y) = 0.65709484f0 (tracked)
loss(X, Y) = 0.63528615f0 (tracked)
loss(X, Y) = 0.64841133f0 (tracked)
loss(X, Y) = 0.63255847f0 (tracked)
loss(X, Y) = 0.62896f0 (tracked)
loss(X, Y) = 0.6275793f0 (tracked)
loss(X, Y) = 0.6482452f0 (tracked)
loss(X, Y) = 0.63881046f0 (tracked)
loss(X, Y) = 0.6215811f0 (tracked)
loss(X, Y) = 0.63259757f0 (tracked)
loss(X, Y) = 0.6020515f0 (tracked)
loss(X, Y) = 0.64133644f0 (tracked)
loss(X, Y) = 0.6645541f0 (tracked)
loss(X, Y) = 0.65003294f0 (tracked)
loss(X, Y) = 0.62201905f0 (tracked)
loss(X, Y) = 0.6306872f0 (tracked)
loss(X, Y) = 0.6171236f0 (tracked)
loss(X, Y) = 0.67498094f0 (tracked)
loss(X, Y) = 0.6525387f0 (tracked)
loss(X, Y) = 0.5929153f0 (tracked)
loss(X, Y) = 0.6593092f0 (tracked)
loss(X, Y) = 0.6510517f0 (tracked)
loss(X, Y) = 0.6497354f0 (tracked)
loss(X, Y) = 0.61437666f0 (tracked)
loss(X, Y) = 0.61846626f0 (tracked)
loss(X, Y) = 0.6488098f0 (tracked)
loss(X, Y) = 0.6086824f0 (tracked)
loss(X, Y) = 0.64053094f0 (tracked)
loss(X, Y) = 0.6711764f0 (tracked)
loss(X, Y) = 0.6462557f0 (tracked)
loss(X, Y) = 0.644424f0 (tracked)
loss(X, Y) = 0.6387843f0 (tracked)
loss(X, Y) = 0.6799979f0 (tracked)
loss(X, Y) = 0.6429915f0 (tracked)
loss(X, Y) = 0.6289682f0 (tracked)
loss(X, Y) = 0.62865895f0 (tracked)
loss(X, Y) = 0.620585f0 (tracked)
loss(X, Y) = 0.61215854f0 (tracked)
loss(X, Y) = 0.62790906f0 (tracked)
loss(X, Y) = 0.6558285f0 (tracked)
loss(X, Y) = 0.61439544f0 (tracked)
loss(X, Y) = 0.6321304f0 (tracked)
loss(X, Y) = 0.62315625f0 (tracked)
loss(X, Y) = 0.63372874f0 (tracked)
loss(X, Y) = 0.6385917f0 (tracked)
loss(X, Y) = 0.6740109f0 (tracked)
loss(X, Y) = 0.6246187f0 (tracked)
loss(X, Y) = 0.67015004f0 (tracked)
loss(X, Y) = 0.60984266f0 (tracked)
loss(X, Y) = 0.65024006f0 (tracked)
loss(X, Y) = 0.6439636f0 (tracked)
loss(X, Y) = 0.64114517f0 (tracked)
loss(X, Y) = 0.6210557f0 (tracked)
loss(X, Y) = 0.59722596f0 (tracked)
loss(X, Y) = 0.63587385f0 (tracked)
loss(X, Y) = 0.63188124f0 (tracked)
loss(X, Y) = 0.63958734f0 (tracked)
loss(X, Y) = 0.6318565f0 (tracked)
loss(X, Y) = 0.65777236f0 (tracked)
loss(X, Y) = 0.6831807f0 (tracked)
loss(X, Y) = 0.6224663f0 (tracked)
loss(X, Y) = 0.6002245f0 (tracked)
loss(X, Y) = 0.6045171f0 (tracked)
```

```
┌ Info: Epoch 900
└ @ Main /Users/kcft114/.julia/packages/Flux/qXNjB/src/optimise/train.jl:105
┌ Info: Epoch 901
└ @ Main /Users/kcft114/.julia/packages/Flux/qXNjB/src/optimise/train.jl:105
┌ Info: Epoch 902
└ @ Main /Users/kcft114/.julia/packages/Flux/qXNjB/src/optimise/train.jl:105
┌ Info: Epoch 903
└ @ Main /Users/kcft114/.julia/packages/Flux/qXNjB/src/optimise/train.jl:105
┌ Info: Epoch 904
└ @ Main /Users/kcft114/.julia/packages/Flux/qXNjB/src/optimise/train.jl:105
┌ Info: Epoch 905
└ @ Main /Users/kcft114/.julia/packages/Flux/qXNjB/src/optimise/train.jl:105
┌ Info: Epoch 906
└ @ Main /Users/kcft114/.julia/packages/Flux/qXNjB/src/optimise/train.jl:105
┌ Info: Epoch 907
└ @ Main /Users/kcft114/.julia/packages/Flux/qXNjB/src/optimise/train.jl:105
┌ Info: Epoch 908
└ @ Main /Users/kcft114/.julia/packages/Flux/qXNjB/src/optimise/train.jl:105
┌ Info: Epoch 909
└ @ Main /Users/kcft114/.julia/packages/Flux/qXNjB/src/optimise/train.jl:105
┌ Info: Epoch 910
└ @ Main /Users/kcft114/.julia/packages/Flux/qXNjB/src/optimise/train.jl:105
┌ Info: Epoch 911
└ @ Main /Users/kcft114/.julia/packages/Flux/qXNjB/src/optimise/train.jl:105
┌ Info: Epoch 912
└ @ Main /Users/kcft114/.julia/packages/Flux/qXNjB/src/optimise/train.jl:105
┌ Info: Epoch 913
└ @ Main /Users/kcft114/.julia/packages/Flux/qXNjB/src/optimise/train.jl:105
┌ Info: Epoch 914
└ @ Main /Users/kcft114/.julia/packages/Flux/qXNjB/src/optimise/train.jl:105
┌ Info: Epoch 915
└ @ Main /Users/kcft114/.julia/packages/Flux/qXNjB/src/optimise/train.jl:105
┌ Info: Epoch 916
└ @ Main /Users/kcft114/.julia/packages/Flux/qXNjB/src/optimise/train.jl:105
┌ Info: Epoch 917
└ @ Main /Users/kcft114/.julia/packages/Flux/qXNjB/src/optimise/train.jl:105
┌ Info: Epoch 918
└ @ Main /Users/kcft114/.julia/packages/Flux/qXNjB/src/optimise/train.jl:105
┌ Info: Epoch 919
└ @ Main /Users/kcft114/.julia/packages/Flux/qXNjB/src/optimise/train.jl:105
┌ Info: Epoch 920
└ @ Main /Users/kcft114/.julia/packages/Flux/qXNjB/src/optimise/train.jl:105
┌ Info: Epoch 921
└ @ Main /Users/kcft114/.julia/packages/Flux/qXNjB/src/optimise/train.jl:105
┌ Info: Epoch 922
└ @ Main /Users/kcft114/.julia/packages/Flux/qXNjB/src/optimise/train.jl:105
┌ Info: Epoch 923
└ @ Main /Users/kcft114/.julia/packages/Flux/qXNjB/src/optimise/train.jl:105
┌ Info: Epoch 924
└ @ Main /Users/kcft114/.julia/packages/Flux/qXNjB/src/optimise/train.jl:105
┌ Info: Epoch 925
└ @ Main /Users/kcft114/.julia/packages/Flux/qXNjB/src/optimise/train.jl:105
┌ Info: Epoch 926
└ @ Main /Users/kcft114/.julia/packages/Flux/qXNjB/src/optimise/train.jl:105
┌ Info: Epoch 927
└ @ Main /Users/kcft114/.julia/packages/Flux/qXNjB/src/optimise/train.jl:105
┌ Info: Epoch 928
└ @ Main /Users/kcft114/.julia/packages/Flux/qXNjB/src/optimise/train.jl:105
┌ Info: Epoch 929
└ @ Main /Users/kcft114/.julia/packages/Flux/qXNjB/src/optimise/train.jl:105
┌ Info: Epoch 930
└ @ Main /Users/kcft114/.julia/packages/Flux/qXNjB/src/optimise/train.jl:105
┌ Info: Epoch 931
└ @ Main /Users/kcft114/.julia/packages/Flux/qXNjB/src/optimise/train.jl:105
┌ Info: Epoch 932
└ @ Main /Users/kcft114/.julia/packages/Flux/qXNjB/src/optimise/train.jl:105
┌ Info: Epoch 933
└ @ Main /Users/kcft114/.julia/packages/Flux/qXNjB/src/optimise/train.jl:105
┌ Info: Epoch 934
└ @ Main /Users/kcft114/.julia/packages/Flux/qXNjB/src/optimise/train.jl:105
┌ Info: Epoch 935
└ @ Main /Users/kcft114/.julia/packages/Flux/qXNjB/src/optimise/train.jl:105
┌ Info: Epoch 936
└ @ Main /Users/kcft114/.julia/packages/Flux/qXNjB/src/optimise/train.jl:105
┌ Info: Epoch 937
└ @ Main /Users/kcft114/.julia/packages/Flux/qXNjB/src/optimise/train.jl:105
┌ Info: Epoch 938
└ @ Main /Users/kcft114/.julia/packages/Flux/qXNjB/src/optimise/train.jl:105
┌ Info: Epoch 939
└ @ Main /Users/kcft114/.julia/packages/Flux/qXNjB/src/optimise/train.jl:105
┌ Info: Epoch 940
└ @ Main /Users/kcft114/.julia/packages/Flux/qXNjB/src/optimise/train.jl:105
┌ Info: Epoch 941
└ @ Main /Users/kcft114/.julia/packages/Flux/qXNjB/src/optimise/train.jl:105
┌ Info: Epoch 942
└ @ Main /Users/kcft114/.julia/packages/Flux/qXNjB/src/optimise/train.jl:105
┌ Info: Epoch 943
└ @ Main /Users/kcft114/.julia/packages/Flux/qXNjB/src/optimise/train.jl:105
┌ Info: Epoch 944
└ @ Main /Users/kcft114/.julia/packages/Flux/qXNjB/src/optimise/train.jl:105
┌ Info: Epoch 945
└ @ Main /Users/kcft114/.julia/packages/Flux/qXNjB/src/optimise/train.jl:105
┌ Info: Epoch 946
└ @ Main /Users/kcft114/.julia/packages/Flux/qXNjB/src/optimise/train.jl:105
┌ Info: Epoch 947
└ @ Main /Users/kcft114/.julia/packages/Flux/qXNjB/src/optimise/train.jl:105
┌ Info: Epoch 948
└ @ Main /Users/kcft114/.julia/packages/Flux/qXNjB/src/optimise/train.jl:105
┌ Info: Epoch 949
└ @ Main /Users/kcft114/.julia/packages/Flux/qXNjB/src/optimise/train.jl:105
┌ Info: Epoch 950
└ @ Main /Users/kcft114/.julia/packages/Flux/qXNjB/src/optimise/train.jl:105
┌ Info: Epoch 951
└ @ Main /Users/kcft114/.julia/packages/Flux/qXNjB/src/optimise/train.jl:105
┌ Info: Epoch 952
└ @ Main /Users/kcft114/.julia/packages/Flux/qXNjB/src/optimise/train.jl:105
┌ Info: Epoch 953
└ @ Main /Users/kcft114/.julia/packages/Flux/qXNjB/src/optimise/train.jl:105
┌ Info: Epoch 954
└ @ Main /Users/kcft114/.julia/packages/Flux/qXNjB/src/optimise/train.jl:105
┌ Info: Epoch 955
└ @ Main /Users/kcft114/.julia/packages/Flux/qXNjB/src/optimise/train.jl:105
┌ Info: Epoch 956
└ @ Main /Users/kcft114/.julia/packages/Flux/qXNjB/src/optimise/train.jl:105
┌ Info: Epoch 957
└ @ Main /Users/kcft114/.julia/packages/Flux/qXNjB/src/optimise/train.jl:105
┌ Info: Epoch 958
└ @ Main /Users/kcft114/.julia/packages/Flux/qXNjB/src/optimise/train.jl:105
┌ Info: Epoch 959
└ @ Main /Users/kcft114/.julia/packages/Flux/qXNjB/src/optimise/train.jl:105
┌ Info: Epoch 960
└ @ Main /Users/kcft114/.julia/packages/Flux/qXNjB/src/optimise/train.jl:105
┌ Info: Epoch 961
└ @ Main /Users/kcft114/.julia/packages/Flux/qXNjB/src/optimise/train.jl:105
┌ Info: Epoch 962
└ @ Main /Users/kcft114/.julia/packages/Flux/qXNjB/src/optimise/train.jl:105
┌ Info: Epoch 963
└ @ Main /Users/kcft114/.julia/packages/Flux/qXNjB/src/optimise/train.jl:105
┌ Info: Epoch 964
└ @ Main /Users/kcft114/.julia/packages/Flux/qXNjB/src/optimise/train.jl:105
┌ Info: Epoch 965
└ @ Main /Users/kcft114/.julia/packages/Flux/qXNjB/src/optimise/train.jl:105
┌ Info: Epoch 966
└ @ Main /Users/kcft114/.julia/packages/Flux/qXNjB/src/optimise/train.jl:105
┌ Info: Epoch 967
└ @ Main /Users/kcft114/.julia/packages/Flux/qXNjB/src/optimise/train.jl:105
┌ Info: Epoch 968
└ @ Main /Users/kcft114/.julia/packages/Flux/qXNjB/src/optimise/train.jl:105
┌ Info: Epoch 969
└ @ Main /Users/kcft114/.julia/packages/Flux/qXNjB/src/optimise/train.jl:105
┌ Info: Epoch 970
└ @ Main /Users/kcft114/.julia/packages/Flux/qXNjB/src/optimise/train.jl:105
┌ Info: Epoch 971
└ @ Main /Users/kcft114/.julia/packages/Flux/qXNjB/src/optimise/train.jl:105
┌ Info: Epoch 972
└ @ Main /Users/kcft114/.julia/packages/Flux/qXNjB/src/optimise/train.jl:105
┌ Info: Epoch 973
└ @ Main /Users/kcft114/.julia/packages/Flux/qXNjB/src/optimise/train.jl:105
┌ Info: Epoch 974
└ @ Main /Users/kcft114/.julia/packages/Flux/qXNjB/src/optimise/train.jl:105
┌ Info: Epoch 975
└ @ Main /Users/kcft114/.julia/packages/Flux/qXNjB/src/optimise/train.jl:105
┌ Info: Epoch 976
└ @ Main /Users/kcft114/.julia/packages/Flux/qXNjB/src/optimise/train.jl:105
┌ Info: Epoch 977
└ @ Main /Users/kcft114/.julia/packages/Flux/qXNjB/src/optimise/train.jl:105
┌ Info: Epoch 978
└ @ Main /Users/kcft114/.julia/packages/Flux/qXNjB/src/optimise/train.jl:105
┌ Info: Epoch 979
└ @ Main /Users/kcft114/.julia/packages/Flux/qXNjB/src/optimise/train.jl:105
┌ Info: Epoch 980
└ @ Main /Users/kcft114/.julia/packages/Flux/qXNjB/src/optimise/train.jl:105
┌ Info: Epoch 981
└ @ Main /Users/kcft114/.julia/packages/Flux/qXNjB/src/optimise/train.jl:105
┌ Info: Epoch 982
└ @ Main /Users/kcft114/.julia/packages/Flux/qXNjB/src/optimise/train.jl:105
┌ Info: Epoch 983
└ @ Main /Users/kcft114/.julia/packages/Flux/qXNjB/src/optimise/train.jl:105
┌ Info: Epoch 984
└ @ Main /Users/kcft114/.julia/packages/Flux/qXNjB/src/optimise/train.jl:105
┌ Info: Epoch 985
└ @ Main /Users/kcft114/.julia/packages/Flux/qXNjB/src/optimise/train.jl:105
┌ Info: Epoch 986
└ @ Main /Users/kcft114/.julia/packages/Flux/qXNjB/src/optimise/train.jl:105
┌ Info: Epoch 987
└ @ Main /Users/kcft114/.julia/packages/Flux/qXNjB/src/optimise/train.jl:105
┌ Info: Epoch 988
└ @ Main /Users/kcft114/.julia/packages/Flux/qXNjB/src/optimise/train.jl:105
┌ Info: Epoch 989
└ @ Main /Users/kcft114/.julia/packages/Flux/qXNjB/src/optimise/train.jl:105
┌ Info: Epoch 990
└ @ Main /Users/kcft114/.julia/packages/Flux/qXNjB/src/optimise/train.jl:105
┌ Info: Epoch 991
└ @ Main /Users/kcft114/.julia/packages/Flux/qXNjB/src/optimise/train.jl:105
┌ Info: Epoch 992
└ @ Main /Users/kcft114/.julia/packages/Flux/qXNjB/src/optimise/train.jl:105
┌ Info: Epoch 993
└ @ Main /Users/kcft114/.julia/packages/Flux/qXNjB/src/optimise/train.jl:105
┌ Info: Epoch 994
└ @ Main /Users/kcft114/.julia/packages/Flux/qXNjB/src/optimise/train.jl:105
┌ Info: Epoch 995
└ @ Main /Users/kcft114/.julia/packages/Flux/qXNjB/src/optimise/train.jl:105
┌ Info: Epoch 996
└ @ Main /Users/kcft114/.julia/packages/Flux/qXNjB/src/optimise/train.jl:105
┌ Info: Epoch 997
└ @ Main /Users/kcft114/.julia/packages/Flux/qXNjB/src/optimise/train.jl:105
┌ Info: Epoch 998
└ @ Main /Users/kcft114/.julia/packages/Flux/qXNjB/src/optimise/train.jl:105
┌ Info: Epoch 999
└ @ Main /Users/kcft114/.julia/packages/Flux/qXNjB/src/optimise/train.jl:105
┌ Info: Epoch 1000
└ @ Main /Users/kcft114/.julia/packages/Flux/qXNjB/src/optimise/train.jl:105
┌ Info: Epoch 1001
└ @ Main /Users/kcft114/.julia/packages/Flux/qXNjB/src/optimise/train.jl:105
┌ Info: Epoch 1002
└ @ Main /Users/kcft114/.julia/packages/Flux/qXNjB/src/optimise/train.jl:105
┌ Info: Epoch 1003
└ @ Main /Users/kcft114/.julia/packages/Flux/qXNjB/src/optimise/train.jl:105
┌ Info: Epoch 1004
└ @ Main /Users/kcft114/.julia/packages/Flux/qXNjB/src/optimise/train.jl:105
┌ Info: Epoch 1005
└ @ Main /Users/kcft114/.julia/packages/Flux/qXNjB/src/optimise/train.jl:105
┌ Info: Epoch 1006
└ @ Main /Users/kcft114/.julia/packages/Flux/qXNjB/src/optimise/train.jl:105
┌ Info: Epoch 1007
└ @ Main /Users/kcft114/.julia/packages/Flux/qXNjB/src/optimise/train.jl:105
```

```
loss(X, Y) = 0.60793173f0 (tracked)
loss(X, Y) = 0.6063072f0 (tracked)
loss(X, Y) = 0.6423209f0 (tracked)
loss(X, Y) = 0.64041156f0 (tracked)
loss(X, Y) = 0.62492746f0 (tracked)
loss(X, Y) = 0.6465782f0 (tracked)
loss(X, Y) = 0.63990504f0 (tracked)
loss(X, Y) = 0.6490135f0 (tracked)
loss(X, Y) = 0.62568295f0 (tracked)
loss(X, Y) = 0.6329541f0 (tracked)
loss(X, Y) = 0.6720444f0 (tracked)
loss(X, Y) = 0.64323676f0 (tracked)
loss(X, Y) = 0.60144293f0 (tracked)
loss(X, Y) = 0.6303693f0 (tracked)
loss(X, Y) = 0.6702721f0 (tracked)
loss(X, Y) = 0.63683784f0 (tracked)
loss(X, Y) = 0.6590518f0 (tracked)
loss(X, Y) = 0.6383255f0 (tracked)
loss(X, Y) = 0.6168021f0 (tracked)
loss(X, Y) = 0.6335767f0 (tracked)
loss(X, Y) = 0.6091538f0 (tracked)
loss(X, Y) = 0.6285477f0 (tracked)
loss(X, Y) = 0.63813174f0 (tracked)
loss(X, Y) = 0.6586439f0 (tracked)
loss(X, Y) = 0.6073172f0 (tracked)
loss(X, Y) = 0.6251194f0 (tracked)
loss(X, Y) = 0.5929964f0 (tracked)
loss(X, Y) = 0.60575473f0 (tracked)
loss(X, Y) = 0.60260606f0 (tracked)
loss(X, Y) = 0.64467084f0 (tracked)
loss(X, Y) = 0.6411121f0 (tracked)
loss(X, Y) = 0.64564836f0 (tracked)
loss(X, Y) = 0.6172382f0 (tracked)
loss(X, Y) = 0.66373754f0 (tracked)
loss(X, Y) = 0.6546255f0 (tracked)
loss(X, Y) = 0.62810326f0 (tracked)
loss(X, Y) = 0.64117754f0 (tracked)
loss(X, Y) = 0.6149943f0 (tracked)
loss(X, Y) = 0.6216031f0 (tracked)
loss(X, Y) = 0.61649144f0 (tracked)
loss(X, Y) = 0.6367774f0 (tracked)
loss(X, Y) = 0.597297f0 (tracked)
loss(X, Y) = 0.62856424f0 (tracked)
loss(X, Y) = 0.6367925f0 (tracked)
loss(X, Y) = 0.5897342f0 (tracked)
loss(X, Y) = 0.61956596f0 (tracked)
loss(X, Y) = 0.597458f0 (tracked)
loss(X, Y) = 0.6051173f0 (tracked)
loss(X, Y) = 0.66144115f0 (tracked)
loss(X, Y) = 0.63185716f0 (tracked)
loss(X, Y) = 0.6289252f0 (tracked)
loss(X, Y) = 0.6339841f0 (tracked)
loss(X, Y) = 0.62994224f0 (tracked)
loss(X, Y) = 0.6003101f0 (tracked)
loss(X, Y) = 0.65939355f0 (tracked)
loss(X, Y) = 0.65175176f0 (tracked)
loss(X, Y) = 0.6407449f0 (tracked)
loss(X, Y) = 0.64739835f0 (tracked)
loss(X, Y) = 0.6077424f0 (tracked)
loss(X, Y) = 0.62903315f0 (tracked)
loss(X, Y) = 0.5957471f0 (tracked)
loss(X, Y) = 0.61763f0 (tracked)
loss(X, Y) = 0.6547045f0 (tracked)
loss(X, Y) = 0.63497794f0 (tracked)
loss(X, Y) = 0.62146f0 (tracked)
loss(X, Y) = 0.6352012f0 (tracked)
loss(X, Y) = 0.5945885f0 (tracked)
loss(X, Y) = 0.6019197f0 (tracked)
loss(X, Y) = 0.6331294f0 (tracked)
loss(X, Y) = 0.6108676f0 (tracked)
loss(X, Y) = 0.6327137f0 (tracked)
loss(X, Y) = 0.5850024f0 (tracked)
loss(X, Y) = 0.61948115f0 (tracked)
loss(X, Y) = 0.5989253f0 (tracked)
loss(X, Y) = 0.6454108f0 (tracked)
loss(X, Y) = 0.6402918f0 (tracked)
loss(X, Y) = 0.6228538f0 (tracked)
loss(X, Y) = 0.6182785f0 (tracked)
loss(X, Y) = 0.6128051f0 (tracked)
loss(X, Y) = 0.5947555f0 (tracked)
loss(X, Y) = 0.6259745f0 (tracked)
loss(X, Y) = 0.6432255f0 (tracked)
loss(X, Y) = 0.57703876f0 (tracked)
loss(X, Y) = 0.6039409f0 (tracked)
loss(X, Y) = 0.648734f0 (tracked)
loss(X, Y) = 0.6010152f0 (tracked)
loss(X, Y) = 0.66257983f0 (tracked)
loss(X, Y) = 0.6490202f0 (tracked)
loss(X, Y) = 0.64666635f0 (tracked)
loss(X, Y) = 0.62766236f0 (tracked)
loss(X, Y) = 0.5973399f0 (tracked)
loss(X, Y) = 0.621776f0 (tracked)
loss(X, Y) = 0.63313437f0 (tracked)
loss(X, Y) = 0.62791646f0 (tracked)
loss(X, Y) = 0.6401814f0 (tracked)
loss(X, Y) = 0.640098f0 (tracked)
loss(X, Y) = 0.6375518f0 (tracked)
loss(X, Y) = 0.62495077f0 (tracked)
loss(X, Y) = 0.63794833f0 (tracked)
loss(X, Y) = 0.5981404f0 (tracked)
loss(X, Y) = 0.61519635f0 (tracked)
loss(X, Y) = 0.6301696f0 (tracked)
loss(X, Y) = 0.59275526f0 (tracked)
loss(X, Y) = 0.6331097f0 (tracked)
loss(X, Y) = 0.61993814f0 (tracked)
loss(X, Y) = 0.6075529f0 (tracked)
loss(X, Y) = 0.6193827f0 (tracked)
loss(X, Y) = 0.60586977f0 (tracked)
loss(X, Y) = 0.6108007f0 (tracked)
loss(X, Y) = 0.6051147f0 (tracked)
loss(X, Y) = 0.6186815f0 (tracked)
loss(X, Y) = 0.635349f0 (tracked)
loss(X, Y) = 0.6011753f0 (tracked)
loss(X, Y) = 0.6201824f0 (tracked)
```

```
┌ Info: Epoch 1008
└ @ Main /Users/kcft114/.julia/packages/Flux/qXNjB/src/optimise/train.jl:105
┌ Info: Epoch 1009
└ @ Main /Users/kcft114/.julia/packages/Flux/qXNjB/src/optimise/train.jl:105
┌ Info: Epoch 1010
└ @ Main /Users/kcft114/.julia/packages/Flux/qXNjB/src/optimise/train.jl:105
┌ Info: Epoch 1011
└ @ Main /Users/kcft114/.julia/packages/Flux/qXNjB/src/optimise/train.jl:105
┌ Info: Epoch 1012
└ @ Main /Users/kcft114/.julia/packages/Flux/qXNjB/src/optimise/train.jl:105
┌ Info: Epoch 1013
└ @ Main /Users/kcft114/.julia/packages/Flux/qXNjB/src/optimise/train.jl:105
┌ Info: Epoch 1014
└ @ Main /Users/kcft114/.julia/packages/Flux/qXNjB/src/optimise/train.jl:105
┌ Info: Epoch 1015
└ @ Main /Users/kcft114/.julia/packages/Flux/qXNjB/src/optimise/train.jl:105
┌ Info: Epoch 1016
└ @ Main /Users/kcft114/.julia/packages/Flux/qXNjB/src/optimise/train.jl:105
┌ Info: Epoch 1017
└ @ Main /Users/kcft114/.julia/packages/Flux/qXNjB/src/optimise/train.jl:105
```

```
loss(X, Y) = 0.5680728f0 (tracked)
loss(X, Y) = 0.6165956f0 (tracked)
loss(X, Y) = 0.60856885f0 (tracked)
loss(X, Y) = 0.6161308f0 (tracked)
loss(X, Y) = 0.6360299f0 (tracked)
loss(X, Y) = 0.62404627f0 (tracked)
loss(X, Y) = 0.6095176f0 (tracked)
loss(X, Y) = 0.59207207f0 (tracked)
loss(X, Y) = 0.5627891f0 (tracked)
loss(X, Y) = 0.58521724f0 (tracked)
loss(X, Y) = 0.6385483f0 (tracked)
loss(X, Y) = 0.6386978f0 (tracked)
loss(X, Y) = 0.6015052f0 (tracked)
loss(X, Y) = 0.6605864f0 (tracked)
loss(X, Y) = 0.5993499f0 (tracked)
loss(X, Y) = 0.64790314f0 (tracked)
loss(X, Y) = 0.6317379f0 (tracked)
loss(X, Y) = 0.60657597f0 (tracked)
loss(X, Y) = 0.5963951f0 (tracked)
loss(X, Y) = 0.54575074f0 (tracked)
loss(X, Y) = 0.6403069f0 (tracked)
loss(X, Y) = 0.58062756f0 (tracked)
loss(X, Y) = 0.6027597f0 (tracked)
loss(X, Y) = 0.6381513f0 (tracked)
loss(X, Y) = 0.61357504f0 (tracked)
loss(X, Y) = 0.6127939f0 (tracked)
loss(X, Y) = 0.6054067f0 (tracked)
loss(X, Y) = 0.6193082f0 (tracked)
loss(X, Y) = 0.63072944f0 (tracked)
loss(X, Y) = 0.6065122f0 (tracked)
loss(X, Y) = 0.5826374f0 (tracked)
loss(X, Y) = 0.6054335f0 (tracked)
loss(X, Y) = 0.6368483f0 (tracked)
loss(X, Y) = 0.5926273f0 (tracked)
loss(X, Y) = 0.5987484f0 (tracked)
loss(X, Y) = 0.62023294f0 (tracked)
loss(X, Y) = 0.62547076f0 (tracked)
loss(X, Y) = 0.58908445f0 (tracked)
loss(X, Y) = 0.61325085f0 (tracked)
loss(X, Y) = 0.653885f0 (tracked)
loss(X, Y) = 0.6252448f0 (tracked)
loss(X, Y) = 0.6244537f0 (tracked)
loss(X, Y) = 0.6020329f0 (tracked)
loss(X, Y) = 0.5733493f0 (tracked)
loss(X, Y) = 0.61399394f0 (tracked)
loss(X, Y) = 0.6115663f0 (tracked)
loss(X, Y) = 0.6237954f0 (tracked)
loss(X, Y) = 0.58417124f0 (tracked)
loss(X, Y) = 0.60966253f0 (tracked)
loss(X, Y) = 0.6049778f0 (tracked)
loss(X, Y) = 0.6174739f0 (tracked)
loss(X, Y) = 0.6039012f0 (tracked)
loss(X, Y) = 0.668107f0 (tracked)
loss(X, Y) = 0.58626133f0 (tracked)
loss(X, Y) = 0.5967062f0 (tracked)
loss(X, Y) = 0.62891763f0 (tracked)
loss(X, Y) = 0.61777675f0 (tracked)
loss(X, Y) = 0.60215163f0 (tracked)
loss(X, Y) = 0.5782227f0 (tracked)
loss(X, Y) = 0.63024664f0 (tracked)
loss(X, Y) = 0.5956474f0 (tracked)
loss(X, Y) = 0.625594f0 (tracked)
loss(X, Y) = 0.62954843f0 (tracked)
loss(X, Y) = 0.6395541f0 (tracked)
loss(X, Y) = 0.6121223f0 (tracked)
loss(X, Y) = 0.60490805f0 (tracked)
loss(X, Y) = 0.6187581f0 (tracked)
loss(X, Y) = 0.60337275f0 (tracked)
loss(X, Y) = 0.6283455f0 (tracked)
loss(X, Y) = 0.6204655f0 (tracked)
loss(X, Y) = 0.58726186f0 (tracked)
loss(X, Y) = 0.618399f0 (tracked)
loss(X, Y) = 0.59675974f0 (tracked)
loss(X, Y) = 0.6062246f0 (tracked)
loss(X, Y) = 0.61061895f0 (tracked)
loss(X, Y) = 0.6341682f0 (tracked)
loss(X, Y) = 0.57816166f0 (tracked)
loss(X, Y) = 0.61187565f0 (tracked)
loss(X, Y) = 0.6598661f0 (tracked)
loss(X, Y) = 0.6470721f0 (tracked)
loss(X, Y) = 0.6687894f0 (tracked)
loss(X, Y) = 0.65604717f0 (tracked)
loss(X, Y) = 0.60747015f0 (tracked)
loss(X, Y) = 0.62102276f0 (tracked)
loss(X, Y) = 0.5893314f0 (tracked)
loss(X, Y) = 0.57333946f0 (tracked)
loss(X, Y) = 0.6108356f0 (tracked)
loss(X, Y) = 0.60629326f0 (tracked)
loss(X, Y) = 0.5910939f0 (tracked)
loss(X, Y) = 0.59164035f0 (tracked)
loss(X, Y) = 0.59826934f0 (tracked)
loss(X, Y) = 0.61699736f0 (tracked)
loss(X, Y) = 0.60142744f0 (tracked)
```

```
┌ Info: Epoch 1018
└ @ Main /Users/kcft114/.julia/packages/Flux/qXNjB/src/optimise/train.jl:105
┌ Info: Epoch 1019
└ @ Main /Users/kcft114/.julia/packages/Flux/qXNjB/src/optimise/train.jl:105
┌ Info: Epoch 1020
└ @ Main /Users/kcft114/.julia/packages/Flux/qXNjB/src/optimise/train.jl:105
┌ Info: Epoch 1021
└ @ Main /Users/kcft114/.julia/packages/Flux/qXNjB/src/optimise/train.jl:105
┌ Info: Epoch 1022
└ @ Main /Users/kcft114/.julia/packages/Flux/qXNjB/src/optimise/train.jl:105
┌ Info: Epoch 1023
└ @ Main /Users/kcft114/.julia/packages/Flux/qXNjB/src/optimise/train.jl:105
┌ Info: Epoch 1024
└ @ Main /Users/kcft114/.julia/packages/Flux/qXNjB/src/optimise/train.jl:105
┌ Info: Epoch 1025
└ @ Main /Users/kcft114/.julia/packages/Flux/qXNjB/src/optimise/train.jl:105
┌ Info: Epoch 1026
└ @ Main /Users/kcft114/.julia/packages/Flux/qXNjB/src/optimise/train.jl:105
┌ Info: Epoch 1027
└ @ Main /Users/kcft114/.julia/packages/Flux/qXNjB/src/optimise/train.jl:105
┌ Info: Epoch 1028
└ @ Main /Users/kcft114/.julia/packages/Flux/qXNjB/src/optimise/train.jl:105
┌ Info: Epoch 1029
└ @ Main /Users/kcft114/.julia/packages/Flux/qXNjB/src/optimise/train.jl:105
┌ Info: Epoch 1030
└ @ Main /Users/kcft114/.julia/packages/Flux/qXNjB/src/optimise/train.jl:105
┌ Info: Epoch 1031
└ @ Main /Users/kcft114/.julia/packages/Flux/qXNjB/src/optimise/train.jl:105
┌ Info: Epoch 1032
└ @ Main /Users/kcft114/.julia/packages/Flux/qXNjB/src/optimise/train.jl:105
┌ Info: Epoch 1033
└ @ Main /Users/kcft114/.julia/packages/Flux/qXNjB/src/optimise/train.jl:105
┌ Info: Epoch 1034
└ @ Main /Users/kcft114/.julia/packages/Flux/qXNjB/src/optimise/train.jl:105
┌ Info: Epoch 1035
└ @ Main /Users/kcft114/.julia/packages/Flux/qXNjB/src/optimise/train.jl:105
┌ Info: Epoch 1036
└ @ Main /Users/kcft114/.julia/packages/Flux/qXNjB/src/optimise/train.jl:105
┌ Info: Epoch 1037
└ @ Main /Users/kcft114/.julia/packages/Flux/qXNjB/src/optimise/train.jl:105
┌ Info: Epoch 1038
└ @ Main /Users/kcft114/.julia/packages/Flux/qXNjB/src/optimise/train.jl:105
┌ Info: Epoch 1039
└ @ Main /Users/kcft114/.julia/packages/Flux/qXNjB/src/optimise/train.jl:105
┌ Info: Epoch 1040
└ @ Main /Users/kcft114/.julia/packages/Flux/qXNjB/src/optimise/train.jl:105
┌ Info: Epoch 1041
└ @ Main /Users/kcft114/.julia/packages/Flux/qXNjB/src/optimise/train.jl:105
┌ Info: Epoch 1042
└ @ Main /Users/kcft114/.julia/packages/Flux/qXNjB/src/optimise/train.jl:105
┌ Info: Epoch 1043
└ @ Main /Users/kcft114/.julia/packages/Flux/qXNjB/src/optimise/train.jl:105
┌ Info: Epoch 1044
└ @ Main /Users/kcft114/.julia/packages/Flux/qXNjB/src/optimise/train.jl:105
┌ Info: Epoch 1045
└ @ Main /Users/kcft114/.julia/packages/Flux/qXNjB/src/optimise/train.jl:105
┌ Info: Epoch 1046
└ @ Main /Users/kcft114/.julia/packages/Flux/qXNjB/src/optimise/train.jl:105
┌ Info: Epoch 1047
└ @ Main /Users/kcft114/.julia/packages/Flux/qXNjB/src/optimise/train.jl:105
┌ Info: Epoch 1048
└ @ Main /Users/kcft114/.julia/packages/Flux/qXNjB/src/optimise/train.jl:105
┌ Info: Epoch 1049
└ @ Main /Users/kcft114/.julia/packages/Flux/qXNjB/src/optimise/train.jl:105
┌ Info: Epoch 1050
└ @ Main /Users/kcft114/.julia/packages/Flux/qXNjB/src/optimise/train.jl:105
┌ Info: Epoch 1051
└ @ Main /Users/kcft114/.julia/packages/Flux/qXNjB/src/optimise/train.jl:105
┌ Info: Epoch 1052
└ @ Main /Users/kcft114/.julia/packages/Flux/qXNjB/src/optimise/train.jl:105
┌ Info: Epoch 1053
└ @ Main /Users/kcft114/.julia/packages/Flux/qXNjB/src/optimise/train.jl:105
┌ Info: Epoch 1054
└ @ Main /Users/kcft114/.julia/packages/Flux/qXNjB/src/optimise/train.jl:105
┌ Info: Epoch 1055
└ @ Main /Users/kcft114/.julia/packages/Flux/qXNjB/src/optimise/train.jl:105
┌ Info: Epoch 1056
└ @ Main /Users/kcft114/.julia/packages/Flux/qXNjB/src/optimise/train.jl:105
┌ Info: Epoch 1057
└ @ Main /Users/kcft114/.julia/packages/Flux/qXNjB/src/optimise/train.jl:105
┌ Info: Epoch 1058
└ @ Main /Users/kcft114/.julia/packages/Flux/qXNjB/src/optimise/train.jl:105
┌ Info: Epoch 1059
└ @ Main /Users/kcft114/.julia/packages/Flux/qXNjB/src/optimise/train.jl:105
┌ Info: Epoch 1060
└ @ Main /Users/kcft114/.julia/packages/Flux/qXNjB/src/optimise/train.jl:105
┌ Info: Epoch 1061
└ @ Main /Users/kcft114/.julia/packages/Flux/qXNjB/src/optimise/train.jl:105
┌ Info: Epoch 1062
└ @ Main /Users/kcft114/.julia/packages/Flux/qXNjB/src/optimise/train.jl:105
┌ Info: Epoch 1063
└ @ Main /Users/kcft114/.julia/packages/Flux/qXNjB/src/optimise/train.jl:105
┌ Info: Epoch 1064
└ @ Main /Users/kcft114/.julia/packages/Flux/qXNjB/src/optimise/train.jl:105
┌ Info: Epoch 1065
└ @ Main /Users/kcft114/.julia/packages/Flux/qXNjB/src/optimise/train.jl:105
┌ Info: Epoch 1066
└ @ Main /Users/kcft114/.julia/packages/Flux/qXNjB/src/optimise/train.jl:105
┌ Info: Epoch 1067
└ @ Main /Users/kcft114/.julia/packages/Flux/qXNjB/src/optimise/train.jl:105
┌ Info: Epoch 1068
└ @ Main /Users/kcft114/.julia/packages/Flux/qXNjB/src/optimise/train.jl:105
┌ Info: Epoch 1069
└ @ Main /Users/kcft114/.julia/packages/Flux/qXNjB/src/optimise/train.jl:105
┌ Info: Epoch 1070
└ @ Main /Users/kcft114/.julia/packages/Flux/qXNjB/src/optimise/train.jl:105
┌ Info: Epoch 1071
└ @ Main /Users/kcft114/.julia/packages/Flux/qXNjB/src/optimise/train.jl:105
┌ Info: Epoch 1072
└ @ Main /Users/kcft114/.julia/packages/Flux/qXNjB/src/optimise/train.jl:105
┌ Info: Epoch 1073
└ @ Main /Users/kcft114/.julia/packages/Flux/qXNjB/src/optimise/train.jl:105
┌ Info: Epoch 1074
└ @ Main /Users/kcft114/.julia/packages/Flux/qXNjB/src/optimise/train.jl:105
┌ Info: Epoch 1075
└ @ Main /Users/kcft114/.julia/packages/Flux/qXNjB/src/optimise/train.jl:105
┌ Info: Epoch 1076
└ @ Main /Users/kcft114/.julia/packages/Flux/qXNjB/src/optimise/train.jl:105
┌ Info: Epoch 1077
└ @ Main /Users/kcft114/.julia/packages/Flux/qXNjB/src/optimise/train.jl:105
┌ Info: Epoch 1078
└ @ Main /Users/kcft114/.julia/packages/Flux/qXNjB/src/optimise/train.jl:105
┌ Info: Epoch 1079
└ @ Main /Users/kcft114/.julia/packages/Flux/qXNjB/src/optimise/train.jl:105
┌ Info: Epoch 1080
└ @ Main /Users/kcft114/.julia/packages/Flux/qXNjB/src/optimise/train.jl:105
┌ Info: Epoch 1081
└ @ Main /Users/kcft114/.julia/packages/Flux/qXNjB/src/optimise/train.jl:105
┌ Info: Epoch 1082
└ @ Main /Users/kcft114/.julia/packages/Flux/qXNjB/src/optimise/train.jl:105
┌ Info: Epoch 1083
└ @ Main /Users/kcft114/.julia/packages/Flux/qXNjB/src/optimise/train.jl:105
┌ Info: Epoch 1084
└ @ Main /Users/kcft114/.julia/packages/Flux/qXNjB/src/optimise/train.jl:105
┌ Info: Epoch 1085
└ @ Main /Users/kcft114/.julia/packages/Flux/qXNjB/src/optimise/train.jl:105
┌ Info: Epoch 1086
└ @ Main /Users/kcft114/.julia/packages/Flux/qXNjB/src/optimise/train.jl:105
┌ Info: Epoch 1087
└ @ Main /Users/kcft114/.julia/packages/Flux/qXNjB/src/optimise/train.jl:105
┌ Info: Epoch 1088
└ @ Main /Users/kcft114/.julia/packages/Flux/qXNjB/src/optimise/train.jl:105
┌ Info: Epoch 1089
└ @ Main /Users/kcft114/.julia/packages/Flux/qXNjB/src/optimise/train.jl:105
┌ Info: Epoch 1090
└ @ Main /Users/kcft114/.julia/packages/Flux/qXNjB/src/optimise/train.jl:105
┌ Info: Epoch 1091
└ @ Main /Users/kcft114/.julia/packages/Flux/qXNjB/src/optimise/train.jl:105
┌ Info: Epoch 1092
└ @ Main /Users/kcft114/.julia/packages/Flux/qXNjB/src/optimise/train.jl:105
┌ Info: Epoch 1093
└ @ Main /Users/kcft114/.julia/packages/Flux/qXNjB/src/optimise/train.jl:105
┌ Info: Epoch 1094
└ @ Main /Users/kcft114/.julia/packages/Flux/qXNjB/src/optimise/train.jl:105
┌ Info: Epoch 1095
└ @ Main /Users/kcft114/.julia/packages/Flux/qXNjB/src/optimise/train.jl:105
┌ Info: Epoch 1096
└ @ Main /Users/kcft114/.julia/packages/Flux/qXNjB/src/optimise/train.jl:105
┌ Info: Epoch 1097
└ @ Main /Users/kcft114/.julia/packages/Flux/qXNjB/src/optimise/train.jl:105
┌ Info: Epoch 1098
└ @ Main /Users/kcft114/.julia/packages/Flux/qXNjB/src/optimise/train.jl:105
┌ Info: Epoch 1099
└ @ Main /Users/kcft114/.julia/packages/Flux/qXNjB/src/optimise/train.jl:105
┌ Info: Epoch 1100
└ @ Main /Users/kcft114/.julia/packages/Flux/qXNjB/src/optimise/train.jl:105
┌ Info: Epoch 1101
└ @ Main /Users/kcft114/.julia/packages/Flux/qXNjB/src/optimise/train.jl:105
┌ Info: Epoch 1102
└ @ Main /Users/kcft114/.julia/packages/Flux/qXNjB/src/optimise/train.jl:105
┌ Info: Epoch 1103
└ @ Main /Users/kcft114/.julia/packages/Flux/qXNjB/src/optimise/train.jl:105
┌ Info: Epoch 1104
└ @ Main /Users/kcft114/.julia/packages/Flux/qXNjB/src/optimise/train.jl:105
┌ Info: Epoch 1105
└ @ Main /Users/kcft114/.julia/packages/Flux/qXNjB/src/optimise/train.jl:105
┌ Info: Epoch 1106
└ @ Main /Users/kcft114/.julia/packages/Flux/qXNjB/src/optimise/train.jl:105
┌ Info: Epoch 1107
└ @ Main /Users/kcft114/.julia/packages/Flux/qXNjB/src/optimise/train.jl:105
┌ Info: Epoch 1108
└ @ Main /Users/kcft114/.julia/packages/Flux/qXNjB/src/optimise/train.jl:105
┌ Info: Epoch 1109
└ @ Main /Users/kcft114/.julia/packages/Flux/qXNjB/src/optimise/train.jl:105
┌ Info: Epoch 1110
└ @ Main /Users/kcft114/.julia/packages/Flux/qXNjB/src/optimise/train.jl:105
┌ Info: Epoch 1111
└ @ Main /Users/kcft114/.julia/packages/Flux/qXNjB/src/optimise/train.jl:105
┌ Info: Epoch 1112
└ @ Main /Users/kcft114/.julia/packages/Flux/qXNjB/src/optimise/train.jl:105
┌ Info: Epoch 1113
└ @ Main /Users/kcft114/.julia/packages/Flux/qXNjB/src/optimise/train.jl:105
┌ Info: Epoch 1114
└ @ Main /Users/kcft114/.julia/packages/Flux/qXNjB/src/optimise/train.jl:105
┌ Info: Epoch 1115
└ @ Main /Users/kcft114/.julia/packages/Flux/qXNjB/src/optimise/train.jl:105
┌ Info: Epoch 1116
└ @ Main /Users/kcft114/.julia/packages/Flux/qXNjB/src/optimise/train.jl:105
┌ Info: Epoch 1117
└ @ Main /Users/kcft114/.julia/packages/Flux/qXNjB/src/optimise/train.jl:105
┌ Info: Epoch 1118
└ @ Main /Users/kcft114/.julia/packages/Flux/qXNjB/src/optimise/train.jl:105
┌ Info: Epoch 1119
└ @ Main /Users/kcft114/.julia/packages/Flux/qXNjB/src/optimise/train.jl:105
┌ Info: Epoch 1120
└ @ Main /Users/kcft114/.julia/packages/Flux/qXNjB/src/optimise/train.jl:105
┌ Info: Epoch 1121
└ @ Main /Users/kcft114/.julia/packages/Flux/qXNjB/src/optimise/train.jl:105
┌ Info: Epoch 1122
└ @ Main /Users/kcft114/.julia/packages/Flux/qXNjB/src/optimise/train.jl:105
┌ Info: Epoch 1123
└ @ Main /Users/kcft114/.julia/packages/Flux/qXNjB/src/optimise/train.jl:105
┌ Info: Epoch 1124
└ @ Main /Users/kcft114/.julia/packages/Flux/qXNjB/src/optimise/train.jl:105
┌ Info: Epoch 1125
└ @ Main /Users/kcft114/.julia/packages/Flux/qXNjB/src/optimise/train.jl:105
┌ Info: Epoch 1126
└ @ Main /Users/kcft114/.julia/packages/Flux/qXNjB/src/optimise/train.jl:105
┌ Info: Epoch 1127
└ @ Main /Users/kcft114/.julia/packages/Flux/qXNjB/src/optimise/train.jl:105
┌ Info: Epoch 1128
└ @ Main /Users/kcft114/.julia/packages/Flux/qXNjB/src/optimise/train.jl:105
┌ Info: Epoch 1129
└ @ Main /Users/kcft114/.julia/packages/Flux/qXNjB/src/optimise/train.jl:105
┌ Info: Epoch 1130
└ @ Main /Users/kcft114/.julia/packages/Flux/qXNjB/src/optimise/train.jl:105
┌ Info: Epoch 1131
└ @ Main /Users/kcft114/.julia/packages/Flux/qXNjB/src/optimise/train.jl:105
┌ Info: Epoch 1132
└ @ Main /Users/kcft114/.julia/packages/Flux/qXNjB/src/optimise/train.jl:105
┌ Info: Epoch 1133
└ @ Main /Users/kcft114/.julia/packages/Flux/qXNjB/src/optimise/train.jl:105
┌ Info: Epoch 1134
└ @ Main /Users/kcft114/.julia/packages/Flux/qXNjB/src/optimise/train.jl:105
┌ Info: Epoch 1135
└ @ Main /Users/kcft114/.julia/packages/Flux/qXNjB/src/optimise/train.jl:105
┌ Info: Epoch 1136
└ @ Main /Users/kcft114/.julia/packages/Flux/qXNjB/src/optimise/train.jl:105
┌ Info: Epoch 1137
└ @ Main /Users/kcft114/.julia/packages/Flux/qXNjB/src/optimise/train.jl:105
┌ Info: Epoch 1138
└ @ Main /Users/kcft114/.julia/packages/Flux/qXNjB/src/optimise/train.jl:105
┌ Info: Epoch 1139
└ @ Main /Users/kcft114/.julia/packages/Flux/qXNjB/src/optimise/train.jl:105
┌ Info: Epoch 1140
└ @ Main /Users/kcft114/.julia/packages/Flux/qXNjB/src/optimise/train.jl:105
┌ Info: Epoch 1141
└ @ Main /Users/kcft114/.julia/packages/Flux/qXNjB/src/optimise/train.jl:105
┌ Info: Epoch 1142
└ @ Main /Users/kcft114/.julia/packages/Flux/qXNjB/src/optimise/train.jl:105
┌ Info: Epoch 1143
└ @ Main /Users/kcft114/.julia/packages/Flux/qXNjB/src/optimise/train.jl:105
┌ Info: Epoch 1144
└ @ Main /Users/kcft114/.julia/packages/Flux/qXNjB/src/optimise/train.jl:105
┌ Info: Epoch 1145
└ @ Main /Users/kcft114/.julia/packages/Flux/qXNjB/src/optimise/train.jl:105
┌ Info: Epoch 1146
└ @ Main /Users/kcft114/.julia/packages/Flux/qXNjB/src/optimise/train.jl:105
┌ Info: Epoch 1147
└ @ Main /Users/kcft114/.julia/packages/Flux/qXNjB/src/optimise/train.jl:105
┌ Info: Epoch 1148
└ @ Main /Users/kcft114/.julia/packages/Flux/qXNjB/src/optimise/train.jl:105
┌ Info: Epoch 1149
└ @ Main /Users/kcft114/.julia/packages/Flux/qXNjB/src/optimise/train.jl:105
┌ Info: Epoch 1150
└ @ Main /Users/kcft114/.julia/packages/Flux/qXNjB/src/optimise/train.jl:105
┌ Info: Epoch 1151
└ @ Main /Users/kcft114/.julia/packages/Flux/qXNjB/src/optimise/train.jl:105
┌ Info: Epoch 1152
└ @ Main /Users/kcft114/.julia/packages/Flux/qXNjB/src/optimise/train.jl:105
┌ Info: Epoch 1153
└ @ Main /Users/kcft114/.julia/packages/Flux/qXNjB/src/optimise/train.jl:105
┌ Info: Epoch 1154
└ @ Main /Users/kcft114/.julia/packages/Flux/qXNjB/src/optimise/train.jl:105
┌ Info: Epoch 1155
└ @ Main /Users/kcft114/.julia/packages/Flux/qXNjB/src/optimise/train.jl:105
┌ Info: Epoch 1156
└ @ Main /Users/kcft114/.julia/packages/Flux/qXNjB/src/optimise/train.jl:105
┌ Info: Epoch 1157
└ @ Main /Users/kcft114/.julia/packages/Flux/qXNjB/src/optimise/train.jl:105
┌ Info: Epoch 1158
└ @ Main /Users/kcft114/.julia/packages/Flux/qXNjB/src/optimise/train.jl:105
┌ Info: Epoch 1159
└ @ Main /Users/kcft114/.julia/packages/Flux/qXNjB/src/optimise/train.jl:105
┌ Info: Epoch 1160
└ @ Main /Users/kcft114/.julia/packages/Flux/qXNjB/src/optimise/train.jl:105
┌ Info: Epoch 1161
└ @ Main /Users/kcft114/.julia/packages/Flux/qXNjB/src/optimise/train.jl:105
┌ Info: Epoch 1162
└ @ Main /Users/kcft114/.julia/packages/Flux/qXNjB/src/optimise/train.jl:105
┌ Info: Epoch 1163
└ @ Main /Users/kcft114/.julia/packages/Flux/qXNjB/src/optimise/train.jl:105
┌ Info: Epoch 1164
└ @ Main /Users/kcft114/.julia/packages/Flux/qXNjB/src/optimise/train.jl:105
┌ Info: Epoch 1165
└ @ Main /Users/kcft114/.julia/packages/Flux/qXNjB/src/optimise/train.jl:105
┌ Info: Epoch 1166
└ @ Main /Users/kcft114/.julia/packages/Flux/qXNjB/src/optimise/train.jl:105
┌ Info: Epoch 1167
└ @ Main /Users/kcft114/.julia/packages/Flux/qXNjB/src/optimise/train.jl:105
┌ Info: Epoch 1168
└ @ Main /Users/kcft114/.julia/packages/Flux/qXNjB/src/optimise/train.jl:105
┌ Info: Epoch 1169
└ @ Main /Users/kcft114/.julia/packages/Flux/qXNjB/src/optimise/train.jl:105
┌ Info: Epoch 1170
└ @ Main /Users/kcft114/.julia/packages/Flux/qXNjB/src/optimise/train.jl:105
┌ Info: Epoch 1171
└ @ Main /Users/kcft114/.julia/packages/Flux/qXNjB/src/optimise/train.jl:105
┌ Info: Epoch 1172
└ @ Main /Users/kcft114/.julia/packages/Flux/qXNjB/src/optimise/train.jl:105
┌ Info: Epoch 1173
└ @ Main /Users/kcft114/.julia/packages/Flux/qXNjB/src/optimise/train.jl:105
┌ Info: Epoch 1174
└ @ Main /Users/kcft114/.julia/packages/Flux/qXNjB/src/optimise/train.jl:105
┌ Info: Epoch 1175
└ @ Main /Users/kcft114/.julia/packages/Flux/qXNjB/src/optimise/train.jl:105
┌ Info: Epoch 1176
└ @ Main /Users/kcft114/.julia/packages/Flux/qXNjB/src/optimise/train.jl:105
┌ Info: Epoch 1177
└ @ Main /Users/kcft114/.julia/packages/Flux/qXNjB/src/optimise/train.jl:105
┌ Info: Epoch 1178
└ @ Main /Users/kcft114/.julia/packages/Flux/qXNjB/src/optimise/train.jl:105
┌ Info: Epoch 1179
└ @ Main /Users/kcft114/.julia/packages/Flux/qXNjB/src/optimise/train.jl:105
┌ Info: Epoch 1180
└ @ Main /Users/kcft114/.julia/packages/Flux/qXNjB/src/optimise/train.jl:105
┌ Info: Epoch 1181
└ @ Main /Users/kcft114/.julia/packages/Flux/qXNjB/src/optimise/train.jl:105
┌ Info: Epoch 1182
└ @ Main /Users/kcft114/.julia/packages/Flux/qXNjB/src/optimise/train.jl:105
┌ Info: Epoch 1183
└ @ Main /Users/kcft114/.julia/packages/Flux/qXNjB/src/optimise/train.jl:105
┌ Info: Epoch 1184
└ @ Main /Users/kcft114/.julia/packages/Flux/qXNjB/src/optimise/train.jl:105
┌ Info: Epoch 1185
└ @ Main /Users/kcft114/.julia/packages/Flux/qXNjB/src/optimise/train.jl:105
┌ Info: Epoch 1186
└ @ Main /Users/kcft114/.julia/packages/Flux/qXNjB/src/optimise/train.jl:105
┌ Info: Epoch 1187
└ @ Main /Users/kcft114/.julia/packages/Flux/qXNjB/src/optimise/train.jl:105
┌ Info: Epoch 1188
└ @ Main /Users/kcft114/.julia/packages/Flux/qXNjB/src/optimise/train.jl:105
┌ Info: Epoch 1189
└ @ Main /Users/kcft114/.julia/packages/Flux/qXNjB/src/optimise/train.jl:105
┌ Info: Epoch 1190
└ @ Main /Users/kcft114/.julia/packages/Flux/qXNjB/src/optimise/train.jl:105
┌ Info: Epoch 1191
└ @ Main /Users/kcft114/.julia/packages/Flux/qXNjB/src/optimise/train.jl:105
┌ Info: Epoch 1192
└ @ Main /Users/kcft114/.julia/packages/Flux/qXNjB/src/optimise/train.jl:105
┌ Info: Epoch 1193
└ @ Main /Users/kcft114/.julia/packages/Flux/qXNjB/src/optimise/train.jl:105
┌ Info: Epoch 1194
└ @ Main /Users/kcft114/.julia/packages/Flux/qXNjB/src/optimise/train.jl:105
┌ Info: Epoch 1195
└ @ Main /Users/kcft114/.julia/packages/Flux/qXNjB/src/optimise/train.jl:105
┌ Info: Epoch 1196
└ @ Main /Users/kcft114/.julia/packages/Flux/qXNjB/src/optimise/train.jl:105
┌ Info: Epoch 1197
└ @ Main /Users/kcft114/.julia/packages/Flux/qXNjB/src/optimise/train.jl:105
┌ Info: Epoch 1198
└ @ Main /Users/kcft114/.julia/packages/Flux/qXNjB/src/optimise/train.jl:105
┌ Info: Epoch 1199
└ @ Main /Users/kcft114/.julia/packages/Flux/qXNjB/src/optimise/train.jl:105
┌ Info: Epoch 1200
└ @ Main /Users/kcft114/.julia/packages/Flux/qXNjB/src/optimise/train.jl:105
┌ Info: Epoch 1201
└ @ Main /Users/kcft114/.julia/packages/Flux/qXNjB/src/optimise/train.jl:105
┌ Info: Epoch 1202
└ @ Main /Users/kcft114/.julia/packages/Flux/qXNjB/src/optimise/train.jl:105
┌ Info: Epoch 1203
└ @ Main /Users/kcft114/.julia/packages/Flux/qXNjB/src/optimise/train.jl:105
┌ Info: Epoch 1204
└ @ Main /Users/kcft114/.julia/packages/Flux/qXNjB/src/optimise/train.jl:105
┌ Info: Epoch 1205
└ @ Main /Users/kcft114/.julia/packages/Flux/qXNjB/src/optimise/train.jl:105
┌ Info: Epoch 1206
└ @ Main /Users/kcft114/.julia/packages/Flux/qXNjB/src/optimise/train.jl:105
┌ Info: Epoch 1207
└ @ Main /Users/kcft114/.julia/packages/Flux/qXNjB/src/optimise/train.jl:105
┌ Info: Epoch 1208
└ @ Main /Users/kcft114/.julia/packages/Flux/qXNjB/src/optimise/train.jl:105
┌ Info: Epoch 1209
└ @ Main /Users/kcft114/.julia/packages/Flux/qXNjB/src/optimise/train.jl:105
┌ Info: Epoch 1210
└ @ Main /Users/kcft114/.julia/packages/Flux/qXNjB/src/optimise/train.jl:105
┌ Info: Epoch 1211
└ @ Main /Users/kcft114/.julia/packages/Flux/qXNjB/src/optimise/train.jl:105
┌ Info: Epoch 1212
└ @ Main /Users/kcft114/.julia/packages/Flux/qXNjB/src/optimise/train.jl:105
┌ Info: Epoch 1213
└ @ Main /Users/kcft114/.julia/packages/Flux/qXNjB/src/optimise/train.jl:105
```

```
loss(X, Y) = 0.6075351f0 (tracked)
loss(X, Y) = 0.6286875f0 (tracked)
loss(X, Y) = 0.6102109f0 (tracked)
loss(X, Y) = 0.6493573f0 (tracked)
loss(X, Y) = 0.632143f0 (tracked)
loss(X, Y) = 0.5659929f0 (tracked)
loss(X, Y) = 0.58544356f0 (tracked)
loss(X, Y) = 0.60152394f0 (tracked)
loss(X, Y) = 0.646009f0 (tracked)
loss(X, Y) = 0.5826686f0 (tracked)
loss(X, Y) = 0.572798f0 (tracked)
loss(X, Y) = 0.6094796f0 (tracked)
loss(X, Y) = 0.60307026f0 (tracked)
loss(X, Y) = 0.64466363f0 (tracked)
loss(X, Y) = 0.60310787f0 (tracked)
loss(X, Y) = 0.6296621f0 (tracked)
loss(X, Y) = 0.58659804f0 (tracked)
loss(X, Y) = 0.5710502f0 (tracked)
loss(X, Y) = 0.58421606f0 (tracked)
loss(X, Y) = 0.62596536f0 (tracked)
loss(X, Y) = 0.5742531f0 (tracked)
loss(X, Y) = 0.61834663f0 (tracked)
loss(X, Y) = 0.6238245f0 (tracked)
loss(X, Y) = 0.61250377f0 (tracked)
loss(X, Y) = 0.5868619f0 (tracked)
loss(X, Y) = 0.6464141f0 (tracked)
loss(X, Y) = 0.6091563f0 (tracked)
loss(X, Y) = 0.61494553f0 (tracked)
loss(X, Y) = 0.6049501f0 (tracked)
loss(X, Y) = 0.6313121f0 (tracked)
loss(X, Y) = 0.5703351f0 (tracked)
loss(X, Y) = 0.6247158f0 (tracked)
loss(X, Y) = 0.61506367f0 (tracked)
loss(X, Y) = 0.57670844f0 (tracked)
loss(X, Y) = 0.63519263f0 (tracked)
loss(X, Y) = 0.5643315f0 (tracked)
loss(X, Y) = 0.5765141f0 (tracked)
loss(X, Y) = 0.59277827f0 (tracked)
loss(X, Y) = 0.61980987f0 (tracked)
loss(X, Y) = 0.5713065f0 (tracked)
loss(X, Y) = 0.6010057f0 (tracked)
loss(X, Y) = 0.5756603f0 (tracked)
loss(X, Y) = 0.60017204f0 (tracked)
loss(X, Y) = 0.61659396f0 (tracked)
loss(X, Y) = 0.6142395f0 (tracked)
loss(X, Y) = 0.6285909f0 (tracked)
loss(X, Y) = 0.6242602f0 (tracked)
loss(X, Y) = 0.60018533f0 (tracked)
loss(X, Y) = 0.65508014f0 (tracked)
loss(X, Y) = 0.5706497f0 (tracked)
loss(X, Y) = 0.57330894f0 (tracked)
loss(X, Y) = 0.6298373f0 (tracked)
loss(X, Y) = 0.5882753f0 (tracked)
loss(X, Y) = 0.58167666f0 (tracked)
loss(X, Y) = 0.585854f0 (tracked)
loss(X, Y) = 0.59047574f0 (tracked)
loss(X, Y) = 0.6016257f0 (tracked)
loss(X, Y) = 0.6058115f0 (tracked)
loss(X, Y) = 0.6181979f0 (tracked)
loss(X, Y) = 0.62663484f0 (tracked)
loss(X, Y) = 0.59889233f0 (tracked)
loss(X, Y) = 0.6311005f0 (tracked)
loss(X, Y) = 0.58609015f0 (tracked)
loss(X, Y) = 0.5794827f0 (tracked)
loss(X, Y) = 0.6184779f0 (tracked)
loss(X, Y) = 0.61659443f0 (tracked)
loss(X, Y) = 0.60170746f0 (tracked)
loss(X, Y) = 0.6071121f0 (tracked)
loss(X, Y) = 0.58004814f0 (tracked)
loss(X, Y) = 0.62504786f0 (tracked)
loss(X, Y) = 0.6023545f0 (tracked)
loss(X, Y) = 0.5753755f0 (tracked)
loss(X, Y) = 0.6443475f0 (tracked)
loss(X, Y) = 0.56863135f0 (tracked)
loss(X, Y) = 0.61203796f0 (tracked)
loss(X, Y) = 0.6043412f0 (tracked)
loss(X, Y) = 0.57193714f0 (tracked)
loss(X, Y) = 0.5799848f0 (tracked)
loss(X, Y) = 0.580765f0 (tracked)
loss(X, Y) = 0.57742935f0 (tracked)
loss(X, Y) = 0.619712f0 (tracked)
loss(X, Y) = 0.5926302f0 (tracked)
loss(X, Y) = 0.62476635f0 (tracked)
loss(X, Y) = 0.6001239f0 (tracked)
loss(X, Y) = 0.6088475f0 (tracked)
loss(X, Y) = 0.5936433f0 (tracked)
loss(X, Y) = 0.58836776f0 (tracked)
loss(X, Y) = 0.59156257f0 (tracked)
loss(X, Y) = 0.57480365f0 (tracked)
loss(X, Y) = 0.5944196f0 (tracked)
loss(X, Y) = 0.5704235f0 (tracked)
loss(X, Y) = 0.6115667f0 (tracked)
loss(X, Y) = 0.5708255f0 (tracked)
loss(X, Y) = 0.6151224f0 (tracked)
loss(X, Y) = 0.57445824f0 (tracked)
loss(X, Y) = 0.57396203f0 (tracked)
loss(X, Y) = 0.58698946f0 (tracked)
loss(X, Y) = 0.5946425f0 (tracked)
loss(X, Y) = 0.6079964f0 (tracked)
loss(X, Y) = 0.5926097f0 (tracked)
loss(X, Y) = 0.58016706f0 (tracked)
loss(X, Y) = 0.58994704f0 (tracked)
loss(X, Y) = 0.57461876f0 (tracked)
loss(X, Y) = 0.5818545f0 (tracked)
loss(X, Y) = 0.6330229f0 (tracked)
loss(X, Y) = 0.55881083f0 (tracked)
loss(X, Y) = 0.6258518f0 (tracked)
loss(X, Y) = 0.582665f0 (tracked)
loss(X, Y) = 0.65089476f0 (tracked)
loss(X, Y) = 0.6180446f0 (tracked)
loss(X, Y) = 0.5790353f0 (tracked)
loss(X, Y) = 0.5820663f0 (tracked)
loss(X, Y) = 0.58292586f0 (tracked)
```

```
┌ Info: Epoch 1214
└ @ Main /Users/kcft114/.julia/packages/Flux/qXNjB/src/optimise/train.jl:105
┌ Info: Epoch 1215
└ @ Main /Users/kcft114/.julia/packages/Flux/qXNjB/src/optimise/train.jl:105
┌ Info: Epoch 1216
└ @ Main /Users/kcft114/.julia/packages/Flux/qXNjB/src/optimise/train.jl:105
┌ Info: Epoch 1217
└ @ Main /Users/kcft114/.julia/packages/Flux/qXNjB/src/optimise/train.jl:105
┌ Info: Epoch 1218
└ @ Main /Users/kcft114/.julia/packages/Flux/qXNjB/src/optimise/train.jl:105
┌ Info: Epoch 1219
└ @ Main /Users/kcft114/.julia/packages/Flux/qXNjB/src/optimise/train.jl:105
┌ Info: Epoch 1220
└ @ Main /Users/kcft114/.julia/packages/Flux/qXNjB/src/optimise/train.jl:105
┌ Info: Epoch 1221
└ @ Main /Users/kcft114/.julia/packages/Flux/qXNjB/src/optimise/train.jl:105
┌ Info: Epoch 1222
└ @ Main /Users/kcft114/.julia/packages/Flux/qXNjB/src/optimise/train.jl:105
┌ Info: Epoch 1223
└ @ Main /Users/kcft114/.julia/packages/Flux/qXNjB/src/optimise/train.jl:105
┌ Info: Epoch 1224
└ @ Main /Users/kcft114/.julia/packages/Flux/qXNjB/src/optimise/train.jl:105
┌ Info: Epoch 1225
└ @ Main /Users/kcft114/.julia/packages/Flux/qXNjB/src/optimise/train.jl:105
┌ Info: Epoch 1226
└ @ Main /Users/kcft114/.julia/packages/Flux/qXNjB/src/optimise/train.jl:105
┌ Info: Epoch 1227
└ @ Main /Users/kcft114/.julia/packages/Flux/qXNjB/src/optimise/train.jl:105
┌ Info: Epoch 1228
└ @ Main /Users/kcft114/.julia/packages/Flux/qXNjB/src/optimise/train.jl:105
┌ Info: Epoch 1229
└ @ Main /Users/kcft114/.julia/packages/Flux/qXNjB/src/optimise/train.jl:105
┌ Info: Epoch 1230
└ @ Main /Users/kcft114/.julia/packages/Flux/qXNjB/src/optimise/train.jl:105
┌ Info: Epoch 1231
└ @ Main /Users/kcft114/.julia/packages/Flux/qXNjB/src/optimise/train.jl:105
┌ Info: Epoch 1232
└ @ Main /Users/kcft114/.julia/packages/Flux/qXNjB/src/optimise/train.jl:105
┌ Info: Epoch 1233
└ @ Main /Users/kcft114/.julia/packages/Flux/qXNjB/src/optimise/train.jl:105
┌ Info: Epoch 1234
└ @ Main /Users/kcft114/.julia/packages/Flux/qXNjB/src/optimise/train.jl:105
┌ Info: Epoch 1235
└ @ Main /Users/kcft114/.julia/packages/Flux/qXNjB/src/optimise/train.jl:105
┌ Info: Epoch 1236
└ @ Main /Users/kcft114/.julia/packages/Flux/qXNjB/src/optimise/train.jl:105
┌ Info: Epoch 1237
└ @ Main /Users/kcft114/.julia/packages/Flux/qXNjB/src/optimise/train.jl:105
┌ Info: Epoch 1238
└ @ Main /Users/kcft114/.julia/packages/Flux/qXNjB/src/optimise/train.jl:105
┌ Info: Epoch 1239
└ @ Main /Users/kcft114/.julia/packages/Flux/qXNjB/src/optimise/train.jl:105
┌ Info: Epoch 1240
└ @ Main /Users/kcft114/.julia/packages/Flux/qXNjB/src/optimise/train.jl:105
┌ Info: Epoch 1241
└ @ Main /Users/kcft114/.julia/packages/Flux/qXNjB/src/optimise/train.jl:105
┌ Info: Epoch 1242
└ @ Main /Users/kcft114/.julia/packages/Flux/qXNjB/src/optimise/train.jl:105
┌ Info: Epoch 1243
└ @ Main /Users/kcft114/.julia/packages/Flux/qXNjB/src/optimise/train.jl:105
┌ Info: Epoch 1244
└ @ Main /Users/kcft114/.julia/packages/Flux/qXNjB/src/optimise/train.jl:105
┌ Info: Epoch 1245
└ @ Main /Users/kcft114/.julia/packages/Flux/qXNjB/src/optimise/train.jl:105
┌ Info: Epoch 1246
└ @ Main /Users/kcft114/.julia/packages/Flux/qXNjB/src/optimise/train.jl:105
┌ Info: Epoch 1247
└ @ Main /Users/kcft114/.julia/packages/Flux/qXNjB/src/optimise/train.jl:105
┌ Info: Epoch 1248
└ @ Main /Users/kcft114/.julia/packages/Flux/qXNjB/src/optimise/train.jl:105
┌ Info: Epoch 1249
└ @ Main /Users/kcft114/.julia/packages/Flux/qXNjB/src/optimise/train.jl:105
┌ Info: Epoch 1250
└ @ Main /Users/kcft114/.julia/packages/Flux/qXNjB/src/optimise/train.jl:105
┌ Info: Epoch 1251
└ @ Main /Users/kcft114/.julia/packages/Flux/qXNjB/src/optimise/train.jl:105
┌ Info: Epoch 1252
└ @ Main /Users/kcft114/.julia/packages/Flux/qXNjB/src/optimise/train.jl:105
┌ Info: Epoch 1253
└ @ Main /Users/kcft114/.julia/packages/Flux/qXNjB/src/optimise/train.jl:105
┌ Info: Epoch 1254
└ @ Main /Users/kcft114/.julia/packages/Flux/qXNjB/src/optimise/train.jl:105
┌ Info: Epoch 1255
└ @ Main /Users/kcft114/.julia/packages/Flux/qXNjB/src/optimise/train.jl:105
┌ Info: Epoch 1256
└ @ Main /Users/kcft114/.julia/packages/Flux/qXNjB/src/optimise/train.jl:105
┌ Info: Epoch 1257
└ @ Main /Users/kcft114/.julia/packages/Flux/qXNjB/src/optimise/train.jl:105
┌ Info: Epoch 1258
└ @ Main /Users/kcft114/.julia/packages/Flux/qXNjB/src/optimise/train.jl:105
┌ Info: Epoch 1259
└ @ Main /Users/kcft114/.julia/packages/Flux/qXNjB/src/optimise/train.jl:105
┌ Info: Epoch 1260
└ @ Main /Users/kcft114/.julia/packages/Flux/qXNjB/src/optimise/train.jl:105
┌ Info: Epoch 1261
└ @ Main /Users/kcft114/.julia/packages/Flux/qXNjB/src/optimise/train.jl:105
┌ Info: Epoch 1262
└ @ Main /Users/kcft114/.julia/packages/Flux/qXNjB/src/optimise/train.jl:105
┌ Info: Epoch 1263
└ @ Main /Users/kcft114/.julia/packages/Flux/qXNjB/src/optimise/train.jl:105
┌ Info: Epoch 1264
└ @ Main /Users/kcft114/.julia/packages/Flux/qXNjB/src/optimise/train.jl:105
┌ Info: Epoch 1265
└ @ Main /Users/kcft114/.julia/packages/Flux/qXNjB/src/optimise/train.jl:105
┌ Info: Epoch 1266
└ @ Main /Users/kcft114/.julia/packages/Flux/qXNjB/src/optimise/train.jl:105
┌ Info: Epoch 1267
└ @ Main /Users/kcft114/.julia/packages/Flux/qXNjB/src/optimise/train.jl:105
┌ Info: Epoch 1268
└ @ Main /Users/kcft114/.julia/packages/Flux/qXNjB/src/optimise/train.jl:105
┌ Info: Epoch 1269
└ @ Main /Users/kcft114/.julia/packages/Flux/qXNjB/src/optimise/train.jl:105
┌ Info: Epoch 1270
└ @ Main /Users/kcft114/.julia/packages/Flux/qXNjB/src/optimise/train.jl:105
┌ Info: Epoch 1271
└ @ Main /Users/kcft114/.julia/packages/Flux/qXNjB/src/optimise/train.jl:105
┌ Info: Epoch 1272
└ @ Main /Users/kcft114/.julia/packages/Flux/qXNjB/src/optimise/train.jl:105
┌ Info: Epoch 1273
└ @ Main /Users/kcft114/.julia/packages/Flux/qXNjB/src/optimise/train.jl:105
┌ Info: Epoch 1274
└ @ Main /Users/kcft114/.julia/packages/Flux/qXNjB/src/optimise/train.jl:105
┌ Info: Epoch 1275
└ @ Main /Users/kcft114/.julia/packages/Flux/qXNjB/src/optimise/train.jl:105
┌ Info: Epoch 1276
└ @ Main /Users/kcft114/.julia/packages/Flux/qXNjB/src/optimise/train.jl:105
┌ Info: Epoch 1277
└ @ Main /Users/kcft114/.julia/packages/Flux/qXNjB/src/optimise/train.jl:105
┌ Info: Epoch 1278
└ @ Main /Users/kcft114/.julia/packages/Flux/qXNjB/src/optimise/train.jl:105
┌ Info: Epoch 1279
└ @ Main /Users/kcft114/.julia/packages/Flux/qXNjB/src/optimise/train.jl:105
┌ Info: Epoch 1280
└ @ Main /Users/kcft114/.julia/packages/Flux/qXNjB/src/optimise/train.jl:105
┌ Info: Epoch 1281
└ @ Main /Users/kcft114/.julia/packages/Flux/qXNjB/src/optimise/train.jl:105
┌ Info: Epoch 1282
└ @ Main /Users/kcft114/.julia/packages/Flux/qXNjB/src/optimise/train.jl:105
┌ Info: Epoch 1283
└ @ Main /Users/kcft114/.julia/packages/Flux/qXNjB/src/optimise/train.jl:105
┌ Info: Epoch 1284
└ @ Main /Users/kcft114/.julia/packages/Flux/qXNjB/src/optimise/train.jl:105
┌ Info: Epoch 1285
└ @ Main /Users/kcft114/.julia/packages/Flux/qXNjB/src/optimise/train.jl:105
┌ Info: Epoch 1286
└ @ Main /Users/kcft114/.julia/packages/Flux/qXNjB/src/optimise/train.jl:105
┌ Info: Epoch 1287
└ @ Main /Users/kcft114/.julia/packages/Flux/qXNjB/src/optimise/train.jl:105
┌ Info: Epoch 1288
└ @ Main /Users/kcft114/.julia/packages/Flux/qXNjB/src/optimise/train.jl:105
┌ Info: Epoch 1289
└ @ Main /Users/kcft114/.julia/packages/Flux/qXNjB/src/optimise/train.jl:105
┌ Info: Epoch 1290
└ @ Main /Users/kcft114/.julia/packages/Flux/qXNjB/src/optimise/train.jl:105
┌ Info: Epoch 1291
└ @ Main /Users/kcft114/.julia/packages/Flux/qXNjB/src/optimise/train.jl:105
┌ Info: Epoch 1292
└ @ Main /Users/kcft114/.julia/packages/Flux/qXNjB/src/optimise/train.jl:105
┌ Info: Epoch 1293
└ @ Main /Users/kcft114/.julia/packages/Flux/qXNjB/src/optimise/train.jl:105
┌ Info: Epoch 1294
└ @ Main /Users/kcft114/.julia/packages/Flux/qXNjB/src/optimise/train.jl:105
┌ Info: Epoch 1295
└ @ Main /Users/kcft114/.julia/packages/Flux/qXNjB/src/optimise/train.jl:105
┌ Info: Epoch 1296
└ @ Main /Users/kcft114/.julia/packages/Flux/qXNjB/src/optimise/train.jl:105
┌ Info: Epoch 1297
└ @ Main /Users/kcft114/.julia/packages/Flux/qXNjB/src/optimise/train.jl:105
┌ Info: Epoch 1298
└ @ Main /Users/kcft114/.julia/packages/Flux/qXNjB/src/optimise/train.jl:105
┌ Info: Epoch 1299
└ @ Main /Users/kcft114/.julia/packages/Flux/qXNjB/src/optimise/train.jl:105
┌ Info: Epoch 1300
└ @ Main /Users/kcft114/.julia/packages/Flux/qXNjB/src/optimise/train.jl:105
┌ Info: Epoch 1301
└ @ Main /Users/kcft114/.julia/packages/Flux/qXNjB/src/optimise/train.jl:105
┌ Info: Epoch 1302
└ @ Main /Users/kcft114/.julia/packages/Flux/qXNjB/src/optimise/train.jl:105
┌ Info: Epoch 1303
└ @ Main /Users/kcft114/.julia/packages/Flux/qXNjB/src/optimise/train.jl:105
┌ Info: Epoch 1304
└ @ Main /Users/kcft114/.julia/packages/Flux/qXNjB/src/optimise/train.jl:105
┌ Info: Epoch 1305
└ @ Main /Users/kcft114/.julia/packages/Flux/qXNjB/src/optimise/train.jl:105
┌ Info: Epoch 1306
└ @ Main /Users/kcft114/.julia/packages/Flux/qXNjB/src/optimise/train.jl:105
┌ Info: Epoch 1307
└ @ Main /Users/kcft114/.julia/packages/Flux/qXNjB/src/optimise/train.jl:105
┌ Info: Epoch 1308
└ @ Main /Users/kcft114/.julia/packages/Flux/qXNjB/src/optimise/train.jl:105
┌ Info: Epoch 1309
└ @ Main /Users/kcft114/.julia/packages/Flux/qXNjB/src/optimise/train.jl:105
┌ Info: Epoch 1310
└ @ Main /Users/kcft114/.julia/packages/Flux/qXNjB/src/optimise/train.jl:105
┌ Info: Epoch 1311
└ @ Main /Users/kcft114/.julia/packages/Flux/qXNjB/src/optimise/train.jl:105
┌ Info: Epoch 1312
└ @ Main /Users/kcft114/.julia/packages/Flux/qXNjB/src/optimise/train.jl:105
┌ Info: Epoch 1313
└ @ Main /Users/kcft114/.julia/packages/Flux/qXNjB/src/optimise/train.jl:105
┌ Info: Epoch 1314
└ @ Main /Users/kcft114/.julia/packages/Flux/qXNjB/src/optimise/train.jl:105
┌ Info: Epoch 1315
└ @ Main /Users/kcft114/.julia/packages/Flux/qXNjB/src/optimise/train.jl:105
┌ Info: Epoch 1316
└ @ Main /Users/kcft114/.julia/packages/Flux/qXNjB/src/optimise/train.jl:105
┌ Info: Epoch 1317
└ @ Main /Users/kcft114/.julia/packages/Flux/qXNjB/src/optimise/train.jl:105
┌ Info: Epoch 1318
└ @ Main /Users/kcft114/.julia/packages/Flux/qXNjB/src/optimise/train.jl:105
┌ Info: Epoch 1319
└ @ Main /Users/kcft114/.julia/packages/Flux/qXNjB/src/optimise/train.jl:105
┌ Info: Epoch 1320
└ @ Main /Users/kcft114/.julia/packages/Flux/qXNjB/src/optimise/train.jl:105
┌ Info: Epoch 1321
└ @ Main /Users/kcft114/.julia/packages/Flux/qXNjB/src/optimise/train.jl:105
┌ Info: Epoch 1322
└ @ Main /Users/kcft114/.julia/packages/Flux/qXNjB/src/optimise/train.jl:105
┌ Info: Epoch 1323
└ @ Main /Users/kcft114/.julia/packages/Flux/qXNjB/src/optimise/train.jl:105
┌ Info: Epoch 1324
└ @ Main /Users/kcft114/.julia/packages/Flux/qXNjB/src/optimise/train.jl:105
┌ Info: Epoch 1325
└ @ Main /Users/kcft114/.julia/packages/Flux/qXNjB/src/optimise/train.jl:105
┌ Info: Epoch 1326
└ @ Main /Users/kcft114/.julia/packages/Flux/qXNjB/src/optimise/train.jl:105
```

```
loss(X, Y) = 0.55733085f0 (tracked)
loss(X, Y) = 0.55034816f0 (tracked)
loss(X, Y) = 0.60768515f0 (tracked)
loss(X, Y) = 0.60037273f0 (tracked)
loss(X, Y) = 0.57425636f0 (tracked)
loss(X, Y) = 0.58563805f0 (tracked)
loss(X, Y) = 0.6230008f0 (tracked)
loss(X, Y) = 0.5964541f0 (tracked)
loss(X, Y) = 0.59633106f0 (tracked)
loss(X, Y) = 0.6159092f0 (tracked)
loss(X, Y) = 0.5858916f0 (tracked)
loss(X, Y) = 0.67316276f0 (tracked)
loss(X, Y) = 0.57058245f0 (tracked)
loss(X, Y) = 0.59897697f0 (tracked)
loss(X, Y) = 0.5832638f0 (tracked)
loss(X, Y) = 0.59136534f0 (tracked)
loss(X, Y) = 0.5804756f0 (tracked)
loss(X, Y) = 0.6409319f0 (tracked)
loss(X, Y) = 0.603554f0 (tracked)
loss(X, Y) = 0.6069946f0 (tracked)
loss(X, Y) = 0.58728296f0 (tracked)
loss(X, Y) = 0.60011053f0 (tracked)
loss(X, Y) = 0.5796108f0 (tracked)
loss(X, Y) = 0.5960717f0 (tracked)
loss(X, Y) = 0.563681f0 (tracked)
loss(X, Y) = 0.57598156f0 (tracked)
loss(X, Y) = 0.5883502f0 (tracked)
loss(X, Y) = 0.5953983f0 (tracked)
loss(X, Y) = 0.56408155f0 (tracked)
loss(X, Y) = 0.6157926f0 (tracked)
loss(X, Y) = 0.585754f0 (tracked)
loss(X, Y) = 0.5942962f0 (tracked)
loss(X, Y) = 0.6442959f0 (tracked)
loss(X, Y) = 0.581715f0 (tracked)
loss(X, Y) = 0.55488807f0 (tracked)
loss(X, Y) = 0.58826536f0 (tracked)
loss(X, Y) = 0.6034508f0 (tracked)
loss(X, Y) = 0.5748235f0 (tracked)
loss(X, Y) = 0.5566685f0 (tracked)
loss(X, Y) = 0.61748517f0 (tracked)
loss(X, Y) = 0.63569397f0 (tracked)
loss(X, Y) = 0.56170547f0 (tracked)
loss(X, Y) = 0.6018256f0 (tracked)
loss(X, Y) = 0.61886287f0 (tracked)
loss(X, Y) = 0.5827662f0 (tracked)
loss(X, Y) = 0.5462585f0 (tracked)
loss(X, Y) = 0.6055061f0 (tracked)
loss(X, Y) = 0.5924961f0 (tracked)
loss(X, Y) = 0.59562325f0 (tracked)
loss(X, Y) = 0.6088742f0 (tracked)
loss(X, Y) = 0.5869417f0 (tracked)
loss(X, Y) = 0.5562113f0 (tracked)
loss(X, Y) = 0.61615604f0 (tracked)
loss(X, Y) = 0.55748165f0 (tracked)
loss(X, Y) = 0.5885746f0 (tracked)
loss(X, Y) = 0.5802818f0 (tracked)
loss(X, Y) = 0.6181514f0 (tracked)
loss(X, Y) = 0.57899886f0 (tracked)
loss(X, Y) = 0.6047908f0 (tracked)
loss(X, Y) = 0.5968272f0 (tracked)
loss(X, Y) = 0.5387546f0 (tracked)
loss(X, Y) = 0.5933943f0 (tracked)
loss(X, Y) = 0.58643186f0 (tracked)
loss(X, Y) = 0.5912065f0 (tracked)
loss(X, Y) = 0.5761832f0 (tracked)
loss(X, Y) = 0.5976979f0 (tracked)
loss(X, Y) = 0.5618882f0 (tracked)
loss(X, Y) = 0.57724035f0 (tracked)
loss(X, Y) = 0.5868513f0 (tracked)
loss(X, Y) = 0.59751856f0 (tracked)
loss(X, Y) = 0.607359f0 (tracked)
loss(X, Y) = 0.59860873f0 (tracked)
loss(X, Y) = 0.6088637f0 (tracked)
loss(X, Y) = 0.5820377f0 (tracked)
loss(X, Y) = 0.6143785f0 (tracked)
loss(X, Y) = 0.5529164f0 (tracked)
loss(X, Y) = 0.572202f0 (tracked)
loss(X, Y) = 0.55991113f0 (tracked)
loss(X, Y) = 0.5858557f0 (tracked)
loss(X, Y) = 0.61262035f0 (tracked)
loss(X, Y) = 0.55266064f0 (tracked)
loss(X, Y) = 0.565831f0 (tracked)
loss(X, Y) = 0.62610877f0 (tracked)
loss(X, Y) = 0.58719087f0 (tracked)
loss(X, Y) = 0.58223933f0 (tracked)
loss(X, Y) = 0.572183f0 (tracked)
loss(X, Y) = 0.58664674f0 (tracked)
loss(X, Y) = 0.61301774f0 (tracked)
loss(X, Y) = 0.58048874f0 (tracked)
loss(X, Y) = 0.5959046f0 (tracked)
loss(X, Y) = 0.5648626f0 (tracked)
loss(X, Y) = 0.60201794f0 (tracked)
loss(X, Y) = 0.567556f0 (tracked)
loss(X, Y) = 0.6179512f0 (tracked)
loss(X, Y) = 0.5971592f0 (tracked)
loss(X, Y) = 0.5948965f0 (tracked)
loss(X, Y) = 0.5844192f0 (tracked)
loss(X, Y) = 0.573622f0 (tracked)
loss(X, Y) = 0.6198337f0 (tracked)
loss(X, Y) = 0.56596637f0 (tracked)
loss(X, Y) = 0.5478051f0 (tracked)
loss(X, Y) = 0.5922326f0 (tracked)
loss(X, Y) = 0.5618539f0 (tracked)
loss(X, Y) = 0.5980137f0 (tracked)
loss(X, Y) = 0.6057039f0 (tracked)
loss(X, Y) = 0.564946f0 (tracked)
loss(X, Y) = 0.55755126f0 (tracked)
loss(X, Y) = 0.52249956f0 (tracked)
loss(X, Y) = 0.56973803f0 (tracked)
loss(X, Y) = 0.58058506f0 (tracked)
loss(X, Y) = 0.5389117f0 (tracked)
loss(X, Y) = 0.60442084f0 (tracked)
loss(X, Y) = 0.5732944f0 (tracked)
loss(X, Y) = 0.60572624f0 (tracked)
loss(X, Y) = 0.547878f0 (tracked)
loss(X, Y) = 0.6177001f0 (tracked)
loss(X, Y) = 0.5376921f0 (tracked)
loss(X, Y) = 0.5479858f0 (tracked)
loss(X, Y) = 0.5784174f0 (tracked)
loss(X, Y) = 0.555604f0 (tracked)
loss(X, Y) = 0.5446847f0 (tracked)
loss(X, Y) = 0.5455998f0 (tracked)
loss(X, Y) = 0.60558325f0 (tracked)
loss(X, Y) = 0.57978874f0 (tracked)
loss(X, Y) = 0.5507627f0 (tracked)
```

```
┌ Info: Epoch 1327
└ @ Main /Users/kcft114/.julia/packages/Flux/qXNjB/src/optimise/train.jl:105
┌ Info: Epoch 1328
└ @ Main /Users/kcft114/.julia/packages/Flux/qXNjB/src/optimise/train.jl:105
┌ Info: Epoch 1329
└ @ Main /Users/kcft114/.julia/packages/Flux/qXNjB/src/optimise/train.jl:105
┌ Info: Epoch 1330
└ @ Main /Users/kcft114/.julia/packages/Flux/qXNjB/src/optimise/train.jl:105
┌ Info: Epoch 1331
└ @ Main /Users/kcft114/.julia/packages/Flux/qXNjB/src/optimise/train.jl:105
┌ Info: Epoch 1332
└ @ Main /Users/kcft114/.julia/packages/Flux/qXNjB/src/optimise/train.jl:105
┌ Info: Epoch 1333
└ @ Main /Users/kcft114/.julia/packages/Flux/qXNjB/src/optimise/train.jl:105
┌ Info: Epoch 1334
└ @ Main /Users/kcft114/.julia/packages/Flux/qXNjB/src/optimise/train.jl:105
┌ Info: Epoch 1335
└ @ Main /Users/kcft114/.julia/packages/Flux/qXNjB/src/optimise/train.jl:105
┌ Info: Epoch 1336
└ @ Main /Users/kcft114/.julia/packages/Flux/qXNjB/src/optimise/train.jl:105
┌ Info: Epoch 1337
└ @ Main /Users/kcft114/.julia/packages/Flux/qXNjB/src/optimise/train.jl:105
┌ Info: Epoch 1338
└ @ Main /Users/kcft114/.julia/packages/Flux/qXNjB/src/optimise/train.jl:105
┌ Info: Epoch 1339
└ @ Main /Users/kcft114/.julia/packages/Flux/qXNjB/src/optimise/train.jl:105
┌ Info: Epoch 1340
└ @ Main /Users/kcft114/.julia/packages/Flux/qXNjB/src/optimise/train.jl:105
┌ Info: Epoch 1341
└ @ Main /Users/kcft114/.julia/packages/Flux/qXNjB/src/optimise/train.jl:105
┌ Info: Epoch 1342
└ @ Main /Users/kcft114/.julia/packages/Flux/qXNjB/src/optimise/train.jl:105
┌ Info: Epoch 1343
└ @ Main /Users/kcft114/.julia/packages/Flux/qXNjB/src/optimise/train.jl:105
┌ Info: Epoch 1344
└ @ Main /Users/kcft114/.julia/packages/Flux/qXNjB/src/optimise/train.jl:105
┌ Info: Epoch 1345
└ @ Main /Users/kcft114/.julia/packages/Flux/qXNjB/src/optimise/train.jl:105
┌ Info: Epoch 1346
└ @ Main /Users/kcft114/.julia/packages/Flux/qXNjB/src/optimise/train.jl:105
┌ Info: Epoch 1347
└ @ Main /Users/kcft114/.julia/packages/Flux/qXNjB/src/optimise/train.jl:105
┌ Info: Epoch 1348
└ @ Main /Users/kcft114/.julia/packages/Flux/qXNjB/src/optimise/train.jl:105
```

```
loss(X, Y) = 0.61120313f0 (tracked)
loss(X, Y) = 0.5804478f0 (tracked)
loss(X, Y) = 0.6077153f0 (tracked)
loss(X, Y) = 0.57060987f0 (tracked)
loss(X, Y) = 0.5862369f0 (tracked)
loss(X, Y) = 0.59937763f0 (tracked)
loss(X, Y) = 0.59976256f0 (tracked)
loss(X, Y) = 0.5561027f0 (tracked)
loss(X, Y) = 0.5927012f0 (tracked)
loss(X, Y) = 0.59137976f0 (tracked)
loss(X, Y) = 0.58849317f0 (tracked)
loss(X, Y) = 0.5966612f0 (tracked)
loss(X, Y) = 0.5511516f0 (tracked)
loss(X, Y) = 0.5187056f0 (tracked)
loss(X, Y) = 0.5462923f0 (tracked)
loss(X, Y) = 0.58523047f0 (tracked)
loss(X, Y) = 0.5900871f0 (tracked)
loss(X, Y) = 0.6064775f0 (tracked)
loss(X, Y) = 0.5550502f0 (tracked)
loss(X, Y) = 0.5641582f0 (tracked)
loss(X, Y) = 0.5948051f0 (tracked)
loss(X, Y) = 0.6263272f0 (tracked)
loss(X, Y) = 0.55566156f0 (tracked)
loss(X, Y) = 0.6067797f0 (tracked)
loss(X, Y) = 0.51969606f0 (tracked)
loss(X, Y) = 0.56463474f0 (tracked)
loss(X, Y) = 0.5362081f0 (tracked)
loss(X, Y) = 0.606618f0 (tracked)
loss(X, Y) = 0.6048938f0 (tracked)
loss(X, Y) = 0.5499413f0 (tracked)
loss(X, Y) = 0.59750605f0 (tracked)
loss(X, Y) = 0.56430644f0 (tracked)
loss(X, Y) = 0.5815154f0 (tracked)
loss(X, Y) = 0.56588995f0 (tracked)
loss(X, Y) = 0.5793356f0 (tracked)
loss(X, Y) = 0.55770063f0 (tracked)
loss(X, Y) = 0.5826711f0 (tracked)
loss(X, Y) = 0.57180035f0 (tracked)
loss(X, Y) = 0.577996f0 (tracked)
loss(X, Y) = 0.57025534f0 (tracked)
loss(X, Y) = 0.5876529f0 (tracked)
loss(X, Y) = 0.58196396f0 (tracked)
loss(X, Y) = 0.5991867f0 (tracked)
loss(X, Y) = 0.586361f0 (tracked)
loss(X, Y) = 0.5668733f0 (tracked)
loss(X, Y) = 0.5573244f0 (tracked)
loss(X, Y) = 0.5456567f0 (tracked)
loss(X, Y) = 0.5957703f0 (tracked)
loss(X, Y) = 0.59310967f0 (tracked)
loss(X, Y) = 0.5758276f0 (tracked)
loss(X, Y) = 0.54306185f0 (tracked)
loss(X, Y) = 0.52378327f0 (tracked)
loss(X, Y) = 0.6008214f0 (tracked)
loss(X, Y) = 0.5722469f0 (tracked)
loss(X, Y) = 0.53527015f0 (tracked)
loss(X, Y) = 0.6296538f0 (tracked)
loss(X, Y) = 0.59274197f0 (tracked)
loss(X, Y) = 0.60437953f0 (tracked)
loss(X, Y) = 0.5622834f0 (tracked)
loss(X, Y) = 0.55727005f0 (tracked)
loss(X, Y) = 0.5606086f0 (tracked)
loss(X, Y) = 0.5450919f0 (tracked)
loss(X, Y) = 0.5740725f0 (tracked)
loss(X, Y) = 0.55431026f0 (tracked)
loss(X, Y) = 0.5469574f0 (tracked)
loss(X, Y) = 0.5865655f0 (tracked)
loss(X, Y) = 0.5530399f0 (tracked)
loss(X, Y) = 0.5563933f0 (tracked)
loss(X, Y) = 0.5981317f0 (tracked)
loss(X, Y) = 0.5612751f0 (tracked)
loss(X, Y) = 0.5731148f0 (tracked)
loss(X, Y) = 0.56276315f0 (tracked)
loss(X, Y) = 0.5291644f0 (tracked)
loss(X, Y) = 0.5403445f0 (tracked)
loss(X, Y) = 0.5703942f0 (tracked)
loss(X, Y) = 0.5548976f0 (tracked)
loss(X, Y) = 0.543404f0 (tracked)
loss(X, Y) = 0.5546706f0 (tracked)
loss(X, Y) = 0.6196043f0 (tracked)
loss(X, Y) = 0.5509681f0 (tracked)
loss(X, Y) = 0.5658166f0 (tracked)
loss(X, Y) = 0.55217797f0 (tracked)
loss(X, Y) = 0.54956263f0 (tracked)
loss(X, Y) = 0.55375147f0 (tracked)
loss(X, Y) = 0.582222f0 (tracked)
loss(X, Y) = 0.5407812f0 (tracked)
```

```
┌ Info: Epoch 1349
└ @ Main /Users/kcft114/.julia/packages/Flux/qXNjB/src/optimise/train.jl:105
┌ Info: Epoch 1350
└ @ Main /Users/kcft114/.julia/packages/Flux/qXNjB/src/optimise/train.jl:105
┌ Info: Epoch 1351
└ @ Main /Users/kcft114/.julia/packages/Flux/qXNjB/src/optimise/train.jl:105
┌ Info: Epoch 1352
└ @ Main /Users/kcft114/.julia/packages/Flux/qXNjB/src/optimise/train.jl:105
┌ Info: Epoch 1353
└ @ Main /Users/kcft114/.julia/packages/Flux/qXNjB/src/optimise/train.jl:105
┌ Info: Epoch 1354
└ @ Main /Users/kcft114/.julia/packages/Flux/qXNjB/src/optimise/train.jl:105
┌ Info: Epoch 1355
└ @ Main /Users/kcft114/.julia/packages/Flux/qXNjB/src/optimise/train.jl:105
┌ Info: Epoch 1356
└ @ Main /Users/kcft114/.julia/packages/Flux/qXNjB/src/optimise/train.jl:105
┌ Info: Epoch 1357
└ @ Main /Users/kcft114/.julia/packages/Flux/qXNjB/src/optimise/train.jl:105
┌ Info: Epoch 1358
└ @ Main /Users/kcft114/.julia/packages/Flux/qXNjB/src/optimise/train.jl:105
┌ Info: Epoch 1359
└ @ Main /Users/kcft114/.julia/packages/Flux/qXNjB/src/optimise/train.jl:105
┌ Info: Epoch 1360
└ @ Main /Users/kcft114/.julia/packages/Flux/qXNjB/src/optimise/train.jl:105
┌ Info: Epoch 1361
└ @ Main /Users/kcft114/.julia/packages/Flux/qXNjB/src/optimise/train.jl:105
┌ Info: Epoch 1362
└ @ Main /Users/kcft114/.julia/packages/Flux/qXNjB/src/optimise/train.jl:105
┌ Info: Epoch 1363
└ @ Main /Users/kcft114/.julia/packages/Flux/qXNjB/src/optimise/train.jl:105
┌ Info: Epoch 1364
└ @ Main /Users/kcft114/.julia/packages/Flux/qXNjB/src/optimise/train.jl:105
┌ Info: Epoch 1365
└ @ Main /Users/kcft114/.julia/packages/Flux/qXNjB/src/optimise/train.jl:105
┌ Info: Epoch 1366
└ @ Main /Users/kcft114/.julia/packages/Flux/qXNjB/src/optimise/train.jl:105
┌ Info: Epoch 1367
└ @ Main /Users/kcft114/.julia/packages/Flux/qXNjB/src/optimise/train.jl:105
┌ Info: Epoch 1368
└ @ Main /Users/kcft114/.julia/packages/Flux/qXNjB/src/optimise/train.jl:105
┌ Info: Epoch 1369
└ @ Main /Users/kcft114/.julia/packages/Flux/qXNjB/src/optimise/train.jl:105
┌ Info: Epoch 1370
└ @ Main /Users/kcft114/.julia/packages/Flux/qXNjB/src/optimise/train.jl:105
┌ Info: Epoch 1371
└ @ Main /Users/kcft114/.julia/packages/Flux/qXNjB/src/optimise/train.jl:105
┌ Info: Epoch 1372
└ @ Main /Users/kcft114/.julia/packages/Flux/qXNjB/src/optimise/train.jl:105
┌ Info: Epoch 1373
└ @ Main /Users/kcft114/.julia/packages/Flux/qXNjB/src/optimise/train.jl:105
┌ Info: Epoch 1374
└ @ Main /Users/kcft114/.julia/packages/Flux/qXNjB/src/optimise/train.jl:105
┌ Info: Epoch 1375
└ @ Main /Users/kcft114/.julia/packages/Flux/qXNjB/src/optimise/train.jl:105
┌ Info: Epoch 1376
└ @ Main /Users/kcft114/.julia/packages/Flux/qXNjB/src/optimise/train.jl:105
┌ Info: Epoch 1377
└ @ Main /Users/kcft114/.julia/packages/Flux/qXNjB/src/optimise/train.jl:105
┌ Info: Epoch 1378
└ @ Main /Users/kcft114/.julia/packages/Flux/qXNjB/src/optimise/train.jl:105
┌ Info: Epoch 1379
└ @ Main /Users/kcft114/.julia/packages/Flux/qXNjB/src/optimise/train.jl:105
┌ Info: Epoch 1380
└ @ Main /Users/kcft114/.julia/packages/Flux/qXNjB/src/optimise/train.jl:105
┌ Info: Epoch 1381
└ @ Main /Users/kcft114/.julia/packages/Flux/qXNjB/src/optimise/train.jl:105
┌ Info: Epoch 1382
└ @ Main /Users/kcft114/.julia/packages/Flux/qXNjB/src/optimise/train.jl:105
┌ Info: Epoch 1383
└ @ Main /Users/kcft114/.julia/packages/Flux/qXNjB/src/optimise/train.jl:105
┌ Info: Epoch 1384
└ @ Main /Users/kcft114/.julia/packages/Flux/qXNjB/src/optimise/train.jl:105
┌ Info: Epoch 1385
└ @ Main /Users/kcft114/.julia/packages/Flux/qXNjB/src/optimise/train.jl:105
┌ Info: Epoch 1386
└ @ Main /Users/kcft114/.julia/packages/Flux/qXNjB/src/optimise/train.jl:105
┌ Info: Epoch 1387
└ @ Main /Users/kcft114/.julia/packages/Flux/qXNjB/src/optimise/train.jl:105
┌ Info: Epoch 1388
└ @ Main /Users/kcft114/.julia/packages/Flux/qXNjB/src/optimise/train.jl:105
┌ Info: Epoch 1389
└ @ Main /Users/kcft114/.julia/packages/Flux/qXNjB/src/optimise/train.jl:105
┌ Info: Epoch 1390
└ @ Main /Users/kcft114/.julia/packages/Flux/qXNjB/src/optimise/train.jl:105
┌ Info: Epoch 1391
└ @ Main /Users/kcft114/.julia/packages/Flux/qXNjB/src/optimise/train.jl:105
┌ Info: Epoch 1392
└ @ Main /Users/kcft114/.julia/packages/Flux/qXNjB/src/optimise/train.jl:105
┌ Info: Epoch 1393
└ @ Main /Users/kcft114/.julia/packages/Flux/qXNjB/src/optimise/train.jl:105
┌ Info: Epoch 1394
└ @ Main /Users/kcft114/.julia/packages/Flux/qXNjB/src/optimise/train.jl:105
┌ Info: Epoch 1395
└ @ Main /Users/kcft114/.julia/packages/Flux/qXNjB/src/optimise/train.jl:105
┌ Info: Epoch 1396
└ @ Main /Users/kcft114/.julia/packages/Flux/qXNjB/src/optimise/train.jl:105
┌ Info: Epoch 1397
└ @ Main /Users/kcft114/.julia/packages/Flux/qXNjB/src/optimise/train.jl:105
┌ Info: Epoch 1398
└ @ Main /Users/kcft114/.julia/packages/Flux/qXNjB/src/optimise/train.jl:105
┌ Info: Epoch 1399
└ @ Main /Users/kcft114/.julia/packages/Flux/qXNjB/src/optimise/train.jl:105
┌ Info: Epoch 1400
└ @ Main /Users/kcft114/.julia/packages/Flux/qXNjB/src/optimise/train.jl:105
┌ Info: Epoch 1401
└ @ Main /Users/kcft114/.julia/packages/Flux/qXNjB/src/optimise/train.jl:105
┌ Info: Epoch 1402
└ @ Main /Users/kcft114/.julia/packages/Flux/qXNjB/src/optimise/train.jl:105
┌ Info: Epoch 1403
└ @ Main /Users/kcft114/.julia/packages/Flux/qXNjB/src/optimise/train.jl:105
┌ Info: Epoch 1404
└ @ Main /Users/kcft114/.julia/packages/Flux/qXNjB/src/optimise/train.jl:105
┌ Info: Epoch 1405
└ @ Main /Users/kcft114/.julia/packages/Flux/qXNjB/src/optimise/train.jl:105
┌ Info: Epoch 1406
└ @ Main /Users/kcft114/.julia/packages/Flux/qXNjB/src/optimise/train.jl:105
┌ Info: Epoch 1407
└ @ Main /Users/kcft114/.julia/packages/Flux/qXNjB/src/optimise/train.jl:105
┌ Info: Epoch 1408
└ @ Main /Users/kcft114/.julia/packages/Flux/qXNjB/src/optimise/train.jl:105
┌ Info: Epoch 1409
└ @ Main /Users/kcft114/.julia/packages/Flux/qXNjB/src/optimise/train.jl:105
┌ Info: Epoch 1410
└ @ Main /Users/kcft114/.julia/packages/Flux/qXNjB/src/optimise/train.jl:105
┌ Info: Epoch 1411
└ @ Main /Users/kcft114/.julia/packages/Flux/qXNjB/src/optimise/train.jl:105
┌ Info: Epoch 1412
└ @ Main /Users/kcft114/.julia/packages/Flux/qXNjB/src/optimise/train.jl:105
┌ Info: Epoch 1413
└ @ Main /Users/kcft114/.julia/packages/Flux/qXNjB/src/optimise/train.jl:105
┌ Info: Epoch 1414
└ @ Main /Users/kcft114/.julia/packages/Flux/qXNjB/src/optimise/train.jl:105
┌ Info: Epoch 1415
└ @ Main /Users/kcft114/.julia/packages/Flux/qXNjB/src/optimise/train.jl:105
┌ Info: Epoch 1416
└ @ Main /Users/kcft114/.julia/packages/Flux/qXNjB/src/optimise/train.jl:105
┌ Info: Epoch 1417
└ @ Main /Users/kcft114/.julia/packages/Flux/qXNjB/src/optimise/train.jl:105
┌ Info: Epoch 1418
└ @ Main /Users/kcft114/.julia/packages/Flux/qXNjB/src/optimise/train.jl:105
┌ Info: Epoch 1419
└ @ Main /Users/kcft114/.julia/packages/Flux/qXNjB/src/optimise/train.jl:105
┌ Info: Epoch 1420
└ @ Main /Users/kcft114/.julia/packages/Flux/qXNjB/src/optimise/train.jl:105
┌ Info: Epoch 1421
└ @ Main /Users/kcft114/.julia/packages/Flux/qXNjB/src/optimise/train.jl:105
┌ Info: Epoch 1422
└ @ Main /Users/kcft114/.julia/packages/Flux/qXNjB/src/optimise/train.jl:105
┌ Info: Epoch 1423
└ @ Main /Users/kcft114/.julia/packages/Flux/qXNjB/src/optimise/train.jl:105
┌ Info: Epoch 1424
└ @ Main /Users/kcft114/.julia/packages/Flux/qXNjB/src/optimise/train.jl:105
┌ Info: Epoch 1425
└ @ Main /Users/kcft114/.julia/packages/Flux/qXNjB/src/optimise/train.jl:105
┌ Info: Epoch 1426
└ @ Main /Users/kcft114/.julia/packages/Flux/qXNjB/src/optimise/train.jl:105
┌ Info: Epoch 1427
└ @ Main /Users/kcft114/.julia/packages/Flux/qXNjB/src/optimise/train.jl:105
┌ Info: Epoch 1428
└ @ Main /Users/kcft114/.julia/packages/Flux/qXNjB/src/optimise/train.jl:105
┌ Info: Epoch 1429
└ @ Main /Users/kcft114/.julia/packages/Flux/qXNjB/src/optimise/train.jl:105
┌ Info: Epoch 1430
└ @ Main /Users/kcft114/.julia/packages/Flux/qXNjB/src/optimise/train.jl:105
┌ Info: Epoch 1431
└ @ Main /Users/kcft114/.julia/packages/Flux/qXNjB/src/optimise/train.jl:105
┌ Info: Epoch 1432
└ @ Main /Users/kcft114/.julia/packages/Flux/qXNjB/src/optimise/train.jl:105
┌ Info: Epoch 1433
└ @ Main /Users/kcft114/.julia/packages/Flux/qXNjB/src/optimise/train.jl:105
┌ Info: Epoch 1434
└ @ Main /Users/kcft114/.julia/packages/Flux/qXNjB/src/optimise/train.jl:105
```

```
loss(X, Y) = 0.5872056f0 (tracked)
loss(X, Y) = 0.5981115f0 (tracked)
loss(X, Y) = 0.6079204f0 (tracked)
loss(X, Y) = 0.59479415f0 (tracked)
loss(X, Y) = 0.56102884f0 (tracked)
loss(X, Y) = 0.58001333f0 (tracked)
loss(X, Y) = 0.5742682f0 (tracked)
loss(X, Y) = 0.6284309f0 (tracked)
loss(X, Y) = 0.5703047f0 (tracked)
loss(X, Y) = 0.55527306f0 (tracked)
loss(X, Y) = 0.59614366f0 (tracked)
loss(X, Y) = 0.588309f0 (tracked)
loss(X, Y) = 0.5728417f0 (tracked)
loss(X, Y) = 0.6010231f0 (tracked)
loss(X, Y) = 0.5733168f0 (tracked)
loss(X, Y) = 0.5833456f0 (tracked)
loss(X, Y) = 0.5643175f0 (tracked)
loss(X, Y) = 0.608491f0 (tracked)
loss(X, Y) = 0.5645901f0 (tracked)
loss(X, Y) = 0.60666966f0 (tracked)
loss(X, Y) = 0.58083427f0 (tracked)
loss(X, Y) = 0.5273523f0 (tracked)
loss(X, Y) = 0.5596379f0 (tracked)
loss(X, Y) = 0.5606582f0 (tracked)
loss(X, Y) = 0.5352532f0 (tracked)
loss(X, Y) = 0.54319096f0 (tracked)
loss(X, Y) = 0.56703204f0 (tracked)
loss(X, Y) = 0.58998793f0 (tracked)
loss(X, Y) = 0.5881896f0 (tracked)
loss(X, Y) = 0.57451135f0 (tracked)
loss(X, Y) = 0.58178043f0 (tracked)
loss(X, Y) = 0.5639603f0 (tracked)
loss(X, Y) = 0.55322725f0 (tracked)
loss(X, Y) = 0.5956825f0 (tracked)
loss(X, Y) = 0.6019256f0 (tracked)
loss(X, Y) = 0.54925627f0 (tracked)
loss(X, Y) = 0.54317397f0 (tracked)
loss(X, Y) = 0.56448185f0 (tracked)
loss(X, Y) = 0.56540406f0 (tracked)
loss(X, Y) = 0.5867327f0 (tracked)
loss(X, Y) = 0.5587326f0 (tracked)
loss(X, Y) = 0.5595006f0 (tracked)
loss(X, Y) = 0.6047236f0 (tracked)
loss(X, Y) = 0.56709284f0 (tracked)
loss(X, Y) = 0.561222f0 (tracked)
loss(X, Y) = 0.5464757f0 (tracked)
loss(X, Y) = 0.6062224f0 (tracked)
loss(X, Y) = 0.60274285f0 (tracked)
loss(X, Y) = 0.60684955f0 (tracked)
loss(X, Y) = 0.56735444f0 (tracked)
loss(X, Y) = 0.5507968f0 (tracked)
loss(X, Y) = 0.5444205f0 (tracked)
loss(X, Y) = 0.5515112f0 (tracked)
loss(X, Y) = 0.5993484f0 (tracked)
loss(X, Y) = 0.5219103f0 (tracked)
loss(X, Y) = 0.5470399f0 (tracked)
loss(X, Y) = 0.5629419f0 (tracked)
loss(X, Y) = 0.55036867f0 (tracked)
loss(X, Y) = 0.55046296f0 (tracked)
loss(X, Y) = 0.57521856f0 (tracked)
loss(X, Y) = 0.57171226f0 (tracked)
loss(X, Y) = 0.58009475f0 (tracked)
loss(X, Y) = 0.5509036f0 (tracked)
loss(X, Y) = 0.48945457f0 (tracked)
loss(X, Y) = 0.61934f0 (tracked)
loss(X, Y) = 0.5497217f0 (tracked)
loss(X, Y) = 0.5578594f0 (tracked)
loss(X, Y) = 0.56427807f0 (tracked)
loss(X, Y) = 0.5672386f0 (tracked)
loss(X, Y) = 0.5534545f0 (tracked)
loss(X, Y) = 0.5191004f0 (tracked)
loss(X, Y) = 0.5437922f0 (tracked)
loss(X, Y) = 0.5293231f0 (tracked)
loss(X, Y) = 0.55595386f0 (tracked)
loss(X, Y) = 0.5573351f0 (tracked)
loss(X, Y) = 0.5493013f0 (tracked)
loss(X, Y) = 0.5523169f0 (tracked)
loss(X, Y) = 0.54492915f0 (tracked)
loss(X, Y) = 0.56646025f0 (tracked)
loss(X, Y) = 0.5308076f0 (tracked)
loss(X, Y) = 0.5829514f0 (tracked)
loss(X, Y) = 0.5686526f0 (tracked)
loss(X, Y) = 0.57537514f0 (tracked)
loss(X, Y) = 0.52120703f0 (tracked)
loss(X, Y) = 0.5566956f0 (tracked)
loss(X, Y) = 0.53679067f0 (tracked)
loss(X, Y) = 0.5322926f0 (tracked)
loss(X, Y) = 0.6093437f0 (tracked)
loss(X, Y) = 0.53557456f0 (tracked)
loss(X, Y) = 0.5658169f0 (tracked)
loss(X, Y) = 0.575334f0 (tracked)
loss(X, Y) = 0.55110765f0 (tracked)
loss(X, Y) = 0.5705798f0 (tracked)
loss(X, Y) = 0.5620761f0 (tracked)
```

```
┌ Info: Epoch 1435
└ @ Main /Users/kcft114/.julia/packages/Flux/qXNjB/src/optimise/train.jl:105
┌ Info: Epoch 1436
└ @ Main /Users/kcft114/.julia/packages/Flux/qXNjB/src/optimise/train.jl:105
┌ Info: Epoch 1437
└ @ Main /Users/kcft114/.julia/packages/Flux/qXNjB/src/optimise/train.jl:105
┌ Info: Epoch 1438
└ @ Main /Users/kcft114/.julia/packages/Flux/qXNjB/src/optimise/train.jl:105
┌ Info: Epoch 1439
└ @ Main /Users/kcft114/.julia/packages/Flux/qXNjB/src/optimise/train.jl:105
┌ Info: Epoch 1440
└ @ Main /Users/kcft114/.julia/packages/Flux/qXNjB/src/optimise/train.jl:105
┌ Info: Epoch 1441
└ @ Main /Users/kcft114/.julia/packages/Flux/qXNjB/src/optimise/train.jl:105
┌ Info: Epoch 1442
└ @ Main /Users/kcft114/.julia/packages/Flux/qXNjB/src/optimise/train.jl:105
┌ Info: Epoch 1443
└ @ Main /Users/kcft114/.julia/packages/Flux/qXNjB/src/optimise/train.jl:105
┌ Info: Epoch 1444
└ @ Main /Users/kcft114/.julia/packages/Flux/qXNjB/src/optimise/train.jl:105
┌ Info: Epoch 1445
└ @ Main /Users/kcft114/.julia/packages/Flux/qXNjB/src/optimise/train.jl:105
┌ Info: Epoch 1446
└ @ Main /Users/kcft114/.julia/packages/Flux/qXNjB/src/optimise/train.jl:105
┌ Info: Epoch 1447
└ @ Main /Users/kcft114/.julia/packages/Flux/qXNjB/src/optimise/train.jl:105
┌ Info: Epoch 1448
└ @ Main /Users/kcft114/.julia/packages/Flux/qXNjB/src/optimise/train.jl:105
┌ Info: Epoch 1449
└ @ Main /Users/kcft114/.julia/packages/Flux/qXNjB/src/optimise/train.jl:105
┌ Info: Epoch 1450
└ @ Main /Users/kcft114/.julia/packages/Flux/qXNjB/src/optimise/train.jl:105
┌ Info: Epoch 1451
└ @ Main /Users/kcft114/.julia/packages/Flux/qXNjB/src/optimise/train.jl:105
┌ Info: Epoch 1452
└ @ Main /Users/kcft114/.julia/packages/Flux/qXNjB/src/optimise/train.jl:105
┌ Info: Epoch 1453
└ @ Main /Users/kcft114/.julia/packages/Flux/qXNjB/src/optimise/train.jl:105
┌ Info: Epoch 1454
└ @ Main /Users/kcft114/.julia/packages/Flux/qXNjB/src/optimise/train.jl:105
┌ Info: Epoch 1455
└ @ Main /Users/kcft114/.julia/packages/Flux/qXNjB/src/optimise/train.jl:105
┌ Info: Epoch 1456
└ @ Main /Users/kcft114/.julia/packages/Flux/qXNjB/src/optimise/train.jl:105
┌ Info: Epoch 1457
└ @ Main /Users/kcft114/.julia/packages/Flux/qXNjB/src/optimise/train.jl:105
┌ Info: Epoch 1458
└ @ Main /Users/kcft114/.julia/packages/Flux/qXNjB/src/optimise/train.jl:105
┌ Info: Epoch 1459
└ @ Main /Users/kcft114/.julia/packages/Flux/qXNjB/src/optimise/train.jl:105
┌ Info: Epoch 1460
└ @ Main /Users/kcft114/.julia/packages/Flux/qXNjB/src/optimise/train.jl:105
┌ Info: Epoch 1461
└ @ Main /Users/kcft114/.julia/packages/Flux/qXNjB/src/optimise/train.jl:105
┌ Info: Epoch 1462
└ @ Main /Users/kcft114/.julia/packages/Flux/qXNjB/src/optimise/train.jl:105
┌ Info: Epoch 1463
└ @ Main /Users/kcft114/.julia/packages/Flux/qXNjB/src/optimise/train.jl:105
┌ Info: Epoch 1464
└ @ Main /Users/kcft114/.julia/packages/Flux/qXNjB/src/optimise/train.jl:105
┌ Info: Epoch 1465
└ @ Main /Users/kcft114/.julia/packages/Flux/qXNjB/src/optimise/train.jl:105
┌ Info: Epoch 1466
└ @ Main /Users/kcft114/.julia/packages/Flux/qXNjB/src/optimise/train.jl:105
┌ Info: Epoch 1467
└ @ Main /Users/kcft114/.julia/packages/Flux/qXNjB/src/optimise/train.jl:105
┌ Info: Epoch 1468
└ @ Main /Users/kcft114/.julia/packages/Flux/qXNjB/src/optimise/train.jl:105
┌ Info: Epoch 1469
└ @ Main /Users/kcft114/.julia/packages/Flux/qXNjB/src/optimise/train.jl:105
┌ Info: Epoch 1470
└ @ Main /Users/kcft114/.julia/packages/Flux/qXNjB/src/optimise/train.jl:105
┌ Info: Epoch 1471
└ @ Main /Users/kcft114/.julia/packages/Flux/qXNjB/src/optimise/train.jl:105
┌ Info: Epoch 1472
└ @ Main /Users/kcft114/.julia/packages/Flux/qXNjB/src/optimise/train.jl:105
┌ Info: Epoch 1473
└ @ Main /Users/kcft114/.julia/packages/Flux/qXNjB/src/optimise/train.jl:105
┌ Info: Epoch 1474
└ @ Main /Users/kcft114/.julia/packages/Flux/qXNjB/src/optimise/train.jl:105
┌ Info: Epoch 1475
└ @ Main /Users/kcft114/.julia/packages/Flux/qXNjB/src/optimise/train.jl:105
┌ Info: Epoch 1476
└ @ Main /Users/kcft114/.julia/packages/Flux/qXNjB/src/optimise/train.jl:105
┌ Info: Epoch 1477
└ @ Main /Users/kcft114/.julia/packages/Flux/qXNjB/src/optimise/train.jl:105
┌ Info: Epoch 1478
└ @ Main /Users/kcft114/.julia/packages/Flux/qXNjB/src/optimise/train.jl:105
┌ Info: Epoch 1479
└ @ Main /Users/kcft114/.julia/packages/Flux/qXNjB/src/optimise/train.jl:105
┌ Info: Epoch 1480
└ @ Main /Users/kcft114/.julia/packages/Flux/qXNjB/src/optimise/train.jl:105
┌ Info: Epoch 1481
└ @ Main /Users/kcft114/.julia/packages/Flux/qXNjB/src/optimise/train.jl:105
┌ Info: Epoch 1482
└ @ Main /Users/kcft114/.julia/packages/Flux/qXNjB/src/optimise/train.jl:105
┌ Info: Epoch 1483
└ @ Main /Users/kcft114/.julia/packages/Flux/qXNjB/src/optimise/train.jl:105
┌ Info: Epoch 1484
└ @ Main /Users/kcft114/.julia/packages/Flux/qXNjB/src/optimise/train.jl:105
┌ Info: Epoch 1485
└ @ Main /Users/kcft114/.julia/packages/Flux/qXNjB/src/optimise/train.jl:105
┌ Info: Epoch 1486
└ @ Main /Users/kcft114/.julia/packages/Flux/qXNjB/src/optimise/train.jl:105
┌ Info: Epoch 1487
└ @ Main /Users/kcft114/.julia/packages/Flux/qXNjB/src/optimise/train.jl:105
┌ Info: Epoch 1488
└ @ Main /Users/kcft114/.julia/packages/Flux/qXNjB/src/optimise/train.jl:105
┌ Info: Epoch 1489
└ @ Main /Users/kcft114/.julia/packages/Flux/qXNjB/src/optimise/train.jl:105
┌ Info: Epoch 1490
└ @ Main /Users/kcft114/.julia/packages/Flux/qXNjB/src/optimise/train.jl:105
┌ Info: Epoch 1491
└ @ Main /Users/kcft114/.julia/packages/Flux/qXNjB/src/optimise/train.jl:105
┌ Info: Epoch 1492
└ @ Main /Users/kcft114/.julia/packages/Flux/qXNjB/src/optimise/train.jl:105
┌ Info: Epoch 1493
└ @ Main /Users/kcft114/.julia/packages/Flux/qXNjB/src/optimise/train.jl:105
┌ Info: Epoch 1494
└ @ Main /Users/kcft114/.julia/packages/Flux/qXNjB/src/optimise/train.jl:105
┌ Info: Epoch 1495
└ @ Main /Users/kcft114/.julia/packages/Flux/qXNjB/src/optimise/train.jl:105
┌ Info: Epoch 1496
└ @ Main /Users/kcft114/.julia/packages/Flux/qXNjB/src/optimise/train.jl:105
┌ Info: Epoch 1497
└ @ Main /Users/kcft114/.julia/packages/Flux/qXNjB/src/optimise/train.jl:105
┌ Info: Epoch 1498
└ @ Main /Users/kcft114/.julia/packages/Flux/qXNjB/src/optimise/train.jl:105
┌ Info: Epoch 1499
└ @ Main /Users/kcft114/.julia/packages/Flux/qXNjB/src/optimise/train.jl:105
┌ Info: Epoch 1500
└ @ Main /Users/kcft114/.julia/packages/Flux/qXNjB/src/optimise/train.jl:105
┌ Info: Epoch 1501
└ @ Main /Users/kcft114/.julia/packages/Flux/qXNjB/src/optimise/train.jl:105
┌ Info: Epoch 1502
└ @ Main /Users/kcft114/.julia/packages/Flux/qXNjB/src/optimise/train.jl:105
┌ Info: Epoch 1503
└ @ Main /Users/kcft114/.julia/packages/Flux/qXNjB/src/optimise/train.jl:105
┌ Info: Epoch 1504
└ @ Main /Users/kcft114/.julia/packages/Flux/qXNjB/src/optimise/train.jl:105
┌ Info: Epoch 1505
└ @ Main /Users/kcft114/.julia/packages/Flux/qXNjB/src/optimise/train.jl:105
┌ Info: Epoch 1506
└ @ Main /Users/kcft114/.julia/packages/Flux/qXNjB/src/optimise/train.jl:105
┌ Info: Epoch 1507
└ @ Main /Users/kcft114/.julia/packages/Flux/qXNjB/src/optimise/train.jl:105
┌ Info: Epoch 1508
└ @ Main /Users/kcft114/.julia/packages/Flux/qXNjB/src/optimise/train.jl:105
┌ Info: Epoch 1509
└ @ Main /Users/kcft114/.julia/packages/Flux/qXNjB/src/optimise/train.jl:105
┌ Info: Epoch 1510
└ @ Main /Users/kcft114/.julia/packages/Flux/qXNjB/src/optimise/train.jl:105
┌ Info: Epoch 1511
└ @ Main /Users/kcft114/.julia/packages/Flux/qXNjB/src/optimise/train.jl:105
┌ Info: Epoch 1512
└ @ Main /Users/kcft114/.julia/packages/Flux/qXNjB/src/optimise/train.jl:105
┌ Info: Epoch 1513
└ @ Main /Users/kcft114/.julia/packages/Flux/qXNjB/src/optimise/train.jl:105
┌ Info: Epoch 1514
└ @ Main /Users/kcft114/.julia/packages/Flux/qXNjB/src/optimise/train.jl:105
┌ Info: Epoch 1515
└ @ Main /Users/kcft114/.julia/packages/Flux/qXNjB/src/optimise/train.jl:105
┌ Info: Epoch 1516
└ @ Main /Users/kcft114/.julia/packages/Flux/qXNjB/src/optimise/train.jl:105
┌ Info: Epoch 1517
└ @ Main /Users/kcft114/.julia/packages/Flux/qXNjB/src/optimise/train.jl:105
┌ Info: Epoch 1518
└ @ Main /Users/kcft114/.julia/packages/Flux/qXNjB/src/optimise/train.jl:105
┌ Info: Epoch 1519
└ @ Main /Users/kcft114/.julia/packages/Flux/qXNjB/src/optimise/train.jl:105
┌ Info: Epoch 1520
└ @ Main /Users/kcft114/.julia/packages/Flux/qXNjB/src/optimise/train.jl:105
┌ Info: Epoch 1521
└ @ Main /Users/kcft114/.julia/packages/Flux/qXNjB/src/optimise/train.jl:105
┌ Info: Epoch 1522
└ @ Main /Users/kcft114/.julia/packages/Flux/qXNjB/src/optimise/train.jl:105
┌ Info: Epoch 1523
└ @ Main /Users/kcft114/.julia/packages/Flux/qXNjB/src/optimise/train.jl:105
┌ Info: Epoch 1524
└ @ Main /Users/kcft114/.julia/packages/Flux/qXNjB/src/optimise/train.jl:105
┌ Info: Epoch 1525
└ @ Main /Users/kcft114/.julia/packages/Flux/qXNjB/src/optimise/train.jl:105
┌ Info: Epoch 1526
└ @ Main /Users/kcft114/.julia/packages/Flux/qXNjB/src/optimise/train.jl:105
┌ Info: Epoch 1527
└ @ Main /Users/kcft114/.julia/packages/Flux/qXNjB/src/optimise/train.jl:105
┌ Info: Epoch 1528
└ @ Main /Users/kcft114/.julia/packages/Flux/qXNjB/src/optimise/train.jl:105
```

```
loss(X, Y) = 0.49946618f0 (tracked)
loss(X, Y) = 0.51886064f0 (tracked)
loss(X, Y) = 0.553418f0 (tracked)
loss(X, Y) = 0.5775767f0 (tracked)
loss(X, Y) = 0.5742577f0 (tracked)
loss(X, Y) = 0.55700666f0 (tracked)
loss(X, Y) = 0.5732626f0 (tracked)
loss(X, Y) = 0.54854685f0 (tracked)
loss(X, Y) = 0.56171435f0 (tracked)
loss(X, Y) = 0.563417f0 (tracked)
loss(X, Y) = 0.54885286f0 (tracked)
loss(X, Y) = 0.57717085f0 (tracked)
loss(X, Y) = 0.58472437f0 (tracked)
loss(X, Y) = 0.5453922f0 (tracked)
loss(X, Y) = 0.54673916f0 (tracked)
loss(X, Y) = 0.5745011f0 (tracked)
loss(X, Y) = 0.56290984f0 (tracked)
loss(X, Y) = 0.5432296f0 (tracked)
loss(X, Y) = 0.5372533f0 (tracked)
loss(X, Y) = 0.55328876f0 (tracked)
loss(X, Y) = 0.52554893f0 (tracked)
loss(X, Y) = 0.5762656f0 (tracked)
loss(X, Y) = 0.59414417f0 (tracked)
loss(X, Y) = 0.5758778f0 (tracked)
loss(X, Y) = 0.5475737f0 (tracked)
loss(X, Y) = 0.6105604f0 (tracked)
loss(X, Y) = 0.5865226f0 (tracked)
loss(X, Y) = 0.5794362f0 (tracked)
loss(X, Y) = 0.58071154f0 (tracked)
loss(X, Y) = 0.55292404f0 (tracked)
loss(X, Y) = 0.60803205f0 (tracked)
loss(X, Y) = 0.579504f0 (tracked)
loss(X, Y) = 0.5884834f0 (tracked)
loss(X, Y) = 0.55048794f0 (tracked)
loss(X, Y) = 0.5759863f0 (tracked)
loss(X, Y) = 0.5780431f0 (tracked)
loss(X, Y) = 0.5897811f0 (tracked)
loss(X, Y) = 0.55134934f0 (tracked)
loss(X, Y) = 0.5939312f0 (tracked)
loss(X, Y) = 0.56264424f0 (tracked)
loss(X, Y) = 0.58179945f0 (tracked)
loss(X, Y) = 0.55944645f0 (tracked)
loss(X, Y) = 0.5567013f0 (tracked)
loss(X, Y) = 0.5904971f0 (tracked)
loss(X, Y) = 0.5590393f0 (tracked)
loss(X, Y) = 0.5607428f0 (tracked)
loss(X, Y) = 0.5372155f0 (tracked)
loss(X, Y) = 0.51098084f0 (tracked)
loss(X, Y) = 0.58579105f0 (tracked)
loss(X, Y) = 0.5642481f0 (tracked)
loss(X, Y) = 0.59917194f0 (tracked)
loss(X, Y) = 0.55534786f0 (tracked)
loss(X, Y) = 0.5832294f0 (tracked)
loss(X, Y) = 0.56907773f0 (tracked)
loss(X, Y) = 0.570431f0 (tracked)
loss(X, Y) = 0.53984475f0 (tracked)
loss(X, Y) = 0.5381166f0 (tracked)
loss(X, Y) = 0.58951265f0 (tracked)
loss(X, Y) = 0.53521454f0 (tracked)
loss(X, Y) = 0.55797404f0 (tracked)
loss(X, Y) = 0.55762976f0 (tracked)
loss(X, Y) = 0.563826f0 (tracked)
loss(X, Y) = 0.55525935f0 (tracked)
loss(X, Y) = 0.5451111f0 (tracked)
loss(X, Y) = 0.5071616f0 (tracked)
loss(X, Y) = 0.6029993f0 (tracked)
loss(X, Y) = 0.54009914f0 (tracked)
loss(X, Y) = 0.5648961f0 (tracked)
loss(X, Y) = 0.54806f0 (tracked)
loss(X, Y) = 0.5634597f0 (tracked)
loss(X, Y) = 0.56416065f0 (tracked)
loss(X, Y) = 0.5528649f0 (tracked)
loss(X, Y) = 0.5622601f0 (tracked)
loss(X, Y) = 0.5418642f0 (tracked)
loss(X, Y) = 0.5414009f0 (tracked)
loss(X, Y) = 0.52719045f0 (tracked)
loss(X, Y) = 0.5435355f0 (tracked)
loss(X, Y) = 0.6102842f0 (tracked)
loss(X, Y) = 0.5544733f0 (tracked)
loss(X, Y) = 0.5242769f0 (tracked)
loss(X, Y) = 0.53556454f0 (tracked)
loss(X, Y) = 0.5325201f0 (tracked)
loss(X, Y) = 0.60533f0 (tracked)
loss(X, Y) = 0.5427863f0 (tracked)
loss(X, Y) = 0.5461134f0 (tracked)
loss(X, Y) = 0.563057f0 (tracked)
loss(X, Y) = 0.58546084f0 (tracked)
loss(X, Y) = 0.5331967f0 (tracked)
loss(X, Y) = 0.6145049f0 (tracked)
loss(X, Y) = 0.5403044f0 (tracked)
loss(X, Y) = 0.53505653f0 (tracked)
loss(X, Y) = 0.5464365f0 (tracked)
loss(X, Y) = 0.55502594f0 (tracked)
loss(X, Y) = 0.5290094f0 (tracked)
loss(X, Y) = 0.6057042f0 (tracked)
loss(X, Y) = 0.5772953f0 (tracked)
loss(X, Y) = 0.56026447f0 (tracked)
loss(X, Y) = 0.5882549f0 (tracked)
loss(X, Y) = 0.5272609f0 (tracked)
loss(X, Y) = 0.53239197f0 (tracked)
loss(X, Y) = 0.53504384f0 (tracked)
loss(X, Y) = 0.552059f0 (tracked)
loss(X, Y) = 0.515501f0 (tracked)
```

```
┌ Info: Epoch 1529
└ @ Main /Users/kcft114/.julia/packages/Flux/qXNjB/src/optimise/train.jl:105
┌ Info: Epoch 1530
└ @ Main /Users/kcft114/.julia/packages/Flux/qXNjB/src/optimise/train.jl:105
┌ Info: Epoch 1531
└ @ Main /Users/kcft114/.julia/packages/Flux/qXNjB/src/optimise/train.jl:105
┌ Info: Epoch 1532
└ @ Main /Users/kcft114/.julia/packages/Flux/qXNjB/src/optimise/train.jl:105
┌ Info: Epoch 1533
└ @ Main /Users/kcft114/.julia/packages/Flux/qXNjB/src/optimise/train.jl:105
┌ Info: Epoch 1534
└ @ Main /Users/kcft114/.julia/packages/Flux/qXNjB/src/optimise/train.jl:105
┌ Info: Epoch 1535
└ @ Main /Users/kcft114/.julia/packages/Flux/qXNjB/src/optimise/train.jl:105
┌ Info: Epoch 1536
└ @ Main /Users/kcft114/.julia/packages/Flux/qXNjB/src/optimise/train.jl:105
┌ Info: Epoch 1537
└ @ Main /Users/kcft114/.julia/packages/Flux/qXNjB/src/optimise/train.jl:105
┌ Info: Epoch 1538
└ @ Main /Users/kcft114/.julia/packages/Flux/qXNjB/src/optimise/train.jl:105
┌ Info: Epoch 1539
└ @ Main /Users/kcft114/.julia/packages/Flux/qXNjB/src/optimise/train.jl:105
┌ Info: Epoch 1540
└ @ Main /Users/kcft114/.julia/packages/Flux/qXNjB/src/optimise/train.jl:105
┌ Info: Epoch 1541
└ @ Main /Users/kcft114/.julia/packages/Flux/qXNjB/src/optimise/train.jl:105
┌ Info: Epoch 1542
└ @ Main /Users/kcft114/.julia/packages/Flux/qXNjB/src/optimise/train.jl:105
┌ Info: Epoch 1543
└ @ Main /Users/kcft114/.julia/packages/Flux/qXNjB/src/optimise/train.jl:105
┌ Info: Epoch 1544
└ @ Main /Users/kcft114/.julia/packages/Flux/qXNjB/src/optimise/train.jl:105
┌ Info: Epoch 1545
└ @ Main /Users/kcft114/.julia/packages/Flux/qXNjB/src/optimise/train.jl:105
┌ Info: Epoch 1546
└ @ Main /Users/kcft114/.julia/packages/Flux/qXNjB/src/optimise/train.jl:105
┌ Info: Epoch 1547
└ @ Main /Users/kcft114/.julia/packages/Flux/qXNjB/src/optimise/train.jl:105
┌ Info: Epoch 1548
└ @ Main /Users/kcft114/.julia/packages/Flux/qXNjB/src/optimise/train.jl:105
┌ Info: Epoch 1549
└ @ Main /Users/kcft114/.julia/packages/Flux/qXNjB/src/optimise/train.jl:105
┌ Info: Epoch 1550
└ @ Main /Users/kcft114/.julia/packages/Flux/qXNjB/src/optimise/train.jl:105
┌ Info: Epoch 1551
└ @ Main /Users/kcft114/.julia/packages/Flux/qXNjB/src/optimise/train.jl:105
┌ Info: Epoch 1552
└ @ Main /Users/kcft114/.julia/packages/Flux/qXNjB/src/optimise/train.jl:105
┌ Info: Epoch 1553
└ @ Main /Users/kcft114/.julia/packages/Flux/qXNjB/src/optimise/train.jl:105
┌ Info: Epoch 1554
└ @ Main /Users/kcft114/.julia/packages/Flux/qXNjB/src/optimise/train.jl:105
┌ Info: Epoch 1555
└ @ Main /Users/kcft114/.julia/packages/Flux/qXNjB/src/optimise/train.jl:105
┌ Info: Epoch 1556
└ @ Main /Users/kcft114/.julia/packages/Flux/qXNjB/src/optimise/train.jl:105
┌ Info: Epoch 1557
└ @ Main /Users/kcft114/.julia/packages/Flux/qXNjB/src/optimise/train.jl:105
┌ Info: Epoch 1558
└ @ Main /Users/kcft114/.julia/packages/Flux/qXNjB/src/optimise/train.jl:105
┌ Info: Epoch 1559
└ @ Main /Users/kcft114/.julia/packages/Flux/qXNjB/src/optimise/train.jl:105
┌ Info: Epoch 1560
└ @ Main /Users/kcft114/.julia/packages/Flux/qXNjB/src/optimise/train.jl:105
┌ Info: Epoch 1561
└ @ Main /Users/kcft114/.julia/packages/Flux/qXNjB/src/optimise/train.jl:105
┌ Info: Epoch 1562
└ @ Main /Users/kcft114/.julia/packages/Flux/qXNjB/src/optimise/train.jl:105
┌ Info: Epoch 1563
└ @ Main /Users/kcft114/.julia/packages/Flux/qXNjB/src/optimise/train.jl:105
┌ Info: Epoch 1564
└ @ Main /Users/kcft114/.julia/packages/Flux/qXNjB/src/optimise/train.jl:105
┌ Info: Epoch 1565
└ @ Main /Users/kcft114/.julia/packages/Flux/qXNjB/src/optimise/train.jl:105
┌ Info: Epoch 1566
└ @ Main /Users/kcft114/.julia/packages/Flux/qXNjB/src/optimise/train.jl:105
┌ Info: Epoch 1567
└ @ Main /Users/kcft114/.julia/packages/Flux/qXNjB/src/optimise/train.jl:105
┌ Info: Epoch 1568
└ @ Main /Users/kcft114/.julia/packages/Flux/qXNjB/src/optimise/train.jl:105
┌ Info: Epoch 1569
└ @ Main /Users/kcft114/.julia/packages/Flux/qXNjB/src/optimise/train.jl:105
┌ Info: Epoch 1570
└ @ Main /Users/kcft114/.julia/packages/Flux/qXNjB/src/optimise/train.jl:105
┌ Info: Epoch 1571
└ @ Main /Users/kcft114/.julia/packages/Flux/qXNjB/src/optimise/train.jl:105
┌ Info: Epoch 1572
└ @ Main /Users/kcft114/.julia/packages/Flux/qXNjB/src/optimise/train.jl:105
┌ Info: Epoch 1573
└ @ Main /Users/kcft114/.julia/packages/Flux/qXNjB/src/optimise/train.jl:105
┌ Info: Epoch 1574
└ @ Main /Users/kcft114/.julia/packages/Flux/qXNjB/src/optimise/train.jl:105
┌ Info: Epoch 1575
└ @ Main /Users/kcft114/.julia/packages/Flux/qXNjB/src/optimise/train.jl:105
┌ Info: Epoch 1576
└ @ Main /Users/kcft114/.julia/packages/Flux/qXNjB/src/optimise/train.jl:105
┌ Info: Epoch 1577
└ @ Main /Users/kcft114/.julia/packages/Flux/qXNjB/src/optimise/train.jl:105
┌ Info: Epoch 1578
└ @ Main /Users/kcft114/.julia/packages/Flux/qXNjB/src/optimise/train.jl:105
┌ Info: Epoch 1579
└ @ Main /Users/kcft114/.julia/packages/Flux/qXNjB/src/optimise/train.jl:105
┌ Info: Epoch 1580
└ @ Main /Users/kcft114/.julia/packages/Flux/qXNjB/src/optimise/train.jl:105
┌ Info: Epoch 1581
└ @ Main /Users/kcft114/.julia/packages/Flux/qXNjB/src/optimise/train.jl:105
┌ Info: Epoch 1582
└ @ Main /Users/kcft114/.julia/packages/Flux/qXNjB/src/optimise/train.jl:105
┌ Info: Epoch 1583
└ @ Main /Users/kcft114/.julia/packages/Flux/qXNjB/src/optimise/train.jl:105
┌ Info: Epoch 1584
└ @ Main /Users/kcft114/.julia/packages/Flux/qXNjB/src/optimise/train.jl:105
┌ Info: Epoch 1585
└ @ Main /Users/kcft114/.julia/packages/Flux/qXNjB/src/optimise/train.jl:105
┌ Info: Epoch 1586
└ @ Main /Users/kcft114/.julia/packages/Flux/qXNjB/src/optimise/train.jl:105
┌ Info: Epoch 1587
└ @ Main /Users/kcft114/.julia/packages/Flux/qXNjB/src/optimise/train.jl:105
┌ Info: Epoch 1588
└ @ Main /Users/kcft114/.julia/packages/Flux/qXNjB/src/optimise/train.jl:105
┌ Info: Epoch 1589
└ @ Main /Users/kcft114/.julia/packages/Flux/qXNjB/src/optimise/train.jl:105
┌ Info: Epoch 1590
└ @ Main /Users/kcft114/.julia/packages/Flux/qXNjB/src/optimise/train.jl:105
┌ Info: Epoch 1591
└ @ Main /Users/kcft114/.julia/packages/Flux/qXNjB/src/optimise/train.jl:105
┌ Info: Epoch 1592
└ @ Main /Users/kcft114/.julia/packages/Flux/qXNjB/src/optimise/train.jl:105
┌ Info: Epoch 1593
└ @ Main /Users/kcft114/.julia/packages/Flux/qXNjB/src/optimise/train.jl:105
┌ Info: Epoch 1594
└ @ Main /Users/kcft114/.julia/packages/Flux/qXNjB/src/optimise/train.jl:105
┌ Info: Epoch 1595
└ @ Main /Users/kcft114/.julia/packages/Flux/qXNjB/src/optimise/train.jl:105
┌ Info: Epoch 1596
└ @ Main /Users/kcft114/.julia/packages/Flux/qXNjB/src/optimise/train.jl:105
┌ Info: Epoch 1597
└ @ Main /Users/kcft114/.julia/packages/Flux/qXNjB/src/optimise/train.jl:105
┌ Info: Epoch 1598
└ @ Main /Users/kcft114/.julia/packages/Flux/qXNjB/src/optimise/train.jl:105
┌ Info: Epoch 1599
└ @ Main /Users/kcft114/.julia/packages/Flux/qXNjB/src/optimise/train.jl:105
┌ Info: Epoch 1600
└ @ Main /Users/kcft114/.julia/packages/Flux/qXNjB/src/optimise/train.jl:105
┌ Info: Epoch 1601
└ @ Main /Users/kcft114/.julia/packages/Flux/qXNjB/src/optimise/train.jl:105
┌ Info: Epoch 1602
└ @ Main /Users/kcft114/.julia/packages/Flux/qXNjB/src/optimise/train.jl:105
┌ Info: Epoch 1603
└ @ Main /Users/kcft114/.julia/packages/Flux/qXNjB/src/optimise/train.jl:105
┌ Info: Epoch 1604
└ @ Main /Users/kcft114/.julia/packages/Flux/qXNjB/src/optimise/train.jl:105
┌ Info: Epoch 1605
└ @ Main /Users/kcft114/.julia/packages/Flux/qXNjB/src/optimise/train.jl:105
┌ Info: Epoch 1606
└ @ Main /Users/kcft114/.julia/packages/Flux/qXNjB/src/optimise/train.jl:105
┌ Info: Epoch 1607
└ @ Main /Users/kcft114/.julia/packages/Flux/qXNjB/src/optimise/train.jl:105
┌ Info: Epoch 1608
└ @ Main /Users/kcft114/.julia/packages/Flux/qXNjB/src/optimise/train.jl:105
┌ Info: Epoch 1609
└ @ Main /Users/kcft114/.julia/packages/Flux/qXNjB/src/optimise/train.jl:105
┌ Info: Epoch 1610
└ @ Main /Users/kcft114/.julia/packages/Flux/qXNjB/src/optimise/train.jl:105
┌ Info: Epoch 1611
└ @ Main /Users/kcft114/.julia/packages/Flux/qXNjB/src/optimise/train.jl:105
┌ Info: Epoch 1612
└ @ Main /Users/kcft114/.julia/packages/Flux/qXNjB/src/optimise/train.jl:105
┌ Info: Epoch 1613
└ @ Main /Users/kcft114/.julia/packages/Flux/qXNjB/src/optimise/train.jl:105
┌ Info: Epoch 1614
└ @ Main /Users/kcft114/.julia/packages/Flux/qXNjB/src/optimise/train.jl:105
┌ Info: Epoch 1615
└ @ Main /Users/kcft114/.julia/packages/Flux/qXNjB/src/optimise/train.jl:105
┌ Info: Epoch 1616
└ @ Main /Users/kcft114/.julia/packages/Flux/qXNjB/src/optimise/train.jl:105
┌ Info: Epoch 1617
└ @ Main /Users/kcft114/.julia/packages/Flux/qXNjB/src/optimise/train.jl:105
┌ Info: Epoch 1618
└ @ Main /Users/kcft114/.julia/packages/Flux/qXNjB/src/optimise/train.jl:105
┌ Info: Epoch 1619
└ @ Main /Users/kcft114/.julia/packages/Flux/qXNjB/src/optimise/train.jl:105
┌ Info: Epoch 1620
└ @ Main /Users/kcft114/.julia/packages/Flux/qXNjB/src/optimise/train.jl:105
┌ Info: Epoch 1621
└ @ Main /Users/kcft114/.julia/packages/Flux/qXNjB/src/optimise/train.jl:105
┌ Info: Epoch 1622
└ @ Main /Users/kcft114/.julia/packages/Flux/qXNjB/src/optimise/train.jl:105
┌ Info: Epoch 1623
└ @ Main /Users/kcft114/.julia/packages/Flux/qXNjB/src/optimise/train.jl:105
┌ Info: Epoch 1624
└ @ Main /Users/kcft114/.julia/packages/Flux/qXNjB/src/optimise/train.jl:105
┌ Info: Epoch 1625
└ @ Main /Users/kcft114/.julia/packages/Flux/qXNjB/src/optimise/train.jl:105
┌ Info: Epoch 1626
└ @ Main /Users/kcft114/.julia/packages/Flux/qXNjB/src/optimise/train.jl:105
┌ Info: Epoch 1627
└ @ Main /Users/kcft114/.julia/packages/Flux/qXNjB/src/optimise/train.jl:105
┌ Info: Epoch 1628
└ @ Main /Users/kcft114/.julia/packages/Flux/qXNjB/src/optimise/train.jl:105
┌ Info: Epoch 1629
└ @ Main /Users/kcft114/.julia/packages/Flux/qXNjB/src/optimise/train.jl:105
┌ Info: Epoch 1630
└ @ Main /Users/kcft114/.julia/packages/Flux/qXNjB/src/optimise/train.jl:105
```

```
loss(X, Y) = 0.5268678f0 (tracked)
loss(X, Y) = 0.59836096f0 (tracked)
loss(X, Y) = 0.5217467f0 (tracked)
loss(X, Y) = 0.60797095f0 (tracked)
loss(X, Y) = 0.5765486f0 (tracked)
loss(X, Y) = 0.5132886f0 (tracked)
loss(X, Y) = 0.54431874f0 (tracked)
loss(X, Y) = 0.5913559f0 (tracked)
loss(X, Y) = 0.5273861f0 (tracked)
loss(X, Y) = 0.52789277f0 (tracked)
loss(X, Y) = 0.5264673f0 (tracked)
loss(X, Y) = 0.527552f0 (tracked)
loss(X, Y) = 0.5680058f0 (tracked)
loss(X, Y) = 0.59911615f0 (tracked)
loss(X, Y) = 0.57512474f0 (tracked)
loss(X, Y) = 0.5626529f0 (tracked)
loss(X, Y) = 0.5351677f0 (tracked)
loss(X, Y) = 0.5425221f0 (tracked)
loss(X, Y) = 0.610958f0 (tracked)
loss(X, Y) = 0.54486537f0 (tracked)
loss(X, Y) = 0.56564826f0 (tracked)
loss(X, Y) = 0.53813773f0 (tracked)
loss(X, Y) = 0.49973872f0 (tracked)
loss(X, Y) = 0.5510776f0 (tracked)
loss(X, Y) = 0.5491359f0 (tracked)
loss(X, Y) = 0.55009425f0 (tracked)
loss(X, Y) = 0.54155946f0 (tracked)
loss(X, Y) = 0.5593633f0 (tracked)
loss(X, Y) = 0.5535161f0 (tracked)
loss(X, Y) = 0.5682498f0 (tracked)
loss(X, Y) = 0.6207022f0 (tracked)
loss(X, Y) = 0.55288553f0 (tracked)
loss(X, Y) = 0.5649689f0 (tracked)
loss(X, Y) = 0.5719933f0 (tracked)
loss(X, Y) = 0.60865194f0 (tracked)
loss(X, Y) = 0.5270776f0 (tracked)
loss(X, Y) = 0.5358741f0 (tracked)
loss(X, Y) = 0.531377f0 (tracked)
loss(X, Y) = 0.5504799f0 (tracked)
loss(X, Y) = 0.5856249f0 (tracked)
loss(X, Y) = 0.5775031f0 (tracked)
loss(X, Y) = 0.55902815f0 (tracked)
loss(X, Y) = 0.5668751f0 (tracked)
loss(X, Y) = 0.5295308f0 (tracked)
loss(X, Y) = 0.5737778f0 (tracked)
loss(X, Y) = 0.588193f0 (tracked)
loss(X, Y) = 0.5618905f0 (tracked)
loss(X, Y) = 0.5322411f0 (tracked)
loss(X, Y) = 0.49850205f0 (tracked)
loss(X, Y) = 0.51030624f0 (tracked)
loss(X, Y) = 0.56854826f0 (tracked)
loss(X, Y) = 0.50128525f0 (tracked)
loss(X, Y) = 0.57561284f0 (tracked)
loss(X, Y) = 0.52766556f0 (tracked)
loss(X, Y) = 0.55730367f0 (tracked)
loss(X, Y) = 0.54890096f0 (tracked)
loss(X, Y) = 0.5226309f0 (tracked)
loss(X, Y) = 0.5305829f0 (tracked)
loss(X, Y) = 0.56642205f0 (tracked)
loss(X, Y) = 0.5253218f0 (tracked)
loss(X, Y) = 0.53049666f0 (tracked)
loss(X, Y) = 0.5406304f0 (tracked)
loss(X, Y) = 0.5005875f0 (tracked)
loss(X, Y) = 0.52996004f0 (tracked)
loss(X, Y) = 0.50587434f0 (tracked)
loss(X, Y) = 0.500908f0 (tracked)
loss(X, Y) = 0.5607283f0 (tracked)
loss(X, Y) = 0.55508745f0 (tracked)
loss(X, Y) = 0.5714572f0 (tracked)
loss(X, Y) = 0.5318408f0 (tracked)
loss(X, Y) = 0.55439144f0 (tracked)
loss(X, Y) = 0.56320673f0 (tracked)
loss(X, Y) = 0.604806f0 (tracked)
loss(X, Y) = 0.53126687f0 (tracked)
loss(X, Y) = 0.59155643f0 (tracked)
loss(X, Y) = 0.5052094f0 (tracked)
loss(X, Y) = 0.57188606f0 (tracked)
loss(X, Y) = 0.5302607f0 (tracked)
loss(X, Y) = 0.52343637f0 (tracked)
loss(X, Y) = 0.5028476f0 (tracked)
loss(X, Y) = 0.54218274f0 (tracked)
loss(X, Y) = 0.53776646f0 (tracked)
loss(X, Y) = 0.5570664f0 (tracked)
loss(X, Y) = 0.5314999f0 (tracked)
loss(X, Y) = 0.5428793f0 (tracked)
loss(X, Y) = 0.5234865f0 (tracked)
loss(X, Y) = 0.5547617f0 (tracked)
loss(X, Y) = 0.522447f0 (tracked)
loss(X, Y) = 0.52942306f0 (tracked)
loss(X, Y) = 0.5779274f0 (tracked)
loss(X, Y) = 0.61520237f0 (tracked)
loss(X, Y) = 0.5305041f0 (tracked)
loss(X, Y) = 0.54684895f0 (tracked)
loss(X, Y) = 0.518372f0 (tracked)
loss(X, Y) = 0.5955378f0 (tracked)
loss(X, Y) = 0.5421176f0 (tracked)
loss(X, Y) = 0.5271924f0 (tracked)
loss(X, Y) = 0.52481693f0 (tracked)
loss(X, Y) = 0.5037967f0 (tracked)
```

```
┌ Info: Epoch 1631
└ @ Main /Users/kcft114/.julia/packages/Flux/qXNjB/src/optimise/train.jl:105
┌ Info: Epoch 1632
└ @ Main /Users/kcft114/.julia/packages/Flux/qXNjB/src/optimise/train.jl:105
┌ Info: Epoch 1633
└ @ Main /Users/kcft114/.julia/packages/Flux/qXNjB/src/optimise/train.jl:105
┌ Info: Epoch 1634
└ @ Main /Users/kcft114/.julia/packages/Flux/qXNjB/src/optimise/train.jl:105
┌ Info: Epoch 1635
└ @ Main /Users/kcft114/.julia/packages/Flux/qXNjB/src/optimise/train.jl:105
┌ Info: Epoch 1636
└ @ Main /Users/kcft114/.julia/packages/Flux/qXNjB/src/optimise/train.jl:105
┌ Info: Epoch 1637
└ @ Main /Users/kcft114/.julia/packages/Flux/qXNjB/src/optimise/train.jl:105
┌ Info: Epoch 1638
└ @ Main /Users/kcft114/.julia/packages/Flux/qXNjB/src/optimise/train.jl:105
┌ Info: Epoch 1639
└ @ Main /Users/kcft114/.julia/packages/Flux/qXNjB/src/optimise/train.jl:105
┌ Info: Epoch 1640
└ @ Main /Users/kcft114/.julia/packages/Flux/qXNjB/src/optimise/train.jl:105
┌ Info: Epoch 1641
└ @ Main /Users/kcft114/.julia/packages/Flux/qXNjB/src/optimise/train.jl:105
┌ Info: Epoch 1642
└ @ Main /Users/kcft114/.julia/packages/Flux/qXNjB/src/optimise/train.jl:105
┌ Info: Epoch 1643
└ @ Main /Users/kcft114/.julia/packages/Flux/qXNjB/src/optimise/train.jl:105
┌ Info: Epoch 1644
└ @ Main /Users/kcft114/.julia/packages/Flux/qXNjB/src/optimise/train.jl:105
┌ Info: Epoch 1645
└ @ Main /Users/kcft114/.julia/packages/Flux/qXNjB/src/optimise/train.jl:105
┌ Info: Epoch 1646
└ @ Main /Users/kcft114/.julia/packages/Flux/qXNjB/src/optimise/train.jl:105
┌ Info: Epoch 1647
└ @ Main /Users/kcft114/.julia/packages/Flux/qXNjB/src/optimise/train.jl:105
┌ Info: Epoch 1648
└ @ Main /Users/kcft114/.julia/packages/Flux/qXNjB/src/optimise/train.jl:105
┌ Info: Epoch 1649
└ @ Main /Users/kcft114/.julia/packages/Flux/qXNjB/src/optimise/train.jl:105
┌ Info: Epoch 1650
└ @ Main /Users/kcft114/.julia/packages/Flux/qXNjB/src/optimise/train.jl:105
┌ Info: Epoch 1651
└ @ Main /Users/kcft114/.julia/packages/Flux/qXNjB/src/optimise/train.jl:105
┌ Info: Epoch 1652
└ @ Main /Users/kcft114/.julia/packages/Flux/qXNjB/src/optimise/train.jl:105
┌ Info: Epoch 1653
└ @ Main /Users/kcft114/.julia/packages/Flux/qXNjB/src/optimise/train.jl:105
┌ Info: Epoch 1654
└ @ Main /Users/kcft114/.julia/packages/Flux/qXNjB/src/optimise/train.jl:105
┌ Info: Epoch 1655
└ @ Main /Users/kcft114/.julia/packages/Flux/qXNjB/src/optimise/train.jl:105
┌ Info: Epoch 1656
└ @ Main /Users/kcft114/.julia/packages/Flux/qXNjB/src/optimise/train.jl:105
┌ Info: Epoch 1657
└ @ Main /Users/kcft114/.julia/packages/Flux/qXNjB/src/optimise/train.jl:105
┌ Info: Epoch 1658
└ @ Main /Users/kcft114/.julia/packages/Flux/qXNjB/src/optimise/train.jl:105
┌ Info: Epoch 1659
└ @ Main /Users/kcft114/.julia/packages/Flux/qXNjB/src/optimise/train.jl:105
┌ Info: Epoch 1660
└ @ Main /Users/kcft114/.julia/packages/Flux/qXNjB/src/optimise/train.jl:105
┌ Info: Epoch 1661
└ @ Main /Users/kcft114/.julia/packages/Flux/qXNjB/src/optimise/train.jl:105
┌ Info: Epoch 1662
└ @ Main /Users/kcft114/.julia/packages/Flux/qXNjB/src/optimise/train.jl:105
┌ Info: Epoch 1663
└ @ Main /Users/kcft114/.julia/packages/Flux/qXNjB/src/optimise/train.jl:105
┌ Info: Epoch 1664
└ @ Main /Users/kcft114/.julia/packages/Flux/qXNjB/src/optimise/train.jl:105
┌ Info: Epoch 1665
└ @ Main /Users/kcft114/.julia/packages/Flux/qXNjB/src/optimise/train.jl:105
┌ Info: Epoch 1666
└ @ Main /Users/kcft114/.julia/packages/Flux/qXNjB/src/optimise/train.jl:105
┌ Info: Epoch 1667
└ @ Main /Users/kcft114/.julia/packages/Flux/qXNjB/src/optimise/train.jl:105
┌ Info: Epoch 1668
└ @ Main /Users/kcft114/.julia/packages/Flux/qXNjB/src/optimise/train.jl:105
┌ Info: Epoch 1669
└ @ Main /Users/kcft114/.julia/packages/Flux/qXNjB/src/optimise/train.jl:105
┌ Info: Epoch 1670
└ @ Main /Users/kcft114/.julia/packages/Flux/qXNjB/src/optimise/train.jl:105
┌ Info: Epoch 1671
└ @ Main /Users/kcft114/.julia/packages/Flux/qXNjB/src/optimise/train.jl:105
┌ Info: Epoch 1672
└ @ Main /Users/kcft114/.julia/packages/Flux/qXNjB/src/optimise/train.jl:105
┌ Info: Epoch 1673
└ @ Main /Users/kcft114/.julia/packages/Flux/qXNjB/src/optimise/train.jl:105
┌ Info: Epoch 1674
└ @ Main /Users/kcft114/.julia/packages/Flux/qXNjB/src/optimise/train.jl:105
┌ Info: Epoch 1675
└ @ Main /Users/kcft114/.julia/packages/Flux/qXNjB/src/optimise/train.jl:105
┌ Info: Epoch 1676
└ @ Main /Users/kcft114/.julia/packages/Flux/qXNjB/src/optimise/train.jl:105
┌ Info: Epoch 1677
└ @ Main /Users/kcft114/.julia/packages/Flux/qXNjB/src/optimise/train.jl:105
┌ Info: Epoch 1678
└ @ Main /Users/kcft114/.julia/packages/Flux/qXNjB/src/optimise/train.jl:105
┌ Info: Epoch 1679
└ @ Main /Users/kcft114/.julia/packages/Flux/qXNjB/src/optimise/train.jl:105
┌ Info: Epoch 1680
└ @ Main /Users/kcft114/.julia/packages/Flux/qXNjB/src/optimise/train.jl:105
┌ Info: Epoch 1681
└ @ Main /Users/kcft114/.julia/packages/Flux/qXNjB/src/optimise/train.jl:105
┌ Info: Epoch 1682
└ @ Main /Users/kcft114/.julia/packages/Flux/qXNjB/src/optimise/train.jl:105
┌ Info: Epoch 1683
└ @ Main /Users/kcft114/.julia/packages/Flux/qXNjB/src/optimise/train.jl:105
┌ Info: Epoch 1684
└ @ Main /Users/kcft114/.julia/packages/Flux/qXNjB/src/optimise/train.jl:105
┌ Info: Epoch 1685
└ @ Main /Users/kcft114/.julia/packages/Flux/qXNjB/src/optimise/train.jl:105
┌ Info: Epoch 1686
└ @ Main /Users/kcft114/.julia/packages/Flux/qXNjB/src/optimise/train.jl:105
┌ Info: Epoch 1687
└ @ Main /Users/kcft114/.julia/packages/Flux/qXNjB/src/optimise/train.jl:105
┌ Info: Epoch 1688
└ @ Main /Users/kcft114/.julia/packages/Flux/qXNjB/src/optimise/train.jl:105
┌ Info: Epoch 1689
└ @ Main /Users/kcft114/.julia/packages/Flux/qXNjB/src/optimise/train.jl:105
┌ Info: Epoch 1690
└ @ Main /Users/kcft114/.julia/packages/Flux/qXNjB/src/optimise/train.jl:105
┌ Info: Epoch 1691
└ @ Main /Users/kcft114/.julia/packages/Flux/qXNjB/src/optimise/train.jl:105
┌ Info: Epoch 1692
└ @ Main /Users/kcft114/.julia/packages/Flux/qXNjB/src/optimise/train.jl:105
┌ Info: Epoch 1693
└ @ Main /Users/kcft114/.julia/packages/Flux/qXNjB/src/optimise/train.jl:105
┌ Info: Epoch 1694
└ @ Main /Users/kcft114/.julia/packages/Flux/qXNjB/src/optimise/train.jl:105
┌ Info: Epoch 1695
└ @ Main /Users/kcft114/.julia/packages/Flux/qXNjB/src/optimise/train.jl:105
┌ Info: Epoch 1696
└ @ Main /Users/kcft114/.julia/packages/Flux/qXNjB/src/optimise/train.jl:105
┌ Info: Epoch 1697
└ @ Main /Users/kcft114/.julia/packages/Flux/qXNjB/src/optimise/train.jl:105
┌ Info: Epoch 1698
└ @ Main /Users/kcft114/.julia/packages/Flux/qXNjB/src/optimise/train.jl:105
┌ Info: Epoch 1699
└ @ Main /Users/kcft114/.julia/packages/Flux/qXNjB/src/optimise/train.jl:105
┌ Info: Epoch 1700
└ @ Main /Users/kcft114/.julia/packages/Flux/qXNjB/src/optimise/train.jl:105
┌ Info: Epoch 1701
└ @ Main /Users/kcft114/.julia/packages/Flux/qXNjB/src/optimise/train.jl:105
┌ Info: Epoch 1702
└ @ Main /Users/kcft114/.julia/packages/Flux/qXNjB/src/optimise/train.jl:105
┌ Info: Epoch 1703
└ @ Main /Users/kcft114/.julia/packages/Flux/qXNjB/src/optimise/train.jl:105
┌ Info: Epoch 1704
└ @ Main /Users/kcft114/.julia/packages/Flux/qXNjB/src/optimise/train.jl:105
┌ Info: Epoch 1705
└ @ Main /Users/kcft114/.julia/packages/Flux/qXNjB/src/optimise/train.jl:105
┌ Info: Epoch 1706
└ @ Main /Users/kcft114/.julia/packages/Flux/qXNjB/src/optimise/train.jl:105
┌ Info: Epoch 1707
└ @ Main /Users/kcft114/.julia/packages/Flux/qXNjB/src/optimise/train.jl:105
┌ Info: Epoch 1708
└ @ Main /Users/kcft114/.julia/packages/Flux/qXNjB/src/optimise/train.jl:105
┌ Info: Epoch 1709
└ @ Main /Users/kcft114/.julia/packages/Flux/qXNjB/src/optimise/train.jl:105
┌ Info: Epoch 1710
└ @ Main /Users/kcft114/.julia/packages/Flux/qXNjB/src/optimise/train.jl:105
┌ Info: Epoch 1711
└ @ Main /Users/kcft114/.julia/packages/Flux/qXNjB/src/optimise/train.jl:105
┌ Info: Epoch 1712
└ @ Main /Users/kcft114/.julia/packages/Flux/qXNjB/src/optimise/train.jl:105
┌ Info: Epoch 1713
└ @ Main /Users/kcft114/.julia/packages/Flux/qXNjB/src/optimise/train.jl:105
┌ Info: Epoch 1714
└ @ Main /Users/kcft114/.julia/packages/Flux/qXNjB/src/optimise/train.jl:105
┌ Info: Epoch 1715
└ @ Main /Users/kcft114/.julia/packages/Flux/qXNjB/src/optimise/train.jl:105
┌ Info: Epoch 1716
└ @ Main /Users/kcft114/.julia/packages/Flux/qXNjB/src/optimise/train.jl:105
┌ Info: Epoch 1717
└ @ Main /Users/kcft114/.julia/packages/Flux/qXNjB/src/optimise/train.jl:105
┌ Info: Epoch 1718
└ @ Main /Users/kcft114/.julia/packages/Flux/qXNjB/src/optimise/train.jl:105
┌ Info: Epoch 1719
└ @ Main /Users/kcft114/.julia/packages/Flux/qXNjB/src/optimise/train.jl:105
┌ Info: Epoch 1720
└ @ Main /Users/kcft114/.julia/packages/Flux/qXNjB/src/optimise/train.jl:105
┌ Info: Epoch 1721
└ @ Main /Users/kcft114/.julia/packages/Flux/qXNjB/src/optimise/train.jl:105
┌ Info: Epoch 1722
└ @ Main /Users/kcft114/.julia/packages/Flux/qXNjB/src/optimise/train.jl:105
┌ Info: Epoch 1723
└ @ Main /Users/kcft114/.julia/packages/Flux/qXNjB/src/optimise/train.jl:105
┌ Info: Epoch 1724
└ @ Main /Users/kcft114/.julia/packages/Flux/qXNjB/src/optimise/train.jl:105
┌ Info: Epoch 1725
└ @ Main /Users/kcft114/.julia/packages/Flux/qXNjB/src/optimise/train.jl:105
┌ Info: Epoch 1726
└ @ Main /Users/kcft114/.julia/packages/Flux/qXNjB/src/optimise/train.jl:105
┌ Info: Epoch 1727
└ @ Main /Users/kcft114/.julia/packages/Flux/qXNjB/src/optimise/train.jl:105
┌ Info: Epoch 1728
└ @ Main /Users/kcft114/.julia/packages/Flux/qXNjB/src/optimise/train.jl:105
┌ Info: Epoch 1729
└ @ Main /Users/kcft114/.julia/packages/Flux/qXNjB/src/optimise/train.jl:105
┌ Info: Epoch 1730
└ @ Main /Users/kcft114/.julia/packages/Flux/qXNjB/src/optimise/train.jl:105
```

```
loss(X, Y) = 0.5292207f0 (tracked)
loss(X, Y) = 0.5089128f0 (tracked)
loss(X, Y) = 0.5063991f0 (tracked)
loss(X, Y) = 0.5016215f0 (tracked)
loss(X, Y) = 0.5667775f0 (tracked)
loss(X, Y) = 0.58505976f0 (tracked)
loss(X, Y) = 0.59936917f0 (tracked)
loss(X, Y) = 0.52164763f0 (tracked)
loss(X, Y) = 0.5004062f0 (tracked)
loss(X, Y) = 0.5315197f0 (tracked)
loss(X, Y) = 0.61700237f0 (tracked)
loss(X, Y) = 0.5004513f0 (tracked)
loss(X, Y) = 0.49816152f0 (tracked)
loss(X, Y) = 0.5341637f0 (tracked)
loss(X, Y) = 0.54208004f0 (tracked)
loss(X, Y) = 0.49803358f0 (tracked)
loss(X, Y) = 0.56928575f0 (tracked)
loss(X, Y) = 0.5439523f0 (tracked)
loss(X, Y) = 0.5449157f0 (tracked)
loss(X, Y) = 0.5267422f0 (tracked)
loss(X, Y) = 0.49432725f0 (tracked)
loss(X, Y) = 0.59515345f0 (tracked)
loss(X, Y) = 0.5500599f0 (tracked)
loss(X, Y) = 0.5118658f0 (tracked)
loss(X, Y) = 0.5492384f0 (tracked)
loss(X, Y) = 0.4684284f0 (tracked)
loss(X, Y) = 0.5307101f0 (tracked)
loss(X, Y) = 0.4865106f0 (tracked)
loss(X, Y) = 0.5310927f0 (tracked)
loss(X, Y) = 0.55773413f0 (tracked)
loss(X, Y) = 0.55052537f0 (tracked)
loss(X, Y) = 0.58839107f0 (tracked)
loss(X, Y) = 0.53011566f0 (tracked)
loss(X, Y) = 0.5242881f0 (tracked)
loss(X, Y) = 0.52833474f0 (tracked)
loss(X, Y) = 0.5520381f0 (tracked)
loss(X, Y) = 0.56166005f0 (tracked)
loss(X, Y) = 0.51222616f0 (tracked)
loss(X, Y) = 0.5203462f0 (tracked)
loss(X, Y) = 0.5791419f0 (tracked)
loss(X, Y) = 0.5684855f0 (tracked)
loss(X, Y) = 0.56393224f0 (tracked)
loss(X, Y) = 0.52555346f0 (tracked)
loss(X, Y) = 0.58420414f0 (tracked)
loss(X, Y) = 0.5519081f0 (tracked)
loss(X, Y) = 0.5305409f0 (tracked)
loss(X, Y) = 0.5218871f0 (tracked)
loss(X, Y) = 0.5645043f0 (tracked)
loss(X, Y) = 0.45533258f0 (tracked)
loss(X, Y) = 0.509243f0 (tracked)
loss(X, Y) = 0.51424944f0 (tracked)
loss(X, Y) = 0.5672799f0 (tracked)
loss(X, Y) = 0.5475983f0 (tracked)
loss(X, Y) = 0.49977273f0 (tracked)
loss(X, Y) = 0.55203366f0 (tracked)
loss(X, Y) = 0.5515814f0 (tracked)
loss(X, Y) = 0.53603995f0 (tracked)
loss(X, Y) = 0.5098086f0 (tracked)
loss(X, Y) = 0.50954056f0 (tracked)
loss(X, Y) = 0.5441899f0 (tracked)
loss(X, Y) = 0.5521936f0 (tracked)
loss(X, Y) = 0.5707014f0 (tracked)
loss(X, Y) = 0.5199535f0 (tracked)
loss(X, Y) = 0.5490095f0 (tracked)
loss(X, Y) = 0.51593965f0 (tracked)
loss(X, Y) = 0.49848616f0 (tracked)
loss(X, Y) = 0.49985126f0 (tracked)
loss(X, Y) = 0.5026869f0 (tracked)
loss(X, Y) = 0.5499277f0 (tracked)
loss(X, Y) = 0.51011777f0 (tracked)
loss(X, Y) = 0.57388663f0 (tracked)
loss(X, Y) = 0.5067123f0 (tracked)
loss(X, Y) = 0.53657055f0 (tracked)
loss(X, Y) = 0.50100935f0 (tracked)
loss(X, Y) = 0.5210158f0 (tracked)
loss(X, Y) = 0.5319103f0 (tracked)
loss(X, Y) = 0.51826835f0 (tracked)
loss(X, Y) = 0.5436712f0 (tracked)
loss(X, Y) = 0.56395817f0 (tracked)
loss(X, Y) = 0.53284186f0 (tracked)
loss(X, Y) = 0.5610746f0 (tracked)
loss(X, Y) = 0.5573498f0 (tracked)
loss(X, Y) = 0.5136747f0 (tracked)
loss(X, Y) = 0.56544584f0 (tracked)
loss(X, Y) = 0.47318453f0 (tracked)
loss(X, Y) = 0.5455121f0 (tracked)
loss(X, Y) = 0.53505117f0 (tracked)
loss(X, Y) = 0.5015188f0 (tracked)
loss(X, Y) = 0.5925925f0 (tracked)
```

```
┌ Info: Epoch 1731
└ @ Main /Users/kcft114/.julia/packages/Flux/qXNjB/src/optimise/train.jl:105
┌ Info: Epoch 1732
└ @ Main /Users/kcft114/.julia/packages/Flux/qXNjB/src/optimise/train.jl:105
┌ Info: Epoch 1733
└ @ Main /Users/kcft114/.julia/packages/Flux/qXNjB/src/optimise/train.jl:105
┌ Info: Epoch 1734
└ @ Main /Users/kcft114/.julia/packages/Flux/qXNjB/src/optimise/train.jl:105
┌ Info: Epoch 1735
└ @ Main /Users/kcft114/.julia/packages/Flux/qXNjB/src/optimise/train.jl:105
┌ Info: Epoch 1736
└ @ Main /Users/kcft114/.julia/packages/Flux/qXNjB/src/optimise/train.jl:105
┌ Info: Epoch 1737
└ @ Main /Users/kcft114/.julia/packages/Flux/qXNjB/src/optimise/train.jl:105
┌ Info: Epoch 1738
└ @ Main /Users/kcft114/.julia/packages/Flux/qXNjB/src/optimise/train.jl:105
┌ Info: Epoch 1739
└ @ Main /Users/kcft114/.julia/packages/Flux/qXNjB/src/optimise/train.jl:105
┌ Info: Epoch 1740
└ @ Main /Users/kcft114/.julia/packages/Flux/qXNjB/src/optimise/train.jl:105
┌ Info: Epoch 1741
└ @ Main /Users/kcft114/.julia/packages/Flux/qXNjB/src/optimise/train.jl:105
┌ Info: Epoch 1742
└ @ Main /Users/kcft114/.julia/packages/Flux/qXNjB/src/optimise/train.jl:105
┌ Info: Epoch 1743
└ @ Main /Users/kcft114/.julia/packages/Flux/qXNjB/src/optimise/train.jl:105
┌ Info: Epoch 1744
└ @ Main /Users/kcft114/.julia/packages/Flux/qXNjB/src/optimise/train.jl:105
┌ Info: Epoch 1745
└ @ Main /Users/kcft114/.julia/packages/Flux/qXNjB/src/optimise/train.jl:105
┌ Info: Epoch 1746
└ @ Main /Users/kcft114/.julia/packages/Flux/qXNjB/src/optimise/train.jl:105
┌ Info: Epoch 1747
└ @ Main /Users/kcft114/.julia/packages/Flux/qXNjB/src/optimise/train.jl:105
┌ Info: Epoch 1748
└ @ Main /Users/kcft114/.julia/packages/Flux/qXNjB/src/optimise/train.jl:105
┌ Info: Epoch 1749
└ @ Main /Users/kcft114/.julia/packages/Flux/qXNjB/src/optimise/train.jl:105
┌ Info: Epoch 1750
└ @ Main /Users/kcft114/.julia/packages/Flux/qXNjB/src/optimise/train.jl:105
┌ Info: Epoch 1751
└ @ Main /Users/kcft114/.julia/packages/Flux/qXNjB/src/optimise/train.jl:105
┌ Info: Epoch 1752
└ @ Main /Users/kcft114/.julia/packages/Flux/qXNjB/src/optimise/train.jl:105
┌ Info: Epoch 1753
└ @ Main /Users/kcft114/.julia/packages/Flux/qXNjB/src/optimise/train.jl:105
┌ Info: Epoch 1754
└ @ Main /Users/kcft114/.julia/packages/Flux/qXNjB/src/optimise/train.jl:105
┌ Info: Epoch 1755
└ @ Main /Users/kcft114/.julia/packages/Flux/qXNjB/src/optimise/train.jl:105
┌ Info: Epoch 1756
└ @ Main /Users/kcft114/.julia/packages/Flux/qXNjB/src/optimise/train.jl:105
┌ Info: Epoch 1757
└ @ Main /Users/kcft114/.julia/packages/Flux/qXNjB/src/optimise/train.jl:105
┌ Info: Epoch 1758
└ @ Main /Users/kcft114/.julia/packages/Flux/qXNjB/src/optimise/train.jl:105
┌ Info: Epoch 1759
└ @ Main /Users/kcft114/.julia/packages/Flux/qXNjB/src/optimise/train.jl:105
┌ Info: Epoch 1760
└ @ Main /Users/kcft114/.julia/packages/Flux/qXNjB/src/optimise/train.jl:105
┌ Info: Epoch 1761
└ @ Main /Users/kcft114/.julia/packages/Flux/qXNjB/src/optimise/train.jl:105
┌ Info: Epoch 1762
└ @ Main /Users/kcft114/.julia/packages/Flux/qXNjB/src/optimise/train.jl:105
┌ Info: Epoch 1763
└ @ Main /Users/kcft114/.julia/packages/Flux/qXNjB/src/optimise/train.jl:105
┌ Info: Epoch 1764
└ @ Main /Users/kcft114/.julia/packages/Flux/qXNjB/src/optimise/train.jl:105
┌ Info: Epoch 1765
└ @ Main /Users/kcft114/.julia/packages/Flux/qXNjB/src/optimise/train.jl:105
┌ Info: Epoch 1766
└ @ Main /Users/kcft114/.julia/packages/Flux/qXNjB/src/optimise/train.jl:105
┌ Info: Epoch 1767
└ @ Main /Users/kcft114/.julia/packages/Flux/qXNjB/src/optimise/train.jl:105
┌ Info: Epoch 1768
└ @ Main /Users/kcft114/.julia/packages/Flux/qXNjB/src/optimise/train.jl:105
┌ Info: Epoch 1769
└ @ Main /Users/kcft114/.julia/packages/Flux/qXNjB/src/optimise/train.jl:105
┌ Info: Epoch 1770
└ @ Main /Users/kcft114/.julia/packages/Flux/qXNjB/src/optimise/train.jl:105
┌ Info: Epoch 1771
└ @ Main /Users/kcft114/.julia/packages/Flux/qXNjB/src/optimise/train.jl:105
┌ Info: Epoch 1772
└ @ Main /Users/kcft114/.julia/packages/Flux/qXNjB/src/optimise/train.jl:105
┌ Info: Epoch 1773
└ @ Main /Users/kcft114/.julia/packages/Flux/qXNjB/src/optimise/train.jl:105
┌ Info: Epoch 1774
└ @ Main /Users/kcft114/.julia/packages/Flux/qXNjB/src/optimise/train.jl:105
┌ Info: Epoch 1775
└ @ Main /Users/kcft114/.julia/packages/Flux/qXNjB/src/optimise/train.jl:105
┌ Info: Epoch 1776
└ @ Main /Users/kcft114/.julia/packages/Flux/qXNjB/src/optimise/train.jl:105
┌ Info: Epoch 1777
└ @ Main /Users/kcft114/.julia/packages/Flux/qXNjB/src/optimise/train.jl:105
┌ Info: Epoch 1778
└ @ Main /Users/kcft114/.julia/packages/Flux/qXNjB/src/optimise/train.jl:105
┌ Info: Epoch 1779
└ @ Main /Users/kcft114/.julia/packages/Flux/qXNjB/src/optimise/train.jl:105
┌ Info: Epoch 1780
└ @ Main /Users/kcft114/.julia/packages/Flux/qXNjB/src/optimise/train.jl:105
┌ Info: Epoch 1781
└ @ Main /Users/kcft114/.julia/packages/Flux/qXNjB/src/optimise/train.jl:105
┌ Info: Epoch 1782
└ @ Main /Users/kcft114/.julia/packages/Flux/qXNjB/src/optimise/train.jl:105
┌ Info: Epoch 1783
└ @ Main /Users/kcft114/.julia/packages/Flux/qXNjB/src/optimise/train.jl:105
┌ Info: Epoch 1784
└ @ Main /Users/kcft114/.julia/packages/Flux/qXNjB/src/optimise/train.jl:105
┌ Info: Epoch 1785
└ @ Main /Users/kcft114/.julia/packages/Flux/qXNjB/src/optimise/train.jl:105
┌ Info: Epoch 1786
└ @ Main /Users/kcft114/.julia/packages/Flux/qXNjB/src/optimise/train.jl:105
┌ Info: Epoch 1787
└ @ Main /Users/kcft114/.julia/packages/Flux/qXNjB/src/optimise/train.jl:105
┌ Info: Epoch 1788
└ @ Main /Users/kcft114/.julia/packages/Flux/qXNjB/src/optimise/train.jl:105
┌ Info: Epoch 1789
└ @ Main /Users/kcft114/.julia/packages/Flux/qXNjB/src/optimise/train.jl:105
┌ Info: Epoch 1790
└ @ Main /Users/kcft114/.julia/packages/Flux/qXNjB/src/optimise/train.jl:105
┌ Info: Epoch 1791
└ @ Main /Users/kcft114/.julia/packages/Flux/qXNjB/src/optimise/train.jl:105
┌ Info: Epoch 1792
└ @ Main /Users/kcft114/.julia/packages/Flux/qXNjB/src/optimise/train.jl:105
┌ Info: Epoch 1793
└ @ Main /Users/kcft114/.julia/packages/Flux/qXNjB/src/optimise/train.jl:105
┌ Info: Epoch 1794
└ @ Main /Users/kcft114/.julia/packages/Flux/qXNjB/src/optimise/train.jl:105
┌ Info: Epoch 1795
└ @ Main /Users/kcft114/.julia/packages/Flux/qXNjB/src/optimise/train.jl:105
┌ Info: Epoch 1796
└ @ Main /Users/kcft114/.julia/packages/Flux/qXNjB/src/optimise/train.jl:105
┌ Info: Epoch 1797
└ @ Main /Users/kcft114/.julia/packages/Flux/qXNjB/src/optimise/train.jl:105
┌ Info: Epoch 1798
└ @ Main /Users/kcft114/.julia/packages/Flux/qXNjB/src/optimise/train.jl:105
┌ Info: Epoch 1799
└ @ Main /Users/kcft114/.julia/packages/Flux/qXNjB/src/optimise/train.jl:105
┌ Info: Epoch 1800
└ @ Main /Users/kcft114/.julia/packages/Flux/qXNjB/src/optimise/train.jl:105
┌ Info: Epoch 1801
└ @ Main /Users/kcft114/.julia/packages/Flux/qXNjB/src/optimise/train.jl:105
┌ Info: Epoch 1802
└ @ Main /Users/kcft114/.julia/packages/Flux/qXNjB/src/optimise/train.jl:105
┌ Info: Epoch 1803
└ @ Main /Users/kcft114/.julia/packages/Flux/qXNjB/src/optimise/train.jl:105
┌ Info: Epoch 1804
└ @ Main /Users/kcft114/.julia/packages/Flux/qXNjB/src/optimise/train.jl:105
┌ Info: Epoch 1805
└ @ Main /Users/kcft114/.julia/packages/Flux/qXNjB/src/optimise/train.jl:105
┌ Info: Epoch 1806
└ @ Main /Users/kcft114/.julia/packages/Flux/qXNjB/src/optimise/train.jl:105
┌ Info: Epoch 1807
└ @ Main /Users/kcft114/.julia/packages/Flux/qXNjB/src/optimise/train.jl:105
┌ Info: Epoch 1808
└ @ Main /Users/kcft114/.julia/packages/Flux/qXNjB/src/optimise/train.jl:105
┌ Info: Epoch 1809
└ @ Main /Users/kcft114/.julia/packages/Flux/qXNjB/src/optimise/train.jl:105
┌ Info: Epoch 1810
└ @ Main /Users/kcft114/.julia/packages/Flux/qXNjB/src/optimise/train.jl:105
┌ Info: Epoch 1811
└ @ Main /Users/kcft114/.julia/packages/Flux/qXNjB/src/optimise/train.jl:105
┌ Info: Epoch 1812
└ @ Main /Users/kcft114/.julia/packages/Flux/qXNjB/src/optimise/train.jl:105
┌ Info: Epoch 1813
└ @ Main /Users/kcft114/.julia/packages/Flux/qXNjB/src/optimise/train.jl:105
┌ Info: Epoch 1814
└ @ Main /Users/kcft114/.julia/packages/Flux/qXNjB/src/optimise/train.jl:105
┌ Info: Epoch 1815
└ @ Main /Users/kcft114/.julia/packages/Flux/qXNjB/src/optimise/train.jl:105
┌ Info: Epoch 1816
└ @ Main /Users/kcft114/.julia/packages/Flux/qXNjB/src/optimise/train.jl:105
┌ Info: Epoch 1817
└ @ Main /Users/kcft114/.julia/packages/Flux/qXNjB/src/optimise/train.jl:105
┌ Info: Epoch 1818
└ @ Main /Users/kcft114/.julia/packages/Flux/qXNjB/src/optimise/train.jl:105
┌ Info: Epoch 1819
└ @ Main /Users/kcft114/.julia/packages/Flux/qXNjB/src/optimise/train.jl:105
```

```
loss(X, Y) = 0.53737205f0 (tracked)
loss(X, Y) = 0.5850337f0 (tracked)
loss(X, Y) = 0.51238453f0 (tracked)
loss(X, Y) = 0.5239811f0 (tracked)
loss(X, Y) = 0.5108983f0 (tracked)
loss(X, Y) = 0.5022948f0 (tracked)
loss(X, Y) = 0.48988742f0 (tracked)
loss(X, Y) = 0.524208f0 (tracked)
loss(X, Y) = 0.53184724f0 (tracked)
loss(X, Y) = 0.56212646f0 (tracked)
loss(X, Y) = 0.543438f0 (tracked)
loss(X, Y) = 0.54889685f0 (tracked)
loss(X, Y) = 0.5404137f0 (tracked)
loss(X, Y) = 0.50972486f0 (tracked)
loss(X, Y) = 0.5217216f0 (tracked)
loss(X, Y) = 0.5412606f0 (tracked)
loss(X, Y) = 0.5281941f0 (tracked)
loss(X, Y) = 0.52566355f0 (tracked)
loss(X, Y) = 0.5457087f0 (tracked)
loss(X, Y) = 0.54021347f0 (tracked)
loss(X, Y) = 0.5988979f0 (tracked)
loss(X, Y) = 0.5433061f0 (tracked)
loss(X, Y) = 0.5346193f0 (tracked)
loss(X, Y) = 0.51730907f0 (tracked)
loss(X, Y) = 0.530571f0 (tracked)
loss(X, Y) = 0.5307566f0 (tracked)
loss(X, Y) = 0.50018865f0 (tracked)
loss(X, Y) = 0.50528514f0 (tracked)
loss(X, Y) = 0.5279745f0 (tracked)
loss(X, Y) = 0.5107784f0 (tracked)
loss(X, Y) = 0.54254854f0 (tracked)
loss(X, Y) = 0.5155395f0 (tracked)
loss(X, Y) = 0.56368256f0 (tracked)
loss(X, Y) = 0.54854834f0 (tracked)
loss(X, Y) = 0.53102845f0 (tracked)
loss(X, Y) = 0.53889954f0 (tracked)
loss(X, Y) = 0.4886394f0 (tracked)
loss(X, Y) = 0.60406023f0 (tracked)
loss(X, Y) = 0.4972647f0 (tracked)
loss(X, Y) = 0.5052632f0 (tracked)
loss(X, Y) = 0.53251696f0 (tracked)
loss(X, Y) = 0.50783277f0 (tracked)
loss(X, Y) = 0.50359035f0 (tracked)
loss(X, Y) = 0.5225068f0 (tracked)
loss(X, Y) = 0.529095f0 (tracked)
loss(X, Y) = 0.5425661f0 (tracked)
loss(X, Y) = 0.5165156f0 (tracked)
loss(X, Y) = 0.5974118f0 (tracked)
loss(X, Y) = 0.49648228f0 (tracked)
loss(X, Y) = 0.48921448f0 (tracked)
loss(X, Y) = 0.5439531f0 (tracked)
loss(X, Y) = 0.5339454f0 (tracked)
loss(X, Y) = 0.523035f0 (tracked)
loss(X, Y) = 0.52324975f0 (tracked)
loss(X, Y) = 0.53436714f0 (tracked)
loss(X, Y) = 0.5329549f0 (tracked)
loss(X, Y) = 0.58006305f0 (tracked)
loss(X, Y) = 0.54699427f0 (tracked)
loss(X, Y) = 0.5355494f0 (tracked)
loss(X, Y) = 0.59069246f0 (tracked)
loss(X, Y) = 0.5260272f0 (tracked)
loss(X, Y) = 0.5430975f0 (tracked)
loss(X, Y) = 0.52114433f0 (tracked)
loss(X, Y) = 0.5537544f0 (tracked)
loss(X, Y) = 0.55025154f0 (tracked)
loss(X, Y) = 0.48508406f0 (tracked)
loss(X, Y) = 0.5545435f0 (tracked)
loss(X, Y) = 0.5673191f0 (tracked)
loss(X, Y) = 0.5712421f0 (tracked)
loss(X, Y) = 0.5433599f0 (tracked)
loss(X, Y) = 0.49906904f0 (tracked)
loss(X, Y) = 0.5283878f0 (tracked)
loss(X, Y) = 0.54388726f0 (tracked)
loss(X, Y) = 0.5657106f0 (tracked)
loss(X, Y) = 0.5470395f0 (tracked)
loss(X, Y) = 0.4975465f0 (tracked)
loss(X, Y) = 0.5693742f0 (tracked)
loss(X, Y) = 0.56024945f0 (tracked)
loss(X, Y) = 0.5354072f0 (tracked)
loss(X, Y) = 0.4966412f0 (tracked)
loss(X, Y) = 0.5433077f0 (tracked)
loss(X, Y) = 0.51655066f0 (tracked)
loss(X, Y) = 0.55533546f0 (tracked)
loss(X, Y) = 0.5540223f0 (tracked)
loss(X, Y) = 0.52658397f0 (tracked)
loss(X, Y) = 0.60532993f0 (tracked)
loss(X, Y) = 0.492376f0 (tracked)
loss(X, Y) = 0.52150744f0 (tracked)
```

```
┌ Info: Epoch 1820
└ @ Main /Users/kcft114/.julia/packages/Flux/qXNjB/src/optimise/train.jl:105
┌ Info: Epoch 1821
└ @ Main /Users/kcft114/.julia/packages/Flux/qXNjB/src/optimise/train.jl:105
┌ Info: Epoch 1822
└ @ Main /Users/kcft114/.julia/packages/Flux/qXNjB/src/optimise/train.jl:105
┌ Info: Epoch 1823
└ @ Main /Users/kcft114/.julia/packages/Flux/qXNjB/src/optimise/train.jl:105
┌ Info: Epoch 1824
└ @ Main /Users/kcft114/.julia/packages/Flux/qXNjB/src/optimise/train.jl:105
┌ Info: Epoch 1825
└ @ Main /Users/kcft114/.julia/packages/Flux/qXNjB/src/optimise/train.jl:105
┌ Info: Epoch 1826
└ @ Main /Users/kcft114/.julia/packages/Flux/qXNjB/src/optimise/train.jl:105
┌ Info: Epoch 1827
└ @ Main /Users/kcft114/.julia/packages/Flux/qXNjB/src/optimise/train.jl:105
┌ Info: Epoch 1828
└ @ Main /Users/kcft114/.julia/packages/Flux/qXNjB/src/optimise/train.jl:105
┌ Info: Epoch 1829
└ @ Main /Users/kcft114/.julia/packages/Flux/qXNjB/src/optimise/train.jl:105
┌ Info: Epoch 1830
└ @ Main /Users/kcft114/.julia/packages/Flux/qXNjB/src/optimise/train.jl:105
┌ Info: Epoch 1831
└ @ Main /Users/kcft114/.julia/packages/Flux/qXNjB/src/optimise/train.jl:105
┌ Info: Epoch 1832
└ @ Main /Users/kcft114/.julia/packages/Flux/qXNjB/src/optimise/train.jl:105
┌ Info: Epoch 1833
└ @ Main /Users/kcft114/.julia/packages/Flux/qXNjB/src/optimise/train.jl:105
┌ Info: Epoch 1834
└ @ Main /Users/kcft114/.julia/packages/Flux/qXNjB/src/optimise/train.jl:105
┌ Info: Epoch 1835
└ @ Main /Users/kcft114/.julia/packages/Flux/qXNjB/src/optimise/train.jl:105
┌ Info: Epoch 1836
└ @ Main /Users/kcft114/.julia/packages/Flux/qXNjB/src/optimise/train.jl:105
┌ Info: Epoch 1837
└ @ Main /Users/kcft114/.julia/packages/Flux/qXNjB/src/optimise/train.jl:105
┌ Info: Epoch 1838
└ @ Main /Users/kcft114/.julia/packages/Flux/qXNjB/src/optimise/train.jl:105
┌ Info: Epoch 1839
└ @ Main /Users/kcft114/.julia/packages/Flux/qXNjB/src/optimise/train.jl:105
┌ Info: Epoch 1840
└ @ Main /Users/kcft114/.julia/packages/Flux/qXNjB/src/optimise/train.jl:105
┌ Info: Epoch 1841
└ @ Main /Users/kcft114/.julia/packages/Flux/qXNjB/src/optimise/train.jl:105
┌ Info: Epoch 1842
└ @ Main /Users/kcft114/.julia/packages/Flux/qXNjB/src/optimise/train.jl:105
┌ Info: Epoch 1843
└ @ Main /Users/kcft114/.julia/packages/Flux/qXNjB/src/optimise/train.jl:105
┌ Info: Epoch 1844
└ @ Main /Users/kcft114/.julia/packages/Flux/qXNjB/src/optimise/train.jl:105
┌ Info: Epoch 1845
└ @ Main /Users/kcft114/.julia/packages/Flux/qXNjB/src/optimise/train.jl:105
┌ Info: Epoch 1846
└ @ Main /Users/kcft114/.julia/packages/Flux/qXNjB/src/optimise/train.jl:105
┌ Info: Epoch 1847
└ @ Main /Users/kcft114/.julia/packages/Flux/qXNjB/src/optimise/train.jl:105
┌ Info: Epoch 1848
└ @ Main /Users/kcft114/.julia/packages/Flux/qXNjB/src/optimise/train.jl:105
┌ Info: Epoch 1849
└ @ Main /Users/kcft114/.julia/packages/Flux/qXNjB/src/optimise/train.jl:105
┌ Info: Epoch 1850
└ @ Main /Users/kcft114/.julia/packages/Flux/qXNjB/src/optimise/train.jl:105
┌ Info: Epoch 1851
└ @ Main /Users/kcft114/.julia/packages/Flux/qXNjB/src/optimise/train.jl:105
┌ Info: Epoch 1852
└ @ Main /Users/kcft114/.julia/packages/Flux/qXNjB/src/optimise/train.jl:105
┌ Info: Epoch 1853
└ @ Main /Users/kcft114/.julia/packages/Flux/qXNjB/src/optimise/train.jl:105
┌ Info: Epoch 1854
└ @ Main /Users/kcft114/.julia/packages/Flux/qXNjB/src/optimise/train.jl:105
┌ Info: Epoch 1855
└ @ Main /Users/kcft114/.julia/packages/Flux/qXNjB/src/optimise/train.jl:105
┌ Info: Epoch 1856
└ @ Main /Users/kcft114/.julia/packages/Flux/qXNjB/src/optimise/train.jl:105
┌ Info: Epoch 1857
└ @ Main /Users/kcft114/.julia/packages/Flux/qXNjB/src/optimise/train.jl:105
┌ Info: Epoch 1858
└ @ Main /Users/kcft114/.julia/packages/Flux/qXNjB/src/optimise/train.jl:105
┌ Info: Epoch 1859
└ @ Main /Users/kcft114/.julia/packages/Flux/qXNjB/src/optimise/train.jl:105
┌ Info: Epoch 1860
└ @ Main /Users/kcft114/.julia/packages/Flux/qXNjB/src/optimise/train.jl:105
┌ Info: Epoch 1861
└ @ Main /Users/kcft114/.julia/packages/Flux/qXNjB/src/optimise/train.jl:105
┌ Info: Epoch 1862
└ @ Main /Users/kcft114/.julia/packages/Flux/qXNjB/src/optimise/train.jl:105
┌ Info: Epoch 1863
└ @ Main /Users/kcft114/.julia/packages/Flux/qXNjB/src/optimise/train.jl:105
┌ Info: Epoch 1864
└ @ Main /Users/kcft114/.julia/packages/Flux/qXNjB/src/optimise/train.jl:105
┌ Info: Epoch 1865
└ @ Main /Users/kcft114/.julia/packages/Flux/qXNjB/src/optimise/train.jl:105
┌ Info: Epoch 1866
└ @ Main /Users/kcft114/.julia/packages/Flux/qXNjB/src/optimise/train.jl:105
┌ Info: Epoch 1867
└ @ Main /Users/kcft114/.julia/packages/Flux/qXNjB/src/optimise/train.jl:105
┌ Info: Epoch 1868
└ @ Main /Users/kcft114/.julia/packages/Flux/qXNjB/src/optimise/train.jl:105
┌ Info: Epoch 1869
└ @ Main /Users/kcft114/.julia/packages/Flux/qXNjB/src/optimise/train.jl:105
┌ Info: Epoch 1870
└ @ Main /Users/kcft114/.julia/packages/Flux/qXNjB/src/optimise/train.jl:105
┌ Info: Epoch 1871
└ @ Main /Users/kcft114/.julia/packages/Flux/qXNjB/src/optimise/train.jl:105
┌ Info: Epoch 1872
└ @ Main /Users/kcft114/.julia/packages/Flux/qXNjB/src/optimise/train.jl:105
┌ Info: Epoch 1873
└ @ Main /Users/kcft114/.julia/packages/Flux/qXNjB/src/optimise/train.jl:105
┌ Info: Epoch 1874
└ @ Main /Users/kcft114/.julia/packages/Flux/qXNjB/src/optimise/train.jl:105
┌ Info: Epoch 1875
└ @ Main /Users/kcft114/.julia/packages/Flux/qXNjB/src/optimise/train.jl:105
┌ Info: Epoch 1876
└ @ Main /Users/kcft114/.julia/packages/Flux/qXNjB/src/optimise/train.jl:105
┌ Info: Epoch 1877
└ @ Main /Users/kcft114/.julia/packages/Flux/qXNjB/src/optimise/train.jl:105
┌ Info: Epoch 1878
└ @ Main /Users/kcft114/.julia/packages/Flux/qXNjB/src/optimise/train.jl:105
┌ Info: Epoch 1879
└ @ Main /Users/kcft114/.julia/packages/Flux/qXNjB/src/optimise/train.jl:105
┌ Info: Epoch 1880
└ @ Main /Users/kcft114/.julia/packages/Flux/qXNjB/src/optimise/train.jl:105
┌ Info: Epoch 1881
└ @ Main /Users/kcft114/.julia/packages/Flux/qXNjB/src/optimise/train.jl:105
┌ Info: Epoch 1882
└ @ Main /Users/kcft114/.julia/packages/Flux/qXNjB/src/optimise/train.jl:105
┌ Info: Epoch 1883
└ @ Main /Users/kcft114/.julia/packages/Flux/qXNjB/src/optimise/train.jl:105
┌ Info: Epoch 1884
└ @ Main /Users/kcft114/.julia/packages/Flux/qXNjB/src/optimise/train.jl:105
┌ Info: Epoch 1885
└ @ Main /Users/kcft114/.julia/packages/Flux/qXNjB/src/optimise/train.jl:105
┌ Info: Epoch 1886
└ @ Main /Users/kcft114/.julia/packages/Flux/qXNjB/src/optimise/train.jl:105
┌ Info: Epoch 1887
└ @ Main /Users/kcft114/.julia/packages/Flux/qXNjB/src/optimise/train.jl:105
┌ Info: Epoch 1888
└ @ Main /Users/kcft114/.julia/packages/Flux/qXNjB/src/optimise/train.jl:105
┌ Info: Epoch 1889
└ @ Main /Users/kcft114/.julia/packages/Flux/qXNjB/src/optimise/train.jl:105
┌ Info: Epoch 1890
└ @ Main /Users/kcft114/.julia/packages/Flux/qXNjB/src/optimise/train.jl:105
┌ Info: Epoch 1891
└ @ Main /Users/kcft114/.julia/packages/Flux/qXNjB/src/optimise/train.jl:105
┌ Info: Epoch 1892
└ @ Main /Users/kcft114/.julia/packages/Flux/qXNjB/src/optimise/train.jl:105
┌ Info: Epoch 1893
└ @ Main /Users/kcft114/.julia/packages/Flux/qXNjB/src/optimise/train.jl:105
┌ Info: Epoch 1894
└ @ Main /Users/kcft114/.julia/packages/Flux/qXNjB/src/optimise/train.jl:105
┌ Info: Epoch 1895
└ @ Main /Users/kcft114/.julia/packages/Flux/qXNjB/src/optimise/train.jl:105
┌ Info: Epoch 1896
└ @ Main /Users/kcft114/.julia/packages/Flux/qXNjB/src/optimise/train.jl:105
┌ Info: Epoch 1897
└ @ Main /Users/kcft114/.julia/packages/Flux/qXNjB/src/optimise/train.jl:105
┌ Info: Epoch 1898
└ @ Main /Users/kcft114/.julia/packages/Flux/qXNjB/src/optimise/train.jl:105
┌ Info: Epoch 1899
└ @ Main /Users/kcft114/.julia/packages/Flux/qXNjB/src/optimise/train.jl:105
┌ Info: Epoch 1900
└ @ Main /Users/kcft114/.julia/packages/Flux/qXNjB/src/optimise/train.jl:105
┌ Info: Epoch 1901
└ @ Main /Users/kcft114/.julia/packages/Flux/qXNjB/src/optimise/train.jl:105
┌ Info: Epoch 1902
└ @ Main /Users/kcft114/.julia/packages/Flux/qXNjB/src/optimise/train.jl:105
┌ Info: Epoch 1903
└ @ Main /Users/kcft114/.julia/packages/Flux/qXNjB/src/optimise/train.jl:105
┌ Info: Epoch 1904
└ @ Main /Users/kcft114/.julia/packages/Flux/qXNjB/src/optimise/train.jl:105
┌ Info: Epoch 1905
└ @ Main /Users/kcft114/.julia/packages/Flux/qXNjB/src/optimise/train.jl:105
┌ Info: Epoch 1906
└ @ Main /Users/kcft114/.julia/packages/Flux/qXNjB/src/optimise/train.jl:105
┌ Info: Epoch 1907
└ @ Main /Users/kcft114/.julia/packages/Flux/qXNjB/src/optimise/train.jl:105
```

```
loss(X, Y) = 0.5546622f0 (tracked)
loss(X, Y) = 0.480631f0 (tracked)
loss(X, Y) = 0.54883265f0 (tracked)
loss(X, Y) = 0.56895167f0 (tracked)
loss(X, Y) = 0.57074195f0 (tracked)
loss(X, Y) = 0.51515794f0 (tracked)
loss(X, Y) = 0.5409047f0 (tracked)
loss(X, Y) = 0.47560477f0 (tracked)
loss(X, Y) = 0.5244418f0 (tracked)
loss(X, Y) = 0.51409966f0 (tracked)
loss(X, Y) = 0.54662776f0 (tracked)
loss(X, Y) = 0.5182299f0 (tracked)
loss(X, Y) = 0.5241443f0 (tracked)
loss(X, Y) = 0.53954f0 (tracked)
loss(X, Y) = 0.52972543f0 (tracked)
loss(X, Y) = 0.5544001f0 (tracked)
loss(X, Y) = 0.48673356f0 (tracked)
loss(X, Y) = 0.46175987f0 (tracked)
loss(X, Y) = 0.50920165f0 (tracked)
loss(X, Y) = 0.5379316f0 (tracked)
loss(X, Y) = 0.51355404f0 (tracked)
loss(X, Y) = 0.5198704f0 (tracked)
loss(X, Y) = 0.555852f0 (tracked)
loss(X, Y) = 0.5853574f0 (tracked)
loss(X, Y) = 0.54762274f0 (tracked)
loss(X, Y) = 0.5256122f0 (tracked)
loss(X, Y) = 0.4862132f0 (tracked)
loss(X, Y) = 0.5856586f0 (tracked)
loss(X, Y) = 0.59551984f0 (tracked)
loss(X, Y) = 0.48716477f0 (tracked)
loss(X, Y) = 0.50055754f0 (tracked)
loss(X, Y) = 0.51351964f0 (tracked)
loss(X, Y) = 0.49796554f0 (tracked)
loss(X, Y) = 0.5774842f0 (tracked)
loss(X, Y) = 0.56313175f0 (tracked)
loss(X, Y) = 0.4938002f0 (tracked)
loss(X, Y) = 0.53668916f0 (tracked)
loss(X, Y) = 0.52852046f0 (tracked)
loss(X, Y) = 0.5443239f0 (tracked)
loss(X, Y) = 0.47583115f0 (tracked)
loss(X, Y) = 0.56455606f0 (tracked)
loss(X, Y) = 0.6152415f0 (tracked)
loss(X, Y) = 0.5414169f0 (tracked)
loss(X, Y) = 0.53247684f0 (tracked)
loss(X, Y) = 0.54274976f0 (tracked)
loss(X, Y) = 0.5097327f0 (tracked)
loss(X, Y) = 0.4940477f0 (tracked)
loss(X, Y) = 0.60816133f0 (tracked)
loss(X, Y) = 0.5523155f0 (tracked)
loss(X, Y) = 0.5169554f0 (tracked)
loss(X, Y) = 0.5582526f0 (tracked)
loss(X, Y) = 0.52691734f0 (tracked)
loss(X, Y) = 0.56295896f0 (tracked)
loss(X, Y) = 0.48978958f0 (tracked)
loss(X, Y) = 0.5395357f0 (tracked)
loss(X, Y) = 0.48925832f0 (tracked)
loss(X, Y) = 0.47205347f0 (tracked)
loss(X, Y) = 0.5646951f0 (tracked)
loss(X, Y) = 0.5617492f0 (tracked)
loss(X, Y) = 0.5430844f0 (tracked)
loss(X, Y) = 0.5159319f0 (tracked)
loss(X, Y) = 0.519831f0 (tracked)
loss(X, Y) = 0.56468254f0 (tracked)
loss(X, Y) = 0.48351288f0 (tracked)
loss(X, Y) = 0.5417003f0 (tracked)
loss(X, Y) = 0.5407022f0 (tracked)
loss(X, Y) = 0.5083993f0 (tracked)
loss(X, Y) = 0.45167446f0 (tracked)
loss(X, Y) = 0.5193101f0 (tracked)
loss(X, Y) = 0.53083634f0 (tracked)
loss(X, Y) = 0.5748156f0 (tracked)
loss(X, Y) = 0.50120604f0 (tracked)
loss(X, Y) = 0.54719895f0 (tracked)
loss(X, Y) = 0.5232758f0 (tracked)
loss(X, Y) = 0.51511234f0 (tracked)
loss(X, Y) = 0.5490587f0 (tracked)
loss(X, Y) = 0.52862847f0 (tracked)
loss(X, Y) = 0.43384162f0 (tracked)
loss(X, Y) = 0.49464118f0 (tracked)
loss(X, Y) = 0.5509336f0 (tracked)
loss(X, Y) = 0.55945385f0 (tracked)
loss(X, Y) = 0.531437f0 (tracked)
loss(X, Y) = 0.4914446f0 (tracked)
loss(X, Y) = 0.5069249f0 (tracked)
loss(X, Y) = 0.516135f0 (tracked)
loss(X, Y) = 0.56117356f0 (tracked)
loss(X, Y) = 0.58962303f0 (tracked)
loss(X, Y) = 0.5147554f0 (tracked)
loss(X, Y) = 0.4977572f0 (tracked)
loss(X, Y) = 0.49891546f0 (tracked)
loss(X, Y) = 0.52702737f0 (tracked)
loss(X, Y) = 0.5314071f0 (tracked)
loss(X, Y) = 0.5260973f0 (tracked)
loss(X, Y) = 0.508558f0 (tracked)
train accuracy = 0.8027210884353742
test accuracy = 0.5135135135135135
train BA = 0.78
test BA = 0.55
loss(X, Y) = 1.333531f0 (tracked)
loss(X, Y) = 1.2854184f0 (tracked)
loss(X, Y) = 1.3101898f0 (tracked)
loss(X, Y) = 1.2622038f0 (tracked)
```

```
┌ Info: Epoch 1908
└ @ Main /Users/kcft114/.julia/packages/Flux/qXNjB/src/optimise/train.jl:105
┌ Info: Epoch 1909
└ @ Main /Users/kcft114/.julia/packages/Flux/qXNjB/src/optimise/train.jl:105
┌ Info: Epoch 1910
└ @ Main /Users/kcft114/.julia/packages/Flux/qXNjB/src/optimise/train.jl:105
┌ Info: Epoch 1911
└ @ Main /Users/kcft114/.julia/packages/Flux/qXNjB/src/optimise/train.jl:105
┌ Info: Epoch 1912
└ @ Main /Users/kcft114/.julia/packages/Flux/qXNjB/src/optimise/train.jl:105
┌ Info: Epoch 1913
└ @ Main /Users/kcft114/.julia/packages/Flux/qXNjB/src/optimise/train.jl:105
┌ Info: Epoch 1914
└ @ Main /Users/kcft114/.julia/packages/Flux/qXNjB/src/optimise/train.jl:105
┌ Info: Epoch 1915
└ @ Main /Users/kcft114/.julia/packages/Flux/qXNjB/src/optimise/train.jl:105
┌ Info: Epoch 1916
└ @ Main /Users/kcft114/.julia/packages/Flux/qXNjB/src/optimise/train.jl:105
┌ Info: Epoch 1917
└ @ Main /Users/kcft114/.julia/packages/Flux/qXNjB/src/optimise/train.jl:105
┌ Info: Epoch 1918
└ @ Main /Users/kcft114/.julia/packages/Flux/qXNjB/src/optimise/train.jl:105
┌ Info: Epoch 1919
└ @ Main /Users/kcft114/.julia/packages/Flux/qXNjB/src/optimise/train.jl:105
┌ Info: Epoch 1920
└ @ Main /Users/kcft114/.julia/packages/Flux/qXNjB/src/optimise/train.jl:105
┌ Info: Epoch 1921
└ @ Main /Users/kcft114/.julia/packages/Flux/qXNjB/src/optimise/train.jl:105
┌ Info: Epoch 1922
└ @ Main /Users/kcft114/.julia/packages/Flux/qXNjB/src/optimise/train.jl:105
┌ Info: Epoch 1923
└ @ Main /Users/kcft114/.julia/packages/Flux/qXNjB/src/optimise/train.jl:105
┌ Info: Epoch 1924
└ @ Main /Users/kcft114/.julia/packages/Flux/qXNjB/src/optimise/train.jl:105
┌ Info: Epoch 1925
└ @ Main /Users/kcft114/.julia/packages/Flux/qXNjB/src/optimise/train.jl:105
┌ Info: Epoch 1926
└ @ Main /Users/kcft114/.julia/packages/Flux/qXNjB/src/optimise/train.jl:105
┌ Info: Epoch 1927
└ @ Main /Users/kcft114/.julia/packages/Flux/qXNjB/src/optimise/train.jl:105
┌ Info: Epoch 1928
└ @ Main /Users/kcft114/.julia/packages/Flux/qXNjB/src/optimise/train.jl:105
┌ Info: Epoch 1929
└ @ Main /Users/kcft114/.julia/packages/Flux/qXNjB/src/optimise/train.jl:105
┌ Info: Epoch 1930
└ @ Main /Users/kcft114/.julia/packages/Flux/qXNjB/src/optimise/train.jl:105
┌ Info: Epoch 1931
└ @ Main /Users/kcft114/.julia/packages/Flux/qXNjB/src/optimise/train.jl:105
┌ Info: Epoch 1932
└ @ Main /Users/kcft114/.julia/packages/Flux/qXNjB/src/optimise/train.jl:105
┌ Info: Epoch 1933
└ @ Main /Users/kcft114/.julia/packages/Flux/qXNjB/src/optimise/train.jl:105
┌ Info: Epoch 1934
└ @ Main /Users/kcft114/.julia/packages/Flux/qXNjB/src/optimise/train.jl:105
┌ Info: Epoch 1935
└ @ Main /Users/kcft114/.julia/packages/Flux/qXNjB/src/optimise/train.jl:105
┌ Info: Epoch 1936
└ @ Main /Users/kcft114/.julia/packages/Flux/qXNjB/src/optimise/train.jl:105
┌ Info: Epoch 1937
└ @ Main /Users/kcft114/.julia/packages/Flux/qXNjB/src/optimise/train.jl:105
┌ Info: Epoch 1938
└ @ Main /Users/kcft114/.julia/packages/Flux/qXNjB/src/optimise/train.jl:105
┌ Info: Epoch 1939
└ @ Main /Users/kcft114/.julia/packages/Flux/qXNjB/src/optimise/train.jl:105
┌ Info: Epoch 1940
└ @ Main /Users/kcft114/.julia/packages/Flux/qXNjB/src/optimise/train.jl:105
┌ Info: Epoch 1941
└ @ Main /Users/kcft114/.julia/packages/Flux/qXNjB/src/optimise/train.jl:105
┌ Info: Epoch 1942
└ @ Main /Users/kcft114/.julia/packages/Flux/qXNjB/src/optimise/train.jl:105
┌ Info: Epoch 1943
└ @ Main /Users/kcft114/.julia/packages/Flux/qXNjB/src/optimise/train.jl:105
┌ Info: Epoch 1944
└ @ Main /Users/kcft114/.julia/packages/Flux/qXNjB/src/optimise/train.jl:105
┌ Info: Epoch 1945
└ @ Main /Users/kcft114/.julia/packages/Flux/qXNjB/src/optimise/train.jl:105
┌ Info: Epoch 1946
└ @ Main /Users/kcft114/.julia/packages/Flux/qXNjB/src/optimise/train.jl:105
┌ Info: Epoch 1947
└ @ Main /Users/kcft114/.julia/packages/Flux/qXNjB/src/optimise/train.jl:105
┌ Info: Epoch 1948
└ @ Main /Users/kcft114/.julia/packages/Flux/qXNjB/src/optimise/train.jl:105
┌ Info: Epoch 1949
└ @ Main /Users/kcft114/.julia/packages/Flux/qXNjB/src/optimise/train.jl:105
┌ Info: Epoch 1950
└ @ Main /Users/kcft114/.julia/packages/Flux/qXNjB/src/optimise/train.jl:105
┌ Info: Epoch 1951
└ @ Main /Users/kcft114/.julia/packages/Flux/qXNjB/src/optimise/train.jl:105
┌ Info: Epoch 1952
└ @ Main /Users/kcft114/.julia/packages/Flux/qXNjB/src/optimise/train.jl:105
┌ Info: Epoch 1953
└ @ Main /Users/kcft114/.julia/packages/Flux/qXNjB/src/optimise/train.jl:105
┌ Info: Epoch 1954
└ @ Main /Users/kcft114/.julia/packages/Flux/qXNjB/src/optimise/train.jl:105
┌ Info: Epoch 1955
└ @ Main /Users/kcft114/.julia/packages/Flux/qXNjB/src/optimise/train.jl:105
┌ Info: Epoch 1956
└ @ Main /Users/kcft114/.julia/packages/Flux/qXNjB/src/optimise/train.jl:105
┌ Info: Epoch 1957
└ @ Main /Users/kcft114/.julia/packages/Flux/qXNjB/src/optimise/train.jl:105
┌ Info: Epoch 1958
└ @ Main /Users/kcft114/.julia/packages/Flux/qXNjB/src/optimise/train.jl:105
┌ Info: Epoch 1959
└ @ Main /Users/kcft114/.julia/packages/Flux/qXNjB/src/optimise/train.jl:105
┌ Info: Epoch 1960
└ @ Main /Users/kcft114/.julia/packages/Flux/qXNjB/src/optimise/train.jl:105
┌ Info: Epoch 1961
└ @ Main /Users/kcft114/.julia/packages/Flux/qXNjB/src/optimise/train.jl:105
┌ Info: Epoch 1962
└ @ Main /Users/kcft114/.julia/packages/Flux/qXNjB/src/optimise/train.jl:105
┌ Info: Epoch 1963
└ @ Main /Users/kcft114/.julia/packages/Flux/qXNjB/src/optimise/train.jl:105
┌ Info: Epoch 1964
└ @ Main /Users/kcft114/.julia/packages/Flux/qXNjB/src/optimise/train.jl:105
┌ Info: Epoch 1965
└ @ Main /Users/kcft114/.julia/packages/Flux/qXNjB/src/optimise/train.jl:105
┌ Info: Epoch 1966
└ @ Main /Users/kcft114/.julia/packages/Flux/qXNjB/src/optimise/train.jl:105
┌ Info: Epoch 1967
└ @ Main /Users/kcft114/.julia/packages/Flux/qXNjB/src/optimise/train.jl:105
┌ Info: Epoch 1968
└ @ Main /Users/kcft114/.julia/packages/Flux/qXNjB/src/optimise/train.jl:105
┌ Info: Epoch 1969
└ @ Main /Users/kcft114/.julia/packages/Flux/qXNjB/src/optimise/train.jl:105
┌ Info: Epoch 1970
└ @ Main /Users/kcft114/.julia/packages/Flux/qXNjB/src/optimise/train.jl:105
┌ Info: Epoch 1971
└ @ Main /Users/kcft114/.julia/packages/Flux/qXNjB/src/optimise/train.jl:105
┌ Info: Epoch 1972
└ @ Main /Users/kcft114/.julia/packages/Flux/qXNjB/src/optimise/train.jl:105
┌ Info: Epoch 1973
└ @ Main /Users/kcft114/.julia/packages/Flux/qXNjB/src/optimise/train.jl:105
┌ Info: Epoch 1974
└ @ Main /Users/kcft114/.julia/packages/Flux/qXNjB/src/optimise/train.jl:105
┌ Info: Epoch 1975
└ @ Main /Users/kcft114/.julia/packages/Flux/qXNjB/src/optimise/train.jl:105
┌ Info: Epoch 1976
└ @ Main /Users/kcft114/.julia/packages/Flux/qXNjB/src/optimise/train.jl:105
┌ Info: Epoch 1977
└ @ Main /Users/kcft114/.julia/packages/Flux/qXNjB/src/optimise/train.jl:105
┌ Info: Epoch 1978
└ @ Main /Users/kcft114/.julia/packages/Flux/qXNjB/src/optimise/train.jl:105
┌ Info: Epoch 1979
└ @ Main /Users/kcft114/.julia/packages/Flux/qXNjB/src/optimise/train.jl:105
┌ Info: Epoch 1980
└ @ Main /Users/kcft114/.julia/packages/Flux/qXNjB/src/optimise/train.jl:105
┌ Info: Epoch 1981
└ @ Main /Users/kcft114/.julia/packages/Flux/qXNjB/src/optimise/train.jl:105
┌ Info: Epoch 1982
└ @ Main /Users/kcft114/.julia/packages/Flux/qXNjB/src/optimise/train.jl:105
┌ Info: Epoch 1983
└ @ Main /Users/kcft114/.julia/packages/Flux/qXNjB/src/optimise/train.jl:105
┌ Info: Epoch 1984
└ @ Main /Users/kcft114/.julia/packages/Flux/qXNjB/src/optimise/train.jl:105
┌ Info: Epoch 1985
└ @ Main /Users/kcft114/.julia/packages/Flux/qXNjB/src/optimise/train.jl:105
┌ Info: Epoch 1986
└ @ Main /Users/kcft114/.julia/packages/Flux/qXNjB/src/optimise/train.jl:105
┌ Info: Epoch 1987
└ @ Main /Users/kcft114/.julia/packages/Flux/qXNjB/src/optimise/train.jl:105
┌ Info: Epoch 1988
└ @ Main /Users/kcft114/.julia/packages/Flux/qXNjB/src/optimise/train.jl:105
┌ Info: Epoch 1989
└ @ Main /Users/kcft114/.julia/packages/Flux/qXNjB/src/optimise/train.jl:105
┌ Info: Epoch 1990
└ @ Main /Users/kcft114/.julia/packages/Flux/qXNjB/src/optimise/train.jl:105
┌ Info: Epoch 1991
└ @ Main /Users/kcft114/.julia/packages/Flux/qXNjB/src/optimise/train.jl:105
┌ Info: Epoch 1992
└ @ Main /Users/kcft114/.julia/packages/Flux/qXNjB/src/optimise/train.jl:105
┌ Info: Epoch 1993
└ @ Main /Users/kcft114/.julia/packages/Flux/qXNjB/src/optimise/train.jl:105
┌ Info: Epoch 1994
└ @ Main /Users/kcft114/.julia/packages/Flux/qXNjB/src/optimise/train.jl:105
┌ Info: Epoch 1995
└ @ Main /Users/kcft114/.julia/packages/Flux/qXNjB/src/optimise/train.jl:105
┌ Info: Epoch 1996
└ @ Main /Users/kcft114/.julia/packages/Flux/qXNjB/src/optimise/train.jl:105
┌ Info: Epoch 1997
└ @ Main /Users/kcft114/.julia/packages/Flux/qXNjB/src/optimise/train.jl:105
┌ Info: Epoch 1998
└ @ Main /Users/kcft114/.julia/packages/Flux/qXNjB/src/optimise/train.jl:105
┌ Info: Epoch 1999
└ @ Main /Users/kcft114/.julia/packages/Flux/qXNjB/src/optimise/train.jl:105
┌ Info: Epoch 2000
└ @ Main /Users/kcft114/.julia/packages/Flux/qXNjB/src/optimise/train.jl:105
┌ Info: Epoch 1
└ @ Main /Users/kcft114/.julia/packages/Flux/qXNjB/src/optimise/train.jl:105
┌ Info: Epoch 2
└ @ Main /Users/kcft114/.julia/packages/Flux/qXNjB/src/optimise/train.jl:105
┌ Info: Epoch 3
└ @ Main /Users/kcft114/.julia/packages/Flux/qXNjB/src/optimise/train.jl:105
┌ Info: Epoch 4
└ @ Main /Users/kcft114/.julia/packages/Flux/qXNjB/src/optimise/train.jl:105
┌ Info: Epoch 5
└ @ Main /Users/kcft114/.julia/packages/Flux/qXNjB/src/optimise/train.jl:105
┌ Info: Epoch 6
└ @ Main /Users/kcft114/.julia/packages/Flux/qXNjB/src/optimise/train.jl:105
┌ Info: Epoch 7
└ @ Main /Users/kcft114/.julia/packages/Flux/qXNjB/src/optimise/train.jl:105
┌ Info: Epoch 8
└ @ Main /Users/kcft114/.julia/packages/Flux/qXNjB/src/optimise/train.jl:105
┌ Info: Epoch 9
└ @ Main /Users/kcft114/.julia/packages/Flux/qXNjB/src/optimise/train.jl:105
┌ Info: Epoch 10
└ @ Main /Users/kcft114/.julia/packages/Flux/qXNjB/src/optimise/train.jl:105
┌ Info: Epoch 11
└ @ Main /Users/kcft114/.julia/packages/Flux/qXNjB/src/optimise/train.jl:105
┌ Info: Epoch 12
└ @ Main /Users/kcft114/.julia/packages/Flux/qXNjB/src/optimise/train.jl:105
┌ Info: Epoch 13
└ @ Main /Users/kcft114/.julia/packages/Flux/qXNjB/src/optimise/train.jl:105
┌ Info: Epoch 14
└ @ Main /Users/kcft114/.julia/packages/Flux/qXNjB/src/optimise/train.jl:105
┌ Info: Epoch 15
└ @ Main /Users/kcft114/.julia/packages/Flux/qXNjB/src/optimise/train.jl:105
┌ Info: Epoch 16
└ @ Main /Users/kcft114/.julia/packages/Flux/qXNjB/src/optimise/train.jl:105
┌ Info: Epoch 17
└ @ Main /Users/kcft114/.julia/packages/Flux/qXNjB/src/optimise/train.jl:105
┌ Info: Epoch 18
└ @ Main /Users/kcft114/.julia/packages/Flux/qXNjB/src/optimise/train.jl:105
┌ Info: Epoch 19
└ @ Main /Users/kcft114/.julia/packages/Flux/qXNjB/src/optimise/train.jl:105
┌ Info: Epoch 20
└ @ Main /Users/kcft114/.julia/packages/Flux/qXNjB/src/optimise/train.jl:105
┌ Info: Epoch 21
└ @ Main /Users/kcft114/.julia/packages/Flux/qXNjB/src/optimise/train.jl:105
┌ Info: Epoch 22
└ @ Main /Users/kcft114/.julia/packages/Flux/qXNjB/src/optimise/train.jl:105
┌ Info: Epoch 23
└ @ Main /Users/kcft114/.julia/packages/Flux/qXNjB/src/optimise/train.jl:105
┌ Info: Epoch 24
└ @ Main /Users/kcft114/.julia/packages/Flux/qXNjB/src/optimise/train.jl:105
┌ Info: Epoch 25
└ @ Main /Users/kcft114/.julia/packages/Flux/qXNjB/src/optimise/train.jl:105
┌ Info: Epoch 26
└ @ Main /Users/kcft114/.julia/packages/Flux/qXNjB/src/optimise/train.jl:105
┌ Info: Epoch 27
└ @ Main /Users/kcft114/.julia/packages/Flux/qXNjB/src/optimise/train.jl:105
┌ Info: Epoch 28
└ @ Main /Users/kcft114/.julia/packages/Flux/qXNjB/src/optimise/train.jl:105
┌ Info: Epoch 29
└ @ Main /Users/kcft114/.julia/packages/Flux/qXNjB/src/optimise/train.jl:105
┌ Info: Epoch 30
└ @ Main /Users/kcft114/.julia/packages/Flux/qXNjB/src/optimise/train.jl:105
┌ Info: Epoch 31
└ @ Main /Users/kcft114/.julia/packages/Flux/qXNjB/src/optimise/train.jl:105
┌ Info: Epoch 32
└ @ Main /Users/kcft114/.julia/packages/Flux/qXNjB/src/optimise/train.jl:105
┌ Info: Epoch 33
└ @ Main /Users/kcft114/.julia/packages/Flux/qXNjB/src/optimise/train.jl:105
┌ Info: Epoch 34
└ @ Main /Users/kcft114/.julia/packages/Flux/qXNjB/src/optimise/train.jl:105
┌ Info: Epoch 35
└ @ Main /Users/kcft114/.julia/packages/Flux/qXNjB/src/optimise/train.jl:105
┌ Info: Epoch 36
└ @ Main /Users/kcft114/.julia/packages/Flux/qXNjB/src/optimise/train.jl:105
┌ Info: Epoch 37
└ @ Main /Users/kcft114/.julia/packages/Flux/qXNjB/src/optimise/train.jl:105
┌ Info: Epoch 38
└ @ Main /Users/kcft114/.julia/packages/Flux/qXNjB/src/optimise/train.jl:105
┌ Info: Epoch 39
└ @ Main /Users/kcft114/.julia/packages/Flux/qXNjB/src/optimise/train.jl:105
┌ Info: Epoch 40
└ @ Main /Users/kcft114/.julia/packages/Flux/qXNjB/src/optimise/train.jl:105
┌ Info: Epoch 41
└ @ Main /Users/kcft114/.julia/packages/Flux/qXNjB/src/optimise/train.jl:105
┌ Info: Epoch 42
└ @ Main /Users/kcft114/.julia/packages/Flux/qXNjB/src/optimise/train.jl:105
┌ Info: Epoch 43
└ @ Main /Users/kcft114/.julia/packages/Flux/qXNjB/src/optimise/train.jl:105
┌ Info: Epoch 44
└ @ Main /Users/kcft114/.julia/packages/Flux/qXNjB/src/optimise/train.jl:105
┌ Info: Epoch 45
└ @ Main /Users/kcft114/.julia/packages/Flux/qXNjB/src/optimise/train.jl:105
┌ Info: Epoch 46
└ @ Main /Users/kcft114/.julia/packages/Flux/qXNjB/src/optimise/train.jl:105
┌ Info: Epoch 47
└ @ Main /Users/kcft114/.julia/packages/Flux/qXNjB/src/optimise/train.jl:105
┌ Info: Epoch 48
└ @ Main /Users/kcft114/.julia/packages/Flux/qXNjB/src/optimise/train.jl:105
┌ Info: Epoch 49
└ @ Main /Users/kcft114/.julia/packages/Flux/qXNjB/src/optimise/train.jl:105
┌ Info: Epoch 50
└ @ Main /Users/kcft114/.julia/packages/Flux/qXNjB/src/optimise/train.jl:105
┌ Info: Epoch 51
└ @ Main /Users/kcft114/.julia/packages/Flux/qXNjB/src/optimise/train.jl:105
┌ Info: Epoch 52
└ @ Main /Users/kcft114/.julia/packages/Flux/qXNjB/src/optimise/train.jl:105
┌ Info: Epoch 53
└ @ Main /Users/kcft114/.julia/packages/Flux/qXNjB/src/optimise/train.jl:105
┌ Info: Epoch 54
└ @ Main /Users/kcft114/.julia/packages/Flux/qXNjB/src/optimise/train.jl:105
┌ Info: Epoch 55
└ @ Main /Users/kcft114/.julia/packages/Flux/qXNjB/src/optimise/train.jl:105
┌ Info: Epoch 56
└ @ Main /Users/kcft114/.julia/packages/Flux/qXNjB/src/optimise/train.jl:105
┌ Info: Epoch 57
└ @ Main /Users/kcft114/.julia/packages/Flux/qXNjB/src/optimise/train.jl:105
┌ Info: Epoch 58
└ @ Main /Users/kcft114/.julia/packages/Flux/qXNjB/src/optimise/train.jl:105
┌ Info: Epoch 59
└ @ Main /Users/kcft114/.julia/packages/Flux/qXNjB/src/optimise/train.jl:105
┌ Info: Epoch 60
└ @ Main /Users/kcft114/.julia/packages/Flux/qXNjB/src/optimise/train.jl:105
┌ Info: Epoch 61
└ @ Main /Users/kcft114/.julia/packages/Flux/qXNjB/src/optimise/train.jl:105
┌ Info: Epoch 62
└ @ Main /Users/kcft114/.julia/packages/Flux/qXNjB/src/optimise/train.jl:105
┌ Info: Epoch 63
└ @ Main /Users/kcft114/.julia/packages/Flux/qXNjB/src/optimise/train.jl:105
┌ Info: Epoch 64
└ @ Main /Users/kcft114/.julia/packages/Flux/qXNjB/src/optimise/train.jl:105
┌ Info: Epoch 65
└ @ Main /Users/kcft114/.julia/packages/Flux/qXNjB/src/optimise/train.jl:105
┌ Info: Epoch 66
└ @ Main /Users/kcft114/.julia/packages/Flux/qXNjB/src/optimise/train.jl:105
┌ Info: Epoch 67
└ @ Main /Users/kcft114/.julia/packages/Flux/qXNjB/src/optimise/train.jl:105
┌ Info: Epoch 68
└ @ Main /Users/kcft114/.julia/packages/Flux/qXNjB/src/optimise/train.jl:105
┌ Info: Epoch 69
└ @ Main /Users/kcft114/.julia/packages/Flux/qXNjB/src/optimise/train.jl:105
┌ Info: Epoch 70
└ @ Main /Users/kcft114/.julia/packages/Flux/qXNjB/src/optimise/train.jl:105
┌ Info: Epoch 71
└ @ Main /Users/kcft114/.julia/packages/Flux/qXNjB/src/optimise/train.jl:105
┌ Info: Epoch 72
└ @ Main /Users/kcft114/.julia/packages/Flux/qXNjB/src/optimise/train.jl:105
┌ Info: Epoch 73
└ @ Main /Users/kcft114/.julia/packages/Flux/qXNjB/src/optimise/train.jl:105
┌ Info: Epoch 74
└ @ Main /Users/kcft114/.julia/packages/Flux/qXNjB/src/optimise/train.jl:105
┌ Info: Epoch 75
└ @ Main /Users/kcft114/.julia/packages/Flux/qXNjB/src/optimise/train.jl:105
┌ Info: Epoch 76
└ @ Main /Users/kcft114/.julia/packages/Flux/qXNjB/src/optimise/train.jl:105
┌ Info: Epoch 77
└ @ Main /Users/kcft114/.julia/packages/Flux/qXNjB/src/optimise/train.jl:105
┌ Info: Epoch 78
└ @ Main /Users/kcft114/.julia/packages/Flux/qXNjB/src/optimise/train.jl:105
┌ Info: Epoch 79
└ @ Main /Users/kcft114/.julia/packages/Flux/qXNjB/src/optimise/train.jl:105
┌ Info: Epoch 80
└ @ Main /Users/kcft114/.julia/packages/Flux/qXNjB/src/optimise/train.jl:105
┌ Info: Epoch 81
└ @ Main /Users/kcft114/.julia/packages/Flux/qXNjB/src/optimise/train.jl:105
┌ Info: Epoch 82
└ @ Main /Users/kcft114/.julia/packages/Flux/qXNjB/src/optimise/train.jl:105
┌ Info: Epoch 83
└ @ Main /Users/kcft114/.julia/packages/Flux/qXNjB/src/optimise/train.jl:105
┌ Info: Epoch 84
└ @ Main /Users/kcft114/.julia/packages/Flux/qXNjB/src/optimise/train.jl:105
┌ Info: Epoch 85
└ @ Main /Users/kcft114/.julia/packages/Flux/qXNjB/src/optimise/train.jl:105
┌ Info: Epoch 86
└ @ Main /Users/kcft114/.julia/packages/Flux/qXNjB/src/optimise/train.jl:105
┌ Info: Epoch 87
└ @ Main /Users/kcft114/.julia/packages/Flux/qXNjB/src/optimise/train.jl:105
┌ Info: Epoch 88
└ @ Main /Users/kcft114/.julia/packages/Flux/qXNjB/src/optimise/train.jl:105
┌ Info: Epoch 89
└ @ Main /Users/kcft114/.julia/packages/Flux/qXNjB/src/optimise/train.jl:105
┌ Info: Epoch 90
└ @ Main /Users/kcft114/.julia/packages/Flux/qXNjB/src/optimise/train.jl:105
┌ Info: Epoch 91
└ @ Main /Users/kcft114/.julia/packages/Flux/qXNjB/src/optimise/train.jl:105
┌ Info: Epoch 92
└ @ Main /Users/kcft114/.julia/packages/Flux/qXNjB/src/optimise/train.jl:105
┌ Info: Epoch 93
└ @ Main /Users/kcft114/.julia/packages/Flux/qXNjB/src/optimise/train.jl:105
┌ Info: Epoch 94
└ @ Main /Users/kcft114/.julia/packages/Flux/qXNjB/src/optimise/train.jl:105
┌ Info: Epoch 95
└ @ Main /Users/kcft114/.julia/packages/Flux/qXNjB/src/optimise/train.jl:105
┌ Info: Epoch 96
└ @ Main /Users/kcft114/.julia/packages/Flux/qXNjB/src/optimise/train.jl:105
┌ Info: Epoch 97
└ @ Main /Users/kcft114/.julia/packages/Flux/qXNjB/src/optimise/train.jl:105
┌ Info: Epoch 98
└ @ Main /Users/kcft114/.julia/packages/Flux/qXNjB/src/optimise/train.jl:105
┌ Info: Epoch 99
└ @ Main /Users/kcft114/.julia/packages/Flux/qXNjB/src/optimise/train.jl:105
┌ Info: Epoch 100
└ @ Main /Users/kcft114/.julia/packages/Flux/qXNjB/src/optimise/train.jl:105
┌ Info: Epoch 101
└ @ Main /Users/kcft114/.julia/packages/Flux/qXNjB/src/optimise/train.jl:105
┌ Info: Epoch 102
└ @ Main /Users/kcft114/.julia/packages/Flux/qXNjB/src/optimise/train.jl:105
┌ Info: Epoch 103
└ @ Main /Users/kcft114/.julia/packages/Flux/qXNjB/src/optimise/train.jl:105
┌ Info: Epoch 104
└ @ Main /Users/kcft114/.julia/packages/Flux/qXNjB/src/optimise/train.jl:105
┌ Info: Epoch 105
└ @ Main /Users/kcft114/.julia/packages/Flux/qXNjB/src/optimise/train.jl:105
┌ Info: Epoch 106
└ @ Main /Users/kcft114/.julia/packages/Flux/qXNjB/src/optimise/train.jl:105
┌ Info: Epoch 107
└ @ Main /Users/kcft114/.julia/packages/Flux/qXNjB/src/optimise/train.jl:105
┌ Info: Epoch 108
└ @ Main /Users/kcft114/.julia/packages/Flux/qXNjB/src/optimise/train.jl:105
┌ Info: Epoch 109
└ @ Main /Users/kcft114/.julia/packages/Flux/qXNjB/src/optimise/train.jl:105
┌ Info: Epoch 110
└ @ Main /Users/kcft114/.julia/packages/Flux/qXNjB/src/optimise/train.jl:105
```

```
loss(X, Y) = 1.3467125f0 (tracked)
loss(X, Y) = 1.3047087f0 (tracked)
loss(X, Y) = 1.3312677f0 (tracked)
loss(X, Y) = 1.2769516f0 (tracked)
loss(X, Y) = 1.2984444f0 (tracked)
loss(X, Y) = 1.2669388f0 (tracked)
loss(X, Y) = 1.3043057f0 (tracked)
loss(X, Y) = 1.2662073f0 (tracked)
loss(X, Y) = 1.3154396f0 (tracked)
loss(X, Y) = 1.290856f0 (tracked)
loss(X, Y) = 1.2551658f0 (tracked)
loss(X, Y) = 1.2386734f0 (tracked)
loss(X, Y) = 1.2204632f0 (tracked)
loss(X, Y) = 1.2175406f0 (tracked)
loss(X, Y) = 1.228545f0 (tracked)
loss(X, Y) = 1.2437586f0 (tracked)
loss(X, Y) = 1.2038511f0 (tracked)
loss(X, Y) = 1.216f0 (tracked)
loss(X, Y) = 1.225382f0 (tracked)
loss(X, Y) = 1.1653671f0 (tracked)
loss(X, Y) = 1.1597867f0 (tracked)
loss(X, Y) = 1.1930068f0 (tracked)
loss(X, Y) = 1.1875393f0 (tracked)
loss(X, Y) = 1.2402166f0 (tracked)
loss(X, Y) = 1.1792388f0 (tracked)
loss(X, Y) = 1.1611946f0 (tracked)
loss(X, Y) = 1.1977848f0 (tracked)
loss(X, Y) = 1.1838388f0 (tracked)
loss(X, Y) = 1.2040635f0 (tracked)
loss(X, Y) = 1.1631953f0 (tracked)
loss(X, Y) = 1.1820755f0 (tracked)
loss(X, Y) = 1.1527005f0 (tracked)
loss(X, Y) = 1.2093433f0 (tracked)
loss(X, Y) = 1.157059f0 (tracked)
loss(X, Y) = 1.1252534f0 (tracked)
loss(X, Y) = 1.1226615f0 (tracked)
loss(X, Y) = 1.1241466f0 (tracked)
loss(X, Y) = 1.1437389f0 (tracked)
loss(X, Y) = 1.1142156f0 (tracked)
loss(X, Y) = 1.0946305f0 (tracked)
loss(X, Y) = 1.085465f0 (tracked)
loss(X, Y) = 1.1147543f0 (tracked)
loss(X, Y) = 1.135716f0 (tracked)
loss(X, Y) = 1.0871335f0 (tracked)
loss(X, Y) = 1.089532f0 (tracked)
loss(X, Y) = 1.1146936f0 (tracked)
loss(X, Y) = 1.1097784f0 (tracked)
loss(X, Y) = 1.1416179f0 (tracked)
loss(X, Y) = 1.1174663f0 (tracked)
loss(X, Y) = 1.1093271f0 (tracked)
loss(X, Y) = 1.0824763f0 (tracked)
loss(X, Y) = 1.1080903f0 (tracked)
loss(X, Y) = 1.0973288f0 (tracked)
loss(X, Y) = 1.0524118f0 (tracked)
loss(X, Y) = 1.0606697f0 (tracked)
loss(X, Y) = 1.1223357f0 (tracked)
loss(X, Y) = 1.084105f0 (tracked)
loss(X, Y) = 1.1162953f0 (tracked)
loss(X, Y) = 1.1007217f0 (tracked)
loss(X, Y) = 1.0604643f0 (tracked)
loss(X, Y) = 1.0918428f0 (tracked)
loss(X, Y) = 1.0655751f0 (tracked)
loss(X, Y) = 1.0473906f0 (tracked)
loss(X, Y) = 1.0638561f0 (tracked)
loss(X, Y) = 1.0627724f0 (tracked)
loss(X, Y) = 1.0484501f0 (tracked)
loss(X, Y) = 1.0437859f0 (tracked)
loss(X, Y) = 1.0799747f0 (tracked)
loss(X, Y) = 1.0793113f0 (tracked)
loss(X, Y) = 1.0299392f0 (tracked)
loss(X, Y) = 1.0312334f0 (tracked)
loss(X, Y) = 1.048593f0 (tracked)
loss(X, Y) = 1.0553751f0 (tracked)
loss(X, Y) = 1.0523022f0 (tracked)
loss(X, Y) = 1.0353171f0 (tracked)
loss(X, Y) = 1.0765704f0 (tracked)
loss(X, Y) = 0.99791586f0 (tracked)
loss(X, Y) = 1.0265497f0 (tracked)
loss(X, Y) = 1.038456f0 (tracked)
loss(X, Y) = 1.0340308f0 (tracked)
loss(X, Y) = 0.98968285f0 (tracked)
loss(X, Y) = 1.0459561f0 (tracked)
loss(X, Y) = 1.0289427f0 (tracked)
loss(X, Y) = 0.9913067f0 (tracked)
loss(X, Y) = 0.9755689f0 (tracked)
loss(X, Y) = 1.0352666f0 (tracked)
loss(X, Y) = 0.9944549f0 (tracked)
loss(X, Y) = 1.0057347f0 (tracked)
loss(X, Y) = 1.0133622f0 (tracked)
loss(X, Y) = 1.0328121f0 (tracked)
loss(X, Y) = 0.9936522f0 (tracked)
loss(X, Y) = 0.97716f0 (tracked)
loss(X, Y) = 1.0565579f0 (tracked)
loss(X, Y) = 0.9754034f0 (tracked)
loss(X, Y) = 1.0299287f0 (tracked)
loss(X, Y) = 1.0103786f0 (tracked)
loss(X, Y) = 0.97637904f0 (tracked)
loss(X, Y) = 1.0166289f0 (tracked)
loss(X, Y) = 1.0008407f0 (tracked)
loss(X, Y) = 0.97993f0 (tracked)
loss(X, Y) = 0.9849493f0 (tracked)
loss(X, Y) = 0.96311545f0 (tracked)
loss(X, Y) = 0.9717797f0 (tracked)
loss(X, Y) = 0.97244126f0 (tracked)
loss(X, Y) = 0.99078095f0 (tracked)
loss(X, Y) = 0.96367246f0 (tracked)
loss(X, Y) = 0.93668836f0 (tracked)
loss(X, Y) = 0.9565912f0 (tracked)
loss(X, Y) = 0.948128f0 (tracked)
```

```
┌ Info: Epoch 111
└ @ Main /Users/kcft114/.julia/packages/Flux/qXNjB/src/optimise/train.jl:105
┌ Info: Epoch 112
└ @ Main /Users/kcft114/.julia/packages/Flux/qXNjB/src/optimise/train.jl:105
┌ Info: Epoch 113
└ @ Main /Users/kcft114/.julia/packages/Flux/qXNjB/src/optimise/train.jl:105
┌ Info: Epoch 114
└ @ Main /Users/kcft114/.julia/packages/Flux/qXNjB/src/optimise/train.jl:105
```

```
loss(X, Y) = 0.9839686f0 (tracked)
loss(X, Y) = 0.9411767f0 (tracked)
loss(X, Y) = 0.96405774f0 (tracked)
loss(X, Y) = 0.9257561f0 (tracked)
loss(X, Y) = 0.95268255f0 (tracked)
loss(X, Y) = 0.9891264f0 (tracked)
loss(X, Y) = 0.976072f0 (tracked)
loss(X, Y) = 0.97650737f0 (tracked)
loss(X, Y) = 0.9724991f0 (tracked)
loss(X, Y) = 0.9577151f0 (tracked)
loss(X, Y) = 0.97228366f0 (tracked)
loss(X, Y) = 0.98017496f0 (tracked)
loss(X, Y) = 0.9470698f0 (tracked)
loss(X, Y) = 0.9409116f0 (tracked)
loss(X, Y) = 0.9300347f0 (tracked)
loss(X, Y) = 0.94383484f0 (tracked)
loss(X, Y) = 0.95992386f0 (tracked)
loss(X, Y) = 0.93809724f0 (tracked)
loss(X, Y) = 0.8878861f0 (tracked)
loss(X, Y) = 0.93080366f0 (tracked)
loss(X, Y) = 0.92006487f0 (tracked)
loss(X, Y) = 0.955726f0 (tracked)
loss(X, Y) = 0.93053406f0 (tracked)
loss(X, Y) = 0.9553296f0 (tracked)
loss(X, Y) = 0.9556966f0 (tracked)
loss(X, Y) = 0.95113397f0 (tracked)
loss(X, Y) = 0.93613356f0 (tracked)
loss(X, Y) = 0.9219866f0 (tracked)
loss(X, Y) = 0.9351169f0 (tracked)
loss(X, Y) = 0.9066087f0 (tracked)
loss(X, Y) = 0.9471087f0 (tracked)
loss(X, Y) = 0.9201276f0 (tracked)
loss(X, Y) = 0.9225372f0 (tracked)
loss(X, Y) = 0.92714137f0 (tracked)
loss(X, Y) = 0.9177879f0 (tracked)
loss(X, Y) = 0.92457986f0 (tracked)
loss(X, Y) = 0.9104595f0 (tracked)
loss(X, Y) = 0.9164769f0 (tracked)
loss(X, Y) = 0.9239802f0 (tracked)
loss(X, Y) = 0.8936834f0 (tracked)
loss(X, Y) = 0.8893945f0 (tracked)
loss(X, Y) = 0.9275367f0 (tracked)
loss(X, Y) = 0.8857354f0 (tracked)
loss(X, Y) = 0.89744264f0 (tracked)
loss(X, Y) = 0.91361696f0 (tracked)
loss(X, Y) = 0.91840345f0 (tracked)
loss(X, Y) = 0.9024936f0 (tracked)
loss(X, Y) = 0.9133095f0 (tracked)
loss(X, Y) = 0.91450965f0 (tracked)
loss(X, Y) = 0.9393171f0 (tracked)
loss(X, Y) = 0.9095812f0 (tracked)
loss(X, Y) = 0.89218044f0 (tracked)
loss(X, Y) = 0.9234688f0 (tracked)
loss(X, Y) = 0.9194043f0 (tracked)
loss(X, Y) = 0.9215759f0 (tracked)
loss(X, Y) = 0.9252668f0 (tracked)
loss(X, Y) = 0.9213628f0 (tracked)
loss(X, Y) = 0.93098646f0 (tracked)
loss(X, Y) = 0.91015935f0 (tracked)
loss(X, Y) = 0.881624f0 (tracked)
loss(X, Y) = 0.91816545f0 (tracked)
loss(X, Y) = 0.8864144f0 (tracked)
loss(X, Y) = 0.8764293f0 (tracked)
loss(X, Y) = 0.8804156f0 (tracked)
loss(X, Y) = 0.8693856f0 (tracked)
loss(X, Y) = 0.8993589f0 (tracked)
loss(X, Y) = 0.8954014f0 (tracked)
loss(X, Y) = 0.8735167f0 (tracked)
loss(X, Y) = 0.8634664f0 (tracked)
loss(X, Y) = 0.8970945f0 (tracked)
loss(X, Y) = 0.8633842f0 (tracked)
loss(X, Y) = 0.879129f0 (tracked)
loss(X, Y) = 0.88793665f0 (tracked)
loss(X, Y) = 0.87114066f0 (tracked)
loss(X, Y) = 0.87879306f0 (tracked)
loss(X, Y) = 0.8766481f0 (tracked)
loss(X, Y) = 0.854134f0 (tracked)
loss(X, Y) = 0.8632811f0 (tracked)
loss(X, Y) = 0.89980775f0 (tracked)
loss(X, Y) = 0.8610014f0 (tracked)
loss(X, Y) = 0.9196308f0 (tracked)
loss(X, Y) = 0.8616388f0 (tracked)
loss(X, Y) = 0.8690876f0 (tracked)
loss(X, Y) = 0.8591752f0 (tracked)
loss(X, Y) = 0.8746648f0 (tracked)
loss(X, Y) = 0.87761974f0 (tracked)
loss(X, Y) = 0.8871033f0 (tracked)
loss(X, Y) = 0.85495716f0 (tracked)
loss(X, Y) = 0.83289796f0 (tracked)
loss(X, Y) = 0.88325953f0 (tracked)
loss(X, Y) = 0.8413782f0 (tracked)
loss(X, Y) = 0.85229176f0 (tracked)
loss(X, Y) = 0.87521815f0 (tracked)
loss(X, Y) = 0.8722207f0 (tracked)
loss(X, Y) = 0.8585913f0 (tracked)
loss(X, Y) = 0.8918467f0 (tracked)
loss(X, Y) = 0.8567065f0 (tracked)
loss(X, Y) = 0.8782688f0 (tracked)
loss(X, Y) = 0.8507946f0 (tracked)
```

```
┌ Info: Epoch 115
└ @ Main /Users/kcft114/.julia/packages/Flux/qXNjB/src/optimise/train.jl:105
┌ Info: Epoch 116
└ @ Main /Users/kcft114/.julia/packages/Flux/qXNjB/src/optimise/train.jl:105
┌ Info: Epoch 117
└ @ Main /Users/kcft114/.julia/packages/Flux/qXNjB/src/optimise/train.jl:105
┌ Info: Epoch 118
└ @ Main /Users/kcft114/.julia/packages/Flux/qXNjB/src/optimise/train.jl:105
┌ Info: Epoch 119
└ @ Main /Users/kcft114/.julia/packages/Flux/qXNjB/src/optimise/train.jl:105
┌ Info: Epoch 120
└ @ Main /Users/kcft114/.julia/packages/Flux/qXNjB/src/optimise/train.jl:105
┌ Info: Epoch 121
└ @ Main /Users/kcft114/.julia/packages/Flux/qXNjB/src/optimise/train.jl:105
┌ Info: Epoch 122
└ @ Main /Users/kcft114/.julia/packages/Flux/qXNjB/src/optimise/train.jl:105
┌ Info: Epoch 123
└ @ Main /Users/kcft114/.julia/packages/Flux/qXNjB/src/optimise/train.jl:105
┌ Info: Epoch 124
└ @ Main /Users/kcft114/.julia/packages/Flux/qXNjB/src/optimise/train.jl:105
┌ Info: Epoch 125
└ @ Main /Users/kcft114/.julia/packages/Flux/qXNjB/src/optimise/train.jl:105
┌ Info: Epoch 126
└ @ Main /Users/kcft114/.julia/packages/Flux/qXNjB/src/optimise/train.jl:105
┌ Info: Epoch 127
└ @ Main /Users/kcft114/.julia/packages/Flux/qXNjB/src/optimise/train.jl:105
┌ Info: Epoch 128
└ @ Main /Users/kcft114/.julia/packages/Flux/qXNjB/src/optimise/train.jl:105
┌ Info: Epoch 129
└ @ Main /Users/kcft114/.julia/packages/Flux/qXNjB/src/optimise/train.jl:105
┌ Info: Epoch 130
└ @ Main /Users/kcft114/.julia/packages/Flux/qXNjB/src/optimise/train.jl:105
┌ Info: Epoch 131
└ @ Main /Users/kcft114/.julia/packages/Flux/qXNjB/src/optimise/train.jl:105
┌ Info: Epoch 132
└ @ Main /Users/kcft114/.julia/packages/Flux/qXNjB/src/optimise/train.jl:105
┌ Info: Epoch 133
└ @ Main /Users/kcft114/.julia/packages/Flux/qXNjB/src/optimise/train.jl:105
┌ Info: Epoch 134
└ @ Main /Users/kcft114/.julia/packages/Flux/qXNjB/src/optimise/train.jl:105
┌ Info: Epoch 135
└ @ Main /Users/kcft114/.julia/packages/Flux/qXNjB/src/optimise/train.jl:105
┌ Info: Epoch 136
└ @ Main /Users/kcft114/.julia/packages/Flux/qXNjB/src/optimise/train.jl:105
┌ Info: Epoch 137
└ @ Main /Users/kcft114/.julia/packages/Flux/qXNjB/src/optimise/train.jl:105
┌ Info: Epoch 138
└ @ Main /Users/kcft114/.julia/packages/Flux/qXNjB/src/optimise/train.jl:105
┌ Info: Epoch 139
└ @ Main /Users/kcft114/.julia/packages/Flux/qXNjB/src/optimise/train.jl:105
┌ Info: Epoch 140
└ @ Main /Users/kcft114/.julia/packages/Flux/qXNjB/src/optimise/train.jl:105
┌ Info: Epoch 141
└ @ Main /Users/kcft114/.julia/packages/Flux/qXNjB/src/optimise/train.jl:105
┌ Info: Epoch 142
└ @ Main /Users/kcft114/.julia/packages/Flux/qXNjB/src/optimise/train.jl:105
┌ Info: Epoch 143
└ @ Main /Users/kcft114/.julia/packages/Flux/qXNjB/src/optimise/train.jl:105
┌ Info: Epoch 144
└ @ Main /Users/kcft114/.julia/packages/Flux/qXNjB/src/optimise/train.jl:105
┌ Info: Epoch 145
└ @ Main /Users/kcft114/.julia/packages/Flux/qXNjB/src/optimise/train.jl:105
┌ Info: Epoch 146
└ @ Main /Users/kcft114/.julia/packages/Flux/qXNjB/src/optimise/train.jl:105
┌ Info: Epoch 147
└ @ Main /Users/kcft114/.julia/packages/Flux/qXNjB/src/optimise/train.jl:105
┌ Info: Epoch 148
└ @ Main /Users/kcft114/.julia/packages/Flux/qXNjB/src/optimise/train.jl:105
┌ Info: Epoch 149
└ @ Main /Users/kcft114/.julia/packages/Flux/qXNjB/src/optimise/train.jl:105
┌ Info: Epoch 150
└ @ Main /Users/kcft114/.julia/packages/Flux/qXNjB/src/optimise/train.jl:105
┌ Info: Epoch 151
└ @ Main /Users/kcft114/.julia/packages/Flux/qXNjB/src/optimise/train.jl:105
┌ Info: Epoch 152
└ @ Main /Users/kcft114/.julia/packages/Flux/qXNjB/src/optimise/train.jl:105
┌ Info: Epoch 153
└ @ Main /Users/kcft114/.julia/packages/Flux/qXNjB/src/optimise/train.jl:105
┌ Info: Epoch 154
└ @ Main /Users/kcft114/.julia/packages/Flux/qXNjB/src/optimise/train.jl:105
┌ Info: Epoch 155
└ @ Main /Users/kcft114/.julia/packages/Flux/qXNjB/src/optimise/train.jl:105
┌ Info: Epoch 156
└ @ Main /Users/kcft114/.julia/packages/Flux/qXNjB/src/optimise/train.jl:105
┌ Info: Epoch 157
└ @ Main /Users/kcft114/.julia/packages/Flux/qXNjB/src/optimise/train.jl:105
┌ Info: Epoch 158
└ @ Main /Users/kcft114/.julia/packages/Flux/qXNjB/src/optimise/train.jl:105
┌ Info: Epoch 159
└ @ Main /Users/kcft114/.julia/packages/Flux/qXNjB/src/optimise/train.jl:105
┌ Info: Epoch 160
└ @ Main /Users/kcft114/.julia/packages/Flux/qXNjB/src/optimise/train.jl:105
┌ Info: Epoch 161
└ @ Main /Users/kcft114/.julia/packages/Flux/qXNjB/src/optimise/train.jl:105
┌ Info: Epoch 162
└ @ Main /Users/kcft114/.julia/packages/Flux/qXNjB/src/optimise/train.jl:105
┌ Info: Epoch 163
└ @ Main /Users/kcft114/.julia/packages/Flux/qXNjB/src/optimise/train.jl:105
┌ Info: Epoch 164
└ @ Main /Users/kcft114/.julia/packages/Flux/qXNjB/src/optimise/train.jl:105
┌ Info: Epoch 165
└ @ Main /Users/kcft114/.julia/packages/Flux/qXNjB/src/optimise/train.jl:105
┌ Info: Epoch 166
└ @ Main /Users/kcft114/.julia/packages/Flux/qXNjB/src/optimise/train.jl:105
┌ Info: Epoch 167
└ @ Main /Users/kcft114/.julia/packages/Flux/qXNjB/src/optimise/train.jl:105
┌ Info: Epoch 168
└ @ Main /Users/kcft114/.julia/packages/Flux/qXNjB/src/optimise/train.jl:105
┌ Info: Epoch 169
└ @ Main /Users/kcft114/.julia/packages/Flux/qXNjB/src/optimise/train.jl:105
┌ Info: Epoch 170
└ @ Main /Users/kcft114/.julia/packages/Flux/qXNjB/src/optimise/train.jl:105
┌ Info: Epoch 171
└ @ Main /Users/kcft114/.julia/packages/Flux/qXNjB/src/optimise/train.jl:105
┌ Info: Epoch 172
└ @ Main /Users/kcft114/.julia/packages/Flux/qXNjB/src/optimise/train.jl:105
┌ Info: Epoch 173
└ @ Main /Users/kcft114/.julia/packages/Flux/qXNjB/src/optimise/train.jl:105
┌ Info: Epoch 174
└ @ Main /Users/kcft114/.julia/packages/Flux/qXNjB/src/optimise/train.jl:105
┌ Info: Epoch 175
└ @ Main /Users/kcft114/.julia/packages/Flux/qXNjB/src/optimise/train.jl:105
┌ Info: Epoch 176
└ @ Main /Users/kcft114/.julia/packages/Flux/qXNjB/src/optimise/train.jl:105
┌ Info: Epoch 177
└ @ Main /Users/kcft114/.julia/packages/Flux/qXNjB/src/optimise/train.jl:105
┌ Info: Epoch 178
└ @ Main /Users/kcft114/.julia/packages/Flux/qXNjB/src/optimise/train.jl:105
┌ Info: Epoch 179
└ @ Main /Users/kcft114/.julia/packages/Flux/qXNjB/src/optimise/train.jl:105
┌ Info: Epoch 180
└ @ Main /Users/kcft114/.julia/packages/Flux/qXNjB/src/optimise/train.jl:105
┌ Info: Epoch 181
└ @ Main /Users/kcft114/.julia/packages/Flux/qXNjB/src/optimise/train.jl:105
┌ Info: Epoch 182
└ @ Main /Users/kcft114/.julia/packages/Flux/qXNjB/src/optimise/train.jl:105
┌ Info: Epoch 183
└ @ Main /Users/kcft114/.julia/packages/Flux/qXNjB/src/optimise/train.jl:105
┌ Info: Epoch 184
└ @ Main /Users/kcft114/.julia/packages/Flux/qXNjB/src/optimise/train.jl:105
┌ Info: Epoch 185
└ @ Main /Users/kcft114/.julia/packages/Flux/qXNjB/src/optimise/train.jl:105
┌ Info: Epoch 186
└ @ Main /Users/kcft114/.julia/packages/Flux/qXNjB/src/optimise/train.jl:105
┌ Info: Epoch 187
└ @ Main /Users/kcft114/.julia/packages/Flux/qXNjB/src/optimise/train.jl:105
┌ Info: Epoch 188
└ @ Main /Users/kcft114/.julia/packages/Flux/qXNjB/src/optimise/train.jl:105
┌ Info: Epoch 189
└ @ Main /Users/kcft114/.julia/packages/Flux/qXNjB/src/optimise/train.jl:105
┌ Info: Epoch 190
└ @ Main /Users/kcft114/.julia/packages/Flux/qXNjB/src/optimise/train.jl:105
┌ Info: Epoch 191
└ @ Main /Users/kcft114/.julia/packages/Flux/qXNjB/src/optimise/train.jl:105
┌ Info: Epoch 192
└ @ Main /Users/kcft114/.julia/packages/Flux/qXNjB/src/optimise/train.jl:105
┌ Info: Epoch 193
└ @ Main /Users/kcft114/.julia/packages/Flux/qXNjB/src/optimise/train.jl:105
┌ Info: Epoch 194
└ @ Main /Users/kcft114/.julia/packages/Flux/qXNjB/src/optimise/train.jl:105
┌ Info: Epoch 195
└ @ Main /Users/kcft114/.julia/packages/Flux/qXNjB/src/optimise/train.jl:105
┌ Info: Epoch 196
└ @ Main /Users/kcft114/.julia/packages/Flux/qXNjB/src/optimise/train.jl:105
┌ Info: Epoch 197
└ @ Main /Users/kcft114/.julia/packages/Flux/qXNjB/src/optimise/train.jl:105
┌ Info: Epoch 198
└ @ Main /Users/kcft114/.julia/packages/Flux/qXNjB/src/optimise/train.jl:105
┌ Info: Epoch 199
└ @ Main /Users/kcft114/.julia/packages/Flux/qXNjB/src/optimise/train.jl:105
┌ Info: Epoch 200
└ @ Main /Users/kcft114/.julia/packages/Flux/qXNjB/src/optimise/train.jl:105
┌ Info: Epoch 201
└ @ Main /Users/kcft114/.julia/packages/Flux/qXNjB/src/optimise/train.jl:105
┌ Info: Epoch 202
└ @ Main /Users/kcft114/.julia/packages/Flux/qXNjB/src/optimise/train.jl:105
┌ Info: Epoch 203
└ @ Main /Users/kcft114/.julia/packages/Flux/qXNjB/src/optimise/train.jl:105
┌ Info: Epoch 204
└ @ Main /Users/kcft114/.julia/packages/Flux/qXNjB/src/optimise/train.jl:105
┌ Info: Epoch 205
└ @ Main /Users/kcft114/.julia/packages/Flux/qXNjB/src/optimise/train.jl:105
┌ Info: Epoch 206
└ @ Main /Users/kcft114/.julia/packages/Flux/qXNjB/src/optimise/train.jl:105
┌ Info: Epoch 207
└ @ Main /Users/kcft114/.julia/packages/Flux/qXNjB/src/optimise/train.jl:105
┌ Info: Epoch 208
└ @ Main /Users/kcft114/.julia/packages/Flux/qXNjB/src/optimise/train.jl:105
┌ Info: Epoch 209
└ @ Main /Users/kcft114/.julia/packages/Flux/qXNjB/src/optimise/train.jl:105
┌ Info: Epoch 210
└ @ Main /Users/kcft114/.julia/packages/Flux/qXNjB/src/optimise/train.jl:105
┌ Info: Epoch 211
└ @ Main /Users/kcft114/.julia/packages/Flux/qXNjB/src/optimise/train.jl:105
┌ Info: Epoch 212
└ @ Main /Users/kcft114/.julia/packages/Flux/qXNjB/src/optimise/train.jl:105
┌ Info: Epoch 213
└ @ Main /Users/kcft114/.julia/packages/Flux/qXNjB/src/optimise/train.jl:105
```

```
loss(X, Y) = 0.8860216f0 (tracked)
loss(X, Y) = 0.8609177f0 (tracked)
loss(X, Y) = 0.8594374f0 (tracked)
loss(X, Y) = 0.8205869f0 (tracked)
loss(X, Y) = 0.84911f0 (tracked)
loss(X, Y) = 0.84408474f0 (tracked)
loss(X, Y) = 0.8429455f0 (tracked)
loss(X, Y) = 0.84232336f0 (tracked)
loss(X, Y) = 0.8496332f0 (tracked)
loss(X, Y) = 0.8742406f0 (tracked)
loss(X, Y) = 0.8688019f0 (tracked)
loss(X, Y) = 0.8725018f0 (tracked)
loss(X, Y) = 0.8730654f0 (tracked)
loss(X, Y) = 0.83709425f0 (tracked)
loss(X, Y) = 0.83062196f0 (tracked)
loss(X, Y) = 0.84921414f0 (tracked)
loss(X, Y) = 0.89192104f0 (tracked)
loss(X, Y) = 0.8403022f0 (tracked)
loss(X, Y) = 0.8532394f0 (tracked)
loss(X, Y) = 0.8682464f0 (tracked)
loss(X, Y) = 0.84390706f0 (tracked)
loss(X, Y) = 0.854736f0 (tracked)
loss(X, Y) = 0.88314515f0 (tracked)
loss(X, Y) = 0.85414165f0 (tracked)
loss(X, Y) = 0.8204616f0 (tracked)
loss(X, Y) = 0.83599013f0 (tracked)
loss(X, Y) = 0.8723713f0 (tracked)
loss(X, Y) = 0.8714295f0 (tracked)
loss(X, Y) = 0.8404717f0 (tracked)
loss(X, Y) = 0.86368924f0 (tracked)
loss(X, Y) = 0.84366924f0 (tracked)
loss(X, Y) = 0.85056514f0 (tracked)
loss(X, Y) = 0.8628842f0 (tracked)
loss(X, Y) = 0.85738426f0 (tracked)
loss(X, Y) = 0.8894725f0 (tracked)
loss(X, Y) = 0.8415478f0 (tracked)
loss(X, Y) = 0.83079505f0 (tracked)
loss(X, Y) = 0.8403887f0 (tracked)
loss(X, Y) = 0.8517525f0 (tracked)
loss(X, Y) = 0.8476496f0 (tracked)
loss(X, Y) = 0.84388363f0 (tracked)
loss(X, Y) = 0.8408921f0 (tracked)
loss(X, Y) = 0.88579005f0 (tracked)
loss(X, Y) = 0.8390226f0 (tracked)
loss(X, Y) = 0.8206099f0 (tracked)
loss(X, Y) = 0.8286992f0 (tracked)
loss(X, Y) = 0.8240276f0 (tracked)
loss(X, Y) = 0.7976628f0 (tracked)
loss(X, Y) = 0.8425422f0 (tracked)
loss(X, Y) = 0.8208648f0 (tracked)
loss(X, Y) = 0.86873275f0 (tracked)
loss(X, Y) = 0.8456189f0 (tracked)
loss(X, Y) = 0.8342966f0 (tracked)
loss(X, Y) = 0.82279617f0 (tracked)
loss(X, Y) = 0.8455843f0 (tracked)
loss(X, Y) = 0.8103625f0 (tracked)
loss(X, Y) = 0.80683345f0 (tracked)
loss(X, Y) = 0.86179f0 (tracked)
loss(X, Y) = 0.8374699f0 (tracked)
loss(X, Y) = 0.8100735f0 (tracked)
loss(X, Y) = 0.8040225f0 (tracked)
loss(X, Y) = 0.8331151f0 (tracked)
loss(X, Y) = 0.8090667f0 (tracked)
loss(X, Y) = 0.7907094f0 (tracked)
loss(X, Y) = 0.8526385f0 (tracked)
loss(X, Y) = 0.8269325f0 (tracked)
loss(X, Y) = 0.7963861f0 (tracked)
loss(X, Y) = 0.8276748f0 (tracked)
loss(X, Y) = 0.86182916f0 (tracked)
loss(X, Y) = 0.88395905f0 (tracked)
loss(X, Y) = 0.83950007f0 (tracked)
loss(X, Y) = 0.8014383f0 (tracked)
loss(X, Y) = 0.8110664f0 (tracked)
loss(X, Y) = 0.8173764f0 (tracked)
loss(X, Y) = 0.8053947f0 (tracked)
loss(X, Y) = 0.84283614f0 (tracked)
loss(X, Y) = 0.818394f0 (tracked)
loss(X, Y) = 0.77193075f0 (tracked)
loss(X, Y) = 0.7881768f0 (tracked)
loss(X, Y) = 0.8317665f0 (tracked)
loss(X, Y) = 0.8126892f0 (tracked)
loss(X, Y) = 0.815563f0 (tracked)
loss(X, Y) = 0.8131646f0 (tracked)
loss(X, Y) = 0.8062998f0 (tracked)
loss(X, Y) = 0.8543282f0 (tracked)
loss(X, Y) = 0.80934286f0 (tracked)
loss(X, Y) = 0.8228883f0 (tracked)
loss(X, Y) = 0.8322991f0 (tracked)
loss(X, Y) = 0.8257181f0 (tracked)
loss(X, Y) = 0.79811865f0 (tracked)
loss(X, Y) = 0.827395f0 (tracked)
loss(X, Y) = 0.79926544f0 (tracked)
loss(X, Y) = 0.8099024f0 (tracked)
loss(X, Y) = 0.8115616f0 (tracked)
loss(X, Y) = 0.8029746f0 (tracked)
loss(X, Y) = 0.79434454f0 (tracked)
loss(X, Y) = 0.7951581f0 (tracked)
loss(X, Y) = 0.8362379f0 (tracked)
loss(X, Y) = 0.78495014f0 (tracked)
loss(X, Y) = 0.8180045f0 (tracked)
loss(X, Y) = 0.79855794f0 (tracked)
loss(X, Y) = 0.82884467f0 (tracked)
```

```
┌ Info: Epoch 214
└ @ Main /Users/kcft114/.julia/packages/Flux/qXNjB/src/optimise/train.jl:105
┌ Info: Epoch 215
└ @ Main /Users/kcft114/.julia/packages/Flux/qXNjB/src/optimise/train.jl:105
┌ Info: Epoch 216
└ @ Main /Users/kcft114/.julia/packages/Flux/qXNjB/src/optimise/train.jl:105
┌ Info: Epoch 217
└ @ Main /Users/kcft114/.julia/packages/Flux/qXNjB/src/optimise/train.jl:105
┌ Info: Epoch 218
└ @ Main /Users/kcft114/.julia/packages/Flux/qXNjB/src/optimise/train.jl:105
┌ Info: Epoch 219
└ @ Main /Users/kcft114/.julia/packages/Flux/qXNjB/src/optimise/train.jl:105
┌ Info: Epoch 220
└ @ Main /Users/kcft114/.julia/packages/Flux/qXNjB/src/optimise/train.jl:105
┌ Info: Epoch 221
└ @ Main /Users/kcft114/.julia/packages/Flux/qXNjB/src/optimise/train.jl:105
┌ Info: Epoch 222
└ @ Main /Users/kcft114/.julia/packages/Flux/qXNjB/src/optimise/train.jl:105
┌ Info: Epoch 223
└ @ Main /Users/kcft114/.julia/packages/Flux/qXNjB/src/optimise/train.jl:105
┌ Info: Epoch 224
└ @ Main /Users/kcft114/.julia/packages/Flux/qXNjB/src/optimise/train.jl:105
┌ Info: Epoch 225
└ @ Main /Users/kcft114/.julia/packages/Flux/qXNjB/src/optimise/train.jl:105
┌ Info: Epoch 226
└ @ Main /Users/kcft114/.julia/packages/Flux/qXNjB/src/optimise/train.jl:105
┌ Info: Epoch 227
└ @ Main /Users/kcft114/.julia/packages/Flux/qXNjB/src/optimise/train.jl:105
┌ Info: Epoch 228
└ @ Main /Users/kcft114/.julia/packages/Flux/qXNjB/src/optimise/train.jl:105
┌ Info: Epoch 229
└ @ Main /Users/kcft114/.julia/packages/Flux/qXNjB/src/optimise/train.jl:105
┌ Info: Epoch 230
└ @ Main /Users/kcft114/.julia/packages/Flux/qXNjB/src/optimise/train.jl:105
┌ Info: Epoch 231
└ @ Main /Users/kcft114/.julia/packages/Flux/qXNjB/src/optimise/train.jl:105
┌ Info: Epoch 232
└ @ Main /Users/kcft114/.julia/packages/Flux/qXNjB/src/optimise/train.jl:105
┌ Info: Epoch 233
└ @ Main /Users/kcft114/.julia/packages/Flux/qXNjB/src/optimise/train.jl:105
┌ Info: Epoch 234
└ @ Main /Users/kcft114/.julia/packages/Flux/qXNjB/src/optimise/train.jl:105
┌ Info: Epoch 235
└ @ Main /Users/kcft114/.julia/packages/Flux/qXNjB/src/optimise/train.jl:105
┌ Info: Epoch 236
└ @ Main /Users/kcft114/.julia/packages/Flux/qXNjB/src/optimise/train.jl:105
┌ Info: Epoch 237
└ @ Main /Users/kcft114/.julia/packages/Flux/qXNjB/src/optimise/train.jl:105
┌ Info: Epoch 238
└ @ Main /Users/kcft114/.julia/packages/Flux/qXNjB/src/optimise/train.jl:105
┌ Info: Epoch 239
└ @ Main /Users/kcft114/.julia/packages/Flux/qXNjB/src/optimise/train.jl:105
┌ Info: Epoch 240
└ @ Main /Users/kcft114/.julia/packages/Flux/qXNjB/src/optimise/train.jl:105
┌ Info: Epoch 241
└ @ Main /Users/kcft114/.julia/packages/Flux/qXNjB/src/optimise/train.jl:105
┌ Info: Epoch 242
└ @ Main /Users/kcft114/.julia/packages/Flux/qXNjB/src/optimise/train.jl:105
┌ Info: Epoch 243
└ @ Main /Users/kcft114/.julia/packages/Flux/qXNjB/src/optimise/train.jl:105
┌ Info: Epoch 244
└ @ Main /Users/kcft114/.julia/packages/Flux/qXNjB/src/optimise/train.jl:105
┌ Info: Epoch 245
└ @ Main /Users/kcft114/.julia/packages/Flux/qXNjB/src/optimise/train.jl:105
┌ Info: Epoch 246
└ @ Main /Users/kcft114/.julia/packages/Flux/qXNjB/src/optimise/train.jl:105
┌ Info: Epoch 247
└ @ Main /Users/kcft114/.julia/packages/Flux/qXNjB/src/optimise/train.jl:105
┌ Info: Epoch 248
└ @ Main /Users/kcft114/.julia/packages/Flux/qXNjB/src/optimise/train.jl:105
┌ Info: Epoch 249
└ @ Main /Users/kcft114/.julia/packages/Flux/qXNjB/src/optimise/train.jl:105
┌ Info: Epoch 250
└ @ Main /Users/kcft114/.julia/packages/Flux/qXNjB/src/optimise/train.jl:105
┌ Info: Epoch 251
└ @ Main /Users/kcft114/.julia/packages/Flux/qXNjB/src/optimise/train.jl:105
┌ Info: Epoch 252
└ @ Main /Users/kcft114/.julia/packages/Flux/qXNjB/src/optimise/train.jl:105
┌ Info: Epoch 253
└ @ Main /Users/kcft114/.julia/packages/Flux/qXNjB/src/optimise/train.jl:105
┌ Info: Epoch 254
└ @ Main /Users/kcft114/.julia/packages/Flux/qXNjB/src/optimise/train.jl:105
┌ Info: Epoch 255
└ @ Main /Users/kcft114/.julia/packages/Flux/qXNjB/src/optimise/train.jl:105
┌ Info: Epoch 256
└ @ Main /Users/kcft114/.julia/packages/Flux/qXNjB/src/optimise/train.jl:105
┌ Info: Epoch 257
└ @ Main /Users/kcft114/.julia/packages/Flux/qXNjB/src/optimise/train.jl:105
┌ Info: Epoch 258
└ @ Main /Users/kcft114/.julia/packages/Flux/qXNjB/src/optimise/train.jl:105
┌ Info: Epoch 259
└ @ Main /Users/kcft114/.julia/packages/Flux/qXNjB/src/optimise/train.jl:105
┌ Info: Epoch 260
└ @ Main /Users/kcft114/.julia/packages/Flux/qXNjB/src/optimise/train.jl:105
┌ Info: Epoch 261
└ @ Main /Users/kcft114/.julia/packages/Flux/qXNjB/src/optimise/train.jl:105
┌ Info: Epoch 262
└ @ Main /Users/kcft114/.julia/packages/Flux/qXNjB/src/optimise/train.jl:105
┌ Info: Epoch 263
└ @ Main /Users/kcft114/.julia/packages/Flux/qXNjB/src/optimise/train.jl:105
┌ Info: Epoch 264
└ @ Main /Users/kcft114/.julia/packages/Flux/qXNjB/src/optimise/train.jl:105
┌ Info: Epoch 265
└ @ Main /Users/kcft114/.julia/packages/Flux/qXNjB/src/optimise/train.jl:105
┌ Info: Epoch 266
└ @ Main /Users/kcft114/.julia/packages/Flux/qXNjB/src/optimise/train.jl:105
┌ Info: Epoch 267
└ @ Main /Users/kcft114/.julia/packages/Flux/qXNjB/src/optimise/train.jl:105
┌ Info: Epoch 268
└ @ Main /Users/kcft114/.julia/packages/Flux/qXNjB/src/optimise/train.jl:105
┌ Info: Epoch 269
└ @ Main /Users/kcft114/.julia/packages/Flux/qXNjB/src/optimise/train.jl:105
┌ Info: Epoch 270
└ @ Main /Users/kcft114/.julia/packages/Flux/qXNjB/src/optimise/train.jl:105
┌ Info: Epoch 271
└ @ Main /Users/kcft114/.julia/packages/Flux/qXNjB/src/optimise/train.jl:105
┌ Info: Epoch 272
└ @ Main /Users/kcft114/.julia/packages/Flux/qXNjB/src/optimise/train.jl:105
┌ Info: Epoch 273
└ @ Main /Users/kcft114/.julia/packages/Flux/qXNjB/src/optimise/train.jl:105
┌ Info: Epoch 274
└ @ Main /Users/kcft114/.julia/packages/Flux/qXNjB/src/optimise/train.jl:105
┌ Info: Epoch 275
└ @ Main /Users/kcft114/.julia/packages/Flux/qXNjB/src/optimise/train.jl:105
┌ Info: Epoch 276
└ @ Main /Users/kcft114/.julia/packages/Flux/qXNjB/src/optimise/train.jl:105
┌ Info: Epoch 277
└ @ Main /Users/kcft114/.julia/packages/Flux/qXNjB/src/optimise/train.jl:105
┌ Info: Epoch 278
└ @ Main /Users/kcft114/.julia/packages/Flux/qXNjB/src/optimise/train.jl:105
┌ Info: Epoch 279
└ @ Main /Users/kcft114/.julia/packages/Flux/qXNjB/src/optimise/train.jl:105
┌ Info: Epoch 280
└ @ Main /Users/kcft114/.julia/packages/Flux/qXNjB/src/optimise/train.jl:105
┌ Info: Epoch 281
└ @ Main /Users/kcft114/.julia/packages/Flux/qXNjB/src/optimise/train.jl:105
┌ Info: Epoch 282
└ @ Main /Users/kcft114/.julia/packages/Flux/qXNjB/src/optimise/train.jl:105
┌ Info: Epoch 283
└ @ Main /Users/kcft114/.julia/packages/Flux/qXNjB/src/optimise/train.jl:105
┌ Info: Epoch 284
└ @ Main /Users/kcft114/.julia/packages/Flux/qXNjB/src/optimise/train.jl:105
┌ Info: Epoch 285
└ @ Main /Users/kcft114/.julia/packages/Flux/qXNjB/src/optimise/train.jl:105
┌ Info: Epoch 286
└ @ Main /Users/kcft114/.julia/packages/Flux/qXNjB/src/optimise/train.jl:105
┌ Info: Epoch 287
└ @ Main /Users/kcft114/.julia/packages/Flux/qXNjB/src/optimise/train.jl:105
┌ Info: Epoch 288
└ @ Main /Users/kcft114/.julia/packages/Flux/qXNjB/src/optimise/train.jl:105
┌ Info: Epoch 289
└ @ Main /Users/kcft114/.julia/packages/Flux/qXNjB/src/optimise/train.jl:105
┌ Info: Epoch 290
└ @ Main /Users/kcft114/.julia/packages/Flux/qXNjB/src/optimise/train.jl:105
┌ Info: Epoch 291
└ @ Main /Users/kcft114/.julia/packages/Flux/qXNjB/src/optimise/train.jl:105
┌ Info: Epoch 292
└ @ Main /Users/kcft114/.julia/packages/Flux/qXNjB/src/optimise/train.jl:105
┌ Info: Epoch 293
└ @ Main /Users/kcft114/.julia/packages/Flux/qXNjB/src/optimise/train.jl:105
┌ Info: Epoch 294
└ @ Main /Users/kcft114/.julia/packages/Flux/qXNjB/src/optimise/train.jl:105
┌ Info: Epoch 295
└ @ Main /Users/kcft114/.julia/packages/Flux/qXNjB/src/optimise/train.jl:105
┌ Info: Epoch 296
└ @ Main /Users/kcft114/.julia/packages/Flux/qXNjB/src/optimise/train.jl:105
┌ Info: Epoch 297
└ @ Main /Users/kcft114/.julia/packages/Flux/qXNjB/src/optimise/train.jl:105
┌ Info: Epoch 298
└ @ Main /Users/kcft114/.julia/packages/Flux/qXNjB/src/optimise/train.jl:105
┌ Info: Epoch 299
└ @ Main /Users/kcft114/.julia/packages/Flux/qXNjB/src/optimise/train.jl:105
┌ Info: Epoch 300
└ @ Main /Users/kcft114/.julia/packages/Flux/qXNjB/src/optimise/train.jl:105
┌ Info: Epoch 301
└ @ Main /Users/kcft114/.julia/packages/Flux/qXNjB/src/optimise/train.jl:105
┌ Info: Epoch 302
└ @ Main /Users/kcft114/.julia/packages/Flux/qXNjB/src/optimise/train.jl:105
┌ Info: Epoch 303
└ @ Main /Users/kcft114/.julia/packages/Flux/qXNjB/src/optimise/train.jl:105
┌ Info: Epoch 304
└ @ Main /Users/kcft114/.julia/packages/Flux/qXNjB/src/optimise/train.jl:105
┌ Info: Epoch 305
└ @ Main /Users/kcft114/.julia/packages/Flux/qXNjB/src/optimise/train.jl:105
┌ Info: Epoch 306
└ @ Main /Users/kcft114/.julia/packages/Flux/qXNjB/src/optimise/train.jl:105
┌ Info: Epoch 307
└ @ Main /Users/kcft114/.julia/packages/Flux/qXNjB/src/optimise/train.jl:105
┌ Info: Epoch 308
└ @ Main /Users/kcft114/.julia/packages/Flux/qXNjB/src/optimise/train.jl:105
┌ Info: Epoch 309
└ @ Main /Users/kcft114/.julia/packages/Flux/qXNjB/src/optimise/train.jl:105
┌ Info: Epoch 310
└ @ Main /Users/kcft114/.julia/packages/Flux/qXNjB/src/optimise/train.jl:105
┌ Info: Epoch 311
└ @ Main /Users/kcft114/.julia/packages/Flux/qXNjB/src/optimise/train.jl:105
┌ Info: Epoch 312
└ @ Main /Users/kcft114/.julia/packages/Flux/qXNjB/src/optimise/train.jl:105
┌ Info: Epoch 313
└ @ Main /Users/kcft114/.julia/packages/Flux/qXNjB/src/optimise/train.jl:105
┌ Info: Epoch 314
└ @ Main /Users/kcft114/.julia/packages/Flux/qXNjB/src/optimise/train.jl:105
┌ Info: Epoch 315
└ @ Main /Users/kcft114/.julia/packages/Flux/qXNjB/src/optimise/train.jl:105
```

```
loss(X, Y) = 0.83246857f0 (tracked)
loss(X, Y) = 0.8019063f0 (tracked)
loss(X, Y) = 0.8167447f0 (tracked)
loss(X, Y) = 0.81687295f0 (tracked)
loss(X, Y) = 0.79665357f0 (tracked)
loss(X, Y) = 0.7890146f0 (tracked)
loss(X, Y) = 0.779886f0 (tracked)
loss(X, Y) = 0.82075477f0 (tracked)
loss(X, Y) = 0.7997042f0 (tracked)
loss(X, Y) = 0.7867913f0 (tracked)
loss(X, Y) = 0.81163853f0 (tracked)
loss(X, Y) = 0.7595922f0 (tracked)
loss(X, Y) = 0.7965541f0 (tracked)
loss(X, Y) = 0.77363986f0 (tracked)
loss(X, Y) = 0.8004299f0 (tracked)
loss(X, Y) = 0.80332255f0 (tracked)
loss(X, Y) = 0.77309185f0 (tracked)
loss(X, Y) = 0.7830155f0 (tracked)
loss(X, Y) = 0.8276161f0 (tracked)
loss(X, Y) = 0.7893118f0 (tracked)
loss(X, Y) = 0.81413925f0 (tracked)
loss(X, Y) = 0.8037464f0 (tracked)
loss(X, Y) = 0.791125f0 (tracked)
loss(X, Y) = 0.79280055f0 (tracked)
loss(X, Y) = 0.7727652f0 (tracked)
loss(X, Y) = 0.8022267f0 (tracked)
loss(X, Y) = 0.7813553f0 (tracked)
loss(X, Y) = 0.84987646f0 (tracked)
loss(X, Y) = 0.7667703f0 (tracked)
loss(X, Y) = 0.7944592f0 (tracked)
loss(X, Y) = 0.78480494f0 (tracked)
loss(X, Y) = 0.8343045f0 (tracked)
loss(X, Y) = 0.81072414f0 (tracked)
loss(X, Y) = 0.7538656f0 (tracked)
loss(X, Y) = 0.7872334f0 (tracked)
loss(X, Y) = 0.7581188f0 (tracked)
loss(X, Y) = 0.794513f0 (tracked)
loss(X, Y) = 0.7898002f0 (tracked)
loss(X, Y) = 0.7734566f0 (tracked)
loss(X, Y) = 0.7802016f0 (tracked)
loss(X, Y) = 0.7945919f0 (tracked)
loss(X, Y) = 0.7702934f0 (tracked)
loss(X, Y) = 0.799928f0 (tracked)
loss(X, Y) = 0.8061271f0 (tracked)
loss(X, Y) = 0.7772341f0 (tracked)
loss(X, Y) = 0.7564092f0 (tracked)
loss(X, Y) = 0.78624624f0 (tracked)
loss(X, Y) = 0.7516837f0 (tracked)
loss(X, Y) = 0.7412778f0 (tracked)
loss(X, Y) = 0.7962636f0 (tracked)
loss(X, Y) = 0.7979005f0 (tracked)
loss(X, Y) = 0.7905396f0 (tracked)
loss(X, Y) = 0.76558983f0 (tracked)
loss(X, Y) = 0.8246294f0 (tracked)
loss(X, Y) = 0.8046663f0 (tracked)
loss(X, Y) = 0.7463221f0 (tracked)
loss(X, Y) = 0.782897f0 (tracked)
loss(X, Y) = 0.75859565f0 (tracked)
loss(X, Y) = 0.7516325f0 (tracked)
loss(X, Y) = 0.75876385f0 (tracked)
loss(X, Y) = 0.8039583f0 (tracked)
loss(X, Y) = 0.7501536f0 (tracked)
loss(X, Y) = 0.8167237f0 (tracked)
loss(X, Y) = 0.78706974f0 (tracked)
loss(X, Y) = 0.7620924f0 (tracked)
loss(X, Y) = 0.75583225f0 (tracked)
loss(X, Y) = 0.78817064f0 (tracked)
loss(X, Y) = 0.7862896f0 (tracked)
loss(X, Y) = 0.7719802f0 (tracked)
loss(X, Y) = 0.7701486f0 (tracked)
loss(X, Y) = 0.77370316f0 (tracked)
loss(X, Y) = 0.79238236f0 (tracked)
loss(X, Y) = 0.7832301f0 (tracked)
loss(X, Y) = 0.81409997f0 (tracked)
loss(X, Y) = 0.72556496f0 (tracked)
loss(X, Y) = 0.79146665f0 (tracked)
loss(X, Y) = 0.7411251f0 (tracked)
loss(X, Y) = 0.77164876f0 (tracked)
loss(X, Y) = 0.74900436f0 (tracked)
loss(X, Y) = 0.7909855f0 (tracked)
loss(X, Y) = 0.7781157f0 (tracked)
loss(X, Y) = 0.76402795f0 (tracked)
loss(X, Y) = 0.7957282f0 (tracked)
loss(X, Y) = 0.75543255f0 (tracked)
loss(X, Y) = 0.7383021f0 (tracked)
loss(X, Y) = 0.77921796f0 (tracked)
loss(X, Y) = 0.77958566f0 (tracked)
loss(X, Y) = 0.78192836f0 (tracked)
```

```
┌ Info: Epoch 316
└ @ Main /Users/kcft114/.julia/packages/Flux/qXNjB/src/optimise/train.jl:105
┌ Info: Epoch 317
└ @ Main /Users/kcft114/.julia/packages/Flux/qXNjB/src/optimise/train.jl:105
┌ Info: Epoch 318
└ @ Main /Users/kcft114/.julia/packages/Flux/qXNjB/src/optimise/train.jl:105
┌ Info: Epoch 319
└ @ Main /Users/kcft114/.julia/packages/Flux/qXNjB/src/optimise/train.jl:105
┌ Info: Epoch 320
└ @ Main /Users/kcft114/.julia/packages/Flux/qXNjB/src/optimise/train.jl:105
┌ Info: Epoch 321
└ @ Main /Users/kcft114/.julia/packages/Flux/qXNjB/src/optimise/train.jl:105
┌ Info: Epoch 322
└ @ Main /Users/kcft114/.julia/packages/Flux/qXNjB/src/optimise/train.jl:105
┌ Info: Epoch 323
└ @ Main /Users/kcft114/.julia/packages/Flux/qXNjB/src/optimise/train.jl:105
┌ Info: Epoch 324
└ @ Main /Users/kcft114/.julia/packages/Flux/qXNjB/src/optimise/train.jl:105
┌ Info: Epoch 325
└ @ Main /Users/kcft114/.julia/packages/Flux/qXNjB/src/optimise/train.jl:105
┌ Info: Epoch 326
└ @ Main /Users/kcft114/.julia/packages/Flux/qXNjB/src/optimise/train.jl:105
┌ Info: Epoch 327
└ @ Main /Users/kcft114/.julia/packages/Flux/qXNjB/src/optimise/train.jl:105
┌ Info: Epoch 328
└ @ Main /Users/kcft114/.julia/packages/Flux/qXNjB/src/optimise/train.jl:105
┌ Info: Epoch 329
└ @ Main /Users/kcft114/.julia/packages/Flux/qXNjB/src/optimise/train.jl:105
┌ Info: Epoch 330
└ @ Main /Users/kcft114/.julia/packages/Flux/qXNjB/src/optimise/train.jl:105
┌ Info: Epoch 331
└ @ Main /Users/kcft114/.julia/packages/Flux/qXNjB/src/optimise/train.jl:105
┌ Info: Epoch 332
└ @ Main /Users/kcft114/.julia/packages/Flux/qXNjB/src/optimise/train.jl:105
┌ Info: Epoch 333
└ @ Main /Users/kcft114/.julia/packages/Flux/qXNjB/src/optimise/train.jl:105
┌ Info: Epoch 334
└ @ Main /Users/kcft114/.julia/packages/Flux/qXNjB/src/optimise/train.jl:105
┌ Info: Epoch 335
└ @ Main /Users/kcft114/.julia/packages/Flux/qXNjB/src/optimise/train.jl:105
┌ Info: Epoch 336
└ @ Main /Users/kcft114/.julia/packages/Flux/qXNjB/src/optimise/train.jl:105
┌ Info: Epoch 337
└ @ Main /Users/kcft114/.julia/packages/Flux/qXNjB/src/optimise/train.jl:105
┌ Info: Epoch 338
└ @ Main /Users/kcft114/.julia/packages/Flux/qXNjB/src/optimise/train.jl:105
┌ Info: Epoch 339
└ @ Main /Users/kcft114/.julia/packages/Flux/qXNjB/src/optimise/train.jl:105
┌ Info: Epoch 340
└ @ Main /Users/kcft114/.julia/packages/Flux/qXNjB/src/optimise/train.jl:105
┌ Info: Epoch 341
└ @ Main /Users/kcft114/.julia/packages/Flux/qXNjB/src/optimise/train.jl:105
┌ Info: Epoch 342
└ @ Main /Users/kcft114/.julia/packages/Flux/qXNjB/src/optimise/train.jl:105
┌ Info: Epoch 343
└ @ Main /Users/kcft114/.julia/packages/Flux/qXNjB/src/optimise/train.jl:105
┌ Info: Epoch 344
└ @ Main /Users/kcft114/.julia/packages/Flux/qXNjB/src/optimise/train.jl:105
┌ Info: Epoch 345
└ @ Main /Users/kcft114/.julia/packages/Flux/qXNjB/src/optimise/train.jl:105
┌ Info: Epoch 346
└ @ Main /Users/kcft114/.julia/packages/Flux/qXNjB/src/optimise/train.jl:105
┌ Info: Epoch 347
└ @ Main /Users/kcft114/.julia/packages/Flux/qXNjB/src/optimise/train.jl:105
┌ Info: Epoch 348
└ @ Main /Users/kcft114/.julia/packages/Flux/qXNjB/src/optimise/train.jl:105
┌ Info: Epoch 349
└ @ Main /Users/kcft114/.julia/packages/Flux/qXNjB/src/optimise/train.jl:105
┌ Info: Epoch 350
└ @ Main /Users/kcft114/.julia/packages/Flux/qXNjB/src/optimise/train.jl:105
┌ Info: Epoch 351
└ @ Main /Users/kcft114/.julia/packages/Flux/qXNjB/src/optimise/train.jl:105
┌ Info: Epoch 352
└ @ Main /Users/kcft114/.julia/packages/Flux/qXNjB/src/optimise/train.jl:105
┌ Info: Epoch 353
└ @ Main /Users/kcft114/.julia/packages/Flux/qXNjB/src/optimise/train.jl:105
┌ Info: Epoch 354
└ @ Main /Users/kcft114/.julia/packages/Flux/qXNjB/src/optimise/train.jl:105
┌ Info: Epoch 355
└ @ Main /Users/kcft114/.julia/packages/Flux/qXNjB/src/optimise/train.jl:105
┌ Info: Epoch 356
└ @ Main /Users/kcft114/.julia/packages/Flux/qXNjB/src/optimise/train.jl:105
┌ Info: Epoch 357
└ @ Main /Users/kcft114/.julia/packages/Flux/qXNjB/src/optimise/train.jl:105
┌ Info: Epoch 358
└ @ Main /Users/kcft114/.julia/packages/Flux/qXNjB/src/optimise/train.jl:105
┌ Info: Epoch 359
└ @ Main /Users/kcft114/.julia/packages/Flux/qXNjB/src/optimise/train.jl:105
┌ Info: Epoch 360
└ @ Main /Users/kcft114/.julia/packages/Flux/qXNjB/src/optimise/train.jl:105
┌ Info: Epoch 361
└ @ Main /Users/kcft114/.julia/packages/Flux/qXNjB/src/optimise/train.jl:105
┌ Info: Epoch 362
└ @ Main /Users/kcft114/.julia/packages/Flux/qXNjB/src/optimise/train.jl:105
┌ Info: Epoch 363
└ @ Main /Users/kcft114/.julia/packages/Flux/qXNjB/src/optimise/train.jl:105
┌ Info: Epoch 364
└ @ Main /Users/kcft114/.julia/packages/Flux/qXNjB/src/optimise/train.jl:105
┌ Info: Epoch 365
└ @ Main /Users/kcft114/.julia/packages/Flux/qXNjB/src/optimise/train.jl:105
┌ Info: Epoch 366
└ @ Main /Users/kcft114/.julia/packages/Flux/qXNjB/src/optimise/train.jl:105
┌ Info: Epoch 367
└ @ Main /Users/kcft114/.julia/packages/Flux/qXNjB/src/optimise/train.jl:105
┌ Info: Epoch 368
└ @ Main /Users/kcft114/.julia/packages/Flux/qXNjB/src/optimise/train.jl:105
┌ Info: Epoch 369
└ @ Main /Users/kcft114/.julia/packages/Flux/qXNjB/src/optimise/train.jl:105
┌ Info: Epoch 370
└ @ Main /Users/kcft114/.julia/packages/Flux/qXNjB/src/optimise/train.jl:105
┌ Info: Epoch 371
└ @ Main /Users/kcft114/.julia/packages/Flux/qXNjB/src/optimise/train.jl:105
┌ Info: Epoch 372
└ @ Main /Users/kcft114/.julia/packages/Flux/qXNjB/src/optimise/train.jl:105
┌ Info: Epoch 373
└ @ Main /Users/kcft114/.julia/packages/Flux/qXNjB/src/optimise/train.jl:105
┌ Info: Epoch 374
└ @ Main /Users/kcft114/.julia/packages/Flux/qXNjB/src/optimise/train.jl:105
┌ Info: Epoch 375
└ @ Main /Users/kcft114/.julia/packages/Flux/qXNjB/src/optimise/train.jl:105
┌ Info: Epoch 376
└ @ Main /Users/kcft114/.julia/packages/Flux/qXNjB/src/optimise/train.jl:105
┌ Info: Epoch 377
└ @ Main /Users/kcft114/.julia/packages/Flux/qXNjB/src/optimise/train.jl:105
┌ Info: Epoch 378
└ @ Main /Users/kcft114/.julia/packages/Flux/qXNjB/src/optimise/train.jl:105
┌ Info: Epoch 379
└ @ Main /Users/kcft114/.julia/packages/Flux/qXNjB/src/optimise/train.jl:105
┌ Info: Epoch 380
└ @ Main /Users/kcft114/.julia/packages/Flux/qXNjB/src/optimise/train.jl:105
┌ Info: Epoch 381
└ @ Main /Users/kcft114/.julia/packages/Flux/qXNjB/src/optimise/train.jl:105
┌ Info: Epoch 382
└ @ Main /Users/kcft114/.julia/packages/Flux/qXNjB/src/optimise/train.jl:105
┌ Info: Epoch 383
└ @ Main /Users/kcft114/.julia/packages/Flux/qXNjB/src/optimise/train.jl:105
┌ Info: Epoch 384
└ @ Main /Users/kcft114/.julia/packages/Flux/qXNjB/src/optimise/train.jl:105
┌ Info: Epoch 385
└ @ Main /Users/kcft114/.julia/packages/Flux/qXNjB/src/optimise/train.jl:105
┌ Info: Epoch 386
└ @ Main /Users/kcft114/.julia/packages/Flux/qXNjB/src/optimise/train.jl:105
┌ Info: Epoch 387
└ @ Main /Users/kcft114/.julia/packages/Flux/qXNjB/src/optimise/train.jl:105
┌ Info: Epoch 388
└ @ Main /Users/kcft114/.julia/packages/Flux/qXNjB/src/optimise/train.jl:105
┌ Info: Epoch 389
└ @ Main /Users/kcft114/.julia/packages/Flux/qXNjB/src/optimise/train.jl:105
┌ Info: Epoch 390
└ @ Main /Users/kcft114/.julia/packages/Flux/qXNjB/src/optimise/train.jl:105
┌ Info: Epoch 391
└ @ Main /Users/kcft114/.julia/packages/Flux/qXNjB/src/optimise/train.jl:105
┌ Info: Epoch 392
└ @ Main /Users/kcft114/.julia/packages/Flux/qXNjB/src/optimise/train.jl:105
┌ Info: Epoch 393
└ @ Main /Users/kcft114/.julia/packages/Flux/qXNjB/src/optimise/train.jl:105
┌ Info: Epoch 394
└ @ Main /Users/kcft114/.julia/packages/Flux/qXNjB/src/optimise/train.jl:105
┌ Info: Epoch 395
└ @ Main /Users/kcft114/.julia/packages/Flux/qXNjB/src/optimise/train.jl:105
┌ Info: Epoch 396
└ @ Main /Users/kcft114/.julia/packages/Flux/qXNjB/src/optimise/train.jl:105
┌ Info: Epoch 397
└ @ Main /Users/kcft114/.julia/packages/Flux/qXNjB/src/optimise/train.jl:105
┌ Info: Epoch 398
└ @ Main /Users/kcft114/.julia/packages/Flux/qXNjB/src/optimise/train.jl:105
┌ Info: Epoch 399
└ @ Main /Users/kcft114/.julia/packages/Flux/qXNjB/src/optimise/train.jl:105
┌ Info: Epoch 400
└ @ Main /Users/kcft114/.julia/packages/Flux/qXNjB/src/optimise/train.jl:105
┌ Info: Epoch 401
└ @ Main /Users/kcft114/.julia/packages/Flux/qXNjB/src/optimise/train.jl:105
┌ Info: Epoch 402
└ @ Main /Users/kcft114/.julia/packages/Flux/qXNjB/src/optimise/train.jl:105
┌ Info: Epoch 403
└ @ Main /Users/kcft114/.julia/packages/Flux/qXNjB/src/optimise/train.jl:105
```

```
loss(X, Y) = 0.7817463f0 (tracked)
loss(X, Y) = 0.771058f0 (tracked)
loss(X, Y) = 0.7685816f0 (tracked)
loss(X, Y) = 0.7989939f0 (tracked)
loss(X, Y) = 0.7412211f0 (tracked)
loss(X, Y) = 0.77366f0 (tracked)
loss(X, Y) = 0.7497833f0 (tracked)
loss(X, Y) = 0.75111973f0 (tracked)
loss(X, Y) = 0.7756746f0 (tracked)
loss(X, Y) = 0.7455674f0 (tracked)
loss(X, Y) = 0.7416666f0 (tracked)
loss(X, Y) = 0.7517082f0 (tracked)
loss(X, Y) = 0.7916704f0 (tracked)
loss(X, Y) = 0.7808445f0 (tracked)
loss(X, Y) = 0.7351819f0 (tracked)
loss(X, Y) = 0.796162f0 (tracked)
loss(X, Y) = 0.8127305f0 (tracked)
loss(X, Y) = 0.7500528f0 (tracked)
loss(X, Y) = 0.7583486f0 (tracked)
loss(X, Y) = 0.7734893f0 (tracked)
loss(X, Y) = 0.7641875f0 (tracked)
loss(X, Y) = 0.759271f0 (tracked)
loss(X, Y) = 0.7309843f0 (tracked)
loss(X, Y) = 0.7575377f0 (tracked)
loss(X, Y) = 0.74290717f0 (tracked)
loss(X, Y) = 0.7312618f0 (tracked)
loss(X, Y) = 0.81692314f0 (tracked)
loss(X, Y) = 0.7913253f0 (tracked)
loss(X, Y) = 0.77931774f0 (tracked)
loss(X, Y) = 0.78239447f0 (tracked)
loss(X, Y) = 0.77525204f0 (tracked)
loss(X, Y) = 0.7463731f0 (tracked)
loss(X, Y) = 0.76336557f0 (tracked)
loss(X, Y) = 0.74982893f0 (tracked)
loss(X, Y) = 0.77536845f0 (tracked)
loss(X, Y) = 0.7291404f0 (tracked)
loss(X, Y) = 0.7318591f0 (tracked)
loss(X, Y) = 0.75129426f0 (tracked)
loss(X, Y) = 0.77770483f0 (tracked)
loss(X, Y) = 0.755878f0 (tracked)
loss(X, Y) = 0.7963867f0 (tracked)
loss(X, Y) = 0.7437519f0 (tracked)
loss(X, Y) = 0.7697099f0 (tracked)
loss(X, Y) = 0.75981903f0 (tracked)
loss(X, Y) = 0.78052413f0 (tracked)
loss(X, Y) = 0.7601094f0 (tracked)
loss(X, Y) = 0.74126315f0 (tracked)
loss(X, Y) = 0.72030324f0 (tracked)
loss(X, Y) = 0.73321176f0 (tracked)
loss(X, Y) = 0.7363602f0 (tracked)
loss(X, Y) = 0.7330205f0 (tracked)
loss(X, Y) = 0.7402575f0 (tracked)
loss(X, Y) = 0.7428928f0 (tracked)
loss(X, Y) = 0.7247577f0 (tracked)
loss(X, Y) = 0.77740306f0 (tracked)
loss(X, Y) = 0.7858582f0 (tracked)
loss(X, Y) = 0.7741117f0 (tracked)
loss(X, Y) = 0.7668387f0 (tracked)
loss(X, Y) = 0.75445753f0 (tracked)
loss(X, Y) = 0.77636606f0 (tracked)
loss(X, Y) = 0.72910136f0 (tracked)
loss(X, Y) = 0.73195887f0 (tracked)
loss(X, Y) = 0.750523f0 (tracked)
loss(X, Y) = 0.7459136f0 (tracked)
loss(X, Y) = 0.7483987f0 (tracked)
loss(X, Y) = 0.7471447f0 (tracked)
loss(X, Y) = 0.7110078f0 (tracked)
loss(X, Y) = 0.72805715f0 (tracked)
loss(X, Y) = 0.7627402f0 (tracked)
loss(X, Y) = 0.7507442f0 (tracked)
loss(X, Y) = 0.73824185f0 (tracked)
loss(X, Y) = 0.74632597f0 (tracked)
loss(X, Y) = 0.672907f0 (tracked)
loss(X, Y) = 0.77334845f0 (tracked)
loss(X, Y) = 0.73934287f0 (tracked)
loss(X, Y) = 0.7407889f0 (tracked)
loss(X, Y) = 0.76572007f0 (tracked)
loss(X, Y) = 0.7428985f0 (tracked)
loss(X, Y) = 0.7584321f0 (tracked)
loss(X, Y) = 0.7385346f0 (tracked)
loss(X, Y) = 0.7616864f0 (tracked)
loss(X, Y) = 0.72663534f0 (tracked)
loss(X, Y) = 0.7095579f0 (tracked)
loss(X, Y) = 0.70315045f0 (tracked)
loss(X, Y) = 0.7440498f0 (tracked)
loss(X, Y) = 0.7424984f0 (tracked)
loss(X, Y) = 0.73047984f0 (tracked)
loss(X, Y) = 0.7310427f0 (tracked)
loss(X, Y) = 0.72759175f0 (tracked)
loss(X, Y) = 0.731241f0 (tracked)
```

```
┌ Info: Epoch 404
└ @ Main /Users/kcft114/.julia/packages/Flux/qXNjB/src/optimise/train.jl:105
┌ Info: Epoch 405
└ @ Main /Users/kcft114/.julia/packages/Flux/qXNjB/src/optimise/train.jl:105
┌ Info: Epoch 406
└ @ Main /Users/kcft114/.julia/packages/Flux/qXNjB/src/optimise/train.jl:105
┌ Info: Epoch 407
└ @ Main /Users/kcft114/.julia/packages/Flux/qXNjB/src/optimise/train.jl:105
┌ Info: Epoch 408
└ @ Main /Users/kcft114/.julia/packages/Flux/qXNjB/src/optimise/train.jl:105
┌ Info: Epoch 409
└ @ Main /Users/kcft114/.julia/packages/Flux/qXNjB/src/optimise/train.jl:105
┌ Info: Epoch 410
└ @ Main /Users/kcft114/.julia/packages/Flux/qXNjB/src/optimise/train.jl:105
┌ Info: Epoch 411
└ @ Main /Users/kcft114/.julia/packages/Flux/qXNjB/src/optimise/train.jl:105
┌ Info: Epoch 412
└ @ Main /Users/kcft114/.julia/packages/Flux/qXNjB/src/optimise/train.jl:105
┌ Info: Epoch 413
└ @ Main /Users/kcft114/.julia/packages/Flux/qXNjB/src/optimise/train.jl:105
┌ Info: Epoch 414
└ @ Main /Users/kcft114/.julia/packages/Flux/qXNjB/src/optimise/train.jl:105
┌ Info: Epoch 415
└ @ Main /Users/kcft114/.julia/packages/Flux/qXNjB/src/optimise/train.jl:105
┌ Info: Epoch 416
└ @ Main /Users/kcft114/.julia/packages/Flux/qXNjB/src/optimise/train.jl:105
┌ Info: Epoch 417
└ @ Main /Users/kcft114/.julia/packages/Flux/qXNjB/src/optimise/train.jl:105
┌ Info: Epoch 418
└ @ Main /Users/kcft114/.julia/packages/Flux/qXNjB/src/optimise/train.jl:105
┌ Info: Epoch 419
└ @ Main /Users/kcft114/.julia/packages/Flux/qXNjB/src/optimise/train.jl:105
┌ Info: Epoch 420
└ @ Main /Users/kcft114/.julia/packages/Flux/qXNjB/src/optimise/train.jl:105
┌ Info: Epoch 421
└ @ Main /Users/kcft114/.julia/packages/Flux/qXNjB/src/optimise/train.jl:105
┌ Info: Epoch 422
└ @ Main /Users/kcft114/.julia/packages/Flux/qXNjB/src/optimise/train.jl:105
┌ Info: Epoch 423
└ @ Main /Users/kcft114/.julia/packages/Flux/qXNjB/src/optimise/train.jl:105
┌ Info: Epoch 424
└ @ Main /Users/kcft114/.julia/packages/Flux/qXNjB/src/optimise/train.jl:105
┌ Info: Epoch 425
└ @ Main /Users/kcft114/.julia/packages/Flux/qXNjB/src/optimise/train.jl:105
┌ Info: Epoch 426
└ @ Main /Users/kcft114/.julia/packages/Flux/qXNjB/src/optimise/train.jl:105
┌ Info: Epoch 427
└ @ Main /Users/kcft114/.julia/packages/Flux/qXNjB/src/optimise/train.jl:105
┌ Info: Epoch 428
└ @ Main /Users/kcft114/.julia/packages/Flux/qXNjB/src/optimise/train.jl:105
┌ Info: Epoch 429
└ @ Main /Users/kcft114/.julia/packages/Flux/qXNjB/src/optimise/train.jl:105
┌ Info: Epoch 430
└ @ Main /Users/kcft114/.julia/packages/Flux/qXNjB/src/optimise/train.jl:105
┌ Info: Epoch 431
└ @ Main /Users/kcft114/.julia/packages/Flux/qXNjB/src/optimise/train.jl:105
┌ Info: Epoch 432
└ @ Main /Users/kcft114/.julia/packages/Flux/qXNjB/src/optimise/train.jl:105
┌ Info: Epoch 433
└ @ Main /Users/kcft114/.julia/packages/Flux/qXNjB/src/optimise/train.jl:105
┌ Info: Epoch 434
└ @ Main /Users/kcft114/.julia/packages/Flux/qXNjB/src/optimise/train.jl:105
┌ Info: Epoch 435
└ @ Main /Users/kcft114/.julia/packages/Flux/qXNjB/src/optimise/train.jl:105
┌ Info: Epoch 436
└ @ Main /Users/kcft114/.julia/packages/Flux/qXNjB/src/optimise/train.jl:105
┌ Info: Epoch 437
└ @ Main /Users/kcft114/.julia/packages/Flux/qXNjB/src/optimise/train.jl:105
┌ Info: Epoch 438
└ @ Main /Users/kcft114/.julia/packages/Flux/qXNjB/src/optimise/train.jl:105
┌ Info: Epoch 439
└ @ Main /Users/kcft114/.julia/packages/Flux/qXNjB/src/optimise/train.jl:105
┌ Info: Epoch 440
└ @ Main /Users/kcft114/.julia/packages/Flux/qXNjB/src/optimise/train.jl:105
┌ Info: Epoch 441
└ @ Main /Users/kcft114/.julia/packages/Flux/qXNjB/src/optimise/train.jl:105
┌ Info: Epoch 442
└ @ Main /Users/kcft114/.julia/packages/Flux/qXNjB/src/optimise/train.jl:105
┌ Info: Epoch 443
└ @ Main /Users/kcft114/.julia/packages/Flux/qXNjB/src/optimise/train.jl:105
┌ Info: Epoch 444
└ @ Main /Users/kcft114/.julia/packages/Flux/qXNjB/src/optimise/train.jl:105
┌ Info: Epoch 445
└ @ Main /Users/kcft114/.julia/packages/Flux/qXNjB/src/optimise/train.jl:105
┌ Info: Epoch 446
└ @ Main /Users/kcft114/.julia/packages/Flux/qXNjB/src/optimise/train.jl:105
┌ Info: Epoch 447
└ @ Main /Users/kcft114/.julia/packages/Flux/qXNjB/src/optimise/train.jl:105
┌ Info: Epoch 448
└ @ Main /Users/kcft114/.julia/packages/Flux/qXNjB/src/optimise/train.jl:105
┌ Info: Epoch 449
└ @ Main /Users/kcft114/.julia/packages/Flux/qXNjB/src/optimise/train.jl:105
┌ Info: Epoch 450
└ @ Main /Users/kcft114/.julia/packages/Flux/qXNjB/src/optimise/train.jl:105
┌ Info: Epoch 451
└ @ Main /Users/kcft114/.julia/packages/Flux/qXNjB/src/optimise/train.jl:105
┌ Info: Epoch 452
└ @ Main /Users/kcft114/.julia/packages/Flux/qXNjB/src/optimise/train.jl:105
┌ Info: Epoch 453
└ @ Main /Users/kcft114/.julia/packages/Flux/qXNjB/src/optimise/train.jl:105
┌ Info: Epoch 454
└ @ Main /Users/kcft114/.julia/packages/Flux/qXNjB/src/optimise/train.jl:105
┌ Info: Epoch 455
└ @ Main /Users/kcft114/.julia/packages/Flux/qXNjB/src/optimise/train.jl:105
┌ Info: Epoch 456
└ @ Main /Users/kcft114/.julia/packages/Flux/qXNjB/src/optimise/train.jl:105
┌ Info: Epoch 457
└ @ Main /Users/kcft114/.julia/packages/Flux/qXNjB/src/optimise/train.jl:105
┌ Info: Epoch 458
└ @ Main /Users/kcft114/.julia/packages/Flux/qXNjB/src/optimise/train.jl:105
┌ Info: Epoch 459
└ @ Main /Users/kcft114/.julia/packages/Flux/qXNjB/src/optimise/train.jl:105
┌ Info: Epoch 460
└ @ Main /Users/kcft114/.julia/packages/Flux/qXNjB/src/optimise/train.jl:105
┌ Info: Epoch 461
└ @ Main /Users/kcft114/.julia/packages/Flux/qXNjB/src/optimise/train.jl:105
┌ Info: Epoch 462
└ @ Main /Users/kcft114/.julia/packages/Flux/qXNjB/src/optimise/train.jl:105
┌ Info: Epoch 463
└ @ Main /Users/kcft114/.julia/packages/Flux/qXNjB/src/optimise/train.jl:105
┌ Info: Epoch 464
└ @ Main /Users/kcft114/.julia/packages/Flux/qXNjB/src/optimise/train.jl:105
┌ Info: Epoch 465
└ @ Main /Users/kcft114/.julia/packages/Flux/qXNjB/src/optimise/train.jl:105
┌ Info: Epoch 466
└ @ Main /Users/kcft114/.julia/packages/Flux/qXNjB/src/optimise/train.jl:105
┌ Info: Epoch 467
└ @ Main /Users/kcft114/.julia/packages/Flux/qXNjB/src/optimise/train.jl:105
┌ Info: Epoch 468
└ @ Main /Users/kcft114/.julia/packages/Flux/qXNjB/src/optimise/train.jl:105
┌ Info: Epoch 469
└ @ Main /Users/kcft114/.julia/packages/Flux/qXNjB/src/optimise/train.jl:105
┌ Info: Epoch 470
└ @ Main /Users/kcft114/.julia/packages/Flux/qXNjB/src/optimise/train.jl:105
┌ Info: Epoch 471
└ @ Main /Users/kcft114/.julia/packages/Flux/qXNjB/src/optimise/train.jl:105
┌ Info: Epoch 472
└ @ Main /Users/kcft114/.julia/packages/Flux/qXNjB/src/optimise/train.jl:105
┌ Info: Epoch 473
└ @ Main /Users/kcft114/.julia/packages/Flux/qXNjB/src/optimise/train.jl:105
┌ Info: Epoch 474
└ @ Main /Users/kcft114/.julia/packages/Flux/qXNjB/src/optimise/train.jl:105
┌ Info: Epoch 475
└ @ Main /Users/kcft114/.julia/packages/Flux/qXNjB/src/optimise/train.jl:105
┌ Info: Epoch 476
└ @ Main /Users/kcft114/.julia/packages/Flux/qXNjB/src/optimise/train.jl:105
┌ Info: Epoch 477
└ @ Main /Users/kcft114/.julia/packages/Flux/qXNjB/src/optimise/train.jl:105
┌ Info: Epoch 478
└ @ Main /Users/kcft114/.julia/packages/Flux/qXNjB/src/optimise/train.jl:105
┌ Info: Epoch 479
└ @ Main /Users/kcft114/.julia/packages/Flux/qXNjB/src/optimise/train.jl:105
┌ Info: Epoch 480
└ @ Main /Users/kcft114/.julia/packages/Flux/qXNjB/src/optimise/train.jl:105
┌ Info: Epoch 481
└ @ Main /Users/kcft114/.julia/packages/Flux/qXNjB/src/optimise/train.jl:105
┌ Info: Epoch 482
└ @ Main /Users/kcft114/.julia/packages/Flux/qXNjB/src/optimise/train.jl:105
┌ Info: Epoch 483
└ @ Main /Users/kcft114/.julia/packages/Flux/qXNjB/src/optimise/train.jl:105
┌ Info: Epoch 484
└ @ Main /Users/kcft114/.julia/packages/Flux/qXNjB/src/optimise/train.jl:105
┌ Info: Epoch 485
└ @ Main /Users/kcft114/.julia/packages/Flux/qXNjB/src/optimise/train.jl:105
┌ Info: Epoch 486
└ @ Main /Users/kcft114/.julia/packages/Flux/qXNjB/src/optimise/train.jl:105
┌ Info: Epoch 487
└ @ Main /Users/kcft114/.julia/packages/Flux/qXNjB/src/optimise/train.jl:105
┌ Info: Epoch 488
└ @ Main /Users/kcft114/.julia/packages/Flux/qXNjB/src/optimise/train.jl:105
┌ Info: Epoch 489
└ @ Main /Users/kcft114/.julia/packages/Flux/qXNjB/src/optimise/train.jl:105
┌ Info: Epoch 490
└ @ Main /Users/kcft114/.julia/packages/Flux/qXNjB/src/optimise/train.jl:105
┌ Info: Epoch 491
└ @ Main /Users/kcft114/.julia/packages/Flux/qXNjB/src/optimise/train.jl:105
┌ Info: Epoch 492
└ @ Main /Users/kcft114/.julia/packages/Flux/qXNjB/src/optimise/train.jl:105
┌ Info: Epoch 493
└ @ Main /Users/kcft114/.julia/packages/Flux/qXNjB/src/optimise/train.jl:105
```

```
loss(X, Y) = 0.7379321f0 (tracked)
loss(X, Y) = 0.7426964f0 (tracked)
loss(X, Y) = 0.6817937f0 (tracked)
loss(X, Y) = 0.7195329f0 (tracked)
loss(X, Y) = 0.74803007f0 (tracked)
loss(X, Y) = 0.7500099f0 (tracked)
loss(X, Y) = 0.7529191f0 (tracked)
loss(X, Y) = 0.76480067f0 (tracked)
loss(X, Y) = 0.72040325f0 (tracked)
loss(X, Y) = 0.7429104f0 (tracked)
loss(X, Y) = 0.75204545f0 (tracked)
loss(X, Y) = 0.69609565f0 (tracked)
loss(X, Y) = 0.7330183f0 (tracked)
loss(X, Y) = 0.77070147f0 (tracked)
loss(X, Y) = 0.7325018f0 (tracked)
loss(X, Y) = 0.7245119f0 (tracked)
loss(X, Y) = 0.7239136f0 (tracked)
loss(X, Y) = 0.7315253f0 (tracked)
loss(X, Y) = 0.7418457f0 (tracked)
loss(X, Y) = 0.7616962f0 (tracked)
loss(X, Y) = 0.7502587f0 (tracked)
loss(X, Y) = 0.7290058f0 (tracked)
loss(X, Y) = 0.73061496f0 (tracked)
loss(X, Y) = 0.7499862f0 (tracked)
loss(X, Y) = 0.7449072f0 (tracked)
loss(X, Y) = 0.71739924f0 (tracked)
loss(X, Y) = 0.70763254f0 (tracked)
loss(X, Y) = 0.7197792f0 (tracked)
loss(X, Y) = 0.7254062f0 (tracked)
loss(X, Y) = 0.7461912f0 (tracked)
loss(X, Y) = 0.7513179f0 (tracked)
loss(X, Y) = 0.73959184f0 (tracked)
loss(X, Y) = 0.6808501f0 (tracked)
loss(X, Y) = 0.7368101f0 (tracked)
loss(X, Y) = 0.7512724f0 (tracked)
loss(X, Y) = 0.7191938f0 (tracked)
loss(X, Y) = 0.7342348f0 (tracked)
loss(X, Y) = 0.7290931f0 (tracked)
loss(X, Y) = 0.7467278f0 (tracked)
loss(X, Y) = 0.73765975f0 (tracked)
loss(X, Y) = 0.73344344f0 (tracked)
loss(X, Y) = 0.7372976f0 (tracked)
loss(X, Y) = 0.6808139f0 (tracked)
loss(X, Y) = 0.71463764f0 (tracked)
loss(X, Y) = 0.7135047f0 (tracked)
loss(X, Y) = 0.75548995f0 (tracked)
loss(X, Y) = 0.6921534f0 (tracked)
loss(X, Y) = 0.7090434f0 (tracked)
loss(X, Y) = 0.6957765f0 (tracked)
loss(X, Y) = 0.74005276f0 (tracked)
loss(X, Y) = 0.7330553f0 (tracked)
loss(X, Y) = 0.7333525f0 (tracked)
loss(X, Y) = 0.7187067f0 (tracked)
loss(X, Y) = 0.7431949f0 (tracked)
loss(X, Y) = 0.71582353f0 (tracked)
loss(X, Y) = 0.7475445f0 (tracked)
loss(X, Y) = 0.7172442f0 (tracked)
loss(X, Y) = 0.6986778f0 (tracked)
loss(X, Y) = 0.7405002f0 (tracked)
loss(X, Y) = 0.7002415f0 (tracked)
loss(X, Y) = 0.7055885f0 (tracked)
loss(X, Y) = 0.711549f0 (tracked)
loss(X, Y) = 0.7261648f0 (tracked)
loss(X, Y) = 0.7050888f0 (tracked)
loss(X, Y) = 0.73839617f0 (tracked)
loss(X, Y) = 0.70630115f0 (tracked)
loss(X, Y) = 0.70400834f0 (tracked)
loss(X, Y) = 0.71470225f0 (tracked)
loss(X, Y) = 0.717468f0 (tracked)
loss(X, Y) = 0.72392315f0 (tracked)
loss(X, Y) = 0.7033103f0 (tracked)
loss(X, Y) = 0.7328105f0 (tracked)
loss(X, Y) = 0.70702577f0 (tracked)
loss(X, Y) = 0.71912366f0 (tracked)
loss(X, Y) = 0.73353726f0 (tracked)
loss(X, Y) = 0.73435473f0 (tracked)
loss(X, Y) = 0.7257317f0 (tracked)
loss(X, Y) = 0.7241476f0 (tracked)
loss(X, Y) = 0.6844202f0 (tracked)
loss(X, Y) = 0.695444f0 (tracked)
loss(X, Y) = 0.71073174f0 (tracked)
loss(X, Y) = 0.69614935f0 (tracked)
loss(X, Y) = 0.69490445f0 (tracked)
loss(X, Y) = 0.70866203f0 (tracked)
loss(X, Y) = 0.7746962f0 (tracked)
loss(X, Y) = 0.7257818f0 (tracked)
loss(X, Y) = 0.72484034f0 (tracked)
loss(X, Y) = 0.6962749f0 (tracked)
loss(X, Y) = 0.6995131f0 (tracked)
loss(X, Y) = 0.70548654f0 (tracked)
loss(X, Y) = 0.736215f0 (tracked)
loss(X, Y) = 0.71154135f0 (tracked)
loss(X, Y) = 0.67855054f0 (tracked)
loss(X, Y) = 0.69358486f0 (tracked)
loss(X, Y) = 0.72032213f0 (tracked)
loss(X, Y) = 0.6808846f0 (tracked)
loss(X, Y) = 0.70306396f0 (tracked)
loss(X, Y) = 0.6908724f0 (tracked)
loss(X, Y) = 0.72686094f0 (tracked)
loss(X, Y) = 0.70346653f0 (tracked)
loss(X, Y) = 0.6781376f0 (tracked)
loss(X, Y) = 0.7180741f0 (tracked)
loss(X, Y) = 0.70265007f0 (tracked)
```

```
┌ Info: Epoch 494
└ @ Main /Users/kcft114/.julia/packages/Flux/qXNjB/src/optimise/train.jl:105
┌ Info: Epoch 495
└ @ Main /Users/kcft114/.julia/packages/Flux/qXNjB/src/optimise/train.jl:105
┌ Info: Epoch 496
└ @ Main /Users/kcft114/.julia/packages/Flux/qXNjB/src/optimise/train.jl:105
┌ Info: Epoch 497
└ @ Main /Users/kcft114/.julia/packages/Flux/qXNjB/src/optimise/train.jl:105
┌ Info: Epoch 498
└ @ Main /Users/kcft114/.julia/packages/Flux/qXNjB/src/optimise/train.jl:105
┌ Info: Epoch 499
└ @ Main /Users/kcft114/.julia/packages/Flux/qXNjB/src/optimise/train.jl:105
┌ Info: Epoch 500
└ @ Main /Users/kcft114/.julia/packages/Flux/qXNjB/src/optimise/train.jl:105
┌ Info: Epoch 501
└ @ Main /Users/kcft114/.julia/packages/Flux/qXNjB/src/optimise/train.jl:105
┌ Info: Epoch 502
└ @ Main /Users/kcft114/.julia/packages/Flux/qXNjB/src/optimise/train.jl:105
┌ Info: Epoch 503
└ @ Main /Users/kcft114/.julia/packages/Flux/qXNjB/src/optimise/train.jl:105
┌ Info: Epoch 504
└ @ Main /Users/kcft114/.julia/packages/Flux/qXNjB/src/optimise/train.jl:105
┌ Info: Epoch 505
└ @ Main /Users/kcft114/.julia/packages/Flux/qXNjB/src/optimise/train.jl:105
┌ Info: Epoch 506
└ @ Main /Users/kcft114/.julia/packages/Flux/qXNjB/src/optimise/train.jl:105
┌ Info: Epoch 507
└ @ Main /Users/kcft114/.julia/packages/Flux/qXNjB/src/optimise/train.jl:105
┌ Info: Epoch 508
└ @ Main /Users/kcft114/.julia/packages/Flux/qXNjB/src/optimise/train.jl:105
┌ Info: Epoch 509
└ @ Main /Users/kcft114/.julia/packages/Flux/qXNjB/src/optimise/train.jl:105
┌ Info: Epoch 510
└ @ Main /Users/kcft114/.julia/packages/Flux/qXNjB/src/optimise/train.jl:105
┌ Info: Epoch 511
└ @ Main /Users/kcft114/.julia/packages/Flux/qXNjB/src/optimise/train.jl:105
┌ Info: Epoch 512
└ @ Main /Users/kcft114/.julia/packages/Flux/qXNjB/src/optimise/train.jl:105
┌ Info: Epoch 513
└ @ Main /Users/kcft114/.julia/packages/Flux/qXNjB/src/optimise/train.jl:105
┌ Info: Epoch 514
└ @ Main /Users/kcft114/.julia/packages/Flux/qXNjB/src/optimise/train.jl:105
┌ Info: Epoch 515
└ @ Main /Users/kcft114/.julia/packages/Flux/qXNjB/src/optimise/train.jl:105
┌ Info: Epoch 516
└ @ Main /Users/kcft114/.julia/packages/Flux/qXNjB/src/optimise/train.jl:105
┌ Info: Epoch 517
└ @ Main /Users/kcft114/.julia/packages/Flux/qXNjB/src/optimise/train.jl:105
┌ Info: Epoch 518
└ @ Main /Users/kcft114/.julia/packages/Flux/qXNjB/src/optimise/train.jl:105
┌ Info: Epoch 519
└ @ Main /Users/kcft114/.julia/packages/Flux/qXNjB/src/optimise/train.jl:105
┌ Info: Epoch 520
└ @ Main /Users/kcft114/.julia/packages/Flux/qXNjB/src/optimise/train.jl:105
┌ Info: Epoch 521
└ @ Main /Users/kcft114/.julia/packages/Flux/qXNjB/src/optimise/train.jl:105
┌ Info: Epoch 522
└ @ Main /Users/kcft114/.julia/packages/Flux/qXNjB/src/optimise/train.jl:105
┌ Info: Epoch 523
└ @ Main /Users/kcft114/.julia/packages/Flux/qXNjB/src/optimise/train.jl:105
┌ Info: Epoch 524
└ @ Main /Users/kcft114/.julia/packages/Flux/qXNjB/src/optimise/train.jl:105
┌ Info: Epoch 525
└ @ Main /Users/kcft114/.julia/packages/Flux/qXNjB/src/optimise/train.jl:105
┌ Info: Epoch 526
└ @ Main /Users/kcft114/.julia/packages/Flux/qXNjB/src/optimise/train.jl:105
┌ Info: Epoch 527
└ @ Main /Users/kcft114/.julia/packages/Flux/qXNjB/src/optimise/train.jl:105
┌ Info: Epoch 528
└ @ Main /Users/kcft114/.julia/packages/Flux/qXNjB/src/optimise/train.jl:105
┌ Info: Epoch 529
└ @ Main /Users/kcft114/.julia/packages/Flux/qXNjB/src/optimise/train.jl:105
┌ Info: Epoch 530
└ @ Main /Users/kcft114/.julia/packages/Flux/qXNjB/src/optimise/train.jl:105
┌ Info: Epoch 531
└ @ Main /Users/kcft114/.julia/packages/Flux/qXNjB/src/optimise/train.jl:105
┌ Info: Epoch 532
└ @ Main /Users/kcft114/.julia/packages/Flux/qXNjB/src/optimise/train.jl:105
┌ Info: Epoch 533
└ @ Main /Users/kcft114/.julia/packages/Flux/qXNjB/src/optimise/train.jl:105
┌ Info: Epoch 534
└ @ Main /Users/kcft114/.julia/packages/Flux/qXNjB/src/optimise/train.jl:105
┌ Info: Epoch 535
└ @ Main /Users/kcft114/.julia/packages/Flux/qXNjB/src/optimise/train.jl:105
┌ Info: Epoch 536
└ @ Main /Users/kcft114/.julia/packages/Flux/qXNjB/src/optimise/train.jl:105
┌ Info: Epoch 537
└ @ Main /Users/kcft114/.julia/packages/Flux/qXNjB/src/optimise/train.jl:105
┌ Info: Epoch 538
└ @ Main /Users/kcft114/.julia/packages/Flux/qXNjB/src/optimise/train.jl:105
┌ Info: Epoch 539
└ @ Main /Users/kcft114/.julia/packages/Flux/qXNjB/src/optimise/train.jl:105
┌ Info: Epoch 540
└ @ Main /Users/kcft114/.julia/packages/Flux/qXNjB/src/optimise/train.jl:105
┌ Info: Epoch 541
└ @ Main /Users/kcft114/.julia/packages/Flux/qXNjB/src/optimise/train.jl:105
┌ Info: Epoch 542
└ @ Main /Users/kcft114/.julia/packages/Flux/qXNjB/src/optimise/train.jl:105
┌ Info: Epoch 543
└ @ Main /Users/kcft114/.julia/packages/Flux/qXNjB/src/optimise/train.jl:105
┌ Info: Epoch 544
└ @ Main /Users/kcft114/.julia/packages/Flux/qXNjB/src/optimise/train.jl:105
┌ Info: Epoch 545
└ @ Main /Users/kcft114/.julia/packages/Flux/qXNjB/src/optimise/train.jl:105
┌ Info: Epoch 546
└ @ Main /Users/kcft114/.julia/packages/Flux/qXNjB/src/optimise/train.jl:105
┌ Info: Epoch 547
└ @ Main /Users/kcft114/.julia/packages/Flux/qXNjB/src/optimise/train.jl:105
┌ Info: Epoch 548
└ @ Main /Users/kcft114/.julia/packages/Flux/qXNjB/src/optimise/train.jl:105
┌ Info: Epoch 549
└ @ Main /Users/kcft114/.julia/packages/Flux/qXNjB/src/optimise/train.jl:105
┌ Info: Epoch 550
└ @ Main /Users/kcft114/.julia/packages/Flux/qXNjB/src/optimise/train.jl:105
┌ Info: Epoch 551
└ @ Main /Users/kcft114/.julia/packages/Flux/qXNjB/src/optimise/train.jl:105
┌ Info: Epoch 552
└ @ Main /Users/kcft114/.julia/packages/Flux/qXNjB/src/optimise/train.jl:105
┌ Info: Epoch 553
└ @ Main /Users/kcft114/.julia/packages/Flux/qXNjB/src/optimise/train.jl:105
┌ Info: Epoch 554
└ @ Main /Users/kcft114/.julia/packages/Flux/qXNjB/src/optimise/train.jl:105
┌ Info: Epoch 555
└ @ Main /Users/kcft114/.julia/packages/Flux/qXNjB/src/optimise/train.jl:105
┌ Info: Epoch 556
└ @ Main /Users/kcft114/.julia/packages/Flux/qXNjB/src/optimise/train.jl:105
┌ Info: Epoch 557
└ @ Main /Users/kcft114/.julia/packages/Flux/qXNjB/src/optimise/train.jl:105
┌ Info: Epoch 558
└ @ Main /Users/kcft114/.julia/packages/Flux/qXNjB/src/optimise/train.jl:105
┌ Info: Epoch 559
└ @ Main /Users/kcft114/.julia/packages/Flux/qXNjB/src/optimise/train.jl:105
┌ Info: Epoch 560
└ @ Main /Users/kcft114/.julia/packages/Flux/qXNjB/src/optimise/train.jl:105
┌ Info: Epoch 561
└ @ Main /Users/kcft114/.julia/packages/Flux/qXNjB/src/optimise/train.jl:105
┌ Info: Epoch 562
└ @ Main /Users/kcft114/.julia/packages/Flux/qXNjB/src/optimise/train.jl:105
┌ Info: Epoch 563
└ @ Main /Users/kcft114/.julia/packages/Flux/qXNjB/src/optimise/train.jl:105
┌ Info: Epoch 564
└ @ Main /Users/kcft114/.julia/packages/Flux/qXNjB/src/optimise/train.jl:105
┌ Info: Epoch 565
└ @ Main /Users/kcft114/.julia/packages/Flux/qXNjB/src/optimise/train.jl:105
┌ Info: Epoch 566
└ @ Main /Users/kcft114/.julia/packages/Flux/qXNjB/src/optimise/train.jl:105
┌ Info: Epoch 567
└ @ Main /Users/kcft114/.julia/packages/Flux/qXNjB/src/optimise/train.jl:105
┌ Info: Epoch 568
└ @ Main /Users/kcft114/.julia/packages/Flux/qXNjB/src/optimise/train.jl:105
┌ Info: Epoch 569
└ @ Main /Users/kcft114/.julia/packages/Flux/qXNjB/src/optimise/train.jl:105
┌ Info: Epoch 570
└ @ Main /Users/kcft114/.julia/packages/Flux/qXNjB/src/optimise/train.jl:105
┌ Info: Epoch 571
└ @ Main /Users/kcft114/.julia/packages/Flux/qXNjB/src/optimise/train.jl:105
┌ Info: Epoch 572
└ @ Main /Users/kcft114/.julia/packages/Flux/qXNjB/src/optimise/train.jl:105
┌ Info: Epoch 573
└ @ Main /Users/kcft114/.julia/packages/Flux/qXNjB/src/optimise/train.jl:105
┌ Info: Epoch 574
└ @ Main /Users/kcft114/.julia/packages/Flux/qXNjB/src/optimise/train.jl:105
┌ Info: Epoch 575
└ @ Main /Users/kcft114/.julia/packages/Flux/qXNjB/src/optimise/train.jl:105
┌ Info: Epoch 576
└ @ Main /Users/kcft114/.julia/packages/Flux/qXNjB/src/optimise/train.jl:105
┌ Info: Epoch 577
└ @ Main /Users/kcft114/.julia/packages/Flux/qXNjB/src/optimise/train.jl:105
┌ Info: Epoch 578
└ @ Main /Users/kcft114/.julia/packages/Flux/qXNjB/src/optimise/train.jl:105
┌ Info: Epoch 579
└ @ Main /Users/kcft114/.julia/packages/Flux/qXNjB/src/optimise/train.jl:105
┌ Info: Epoch 580
└ @ Main /Users/kcft114/.julia/packages/Flux/qXNjB/src/optimise/train.jl:105
┌ Info: Epoch 581
└ @ Main /Users/kcft114/.julia/packages/Flux/qXNjB/src/optimise/train.jl:105
┌ Info: Epoch 582
└ @ Main /Users/kcft114/.julia/packages/Flux/qXNjB/src/optimise/train.jl:105
┌ Info: Epoch 583
└ @ Main /Users/kcft114/.julia/packages/Flux/qXNjB/src/optimise/train.jl:105
┌ Info: Epoch 584
└ @ Main /Users/kcft114/.julia/packages/Flux/qXNjB/src/optimise/train.jl:105
┌ Info: Epoch 585
└ @ Main /Users/kcft114/.julia/packages/Flux/qXNjB/src/optimise/train.jl:105
┌ Info: Epoch 586
└ @ Main /Users/kcft114/.julia/packages/Flux/qXNjB/src/optimise/train.jl:105
┌ Info: Epoch 587
└ @ Main /Users/kcft114/.julia/packages/Flux/qXNjB/src/optimise/train.jl:105
┌ Info: Epoch 588
└ @ Main /Users/kcft114/.julia/packages/Flux/qXNjB/src/optimise/train.jl:105
┌ Info: Epoch 589
└ @ Main /Users/kcft114/.julia/packages/Flux/qXNjB/src/optimise/train.jl:105
┌ Info: Epoch 590
└ @ Main /Users/kcft114/.julia/packages/Flux/qXNjB/src/optimise/train.jl:105
┌ Info: Epoch 591
└ @ Main /Users/kcft114/.julia/packages/Flux/qXNjB/src/optimise/train.jl:105
┌ Info: Epoch 592
└ @ Main /Users/kcft114/.julia/packages/Flux/qXNjB/src/optimise/train.jl:105
┌ Info: Epoch 593
└ @ Main /Users/kcft114/.julia/packages/Flux/qXNjB/src/optimise/train.jl:105
┌ Info: Epoch 594
└ @ Main /Users/kcft114/.julia/packages/Flux/qXNjB/src/optimise/train.jl:105
┌ Info: Epoch 595
└ @ Main /Users/kcft114/.julia/packages/Flux/qXNjB/src/optimise/train.jl:105
┌ Info: Epoch 596
└ @ Main /Users/kcft114/.julia/packages/Flux/qXNjB/src/optimise/train.jl:105
```

```
loss(X, Y) = 0.75190467f0 (tracked)
loss(X, Y) = 0.73761916f0 (tracked)
loss(X, Y) = 0.72725487f0 (tracked)
loss(X, Y) = 0.6801725f0 (tracked)
loss(X, Y) = 0.7141217f0 (tracked)
loss(X, Y) = 0.73427266f0 (tracked)
loss(X, Y) = 0.72598135f0 (tracked)
loss(X, Y) = 0.67898613f0 (tracked)
loss(X, Y) = 0.7409581f0 (tracked)
loss(X, Y) = 0.69008267f0 (tracked)
loss(X, Y) = 0.6632458f0 (tracked)
loss(X, Y) = 0.70038176f0 (tracked)
loss(X, Y) = 0.71776384f0 (tracked)
loss(X, Y) = 0.6915665f0 (tracked)
loss(X, Y) = 0.6947145f0 (tracked)
loss(X, Y) = 0.7300155f0 (tracked)
loss(X, Y) = 0.6864665f0 (tracked)
loss(X, Y) = 0.69461656f0 (tracked)
loss(X, Y) = 0.6844215f0 (tracked)
loss(X, Y) = 0.7135685f0 (tracked)
loss(X, Y) = 0.68195564f0 (tracked)
loss(X, Y) = 0.72684526f0 (tracked)
loss(X, Y) = 0.74407774f0 (tracked)
loss(X, Y) = 0.6900433f0 (tracked)
loss(X, Y) = 0.6632197f0 (tracked)
loss(X, Y) = 0.7057307f0 (tracked)
loss(X, Y) = 0.69842273f0 (tracked)
loss(X, Y) = 0.6948004f0 (tracked)
loss(X, Y) = 0.7139594f0 (tracked)
loss(X, Y) = 0.73458713f0 (tracked)
loss(X, Y) = 0.750081f0 (tracked)
loss(X, Y) = 0.7303361f0 (tracked)
loss(X, Y) = 0.7006592f0 (tracked)
loss(X, Y) = 0.693133f0 (tracked)
loss(X, Y) = 0.70222455f0 (tracked)
loss(X, Y) = 0.74639034f0 (tracked)
loss(X, Y) = 0.6414073f0 (tracked)
loss(X, Y) = 0.6962647f0 (tracked)
loss(X, Y) = 0.72828364f0 (tracked)
loss(X, Y) = 0.7267945f0 (tracked)
loss(X, Y) = 0.68901956f0 (tracked)
loss(X, Y) = 0.7047121f0 (tracked)
loss(X, Y) = 0.70850533f0 (tracked)
loss(X, Y) = 0.74512875f0 (tracked)
loss(X, Y) = 0.69012856f0 (tracked)
loss(X, Y) = 0.72702557f0 (tracked)
loss(X, Y) = 0.69472593f0 (tracked)
loss(X, Y) = 0.7199358f0 (tracked)
loss(X, Y) = 0.7010459f0 (tracked)
loss(X, Y) = 0.6975255f0 (tracked)
loss(X, Y) = 0.6967493f0 (tracked)
loss(X, Y) = 0.71459997f0 (tracked)
loss(X, Y) = 0.698681f0 (tracked)
loss(X, Y) = 0.7327166f0 (tracked)
loss(X, Y) = 0.6977215f0 (tracked)
loss(X, Y) = 0.7137824f0 (tracked)
loss(X, Y) = 0.6494397f0 (tracked)
loss(X, Y) = 0.7130699f0 (tracked)
loss(X, Y) = 0.6639482f0 (tracked)
loss(X, Y) = 0.73208755f0 (tracked)
loss(X, Y) = 0.6852256f0 (tracked)
loss(X, Y) = 0.69782907f0 (tracked)
loss(X, Y) = 0.7236633f0 (tracked)
loss(X, Y) = 0.7156138f0 (tracked)
loss(X, Y) = 0.70345014f0 (tracked)
loss(X, Y) = 0.71077245f0 (tracked)
loss(X, Y) = 0.65580726f0 (tracked)
loss(X, Y) = 0.6981912f0 (tracked)
loss(X, Y) = 0.6741312f0 (tracked)
loss(X, Y) = 0.6908085f0 (tracked)
loss(X, Y) = 0.686179f0 (tracked)
loss(X, Y) = 0.7380404f0 (tracked)
loss(X, Y) = 0.676912f0 (tracked)
loss(X, Y) = 0.67040676f0 (tracked)
loss(X, Y) = 0.71439534f0 (tracked)
loss(X, Y) = 0.6899384f0 (tracked)
loss(X, Y) = 0.69529223f0 (tracked)
loss(X, Y) = 0.6521206f0 (tracked)
loss(X, Y) = 0.7346227f0 (tracked)
loss(X, Y) = 0.6810336f0 (tracked)
loss(X, Y) = 0.698119f0 (tracked)
loss(X, Y) = 0.7080441f0 (tracked)
loss(X, Y) = 0.69931465f0 (tracked)
loss(X, Y) = 0.7082349f0 (tracked)
loss(X, Y) = 0.672103f0 (tracked)
loss(X, Y) = 0.6792659f0 (tracked)
loss(X, Y) = 0.72471803f0 (tracked)
loss(X, Y) = 0.7149587f0 (tracked)
loss(X, Y) = 0.6900415f0 (tracked)
loss(X, Y) = 0.72085553f0 (tracked)
loss(X, Y) = 0.69564813f0 (tracked)
loss(X, Y) = 0.67797774f0 (tracked)
loss(X, Y) = 0.6743399f0 (tracked)
loss(X, Y) = 0.6858558f0 (tracked)
loss(X, Y) = 0.72442234f0 (tracked)
loss(X, Y) = 0.66632426f0 (tracked)
loss(X, Y) = 0.72304577f0 (tracked)
```

```
┌ Info: Epoch 597
└ @ Main /Users/kcft114/.julia/packages/Flux/qXNjB/src/optimise/train.jl:105
┌ Info: Epoch 598
└ @ Main /Users/kcft114/.julia/packages/Flux/qXNjB/src/optimise/train.jl:105
┌ Info: Epoch 599
└ @ Main /Users/kcft114/.julia/packages/Flux/qXNjB/src/optimise/train.jl:105
┌ Info: Epoch 600
└ @ Main /Users/kcft114/.julia/packages/Flux/qXNjB/src/optimise/train.jl:105
┌ Info: Epoch 601
└ @ Main /Users/kcft114/.julia/packages/Flux/qXNjB/src/optimise/train.jl:105
┌ Info: Epoch 602
└ @ Main /Users/kcft114/.julia/packages/Flux/qXNjB/src/optimise/train.jl:105
┌ Info: Epoch 603
└ @ Main /Users/kcft114/.julia/packages/Flux/qXNjB/src/optimise/train.jl:105
┌ Info: Epoch 604
└ @ Main /Users/kcft114/.julia/packages/Flux/qXNjB/src/optimise/train.jl:105
┌ Info: Epoch 605
└ @ Main /Users/kcft114/.julia/packages/Flux/qXNjB/src/optimise/train.jl:105
┌ Info: Epoch 606
└ @ Main /Users/kcft114/.julia/packages/Flux/qXNjB/src/optimise/train.jl:105
┌ Info: Epoch 607
└ @ Main /Users/kcft114/.julia/packages/Flux/qXNjB/src/optimise/train.jl:105
┌ Info: Epoch 608
└ @ Main /Users/kcft114/.julia/packages/Flux/qXNjB/src/optimise/train.jl:105
┌ Info: Epoch 609
└ @ Main /Users/kcft114/.julia/packages/Flux/qXNjB/src/optimise/train.jl:105
┌ Info: Epoch 610
└ @ Main /Users/kcft114/.julia/packages/Flux/qXNjB/src/optimise/train.jl:105
┌ Info: Epoch 611
└ @ Main /Users/kcft114/.julia/packages/Flux/qXNjB/src/optimise/train.jl:105
┌ Info: Epoch 612
└ @ Main /Users/kcft114/.julia/packages/Flux/qXNjB/src/optimise/train.jl:105
┌ Info: Epoch 613
└ @ Main /Users/kcft114/.julia/packages/Flux/qXNjB/src/optimise/train.jl:105
┌ Info: Epoch 614
└ @ Main /Users/kcft114/.julia/packages/Flux/qXNjB/src/optimise/train.jl:105
┌ Info: Epoch 615
└ @ Main /Users/kcft114/.julia/packages/Flux/qXNjB/src/optimise/train.jl:105
┌ Info: Epoch 616
└ @ Main /Users/kcft114/.julia/packages/Flux/qXNjB/src/optimise/train.jl:105
┌ Info: Epoch 617
└ @ Main /Users/kcft114/.julia/packages/Flux/qXNjB/src/optimise/train.jl:105
┌ Info: Epoch 618
└ @ Main /Users/kcft114/.julia/packages/Flux/qXNjB/src/optimise/train.jl:105
┌ Info: Epoch 619
└ @ Main /Users/kcft114/.julia/packages/Flux/qXNjB/src/optimise/train.jl:105
┌ Info: Epoch 620
└ @ Main /Users/kcft114/.julia/packages/Flux/qXNjB/src/optimise/train.jl:105
┌ Info: Epoch 621
└ @ Main /Users/kcft114/.julia/packages/Flux/qXNjB/src/optimise/train.jl:105
┌ Info: Epoch 622
└ @ Main /Users/kcft114/.julia/packages/Flux/qXNjB/src/optimise/train.jl:105
┌ Info: Epoch 623
└ @ Main /Users/kcft114/.julia/packages/Flux/qXNjB/src/optimise/train.jl:105
┌ Info: Epoch 624
└ @ Main /Users/kcft114/.julia/packages/Flux/qXNjB/src/optimise/train.jl:105
┌ Info: Epoch 625
└ @ Main /Users/kcft114/.julia/packages/Flux/qXNjB/src/optimise/train.jl:105
┌ Info: Epoch 626
└ @ Main /Users/kcft114/.julia/packages/Flux/qXNjB/src/optimise/train.jl:105
┌ Info: Epoch 627
└ @ Main /Users/kcft114/.julia/packages/Flux/qXNjB/src/optimise/train.jl:105
┌ Info: Epoch 628
└ @ Main /Users/kcft114/.julia/packages/Flux/qXNjB/src/optimise/train.jl:105
┌ Info: Epoch 629
└ @ Main /Users/kcft114/.julia/packages/Flux/qXNjB/src/optimise/train.jl:105
┌ Info: Epoch 630
└ @ Main /Users/kcft114/.julia/packages/Flux/qXNjB/src/optimise/train.jl:105
┌ Info: Epoch 631
└ @ Main /Users/kcft114/.julia/packages/Flux/qXNjB/src/optimise/train.jl:105
┌ Info: Epoch 632
└ @ Main /Users/kcft114/.julia/packages/Flux/qXNjB/src/optimise/train.jl:105
┌ Info: Epoch 633
└ @ Main /Users/kcft114/.julia/packages/Flux/qXNjB/src/optimise/train.jl:105
┌ Info: Epoch 634
└ @ Main /Users/kcft114/.julia/packages/Flux/qXNjB/src/optimise/train.jl:105
┌ Info: Epoch 635
└ @ Main /Users/kcft114/.julia/packages/Flux/qXNjB/src/optimise/train.jl:105
┌ Info: Epoch 636
└ @ Main /Users/kcft114/.julia/packages/Flux/qXNjB/src/optimise/train.jl:105
┌ Info: Epoch 637
└ @ Main /Users/kcft114/.julia/packages/Flux/qXNjB/src/optimise/train.jl:105
┌ Info: Epoch 638
└ @ Main /Users/kcft114/.julia/packages/Flux/qXNjB/src/optimise/train.jl:105
┌ Info: Epoch 639
└ @ Main /Users/kcft114/.julia/packages/Flux/qXNjB/src/optimise/train.jl:105
┌ Info: Epoch 640
└ @ Main /Users/kcft114/.julia/packages/Flux/qXNjB/src/optimise/train.jl:105
┌ Info: Epoch 641
└ @ Main /Users/kcft114/.julia/packages/Flux/qXNjB/src/optimise/train.jl:105
┌ Info: Epoch 642
└ @ Main /Users/kcft114/.julia/packages/Flux/qXNjB/src/optimise/train.jl:105
┌ Info: Epoch 643
└ @ Main /Users/kcft114/.julia/packages/Flux/qXNjB/src/optimise/train.jl:105
┌ Info: Epoch 644
└ @ Main /Users/kcft114/.julia/packages/Flux/qXNjB/src/optimise/train.jl:105
┌ Info: Epoch 645
└ @ Main /Users/kcft114/.julia/packages/Flux/qXNjB/src/optimise/train.jl:105
┌ Info: Epoch 646
└ @ Main /Users/kcft114/.julia/packages/Flux/qXNjB/src/optimise/train.jl:105
┌ Info: Epoch 647
└ @ Main /Users/kcft114/.julia/packages/Flux/qXNjB/src/optimise/train.jl:105
┌ Info: Epoch 648
└ @ Main /Users/kcft114/.julia/packages/Flux/qXNjB/src/optimise/train.jl:105
┌ Info: Epoch 649
└ @ Main /Users/kcft114/.julia/packages/Flux/qXNjB/src/optimise/train.jl:105
┌ Info: Epoch 650
└ @ Main /Users/kcft114/.julia/packages/Flux/qXNjB/src/optimise/train.jl:105
┌ Info: Epoch 651
└ @ Main /Users/kcft114/.julia/packages/Flux/qXNjB/src/optimise/train.jl:105
┌ Info: Epoch 652
└ @ Main /Users/kcft114/.julia/packages/Flux/qXNjB/src/optimise/train.jl:105
┌ Info: Epoch 653
└ @ Main /Users/kcft114/.julia/packages/Flux/qXNjB/src/optimise/train.jl:105
┌ Info: Epoch 654
└ @ Main /Users/kcft114/.julia/packages/Flux/qXNjB/src/optimise/train.jl:105
┌ Info: Epoch 655
└ @ Main /Users/kcft114/.julia/packages/Flux/qXNjB/src/optimise/train.jl:105
┌ Info: Epoch 656
└ @ Main /Users/kcft114/.julia/packages/Flux/qXNjB/src/optimise/train.jl:105
┌ Info: Epoch 657
└ @ Main /Users/kcft114/.julia/packages/Flux/qXNjB/src/optimise/train.jl:105
┌ Info: Epoch 658
└ @ Main /Users/kcft114/.julia/packages/Flux/qXNjB/src/optimise/train.jl:105
┌ Info: Epoch 659
└ @ Main /Users/kcft114/.julia/packages/Flux/qXNjB/src/optimise/train.jl:105
┌ Info: Epoch 660
└ @ Main /Users/kcft114/.julia/packages/Flux/qXNjB/src/optimise/train.jl:105
┌ Info: Epoch 661
└ @ Main /Users/kcft114/.julia/packages/Flux/qXNjB/src/optimise/train.jl:105
┌ Info: Epoch 662
└ @ Main /Users/kcft114/.julia/packages/Flux/qXNjB/src/optimise/train.jl:105
┌ Info: Epoch 663
└ @ Main /Users/kcft114/.julia/packages/Flux/qXNjB/src/optimise/train.jl:105
┌ Info: Epoch 664
└ @ Main /Users/kcft114/.julia/packages/Flux/qXNjB/src/optimise/train.jl:105
┌ Info: Epoch 665
└ @ Main /Users/kcft114/.julia/packages/Flux/qXNjB/src/optimise/train.jl:105
┌ Info: Epoch 666
└ @ Main /Users/kcft114/.julia/packages/Flux/qXNjB/src/optimise/train.jl:105
┌ Info: Epoch 667
└ @ Main /Users/kcft114/.julia/packages/Flux/qXNjB/src/optimise/train.jl:105
┌ Info: Epoch 668
└ @ Main /Users/kcft114/.julia/packages/Flux/qXNjB/src/optimise/train.jl:105
┌ Info: Epoch 669
└ @ Main /Users/kcft114/.julia/packages/Flux/qXNjB/src/optimise/train.jl:105
┌ Info: Epoch 670
└ @ Main /Users/kcft114/.julia/packages/Flux/qXNjB/src/optimise/train.jl:105
┌ Info: Epoch 671
└ @ Main /Users/kcft114/.julia/packages/Flux/qXNjB/src/optimise/train.jl:105
┌ Info: Epoch 672
└ @ Main /Users/kcft114/.julia/packages/Flux/qXNjB/src/optimise/train.jl:105
┌ Info: Epoch 673
└ @ Main /Users/kcft114/.julia/packages/Flux/qXNjB/src/optimise/train.jl:105
┌ Info: Epoch 674
└ @ Main /Users/kcft114/.julia/packages/Flux/qXNjB/src/optimise/train.jl:105
┌ Info: Epoch 675
└ @ Main /Users/kcft114/.julia/packages/Flux/qXNjB/src/optimise/train.jl:105
┌ Info: Epoch 676
└ @ Main /Users/kcft114/.julia/packages/Flux/qXNjB/src/optimise/train.jl:105
┌ Info: Epoch 677
└ @ Main /Users/kcft114/.julia/packages/Flux/qXNjB/src/optimise/train.jl:105
┌ Info: Epoch 678
└ @ Main /Users/kcft114/.julia/packages/Flux/qXNjB/src/optimise/train.jl:105
┌ Info: Epoch 679
└ @ Main /Users/kcft114/.julia/packages/Flux/qXNjB/src/optimise/train.jl:105
┌ Info: Epoch 680
└ @ Main /Users/kcft114/.julia/packages/Flux/qXNjB/src/optimise/train.jl:105
┌ Info: Epoch 681
└ @ Main /Users/kcft114/.julia/packages/Flux/qXNjB/src/optimise/train.jl:105
┌ Info: Epoch 682
└ @ Main /Users/kcft114/.julia/packages/Flux/qXNjB/src/optimise/train.jl:105
┌ Info: Epoch 683
└ @ Main /Users/kcft114/.julia/packages/Flux/qXNjB/src/optimise/train.jl:105
┌ Info: Epoch 684
└ @ Main /Users/kcft114/.julia/packages/Flux/qXNjB/src/optimise/train.jl:105
┌ Info: Epoch 685
└ @ Main /Users/kcft114/.julia/packages/Flux/qXNjB/src/optimise/train.jl:105
┌ Info: Epoch 686
└ @ Main /Users/kcft114/.julia/packages/Flux/qXNjB/src/optimise/train.jl:105
┌ Info: Epoch 687
└ @ Main /Users/kcft114/.julia/packages/Flux/qXNjB/src/optimise/train.jl:105
┌ Info: Epoch 688
└ @ Main /Users/kcft114/.julia/packages/Flux/qXNjB/src/optimise/train.jl:105
┌ Info: Epoch 689
└ @ Main /Users/kcft114/.julia/packages/Flux/qXNjB/src/optimise/train.jl:105
┌ Info: Epoch 690
└ @ Main /Users/kcft114/.julia/packages/Flux/qXNjB/src/optimise/train.jl:105
┌ Info: Epoch 691
└ @ Main /Users/kcft114/.julia/packages/Flux/qXNjB/src/optimise/train.jl:105
┌ Info: Epoch 692
└ @ Main /Users/kcft114/.julia/packages/Flux/qXNjB/src/optimise/train.jl:105
┌ Info: Epoch 693
└ @ Main /Users/kcft114/.julia/packages/Flux/qXNjB/src/optimise/train.jl:105
```

```
loss(X, Y) = 0.7055934f0 (tracked)
loss(X, Y) = 0.699649f0 (tracked)
loss(X, Y) = 0.7012699f0 (tracked)
loss(X, Y) = 0.6675111f0 (tracked)
loss(X, Y) = 0.6749224f0 (tracked)
loss(X, Y) = 0.6954035f0 (tracked)
loss(X, Y) = 0.7059613f0 (tracked)
loss(X, Y) = 0.6622683f0 (tracked)
loss(X, Y) = 0.6905897f0 (tracked)
loss(X, Y) = 0.6884678f0 (tracked)
loss(X, Y) = 0.69129425f0 (tracked)
loss(X, Y) = 0.69586104f0 (tracked)
loss(X, Y) = 0.6877215f0 (tracked)
loss(X, Y) = 0.70434225f0 (tracked)
loss(X, Y) = 0.69703406f0 (tracked)
loss(X, Y) = 0.6790692f0 (tracked)
loss(X, Y) = 0.69012463f0 (tracked)
loss(X, Y) = 0.69512224f0 (tracked)
loss(X, Y) = 0.68063116f0 (tracked)
loss(X, Y) = 0.67920613f0 (tracked)
loss(X, Y) = 0.6680033f0 (tracked)
loss(X, Y) = 0.69472665f0 (tracked)
loss(X, Y) = 0.7012124f0 (tracked)
loss(X, Y) = 0.6846852f0 (tracked)
loss(X, Y) = 0.67636955f0 (tracked)
loss(X, Y) = 0.685309f0 (tracked)
loss(X, Y) = 0.7037586f0 (tracked)
loss(X, Y) = 0.6872026f0 (tracked)
loss(X, Y) = 0.7202507f0 (tracked)
loss(X, Y) = 0.6962537f0 (tracked)
loss(X, Y) = 0.72549987f0 (tracked)
loss(X, Y) = 0.6921462f0 (tracked)
loss(X, Y) = 0.6626569f0 (tracked)
loss(X, Y) = 0.68225396f0 (tracked)
loss(X, Y) = 0.68253493f0 (tracked)
loss(X, Y) = 0.698693f0 (tracked)
loss(X, Y) = 0.6924535f0 (tracked)
loss(X, Y) = 0.6971162f0 (tracked)
loss(X, Y) = 0.70779026f0 (tracked)
loss(X, Y) = 0.6666872f0 (tracked)
loss(X, Y) = 0.668002f0 (tracked)
loss(X, Y) = 0.69161534f0 (tracked)
loss(X, Y) = 0.6753659f0 (tracked)
loss(X, Y) = 0.6912782f0 (tracked)
loss(X, Y) = 0.7082317f0 (tracked)
loss(X, Y) = 0.649907f0 (tracked)
loss(X, Y) = 0.64683783f0 (tracked)
loss(X, Y) = 0.7097732f0 (tracked)
loss(X, Y) = 0.6527807f0 (tracked)
loss(X, Y) = 0.7033601f0 (tracked)
loss(X, Y) = 0.68801427f0 (tracked)
loss(X, Y) = 0.70758605f0 (tracked)
loss(X, Y) = 0.67575f0 (tracked)
loss(X, Y) = 0.70181227f0 (tracked)
loss(X, Y) = 0.7106234f0 (tracked)
loss(X, Y) = 0.71640766f0 (tracked)
loss(X, Y) = 0.7162051f0 (tracked)
loss(X, Y) = 0.74148977f0 (tracked)
loss(X, Y) = 0.7063311f0 (tracked)
loss(X, Y) = 0.7165143f0 (tracked)
loss(X, Y) = 0.6844802f0 (tracked)
loss(X, Y) = 0.6900535f0 (tracked)
loss(X, Y) = 0.6810328f0 (tracked)
loss(X, Y) = 0.6758583f0 (tracked)
loss(X, Y) = 0.70045763f0 (tracked)
loss(X, Y) = 0.7231896f0 (tracked)
loss(X, Y) = 0.6893898f0 (tracked)
loss(X, Y) = 0.67323774f0 (tracked)
loss(X, Y) = 0.7324545f0 (tracked)
loss(X, Y) = 0.68426895f0 (tracked)
loss(X, Y) = 0.7100366f0 (tracked)
loss(X, Y) = 0.68158114f0 (tracked)
loss(X, Y) = 0.6640649f0 (tracked)
loss(X, Y) = 0.71757406f0 (tracked)
loss(X, Y) = 0.6717794f0 (tracked)
loss(X, Y) = 0.7067284f0 (tracked)
loss(X, Y) = 0.7022853f0 (tracked)
loss(X, Y) = 0.68704396f0 (tracked)
loss(X, Y) = 0.6708038f0 (tracked)
loss(X, Y) = 0.7010561f0 (tracked)
loss(X, Y) = 0.6748065f0 (tracked)
loss(X, Y) = 0.69151235f0 (tracked)
loss(X, Y) = 0.6659712f0 (tracked)
loss(X, Y) = 0.68626857f0 (tracked)
loss(X, Y) = 0.6699607f0 (tracked)
loss(X, Y) = 0.6727131f0 (tracked)
loss(X, Y) = 0.68483603f0 (tracked)
loss(X, Y) = 0.64837f0 (tracked)
loss(X, Y) = 0.7041512f0 (tracked)
loss(X, Y) = 0.72727704f0 (tracked)
```

```
┌ Info: Epoch 694
└ @ Main /Users/kcft114/.julia/packages/Flux/qXNjB/src/optimise/train.jl:105
┌ Info: Epoch 695
└ @ Main /Users/kcft114/.julia/packages/Flux/qXNjB/src/optimise/train.jl:105
┌ Info: Epoch 696
└ @ Main /Users/kcft114/.julia/packages/Flux/qXNjB/src/optimise/train.jl:105
┌ Info: Epoch 697
└ @ Main /Users/kcft114/.julia/packages/Flux/qXNjB/src/optimise/train.jl:105
┌ Info: Epoch 698
└ @ Main /Users/kcft114/.julia/packages/Flux/qXNjB/src/optimise/train.jl:105
┌ Info: Epoch 699
└ @ Main /Users/kcft114/.julia/packages/Flux/qXNjB/src/optimise/train.jl:105
┌ Info: Epoch 700
└ @ Main /Users/kcft114/.julia/packages/Flux/qXNjB/src/optimise/train.jl:105
┌ Info: Epoch 701
└ @ Main /Users/kcft114/.julia/packages/Flux/qXNjB/src/optimise/train.jl:105
┌ Info: Epoch 702
└ @ Main /Users/kcft114/.julia/packages/Flux/qXNjB/src/optimise/train.jl:105
┌ Info: Epoch 703
└ @ Main /Users/kcft114/.julia/packages/Flux/qXNjB/src/optimise/train.jl:105
┌ Info: Epoch 704
└ @ Main /Users/kcft114/.julia/packages/Flux/qXNjB/src/optimise/train.jl:105
┌ Info: Epoch 705
└ @ Main /Users/kcft114/.julia/packages/Flux/qXNjB/src/optimise/train.jl:105
┌ Info: Epoch 706
└ @ Main /Users/kcft114/.julia/packages/Flux/qXNjB/src/optimise/train.jl:105
┌ Info: Epoch 707
└ @ Main /Users/kcft114/.julia/packages/Flux/qXNjB/src/optimise/train.jl:105
┌ Info: Epoch 708
└ @ Main /Users/kcft114/.julia/packages/Flux/qXNjB/src/optimise/train.jl:105
┌ Info: Epoch 709
└ @ Main /Users/kcft114/.julia/packages/Flux/qXNjB/src/optimise/train.jl:105
┌ Info: Epoch 710
└ @ Main /Users/kcft114/.julia/packages/Flux/qXNjB/src/optimise/train.jl:105
┌ Info: Epoch 711
└ @ Main /Users/kcft114/.julia/packages/Flux/qXNjB/src/optimise/train.jl:105
┌ Info: Epoch 712
└ @ Main /Users/kcft114/.julia/packages/Flux/qXNjB/src/optimise/train.jl:105
┌ Info: Epoch 713
└ @ Main /Users/kcft114/.julia/packages/Flux/qXNjB/src/optimise/train.jl:105
┌ Info: Epoch 714
└ @ Main /Users/kcft114/.julia/packages/Flux/qXNjB/src/optimise/train.jl:105
┌ Info: Epoch 715
└ @ Main /Users/kcft114/.julia/packages/Flux/qXNjB/src/optimise/train.jl:105
┌ Info: Epoch 716
└ @ Main /Users/kcft114/.julia/packages/Flux/qXNjB/src/optimise/train.jl:105
┌ Info: Epoch 717
└ @ Main /Users/kcft114/.julia/packages/Flux/qXNjB/src/optimise/train.jl:105
┌ Info: Epoch 718
└ @ Main /Users/kcft114/.julia/packages/Flux/qXNjB/src/optimise/train.jl:105
┌ Info: Epoch 719
└ @ Main /Users/kcft114/.julia/packages/Flux/qXNjB/src/optimise/train.jl:105
┌ Info: Epoch 720
└ @ Main /Users/kcft114/.julia/packages/Flux/qXNjB/src/optimise/train.jl:105
┌ Info: Epoch 721
└ @ Main /Users/kcft114/.julia/packages/Flux/qXNjB/src/optimise/train.jl:105
┌ Info: Epoch 722
└ @ Main /Users/kcft114/.julia/packages/Flux/qXNjB/src/optimise/train.jl:105
┌ Info: Epoch 723
└ @ Main /Users/kcft114/.julia/packages/Flux/qXNjB/src/optimise/train.jl:105
┌ Info: Epoch 724
└ @ Main /Users/kcft114/.julia/packages/Flux/qXNjB/src/optimise/train.jl:105
┌ Info: Epoch 725
└ @ Main /Users/kcft114/.julia/packages/Flux/qXNjB/src/optimise/train.jl:105
┌ Info: Epoch 726
└ @ Main /Users/kcft114/.julia/packages/Flux/qXNjB/src/optimise/train.jl:105
┌ Info: Epoch 727
└ @ Main /Users/kcft114/.julia/packages/Flux/qXNjB/src/optimise/train.jl:105
┌ Info: Epoch 728
└ @ Main /Users/kcft114/.julia/packages/Flux/qXNjB/src/optimise/train.jl:105
┌ Info: Epoch 729
└ @ Main /Users/kcft114/.julia/packages/Flux/qXNjB/src/optimise/train.jl:105
┌ Info: Epoch 730
└ @ Main /Users/kcft114/.julia/packages/Flux/qXNjB/src/optimise/train.jl:105
┌ Info: Epoch 731
└ @ Main /Users/kcft114/.julia/packages/Flux/qXNjB/src/optimise/train.jl:105
┌ Info: Epoch 732
└ @ Main /Users/kcft114/.julia/packages/Flux/qXNjB/src/optimise/train.jl:105
┌ Info: Epoch 733
└ @ Main /Users/kcft114/.julia/packages/Flux/qXNjB/src/optimise/train.jl:105
┌ Info: Epoch 734
└ @ Main /Users/kcft114/.julia/packages/Flux/qXNjB/src/optimise/train.jl:105
┌ Info: Epoch 735
└ @ Main /Users/kcft114/.julia/packages/Flux/qXNjB/src/optimise/train.jl:105
┌ Info: Epoch 736
└ @ Main /Users/kcft114/.julia/packages/Flux/qXNjB/src/optimise/train.jl:105
┌ Info: Epoch 737
└ @ Main /Users/kcft114/.julia/packages/Flux/qXNjB/src/optimise/train.jl:105
┌ Info: Epoch 738
└ @ Main /Users/kcft114/.julia/packages/Flux/qXNjB/src/optimise/train.jl:105
┌ Info: Epoch 739
└ @ Main /Users/kcft114/.julia/packages/Flux/qXNjB/src/optimise/train.jl:105
┌ Info: Epoch 740
└ @ Main /Users/kcft114/.julia/packages/Flux/qXNjB/src/optimise/train.jl:105
┌ Info: Epoch 741
└ @ Main /Users/kcft114/.julia/packages/Flux/qXNjB/src/optimise/train.jl:105
┌ Info: Epoch 742
└ @ Main /Users/kcft114/.julia/packages/Flux/qXNjB/src/optimise/train.jl:105
┌ Info: Epoch 743
└ @ Main /Users/kcft114/.julia/packages/Flux/qXNjB/src/optimise/train.jl:105
┌ Info: Epoch 744
└ @ Main /Users/kcft114/.julia/packages/Flux/qXNjB/src/optimise/train.jl:105
┌ Info: Epoch 745
└ @ Main /Users/kcft114/.julia/packages/Flux/qXNjB/src/optimise/train.jl:105
┌ Info: Epoch 746
└ @ Main /Users/kcft114/.julia/packages/Flux/qXNjB/src/optimise/train.jl:105
┌ Info: Epoch 747
└ @ Main /Users/kcft114/.julia/packages/Flux/qXNjB/src/optimise/train.jl:105
┌ Info: Epoch 748
└ @ Main /Users/kcft114/.julia/packages/Flux/qXNjB/src/optimise/train.jl:105
┌ Info: Epoch 749
└ @ Main /Users/kcft114/.julia/packages/Flux/qXNjB/src/optimise/train.jl:105
┌ Info: Epoch 750
└ @ Main /Users/kcft114/.julia/packages/Flux/qXNjB/src/optimise/train.jl:105
┌ Info: Epoch 751
└ @ Main /Users/kcft114/.julia/packages/Flux/qXNjB/src/optimise/train.jl:105
┌ Info: Epoch 752
└ @ Main /Users/kcft114/.julia/packages/Flux/qXNjB/src/optimise/train.jl:105
┌ Info: Epoch 753
└ @ Main /Users/kcft114/.julia/packages/Flux/qXNjB/src/optimise/train.jl:105
┌ Info: Epoch 754
└ @ Main /Users/kcft114/.julia/packages/Flux/qXNjB/src/optimise/train.jl:105
┌ Info: Epoch 755
└ @ Main /Users/kcft114/.julia/packages/Flux/qXNjB/src/optimise/train.jl:105
┌ Info: Epoch 756
└ @ Main /Users/kcft114/.julia/packages/Flux/qXNjB/src/optimise/train.jl:105
┌ Info: Epoch 757
└ @ Main /Users/kcft114/.julia/packages/Flux/qXNjB/src/optimise/train.jl:105
┌ Info: Epoch 758
└ @ Main /Users/kcft114/.julia/packages/Flux/qXNjB/src/optimise/train.jl:105
┌ Info: Epoch 759
└ @ Main /Users/kcft114/.julia/packages/Flux/qXNjB/src/optimise/train.jl:105
┌ Info: Epoch 760
└ @ Main /Users/kcft114/.julia/packages/Flux/qXNjB/src/optimise/train.jl:105
┌ Info: Epoch 761
└ @ Main /Users/kcft114/.julia/packages/Flux/qXNjB/src/optimise/train.jl:105
┌ Info: Epoch 762
└ @ Main /Users/kcft114/.julia/packages/Flux/qXNjB/src/optimise/train.jl:105
┌ Info: Epoch 763
└ @ Main /Users/kcft114/.julia/packages/Flux/qXNjB/src/optimise/train.jl:105
┌ Info: Epoch 764
└ @ Main /Users/kcft114/.julia/packages/Flux/qXNjB/src/optimise/train.jl:105
┌ Info: Epoch 765
└ @ Main /Users/kcft114/.julia/packages/Flux/qXNjB/src/optimise/train.jl:105
┌ Info: Epoch 766
└ @ Main /Users/kcft114/.julia/packages/Flux/qXNjB/src/optimise/train.jl:105
┌ Info: Epoch 767
└ @ Main /Users/kcft114/.julia/packages/Flux/qXNjB/src/optimise/train.jl:105
┌ Info: Epoch 768
└ @ Main /Users/kcft114/.julia/packages/Flux/qXNjB/src/optimise/train.jl:105
┌ Info: Epoch 769
└ @ Main /Users/kcft114/.julia/packages/Flux/qXNjB/src/optimise/train.jl:105
┌ Info: Epoch 770
└ @ Main /Users/kcft114/.julia/packages/Flux/qXNjB/src/optimise/train.jl:105
┌ Info: Epoch 771
└ @ Main /Users/kcft114/.julia/packages/Flux/qXNjB/src/optimise/train.jl:105
┌ Info: Epoch 772
└ @ Main /Users/kcft114/.julia/packages/Flux/qXNjB/src/optimise/train.jl:105
┌ Info: Epoch 773
└ @ Main /Users/kcft114/.julia/packages/Flux/qXNjB/src/optimise/train.jl:105
┌ Info: Epoch 774
└ @ Main /Users/kcft114/.julia/packages/Flux/qXNjB/src/optimise/train.jl:105
┌ Info: Epoch 775
└ @ Main /Users/kcft114/.julia/packages/Flux/qXNjB/src/optimise/train.jl:105
┌ Info: Epoch 776
└ @ Main /Users/kcft114/.julia/packages/Flux/qXNjB/src/optimise/train.jl:105
┌ Info: Epoch 777
└ @ Main /Users/kcft114/.julia/packages/Flux/qXNjB/src/optimise/train.jl:105
┌ Info: Epoch 778
└ @ Main /Users/kcft114/.julia/packages/Flux/qXNjB/src/optimise/train.jl:105
┌ Info: Epoch 779
└ @ Main /Users/kcft114/.julia/packages/Flux/qXNjB/src/optimise/train.jl:105
┌ Info: Epoch 780
└ @ Main /Users/kcft114/.julia/packages/Flux/qXNjB/src/optimise/train.jl:105
┌ Info: Epoch 781
└ @ Main /Users/kcft114/.julia/packages/Flux/qXNjB/src/optimise/train.jl:105
┌ Info: Epoch 782
└ @ Main /Users/kcft114/.julia/packages/Flux/qXNjB/src/optimise/train.jl:105
┌ Info: Epoch 783
└ @ Main /Users/kcft114/.julia/packages/Flux/qXNjB/src/optimise/train.jl:105
```

```
loss(X, Y) = 0.7195205f0 (tracked)
loss(X, Y) = 0.66725373f0 (tracked)
loss(X, Y) = 0.6457521f0 (tracked)
loss(X, Y) = 0.6844163f0 (tracked)
loss(X, Y) = 0.70795906f0 (tracked)
loss(X, Y) = 0.703994f0 (tracked)
loss(X, Y) = 0.69779867f0 (tracked)
loss(X, Y) = 0.6691406f0 (tracked)
loss(X, Y) = 0.6997587f0 (tracked)
loss(X, Y) = 0.63613784f0 (tracked)
loss(X, Y) = 0.6541954f0 (tracked)
loss(X, Y) = 0.68679297f0 (tracked)
loss(X, Y) = 0.6693212f0 (tracked)
loss(X, Y) = 0.69165486f0 (tracked)
loss(X, Y) = 0.73390865f0 (tracked)
loss(X, Y) = 0.6557339f0 (tracked)
loss(X, Y) = 0.6583098f0 (tracked)
loss(X, Y) = 0.67023f0 (tracked)
loss(X, Y) = 0.7039299f0 (tracked)
loss(X, Y) = 0.6576727f0 (tracked)
loss(X, Y) = 0.646566f0 (tracked)
loss(X, Y) = 0.6948965f0 (tracked)
loss(X, Y) = 0.6808419f0 (tracked)
loss(X, Y) = 0.6574098f0 (tracked)
loss(X, Y) = 0.6480497f0 (tracked)
loss(X, Y) = 0.6762754f0 (tracked)
loss(X, Y) = 0.69416296f0 (tracked)
loss(X, Y) = 0.6083664f0 (tracked)
loss(X, Y) = 0.69158244f0 (tracked)
loss(X, Y) = 0.69179183f0 (tracked)
loss(X, Y) = 0.6678556f0 (tracked)
loss(X, Y) = 0.69380563f0 (tracked)
loss(X, Y) = 0.6802714f0 (tracked)
loss(X, Y) = 0.7062808f0 (tracked)
loss(X, Y) = 0.66685873f0 (tracked)
loss(X, Y) = 0.6786481f0 (tracked)
loss(X, Y) = 0.67989385f0 (tracked)
loss(X, Y) = 0.6678381f0 (tracked)
loss(X, Y) = 0.7091412f0 (tracked)
loss(X, Y) = 0.61341834f0 (tracked)
loss(X, Y) = 0.67806786f0 (tracked)
loss(X, Y) = 0.6689573f0 (tracked)
loss(X, Y) = 0.67428726f0 (tracked)
loss(X, Y) = 0.6506438f0 (tracked)
loss(X, Y) = 0.6948227f0 (tracked)
loss(X, Y) = 0.69588345f0 (tracked)
loss(X, Y) = 0.6788713f0 (tracked)
loss(X, Y) = 0.6531901f0 (tracked)
loss(X, Y) = 0.68700665f0 (tracked)
loss(X, Y) = 0.7001625f0 (tracked)
loss(X, Y) = 0.6597629f0 (tracked)
loss(X, Y) = 0.6502764f0 (tracked)
loss(X, Y) = 0.65643096f0 (tracked)
loss(X, Y) = 0.6800546f0 (tracked)
loss(X, Y) = 0.68498826f0 (tracked)
loss(X, Y) = 0.633734f0 (tracked)
loss(X, Y) = 0.65802175f0 (tracked)
loss(X, Y) = 0.6449654f0 (tracked)
loss(X, Y) = 0.7041659f0 (tracked)
loss(X, Y) = 0.6570617f0 (tracked)
loss(X, Y) = 0.6921433f0 (tracked)
loss(X, Y) = 0.669958f0 (tracked)
loss(X, Y) = 0.69635904f0 (tracked)
loss(X, Y) = 0.67560345f0 (tracked)
loss(X, Y) = 0.6702488f0 (tracked)
```

```
┌ Info: Epoch 784
└ @ Main /Users/kcft114/.julia/packages/Flux/qXNjB/src/optimise/train.jl:105
┌ Info: Epoch 785
└ @ Main /Users/kcft114/.julia/packages/Flux/qXNjB/src/optimise/train.jl:105
┌ Info: Epoch 786
└ @ Main /Users/kcft114/.julia/packages/Flux/qXNjB/src/optimise/train.jl:105
┌ Info: Epoch 787
└ @ Main /Users/kcft114/.julia/packages/Flux/qXNjB/src/optimise/train.jl:105
┌ Info: Epoch 788
└ @ Main /Users/kcft114/.julia/packages/Flux/qXNjB/src/optimise/train.jl:105
┌ Info: Epoch 789
└ @ Main /Users/kcft114/.julia/packages/Flux/qXNjB/src/optimise/train.jl:105
┌ Info: Epoch 790
└ @ Main /Users/kcft114/.julia/packages/Flux/qXNjB/src/optimise/train.jl:105
┌ Info: Epoch 791
└ @ Main /Users/kcft114/.julia/packages/Flux/qXNjB/src/optimise/train.jl:105
┌ Info: Epoch 792
└ @ Main /Users/kcft114/.julia/packages/Flux/qXNjB/src/optimise/train.jl:105
┌ Info: Epoch 793
└ @ Main /Users/kcft114/.julia/packages/Flux/qXNjB/src/optimise/train.jl:105
┌ Info: Epoch 794
└ @ Main /Users/kcft114/.julia/packages/Flux/qXNjB/src/optimise/train.jl:105
┌ Info: Epoch 795
└ @ Main /Users/kcft114/.julia/packages/Flux/qXNjB/src/optimise/train.jl:105
┌ Info: Epoch 796
└ @ Main /Users/kcft114/.julia/packages/Flux/qXNjB/src/optimise/train.jl:105
┌ Info: Epoch 797
└ @ Main /Users/kcft114/.julia/packages/Flux/qXNjB/src/optimise/train.jl:105
┌ Info: Epoch 798
└ @ Main /Users/kcft114/.julia/packages/Flux/qXNjB/src/optimise/train.jl:105
┌ Info: Epoch 799
└ @ Main /Users/kcft114/.julia/packages/Flux/qXNjB/src/optimise/train.jl:105
┌ Info: Epoch 800
└ @ Main /Users/kcft114/.julia/packages/Flux/qXNjB/src/optimise/train.jl:105
┌ Info: Epoch 801
└ @ Main /Users/kcft114/.julia/packages/Flux/qXNjB/src/optimise/train.jl:105
┌ Info: Epoch 802
└ @ Main /Users/kcft114/.julia/packages/Flux/qXNjB/src/optimise/train.jl:105
┌ Info: Epoch 803
└ @ Main /Users/kcft114/.julia/packages/Flux/qXNjB/src/optimise/train.jl:105
┌ Info: Epoch 804
└ @ Main /Users/kcft114/.julia/packages/Flux/qXNjB/src/optimise/train.jl:105
┌ Info: Epoch 805
└ @ Main /Users/kcft114/.julia/packages/Flux/qXNjB/src/optimise/train.jl:105
┌ Info: Epoch 806
└ @ Main /Users/kcft114/.julia/packages/Flux/qXNjB/src/optimise/train.jl:105
┌ Info: Epoch 807
└ @ Main /Users/kcft114/.julia/packages/Flux/qXNjB/src/optimise/train.jl:105
┌ Info: Epoch 808
└ @ Main /Users/kcft114/.julia/packages/Flux/qXNjB/src/optimise/train.jl:105
┌ Info: Epoch 809
└ @ Main /Users/kcft114/.julia/packages/Flux/qXNjB/src/optimise/train.jl:105
┌ Info: Epoch 810
└ @ Main /Users/kcft114/.julia/packages/Flux/qXNjB/src/optimise/train.jl:105
┌ Info: Epoch 811
└ @ Main /Users/kcft114/.julia/packages/Flux/qXNjB/src/optimise/train.jl:105
┌ Info: Epoch 812
└ @ Main /Users/kcft114/.julia/packages/Flux/qXNjB/src/optimise/train.jl:105
┌ Info: Epoch 813
└ @ Main /Users/kcft114/.julia/packages/Flux/qXNjB/src/optimise/train.jl:105
┌ Info: Epoch 814
└ @ Main /Users/kcft114/.julia/packages/Flux/qXNjB/src/optimise/train.jl:105
┌ Info: Epoch 815
└ @ Main /Users/kcft114/.julia/packages/Flux/qXNjB/src/optimise/train.jl:105
┌ Info: Epoch 816
└ @ Main /Users/kcft114/.julia/packages/Flux/qXNjB/src/optimise/train.jl:105
┌ Info: Epoch 817
└ @ Main /Users/kcft114/.julia/packages/Flux/qXNjB/src/optimise/train.jl:105
┌ Info: Epoch 818
└ @ Main /Users/kcft114/.julia/packages/Flux/qXNjB/src/optimise/train.jl:105
┌ Info: Epoch 819
└ @ Main /Users/kcft114/.julia/packages/Flux/qXNjB/src/optimise/train.jl:105
┌ Info: Epoch 820
└ @ Main /Users/kcft114/.julia/packages/Flux/qXNjB/src/optimise/train.jl:105
┌ Info: Epoch 821
└ @ Main /Users/kcft114/.julia/packages/Flux/qXNjB/src/optimise/train.jl:105
┌ Info: Epoch 822
└ @ Main /Users/kcft114/.julia/packages/Flux/qXNjB/src/optimise/train.jl:105
┌ Info: Epoch 823
└ @ Main /Users/kcft114/.julia/packages/Flux/qXNjB/src/optimise/train.jl:105
┌ Info: Epoch 824
└ @ Main /Users/kcft114/.julia/packages/Flux/qXNjB/src/optimise/train.jl:105
┌ Info: Epoch 825
└ @ Main /Users/kcft114/.julia/packages/Flux/qXNjB/src/optimise/train.jl:105
┌ Info: Epoch 826
└ @ Main /Users/kcft114/.julia/packages/Flux/qXNjB/src/optimise/train.jl:105
┌ Info: Epoch 827
└ @ Main /Users/kcft114/.julia/packages/Flux/qXNjB/src/optimise/train.jl:105
┌ Info: Epoch 828
└ @ Main /Users/kcft114/.julia/packages/Flux/qXNjB/src/optimise/train.jl:105
┌ Info: Epoch 829
└ @ Main /Users/kcft114/.julia/packages/Flux/qXNjB/src/optimise/train.jl:105
┌ Info: Epoch 830
└ @ Main /Users/kcft114/.julia/packages/Flux/qXNjB/src/optimise/train.jl:105
┌ Info: Epoch 831
└ @ Main /Users/kcft114/.julia/packages/Flux/qXNjB/src/optimise/train.jl:105
┌ Info: Epoch 832
└ @ Main /Users/kcft114/.julia/packages/Flux/qXNjB/src/optimise/train.jl:105
┌ Info: Epoch 833
└ @ Main /Users/kcft114/.julia/packages/Flux/qXNjB/src/optimise/train.jl:105
┌ Info: Epoch 834
└ @ Main /Users/kcft114/.julia/packages/Flux/qXNjB/src/optimise/train.jl:105
┌ Info: Epoch 835
└ @ Main /Users/kcft114/.julia/packages/Flux/qXNjB/src/optimise/train.jl:105
┌ Info: Epoch 836
└ @ Main /Users/kcft114/.julia/packages/Flux/qXNjB/src/optimise/train.jl:105
┌ Info: Epoch 837
└ @ Main /Users/kcft114/.julia/packages/Flux/qXNjB/src/optimise/train.jl:105
┌ Info: Epoch 838
└ @ Main /Users/kcft114/.julia/packages/Flux/qXNjB/src/optimise/train.jl:105
┌ Info: Epoch 839
└ @ Main /Users/kcft114/.julia/packages/Flux/qXNjB/src/optimise/train.jl:105
┌ Info: Epoch 840
└ @ Main /Users/kcft114/.julia/packages/Flux/qXNjB/src/optimise/train.jl:105
┌ Info: Epoch 841
└ @ Main /Users/kcft114/.julia/packages/Flux/qXNjB/src/optimise/train.jl:105
┌ Info: Epoch 842
└ @ Main /Users/kcft114/.julia/packages/Flux/qXNjB/src/optimise/train.jl:105
┌ Info: Epoch 843
└ @ Main /Users/kcft114/.julia/packages/Flux/qXNjB/src/optimise/train.jl:105
┌ Info: Epoch 844
└ @ Main /Users/kcft114/.julia/packages/Flux/qXNjB/src/optimise/train.jl:105
┌ Info: Epoch 845
└ @ Main /Users/kcft114/.julia/packages/Flux/qXNjB/src/optimise/train.jl:105
┌ Info: Epoch 846
└ @ Main /Users/kcft114/.julia/packages/Flux/qXNjB/src/optimise/train.jl:105
┌ Info: Epoch 847
└ @ Main /Users/kcft114/.julia/packages/Flux/qXNjB/src/optimise/train.jl:105
┌ Info: Epoch 848
└ @ Main /Users/kcft114/.julia/packages/Flux/qXNjB/src/optimise/train.jl:105
┌ Info: Epoch 849
└ @ Main /Users/kcft114/.julia/packages/Flux/qXNjB/src/optimise/train.jl:105
┌ Info: Epoch 850
└ @ Main /Users/kcft114/.julia/packages/Flux/qXNjB/src/optimise/train.jl:105
┌ Info: Epoch 851
└ @ Main /Users/kcft114/.julia/packages/Flux/qXNjB/src/optimise/train.jl:105
┌ Info: Epoch 852
└ @ Main /Users/kcft114/.julia/packages/Flux/qXNjB/src/optimise/train.jl:105
┌ Info: Epoch 853
└ @ Main /Users/kcft114/.julia/packages/Flux/qXNjB/src/optimise/train.jl:105
┌ Info: Epoch 854
└ @ Main /Users/kcft114/.julia/packages/Flux/qXNjB/src/optimise/train.jl:105
┌ Info: Epoch 855
└ @ Main /Users/kcft114/.julia/packages/Flux/qXNjB/src/optimise/train.jl:105
┌ Info: Epoch 856
└ @ Main /Users/kcft114/.julia/packages/Flux/qXNjB/src/optimise/train.jl:105
┌ Info: Epoch 857
└ @ Main /Users/kcft114/.julia/packages/Flux/qXNjB/src/optimise/train.jl:105
┌ Info: Epoch 858
└ @ Main /Users/kcft114/.julia/packages/Flux/qXNjB/src/optimise/train.jl:105
┌ Info: Epoch 859
└ @ Main /Users/kcft114/.julia/packages/Flux/qXNjB/src/optimise/train.jl:105
┌ Info: Epoch 860
└ @ Main /Users/kcft114/.julia/packages/Flux/qXNjB/src/optimise/train.jl:105
┌ Info: Epoch 861
└ @ Main /Users/kcft114/.julia/packages/Flux/qXNjB/src/optimise/train.jl:105
┌ Info: Epoch 862
└ @ Main /Users/kcft114/.julia/packages/Flux/qXNjB/src/optimise/train.jl:105
┌ Info: Epoch 863
└ @ Main /Users/kcft114/.julia/packages/Flux/qXNjB/src/optimise/train.jl:105
┌ Info: Epoch 864
└ @ Main /Users/kcft114/.julia/packages/Flux/qXNjB/src/optimise/train.jl:105
┌ Info: Epoch 865
└ @ Main /Users/kcft114/.julia/packages/Flux/qXNjB/src/optimise/train.jl:105
┌ Info: Epoch 866
└ @ Main /Users/kcft114/.julia/packages/Flux/qXNjB/src/optimise/train.jl:105
┌ Info: Epoch 867
└ @ Main /Users/kcft114/.julia/packages/Flux/qXNjB/src/optimise/train.jl:105
┌ Info: Epoch 868
└ @ Main /Users/kcft114/.julia/packages/Flux/qXNjB/src/optimise/train.jl:105
┌ Info: Epoch 869
└ @ Main /Users/kcft114/.julia/packages/Flux/qXNjB/src/optimise/train.jl:105
┌ Info: Epoch 870
└ @ Main /Users/kcft114/.julia/packages/Flux/qXNjB/src/optimise/train.jl:105
┌ Info: Epoch 871
└ @ Main /Users/kcft114/.julia/packages/Flux/qXNjB/src/optimise/train.jl:105
┌ Info: Epoch 872
└ @ Main /Users/kcft114/.julia/packages/Flux/qXNjB/src/optimise/train.jl:105
┌ Info: Epoch 873
└ @ Main /Users/kcft114/.julia/packages/Flux/qXNjB/src/optimise/train.jl:105
┌ Info: Epoch 874
└ @ Main /Users/kcft114/.julia/packages/Flux/qXNjB/src/optimise/train.jl:105
┌ Info: Epoch 875
└ @ Main /Users/kcft114/.julia/packages/Flux/qXNjB/src/optimise/train.jl:105
┌ Info: Epoch 876
└ @ Main /Users/kcft114/.julia/packages/Flux/qXNjB/src/optimise/train.jl:105
┌ Info: Epoch 877
└ @ Main /Users/kcft114/.julia/packages/Flux/qXNjB/src/optimise/train.jl:105
┌ Info: Epoch 878
└ @ Main /Users/kcft114/.julia/packages/Flux/qXNjB/src/optimise/train.jl:105
┌ Info: Epoch 879
└ @ Main /Users/kcft114/.julia/packages/Flux/qXNjB/src/optimise/train.jl:105
┌ Info: Epoch 880
└ @ Main /Users/kcft114/.julia/packages/Flux/qXNjB/src/optimise/train.jl:105
┌ Info: Epoch 881
└ @ Main /Users/kcft114/.julia/packages/Flux/qXNjB/src/optimise/train.jl:105
┌ Info: Epoch 882
└ @ Main /Users/kcft114/.julia/packages/Flux/qXNjB/src/optimise/train.jl:105
┌ Info: Epoch 883
└ @ Main /Users/kcft114/.julia/packages/Flux/qXNjB/src/optimise/train.jl:105
┌ Info: Epoch 884
└ @ Main /Users/kcft114/.julia/packages/Flux/qXNjB/src/optimise/train.jl:105
┌ Info: Epoch 885
└ @ Main /Users/kcft114/.julia/packages/Flux/qXNjB/src/optimise/train.jl:105
┌ Info: Epoch 886
└ @ Main /Users/kcft114/.julia/packages/Flux/qXNjB/src/optimise/train.jl:105
┌ Info: Epoch 887
└ @ Main /Users/kcft114/.julia/packages/Flux/qXNjB/src/optimise/train.jl:105
┌ Info: Epoch 888
└ @ Main /Users/kcft114/.julia/packages/Flux/qXNjB/src/optimise/train.jl:105
┌ Info: Epoch 889
└ @ Main /Users/kcft114/.julia/packages/Flux/qXNjB/src/optimise/train.jl:105
┌ Info: Epoch 890
└ @ Main /Users/kcft114/.julia/packages/Flux/qXNjB/src/optimise/train.jl:105
┌ Info: Epoch 891
└ @ Main /Users/kcft114/.julia/packages/Flux/qXNjB/src/optimise/train.jl:105
┌ Info: Epoch 892
└ @ Main /Users/kcft114/.julia/packages/Flux/qXNjB/src/optimise/train.jl:105
┌ Info: Epoch 893
└ @ Main /Users/kcft114/.julia/packages/Flux/qXNjB/src/optimise/train.jl:105
┌ Info: Epoch 894
└ @ Main /Users/kcft114/.julia/packages/Flux/qXNjB/src/optimise/train.jl:105
┌ Info: Epoch 895
└ @ Main /Users/kcft114/.julia/packages/Flux/qXNjB/src/optimise/train.jl:105
┌ Info: Epoch 896
└ @ Main /Users/kcft114/.julia/packages/Flux/qXNjB/src/optimise/train.jl:105
┌ Info: Epoch 897
└ @ Main /Users/kcft114/.julia/packages/Flux/qXNjB/src/optimise/train.jl:105
┌ Info: Epoch 898
└ @ Main /Users/kcft114/.julia/packages/Flux/qXNjB/src/optimise/train.jl:105
┌ Info: Epoch 899
└ @ Main /Users/kcft114/.julia/packages/Flux/qXNjB/src/optimise/train.jl:105
┌ Info: Epoch 900
└ @ Main /Users/kcft114/.julia/packages/Flux/qXNjB/src/optimise/train.jl:105
┌ Info: Epoch 901
└ @ Main /Users/kcft114/.julia/packages/Flux/qXNjB/src/optimise/train.jl:105
┌ Info: Epoch 902
└ @ Main /Users/kcft114/.julia/packages/Flux/qXNjB/src/optimise/train.jl:105
┌ Info: Epoch 903
└ @ Main /Users/kcft114/.julia/packages/Flux/qXNjB/src/optimise/train.jl:105
┌ Info: Epoch 904
└ @ Main /Users/kcft114/.julia/packages/Flux/qXNjB/src/optimise/train.jl:105
┌ Info: Epoch 905
└ @ Main /Users/kcft114/.julia/packages/Flux/qXNjB/src/optimise/train.jl:105
┌ Info: Epoch 906
└ @ Main /Users/kcft114/.julia/packages/Flux/qXNjB/src/optimise/train.jl:105
┌ Info: Epoch 907
└ @ Main /Users/kcft114/.julia/packages/Flux/qXNjB/src/optimise/train.jl:105
┌ Info: Epoch 908
└ @ Main /Users/kcft114/.julia/packages/Flux/qXNjB/src/optimise/train.jl:105
┌ Info: Epoch 909
└ @ Main /Users/kcft114/.julia/packages/Flux/qXNjB/src/optimise/train.jl:105
┌ Info: Epoch 910
└ @ Main /Users/kcft114/.julia/packages/Flux/qXNjB/src/optimise/train.jl:105
┌ Info: Epoch 911
└ @ Main /Users/kcft114/.julia/packages/Flux/qXNjB/src/optimise/train.jl:105
┌ Info: Epoch 912
└ @ Main /Users/kcft114/.julia/packages/Flux/qXNjB/src/optimise/train.jl:105
┌ Info: Epoch 913
└ @ Main /Users/kcft114/.julia/packages/Flux/qXNjB/src/optimise/train.jl:105
┌ Info: Epoch 914
└ @ Main /Users/kcft114/.julia/packages/Flux/qXNjB/src/optimise/train.jl:105
┌ Info: Epoch 915
└ @ Main /Users/kcft114/.julia/packages/Flux/qXNjB/src/optimise/train.jl:105
┌ Info: Epoch 916
└ @ Main /Users/kcft114/.julia/packages/Flux/qXNjB/src/optimise/train.jl:105
┌ Info: Epoch 917
└ @ Main /Users/kcft114/.julia/packages/Flux/qXNjB/src/optimise/train.jl:105
┌ Info: Epoch 918
└ @ Main /Users/kcft114/.julia/packages/Flux/qXNjB/src/optimise/train.jl:105
┌ Info: Epoch 919
└ @ Main /Users/kcft114/.julia/packages/Flux/qXNjB/src/optimise/train.jl:105
┌ Info: Epoch 920
└ @ Main /Users/kcft114/.julia/packages/Flux/qXNjB/src/optimise/train.jl:105
┌ Info: Epoch 921
└ @ Main /Users/kcft114/.julia/packages/Flux/qXNjB/src/optimise/train.jl:105
┌ Info: Epoch 922
└ @ Main /Users/kcft114/.julia/packages/Flux/qXNjB/src/optimise/train.jl:105
┌ Info: Epoch 923
└ @ Main /Users/kcft114/.julia/packages/Flux/qXNjB/src/optimise/train.jl:105
┌ Info: Epoch 924
└ @ Main /Users/kcft114/.julia/packages/Flux/qXNjB/src/optimise/train.jl:105
┌ Info: Epoch 925
└ @ Main /Users/kcft114/.julia/packages/Flux/qXNjB/src/optimise/train.jl:105
┌ Info: Epoch 926
└ @ Main /Users/kcft114/.julia/packages/Flux/qXNjB/src/optimise/train.jl:105
┌ Info: Epoch 927
└ @ Main /Users/kcft114/.julia/packages/Flux/qXNjB/src/optimise/train.jl:105
┌ Info: Epoch 928
└ @ Main /Users/kcft114/.julia/packages/Flux/qXNjB/src/optimise/train.jl:105
┌ Info: Epoch 929
└ @ Main /Users/kcft114/.julia/packages/Flux/qXNjB/src/optimise/train.jl:105
┌ Info: Epoch 930
└ @ Main /Users/kcft114/.julia/packages/Flux/qXNjB/src/optimise/train.jl:105
┌ Info: Epoch 931
└ @ Main /Users/kcft114/.julia/packages/Flux/qXNjB/src/optimise/train.jl:105
┌ Info: Epoch 932
└ @ Main /Users/kcft114/.julia/packages/Flux/qXNjB/src/optimise/train.jl:105
┌ Info: Epoch 933
└ @ Main /Users/kcft114/.julia/packages/Flux/qXNjB/src/optimise/train.jl:105
┌ Info: Epoch 934
└ @ Main /Users/kcft114/.julia/packages/Flux/qXNjB/src/optimise/train.jl:105
┌ Info: Epoch 935
└ @ Main /Users/kcft114/.julia/packages/Flux/qXNjB/src/optimise/train.jl:105
┌ Info: Epoch 936
└ @ Main /Users/kcft114/.julia/packages/Flux/qXNjB/src/optimise/train.jl:105
┌ Info: Epoch 937
└ @ Main /Users/kcft114/.julia/packages/Flux/qXNjB/src/optimise/train.jl:105
┌ Info: Epoch 938
└ @ Main /Users/kcft114/.julia/packages/Flux/qXNjB/src/optimise/train.jl:105
┌ Info: Epoch 939
└ @ Main /Users/kcft114/.julia/packages/Flux/qXNjB/src/optimise/train.jl:105
┌ Info: Epoch 940
└ @ Main /Users/kcft114/.julia/packages/Flux/qXNjB/src/optimise/train.jl:105
┌ Info: Epoch 941
└ @ Main /Users/kcft114/.julia/packages/Flux/qXNjB/src/optimise/train.jl:105
┌ Info: Epoch 942
└ @ Main /Users/kcft114/.julia/packages/Flux/qXNjB/src/optimise/train.jl:105
┌ Info: Epoch 943
└ @ Main /Users/kcft114/.julia/packages/Flux/qXNjB/src/optimise/train.jl:105
┌ Info: Epoch 944
└ @ Main /Users/kcft114/.julia/packages/Flux/qXNjB/src/optimise/train.jl:105
┌ Info: Epoch 945
└ @ Main /Users/kcft114/.julia/packages/Flux/qXNjB/src/optimise/train.jl:105
┌ Info: Epoch 946
└ @ Main /Users/kcft114/.julia/packages/Flux/qXNjB/src/optimise/train.jl:105
┌ Info: Epoch 947
└ @ Main /Users/kcft114/.julia/packages/Flux/qXNjB/src/optimise/train.jl:105
┌ Info: Epoch 948
└ @ Main /Users/kcft114/.julia/packages/Flux/qXNjB/src/optimise/train.jl:105
┌ Info: Epoch 949
└ @ Main /Users/kcft114/.julia/packages/Flux/qXNjB/src/optimise/train.jl:105
┌ Info: Epoch 950
└ @ Main /Users/kcft114/.julia/packages/Flux/qXNjB/src/optimise/train.jl:105
┌ Info: Epoch 951
└ @ Main /Users/kcft114/.julia/packages/Flux/qXNjB/src/optimise/train.jl:105
┌ Info: Epoch 952
└ @ Main /Users/kcft114/.julia/packages/Flux/qXNjB/src/optimise/train.jl:105
```

```
loss(X, Y) = 0.6671838f0 (tracked)
loss(X, Y) = 0.6841746f0 (tracked)
loss(X, Y) = 0.64192164f0 (tracked)
loss(X, Y) = 0.6525643f0 (tracked)
loss(X, Y) = 0.7022214f0 (tracked)
loss(X, Y) = 0.67699194f0 (tracked)
loss(X, Y) = 0.6744363f0 (tracked)
loss(X, Y) = 0.69266474f0 (tracked)
loss(X, Y) = 0.67911303f0 (tracked)
loss(X, Y) = 0.69073343f0 (tracked)
loss(X, Y) = 0.6561334f0 (tracked)
loss(X, Y) = 0.64704204f0 (tracked)
loss(X, Y) = 0.7204349f0 (tracked)
loss(X, Y) = 0.64669544f0 (tracked)
loss(X, Y) = 0.64767843f0 (tracked)
loss(X, Y) = 0.6810389f0 (tracked)
loss(X, Y) = 0.6593967f0 (tracked)
loss(X, Y) = 0.666346f0 (tracked)
loss(X, Y) = 0.67135435f0 (tracked)
loss(X, Y) = 0.646866f0 (tracked)
loss(X, Y) = 0.6927729f0 (tracked)
loss(X, Y) = 0.67230994f0 (tracked)
loss(X, Y) = 0.6623483f0 (tracked)
loss(X, Y) = 0.6656063f0 (tracked)
loss(X, Y) = 0.638704f0 (tracked)
loss(X, Y) = 0.6682279f0 (tracked)
loss(X, Y) = 0.66512406f0 (tracked)
loss(X, Y) = 0.67116964f0 (tracked)
loss(X, Y) = 0.6606948f0 (tracked)
loss(X, Y) = 0.67555577f0 (tracked)
loss(X, Y) = 0.63331443f0 (tracked)
loss(X, Y) = 0.65619093f0 (tracked)
loss(X, Y) = 0.647464f0 (tracked)
loss(X, Y) = 0.6677524f0 (tracked)
loss(X, Y) = 0.6593078f0 (tracked)
loss(X, Y) = 0.6754581f0 (tracked)
loss(X, Y) = 0.6542649f0 (tracked)
loss(X, Y) = 0.63884795f0 (tracked)
loss(X, Y) = 0.643803f0 (tracked)
loss(X, Y) = 0.6670376f0 (tracked)
loss(X, Y) = 0.6685368f0 (tracked)
loss(X, Y) = 0.6440031f0 (tracked)
loss(X, Y) = 0.64561206f0 (tracked)
loss(X, Y) = 0.690761f0 (tracked)
loss(X, Y) = 0.66420096f0 (tracked)
loss(X, Y) = 0.65595216f0 (tracked)
loss(X, Y) = 0.63526624f0 (tracked)
loss(X, Y) = 0.6262139f0 (tracked)
loss(X, Y) = 0.65766436f0 (tracked)
loss(X, Y) = 0.63342893f0 (tracked)
loss(X, Y) = 0.6396573f0 (tracked)
loss(X, Y) = 0.6749461f0 (tracked)
loss(X, Y) = 0.64899105f0 (tracked)
loss(X, Y) = 0.6499046f0 (tracked)
loss(X, Y) = 0.6584036f0 (tracked)
loss(X, Y) = 0.6599826f0 (tracked)
loss(X, Y) = 0.6286549f0 (tracked)
loss(X, Y) = 0.6582334f0 (tracked)
loss(X, Y) = 0.68676233f0 (tracked)
loss(X, Y) = 0.64581954f0 (tracked)
loss(X, Y) = 0.6929414f0 (tracked)
loss(X, Y) = 0.68261486f0 (tracked)
loss(X, Y) = 0.6478984f0 (tracked)
loss(X, Y) = 0.6893622f0 (tracked)
loss(X, Y) = 0.6684013f0 (tracked)
loss(X, Y) = 0.66696733f0 (tracked)
loss(X, Y) = 0.64228064f0 (tracked)
loss(X, Y) = 0.71711266f0 (tracked)
loss(X, Y) = 0.7017198f0 (tracked)
loss(X, Y) = 0.6948995f0 (tracked)
loss(X, Y) = 0.67274773f0 (tracked)
loss(X, Y) = 0.6820771f0 (tracked)
loss(X, Y) = 0.6657966f0 (tracked)
loss(X, Y) = 0.67506886f0 (tracked)
loss(X, Y) = 0.68224305f0 (tracked)
loss(X, Y) = 0.64385825f0 (tracked)
loss(X, Y) = 0.6669282f0 (tracked)
loss(X, Y) = 0.6583961f0 (tracked)
loss(X, Y) = 0.6318741f0 (tracked)
loss(X, Y) = 0.64539164f0 (tracked)
loss(X, Y) = 0.6532544f0 (tracked)
loss(X, Y) = 0.665437f0 (tracked)
loss(X, Y) = 0.6320447f0 (tracked)
loss(X, Y) = 0.6460534f0 (tracked)
loss(X, Y) = 0.657612f0 (tracked)
loss(X, Y) = 0.6838117f0 (tracked)
loss(X, Y) = 0.6538039f0 (tracked)
loss(X, Y) = 0.6478201f0 (tracked)
loss(X, Y) = 0.6230047f0 (tracked)
loss(X, Y) = 0.64436406f0 (tracked)
loss(X, Y) = 0.6716414f0 (tracked)
loss(X, Y) = 0.6940584f0 (tracked)
loss(X, Y) = 0.670569f0 (tracked)
loss(X, Y) = 0.6430569f0 (tracked)
loss(X, Y) = 0.63262355f0 (tracked)
loss(X, Y) = 0.64480174f0 (tracked)
loss(X, Y) = 0.68139297f0 (tracked)
loss(X, Y) = 0.6632412f0 (tracked)
loss(X, Y) = 0.68888956f0 (tracked)
loss(X, Y) = 0.64061743f0 (tracked)
loss(X, Y) = 0.65730727f0 (tracked)
loss(X, Y) = 0.65562224f0 (tracked)
loss(X, Y) = 0.67598516f0 (tracked)
loss(X, Y) = 0.63861763f0 (tracked)
loss(X, Y) = 0.67218965f0 (tracked)
loss(X, Y) = 0.67550665f0 (tracked)
loss(X, Y) = 0.6823026f0 (tracked)
loss(X, Y) = 0.683166f0 (tracked)
```

```
┌ Info: Epoch 953
└ @ Main /Users/kcft114/.julia/packages/Flux/qXNjB/src/optimise/train.jl:105
┌ Info: Epoch 954
└ @ Main /Users/kcft114/.julia/packages/Flux/qXNjB/src/optimise/train.jl:105
┌ Info: Epoch 955
└ @ Main /Users/kcft114/.julia/packages/Flux/qXNjB/src/optimise/train.jl:105
┌ Info: Epoch 956
└ @ Main /Users/kcft114/.julia/packages/Flux/qXNjB/src/optimise/train.jl:105
```

```
loss(X, Y) = 0.61935663f0 (tracked)
loss(X, Y) = 0.64460266f0 (tracked)
loss(X, Y) = 0.6532207f0 (tracked)
loss(X, Y) = 0.68134886f0 (tracked)
loss(X, Y) = 0.6519798f0 (tracked)
loss(X, Y) = 0.6556974f0 (tracked)
loss(X, Y) = 0.6629607f0 (tracked)
loss(X, Y) = 0.63342106f0 (tracked)
loss(X, Y) = 0.6679827f0 (tracked)
loss(X, Y) = 0.69018334f0 (tracked)
loss(X, Y) = 0.6666238f0 (tracked)
loss(X, Y) = 0.6185691f0 (tracked)
loss(X, Y) = 0.6233088f0 (tracked)
loss(X, Y) = 0.6715278f0 (tracked)
loss(X, Y) = 0.62451154f0 (tracked)
loss(X, Y) = 0.65241534f0 (tracked)
loss(X, Y) = 0.6634822f0 (tracked)
loss(X, Y) = 0.6475164f0 (tracked)
loss(X, Y) = 0.67379445f0 (tracked)
loss(X, Y) = 0.6438339f0 (tracked)
loss(X, Y) = 0.66007006f0 (tracked)
loss(X, Y) = 0.62721837f0 (tracked)
loss(X, Y) = 0.68052846f0 (tracked)
loss(X, Y) = 0.6521371f0 (tracked)
loss(X, Y) = 0.7008917f0 (tracked)
loss(X, Y) = 0.6153885f0 (tracked)
loss(X, Y) = 0.6499335f0 (tracked)
loss(X, Y) = 0.65977097f0 (tracked)
loss(X, Y) = 0.6351008f0 (tracked)
loss(X, Y) = 0.64755595f0 (tracked)
loss(X, Y) = 0.6469627f0 (tracked)
loss(X, Y) = 0.70673615f0 (tracked)
loss(X, Y) = 0.63079697f0 (tracked)
loss(X, Y) = 0.64128655f0 (tracked)
loss(X, Y) = 0.67120004f0 (tracked)
loss(X, Y) = 0.64961225f0 (tracked)
loss(X, Y) = 0.60824525f0 (tracked)
loss(X, Y) = 0.6511185f0 (tracked)
loss(X, Y) = 0.6574531f0 (tracked)
loss(X, Y) = 0.64467597f0 (tracked)
loss(X, Y) = 0.60814446f0 (tracked)
loss(X, Y) = 0.65094006f0 (tracked)
loss(X, Y) = 0.67883724f0 (tracked)
loss(X, Y) = 0.6004928f0 (tracked)
loss(X, Y) = 0.61470497f0 (tracked)
loss(X, Y) = 0.62750936f0 (tracked)
loss(X, Y) = 0.6456348f0 (tracked)
loss(X, Y) = 0.65042853f0 (tracked)
loss(X, Y) = 0.6662066f0 (tracked)
loss(X, Y) = 0.66983634f0 (tracked)
loss(X, Y) = 0.6269243f0 (tracked)
loss(X, Y) = 0.6510842f0 (tracked)
loss(X, Y) = 0.6565717f0 (tracked)
loss(X, Y) = 0.6720992f0 (tracked)
loss(X, Y) = 0.62722224f0 (tracked)
loss(X, Y) = 0.61407554f0 (tracked)
loss(X, Y) = 0.6632952f0 (tracked)
loss(X, Y) = 0.6889738f0 (tracked)
loss(X, Y) = 0.6386887f0 (tracked)
loss(X, Y) = 0.66336036f0 (tracked)
loss(X, Y) = 0.6559141f0 (tracked)
loss(X, Y) = 0.65082896f0 (tracked)
loss(X, Y) = 0.60981315f0 (tracked)
loss(X, Y) = 0.6186922f0 (tracked)
loss(X, Y) = 0.61150146f0 (tracked)
loss(X, Y) = 0.67120785f0 (tracked)
loss(X, Y) = 0.6267077f0 (tracked)
loss(X, Y) = 0.6351592f0 (tracked)
loss(X, Y) = 0.6599216f0 (tracked)
loss(X, Y) = 0.67075336f0 (tracked)
loss(X, Y) = 0.66791505f0 (tracked)
loss(X, Y) = 0.63932747f0 (tracked)
loss(X, Y) = 0.6693566f0 (tracked)
loss(X, Y) = 0.64593154f0 (tracked)
loss(X, Y) = 0.67147756f0 (tracked)
loss(X, Y) = 0.673299f0 (tracked)
loss(X, Y) = 0.63746333f0 (tracked)
loss(X, Y) = 0.64072067f0 (tracked)
loss(X, Y) = 0.6469931f0 (tracked)
loss(X, Y) = 0.6079817f0 (tracked)
loss(X, Y) = 0.6803706f0 (tracked)
loss(X, Y) = 0.6718687f0 (tracked)
loss(X, Y) = 0.6164563f0 (tracked)
loss(X, Y) = 0.65725f0 (tracked)
loss(X, Y) = 0.5998259f0 (tracked)
loss(X, Y) = 0.65219045f0 (tracked)
loss(X, Y) = 0.6502825f0 (tracked)
loss(X, Y) = 0.6914232f0 (tracked)
loss(X, Y) = 0.6397973f0 (tracked)
loss(X, Y) = 0.68723017f0 (tracked)
loss(X, Y) = 0.6299858f0 (tracked)
loss(X, Y) = 0.6922147f0 (tracked)
loss(X, Y) = 0.61831844f0 (tracked)
loss(X, Y) = 0.68886894f0 (tracked)
loss(X, Y) = 0.6567727f0 (tracked)
loss(X, Y) = 0.6225352f0 (tracked)
loss(X, Y) = 0.6509442f0 (tracked)
loss(X, Y) = 0.6353344f0 (tracked)
loss(X, Y) = 0.65574867f0 (tracked)
loss(X, Y) = 0.6225042f0 (tracked)
```

```
┌ Info: Epoch 957
└ @ Main /Users/kcft114/.julia/packages/Flux/qXNjB/src/optimise/train.jl:105
┌ Info: Epoch 958
└ @ Main /Users/kcft114/.julia/packages/Flux/qXNjB/src/optimise/train.jl:105
┌ Info: Epoch 959
└ @ Main /Users/kcft114/.julia/packages/Flux/qXNjB/src/optimise/train.jl:105
┌ Info: Epoch 960
└ @ Main /Users/kcft114/.julia/packages/Flux/qXNjB/src/optimise/train.jl:105
┌ Info: Epoch 961
└ @ Main /Users/kcft114/.julia/packages/Flux/qXNjB/src/optimise/train.jl:105
┌ Info: Epoch 962
└ @ Main /Users/kcft114/.julia/packages/Flux/qXNjB/src/optimise/train.jl:105
┌ Info: Epoch 963
└ @ Main /Users/kcft114/.julia/packages/Flux/qXNjB/src/optimise/train.jl:105
┌ Info: Epoch 964
└ @ Main /Users/kcft114/.julia/packages/Flux/qXNjB/src/optimise/train.jl:105
┌ Info: Epoch 965
└ @ Main /Users/kcft114/.julia/packages/Flux/qXNjB/src/optimise/train.jl:105
┌ Info: Epoch 966
└ @ Main /Users/kcft114/.julia/packages/Flux/qXNjB/src/optimise/train.jl:105
┌ Info: Epoch 967
└ @ Main /Users/kcft114/.julia/packages/Flux/qXNjB/src/optimise/train.jl:105
┌ Info: Epoch 968
└ @ Main /Users/kcft114/.julia/packages/Flux/qXNjB/src/optimise/train.jl:105
┌ Info: Epoch 969
└ @ Main /Users/kcft114/.julia/packages/Flux/qXNjB/src/optimise/train.jl:105
┌ Info: Epoch 970
└ @ Main /Users/kcft114/.julia/packages/Flux/qXNjB/src/optimise/train.jl:105
┌ Info: Epoch 971
└ @ Main /Users/kcft114/.julia/packages/Flux/qXNjB/src/optimise/train.jl:105
┌ Info: Epoch 972
└ @ Main /Users/kcft114/.julia/packages/Flux/qXNjB/src/optimise/train.jl:105
┌ Info: Epoch 973
└ @ Main /Users/kcft114/.julia/packages/Flux/qXNjB/src/optimise/train.jl:105
┌ Info: Epoch 974
└ @ Main /Users/kcft114/.julia/packages/Flux/qXNjB/src/optimise/train.jl:105
┌ Info: Epoch 975
└ @ Main /Users/kcft114/.julia/packages/Flux/qXNjB/src/optimise/train.jl:105
┌ Info: Epoch 976
└ @ Main /Users/kcft114/.julia/packages/Flux/qXNjB/src/optimise/train.jl:105
┌ Info: Epoch 977
└ @ Main /Users/kcft114/.julia/packages/Flux/qXNjB/src/optimise/train.jl:105
┌ Info: Epoch 978
└ @ Main /Users/kcft114/.julia/packages/Flux/qXNjB/src/optimise/train.jl:105
┌ Info: Epoch 979
└ @ Main /Users/kcft114/.julia/packages/Flux/qXNjB/src/optimise/train.jl:105
┌ Info: Epoch 980
└ @ Main /Users/kcft114/.julia/packages/Flux/qXNjB/src/optimise/train.jl:105
┌ Info: Epoch 981
└ @ Main /Users/kcft114/.julia/packages/Flux/qXNjB/src/optimise/train.jl:105
┌ Info: Epoch 982
└ @ Main /Users/kcft114/.julia/packages/Flux/qXNjB/src/optimise/train.jl:105
┌ Info: Epoch 983
└ @ Main /Users/kcft114/.julia/packages/Flux/qXNjB/src/optimise/train.jl:105
┌ Info: Epoch 984
└ @ Main /Users/kcft114/.julia/packages/Flux/qXNjB/src/optimise/train.jl:105
┌ Info: Epoch 985
└ @ Main /Users/kcft114/.julia/packages/Flux/qXNjB/src/optimise/train.jl:105
┌ Info: Epoch 986
└ @ Main /Users/kcft114/.julia/packages/Flux/qXNjB/src/optimise/train.jl:105
┌ Info: Epoch 987
└ @ Main /Users/kcft114/.julia/packages/Flux/qXNjB/src/optimise/train.jl:105
┌ Info: Epoch 988
└ @ Main /Users/kcft114/.julia/packages/Flux/qXNjB/src/optimise/train.jl:105
┌ Info: Epoch 989
└ @ Main /Users/kcft114/.julia/packages/Flux/qXNjB/src/optimise/train.jl:105
┌ Info: Epoch 990
└ @ Main /Users/kcft114/.julia/packages/Flux/qXNjB/src/optimise/train.jl:105
┌ Info: Epoch 991
└ @ Main /Users/kcft114/.julia/packages/Flux/qXNjB/src/optimise/train.jl:105
┌ Info: Epoch 992
└ @ Main /Users/kcft114/.julia/packages/Flux/qXNjB/src/optimise/train.jl:105
┌ Info: Epoch 993
└ @ Main /Users/kcft114/.julia/packages/Flux/qXNjB/src/optimise/train.jl:105
┌ Info: Epoch 994
└ @ Main /Users/kcft114/.julia/packages/Flux/qXNjB/src/optimise/train.jl:105
┌ Info: Epoch 995
└ @ Main /Users/kcft114/.julia/packages/Flux/qXNjB/src/optimise/train.jl:105
┌ Info: Epoch 996
└ @ Main /Users/kcft114/.julia/packages/Flux/qXNjB/src/optimise/train.jl:105
┌ Info: Epoch 997
└ @ Main /Users/kcft114/.julia/packages/Flux/qXNjB/src/optimise/train.jl:105
┌ Info: Epoch 998
└ @ Main /Users/kcft114/.julia/packages/Flux/qXNjB/src/optimise/train.jl:105
┌ Info: Epoch 999
└ @ Main /Users/kcft114/.julia/packages/Flux/qXNjB/src/optimise/train.jl:105
┌ Info: Epoch 1000
└ @ Main /Users/kcft114/.julia/packages/Flux/qXNjB/src/optimise/train.jl:105
┌ Info: Epoch 1001
└ @ Main /Users/kcft114/.julia/packages/Flux/qXNjB/src/optimise/train.jl:105
┌ Info: Epoch 1002
└ @ Main /Users/kcft114/.julia/packages/Flux/qXNjB/src/optimise/train.jl:105
┌ Info: Epoch 1003
└ @ Main /Users/kcft114/.julia/packages/Flux/qXNjB/src/optimise/train.jl:105
┌ Info: Epoch 1004
└ @ Main /Users/kcft114/.julia/packages/Flux/qXNjB/src/optimise/train.jl:105
┌ Info: Epoch 1005
└ @ Main /Users/kcft114/.julia/packages/Flux/qXNjB/src/optimise/train.jl:105
┌ Info: Epoch 1006
└ @ Main /Users/kcft114/.julia/packages/Flux/qXNjB/src/optimise/train.jl:105
┌ Info: Epoch 1007
└ @ Main /Users/kcft114/.julia/packages/Flux/qXNjB/src/optimise/train.jl:105
┌ Info: Epoch 1008
└ @ Main /Users/kcft114/.julia/packages/Flux/qXNjB/src/optimise/train.jl:105
┌ Info: Epoch 1009
└ @ Main /Users/kcft114/.julia/packages/Flux/qXNjB/src/optimise/train.jl:105
┌ Info: Epoch 1010
└ @ Main /Users/kcft114/.julia/packages/Flux/qXNjB/src/optimise/train.jl:105
┌ Info: Epoch 1011
└ @ Main /Users/kcft114/.julia/packages/Flux/qXNjB/src/optimise/train.jl:105
┌ Info: Epoch 1012
└ @ Main /Users/kcft114/.julia/packages/Flux/qXNjB/src/optimise/train.jl:105
┌ Info: Epoch 1013
└ @ Main /Users/kcft114/.julia/packages/Flux/qXNjB/src/optimise/train.jl:105
┌ Info: Epoch 1014
└ @ Main /Users/kcft114/.julia/packages/Flux/qXNjB/src/optimise/train.jl:105
┌ Info: Epoch 1015
└ @ Main /Users/kcft114/.julia/packages/Flux/qXNjB/src/optimise/train.jl:105
┌ Info: Epoch 1016
└ @ Main /Users/kcft114/.julia/packages/Flux/qXNjB/src/optimise/train.jl:105
┌ Info: Epoch 1017
└ @ Main /Users/kcft114/.julia/packages/Flux/qXNjB/src/optimise/train.jl:105
┌ Info: Epoch 1018
└ @ Main /Users/kcft114/.julia/packages/Flux/qXNjB/src/optimise/train.jl:105
┌ Info: Epoch 1019
└ @ Main /Users/kcft114/.julia/packages/Flux/qXNjB/src/optimise/train.jl:105
┌ Info: Epoch 1020
└ @ Main /Users/kcft114/.julia/packages/Flux/qXNjB/src/optimise/train.jl:105
┌ Info: Epoch 1021
└ @ Main /Users/kcft114/.julia/packages/Flux/qXNjB/src/optimise/train.jl:105
┌ Info: Epoch 1022
└ @ Main /Users/kcft114/.julia/packages/Flux/qXNjB/src/optimise/train.jl:105
┌ Info: Epoch 1023
└ @ Main /Users/kcft114/.julia/packages/Flux/qXNjB/src/optimise/train.jl:105
┌ Info: Epoch 1024
└ @ Main /Users/kcft114/.julia/packages/Flux/qXNjB/src/optimise/train.jl:105
┌ Info: Epoch 1025
└ @ Main /Users/kcft114/.julia/packages/Flux/qXNjB/src/optimise/train.jl:105
┌ Info: Epoch 1026
└ @ Main /Users/kcft114/.julia/packages/Flux/qXNjB/src/optimise/train.jl:105
┌ Info: Epoch 1027
└ @ Main /Users/kcft114/.julia/packages/Flux/qXNjB/src/optimise/train.jl:105
┌ Info: Epoch 1028
└ @ Main /Users/kcft114/.julia/packages/Flux/qXNjB/src/optimise/train.jl:105
┌ Info: Epoch 1029
└ @ Main /Users/kcft114/.julia/packages/Flux/qXNjB/src/optimise/train.jl:105
┌ Info: Epoch 1030
└ @ Main /Users/kcft114/.julia/packages/Flux/qXNjB/src/optimise/train.jl:105
┌ Info: Epoch 1031
└ @ Main /Users/kcft114/.julia/packages/Flux/qXNjB/src/optimise/train.jl:105
┌ Info: Epoch 1032
└ @ Main /Users/kcft114/.julia/packages/Flux/qXNjB/src/optimise/train.jl:105
┌ Info: Epoch 1033
└ @ Main /Users/kcft114/.julia/packages/Flux/qXNjB/src/optimise/train.jl:105
┌ Info: Epoch 1034
└ @ Main /Users/kcft114/.julia/packages/Flux/qXNjB/src/optimise/train.jl:105
┌ Info: Epoch 1035
└ @ Main /Users/kcft114/.julia/packages/Flux/qXNjB/src/optimise/train.jl:105
┌ Info: Epoch 1036
└ @ Main /Users/kcft114/.julia/packages/Flux/qXNjB/src/optimise/train.jl:105
┌ Info: Epoch 1037
└ @ Main /Users/kcft114/.julia/packages/Flux/qXNjB/src/optimise/train.jl:105
┌ Info: Epoch 1038
└ @ Main /Users/kcft114/.julia/packages/Flux/qXNjB/src/optimise/train.jl:105
┌ Info: Epoch 1039
└ @ Main /Users/kcft114/.julia/packages/Flux/qXNjB/src/optimise/train.jl:105
┌ Info: Epoch 1040
└ @ Main /Users/kcft114/.julia/packages/Flux/qXNjB/src/optimise/train.jl:105
┌ Info: Epoch 1041
└ @ Main /Users/kcft114/.julia/packages/Flux/qXNjB/src/optimise/train.jl:105
┌ Info: Epoch 1042
└ @ Main /Users/kcft114/.julia/packages/Flux/qXNjB/src/optimise/train.jl:105
┌ Info: Epoch 1043
└ @ Main /Users/kcft114/.julia/packages/Flux/qXNjB/src/optimise/train.jl:105
┌ Info: Epoch 1044
└ @ Main /Users/kcft114/.julia/packages/Flux/qXNjB/src/optimise/train.jl:105
┌ Info: Epoch 1045
└ @ Main /Users/kcft114/.julia/packages/Flux/qXNjB/src/optimise/train.jl:105
┌ Info: Epoch 1046
└ @ Main /Users/kcft114/.julia/packages/Flux/qXNjB/src/optimise/train.jl:105
┌ Info: Epoch 1047
└ @ Main /Users/kcft114/.julia/packages/Flux/qXNjB/src/optimise/train.jl:105
┌ Info: Epoch 1048
└ @ Main /Users/kcft114/.julia/packages/Flux/qXNjB/src/optimise/train.jl:105
┌ Info: Epoch 1049
└ @ Main /Users/kcft114/.julia/packages/Flux/qXNjB/src/optimise/train.jl:105
┌ Info: Epoch 1050
└ @ Main /Users/kcft114/.julia/packages/Flux/qXNjB/src/optimise/train.jl:105
┌ Info: Epoch 1051
└ @ Main /Users/kcft114/.julia/packages/Flux/qXNjB/src/optimise/train.jl:105
┌ Info: Epoch 1052
└ @ Main /Users/kcft114/.julia/packages/Flux/qXNjB/src/optimise/train.jl:105
┌ Info: Epoch 1053
└ @ Main /Users/kcft114/.julia/packages/Flux/qXNjB/src/optimise/train.jl:105
┌ Info: Epoch 1054
└ @ Main /Users/kcft114/.julia/packages/Flux/qXNjB/src/optimise/train.jl:105
┌ Info: Epoch 1055
└ @ Main /Users/kcft114/.julia/packages/Flux/qXNjB/src/optimise/train.jl:105
┌ Info: Epoch 1056
└ @ Main /Users/kcft114/.julia/packages/Flux/qXNjB/src/optimise/train.jl:105
┌ Info: Epoch 1057
└ @ Main /Users/kcft114/.julia/packages/Flux/qXNjB/src/optimise/train.jl:105
┌ Info: Epoch 1058
└ @ Main /Users/kcft114/.julia/packages/Flux/qXNjB/src/optimise/train.jl:105
┌ Info: Epoch 1059
└ @ Main /Users/kcft114/.julia/packages/Flux/qXNjB/src/optimise/train.jl:105
┌ Info: Epoch 1060
└ @ Main /Users/kcft114/.julia/packages/Flux/qXNjB/src/optimise/train.jl:105
┌ Info: Epoch 1061
└ @ Main /Users/kcft114/.julia/packages/Flux/qXNjB/src/optimise/train.jl:105
┌ Info: Epoch 1062
└ @ Main /Users/kcft114/.julia/packages/Flux/qXNjB/src/optimise/train.jl:105
┌ Info: Epoch 1063
└ @ Main /Users/kcft114/.julia/packages/Flux/qXNjB/src/optimise/train.jl:105
┌ Info: Epoch 1064
└ @ Main /Users/kcft114/.julia/packages/Flux/qXNjB/src/optimise/train.jl:105
┌ Info: Epoch 1065
└ @ Main /Users/kcft114/.julia/packages/Flux/qXNjB/src/optimise/train.jl:105
┌ Info: Epoch 1066
└ @ Main /Users/kcft114/.julia/packages/Flux/qXNjB/src/optimise/train.jl:105
┌ Info: Epoch 1067
└ @ Main /Users/kcft114/.julia/packages/Flux/qXNjB/src/optimise/train.jl:105
┌ Info: Epoch 1068
└ @ Main /Users/kcft114/.julia/packages/Flux/qXNjB/src/optimise/train.jl:105
┌ Info: Epoch 1069
└ @ Main /Users/kcft114/.julia/packages/Flux/qXNjB/src/optimise/train.jl:105
┌ Info: Epoch 1070
└ @ Main /Users/kcft114/.julia/packages/Flux/qXNjB/src/optimise/train.jl:105
┌ Info: Epoch 1071
└ @ Main /Users/kcft114/.julia/packages/Flux/qXNjB/src/optimise/train.jl:105
┌ Info: Epoch 1072
└ @ Main /Users/kcft114/.julia/packages/Flux/qXNjB/src/optimise/train.jl:105
┌ Info: Epoch 1073
└ @ Main /Users/kcft114/.julia/packages/Flux/qXNjB/src/optimise/train.jl:105
┌ Info: Epoch 1074
└ @ Main /Users/kcft114/.julia/packages/Flux/qXNjB/src/optimise/train.jl:105
┌ Info: Epoch 1075
└ @ Main /Users/kcft114/.julia/packages/Flux/qXNjB/src/optimise/train.jl:105
┌ Info: Epoch 1076
└ @ Main /Users/kcft114/.julia/packages/Flux/qXNjB/src/optimise/train.jl:105
┌ Info: Epoch 1077
└ @ Main /Users/kcft114/.julia/packages/Flux/qXNjB/src/optimise/train.jl:105
┌ Info: Epoch 1078
└ @ Main /Users/kcft114/.julia/packages/Flux/qXNjB/src/optimise/train.jl:105
┌ Info: Epoch 1079
└ @ Main /Users/kcft114/.julia/packages/Flux/qXNjB/src/optimise/train.jl:105
┌ Info: Epoch 1080
└ @ Main /Users/kcft114/.julia/packages/Flux/qXNjB/src/optimise/train.jl:105
┌ Info: Epoch 1081
└ @ Main /Users/kcft114/.julia/packages/Flux/qXNjB/src/optimise/train.jl:105
┌ Info: Epoch 1082
└ @ Main /Users/kcft114/.julia/packages/Flux/qXNjB/src/optimise/train.jl:105
┌ Info: Epoch 1083
└ @ Main /Users/kcft114/.julia/packages/Flux/qXNjB/src/optimise/train.jl:105
┌ Info: Epoch 1084
└ @ Main /Users/kcft114/.julia/packages/Flux/qXNjB/src/optimise/train.jl:105
┌ Info: Epoch 1085
└ @ Main /Users/kcft114/.julia/packages/Flux/qXNjB/src/optimise/train.jl:105
┌ Info: Epoch 1086
└ @ Main /Users/kcft114/.julia/packages/Flux/qXNjB/src/optimise/train.jl:105
┌ Info: Epoch 1087
└ @ Main /Users/kcft114/.julia/packages/Flux/qXNjB/src/optimise/train.jl:105
┌ Info: Epoch 1088
└ @ Main /Users/kcft114/.julia/packages/Flux/qXNjB/src/optimise/train.jl:105
┌ Info: Epoch 1089
└ @ Main /Users/kcft114/.julia/packages/Flux/qXNjB/src/optimise/train.jl:105
┌ Info: Epoch 1090
└ @ Main /Users/kcft114/.julia/packages/Flux/qXNjB/src/optimise/train.jl:105
┌ Info: Epoch 1091
└ @ Main /Users/kcft114/.julia/packages/Flux/qXNjB/src/optimise/train.jl:105
┌ Info: Epoch 1092
└ @ Main /Users/kcft114/.julia/packages/Flux/qXNjB/src/optimise/train.jl:105
┌ Info: Epoch 1093
└ @ Main /Users/kcft114/.julia/packages/Flux/qXNjB/src/optimise/train.jl:105
┌ Info: Epoch 1094
└ @ Main /Users/kcft114/.julia/packages/Flux/qXNjB/src/optimise/train.jl:105
┌ Info: Epoch 1095
└ @ Main /Users/kcft114/.julia/packages/Flux/qXNjB/src/optimise/train.jl:105
┌ Info: Epoch 1096
└ @ Main /Users/kcft114/.julia/packages/Flux/qXNjB/src/optimise/train.jl:105
┌ Info: Epoch 1097
└ @ Main /Users/kcft114/.julia/packages/Flux/qXNjB/src/optimise/train.jl:105
┌ Info: Epoch 1098
└ @ Main /Users/kcft114/.julia/packages/Flux/qXNjB/src/optimise/train.jl:105
┌ Info: Epoch 1099
└ @ Main /Users/kcft114/.julia/packages/Flux/qXNjB/src/optimise/train.jl:105
┌ Info: Epoch 1100
└ @ Main /Users/kcft114/.julia/packages/Flux/qXNjB/src/optimise/train.jl:105
┌ Info: Epoch 1101
└ @ Main /Users/kcft114/.julia/packages/Flux/qXNjB/src/optimise/train.jl:105
┌ Info: Epoch 1102
└ @ Main /Users/kcft114/.julia/packages/Flux/qXNjB/src/optimise/train.jl:105
┌ Info: Epoch 1103
└ @ Main /Users/kcft114/.julia/packages/Flux/qXNjB/src/optimise/train.jl:105
┌ Info: Epoch 1104
└ @ Main /Users/kcft114/.julia/packages/Flux/qXNjB/src/optimise/train.jl:105
┌ Info: Epoch 1105
└ @ Main /Users/kcft114/.julia/packages/Flux/qXNjB/src/optimise/train.jl:105
┌ Info: Epoch 1106
└ @ Main /Users/kcft114/.julia/packages/Flux/qXNjB/src/optimise/train.jl:105
┌ Info: Epoch 1107
└ @ Main /Users/kcft114/.julia/packages/Flux/qXNjB/src/optimise/train.jl:105
┌ Info: Epoch 1108
└ @ Main /Users/kcft114/.julia/packages/Flux/qXNjB/src/optimise/train.jl:105
┌ Info: Epoch 1109
└ @ Main /Users/kcft114/.julia/packages/Flux/qXNjB/src/optimise/train.jl:105
┌ Info: Epoch 1110
└ @ Main /Users/kcft114/.julia/packages/Flux/qXNjB/src/optimise/train.jl:105
┌ Info: Epoch 1111
└ @ Main /Users/kcft114/.julia/packages/Flux/qXNjB/src/optimise/train.jl:105
┌ Info: Epoch 1112
└ @ Main /Users/kcft114/.julia/packages/Flux/qXNjB/src/optimise/train.jl:105
┌ Info: Epoch 1113
└ @ Main /Users/kcft114/.julia/packages/Flux/qXNjB/src/optimise/train.jl:105
┌ Info: Epoch 1114
└ @ Main /Users/kcft114/.julia/packages/Flux/qXNjB/src/optimise/train.jl:105
┌ Info: Epoch 1115
└ @ Main /Users/kcft114/.julia/packages/Flux/qXNjB/src/optimise/train.jl:105
┌ Info: Epoch 1116
└ @ Main /Users/kcft114/.julia/packages/Flux/qXNjB/src/optimise/train.jl:105
┌ Info: Epoch 1117
└ @ Main /Users/kcft114/.julia/packages/Flux/qXNjB/src/optimise/train.jl:105
┌ Info: Epoch 1118
└ @ Main /Users/kcft114/.julia/packages/Flux/qXNjB/src/optimise/train.jl:105
┌ Info: Epoch 1119
└ @ Main /Users/kcft114/.julia/packages/Flux/qXNjB/src/optimise/train.jl:105
┌ Info: Epoch 1120
└ @ Main /Users/kcft114/.julia/packages/Flux/qXNjB/src/optimise/train.jl:105
┌ Info: Epoch 1121
└ @ Main /Users/kcft114/.julia/packages/Flux/qXNjB/src/optimise/train.jl:105
┌ Info: Epoch 1122
└ @ Main /Users/kcft114/.julia/packages/Flux/qXNjB/src/optimise/train.jl:105
┌ Info: Epoch 1123
└ @ Main /Users/kcft114/.julia/packages/Flux/qXNjB/src/optimise/train.jl:105
┌ Info: Epoch 1124
└ @ Main /Users/kcft114/.julia/packages/Flux/qXNjB/src/optimise/train.jl:105
┌ Info: Epoch 1125
└ @ Main /Users/kcft114/.julia/packages/Flux/qXNjB/src/optimise/train.jl:105
┌ Info: Epoch 1126
└ @ Main /Users/kcft114/.julia/packages/Flux/qXNjB/src/optimise/train.jl:105
┌ Info: Epoch 1127
└ @ Main /Users/kcft114/.julia/packages/Flux/qXNjB/src/optimise/train.jl:105
┌ Info: Epoch 1128
└ @ Main /Users/kcft114/.julia/packages/Flux/qXNjB/src/optimise/train.jl:105
┌ Info: Epoch 1129
└ @ Main /Users/kcft114/.julia/packages/Flux/qXNjB/src/optimise/train.jl:105
┌ Info: Epoch 1130
└ @ Main /Users/kcft114/.julia/packages/Flux/qXNjB/src/optimise/train.jl:105
┌ Info: Epoch 1131
└ @ Main /Users/kcft114/.julia/packages/Flux/qXNjB/src/optimise/train.jl:105
┌ Info: Epoch 1132
└ @ Main /Users/kcft114/.julia/packages/Flux/qXNjB/src/optimise/train.jl:105
┌ Info: Epoch 1133
└ @ Main /Users/kcft114/.julia/packages/Flux/qXNjB/src/optimise/train.jl:105
┌ Info: Epoch 1134
└ @ Main /Users/kcft114/.julia/packages/Flux/qXNjB/src/optimise/train.jl:105
┌ Info: Epoch 1135
└ @ Main /Users/kcft114/.julia/packages/Flux/qXNjB/src/optimise/train.jl:105
┌ Info: Epoch 1136
└ @ Main /Users/kcft114/.julia/packages/Flux/qXNjB/src/optimise/train.jl:105
┌ Info: Epoch 1137
└ @ Main /Users/kcft114/.julia/packages/Flux/qXNjB/src/optimise/train.jl:105
┌ Info: Epoch 1138
└ @ Main /Users/kcft114/.julia/packages/Flux/qXNjB/src/optimise/train.jl:105
┌ Info: Epoch 1139
└ @ Main /Users/kcft114/.julia/packages/Flux/qXNjB/src/optimise/train.jl:105
┌ Info: Epoch 1140
└ @ Main /Users/kcft114/.julia/packages/Flux/qXNjB/src/optimise/train.jl:105
┌ Info: Epoch 1141
└ @ Main /Users/kcft114/.julia/packages/Flux/qXNjB/src/optimise/train.jl:105
┌ Info: Epoch 1142
└ @ Main /Users/kcft114/.julia/packages/Flux/qXNjB/src/optimise/train.jl:105
┌ Info: Epoch 1143
└ @ Main /Users/kcft114/.julia/packages/Flux/qXNjB/src/optimise/train.jl:105
┌ Info: Epoch 1144
└ @ Main /Users/kcft114/.julia/packages/Flux/qXNjB/src/optimise/train.jl:105
┌ Info: Epoch 1145
└ @ Main /Users/kcft114/.julia/packages/Flux/qXNjB/src/optimise/train.jl:105
┌ Info: Epoch 1146
└ @ Main /Users/kcft114/.julia/packages/Flux/qXNjB/src/optimise/train.jl:105
┌ Info: Epoch 1147
└ @ Main /Users/kcft114/.julia/packages/Flux/qXNjB/src/optimise/train.jl:105
┌ Info: Epoch 1148
└ @ Main /Users/kcft114/.julia/packages/Flux/qXNjB/src/optimise/train.jl:105
┌ Info: Epoch 1149
└ @ Main /Users/kcft114/.julia/packages/Flux/qXNjB/src/optimise/train.jl:105
┌ Info: Epoch 1150
└ @ Main /Users/kcft114/.julia/packages/Flux/qXNjB/src/optimise/train.jl:105
┌ Info: Epoch 1151
└ @ Main /Users/kcft114/.julia/packages/Flux/qXNjB/src/optimise/train.jl:105
┌ Info: Epoch 1152
└ @ Main /Users/kcft114/.julia/packages/Flux/qXNjB/src/optimise/train.jl:105
┌ Info: Epoch 1153
└ @ Main /Users/kcft114/.julia/packages/Flux/qXNjB/src/optimise/train.jl:105
┌ Info: Epoch 1154
└ @ Main /Users/kcft114/.julia/packages/Flux/qXNjB/src/optimise/train.jl:105
┌ Info: Epoch 1155
└ @ Main /Users/kcft114/.julia/packages/Flux/qXNjB/src/optimise/train.jl:105
┌ Info: Epoch 1156
└ @ Main /Users/kcft114/.julia/packages/Flux/qXNjB/src/optimise/train.jl:105
┌ Info: Epoch 1157
└ @ Main /Users/kcft114/.julia/packages/Flux/qXNjB/src/optimise/train.jl:105
┌ Info: Epoch 1158
└ @ Main /Users/kcft114/.julia/packages/Flux/qXNjB/src/optimise/train.jl:105
┌ Info: Epoch 1159
└ @ Main /Users/kcft114/.julia/packages/Flux/qXNjB/src/optimise/train.jl:105
```

```
loss(X, Y) = 0.71663797f0 (tracked)
loss(X, Y) = 0.6488766f0 (tracked)
loss(X, Y) = 0.67830724f0 (tracked)
loss(X, Y) = 0.67146504f0 (tracked)
loss(X, Y) = 0.6790889f0 (tracked)
loss(X, Y) = 0.6636226f0 (tracked)
loss(X, Y) = 0.6149886f0 (tracked)
loss(X, Y) = 0.61344737f0 (tracked)
loss(X, Y) = 0.6545407f0 (tracked)
loss(X, Y) = 0.6386875f0 (tracked)
loss(X, Y) = 0.66448224f0 (tracked)
loss(X, Y) = 0.6476728f0 (tracked)
loss(X, Y) = 0.6644372f0 (tracked)
loss(X, Y) = 0.6147415f0 (tracked)
loss(X, Y) = 0.6641573f0 (tracked)
loss(X, Y) = 0.6605932f0 (tracked)
loss(X, Y) = 0.6816602f0 (tracked)
loss(X, Y) = 0.65697706f0 (tracked)
loss(X, Y) = 0.640668f0 (tracked)
loss(X, Y) = 0.6557974f0 (tracked)
loss(X, Y) = 0.6310639f0 (tracked)
loss(X, Y) = 0.6727289f0 (tracked)
loss(X, Y) = 0.6200759f0 (tracked)
loss(X, Y) = 0.64335936f0 (tracked)
loss(X, Y) = 0.6328566f0 (tracked)
loss(X, Y) = 0.64272594f0 (tracked)
loss(X, Y) = 0.6451187f0 (tracked)
loss(X, Y) = 0.60523075f0 (tracked)
loss(X, Y) = 0.6541132f0 (tracked)
loss(X, Y) = 0.6087681f0 (tracked)
loss(X, Y) = 0.64279675f0 (tracked)
loss(X, Y) = 0.61273044f0 (tracked)
loss(X, Y) = 0.61727273f0 (tracked)
loss(X, Y) = 0.6467625f0 (tracked)
loss(X, Y) = 0.6160432f0 (tracked)
loss(X, Y) = 0.6687918f0 (tracked)
loss(X, Y) = 0.62947416f0 (tracked)
loss(X, Y) = 0.6232027f0 (tracked)
loss(X, Y) = 0.64016914f0 (tracked)
loss(X, Y) = 0.5975949f0 (tracked)
loss(X, Y) = 0.6975129f0 (tracked)
loss(X, Y) = 0.6236321f0 (tracked)
loss(X, Y) = 0.5871613f0 (tracked)
loss(X, Y) = 0.6843003f0 (tracked)
loss(X, Y) = 0.64775455f0 (tracked)
loss(X, Y) = 0.662617f0 (tracked)
loss(X, Y) = 0.64235324f0 (tracked)
loss(X, Y) = 0.6011195f0 (tracked)
loss(X, Y) = 0.6800201f0 (tracked)
loss(X, Y) = 0.6352203f0 (tracked)
loss(X, Y) = 0.6284643f0 (tracked)
loss(X, Y) = 0.65483665f0 (tracked)
loss(X, Y) = 0.67133206f0 (tracked)
loss(X, Y) = 0.6241156f0 (tracked)
loss(X, Y) = 0.62277365f0 (tracked)
loss(X, Y) = 0.57871574f0 (tracked)
loss(X, Y) = 0.64859676f0 (tracked)
loss(X, Y) = 0.6626723f0 (tracked)
loss(X, Y) = 0.66448146f0 (tracked)
loss(X, Y) = 0.648077f0 (tracked)
loss(X, Y) = 0.6642463f0 (tracked)
loss(X, Y) = 0.66324264f0 (tracked)
loss(X, Y) = 0.6029891f0 (tracked)
loss(X, Y) = 0.6160023f0 (tracked)
loss(X, Y) = 0.65621436f0 (tracked)
loss(X, Y) = 0.6435348f0 (tracked)
loss(X, Y) = 0.6149486f0 (tracked)
loss(X, Y) = 0.6332762f0 (tracked)
loss(X, Y) = 0.6332153f0 (tracked)
loss(X, Y) = 0.6694901f0 (tracked)
loss(X, Y) = 0.5835621f0 (tracked)
loss(X, Y) = 0.6539594f0 (tracked)
loss(X, Y) = 0.6302168f0 (tracked)
loss(X, Y) = 0.6420661f0 (tracked)
loss(X, Y) = 0.64767736f0 (tracked)
loss(X, Y) = 0.6416611f0 (tracked)
loss(X, Y) = 0.634001f0 (tracked)
loss(X, Y) = 0.64844894f0 (tracked)
loss(X, Y) = 0.66095805f0 (tracked)
loss(X, Y) = 0.6013113f0 (tracked)
loss(X, Y) = 0.62785155f0 (tracked)
loss(X, Y) = 0.63771063f0 (tracked)
loss(X, Y) = 0.6367136f0 (tracked)
loss(X, Y) = 0.64153194f0 (tracked)
loss(X, Y) = 0.61954975f0 (tracked)
loss(X, Y) = 0.6461648f0 (tracked)
loss(X, Y) = 0.66970366f0 (tracked)
loss(X, Y) = 0.6155341f0 (tracked)
loss(X, Y) = 0.6400267f0 (tracked)
loss(X, Y) = 0.63433486f0 (tracked)
loss(X, Y) = 0.6257223f0 (tracked)
loss(X, Y) = 0.6277698f0 (tracked)
loss(X, Y) = 0.66471064f0 (tracked)
loss(X, Y) = 0.66231203f0 (tracked)
loss(X, Y) = 0.68069476f0 (tracked)
loss(X, Y) = 0.65125465f0 (tracked)
loss(X, Y) = 0.61924905f0 (tracked)
loss(X, Y) = 0.6275967f0 (tracked)
loss(X, Y) = 0.6317573f0 (tracked)
loss(X, Y) = 0.60801625f0 (tracked)
loss(X, Y) = 0.6548127f0 (tracked)
loss(X, Y) = 0.6133977f0 (tracked)
loss(X, Y) = 0.6277277f0 (tracked)
loss(X, Y) = 0.6228307f0 (tracked)
loss(X, Y) = 0.6422386f0 (tracked)
loss(X, Y) = 0.5740463f0 (tracked)
loss(X, Y) = 0.63048923f0 (tracked)
loss(X, Y) = 0.6515221f0 (tracked)
loss(X, Y) = 0.6589012f0 (tracked)
loss(X, Y) = 0.61626494f0 (tracked)
loss(X, Y) = 0.63010687f0 (tracked)
loss(X, Y) = 0.60252076f0 (tracked)
loss(X, Y) = 0.61866915f0 (tracked)
loss(X, Y) = 0.6160215f0 (tracked)
loss(X, Y) = 0.64333916f0 (tracked)
loss(X, Y) = 0.6194178f0 (tracked)
loss(X, Y) = 0.6613978f0 (tracked)
loss(X, Y) = 0.62129694f0 (tracked)
loss(X, Y) = 0.6305857f0 (tracked)
loss(X, Y) = 0.6198785f0 (tracked)
loss(X, Y) = 0.64134336f0 (tracked)
loss(X, Y) = 0.6197618f0 (tracked)
loss(X, Y) = 0.65698797f0 (tracked)
loss(X, Y) = 0.61271954f0 (tracked)
```

```
┌ Info: Epoch 1160
└ @ Main /Users/kcft114/.julia/packages/Flux/qXNjB/src/optimise/train.jl:105
┌ Info: Epoch 1161
└ @ Main /Users/kcft114/.julia/packages/Flux/qXNjB/src/optimise/train.jl:105
┌ Info: Epoch 1162
└ @ Main /Users/kcft114/.julia/packages/Flux/qXNjB/src/optimise/train.jl:105
┌ Info: Epoch 1163
└ @ Main /Users/kcft114/.julia/packages/Flux/qXNjB/src/optimise/train.jl:105
┌ Info: Epoch 1164
└ @ Main /Users/kcft114/.julia/packages/Flux/qXNjB/src/optimise/train.jl:105
┌ Info: Epoch 1165
└ @ Main /Users/kcft114/.julia/packages/Flux/qXNjB/src/optimise/train.jl:105
┌ Info: Epoch 1166
└ @ Main /Users/kcft114/.julia/packages/Flux/qXNjB/src/optimise/train.jl:105
┌ Info: Epoch 1167
└ @ Main /Users/kcft114/.julia/packages/Flux/qXNjB/src/optimise/train.jl:105
┌ Info: Epoch 1168
└ @ Main /Users/kcft114/.julia/packages/Flux/qXNjB/src/optimise/train.jl:105
┌ Info: Epoch 1169
└ @ Main /Users/kcft114/.julia/packages/Flux/qXNjB/src/optimise/train.jl:105
┌ Info: Epoch 1170
└ @ Main /Users/kcft114/.julia/packages/Flux/qXNjB/src/optimise/train.jl:105
┌ Info: Epoch 1171
└ @ Main /Users/kcft114/.julia/packages/Flux/qXNjB/src/optimise/train.jl:105
┌ Info: Epoch 1172
└ @ Main /Users/kcft114/.julia/packages/Flux/qXNjB/src/optimise/train.jl:105
┌ Info: Epoch 1173
└ @ Main /Users/kcft114/.julia/packages/Flux/qXNjB/src/optimise/train.jl:105
┌ Info: Epoch 1174
└ @ Main /Users/kcft114/.julia/packages/Flux/qXNjB/src/optimise/train.jl:105
┌ Info: Epoch 1175
└ @ Main /Users/kcft114/.julia/packages/Flux/qXNjB/src/optimise/train.jl:105
┌ Info: Epoch 1176
└ @ Main /Users/kcft114/.julia/packages/Flux/qXNjB/src/optimise/train.jl:105
┌ Info: Epoch 1177
└ @ Main /Users/kcft114/.julia/packages/Flux/qXNjB/src/optimise/train.jl:105
┌ Info: Epoch 1178
└ @ Main /Users/kcft114/.julia/packages/Flux/qXNjB/src/optimise/train.jl:105
┌ Info: Epoch 1179
└ @ Main /Users/kcft114/.julia/packages/Flux/qXNjB/src/optimise/train.jl:105
┌ Info: Epoch 1180
└ @ Main /Users/kcft114/.julia/packages/Flux/qXNjB/src/optimise/train.jl:105
┌ Info: Epoch 1181
└ @ Main /Users/kcft114/.julia/packages/Flux/qXNjB/src/optimise/train.jl:105
┌ Info: Epoch 1182
└ @ Main /Users/kcft114/.julia/packages/Flux/qXNjB/src/optimise/train.jl:105
┌ Info: Epoch 1183
└ @ Main /Users/kcft114/.julia/packages/Flux/qXNjB/src/optimise/train.jl:105
┌ Info: Epoch 1184
└ @ Main /Users/kcft114/.julia/packages/Flux/qXNjB/src/optimise/train.jl:105
┌ Info: Epoch 1185
└ @ Main /Users/kcft114/.julia/packages/Flux/qXNjB/src/optimise/train.jl:105
┌ Info: Epoch 1186
└ @ Main /Users/kcft114/.julia/packages/Flux/qXNjB/src/optimise/train.jl:105
┌ Info: Epoch 1187
└ @ Main /Users/kcft114/.julia/packages/Flux/qXNjB/src/optimise/train.jl:105
┌ Info: Epoch 1188
└ @ Main /Users/kcft114/.julia/packages/Flux/qXNjB/src/optimise/train.jl:105
┌ Info: Epoch 1189
└ @ Main /Users/kcft114/.julia/packages/Flux/qXNjB/src/optimise/train.jl:105
┌ Info: Epoch 1190
└ @ Main /Users/kcft114/.julia/packages/Flux/qXNjB/src/optimise/train.jl:105
┌ Info: Epoch 1191
└ @ Main /Users/kcft114/.julia/packages/Flux/qXNjB/src/optimise/train.jl:105
┌ Info: Epoch 1192
└ @ Main /Users/kcft114/.julia/packages/Flux/qXNjB/src/optimise/train.jl:105
┌ Info: Epoch 1193
└ @ Main /Users/kcft114/.julia/packages/Flux/qXNjB/src/optimise/train.jl:105
┌ Info: Epoch 1194
└ @ Main /Users/kcft114/.julia/packages/Flux/qXNjB/src/optimise/train.jl:105
┌ Info: Epoch 1195
└ @ Main /Users/kcft114/.julia/packages/Flux/qXNjB/src/optimise/train.jl:105
┌ Info: Epoch 1196
└ @ Main /Users/kcft114/.julia/packages/Flux/qXNjB/src/optimise/train.jl:105
┌ Info: Epoch 1197
└ @ Main /Users/kcft114/.julia/packages/Flux/qXNjB/src/optimise/train.jl:105
┌ Info: Epoch 1198
└ @ Main /Users/kcft114/.julia/packages/Flux/qXNjB/src/optimise/train.jl:105
┌ Info: Epoch 1199
└ @ Main /Users/kcft114/.julia/packages/Flux/qXNjB/src/optimise/train.jl:105
┌ Info: Epoch 1200
└ @ Main /Users/kcft114/.julia/packages/Flux/qXNjB/src/optimise/train.jl:105
┌ Info: Epoch 1201
└ @ Main /Users/kcft114/.julia/packages/Flux/qXNjB/src/optimise/train.jl:105
┌ Info: Epoch 1202
└ @ Main /Users/kcft114/.julia/packages/Flux/qXNjB/src/optimise/train.jl:105
┌ Info: Epoch 1203
└ @ Main /Users/kcft114/.julia/packages/Flux/qXNjB/src/optimise/train.jl:105
┌ Info: Epoch 1204
└ @ Main /Users/kcft114/.julia/packages/Flux/qXNjB/src/optimise/train.jl:105
┌ Info: Epoch 1205
└ @ Main /Users/kcft114/.julia/packages/Flux/qXNjB/src/optimise/train.jl:105
┌ Info: Epoch 1206
└ @ Main /Users/kcft114/.julia/packages/Flux/qXNjB/src/optimise/train.jl:105
┌ Info: Epoch 1207
└ @ Main /Users/kcft114/.julia/packages/Flux/qXNjB/src/optimise/train.jl:105
┌ Info: Epoch 1208
└ @ Main /Users/kcft114/.julia/packages/Flux/qXNjB/src/optimise/train.jl:105
┌ Info: Epoch 1209
└ @ Main /Users/kcft114/.julia/packages/Flux/qXNjB/src/optimise/train.jl:105
┌ Info: Epoch 1210
└ @ Main /Users/kcft114/.julia/packages/Flux/qXNjB/src/optimise/train.jl:105
┌ Info: Epoch 1211
└ @ Main /Users/kcft114/.julia/packages/Flux/qXNjB/src/optimise/train.jl:105
┌ Info: Epoch 1212
└ @ Main /Users/kcft114/.julia/packages/Flux/qXNjB/src/optimise/train.jl:105
┌ Info: Epoch 1213
└ @ Main /Users/kcft114/.julia/packages/Flux/qXNjB/src/optimise/train.jl:105
┌ Info: Epoch 1214
└ @ Main /Users/kcft114/.julia/packages/Flux/qXNjB/src/optimise/train.jl:105
┌ Info: Epoch 1215
└ @ Main /Users/kcft114/.julia/packages/Flux/qXNjB/src/optimise/train.jl:105
┌ Info: Epoch 1216
└ @ Main /Users/kcft114/.julia/packages/Flux/qXNjB/src/optimise/train.jl:105
┌ Info: Epoch 1217
└ @ Main /Users/kcft114/.julia/packages/Flux/qXNjB/src/optimise/train.jl:105
┌ Info: Epoch 1218
└ @ Main /Users/kcft114/.julia/packages/Flux/qXNjB/src/optimise/train.jl:105
┌ Info: Epoch 1219
└ @ Main /Users/kcft114/.julia/packages/Flux/qXNjB/src/optimise/train.jl:105
┌ Info: Epoch 1220
└ @ Main /Users/kcft114/.julia/packages/Flux/qXNjB/src/optimise/train.jl:105
┌ Info: Epoch 1221
└ @ Main /Users/kcft114/.julia/packages/Flux/qXNjB/src/optimise/train.jl:105
┌ Info: Epoch 1222
└ @ Main /Users/kcft114/.julia/packages/Flux/qXNjB/src/optimise/train.jl:105
┌ Info: Epoch 1223
└ @ Main /Users/kcft114/.julia/packages/Flux/qXNjB/src/optimise/train.jl:105
┌ Info: Epoch 1224
└ @ Main /Users/kcft114/.julia/packages/Flux/qXNjB/src/optimise/train.jl:105
┌ Info: Epoch 1225
└ @ Main /Users/kcft114/.julia/packages/Flux/qXNjB/src/optimise/train.jl:105
┌ Info: Epoch 1226
└ @ Main /Users/kcft114/.julia/packages/Flux/qXNjB/src/optimise/train.jl:105
┌ Info: Epoch 1227
└ @ Main /Users/kcft114/.julia/packages/Flux/qXNjB/src/optimise/train.jl:105
┌ Info: Epoch 1228
└ @ Main /Users/kcft114/.julia/packages/Flux/qXNjB/src/optimise/train.jl:105
┌ Info: Epoch 1229
└ @ Main /Users/kcft114/.julia/packages/Flux/qXNjB/src/optimise/train.jl:105
┌ Info: Epoch 1230
└ @ Main /Users/kcft114/.julia/packages/Flux/qXNjB/src/optimise/train.jl:105
┌ Info: Epoch 1231
└ @ Main /Users/kcft114/.julia/packages/Flux/qXNjB/src/optimise/train.jl:105
┌ Info: Epoch 1232
└ @ Main /Users/kcft114/.julia/packages/Flux/qXNjB/src/optimise/train.jl:105
┌ Info: Epoch 1233
└ @ Main /Users/kcft114/.julia/packages/Flux/qXNjB/src/optimise/train.jl:105
┌ Info: Epoch 1234
└ @ Main /Users/kcft114/.julia/packages/Flux/qXNjB/src/optimise/train.jl:105
┌ Info: Epoch 1235
└ @ Main /Users/kcft114/.julia/packages/Flux/qXNjB/src/optimise/train.jl:105
┌ Info: Epoch 1236
└ @ Main /Users/kcft114/.julia/packages/Flux/qXNjB/src/optimise/train.jl:105
┌ Info: Epoch 1237
└ @ Main /Users/kcft114/.julia/packages/Flux/qXNjB/src/optimise/train.jl:105
┌ Info: Epoch 1238
└ @ Main /Users/kcft114/.julia/packages/Flux/qXNjB/src/optimise/train.jl:105
┌ Info: Epoch 1239
└ @ Main /Users/kcft114/.julia/packages/Flux/qXNjB/src/optimise/train.jl:105
┌ Info: Epoch 1240
└ @ Main /Users/kcft114/.julia/packages/Flux/qXNjB/src/optimise/train.jl:105
┌ Info: Epoch 1241
└ @ Main /Users/kcft114/.julia/packages/Flux/qXNjB/src/optimise/train.jl:105
┌ Info: Epoch 1242
└ @ Main /Users/kcft114/.julia/packages/Flux/qXNjB/src/optimise/train.jl:105
┌ Info: Epoch 1243
└ @ Main /Users/kcft114/.julia/packages/Flux/qXNjB/src/optimise/train.jl:105
┌ Info: Epoch 1244
└ @ Main /Users/kcft114/.julia/packages/Flux/qXNjB/src/optimise/train.jl:105
┌ Info: Epoch 1245
└ @ Main /Users/kcft114/.julia/packages/Flux/qXNjB/src/optimise/train.jl:105
┌ Info: Epoch 1246
└ @ Main /Users/kcft114/.julia/packages/Flux/qXNjB/src/optimise/train.jl:105
┌ Info: Epoch 1247
└ @ Main /Users/kcft114/.julia/packages/Flux/qXNjB/src/optimise/train.jl:105
┌ Info: Epoch 1248
└ @ Main /Users/kcft114/.julia/packages/Flux/qXNjB/src/optimise/train.jl:105
┌ Info: Epoch 1249
└ @ Main /Users/kcft114/.julia/packages/Flux/qXNjB/src/optimise/train.jl:105
┌ Info: Epoch 1250
└ @ Main /Users/kcft114/.julia/packages/Flux/qXNjB/src/optimise/train.jl:105
┌ Info: Epoch 1251
└ @ Main /Users/kcft114/.julia/packages/Flux/qXNjB/src/optimise/train.jl:105
┌ Info: Epoch 1252
└ @ Main /Users/kcft114/.julia/packages/Flux/qXNjB/src/optimise/train.jl:105
┌ Info: Epoch 1253
└ @ Main /Users/kcft114/.julia/packages/Flux/qXNjB/src/optimise/train.jl:105
┌ Info: Epoch 1254
└ @ Main /Users/kcft114/.julia/packages/Flux/qXNjB/src/optimise/train.jl:105
┌ Info: Epoch 1255
└ @ Main /Users/kcft114/.julia/packages/Flux/qXNjB/src/optimise/train.jl:105
┌ Info: Epoch 1256
└ @ Main /Users/kcft114/.julia/packages/Flux/qXNjB/src/optimise/train.jl:105
┌ Info: Epoch 1257
└ @ Main /Users/kcft114/.julia/packages/Flux/qXNjB/src/optimise/train.jl:105
┌ Info: Epoch 1258
└ @ Main /Users/kcft114/.julia/packages/Flux/qXNjB/src/optimise/train.jl:105
┌ Info: Epoch 1259
└ @ Main /Users/kcft114/.julia/packages/Flux/qXNjB/src/optimise/train.jl:105
┌ Info: Epoch 1260
└ @ Main /Users/kcft114/.julia/packages/Flux/qXNjB/src/optimise/train.jl:105
┌ Info: Epoch 1261
└ @ Main /Users/kcft114/.julia/packages/Flux/qXNjB/src/optimise/train.jl:105
┌ Info: Epoch 1262
└ @ Main /Users/kcft114/.julia/packages/Flux/qXNjB/src/optimise/train.jl:105
┌ Info: Epoch 1263
└ @ Main /Users/kcft114/.julia/packages/Flux/qXNjB/src/optimise/train.jl:105
┌ Info: Epoch 1264
└ @ Main /Users/kcft114/.julia/packages/Flux/qXNjB/src/optimise/train.jl:105
┌ Info: Epoch 1265
└ @ Main /Users/kcft114/.julia/packages/Flux/qXNjB/src/optimise/train.jl:105
┌ Info: Epoch 1266
└ @ Main /Users/kcft114/.julia/packages/Flux/qXNjB/src/optimise/train.jl:105
┌ Info: Epoch 1267
└ @ Main /Users/kcft114/.julia/packages/Flux/qXNjB/src/optimise/train.jl:105
┌ Info: Epoch 1268
└ @ Main /Users/kcft114/.julia/packages/Flux/qXNjB/src/optimise/train.jl:105
┌ Info: Epoch 1269
└ @ Main /Users/kcft114/.julia/packages/Flux/qXNjB/src/optimise/train.jl:105
┌ Info: Epoch 1270
└ @ Main /Users/kcft114/.julia/packages/Flux/qXNjB/src/optimise/train.jl:105
┌ Info: Epoch 1271
└ @ Main /Users/kcft114/.julia/packages/Flux/qXNjB/src/optimise/train.jl:105
┌ Info: Epoch 1272
└ @ Main /Users/kcft114/.julia/packages/Flux/qXNjB/src/optimise/train.jl:105
┌ Info: Epoch 1273
└ @ Main /Users/kcft114/.julia/packages/Flux/qXNjB/src/optimise/train.jl:105
┌ Info: Epoch 1274
└ @ Main /Users/kcft114/.julia/packages/Flux/qXNjB/src/optimise/train.jl:105
┌ Info: Epoch 1275
└ @ Main /Users/kcft114/.julia/packages/Flux/qXNjB/src/optimise/train.jl:105
┌ Info: Epoch 1276
└ @ Main /Users/kcft114/.julia/packages/Flux/qXNjB/src/optimise/train.jl:105
┌ Info: Epoch 1277
└ @ Main /Users/kcft114/.julia/packages/Flux/qXNjB/src/optimise/train.jl:105
┌ Info: Epoch 1278
└ @ Main /Users/kcft114/.julia/packages/Flux/qXNjB/src/optimise/train.jl:105
┌ Info: Epoch 1279
└ @ Main /Users/kcft114/.julia/packages/Flux/qXNjB/src/optimise/train.jl:105
┌ Info: Epoch 1280
└ @ Main /Users/kcft114/.julia/packages/Flux/qXNjB/src/optimise/train.jl:105
┌ Info: Epoch 1281
└ @ Main /Users/kcft114/.julia/packages/Flux/qXNjB/src/optimise/train.jl:105
┌ Info: Epoch 1282
└ @ Main /Users/kcft114/.julia/packages/Flux/qXNjB/src/optimise/train.jl:105
┌ Info: Epoch 1283
└ @ Main /Users/kcft114/.julia/packages/Flux/qXNjB/src/optimise/train.jl:105
```

```
loss(X, Y) = 0.6218399f0 (tracked)
loss(X, Y) = 0.65260935f0 (tracked)
loss(X, Y) = 0.625877f0 (tracked)
loss(X, Y) = 0.6660999f0 (tracked)
loss(X, Y) = 0.6278848f0 (tracked)
loss(X, Y) = 0.60648876f0 (tracked)
loss(X, Y) = 0.6419599f0 (tracked)
loss(X, Y) = 0.64708596f0 (tracked)
loss(X, Y) = 0.6106237f0 (tracked)
loss(X, Y) = 0.6659503f0 (tracked)
loss(X, Y) = 0.62869877f0 (tracked)
loss(X, Y) = 0.6790052f0 (tracked)
loss(X, Y) = 0.6900519f0 (tracked)
loss(X, Y) = 0.5961804f0 (tracked)
loss(X, Y) = 0.67167825f0 (tracked)
loss(X, Y) = 0.60255885f0 (tracked)
loss(X, Y) = 0.6369214f0 (tracked)
loss(X, Y) = 0.6185106f0 (tracked)
loss(X, Y) = 0.63261896f0 (tracked)
loss(X, Y) = 0.6653484f0 (tracked)
loss(X, Y) = 0.64047533f0 (tracked)
loss(X, Y) = 0.6571294f0 (tracked)
loss(X, Y) = 0.6203486f0 (tracked)
loss(X, Y) = 0.66616255f0 (tracked)
loss(X, Y) = 0.5896824f0 (tracked)
loss(X, Y) = 0.5984948f0 (tracked)
loss(X, Y) = 0.595751f0 (tracked)
loss(X, Y) = 0.65758944f0 (tracked)
loss(X, Y) = 0.62064195f0 (tracked)
loss(X, Y) = 0.6238625f0 (tracked)
loss(X, Y) = 0.6496383f0 (tracked)
loss(X, Y) = 0.6712444f0 (tracked)
loss(X, Y) = 0.62800926f0 (tracked)
loss(X, Y) = 0.65714103f0 (tracked)
loss(X, Y) = 0.60279095f0 (tracked)
loss(X, Y) = 0.60133785f0 (tracked)
loss(X, Y) = 0.60806113f0 (tracked)
loss(X, Y) = 0.6593898f0 (tracked)
loss(X, Y) = 0.5953912f0 (tracked)
loss(X, Y) = 0.62955403f0 (tracked)
loss(X, Y) = 0.613435f0 (tracked)
loss(X, Y) = 0.6299619f0 (tracked)
loss(X, Y) = 0.6310796f0 (tracked)
loss(X, Y) = 0.60146224f0 (tracked)
loss(X, Y) = 0.659257f0 (tracked)
loss(X, Y) = 0.60895693f0 (tracked)
loss(X, Y) = 0.6317545f0 (tracked)
loss(X, Y) = 0.6267933f0 (tracked)
loss(X, Y) = 0.6693763f0 (tracked)
loss(X, Y) = 0.6480699f0 (tracked)
loss(X, Y) = 0.661617f0 (tracked)
loss(X, Y) = 0.63976496f0 (tracked)
loss(X, Y) = 0.6158444f0 (tracked)
loss(X, Y) = 0.59838134f0 (tracked)
loss(X, Y) = 0.6243677f0 (tracked)
loss(X, Y) = 0.6338309f0 (tracked)
loss(X, Y) = 0.62505424f0 (tracked)
loss(X, Y) = 0.62322086f0 (tracked)
loss(X, Y) = 0.6427446f0 (tracked)
loss(X, Y) = 0.645981f0 (tracked)
loss(X, Y) = 0.6189468f0 (tracked)
loss(X, Y) = 0.63415587f0 (tracked)
loss(X, Y) = 0.5947249f0 (tracked)
loss(X, Y) = 0.62113655f0 (tracked)
loss(X, Y) = 0.60868603f0 (tracked)
loss(X, Y) = 0.65801054f0 (tracked)
loss(X, Y) = 0.6349294f0 (tracked)
loss(X, Y) = 0.640145f0 (tracked)
loss(X, Y) = 0.6063215f0 (tracked)
loss(X, Y) = 0.5935204f0 (tracked)
loss(X, Y) = 0.6327011f0 (tracked)
loss(X, Y) = 0.6106438f0 (tracked)
loss(X, Y) = 0.6298187f0 (tracked)
loss(X, Y) = 0.6069974f0 (tracked)
loss(X, Y) = 0.6616681f0 (tracked)
loss(X, Y) = 0.65383536f0 (tracked)
loss(X, Y) = 0.6200465f0 (tracked)
loss(X, Y) = 0.61743444f0 (tracked)
loss(X, Y) = 0.6314425f0 (tracked)
loss(X, Y) = 0.6521996f0 (tracked)
loss(X, Y) = 0.6500195f0 (tracked)
loss(X, Y) = 0.6463038f0 (tracked)
loss(X, Y) = 0.5778621f0 (tracked)
loss(X, Y) = 0.6314115f0 (tracked)
loss(X, Y) = 0.58887196f0 (tracked)
loss(X, Y) = 0.6089532f0 (tracked)
loss(X, Y) = 0.6188347f0 (tracked)
loss(X, Y) = 0.59570545f0 (tracked)
loss(X, Y) = 0.6015903f0 (tracked)
loss(X, Y) = 0.6282968f0 (tracked)
loss(X, Y) = 0.5950037f0 (tracked)
loss(X, Y) = 0.6235613f0 (tracked)
loss(X, Y) = 0.604235f0 (tracked)
loss(X, Y) = 0.61147505f0 (tracked)
loss(X, Y) = 0.5685922f0 (tracked)
loss(X, Y) = 0.6252985f0 (tracked)
loss(X, Y) = 0.6363887f0 (tracked)
loss(X, Y) = 0.63080984f0 (tracked)
loss(X, Y) = 0.6240838f0 (tracked)
loss(X, Y) = 0.6503475f0 (tracked)
loss(X, Y) = 0.6075508f0 (tracked)
loss(X, Y) = 0.6868223f0 (tracked)
loss(X, Y) = 0.59735715f0 (tracked)
loss(X, Y) = 0.5991147f0 (tracked)
loss(X, Y) = 0.61571026f0 (tracked)
loss(X, Y) = 0.61915916f0 (tracked)
loss(X, Y) = 0.6353086f0 (tracked)
loss(X, Y) = 0.626602f0 (tracked)
loss(X, Y) = 0.67258704f0 (tracked)
loss(X, Y) = 0.61666f0 (tracked)
loss(X, Y) = 0.6168443f0 (tracked)
loss(X, Y) = 0.6118459f0 (tracked)
loss(X, Y) = 0.6191014f0 (tracked)
loss(X, Y) = 0.59591967f0 (tracked)
loss(X, Y) = 0.6403572f0 (tracked)
loss(X, Y) = 0.64526963f0 (tracked)
loss(X, Y) = 0.6245734f0 (tracked)
loss(X, Y) = 0.59318787f0 (tracked)
loss(X, Y) = 0.62040913f0 (tracked)
loss(X, Y) = 0.62916416f0 (tracked)
loss(X, Y) = 0.5959814f0 (tracked)
loss(X, Y) = 0.643995f0 (tracked)
loss(X, Y) = 0.6155773f0 (tracked)
loss(X, Y) = 0.5978483f0 (tracked)
loss(X, Y) = 0.5819404f0 (tracked)
loss(X, Y) = 0.59228146f0 (tracked)
loss(X, Y) = 0.6251637f0 (tracked)
loss(X, Y) = 0.6038252f0 (tracked)
loss(X, Y) = 0.6372946f0 (tracked)
loss(X, Y) = 0.631019f0 (tracked)
loss(X, Y) = 0.56991225f0 (tracked)
loss(X, Y) = 0.6554759f0 (tracked)
```

```
┌ Info: Epoch 1284
└ @ Main /Users/kcft114/.julia/packages/Flux/qXNjB/src/optimise/train.jl:105
┌ Info: Epoch 1285
└ @ Main /Users/kcft114/.julia/packages/Flux/qXNjB/src/optimise/train.jl:105
┌ Info: Epoch 1286
└ @ Main /Users/kcft114/.julia/packages/Flux/qXNjB/src/optimise/train.jl:105
┌ Info: Epoch 1287
└ @ Main /Users/kcft114/.julia/packages/Flux/qXNjB/src/optimise/train.jl:105
┌ Info: Epoch 1288
└ @ Main /Users/kcft114/.julia/packages/Flux/qXNjB/src/optimise/train.jl:105
┌ Info: Epoch 1289
└ @ Main /Users/kcft114/.julia/packages/Flux/qXNjB/src/optimise/train.jl:105
┌ Info: Epoch 1290
└ @ Main /Users/kcft114/.julia/packages/Flux/qXNjB/src/optimise/train.jl:105
┌ Info: Epoch 1291
└ @ Main /Users/kcft114/.julia/packages/Flux/qXNjB/src/optimise/train.jl:105
┌ Info: Epoch 1292
└ @ Main /Users/kcft114/.julia/packages/Flux/qXNjB/src/optimise/train.jl:105
┌ Info: Epoch 1293
└ @ Main /Users/kcft114/.julia/packages/Flux/qXNjB/src/optimise/train.jl:105
┌ Info: Epoch 1294
└ @ Main /Users/kcft114/.julia/packages/Flux/qXNjB/src/optimise/train.jl:105
┌ Info: Epoch 1295
└ @ Main /Users/kcft114/.julia/packages/Flux/qXNjB/src/optimise/train.jl:105
┌ Info: Epoch 1296
└ @ Main /Users/kcft114/.julia/packages/Flux/qXNjB/src/optimise/train.jl:105
┌ Info: Epoch 1297
└ @ Main /Users/kcft114/.julia/packages/Flux/qXNjB/src/optimise/train.jl:105
┌ Info: Epoch 1298
└ @ Main /Users/kcft114/.julia/packages/Flux/qXNjB/src/optimise/train.jl:105
┌ Info: Epoch 1299
└ @ Main /Users/kcft114/.julia/packages/Flux/qXNjB/src/optimise/train.jl:105
┌ Info: Epoch 1300
└ @ Main /Users/kcft114/.julia/packages/Flux/qXNjB/src/optimise/train.jl:105
┌ Info: Epoch 1301
└ @ Main /Users/kcft114/.julia/packages/Flux/qXNjB/src/optimise/train.jl:105
┌ Info: Epoch 1302
└ @ Main /Users/kcft114/.julia/packages/Flux/qXNjB/src/optimise/train.jl:105
┌ Info: Epoch 1303
└ @ Main /Users/kcft114/.julia/packages/Flux/qXNjB/src/optimise/train.jl:105
┌ Info: Epoch 1304
└ @ Main /Users/kcft114/.julia/packages/Flux/qXNjB/src/optimise/train.jl:105
┌ Info: Epoch 1305
└ @ Main /Users/kcft114/.julia/packages/Flux/qXNjB/src/optimise/train.jl:105
┌ Info: Epoch 1306
└ @ Main /Users/kcft114/.julia/packages/Flux/qXNjB/src/optimise/train.jl:105
┌ Info: Epoch 1307
└ @ Main /Users/kcft114/.julia/packages/Flux/qXNjB/src/optimise/train.jl:105
┌ Info: Epoch 1308
└ @ Main /Users/kcft114/.julia/packages/Flux/qXNjB/src/optimise/train.jl:105
┌ Info: Epoch 1309
└ @ Main /Users/kcft114/.julia/packages/Flux/qXNjB/src/optimise/train.jl:105
┌ Info: Epoch 1310
└ @ Main /Users/kcft114/.julia/packages/Flux/qXNjB/src/optimise/train.jl:105
┌ Info: Epoch 1311
└ @ Main /Users/kcft114/.julia/packages/Flux/qXNjB/src/optimise/train.jl:105
┌ Info: Epoch 1312
└ @ Main /Users/kcft114/.julia/packages/Flux/qXNjB/src/optimise/train.jl:105
┌ Info: Epoch 1313
└ @ Main /Users/kcft114/.julia/packages/Flux/qXNjB/src/optimise/train.jl:105
┌ Info: Epoch 1314
└ @ Main /Users/kcft114/.julia/packages/Flux/qXNjB/src/optimise/train.jl:105
┌ Info: Epoch 1315
└ @ Main /Users/kcft114/.julia/packages/Flux/qXNjB/src/optimise/train.jl:105
┌ Info: Epoch 1316
└ @ Main /Users/kcft114/.julia/packages/Flux/qXNjB/src/optimise/train.jl:105
┌ Info: Epoch 1317
└ @ Main /Users/kcft114/.julia/packages/Flux/qXNjB/src/optimise/train.jl:105
┌ Info: Epoch 1318
└ @ Main /Users/kcft114/.julia/packages/Flux/qXNjB/src/optimise/train.jl:105
┌ Info: Epoch 1319
└ @ Main /Users/kcft114/.julia/packages/Flux/qXNjB/src/optimise/train.jl:105
┌ Info: Epoch 1320
└ @ Main /Users/kcft114/.julia/packages/Flux/qXNjB/src/optimise/train.jl:105
┌ Info: Epoch 1321
└ @ Main /Users/kcft114/.julia/packages/Flux/qXNjB/src/optimise/train.jl:105
┌ Info: Epoch 1322
└ @ Main /Users/kcft114/.julia/packages/Flux/qXNjB/src/optimise/train.jl:105
┌ Info: Epoch 1323
└ @ Main /Users/kcft114/.julia/packages/Flux/qXNjB/src/optimise/train.jl:105
┌ Info: Epoch 1324
└ @ Main /Users/kcft114/.julia/packages/Flux/qXNjB/src/optimise/train.jl:105
┌ Info: Epoch 1325
└ @ Main /Users/kcft114/.julia/packages/Flux/qXNjB/src/optimise/train.jl:105
┌ Info: Epoch 1326
└ @ Main /Users/kcft114/.julia/packages/Flux/qXNjB/src/optimise/train.jl:105
┌ Info: Epoch 1327
└ @ Main /Users/kcft114/.julia/packages/Flux/qXNjB/src/optimise/train.jl:105
┌ Info: Epoch 1328
└ @ Main /Users/kcft114/.julia/packages/Flux/qXNjB/src/optimise/train.jl:105
┌ Info: Epoch 1329
└ @ Main /Users/kcft114/.julia/packages/Flux/qXNjB/src/optimise/train.jl:105
┌ Info: Epoch 1330
└ @ Main /Users/kcft114/.julia/packages/Flux/qXNjB/src/optimise/train.jl:105
┌ Info: Epoch 1331
└ @ Main /Users/kcft114/.julia/packages/Flux/qXNjB/src/optimise/train.jl:105
┌ Info: Epoch 1332
└ @ Main /Users/kcft114/.julia/packages/Flux/qXNjB/src/optimise/train.jl:105
┌ Info: Epoch 1333
└ @ Main /Users/kcft114/.julia/packages/Flux/qXNjB/src/optimise/train.jl:105
┌ Info: Epoch 1334
└ @ Main /Users/kcft114/.julia/packages/Flux/qXNjB/src/optimise/train.jl:105
┌ Info: Epoch 1335
└ @ Main /Users/kcft114/.julia/packages/Flux/qXNjB/src/optimise/train.jl:105
┌ Info: Epoch 1336
└ @ Main /Users/kcft114/.julia/packages/Flux/qXNjB/src/optimise/train.jl:105
┌ Info: Epoch 1337
└ @ Main /Users/kcft114/.julia/packages/Flux/qXNjB/src/optimise/train.jl:105
┌ Info: Epoch 1338
└ @ Main /Users/kcft114/.julia/packages/Flux/qXNjB/src/optimise/train.jl:105
┌ Info: Epoch 1339
└ @ Main /Users/kcft114/.julia/packages/Flux/qXNjB/src/optimise/train.jl:105
┌ Info: Epoch 1340
└ @ Main /Users/kcft114/.julia/packages/Flux/qXNjB/src/optimise/train.jl:105
┌ Info: Epoch 1341
└ @ Main /Users/kcft114/.julia/packages/Flux/qXNjB/src/optimise/train.jl:105
┌ Info: Epoch 1342
└ @ Main /Users/kcft114/.julia/packages/Flux/qXNjB/src/optimise/train.jl:105
┌ Info: Epoch 1343
└ @ Main /Users/kcft114/.julia/packages/Flux/qXNjB/src/optimise/train.jl:105
┌ Info: Epoch 1344
└ @ Main /Users/kcft114/.julia/packages/Flux/qXNjB/src/optimise/train.jl:105
┌ Info: Epoch 1345
└ @ Main /Users/kcft114/.julia/packages/Flux/qXNjB/src/optimise/train.jl:105
┌ Info: Epoch 1346
└ @ Main /Users/kcft114/.julia/packages/Flux/qXNjB/src/optimise/train.jl:105
┌ Info: Epoch 1347
└ @ Main /Users/kcft114/.julia/packages/Flux/qXNjB/src/optimise/train.jl:105
┌ Info: Epoch 1348
└ @ Main /Users/kcft114/.julia/packages/Flux/qXNjB/src/optimise/train.jl:105
┌ Info: Epoch 1349
└ @ Main /Users/kcft114/.julia/packages/Flux/qXNjB/src/optimise/train.jl:105
┌ Info: Epoch 1350
└ @ Main /Users/kcft114/.julia/packages/Flux/qXNjB/src/optimise/train.jl:105
┌ Info: Epoch 1351
└ @ Main /Users/kcft114/.julia/packages/Flux/qXNjB/src/optimise/train.jl:105
┌ Info: Epoch 1352
└ @ Main /Users/kcft114/.julia/packages/Flux/qXNjB/src/optimise/train.jl:105
┌ Info: Epoch 1353
└ @ Main /Users/kcft114/.julia/packages/Flux/qXNjB/src/optimise/train.jl:105
┌ Info: Epoch 1354
└ @ Main /Users/kcft114/.julia/packages/Flux/qXNjB/src/optimise/train.jl:105
┌ Info: Epoch 1355
└ @ Main /Users/kcft114/.julia/packages/Flux/qXNjB/src/optimise/train.jl:105
┌ Info: Epoch 1356
└ @ Main /Users/kcft114/.julia/packages/Flux/qXNjB/src/optimise/train.jl:105
┌ Info: Epoch 1357
└ @ Main /Users/kcft114/.julia/packages/Flux/qXNjB/src/optimise/train.jl:105
┌ Info: Epoch 1358
└ @ Main /Users/kcft114/.julia/packages/Flux/qXNjB/src/optimise/train.jl:105
┌ Info: Epoch 1359
└ @ Main /Users/kcft114/.julia/packages/Flux/qXNjB/src/optimise/train.jl:105
┌ Info: Epoch 1360
└ @ Main /Users/kcft114/.julia/packages/Flux/qXNjB/src/optimise/train.jl:105
┌ Info: Epoch 1361
└ @ Main /Users/kcft114/.julia/packages/Flux/qXNjB/src/optimise/train.jl:105
┌ Info: Epoch 1362
└ @ Main /Users/kcft114/.julia/packages/Flux/qXNjB/src/optimise/train.jl:105
┌ Info: Epoch 1363
└ @ Main /Users/kcft114/.julia/packages/Flux/qXNjB/src/optimise/train.jl:105
┌ Info: Epoch 1364
└ @ Main /Users/kcft114/.julia/packages/Flux/qXNjB/src/optimise/train.jl:105
┌ Info: Epoch 1365
└ @ Main /Users/kcft114/.julia/packages/Flux/qXNjB/src/optimise/train.jl:105
┌ Info: Epoch 1366
└ @ Main /Users/kcft114/.julia/packages/Flux/qXNjB/src/optimise/train.jl:105
┌ Info: Epoch 1367
└ @ Main /Users/kcft114/.julia/packages/Flux/qXNjB/src/optimise/train.jl:105
┌ Info: Epoch 1368
└ @ Main /Users/kcft114/.julia/packages/Flux/qXNjB/src/optimise/train.jl:105
┌ Info: Epoch 1369
└ @ Main /Users/kcft114/.julia/packages/Flux/qXNjB/src/optimise/train.jl:105
┌ Info: Epoch 1370
└ @ Main /Users/kcft114/.julia/packages/Flux/qXNjB/src/optimise/train.jl:105
┌ Info: Epoch 1371
└ @ Main /Users/kcft114/.julia/packages/Flux/qXNjB/src/optimise/train.jl:105
┌ Info: Epoch 1372
└ @ Main /Users/kcft114/.julia/packages/Flux/qXNjB/src/optimise/train.jl:105
┌ Info: Epoch 1373
└ @ Main /Users/kcft114/.julia/packages/Flux/qXNjB/src/optimise/train.jl:105
┌ Info: Epoch 1374
└ @ Main /Users/kcft114/.julia/packages/Flux/qXNjB/src/optimise/train.jl:105
┌ Info: Epoch 1375
└ @ Main /Users/kcft114/.julia/packages/Flux/qXNjB/src/optimise/train.jl:105
┌ Info: Epoch 1376
└ @ Main /Users/kcft114/.julia/packages/Flux/qXNjB/src/optimise/train.jl:105
┌ Info: Epoch 1377
└ @ Main /Users/kcft114/.julia/packages/Flux/qXNjB/src/optimise/train.jl:105
┌ Info: Epoch 1378
└ @ Main /Users/kcft114/.julia/packages/Flux/qXNjB/src/optimise/train.jl:105
┌ Info: Epoch 1379
└ @ Main /Users/kcft114/.julia/packages/Flux/qXNjB/src/optimise/train.jl:105
┌ Info: Epoch 1380
└ @ Main /Users/kcft114/.julia/packages/Flux/qXNjB/src/optimise/train.jl:105
┌ Info: Epoch 1381
└ @ Main /Users/kcft114/.julia/packages/Flux/qXNjB/src/optimise/train.jl:105
┌ Info: Epoch 1382
└ @ Main /Users/kcft114/.julia/packages/Flux/qXNjB/src/optimise/train.jl:105
┌ Info: Epoch 1383
└ @ Main /Users/kcft114/.julia/packages/Flux/qXNjB/src/optimise/train.jl:105
┌ Info: Epoch 1384
└ @ Main /Users/kcft114/.julia/packages/Flux/qXNjB/src/optimise/train.jl:105
┌ Info: Epoch 1385
└ @ Main /Users/kcft114/.julia/packages/Flux/qXNjB/src/optimise/train.jl:105
┌ Info: Epoch 1386
└ @ Main /Users/kcft114/.julia/packages/Flux/qXNjB/src/optimise/train.jl:105
┌ Info: Epoch 1387
└ @ Main /Users/kcft114/.julia/packages/Flux/qXNjB/src/optimise/train.jl:105
┌ Info: Epoch 1388
└ @ Main /Users/kcft114/.julia/packages/Flux/qXNjB/src/optimise/train.jl:105
┌ Info: Epoch 1389
└ @ Main /Users/kcft114/.julia/packages/Flux/qXNjB/src/optimise/train.jl:105
┌ Info: Epoch 1390
└ @ Main /Users/kcft114/.julia/packages/Flux/qXNjB/src/optimise/train.jl:105
┌ Info: Epoch 1391
└ @ Main /Users/kcft114/.julia/packages/Flux/qXNjB/src/optimise/train.jl:105
┌ Info: Epoch 1392
└ @ Main /Users/kcft114/.julia/packages/Flux/qXNjB/src/optimise/train.jl:105
┌ Info: Epoch 1393
└ @ Main /Users/kcft114/.julia/packages/Flux/qXNjB/src/optimise/train.jl:105
┌ Info: Epoch 1394
└ @ Main /Users/kcft114/.julia/packages/Flux/qXNjB/src/optimise/train.jl:105
┌ Info: Epoch 1395
└ @ Main /Users/kcft114/.julia/packages/Flux/qXNjB/src/optimise/train.jl:105
┌ Info: Epoch 1396
└ @ Main /Users/kcft114/.julia/packages/Flux/qXNjB/src/optimise/train.jl:105
┌ Info: Epoch 1397
└ @ Main /Users/kcft114/.julia/packages/Flux/qXNjB/src/optimise/train.jl:105
┌ Info: Epoch 1398
└ @ Main /Users/kcft114/.julia/packages/Flux/qXNjB/src/optimise/train.jl:105
┌ Info: Epoch 1399
└ @ Main /Users/kcft114/.julia/packages/Flux/qXNjB/src/optimise/train.jl:105
┌ Info: Epoch 1400
└ @ Main /Users/kcft114/.julia/packages/Flux/qXNjB/src/optimise/train.jl:105
┌ Info: Epoch 1401
└ @ Main /Users/kcft114/.julia/packages/Flux/qXNjB/src/optimise/train.jl:105
┌ Info: Epoch 1402
└ @ Main /Users/kcft114/.julia/packages/Flux/qXNjB/src/optimise/train.jl:105
┌ Info: Epoch 1403
└ @ Main /Users/kcft114/.julia/packages/Flux/qXNjB/src/optimise/train.jl:105
┌ Info: Epoch 1404
└ @ Main /Users/kcft114/.julia/packages/Flux/qXNjB/src/optimise/train.jl:105
┌ Info: Epoch 1405
└ @ Main /Users/kcft114/.julia/packages/Flux/qXNjB/src/optimise/train.jl:105
┌ Info: Epoch 1406
└ @ Main /Users/kcft114/.julia/packages/Flux/qXNjB/src/optimise/train.jl:105
┌ Info: Epoch 1407
└ @ Main /Users/kcft114/.julia/packages/Flux/qXNjB/src/optimise/train.jl:105
┌ Info: Epoch 1408
└ @ Main /Users/kcft114/.julia/packages/Flux/qXNjB/src/optimise/train.jl:105
┌ Info: Epoch 1409
└ @ Main /Users/kcft114/.julia/packages/Flux/qXNjB/src/optimise/train.jl:105
┌ Info: Epoch 1410
└ @ Main /Users/kcft114/.julia/packages/Flux/qXNjB/src/optimise/train.jl:105
┌ Info: Epoch 1411
└ @ Main /Users/kcft114/.julia/packages/Flux/qXNjB/src/optimise/train.jl:105
┌ Info: Epoch 1412
└ @ Main /Users/kcft114/.julia/packages/Flux/qXNjB/src/optimise/train.jl:105
┌ Info: Epoch 1413
└ @ Main /Users/kcft114/.julia/packages/Flux/qXNjB/src/optimise/train.jl:105
┌ Info: Epoch 1414
└ @ Main /Users/kcft114/.julia/packages/Flux/qXNjB/src/optimise/train.jl:105
┌ Info: Epoch 1415
└ @ Main /Users/kcft114/.julia/packages/Flux/qXNjB/src/optimise/train.jl:105
```

```
loss(X, Y) = 0.59091496f0 (tracked)
loss(X, Y) = 0.60670865f0 (tracked)
loss(X, Y) = 0.601181f0 (tracked)
loss(X, Y) = 0.5988634f0 (tracked)
loss(X, Y) = 0.6395702f0 (tracked)
loss(X, Y) = 0.67590255f0 (tracked)
loss(X, Y) = 0.60094756f0 (tracked)
loss(X, Y) = 0.6023563f0 (tracked)
loss(X, Y) = 0.60090274f0 (tracked)
loss(X, Y) = 0.6151663f0 (tracked)
loss(X, Y) = 0.601555f0 (tracked)
loss(X, Y) = 0.65734875f0 (tracked)
loss(X, Y) = 0.59961283f0 (tracked)
loss(X, Y) = 0.5850407f0 (tracked)
loss(X, Y) = 0.6335523f0 (tracked)
loss(X, Y) = 0.6283254f0 (tracked)
loss(X, Y) = 0.6193323f0 (tracked)
loss(X, Y) = 0.6304408f0 (tracked)
loss(X, Y) = 0.6417276f0 (tracked)
loss(X, Y) = 0.6120141f0 (tracked)
loss(X, Y) = 0.63428265f0 (tracked)
loss(X, Y) = 0.62747484f0 (tracked)
loss(X, Y) = 0.61044043f0 (tracked)
loss(X, Y) = 0.6395432f0 (tracked)
loss(X, Y) = 0.5972257f0 (tracked)
loss(X, Y) = 0.623217f0 (tracked)
loss(X, Y) = 0.65538484f0 (tracked)
loss(X, Y) = 0.5937715f0 (tracked)
loss(X, Y) = 0.6066198f0 (tracked)
loss(X, Y) = 0.6521239f0 (tracked)
loss(X, Y) = 0.6711213f0 (tracked)
loss(X, Y) = 0.58680964f0 (tracked)
loss(X, Y) = 0.5834554f0 (tracked)
loss(X, Y) = 0.6332384f0 (tracked)
loss(X, Y) = 0.6690228f0 (tracked)
loss(X, Y) = 0.55551356f0 (tracked)
loss(X, Y) = 0.61836576f0 (tracked)
loss(X, Y) = 0.61485755f0 (tracked)
loss(X, Y) = 0.63158035f0 (tracked)
loss(X, Y) = 0.62958807f0 (tracked)
loss(X, Y) = 0.62179697f0 (tracked)
loss(X, Y) = 0.60974383f0 (tracked)
loss(X, Y) = 0.62735265f0 (tracked)
loss(X, Y) = 0.6405551f0 (tracked)
loss(X, Y) = 0.60521185f0 (tracked)
loss(X, Y) = 0.60868573f0 (tracked)
loss(X, Y) = 0.6425455f0 (tracked)
loss(X, Y) = 0.5734903f0 (tracked)
loss(X, Y) = 0.6106523f0 (tracked)
loss(X, Y) = 0.5837751f0 (tracked)
loss(X, Y) = 0.5746031f0 (tracked)
loss(X, Y) = 0.6052302f0 (tracked)
loss(X, Y) = 0.62887424f0 (tracked)
loss(X, Y) = 0.64170593f0 (tracked)
loss(X, Y) = 0.58259726f0 (tracked)
loss(X, Y) = 0.5943821f0 (tracked)
loss(X, Y) = 0.63145095f0 (tracked)
loss(X, Y) = 0.58857757f0 (tracked)
loss(X, Y) = 0.6140836f0 (tracked)
loss(X, Y) = 0.6180825f0 (tracked)
loss(X, Y) = 0.5862258f0 (tracked)
loss(X, Y) = 0.6392244f0 (tracked)
loss(X, Y) = 0.622398f0 (tracked)
loss(X, Y) = 0.6160064f0 (tracked)
loss(X, Y) = 0.5947708f0 (tracked)
loss(X, Y) = 0.5860163f0 (tracked)
loss(X, Y) = 0.61743873f0 (tracked)
loss(X, Y) = 0.59850985f0 (tracked)
loss(X, Y) = 0.616492f0 (tracked)
loss(X, Y) = 0.57928014f0 (tracked)
loss(X, Y) = 0.6388621f0 (tracked)
loss(X, Y) = 0.63012004f0 (tracked)
loss(X, Y) = 0.64058757f0 (tracked)
loss(X, Y) = 0.5919291f0 (tracked)
loss(X, Y) = 0.62092245f0 (tracked)
loss(X, Y) = 0.5715772f0 (tracked)
loss(X, Y) = 0.60128844f0 (tracked)
loss(X, Y) = 0.59328634f0 (tracked)
loss(X, Y) = 0.58552414f0 (tracked)
loss(X, Y) = 0.62003106f0 (tracked)
loss(X, Y) = 0.6601665f0 (tracked)
loss(X, Y) = 0.59453875f0 (tracked)
loss(X, Y) = 0.6070547f0 (tracked)
loss(X, Y) = 0.6162787f0 (tracked)
loss(X, Y) = 0.5928559f0 (tracked)
loss(X, Y) = 0.6352895f0 (tracked)
loss(X, Y) = 0.60741127f0 (tracked)
loss(X, Y) = 0.62550724f0 (tracked)
loss(X, Y) = 0.58711755f0 (tracked)
loss(X, Y) = 0.5551493f0 (tracked)
loss(X, Y) = 0.6284844f0 (tracked)
loss(X, Y) = 0.6035333f0 (tracked)
loss(X, Y) = 0.58848417f0 (tracked)
loss(X, Y) = 0.57836217f0 (tracked)
loss(X, Y) = 0.59359926f0 (tracked)
loss(X, Y) = 0.5613754f0 (tracked)
loss(X, Y) = 0.5700846f0 (tracked)
loss(X, Y) = 0.59026754f0 (tracked)
loss(X, Y) = 0.5908896f0 (tracked)
loss(X, Y) = 0.5933106f0 (tracked)
loss(X, Y) = 0.5879881f0 (tracked)
loss(X, Y) = 0.65452194f0 (tracked)
loss(X, Y) = 0.633935f0 (tracked)
loss(X, Y) = 0.6199904f0 (tracked)
loss(X, Y) = 0.6316927f0 (tracked)
loss(X, Y) = 0.59679294f0 (tracked)
loss(X, Y) = 0.5945174f0 (tracked)
loss(X, Y) = 0.6085847f0 (tracked)
loss(X, Y) = 0.59990776f0 (tracked)
loss(X, Y) = 0.5681915f0 (tracked)
loss(X, Y) = 0.61458373f0 (tracked)
loss(X, Y) = 0.6255189f0 (tracked)
loss(X, Y) = 0.57800084f0 (tracked)
loss(X, Y) = 0.62511975f0 (tracked)
loss(X, Y) = 0.5897507f0 (tracked)
loss(X, Y) = 0.6081103f0 (tracked)
loss(X, Y) = 0.5817309f0 (tracked)
loss(X, Y) = 0.63040644f0 (tracked)
loss(X, Y) = 0.5944788f0 (tracked)
loss(X, Y) = 0.618949f0 (tracked)
loss(X, Y) = 0.5768233f0 (tracked)
loss(X, Y) = 0.63749504f0 (tracked)
loss(X, Y) = 0.59141076f0 (tracked)
loss(X, Y) = 0.66025865f0 (tracked)
loss(X, Y) = 0.5752598f0 (tracked)
loss(X, Y) = 0.5800357f0 (tracked)
loss(X, Y) = 0.57229966f0 (tracked)
loss(X, Y) = 0.632325f0 (tracked)
loss(X, Y) = 0.6233144f0 (tracked)
loss(X, Y) = 0.56245154f0 (tracked)
loss(X, Y) = 0.6117411f0 (tracked)
loss(X, Y) = 0.61006457f0 (tracked)
loss(X, Y) = 0.5933698f0 (tracked)
loss(X, Y) = 0.6086571f0 (tracked)
loss(X, Y) = 0.6120535f0 (tracked)
loss(X, Y) = 0.61296207f0 (tracked)
loss(X, Y) = 0.59774435f0 (tracked)
loss(X, Y) = 0.6327109f0 (tracked)
loss(X, Y) = 0.59253234f0 (tracked)
loss(X, Y) = 0.619497f0 (tracked)
```

```
┌ Info: Epoch 1416
└ @ Main /Users/kcft114/.julia/packages/Flux/qXNjB/src/optimise/train.jl:105
┌ Info: Epoch 1417
└ @ Main /Users/kcft114/.julia/packages/Flux/qXNjB/src/optimise/train.jl:105
┌ Info: Epoch 1418
└ @ Main /Users/kcft114/.julia/packages/Flux/qXNjB/src/optimise/train.jl:105
┌ Info: Epoch 1419
└ @ Main /Users/kcft114/.julia/packages/Flux/qXNjB/src/optimise/train.jl:105
┌ Info: Epoch 1420
└ @ Main /Users/kcft114/.julia/packages/Flux/qXNjB/src/optimise/train.jl:105
┌ Info: Epoch 1421
└ @ Main /Users/kcft114/.julia/packages/Flux/qXNjB/src/optimise/train.jl:105
┌ Info: Epoch 1422
└ @ Main /Users/kcft114/.julia/packages/Flux/qXNjB/src/optimise/train.jl:105
┌ Info: Epoch 1423
└ @ Main /Users/kcft114/.julia/packages/Flux/qXNjB/src/optimise/train.jl:105
┌ Info: Epoch 1424
└ @ Main /Users/kcft114/.julia/packages/Flux/qXNjB/src/optimise/train.jl:105
┌ Info: Epoch 1425
└ @ Main /Users/kcft114/.julia/packages/Flux/qXNjB/src/optimise/train.jl:105
┌ Info: Epoch 1426
└ @ Main /Users/kcft114/.julia/packages/Flux/qXNjB/src/optimise/train.jl:105
┌ Info: Epoch 1427
└ @ Main /Users/kcft114/.julia/packages/Flux/qXNjB/src/optimise/train.jl:105
┌ Info: Epoch 1428
└ @ Main /Users/kcft114/.julia/packages/Flux/qXNjB/src/optimise/train.jl:105
┌ Info: Epoch 1429
└ @ Main /Users/kcft114/.julia/packages/Flux/qXNjB/src/optimise/train.jl:105
┌ Info: Epoch 1430
└ @ Main /Users/kcft114/.julia/packages/Flux/qXNjB/src/optimise/train.jl:105
┌ Info: Epoch 1431
└ @ Main /Users/kcft114/.julia/packages/Flux/qXNjB/src/optimise/train.jl:105
┌ Info: Epoch 1432
└ @ Main /Users/kcft114/.julia/packages/Flux/qXNjB/src/optimise/train.jl:105
┌ Info: Epoch 1433
└ @ Main /Users/kcft114/.julia/packages/Flux/qXNjB/src/optimise/train.jl:105
┌ Info: Epoch 1434
└ @ Main /Users/kcft114/.julia/packages/Flux/qXNjB/src/optimise/train.jl:105
┌ Info: Epoch 1435
└ @ Main /Users/kcft114/.julia/packages/Flux/qXNjB/src/optimise/train.jl:105
┌ Info: Epoch 1436
└ @ Main /Users/kcft114/.julia/packages/Flux/qXNjB/src/optimise/train.jl:105
┌ Info: Epoch 1437
└ @ Main /Users/kcft114/.julia/packages/Flux/qXNjB/src/optimise/train.jl:105
┌ Info: Epoch 1438
└ @ Main /Users/kcft114/.julia/packages/Flux/qXNjB/src/optimise/train.jl:105
┌ Info: Epoch 1439
└ @ Main /Users/kcft114/.julia/packages/Flux/qXNjB/src/optimise/train.jl:105
┌ Info: Epoch 1440
└ @ Main /Users/kcft114/.julia/packages/Flux/qXNjB/src/optimise/train.jl:105
┌ Info: Epoch 1441
└ @ Main /Users/kcft114/.julia/packages/Flux/qXNjB/src/optimise/train.jl:105
┌ Info: Epoch 1442
└ @ Main /Users/kcft114/.julia/packages/Flux/qXNjB/src/optimise/train.jl:105
┌ Info: Epoch 1443
└ @ Main /Users/kcft114/.julia/packages/Flux/qXNjB/src/optimise/train.jl:105
┌ Info: Epoch 1444
└ @ Main /Users/kcft114/.julia/packages/Flux/qXNjB/src/optimise/train.jl:105
┌ Info: Epoch 1445
└ @ Main /Users/kcft114/.julia/packages/Flux/qXNjB/src/optimise/train.jl:105
┌ Info: Epoch 1446
└ @ Main /Users/kcft114/.julia/packages/Flux/qXNjB/src/optimise/train.jl:105
┌ Info: Epoch 1447
└ @ Main /Users/kcft114/.julia/packages/Flux/qXNjB/src/optimise/train.jl:105
┌ Info: Epoch 1448
└ @ Main /Users/kcft114/.julia/packages/Flux/qXNjB/src/optimise/train.jl:105
┌ Info: Epoch 1449
└ @ Main /Users/kcft114/.julia/packages/Flux/qXNjB/src/optimise/train.jl:105
┌ Info: Epoch 1450
└ @ Main /Users/kcft114/.julia/packages/Flux/qXNjB/src/optimise/train.jl:105
┌ Info: Epoch 1451
└ @ Main /Users/kcft114/.julia/packages/Flux/qXNjB/src/optimise/train.jl:105
┌ Info: Epoch 1452
└ @ Main /Users/kcft114/.julia/packages/Flux/qXNjB/src/optimise/train.jl:105
┌ Info: Epoch 1453
└ @ Main /Users/kcft114/.julia/packages/Flux/qXNjB/src/optimise/train.jl:105
┌ Info: Epoch 1454
└ @ Main /Users/kcft114/.julia/packages/Flux/qXNjB/src/optimise/train.jl:105
┌ Info: Epoch 1455
└ @ Main /Users/kcft114/.julia/packages/Flux/qXNjB/src/optimise/train.jl:105
┌ Info: Epoch 1456
└ @ Main /Users/kcft114/.julia/packages/Flux/qXNjB/src/optimise/train.jl:105
┌ Info: Epoch 1457
└ @ Main /Users/kcft114/.julia/packages/Flux/qXNjB/src/optimise/train.jl:105
┌ Info: Epoch 1458
└ @ Main /Users/kcft114/.julia/packages/Flux/qXNjB/src/optimise/train.jl:105
┌ Info: Epoch 1459
└ @ Main /Users/kcft114/.julia/packages/Flux/qXNjB/src/optimise/train.jl:105
┌ Info: Epoch 1460
└ @ Main /Users/kcft114/.julia/packages/Flux/qXNjB/src/optimise/train.jl:105
┌ Info: Epoch 1461
└ @ Main /Users/kcft114/.julia/packages/Flux/qXNjB/src/optimise/train.jl:105
┌ Info: Epoch 1462
└ @ Main /Users/kcft114/.julia/packages/Flux/qXNjB/src/optimise/train.jl:105
┌ Info: Epoch 1463
└ @ Main /Users/kcft114/.julia/packages/Flux/qXNjB/src/optimise/train.jl:105
┌ Info: Epoch 1464
└ @ Main /Users/kcft114/.julia/packages/Flux/qXNjB/src/optimise/train.jl:105
┌ Info: Epoch 1465
└ @ Main /Users/kcft114/.julia/packages/Flux/qXNjB/src/optimise/train.jl:105
┌ Info: Epoch 1466
└ @ Main /Users/kcft114/.julia/packages/Flux/qXNjB/src/optimise/train.jl:105
┌ Info: Epoch 1467
└ @ Main /Users/kcft114/.julia/packages/Flux/qXNjB/src/optimise/train.jl:105
┌ Info: Epoch 1468
└ @ Main /Users/kcft114/.julia/packages/Flux/qXNjB/src/optimise/train.jl:105
┌ Info: Epoch 1469
└ @ Main /Users/kcft114/.julia/packages/Flux/qXNjB/src/optimise/train.jl:105
┌ Info: Epoch 1470
└ @ Main /Users/kcft114/.julia/packages/Flux/qXNjB/src/optimise/train.jl:105
┌ Info: Epoch 1471
└ @ Main /Users/kcft114/.julia/packages/Flux/qXNjB/src/optimise/train.jl:105
┌ Info: Epoch 1472
└ @ Main /Users/kcft114/.julia/packages/Flux/qXNjB/src/optimise/train.jl:105
┌ Info: Epoch 1473
└ @ Main /Users/kcft114/.julia/packages/Flux/qXNjB/src/optimise/train.jl:105
┌ Info: Epoch 1474
└ @ Main /Users/kcft114/.julia/packages/Flux/qXNjB/src/optimise/train.jl:105
┌ Info: Epoch 1475
└ @ Main /Users/kcft114/.julia/packages/Flux/qXNjB/src/optimise/train.jl:105
┌ Info: Epoch 1476
└ @ Main /Users/kcft114/.julia/packages/Flux/qXNjB/src/optimise/train.jl:105
┌ Info: Epoch 1477
└ @ Main /Users/kcft114/.julia/packages/Flux/qXNjB/src/optimise/train.jl:105
┌ Info: Epoch 1478
└ @ Main /Users/kcft114/.julia/packages/Flux/qXNjB/src/optimise/train.jl:105
┌ Info: Epoch 1479
└ @ Main /Users/kcft114/.julia/packages/Flux/qXNjB/src/optimise/train.jl:105
┌ Info: Epoch 1480
└ @ Main /Users/kcft114/.julia/packages/Flux/qXNjB/src/optimise/train.jl:105
┌ Info: Epoch 1481
└ @ Main /Users/kcft114/.julia/packages/Flux/qXNjB/src/optimise/train.jl:105
┌ Info: Epoch 1482
└ @ Main /Users/kcft114/.julia/packages/Flux/qXNjB/src/optimise/train.jl:105
┌ Info: Epoch 1483
└ @ Main /Users/kcft114/.julia/packages/Flux/qXNjB/src/optimise/train.jl:105
┌ Info: Epoch 1484
└ @ Main /Users/kcft114/.julia/packages/Flux/qXNjB/src/optimise/train.jl:105
┌ Info: Epoch 1485
└ @ Main /Users/kcft114/.julia/packages/Flux/qXNjB/src/optimise/train.jl:105
┌ Info: Epoch 1486
└ @ Main /Users/kcft114/.julia/packages/Flux/qXNjB/src/optimise/train.jl:105
┌ Info: Epoch 1487
└ @ Main /Users/kcft114/.julia/packages/Flux/qXNjB/src/optimise/train.jl:105
┌ Info: Epoch 1488
└ @ Main /Users/kcft114/.julia/packages/Flux/qXNjB/src/optimise/train.jl:105
┌ Info: Epoch 1489
└ @ Main /Users/kcft114/.julia/packages/Flux/qXNjB/src/optimise/train.jl:105
┌ Info: Epoch 1490
└ @ Main /Users/kcft114/.julia/packages/Flux/qXNjB/src/optimise/train.jl:105
┌ Info: Epoch 1491
└ @ Main /Users/kcft114/.julia/packages/Flux/qXNjB/src/optimise/train.jl:105
┌ Info: Epoch 1492
└ @ Main /Users/kcft114/.julia/packages/Flux/qXNjB/src/optimise/train.jl:105
┌ Info: Epoch 1493
└ @ Main /Users/kcft114/.julia/packages/Flux/qXNjB/src/optimise/train.jl:105
┌ Info: Epoch 1494
└ @ Main /Users/kcft114/.julia/packages/Flux/qXNjB/src/optimise/train.jl:105
┌ Info: Epoch 1495
└ @ Main /Users/kcft114/.julia/packages/Flux/qXNjB/src/optimise/train.jl:105
┌ Info: Epoch 1496
└ @ Main /Users/kcft114/.julia/packages/Flux/qXNjB/src/optimise/train.jl:105
┌ Info: Epoch 1497
└ @ Main /Users/kcft114/.julia/packages/Flux/qXNjB/src/optimise/train.jl:105
┌ Info: Epoch 1498
└ @ Main /Users/kcft114/.julia/packages/Flux/qXNjB/src/optimise/train.jl:105
┌ Info: Epoch 1499
└ @ Main /Users/kcft114/.julia/packages/Flux/qXNjB/src/optimise/train.jl:105
┌ Info: Epoch 1500
└ @ Main /Users/kcft114/.julia/packages/Flux/qXNjB/src/optimise/train.jl:105
┌ Info: Epoch 1501
└ @ Main /Users/kcft114/.julia/packages/Flux/qXNjB/src/optimise/train.jl:105
┌ Info: Epoch 1502
└ @ Main /Users/kcft114/.julia/packages/Flux/qXNjB/src/optimise/train.jl:105
┌ Info: Epoch 1503
└ @ Main /Users/kcft114/.julia/packages/Flux/qXNjB/src/optimise/train.jl:105
┌ Info: Epoch 1504
└ @ Main /Users/kcft114/.julia/packages/Flux/qXNjB/src/optimise/train.jl:105
┌ Info: Epoch 1505
└ @ Main /Users/kcft114/.julia/packages/Flux/qXNjB/src/optimise/train.jl:105
┌ Info: Epoch 1506
└ @ Main /Users/kcft114/.julia/packages/Flux/qXNjB/src/optimise/train.jl:105
┌ Info: Epoch 1507
└ @ Main /Users/kcft114/.julia/packages/Flux/qXNjB/src/optimise/train.jl:105
┌ Info: Epoch 1508
└ @ Main /Users/kcft114/.julia/packages/Flux/qXNjB/src/optimise/train.jl:105
┌ Info: Epoch 1509
└ @ Main /Users/kcft114/.julia/packages/Flux/qXNjB/src/optimise/train.jl:105
┌ Info: Epoch 1510
└ @ Main /Users/kcft114/.julia/packages/Flux/qXNjB/src/optimise/train.jl:105
┌ Info: Epoch 1511
└ @ Main /Users/kcft114/.julia/packages/Flux/qXNjB/src/optimise/train.jl:105
┌ Info: Epoch 1512
└ @ Main /Users/kcft114/.julia/packages/Flux/qXNjB/src/optimise/train.jl:105
┌ Info: Epoch 1513
└ @ Main /Users/kcft114/.julia/packages/Flux/qXNjB/src/optimise/train.jl:105
┌ Info: Epoch 1514
└ @ Main /Users/kcft114/.julia/packages/Flux/qXNjB/src/optimise/train.jl:105
┌ Info: Epoch 1515
└ @ Main /Users/kcft114/.julia/packages/Flux/qXNjB/src/optimise/train.jl:105
┌ Info: Epoch 1516
└ @ Main /Users/kcft114/.julia/packages/Flux/qXNjB/src/optimise/train.jl:105
┌ Info: Epoch 1517
└ @ Main /Users/kcft114/.julia/packages/Flux/qXNjB/src/optimise/train.jl:105
┌ Info: Epoch 1518
└ @ Main /Users/kcft114/.julia/packages/Flux/qXNjB/src/optimise/train.jl:105
┌ Info: Epoch 1519
└ @ Main /Users/kcft114/.julia/packages/Flux/qXNjB/src/optimise/train.jl:105
┌ Info: Epoch 1520
└ @ Main /Users/kcft114/.julia/packages/Flux/qXNjB/src/optimise/train.jl:105
┌ Info: Epoch 1521
└ @ Main /Users/kcft114/.julia/packages/Flux/qXNjB/src/optimise/train.jl:105
┌ Info: Epoch 1522
└ @ Main /Users/kcft114/.julia/packages/Flux/qXNjB/src/optimise/train.jl:105
┌ Info: Epoch 1523
└ @ Main /Users/kcft114/.julia/packages/Flux/qXNjB/src/optimise/train.jl:105
┌ Info: Epoch 1524
└ @ Main /Users/kcft114/.julia/packages/Flux/qXNjB/src/optimise/train.jl:105
┌ Info: Epoch 1525
└ @ Main /Users/kcft114/.julia/packages/Flux/qXNjB/src/optimise/train.jl:105
┌ Info: Epoch 1526
└ @ Main /Users/kcft114/.julia/packages/Flux/qXNjB/src/optimise/train.jl:105
┌ Info: Epoch 1527
└ @ Main /Users/kcft114/.julia/packages/Flux/qXNjB/src/optimise/train.jl:105
┌ Info: Epoch 1528
└ @ Main /Users/kcft114/.julia/packages/Flux/qXNjB/src/optimise/train.jl:105
┌ Info: Epoch 1529
└ @ Main /Users/kcft114/.julia/packages/Flux/qXNjB/src/optimise/train.jl:105
┌ Info: Epoch 1530
└ @ Main /Users/kcft114/.julia/packages/Flux/qXNjB/src/optimise/train.jl:105
┌ Info: Epoch 1531
└ @ Main /Users/kcft114/.julia/packages/Flux/qXNjB/src/optimise/train.jl:105
┌ Info: Epoch 1532
└ @ Main /Users/kcft114/.julia/packages/Flux/qXNjB/src/optimise/train.jl:105
┌ Info: Epoch 1533
└ @ Main /Users/kcft114/.julia/packages/Flux/qXNjB/src/optimise/train.jl:105
┌ Info: Epoch 1534
└ @ Main /Users/kcft114/.julia/packages/Flux/qXNjB/src/optimise/train.jl:105
┌ Info: Epoch 1535
└ @ Main /Users/kcft114/.julia/packages/Flux/qXNjB/src/optimise/train.jl:105
┌ Info: Epoch 1536
└ @ Main /Users/kcft114/.julia/packages/Flux/qXNjB/src/optimise/train.jl:105
┌ Info: Epoch 1537
└ @ Main /Users/kcft114/.julia/packages/Flux/qXNjB/src/optimise/train.jl:105
┌ Info: Epoch 1538
└ @ Main /Users/kcft114/.julia/packages/Flux/qXNjB/src/optimise/train.jl:105
┌ Info: Epoch 1539
└ @ Main /Users/kcft114/.julia/packages/Flux/qXNjB/src/optimise/train.jl:105
┌ Info: Epoch 1540
└ @ Main /Users/kcft114/.julia/packages/Flux/qXNjB/src/optimise/train.jl:105
┌ Info: Epoch 1541
└ @ Main /Users/kcft114/.julia/packages/Flux/qXNjB/src/optimise/train.jl:105
┌ Info: Epoch 1542
└ @ Main /Users/kcft114/.julia/packages/Flux/qXNjB/src/optimise/train.jl:105
┌ Info: Epoch 1543
└ @ Main /Users/kcft114/.julia/packages/Flux/qXNjB/src/optimise/train.jl:105
┌ Info: Epoch 1544
└ @ Main /Users/kcft114/.julia/packages/Flux/qXNjB/src/optimise/train.jl:105
┌ Info: Epoch 1545
└ @ Main /Users/kcft114/.julia/packages/Flux/qXNjB/src/optimise/train.jl:105
┌ Info: Epoch 1546
└ @ Main /Users/kcft114/.julia/packages/Flux/qXNjB/src/optimise/train.jl:105
┌ Info: Epoch 1547
└ @ Main /Users/kcft114/.julia/packages/Flux/qXNjB/src/optimise/train.jl:105
┌ Info: Epoch 1548
└ @ Main /Users/kcft114/.julia/packages/Flux/qXNjB/src/optimise/train.jl:105
┌ Info: Epoch 1549
└ @ Main /Users/kcft114/.julia/packages/Flux/qXNjB/src/optimise/train.jl:105
┌ Info: Epoch 1550
└ @ Main /Users/kcft114/.julia/packages/Flux/qXNjB/src/optimise/train.jl:105
┌ Info: Epoch 1551
└ @ Main /Users/kcft114/.julia/packages/Flux/qXNjB/src/optimise/train.jl:105
┌ Info: Epoch 1552
└ @ Main /Users/kcft114/.julia/packages/Flux/qXNjB/src/optimise/train.jl:105
┌ Info: Epoch 1553
└ @ Main /Users/kcft114/.julia/packages/Flux/qXNjB/src/optimise/train.jl:105
┌ Info: Epoch 1554
└ @ Main /Users/kcft114/.julia/packages/Flux/qXNjB/src/optimise/train.jl:105
┌ Info: Epoch 1555
└ @ Main /Users/kcft114/.julia/packages/Flux/qXNjB/src/optimise/train.jl:105
```

```
loss(X, Y) = 0.64419234f0 (tracked)
loss(X, Y) = 0.6024491f0 (tracked)
loss(X, Y) = 0.6001922f0 (tracked)
loss(X, Y) = 0.59617025f0 (tracked)
loss(X, Y) = 0.5729482f0 (tracked)
loss(X, Y) = 0.6551821f0 (tracked)
loss(X, Y) = 0.5446839f0 (tracked)
loss(X, Y) = 0.6415595f0 (tracked)
loss(X, Y) = 0.58512497f0 (tracked)
loss(X, Y) = 0.6124457f0 (tracked)
loss(X, Y) = 0.6243346f0 (tracked)
loss(X, Y) = 0.57899195f0 (tracked)
loss(X, Y) = 0.5976308f0 (tracked)
loss(X, Y) = 0.6038144f0 (tracked)
loss(X, Y) = 0.57946f0 (tracked)
loss(X, Y) = 0.58122224f0 (tracked)
loss(X, Y) = 0.6154147f0 (tracked)
loss(X, Y) = 0.5975482f0 (tracked)
loss(X, Y) = 0.5901907f0 (tracked)
loss(X, Y) = 0.5899133f0 (tracked)
loss(X, Y) = 0.6482003f0 (tracked)
loss(X, Y) = 0.61246145f0 (tracked)
loss(X, Y) = 0.5872698f0 (tracked)
loss(X, Y) = 0.59977907f0 (tracked)
loss(X, Y) = 0.5897221f0 (tracked)
loss(X, Y) = 0.596882f0 (tracked)
loss(X, Y) = 0.6242709f0 (tracked)
loss(X, Y) = 0.58377063f0 (tracked)
loss(X, Y) = 0.5499279f0 (tracked)
loss(X, Y) = 0.5837796f0 (tracked)
loss(X, Y) = 0.62257564f0 (tracked)
loss(X, Y) = 0.6001522f0 (tracked)
loss(X, Y) = 0.5996005f0 (tracked)
loss(X, Y) = 0.5683668f0 (tracked)
loss(X, Y) = 0.60862744f0 (tracked)
loss(X, Y) = 0.6175161f0 (tracked)
loss(X, Y) = 0.5976075f0 (tracked)
loss(X, Y) = 0.55998445f0 (tracked)
loss(X, Y) = 0.596443f0 (tracked)
loss(X, Y) = 0.5905653f0 (tracked)
loss(X, Y) = 0.60749567f0 (tracked)
loss(X, Y) = 0.6089612f0 (tracked)
loss(X, Y) = 0.60287243f0 (tracked)
loss(X, Y) = 0.60734135f0 (tracked)
loss(X, Y) = 0.61831015f0 (tracked)
loss(X, Y) = 0.60017943f0 (tracked)
loss(X, Y) = 0.5976696f0 (tracked)
loss(X, Y) = 0.5928406f0 (tracked)
loss(X, Y) = 0.64141965f0 (tracked)
loss(X, Y) = 0.5645525f0 (tracked)
loss(X, Y) = 0.60185117f0 (tracked)
loss(X, Y) = 0.5608508f0 (tracked)
loss(X, Y) = 0.62405425f0 (tracked)
loss(X, Y) = 0.5818948f0 (tracked)
loss(X, Y) = 0.5659576f0 (tracked)
loss(X, Y) = 0.5875023f0 (tracked)
loss(X, Y) = 0.5959815f0 (tracked)
loss(X, Y) = 0.5896125f0 (tracked)
loss(X, Y) = 0.6005078f0 (tracked)
loss(X, Y) = 0.56159234f0 (tracked)
loss(X, Y) = 0.6047096f0 (tracked)
loss(X, Y) = 0.6154098f0 (tracked)
loss(X, Y) = 0.59120584f0 (tracked)
loss(X, Y) = 0.6053341f0 (tracked)
loss(X, Y) = 0.6012075f0 (tracked)
loss(X, Y) = 0.5974082f0 (tracked)
loss(X, Y) = 0.5800778f0 (tracked)
loss(X, Y) = 0.57635766f0 (tracked)
loss(X, Y) = 0.5653049f0 (tracked)
loss(X, Y) = 0.59522045f0 (tracked)
loss(X, Y) = 0.61238474f0 (tracked)
loss(X, Y) = 0.60274565f0 (tracked)
loss(X, Y) = 0.6083589f0 (tracked)
loss(X, Y) = 0.566541f0 (tracked)
loss(X, Y) = 0.59475034f0 (tracked)
loss(X, Y) = 0.5690452f0 (tracked)
loss(X, Y) = 0.57209986f0 (tracked)
loss(X, Y) = 0.61305004f0 (tracked)
loss(X, Y) = 0.5869542f0 (tracked)
loss(X, Y) = 0.63012815f0 (tracked)
loss(X, Y) = 0.6095201f0 (tracked)
loss(X, Y) = 0.57739973f0 (tracked)
loss(X, Y) = 0.5325065f0 (tracked)
loss(X, Y) = 0.59041864f0 (tracked)
loss(X, Y) = 0.5688756f0 (tracked)
loss(X, Y) = 0.6103827f0 (tracked)
loss(X, Y) = 0.56630117f0 (tracked)
loss(X, Y) = 0.56198466f0 (tracked)
loss(X, Y) = 0.5752691f0 (tracked)
loss(X, Y) = 0.5703543f0 (tracked)
loss(X, Y) = 0.59332275f0 (tracked)
loss(X, Y) = 0.5576941f0 (tracked)
loss(X, Y) = 0.5927703f0 (tracked)
loss(X, Y) = 0.5970319f0 (tracked)
loss(X, Y) = 0.5534188f0 (tracked)
loss(X, Y) = 0.53878057f0 (tracked)
loss(X, Y) = 0.60122514f0 (tracked)
loss(X, Y) = 0.58153325f0 (tracked)
loss(X, Y) = 0.5774822f0 (tracked)
loss(X, Y) = 0.5928292f0 (tracked)
loss(X, Y) = 0.577775f0 (tracked)
loss(X, Y) = 0.5635516f0 (tracked)
loss(X, Y) = 0.582151f0 (tracked)
loss(X, Y) = 0.5975529f0 (tracked)
loss(X, Y) = 0.5866114f0 (tracked)
loss(X, Y) = 0.59944606f0 (tracked)
loss(X, Y) = 0.5839187f0 (tracked)
loss(X, Y) = 0.57263315f0 (tracked)
loss(X, Y) = 0.5916532f0 (tracked)
loss(X, Y) = 0.5897748f0 (tracked)
loss(X, Y) = 0.5993745f0 (tracked)
loss(X, Y) = 0.5912861f0 (tracked)
loss(X, Y) = 0.6052588f0 (tracked)
loss(X, Y) = 0.630211f0 (tracked)
loss(X, Y) = 0.5744543f0 (tracked)
loss(X, Y) = 0.57963187f0 (tracked)
loss(X, Y) = 0.6042212f0 (tracked)
loss(X, Y) = 0.5560711f0 (tracked)
loss(X, Y) = 0.5481517f0 (tracked)
loss(X, Y) = 0.59543926f0 (tracked)
loss(X, Y) = 0.57372785f0 (tracked)
loss(X, Y) = 0.5956538f0 (tracked)
loss(X, Y) = 0.59829146f0 (tracked)
loss(X, Y) = 0.57295007f0 (tracked)
```

```
┌ Info: Epoch 1556
└ @ Main /Users/kcft114/.julia/packages/Flux/qXNjB/src/optimise/train.jl:105
┌ Info: Epoch 1557
└ @ Main /Users/kcft114/.julia/packages/Flux/qXNjB/src/optimise/train.jl:105
┌ Info: Epoch 1558
└ @ Main /Users/kcft114/.julia/packages/Flux/qXNjB/src/optimise/train.jl:105
┌ Info: Epoch 1559
└ @ Main /Users/kcft114/.julia/packages/Flux/qXNjB/src/optimise/train.jl:105
┌ Info: Epoch 1560
└ @ Main /Users/kcft114/.julia/packages/Flux/qXNjB/src/optimise/train.jl:105
┌ Info: Epoch 1561
└ @ Main /Users/kcft114/.julia/packages/Flux/qXNjB/src/optimise/train.jl:105
┌ Info: Epoch 1562
└ @ Main /Users/kcft114/.julia/packages/Flux/qXNjB/src/optimise/train.jl:105
┌ Info: Epoch 1563
└ @ Main /Users/kcft114/.julia/packages/Flux/qXNjB/src/optimise/train.jl:105
┌ Info: Epoch 1564
└ @ Main /Users/kcft114/.julia/packages/Flux/qXNjB/src/optimise/train.jl:105
┌ Info: Epoch 1565
└ @ Main /Users/kcft114/.julia/packages/Flux/qXNjB/src/optimise/train.jl:105
┌ Info: Epoch 1566
└ @ Main /Users/kcft114/.julia/packages/Flux/qXNjB/src/optimise/train.jl:105
┌ Info: Epoch 1567
└ @ Main /Users/kcft114/.julia/packages/Flux/qXNjB/src/optimise/train.jl:105
┌ Info: Epoch 1568
└ @ Main /Users/kcft114/.julia/packages/Flux/qXNjB/src/optimise/train.jl:105
┌ Info: Epoch 1569
└ @ Main /Users/kcft114/.julia/packages/Flux/qXNjB/src/optimise/train.jl:105
┌ Info: Epoch 1570
└ @ Main /Users/kcft114/.julia/packages/Flux/qXNjB/src/optimise/train.jl:105
┌ Info: Epoch 1571
└ @ Main /Users/kcft114/.julia/packages/Flux/qXNjB/src/optimise/train.jl:105
┌ Info: Epoch 1572
└ @ Main /Users/kcft114/.julia/packages/Flux/qXNjB/src/optimise/train.jl:105
┌ Info: Epoch 1573
└ @ Main /Users/kcft114/.julia/packages/Flux/qXNjB/src/optimise/train.jl:105
┌ Info: Epoch 1574
└ @ Main /Users/kcft114/.julia/packages/Flux/qXNjB/src/optimise/train.jl:105
┌ Info: Epoch 1575
└ @ Main /Users/kcft114/.julia/packages/Flux/qXNjB/src/optimise/train.jl:105
┌ Info: Epoch 1576
└ @ Main /Users/kcft114/.julia/packages/Flux/qXNjB/src/optimise/train.jl:105
┌ Info: Epoch 1577
└ @ Main /Users/kcft114/.julia/packages/Flux/qXNjB/src/optimise/train.jl:105
┌ Info: Epoch 1578
└ @ Main /Users/kcft114/.julia/packages/Flux/qXNjB/src/optimise/train.jl:105
┌ Info: Epoch 1579
└ @ Main /Users/kcft114/.julia/packages/Flux/qXNjB/src/optimise/train.jl:105
┌ Info: Epoch 1580
└ @ Main /Users/kcft114/.julia/packages/Flux/qXNjB/src/optimise/train.jl:105
┌ Info: Epoch 1581
└ @ Main /Users/kcft114/.julia/packages/Flux/qXNjB/src/optimise/train.jl:105
┌ Info: Epoch 1582
└ @ Main /Users/kcft114/.julia/packages/Flux/qXNjB/src/optimise/train.jl:105
┌ Info: Epoch 1583
└ @ Main /Users/kcft114/.julia/packages/Flux/qXNjB/src/optimise/train.jl:105
┌ Info: Epoch 1584
└ @ Main /Users/kcft114/.julia/packages/Flux/qXNjB/src/optimise/train.jl:105
┌ Info: Epoch 1585
└ @ Main /Users/kcft114/.julia/packages/Flux/qXNjB/src/optimise/train.jl:105
┌ Info: Epoch 1586
└ @ Main /Users/kcft114/.julia/packages/Flux/qXNjB/src/optimise/train.jl:105
┌ Info: Epoch 1587
└ @ Main /Users/kcft114/.julia/packages/Flux/qXNjB/src/optimise/train.jl:105
┌ Info: Epoch 1588
└ @ Main /Users/kcft114/.julia/packages/Flux/qXNjB/src/optimise/train.jl:105
┌ Info: Epoch 1589
└ @ Main /Users/kcft114/.julia/packages/Flux/qXNjB/src/optimise/train.jl:105
┌ Info: Epoch 1590
└ @ Main /Users/kcft114/.julia/packages/Flux/qXNjB/src/optimise/train.jl:105
┌ Info: Epoch 1591
└ @ Main /Users/kcft114/.julia/packages/Flux/qXNjB/src/optimise/train.jl:105
┌ Info: Epoch 1592
└ @ Main /Users/kcft114/.julia/packages/Flux/qXNjB/src/optimise/train.jl:105
┌ Info: Epoch 1593
└ @ Main /Users/kcft114/.julia/packages/Flux/qXNjB/src/optimise/train.jl:105
┌ Info: Epoch 1594
└ @ Main /Users/kcft114/.julia/packages/Flux/qXNjB/src/optimise/train.jl:105
┌ Info: Epoch 1595
└ @ Main /Users/kcft114/.julia/packages/Flux/qXNjB/src/optimise/train.jl:105
┌ Info: Epoch 1596
└ @ Main /Users/kcft114/.julia/packages/Flux/qXNjB/src/optimise/train.jl:105
┌ Info: Epoch 1597
└ @ Main /Users/kcft114/.julia/packages/Flux/qXNjB/src/optimise/train.jl:105
┌ Info: Epoch 1598
└ @ Main /Users/kcft114/.julia/packages/Flux/qXNjB/src/optimise/train.jl:105
┌ Info: Epoch 1599
└ @ Main /Users/kcft114/.julia/packages/Flux/qXNjB/src/optimise/train.jl:105
┌ Info: Epoch 1600
└ @ Main /Users/kcft114/.julia/packages/Flux/qXNjB/src/optimise/train.jl:105
┌ Info: Epoch 1601
└ @ Main /Users/kcft114/.julia/packages/Flux/qXNjB/src/optimise/train.jl:105
┌ Info: Epoch 1602
└ @ Main /Users/kcft114/.julia/packages/Flux/qXNjB/src/optimise/train.jl:105
┌ Info: Epoch 1603
└ @ Main /Users/kcft114/.julia/packages/Flux/qXNjB/src/optimise/train.jl:105
┌ Info: Epoch 1604
└ @ Main /Users/kcft114/.julia/packages/Flux/qXNjB/src/optimise/train.jl:105
┌ Info: Epoch 1605
└ @ Main /Users/kcft114/.julia/packages/Flux/qXNjB/src/optimise/train.jl:105
┌ Info: Epoch 1606
└ @ Main /Users/kcft114/.julia/packages/Flux/qXNjB/src/optimise/train.jl:105
┌ Info: Epoch 1607
└ @ Main /Users/kcft114/.julia/packages/Flux/qXNjB/src/optimise/train.jl:105
┌ Info: Epoch 1608
└ @ Main /Users/kcft114/.julia/packages/Flux/qXNjB/src/optimise/train.jl:105
┌ Info: Epoch 1609
└ @ Main /Users/kcft114/.julia/packages/Flux/qXNjB/src/optimise/train.jl:105
┌ Info: Epoch 1610
└ @ Main /Users/kcft114/.julia/packages/Flux/qXNjB/src/optimise/train.jl:105
┌ Info: Epoch 1611
└ @ Main /Users/kcft114/.julia/packages/Flux/qXNjB/src/optimise/train.jl:105
┌ Info: Epoch 1612
└ @ Main /Users/kcft114/.julia/packages/Flux/qXNjB/src/optimise/train.jl:105
┌ Info: Epoch 1613
└ @ Main /Users/kcft114/.julia/packages/Flux/qXNjB/src/optimise/train.jl:105
┌ Info: Epoch 1614
└ @ Main /Users/kcft114/.julia/packages/Flux/qXNjB/src/optimise/train.jl:105
┌ Info: Epoch 1615
└ @ Main /Users/kcft114/.julia/packages/Flux/qXNjB/src/optimise/train.jl:105
┌ Info: Epoch 1616
└ @ Main /Users/kcft114/.julia/packages/Flux/qXNjB/src/optimise/train.jl:105
┌ Info: Epoch 1617
└ @ Main /Users/kcft114/.julia/packages/Flux/qXNjB/src/optimise/train.jl:105
┌ Info: Epoch 1618
└ @ Main /Users/kcft114/.julia/packages/Flux/qXNjB/src/optimise/train.jl:105
┌ Info: Epoch 1619
└ @ Main /Users/kcft114/.julia/packages/Flux/qXNjB/src/optimise/train.jl:105
┌ Info: Epoch 1620
└ @ Main /Users/kcft114/.julia/packages/Flux/qXNjB/src/optimise/train.jl:105
┌ Info: Epoch 1621
└ @ Main /Users/kcft114/.julia/packages/Flux/qXNjB/src/optimise/train.jl:105
┌ Info: Epoch 1622
└ @ Main /Users/kcft114/.julia/packages/Flux/qXNjB/src/optimise/train.jl:105
┌ Info: Epoch 1623
└ @ Main /Users/kcft114/.julia/packages/Flux/qXNjB/src/optimise/train.jl:105
┌ Info: Epoch 1624
└ @ Main /Users/kcft114/.julia/packages/Flux/qXNjB/src/optimise/train.jl:105
┌ Info: Epoch 1625
└ @ Main /Users/kcft114/.julia/packages/Flux/qXNjB/src/optimise/train.jl:105
┌ Info: Epoch 1626
└ @ Main /Users/kcft114/.julia/packages/Flux/qXNjB/src/optimise/train.jl:105
┌ Info: Epoch 1627
└ @ Main /Users/kcft114/.julia/packages/Flux/qXNjB/src/optimise/train.jl:105
┌ Info: Epoch 1628
└ @ Main /Users/kcft114/.julia/packages/Flux/qXNjB/src/optimise/train.jl:105
┌ Info: Epoch 1629
└ @ Main /Users/kcft114/.julia/packages/Flux/qXNjB/src/optimise/train.jl:105
┌ Info: Epoch 1630
└ @ Main /Users/kcft114/.julia/packages/Flux/qXNjB/src/optimise/train.jl:105
┌ Info: Epoch 1631
└ @ Main /Users/kcft114/.julia/packages/Flux/qXNjB/src/optimise/train.jl:105
┌ Info: Epoch 1632
└ @ Main /Users/kcft114/.julia/packages/Flux/qXNjB/src/optimise/train.jl:105
┌ Info: Epoch 1633
└ @ Main /Users/kcft114/.julia/packages/Flux/qXNjB/src/optimise/train.jl:105
┌ Info: Epoch 1634
└ @ Main /Users/kcft114/.julia/packages/Flux/qXNjB/src/optimise/train.jl:105
┌ Info: Epoch 1635
└ @ Main /Users/kcft114/.julia/packages/Flux/qXNjB/src/optimise/train.jl:105
┌ Info: Epoch 1636
└ @ Main /Users/kcft114/.julia/packages/Flux/qXNjB/src/optimise/train.jl:105
┌ Info: Epoch 1637
└ @ Main /Users/kcft114/.julia/packages/Flux/qXNjB/src/optimise/train.jl:105
┌ Info: Epoch 1638
└ @ Main /Users/kcft114/.julia/packages/Flux/qXNjB/src/optimise/train.jl:105
┌ Info: Epoch 1639
└ @ Main /Users/kcft114/.julia/packages/Flux/qXNjB/src/optimise/train.jl:105
┌ Info: Epoch 1640
└ @ Main /Users/kcft114/.julia/packages/Flux/qXNjB/src/optimise/train.jl:105
┌ Info: Epoch 1641
└ @ Main /Users/kcft114/.julia/packages/Flux/qXNjB/src/optimise/train.jl:105
┌ Info: Epoch 1642
└ @ Main /Users/kcft114/.julia/packages/Flux/qXNjB/src/optimise/train.jl:105
┌ Info: Epoch 1643
└ @ Main /Users/kcft114/.julia/packages/Flux/qXNjB/src/optimise/train.jl:105
┌ Info: Epoch 1644
└ @ Main /Users/kcft114/.julia/packages/Flux/qXNjB/src/optimise/train.jl:105
┌ Info: Epoch 1645
└ @ Main /Users/kcft114/.julia/packages/Flux/qXNjB/src/optimise/train.jl:105
┌ Info: Epoch 1646
└ @ Main /Users/kcft114/.julia/packages/Flux/qXNjB/src/optimise/train.jl:105
┌ Info: Epoch 1647
└ @ Main /Users/kcft114/.julia/packages/Flux/qXNjB/src/optimise/train.jl:105
┌ Info: Epoch 1648
└ @ Main /Users/kcft114/.julia/packages/Flux/qXNjB/src/optimise/train.jl:105
┌ Info: Epoch 1649
└ @ Main /Users/kcft114/.julia/packages/Flux/qXNjB/src/optimise/train.jl:105
┌ Info: Epoch 1650
└ @ Main /Users/kcft114/.julia/packages/Flux/qXNjB/src/optimise/train.jl:105
┌ Info: Epoch 1651
└ @ Main /Users/kcft114/.julia/packages/Flux/qXNjB/src/optimise/train.jl:105
┌ Info: Epoch 1652
└ @ Main /Users/kcft114/.julia/packages/Flux/qXNjB/src/optimise/train.jl:105
┌ Info: Epoch 1653
└ @ Main /Users/kcft114/.julia/packages/Flux/qXNjB/src/optimise/train.jl:105
┌ Info: Epoch 1654
└ @ Main /Users/kcft114/.julia/packages/Flux/qXNjB/src/optimise/train.jl:105
┌ Info: Epoch 1655
└ @ Main /Users/kcft114/.julia/packages/Flux/qXNjB/src/optimise/train.jl:105
┌ Info: Epoch 1656
└ @ Main /Users/kcft114/.julia/packages/Flux/qXNjB/src/optimise/train.jl:105
┌ Info: Epoch 1657
└ @ Main /Users/kcft114/.julia/packages/Flux/qXNjB/src/optimise/train.jl:105
┌ Info: Epoch 1658
└ @ Main /Users/kcft114/.julia/packages/Flux/qXNjB/src/optimise/train.jl:105
┌ Info: Epoch 1659
└ @ Main /Users/kcft114/.julia/packages/Flux/qXNjB/src/optimise/train.jl:105
┌ Info: Epoch 1660
└ @ Main /Users/kcft114/.julia/packages/Flux/qXNjB/src/optimise/train.jl:105
┌ Info: Epoch 1661
└ @ Main /Users/kcft114/.julia/packages/Flux/qXNjB/src/optimise/train.jl:105
┌ Info: Epoch 1662
└ @ Main /Users/kcft114/.julia/packages/Flux/qXNjB/src/optimise/train.jl:105
┌ Info: Epoch 1663
└ @ Main /Users/kcft114/.julia/packages/Flux/qXNjB/src/optimise/train.jl:105
┌ Info: Epoch 1664
└ @ Main /Users/kcft114/.julia/packages/Flux/qXNjB/src/optimise/train.jl:105
┌ Info: Epoch 1665
└ @ Main /Users/kcft114/.julia/packages/Flux/qXNjB/src/optimise/train.jl:105
┌ Info: Epoch 1666
└ @ Main /Users/kcft114/.julia/packages/Flux/qXNjB/src/optimise/train.jl:105
┌ Info: Epoch 1667
└ @ Main /Users/kcft114/.julia/packages/Flux/qXNjB/src/optimise/train.jl:105
┌ Info: Epoch 1668
└ @ Main /Users/kcft114/.julia/packages/Flux/qXNjB/src/optimise/train.jl:105
┌ Info: Epoch 1669
└ @ Main /Users/kcft114/.julia/packages/Flux/qXNjB/src/optimise/train.jl:105
┌ Info: Epoch 1670
└ @ Main /Users/kcft114/.julia/packages/Flux/qXNjB/src/optimise/train.jl:105
┌ Info: Epoch 1671
└ @ Main /Users/kcft114/.julia/packages/Flux/qXNjB/src/optimise/train.jl:105
┌ Info: Epoch 1672
└ @ Main /Users/kcft114/.julia/packages/Flux/qXNjB/src/optimise/train.jl:105
┌ Info: Epoch 1673
└ @ Main /Users/kcft114/.julia/packages/Flux/qXNjB/src/optimise/train.jl:105
┌ Info: Epoch 1674
└ @ Main /Users/kcft114/.julia/packages/Flux/qXNjB/src/optimise/train.jl:105
┌ Info: Epoch 1675
└ @ Main /Users/kcft114/.julia/packages/Flux/qXNjB/src/optimise/train.jl:105
┌ Info: Epoch 1676
└ @ Main /Users/kcft114/.julia/packages/Flux/qXNjB/src/optimise/train.jl:105
┌ Info: Epoch 1677
└ @ Main /Users/kcft114/.julia/packages/Flux/qXNjB/src/optimise/train.jl:105
┌ Info: Epoch 1678
└ @ Main /Users/kcft114/.julia/packages/Flux/qXNjB/src/optimise/train.jl:105
┌ Info: Epoch 1679
└ @ Main /Users/kcft114/.julia/packages/Flux/qXNjB/src/optimise/train.jl:105
```

```
loss(X, Y) = 0.6122293f0 (tracked)
loss(X, Y) = 0.61313903f0 (tracked)
loss(X, Y) = 0.5716195f0 (tracked)
loss(X, Y) = 0.58293337f0 (tracked)
loss(X, Y) = 0.5595803f0 (tracked)
loss(X, Y) = 0.57201904f0 (tracked)
loss(X, Y) = 0.5765407f0 (tracked)
loss(X, Y) = 0.5682413f0 (tracked)
loss(X, Y) = 0.5809502f0 (tracked)
loss(X, Y) = 0.5768004f0 (tracked)
loss(X, Y) = 0.5691236f0 (tracked)
loss(X, Y) = 0.57014877f0 (tracked)
loss(X, Y) = 0.6078463f0 (tracked)
loss(X, Y) = 0.6168567f0 (tracked)
loss(X, Y) = 0.6351762f0 (tracked)
loss(X, Y) = 0.6021023f0 (tracked)
loss(X, Y) = 0.6217576f0 (tracked)
loss(X, Y) = 0.6025807f0 (tracked)
loss(X, Y) = 0.57461923f0 (tracked)
loss(X, Y) = 0.5789958f0 (tracked)
loss(X, Y) = 0.5563594f0 (tracked)
loss(X, Y) = 0.6150642f0 (tracked)
loss(X, Y) = 0.5487784f0 (tracked)
loss(X, Y) = 0.6124406f0 (tracked)
loss(X, Y) = 0.602256f0 (tracked)
loss(X, Y) = 0.54991883f0 (tracked)
loss(X, Y) = 0.63324195f0 (tracked)
loss(X, Y) = 0.59845877f0 (tracked)
loss(X, Y) = 0.6016636f0 (tracked)
loss(X, Y) = 0.56027645f0 (tracked)
loss(X, Y) = 0.5945918f0 (tracked)
loss(X, Y) = 0.5735155f0 (tracked)
loss(X, Y) = 0.60034674f0 (tracked)
loss(X, Y) = 0.5850193f0 (tracked)
loss(X, Y) = 0.60778135f0 (tracked)
loss(X, Y) = 0.57182795f0 (tracked)
loss(X, Y) = 0.5999773f0 (tracked)
loss(X, Y) = 0.5899615f0 (tracked)
loss(X, Y) = 0.5523743f0 (tracked)
loss(X, Y) = 0.59528446f0 (tracked)
loss(X, Y) = 0.5810351f0 (tracked)
loss(X, Y) = 0.58949965f0 (tracked)
loss(X, Y) = 0.57519126f0 (tracked)
loss(X, Y) = 0.5909097f0 (tracked)
loss(X, Y) = 0.5595017f0 (tracked)
loss(X, Y) = 0.5864904f0 (tracked)
loss(X, Y) = 0.5616782f0 (tracked)
loss(X, Y) = 0.54475653f0 (tracked)
loss(X, Y) = 0.5600272f0 (tracked)
loss(X, Y) = 0.56674707f0 (tracked)
loss(X, Y) = 0.61999655f0 (tracked)
loss(X, Y) = 0.5507797f0 (tracked)
loss(X, Y) = 0.5741348f0 (tracked)
loss(X, Y) = 0.5805508f0 (tracked)
loss(X, Y) = 0.6230081f0 (tracked)
loss(X, Y) = 0.5734544f0 (tracked)
loss(X, Y) = 0.589037f0 (tracked)
loss(X, Y) = 0.65263426f0 (tracked)
loss(X, Y) = 0.57135826f0 (tracked)
loss(X, Y) = 0.57593185f0 (tracked)
loss(X, Y) = 0.57167685f0 (tracked)
loss(X, Y) = 0.5772657f0 (tracked)
loss(X, Y) = 0.58728987f0 (tracked)
loss(X, Y) = 0.5766025f0 (tracked)
loss(X, Y) = 0.5579392f0 (tracked)
loss(X, Y) = 0.62133795f0 (tracked)
loss(X, Y) = 0.5783697f0 (tracked)
loss(X, Y) = 0.5807509f0 (tracked)
loss(X, Y) = 0.564285f0 (tracked)
loss(X, Y) = 0.5839893f0 (tracked)
loss(X, Y) = 0.558118f0 (tracked)
loss(X, Y) = 0.54874825f0 (tracked)
loss(X, Y) = 0.6514926f0 (tracked)
loss(X, Y) = 0.55948955f0 (tracked)
loss(X, Y) = 0.60274315f0 (tracked)
loss(X, Y) = 0.59180266f0 (tracked)
loss(X, Y) = 0.5504696f0 (tracked)
loss(X, Y) = 0.56178033f0 (tracked)
loss(X, Y) = 0.59248227f0 (tracked)
loss(X, Y) = 0.6064298f0 (tracked)
loss(X, Y) = 0.6113405f0 (tracked)
loss(X, Y) = 0.58342326f0 (tracked)
loss(X, Y) = 0.5434726f0 (tracked)
loss(X, Y) = 0.61018836f0 (tracked)
loss(X, Y) = 0.54424924f0 (tracked)
loss(X, Y) = 0.576647f0 (tracked)
loss(X, Y) = 0.553283f0 (tracked)
loss(X, Y) = 0.6008659f0 (tracked)
loss(X, Y) = 0.57297015f0 (tracked)
loss(X, Y) = 0.6197977f0 (tracked)
loss(X, Y) = 0.57422423f0 (tracked)
loss(X, Y) = 0.581446f0 (tracked)
loss(X, Y) = 0.5864971f0 (tracked)
loss(X, Y) = 0.56549203f0 (tracked)
loss(X, Y) = 0.5850217f0 (tracked)
loss(X, Y) = 0.62801373f0 (tracked)
loss(X, Y) = 0.55068505f0 (tracked)
loss(X, Y) = 0.5437788f0 (tracked)
loss(X, Y) = 0.5614204f0 (tracked)
loss(X, Y) = 0.5954191f0 (tracked)
loss(X, Y) = 0.5398421f0 (tracked)
loss(X, Y) = 0.6012662f0 (tracked)
loss(X, Y) = 0.5876521f0 (tracked)
loss(X, Y) = 0.58171713f0 (tracked)
loss(X, Y) = 0.63670444f0 (tracked)
```

```
┌ Info: Epoch 1680
└ @ Main /Users/kcft114/.julia/packages/Flux/qXNjB/src/optimise/train.jl:105
┌ Info: Epoch 1681
└ @ Main /Users/kcft114/.julia/packages/Flux/qXNjB/src/optimise/train.jl:105
┌ Info: Epoch 1682
└ @ Main /Users/kcft114/.julia/packages/Flux/qXNjB/src/optimise/train.jl:105
┌ Info: Epoch 1683
└ @ Main /Users/kcft114/.julia/packages/Flux/qXNjB/src/optimise/train.jl:105
┌ Info: Epoch 1684
└ @ Main /Users/kcft114/.julia/packages/Flux/qXNjB/src/optimise/train.jl:105
┌ Info: Epoch 1685
└ @ Main /Users/kcft114/.julia/packages/Flux/qXNjB/src/optimise/train.jl:105
┌ Info: Epoch 1686
└ @ Main /Users/kcft114/.julia/packages/Flux/qXNjB/src/optimise/train.jl:105
┌ Info: Epoch 1687
└ @ Main /Users/kcft114/.julia/packages/Flux/qXNjB/src/optimise/train.jl:105
┌ Info: Epoch 1688
└ @ Main /Users/kcft114/.julia/packages/Flux/qXNjB/src/optimise/train.jl:105
┌ Info: Epoch 1689
└ @ Main /Users/kcft114/.julia/packages/Flux/qXNjB/src/optimise/train.jl:105
┌ Info: Epoch 1690
└ @ Main /Users/kcft114/.julia/packages/Flux/qXNjB/src/optimise/train.jl:105
┌ Info: Epoch 1691
└ @ Main /Users/kcft114/.julia/packages/Flux/qXNjB/src/optimise/train.jl:105
┌ Info: Epoch 1692
└ @ Main /Users/kcft114/.julia/packages/Flux/qXNjB/src/optimise/train.jl:105
┌ Info: Epoch 1693
└ @ Main /Users/kcft114/.julia/packages/Flux/qXNjB/src/optimise/train.jl:105
┌ Info: Epoch 1694
└ @ Main /Users/kcft114/.julia/packages/Flux/qXNjB/src/optimise/train.jl:105
┌ Info: Epoch 1695
└ @ Main /Users/kcft114/.julia/packages/Flux/qXNjB/src/optimise/train.jl:105
┌ Info: Epoch 1696
└ @ Main /Users/kcft114/.julia/packages/Flux/qXNjB/src/optimise/train.jl:105
┌ Info: Epoch 1697
└ @ Main /Users/kcft114/.julia/packages/Flux/qXNjB/src/optimise/train.jl:105
┌ Info: Epoch 1698
└ @ Main /Users/kcft114/.julia/packages/Flux/qXNjB/src/optimise/train.jl:105
┌ Info: Epoch 1699
└ @ Main /Users/kcft114/.julia/packages/Flux/qXNjB/src/optimise/train.jl:105
┌ Info: Epoch 1700
└ @ Main /Users/kcft114/.julia/packages/Flux/qXNjB/src/optimise/train.jl:105
┌ Info: Epoch 1701
└ @ Main /Users/kcft114/.julia/packages/Flux/qXNjB/src/optimise/train.jl:105
┌ Info: Epoch 1702
└ @ Main /Users/kcft114/.julia/packages/Flux/qXNjB/src/optimise/train.jl:105
┌ Info: Epoch 1703
└ @ Main /Users/kcft114/.julia/packages/Flux/qXNjB/src/optimise/train.jl:105
┌ Info: Epoch 1704
└ @ Main /Users/kcft114/.julia/packages/Flux/qXNjB/src/optimise/train.jl:105
┌ Info: Epoch 1705
└ @ Main /Users/kcft114/.julia/packages/Flux/qXNjB/src/optimise/train.jl:105
┌ Info: Epoch 1706
└ @ Main /Users/kcft114/.julia/packages/Flux/qXNjB/src/optimise/train.jl:105
┌ Info: Epoch 1707
└ @ Main /Users/kcft114/.julia/packages/Flux/qXNjB/src/optimise/train.jl:105
┌ Info: Epoch 1708
└ @ Main /Users/kcft114/.julia/packages/Flux/qXNjB/src/optimise/train.jl:105
┌ Info: Epoch 1709
└ @ Main /Users/kcft114/.julia/packages/Flux/qXNjB/src/optimise/train.jl:105
┌ Info: Epoch 1710
└ @ Main /Users/kcft114/.julia/packages/Flux/qXNjB/src/optimise/train.jl:105
┌ Info: Epoch 1711
└ @ Main /Users/kcft114/.julia/packages/Flux/qXNjB/src/optimise/train.jl:105
┌ Info: Epoch 1712
└ @ Main /Users/kcft114/.julia/packages/Flux/qXNjB/src/optimise/train.jl:105
┌ Info: Epoch 1713
└ @ Main /Users/kcft114/.julia/packages/Flux/qXNjB/src/optimise/train.jl:105
┌ Info: Epoch 1714
└ @ Main /Users/kcft114/.julia/packages/Flux/qXNjB/src/optimise/train.jl:105
┌ Info: Epoch 1715
└ @ Main /Users/kcft114/.julia/packages/Flux/qXNjB/src/optimise/train.jl:105
┌ Info: Epoch 1716
└ @ Main /Users/kcft114/.julia/packages/Flux/qXNjB/src/optimise/train.jl:105
┌ Info: Epoch 1717
└ @ Main /Users/kcft114/.julia/packages/Flux/qXNjB/src/optimise/train.jl:105
┌ Info: Epoch 1718
└ @ Main /Users/kcft114/.julia/packages/Flux/qXNjB/src/optimise/train.jl:105
┌ Info: Epoch 1719
└ @ Main /Users/kcft114/.julia/packages/Flux/qXNjB/src/optimise/train.jl:105
┌ Info: Epoch 1720
└ @ Main /Users/kcft114/.julia/packages/Flux/qXNjB/src/optimise/train.jl:105
┌ Info: Epoch 1721
└ @ Main /Users/kcft114/.julia/packages/Flux/qXNjB/src/optimise/train.jl:105
┌ Info: Epoch 1722
└ @ Main /Users/kcft114/.julia/packages/Flux/qXNjB/src/optimise/train.jl:105
┌ Info: Epoch 1723
└ @ Main /Users/kcft114/.julia/packages/Flux/qXNjB/src/optimise/train.jl:105
┌ Info: Epoch 1724
└ @ Main /Users/kcft114/.julia/packages/Flux/qXNjB/src/optimise/train.jl:105
┌ Info: Epoch 1725
└ @ Main /Users/kcft114/.julia/packages/Flux/qXNjB/src/optimise/train.jl:105
┌ Info: Epoch 1726
└ @ Main /Users/kcft114/.julia/packages/Flux/qXNjB/src/optimise/train.jl:105
┌ Info: Epoch 1727
└ @ Main /Users/kcft114/.julia/packages/Flux/qXNjB/src/optimise/train.jl:105
┌ Info: Epoch 1728
└ @ Main /Users/kcft114/.julia/packages/Flux/qXNjB/src/optimise/train.jl:105
┌ Info: Epoch 1729
└ @ Main /Users/kcft114/.julia/packages/Flux/qXNjB/src/optimise/train.jl:105
┌ Info: Epoch 1730
└ @ Main /Users/kcft114/.julia/packages/Flux/qXNjB/src/optimise/train.jl:105
┌ Info: Epoch 1731
└ @ Main /Users/kcft114/.julia/packages/Flux/qXNjB/src/optimise/train.jl:105
┌ Info: Epoch 1732
└ @ Main /Users/kcft114/.julia/packages/Flux/qXNjB/src/optimise/train.jl:105
┌ Info: Epoch 1733
└ @ Main /Users/kcft114/.julia/packages/Flux/qXNjB/src/optimise/train.jl:105
┌ Info: Epoch 1734
└ @ Main /Users/kcft114/.julia/packages/Flux/qXNjB/src/optimise/train.jl:105
┌ Info: Epoch 1735
└ @ Main /Users/kcft114/.julia/packages/Flux/qXNjB/src/optimise/train.jl:105
┌ Info: Epoch 1736
└ @ Main /Users/kcft114/.julia/packages/Flux/qXNjB/src/optimise/train.jl:105
┌ Info: Epoch 1737
└ @ Main /Users/kcft114/.julia/packages/Flux/qXNjB/src/optimise/train.jl:105
┌ Info: Epoch 1738
└ @ Main /Users/kcft114/.julia/packages/Flux/qXNjB/src/optimise/train.jl:105
┌ Info: Epoch 1739
└ @ Main /Users/kcft114/.julia/packages/Flux/qXNjB/src/optimise/train.jl:105
┌ Info: Epoch 1740
└ @ Main /Users/kcft114/.julia/packages/Flux/qXNjB/src/optimise/train.jl:105
┌ Info: Epoch 1741
└ @ Main /Users/kcft114/.julia/packages/Flux/qXNjB/src/optimise/train.jl:105
┌ Info: Epoch 1742
└ @ Main /Users/kcft114/.julia/packages/Flux/qXNjB/src/optimise/train.jl:105
┌ Info: Epoch 1743
└ @ Main /Users/kcft114/.julia/packages/Flux/qXNjB/src/optimise/train.jl:105
┌ Info: Epoch 1744
└ @ Main /Users/kcft114/.julia/packages/Flux/qXNjB/src/optimise/train.jl:105
┌ Info: Epoch 1745
└ @ Main /Users/kcft114/.julia/packages/Flux/qXNjB/src/optimise/train.jl:105
┌ Info: Epoch 1746
└ @ Main /Users/kcft114/.julia/packages/Flux/qXNjB/src/optimise/train.jl:105
┌ Info: Epoch 1747
└ @ Main /Users/kcft114/.julia/packages/Flux/qXNjB/src/optimise/train.jl:105
┌ Info: Epoch 1748
└ @ Main /Users/kcft114/.julia/packages/Flux/qXNjB/src/optimise/train.jl:105
┌ Info: Epoch 1749
└ @ Main /Users/kcft114/.julia/packages/Flux/qXNjB/src/optimise/train.jl:105
┌ Info: Epoch 1750
└ @ Main /Users/kcft114/.julia/packages/Flux/qXNjB/src/optimise/train.jl:105
┌ Info: Epoch 1751
└ @ Main /Users/kcft114/.julia/packages/Flux/qXNjB/src/optimise/train.jl:105
┌ Info: Epoch 1752
└ @ Main /Users/kcft114/.julia/packages/Flux/qXNjB/src/optimise/train.jl:105
┌ Info: Epoch 1753
└ @ Main /Users/kcft114/.julia/packages/Flux/qXNjB/src/optimise/train.jl:105
┌ Info: Epoch 1754
└ @ Main /Users/kcft114/.julia/packages/Flux/qXNjB/src/optimise/train.jl:105
┌ Info: Epoch 1755
└ @ Main /Users/kcft114/.julia/packages/Flux/qXNjB/src/optimise/train.jl:105
┌ Info: Epoch 1756
└ @ Main /Users/kcft114/.julia/packages/Flux/qXNjB/src/optimise/train.jl:105
┌ Info: Epoch 1757
└ @ Main /Users/kcft114/.julia/packages/Flux/qXNjB/src/optimise/train.jl:105
┌ Info: Epoch 1758
└ @ Main /Users/kcft114/.julia/packages/Flux/qXNjB/src/optimise/train.jl:105
┌ Info: Epoch 1759
└ @ Main /Users/kcft114/.julia/packages/Flux/qXNjB/src/optimise/train.jl:105
┌ Info: Epoch 1760
└ @ Main /Users/kcft114/.julia/packages/Flux/qXNjB/src/optimise/train.jl:105
┌ Info: Epoch 1761
└ @ Main /Users/kcft114/.julia/packages/Flux/qXNjB/src/optimise/train.jl:105
┌ Info: Epoch 1762
└ @ Main /Users/kcft114/.julia/packages/Flux/qXNjB/src/optimise/train.jl:105
┌ Info: Epoch 1763
└ @ Main /Users/kcft114/.julia/packages/Flux/qXNjB/src/optimise/train.jl:105
┌ Info: Epoch 1764
└ @ Main /Users/kcft114/.julia/packages/Flux/qXNjB/src/optimise/train.jl:105
┌ Info: Epoch 1765
└ @ Main /Users/kcft114/.julia/packages/Flux/qXNjB/src/optimise/train.jl:105
┌ Info: Epoch 1766
└ @ Main /Users/kcft114/.julia/packages/Flux/qXNjB/src/optimise/train.jl:105
┌ Info: Epoch 1767
└ @ Main /Users/kcft114/.julia/packages/Flux/qXNjB/src/optimise/train.jl:105
┌ Info: Epoch 1768
└ @ Main /Users/kcft114/.julia/packages/Flux/qXNjB/src/optimise/train.jl:105
┌ Info: Epoch 1769
└ @ Main /Users/kcft114/.julia/packages/Flux/qXNjB/src/optimise/train.jl:105
┌ Info: Epoch 1770
└ @ Main /Users/kcft114/.julia/packages/Flux/qXNjB/src/optimise/train.jl:105
┌ Info: Epoch 1771
└ @ Main /Users/kcft114/.julia/packages/Flux/qXNjB/src/optimise/train.jl:105
┌ Info: Epoch 1772
└ @ Main /Users/kcft114/.julia/packages/Flux/qXNjB/src/optimise/train.jl:105
┌ Info: Epoch 1773
└ @ Main /Users/kcft114/.julia/packages/Flux/qXNjB/src/optimise/train.jl:105
┌ Info: Epoch 1774
└ @ Main /Users/kcft114/.julia/packages/Flux/qXNjB/src/optimise/train.jl:105
┌ Info: Epoch 1775
└ @ Main /Users/kcft114/.julia/packages/Flux/qXNjB/src/optimise/train.jl:105
┌ Info: Epoch 1776
└ @ Main /Users/kcft114/.julia/packages/Flux/qXNjB/src/optimise/train.jl:105
┌ Info: Epoch 1777
└ @ Main /Users/kcft114/.julia/packages/Flux/qXNjB/src/optimise/train.jl:105
┌ Info: Epoch 1778
└ @ Main /Users/kcft114/.julia/packages/Flux/qXNjB/src/optimise/train.jl:105
┌ Info: Epoch 1779
└ @ Main /Users/kcft114/.julia/packages/Flux/qXNjB/src/optimise/train.jl:105
┌ Info: Epoch 1780
└ @ Main /Users/kcft114/.julia/packages/Flux/qXNjB/src/optimise/train.jl:105
┌ Info: Epoch 1781
└ @ Main /Users/kcft114/.julia/packages/Flux/qXNjB/src/optimise/train.jl:105
┌ Info: Epoch 1782
└ @ Main /Users/kcft114/.julia/packages/Flux/qXNjB/src/optimise/train.jl:105
┌ Info: Epoch 1783
└ @ Main /Users/kcft114/.julia/packages/Flux/qXNjB/src/optimise/train.jl:105
┌ Info: Epoch 1784
└ @ Main /Users/kcft114/.julia/packages/Flux/qXNjB/src/optimise/train.jl:105
```

```
loss(X, Y) = 0.6087769f0 (tracked)
loss(X, Y) = 0.57817787f0 (tracked)
loss(X, Y) = 0.56418145f0 (tracked)
loss(X, Y) = 0.5698657f0 (tracked)
loss(X, Y) = 0.55719525f0 (tracked)
loss(X, Y) = 0.5826232f0 (tracked)
loss(X, Y) = 0.56152755f0 (tracked)
loss(X, Y) = 0.550301f0 (tracked)
loss(X, Y) = 0.5747475f0 (tracked)
loss(X, Y) = 0.5812243f0 (tracked)
loss(X, Y) = 0.55625665f0 (tracked)
loss(X, Y) = 0.58164936f0 (tracked)
loss(X, Y) = 0.5748991f0 (tracked)
loss(X, Y) = 0.56106526f0 (tracked)
loss(X, Y) = 0.5850826f0 (tracked)
loss(X, Y) = 0.557883f0 (tracked)
loss(X, Y) = 0.62918246f0 (tracked)
loss(X, Y) = 0.572292f0 (tracked)
loss(X, Y) = 0.5832315f0 (tracked)
loss(X, Y) = 0.5510777f0 (tracked)
loss(X, Y) = 0.55407083f0 (tracked)
loss(X, Y) = 0.5716451f0 (tracked)
loss(X, Y) = 0.60161877f0 (tracked)
loss(X, Y) = 0.5965039f0 (tracked)
loss(X, Y) = 0.57607913f0 (tracked)
loss(X, Y) = 0.60864633f0 (tracked)
loss(X, Y) = 0.5522997f0 (tracked)
loss(X, Y) = 0.5582816f0 (tracked)
loss(X, Y) = 0.5846598f0 (tracked)
loss(X, Y) = 0.57613856f0 (tracked)
loss(X, Y) = 0.5696285f0 (tracked)
loss(X, Y) = 0.60838234f0 (tracked)
loss(X, Y) = 0.5853387f0 (tracked)
loss(X, Y) = 0.59633845f0 (tracked)
loss(X, Y) = 0.5574718f0 (tracked)
loss(X, Y) = 0.53505874f0 (tracked)
loss(X, Y) = 0.5616112f0 (tracked)
loss(X, Y) = 0.5719944f0 (tracked)
loss(X, Y) = 0.5605314f0 (tracked)
loss(X, Y) = 0.5713115f0 (tracked)
loss(X, Y) = 0.5825594f0 (tracked)
loss(X, Y) = 0.6123417f0 (tracked)
loss(X, Y) = 0.5990066f0 (tracked)
loss(X, Y) = 0.545265f0 (tracked)
loss(X, Y) = 0.5771256f0 (tracked)
loss(X, Y) = 0.5970912f0 (tracked)
loss(X, Y) = 0.563134f0 (tracked)
loss(X, Y) = 0.5573326f0 (tracked)
loss(X, Y) = 0.5871947f0 (tracked)
loss(X, Y) = 0.5579035f0 (tracked)
loss(X, Y) = 0.5890942f0 (tracked)
loss(X, Y) = 0.5659854f0 (tracked)
loss(X, Y) = 0.59079313f0 (tracked)
loss(X, Y) = 0.5819968f0 (tracked)
loss(X, Y) = 0.5406468f0 (tracked)
loss(X, Y) = 0.5670439f0 (tracked)
loss(X, Y) = 0.5649629f0 (tracked)
loss(X, Y) = 0.54736644f0 (tracked)
loss(X, Y) = 0.6160095f0 (tracked)
loss(X, Y) = 0.5226951f0 (tracked)
loss(X, Y) = 0.59508973f0 (tracked)
loss(X, Y) = 0.5858697f0 (tracked)
loss(X, Y) = 0.5857252f0 (tracked)
loss(X, Y) = 0.54463893f0 (tracked)
loss(X, Y) = 0.5659421f0 (tracked)
loss(X, Y) = 0.5876072f0 (tracked)
loss(X, Y) = 0.59968174f0 (tracked)
loss(X, Y) = 0.5485953f0 (tracked)
loss(X, Y) = 0.5899976f0 (tracked)
loss(X, Y) = 0.5554502f0 (tracked)
loss(X, Y) = 0.55735284f0 (tracked)
loss(X, Y) = 0.54236555f0 (tracked)
loss(X, Y) = 0.6416947f0 (tracked)
loss(X, Y) = 0.5429419f0 (tracked)
loss(X, Y) = 0.5190532f0 (tracked)
loss(X, Y) = 0.5857589f0 (tracked)
loss(X, Y) = 0.588074f0 (tracked)
loss(X, Y) = 0.5365791f0 (tracked)
loss(X, Y) = 0.5729009f0 (tracked)
loss(X, Y) = 0.59439445f0 (tracked)
loss(X, Y) = 0.55788094f0 (tracked)
loss(X, Y) = 0.5375739f0 (tracked)
loss(X, Y) = 0.5691319f0 (tracked)
loss(X, Y) = 0.5795059f0 (tracked)
loss(X, Y) = 0.6029161f0 (tracked)
loss(X, Y) = 0.58543205f0 (tracked)
loss(X, Y) = 0.5430142f0 (tracked)
loss(X, Y) = 0.5421662f0 (tracked)
loss(X, Y) = 0.6177355f0 (tracked)
loss(X, Y) = 0.5889621f0 (tracked)
loss(X, Y) = 0.595718f0 (tracked)
loss(X, Y) = 0.6128009f0 (tracked)
loss(X, Y) = 0.61008364f0 (tracked)
loss(X, Y) = 0.60008556f0 (tracked)
loss(X, Y) = 0.5677688f0 (tracked)
loss(X, Y) = 0.57292306f0 (tracked)
loss(X, Y) = 0.6360126f0 (tracked)
loss(X, Y) = 0.5776093f0 (tracked)
loss(X, Y) = 0.59821624f0 (tracked)
loss(X, Y) = 0.5688033f0 (tracked)
loss(X, Y) = 0.5818693f0 (tracked)
loss(X, Y) = 0.5794205f0 (tracked)
loss(X, Y) = 0.6180373f0 (tracked)
loss(X, Y) = 0.5719078f0 (tracked)
loss(X, Y) = 0.5562393f0 (tracked)
loss(X, Y) = 0.55684865f0 (tracked)
loss(X, Y) = 0.53954273f0 (tracked)
loss(X, Y) = 0.5778591f0 (tracked)
```

```
┌ Info: Epoch 1785
└ @ Main /Users/kcft114/.julia/packages/Flux/qXNjB/src/optimise/train.jl:105
┌ Info: Epoch 1786
└ @ Main /Users/kcft114/.julia/packages/Flux/qXNjB/src/optimise/train.jl:105
┌ Info: Epoch 1787
└ @ Main /Users/kcft114/.julia/packages/Flux/qXNjB/src/optimise/train.jl:105
┌ Info: Epoch 1788
└ @ Main /Users/kcft114/.julia/packages/Flux/qXNjB/src/optimise/train.jl:105
┌ Info: Epoch 1789
└ @ Main /Users/kcft114/.julia/packages/Flux/qXNjB/src/optimise/train.jl:105
Excessive output truncated after 524301 bytes.
```

```
loss(X, Y) = 0.5960616f0 (tracked)
loss(X, Y) = 0.55993843f0 (tracked)
loss(X, Y) = 0.5712862f0 (tracked)
loss(X, Y) = 0.56111336f0 (tracked)
loss(X, Y) = 0.57090145f0 (tracked)
loss(X, Y) = 0.5803066f0 (tracked)
loss(X, Y) = 0.5156924f0 (tracked)
loss(X, Y) = 0.55351335f0 (tracked)
loss(X, Y) = 0.5479171f0 (tracked)
loss(X, Y) = 0.521396f0 (tracked)
loss(X, Y) = 0.56771517f0 (tracked)
loss(X, Y) = 0.5995986f0 (tracked)
loss(X, Y) = 0.53819674f0 (tracked)
loss(X, Y) = 0.54017943f0 (tracked)
loss(X, Y) = 0.5748983f0 (tracked)
loss(X, Y) = 0.60641795f0 (tracked)
loss(X, Y) = 0.582727f0 (tracked)
loss(X, Y) = 0.55142754f0 (tracked)
loss(X, Y) = 0.5825109f0 (tracked)
loss(X, Y) = 0.5706214f0 (tracked)
loss(X, Y) = 0.52800757f0 (tracked)
loss(X, Y) = 0.56999755f0 (tracked)
loss(X, Y) = 0.60998744f0 (tracked)
loss(X, Y) = 0.56740785f0 (tracked)
loss(X, Y) = 0.58124775f0 (tracked)
loss(X, Y) = 0.5797334f0 (tracked)
loss(X, Y) = 0.5583267f0 (tracked)
loss(X, Y) = 0.5724017f0 (tracked)
loss(X, Y) = 0.5811451f0 (tracked)
loss(X, Y) = 0.5633649f0 (tracked)
loss(X, Y) = 0.5570208f0 (tracked)
loss(X, Y) = 0.5753409f0 (tracked)
loss(X, Y) = 0.5700249f0 (tracked)
loss(X, Y) = 0.57959855f0 (tracked)
loss(X, Y) = 0.5841565f0 (tracked)
loss(X, Y) = 0.5089195f0 (tracked)
loss(X, Y) = 0.547034f0 (tracked)
loss(X, Y) = 0.5655582f0 (tracked)
loss(X, Y) = 0.58361095f0 (tracked)
loss(X, Y) = 0.58106375f0 (tracked)
loss(X, Y) = 0.5631057f0 (tracked)
loss(X, Y) = 0.5602211f0 (tracked)
loss(X, Y) = 0.56315786f0 (tracked)
loss(X, Y) = 0.6178744f0 (tracked)
loss(X, Y) = 0.6012879f0 (tracked)
loss(X, Y) = 0.54092455f0 (tracked)
loss(X, Y) = 0.58230895f0 (tracked)
loss(X, Y) = 0.5586653f0 (tracked)
loss(X, Y) = 0.5636808f0 (tracked)
loss(X, Y) = 0.63843226f0 (tracked)
loss(X, Y) = 0.54798573f0 (tracked)
loss(X, Y) = 0.5841104f0 (tracked)
loss(X, Y) = 0.5853926f0 (tracked)
loss(X, Y) = 0.5352356f0 (tracked)
loss(X, Y) = 0.5563442f0 (tracked)
loss(X, Y) = 0.56778866f0 (tracked)
loss(X, Y) = 0.58771527f0 (tracked)
loss(X, Y) = 0.5465077f0 (tracked)
loss(X, Y) = 0.54295456f0 (tracked)
loss(X, Y) = 0.53277105f0 (tracked)
loss(X, Y) = 0.5358535f0 (tracked)
loss(X, Y) = 0.559613f0 (tracked)
loss(X, Y) = 0.592009f0 (tracked)
loss(X, Y) = 0.60132337f0 (tracked)
loss(X, Y) = 0.5500893f0 (tracked)
loss(X, Y) = 0.5254765f0 (tracked)
loss(X, Y) = 0.5672923f0 (tracked)
loss(X, Y) = 0.5732325f0 (tracked)
loss(X, Y) = 0.58006036f0 (tracked)
loss(X, Y) = 0.5361335f0 (tracked)
loss(X, Y) = 0.531928f0 (tracked)
loss(X, Y) = 0.5235769f0 (tracked)
loss(X, Y) = 0.5577219f0 (tracked)
loss(X, Y) = 0.5819551f0 (tracked)
loss(X, Y) = 0.5449086f0 (tracked)
loss(X, Y) = 0.54256403f0 (tracked)
loss(X, Y) = 0.58178514f0 (tracked)
loss(X, Y) = 0.49799025f0 (tracked)
loss(X, Y) = 0.5639648f0 (tracked)
loss(X, Y) = 0.5865064f0 (tracked)
loss(X, Y) = 0.56966937f0 (tracked)
loss(X, Y) = 0.5772064f0 (tracked)
loss(X, Y) = 0.5879902f0 (tracked)
loss(X, Y) = 0.582337f0 (tracked)
loss(X, Y) = 0.58467025f0 (tracked)
loss(X, Y) = 0.56690073f0 (tracked)
loss(X, Y) = 0.57402986f0 (tracked)
loss(X, Y) = 0.5982362f0 (tracked)
loss(X, Y) = 0.55619377f0 (tracked)
loss(X, Y) = 0.537548f0 (tracked)
loss(X, Y) = 0.5407651f0 (tracked)
loss(X, Y) = 0.5643661f0 (tracked)
loss(X, Y) = 0.5793595f0 (tracked)
loss(X, Y) = 0.5784386f0 (tracked)
loss(X, Y) = 0.54244983f0 (tracked)
loss(X, Y) = 0.57728124f0 (tracked)
loss(X, Y) = 0.5452228f0 (tracked)
loss(X, Y) = 0.53987014f0 (tracked)
loss(X, Y) = 0.592807f0 (tracked)
loss(X, Y) = 0.55019706f0 (tracked)
loss(X, Y) = 0.60871255f0 (tracked)
loss(X, Y) = 0.50114965f0 (tracked)
```

```
┌ Info: Epoch 1790
└ @ Main /Users/kcft114/.julia/packages/Flux/qXNjB/src/optimise/train.jl:105
┌ Info: Epoch 1791
└ @ Main /Users/kcft114/.julia/packages/Flux/qXNjB/src/optimise/train.jl:105
┌ Info: Epoch 1792
└ @ Main /Users/kcft114/.julia/packages/Flux/qXNjB/src/optimise/train.jl:105
┌ Info: Epoch 1793
└ @ Main /Users/kcft114/.julia/packages/Flux/qXNjB/src/optimise/train.jl:105
┌ Info: Epoch 1794
└ @ Main /Users/kcft114/.julia/packages/Flux/qXNjB/src/optimise/train.jl:105
┌ Info: Epoch 1795
└ @ Main /Users/kcft114/.julia/packages/Flux/qXNjB/src/optimise/train.jl:105
┌ Info: Epoch 1796
└ @ Main /Users/kcft114/.julia/packages/Flux/qXNjB/src/optimise/train.jl:105
┌ Info: Epoch 1797
└ @ Main /Users/kcft114/.julia/packages/Flux/qXNjB/src/optimise/train.jl:105
┌ Info: Epoch 1798
└ @ Main /Users/kcft114/.julia/packages/Flux/qXNjB/src/optimise/train.jl:105
┌ Info: Epoch 1799
└ @ Main /Users/kcft114/.julia/packages/Flux/qXNjB/src/optimise/train.jl:105
┌ Info: Epoch 1800
└ @ Main /Users/kcft114/.julia/packages/Flux/qXNjB/src/optimise/train.jl:105
┌ Info: Epoch 1801
└ @ Main /Users/kcft114/.julia/packages/Flux/qXNjB/src/optimise/train.jl:105
┌ Info: Epoch 1802
└ @ Main /Users/kcft114/.julia/packages/Flux/qXNjB/src/optimise/train.jl:105
┌ Info: Epoch 1803
└ @ Main /Users/kcft114/.julia/packages/Flux/qXNjB/src/optimise/train.jl:105
┌ Info: Epoch 1804
└ @ Main /Users/kcft114/.julia/packages/Flux/qXNjB/src/optimise/train.jl:105
┌ Info: Epoch 1805
└ @ Main /Users/kcft114/.julia/packages/Flux/qXNjB/src/optimise/train.jl:105
┌ Info: Epoch 1806
└ @ Main /Users/kcft114/.julia/packages/Flux/qXNjB/src/optimise/train.jl:105
┌ Info: Epoch 1807
└ @ Main /Users/kcft114/.julia/packages/Flux/qXNjB/src/optimise/train.jl:105
┌ Info: Epoch 1808
└ @ Main /Users/kcft114/.julia/packages/Flux/qXNjB/src/optimise/train.jl:105
┌ Info: Epoch 1809
└ @ Main /Users/kcft114/.julia/packages/Flux/qXNjB/src/optimise/train.jl:105
┌ Info: Epoch 1810
└ @ Main /Users/kcft114/.julia/packages/Flux/qXNjB/src/optimise/train.jl:105
┌ Info: Epoch 1811
└ @ Main /Users/kcft114/.julia/packages/Flux/qXNjB/src/optimise/train.jl:105
┌ Info: Epoch 1812
└ @ Main /Users/kcft114/.julia/packages/Flux/qXNjB/src/optimise/train.jl:105
┌ Info: Epoch 1813
└ @ Main /Users/kcft114/.julia/packages/Flux/qXNjB/src/optimise/train.jl:105
┌ Info: Epoch 1814
└ @ Main /Users/kcft114/.julia/packages/Flux/qXNjB/src/optimise/train.jl:105
┌ Info: Epoch 1815
└ @ Main /Users/kcft114/.julia/packages/Flux/qXNjB/src/optimise/train.jl:105
┌ Info: Epoch 1816
└ @ Main /Users/kcft114/.julia/packages/Flux/qXNjB/src/optimise/train.jl:105
┌ Info: Epoch 1817
└ @ Main /Users/kcft114/.julia/packages/Flux/qXNjB/src/optimise/train.jl:105
┌ Info: Epoch 1818
└ @ Main /Users/kcft114/.julia/packages/Flux/qXNjB/src/optimise/train.jl:105
┌ Info: Epoch 1819
└ @ Main /Users/kcft114/.julia/packages/Flux/qXNjB/src/optimise/train.jl:105
┌ Info: Epoch 1820
└ @ Main /Users/kcft114/.julia/packages/Flux/qXNjB/src/optimise/train.jl:105
┌ Info: Epoch 1821
└ @ Main /Users/kcft114/.julia/packages/Flux/qXNjB/src/optimise/train.jl:105
┌ Info: Epoch 1822
└ @ Main /Users/kcft114/.julia/packages/Flux/qXNjB/src/optimise/train.jl:105
┌ Info: Epoch 1823
└ @ Main /Users/kcft114/.julia/packages/Flux/qXNjB/src/optimise/train.jl:105
┌ Info: Epoch 1824
└ @ Main /Users/kcft114/.julia/packages/Flux/qXNjB/src/optimise/train.jl:105
┌ Info: Epoch 1825
└ @ Main /Users/kcft114/.julia/packages/Flux/qXNjB/src/optimise/train.jl:105
┌ Info: Epoch 1826
└ @ Main /Users/kcft114/.julia/packages/Flux/qXNjB/src/optimise/train.jl:105
┌ Info: Epoch 1827
└ @ Main /Users/kcft114/.julia/packages/Flux/qXNjB/src/optimise/train.jl:105
┌ Info: Epoch 1828
└ @ Main /Users/kcft114/.julia/packages/Flux/qXNjB/src/optimise/train.jl:105
┌ Info: Epoch 1829
└ @ Main /Users/kcft114/.julia/packages/Flux/qXNjB/src/optimise/train.jl:105
┌ Info: Epoch 1830
└ @ Main /Users/kcft114/.julia/packages/Flux/qXNjB/src/optimise/train.jl:105
┌ Info: Epoch 1831
└ @ Main /Users/kcft114/.julia/packages/Flux/qXNjB/src/optimise/train.jl:105
┌ Info: Epoch 1832
└ @ Main /Users/kcft114/.julia/packages/Flux/qXNjB/src/optimise/train.jl:105
┌ Info: Epoch 1833
└ @ Main /Users/kcft114/.julia/packages/Flux/qXNjB/src/optimise/train.jl:105
┌ Info: Epoch 1834
└ @ Main /Users/kcft114/.julia/packages/Flux/qXNjB/src/optimise/train.jl:105
┌ Info: Epoch 1835
└ @ Main /Users/kcft114/.julia/packages/Flux/qXNjB/src/optimise/train.jl:105
┌ Info: Epoch 1836
└ @ Main /Users/kcft114/.julia/packages/Flux/qXNjB/src/optimise/train.jl:105
┌ Info: Epoch 1837
└ @ Main /Users/kcft114/.julia/packages/Flux/qXNjB/src/optimise/train.jl:105
┌ Info: Epoch 1838
└ @ Main /Users/kcft114/.julia/packages/Flux/qXNjB/src/optimise/train.jl:105
┌ Info: Epoch 1839
└ @ Main /Users/kcft114/.julia/packages/Flux/qXNjB/src/optimise/train.jl:105
┌ Info: Epoch 1840
└ @ Main /Users/kcft114/.julia/packages/Flux/qXNjB/src/optimise/train.jl:105
┌ Info: Epoch 1841
└ @ Main /Users/kcft114/.julia/packages/Flux/qXNjB/src/optimise/train.jl:105
┌ Info: Epoch 1842
└ @ Main /Users/kcft114/.julia/packages/Flux/qXNjB/src/optimise/train.jl:105
┌ Info: Epoch 1843
└ @ Main /Users/kcft114/.julia/packages/Flux/qXNjB/src/optimise/train.jl:105
┌ Info: Epoch 1844
└ @ Main /Users/kcft114/.julia/packages/Flux/qXNjB/src/optimise/train.jl:105
┌ Info: Epoch 1845
└ @ Main /Users/kcft114/.julia/packages/Flux/qXNjB/src/optimise/train.jl:105
┌ Info: Epoch 1846
└ @ Main /Users/kcft114/.julia/packages/Flux/qXNjB/src/optimise/train.jl:105
┌ Info: Epoch 1847
└ @ Main /Users/kcft114/.julia/packages/Flux/qXNjB/src/optimise/train.jl:105
┌ Info: Epoch 1848
└ @ Main /Users/kcft114/.julia/packages/Flux/qXNjB/src/optimise/train.jl:105
┌ Info: Epoch 1849
└ @ Main /Users/kcft114/.julia/packages/Flux/qXNjB/src/optimise/train.jl:105
┌ Info: Epoch 1850
└ @ Main /Users/kcft114/.julia/packages/Flux/qXNjB/src/optimise/train.jl:105
┌ Info: Epoch 1851
└ @ Main /Users/kcft114/.julia/packages/Flux/qXNjB/src/optimise/train.jl:105
┌ Info: Epoch 1852
└ @ Main /Users/kcft114/.julia/packages/Flux/qXNjB/src/optimise/train.jl:105
┌ Info: Epoch 1853
└ @ Main /Users/kcft114/.julia/packages/Flux/qXNjB/src/optimise/train.jl:105
┌ Info: Epoch 1854
└ @ Main /Users/kcft114/.julia/packages/Flux/qXNjB/src/optimise/train.jl:105
┌ Info: Epoch 1855
└ @ Main /Users/kcft114/.julia/packages/Flux/qXNjB/src/optimise/train.jl:105
┌ Info: Epoch 1856
└ @ Main /Users/kcft114/.julia/packages/Flux/qXNjB/src/optimise/train.jl:105
┌ Info: Epoch 1857
└ @ Main /Users/kcft114/.julia/packages/Flux/qXNjB/src/optimise/train.jl:105
┌ Info: Epoch 1858
└ @ Main /Users/kcft114/.julia/packages/Flux/qXNjB/src/optimise/train.jl:105
┌ Info: Epoch 1859
└ @ Main /Users/kcft114/.julia/packages/Flux/qXNjB/src/optimise/train.jl:105
┌ Info: Epoch 1860
└ @ Main /Users/kcft114/.julia/packages/Flux/qXNjB/src/optimise/train.jl:105
┌ Info: Epoch 1861
└ @ Main /Users/kcft114/.julia/packages/Flux/qXNjB/src/optimise/train.jl:105
┌ Info: Epoch 1862
└ @ Main /Users/kcft114/.julia/packages/Flux/qXNjB/src/optimise/train.jl:105
┌ Info: Epoch 1863
└ @ Main /Users/kcft114/.julia/packages/Flux/qXNjB/src/optimise/train.jl:105
┌ Info: Epoch 1864
└ @ Main /Users/kcft114/.julia/packages/Flux/qXNjB/src/optimise/train.jl:105
┌ Info: Epoch 1865
└ @ Main /Users/kcft114/.julia/packages/Flux/qXNjB/src/optimise/train.jl:105
┌ Info: Epoch 1866
└ @ Main /Users/kcft114/.julia/packages/Flux/qXNjB/src/optimise/train.jl:105
┌ Info: Epoch 1867
└ @ Main /Users/kcft114/.julia/packages/Flux/qXNjB/src/optimise/train.jl:105
┌ Info: Epoch 1868
└ @ Main /Users/kcft114/.julia/packages/Flux/qXNjB/src/optimise/train.jl:105
┌ Info: Epoch 1869
└ @ Main /Users/kcft114/.julia/packages/Flux/qXNjB/src/optimise/train.jl:105
┌ Info: Epoch 1870
└ @ Main /Users/kcft114/.julia/packages/Flux/qXNjB/src/optimise/train.jl:105
┌ Info: Epoch 1871
└ @ Main /Users/kcft114/.julia/packages/Flux/qXNjB/src/optimise/train.jl:105
┌ Info: Epoch 1872
└ @ Main /Users/kcft114/.julia/packages/Flux/qXNjB/src/optimise/train.jl:105
┌ Info: Epoch 1873
└ @ Main /Users/kcft114/.julia/packages/Flux/qXNjB/src/optimise/train.jl:105
┌ Info: Epoch 1874
└ @ Main /Users/kcft114/.julia/packages/Flux/qXNjB/src/optimise/train.jl:105
┌ Info: Epoch 1875
└ @ Main /Users/kcft114/.julia/packages/Flux/qXNjB/src/optimise/train.jl:105
┌ Info: Epoch 1876
└ @ Main /Users/kcft114/.julia/packages/Flux/qXNjB/src/optimise/train.jl:105
┌ Info: Epoch 1877
└ @ Main /Users/kcft114/.julia/packages/Flux/qXNjB/src/optimise/train.jl:105
┌ Info: Epoch 1878
└ @ Main /Users/kcft114/.julia/packages/Flux/qXNjB/src/optimise/train.jl:105
┌ Info: Epoch 1879
└ @ Main /Users/kcft114/.julia/packages/Flux/qXNjB/src/optimise/train.jl:105
┌ Info: Epoch 1880
└ @ Main /Users/kcft114/.julia/packages/Flux/qXNjB/src/optimise/train.jl:105
┌ Info: Epoch 1881
└ @ Main /Users/kcft114/.julia/packages/Flux/qXNjB/src/optimise/train.jl:105
┌ Info: Epoch 1882
└ @ Main /Users/kcft114/.julia/packages/Flux/qXNjB/src/optimise/train.jl:105
┌ Info: Epoch 1883
└ @ Main /Users/kcft114/.julia/packages/Flux/qXNjB/src/optimise/train.jl:105
┌ Info: Epoch 1884
└ @ Main /Users/kcft114/.julia/packages/Flux/qXNjB/src/optimise/train.jl:105
┌ Info: Epoch 1885
└ @ Main /Users/kcft114/.julia/packages/Flux/qXNjB/src/optimise/train.jl:105
┌ Info: Epoch 1886
└ @ Main /Users/kcft114/.julia/packages/Flux/qXNjB/src/optimise/train.jl:105
┌ Info: Epoch 1887
└ @ Main /Users/kcft114/.julia/packages/Flux/qXNjB/src/optimise/train.jl:105
┌ Info: Epoch 1888
└ @ Main /Users/kcft114/.julia/packages/Flux/qXNjB/src/optimise/train.jl:105
┌ Info: Epoch 1889
└ @ Main /Users/kcft114/.julia/packages/Flux/qXNjB/src/optimise/train.jl:105
┌ Info: Epoch 1890
└ @ Main /Users/kcft114/.julia/packages/Flux/qXNjB/src/optimise/train.jl:105
```

In [15]:

```
p_dropout=0.3

n_exper = 100
experiments_03 = zeros(n_exper, 4)

for i=1:n_exper
    
    nn = Chain(
              Dense(8, n_h, relu),
              Dropout(p_dropout),
              Dense(n_h, 3),
              softmax)
    
    loss(x, y) = crossentropy(nn(x), y)
    accuracy(x, y) = mean(onecold(nn(x)) .== onecold(y))

    dataset = [(X, Y)]
    evalcb = () -> @show(loss(X, Y))
    
    opt = ADAM(0.001)
    
    @epochs 2000 Flux.train!(loss, Flux.params(nn), dataset, opt, cb = throttle(evalcb, 10));
    
    ac_train, ac_test, ba_train, ba_test =  res(X, Y, Xt, Yt);
    experiments_03[i,:] = [ac_train, ac_test, ba_train, ba_test]
end
```

```
┌ Info: Epoch 1
└ @ Main /Users/kcft114/.julia/packages/Flux/qXNjB/src/optimise/train.jl:105
```

```
loss(X, Y) = 1.4015378f0 (tracked)
loss(X, Y) = 1.4150095f0 (tracked)
loss(X, Y) = 1.39594f0 (tracked)
loss(X, Y) = 1.4572552f0 (tracked)
loss(X, Y) = 1.3832972f0 (tracked)
loss(X, Y) = 1.3883202f0 (tracked)
loss(X, Y) = 1.3929083f0 (tracked)
loss(X, Y) = 1.4198076f0 (tracked)
loss(X, Y) = 1.3640277f0 (tracked)
loss(X, Y) = 1.393755f0 (tracked)
loss(X, Y) = 1.383044f0 (tracked)
loss(X, Y) = 1.3541162f0 (tracked)
loss(X, Y) = 1.3175077f0 (tracked)
loss(X, Y) = 1.3832573f0 (tracked)
loss(X, Y) = 1.3374555f0 (tracked)
loss(X, Y) = 1.3434267f0 (tracked)
loss(X, Y) = 1.286337f0 (tracked)
loss(X, Y) = 1.358889f0 (tracked)
loss(X, Y) = 1.3483764f0 (tracked)
loss(X, Y) = 1.3131049f0 (tracked)
loss(X, Y) = 1.3057462f0 (tracked)
loss(X, Y) = 1.2772628f0 (tracked)
loss(X, Y) = 1.2866695f0 (tracked)
loss(X, Y) = 1.252662f0 (tracked)
loss(X, Y) = 1.2530156f0 (tracked)
loss(X, Y) = 1.2982914f0 (tracked)
loss(X, Y) = 1.2626579f0 (tracked)
loss(X, Y) = 1.2717128f0 (tracked)
loss(X, Y) = 1.2690954f0 (tracked)
loss(X, Y) = 1.3182354f0 (tracked)
loss(X, Y) = 1.2250469f0 (tracked)
loss(X, Y) = 1.2371523f0 (tracked)
loss(X, Y) = 1.2862895f0 (tracked)
loss(X, Y) = 1.2044495f0 (tracked)
loss(X, Y) = 1.2486861f0 (tracked)
loss(X, Y) = 1.2404298f0 (tracked)
loss(X, Y) = 1.2226276f0 (tracked)
loss(X, Y) = 1.2123513f0 (tracked)
loss(X, Y) = 1.2271961f0 (tracked)
loss(X, Y) = 1.2360799f0 (tracked)
loss(X, Y) = 1.2071867f0 (tracked)
loss(X, Y) = 1.1871043f0 (tracked)
loss(X, Y) = 1.2046088f0 (tracked)
loss(X, Y) = 1.1556908f0 (tracked)
loss(X, Y) = 1.1770164f0 (tracked)
loss(X, Y) = 1.1760759f0 (tracked)
loss(X, Y) = 1.1475737f0 (tracked)
loss(X, Y) = 1.1585138f0 (tracked)
loss(X, Y) = 1.1760298f0 (tracked)
loss(X, Y) = 1.1781244f0 (tracked)
loss(X, Y) = 1.1477431f0 (tracked)
loss(X, Y) = 1.161496f0 (tracked)
loss(X, Y) = 1.2213855f0 (tracked)
loss(X, Y) = 1.1383508f0 (tracked)
loss(X, Y) = 1.172035f0 (tracked)
loss(X, Y) = 1.1527739f0 (tracked)
loss(X, Y) = 1.1253704f0 (tracked)
loss(X, Y) = 1.1464657f0 (tracked)
loss(X, Y) = 1.1384068f0 (tracked)
loss(X, Y) = 1.1420487f0 (tracked)
loss(X, Y) = 1.1588707f0 (tracked)
loss(X, Y) = 1.117178f0 (tracked)
loss(X, Y) = 1.125533f0 (tracked)
loss(X, Y) = 1.160956f0 (tracked)
loss(X, Y) = 1.0933455f0 (tracked)
loss(X, Y) = 1.0996755f0 (tracked)
loss(X, Y) = 1.1206036f0 (tracked)
loss(X, Y) = 1.1143405f0 (tracked)
loss(X, Y) = 1.1615052f0 (tracked)
loss(X, Y) = 1.0728835f0 (tracked)
loss(X, Y) = 1.1151575f0 (tracked)
loss(X, Y) = 1.0703918f0 (tracked)
loss(X, Y) = 1.1037776f0 (tracked)
loss(X, Y) = 1.1291155f0 (tracked)
loss(X, Y) = 1.0941086f0 (tracked)
loss(X, Y) = 1.0872505f0 (tracked)
loss(X, Y) = 1.0483531f0 (tracked)
loss(X, Y) = 1.0642141f0 (tracked)
loss(X, Y) = 1.1068438f0 (tracked)
loss(X, Y) = 1.0850729f0 (tracked)
loss(X, Y) = 1.0677036f0 (tracked)
loss(X, Y) = 1.0960513f0 (tracked)
loss(X, Y) = 1.1036851f0 (tracked)
loss(X, Y) = 1.1091121f0 (tracked)
loss(X, Y) = 1.0769372f0 (tracked)
loss(X, Y) = 1.0825312f0 (tracked)
loss(X, Y) = 1.0981294f0 (tracked)
loss(X, Y) = 1.0499734f0 (tracked)
loss(X, Y) = 1.0671715f0 (tracked)
loss(X, Y) = 1.0282348f0 (tracked)
loss(X, Y) = 1.1130214f0 (tracked)
loss(X, Y) = 1.0590674f0 (tracked)
loss(X, Y) = 1.0093794f0 (tracked)
loss(X, Y) = 1.04696f0 (tracked)
loss(X, Y) = 1.0547296f0 (tracked)
loss(X, Y) = 1.0357682f0 (tracked)
loss(X, Y) = 1.0320774f0 (tracked)
loss(X, Y) = 1.0542581f0 (tracked)
loss(X, Y) = 1.0617315f0 (tracked)
```

```
┌ Info: Epoch 2
└ @ Main /Users/kcft114/.julia/packages/Flux/qXNjB/src/optimise/train.jl:105
┌ Info: Epoch 3
└ @ Main /Users/kcft114/.julia/packages/Flux/qXNjB/src/optimise/train.jl:105
┌ Info: Epoch 4
└ @ Main /Users/kcft114/.julia/packages/Flux/qXNjB/src/optimise/train.jl:105
┌ Info: Epoch 5
└ @ Main /Users/kcft114/.julia/packages/Flux/qXNjB/src/optimise/train.jl:105
┌ Info: Epoch 6
└ @ Main /Users/kcft114/.julia/packages/Flux/qXNjB/src/optimise/train.jl:105
┌ Info: Epoch 7
└ @ Main /Users/kcft114/.julia/packages/Flux/qXNjB/src/optimise/train.jl:105
┌ Info: Epoch 8
└ @ Main /Users/kcft114/.julia/packages/Flux/qXNjB/src/optimise/train.jl:105
┌ Info: Epoch 9
└ @ Main /Users/kcft114/.julia/packages/Flux/qXNjB/src/optimise/train.jl:105
┌ Info: Epoch 10
└ @ Main /Users/kcft114/.julia/packages/Flux/qXNjB/src/optimise/train.jl:105
┌ Info: Epoch 11
└ @ Main /Users/kcft114/.julia/packages/Flux/qXNjB/src/optimise/train.jl:105
┌ Info: Epoch 12
└ @ Main /Users/kcft114/.julia/packages/Flux/qXNjB/src/optimise/train.jl:105
┌ Info: Epoch 13
└ @ Main /Users/kcft114/.julia/packages/Flux/qXNjB/src/optimise/train.jl:105
┌ Info: Epoch 14
└ @ Main /Users/kcft114/.julia/packages/Flux/qXNjB/src/optimise/train.jl:105
┌ Info: Epoch 15
└ @ Main /Users/kcft114/.julia/packages/Flux/qXNjB/src/optimise/train.jl:105
┌ Info: Epoch 16
└ @ Main /Users/kcft114/.julia/packages/Flux/qXNjB/src/optimise/train.jl:105
┌ Info: Epoch 17
└ @ Main /Users/kcft114/.julia/packages/Flux/qXNjB/src/optimise/train.jl:105
┌ Info: Epoch 18
└ @ Main /Users/kcft114/.julia/packages/Flux/qXNjB/src/optimise/train.jl:105
┌ Info: Epoch 19
└ @ Main /Users/kcft114/.julia/packages/Flux/qXNjB/src/optimise/train.jl:105
┌ Info: Epoch 20
└ @ Main /Users/kcft114/.julia/packages/Flux/qXNjB/src/optimise/train.jl:105
┌ Info: Epoch 21
└ @ Main /Users/kcft114/.julia/packages/Flux/qXNjB/src/optimise/train.jl:105
┌ Info: Epoch 22
└ @ Main /Users/kcft114/.julia/packages/Flux/qXNjB/src/optimise/train.jl:105
┌ Info: Epoch 23
└ @ Main /Users/kcft114/.julia/packages/Flux/qXNjB/src/optimise/train.jl:105
┌ Info: Epoch 24
└ @ Main /Users/kcft114/.julia/packages/Flux/qXNjB/src/optimise/train.jl:105
┌ Info: Epoch 25
└ @ Main /Users/kcft114/.julia/packages/Flux/qXNjB/src/optimise/train.jl:105
┌ Info: Epoch 26
└ @ Main /Users/kcft114/.julia/packages/Flux/qXNjB/src/optimise/train.jl:105
┌ Info: Epoch 27
└ @ Main /Users/kcft114/.julia/packages/Flux/qXNjB/src/optimise/train.jl:105
┌ Info: Epoch 28
└ @ Main /Users/kcft114/.julia/packages/Flux/qXNjB/src/optimise/train.jl:105
┌ Info: Epoch 29
└ @ Main /Users/kcft114/.julia/packages/Flux/qXNjB/src/optimise/train.jl:105
┌ Info: Epoch 30
└ @ Main /Users/kcft114/.julia/packages/Flux/qXNjB/src/optimise/train.jl:105
┌ Info: Epoch 31
└ @ Main /Users/kcft114/.julia/packages/Flux/qXNjB/src/optimise/train.jl:105
┌ Info: Epoch 32
└ @ Main /Users/kcft114/.julia/packages/Flux/qXNjB/src/optimise/train.jl:105
┌ Info: Epoch 33
└ @ Main /Users/kcft114/.julia/packages/Flux/qXNjB/src/optimise/train.jl:105
┌ Info: Epoch 34
└ @ Main /Users/kcft114/.julia/packages/Flux/qXNjB/src/optimise/train.jl:105
┌ Info: Epoch 35
└ @ Main /Users/kcft114/.julia/packages/Flux/qXNjB/src/optimise/train.jl:105
┌ Info: Epoch 36
└ @ Main /Users/kcft114/.julia/packages/Flux/qXNjB/src/optimise/train.jl:105
┌ Info: Epoch 37
└ @ Main /Users/kcft114/.julia/packages/Flux/qXNjB/src/optimise/train.jl:105
┌ Info: Epoch 38
└ @ Main /Users/kcft114/.julia/packages/Flux/qXNjB/src/optimise/train.jl:105
┌ Info: Epoch 39
└ @ Main /Users/kcft114/.julia/packages/Flux/qXNjB/src/optimise/train.jl:105
┌ Info: Epoch 40
└ @ Main /Users/kcft114/.julia/packages/Flux/qXNjB/src/optimise/train.jl:105
┌ Info: Epoch 41
└ @ Main /Users/kcft114/.julia/packages/Flux/qXNjB/src/optimise/train.jl:105
┌ Info: Epoch 42
└ @ Main /Users/kcft114/.julia/packages/Flux/qXNjB/src/optimise/train.jl:105
┌ Info: Epoch 43
└ @ Main /Users/kcft114/.julia/packages/Flux/qXNjB/src/optimise/train.jl:105
┌ Info: Epoch 44
└ @ Main /Users/kcft114/.julia/packages/Flux/qXNjB/src/optimise/train.jl:105
┌ Info: Epoch 45
└ @ Main /Users/kcft114/.julia/packages/Flux/qXNjB/src/optimise/train.jl:105
┌ Info: Epoch 46
└ @ Main /Users/kcft114/.julia/packages/Flux/qXNjB/src/optimise/train.jl:105
┌ Info: Epoch 47
└ @ Main /Users/kcft114/.julia/packages/Flux/qXNjB/src/optimise/train.jl:105
┌ Info: Epoch 48
└ @ Main /Users/kcft114/.julia/packages/Flux/qXNjB/src/optimise/train.jl:105
┌ Info: Epoch 49
└ @ Main /Users/kcft114/.julia/packages/Flux/qXNjB/src/optimise/train.jl:105
┌ Info: Epoch 50
└ @ Main /Users/kcft114/.julia/packages/Flux/qXNjB/src/optimise/train.jl:105
┌ Info: Epoch 51
└ @ Main /Users/kcft114/.julia/packages/Flux/qXNjB/src/optimise/train.jl:105
┌ Info: Epoch 52
└ @ Main /Users/kcft114/.julia/packages/Flux/qXNjB/src/optimise/train.jl:105
┌ Info: Epoch 53
└ @ Main /Users/kcft114/.julia/packages/Flux/qXNjB/src/optimise/train.jl:105
┌ Info: Epoch 54
└ @ Main /Users/kcft114/.julia/packages/Flux/qXNjB/src/optimise/train.jl:105
┌ Info: Epoch 55
└ @ Main /Users/kcft114/.julia/packages/Flux/qXNjB/src/optimise/train.jl:105
┌ Info: Epoch 56
└ @ Main /Users/kcft114/.julia/packages/Flux/qXNjB/src/optimise/train.jl:105
┌ Info: Epoch 57
└ @ Main /Users/kcft114/.julia/packages/Flux/qXNjB/src/optimise/train.jl:105
┌ Info: Epoch 58
└ @ Main /Users/kcft114/.julia/packages/Flux/qXNjB/src/optimise/train.jl:105
┌ Info: Epoch 59
└ @ Main /Users/kcft114/.julia/packages/Flux/qXNjB/src/optimise/train.jl:105
┌ Info: Epoch 60
└ @ Main /Users/kcft114/.julia/packages/Flux/qXNjB/src/optimise/train.jl:105
┌ Info: Epoch 61
└ @ Main /Users/kcft114/.julia/packages/Flux/qXNjB/src/optimise/train.jl:105
┌ Info: Epoch 62
└ @ Main /Users/kcft114/.julia/packages/Flux/qXNjB/src/optimise/train.jl:105
┌ Info: Epoch 63
└ @ Main /Users/kcft114/.julia/packages/Flux/qXNjB/src/optimise/train.jl:105
┌ Info: Epoch 64
└ @ Main /Users/kcft114/.julia/packages/Flux/qXNjB/src/optimise/train.jl:105
┌ Info: Epoch 65
└ @ Main /Users/kcft114/.julia/packages/Flux/qXNjB/src/optimise/train.jl:105
┌ Info: Epoch 66
└ @ Main /Users/kcft114/.julia/packages/Flux/qXNjB/src/optimise/train.jl:105
┌ Info: Epoch 67
└ @ Main /Users/kcft114/.julia/packages/Flux/qXNjB/src/optimise/train.jl:105
┌ Info: Epoch 68
└ @ Main /Users/kcft114/.julia/packages/Flux/qXNjB/src/optimise/train.jl:105
┌ Info: Epoch 69
└ @ Main /Users/kcft114/.julia/packages/Flux/qXNjB/src/optimise/train.jl:105
┌ Info: Epoch 70
└ @ Main /Users/kcft114/.julia/packages/Flux/qXNjB/src/optimise/train.jl:105
┌ Info: Epoch 71
└ @ Main /Users/kcft114/.julia/packages/Flux/qXNjB/src/optimise/train.jl:105
┌ Info: Epoch 72
└ @ Main /Users/kcft114/.julia/packages/Flux/qXNjB/src/optimise/train.jl:105
┌ Info: Epoch 73
└ @ Main /Users/kcft114/.julia/packages/Flux/qXNjB/src/optimise/train.jl:105
┌ Info: Epoch 74
└ @ Main /Users/kcft114/.julia/packages/Flux/qXNjB/src/optimise/train.jl:105
┌ Info: Epoch 75
└ @ Main /Users/kcft114/.julia/packages/Flux/qXNjB/src/optimise/train.jl:105
┌ Info: Epoch 76
└ @ Main /Users/kcft114/.julia/packages/Flux/qXNjB/src/optimise/train.jl:105
┌ Info: Epoch 77
└ @ Main /Users/kcft114/.julia/packages/Flux/qXNjB/src/optimise/train.jl:105
┌ Info: Epoch 78
└ @ Main /Users/kcft114/.julia/packages/Flux/qXNjB/src/optimise/train.jl:105
┌ Info: Epoch 79
└ @ Main /Users/kcft114/.julia/packages/Flux/qXNjB/src/optimise/train.jl:105
┌ Info: Epoch 80
└ @ Main /Users/kcft114/.julia/packages/Flux/qXNjB/src/optimise/train.jl:105
┌ Info: Epoch 81
└ @ Main /Users/kcft114/.julia/packages/Flux/qXNjB/src/optimise/train.jl:105
┌ Info: Epoch 82
└ @ Main /Users/kcft114/.julia/packages/Flux/qXNjB/src/optimise/train.jl:105
┌ Info: Epoch 83
└ @ Main /Users/kcft114/.julia/packages/Flux/qXNjB/src/optimise/train.jl:105
┌ Info: Epoch 84
└ @ Main /Users/kcft114/.julia/packages/Flux/qXNjB/src/optimise/train.jl:105
┌ Info: Epoch 85
└ @ Main /Users/kcft114/.julia/packages/Flux/qXNjB/src/optimise/train.jl:105
┌ Info: Epoch 86
└ @ Main /Users/kcft114/.julia/packages/Flux/qXNjB/src/optimise/train.jl:105
┌ Info: Epoch 87
└ @ Main /Users/kcft114/.julia/packages/Flux/qXNjB/src/optimise/train.jl:105
┌ Info: Epoch 88
└ @ Main /Users/kcft114/.julia/packages/Flux/qXNjB/src/optimise/train.jl:105
┌ Info: Epoch 89
└ @ Main /Users/kcft114/.julia/packages/Flux/qXNjB/src/optimise/train.jl:105
┌ Info: Epoch 90
└ @ Main /Users/kcft114/.julia/packages/Flux/qXNjB/src/optimise/train.jl:105
┌ Info: Epoch 91
└ @ Main /Users/kcft114/.julia/packages/Flux/qXNjB/src/optimise/train.jl:105
┌ Info: Epoch 92
└ @ Main /Users/kcft114/.julia/packages/Flux/qXNjB/src/optimise/train.jl:105
┌ Info: Epoch 93
└ @ Main /Users/kcft114/.julia/packages/Flux/qXNjB/src/optimise/train.jl:105
┌ Info: Epoch 94
└ @ Main /Users/kcft114/.julia/packages/Flux/qXNjB/src/optimise/train.jl:105
┌ Info: Epoch 95
└ @ Main /Users/kcft114/.julia/packages/Flux/qXNjB/src/optimise/train.jl:105
┌ Info: Epoch 96
└ @ Main /Users/kcft114/.julia/packages/Flux/qXNjB/src/optimise/train.jl:105
┌ Info: Epoch 97
└ @ Main /Users/kcft114/.julia/packages/Flux/qXNjB/src/optimise/train.jl:105
┌ Info: Epoch 98
└ @ Main /Users/kcft114/.julia/packages/Flux/qXNjB/src/optimise/train.jl:105
┌ Info: Epoch 99
└ @ Main /Users/kcft114/.julia/packages/Flux/qXNjB/src/optimise/train.jl:105
┌ Info: Epoch 100
└ @ Main /Users/kcft114/.julia/packages/Flux/qXNjB/src/optimise/train.jl:105
```

```
loss(X, Y) = 1.0426769f0 (tracked)
loss(X, Y) = 1.052208f0 (tracked)
loss(X, Y) = 1.0104861f0 (tracked)
loss(X, Y) = 1.0717872f0 (tracked)
loss(X, Y) = 1.0628359f0 (tracked)
loss(X, Y) = 1.0331645f0 (tracked)
loss(X, Y) = 1.0798447f0 (tracked)
loss(X, Y) = 1.0796754f0 (tracked)
loss(X, Y) = 1.0101264f0 (tracked)
loss(X, Y) = 1.0197371f0 (tracked)
loss(X, Y) = 0.99604505f0 (tracked)
loss(X, Y) = 1.0413122f0 (tracked)
loss(X, Y) = 1.0348955f0 (tracked)
loss(X, Y) = 1.0444361f0 (tracked)
loss(X, Y) = 1.0277448f0 (tracked)
loss(X, Y) = 1.0043896f0 (tracked)
loss(X, Y) = 0.9974853f0 (tracked)
loss(X, Y) = 1.023557f0 (tracked)
loss(X, Y) = 0.9889581f0 (tracked)
loss(X, Y) = 0.999776f0 (tracked)
loss(X, Y) = 0.99360335f0 (tracked)
loss(X, Y) = 1.0150256f0 (tracked)
loss(X, Y) = 1.0272121f0 (tracked)
loss(X, Y) = 1.008893f0 (tracked)
loss(X, Y) = 0.96097153f0 (tracked)
loss(X, Y) = 0.98541826f0 (tracked)
loss(X, Y) = 1.0201582f0 (tracked)
loss(X, Y) = 0.9742531f0 (tracked)
loss(X, Y) = 1.0083301f0 (tracked)
loss(X, Y) = 0.9708524f0 (tracked)
loss(X, Y) = 0.9743122f0 (tracked)
loss(X, Y) = 0.9422764f0 (tracked)
loss(X, Y) = 0.9776928f0 (tracked)
loss(X, Y) = 0.9766117f0 (tracked)
loss(X, Y) = 0.9954465f0 (tracked)
loss(X, Y) = 0.9905516f0 (tracked)
loss(X, Y) = 0.94867086f0 (tracked)
loss(X, Y) = 0.980568f0 (tracked)
loss(X, Y) = 0.9797987f0 (tracked)
loss(X, Y) = 0.98623604f0 (tracked)
loss(X, Y) = 0.9909324f0 (tracked)
loss(X, Y) = 0.9522067f0 (tracked)
loss(X, Y) = 1.005724f0 (tracked)
loss(X, Y) = 1.0225908f0 (tracked)
loss(X, Y) = 0.95729864f0 (tracked)
loss(X, Y) = 0.9871408f0 (tracked)
loss(X, Y) = 1.0164142f0 (tracked)
loss(X, Y) = 1.0297016f0 (tracked)
loss(X, Y) = 0.935324f0 (tracked)
loss(X, Y) = 0.99795747f0 (tracked)
loss(X, Y) = 1.0143099f0 (tracked)
loss(X, Y) = 1.0044863f0 (tracked)
loss(X, Y) = 0.94431144f0 (tracked)
loss(X, Y) = 0.9542808f0 (tracked)
loss(X, Y) = 0.952812f0 (tracked)
loss(X, Y) = 0.9604814f0 (tracked)
loss(X, Y) = 0.9742621f0 (tracked)
loss(X, Y) = 0.92424965f0 (tracked)
loss(X, Y) = 0.946916f0 (tracked)
loss(X, Y) = 0.9344596f0 (tracked)
loss(X, Y) = 0.96294457f0 (tracked)
loss(X, Y) = 0.9787427f0 (tracked)
loss(X, Y) = 0.9686918f0 (tracked)
loss(X, Y) = 0.9448969f0 (tracked)
loss(X, Y) = 0.99337995f0 (tracked)
loss(X, Y) = 0.92758113f0 (tracked)
loss(X, Y) = 0.93516517f0 (tracked)
loss(X, Y) = 0.94423336f0 (tracked)
loss(X, Y) = 0.95680857f0 (tracked)
loss(X, Y) = 0.9374736f0 (tracked)
loss(X, Y) = 0.91867f0 (tracked)
loss(X, Y) = 0.9697691f0 (tracked)
loss(X, Y) = 0.9806023f0 (tracked)
loss(X, Y) = 0.91937524f0 (tracked)
loss(X, Y) = 0.92711186f0 (tracked)
loss(X, Y) = 0.926498f0 (tracked)
loss(X, Y) = 0.957151f0 (tracked)
loss(X, Y) = 0.94757324f0 (tracked)
loss(X, Y) = 0.92880577f0 (tracked)
loss(X, Y) = 0.9594909f0 (tracked)
loss(X, Y) = 0.94247425f0 (tracked)
loss(X, Y) = 0.93399596f0 (tracked)
loss(X, Y) = 0.91138864f0 (tracked)
loss(X, Y) = 0.94118565f0 (tracked)
loss(X, Y) = 0.9345464f0 (tracked)
loss(X, Y) = 0.94063914f0 (tracked)
loss(X, Y) = 0.8960351f0 (tracked)
loss(X, Y) = 0.9308087f0 (tracked)
loss(X, Y) = 0.92805594f0 (tracked)
loss(X, Y) = 0.9021842f0 (tracked)
loss(X, Y) = 0.9154246f0 (tracked)
loss(X, Y) = 0.94456637f0 (tracked)
loss(X, Y) = 0.9432428f0 (tracked)
loss(X, Y) = 0.92999953f0 (tracked)
loss(X, Y) = 0.93090683f0 (tracked)
```

```
┌ Info: Epoch 101
└ @ Main /Users/kcft114/.julia/packages/Flux/qXNjB/src/optimise/train.jl:105
┌ Info: Epoch 102
└ @ Main /Users/kcft114/.julia/packages/Flux/qXNjB/src/optimise/train.jl:105
┌ Info: Epoch 103
└ @ Main /Users/kcft114/.julia/packages/Flux/qXNjB/src/optimise/train.jl:105
┌ Info: Epoch 104
└ @ Main /Users/kcft114/.julia/packages/Flux/qXNjB/src/optimise/train.jl:105
┌ Info: Epoch 105
└ @ Main /Users/kcft114/.julia/packages/Flux/qXNjB/src/optimise/train.jl:105
┌ Info: Epoch 106
└ @ Main /Users/kcft114/.julia/packages/Flux/qXNjB/src/optimise/train.jl:105
┌ Info: Epoch 107
└ @ Main /Users/kcft114/.julia/packages/Flux/qXNjB/src/optimise/train.jl:105
┌ Info: Epoch 108
└ @ Main /Users/kcft114/.julia/packages/Flux/qXNjB/src/optimise/train.jl:105
┌ Info: Epoch 109
└ @ Main /Users/kcft114/.julia/packages/Flux/qXNjB/src/optimise/train.jl:105
┌ Info: Epoch 110
└ @ Main /Users/kcft114/.julia/packages/Flux/qXNjB/src/optimise/train.jl:105
┌ Info: Epoch 111
└ @ Main /Users/kcft114/.julia/packages/Flux/qXNjB/src/optimise/train.jl:105
┌ Info: Epoch 112
└ @ Main /Users/kcft114/.julia/packages/Flux/qXNjB/src/optimise/train.jl:105
┌ Info: Epoch 113
└ @ Main /Users/kcft114/.julia/packages/Flux/qXNjB/src/optimise/train.jl:105
┌ Info: Epoch 114
└ @ Main /Users/kcft114/.julia/packages/Flux/qXNjB/src/optimise/train.jl:105
┌ Info: Epoch 115
└ @ Main /Users/kcft114/.julia/packages/Flux/qXNjB/src/optimise/train.jl:105
┌ Info: Epoch 116
└ @ Main /Users/kcft114/.julia/packages/Flux/qXNjB/src/optimise/train.jl:105
┌ Info: Epoch 117
└ @ Main /Users/kcft114/.julia/packages/Flux/qXNjB/src/optimise/train.jl:105
┌ Info: Epoch 118
└ @ Main /Users/kcft114/.julia/packages/Flux/qXNjB/src/optimise/train.jl:105
┌ Info: Epoch 119
└ @ Main /Users/kcft114/.julia/packages/Flux/qXNjB/src/optimise/train.jl:105
┌ Info: Epoch 120
└ @ Main /Users/kcft114/.julia/packages/Flux/qXNjB/src/optimise/train.jl:105
┌ Info: Epoch 121
└ @ Main /Users/kcft114/.julia/packages/Flux/qXNjB/src/optimise/train.jl:105
┌ Info: Epoch 122
└ @ Main /Users/kcft114/.julia/packages/Flux/qXNjB/src/optimise/train.jl:105
┌ Info: Epoch 123
└ @ Main /Users/kcft114/.julia/packages/Flux/qXNjB/src/optimise/train.jl:105
┌ Info: Epoch 124
└ @ Main /Users/kcft114/.julia/packages/Flux/qXNjB/src/optimise/train.jl:105
┌ Info: Epoch 125
└ @ Main /Users/kcft114/.julia/packages/Flux/qXNjB/src/optimise/train.jl:105
┌ Info: Epoch 126
└ @ Main /Users/kcft114/.julia/packages/Flux/qXNjB/src/optimise/train.jl:105
┌ Info: Epoch 127
└ @ Main /Users/kcft114/.julia/packages/Flux/qXNjB/src/optimise/train.jl:105
┌ Info: Epoch 128
└ @ Main /Users/kcft114/.julia/packages/Flux/qXNjB/src/optimise/train.jl:105
┌ Info: Epoch 129
└ @ Main /Users/kcft114/.julia/packages/Flux/qXNjB/src/optimise/train.jl:105
┌ Info: Epoch 130
└ @ Main /Users/kcft114/.julia/packages/Flux/qXNjB/src/optimise/train.jl:105
┌ Info: Epoch 131
└ @ Main /Users/kcft114/.julia/packages/Flux/qXNjB/src/optimise/train.jl:105
┌ Info: Epoch 132
└ @ Main /Users/kcft114/.julia/packages/Flux/qXNjB/src/optimise/train.jl:105
┌ Info: Epoch 133
└ @ Main /Users/kcft114/.julia/packages/Flux/qXNjB/src/optimise/train.jl:105
┌ Info: Epoch 134
└ @ Main /Users/kcft114/.julia/packages/Flux/qXNjB/src/optimise/train.jl:105
┌ Info: Epoch 135
└ @ Main /Users/kcft114/.julia/packages/Flux/qXNjB/src/optimise/train.jl:105
┌ Info: Epoch 136
└ @ Main /Users/kcft114/.julia/packages/Flux/qXNjB/src/optimise/train.jl:105
┌ Info: Epoch 137
└ @ Main /Users/kcft114/.julia/packages/Flux/qXNjB/src/optimise/train.jl:105
┌ Info: Epoch 138
└ @ Main /Users/kcft114/.julia/packages/Flux/qXNjB/src/optimise/train.jl:105
┌ Info: Epoch 139
└ @ Main /Users/kcft114/.julia/packages/Flux/qXNjB/src/optimise/train.jl:105
┌ Info: Epoch 140
└ @ Main /Users/kcft114/.julia/packages/Flux/qXNjB/src/optimise/train.jl:105
┌ Info: Epoch 141
└ @ Main /Users/kcft114/.julia/packages/Flux/qXNjB/src/optimise/train.jl:105
┌ Info: Epoch 142
└ @ Main /Users/kcft114/.julia/packages/Flux/qXNjB/src/optimise/train.jl:105
┌ Info: Epoch 143
└ @ Main /Users/kcft114/.julia/packages/Flux/qXNjB/src/optimise/train.jl:105
┌ Info: Epoch 144
└ @ Main /Users/kcft114/.julia/packages/Flux/qXNjB/src/optimise/train.jl:105
┌ Info: Epoch 145
└ @ Main /Users/kcft114/.julia/packages/Flux/qXNjB/src/optimise/train.jl:105
┌ Info: Epoch 146
└ @ Main /Users/kcft114/.julia/packages/Flux/qXNjB/src/optimise/train.jl:105
┌ Info: Epoch 147
└ @ Main /Users/kcft114/.julia/packages/Flux/qXNjB/src/optimise/train.jl:105
┌ Info: Epoch 148
└ @ Main /Users/kcft114/.julia/packages/Flux/qXNjB/src/optimise/train.jl:105
┌ Info: Epoch 149
└ @ Main /Users/kcft114/.julia/packages/Flux/qXNjB/src/optimise/train.jl:105
┌ Info: Epoch 150
└ @ Main /Users/kcft114/.julia/packages/Flux/qXNjB/src/optimise/train.jl:105
┌ Info: Epoch 151
└ @ Main /Users/kcft114/.julia/packages/Flux/qXNjB/src/optimise/train.jl:105
┌ Info: Epoch 152
└ @ Main /Users/kcft114/.julia/packages/Flux/qXNjB/src/optimise/train.jl:105
┌ Info: Epoch 153
└ @ Main /Users/kcft114/.julia/packages/Flux/qXNjB/src/optimise/train.jl:105
┌ Info: Epoch 154
└ @ Main /Users/kcft114/.julia/packages/Flux/qXNjB/src/optimise/train.jl:105
┌ Info: Epoch 155
└ @ Main /Users/kcft114/.julia/packages/Flux/qXNjB/src/optimise/train.jl:105
┌ Info: Epoch 156
└ @ Main /Users/kcft114/.julia/packages/Flux/qXNjB/src/optimise/train.jl:105
┌ Info: Epoch 157
└ @ Main /Users/kcft114/.julia/packages/Flux/qXNjB/src/optimise/train.jl:105
┌ Info: Epoch 158
└ @ Main /Users/kcft114/.julia/packages/Flux/qXNjB/src/optimise/train.jl:105
┌ Info: Epoch 159
└ @ Main /Users/kcft114/.julia/packages/Flux/qXNjB/src/optimise/train.jl:105
┌ Info: Epoch 160
└ @ Main /Users/kcft114/.julia/packages/Flux/qXNjB/src/optimise/train.jl:105
┌ Info: Epoch 161
└ @ Main /Users/kcft114/.julia/packages/Flux/qXNjB/src/optimise/train.jl:105
┌ Info: Epoch 162
└ @ Main /Users/kcft114/.julia/packages/Flux/qXNjB/src/optimise/train.jl:105
┌ Info: Epoch 163
└ @ Main /Users/kcft114/.julia/packages/Flux/qXNjB/src/optimise/train.jl:105
┌ Info: Epoch 164
└ @ Main /Users/kcft114/.julia/packages/Flux/qXNjB/src/optimise/train.jl:105
┌ Info: Epoch 165
└ @ Main /Users/kcft114/.julia/packages/Flux/qXNjB/src/optimise/train.jl:105
┌ Info: Epoch 166
└ @ Main /Users/kcft114/.julia/packages/Flux/qXNjB/src/optimise/train.jl:105
┌ Info: Epoch 167
└ @ Main /Users/kcft114/.julia/packages/Flux/qXNjB/src/optimise/train.jl:105
┌ Info: Epoch 168
└ @ Main /Users/kcft114/.julia/packages/Flux/qXNjB/src/optimise/train.jl:105
┌ Info: Epoch 169
└ @ Main /Users/kcft114/.julia/packages/Flux/qXNjB/src/optimise/train.jl:105
┌ Info: Epoch 170
└ @ Main /Users/kcft114/.julia/packages/Flux/qXNjB/src/optimise/train.jl:105
┌ Info: Epoch 171
└ @ Main /Users/kcft114/.julia/packages/Flux/qXNjB/src/optimise/train.jl:105
┌ Info: Epoch 172
└ @ Main /Users/kcft114/.julia/packages/Flux/qXNjB/src/optimise/train.jl:105
┌ Info: Epoch 173
└ @ Main /Users/kcft114/.julia/packages/Flux/qXNjB/src/optimise/train.jl:105
┌ Info: Epoch 174
└ @ Main /Users/kcft114/.julia/packages/Flux/qXNjB/src/optimise/train.jl:105
┌ Info: Epoch 175
└ @ Main /Users/kcft114/.julia/packages/Flux/qXNjB/src/optimise/train.jl:105
┌ Info: Epoch 176
└ @ Main /Users/kcft114/.julia/packages/Flux/qXNjB/src/optimise/train.jl:105
┌ Info: Epoch 177
└ @ Main /Users/kcft114/.julia/packages/Flux/qXNjB/src/optimise/train.jl:105
┌ Info: Epoch 178
└ @ Main /Users/kcft114/.julia/packages/Flux/qXNjB/src/optimise/train.jl:105
┌ Info: Epoch 179
└ @ Main /Users/kcft114/.julia/packages/Flux/qXNjB/src/optimise/train.jl:105
┌ Info: Epoch 180
└ @ Main /Users/kcft114/.julia/packages/Flux/qXNjB/src/optimise/train.jl:105
┌ Info: Epoch 181
└ @ Main /Users/kcft114/.julia/packages/Flux/qXNjB/src/optimise/train.jl:105
┌ Info: Epoch 182
└ @ Main /Users/kcft114/.julia/packages/Flux/qXNjB/src/optimise/train.jl:105
┌ Info: Epoch 183
└ @ Main /Users/kcft114/.julia/packages/Flux/qXNjB/src/optimise/train.jl:105
┌ Info: Epoch 184
└ @ Main /Users/kcft114/.julia/packages/Flux/qXNjB/src/optimise/train.jl:105
┌ Info: Epoch 185
└ @ Main /Users/kcft114/.julia/packages/Flux/qXNjB/src/optimise/train.jl:105
┌ Info: Epoch 186
└ @ Main /Users/kcft114/.julia/packages/Flux/qXNjB/src/optimise/train.jl:105
┌ Info: Epoch 187
└ @ Main /Users/kcft114/.julia/packages/Flux/qXNjB/src/optimise/train.jl:105
┌ Info: Epoch 188
└ @ Main /Users/kcft114/.julia/packages/Flux/qXNjB/src/optimise/train.jl:105
┌ Info: Epoch 189
└ @ Main /Users/kcft114/.julia/packages/Flux/qXNjB/src/optimise/train.jl:105
┌ Info: Epoch 190
└ @ Main /Users/kcft114/.julia/packages/Flux/qXNjB/src/optimise/train.jl:105
┌ Info: Epoch 191
└ @ Main /Users/kcft114/.julia/packages/Flux/qXNjB/src/optimise/train.jl:105
┌ Info: Epoch 192
└ @ Main /Users/kcft114/.julia/packages/Flux/qXNjB/src/optimise/train.jl:105
┌ Info: Epoch 193
└ @ Main /Users/kcft114/.julia/packages/Flux/qXNjB/src/optimise/train.jl:105
┌ Info: Epoch 194
└ @ Main /Users/kcft114/.julia/packages/Flux/qXNjB/src/optimise/train.jl:105
┌ Info: Epoch 195
└ @ Main /Users/kcft114/.julia/packages/Flux/qXNjB/src/optimise/train.jl:105
┌ Info: Epoch 196
└ @ Main /Users/kcft114/.julia/packages/Flux/qXNjB/src/optimise/train.jl:105
┌ Info: Epoch 197
└ @ Main /Users/kcft114/.julia/packages/Flux/qXNjB/src/optimise/train.jl:105
┌ Info: Epoch 198
└ @ Main /Users/kcft114/.julia/packages/Flux/qXNjB/src/optimise/train.jl:105
┌ Info: Epoch 199
└ @ Main /Users/kcft114/.julia/packages/Flux/qXNjB/src/optimise/train.jl:105
┌ Info: Epoch 200
└ @ Main /Users/kcft114/.julia/packages/Flux/qXNjB/src/optimise/train.jl:105
┌ Info: Epoch 201
└ @ Main /Users/kcft114/.julia/packages/Flux/qXNjB/src/optimise/train.jl:105
┌ Info: Epoch 202
└ @ Main /Users/kcft114/.julia/packages/Flux/qXNjB/src/optimise/train.jl:105
┌ Info: Epoch 203
└ @ Main /Users/kcft114/.julia/packages/Flux/qXNjB/src/optimise/train.jl:105
┌ Info: Epoch 204
└ @ Main /Users/kcft114/.julia/packages/Flux/qXNjB/src/optimise/train.jl:105
┌ Info: Epoch 205
└ @ Main /Users/kcft114/.julia/packages/Flux/qXNjB/src/optimise/train.jl:105
┌ Info: Epoch 206
└ @ Main /Users/kcft114/.julia/packages/Flux/qXNjB/src/optimise/train.jl:105
┌ Info: Epoch 207
└ @ Main /Users/kcft114/.julia/packages/Flux/qXNjB/src/optimise/train.jl:105
┌ Info: Epoch 208
└ @ Main /Users/kcft114/.julia/packages/Flux/qXNjB/src/optimise/train.jl:105
┌ Info: Epoch 209
└ @ Main /Users/kcft114/.julia/packages/Flux/qXNjB/src/optimise/train.jl:105
┌ Info: Epoch 210
└ @ Main /Users/kcft114/.julia/packages/Flux/qXNjB/src/optimise/train.jl:105
┌ Info: Epoch 211
└ @ Main /Users/kcft114/.julia/packages/Flux/qXNjB/src/optimise/train.jl:105
┌ Info: Epoch 212
└ @ Main /Users/kcft114/.julia/packages/Flux/qXNjB/src/optimise/train.jl:105
┌ Info: Epoch 213
└ @ Main /Users/kcft114/.julia/packages/Flux/qXNjB/src/optimise/train.jl:105
┌ Info: Epoch 214
└ @ Main /Users/kcft114/.julia/packages/Flux/qXNjB/src/optimise/train.jl:105
┌ Info: Epoch 215
└ @ Main /Users/kcft114/.julia/packages/Flux/qXNjB/src/optimise/train.jl:105
┌ Info: Epoch 216
└ @ Main /Users/kcft114/.julia/packages/Flux/qXNjB/src/optimise/train.jl:105
┌ Info: Epoch 217
└ @ Main /Users/kcft114/.julia/packages/Flux/qXNjB/src/optimise/train.jl:105
┌ Info: Epoch 218
└ @ Main /Users/kcft114/.julia/packages/Flux/qXNjB/src/optimise/train.jl:105
┌ Info: Epoch 219
└ @ Main /Users/kcft114/.julia/packages/Flux/qXNjB/src/optimise/train.jl:105
┌ Info: Epoch 220
└ @ Main /Users/kcft114/.julia/packages/Flux/qXNjB/src/optimise/train.jl:105
┌ Info: Epoch 221
└ @ Main /Users/kcft114/.julia/packages/Flux/qXNjB/src/optimise/train.jl:105
┌ Info: Epoch 222
└ @ Main /Users/kcft114/.julia/packages/Flux/qXNjB/src/optimise/train.jl:105
┌ Info: Epoch 223
└ @ Main /Users/kcft114/.julia/packages/Flux/qXNjB/src/optimise/train.jl:105
┌ Info: Epoch 224
└ @ Main /Users/kcft114/.julia/packages/Flux/qXNjB/src/optimise/train.jl:105
┌ Info: Epoch 225
└ @ Main /Users/kcft114/.julia/packages/Flux/qXNjB/src/optimise/train.jl:105
┌ Info: Epoch 226
└ @ Main /Users/kcft114/.julia/packages/Flux/qXNjB/src/optimise/train.jl:105
┌ Info: Epoch 227
└ @ Main /Users/kcft114/.julia/packages/Flux/qXNjB/src/optimise/train.jl:105
┌ Info: Epoch 228
└ @ Main /Users/kcft114/.julia/packages/Flux/qXNjB/src/optimise/train.jl:105
┌ Info: Epoch 229
└ @ Main /Users/kcft114/.julia/packages/Flux/qXNjB/src/optimise/train.jl:105
┌ Info: Epoch 230
└ @ Main /Users/kcft114/.julia/packages/Flux/qXNjB/src/optimise/train.jl:105
┌ Info: Epoch 231
└ @ Main /Users/kcft114/.julia/packages/Flux/qXNjB/src/optimise/train.jl:105
┌ Info: Epoch 232
└ @ Main /Users/kcft114/.julia/packages/Flux/qXNjB/src/optimise/train.jl:105
┌ Info: Epoch 233
└ @ Main /Users/kcft114/.julia/packages/Flux/qXNjB/src/optimise/train.jl:105
┌ Info: Epoch 234
└ @ Main /Users/kcft114/.julia/packages/Flux/qXNjB/src/optimise/train.jl:105
┌ Info: Epoch 235
└ @ Main /Users/kcft114/.julia/packages/Flux/qXNjB/src/optimise/train.jl:105
┌ Info: Epoch 236
└ @ Main /Users/kcft114/.julia/packages/Flux/qXNjB/src/optimise/train.jl:105
┌ Info: Epoch 237
└ @ Main /Users/kcft114/.julia/packages/Flux/qXNjB/src/optimise/train.jl:105
┌ Info: Epoch 238
└ @ Main /Users/kcft114/.julia/packages/Flux/qXNjB/src/optimise/train.jl:105
┌ Info: Epoch 239
└ @ Main /Users/kcft114/.julia/packages/Flux/qXNjB/src/optimise/train.jl:105
┌ Info: Epoch 240
└ @ Main /Users/kcft114/.julia/packages/Flux/qXNjB/src/optimise/train.jl:105
┌ Info: Epoch 241
└ @ Main /Users/kcft114/.julia/packages/Flux/qXNjB/src/optimise/train.jl:105
┌ Info: Epoch 242
└ @ Main /Users/kcft114/.julia/packages/Flux/qXNjB/src/optimise/train.jl:105
┌ Info: Epoch 243
└ @ Main /Users/kcft114/.julia/packages/Flux/qXNjB/src/optimise/train.jl:105
┌ Info: Epoch 244
└ @ Main /Users/kcft114/.julia/packages/Flux/qXNjB/src/optimise/train.jl:105
┌ Info: Epoch 245
└ @ Main /Users/kcft114/.julia/packages/Flux/qXNjB/src/optimise/train.jl:105
┌ Info: Epoch 246
└ @ Main /Users/kcft114/.julia/packages/Flux/qXNjB/src/optimise/train.jl:105
┌ Info: Epoch 247
└ @ Main /Users/kcft114/.julia/packages/Flux/qXNjB/src/optimise/train.jl:105
┌ Info: Epoch 248
└ @ Main /Users/kcft114/.julia/packages/Flux/qXNjB/src/optimise/train.jl:105
┌ Info: Epoch 249
└ @ Main /Users/kcft114/.julia/packages/Flux/qXNjB/src/optimise/train.jl:105
┌ Info: Epoch 250
└ @ Main /Users/kcft114/.julia/packages/Flux/qXNjB/src/optimise/train.jl:105
┌ Info: Epoch 251
└ @ Main /Users/kcft114/.julia/packages/Flux/qXNjB/src/optimise/train.jl:105
┌ Info: Epoch 252
└ @ Main /Users/kcft114/.julia/packages/Flux/qXNjB/src/optimise/train.jl:105
┌ Info: Epoch 253
└ @ Main /Users/kcft114/.julia/packages/Flux/qXNjB/src/optimise/train.jl:105
┌ Info: Epoch 254
└ @ Main /Users/kcft114/.julia/packages/Flux/qXNjB/src/optimise/train.jl:105
┌ Info: Epoch 255
└ @ Main /Users/kcft114/.julia/packages/Flux/qXNjB/src/optimise/train.jl:105
┌ Info: Epoch 256
└ @ Main /Users/kcft114/.julia/packages/Flux/qXNjB/src/optimise/train.jl:105
┌ Info: Epoch 257
└ @ Main /Users/kcft114/.julia/packages/Flux/qXNjB/src/optimise/train.jl:105
┌ Info: Epoch 258
└ @ Main /Users/kcft114/.julia/packages/Flux/qXNjB/src/optimise/train.jl:105
┌ Info: Epoch 259
└ @ Main /Users/kcft114/.julia/packages/Flux/qXNjB/src/optimise/train.jl:105
┌ Info: Epoch 260
└ @ Main /Users/kcft114/.julia/packages/Flux/qXNjB/src/optimise/train.jl:105
┌ Info: Epoch 261
└ @ Main /Users/kcft114/.julia/packages/Flux/qXNjB/src/optimise/train.jl:105
┌ Info: Epoch 262
└ @ Main /Users/kcft114/.julia/packages/Flux/qXNjB/src/optimise/train.jl:105
┌ Info: Epoch 263
└ @ Main /Users/kcft114/.julia/packages/Flux/qXNjB/src/optimise/train.jl:105
┌ Info: Epoch 264
└ @ Main /Users/kcft114/.julia/packages/Flux/qXNjB/src/optimise/train.jl:105
┌ Info: Epoch 265
└ @ Main /Users/kcft114/.julia/packages/Flux/qXNjB/src/optimise/train.jl:105
┌ Info: Epoch 266
└ @ Main /Users/kcft114/.julia/packages/Flux/qXNjB/src/optimise/train.jl:105
┌ Info: Epoch 267
└ @ Main /Users/kcft114/.julia/packages/Flux/qXNjB/src/optimise/train.jl:105
┌ Info: Epoch 268
└ @ Main /Users/kcft114/.julia/packages/Flux/qXNjB/src/optimise/train.jl:105
┌ Info: Epoch 269
└ @ Main /Users/kcft114/.julia/packages/Flux/qXNjB/src/optimise/train.jl:105
┌ Info: Epoch 270
└ @ Main /Users/kcft114/.julia/packages/Flux/qXNjB/src/optimise/train.jl:105
┌ Info: Epoch 271
└ @ Main /Users/kcft114/.julia/packages/Flux/qXNjB/src/optimise/train.jl:105
┌ Info: Epoch 272
└ @ Main /Users/kcft114/.julia/packages/Flux/qXNjB/src/optimise/train.jl:105
┌ Info: Epoch 273
└ @ Main /Users/kcft114/.julia/packages/Flux/qXNjB/src/optimise/train.jl:105
┌ Info: Epoch 274
└ @ Main /Users/kcft114/.julia/packages/Flux/qXNjB/src/optimise/train.jl:105
┌ Info: Epoch 275
└ @ Main /Users/kcft114/.julia/packages/Flux/qXNjB/src/optimise/train.jl:105
┌ Info: Epoch 276
└ @ Main /Users/kcft114/.julia/packages/Flux/qXNjB/src/optimise/train.jl:105
┌ Info: Epoch 277
└ @ Main /Users/kcft114/.julia/packages/Flux/qXNjB/src/optimise/train.jl:105
┌ Info: Epoch 278
└ @ Main /Users/kcft114/.julia/packages/Flux/qXNjB/src/optimise/train.jl:105
┌ Info: Epoch 279
└ @ Main /Users/kcft114/.julia/packages/Flux/qXNjB/src/optimise/train.jl:105
┌ Info: Epoch 280
└ @ Main /Users/kcft114/.julia/packages/Flux/qXNjB/src/optimise/train.jl:105
┌ Info: Epoch 281
└ @ Main /Users/kcft114/.julia/packages/Flux/qXNjB/src/optimise/train.jl:105
┌ Info: Epoch 282
└ @ Main /Users/kcft114/.julia/packages/Flux/qXNjB/src/optimise/train.jl:105
┌ Info: Epoch 283
└ @ Main /Users/kcft114/.julia/packages/Flux/qXNjB/src/optimise/train.jl:105
┌ Info: Epoch 284
└ @ Main /Users/kcft114/.julia/packages/Flux/qXNjB/src/optimise/train.jl:105
┌ Info: Epoch 285
└ @ Main /Users/kcft114/.julia/packages/Flux/qXNjB/src/optimise/train.jl:105
┌ Info: Epoch 286
└ @ Main /Users/kcft114/.julia/packages/Flux/qXNjB/src/optimise/train.jl:105
┌ Info: Epoch 287
└ @ Main /Users/kcft114/.julia/packages/Flux/qXNjB/src/optimise/train.jl:105
┌ Info: Epoch 288
└ @ Main /Users/kcft114/.julia/packages/Flux/qXNjB/src/optimise/train.jl:105
┌ Info: Epoch 289
└ @ Main /Users/kcft114/.julia/packages/Flux/qXNjB/src/optimise/train.jl:105
┌ Info: Epoch 290
└ @ Main /Users/kcft114/.julia/packages/Flux/qXNjB/src/optimise/train.jl:105
┌ Info: Epoch 291
└ @ Main /Users/kcft114/.julia/packages/Flux/qXNjB/src/optimise/train.jl:105
┌ Info: Epoch 292
└ @ Main /Users/kcft114/.julia/packages/Flux/qXNjB/src/optimise/train.jl:105
┌ Info: Epoch 293
└ @ Main /Users/kcft114/.julia/packages/Flux/qXNjB/src/optimise/train.jl:105
┌ Info: Epoch 294
└ @ Main /Users/kcft114/.julia/packages/Flux/qXNjB/src/optimise/train.jl:105
┌ Info: Epoch 295
└ @ Main /Users/kcft114/.julia/packages/Flux/qXNjB/src/optimise/train.jl:105
┌ Info: Epoch 296
└ @ Main /Users/kcft114/.julia/packages/Flux/qXNjB/src/optimise/train.jl:105
┌ Info: Epoch 297
└ @ Main /Users/kcft114/.julia/packages/Flux/qXNjB/src/optimise/train.jl:105
┌ Info: Epoch 298
└ @ Main /Users/kcft114/.julia/packages/Flux/qXNjB/src/optimise/train.jl:105
```

```
loss(X, Y) = 0.88790643f0 (tracked)
loss(X, Y) = 0.9286911f0 (tracked)
loss(X, Y) = 0.92944187f0 (tracked)
loss(X, Y) = 0.9287223f0 (tracked)
loss(X, Y) = 0.8905263f0 (tracked)
loss(X, Y) = 0.9153553f0 (tracked)
loss(X, Y) = 0.910957f0 (tracked)
loss(X, Y) = 0.8995635f0 (tracked)
loss(X, Y) = 0.9202243f0 (tracked)
loss(X, Y) = 0.9325927f0 (tracked)
loss(X, Y) = 0.9232343f0 (tracked)
loss(X, Y) = 0.8778821f0 (tracked)
loss(X, Y) = 0.90124124f0 (tracked)
loss(X, Y) = 0.8965982f0 (tracked)
loss(X, Y) = 0.9335777f0 (tracked)
loss(X, Y) = 0.89111274f0 (tracked)
loss(X, Y) = 0.89780754f0 (tracked)
loss(X, Y) = 0.9587792f0 (tracked)
loss(X, Y) = 0.89368796f0 (tracked)
loss(X, Y) = 0.8573135f0 (tracked)
loss(X, Y) = 0.87292635f0 (tracked)
loss(X, Y) = 0.937667f0 (tracked)
loss(X, Y) = 0.90031195f0 (tracked)
loss(X, Y) = 0.84533566f0 (tracked)
loss(X, Y) = 0.9133903f0 (tracked)
loss(X, Y) = 0.913899f0 (tracked)
loss(X, Y) = 0.8939537f0 (tracked)
loss(X, Y) = 0.88805664f0 (tracked)
loss(X, Y) = 0.89258265f0 (tracked)
loss(X, Y) = 0.9204196f0 (tracked)
loss(X, Y) = 0.91185015f0 (tracked)
loss(X, Y) = 0.92145604f0 (tracked)
loss(X, Y) = 0.88126624f0 (tracked)
loss(X, Y) = 0.9194093f0 (tracked)
loss(X, Y) = 0.8782507f0 (tracked)
loss(X, Y) = 0.87737995f0 (tracked)
loss(X, Y) = 0.9057361f0 (tracked)
loss(X, Y) = 0.9057896f0 (tracked)
loss(X, Y) = 0.88578534f0 (tracked)
loss(X, Y) = 0.9101969f0 (tracked)
loss(X, Y) = 0.8716062f0 (tracked)
loss(X, Y) = 0.887712f0 (tracked)
loss(X, Y) = 0.87261546f0 (tracked)
loss(X, Y) = 0.853206f0 (tracked)
loss(X, Y) = 0.9062352f0 (tracked)
loss(X, Y) = 0.8916817f0 (tracked)
loss(X, Y) = 0.87655765f0 (tracked)
loss(X, Y) = 0.9175773f0 (tracked)
loss(X, Y) = 0.8848004f0 (tracked)
loss(X, Y) = 0.863041f0 (tracked)
loss(X, Y) = 0.9342768f0 (tracked)
loss(X, Y) = 0.84446174f0 (tracked)
loss(X, Y) = 0.90235835f0 (tracked)
loss(X, Y) = 0.8989537f0 (tracked)
loss(X, Y) = 0.85278875f0 (tracked)
loss(X, Y) = 0.876268f0 (tracked)
loss(X, Y) = 0.89824057f0 (tracked)
loss(X, Y) = 0.8975349f0 (tracked)
loss(X, Y) = 0.8881649f0 (tracked)
loss(X, Y) = 0.8679238f0 (tracked)
loss(X, Y) = 0.85853934f0 (tracked)
loss(X, Y) = 0.88619757f0 (tracked)
loss(X, Y) = 0.8639636f0 (tracked)
loss(X, Y) = 0.8389434f0 (tracked)
loss(X, Y) = 0.85149354f0 (tracked)
loss(X, Y) = 0.8791238f0 (tracked)
loss(X, Y) = 0.8959718f0 (tracked)
loss(X, Y) = 0.9023023f0 (tracked)
loss(X, Y) = 0.8703662f0 (tracked)
loss(X, Y) = 0.8812697f0 (tracked)
loss(X, Y) = 0.8907325f0 (tracked)
loss(X, Y) = 0.87691015f0 (tracked)
loss(X, Y) = 0.9305509f0 (tracked)
loss(X, Y) = 0.8818328f0 (tracked)
loss(X, Y) = 0.8440602f0 (tracked)
loss(X, Y) = 0.8527147f0 (tracked)
loss(X, Y) = 0.8743996f0 (tracked)
loss(X, Y) = 0.85812646f0 (tracked)
loss(X, Y) = 0.87494135f0 (tracked)
loss(X, Y) = 0.8911969f0 (tracked)
loss(X, Y) = 0.86487204f0 (tracked)
loss(X, Y) = 0.85958636f0 (tracked)
loss(X, Y) = 0.8412272f0 (tracked)
loss(X, Y) = 0.85209477f0 (tracked)
loss(X, Y) = 0.8326707f0 (tracked)
loss(X, Y) = 0.8618236f0 (tracked)
loss(X, Y) = 0.8756697f0 (tracked)
loss(X, Y) = 0.8463271f0 (tracked)
loss(X, Y) = 0.84580886f0 (tracked)
loss(X, Y) = 0.83948225f0 (tracked)
loss(X, Y) = 0.86566406f0 (tracked)
loss(X, Y) = 0.85640323f0 (tracked)
loss(X, Y) = 0.8716936f0 (tracked)
loss(X, Y) = 0.83407587f0 (tracked)
loss(X, Y) = 0.82902014f0 (tracked)
loss(X, Y) = 0.82384074f0 (tracked)
loss(X, Y) = 0.8399188f0 (tracked)
loss(X, Y) = 0.875844f0 (tracked)
loss(X, Y) = 0.83422005f0 (tracked)
loss(X, Y) = 0.8314742f0 (tracked)
loss(X, Y) = 0.87907696f0 (tracked)
loss(X, Y) = 0.82709986f0 (tracked)
loss(X, Y) = 0.84948933f0 (tracked)
```

```
┌ Info: Epoch 299
└ @ Main /Users/kcft114/.julia/packages/Flux/qXNjB/src/optimise/train.jl:105
┌ Info: Epoch 300
└ @ Main /Users/kcft114/.julia/packages/Flux/qXNjB/src/optimise/train.jl:105
┌ Info: Epoch 301
└ @ Main /Users/kcft114/.julia/packages/Flux/qXNjB/src/optimise/train.jl:105
┌ Info: Epoch 302
└ @ Main /Users/kcft114/.julia/packages/Flux/qXNjB/src/optimise/train.jl:105
┌ Info: Epoch 303
└ @ Main /Users/kcft114/.julia/packages/Flux/qXNjB/src/optimise/train.jl:105
┌ Info: Epoch 304
└ @ Main /Users/kcft114/.julia/packages/Flux/qXNjB/src/optimise/train.jl:105
┌ Info: Epoch 305
└ @ Main /Users/kcft114/.julia/packages/Flux/qXNjB/src/optimise/train.jl:105
┌ Info: Epoch 306
└ @ Main /Users/kcft114/.julia/packages/Flux/qXNjB/src/optimise/train.jl:105
┌ Info: Epoch 307
└ @ Main /Users/kcft114/.julia/packages/Flux/qXNjB/src/optimise/train.jl:105
┌ Info: Epoch 308
└ @ Main /Users/kcft114/.julia/packages/Flux/qXNjB/src/optimise/train.jl:105
┌ Info: Epoch 309
└ @ Main /Users/kcft114/.julia/packages/Flux/qXNjB/src/optimise/train.jl:105
┌ Info: Epoch 310
└ @ Main /Users/kcft114/.julia/packages/Flux/qXNjB/src/optimise/train.jl:105
┌ Info: Epoch 311
└ @ Main /Users/kcft114/.julia/packages/Flux/qXNjB/src/optimise/train.jl:105
┌ Info: Epoch 312
└ @ Main /Users/kcft114/.julia/packages/Flux/qXNjB/src/optimise/train.jl:105
┌ Info: Epoch 313
└ @ Main /Users/kcft114/.julia/packages/Flux/qXNjB/src/optimise/train.jl:105
┌ Info: Epoch 314
└ @ Main /Users/kcft114/.julia/packages/Flux/qXNjB/src/optimise/train.jl:105
┌ Info: Epoch 315
└ @ Main /Users/kcft114/.julia/packages/Flux/qXNjB/src/optimise/train.jl:105
┌ Info: Epoch 316
└ @ Main /Users/kcft114/.julia/packages/Flux/qXNjB/src/optimise/train.jl:105
┌ Info: Epoch 317
└ @ Main /Users/kcft114/.julia/packages/Flux/qXNjB/src/optimise/train.jl:105
┌ Info: Epoch 318
└ @ Main /Users/kcft114/.julia/packages/Flux/qXNjB/src/optimise/train.jl:105
┌ Info: Epoch 319
└ @ Main /Users/kcft114/.julia/packages/Flux/qXNjB/src/optimise/train.jl:105
┌ Info: Epoch 320
└ @ Main /Users/kcft114/.julia/packages/Flux/qXNjB/src/optimise/train.jl:105
┌ Info: Epoch 321
└ @ Main /Users/kcft114/.julia/packages/Flux/qXNjB/src/optimise/train.jl:105
┌ Info: Epoch 322
└ @ Main /Users/kcft114/.julia/packages/Flux/qXNjB/src/optimise/train.jl:105
┌ Info: Epoch 323
└ @ Main /Users/kcft114/.julia/packages/Flux/qXNjB/src/optimise/train.jl:105
┌ Info: Epoch 324
└ @ Main /Users/kcft114/.julia/packages/Flux/qXNjB/src/optimise/train.jl:105
┌ Info: Epoch 325
└ @ Main /Users/kcft114/.julia/packages/Flux/qXNjB/src/optimise/train.jl:105
┌ Info: Epoch 326
└ @ Main /Users/kcft114/.julia/packages/Flux/qXNjB/src/optimise/train.jl:105
┌ Info: Epoch 327
└ @ Main /Users/kcft114/.julia/packages/Flux/qXNjB/src/optimise/train.jl:105
┌ Info: Epoch 328
└ @ Main /Users/kcft114/.julia/packages/Flux/qXNjB/src/optimise/train.jl:105
┌ Info: Epoch 329
└ @ Main /Users/kcft114/.julia/packages/Flux/qXNjB/src/optimise/train.jl:105
┌ Info: Epoch 330
└ @ Main /Users/kcft114/.julia/packages/Flux/qXNjB/src/optimise/train.jl:105
┌ Info: Epoch 331
└ @ Main /Users/kcft114/.julia/packages/Flux/qXNjB/src/optimise/train.jl:105
┌ Info: Epoch 332
└ @ Main /Users/kcft114/.julia/packages/Flux/qXNjB/src/optimise/train.jl:105
┌ Info: Epoch 333
└ @ Main /Users/kcft114/.julia/packages/Flux/qXNjB/src/optimise/train.jl:105
┌ Info: Epoch 334
└ @ Main /Users/kcft114/.julia/packages/Flux/qXNjB/src/optimise/train.jl:105
┌ Info: Epoch 335
└ @ Main /Users/kcft114/.julia/packages/Flux/qXNjB/src/optimise/train.jl:105
┌ Info: Epoch 336
└ @ Main /Users/kcft114/.julia/packages/Flux/qXNjB/src/optimise/train.jl:105
┌ Info: Epoch 337
└ @ Main /Users/kcft114/.julia/packages/Flux/qXNjB/src/optimise/train.jl:105
┌ Info: Epoch 338
└ @ Main /Users/kcft114/.julia/packages/Flux/qXNjB/src/optimise/train.jl:105
┌ Info: Epoch 339
└ @ Main /Users/kcft114/.julia/packages/Flux/qXNjB/src/optimise/train.jl:105
┌ Info: Epoch 340
└ @ Main /Users/kcft114/.julia/packages/Flux/qXNjB/src/optimise/train.jl:105
┌ Info: Epoch 341
└ @ Main /Users/kcft114/.julia/packages/Flux/qXNjB/src/optimise/train.jl:105
┌ Info: Epoch 342
└ @ Main /Users/kcft114/.julia/packages/Flux/qXNjB/src/optimise/train.jl:105
┌ Info: Epoch 343
└ @ Main /Users/kcft114/.julia/packages/Flux/qXNjB/src/optimise/train.jl:105
┌ Info: Epoch 344
└ @ Main /Users/kcft114/.julia/packages/Flux/qXNjB/src/optimise/train.jl:105
┌ Info: Epoch 345
└ @ Main /Users/kcft114/.julia/packages/Flux/qXNjB/src/optimise/train.jl:105
┌ Info: Epoch 346
└ @ Main /Users/kcft114/.julia/packages/Flux/qXNjB/src/optimise/train.jl:105
┌ Info: Epoch 347
└ @ Main /Users/kcft114/.julia/packages/Flux/qXNjB/src/optimise/train.jl:105
┌ Info: Epoch 348
└ @ Main /Users/kcft114/.julia/packages/Flux/qXNjB/src/optimise/train.jl:105
┌ Info: Epoch 349
└ @ Main /Users/kcft114/.julia/packages/Flux/qXNjB/src/optimise/train.jl:105
┌ Info: Epoch 350
└ @ Main /Users/kcft114/.julia/packages/Flux/qXNjB/src/optimise/train.jl:105
┌ Info: Epoch 351
└ @ Main /Users/kcft114/.julia/packages/Flux/qXNjB/src/optimise/train.jl:105
┌ Info: Epoch 352
└ @ Main /Users/kcft114/.julia/packages/Flux/qXNjB/src/optimise/train.jl:105
┌ Info: Epoch 353
└ @ Main /Users/kcft114/.julia/packages/Flux/qXNjB/src/optimise/train.jl:105
┌ Info: Epoch 354
└ @ Main /Users/kcft114/.julia/packages/Flux/qXNjB/src/optimise/train.jl:105
┌ Info: Epoch 355
└ @ Main /Users/kcft114/.julia/packages/Flux/qXNjB/src/optimise/train.jl:105
┌ Info: Epoch 356
└ @ Main /Users/kcft114/.julia/packages/Flux/qXNjB/src/optimise/train.jl:105
┌ Info: Epoch 357
└ @ Main /Users/kcft114/.julia/packages/Flux/qXNjB/src/optimise/train.jl:105
┌ Info: Epoch 358
└ @ Main /Users/kcft114/.julia/packages/Flux/qXNjB/src/optimise/train.jl:105
┌ Info: Epoch 359
└ @ Main /Users/kcft114/.julia/packages/Flux/qXNjB/src/optimise/train.jl:105
┌ Info: Epoch 360
└ @ Main /Users/kcft114/.julia/packages/Flux/qXNjB/src/optimise/train.jl:105
┌ Info: Epoch 361
└ @ Main /Users/kcft114/.julia/packages/Flux/qXNjB/src/optimise/train.jl:105
┌ Info: Epoch 362
└ @ Main /Users/kcft114/.julia/packages/Flux/qXNjB/src/optimise/train.jl:105
┌ Info: Epoch 363
└ @ Main /Users/kcft114/.julia/packages/Flux/qXNjB/src/optimise/train.jl:105
┌ Info: Epoch 364
└ @ Main /Users/kcft114/.julia/packages/Flux/qXNjB/src/optimise/train.jl:105
┌ Info: Epoch 365
└ @ Main /Users/kcft114/.julia/packages/Flux/qXNjB/src/optimise/train.jl:105
┌ Info: Epoch 366
└ @ Main /Users/kcft114/.julia/packages/Flux/qXNjB/src/optimise/train.jl:105
┌ Info: Epoch 367
└ @ Main /Users/kcft114/.julia/packages/Flux/qXNjB/src/optimise/train.jl:105
┌ Info: Epoch 368
└ @ Main /Users/kcft114/.julia/packages/Flux/qXNjB/src/optimise/train.jl:105
┌ Info: Epoch 369
└ @ Main /Users/kcft114/.julia/packages/Flux/qXNjB/src/optimise/train.jl:105
┌ Info: Epoch 370
└ @ Main /Users/kcft114/.julia/packages/Flux/qXNjB/src/optimise/train.jl:105
┌ Info: Epoch 371
└ @ Main /Users/kcft114/.julia/packages/Flux/qXNjB/src/optimise/train.jl:105
┌ Info: Epoch 372
└ @ Main /Users/kcft114/.julia/packages/Flux/qXNjB/src/optimise/train.jl:105
┌ Info: Epoch 373
└ @ Main /Users/kcft114/.julia/packages/Flux/qXNjB/src/optimise/train.jl:105
┌ Info: Epoch 374
└ @ Main /Users/kcft114/.julia/packages/Flux/qXNjB/src/optimise/train.jl:105
┌ Info: Epoch 375
└ @ Main /Users/kcft114/.julia/packages/Flux/qXNjB/src/optimise/train.jl:105
┌ Info: Epoch 376
└ @ Main /Users/kcft114/.julia/packages/Flux/qXNjB/src/optimise/train.jl:105
┌ Info: Epoch 377
└ @ Main /Users/kcft114/.julia/packages/Flux/qXNjB/src/optimise/train.jl:105
┌ Info: Epoch 378
└ @ Main /Users/kcft114/.julia/packages/Flux/qXNjB/src/optimise/train.jl:105
┌ Info: Epoch 379
└ @ Main /Users/kcft114/.julia/packages/Flux/qXNjB/src/optimise/train.jl:105
┌ Info: Epoch 380
└ @ Main /Users/kcft114/.julia/packages/Flux/qXNjB/src/optimise/train.jl:105
┌ Info: Epoch 381
└ @ Main /Users/kcft114/.julia/packages/Flux/qXNjB/src/optimise/train.jl:105
┌ Info: Epoch 382
└ @ Main /Users/kcft114/.julia/packages/Flux/qXNjB/src/optimise/train.jl:105
┌ Info: Epoch 383
└ @ Main /Users/kcft114/.julia/packages/Flux/qXNjB/src/optimise/train.jl:105
┌ Info: Epoch 384
└ @ Main /Users/kcft114/.julia/packages/Flux/qXNjB/src/optimise/train.jl:105
┌ Info: Epoch 385
└ @ Main /Users/kcft114/.julia/packages/Flux/qXNjB/src/optimise/train.jl:105
┌ Info: Epoch 386
└ @ Main /Users/kcft114/.julia/packages/Flux/qXNjB/src/optimise/train.jl:105
┌ Info: Epoch 387
└ @ Main /Users/kcft114/.julia/packages/Flux/qXNjB/src/optimise/train.jl:105
┌ Info: Epoch 388
└ @ Main /Users/kcft114/.julia/packages/Flux/qXNjB/src/optimise/train.jl:105
┌ Info: Epoch 389
└ @ Main /Users/kcft114/.julia/packages/Flux/qXNjB/src/optimise/train.jl:105
┌ Info: Epoch 390
└ @ Main /Users/kcft114/.julia/packages/Flux/qXNjB/src/optimise/train.jl:105
┌ Info: Epoch 391
└ @ Main /Users/kcft114/.julia/packages/Flux/qXNjB/src/optimise/train.jl:105
┌ Info: Epoch 392
└ @ Main /Users/kcft114/.julia/packages/Flux/qXNjB/src/optimise/train.jl:105
┌ Info: Epoch 393
└ @ Main /Users/kcft114/.julia/packages/Flux/qXNjB/src/optimise/train.jl:105
┌ Info: Epoch 394
└ @ Main /Users/kcft114/.julia/packages/Flux/qXNjB/src/optimise/train.jl:105
┌ Info: Epoch 395
└ @ Main /Users/kcft114/.julia/packages/Flux/qXNjB/src/optimise/train.jl:105
┌ Info: Epoch 396
└ @ Main /Users/kcft114/.julia/packages/Flux/qXNjB/src/optimise/train.jl:105
┌ Info: Epoch 397
└ @ Main /Users/kcft114/.julia/packages/Flux/qXNjB/src/optimise/train.jl:105
┌ Info: Epoch 398
└ @ Main /Users/kcft114/.julia/packages/Flux/qXNjB/src/optimise/train.jl:105
┌ Info: Epoch 399
└ @ Main /Users/kcft114/.julia/packages/Flux/qXNjB/src/optimise/train.jl:105
┌ Info: Epoch 400
└ @ Main /Users/kcft114/.julia/packages/Flux/qXNjB/src/optimise/train.jl:105
┌ Info: Epoch 401
└ @ Main /Users/kcft114/.julia/packages/Flux/qXNjB/src/optimise/train.jl:105
┌ Info: Epoch 402
└ @ Main /Users/kcft114/.julia/packages/Flux/qXNjB/src/optimise/train.jl:105
```

```
loss(X, Y) = 0.86344635f0 (tracked)
loss(X, Y) = 0.86225414f0 (tracked)
loss(X, Y) = 0.8588022f0 (tracked)
loss(X, Y) = 0.83230627f0 (tracked)
loss(X, Y) = 0.84258133f0 (tracked)
loss(X, Y) = 0.8426161f0 (tracked)
loss(X, Y) = 0.84106505f0 (tracked)
loss(X, Y) = 0.8410611f0 (tracked)
loss(X, Y) = 0.8705449f0 (tracked)
loss(X, Y) = 0.84748703f0 (tracked)
loss(X, Y) = 0.85165966f0 (tracked)
loss(X, Y) = 0.85785145f0 (tracked)
loss(X, Y) = 0.8747703f0 (tracked)
loss(X, Y) = 0.8369053f0 (tracked)
loss(X, Y) = 0.8476479f0 (tracked)
loss(X, Y) = 0.86867857f0 (tracked)
loss(X, Y) = 0.83247995f0 (tracked)
loss(X, Y) = 0.8325211f0 (tracked)
loss(X, Y) = 0.8013926f0 (tracked)
loss(X, Y) = 0.8132082f0 (tracked)
loss(X, Y) = 0.83075374f0 (tracked)
loss(X, Y) = 0.8509131f0 (tracked)
loss(X, Y) = 0.85460204f0 (tracked)
loss(X, Y) = 0.82951915f0 (tracked)
loss(X, Y) = 0.815464f0 (tracked)
loss(X, Y) = 0.8541885f0 (tracked)
loss(X, Y) = 0.80340785f0 (tracked)
loss(X, Y) = 0.7992977f0 (tracked)
loss(X, Y) = 0.8415171f0 (tracked)
loss(X, Y) = 0.8413066f0 (tracked)
loss(X, Y) = 0.8168176f0 (tracked)
loss(X, Y) = 0.8371206f0 (tracked)
loss(X, Y) = 0.8328261f0 (tracked)
loss(X, Y) = 0.839933f0 (tracked)
loss(X, Y) = 0.79159313f0 (tracked)
loss(X, Y) = 0.80421543f0 (tracked)
loss(X, Y) = 0.81811f0 (tracked)
loss(X, Y) = 0.8387244f0 (tracked)
loss(X, Y) = 0.86280024f0 (tracked)
loss(X, Y) = 0.81813884f0 (tracked)
loss(X, Y) = 0.8016678f0 (tracked)
loss(X, Y) = 0.834359f0 (tracked)
loss(X, Y) = 0.83953404f0 (tracked)
loss(X, Y) = 0.85046285f0 (tracked)
loss(X, Y) = 0.80221695f0 (tracked)
loss(X, Y) = 0.8534946f0 (tracked)
loss(X, Y) = 0.8340242f0 (tracked)
loss(X, Y) = 0.79579544f0 (tracked)
loss(X, Y) = 0.79228264f0 (tracked)
loss(X, Y) = 0.81045455f0 (tracked)
loss(X, Y) = 0.8431229f0 (tracked)
loss(X, Y) = 0.8212987f0 (tracked)
loss(X, Y) = 0.8455129f0 (tracked)
loss(X, Y) = 0.81695f0 (tracked)
loss(X, Y) = 0.8080729f0 (tracked)
loss(X, Y) = 0.8240646f0 (tracked)
loss(X, Y) = 0.8560898f0 (tracked)
loss(X, Y) = 0.8192509f0 (tracked)
loss(X, Y) = 0.8061423f0 (tracked)
loss(X, Y) = 0.8333934f0 (tracked)
loss(X, Y) = 0.8352553f0 (tracked)
loss(X, Y) = 0.81054777f0 (tracked)
loss(X, Y) = 0.807786f0 (tracked)
loss(X, Y) = 0.8118674f0 (tracked)
loss(X, Y) = 0.842276f0 (tracked)
loss(X, Y) = 0.80674034f0 (tracked)
loss(X, Y) = 0.82724625f0 (tracked)
loss(X, Y) = 0.7905523f0 (tracked)
loss(X, Y) = 0.85005736f0 (tracked)
loss(X, Y) = 0.8475037f0 (tracked)
loss(X, Y) = 0.82774615f0 (tracked)
loss(X, Y) = 0.80599034f0 (tracked)
loss(X, Y) = 0.82872915f0 (tracked)
loss(X, Y) = 0.80525917f0 (tracked)
loss(X, Y) = 0.8024046f0 (tracked)
loss(X, Y) = 0.7990033f0 (tracked)
loss(X, Y) = 0.8287238f0 (tracked)
loss(X, Y) = 0.7776839f0 (tracked)
loss(X, Y) = 0.7966531f0 (tracked)
loss(X, Y) = 0.79551566f0 (tracked)
loss(X, Y) = 0.83996654f0 (tracked)
loss(X, Y) = 0.78248644f0 (tracked)
loss(X, Y) = 0.7919257f0 (tracked)
loss(X, Y) = 0.8092085f0 (tracked)
loss(X, Y) = 0.8319242f0 (tracked)
loss(X, Y) = 0.8111164f0 (tracked)
loss(X, Y) = 0.81892014f0 (tracked)
loss(X, Y) = 0.80995375f0 (tracked)
loss(X, Y) = 0.76956826f0 (tracked)
loss(X, Y) = 0.82592714f0 (tracked)
loss(X, Y) = 0.79111165f0 (tracked)
loss(X, Y) = 0.8166816f0 (tracked)
loss(X, Y) = 0.8125009f0 (tracked)
loss(X, Y) = 0.792382f0 (tracked)
loss(X, Y) = 0.8112472f0 (tracked)
loss(X, Y) = 0.77497536f0 (tracked)
loss(X, Y) = 0.80952126f0 (tracked)
loss(X, Y) = 0.8067982f0 (tracked)
loss(X, Y) = 0.82489675f0 (tracked)
loss(X, Y) = 0.78897685f0 (tracked)
loss(X, Y) = 0.82969695f0 (tracked)
loss(X, Y) = 0.77619284f0 (tracked)
loss(X, Y) = 0.79684734f0 (tracked)
loss(X, Y) = 0.796362f0 (tracked)
```

```
┌ Info: Epoch 403
└ @ Main /Users/kcft114/.julia/packages/Flux/qXNjB/src/optimise/train.jl:105
┌ Info: Epoch 404
└ @ Main /Users/kcft114/.julia/packages/Flux/qXNjB/src/optimise/train.jl:105
┌ Info: Epoch 405
└ @ Main /Users/kcft114/.julia/packages/Flux/qXNjB/src/optimise/train.jl:105
┌ Info: Epoch 406
└ @ Main /Users/kcft114/.julia/packages/Flux/qXNjB/src/optimise/train.jl:105
┌ Info: Epoch 407
└ @ Main /Users/kcft114/.julia/packages/Flux/qXNjB/src/optimise/train.jl:105
┌ Info: Epoch 408
└ @ Main /Users/kcft114/.julia/packages/Flux/qXNjB/src/optimise/train.jl:105
┌ Info: Epoch 409
└ @ Main /Users/kcft114/.julia/packages/Flux/qXNjB/src/optimise/train.jl:105
┌ Info: Epoch 410
└ @ Main /Users/kcft114/.julia/packages/Flux/qXNjB/src/optimise/train.jl:105
┌ Info: Epoch 411
└ @ Main /Users/kcft114/.julia/packages/Flux/qXNjB/src/optimise/train.jl:105
┌ Info: Epoch 412
└ @ Main /Users/kcft114/.julia/packages/Flux/qXNjB/src/optimise/train.jl:105
┌ Info: Epoch 413
└ @ Main /Users/kcft114/.julia/packages/Flux/qXNjB/src/optimise/train.jl:105
┌ Info: Epoch 414
└ @ Main /Users/kcft114/.julia/packages/Flux/qXNjB/src/optimise/train.jl:105
┌ Info: Epoch 415
└ @ Main /Users/kcft114/.julia/packages/Flux/qXNjB/src/optimise/train.jl:105
┌ Info: Epoch 416
└ @ Main /Users/kcft114/.julia/packages/Flux/qXNjB/src/optimise/train.jl:105
┌ Info: Epoch 417
└ @ Main /Users/kcft114/.julia/packages/Flux/qXNjB/src/optimise/train.jl:105
┌ Info: Epoch 418
└ @ Main /Users/kcft114/.julia/packages/Flux/qXNjB/src/optimise/train.jl:105
┌ Info: Epoch 419
└ @ Main /Users/kcft114/.julia/packages/Flux/qXNjB/src/optimise/train.jl:105
┌ Info: Epoch 420
└ @ Main /Users/kcft114/.julia/packages/Flux/qXNjB/src/optimise/train.jl:105
┌ Info: Epoch 421
└ @ Main /Users/kcft114/.julia/packages/Flux/qXNjB/src/optimise/train.jl:105
┌ Info: Epoch 422
└ @ Main /Users/kcft114/.julia/packages/Flux/qXNjB/src/optimise/train.jl:105
┌ Info: Epoch 423
└ @ Main /Users/kcft114/.julia/packages/Flux/qXNjB/src/optimise/train.jl:105
┌ Info: Epoch 424
└ @ Main /Users/kcft114/.julia/packages/Flux/qXNjB/src/optimise/train.jl:105
┌ Info: Epoch 425
└ @ Main /Users/kcft114/.julia/packages/Flux/qXNjB/src/optimise/train.jl:105
┌ Info: Epoch 426
└ @ Main /Users/kcft114/.julia/packages/Flux/qXNjB/src/optimise/train.jl:105
┌ Info: Epoch 427
└ @ Main /Users/kcft114/.julia/packages/Flux/qXNjB/src/optimise/train.jl:105
┌ Info: Epoch 428
└ @ Main /Users/kcft114/.julia/packages/Flux/qXNjB/src/optimise/train.jl:105
┌ Info: Epoch 429
└ @ Main /Users/kcft114/.julia/packages/Flux/qXNjB/src/optimise/train.jl:105
┌ Info: Epoch 430
└ @ Main /Users/kcft114/.julia/packages/Flux/qXNjB/src/optimise/train.jl:105
┌ Info: Epoch 431
└ @ Main /Users/kcft114/.julia/packages/Flux/qXNjB/src/optimise/train.jl:105
┌ Info: Epoch 432
└ @ Main /Users/kcft114/.julia/packages/Flux/qXNjB/src/optimise/train.jl:105
┌ Info: Epoch 433
└ @ Main /Users/kcft114/.julia/packages/Flux/qXNjB/src/optimise/train.jl:105
┌ Info: Epoch 434
└ @ Main /Users/kcft114/.julia/packages/Flux/qXNjB/src/optimise/train.jl:105
┌ Info: Epoch 435
└ @ Main /Users/kcft114/.julia/packages/Flux/qXNjB/src/optimise/train.jl:105
┌ Info: Epoch 436
└ @ Main /Users/kcft114/.julia/packages/Flux/qXNjB/src/optimise/train.jl:105
┌ Info: Epoch 437
└ @ Main /Users/kcft114/.julia/packages/Flux/qXNjB/src/optimise/train.jl:105
┌ Info: Epoch 438
└ @ Main /Users/kcft114/.julia/packages/Flux/qXNjB/src/optimise/train.jl:105
┌ Info: Epoch 439
└ @ Main /Users/kcft114/.julia/packages/Flux/qXNjB/src/optimise/train.jl:105
┌ Info: Epoch 440
└ @ Main /Users/kcft114/.julia/packages/Flux/qXNjB/src/optimise/train.jl:105
┌ Info: Epoch 441
└ @ Main /Users/kcft114/.julia/packages/Flux/qXNjB/src/optimise/train.jl:105
┌ Info: Epoch 442
└ @ Main /Users/kcft114/.julia/packages/Flux/qXNjB/src/optimise/train.jl:105
┌ Info: Epoch 443
└ @ Main /Users/kcft114/.julia/packages/Flux/qXNjB/src/optimise/train.jl:105
┌ Info: Epoch 444
└ @ Main /Users/kcft114/.julia/packages/Flux/qXNjB/src/optimise/train.jl:105
┌ Info: Epoch 445
└ @ Main /Users/kcft114/.julia/packages/Flux/qXNjB/src/optimise/train.jl:105
┌ Info: Epoch 446
└ @ Main /Users/kcft114/.julia/packages/Flux/qXNjB/src/optimise/train.jl:105
┌ Info: Epoch 447
└ @ Main /Users/kcft114/.julia/packages/Flux/qXNjB/src/optimise/train.jl:105
┌ Info: Epoch 448
└ @ Main /Users/kcft114/.julia/packages/Flux/qXNjB/src/optimise/train.jl:105
┌ Info: Epoch 449
└ @ Main /Users/kcft114/.julia/packages/Flux/qXNjB/src/optimise/train.jl:105
┌ Info: Epoch 450
└ @ Main /Users/kcft114/.julia/packages/Flux/qXNjB/src/optimise/train.jl:105
┌ Info: Epoch 451
└ @ Main /Users/kcft114/.julia/packages/Flux/qXNjB/src/optimise/train.jl:105
┌ Info: Epoch 452
└ @ Main /Users/kcft114/.julia/packages/Flux/qXNjB/src/optimise/train.jl:105
┌ Info: Epoch 453
└ @ Main /Users/kcft114/.julia/packages/Flux/qXNjB/src/optimise/train.jl:105
┌ Info: Epoch 454
└ @ Main /Users/kcft114/.julia/packages/Flux/qXNjB/src/optimise/train.jl:105
┌ Info: Epoch 455
└ @ Main /Users/kcft114/.julia/packages/Flux/qXNjB/src/optimise/train.jl:105
┌ Info: Epoch 456
└ @ Main /Users/kcft114/.julia/packages/Flux/qXNjB/src/optimise/train.jl:105
┌ Info: Epoch 457
└ @ Main /Users/kcft114/.julia/packages/Flux/qXNjB/src/optimise/train.jl:105
┌ Info: Epoch 458
└ @ Main /Users/kcft114/.julia/packages/Flux/qXNjB/src/optimise/train.jl:105
┌ Info: Epoch 459
└ @ Main /Users/kcft114/.julia/packages/Flux/qXNjB/src/optimise/train.jl:105
┌ Info: Epoch 460
└ @ Main /Users/kcft114/.julia/packages/Flux/qXNjB/src/optimise/train.jl:105
┌ Info: Epoch 461
└ @ Main /Users/kcft114/.julia/packages/Flux/qXNjB/src/optimise/train.jl:105
┌ Info: Epoch 462
└ @ Main /Users/kcft114/.julia/packages/Flux/qXNjB/src/optimise/train.jl:105
┌ Info: Epoch 463
└ @ Main /Users/kcft114/.julia/packages/Flux/qXNjB/src/optimise/train.jl:105
┌ Info: Epoch 464
└ @ Main /Users/kcft114/.julia/packages/Flux/qXNjB/src/optimise/train.jl:105
┌ Info: Epoch 465
└ @ Main /Users/kcft114/.julia/packages/Flux/qXNjB/src/optimise/train.jl:105
┌ Info: Epoch 466
└ @ Main /Users/kcft114/.julia/packages/Flux/qXNjB/src/optimise/train.jl:105
┌ Info: Epoch 467
└ @ Main /Users/kcft114/.julia/packages/Flux/qXNjB/src/optimise/train.jl:105
┌ Info: Epoch 468
└ @ Main /Users/kcft114/.julia/packages/Flux/qXNjB/src/optimise/train.jl:105
┌ Info: Epoch 469
└ @ Main /Users/kcft114/.julia/packages/Flux/qXNjB/src/optimise/train.jl:105
┌ Info: Epoch 470
└ @ Main /Users/kcft114/.julia/packages/Flux/qXNjB/src/optimise/train.jl:105
┌ Info: Epoch 471
└ @ Main /Users/kcft114/.julia/packages/Flux/qXNjB/src/optimise/train.jl:105
┌ Info: Epoch 472
└ @ Main /Users/kcft114/.julia/packages/Flux/qXNjB/src/optimise/train.jl:105
┌ Info: Epoch 473
└ @ Main /Users/kcft114/.julia/packages/Flux/qXNjB/src/optimise/train.jl:105
┌ Info: Epoch 474
└ @ Main /Users/kcft114/.julia/packages/Flux/qXNjB/src/optimise/train.jl:105
┌ Info: Epoch 475
└ @ Main /Users/kcft114/.julia/packages/Flux/qXNjB/src/optimise/train.jl:105
┌ Info: Epoch 476
└ @ Main /Users/kcft114/.julia/packages/Flux/qXNjB/src/optimise/train.jl:105
┌ Info: Epoch 477
└ @ Main /Users/kcft114/.julia/packages/Flux/qXNjB/src/optimise/train.jl:105
┌ Info: Epoch 478
└ @ Main /Users/kcft114/.julia/packages/Flux/qXNjB/src/optimise/train.jl:105
┌ Info: Epoch 479
└ @ Main /Users/kcft114/.julia/packages/Flux/qXNjB/src/optimise/train.jl:105
┌ Info: Epoch 480
└ @ Main /Users/kcft114/.julia/packages/Flux/qXNjB/src/optimise/train.jl:105
┌ Info: Epoch 481
└ @ Main /Users/kcft114/.julia/packages/Flux/qXNjB/src/optimise/train.jl:105
┌ Info: Epoch 482
└ @ Main /Users/kcft114/.julia/packages/Flux/qXNjB/src/optimise/train.jl:105
┌ Info: Epoch 483
└ @ Main /Users/kcft114/.julia/packages/Flux/qXNjB/src/optimise/train.jl:105
┌ Info: Epoch 484
└ @ Main /Users/kcft114/.julia/packages/Flux/qXNjB/src/optimise/train.jl:105
┌ Info: Epoch 485
└ @ Main /Users/kcft114/.julia/packages/Flux/qXNjB/src/optimise/train.jl:105
┌ Info: Epoch 486
└ @ Main /Users/kcft114/.julia/packages/Flux/qXNjB/src/optimise/train.jl:105
┌ Info: Epoch 487
└ @ Main /Users/kcft114/.julia/packages/Flux/qXNjB/src/optimise/train.jl:105
┌ Info: Epoch 488
└ @ Main /Users/kcft114/.julia/packages/Flux/qXNjB/src/optimise/train.jl:105
┌ Info: Epoch 489
└ @ Main /Users/kcft114/.julia/packages/Flux/qXNjB/src/optimise/train.jl:105
┌ Info: Epoch 490
└ @ Main /Users/kcft114/.julia/packages/Flux/qXNjB/src/optimise/train.jl:105
┌ Info: Epoch 491
└ @ Main /Users/kcft114/.julia/packages/Flux/qXNjB/src/optimise/train.jl:105
┌ Info: Epoch 492
└ @ Main /Users/kcft114/.julia/packages/Flux/qXNjB/src/optimise/train.jl:105
┌ Info: Epoch 493
└ @ Main /Users/kcft114/.julia/packages/Flux/qXNjB/src/optimise/train.jl:105
┌ Info: Epoch 494
└ @ Main /Users/kcft114/.julia/packages/Flux/qXNjB/src/optimise/train.jl:105
┌ Info: Epoch 495
└ @ Main /Users/kcft114/.julia/packages/Flux/qXNjB/src/optimise/train.jl:105
┌ Info: Epoch 496
└ @ Main /Users/kcft114/.julia/packages/Flux/qXNjB/src/optimise/train.jl:105
┌ Info: Epoch 497
└ @ Main /Users/kcft114/.julia/packages/Flux/qXNjB/src/optimise/train.jl:105
┌ Info: Epoch 498
└ @ Main /Users/kcft114/.julia/packages/Flux/qXNjB/src/optimise/train.jl:105
┌ Info: Epoch 499
└ @ Main /Users/kcft114/.julia/packages/Flux/qXNjB/src/optimise/train.jl:105
┌ Info: Epoch 500
└ @ Main /Users/kcft114/.julia/packages/Flux/qXNjB/src/optimise/train.jl:105
┌ Info: Epoch 501
└ @ Main /Users/kcft114/.julia/packages/Flux/qXNjB/src/optimise/train.jl:105
┌ Info: Epoch 502
└ @ Main /Users/kcft114/.julia/packages/Flux/qXNjB/src/optimise/train.jl:105
┌ Info: Epoch 503
└ @ Main /Users/kcft114/.julia/packages/Flux/qXNjB/src/optimise/train.jl:105
┌ Info: Epoch 504
└ @ Main /Users/kcft114/.julia/packages/Flux/qXNjB/src/optimise/train.jl:105
┌ Info: Epoch 505
└ @ Main /Users/kcft114/.julia/packages/Flux/qXNjB/src/optimise/train.jl:105
┌ Info: Epoch 506
└ @ Main /Users/kcft114/.julia/packages/Flux/qXNjB/src/optimise/train.jl:105
```

```
loss(X, Y) = 0.7932901f0 (tracked)
loss(X, Y) = 0.7726076f0 (tracked)
loss(X, Y) = 0.80511516f0 (tracked)
loss(X, Y) = 0.7826729f0 (tracked)
loss(X, Y) = 0.7676051f0 (tracked)
loss(X, Y) = 0.7955329f0 (tracked)
loss(X, Y) = 0.8119369f0 (tracked)
loss(X, Y) = 0.772674f0 (tracked)
loss(X, Y) = 0.759455f0 (tracked)
loss(X, Y) = 0.7931981f0 (tracked)
loss(X, Y) = 0.7857211f0 (tracked)
loss(X, Y) = 0.7660744f0 (tracked)
loss(X, Y) = 0.8206698f0 (tracked)
loss(X, Y) = 0.7771679f0 (tracked)
loss(X, Y) = 0.7651627f0 (tracked)
loss(X, Y) = 0.81146693f0 (tracked)
loss(X, Y) = 0.81643564f0 (tracked)
loss(X, Y) = 0.772239f0 (tracked)
loss(X, Y) = 0.7714324f0 (tracked)
loss(X, Y) = 0.7907431f0 (tracked)
loss(X, Y) = 0.79079175f0 (tracked)
loss(X, Y) = 0.8050814f0 (tracked)
loss(X, Y) = 0.7964571f0 (tracked)
loss(X, Y) = 0.7991944f0 (tracked)
loss(X, Y) = 0.7910977f0 (tracked)
loss(X, Y) = 0.7847078f0 (tracked)
loss(X, Y) = 0.7621396f0 (tracked)
loss(X, Y) = 0.77758634f0 (tracked)
loss(X, Y) = 0.77256495f0 (tracked)
loss(X, Y) = 0.7914907f0 (tracked)
loss(X, Y) = 0.778197f0 (tracked)
loss(X, Y) = 0.8008472f0 (tracked)
loss(X, Y) = 0.76652104f0 (tracked)
loss(X, Y) = 0.7870799f0 (tracked)
loss(X, Y) = 0.8110459f0 (tracked)
loss(X, Y) = 0.7834569f0 (tracked)
loss(X, Y) = 0.80108595f0 (tracked)
loss(X, Y) = 0.7972263f0 (tracked)
loss(X, Y) = 0.79169697f0 (tracked)
loss(X, Y) = 0.77145034f0 (tracked)
loss(X, Y) = 0.7992806f0 (tracked)
loss(X, Y) = 0.79691565f0 (tracked)
loss(X, Y) = 0.8017893f0 (tracked)
loss(X, Y) = 0.7771863f0 (tracked)
loss(X, Y) = 0.79257876f0 (tracked)
loss(X, Y) = 0.7955068f0 (tracked)
loss(X, Y) = 0.76340795f0 (tracked)
loss(X, Y) = 0.77451265f0 (tracked)
loss(X, Y) = 0.76421165f0 (tracked)
loss(X, Y) = 0.7537883f0 (tracked)
loss(X, Y) = 0.79372764f0 (tracked)
loss(X, Y) = 0.75919825f0 (tracked)
loss(X, Y) = 0.7449884f0 (tracked)
loss(X, Y) = 0.7284684f0 (tracked)
loss(X, Y) = 0.75822556f0 (tracked)
loss(X, Y) = 0.7784279f0 (tracked)
loss(X, Y) = 0.7588139f0 (tracked)
loss(X, Y) = 0.7667141f0 (tracked)
loss(X, Y) = 0.7692669f0 (tracked)
loss(X, Y) = 0.7590804f0 (tracked)
loss(X, Y) = 0.78065443f0 (tracked)
loss(X, Y) = 0.7493476f0 (tracked)
loss(X, Y) = 0.7697596f0 (tracked)
loss(X, Y) = 0.7396147f0 (tracked)
loss(X, Y) = 0.7977597f0 (tracked)
loss(X, Y) = 0.7814123f0 (tracked)
loss(X, Y) = 0.7756565f0 (tracked)
loss(X, Y) = 0.763024f0 (tracked)
loss(X, Y) = 0.7900003f0 (tracked)
loss(X, Y) = 0.80385476f0 (tracked)
loss(X, Y) = 0.7257791f0 (tracked)
loss(X, Y) = 0.7819589f0 (tracked)
loss(X, Y) = 0.7587526f0 (tracked)
loss(X, Y) = 0.78802055f0 (tracked)
loss(X, Y) = 0.745415f0 (tracked)
loss(X, Y) = 0.7743769f0 (tracked)
loss(X, Y) = 0.7703544f0 (tracked)
loss(X, Y) = 0.78825706f0 (tracked)
loss(X, Y) = 0.7694886f0 (tracked)
loss(X, Y) = 0.7958584f0 (tracked)
loss(X, Y) = 0.75725955f0 (tracked)
loss(X, Y) = 0.7814585f0 (tracked)
loss(X, Y) = 0.75336623f0 (tracked)
loss(X, Y) = 0.76728827f0 (tracked)
loss(X, Y) = 0.7701327f0 (tracked)
loss(X, Y) = 0.76699156f0 (tracked)
loss(X, Y) = 0.78831214f0 (tracked)
loss(X, Y) = 0.7297902f0 (tracked)
loss(X, Y) = 0.75468725f0 (tracked)
loss(X, Y) = 0.7765387f0 (tracked)
loss(X, Y) = 0.7824793f0 (tracked)
loss(X, Y) = 0.75055146f0 (tracked)
loss(X, Y) = 0.7438659f0 (tracked)
loss(X, Y) = 0.79263914f0 (tracked)
loss(X, Y) = 0.794572f0 (tracked)
loss(X, Y) = 0.8043518f0 (tracked)
loss(X, Y) = 0.7618205f0 (tracked)
loss(X, Y) = 0.76163363f0 (tracked)
loss(X, Y) = 0.80050975f0 (tracked)
loss(X, Y) = 0.77449137f0 (tracked)
loss(X, Y) = 0.7363064f0 (tracked)
loss(X, Y) = 0.7676794f0 (tracked)
loss(X, Y) = 0.70255256f0 (tracked)
loss(X, Y) = 0.8017512f0 (tracked)
loss(X, Y) = 0.7527331f0 (tracked)
loss(X, Y) = 0.7961507f0 (tracked)
loss(X, Y) = 0.7362218f0 (tracked)
```

```
┌ Info: Epoch 507
└ @ Main /Users/kcft114/.julia/packages/Flux/qXNjB/src/optimise/train.jl:105
┌ Info: Epoch 508
└ @ Main /Users/kcft114/.julia/packages/Flux/qXNjB/src/optimise/train.jl:105
┌ Info: Epoch 509
└ @ Main /Users/kcft114/.julia/packages/Flux/qXNjB/src/optimise/train.jl:105
┌ Info: Epoch 510
└ @ Main /Users/kcft114/.julia/packages/Flux/qXNjB/src/optimise/train.jl:105
┌ Info: Epoch 511
└ @ Main /Users/kcft114/.julia/packages/Flux/qXNjB/src/optimise/train.jl:105
┌ Info: Epoch 512
└ @ Main /Users/kcft114/.julia/packages/Flux/qXNjB/src/optimise/train.jl:105
┌ Info: Epoch 513
└ @ Main /Users/kcft114/.julia/packages/Flux/qXNjB/src/optimise/train.jl:105
┌ Info: Epoch 514
└ @ Main /Users/kcft114/.julia/packages/Flux/qXNjB/src/optimise/train.jl:105
┌ Info: Epoch 515
└ @ Main /Users/kcft114/.julia/packages/Flux/qXNjB/src/optimise/train.jl:105
┌ Info: Epoch 516
└ @ Main /Users/kcft114/.julia/packages/Flux/qXNjB/src/optimise/train.jl:105
┌ Info: Epoch 517
└ @ Main /Users/kcft114/.julia/packages/Flux/qXNjB/src/optimise/train.jl:105
┌ Info: Epoch 518
└ @ Main /Users/kcft114/.julia/packages/Flux/qXNjB/src/optimise/train.jl:105
┌ Info: Epoch 519
└ @ Main /Users/kcft114/.julia/packages/Flux/qXNjB/src/optimise/train.jl:105
┌ Info: Epoch 520
└ @ Main /Users/kcft114/.julia/packages/Flux/qXNjB/src/optimise/train.jl:105
┌ Info: Epoch 521
└ @ Main /Users/kcft114/.julia/packages/Flux/qXNjB/src/optimise/train.jl:105
┌ Info: Epoch 522
└ @ Main /Users/kcft114/.julia/packages/Flux/qXNjB/src/optimise/train.jl:105
┌ Info: Epoch 523
└ @ Main /Users/kcft114/.julia/packages/Flux/qXNjB/src/optimise/train.jl:105
┌ Info: Epoch 524
└ @ Main /Users/kcft114/.julia/packages/Flux/qXNjB/src/optimise/train.jl:105
┌ Info: Epoch 525
└ @ Main /Users/kcft114/.julia/packages/Flux/qXNjB/src/optimise/train.jl:105
┌ Info: Epoch 526
└ @ Main /Users/kcft114/.julia/packages/Flux/qXNjB/src/optimise/train.jl:105
┌ Info: Epoch 527
└ @ Main /Users/kcft114/.julia/packages/Flux/qXNjB/src/optimise/train.jl:105
┌ Info: Epoch 528
└ @ Main /Users/kcft114/.julia/packages/Flux/qXNjB/src/optimise/train.jl:105
┌ Info: Epoch 529
└ @ Main /Users/kcft114/.julia/packages/Flux/qXNjB/src/optimise/train.jl:105
┌ Info: Epoch 530
└ @ Main /Users/kcft114/.julia/packages/Flux/qXNjB/src/optimise/train.jl:105
┌ Info: Epoch 531
└ @ Main /Users/kcft114/.julia/packages/Flux/qXNjB/src/optimise/train.jl:105
┌ Info: Epoch 532
└ @ Main /Users/kcft114/.julia/packages/Flux/qXNjB/src/optimise/train.jl:105
┌ Info: Epoch 533
└ @ Main /Users/kcft114/.julia/packages/Flux/qXNjB/src/optimise/train.jl:105
┌ Info: Epoch 534
└ @ Main /Users/kcft114/.julia/packages/Flux/qXNjB/src/optimise/train.jl:105
┌ Info: Epoch 535
└ @ Main /Users/kcft114/.julia/packages/Flux/qXNjB/src/optimise/train.jl:105
┌ Info: Epoch 536
└ @ Main /Users/kcft114/.julia/packages/Flux/qXNjB/src/optimise/train.jl:105
┌ Info: Epoch 537
└ @ Main /Users/kcft114/.julia/packages/Flux/qXNjB/src/optimise/train.jl:105
┌ Info: Epoch 538
└ @ Main /Users/kcft114/.julia/packages/Flux/qXNjB/src/optimise/train.jl:105
┌ Info: Epoch 539
└ @ Main /Users/kcft114/.julia/packages/Flux/qXNjB/src/optimise/train.jl:105
┌ Info: Epoch 540
└ @ Main /Users/kcft114/.julia/packages/Flux/qXNjB/src/optimise/train.jl:105
┌ Info: Epoch 541
└ @ Main /Users/kcft114/.julia/packages/Flux/qXNjB/src/optimise/train.jl:105
┌ Info: Epoch 542
└ @ Main /Users/kcft114/.julia/packages/Flux/qXNjB/src/optimise/train.jl:105
┌ Info: Epoch 543
└ @ Main /Users/kcft114/.julia/packages/Flux/qXNjB/src/optimise/train.jl:105
┌ Info: Epoch 544
└ @ Main /Users/kcft114/.julia/packages/Flux/qXNjB/src/optimise/train.jl:105
┌ Info: Epoch 545
└ @ Main /Users/kcft114/.julia/packages/Flux/qXNjB/src/optimise/train.jl:105
┌ Info: Epoch 546
└ @ Main /Users/kcft114/.julia/packages/Flux/qXNjB/src/optimise/train.jl:105
┌ Info: Epoch 547
└ @ Main /Users/kcft114/.julia/packages/Flux/qXNjB/src/optimise/train.jl:105
┌ Info: Epoch 548
└ @ Main /Users/kcft114/.julia/packages/Flux/qXNjB/src/optimise/train.jl:105
┌ Info: Epoch 549
└ @ Main /Users/kcft114/.julia/packages/Flux/qXNjB/src/optimise/train.jl:105
┌ Info: Epoch 550
└ @ Main /Users/kcft114/.julia/packages/Flux/qXNjB/src/optimise/train.jl:105
┌ Info: Epoch 551
└ @ Main /Users/kcft114/.julia/packages/Flux/qXNjB/src/optimise/train.jl:105
┌ Info: Epoch 552
└ @ Main /Users/kcft114/.julia/packages/Flux/qXNjB/src/optimise/train.jl:105
┌ Info: Epoch 553
└ @ Main /Users/kcft114/.julia/packages/Flux/qXNjB/src/optimise/train.jl:105
┌ Info: Epoch 554
└ @ Main /Users/kcft114/.julia/packages/Flux/qXNjB/src/optimise/train.jl:105
┌ Info: Epoch 555
└ @ Main /Users/kcft114/.julia/packages/Flux/qXNjB/src/optimise/train.jl:105
┌ Info: Epoch 556
└ @ Main /Users/kcft114/.julia/packages/Flux/qXNjB/src/optimise/train.jl:105
┌ Info: Epoch 557
└ @ Main /Users/kcft114/.julia/packages/Flux/qXNjB/src/optimise/train.jl:105
┌ Info: Epoch 558
└ @ Main /Users/kcft114/.julia/packages/Flux/qXNjB/src/optimise/train.jl:105
┌ Info: Epoch 559
└ @ Main /Users/kcft114/.julia/packages/Flux/qXNjB/src/optimise/train.jl:105
┌ Info: Epoch 560
└ @ Main /Users/kcft114/.julia/packages/Flux/qXNjB/src/optimise/train.jl:105
┌ Info: Epoch 561
└ @ Main /Users/kcft114/.julia/packages/Flux/qXNjB/src/optimise/train.jl:105
┌ Info: Epoch 562
└ @ Main /Users/kcft114/.julia/packages/Flux/qXNjB/src/optimise/train.jl:105
┌ Info: Epoch 563
└ @ Main /Users/kcft114/.julia/packages/Flux/qXNjB/src/optimise/train.jl:105
┌ Info: Epoch 564
└ @ Main /Users/kcft114/.julia/packages/Flux/qXNjB/src/optimise/train.jl:105
┌ Info: Epoch 565
└ @ Main /Users/kcft114/.julia/packages/Flux/qXNjB/src/optimise/train.jl:105
┌ Info: Epoch 566
└ @ Main /Users/kcft114/.julia/packages/Flux/qXNjB/src/optimise/train.jl:105
┌ Info: Epoch 567
└ @ Main /Users/kcft114/.julia/packages/Flux/qXNjB/src/optimise/train.jl:105
┌ Info: Epoch 568
└ @ Main /Users/kcft114/.julia/packages/Flux/qXNjB/src/optimise/train.jl:105
┌ Info: Epoch 569
└ @ Main /Users/kcft114/.julia/packages/Flux/qXNjB/src/optimise/train.jl:105
┌ Info: Epoch 570
└ @ Main /Users/kcft114/.julia/packages/Flux/qXNjB/src/optimise/train.jl:105
┌ Info: Epoch 571
└ @ Main /Users/kcft114/.julia/packages/Flux/qXNjB/src/optimise/train.jl:105
┌ Info: Epoch 572
└ @ Main /Users/kcft114/.julia/packages/Flux/qXNjB/src/optimise/train.jl:105
┌ Info: Epoch 573
└ @ Main /Users/kcft114/.julia/packages/Flux/qXNjB/src/optimise/train.jl:105
┌ Info: Epoch 574
└ @ Main /Users/kcft114/.julia/packages/Flux/qXNjB/src/optimise/train.jl:105
┌ Info: Epoch 575
└ @ Main /Users/kcft114/.julia/packages/Flux/qXNjB/src/optimise/train.jl:105
┌ Info: Epoch 576
└ @ Main /Users/kcft114/.julia/packages/Flux/qXNjB/src/optimise/train.jl:105
┌ Info: Epoch 577
└ @ Main /Users/kcft114/.julia/packages/Flux/qXNjB/src/optimise/train.jl:105
┌ Info: Epoch 578
└ @ Main /Users/kcft114/.julia/packages/Flux/qXNjB/src/optimise/train.jl:105
┌ Info: Epoch 579
└ @ Main /Users/kcft114/.julia/packages/Flux/qXNjB/src/optimise/train.jl:105
┌ Info: Epoch 580
└ @ Main /Users/kcft114/.julia/packages/Flux/qXNjB/src/optimise/train.jl:105
┌ Info: Epoch 581
└ @ Main /Users/kcft114/.julia/packages/Flux/qXNjB/src/optimise/train.jl:105
┌ Info: Epoch 582
└ @ Main /Users/kcft114/.julia/packages/Flux/qXNjB/src/optimise/train.jl:105
┌ Info: Epoch 583
└ @ Main /Users/kcft114/.julia/packages/Flux/qXNjB/src/optimise/train.jl:105
┌ Info: Epoch 584
└ @ Main /Users/kcft114/.julia/packages/Flux/qXNjB/src/optimise/train.jl:105
┌ Info: Epoch 585
└ @ Main /Users/kcft114/.julia/packages/Flux/qXNjB/src/optimise/train.jl:105
┌ Info: Epoch 586
└ @ Main /Users/kcft114/.julia/packages/Flux/qXNjB/src/optimise/train.jl:105
┌ Info: Epoch 587
└ @ Main /Users/kcft114/.julia/packages/Flux/qXNjB/src/optimise/train.jl:105
┌ Info: Epoch 588
└ @ Main /Users/kcft114/.julia/packages/Flux/qXNjB/src/optimise/train.jl:105
┌ Info: Epoch 589
└ @ Main /Users/kcft114/.julia/packages/Flux/qXNjB/src/optimise/train.jl:105
┌ Info: Epoch 590
└ @ Main /Users/kcft114/.julia/packages/Flux/qXNjB/src/optimise/train.jl:105
┌ Info: Epoch 591
└ @ Main /Users/kcft114/.julia/packages/Flux/qXNjB/src/optimise/train.jl:105
┌ Info: Epoch 592
└ @ Main /Users/kcft114/.julia/packages/Flux/qXNjB/src/optimise/train.jl:105
┌ Info: Epoch 593
└ @ Main /Users/kcft114/.julia/packages/Flux/qXNjB/src/optimise/train.jl:105
┌ Info: Epoch 594
└ @ Main /Users/kcft114/.julia/packages/Flux/qXNjB/src/optimise/train.jl:105
┌ Info: Epoch 595
└ @ Main /Users/kcft114/.julia/packages/Flux/qXNjB/src/optimise/train.jl:105
┌ Info: Epoch 596
└ @ Main /Users/kcft114/.julia/packages/Flux/qXNjB/src/optimise/train.jl:105
┌ Info: Epoch 597
└ @ Main /Users/kcft114/.julia/packages/Flux/qXNjB/src/optimise/train.jl:105
┌ Info: Epoch 598
└ @ Main /Users/kcft114/.julia/packages/Flux/qXNjB/src/optimise/train.jl:105
┌ Info: Epoch 599
└ @ Main /Users/kcft114/.julia/packages/Flux/qXNjB/src/optimise/train.jl:105
┌ Info: Epoch 600
└ @ Main /Users/kcft114/.julia/packages/Flux/qXNjB/src/optimise/train.jl:105
┌ Info: Epoch 601
└ @ Main /Users/kcft114/.julia/packages/Flux/qXNjB/src/optimise/train.jl:105
┌ Info: Epoch 602
└ @ Main /Users/kcft114/.julia/packages/Flux/qXNjB/src/optimise/train.jl:105
┌ Info: Epoch 603
└ @ Main /Users/kcft114/.julia/packages/Flux/qXNjB/src/optimise/train.jl:105
┌ Info: Epoch 604
└ @ Main /Users/kcft114/.julia/packages/Flux/qXNjB/src/optimise/train.jl:105
┌ Info: Epoch 605
└ @ Main /Users/kcft114/.julia/packages/Flux/qXNjB/src/optimise/train.jl:105
┌ Info: Epoch 606
└ @ Main /Users/kcft114/.julia/packages/Flux/qXNjB/src/optimise/train.jl:105
┌ Info: Epoch 607
└ @ Main /Users/kcft114/.julia/packages/Flux/qXNjB/src/optimise/train.jl:105
┌ Info: Epoch 608
└ @ Main /Users/kcft114/.julia/packages/Flux/qXNjB/src/optimise/train.jl:105
┌ Info: Epoch 609
└ @ Main /Users/kcft114/.julia/packages/Flux/qXNjB/src/optimise/train.jl:105
┌ Info: Epoch 610
└ @ Main /Users/kcft114/.julia/packages/Flux/qXNjB/src/optimise/train.jl:105
┌ Info: Epoch 611
└ @ Main /Users/kcft114/.julia/packages/Flux/qXNjB/src/optimise/train.jl:105
┌ Info: Epoch 612
└ @ Main /Users/kcft114/.julia/packages/Flux/qXNjB/src/optimise/train.jl:105
┌ Info: Epoch 613
└ @ Main /Users/kcft114/.julia/packages/Flux/qXNjB/src/optimise/train.jl:105
```

```
loss(X, Y) = 0.7492642f0 (tracked)
loss(X, Y) = 0.73979914f0 (tracked)
loss(X, Y) = 0.7479855f0 (tracked)
loss(X, Y) = 0.7618842f0 (tracked)
loss(X, Y) = 0.7908872f0 (tracked)
loss(X, Y) = 0.7750089f0 (tracked)
loss(X, Y) = 0.72769064f0 (tracked)
loss(X, Y) = 0.7547323f0 (tracked)
loss(X, Y) = 0.76770014f0 (tracked)
loss(X, Y) = 0.77933204f0 (tracked)
loss(X, Y) = 0.74718016f0 (tracked)
loss(X, Y) = 0.7199362f0 (tracked)
loss(X, Y) = 0.7976301f0 (tracked)
loss(X, Y) = 0.77669144f0 (tracked)
loss(X, Y) = 0.773267f0 (tracked)
loss(X, Y) = 0.7898686f0 (tracked)
loss(X, Y) = 0.80281436f0 (tracked)
loss(X, Y) = 0.7797168f0 (tracked)
loss(X, Y) = 0.7572541f0 (tracked)
loss(X, Y) = 0.7592449f0 (tracked)
loss(X, Y) = 0.77001524f0 (tracked)
loss(X, Y) = 0.74159306f0 (tracked)
loss(X, Y) = 0.7410458f0 (tracked)
loss(X, Y) = 0.7488606f0 (tracked)
loss(X, Y) = 0.73112094f0 (tracked)
loss(X, Y) = 0.74929225f0 (tracked)
loss(X, Y) = 0.75655574f0 (tracked)
loss(X, Y) = 0.77102625f0 (tracked)
loss(X, Y) = 0.7631667f0 (tracked)
loss(X, Y) = 0.7445836f0 (tracked)
loss(X, Y) = 0.78620416f0 (tracked)
loss(X, Y) = 0.741381f0 (tracked)
loss(X, Y) = 0.7747346f0 (tracked)
loss(X, Y) = 0.7695766f0 (tracked)
loss(X, Y) = 0.7402482f0 (tracked)
loss(X, Y) = 0.7375736f0 (tracked)
loss(X, Y) = 0.77723294f0 (tracked)
loss(X, Y) = 0.7641498f0 (tracked)
loss(X, Y) = 0.74534327f0 (tracked)
loss(X, Y) = 0.7629425f0 (tracked)
loss(X, Y) = 0.7655783f0 (tracked)
loss(X, Y) = 0.7744168f0 (tracked)
loss(X, Y) = 0.7731227f0 (tracked)
loss(X, Y) = 0.74720865f0 (tracked)
loss(X, Y) = 0.7412424f0 (tracked)
loss(X, Y) = 0.73158765f0 (tracked)
loss(X, Y) = 0.717054f0 (tracked)
loss(X, Y) = 0.73398006f0 (tracked)
loss(X, Y) = 0.7562152f0 (tracked)
loss(X, Y) = 0.7429742f0 (tracked)
loss(X, Y) = 0.70352024f0 (tracked)
loss(X, Y) = 0.7568818f0 (tracked)
loss(X, Y) = 0.7258394f0 (tracked)
loss(X, Y) = 0.73988783f0 (tracked)
loss(X, Y) = 0.75724757f0 (tracked)
loss(X, Y) = 0.72980255f0 (tracked)
loss(X, Y) = 0.72615963f0 (tracked)
loss(X, Y) = 0.75029194f0 (tracked)
loss(X, Y) = 0.7650335f0 (tracked)
loss(X, Y) = 0.7581923f0 (tracked)
loss(X, Y) = 0.73938245f0 (tracked)
loss(X, Y) = 0.76504636f0 (tracked)
loss(X, Y) = 0.73823595f0 (tracked)
loss(X, Y) = 0.73063713f0 (tracked)
loss(X, Y) = 0.766849f0 (tracked)
loss(X, Y) = 0.7406775f0 (tracked)
loss(X, Y) = 0.7452379f0 (tracked)
loss(X, Y) = 0.7051581f0 (tracked)
loss(X, Y) = 0.73653024f0 (tracked)
loss(X, Y) = 0.773958f0 (tracked)
loss(X, Y) = 0.75759864f0 (tracked)
loss(X, Y) = 0.732686f0 (tracked)
loss(X, Y) = 0.7625657f0 (tracked)
loss(X, Y) = 0.74418336f0 (tracked)
loss(X, Y) = 0.75245816f0 (tracked)
loss(X, Y) = 0.7154732f0 (tracked)
loss(X, Y) = 0.802215f0 (tracked)
loss(X, Y) = 0.7697058f0 (tracked)
loss(X, Y) = 0.7123144f0 (tracked)
loss(X, Y) = 0.7508701f0 (tracked)
loss(X, Y) = 0.7543895f0 (tracked)
loss(X, Y) = 0.7618959f0 (tracked)
loss(X, Y) = 0.69272184f0 (tracked)
loss(X, Y) = 0.7506181f0 (tracked)
loss(X, Y) = 0.7456382f0 (tracked)
loss(X, Y) = 0.7544996f0 (tracked)
loss(X, Y) = 0.75575227f0 (tracked)
loss(X, Y) = 0.7521175f0 (tracked)
loss(X, Y) = 0.74570286f0 (tracked)
loss(X, Y) = 0.7456896f0 (tracked)
loss(X, Y) = 0.7548668f0 (tracked)
loss(X, Y) = 0.72851557f0 (tracked)
loss(X, Y) = 0.74550074f0 (tracked)
loss(X, Y) = 0.7631668f0 (tracked)
loss(X, Y) = 0.7481513f0 (tracked)
loss(X, Y) = 0.75505877f0 (tracked)
loss(X, Y) = 0.75655824f0 (tracked)
loss(X, Y) = 0.71642554f0 (tracked)
loss(X, Y) = 0.76925147f0 (tracked)
loss(X, Y) = 0.7069647f0 (tracked)
loss(X, Y) = 0.7282737f0 (tracked)
loss(X, Y) = 0.7627425f0 (tracked)
loss(X, Y) = 0.77748555f0 (tracked)
loss(X, Y) = 0.73620224f0 (tracked)
loss(X, Y) = 0.77735f0 (tracked)
```

```
┌ Info: Epoch 614
└ @ Main /Users/kcft114/.julia/packages/Flux/qXNjB/src/optimise/train.jl:105
```

```
loss(X, Y) = 0.76704735f0 (tracked)
loss(X, Y) = 0.73893946f0 (tracked)
loss(X, Y) = 0.72116745f0 (tracked)
loss(X, Y) = 0.7584146f0 (tracked)
loss(X, Y) = 0.7647898f0 (tracked)
loss(X, Y) = 0.73164874f0 (tracked)
loss(X, Y) = 0.7268402f0 (tracked)
loss(X, Y) = 0.7244966f0 (tracked)
loss(X, Y) = 0.73856264f0 (tracked)
loss(X, Y) = 0.73068845f0 (tracked)
loss(X, Y) = 0.6934631f0 (tracked)
loss(X, Y) = 0.7174833f0 (tracked)
loss(X, Y) = 0.74291104f0 (tracked)
loss(X, Y) = 0.7177957f0 (tracked)
loss(X, Y) = 0.74340296f0 (tracked)
loss(X, Y) = 0.75527686f0 (tracked)
loss(X, Y) = 0.70718616f0 (tracked)
loss(X, Y) = 0.7518172f0 (tracked)
loss(X, Y) = 0.7713889f0 (tracked)
loss(X, Y) = 0.7457037f0 (tracked)
loss(X, Y) = 0.71911776f0 (tracked)
loss(X, Y) = 0.6850661f0 (tracked)
loss(X, Y) = 0.7212687f0 (tracked)
loss(X, Y) = 0.7272428f0 (tracked)
loss(X, Y) = 0.7610036f0 (tracked)
loss(X, Y) = 0.7681496f0 (tracked)
loss(X, Y) = 0.70573604f0 (tracked)
loss(X, Y) = 0.7380822f0 (tracked)
loss(X, Y) = 0.71804744f0 (tracked)
loss(X, Y) = 0.7537408f0 (tracked)
loss(X, Y) = 0.72895986f0 (tracked)
loss(X, Y) = 0.7658888f0 (tracked)
loss(X, Y) = 0.6765052f0 (tracked)
loss(X, Y) = 0.7131405f0 (tracked)
loss(X, Y) = 0.7627419f0 (tracked)
loss(X, Y) = 0.75484115f0 (tracked)
loss(X, Y) = 0.70537925f0 (tracked)
loss(X, Y) = 0.7617015f0 (tracked)
loss(X, Y) = 0.73713547f0 (tracked)
loss(X, Y) = 0.71265614f0 (tracked)
loss(X, Y) = 0.73011315f0 (tracked)
loss(X, Y) = 0.7096587f0 (tracked)
loss(X, Y) = 0.70729554f0 (tracked)
loss(X, Y) = 0.7269213f0 (tracked)
loss(X, Y) = 0.7333256f0 (tracked)
loss(X, Y) = 0.7285676f0 (tracked)
loss(X, Y) = 0.746601f0 (tracked)
loss(X, Y) = 0.7254054f0 (tracked)
loss(X, Y) = 0.7065961f0 (tracked)
loss(X, Y) = 0.70142525f0 (tracked)
loss(X, Y) = 0.76985717f0 (tracked)
loss(X, Y) = 0.72787595f0 (tracked)
loss(X, Y) = 0.74965477f0 (tracked)
loss(X, Y) = 0.6901199f0 (tracked)
loss(X, Y) = 0.7161253f0 (tracked)
loss(X, Y) = 0.7579199f0 (tracked)
loss(X, Y) = 0.7245859f0 (tracked)
loss(X, Y) = 0.71649456f0 (tracked)
loss(X, Y) = 0.69521135f0 (tracked)
loss(X, Y) = 0.74362f0 (tracked)
loss(X, Y) = 0.73039687f0 (tracked)
loss(X, Y) = 0.71995354f0 (tracked)
loss(X, Y) = 0.7029117f0 (tracked)
loss(X, Y) = 0.7143169f0 (tracked)
loss(X, Y) = 0.7088924f0 (tracked)
loss(X, Y) = 0.73568857f0 (tracked)
loss(X, Y) = 0.726342f0 (tracked)
loss(X, Y) = 0.7542218f0 (tracked)
loss(X, Y) = 0.7644404f0 (tracked)
loss(X, Y) = 0.7324933f0 (tracked)
loss(X, Y) = 0.72836125f0 (tracked)
loss(X, Y) = 0.68689454f0 (tracked)
loss(X, Y) = 0.74413556f0 (tracked)
loss(X, Y) = 0.719356f0 (tracked)
loss(X, Y) = 0.7094047f0 (tracked)
loss(X, Y) = 0.74616325f0 (tracked)
loss(X, Y) = 0.70834213f0 (tracked)
loss(X, Y) = 0.7280569f0 (tracked)
loss(X, Y) = 0.72382647f0 (tracked)
loss(X, Y) = 0.7141946f0 (tracked)
loss(X, Y) = 0.71253854f0 (tracked)
loss(X, Y) = 0.7120496f0 (tracked)
loss(X, Y) = 0.74203074f0 (tracked)
loss(X, Y) = 0.7580944f0 (tracked)
loss(X, Y) = 0.74381083f0 (tracked)
loss(X, Y) = 0.7432939f0 (tracked)
loss(X, Y) = 0.7057753f0 (tracked)
loss(X, Y) = 0.7767504f0 (tracked)
loss(X, Y) = 0.72341603f0 (tracked)
loss(X, Y) = 0.6900688f0 (tracked)
loss(X, Y) = 0.72917044f0 (tracked)
loss(X, Y) = 0.7073558f0 (tracked)
loss(X, Y) = 0.7369823f0 (tracked)
loss(X, Y) = 0.7451153f0 (tracked)
loss(X, Y) = 0.6924271f0 (tracked)
loss(X, Y) = 0.7491702f0 (tracked)
```

```
┌ Info: Epoch 615
└ @ Main /Users/kcft114/.julia/packages/Flux/qXNjB/src/optimise/train.jl:105
┌ Info: Epoch 616
└ @ Main /Users/kcft114/.julia/packages/Flux/qXNjB/src/optimise/train.jl:105
┌ Info: Epoch 617
└ @ Main /Users/kcft114/.julia/packages/Flux/qXNjB/src/optimise/train.jl:105
┌ Info: Epoch 618
└ @ Main /Users/kcft114/.julia/packages/Flux/qXNjB/src/optimise/train.jl:105
┌ Info: Epoch 619
└ @ Main /Users/kcft114/.julia/packages/Flux/qXNjB/src/optimise/train.jl:105
┌ Info: Epoch 620
└ @ Main /Users/kcft114/.julia/packages/Flux/qXNjB/src/optimise/train.jl:105
┌ Info: Epoch 621
└ @ Main /Users/kcft114/.julia/packages/Flux/qXNjB/src/optimise/train.jl:105
┌ Info: Epoch 622
└ @ Main /Users/kcft114/.julia/packages/Flux/qXNjB/src/optimise/train.jl:105
┌ Info: Epoch 623
└ @ Main /Users/kcft114/.julia/packages/Flux/qXNjB/src/optimise/train.jl:105
┌ Info: Epoch 624
└ @ Main /Users/kcft114/.julia/packages/Flux/qXNjB/src/optimise/train.jl:105
┌ Info: Epoch 625
└ @ Main /Users/kcft114/.julia/packages/Flux/qXNjB/src/optimise/train.jl:105
┌ Info: Epoch 626
└ @ Main /Users/kcft114/.julia/packages/Flux/qXNjB/src/optimise/train.jl:105
┌ Info: Epoch 627
└ @ Main /Users/kcft114/.julia/packages/Flux/qXNjB/src/optimise/train.jl:105
┌ Info: Epoch 628
└ @ Main /Users/kcft114/.julia/packages/Flux/qXNjB/src/optimise/train.jl:105
┌ Info: Epoch 629
└ @ Main /Users/kcft114/.julia/packages/Flux/qXNjB/src/optimise/train.jl:105
┌ Info: Epoch 630
└ @ Main /Users/kcft114/.julia/packages/Flux/qXNjB/src/optimise/train.jl:105
┌ Info: Epoch 631
└ @ Main /Users/kcft114/.julia/packages/Flux/qXNjB/src/optimise/train.jl:105
┌ Info: Epoch 632
└ @ Main /Users/kcft114/.julia/packages/Flux/qXNjB/src/optimise/train.jl:105
┌ Info: Epoch 633
└ @ Main /Users/kcft114/.julia/packages/Flux/qXNjB/src/optimise/train.jl:105
┌ Info: Epoch 634
└ @ Main /Users/kcft114/.julia/packages/Flux/qXNjB/src/optimise/train.jl:105
┌ Info: Epoch 635
└ @ Main /Users/kcft114/.julia/packages/Flux/qXNjB/src/optimise/train.jl:105
┌ Info: Epoch 636
└ @ Main /Users/kcft114/.julia/packages/Flux/qXNjB/src/optimise/train.jl:105
┌ Info: Epoch 637
└ @ Main /Users/kcft114/.julia/packages/Flux/qXNjB/src/optimise/train.jl:105
┌ Info: Epoch 638
└ @ Main /Users/kcft114/.julia/packages/Flux/qXNjB/src/optimise/train.jl:105
┌ Info: Epoch 639
└ @ Main /Users/kcft114/.julia/packages/Flux/qXNjB/src/optimise/train.jl:105
┌ Info: Epoch 640
└ @ Main /Users/kcft114/.julia/packages/Flux/qXNjB/src/optimise/train.jl:105
┌ Info: Epoch 641
└ @ Main /Users/kcft114/.julia/packages/Flux/qXNjB/src/optimise/train.jl:105
┌ Info: Epoch 642
└ @ Main /Users/kcft114/.julia/packages/Flux/qXNjB/src/optimise/train.jl:105
┌ Info: Epoch 643
└ @ Main /Users/kcft114/.julia/packages/Flux/qXNjB/src/optimise/train.jl:105
┌ Info: Epoch 644
└ @ Main /Users/kcft114/.julia/packages/Flux/qXNjB/src/optimise/train.jl:105
┌ Info: Epoch 645
└ @ Main /Users/kcft114/.julia/packages/Flux/qXNjB/src/optimise/train.jl:105
┌ Info: Epoch 646
└ @ Main /Users/kcft114/.julia/packages/Flux/qXNjB/src/optimise/train.jl:105
┌ Info: Epoch 647
└ @ Main /Users/kcft114/.julia/packages/Flux/qXNjB/src/optimise/train.jl:105
┌ Info: Epoch 648
└ @ Main /Users/kcft114/.julia/packages/Flux/qXNjB/src/optimise/train.jl:105
┌ Info: Epoch 649
└ @ Main /Users/kcft114/.julia/packages/Flux/qXNjB/src/optimise/train.jl:105
┌ Info: Epoch 650
└ @ Main /Users/kcft114/.julia/packages/Flux/qXNjB/src/optimise/train.jl:105
┌ Info: Epoch 651
└ @ Main /Users/kcft114/.julia/packages/Flux/qXNjB/src/optimise/train.jl:105
┌ Info: Epoch 652
└ @ Main /Users/kcft114/.julia/packages/Flux/qXNjB/src/optimise/train.jl:105
┌ Info: Epoch 653
└ @ Main /Users/kcft114/.julia/packages/Flux/qXNjB/src/optimise/train.jl:105
┌ Info: Epoch 654
└ @ Main /Users/kcft114/.julia/packages/Flux/qXNjB/src/optimise/train.jl:105
┌ Info: Epoch 655
└ @ Main /Users/kcft114/.julia/packages/Flux/qXNjB/src/optimise/train.jl:105
┌ Info: Epoch 656
└ @ Main /Users/kcft114/.julia/packages/Flux/qXNjB/src/optimise/train.jl:105
┌ Info: Epoch 657
└ @ Main /Users/kcft114/.julia/packages/Flux/qXNjB/src/optimise/train.jl:105
┌ Info: Epoch 658
└ @ Main /Users/kcft114/.julia/packages/Flux/qXNjB/src/optimise/train.jl:105
┌ Info: Epoch 659
└ @ Main /Users/kcft114/.julia/packages/Flux/qXNjB/src/optimise/train.jl:105
┌ Info: Epoch 660
└ @ Main /Users/kcft114/.julia/packages/Flux/qXNjB/src/optimise/train.jl:105
┌ Info: Epoch 661
└ @ Main /Users/kcft114/.julia/packages/Flux/qXNjB/src/optimise/train.jl:105
┌ Info: Epoch 662
└ @ Main /Users/kcft114/.julia/packages/Flux/qXNjB/src/optimise/train.jl:105
┌ Info: Epoch 663
└ @ Main /Users/kcft114/.julia/packages/Flux/qXNjB/src/optimise/train.jl:105
┌ Info: Epoch 664
└ @ Main /Users/kcft114/.julia/packages/Flux/qXNjB/src/optimise/train.jl:105
┌ Info: Epoch 665
└ @ Main /Users/kcft114/.julia/packages/Flux/qXNjB/src/optimise/train.jl:105
┌ Info: Epoch 666
└ @ Main /Users/kcft114/.julia/packages/Flux/qXNjB/src/optimise/train.jl:105
┌ Info: Epoch 667
└ @ Main /Users/kcft114/.julia/packages/Flux/qXNjB/src/optimise/train.jl:105
┌ Info: Epoch 668
└ @ Main /Users/kcft114/.julia/packages/Flux/qXNjB/src/optimise/train.jl:105
┌ Info: Epoch 669
└ @ Main /Users/kcft114/.julia/packages/Flux/qXNjB/src/optimise/train.jl:105
┌ Info: Epoch 670
└ @ Main /Users/kcft114/.julia/packages/Flux/qXNjB/src/optimise/train.jl:105
┌ Info: Epoch 671
└ @ Main /Users/kcft114/.julia/packages/Flux/qXNjB/src/optimise/train.jl:105
┌ Info: Epoch 672
└ @ Main /Users/kcft114/.julia/packages/Flux/qXNjB/src/optimise/train.jl:105
┌ Info: Epoch 673
└ @ Main /Users/kcft114/.julia/packages/Flux/qXNjB/src/optimise/train.jl:105
┌ Info: Epoch 674
└ @ Main /Users/kcft114/.julia/packages/Flux/qXNjB/src/optimise/train.jl:105
┌ Info: Epoch 675
└ @ Main /Users/kcft114/.julia/packages/Flux/qXNjB/src/optimise/train.jl:105
┌ Info: Epoch 676
└ @ Main /Users/kcft114/.julia/packages/Flux/qXNjB/src/optimise/train.jl:105
┌ Info: Epoch 677
└ @ Main /Users/kcft114/.julia/packages/Flux/qXNjB/src/optimise/train.jl:105
┌ Info: Epoch 678
└ @ Main /Users/kcft114/.julia/packages/Flux/qXNjB/src/optimise/train.jl:105
┌ Info: Epoch 679
└ @ Main /Users/kcft114/.julia/packages/Flux/qXNjB/src/optimise/train.jl:105
┌ Info: Epoch 680
└ @ Main /Users/kcft114/.julia/packages/Flux/qXNjB/src/optimise/train.jl:105
┌ Info: Epoch 681
└ @ Main /Users/kcft114/.julia/packages/Flux/qXNjB/src/optimise/train.jl:105
┌ Info: Epoch 682
└ @ Main /Users/kcft114/.julia/packages/Flux/qXNjB/src/optimise/train.jl:105
┌ Info: Epoch 683
└ @ Main /Users/kcft114/.julia/packages/Flux/qXNjB/src/optimise/train.jl:105
┌ Info: Epoch 684
└ @ Main /Users/kcft114/.julia/packages/Flux/qXNjB/src/optimise/train.jl:105
┌ Info: Epoch 685
└ @ Main /Users/kcft114/.julia/packages/Flux/qXNjB/src/optimise/train.jl:105
┌ Info: Epoch 686
└ @ Main /Users/kcft114/.julia/packages/Flux/qXNjB/src/optimise/train.jl:105
┌ Info: Epoch 687
└ @ Main /Users/kcft114/.julia/packages/Flux/qXNjB/src/optimise/train.jl:105
┌ Info: Epoch 688
└ @ Main /Users/kcft114/.julia/packages/Flux/qXNjB/src/optimise/train.jl:105
┌ Info: Epoch 689
└ @ Main /Users/kcft114/.julia/packages/Flux/qXNjB/src/optimise/train.jl:105
┌ Info: Epoch 690
└ @ Main /Users/kcft114/.julia/packages/Flux/qXNjB/src/optimise/train.jl:105
┌ Info: Epoch 691
└ @ Main /Users/kcft114/.julia/packages/Flux/qXNjB/src/optimise/train.jl:105
┌ Info: Epoch 692
└ @ Main /Users/kcft114/.julia/packages/Flux/qXNjB/src/optimise/train.jl:105
┌ Info: Epoch 693
└ @ Main /Users/kcft114/.julia/packages/Flux/qXNjB/src/optimise/train.jl:105
┌ Info: Epoch 694
└ @ Main /Users/kcft114/.julia/packages/Flux/qXNjB/src/optimise/train.jl:105
┌ Info: Epoch 695
└ @ Main /Users/kcft114/.julia/packages/Flux/qXNjB/src/optimise/train.jl:105
┌ Info: Epoch 696
└ @ Main /Users/kcft114/.julia/packages/Flux/qXNjB/src/optimise/train.jl:105
┌ Info: Epoch 697
└ @ Main /Users/kcft114/.julia/packages/Flux/qXNjB/src/optimise/train.jl:105
┌ Info: Epoch 698
└ @ Main /Users/kcft114/.julia/packages/Flux/qXNjB/src/optimise/train.jl:105
┌ Info: Epoch 699
└ @ Main /Users/kcft114/.julia/packages/Flux/qXNjB/src/optimise/train.jl:105
┌ Info: Epoch 700
└ @ Main /Users/kcft114/.julia/packages/Flux/qXNjB/src/optimise/train.jl:105
┌ Info: Epoch 701
└ @ Main /Users/kcft114/.julia/packages/Flux/qXNjB/src/optimise/train.jl:105
┌ Info: Epoch 702
└ @ Main /Users/kcft114/.julia/packages/Flux/qXNjB/src/optimise/train.jl:105
┌ Info: Epoch 703
└ @ Main /Users/kcft114/.julia/packages/Flux/qXNjB/src/optimise/train.jl:105
┌ Info: Epoch 704
└ @ Main /Users/kcft114/.julia/packages/Flux/qXNjB/src/optimise/train.jl:105
┌ Info: Epoch 705
└ @ Main /Users/kcft114/.julia/packages/Flux/qXNjB/src/optimise/train.jl:105
┌ Info: Epoch 706
└ @ Main /Users/kcft114/.julia/packages/Flux/qXNjB/src/optimise/train.jl:105
┌ Info: Epoch 707
└ @ Main /Users/kcft114/.julia/packages/Flux/qXNjB/src/optimise/train.jl:105
┌ Info: Epoch 708
└ @ Main /Users/kcft114/.julia/packages/Flux/qXNjB/src/optimise/train.jl:105
┌ Info: Epoch 709
└ @ Main /Users/kcft114/.julia/packages/Flux/qXNjB/src/optimise/train.jl:105
┌ Info: Epoch 710
└ @ Main /Users/kcft114/.julia/packages/Flux/qXNjB/src/optimise/train.jl:105
```

```
loss(X, Y) = 0.7624088f0 (tracked)
loss(X, Y) = 0.70465815f0 (tracked)
loss(X, Y) = 0.7045377f0 (tracked)
loss(X, Y) = 0.6898379f0 (tracked)
loss(X, Y) = 0.72161305f0 (tracked)
loss(X, Y) = 0.7597183f0 (tracked)
loss(X, Y) = 0.7137113f0 (tracked)
loss(X, Y) = 0.710532f0 (tracked)
loss(X, Y) = 0.72440326f0 (tracked)
loss(X, Y) = 0.7503361f0 (tracked)
loss(X, Y) = 0.74908364f0 (tracked)
loss(X, Y) = 0.7090278f0 (tracked)
loss(X, Y) = 0.7177918f0 (tracked)
loss(X, Y) = 0.7102163f0 (tracked)
loss(X, Y) = 0.70971406f0 (tracked)
loss(X, Y) = 0.7417348f0 (tracked)
loss(X, Y) = 0.6906067f0 (tracked)
loss(X, Y) = 0.72205013f0 (tracked)
loss(X, Y) = 0.7140626f0 (tracked)
loss(X, Y) = 0.75302076f0 (tracked)
loss(X, Y) = 0.6485054f0 (tracked)
loss(X, Y) = 0.73091376f0 (tracked)
loss(X, Y) = 0.7218902f0 (tracked)
loss(X, Y) = 0.7473136f0 (tracked)
loss(X, Y) = 0.7287833f0 (tracked)
loss(X, Y) = 0.71361786f0 (tracked)
loss(X, Y) = 0.71408576f0 (tracked)
loss(X, Y) = 0.776506f0 (tracked)
loss(X, Y) = 0.7060277f0 (tracked)
loss(X, Y) = 0.7550056f0 (tracked)
loss(X, Y) = 0.7020893f0 (tracked)
loss(X, Y) = 0.72746974f0 (tracked)
loss(X, Y) = 0.7340105f0 (tracked)
loss(X, Y) = 0.74197805f0 (tracked)
loss(X, Y) = 0.6651162f0 (tracked)
loss(X, Y) = 0.7396036f0 (tracked)
loss(X, Y) = 0.72045004f0 (tracked)
loss(X, Y) = 0.7081742f0 (tracked)
loss(X, Y) = 0.7209229f0 (tracked)
loss(X, Y) = 0.6869993f0 (tracked)
loss(X, Y) = 0.7208205f0 (tracked)
loss(X, Y) = 0.70584196f0 (tracked)
loss(X, Y) = 0.66061866f0 (tracked)
loss(X, Y) = 0.73413014f0 (tracked)
loss(X, Y) = 0.72155255f0 (tracked)
loss(X, Y) = 0.6947195f0 (tracked)
loss(X, Y) = 0.67224365f0 (tracked)
loss(X, Y) = 0.7162396f0 (tracked)
loss(X, Y) = 0.74082637f0 (tracked)
loss(X, Y) = 0.70519584f0 (tracked)
loss(X, Y) = 0.76452667f0 (tracked)
loss(X, Y) = 0.67613506f0 (tracked)
loss(X, Y) = 0.74213064f0 (tracked)
loss(X, Y) = 0.77123743f0 (tracked)
loss(X, Y) = 0.757242f0 (tracked)
loss(X, Y) = 0.7180773f0 (tracked)
loss(X, Y) = 0.7200587f0 (tracked)
loss(X, Y) = 0.7462801f0 (tracked)
loss(X, Y) = 0.7069417f0 (tracked)
loss(X, Y) = 0.7258382f0 (tracked)
loss(X, Y) = 0.76809365f0 (tracked)
loss(X, Y) = 0.70545363f0 (tracked)
loss(X, Y) = 0.7100269f0 (tracked)
loss(X, Y) = 0.7214894f0 (tracked)
loss(X, Y) = 0.73400646f0 (tracked)
loss(X, Y) = 0.7004624f0 (tracked)
loss(X, Y) = 0.7102445f0 (tracked)
loss(X, Y) = 0.718355f0 (tracked)
loss(X, Y) = 0.6936366f0 (tracked)
loss(X, Y) = 0.6632039f0 (tracked)
loss(X, Y) = 0.68442196f0 (tracked)
loss(X, Y) = 0.7186361f0 (tracked)
loss(X, Y) = 0.6957198f0 (tracked)
loss(X, Y) = 0.7105029f0 (tracked)
loss(X, Y) = 0.7072867f0 (tracked)
loss(X, Y) = 0.7467813f0 (tracked)
loss(X, Y) = 0.68205106f0 (tracked)
loss(X, Y) = 0.69839954f0 (tracked)
loss(X, Y) = 0.73567307f0 (tracked)
loss(X, Y) = 0.7046206f0 (tracked)
loss(X, Y) = 0.75119257f0 (tracked)
```

```
┌ Info: Epoch 711
└ @ Main /Users/kcft114/.julia/packages/Flux/qXNjB/src/optimise/train.jl:105
┌ Info: Epoch 712
└ @ Main /Users/kcft114/.julia/packages/Flux/qXNjB/src/optimise/train.jl:105
┌ Info: Epoch 713
└ @ Main /Users/kcft114/.julia/packages/Flux/qXNjB/src/optimise/train.jl:105
┌ Info: Epoch 714
└ @ Main /Users/kcft114/.julia/packages/Flux/qXNjB/src/optimise/train.jl:105
┌ Info: Epoch 715
└ @ Main /Users/kcft114/.julia/packages/Flux/qXNjB/src/optimise/train.jl:105
┌ Info: Epoch 716
└ @ Main /Users/kcft114/.julia/packages/Flux/qXNjB/src/optimise/train.jl:105
┌ Info: Epoch 717
└ @ Main /Users/kcft114/.julia/packages/Flux/qXNjB/src/optimise/train.jl:105
┌ Info: Epoch 718
└ @ Main /Users/kcft114/.julia/packages/Flux/qXNjB/src/optimise/train.jl:105
┌ Info: Epoch 719
└ @ Main /Users/kcft114/.julia/packages/Flux/qXNjB/src/optimise/train.jl:105
┌ Info: Epoch 720
└ @ Main /Users/kcft114/.julia/packages/Flux/qXNjB/src/optimise/train.jl:105
┌ Info: Epoch 721
└ @ Main /Users/kcft114/.julia/packages/Flux/qXNjB/src/optimise/train.jl:105
┌ Info: Epoch 722
└ @ Main /Users/kcft114/.julia/packages/Flux/qXNjB/src/optimise/train.jl:105
┌ Info: Epoch 723
└ @ Main /Users/kcft114/.julia/packages/Flux/qXNjB/src/optimise/train.jl:105
┌ Info: Epoch 724
└ @ Main /Users/kcft114/.julia/packages/Flux/qXNjB/src/optimise/train.jl:105
┌ Info: Epoch 725
└ @ Main /Users/kcft114/.julia/packages/Flux/qXNjB/src/optimise/train.jl:105
┌ Info: Epoch 726
└ @ Main /Users/kcft114/.julia/packages/Flux/qXNjB/src/optimise/train.jl:105
┌ Info: Epoch 727
└ @ Main /Users/kcft114/.julia/packages/Flux/qXNjB/src/optimise/train.jl:105
┌ Info: Epoch 728
└ @ Main /Users/kcft114/.julia/packages/Flux/qXNjB/src/optimise/train.jl:105
┌ Info: Epoch 729
└ @ Main /Users/kcft114/.julia/packages/Flux/qXNjB/src/optimise/train.jl:105
┌ Info: Epoch 730
└ @ Main /Users/kcft114/.julia/packages/Flux/qXNjB/src/optimise/train.jl:105
┌ Info: Epoch 731
└ @ Main /Users/kcft114/.julia/packages/Flux/qXNjB/src/optimise/train.jl:105
┌ Info: Epoch 732
└ @ Main /Users/kcft114/.julia/packages/Flux/qXNjB/src/optimise/train.jl:105
┌ Info: Epoch 733
└ @ Main /Users/kcft114/.julia/packages/Flux/qXNjB/src/optimise/train.jl:105
┌ Info: Epoch 734
└ @ Main /Users/kcft114/.julia/packages/Flux/qXNjB/src/optimise/train.jl:105
┌ Info: Epoch 735
└ @ Main /Users/kcft114/.julia/packages/Flux/qXNjB/src/optimise/train.jl:105
┌ Info: Epoch 736
└ @ Main /Users/kcft114/.julia/packages/Flux/qXNjB/src/optimise/train.jl:105
┌ Info: Epoch 737
└ @ Main /Users/kcft114/.julia/packages/Flux/qXNjB/src/optimise/train.jl:105
┌ Info: Epoch 738
└ @ Main /Users/kcft114/.julia/packages/Flux/qXNjB/src/optimise/train.jl:105
┌ Info: Epoch 739
└ @ Main /Users/kcft114/.julia/packages/Flux/qXNjB/src/optimise/train.jl:105
┌ Info: Epoch 740
└ @ Main /Users/kcft114/.julia/packages/Flux/qXNjB/src/optimise/train.jl:105
┌ Info: Epoch 741
└ @ Main /Users/kcft114/.julia/packages/Flux/qXNjB/src/optimise/train.jl:105
┌ Info: Epoch 742
└ @ Main /Users/kcft114/.julia/packages/Flux/qXNjB/src/optimise/train.jl:105
┌ Info: Epoch 743
└ @ Main /Users/kcft114/.julia/packages/Flux/qXNjB/src/optimise/train.jl:105
┌ Info: Epoch 744
└ @ Main /Users/kcft114/.julia/packages/Flux/qXNjB/src/optimise/train.jl:105
┌ Info: Epoch 745
└ @ Main /Users/kcft114/.julia/packages/Flux/qXNjB/src/optimise/train.jl:105
┌ Info: Epoch 746
└ @ Main /Users/kcft114/.julia/packages/Flux/qXNjB/src/optimise/train.jl:105
┌ Info: Epoch 747
└ @ Main /Users/kcft114/.julia/packages/Flux/qXNjB/src/optimise/train.jl:105
┌ Info: Epoch 748
└ @ Main /Users/kcft114/.julia/packages/Flux/qXNjB/src/optimise/train.jl:105
┌ Info: Epoch 749
└ @ Main /Users/kcft114/.julia/packages/Flux/qXNjB/src/optimise/train.jl:105
┌ Info: Epoch 750
└ @ Main /Users/kcft114/.julia/packages/Flux/qXNjB/src/optimise/train.jl:105
┌ Info: Epoch 751
└ @ Main /Users/kcft114/.julia/packages/Flux/qXNjB/src/optimise/train.jl:105
┌ Info: Epoch 752
└ @ Main /Users/kcft114/.julia/packages/Flux/qXNjB/src/optimise/train.jl:105
┌ Info: Epoch 753
└ @ Main /Users/kcft114/.julia/packages/Flux/qXNjB/src/optimise/train.jl:105
┌ Info: Epoch 754
└ @ Main /Users/kcft114/.julia/packages/Flux/qXNjB/src/optimise/train.jl:105
┌ Info: Epoch 755
└ @ Main /Users/kcft114/.julia/packages/Flux/qXNjB/src/optimise/train.jl:105
┌ Info: Epoch 756
└ @ Main /Users/kcft114/.julia/packages/Flux/qXNjB/src/optimise/train.jl:105
┌ Info: Epoch 757
└ @ Main /Users/kcft114/.julia/packages/Flux/qXNjB/src/optimise/train.jl:105
┌ Info: Epoch 758
└ @ Main /Users/kcft114/.julia/packages/Flux/qXNjB/src/optimise/train.jl:105
┌ Info: Epoch 759
└ @ Main /Users/kcft114/.julia/packages/Flux/qXNjB/src/optimise/train.jl:105
┌ Info: Epoch 760
└ @ Main /Users/kcft114/.julia/packages/Flux/qXNjB/src/optimise/train.jl:105
┌ Info: Epoch 761
└ @ Main /Users/kcft114/.julia/packages/Flux/qXNjB/src/optimise/train.jl:105
┌ Info: Epoch 762
└ @ Main /Users/kcft114/.julia/packages/Flux/qXNjB/src/optimise/train.jl:105
┌ Info: Epoch 763
└ @ Main /Users/kcft114/.julia/packages/Flux/qXNjB/src/optimise/train.jl:105
┌ Info: Epoch 764
└ @ Main /Users/kcft114/.julia/packages/Flux/qXNjB/src/optimise/train.jl:105
┌ Info: Epoch 765
└ @ Main /Users/kcft114/.julia/packages/Flux/qXNjB/src/optimise/train.jl:105
┌ Info: Epoch 766
└ @ Main /Users/kcft114/.julia/packages/Flux/qXNjB/src/optimise/train.jl:105
┌ Info: Epoch 767
└ @ Main /Users/kcft114/.julia/packages/Flux/qXNjB/src/optimise/train.jl:105
┌ Info: Epoch 768
└ @ Main /Users/kcft114/.julia/packages/Flux/qXNjB/src/optimise/train.jl:105
┌ Info: Epoch 769
└ @ Main /Users/kcft114/.julia/packages/Flux/qXNjB/src/optimise/train.jl:105
┌ Info: Epoch 770
└ @ Main /Users/kcft114/.julia/packages/Flux/qXNjB/src/optimise/train.jl:105
┌ Info: Epoch 771
└ @ Main /Users/kcft114/.julia/packages/Flux/qXNjB/src/optimise/train.jl:105
┌ Info: Epoch 772
└ @ Main /Users/kcft114/.julia/packages/Flux/qXNjB/src/optimise/train.jl:105
┌ Info: Epoch 773
└ @ Main /Users/kcft114/.julia/packages/Flux/qXNjB/src/optimise/train.jl:105
┌ Info: Epoch 774
└ @ Main /Users/kcft114/.julia/packages/Flux/qXNjB/src/optimise/train.jl:105
┌ Info: Epoch 775
└ @ Main /Users/kcft114/.julia/packages/Flux/qXNjB/src/optimise/train.jl:105
┌ Info: Epoch 776
└ @ Main /Users/kcft114/.julia/packages/Flux/qXNjB/src/optimise/train.jl:105
┌ Info: Epoch 777
└ @ Main /Users/kcft114/.julia/packages/Flux/qXNjB/src/optimise/train.jl:105
┌ Info: Epoch 778
└ @ Main /Users/kcft114/.julia/packages/Flux/qXNjB/src/optimise/train.jl:105
┌ Info: Epoch 779
└ @ Main /Users/kcft114/.julia/packages/Flux/qXNjB/src/optimise/train.jl:105
┌ Info: Epoch 780
└ @ Main /Users/kcft114/.julia/packages/Flux/qXNjB/src/optimise/train.jl:105
┌ Info: Epoch 781
└ @ Main /Users/kcft114/.julia/packages/Flux/qXNjB/src/optimise/train.jl:105
┌ Info: Epoch 782
└ @ Main /Users/kcft114/.julia/packages/Flux/qXNjB/src/optimise/train.jl:105
┌ Info: Epoch 783
└ @ Main /Users/kcft114/.julia/packages/Flux/qXNjB/src/optimise/train.jl:105
┌ Info: Epoch 784
└ @ Main /Users/kcft114/.julia/packages/Flux/qXNjB/src/optimise/train.jl:105
┌ Info: Epoch 785
└ @ Main /Users/kcft114/.julia/packages/Flux/qXNjB/src/optimise/train.jl:105
┌ Info: Epoch 786
└ @ Main /Users/kcft114/.julia/packages/Flux/qXNjB/src/optimise/train.jl:105
┌ Info: Epoch 787
└ @ Main /Users/kcft114/.julia/packages/Flux/qXNjB/src/optimise/train.jl:105
┌ Info: Epoch 788
└ @ Main /Users/kcft114/.julia/packages/Flux/qXNjB/src/optimise/train.jl:105
┌ Info: Epoch 789
└ @ Main /Users/kcft114/.julia/packages/Flux/qXNjB/src/optimise/train.jl:105
┌ Info: Epoch 790
└ @ Main /Users/kcft114/.julia/packages/Flux/qXNjB/src/optimise/train.jl:105
┌ Info: Epoch 791
└ @ Main /Users/kcft114/.julia/packages/Flux/qXNjB/src/optimise/train.jl:105
```

```
loss(X, Y) = 0.7119716f0 (tracked)
loss(X, Y) = 0.7766874f0 (tracked)
loss(X, Y) = 0.7320691f0 (tracked)
loss(X, Y) = 0.71538f0 (tracked)
loss(X, Y) = 0.7520833f0 (tracked)
loss(X, Y) = 0.66008604f0 (tracked)
loss(X, Y) = 0.7126327f0 (tracked)
loss(X, Y) = 0.7043376f0 (tracked)
loss(X, Y) = 0.7023567f0 (tracked)
loss(X, Y) = 0.76381725f0 (tracked)
loss(X, Y) = 0.6808479f0 (tracked)
loss(X, Y) = 0.6986334f0 (tracked)
loss(X, Y) = 0.7355853f0 (tracked)
loss(X, Y) = 0.7399476f0 (tracked)
loss(X, Y) = 0.7804803f0 (tracked)
loss(X, Y) = 0.6666032f0 (tracked)
loss(X, Y) = 0.73758215f0 (tracked)
loss(X, Y) = 0.67337084f0 (tracked)
loss(X, Y) = 0.709966f0 (tracked)
loss(X, Y) = 0.706935f0 (tracked)
loss(X, Y) = 0.6841793f0 (tracked)
loss(X, Y) = 0.66881436f0 (tracked)
loss(X, Y) = 0.7355023f0 (tracked)
loss(X, Y) = 0.7371907f0 (tracked)
loss(X, Y) = 0.7410555f0 (tracked)
loss(X, Y) = 0.72842497f0 (tracked)
loss(X, Y) = 0.71209604f0 (tracked)
loss(X, Y) = 0.6817917f0 (tracked)
loss(X, Y) = 0.7154079f0 (tracked)
loss(X, Y) = 0.645933f0 (tracked)
loss(X, Y) = 0.68363154f0 (tracked)
loss(X, Y) = 0.7241707f0 (tracked)
loss(X, Y) = 0.67259747f0 (tracked)
loss(X, Y) = 0.71457684f0 (tracked)
loss(X, Y) = 0.69414425f0 (tracked)
loss(X, Y) = 0.6692786f0 (tracked)
loss(X, Y) = 0.73218304f0 (tracked)
loss(X, Y) = 0.7380397f0 (tracked)
loss(X, Y) = 0.687688f0 (tracked)
loss(X, Y) = 0.6863628f0 (tracked)
loss(X, Y) = 0.7321916f0 (tracked)
loss(X, Y) = 0.6943654f0 (tracked)
loss(X, Y) = 0.7108276f0 (tracked)
loss(X, Y) = 0.764429f0 (tracked)
loss(X, Y) = 0.7210932f0 (tracked)
loss(X, Y) = 0.743138f0 (tracked)
loss(X, Y) = 0.6702884f0 (tracked)
loss(X, Y) = 0.6755032f0 (tracked)
loss(X, Y) = 0.7404949f0 (tracked)
loss(X, Y) = 0.7373663f0 (tracked)
loss(X, Y) = 0.71392834f0 (tracked)
loss(X, Y) = 0.70701903f0 (tracked)
loss(X, Y) = 0.70079505f0 (tracked)
loss(X, Y) = 0.709598f0 (tracked)
loss(X, Y) = 0.68053025f0 (tracked)
loss(X, Y) = 0.69035155f0 (tracked)
loss(X, Y) = 0.6654778f0 (tracked)
loss(X, Y) = 0.7819221f0 (tracked)
loss(X, Y) = 0.7084835f0 (tracked)
loss(X, Y) = 0.66017514f0 (tracked)
loss(X, Y) = 0.74539584f0 (tracked)
loss(X, Y) = 0.7461001f0 (tracked)
loss(X, Y) = 0.70667225f0 (tracked)
loss(X, Y) = 0.702872f0 (tracked)
loss(X, Y) = 0.67073697f0 (tracked)
loss(X, Y) = 0.6844779f0 (tracked)
loss(X, Y) = 0.68626136f0 (tracked)
loss(X, Y) = 0.76994574f0 (tracked)
loss(X, Y) = 0.72684085f0 (tracked)
loss(X, Y) = 0.7365808f0 (tracked)
loss(X, Y) = 0.73322093f0 (tracked)
loss(X, Y) = 0.69672644f0 (tracked)
loss(X, Y) = 0.7289116f0 (tracked)
loss(X, Y) = 0.7014691f0 (tracked)
loss(X, Y) = 0.687638f0 (tracked)
loss(X, Y) = 0.7076198f0 (tracked)
loss(X, Y) = 0.7216815f0 (tracked)
loss(X, Y) = 0.74557024f0 (tracked)
```

```
┌ Info: Epoch 792
└ @ Main /Users/kcft114/.julia/packages/Flux/qXNjB/src/optimise/train.jl:105
┌ Info: Epoch 793
└ @ Main /Users/kcft114/.julia/packages/Flux/qXNjB/src/optimise/train.jl:105
┌ Info: Epoch 794
└ @ Main /Users/kcft114/.julia/packages/Flux/qXNjB/src/optimise/train.jl:105
┌ Info: Epoch 795
└ @ Main /Users/kcft114/.julia/packages/Flux/qXNjB/src/optimise/train.jl:105
┌ Info: Epoch 796
└ @ Main /Users/kcft114/.julia/packages/Flux/qXNjB/src/optimise/train.jl:105
┌ Info: Epoch 797
└ @ Main /Users/kcft114/.julia/packages/Flux/qXNjB/src/optimise/train.jl:105
┌ Info: Epoch 798
└ @ Main /Users/kcft114/.julia/packages/Flux/qXNjB/src/optimise/train.jl:105
┌ Info: Epoch 799
└ @ Main /Users/kcft114/.julia/packages/Flux/qXNjB/src/optimise/train.jl:105
┌ Info: Epoch 800
└ @ Main /Users/kcft114/.julia/packages/Flux/qXNjB/src/optimise/train.jl:105
┌ Info: Epoch 801
└ @ Main /Users/kcft114/.julia/packages/Flux/qXNjB/src/optimise/train.jl:105
┌ Info: Epoch 802
└ @ Main /Users/kcft114/.julia/packages/Flux/qXNjB/src/optimise/train.jl:105
┌ Info: Epoch 803
└ @ Main /Users/kcft114/.julia/packages/Flux/qXNjB/src/optimise/train.jl:105
┌ Info: Epoch 804
└ @ Main /Users/kcft114/.julia/packages/Flux/qXNjB/src/optimise/train.jl:105
┌ Info: Epoch 805
└ @ Main /Users/kcft114/.julia/packages/Flux/qXNjB/src/optimise/train.jl:105
┌ Info: Epoch 806
└ @ Main /Users/kcft114/.julia/packages/Flux/qXNjB/src/optimise/train.jl:105
┌ Info: Epoch 807
└ @ Main /Users/kcft114/.julia/packages/Flux/qXNjB/src/optimise/train.jl:105
┌ Info: Epoch 808
└ @ Main /Users/kcft114/.julia/packages/Flux/qXNjB/src/optimise/train.jl:105
┌ Info: Epoch 809
└ @ Main /Users/kcft114/.julia/packages/Flux/qXNjB/src/optimise/train.jl:105
┌ Info: Epoch 810
└ @ Main /Users/kcft114/.julia/packages/Flux/qXNjB/src/optimise/train.jl:105
┌ Info: Epoch 811
└ @ Main /Users/kcft114/.julia/packages/Flux/qXNjB/src/optimise/train.jl:105
┌ Info: Epoch 812
└ @ Main /Users/kcft114/.julia/packages/Flux/qXNjB/src/optimise/train.jl:105
┌ Info: Epoch 813
└ @ Main /Users/kcft114/.julia/packages/Flux/qXNjB/src/optimise/train.jl:105
┌ Info: Epoch 814
└ @ Main /Users/kcft114/.julia/packages/Flux/qXNjB/src/optimise/train.jl:105
┌ Info: Epoch 815
└ @ Main /Users/kcft114/.julia/packages/Flux/qXNjB/src/optimise/train.jl:105
┌ Info: Epoch 816
└ @ Main /Users/kcft114/.julia/packages/Flux/qXNjB/src/optimise/train.jl:105
┌ Info: Epoch 817
└ @ Main /Users/kcft114/.julia/packages/Flux/qXNjB/src/optimise/train.jl:105
┌ Info: Epoch 818
└ @ Main /Users/kcft114/.julia/packages/Flux/qXNjB/src/optimise/train.jl:105
┌ Info: Epoch 819
└ @ Main /Users/kcft114/.julia/packages/Flux/qXNjB/src/optimise/train.jl:105
┌ Info: Epoch 820
└ @ Main /Users/kcft114/.julia/packages/Flux/qXNjB/src/optimise/train.jl:105
┌ Info: Epoch 821
└ @ Main /Users/kcft114/.julia/packages/Flux/qXNjB/src/optimise/train.jl:105
┌ Info: Epoch 822
└ @ Main /Users/kcft114/.julia/packages/Flux/qXNjB/src/optimise/train.jl:105
┌ Info: Epoch 823
└ @ Main /Users/kcft114/.julia/packages/Flux/qXNjB/src/optimise/train.jl:105
┌ Info: Epoch 824
└ @ Main /Users/kcft114/.julia/packages/Flux/qXNjB/src/optimise/train.jl:105
┌ Info: Epoch 825
└ @ Main /Users/kcft114/.julia/packages/Flux/qXNjB/src/optimise/train.jl:105
┌ Info: Epoch 826
└ @ Main /Users/kcft114/.julia/packages/Flux/qXNjB/src/optimise/train.jl:105
┌ Info: Epoch 827
└ @ Main /Users/kcft114/.julia/packages/Flux/qXNjB/src/optimise/train.jl:105
┌ Info: Epoch 828
└ @ Main /Users/kcft114/.julia/packages/Flux/qXNjB/src/optimise/train.jl:105
┌ Info: Epoch 829
└ @ Main /Users/kcft114/.julia/packages/Flux/qXNjB/src/optimise/train.jl:105
┌ Info: Epoch 830
└ @ Main /Users/kcft114/.julia/packages/Flux/qXNjB/src/optimise/train.jl:105
┌ Info: Epoch 831
└ @ Main /Users/kcft114/.julia/packages/Flux/qXNjB/src/optimise/train.jl:105
┌ Info: Epoch 832
└ @ Main /Users/kcft114/.julia/packages/Flux/qXNjB/src/optimise/train.jl:105
┌ Info: Epoch 833
└ @ Main /Users/kcft114/.julia/packages/Flux/qXNjB/src/optimise/train.jl:105
┌ Info: Epoch 834
└ @ Main /Users/kcft114/.julia/packages/Flux/qXNjB/src/optimise/train.jl:105
┌ Info: Epoch 835
└ @ Main /Users/kcft114/.julia/packages/Flux/qXNjB/src/optimise/train.jl:105
┌ Info: Epoch 836
└ @ Main /Users/kcft114/.julia/packages/Flux/qXNjB/src/optimise/train.jl:105
┌ Info: Epoch 837
└ @ Main /Users/kcft114/.julia/packages/Flux/qXNjB/src/optimise/train.jl:105
┌ Info: Epoch 838
└ @ Main /Users/kcft114/.julia/packages/Flux/qXNjB/src/optimise/train.jl:105
┌ Info: Epoch 839
└ @ Main /Users/kcft114/.julia/packages/Flux/qXNjB/src/optimise/train.jl:105
┌ Info: Epoch 840
└ @ Main /Users/kcft114/.julia/packages/Flux/qXNjB/src/optimise/train.jl:105
┌ Info: Epoch 841
└ @ Main /Users/kcft114/.julia/packages/Flux/qXNjB/src/optimise/train.jl:105
┌ Info: Epoch 842
└ @ Main /Users/kcft114/.julia/packages/Flux/qXNjB/src/optimise/train.jl:105
┌ Info: Epoch 843
└ @ Main /Users/kcft114/.julia/packages/Flux/qXNjB/src/optimise/train.jl:105
┌ Info: Epoch 844
└ @ Main /Users/kcft114/.julia/packages/Flux/qXNjB/src/optimise/train.jl:105
┌ Info: Epoch 845
└ @ Main /Users/kcft114/.julia/packages/Flux/qXNjB/src/optimise/train.jl:105
┌ Info: Epoch 846
└ @ Main /Users/kcft114/.julia/packages/Flux/qXNjB/src/optimise/train.jl:105
┌ Info: Epoch 847
└ @ Main /Users/kcft114/.julia/packages/Flux/qXNjB/src/optimise/train.jl:105
┌ Info: Epoch 848
└ @ Main /Users/kcft114/.julia/packages/Flux/qXNjB/src/optimise/train.jl:105
┌ Info: Epoch 849
└ @ Main /Users/kcft114/.julia/packages/Flux/qXNjB/src/optimise/train.jl:105
┌ Info: Epoch 850
└ @ Main /Users/kcft114/.julia/packages/Flux/qXNjB/src/optimise/train.jl:105
┌ Info: Epoch 851
└ @ Main /Users/kcft114/.julia/packages/Flux/qXNjB/src/optimise/train.jl:105
┌ Info: Epoch 852
└ @ Main /Users/kcft114/.julia/packages/Flux/qXNjB/src/optimise/train.jl:105
┌ Info: Epoch 853
└ @ Main /Users/kcft114/.julia/packages/Flux/qXNjB/src/optimise/train.jl:105
┌ Info: Epoch 854
└ @ Main /Users/kcft114/.julia/packages/Flux/qXNjB/src/optimise/train.jl:105
┌ Info: Epoch 855
└ @ Main /Users/kcft114/.julia/packages/Flux/qXNjB/src/optimise/train.jl:105
┌ Info: Epoch 856
└ @ Main /Users/kcft114/.julia/packages/Flux/qXNjB/src/optimise/train.jl:105
┌ Info: Epoch 857
└ @ Main /Users/kcft114/.julia/packages/Flux/qXNjB/src/optimise/train.jl:105
┌ Info: Epoch 858
└ @ Main /Users/kcft114/.julia/packages/Flux/qXNjB/src/optimise/train.jl:105
┌ Info: Epoch 859
└ @ Main /Users/kcft114/.julia/packages/Flux/qXNjB/src/optimise/train.jl:105
┌ Info: Epoch 860
└ @ Main /Users/kcft114/.julia/packages/Flux/qXNjB/src/optimise/train.jl:105
┌ Info: Epoch 861
└ @ Main /Users/kcft114/.julia/packages/Flux/qXNjB/src/optimise/train.jl:105
┌ Info: Epoch 862
└ @ Main /Users/kcft114/.julia/packages/Flux/qXNjB/src/optimise/train.jl:105
┌ Info: Epoch 863
└ @ Main /Users/kcft114/.julia/packages/Flux/qXNjB/src/optimise/train.jl:105
┌ Info: Epoch 864
└ @ Main /Users/kcft114/.julia/packages/Flux/qXNjB/src/optimise/train.jl:105
┌ Info: Epoch 865
└ @ Main /Users/kcft114/.julia/packages/Flux/qXNjB/src/optimise/train.jl:105
┌ Info: Epoch 866
└ @ Main /Users/kcft114/.julia/packages/Flux/qXNjB/src/optimise/train.jl:105
┌ Info: Epoch 867
└ @ Main /Users/kcft114/.julia/packages/Flux/qXNjB/src/optimise/train.jl:105
┌ Info: Epoch 868
└ @ Main /Users/kcft114/.julia/packages/Flux/qXNjB/src/optimise/train.jl:105
┌ Info: Epoch 869
└ @ Main /Users/kcft114/.julia/packages/Flux/qXNjB/src/optimise/train.jl:105
┌ Info: Epoch 870
└ @ Main /Users/kcft114/.julia/packages/Flux/qXNjB/src/optimise/train.jl:105
┌ Info: Epoch 871
└ @ Main /Users/kcft114/.julia/packages/Flux/qXNjB/src/optimise/train.jl:105
┌ Info: Epoch 872
└ @ Main /Users/kcft114/.julia/packages/Flux/qXNjB/src/optimise/train.jl:105
┌ Info: Epoch 873
└ @ Main /Users/kcft114/.julia/packages/Flux/qXNjB/src/optimise/train.jl:105
┌ Info: Epoch 874
└ @ Main /Users/kcft114/.julia/packages/Flux/qXNjB/src/optimise/train.jl:105
┌ Info: Epoch 875
└ @ Main /Users/kcft114/.julia/packages/Flux/qXNjB/src/optimise/train.jl:105
┌ Info: Epoch 876
└ @ Main /Users/kcft114/.julia/packages/Flux/qXNjB/src/optimise/train.jl:105
┌ Info: Epoch 877
└ @ Main /Users/kcft114/.julia/packages/Flux/qXNjB/src/optimise/train.jl:105
┌ Info: Epoch 878
└ @ Main /Users/kcft114/.julia/packages/Flux/qXNjB/src/optimise/train.jl:105
┌ Info: Epoch 879
└ @ Main /Users/kcft114/.julia/packages/Flux/qXNjB/src/optimise/train.jl:105
┌ Info: Epoch 880
└ @ Main /Users/kcft114/.julia/packages/Flux/qXNjB/src/optimise/train.jl:105
┌ Info: Epoch 881
└ @ Main /Users/kcft114/.julia/packages/Flux/qXNjB/src/optimise/train.jl:105
┌ Info: Epoch 882
└ @ Main /Users/kcft114/.julia/packages/Flux/qXNjB/src/optimise/train.jl:105
┌ Info: Epoch 883
└ @ Main /Users/kcft114/.julia/packages/Flux/qXNjB/src/optimise/train.jl:105
┌ Info: Epoch 884
└ @ Main /Users/kcft114/.julia/packages/Flux/qXNjB/src/optimise/train.jl:105
┌ Info: Epoch 885
└ @ Main /Users/kcft114/.julia/packages/Flux/qXNjB/src/optimise/train.jl:105
┌ Info: Epoch 886
└ @ Main /Users/kcft114/.julia/packages/Flux/qXNjB/src/optimise/train.jl:105
┌ Info: Epoch 887
└ @ Main /Users/kcft114/.julia/packages/Flux/qXNjB/src/optimise/train.jl:105
┌ Info: Epoch 888
└ @ Main /Users/kcft114/.julia/packages/Flux/qXNjB/src/optimise/train.jl:105
┌ Info: Epoch 889
└ @ Main /Users/kcft114/.julia/packages/Flux/qXNjB/src/optimise/train.jl:105
┌ Info: Epoch 890
└ @ Main /Users/kcft114/.julia/packages/Flux/qXNjB/src/optimise/train.jl:105
┌ Info: Epoch 891
└ @ Main /Users/kcft114/.julia/packages/Flux/qXNjB/src/optimise/train.jl:105
┌ Info: Epoch 892
└ @ Main /Users/kcft114/.julia/packages/Flux/qXNjB/src/optimise/train.jl:105
┌ Info: Epoch 893
└ @ Main /Users/kcft114/.julia/packages/Flux/qXNjB/src/optimise/train.jl:105
┌ Info: Epoch 894
└ @ Main /Users/kcft114/.julia/packages/Flux/qXNjB/src/optimise/train.jl:105
┌ Info: Epoch 895
└ @ Main /Users/kcft114/.julia/packages/Flux/qXNjB/src/optimise/train.jl:105
┌ Info: Epoch 896
└ @ Main /Users/kcft114/.julia/packages/Flux/qXNjB/src/optimise/train.jl:105
┌ Info: Epoch 897
└ @ Main /Users/kcft114/.julia/packages/Flux/qXNjB/src/optimise/train.jl:105
┌ Info: Epoch 898
└ @ Main /Users/kcft114/.julia/packages/Flux/qXNjB/src/optimise/train.jl:105
┌ Info: Epoch 899
└ @ Main /Users/kcft114/.julia/packages/Flux/qXNjB/src/optimise/train.jl:105
┌ Info: Epoch 900
└ @ Main /Users/kcft114/.julia/packages/Flux/qXNjB/src/optimise/train.jl:105
┌ Info: Epoch 901
└ @ Main /Users/kcft114/.julia/packages/Flux/qXNjB/src/optimise/train.jl:105
┌ Info: Epoch 902
└ @ Main /Users/kcft114/.julia/packages/Flux/qXNjB/src/optimise/train.jl:105
┌ Info: Epoch 903
└ @ Main /Users/kcft114/.julia/packages/Flux/qXNjB/src/optimise/train.jl:105
┌ Info: Epoch 904
└ @ Main /Users/kcft114/.julia/packages/Flux/qXNjB/src/optimise/train.jl:105
┌ Info: Epoch 905
└ @ Main /Users/kcft114/.julia/packages/Flux/qXNjB/src/optimise/train.jl:105
┌ Info: Epoch 906
└ @ Main /Users/kcft114/.julia/packages/Flux/qXNjB/src/optimise/train.jl:105
┌ Info: Epoch 907
└ @ Main /Users/kcft114/.julia/packages/Flux/qXNjB/src/optimise/train.jl:105
┌ Info: Epoch 908
└ @ Main /Users/kcft114/.julia/packages/Flux/qXNjB/src/optimise/train.jl:105
┌ Info: Epoch 909
└ @ Main /Users/kcft114/.julia/packages/Flux/qXNjB/src/optimise/train.jl:105
┌ Info: Epoch 910
└ @ Main /Users/kcft114/.julia/packages/Flux/qXNjB/src/optimise/train.jl:105
┌ Info: Epoch 911
└ @ Main /Users/kcft114/.julia/packages/Flux/qXNjB/src/optimise/train.jl:105
┌ Info: Epoch 912
└ @ Main /Users/kcft114/.julia/packages/Flux/qXNjB/src/optimise/train.jl:105
┌ Info: Epoch 913
└ @ Main /Users/kcft114/.julia/packages/Flux/qXNjB/src/optimise/train.jl:105
┌ Info: Epoch 914
└ @ Main /Users/kcft114/.julia/packages/Flux/qXNjB/src/optimise/train.jl:105
┌ Info: Epoch 915
└ @ Main /Users/kcft114/.julia/packages/Flux/qXNjB/src/optimise/train.jl:105
┌ Info: Epoch 916
└ @ Main /Users/kcft114/.julia/packages/Flux/qXNjB/src/optimise/train.jl:105
┌ Info: Epoch 917
└ @ Main /Users/kcft114/.julia/packages/Flux/qXNjB/src/optimise/train.jl:105
┌ Info: Epoch 918
└ @ Main /Users/kcft114/.julia/packages/Flux/qXNjB/src/optimise/train.jl:105
┌ Info: Epoch 919
└ @ Main /Users/kcft114/.julia/packages/Flux/qXNjB/src/optimise/train.jl:105
┌ Info: Epoch 920
└ @ Main /Users/kcft114/.julia/packages/Flux/qXNjB/src/optimise/train.jl:105
┌ Info: Epoch 921
└ @ Main /Users/kcft114/.julia/packages/Flux/qXNjB/src/optimise/train.jl:105
┌ Info: Epoch 922
└ @ Main /Users/kcft114/.julia/packages/Flux/qXNjB/src/optimise/train.jl:105
┌ Info: Epoch 923
└ @ Main /Users/kcft114/.julia/packages/Flux/qXNjB/src/optimise/train.jl:105
┌ Info: Epoch 924
└ @ Main /Users/kcft114/.julia/packages/Flux/qXNjB/src/optimise/train.jl:105
┌ Info: Epoch 925
└ @ Main /Users/kcft114/.julia/packages/Flux/qXNjB/src/optimise/train.jl:105
┌ Info: Epoch 926
└ @ Main /Users/kcft114/.julia/packages/Flux/qXNjB/src/optimise/train.jl:105
┌ Info: Epoch 927
└ @ Main /Users/kcft114/.julia/packages/Flux/qXNjB/src/optimise/train.jl:105
┌ Info: Epoch 928
└ @ Main /Users/kcft114/.julia/packages/Flux/qXNjB/src/optimise/train.jl:105
┌ Info: Epoch 929
└ @ Main /Users/kcft114/.julia/packages/Flux/qXNjB/src/optimise/train.jl:105
┌ Info: Epoch 930
└ @ Main /Users/kcft114/.julia/packages/Flux/qXNjB/src/optimise/train.jl:105
┌ Info: Epoch 931
└ @ Main /Users/kcft114/.julia/packages/Flux/qXNjB/src/optimise/train.jl:105
┌ Info: Epoch 932
└ @ Main /Users/kcft114/.julia/packages/Flux/qXNjB/src/optimise/train.jl:105
┌ Info: Epoch 933
└ @ Main /Users/kcft114/.julia/packages/Flux/qXNjB/src/optimise/train.jl:105
┌ Info: Epoch 934
└ @ Main /Users/kcft114/.julia/packages/Flux/qXNjB/src/optimise/train.jl:105
┌ Info: Epoch 935
└ @ Main /Users/kcft114/.julia/packages/Flux/qXNjB/src/optimise/train.jl:105
┌ Info: Epoch 936
└ @ Main /Users/kcft114/.julia/packages/Flux/qXNjB/src/optimise/train.jl:105
┌ Info: Epoch 937
└ @ Main /Users/kcft114/.julia/packages/Flux/qXNjB/src/optimise/train.jl:105
┌ Info: Epoch 938
└ @ Main /Users/kcft114/.julia/packages/Flux/qXNjB/src/optimise/train.jl:105
┌ Info: Epoch 939
└ @ Main /Users/kcft114/.julia/packages/Flux/qXNjB/src/optimise/train.jl:105
┌ Info: Epoch 940
└ @ Main /Users/kcft114/.julia/packages/Flux/qXNjB/src/optimise/train.jl:105
┌ Info: Epoch 941
└ @ Main /Users/kcft114/.julia/packages/Flux/qXNjB/src/optimise/train.jl:105
┌ Info: Epoch 942
└ @ Main /Users/kcft114/.julia/packages/Flux/qXNjB/src/optimise/train.jl:105
┌ Info: Epoch 943
└ @ Main /Users/kcft114/.julia/packages/Flux/qXNjB/src/optimise/train.jl:105
┌ Info: Epoch 944
└ @ Main /Users/kcft114/.julia/packages/Flux/qXNjB/src/optimise/train.jl:105
┌ Info: Epoch 945
└ @ Main /Users/kcft114/.julia/packages/Flux/qXNjB/src/optimise/train.jl:105
┌ Info: Epoch 946
└ @ Main /Users/kcft114/.julia/packages/Flux/qXNjB/src/optimise/train.jl:105
┌ Info: Epoch 947
└ @ Main /Users/kcft114/.julia/packages/Flux/qXNjB/src/optimise/train.jl:105
┌ Info: Epoch 948
└ @ Main /Users/kcft114/.julia/packages/Flux/qXNjB/src/optimise/train.jl:105
┌ Info: Epoch 949
└ @ Main /Users/kcft114/.julia/packages/Flux/qXNjB/src/optimise/train.jl:105
┌ Info: Epoch 950
└ @ Main /Users/kcft114/.julia/packages/Flux/qXNjB/src/optimise/train.jl:105
┌ Info: Epoch 951
└ @ Main /Users/kcft114/.julia/packages/Flux/qXNjB/src/optimise/train.jl:105
┌ Info: Epoch 952
└ @ Main /Users/kcft114/.julia/packages/Flux/qXNjB/src/optimise/train.jl:105
┌ Info: Epoch 953
└ @ Main /Users/kcft114/.julia/packages/Flux/qXNjB/src/optimise/train.jl:105
┌ Info: Epoch 954
└ @ Main /Users/kcft114/.julia/packages/Flux/qXNjB/src/optimise/train.jl:105
┌ Info: Epoch 955
└ @ Main /Users/kcft114/.julia/packages/Flux/qXNjB/src/optimise/train.jl:105
┌ Info: Epoch 956
└ @ Main /Users/kcft114/.julia/packages/Flux/qXNjB/src/optimise/train.jl:105
┌ Info: Epoch 957
└ @ Main /Users/kcft114/.julia/packages/Flux/qXNjB/src/optimise/train.jl:105
┌ Info: Epoch 958
└ @ Main /Users/kcft114/.julia/packages/Flux/qXNjB/src/optimise/train.jl:105
┌ Info: Epoch 959
└ @ Main /Users/kcft114/.julia/packages/Flux/qXNjB/src/optimise/train.jl:105
┌ Info: Epoch 960
└ @ Main /Users/kcft114/.julia/packages/Flux/qXNjB/src/optimise/train.jl:105
┌ Info: Epoch 961
└ @ Main /Users/kcft114/.julia/packages/Flux/qXNjB/src/optimise/train.jl:105
┌ Info: Epoch 962
└ @ Main /Users/kcft114/.julia/packages/Flux/qXNjB/src/optimise/train.jl:105
┌ Info: Epoch 963
└ @ Main /Users/kcft114/.julia/packages/Flux/qXNjB/src/optimise/train.jl:105
┌ Info: Epoch 964
└ @ Main /Users/kcft114/.julia/packages/Flux/qXNjB/src/optimise/train.jl:105
┌ Info: Epoch 965
└ @ Main /Users/kcft114/.julia/packages/Flux/qXNjB/src/optimise/train.jl:105
┌ Info: Epoch 966
└ @ Main /Users/kcft114/.julia/packages/Flux/qXNjB/src/optimise/train.jl:105
┌ Info: Epoch 967
└ @ Main /Users/kcft114/.julia/packages/Flux/qXNjB/src/optimise/train.jl:105
┌ Info: Epoch 968
└ @ Main /Users/kcft114/.julia/packages/Flux/qXNjB/src/optimise/train.jl:105
┌ Info: Epoch 969
└ @ Main /Users/kcft114/.julia/packages/Flux/qXNjB/src/optimise/train.jl:105
┌ Info: Epoch 970
└ @ Main /Users/kcft114/.julia/packages/Flux/qXNjB/src/optimise/train.jl:105
┌ Info: Epoch 971
└ @ Main /Users/kcft114/.julia/packages/Flux/qXNjB/src/optimise/train.jl:105
┌ Info: Epoch 972
└ @ Main /Users/kcft114/.julia/packages/Flux/qXNjB/src/optimise/train.jl:105
┌ Info: Epoch 973
└ @ Main /Users/kcft114/.julia/packages/Flux/qXNjB/src/optimise/train.jl:105
```

```
loss(X, Y) = 0.682411f0 (tracked)
loss(X, Y) = 0.7180946f0 (tracked)
loss(X, Y) = 0.6843733f0 (tracked)
loss(X, Y) = 0.7834336f0 (tracked)
loss(X, Y) = 0.69214016f0 (tracked)
loss(X, Y) = 0.69555026f0 (tracked)
loss(X, Y) = 0.75220466f0 (tracked)
loss(X, Y) = 0.664167f0 (tracked)
loss(X, Y) = 0.67442757f0 (tracked)
loss(X, Y) = 0.7361967f0 (tracked)
loss(X, Y) = 0.70052505f0 (tracked)
loss(X, Y) = 0.7170178f0 (tracked)
loss(X, Y) = 0.706131f0 (tracked)
loss(X, Y) = 0.68404627f0 (tracked)
loss(X, Y) = 0.72323185f0 (tracked)
loss(X, Y) = 0.662059f0 (tracked)
loss(X, Y) = 0.66919816f0 (tracked)
loss(X, Y) = 0.6780283f0 (tracked)
loss(X, Y) = 0.7520018f0 (tracked)
loss(X, Y) = 0.7135429f0 (tracked)
loss(X, Y) = 0.6863023f0 (tracked)
loss(X, Y) = 0.6919742f0 (tracked)
loss(X, Y) = 0.7206456f0 (tracked)
loss(X, Y) = 0.73409593f0 (tracked)
loss(X, Y) = 0.69943446f0 (tracked)
loss(X, Y) = 0.7300585f0 (tracked)
loss(X, Y) = 0.70576537f0 (tracked)
loss(X, Y) = 0.7278941f0 (tracked)
loss(X, Y) = 0.6751521f0 (tracked)
loss(X, Y) = 0.68081313f0 (tracked)
loss(X, Y) = 0.7072225f0 (tracked)
loss(X, Y) = 0.69669324f0 (tracked)
loss(X, Y) = 0.7353705f0 (tracked)
loss(X, Y) = 0.73397017f0 (tracked)
loss(X, Y) = 0.7286543f0 (tracked)
loss(X, Y) = 0.70915306f0 (tracked)
loss(X, Y) = 0.66486084f0 (tracked)
loss(X, Y) = 0.74751353f0 (tracked)
loss(X, Y) = 0.6833486f0 (tracked)
loss(X, Y) = 0.6836099f0 (tracked)
loss(X, Y) = 0.69403964f0 (tracked)
loss(X, Y) = 0.66686034f0 (tracked)
loss(X, Y) = 0.6934331f0 (tracked)
loss(X, Y) = 0.6940967f0 (tracked)
loss(X, Y) = 0.6833016f0 (tracked)
loss(X, Y) = 0.6763581f0 (tracked)
loss(X, Y) = 0.7042978f0 (tracked)
loss(X, Y) = 0.73882794f0 (tracked)
loss(X, Y) = 0.6243916f0 (tracked)
loss(X, Y) = 0.7225308f0 (tracked)
loss(X, Y) = 0.70890635f0 (tracked)
loss(X, Y) = 0.6963944f0 (tracked)
loss(X, Y) = 0.7315385f0 (tracked)
loss(X, Y) = 0.68078965f0 (tracked)
loss(X, Y) = 0.6987438f0 (tracked)
loss(X, Y) = 0.6885959f0 (tracked)
loss(X, Y) = 0.7084215f0 (tracked)
loss(X, Y) = 0.71097124f0 (tracked)
loss(X, Y) = 0.6895757f0 (tracked)
loss(X, Y) = 0.71774733f0 (tracked)
loss(X, Y) = 0.7643937f0 (tracked)
loss(X, Y) = 0.69668424f0 (tracked)
loss(X, Y) = 0.6866102f0 (tracked)
loss(X, Y) = 0.7090309f0 (tracked)
loss(X, Y) = 0.711933f0 (tracked)
loss(X, Y) = 0.6906861f0 (tracked)
loss(X, Y) = 0.6907204f0 (tracked)
loss(X, Y) = 0.69767046f0 (tracked)
loss(X, Y) = 0.7194098f0 (tracked)
loss(X, Y) = 0.6760449f0 (tracked)
loss(X, Y) = 0.6975015f0 (tracked)
loss(X, Y) = 0.6996736f0 (tracked)
loss(X, Y) = 0.6876031f0 (tracked)
loss(X, Y) = 0.6940381f0 (tracked)
loss(X, Y) = 0.6940019f0 (tracked)
loss(X, Y) = 0.6806128f0 (tracked)
loss(X, Y) = 0.754417f0 (tracked)
loss(X, Y) = 0.64789987f0 (tracked)
loss(X, Y) = 0.7099273f0 (tracked)
loss(X, Y) = 0.6914092f0 (tracked)
loss(X, Y) = 0.724336f0 (tracked)
loss(X, Y) = 0.7326263f0 (tracked)
loss(X, Y) = 0.7253736f0 (tracked)
loss(X, Y) = 0.663991f0 (tracked)
loss(X, Y) = 0.69819766f0 (tracked)
loss(X, Y) = 0.70349574f0 (tracked)
loss(X, Y) = 0.70838857f0 (tracked)
loss(X, Y) = 0.7019772f0 (tracked)
loss(X, Y) = 0.6881237f0 (tracked)
loss(X, Y) = 0.7222554f0 (tracked)
loss(X, Y) = 0.6767961f0 (tracked)
loss(X, Y) = 0.6868342f0 (tracked)
loss(X, Y) = 0.6753985f0 (tracked)
loss(X, Y) = 0.68959814f0 (tracked)
loss(X, Y) = 0.7233196f0 (tracked)
loss(X, Y) = 0.683153f0 (tracked)
loss(X, Y) = 0.68152493f0 (tracked)
loss(X, Y) = 0.66515934f0 (tracked)
loss(X, Y) = 0.68585426f0 (tracked)
loss(X, Y) = 0.7210099f0 (tracked)
loss(X, Y) = 0.7012255f0 (tracked)
loss(X, Y) = 0.7094808f0 (tracked)
loss(X, Y) = 0.72254837f0 (tracked)
loss(X, Y) = 0.6669809f0 (tracked)
loss(X, Y) = 0.74629706f0 (tracked)
loss(X, Y) = 0.699216f0 (tracked)
loss(X, Y) = 0.69070345f0 (tracked)
loss(X, Y) = 0.6932285f0 (tracked)
loss(X, Y) = 0.6749637f0 (tracked)
loss(X, Y) = 0.62749654f0 (tracked)
loss(X, Y) = 0.66145223f0 (tracked)
```

```
┌ Info: Epoch 974
└ @ Main /Users/kcft114/.julia/packages/Flux/qXNjB/src/optimise/train.jl:105
┌ Info: Epoch 975
└ @ Main /Users/kcft114/.julia/packages/Flux/qXNjB/src/optimise/train.jl:105
┌ Info: Epoch 976
└ @ Main /Users/kcft114/.julia/packages/Flux/qXNjB/src/optimise/train.jl:105
┌ Info: Epoch 977
└ @ Main /Users/kcft114/.julia/packages/Flux/qXNjB/src/optimise/train.jl:105
┌ Info: Epoch 978
└ @ Main /Users/kcft114/.julia/packages/Flux/qXNjB/src/optimise/train.jl:105
┌ Info: Epoch 979
└ @ Main /Users/kcft114/.julia/packages/Flux/qXNjB/src/optimise/train.jl:105
┌ Info: Epoch 980
└ @ Main /Users/kcft114/.julia/packages/Flux/qXNjB/src/optimise/train.jl:105
┌ Info: Epoch 981
└ @ Main /Users/kcft114/.julia/packages/Flux/qXNjB/src/optimise/train.jl:105
┌ Info: Epoch 982
└ @ Main /Users/kcft114/.julia/packages/Flux/qXNjB/src/optimise/train.jl:105
┌ Info: Epoch 983
└ @ Main /Users/kcft114/.julia/packages/Flux/qXNjB/src/optimise/train.jl:105
┌ Info: Epoch 984
└ @ Main /Users/kcft114/.julia/packages/Flux/qXNjB/src/optimise/train.jl:105
┌ Info: Epoch 985
└ @ Main /Users/kcft114/.julia/packages/Flux/qXNjB/src/optimise/train.jl:105
┌ Info: Epoch 986
└ @ Main /Users/kcft114/.julia/packages/Flux/qXNjB/src/optimise/train.jl:105
┌ Info: Epoch 987
└ @ Main /Users/kcft114/.julia/packages/Flux/qXNjB/src/optimise/train.jl:105
┌ Info: Epoch 988
└ @ Main /Users/kcft114/.julia/packages/Flux/qXNjB/src/optimise/train.jl:105
┌ Info: Epoch 989
└ @ Main /Users/kcft114/.julia/packages/Flux/qXNjB/src/optimise/train.jl:105
┌ Info: Epoch 990
└ @ Main /Users/kcft114/.julia/packages/Flux/qXNjB/src/optimise/train.jl:105
┌ Info: Epoch 991
└ @ Main /Users/kcft114/.julia/packages/Flux/qXNjB/src/optimise/train.jl:105
┌ Info: Epoch 992
└ @ Main /Users/kcft114/.julia/packages/Flux/qXNjB/src/optimise/train.jl:105
┌ Info: Epoch 993
└ @ Main /Users/kcft114/.julia/packages/Flux/qXNjB/src/optimise/train.jl:105
┌ Info: Epoch 994
└ @ Main /Users/kcft114/.julia/packages/Flux/qXNjB/src/optimise/train.jl:105
┌ Info: Epoch 995
└ @ Main /Users/kcft114/.julia/packages/Flux/qXNjB/src/optimise/train.jl:105
┌ Info: Epoch 996
└ @ Main /Users/kcft114/.julia/packages/Flux/qXNjB/src/optimise/train.jl:105
┌ Info: Epoch 997
└ @ Main /Users/kcft114/.julia/packages/Flux/qXNjB/src/optimise/train.jl:105
┌ Info: Epoch 998
└ @ Main /Users/kcft114/.julia/packages/Flux/qXNjB/src/optimise/train.jl:105
┌ Info: Epoch 999
└ @ Main /Users/kcft114/.julia/packages/Flux/qXNjB/src/optimise/train.jl:105
┌ Info: Epoch 1000
└ @ Main /Users/kcft114/.julia/packages/Flux/qXNjB/src/optimise/train.jl:105
┌ Info: Epoch 1001
└ @ Main /Users/kcft114/.julia/packages/Flux/qXNjB/src/optimise/train.jl:105
┌ Info: Epoch 1002
└ @ Main /Users/kcft114/.julia/packages/Flux/qXNjB/src/optimise/train.jl:105
┌ Info: Epoch 1003
└ @ Main /Users/kcft114/.julia/packages/Flux/qXNjB/src/optimise/train.jl:105
┌ Info: Epoch 1004
└ @ Main /Users/kcft114/.julia/packages/Flux/qXNjB/src/optimise/train.jl:105
┌ Info: Epoch 1005
└ @ Main /Users/kcft114/.julia/packages/Flux/qXNjB/src/optimise/train.jl:105
┌ Info: Epoch 1006
└ @ Main /Users/kcft114/.julia/packages/Flux/qXNjB/src/optimise/train.jl:105
┌ Info: Epoch 1007
└ @ Main /Users/kcft114/.julia/packages/Flux/qXNjB/src/optimise/train.jl:105
┌ Info: Epoch 1008
└ @ Main /Users/kcft114/.julia/packages/Flux/qXNjB/src/optimise/train.jl:105
┌ Info: Epoch 1009
└ @ Main /Users/kcft114/.julia/packages/Flux/qXNjB/src/optimise/train.jl:105
┌ Info: Epoch 1010
└ @ Main /Users/kcft114/.julia/packages/Flux/qXNjB/src/optimise/train.jl:105
┌ Info: Epoch 1011
└ @ Main /Users/kcft114/.julia/packages/Flux/qXNjB/src/optimise/train.jl:105
┌ Info: Epoch 1012
└ @ Main /Users/kcft114/.julia/packages/Flux/qXNjB/src/optimise/train.jl:105
┌ Info: Epoch 1013
└ @ Main /Users/kcft114/.julia/packages/Flux/qXNjB/src/optimise/train.jl:105
┌ Info: Epoch 1014
└ @ Main /Users/kcft114/.julia/packages/Flux/qXNjB/src/optimise/train.jl:105
┌ Info: Epoch 1015
└ @ Main /Users/kcft114/.julia/packages/Flux/qXNjB/src/optimise/train.jl:105
┌ Info: Epoch 1016
└ @ Main /Users/kcft114/.julia/packages/Flux/qXNjB/src/optimise/train.jl:105
┌ Info: Epoch 1017
└ @ Main /Users/kcft114/.julia/packages/Flux/qXNjB/src/optimise/train.jl:105
┌ Info: Epoch 1018
└ @ Main /Users/kcft114/.julia/packages/Flux/qXNjB/src/optimise/train.jl:105
┌ Info: Epoch 1019
└ @ Main /Users/kcft114/.julia/packages/Flux/qXNjB/src/optimise/train.jl:105
┌ Info: Epoch 1020
└ @ Main /Users/kcft114/.julia/packages/Flux/qXNjB/src/optimise/train.jl:105
┌ Info: Epoch 1021
└ @ Main /Users/kcft114/.julia/packages/Flux/qXNjB/src/optimise/train.jl:105
┌ Info: Epoch 1022
└ @ Main /Users/kcft114/.julia/packages/Flux/qXNjB/src/optimise/train.jl:105
┌ Info: Epoch 1023
└ @ Main /Users/kcft114/.julia/packages/Flux/qXNjB/src/optimise/train.jl:105
┌ Info: Epoch 1024
└ @ Main /Users/kcft114/.julia/packages/Flux/qXNjB/src/optimise/train.jl:105
┌ Info: Epoch 1025
└ @ Main /Users/kcft114/.julia/packages/Flux/qXNjB/src/optimise/train.jl:105
┌ Info: Epoch 1026
└ @ Main /Users/kcft114/.julia/packages/Flux/qXNjB/src/optimise/train.jl:105
┌ Info: Epoch 1027
└ @ Main /Users/kcft114/.julia/packages/Flux/qXNjB/src/optimise/train.jl:105
┌ Info: Epoch 1028
└ @ Main /Users/kcft114/.julia/packages/Flux/qXNjB/src/optimise/train.jl:105
┌ Info: Epoch 1029
└ @ Main /Users/kcft114/.julia/packages/Flux/qXNjB/src/optimise/train.jl:105
┌ Info: Epoch 1030
└ @ Main /Users/kcft114/.julia/packages/Flux/qXNjB/src/optimise/train.jl:105
┌ Info: Epoch 1031
└ @ Main /Users/kcft114/.julia/packages/Flux/qXNjB/src/optimise/train.jl:105
┌ Info: Epoch 1032
└ @ Main /Users/kcft114/.julia/packages/Flux/qXNjB/src/optimise/train.jl:105
┌ Info: Epoch 1033
└ @ Main /Users/kcft114/.julia/packages/Flux/qXNjB/src/optimise/train.jl:105
┌ Info: Epoch 1034
└ @ Main /Users/kcft114/.julia/packages/Flux/qXNjB/src/optimise/train.jl:105
┌ Info: Epoch 1035
└ @ Main /Users/kcft114/.julia/packages/Flux/qXNjB/src/optimise/train.jl:105
┌ Info: Epoch 1036
└ @ Main /Users/kcft114/.julia/packages/Flux/qXNjB/src/optimise/train.jl:105
┌ Info: Epoch 1037
└ @ Main /Users/kcft114/.julia/packages/Flux/qXNjB/src/optimise/train.jl:105
┌ Info: Epoch 1038
└ @ Main /Users/kcft114/.julia/packages/Flux/qXNjB/src/optimise/train.jl:105
┌ Info: Epoch 1039
└ @ Main /Users/kcft114/.julia/packages/Flux/qXNjB/src/optimise/train.jl:105
┌ Info: Epoch 1040
└ @ Main /Users/kcft114/.julia/packages/Flux/qXNjB/src/optimise/train.jl:105
┌ Info: Epoch 1041
└ @ Main /Users/kcft114/.julia/packages/Flux/qXNjB/src/optimise/train.jl:105
┌ Info: Epoch 1042
└ @ Main /Users/kcft114/.julia/packages/Flux/qXNjB/src/optimise/train.jl:105
┌ Info: Epoch 1043
└ @ Main /Users/kcft114/.julia/packages/Flux/qXNjB/src/optimise/train.jl:105
┌ Info: Epoch 1044
└ @ Main /Users/kcft114/.julia/packages/Flux/qXNjB/src/optimise/train.jl:105
┌ Info: Epoch 1045
└ @ Main /Users/kcft114/.julia/packages/Flux/qXNjB/src/optimise/train.jl:105
┌ Info: Epoch 1046
└ @ Main /Users/kcft114/.julia/packages/Flux/qXNjB/src/optimise/train.jl:105
┌ Info: Epoch 1047
└ @ Main /Users/kcft114/.julia/packages/Flux/qXNjB/src/optimise/train.jl:105
┌ Info: Epoch 1048
└ @ Main /Users/kcft114/.julia/packages/Flux/qXNjB/src/optimise/train.jl:105
┌ Info: Epoch 1049
└ @ Main /Users/kcft114/.julia/packages/Flux/qXNjB/src/optimise/train.jl:105
┌ Info: Epoch 1050
└ @ Main /Users/kcft114/.julia/packages/Flux/qXNjB/src/optimise/train.jl:105
┌ Info: Epoch 1051
└ @ Main /Users/kcft114/.julia/packages/Flux/qXNjB/src/optimise/train.jl:105
┌ Info: Epoch 1052
└ @ Main /Users/kcft114/.julia/packages/Flux/qXNjB/src/optimise/train.jl:105
┌ Info: Epoch 1053
└ @ Main /Users/kcft114/.julia/packages/Flux/qXNjB/src/optimise/train.jl:105
┌ Info: Epoch 1054
└ @ Main /Users/kcft114/.julia/packages/Flux/qXNjB/src/optimise/train.jl:105
┌ Info: Epoch 1055
└ @ Main /Users/kcft114/.julia/packages/Flux/qXNjB/src/optimise/train.jl:105
┌ Info: Epoch 1056
└ @ Main /Users/kcft114/.julia/packages/Flux/qXNjB/src/optimise/train.jl:105
┌ Info: Epoch 1057
└ @ Main /Users/kcft114/.julia/packages/Flux/qXNjB/src/optimise/train.jl:105
┌ Info: Epoch 1058
└ @ Main /Users/kcft114/.julia/packages/Flux/qXNjB/src/optimise/train.jl:105
┌ Info: Epoch 1059
└ @ Main /Users/kcft114/.julia/packages/Flux/qXNjB/src/optimise/train.jl:105
┌ Info: Epoch 1060
└ @ Main /Users/kcft114/.julia/packages/Flux/qXNjB/src/optimise/train.jl:105
┌ Info: Epoch 1061
└ @ Main /Users/kcft114/.julia/packages/Flux/qXNjB/src/optimise/train.jl:105
┌ Info: Epoch 1062
└ @ Main /Users/kcft114/.julia/packages/Flux/qXNjB/src/optimise/train.jl:105
┌ Info: Epoch 1063
└ @ Main /Users/kcft114/.julia/packages/Flux/qXNjB/src/optimise/train.jl:105
┌ Info: Epoch 1064
└ @ Main /Users/kcft114/.julia/packages/Flux/qXNjB/src/optimise/train.jl:105
┌ Info: Epoch 1065
└ @ Main /Users/kcft114/.julia/packages/Flux/qXNjB/src/optimise/train.jl:105
┌ Info: Epoch 1066
└ @ Main /Users/kcft114/.julia/packages/Flux/qXNjB/src/optimise/train.jl:105
┌ Info: Epoch 1067
└ @ Main /Users/kcft114/.julia/packages/Flux/qXNjB/src/optimise/train.jl:105
┌ Info: Epoch 1068
└ @ Main /Users/kcft114/.julia/packages/Flux/qXNjB/src/optimise/train.jl:105
┌ Info: Epoch 1069
└ @ Main /Users/kcft114/.julia/packages/Flux/qXNjB/src/optimise/train.jl:105
┌ Info: Epoch 1070
└ @ Main /Users/kcft114/.julia/packages/Flux/qXNjB/src/optimise/train.jl:105
┌ Info: Epoch 1071
└ @ Main /Users/kcft114/.julia/packages/Flux/qXNjB/src/optimise/train.jl:105
┌ Info: Epoch 1072
└ @ Main /Users/kcft114/.julia/packages/Flux/qXNjB/src/optimise/train.jl:105
┌ Info: Epoch 1073
└ @ Main /Users/kcft114/.julia/packages/Flux/qXNjB/src/optimise/train.jl:105
┌ Info: Epoch 1074
└ @ Main /Users/kcft114/.julia/packages/Flux/qXNjB/src/optimise/train.jl:105
┌ Info: Epoch 1075
└ @ Main /Users/kcft114/.julia/packages/Flux/qXNjB/src/optimise/train.jl:105
┌ Info: Epoch 1076
└ @ Main /Users/kcft114/.julia/packages/Flux/qXNjB/src/optimise/train.jl:105
┌ Info: Epoch 1077
└ @ Main /Users/kcft114/.julia/packages/Flux/qXNjB/src/optimise/train.jl:105
┌ Info: Epoch 1078
└ @ Main /Users/kcft114/.julia/packages/Flux/qXNjB/src/optimise/train.jl:105
┌ Info: Epoch 1079
└ @ Main /Users/kcft114/.julia/packages/Flux/qXNjB/src/optimise/train.jl:105
┌ Info: Epoch 1080
└ @ Main /Users/kcft114/.julia/packages/Flux/qXNjB/src/optimise/train.jl:105
┌ Info: Epoch 1081
└ @ Main /Users/kcft114/.julia/packages/Flux/qXNjB/src/optimise/train.jl:105
┌ Info: Epoch 1082
└ @ Main /Users/kcft114/.julia/packages/Flux/qXNjB/src/optimise/train.jl:105
```

```
loss(X, Y) = 0.6936304f0 (tracked)
loss(X, Y) = 0.70037895f0 (tracked)
loss(X, Y) = 0.6785544f0 (tracked)
loss(X, Y) = 0.64313275f0 (tracked)
loss(X, Y) = 0.7000257f0 (tracked)
loss(X, Y) = 0.710205f0 (tracked)
loss(X, Y) = 0.64600813f0 (tracked)
loss(X, Y) = 0.67955667f0 (tracked)
loss(X, Y) = 0.67777556f0 (tracked)
loss(X, Y) = 0.7075046f0 (tracked)
loss(X, Y) = 0.7428704f0 (tracked)
loss(X, Y) = 0.70675063f0 (tracked)
loss(X, Y) = 0.6926712f0 (tracked)
loss(X, Y) = 0.709556f0 (tracked)
loss(X, Y) = 0.7130914f0 (tracked)
loss(X, Y) = 0.71285623f0 (tracked)
loss(X, Y) = 0.66888237f0 (tracked)
loss(X, Y) = 0.66927636f0 (tracked)
loss(X, Y) = 0.6774139f0 (tracked)
loss(X, Y) = 0.7160972f0 (tracked)
loss(X, Y) = 0.7325412f0 (tracked)
loss(X, Y) = 0.69695294f0 (tracked)
loss(X, Y) = 0.6813395f0 (tracked)
loss(X, Y) = 0.68256193f0 (tracked)
loss(X, Y) = 0.6691176f0 (tracked)
loss(X, Y) = 0.6319757f0 (tracked)
loss(X, Y) = 0.6764093f0 (tracked)
loss(X, Y) = 0.6990234f0 (tracked)
loss(X, Y) = 0.6903091f0 (tracked)
loss(X, Y) = 0.7636559f0 (tracked)
loss(X, Y) = 0.66324866f0 (tracked)
loss(X, Y) = 0.6721868f0 (tracked)
loss(X, Y) = 0.6982198f0 (tracked)
loss(X, Y) = 0.68026173f0 (tracked)
loss(X, Y) = 0.69009745f0 (tracked)
loss(X, Y) = 0.68847114f0 (tracked)
loss(X, Y) = 0.7253738f0 (tracked)
loss(X, Y) = 0.6612879f0 (tracked)
loss(X, Y) = 0.71758276f0 (tracked)
loss(X, Y) = 0.6688177f0 (tracked)
loss(X, Y) = 0.7250359f0 (tracked)
loss(X, Y) = 0.6381049f0 (tracked)
loss(X, Y) = 0.66990435f0 (tracked)
loss(X, Y) = 0.72255945f0 (tracked)
loss(X, Y) = 0.721212f0 (tracked)
loss(X, Y) = 0.71423155f0 (tracked)
loss(X, Y) = 0.70097595f0 (tracked)
loss(X, Y) = 0.69080335f0 (tracked)
loss(X, Y) = 0.7104102f0 (tracked)
loss(X, Y) = 0.6973223f0 (tracked)
loss(X, Y) = 0.71721655f0 (tracked)
loss(X, Y) = 0.70636034f0 (tracked)
loss(X, Y) = 0.70395696f0 (tracked)
loss(X, Y) = 0.66052634f0 (tracked)
loss(X, Y) = 0.668006f0 (tracked)
loss(X, Y) = 0.71468526f0 (tracked)
loss(X, Y) = 0.71314967f0 (tracked)
loss(X, Y) = 0.68808573f0 (tracked)
loss(X, Y) = 0.72185117f0 (tracked)
loss(X, Y) = 0.68903184f0 (tracked)
loss(X, Y) = 0.64671993f0 (tracked)
loss(X, Y) = 0.6581924f0 (tracked)
loss(X, Y) = 0.6997162f0 (tracked)
loss(X, Y) = 0.6645505f0 (tracked)
loss(X, Y) = 0.67549664f0 (tracked)
loss(X, Y) = 0.64725924f0 (tracked)
loss(X, Y) = 0.6904017f0 (tracked)
loss(X, Y) = 0.70576525f0 (tracked)
loss(X, Y) = 0.67661375f0 (tracked)
loss(X, Y) = 0.6715332f0 (tracked)
loss(X, Y) = 0.6621552f0 (tracked)
loss(X, Y) = 0.68898886f0 (tracked)
loss(X, Y) = 0.70596033f0 (tracked)
loss(X, Y) = 0.7105352f0 (tracked)
loss(X, Y) = 0.69626963f0 (tracked)
loss(X, Y) = 0.69083196f0 (tracked)
loss(X, Y) = 0.64669687f0 (tracked)
loss(X, Y) = 0.6892454f0 (tracked)
loss(X, Y) = 0.6920908f0 (tracked)
loss(X, Y) = 0.6939998f0 (tracked)
loss(X, Y) = 0.6822931f0 (tracked)
loss(X, Y) = 0.6422962f0 (tracked)
loss(X, Y) = 0.69492364f0 (tracked)
loss(X, Y) = 0.65496653f0 (tracked)
loss(X, Y) = 0.7267519f0 (tracked)
loss(X, Y) = 0.7241204f0 (tracked)
loss(X, Y) = 0.6774542f0 (tracked)
loss(X, Y) = 0.6716914f0 (tracked)
loss(X, Y) = 0.7171605f0 (tracked)
loss(X, Y) = 0.72760797f0 (tracked)
loss(X, Y) = 0.6642383f0 (tracked)
loss(X, Y) = 0.69615364f0 (tracked)
loss(X, Y) = 0.69233096f0 (tracked)
loss(X, Y) = 0.6887259f0 (tracked)
loss(X, Y) = 0.6952232f0 (tracked)
loss(X, Y) = 0.66579556f0 (tracked)
loss(X, Y) = 0.69325244f0 (tracked)
loss(X, Y) = 0.6830598f0 (tracked)
loss(X, Y) = 0.70192146f0 (tracked)
loss(X, Y) = 0.65909123f0 (tracked)
loss(X, Y) = 0.64879215f0 (tracked)
loss(X, Y) = 0.6814489f0 (tracked)
loss(X, Y) = 0.68627656f0 (tracked)
loss(X, Y) = 0.65116984f0 (tracked)
loss(X, Y) = 0.6460765f0 (tracked)
loss(X, Y) = 0.7610257f0 (tracked)
loss(X, Y) = 0.6758666f0 (tracked)
loss(X, Y) = 0.7387963f0 (tracked)
loss(X, Y) = 0.69045115f0 (tracked)
loss(X, Y) = 0.68600583f0 (tracked)
loss(X, Y) = 0.7153495f0 (tracked)
loss(X, Y) = 0.7189269f0 (tracked)
loss(X, Y) = 0.6532331f0 (tracked)
loss(X, Y) = 0.670089f0 (tracked)
loss(X, Y) = 0.6957624f0 (tracked)
loss(X, Y) = 0.6664974f0 (tracked)
loss(X, Y) = 0.6246372f0 (tracked)
loss(X, Y) = 0.71189976f0 (tracked)
```

```
┌ Info: Epoch 1083
└ @ Main /Users/kcft114/.julia/packages/Flux/qXNjB/src/optimise/train.jl:105
┌ Info: Epoch 1084
└ @ Main /Users/kcft114/.julia/packages/Flux/qXNjB/src/optimise/train.jl:105
┌ Info: Epoch 1085
└ @ Main /Users/kcft114/.julia/packages/Flux/qXNjB/src/optimise/train.jl:105
┌ Info: Epoch 1086
└ @ Main /Users/kcft114/.julia/packages/Flux/qXNjB/src/optimise/train.jl:105
┌ Info: Epoch 1087
└ @ Main /Users/kcft114/.julia/packages/Flux/qXNjB/src/optimise/train.jl:105
┌ Info: Epoch 1088
└ @ Main /Users/kcft114/.julia/packages/Flux/qXNjB/src/optimise/train.jl:105
┌ Info: Epoch 1089
└ @ Main /Users/kcft114/.julia/packages/Flux/qXNjB/src/optimise/train.jl:105
┌ Info: Epoch 1090
└ @ Main /Users/kcft114/.julia/packages/Flux/qXNjB/src/optimise/train.jl:105
┌ Info: Epoch 1091
└ @ Main /Users/kcft114/.julia/packages/Flux/qXNjB/src/optimise/train.jl:105
┌ Info: Epoch 1092
└ @ Main /Users/kcft114/.julia/packages/Flux/qXNjB/src/optimise/train.jl:105
┌ Info: Epoch 1093
└ @ Main /Users/kcft114/.julia/packages/Flux/qXNjB/src/optimise/train.jl:105
┌ Info: Epoch 1094
└ @ Main /Users/kcft114/.julia/packages/Flux/qXNjB/src/optimise/train.jl:105
┌ Info: Epoch 1095
└ @ Main /Users/kcft114/.julia/packages/Flux/qXNjB/src/optimise/train.jl:105
┌ Info: Epoch 1096
└ @ Main /Users/kcft114/.julia/packages/Flux/qXNjB/src/optimise/train.jl:105
┌ Info: Epoch 1097
└ @ Main /Users/kcft114/.julia/packages/Flux/qXNjB/src/optimise/train.jl:105
┌ Info: Epoch 1098
└ @ Main /Users/kcft114/.julia/packages/Flux/qXNjB/src/optimise/train.jl:105
┌ Info: Epoch 1099
└ @ Main /Users/kcft114/.julia/packages/Flux/qXNjB/src/optimise/train.jl:105
┌ Info: Epoch 1100
└ @ Main /Users/kcft114/.julia/packages/Flux/qXNjB/src/optimise/train.jl:105
┌ Info: Epoch 1101
└ @ Main /Users/kcft114/.julia/packages/Flux/qXNjB/src/optimise/train.jl:105
┌ Info: Epoch 1102
└ @ Main /Users/kcft114/.julia/packages/Flux/qXNjB/src/optimise/train.jl:105
┌ Info: Epoch 1103
└ @ Main /Users/kcft114/.julia/packages/Flux/qXNjB/src/optimise/train.jl:105
┌ Info: Epoch 1104
└ @ Main /Users/kcft114/.julia/packages/Flux/qXNjB/src/optimise/train.jl:105
┌ Info: Epoch 1105
└ @ Main /Users/kcft114/.julia/packages/Flux/qXNjB/src/optimise/train.jl:105
┌ Info: Epoch 1106
└ @ Main /Users/kcft114/.julia/packages/Flux/qXNjB/src/optimise/train.jl:105
┌ Info: Epoch 1107
└ @ Main /Users/kcft114/.julia/packages/Flux/qXNjB/src/optimise/train.jl:105
┌ Info: Epoch 1108
└ @ Main /Users/kcft114/.julia/packages/Flux/qXNjB/src/optimise/train.jl:105
┌ Info: Epoch 1109
└ @ Main /Users/kcft114/.julia/packages/Flux/qXNjB/src/optimise/train.jl:105
┌ Info: Epoch 1110
└ @ Main /Users/kcft114/.julia/packages/Flux/qXNjB/src/optimise/train.jl:105
┌ Info: Epoch 1111
└ @ Main /Users/kcft114/.julia/packages/Flux/qXNjB/src/optimise/train.jl:105
┌ Info: Epoch 1112
└ @ Main /Users/kcft114/.julia/packages/Flux/qXNjB/src/optimise/train.jl:105
┌ Info: Epoch 1113
└ @ Main /Users/kcft114/.julia/packages/Flux/qXNjB/src/optimise/train.jl:105
┌ Info: Epoch 1114
└ @ Main /Users/kcft114/.julia/packages/Flux/qXNjB/src/optimise/train.jl:105
┌ Info: Epoch 1115
└ @ Main /Users/kcft114/.julia/packages/Flux/qXNjB/src/optimise/train.jl:105
┌ Info: Epoch 1116
└ @ Main /Users/kcft114/.julia/packages/Flux/qXNjB/src/optimise/train.jl:105
┌ Info: Epoch 1117
└ @ Main /Users/kcft114/.julia/packages/Flux/qXNjB/src/optimise/train.jl:105
┌ Info: Epoch 1118
└ @ Main /Users/kcft114/.julia/packages/Flux/qXNjB/src/optimise/train.jl:105
┌ Info: Epoch 1119
└ @ Main /Users/kcft114/.julia/packages/Flux/qXNjB/src/optimise/train.jl:105
┌ Info: Epoch 1120
└ @ Main /Users/kcft114/.julia/packages/Flux/qXNjB/src/optimise/train.jl:105
┌ Info: Epoch 1121
└ @ Main /Users/kcft114/.julia/packages/Flux/qXNjB/src/optimise/train.jl:105
┌ Info: Epoch 1122
└ @ Main /Users/kcft114/.julia/packages/Flux/qXNjB/src/optimise/train.jl:105
┌ Info: Epoch 1123
└ @ Main /Users/kcft114/.julia/packages/Flux/qXNjB/src/optimise/train.jl:105
┌ Info: Epoch 1124
└ @ Main /Users/kcft114/.julia/packages/Flux/qXNjB/src/optimise/train.jl:105
┌ Info: Epoch 1125
└ @ Main /Users/kcft114/.julia/packages/Flux/qXNjB/src/optimise/train.jl:105
┌ Info: Epoch 1126
└ @ Main /Users/kcft114/.julia/packages/Flux/qXNjB/src/optimise/train.jl:105
┌ Info: Epoch 1127
└ @ Main /Users/kcft114/.julia/packages/Flux/qXNjB/src/optimise/train.jl:105
┌ Info: Epoch 1128
└ @ Main /Users/kcft114/.julia/packages/Flux/qXNjB/src/optimise/train.jl:105
┌ Info: Epoch 1129
└ @ Main /Users/kcft114/.julia/packages/Flux/qXNjB/src/optimise/train.jl:105
┌ Info: Epoch 1130
└ @ Main /Users/kcft114/.julia/packages/Flux/qXNjB/src/optimise/train.jl:105
┌ Info: Epoch 1131
└ @ Main /Users/kcft114/.julia/packages/Flux/qXNjB/src/optimise/train.jl:105
┌ Info: Epoch 1132
└ @ Main /Users/kcft114/.julia/packages/Flux/qXNjB/src/optimise/train.jl:105
┌ Info: Epoch 1133
└ @ Main /Users/kcft114/.julia/packages/Flux/qXNjB/src/optimise/train.jl:105
┌ Info: Epoch 1134
└ @ Main /Users/kcft114/.julia/packages/Flux/qXNjB/src/optimise/train.jl:105
┌ Info: Epoch 1135
└ @ Main /Users/kcft114/.julia/packages/Flux/qXNjB/src/optimise/train.jl:105
┌ Info: Epoch 1136
└ @ Main /Users/kcft114/.julia/packages/Flux/qXNjB/src/optimise/train.jl:105
┌ Info: Epoch 1137
└ @ Main /Users/kcft114/.julia/packages/Flux/qXNjB/src/optimise/train.jl:105
┌ Info: Epoch 1138
└ @ Main /Users/kcft114/.julia/packages/Flux/qXNjB/src/optimise/train.jl:105
┌ Info: Epoch 1139
└ @ Main /Users/kcft114/.julia/packages/Flux/qXNjB/src/optimise/train.jl:105
┌ Info: Epoch 1140
└ @ Main /Users/kcft114/.julia/packages/Flux/qXNjB/src/optimise/train.jl:105
┌ Info: Epoch 1141
└ @ Main /Users/kcft114/.julia/packages/Flux/qXNjB/src/optimise/train.jl:105
┌ Info: Epoch 1142
└ @ Main /Users/kcft114/.julia/packages/Flux/qXNjB/src/optimise/train.jl:105
┌ Info: Epoch 1143
└ @ Main /Users/kcft114/.julia/packages/Flux/qXNjB/src/optimise/train.jl:105
┌ Info: Epoch 1144
└ @ Main /Users/kcft114/.julia/packages/Flux/qXNjB/src/optimise/train.jl:105
┌ Info: Epoch 1145
└ @ Main /Users/kcft114/.julia/packages/Flux/qXNjB/src/optimise/train.jl:105
┌ Info: Epoch 1146
└ @ Main /Users/kcft114/.julia/packages/Flux/qXNjB/src/optimise/train.jl:105
┌ Info: Epoch 1147
└ @ Main /Users/kcft114/.julia/packages/Flux/qXNjB/src/optimise/train.jl:105
┌ Info: Epoch 1148
└ @ Main /Users/kcft114/.julia/packages/Flux/qXNjB/src/optimise/train.jl:105
┌ Info: Epoch 1149
└ @ Main /Users/kcft114/.julia/packages/Flux/qXNjB/src/optimise/train.jl:105
┌ Info: Epoch 1150
└ @ Main /Users/kcft114/.julia/packages/Flux/qXNjB/src/optimise/train.jl:105
┌ Info: Epoch 1151
└ @ Main /Users/kcft114/.julia/packages/Flux/qXNjB/src/optimise/train.jl:105
┌ Info: Epoch 1152
└ @ Main /Users/kcft114/.julia/packages/Flux/qXNjB/src/optimise/train.jl:105
┌ Info: Epoch 1153
└ @ Main /Users/kcft114/.julia/packages/Flux/qXNjB/src/optimise/train.jl:105
┌ Info: Epoch 1154
└ @ Main /Users/kcft114/.julia/packages/Flux/qXNjB/src/optimise/train.jl:105
┌ Info: Epoch 1155
└ @ Main /Users/kcft114/.julia/packages/Flux/qXNjB/src/optimise/train.jl:105
┌ Info: Epoch 1156
└ @ Main /Users/kcft114/.julia/packages/Flux/qXNjB/src/optimise/train.jl:105
┌ Info: Epoch 1157
└ @ Main /Users/kcft114/.julia/packages/Flux/qXNjB/src/optimise/train.jl:105
┌ Info: Epoch 1158
└ @ Main /Users/kcft114/.julia/packages/Flux/qXNjB/src/optimise/train.jl:105
┌ Info: Epoch 1159
└ @ Main /Users/kcft114/.julia/packages/Flux/qXNjB/src/optimise/train.jl:105
┌ Info: Epoch 1160
└ @ Main /Users/kcft114/.julia/packages/Flux/qXNjB/src/optimise/train.jl:105
┌ Info: Epoch 1161
└ @ Main /Users/kcft114/.julia/packages/Flux/qXNjB/src/optimise/train.jl:105
┌ Info: Epoch 1162
└ @ Main /Users/kcft114/.julia/packages/Flux/qXNjB/src/optimise/train.jl:105
┌ Info: Epoch 1163
└ @ Main /Users/kcft114/.julia/packages/Flux/qXNjB/src/optimise/train.jl:105
┌ Info: Epoch 1164
└ @ Main /Users/kcft114/.julia/packages/Flux/qXNjB/src/optimise/train.jl:105
┌ Info: Epoch 1165
└ @ Main /Users/kcft114/.julia/packages/Flux/qXNjB/src/optimise/train.jl:105
┌ Info: Epoch 1166
└ @ Main /Users/kcft114/.julia/packages/Flux/qXNjB/src/optimise/train.jl:105
┌ Info: Epoch 1167
└ @ Main /Users/kcft114/.julia/packages/Flux/qXNjB/src/optimise/train.jl:105
┌ Info: Epoch 1168
└ @ Main /Users/kcft114/.julia/packages/Flux/qXNjB/src/optimise/train.jl:105
┌ Info: Epoch 1169
└ @ Main /Users/kcft114/.julia/packages/Flux/qXNjB/src/optimise/train.jl:105
┌ Info: Epoch 1170
└ @ Main /Users/kcft114/.julia/packages/Flux/qXNjB/src/optimise/train.jl:105
┌ Info: Epoch 1171
└ @ Main /Users/kcft114/.julia/packages/Flux/qXNjB/src/optimise/train.jl:105
┌ Info: Epoch 1172
└ @ Main /Users/kcft114/.julia/packages/Flux/qXNjB/src/optimise/train.jl:105
┌ Info: Epoch 1173
└ @ Main /Users/kcft114/.julia/packages/Flux/qXNjB/src/optimise/train.jl:105
┌ Info: Epoch 1174
└ @ Main /Users/kcft114/.julia/packages/Flux/qXNjB/src/optimise/train.jl:105
┌ Info: Epoch 1175
└ @ Main /Users/kcft114/.julia/packages/Flux/qXNjB/src/optimise/train.jl:105
┌ Info: Epoch 1176
└ @ Main /Users/kcft114/.julia/packages/Flux/qXNjB/src/optimise/train.jl:105
┌ Info: Epoch 1177
└ @ Main /Users/kcft114/.julia/packages/Flux/qXNjB/src/optimise/train.jl:105
┌ Info: Epoch 1178
└ @ Main /Users/kcft114/.julia/packages/Flux/qXNjB/src/optimise/train.jl:105
┌ Info: Epoch 1179
└ @ Main /Users/kcft114/.julia/packages/Flux/qXNjB/src/optimise/train.jl:105
┌ Info: Epoch 1180
└ @ Main /Users/kcft114/.julia/packages/Flux/qXNjB/src/optimise/train.jl:105
┌ Info: Epoch 1181
└ @ Main /Users/kcft114/.julia/packages/Flux/qXNjB/src/optimise/train.jl:105
┌ Info: Epoch 1182
└ @ Main /Users/kcft114/.julia/packages/Flux/qXNjB/src/optimise/train.jl:105
┌ Info: Epoch 1183
└ @ Main /Users/kcft114/.julia/packages/Flux/qXNjB/src/optimise/train.jl:105
┌ Info: Epoch 1184
└ @ Main /Users/kcft114/.julia/packages/Flux/qXNjB/src/optimise/train.jl:105
┌ Info: Epoch 1185
└ @ Main /Users/kcft114/.julia/packages/Flux/qXNjB/src/optimise/train.jl:105
┌ Info: Epoch 1186
└ @ Main /Users/kcft114/.julia/packages/Flux/qXNjB/src/optimise/train.jl:105
┌ Info: Epoch 1187
└ @ Main /Users/kcft114/.julia/packages/Flux/qXNjB/src/optimise/train.jl:105
┌ Info: Epoch 1188
└ @ Main /Users/kcft114/.julia/packages/Flux/qXNjB/src/optimise/train.jl:105
┌ Info: Epoch 1189
└ @ Main /Users/kcft114/.julia/packages/Flux/qXNjB/src/optimise/train.jl:105
┌ Info: Epoch 1190
└ @ Main /Users/kcft114/.julia/packages/Flux/qXNjB/src/optimise/train.jl:105
┌ Info: Epoch 1191
└ @ Main /Users/kcft114/.julia/packages/Flux/qXNjB/src/optimise/train.jl:105
┌ Info: Epoch 1192
└ @ Main /Users/kcft114/.julia/packages/Flux/qXNjB/src/optimise/train.jl:105
┌ Info: Epoch 1193
└ @ Main /Users/kcft114/.julia/packages/Flux/qXNjB/src/optimise/train.jl:105
┌ Info: Epoch 1194
└ @ Main /Users/kcft114/.julia/packages/Flux/qXNjB/src/optimise/train.jl:105
┌ Info: Epoch 1195
└ @ Main /Users/kcft114/.julia/packages/Flux/qXNjB/src/optimise/train.jl:105
┌ Info: Epoch 1196
└ @ Main /Users/kcft114/.julia/packages/Flux/qXNjB/src/optimise/train.jl:105
┌ Info: Epoch 1197
└ @ Main /Users/kcft114/.julia/packages/Flux/qXNjB/src/optimise/train.jl:105
┌ Info: Epoch 1198
└ @ Main /Users/kcft114/.julia/packages/Flux/qXNjB/src/optimise/train.jl:105
┌ Info: Epoch 1199
└ @ Main /Users/kcft114/.julia/packages/Flux/qXNjB/src/optimise/train.jl:105
┌ Info: Epoch 1200
└ @ Main /Users/kcft114/.julia/packages/Flux/qXNjB/src/optimise/train.jl:105
┌ Info: Epoch 1201
└ @ Main /Users/kcft114/.julia/packages/Flux/qXNjB/src/optimise/train.jl:105
```

```
loss(X, Y) = 0.6342023f0 (tracked)
loss(X, Y) = 0.7093195f0 (tracked)
loss(X, Y) = 0.6395287f0 (tracked)
loss(X, Y) = 0.7153815f0 (tracked)
loss(X, Y) = 0.6545809f0 (tracked)
loss(X, Y) = 0.68473905f0 (tracked)
loss(X, Y) = 0.7134327f0 (tracked)
loss(X, Y) = 0.7096374f0 (tracked)
loss(X, Y) = 0.689814f0 (tracked)
loss(X, Y) = 0.68655926f0 (tracked)
loss(X, Y) = 0.6749837f0 (tracked)
loss(X, Y) = 0.689297f0 (tracked)
loss(X, Y) = 0.69848657f0 (tracked)
loss(X, Y) = 0.6500114f0 (tracked)
loss(X, Y) = 0.66133237f0 (tracked)
loss(X, Y) = 0.6243066f0 (tracked)
loss(X, Y) = 0.693343f0 (tracked)
loss(X, Y) = 0.6788756f0 (tracked)
loss(X, Y) = 0.69033116f0 (tracked)
loss(X, Y) = 0.68866014f0 (tracked)
loss(X, Y) = 0.6418316f0 (tracked)
loss(X, Y) = 0.6725401f0 (tracked)
loss(X, Y) = 0.7292098f0 (tracked)
loss(X, Y) = 0.6834111f0 (tracked)
loss(X, Y) = 0.7273809f0 (tracked)
loss(X, Y) = 0.7054128f0 (tracked)
loss(X, Y) = 0.6593441f0 (tracked)
loss(X, Y) = 0.64351f0 (tracked)
loss(X, Y) = 0.6338984f0 (tracked)
loss(X, Y) = 0.72741544f0 (tracked)
loss(X, Y) = 0.6827716f0 (tracked)
loss(X, Y) = 0.6880737f0 (tracked)
loss(X, Y) = 0.6784719f0 (tracked)
loss(X, Y) = 0.6330573f0 (tracked)
loss(X, Y) = 0.68422675f0 (tracked)
loss(X, Y) = 0.7068919f0 (tracked)
loss(X, Y) = 0.6709929f0 (tracked)
loss(X, Y) = 0.6768805f0 (tracked)
loss(X, Y) = 0.74458253f0 (tracked)
loss(X, Y) = 0.68304753f0 (tracked)
loss(X, Y) = 0.6675152f0 (tracked)
loss(X, Y) = 0.6812478f0 (tracked)
loss(X, Y) = 0.67225146f0 (tracked)
loss(X, Y) = 0.67378086f0 (tracked)
loss(X, Y) = 0.70624584f0 (tracked)
loss(X, Y) = 0.69784814f0 (tracked)
loss(X, Y) = 0.7059017f0 (tracked)
loss(X, Y) = 0.6653181f0 (tracked)
loss(X, Y) = 0.706143f0 (tracked)
loss(X, Y) = 0.6590497f0 (tracked)
loss(X, Y) = 0.6669431f0 (tracked)
loss(X, Y) = 0.71014476f0 (tracked)
loss(X, Y) = 0.6776856f0 (tracked)
loss(X, Y) = 0.6823215f0 (tracked)
loss(X, Y) = 0.68402785f0 (tracked)
loss(X, Y) = 0.66978705f0 (tracked)
loss(X, Y) = 0.6577397f0 (tracked)
loss(X, Y) = 0.65926534f0 (tracked)
loss(X, Y) = 0.6736515f0 (tracked)
loss(X, Y) = 0.62487197f0 (tracked)
loss(X, Y) = 0.67917645f0 (tracked)
loss(X, Y) = 0.71261805f0 (tracked)
loss(X, Y) = 0.6762094f0 (tracked)
loss(X, Y) = 0.6472725f0 (tracked)
loss(X, Y) = 0.69074994f0 (tracked)
loss(X, Y) = 0.66735315f0 (tracked)
loss(X, Y) = 0.637026f0 (tracked)
loss(X, Y) = 0.6933585f0 (tracked)
loss(X, Y) = 0.64486223f0 (tracked)
loss(X, Y) = 0.69249374f0 (tracked)
loss(X, Y) = 0.71436745f0 (tracked)
loss(X, Y) = 0.6850248f0 (tracked)
loss(X, Y) = 0.67271715f0 (tracked)
loss(X, Y) = 0.6444794f0 (tracked)
loss(X, Y) = 0.6545587f0 (tracked)
loss(X, Y) = 0.6874479f0 (tracked)
loss(X, Y) = 0.63568294f0 (tracked)
loss(X, Y) = 0.6950766f0 (tracked)
loss(X, Y) = 0.6919536f0 (tracked)
loss(X, Y) = 0.63043934f0 (tracked)
loss(X, Y) = 0.7027645f0 (tracked)
loss(X, Y) = 0.6781396f0 (tracked)
loss(X, Y) = 0.6912108f0 (tracked)
loss(X, Y) = 0.6752447f0 (tracked)
loss(X, Y) = 0.6913357f0 (tracked)
loss(X, Y) = 0.63648087f0 (tracked)
loss(X, Y) = 0.69451374f0 (tracked)
loss(X, Y) = 0.65898174f0 (tracked)
loss(X, Y) = 0.6639598f0 (tracked)
loss(X, Y) = 0.61907077f0 (tracked)
loss(X, Y) = 0.66171587f0 (tracked)
loss(X, Y) = 0.6658953f0 (tracked)
loss(X, Y) = 0.697267f0 (tracked)
loss(X, Y) = 0.681913f0 (tracked)
loss(X, Y) = 0.7074685f0 (tracked)
loss(X, Y) = 0.63839f0 (tracked)
loss(X, Y) = 0.67797214f0 (tracked)
loss(X, Y) = 0.6689722f0 (tracked)
loss(X, Y) = 0.6532903f0 (tracked)
loss(X, Y) = 0.6624873f0 (tracked)
loss(X, Y) = 0.6887596f0 (tracked)
loss(X, Y) = 0.6892155f0 (tracked)
loss(X, Y) = 0.6840002f0 (tracked)
loss(X, Y) = 0.67713827f0 (tracked)
loss(X, Y) = 0.65462923f0 (tracked)
loss(X, Y) = 0.6525042f0 (tracked)
```

```
┌ Info: Epoch 1202
└ @ Main /Users/kcft114/.julia/packages/Flux/qXNjB/src/optimise/train.jl:105
┌ Info: Epoch 1203
└ @ Main /Users/kcft114/.julia/packages/Flux/qXNjB/src/optimise/train.jl:105
┌ Info: Epoch 1204
└ @ Main /Users/kcft114/.julia/packages/Flux/qXNjB/src/optimise/train.jl:105
```

```
loss(X, Y) = 0.6852579f0 (tracked)
loss(X, Y) = 0.6815297f0 (tracked)
loss(X, Y) = 0.71649194f0 (tracked)
loss(X, Y) = 0.68136793f0 (tracked)
loss(X, Y) = 0.6377204f0 (tracked)
loss(X, Y) = 0.6491617f0 (tracked)
loss(X, Y) = 0.67729133f0 (tracked)
loss(X, Y) = 0.65339893f0 (tracked)
loss(X, Y) = 0.73092926f0 (tracked)
loss(X, Y) = 0.68339384f0 (tracked)
loss(X, Y) = 0.71496916f0 (tracked)
loss(X, Y) = 0.74130595f0 (tracked)
loss(X, Y) = 0.6355944f0 (tracked)
loss(X, Y) = 0.711003f0 (tracked)
loss(X, Y) = 0.61913824f0 (tracked)
loss(X, Y) = 0.64442426f0 (tracked)
loss(X, Y) = 0.6797327f0 (tracked)
loss(X, Y) = 0.6767847f0 (tracked)
loss(X, Y) = 0.66356117f0 (tracked)
loss(X, Y) = 0.6868195f0 (tracked)
loss(X, Y) = 0.6847917f0 (tracked)
loss(X, Y) = 0.6616465f0 (tracked)
loss(X, Y) = 0.64710015f0 (tracked)
loss(X, Y) = 0.6517122f0 (tracked)
loss(X, Y) = 0.6471298f0 (tracked)
loss(X, Y) = 0.65506744f0 (tracked)
loss(X, Y) = 0.6464066f0 (tracked)
loss(X, Y) = 0.67776537f0 (tracked)
loss(X, Y) = 0.6680802f0 (tracked)
loss(X, Y) = 0.69760615f0 (tracked)
loss(X, Y) = 0.6648627f0 (tracked)
loss(X, Y) = 0.66454077f0 (tracked)
loss(X, Y) = 0.68037736f0 (tracked)
loss(X, Y) = 0.63622814f0 (tracked)
loss(X, Y) = 0.64426416f0 (tracked)
loss(X, Y) = 0.6761191f0 (tracked)
loss(X, Y) = 0.63654184f0 (tracked)
loss(X, Y) = 0.6897423f0 (tracked)
loss(X, Y) = 0.71914935f0 (tracked)
loss(X, Y) = 0.6861103f0 (tracked)
loss(X, Y) = 0.68468523f0 (tracked)
loss(X, Y) = 0.6594952f0 (tracked)
loss(X, Y) = 0.6542608f0 (tracked)
loss(X, Y) = 0.6669708f0 (tracked)
loss(X, Y) = 0.6333629f0 (tracked)
loss(X, Y) = 0.64406407f0 (tracked)
loss(X, Y) = 0.68430877f0 (tracked)
loss(X, Y) = 0.7087895f0 (tracked)
loss(X, Y) = 0.6601346f0 (tracked)
loss(X, Y) = 0.6564464f0 (tracked)
loss(X, Y) = 0.70302236f0 (tracked)
loss(X, Y) = 0.7013114f0 (tracked)
loss(X, Y) = 0.6515772f0 (tracked)
loss(X, Y) = 0.68510395f0 (tracked)
loss(X, Y) = 0.72276837f0 (tracked)
loss(X, Y) = 0.69378006f0 (tracked)
loss(X, Y) = 0.7172118f0 (tracked)
loss(X, Y) = 0.672392f0 (tracked)
loss(X, Y) = 0.64319795f0 (tracked)
loss(X, Y) = 0.7119784f0 (tracked)
loss(X, Y) = 0.7340362f0 (tracked)
loss(X, Y) = 0.628783f0 (tracked)
loss(X, Y) = 0.6862802f0 (tracked)
loss(X, Y) = 0.7388174f0 (tracked)
loss(X, Y) = 0.7232539f0 (tracked)
loss(X, Y) = 0.66773635f0 (tracked)
loss(X, Y) = 0.68709314f0 (tracked)
loss(X, Y) = 0.6395298f0 (tracked)
loss(X, Y) = 0.69613576f0 (tracked)
loss(X, Y) = 0.65699357f0 (tracked)
loss(X, Y) = 0.67732424f0 (tracked)
loss(X, Y) = 0.6858492f0 (tracked)
loss(X, Y) = 0.65598f0 (tracked)
loss(X, Y) = 0.7014423f0 (tracked)
loss(X, Y) = 0.66167426f0 (tracked)
loss(X, Y) = 0.65571725f0 (tracked)
loss(X, Y) = 0.66905093f0 (tracked)
loss(X, Y) = 0.670471f0 (tracked)
loss(X, Y) = 0.6813998f0 (tracked)
loss(X, Y) = 0.6205204f0 (tracked)
loss(X, Y) = 0.67490846f0 (tracked)
loss(X, Y) = 0.648592f0 (tracked)
loss(X, Y) = 0.6353477f0 (tracked)
loss(X, Y) = 0.7320204f0 (tracked)
loss(X, Y) = 0.69658685f0 (tracked)
loss(X, Y) = 0.673018f0 (tracked)
loss(X, Y) = 0.70244426f0 (tracked)
loss(X, Y) = 0.6477783f0 (tracked)
loss(X, Y) = 0.67263657f0 (tracked)
loss(X, Y) = 0.6665536f0 (tracked)
loss(X, Y) = 0.69258434f0 (tracked)
loss(X, Y) = 0.6960204f0 (tracked)
loss(X, Y) = 0.70434296f0 (tracked)
loss(X, Y) = 0.71952915f0 (tracked)
loss(X, Y) = 0.6611061f0 (tracked)
loss(X, Y) = 0.6589516f0 (tracked)
loss(X, Y) = 0.63675094f0 (tracked)
loss(X, Y) = 0.68481773f0 (tracked)
loss(X, Y) = 0.6372627f0 (tracked)
loss(X, Y) = 0.6534951f0 (tracked)
loss(X, Y) = 0.65206826f0 (tracked)
loss(X, Y) = 0.66116756f0 (tracked)
```

```
┌ Info: Epoch 1205
└ @ Main /Users/kcft114/.julia/packages/Flux/qXNjB/src/optimise/train.jl:105
┌ Info: Epoch 1206
└ @ Main /Users/kcft114/.julia/packages/Flux/qXNjB/src/optimise/train.jl:105
┌ Info: Epoch 1207
└ @ Main /Users/kcft114/.julia/packages/Flux/qXNjB/src/optimise/train.jl:105
┌ Info: Epoch 1208
└ @ Main /Users/kcft114/.julia/packages/Flux/qXNjB/src/optimise/train.jl:105
┌ Info: Epoch 1209
└ @ Main /Users/kcft114/.julia/packages/Flux/qXNjB/src/optimise/train.jl:105
┌ Info: Epoch 1210
└ @ Main /Users/kcft114/.julia/packages/Flux/qXNjB/src/optimise/train.jl:105
┌ Info: Epoch 1211
└ @ Main /Users/kcft114/.julia/packages/Flux/qXNjB/src/optimise/train.jl:105
┌ Info: Epoch 1212
└ @ Main /Users/kcft114/.julia/packages/Flux/qXNjB/src/optimise/train.jl:105
┌ Info: Epoch 1213
└ @ Main /Users/kcft114/.julia/packages/Flux/qXNjB/src/optimise/train.jl:105
┌ Info: Epoch 1214
└ @ Main /Users/kcft114/.julia/packages/Flux/qXNjB/src/optimise/train.jl:105
┌ Info: Epoch 1215
└ @ Main /Users/kcft114/.julia/packages/Flux/qXNjB/src/optimise/train.jl:105
┌ Info: Epoch 1216
└ @ Main /Users/kcft114/.julia/packages/Flux/qXNjB/src/optimise/train.jl:105
┌ Info: Epoch 1217
└ @ Main /Users/kcft114/.julia/packages/Flux/qXNjB/src/optimise/train.jl:105
┌ Info: Epoch 1218
└ @ Main /Users/kcft114/.julia/packages/Flux/qXNjB/src/optimise/train.jl:105
┌ Info: Epoch 1219
└ @ Main /Users/kcft114/.julia/packages/Flux/qXNjB/src/optimise/train.jl:105
┌ Info: Epoch 1220
└ @ Main /Users/kcft114/.julia/packages/Flux/qXNjB/src/optimise/train.jl:105
┌ Info: Epoch 1221
└ @ Main /Users/kcft114/.julia/packages/Flux/qXNjB/src/optimise/train.jl:105
┌ Info: Epoch 1222
└ @ Main /Users/kcft114/.julia/packages/Flux/qXNjB/src/optimise/train.jl:105
┌ Info: Epoch 1223
└ @ Main /Users/kcft114/.julia/packages/Flux/qXNjB/src/optimise/train.jl:105
┌ Info: Epoch 1224
└ @ Main /Users/kcft114/.julia/packages/Flux/qXNjB/src/optimise/train.jl:105
┌ Info: Epoch 1225
└ @ Main /Users/kcft114/.julia/packages/Flux/qXNjB/src/optimise/train.jl:105
┌ Info: Epoch 1226
└ @ Main /Users/kcft114/.julia/packages/Flux/qXNjB/src/optimise/train.jl:105
┌ Info: Epoch 1227
└ @ Main /Users/kcft114/.julia/packages/Flux/qXNjB/src/optimise/train.jl:105
┌ Info: Epoch 1228
└ @ Main /Users/kcft114/.julia/packages/Flux/qXNjB/src/optimise/train.jl:105
┌ Info: Epoch 1229
└ @ Main /Users/kcft114/.julia/packages/Flux/qXNjB/src/optimise/train.jl:105
┌ Info: Epoch 1230
└ @ Main /Users/kcft114/.julia/packages/Flux/qXNjB/src/optimise/train.jl:105
┌ Info: Epoch 1231
└ @ Main /Users/kcft114/.julia/packages/Flux/qXNjB/src/optimise/train.jl:105
┌ Info: Epoch 1232
└ @ Main /Users/kcft114/.julia/packages/Flux/qXNjB/src/optimise/train.jl:105
┌ Info: Epoch 1233
└ @ Main /Users/kcft114/.julia/packages/Flux/qXNjB/src/optimise/train.jl:105
┌ Info: Epoch 1234
└ @ Main /Users/kcft114/.julia/packages/Flux/qXNjB/src/optimise/train.jl:105
┌ Info: Epoch 1235
└ @ Main /Users/kcft114/.julia/packages/Flux/qXNjB/src/optimise/train.jl:105
┌ Info: Epoch 1236
└ @ Main /Users/kcft114/.julia/packages/Flux/qXNjB/src/optimise/train.jl:105
┌ Info: Epoch 1237
└ @ Main /Users/kcft114/.julia/packages/Flux/qXNjB/src/optimise/train.jl:105
┌ Info: Epoch 1238
└ @ Main /Users/kcft114/.julia/packages/Flux/qXNjB/src/optimise/train.jl:105
┌ Info: Epoch 1239
└ @ Main /Users/kcft114/.julia/packages/Flux/qXNjB/src/optimise/train.jl:105
┌ Info: Epoch 1240
└ @ Main /Users/kcft114/.julia/packages/Flux/qXNjB/src/optimise/train.jl:105
┌ Info: Epoch 1241
└ @ Main /Users/kcft114/.julia/packages/Flux/qXNjB/src/optimise/train.jl:105
┌ Info: Epoch 1242
└ @ Main /Users/kcft114/.julia/packages/Flux/qXNjB/src/optimise/train.jl:105
┌ Info: Epoch 1243
└ @ Main /Users/kcft114/.julia/packages/Flux/qXNjB/src/optimise/train.jl:105
┌ Info: Epoch 1244
└ @ Main /Users/kcft114/.julia/packages/Flux/qXNjB/src/optimise/train.jl:105
┌ Info: Epoch 1245
└ @ Main /Users/kcft114/.julia/packages/Flux/qXNjB/src/optimise/train.jl:105
┌ Info: Epoch 1246
└ @ Main /Users/kcft114/.julia/packages/Flux/qXNjB/src/optimise/train.jl:105
┌ Info: Epoch 1247
└ @ Main /Users/kcft114/.julia/packages/Flux/qXNjB/src/optimise/train.jl:105
┌ Info: Epoch 1248
└ @ Main /Users/kcft114/.julia/packages/Flux/qXNjB/src/optimise/train.jl:105
┌ Info: Epoch 1249
└ @ Main /Users/kcft114/.julia/packages/Flux/qXNjB/src/optimise/train.jl:105
┌ Info: Epoch 1250
└ @ Main /Users/kcft114/.julia/packages/Flux/qXNjB/src/optimise/train.jl:105
┌ Info: Epoch 1251
└ @ Main /Users/kcft114/.julia/packages/Flux/qXNjB/src/optimise/train.jl:105
┌ Info: Epoch 1252
└ @ Main /Users/kcft114/.julia/packages/Flux/qXNjB/src/optimise/train.jl:105
┌ Info: Epoch 1253
└ @ Main /Users/kcft114/.julia/packages/Flux/qXNjB/src/optimise/train.jl:105
┌ Info: Epoch 1254
└ @ Main /Users/kcft114/.julia/packages/Flux/qXNjB/src/optimise/train.jl:105
┌ Info: Epoch 1255
└ @ Main /Users/kcft114/.julia/packages/Flux/qXNjB/src/optimise/train.jl:105
┌ Info: Epoch 1256
└ @ Main /Users/kcft114/.julia/packages/Flux/qXNjB/src/optimise/train.jl:105
┌ Info: Epoch 1257
└ @ Main /Users/kcft114/.julia/packages/Flux/qXNjB/src/optimise/train.jl:105
┌ Info: Epoch 1258
└ @ Main /Users/kcft114/.julia/packages/Flux/qXNjB/src/optimise/train.jl:105
┌ Info: Epoch 1259
└ @ Main /Users/kcft114/.julia/packages/Flux/qXNjB/src/optimise/train.jl:105
┌ Info: Epoch 1260
└ @ Main /Users/kcft114/.julia/packages/Flux/qXNjB/src/optimise/train.jl:105
┌ Info: Epoch 1261
└ @ Main /Users/kcft114/.julia/packages/Flux/qXNjB/src/optimise/train.jl:105
┌ Info: Epoch 1262
└ @ Main /Users/kcft114/.julia/packages/Flux/qXNjB/src/optimise/train.jl:105
┌ Info: Epoch 1263
└ @ Main /Users/kcft114/.julia/packages/Flux/qXNjB/src/optimise/train.jl:105
┌ Info: Epoch 1264
└ @ Main /Users/kcft114/.julia/packages/Flux/qXNjB/src/optimise/train.jl:105
┌ Info: Epoch 1265
└ @ Main /Users/kcft114/.julia/packages/Flux/qXNjB/src/optimise/train.jl:105
┌ Info: Epoch 1266
└ @ Main /Users/kcft114/.julia/packages/Flux/qXNjB/src/optimise/train.jl:105
┌ Info: Epoch 1267
└ @ Main /Users/kcft114/.julia/packages/Flux/qXNjB/src/optimise/train.jl:105
┌ Info: Epoch 1268
└ @ Main /Users/kcft114/.julia/packages/Flux/qXNjB/src/optimise/train.jl:105
┌ Info: Epoch 1269
└ @ Main /Users/kcft114/.julia/packages/Flux/qXNjB/src/optimise/train.jl:105
┌ Info: Epoch 1270
└ @ Main /Users/kcft114/.julia/packages/Flux/qXNjB/src/optimise/train.jl:105
┌ Info: Epoch 1271
└ @ Main /Users/kcft114/.julia/packages/Flux/qXNjB/src/optimise/train.jl:105
┌ Info: Epoch 1272
└ @ Main /Users/kcft114/.julia/packages/Flux/qXNjB/src/optimise/train.jl:105
┌ Info: Epoch 1273
└ @ Main /Users/kcft114/.julia/packages/Flux/qXNjB/src/optimise/train.jl:105
┌ Info: Epoch 1274
└ @ Main /Users/kcft114/.julia/packages/Flux/qXNjB/src/optimise/train.jl:105
┌ Info: Epoch 1275
└ @ Main /Users/kcft114/.julia/packages/Flux/qXNjB/src/optimise/train.jl:105
┌ Info: Epoch 1276
└ @ Main /Users/kcft114/.julia/packages/Flux/qXNjB/src/optimise/train.jl:105
┌ Info: Epoch 1277
└ @ Main /Users/kcft114/.julia/packages/Flux/qXNjB/src/optimise/train.jl:105
┌ Info: Epoch 1278
└ @ Main /Users/kcft114/.julia/packages/Flux/qXNjB/src/optimise/train.jl:105
┌ Info: Epoch 1279
└ @ Main /Users/kcft114/.julia/packages/Flux/qXNjB/src/optimise/train.jl:105
┌ Info: Epoch 1280
└ @ Main /Users/kcft114/.julia/packages/Flux/qXNjB/src/optimise/train.jl:105
┌ Info: Epoch 1281
└ @ Main /Users/kcft114/.julia/packages/Flux/qXNjB/src/optimise/train.jl:105
┌ Info: Epoch 1282
└ @ Main /Users/kcft114/.julia/packages/Flux/qXNjB/src/optimise/train.jl:105
┌ Info: Epoch 1283
└ @ Main /Users/kcft114/.julia/packages/Flux/qXNjB/src/optimise/train.jl:105
┌ Info: Epoch 1284
└ @ Main /Users/kcft114/.julia/packages/Flux/qXNjB/src/optimise/train.jl:105
┌ Info: Epoch 1285
└ @ Main /Users/kcft114/.julia/packages/Flux/qXNjB/src/optimise/train.jl:105
┌ Info: Epoch 1286
└ @ Main /Users/kcft114/.julia/packages/Flux/qXNjB/src/optimise/train.jl:105
┌ Info: Epoch 1287
└ @ Main /Users/kcft114/.julia/packages/Flux/qXNjB/src/optimise/train.jl:105
┌ Info: Epoch 1288
└ @ Main /Users/kcft114/.julia/packages/Flux/qXNjB/src/optimise/train.jl:105
┌ Info: Epoch 1289
└ @ Main /Users/kcft114/.julia/packages/Flux/qXNjB/src/optimise/train.jl:105
┌ Info: Epoch 1290
└ @ Main /Users/kcft114/.julia/packages/Flux/qXNjB/src/optimise/train.jl:105
┌ Info: Epoch 1291
└ @ Main /Users/kcft114/.julia/packages/Flux/qXNjB/src/optimise/train.jl:105
┌ Info: Epoch 1292
└ @ Main /Users/kcft114/.julia/packages/Flux/qXNjB/src/optimise/train.jl:105
┌ Info: Epoch 1293
└ @ Main /Users/kcft114/.julia/packages/Flux/qXNjB/src/optimise/train.jl:105
┌ Info: Epoch 1294
└ @ Main /Users/kcft114/.julia/packages/Flux/qXNjB/src/optimise/train.jl:105
┌ Info: Epoch 1295
└ @ Main /Users/kcft114/.julia/packages/Flux/qXNjB/src/optimise/train.jl:105
┌ Info: Epoch 1296
└ @ Main /Users/kcft114/.julia/packages/Flux/qXNjB/src/optimise/train.jl:105
┌ Info: Epoch 1297
└ @ Main /Users/kcft114/.julia/packages/Flux/qXNjB/src/optimise/train.jl:105
┌ Info: Epoch 1298
└ @ Main /Users/kcft114/.julia/packages/Flux/qXNjB/src/optimise/train.jl:105
┌ Info: Epoch 1299
└ @ Main /Users/kcft114/.julia/packages/Flux/qXNjB/src/optimise/train.jl:105
┌ Info: Epoch 1300
└ @ Main /Users/kcft114/.julia/packages/Flux/qXNjB/src/optimise/train.jl:105
┌ Info: Epoch 1301
└ @ Main /Users/kcft114/.julia/packages/Flux/qXNjB/src/optimise/train.jl:105
┌ Info: Epoch 1302
└ @ Main /Users/kcft114/.julia/packages/Flux/qXNjB/src/optimise/train.jl:105
┌ Info: Epoch 1303
└ @ Main /Users/kcft114/.julia/packages/Flux/qXNjB/src/optimise/train.jl:105
┌ Info: Epoch 1304
└ @ Main /Users/kcft114/.julia/packages/Flux/qXNjB/src/optimise/train.jl:105
┌ Info: Epoch 1305
└ @ Main /Users/kcft114/.julia/packages/Flux/qXNjB/src/optimise/train.jl:105
┌ Info: Epoch 1306
└ @ Main /Users/kcft114/.julia/packages/Flux/qXNjB/src/optimise/train.jl:105
```

```
loss(X, Y) = 0.6534927f0 (tracked)
loss(X, Y) = 0.6382581f0 (tracked)
loss(X, Y) = 0.64687943f0 (tracked)
loss(X, Y) = 0.63563746f0 (tracked)
loss(X, Y) = 0.66766244f0 (tracked)
loss(X, Y) = 0.6441209f0 (tracked)
loss(X, Y) = 0.6987818f0 (tracked)
loss(X, Y) = 0.64664185f0 (tracked)
loss(X, Y) = 0.643846f0 (tracked)
loss(X, Y) = 0.66940826f0 (tracked)
loss(X, Y) = 0.6731411f0 (tracked)
loss(X, Y) = 0.65261954f0 (tracked)
loss(X, Y) = 0.63389194f0 (tracked)
loss(X, Y) = 0.65031517f0 (tracked)
loss(X, Y) = 0.6680568f0 (tracked)
loss(X, Y) = 0.6757077f0 (tracked)
loss(X, Y) = 0.69482565f0 (tracked)
loss(X, Y) = 0.6606259f0 (tracked)
loss(X, Y) = 0.69337964f0 (tracked)
loss(X, Y) = 0.6834681f0 (tracked)
loss(X, Y) = 0.7324225f0 (tracked)
loss(X, Y) = 0.6164684f0 (tracked)
loss(X, Y) = 0.68250364f0 (tracked)
loss(X, Y) = 0.6486711f0 (tracked)
loss(X, Y) = 0.6635099f0 (tracked)
loss(X, Y) = 0.7033808f0 (tracked)
loss(X, Y) = 0.6392321f0 (tracked)
loss(X, Y) = 0.65720344f0 (tracked)
loss(X, Y) = 0.69175166f0 (tracked)
loss(X, Y) = 0.6842302f0 (tracked)
loss(X, Y) = 0.6573821f0 (tracked)
loss(X, Y) = 0.6563909f0 (tracked)
loss(X, Y) = 0.6065194f0 (tracked)
loss(X, Y) = 0.6270826f0 (tracked)
loss(X, Y) = 0.64759135f0 (tracked)
loss(X, Y) = 0.66640747f0 (tracked)
loss(X, Y) = 0.65055156f0 (tracked)
loss(X, Y) = 0.65818423f0 (tracked)
loss(X, Y) = 0.747412f0 (tracked)
loss(X, Y) = 0.6182108f0 (tracked)
loss(X, Y) = 0.6503543f0 (tracked)
loss(X, Y) = 0.6433886f0 (tracked)
loss(X, Y) = 0.6594179f0 (tracked)
loss(X, Y) = 0.6965361f0 (tracked)
loss(X, Y) = 0.65179485f0 (tracked)
loss(X, Y) = 0.6600359f0 (tracked)
loss(X, Y) = 0.66232485f0 (tracked)
loss(X, Y) = 0.6576227f0 (tracked)
loss(X, Y) = 0.673043f0 (tracked)
loss(X, Y) = 0.6968394f0 (tracked)
loss(X, Y) = 0.69503725f0 (tracked)
loss(X, Y) = 0.6480979f0 (tracked)
loss(X, Y) = 0.6491667f0 (tracked)
loss(X, Y) = 0.6866962f0 (tracked)
loss(X, Y) = 0.66502815f0 (tracked)
loss(X, Y) = 0.6882442f0 (tracked)
loss(X, Y) = 0.666071f0 (tracked)
loss(X, Y) = 0.69358385f0 (tracked)
loss(X, Y) = 0.6036964f0 (tracked)
loss(X, Y) = 0.64616275f0 (tracked)
loss(X, Y) = 0.6622785f0 (tracked)
loss(X, Y) = 0.68390733f0 (tracked)
loss(X, Y) = 0.67541355f0 (tracked)
loss(X, Y) = 0.63590735f0 (tracked)
loss(X, Y) = 0.6889175f0 (tracked)
loss(X, Y) = 0.6585421f0 (tracked)
loss(X, Y) = 0.66215324f0 (tracked)
loss(X, Y) = 0.65426564f0 (tracked)
loss(X, Y) = 0.68065536f0 (tracked)
loss(X, Y) = 0.6492629f0 (tracked)
loss(X, Y) = 0.6319065f0 (tracked)
loss(X, Y) = 0.6529908f0 (tracked)
loss(X, Y) = 0.6922926f0 (tracked)
loss(X, Y) = 0.68995744f0 (tracked)
loss(X, Y) = 0.6313281f0 (tracked)
loss(X, Y) = 0.6784304f0 (tracked)
loss(X, Y) = 0.65382075f0 (tracked)
loss(X, Y) = 0.6412715f0 (tracked)
loss(X, Y) = 0.6993496f0 (tracked)
loss(X, Y) = 0.67600054f0 (tracked)
loss(X, Y) = 0.6848031f0 (tracked)
loss(X, Y) = 0.68887055f0 (tracked)
loss(X, Y) = 0.67243874f0 (tracked)
loss(X, Y) = 0.6699682f0 (tracked)
loss(X, Y) = 0.64410144f0 (tracked)
loss(X, Y) = 0.58822167f0 (tracked)
```

```
┌ Info: Epoch 1307
└ @ Main /Users/kcft114/.julia/packages/Flux/qXNjB/src/optimise/train.jl:105
┌ Info: Epoch 1308
└ @ Main /Users/kcft114/.julia/packages/Flux/qXNjB/src/optimise/train.jl:105
┌ Info: Epoch 1309
└ @ Main /Users/kcft114/.julia/packages/Flux/qXNjB/src/optimise/train.jl:105
┌ Info: Epoch 1310
└ @ Main /Users/kcft114/.julia/packages/Flux/qXNjB/src/optimise/train.jl:105
┌ Info: Epoch 1311
└ @ Main /Users/kcft114/.julia/packages/Flux/qXNjB/src/optimise/train.jl:105
┌ Info: Epoch 1312
└ @ Main /Users/kcft114/.julia/packages/Flux/qXNjB/src/optimise/train.jl:105
┌ Info: Epoch 1313
└ @ Main /Users/kcft114/.julia/packages/Flux/qXNjB/src/optimise/train.jl:105
┌ Info: Epoch 1314
└ @ Main /Users/kcft114/.julia/packages/Flux/qXNjB/src/optimise/train.jl:105
┌ Info: Epoch 1315
└ @ Main /Users/kcft114/.julia/packages/Flux/qXNjB/src/optimise/train.jl:105
┌ Info: Epoch 1316
└ @ Main /Users/kcft114/.julia/packages/Flux/qXNjB/src/optimise/train.jl:105
┌ Info: Epoch 1317
└ @ Main /Users/kcft114/.julia/packages/Flux/qXNjB/src/optimise/train.jl:105
┌ Info: Epoch 1318
└ @ Main /Users/kcft114/.julia/packages/Flux/qXNjB/src/optimise/train.jl:105
┌ Info: Epoch 1319
└ @ Main /Users/kcft114/.julia/packages/Flux/qXNjB/src/optimise/train.jl:105
┌ Info: Epoch 1320
└ @ Main /Users/kcft114/.julia/packages/Flux/qXNjB/src/optimise/train.jl:105
┌ Info: Epoch 1321
└ @ Main /Users/kcft114/.julia/packages/Flux/qXNjB/src/optimise/train.jl:105
┌ Info: Epoch 1322
└ @ Main /Users/kcft114/.julia/packages/Flux/qXNjB/src/optimise/train.jl:105
┌ Info: Epoch 1323
└ @ Main /Users/kcft114/.julia/packages/Flux/qXNjB/src/optimise/train.jl:105
┌ Info: Epoch 1324
└ @ Main /Users/kcft114/.julia/packages/Flux/qXNjB/src/optimise/train.jl:105
┌ Info: Epoch 1325
└ @ Main /Users/kcft114/.julia/packages/Flux/qXNjB/src/optimise/train.jl:105
┌ Info: Epoch 1326
└ @ Main /Users/kcft114/.julia/packages/Flux/qXNjB/src/optimise/train.jl:105
┌ Info: Epoch 1327
└ @ Main /Users/kcft114/.julia/packages/Flux/qXNjB/src/optimise/train.jl:105
┌ Info: Epoch 1328
└ @ Main /Users/kcft114/.julia/packages/Flux/qXNjB/src/optimise/train.jl:105
┌ Info: Epoch 1329
└ @ Main /Users/kcft114/.julia/packages/Flux/qXNjB/src/optimise/train.jl:105
┌ Info: Epoch 1330
└ @ Main /Users/kcft114/.julia/packages/Flux/qXNjB/src/optimise/train.jl:105
┌ Info: Epoch 1331
└ @ Main /Users/kcft114/.julia/packages/Flux/qXNjB/src/optimise/train.jl:105
┌ Info: Epoch 1332
└ @ Main /Users/kcft114/.julia/packages/Flux/qXNjB/src/optimise/train.jl:105
┌ Info: Epoch 1333
└ @ Main /Users/kcft114/.julia/packages/Flux/qXNjB/src/optimise/train.jl:105
┌ Info: Epoch 1334
└ @ Main /Users/kcft114/.julia/packages/Flux/qXNjB/src/optimise/train.jl:105
┌ Info: Epoch 1335
└ @ Main /Users/kcft114/.julia/packages/Flux/qXNjB/src/optimise/train.jl:105
┌ Info: Epoch 1336
└ @ Main /Users/kcft114/.julia/packages/Flux/qXNjB/src/optimise/train.jl:105
┌ Info: Epoch 1337
└ @ Main /Users/kcft114/.julia/packages/Flux/qXNjB/src/optimise/train.jl:105
┌ Info: Epoch 1338
└ @ Main /Users/kcft114/.julia/packages/Flux/qXNjB/src/optimise/train.jl:105
┌ Info: Epoch 1339
└ @ Main /Users/kcft114/.julia/packages/Flux/qXNjB/src/optimise/train.jl:105
┌ Info: Epoch 1340
└ @ Main /Users/kcft114/.julia/packages/Flux/qXNjB/src/optimise/train.jl:105
┌ Info: Epoch 1341
└ @ Main /Users/kcft114/.julia/packages/Flux/qXNjB/src/optimise/train.jl:105
┌ Info: Epoch 1342
└ @ Main /Users/kcft114/.julia/packages/Flux/qXNjB/src/optimise/train.jl:105
┌ Info: Epoch 1343
└ @ Main /Users/kcft114/.julia/packages/Flux/qXNjB/src/optimise/train.jl:105
┌ Info: Epoch 1344
└ @ Main /Users/kcft114/.julia/packages/Flux/qXNjB/src/optimise/train.jl:105
┌ Info: Epoch 1345
└ @ Main /Users/kcft114/.julia/packages/Flux/qXNjB/src/optimise/train.jl:105
┌ Info: Epoch 1346
└ @ Main /Users/kcft114/.julia/packages/Flux/qXNjB/src/optimise/train.jl:105
┌ Info: Epoch 1347
└ @ Main /Users/kcft114/.julia/packages/Flux/qXNjB/src/optimise/train.jl:105
┌ Info: Epoch 1348
└ @ Main /Users/kcft114/.julia/packages/Flux/qXNjB/src/optimise/train.jl:105
┌ Info: Epoch 1349
└ @ Main /Users/kcft114/.julia/packages/Flux/qXNjB/src/optimise/train.jl:105
┌ Info: Epoch 1350
└ @ Main /Users/kcft114/.julia/packages/Flux/qXNjB/src/optimise/train.jl:105
┌ Info: Epoch 1351
└ @ Main /Users/kcft114/.julia/packages/Flux/qXNjB/src/optimise/train.jl:105
┌ Info: Epoch 1352
└ @ Main /Users/kcft114/.julia/packages/Flux/qXNjB/src/optimise/train.jl:105
┌ Info: Epoch 1353
└ @ Main /Users/kcft114/.julia/packages/Flux/qXNjB/src/optimise/train.jl:105
┌ Info: Epoch 1354
└ @ Main /Users/kcft114/.julia/packages/Flux/qXNjB/src/optimise/train.jl:105
┌ Info: Epoch 1355
└ @ Main /Users/kcft114/.julia/packages/Flux/qXNjB/src/optimise/train.jl:105
┌ Info: Epoch 1356
└ @ Main /Users/kcft114/.julia/packages/Flux/qXNjB/src/optimise/train.jl:105
┌ Info: Epoch 1357
└ @ Main /Users/kcft114/.julia/packages/Flux/qXNjB/src/optimise/train.jl:105
┌ Info: Epoch 1358
└ @ Main /Users/kcft114/.julia/packages/Flux/qXNjB/src/optimise/train.jl:105
┌ Info: Epoch 1359
└ @ Main /Users/kcft114/.julia/packages/Flux/qXNjB/src/optimise/train.jl:105
┌ Info: Epoch 1360
└ @ Main /Users/kcft114/.julia/packages/Flux/qXNjB/src/optimise/train.jl:105
┌ Info: Epoch 1361
└ @ Main /Users/kcft114/.julia/packages/Flux/qXNjB/src/optimise/train.jl:105
┌ Info: Epoch 1362
└ @ Main /Users/kcft114/.julia/packages/Flux/qXNjB/src/optimise/train.jl:105
┌ Info: Epoch 1363
└ @ Main /Users/kcft114/.julia/packages/Flux/qXNjB/src/optimise/train.jl:105
┌ Info: Epoch 1364
└ @ Main /Users/kcft114/.julia/packages/Flux/qXNjB/src/optimise/train.jl:105
┌ Info: Epoch 1365
└ @ Main /Users/kcft114/.julia/packages/Flux/qXNjB/src/optimise/train.jl:105
┌ Info: Epoch 1366
└ @ Main /Users/kcft114/.julia/packages/Flux/qXNjB/src/optimise/train.jl:105
┌ Info: Epoch 1367
└ @ Main /Users/kcft114/.julia/packages/Flux/qXNjB/src/optimise/train.jl:105
┌ Info: Epoch 1368
└ @ Main /Users/kcft114/.julia/packages/Flux/qXNjB/src/optimise/train.jl:105
┌ Info: Epoch 1369
└ @ Main /Users/kcft114/.julia/packages/Flux/qXNjB/src/optimise/train.jl:105
┌ Info: Epoch 1370
└ @ Main /Users/kcft114/.julia/packages/Flux/qXNjB/src/optimise/train.jl:105
┌ Info: Epoch 1371
└ @ Main /Users/kcft114/.julia/packages/Flux/qXNjB/src/optimise/train.jl:105
┌ Info: Epoch 1372
└ @ Main /Users/kcft114/.julia/packages/Flux/qXNjB/src/optimise/train.jl:105
┌ Info: Epoch 1373
└ @ Main /Users/kcft114/.julia/packages/Flux/qXNjB/src/optimise/train.jl:105
┌ Info: Epoch 1374
└ @ Main /Users/kcft114/.julia/packages/Flux/qXNjB/src/optimise/train.jl:105
┌ Info: Epoch 1375
└ @ Main /Users/kcft114/.julia/packages/Flux/qXNjB/src/optimise/train.jl:105
┌ Info: Epoch 1376
└ @ Main /Users/kcft114/.julia/packages/Flux/qXNjB/src/optimise/train.jl:105
┌ Info: Epoch 1377
└ @ Main /Users/kcft114/.julia/packages/Flux/qXNjB/src/optimise/train.jl:105
┌ Info: Epoch 1378
└ @ Main /Users/kcft114/.julia/packages/Flux/qXNjB/src/optimise/train.jl:105
┌ Info: Epoch 1379
└ @ Main /Users/kcft114/.julia/packages/Flux/qXNjB/src/optimise/train.jl:105
┌ Info: Epoch 1380
└ @ Main /Users/kcft114/.julia/packages/Flux/qXNjB/src/optimise/train.jl:105
┌ Info: Epoch 1381
└ @ Main /Users/kcft114/.julia/packages/Flux/qXNjB/src/optimise/train.jl:105
┌ Info: Epoch 1382
└ @ Main /Users/kcft114/.julia/packages/Flux/qXNjB/src/optimise/train.jl:105
┌ Info: Epoch 1383
└ @ Main /Users/kcft114/.julia/packages/Flux/qXNjB/src/optimise/train.jl:105
┌ Info: Epoch 1384
└ @ Main /Users/kcft114/.julia/packages/Flux/qXNjB/src/optimise/train.jl:105
┌ Info: Epoch 1385
└ @ Main /Users/kcft114/.julia/packages/Flux/qXNjB/src/optimise/train.jl:105
┌ Info: Epoch 1386
└ @ Main /Users/kcft114/.julia/packages/Flux/qXNjB/src/optimise/train.jl:105
┌ Info: Epoch 1387
└ @ Main /Users/kcft114/.julia/packages/Flux/qXNjB/src/optimise/train.jl:105
┌ Info: Epoch 1388
└ @ Main /Users/kcft114/.julia/packages/Flux/qXNjB/src/optimise/train.jl:105
┌ Info: Epoch 1389
└ @ Main /Users/kcft114/.julia/packages/Flux/qXNjB/src/optimise/train.jl:105
┌ Info: Epoch 1390
└ @ Main /Users/kcft114/.julia/packages/Flux/qXNjB/src/optimise/train.jl:105
┌ Info: Epoch 1391
└ @ Main /Users/kcft114/.julia/packages/Flux/qXNjB/src/optimise/train.jl:105
┌ Info: Epoch 1392
└ @ Main /Users/kcft114/.julia/packages/Flux/qXNjB/src/optimise/train.jl:105
```

```
loss(X, Y) = 0.6597326f0 (tracked)
loss(X, Y) = 0.65700775f0 (tracked)
loss(X, Y) = 0.70851225f0 (tracked)
loss(X, Y) = 0.6286214f0 (tracked)
loss(X, Y) = 0.6128145f0 (tracked)
loss(X, Y) = 0.67416817f0 (tracked)
loss(X, Y) = 0.68884027f0 (tracked)
loss(X, Y) = 0.6669521f0 (tracked)
loss(X, Y) = 0.65002257f0 (tracked)
loss(X, Y) = 0.672754f0 (tracked)
loss(X, Y) = 0.7063492f0 (tracked)
loss(X, Y) = 0.6570394f0 (tracked)
loss(X, Y) = 0.6504315f0 (tracked)
loss(X, Y) = 0.7005543f0 (tracked)
loss(X, Y) = 0.688511f0 (tracked)
loss(X, Y) = 0.6733986f0 (tracked)
loss(X, Y) = 0.6559168f0 (tracked)
loss(X, Y) = 0.6732764f0 (tracked)
loss(X, Y) = 0.6375938f0 (tracked)
loss(X, Y) = 0.6669925f0 (tracked)
loss(X, Y) = 0.6268444f0 (tracked)
loss(X, Y) = 0.66034013f0 (tracked)
loss(X, Y) = 0.6522437f0 (tracked)
loss(X, Y) = 0.6998942f0 (tracked)
loss(X, Y) = 0.6515588f0 (tracked)
loss(X, Y) = 0.6561513f0 (tracked)
loss(X, Y) = 0.6529107f0 (tracked)
loss(X, Y) = 0.6538222f0 (tracked)
loss(X, Y) = 0.67024416f0 (tracked)
loss(X, Y) = 0.6384957f0 (tracked)
loss(X, Y) = 0.64609075f0 (tracked)
loss(X, Y) = 0.69564116f0 (tracked)
loss(X, Y) = 0.67059976f0 (tracked)
loss(X, Y) = 0.6979861f0 (tracked)
loss(X, Y) = 0.6567978f0 (tracked)
loss(X, Y) = 0.6387662f0 (tracked)
loss(X, Y) = 0.6984767f0 (tracked)
loss(X, Y) = 0.6548577f0 (tracked)
loss(X, Y) = 0.6731368f0 (tracked)
loss(X, Y) = 0.6232724f0 (tracked)
loss(X, Y) = 0.64905226f0 (tracked)
loss(X, Y) = 0.6897416f0 (tracked)
loss(X, Y) = 0.6367858f0 (tracked)
loss(X, Y) = 0.6397547f0 (tracked)
loss(X, Y) = 0.64486593f0 (tracked)
loss(X, Y) = 0.6191149f0 (tracked)
loss(X, Y) = 0.68767536f0 (tracked)
loss(X, Y) = 0.67727196f0 (tracked)
loss(X, Y) = 0.6294089f0 (tracked)
loss(X, Y) = 0.6808517f0 (tracked)
loss(X, Y) = 0.69478303f0 (tracked)
loss(X, Y) = 0.6255055f0 (tracked)
loss(X, Y) = 0.7019314f0 (tracked)
loss(X, Y) = 0.65219444f0 (tracked)
loss(X, Y) = 0.63088286f0 (tracked)
loss(X, Y) = 0.6482323f0 (tracked)
loss(X, Y) = 0.63600445f0 (tracked)
loss(X, Y) = 0.65388596f0 (tracked)
loss(X, Y) = 0.6796197f0 (tracked)
loss(X, Y) = 0.63050383f0 (tracked)
loss(X, Y) = 0.6401099f0 (tracked)
loss(X, Y) = 0.6167498f0 (tracked)
loss(X, Y) = 0.6642481f0 (tracked)
loss(X, Y) = 0.6391478f0 (tracked)
loss(X, Y) = 0.65067863f0 (tracked)
loss(X, Y) = 0.59448725f0 (tracked)
loss(X, Y) = 0.7043593f0 (tracked)
loss(X, Y) = 0.63208055f0 (tracked)
loss(X, Y) = 0.6539814f0 (tracked)
loss(X, Y) = 0.6222352f0 (tracked)
loss(X, Y) = 0.61518687f0 (tracked)
loss(X, Y) = 0.65624326f0 (tracked)
loss(X, Y) = 0.62686396f0 (tracked)
loss(X, Y) = 0.65024424f0 (tracked)
loss(X, Y) = 0.6334348f0 (tracked)
loss(X, Y) = 0.62500495f0 (tracked)
loss(X, Y) = 0.6312349f0 (tracked)
loss(X, Y) = 0.62217367f0 (tracked)
loss(X, Y) = 0.65334845f0 (tracked)
loss(X, Y) = 0.6321011f0 (tracked)
loss(X, Y) = 0.687434f0 (tracked)
loss(X, Y) = 0.618666f0 (tracked)
loss(X, Y) = 0.6313139f0 (tracked)
loss(X, Y) = 0.6214606f0 (tracked)
loss(X, Y) = 0.6722478f0 (tracked)
loss(X, Y) = 0.68595624f0 (tracked)
loss(X, Y) = 0.6595492f0 (tracked)
loss(X, Y) = 0.66640717f0 (tracked)
loss(X, Y) = 0.6196415f0 (tracked)
loss(X, Y) = 0.66541815f0 (tracked)
loss(X, Y) = 0.7081881f0 (tracked)
loss(X, Y) = 0.66744584f0 (tracked)
loss(X, Y) = 0.6363587f0 (tracked)
loss(X, Y) = 0.62415767f0 (tracked)
loss(X, Y) = 0.6128806f0 (tracked)
loss(X, Y) = 0.67744976f0 (tracked)
loss(X, Y) = 0.6283347f0 (tracked)
```

```
┌ Info: Epoch 1393
└ @ Main /Users/kcft114/.julia/packages/Flux/qXNjB/src/optimise/train.jl:105
┌ Info: Epoch 1394
└ @ Main /Users/kcft114/.julia/packages/Flux/qXNjB/src/optimise/train.jl:105
┌ Info: Epoch 1395
└ @ Main /Users/kcft114/.julia/packages/Flux/qXNjB/src/optimise/train.jl:105
┌ Info: Epoch 1396
└ @ Main /Users/kcft114/.julia/packages/Flux/qXNjB/src/optimise/train.jl:105
┌ Info: Epoch 1397
└ @ Main /Users/kcft114/.julia/packages/Flux/qXNjB/src/optimise/train.jl:105
┌ Info: Epoch 1398
└ @ Main /Users/kcft114/.julia/packages/Flux/qXNjB/src/optimise/train.jl:105
┌ Info: Epoch 1399
└ @ Main /Users/kcft114/.julia/packages/Flux/qXNjB/src/optimise/train.jl:105
┌ Info: Epoch 1400
└ @ Main /Users/kcft114/.julia/packages/Flux/qXNjB/src/optimise/train.jl:105
┌ Info: Epoch 1401
└ @ Main /Users/kcft114/.julia/packages/Flux/qXNjB/src/optimise/train.jl:105
┌ Info: Epoch 1402
└ @ Main /Users/kcft114/.julia/packages/Flux/qXNjB/src/optimise/train.jl:105
┌ Info: Epoch 1403
└ @ Main /Users/kcft114/.julia/packages/Flux/qXNjB/src/optimise/train.jl:105
┌ Info: Epoch 1404
└ @ Main /Users/kcft114/.julia/packages/Flux/qXNjB/src/optimise/train.jl:105
┌ Info: Epoch 1405
└ @ Main /Users/kcft114/.julia/packages/Flux/qXNjB/src/optimise/train.jl:105
┌ Info: Epoch 1406
└ @ Main /Users/kcft114/.julia/packages/Flux/qXNjB/src/optimise/train.jl:105
┌ Info: Epoch 1407
└ @ Main /Users/kcft114/.julia/packages/Flux/qXNjB/src/optimise/train.jl:105
┌ Info: Epoch 1408
└ @ Main /Users/kcft114/.julia/packages/Flux/qXNjB/src/optimise/train.jl:105
┌ Info: Epoch 1409
└ @ Main /Users/kcft114/.julia/packages/Flux/qXNjB/src/optimise/train.jl:105
┌ Info: Epoch 1410
└ @ Main /Users/kcft114/.julia/packages/Flux/qXNjB/src/optimise/train.jl:105
┌ Info: Epoch 1411
└ @ Main /Users/kcft114/.julia/packages/Flux/qXNjB/src/optimise/train.jl:105
┌ Info: Epoch 1412
└ @ Main /Users/kcft114/.julia/packages/Flux/qXNjB/src/optimise/train.jl:105
┌ Info: Epoch 1413
└ @ Main /Users/kcft114/.julia/packages/Flux/qXNjB/src/optimise/train.jl:105
┌ Info: Epoch 1414
└ @ Main /Users/kcft114/.julia/packages/Flux/qXNjB/src/optimise/train.jl:105
┌ Info: Epoch 1415
└ @ Main /Users/kcft114/.julia/packages/Flux/qXNjB/src/optimise/train.jl:105
┌ Info: Epoch 1416
└ @ Main /Users/kcft114/.julia/packages/Flux/qXNjB/src/optimise/train.jl:105
┌ Info: Epoch 1417
└ @ Main /Users/kcft114/.julia/packages/Flux/qXNjB/src/optimise/train.jl:105
┌ Info: Epoch 1418
└ @ Main /Users/kcft114/.julia/packages/Flux/qXNjB/src/optimise/train.jl:105
┌ Info: Epoch 1419
└ @ Main /Users/kcft114/.julia/packages/Flux/qXNjB/src/optimise/train.jl:105
┌ Info: Epoch 1420
└ @ Main /Users/kcft114/.julia/packages/Flux/qXNjB/src/optimise/train.jl:105
┌ Info: Epoch 1421
└ @ Main /Users/kcft114/.julia/packages/Flux/qXNjB/src/optimise/train.jl:105
┌ Info: Epoch 1422
└ @ Main /Users/kcft114/.julia/packages/Flux/qXNjB/src/optimise/train.jl:105
┌ Info: Epoch 1423
└ @ Main /Users/kcft114/.julia/packages/Flux/qXNjB/src/optimise/train.jl:105
┌ Info: Epoch 1424
└ @ Main /Users/kcft114/.julia/packages/Flux/qXNjB/src/optimise/train.jl:105
┌ Info: Epoch 1425
└ @ Main /Users/kcft114/.julia/packages/Flux/qXNjB/src/optimise/train.jl:105
┌ Info: Epoch 1426
└ @ Main /Users/kcft114/.julia/packages/Flux/qXNjB/src/optimise/train.jl:105
┌ Info: Epoch 1427
└ @ Main /Users/kcft114/.julia/packages/Flux/qXNjB/src/optimise/train.jl:105
┌ Info: Epoch 1428
└ @ Main /Users/kcft114/.julia/packages/Flux/qXNjB/src/optimise/train.jl:105
┌ Info: Epoch 1429
└ @ Main /Users/kcft114/.julia/packages/Flux/qXNjB/src/optimise/train.jl:105
┌ Info: Epoch 1430
└ @ Main /Users/kcft114/.julia/packages/Flux/qXNjB/src/optimise/train.jl:105
┌ Info: Epoch 1431
└ @ Main /Users/kcft114/.julia/packages/Flux/qXNjB/src/optimise/train.jl:105
┌ Info: Epoch 1432
└ @ Main /Users/kcft114/.julia/packages/Flux/qXNjB/src/optimise/train.jl:105
┌ Info: Epoch 1433
└ @ Main /Users/kcft114/.julia/packages/Flux/qXNjB/src/optimise/train.jl:105
┌ Info: Epoch 1434
└ @ Main /Users/kcft114/.julia/packages/Flux/qXNjB/src/optimise/train.jl:105
┌ Info: Epoch 1435
└ @ Main /Users/kcft114/.julia/packages/Flux/qXNjB/src/optimise/train.jl:105
┌ Info: Epoch 1436
└ @ Main /Users/kcft114/.julia/packages/Flux/qXNjB/src/optimise/train.jl:105
┌ Info: Epoch 1437
└ @ Main /Users/kcft114/.julia/packages/Flux/qXNjB/src/optimise/train.jl:105
┌ Info: Epoch 1438
└ @ Main /Users/kcft114/.julia/packages/Flux/qXNjB/src/optimise/train.jl:105
┌ Info: Epoch 1439
└ @ Main /Users/kcft114/.julia/packages/Flux/qXNjB/src/optimise/train.jl:105
┌ Info: Epoch 1440
└ @ Main /Users/kcft114/.julia/packages/Flux/qXNjB/src/optimise/train.jl:105
┌ Info: Epoch 1441
└ @ Main /Users/kcft114/.julia/packages/Flux/qXNjB/src/optimise/train.jl:105
┌ Info: Epoch 1442
└ @ Main /Users/kcft114/.julia/packages/Flux/qXNjB/src/optimise/train.jl:105
┌ Info: Epoch 1443
└ @ Main /Users/kcft114/.julia/packages/Flux/qXNjB/src/optimise/train.jl:105
┌ Info: Epoch 1444
└ @ Main /Users/kcft114/.julia/packages/Flux/qXNjB/src/optimise/train.jl:105
┌ Info: Epoch 1445
└ @ Main /Users/kcft114/.julia/packages/Flux/qXNjB/src/optimise/train.jl:105
┌ Info: Epoch 1446
└ @ Main /Users/kcft114/.julia/packages/Flux/qXNjB/src/optimise/train.jl:105
┌ Info: Epoch 1447
└ @ Main /Users/kcft114/.julia/packages/Flux/qXNjB/src/optimise/train.jl:105
┌ Info: Epoch 1448
└ @ Main /Users/kcft114/.julia/packages/Flux/qXNjB/src/optimise/train.jl:105
┌ Info: Epoch 1449
└ @ Main /Users/kcft114/.julia/packages/Flux/qXNjB/src/optimise/train.jl:105
┌ Info: Epoch 1450
└ @ Main /Users/kcft114/.julia/packages/Flux/qXNjB/src/optimise/train.jl:105
┌ Info: Epoch 1451
└ @ Main /Users/kcft114/.julia/packages/Flux/qXNjB/src/optimise/train.jl:105
┌ Info: Epoch 1452
└ @ Main /Users/kcft114/.julia/packages/Flux/qXNjB/src/optimise/train.jl:105
┌ Info: Epoch 1453
└ @ Main /Users/kcft114/.julia/packages/Flux/qXNjB/src/optimise/train.jl:105
┌ Info: Epoch 1454
└ @ Main /Users/kcft114/.julia/packages/Flux/qXNjB/src/optimise/train.jl:105
┌ Info: Epoch 1455
└ @ Main /Users/kcft114/.julia/packages/Flux/qXNjB/src/optimise/train.jl:105
┌ Info: Epoch 1456
└ @ Main /Users/kcft114/.julia/packages/Flux/qXNjB/src/optimise/train.jl:105
┌ Info: Epoch 1457
└ @ Main /Users/kcft114/.julia/packages/Flux/qXNjB/src/optimise/train.jl:105
┌ Info: Epoch 1458
└ @ Main /Users/kcft114/.julia/packages/Flux/qXNjB/src/optimise/train.jl:105
┌ Info: Epoch 1459
└ @ Main /Users/kcft114/.julia/packages/Flux/qXNjB/src/optimise/train.jl:105
┌ Info: Epoch 1460
└ @ Main /Users/kcft114/.julia/packages/Flux/qXNjB/src/optimise/train.jl:105
┌ Info: Epoch 1461
└ @ Main /Users/kcft114/.julia/packages/Flux/qXNjB/src/optimise/train.jl:105
┌ Info: Epoch 1462
└ @ Main /Users/kcft114/.julia/packages/Flux/qXNjB/src/optimise/train.jl:105
┌ Info: Epoch 1463
└ @ Main /Users/kcft114/.julia/packages/Flux/qXNjB/src/optimise/train.jl:105
┌ Info: Epoch 1464
└ @ Main /Users/kcft114/.julia/packages/Flux/qXNjB/src/optimise/train.jl:105
┌ Info: Epoch 1465
└ @ Main /Users/kcft114/.julia/packages/Flux/qXNjB/src/optimise/train.jl:105
┌ Info: Epoch 1466
└ @ Main /Users/kcft114/.julia/packages/Flux/qXNjB/src/optimise/train.jl:105
┌ Info: Epoch 1467
└ @ Main /Users/kcft114/.julia/packages/Flux/qXNjB/src/optimise/train.jl:105
┌ Info: Epoch 1468
└ @ Main /Users/kcft114/.julia/packages/Flux/qXNjB/src/optimise/train.jl:105
┌ Info: Epoch 1469
└ @ Main /Users/kcft114/.julia/packages/Flux/qXNjB/src/optimise/train.jl:105
┌ Info: Epoch 1470
└ @ Main /Users/kcft114/.julia/packages/Flux/qXNjB/src/optimise/train.jl:105
┌ Info: Epoch 1471
└ @ Main /Users/kcft114/.julia/packages/Flux/qXNjB/src/optimise/train.jl:105
┌ Info: Epoch 1472
└ @ Main /Users/kcft114/.julia/packages/Flux/qXNjB/src/optimise/train.jl:105
┌ Info: Epoch 1473
└ @ Main /Users/kcft114/.julia/packages/Flux/qXNjB/src/optimise/train.jl:105
┌ Info: Epoch 1474
└ @ Main /Users/kcft114/.julia/packages/Flux/qXNjB/src/optimise/train.jl:105
┌ Info: Epoch 1475
└ @ Main /Users/kcft114/.julia/packages/Flux/qXNjB/src/optimise/train.jl:105
┌ Info: Epoch 1476
└ @ Main /Users/kcft114/.julia/packages/Flux/qXNjB/src/optimise/train.jl:105
┌ Info: Epoch 1477
└ @ Main /Users/kcft114/.julia/packages/Flux/qXNjB/src/optimise/train.jl:105
┌ Info: Epoch 1478
└ @ Main /Users/kcft114/.julia/packages/Flux/qXNjB/src/optimise/train.jl:105
┌ Info: Epoch 1479
└ @ Main /Users/kcft114/.julia/packages/Flux/qXNjB/src/optimise/train.jl:105
┌ Info: Epoch 1480
└ @ Main /Users/kcft114/.julia/packages/Flux/qXNjB/src/optimise/train.jl:105
┌ Info: Epoch 1481
└ @ Main /Users/kcft114/.julia/packages/Flux/qXNjB/src/optimise/train.jl:105
┌ Info: Epoch 1482
└ @ Main /Users/kcft114/.julia/packages/Flux/qXNjB/src/optimise/train.jl:105
┌ Info: Epoch 1483
└ @ Main /Users/kcft114/.julia/packages/Flux/qXNjB/src/optimise/train.jl:105
┌ Info: Epoch 1484
└ @ Main /Users/kcft114/.julia/packages/Flux/qXNjB/src/optimise/train.jl:105
┌ Info: Epoch 1485
└ @ Main /Users/kcft114/.julia/packages/Flux/qXNjB/src/optimise/train.jl:105
┌ Info: Epoch 1486
└ @ Main /Users/kcft114/.julia/packages/Flux/qXNjB/src/optimise/train.jl:105
┌ Info: Epoch 1487
└ @ Main /Users/kcft114/.julia/packages/Flux/qXNjB/src/optimise/train.jl:105
┌ Info: Epoch 1488
└ @ Main /Users/kcft114/.julia/packages/Flux/qXNjB/src/optimise/train.jl:105
┌ Info: Epoch 1489
└ @ Main /Users/kcft114/.julia/packages/Flux/qXNjB/src/optimise/train.jl:105
```

```
loss(X, Y) = 0.6764195f0 (tracked)
loss(X, Y) = 0.6546653f0 (tracked)
loss(X, Y) = 0.6292179f0 (tracked)
loss(X, Y) = 0.6817438f0 (tracked)
loss(X, Y) = 0.6099831f0 (tracked)
loss(X, Y) = 0.6123422f0 (tracked)
loss(X, Y) = 0.63874334f0 (tracked)
loss(X, Y) = 0.63543963f0 (tracked)
loss(X, Y) = 0.58599555f0 (tracked)
loss(X, Y) = 0.65844274f0 (tracked)
loss(X, Y) = 0.6864269f0 (tracked)
loss(X, Y) = 0.63847363f0 (tracked)
loss(X, Y) = 0.6287334f0 (tracked)
loss(X, Y) = 0.604869f0 (tracked)
loss(X, Y) = 0.65039635f0 (tracked)
loss(X, Y) = 0.6495059f0 (tracked)
loss(X, Y) = 0.61603194f0 (tracked)
loss(X, Y) = 0.6750672f0 (tracked)
loss(X, Y) = 0.66994447f0 (tracked)
loss(X, Y) = 0.69157314f0 (tracked)
loss(X, Y) = 0.63131213f0 (tracked)
loss(X, Y) = 0.68224066f0 (tracked)
loss(X, Y) = 0.60577285f0 (tracked)
loss(X, Y) = 0.6651509f0 (tracked)
loss(X, Y) = 0.6342928f0 (tracked)
loss(X, Y) = 0.6404424f0 (tracked)
loss(X, Y) = 0.64677763f0 (tracked)
loss(X, Y) = 0.6071711f0 (tracked)
loss(X, Y) = 0.6388054f0 (tracked)
loss(X, Y) = 0.67392236f0 (tracked)
loss(X, Y) = 0.6507409f0 (tracked)
loss(X, Y) = 0.6325811f0 (tracked)
loss(X, Y) = 0.6530342f0 (tracked)
loss(X, Y) = 0.59239525f0 (tracked)
loss(X, Y) = 0.62015104f0 (tracked)
loss(X, Y) = 0.706327f0 (tracked)
loss(X, Y) = 0.6252258f0 (tracked)
loss(X, Y) = 0.6346023f0 (tracked)
loss(X, Y) = 0.6690571f0 (tracked)
loss(X, Y) = 0.6526177f0 (tracked)
loss(X, Y) = 0.63281584f0 (tracked)
loss(X, Y) = 0.624864f0 (tracked)
loss(X, Y) = 0.62846786f0 (tracked)
loss(X, Y) = 0.6820903f0 (tracked)
loss(X, Y) = 0.6461117f0 (tracked)
loss(X, Y) = 0.6247505f0 (tracked)
loss(X, Y) = 0.6746108f0 (tracked)
loss(X, Y) = 0.64650804f0 (tracked)
loss(X, Y) = 0.63136023f0 (tracked)
loss(X, Y) = 0.67411906f0 (tracked)
loss(X, Y) = 0.67223495f0 (tracked)
loss(X, Y) = 0.62320125f0 (tracked)
loss(X, Y) = 0.65855974f0 (tracked)
loss(X, Y) = 0.71357375f0 (tracked)
loss(X, Y) = 0.6300557f0 (tracked)
loss(X, Y) = 0.62527233f0 (tracked)
loss(X, Y) = 0.6550604f0 (tracked)
loss(X, Y) = 0.68419874f0 (tracked)
loss(X, Y) = 0.69268596f0 (tracked)
loss(X, Y) = 0.6758319f0 (tracked)
loss(X, Y) = 0.67102903f0 (tracked)
loss(X, Y) = 0.68008536f0 (tracked)
loss(X, Y) = 0.613136f0 (tracked)
loss(X, Y) = 0.6193403f0 (tracked)
loss(X, Y) = 0.6142371f0 (tracked)
loss(X, Y) = 0.6684183f0 (tracked)
loss(X, Y) = 0.66614914f0 (tracked)
loss(X, Y) = 0.6508379f0 (tracked)
loss(X, Y) = 0.61209196f0 (tracked)
loss(X, Y) = 0.68361306f0 (tracked)
loss(X, Y) = 0.67074835f0 (tracked)
loss(X, Y) = 0.66886324f0 (tracked)
loss(X, Y) = 0.67859805f0 (tracked)
loss(X, Y) = 0.6682615f0 (tracked)
loss(X, Y) = 0.65015f0 (tracked)
loss(X, Y) = 0.6117866f0 (tracked)
loss(X, Y) = 0.66469413f0 (tracked)
loss(X, Y) = 0.6083329f0 (tracked)
loss(X, Y) = 0.6605795f0 (tracked)
loss(X, Y) = 0.67618823f0 (tracked)
loss(X, Y) = 0.61304456f0 (tracked)
loss(X, Y) = 0.6690363f0 (tracked)
loss(X, Y) = 0.6352091f0 (tracked)
loss(X, Y) = 0.6270118f0 (tracked)
loss(X, Y) = 0.627662f0 (tracked)
loss(X, Y) = 0.68302834f0 (tracked)
loss(X, Y) = 0.6857926f0 (tracked)
loss(X, Y) = 0.7005788f0 (tracked)
loss(X, Y) = 0.6611881f0 (tracked)
loss(X, Y) = 0.6776063f0 (tracked)
loss(X, Y) = 0.62317973f0 (tracked)
loss(X, Y) = 0.67424643f0 (tracked)
```

```
┌ Info: Epoch 1490
└ @ Main /Users/kcft114/.julia/packages/Flux/qXNjB/src/optimise/train.jl:105
┌ Info: Epoch 1491
└ @ Main /Users/kcft114/.julia/packages/Flux/qXNjB/src/optimise/train.jl:105
┌ Info: Epoch 1492
└ @ Main /Users/kcft114/.julia/packages/Flux/qXNjB/src/optimise/train.jl:105
┌ Info: Epoch 1493
└ @ Main /Users/kcft114/.julia/packages/Flux/qXNjB/src/optimise/train.jl:105
┌ Info: Epoch 1494
└ @ Main /Users/kcft114/.julia/packages/Flux/qXNjB/src/optimise/train.jl:105
┌ Info: Epoch 1495
└ @ Main /Users/kcft114/.julia/packages/Flux/qXNjB/src/optimise/train.jl:105
┌ Info: Epoch 1496
└ @ Main /Users/kcft114/.julia/packages/Flux/qXNjB/src/optimise/train.jl:105
┌ Info: Epoch 1497
└ @ Main /Users/kcft114/.julia/packages/Flux/qXNjB/src/optimise/train.jl:105
┌ Info: Epoch 1498
└ @ Main /Users/kcft114/.julia/packages/Flux/qXNjB/src/optimise/train.jl:105
┌ Info: Epoch 1499
└ @ Main /Users/kcft114/.julia/packages/Flux/qXNjB/src/optimise/train.jl:105
┌ Info: Epoch 1500
└ @ Main /Users/kcft114/.julia/packages/Flux/qXNjB/src/optimise/train.jl:105
┌ Info: Epoch 1501
└ @ Main /Users/kcft114/.julia/packages/Flux/qXNjB/src/optimise/train.jl:105
┌ Info: Epoch 1502
└ @ Main /Users/kcft114/.julia/packages/Flux/qXNjB/src/optimise/train.jl:105
┌ Info: Epoch 1503
└ @ Main /Users/kcft114/.julia/packages/Flux/qXNjB/src/optimise/train.jl:105
┌ Info: Epoch 1504
└ @ Main /Users/kcft114/.julia/packages/Flux/qXNjB/src/optimise/train.jl:105
┌ Info: Epoch 1505
└ @ Main /Users/kcft114/.julia/packages/Flux/qXNjB/src/optimise/train.jl:105
┌ Info: Epoch 1506
└ @ Main /Users/kcft114/.julia/packages/Flux/qXNjB/src/optimise/train.jl:105
┌ Info: Epoch 1507
└ @ Main /Users/kcft114/.julia/packages/Flux/qXNjB/src/optimise/train.jl:105
┌ Info: Epoch 1508
└ @ Main /Users/kcft114/.julia/packages/Flux/qXNjB/src/optimise/train.jl:105
┌ Info: Epoch 1509
└ @ Main /Users/kcft114/.julia/packages/Flux/qXNjB/src/optimise/train.jl:105
┌ Info: Epoch 1510
└ @ Main /Users/kcft114/.julia/packages/Flux/qXNjB/src/optimise/train.jl:105
┌ Info: Epoch 1511
└ @ Main /Users/kcft114/.julia/packages/Flux/qXNjB/src/optimise/train.jl:105
┌ Info: Epoch 1512
└ @ Main /Users/kcft114/.julia/packages/Flux/qXNjB/src/optimise/train.jl:105
┌ Info: Epoch 1513
└ @ Main /Users/kcft114/.julia/packages/Flux/qXNjB/src/optimise/train.jl:105
┌ Info: Epoch 1514
└ @ Main /Users/kcft114/.julia/packages/Flux/qXNjB/src/optimise/train.jl:105
┌ Info: Epoch 1515
└ @ Main /Users/kcft114/.julia/packages/Flux/qXNjB/src/optimise/train.jl:105
┌ Info: Epoch 1516
└ @ Main /Users/kcft114/.julia/packages/Flux/qXNjB/src/optimise/train.jl:105
┌ Info: Epoch 1517
└ @ Main /Users/kcft114/.julia/packages/Flux/qXNjB/src/optimise/train.jl:105
┌ Info: Epoch 1518
└ @ Main /Users/kcft114/.julia/packages/Flux/qXNjB/src/optimise/train.jl:105
┌ Info: Epoch 1519
└ @ Main /Users/kcft114/.julia/packages/Flux/qXNjB/src/optimise/train.jl:105
┌ Info: Epoch 1520
└ @ Main /Users/kcft114/.julia/packages/Flux/qXNjB/src/optimise/train.jl:105
┌ Info: Epoch 1521
└ @ Main /Users/kcft114/.julia/packages/Flux/qXNjB/src/optimise/train.jl:105
┌ Info: Epoch 1522
└ @ Main /Users/kcft114/.julia/packages/Flux/qXNjB/src/optimise/train.jl:105
┌ Info: Epoch 1523
└ @ Main /Users/kcft114/.julia/packages/Flux/qXNjB/src/optimise/train.jl:105
┌ Info: Epoch 1524
└ @ Main /Users/kcft114/.julia/packages/Flux/qXNjB/src/optimise/train.jl:105
┌ Info: Epoch 1525
└ @ Main /Users/kcft114/.julia/packages/Flux/qXNjB/src/optimise/train.jl:105
┌ Info: Epoch 1526
└ @ Main /Users/kcft114/.julia/packages/Flux/qXNjB/src/optimise/train.jl:105
┌ Info: Epoch 1527
└ @ Main /Users/kcft114/.julia/packages/Flux/qXNjB/src/optimise/train.jl:105
┌ Info: Epoch 1528
└ @ Main /Users/kcft114/.julia/packages/Flux/qXNjB/src/optimise/train.jl:105
┌ Info: Epoch 1529
└ @ Main /Users/kcft114/.julia/packages/Flux/qXNjB/src/optimise/train.jl:105
┌ Info: Epoch 1530
└ @ Main /Users/kcft114/.julia/packages/Flux/qXNjB/src/optimise/train.jl:105
┌ Info: Epoch 1531
└ @ Main /Users/kcft114/.julia/packages/Flux/qXNjB/src/optimise/train.jl:105
┌ Info: Epoch 1532
└ @ Main /Users/kcft114/.julia/packages/Flux/qXNjB/src/optimise/train.jl:105
┌ Info: Epoch 1533
└ @ Main /Users/kcft114/.julia/packages/Flux/qXNjB/src/optimise/train.jl:105
┌ Info: Epoch 1534
└ @ Main /Users/kcft114/.julia/packages/Flux/qXNjB/src/optimise/train.jl:105
┌ Info: Epoch 1535
└ @ Main /Users/kcft114/.julia/packages/Flux/qXNjB/src/optimise/train.jl:105
┌ Info: Epoch 1536
└ @ Main /Users/kcft114/.julia/packages/Flux/qXNjB/src/optimise/train.jl:105
┌ Info: Epoch 1537
└ @ Main /Users/kcft114/.julia/packages/Flux/qXNjB/src/optimise/train.jl:105
┌ Info: Epoch 1538
└ @ Main /Users/kcft114/.julia/packages/Flux/qXNjB/src/optimise/train.jl:105
┌ Info: Epoch 1539
└ @ Main /Users/kcft114/.julia/packages/Flux/qXNjB/src/optimise/train.jl:105
┌ Info: Epoch 1540
└ @ Main /Users/kcft114/.julia/packages/Flux/qXNjB/src/optimise/train.jl:105
┌ Info: Epoch 1541
└ @ Main /Users/kcft114/.julia/packages/Flux/qXNjB/src/optimise/train.jl:105
┌ Info: Epoch 1542
└ @ Main /Users/kcft114/.julia/packages/Flux/qXNjB/src/optimise/train.jl:105
┌ Info: Epoch 1543
└ @ Main /Users/kcft114/.julia/packages/Flux/qXNjB/src/optimise/train.jl:105
┌ Info: Epoch 1544
└ @ Main /Users/kcft114/.julia/packages/Flux/qXNjB/src/optimise/train.jl:105
┌ Info: Epoch 1545
└ @ Main /Users/kcft114/.julia/packages/Flux/qXNjB/src/optimise/train.jl:105
┌ Info: Epoch 1546
└ @ Main /Users/kcft114/.julia/packages/Flux/qXNjB/src/optimise/train.jl:105
┌ Info: Epoch 1547
└ @ Main /Users/kcft114/.julia/packages/Flux/qXNjB/src/optimise/train.jl:105
┌ Info: Epoch 1548
└ @ Main /Users/kcft114/.julia/packages/Flux/qXNjB/src/optimise/train.jl:105
┌ Info: Epoch 1549
└ @ Main /Users/kcft114/.julia/packages/Flux/qXNjB/src/optimise/train.jl:105
┌ Info: Epoch 1550
└ @ Main /Users/kcft114/.julia/packages/Flux/qXNjB/src/optimise/train.jl:105
┌ Info: Epoch 1551
└ @ Main /Users/kcft114/.julia/packages/Flux/qXNjB/src/optimise/train.jl:105
┌ Info: Epoch 1552
└ @ Main /Users/kcft114/.julia/packages/Flux/qXNjB/src/optimise/train.jl:105
┌ Info: Epoch 1553
└ @ Main /Users/kcft114/.julia/packages/Flux/qXNjB/src/optimise/train.jl:105
┌ Info: Epoch 1554
└ @ Main /Users/kcft114/.julia/packages/Flux/qXNjB/src/optimise/train.jl:105
┌ Info: Epoch 1555
└ @ Main /Users/kcft114/.julia/packages/Flux/qXNjB/src/optimise/train.jl:105
┌ Info: Epoch 1556
└ @ Main /Users/kcft114/.julia/packages/Flux/qXNjB/src/optimise/train.jl:105
┌ Info: Epoch 1557
└ @ Main /Users/kcft114/.julia/packages/Flux/qXNjB/src/optimise/train.jl:105
┌ Info: Epoch 1558
└ @ Main /Users/kcft114/.julia/packages/Flux/qXNjB/src/optimise/train.jl:105
┌ Info: Epoch 1559
└ @ Main /Users/kcft114/.julia/packages/Flux/qXNjB/src/optimise/train.jl:105
┌ Info: Epoch 1560
└ @ Main /Users/kcft114/.julia/packages/Flux/qXNjB/src/optimise/train.jl:105
┌ Info: Epoch 1561
└ @ Main /Users/kcft114/.julia/packages/Flux/qXNjB/src/optimise/train.jl:105
┌ Info: Epoch 1562
└ @ Main /Users/kcft114/.julia/packages/Flux/qXNjB/src/optimise/train.jl:105
┌ Info: Epoch 1563
└ @ Main /Users/kcft114/.julia/packages/Flux/qXNjB/src/optimise/train.jl:105
┌ Info: Epoch 1564
└ @ Main /Users/kcft114/.julia/packages/Flux/qXNjB/src/optimise/train.jl:105
┌ Info: Epoch 1565
└ @ Main /Users/kcft114/.julia/packages/Flux/qXNjB/src/optimise/train.jl:105
┌ Info: Epoch 1566
└ @ Main /Users/kcft114/.julia/packages/Flux/qXNjB/src/optimise/train.jl:105
┌ Info: Epoch 1567
└ @ Main /Users/kcft114/.julia/packages/Flux/qXNjB/src/optimise/train.jl:105
┌ Info: Epoch 1568
└ @ Main /Users/kcft114/.julia/packages/Flux/qXNjB/src/optimise/train.jl:105
┌ Info: Epoch 1569
└ @ Main /Users/kcft114/.julia/packages/Flux/qXNjB/src/optimise/train.jl:105
┌ Info: Epoch 1570
└ @ Main /Users/kcft114/.julia/packages/Flux/qXNjB/src/optimise/train.jl:105
┌ Info: Epoch 1571
└ @ Main /Users/kcft114/.julia/packages/Flux/qXNjB/src/optimise/train.jl:105
┌ Info: Epoch 1572
└ @ Main /Users/kcft114/.julia/packages/Flux/qXNjB/src/optimise/train.jl:105
┌ Info: Epoch 1573
└ @ Main /Users/kcft114/.julia/packages/Flux/qXNjB/src/optimise/train.jl:105
┌ Info: Epoch 1574
└ @ Main /Users/kcft114/.julia/packages/Flux/qXNjB/src/optimise/train.jl:105
┌ Info: Epoch 1575
└ @ Main /Users/kcft114/.julia/packages/Flux/qXNjB/src/optimise/train.jl:105
┌ Info: Epoch 1576
└ @ Main /Users/kcft114/.julia/packages/Flux/qXNjB/src/optimise/train.jl:105
┌ Info: Epoch 1577
└ @ Main /Users/kcft114/.julia/packages/Flux/qXNjB/src/optimise/train.jl:105
┌ Info: Epoch 1578
└ @ Main /Users/kcft114/.julia/packages/Flux/qXNjB/src/optimise/train.jl:105
┌ Info: Epoch 1579
└ @ Main /Users/kcft114/.julia/packages/Flux/qXNjB/src/optimise/train.jl:105
┌ Info: Epoch 1580
└ @ Main /Users/kcft114/.julia/packages/Flux/qXNjB/src/optimise/train.jl:105
┌ Info: Epoch 1581
└ @ Main /Users/kcft114/.julia/packages/Flux/qXNjB/src/optimise/train.jl:105
```

```
loss(X, Y) = 0.644758f0 (tracked)
loss(X, Y) = 0.60574603f0 (tracked)
loss(X, Y) = 0.6787266f0 (tracked)
loss(X, Y) = 0.6543716f0 (tracked)
loss(X, Y) = 0.62627894f0 (tracked)
loss(X, Y) = 0.66617715f0 (tracked)
loss(X, Y) = 0.6889024f0 (tracked)
loss(X, Y) = 0.6661057f0 (tracked)
loss(X, Y) = 0.64056474f0 (tracked)
loss(X, Y) = 0.61301464f0 (tracked)
loss(X, Y) = 0.6648921f0 (tracked)
loss(X, Y) = 0.6492206f0 (tracked)
loss(X, Y) = 0.6265632f0 (tracked)
loss(X, Y) = 0.6080514f0 (tracked)
loss(X, Y) = 0.6968759f0 (tracked)
loss(X, Y) = 0.60652614f0 (tracked)
loss(X, Y) = 0.62427104f0 (tracked)
loss(X, Y) = 0.6223374f0 (tracked)
loss(X, Y) = 0.59098583f0 (tracked)
loss(X, Y) = 0.60462356f0 (tracked)
loss(X, Y) = 0.6635046f0 (tracked)
loss(X, Y) = 0.70912653f0 (tracked)
loss(X, Y) = 0.67627096f0 (tracked)
loss(X, Y) = 0.63066393f0 (tracked)
loss(X, Y) = 0.663024f0 (tracked)
loss(X, Y) = 0.61439174f0 (tracked)
loss(X, Y) = 0.64784276f0 (tracked)
loss(X, Y) = 0.6719536f0 (tracked)
loss(X, Y) = 0.6427986f0 (tracked)
loss(X, Y) = 0.67834747f0 (tracked)
loss(X, Y) = 0.63190633f0 (tracked)
loss(X, Y) = 0.6383498f0 (tracked)
loss(X, Y) = 0.6389928f0 (tracked)
loss(X, Y) = 0.6253382f0 (tracked)
loss(X, Y) = 0.68020433f0 (tracked)
loss(X, Y) = 0.6525936f0 (tracked)
loss(X, Y) = 0.66959673f0 (tracked)
loss(X, Y) = 0.6405792f0 (tracked)
loss(X, Y) = 0.6588123f0 (tracked)
loss(X, Y) = 0.6128562f0 (tracked)
loss(X, Y) = 0.61361647f0 (tracked)
loss(X, Y) = 0.63872075f0 (tracked)
loss(X, Y) = 0.6252902f0 (tracked)
loss(X, Y) = 0.60887235f0 (tracked)
loss(X, Y) = 0.6327966f0 (tracked)
loss(X, Y) = 0.62330836f0 (tracked)
loss(X, Y) = 0.6341285f0 (tracked)
loss(X, Y) = 0.6393973f0 (tracked)
loss(X, Y) = 0.64264953f0 (tracked)
loss(X, Y) = 0.6539547f0 (tracked)
loss(X, Y) = 0.6310384f0 (tracked)
loss(X, Y) = 0.63477737f0 (tracked)
loss(X, Y) = 0.58370745f0 (tracked)
loss(X, Y) = 0.64614797f0 (tracked)
loss(X, Y) = 0.6177799f0 (tracked)
loss(X, Y) = 0.6581924f0 (tracked)
loss(X, Y) = 0.6655234f0 (tracked)
loss(X, Y) = 0.64274627f0 (tracked)
loss(X, Y) = 0.6311921f0 (tracked)
loss(X, Y) = 0.65236926f0 (tracked)
loss(X, Y) = 0.68886256f0 (tracked)
loss(X, Y) = 0.58882236f0 (tracked)
loss(X, Y) = 0.64576787f0 (tracked)
loss(X, Y) = 0.62275326f0 (tracked)
loss(X, Y) = 0.5940406f0 (tracked)
loss(X, Y) = 0.6833065f0 (tracked)
loss(X, Y) = 0.642017f0 (tracked)
loss(X, Y) = 0.6689957f0 (tracked)
loss(X, Y) = 0.6718996f0 (tracked)
loss(X, Y) = 0.6167113f0 (tracked)
loss(X, Y) = 0.61681014f0 (tracked)
loss(X, Y) = 0.62875676f0 (tracked)
loss(X, Y) = 0.6118279f0 (tracked)
loss(X, Y) = 0.6638719f0 (tracked)
loss(X, Y) = 0.63718367f0 (tracked)
loss(X, Y) = 0.6779795f0 (tracked)
loss(X, Y) = 0.6748627f0 (tracked)
loss(X, Y) = 0.6372499f0 (tracked)
loss(X, Y) = 0.62984854f0 (tracked)
loss(X, Y) = 0.6462485f0 (tracked)
loss(X, Y) = 0.6294144f0 (tracked)
loss(X, Y) = 0.6480343f0 (tracked)
```

```
┌ Info: Epoch 1582
└ @ Main /Users/kcft114/.julia/packages/Flux/qXNjB/src/optimise/train.jl:105
┌ Info: Epoch 1583
└ @ Main /Users/kcft114/.julia/packages/Flux/qXNjB/src/optimise/train.jl:105
┌ Info: Epoch 1584
└ @ Main /Users/kcft114/.julia/packages/Flux/qXNjB/src/optimise/train.jl:105
┌ Info: Epoch 1585
└ @ Main /Users/kcft114/.julia/packages/Flux/qXNjB/src/optimise/train.jl:105
┌ Info: Epoch 1586
└ @ Main /Users/kcft114/.julia/packages/Flux/qXNjB/src/optimise/train.jl:105
┌ Info: Epoch 1587
└ @ Main /Users/kcft114/.julia/packages/Flux/qXNjB/src/optimise/train.jl:105
┌ Info: Epoch 1588
└ @ Main /Users/kcft114/.julia/packages/Flux/qXNjB/src/optimise/train.jl:105
┌ Info: Epoch 1589
└ @ Main /Users/kcft114/.julia/packages/Flux/qXNjB/src/optimise/train.jl:105
┌ Info: Epoch 1590
└ @ Main /Users/kcft114/.julia/packages/Flux/qXNjB/src/optimise/train.jl:105
┌ Info: Epoch 1591
└ @ Main /Users/kcft114/.julia/packages/Flux/qXNjB/src/optimise/train.jl:105
┌ Info: Epoch 1592
└ @ Main /Users/kcft114/.julia/packages/Flux/qXNjB/src/optimise/train.jl:105
┌ Info: Epoch 1593
└ @ Main /Users/kcft114/.julia/packages/Flux/qXNjB/src/optimise/train.jl:105
┌ Info: Epoch 1594
└ @ Main /Users/kcft114/.julia/packages/Flux/qXNjB/src/optimise/train.jl:105
┌ Info: Epoch 1595
└ @ Main /Users/kcft114/.julia/packages/Flux/qXNjB/src/optimise/train.jl:105
┌ Info: Epoch 1596
└ @ Main /Users/kcft114/.julia/packages/Flux/qXNjB/src/optimise/train.jl:105
┌ Info: Epoch 1597
└ @ Main /Users/kcft114/.julia/packages/Flux/qXNjB/src/optimise/train.jl:105
┌ Info: Epoch 1598
└ @ Main /Users/kcft114/.julia/packages/Flux/qXNjB/src/optimise/train.jl:105
┌ Info: Epoch 1599
└ @ Main /Users/kcft114/.julia/packages/Flux/qXNjB/src/optimise/train.jl:105
┌ Info: Epoch 1600
└ @ Main /Users/kcft114/.julia/packages/Flux/qXNjB/src/optimise/train.jl:105
┌ Info: Epoch 1601
└ @ Main /Users/kcft114/.julia/packages/Flux/qXNjB/src/optimise/train.jl:105
┌ Info: Epoch 1602
└ @ Main /Users/kcft114/.julia/packages/Flux/qXNjB/src/optimise/train.jl:105
┌ Info: Epoch 1603
└ @ Main /Users/kcft114/.julia/packages/Flux/qXNjB/src/optimise/train.jl:105
┌ Info: Epoch 1604
└ @ Main /Users/kcft114/.julia/packages/Flux/qXNjB/src/optimise/train.jl:105
┌ Info: Epoch 1605
└ @ Main /Users/kcft114/.julia/packages/Flux/qXNjB/src/optimise/train.jl:105
┌ Info: Epoch 1606
└ @ Main /Users/kcft114/.julia/packages/Flux/qXNjB/src/optimise/train.jl:105
┌ Info: Epoch 1607
└ @ Main /Users/kcft114/.julia/packages/Flux/qXNjB/src/optimise/train.jl:105
┌ Info: Epoch 1608
└ @ Main /Users/kcft114/.julia/packages/Flux/qXNjB/src/optimise/train.jl:105
┌ Info: Epoch 1609
└ @ Main /Users/kcft114/.julia/packages/Flux/qXNjB/src/optimise/train.jl:105
┌ Info: Epoch 1610
└ @ Main /Users/kcft114/.julia/packages/Flux/qXNjB/src/optimise/train.jl:105
┌ Info: Epoch 1611
└ @ Main /Users/kcft114/.julia/packages/Flux/qXNjB/src/optimise/train.jl:105
┌ Info: Epoch 1612
└ @ Main /Users/kcft114/.julia/packages/Flux/qXNjB/src/optimise/train.jl:105
┌ Info: Epoch 1613
└ @ Main /Users/kcft114/.julia/packages/Flux/qXNjB/src/optimise/train.jl:105
┌ Info: Epoch 1614
└ @ Main /Users/kcft114/.julia/packages/Flux/qXNjB/src/optimise/train.jl:105
┌ Info: Epoch 1615
└ @ Main /Users/kcft114/.julia/packages/Flux/qXNjB/src/optimise/train.jl:105
┌ Info: Epoch 1616
└ @ Main /Users/kcft114/.julia/packages/Flux/qXNjB/src/optimise/train.jl:105
┌ Info: Epoch 1617
└ @ Main /Users/kcft114/.julia/packages/Flux/qXNjB/src/optimise/train.jl:105
┌ Info: Epoch 1618
└ @ Main /Users/kcft114/.julia/packages/Flux/qXNjB/src/optimise/train.jl:105
┌ Info: Epoch 1619
└ @ Main /Users/kcft114/.julia/packages/Flux/qXNjB/src/optimise/train.jl:105
┌ Info: Epoch 1620
└ @ Main /Users/kcft114/.julia/packages/Flux/qXNjB/src/optimise/train.jl:105
┌ Info: Epoch 1621
└ @ Main /Users/kcft114/.julia/packages/Flux/qXNjB/src/optimise/train.jl:105
┌ Info: Epoch 1622
└ @ Main /Users/kcft114/.julia/packages/Flux/qXNjB/src/optimise/train.jl:105
┌ Info: Epoch 1623
└ @ Main /Users/kcft114/.julia/packages/Flux/qXNjB/src/optimise/train.jl:105
┌ Info: Epoch 1624
└ @ Main /Users/kcft114/.julia/packages/Flux/qXNjB/src/optimise/train.jl:105
┌ Info: Epoch 1625
└ @ Main /Users/kcft114/.julia/packages/Flux/qXNjB/src/optimise/train.jl:105
┌ Info: Epoch 1626
└ @ Main /Users/kcft114/.julia/packages/Flux/qXNjB/src/optimise/train.jl:105
┌ Info: Epoch 1627
└ @ Main /Users/kcft114/.julia/packages/Flux/qXNjB/src/optimise/train.jl:105
┌ Info: Epoch 1628
└ @ Main /Users/kcft114/.julia/packages/Flux/qXNjB/src/optimise/train.jl:105
┌ Info: Epoch 1629
└ @ Main /Users/kcft114/.julia/packages/Flux/qXNjB/src/optimise/train.jl:105
┌ Info: Epoch 1630
└ @ Main /Users/kcft114/.julia/packages/Flux/qXNjB/src/optimise/train.jl:105
┌ Info: Epoch 1631
└ @ Main /Users/kcft114/.julia/packages/Flux/qXNjB/src/optimise/train.jl:105
┌ Info: Epoch 1632
└ @ Main /Users/kcft114/.julia/packages/Flux/qXNjB/src/optimise/train.jl:105
┌ Info: Epoch 1633
└ @ Main /Users/kcft114/.julia/packages/Flux/qXNjB/src/optimise/train.jl:105
┌ Info: Epoch 1634
└ @ Main /Users/kcft114/.julia/packages/Flux/qXNjB/src/optimise/train.jl:105
┌ Info: Epoch 1635
└ @ Main /Users/kcft114/.julia/packages/Flux/qXNjB/src/optimise/train.jl:105
┌ Info: Epoch 1636
└ @ Main /Users/kcft114/.julia/packages/Flux/qXNjB/src/optimise/train.jl:105
┌ Info: Epoch 1637
└ @ Main /Users/kcft114/.julia/packages/Flux/qXNjB/src/optimise/train.jl:105
┌ Info: Epoch 1638
└ @ Main /Users/kcft114/.julia/packages/Flux/qXNjB/src/optimise/train.jl:105
┌ Info: Epoch 1639
└ @ Main /Users/kcft114/.julia/packages/Flux/qXNjB/src/optimise/train.jl:105
┌ Info: Epoch 1640
└ @ Main /Users/kcft114/.julia/packages/Flux/qXNjB/src/optimise/train.jl:105
┌ Info: Epoch 1641
└ @ Main /Users/kcft114/.julia/packages/Flux/qXNjB/src/optimise/train.jl:105
┌ Info: Epoch 1642
└ @ Main /Users/kcft114/.julia/packages/Flux/qXNjB/src/optimise/train.jl:105
┌ Info: Epoch 1643
└ @ Main /Users/kcft114/.julia/packages/Flux/qXNjB/src/optimise/train.jl:105
┌ Info: Epoch 1644
└ @ Main /Users/kcft114/.julia/packages/Flux/qXNjB/src/optimise/train.jl:105
┌ Info: Epoch 1645
└ @ Main /Users/kcft114/.julia/packages/Flux/qXNjB/src/optimise/train.jl:105
┌ Info: Epoch 1646
└ @ Main /Users/kcft114/.julia/packages/Flux/qXNjB/src/optimise/train.jl:105
┌ Info: Epoch 1647
└ @ Main /Users/kcft114/.julia/packages/Flux/qXNjB/src/optimise/train.jl:105
┌ Info: Epoch 1648
└ @ Main /Users/kcft114/.julia/packages/Flux/qXNjB/src/optimise/train.jl:105
┌ Info: Epoch 1649
└ @ Main /Users/kcft114/.julia/packages/Flux/qXNjB/src/optimise/train.jl:105
┌ Info: Epoch 1650
└ @ Main /Users/kcft114/.julia/packages/Flux/qXNjB/src/optimise/train.jl:105
┌ Info: Epoch 1651
└ @ Main /Users/kcft114/.julia/packages/Flux/qXNjB/src/optimise/train.jl:105
┌ Info: Epoch 1652
└ @ Main /Users/kcft114/.julia/packages/Flux/qXNjB/src/optimise/train.jl:105
┌ Info: Epoch 1653
└ @ Main /Users/kcft114/.julia/packages/Flux/qXNjB/src/optimise/train.jl:105
┌ Info: Epoch 1654
└ @ Main /Users/kcft114/.julia/packages/Flux/qXNjB/src/optimise/train.jl:105
┌ Info: Epoch 1655
└ @ Main /Users/kcft114/.julia/packages/Flux/qXNjB/src/optimise/train.jl:105
┌ Info: Epoch 1656
└ @ Main /Users/kcft114/.julia/packages/Flux/qXNjB/src/optimise/train.jl:105
┌ Info: Epoch 1657
└ @ Main /Users/kcft114/.julia/packages/Flux/qXNjB/src/optimise/train.jl:105
┌ Info: Epoch 1658
└ @ Main /Users/kcft114/.julia/packages/Flux/qXNjB/src/optimise/train.jl:105
┌ Info: Epoch 1659
└ @ Main /Users/kcft114/.julia/packages/Flux/qXNjB/src/optimise/train.jl:105
┌ Info: Epoch 1660
└ @ Main /Users/kcft114/.julia/packages/Flux/qXNjB/src/optimise/train.jl:105
┌ Info: Epoch 1661
└ @ Main /Users/kcft114/.julia/packages/Flux/qXNjB/src/optimise/train.jl:105
┌ Info: Epoch 1662
└ @ Main /Users/kcft114/.julia/packages/Flux/qXNjB/src/optimise/train.jl:105
┌ Info: Epoch 1663
└ @ Main /Users/kcft114/.julia/packages/Flux/qXNjB/src/optimise/train.jl:105
```

```
loss(X, Y) = 0.657114f0 (tracked)
loss(X, Y) = 0.6151941f0 (tracked)
loss(X, Y) = 0.6577829f0 (tracked)
loss(X, Y) = 0.6509336f0 (tracked)
loss(X, Y) = 0.6690123f0 (tracked)
loss(X, Y) = 0.64622426f0 (tracked)
loss(X, Y) = 0.6667024f0 (tracked)
loss(X, Y) = 0.6470658f0 (tracked)
loss(X, Y) = 0.65944934f0 (tracked)
loss(X, Y) = 0.6678487f0 (tracked)
loss(X, Y) = 0.6415548f0 (tracked)
loss(X, Y) = 0.6204887f0 (tracked)
loss(X, Y) = 0.6084731f0 (tracked)
loss(X, Y) = 0.5753247f0 (tracked)
loss(X, Y) = 0.665178f0 (tracked)
loss(X, Y) = 0.5980356f0 (tracked)
loss(X, Y) = 0.64094496f0 (tracked)
loss(X, Y) = 0.6312919f0 (tracked)
loss(X, Y) = 0.66584045f0 (tracked)
loss(X, Y) = 0.65870947f0 (tracked)
loss(X, Y) = 0.6544794f0 (tracked)
loss(X, Y) = 0.6675962f0 (tracked)
loss(X, Y) = 0.65485877f0 (tracked)
loss(X, Y) = 0.605903f0 (tracked)
loss(X, Y) = 0.6208688f0 (tracked)
loss(X, Y) = 0.63036567f0 (tracked)
loss(X, Y) = 0.64858896f0 (tracked)
loss(X, Y) = 0.6294745f0 (tracked)
loss(X, Y) = 0.69753397f0 (tracked)
loss(X, Y) = 0.65160173f0 (tracked)
loss(X, Y) = 0.6522489f0 (tracked)
loss(X, Y) = 0.65113115f0 (tracked)
loss(X, Y) = 0.64577436f0 (tracked)
loss(X, Y) = 0.62927693f0 (tracked)
loss(X, Y) = 0.58560485f0 (tracked)
loss(X, Y) = 0.65795934f0 (tracked)
loss(X, Y) = 0.66291916f0 (tracked)
loss(X, Y) = 0.6160067f0 (tracked)
loss(X, Y) = 0.6424431f0 (tracked)
loss(X, Y) = 0.61829144f0 (tracked)
loss(X, Y) = 0.65363127f0 (tracked)
loss(X, Y) = 0.638464f0 (tracked)
loss(X, Y) = 0.6157034f0 (tracked)
loss(X, Y) = 0.6100794f0 (tracked)
loss(X, Y) = 0.6686302f0 (tracked)
loss(X, Y) = 0.6442752f0 (tracked)
loss(X, Y) = 0.6549493f0 (tracked)
loss(X, Y) = 0.6900487f0 (tracked)
loss(X, Y) = 0.62855864f0 (tracked)
loss(X, Y) = 0.65128255f0 (tracked)
loss(X, Y) = 0.6323103f0 (tracked)
loss(X, Y) = 0.6634602f0 (tracked)
loss(X, Y) = 0.5876863f0 (tracked)
loss(X, Y) = 0.70675385f0 (tracked)
loss(X, Y) = 0.6556542f0 (tracked)
loss(X, Y) = 0.6517337f0 (tracked)
loss(X, Y) = 0.6543001f0 (tracked)
loss(X, Y) = 0.65429777f0 (tracked)
loss(X, Y) = 0.677972f0 (tracked)
loss(X, Y) = 0.633483f0 (tracked)
loss(X, Y) = 0.65123904f0 (tracked)
loss(X, Y) = 0.63604444f0 (tracked)
loss(X, Y) = 0.5937298f0 (tracked)
loss(X, Y) = 0.61056566f0 (tracked)
loss(X, Y) = 0.65980387f0 (tracked)
loss(X, Y) = 0.6644056f0 (tracked)
loss(X, Y) = 0.62604713f0 (tracked)
loss(X, Y) = 0.6069375f0 (tracked)
loss(X, Y) = 0.644539f0 (tracked)
loss(X, Y) = 0.6219262f0 (tracked)
loss(X, Y) = 0.6340136f0 (tracked)
loss(X, Y) = 0.6687287f0 (tracked)
loss(X, Y) = 0.62424153f0 (tracked)
loss(X, Y) = 0.6661876f0 (tracked)
```

```
┌ Info: Epoch 1664
└ @ Main /Users/kcft114/.julia/packages/Flux/qXNjB/src/optimise/train.jl:105
┌ Info: Epoch 1665
└ @ Main /Users/kcft114/.julia/packages/Flux/qXNjB/src/optimise/train.jl:105
┌ Info: Epoch 1666
└ @ Main /Users/kcft114/.julia/packages/Flux/qXNjB/src/optimise/train.jl:105
┌ Info: Epoch 1667
└ @ Main /Users/kcft114/.julia/packages/Flux/qXNjB/src/optimise/train.jl:105
┌ Info: Epoch 1668
└ @ Main /Users/kcft114/.julia/packages/Flux/qXNjB/src/optimise/train.jl:105
┌ Info: Epoch 1669
└ @ Main /Users/kcft114/.julia/packages/Flux/qXNjB/src/optimise/train.jl:105
┌ Info: Epoch 1670
└ @ Main /Users/kcft114/.julia/packages/Flux/qXNjB/src/optimise/train.jl:105
┌ Info: Epoch 1671
└ @ Main /Users/kcft114/.julia/packages/Flux/qXNjB/src/optimise/train.jl:105
┌ Info: Epoch 1672
└ @ Main /Users/kcft114/.julia/packages/Flux/qXNjB/src/optimise/train.jl:105
┌ Info: Epoch 1673
└ @ Main /Users/kcft114/.julia/packages/Flux/qXNjB/src/optimise/train.jl:105
┌ Info: Epoch 1674
└ @ Main /Users/kcft114/.julia/packages/Flux/qXNjB/src/optimise/train.jl:105
┌ Info: Epoch 1675
└ @ Main /Users/kcft114/.julia/packages/Flux/qXNjB/src/optimise/train.jl:105
┌ Info: Epoch 1676
└ @ Main /Users/kcft114/.julia/packages/Flux/qXNjB/src/optimise/train.jl:105
┌ Info: Epoch 1677
└ @ Main /Users/kcft114/.julia/packages/Flux/qXNjB/src/optimise/train.jl:105
┌ Info: Epoch 1678
└ @ Main /Users/kcft114/.julia/packages/Flux/qXNjB/src/optimise/train.jl:105
┌ Info: Epoch 1679
└ @ Main /Users/kcft114/.julia/packages/Flux/qXNjB/src/optimise/train.jl:105
┌ Info: Epoch 1680
└ @ Main /Users/kcft114/.julia/packages/Flux/qXNjB/src/optimise/train.jl:105
┌ Info: Epoch 1681
└ @ Main /Users/kcft114/.julia/packages/Flux/qXNjB/src/optimise/train.jl:105
┌ Info: Epoch 1682
└ @ Main /Users/kcft114/.julia/packages/Flux/qXNjB/src/optimise/train.jl:105
┌ Info: Epoch 1683
└ @ Main /Users/kcft114/.julia/packages/Flux/qXNjB/src/optimise/train.jl:105
┌ Info: Epoch 1684
└ @ Main /Users/kcft114/.julia/packages/Flux/qXNjB/src/optimise/train.jl:105
┌ Info: Epoch 1685
└ @ Main /Users/kcft114/.julia/packages/Flux/qXNjB/src/optimise/train.jl:105
┌ Info: Epoch 1686
└ @ Main /Users/kcft114/.julia/packages/Flux/qXNjB/src/optimise/train.jl:105
┌ Info: Epoch 1687
└ @ Main /Users/kcft114/.julia/packages/Flux/qXNjB/src/optimise/train.jl:105
┌ Info: Epoch 1688
└ @ Main /Users/kcft114/.julia/packages/Flux/qXNjB/src/optimise/train.jl:105
┌ Info: Epoch 1689
└ @ Main /Users/kcft114/.julia/packages/Flux/qXNjB/src/optimise/train.jl:105
┌ Info: Epoch 1690
└ @ Main /Users/kcft114/.julia/packages/Flux/qXNjB/src/optimise/train.jl:105
┌ Info: Epoch 1691
└ @ Main /Users/kcft114/.julia/packages/Flux/qXNjB/src/optimise/train.jl:105
┌ Info: Epoch 1692
└ @ Main /Users/kcft114/.julia/packages/Flux/qXNjB/src/optimise/train.jl:105
┌ Info: Epoch 1693
└ @ Main /Users/kcft114/.julia/packages/Flux/qXNjB/src/optimise/train.jl:105
┌ Info: Epoch 1694
└ @ Main /Users/kcft114/.julia/packages/Flux/qXNjB/src/optimise/train.jl:105
┌ Info: Epoch 1695
└ @ Main /Users/kcft114/.julia/packages/Flux/qXNjB/src/optimise/train.jl:105
┌ Info: Epoch 1696
└ @ Main /Users/kcft114/.julia/packages/Flux/qXNjB/src/optimise/train.jl:105
┌ Info: Epoch 1697
└ @ Main /Users/kcft114/.julia/packages/Flux/qXNjB/src/optimise/train.jl:105
┌ Info: Epoch 1698
└ @ Main /Users/kcft114/.julia/packages/Flux/qXNjB/src/optimise/train.jl:105
┌ Info: Epoch 1699
└ @ Main /Users/kcft114/.julia/packages/Flux/qXNjB/src/optimise/train.jl:105
┌ Info: Epoch 1700
└ @ Main /Users/kcft114/.julia/packages/Flux/qXNjB/src/optimise/train.jl:105
┌ Info: Epoch 1701
└ @ Main /Users/kcft114/.julia/packages/Flux/qXNjB/src/optimise/train.jl:105
┌ Info: Epoch 1702
└ @ Main /Users/kcft114/.julia/packages/Flux/qXNjB/src/optimise/train.jl:105
┌ Info: Epoch 1703
└ @ Main /Users/kcft114/.julia/packages/Flux/qXNjB/src/optimise/train.jl:105
┌ Info: Epoch 1704
└ @ Main /Users/kcft114/.julia/packages/Flux/qXNjB/src/optimise/train.jl:105
┌ Info: Epoch 1705
└ @ Main /Users/kcft114/.julia/packages/Flux/qXNjB/src/optimise/train.jl:105
┌ Info: Epoch 1706
└ @ Main /Users/kcft114/.julia/packages/Flux/qXNjB/src/optimise/train.jl:105
┌ Info: Epoch 1707
└ @ Main /Users/kcft114/.julia/packages/Flux/qXNjB/src/optimise/train.jl:105
┌ Info: Epoch 1708
└ @ Main /Users/kcft114/.julia/packages/Flux/qXNjB/src/optimise/train.jl:105
┌ Info: Epoch 1709
└ @ Main /Users/kcft114/.julia/packages/Flux/qXNjB/src/optimise/train.jl:105
┌ Info: Epoch 1710
└ @ Main /Users/kcft114/.julia/packages/Flux/qXNjB/src/optimise/train.jl:105
┌ Info: Epoch 1711
└ @ Main /Users/kcft114/.julia/packages/Flux/qXNjB/src/optimise/train.jl:105
┌ Info: Epoch 1712
└ @ Main /Users/kcft114/.julia/packages/Flux/qXNjB/src/optimise/train.jl:105
┌ Info: Epoch 1713
└ @ Main /Users/kcft114/.julia/packages/Flux/qXNjB/src/optimise/train.jl:105
┌ Info: Epoch 1714
└ @ Main /Users/kcft114/.julia/packages/Flux/qXNjB/src/optimise/train.jl:105
┌ Info: Epoch 1715
└ @ Main /Users/kcft114/.julia/packages/Flux/qXNjB/src/optimise/train.jl:105
┌ Info: Epoch 1716
└ @ Main /Users/kcft114/.julia/packages/Flux/qXNjB/src/optimise/train.jl:105
┌ Info: Epoch 1717
└ @ Main /Users/kcft114/.julia/packages/Flux/qXNjB/src/optimise/train.jl:105
┌ Info: Epoch 1718
└ @ Main /Users/kcft114/.julia/packages/Flux/qXNjB/src/optimise/train.jl:105
┌ Info: Epoch 1719
└ @ Main /Users/kcft114/.julia/packages/Flux/qXNjB/src/optimise/train.jl:105
┌ Info: Epoch 1720
└ @ Main /Users/kcft114/.julia/packages/Flux/qXNjB/src/optimise/train.jl:105
┌ Info: Epoch 1721
└ @ Main /Users/kcft114/.julia/packages/Flux/qXNjB/src/optimise/train.jl:105
┌ Info: Epoch 1722
└ @ Main /Users/kcft114/.julia/packages/Flux/qXNjB/src/optimise/train.jl:105
┌ Info: Epoch 1723
└ @ Main /Users/kcft114/.julia/packages/Flux/qXNjB/src/optimise/train.jl:105
┌ Info: Epoch 1724
└ @ Main /Users/kcft114/.julia/packages/Flux/qXNjB/src/optimise/train.jl:105
┌ Info: Epoch 1725
└ @ Main /Users/kcft114/.julia/packages/Flux/qXNjB/src/optimise/train.jl:105
┌ Info: Epoch 1726
└ @ Main /Users/kcft114/.julia/packages/Flux/qXNjB/src/optimise/train.jl:105
┌ Info: Epoch 1727
└ @ Main /Users/kcft114/.julia/packages/Flux/qXNjB/src/optimise/train.jl:105
┌ Info: Epoch 1728
└ @ Main /Users/kcft114/.julia/packages/Flux/qXNjB/src/optimise/train.jl:105
┌ Info: Epoch 1729
└ @ Main /Users/kcft114/.julia/packages/Flux/qXNjB/src/optimise/train.jl:105
┌ Info: Epoch 1730
└ @ Main /Users/kcft114/.julia/packages/Flux/qXNjB/src/optimise/train.jl:105
┌ Info: Epoch 1731
└ @ Main /Users/kcft114/.julia/packages/Flux/qXNjB/src/optimise/train.jl:105
┌ Info: Epoch 1732
└ @ Main /Users/kcft114/.julia/packages/Flux/qXNjB/src/optimise/train.jl:105
┌ Info: Epoch 1733
└ @ Main /Users/kcft114/.julia/packages/Flux/qXNjB/src/optimise/train.jl:105
┌ Info: Epoch 1734
└ @ Main /Users/kcft114/.julia/packages/Flux/qXNjB/src/optimise/train.jl:105
┌ Info: Epoch 1735
└ @ Main /Users/kcft114/.julia/packages/Flux/qXNjB/src/optimise/train.jl:105
┌ Info: Epoch 1736
└ @ Main /Users/kcft114/.julia/packages/Flux/qXNjB/src/optimise/train.jl:105
```

```
loss(X, Y) = 0.66782266f0 (tracked)
loss(X, Y) = 0.63757694f0 (tracked)
loss(X, Y) = 0.63132155f0 (tracked)
loss(X, Y) = 0.6363062f0 (tracked)
loss(X, Y) = 0.6194915f0 (tracked)
loss(X, Y) = 0.64897203f0 (tracked)
loss(X, Y) = 0.65347046f0 (tracked)
loss(X, Y) = 0.6148855f0 (tracked)
loss(X, Y) = 0.63988024f0 (tracked)
loss(X, Y) = 0.64830166f0 (tracked)
loss(X, Y) = 0.66076285f0 (tracked)
loss(X, Y) = 0.6131661f0 (tracked)
loss(X, Y) = 0.70598406f0 (tracked)
loss(X, Y) = 0.60872096f0 (tracked)
loss(X, Y) = 0.5868491f0 (tracked)
loss(X, Y) = 0.64582914f0 (tracked)
loss(X, Y) = 0.62257683f0 (tracked)
loss(X, Y) = 0.6174172f0 (tracked)
loss(X, Y) = 0.67186856f0 (tracked)
loss(X, Y) = 0.6778028f0 (tracked)
loss(X, Y) = 0.66380674f0 (tracked)
loss(X, Y) = 0.617078f0 (tracked)
loss(X, Y) = 0.6368944f0 (tracked)
loss(X, Y) = 0.6167321f0 (tracked)
loss(X, Y) = 0.6613857f0 (tracked)
loss(X, Y) = 0.62482464f0 (tracked)
loss(X, Y) = 0.6289133f0 (tracked)
loss(X, Y) = 0.67457354f0 (tracked)
loss(X, Y) = 0.609319f0 (tracked)
loss(X, Y) = 0.6563459f0 (tracked)
loss(X, Y) = 0.6059859f0 (tracked)
loss(X, Y) = 0.5923609f0 (tracked)
loss(X, Y) = 0.61862206f0 (tracked)
loss(X, Y) = 0.58407164f0 (tracked)
loss(X, Y) = 0.6297339f0 (tracked)
loss(X, Y) = 0.6323107f0 (tracked)
loss(X, Y) = 0.59723324f0 (tracked)
loss(X, Y) = 0.65243316f0 (tracked)
loss(X, Y) = 0.6656515f0 (tracked)
loss(X, Y) = 0.6267932f0 (tracked)
loss(X, Y) = 0.6031629f0 (tracked)
loss(X, Y) = 0.64422405f0 (tracked)
loss(X, Y) = 0.6667651f0 (tracked)
loss(X, Y) = 0.61817044f0 (tracked)
loss(X, Y) = 0.68404573f0 (tracked)
loss(X, Y) = 0.6394195f0 (tracked)
loss(X, Y) = 0.64628637f0 (tracked)
loss(X, Y) = 0.66935855f0 (tracked)
loss(X, Y) = 0.6312175f0 (tracked)
loss(X, Y) = 0.6606559f0 (tracked)
loss(X, Y) = 0.63791555f0 (tracked)
loss(X, Y) = 0.66063243f0 (tracked)
loss(X, Y) = 0.66370475f0 (tracked)
loss(X, Y) = 0.60543156f0 (tracked)
loss(X, Y) = 0.6686704f0 (tracked)
loss(X, Y) = 0.62924755f0 (tracked)
loss(X, Y) = 0.6733442f0 (tracked)
loss(X, Y) = 0.6194672f0 (tracked)
loss(X, Y) = 0.6389753f0 (tracked)
loss(X, Y) = 0.6324817f0 (tracked)
loss(X, Y) = 0.6359898f0 (tracked)
loss(X, Y) = 0.68027f0 (tracked)
loss(X, Y) = 0.6271817f0 (tracked)
loss(X, Y) = 0.63048846f0 (tracked)
loss(X, Y) = 0.6570161f0 (tracked)
loss(X, Y) = 0.64128685f0 (tracked)
loss(X, Y) = 0.5999285f0 (tracked)
loss(X, Y) = 0.6406757f0 (tracked)
loss(X, Y) = 0.62836564f0 (tracked)
loss(X, Y) = 0.64507014f0 (tracked)
loss(X, Y) = 0.6733851f0 (tracked)
loss(X, Y) = 0.61453426f0 (tracked)
loss(X, Y) = 0.66585946f0 (tracked)
loss(X, Y) = 0.65722406f0 (tracked)
```

```
┌ Info: Epoch 1737
└ @ Main /Users/kcft114/.julia/packages/Flux/qXNjB/src/optimise/train.jl:105
┌ Info: Epoch 1738
└ @ Main /Users/kcft114/.julia/packages/Flux/qXNjB/src/optimise/train.jl:105
┌ Info: Epoch 1739
└ @ Main /Users/kcft114/.julia/packages/Flux/qXNjB/src/optimise/train.jl:105
┌ Info: Epoch 1740
└ @ Main /Users/kcft114/.julia/packages/Flux/qXNjB/src/optimise/train.jl:105
┌ Info: Epoch 1741
└ @ Main /Users/kcft114/.julia/packages/Flux/qXNjB/src/optimise/train.jl:105
┌ Info: Epoch 1742
└ @ Main /Users/kcft114/.julia/packages/Flux/qXNjB/src/optimise/train.jl:105
┌ Info: Epoch 1743
└ @ Main /Users/kcft114/.julia/packages/Flux/qXNjB/src/optimise/train.jl:105
┌ Info: Epoch 1744
└ @ Main /Users/kcft114/.julia/packages/Flux/qXNjB/src/optimise/train.jl:105
┌ Info: Epoch 1745
└ @ Main /Users/kcft114/.julia/packages/Flux/qXNjB/src/optimise/train.jl:105
┌ Info: Epoch 1746
└ @ Main /Users/kcft114/.julia/packages/Flux/qXNjB/src/optimise/train.jl:105
┌ Info: Epoch 1747
└ @ Main /Users/kcft114/.julia/packages/Flux/qXNjB/src/optimise/train.jl:105
┌ Info: Epoch 1748
└ @ Main /Users/kcft114/.julia/packages/Flux/qXNjB/src/optimise/train.jl:105
┌ Info: Epoch 1749
└ @ Main /Users/kcft114/.julia/packages/Flux/qXNjB/src/optimise/train.jl:105
┌ Info: Epoch 1750
└ @ Main /Users/kcft114/.julia/packages/Flux/qXNjB/src/optimise/train.jl:105
┌ Info: Epoch 1751
└ @ Main /Users/kcft114/.julia/packages/Flux/qXNjB/src/optimise/train.jl:105
┌ Info: Epoch 1752
└ @ Main /Users/kcft114/.julia/packages/Flux/qXNjB/src/optimise/train.jl:105
┌ Info: Epoch 1753
└ @ Main /Users/kcft114/.julia/packages/Flux/qXNjB/src/optimise/train.jl:105
┌ Info: Epoch 1754
└ @ Main /Users/kcft114/.julia/packages/Flux/qXNjB/src/optimise/train.jl:105
┌ Info: Epoch 1755
└ @ Main /Users/kcft114/.julia/packages/Flux/qXNjB/src/optimise/train.jl:105
┌ Info: Epoch 1756
└ @ Main /Users/kcft114/.julia/packages/Flux/qXNjB/src/optimise/train.jl:105
┌ Info: Epoch 1757
└ @ Main /Users/kcft114/.julia/packages/Flux/qXNjB/src/optimise/train.jl:105
┌ Info: Epoch 1758
└ @ Main /Users/kcft114/.julia/packages/Flux/qXNjB/src/optimise/train.jl:105
┌ Info: Epoch 1759
└ @ Main /Users/kcft114/.julia/packages/Flux/qXNjB/src/optimise/train.jl:105
┌ Info: Epoch 1760
└ @ Main /Users/kcft114/.julia/packages/Flux/qXNjB/src/optimise/train.jl:105
┌ Info: Epoch 1761
└ @ Main /Users/kcft114/.julia/packages/Flux/qXNjB/src/optimise/train.jl:105
┌ Info: Epoch 1762
└ @ Main /Users/kcft114/.julia/packages/Flux/qXNjB/src/optimise/train.jl:105
┌ Info: Epoch 1763
└ @ Main /Users/kcft114/.julia/packages/Flux/qXNjB/src/optimise/train.jl:105
┌ Info: Epoch 1764
└ @ Main /Users/kcft114/.julia/packages/Flux/qXNjB/src/optimise/train.jl:105
┌ Info: Epoch 1765
└ @ Main /Users/kcft114/.julia/packages/Flux/qXNjB/src/optimise/train.jl:105
┌ Info: Epoch 1766
└ @ Main /Users/kcft114/.julia/packages/Flux/qXNjB/src/optimise/train.jl:105
┌ Info: Epoch 1767
└ @ Main /Users/kcft114/.julia/packages/Flux/qXNjB/src/optimise/train.jl:105
┌ Info: Epoch 1768
└ @ Main /Users/kcft114/.julia/packages/Flux/qXNjB/src/optimise/train.jl:105
┌ Info: Epoch 1769
└ @ Main /Users/kcft114/.julia/packages/Flux/qXNjB/src/optimise/train.jl:105
┌ Info: Epoch 1770
└ @ Main /Users/kcft114/.julia/packages/Flux/qXNjB/src/optimise/train.jl:105
┌ Info: Epoch 1771
└ @ Main /Users/kcft114/.julia/packages/Flux/qXNjB/src/optimise/train.jl:105
┌ Info: Epoch 1772
└ @ Main /Users/kcft114/.julia/packages/Flux/qXNjB/src/optimise/train.jl:105
┌ Info: Epoch 1773
└ @ Main /Users/kcft114/.julia/packages/Flux/qXNjB/src/optimise/train.jl:105
┌ Info: Epoch 1774
└ @ Main /Users/kcft114/.julia/packages/Flux/qXNjB/src/optimise/train.jl:105
┌ Info: Epoch 1775
└ @ Main /Users/kcft114/.julia/packages/Flux/qXNjB/src/optimise/train.jl:105
┌ Info: Epoch 1776
└ @ Main /Users/kcft114/.julia/packages/Flux/qXNjB/src/optimise/train.jl:105
┌ Info: Epoch 1777
└ @ Main /Users/kcft114/.julia/packages/Flux/qXNjB/src/optimise/train.jl:105
┌ Info: Epoch 1778
└ @ Main /Users/kcft114/.julia/packages/Flux/qXNjB/src/optimise/train.jl:105
┌ Info: Epoch 1779
└ @ Main /Users/kcft114/.julia/packages/Flux/qXNjB/src/optimise/train.jl:105
┌ Info: Epoch 1780
└ @ Main /Users/kcft114/.julia/packages/Flux/qXNjB/src/optimise/train.jl:105
┌ Info: Epoch 1781
└ @ Main /Users/kcft114/.julia/packages/Flux/qXNjB/src/optimise/train.jl:105
┌ Info: Epoch 1782
└ @ Main /Users/kcft114/.julia/packages/Flux/qXNjB/src/optimise/train.jl:105
┌ Info: Epoch 1783
└ @ Main /Users/kcft114/.julia/packages/Flux/qXNjB/src/optimise/train.jl:105
┌ Info: Epoch 1784
└ @ Main /Users/kcft114/.julia/packages/Flux/qXNjB/src/optimise/train.jl:105
┌ Info: Epoch 1785
└ @ Main /Users/kcft114/.julia/packages/Flux/qXNjB/src/optimise/train.jl:105
┌ Info: Epoch 1786
└ @ Main /Users/kcft114/.julia/packages/Flux/qXNjB/src/optimise/train.jl:105
┌ Info: Epoch 1787
└ @ Main /Users/kcft114/.julia/packages/Flux/qXNjB/src/optimise/train.jl:105
┌ Info: Epoch 1788
└ @ Main /Users/kcft114/.julia/packages/Flux/qXNjB/src/optimise/train.jl:105
┌ Info: Epoch 1789
└ @ Main /Users/kcft114/.julia/packages/Flux/qXNjB/src/optimise/train.jl:105
┌ Info: Epoch 1790
└ @ Main /Users/kcft114/.julia/packages/Flux/qXNjB/src/optimise/train.jl:105
┌ Info: Epoch 1791
└ @ Main /Users/kcft114/.julia/packages/Flux/qXNjB/src/optimise/train.jl:105
┌ Info: Epoch 1792
└ @ Main /Users/kcft114/.julia/packages/Flux/qXNjB/src/optimise/train.jl:105
┌ Info: Epoch 1793
└ @ Main /Users/kcft114/.julia/packages/Flux/qXNjB/src/optimise/train.jl:105
┌ Info: Epoch 1794
└ @ Main /Users/kcft114/.julia/packages/Flux/qXNjB/src/optimise/train.jl:105
┌ Info: Epoch 1795
└ @ Main /Users/kcft114/.julia/packages/Flux/qXNjB/src/optimise/train.jl:105
┌ Info: Epoch 1796
└ @ Main /Users/kcft114/.julia/packages/Flux/qXNjB/src/optimise/train.jl:105
┌ Info: Epoch 1797
└ @ Main /Users/kcft114/.julia/packages/Flux/qXNjB/src/optimise/train.jl:105
┌ Info: Epoch 1798
└ @ Main /Users/kcft114/.julia/packages/Flux/qXNjB/src/optimise/train.jl:105
┌ Info: Epoch 1799
└ @ Main /Users/kcft114/.julia/packages/Flux/qXNjB/src/optimise/train.jl:105
┌ Info: Epoch 1800
└ @ Main /Users/kcft114/.julia/packages/Flux/qXNjB/src/optimise/train.jl:105
┌ Info: Epoch 1801
└ @ Main /Users/kcft114/.julia/packages/Flux/qXNjB/src/optimise/train.jl:105
┌ Info: Epoch 1802
└ @ Main /Users/kcft114/.julia/packages/Flux/qXNjB/src/optimise/train.jl:105
┌ Info: Epoch 1803
└ @ Main /Users/kcft114/.julia/packages/Flux/qXNjB/src/optimise/train.jl:105
┌ Info: Epoch 1804
└ @ Main /Users/kcft114/.julia/packages/Flux/qXNjB/src/optimise/train.jl:105
┌ Info: Epoch 1805
└ @ Main /Users/kcft114/.julia/packages/Flux/qXNjB/src/optimise/train.jl:105
┌ Info: Epoch 1806
└ @ Main /Users/kcft114/.julia/packages/Flux/qXNjB/src/optimise/train.jl:105
┌ Info: Epoch 1807
└ @ Main /Users/kcft114/.julia/packages/Flux/qXNjB/src/optimise/train.jl:105
┌ Info: Epoch 1808
└ @ Main /Users/kcft114/.julia/packages/Flux/qXNjB/src/optimise/train.jl:105
┌ Info: Epoch 1809
└ @ Main /Users/kcft114/.julia/packages/Flux/qXNjB/src/optimise/train.jl:105
┌ Info: Epoch 1810
└ @ Main /Users/kcft114/.julia/packages/Flux/qXNjB/src/optimise/train.jl:105
┌ Info: Epoch 1811
└ @ Main /Users/kcft114/.julia/packages/Flux/qXNjB/src/optimise/train.jl:105
```

```
loss(X, Y) = 0.65944f0 (tracked)
loss(X, Y) = 0.59645236f0 (tracked)
loss(X, Y) = 0.65685886f0 (tracked)
loss(X, Y) = 0.6097052f0 (tracked)
loss(X, Y) = 0.6374205f0 (tracked)
loss(X, Y) = 0.6443402f0 (tracked)
loss(X, Y) = 0.63929576f0 (tracked)
loss(X, Y) = 0.6466576f0 (tracked)
loss(X, Y) = 0.6551776f0 (tracked)
loss(X, Y) = 0.61402655f0 (tracked)
loss(X, Y) = 0.6123994f0 (tracked)
loss(X, Y) = 0.57711655f0 (tracked)
loss(X, Y) = 0.60567445f0 (tracked)
loss(X, Y) = 0.6280725f0 (tracked)
loss(X, Y) = 0.6333788f0 (tracked)
loss(X, Y) = 0.618852f0 (tracked)
loss(X, Y) = 0.5921403f0 (tracked)
loss(X, Y) = 0.5912162f0 (tracked)
loss(X, Y) = 0.601928f0 (tracked)
loss(X, Y) = 0.5617615f0 (tracked)
loss(X, Y) = 0.64007604f0 (tracked)
loss(X, Y) = 0.6388574f0 (tracked)
loss(X, Y) = 0.5835199f0 (tracked)
loss(X, Y) = 0.6051281f0 (tracked)
loss(X, Y) = 0.6421856f0 (tracked)
loss(X, Y) = 0.641707f0 (tracked)
loss(X, Y) = 0.6060694f0 (tracked)
loss(X, Y) = 0.6146595f0 (tracked)
loss(X, Y) = 0.6075559f0 (tracked)
loss(X, Y) = 0.6661756f0 (tracked)
loss(X, Y) = 0.6254025f0 (tracked)
loss(X, Y) = 0.7074504f0 (tracked)
loss(X, Y) = 0.61495227f0 (tracked)
loss(X, Y) = 0.62525445f0 (tracked)
loss(X, Y) = 0.6275575f0 (tracked)
loss(X, Y) = 0.6563438f0 (tracked)
loss(X, Y) = 0.64095336f0 (tracked)
loss(X, Y) = 0.64938563f0 (tracked)
loss(X, Y) = 0.6538995f0 (tracked)
loss(X, Y) = 0.5911316f0 (tracked)
loss(X, Y) = 0.646446f0 (tracked)
loss(X, Y) = 0.60341394f0 (tracked)
loss(X, Y) = 0.64601105f0 (tracked)
loss(X, Y) = 0.5936107f0 (tracked)
loss(X, Y) = 0.6458488f0 (tracked)
loss(X, Y) = 0.6362863f0 (tracked)
loss(X, Y) = 0.62410074f0 (tracked)
loss(X, Y) = 0.5910361f0 (tracked)
loss(X, Y) = 0.60086197f0 (tracked)
loss(X, Y) = 0.67026126f0 (tracked)
loss(X, Y) = 0.640184f0 (tracked)
loss(X, Y) = 0.6201023f0 (tracked)
loss(X, Y) = 0.66991436f0 (tracked)
loss(X, Y) = 0.6416015f0 (tracked)
loss(X, Y) = 0.63790196f0 (tracked)
loss(X, Y) = 0.6029853f0 (tracked)
loss(X, Y) = 0.641172f0 (tracked)
loss(X, Y) = 0.66983664f0 (tracked)
loss(X, Y) = 0.64579946f0 (tracked)
loss(X, Y) = 0.62625414f0 (tracked)
loss(X, Y) = 0.5965335f0 (tracked)
loss(X, Y) = 0.6285444f0 (tracked)
loss(X, Y) = 0.5965808f0 (tracked)
loss(X, Y) = 0.638183f0 (tracked)
loss(X, Y) = 0.6289893f0 (tracked)
loss(X, Y) = 0.6377094f0 (tracked)
loss(X, Y) = 0.6196705f0 (tracked)
loss(X, Y) = 0.6124847f0 (tracked)
loss(X, Y) = 0.6287056f0 (tracked)
loss(X, Y) = 0.60782313f0 (tracked)
loss(X, Y) = 0.63079727f0 (tracked)
loss(X, Y) = 0.65371114f0 (tracked)
loss(X, Y) = 0.66690755f0 (tracked)
loss(X, Y) = 0.60166276f0 (tracked)
loss(X, Y) = 0.643429f0 (tracked)
loss(X, Y) = 0.6251985f0 (tracked)
loss(X, Y) = 0.6535734f0 (tracked)
loss(X, Y) = 0.65374744f0 (tracked)
loss(X, Y) = 0.6018902f0 (tracked)
loss(X, Y) = 0.6196172f0 (tracked)
loss(X, Y) = 0.641336f0 (tracked)
loss(X, Y) = 0.59489733f0 (tracked)
loss(X, Y) = 0.58486146f0 (tracked)
loss(X, Y) = 0.6127011f0 (tracked)
loss(X, Y) = 0.64984584f0 (tracked)
```

```
┌ Info: Epoch 1812
└ @ Main /Users/kcft114/.julia/packages/Flux/qXNjB/src/optimise/train.jl:105
┌ Info: Epoch 1813
└ @ Main /Users/kcft114/.julia/packages/Flux/qXNjB/src/optimise/train.jl:105
┌ Info: Epoch 1814
└ @ Main /Users/kcft114/.julia/packages/Flux/qXNjB/src/optimise/train.jl:105
┌ Info: Epoch 1815
└ @ Main /Users/kcft114/.julia/packages/Flux/qXNjB/src/optimise/train.jl:105
┌ Info: Epoch 1816
└ @ Main /Users/kcft114/.julia/packages/Flux/qXNjB/src/optimise/train.jl:105
┌ Info: Epoch 1817
└ @ Main /Users/kcft114/.julia/packages/Flux/qXNjB/src/optimise/train.jl:105
┌ Info: Epoch 1818
└ @ Main /Users/kcft114/.julia/packages/Flux/qXNjB/src/optimise/train.jl:105
┌ Info: Epoch 1819
└ @ Main /Users/kcft114/.julia/packages/Flux/qXNjB/src/optimise/train.jl:105
┌ Info: Epoch 1820
└ @ Main /Users/kcft114/.julia/packages/Flux/qXNjB/src/optimise/train.jl:105
┌ Info: Epoch 1821
└ @ Main /Users/kcft114/.julia/packages/Flux/qXNjB/src/optimise/train.jl:105
┌ Info: Epoch 1822
└ @ Main /Users/kcft114/.julia/packages/Flux/qXNjB/src/optimise/train.jl:105
┌ Info: Epoch 1823
└ @ Main /Users/kcft114/.julia/packages/Flux/qXNjB/src/optimise/train.jl:105
┌ Info: Epoch 1824
└ @ Main /Users/kcft114/.julia/packages/Flux/qXNjB/src/optimise/train.jl:105
┌ Info: Epoch 1825
└ @ Main /Users/kcft114/.julia/packages/Flux/qXNjB/src/optimise/train.jl:105
┌ Info: Epoch 1826
└ @ Main /Users/kcft114/.julia/packages/Flux/qXNjB/src/optimise/train.jl:105
┌ Info: Epoch 1827
└ @ Main /Users/kcft114/.julia/packages/Flux/qXNjB/src/optimise/train.jl:105
┌ Info: Epoch 1828
└ @ Main /Users/kcft114/.julia/packages/Flux/qXNjB/src/optimise/train.jl:105
┌ Info: Epoch 1829
└ @ Main /Users/kcft114/.julia/packages/Flux/qXNjB/src/optimise/train.jl:105
┌ Info: Epoch 1830
└ @ Main /Users/kcft114/.julia/packages/Flux/qXNjB/src/optimise/train.jl:105
┌ Info: Epoch 1831
└ @ Main /Users/kcft114/.julia/packages/Flux/qXNjB/src/optimise/train.jl:105
┌ Info: Epoch 1832
└ @ Main /Users/kcft114/.julia/packages/Flux/qXNjB/src/optimise/train.jl:105
┌ Info: Epoch 1833
└ @ Main /Users/kcft114/.julia/packages/Flux/qXNjB/src/optimise/train.jl:105
┌ Info: Epoch 1834
└ @ Main /Users/kcft114/.julia/packages/Flux/qXNjB/src/optimise/train.jl:105
┌ Info: Epoch 1835
└ @ Main /Users/kcft114/.julia/packages/Flux/qXNjB/src/optimise/train.jl:105
┌ Info: Epoch 1836
└ @ Main /Users/kcft114/.julia/packages/Flux/qXNjB/src/optimise/train.jl:105
┌ Info: Epoch 1837
└ @ Main /Users/kcft114/.julia/packages/Flux/qXNjB/src/optimise/train.jl:105
┌ Info: Epoch 1838
└ @ Main /Users/kcft114/.julia/packages/Flux/qXNjB/src/optimise/train.jl:105
┌ Info: Epoch 1839
└ @ Main /Users/kcft114/.julia/packages/Flux/qXNjB/src/optimise/train.jl:105
┌ Info: Epoch 1840
└ @ Main /Users/kcft114/.julia/packages/Flux/qXNjB/src/optimise/train.jl:105
┌ Info: Epoch 1841
└ @ Main /Users/kcft114/.julia/packages/Flux/qXNjB/src/optimise/train.jl:105
┌ Info: Epoch 1842
└ @ Main /Users/kcft114/.julia/packages/Flux/qXNjB/src/optimise/train.jl:105
┌ Info: Epoch 1843
└ @ Main /Users/kcft114/.julia/packages/Flux/qXNjB/src/optimise/train.jl:105
┌ Info: Epoch 1844
└ @ Main /Users/kcft114/.julia/packages/Flux/qXNjB/src/optimise/train.jl:105
┌ Info: Epoch 1845
└ @ Main /Users/kcft114/.julia/packages/Flux/qXNjB/src/optimise/train.jl:105
┌ Info: Epoch 1846
└ @ Main /Users/kcft114/.julia/packages/Flux/qXNjB/src/optimise/train.jl:105
┌ Info: Epoch 1847
└ @ Main /Users/kcft114/.julia/packages/Flux/qXNjB/src/optimise/train.jl:105
┌ Info: Epoch 1848
└ @ Main /Users/kcft114/.julia/packages/Flux/qXNjB/src/optimise/train.jl:105
┌ Info: Epoch 1849
└ @ Main /Users/kcft114/.julia/packages/Flux/qXNjB/src/optimise/train.jl:105
┌ Info: Epoch 1850
└ @ Main /Users/kcft114/.julia/packages/Flux/qXNjB/src/optimise/train.jl:105
┌ Info: Epoch 1851
└ @ Main /Users/kcft114/.julia/packages/Flux/qXNjB/src/optimise/train.jl:105
┌ Info: Epoch 1852
└ @ Main /Users/kcft114/.julia/packages/Flux/qXNjB/src/optimise/train.jl:105
┌ Info: Epoch 1853
└ @ Main /Users/kcft114/.julia/packages/Flux/qXNjB/src/optimise/train.jl:105
┌ Info: Epoch 1854
└ @ Main /Users/kcft114/.julia/packages/Flux/qXNjB/src/optimise/train.jl:105
┌ Info: Epoch 1855
└ @ Main /Users/kcft114/.julia/packages/Flux/qXNjB/src/optimise/train.jl:105
┌ Info: Epoch 1856
└ @ Main /Users/kcft114/.julia/packages/Flux/qXNjB/src/optimise/train.jl:105
┌ Info: Epoch 1857
└ @ Main /Users/kcft114/.julia/packages/Flux/qXNjB/src/optimise/train.jl:105
┌ Info: Epoch 1858
└ @ Main /Users/kcft114/.julia/packages/Flux/qXNjB/src/optimise/train.jl:105
┌ Info: Epoch 1859
└ @ Main /Users/kcft114/.julia/packages/Flux/qXNjB/src/optimise/train.jl:105
┌ Info: Epoch 1860
└ @ Main /Users/kcft114/.julia/packages/Flux/qXNjB/src/optimise/train.jl:105
┌ Info: Epoch 1861
└ @ Main /Users/kcft114/.julia/packages/Flux/qXNjB/src/optimise/train.jl:105
┌ Info: Epoch 1862
└ @ Main /Users/kcft114/.julia/packages/Flux/qXNjB/src/optimise/train.jl:105
┌ Info: Epoch 1863
└ @ Main /Users/kcft114/.julia/packages/Flux/qXNjB/src/optimise/train.jl:105
┌ Info: Epoch 1864
└ @ Main /Users/kcft114/.julia/packages/Flux/qXNjB/src/optimise/train.jl:105
┌ Info: Epoch 1865
└ @ Main /Users/kcft114/.julia/packages/Flux/qXNjB/src/optimise/train.jl:105
┌ Info: Epoch 1866
└ @ Main /Users/kcft114/.julia/packages/Flux/qXNjB/src/optimise/train.jl:105
┌ Info: Epoch 1867
└ @ Main /Users/kcft114/.julia/packages/Flux/qXNjB/src/optimise/train.jl:105
┌ Info: Epoch 1868
└ @ Main /Users/kcft114/.julia/packages/Flux/qXNjB/src/optimise/train.jl:105
┌ Info: Epoch 1869
└ @ Main /Users/kcft114/.julia/packages/Flux/qXNjB/src/optimise/train.jl:105
┌ Info: Epoch 1870
└ @ Main /Users/kcft114/.julia/packages/Flux/qXNjB/src/optimise/train.jl:105
┌ Info: Epoch 1871
└ @ Main /Users/kcft114/.julia/packages/Flux/qXNjB/src/optimise/train.jl:105
┌ Info: Epoch 1872
└ @ Main /Users/kcft114/.julia/packages/Flux/qXNjB/src/optimise/train.jl:105
┌ Info: Epoch 1873
└ @ Main /Users/kcft114/.julia/packages/Flux/qXNjB/src/optimise/train.jl:105
┌ Info: Epoch 1874
└ @ Main /Users/kcft114/.julia/packages/Flux/qXNjB/src/optimise/train.jl:105
┌ Info: Epoch 1875
└ @ Main /Users/kcft114/.julia/packages/Flux/qXNjB/src/optimise/train.jl:105
┌ Info: Epoch 1876
└ @ Main /Users/kcft114/.julia/packages/Flux/qXNjB/src/optimise/train.jl:105
┌ Info: Epoch 1877
└ @ Main /Users/kcft114/.julia/packages/Flux/qXNjB/src/optimise/train.jl:105
┌ Info: Epoch 1878
└ @ Main /Users/kcft114/.julia/packages/Flux/qXNjB/src/optimise/train.jl:105
┌ Info: Epoch 1879
└ @ Main /Users/kcft114/.julia/packages/Flux/qXNjB/src/optimise/train.jl:105
┌ Info: Epoch 1880
└ @ Main /Users/kcft114/.julia/packages/Flux/qXNjB/src/optimise/train.jl:105
┌ Info: Epoch 1881
└ @ Main /Users/kcft114/.julia/packages/Flux/qXNjB/src/optimise/train.jl:105
┌ Info: Epoch 1882
└ @ Main /Users/kcft114/.julia/packages/Flux/qXNjB/src/optimise/train.jl:105
┌ Info: Epoch 1883
└ @ Main /Users/kcft114/.julia/packages/Flux/qXNjB/src/optimise/train.jl:105
┌ Info: Epoch 1884
└ @ Main /Users/kcft114/.julia/packages/Flux/qXNjB/src/optimise/train.jl:105
┌ Info: Epoch 1885
└ @ Main /Users/kcft114/.julia/packages/Flux/qXNjB/src/optimise/train.jl:105
┌ Info: Epoch 1886
└ @ Main /Users/kcft114/.julia/packages/Flux/qXNjB/src/optimise/train.jl:105
┌ Info: Epoch 1887
└ @ Main /Users/kcft114/.julia/packages/Flux/qXNjB/src/optimise/train.jl:105
┌ Info: Epoch 1888
└ @ Main /Users/kcft114/.julia/packages/Flux/qXNjB/src/optimise/train.jl:105
┌ Info: Epoch 1889
└ @ Main /Users/kcft114/.julia/packages/Flux/qXNjB/src/optimise/train.jl:105
┌ Info: Epoch 1890
└ @ Main /Users/kcft114/.julia/packages/Flux/qXNjB/src/optimise/train.jl:105
┌ Info: Epoch 1891
└ @ Main /Users/kcft114/.julia/packages/Flux/qXNjB/src/optimise/train.jl:105
┌ Info: Epoch 1892
└ @ Main /Users/kcft114/.julia/packages/Flux/qXNjB/src/optimise/train.jl:105
┌ Info: Epoch 1893
└ @ Main /Users/kcft114/.julia/packages/Flux/qXNjB/src/optimise/train.jl:105
┌ Info: Epoch 1894
└ @ Main /Users/kcft114/.julia/packages/Flux/qXNjB/src/optimise/train.jl:105
┌ Info: Epoch 1895
└ @ Main /Users/kcft114/.julia/packages/Flux/qXNjB/src/optimise/train.jl:105
┌ Info: Epoch 1896
└ @ Main /Users/kcft114/.julia/packages/Flux/qXNjB/src/optimise/train.jl:105
```

```
loss(X, Y) = 0.6337289f0 (tracked)
loss(X, Y) = 0.635329f0 (tracked)
loss(X, Y) = 0.62219435f0 (tracked)
loss(X, Y) = 0.62310624f0 (tracked)
loss(X, Y) = 0.66152966f0 (tracked)
loss(X, Y) = 0.64648676f0 (tracked)
loss(X, Y) = 0.65963626f0 (tracked)
loss(X, Y) = 0.6229478f0 (tracked)
loss(X, Y) = 0.6334693f0 (tracked)
loss(X, Y) = 0.6520282f0 (tracked)
loss(X, Y) = 0.6626635f0 (tracked)
loss(X, Y) = 0.6262247f0 (tracked)
loss(X, Y) = 0.6460532f0 (tracked)
loss(X, Y) = 0.6618825f0 (tracked)
loss(X, Y) = 0.6715101f0 (tracked)
loss(X, Y) = 0.6176911f0 (tracked)
loss(X, Y) = 0.6285217f0 (tracked)
loss(X, Y) = 0.6433223f0 (tracked)
loss(X, Y) = 0.57812595f0 (tracked)
loss(X, Y) = 0.6391979f0 (tracked)
loss(X, Y) = 0.6240054f0 (tracked)
loss(X, Y) = 0.64610016f0 (tracked)
loss(X, Y) = 0.63141346f0 (tracked)
loss(X, Y) = 0.5561694f0 (tracked)
loss(X, Y) = 0.6081138f0 (tracked)
loss(X, Y) = 0.61775076f0 (tracked)
loss(X, Y) = 0.640878f0 (tracked)
loss(X, Y) = 0.61118096f0 (tracked)
loss(X, Y) = 0.6749254f0 (tracked)
loss(X, Y) = 0.6030777f0 (tracked)
loss(X, Y) = 0.6733647f0 (tracked)
loss(X, Y) = 0.6404432f0 (tracked)
loss(X, Y) = 0.6025025f0 (tracked)
loss(X, Y) = 0.6343694f0 (tracked)
loss(X, Y) = 0.6316181f0 (tracked)
loss(X, Y) = 0.6073894f0 (tracked)
loss(X, Y) = 0.6128019f0 (tracked)
loss(X, Y) = 0.6630304f0 (tracked)
loss(X, Y) = 0.60822517f0 (tracked)
loss(X, Y) = 0.5942634f0 (tracked)
loss(X, Y) = 0.57467175f0 (tracked)
loss(X, Y) = 0.6562902f0 (tracked)
loss(X, Y) = 0.6187325f0 (tracked)
loss(X, Y) = 0.7018032f0 (tracked)
loss(X, Y) = 0.6448602f0 (tracked)
loss(X, Y) = 0.67109f0 (tracked)
loss(X, Y) = 0.6856441f0 (tracked)
loss(X, Y) = 0.6894942f0 (tracked)
loss(X, Y) = 0.6079464f0 (tracked)
loss(X, Y) = 0.59303623f0 (tracked)
loss(X, Y) = 0.59607804f0 (tracked)
loss(X, Y) = 0.6170647f0 (tracked)
loss(X, Y) = 0.64101416f0 (tracked)
loss(X, Y) = 0.62971425f0 (tracked)
loss(X, Y) = 0.5731932f0 (tracked)
loss(X, Y) = 0.6476189f0 (tracked)
loss(X, Y) = 0.68732965f0 (tracked)
loss(X, Y) = 0.65336436f0 (tracked)
loss(X, Y) = 0.6257293f0 (tracked)
loss(X, Y) = 0.6878567f0 (tracked)
loss(X, Y) = 0.6525229f0 (tracked)
loss(X, Y) = 0.60134244f0 (tracked)
loss(X, Y) = 0.6472801f0 (tracked)
loss(X, Y) = 0.5892767f0 (tracked)
loss(X, Y) = 0.63882333f0 (tracked)
loss(X, Y) = 0.5886983f0 (tracked)
loss(X, Y) = 0.6182731f0 (tracked)
loss(X, Y) = 0.65370786f0 (tracked)
loss(X, Y) = 0.6391635f0 (tracked)
loss(X, Y) = 0.681659f0 (tracked)
loss(X, Y) = 0.5877134f0 (tracked)
loss(X, Y) = 0.5942362f0 (tracked)
loss(X, Y) = 0.6283484f0 (tracked)
loss(X, Y) = 0.6419956f0 (tracked)
loss(X, Y) = 0.5782308f0 (tracked)
loss(X, Y) = 0.60508937f0 (tracked)
loss(X, Y) = 0.6050126f0 (tracked)
loss(X, Y) = 0.60176176f0 (tracked)
loss(X, Y) = 0.6291244f0 (tracked)
loss(X, Y) = 0.5766114f0 (tracked)
loss(X, Y) = 0.6013237f0 (tracked)
loss(X, Y) = 0.62196743f0 (tracked)
loss(X, Y) = 0.61812943f0 (tracked)
loss(X, Y) = 0.6347674f0 (tracked)
loss(X, Y) = 0.6148805f0 (tracked)
```

```
┌ Info: Epoch 1897
└ @ Main /Users/kcft114/.julia/packages/Flux/qXNjB/src/optimise/train.jl:105
┌ Info: Epoch 1898
└ @ Main /Users/kcft114/.julia/packages/Flux/qXNjB/src/optimise/train.jl:105
┌ Info: Epoch 1899
└ @ Main /Users/kcft114/.julia/packages/Flux/qXNjB/src/optimise/train.jl:105
┌ Info: Epoch 1900
└ @ Main /Users/kcft114/.julia/packages/Flux/qXNjB/src/optimise/train.jl:105
┌ Info: Epoch 1901
└ @ Main /Users/kcft114/.julia/packages/Flux/qXNjB/src/optimise/train.jl:105
┌ Info: Epoch 1902
└ @ Main /Users/kcft114/.julia/packages/Flux/qXNjB/src/optimise/train.jl:105
┌ Info: Epoch 1903
└ @ Main /Users/kcft114/.julia/packages/Flux/qXNjB/src/optimise/train.jl:105
┌ Info: Epoch 1904
└ @ Main /Users/kcft114/.julia/packages/Flux/qXNjB/src/optimise/train.jl:105
┌ Info: Epoch 1905
└ @ Main /Users/kcft114/.julia/packages/Flux/qXNjB/src/optimise/train.jl:105
┌ Info: Epoch 1906
└ @ Main /Users/kcft114/.julia/packages/Flux/qXNjB/src/optimise/train.jl:105
┌ Info: Epoch 1907
└ @ Main /Users/kcft114/.julia/packages/Flux/qXNjB/src/optimise/train.jl:105
┌ Info: Epoch 1908
└ @ Main /Users/kcft114/.julia/packages/Flux/qXNjB/src/optimise/train.jl:105
┌ Info: Epoch 1909
└ @ Main /Users/kcft114/.julia/packages/Flux/qXNjB/src/optimise/train.jl:105
┌ Info: Epoch 1910
└ @ Main /Users/kcft114/.julia/packages/Flux/qXNjB/src/optimise/train.jl:105
┌ Info: Epoch 1911
└ @ Main /Users/kcft114/.julia/packages/Flux/qXNjB/src/optimise/train.jl:105
┌ Info: Epoch 1912
└ @ Main /Users/kcft114/.julia/packages/Flux/qXNjB/src/optimise/train.jl:105
┌ Info: Epoch 1913
└ @ Main /Users/kcft114/.julia/packages/Flux/qXNjB/src/optimise/train.jl:105
┌ Info: Epoch 1914
└ @ Main /Users/kcft114/.julia/packages/Flux/qXNjB/src/optimise/train.jl:105
┌ Info: Epoch 1915
└ @ Main /Users/kcft114/.julia/packages/Flux/qXNjB/src/optimise/train.jl:105
┌ Info: Epoch 1916
└ @ Main /Users/kcft114/.julia/packages/Flux/qXNjB/src/optimise/train.jl:105
┌ Info: Epoch 1917
└ @ Main /Users/kcft114/.julia/packages/Flux/qXNjB/src/optimise/train.jl:105
┌ Info: Epoch 1918
└ @ Main /Users/kcft114/.julia/packages/Flux/qXNjB/src/optimise/train.jl:105
┌ Info: Epoch 1919
└ @ Main /Users/kcft114/.julia/packages/Flux/qXNjB/src/optimise/train.jl:105
┌ Info: Epoch 1920
└ @ Main /Users/kcft114/.julia/packages/Flux/qXNjB/src/optimise/train.jl:105
┌ Info: Epoch 1921
└ @ Main /Users/kcft114/.julia/packages/Flux/qXNjB/src/optimise/train.jl:105
┌ Info: Epoch 1922
└ @ Main /Users/kcft114/.julia/packages/Flux/qXNjB/src/optimise/train.jl:105
┌ Info: Epoch 1923
└ @ Main /Users/kcft114/.julia/packages/Flux/qXNjB/src/optimise/train.jl:105
┌ Info: Epoch 1924
└ @ Main /Users/kcft114/.julia/packages/Flux/qXNjB/src/optimise/train.jl:105
┌ Info: Epoch 1925
└ @ Main /Users/kcft114/.julia/packages/Flux/qXNjB/src/optimise/train.jl:105
┌ Info: Epoch 1926
└ @ Main /Users/kcft114/.julia/packages/Flux/qXNjB/src/optimise/train.jl:105
┌ Info: Epoch 1927
└ @ Main /Users/kcft114/.julia/packages/Flux/qXNjB/src/optimise/train.jl:105
┌ Info: Epoch 1928
└ @ Main /Users/kcft114/.julia/packages/Flux/qXNjB/src/optimise/train.jl:105
┌ Info: Epoch 1929
└ @ Main /Users/kcft114/.julia/packages/Flux/qXNjB/src/optimise/train.jl:105
┌ Info: Epoch 1930
└ @ Main /Users/kcft114/.julia/packages/Flux/qXNjB/src/optimise/train.jl:105
┌ Info: Epoch 1931
└ @ Main /Users/kcft114/.julia/packages/Flux/qXNjB/src/optimise/train.jl:105
┌ Info: Epoch 1932
└ @ Main /Users/kcft114/.julia/packages/Flux/qXNjB/src/optimise/train.jl:105
┌ Info: Epoch 1933
└ @ Main /Users/kcft114/.julia/packages/Flux/qXNjB/src/optimise/train.jl:105
┌ Info: Epoch 1934
└ @ Main /Users/kcft114/.julia/packages/Flux/qXNjB/src/optimise/train.jl:105
┌ Info: Epoch 1935
└ @ Main /Users/kcft114/.julia/packages/Flux/qXNjB/src/optimise/train.jl:105
┌ Info: Epoch 1936
└ @ Main /Users/kcft114/.julia/packages/Flux/qXNjB/src/optimise/train.jl:105
┌ Info: Epoch 1937
└ @ Main /Users/kcft114/.julia/packages/Flux/qXNjB/src/optimise/train.jl:105
┌ Info: Epoch 1938
└ @ Main /Users/kcft114/.julia/packages/Flux/qXNjB/src/optimise/train.jl:105
┌ Info: Epoch 1939
└ @ Main /Users/kcft114/.julia/packages/Flux/qXNjB/src/optimise/train.jl:105
┌ Info: Epoch 1940
└ @ Main /Users/kcft114/.julia/packages/Flux/qXNjB/src/optimise/train.jl:105
┌ Info: Epoch 1941
└ @ Main /Users/kcft114/.julia/packages/Flux/qXNjB/src/optimise/train.jl:105
┌ Info: Epoch 1942
└ @ Main /Users/kcft114/.julia/packages/Flux/qXNjB/src/optimise/train.jl:105
┌ Info: Epoch 1943
└ @ Main /Users/kcft114/.julia/packages/Flux/qXNjB/src/optimise/train.jl:105
┌ Info: Epoch 1944
└ @ Main /Users/kcft114/.julia/packages/Flux/qXNjB/src/optimise/train.jl:105
┌ Info: Epoch 1945
└ @ Main /Users/kcft114/.julia/packages/Flux/qXNjB/src/optimise/train.jl:105
┌ Info: Epoch 1946
└ @ Main /Users/kcft114/.julia/packages/Flux/qXNjB/src/optimise/train.jl:105
┌ Info: Epoch 1947
└ @ Main /Users/kcft114/.julia/packages/Flux/qXNjB/src/optimise/train.jl:105
┌ Info: Epoch 1948
└ @ Main /Users/kcft114/.julia/packages/Flux/qXNjB/src/optimise/train.jl:105
┌ Info: Epoch 1949
└ @ Main /Users/kcft114/.julia/packages/Flux/qXNjB/src/optimise/train.jl:105
┌ Info: Epoch 1950
└ @ Main /Users/kcft114/.julia/packages/Flux/qXNjB/src/optimise/train.jl:105
┌ Info: Epoch 1951
└ @ Main /Users/kcft114/.julia/packages/Flux/qXNjB/src/optimise/train.jl:105
┌ Info: Epoch 1952
└ @ Main /Users/kcft114/.julia/packages/Flux/qXNjB/src/optimise/train.jl:105
┌ Info: Epoch 1953
└ @ Main /Users/kcft114/.julia/packages/Flux/qXNjB/src/optimise/train.jl:105
┌ Info: Epoch 1954
└ @ Main /Users/kcft114/.julia/packages/Flux/qXNjB/src/optimise/train.jl:105
┌ Info: Epoch 1955
└ @ Main /Users/kcft114/.julia/packages/Flux/qXNjB/src/optimise/train.jl:105
┌ Info: Epoch 1956
└ @ Main /Users/kcft114/.julia/packages/Flux/qXNjB/src/optimise/train.jl:105
┌ Info: Epoch 1957
└ @ Main /Users/kcft114/.julia/packages/Flux/qXNjB/src/optimise/train.jl:105
┌ Info: Epoch 1958
└ @ Main /Users/kcft114/.julia/packages/Flux/qXNjB/src/optimise/train.jl:105
┌ Info: Epoch 1959
└ @ Main /Users/kcft114/.julia/packages/Flux/qXNjB/src/optimise/train.jl:105
┌ Info: Epoch 1960
└ @ Main /Users/kcft114/.julia/packages/Flux/qXNjB/src/optimise/train.jl:105
┌ Info: Epoch 1961
└ @ Main /Users/kcft114/.julia/packages/Flux/qXNjB/src/optimise/train.jl:105
┌ Info: Epoch 1962
└ @ Main /Users/kcft114/.julia/packages/Flux/qXNjB/src/optimise/train.jl:105
┌ Info: Epoch 1963
└ @ Main /Users/kcft114/.julia/packages/Flux/qXNjB/src/optimise/train.jl:105
┌ Info: Epoch 1964
└ @ Main /Users/kcft114/.julia/packages/Flux/qXNjB/src/optimise/train.jl:105
┌ Info: Epoch 1965
└ @ Main /Users/kcft114/.julia/packages/Flux/qXNjB/src/optimise/train.jl:105
┌ Info: Epoch 1966
└ @ Main /Users/kcft114/.julia/packages/Flux/qXNjB/src/optimise/train.jl:105
┌ Info: Epoch 1967
└ @ Main /Users/kcft114/.julia/packages/Flux/qXNjB/src/optimise/train.jl:105
┌ Info: Epoch 1968
└ @ Main /Users/kcft114/.julia/packages/Flux/qXNjB/src/optimise/train.jl:105
┌ Info: Epoch 1969
└ @ Main /Users/kcft114/.julia/packages/Flux/qXNjB/src/optimise/train.jl:105
┌ Info: Epoch 1970
└ @ Main /Users/kcft114/.julia/packages/Flux/qXNjB/src/optimise/train.jl:105
┌ Info: Epoch 1971
└ @ Main /Users/kcft114/.julia/packages/Flux/qXNjB/src/optimise/train.jl:105
┌ Info: Epoch 1972
└ @ Main /Users/kcft114/.julia/packages/Flux/qXNjB/src/optimise/train.jl:105
┌ Info: Epoch 1973
└ @ Main /Users/kcft114/.julia/packages/Flux/qXNjB/src/optimise/train.jl:105
┌ Info: Epoch 1974
└ @ Main /Users/kcft114/.julia/packages/Flux/qXNjB/src/optimise/train.jl:105
┌ Info: Epoch 1975
└ @ Main /Users/kcft114/.julia/packages/Flux/qXNjB/src/optimise/train.jl:105
┌ Info: Epoch 1976
└ @ Main /Users/kcft114/.julia/packages/Flux/qXNjB/src/optimise/train.jl:105
┌ Info: Epoch 1977
└ @ Main /Users/kcft114/.julia/packages/Flux/qXNjB/src/optimise/train.jl:105
┌ Info: Epoch 1978
└ @ Main /Users/kcft114/.julia/packages/Flux/qXNjB/src/optimise/train.jl:105
┌ Info: Epoch 1979
└ @ Main /Users/kcft114/.julia/packages/Flux/qXNjB/src/optimise/train.jl:105
┌ Info: Epoch 1980
└ @ Main /Users/kcft114/.julia/packages/Flux/qXNjB/src/optimise/train.jl:105
```

```
loss(X, Y) = 0.64158624f0 (tracked)
loss(X, Y) = 0.64521545f0 (tracked)
loss(X, Y) = 0.62413615f0 (tracked)
loss(X, Y) = 0.5606213f0 (tracked)
loss(X, Y) = 0.6348677f0 (tracked)
loss(X, Y) = 0.65725636f0 (tracked)
loss(X, Y) = 0.6101086f0 (tracked)
loss(X, Y) = 0.6440645f0 (tracked)
loss(X, Y) = 0.600159f0 (tracked)
loss(X, Y) = 0.65581733f0 (tracked)
loss(X, Y) = 0.6476257f0 (tracked)
loss(X, Y) = 0.6527225f0 (tracked)
loss(X, Y) = 0.6395538f0 (tracked)
loss(X, Y) = 0.58349884f0 (tracked)
loss(X, Y) = 0.6071758f0 (tracked)
loss(X, Y) = 0.6390331f0 (tracked)
loss(X, Y) = 0.6390869f0 (tracked)
loss(X, Y) = 0.619683f0 (tracked)
loss(X, Y) = 0.69273436f0 (tracked)
loss(X, Y) = 0.6265303f0 (tracked)
train accuracy = 0.7346938775510204
test accuracy = 0.5675675675675675
train BA = 0.72
test BA = 0.7
loss(X, Y) = 1.2443976f0 (tracked)
loss(X, Y) = 1.2292174f0 (tracked)
loss(X, Y) = 1.2368766f0 (tracked)
loss(X, Y) = 1.2741996f0 (tracked)
loss(X, Y) = 1.2369813f0 (tracked)
loss(X, Y) = 1.1361145f0 (tracked)
loss(X, Y) = 1.1671224f0 (tracked)
loss(X, Y) = 1.1832021f0 (tracked)
loss(X, Y) = 1.1907928f0 (tracked)
loss(X, Y) = 1.2134501f0 (tracked)
loss(X, Y) = 1.1112274f0 (tracked)
loss(X, Y) = 1.1736993f0 (tracked)
loss(X, Y) = 1.1703459f0 (tracked)
loss(X, Y) = 1.1776077f0 (tracked)
loss(X, Y) = 1.104146f0 (tracked)
loss(X, Y) = 1.1303027f0 (tracked)
loss(X, Y) = 1.1359757f0 (tracked)
loss(X, Y) = 1.1405015f0 (tracked)
loss(X, Y) = 1.096589f0 (tracked)
loss(X, Y) = 1.1613358f0 (tracked)
loss(X, Y) = 1.169162f0 (tracked)
loss(X, Y) = 1.1090144f0 (tracked)
loss(X, Y) = 1.1392947f0 (tracked)
loss(X, Y) = 1.1044648f0 (tracked)
loss(X, Y) = 1.0519712f0 (tracked)
loss(X, Y) = 1.0691546f0 (tracked)
loss(X, Y) = 1.137708f0 (tracked)
loss(X, Y) = 1.0838076f0 (tracked)
loss(X, Y) = 1.0409431f0 (tracked)
loss(X, Y) = 1.0574038f0 (tracked)
loss(X, Y) = 1.0932288f0 (tracked)
loss(X, Y) = 1.0988727f0 (tracked)
loss(X, Y) = 1.0771126f0 (tracked)
loss(X, Y) = 1.0798501f0 (tracked)
loss(X, Y) = 1.0495391f0 (tracked)
loss(X, Y) = 1.0869085f0 (tracked)
loss(X, Y) = 1.031274f0 (tracked)
loss(X, Y) = 1.0238745f0 (tracked)
loss(X, Y) = 1.1114655f0 (tracked)
loss(X, Y) = 1.0840786f0 (tracked)
loss(X, Y) = 1.0390915f0 (tracked)
loss(X, Y) = 1.074995f0 (tracked)
loss(X, Y) = 1.0446953f0 (tracked)
loss(X, Y) = 1.0212973f0 (tracked)
loss(X, Y) = 0.99588853f0 (tracked)
loss(X, Y) = 1.0285969f0 (tracked)
loss(X, Y) = 0.9667511f0 (tracked)
loss(X, Y) = 1.0655057f0 (tracked)
loss(X, Y) = 1.0849724f0 (tracked)
loss(X, Y) = 0.9957508f0 (tracked)
loss(X, Y) = 0.9970628f0 (tracked)
loss(X, Y) = 1.0845549f0 (tracked)
loss(X, Y) = 1.0715272f0 (tracked)
loss(X, Y) = 1.0101036f0 (tracked)
loss(X, Y) = 1.035316f0 (tracked)
loss(X, Y) = 1.0106344f0 (tracked)
loss(X, Y) = 1.0173042f0 (tracked)
loss(X, Y) = 0.96101606f0 (tracked)
loss(X, Y) = 1.0118579f0 (tracked)
loss(X, Y) = 1.0133506f0 (tracked)
loss(X, Y) = 0.98635167f0 (tracked)
loss(X, Y) = 1.0016145f0 (tracked)
loss(X, Y) = 1.0290744f0 (tracked)
loss(X, Y) = 1.0156257f0 (tracked)
```

```
┌ Info: Epoch 1981
└ @ Main /Users/kcft114/.julia/packages/Flux/qXNjB/src/optimise/train.jl:105
┌ Info: Epoch 1982
└ @ Main /Users/kcft114/.julia/packages/Flux/qXNjB/src/optimise/train.jl:105
┌ Info: Epoch 1983
└ @ Main /Users/kcft114/.julia/packages/Flux/qXNjB/src/optimise/train.jl:105
┌ Info: Epoch 1984
└ @ Main /Users/kcft114/.julia/packages/Flux/qXNjB/src/optimise/train.jl:105
┌ Info: Epoch 1985
└ @ Main /Users/kcft114/.julia/packages/Flux/qXNjB/src/optimise/train.jl:105
┌ Info: Epoch 1986
└ @ Main /Users/kcft114/.julia/packages/Flux/qXNjB/src/optimise/train.jl:105
┌ Info: Epoch 1987
└ @ Main /Users/kcft114/.julia/packages/Flux/qXNjB/src/optimise/train.jl:105
┌ Info: Epoch 1988
└ @ Main /Users/kcft114/.julia/packages/Flux/qXNjB/src/optimise/train.jl:105
┌ Info: Epoch 1989
└ @ Main /Users/kcft114/.julia/packages/Flux/qXNjB/src/optimise/train.jl:105
┌ Info: Epoch 1990
└ @ Main /Users/kcft114/.julia/packages/Flux/qXNjB/src/optimise/train.jl:105
┌ Info: Epoch 1991
└ @ Main /Users/kcft114/.julia/packages/Flux/qXNjB/src/optimise/train.jl:105
┌ Info: Epoch 1992
└ @ Main /Users/kcft114/.julia/packages/Flux/qXNjB/src/optimise/train.jl:105
┌ Info: Epoch 1993
└ @ Main /Users/kcft114/.julia/packages/Flux/qXNjB/src/optimise/train.jl:105
┌ Info: Epoch 1994
└ @ Main /Users/kcft114/.julia/packages/Flux/qXNjB/src/optimise/train.jl:105
┌ Info: Epoch 1995
└ @ Main /Users/kcft114/.julia/packages/Flux/qXNjB/src/optimise/train.jl:105
┌ Info: Epoch 1996
└ @ Main /Users/kcft114/.julia/packages/Flux/qXNjB/src/optimise/train.jl:105
┌ Info: Epoch 1997
└ @ Main /Users/kcft114/.julia/packages/Flux/qXNjB/src/optimise/train.jl:105
┌ Info: Epoch 1998
└ @ Main /Users/kcft114/.julia/packages/Flux/qXNjB/src/optimise/train.jl:105
┌ Info: Epoch 1999
└ @ Main /Users/kcft114/.julia/packages/Flux/qXNjB/src/optimise/train.jl:105
┌ Info: Epoch 2000
└ @ Main /Users/kcft114/.julia/packages/Flux/qXNjB/src/optimise/train.jl:105
┌ Info: Epoch 1
└ @ Main /Users/kcft114/.julia/packages/Flux/qXNjB/src/optimise/train.jl:105
┌ Info: Epoch 2
└ @ Main /Users/kcft114/.julia/packages/Flux/qXNjB/src/optimise/train.jl:105
┌ Info: Epoch 3
└ @ Main /Users/kcft114/.julia/packages/Flux/qXNjB/src/optimise/train.jl:105
┌ Info: Epoch 4
└ @ Main /Users/kcft114/.julia/packages/Flux/qXNjB/src/optimise/train.jl:105
┌ Info: Epoch 5
└ @ Main /Users/kcft114/.julia/packages/Flux/qXNjB/src/optimise/train.jl:105
┌ Info: Epoch 6
└ @ Main /Users/kcft114/.julia/packages/Flux/qXNjB/src/optimise/train.jl:105
┌ Info: Epoch 7
└ @ Main /Users/kcft114/.julia/packages/Flux/qXNjB/src/optimise/train.jl:105
┌ Info: Epoch 8
└ @ Main /Users/kcft114/.julia/packages/Flux/qXNjB/src/optimise/train.jl:105
┌ Info: Epoch 9
└ @ Main /Users/kcft114/.julia/packages/Flux/qXNjB/src/optimise/train.jl:105
┌ Info: Epoch 10
└ @ Main /Users/kcft114/.julia/packages/Flux/qXNjB/src/optimise/train.jl:105
┌ Info: Epoch 11
└ @ Main /Users/kcft114/.julia/packages/Flux/qXNjB/src/optimise/train.jl:105
┌ Info: Epoch 12
└ @ Main /Users/kcft114/.julia/packages/Flux/qXNjB/src/optimise/train.jl:105
┌ Info: Epoch 13
└ @ Main /Users/kcft114/.julia/packages/Flux/qXNjB/src/optimise/train.jl:105
┌ Info: Epoch 14
└ @ Main /Users/kcft114/.julia/packages/Flux/qXNjB/src/optimise/train.jl:105
┌ Info: Epoch 15
└ @ Main /Users/kcft114/.julia/packages/Flux/qXNjB/src/optimise/train.jl:105
┌ Info: Epoch 16
└ @ Main /Users/kcft114/.julia/packages/Flux/qXNjB/src/optimise/train.jl:105
┌ Info: Epoch 17
└ @ Main /Users/kcft114/.julia/packages/Flux/qXNjB/src/optimise/train.jl:105
┌ Info: Epoch 18
└ @ Main /Users/kcft114/.julia/packages/Flux/qXNjB/src/optimise/train.jl:105
┌ Info: Epoch 19
└ @ Main /Users/kcft114/.julia/packages/Flux/qXNjB/src/optimise/train.jl:105
┌ Info: Epoch 20
└ @ Main /Users/kcft114/.julia/packages/Flux/qXNjB/src/optimise/train.jl:105
┌ Info: Epoch 21
└ @ Main /Users/kcft114/.julia/packages/Flux/qXNjB/src/optimise/train.jl:105
┌ Info: Epoch 22
└ @ Main /Users/kcft114/.julia/packages/Flux/qXNjB/src/optimise/train.jl:105
┌ Info: Epoch 23
└ @ Main /Users/kcft114/.julia/packages/Flux/qXNjB/src/optimise/train.jl:105
┌ Info: Epoch 24
└ @ Main /Users/kcft114/.julia/packages/Flux/qXNjB/src/optimise/train.jl:105
┌ Info: Epoch 25
└ @ Main /Users/kcft114/.julia/packages/Flux/qXNjB/src/optimise/train.jl:105
┌ Info: Epoch 26
└ @ Main /Users/kcft114/.julia/packages/Flux/qXNjB/src/optimise/train.jl:105
┌ Info: Epoch 27
└ @ Main /Users/kcft114/.julia/packages/Flux/qXNjB/src/optimise/train.jl:105
┌ Info: Epoch 28
└ @ Main /Users/kcft114/.julia/packages/Flux/qXNjB/src/optimise/train.jl:105
┌ Info: Epoch 29
└ @ Main /Users/kcft114/.julia/packages/Flux/qXNjB/src/optimise/train.jl:105
┌ Info: Epoch 30
└ @ Main /Users/kcft114/.julia/packages/Flux/qXNjB/src/optimise/train.jl:105
┌ Info: Epoch 31
└ @ Main /Users/kcft114/.julia/packages/Flux/qXNjB/src/optimise/train.jl:105
┌ Info: Epoch 32
└ @ Main /Users/kcft114/.julia/packages/Flux/qXNjB/src/optimise/train.jl:105
┌ Info: Epoch 33
└ @ Main /Users/kcft114/.julia/packages/Flux/qXNjB/src/optimise/train.jl:105
┌ Info: Epoch 34
└ @ Main /Users/kcft114/.julia/packages/Flux/qXNjB/src/optimise/train.jl:105
┌ Info: Epoch 35
└ @ Main /Users/kcft114/.julia/packages/Flux/qXNjB/src/optimise/train.jl:105
┌ Info: Epoch 36
└ @ Main /Users/kcft114/.julia/packages/Flux/qXNjB/src/optimise/train.jl:105
┌ Info: Epoch 37
└ @ Main /Users/kcft114/.julia/packages/Flux/qXNjB/src/optimise/train.jl:105
┌ Info: Epoch 38
└ @ Main /Users/kcft114/.julia/packages/Flux/qXNjB/src/optimise/train.jl:105
┌ Info: Epoch 39
└ @ Main /Users/kcft114/.julia/packages/Flux/qXNjB/src/optimise/train.jl:105
┌ Info: Epoch 40
└ @ Main /Users/kcft114/.julia/packages/Flux/qXNjB/src/optimise/train.jl:105
┌ Info: Epoch 41
└ @ Main /Users/kcft114/.julia/packages/Flux/qXNjB/src/optimise/train.jl:105
┌ Info: Epoch 42
└ @ Main /Users/kcft114/.julia/packages/Flux/qXNjB/src/optimise/train.jl:105
┌ Info: Epoch 43
└ @ Main /Users/kcft114/.julia/packages/Flux/qXNjB/src/optimise/train.jl:105
┌ Info: Epoch 44
└ @ Main /Users/kcft114/.julia/packages/Flux/qXNjB/src/optimise/train.jl:105
┌ Info: Epoch 45
└ @ Main /Users/kcft114/.julia/packages/Flux/qXNjB/src/optimise/train.jl:105
┌ Info: Epoch 46
└ @ Main /Users/kcft114/.julia/packages/Flux/qXNjB/src/optimise/train.jl:105
┌ Info: Epoch 47
└ @ Main /Users/kcft114/.julia/packages/Flux/qXNjB/src/optimise/train.jl:105
┌ Info: Epoch 48
└ @ Main /Users/kcft114/.julia/packages/Flux/qXNjB/src/optimise/train.jl:105
┌ Info: Epoch 49
└ @ Main /Users/kcft114/.julia/packages/Flux/qXNjB/src/optimise/train.jl:105
┌ Info: Epoch 50
└ @ Main /Users/kcft114/.julia/packages/Flux/qXNjB/src/optimise/train.jl:105
┌ Info: Epoch 51
└ @ Main /Users/kcft114/.julia/packages/Flux/qXNjB/src/optimise/train.jl:105
┌ Info: Epoch 52
└ @ Main /Users/kcft114/.julia/packages/Flux/qXNjB/src/optimise/train.jl:105
┌ Info: Epoch 53
└ @ Main /Users/kcft114/.julia/packages/Flux/qXNjB/src/optimise/train.jl:105
┌ Info: Epoch 54
└ @ Main /Users/kcft114/.julia/packages/Flux/qXNjB/src/optimise/train.jl:105
┌ Info: Epoch 55
└ @ Main /Users/kcft114/.julia/packages/Flux/qXNjB/src/optimise/train.jl:105
┌ Info: Epoch 56
└ @ Main /Users/kcft114/.julia/packages/Flux/qXNjB/src/optimise/train.jl:105
┌ Info: Epoch 57
└ @ Main /Users/kcft114/.julia/packages/Flux/qXNjB/src/optimise/train.jl:105
┌ Info: Epoch 58
└ @ Main /Users/kcft114/.julia/packages/Flux/qXNjB/src/optimise/train.jl:105
┌ Info: Epoch 59
└ @ Main /Users/kcft114/.julia/packages/Flux/qXNjB/src/optimise/train.jl:105
┌ Info: Epoch 60
└ @ Main /Users/kcft114/.julia/packages/Flux/qXNjB/src/optimise/train.jl:105
┌ Info: Epoch 61
└ @ Main /Users/kcft114/.julia/packages/Flux/qXNjB/src/optimise/train.jl:105
┌ Info: Epoch 62
└ @ Main /Users/kcft114/.julia/packages/Flux/qXNjB/src/optimise/train.jl:105
┌ Info: Epoch 63
└ @ Main /Users/kcft114/.julia/packages/Flux/qXNjB/src/optimise/train.jl:105
┌ Info: Epoch 64
└ @ Main /Users/kcft114/.julia/packages/Flux/qXNjB/src/optimise/train.jl:105
┌ Info: Epoch 65
└ @ Main /Users/kcft114/.julia/packages/Flux/qXNjB/src/optimise/train.jl:105
```

```
loss(X, Y) = 0.9497451f0 (tracked)
loss(X, Y) = 0.94427884f0 (tracked)
loss(X, Y) = 0.9695559f0 (tracked)
loss(X, Y) = 1.0120207f0 (tracked)
loss(X, Y) = 0.9563005f0 (tracked)
loss(X, Y) = 0.9626469f0 (tracked)
loss(X, Y) = 0.97485244f0 (tracked)
loss(X, Y) = 0.9666041f0 (tracked)
loss(X, Y) = 0.98447454f0 (tracked)
loss(X, Y) = 0.9607599f0 (tracked)
loss(X, Y) = 0.9374058f0 (tracked)
loss(X, Y) = 0.9448182f0 (tracked)
loss(X, Y) = 0.98502433f0 (tracked)
loss(X, Y) = 0.98629117f0 (tracked)
loss(X, Y) = 0.9837909f0 (tracked)
loss(X, Y) = 0.93073356f0 (tracked)
loss(X, Y) = 0.99434394f0 (tracked)
loss(X, Y) = 0.9767252f0 (tracked)
loss(X, Y) = 0.9428346f0 (tracked)
loss(X, Y) = 0.9448683f0 (tracked)
loss(X, Y) = 0.9419109f0 (tracked)
loss(X, Y) = 0.9596738f0 (tracked)
loss(X, Y) = 0.9697635f0 (tracked)
loss(X, Y) = 0.943238f0 (tracked)
loss(X, Y) = 0.94123423f0 (tracked)
loss(X, Y) = 0.94096506f0 (tracked)
loss(X, Y) = 0.97007f0 (tracked)
loss(X, Y) = 0.95462614f0 (tracked)
loss(X, Y) = 0.95066303f0 (tracked)
loss(X, Y) = 0.9190356f0 (tracked)
loss(X, Y) = 0.924714f0 (tracked)
loss(X, Y) = 0.9865251f0 (tracked)
loss(X, Y) = 0.98967403f0 (tracked)
loss(X, Y) = 0.9412464f0 (tracked)
loss(X, Y) = 0.9402161f0 (tracked)
loss(X, Y) = 0.9596682f0 (tracked)
loss(X, Y) = 0.93741363f0 (tracked)
loss(X, Y) = 0.9465758f0 (tracked)
loss(X, Y) = 1.0093291f0 (tracked)
loss(X, Y) = 0.9182328f0 (tracked)
loss(X, Y) = 0.88304996f0 (tracked)
loss(X, Y) = 0.92116374f0 (tracked)
loss(X, Y) = 0.9404692f0 (tracked)
loss(X, Y) = 0.9258454f0 (tracked)
loss(X, Y) = 0.95137376f0 (tracked)
loss(X, Y) = 0.85321933f0 (tracked)
loss(X, Y) = 0.90341216f0 (tracked)
loss(X, Y) = 0.8684617f0 (tracked)
loss(X, Y) = 0.95561624f0 (tracked)
loss(X, Y) = 0.89610183f0 (tracked)
loss(X, Y) = 0.92908263f0 (tracked)
loss(X, Y) = 0.9499358f0 (tracked)
loss(X, Y) = 0.8944508f0 (tracked)
loss(X, Y) = 0.87999845f0 (tracked)
loss(X, Y) = 0.9469507f0 (tracked)
loss(X, Y) = 0.87968796f0 (tracked)
loss(X, Y) = 0.97039276f0 (tracked)
loss(X, Y) = 0.9212363f0 (tracked)
loss(X, Y) = 0.8840013f0 (tracked)
loss(X, Y) = 0.8526903f0 (tracked)
loss(X, Y) = 0.934335f0 (tracked)
loss(X, Y) = 0.9055735f0 (tracked)
loss(X, Y) = 0.8997711f0 (tracked)
loss(X, Y) = 0.9011472f0 (tracked)
loss(X, Y) = 0.85034925f0 (tracked)
loss(X, Y) = 0.91609436f0 (tracked)
loss(X, Y) = 0.87688494f0 (tracked)
loss(X, Y) = 0.9214629f0 (tracked)
loss(X, Y) = 0.9418016f0 (tracked)
loss(X, Y) = 0.9516644f0 (tracked)
loss(X, Y) = 0.8849272f0 (tracked)
loss(X, Y) = 0.8840609f0 (tracked)
loss(X, Y) = 0.8730249f0 (tracked)
loss(X, Y) = 0.8944421f0 (tracked)
loss(X, Y) = 0.8999124f0 (tracked)
loss(X, Y) = 0.8951156f0 (tracked)
loss(X, Y) = 0.8930753f0 (tracked)
loss(X, Y) = 0.9338351f0 (tracked)
```

```
┌ Info: Epoch 66
└ @ Main /Users/kcft114/.julia/packages/Flux/qXNjB/src/optimise/train.jl:105
┌ Info: Epoch 67
└ @ Main /Users/kcft114/.julia/packages/Flux/qXNjB/src/optimise/train.jl:105
┌ Info: Epoch 68
└ @ Main /Users/kcft114/.julia/packages/Flux/qXNjB/src/optimise/train.jl:105
┌ Info: Epoch 69
└ @ Main /Users/kcft114/.julia/packages/Flux/qXNjB/src/optimise/train.jl:105
┌ Info: Epoch 70
└ @ Main /Users/kcft114/.julia/packages/Flux/qXNjB/src/optimise/train.jl:105
┌ Info: Epoch 71
└ @ Main /Users/kcft114/.julia/packages/Flux/qXNjB/src/optimise/train.jl:105
┌ Info: Epoch 72
└ @ Main /Users/kcft114/.julia/packages/Flux/qXNjB/src/optimise/train.jl:105
┌ Info: Epoch 73
└ @ Main /Users/kcft114/.julia/packages/Flux/qXNjB/src/optimise/train.jl:105
┌ Info: Epoch 74
└ @ Main /Users/kcft114/.julia/packages/Flux/qXNjB/src/optimise/train.jl:105
┌ Info: Epoch 75
└ @ Main /Users/kcft114/.julia/packages/Flux/qXNjB/src/optimise/train.jl:105
┌ Info: Epoch 76
└ @ Main /Users/kcft114/.julia/packages/Flux/qXNjB/src/optimise/train.jl:105
┌ Info: Epoch 77
└ @ Main /Users/kcft114/.julia/packages/Flux/qXNjB/src/optimise/train.jl:105
┌ Info: Epoch 78
└ @ Main /Users/kcft114/.julia/packages/Flux/qXNjB/src/optimise/train.jl:105
┌ Info: Epoch 79
└ @ Main /Users/kcft114/.julia/packages/Flux/qXNjB/src/optimise/train.jl:105
┌ Info: Epoch 80
└ @ Main /Users/kcft114/.julia/packages/Flux/qXNjB/src/optimise/train.jl:105
┌ Info: Epoch 81
└ @ Main /Users/kcft114/.julia/packages/Flux/qXNjB/src/optimise/train.jl:105
┌ Info: Epoch 82
└ @ Main /Users/kcft114/.julia/packages/Flux/qXNjB/src/optimise/train.jl:105
┌ Info: Epoch 83
└ @ Main /Users/kcft114/.julia/packages/Flux/qXNjB/src/optimise/train.jl:105
┌ Info: Epoch 84
└ @ Main /Users/kcft114/.julia/packages/Flux/qXNjB/src/optimise/train.jl:105
┌ Info: Epoch 85
└ @ Main /Users/kcft114/.julia/packages/Flux/qXNjB/src/optimise/train.jl:105
┌ Info: Epoch 86
└ @ Main /Users/kcft114/.julia/packages/Flux/qXNjB/src/optimise/train.jl:105
┌ Info: Epoch 87
└ @ Main /Users/kcft114/.julia/packages/Flux/qXNjB/src/optimise/train.jl:105
┌ Info: Epoch 88
└ @ Main /Users/kcft114/.julia/packages/Flux/qXNjB/src/optimise/train.jl:105
┌ Info: Epoch 89
└ @ Main /Users/kcft114/.julia/packages/Flux/qXNjB/src/optimise/train.jl:105
┌ Info: Epoch 90
└ @ Main /Users/kcft114/.julia/packages/Flux/qXNjB/src/optimise/train.jl:105
┌ Info: Epoch 91
└ @ Main /Users/kcft114/.julia/packages/Flux/qXNjB/src/optimise/train.jl:105
┌ Info: Epoch 92
└ @ Main /Users/kcft114/.julia/packages/Flux/qXNjB/src/optimise/train.jl:105
┌ Info: Epoch 93
└ @ Main /Users/kcft114/.julia/packages/Flux/qXNjB/src/optimise/train.jl:105
┌ Info: Epoch 94
└ @ Main /Users/kcft114/.julia/packages/Flux/qXNjB/src/optimise/train.jl:105
┌ Info: Epoch 95
└ @ Main /Users/kcft114/.julia/packages/Flux/qXNjB/src/optimise/train.jl:105
┌ Info: Epoch 96
└ @ Main /Users/kcft114/.julia/packages/Flux/qXNjB/src/optimise/train.jl:105
┌ Info: Epoch 97
└ @ Main /Users/kcft114/.julia/packages/Flux/qXNjB/src/optimise/train.jl:105
┌ Info: Epoch 98
└ @ Main /Users/kcft114/.julia/packages/Flux/qXNjB/src/optimise/train.jl:105
┌ Info: Epoch 99
└ @ Main /Users/kcft114/.julia/packages/Flux/qXNjB/src/optimise/train.jl:105
┌ Info: Epoch 100
└ @ Main /Users/kcft114/.julia/packages/Flux/qXNjB/src/optimise/train.jl:105
┌ Info: Epoch 101
└ @ Main /Users/kcft114/.julia/packages/Flux/qXNjB/src/optimise/train.jl:105
┌ Info: Epoch 102
└ @ Main /Users/kcft114/.julia/packages/Flux/qXNjB/src/optimise/train.jl:105
┌ Info: Epoch 103
└ @ Main /Users/kcft114/.julia/packages/Flux/qXNjB/src/optimise/train.jl:105
┌ Info: Epoch 104
└ @ Main /Users/kcft114/.julia/packages/Flux/qXNjB/src/optimise/train.jl:105
┌ Info: Epoch 105
└ @ Main /Users/kcft114/.julia/packages/Flux/qXNjB/src/optimise/train.jl:105
┌ Info: Epoch 106
└ @ Main /Users/kcft114/.julia/packages/Flux/qXNjB/src/optimise/train.jl:105
┌ Info: Epoch 107
└ @ Main /Users/kcft114/.julia/packages/Flux/qXNjB/src/optimise/train.jl:105
┌ Info: Epoch 108
└ @ Main /Users/kcft114/.julia/packages/Flux/qXNjB/src/optimise/train.jl:105
┌ Info: Epoch 109
└ @ Main /Users/kcft114/.julia/packages/Flux/qXNjB/src/optimise/train.jl:105
┌ Info: Epoch 110
└ @ Main /Users/kcft114/.julia/packages/Flux/qXNjB/src/optimise/train.jl:105
┌ Info: Epoch 111
└ @ Main /Users/kcft114/.julia/packages/Flux/qXNjB/src/optimise/train.jl:105
┌ Info: Epoch 112
└ @ Main /Users/kcft114/.julia/packages/Flux/qXNjB/src/optimise/train.jl:105
┌ Info: Epoch 113
└ @ Main /Users/kcft114/.julia/packages/Flux/qXNjB/src/optimise/train.jl:105
┌ Info: Epoch 114
└ @ Main /Users/kcft114/.julia/packages/Flux/qXNjB/src/optimise/train.jl:105
┌ Info: Epoch 115
└ @ Main /Users/kcft114/.julia/packages/Flux/qXNjB/src/optimise/train.jl:105
┌ Info: Epoch 116
└ @ Main /Users/kcft114/.julia/packages/Flux/qXNjB/src/optimise/train.jl:105
┌ Info: Epoch 117
└ @ Main /Users/kcft114/.julia/packages/Flux/qXNjB/src/optimise/train.jl:105
┌ Info: Epoch 118
└ @ Main /Users/kcft114/.julia/packages/Flux/qXNjB/src/optimise/train.jl:105
┌ Info: Epoch 119
└ @ Main /Users/kcft114/.julia/packages/Flux/qXNjB/src/optimise/train.jl:105
┌ Info: Epoch 120
└ @ Main /Users/kcft114/.julia/packages/Flux/qXNjB/src/optimise/train.jl:105
┌ Info: Epoch 121
└ @ Main /Users/kcft114/.julia/packages/Flux/qXNjB/src/optimise/train.jl:105
┌ Info: Epoch 122
└ @ Main /Users/kcft114/.julia/packages/Flux/qXNjB/src/optimise/train.jl:105
┌ Info: Epoch 123
└ @ Main /Users/kcft114/.julia/packages/Flux/qXNjB/src/optimise/train.jl:105
┌ Info: Epoch 124
└ @ Main /Users/kcft114/.julia/packages/Flux/qXNjB/src/optimise/train.jl:105
┌ Info: Epoch 125
└ @ Main /Users/kcft114/.julia/packages/Flux/qXNjB/src/optimise/train.jl:105
┌ Info: Epoch 126
└ @ Main /Users/kcft114/.julia/packages/Flux/qXNjB/src/optimise/train.jl:105
┌ Info: Epoch 127
└ @ Main /Users/kcft114/.julia/packages/Flux/qXNjB/src/optimise/train.jl:105
┌ Info: Epoch 128
└ @ Main /Users/kcft114/.julia/packages/Flux/qXNjB/src/optimise/train.jl:105
┌ Info: Epoch 129
└ @ Main /Users/kcft114/.julia/packages/Flux/qXNjB/src/optimise/train.jl:105
┌ Info: Epoch 130
└ @ Main /Users/kcft114/.julia/packages/Flux/qXNjB/src/optimise/train.jl:105
┌ Info: Epoch 131
└ @ Main /Users/kcft114/.julia/packages/Flux/qXNjB/src/optimise/train.jl:105
┌ Info: Epoch 132
└ @ Main /Users/kcft114/.julia/packages/Flux/qXNjB/src/optimise/train.jl:105
┌ Info: Epoch 133
└ @ Main /Users/kcft114/.julia/packages/Flux/qXNjB/src/optimise/train.jl:105
┌ Info: Epoch 134
└ @ Main /Users/kcft114/.julia/packages/Flux/qXNjB/src/optimise/train.jl:105
┌ Info: Epoch 135
└ @ Main /Users/kcft114/.julia/packages/Flux/qXNjB/src/optimise/train.jl:105
┌ Info: Epoch 136
└ @ Main /Users/kcft114/.julia/packages/Flux/qXNjB/src/optimise/train.jl:105
┌ Info: Epoch 137
└ @ Main /Users/kcft114/.julia/packages/Flux/qXNjB/src/optimise/train.jl:105
┌ Info: Epoch 138
└ @ Main /Users/kcft114/.julia/packages/Flux/qXNjB/src/optimise/train.jl:105
┌ Info: Epoch 139
└ @ Main /Users/kcft114/.julia/packages/Flux/qXNjB/src/optimise/train.jl:105
┌ Info: Epoch 140
└ @ Main /Users/kcft114/.julia/packages/Flux/qXNjB/src/optimise/train.jl:105
┌ Info: Epoch 141
└ @ Main /Users/kcft114/.julia/packages/Flux/qXNjB/src/optimise/train.jl:105
┌ Info: Epoch 142
└ @ Main /Users/kcft114/.julia/packages/Flux/qXNjB/src/optimise/train.jl:105
┌ Info: Epoch 143
└ @ Main /Users/kcft114/.julia/packages/Flux/qXNjB/src/optimise/train.jl:105
```

```
loss(X, Y) = 0.8870154f0 (tracked)
loss(X, Y) = 0.9150373f0 (tracked)
loss(X, Y) = 0.8593738f0 (tracked)
loss(X, Y) = 0.9049018f0 (tracked)
loss(X, Y) = 0.93948126f0 (tracked)
loss(X, Y) = 0.8999963f0 (tracked)
loss(X, Y) = 0.8939562f0 (tracked)
loss(X, Y) = 0.86704016f0 (tracked)
loss(X, Y) = 0.8975175f0 (tracked)
loss(X, Y) = 0.9465345f0 (tracked)
loss(X, Y) = 0.9066081f0 (tracked)
loss(X, Y) = 0.8966528f0 (tracked)
loss(X, Y) = 0.85192364f0 (tracked)
loss(X, Y) = 0.8464648f0 (tracked)
loss(X, Y) = 0.86607164f0 (tracked)
loss(X, Y) = 0.85108906f0 (tracked)
loss(X, Y) = 0.87136817f0 (tracked)
loss(X, Y) = 0.89522517f0 (tracked)
loss(X, Y) = 0.91759735f0 (tracked)
loss(X, Y) = 0.8666294f0 (tracked)
loss(X, Y) = 0.9016882f0 (tracked)
loss(X, Y) = 0.86922914f0 (tracked)
loss(X, Y) = 0.8935455f0 (tracked)
loss(X, Y) = 0.8695436f0 (tracked)
loss(X, Y) = 0.83998156f0 (tracked)
loss(X, Y) = 0.86254317f0 (tracked)
loss(X, Y) = 0.8497433f0 (tracked)
loss(X, Y) = 0.88254094f0 (tracked)
loss(X, Y) = 0.8827177f0 (tracked)
loss(X, Y) = 0.87649214f0 (tracked)
loss(X, Y) = 0.9278769f0 (tracked)
loss(X, Y) = 0.9077661f0 (tracked)
loss(X, Y) = 0.81498003f0 (tracked)
loss(X, Y) = 0.8516158f0 (tracked)
loss(X, Y) = 0.8409706f0 (tracked)
loss(X, Y) = 0.844396f0 (tracked)
loss(X, Y) = 0.8032423f0 (tracked)
loss(X, Y) = 0.87843454f0 (tracked)
loss(X, Y) = 0.8589989f0 (tracked)
loss(X, Y) = 0.85062337f0 (tracked)
loss(X, Y) = 0.8484891f0 (tracked)
loss(X, Y) = 0.87685823f0 (tracked)
loss(X, Y) = 0.8370744f0 (tracked)
loss(X, Y) = 0.8468203f0 (tracked)
loss(X, Y) = 0.8836431f0 (tracked)
loss(X, Y) = 0.8948309f0 (tracked)
loss(X, Y) = 0.8584114f0 (tracked)
loss(X, Y) = 0.8902958f0 (tracked)
loss(X, Y) = 0.84273714f0 (tracked)
loss(X, Y) = 0.8228445f0 (tracked)
loss(X, Y) = 0.8303025f0 (tracked)
loss(X, Y) = 0.857155f0 (tracked)
loss(X, Y) = 0.85169065f0 (tracked)
loss(X, Y) = 0.85581887f0 (tracked)
loss(X, Y) = 0.8312594f0 (tracked)
loss(X, Y) = 0.86449575f0 (tracked)
loss(X, Y) = 0.8311636f0 (tracked)
loss(X, Y) = 0.81745887f0 (tracked)
loss(X, Y) = 0.8351815f0 (tracked)
loss(X, Y) = 0.87061274f0 (tracked)
loss(X, Y) = 0.8220462f0 (tracked)
loss(X, Y) = 0.83009624f0 (tracked)
loss(X, Y) = 0.82091194f0 (tracked)
loss(X, Y) = 0.84782135f0 (tracked)
loss(X, Y) = 0.82636064f0 (tracked)
loss(X, Y) = 0.8596973f0 (tracked)
loss(X, Y) = 0.8620874f0 (tracked)
loss(X, Y) = 0.8504708f0 (tracked)
loss(X, Y) = 0.83866537f0 (tracked)
loss(X, Y) = 0.839313f0 (tracked)
loss(X, Y) = 0.81971407f0 (tracked)
loss(X, Y) = 0.8187866f0 (tracked)
loss(X, Y) = 0.82899565f0 (tracked)
loss(X, Y) = 0.84740746f0 (tracked)
loss(X, Y) = 0.87150705f0 (tracked)
loss(X, Y) = 0.81679744f0 (tracked)
loss(X, Y) = 0.84586895f0 (tracked)
loss(X, Y) = 0.78860533f0 (tracked)
loss(X, Y) = 0.8408203f0 (tracked)
loss(X, Y) = 0.79887503f0 (tracked)
loss(X, Y) = 0.88467187f0 (tracked)
loss(X, Y) = 0.84681773f0 (tracked)
loss(X, Y) = 0.7931957f0 (tracked)
loss(X, Y) = 0.8124792f0 (tracked)
loss(X, Y) = 0.7989896f0 (tracked)
loss(X, Y) = 0.8431828f0 (tracked)
loss(X, Y) = 0.8746423f0 (tracked)
loss(X, Y) = 0.7993517f0 (tracked)
loss(X, Y) = 0.81941205f0 (tracked)
loss(X, Y) = 0.8552527f0 (tracked)
loss(X, Y) = 0.80508006f0 (tracked)
loss(X, Y) = 0.8599043f0 (tracked)
loss(X, Y) = 0.81718653f0 (tracked)
loss(X, Y) = 0.798484f0 (tracked)
loss(X, Y) = 0.8315489f0 (tracked)
loss(X, Y) = 0.8286657f0 (tracked)
loss(X, Y) = 0.82157874f0 (tracked)
loss(X, Y) = 0.8078269f0 (tracked)
loss(X, Y) = 0.86382806f0 (tracked)
loss(X, Y) = 0.78116703f0 (tracked)
```

```
┌ Info: Epoch 144
└ @ Main /Users/kcft114/.julia/packages/Flux/qXNjB/src/optimise/train.jl:105
┌ Info: Epoch 145
└ @ Main /Users/kcft114/.julia/packages/Flux/qXNjB/src/optimise/train.jl:105
┌ Info: Epoch 146
└ @ Main /Users/kcft114/.julia/packages/Flux/qXNjB/src/optimise/train.jl:105
┌ Info: Epoch 147
└ @ Main /Users/kcft114/.julia/packages/Flux/qXNjB/src/optimise/train.jl:105
┌ Info: Epoch 148
└ @ Main /Users/kcft114/.julia/packages/Flux/qXNjB/src/optimise/train.jl:105
┌ Info: Epoch 149
└ @ Main /Users/kcft114/.julia/packages/Flux/qXNjB/src/optimise/train.jl:105
┌ Info: Epoch 150
└ @ Main /Users/kcft114/.julia/packages/Flux/qXNjB/src/optimise/train.jl:105
┌ Info: Epoch 151
└ @ Main /Users/kcft114/.julia/packages/Flux/qXNjB/src/optimise/train.jl:105
┌ Info: Epoch 152
└ @ Main /Users/kcft114/.julia/packages/Flux/qXNjB/src/optimise/train.jl:105
┌ Info: Epoch 153
└ @ Main /Users/kcft114/.julia/packages/Flux/qXNjB/src/optimise/train.jl:105
┌ Info: Epoch 154
└ @ Main /Users/kcft114/.julia/packages/Flux/qXNjB/src/optimise/train.jl:105
┌ Info: Epoch 155
└ @ Main /Users/kcft114/.julia/packages/Flux/qXNjB/src/optimise/train.jl:105
┌ Info: Epoch 156
└ @ Main /Users/kcft114/.julia/packages/Flux/qXNjB/src/optimise/train.jl:105
┌ Info: Epoch 157
└ @ Main /Users/kcft114/.julia/packages/Flux/qXNjB/src/optimise/train.jl:105
┌ Info: Epoch 158
└ @ Main /Users/kcft114/.julia/packages/Flux/qXNjB/src/optimise/train.jl:105
┌ Info: Epoch 159
└ @ Main /Users/kcft114/.julia/packages/Flux/qXNjB/src/optimise/train.jl:105
┌ Info: Epoch 160
└ @ Main /Users/kcft114/.julia/packages/Flux/qXNjB/src/optimise/train.jl:105
┌ Info: Epoch 161
└ @ Main /Users/kcft114/.julia/packages/Flux/qXNjB/src/optimise/train.jl:105
┌ Info: Epoch 162
└ @ Main /Users/kcft114/.julia/packages/Flux/qXNjB/src/optimise/train.jl:105
┌ Info: Epoch 163
└ @ Main /Users/kcft114/.julia/packages/Flux/qXNjB/src/optimise/train.jl:105
┌ Info: Epoch 164
└ @ Main /Users/kcft114/.julia/packages/Flux/qXNjB/src/optimise/train.jl:105
┌ Info: Epoch 165
└ @ Main /Users/kcft114/.julia/packages/Flux/qXNjB/src/optimise/train.jl:105
┌ Info: Epoch 166
└ @ Main /Users/kcft114/.julia/packages/Flux/qXNjB/src/optimise/train.jl:105
┌ Info: Epoch 167
└ @ Main /Users/kcft114/.julia/packages/Flux/qXNjB/src/optimise/train.jl:105
┌ Info: Epoch 168
└ @ Main /Users/kcft114/.julia/packages/Flux/qXNjB/src/optimise/train.jl:105
┌ Info: Epoch 169
└ @ Main /Users/kcft114/.julia/packages/Flux/qXNjB/src/optimise/train.jl:105
┌ Info: Epoch 170
└ @ Main /Users/kcft114/.julia/packages/Flux/qXNjB/src/optimise/train.jl:105
┌ Info: Epoch 171
└ @ Main /Users/kcft114/.julia/packages/Flux/qXNjB/src/optimise/train.jl:105
┌ Info: Epoch 172
└ @ Main /Users/kcft114/.julia/packages/Flux/qXNjB/src/optimise/train.jl:105
┌ Info: Epoch 173
└ @ Main /Users/kcft114/.julia/packages/Flux/qXNjB/src/optimise/train.jl:105
┌ Info: Epoch 174
└ @ Main /Users/kcft114/.julia/packages/Flux/qXNjB/src/optimise/train.jl:105
┌ Info: Epoch 175
└ @ Main /Users/kcft114/.julia/packages/Flux/qXNjB/src/optimise/train.jl:105
┌ Info: Epoch 176
└ @ Main /Users/kcft114/.julia/packages/Flux/qXNjB/src/optimise/train.jl:105
┌ Info: Epoch 177
└ @ Main /Users/kcft114/.julia/packages/Flux/qXNjB/src/optimise/train.jl:105
┌ Info: Epoch 178
└ @ Main /Users/kcft114/.julia/packages/Flux/qXNjB/src/optimise/train.jl:105
┌ Info: Epoch 179
└ @ Main /Users/kcft114/.julia/packages/Flux/qXNjB/src/optimise/train.jl:105
┌ Info: Epoch 180
└ @ Main /Users/kcft114/.julia/packages/Flux/qXNjB/src/optimise/train.jl:105
┌ Info: Epoch 181
└ @ Main /Users/kcft114/.julia/packages/Flux/qXNjB/src/optimise/train.jl:105
┌ Info: Epoch 182
└ @ Main /Users/kcft114/.julia/packages/Flux/qXNjB/src/optimise/train.jl:105
┌ Info: Epoch 183
└ @ Main /Users/kcft114/.julia/packages/Flux/qXNjB/src/optimise/train.jl:105
┌ Info: Epoch 184
└ @ Main /Users/kcft114/.julia/packages/Flux/qXNjB/src/optimise/train.jl:105
┌ Info: Epoch 185
└ @ Main /Users/kcft114/.julia/packages/Flux/qXNjB/src/optimise/train.jl:105
┌ Info: Epoch 186
└ @ Main /Users/kcft114/.julia/packages/Flux/qXNjB/src/optimise/train.jl:105
┌ Info: Epoch 187
└ @ Main /Users/kcft114/.julia/packages/Flux/qXNjB/src/optimise/train.jl:105
┌ Info: Epoch 188
└ @ Main /Users/kcft114/.julia/packages/Flux/qXNjB/src/optimise/train.jl:105
┌ Info: Epoch 189
└ @ Main /Users/kcft114/.julia/packages/Flux/qXNjB/src/optimise/train.jl:105
┌ Info: Epoch 190
└ @ Main /Users/kcft114/.julia/packages/Flux/qXNjB/src/optimise/train.jl:105
┌ Info: Epoch 191
└ @ Main /Users/kcft114/.julia/packages/Flux/qXNjB/src/optimise/train.jl:105
┌ Info: Epoch 192
└ @ Main /Users/kcft114/.julia/packages/Flux/qXNjB/src/optimise/train.jl:105
┌ Info: Epoch 193
└ @ Main /Users/kcft114/.julia/packages/Flux/qXNjB/src/optimise/train.jl:105
┌ Info: Epoch 194
└ @ Main /Users/kcft114/.julia/packages/Flux/qXNjB/src/optimise/train.jl:105
┌ Info: Epoch 195
└ @ Main /Users/kcft114/.julia/packages/Flux/qXNjB/src/optimise/train.jl:105
┌ Info: Epoch 196
└ @ Main /Users/kcft114/.julia/packages/Flux/qXNjB/src/optimise/train.jl:105
┌ Info: Epoch 197
└ @ Main /Users/kcft114/.julia/packages/Flux/qXNjB/src/optimise/train.jl:105
┌ Info: Epoch 198
└ @ Main /Users/kcft114/.julia/packages/Flux/qXNjB/src/optimise/train.jl:105
┌ Info: Epoch 199
└ @ Main /Users/kcft114/.julia/packages/Flux/qXNjB/src/optimise/train.jl:105
┌ Info: Epoch 200
└ @ Main /Users/kcft114/.julia/packages/Flux/qXNjB/src/optimise/train.jl:105
┌ Info: Epoch 201
└ @ Main /Users/kcft114/.julia/packages/Flux/qXNjB/src/optimise/train.jl:105
┌ Info: Epoch 202
└ @ Main /Users/kcft114/.julia/packages/Flux/qXNjB/src/optimise/train.jl:105
┌ Info: Epoch 203
└ @ Main /Users/kcft114/.julia/packages/Flux/qXNjB/src/optimise/train.jl:105
┌ Info: Epoch 204
└ @ Main /Users/kcft114/.julia/packages/Flux/qXNjB/src/optimise/train.jl:105
┌ Info: Epoch 205
└ @ Main /Users/kcft114/.julia/packages/Flux/qXNjB/src/optimise/train.jl:105
┌ Info: Epoch 206
└ @ Main /Users/kcft114/.julia/packages/Flux/qXNjB/src/optimise/train.jl:105
┌ Info: Epoch 207
└ @ Main /Users/kcft114/.julia/packages/Flux/qXNjB/src/optimise/train.jl:105
┌ Info: Epoch 208
└ @ Main /Users/kcft114/.julia/packages/Flux/qXNjB/src/optimise/train.jl:105
┌ Info: Epoch 209
└ @ Main /Users/kcft114/.julia/packages/Flux/qXNjB/src/optimise/train.jl:105
┌ Info: Epoch 210
└ @ Main /Users/kcft114/.julia/packages/Flux/qXNjB/src/optimise/train.jl:105
┌ Info: Epoch 211
└ @ Main /Users/kcft114/.julia/packages/Flux/qXNjB/src/optimise/train.jl:105
┌ Info: Epoch 212
└ @ Main /Users/kcft114/.julia/packages/Flux/qXNjB/src/optimise/train.jl:105
┌ Info: Epoch 213
└ @ Main /Users/kcft114/.julia/packages/Flux/qXNjB/src/optimise/train.jl:105
┌ Info: Epoch 214
└ @ Main /Users/kcft114/.julia/packages/Flux/qXNjB/src/optimise/train.jl:105
┌ Info: Epoch 215
└ @ Main /Users/kcft114/.julia/packages/Flux/qXNjB/src/optimise/train.jl:105
┌ Info: Epoch 216
└ @ Main /Users/kcft114/.julia/packages/Flux/qXNjB/src/optimise/train.jl:105
┌ Info: Epoch 217
└ @ Main /Users/kcft114/.julia/packages/Flux/qXNjB/src/optimise/train.jl:105
┌ Info: Epoch 218
└ @ Main /Users/kcft114/.julia/packages/Flux/qXNjB/src/optimise/train.jl:105
┌ Info: Epoch 219
└ @ Main /Users/kcft114/.julia/packages/Flux/qXNjB/src/optimise/train.jl:105
┌ Info: Epoch 220
└ @ Main /Users/kcft114/.julia/packages/Flux/qXNjB/src/optimise/train.jl:105
┌ Info: Epoch 221
└ @ Main /Users/kcft114/.julia/packages/Flux/qXNjB/src/optimise/train.jl:105
┌ Info: Epoch 222
└ @ Main /Users/kcft114/.julia/packages/Flux/qXNjB/src/optimise/train.jl:105
┌ Info: Epoch 223
└ @ Main /Users/kcft114/.julia/packages/Flux/qXNjB/src/optimise/train.jl:105
┌ Info: Epoch 224
└ @ Main /Users/kcft114/.julia/packages/Flux/qXNjB/src/optimise/train.jl:105
┌ Info: Epoch 225
└ @ Main /Users/kcft114/.julia/packages/Flux/qXNjB/src/optimise/train.jl:105
┌ Info: Epoch 226
└ @ Main /Users/kcft114/.julia/packages/Flux/qXNjB/src/optimise/train.jl:105
┌ Info: Epoch 227
└ @ Main /Users/kcft114/.julia/packages/Flux/qXNjB/src/optimise/train.jl:105
┌ Info: Epoch 228
└ @ Main /Users/kcft114/.julia/packages/Flux/qXNjB/src/optimise/train.jl:105
┌ Info: Epoch 229
└ @ Main /Users/kcft114/.julia/packages/Flux/qXNjB/src/optimise/train.jl:105
┌ Info: Epoch 230
└ @ Main /Users/kcft114/.julia/packages/Flux/qXNjB/src/optimise/train.jl:105
┌ Info: Epoch 231
└ @ Main /Users/kcft114/.julia/packages/Flux/qXNjB/src/optimise/train.jl:105
┌ Info: Epoch 232
└ @ Main /Users/kcft114/.julia/packages/Flux/qXNjB/src/optimise/train.jl:105
┌ Info: Epoch 233
└ @ Main /Users/kcft114/.julia/packages/Flux/qXNjB/src/optimise/train.jl:105
┌ Info: Epoch 234
└ @ Main /Users/kcft114/.julia/packages/Flux/qXNjB/src/optimise/train.jl:105
┌ Info: Epoch 235
└ @ Main /Users/kcft114/.julia/packages/Flux/qXNjB/src/optimise/train.jl:105
┌ Info: Epoch 236
└ @ Main /Users/kcft114/.julia/packages/Flux/qXNjB/src/optimise/train.jl:105
┌ Info: Epoch 237
└ @ Main /Users/kcft114/.julia/packages/Flux/qXNjB/src/optimise/train.jl:105
┌ Info: Epoch 238
└ @ Main /Users/kcft114/.julia/packages/Flux/qXNjB/src/optimise/train.jl:105
┌ Info: Epoch 239
└ @ Main /Users/kcft114/.julia/packages/Flux/qXNjB/src/optimise/train.jl:105
┌ Info: Epoch 240
└ @ Main /Users/kcft114/.julia/packages/Flux/qXNjB/src/optimise/train.jl:105
┌ Info: Epoch 241
└ @ Main /Users/kcft114/.julia/packages/Flux/qXNjB/src/optimise/train.jl:105
┌ Info: Epoch 242
└ @ Main /Users/kcft114/.julia/packages/Flux/qXNjB/src/optimise/train.jl:105
┌ Info: Epoch 243
└ @ Main /Users/kcft114/.julia/packages/Flux/qXNjB/src/optimise/train.jl:105
┌ Info: Epoch 244
└ @ Main /Users/kcft114/.julia/packages/Flux/qXNjB/src/optimise/train.jl:105
┌ Info: Epoch 245
└ @ Main /Users/kcft114/.julia/packages/Flux/qXNjB/src/optimise/train.jl:105
┌ Info: Epoch 246
└ @ Main /Users/kcft114/.julia/packages/Flux/qXNjB/src/optimise/train.jl:105
┌ Info: Epoch 247
└ @ Main /Users/kcft114/.julia/packages/Flux/qXNjB/src/optimise/train.jl:105
┌ Info: Epoch 248
└ @ Main /Users/kcft114/.julia/packages/Flux/qXNjB/src/optimise/train.jl:105
┌ Info: Epoch 249
└ @ Main /Users/kcft114/.julia/packages/Flux/qXNjB/src/optimise/train.jl:105
┌ Info: Epoch 250
└ @ Main /Users/kcft114/.julia/packages/Flux/qXNjB/src/optimise/train.jl:105
┌ Info: Epoch 251
└ @ Main /Users/kcft114/.julia/packages/Flux/qXNjB/src/optimise/train.jl:105
┌ Info: Epoch 252
└ @ Main /Users/kcft114/.julia/packages/Flux/qXNjB/src/optimise/train.jl:105
┌ Info: Epoch 253
└ @ Main /Users/kcft114/.julia/packages/Flux/qXNjB/src/optimise/train.jl:105
┌ Info: Epoch 254
└ @ Main /Users/kcft114/.julia/packages/Flux/qXNjB/src/optimise/train.jl:105
┌ Info: Epoch 255
└ @ Main /Users/kcft114/.julia/packages/Flux/qXNjB/src/optimise/train.jl:105
┌ Info: Epoch 256
└ @ Main /Users/kcft114/.julia/packages/Flux/qXNjB/src/optimise/train.jl:105
┌ Info: Epoch 257
└ @ Main /Users/kcft114/.julia/packages/Flux/qXNjB/src/optimise/train.jl:105
┌ Info: Epoch 258
└ @ Main /Users/kcft114/.julia/packages/Flux/qXNjB/src/optimise/train.jl:105
┌ Info: Epoch 259
└ @ Main /Users/kcft114/.julia/packages/Flux/qXNjB/src/optimise/train.jl:105
┌ Info: Epoch 260
└ @ Main /Users/kcft114/.julia/packages/Flux/qXNjB/src/optimise/train.jl:105
┌ Info: Epoch 261
└ @ Main /Users/kcft114/.julia/packages/Flux/qXNjB/src/optimise/train.jl:105
┌ Info: Epoch 262
└ @ Main /Users/kcft114/.julia/packages/Flux/qXNjB/src/optimise/train.jl:105
┌ Info: Epoch 263
└ @ Main /Users/kcft114/.julia/packages/Flux/qXNjB/src/optimise/train.jl:105
┌ Info: Epoch 264
└ @ Main /Users/kcft114/.julia/packages/Flux/qXNjB/src/optimise/train.jl:105
┌ Info: Epoch 265
└ @ Main /Users/kcft114/.julia/packages/Flux/qXNjB/src/optimise/train.jl:105
┌ Info: Epoch 266
└ @ Main /Users/kcft114/.julia/packages/Flux/qXNjB/src/optimise/train.jl:105
┌ Info: Epoch 267
└ @ Main /Users/kcft114/.julia/packages/Flux/qXNjB/src/optimise/train.jl:105
┌ Info: Epoch 268
└ @ Main /Users/kcft114/.julia/packages/Flux/qXNjB/src/optimise/train.jl:105
┌ Info: Epoch 269
└ @ Main /Users/kcft114/.julia/packages/Flux/qXNjB/src/optimise/train.jl:105
┌ Info: Epoch 270
└ @ Main /Users/kcft114/.julia/packages/Flux/qXNjB/src/optimise/train.jl:105
┌ Info: Epoch 271
└ @ Main /Users/kcft114/.julia/packages/Flux/qXNjB/src/optimise/train.jl:105
┌ Info: Epoch 272
└ @ Main /Users/kcft114/.julia/packages/Flux/qXNjB/src/optimise/train.jl:105
┌ Info: Epoch 273
└ @ Main /Users/kcft114/.julia/packages/Flux/qXNjB/src/optimise/train.jl:105
┌ Info: Epoch 274
└ @ Main /Users/kcft114/.julia/packages/Flux/qXNjB/src/optimise/train.jl:105
┌ Info: Epoch 275
└ @ Main /Users/kcft114/.julia/packages/Flux/qXNjB/src/optimise/train.jl:105
┌ Info: Epoch 276
└ @ Main /Users/kcft114/.julia/packages/Flux/qXNjB/src/optimise/train.jl:105
┌ Info: Epoch 277
└ @ Main /Users/kcft114/.julia/packages/Flux/qXNjB/src/optimise/train.jl:105
┌ Info: Epoch 278
└ @ Main /Users/kcft114/.julia/packages/Flux/qXNjB/src/optimise/train.jl:105
┌ Info: Epoch 279
└ @ Main /Users/kcft114/.julia/packages/Flux/qXNjB/src/optimise/train.jl:105
┌ Info: Epoch 280
└ @ Main /Users/kcft114/.julia/packages/Flux/qXNjB/src/optimise/train.jl:105
┌ Info: Epoch 281
└ @ Main /Users/kcft114/.julia/packages/Flux/qXNjB/src/optimise/train.jl:105
┌ Info: Epoch 282
└ @ Main /Users/kcft114/.julia/packages/Flux/qXNjB/src/optimise/train.jl:105
┌ Info: Epoch 283
└ @ Main /Users/kcft114/.julia/packages/Flux/qXNjB/src/optimise/train.jl:105
┌ Info: Epoch 284
└ @ Main /Users/kcft114/.julia/packages/Flux/qXNjB/src/optimise/train.jl:105
┌ Info: Epoch 285
└ @ Main /Users/kcft114/.julia/packages/Flux/qXNjB/src/optimise/train.jl:105
┌ Info: Epoch 286
└ @ Main /Users/kcft114/.julia/packages/Flux/qXNjB/src/optimise/train.jl:105
┌ Info: Epoch 287
└ @ Main /Users/kcft114/.julia/packages/Flux/qXNjB/src/optimise/train.jl:105
┌ Info: Epoch 288
└ @ Main /Users/kcft114/.julia/packages/Flux/qXNjB/src/optimise/train.jl:105
┌ Info: Epoch 289
└ @ Main /Users/kcft114/.julia/packages/Flux/qXNjB/src/optimise/train.jl:105
┌ Info: Epoch 290
└ @ Main /Users/kcft114/.julia/packages/Flux/qXNjB/src/optimise/train.jl:105
┌ Info: Epoch 291
└ @ Main /Users/kcft114/.julia/packages/Flux/qXNjB/src/optimise/train.jl:105
┌ Info: Epoch 292
└ @ Main /Users/kcft114/.julia/packages/Flux/qXNjB/src/optimise/train.jl:105
┌ Info: Epoch 293
└ @ Main /Users/kcft114/.julia/packages/Flux/qXNjB/src/optimise/train.jl:105
┌ Info: Epoch 294
└ @ Main /Users/kcft114/.julia/packages/Flux/qXNjB/src/optimise/train.jl:105
┌ Info: Epoch 295
└ @ Main /Users/kcft114/.julia/packages/Flux/qXNjB/src/optimise/train.jl:105
┌ Info: Epoch 296
└ @ Main /Users/kcft114/.julia/packages/Flux/qXNjB/src/optimise/train.jl:105
┌ Info: Epoch 297
└ @ Main /Users/kcft114/.julia/packages/Flux/qXNjB/src/optimise/train.jl:105
┌ Info: Epoch 298
└ @ Main /Users/kcft114/.julia/packages/Flux/qXNjB/src/optimise/train.jl:105
┌ Info: Epoch 299
└ @ Main /Users/kcft114/.julia/packages/Flux/qXNjB/src/optimise/train.jl:105
┌ Info: Epoch 300
└ @ Main /Users/kcft114/.julia/packages/Flux/qXNjB/src/optimise/train.jl:105
┌ Info: Epoch 301
└ @ Main /Users/kcft114/.julia/packages/Flux/qXNjB/src/optimise/train.jl:105
┌ Info: Epoch 302
└ @ Main /Users/kcft114/.julia/packages/Flux/qXNjB/src/optimise/train.jl:105
┌ Info: Epoch 303
└ @ Main /Users/kcft114/.julia/packages/Flux/qXNjB/src/optimise/train.jl:105
┌ Info: Epoch 304
└ @ Main /Users/kcft114/.julia/packages/Flux/qXNjB/src/optimise/train.jl:105
┌ Info: Epoch 305
└ @ Main /Users/kcft114/.julia/packages/Flux/qXNjB/src/optimise/train.jl:105
┌ Info: Epoch 306
└ @ Main /Users/kcft114/.julia/packages/Flux/qXNjB/src/optimise/train.jl:105
┌ Info: Epoch 307
└ @ Main /Users/kcft114/.julia/packages/Flux/qXNjB/src/optimise/train.jl:105
┌ Info: Epoch 308
└ @ Main /Users/kcft114/.julia/packages/Flux/qXNjB/src/optimise/train.jl:105
┌ Info: Epoch 309
└ @ Main /Users/kcft114/.julia/packages/Flux/qXNjB/src/optimise/train.jl:105
┌ Info: Epoch 310
└ @ Main /Users/kcft114/.julia/packages/Flux/qXNjB/src/optimise/train.jl:105
┌ Info: Epoch 311
└ @ Main /Users/kcft114/.julia/packages/Flux/qXNjB/src/optimise/train.jl:105
┌ Info: Epoch 312
└ @ Main /Users/kcft114/.julia/packages/Flux/qXNjB/src/optimise/train.jl:105
┌ Info: Epoch 313
└ @ Main /Users/kcft114/.julia/packages/Flux/qXNjB/src/optimise/train.jl:105
┌ Info: Epoch 314
└ @ Main /Users/kcft114/.julia/packages/Flux/qXNjB/src/optimise/train.jl:105
┌ Info: Epoch 315
└ @ Main /Users/kcft114/.julia/packages/Flux/qXNjB/src/optimise/train.jl:105
┌ Info: Epoch 316
└ @ Main /Users/kcft114/.julia/packages/Flux/qXNjB/src/optimise/train.jl:105
┌ Info: Epoch 317
└ @ Main /Users/kcft114/.julia/packages/Flux/qXNjB/src/optimise/train.jl:105
┌ Info: Epoch 318
└ @ Main /Users/kcft114/.julia/packages/Flux/qXNjB/src/optimise/train.jl:105
┌ Info: Epoch 319
└ @ Main /Users/kcft114/.julia/packages/Flux/qXNjB/src/optimise/train.jl:105
┌ Info: Epoch 320
└ @ Main /Users/kcft114/.julia/packages/Flux/qXNjB/src/optimise/train.jl:105
┌ Info: Epoch 321
└ @ Main /Users/kcft114/.julia/packages/Flux/qXNjB/src/optimise/train.jl:105
┌ Info: Epoch 322
└ @ Main /Users/kcft114/.julia/packages/Flux/qXNjB/src/optimise/train.jl:105
┌ Info: Epoch 323
└ @ Main /Users/kcft114/.julia/packages/Flux/qXNjB/src/optimise/train.jl:105
┌ Info: Epoch 324
└ @ Main /Users/kcft114/.julia/packages/Flux/qXNjB/src/optimise/train.jl:105
┌ Info: Epoch 325
└ @ Main /Users/kcft114/.julia/packages/Flux/qXNjB/src/optimise/train.jl:105
┌ Info: Epoch 326
└ @ Main /Users/kcft114/.julia/packages/Flux/qXNjB/src/optimise/train.jl:105
┌ Info: Epoch 327
└ @ Main /Users/kcft114/.julia/packages/Flux/qXNjB/src/optimise/train.jl:105
┌ Info: Epoch 328
└ @ Main /Users/kcft114/.julia/packages/Flux/qXNjB/src/optimise/train.jl:105
┌ Info: Epoch 329
└ @ Main /Users/kcft114/.julia/packages/Flux/qXNjB/src/optimise/train.jl:105
┌ Info: Epoch 330
└ @ Main /Users/kcft114/.julia/packages/Flux/qXNjB/src/optimise/train.jl:105
┌ Info: Epoch 331
└ @ Main /Users/kcft114/.julia/packages/Flux/qXNjB/src/optimise/train.jl:105
┌ Info: Epoch 332
└ @ Main /Users/kcft114/.julia/packages/Flux/qXNjB/src/optimise/train.jl:105
┌ Info: Epoch 333
└ @ Main /Users/kcft114/.julia/packages/Flux/qXNjB/src/optimise/train.jl:105
┌ Info: Epoch 334
└ @ Main /Users/kcft114/.julia/packages/Flux/qXNjB/src/optimise/train.jl:105
┌ Info: Epoch 335
└ @ Main /Users/kcft114/.julia/packages/Flux/qXNjB/src/optimise/train.jl:105
┌ Info: Epoch 336
└ @ Main /Users/kcft114/.julia/packages/Flux/qXNjB/src/optimise/train.jl:105
┌ Info: Epoch 337
└ @ Main /Users/kcft114/.julia/packages/Flux/qXNjB/src/optimise/train.jl:105
┌ Info: Epoch 338
└ @ Main /Users/kcft114/.julia/packages/Flux/qXNjB/src/optimise/train.jl:105
┌ Info: Epoch 339
└ @ Main /Users/kcft114/.julia/packages/Flux/qXNjB/src/optimise/train.jl:105
┌ Info: Epoch 340
└ @ Main /Users/kcft114/.julia/packages/Flux/qXNjB/src/optimise/train.jl:105
┌ Info: Epoch 341
└ @ Main /Users/kcft114/.julia/packages/Flux/qXNjB/src/optimise/train.jl:105
┌ Info: Epoch 342
└ @ Main /Users/kcft114/.julia/packages/Flux/qXNjB/src/optimise/train.jl:105
┌ Info: Epoch 343
└ @ Main /Users/kcft114/.julia/packages/Flux/qXNjB/src/optimise/train.jl:105
┌ Info: Epoch 344
└ @ Main /Users/kcft114/.julia/packages/Flux/qXNjB/src/optimise/train.jl:105
┌ Info: Epoch 345
└ @ Main /Users/kcft114/.julia/packages/Flux/qXNjB/src/optimise/train.jl:105
┌ Info: Epoch 346
└ @ Main /Users/kcft114/.julia/packages/Flux/qXNjB/src/optimise/train.jl:105
┌ Info: Epoch 347
└ @ Main /Users/kcft114/.julia/packages/Flux/qXNjB/src/optimise/train.jl:105
```

```
loss(X, Y) = 0.84117687f0 (tracked)
loss(X, Y) = 0.8438544f0 (tracked)
loss(X, Y) = 0.8061088f0 (tracked)
loss(X, Y) = 0.825175f0 (tracked)
loss(X, Y) = 0.84221303f0 (tracked)
loss(X, Y) = 0.8080887f0 (tracked)
loss(X, Y) = 0.85795796f0 (tracked)
loss(X, Y) = 0.81222975f0 (tracked)
loss(X, Y) = 0.813777f0 (tracked)
loss(X, Y) = 0.79057366f0 (tracked)
loss(X, Y) = 0.78214115f0 (tracked)
loss(X, Y) = 0.81286305f0 (tracked)
loss(X, Y) = 0.80297583f0 (tracked)
loss(X, Y) = 0.7888559f0 (tracked)
loss(X, Y) = 0.81840384f0 (tracked)
loss(X, Y) = 0.76677984f0 (tracked)
loss(X, Y) = 0.8541526f0 (tracked)
loss(X, Y) = 0.8011451f0 (tracked)
loss(X, Y) = 0.8321096f0 (tracked)
loss(X, Y) = 0.7978486f0 (tracked)
loss(X, Y) = 0.8405903f0 (tracked)
loss(X, Y) = 0.8117591f0 (tracked)
loss(X, Y) = 0.83430207f0 (tracked)
loss(X, Y) = 0.8169931f0 (tracked)
loss(X, Y) = 0.7932198f0 (tracked)
loss(X, Y) = 0.80760896f0 (tracked)
loss(X, Y) = 0.83401823f0 (tracked)
loss(X, Y) = 0.84582293f0 (tracked)
loss(X, Y) = 0.8150649f0 (tracked)
loss(X, Y) = 0.84480363f0 (tracked)
loss(X, Y) = 0.8660638f0 (tracked)
loss(X, Y) = 0.80538166f0 (tracked)
loss(X, Y) = 0.7701956f0 (tracked)
loss(X, Y) = 0.7881318f0 (tracked)
loss(X, Y) = 0.85840726f0 (tracked)
loss(X, Y) = 0.8303439f0 (tracked)
loss(X, Y) = 0.7754598f0 (tracked)
loss(X, Y) = 0.8561206f0 (tracked)
loss(X, Y) = 0.792406f0 (tracked)
loss(X, Y) = 0.7903921f0 (tracked)
loss(X, Y) = 0.8116956f0 (tracked)
loss(X, Y) = 0.8276532f0 (tracked)
loss(X, Y) = 0.7882154f0 (tracked)
loss(X, Y) = 0.79824126f0 (tracked)
loss(X, Y) = 0.82909685f0 (tracked)
loss(X, Y) = 0.8090938f0 (tracked)
loss(X, Y) = 0.80035716f0 (tracked)
loss(X, Y) = 0.8146396f0 (tracked)
loss(X, Y) = 0.8342784f0 (tracked)
loss(X, Y) = 0.8372273f0 (tracked)
loss(X, Y) = 0.78660554f0 (tracked)
loss(X, Y) = 0.8245268f0 (tracked)
loss(X, Y) = 0.79888713f0 (tracked)
loss(X, Y) = 0.78311205f0 (tracked)
loss(X, Y) = 0.8251619f0 (tracked)
loss(X, Y) = 0.84401673f0 (tracked)
loss(X, Y) = 0.78878534f0 (tracked)
loss(X, Y) = 0.76142526f0 (tracked)
loss(X, Y) = 0.7830081f0 (tracked)
loss(X, Y) = 0.8468484f0 (tracked)
loss(X, Y) = 0.8221331f0 (tracked)
loss(X, Y) = 0.8089215f0 (tracked)
loss(X, Y) = 0.83232737f0 (tracked)
loss(X, Y) = 0.8136237f0 (tracked)
loss(X, Y) = 0.7815425f0 (tracked)
loss(X, Y) = 0.7782978f0 (tracked)
loss(X, Y) = 0.7882758f0 (tracked)
loss(X, Y) = 0.7793027f0 (tracked)
loss(X, Y) = 0.8041767f0 (tracked)
loss(X, Y) = 0.8407127f0 (tracked)
loss(X, Y) = 0.81995684f0 (tracked)
loss(X, Y) = 0.82684076f0 (tracked)
loss(X, Y) = 0.85357744f0 (tracked)
loss(X, Y) = 0.77313787f0 (tracked)
loss(X, Y) = 0.7630182f0 (tracked)
loss(X, Y) = 0.81418896f0 (tracked)
loss(X, Y) = 0.8004599f0 (tracked)
loss(X, Y) = 0.76574063f0 (tracked)
loss(X, Y) = 0.7870877f0 (tracked)
loss(X, Y) = 0.7856661f0 (tracked)
loss(X, Y) = 0.8185299f0 (tracked)
loss(X, Y) = 0.7967444f0 (tracked)
loss(X, Y) = 0.76997143f0 (tracked)
loss(X, Y) = 0.7957077f0 (tracked)
loss(X, Y) = 0.77477837f0 (tracked)
loss(X, Y) = 0.80786234f0 (tracked)
loss(X, Y) = 0.8136959f0 (tracked)
loss(X, Y) = 0.7809512f0 (tracked)
loss(X, Y) = 0.7837749f0 (tracked)
loss(X, Y) = 0.79504836f0 (tracked)
loss(X, Y) = 0.7973735f0 (tracked)
loss(X, Y) = 0.7753117f0 (tracked)
loss(X, Y) = 0.76596504f0 (tracked)
loss(X, Y) = 0.79471904f0 (tracked)
loss(X, Y) = 0.76247776f0 (tracked)
loss(X, Y) = 0.8061022f0 (tracked)
loss(X, Y) = 0.8311222f0 (tracked)
loss(X, Y) = 0.81144494f0 (tracked)
loss(X, Y) = 0.77291316f0 (tracked)
loss(X, Y) = 0.7942616f0 (tracked)
loss(X, Y) = 0.82278126f0 (tracked)
loss(X, Y) = 0.7832268f0 (tracked)
loss(X, Y) = 0.79920363f0 (tracked)
loss(X, Y) = 0.7887807f0 (tracked)
loss(X, Y) = 0.7746326f0 (tracked)
loss(X, Y) = 0.7827619f0 (tracked)
loss(X, Y) = 0.7981733f0 (tracked)
```

```
┌ Info: Epoch 348
└ @ Main /Users/kcft114/.julia/packages/Flux/qXNjB/src/optimise/train.jl:105
┌ Info: Epoch 349
└ @ Main /Users/kcft114/.julia/packages/Flux/qXNjB/src/optimise/train.jl:105
┌ Info: Epoch 350
└ @ Main /Users/kcft114/.julia/packages/Flux/qXNjB/src/optimise/train.jl:105
```

```
loss(X, Y) = 0.79081976f0 (tracked)
loss(X, Y) = 0.76280457f0 (tracked)
loss(X, Y) = 0.7862827f0 (tracked)
loss(X, Y) = 0.76477486f0 (tracked)
loss(X, Y) = 0.77507776f0 (tracked)
loss(X, Y) = 0.7883855f0 (tracked)
loss(X, Y) = 0.8006591f0 (tracked)
loss(X, Y) = 0.79431516f0 (tracked)
loss(X, Y) = 0.7709993f0 (tracked)
loss(X, Y) = 0.7821403f0 (tracked)
loss(X, Y) = 0.7609145f0 (tracked)
loss(X, Y) = 0.7608704f0 (tracked)
loss(X, Y) = 0.8238525f0 (tracked)
loss(X, Y) = 0.76651293f0 (tracked)
loss(X, Y) = 0.79552627f0 (tracked)
loss(X, Y) = 0.79934764f0 (tracked)
loss(X, Y) = 0.803073f0 (tracked)
loss(X, Y) = 0.7792558f0 (tracked)
loss(X, Y) = 0.7974174f0 (tracked)
loss(X, Y) = 0.7550437f0 (tracked)
loss(X, Y) = 0.7685101f0 (tracked)
loss(X, Y) = 0.7857062f0 (tracked)
loss(X, Y) = 0.772215f0 (tracked)
loss(X, Y) = 0.76532435f0 (tracked)
loss(X, Y) = 0.7801686f0 (tracked)
loss(X, Y) = 0.7439239f0 (tracked)
loss(X, Y) = 0.82695943f0 (tracked)
loss(X, Y) = 0.7523653f0 (tracked)
loss(X, Y) = 0.7630238f0 (tracked)
loss(X, Y) = 0.80530643f0 (tracked)
loss(X, Y) = 0.76062953f0 (tracked)
loss(X, Y) = 0.7997552f0 (tracked)
loss(X, Y) = 0.8361595f0 (tracked)
loss(X, Y) = 0.7599877f0 (tracked)
loss(X, Y) = 0.8186188f0 (tracked)
loss(X, Y) = 0.7644794f0 (tracked)
loss(X, Y) = 0.7669887f0 (tracked)
loss(X, Y) = 0.7728953f0 (tracked)
loss(X, Y) = 0.7986659f0 (tracked)
loss(X, Y) = 0.71948785f0 (tracked)
loss(X, Y) = 0.77906966f0 (tracked)
loss(X, Y) = 0.78712994f0 (tracked)
loss(X, Y) = 0.8167696f0 (tracked)
loss(X, Y) = 0.80042064f0 (tracked)
loss(X, Y) = 0.79053515f0 (tracked)
loss(X, Y) = 0.81388795f0 (tracked)
loss(X, Y) = 0.77239513f0 (tracked)
loss(X, Y) = 0.7261336f0 (tracked)
loss(X, Y) = 0.7422736f0 (tracked)
loss(X, Y) = 0.7971348f0 (tracked)
loss(X, Y) = 0.7869447f0 (tracked)
loss(X, Y) = 0.7598878f0 (tracked)
loss(X, Y) = 0.748417f0 (tracked)
loss(X, Y) = 0.7838186f0 (tracked)
loss(X, Y) = 0.7769732f0 (tracked)
loss(X, Y) = 0.7909031f0 (tracked)
loss(X, Y) = 0.7553956f0 (tracked)
loss(X, Y) = 0.7918248f0 (tracked)
loss(X, Y) = 0.7804014f0 (tracked)
loss(X, Y) = 0.7983215f0 (tracked)
loss(X, Y) = 0.7827771f0 (tracked)
loss(X, Y) = 0.79180604f0 (tracked)
loss(X, Y) = 0.7951127f0 (tracked)
loss(X, Y) = 0.7704822f0 (tracked)
loss(X, Y) = 0.7812029f0 (tracked)
loss(X, Y) = 0.78319526f0 (tracked)
loss(X, Y) = 0.79686725f0 (tracked)
loss(X, Y) = 0.7864942f0 (tracked)
loss(X, Y) = 0.7490713f0 (tracked)
loss(X, Y) = 0.7614974f0 (tracked)
loss(X, Y) = 0.7789283f0 (tracked)
loss(X, Y) = 0.77151215f0 (tracked)
loss(X, Y) = 0.8092371f0 (tracked)
loss(X, Y) = 0.78401893f0 (tracked)
loss(X, Y) = 0.77248514f0 (tracked)
loss(X, Y) = 0.7825707f0 (tracked)
loss(X, Y) = 0.7816429f0 (tracked)
loss(X, Y) = 0.7968442f0 (tracked)
loss(X, Y) = 0.802255f0 (tracked)
loss(X, Y) = 0.7263225f0 (tracked)
loss(X, Y) = 0.74846685f0 (tracked)
loss(X, Y) = 0.769853f0 (tracked)
loss(X, Y) = 0.78153944f0 (tracked)
loss(X, Y) = 0.7592337f0 (tracked)
loss(X, Y) = 0.7964479f0 (tracked)
loss(X, Y) = 0.748965f0 (tracked)
loss(X, Y) = 0.8237157f0 (tracked)
```

```
┌ Info: Epoch 351
└ @ Main /Users/kcft114/.julia/packages/Flux/qXNjB/src/optimise/train.jl:105
┌ Info: Epoch 352
└ @ Main /Users/kcft114/.julia/packages/Flux/qXNjB/src/optimise/train.jl:105
┌ Info: Epoch 353
└ @ Main /Users/kcft114/.julia/packages/Flux/qXNjB/src/optimise/train.jl:105
┌ Info: Epoch 354
└ @ Main /Users/kcft114/.julia/packages/Flux/qXNjB/src/optimise/train.jl:105
┌ Info: Epoch 355
└ @ Main /Users/kcft114/.julia/packages/Flux/qXNjB/src/optimise/train.jl:105
┌ Info: Epoch 356
└ @ Main /Users/kcft114/.julia/packages/Flux/qXNjB/src/optimise/train.jl:105
┌ Info: Epoch 357
└ @ Main /Users/kcft114/.julia/packages/Flux/qXNjB/src/optimise/train.jl:105
┌ Info: Epoch 358
└ @ Main /Users/kcft114/.julia/packages/Flux/qXNjB/src/optimise/train.jl:105
┌ Info: Epoch 359
└ @ Main /Users/kcft114/.julia/packages/Flux/qXNjB/src/optimise/train.jl:105
┌ Info: Epoch 360
└ @ Main /Users/kcft114/.julia/packages/Flux/qXNjB/src/optimise/train.jl:105
┌ Info: Epoch 361
└ @ Main /Users/kcft114/.julia/packages/Flux/qXNjB/src/optimise/train.jl:105
┌ Info: Epoch 362
└ @ Main /Users/kcft114/.julia/packages/Flux/qXNjB/src/optimise/train.jl:105
┌ Info: Epoch 363
└ @ Main /Users/kcft114/.julia/packages/Flux/qXNjB/src/optimise/train.jl:105
┌ Info: Epoch 364
└ @ Main /Users/kcft114/.julia/packages/Flux/qXNjB/src/optimise/train.jl:105
┌ Info: Epoch 365
└ @ Main /Users/kcft114/.julia/packages/Flux/qXNjB/src/optimise/train.jl:105
┌ Info: Epoch 366
└ @ Main /Users/kcft114/.julia/packages/Flux/qXNjB/src/optimise/train.jl:105
┌ Info: Epoch 367
└ @ Main /Users/kcft114/.julia/packages/Flux/qXNjB/src/optimise/train.jl:105
┌ Info: Epoch 368
└ @ Main /Users/kcft114/.julia/packages/Flux/qXNjB/src/optimise/train.jl:105
┌ Info: Epoch 369
└ @ Main /Users/kcft114/.julia/packages/Flux/qXNjB/src/optimise/train.jl:105
┌ Info: Epoch 370
└ @ Main /Users/kcft114/.julia/packages/Flux/qXNjB/src/optimise/train.jl:105
┌ Info: Epoch 371
└ @ Main /Users/kcft114/.julia/packages/Flux/qXNjB/src/optimise/train.jl:105
┌ Info: Epoch 372
└ @ Main /Users/kcft114/.julia/packages/Flux/qXNjB/src/optimise/train.jl:105
┌ Info: Epoch 373
└ @ Main /Users/kcft114/.julia/packages/Flux/qXNjB/src/optimise/train.jl:105
┌ Info: Epoch 374
└ @ Main /Users/kcft114/.julia/packages/Flux/qXNjB/src/optimise/train.jl:105
┌ Info: Epoch 375
└ @ Main /Users/kcft114/.julia/packages/Flux/qXNjB/src/optimise/train.jl:105
┌ Info: Epoch 376
└ @ Main /Users/kcft114/.julia/packages/Flux/qXNjB/src/optimise/train.jl:105
┌ Info: Epoch 377
└ @ Main /Users/kcft114/.julia/packages/Flux/qXNjB/src/optimise/train.jl:105
┌ Info: Epoch 378
└ @ Main /Users/kcft114/.julia/packages/Flux/qXNjB/src/optimise/train.jl:105
┌ Info: Epoch 379
└ @ Main /Users/kcft114/.julia/packages/Flux/qXNjB/src/optimise/train.jl:105
┌ Info: Epoch 380
└ @ Main /Users/kcft114/.julia/packages/Flux/qXNjB/src/optimise/train.jl:105
┌ Info: Epoch 381
└ @ Main /Users/kcft114/.julia/packages/Flux/qXNjB/src/optimise/train.jl:105
┌ Info: Epoch 382
└ @ Main /Users/kcft114/.julia/packages/Flux/qXNjB/src/optimise/train.jl:105
┌ Info: Epoch 383
└ @ Main /Users/kcft114/.julia/packages/Flux/qXNjB/src/optimise/train.jl:105
┌ Info: Epoch 384
└ @ Main /Users/kcft114/.julia/packages/Flux/qXNjB/src/optimise/train.jl:105
┌ Info: Epoch 385
└ @ Main /Users/kcft114/.julia/packages/Flux/qXNjB/src/optimise/train.jl:105
┌ Info: Epoch 386
└ @ Main /Users/kcft114/.julia/packages/Flux/qXNjB/src/optimise/train.jl:105
┌ Info: Epoch 387
└ @ Main /Users/kcft114/.julia/packages/Flux/qXNjB/src/optimise/train.jl:105
┌ Info: Epoch 388
└ @ Main /Users/kcft114/.julia/packages/Flux/qXNjB/src/optimise/train.jl:105
┌ Info: Epoch 389
└ @ Main /Users/kcft114/.julia/packages/Flux/qXNjB/src/optimise/train.jl:105
┌ Info: Epoch 390
└ @ Main /Users/kcft114/.julia/packages/Flux/qXNjB/src/optimise/train.jl:105
┌ Info: Epoch 391
└ @ Main /Users/kcft114/.julia/packages/Flux/qXNjB/src/optimise/train.jl:105
┌ Info: Epoch 392
└ @ Main /Users/kcft114/.julia/packages/Flux/qXNjB/src/optimise/train.jl:105
┌ Info: Epoch 393
└ @ Main /Users/kcft114/.julia/packages/Flux/qXNjB/src/optimise/train.jl:105
┌ Info: Epoch 394
└ @ Main /Users/kcft114/.julia/packages/Flux/qXNjB/src/optimise/train.jl:105
┌ Info: Epoch 395
└ @ Main /Users/kcft114/.julia/packages/Flux/qXNjB/src/optimise/train.jl:105
┌ Info: Epoch 396
└ @ Main /Users/kcft114/.julia/packages/Flux/qXNjB/src/optimise/train.jl:105
┌ Info: Epoch 397
└ @ Main /Users/kcft114/.julia/packages/Flux/qXNjB/src/optimise/train.jl:105
┌ Info: Epoch 398
└ @ Main /Users/kcft114/.julia/packages/Flux/qXNjB/src/optimise/train.jl:105
┌ Info: Epoch 399
└ @ Main /Users/kcft114/.julia/packages/Flux/qXNjB/src/optimise/train.jl:105
┌ Info: Epoch 400
└ @ Main /Users/kcft114/.julia/packages/Flux/qXNjB/src/optimise/train.jl:105
┌ Info: Epoch 401
└ @ Main /Users/kcft114/.julia/packages/Flux/qXNjB/src/optimise/train.jl:105
┌ Info: Epoch 402
└ @ Main /Users/kcft114/.julia/packages/Flux/qXNjB/src/optimise/train.jl:105
┌ Info: Epoch 403
└ @ Main /Users/kcft114/.julia/packages/Flux/qXNjB/src/optimise/train.jl:105
┌ Info: Epoch 404
└ @ Main /Users/kcft114/.julia/packages/Flux/qXNjB/src/optimise/train.jl:105
┌ Info: Epoch 405
└ @ Main /Users/kcft114/.julia/packages/Flux/qXNjB/src/optimise/train.jl:105
┌ Info: Epoch 406
└ @ Main /Users/kcft114/.julia/packages/Flux/qXNjB/src/optimise/train.jl:105
┌ Info: Epoch 407
└ @ Main /Users/kcft114/.julia/packages/Flux/qXNjB/src/optimise/train.jl:105
┌ Info: Epoch 408
└ @ Main /Users/kcft114/.julia/packages/Flux/qXNjB/src/optimise/train.jl:105
┌ Info: Epoch 409
└ @ Main /Users/kcft114/.julia/packages/Flux/qXNjB/src/optimise/train.jl:105
┌ Info: Epoch 410
└ @ Main /Users/kcft114/.julia/packages/Flux/qXNjB/src/optimise/train.jl:105
┌ Info: Epoch 411
└ @ Main /Users/kcft114/.julia/packages/Flux/qXNjB/src/optimise/train.jl:105
┌ Info: Epoch 412
└ @ Main /Users/kcft114/.julia/packages/Flux/qXNjB/src/optimise/train.jl:105
┌ Info: Epoch 413
└ @ Main /Users/kcft114/.julia/packages/Flux/qXNjB/src/optimise/train.jl:105
┌ Info: Epoch 414
└ @ Main /Users/kcft114/.julia/packages/Flux/qXNjB/src/optimise/train.jl:105
┌ Info: Epoch 415
└ @ Main /Users/kcft114/.julia/packages/Flux/qXNjB/src/optimise/train.jl:105
┌ Info: Epoch 416
└ @ Main /Users/kcft114/.julia/packages/Flux/qXNjB/src/optimise/train.jl:105
┌ Info: Epoch 417
└ @ Main /Users/kcft114/.julia/packages/Flux/qXNjB/src/optimise/train.jl:105
┌ Info: Epoch 418
└ @ Main /Users/kcft114/.julia/packages/Flux/qXNjB/src/optimise/train.jl:105
┌ Info: Epoch 419
└ @ Main /Users/kcft114/.julia/packages/Flux/qXNjB/src/optimise/train.jl:105
┌ Info: Epoch 420
└ @ Main /Users/kcft114/.julia/packages/Flux/qXNjB/src/optimise/train.jl:105
┌ Info: Epoch 421
└ @ Main /Users/kcft114/.julia/packages/Flux/qXNjB/src/optimise/train.jl:105
┌ Info: Epoch 422
└ @ Main /Users/kcft114/.julia/packages/Flux/qXNjB/src/optimise/train.jl:105
┌ Info: Epoch 423
└ @ Main /Users/kcft114/.julia/packages/Flux/qXNjB/src/optimise/train.jl:105
┌ Info: Epoch 424
└ @ Main /Users/kcft114/.julia/packages/Flux/qXNjB/src/optimise/train.jl:105
┌ Info: Epoch 425
└ @ Main /Users/kcft114/.julia/packages/Flux/qXNjB/src/optimise/train.jl:105
┌ Info: Epoch 426
└ @ Main /Users/kcft114/.julia/packages/Flux/qXNjB/src/optimise/train.jl:105
┌ Info: Epoch 427
└ @ Main /Users/kcft114/.julia/packages/Flux/qXNjB/src/optimise/train.jl:105
┌ Info: Epoch 428
└ @ Main /Users/kcft114/.julia/packages/Flux/qXNjB/src/optimise/train.jl:105
┌ Info: Epoch 429
└ @ Main /Users/kcft114/.julia/packages/Flux/qXNjB/src/optimise/train.jl:105
┌ Info: Epoch 430
└ @ Main /Users/kcft114/.julia/packages/Flux/qXNjB/src/optimise/train.jl:105
┌ Info: Epoch 431
└ @ Main /Users/kcft114/.julia/packages/Flux/qXNjB/src/optimise/train.jl:105
┌ Info: Epoch 432
└ @ Main /Users/kcft114/.julia/packages/Flux/qXNjB/src/optimise/train.jl:105
┌ Info: Epoch 433
└ @ Main /Users/kcft114/.julia/packages/Flux/qXNjB/src/optimise/train.jl:105
┌ Info: Epoch 434
└ @ Main /Users/kcft114/.julia/packages/Flux/qXNjB/src/optimise/train.jl:105
┌ Info: Epoch 435
└ @ Main /Users/kcft114/.julia/packages/Flux/qXNjB/src/optimise/train.jl:105
┌ Info: Epoch 436
└ @ Main /Users/kcft114/.julia/packages/Flux/qXNjB/src/optimise/train.jl:105
┌ Info: Epoch 437
└ @ Main /Users/kcft114/.julia/packages/Flux/qXNjB/src/optimise/train.jl:105
```

```
loss(X, Y) = 0.75432193f0 (tracked)
loss(X, Y) = 0.76627094f0 (tracked)
loss(X, Y) = 0.7532768f0 (tracked)
loss(X, Y) = 0.71886957f0 (tracked)
loss(X, Y) = 0.7740851f0 (tracked)
loss(X, Y) = 0.7167764f0 (tracked)
loss(X, Y) = 0.74433696f0 (tracked)
loss(X, Y) = 0.7766697f0 (tracked)
loss(X, Y) = 0.7677466f0 (tracked)
loss(X, Y) = 0.7888645f0 (tracked)
loss(X, Y) = 0.78386813f0 (tracked)
loss(X, Y) = 0.7569144f0 (tracked)
loss(X, Y) = 0.77055866f0 (tracked)
loss(X, Y) = 0.7645961f0 (tracked)
loss(X, Y) = 0.79600775f0 (tracked)
loss(X, Y) = 0.7792018f0 (tracked)
loss(X, Y) = 0.7858327f0 (tracked)
loss(X, Y) = 0.79004395f0 (tracked)
loss(X, Y) = 0.7611087f0 (tracked)
loss(X, Y) = 0.8044269f0 (tracked)
loss(X, Y) = 0.75565535f0 (tracked)
loss(X, Y) = 0.7390081f0 (tracked)
loss(X, Y) = 0.7368619f0 (tracked)
loss(X, Y) = 0.76894134f0 (tracked)
loss(X, Y) = 0.7580368f0 (tracked)
loss(X, Y) = 0.7302556f0 (tracked)
loss(X, Y) = 0.760076f0 (tracked)
loss(X, Y) = 0.74317807f0 (tracked)
loss(X, Y) = 0.7215585f0 (tracked)
loss(X, Y) = 0.75073236f0 (tracked)
loss(X, Y) = 0.7801718f0 (tracked)
loss(X, Y) = 0.71489155f0 (tracked)
loss(X, Y) = 0.80357015f0 (tracked)
loss(X, Y) = 0.7448495f0 (tracked)
loss(X, Y) = 0.7448198f0 (tracked)
loss(X, Y) = 0.7569419f0 (tracked)
loss(X, Y) = 0.7525051f0 (tracked)
loss(X, Y) = 0.797273f0 (tracked)
loss(X, Y) = 0.73908913f0 (tracked)
loss(X, Y) = 0.8013363f0 (tracked)
loss(X, Y) = 0.76935095f0 (tracked)
loss(X, Y) = 0.7516254f0 (tracked)
loss(X, Y) = 0.76135784f0 (tracked)
loss(X, Y) = 0.75758624f0 (tracked)
loss(X, Y) = 0.7637363f0 (tracked)
loss(X, Y) = 0.7849119f0 (tracked)
loss(X, Y) = 0.78013206f0 (tracked)
loss(X, Y) = 0.75613844f0 (tracked)
loss(X, Y) = 0.7611557f0 (tracked)
loss(X, Y) = 0.7389563f0 (tracked)
loss(X, Y) = 0.72775936f0 (tracked)
loss(X, Y) = 0.765957f0 (tracked)
loss(X, Y) = 0.71530765f0 (tracked)
loss(X, Y) = 0.7694842f0 (tracked)
loss(X, Y) = 0.7525191f0 (tracked)
loss(X, Y) = 0.7418156f0 (tracked)
loss(X, Y) = 0.7437229f0 (tracked)
loss(X, Y) = 0.7779852f0 (tracked)
loss(X, Y) = 0.73223317f0 (tracked)
loss(X, Y) = 0.75658715f0 (tracked)
loss(X, Y) = 0.7477881f0 (tracked)
loss(X, Y) = 0.7719633f0 (tracked)
loss(X, Y) = 0.78089726f0 (tracked)
loss(X, Y) = 0.74898314f0 (tracked)
loss(X, Y) = 0.7587797f0 (tracked)
loss(X, Y) = 0.7476977f0 (tracked)
loss(X, Y) = 0.7400585f0 (tracked)
loss(X, Y) = 0.75047266f0 (tracked)
loss(X, Y) = 0.7357777f0 (tracked)
loss(X, Y) = 0.7354299f0 (tracked)
loss(X, Y) = 0.7256697f0 (tracked)
loss(X, Y) = 0.7761278f0 (tracked)
loss(X, Y) = 0.7325568f0 (tracked)
loss(X, Y) = 0.7861229f0 (tracked)
loss(X, Y) = 0.73570865f0 (tracked)
loss(X, Y) = 0.74500525f0 (tracked)
loss(X, Y) = 0.7348451f0 (tracked)
loss(X, Y) = 0.7561309f0 (tracked)
loss(X, Y) = 0.75913507f0 (tracked)
loss(X, Y) = 0.7581595f0 (tracked)
loss(X, Y) = 0.74138373f0 (tracked)
loss(X, Y) = 0.7315237f0 (tracked)
loss(X, Y) = 0.79060864f0 (tracked)
loss(X, Y) = 0.76717216f0 (tracked)
loss(X, Y) = 0.74276173f0 (tracked)
loss(X, Y) = 0.7371859f0 (tracked)
loss(X, Y) = 0.7436273f0 (tracked)
loss(X, Y) = 0.750863f0 (tracked)
loss(X, Y) = 0.7330692f0 (tracked)
loss(X, Y) = 0.78198475f0 (tracked)
```

```
┌ Info: Epoch 438
└ @ Main /Users/kcft114/.julia/packages/Flux/qXNjB/src/optimise/train.jl:105
┌ Info: Epoch 439
└ @ Main /Users/kcft114/.julia/packages/Flux/qXNjB/src/optimise/train.jl:105
┌ Info: Epoch 440
└ @ Main /Users/kcft114/.julia/packages/Flux/qXNjB/src/optimise/train.jl:105
┌ Info: Epoch 441
└ @ Main /Users/kcft114/.julia/packages/Flux/qXNjB/src/optimise/train.jl:105
┌ Info: Epoch 442
└ @ Main /Users/kcft114/.julia/packages/Flux/qXNjB/src/optimise/train.jl:105
┌ Info: Epoch 443
└ @ Main /Users/kcft114/.julia/packages/Flux/qXNjB/src/optimise/train.jl:105
┌ Info: Epoch 444
└ @ Main /Users/kcft114/.julia/packages/Flux/qXNjB/src/optimise/train.jl:105
┌ Info: Epoch 445
└ @ Main /Users/kcft114/.julia/packages/Flux/qXNjB/src/optimise/train.jl:105
┌ Info: Epoch 446
└ @ Main /Users/kcft114/.julia/packages/Flux/qXNjB/src/optimise/train.jl:105
┌ Info: Epoch 447
└ @ Main /Users/kcft114/.julia/packages/Flux/qXNjB/src/optimise/train.jl:105
┌ Info: Epoch 448
└ @ Main /Users/kcft114/.julia/packages/Flux/qXNjB/src/optimise/train.jl:105
┌ Info: Epoch 449
└ @ Main /Users/kcft114/.julia/packages/Flux/qXNjB/src/optimise/train.jl:105
┌ Info: Epoch 450
└ @ Main /Users/kcft114/.julia/packages/Flux/qXNjB/src/optimise/train.jl:105
┌ Info: Epoch 451
└ @ Main /Users/kcft114/.julia/packages/Flux/qXNjB/src/optimise/train.jl:105
┌ Info: Epoch 452
└ @ Main /Users/kcft114/.julia/packages/Flux/qXNjB/src/optimise/train.jl:105
┌ Info: Epoch 453
└ @ Main /Users/kcft114/.julia/packages/Flux/qXNjB/src/optimise/train.jl:105
┌ Info: Epoch 454
└ @ Main /Users/kcft114/.julia/packages/Flux/qXNjB/src/optimise/train.jl:105
┌ Info: Epoch 455
└ @ Main /Users/kcft114/.julia/packages/Flux/qXNjB/src/optimise/train.jl:105
┌ Info: Epoch 456
└ @ Main /Users/kcft114/.julia/packages/Flux/qXNjB/src/optimise/train.jl:105
┌ Info: Epoch 457
└ @ Main /Users/kcft114/.julia/packages/Flux/qXNjB/src/optimise/train.jl:105
┌ Info: Epoch 458
└ @ Main /Users/kcft114/.julia/packages/Flux/qXNjB/src/optimise/train.jl:105
┌ Info: Epoch 459
└ @ Main /Users/kcft114/.julia/packages/Flux/qXNjB/src/optimise/train.jl:105
┌ Info: Epoch 460
└ @ Main /Users/kcft114/.julia/packages/Flux/qXNjB/src/optimise/train.jl:105
┌ Info: Epoch 461
└ @ Main /Users/kcft114/.julia/packages/Flux/qXNjB/src/optimise/train.jl:105
┌ Info: Epoch 462
└ @ Main /Users/kcft114/.julia/packages/Flux/qXNjB/src/optimise/train.jl:105
┌ Info: Epoch 463
└ @ Main /Users/kcft114/.julia/packages/Flux/qXNjB/src/optimise/train.jl:105
┌ Info: Epoch 464
└ @ Main /Users/kcft114/.julia/packages/Flux/qXNjB/src/optimise/train.jl:105
┌ Info: Epoch 465
└ @ Main /Users/kcft114/.julia/packages/Flux/qXNjB/src/optimise/train.jl:105
┌ Info: Epoch 466
└ @ Main /Users/kcft114/.julia/packages/Flux/qXNjB/src/optimise/train.jl:105
┌ Info: Epoch 467
└ @ Main /Users/kcft114/.julia/packages/Flux/qXNjB/src/optimise/train.jl:105
┌ Info: Epoch 468
└ @ Main /Users/kcft114/.julia/packages/Flux/qXNjB/src/optimise/train.jl:105
┌ Info: Epoch 469
└ @ Main /Users/kcft114/.julia/packages/Flux/qXNjB/src/optimise/train.jl:105
┌ Info: Epoch 470
└ @ Main /Users/kcft114/.julia/packages/Flux/qXNjB/src/optimise/train.jl:105
┌ Info: Epoch 471
└ @ Main /Users/kcft114/.julia/packages/Flux/qXNjB/src/optimise/train.jl:105
┌ Info: Epoch 472
└ @ Main /Users/kcft114/.julia/packages/Flux/qXNjB/src/optimise/train.jl:105
┌ Info: Epoch 473
└ @ Main /Users/kcft114/.julia/packages/Flux/qXNjB/src/optimise/train.jl:105
┌ Info: Epoch 474
└ @ Main /Users/kcft114/.julia/packages/Flux/qXNjB/src/optimise/train.jl:105
┌ Info: Epoch 475
└ @ Main /Users/kcft114/.julia/packages/Flux/qXNjB/src/optimise/train.jl:105
┌ Info: Epoch 476
└ @ Main /Users/kcft114/.julia/packages/Flux/qXNjB/src/optimise/train.jl:105
┌ Info: Epoch 477
└ @ Main /Users/kcft114/.julia/packages/Flux/qXNjB/src/optimise/train.jl:105
┌ Info: Epoch 478
└ @ Main /Users/kcft114/.julia/packages/Flux/qXNjB/src/optimise/train.jl:105
┌ Info: Epoch 479
└ @ Main /Users/kcft114/.julia/packages/Flux/qXNjB/src/optimise/train.jl:105
┌ Info: Epoch 480
└ @ Main /Users/kcft114/.julia/packages/Flux/qXNjB/src/optimise/train.jl:105
┌ Info: Epoch 481
└ @ Main /Users/kcft114/.julia/packages/Flux/qXNjB/src/optimise/train.jl:105
┌ Info: Epoch 482
└ @ Main /Users/kcft114/.julia/packages/Flux/qXNjB/src/optimise/train.jl:105
┌ Info: Epoch 483
└ @ Main /Users/kcft114/.julia/packages/Flux/qXNjB/src/optimise/train.jl:105
┌ Info: Epoch 484
└ @ Main /Users/kcft114/.julia/packages/Flux/qXNjB/src/optimise/train.jl:105
┌ Info: Epoch 485
└ @ Main /Users/kcft114/.julia/packages/Flux/qXNjB/src/optimise/train.jl:105
┌ Info: Epoch 486
└ @ Main /Users/kcft114/.julia/packages/Flux/qXNjB/src/optimise/train.jl:105
┌ Info: Epoch 487
└ @ Main /Users/kcft114/.julia/packages/Flux/qXNjB/src/optimise/train.jl:105
┌ Info: Epoch 488
└ @ Main /Users/kcft114/.julia/packages/Flux/qXNjB/src/optimise/train.jl:105
┌ Info: Epoch 489
└ @ Main /Users/kcft114/.julia/packages/Flux/qXNjB/src/optimise/train.jl:105
┌ Info: Epoch 490
└ @ Main /Users/kcft114/.julia/packages/Flux/qXNjB/src/optimise/train.jl:105
┌ Info: Epoch 491
└ @ Main /Users/kcft114/.julia/packages/Flux/qXNjB/src/optimise/train.jl:105
┌ Info: Epoch 492
└ @ Main /Users/kcft114/.julia/packages/Flux/qXNjB/src/optimise/train.jl:105
┌ Info: Epoch 493
└ @ Main /Users/kcft114/.julia/packages/Flux/qXNjB/src/optimise/train.jl:105
┌ Info: Epoch 494
└ @ Main /Users/kcft114/.julia/packages/Flux/qXNjB/src/optimise/train.jl:105
┌ Info: Epoch 495
└ @ Main /Users/kcft114/.julia/packages/Flux/qXNjB/src/optimise/train.jl:105
┌ Info: Epoch 496
└ @ Main /Users/kcft114/.julia/packages/Flux/qXNjB/src/optimise/train.jl:105
┌ Info: Epoch 497
└ @ Main /Users/kcft114/.julia/packages/Flux/qXNjB/src/optimise/train.jl:105
┌ Info: Epoch 498
└ @ Main /Users/kcft114/.julia/packages/Flux/qXNjB/src/optimise/train.jl:105
┌ Info: Epoch 499
└ @ Main /Users/kcft114/.julia/packages/Flux/qXNjB/src/optimise/train.jl:105
┌ Info: Epoch 500
└ @ Main /Users/kcft114/.julia/packages/Flux/qXNjB/src/optimise/train.jl:105
┌ Info: Epoch 501
└ @ Main /Users/kcft114/.julia/packages/Flux/qXNjB/src/optimise/train.jl:105
┌ Info: Epoch 502
└ @ Main /Users/kcft114/.julia/packages/Flux/qXNjB/src/optimise/train.jl:105
┌ Info: Epoch 503
└ @ Main /Users/kcft114/.julia/packages/Flux/qXNjB/src/optimise/train.jl:105
┌ Info: Epoch 504
└ @ Main /Users/kcft114/.julia/packages/Flux/qXNjB/src/optimise/train.jl:105
┌ Info: Epoch 505
└ @ Main /Users/kcft114/.julia/packages/Flux/qXNjB/src/optimise/train.jl:105
┌ Info: Epoch 506
└ @ Main /Users/kcft114/.julia/packages/Flux/qXNjB/src/optimise/train.jl:105
┌ Info: Epoch 507
└ @ Main /Users/kcft114/.julia/packages/Flux/qXNjB/src/optimise/train.jl:105
┌ Info: Epoch 508
└ @ Main /Users/kcft114/.julia/packages/Flux/qXNjB/src/optimise/train.jl:105
┌ Info: Epoch 509
└ @ Main /Users/kcft114/.julia/packages/Flux/qXNjB/src/optimise/train.jl:105
┌ Info: Epoch 510
└ @ Main /Users/kcft114/.julia/packages/Flux/qXNjB/src/optimise/train.jl:105
┌ Info: Epoch 511
└ @ Main /Users/kcft114/.julia/packages/Flux/qXNjB/src/optimise/train.jl:105
┌ Info: Epoch 512
└ @ Main /Users/kcft114/.julia/packages/Flux/qXNjB/src/optimise/train.jl:105
┌ Info: Epoch 513
└ @ Main /Users/kcft114/.julia/packages/Flux/qXNjB/src/optimise/train.jl:105
┌ Info: Epoch 514
└ @ Main /Users/kcft114/.julia/packages/Flux/qXNjB/src/optimise/train.jl:105
┌ Info: Epoch 515
└ @ Main /Users/kcft114/.julia/packages/Flux/qXNjB/src/optimise/train.jl:105
┌ Info: Epoch 516
└ @ Main /Users/kcft114/.julia/packages/Flux/qXNjB/src/optimise/train.jl:105
┌ Info: Epoch 517
└ @ Main /Users/kcft114/.julia/packages/Flux/qXNjB/src/optimise/train.jl:105
┌ Info: Epoch 518
└ @ Main /Users/kcft114/.julia/packages/Flux/qXNjB/src/optimise/train.jl:105
┌ Info: Epoch 519
└ @ Main /Users/kcft114/.julia/packages/Flux/qXNjB/src/optimise/train.jl:105
┌ Info: Epoch 520
└ @ Main /Users/kcft114/.julia/packages/Flux/qXNjB/src/optimise/train.jl:105
┌ Info: Epoch 521
└ @ Main /Users/kcft114/.julia/packages/Flux/qXNjB/src/optimise/train.jl:105
┌ Info: Epoch 522
└ @ Main /Users/kcft114/.julia/packages/Flux/qXNjB/src/optimise/train.jl:105
┌ Info: Epoch 523
└ @ Main /Users/kcft114/.julia/packages/Flux/qXNjB/src/optimise/train.jl:105
┌ Info: Epoch 524
└ @ Main /Users/kcft114/.julia/packages/Flux/qXNjB/src/optimise/train.jl:105
┌ Info: Epoch 525
└ @ Main /Users/kcft114/.julia/packages/Flux/qXNjB/src/optimise/train.jl:105
┌ Info: Epoch 526
└ @ Main /Users/kcft114/.julia/packages/Flux/qXNjB/src/optimise/train.jl:105
┌ Info: Epoch 527
└ @ Main /Users/kcft114/.julia/packages/Flux/qXNjB/src/optimise/train.jl:105
```

```
loss(X, Y) = 0.75980794f0 (tracked)
loss(X, Y) = 0.76385844f0 (tracked)
loss(X, Y) = 0.775675f0 (tracked)
loss(X, Y) = 0.7551488f0 (tracked)
loss(X, Y) = 0.72274476f0 (tracked)
loss(X, Y) = 0.7549279f0 (tracked)
loss(X, Y) = 0.7340742f0 (tracked)
loss(X, Y) = 0.7897401f0 (tracked)
loss(X, Y) = 0.7453581f0 (tracked)
loss(X, Y) = 0.7455767f0 (tracked)
loss(X, Y) = 0.7323263f0 (tracked)
loss(X, Y) = 0.7392322f0 (tracked)
loss(X, Y) = 0.7563847f0 (tracked)
loss(X, Y) = 0.7273271f0 (tracked)
loss(X, Y) = 0.7566583f0 (tracked)
loss(X, Y) = 0.7398385f0 (tracked)
loss(X, Y) = 0.7290916f0 (tracked)
loss(X, Y) = 0.7433956f0 (tracked)
loss(X, Y) = 0.71551f0 (tracked)
loss(X, Y) = 0.7827436f0 (tracked)
loss(X, Y) = 0.71132725f0 (tracked)
loss(X, Y) = 0.7697054f0 (tracked)
loss(X, Y) = 0.7432711f0 (tracked)
loss(X, Y) = 0.71374345f0 (tracked)
loss(X, Y) = 0.7570431f0 (tracked)
loss(X, Y) = 0.75523216f0 (tracked)
loss(X, Y) = 0.76335067f0 (tracked)
loss(X, Y) = 0.75219685f0 (tracked)
loss(X, Y) = 0.75572896f0 (tracked)
loss(X, Y) = 0.7038669f0 (tracked)
loss(X, Y) = 0.73188245f0 (tracked)
loss(X, Y) = 0.7361507f0 (tracked)
loss(X, Y) = 0.7150385f0 (tracked)
loss(X, Y) = 0.7268066f0 (tracked)
loss(X, Y) = 0.74298215f0 (tracked)
loss(X, Y) = 0.73003787f0 (tracked)
loss(X, Y) = 0.7429833f0 (tracked)
loss(X, Y) = 0.737121f0 (tracked)
loss(X, Y) = 0.74137247f0 (tracked)
loss(X, Y) = 0.74519575f0 (tracked)
loss(X, Y) = 0.7374104f0 (tracked)
loss(X, Y) = 0.7338884f0 (tracked)
loss(X, Y) = 0.72864753f0 (tracked)
loss(X, Y) = 0.7338339f0 (tracked)
loss(X, Y) = 0.75646955f0 (tracked)
loss(X, Y) = 0.76877904f0 (tracked)
loss(X, Y) = 0.7326942f0 (tracked)
loss(X, Y) = 0.7653967f0 (tracked)
loss(X, Y) = 0.7488681f0 (tracked)
loss(X, Y) = 0.7515241f0 (tracked)
loss(X, Y) = 0.7618703f0 (tracked)
loss(X, Y) = 0.7017947f0 (tracked)
loss(X, Y) = 0.7295883f0 (tracked)
loss(X, Y) = 0.71439433f0 (tracked)
loss(X, Y) = 0.7615108f0 (tracked)
loss(X, Y) = 0.7527541f0 (tracked)
loss(X, Y) = 0.7376375f0 (tracked)
loss(X, Y) = 0.75613075f0 (tracked)
loss(X, Y) = 0.73523325f0 (tracked)
loss(X, Y) = 0.7304719f0 (tracked)
loss(X, Y) = 0.73056227f0 (tracked)
loss(X, Y) = 0.722101f0 (tracked)
loss(X, Y) = 0.77372503f0 (tracked)
loss(X, Y) = 0.73546195f0 (tracked)
loss(X, Y) = 0.80458194f0 (tracked)
loss(X, Y) = 0.73708063f0 (tracked)
loss(X, Y) = 0.77611214f0 (tracked)
loss(X, Y) = 0.74830735f0 (tracked)
loss(X, Y) = 0.7152305f0 (tracked)
loss(X, Y) = 0.7125095f0 (tracked)
loss(X, Y) = 0.71941626f0 (tracked)
loss(X, Y) = 0.7273809f0 (tracked)
loss(X, Y) = 0.7537164f0 (tracked)
loss(X, Y) = 0.7504297f0 (tracked)
loss(X, Y) = 0.73555243f0 (tracked)
loss(X, Y) = 0.71781033f0 (tracked)
loss(X, Y) = 0.74129134f0 (tracked)
loss(X, Y) = 0.77056265f0 (tracked)
loss(X, Y) = 0.7391844f0 (tracked)
loss(X, Y) = 0.7201659f0 (tracked)
```

```
┌ Info: Epoch 528
└ @ Main /Users/kcft114/.julia/packages/Flux/qXNjB/src/optimise/train.jl:105
┌ Info: Epoch 529
└ @ Main /Users/kcft114/.julia/packages/Flux/qXNjB/src/optimise/train.jl:105
┌ Info: Epoch 530
└ @ Main /Users/kcft114/.julia/packages/Flux/qXNjB/src/optimise/train.jl:105
┌ Info: Epoch 531
└ @ Main /Users/kcft114/.julia/packages/Flux/qXNjB/src/optimise/train.jl:105
┌ Info: Epoch 532
└ @ Main /Users/kcft114/.julia/packages/Flux/qXNjB/src/optimise/train.jl:105
┌ Info: Epoch 533
└ @ Main /Users/kcft114/.julia/packages/Flux/qXNjB/src/optimise/train.jl:105
┌ Info: Epoch 534
└ @ Main /Users/kcft114/.julia/packages/Flux/qXNjB/src/optimise/train.jl:105
┌ Info: Epoch 535
└ @ Main /Users/kcft114/.julia/packages/Flux/qXNjB/src/optimise/train.jl:105
┌ Info: Epoch 536
└ @ Main /Users/kcft114/.julia/packages/Flux/qXNjB/src/optimise/train.jl:105
┌ Info: Epoch 537
└ @ Main /Users/kcft114/.julia/packages/Flux/qXNjB/src/optimise/train.jl:105
┌ Info: Epoch 538
└ @ Main /Users/kcft114/.julia/packages/Flux/qXNjB/src/optimise/train.jl:105
┌ Info: Epoch 539
└ @ Main /Users/kcft114/.julia/packages/Flux/qXNjB/src/optimise/train.jl:105
┌ Info: Epoch 540
└ @ Main /Users/kcft114/.julia/packages/Flux/qXNjB/src/optimise/train.jl:105
┌ Info: Epoch 541
└ @ Main /Users/kcft114/.julia/packages/Flux/qXNjB/src/optimise/train.jl:105
┌ Info: Epoch 542
└ @ Main /Users/kcft114/.julia/packages/Flux/qXNjB/src/optimise/train.jl:105
┌ Info: Epoch 543
└ @ Main /Users/kcft114/.julia/packages/Flux/qXNjB/src/optimise/train.jl:105
┌ Info: Epoch 544
└ @ Main /Users/kcft114/.julia/packages/Flux/qXNjB/src/optimise/train.jl:105
┌ Info: Epoch 545
└ @ Main /Users/kcft114/.julia/packages/Flux/qXNjB/src/optimise/train.jl:105
┌ Info: Epoch 546
└ @ Main /Users/kcft114/.julia/packages/Flux/qXNjB/src/optimise/train.jl:105
┌ Info: Epoch 547
└ @ Main /Users/kcft114/.julia/packages/Flux/qXNjB/src/optimise/train.jl:105
┌ Info: Epoch 548
└ @ Main /Users/kcft114/.julia/packages/Flux/qXNjB/src/optimise/train.jl:105
┌ Info: Epoch 549
└ @ Main /Users/kcft114/.julia/packages/Flux/qXNjB/src/optimise/train.jl:105
┌ Info: Epoch 550
└ @ Main /Users/kcft114/.julia/packages/Flux/qXNjB/src/optimise/train.jl:105
┌ Info: Epoch 551
└ @ Main /Users/kcft114/.julia/packages/Flux/qXNjB/src/optimise/train.jl:105
┌ Info: Epoch 552
└ @ Main /Users/kcft114/.julia/packages/Flux/qXNjB/src/optimise/train.jl:105
┌ Info: Epoch 553
└ @ Main /Users/kcft114/.julia/packages/Flux/qXNjB/src/optimise/train.jl:105
┌ Info: Epoch 554
└ @ Main /Users/kcft114/.julia/packages/Flux/qXNjB/src/optimise/train.jl:105
┌ Info: Epoch 555
└ @ Main /Users/kcft114/.julia/packages/Flux/qXNjB/src/optimise/train.jl:105
┌ Info: Epoch 556
└ @ Main /Users/kcft114/.julia/packages/Flux/qXNjB/src/optimise/train.jl:105
┌ Info: Epoch 557
└ @ Main /Users/kcft114/.julia/packages/Flux/qXNjB/src/optimise/train.jl:105
┌ Info: Epoch 558
└ @ Main /Users/kcft114/.julia/packages/Flux/qXNjB/src/optimise/train.jl:105
┌ Info: Epoch 559
└ @ Main /Users/kcft114/.julia/packages/Flux/qXNjB/src/optimise/train.jl:105
┌ Info: Epoch 560
└ @ Main /Users/kcft114/.julia/packages/Flux/qXNjB/src/optimise/train.jl:105
┌ Info: Epoch 561
└ @ Main /Users/kcft114/.julia/packages/Flux/qXNjB/src/optimise/train.jl:105
┌ Info: Epoch 562
└ @ Main /Users/kcft114/.julia/packages/Flux/qXNjB/src/optimise/train.jl:105
┌ Info: Epoch 563
└ @ Main /Users/kcft114/.julia/packages/Flux/qXNjB/src/optimise/train.jl:105
┌ Info: Epoch 564
└ @ Main /Users/kcft114/.julia/packages/Flux/qXNjB/src/optimise/train.jl:105
┌ Info: Epoch 565
└ @ Main /Users/kcft114/.julia/packages/Flux/qXNjB/src/optimise/train.jl:105
┌ Info: Epoch 566
└ @ Main /Users/kcft114/.julia/packages/Flux/qXNjB/src/optimise/train.jl:105
┌ Info: Epoch 567
└ @ Main /Users/kcft114/.julia/packages/Flux/qXNjB/src/optimise/train.jl:105
┌ Info: Epoch 568
└ @ Main /Users/kcft114/.julia/packages/Flux/qXNjB/src/optimise/train.jl:105
┌ Info: Epoch 569
└ @ Main /Users/kcft114/.julia/packages/Flux/qXNjB/src/optimise/train.jl:105
┌ Info: Epoch 570
└ @ Main /Users/kcft114/.julia/packages/Flux/qXNjB/src/optimise/train.jl:105
┌ Info: Epoch 571
└ @ Main /Users/kcft114/.julia/packages/Flux/qXNjB/src/optimise/train.jl:105
┌ Info: Epoch 572
└ @ Main /Users/kcft114/.julia/packages/Flux/qXNjB/src/optimise/train.jl:105
┌ Info: Epoch 573
└ @ Main /Users/kcft114/.julia/packages/Flux/qXNjB/src/optimise/train.jl:105
┌ Info: Epoch 574
└ @ Main /Users/kcft114/.julia/packages/Flux/qXNjB/src/optimise/train.jl:105
┌ Info: Epoch 575
└ @ Main /Users/kcft114/.julia/packages/Flux/qXNjB/src/optimise/train.jl:105
┌ Info: Epoch 576
└ @ Main /Users/kcft114/.julia/packages/Flux/qXNjB/src/optimise/train.jl:105
┌ Info: Epoch 577
└ @ Main /Users/kcft114/.julia/packages/Flux/qXNjB/src/optimise/train.jl:105
┌ Info: Epoch 578
└ @ Main /Users/kcft114/.julia/packages/Flux/qXNjB/src/optimise/train.jl:105
┌ Info: Epoch 579
└ @ Main /Users/kcft114/.julia/packages/Flux/qXNjB/src/optimise/train.jl:105
┌ Info: Epoch 580
└ @ Main /Users/kcft114/.julia/packages/Flux/qXNjB/src/optimise/train.jl:105
┌ Info: Epoch 581
└ @ Main /Users/kcft114/.julia/packages/Flux/qXNjB/src/optimise/train.jl:105
┌ Info: Epoch 582
└ @ Main /Users/kcft114/.julia/packages/Flux/qXNjB/src/optimise/train.jl:105
┌ Info: Epoch 583
└ @ Main /Users/kcft114/.julia/packages/Flux/qXNjB/src/optimise/train.jl:105
┌ Info: Epoch 584
└ @ Main /Users/kcft114/.julia/packages/Flux/qXNjB/src/optimise/train.jl:105
┌ Info: Epoch 585
└ @ Main /Users/kcft114/.julia/packages/Flux/qXNjB/src/optimise/train.jl:105
┌ Info: Epoch 586
└ @ Main /Users/kcft114/.julia/packages/Flux/qXNjB/src/optimise/train.jl:105
┌ Info: Epoch 587
└ @ Main /Users/kcft114/.julia/packages/Flux/qXNjB/src/optimise/train.jl:105
┌ Info: Epoch 588
└ @ Main /Users/kcft114/.julia/packages/Flux/qXNjB/src/optimise/train.jl:105
┌ Info: Epoch 589
└ @ Main /Users/kcft114/.julia/packages/Flux/qXNjB/src/optimise/train.jl:105
┌ Info: Epoch 590
└ @ Main /Users/kcft114/.julia/packages/Flux/qXNjB/src/optimise/train.jl:105
┌ Info: Epoch 591
└ @ Main /Users/kcft114/.julia/packages/Flux/qXNjB/src/optimise/train.jl:105
┌ Info: Epoch 592
└ @ Main /Users/kcft114/.julia/packages/Flux/qXNjB/src/optimise/train.jl:105
┌ Info: Epoch 593
└ @ Main /Users/kcft114/.julia/packages/Flux/qXNjB/src/optimise/train.jl:105
┌ Info: Epoch 594
└ @ Main /Users/kcft114/.julia/packages/Flux/qXNjB/src/optimise/train.jl:105
┌ Info: Epoch 595
└ @ Main /Users/kcft114/.julia/packages/Flux/qXNjB/src/optimise/train.jl:105
┌ Info: Epoch 596
└ @ Main /Users/kcft114/.julia/packages/Flux/qXNjB/src/optimise/train.jl:105
┌ Info: Epoch 597
└ @ Main /Users/kcft114/.julia/packages/Flux/qXNjB/src/optimise/train.jl:105
┌ Info: Epoch 598
└ @ Main /Users/kcft114/.julia/packages/Flux/qXNjB/src/optimise/train.jl:105
┌ Info: Epoch 599
└ @ Main /Users/kcft114/.julia/packages/Flux/qXNjB/src/optimise/train.jl:105
┌ Info: Epoch 600
└ @ Main /Users/kcft114/.julia/packages/Flux/qXNjB/src/optimise/train.jl:105
┌ Info: Epoch 601
└ @ Main /Users/kcft114/.julia/packages/Flux/qXNjB/src/optimise/train.jl:105
┌ Info: Epoch 602
└ @ Main /Users/kcft114/.julia/packages/Flux/qXNjB/src/optimise/train.jl:105
┌ Info: Epoch 603
└ @ Main /Users/kcft114/.julia/packages/Flux/qXNjB/src/optimise/train.jl:105
┌ Info: Epoch 604
└ @ Main /Users/kcft114/.julia/packages/Flux/qXNjB/src/optimise/train.jl:105
┌ Info: Epoch 605
└ @ Main /Users/kcft114/.julia/packages/Flux/qXNjB/src/optimise/train.jl:105
┌ Info: Epoch 606
└ @ Main /Users/kcft114/.julia/packages/Flux/qXNjB/src/optimise/train.jl:105
┌ Info: Epoch 607
└ @ Main /Users/kcft114/.julia/packages/Flux/qXNjB/src/optimise/train.jl:105
┌ Info: Epoch 608
└ @ Main /Users/kcft114/.julia/packages/Flux/qXNjB/src/optimise/train.jl:105
┌ Info: Epoch 609
└ @ Main /Users/kcft114/.julia/packages/Flux/qXNjB/src/optimise/train.jl:105
┌ Info: Epoch 610
└ @ Main /Users/kcft114/.julia/packages/Flux/qXNjB/src/optimise/train.jl:105
┌ Info: Epoch 611
└ @ Main /Users/kcft114/.julia/packages/Flux/qXNjB/src/optimise/train.jl:105
┌ Info: Epoch 612
└ @ Main /Users/kcft114/.julia/packages/Flux/qXNjB/src/optimise/train.jl:105
┌ Info: Epoch 613
└ @ Main /Users/kcft114/.julia/packages/Flux/qXNjB/src/optimise/train.jl:105
┌ Info: Epoch 614
└ @ Main /Users/kcft114/.julia/packages/Flux/qXNjB/src/optimise/train.jl:105
┌ Info: Epoch 615
└ @ Main /Users/kcft114/.julia/packages/Flux/qXNjB/src/optimise/train.jl:105
┌ Info: Epoch 616
└ @ Main /Users/kcft114/.julia/packages/Flux/qXNjB/src/optimise/train.jl:105
┌ Info: Epoch 617
└ @ Main /Users/kcft114/.julia/packages/Flux/qXNjB/src/optimise/train.jl:105
┌ Info: Epoch 618
└ @ Main /Users/kcft114/.julia/packages/Flux/qXNjB/src/optimise/train.jl:105
┌ Info: Epoch 619
└ @ Main /Users/kcft114/.julia/packages/Flux/qXNjB/src/optimise/train.jl:105
┌ Info: Epoch 620
└ @ Main /Users/kcft114/.julia/packages/Flux/qXNjB/src/optimise/train.jl:105
┌ Info: Epoch 621
└ @ Main /Users/kcft114/.julia/packages/Flux/qXNjB/src/optimise/train.jl:105
┌ Info: Epoch 622
└ @ Main /Users/kcft114/.julia/packages/Flux/qXNjB/src/optimise/train.jl:105
┌ Info: Epoch 623
└ @ Main /Users/kcft114/.julia/packages/Flux/qXNjB/src/optimise/train.jl:105
┌ Info: Epoch 624
└ @ Main /Users/kcft114/.julia/packages/Flux/qXNjB/src/optimise/train.jl:105
┌ Info: Epoch 625
└ @ Main /Users/kcft114/.julia/packages/Flux/qXNjB/src/optimise/train.jl:105
┌ Info: Epoch 626
└ @ Main /Users/kcft114/.julia/packages/Flux/qXNjB/src/optimise/train.jl:105
┌ Info: Epoch 627
└ @ Main /Users/kcft114/.julia/packages/Flux/qXNjB/src/optimise/train.jl:105
┌ Info: Epoch 628
└ @ Main /Users/kcft114/.julia/packages/Flux/qXNjB/src/optimise/train.jl:105
┌ Info: Epoch 629
└ @ Main /Users/kcft114/.julia/packages/Flux/qXNjB/src/optimise/train.jl:105
┌ Info: Epoch 630
└ @ Main /Users/kcft114/.julia/packages/Flux/qXNjB/src/optimise/train.jl:105
┌ Info: Epoch 631
└ @ Main /Users/kcft114/.julia/packages/Flux/qXNjB/src/optimise/train.jl:105
┌ Info: Epoch 632
└ @ Main /Users/kcft114/.julia/packages/Flux/qXNjB/src/optimise/train.jl:105
┌ Info: Epoch 633
└ @ Main /Users/kcft114/.julia/packages/Flux/qXNjB/src/optimise/train.jl:105
┌ Info: Epoch 634
└ @ Main /Users/kcft114/.julia/packages/Flux/qXNjB/src/optimise/train.jl:105
┌ Info: Epoch 635
└ @ Main /Users/kcft114/.julia/packages/Flux/qXNjB/src/optimise/train.jl:105
┌ Info: Epoch 636
└ @ Main /Users/kcft114/.julia/packages/Flux/qXNjB/src/optimise/train.jl:105
┌ Info: Epoch 637
└ @ Main /Users/kcft114/.julia/packages/Flux/qXNjB/src/optimise/train.jl:105
┌ Info: Epoch 638
└ @ Main /Users/kcft114/.julia/packages/Flux/qXNjB/src/optimise/train.jl:105
┌ Info: Epoch 639
└ @ Main /Users/kcft114/.julia/packages/Flux/qXNjB/src/optimise/train.jl:105
┌ Info: Epoch 640
└ @ Main /Users/kcft114/.julia/packages/Flux/qXNjB/src/optimise/train.jl:105
┌ Info: Epoch 641
└ @ Main /Users/kcft114/.julia/packages/Flux/qXNjB/src/optimise/train.jl:105
┌ Info: Epoch 642
└ @ Main /Users/kcft114/.julia/packages/Flux/qXNjB/src/optimise/train.jl:105
┌ Info: Epoch 643
└ @ Main /Users/kcft114/.julia/packages/Flux/qXNjB/src/optimise/train.jl:105
┌ Info: Epoch 644
└ @ Main /Users/kcft114/.julia/packages/Flux/qXNjB/src/optimise/train.jl:105
┌ Info: Epoch 645
└ @ Main /Users/kcft114/.julia/packages/Flux/qXNjB/src/optimise/train.jl:105
┌ Info: Epoch 646
└ @ Main /Users/kcft114/.julia/packages/Flux/qXNjB/src/optimise/train.jl:105
┌ Info: Epoch 647
└ @ Main /Users/kcft114/.julia/packages/Flux/qXNjB/src/optimise/train.jl:105
┌ Info: Epoch 648
└ @ Main /Users/kcft114/.julia/packages/Flux/qXNjB/src/optimise/train.jl:105
┌ Info: Epoch 649
└ @ Main /Users/kcft114/.julia/packages/Flux/qXNjB/src/optimise/train.jl:105
┌ Info: Epoch 650
└ @ Main /Users/kcft114/.julia/packages/Flux/qXNjB/src/optimise/train.jl:105
┌ Info: Epoch 651
└ @ Main /Users/kcft114/.julia/packages/Flux/qXNjB/src/optimise/train.jl:105
┌ Info: Epoch 652
└ @ Main /Users/kcft114/.julia/packages/Flux/qXNjB/src/optimise/train.jl:105
┌ Info: Epoch 653
└ @ Main /Users/kcft114/.julia/packages/Flux/qXNjB/src/optimise/train.jl:105
┌ Info: Epoch 654
└ @ Main /Users/kcft114/.julia/packages/Flux/qXNjB/src/optimise/train.jl:105
┌ Info: Epoch 655
└ @ Main /Users/kcft114/.julia/packages/Flux/qXNjB/src/optimise/train.jl:105
┌ Info: Epoch 656
└ @ Main /Users/kcft114/.julia/packages/Flux/qXNjB/src/optimise/train.jl:105
┌ Info: Epoch 657
└ @ Main /Users/kcft114/.julia/packages/Flux/qXNjB/src/optimise/train.jl:105
┌ Info: Epoch 658
└ @ Main /Users/kcft114/.julia/packages/Flux/qXNjB/src/optimise/train.jl:105
┌ Info: Epoch 659
└ @ Main /Users/kcft114/.julia/packages/Flux/qXNjB/src/optimise/train.jl:105
┌ Info: Epoch 660
└ @ Main /Users/kcft114/.julia/packages/Flux/qXNjB/src/optimise/train.jl:105
┌ Info: Epoch 661
└ @ Main /Users/kcft114/.julia/packages/Flux/qXNjB/src/optimise/train.jl:105
┌ Info: Epoch 662
└ @ Main /Users/kcft114/.julia/packages/Flux/qXNjB/src/optimise/train.jl:105
┌ Info: Epoch 663
└ @ Main /Users/kcft114/.julia/packages/Flux/qXNjB/src/optimise/train.jl:105
┌ Info: Epoch 664
└ @ Main /Users/kcft114/.julia/packages/Flux/qXNjB/src/optimise/train.jl:105
┌ Info: Epoch 665
└ @ Main /Users/kcft114/.julia/packages/Flux/qXNjB/src/optimise/train.jl:105
┌ Info: Epoch 666
└ @ Main /Users/kcft114/.julia/packages/Flux/qXNjB/src/optimise/train.jl:105
┌ Info: Epoch 667
└ @ Main /Users/kcft114/.julia/packages/Flux/qXNjB/src/optimise/train.jl:105
┌ Info: Epoch 668
└ @ Main /Users/kcft114/.julia/packages/Flux/qXNjB/src/optimise/train.jl:105
┌ Info: Epoch 669
└ @ Main /Users/kcft114/.julia/packages/Flux/qXNjB/src/optimise/train.jl:105
┌ Info: Epoch 670
└ @ Main /Users/kcft114/.julia/packages/Flux/qXNjB/src/optimise/train.jl:105
┌ Info: Epoch 671
└ @ Main /Users/kcft114/.julia/packages/Flux/qXNjB/src/optimise/train.jl:105
┌ Info: Epoch 672
└ @ Main /Users/kcft114/.julia/packages/Flux/qXNjB/src/optimise/train.jl:105
┌ Info: Epoch 673
└ @ Main /Users/kcft114/.julia/packages/Flux/qXNjB/src/optimise/train.jl:105
┌ Info: Epoch 674
└ @ Main /Users/kcft114/.julia/packages/Flux/qXNjB/src/optimise/train.jl:105
┌ Info: Epoch 675
└ @ Main /Users/kcft114/.julia/packages/Flux/qXNjB/src/optimise/train.jl:105
┌ Info: Epoch 676
└ @ Main /Users/kcft114/.julia/packages/Flux/qXNjB/src/optimise/train.jl:105
┌ Info: Epoch 677
└ @ Main /Users/kcft114/.julia/packages/Flux/qXNjB/src/optimise/train.jl:105
┌ Info: Epoch 678
└ @ Main /Users/kcft114/.julia/packages/Flux/qXNjB/src/optimise/train.jl:105
┌ Info: Epoch 679
└ @ Main /Users/kcft114/.julia/packages/Flux/qXNjB/src/optimise/train.jl:105
┌ Info: Epoch 680
└ @ Main /Users/kcft114/.julia/packages/Flux/qXNjB/src/optimise/train.jl:105
┌ Info: Epoch 681
└ @ Main /Users/kcft114/.julia/packages/Flux/qXNjB/src/optimise/train.jl:105
┌ Info: Epoch 682
└ @ Main /Users/kcft114/.julia/packages/Flux/qXNjB/src/optimise/train.jl:105
┌ Info: Epoch 683
└ @ Main /Users/kcft114/.julia/packages/Flux/qXNjB/src/optimise/train.jl:105
┌ Info: Epoch 684
└ @ Main /Users/kcft114/.julia/packages/Flux/qXNjB/src/optimise/train.jl:105
┌ Info: Epoch 685
└ @ Main /Users/kcft114/.julia/packages/Flux/qXNjB/src/optimise/train.jl:105
┌ Info: Epoch 686
└ @ Main /Users/kcft114/.julia/packages/Flux/qXNjB/src/optimise/train.jl:105
┌ Info: Epoch 687
└ @ Main /Users/kcft114/.julia/packages/Flux/qXNjB/src/optimise/train.jl:105
┌ Info: Epoch 688
└ @ Main /Users/kcft114/.julia/packages/Flux/qXNjB/src/optimise/train.jl:105
┌ Info: Epoch 689
└ @ Main /Users/kcft114/.julia/packages/Flux/qXNjB/src/optimise/train.jl:105
┌ Info: Epoch 690
└ @ Main /Users/kcft114/.julia/packages/Flux/qXNjB/src/optimise/train.jl:105
┌ Info: Epoch 691
└ @ Main /Users/kcft114/.julia/packages/Flux/qXNjB/src/optimise/train.jl:105
┌ Info: Epoch 692
└ @ Main /Users/kcft114/.julia/packages/Flux/qXNjB/src/optimise/train.jl:105
┌ Info: Epoch 693
└ @ Main /Users/kcft114/.julia/packages/Flux/qXNjB/src/optimise/train.jl:105
┌ Info: Epoch 694
└ @ Main /Users/kcft114/.julia/packages/Flux/qXNjB/src/optimise/train.jl:105
┌ Info: Epoch 695
└ @ Main /Users/kcft114/.julia/packages/Flux/qXNjB/src/optimise/train.jl:105
┌ Info: Epoch 696
└ @ Main /Users/kcft114/.julia/packages/Flux/qXNjB/src/optimise/train.jl:105
┌ Info: Epoch 697
└ @ Main /Users/kcft114/.julia/packages/Flux/qXNjB/src/optimise/train.jl:105
┌ Info: Epoch 698
└ @ Main /Users/kcft114/.julia/packages/Flux/qXNjB/src/optimise/train.jl:105
┌ Info: Epoch 699
└ @ Main /Users/kcft114/.julia/packages/Flux/qXNjB/src/optimise/train.jl:105
┌ Info: Epoch 700
└ @ Main /Users/kcft114/.julia/packages/Flux/qXNjB/src/optimise/train.jl:105
┌ Info: Epoch 701
└ @ Main /Users/kcft114/.julia/packages/Flux/qXNjB/src/optimise/train.jl:105
┌ Info: Epoch 702
└ @ Main /Users/kcft114/.julia/packages/Flux/qXNjB/src/optimise/train.jl:105
┌ Info: Epoch 703
└ @ Main /Users/kcft114/.julia/packages/Flux/qXNjB/src/optimise/train.jl:105
┌ Info: Epoch 704
└ @ Main /Users/kcft114/.julia/packages/Flux/qXNjB/src/optimise/train.jl:105
┌ Info: Epoch 705
└ @ Main /Users/kcft114/.julia/packages/Flux/qXNjB/src/optimise/train.jl:105
┌ Info: Epoch 706
└ @ Main /Users/kcft114/.julia/packages/Flux/qXNjB/src/optimise/train.jl:105
┌ Info: Epoch 707
└ @ Main /Users/kcft114/.julia/packages/Flux/qXNjB/src/optimise/train.jl:105
┌ Info: Epoch 708
└ @ Main /Users/kcft114/.julia/packages/Flux/qXNjB/src/optimise/train.jl:105
┌ Info: Epoch 709
└ @ Main /Users/kcft114/.julia/packages/Flux/qXNjB/src/optimise/train.jl:105
┌ Info: Epoch 710
└ @ Main /Users/kcft114/.julia/packages/Flux/qXNjB/src/optimise/train.jl:105
┌ Info: Epoch 711
└ @ Main /Users/kcft114/.julia/packages/Flux/qXNjB/src/optimise/train.jl:105
```

```
loss(X, Y) = 0.7733187f0 (tracked)
loss(X, Y) = 0.7039687f0 (tracked)
loss(X, Y) = 0.7515414f0 (tracked)
loss(X, Y) = 0.73083895f0 (tracked)
loss(X, Y) = 0.7665408f0 (tracked)
loss(X, Y) = 0.75864565f0 (tracked)
loss(X, Y) = 0.7565176f0 (tracked)
loss(X, Y) = 0.740115f0 (tracked)
loss(X, Y) = 0.7095126f0 (tracked)
loss(X, Y) = 0.7225706f0 (tracked)
loss(X, Y) = 0.7600797f0 (tracked)
loss(X, Y) = 0.72207844f0 (tracked)
loss(X, Y) = 0.70342356f0 (tracked)
loss(X, Y) = 0.73068035f0 (tracked)
loss(X, Y) = 0.7400701f0 (tracked)
loss(X, Y) = 0.7157904f0 (tracked)
loss(X, Y) = 0.7427011f0 (tracked)
loss(X, Y) = 0.77508223f0 (tracked)
loss(X, Y) = 0.74357027f0 (tracked)
loss(X, Y) = 0.7144254f0 (tracked)
loss(X, Y) = 0.7442501f0 (tracked)
loss(X, Y) = 0.7406966f0 (tracked)
loss(X, Y) = 0.70667017f0 (tracked)
loss(X, Y) = 0.7684857f0 (tracked)
loss(X, Y) = 0.7273827f0 (tracked)
loss(X, Y) = 0.7440459f0 (tracked)
loss(X, Y) = 0.7247026f0 (tracked)
loss(X, Y) = 0.7338699f0 (tracked)
loss(X, Y) = 0.72227085f0 (tracked)
loss(X, Y) = 0.7082627f0 (tracked)
loss(X, Y) = 0.7528664f0 (tracked)
loss(X, Y) = 0.71666336f0 (tracked)
loss(X, Y) = 0.75377464f0 (tracked)
loss(X, Y) = 0.7145989f0 (tracked)
loss(X, Y) = 0.74024206f0 (tracked)
loss(X, Y) = 0.7088094f0 (tracked)
loss(X, Y) = 0.743436f0 (tracked)
loss(X, Y) = 0.7321192f0 (tracked)
loss(X, Y) = 0.72824633f0 (tracked)
loss(X, Y) = 0.7302763f0 (tracked)
loss(X, Y) = 0.72020185f0 (tracked)
loss(X, Y) = 0.7329098f0 (tracked)
loss(X, Y) = 0.72347575f0 (tracked)
loss(X, Y) = 0.7011565f0 (tracked)
loss(X, Y) = 0.6811183f0 (tracked)
loss(X, Y) = 0.7312149f0 (tracked)
loss(X, Y) = 0.7461938f0 (tracked)
loss(X, Y) = 0.7646711f0 (tracked)
loss(X, Y) = 0.7841954f0 (tracked)
loss(X, Y) = 0.7077831f0 (tracked)
loss(X, Y) = 0.71341175f0 (tracked)
loss(X, Y) = 0.68391246f0 (tracked)
loss(X, Y) = 0.7681564f0 (tracked)
loss(X, Y) = 0.73760366f0 (tracked)
loss(X, Y) = 0.73895675f0 (tracked)
loss(X, Y) = 0.7088361f0 (tracked)
loss(X, Y) = 0.70196545f0 (tracked)
loss(X, Y) = 0.7651288f0 (tracked)
loss(X, Y) = 0.74042463f0 (tracked)
loss(X, Y) = 0.73337847f0 (tracked)
loss(X, Y) = 0.713479f0 (tracked)
loss(X, Y) = 0.72417027f0 (tracked)
loss(X, Y) = 0.7000363f0 (tracked)
loss(X, Y) = 0.7181434f0 (tracked)
loss(X, Y) = 0.7866115f0 (tracked)
loss(X, Y) = 0.7384211f0 (tracked)
loss(X, Y) = 0.7218398f0 (tracked)
loss(X, Y) = 0.73533064f0 (tracked)
loss(X, Y) = 0.70769995f0 (tracked)
loss(X, Y) = 0.713083f0 (tracked)
loss(X, Y) = 0.70625633f0 (tracked)
loss(X, Y) = 0.73935395f0 (tracked)
loss(X, Y) = 0.721373f0 (tracked)
loss(X, Y) = 0.7692492f0 (tracked)
loss(X, Y) = 0.7277862f0 (tracked)
loss(X, Y) = 0.73637056f0 (tracked)
loss(X, Y) = 0.73405474f0 (tracked)
loss(X, Y) = 0.7342775f0 (tracked)
loss(X, Y) = 0.7414967f0 (tracked)
loss(X, Y) = 0.7005826f0 (tracked)
loss(X, Y) = 0.71261144f0 (tracked)
loss(X, Y) = 0.76091206f0 (tracked)
loss(X, Y) = 0.7604916f0 (tracked)
loss(X, Y) = 0.7032497f0 (tracked)
loss(X, Y) = 0.71181285f0 (tracked)
loss(X, Y) = 0.7320068f0 (tracked)
loss(X, Y) = 0.7286856f0 (tracked)
loss(X, Y) = 0.718248f0 (tracked)
loss(X, Y) = 0.7522193f0 (tracked)
loss(X, Y) = 0.6949429f0 (tracked)
loss(X, Y) = 0.7299261f0 (tracked)
loss(X, Y) = 0.72835124f0 (tracked)
loss(X, Y) = 0.7067801f0 (tracked)
loss(X, Y) = 0.7474545f0 (tracked)
loss(X, Y) = 0.78239065f0 (tracked)
loss(X, Y) = 0.7537122f0 (tracked)
loss(X, Y) = 0.6787727f0 (tracked)
loss(X, Y) = 0.7470605f0 (tracked)
loss(X, Y) = 0.75448847f0 (tracked)
loss(X, Y) = 0.7344839f0 (tracked)
loss(X, Y) = 0.6761012f0 (tracked)
loss(X, Y) = 0.74980044f0 (tracked)
loss(X, Y) = 0.70945275f0 (tracked)
loss(X, Y) = 0.67769754f0 (tracked)
loss(X, Y) = 0.7607836f0 (tracked)
```

```
┌ Info: Epoch 712
└ @ Main /Users/kcft114/.julia/packages/Flux/qXNjB/src/optimise/train.jl:105
┌ Info: Epoch 713
└ @ Main /Users/kcft114/.julia/packages/Flux/qXNjB/src/optimise/train.jl:105
┌ Info: Epoch 714
└ @ Main /Users/kcft114/.julia/packages/Flux/qXNjB/src/optimise/train.jl:105
┌ Info: Epoch 715
└ @ Main /Users/kcft114/.julia/packages/Flux/qXNjB/src/optimise/train.jl:105
┌ Info: Epoch 716
└ @ Main /Users/kcft114/.julia/packages/Flux/qXNjB/src/optimise/train.jl:105
┌ Info: Epoch 717
└ @ Main /Users/kcft114/.julia/packages/Flux/qXNjB/src/optimise/train.jl:105
┌ Info: Epoch 718
└ @ Main /Users/kcft114/.julia/packages/Flux/qXNjB/src/optimise/train.jl:105
┌ Info: Epoch 719
└ @ Main /Users/kcft114/.julia/packages/Flux/qXNjB/src/optimise/train.jl:105
┌ Info: Epoch 720
└ @ Main /Users/kcft114/.julia/packages/Flux/qXNjB/src/optimise/train.jl:105
┌ Info: Epoch 721
└ @ Main /Users/kcft114/.julia/packages/Flux/qXNjB/src/optimise/train.jl:105
┌ Info: Epoch 722
└ @ Main /Users/kcft114/.julia/packages/Flux/qXNjB/src/optimise/train.jl:105
┌ Info: Epoch 723
└ @ Main /Users/kcft114/.julia/packages/Flux/qXNjB/src/optimise/train.jl:105
┌ Info: Epoch 724
└ @ Main /Users/kcft114/.julia/packages/Flux/qXNjB/src/optimise/train.jl:105
┌ Info: Epoch 725
└ @ Main /Users/kcft114/.julia/packages/Flux/qXNjB/src/optimise/train.jl:105
┌ Info: Epoch 726
└ @ Main /Users/kcft114/.julia/packages/Flux/qXNjB/src/optimise/train.jl:105
┌ Info: Epoch 727
└ @ Main /Users/kcft114/.julia/packages/Flux/qXNjB/src/optimise/train.jl:105
┌ Info: Epoch 728
└ @ Main /Users/kcft114/.julia/packages/Flux/qXNjB/src/optimise/train.jl:105
┌ Info: Epoch 729
└ @ Main /Users/kcft114/.julia/packages/Flux/qXNjB/src/optimise/train.jl:105
┌ Info: Epoch 730
└ @ Main /Users/kcft114/.julia/packages/Flux/qXNjB/src/optimise/train.jl:105
┌ Info: Epoch 731
└ @ Main /Users/kcft114/.julia/packages/Flux/qXNjB/src/optimise/train.jl:105
┌ Info: Epoch 732
└ @ Main /Users/kcft114/.julia/packages/Flux/qXNjB/src/optimise/train.jl:105
┌ Info: Epoch 733
└ @ Main /Users/kcft114/.julia/packages/Flux/qXNjB/src/optimise/train.jl:105
┌ Info: Epoch 734
└ @ Main /Users/kcft114/.julia/packages/Flux/qXNjB/src/optimise/train.jl:105
┌ Info: Epoch 735
└ @ Main /Users/kcft114/.julia/packages/Flux/qXNjB/src/optimise/train.jl:105
┌ Info: Epoch 736
└ @ Main /Users/kcft114/.julia/packages/Flux/qXNjB/src/optimise/train.jl:105
┌ Info: Epoch 737
└ @ Main /Users/kcft114/.julia/packages/Flux/qXNjB/src/optimise/train.jl:105
┌ Info: Epoch 738
└ @ Main /Users/kcft114/.julia/packages/Flux/qXNjB/src/optimise/train.jl:105
┌ Info: Epoch 739
└ @ Main /Users/kcft114/.julia/packages/Flux/qXNjB/src/optimise/train.jl:105
┌ Info: Epoch 740
└ @ Main /Users/kcft114/.julia/packages/Flux/qXNjB/src/optimise/train.jl:105
┌ Info: Epoch 741
└ @ Main /Users/kcft114/.julia/packages/Flux/qXNjB/src/optimise/train.jl:105
┌ Info: Epoch 742
└ @ Main /Users/kcft114/.julia/packages/Flux/qXNjB/src/optimise/train.jl:105
┌ Info: Epoch 743
└ @ Main /Users/kcft114/.julia/packages/Flux/qXNjB/src/optimise/train.jl:105
┌ Info: Epoch 744
└ @ Main /Users/kcft114/.julia/packages/Flux/qXNjB/src/optimise/train.jl:105
┌ Info: Epoch 745
└ @ Main /Users/kcft114/.julia/packages/Flux/qXNjB/src/optimise/train.jl:105
┌ Info: Epoch 746
└ @ Main /Users/kcft114/.julia/packages/Flux/qXNjB/src/optimise/train.jl:105
┌ Info: Epoch 747
└ @ Main /Users/kcft114/.julia/packages/Flux/qXNjB/src/optimise/train.jl:105
┌ Info: Epoch 748
└ @ Main /Users/kcft114/.julia/packages/Flux/qXNjB/src/optimise/train.jl:105
┌ Info: Epoch 749
└ @ Main /Users/kcft114/.julia/packages/Flux/qXNjB/src/optimise/train.jl:105
┌ Info: Epoch 750
└ @ Main /Users/kcft114/.julia/packages/Flux/qXNjB/src/optimise/train.jl:105
┌ Info: Epoch 751
└ @ Main /Users/kcft114/.julia/packages/Flux/qXNjB/src/optimise/train.jl:105
┌ Info: Epoch 752
└ @ Main /Users/kcft114/.julia/packages/Flux/qXNjB/src/optimise/train.jl:105
┌ Info: Epoch 753
└ @ Main /Users/kcft114/.julia/packages/Flux/qXNjB/src/optimise/train.jl:105
┌ Info: Epoch 754
└ @ Main /Users/kcft114/.julia/packages/Flux/qXNjB/src/optimise/train.jl:105
┌ Info: Epoch 755
└ @ Main /Users/kcft114/.julia/packages/Flux/qXNjB/src/optimise/train.jl:105
┌ Info: Epoch 756
└ @ Main /Users/kcft114/.julia/packages/Flux/qXNjB/src/optimise/train.jl:105
┌ Info: Epoch 757
└ @ Main /Users/kcft114/.julia/packages/Flux/qXNjB/src/optimise/train.jl:105
┌ Info: Epoch 758
└ @ Main /Users/kcft114/.julia/packages/Flux/qXNjB/src/optimise/train.jl:105
┌ Info: Epoch 759
└ @ Main /Users/kcft114/.julia/packages/Flux/qXNjB/src/optimise/train.jl:105
┌ Info: Epoch 760
└ @ Main /Users/kcft114/.julia/packages/Flux/qXNjB/src/optimise/train.jl:105
┌ Info: Epoch 761
└ @ Main /Users/kcft114/.julia/packages/Flux/qXNjB/src/optimise/train.jl:105
┌ Info: Epoch 762
└ @ Main /Users/kcft114/.julia/packages/Flux/qXNjB/src/optimise/train.jl:105
┌ Info: Epoch 763
└ @ Main /Users/kcft114/.julia/packages/Flux/qXNjB/src/optimise/train.jl:105
┌ Info: Epoch 764
└ @ Main /Users/kcft114/.julia/packages/Flux/qXNjB/src/optimise/train.jl:105
┌ Info: Epoch 765
└ @ Main /Users/kcft114/.julia/packages/Flux/qXNjB/src/optimise/train.jl:105
┌ Info: Epoch 766
└ @ Main /Users/kcft114/.julia/packages/Flux/qXNjB/src/optimise/train.jl:105
┌ Info: Epoch 767
└ @ Main /Users/kcft114/.julia/packages/Flux/qXNjB/src/optimise/train.jl:105
┌ Info: Epoch 768
└ @ Main /Users/kcft114/.julia/packages/Flux/qXNjB/src/optimise/train.jl:105
┌ Info: Epoch 769
└ @ Main /Users/kcft114/.julia/packages/Flux/qXNjB/src/optimise/train.jl:105
┌ Info: Epoch 770
└ @ Main /Users/kcft114/.julia/packages/Flux/qXNjB/src/optimise/train.jl:105
┌ Info: Epoch 771
└ @ Main /Users/kcft114/.julia/packages/Flux/qXNjB/src/optimise/train.jl:105
┌ Info: Epoch 772
└ @ Main /Users/kcft114/.julia/packages/Flux/qXNjB/src/optimise/train.jl:105
┌ Info: Epoch 773
└ @ Main /Users/kcft114/.julia/packages/Flux/qXNjB/src/optimise/train.jl:105
┌ Info: Epoch 774
└ @ Main /Users/kcft114/.julia/packages/Flux/qXNjB/src/optimise/train.jl:105
┌ Info: Epoch 775
└ @ Main /Users/kcft114/.julia/packages/Flux/qXNjB/src/optimise/train.jl:105
┌ Info: Epoch 776
└ @ Main /Users/kcft114/.julia/packages/Flux/qXNjB/src/optimise/train.jl:105
┌ Info: Epoch 777
└ @ Main /Users/kcft114/.julia/packages/Flux/qXNjB/src/optimise/train.jl:105
┌ Info: Epoch 778
└ @ Main /Users/kcft114/.julia/packages/Flux/qXNjB/src/optimise/train.jl:105
┌ Info: Epoch 779
└ @ Main /Users/kcft114/.julia/packages/Flux/qXNjB/src/optimise/train.jl:105
┌ Info: Epoch 780
└ @ Main /Users/kcft114/.julia/packages/Flux/qXNjB/src/optimise/train.jl:105
┌ Info: Epoch 781
└ @ Main /Users/kcft114/.julia/packages/Flux/qXNjB/src/optimise/train.jl:105
┌ Info: Epoch 782
└ @ Main /Users/kcft114/.julia/packages/Flux/qXNjB/src/optimise/train.jl:105
┌ Info: Epoch 783
└ @ Main /Users/kcft114/.julia/packages/Flux/qXNjB/src/optimise/train.jl:105
┌ Info: Epoch 784
└ @ Main /Users/kcft114/.julia/packages/Flux/qXNjB/src/optimise/train.jl:105
┌ Info: Epoch 785
└ @ Main /Users/kcft114/.julia/packages/Flux/qXNjB/src/optimise/train.jl:105
┌ Info: Epoch 786
└ @ Main /Users/kcft114/.julia/packages/Flux/qXNjB/src/optimise/train.jl:105
┌ Info: Epoch 787
└ @ Main /Users/kcft114/.julia/packages/Flux/qXNjB/src/optimise/train.jl:105
┌ Info: Epoch 788
└ @ Main /Users/kcft114/.julia/packages/Flux/qXNjB/src/optimise/train.jl:105
┌ Info: Epoch 789
└ @ Main /Users/kcft114/.julia/packages/Flux/qXNjB/src/optimise/train.jl:105
┌ Info: Epoch 790
└ @ Main /Users/kcft114/.julia/packages/Flux/qXNjB/src/optimise/train.jl:105
┌ Info: Epoch 791
└ @ Main /Users/kcft114/.julia/packages/Flux/qXNjB/src/optimise/train.jl:105
┌ Info: Epoch 792
└ @ Main /Users/kcft114/.julia/packages/Flux/qXNjB/src/optimise/train.jl:105
┌ Info: Epoch 793
└ @ Main /Users/kcft114/.julia/packages/Flux/qXNjB/src/optimise/train.jl:105
┌ Info: Epoch 794
└ @ Main /Users/kcft114/.julia/packages/Flux/qXNjB/src/optimise/train.jl:105
┌ Info: Epoch 795
└ @ Main /Users/kcft114/.julia/packages/Flux/qXNjB/src/optimise/train.jl:105
┌ Info: Epoch 796
└ @ Main /Users/kcft114/.julia/packages/Flux/qXNjB/src/optimise/train.jl:105
┌ Info: Epoch 797
└ @ Main /Users/kcft114/.julia/packages/Flux/qXNjB/src/optimise/train.jl:105
┌ Info: Epoch 798
└ @ Main /Users/kcft114/.julia/packages/Flux/qXNjB/src/optimise/train.jl:105
┌ Info: Epoch 799
└ @ Main /Users/kcft114/.julia/packages/Flux/qXNjB/src/optimise/train.jl:105
┌ Info: Epoch 800
└ @ Main /Users/kcft114/.julia/packages/Flux/qXNjB/src/optimise/train.jl:105
┌ Info: Epoch 801
└ @ Main /Users/kcft114/.julia/packages/Flux/qXNjB/src/optimise/train.jl:105
┌ Info: Epoch 802
└ @ Main /Users/kcft114/.julia/packages/Flux/qXNjB/src/optimise/train.jl:105
┌ Info: Epoch 803
└ @ Main /Users/kcft114/.julia/packages/Flux/qXNjB/src/optimise/train.jl:105
┌ Info: Epoch 804
└ @ Main /Users/kcft114/.julia/packages/Flux/qXNjB/src/optimise/train.jl:105
┌ Info: Epoch 805
└ @ Main /Users/kcft114/.julia/packages/Flux/qXNjB/src/optimise/train.jl:105
┌ Info: Epoch 806
└ @ Main /Users/kcft114/.julia/packages/Flux/qXNjB/src/optimise/train.jl:105
┌ Info: Epoch 807
└ @ Main /Users/kcft114/.julia/packages/Flux/qXNjB/src/optimise/train.jl:105
┌ Info: Epoch 808
└ @ Main /Users/kcft114/.julia/packages/Flux/qXNjB/src/optimise/train.jl:105
┌ Info: Epoch 809
└ @ Main /Users/kcft114/.julia/packages/Flux/qXNjB/src/optimise/train.jl:105
┌ Info: Epoch 810
└ @ Main /Users/kcft114/.julia/packages/Flux/qXNjB/src/optimise/train.jl:105
┌ Info: Epoch 811
└ @ Main /Users/kcft114/.julia/packages/Flux/qXNjB/src/optimise/train.jl:105
┌ Info: Epoch 812
└ @ Main /Users/kcft114/.julia/packages/Flux/qXNjB/src/optimise/train.jl:105
┌ Info: Epoch 813
└ @ Main /Users/kcft114/.julia/packages/Flux/qXNjB/src/optimise/train.jl:105
┌ Info: Epoch 814
└ @ Main /Users/kcft114/.julia/packages/Flux/qXNjB/src/optimise/train.jl:105
┌ Info: Epoch 815
└ @ Main /Users/kcft114/.julia/packages/Flux/qXNjB/src/optimise/train.jl:105
┌ Info: Epoch 816
└ @ Main /Users/kcft114/.julia/packages/Flux/qXNjB/src/optimise/train.jl:105
```

```
loss(X, Y) = 0.6922104f0 (tracked)
loss(X, Y) = 0.7473677f0 (tracked)
loss(X, Y) = 0.71103126f0 (tracked)
loss(X, Y) = 0.68093365f0 (tracked)
loss(X, Y) = 0.72543275f0 (tracked)
loss(X, Y) = 0.72837174f0 (tracked)
loss(X, Y) = 0.74800116f0 (tracked)
loss(X, Y) = 0.76573056f0 (tracked)
loss(X, Y) = 0.7367984f0 (tracked)
loss(X, Y) = 0.68340766f0 (tracked)
loss(X, Y) = 0.7358576f0 (tracked)
loss(X, Y) = 0.7217064f0 (tracked)
loss(X, Y) = 0.70777905f0 (tracked)
loss(X, Y) = 0.7124905f0 (tracked)
loss(X, Y) = 0.73052835f0 (tracked)
loss(X, Y) = 0.7095141f0 (tracked)
loss(X, Y) = 0.7257066f0 (tracked)
loss(X, Y) = 0.72542894f0 (tracked)
loss(X, Y) = 0.76194006f0 (tracked)
loss(X, Y) = 0.6892919f0 (tracked)
loss(X, Y) = 0.71858895f0 (tracked)
loss(X, Y) = 0.7235664f0 (tracked)
loss(X, Y) = 0.7268888f0 (tracked)
loss(X, Y) = 0.67814153f0 (tracked)
loss(X, Y) = 0.71437246f0 (tracked)
loss(X, Y) = 0.70541954f0 (tracked)
loss(X, Y) = 0.73448807f0 (tracked)
loss(X, Y) = 0.6772947f0 (tracked)
loss(X, Y) = 0.7202785f0 (tracked)
loss(X, Y) = 0.7347961f0 (tracked)
loss(X, Y) = 0.7025054f0 (tracked)
loss(X, Y) = 0.73656714f0 (tracked)
loss(X, Y) = 0.7138653f0 (tracked)
loss(X, Y) = 0.69457453f0 (tracked)
loss(X, Y) = 0.7094671f0 (tracked)
loss(X, Y) = 0.7384987f0 (tracked)
loss(X, Y) = 0.76758724f0 (tracked)
loss(X, Y) = 0.6971399f0 (tracked)
loss(X, Y) = 0.7092683f0 (tracked)
loss(X, Y) = 0.7217919f0 (tracked)
loss(X, Y) = 0.71540517f0 (tracked)
loss(X, Y) = 0.71751976f0 (tracked)
loss(X, Y) = 0.7263364f0 (tracked)
loss(X, Y) = 0.73234147f0 (tracked)
loss(X, Y) = 0.70991266f0 (tracked)
loss(X, Y) = 0.70873004f0 (tracked)
loss(X, Y) = 0.7336822f0 (tracked)
loss(X, Y) = 0.713706f0 (tracked)
loss(X, Y) = 0.7536578f0 (tracked)
loss(X, Y) = 0.70653063f0 (tracked)
loss(X, Y) = 0.70677006f0 (tracked)
loss(X, Y) = 0.72858924f0 (tracked)
loss(X, Y) = 0.68342197f0 (tracked)
loss(X, Y) = 0.70033395f0 (tracked)
loss(X, Y) = 0.7250937f0 (tracked)
loss(X, Y) = 0.68512285f0 (tracked)
loss(X, Y) = 0.75631917f0 (tracked)
loss(X, Y) = 0.73532623f0 (tracked)
loss(X, Y) = 0.66119814f0 (tracked)
loss(X, Y) = 0.7346773f0 (tracked)
loss(X, Y) = 0.749409f0 (tracked)
loss(X, Y) = 0.68341744f0 (tracked)
loss(X, Y) = 0.7276292f0 (tracked)
loss(X, Y) = 0.67245364f0 (tracked)
loss(X, Y) = 0.7112132f0 (tracked)
loss(X, Y) = 0.7324226f0 (tracked)
loss(X, Y) = 0.68268174f0 (tracked)
loss(X, Y) = 0.7026228f0 (tracked)
loss(X, Y) = 0.7176752f0 (tracked)
loss(X, Y) = 0.67181945f0 (tracked)
loss(X, Y) = 0.71901315f0 (tracked)
loss(X, Y) = 0.6837347f0 (tracked)
loss(X, Y) = 0.73547256f0 (tracked)
loss(X, Y) = 0.7010715f0 (tracked)
loss(X, Y) = 0.73292196f0 (tracked)
loss(X, Y) = 0.68408835f0 (tracked)
loss(X, Y) = 0.7090739f0 (tracked)
loss(X, Y) = 0.7247579f0 (tracked)
loss(X, Y) = 0.7286262f0 (tracked)
loss(X, Y) = 0.7138246f0 (tracked)
loss(X, Y) = 0.7104435f0 (tracked)
loss(X, Y) = 0.7167241f0 (tracked)
loss(X, Y) = 0.68698704f0 (tracked)
loss(X, Y) = 0.73183286f0 (tracked)
loss(X, Y) = 0.7480545f0 (tracked)
loss(X, Y) = 0.6900343f0 (tracked)
loss(X, Y) = 0.70385945f0 (tracked)
loss(X, Y) = 0.73436683f0 (tracked)
loss(X, Y) = 0.7270227f0 (tracked)
loss(X, Y) = 0.7273345f0 (tracked)
loss(X, Y) = 0.72931296f0 (tracked)
loss(X, Y) = 0.7150417f0 (tracked)
loss(X, Y) = 0.7162678f0 (tracked)
loss(X, Y) = 0.71413535f0 (tracked)
loss(X, Y) = 0.6993854f0 (tracked)
loss(X, Y) = 0.73226714f0 (tracked)
loss(X, Y) = 0.69950306f0 (tracked)
loss(X, Y) = 0.7001716f0 (tracked)
loss(X, Y) = 0.7459269f0 (tracked)
loss(X, Y) = 0.7372564f0 (tracked)
loss(X, Y) = 0.6878915f0 (tracked)
loss(X, Y) = 0.74370676f0 (tracked)
loss(X, Y) = 0.7318694f0 (tracked)
loss(X, Y) = 0.6858907f0 (tracked)
loss(X, Y) = 0.71278614f0 (tracked)
```

```
┌ Info: Epoch 817
└ @ Main /Users/kcft114/.julia/packages/Flux/qXNjB/src/optimise/train.jl:105
┌ Info: Epoch 818
└ @ Main /Users/kcft114/.julia/packages/Flux/qXNjB/src/optimise/train.jl:105
┌ Info: Epoch 819
└ @ Main /Users/kcft114/.julia/packages/Flux/qXNjB/src/optimise/train.jl:105
┌ Info: Epoch 820
└ @ Main /Users/kcft114/.julia/packages/Flux/qXNjB/src/optimise/train.jl:105
┌ Info: Epoch 821
└ @ Main /Users/kcft114/.julia/packages/Flux/qXNjB/src/optimise/train.jl:105
┌ Info: Epoch 822
└ @ Main /Users/kcft114/.julia/packages/Flux/qXNjB/src/optimise/train.jl:105
┌ Info: Epoch 823
└ @ Main /Users/kcft114/.julia/packages/Flux/qXNjB/src/optimise/train.jl:105
┌ Info: Epoch 824
└ @ Main /Users/kcft114/.julia/packages/Flux/qXNjB/src/optimise/train.jl:105
┌ Info: Epoch 825
└ @ Main /Users/kcft114/.julia/packages/Flux/qXNjB/src/optimise/train.jl:105
┌ Info: Epoch 826
└ @ Main /Users/kcft114/.julia/packages/Flux/qXNjB/src/optimise/train.jl:105
┌ Info: Epoch 827
└ @ Main /Users/kcft114/.julia/packages/Flux/qXNjB/src/optimise/train.jl:105
┌ Info: Epoch 828
└ @ Main /Users/kcft114/.julia/packages/Flux/qXNjB/src/optimise/train.jl:105
┌ Info: Epoch 829
└ @ Main /Users/kcft114/.julia/packages/Flux/qXNjB/src/optimise/train.jl:105
┌ Info: Epoch 830
└ @ Main /Users/kcft114/.julia/packages/Flux/qXNjB/src/optimise/train.jl:105
┌ Info: Epoch 831
└ @ Main /Users/kcft114/.julia/packages/Flux/qXNjB/src/optimise/train.jl:105
┌ Info: Epoch 832
└ @ Main /Users/kcft114/.julia/packages/Flux/qXNjB/src/optimise/train.jl:105
┌ Info: Epoch 833
└ @ Main /Users/kcft114/.julia/packages/Flux/qXNjB/src/optimise/train.jl:105
┌ Info: Epoch 834
└ @ Main /Users/kcft114/.julia/packages/Flux/qXNjB/src/optimise/train.jl:105
┌ Info: Epoch 835
└ @ Main /Users/kcft114/.julia/packages/Flux/qXNjB/src/optimise/train.jl:105
┌ Info: Epoch 836
└ @ Main /Users/kcft114/.julia/packages/Flux/qXNjB/src/optimise/train.jl:105
┌ Info: Epoch 837
└ @ Main /Users/kcft114/.julia/packages/Flux/qXNjB/src/optimise/train.jl:105
┌ Info: Epoch 838
└ @ Main /Users/kcft114/.julia/packages/Flux/qXNjB/src/optimise/train.jl:105
┌ Info: Epoch 839
└ @ Main /Users/kcft114/.julia/packages/Flux/qXNjB/src/optimise/train.jl:105
┌ Info: Epoch 840
└ @ Main /Users/kcft114/.julia/packages/Flux/qXNjB/src/optimise/train.jl:105
┌ Info: Epoch 841
└ @ Main /Users/kcft114/.julia/packages/Flux/qXNjB/src/optimise/train.jl:105
┌ Info: Epoch 842
└ @ Main /Users/kcft114/.julia/packages/Flux/qXNjB/src/optimise/train.jl:105
┌ Info: Epoch 843
└ @ Main /Users/kcft114/.julia/packages/Flux/qXNjB/src/optimise/train.jl:105
┌ Info: Epoch 844
└ @ Main /Users/kcft114/.julia/packages/Flux/qXNjB/src/optimise/train.jl:105
┌ Info: Epoch 845
└ @ Main /Users/kcft114/.julia/packages/Flux/qXNjB/src/optimise/train.jl:105
┌ Info: Epoch 846
└ @ Main /Users/kcft114/.julia/packages/Flux/qXNjB/src/optimise/train.jl:105
┌ Info: Epoch 847
└ @ Main /Users/kcft114/.julia/packages/Flux/qXNjB/src/optimise/train.jl:105
┌ Info: Epoch 848
└ @ Main /Users/kcft114/.julia/packages/Flux/qXNjB/src/optimise/train.jl:105
┌ Info: Epoch 849
└ @ Main /Users/kcft114/.julia/packages/Flux/qXNjB/src/optimise/train.jl:105
┌ Info: Epoch 850
└ @ Main /Users/kcft114/.julia/packages/Flux/qXNjB/src/optimise/train.jl:105
┌ Info: Epoch 851
└ @ Main /Users/kcft114/.julia/packages/Flux/qXNjB/src/optimise/train.jl:105
┌ Info: Epoch 852
└ @ Main /Users/kcft114/.julia/packages/Flux/qXNjB/src/optimise/train.jl:105
┌ Info: Epoch 853
└ @ Main /Users/kcft114/.julia/packages/Flux/qXNjB/src/optimise/train.jl:105
┌ Info: Epoch 854
└ @ Main /Users/kcft114/.julia/packages/Flux/qXNjB/src/optimise/train.jl:105
┌ Info: Epoch 855
└ @ Main /Users/kcft114/.julia/packages/Flux/qXNjB/src/optimise/train.jl:105
┌ Info: Epoch 856
└ @ Main /Users/kcft114/.julia/packages/Flux/qXNjB/src/optimise/train.jl:105
┌ Info: Epoch 857
└ @ Main /Users/kcft114/.julia/packages/Flux/qXNjB/src/optimise/train.jl:105
┌ Info: Epoch 858
└ @ Main /Users/kcft114/.julia/packages/Flux/qXNjB/src/optimise/train.jl:105
┌ Info: Epoch 859
└ @ Main /Users/kcft114/.julia/packages/Flux/qXNjB/src/optimise/train.jl:105
┌ Info: Epoch 860
└ @ Main /Users/kcft114/.julia/packages/Flux/qXNjB/src/optimise/train.jl:105
┌ Info: Epoch 861
└ @ Main /Users/kcft114/.julia/packages/Flux/qXNjB/src/optimise/train.jl:105
┌ Info: Epoch 862
└ @ Main /Users/kcft114/.julia/packages/Flux/qXNjB/src/optimise/train.jl:105
┌ Info: Epoch 863
└ @ Main /Users/kcft114/.julia/packages/Flux/qXNjB/src/optimise/train.jl:105
┌ Info: Epoch 864
└ @ Main /Users/kcft114/.julia/packages/Flux/qXNjB/src/optimise/train.jl:105
┌ Info: Epoch 865
└ @ Main /Users/kcft114/.julia/packages/Flux/qXNjB/src/optimise/train.jl:105
┌ Info: Epoch 866
└ @ Main /Users/kcft114/.julia/packages/Flux/qXNjB/src/optimise/train.jl:105
┌ Info: Epoch 867
└ @ Main /Users/kcft114/.julia/packages/Flux/qXNjB/src/optimise/train.jl:105
┌ Info: Epoch 868
└ @ Main /Users/kcft114/.julia/packages/Flux/qXNjB/src/optimise/train.jl:105
┌ Info: Epoch 869
└ @ Main /Users/kcft114/.julia/packages/Flux/qXNjB/src/optimise/train.jl:105
┌ Info: Epoch 870
└ @ Main /Users/kcft114/.julia/packages/Flux/qXNjB/src/optimise/train.jl:105
┌ Info: Epoch 871
└ @ Main /Users/kcft114/.julia/packages/Flux/qXNjB/src/optimise/train.jl:105
┌ Info: Epoch 872
└ @ Main /Users/kcft114/.julia/packages/Flux/qXNjB/src/optimise/train.jl:105
┌ Info: Epoch 873
└ @ Main /Users/kcft114/.julia/packages/Flux/qXNjB/src/optimise/train.jl:105
┌ Info: Epoch 874
└ @ Main /Users/kcft114/.julia/packages/Flux/qXNjB/src/optimise/train.jl:105
┌ Info: Epoch 875
└ @ Main /Users/kcft114/.julia/packages/Flux/qXNjB/src/optimise/train.jl:105
┌ Info: Epoch 876
└ @ Main /Users/kcft114/.julia/packages/Flux/qXNjB/src/optimise/train.jl:105
┌ Info: Epoch 877
└ @ Main /Users/kcft114/.julia/packages/Flux/qXNjB/src/optimise/train.jl:105
┌ Info: Epoch 878
└ @ Main /Users/kcft114/.julia/packages/Flux/qXNjB/src/optimise/train.jl:105
┌ Info: Epoch 879
└ @ Main /Users/kcft114/.julia/packages/Flux/qXNjB/src/optimise/train.jl:105
┌ Info: Epoch 880
└ @ Main /Users/kcft114/.julia/packages/Flux/qXNjB/src/optimise/train.jl:105
┌ Info: Epoch 881
└ @ Main /Users/kcft114/.julia/packages/Flux/qXNjB/src/optimise/train.jl:105
┌ Info: Epoch 882
└ @ Main /Users/kcft114/.julia/packages/Flux/qXNjB/src/optimise/train.jl:105
┌ Info: Epoch 883
└ @ Main /Users/kcft114/.julia/packages/Flux/qXNjB/src/optimise/train.jl:105
┌ Info: Epoch 884
└ @ Main /Users/kcft114/.julia/packages/Flux/qXNjB/src/optimise/train.jl:105
┌ Info: Epoch 885
└ @ Main /Users/kcft114/.julia/packages/Flux/qXNjB/src/optimise/train.jl:105
┌ Info: Epoch 886
└ @ Main /Users/kcft114/.julia/packages/Flux/qXNjB/src/optimise/train.jl:105
┌ Info: Epoch 887
└ @ Main /Users/kcft114/.julia/packages/Flux/qXNjB/src/optimise/train.jl:105
┌ Info: Epoch 888
└ @ Main /Users/kcft114/.julia/packages/Flux/qXNjB/src/optimise/train.jl:105
┌ Info: Epoch 889
└ @ Main /Users/kcft114/.julia/packages/Flux/qXNjB/src/optimise/train.jl:105
┌ Info: Epoch 890
└ @ Main /Users/kcft114/.julia/packages/Flux/qXNjB/src/optimise/train.jl:105
┌ Info: Epoch 891
└ @ Main /Users/kcft114/.julia/packages/Flux/qXNjB/src/optimise/train.jl:105
┌ Info: Epoch 892
└ @ Main /Users/kcft114/.julia/packages/Flux/qXNjB/src/optimise/train.jl:105
┌ Info: Epoch 893
└ @ Main /Users/kcft114/.julia/packages/Flux/qXNjB/src/optimise/train.jl:105
┌ Info: Epoch 894
└ @ Main /Users/kcft114/.julia/packages/Flux/qXNjB/src/optimise/train.jl:105
┌ Info: Epoch 895
└ @ Main /Users/kcft114/.julia/packages/Flux/qXNjB/src/optimise/train.jl:105
┌ Info: Epoch 896
└ @ Main /Users/kcft114/.julia/packages/Flux/qXNjB/src/optimise/train.jl:105
┌ Info: Epoch 897
└ @ Main /Users/kcft114/.julia/packages/Flux/qXNjB/src/optimise/train.jl:105
┌ Info: Epoch 898
└ @ Main /Users/kcft114/.julia/packages/Flux/qXNjB/src/optimise/train.jl:105
┌ Info: Epoch 899
└ @ Main /Users/kcft114/.julia/packages/Flux/qXNjB/src/optimise/train.jl:105
┌ Info: Epoch 900
└ @ Main /Users/kcft114/.julia/packages/Flux/qXNjB/src/optimise/train.jl:105
┌ Info: Epoch 901
└ @ Main /Users/kcft114/.julia/packages/Flux/qXNjB/src/optimise/train.jl:105
┌ Info: Epoch 902
└ @ Main /Users/kcft114/.julia/packages/Flux/qXNjB/src/optimise/train.jl:105
┌ Info: Epoch 903
└ @ Main /Users/kcft114/.julia/packages/Flux/qXNjB/src/optimise/train.jl:105
┌ Info: Epoch 904
└ @ Main /Users/kcft114/.julia/packages/Flux/qXNjB/src/optimise/train.jl:105
┌ Info: Epoch 905
└ @ Main /Users/kcft114/.julia/packages/Flux/qXNjB/src/optimise/train.jl:105
┌ Info: Epoch 906
└ @ Main /Users/kcft114/.julia/packages/Flux/qXNjB/src/optimise/train.jl:105
┌ Info: Epoch 907
└ @ Main /Users/kcft114/.julia/packages/Flux/qXNjB/src/optimise/train.jl:105
┌ Info: Epoch 908
└ @ Main /Users/kcft114/.julia/packages/Flux/qXNjB/src/optimise/train.jl:105
┌ Info: Epoch 909
└ @ Main /Users/kcft114/.julia/packages/Flux/qXNjB/src/optimise/train.jl:105
┌ Info: Epoch 910
└ @ Main /Users/kcft114/.julia/packages/Flux/qXNjB/src/optimise/train.jl:105
┌ Info: Epoch 911
└ @ Main /Users/kcft114/.julia/packages/Flux/qXNjB/src/optimise/train.jl:105
┌ Info: Epoch 912
└ @ Main /Users/kcft114/.julia/packages/Flux/qXNjB/src/optimise/train.jl:105
┌ Info: Epoch 913
└ @ Main /Users/kcft114/.julia/packages/Flux/qXNjB/src/optimise/train.jl:105
┌ Info: Epoch 914
└ @ Main /Users/kcft114/.julia/packages/Flux/qXNjB/src/optimise/train.jl:105
┌ Info: Epoch 915
└ @ Main /Users/kcft114/.julia/packages/Flux/qXNjB/src/optimise/train.jl:105
┌ Info: Epoch 916
└ @ Main /Users/kcft114/.julia/packages/Flux/qXNjB/src/optimise/train.jl:105
┌ Info: Epoch 917
└ @ Main /Users/kcft114/.julia/packages/Flux/qXNjB/src/optimise/train.jl:105
┌ Info: Epoch 918
└ @ Main /Users/kcft114/.julia/packages/Flux/qXNjB/src/optimise/train.jl:105
┌ Info: Epoch 919
└ @ Main /Users/kcft114/.julia/packages/Flux/qXNjB/src/optimise/train.jl:105
┌ Info: Epoch 920
└ @ Main /Users/kcft114/.julia/packages/Flux/qXNjB/src/optimise/train.jl:105
┌ Info: Epoch 921
└ @ Main /Users/kcft114/.julia/packages/Flux/qXNjB/src/optimise/train.jl:105
```

```
loss(X, Y) = 0.75883776f0 (tracked)
loss(X, Y) = 0.72570086f0 (tracked)
loss(X, Y) = 0.71485525f0 (tracked)
loss(X, Y) = 0.6968212f0 (tracked)
loss(X, Y) = 0.70334446f0 (tracked)
loss(X, Y) = 0.6706896f0 (tracked)
loss(X, Y) = 0.6877144f0 (tracked)
loss(X, Y) = 0.711464f0 (tracked)
loss(X, Y) = 0.6923495f0 (tracked)
loss(X, Y) = 0.7351494f0 (tracked)
loss(X, Y) = 0.7307077f0 (tracked)
loss(X, Y) = 0.66625816f0 (tracked)
loss(X, Y) = 0.73522294f0 (tracked)
loss(X, Y) = 0.7150659f0 (tracked)
loss(X, Y) = 0.7112949f0 (tracked)
loss(X, Y) = 0.68689257f0 (tracked)
loss(X, Y) = 0.7495173f0 (tracked)
loss(X, Y) = 0.6926665f0 (tracked)
loss(X, Y) = 0.7006073f0 (tracked)
loss(X, Y) = 0.69800895f0 (tracked)
loss(X, Y) = 0.72881025f0 (tracked)
loss(X, Y) = 0.7398764f0 (tracked)
loss(X, Y) = 0.7169691f0 (tracked)
loss(X, Y) = 0.72438085f0 (tracked)
loss(X, Y) = 0.7110641f0 (tracked)
loss(X, Y) = 0.6808945f0 (tracked)
loss(X, Y) = 0.6803195f0 (tracked)
loss(X, Y) = 0.69081116f0 (tracked)
loss(X, Y) = 0.6940091f0 (tracked)
loss(X, Y) = 0.7289362f0 (tracked)
loss(X, Y) = 0.7444154f0 (tracked)
loss(X, Y) = 0.72711563f0 (tracked)
loss(X, Y) = 0.730314f0 (tracked)
loss(X, Y) = 0.68708414f0 (tracked)
loss(X, Y) = 0.71014106f0 (tracked)
loss(X, Y) = 0.71175826f0 (tracked)
loss(X, Y) = 0.73348826f0 (tracked)
loss(X, Y) = 0.76548856f0 (tracked)
loss(X, Y) = 0.6928718f0 (tracked)
loss(X, Y) = 0.69542456f0 (tracked)
loss(X, Y) = 0.6853662f0 (tracked)
loss(X, Y) = 0.72453576f0 (tracked)
loss(X, Y) = 0.6894609f0 (tracked)
loss(X, Y) = 0.747375f0 (tracked)
loss(X, Y) = 0.71158147f0 (tracked)
loss(X, Y) = 0.69655365f0 (tracked)
loss(X, Y) = 0.65436935f0 (tracked)
loss(X, Y) = 0.7054951f0 (tracked)
loss(X, Y) = 0.69328f0 (tracked)
loss(X, Y) = 0.70199364f0 (tracked)
loss(X, Y) = 0.7518067f0 (tracked)
loss(X, Y) = 0.7254518f0 (tracked)
loss(X, Y) = 0.6662818f0 (tracked)
loss(X, Y) = 0.6691554f0 (tracked)
loss(X, Y) = 0.70891595f0 (tracked)
loss(X, Y) = 0.74972355f0 (tracked)
loss(X, Y) = 0.6793388f0 (tracked)
loss(X, Y) = 0.6897884f0 (tracked)
loss(X, Y) = 0.6931948f0 (tracked)
loss(X, Y) = 0.7265213f0 (tracked)
loss(X, Y) = 0.70535433f0 (tracked)
loss(X, Y) = 0.7160039f0 (tracked)
loss(X, Y) = 0.6825211f0 (tracked)
loss(X, Y) = 0.72929674f0 (tracked)
loss(X, Y) = 0.71191376f0 (tracked)
loss(X, Y) = 0.70807403f0 (tracked)
loss(X, Y) = 0.7262443f0 (tracked)
loss(X, Y) = 0.64582026f0 (tracked)
loss(X, Y) = 0.7303186f0 (tracked)
loss(X, Y) = 0.74089074f0 (tracked)
loss(X, Y) = 0.70170474f0 (tracked)
loss(X, Y) = 0.69651026f0 (tracked)
loss(X, Y) = 0.65704566f0 (tracked)
loss(X, Y) = 0.7221996f0 (tracked)
loss(X, Y) = 0.71676844f0 (tracked)
loss(X, Y) = 0.68702185f0 (tracked)
loss(X, Y) = 0.65819013f0 (tracked)
loss(X, Y) = 0.7278062f0 (tracked)
loss(X, Y) = 0.70790505f0 (tracked)
loss(X, Y) = 0.73823f0 (tracked)
loss(X, Y) = 0.6766274f0 (tracked)
loss(X, Y) = 0.6989436f0 (tracked)
loss(X, Y) = 0.7247779f0 (tracked)
loss(X, Y) = 0.68107945f0 (tracked)
loss(X, Y) = 0.7258583f0 (tracked)
loss(X, Y) = 0.7270309f0 (tracked)
loss(X, Y) = 0.6986252f0 (tracked)
loss(X, Y) = 0.69854945f0 (tracked)
loss(X, Y) = 0.70288056f0 (tracked)
loss(X, Y) = 0.67136395f0 (tracked)
loss(X, Y) = 0.66308224f0 (tracked)
loss(X, Y) = 0.71071106f0 (tracked)
loss(X, Y) = 0.70126355f0 (tracked)
loss(X, Y) = 0.66049486f0 (tracked)
loss(X, Y) = 0.6810158f0 (tracked)
loss(X, Y) = 0.681141f0 (tracked)
loss(X, Y) = 0.666336f0 (tracked)
loss(X, Y) = 0.72453314f0 (tracked)
loss(X, Y) = 0.71698564f0 (tracked)
loss(X, Y) = 0.7142493f0 (tracked)
loss(X, Y) = 0.6783712f0 (tracked)
loss(X, Y) = 0.6728902f0 (tracked)
loss(X, Y) = 0.6872576f0 (tracked)
loss(X, Y) = 0.6961124f0 (tracked)
loss(X, Y) = 0.7126976f0 (tracked)
loss(X, Y) = 0.67372143f0 (tracked)
loss(X, Y) = 0.6556979f0 (tracked)
loss(X, Y) = 0.7110565f0 (tracked)
loss(X, Y) = 0.725835f0 (tracked)
loss(X, Y) = 0.6763212f0 (tracked)
loss(X, Y) = 0.68695754f0 (tracked)
loss(X, Y) = 0.6597861f0 (tracked)
loss(X, Y) = 0.71537596f0 (tracked)
loss(X, Y) = 0.72663254f0 (tracked)
loss(X, Y) = 0.6986312f0 (tracked)
loss(X, Y) = 0.7045031f0 (tracked)
loss(X, Y) = 0.71135473f0 (tracked)
loss(X, Y) = 0.7075517f0 (tracked)
loss(X, Y) = 0.67857057f0 (tracked)
loss(X, Y) = 0.7127943f0 (tracked)
loss(X, Y) = 0.710931f0 (tracked)
loss(X, Y) = 0.7010061f0 (tracked)
loss(X, Y) = 0.7169499f0 (tracked)
loss(X, Y) = 0.7045616f0 (tracked)
loss(X, Y) = 0.7127537f0 (tracked)
loss(X, Y) = 0.7219007f0 (tracked)
loss(X, Y) = 0.71793455f0 (tracked)
loss(X, Y) = 0.7155184f0 (tracked)
loss(X, Y) = 0.69881225f0 (tracked)
loss(X, Y) = 0.70449495f0 (tracked)
loss(X, Y) = 0.6841485f0 (tracked)
loss(X, Y) = 0.7228137f0 (tracked)
loss(X, Y) = 0.64708525f0 (tracked)
loss(X, Y) = 0.662758f0 (tracked)
loss(X, Y) = 0.682634f0 (tracked)
loss(X, Y) = 0.6961398f0 (tracked)
loss(X, Y) = 0.7201903f0 (tracked)
loss(X, Y) = 0.66032505f0 (tracked)
loss(X, Y) = 0.71415555f0 (tracked)
loss(X, Y) = 0.6303857f0 (tracked)
loss(X, Y) = 0.6861497f0 (tracked)
loss(X, Y) = 0.70071626f0 (tracked)
loss(X, Y) = 0.6836914f0 (tracked)
loss(X, Y) = 0.6587747f0 (tracked)
loss(X, Y) = 0.68907654f0 (tracked)
loss(X, Y) = 0.6941822f0 (tracked)
loss(X, Y) = 0.6741439f0 (tracked)
loss(X, Y) = 0.70234865f0 (tracked)
loss(X, Y) = 0.68713856f0 (tracked)
loss(X, Y) = 0.72878844f0 (tracked)
loss(X, Y) = 0.7018873f0 (tracked)
loss(X, Y) = 0.71646345f0 (tracked)
loss(X, Y) = 0.6919575f0 (tracked)
loss(X, Y) = 0.68772095f0 (tracked)
loss(X, Y) = 0.6698317f0 (tracked)
loss(X, Y) = 0.7353323f0 (tracked)
loss(X, Y) = 0.69434476f0 (tracked)
loss(X, Y) = 0.7318185f0 (tracked)
loss(X, Y) = 0.66780984f0 (tracked)
loss(X, Y) = 0.71204317f0 (tracked)
loss(X, Y) = 0.6836856f0 (tracked)
loss(X, Y) = 0.69212866f0 (tracked)
loss(X, Y) = 0.6942358f0 (tracked)
loss(X, Y) = 0.7123323f0 (tracked)
loss(X, Y) = 0.70697385f0 (tracked)
loss(X, Y) = 0.69439346f0 (tracked)
loss(X, Y) = 0.6630729f0 (tracked)
loss(X, Y) = 0.72195154f0 (tracked)
loss(X, Y) = 0.65047824f0 (tracked)
loss(X, Y) = 0.6984784f0 (tracked)
loss(X, Y) = 0.67807806f0 (tracked)
loss(X, Y) = 0.70815426f0 (tracked)
loss(X, Y) = 0.6892966f0 (tracked)
loss(X, Y) = 0.70570993f0 (tracked)
loss(X, Y) = 0.71375686f0 (tracked)
loss(X, Y) = 0.656118f0 (tracked)
loss(X, Y) = 0.7017272f0 (tracked)
loss(X, Y) = 0.6993947f0 (tracked)
loss(X, Y) = 0.7074386f0 (tracked)
loss(X, Y) = 0.6601378f0 (tracked)
loss(X, Y) = 0.69423443f0 (tracked)
loss(X, Y) = 0.674727f0 (tracked)
loss(X, Y) = 0.68649065f0 (tracked)
loss(X, Y) = 0.7152237f0 (tracked)
loss(X, Y) = 0.69924194f0 (tracked)
loss(X, Y) = 0.6642384f0 (tracked)
loss(X, Y) = 0.6793967f0 (tracked)
```

```
┌ Info: Epoch 922
└ @ Main /Users/kcft114/.julia/packages/Flux/qXNjB/src/optimise/train.jl:105
┌ Info: Epoch 923
└ @ Main /Users/kcft114/.julia/packages/Flux/qXNjB/src/optimise/train.jl:105
┌ Info: Epoch 924
└ @ Main /Users/kcft114/.julia/packages/Flux/qXNjB/src/optimise/train.jl:105
┌ Info: Epoch 925
└ @ Main /Users/kcft114/.julia/packages/Flux/qXNjB/src/optimise/train.jl:105
┌ Info: Epoch 926
└ @ Main /Users/kcft114/.julia/packages/Flux/qXNjB/src/optimise/train.jl:105
┌ Info: Epoch 927
└ @ Main /Users/kcft114/.julia/packages/Flux/qXNjB/src/optimise/train.jl:105
┌ Info: Epoch 928
└ @ Main /Users/kcft114/.julia/packages/Flux/qXNjB/src/optimise/train.jl:105
┌ Info: Epoch 929
└ @ Main /Users/kcft114/.julia/packages/Flux/qXNjB/src/optimise/train.jl:105
┌ Info: Epoch 930
└ @ Main /Users/kcft114/.julia/packages/Flux/qXNjB/src/optimise/train.jl:105
┌ Info: Epoch 931
└ @ Main /Users/kcft114/.julia/packages/Flux/qXNjB/src/optimise/train.jl:105
┌ Info: Epoch 932
└ @ Main /Users/kcft114/.julia/packages/Flux/qXNjB/src/optimise/train.jl:105
┌ Info: Epoch 933
└ @ Main /Users/kcft114/.julia/packages/Flux/qXNjB/src/optimise/train.jl:105
┌ Info: Epoch 934
└ @ Main /Users/kcft114/.julia/packages/Flux/qXNjB/src/optimise/train.jl:105
┌ Info: Epoch 935
└ @ Main /Users/kcft114/.julia/packages/Flux/qXNjB/src/optimise/train.jl:105
┌ Info: Epoch 936
└ @ Main /Users/kcft114/.julia/packages/Flux/qXNjB/src/optimise/train.jl:105
┌ Info: Epoch 937
└ @ Main /Users/kcft114/.julia/packages/Flux/qXNjB/src/optimise/train.jl:105
┌ Info: Epoch 938
└ @ Main /Users/kcft114/.julia/packages/Flux/qXNjB/src/optimise/train.jl:105
┌ Info: Epoch 939
└ @ Main /Users/kcft114/.julia/packages/Flux/qXNjB/src/optimise/train.jl:105
┌ Info: Epoch 940
└ @ Main /Users/kcft114/.julia/packages/Flux/qXNjB/src/optimise/train.jl:105
┌ Info: Epoch 941
└ @ Main /Users/kcft114/.julia/packages/Flux/qXNjB/src/optimise/train.jl:105
┌ Info: Epoch 942
└ @ Main /Users/kcft114/.julia/packages/Flux/qXNjB/src/optimise/train.jl:105
┌ Info: Epoch 943
└ @ Main /Users/kcft114/.julia/packages/Flux/qXNjB/src/optimise/train.jl:105
┌ Info: Epoch 944
└ @ Main /Users/kcft114/.julia/packages/Flux/qXNjB/src/optimise/train.jl:105
┌ Info: Epoch 945
└ @ Main /Users/kcft114/.julia/packages/Flux/qXNjB/src/optimise/train.jl:105
┌ Info: Epoch 946
└ @ Main /Users/kcft114/.julia/packages/Flux/qXNjB/src/optimise/train.jl:105
┌ Info: Epoch 947
└ @ Main /Users/kcft114/.julia/packages/Flux/qXNjB/src/optimise/train.jl:105
┌ Info: Epoch 948
└ @ Main /Users/kcft114/.julia/packages/Flux/qXNjB/src/optimise/train.jl:105
┌ Info: Epoch 949
└ @ Main /Users/kcft114/.julia/packages/Flux/qXNjB/src/optimise/train.jl:105
┌ Info: Epoch 950
└ @ Main /Users/kcft114/.julia/packages/Flux/qXNjB/src/optimise/train.jl:105
┌ Info: Epoch 951
└ @ Main /Users/kcft114/.julia/packages/Flux/qXNjB/src/optimise/train.jl:105
┌ Info: Epoch 952
└ @ Main /Users/kcft114/.julia/packages/Flux/qXNjB/src/optimise/train.jl:105
┌ Info: Epoch 953
└ @ Main /Users/kcft114/.julia/packages/Flux/qXNjB/src/optimise/train.jl:105
┌ Info: Epoch 954
└ @ Main /Users/kcft114/.julia/packages/Flux/qXNjB/src/optimise/train.jl:105
┌ Info: Epoch 955
└ @ Main /Users/kcft114/.julia/packages/Flux/qXNjB/src/optimise/train.jl:105
┌ Info: Epoch 956
└ @ Main /Users/kcft114/.julia/packages/Flux/qXNjB/src/optimise/train.jl:105
┌ Info: Epoch 957
└ @ Main /Users/kcft114/.julia/packages/Flux/qXNjB/src/optimise/train.jl:105
┌ Info: Epoch 958
└ @ Main /Users/kcft114/.julia/packages/Flux/qXNjB/src/optimise/train.jl:105
┌ Info: Epoch 959
└ @ Main /Users/kcft114/.julia/packages/Flux/qXNjB/src/optimise/train.jl:105
┌ Info: Epoch 960
└ @ Main /Users/kcft114/.julia/packages/Flux/qXNjB/src/optimise/train.jl:105
┌ Info: Epoch 961
└ @ Main /Users/kcft114/.julia/packages/Flux/qXNjB/src/optimise/train.jl:105
┌ Info: Epoch 962
└ @ Main /Users/kcft114/.julia/packages/Flux/qXNjB/src/optimise/train.jl:105
┌ Info: Epoch 963
└ @ Main /Users/kcft114/.julia/packages/Flux/qXNjB/src/optimise/train.jl:105
┌ Info: Epoch 964
└ @ Main /Users/kcft114/.julia/packages/Flux/qXNjB/src/optimise/train.jl:105
┌ Info: Epoch 965
└ @ Main /Users/kcft114/.julia/packages/Flux/qXNjB/src/optimise/train.jl:105
┌ Info: Epoch 966
└ @ Main /Users/kcft114/.julia/packages/Flux/qXNjB/src/optimise/train.jl:105
┌ Info: Epoch 967
└ @ Main /Users/kcft114/.julia/packages/Flux/qXNjB/src/optimise/train.jl:105
┌ Info: Epoch 968
└ @ Main /Users/kcft114/.julia/packages/Flux/qXNjB/src/optimise/train.jl:105
┌ Info: Epoch 969
└ @ Main /Users/kcft114/.julia/packages/Flux/qXNjB/src/optimise/train.jl:105
┌ Info: Epoch 970
└ @ Main /Users/kcft114/.julia/packages/Flux/qXNjB/src/optimise/train.jl:105
┌ Info: Epoch 971
└ @ Main /Users/kcft114/.julia/packages/Flux/qXNjB/src/optimise/train.jl:105
┌ Info: Epoch 972
└ @ Main /Users/kcft114/.julia/packages/Flux/qXNjB/src/optimise/train.jl:105
┌ Info: Epoch 973
└ @ Main /Users/kcft114/.julia/packages/Flux/qXNjB/src/optimise/train.jl:105
┌ Info: Epoch 974
└ @ Main /Users/kcft114/.julia/packages/Flux/qXNjB/src/optimise/train.jl:105
┌ Info: Epoch 975
└ @ Main /Users/kcft114/.julia/packages/Flux/qXNjB/src/optimise/train.jl:105
┌ Info: Epoch 976
└ @ Main /Users/kcft114/.julia/packages/Flux/qXNjB/src/optimise/train.jl:105
┌ Info: Epoch 977
└ @ Main /Users/kcft114/.julia/packages/Flux/qXNjB/src/optimise/train.jl:105
┌ Info: Epoch 978
└ @ Main /Users/kcft114/.julia/packages/Flux/qXNjB/src/optimise/train.jl:105
┌ Info: Epoch 979
└ @ Main /Users/kcft114/.julia/packages/Flux/qXNjB/src/optimise/train.jl:105
┌ Info: Epoch 980
└ @ Main /Users/kcft114/.julia/packages/Flux/qXNjB/src/optimise/train.jl:105
┌ Info: Epoch 981
└ @ Main /Users/kcft114/.julia/packages/Flux/qXNjB/src/optimise/train.jl:105
┌ Info: Epoch 982
└ @ Main /Users/kcft114/.julia/packages/Flux/qXNjB/src/optimise/train.jl:105
┌ Info: Epoch 983
└ @ Main /Users/kcft114/.julia/packages/Flux/qXNjB/src/optimise/train.jl:105
┌ Info: Epoch 984
└ @ Main /Users/kcft114/.julia/packages/Flux/qXNjB/src/optimise/train.jl:105
┌ Info: Epoch 985
└ @ Main /Users/kcft114/.julia/packages/Flux/qXNjB/src/optimise/train.jl:105
┌ Info: Epoch 986
└ @ Main /Users/kcft114/.julia/packages/Flux/qXNjB/src/optimise/train.jl:105
┌ Info: Epoch 987
└ @ Main /Users/kcft114/.julia/packages/Flux/qXNjB/src/optimise/train.jl:105
┌ Info: Epoch 988
└ @ Main /Users/kcft114/.julia/packages/Flux/qXNjB/src/optimise/train.jl:105
┌ Info: Epoch 989
└ @ Main /Users/kcft114/.julia/packages/Flux/qXNjB/src/optimise/train.jl:105
┌ Info: Epoch 990
└ @ Main /Users/kcft114/.julia/packages/Flux/qXNjB/src/optimise/train.jl:105
┌ Info: Epoch 991
└ @ Main /Users/kcft114/.julia/packages/Flux/qXNjB/src/optimise/train.jl:105
┌ Info: Epoch 992
└ @ Main /Users/kcft114/.julia/packages/Flux/qXNjB/src/optimise/train.jl:105
┌ Info: Epoch 993
└ @ Main /Users/kcft114/.julia/packages/Flux/qXNjB/src/optimise/train.jl:105
┌ Info: Epoch 994
└ @ Main /Users/kcft114/.julia/packages/Flux/qXNjB/src/optimise/train.jl:105
┌ Info: Epoch 995
└ @ Main /Users/kcft114/.julia/packages/Flux/qXNjB/src/optimise/train.jl:105
┌ Info: Epoch 996
└ @ Main /Users/kcft114/.julia/packages/Flux/qXNjB/src/optimise/train.jl:105
┌ Info: Epoch 997
└ @ Main /Users/kcft114/.julia/packages/Flux/qXNjB/src/optimise/train.jl:105
┌ Info: Epoch 998
└ @ Main /Users/kcft114/.julia/packages/Flux/qXNjB/src/optimise/train.jl:105
┌ Info: Epoch 999
└ @ Main /Users/kcft114/.julia/packages/Flux/qXNjB/src/optimise/train.jl:105
┌ Info: Epoch 1000
└ @ Main /Users/kcft114/.julia/packages/Flux/qXNjB/src/optimise/train.jl:105
┌ Info: Epoch 1001
└ @ Main /Users/kcft114/.julia/packages/Flux/qXNjB/src/optimise/train.jl:105
┌ Info: Epoch 1002
└ @ Main /Users/kcft114/.julia/packages/Flux/qXNjB/src/optimise/train.jl:105
┌ Info: Epoch 1003
└ @ Main /Users/kcft114/.julia/packages/Flux/qXNjB/src/optimise/train.jl:105
┌ Info: Epoch 1004
└ @ Main /Users/kcft114/.julia/packages/Flux/qXNjB/src/optimise/train.jl:105
```

```
loss(X, Y) = 0.67595094f0 (tracked)
loss(X, Y) = 0.6426698f0 (tracked)
loss(X, Y) = 0.71189266f0 (tracked)
loss(X, Y) = 0.73451525f0 (tracked)
loss(X, Y) = 0.6891168f0 (tracked)
loss(X, Y) = 0.70423394f0 (tracked)
loss(X, Y) = 0.6879899f0 (tracked)
loss(X, Y) = 0.7028272f0 (tracked)
loss(X, Y) = 0.70322263f0 (tracked)
loss(X, Y) = 0.69343305f0 (tracked)
loss(X, Y) = 0.67835534f0 (tracked)
loss(X, Y) = 0.7070978f0 (tracked)
loss(X, Y) = 0.6552592f0 (tracked)
loss(X, Y) = 0.6892364f0 (tracked)
loss(X, Y) = 0.6948683f0 (tracked)
loss(X, Y) = 0.71286076f0 (tracked)
loss(X, Y) = 0.68415767f0 (tracked)
loss(X, Y) = 0.6699308f0 (tracked)
loss(X, Y) = 0.6763815f0 (tracked)
loss(X, Y) = 0.66815114f0 (tracked)
loss(X, Y) = 0.71366435f0 (tracked)
loss(X, Y) = 0.7151708f0 (tracked)
loss(X, Y) = 0.67802954f0 (tracked)
loss(X, Y) = 0.7028352f0 (tracked)
loss(X, Y) = 0.67310333f0 (tracked)
loss(X, Y) = 0.6563865f0 (tracked)
loss(X, Y) = 0.69165236f0 (tracked)
loss(X, Y) = 0.667946f0 (tracked)
loss(X, Y) = 0.6990464f0 (tracked)
loss(X, Y) = 0.6858905f0 (tracked)
loss(X, Y) = 0.7072953f0 (tracked)
loss(X, Y) = 0.7054481f0 (tracked)
loss(X, Y) = 0.6792316f0 (tracked)
loss(X, Y) = 0.6667893f0 (tracked)
loss(X, Y) = 0.71663624f0 (tracked)
loss(X, Y) = 0.6545974f0 (tracked)
loss(X, Y) = 0.7117643f0 (tracked)
loss(X, Y) = 0.7026523f0 (tracked)
loss(X, Y) = 0.6814642f0 (tracked)
loss(X, Y) = 0.6711185f0 (tracked)
loss(X, Y) = 0.6486459f0 (tracked)
loss(X, Y) = 0.6683435f0 (tracked)
loss(X, Y) = 0.69707686f0 (tracked)
loss(X, Y) = 0.6907591f0 (tracked)
loss(X, Y) = 0.693238f0 (tracked)
loss(X, Y) = 0.71681195f0 (tracked)
loss(X, Y) = 0.7200767f0 (tracked)
loss(X, Y) = 0.7064458f0 (tracked)
loss(X, Y) = 0.7002218f0 (tracked)
loss(X, Y) = 0.6850381f0 (tracked)
loss(X, Y) = 0.7430745f0 (tracked)
loss(X, Y) = 0.69393843f0 (tracked)
loss(X, Y) = 0.65876377f0 (tracked)
loss(X, Y) = 0.6882549f0 (tracked)
loss(X, Y) = 0.6734336f0 (tracked)
loss(X, Y) = 0.689227f0 (tracked)
loss(X, Y) = 0.70016587f0 (tracked)
loss(X, Y) = 0.6899596f0 (tracked)
loss(X, Y) = 0.71117216f0 (tracked)
loss(X, Y) = 0.69133663f0 (tracked)
loss(X, Y) = 0.7075567f0 (tracked)
loss(X, Y) = 0.7144218f0 (tracked)
loss(X, Y) = 0.682909f0 (tracked)
loss(X, Y) = 0.6631559f0 (tracked)
loss(X, Y) = 0.7355378f0 (tracked)
loss(X, Y) = 0.6483132f0 (tracked)
loss(X, Y) = 0.6808718f0 (tracked)
loss(X, Y) = 0.6865473f0 (tracked)
loss(X, Y) = 0.7092115f0 (tracked)
loss(X, Y) = 0.76189256f0 (tracked)
loss(X, Y) = 0.6690209f0 (tracked)
loss(X, Y) = 0.67438877f0 (tracked)
loss(X, Y) = 0.68810153f0 (tracked)
loss(X, Y) = 0.66864866f0 (tracked)
loss(X, Y) = 0.66516405f0 (tracked)
loss(X, Y) = 0.7018896f0 (tracked)
loss(X, Y) = 0.67651576f0 (tracked)
loss(X, Y) = 0.6639716f0 (tracked)
loss(X, Y) = 0.70790845f0 (tracked)
loss(X, Y) = 0.64809453f0 (tracked)
loss(X, Y) = 0.6914787f0 (tracked)
loss(X, Y) = 0.7151507f0 (tracked)
loss(X, Y) = 0.66682595f0 (tracked)
loss(X, Y) = 0.6685988f0 (tracked)
loss(X, Y) = 0.7019008f0 (tracked)
loss(X, Y) = 0.63253516f0 (tracked)
loss(X, Y) = 0.6828697f0 (tracked)
loss(X, Y) = 0.6510263f0 (tracked)
loss(X, Y) = 0.6767379f0 (tracked)
loss(X, Y) = 0.7070169f0 (tracked)
loss(X, Y) = 0.7198915f0 (tracked)
loss(X, Y) = 0.7111322f0 (tracked)
loss(X, Y) = 0.67073494f0 (tracked)
loss(X, Y) = 0.658716f0 (tracked)
loss(X, Y) = 0.6948668f0 (tracked)
loss(X, Y) = 0.6644087f0 (tracked)
loss(X, Y) = 0.6629499f0 (tracked)
loss(X, Y) = 0.66943747f0 (tracked)
loss(X, Y) = 0.7235976f0 (tracked)
loss(X, Y) = 0.68035185f0 (tracked)
loss(X, Y) = 0.67064834f0 (tracked)
```

```
┌ Info: Epoch 1005
└ @ Main /Users/kcft114/.julia/packages/Flux/qXNjB/src/optimise/train.jl:105
┌ Info: Epoch 1006
└ @ Main /Users/kcft114/.julia/packages/Flux/qXNjB/src/optimise/train.jl:105
┌ Info: Epoch 1007
└ @ Main /Users/kcft114/.julia/packages/Flux/qXNjB/src/optimise/train.jl:105
┌ Info: Epoch 1008
└ @ Main /Users/kcft114/.julia/packages/Flux/qXNjB/src/optimise/train.jl:105
┌ Info: Epoch 1009
└ @ Main /Users/kcft114/.julia/packages/Flux/qXNjB/src/optimise/train.jl:105
┌ Info: Epoch 1010
└ @ Main /Users/kcft114/.julia/packages/Flux/qXNjB/src/optimise/train.jl:105
┌ Info: Epoch 1011
└ @ Main /Users/kcft114/.julia/packages/Flux/qXNjB/src/optimise/train.jl:105
┌ Info: Epoch 1012
└ @ Main /Users/kcft114/.julia/packages/Flux/qXNjB/src/optimise/train.jl:105
┌ Info: Epoch 1013
└ @ Main /Users/kcft114/.julia/packages/Flux/qXNjB/src/optimise/train.jl:105
┌ Info: Epoch 1014
└ @ Main /Users/kcft114/.julia/packages/Flux/qXNjB/src/optimise/train.jl:105
┌ Info: Epoch 1015
└ @ Main /Users/kcft114/.julia/packages/Flux/qXNjB/src/optimise/train.jl:105
┌ Info: Epoch 1016
└ @ Main /Users/kcft114/.julia/packages/Flux/qXNjB/src/optimise/train.jl:105
┌ Info: Epoch 1017
└ @ Main /Users/kcft114/.julia/packages/Flux/qXNjB/src/optimise/train.jl:105
┌ Info: Epoch 1018
└ @ Main /Users/kcft114/.julia/packages/Flux/qXNjB/src/optimise/train.jl:105
┌ Info: Epoch 1019
└ @ Main /Users/kcft114/.julia/packages/Flux/qXNjB/src/optimise/train.jl:105
┌ Info: Epoch 1020
└ @ Main /Users/kcft114/.julia/packages/Flux/qXNjB/src/optimise/train.jl:105
┌ Info: Epoch 1021
└ @ Main /Users/kcft114/.julia/packages/Flux/qXNjB/src/optimise/train.jl:105
┌ Info: Epoch 1022
└ @ Main /Users/kcft114/.julia/packages/Flux/qXNjB/src/optimise/train.jl:105
┌ Info: Epoch 1023
└ @ Main /Users/kcft114/.julia/packages/Flux/qXNjB/src/optimise/train.jl:105
┌ Info: Epoch 1024
└ @ Main /Users/kcft114/.julia/packages/Flux/qXNjB/src/optimise/train.jl:105
┌ Info: Epoch 1025
└ @ Main /Users/kcft114/.julia/packages/Flux/qXNjB/src/optimise/train.jl:105
┌ Info: Epoch 1026
└ @ Main /Users/kcft114/.julia/packages/Flux/qXNjB/src/optimise/train.jl:105
┌ Info: Epoch 1027
└ @ Main /Users/kcft114/.julia/packages/Flux/qXNjB/src/optimise/train.jl:105
┌ Info: Epoch 1028
└ @ Main /Users/kcft114/.julia/packages/Flux/qXNjB/src/optimise/train.jl:105
┌ Info: Epoch 1029
└ @ Main /Users/kcft114/.julia/packages/Flux/qXNjB/src/optimise/train.jl:105
┌ Info: Epoch 1030
└ @ Main /Users/kcft114/.julia/packages/Flux/qXNjB/src/optimise/train.jl:105
┌ Info: Epoch 1031
└ @ Main /Users/kcft114/.julia/packages/Flux/qXNjB/src/optimise/train.jl:105
┌ Info: Epoch 1032
└ @ Main /Users/kcft114/.julia/packages/Flux/qXNjB/src/optimise/train.jl:105
┌ Info: Epoch 1033
└ @ Main /Users/kcft114/.julia/packages/Flux/qXNjB/src/optimise/train.jl:105
┌ Info: Epoch 1034
└ @ Main /Users/kcft114/.julia/packages/Flux/qXNjB/src/optimise/train.jl:105
┌ Info: Epoch 1035
└ @ Main /Users/kcft114/.julia/packages/Flux/qXNjB/src/optimise/train.jl:105
┌ Info: Epoch 1036
└ @ Main /Users/kcft114/.julia/packages/Flux/qXNjB/src/optimise/train.jl:105
┌ Info: Epoch 1037
└ @ Main /Users/kcft114/.julia/packages/Flux/qXNjB/src/optimise/train.jl:105
┌ Info: Epoch 1038
└ @ Main /Users/kcft114/.julia/packages/Flux/qXNjB/src/optimise/train.jl:105
┌ Info: Epoch 1039
└ @ Main /Users/kcft114/.julia/packages/Flux/qXNjB/src/optimise/train.jl:105
┌ Info: Epoch 1040
└ @ Main /Users/kcft114/.julia/packages/Flux/qXNjB/src/optimise/train.jl:105
┌ Info: Epoch 1041
└ @ Main /Users/kcft114/.julia/packages/Flux/qXNjB/src/optimise/train.jl:105
┌ Info: Epoch 1042
└ @ Main /Users/kcft114/.julia/packages/Flux/qXNjB/src/optimise/train.jl:105
┌ Info: Epoch 1043
└ @ Main /Users/kcft114/.julia/packages/Flux/qXNjB/src/optimise/train.jl:105
┌ Info: Epoch 1044
└ @ Main /Users/kcft114/.julia/packages/Flux/qXNjB/src/optimise/train.jl:105
┌ Info: Epoch 1045
└ @ Main /Users/kcft114/.julia/packages/Flux/qXNjB/src/optimise/train.jl:105
┌ Info: Epoch 1046
└ @ Main /Users/kcft114/.julia/packages/Flux/qXNjB/src/optimise/train.jl:105
┌ Info: Epoch 1047
└ @ Main /Users/kcft114/.julia/packages/Flux/qXNjB/src/optimise/train.jl:105
┌ Info: Epoch 1048
└ @ Main /Users/kcft114/.julia/packages/Flux/qXNjB/src/optimise/train.jl:105
┌ Info: Epoch 1049
└ @ Main /Users/kcft114/.julia/packages/Flux/qXNjB/src/optimise/train.jl:105
┌ Info: Epoch 1050
└ @ Main /Users/kcft114/.julia/packages/Flux/qXNjB/src/optimise/train.jl:105
┌ Info: Epoch 1051
└ @ Main /Users/kcft114/.julia/packages/Flux/qXNjB/src/optimise/train.jl:105
┌ Info: Epoch 1052
└ @ Main /Users/kcft114/.julia/packages/Flux/qXNjB/src/optimise/train.jl:105
┌ Info: Epoch 1053
└ @ Main /Users/kcft114/.julia/packages/Flux/qXNjB/src/optimise/train.jl:105
┌ Info: Epoch 1054
└ @ Main /Users/kcft114/.julia/packages/Flux/qXNjB/src/optimise/train.jl:105
┌ Info: Epoch 1055
└ @ Main /Users/kcft114/.julia/packages/Flux/qXNjB/src/optimise/train.jl:105
┌ Info: Epoch 1056
└ @ Main /Users/kcft114/.julia/packages/Flux/qXNjB/src/optimise/train.jl:105
┌ Info: Epoch 1057
└ @ Main /Users/kcft114/.julia/packages/Flux/qXNjB/src/optimise/train.jl:105
┌ Info: Epoch 1058
└ @ Main /Users/kcft114/.julia/packages/Flux/qXNjB/src/optimise/train.jl:105
┌ Info: Epoch 1059
└ @ Main /Users/kcft114/.julia/packages/Flux/qXNjB/src/optimise/train.jl:105
┌ Info: Epoch 1060
└ @ Main /Users/kcft114/.julia/packages/Flux/qXNjB/src/optimise/train.jl:105
┌ Info: Epoch 1061
└ @ Main /Users/kcft114/.julia/packages/Flux/qXNjB/src/optimise/train.jl:105
┌ Info: Epoch 1062
└ @ Main /Users/kcft114/.julia/packages/Flux/qXNjB/src/optimise/train.jl:105
┌ Info: Epoch 1063
└ @ Main /Users/kcft114/.julia/packages/Flux/qXNjB/src/optimise/train.jl:105
┌ Info: Epoch 1064
└ @ Main /Users/kcft114/.julia/packages/Flux/qXNjB/src/optimise/train.jl:105
┌ Info: Epoch 1065
└ @ Main /Users/kcft114/.julia/packages/Flux/qXNjB/src/optimise/train.jl:105
┌ Info: Epoch 1066
└ @ Main /Users/kcft114/.julia/packages/Flux/qXNjB/src/optimise/train.jl:105
┌ Info: Epoch 1067
└ @ Main /Users/kcft114/.julia/packages/Flux/qXNjB/src/optimise/train.jl:105
┌ Info: Epoch 1068
└ @ Main /Users/kcft114/.julia/packages/Flux/qXNjB/src/optimise/train.jl:105
┌ Info: Epoch 1069
└ @ Main /Users/kcft114/.julia/packages/Flux/qXNjB/src/optimise/train.jl:105
┌ Info: Epoch 1070
└ @ Main /Users/kcft114/.julia/packages/Flux/qXNjB/src/optimise/train.jl:105
┌ Info: Epoch 1071
└ @ Main /Users/kcft114/.julia/packages/Flux/qXNjB/src/optimise/train.jl:105
┌ Info: Epoch 1072
└ @ Main /Users/kcft114/.julia/packages/Flux/qXNjB/src/optimise/train.jl:105
┌ Info: Epoch 1073
└ @ Main /Users/kcft114/.julia/packages/Flux/qXNjB/src/optimise/train.jl:105
┌ Info: Epoch 1074
└ @ Main /Users/kcft114/.julia/packages/Flux/qXNjB/src/optimise/train.jl:105
┌ Info: Epoch 1075
└ @ Main /Users/kcft114/.julia/packages/Flux/qXNjB/src/optimise/train.jl:105
┌ Info: Epoch 1076
└ @ Main /Users/kcft114/.julia/packages/Flux/qXNjB/src/optimise/train.jl:105
┌ Info: Epoch 1077
└ @ Main /Users/kcft114/.julia/packages/Flux/qXNjB/src/optimise/train.jl:105
┌ Info: Epoch 1078
└ @ Main /Users/kcft114/.julia/packages/Flux/qXNjB/src/optimise/train.jl:105
┌ Info: Epoch 1079
└ @ Main /Users/kcft114/.julia/packages/Flux/qXNjB/src/optimise/train.jl:105
┌ Info: Epoch 1080
└ @ Main /Users/kcft114/.julia/packages/Flux/qXNjB/src/optimise/train.jl:105
┌ Info: Epoch 1081
└ @ Main /Users/kcft114/.julia/packages/Flux/qXNjB/src/optimise/train.jl:105
┌ Info: Epoch 1082
└ @ Main /Users/kcft114/.julia/packages/Flux/qXNjB/src/optimise/train.jl:105
┌ Info: Epoch 1083
└ @ Main /Users/kcft114/.julia/packages/Flux/qXNjB/src/optimise/train.jl:105
┌ Info: Epoch 1084
└ @ Main /Users/kcft114/.julia/packages/Flux/qXNjB/src/optimise/train.jl:105
┌ Info: Epoch 1085
└ @ Main /Users/kcft114/.julia/packages/Flux/qXNjB/src/optimise/train.jl:105
┌ Info: Epoch 1086
└ @ Main /Users/kcft114/.julia/packages/Flux/qXNjB/src/optimise/train.jl:105
┌ Info: Epoch 1087
└ @ Main /Users/kcft114/.julia/packages/Flux/qXNjB/src/optimise/train.jl:105
┌ Info: Epoch 1088
└ @ Main /Users/kcft114/.julia/packages/Flux/qXNjB/src/optimise/train.jl:105
┌ Info: Epoch 1089
└ @ Main /Users/kcft114/.julia/packages/Flux/qXNjB/src/optimise/train.jl:105
┌ Info: Epoch 1090
└ @ Main /Users/kcft114/.julia/packages/Flux/qXNjB/src/optimise/train.jl:105
┌ Info: Epoch 1091
└ @ Main /Users/kcft114/.julia/packages/Flux/qXNjB/src/optimise/train.jl:105
┌ Info: Epoch 1092
└ @ Main /Users/kcft114/.julia/packages/Flux/qXNjB/src/optimise/train.jl:105
┌ Info: Epoch 1093
└ @ Main /Users/kcft114/.julia/packages/Flux/qXNjB/src/optimise/train.jl:105
┌ Info: Epoch 1094
└ @ Main /Users/kcft114/.julia/packages/Flux/qXNjB/src/optimise/train.jl:105
┌ Info: Epoch 1095
└ @ Main /Users/kcft114/.julia/packages/Flux/qXNjB/src/optimise/train.jl:105
┌ Info: Epoch 1096
└ @ Main /Users/kcft114/.julia/packages/Flux/qXNjB/src/optimise/train.jl:105
┌ Info: Epoch 1097
└ @ Main /Users/kcft114/.julia/packages/Flux/qXNjB/src/optimise/train.jl:105
┌ Info: Epoch 1098
└ @ Main /Users/kcft114/.julia/packages/Flux/qXNjB/src/optimise/train.jl:105
┌ Info: Epoch 1099
└ @ Main /Users/kcft114/.julia/packages/Flux/qXNjB/src/optimise/train.jl:105
┌ Info: Epoch 1100
└ @ Main /Users/kcft114/.julia/packages/Flux/qXNjB/src/optimise/train.jl:105
┌ Info: Epoch 1101
└ @ Main /Users/kcft114/.julia/packages/Flux/qXNjB/src/optimise/train.jl:105
┌ Info: Epoch 1102
└ @ Main /Users/kcft114/.julia/packages/Flux/qXNjB/src/optimise/train.jl:105
┌ Info: Epoch 1103
└ @ Main /Users/kcft114/.julia/packages/Flux/qXNjB/src/optimise/train.jl:105
┌ Info: Epoch 1104
└ @ Main /Users/kcft114/.julia/packages/Flux/qXNjB/src/optimise/train.jl:105
┌ Info: Epoch 1105
└ @ Main /Users/kcft114/.julia/packages/Flux/qXNjB/src/optimise/train.jl:105
┌ Info: Epoch 1106
└ @ Main /Users/kcft114/.julia/packages/Flux/qXNjB/src/optimise/train.jl:105
┌ Info: Epoch 1107
└ @ Main /Users/kcft114/.julia/packages/Flux/qXNjB/src/optimise/train.jl:105
┌ Info: Epoch 1108
└ @ Main /Users/kcft114/.julia/packages/Flux/qXNjB/src/optimise/train.jl:105
┌ Info: Epoch 1109
└ @ Main /Users/kcft114/.julia/packages/Flux/qXNjB/src/optimise/train.jl:105
┌ Info: Epoch 1110
└ @ Main /Users/kcft114/.julia/packages/Flux/qXNjB/src/optimise/train.jl:105
┌ Info: Epoch 1111
└ @ Main /Users/kcft114/.julia/packages/Flux/qXNjB/src/optimise/train.jl:105
┌ Info: Epoch 1112
└ @ Main /Users/kcft114/.julia/packages/Flux/qXNjB/src/optimise/train.jl:105
┌ Info: Epoch 1113
└ @ Main /Users/kcft114/.julia/packages/Flux/qXNjB/src/optimise/train.jl:105
┌ Info: Epoch 1114
└ @ Main /Users/kcft114/.julia/packages/Flux/qXNjB/src/optimise/train.jl:105
┌ Info: Epoch 1115
└ @ Main /Users/kcft114/.julia/packages/Flux/qXNjB/src/optimise/train.jl:105
┌ Info: Epoch 1116
└ @ Main /Users/kcft114/.julia/packages/Flux/qXNjB/src/optimise/train.jl:105
┌ Info: Epoch 1117
└ @ Main /Users/kcft114/.julia/packages/Flux/qXNjB/src/optimise/train.jl:105
┌ Info: Epoch 1118
└ @ Main /Users/kcft114/.julia/packages/Flux/qXNjB/src/optimise/train.jl:105
┌ Info: Epoch 1119
└ @ Main /Users/kcft114/.julia/packages/Flux/qXNjB/src/optimise/train.jl:105
┌ Info: Epoch 1120
└ @ Main /Users/kcft114/.julia/packages/Flux/qXNjB/src/optimise/train.jl:105
┌ Info: Epoch 1121
└ @ Main /Users/kcft114/.julia/packages/Flux/qXNjB/src/optimise/train.jl:105
┌ Info: Epoch 1122
└ @ Main /Users/kcft114/.julia/packages/Flux/qXNjB/src/optimise/train.jl:105
┌ Info: Epoch 1123
└ @ Main /Users/kcft114/.julia/packages/Flux/qXNjB/src/optimise/train.jl:105
┌ Info: Epoch 1124
└ @ Main /Users/kcft114/.julia/packages/Flux/qXNjB/src/optimise/train.jl:105
┌ Info: Epoch 1125
└ @ Main /Users/kcft114/.julia/packages/Flux/qXNjB/src/optimise/train.jl:105
┌ Info: Epoch 1126
└ @ Main /Users/kcft114/.julia/packages/Flux/qXNjB/src/optimise/train.jl:105
┌ Info: Epoch 1127
└ @ Main /Users/kcft114/.julia/packages/Flux/qXNjB/src/optimise/train.jl:105
┌ Info: Epoch 1128
└ @ Main /Users/kcft114/.julia/packages/Flux/qXNjB/src/optimise/train.jl:105
┌ Info: Epoch 1129
└ @ Main /Users/kcft114/.julia/packages/Flux/qXNjB/src/optimise/train.jl:105
┌ Info: Epoch 1130
└ @ Main /Users/kcft114/.julia/packages/Flux/qXNjB/src/optimise/train.jl:105
┌ Info: Epoch 1131
└ @ Main /Users/kcft114/.julia/packages/Flux/qXNjB/src/optimise/train.jl:105
┌ Info: Epoch 1132
└ @ Main /Users/kcft114/.julia/packages/Flux/qXNjB/src/optimise/train.jl:105
┌ Info: Epoch 1133
└ @ Main /Users/kcft114/.julia/packages/Flux/qXNjB/src/optimise/train.jl:105
┌ Info: Epoch 1134
└ @ Main /Users/kcft114/.julia/packages/Flux/qXNjB/src/optimise/train.jl:105
┌ Info: Epoch 1135
└ @ Main /Users/kcft114/.julia/packages/Flux/qXNjB/src/optimise/train.jl:105
┌ Info: Epoch 1136
└ @ Main /Users/kcft114/.julia/packages/Flux/qXNjB/src/optimise/train.jl:105
┌ Info: Epoch 1137
└ @ Main /Users/kcft114/.julia/packages/Flux/qXNjB/src/optimise/train.jl:105
┌ Info: Epoch 1138
└ @ Main /Users/kcft114/.julia/packages/Flux/qXNjB/src/optimise/train.jl:105
┌ Info: Epoch 1139
└ @ Main /Users/kcft114/.julia/packages/Flux/qXNjB/src/optimise/train.jl:105
┌ Info: Epoch 1140
└ @ Main /Users/kcft114/.julia/packages/Flux/qXNjB/src/optimise/train.jl:105
┌ Info: Epoch 1141
└ @ Main /Users/kcft114/.julia/packages/Flux/qXNjB/src/optimise/train.jl:105
┌ Info: Epoch 1142
└ @ Main /Users/kcft114/.julia/packages/Flux/qXNjB/src/optimise/train.jl:105
┌ Info: Epoch 1143
└ @ Main /Users/kcft114/.julia/packages/Flux/qXNjB/src/optimise/train.jl:105
┌ Info: Epoch 1144
└ @ Main /Users/kcft114/.julia/packages/Flux/qXNjB/src/optimise/train.jl:105
┌ Info: Epoch 1145
└ @ Main /Users/kcft114/.julia/packages/Flux/qXNjB/src/optimise/train.jl:105
┌ Info: Epoch 1146
└ @ Main /Users/kcft114/.julia/packages/Flux/qXNjB/src/optimise/train.jl:105
┌ Info: Epoch 1147
└ @ Main /Users/kcft114/.julia/packages/Flux/qXNjB/src/optimise/train.jl:105
┌ Info: Epoch 1148
└ @ Main /Users/kcft114/.julia/packages/Flux/qXNjB/src/optimise/train.jl:105
┌ Info: Epoch 1149
└ @ Main /Users/kcft114/.julia/packages/Flux/qXNjB/src/optimise/train.jl:105
┌ Info: Epoch 1150
└ @ Main /Users/kcft114/.julia/packages/Flux/qXNjB/src/optimise/train.jl:105
┌ Info: Epoch 1151
└ @ Main /Users/kcft114/.julia/packages/Flux/qXNjB/src/optimise/train.jl:105
┌ Info: Epoch 1152
└ @ Main /Users/kcft114/.julia/packages/Flux/qXNjB/src/optimise/train.jl:105
┌ Info: Epoch 1153
└ @ Main /Users/kcft114/.julia/packages/Flux/qXNjB/src/optimise/train.jl:105
┌ Info: Epoch 1154
└ @ Main /Users/kcft114/.julia/packages/Flux/qXNjB/src/optimise/train.jl:105
┌ Info: Epoch 1155
└ @ Main /Users/kcft114/.julia/packages/Flux/qXNjB/src/optimise/train.jl:105
┌ Info: Epoch 1156
└ @ Main /Users/kcft114/.julia/packages/Flux/qXNjB/src/optimise/train.jl:105
┌ Info: Epoch 1157
└ @ Main /Users/kcft114/.julia/packages/Flux/qXNjB/src/optimise/train.jl:105
┌ Info: Epoch 1158
└ @ Main /Users/kcft114/.julia/packages/Flux/qXNjB/src/optimise/train.jl:105
┌ Info: Epoch 1159
└ @ Main /Users/kcft114/.julia/packages/Flux/qXNjB/src/optimise/train.jl:105
┌ Info: Epoch 1160
└ @ Main /Users/kcft114/.julia/packages/Flux/qXNjB/src/optimise/train.jl:105
┌ Info: Epoch 1161
└ @ Main /Users/kcft114/.julia/packages/Flux/qXNjB/src/optimise/train.jl:105
┌ Info: Epoch 1162
└ @ Main /Users/kcft114/.julia/packages/Flux/qXNjB/src/optimise/train.jl:105
┌ Info: Epoch 1163
└ @ Main /Users/kcft114/.julia/packages/Flux/qXNjB/src/optimise/train.jl:105
┌ Info: Epoch 1164
└ @ Main /Users/kcft114/.julia/packages/Flux/qXNjB/src/optimise/train.jl:105
┌ Info: Epoch 1165
└ @ Main /Users/kcft114/.julia/packages/Flux/qXNjB/src/optimise/train.jl:105
┌ Info: Epoch 1166
└ @ Main /Users/kcft114/.julia/packages/Flux/qXNjB/src/optimise/train.jl:105
┌ Info: Epoch 1167
└ @ Main /Users/kcft114/.julia/packages/Flux/qXNjB/src/optimise/train.jl:105
┌ Info: Epoch 1168
└ @ Main /Users/kcft114/.julia/packages/Flux/qXNjB/src/optimise/train.jl:105
┌ Info: Epoch 1169
└ @ Main /Users/kcft114/.julia/packages/Flux/qXNjB/src/optimise/train.jl:105
┌ Info: Epoch 1170
└ @ Main /Users/kcft114/.julia/packages/Flux/qXNjB/src/optimise/train.jl:105
┌ Info: Epoch 1171
└ @ Main /Users/kcft114/.julia/packages/Flux/qXNjB/src/optimise/train.jl:105
┌ Info: Epoch 1172
└ @ Main /Users/kcft114/.julia/packages/Flux/qXNjB/src/optimise/train.jl:105
┌ Info: Epoch 1173
└ @ Main /Users/kcft114/.julia/packages/Flux/qXNjB/src/optimise/train.jl:105
┌ Info: Epoch 1174
└ @ Main /Users/kcft114/.julia/packages/Flux/qXNjB/src/optimise/train.jl:105
┌ Info: Epoch 1175
└ @ Main /Users/kcft114/.julia/packages/Flux/qXNjB/src/optimise/train.jl:105
┌ Info: Epoch 1176
└ @ Main /Users/kcft114/.julia/packages/Flux/qXNjB/src/optimise/train.jl:105
┌ Info: Epoch 1177
└ @ Main /Users/kcft114/.julia/packages/Flux/qXNjB/src/optimise/train.jl:105
┌ Info: Epoch 1178
└ @ Main /Users/kcft114/.julia/packages/Flux/qXNjB/src/optimise/train.jl:105
┌ Info: Epoch 1179
└ @ Main /Users/kcft114/.julia/packages/Flux/qXNjB/src/optimise/train.jl:105
┌ Info: Epoch 1180
└ @ Main /Users/kcft114/.julia/packages/Flux/qXNjB/src/optimise/train.jl:105
┌ Info: Epoch 1181
└ @ Main /Users/kcft114/.julia/packages/Flux/qXNjB/src/optimise/train.jl:105
┌ Info: Epoch 1182
└ @ Main /Users/kcft114/.julia/packages/Flux/qXNjB/src/optimise/train.jl:105
┌ Info: Epoch 1183
└ @ Main /Users/kcft114/.julia/packages/Flux/qXNjB/src/optimise/train.jl:105
┌ Info: Epoch 1184
└ @ Main /Users/kcft114/.julia/packages/Flux/qXNjB/src/optimise/train.jl:105
┌ Info: Epoch 1185
└ @ Main /Users/kcft114/.julia/packages/Flux/qXNjB/src/optimise/train.jl:105
┌ Info: Epoch 1186
└ @ Main /Users/kcft114/.julia/packages/Flux/qXNjB/src/optimise/train.jl:105
┌ Info: Epoch 1187
└ @ Main /Users/kcft114/.julia/packages/Flux/qXNjB/src/optimise/train.jl:105
┌ Info: Epoch 1188
└ @ Main /Users/kcft114/.julia/packages/Flux/qXNjB/src/optimise/train.jl:105
┌ Info: Epoch 1189
└ @ Main /Users/kcft114/.julia/packages/Flux/qXNjB/src/optimise/train.jl:105
┌ Info: Epoch 1190
└ @ Main /Users/kcft114/.julia/packages/Flux/qXNjB/src/optimise/train.jl:105
┌ Info: Epoch 1191
└ @ Main /Users/kcft114/.julia/packages/Flux/qXNjB/src/optimise/train.jl:105
┌ Info: Epoch 1192
└ @ Main /Users/kcft114/.julia/packages/Flux/qXNjB/src/optimise/train.jl:105
┌ Info: Epoch 1193
└ @ Main /Users/kcft114/.julia/packages/Flux/qXNjB/src/optimise/train.jl:105
┌ Info: Epoch 1194
└ @ Main /Users/kcft114/.julia/packages/Flux/qXNjB/src/optimise/train.jl:105
┌ Info: Epoch 1195
└ @ Main /Users/kcft114/.julia/packages/Flux/qXNjB/src/optimise/train.jl:105
┌ Info: Epoch 1196
└ @ Main /Users/kcft114/.julia/packages/Flux/qXNjB/src/optimise/train.jl:105
┌ Info: Epoch 1197
└ @ Main /Users/kcft114/.julia/packages/Flux/qXNjB/src/optimise/train.jl:105
┌ Info: Epoch 1198
└ @ Main /Users/kcft114/.julia/packages/Flux/qXNjB/src/optimise/train.jl:105
┌ Info: Epoch 1199
└ @ Main /Users/kcft114/.julia/packages/Flux/qXNjB/src/optimise/train.jl:105
┌ Info: Epoch 1200
└ @ Main /Users/kcft114/.julia/packages/Flux/qXNjB/src/optimise/train.jl:105
┌ Info: Epoch 1201
└ @ Main /Users/kcft114/.julia/packages/Flux/qXNjB/src/optimise/train.jl:105
┌ Info: Epoch 1202
└ @ Main /Users/kcft114/.julia/packages/Flux/qXNjB/src/optimise/train.jl:105
┌ Info: Epoch 1203
└ @ Main /Users/kcft114/.julia/packages/Flux/qXNjB/src/optimise/train.jl:105
┌ Info: Epoch 1204
└ @ Main /Users/kcft114/.julia/packages/Flux/qXNjB/src/optimise/train.jl:105
┌ Info: Epoch 1205
└ @ Main /Users/kcft114/.julia/packages/Flux/qXNjB/src/optimise/train.jl:105
┌ Info: Epoch 1206
└ @ Main /Users/kcft114/.julia/packages/Flux/qXNjB/src/optimise/train.jl:105
┌ Info: Epoch 1207
└ @ Main /Users/kcft114/.julia/packages/Flux/qXNjB/src/optimise/train.jl:105
┌ Info: Epoch 1208
└ @ Main /Users/kcft114/.julia/packages/Flux/qXNjB/src/optimise/train.jl:105
```

```
loss(X, Y) = 0.6965608f0 (tracked)
loss(X, Y) = 0.6802687f0 (tracked)
loss(X, Y) = 0.65159947f0 (tracked)
loss(X, Y) = 0.7101655f0 (tracked)
loss(X, Y) = 0.698598f0 (tracked)
loss(X, Y) = 0.7217573f0 (tracked)
loss(X, Y) = 0.68595f0 (tracked)
loss(X, Y) = 0.65557665f0 (tracked)
loss(X, Y) = 0.68220115f0 (tracked)
loss(X, Y) = 0.69954926f0 (tracked)
loss(X, Y) = 0.70626205f0 (tracked)
loss(X, Y) = 0.706102f0 (tracked)
loss(X, Y) = 0.64914995f0 (tracked)
loss(X, Y) = 0.6430018f0 (tracked)
loss(X, Y) = 0.7133351f0 (tracked)
loss(X, Y) = 0.6581708f0 (tracked)
loss(X, Y) = 0.64298284f0 (tracked)
loss(X, Y) = 0.69865113f0 (tracked)
loss(X, Y) = 0.66491306f0 (tracked)
loss(X, Y) = 0.68559504f0 (tracked)
loss(X, Y) = 0.6926861f0 (tracked)
loss(X, Y) = 0.73530424f0 (tracked)
loss(X, Y) = 0.6672139f0 (tracked)
loss(X, Y) = 0.65817225f0 (tracked)
loss(X, Y) = 0.62593985f0 (tracked)
loss(X, Y) = 0.63504523f0 (tracked)
loss(X, Y) = 0.71866053f0 (tracked)
loss(X, Y) = 0.64061826f0 (tracked)
loss(X, Y) = 0.6880855f0 (tracked)
loss(X, Y) = 0.72988594f0 (tracked)
loss(X, Y) = 0.67137986f0 (tracked)
loss(X, Y) = 0.67293787f0 (tracked)
loss(X, Y) = 0.69234294f0 (tracked)
loss(X, Y) = 0.6576415f0 (tracked)
loss(X, Y) = 0.6833068f0 (tracked)
loss(X, Y) = 0.6786404f0 (tracked)
loss(X, Y) = 0.6818827f0 (tracked)
loss(X, Y) = 0.67901075f0 (tracked)
loss(X, Y) = 0.6924675f0 (tracked)
loss(X, Y) = 0.6806448f0 (tracked)
loss(X, Y) = 0.6802397f0 (tracked)
loss(X, Y) = 0.6685333f0 (tracked)
loss(X, Y) = 0.66838133f0 (tracked)
loss(X, Y) = 0.6526884f0 (tracked)
loss(X, Y) = 0.6771602f0 (tracked)
loss(X, Y) = 0.6941834f0 (tracked)
loss(X, Y) = 0.6757105f0 (tracked)
loss(X, Y) = 0.6802244f0 (tracked)
loss(X, Y) = 0.6610573f0 (tracked)
loss(X, Y) = 0.6646411f0 (tracked)
loss(X, Y) = 0.6604653f0 (tracked)
loss(X, Y) = 0.6847386f0 (tracked)
loss(X, Y) = 0.6582564f0 (tracked)
loss(X, Y) = 0.64805907f0 (tracked)
loss(X, Y) = 0.71832085f0 (tracked)
loss(X, Y) = 0.6639067f0 (tracked)
loss(X, Y) = 0.7087463f0 (tracked)
loss(X, Y) = 0.6565331f0 (tracked)
loss(X, Y) = 0.67684406f0 (tracked)
loss(X, Y) = 0.6758634f0 (tracked)
loss(X, Y) = 0.6962008f0 (tracked)
loss(X, Y) = 0.7002502f0 (tracked)
loss(X, Y) = 0.6500984f0 (tracked)
loss(X, Y) = 0.695027f0 (tracked)
loss(X, Y) = 0.6777928f0 (tracked)
loss(X, Y) = 0.6593975f0 (tracked)
loss(X, Y) = 0.6757674f0 (tracked)
loss(X, Y) = 0.66385764f0 (tracked)
loss(X, Y) = 0.7126738f0 (tracked)
loss(X, Y) = 0.72564703f0 (tracked)
loss(X, Y) = 0.69283676f0 (tracked)
loss(X, Y) = 0.68570274f0 (tracked)
loss(X, Y) = 0.642318f0 (tracked)
loss(X, Y) = 0.70713854f0 (tracked)
loss(X, Y) = 0.7148539f0 (tracked)
loss(X, Y) = 0.7072607f0 (tracked)
loss(X, Y) = 0.6621681f0 (tracked)
loss(X, Y) = 0.7029167f0 (tracked)
loss(X, Y) = 0.69864726f0 (tracked)
loss(X, Y) = 0.6667511f0 (tracked)
loss(X, Y) = 0.6180716f0 (tracked)
loss(X, Y) = 0.72252923f0 (tracked)
loss(X, Y) = 0.7021649f0 (tracked)
loss(X, Y) = 0.66779774f0 (tracked)
loss(X, Y) = 0.6499406f0 (tracked)
loss(X, Y) = 0.68133837f0 (tracked)
loss(X, Y) = 0.6944643f0 (tracked)
loss(X, Y) = 0.63885945f0 (tracked)
loss(X, Y) = 0.6249309f0 (tracked)
loss(X, Y) = 0.6610752f0 (tracked)
loss(X, Y) = 0.71879023f0 (tracked)
loss(X, Y) = 0.6890728f0 (tracked)
loss(X, Y) = 0.7020124f0 (tracked)
loss(X, Y) = 0.6460753f0 (tracked)
loss(X, Y) = 0.7186115f0 (tracked)
loss(X, Y) = 0.69836426f0 (tracked)
loss(X, Y) = 0.65317345f0 (tracked)
loss(X, Y) = 0.70160925f0 (tracked)
loss(X, Y) = 0.68258643f0 (tracked)
loss(X, Y) = 0.6711817f0 (tracked)
loss(X, Y) = 0.6733705f0 (tracked)
loss(X, Y) = 0.6533822f0 (tracked)
loss(X, Y) = 0.6830475f0 (tracked)
loss(X, Y) = 0.68697226f0 (tracked)
loss(X, Y) = 0.63650936f0 (tracked)
loss(X, Y) = 0.6288943f0 (tracked)
loss(X, Y) = 0.6651022f0 (tracked)
loss(X, Y) = 0.7279023f0 (tracked)
loss(X, Y) = 0.6449984f0 (tracked)
loss(X, Y) = 0.67972016f0 (tracked)
loss(X, Y) = 0.65115845f0 (tracked)
loss(X, Y) = 0.659006f0 (tracked)
loss(X, Y) = 0.6696141f0 (tracked)
loss(X, Y) = 0.6621102f0 (tracked)
loss(X, Y) = 0.6512224f0 (tracked)
loss(X, Y) = 0.60455316f0 (tracked)
loss(X, Y) = 0.6446687f0 (tracked)
loss(X, Y) = 0.6511131f0 (tracked)
loss(X, Y) = 0.67983514f0 (tracked)
loss(X, Y) = 0.6833304f0 (tracked)
loss(X, Y) = 0.6633861f0 (tracked)
loss(X, Y) = 0.7029793f0 (tracked)
loss(X, Y) = 0.70573616f0 (tracked)
loss(X, Y) = 0.6899373f0 (tracked)
loss(X, Y) = 0.64745474f0 (tracked)
loss(X, Y) = 0.6532623f0 (tracked)
loss(X, Y) = 0.65863025f0 (tracked)
loss(X, Y) = 0.6305327f0 (tracked)
loss(X, Y) = 0.6851355f0 (tracked)
loss(X, Y) = 0.6616162f0 (tracked)
loss(X, Y) = 0.6330949f0 (tracked)
```

```
┌ Info: Epoch 1209
└ @ Main /Users/kcft114/.julia/packages/Flux/qXNjB/src/optimise/train.jl:105
┌ Info: Epoch 1210
└ @ Main /Users/kcft114/.julia/packages/Flux/qXNjB/src/optimise/train.jl:105
┌ Info: Epoch 1211
└ @ Main /Users/kcft114/.julia/packages/Flux/qXNjB/src/optimise/train.jl:105
┌ Info: Epoch 1212
└ @ Main /Users/kcft114/.julia/packages/Flux/qXNjB/src/optimise/train.jl:105
┌ Info: Epoch 1213
└ @ Main /Users/kcft114/.julia/packages/Flux/qXNjB/src/optimise/train.jl:105
┌ Info: Epoch 1214
└ @ Main /Users/kcft114/.julia/packages/Flux/qXNjB/src/optimise/train.jl:105
┌ Info: Epoch 1215
└ @ Main /Users/kcft114/.julia/packages/Flux/qXNjB/src/optimise/train.jl:105
┌ Info: Epoch 1216
└ @ Main /Users/kcft114/.julia/packages/Flux/qXNjB/src/optimise/train.jl:105
┌ Info: Epoch 1217
└ @ Main /Users/kcft114/.julia/packages/Flux/qXNjB/src/optimise/train.jl:105
┌ Info: Epoch 1218
└ @ Main /Users/kcft114/.julia/packages/Flux/qXNjB/src/optimise/train.jl:105
┌ Info: Epoch 1219
└ @ Main /Users/kcft114/.julia/packages/Flux/qXNjB/src/optimise/train.jl:105
┌ Info: Epoch 1220
└ @ Main /Users/kcft114/.julia/packages/Flux/qXNjB/src/optimise/train.jl:105
┌ Info: Epoch 1221
└ @ Main /Users/kcft114/.julia/packages/Flux/qXNjB/src/optimise/train.jl:105
┌ Info: Epoch 1222
└ @ Main /Users/kcft114/.julia/packages/Flux/qXNjB/src/optimise/train.jl:105
┌ Info: Epoch 1223
└ @ Main /Users/kcft114/.julia/packages/Flux/qXNjB/src/optimise/train.jl:105
┌ Info: Epoch 1224
└ @ Main /Users/kcft114/.julia/packages/Flux/qXNjB/src/optimise/train.jl:105
┌ Info: Epoch 1225
└ @ Main /Users/kcft114/.julia/packages/Flux/qXNjB/src/optimise/train.jl:105
┌ Info: Epoch 1226
└ @ Main /Users/kcft114/.julia/packages/Flux/qXNjB/src/optimise/train.jl:105
┌ Info: Epoch 1227
└ @ Main /Users/kcft114/.julia/packages/Flux/qXNjB/src/optimise/train.jl:105
┌ Info: Epoch 1228
└ @ Main /Users/kcft114/.julia/packages/Flux/qXNjB/src/optimise/train.jl:105
┌ Info: Epoch 1229
└ @ Main /Users/kcft114/.julia/packages/Flux/qXNjB/src/optimise/train.jl:105
┌ Info: Epoch 1230
└ @ Main /Users/kcft114/.julia/packages/Flux/qXNjB/src/optimise/train.jl:105
┌ Info: Epoch 1231
└ @ Main /Users/kcft114/.julia/packages/Flux/qXNjB/src/optimise/train.jl:105
┌ Info: Epoch 1232
└ @ Main /Users/kcft114/.julia/packages/Flux/qXNjB/src/optimise/train.jl:105
┌ Info: Epoch 1233
└ @ Main /Users/kcft114/.julia/packages/Flux/qXNjB/src/optimise/train.jl:105
┌ Info: Epoch 1234
└ @ Main /Users/kcft114/.julia/packages/Flux/qXNjB/src/optimise/train.jl:105
┌ Info: Epoch 1235
└ @ Main /Users/kcft114/.julia/packages/Flux/qXNjB/src/optimise/train.jl:105
┌ Info: Epoch 1236
└ @ Main /Users/kcft114/.julia/packages/Flux/qXNjB/src/optimise/train.jl:105
┌ Info: Epoch 1237
└ @ Main /Users/kcft114/.julia/packages/Flux/qXNjB/src/optimise/train.jl:105
┌ Info: Epoch 1238
└ @ Main /Users/kcft114/.julia/packages/Flux/qXNjB/src/optimise/train.jl:105
┌ Info: Epoch 1239
└ @ Main /Users/kcft114/.julia/packages/Flux/qXNjB/src/optimise/train.jl:105
┌ Info: Epoch 1240
└ @ Main /Users/kcft114/.julia/packages/Flux/qXNjB/src/optimise/train.jl:105
┌ Info: Epoch 1241
└ @ Main /Users/kcft114/.julia/packages/Flux/qXNjB/src/optimise/train.jl:105
┌ Info: Epoch 1242
└ @ Main /Users/kcft114/.julia/packages/Flux/qXNjB/src/optimise/train.jl:105
┌ Info: Epoch 1243
└ @ Main /Users/kcft114/.julia/packages/Flux/qXNjB/src/optimise/train.jl:105
┌ Info: Epoch 1244
└ @ Main /Users/kcft114/.julia/packages/Flux/qXNjB/src/optimise/train.jl:105
┌ Info: Epoch 1245
└ @ Main /Users/kcft114/.julia/packages/Flux/qXNjB/src/optimise/train.jl:105
┌ Info: Epoch 1246
└ @ Main /Users/kcft114/.julia/packages/Flux/qXNjB/src/optimise/train.jl:105
┌ Info: Epoch 1247
└ @ Main /Users/kcft114/.julia/packages/Flux/qXNjB/src/optimise/train.jl:105
┌ Info: Epoch 1248
└ @ Main /Users/kcft114/.julia/packages/Flux/qXNjB/src/optimise/train.jl:105
┌ Info: Epoch 1249
└ @ Main /Users/kcft114/.julia/packages/Flux/qXNjB/src/optimise/train.jl:105
┌ Info: Epoch 1250
└ @ Main /Users/kcft114/.julia/packages/Flux/qXNjB/src/optimise/train.jl:105
┌ Info: Epoch 1251
└ @ Main /Users/kcft114/.julia/packages/Flux/qXNjB/src/optimise/train.jl:105
┌ Info: Epoch 1252
└ @ Main /Users/kcft114/.julia/packages/Flux/qXNjB/src/optimise/train.jl:105
┌ Info: Epoch 1253
└ @ Main /Users/kcft114/.julia/packages/Flux/qXNjB/src/optimise/train.jl:105
┌ Info: Epoch 1254
└ @ Main /Users/kcft114/.julia/packages/Flux/qXNjB/src/optimise/train.jl:105
┌ Info: Epoch 1255
└ @ Main /Users/kcft114/.julia/packages/Flux/qXNjB/src/optimise/train.jl:105
┌ Info: Epoch 1256
└ @ Main /Users/kcft114/.julia/packages/Flux/qXNjB/src/optimise/train.jl:105
┌ Info: Epoch 1257
└ @ Main /Users/kcft114/.julia/packages/Flux/qXNjB/src/optimise/train.jl:105
┌ Info: Epoch 1258
└ @ Main /Users/kcft114/.julia/packages/Flux/qXNjB/src/optimise/train.jl:105
┌ Info: Epoch 1259
└ @ Main /Users/kcft114/.julia/packages/Flux/qXNjB/src/optimise/train.jl:105
┌ Info: Epoch 1260
└ @ Main /Users/kcft114/.julia/packages/Flux/qXNjB/src/optimise/train.jl:105
┌ Info: Epoch 1261
└ @ Main /Users/kcft114/.julia/packages/Flux/qXNjB/src/optimise/train.jl:105
┌ Info: Epoch 1262
└ @ Main /Users/kcft114/.julia/packages/Flux/qXNjB/src/optimise/train.jl:105
┌ Info: Epoch 1263
└ @ Main /Users/kcft114/.julia/packages/Flux/qXNjB/src/optimise/train.jl:105
┌ Info: Epoch 1264
└ @ Main /Users/kcft114/.julia/packages/Flux/qXNjB/src/optimise/train.jl:105
┌ Info: Epoch 1265
└ @ Main /Users/kcft114/.julia/packages/Flux/qXNjB/src/optimise/train.jl:105
┌ Info: Epoch 1266
└ @ Main /Users/kcft114/.julia/packages/Flux/qXNjB/src/optimise/train.jl:105
┌ Info: Epoch 1267
└ @ Main /Users/kcft114/.julia/packages/Flux/qXNjB/src/optimise/train.jl:105
┌ Info: Epoch 1268
└ @ Main /Users/kcft114/.julia/packages/Flux/qXNjB/src/optimise/train.jl:105
┌ Info: Epoch 1269
└ @ Main /Users/kcft114/.julia/packages/Flux/qXNjB/src/optimise/train.jl:105
┌ Info: Epoch 1270
└ @ Main /Users/kcft114/.julia/packages/Flux/qXNjB/src/optimise/train.jl:105
┌ Info: Epoch 1271
└ @ Main /Users/kcft114/.julia/packages/Flux/qXNjB/src/optimise/train.jl:105
┌ Info: Epoch 1272
└ @ Main /Users/kcft114/.julia/packages/Flux/qXNjB/src/optimise/train.jl:105
┌ Info: Epoch 1273
└ @ Main /Users/kcft114/.julia/packages/Flux/qXNjB/src/optimise/train.jl:105
┌ Info: Epoch 1274
└ @ Main /Users/kcft114/.julia/packages/Flux/qXNjB/src/optimise/train.jl:105
┌ Info: Epoch 1275
└ @ Main /Users/kcft114/.julia/packages/Flux/qXNjB/src/optimise/train.jl:105
┌ Info: Epoch 1276
└ @ Main /Users/kcft114/.julia/packages/Flux/qXNjB/src/optimise/train.jl:105
┌ Info: Epoch 1277
└ @ Main /Users/kcft114/.julia/packages/Flux/qXNjB/src/optimise/train.jl:105
┌ Info: Epoch 1278
└ @ Main /Users/kcft114/.julia/packages/Flux/qXNjB/src/optimise/train.jl:105
┌ Info: Epoch 1279
└ @ Main /Users/kcft114/.julia/packages/Flux/qXNjB/src/optimise/train.jl:105
┌ Info: Epoch 1280
└ @ Main /Users/kcft114/.julia/packages/Flux/qXNjB/src/optimise/train.jl:105
┌ Info: Epoch 1281
└ @ Main /Users/kcft114/.julia/packages/Flux/qXNjB/src/optimise/train.jl:105
┌ Info: Epoch 1282
└ @ Main /Users/kcft114/.julia/packages/Flux/qXNjB/src/optimise/train.jl:105
┌ Info: Epoch 1283
└ @ Main /Users/kcft114/.julia/packages/Flux/qXNjB/src/optimise/train.jl:105
┌ Info: Epoch 1284
└ @ Main /Users/kcft114/.julia/packages/Flux/qXNjB/src/optimise/train.jl:105
┌ Info: Epoch 1285
└ @ Main /Users/kcft114/.julia/packages/Flux/qXNjB/src/optimise/train.jl:105
┌ Info: Epoch 1286
└ @ Main /Users/kcft114/.julia/packages/Flux/qXNjB/src/optimise/train.jl:105
┌ Info: Epoch 1287
└ @ Main /Users/kcft114/.julia/packages/Flux/qXNjB/src/optimise/train.jl:105
┌ Info: Epoch 1288
└ @ Main /Users/kcft114/.julia/packages/Flux/qXNjB/src/optimise/train.jl:105
┌ Info: Epoch 1289
└ @ Main /Users/kcft114/.julia/packages/Flux/qXNjB/src/optimise/train.jl:105
┌ Info: Epoch 1290
└ @ Main /Users/kcft114/.julia/packages/Flux/qXNjB/src/optimise/train.jl:105
┌ Info: Epoch 1291
└ @ Main /Users/kcft114/.julia/packages/Flux/qXNjB/src/optimise/train.jl:105
┌ Info: Epoch 1292
└ @ Main /Users/kcft114/.julia/packages/Flux/qXNjB/src/optimise/train.jl:105
┌ Info: Epoch 1293
└ @ Main /Users/kcft114/.julia/packages/Flux/qXNjB/src/optimise/train.jl:105
┌ Info: Epoch 1294
└ @ Main /Users/kcft114/.julia/packages/Flux/qXNjB/src/optimise/train.jl:105
┌ Info: Epoch 1295
└ @ Main /Users/kcft114/.julia/packages/Flux/qXNjB/src/optimise/train.jl:105
┌ Info: Epoch 1296
└ @ Main /Users/kcft114/.julia/packages/Flux/qXNjB/src/optimise/train.jl:105
┌ Info: Epoch 1297
└ @ Main /Users/kcft114/.julia/packages/Flux/qXNjB/src/optimise/train.jl:105
┌ Info: Epoch 1298
└ @ Main /Users/kcft114/.julia/packages/Flux/qXNjB/src/optimise/train.jl:105
┌ Info: Epoch 1299
└ @ Main /Users/kcft114/.julia/packages/Flux/qXNjB/src/optimise/train.jl:105
┌ Info: Epoch 1300
└ @ Main /Users/kcft114/.julia/packages/Flux/qXNjB/src/optimise/train.jl:105
┌ Info: Epoch 1301
└ @ Main /Users/kcft114/.julia/packages/Flux/qXNjB/src/optimise/train.jl:105
┌ Info: Epoch 1302
└ @ Main /Users/kcft114/.julia/packages/Flux/qXNjB/src/optimise/train.jl:105
┌ Info: Epoch 1303
└ @ Main /Users/kcft114/.julia/packages/Flux/qXNjB/src/optimise/train.jl:105
┌ Info: Epoch 1304
└ @ Main /Users/kcft114/.julia/packages/Flux/qXNjB/src/optimise/train.jl:105
┌ Info: Epoch 1305
└ @ Main /Users/kcft114/.julia/packages/Flux/qXNjB/src/optimise/train.jl:105
┌ Info: Epoch 1306
└ @ Main /Users/kcft114/.julia/packages/Flux/qXNjB/src/optimise/train.jl:105
┌ Info: Epoch 1307
└ @ Main /Users/kcft114/.julia/packages/Flux/qXNjB/src/optimise/train.jl:105
┌ Info: Epoch 1308
└ @ Main /Users/kcft114/.julia/packages/Flux/qXNjB/src/optimise/train.jl:105
┌ Info: Epoch 1309
└ @ Main /Users/kcft114/.julia/packages/Flux/qXNjB/src/optimise/train.jl:105
┌ Info: Epoch 1310
└ @ Main /Users/kcft114/.julia/packages/Flux/qXNjB/src/optimise/train.jl:105
┌ Info: Epoch 1311
└ @ Main /Users/kcft114/.julia/packages/Flux/qXNjB/src/optimise/train.jl:105
┌ Info: Epoch 1312
└ @ Main /Users/kcft114/.julia/packages/Flux/qXNjB/src/optimise/train.jl:105
┌ Info: Epoch 1313
└ @ Main /Users/kcft114/.julia/packages/Flux/qXNjB/src/optimise/train.jl:105
┌ Info: Epoch 1314
└ @ Main /Users/kcft114/.julia/packages/Flux/qXNjB/src/optimise/train.jl:105
┌ Info: Epoch 1315
└ @ Main /Users/kcft114/.julia/packages/Flux/qXNjB/src/optimise/train.jl:105
┌ Info: Epoch 1316
└ @ Main /Users/kcft114/.julia/packages/Flux/qXNjB/src/optimise/train.jl:105
┌ Info: Epoch 1317
└ @ Main /Users/kcft114/.julia/packages/Flux/qXNjB/src/optimise/train.jl:105
┌ Info: Epoch 1318
└ @ Main /Users/kcft114/.julia/packages/Flux/qXNjB/src/optimise/train.jl:105
┌ Info: Epoch 1319
└ @ Main /Users/kcft114/.julia/packages/Flux/qXNjB/src/optimise/train.jl:105
┌ Info: Epoch 1320
└ @ Main /Users/kcft114/.julia/packages/Flux/qXNjB/src/optimise/train.jl:105
┌ Info: Epoch 1321
└ @ Main /Users/kcft114/.julia/packages/Flux/qXNjB/src/optimise/train.jl:105
┌ Info: Epoch 1322
└ @ Main /Users/kcft114/.julia/packages/Flux/qXNjB/src/optimise/train.jl:105
┌ Info: Epoch 1323
└ @ Main /Users/kcft114/.julia/packages/Flux/qXNjB/src/optimise/train.jl:105
┌ Info: Epoch 1324
└ @ Main /Users/kcft114/.julia/packages/Flux/qXNjB/src/optimise/train.jl:105
┌ Info: Epoch 1325
└ @ Main /Users/kcft114/.julia/packages/Flux/qXNjB/src/optimise/train.jl:105
┌ Info: Epoch 1326
└ @ Main /Users/kcft114/.julia/packages/Flux/qXNjB/src/optimise/train.jl:105
┌ Info: Epoch 1327
└ @ Main /Users/kcft114/.julia/packages/Flux/qXNjB/src/optimise/train.jl:105
┌ Info: Epoch 1328
└ @ Main /Users/kcft114/.julia/packages/Flux/qXNjB/src/optimise/train.jl:105
┌ Info: Epoch 1329
└ @ Main /Users/kcft114/.julia/packages/Flux/qXNjB/src/optimise/train.jl:105
┌ Info: Epoch 1330
└ @ Main /Users/kcft114/.julia/packages/Flux/qXNjB/src/optimise/train.jl:105
┌ Info: Epoch 1331
└ @ Main /Users/kcft114/.julia/packages/Flux/qXNjB/src/optimise/train.jl:105
┌ Info: Epoch 1332
└ @ Main /Users/kcft114/.julia/packages/Flux/qXNjB/src/optimise/train.jl:105
┌ Info: Epoch 1333
└ @ Main /Users/kcft114/.julia/packages/Flux/qXNjB/src/optimise/train.jl:105
┌ Info: Epoch 1334
└ @ Main /Users/kcft114/.julia/packages/Flux/qXNjB/src/optimise/train.jl:105
┌ Info: Epoch 1335
└ @ Main /Users/kcft114/.julia/packages/Flux/qXNjB/src/optimise/train.jl:105
┌ Info: Epoch 1336
└ @ Main /Users/kcft114/.julia/packages/Flux/qXNjB/src/optimise/train.jl:105
┌ Info: Epoch 1337
└ @ Main /Users/kcft114/.julia/packages/Flux/qXNjB/src/optimise/train.jl:105
┌ Info: Epoch 1338
└ @ Main /Users/kcft114/.julia/packages/Flux/qXNjB/src/optimise/train.jl:105
```

```
loss(X, Y) = 0.6825727f0 (tracked)
loss(X, Y) = 0.688727f0 (tracked)
loss(X, Y) = 0.6768936f0 (tracked)
loss(X, Y) = 0.6784141f0 (tracked)
loss(X, Y) = 0.686315f0 (tracked)
loss(X, Y) = 0.6721959f0 (tracked)
loss(X, Y) = 0.6661091f0 (tracked)
loss(X, Y) = 0.68310606f0 (tracked)
loss(X, Y) = 0.7142688f0 (tracked)
loss(X, Y) = 0.68978935f0 (tracked)
loss(X, Y) = 0.7005074f0 (tracked)
loss(X, Y) = 0.69343525f0 (tracked)
loss(X, Y) = 0.6931788f0 (tracked)
loss(X, Y) = 0.70686877f0 (tracked)
loss(X, Y) = 0.68373173f0 (tracked)
loss(X, Y) = 0.6034798f0 (tracked)
loss(X, Y) = 0.6693458f0 (tracked)
loss(X, Y) = 0.6724115f0 (tracked)
loss(X, Y) = 0.67078424f0 (tracked)
loss(X, Y) = 0.7408765f0 (tracked)
loss(X, Y) = 0.6657461f0 (tracked)
loss(X, Y) = 0.6740876f0 (tracked)
loss(X, Y) = 0.7095576f0 (tracked)
loss(X, Y) = 0.7011606f0 (tracked)
loss(X, Y) = 0.6780589f0 (tracked)
loss(X, Y) = 0.68195844f0 (tracked)
loss(X, Y) = 0.6662114f0 (tracked)
loss(X, Y) = 0.61401933f0 (tracked)
loss(X, Y) = 0.6692888f0 (tracked)
loss(X, Y) = 0.6366968f0 (tracked)
loss(X, Y) = 0.6839248f0 (tracked)
loss(X, Y) = 0.6957952f0 (tracked)
loss(X, Y) = 0.66393805f0 (tracked)
loss(X, Y) = 0.6762401f0 (tracked)
loss(X, Y) = 0.6962743f0 (tracked)
loss(X, Y) = 0.6777634f0 (tracked)
loss(X, Y) = 0.6259486f0 (tracked)
loss(X, Y) = 0.65727854f0 (tracked)
loss(X, Y) = 0.7028709f0 (tracked)
loss(X, Y) = 0.6833862f0 (tracked)
loss(X, Y) = 0.64326096f0 (tracked)
loss(X, Y) = 0.6806216f0 (tracked)
loss(X, Y) = 0.6522511f0 (tracked)
loss(X, Y) = 0.67379504f0 (tracked)
loss(X, Y) = 0.68463224f0 (tracked)
loss(X, Y) = 0.64690334f0 (tracked)
loss(X, Y) = 0.6563734f0 (tracked)
loss(X, Y) = 0.6350112f0 (tracked)
loss(X, Y) = 0.6645569f0 (tracked)
loss(X, Y) = 0.7011848f0 (tracked)
loss(X, Y) = 0.59970146f0 (tracked)
loss(X, Y) = 0.64776725f0 (tracked)
loss(X, Y) = 0.6725102f0 (tracked)
loss(X, Y) = 0.61569816f0 (tracked)
loss(X, Y) = 0.6507119f0 (tracked)
loss(X, Y) = 0.669793f0 (tracked)
loss(X, Y) = 0.7017598f0 (tracked)
loss(X, Y) = 0.6827177f0 (tracked)
loss(X, Y) = 0.7063337f0 (tracked)
loss(X, Y) = 0.68901753f0 (tracked)
loss(X, Y) = 0.69181675f0 (tracked)
loss(X, Y) = 0.63317794f0 (tracked)
loss(X, Y) = 0.66718227f0 (tracked)
loss(X, Y) = 0.6926866f0 (tracked)
loss(X, Y) = 0.6613034f0 (tracked)
loss(X, Y) = 0.6622927f0 (tracked)
loss(X, Y) = 0.72102165f0 (tracked)
loss(X, Y) = 0.6889685f0 (tracked)
loss(X, Y) = 0.6954836f0 (tracked)
loss(X, Y) = 0.6337068f0 (tracked)
loss(X, Y) = 0.6630601f0 (tracked)
loss(X, Y) = 0.6127039f0 (tracked)
loss(X, Y) = 0.664256f0 (tracked)
loss(X, Y) = 0.6663506f0 (tracked)
loss(X, Y) = 0.70042527f0 (tracked)
loss(X, Y) = 0.6543753f0 (tracked)
loss(X, Y) = 0.6670537f0 (tracked)
loss(X, Y) = 0.6707404f0 (tracked)
loss(X, Y) = 0.72387993f0 (tracked)
loss(X, Y) = 0.620979f0 (tracked)
loss(X, Y) = 0.6756536f0 (tracked)
loss(X, Y) = 0.6519062f0 (tracked)
loss(X, Y) = 0.6664638f0 (tracked)
loss(X, Y) = 0.6428914f0 (tracked)
loss(X, Y) = 0.6732205f0 (tracked)
loss(X, Y) = 0.69586813f0 (tracked)
loss(X, Y) = 0.6596593f0 (tracked)
loss(X, Y) = 0.68925214f0 (tracked)
loss(X, Y) = 0.6833043f0 (tracked)
loss(X, Y) = 0.6878874f0 (tracked)
loss(X, Y) = 0.695259f0 (tracked)
loss(X, Y) = 0.665226f0 (tracked)
loss(X, Y) = 0.64416355f0 (tracked)
loss(X, Y) = 0.65849286f0 (tracked)
loss(X, Y) = 0.6287957f0 (tracked)
loss(X, Y) = 0.6807318f0 (tracked)
loss(X, Y) = 0.6392945f0 (tracked)
loss(X, Y) = 0.6809157f0 (tracked)
loss(X, Y) = 0.66051906f0 (tracked)
loss(X, Y) = 0.68157804f0 (tracked)
loss(X, Y) = 0.6903622f0 (tracked)
loss(X, Y) = 0.6458303f0 (tracked)
loss(X, Y) = 0.65936154f0 (tracked)
loss(X, Y) = 0.68183196f0 (tracked)
loss(X, Y) = 0.6406925f0 (tracked)
loss(X, Y) = 0.63794464f0 (tracked)
loss(X, Y) = 0.6332186f0 (tracked)
loss(X, Y) = 0.6946383f0 (tracked)
loss(X, Y) = 0.7132806f0 (tracked)
loss(X, Y) = 0.6728592f0 (tracked)
loss(X, Y) = 0.6569104f0 (tracked)
loss(X, Y) = 0.67075366f0 (tracked)
loss(X, Y) = 0.67380583f0 (tracked)
loss(X, Y) = 0.6365691f0 (tracked)
loss(X, Y) = 0.6568996f0 (tracked)
loss(X, Y) = 0.7017641f0 (tracked)
loss(X, Y) = 0.6731014f0 (tracked)
loss(X, Y) = 0.69374365f0 (tracked)
loss(X, Y) = 0.65648615f0 (tracked)
loss(X, Y) = 0.67647254f0 (tracked)
loss(X, Y) = 0.6580716f0 (tracked)
loss(X, Y) = 0.6779391f0 (tracked)
loss(X, Y) = 0.6483516f0 (tracked)
loss(X, Y) = 0.6782299f0 (tracked)
loss(X, Y) = 0.682853f0 (tracked)
loss(X, Y) = 0.6693316f0 (tracked)
loss(X, Y) = 0.59366006f0 (tracked)
loss(X, Y) = 0.64833057f0 (tracked)
loss(X, Y) = 0.6744772f0 (tracked)
loss(X, Y) = 0.6535638f0 (tracked)
loss(X, Y) = 0.66736436f0 (tracked)
loss(X, Y) = 0.688239f0 (tracked)
loss(X, Y) = 0.61790013f0 (tracked)
loss(X, Y) = 0.6580924f0 (tracked)
loss(X, Y) = 0.67417485f0 (tracked)
loss(X, Y) = 0.66816187f0 (tracked)
```

```
┌ Info: Epoch 1339
└ @ Main /Users/kcft114/.julia/packages/Flux/qXNjB/src/optimise/train.jl:105
┌ Info: Epoch 1340
└ @ Main /Users/kcft114/.julia/packages/Flux/qXNjB/src/optimise/train.jl:105
┌ Info: Epoch 1341
└ @ Main /Users/kcft114/.julia/packages/Flux/qXNjB/src/optimise/train.jl:105
┌ Info: Epoch 1342
└ @ Main /Users/kcft114/.julia/packages/Flux/qXNjB/src/optimise/train.jl:105
┌ Info: Epoch 1343
└ @ Main /Users/kcft114/.julia/packages/Flux/qXNjB/src/optimise/train.jl:105
┌ Info: Epoch 1344
└ @ Main /Users/kcft114/.julia/packages/Flux/qXNjB/src/optimise/train.jl:105
┌ Info: Epoch 1345
└ @ Main /Users/kcft114/.julia/packages/Flux/qXNjB/src/optimise/train.jl:105
┌ Info: Epoch 1346
└ @ Main /Users/kcft114/.julia/packages/Flux/qXNjB/src/optimise/train.jl:105
┌ Info: Epoch 1347
└ @ Main /Users/kcft114/.julia/packages/Flux/qXNjB/src/optimise/train.jl:105
┌ Info: Epoch 1348
└ @ Main /Users/kcft114/.julia/packages/Flux/qXNjB/src/optimise/train.jl:105
┌ Info: Epoch 1349
└ @ Main /Users/kcft114/.julia/packages/Flux/qXNjB/src/optimise/train.jl:105
┌ Info: Epoch 1350
└ @ Main /Users/kcft114/.julia/packages/Flux/qXNjB/src/optimise/train.jl:105
┌ Info: Epoch 1351
└ @ Main /Users/kcft114/.julia/packages/Flux/qXNjB/src/optimise/train.jl:105
┌ Info: Epoch 1352
└ @ Main /Users/kcft114/.julia/packages/Flux/qXNjB/src/optimise/train.jl:105
┌ Info: Epoch 1353
└ @ Main /Users/kcft114/.julia/packages/Flux/qXNjB/src/optimise/train.jl:105
┌ Info: Epoch 1354
└ @ Main /Users/kcft114/.julia/packages/Flux/qXNjB/src/optimise/train.jl:105
┌ Info: Epoch 1355
└ @ Main /Users/kcft114/.julia/packages/Flux/qXNjB/src/optimise/train.jl:105
┌ Info: Epoch 1356
└ @ Main /Users/kcft114/.julia/packages/Flux/qXNjB/src/optimise/train.jl:105
┌ Info: Epoch 1357
└ @ Main /Users/kcft114/.julia/packages/Flux/qXNjB/src/optimise/train.jl:105
┌ Info: Epoch 1358
└ @ Main /Users/kcft114/.julia/packages/Flux/qXNjB/src/optimise/train.jl:105
┌ Info: Epoch 1359
└ @ Main /Users/kcft114/.julia/packages/Flux/qXNjB/src/optimise/train.jl:105
┌ Info: Epoch 1360
└ @ Main /Users/kcft114/.julia/packages/Flux/qXNjB/src/optimise/train.jl:105
┌ Info: Epoch 1361
└ @ Main /Users/kcft114/.julia/packages/Flux/qXNjB/src/optimise/train.jl:105
┌ Info: Epoch 1362
└ @ Main /Users/kcft114/.julia/packages/Flux/qXNjB/src/optimise/train.jl:105
┌ Info: Epoch 1363
└ @ Main /Users/kcft114/.julia/packages/Flux/qXNjB/src/optimise/train.jl:105
┌ Info: Epoch 1364
└ @ Main /Users/kcft114/.julia/packages/Flux/qXNjB/src/optimise/train.jl:105
┌ Info: Epoch 1365
└ @ Main /Users/kcft114/.julia/packages/Flux/qXNjB/src/optimise/train.jl:105
┌ Info: Epoch 1366
└ @ Main /Users/kcft114/.julia/packages/Flux/qXNjB/src/optimise/train.jl:105
┌ Info: Epoch 1367
└ @ Main /Users/kcft114/.julia/packages/Flux/qXNjB/src/optimise/train.jl:105
┌ Info: Epoch 1368
└ @ Main /Users/kcft114/.julia/packages/Flux/qXNjB/src/optimise/train.jl:105
┌ Info: Epoch 1369
└ @ Main /Users/kcft114/.julia/packages/Flux/qXNjB/src/optimise/train.jl:105
┌ Info: Epoch 1370
└ @ Main /Users/kcft114/.julia/packages/Flux/qXNjB/src/optimise/train.jl:105
┌ Info: Epoch 1371
└ @ Main /Users/kcft114/.julia/packages/Flux/qXNjB/src/optimise/train.jl:105
┌ Info: Epoch 1372
└ @ Main /Users/kcft114/.julia/packages/Flux/qXNjB/src/optimise/train.jl:105
┌ Info: Epoch 1373
└ @ Main /Users/kcft114/.julia/packages/Flux/qXNjB/src/optimise/train.jl:105
┌ Info: Epoch 1374
└ @ Main /Users/kcft114/.julia/packages/Flux/qXNjB/src/optimise/train.jl:105
┌ Info: Epoch 1375
└ @ Main /Users/kcft114/.julia/packages/Flux/qXNjB/src/optimise/train.jl:105
┌ Info: Epoch 1376
└ @ Main /Users/kcft114/.julia/packages/Flux/qXNjB/src/optimise/train.jl:105
┌ Info: Epoch 1377
└ @ Main /Users/kcft114/.julia/packages/Flux/qXNjB/src/optimise/train.jl:105
┌ Info: Epoch 1378
└ @ Main /Users/kcft114/.julia/packages/Flux/qXNjB/src/optimise/train.jl:105
┌ Info: Epoch 1379
└ @ Main /Users/kcft114/.julia/packages/Flux/qXNjB/src/optimise/train.jl:105
┌ Info: Epoch 1380
└ @ Main /Users/kcft114/.julia/packages/Flux/qXNjB/src/optimise/train.jl:105
┌ Info: Epoch 1381
└ @ Main /Users/kcft114/.julia/packages/Flux/qXNjB/src/optimise/train.jl:105
┌ Info: Epoch 1382
└ @ Main /Users/kcft114/.julia/packages/Flux/qXNjB/src/optimise/train.jl:105
┌ Info: Epoch 1383
└ @ Main /Users/kcft114/.julia/packages/Flux/qXNjB/src/optimise/train.jl:105
┌ Info: Epoch 1384
└ @ Main /Users/kcft114/.julia/packages/Flux/qXNjB/src/optimise/train.jl:105
┌ Info: Epoch 1385
└ @ Main /Users/kcft114/.julia/packages/Flux/qXNjB/src/optimise/train.jl:105
┌ Info: Epoch 1386
└ @ Main /Users/kcft114/.julia/packages/Flux/qXNjB/src/optimise/train.jl:105
┌ Info: Epoch 1387
└ @ Main /Users/kcft114/.julia/packages/Flux/qXNjB/src/optimise/train.jl:105
┌ Info: Epoch 1388
└ @ Main /Users/kcft114/.julia/packages/Flux/qXNjB/src/optimise/train.jl:105
┌ Info: Epoch 1389
└ @ Main /Users/kcft114/.julia/packages/Flux/qXNjB/src/optimise/train.jl:105
┌ Info: Epoch 1390
└ @ Main /Users/kcft114/.julia/packages/Flux/qXNjB/src/optimise/train.jl:105
┌ Info: Epoch 1391
└ @ Main /Users/kcft114/.julia/packages/Flux/qXNjB/src/optimise/train.jl:105
┌ Info: Epoch 1392
└ @ Main /Users/kcft114/.julia/packages/Flux/qXNjB/src/optimise/train.jl:105
┌ Info: Epoch 1393
└ @ Main /Users/kcft114/.julia/packages/Flux/qXNjB/src/optimise/train.jl:105
┌ Info: Epoch 1394
└ @ Main /Users/kcft114/.julia/packages/Flux/qXNjB/src/optimise/train.jl:105
┌ Info: Epoch 1395
└ @ Main /Users/kcft114/.julia/packages/Flux/qXNjB/src/optimise/train.jl:105
┌ Info: Epoch 1396
└ @ Main /Users/kcft114/.julia/packages/Flux/qXNjB/src/optimise/train.jl:105
┌ Info: Epoch 1397
└ @ Main /Users/kcft114/.julia/packages/Flux/qXNjB/src/optimise/train.jl:105
┌ Info: Epoch 1398
└ @ Main /Users/kcft114/.julia/packages/Flux/qXNjB/src/optimise/train.jl:105
┌ Info: Epoch 1399
└ @ Main /Users/kcft114/.julia/packages/Flux/qXNjB/src/optimise/train.jl:105
┌ Info: Epoch 1400
└ @ Main /Users/kcft114/.julia/packages/Flux/qXNjB/src/optimise/train.jl:105
┌ Info: Epoch 1401
└ @ Main /Users/kcft114/.julia/packages/Flux/qXNjB/src/optimise/train.jl:105
┌ Info: Epoch 1402
└ @ Main /Users/kcft114/.julia/packages/Flux/qXNjB/src/optimise/train.jl:105
┌ Info: Epoch 1403
└ @ Main /Users/kcft114/.julia/packages/Flux/qXNjB/src/optimise/train.jl:105
┌ Info: Epoch 1404
└ @ Main /Users/kcft114/.julia/packages/Flux/qXNjB/src/optimise/train.jl:105
┌ Info: Epoch 1405
└ @ Main /Users/kcft114/.julia/packages/Flux/qXNjB/src/optimise/train.jl:105
┌ Info: Epoch 1406
└ @ Main /Users/kcft114/.julia/packages/Flux/qXNjB/src/optimise/train.jl:105
┌ Info: Epoch 1407
└ @ Main /Users/kcft114/.julia/packages/Flux/qXNjB/src/optimise/train.jl:105
┌ Info: Epoch 1408
└ @ Main /Users/kcft114/.julia/packages/Flux/qXNjB/src/optimise/train.jl:105
┌ Info: Epoch 1409
└ @ Main /Users/kcft114/.julia/packages/Flux/qXNjB/src/optimise/train.jl:105
┌ Info: Epoch 1410
└ @ Main /Users/kcft114/.julia/packages/Flux/qXNjB/src/optimise/train.jl:105
┌ Info: Epoch 1411
└ @ Main /Users/kcft114/.julia/packages/Flux/qXNjB/src/optimise/train.jl:105
┌ Info: Epoch 1412
└ @ Main /Users/kcft114/.julia/packages/Flux/qXNjB/src/optimise/train.jl:105
┌ Info: Epoch 1413
└ @ Main /Users/kcft114/.julia/packages/Flux/qXNjB/src/optimise/train.jl:105
┌ Info: Epoch 1414
└ @ Main /Users/kcft114/.julia/packages/Flux/qXNjB/src/optimise/train.jl:105
┌ Info: Epoch 1415
└ @ Main /Users/kcft114/.julia/packages/Flux/qXNjB/src/optimise/train.jl:105
┌ Info: Epoch 1416
└ @ Main /Users/kcft114/.julia/packages/Flux/qXNjB/src/optimise/train.jl:105
┌ Info: Epoch 1417
└ @ Main /Users/kcft114/.julia/packages/Flux/qXNjB/src/optimise/train.jl:105
┌ Info: Epoch 1418
└ @ Main /Users/kcft114/.julia/packages/Flux/qXNjB/src/optimise/train.jl:105
┌ Info: Epoch 1419
└ @ Main /Users/kcft114/.julia/packages/Flux/qXNjB/src/optimise/train.jl:105
┌ Info: Epoch 1420
└ @ Main /Users/kcft114/.julia/packages/Flux/qXNjB/src/optimise/train.jl:105
┌ Info: Epoch 1421
└ @ Main /Users/kcft114/.julia/packages/Flux/qXNjB/src/optimise/train.jl:105
┌ Info: Epoch 1422
└ @ Main /Users/kcft114/.julia/packages/Flux/qXNjB/src/optimise/train.jl:105
┌ Info: Epoch 1423
└ @ Main /Users/kcft114/.julia/packages/Flux/qXNjB/src/optimise/train.jl:105
┌ Info: Epoch 1424
└ @ Main /Users/kcft114/.julia/packages/Flux/qXNjB/src/optimise/train.jl:105
┌ Info: Epoch 1425
└ @ Main /Users/kcft114/.julia/packages/Flux/qXNjB/src/optimise/train.jl:105
┌ Info: Epoch 1426
└ @ Main /Users/kcft114/.julia/packages/Flux/qXNjB/src/optimise/train.jl:105
┌ Info: Epoch 1427
└ @ Main /Users/kcft114/.julia/packages/Flux/qXNjB/src/optimise/train.jl:105
┌ Info: Epoch 1428
└ @ Main /Users/kcft114/.julia/packages/Flux/qXNjB/src/optimise/train.jl:105
┌ Info: Epoch 1429
└ @ Main /Users/kcft114/.julia/packages/Flux/qXNjB/src/optimise/train.jl:105
┌ Info: Epoch 1430
└ @ Main /Users/kcft114/.julia/packages/Flux/qXNjB/src/optimise/train.jl:105
┌ Info: Epoch 1431
└ @ Main /Users/kcft114/.julia/packages/Flux/qXNjB/src/optimise/train.jl:105
┌ Info: Epoch 1432
└ @ Main /Users/kcft114/.julia/packages/Flux/qXNjB/src/optimise/train.jl:105
┌ Info: Epoch 1433
└ @ Main /Users/kcft114/.julia/packages/Flux/qXNjB/src/optimise/train.jl:105
┌ Info: Epoch 1434
└ @ Main /Users/kcft114/.julia/packages/Flux/qXNjB/src/optimise/train.jl:105
┌ Info: Epoch 1435
└ @ Main /Users/kcft114/.julia/packages/Flux/qXNjB/src/optimise/train.jl:105
┌ Info: Epoch 1436
└ @ Main /Users/kcft114/.julia/packages/Flux/qXNjB/src/optimise/train.jl:105
┌ Info: Epoch 1437
└ @ Main /Users/kcft114/.julia/packages/Flux/qXNjB/src/optimise/train.jl:105
┌ Info: Epoch 1438
└ @ Main /Users/kcft114/.julia/packages/Flux/qXNjB/src/optimise/train.jl:105
┌ Info: Epoch 1439
└ @ Main /Users/kcft114/.julia/packages/Flux/qXNjB/src/optimise/train.jl:105
┌ Info: Epoch 1440
└ @ Main /Users/kcft114/.julia/packages/Flux/qXNjB/src/optimise/train.jl:105
┌ Info: Epoch 1441
└ @ Main /Users/kcft114/.julia/packages/Flux/qXNjB/src/optimise/train.jl:105
┌ Info: Epoch 1442
└ @ Main /Users/kcft114/.julia/packages/Flux/qXNjB/src/optimise/train.jl:105
┌ Info: Epoch 1443
└ @ Main /Users/kcft114/.julia/packages/Flux/qXNjB/src/optimise/train.jl:105
┌ Info: Epoch 1444
└ @ Main /Users/kcft114/.julia/packages/Flux/qXNjB/src/optimise/train.jl:105
┌ Info: Epoch 1445
└ @ Main /Users/kcft114/.julia/packages/Flux/qXNjB/src/optimise/train.jl:105
┌ Info: Epoch 1446
└ @ Main /Users/kcft114/.julia/packages/Flux/qXNjB/src/optimise/train.jl:105
┌ Info: Epoch 1447
└ @ Main /Users/kcft114/.julia/packages/Flux/qXNjB/src/optimise/train.jl:105
┌ Info: Epoch 1448
└ @ Main /Users/kcft114/.julia/packages/Flux/qXNjB/src/optimise/train.jl:105
┌ Info: Epoch 1449
└ @ Main /Users/kcft114/.julia/packages/Flux/qXNjB/src/optimise/train.jl:105
┌ Info: Epoch 1450
└ @ Main /Users/kcft114/.julia/packages/Flux/qXNjB/src/optimise/train.jl:105
┌ Info: Epoch 1451
└ @ Main /Users/kcft114/.julia/packages/Flux/qXNjB/src/optimise/train.jl:105
┌ Info: Epoch 1452
└ @ Main /Users/kcft114/.julia/packages/Flux/qXNjB/src/optimise/train.jl:105
┌ Info: Epoch 1453
└ @ Main /Users/kcft114/.julia/packages/Flux/qXNjB/src/optimise/train.jl:105
┌ Info: Epoch 1454
└ @ Main /Users/kcft114/.julia/packages/Flux/qXNjB/src/optimise/train.jl:105
┌ Info: Epoch 1455
└ @ Main /Users/kcft114/.julia/packages/Flux/qXNjB/src/optimise/train.jl:105
┌ Info: Epoch 1456
└ @ Main /Users/kcft114/.julia/packages/Flux/qXNjB/src/optimise/train.jl:105
┌ Info: Epoch 1457
└ @ Main /Users/kcft114/.julia/packages/Flux/qXNjB/src/optimise/train.jl:105
┌ Info: Epoch 1458
└ @ Main /Users/kcft114/.julia/packages/Flux/qXNjB/src/optimise/train.jl:105
┌ Info: Epoch 1459
└ @ Main /Users/kcft114/.julia/packages/Flux/qXNjB/src/optimise/train.jl:105
┌ Info: Epoch 1460
└ @ Main /Users/kcft114/.julia/packages/Flux/qXNjB/src/optimise/train.jl:105
┌ Info: Epoch 1461
└ @ Main /Users/kcft114/.julia/packages/Flux/qXNjB/src/optimise/train.jl:105
┌ Info: Epoch 1462
└ @ Main /Users/kcft114/.julia/packages/Flux/qXNjB/src/optimise/train.jl:105
┌ Info: Epoch 1463
└ @ Main /Users/kcft114/.julia/packages/Flux/qXNjB/src/optimise/train.jl:105
┌ Info: Epoch 1464
└ @ Main /Users/kcft114/.julia/packages/Flux/qXNjB/src/optimise/train.jl:105
┌ Info: Epoch 1465
└ @ Main /Users/kcft114/.julia/packages/Flux/qXNjB/src/optimise/train.jl:105
┌ Info: Epoch 1466
└ @ Main /Users/kcft114/.julia/packages/Flux/qXNjB/src/optimise/train.jl:105
┌ Info: Epoch 1467
└ @ Main /Users/kcft114/.julia/packages/Flux/qXNjB/src/optimise/train.jl:105
┌ Info: Epoch 1468
└ @ Main /Users/kcft114/.julia/packages/Flux/qXNjB/src/optimise/train.jl:105
┌ Info: Epoch 1469
└ @ Main /Users/kcft114/.julia/packages/Flux/qXNjB/src/optimise/train.jl:105
┌ Info: Epoch 1470
└ @ Main /Users/kcft114/.julia/packages/Flux/qXNjB/src/optimise/train.jl:105
┌ Info: Epoch 1471
└ @ Main /Users/kcft114/.julia/packages/Flux/qXNjB/src/optimise/train.jl:105
┌ Info: Epoch 1472
└ @ Main /Users/kcft114/.julia/packages/Flux/qXNjB/src/optimise/train.jl:105
┌ Info: Epoch 1473
└ @ Main /Users/kcft114/.julia/packages/Flux/qXNjB/src/optimise/train.jl:105
┌ Info: Epoch 1474
└ @ Main /Users/kcft114/.julia/packages/Flux/qXNjB/src/optimise/train.jl:105
┌ Info: Epoch 1475
└ @ Main /Users/kcft114/.julia/packages/Flux/qXNjB/src/optimise/train.jl:105
```

```
loss(X, Y) = 0.6959116f0 (tracked)
loss(X, Y) = 0.6678538f0 (tracked)
loss(X, Y) = 0.61296195f0 (tracked)
loss(X, Y) = 0.65493333f0 (tracked)
loss(X, Y) = 0.66609186f0 (tracked)
loss(X, Y) = 0.66309524f0 (tracked)
loss(X, Y) = 0.6598352f0 (tracked)
loss(X, Y) = 0.6695768f0 (tracked)
loss(X, Y) = 0.6818154f0 (tracked)
loss(X, Y) = 0.6568318f0 (tracked)
loss(X, Y) = 0.6458922f0 (tracked)
loss(X, Y) = 0.6421903f0 (tracked)
loss(X, Y) = 0.64641315f0 (tracked)
loss(X, Y) = 0.6958996f0 (tracked)
loss(X, Y) = 0.6542654f0 (tracked)
loss(X, Y) = 0.68100196f0 (tracked)
loss(X, Y) = 0.6795845f0 (tracked)
loss(X, Y) = 0.65962684f0 (tracked)
loss(X, Y) = 0.6535537f0 (tracked)
loss(X, Y) = 0.6874785f0 (tracked)
loss(X, Y) = 0.66340536f0 (tracked)
loss(X, Y) = 0.6723143f0 (tracked)
loss(X, Y) = 0.6446178f0 (tracked)
loss(X, Y) = 0.6538817f0 (tracked)
loss(X, Y) = 0.6888384f0 (tracked)
loss(X, Y) = 0.6530408f0 (tracked)
loss(X, Y) = 0.6647593f0 (tracked)
loss(X, Y) = 0.6751719f0 (tracked)
loss(X, Y) = 0.6711021f0 (tracked)
loss(X, Y) = 0.66386384f0 (tracked)
loss(X, Y) = 0.6270536f0 (tracked)
loss(X, Y) = 0.6617138f0 (tracked)
loss(X, Y) = 0.66805667f0 (tracked)
loss(X, Y) = 0.6599437f0 (tracked)
loss(X, Y) = 0.621299f0 (tracked)
loss(X, Y) = 0.7114187f0 (tracked)
loss(X, Y) = 0.62056065f0 (tracked)
loss(X, Y) = 0.6347945f0 (tracked)
loss(X, Y) = 0.67499185f0 (tracked)
loss(X, Y) = 0.6383294f0 (tracked)
loss(X, Y) = 0.65953463f0 (tracked)
loss(X, Y) = 0.6605863f0 (tracked)
loss(X, Y) = 0.6636992f0 (tracked)
loss(X, Y) = 0.63137406f0 (tracked)
loss(X, Y) = 0.6593082f0 (tracked)
loss(X, Y) = 0.65991f0 (tracked)
loss(X, Y) = 0.6827634f0 (tracked)
loss(X, Y) = 0.63084745f0 (tracked)
loss(X, Y) = 0.69152373f0 (tracked)
loss(X, Y) = 0.67112243f0 (tracked)
loss(X, Y) = 0.64415973f0 (tracked)
loss(X, Y) = 0.6676149f0 (tracked)
loss(X, Y) = 0.6318966f0 (tracked)
loss(X, Y) = 0.6780599f0 (tracked)
loss(X, Y) = 0.6683469f0 (tracked)
loss(X, Y) = 0.6345388f0 (tracked)
loss(X, Y) = 0.6797041f0 (tracked)
loss(X, Y) = 0.64253926f0 (tracked)
loss(X, Y) = 0.70591533f0 (tracked)
loss(X, Y) = 0.6324592f0 (tracked)
loss(X, Y) = 0.6680792f0 (tracked)
loss(X, Y) = 0.6673136f0 (tracked)
loss(X, Y) = 0.6711165f0 (tracked)
loss(X, Y) = 0.66762805f0 (tracked)
loss(X, Y) = 0.66849947f0 (tracked)
loss(X, Y) = 0.6566562f0 (tracked)
loss(X, Y) = 0.63407665f0 (tracked)
loss(X, Y) = 0.65135545f0 (tracked)
loss(X, Y) = 0.6554306f0 (tracked)
loss(X, Y) = 0.6716359f0 (tracked)
loss(X, Y) = 0.6448614f0 (tracked)
loss(X, Y) = 0.69008064f0 (tracked)
loss(X, Y) = 0.638071f0 (tracked)
loss(X, Y) = 0.65534264f0 (tracked)
loss(X, Y) = 0.7088086f0 (tracked)
loss(X, Y) = 0.69971794f0 (tracked)
loss(X, Y) = 0.70153755f0 (tracked)
loss(X, Y) = 0.6738605f0 (tracked)
loss(X, Y) = 0.6497817f0 (tracked)
loss(X, Y) = 0.6676745f0 (tracked)
loss(X, Y) = 0.6081415f0 (tracked)
loss(X, Y) = 0.68392265f0 (tracked)
loss(X, Y) = 0.6614556f0 (tracked)
loss(X, Y) = 0.66021794f0 (tracked)
loss(X, Y) = 0.68779606f0 (tracked)
loss(X, Y) = 0.65620196f0 (tracked)
loss(X, Y) = 0.6770875f0 (tracked)
loss(X, Y) = 0.6273709f0 (tracked)
loss(X, Y) = 0.6399662f0 (tracked)
loss(X, Y) = 0.6524677f0 (tracked)
loss(X, Y) = 0.6863307f0 (tracked)
loss(X, Y) = 0.65878433f0 (tracked)
loss(X, Y) = 0.6364464f0 (tracked)
loss(X, Y) = 0.6794976f0 (tracked)
loss(X, Y) = 0.6396266f0 (tracked)
loss(X, Y) = 0.6390237f0 (tracked)
loss(X, Y) = 0.6553628f0 (tracked)
loss(X, Y) = 0.6669782f0 (tracked)
loss(X, Y) = 0.6064821f0 (tracked)
loss(X, Y) = 0.61636376f0 (tracked)
loss(X, Y) = 0.7042939f0 (tracked)
loss(X, Y) = 0.6478337f0 (tracked)
loss(X, Y) = 0.6708485f0 (tracked)
loss(X, Y) = 0.6476134f0 (tracked)
loss(X, Y) = 0.6122503f0 (tracked)
loss(X, Y) = 0.63146895f0 (tracked)
loss(X, Y) = 0.6629837f0 (tracked)
loss(X, Y) = 0.7258536f0 (tracked)
loss(X, Y) = 0.6480614f0 (tracked)
loss(X, Y) = 0.6748813f0 (tracked)
loss(X, Y) = 0.634684f0 (tracked)
loss(X, Y) = 0.6688082f0 (tracked)
loss(X, Y) = 0.6368485f0 (tracked)
loss(X, Y) = 0.6611579f0 (tracked)
loss(X, Y) = 0.6707258f0 (tracked)
loss(X, Y) = 0.6480164f0 (tracked)
loss(X, Y) = 0.6817952f0 (tracked)
loss(X, Y) = 0.6643745f0 (tracked)
loss(X, Y) = 0.68989974f0 (tracked)
loss(X, Y) = 0.6481807f0 (tracked)
loss(X, Y) = 0.67140317f0 (tracked)
loss(X, Y) = 0.61408013f0 (tracked)
loss(X, Y) = 0.6433641f0 (tracked)
loss(X, Y) = 0.6824888f0 (tracked)
loss(X, Y) = 0.6394039f0 (tracked)
loss(X, Y) = 0.66903096f0 (tracked)
loss(X, Y) = 0.65654916f0 (tracked)
loss(X, Y) = 0.6356459f0 (tracked)
loss(X, Y) = 0.6562736f0 (tracked)
loss(X, Y) = 0.65234077f0 (tracked)
loss(X, Y) = 0.60321414f0 (tracked)
loss(X, Y) = 0.67640907f0 (tracked)
loss(X, Y) = 0.63158655f0 (tracked)
loss(X, Y) = 0.63253176f0 (tracked)
loss(X, Y) = 0.6311949f0 (tracked)
loss(X, Y) = 0.6668478f0 (tracked)
loss(X, Y) = 0.65309f0 (tracked)
```

```
┌ Info: Epoch 1476
└ @ Main /Users/kcft114/.julia/packages/Flux/qXNjB/src/optimise/train.jl:105
┌ Info: Epoch 1477
└ @ Main /Users/kcft114/.julia/packages/Flux/qXNjB/src/optimise/train.jl:105
┌ Info: Epoch 1478
└ @ Main /Users/kcft114/.julia/packages/Flux/qXNjB/src/optimise/train.jl:105
┌ Info: Epoch 1479
└ @ Main /Users/kcft114/.julia/packages/Flux/qXNjB/src/optimise/train.jl:105
┌ Info: Epoch 1480
└ @ Main /Users/kcft114/.julia/packages/Flux/qXNjB/src/optimise/train.jl:105
┌ Info: Epoch 1481
└ @ Main /Users/kcft114/.julia/packages/Flux/qXNjB/src/optimise/train.jl:105
┌ Info: Epoch 1482
└ @ Main /Users/kcft114/.julia/packages/Flux/qXNjB/src/optimise/train.jl:105
┌ Info: Epoch 1483
└ @ Main /Users/kcft114/.julia/packages/Flux/qXNjB/src/optimise/train.jl:105
┌ Info: Epoch 1484
└ @ Main /Users/kcft114/.julia/packages/Flux/qXNjB/src/optimise/train.jl:105
┌ Info: Epoch 1485
└ @ Main /Users/kcft114/.julia/packages/Flux/qXNjB/src/optimise/train.jl:105
┌ Info: Epoch 1486
└ @ Main /Users/kcft114/.julia/packages/Flux/qXNjB/src/optimise/train.jl:105
┌ Info: Epoch 1487
└ @ Main /Users/kcft114/.julia/packages/Flux/qXNjB/src/optimise/train.jl:105
┌ Info: Epoch 1488
└ @ Main /Users/kcft114/.julia/packages/Flux/qXNjB/src/optimise/train.jl:105
┌ Info: Epoch 1489
└ @ Main /Users/kcft114/.julia/packages/Flux/qXNjB/src/optimise/train.jl:105
┌ Info: Epoch 1490
└ @ Main /Users/kcft114/.julia/packages/Flux/qXNjB/src/optimise/train.jl:105
┌ Info: Epoch 1491
└ @ Main /Users/kcft114/.julia/packages/Flux/qXNjB/src/optimise/train.jl:105
┌ Info: Epoch 1492
└ @ Main /Users/kcft114/.julia/packages/Flux/qXNjB/src/optimise/train.jl:105
┌ Info: Epoch 1493
└ @ Main /Users/kcft114/.julia/packages/Flux/qXNjB/src/optimise/train.jl:105
┌ Info: Epoch 1494
└ @ Main /Users/kcft114/.julia/packages/Flux/qXNjB/src/optimise/train.jl:105
┌ Info: Epoch 1495
└ @ Main /Users/kcft114/.julia/packages/Flux/qXNjB/src/optimise/train.jl:105
┌ Info: Epoch 1496
└ @ Main /Users/kcft114/.julia/packages/Flux/qXNjB/src/optimise/train.jl:105
┌ Info: Epoch 1497
└ @ Main /Users/kcft114/.julia/packages/Flux/qXNjB/src/optimise/train.jl:105
┌ Info: Epoch 1498
└ @ Main /Users/kcft114/.julia/packages/Flux/qXNjB/src/optimise/train.jl:105
┌ Info: Epoch 1499
└ @ Main /Users/kcft114/.julia/packages/Flux/qXNjB/src/optimise/train.jl:105
┌ Info: Epoch 1500
└ @ Main /Users/kcft114/.julia/packages/Flux/qXNjB/src/optimise/train.jl:105
┌ Info: Epoch 1501
└ @ Main /Users/kcft114/.julia/packages/Flux/qXNjB/src/optimise/train.jl:105
┌ Info: Epoch 1502
└ @ Main /Users/kcft114/.julia/packages/Flux/qXNjB/src/optimise/train.jl:105
┌ Info: Epoch 1503
└ @ Main /Users/kcft114/.julia/packages/Flux/qXNjB/src/optimise/train.jl:105
┌ Info: Epoch 1504
└ @ Main /Users/kcft114/.julia/packages/Flux/qXNjB/src/optimise/train.jl:105
┌ Info: Epoch 1505
└ @ Main /Users/kcft114/.julia/packages/Flux/qXNjB/src/optimise/train.jl:105
┌ Info: Epoch 1506
└ @ Main /Users/kcft114/.julia/packages/Flux/qXNjB/src/optimise/train.jl:105
┌ Info: Epoch 1507
└ @ Main /Users/kcft114/.julia/packages/Flux/qXNjB/src/optimise/train.jl:105
┌ Info: Epoch 1508
└ @ Main /Users/kcft114/.julia/packages/Flux/qXNjB/src/optimise/train.jl:105
┌ Info: Epoch 1509
└ @ Main /Users/kcft114/.julia/packages/Flux/qXNjB/src/optimise/train.jl:105
┌ Info: Epoch 1510
└ @ Main /Users/kcft114/.julia/packages/Flux/qXNjB/src/optimise/train.jl:105
┌ Info: Epoch 1511
└ @ Main /Users/kcft114/.julia/packages/Flux/qXNjB/src/optimise/train.jl:105
┌ Info: Epoch 1512
└ @ Main /Users/kcft114/.julia/packages/Flux/qXNjB/src/optimise/train.jl:105
┌ Info: Epoch 1513
└ @ Main /Users/kcft114/.julia/packages/Flux/qXNjB/src/optimise/train.jl:105
┌ Info: Epoch 1514
└ @ Main /Users/kcft114/.julia/packages/Flux/qXNjB/src/optimise/train.jl:105
┌ Info: Epoch 1515
└ @ Main /Users/kcft114/.julia/packages/Flux/qXNjB/src/optimise/train.jl:105
┌ Info: Epoch 1516
└ @ Main /Users/kcft114/.julia/packages/Flux/qXNjB/src/optimise/train.jl:105
┌ Info: Epoch 1517
└ @ Main /Users/kcft114/.julia/packages/Flux/qXNjB/src/optimise/train.jl:105
┌ Info: Epoch 1518
└ @ Main /Users/kcft114/.julia/packages/Flux/qXNjB/src/optimise/train.jl:105
┌ Info: Epoch 1519
└ @ Main /Users/kcft114/.julia/packages/Flux/qXNjB/src/optimise/train.jl:105
┌ Info: Epoch 1520
└ @ Main /Users/kcft114/.julia/packages/Flux/qXNjB/src/optimise/train.jl:105
┌ Info: Epoch 1521
└ @ Main /Users/kcft114/.julia/packages/Flux/qXNjB/src/optimise/train.jl:105
┌ Info: Epoch 1522
└ @ Main /Users/kcft114/.julia/packages/Flux/qXNjB/src/optimise/train.jl:105
┌ Info: Epoch 1523
└ @ Main /Users/kcft114/.julia/packages/Flux/qXNjB/src/optimise/train.jl:105
┌ Info: Epoch 1524
└ @ Main /Users/kcft114/.julia/packages/Flux/qXNjB/src/optimise/train.jl:105
┌ Info: Epoch 1525
└ @ Main /Users/kcft114/.julia/packages/Flux/qXNjB/src/optimise/train.jl:105
┌ Info: Epoch 1526
└ @ Main /Users/kcft114/.julia/packages/Flux/qXNjB/src/optimise/train.jl:105
┌ Info: Epoch 1527
└ @ Main /Users/kcft114/.julia/packages/Flux/qXNjB/src/optimise/train.jl:105
┌ Info: Epoch 1528
└ @ Main /Users/kcft114/.julia/packages/Flux/qXNjB/src/optimise/train.jl:105
┌ Info: Epoch 1529
└ @ Main /Users/kcft114/.julia/packages/Flux/qXNjB/src/optimise/train.jl:105
┌ Info: Epoch 1530
└ @ Main /Users/kcft114/.julia/packages/Flux/qXNjB/src/optimise/train.jl:105
┌ Info: Epoch 1531
└ @ Main /Users/kcft114/.julia/packages/Flux/qXNjB/src/optimise/train.jl:105
┌ Info: Epoch 1532
└ @ Main /Users/kcft114/.julia/packages/Flux/qXNjB/src/optimise/train.jl:105
┌ Info: Epoch 1533
└ @ Main /Users/kcft114/.julia/packages/Flux/qXNjB/src/optimise/train.jl:105
┌ Info: Epoch 1534
└ @ Main /Users/kcft114/.julia/packages/Flux/qXNjB/src/optimise/train.jl:105
┌ Info: Epoch 1535
└ @ Main /Users/kcft114/.julia/packages/Flux/qXNjB/src/optimise/train.jl:105
┌ Info: Epoch 1536
└ @ Main /Users/kcft114/.julia/packages/Flux/qXNjB/src/optimise/train.jl:105
┌ Info: Epoch 1537
└ @ Main /Users/kcft114/.julia/packages/Flux/qXNjB/src/optimise/train.jl:105
┌ Info: Epoch 1538
└ @ Main /Users/kcft114/.julia/packages/Flux/qXNjB/src/optimise/train.jl:105
┌ Info: Epoch 1539
└ @ Main /Users/kcft114/.julia/packages/Flux/qXNjB/src/optimise/train.jl:105
┌ Info: Epoch 1540
└ @ Main /Users/kcft114/.julia/packages/Flux/qXNjB/src/optimise/train.jl:105
┌ Info: Epoch 1541
└ @ Main /Users/kcft114/.julia/packages/Flux/qXNjB/src/optimise/train.jl:105
┌ Info: Epoch 1542
└ @ Main /Users/kcft114/.julia/packages/Flux/qXNjB/src/optimise/train.jl:105
┌ Info: Epoch 1543
└ @ Main /Users/kcft114/.julia/packages/Flux/qXNjB/src/optimise/train.jl:105
┌ Info: Epoch 1544
└ @ Main /Users/kcft114/.julia/packages/Flux/qXNjB/src/optimise/train.jl:105
┌ Info: Epoch 1545
└ @ Main /Users/kcft114/.julia/packages/Flux/qXNjB/src/optimise/train.jl:105
┌ Info: Epoch 1546
└ @ Main /Users/kcft114/.julia/packages/Flux/qXNjB/src/optimise/train.jl:105
┌ Info: Epoch 1547
└ @ Main /Users/kcft114/.julia/packages/Flux/qXNjB/src/optimise/train.jl:105
┌ Info: Epoch 1548
└ @ Main /Users/kcft114/.julia/packages/Flux/qXNjB/src/optimise/train.jl:105
┌ Info: Epoch 1549
└ @ Main /Users/kcft114/.julia/packages/Flux/qXNjB/src/optimise/train.jl:105
┌ Info: Epoch 1550
└ @ Main /Users/kcft114/.julia/packages/Flux/qXNjB/src/optimise/train.jl:105
┌ Info: Epoch 1551
└ @ Main /Users/kcft114/.julia/packages/Flux/qXNjB/src/optimise/train.jl:105
┌ Info: Epoch 1552
└ @ Main /Users/kcft114/.julia/packages/Flux/qXNjB/src/optimise/train.jl:105
┌ Info: Epoch 1553
└ @ Main /Users/kcft114/.julia/packages/Flux/qXNjB/src/optimise/train.jl:105
┌ Info: Epoch 1554
└ @ Main /Users/kcft114/.julia/packages/Flux/qXNjB/src/optimise/train.jl:105
┌ Info: Epoch 1555
