## Supplementary material for "A Bayesian neural network for toxicity prediction": 4_NN_multiclass.ipynb

```
# %% [code]
## ---------------------------------------------------------
## Import libraries 
## ---------------------------------------------------------

using CSV
using Statistics
using Flux
using Flux: onehotbatch, throttle, crossentropy, onecold, @epochs
using Base.Iterators: repeated
using MLBase
using LinearAlgebra
using MLPreprocessing

using Random
Random.seed!(1234);

# %% [code]
function res(X, Y, Xt, Yt)
    println("train accuracy = ", accuracy(X, Y))
    println("test accuracy = ", accuracy(Xt, Yt))

    y_pred = onecold(nn(X)) ; y = onecold(Y); C = confusmat(3, y, y_pred)
    bacc_train = 1/3 *(C[1,1] / sum(C[1, :]) + C[2,2] / sum(C[2, :]) + C[3,3] / sum(C[3, :]))
    println("train BA = ", round(bacc_train, digits=2))

    y_pred = onecold(nn(Xt)) ; y = onecold(Yt); C = confusmat(3, y, y_pred)
    bacc_test = 1/3 *(C[1,1] / sum(C[1, :]) + C[2,2] / sum(C[2, :]) + C[3,3] / sum(C[3, :]))
    println("test BA = ", round(bacc_test, digits=2))
    
    return accuracy(X, Y), accuracy(Xt, Yt), bacc_train, bacc_test
end


# %% [code]
## ---------------------------------------------------------
## Read and prepare data
## ---------------------------------------------------------

df_train = CSV.read("/Users/kcft114/Box Sync/27_BNN_DILI/data/Aleo_train_match.csv")
df_test = CSV.read("/Users/kcft114/Box Sync/27_BNN_DILI/data/Aleo_test_match.csv")

df_Xtrain = df_train[!, [:ClogP, :BSEP, :Glu, :Glu_Gal, :THLE, :HepG2, :Fsp3, :log10cmax]]
df_Xtest = df_test[!, [:ClogP, :BSEP, :Glu, :Glu_Gal, :THLE, :HepG2, :Fsp3, :log10cmax]]

scaler = fit(StandardScaler, df_Xtrain)
transform!(df_Xtrain, scaler)
transform!(df_Xtest, scaler)

df_y_train = df_train[!, :dili_sev]
df_y_test = df_test[!, :dili_sev];

df_yXtrain = hcat(df_y_train, df_Xtrain)
df_yXtest = hcat(df_y_test, df_Xtest)

X_train = convert(Matrix, df_Xtrain)
X_test = convert(Matrix, df_Xtest)

df_y_train = df_train[!, :dili_sev]
df_y_test = df_test[!, :dili_sev]

Y_train = Int.(convert(Array, df_y_train))
Y_test = Int.(convert(Array, df_y_test))

Y = onehotbatch(Y_train, 1:3)
X = X_train'

Yt = onehotbatch(Y_test, 1:3)
Xt = X_test'

println(size(X))
println(size(Y))

# %% [code]
n_h = 15

# %% [code]
## ---------------------------------------------------------
## Experiments - no regularization
## ---------------------------------------------------------

n_exper = 100
experiments_none = zeros(n_exper, 4)


for i=1:n_exper
    
    nn = Chain(
          Dense(8, n_h, relu),
          Dense(n_h, 3),
          softmax)
    
    loss(x, y) = crossentropy(nn(x), y)
    accuracy(x, y) = mean(onecold(nn(x)) .== onecold(y))

    dataset = [(X, Y)]
    evalcb = () -> @show(loss(X, Y))
    
    opt = ADAM(0.001)
    
    @epochs 20000 Flux.train!(loss, Flux.params(nn), dataset, opt, cb = throttle(evalcb, 10));
    
    ac_train, ac_test, ba_train, ba_test =  res(X, Y, Xt, Yt);
    global experiments_none[i,:] = [ac_train, ac_test, ba_train, ba_test]
end

# %% [code]
## ---------------------------------------------------------
## Experiments - Regularization via Dropout
## ---------------------------------------------------------


# %% [code]
p_dropout=0.1

n_exper = 100
experiments_01 = zeros(n_exper, 4)


for i=1:n_exper
    
    nn = Chain(
              Dense(8, n_h, relu),
              Dropout(p_dropout),
              Dense(n_h, 3),
              softmax)
    
    loss(x, y) = crossentropy(nn(x), y)
    accuracy(x, y) = mean(
    onecold(nn(x)) .== onecold(y))

    dataset = [(X, Y)]

    evalcb = () -> @show(loss(X, Y))
    
    opt = ADAM(0.001)
    
    @epochs 2000 Flux.train!(loss, Flux.params(nn), dataset, opt, cb = throttle(evalcb, 10));
    
    ac_train, ac_test, ba_train, ba_test =  res(X, Y, Xt, Yt);
    experiments_01[i,:] = [ac_train, ac_test, ba_train, ba_test]
end

# %% [code]
p_dropout=0.2

n_exper = 100
experiments_02 = zeros(n_exper, 4)

for i=1:n_exper
    
    nn = Chain(
              Dense(8, n_h, relu),
              Dropout(p_dropout),
              Dense(n_h, 3),
              softmax)
    
    loss(x, y) = crossentropy(nn(x), y)
    accuracy(x, y) = mean(onecold(nn(x)) .== onecold(y))

    dataset = [(X, Y)]
    evalcb = () -> @show(loss(X, Y))
    
    opt = ADAM(0.001)
    
    @epochs 2000 Flux.train!(loss, Flux.params(nn), dataset, opt, cb = throttle(evalcb, 10));
    
    ac_train, ac_test, ba_train, ba_test =  res(X, Y, Xt, Yt);
    experiments_02[i,:] = [ac_train, ac_test, ba_train, ba_test]
end

# %% [code]
p_dropout=0.3

n_exper = 100
experiments_03 = zeros(n_exper, 4)

for i=1:n_exper
    
    nn = Chain(
              Dense(8, n_h, relu),
              Dropout(p_dropout),
              Dense(n_h, 3),
              softmax)
    
    loss(x, y) = crossentropy(nn(x), y)
    accuracy(x, y) = mean(onecold(nn(x)) .== onecold(y))

    dataset = [(X, Y)]
    evalcb = () -> @show(loss(X, Y))
    
    opt = ADAM(0.001)
    
    @epochs 2000 Flux.train!(loss, Flux.params(nn), dataset, opt, cb = throttle(evalcb, 10));
    
    ac_train, ac_test, ba_train, ba_test =  res(X, Y, Xt, Yt);
    experiments_03[i,:] = [ac_train, ac_test, ba_train, ba_test]
end

# %% [code]
p_dropout=0.4

n_exper = 100
experiments_04 = zeros(n_exper, 4)

for i=1:n_exper
    
    nn = Chain(
              Dense(8, n_h, relu),
              Dropout(p_dropout),
              Dense(n_h, 3),
              softmax)
    
    loss(x, y) = crossentropy(nn(x), y)
    accuracy(x, y) = mean(onecold(nn(x)) .== onecold(y))

    dataset = [(X, Y)]
    evalcb = () -> @show(loss(X, Y))
    
    opt = ADAM(0.001)
    
    @epochs 2000 Flux.train!(loss, Flux.params(nn), dataset, opt, cb = throttle(evalcb, 10));
    
    ac_train, ac_test, ba_train, ba_test =  res(X, Y, Xt, Yt);
    experiments_04[i,:] = [ac_train, ac_test, ba_train, ba_test]
end

# %% [code]
p_dropout=0.5

n_exper = 100
experiments_05 = zeros(n_exper, 4)

for i=1:n_exper
    
    nn = Chain(
              Dense(8, n_h, relu),
              Dropout(p_dropout),
              Dense(n_h, 3),
              softmax)
    
    loss(x, y) = crossentropy(nn(x), y)
    accuracy(x, y) = mean(onecold(nn(x)) .== onecold(y))

    dataset = [(X, Y)]
    evalcb = () -> @show(loss(X, Y))
    
    opt = ADAM(0.001)
    
    @epochs 2000 Flux.train!(loss, Flux.params(nn), dataset, opt, cb = throttle(evalcb, 10));
    
    ac_train, ac_test, ba_train, ba_test =  res(X, Y, Xt, Yt);
    experiments_05[i,:] = [ac_train, ac_test, ba_train, ba_test]
end

# %% [code]
p_dropout=0.6

n_exper = 100
experiments_06 = zeros(n_exper, 4)

for i=1:n_exper
    
    nn = Chain(
              Dense(8, n_h, relu),
              Dropout(p_dropout),
              Dense(n_h, 3),
              softmax)
    
    loss(x, y) = crossentropy(nn(x), y)
    accuracy(x, y) = mean(onecold(nn(x)) .== onecold(y))

    dataset = [(X, Y)]
    evalcb = () -> @show(loss(X, Y))
    
    opt = ADAM(0.001)
    
    @epochs 2000 Flux.train!(loss, Flux.params(nn), dataset, opt, cb = throttle(evalcb, 10));
    
    ac_train, ac_test, ba_train, ba_test =  res(X, Y, Xt, Yt);
    experiments_06[i,:] = [ac_train, ac_test, ba_train, ba_test]
end

# %% [code]
p_dropout=0.7

n_exper = 100
experiments_07 = zeros(n_exper, 4)

for i=1:n_exper
    
    nn = Chain(
              Dense(8, n_h, relu),
              Dropout(p_dropout),
              Dense(n_h, 3),
              softmax)
    
    loss(x, y) = crossentropy(nn(x), y)
    accuracy(x, y) = mean(onecold(nn(x)) .== onecold(y))

    dataset = [(X, Y)]
    evalcb = () -> @show(loss(X, Y))
    
    opt = ADAM(0.001)
    
    @epochs 2000 Flux.train!(loss, Flux.params(nn), dataset, opt, cb = throttle(evalcb, 10));
    
    ac_train, ac_test, ba_train, ba_test =  res(X, Y, Xt, Yt);
    experiments_07[i,:] = [ac_train, ac_test, ba_train, ba_test]
end

# %% [code]
#========================
        Results
=========================#

# %% [code]
println(mean(experiments_none, dims=1))
println(std(experiments_none, dims=1))
experiments_none

# %% [code]
println(mean(experiments_01, dims=1))
println(std(experiments_01, dims=1))
experiments_01

# %% [code]
println(mean(experiments_02, dims=1))
println(std(experiments_02, dims=1))
experiments_02

# %% [code]
println(mean(experiments_03, dims=1))
println(std(experiments_03, dims=1))
experiments_03

# %% [code]
println(mean(experiments_04, dims=1))
println(std(experiments_04, dims=1))
experiments_04

# %% [code]
println(mean(experiments_05, dims=1))
println(std(experiments_05, dims=1))
experiments_05

# %% [code]
println(mean(experiments_06, dims=1))
println(std(experiments_06, dims=1))
experiments_06

# %% [code]
println(mean(experiments_07, dims=1))
println(std(experiments_07, dims=1))
experiments_07

# %% [code]
## ---------------------------------------------------------
## Regularization via penalty - https://fluxml.ai/Flux.jl/stable/models/regularisation/
## ---------------------------------------------------------

# %% [code]
lambda = 0.001

n_exper = 100
experiments_pen_001 = zeros(n_exper, 4)

for i=1:n_exper
    
    nn = Chain(
      Dense(8, n_h, relu),
      Dense(n_h, 3),
      softmax)

    penalty() =  lambda * norm(params(nn)) .* norm(params(nn)) #sum(norm, params(nn))
    #penalty() =  lambda * sum(norm, params(nn))

    loss(x, y) = crossentropy(nn(x), y) + penalty() 
    accuracy(x, y) = mean(onecold(nn(x)) .== onecold(y))

    dataset = [(X, Y)]

    evalcb = () -> @show(loss(X, Y))

    opt = ADAM(0.001)

    @epochs 2000 Flux.train!(loss, Flux.params(nn), dataset, opt, cb = throttle(evalcb, 10));
    
    ac_train, ac_test, ba_train, ba_test =  res(X, Y, Xt, Yt);
    experiments_pen_001[i,:] = [ac_train, ac_test, ba_train, ba_test]
end

# %% [code]
println(mean(experiments_pen_001, dims=1))
println(std(experiments_pen_001, dims=1))
experiments_pen_001

# %% [code]
lambda = 0.01

n_exper = 10
experiments_pen_01 = zeros(n_exper, 4)

for i=1:n_exper
    
    nn = Chain(
      Dense(8, n_h, relu),
      Dense(n_h, 3),
      softmax)

    penalty() =  lambda * norm(params(nn)) .* norm(params(nn)) #sum(norm, params(nn))
    #penalty() =  lambda * sum(norm, params(nn))

    loss(x, y) = crossentropy(nn(x), y) + penalty() 
    accuracy(x, y) = mean(onecold(nn(x)) .== onecold(y))

    dataset = [(X, Y)]

    evalcb = () -> @show(loss(X, Y))

    opt = ADAM(0.001)

    @epochs 2000 Flux.train!(loss, Flux.params(nn), dataset, opt, cb = throttle(evalcb, 10));
    
    ac_train, ac_test, ba_train, ba_test =  res(X, Y, Xt, Yt);
    experiments_pen_01[i,:] = [ac_train, ac_test, ba_train, ba_test]
end

# %% [code]
println(mean(experiments_pen_01, dims=1))
println(std(experiments_pen_01, dims=1))
experiments_pen_01

# %% [code]
lambda = 0.01

n_exper = 100
experiments_pen_01 = zeros(n_exper, 4)

for i=1:n_exper
    
    nn = Chain(
      Dense(8, n_h, relu),
      Dense(n_h, 3),
      softmax)

    penalty() =  lambda * norm(params(nn)) .* norm(params(nn)) #sum(norm, params(nn))
    #penalty() =  lambda * sum(norm, params(nn))

    loss(x, y) = crossentropy(nn(x), y) + penalty() 
    accuracy(x, y) = mean(onecold(nn(x)) .== onecold(y))

    dataset = [(X, Y)]

    evalcb = () -> @show(loss(X, Y))

    opt = ADAM(0.001)

    @epochs 2000 Flux.train!(loss, Flux.params(nn), dataset, opt, cb = throttle(evalcb, 10));
    
    ac_train, ac_test, ba_train, ba_test =  res(X, Y, Xt, Yt);
    experiments_pen_01[i,:] = [ac_train, ac_test, ba_train, ba_test]
end

# %% [code]
println(mean(experiments_pen_01, dims=1))
println(std(experiments_pen_01, dims=1))
experiments_pen_01

# %% [code]
lambda = 0.1

n_exper = 100
experiments_pen_1 = zeros(n_exper, 4)

for i=1:n_exper
    
    nn = Chain(
      Dense(8, n_h, relu),
      Dense(n_h, 3),
      softmax)

    penalty() =  lambda * norm(params(nn)) .* norm(params(nn)) #sum(norm, params(nn))
    #penalty() =  lambda * sum(norm, params(nn))

    loss(x, y) = crossentropy(nn(x), y) + penalty() 
    accuracy(x, y) = mean(onecold(nn(x)) .== onecold(y))

    dataset = [(X, Y)]

    evalcb = () -> @show(loss(X, Y))

    opt = ADAM(0.001)

    @epochs 2000 Flux.train!(loss, Flux.params(nn), dataset, opt, cb = throttle(evalcb, 10));
    
    ac_train, ac_test, ba_train, ba_test =  res(X, Y, Xt, Yt);
    experiments_pen_1[i,:] = [ac_train, ac_test, ba_train, ba_test]
end

# %% [code]
println(mean(experiments_pen_1, dims=1))
println(std(experiments_pen_1, dims=1))
experiments_pen_1

# %% [code]
lambda = 0.007

n_exper = 100
experiments_pen_007 = zeros(n_exper, 4)

for i=1:n_exper
    
    nn = Chain(
      Dense(8, n_h, relu),
      Dense(n_h, 3),
      softmax)

    penalty() =  lambda * norm(params(nn)) .* norm(params(nn)) #sum(norm, params(nn))
    #penalty() =  lambda * sum(norm, params(nn))

    loss(x, y) = crossentropy(nn(x), y) + penalty() 
    accuracy(x, y) = mean(onecold(nn(x)) .== onecold(y))

    dataset = [(X, Y)]

    evalcb = () -> @show(loss(X, Y))

    opt = ADAM(0.001)

    @epochs 2000 Flux.train!(loss, Flux.params(nn), dataset, opt, cb = throttle(evalcb, 10));
    
    ac_train, ac_test, ba_train, ba_test =  res(X, Y, Xt, Yt);
    experiments_pen_007[i,:] = [ac_train, ac_test, ba_train, ba_test]
end

# %% [code]
lambda = 0.05

n_exper = 100
experiments_pen_05 = zeros(n_exper, 4)

for i=1:n_exper
    
    nn = Chain(
      Dense(8, n_h, relu),
      Dense(n_h, 3),
      softmax)

    penalty() =  lambda * norm(params(nn)) .* norm(params(nn)) #sum(norm, params(nn))
    #penalty() =  lambda * sum(norm, params(nn))

    loss(x, y) = crossentropy(nn(x), y) + penalty() 
    accuracy(x, y) = mean(onecold(nn(x)) .== onecold(y))

    dataset = [(X, Y)]

    evalcb = () -> @show(loss(X, Y))

    opt = ADAM(0.001)

    @epochs 2000 Flux.train!(loss, Flux.params(nn), dataset, opt, cb = throttle(evalcb, 10));
    
    ac_train, ac_test, ba_train, ba_test =  res(X, Y, Xt, Yt);
    experiments_pen_05[i,:] = [ac_train, ac_test, ba_train, ba_test]
end

# %% [code]
lambda = 0.002

n_exper = 100
experiments_pen_002 = zeros(n_exper, 4)

for i=1:n_exper
    
    nn = Chain(
      Dense(8, n_h, relu),
      Dense(n_h, 3),
      softmax)

    penalty() =  lambda * norm(params(nn)) .* norm(params(nn)) #sum(norm, params(nn))
    #penalty() =  lambda * sum(norm, params(nn))

    loss(x, y) = crossentropy(nn(x), y) + penalty() 
    accuracy(x, y) = mean(onecold(nn(x)) .== onecold(y))

    dataset = [(X, Y)]

    evalcb = () -> @show(loss(X, Y))

    opt = ADAM(0.001)

    @epochs 2000 Flux.train!(loss, Flux.params(nn), dataset, opt, cb = throttle(evalcb, 10));
    
    ac_train, ac_test, ba_train, ba_test =  res(X, Y, Xt, Yt);
    experiments_pen_002[i,:] = [ac_train, ac_test, ba_train, ba_test]
end

# %% [code]
println(mean(experiments_pen_001, dims=1))
println(std(experiments_pen_001, dims=1))
experiments_pen_001

# %% [code]
println(mean(experiments_pen_002, dims=1))
println(std(experiments_pen_002, dims=1))
experiments_pen_002

# %% [code]
println(mean(experiments_pen_01, dims=1))
println(std(experiments_pen_01, dims=1))
experiments_pen_01

# %% [code]
println(mean(experiments_pen_05, dims=1))
println(std(experiments_pen_05, dims=1))
experiments_pen_01

# %% [code]
println(mean(experiments_pen_1, dims=1))
println(std(experiments_pen_1, dims=1))
experiments_pen_1

# %% [code]
println(mean(experiments_pen_007, dims=1))
println(std(experiments_pen_007, dims=1))
experiments_pen_007

# %% [code]
```
