## Supplementary material for "A Bayesian neural network for toxicity prediction": 5_BNN_multiclass_separate_priors.html


In [1]:

```
## ---------------------------------------------------------
## Import libraries and functions
## ---------------------------------------------------------

using CSV
using DataFrames
using Turing, Flux, Plots, Random
using StatsBase
using StatsFuns: logistic, logsumexp
using MLBase
using StatsModels
using FreqTables
using KernelDensity
using MLPreprocessing

Turing.turnprogress(true);
Turing.setadbackend(:reverse_diff)
```

```
┌ Info: [Turing]: global PROGRESS is set as true
└ @ Turing /Users/kcft114/.julia/packages/Turing/m05p3/src/Turing.jl:24
```

Out[1]:

```
:reverse_diff
```

In [2]:

```
## ---------------------------------------------------------
## Read and prepare data
## ---------------------------------------------------------

df_train = CSV.read("data/Aleo_train_match.csv")
df_test = CSV.read("data/Aleo_test_match.csv")

df_Xtrain = df_train[!, [:ClogP, :BSEP, :Glu, :Glu_Gal, :THLE, :HepG2, :Fsp3, :log10cmax]]
df_Xtest = df_test[!, [:ClogP, :BSEP, :Glu, :Glu_Gal, :THLE, :HepG2, :Fsp3, :log10cmax]]

scaler = fit(StandardScaler, df_Xtrain)
transform!(df_Xtrain, scaler)
transform!(df_Xtest, scaler)

df_y_train = df_train[!, :dili_sev]
df_y_test = df_test[!, :dili_sev];

X_train = convert(Matrix, df_Xtrain)
X_test = convert(Matrix, df_Xtest)

Y_train = Int.(convert(Array, df_y_train)) .- 1
Y_test = Int.(convert(Array, df_y_test)) .- 1

println(sort(countmap(Y_train)))
println(sort(countmap(Y_test)))
```

```
┌ Warning: `getindex(df::DataFrame, col_ind::ColumnIndex)` is deprecated, use `df[!, col_ind]` instead.
│   caller = valid_columns(::DataFrame) at scaleselection.jl:28
└ @ MLPreprocessing /Users/kcft114/.julia/packages/MLPreprocessing/T8hq3/src/scaleselection.jl:28
┌ Warning: `getindex(df::DataFrame, col_ind::ColumnIndex)` is deprecated, use `df[!, col_ind]` instead.
│   caller = valid_columns(::DataFrame) at scaleselection.jl:28
└ @ MLPreprocessing /Users/kcft114/.julia/packages/MLPreprocessing/T8hq3/src/scaleselection.jl:28
┌ Warning: `getindex(df::DataFrame, col_ind::ColumnIndex)` is deprecated, use `df[!, col_ind]` instead.
│   caller = valid_columns(::DataFrame, ::Array{Symbol,1}) at scaleselection.jl:40
└ @ MLPreprocessing /Users/kcft114/.julia/packages/MLPreprocessing/T8hq3/src/scaleselection.jl:40
┌ Warning: `getindex(df::DataFrame, col_ind::ColumnIndex)` is deprecated, use `df[!, col_ind]` instead.
│   caller = valid_columns(::DataFrame, ::Array{Symbol,1}) at scaleselection.jl:40
└ @ MLPreprocessing /Users/kcft114/.julia/packages/MLPreprocessing/T8hq3/src/scaleselection.jl:40
┌ Warning: `getindex(df::DataFrame, col_ind::ColumnIndex)` is deprecated, use `df[!, col_ind]` instead.
│   caller = StandardScaler(::DataFrame, ::Array{Symbol,1}) at standardize.jl:260
└ @ MLPreprocessing /Users/kcft114/.julia/packages/MLPreprocessing/T8hq3/src/standardize.jl:260
┌ Warning: `getindex(df::DataFrame, col_ind::ColumnIndex)` is deprecated, use `df[!, col_ind]` instead.
│   caller = StandardScaler(::DataFrame, ::Array{Symbol,1}) at standardize.jl:261
└ @ MLPreprocessing /Users/kcft114/.julia/packages/MLPreprocessing/T8hq3/src/standardize.jl:261
┌ Warning: `getindex(df::DataFrame, col_ind::ColumnIndex)` is deprecated, use `df[!, col_ind]` instead.
│   caller = standardize!(::DataFrame, ::Float64, ::Float64, ::Symbol) at standardize.jl:156
└ @ MLPreprocessing /Users/kcft114/.julia/packages/MLPreprocessing/T8hq3/src/standardize.jl:156
┌ Warning: `getindex(df::DataFrame, col_ind::ColumnIndex)` is deprecated, use `df[!, col_ind]` instead.
│   caller = standardize!(::DataFrame, ::Float64, ::Float64, ::Symbol) at standardize.jl:156
└ @ MLPreprocessing /Users/kcft114/.julia/packages/MLPreprocessing/T8hq3/src/standardize.jl:156
┌ Warning: `getindex(df::DataFrame, col_ind::ColumnIndex)` is deprecated, use `df[!, col_ind]` instead.
│   caller = standardize!(::DataFrame, ::Float64, ::Float64, ::Symbol) at standardize.jl:159
└ @ MLPreprocessing /Users/kcft114/.julia/packages/MLPreprocessing/T8hq3/src/standardize.jl:159
┌ Warning: `setindex!(df::DataFrame, v::AbstractVector, col_ind::ColumnIndex)` is deprecated, use `begin
│     df[!, col_ind] = v
│     df
│ end` instead.
│   caller = standardize!(::DataFrame, ::Float64, ::Float64, ::Symbol) at standardize.jl:164
└ @ MLPreprocessing /Users/kcft114/.julia/packages/MLPreprocessing/T8hq3/src/standardize.jl:164
```

```
OrderedCollections.OrderedDict(0=>37,1=>45,2=>65)
OrderedCollections.OrderedDict(0=>10,1=>11,2=>16)
```

In [3]:

```
n0 = 8
n1 = 15
K = 3
```

Out[3]:

```
3
```

In [4]:

```
## ---------------------------------------------------------
## Define model
## ---------------------------------------------------------

function weights(theta::AbstractVector)
    W0 = reshape(theta[ 1:(n0*n1)], n1, n0); 
    b0 = reshape(theta[(n0*n1 + 1): (n0*n1 + n1)], n1)
    W1 = reshape(theta[(n0*n1 + n1 + 1): (n1 * (n0 + 1 + K))], K, n1); 
    b1 = reshape(theta[(n1 * (n0 + 1 + K) + 1): (n1 * (n0 + 1 + K) + K)], K)
    return W0, b0, W1, b1
end

function feedforward(inp::AbstractArray, theta::AbstractVector)
    W0, b0, W1, b1 = weights(theta)
    model = Chain(
        Dense(W0, b0, relu),
        Dense(W1, b1)
    )
    return model(inp)
end

# Create `CategoricalLogit` to prevent numerical issues
struct CategoricalLogit <: DiscreteUnivariateDistribution
    logitp
end

# Mainly the `logsumexp` avoids the numerical issue
function Distributions.logpdf(d::CategoricalLogit, k::Int)
    return (d.logitp .- logsumexp(d.logitp))[k+1]    # use k+1 as lab[i] is 0-indexed
end

@model bayesnn_sep_priors_multiclass(inp, lab) = begin
    
    sig_w ~ TruncatedNormal(0, 1, 0, Inf)
    sig_b ~ TruncatedNormal(0, 1, 0, Inf)
    
    W0_v ~ MvNormal(zeros(n0*n1), sig_w .* ones(n0*n1))
    b0 ~ MvNormal(zeros(n1), sig_b .* ones(n1))
    W1_v ~ MvNormal(zeros(n1*K), sig_w .* ones(n1*K))
    b1 ~ MvNormal(zeros(K), sig_b .* ones(K))
        
    theta = vcat(W0_v, b0, W1_v, b1)

    preds = feedforward(inp, theta)
    for i = 1:length(lab)
        lab[i] ~ CategoricalLogit(preds[:,i])
    end
end
```

Out[4]:

```
bayesnn_sep_priors_multiclass (generic function with 3 methods)
```

In [5]:

```
inputs = X_train'
labels = Y_train;
```

In [6]:

```
## ---------------------------------------------------------
## Inference
## ---------------------------------------------------------

steps = 20_000
chain = sample(bayesnn_sep_priors_multiclass(Array(inputs), labels), NUTS(steps, 0.65));
```

```
┌ Info: Found initial step size
│   init_ϵ = 0.05
└ @ Turing.Inference /Users/kcft114/.julia/packages/Turing/m05p3/src/inference/hmc.jl:365
┌ Warning: The current proposal will be rejected due to numerical error(s).
│   isfiniteθ = true
│   isfiniter = false
│   isfiniteℓπ = false
│   isfiniteℓκ = false
└ @ AdvancedHMC /Users/kcft114/.julia/packages/AdvancedHMC/YWXfk/src/hamiltonian.jl:36
┌ Warning: The current proposal will be rejected due to numerical error(s).
│   isfiniteθ = true
│   isfiniter = false
│   isfiniteℓπ = false
│   isfiniteℓκ = false
└ @ AdvancedHMC /Users/kcft114/.julia/packages/AdvancedHMC/YWXfk/src/hamiltonian.jl:36
┌ Info: Finished 1000 adapation steps
│   adaptor = StanHMCAdaptor(n_adapts=1000, pc=DiagPreconditioner, ssa=NesterovDualAveraging(γ=0.05, t_0=10.0, κ=0.75, δ=0.65, state.ϵ=0.06482588669878515), init_buffer=75, term_buffer=50)
│   τ.integrator = Leapfrog(ϵ=0.0648)
│   h.metric = DiagEuclideanMetric([0.032094, 0.767181, 0.2024 ...])
└ @ AdvancedHMC /Users/kcft114/.julia/packages/AdvancedHMC/YWXfk/src/sampler.jl:67
┌ Info: Finished 20000 sampling steps in 20350.053057033 (s)
│   h = Hamiltonian(metric=DiagEuclideanMetric([0.032094, 0.767181, 0.2024 ...]))
│   τ = NUTS{Multinomial}(integrator=Leapfrog(ϵ=0.0648), max_depth=5), Δ_max=1000.0)
│   EBFMI(Hs) = 4125.4146707271875
│   mean(αs) = 0.7975520046907789
└ @ AdvancedHMC /Users/kcft114/.julia/packages/AdvancedHMC/YWXfk/src/sampler.jl:77
```

In [11]:

```
## ---------------------------------------------------------
## Save/read results
## ---------------------------------------------------------

#write("chains/BNN_multiclass_separate_priors.jls", chain)

chain = read("chains/BNN_multiclass_separate_priors.jls", Chains);
```

In [8]:

```
# check convergence

!(sum(isnan.(summarystats(chain)[:, :r_hat]))>0) & !(sum(abs.(summarystats(chain)[:, :r_hat] .- 1) .> 0.1) > 0) & !(sum(summarystats(chain)[:, :std] .< 1e-13) > 0)
```

Out[8]:

```
true
```

In [7]:

```
show(chain)
```

```
Object of type Chains, with data of type 19000×195×1 Array{Union{Missing, Float64},3}

Log evidence      = 0.0
Iterations        = 1:19000
Thinning interval = 1
Chains            = 1
Samples per chain = 19000
internals         = eval_num, lp, acceptance_rate, hamiltonian_energy, is_accept, log_density, n_steps, numerical_error, step_size, tree_depth
parameters        = W1_v[25], W0_v[85], W0_v[7], W0_v[36], W0_v[72], W0_v[26], b1[2], W0_v[47], W1_v[7], W0_v[110], W0_v[112], W1_v[5], b0[4], W0_v[60], b0[15], W0_v[100], b0[12], W0_v[109], W0_v[14], W1_v[10], W0_v[113], W0_v[32], W1_v[3], W0_v[50], W0_v[52], W1_v[14], W0_v[102], W0_v[105], b0[7], W0_v[17], W0_v[8], W0_v[13], W0_v[46], W0_v[31], W0_v[115], W0_v[61], W0_v[117], W1_v[17], W0_v[120], W0_v[97], W0_v[48], W0_v[19], W1_v[18], W0_v[116], W0_v[64], W0_v[54], W0_v[94], W1_v[20], W1_v[32], W1_v[45], W0_v[15], W0_v[101], W1_v[41], sig_w, W0_v[43], W1_v[12], b0[3], W0_v[3], W0_v[35], W0_v[4], W1_v[24], W0_v[20], W0_v[90], b0[5], W0_v[30], W1_v[39], W0_v[68], W0_v[62], W0_v[67], W1_v[42], W1_v[23], W0_v[9], W0_v[70], W0_v[111], W0_v[16], W1_v[44], W0_v[27], W0_v[75], W0_v[53], W0_v[86], W0_v[22], W0_v[59], W0_v[18], W0_v[95], W1_v[2], W0_v[78], W0_v[34], W0_v[37], W0_v[40], W0_v[118], W0_v[51], W0_v[107], b0[8], sig_b, W0_v[1], b0[11], W0_v[77], W0_v[76], W0_v[21], W0_v[38], W0_v[119], W0_v[106], b0[2], W1_v[40], W0_v[57], W0_v[65], W1_v[9], W0_v[96], b0[6], W0_v[87], b0[10], W0_v[103], W0_v[99], W0_v[108], W0_v[6], W1_v[30], W0_v[24], W1_v[43], W0_v[45], W1_v[34], W1_v[16], W0_v[23], W0_v[41], W0_v[69], W0_v[73], W1_v[27], W0_v[114], W0_v[11], W0_v[49], W0_v[33], b1[1], W0_v[29], W0_v[63], W0_v[28], W0_v[74], W1_v[38], b1[3], W0_v[44], W0_v[80], W1_v[1], W1_v[37], W1_v[29], W0_v[55], W0_v[12], W0_v[81], W0_v[104], W0_v[83], W0_v[93], W1_v[19], W0_v[82], W1_v[13], W0_v[39], W0_v[2], W1_v[28], W0_v[91], W0_v[84], W1_v[36], W0_v[58], W0_v[5], W0_v[25], W0_v[56], W1_v[21], W0_v[71], W1_v[4], W1_v[33], W0_v[10], W0_v[79], b0[1], W1_v[11], b0[9], W1_v[6], W0_v[66], W0_v[42], W0_v[98], W1_v[31], W1_v[15], W1_v[22], W1_v[26], W0_v[92], W0_v[88], b0[14], W1_v[35], b0[13], W1_v[8], W0_v[89]

2-element Array{ChainDataFrame,1}

Summary Statistics

│ Row │ parameters │ mean        │ std       │ naive_se    │ mcse       │ ess     │ r_hat    │
│     │ Symbol     │ Float64     │ Float64   │ Float64     │ Float64    │ Any     │ Any      │
├─────┼────────────┼─────────────┼───────────┼─────────────┼────────────┼─────────┼──────────┤
│ 1   │ W0_v[1]    │ 0.0321523   │ 0.450414  │ 0.00326764  │ 0.00371478 │ 12984.4 │ 0.999998 │
│ 2   │ W0_v[2]    │ 0.0243757   │ 0.449902  │ 0.00326393  │ 0.00357065 │ 14183.2 │ 0.999955 │
│ 3   │ W0_v[3]    │ 0.0224524   │ 0.456217  │ 0.00330974  │ 0.00398626 │ 12530.5 │ 1.00007  │
│ 4   │ W0_v[4]    │ 0.0199513   │ 0.460009  │ 0.00333726  │ 0.00398394 │ 14697.2 │ 0.999948 │
│ 5   │ W0_v[5]    │ 0.0254575   │ 0.456407  │ 0.00331113  │ 0.00341593 │ 15459.1 │ 0.999993 │
│ 6   │ W0_v[6]    │ 0.0249979   │ 0.45623   │ 0.00330984  │ 0.00375494 │ 14912.7 │ 1.00025  │
│ 7   │ W0_v[7]    │ 0.023537    │ 0.458445  │ 0.00332591  │ 0.00397152 │ 14390.4 │ 1.00015  │
│ 8   │ W0_v[8]    │ 0.028311    │ 0.455124  │ 0.00330182  │ 0.00383516 │ 13727.8 │ 0.99998  │
│ 9   │ W0_v[9]    │ 0.0257591   │ 0.450334  │ 0.00326707  │ 0.00375726 │ 13679.9 │ 1.00001  │
│ 10  │ W0_v[10]   │ 0.0276999   │ 0.457612  │ 0.00331986  │ 0.0035876  │ 15346.9 │ 0.999949 │
│ 11  │ W0_v[11]   │ 0.0230528   │ 0.456615  │ 0.00331264  │ 0.00385356 │ 14261.4 │ 0.999996 │
│ 12  │ W0_v[12]   │ 0.026686    │ 0.457861  │ 0.00332168  │ 0.00375265 │ 14324.9 │ 0.99999  │
│ 13  │ W0_v[13]   │ 0.0214385   │ 0.455811  │ 0.0033068   │ 0.00390983 │ 14105.8 │ 0.999951 │
│ 14  │ W0_v[14]   │ 0.0205271   │ 0.450286  │ 0.00326672  │ 0.00357367 │ 16110.6 │ 1.00008  │
│ 15  │ W0_v[15]   │ 0.0229797   │ 0.454442  │ 0.00329687  │ 0.00376511 │ 14824.4 │ 1.00002  │
│ 16  │ W0_v[16]   │ -0.0159515  │ 0.513574  │ 0.00372586  │ 0.005262   │ 9635.17 │ 0.999964 │
│ 17  │ W0_v[17]   │ -0.01373    │ 0.508272  │ 0.00368739  │ 0.0052388  │ 9350.85 │ 0.999983 │
│ 18  │ W0_v[18]   │ -0.00264247 │ 0.51314   │ 0.00372271  │ 0.00519899 │ 10572.3 │ 1.00009  │
│ 19  │ W0_v[19]   │ -0.0131384  │ 0.503644  │ 0.00365381  │ 0.00444836 │ 11093.3 │ 0.999977 │
│ 20  │ W0_v[20]   │ -0.016104   │ 0.511662  │ 0.00371198  │ 0.0054198  │ 9628.11 │ 0.99997  │
│ 21  │ W0_v[21]   │ -0.00666653 │ 0.514446  │ 0.00373218  │ 0.00451394 │ 10990.0 │ 1.00017  │
│ 22  │ W0_v[22]   │ -0.0133629  │ 0.508131  │ 0.00368637  │ 0.00447919 │ 10213.2 │ 0.99995  │
│ 23  │ W0_v[23]   │ -0.0083939  │ 0.517336  │ 0.00375315  │ 0.00609386 │ 7043.47 │ 1.00017  │
│ 24  │ W0_v[24]   │ -0.00899534 │ 0.50634   │ 0.00367338  │ 0.00485509 │ 9125.54 │ 0.999949 │
│ 25  │ W0_v[25]   │ -0.0173875  │ 0.507319  │ 0.00368048  │ 0.00523876 │ 9499.87 │ 0.999972 │
│ 26  │ W0_v[26]   │ -0.0227654  │ 0.50627   │ 0.00367287  │ 0.00526806 │ 8786.69 │ 1.00019  │
│ 27  │ W0_v[27]   │ -0.00865392 │ 0.509136  │ 0.00369366  │ 0.00565643 │ 9601.52 │ 0.999948 │
│ 28  │ W0_v[28]   │ -0.0158884  │ 0.511779  │ 0.00371284  │ 0.0049809  │ 9782.32 │ 1.00015  │
│ 29  │ W0_v[29]   │ -0.0183499  │ 0.507101  │ 0.0036789   │ 0.00498634 │ 9516.15 │ 1.00001  │
│ 30  │ W0_v[30]   │ -0.016366   │ 0.510352  │ 0.00370248  │ 0.00510197 │ 10408.4 │ 1.00026  │
│ 31  │ W0_v[31]   │ -0.0291177  │ 0.457572  │ 0.00331958  │ 0.00353384 │ 17493.0 │ 0.999954 │
│ 32  │ W0_v[32]   │ -0.0234506  │ 0.456707  │ 0.0033133   │ 0.00354347 │ 17967.3 │ 1.00004  │
│ 33  │ W0_v[33]   │ -0.0230066  │ 0.465249  │ 0.00337527  │ 0.00355938 │ 15772.5 │ 1.00022  │
│ 34  │ W0_v[34]   │ -0.0221206  │ 0.45954   │ 0.00333385  │ 0.00323813 │ 19000.0 │ 0.999994 │
│ 35  │ W0_v[35]   │ -0.0223897  │ 0.461612  │ 0.00334889  │ 0.00384414 │ 18583.2 │ 0.999948 │
│ 36  │ W0_v[36]   │ -0.0247588  │ 0.459762  │ 0.00333546  │ 0.00375607 │ 16314.9 │ 1.00002  │
│ 37  │ W0_v[37]   │ -0.0201688  │ 0.46043   │ 0.00334031  │ 0.00322193 │ 19000.0 │ 0.999949 │
│ 38  │ W0_v[38]   │ -0.0214791  │ 0.454969  │ 0.00330069  │ 0.00325575 │ 19000.0 │ 0.999955 │
│ 39  │ W0_v[39]   │ -0.0260605  │ 0.464202  │ 0.00336768  │ 0.00381425 │ 16786.0 │ 0.999989 │
│ 40  │ W0_v[40]   │ -0.0171129  │ 0.455989  │ 0.00330809  │ 0.00344455 │ 16968.5 │ 1.00001  │
│ 41  │ W0_v[41]   │ -0.0267193  │ 0.456314  │ 0.00331045  │ 0.00331196 │ 18551.6 │ 1.00004  │
│ 42  │ W0_v[42]   │ -0.0205009  │ 0.449363  │ 0.00326002  │ 0.00354611 │ 18711.7 │ 0.999991 │
│ 43  │ W0_v[43]   │ -0.0273384  │ 0.454957  │ 0.00330061  │ 0.00342336 │ 17080.7 │ 0.99995  │
│ 44  │ W0_v[44]   │ -0.0306693  │ 0.458398  │ 0.00332557  │ 0.0035284  │ 18920.6 │ 0.99995  │
│ 45  │ W0_v[45]   │ -0.0297267  │ 0.454356  │ 0.00329625  │ 0.00361789 │ 16711.8 │ 0.999951 │
│ 46  │ W0_v[46]   │ 0.0287357   │ 0.472459  │ 0.00342758  │ 0.00372872 │ 17916.2 │ 0.999951 │
│ 47  │ W0_v[47]   │ 0.0267056   │ 0.470961  │ 0.00341671  │ 0.00383844 │ 16275.4 │ 1.00022  │
│ 48  │ W0_v[48]   │ 0.0260031   │ 0.472259  │ 0.00342613  │ 0.00365369 │ 18706.0 │ 0.999949 │
│ 49  │ W0_v[49]   │ 0.0272851   │ 0.478376  │ 0.0034705   │ 0.00372401 │ 17739.6 │ 1.00014  │
│ 50  │ W0_v[50]   │ 0.0243619   │ 0.480173  │ 0.00348354  │ 0.00362857 │ 16869.5 │ 0.999948 │
│ 51  │ W0_v[51]   │ 0.0261278   │ 0.468442  │ 0.00339843  │ 0.00376411 │ 15907.7 │ 1.00004  │
│ 52  │ W0_v[52]   │ 0.0253648   │ 0.473036  │ 0.00343176  │ 0.00384527 │ 15612.9 │ 0.999958 │
│ 53  │ W0_v[53]   │ 0.0266892   │ 0.471257  │ 0.00341886  │ 0.00343929 │ 17052.4 │ 0.999987 │
│ 54  │ W0_v[54]   │ 0.0266619   │ 0.475205  │ 0.0034475   │ 0.00384887 │ 15075.8 │ 0.99995  │
│ 55  │ W0_v[55]   │ 0.0245396   │ 0.468036  │ 0.00339549  │ 0.00419734 │ 15404.3 │ 0.999955 │
│ 56  │ W0_v[56]   │ 0.0321173   │ 0.468488  │ 0.00339877  │ 0.00323962 │ 19000.0 │ 1.00001  │
│ 57  │ W0_v[57]   │ 0.0223328   │ 0.470527  │ 0.00341356  │ 0.00401924 │ 16108.6 │ 1.00026  │
│ 58  │ W0_v[58]   │ 0.0346004   │ 0.464193  │ 0.00336761  │ 0.00352811 │ 17342.5 │ 1.00006  │
│ 59  │ W0_v[59]   │ 0.0292955   │ 0.465434  │ 0.00337661  │ 0.003635   │ 16522.2 │ 0.99995  │
│ 60  │ W0_v[60]   │ 0.0285045   │ 0.473718  │ 0.00343671  │ 0.00380895 │ 15740.9 │ 1.00002  │
│ 61  │ W0_v[61]   │ -0.033281   │ 0.450952  │ 0.00327155  │ 0.00354821 │ 16192.0 │ 0.999963 │
│ 62  │ W0_v[62]   │ -0.0180712  │ 0.449424  │ 0.00326046  │ 0.0036769  │ 16516.6 │ 1.00003  │
│ 63  │ W0_v[63]   │ -0.0185295  │ 0.450691  │ 0.00326966  │ 0.00323972 │ 18042.8 │ 1.00012  │
│ 64  │ W0_v[64]   │ -0.0186308  │ 0.450524  │ 0.00326845  │ 0.00339202 │ 17246.3 │ 1.00001  │
│ 65  │ W0_v[65]   │ -0.0234228  │ 0.452408  │ 0.00328211  │ 0.00351661 │ 17076.1 │ 1.00018  │
│ 66  │ W0_v[66]   │ -0.0160867  │ 0.450054  │ 0.00326504  │ 0.00361572 │ 16297.6 │ 1.00058  │
│ 67  │ W0_v[67]   │ -0.0206726  │ 0.449879  │ 0.00326377  │ 0.00312154 │ 19000.0 │ 1.00002  │
│ 68  │ W0_v[68]   │ -0.0253366  │ 0.447458  │ 0.0032462   │ 0.00358065 │ 17482.0 │ 0.999962 │
│ 69  │ W0_v[69]   │ -0.0263828  │ 0.447563  │ 0.00324696  │ 0.00343208 │ 18555.6 │ 0.999982 │
│ 70  │ W0_v[70]   │ -0.0253947  │ 0.452995  │ 0.00328637  │ 0.00348145 │ 16678.6 │ 0.99995  │
│ 71  │ W0_v[71]   │ -0.0242516  │ 0.457389  │ 0.00331825  │ 0.00373474 │ 16626.0 │ 0.999947 │
│ 72  │ W0_v[72]   │ -0.0239997  │ 0.450043  │ 0.00326495  │ 0.00336954 │ 17597.3 │ 0.999948 │
│ 73  │ W0_v[73]   │ -0.0289937  │ 0.459908  │ 0.00333652  │ 0.00358841 │ 17967.8 │ 0.999991 │
│ 74  │ W0_v[74]   │ -0.0241001  │ 0.456262  │ 0.00331007  │ 0.00409079 │ 12413.6 │ 1.00064  │
│ 75  │ W0_v[75]   │ -0.022208   │ 0.454124  │ 0.00329456  │ 0.00385723 │ 15569.2 │ 0.999973 │
│ 76  │ W0_v[76]   │ 0.0202839   │ 0.463787  │ 0.00336466  │ 0.0038598  │ 15813.3 │ 0.999951 │
│ 77  │ W0_v[77]   │ 0.0131385   │ 0.463302  │ 0.00336115  │ 0.00371839 │ 15342.4 │ 0.999955 │
│ 78  │ W0_v[78]   │ 0.0137288   │ 0.462497  │ 0.0033553   │ 0.00404872 │ 13964.0 │ 0.999947 │
│ 79  │ W0_v[79]   │ 0.0244656   │ 0.467191  │ 0.00338936  │ 0.00380075 │ 16265.6 │ 0.999972 │
│ 80  │ W0_v[80]   │ 0.0140998   │ 0.45624   │ 0.00330991  │ 0.00364405 │ 18080.0 │ 1.00004  │
│ 81  │ W0_v[81]   │ 0.0170199   │ 0.460711  │ 0.00334235  │ 0.00340983 │ 16073.8 │ 0.99997  │
│ 82  │ W0_v[82]   │ 0.0185686   │ 0.457306  │ 0.00331765  │ 0.00355068 │ 15299.0 │ 1.00002  │
│ 83  │ W0_v[83]   │ 0.0177278   │ 0.457672  │ 0.0033203   │ 0.00325808 │ 18697.7 │ 0.99996  │
│ 84  │ W0_v[84]   │ 0.0137794   │ 0.455565  │ 0.00330501  │ 0.00354347 │ 18570.9 │ 1.00026  │
│ 85  │ W0_v[85]   │ 0.016467    │ 0.461376  │ 0.00334717  │ 0.00343588 │ 15131.5 │ 0.999949 │
│ 86  │ W0_v[86]   │ 0.0142303   │ 0.463104  │ 0.00335971  │ 0.00351121 │ 15627.3 │ 1.00004  │
│ 87  │ W0_v[87]   │ 0.0168364   │ 0.464841  │ 0.00337231  │ 0.00336715 │ 17151.1 │ 0.999947 │
│ 88  │ W0_v[88]   │ 0.00828623  │ 0.462693  │ 0.00335673  │ 0.00340347 │ 19000.0 │ 1.00017  │
│ 89  │ W0_v[89]   │ 0.0175061   │ 0.458523  │ 0.00332648  │ 0.00364183 │ 16339.9 │ 0.999992 │
│ 90  │ W0_v[90]   │ 0.0164298   │ 0.460242  │ 0.00333894  │ 0.00342328 │ 15228.1 │ 0.999948 │
│ 91  │ W0_v[91]   │ -0.0756048  │ 0.466271  │ 0.00338269  │ 0.00426825 │ 13012.4 │ 0.999947 │
│ 92  │ W0_v[92]   │ -0.0765559  │ 0.463284  │ 0.00336102  │ 0.00466037 │ 11797.0 │ 0.999952 │
│ 93  │ W0_v[93]   │ -0.0678103  │ 0.465424  │ 0.00337654  │ 0.00414564 │ 12592.9 │ 0.999962 │
│ 94  │ W0_v[94]   │ -0.0675345  │ 0.461986  │ 0.0033516   │ 0.00414459 │ 13293.7 │ 0.999947 │
│ 95  │ W0_v[95]   │ -0.0741022  │ 0.461312  │ 0.00334671  │ 0.00417198 │ 11794.6 │ 0.999951 │
│ 96  │ W0_v[96]   │ -0.0671558  │ 0.465175  │ 0.00337474  │ 0.00397788 │ 13090.6 │ 1.00019  │
│ 97  │ W0_v[97]   │ -0.0737511  │ 0.460323  │ 0.00333954  │ 0.00417918 │ 12395.8 │ 0.999961 │
│ 98  │ W0_v[98]   │ -0.0641651  │ 0.459798  │ 0.00333573  │ 0.00402672 │ 13311.7 │ 0.99995  │
│ 99  │ W0_v[99]   │ -0.0758204  │ 0.463421  │ 0.00336201  │ 0.00398051 │ 11633.6 │ 0.999947 │
│ 100 │ W0_v[100]  │ -0.0735417  │ 0.46741   │ 0.00339095  │ 0.00412148 │ 12479.4 │ 0.999974 │
│ 101 │ W0_v[101]  │ -0.073384   │ 0.455538  │ 0.00330482  │ 0.00410716 │ 12123.4 │ 1.00001  │
│ 102 │ W0_v[102]  │ -0.0704332  │ 0.46527   │ 0.00337543  │ 0.00417694 │ 13531.2 │ 1.00005  │
│ 103 │ W0_v[103]  │ -0.0756618  │ 0.460136  │ 0.00333818  │ 0.00403518 │ 12343.8 │ 0.999958 │
│ 104 │ W0_v[104]  │ -0.0761498  │ 0.460554  │ 0.00334121  │ 0.0040995  │ 11660.2 │ 0.999993 │
│ 105 │ W0_v[105]  │ -0.0706223  │ 0.459925  │ 0.00333665  │ 0.00409238 │ 12353.1 │ 0.999966 │
│ 106 │ W0_v[106]  │ 0.0938674   │ 0.532621  │ 0.00386404  │ 0.00536748 │ 8278.76 │ 1.00022  │
│ 107 │ W0_v[107]  │ 0.0854765   │ 0.538945  │ 0.00390991  │ 0.00584044 │ 8743.94 │ 0.999967 │
│ 108 │ W0_v[108]  │ 0.0717528   │ 0.529163  │ 0.00383895  │ 0.00581039 │ 9421.05 │ 0.999958 │
│ 109 │ W0_v[109]  │ 0.0797904   │ 0.525689  │ 0.00381375  │ 0.00578551 │ 9447.38 │ 0.999986 │
│ 110 │ W0_v[110]  │ 0.0768155   │ 0.531188  │ 0.00385364  │ 0.00529164 │ 9098.97 │ 1.00015  │
│ 111 │ W0_v[111]  │ 0.070131    │ 0.534811  │ 0.00387993  │ 0.00517809 │ 10212.2 │ 0.999953 │
│ 112 │ W0_v[112]  │ 0.089385    │ 0.530028  │ 0.00384523  │ 0.00599044 │ 8615.66 │ 1.00005  │
│ 113 │ W0_v[113]  │ 0.0739345   │ 0.52803   │ 0.00383073  │ 0.00538569 │ 9390.33 │ 0.999973 │
│ 114 │ W0_v[114]  │ 0.0793919   │ 0.532554  │ 0.00386355  │ 0.00606634 │ 8824.5  │ 1.00003  │
│ 115 │ W0_v[115]  │ 0.087148    │ 0.538503  │ 0.00390671  │ 0.00534515 │ 8402.82 │ 0.999949 │
│ 116 │ W0_v[116]  │ 0.0705159   │ 0.535872  │ 0.00388763  │ 0.00532656 │ 8446.38 │ 0.999956 │
│ 117 │ W0_v[117]  │ 0.072154    │ 0.537037  │ 0.00389607  │ 0.00590714 │ 9421.91 │ 0.999994 │
│ 118 │ W0_v[118]  │ 0.0869172   │ 0.534637  │ 0.00387866  │ 0.00525493 │ 9085.06 │ 0.999953 │
│ 119 │ W0_v[119]  │ 0.0797283   │ 0.530924  │ 0.00385173  │ 0.00554971 │ 8554.09 │ 1.00021  │
│ 120 │ W0_v[120]  │ 0.0749819   │ 0.536583  │ 0.00389278  │ 0.00559464 │ 8667.03 │ 0.999984 │
│ 121 │ W1_v[1]    │ -0.0463532  │ 0.509014  │ 0.00369278  │ 0.00518258 │ 8316.99 │ 1.00007  │
│ 122 │ W1_v[2]    │ 0.0074551   │ 0.431989  │ 0.00313398  │ 0.00408669 │ 12279.2 │ 0.999994 │
│ 123 │ W1_v[3]    │ 0.0445308   │ 0.489447  │ 0.00355082  │ 0.00522779 │ 8751.67 │ 0.999947 │
│ 124 │ W1_v[4]    │ -0.047475   │ 0.503576  │ 0.00365332  │ 0.00568197 │ 8278.21 │ 0.999975 │
│ 125 │ W1_v[5]    │ 0.0107421   │ 0.423238  │ 0.00307049  │ 0.00350902 │ 14109.2 │ 0.999954 │
│ 126 │ W1_v[6]    │ 0.0300196   │ 0.488923  │ 0.00354702  │ 0.00529146 │ 8310.38 │ 0.999951 │
│ 127 │ W1_v[7]    │ -0.0351243  │ 0.496164  │ 0.00359955  │ 0.00557607 │ 9147.0  │ 1.00002  │
│ 128 │ W1_v[8]    │ 0.00458213  │ 0.425265  │ 0.0030852   │ 0.0033372  │ 15376.6 │ 0.999949 │
│ 129 │ W1_v[9]    │ 0.0216794   │ 0.486395  │ 0.00352868  │ 0.00507575 │ 9859.16 │ 0.999979 │
│ 130 │ W1_v[10]   │ -0.042101   │ 0.502433  │ 0.00364503  │ 0.00555277 │ 8007.89 │ 0.999959 │
│ 131 │ W1_v[11]   │ 0.0089233   │ 0.42136   │ 0.00305687  │ 0.00338385 │ 13728.3 │ 0.99995  │
│ 132 │ W1_v[12]   │ 0.031123    │ 0.482262  │ 0.00349869  │ 0.00493472 │ 9154.18 │ 0.999952 │
│ 133 │ W1_v[13]   │ -0.0405601  │ 0.50078   │ 0.00363304  │ 0.00500223 │ 8838.26 │ 1.00018  │
│ 134 │ W1_v[14]   │ 0.00478547  │ 0.422725  │ 0.00306677  │ 0.00355108 │ 14131.4 │ 1.00003  │
│ 135 │ W1_v[15]   │ 0.0280161   │ 0.4855    │ 0.00352219  │ 0.00484592 │ 9749.68 │ 0.999968 │
│ 136 │ W1_v[16]   │ -0.0354992  │ 0.495099  │ 0.00359183  │ 0.00482912 │ 9545.61 │ 0.999985 │
│ 137 │ W1_v[17]   │ 0.00645782  │ 0.424234  │ 0.00307772  │ 0.00372857 │ 13575.2 │ 1.00003  │
│ 138 │ W1_v[18]   │ 0.0272964   │ 0.484326  │ 0.00351367  │ 0.0045922  │ 9755.47 │ 1.00015  │
│ 139 │ W1_v[19]   │ -0.0425503  │ 0.502787  │ 0.0036476   │ 0.00512493 │ 8056.54 │ 1.00005  │
│ 140 │ W1_v[20]   │ 0.00262421  │ 0.427671  │ 0.00310265  │ 0.00362127 │ 14130.3 │ 0.999993 │
│ 141 │ W1_v[21]   │ 0.0311537   │ 0.483847  │ 0.00351019  │ 0.00474437 │ 9041.55 │ 1.00015  │
│ 142 │ W1_v[22]   │ -0.0308146  │ 0.504825  │ 0.00366239  │ 0.0056461  │ 8062.88 │ 1.00019  │
│ 143 │ W1_v[23]   │ 0.00127137  │ 0.428824  │ 0.00311102  │ 0.00428632 │ 10263.3 │ 1.0004   │
│ 144 │ W1_v[24]   │ 0.0265943   │ 0.488532  │ 0.00354419  │ 0.00511359 │ 9386.46 │ 1.0004   │
│ 145 │ W1_v[25]   │ -0.0427208  │ 0.504303  │ 0.0036586   │ 0.00532903 │ 8928.38 │ 0.999992 │
│ 146 │ W1_v[26]   │ 0.00442513  │ 0.424917  │ 0.00308267  │ 0.00355936 │ 14774.4 │ 1.00006  │
│ 147 │ W1_v[27]   │ 0.0344462   │ 0.489184  │ 0.00354892  │ 0.00535892 │ 8397.59 │ 0.999968 │
│ 148 │ W1_v[28]   │ -0.0459787  │ 0.504037  │ 0.00365667  │ 0.00524148 │ 8213.5  │ 0.999953 │
│ 149 │ W1_v[29]   │ 0.00273668  │ 0.421398  │ 0.00305714  │ 0.00319673 │ 14154.6 │ 0.999949 │
│ 150 │ W1_v[30]   │ 0.0340741   │ 0.488423  │ 0.0035434   │ 0.00486226 │ 9161.75 │ 0.999957 │
│ 151 │ W1_v[31]   │ -0.0396565  │ 0.501077  │ 0.0036352   │ 0.00493315 │ 8713.32 │ 1.00001  │
│ 152 │ W1_v[32]   │ 0.00321648  │ 0.426096  │ 0.00309122  │ 0.00331054 │ 14956.4 │ 0.999948 │
│ 153 │ W1_v[33]   │ 0.0276265   │ 0.482635  │ 0.0035014   │ 0.00471997 │ 9052.29 │ 1.00005  │
│ 154 │ W1_v[34]   │ -0.039768   │ 0.499292  │ 0.00362225  │ 0.00544414 │ 8841.62 │ 1.00008  │
│ 155 │ W1_v[35]   │ 0.00282264  │ 0.424127  │ 0.00307694  │ 0.00354051 │ 14834.3 │ 0.999956 │
│ 156 │ W1_v[36]   │ 0.0286007   │ 0.484633  │ 0.0035159   │ 0.00534778 │ 8620.17 │ 1.00013  │
│ 157 │ W1_v[37]   │ -0.0405884  │ 0.501767  │ 0.0036402   │ 0.00524997 │ 8727.06 │ 0.999968 │
│ 158 │ W1_v[38]   │ 0.00456374  │ 0.421691  │ 0.00305927  │ 0.00366344 │ 14443.9 │ 0.999985 │
│ 159 │ W1_v[39]   │ 0.0344229   │ 0.478136  │ 0.00346876  │ 0.00519819 │ 8671.39 │ 0.99995  │
│ 160 │ W1_v[40]   │ -0.0417355  │ 0.497678  │ 0.00361054  │ 0.00589861 │ 6801.77 │ 1.00026  │
│ 161 │ W1_v[41]   │ 0.00649939  │ 0.421333  │ 0.00305667  │ 0.00360941 │ 15185.2 │ 0.999949 │
│ 162 │ W1_v[42]   │ 0.0354214   │ 0.486519  │ 0.00352958  │ 0.00528821 │ 8733.45 │ 1.00024  │
│ 163 │ W1_v[43]   │ -0.0388029  │ 0.499907  │ 0.0036267   │ 0.00484588 │ 8367.1  │ 0.999948 │
│ 164 │ W1_v[44]   │ 0.00764418  │ 0.422123  │ 0.0030624   │ 0.00355992 │ 14603.4 │ 0.999992 │
│ 165 │ W1_v[45]   │ 0.0238739   │ 0.494184  │ 0.00358519  │ 0.0051414  │ 8848.86 │ 0.999965 │
│ 166 │ b0[1]      │ -0.0407263  │ 0.655278  │ 0.00475388  │ 0.0108198  │ 3517.38 │ 0.999947 │
│ 167 │ b0[2]      │ -0.0342221  │ 0.637253  │ 0.00462312  │ 0.0109936  │ 3763.66 │ 1.0      │
│ 168 │ b0[3]      │ -0.0253737  │ 0.6281    │ 0.00455671  │ 0.00819297 │ 3800.58 │ 1.00014  │
│ 169 │ b0[4]      │ -0.04214    │ 0.632264  │ 0.00458692  │ 0.011523   │ 2950.78 │ 1.00011  │
│ 170 │ b0[5]      │ -0.0489727  │ 0.678414  │ 0.00492173  │ 0.0119612  │ 2761.49 │ 1.00039  │
│ 171 │ b0[6]      │ -0.038801   │ 0.64178   │ 0.00465596  │ 0.0116802  │ 3127.64 │ 0.999947 │
│ 172 │ b0[7]      │ -0.0321498  │ 0.6299    │ 0.00456977  │ 0.00982739 │ 3970.82 │ 0.999948 │
│ 173 │ b0[8]      │ -0.0364933  │ 0.636217  │ 0.0046156   │ 0.00931624 │ 3857.02 │ 1.00004  │
│ 174 │ b0[9]      │ -0.0454393  │ 0.68085   │ 0.0049394   │ 0.0105904  │ 3241.72 │ 1.0003   │
│ 175 │ b0[10]     │ -0.0150773  │ 0.653226  │ 0.004739    │ 0.0127589  │ 2075.27 │ 0.999992 │
│ 176 │ b0[11]     │ -0.0272865  │ 0.671051  │ 0.00486832  │ 0.0109996  │ 3192.46 │ 0.999956 │
│ 177 │ b0[12]     │ -0.0271795  │ 0.657582  │ 0.0047706   │ 0.00956038 │ 3540.77 │ 1.00006  │
│ 178 │ b0[13]     │ -0.0218523  │ 0.623787  │ 0.00452542  │ 0.0111792  │ 3221.85 │ 1.00012  │
│ 179 │ b0[14]     │ -0.0292943  │ 0.639083  │ 0.0046364   │ 0.00932015 │ 4432.53 │ 0.999948 │
│ 180 │ b0[15]     │ -0.035384   │ 0.647875  │ 0.00470018  │ 0.0107247  │ 3849.22 │ 1.00104  │
│ 181 │ b1[1]      │ -0.0216994  │ 0.547463  │ 0.00397171  │ 0.00799579 │ 4323.33 │ 0.999949 │
│ 182 │ b1[2]      │ 0.00240867  │ 0.513659  │ 0.00372647  │ 0.00784905 │ 4324.16 │ 1.00038  │
│ 183 │ b1[3]      │ -0.0115605  │ 0.540646  │ 0.00392226  │ 0.00697793 │ 5705.93 │ 1.00031  │
│ 184 │ sig_b      │ 0.50021     │ 0.417976  │ 0.00303232  │ 0.0174835  │ 502.304 │ 1.00037  │
│ 185 │ sig_w      │ 0.473077    │ 0.0856438 │ 0.000621326 │ 0.00199924 │ 1615.74 │ 0.999984 │

Quantiles

│ Row │ parameters │ 2.5%      │ 25.0%     │ 50.0%       │ 75.0%    │ 97.5%    │
│     │ Symbol     │ Float64   │ Float64   │ Float64     │ Float64  │ Float64  │
├─────┼────────────┼───────────┼───────────┼─────────────┼──────────┼──────────┤
│ 1   │ W0_v[1]    │ -0.868981 │ -0.258174 │ 0.0391126   │ 0.324573 │ 0.905066 │
│ 2   │ W0_v[2]    │ -0.877551 │ -0.26566  │ 0.0270459   │ 0.318265 │ 0.90835  │
│ 3   │ W0_v[3]    │ -0.900729 │ -0.272417 │ 0.0261425   │ 0.321861 │ 0.913871 │
│ 4   │ W0_v[4]    │ -0.907255 │ -0.280394 │ 0.0275826   │ 0.326679 │ 0.912614 │
│ 5   │ W0_v[5]    │ -0.879623 │ -0.271441 │ 0.0285533   │ 0.325256 │ 0.930684 │
│ 6   │ W0_v[6]    │ -0.887809 │ -0.266508 │ 0.0282573   │ 0.326186 │ 0.91792  │
│ 7   │ W0_v[7]    │ -0.909853 │ -0.270692 │ 0.0296331   │ 0.323101 │ 0.926149 │
│ 8   │ W0_v[8]    │ -0.882485 │ -0.268363 │ 0.0349947   │ 0.328985 │ 0.907019 │
│ 9   │ W0_v[9]    │ -0.880739 │ -0.267986 │ 0.0306908   │ 0.32352  │ 0.902501 │
│ 10  │ W0_v[10]   │ -0.896742 │ -0.269168 │ 0.0355923   │ 0.331916 │ 0.923476 │
│ 11  │ W0_v[11]   │ -0.891043 │ -0.275232 │ 0.0271608   │ 0.321923 │ 0.932311 │
│ 12  │ W0_v[12]   │ -0.902963 │ -0.268939 │ 0.0365429   │ 0.322922 │ 0.9168   │
│ 13  │ W0_v[13]   │ -0.884108 │ -0.273917 │ 0.025356    │ 0.314326 │ 0.933286 │
│ 14  │ W0_v[14]   │ -0.878419 │ -0.272089 │ 0.0260703   │ 0.318129 │ 0.882215 │
│ 15  │ W0_v[15]   │ -0.889296 │ -0.273449 │ 0.0320311   │ 0.32148  │ 0.907512 │
│ 16  │ W0_v[16]   │ -0.95905  │ -0.357256 │ -0.0507569  │ 0.293004 │ 1.09735  │
│ 17  │ W0_v[17]   │ -0.933136 │ -0.346612 │ -0.0522963  │ 0.299892 │ 1.07137  │
│ 18  │ W0_v[18]   │ -0.950879 │ -0.341813 │ -0.0352247  │ 0.313419 │ 1.10324  │
│ 19  │ W0_v[19]   │ -0.952971 │ -0.343732 │ -0.0434591  │ 0.295766 │ 1.05729  │
│ 20  │ W0_v[20]   │ -0.990881 │ -0.354278 │ -0.0424297  │ 0.294927 │ 1.05727  │
│ 21  │ W0_v[21]   │ -0.95542  │ -0.342314 │ -0.0426518  │ 0.313365 │ 1.09343  │
│ 22  │ W0_v[22]   │ -0.942172 │ -0.351744 │ -0.0449997  │ 0.290626 │ 1.08318  │
│ 23  │ W0_v[23]   │ -0.961989 │ -0.347453 │ -0.0445168  │ 0.305568 │ 1.1117   │
│ 24  │ W0_v[24]   │ -0.938198 │ -0.34339  │ -0.0462963  │ 0.29501  │ 1.08162  │
│ 25  │ W0_v[25]   │ -0.961421 │ -0.353719 │ -0.047905   │ 0.29206  │ 1.0658   │
│ 26  │ W0_v[26]   │ -0.954685 │ -0.357913 │ -0.0531079  │ 0.285518 │ 1.06917  │
│ 27  │ W0_v[27]   │ -0.966527 │ -0.344672 │ -0.0375302  │ 0.311271 │ 1.05625  │
│ 28  │ W0_v[28]   │ -0.970335 │ -0.351737 │ -0.0411854  │ 0.296965 │ 1.05616  │
│ 29  │ W0_v[29]   │ -0.950533 │ -0.360947 │ -0.051989   │ 0.295152 │ 1.07559  │
│ 30  │ W0_v[30]   │ -0.964937 │ -0.357063 │ -0.0453984  │ 0.300293 │ 1.05943  │
│ 31  │ W0_v[31]   │ -0.939338 │ -0.321471 │ -0.0262558  │ 0.261929 │ 0.895882 │
│ 32  │ W0_v[32]   │ -0.92317  │ -0.322093 │ -0.0285049  │ 0.276076 │ 0.895421 │
│ 33  │ W0_v[33]   │ -0.948938 │ -0.323298 │ -0.0259914  │ 0.276886 │ 0.902888 │
│ 34  │ W0_v[34]   │ -0.928752 │ -0.317683 │ -0.0206732  │ 0.274947 │ 0.883152 │
│ 35  │ W0_v[35]   │ -0.932703 │ -0.327786 │ -0.0217956  │ 0.277814 │ 0.890595 │
│ 36  │ W0_v[36]   │ -0.935248 │ -0.323029 │ -0.0297682  │ 0.273948 │ 0.900663 │
│ 37  │ W0_v[37]   │ -0.924134 │ -0.320165 │ -0.0200678  │ 0.278165 │ 0.891819 │
│ 38  │ W0_v[38]   │ -0.920358 │ -0.318206 │ -0.0224794  │ 0.270448 │ 0.880938 │
│ 39  │ W0_v[39]   │ -0.940794 │ -0.331326 │ -0.0265211  │ 0.277541 │ 0.908165 │
│ 40  │ W0_v[40]   │ -0.913458 │ -0.316217 │ -0.015364   │ 0.278089 │ 0.888473 │
│ 41  │ W0_v[41]   │ -0.913467 │ -0.324595 │ -0.0293269  │ 0.268517 │ 0.885154 │
│ 42  │ W0_v[42]   │ -0.913694 │ -0.310989 │ -0.0189628  │ 0.269178 │ 0.887094 │
│ 43  │ W0_v[43]   │ -0.932538 │ -0.32016  │ -0.0275748  │ 0.259686 │ 0.875128 │
│ 44  │ W0_v[44]   │ -0.927054 │ -0.333978 │ -0.0341004  │ 0.272239 │ 0.878946 │
│ 45  │ W0_v[45]   │ -0.937861 │ -0.320216 │ -0.0298145  │ 0.265874 │ 0.863779 │
│ 46  │ W0_v[46]   │ -0.904933 │ -0.279425 │ 0.0251752   │ 0.341278 │ 0.95907  │
│ 47  │ W0_v[47]   │ -0.917193 │ -0.276143 │ 0.0271656   │ 0.33441  │ 0.957425 │
│ 48  │ W0_v[48]   │ -0.909191 │ -0.277865 │ 0.0233626   │ 0.32982  │ 0.958497 │
│ 49  │ W0_v[49]   │ -0.915789 │ -0.287231 │ 0.0257008   │ 0.336825 │ 0.968066 │
│ 50  │ W0_v[50]   │ -0.937367 │ -0.291463 │ 0.0270819   │ 0.3438   │ 0.952521 │
│ 51  │ W0_v[51]   │ -0.901252 │ -0.280112 │ 0.0308484   │ 0.337925 │ 0.94638  │
│ 52  │ W0_v[52]   │ -0.914423 │ -0.281424 │ 0.026887    │ 0.333658 │ 0.96928  │
│ 53  │ W0_v[53]   │ -0.908783 │ -0.278114 │ 0.0248652   │ 0.334175 │ 0.951337 │
│ 54  │ W0_v[54]   │ -0.903964 │ -0.28928  │ 0.024819    │ 0.338372 │ 0.964437 │
│ 55  │ W0_v[55]   │ -0.89884  │ -0.280799 │ 0.025574    │ 0.33327  │ 0.930986 │
│ 56  │ W0_v[56]   │ -0.896413 │ -0.270839 │ 0.0360476   │ 0.341039 │ 0.947073 │
│ 57  │ W0_v[57]   │ -0.922791 │ -0.287743 │ 0.0276353   │ 0.33161  │ 0.943665 │
│ 58  │ W0_v[58]   │ -0.881806 │ -0.269339 │ 0.0381993   │ 0.339457 │ 0.95368  │
│ 59  │ W0_v[59]   │ -0.8922   │ -0.273231 │ 0.0273571   │ 0.32839  │ 0.961219 │
│ 60  │ W0_v[60]   │ -0.908624 │ -0.277442 │ 0.0264184   │ 0.333547 │ 0.968653 │
│ 61  │ W0_v[61]   │ -0.937064 │ -0.322078 │ -0.0273059  │ 0.256721 │ 0.858801 │
│ 62  │ W0_v[62]   │ -0.912872 │ -0.310095 │ -0.0171236  │ 0.271146 │ 0.878464 │
│ 63  │ W0_v[63]   │ -0.920764 │ -0.309325 │ -0.014521   │ 0.27705  │ 0.868224 │
│ 64  │ W0_v[64]   │ -0.916713 │ -0.312686 │ -0.0147357  │ 0.275736 │ 0.860402 │
│ 65  │ W0_v[65]   │ -0.925193 │ -0.31388  │ -0.0193599  │ 0.271352 │ 0.85926  │
│ 66  │ W0_v[66]   │ -0.90678  │ -0.310309 │ -0.0176734  │ 0.278308 │ 0.87001  │
│ 67  │ W0_v[67]   │ -0.906496 │ -0.31091  │ -0.0248918  │ 0.274715 │ 0.86797  │
│ 68  │ W0_v[68]   │ -0.917958 │ -0.313199 │ -0.023282   │ 0.264376 │ 0.851067 │
│ 69  │ W0_v[69]   │ -0.922717 │ -0.3134   │ -0.0235413  │ 0.264808 │ 0.847491 │
│ 70  │ W0_v[70]   │ -0.933058 │ -0.316474 │ -0.0249148  │ 0.270448 │ 0.874832 │
│ 71  │ W0_v[71]   │ -0.936375 │ -0.321539 │ -0.0199353  │ 0.272642 │ 0.870207 │
│ 72  │ W0_v[72]   │ -0.921344 │ -0.314191 │ -0.0208458  │ 0.264275 │ 0.857442 │
│ 73  │ W0_v[73]   │ -0.955163 │ -0.3252   │ -0.0271828  │ 0.272555 │ 0.874356 │
│ 74  │ W0_v[74]   │ -0.950776 │ -0.313277 │ -0.0209333  │ 0.271928 │ 0.881014 │
│ 75  │ W0_v[75]   │ -0.92653  │ -0.314798 │ -0.0210771  │ 0.27514  │ 0.861468 │
│ 76  │ W0_v[76]   │ -0.886612 │ -0.282969 │ 0.0160505   │ 0.316929 │ 0.943349 │
│ 77  │ W0_v[77]   │ -0.901393 │ -0.289388 │ 0.00723113  │ 0.31729  │ 0.93051  │
│ 78  │ W0_v[78]   │ -0.905198 │ -0.286259 │ 0.0147565   │ 0.310636 │ 0.934716 │
│ 79  │ W0_v[79]   │ -0.893616 │ -0.278426 │ 0.0237365   │ 0.328286 │ 0.956729 │
│ 80  │ W0_v[80]   │ -0.883922 │ -0.283277 │ 0.00847741  │ 0.308513 │ 0.937798 │
│ 81  │ W0_v[81]   │ -0.881616 │ -0.285981 │ 0.0152882   │ 0.310808 │ 0.948217 │
│ 82  │ W0_v[82]   │ -0.889223 │ -0.278472 │ 0.00992292  │ 0.31489  │ 0.930697 │
│ 83  │ W0_v[83]   │ -0.877699 │ -0.285245 │ 0.0146412   │ 0.317581 │ 0.933793 │
│ 84  │ W0_v[84]   │ -0.874794 │ -0.287207 │ 0.00979989  │ 0.310389 │ 0.932987 │
│ 85  │ W0_v[85]   │ -0.891599 │ -0.285372 │ 0.0126733   │ 0.314172 │ 0.942606 │
│ 86  │ W0_v[86]   │ -0.898893 │ -0.288216 │ 0.0148557   │ 0.315911 │ 0.931763 │
│ 87  │ W0_v[87]   │ -0.903145 │ -0.28653  │ 0.0162842   │ 0.321165 │ 0.935251 │
│ 88  │ W0_v[88]   │ -0.896921 │ -0.293673 │ 0.00189332  │ 0.304861 │ 0.944897 │
│ 89  │ W0_v[89]   │ -0.874804 │ -0.28504  │ 0.0124658   │ 0.313537 │ 0.928997 │
│ 90  │ W0_v[90]   │ -0.878054 │ -0.283752 │ 0.012848    │ 0.307585 │ 0.942783 │
│ 91  │ W0_v[91]   │ -1.02099  │ -0.375235 │ -0.0627842  │ 0.231321 │ 0.827668 │
│ 92  │ W0_v[92]   │ -1.00458  │ -0.370739 │ -0.0675911  │ 0.220191 │ 0.830883 │
│ 93  │ W0_v[93]   │ -1.00079  │ -0.368278 │ -0.0630816  │ 0.236253 │ 0.842058 │
│ 94  │ W0_v[94]   │ -0.997451 │ -0.36579  │ -0.0595495  │ 0.239291 │ 0.820854 │
│ 95  │ W0_v[95]   │ -1.00375  │ -0.36841  │ -0.0669159  │ 0.22539  │ 0.829798 │
│ 96  │ W0_v[96]   │ -0.997194 │ -0.363399 │ -0.0668978  │ 0.233046 │ 0.841424 │
│ 97  │ W0_v[97]   │ -1.00326  │ -0.369443 │ -0.0647881  │ 0.226745 │ 0.821219 │
│ 98  │ W0_v[98]   │ -0.995653 │ -0.355998 │ -0.0553313  │ 0.2376   │ 0.821946 │
│ 99  │ W0_v[99]   │ -0.994886 │ -0.374539 │ -0.0682981  │ 0.234253 │ 0.832033 │
│ 100 │ W0_v[100]  │ -1.01496  │ -0.371942 │ -0.0606704  │ 0.227482 │ 0.841509 │
│ 101 │ W0_v[101]  │ -0.977907 │ -0.372782 │ -0.0627221  │ 0.224829 │ 0.809956 │
│ 102 │ W0_v[102]  │ -1.00896  │ -0.368941 │ -0.0600787  │ 0.236687 │ 0.836694 │
│ 103 │ W0_v[103]  │ -1.02035  │ -0.370295 │ -0.0613122  │ 0.229105 │ 0.800596 │
│ 104 │ W0_v[104]  │ -1.0066   │ -0.375448 │ -0.0706892  │ 0.227802 │ 0.823729 │
│ 105 │ W0_v[105]  │ -1.00014  │ -0.36707  │ -0.0616934  │ 0.23035  │ 0.827374 │
│ 106 │ W0_v[106]  │ -0.980269 │ -0.250901 │ 0.105489    │ 0.453989 │ 1.11422  │
│ 107 │ W0_v[107]  │ -1.00159  │ -0.26755  │ 0.0969791   │ 0.445509 │ 1.11959  │
│ 108 │ W0_v[108]  │ -1.01078  │ -0.2655   │ 0.0843049   │ 0.423122 │ 1.08158  │
│ 109 │ W0_v[109]  │ -0.996246 │ -0.263808 │ 0.0900765   │ 0.434179 │ 1.09779  │
│ 110 │ W0_v[110]  │ -1.00966  │ -0.269221 │ 0.0889433   │ 0.432218 │ 1.10286  │
│ 111 │ W0_v[111]  │ -1.01518  │ -0.273835 │ 0.0803719   │ 0.430505 │ 1.08495  │
│ 112 │ W0_v[112]  │ -0.981498 │ -0.253201 │ 0.0992978   │ 0.443544 │ 1.1094   │
│ 113 │ W0_v[113]  │ -0.99455  │ -0.267691 │ 0.0813729   │ 0.427839 │ 1.09334  │
│ 114 │ W0_v[114]  │ -0.994691 │ -0.277103 │ 0.0889361   │ 0.439105 │ 1.08909  │
│ 115 │ W0_v[115]  │ -0.999938 │ -0.264345 │ 0.0964998   │ 0.44308  │ 1.1244   │
│ 116 │ W0_v[116]  │ -1.02728  │ -0.275271 │ 0.0823798   │ 0.433422 │ 1.0863   │
│ 117 │ W0_v[117]  │ -1.01134  │ -0.275426 │ 0.0792584   │ 0.427877 │ 1.10645  │
│ 118 │ W0_v[118]  │ -0.993065 │ -0.257715 │ 0.0952186   │ 0.442464 │ 1.11618  │
│ 119 │ W0_v[119]  │ -0.988673 │ -0.265848 │ 0.0853159   │ 0.441889 │ 1.0839   │
│ 120 │ W0_v[120]  │ -1.03104  │ -0.275092 │ 0.0891599   │ 0.429749 │ 1.09812  │
│ 121 │ W1_v[1]    │ -1.07129  │ -0.375943 │ -0.0361655  │ 0.292751 │ 0.929683 │
│ 122 │ W1_v[2]    │ -0.852419 │ -0.273802 │ 0.004028    │ 0.288651 │ 0.867446 │
│ 123 │ W1_v[3]    │ -0.904939 │ -0.279224 │ 0.0417746   │ 0.370224 │ 1.00511  │
│ 124 │ W1_v[4]    │ -1.05932  │ -0.378134 │ -0.0343764  │ 0.289819 │ 0.919382 │
│ 125 │ W1_v[5]    │ -0.839541 │ -0.259982 │ 0.010866    │ 0.286079 │ 0.833704 │
│ 126 │ W1_v[6]    │ -0.942313 │ -0.291771 │ 0.0273312   │ 0.352234 │ 0.991194 │
│ 127 │ W1_v[7]    │ -1.03851  │ -0.359509 │ -0.0243425  │ 0.292565 │ 0.924192 │
│ 128 │ W1_v[8]    │ -0.845494 │ -0.271096 │ 0.00975352  │ 0.280784 │ 0.838329 │
│ 129 │ W1_v[9]    │ -0.942564 │ -0.298763 │ 0.0235451   │ 0.344459 │ 0.966265 │
│ 130 │ W1_v[10]   │ -1.04179  │ -0.371589 │ -0.0362344  │ 0.28822  │ 0.940258 │
│ 131 │ W1_v[11]   │ -0.821044 │ -0.267687 │ 0.0121341   │ 0.281342 │ 0.845138 │
│ 132 │ W1_v[12]   │ -0.927906 │ -0.286612 │ 0.0333278   │ 0.354188 │ 0.970196 │
│ 133 │ W1_v[13]   │ -1.07061  │ -0.366635 │ -0.0270123  │ 0.296561 │ 0.909973 │
│ 134 │ W1_v[14]   │ -0.828748 │ -0.269479 │ 0.00476403  │ 0.281696 │ 0.833778 │
│ 135 │ W1_v[15]   │ -0.93334  │ -0.295129 │ 0.0313352   │ 0.349078 │ 0.98822  │
│ 136 │ W1_v[16]   │ -1.03624  │ -0.362855 │ -0.0224666  │ 0.296125 │ 0.930736 │
│ 137 │ W1_v[17]   │ -0.838618 │ -0.269158 │ 0.0080396   │ 0.281955 │ 0.847183 │
│ 138 │ W1_v[18]   │ -0.932287 │ -0.289158 │ 0.028883    │ 0.350397 │ 0.985583 │
│ 139 │ W1_v[19]   │ -1.05615  │ -0.369696 │ -0.0360908  │ 0.291501 │ 0.93569  │
│ 140 │ W1_v[20]   │ -0.846894 │ -0.277692 │ 0.00691907  │ 0.284494 │ 0.831633 │
│ 141 │ W1_v[21]   │ -0.92062  │ -0.286869 │ 0.0347591   │ 0.353922 │ 0.969521 │
│ 142 │ W1_v[22]   │ -1.05236  │ -0.357255 │ -0.0271352  │ 0.306415 │ 0.940975 │
│ 143 │ W1_v[23]   │ -0.842594 │ -0.279627 │ 0.0020528   │ 0.280489 │ 0.844383 │
│ 144 │ W1_v[24]   │ -0.932049 │ -0.295995 │ 0.0296794   │ 0.351711 │ 0.991074 │
│ 145 │ W1_v[25]   │ -1.04925  │ -0.374403 │ -0.033351   │ 0.288385 │ 0.948926 │
│ 146 │ W1_v[26]   │ -0.828236 │ -0.274651 │ 0.00473431  │ 0.279252 │ 0.844046 │
│ 147 │ W1_v[27]   │ -0.92195  │ -0.293278 │ 0.0363512   │ 0.357855 │ 1.00909  │
│ 148 │ W1_v[28]   │ -1.05124  │ -0.374113 │ -0.0386497  │ 0.291188 │ 0.932815 │
│ 149 │ W1_v[29]   │ -0.824919 │ -0.275119 │ 0.00475385  │ 0.280596 │ 0.83226  │
│ 150 │ W1_v[30]   │ -0.923334 │ -0.289958 │ 0.0325247   │ 0.36136  │ 0.98617  │
│ 151 │ W1_v[31]   │ -1.06257  │ -0.365276 │ -0.0256236  │ 0.294361 │ 0.912253 │
│ 152 │ W1_v[32]   │ -0.846144 │ -0.272443 │ 0.00615424  │ 0.28668  │ 0.827329 │
│ 153 │ W1_v[33]   │ -0.925864 │ -0.296503 │ 0.0271151   │ 0.347769 │ 0.99008  │
│ 154 │ W1_v[34]   │ -1.04872  │ -0.364919 │ -0.0265945  │ 0.295246 │ 0.92026  │
│ 155 │ W1_v[35]   │ -0.852263 │ -0.271387 │ 0.00893971  │ 0.282245 │ 0.83111  │
│ 156 │ W1_v[36]   │ -0.94167  │ -0.287491 │ 0.0315247   │ 0.348978 │ 0.972264 │
│ 157 │ W1_v[37]   │ -1.0601   │ -0.367878 │ -0.0335812  │ 0.290469 │ 0.931686 │
│ 158 │ W1_v[38]   │ -0.836547 │ -0.268454 │ 0.00804648  │ 0.279678 │ 0.845646 │
│ 159 │ W1_v[39]   │ -0.904901 │ -0.284768 │ 0.036467    │ 0.352145 │ 0.971383 │
│ 160 │ W1_v[40]   │ -1.05367  │ -0.366915 │ -0.0298884  │ 0.29016  │ 0.911355 │
│ 161 │ W1_v[41]   │ -0.831659 │ -0.268641 │ 0.00460723  │ 0.280712 │ 0.821587 │
│ 162 │ W1_v[42]   │ -0.921209 │ -0.285729 │ 0.0294515   │ 0.35684  │ 1.00566  │
│ 163 │ W1_v[43]   │ -1.05029  │ -0.365172 │ -0.0299691  │ 0.298296 │ 0.92374  │
│ 164 │ W1_v[44]   │ -0.832759 │ -0.271493 │ 0.00947347  │ 0.288497 │ 0.833156 │
│ 165 │ W1_v[45]   │ -0.95395  │ -0.300447 │ 0.0231424   │ 0.35302  │ 0.983544 │
│ 166 │ b0[1]      │ -1.53829  │ -0.240923 │ -0.00602439 │ 0.199126 │ 1.2628   │
│ 167 │ b0[2]      │ -1.53617  │ -0.237983 │ -0.00702357 │ 0.203669 │ 1.25299  │
│ 168 │ b0[3]      │ -1.41485  │ -0.238475 │ -0.00759429 │ 0.199858 │ 1.267    │
│ 169 │ b0[4]      │ -1.51605  │ -0.245626 │ -0.00957149 │ 0.201278 │ 1.23553  │
│ 170 │ b0[5]      │ -1.58504  │ -0.236865 │ -0.00543063 │ 0.199245 │ 1.26714  │
│ 171 │ b0[6]      │ -1.45821  │ -0.239693 │ -0.00712823 │ 0.200799 │ 1.21499  │
│ 172 │ b0[7]      │ -1.39519  │ -0.242097 │ -0.00870203 │ 0.197194 │ 1.26371  │
│ 173 │ b0[8]      │ -1.5168   │ -0.241274 │ -0.00488668 │ 0.200445 │ 1.25959  │
│ 174 │ b0[9]      │ -1.55377  │ -0.246532 │ -0.0111446  │ 0.194221 │ 1.23227  │
│ 175 │ b0[10]     │ -1.42389  │ -0.231985 │ -0.00445532 │ 0.210698 │ 1.32526  │
│ 176 │ b0[11]     │ -1.52712  │ -0.231486 │ -0.00373576 │ 0.202539 │ 1.32162  │
│ 177 │ b0[12]     │ -1.47755  │ -0.232708 │ -0.00717565 │ 0.206018 │ 1.29006  │
│ 178 │ b0[13]     │ -1.43747  │ -0.233036 │ -0.00340307 │ 0.207389 │ 1.32408  │
│ 179 │ b0[14]     │ -1.45209  │ -0.23527  │ -0.0050225  │ 0.20411  │ 1.25888  │
│ 180 │ b0[15]     │ -1.46577  │ -0.239984 │ -0.00796984 │ 0.205638 │ 1.25998  │
│ 181 │ b1[1]      │ -1.20429  │ -0.227534 │ -0.0133115  │ 0.174883 │ 1.16863  │
│ 182 │ b1[2]      │ -1.13851  │ -0.185417 │ 0.00366635  │ 0.204254 │ 1.09008  │
│ 183 │ b1[3]      │ -1.24406  │ -0.20011  │ 0.00103772  │ 0.197437 │ 1.10835  │
│ 184 │ sig_b      │ 0.0370193 │ 0.191547  │ 0.389816    │ 0.688069 │ 1.60436  │
│ 185 │ sig_w      │ 0.330451  │ 0.412082  │ 0.465062    │ 0.524651 │ 0.665317 │
```

In [9]:

```
# Extract all weight and bias parameters.

W0_v = chain[:W0_v].value.data[:,:,1]
b0 = chain[:b0].value.data[:,:,1]
W1_v = chain[:W1_v].value.data[:,:,1]
b1 = chain[:b1].value.data[:,:,1]

theta = [W0_v b0 W1_v b1];

niter, _ = size(theta)
```

Out[9]:

```
(19000, 183)
```

In [10]:

```
## ---------------------------------------------------------
## predict for training set
## ---------------------------------------------------------

inputs = X_train'
labels = Y_train;

y_pred_samps = zeros(niter, length(labels))
y_pred = zeros(length(labels))

for j in 1:length(labels)
   for i in 1:niter
        preds = softmax(feedforward(inputs[:,j], theta[i,:]))
        dist = Categorical(preds)
        y_pred_samps[i,j] = rand(dist)
    end
    probs = [mean(y_pred_samps[:,j] .== 1), mean(y_pred_samps[:,j] .== 2), mean(y_pred_samps[:,j] .== 3)]
    y_pred[j] = sum((probs .== maximum(probs)) .* [1, 2, 3])
end

println("train accuracy = ", mean(y_pred .- 1 .== labels))

y_pred = convert(Array{Int64,1}, y_pred);
C = confusmat(3, labels .+ 1, y_pred)
bacc = 1/3 *(C[1,1] / sum(C[1, :]) + C[2,2] / sum(C[2, :]) + C[3,3] / sum(C[3, :]))
println("train BA = ", round(bacc, digits=2))
```

```
train accuracy = 0.7006802721088435
train BA = 0.69
```

In [11]:

```
## ---------------------------------------------------------
## predict for training set
## ---------------------------------------------------------

inputs = X_train'
labels = Y_train;

n_exper = 10
ac_ba_train = zeros(n_exper,2)

for exp=1:n_exper
    
    y_pred_samps = zeros(niter, length(labels))
    y_pred = zeros(length(labels))

    for j in 1:length(labels)
       for i in 1:niter
            preds = softmax(feedforward(inputs[:,j], theta[i,:]))
            dist = Categorical(preds)
            y_pred_samps[i,j] = rand(dist)
        end
        probs = [mean(y_pred_samps[:,j] .== 1), mean(y_pred_samps[:,j] .== 2), mean(y_pred_samps[:,j] .== 3)]
        y_pred[j] = sum((probs .== maximum(probs)) .* [1, 2, 3])
    end

    #println("test accuracy = ", mean(y_pred .- 1 .== labels))

    y_pred = convert(Array{Int64,1}, y_pred);
    C = confusmat(3, labels .+ 1, y_pred)
    bacc = 1/3 *(C[1,1] / sum(C[1, :]) + C[2,2] / sum(C[2, :]) + C[3,3] / sum(C[3, :]))
    #println("train BA = ", round(bacc, digits=2))
    
    ac_ba_train[exp,1] = mean(y_pred .- 1 .== labels)
    ac_ba_train[exp,2] = bacc
    
end

println(round.(mean(ac_ba_train, dims=1), digits=2))
println(round.(std(ac_ba_train, dims=1), digits = 2))
```

```
[0.7 0.69]
[0.01 0.01]
```

In [13]:

```
## ---------------------------------------------------------
## predict for test set
## ---------------------------------------------------------


inputs = X_test'
labels = Y_test

n_exper = 10
ac_ba_test = zeros(n_exper,2)

for exp=1:n_exper
    
    y_pred_samps = zeros(niter, length(labels))
    y_pred = zeros(length(labels))

    for j in 1:length(labels)
       for i in 1:niter
            preds = softmax(feedforward(inputs[:,j], theta[i,:]))
            dist = Categorical(preds)
            y_pred_samps[i,j] = rand(dist)
        end
        probs = [mean(y_pred_samps[:,j] .== 1), mean(y_pred_samps[:,j] .== 2), mean(y_pred_samps[:,j] .== 3)]
        y_pred[j] = sum((probs .== maximum(probs)) .* [1, 2, 3])
    end

    #println("test accuracy = ", mean(y_pred .- 1 .== labels))

    y_pred = convert(Array{Int64,1}, y_pred);
    C = confusmat(3, labels .+ 1, y_pred)
    bacc = 1/3 *(C[1,1] / sum(C[1, :]) + C[2,2] / sum(C[2, :]) + C[3,3] / sum(C[3, :]))
    #println("train BA = ", round(bacc, digits=2))
    
    ac_ba_test[exp,1] = mean(y_pred .- 1 .== labels)
    ac_ba_test[exp,2] = bacc
    
end

println(round.(mean(ac_ba_test, dims=1), digits=2))
println(round.(std(ac_ba_test, dims=1), digits = 2))
```

```
[0.62 0.62]
[0.01 0.01]
```

In [ ]:
