## Supplementary material for "A Bayesian neural network for toxicity prediction": functions.jl

```
function slicematrix(A::AbstractMatrix)
    return [A[i, :] for i in 1:size(A,1)]
end


# This modification of the unpack function generates a series of vectors
# given a network shape.
function unpack(θ::AbstractVector, network_shape::AbstractVector)
    index = 1
    weights = []
    biases = []
    for layer in network_shape
        rows, cols, _, b = layer
        size_w = rows * cols
        last_index_w = size_w + index - 1
        push!(weights, reshape(θ[index:last_index_w], rows, cols))

        if b == 1
            last_index_b = last_index_w + rows
            push!(biases, reshape(θ[last_index_w+1:last_index_b], rows))
            index = last_index_b + 1
        else
            push!(biases, 0.0)
            index = last_index_w + 1
        end

    end
    return weights, biases
end


# Generate an abstract neural network given a shape,
# and return a prediction.
function nn_forward(x, θ::AbstractVector, network_shape::AbstractVector)
    weights, biases = unpack(θ, network_shape)
    layers = []
    for i in eachindex(network_shape)
        push!(layers, Dense(weights[i],
            biases[i],
            eval(network_shape[i][3])))
    end
    nn = Chain(layers...)
    return nn(x)
end

struct OrderedLogistic{T1, T2<:AbstractVector} <: DiscreteUnivariateDistribution
   η::T1
   cutpoints::T2

   function OrderedLogistic(η, cutpoints)
        if !issorted(cutpoints)
            error("cutpoints are not sorted")
        end
        return new{typeof(η), typeof(cutpoints)}(η, cutpoints)
   end

end

function Distributions.logpdf(d::OrderedLogistic, k::Int)
    K = length(d.cutpoints)+1
    c =  d.cutpoints

    if k==1
        logp= -softplus(-(c[k]-d.η))  #logp= log(logistic(c[k]-d.η))
    elseif k<K
        logp= log(logistic(c[k]-d.η) - logistic(c[k-1]-d.η))
    else
        logp= -softplus(c[k-1]-d.η)  #logp= log(1-logistic(c[k-1]-d.η))
    end

    return logp
end

function Distributions.rand(rng::AbstractRNG, d::OrderedLogistic)
    cutpoints = d.cutpoints
    η = d.η

    K = length(cutpoints)+1
    c = vcat(-Inf, cutpoints, Inf)

    ps = [logistic(η - i[1]) - logistic(η - i[2]) for i in zip(c[1:(end-1)],c[2:end])]

    k = rand(rng, Categorical(ps))

    if all(ps.>0)
        return(k)
    else
        return(-Inf)
    end
end


function DIC_logpfd(logpdf_mat)
    Dev = -2 .* sum(logpdf_mat, dims=2)
    DIC = mean(Dev) + var(Dev)/2
    return DIC
end

function WAIC_logpfd(logpdf_mat)
    lppd = sum(log.(mean(exp.(logpdf_mat), dims = 1)))
    pWAIC1 = 2 * sum(log.(mean(exp.(logpdf_mat), dims=1))  - mean(exp.(logpdf_mat), dims = 1))
    pWAIC2 = sum(var(logpdf_mat, dims=1))
    WAIC = -2 * lppd + 2* pWAIC2
    return WAIC
end

function one_hot(x)
 return [x ==1, x ==2, x ==3] *1.0
end

function cumBrier(probs_mat, y)
    K = 3
    cum_probs = cumsum(probs_mat, dims=2)

    RPS = zeros(length(y));

    for i in 1:length(RPS)
        RPS[i] = sum((cum_probs[i, 1:(K-1)] - cumsum(one_hot(y[i]))[1:(K-1)]).^2)
    end

    RPS = RPS/(K-1)

    return RPS
end


function probsplot(drug_names, y_pred_samps, j, y)
    tbl = freqtable(y_pred_samps[:,j])
    ptbl = prop(tbl)

    if typeof(y) == Bool
        title = ""
    else
        title = drug_names[j] * ", true category = " * string(y[j])
    end

    b = bar( ptbl,
             labels="10 sin values",
             size=[250,250],
              xlabel="DILI class",
              ylabel="Predicted probability",
              xticks = 1:1:3,
              legend=false,
              yticks = true,
              framestyle = :box,
              ylimit = (0,1),
              title = title ,
              titlefont = font(7, "Calibri"),
              xguidefontsize=font(7, "Calibri"),
              yguidefontsize=font(7, "Calibri"),
              #fillcolor = [:navy, :purple, :magenta],
              fillcolor = [:green4, :darkgoldenrod, :firebrick],
              #alpha= [0.9, 0.9,0.6],
              minorgrid = true,
              #gridcolor = :black,
              gridopacity = 0.5,
              grid = :xy)
    display(b)

end


## ---------------------------------------------------------
## Plot posterior
## ---------------------------------------------------------

function postplot(drug_names, post, ind, y)
#function postplot(drug_names, post, ind)
    post_ind = post[:,ind]

    post_ind = vcat(0, post_ind, 1)

    kde_npoints = 2048
    #dens = kde(post_ind, npoints=kde_npoints, bandwidth=0.1)
    dens = kde(post_ind, npoints=kde_npoints)

    p1 = plot(dens.x, dens.density,
              fill = (0, 0.2, :blue),
              title = drug_names[ind],
              xlabel="P(DILI)",
              xlims = (0,1.01),
              ylims = (0,6),
              legend=false,
              yticks = false,
              framestyle = :box,
              size=[250,250],
              titlefont = font(7, "Calibri"),
              xguidefontsize=font(7, "Calibri"))
    vline!([c1, c2], color = :black, linestyle = :dash)

    # define a function that returns a Plots.Shape
    rectangle(w, h, x, y) = Shape(x .+ [0,w,w,0], y .+ [0,0,h,h])

    plot!(rectangle(c1+0.05, 0.25, -0.05,0), color = :green4, alpha = 0.9)
    plot!(rectangle(c2-c1,0.25,c1,0), color = :darkgoldenrod, alpha = 0.9)
    plot!(rectangle(1.05-c2,0.25,c2,0), color = :firebrick, alpha = 0.9)

    return p1
end


## ---------------------------------------------------------
## Turing model for BNN
## ---------------------------------------------------------

@model bayes_nn(X, y, network_shape, num_params) = begin

    sig ~ TruncatedNormal(0, 1, 0, Inf)
    θ ~ MvNormal(zeros(num_params), sig .* ones(num_params))

    eta = nn_forward(X, θ, network_shape)

    c1 ~ Normal(0, 20)
    log_diff_c ~ Normal(0, 2)
    c2 = c1 + exp(log_diff_c)
    c = [c1, c2]

    for i = 1:length(y)
        y[i] ~ OrderedLogistic(eta[i], c)
    end
end


function predict(X, y, num_samples)

    n = size(X, 1)

    eta_post = zeros(num_samples, n);
    y_pred_samps = zeros(num_samples, n);
    y_pred = zeros(n);
    logpdf_mat = zeros(num_samples, n);
    probs_mat = zeros(n, 3);

    for j in 1:n
        for i in 1:num_samples

            eta_post[i, j] = nn_forward(X[j], params2[i,:], network_shape)[1]

            c1 = c1_est[i]
            c2 = c2_est[i]
            c = [c1, c2]

            dist = OrderedLogistic(eta_post[i,j], c)

            y_pred_samps[i,j] = rand(dist)

            if !(typeof(y) == Bool)
                logpdf_mat[i, j] = logpdf(dist, y[j])
            end

        end

        probs = [mean(y_pred_samps[:,j] .== 1), mean(y_pred_samps[:,j] .== 2), mean(y_pred_samps[:,j] .== 3)]
        y_pred[j] = sum((probs .== maximum(probs)) .* [1, 2, 3])
        probs_mat[j,:] = probs

    end

    if !(typeof(y) == Bool)
        return probs_mat, y_pred_samps, logpdf_mat, y_pred, eta_post
    else
        return probs_mat, y_pred_samps, false, y_pred, eta_post
    end
end


function res(X, Y, Xt, Yt)
    println("train accuracy = ", accuracy(X, Y))
    println("test accuracy = ", accuracy(Xt, Yt))

    y_pred = onecold(nn(X)) ; y = onecold(Y); C = confusmat(3, y, y_pred)
    bacc = 1/3 *(C[1,1] / sum(C[1, :]) + C[2,2] / sum(C[2, :]) + C[3,3] / sum(C[3, :]))
    println("train BA = ", round(bacc, digits=2))

    y_pred = onecold(nn(Xt)) ; y = onecold(Yt); C = confusmat(3, y, y_pred)
    bacc = 1/3 *(C[1,1] / sum(C[1, :]) + C[2,2] / sum(C[2, :]) + C[3,3] / sum(C[3, :]))
    println("test BA = ", round(bacc, digits=2))
end

## Euclidean distance to cluster centre for each sample
function euc_dist(cluster_centres, x)
    sqrt(sum((cluster_centres - x).^2))
end
```
