## Supplementary material for "A Bayesian neural network for toxicity prediction": Manifest.toml

```
[[AbstractFFTs]]
deps = ["LinearAlgebra"]
git-tree-sha1 = "051c95d6836228d120f5f4b984dd5aba1624f716"
uuid = "621f4979-c628-5d54-868e-fcf4e3e8185c"
version = "0.5.0"

[[AbstractTrees]]
deps = ["Markdown", "Test"]
git-tree-sha1 = "6621d9645702c1c4e6970cc6a3eae440c768000b"
uuid = "1520ce14-60c1-5f80-bbc7-55ef81b5835c"
version = "0.2.1"

[[Adapt]]
deps = ["LinearAlgebra"]
git-tree-sha1 = "82dab828020b872fa9efd3abec1152b075bc7cbf"
uuid = "79e6a3ab-5dfb-504d-930d-738a2a938a0e"
version = "1.0.0"

[[AdvancedHMC]]
deps = ["ArgCheck", "InplaceOps", "LazyArrays", "LinearAlgebra", "Parameters", "ProgressMeter", "Random", "Statistics", "StatsBase", "StatsFuns"]
git-tree-sha1 = "f12a172f2972fa0c2e69505396b77b59e50681a2"
uuid = "0bf59076-c3b1-5ca4-86bd-e02cd72cde3d"
version = "0.2.2"

[[ArgCheck]]
deps = ["Random"]
git-tree-sha1 = "dab25d711a1dedb707a55dbc1eb9fd578f76ff32"
uuid = "dce04be8-c92d-5529-be00-80e4d2c0e197"
version = "1.0.1"

[[Arpack]]
deps = ["BinaryProvider", "Libdl", "LinearAlgebra"]
git-tree-sha1 = "cd64c112638582ba4f0be9c3e20656499c508565"
uuid = "7d9fca2a-8960-54d3-9f78-7d1dccf2cb97"
version = "0.3.2"

[[ArrayInterface]]
deps = ["LinearAlgebra", "Requires", "SparseArrays"]
git-tree-sha1 = "7b26f0015d9459082a16ebe3fe934c4abb8f1327"
uuid = "4fba245c-0d91-5ea0-9b3e-6abc04ee57a9"
version = "2.2.0"

[[AxisAlgorithms]]
deps = ["LinearAlgebra", "Random", "SparseArrays", "WoodburyMatrices"]
git-tree-sha1 = "a4d07a1c313392a77042855df46c5f534076fab9"
uuid = "13072b0f-2c55-5437-9ae7-d433b7a33950"
version = "1.0.0"

[[AxisArrays]]
deps = ["Dates", "IntervalSets", "IterTools", "RangeArrays"]
git-tree-sha1 = "d63ba0315a1d287c9467e61e932578f2fdd048e0"
uuid = "39de3d68-74b9-583c-8d2d-e117c070f3a9"
version = "0.3.3"

[[Base64]]
uuid = "2a0f44e3-6c83-55bd-87e4-b1978d98bd5f"

[[Bijectors]]
deps = ["Distributions", "LinearAlgebra", "MappedArrays", "Reexport", "Requires", "StatsFuns"]
git-tree-sha1 = "8b3920a2fbd1fbb316fd8b6dd71f757f5d929e77"
uuid = "76274a88-744f-5084-9051-94815aaf08c4"
version = "0.3.1"

[[BinDeps]]
deps = ["Compat", "Libdl", "SHA", "URIParser"]
git-tree-sha1 = "12093ca6cdd0ee547c39b1870e0c9c3f154d9ca9"
uuid = "9e28174c-4ba2-5203-b857-d8d62c4213ee"
version = "0.8.10"

[[BinaryProvider]]
deps = ["Libdl", "SHA"]
git-tree-sha1 = "5b08ed6036d9d3f0ee6369410b830f8873d4024c"
uuid = "b99e7846-7c00-51b0-8f62-c81ae34c0232"
version = "0.5.8"

[[Blosc]]
deps = ["BinaryProvider", "CMakeWrapper", "Libdl"]
git-tree-sha1 = "9981f1795919b8f770dc064fe733ba09c2e7c7a9"
uuid = "a74b3585-a348-5f62-a45c-50e91977d574"
version = "0.6.0"

[[CMake]]
deps = ["BinDeps"]
git-tree-sha1 = "c67a8689dc5444adc5eb2be7d837100340ecba11"
uuid = "631607c0-34d2-5d66-819e-eb0f9aa2061a"
version = "1.1.2"

[[CMakeWrapper]]
deps = ["BinDeps", "CMake", "Libdl", "Parameters", "Test"]
git-tree-sha1 = "16d4acb3d37dc05b714977ffefa8890843dc8985"
uuid = "d5fb7624-851a-54ee-a528-d3f3bac0b4a0"
version = "0.2.3"

[[CSV]]
deps = ["CategoricalArrays", "DataFrames", "Dates", "Mmap", "Parsers", "PooledArrays", "Profile", "Tables", "Unicode", "WeakRefStrings"]
git-tree-sha1 = "a7df9250dff3aba96436580dd6ac00d712364cab"
uuid = "336ed68f-0bac-5ca0-87d4-7b16caf5d00b"
version = "0.5.9"

[[Calculus]]
deps = ["LinearAlgebra"]
git-tree-sha1 = "f641eb0a4f00c343bbc32346e1217b86f3ce9dad"
uuid = "49dc2e85-a5d0-5ad3-a950-438e2897f1b9"
version = "0.5.1"

[[CategoricalArrays]]
deps = ["Compat", "DataAPI", "Future", "JSON", "Missings", "Printf", "Reexport"]
git-tree-sha1 = "13240cfcc884837fc1aa89b60d500a652bcc3f10"
uuid = "324d7699-5711-5eae-9e2f-1d82baa6b597"
version = "0.5.5"

[[CodecZlib]]
deps = ["BinaryProvider", "Libdl", "Test", "TranscodingStreams"]
git-tree-sha1 = "36bbf5374c661054d41410dc53ff752972583b9b"
uuid = "944b1d66-785c-5afd-91f1-9de20f533193"
version = "0.5.2"

[[ColorTypes]]
deps = ["FixedPointNumbers", "Random"]
git-tree-sha1 = "b41d0f666d46ba128c6a0fc632d6e33fbfc03604"
uuid = "3da002f7-5984-5a60-b8a6-cbb66c0b333f"
version = "0.9.0"

[[Colors]]
deps = ["ColorTypes", "FixedPointNumbers", "InteractiveUtils", "Printf", "Reexport"]
git-tree-sha1 = "177d8b959d3c103a6d57574c38ee79c81059c31b"
uuid = "5ae59095-9a9b-59fe-a467-6f913c188581"
version = "0.11.2"

[[Combinatorics]]
deps = ["Polynomials"]
git-tree-sha1 = "140cc833258df8e5aafabdb068875ac0c256be23"
uuid = "861a8166-3701-5b0c-9a16-15d98fcdc6aa"
version = "1.0.0"

[[CommonSubexpressions]]
deps = ["Test"]
git-tree-sha1 = "efdaf19ab11c7889334ca247ff4c9f7c322817b0"
uuid = "bbf7d656-a473-5ed7-a52c-81e309532950"
version = "0.2.0"

[[Compat]]
deps = ["Base64", "Dates", "DelimitedFiles", "Distributed", "InteractiveUtils", "LibGit2", "Libdl", "LinearAlgebra", "Markdown", "Mmap", "Pkg", "Printf", "REPL", "Random", "Serialization", "SharedArrays", "Sockets", "SparseArrays", "Statistics", "Test", "UUIDs", "Unicode"]
git-tree-sha1 = "ed2c4abadf84c53d9e58510b5fc48912c2336fbb"
uuid = "34da2185-b29b-5c13-b0c7-acf172513d20"
version = "2.2.0"

[[Conda]]
deps = ["JSON", "VersionParsing"]
git-tree-sha1 = "9a11d428dcdc425072af4aea19ab1e8c3e01c032"
uuid = "8f4d0f93-b110-5947-807f-2305c1781a2d"
version = "1.3.0"

[[Contour]]
deps = ["LinearAlgebra", "StaticArrays", "Test"]
git-tree-sha1 = "b974e164358fea753ef853ce7bad97afec15bb80"
uuid = "d38c429a-6771-53c6-b99e-75d170b6e991"
version = "0.5.1"

[[DataAPI]]
git-tree-sha1 = "674b67f344687a88310213ddfa8a2b3c76cc4252"
uuid = "9a962f9c-6df0-11e9-0e5d-c546b8b5ee8a"
version = "1.1.0"

[[DataFrames]]
deps = ["CategoricalArrays", "Compat", "DataAPI", "InvertedIndices", "IteratorInterfaceExtensions", "Missings", "PooledArrays", "Printf", "REPL", "Reexport", "SortingAlgorithms", "Statistics", "TableTraits", "Tables", "Unicode"]
git-tree-sha1 = "12a96f70fc126f8a308eadd4eda843dfa630bbd4"
uuid = "a93c6f00-e57d-5684-b7b6-d8193f3e46c0"
version = "0.19.1"

[[DataStructures]]
deps = ["InteractiveUtils", "OrderedCollections"]
git-tree-sha1 = "f784254f428fb8fd7ac15982e5862a38a44523d3"
uuid = "864edb3b-99cc-5e75-8d2d-829cb0a9cfe8"
version = "0.17.7"

[[DataValueInterfaces]]
git-tree-sha1 = "bfc1187b79289637fa0ef6d4436ebdfe6905cbd6"
uuid = "e2d170a0-9d28-54be-80f0-106bbe20a464"
version = "1.0.0"

[[Dates]]
deps = ["Printf"]
uuid = "ade2ca70-3891-5945-98fb-dc099432e06a"

[[DelimitedFiles]]
deps = ["Mmap"]
uuid = "8bb1440f-4735-579b-a4ab-409b98df4dab"

[[DiffEqDiffTools]]
deps = ["ArrayInterface", "LinearAlgebra", "Requires", "SparseArrays", "StaticArrays"]
git-tree-sha1 = "951600cc05d9626c3282e2ed125492931596e430"
uuid = "01453d9d-ee7c-5054-8395-0335cb756afa"
version = "1.6.0"

[[DiffResults]]
deps = ["Compat", "StaticArrays"]
git-tree-sha1 = "34a4a1e8be7bc99bc9c611b895b5baf37a80584c"
uuid = "163ba53b-c6d8-5494-b064-1a9d43ac40c5"
version = "0.0.4"

[[DiffRules]]
deps = ["Random", "Test"]
git-tree-sha1 = "dc0869fb2f5b23466b32ea799bd82c76480167f7"
uuid = "b552c78f-8df3-52c6-915a-8e097449b14b"
version = "0.0.10"

[[Distributed]]
deps = ["LinearAlgebra", "Random", "Serialization", "Sockets"]
uuid = "8ba89e20-285c-5b6f-9357-94700520ee1b"

[[Distributions]]
deps = ["LinearAlgebra", "PDMats", "Printf", "QuadGK", "Random", "SpecialFunctions", "Statistics", "StatsBase", "StatsFuns"]
git-tree-sha1 = "baaf9e165ba8a2d11fb4fb3511782ee070ee3694"
uuid = "31c24e10-a181-5473-b8eb-7969acd0382f"
version = "0.21.1"

[[FFTW]]
deps = ["AbstractFFTs", "BinaryProvider", "Conda", "Libdl", "LinearAlgebra", "Reexport"]
git-tree-sha1 = "4cfd3d43819228b9e73ab46600d0af0aa5cedceb"
uuid = "7a1cc6ca-52ef-59f5-83cd-3a7055c09341"
version = "1.1.0"

[[FileIO]]
deps = ["Pkg"]
git-tree-sha1 = "74585bf1f7ed7259e166011e89f49363d7fa89a6"
uuid = "5789e2e9-d7fb-5bc7-8068-2c6fae9b9549"
version = "1.2.1"

[[FileWatching]]
uuid = "7b1f6079-737a-58dc-b8bc-7a2ca5c1b5ee"

[[FillArrays]]
deps = ["LinearAlgebra", "Random", "SparseArrays"]
git-tree-sha1 = "8fba6ddaf66b45dec830233cea0aae43eb1261ad"
uuid = "1a297f60-69ca-5386-bcde-b61e274b549b"
version = "0.6.4"

[[FixedPointNumbers]]
git-tree-sha1 = "bd1386f890e172ef38e1c735cda58cbf004a7c9a"
uuid = "53c48c17-4a7d-5ca2-90c5-79b7896eea93"
version = "0.7.0"

[[Flux]]
deps = ["AbstractTrees", "Adapt", "CodecZlib", "Colors", "DelimitedFiles", "Juno", "LinearAlgebra", "MacroTools", "NNlib", "Pkg", "Printf", "Random", "Reexport", "Requires", "SHA", "Statistics", "StatsBase", "Tracker", "ZipFile"]
git-tree-sha1 = "08212989c2856f95f90709ea5fd824bd27b34514"
uuid = "587475ba-b771-5e3f-ad9e-33799f191a9c"
version = "0.8.3"

[[ForwardDiff]]
deps = ["CommonSubexpressions", "DiffResults", "DiffRules", "NaNMath", "Random", "SpecialFunctions", "StaticArrays"]
git-tree-sha1 = "840700059391d36e2498d89c2e82c08f261f2a2a"
uuid = "f6369f11-7733-5829-9624-2563aa707210"
version = "0.10.8"

[[FreqTables]]
deps = ["CategoricalArrays", "NamedArrays", "Tables", "Test"]
git-tree-sha1 = "93ce75cae27c2d1c6518771d597b6dd0a3d623d3"
uuid = "da1fdf0e-e0ff-5433-a45f-9bb5ff651cb1"
version = "0.3.1"

[[Future]]
deps = ["Random"]
uuid = "9fa8497b-333b-5362-9e8d-4d0656e87820"

[[GR]]
deps = ["Base64", "DelimitedFiles", "LinearAlgebra", "Pkg", "Printf", "Random", "Serialization", "Sockets", "Test"]
git-tree-sha1 = "b4c31b6377b6d51b6c69a3a9737d10c34d43974e"
uuid = "28b8d3ca-fb5f-59d9-8090-bfdbd6d07a71"
version = "0.41.0"

[[GeometryTypes]]
deps = ["ColorTypes", "FixedPointNumbers", "LinearAlgebra", "StaticArrays"]
git-tree-sha1 = "a96baa00f5ac755689c82c29bc3395e161337c69"
uuid = "4d00f742-c7ba-57c2-abde-4428a4b178cb"
version = "0.7.7"

[[HDF5]]
deps = ["BinaryProvider", "Blosc", "CMakeWrapper", "Libdl", "Mmap"]
git-tree-sha1 = "d3ea5532668bf9bdd5e8d5f16571e0b520cbbb9f"
uuid = "f67ccb44-e63f-5c2f-98bd-6dc0ccc4ba2f"
version = "0.12.5"

[[IRTools]]
deps = ["InteractiveUtils", "MacroTools", "Test"]
git-tree-sha1 = "72421971e60917b8cd7737f9577c4f0f87eab306"
uuid = "7869d1d1-7146-5819-86e3-90919afe41df"
version = "0.3.0"

[[InplaceOps]]
deps = ["LinearAlgebra", "Test"]
git-tree-sha1 = "50b41d59e7164ab6fda65e71049fee9d890731ff"
uuid = "505f98c9-085e-5b2c-8e89-488be7bf1f34"
version = "0.3.0"

[[InteractiveUtils]]
deps = ["LinearAlgebra", "Markdown"]
uuid = "b77e0a4c-d291-57a0-90e8-8db25a27a240"

[[Interpolations]]
deps = ["AxisAlgorithms", "LinearAlgebra", "OffsetArrays", "Random", "Ratios", "SharedArrays", "SparseArrays", "StaticArrays", "WoodburyMatrices"]
git-tree-sha1 = "e4a219889f33a8c2435c26bf1bca6f4fbf8c1c0b"
uuid = "a98d9a8b-a2ab-59e6-89dd-64a1c18fca59"
version = "0.12.6"

[[IntervalSets]]
deps = ["Dates", "Statistics"]
git-tree-sha1 = "4214b48a62eb8f2c292b2ee34a508c256c0cdbc9"
uuid = "8197267c-284f-5f27-9208-e0e47529a953"
version = "0.3.2"

[[InvertedIndices]]
deps = ["Test"]
git-tree-sha1 = "15732c475062348b0165684ffe28e85ea8396afc"
uuid = "41ab1584-1d38-5bbf-9106-f11c6c58b48f"
version = "1.0.0"

[[IterTools]]
git-tree-sha1 = "05110a2ab1fc5f932622ffea2a003221f4782c18"
uuid = "c8e1da08-722c-5040-9ed9-7db0dc04731e"
version = "1.3.0"

[[IteratorInterfaceExtensions]]
git-tree-sha1 = "a3f24677c21f5bbe9d2a714f95dcd58337fb2856"
uuid = "82899510-4779-5014-852e-03e436cf321d"
version = "1.0.0"

[[JLD]]
deps = ["Compat", "FileIO", "HDF5", "LegacyStrings"]
git-tree-sha1 = "c8c7a9ca4c17a519961a8ec27e508b07ffc9c1a9"
uuid = "4138dd39-2aa7-5051-a626-17a0bb65d9c8"
version = "0.9.2"

[[JSON]]
deps = ["Dates", "Distributed", "Mmap", "Sockets", "Test", "Unicode"]
git-tree-sha1 = "1f7a25b53ec67f5e9422f1f551ee216503f4a0fa"
uuid = "682c06a0-de6a-54ab-a142-c8b1cf79cde6"
version = "0.20.0"

[[Juno]]
deps = ["Base64", "Logging", "Media", "Profile", "Test"]
git-tree-sha1 = "4e4a8d43aa7ecec66cadaf311fbd1e5c9d7b9175"
uuid = "e5e0dc1b-0480-54bc-9374-aad01c23163d"
version = "0.7.0"

[[KernelDensity]]
deps = ["Distributions", "FFTW", "Interpolations", "Optim", "StatsBase", "Test"]
git-tree-sha1 = "c1048817fe5711f699abc8fabd47b1ac6ba4db04"
uuid = "5ab0869b-81aa-558d-bb23-cbf5423bbe9b"
version = "0.5.1"

[[LazyArrays]]
deps = ["FillArrays", "LinearAlgebra", "MacroTools", "StaticArrays", "Test"]
git-tree-sha1 = "0d9a7174d47407a61953e17c3dcecfb4ca8c5c44"
uuid = "5078a376-72f3-5289-bfd5-ec5146d43c02"
version = "0.9.1"

[[LegacyStrings]]
deps = ["Compat"]
git-tree-sha1 = "d4b9bde2694c552fe579cc4462733f1ce08733fe"
uuid = "1b4a561d-cfcb-5daf-8433-43fcf8b4bea3"
version = "0.4.1"

[[LibGit2]]
uuid = "76f85450-5226-5b5a-8eaa-529ad045b433"

[[Libdl]]
uuid = "8f399da3-3557-5675-b5ff-fb832c97cbdb"

[[Libtask]]
deps = ["BinaryProvider", "Libdl", "Pkg"]
git-tree-sha1 = "4f39d56bf87bf17f5a04370bd036baf0d1fe8cc9"
uuid = "6f1fad26-d15e-5dc8-ae53-837a1d7b8c9f"
version = "0.3.1"

[[LineSearches]]
deps = ["LinearAlgebra", "NLSolversBase", "NaNMath", "Parameters", "Printf", "Test"]
git-tree-sha1 = "54eb90e8dbe745d617c78dee1d6ae95c7f6f5779"
uuid = "d3d80556-e9d4-5f37-9878-2ab0fcc64255"
version = "7.0.1"

[[LinearAlgebra]]
deps = ["Libdl"]
uuid = "37e2e46d-f89d-539d-b4ee-838fcccc9c8e"

[[Logging]]
uuid = "56ddb016-857b-54e1-b83d-db4d58db5568"

[[MCMCChains]]
deps = ["AxisArrays", "DataFrames", "Distributions", "KernelDensity", "LinearAlgebra", "Plots", "Random", "RecipesBase", "Serialization", "Showoff", "SpecialFunctions", "Statistics", "StatsBase"]
git-tree-sha1 = "0f103eec2477f5900fe6ebb291b84ca596e1b33a"
uuid = "c7f686f2-ff18-58e9-bc7b-31028e88f75d"
version = "0.3.11"

[[MLBase]]
deps = ["IterTools", "Random", "Reexport", "StatsBase", "Test"]
git-tree-sha1 = "f63a8d37429568b8c4384d76c4a96fc2897d6ddf"
uuid = "f0e99cf1-93fa-52ec-9ecc-5026115318e0"
version = "0.8.0"

[[MacroTools]]
deps = ["DataStructures", "Markdown", "Random"]
git-tree-sha1 = "e2fc7a55bb2224e203bbd8b59f72b91323233458"
uuid = "1914dd2f-81c6-5fcd-8719-6d5c9610ff09"
version = "0.5.3"

[[MappedArrays]]
git-tree-sha1 = "e2a02fe7ee86a10c707ff1756ab1650b40b140bb"
uuid = "dbb5928d-eab1-5f90-85c2-b9b0edb7c900"
version = "0.2.2"

[[Markdown]]
deps = ["Base64"]
uuid = "d6f4376e-aef5-505a-96c1-9c027394607a"

[[Measures]]
deps = ["Test"]
git-tree-sha1 = "ddfd6d13e330beacdde2c80de27c1c671945e7d9"
uuid = "442fdcdd-2543-5da2-b0f3-8c86c306513e"
version = "0.3.0"

[[Media]]
deps = ["MacroTools", "Test"]
git-tree-sha1 = "75a54abd10709c01f1b86b84ec225d26e840ed58"
uuid = "e89f7d12-3494-54d1-8411-f7d8b9ae1f27"
version = "0.5.0"

[[Missings]]
deps = ["DataAPI"]
git-tree-sha1 = "de0a5ce9e5289f27df672ffabef4d1e5861247d5"
uuid = "e1d29d7a-bbdc-5cf2-9ac0-f12de2c33e28"
version = "0.4.3"

[[Mmap]]
uuid = "a63ad114-7e13-5084-954f-fe012c677804"

[[NLSolversBase]]
deps = ["Calculus", "DiffEqDiffTools", "DiffResults", "Distributed", "ForwardDiff"]
git-tree-sha1 = "f1b8ed89fa332f410cfc7c937682eb4d0b361521"
uuid = "d41bc354-129a-5804-8e4c-c37616107c6c"
version = "7.5.0"

[[NNlib]]
deps = ["BinaryProvider", "Libdl", "LinearAlgebra", "Requires", "Statistics"]
git-tree-sha1 = "135c0de4794d5e214b06f1fb4787af4a72896e61"
uuid = "872c559c-99b0-510c-b3b7-b6c96a88d5cd"
version = "0.6.2"

[[NaNMath]]
git-tree-sha1 = "928b8ca9b2791081dc71a51c55347c27c618760f"
uuid = "77ba4419-2d1f-58cd-9bb1-8ffee604a2e3"
version = "0.3.3"

[[NamedArrays]]
deps = ["Combinatorics", "DataStructures", "DelimitedFiles", "InvertedIndices", "LinearAlgebra", "Random", "Requires", "SparseArrays", "Statistics"]
git-tree-sha1 = "2681361e30e0ee97aad617f1a6a728e53179bd15"
uuid = "86f7a689-2022-50b4-a561-43c23ac3c673"
version = "0.9.3"

[[OffsetArrays]]
git-tree-sha1 = "87d0a91efe29352d5caaa271ae3927083c096e33"
uuid = "6fe1bfb0-de20-5000-8ca7-80f57d26f881"
version = "0.11.4"

[[Optim]]
deps = ["Calculus", "DiffEqDiffTools", "FillArrays", "ForwardDiff", "LineSearches", "LinearAlgebra", "NLSolversBase", "NaNMath", "Parameters", "PositiveFactorizations", "Printf", "Random", "SparseArrays", "StatsBase"]
git-tree-sha1 = "ca8f8917df469d607f8cf116784e58c9c084d0b4"
uuid = "429524aa-4258-5aef-a3af-852621145aeb"
version = "0.19.2"

[[OrderedCollections]]
deps = ["Random", "Serialization", "Test"]
git-tree-sha1 = "c4c13474d23c60d20a67b217f1d7f22a40edf8f1"
uuid = "bac558e1-5e72-5ebc-8fee-abe8a469f55d"
version = "1.1.0"

[[PDMats]]
deps = ["Arpack", "LinearAlgebra", "SparseArrays", "SuiteSparse", "Test"]
git-tree-sha1 = "035f8d60ba2a22cb1d2580b1e0e5ce0cb05e4563"
uuid = "90014a1f-27ba-587c-ab20-58faa44d9150"
version = "0.9.10"

[[Parameters]]
deps = ["Markdown", "OrderedCollections", "REPL", "Test"]
git-tree-sha1 = "70bdbfb2bceabb15345c0b54be4544813b3444e4"
uuid = "d96e819e-fc66-5662-9728-84c9c7592b0a"
version = "0.10.3"

[[Parsers]]
deps = ["Dates", "Test"]
git-tree-sha1 = "0139ba59ce9bc680e2925aec5b7db79065d60556"
uuid = "69de0a69-1ddd-5017-9359-2bf0b02dc9f0"
version = "0.3.10"

[[Pkg]]
deps = ["Dates", "LibGit2", "Markdown", "Printf", "REPL", "Random", "SHA", "UUIDs"]
uuid = "44cfe95a-1eb2-52ea-b672-e2afdf69b78f"

[[PlotThemes]]
deps = ["PlotUtils", "Requires", "Statistics"]
git-tree-sha1 = "df772cc7c78862da96af1ee85cd0111c6640e44e"
uuid = "ccf2f8ad-2431-5c83-bf29-c5338b663b6a"
version = "1.0.1"

[[PlotUtils]]
deps = ["Colors", "Dates", "Printf", "Random", "Reexport"]
git-tree-sha1 = "7622cbde3200a9876a14ba85d66f25d7f4e7a6ca"
uuid = "995b91a9-d308-5afd-9ec6-746e21dbc043"
version = "0.6.1"

[[Plots]]
deps = ["Base64", "Contour", "Dates", "FixedPointNumbers", "GR", "GeometryTypes", "JSON", "LinearAlgebra", "Measures", "NaNMath", "Pkg", "PlotThemes", "PlotUtils", "Printf", "REPL", "Random", "RecipesBase", "Reexport", "Requires", "Showoff", "SparseArrays", "Statistics", "StatsBase", "UUIDs"]
git-tree-sha1 = "a9c011a7176a9010dda4853373e8d89a83f9d69d"
uuid = "91a5bcdd-55d7-5caf-9e0b-520d859cae80"
version = "0.25.3"

[[Polynomials]]
deps = ["LinearAlgebra", "RecipesBase"]
git-tree-sha1 = "f7c0c07e82798aef542d60a6e6e85e39f4590750"
uuid = "f27b6e38-b328-58d1-80ce-0feddd5e7a45"
version = "0.5.3"

[[PooledArrays]]
deps = ["DataAPI"]
git-tree-sha1 = "b1333d4eced1826e15adbdf01a4ecaccca9d353c"
uuid = "2dfb63ee-cc39-5dd5-95bd-886bf059d720"
version = "0.5.3"

[[PositiveFactorizations]]
deps = ["LinearAlgebra", "Test"]
git-tree-sha1 = "127c47b91990c101ee3752291c4f45640eeb03d1"
uuid = "85a6dd25-e78a-55b7-8502-1745935b8125"
version = "0.2.3"

[[Printf]]
deps = ["Unicode"]
uuid = "de0858da-6303-5e67-8744-51eddeeeb8d7"

[[Profile]]
deps = ["Printf"]
uuid = "9abbd945-dff8-562f-b5e8-e1ebf5ef1b79"

[[ProgressMeter]]
deps = ["Distributed", "Printf"]
git-tree-sha1 = "0f08e0e74e5b160ca20d3962a2620038b75881c7"
uuid = "92933f4c-e287-5a05-a399-4b506db050ca"
version = "1.0.0"

[[QuadGK]]
deps = ["DataStructures", "LinearAlgebra"]
git-tree-sha1 = "dc84e810393cfc6294248c9032a9cdacc14a3db4"
uuid = "1fd47b50-473d-5c70-9696-f719f8f3bcdc"
version = "2.3.1"

[[REPL]]
deps = ["InteractiveUtils", "Markdown", "Sockets"]
uuid = "3fa0cd96-eef1-5676-8a61-b3b8758bbffb"

[[Random]]
deps = ["Serialization"]
uuid = "9a3f8284-a2c9-5f02-9a11-845980a1fd5c"

[[RangeArrays]]
git-tree-sha1 = "b9039e93773ddcfc828f12aadf7115b4b4d225f5"
uuid = "b3c3ace0-ae52-54e7-9d0b-2c1406fd6b9d"
version = "0.3.2"

[[Ratios]]
deps = ["Compat"]
git-tree-sha1 = "cdbbe0f350581296f3a2e3e7a91b214121934407"
uuid = "c84ed2f1-dad5-54f0-aa8e-dbefe2724439"
version = "0.3.1"

[[RecipesBase]]
deps = ["Random", "Test"]
git-tree-sha1 = "0b3cb370ee4dc00f47f1193101600949f3dcf884"
uuid = "3cdcf5f2-1ef4-517c-9805-6587b60abb01"
version = "0.6.0"

[[Reexport]]
deps = ["Pkg"]
git-tree-sha1 = "7b1d07f411bc8ddb7977ec7f377b97b158514fe0"
uuid = "189a3867-3050-52da-a836-e630ba90ab69"
version = "0.2.0"

[[Requires]]
deps = ["Test"]
git-tree-sha1 = "f6fbf4ba64d295e146e49e021207993b6b48c7d1"
uuid = "ae029012-a4dd-5104-9daa-d747884805df"
version = "0.5.2"

[[Rmath]]
deps = ["BinaryProvider", "Libdl", "Random", "Statistics"]
git-tree-sha1 = "2bbddcb984a1d08612d0c4abb5b4774883f6fa98"
uuid = "79098fc4-a85e-5d69-aa6a-4863f24498fa"
version = "0.6.0"

[[SHA]]
uuid = "ea8e919c-243c-51af-8825-aaa63cd721ce"

[[Serialization]]
uuid = "9e88b42a-f829-5b0c-bbe9-9e923198166b"

[[SharedArrays]]
deps = ["Distributed", "Mmap", "Random", "Serialization"]
uuid = "1a1011a3-84de-559e-8e89-a11a2f7dc383"

[[ShiftedArrays]]
git-tree-sha1 = "22395afdcf37d6709a5a0766cc4a5ca52cb85ea0"
uuid = "1277b4bf-5013-50f5-be3d-901d8477a67a"
version = "1.0.0"

[[Showoff]]
deps = ["Dates"]
git-tree-sha1 = "e032c9df551fb23c9f98ae1064de074111b7bc39"
uuid = "992d4aef-0814-514b-bc4d-f2e9a6c4116f"
version = "0.3.1"

[[Sockets]]
uuid = "6462fe0b-24de-5631-8697-dd941f90decc"

[[SortingAlgorithms]]
deps = ["DataStructures", "Random", "Test"]
git-tree-sha1 = "03f5898c9959f8115e30bc7226ada7d0df554ddd"
uuid = "a2af1166-a08f-5f64-846c-94a0d3cef48c"
version = "0.3.1"

[[SparseArrays]]
deps = ["LinearAlgebra", "Random"]
uuid = "2f01184e-e22b-5df5-ae63-d93ebab69eaf"

[[SpecialFunctions]]
deps = ["BinDeps", "BinaryProvider", "Libdl", "Test"]
git-tree-sha1 = "0b45dc2e45ed77f445617b99ff2adf0f5b0f23ea"
uuid = "276daf66-3868-5448-9aa4-cd146d93841b"
version = "0.7.2"

[[StaticArrays]]
deps = ["LinearAlgebra", "Random", "Statistics"]
git-tree-sha1 = "5a3bcb6233adabde68ebc97be66e95dcb787424c"
uuid = "90137ffa-7385-5640-81b9-e52037218182"
version = "0.12.1"

[[Statistics]]
deps = ["LinearAlgebra", "SparseArrays"]
uuid = "10745b16-79ce-11e8-11f9-7d13ad32a3b2"

[[StatsBase]]
deps = ["DataAPI", "DataStructures", "LinearAlgebra", "Missings", "Printf", "Random", "SortingAlgorithms", "SparseArrays", "Statistics"]
git-tree-sha1 = "c53e809e63fe5cf5de13632090bc3520649c9950"
uuid = "2913bbd2-ae8a-5f71-8c99-4fb6c76f3a91"
version = "0.32.0"

[[StatsFuns]]
deps = ["Rmath", "SpecialFunctions", "Test"]
git-tree-sha1 = "b3a4e86aa13c732b8a8c0ba0c3d3264f55e6bb3e"
uuid = "4c63d2b9-4356-54db-8cca-17b64c39e42c"
version = "0.8.0"

[[StatsModels]]
deps = ["CategoricalArrays", "DataStructures", "LinearAlgebra", "Missings", "ShiftedArrays", "SparseArrays", "StatsBase", "Tables"]
git-tree-sha1 = "c999ba0765897682e8262323f3bfe806acdf126e"
uuid = "3eaba693-59b7-5ba5-a881-562e759f1c8d"
version = "0.6.2"

[[SuiteSparse]]
deps = ["Libdl", "LinearAlgebra", "Serialization", "SparseArrays"]
uuid = "4607b0f0-06f3-5cda-b6b1-a6196a1729e9"

[[TableTraits]]
deps = ["IteratorInterfaceExtensions"]
git-tree-sha1 = "b1ad568ba658d8cbb3b892ed5380a6f3e781a81e"
uuid = "3783bdb8-4a98-5b6b-af9a-565f29a5fe9c"
version = "1.0.0"

[[Tables]]
deps = ["DataAPI", "DataValueInterfaces", "IteratorInterfaceExtensions", "LinearAlgebra", "TableTraits", "Test"]
git-tree-sha1 = "aaed7b3b00248ff6a794375ad6adf30f30ca5591"
uuid = "bd369af6-aec1-5ad0-b16a-f7cc5008161c"
version = "0.2.11"

[[Test]]
deps = ["Distributed", "InteractiveUtils", "Logging", "Random"]
uuid = "8dfed614-e22c-5e08-85e1-65c5234f0b40"

[[Tracker]]
deps = ["Adapt", "DiffRules", "ForwardDiff", "LinearAlgebra", "MacroTools", "NNlib", "NaNMath", "Printf", "Random", "Requires", "SpecialFunctions", "Statistics", "Test"]
git-tree-sha1 = "327342fec6e09f68ced0c2dc5731ed475e4b696b"
uuid = "9f7883ad-71c0-57eb-9f7f-b5c9e6d3789c"
version = "0.2.2"

[[TranscodingStreams]]
deps = ["Random", "Test"]
git-tree-sha1 = "7c53c35547de1c5b9d46a4797cf6d8253807108c"
uuid = "3bb67fe8-82b1-5028-8e26-92a6c54297fa"
version = "0.9.5"

[[Turing]]
deps = ["AdvancedHMC", "Bijectors", "BinaryProvider", "Distributions", "ForwardDiff", "Libtask", "LinearAlgebra", "MCMCChains", "MacroTools", "Markdown", "PDMats", "ProgressMeter", "Random", "Reexport", "Requires", "SpecialFunctions", "Statistics", "StatsFuns", "Tracker", "Zygote"]
git-tree-sha1 = "dfc4dfdda24f816ee3302957a9e0d9971e961542"
repo-rev = "master"
repo-url = "https://github.com/TuringLang/Turing.jl.git"
uuid = "fce5fe82-541a-59a6-adf8-730c64b5f9a0"
version = "0.6.23"

[[URIParser]]
deps = ["Test", "Unicode"]
git-tree-sha1 = "6ddf8244220dfda2f17539fa8c9de20d6c575b69"
uuid = "30578b45-9adc-5946-b283-645ec420af67"
version = "0.4.0"

[[UUIDs]]
deps = ["Random"]
uuid = "cf7118a7-6976-5b1a-9a39-7adc72f591a4"

[[Unicode]]
uuid = "4ec0a83e-493e-50e2-b9ac-8f72acf5a8f5"

[[VersionParsing]]
git-tree-sha1 = "80229be1f670524750d905f8fc8148e5a8c4537f"
uuid = "81def892-9a0e-5fdd-b105-ffc91e053289"
version = "1.2.0"

[[WeakRefStrings]]
deps = ["DataAPI", "Random", "Test"]
git-tree-sha1 = "28807f85197eaad3cbd2330386fac1dcb9e7e11d"
uuid = "ea10d353-3f73-51f8-a26c-33c1cb351aa5"
version = "0.6.2"

[[WoodburyMatrices]]
deps = ["LinearAlgebra", "Random", "SparseArrays", "Test"]
git-tree-sha1 = "21772c33b447757ec7d3e61fcdfb9ea5c47eedcf"
uuid = "efce3f68-66dc-5838-9240-27a6d6f5f9b6"
version = "0.4.1"

[[ZipFile]]
deps = ["BinaryProvider", "Libdl", "Printf"]
git-tree-sha1 = "580ce62b6c14244916cc28ad54f8a2e2886f843d"
uuid = "a5390f91-8eb1-5f08-bee0-b1d1ffed6cea"
version = "0.8.3"

[[Zygote]]
deps = ["DiffRules", "FFTW", "FillArrays", "ForwardDiff", "IRTools", "InteractiveUtils", "LinearAlgebra", "MacroTools", "NNlib", "NaNMath", "Random", "Requires", "SpecialFunctions", "Statistics", "ZygoteRules"]
git-tree-sha1 = "e4245b9c5362346e154b62842a89a18e0210b92b"
uuid = "e88e6eb3-aa80-5325-afca-941959d7151f"
version = "0.4.1"

[[ZygoteRules]]
deps = ["MacroTools"]
git-tree-sha1 = "b3b4882cc9accf6731a08cc39543fbc6b669dca8"
uuid = "700de1a5-db45-46bc-99cf-38207098b444"
version = "0.2.0"
```
