## Supplementary material for "A Bayesian neural network for toxicity prediction": Project.toml

```
[deps]
CSV = "336ed68f-0bac-5ca0-87d4-7b16caf5d00b"
DataFrames = "a93c6f00-e57d-5684-b7b6-d8193f3e46c0"
Flux = "587475ba-b771-5e3f-ad9e-33799f191a9c"
FreqTables = "da1fdf0e-e0ff-5433-a45f-9bb5ff651cb1"
JLD = "4138dd39-2aa7-5051-a626-17a0bb65d9c8"
KernelDensity = "5ab0869b-81aa-558d-bb23-cbf5423bbe9b"
MLBase = "f0e99cf1-93fa-52ec-9ecc-5026115318e0"
Plots = "91a5bcdd-55d7-5caf-9e0b-520d859cae80"
Random = "9a3f8284-a2c9-5f02-9a11-845980a1fd5c"
StatsBase = "2913bbd2-ae8a-5f71-8c99-4fb6c76f3a91"
StatsFuns = "4c63d2b9-4356-54db-8cca-17b64c39e42c"
StatsModels = "3eaba693-59b7-5ba5-a881-562e759f1c8d"
Turing = "fce5fe82-541a-59a6-adf8-730c64b5f9a0"

[compat]
CSV = "=0.5.9"
DataFrames = "=0.19.1"
Flux = "=0.8.3"
FreqTables = "=0.3.1"
JLD = "=0.9.2"
KernelDensity = "=0.5.1"
MLBase = "=0.8.0"
Plots = "=0.25.3"
StatsBase = "=0.32.0"
StatsFuns = "=0.8.0"
StatsModels = "=0.6.2"
Turing = "=0.6.23"
julia = "=1.0.4"
```
